# Every which way? On predicting tumor evolution using cancer progression models

**DOI:** 10.1101/371039

**Authors:** Ramon Diaz-Uriarte, Claudia Vasallo

**Affiliations:** Dept. Biochemistry, Universidad Autónoma de Madrid Instituto de Investigaciones Biomédicas “Alberto Sols” (UAM-CSIC) Madrid, Spain

## Abstract

Successful prediction of the likely paths of tumor progression is valuable for diagnostic, prognostic, and treatment purposes. Cancer progression models (CPMs) use cross-sectional samples to identify restrictions in the order of accumulation of driver mutations and thus CPMs encode the paths of tumor progression. Here we analyze the performance of four CPMs to examine whether they can be used to predict the true distribution of paths of tumor progression and to estimate evolutionary unpredictability. Employing simulations we show that if fitness landscapes are single peaked (have a single fitness maximum) there is good agreement between true and predicted distributions of paths of tumor progression when sample sizes are large, but performance is poor with the currently common much smaller sample sizes. Under multi-peaked fitness landscapes (i.e., those with multiple fitness maxima), performance is poor and improves only slightly with sample size. In all cases, detection regime (when tumors are sampled) is a key determinant of performance. Estimates of evolutionary unpredictability from the best performing CPM, among the four examined, tend to overestimate the true unpredictability and the bias is affected by detection regime; CPMs could be useful for estimating upper bounds to the true evolutionary unpredictability. Analysis of twenty-two cancer data sets shows low evolutionary unpredictability for several of the data sets. But most of the predictions of paths of tumor progression are very unreliable, and unreliability increases with the number of features analyzed. Our results indicate that CPMs could be valuable tools for predicting cancer progression but that, currently, obtaining useful predictions of paths of tumor progression from CPMs is dubious, and emphasize the need for methodological work that can account for the probably multi-peaked fitness landscapes in cancer.

**Author Summary:** Knowing the likely paths of tumor progression is instrumental for cancer precision medicine as it would allow us to identify genetic targets that block disease progression and to improve therapeutic decisions. Direct information about paths of tumor progression is scarce, but cancer progression models (CPMs), which use as input cross-sectional data on genetic alterations, can be used to predict these paths. CPMs, however, make assumptions about fitness landscapes (genotype-fitness maps) that might not be met in cancer. We examine if four CPMs can be used to predict successfully the distribution of tumor progression paths; we find that some CPMs work well when sample sizes are large and fitness landscapes have a single fitness maximum, but in fitness landscapes with multiple fitness maxima prediction is poor. However, the best performing CPM in our study could be used to estimate evolutionary unpredictability. When we apply the best performing CPM in our study to twenty-two cancer data sets we find that predictions are generally unreliable but that some cancer data sets show low unpredictability. Our results highlight that CPMs could be valuable tools for predicting disease progression, but emphasize the need for methodological work to account for multi-peaked fitness landscapes.

## 1 Introduction

Improving our ability to predict the paths of tumor progression is helpful for diagnostic, prognostic, and treatment purposes as, for example, it would allow us to identify genes that block the most common paths of disease progression (Greaves, 2015; Lipinski *et al*., 2016; McPherson *et al*., 2018; Williams *et al*., 2018). This interest in predicting paths of progression is, of course, not exclusive to cancer (see e.g., reviews in Lässig *et al*., 2017; Losos, 2018). For example, in some cases antibiotic resistance shows parallel evolution with mutations being acquired in a similar order (Toprak *et al*., 2012), and here “Even a modest predictive power might improve therapeutic outcomes by informing the selection of drugs, the preference between monotherapy or combination therapy and the temporal dosing regimen (…)” (Palmer and Kishony, 2013, p. 243i). But detailed information about the paths of tumor evolution and their distribution, obtained from multiple within-patient samples with timing information, is not available.

Cancer progression models (CPMs), such as conjunctive Bayesian networks (CBN) (Gerstung *et al*., 2009, 2011; Montazeri *et al*., 2016), oncogenetic trees (OT) (Desper *et al*., 1999; Szabo and Boucher, 2008), CAncer PRogression Inference (CAPRI) (Caravagna *et al*., 2016; Ramazzotti *et al*., 2015), or CAncer PRogression Extraction with Single Edges (CAPRESE) (Olde Loohuis *et al*., 2014), can be used to predict paths of tumor progression. CPMs were originally developed to identify restrictions in the order of accumulation of mutations during tumor progression from cross-sectional data (Beerenwinkel *et al*., 2015, 2016). But CPMs also encode all the possible mutational paths or trajectories of tumor progression, from the initial genotype to the genotype with all driver genes mutated (see Figure 1); in fact, mutational pathways and evolutionary trajectories are already mentioned in the papers that describe CBN (Gerstung *et al*., 2011), CAPRI (Caravagna *et al*., 2016; Ramazzotti *et al*., 2015) and in general overviews of CPMs (Beerenwinkel *et al*., 2016). Thus, CPMs could improve our ability to predict disease progression by leveraging on the available, and growing, number of cross-sectional data sets.

**Figure 1:**
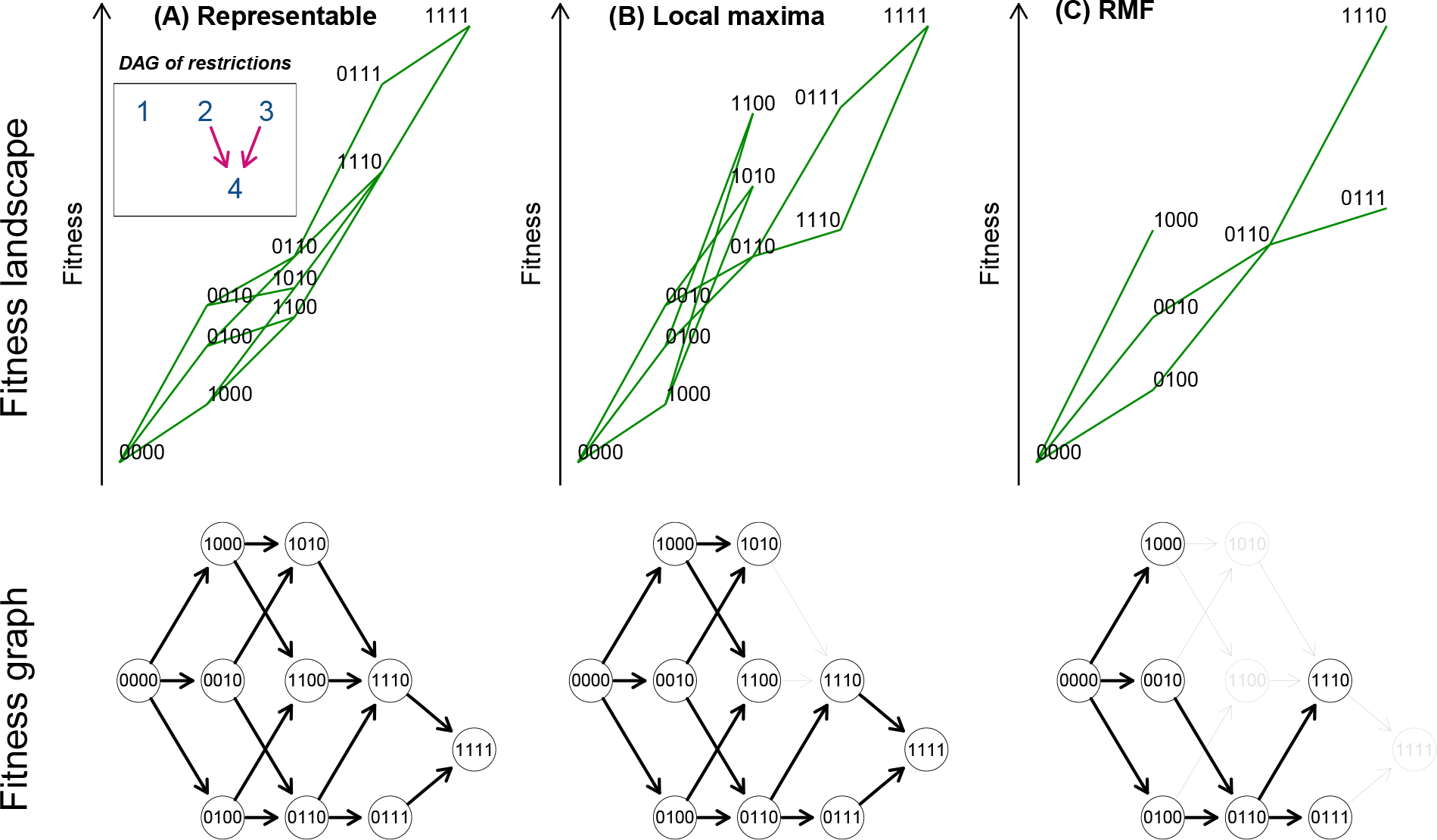
Fitness landscapes, paths of tumor progression, and directed acyclic graphs (DAGs) of restrictions in the order of accumulation of mutations for the three types of landscapes used. (A) Representable; (B) Local maxima; (C) Rough Mount Fuji (RMF). Top row: fitness landscapes (representation based on Brouillet *et al*., 2015); genotypes are shown as sequences of 0s and 1s, where “1010” means a genotype with the first and third genes mutated; the vertical position of a genotype is its fitness; its horizontal position on the x-axis is given by its Hamming distance to the “0000” genotype. Green segments connect mutational neighbors of increasing fitness. The inset in the first row shows the DAG of restrictions in the order of accumulation of mutations that applies to (A) and (B). A DAG of restrictions shows genes (not genotypes) in the nodes; an arrow from gene *i* to gene *j* means that a mutation in *i* must occur before a mutation in *j* can occur; an arrow indicates a direct dependency of a mutation in gene *j* on a mutation in gene *i*. In the figure, a mutation in the fourth gene can be observed only if both the second and third genes are mutated. Note that among the models considered in this paper, CAPRESE and OT can only represent trees (so they can not account for the fourth gene having two, or more, incoming arrows). The absence of an arrow between two genes means that there are no direct dependencies between the two genes. Bottom row: fitness graphs or graphs of mutational paths; in fitness graphs nodes are genotypes and arrows point toward mutational neighbors of higher fitness (i.e., two genotypes connected by an arrow differ in one mutation that increases fitness —Crona *et al*., 2013; de Visser and Krug, 2014; Franke *et al*., 2011). These fitness graphs show all the paths of tumor progression, the set of accessible mutational paths and adaptive walks that, under the restriction that there can be no back mutations, start from the “0000” genotype and end in a fitness maximum. When evolution runs until fixation in a fitness maximum, each path from “0000” to a fitness maximum corresponds to a different Line of Descent (LOD). For (B) and (C), gray edges and nodes in the fitness graphs show edges and nodes that are present in (A) but missing in (B) or (C). Under CPMs, since each new driver mutation with its dependencies satisfied increases fitness, all accessible genotypes that differ by exactly one mutation are connected in the fitness graph, as shown in (A), and the genotype with all genes mutated is the single global fitness maximum. For (B), the fitness landscape —and its fitness graph— has the same accessible genotypes as the fitness landscape in (A). But the fitness landscape in (B) has three maxima and, compared to (A), there are fewer paths to the genotype with all genes mutated, “1111”, and several paths end in the other two maxima (“1100”, “1010”). Thus, the fitness graph of (B) does not fulfill the assumptions of CPMs. Compared to the fitness graph of (A), in the fitness graph of (B) not all accessible genotypes that differ by one mutation are connected —e.g., genotypes “1100” and “1110”. In terms of acquisition of mutations, in (B), and in contrast to (A), we cannot reach genotype “1110” from genotype “1100”, even when a mutation in the third gene does not depend on any previous mutation according to the DAG of restrictions. So if we go from “1000” to “1100”, the acquisition of the second mutation precludes acquiring the third mutation (a violation of an assumption of CPMs), and this creates a local fitness maximum. The fitness landscape in (C) cannot be represented by any DAG of restrictions; e.g., no DAG of restrictions can account at the same time for the presence of genotypes “1000”, “0100”, “0010”, and the absence of every double mutant with the first gene mutated. Relative to (A), the fitness graph in (C) is missing both paths and genotypes (relative to the fitness graph from any possible DAG of restrictions it could either be missing and/or adding genotypes and paths).

The first question we address in this study is whether we can predict the paths of tumor progression using CPMs. To answer this question we will examine how close to the truth are the predictions made by four CPMs (CBN, OT, CAPRI, and CAPRESE) about the distribution of paths of tumor progression. When addressing this question we need to take into account possible deviations from the models assumed by CPMs. In particular, most CPMs assume that the acquisition of a mutation in a driver gene, when all its possible dependencies on other genes are satisfied, does not decrease the probability of gaining a mutation in another driver gene (Misra *et al*., 2014). In other words, acquiring driver mutations (when their dependencies on other genes are satisfied) cannot decrease fitness, which implies that the fitness landscapes assumed by CPMs only have a single global fitness maximum (the genotype with all drivers mutated —see Figure 1). But it is likely that many cancer fitness landscapes have several local fitness maxima (i.e., they are rugged, multi-peaked landscapes): this can happen if there are many combinations of a small number of drivers, out of a larger pool of drivers (Tomasetti *et al*., 2015), that result in the escape genotypes; moreover, synthetic lethality is common in both cancer cells (Beijersbergen *et al*., 2017; O’Neil *et al*., 2017) and the human genome (Blomen *et al*., 2015), and it can lead to local fitness maxima when it affects mutations that individually increase fitness —see also Chiotti *et al*., 2014. Thus, to examine if CPMs can be used to predict paths of tumor progression we will need to assess how the quality of the predictions is affected by multi-peaked fitness landscapes.

The second question addressed in this paper is whether we can we use CPMs to estimate evolutionary unpredictability, regardless of the performance when predicting the actual paths of tumor progression. A model could be useful if it suggests few paths are possible, even if its actual predictions about the distribution of paths are not trustworthy. Conversely, predicting correctly the distribution of paths of tumor progression might be of little importance in scenarios where the true evolutionary unpredictability itself is very large (where disease progression follows a very large number of possible paths); for practical purposes, forecasting here would be useless.

To address the above questions (can we predict the paths of tumor progression using CPMs?; can we estimate evolutionary unpredictability using CPMs?) we use evolutionary simulations on 1260 fitness landscapes that include from none to severe deviations from the assumptions that CPMs make about the structure of fitness landscapes, and we analyze the data with four different CPMs. This paper does not attempt to understand the determinants of evolutionary (un)predictability (see, e.g., Bank *et al*., 2016; de Visser and Krug, 2014; Lässig *et al*., 2017; Losos, 2018; Szendro *et al*., 2013) but, instead, we focus on the effects of evolutionary unpredictability for CPMs. This is why we use variation in key determinants of evolutionary unpredictability (e.g., variation in population sizes and mutation rates) but these factors are only used to generate variability in unpredictability, and not themselves the focus of the study. To better assess the quality of predictions, we use sample sizes that cover the range from what is commonly used to what are much larger sample sizes than currently available. We also include variation in the cancer detection process or detection regime (when cancer samples are taken, or when patients are sampled), since previous studies have shown that it affects the quality of inferences from CPMs (Diaz-Uriarte, 2018).

We have shown before (Diaz-Uriarte, 2018) that the performance of two CPMs (CBN and CAPRI) for predicting accessible genotypes degrades considerably when the fitness landscapes contain reciprocal sign epistasis. That study focused on predicting accessible genotypes and its results cannot provide an answer to the questions about predicting paths of tumor progression and estimating evolutionary unpredictability. To answer whether CPMs can be used to predict paths of progression and to estimate evolutionary unpredictability we need to look directly at the prediction of paths (not genotypes), and compare them with the true paths of progression, as we do in the current work. Thus, the two studies differ in objectives, methods (here we follow evolution until fixation, and we develop procedures to compare predicted with true paths of tumor progression), and scenarios considered (the types fitness landscapes used and the extent of evolutionary unpredictability); see details in S1 Text.

Here we find that the agreement between the predicted and true distributions of paths is generally poor, unless sample sizes are very large and fitness landscapes conform to the assumptions of CPMs. Both detection regime and evolutionary unpredictability itself have major effects on performance. But in spite of the unreliability of the predictions of paths of tumor progression, we find that CPMs can be useful for estimating upper bounds to the true evolutionary unpredictability.

What are the implications of our results for the analysis and interpretation of the use of CPMs with cancer data sets? We analyze twenty-two real cancer data sets with H-CBN, the best performing CPM in the simulations. We cannot examine how close predictions are to the truth, since the truth is unknown; thus, we use bootstrap samples to examine the reliability of the inferences. Many of the cancer data sets reflect conditions where useful predictions could be possible, based on the estimates of evolutionary unpredictability from H-CBN. But for most data sets these results are thwarted by the unreliability of the predictions themselves, which increases with the number of features analyzed. Our results question uncritical use of CPMs for predicting paths of tumor progression, and suggest the need for methodological work that can account for the probably multi-peaked fitness landscapes in cancer.

## 2 Materials and methods

This paper involves both a simulation study where results from four CPMs (CBN —variants H-CBN and MCCBN—, OT, CAPRI —variants CAPRI_AIC and CAPRI_BIC—, and CAPRESE) are compared to the known truth from the simulations, and the analysis of twenty-two cancer data sets using the best performing of the above CPMs (H-CBN). Section 2.1 describes the CPMs used and how predicted paths of tumor progression are obtained from them. Section 2.2 provides an overview of the simulation study. Sections 2.3 to 2.5 provide details on how the simulations were conducted. How the performance of CPMs was assessed is explained in section 2.6. Section 2.7 summarizes the main features of the simulated landscapes and data sets used to evaluate the performance of the CPMs. The cancer data sets and the methods used to analyze them are described in section 2.8.

### 2.1 Cancer Progression Models used and paths of tumor progression

We have compared four distinct CPMs: CBN, OT, CAPRI, and CAPRESE. Two of the models used, CBN and CAPRI, have been used in two variants (H-CBN and MCCBN for CBN, CAPRI_AIC and CAPRI_BIC for CAPRI), yielding a total of six different procedures for obtaining CPMs. Only a brief overview of these CPMs is provided here; detailed descriptions can be found in the original references for each model: H-CBN (Gerstung *et al*., 2009, 2011), MCCBN (Montazeri *et al*., 2016), OT (Desper *et al*., 1999; Szabo and Boucher, 2008), CAPRI (Caravagna *et al*., 2016; Ramazzotti *et al*., 2015), and CAPRESE (Olde Loohuis *et al*., 2014). (Other CPMs exist but they have not been considered here because they are too slow for routine work, have no software available, or have dependencies on non-open source external libraries —see S4 Text).

The CPMs considered try to identify restrictions in the order of accumulation of mutations from cross-sectional data. CPMs assume that the different observations in the cross-sectional data set constitute independent realizations of evolutionary processes where the same constraints hold for all tumors (Beerenwinkel *et al*., 2015, 2016; Gerstung *et al*., 2011). Thus, a data set can be regarded as a set of replicate evolutionary experiments where all individuals are under the same genetic constraints. For the four CPMs considered in this paper, the cross-sectional data is a matrix of subjects (or individuals) by driver alteration events, where each entry in the matrix is binary coded as mutated or not-mutated (or, equivalently, altered or non-altered). CPMs assume there are no back mutations in these events —i.e., once gained, an alteration is not lost. CPMs further assume that the driver genes are known. For the simulations, we will refer to these driver alteration events as “genes”, but they can be individual genes, parts or states of genes, or modules or pathways made from several genes (e.g. Caravagna *et al*., 2016; Gerstung *et al*., 2011). When we analyze the twenty-two cancer data sets (see section 2.8) we will use the generic term “features” as some of those data sets use genes whereas others use pathway or module information. CPMs assume that all tumors start cancer progression without any of the mutations considered in the study (the above matrix of subjects by driver alterations), but other mutations could be present that have caused the initial tumor growth. All these other mutations are absorbed in the root node from which cancer is initiated (Attolini *et al*., 2010); note that the way the data are simulated to generate cross-sectional observations (see section 2.2) is consistent with this assumption.

The above assumptions are common to the CPMs considered. The models examined here differ, however, in the types of restrictions they can represent and on their model fitting procedures. Both OT (Desper *et al*., 1999; Szabo and Boucher, 2008) and CAPRESE (Olde Loohuis *et al*., 2014) describe the accumulation of mutations with order constraints that can be represented as trees. Thus, among the “representable” fitness landscapes used in this paper (section 2.4), OT and CAPRESE can only faithfully model the subset that are trees, those where a gene mutation has a direct dependency on only one other gene’s mutation. A key difference between OT and CAPRESE is that CAPRESE reconstructs these models using a probability raising notion of causation in the framework of Suppes’ probabilistic causation, whereas in OT weights along edges can be directly interpreted as probabilities of transition along the edges by the time of observation (Szabo and Boucher, 2008, p. 4). In contrast to OT and CAPRESE, both CAPRI (Caravagna *et al*., 2016; Ramazzotti *et al*., 2015) and CBN (Gerstung *et al*., 2009, 2011; Montazeri *et al*., 2016) allow modeling the dependence of an event on more than one previous event: the output of the models are directed acyclic graphs (DAGs) where some nodes have multiple parents, instead of a single parent (as in trees). CAPRI tries to identify events (alterations) that constitute “selective advantage relationships”, again using probability raising in the framework of Suppes’ probabilistic causation. We have used two versions of CAPRI, that we will call CAPRI_AIC and CAPRI_BIC, that differ in the penalization used in the maximum likelihood fit, Akaike Information Criterion (AIC), or Bayesian Information Criterion (BIC), respectively. For CBN we have also used two variants, H-CBN, described in Gerstung *et al*. (2009, 2011) that uses simulated annealing with a nested expectation-maximization (EM) algorithm for estimation, and MCCBN, described in Montazeri *et al*. (2016), that uses a Monte-Carlo EM algorithm. Thus, these six procedures can be divided into three groups: models that return trees (OT and CAPRESE) and two families of models that return DAGs, CBN (H-CBN and MCCBN) and CAPRI (CAPRI_AIC and CAPRI_BIC).

Because (the transitive reduction of) a DAG of restrictions determines a fitness graph (see Figure 1 and Diaz-Uriarte, 2018), the set of paths to the maximum encoded by the output from a CPM is obtained from the fitness graph. This we did for all models. From H-CBN and MCCBN we can also obtain the estimated probability of each path of tumor progression to the fitness maximum, since both H-CBN and MCCBN return the parameters of the waiting time to occurrence of each mutation (given its restrictions are satisfied; e.g., p. i729 in Montazeri *et al*., 2016, section 2.2 in Gerstung *et al*., 2009, or Hosseini, 2018; details and example in section 3 of S4 Text). It is also possible to perform a similar operation with the output of OT, and use the edge weights from the fits of OT to obtain the probabilities of transition to each descendant genotype and, from them, the probabilities of the different paths to the global maximum. It must be noted that these probabilities are not really returned by the model, since the OTs used are untimed oncogenetic trees (Desper *et al*., 1999; Szabo and Boucher, 2008). We will refer to paths with probabilities assigned in the above way as **probability-weighted paths**. For CAPRESE and CAPRI, it is not possible to map the output to different probabilities of paths of progression (see section 3 of S4 Text) and in all computations that required probability of paths we assigned the same probability to each path.

### 2.2 Overview of the simulation study

We have used simulations of tumor evolution on fitness landscapes of three different types (see Figure 1), for landscapes of 7 and 10 genes, under different initial population sizes and mutation rates. We have used a total of 1260 fitness landscapes = 35 random fitness landscapes × 2 conditions of numbers of genes × 3 types of fitness landscapes × 3 initial population sizes × 2 mutation regimes. For each one of the 1260 fitness landscapes, we simulated 20000 independent evolutionary processes (with the specified parameters for initial population size and mutation rate) using a logistic-like growth model; each simulated evolutionary process was run until one of the genotypes at the local fitness maxima (or the single global fitness maximum) reached fixation. Each set of 20000 simulated evolutionary processes was then sampled under three detection regimes, so that each fitness landscape generated three sets of 20000 simulated genotypes. From each of these sets, we obtained five different splits of the genotypes for each of three sample sizes (50, 200, 4000); thus a total of 56700 (= 1260 × 3 × 3 × 5 combinations of 1260 fitness landscapes, 3 detection regimes, 3 sample sizes, 5 splits) data sets were produced. Each of these 56700 data sets was analyzed with every one of the CPMs compared (H-CBN, MCCBN, OT, CAPRI_AIC, CAPRI_BIC, and CAPRESE), to obtain predicted paths of tumor progression. These predictions were then compared with the true, recorded, paths of tumor progression from the simulations (see section 2.6). A schematic view of the simulation study is provided in Figure 2.

**Figure 2:**
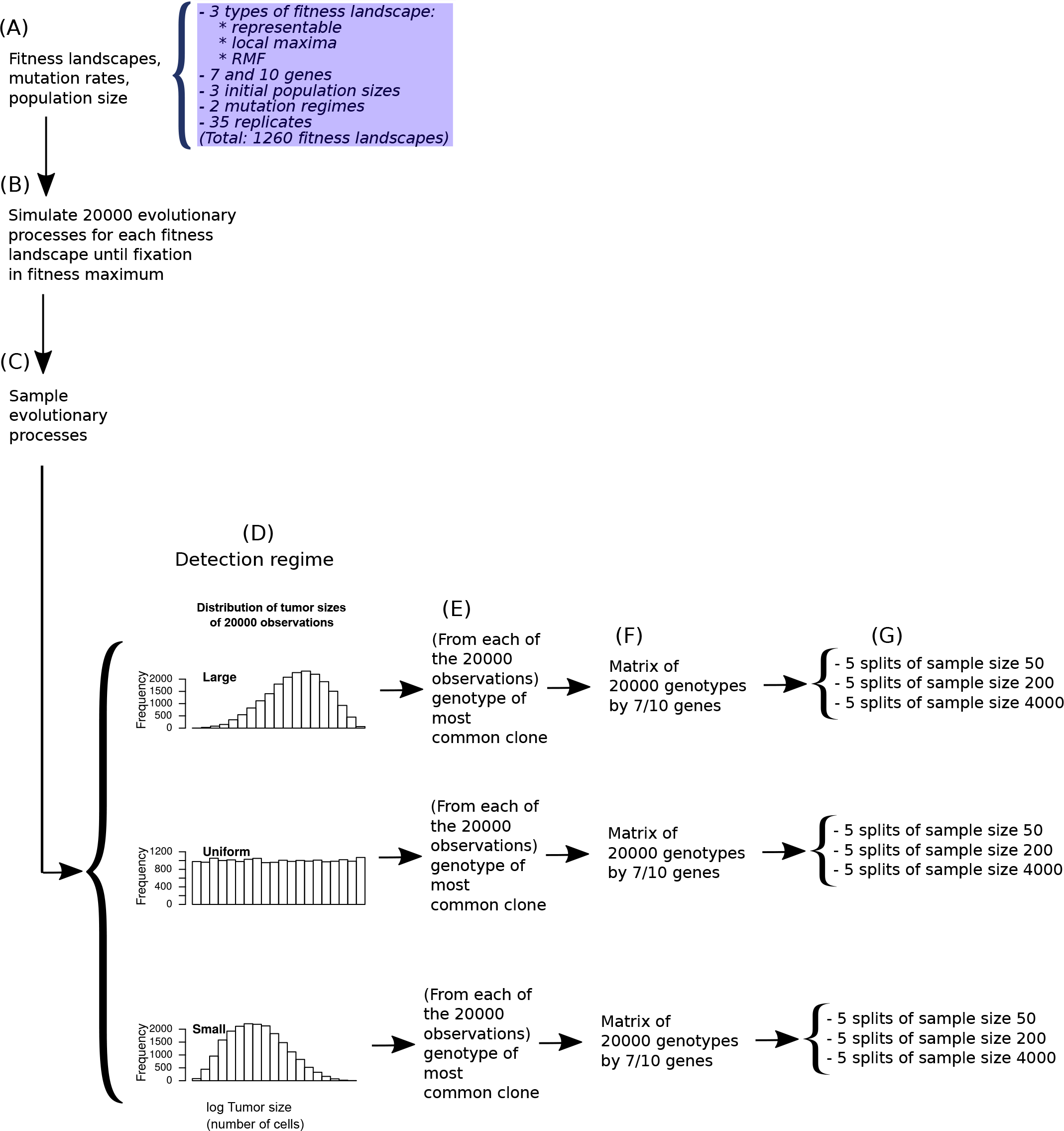
Scheme of the simulation study. (A) On each of the 1260 fitness landscapes, (B) 20000 evolutionary processes were simulated. The set of 20000 evolutionary processes from a fitness landscape were then (C) sampled under three detection regimes, obtaining one observation per simulation under each detection regime, (D) leading to 20000 observations that are enriched in large-sized tumors —large detection regime—, 20000 observations enriched on small-sized tumors —small detection regime—, and 20000 observations with uniform distribution with respect to the logarithm of tumor size (so that the large and small detection regimes emulates cases when cancer tends to be detected at late or early stages, respectively —see text). (E) From each of the individual observations, we obtained the genotype of the most common clone; therefore, (F) each fitness landscape provides 3 sets of 20000 genotypes, one for each detection regime. (G) These sets were split in 5 non-overlapping sets of 50 observations, 5 non-overlapping sets of 200 observations, and 5 non-overlapping sets of 4000 observations. Each of these data sets was analyzed with each of the CPMs considered to obtain the predicted paths of tumor progression.

### 2.3 Fitness landscapes

We have used three different kinds of random fitness landscapes (see Figure 1). **Representable** fitness landscapes are fitness landscapes for which a DAG of restrictions exists with the same accessible genotypes and accessible mutational paths. (Accessible mutational path: a trajectory through a collection of genotypes, where each genotype is separated from the preceding genotype by one mutation, along which fitness increases monotonically —Franke *et al*. (2011); accessible genotypes: genotypes along accessible mutational paths). An example of a representable fitness landscape with its corresponding DAG of restrictions and fitness graph is shown in Figure 1A. A defining characteristic of representable fitness landscapes is that all accessible genotypes that differ by exactly one mutation are connected in the fitness graph; thus, there is a single global fitness maximum, the genotype with all genes mutated, and all accessible mutational paths in the fitness graph end in that single global maximum. For representable fitness landscapes there is a one-to-one correspondence between DAGs of restrictions and fitness graphs.

In **local maxima** fitness landscapes (Figure 1B) the set of accessible genotypes can be represented by a DAG of restrictions, but there are local fitness maxima and the fitness graph has missing paths; the genotype with all genes mutated might or might not be the genotype with largest fitness. In other words, in the local maxima fitness landscapes, the DAG of restrictions and the fitness landscape agree on the genotypes that should and should not be accessible; what the local maxima landscapes are missing are mutational paths to the genotype with all genes mutated, because we have introduced local fitness maxima. Once we introduce local maxima there is no longer a one-to-one correspondence between DAGs of restrictions and fitness graphs (thus, there is no longer a one-to-one correspondence between DAGs of restrictions and sets of tumor progression paths). These local maxima landscapes should not be as challenging to CPMs as the DAG-derived non-representable fitness landscapes used in Diaz-Uriarte (2018), as those also missed some genotypes that should exist under the DAG of restrictions. This is by design: here we want to isolate the effect of multi-peaked landscapes or local maxima (or, equivalently, missing paths), without the additional burden, for the CPMs, of missing genotypes.

The third type of fitness landscapes used are Rough Mount Fuji (**RMF**) fitness landscapes. The RMF model (de Visser and Krug, 2014; Neidhart *et al*., 2014) combines a random House of Cards model (where fitness is assigned to genotypes by independently sampling from a fixed probability distribution) and an additive fitness landscape (where the reference genotype, or the genotype with largest fitness, need not be the one with all genes mutated). The RMF model is a very flexible one, where the ruggedness of the landscape can be modified by changing the ratio of the additive component (the change in fitness per unit increase in Hamming distance from the reference genotype) relative to the variance of the random fitness component (the House of Cards component). The RMF model has been useful to model empirical fitness landscapes (de Visser and Krug, 2014; Franke *et al*., 2011; Neidhart *et al*., 2014). RMF fitness landscapes generally have multiple local fitness maxima and considerable reciprocal sign epistasis and thus not even the set of accessible genotypes can be represented by a DAG of restrictions (see Diaz-Uriarte, 2018, and Figure 1).

We generated the DAG-derived **representable** fitness landscapes by generating a random DAG of restrictions and from it the fitness graph. We then assigned birth rates to genotypes using an iterative procedure on the fitness graph where, starting from the genotype without any driver mutation with a birth rate of 1, the birth rate of each descendant genotype was set equal to the maximum fitness of its parent genotypes times a random uniform variate between 1.01 and 1.19 (*U*(1.01, 1.19)) yielding, therefore, an average multiplicative increase in fitness of 0.1 (which is within values previously used: Bozic *et al*., 2010; McFarland *et al*., 2013; Williams *et al*., 2018). The birth rate of genotypes that were not accessible according to the DAG of restrictions was set to 0. For example, if the DAG of restrictions was the one shown in Figure 1A, a cell with genotype “1001” would have a birth rate of 0, since the dependencies of the DAG of restrictions are not satisfied —mutations in genes 2 and 3 must occur before a mutation in gene 4. Therefore, this simulation scheme strictly adheres to the assumptions about accessible and non-accessible genotypes under the CPM model. (For the growth model used here —see below— birth rates determine fitness at any population size as death rates are identical for all genotypes and depend only on population size. Genotypes with a birth rate of 0 are never added to the population and, thus, they cannot mutate before dying). We generated the DAG-derived **local maxima** fitness landscapes by first generating a random DAG and from it the fitness graph, identically to what was done for representable fitness landscapes. Though in contrast to representable fitness landscapes, before assigning fitness to genotypes a random selection of edges of the fitness graph were removed so that all accessible genotypes remained accessible but now from a possibly much smaller set of parents. Birth rate was then assigned as for the representable fitness landscapes (using the iterative procedure on the fitness graph, where birth rate of descendant genotype = max(birth rate parent genotypes) * *U*(1.01, 1.19), and with all non-accessible genotypes with a birth rate of 0). For each DAG we repeated this procedure 50 times, and kept the one that introduced the largest number of local maxima. Creating local maxima almost always resulted in creating reciprocal sign epistasis (see also section 2.7 and S2 Text). We generated the **RMF** fitness landscapes by randomly choosing the reference genotype (i.e., the genotype with the largest fitness) and the decrease in birth rate of a genotype per each unit increase in Hamming distance from the reference genotype (which affects the ruggedness of the landscape); see details in S2 Text.

### 2.4 Evolutionary simulations

Once a fitness landscape had been generated, we simulated 20000 evolutionary processes (step B in Figure 2). We used the continuous-time, logistic-like model of McFarland *et al*. (2013), in which death rate depends on total population size, as implemented in OncoSimulR (Diaz-Uriarte, 2017), with the specified parameters of initial population size and mutation rate (below). Each individual evolutionary process was run until one of the genotypes at the local fitness maxima (or the single global fitness maximum) reached fixation (see details in S3 Text). We also verified that all 7 or 10 genes had appeared in at least some genotypes, i.e., were part of the paths of tumor progression. If this condition was not fulfilled, a new fitness landscape was generated and the processes started again. This procedure is independent of the detection process that returns the genotypes analyzed by the CPMs (section 2.5).

We used three **initial population sizes**, 2000, 50000, and 1 × 10^6^ cells, for the simulations; these cover a range of population sizes at tumor initiation that have previously been used in the literature (e.g. Beerenwinkel *et al*., 2007; Gerstung *et al*., 2011; McFarland *et al*., 2013; Wodarz and Komarova, 2014). We also used two **mutation regimes**; in the first one, all genes had a common mutation rate of 1 × 10^−5^; in the second, genes had different mutation rates, uniformly distributed in the log scale between (1/5) 1 × 10^−5^ and 5 × 10^−5^ (i.e., the largest ratio between largest and smallest mutation rates was 25), so that the arithmetic mean of mutation rates was 1.5 × 10^−5^ and the geometric mean 1 × 10^−5^. These mutation rates are within ranges previously used in the literature (Bozic *et al*., 2010; McFarland *et al*., 2013; Nowak *et al*., 2004), with a bias towards larger numbers (since we use only 7 or 10 genes relevant for population growth and we could be modeling pathways, not individual genes). Initial population size and mutation rates are not of intrinsic interest here (since our focus are not the determinants of evolutionary predictability *per se*), but are used to generate variability in evolutionary predictability and to allow for deviations from the strong-selection-weak-mutation (SSWM) regime (de Visser and Krug, 2014); see section 2.7.

Only in the representable fitness landscapes are simulations restricted to move uphill in the fitness landscapes. In all three types of fitness landscapes, mutations can lead to either increases or decreases in fitness. In the representable and local maxima fitness landscapes, as explained above, mutation events that do not fulfill the restrictions in the order of accumulation of mutations lead to a birth rate (and, thus, fitness) of 0. Therefore, in the simulations in representable and local maxima fitness landscapes, no path from the “0000” genotype to a fitness maximum can ever go through a non-accessible genotype. This is by design, so that these fitness landscapes strictly adhere to the assumption of CPMs about restrictions in the accumulation of mutations. But in both RMF and local maxima fitness landscapes it is possible to move through a fitness valley (i.e., make moves from ancestor to descendant that are not always monotonically increasing in fitness), phenomena that are more frequent as we deviate from the SSWM assumption (de Visser and Krug, 2014; commented example in Section 5 in S3 Text). (Note that this is possible in local maxima fitness landscapes, even when non-accessible genotypes can never be part of evolutionary paths, because with no back mutations an accessible genotype can be along an uphill path when coming from one ancestor but in a valley when coming from another ancestor; no such genotypes can exist in the representable fitness landscape as in the representable landscapes all accessible genotypes that differ by exactly one mutation are connected in the fitness graph). In addition, in the RMF fitness landscape, we can move through fitness valleys of non-accessible genotypes as non-accessible genotypes need not have a birth rate of 0 in the RMF; see Section 5 in S3 Text).

### 2.5 Detection regimes and obtaining data sets from the simulations

To obtain the genotypes that were analyzed by the CPMs, we first sampled the simulated evolutionary processes, obtaining one observation per evolutionary processes, using three different **detection regimes** (Figure 2, C and D); then, for each observation of each detection regime, we obtained the genotype corresponding to each observation (Figure 2E), which lead to matrices of 20000 genotypes (Figure 2F); finally we split these matrices into non-overlapping subsets to be analyzed with the CPMs (Figure 2G).

The three detection regimes differ in the distribution of sizes of the sampled tumors (Figure 2D). Under the **large** detection regime a large fraction of the samples correspond to large tumors. In contrast, under the **small** detection regime a large fraction of the samples correspond to small tumors. Finally, under the **uniform** detection regime the distribution of sizes of the sampled tumors is approximately uniform. Thus, the large detection regime would emulate scenarios where cancer tends to be detected at late, advanced stages, and the small detection regime would emulate scenarios where cancer tends to be detected at early stages.

To implement these detection regimes, we drew random deviates from beta distributions with parameters B(1, 1), B(5, 3), and B(3, 5) (for uniform, large, and small, respectively), rescaled them to the range of the log-transformed distribution of observed tumor sizes (log of number of cells), and obtained the observation with population size closest to the target (see details in section 2 in S3 Text). (We used the log-scale of tumor size because in the model of McFarland *et al*., 2013 tumor population size increases logarithmically with number of driver mutations; thus, distributions of sampled tumors that are biased towards large sizes in the log scale will mimic sampling of late-stage tumors —tumors with a large number of drivers—, and distributions of sampled tumors that are biased towards small sizes in the log scale will mimic sampling of early-stage tumors, as intended.).

For each observation, the genotype returned was the genotype of the most abundant clone (Figure 2E). Finally, the set of 20000 genotypes (Figure 2F) was then split into five sets of non-overlapping data sets for each of the three **sample sizes** of 50, 200, and 4000 (Figure 2G). These are the data sets that were analyzed with the CPMs.

### 2.6 Measures of performance and predictability

We have characterized evolutionary unpredictability using the diversity of **Lines of Descent (LODs)**. LODs were introduced by Szendro *et al*. (2013) and “(…) represent the lineages that arrive at the most populated genotype at the final time” (p. 572). In other words, in our simulations a LOD is a sequence of parent-child genotypes, from the initial genotype to a local maximum: a LOD is the path that a tumor has taken until fixation. The final genotype in a LOD is a local fitness maximum, but there are no guarantees that any intermediate genotype in the LOD will have been the most common genotype at any time point (especially under deviations from SSWM such as clonal interference and stochastic tunneling —de Visser and Krug, 2014; Sniegowski and Gerrish, 2010; Szendro *et al*., 2013). As in Szendro *et al*. (2013), we can use the entropy of these paths to measure the indeterminism of the paths of evolution, or evolutionary unpredictability, and we will define *S*_*p*_ = − ∑ *p*_*i*_ ln *p*_*i*_, where *p*_*i*_ is the observed probability of each LOD (each path) computed from the 20000 simulations, and the sum is over all paths or LODs. Evolutionary unpredictability, as estimated by the CPMs, will analogously be defined as *S*_*c*_ = − ∑ *q*_*j*_ ln *q*_*j*_, where *q*_*j*_ is the probability of each path to the maximum according to the cancer progression model considered, and the sum is over all paths predicted by the CPMs. (Hosseini, 2018, normalizes predictability by dividing by the maximum entropy, similar to dividing by the prior entropy in the “information gain” statistic in Lässig *et al*., 2017; but the maximum entropy is a constant for each number of genes, i.e., 7! or 10! for our simulations).

To measure how well CPMs predict tumor progression, we used three different statistics. To compare the overall similarity of the distribution of paths predicted by CPMs with the true observed one (i.e., the distribution of LODs) we used the Jensen-Shannon divergence (**JS**) (Crooks, 2017; Lin, 1991), scaled between 0 and 1 (equivalent to using the logarithm of base 2). JS is a symmetrized Kullback-Leibler divergence between two distributions and is defined even if the two distributions do not have the same sample space, i.e., even if *P*(*i*) ≠ 0 and *Q*(*i*) = 0 (or *Q*(*i*) ≠ 0 and *P*(*i*) = 0), as can often be the case for our data. A JS value of 0 means that the distributions are identical, and a value of 1 that they do not overlap. Therefore, predictions of CPMs are closer to the truth the smaller the value of JS. The sum of the probabilities of the paths in the LODs that are not among the paths allowed by the CPMs, *P*(¬*DAG*|*LOD*), is equivalent to **1 - recall**. Larger values of 1-recall mean that the CPM is not capturing a large fraction of the evolutionary paths to the maximum (or maxima). The sum of the predicted probabilities of paths according to the CPMs that are not used by evolution (i.e., that are not LODs), *P*(¬*LOD*|*DAG*), is equivalent to **1 - precision**. Larger values of 1-precision mean that the CPMs predict larger numbers of paths to the maximum that are not used by evolution. In S6 Text we also use as statistic the **probability of recovering the most common LOD**; we will rarely refer to this statistic in the main paper since it follows a pattern very similar to recall (Section 2 in S6 Text). Statistics 1-recall and 1-precision can overestimate performance: they could both have a value of 0, even when JS is very close to 1 (see example in Section 4 in S4 Text). Thus, the main overall performance measure will be JS.

#### 2.6.1 Comparing paths from CPMs with LODs of different lengths

When all paths from the CPM and the LOD have equal length (they end in a genotype with the same number of genes mutated, *K*) computing the above statistics is straightforward. But paths could differ in length. In fitness landscapes with local maxima, LODs can differ in length; some LODs could have a length (or number of mutations of the fixated genotype), *K*_*i*_, shorter than the length of the paths from the CPM, *K*_*C*_ (all paths from a CPM have the same number of mutations, since all arrive at the genotype with all *K*_*C*_ genes mutated). It is also possible that some or all *K*_*i*_ > *K*_*C*_, i.e., some or all LODs have a length larger than the length of the paths from the CPM. This will happen if the CPM has been built from a data set that contains fewer genes than the number of genes in the landscape (e.g., because one or more genes were absent —see Section 2 in S4 Text); if the sampled data set has fewer genes than the landscape in a representable fitness landscape, then all *K*_*i*_ > *K*_*C*_ (as *K*_*i*_ will be equal to either 7 or 10).

To compute JS, 1-recall, and 1-precision that will cover all those cases we used the following procedure (that reduces to the simpler procedure in the previous section when all *K*_*i*_ = *K*_*C*_). Let *i* and *j* denote two paths, one from the LOD and the other from the CPM, with corresponding probabilities *p*_*i*_ and *q*_*j*_; in contrast to the previous section, and to minimize notation, *i*, *j* (and *p*_*i*_, *q*_*j*_) could refer to a path from the LOD and a path from the CPM or, alternatively, a path from the CPM and a path from a LOD. Let *K*_*i*_, *K*_*j*_ denote the length of paths *i* and *j*, respectively. At least one set of either *K*_*i*_s or *K*_*j*_s has all elements identical (e.g., if *j* refers to indices of the paths from the CPM, it is necessarily the case that *K*_1_ = *K*_2_ = … = *K*_*m*_ = *K*_*C*_, with *m* the total number of different paths from the CPM).

Now if *K*_*i*_ > *K*_*j*_ and the path *i* up to *K*_*j*_ mutations (i.e., from the “0000” genotype to the genotype with *K*_*j*_ mutations) is identical to *j*, then path *j* is included in path *i*: all of *q*_*j*_ is accounted for by *i*. This also means that path *i* is partially included in (or accounted for by) path *j*, but a fraction of it, (*K*_*i*_ − *K*_*j*_)/*K*_*i*_, is missing or unaccounted for. The above applies directly to calculations of 1-recall and 1-precision. For computing JS, there will be two entries in the vectors with the probability distributions that will be compared: 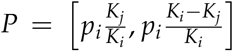, *Q* = [*q*_*i*_, 0]. This procedure can be applied to all elements *i*, *j*, summing all unmatched entries: 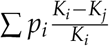 is the total flow in the set of paths *i* that cannot be matched by the *j*s because they are shorter. To simplify computations, that unmatched term can also include ∑ *p*_*u*_, where *u* denotes those paths *i* that do not match any *j*. Conversely, all paths *i* with *K*_*i*_ > *K*_*j*_ such that the paths become indistinguishable up to *K*_*j*_ can be summed in a single entry so that we obtain 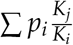 and 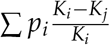 for the matched and unmatched fractions, respectively. All computations have their corresponding counterparts for elements *i*, *j* when *K*_*i*_ < *K*_*j*_. The above procedure is applied at all distinct *k*, the number of mutations of the final genotypes of the true LODs. The final JS (and 1-precision and 1-recall) is the weighted sum of each of those JS (and 1-precision and 1-recall), weighted by *w*_*k*_, the frequency of all paths from the LOD that end at *k* mutations. This procedure results in unique JS (remember the *K* are all the same for at least one of the sets of paths) as well as unique 1-precision and 1-recall, and it reduces to the procedure (see above) when all *K*_*i*_ are equal and equal to all *K*_*j*_. A commented example and further details are provided in the Supporting Information (Section 5.1 in S4 Text).

#### 2.6.2 Statistical modeling of performance

We have used generalized linear mixed-effects models, with a beta model for the dependent variable (Ferrari and Cribari-Neto, 2004; Grün *et al*., 2012; Smithson and Verkuilen, 2006), to model how JS, 1-recall, and 1-precision, are affected by *S*_*p*_, detection regime, sample size, number of genes, type of fitness landscape, and CPM. In all models, the response variable was the average from the five split replicates of each fitness landscape by sample size by detection regime combination, and fitness landscape id (not type) was a random effect. When the dependent variable had values exactly equal to 0 or 1, we used the transformation suggested in Smithson and Verkuilen (2006). Models were fitted using sum-to-zero contrasts (McCullagh and Nelder, 1989) and all regressors were used as discrete regressors, except *S*_*p*_, which has been scaled (mean 0, variance 1) for easier interpretation; the coefficients of the main effect terms of the discrete regressors are the deviations from the average (see further details in Section 6 in S4 Text). We have used the glmmTMB (Brooks *et al*., 2017) and car (Fox and Weisberg, 2011) packages for R (R Core Team, 2018) for statistical model fitting and analysis.

### 2.7 Characteristics of the simulated fitness landscapes and genotypes

All the fitness landscapes used are shown in S1 Figure. We provide here a brief description of the main features of the three different fitness landscapes and the simulated data sets. The three types of fitness landscapes had comparable numbers of accessible genotypes but differed in the number of local fitness maxima and reciprocal sign epistasis, as shown in Figures 1 to 3 in S2 Figure (representable fitness landscapes had a single fitness maximum with no reciprocal sign epistasis, whereas RMF landscapes had the largest of both, and local maxima landscapes were intermediate).

Simulations resulted in varied amounts of clonal interference, as measured by the average frequency of the most common genotype (Figures 4 and 5 in S2 Figure); scenarios where clonal sweeps dominated (i.e., those characterized by the smallest clonal interference) corresponded to initial population sizes of 2000, with clonal interference being much larger at the other population sizes (Figure 4 in S2 Figure).

Simulations resulted in a wide range of numbers of paths to the maximum (number of distinct LODs: Figure 6 in S2 Figure). LOD diversities (*S*_*p*_) ranged from 0.3 to 8.7 (Figure 7 in S2 Figure) with RMF models showing smaller *S*_*p*_; RMF landscapes had the largest number and diversity of observed local fitness maxima (Figures 8 and 9 in S2 Figure and *Sp* was strongly associated to the number of accessible genotypes (Figure 10 in S2 Figure). Of course, the number of mutations of the fitness maxima were 7 and 10 in the representable landscapes, and smaller in the local maxima and RMF landscapes (Figure 11 in S2 Figure).

The number of different sampled genotypes was comparable between detection regimes (Figure 12 in S2 Figure), but diversity differed (Figure 13 in S2 Figure), with the uniform detection regime showing generally larger sampled diversity. The mean and median number of mutations of sampled genotypes (Figures 14 and 15 in S2 Figure) differed between detection regimes, being largest in the large detection regime, and smallest in the small detection regime; the standard deviation and coefficient of variation in the number of mutations (Figures 16 and 17 in S2 Figure) were largest in the uniform detection regime (thus, the uniform detection regime showed both the largest variation in number of mutations of genotypes and the largest diversity of genotypes). Sample characteristics and the difference in sample characteristics between detection regimes were affected by type of fitness landscape (Figures 13 and 16 in S2 Figure).

### 2.8 Cancer data sets

We have used twenty-two cancer data sets (including six different cancer types: breast, glioblastoma, lung, ovarian, colorectal, and pancreatic cancer). All of these data sets, except for the breast cancer data sets BRCA_ba_s and BRCA_he_s (from Cancer Genome Atlas Research Network, 2012b), have been used previously as input for some CPM algorithms in Attolini *et al*. (2010); Caravagna *et al*. (2016); Cheng *et al*. (2012); Gerstung *et al*. (2011); Misra *et al*. (2014); Olde Loohuis *et al*. (2014); Ramazzotti *et al*. (2015), with the original sources of the data being Bamford *et al*. (2004); Brennan *et al*. (2013); Cancer Genome Atlas Research Network (2008, 2011, 2012a); Ding *et al*. (2008); Jones *et al*. (2008); Knutsen *et al*. (2005); Parsons *et al*. (2008); Piazza *et al*. (2013); Wood *et al*. (2007). Details on sources, names, and how the data were obtained and processed are provided in S5 Text.

These data sets vary in sample size (27 to 594 samples), number of features (from 7 to over 100), data types (nonsynonymous somatic mutations and copy number aberrations or both), levels of analysis (genes, modules and pathways, exclusivity groups), patterns of number of mutations per subject and frequency of mutations analyzed, and procedures for driver selection, and restriction of patient subtypes. The data sets, therefore, are a large representative ensemble of data sets to which researchers have previously applied or might apply CPMs.

We have run the CPM analyses three times per data set, limiting the number of features analyzed to the 7, 10, and 12 most common ones, so as to examine how our assessments depend on the number of features analyzed; the first two thresholds use the same number of features as the simulations. (Of course, for data sets with 7 or fewer features, there are no differences in the data sets used under the 7, 10, and 12 thresholds; ditto for data sets with 8 to 10 features with respect to thresholds 10 and 12).

We do not know the true paths of tumor progression, but we can use the bootstrap to assess the robustness or reliability of the inferences. To do so, we repeated the process above with 100 bootstrap samples (Section 1.2 in S5 Text). We computed *JS*_*o*,*b*_, the average JS between the distribution of paths to the maximum from the original data set and each of the bootstrapped samples. Large differences in the distribution of paths between the analyses with the bootstrap samples and the analysis with the original sample (i.e., large *JS*_*o*,*b*_) would suggest that the inferences are unreliable and cannot be trusted (but small differences do not indicate that the inferred paths match the distribution of the true ones).

## 3 Results

### 3.1 Predicting paths of evolution with CPMs

The CPMs used (four, two with two variants, yielding a total of six different procedures for obtaining CPMs: H-CBN, MCCBN, OT, CAPRI_AIC, CAPRI_BIC, and CAPRESE) can be divided into three groups: models that return trees (OT and CAPRESE) and two families of models that return DAGs, CBN (H-CBN and MCCBN) and CAPRI (CAPRI_AIC and CAPRI_BIC). Comparing within groups with respect to JS one member of the pair consistently outperformed the other (see Figure 1 in S6 Text). OT (using probability-weighted paths, see below) was significantly better than CAPRESE (paired *t*-test over all non-missing 56595 pairs of results: *t*_56594_ = −161.1, *P* < 0.0001), H-CBN was significantly better than MCCBN (*t*_56593_ = −42.6, *P* < 0.0001), and CAPRI_AIC was significantly better than CAPRI_BIC (*t*_56594_ = −41.9, *P* < 0.0001). In what follows, therefore, and for the sake of brevity, we will focus on OT, H-CBN, and CAPRI_AIC, since the overall performance of their alternatives is worse.

Figure 3 shows how the performance measures for OT, H-CBN, and CAPRI_AIC change with sample size for all combinations of type of landscape, detection regime, and number of genes (see Figure 2 in S6 Text for the probability of recovering the most common LOD). The measures of JS and 1-precision for OT and H-CBN (and MCCBN) use probability-weighted paths computed as explained in 2.6, because there was strong evidence for all three models that the probability-weighted paths led to better results (JS, paired *t*-test over all pairs: OT, *t*_56594_ = −195.8, *P* < 0.0001; H-CBN: *t*_56594_ = −222.3, *P* < 0.0001; MCCBN: *t*_56593_ = −149.0, *P* < 0.0001; 1-precision: OT: *t*_56594_ = −187.6, *P* < 0.0001; H-CBN: *t*_56594_ = −217.6, *P* < 0.0001; MCCBN: *t*_56593_ = −130.3, *P* < 0.0001). (See also Figures 4 to 6 in S6 Text). Overall, H-CBN was the model with the best performance (*P* < 0.0001 from all pairwise comparisons between the six procedures with Tukey’s contrasts and single-step multiple testing p-value adjustment —Hothorn *et al*., 2008— on linear mixed-effects models with landscape by split replicate as random effect). It must be noted, however, that all CPMs can show large variability in performance (Figure 7 in S6 Text).

**Figure 3:**
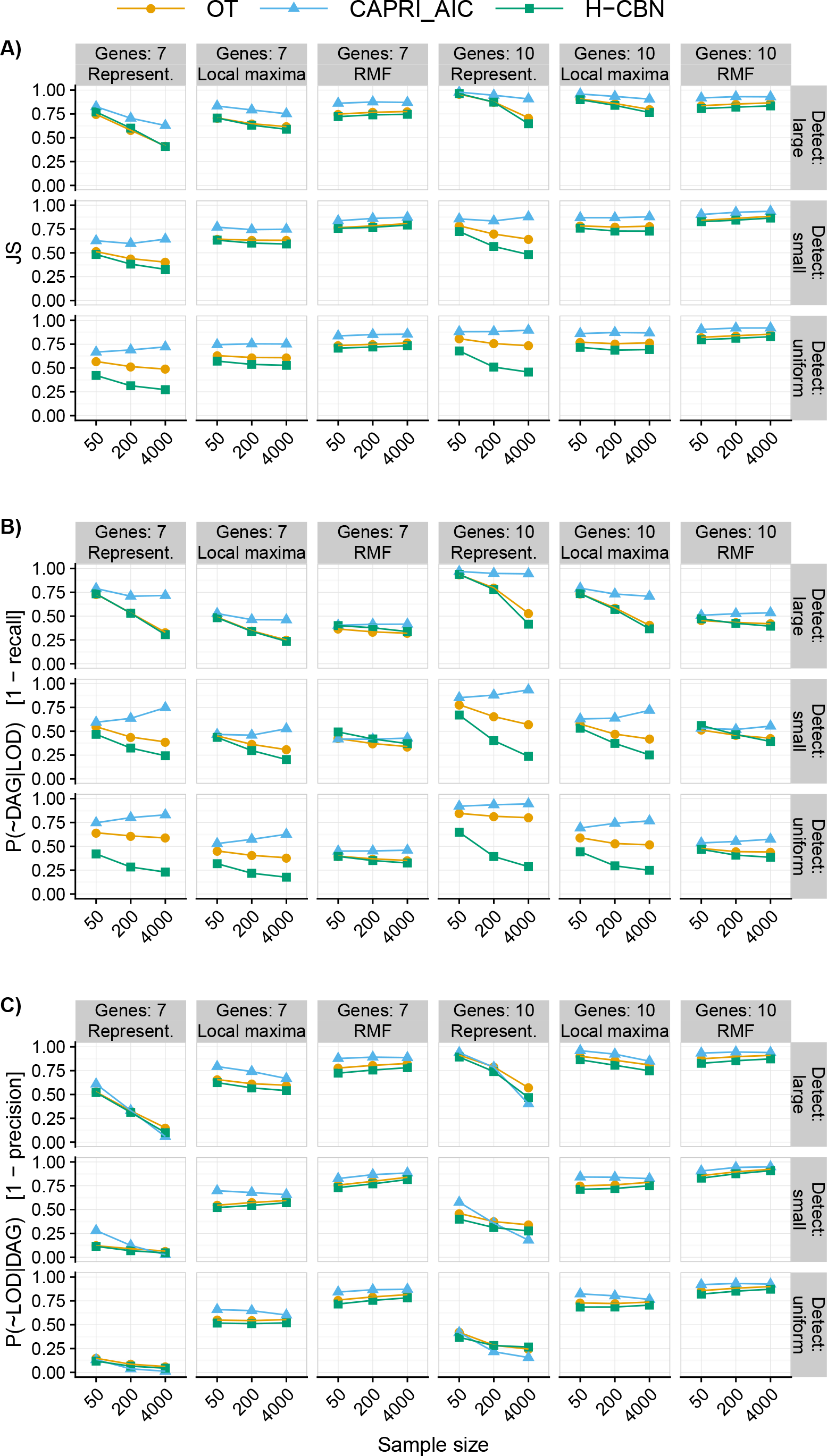
Summary performance measures (see definitions in 2.6) for OT, CAPRI_AIC, and H-CBN for all combinations of sample size, type of landscape, detection regime, and number of genes. (A) Jensen-Shannon divergence (JS); (B) 1 - recall; (C) 1 - precision. For all measures, smaller is better. For OT and H-CBN, JS (panel A) and 1-precision (panel C) use probability-weighted paths (see text). Each point represented is the average of 210 points (35 replicates of each one of the six combinations of 3 initial size by 2 mutation rate regimes —see 2.2); we are thus marginalizing over mutation rate by initial simulation size combinations. Each one of the 210 points is, itself, the average of five runs on different partitions of the simulated data. See Figure 1 in S6 Text for results for all six procedures used (four, two with two variants: H-CBN, MCCBN, OT, CAPRI_AIC, CAPRI_BIC, and CAPRESE).

JS differed between type of landscape, number of genes, detection regime, and sample size, but the magnitude and even direction of effects differed between combinations of those factors, as seen in Figures 3 and 4. Generalized linear mixed-effects models fitted to the complete data set and to the different combinations of CPM and type of landscape (see Section 11 in S6 Text) also showed highly significant (*P* < 0.0001) two-, three-, and four-way interactions between most of the factors, in particular those involving type of landscape and detection regime. Type of landscape and detection regime also had very strong effects in the variability of the estimates, with relative variabilities that could reach 20% with small sample sizes (Figure 7 in S6 Text).

**Figure 4:**
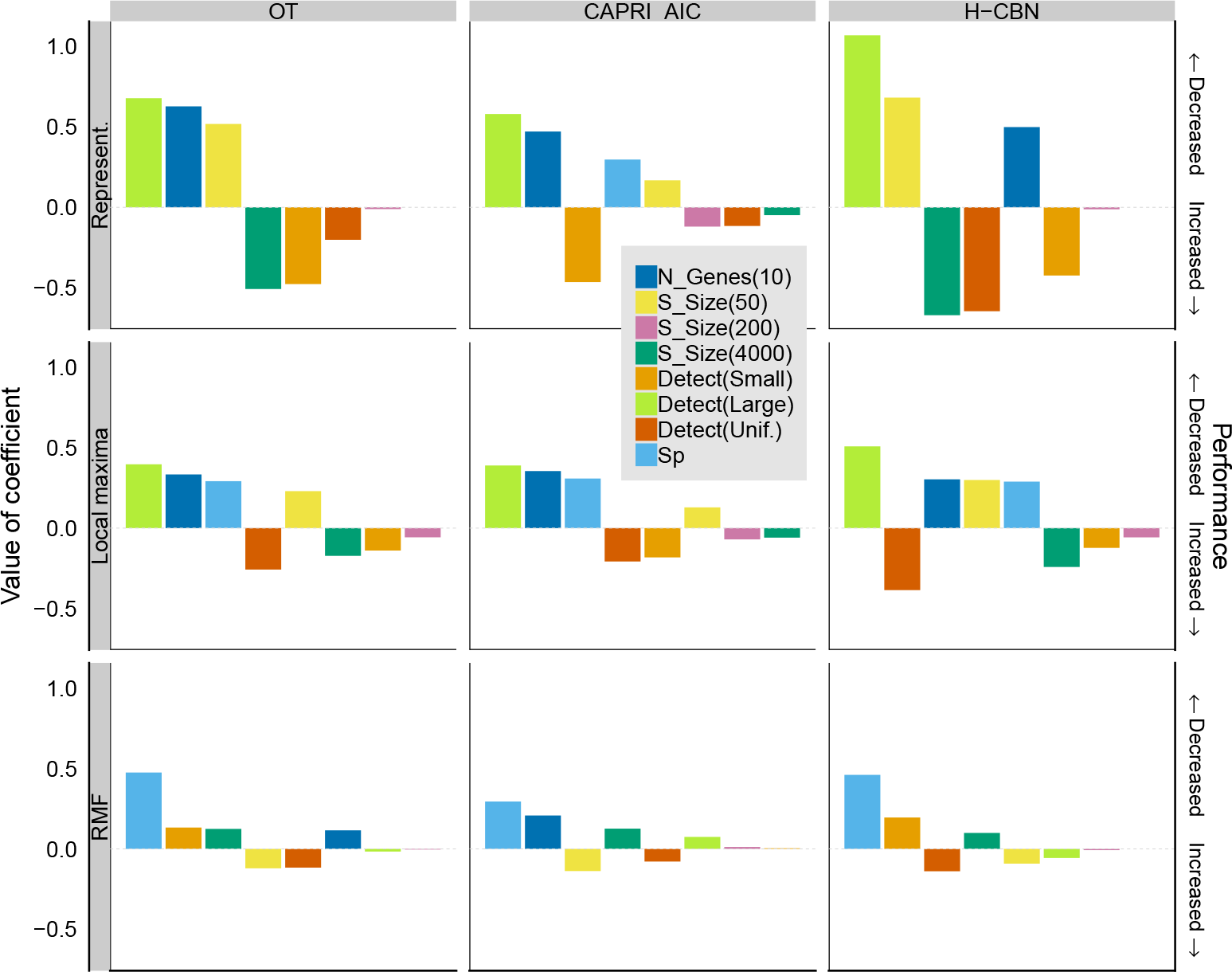
Coefficients from generalized linear mixed-effects models, for JS as dependent variable, with separate models fitted to each combination of CPM and type of fitness landscape. Co-efficients are from models with sum-to-zero contrasts (see text and Section 6 in S4 Text). Within each panel, coefficients have been ordered from left to right according to decreasing absolute value of coefficient. The dotted horizontal gray line indicates 0 (i.e. no effect). Coefficients with a large positive value indicate factors that lead to a large decrease in performance (increase in JS). Only coefficients that correspond to a term with a P-value < 0.05 in Type II Wald chi-square tests are shown. The coefficient that corresponds to Number of genes 7 is not shown (as it is minus the coefficient for 10 genes —from using sum-to-zero contrasts). “N Genes”: number of genes; “S Size”: sample size; “Detect”: detection regime; “Sp”: LOD diversity (*S*_*p*_).

Under representable fitness landscapes, performance improved with increasing sample size and with the uniform detection regime. Performance was worse in fitness landscapes of 10 genes (Figure 3, panel A; Figure 4, top row); the decrease in performance with increasing number of genes is related to CPMs both missing evolutionary paths (Figure 3B), and allowing paths that are not used by evolution (Figure 3C). With CAPRI_AIC the effect of sample size was much weaker and increases in sample size could lead to decreases in performance, specially under the uniform detection regime (highly significant, *P* < 0.0001, interactions of detection and sample size —Section 11 in S6 Text). This is attributable to CAPRI_AIC excluding many paths taken during evolution (Figure 3B). This behavior was caused by CAPRI_AIC sometimes allowing only a few or even just one path to the maximum (Figure 8 in S6 Text); see also next section.

Under the RMF landscape overall performance was worse. Increasing sample size for OT and H-CBN led to minor decreases in performance (Figure 3 and Figure 4 bottom row). CPMs failed to capture about 50% of the evolutionary paths (or fractions of paths) to the local maxima (Figure 3B) and included more than 75% of paths (or fractions of paths) that were never taken by evolution (Figure 3C). The behavior under local maxima was similar to that of representable fitness landscapes in terms of the direction of most effects, but effects were generally weaker, with the exception of evolutionary unpredictability.

Evolutionary unpredictability itself had a strong effect on performance. There were highly significant interactions (*P* < 0.0001) between evolutionary unpredictability (as measured with *S*_*p*_), detection regime, and sample size, within representable and local maxima landscapes, as well as in the overall models (Section 11 in S6 Text). In most scenarios, performance was worse with larger unpredictability (larger *S*_*p*_) as seen by the positive slopes of JS on *S*_*p*_ (Figure 5). But under representable landscapes, in the large detection regime and for sample sizes 50 and 200, larger evolutionary unpredictability was associated with better performance. Under RMF fitness landscapes, large evolutionary unpredictability was associated with poorer performance over all sample sizes. Under local maxima, the effect of evolutionary unpredictability depended strongly on sample size and detection regime, with reversal of effects from sample size of 50 compared to 4000 under the large detection regime, similar to the ones in representable landscapes.

**Figure 5:**
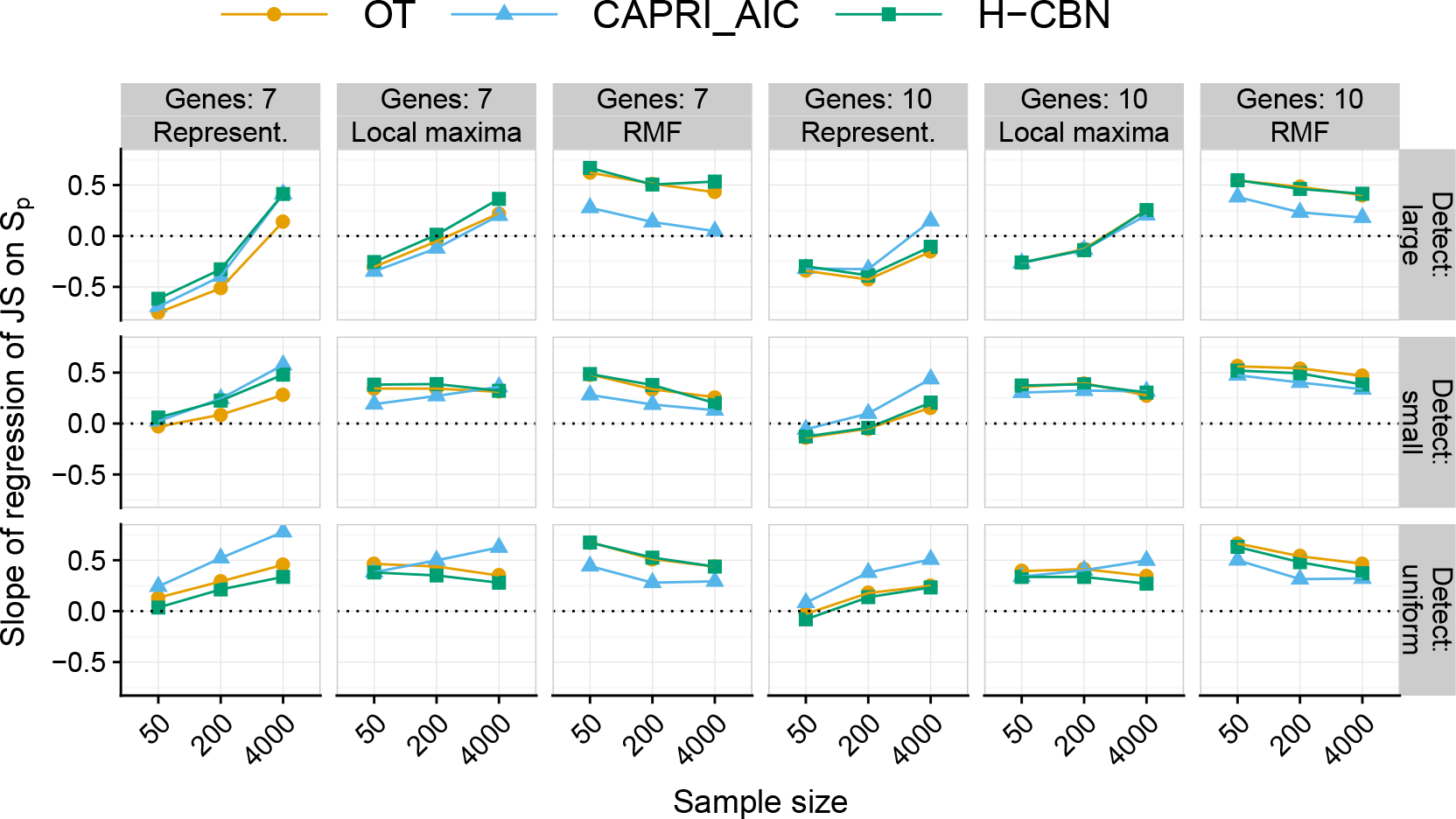
Estimated slopes of the regression of Jensen-Shannon divergence (JS) on LOD diversity (*S*_*p*_) for all combinations of sample size by type of landscape by detection regime by number of genes. A beta regression was fitted to each subset of data. Each regression was fitted to 210 points, each of which is itself the average of five replicates, one for each of the five runs on different partitions of the simulated data.

### 3.2 Inferring evolutionary unpredictability from CPMs

Figure 6A shows the ratio of inferred to true evolutionary unpredictability, *S*_*c*_/*S*_*p*_. Under representable fitness landscapes, for H-CBN this ratio remained close to 1 over all combinations of detection regime, number of genes, and sample size; the values were closest to one with sample size 4000 and under the uniform detection regime. This is in spite of large differences in the ratio of estimated number of paths to the maximum over true number of paths to the maximum (Figure 6B). This good performance is a consequence of both using probability-weighted paths by H-CBN (and OT) and of changes in scale (diversities use logarithms). Patterns for CAPRI_AIC seemed dominated by the tendency of CAPRI_AIC to only allow very few paths as the sample size grows large (see also Figure 8 in S6 Text) and were also the consequence of CAPRI_AIC’s inability to produce probability-weighted paths. For all CPMs type of landscape affected the quality of estimates: under local maxima and specially RMF the number and diversity of paths tended to be overestimated, sometimes by large factors. In summary, and regardless of fitness landscape, the estimates of evolutionary unpredictability from H-CBN (*S*_*c*_) could be used to obtain an upper bound of the true evolutionary unpredictability.

**Figure 6:**
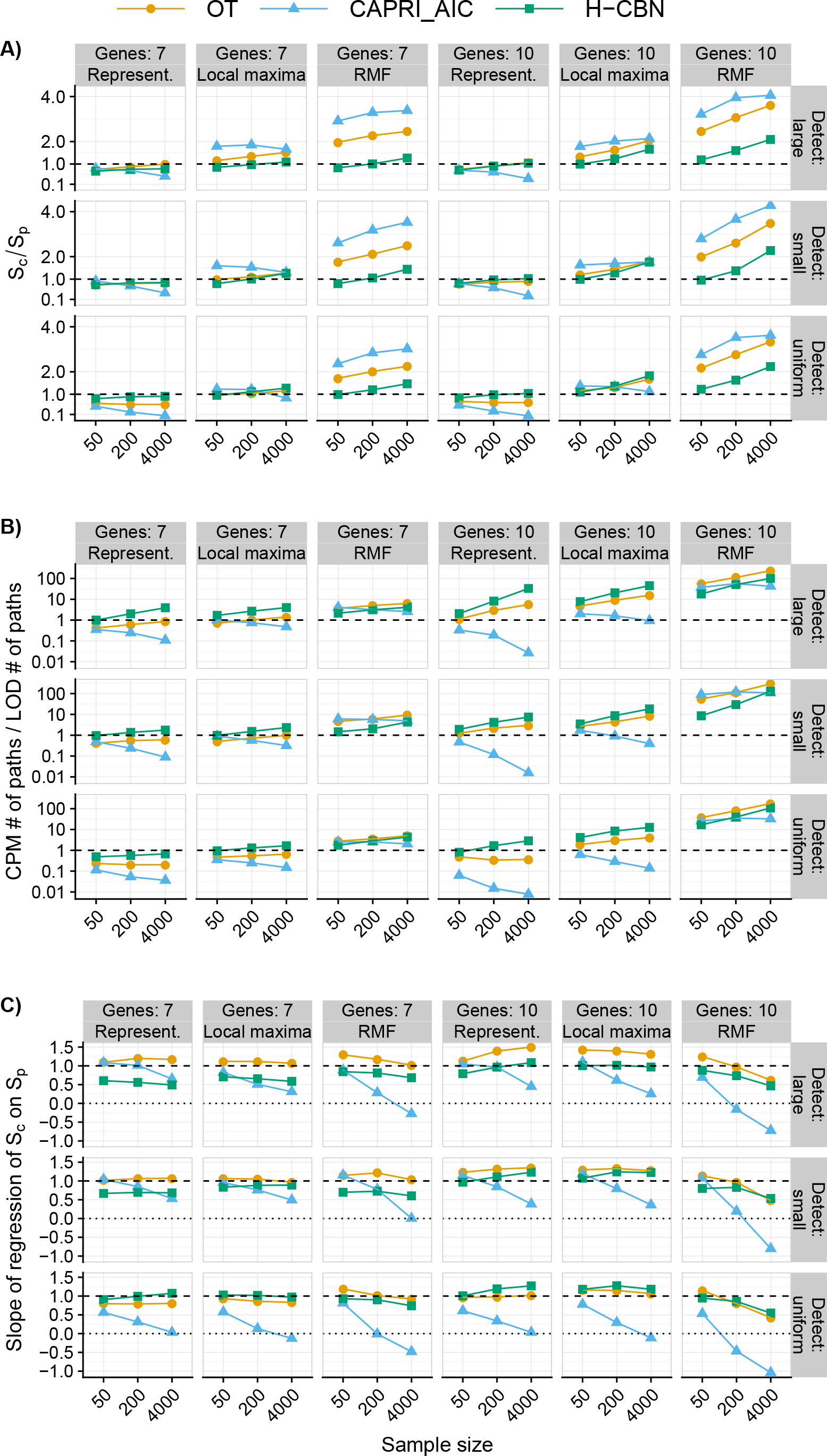
Path diversities and number of paths inferred from CPMs relative to the values from LODs. (A) Average of the ratio of diversity of paths to the maximum inferred by the CPMs (*S*_*c*_) relative to the true LOD diversity (*S*_*p*_), for all combinations of type of landscape by detection regime by number of genes by sample size. (B) Like (A), but for number of paths to the maximum from the CPMs relative to the observed number of distinct LODs As in Figure 3, each point is the average of 210 points. (C) Slope of the regression of *S*_*c*_ on *S*_*p*_; each point is thus a slope from a regression of 210 points, each of which is itself the average of 5 replicates (see Figure 5). (A) shows whether evolutionary unpredictability (*S*_*p*_) tends to be over- or under-estimated by *S*_*c*_; (C) shows how *S*_*c*_ changes with *S*_*p*_ —see Section 13 in S6 Text for an example of positive ratios with negative slopes.

And how does the estimated evolutionary unpredictability change with the true evolutionary unpredictability? Figure 6C shows that the slopes of regressions of estimated unpredictability from CPMs (*S*_*c*_) on true unpredictability (*S*_*p*_) changed depending on fitness landscape, detection regime, and sample size, including slopes over and under 1, and even inversion of signs.

### 3.3 Cancer data sets

We have used H-CBN (the best performing model in the simulations) on twenty-two cancer data sets to examine the estimated evolutionary unpredictability, and assess the reliability of the estimates. The results are shown in Figure 7 (see Figure 1 in S5 Text for ranges of bootstrap runs). Unreliability (*JS*_*o*,*b*_ —section 2.8) was large for most data sets, and very large for some of them. These results would be expected, even if the true fitness landscapes were representable ones, as most of the data sets have small sample sizes (less than 1000), and we have seen that performance is poor (large *JS*) for that range of sample sizes (Figure 3A). For these data sets there was no relationship between *JS*_*o*,*b*_ and sample size (Figure 7A), and when the same data set was analyzed using pathways/modules and genes, performance was generally better using pathways or modules (Pan_pa vs. Pan_ge, Col_pa vs. Col_g, GBM_pa vs. GBM_ge, GBM_mo_vs. GBM_CNA). Within data sets, and for all data sets, as the number of features analyzed increased performance either decreased or stayed the same (i.e., for data sets with more than 7 features, unreliability at the 10 feature threshold, 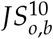, was larger or equal to unreliability at the 7 feature threshold, 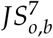; for data sets with more than 10 features, 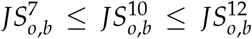: Figure 1A in S5 Text).

**Figure 7:**
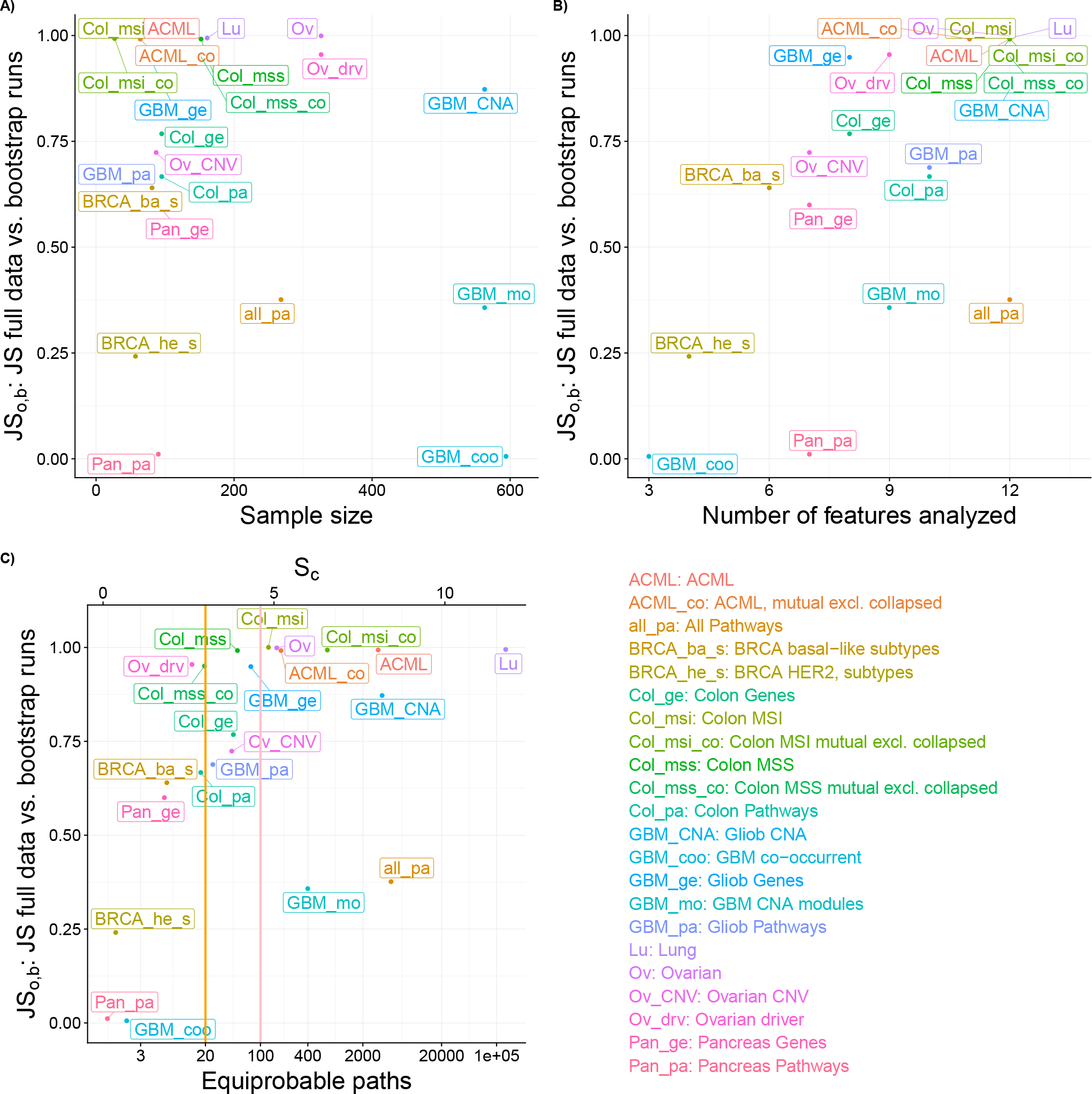
Summary patterns for *JS*_*o*,*b*_ (average JS for the full data compared to the bootstrap runs) and unpredictability for the cancer data sets. (A) *JS*_*o*,*b*_ vs. sample size of data sets. (B) *JS*_*o*,*b*_ vs. number of features analyzed for each data set. (C) *JS*_*o*,*b*_ vs. estimated evolutionary unpredictability; in the bottom x-axis, *S*_*c*_ is shown in terms of number of equiprobable paths; orange and salmon vertical lines indicate 20 and 100 equiprobable paths, respectively. All results shown are from analysis with up to 12 features. Values shown for *JS*_*ob*_ are the average of the 100 bootstrap runs; values for unpredictability (*S*_*c*_ or equiprobable paths) are from the analysis with the original, non-bootstrapped, data.

There were mild trends for an association between smaller *JS*_*o*,*b*_ and smaller numbers of features and smaller *S*_*c*_ (Figure 7B, C), with notable exceptions: the Pancreas Pathways (Pan_pa) data set had very small *JS*_*o*,*b*_ even for moderate number of features, and the All Pathways (all_pa) data set had a relatively small *JS*_*o*,*b*_ even though it used 12 features and had a large *S*_*c*_; the GBM CNA modules (GBM_mo) data set also showed moderate *JS*_*o*,*b*_ in spite of having nine features and relatively large *S*_*c*_. Conversely, some data sets with small *S*_*c*_ had extremely unreliable path predictions (e.g., BRCA_ba_s, Col_mss_co, Col_msi_co, GBM_ge).

Values for *S*_*c*_ were well within the ranges of *S*_*c*_ estimated by H-CBN for the simulated data (Figure 11 in S6 Text). Of course, *S*_*c*_ increased with number of features analyzed (see also Figure 4 in S5 Text). Given the results from section 3.2, where generally *S*_*p*_ < *S*_*c*_, this suggests that the true evolutionary unpredictability (when analyzing up to 12 features) for 13 of the data sets should be less than that corresponding to about 100 equiprobable paths to the maximum, but only eight are below the much more manageable, and useful, 20 equiprobable paths (Figure 7D). The Pan_pa, GBM_coo, and BRCA_he_s show outstanding patterns in Figure 7. Examination of the output from H-CBN revealed that there was one single path with estimated probability > 0.97 for Pan_pa, and two paths to the maximum of about equal probability that together added > 0.95 for GBM_coo. BRCA he s had only four features but mutations in SRPRA and PIK3R1 were present each in only four individuals (different individuals for the two mutations); repeated runs of H-CBN led to different sets of restrictions being inferred which, because there are few paths to the maximum, and some had large probabilities (> 0.5), resulted in large differences in JS statistic between runs.

## 4 Discussion

Can we predict the likely course of tumor progression using CPMs? We have examined the performance of six different procedures for obtaining CPMs (four CPMs, two of them with two variants: H-CBN and MCCBN, OT, CAPRI_AIC and CAPRI_BIC, and CAPRESE). H-CBN was the best performing CPM in our study. Using H-CBN under the representable fitness landscapes (the scenarios that agree with CPMs’ assumptions) returned estimates of the probability of paths of tumor evolution that were not far from the true distribution of paths of evolution (Figure 3A) when sample size was very large. But we find that, even under representable fitness landscapes, performance with moderate (and more realistic) sample sizes was considerably worse and was affected by detection regime. The analysis of the twenty-two cancer data sets revealed that performance (as measured by *JS*_*o*,*b*_, an indicator of unreliability of inferences) was poor or very poor for most data sets. Even data sets with few features and small estimated diversity of paths to the maximum, *S*_*c*_, showed very unreliable predictions.

What factors, and how, affect performance? Under representable fitness landscapes, performance on simulated data was of course affected by the number of features, the dimension of the fitness landscape: JS was worse with 10 than with 7 genes (Figures 3, 4). Increasing sample size improved performance (Figures 3). Detection regime and evolutionary unpredictability, as measured by LOD diversity (*S*_*p*_), affected individually and jointly all performance measures (Figures 3, 5). Increased evolutionary unpredictability hurt performance under most conditions (Figure 5). Detection regime was a key determinant of performance, as already found in previous work (Diaz-Uriarte, 2015, 2018); performance was better under the uniform detection regime and, more importantly, detection regime affected how the rest of the factors (evolutionary unpredictability, sample size, and number of features) impacted on performance (Figures 3 to 5).

The analysis of the twenty-two cancer data sets also indicated number of features as a major determinant of performance. Across data sets, unreliability of inferences (*JS*_*o*,*b*_) increased with number of features (Figure 7). Within data set unreliability also increased as the number of features increased (Figure 1 in S5 Text; note that an increase in the number of features analyzed leads to an increase in the number of features with low frequency events). Interestingly, the driver-selected data sets (Col_mss, Col_msi, BRCA_he_s, BRCA_ba_s) did not perform much better than data sets with a simple frequency-based selection of features (e.g., Lu, Ov, or comparison Ov with Ov_drv). Even data sets with very careful, manually-curated selection of drivers and “exclusivity groups” and where variability due to subtypes has been minimized (Col_msi, Col_msi_co, Col_mss, Col_mss_co, ACML_co, BRCA_he_s and BRCA_ba_s) showed very large *JS*_*o*,*b*_. And BRCA_he_s, with only four features, showed much larger *JS*_*o*,*b*_ than GBM_coo and Pan_pa (with 3 and 7 features, respectively), due to the presence of two low frequency alterations.

These results bring forth the problem of the selection of the relevant features for analysis (Caravagna *et al*., 2016; Cristea *et al*., 2016; Gerstung *et al*., 2011) and whether sample size is large enough relative to the number and frequency of features considered. We have previously shown that feature selection can have a very detrimental impact on the performance of CPMs (Diaz-Uriarte, 2015). Using pathways instead of genes in the analyses (e.g., Cristea *et al*., 2016; Raphael and Vandin, 2015) can alleviate some of the problems of feature selection. Data sets coded as pathways or modules generally reduced the presence of low-frequency alterations (Figures 2 and 3 in S5 Text). Pathways can also improve predictability and how close the estimates of path distributions are to the truth because they are more similar to heritable phenotypes, which often have smoother phenotype-fitness maps and tend to show more repeatable evolution (Lässig *et al*., 2017; see also Wang *et al*., 2015, but also Chebib and Guillaume, 2017; Sailer and Harms, 2017). Gerstung *et al*. (2011) found that analysis using pathways gave stronger evidence for order constraints than analysis using genes, and we also see in Figure 7 that both *S*_*c*_ and *JS*_*o*,*b*_ tend to decrease if we use pathways or modules (Pan_pa vs. Pan_ge, Col_pa vs. Col_g, GBM_ge vs. GBM_pa, GBM_mo vs. GBM_CNA). Using so-called “exclusivity groups” (*sensu* Caravagna *et al*., 2016) to identify “fitness equivalent alterations” is a similar, though not identical, procedure that in this paper showed only modest improvements in *JS*_*o*,*b*_ (Col_mss_co vs. Col_mss, Col_msi_co vs. Col_msi, ACML_co vs. ACML). This can of course be due to particularities of these data sets (e.g., large number of features relative to number of subjects) or the intrinsic difficulties of identifying true fitness equivalent groups via “hard/soft exclusivities”. However, although analysis using pathways/modules/exclusivity groups might lead to more reliable results from the predictability point of view, the identification of paths at the gene level is still the ultimate goal for therapeutic interventions (see Ashworth *et al*., 2011). Regardless of the details of the procedure for collapsing and reducing features, our results suggest that further work on feature selection should consider reduction of variability of estimates of evolutionary paths as a key component.

Hosseini (2018) has reanalized the DAG-derived representable and a subset (those where the fully mutated genotype has the largest fitness) of the DAG-derived non-representable fitness landscapes in Diaz-Uriarte (2018). He finds good agreement between the distributions of paths to the maximum from H-CBN and the fitness landscape-based probability distribution of paths to the maximum computed assuming SSWM. Our results for H-CBN under the best conditions are not as optimistic. Two differences in the studies explain the differences. First, Hosseini (2018) computes the fitness landscape-based probability of paths assuming a SSWM regime and restricting the analysis to fitness landscapes where the fully mutated genotype has the largest fitness, while our analyses directly examine the distribution of the paths to the maximum in each simulation (LODs), without restricting the evolutionary regime and the fitness landscapes; second, he uses H-CBN with the very large sample size of 20000 (the full data sets in Diaz-Uriarte, 2018), while we use a more realistic range of sample sizes.

Even very good performance, though, needs to be interpreted with care. Very good performance simply tells us that the true and estimated probability distributions of the paths to the maximum agree closely. If the true evolutionary unpredictability is large, then for practical purposes our capacity to predict what will happen (in the sense of providing a small set of likely outcomes) is very limited. Ranges of diversities of 3.2 to 6.0, equivalent to 25 to 400 equiprobable paths, were common in the simulated data (Figure 11 in S6 Text) and are comparable to the ranges in most cancer data sets with 7 and 10 feature thresholds (Figure 1 in S5 Text). The inability to narrow down the likely paths to a small set of paths in these cases is, of course, not a limitation of the CPMs, but a problem inherent to the unpredictability of the evolutionary process in many scenarios, which could severely limit the usefulness of even perfect predictions.

The discussion above has centered on representable fitness landscapes. As argued before, fitness landscapes with local fitness maxima are probably common in cancer. With local fitness maxima, achieving good recall involves the relatively easier task of getting right the first part of short paths to the maximum. But good recall was more than offset by low precision and overall predictability was very poor. In fitness landscapes with local maxima, CPMs are fitting models with paths of tumor progression that extend beyond the true end point of the progression. In RMF fitness landscapes, in addition to local peaks, not even the set of accessible genotypes can be represented by DAGs of restrictions (see Diaz-Uriarte, 2018, and Figure 1). The violations of assumptions in RMF and local maxima fitness landscapes explain the decreases in the relevance of sampling regime and why increasing sample size has negligible (or even detrimental) effects in these fitness landscapes (Figures 3 and 4). Remarkably, regardless of type of fitness landscape (i.e., even under violation of assumptions), and for the two tasks considered (prediction of paths and estimating unpredictability) performance of CPMs that could return probability-weighted paths (H-CBN, MCCBN, OT) was better when using probability-weighted paths; thus, further improvement in these CPMs, even under violations of assumptions, might be possible by recalibrating their output.

And we return to our second original question, as even if achieving good performance in predicting the paths of tumor progression is unlikely, inferring evolutionary unpredictability could be an easier task. Can we use inferences of evolutionary unpredictability from CPMs as estimates of the true evolutionary unpredictability? Under representable fitness landscapes, H-CBN, the best performing model also for this task (Figure 6B), returned values of *S*_*c*_ very similar to *S*_*p*_, the evolutionary unpredictability estimated from the diversity of paths, and this held over detection regimes and sample sizes. Hosseini (2018) also finds that the estimates of predictability from H-CBN correlate well with the fitness landscape-based evolutionary predictability (estimated assuming SSWM in fitness landscapes where the fully mutated genotype has largest fitness), with slopes of the regression of CPM-based on landscape-based predictability generally slightly below 1, similar to our findings (Figure 6C). These good results do not hold under the other two types of two fitness landscapes that we analyzed: evolutionary unpredictability is overestimated, and increasing sample sizes made the problems worse and different evolutionary scenarios, sample sizes, and detection regimes have different relationships of estimated and true unpredictability (Figure 6C). But our results indicate that we can use H-CBN to set upper bounds on the true *S*_*p*_; obtaining tighter estimates is an objective for further research to explore. And here our analysis of twenty-two cancer data sets suggests that the true evolutionary unpredictability of at least some cancer scenarios might be reasonably small, specially if *S*_*c*_ is overestimating the true unpredictability.

### 4.1 Conclusion

The answer to the question “can we predict the likely course of tumor progression using CPMs?” is, unfortunately, at least for the models examined, “only with moderate success and only under representable fitness landscapes and with very large sample sizes; but even perfect predictions might be of little use if evolutionary unpredictability is large”. Estimating upper bounds to evolutionary unpredictability is a more modest, though more likely to succeed, use of CPMs. Promisingly, several cancer data sets showed low evolutionary unpredictability. There are three key difficulties for successful prediction: the sheer size of the problem even for moderate numbers of genes, the intrinsic evolutionary unpredictability in many scenarios, and the deviations from the assumptions of CPMs that are likely to hold in most cancer data. Further methodological work to allow CPMs to deal with rugged, multi-peaked, fitness landscapes could improve their usefulness to predict tumor evolution. In addition to the caveat about using these models under scenarios where performance is very poor, this paper raises the general question of what can we really predict about likely paths of tumor progression from cross-sectional data, for instance to guide therapeutic interventions. At a minimum, measures such as *JS*_*o*,*b*_ and *S*_*c*_ with CPMs that return probability-weighted paths should probably become routine as ways of providing a sense of the reliability of predictions and for assessing whether the predictions could be of any practical use.

## Supporting information

S1_Dataset.zip (First part of split zip file)

S2_Dataset.zip (Second part of split zip file: You must rename to S1_Dataset.z01 to uncompress)

## 5 Acknowledgments

N. Beerenwinkel, S. Posada-Céspedes, and G. Caravagna for discussion about progression models or software; S.-R. Hosseini for providing a preprint of his MSc. thesis and for comments that helped us clarify our methods. C. Lázaro-Perea and two anonymous reviewers for comments on the ms.

## 6 Funding

Supported by BFU2015-67302-R (MINECO/FEDER, EU) to RDU. CV supported by PEJD-2016-BMD-2116 from Comunidad de Madrid.

## 7 Data and code availability

All data for this article, as well as source code, is available from the Supporting Information (S7 Text, S1 Dataset, S2 Dataset).

## 9 Supporting information captions

**S1 Figure. Plots of simulated fitness landscapes and fitness graphs.** Plots of the 1260 fitness landscapes (and corresponding fitness graphs) used.

**S2 Figure. Simulated fitness landscapes and fitness graphs: characteristics, evolutionary unpredictability, clonal interference, and sampled genotypes.**

**S1 Text. Differences in fitness landscapes, simulations, methods, and objectives, with Diaz-Uriarte, 2018.**

**S2 Text. Generating random fitness landscapes.**

**S3 Text. Evolutionary simulations.** Runs until fixation; detection regimes and sampling; other parameters of the simulations; number of genes used; LODs through non-accessible genotypes, LODs that go beyond a local maximum, and moving through fitness valleys.

**S4 Text. CPMs: software, probabilities of paths, statistics of performance, linear models.**

**S5 Text. Cancer data sets: sources, characteristics, additional results.**

**S6 Text. Additional results.**

**S7 Text. Data and code availability.**

**S1 Dataset. Compressed file with data and code.** This is the first of a two-part zip file (made up of files S1_Dataset.zip and S2_Dataset.z01). See instructions in S7 Text (briefly: rename S2_Dataset.z01 to S1_Dataset.z01 and uncompress the split archive.)

**S2 Dataset. Compressed file with data and code.** This is the second of a two-part zip file (made up of files S1_Dataset.zip and S2_Dataset.z01). See instructions in S7 Text (briefly: rename S2_Dataset.z01 to S1_Dataset.z01 and uncompress the split archive.)

## References

Ashworth, A., Lord, C.J. and Reis-Filho, J.S. (2011). Genetic interactions in cancer progression and treatment. Cell, 145(1), 30–38.

Attolini, C. et al (2010). A mathematical framework to determine the temporal sequence of somatic genetic events in cancer. Proceedings of the National Academy of Sciences, 107(41), 17604–17609.

Bamford, S. et al (2004). The COSMIC (Catalogue of Somatic Mutations in Cancer) database and website. Br J Cancer, 91(2), 355–358.

Bank, C. et al (2016). On the (un)predictability of a large intragenic fitness landscape. PNAS, 113(49), 14085–14090.

Beerenwinkel, N., Eriksson, N. and Sturmfels, B. (2006). Evolution on distributive lattices. Journal of Theoretical Biology, 242(2), 409–420.

Beerenwinkel, N. et al (2007). Genetic progression and the waiting time to cancer. PLoS computational biology, 3(11), e225.

Beerenwinkel, N. et al (2015). Cancer evolution: Mathematical models and computational inference. Systematic Biology, 64(1), e1–e25.

Beerenwinkel, N., Greenman, C.D. and Lagergren, J. (2016). Computational Cancer Biology: An Evolutionary Perspective. PLoS Comput. Biol., 12(2), e1004717.

Beijersbergen, R.L., Wessels, L.F.A. and Bernards, R. (2017). Synthetic Lethality in Cancer Therapeutics. Annual Review of Cancer Biology, 1(1), 141–161.

Blomen, V.A. et al (2015). Gene essentiality and synthetic lethality in haploid human cells. Science, 350(6264), 1092–1096.

Bozic, I. et al (2010). Accumulation of driver and passenger mutations during tumor progression. Proceedings of the National Academy of Sciences of the United States of America, 107, 18545–18550.

Brennan, C.W. et al (2013). The Somatic Genomic Landscape of Glioblastoma. Cell, 155(2), 462–477.

Brooks, M.E. et al (2017). glmmTMB Balances Speed and Flexibility Among Packages for Zeroinflated Generalized Linear Mixed Modeling. The R Journal, 9(2), 378–400.

Brouillet, S. et al (2015). MAGELLAN: A tool to explore small fitness landscapes. bioRxiv, page 031583.

Cancer Genome Atlas Research Network (2008). Comprehensive genomic characterization defines human glioblastoma genes and core pathways. Nature, 455(7216), 1061–1068.

Cancer Genome Atlas Research Network (2011). Integrated genomic analyses of ovarian carcinoma. Nature, 474(7353), 609–615.

Cancer Genome Atlas Research Network (2012a). Comprehensive molecular characterization of human colon and rectal cancer. Nature, 487(7407), 330–337.

Cancer Genome Atlas Research Network (2012b). Comprehensive molecular portraits of human breast tumours. Nature, 490(7418), 61–70.

Caravagna, G. et al (2016). Algorithmic methods to infer the evolutionary trajectories in cancer progression. PNAS, 113(28), E4025–E4034.

Chebib, J. and Guillaume, F. (2017). What affects the predictability of evolutionary constraints using a G-matrix? The relative effects of modular pleiotropy and mutational correlation. Evolution, 71(10), 2298–2312.

Cheng, Y.K. et al (2012). A mathematical methodology for determining the temporal order of pathway alterations arising during gliomagenesis. PLoS computational biology, 8(1), e1002337.

Chiotti, K.E. et al (2014). The Valley-of-Death: Reciprocal sign epistasis constrains adaptive trajectories in a constant, nutrient limiting environment. Genomics, 104(6, Part A), 431–437.

Cristea, S., Kuipers, J. and Beerenwinkel, N. (2016). pathTiMEx: Joint Inference of Mutually Exclusive Cancer Pathways and Their Progression Dynamics. Journal of Computational Biology.

Crona, K., Greene, D. and Barlow, M. (2013). The peaks and geometry of fitness landscapes. Journal of Theoretical Biology, 317, 1–10.

Crooks, G.E. (2017). On measures of entropy and information. Technical report.

de Visser, J.A.G.M. and Krug, J. (2014). Empirical fitness landscapes and the predictability of evolution. Nat Rev Genet, 15(7), 480–490.

Desper, R. et al (1999). Inferring tree models for oncogenesis from comparative genome hybridization data. J Comput Biol, 6(1), 37–51.

Diaz-Uriarte, R. (2015). Identifying restrictions in the order of accumulation of mutations during tumor progression: Effects of passengers, evolutionary models, and sampling. BMC Bioinformatics, 16(41).

Diaz-Uriarte, R. (2017). OncoSimulR: Genetic simulation with arbitrary epistasis and mutator genes in asexual populations. Bioinformatics, 33(12), 1898–1899.

Diaz-Uriarte, R. (2018). Cancer progression models and fitness landscapes: A many-to-many relationship. Bioinformatics, 34(5), 836–844.

Ding, L. et al (2008). Somatic mutations affect key pathways in lung adenocarcinoma. Nature, 455(7216), 1069–1075.

Ferrari, S. and Cribari-Neto, F. (2004). Beta Regression for Modelling Rates and Proportions. Journal of Applied Statistics, 31(7), 799–815.

Fox, J. and Weisberg, S. (2011). An R Companion to Applied Regression, 2nd Ed. Sage, Thousand Oaks, CA.

Franke, J. et al (2011). Evolutionary Accessibility of Mutational Pathways. PLoS Comput Biol, 7(8), e1002134.

Gerstung, M. and Beerenwinkel, N. (2010). Waiting time models of cancer progression. Mathematical Population Studies, 17, 115–135.

Gerstung, M. et al (2009). Quantifying cancer progression with conjunctive Bayesian networks. Bioinformatics (Oxford, England), 25(21), 2809–2815.

Gerstung, M. et al (2011). The Temporal Order of Genetic and Pathway Alterations in Tumorigenesis. PLoS ONE, 6(11), e27136.

Greaves, M. (2015). Evolutionary Determinants of Cancer. Cancer Discovery, 5(8), 806–820.

Grün, B., Kosmidis, I. and Zeileis, A. (2012). Extended Beta Regression in *R*: Shaken, Stirred, Mixed, and Partitioned. Journal of Statistical Software, 48(11).

Hosseini, S.R. (2018). Quantifying the predictability of cancer progression using Conjunctive Bayesian Networks. M.Sc. Thesis, Swiss Federal Institute of Technology, Zürich.

Hothorn, T., Bretz, F. and Westfall, P. (2008). Simultaneous inference in general parametric models. Biom J, 50(3), 346–363.

Jones, S. et al (2008). Core signaling pathways in human pancreatic cancers revealed by global genomic analyses. Science (New York, N. Y.), 321(5897), 1801–6.

Knutsen, T. et al (2005). The Interactive Online SKY/M-FISH & CGH Database and the Entrez Cancer Chromosomes Search Database: Linkage of Chromosomal Aberrations with the Genome Sequence. Genes, Chromosomes and Cancer, 44(1), 52–64.

Lässig, M., Mustonen, V. and Walczak, A.M. (2017). Predicting evolution. Nature Ecology & Evolution, 1(3), s41559–017–0077–017.

Lin, J. (1991). Divergence measures based on the Shannon entropy. IEEE Transactions on Information theory, 37(1), 145–151.

Lipinski, K.A. et al (2016). Cancer Evolution and the Limits of Predictability in Precision Cancer Medicine. Trends in Cancer, 2(1), 49–63.

Losos, J.B. (2018). Improbable Destinies: Fate, Chance, and the Future of Evolution. Riverhead Books, S.l.

McCullagh, P. and Nelder, J. (1989). Generalized Linear Models, 2nd Ed. Chapman and Hall/CRC, London.

McFarland, C.D. et al (2013). Impact of deleterious passenger mutations on cancer progression. Proceedings of the National Academy of Sciences of the United States of America, 110(8), 2910–5.

McPherson, A.W., Chan, F.C. and Shah, S.P. (2018). Observing Clonal Dynamics across Spatiotemporal Axes: A Prelude to Quantitative Fitness Models for Cancer. Cold Spring Harb Perspect Med, 8(2), a029603.

Misra, N., Szczurek, E. and Vingron, M. (2014). Inferring the paths of somatic evolution in cancer. Bioinformatics (Oxford, England), 30(17), 2456–2463.

Montazeri, H. et al (2016). Large-scale inference of conjunctive Bayesian networks. Bioinformatics, 32(17), i727–i735.

Neidhart, J., Szendro, I.G. and Krug, J. (2014). Adaptation in Tunably Rugged Fitness Landscapes: The Rough Mount Fuji Model. Genetics, 198(2), 699–721.

Nowak, M.A. et al (2004). Evolutionary dynamics of tumor suppressor gene inactivation. PNAS, 101(29), 10635–10638.

Olde Loohuis, L. et al (2014). Inferring Tree Causal Models of Cancer Progression with Probability Raising. PLOS ONE, 9(10), e108358.

O’Neil, N.J., Bailey, M.L. and Hieter, P. (2017). Synthetic lethality and cancer. Nat Rev Genet, 18(10), 613–623.

Palmer, A.C. and Kishony, R. (2013). Understanding, predicting and manipulating the genotypic evolution of antibiotic resistance. Nature Reviews Genetics, 14(4), 243–248.

Parsons, D.W. et al (2008). An Integrated Genomic Analysis of Human Glioblastoma Multiforme. Science, 321(5897), 1807–1812.

Piazza, R. et al (2013). Recurrent SETBP1 mutations in atypical chronic myeloid leukemia. Nature Genetics, 45(1), 18–24.

R Core Team (2018). R: A Language and Environment for Statistical Computing. Vienna, Austria.

Ramazzotti, D. et al (2015). CAPRI: Efficient inference of cancer progression models from cross-sectional data. Bioinformatics, 31(18), 3016–3026.

Raphael, B.J. and Vandin, F. (2015). Simultaneous Inference of Cancer Pathways and Tumor Progression from CrossSectional Mutation Data. Journal of Computational Biology, 22(00), 250–264.

Sailer, Z.R. and Harms, M.J. (2017). Molecular ensembles make evolution unpredictable. PNAS, 114(45), 11938–11943.

Smithson, M. and Verkuilen, J. (2006). A better lemon squeezer? Maximum-likelihood regression with beta-distributed dependent variables. Psychological methods, 11(1), 54–71.

Sniegowski, P.D. and Gerrish, P.J. (2010). Beneficial mutations and the dynamics of adaptation in asexual populations. Philosophical Transactions of the Royal Society of London B: Biological Sciences, 365(1544), 1255–1263.

Szabo, A. and Boucher, K.M. (2008). Oncogenetic trees. In W.-Y. Tan and L. Hanin, editors, Handbook of Cancer Models with Applications, pages 1–24. World Scientific.

Szendro, I.G. et al (2013). Predictability of evolution depends nonmonotonically on population size. PNAS, 110(2), 571–576.

Tomasetti, C. et al (2015). Only three driver gene mutations are required for the development of lung and colorectal cancers. PNAS, 112(1), 118–123.

Toprak, E. et al (2012). Evolutionary paths to antibiotic resistance under dynamically sustained drug selection. Nature Genetics, 44(1), 101–105.

Wang, E. et al (2015). Predictive genomics: A cancer hallmark network framework for predicting tumor clinical phenotypes using genome sequencing data. Seminars in Cancer Biology, 30, 4–12.

Williams, M. J. et al (2018). Quantification of subclonal selection in cancer from bulk sequencing data. Nature Genetics, 50(6), 895–903.

Wodarz, D. and Komarova, N.L. (2014). Dynamics of Cancer: Mathematical Foundations of Oncology.

Wood, L.D. et al (2007). The Genomic Landscapes of Human Breast and Colorectal Cancers. Science, 318(5853), 1108–1113.

