## Supplementary material for "Every which way? On predicting tumor evolution using cancer progression models": S1_Text

### Supporting Information for "Every which way? On predicting tumor evolution using cancer progression models"

2019-04-17

#### S1 Text. Differences in fitness landscapes, simulations, methods, and objectives, with Diaz-Uriarte, 2018

Here we list the key differences between the study in Diaz-Uriarte, R., 2018. "Cancer progression models and fitness landscapes: A many-to- many relationship.", *Bioinformatics*, 34(5):836–844, https://academic.oup.com/bioinformatics/article/34/5/836/4557185, (henceforth RDU18), and the current paper in terms of objectives and methods.

1. **Objectives**

   In RDU18 we focused on predicting genotypes that can and cannot be observed. Here, we focus on predicting paths of tumor progression and estimating evolutionary unpredictability. The questions addressed are, thus, completely different.

   In RDU18 the main source of violations of the CPM assumptions were related to reciprocal sign epistasis. Here, we focus on scenarios with local maxima, not simply reciprocal sign epistasis (see also below).

   (The paper in RDU18 also addressed questions about variability in inferred DAGs of restrictions; these are not relevant, *per se*, in the current paper, as the focus are not DAGs of restrictions as such, but predicting paths of tumor progression. Note, however, that information about variability of the estimated paths and JS divergences is available from Figures 7 and 10, S6 Text).

2. **Methods**: because the objectives are entirely different, so are the methods.

   (a) **Fitness landscapes used**: the focus on RDU18 was reciprocal sign epistasis. The focus here are local fitness maxima. They are related, but are not the same. As we explain in the ms., the "local maxima" fitness landscapes we use here are different from the fitness maxima with reciprocal sign epistasis in RDU18. In fact, all fitness landscapes are different:

      i. **Representable fitness landscapes**: the representable fitness landscapes are generated by a different procedure. Here we focus on respecting the assumption of accessibility and lack of fitness maxima. Thus, as explained in the main manuscript, the birth rate of each descendant genotype was set equal to the maximum fitness of its parent genotypes times a random uniform variate between 1.01 and 1.19. This is done on purpose, so as not to assume any particular model for (lack of) positive/negative epistasis: we simply focus on the key

*, http://ligarto.org/rdiaz

structural features, which are captured via sign/reciprocal sign epistasis (recall that positive/negative/magnitude epistasis are all susceptible to change under monotonic transformations, such as taking the log of birth rates, which is not the case for the structural types, such as sign or reciprocal sign epistasis). Of course, these can no longer be interpreted as per-gene lambdas; again, this is on purpose.

In RDU18 the fitness of children is assigned by using a multiplicative fitness model of the effects of genes (with restrictions satisfied); see Section 2.1, "DAG-derived, representable fitness landscapes" of the Supplementary Material of RDU18.

ii. **Local maxima vs. non-representable fitness landscapes**: as mentioned above, here we focused on creating local maxima. In RDU18 we generated reciprocal sign epistasis by creating synthetic lethals. The local maxima fitness landscapes in the current ms. have no synthetic lethals. As explained in the ms., for the local maxima fitness landscapes the DAG of restrictions and the fitness landscape agree on the genotypes that should and should not be accessible (this was not the case in the non-representable landscapes in RDU18). Here we want to isolate the effect of multi-peaked landscapes or local maxima (or, equivalently, missing paths), without the additional burden for the CPMs of missing genotypes.

iii. **Rough Mount Fuji**: all fitness landscapes were generated anew for this ms.

(b) **Number of genes used**: we consider here scenarios with 7 and 10 genes. Only 7 were used in RDU18.

(c) **Simulations run until fixation**: a major difference here is that simulations in the current ms. are all run until fixation (as we need to record the true, complete, Line of Descent); in contrast, in RDU18 simulations were stopped at sampling (which was adequate for that study, which did not focus on complete tumor trajectories, and thus did not need to follow simulations until fixation).

(d) **Sampling**: the detection regime mechanism is completely different. In the current ms. we sample, with three different regimes, from the completed simulation (the simulation that has run its course until fixation). As explained above, in RDU18 we sampled (and stopped the simulation) while the simulation was running with a mechanism that tended to be enriched in either large- or small-sized tumors. This also highlights another difference: in the current ms. three different detection regimes are used (whereas in RDU18 two were used).

In addition, in RDU18, a bulk sequencing-like procedure was used (see section "5.1 Stopping the simulations: detection" in the Supplementary Material of RDU18), in contrast to the single-cell sampling used in the current ms. (which is a more appropriate procedure for the objective of reconstructing actual paths of tumor progression and comparing with Lines of Descent).

Finally, in RDU18 all analyses use only a single sample size of 1000 in contrast to the 50, 200, and 400 used here.

(e) **Simulations: other parameters**: in RDU18 all simulations were run with **initial population** size of 2000. Here we use 2000, 50000, and 1000000 cells as initial population sizes to provide variability in evolutionary predictability. The **mutation rates** used between the two studies also differ (in particular in the variable mutation rate, as in the current paper we are interested in creating variability in evolutionary predictability).

(f) **Measures of performance**: our focus in the current ms. are paths of tumor progression. We compare against Lines of Descent, and that has also required us to: a) obtain paths of tumor progression from CPMs; b) more importantly, develop new methods to compare these different graphs, as detailed in the ms. (see section 2.6) and "Computing probabilities of paths" in S4 Text .

In RDU18 what were compared where the genotypes predicted to exist under the CPM with the genotypes actually observed during the evolutionary runs.

3. **CPMs compared are different**: in RDU18 we used only CBN (H-CBN in the current ms.) and CAPRI (CAPRI_BIC in the current paper). In the current ms. we have added: MCCBN, OT, CAPRESE, CAPRI_AIC.
