## Supplementary material for "Every which way? On predicting tumor evolution using cancer progression models": S2_Text

### Supporting Information for "Every which way? On predicting tumor evolution using cancer progression models"

2019-04-17

### S2 Text. Generating random fitness landscapes

---

#### Contents

1 Representable fitness landscapes 1

2 Rough Mount Fuji 2

3 Local maxima without reciprocal sign epistasis? 2

4 References 3

---

#### 1. Representable fitness landscapes

For the representable and local maxima fitness landscapes, we started by generating random DAGs. Since no agreed upon model exists for the distribution of DAGs in CPMs, we have used two different procedures, choosing each one randomly with the same probability. One of the procedures uses the function `simOGraph`, in the OncoSimulR package. To generate random DAGs with `simOGraph` for $N$ genes, the genes were first randomly split in a number of levels, where the number of levels used was a randomly chosen integer between 3 and $N-1$, both included. Then, each gene from each level $i$ was randomly connected (as descendant) to randomly chosen genes (the ancestors) from levels $j$, where $j < i$; the number of incoming connections of each gene is a randomly chosen integer between 1 and $maxp$ (both included), where $maxp$ is a randomly chosen integer between 2 and $N-2$ ($maxp$ is common for all genes in a DAG, but can vary between DAGs). The DAG we use is the transitive reduction of the above generated DAG. (Note that this procedure can occasionally result in star DAGs, i.e., DAGs without any dependencies; in such a case, the DAG was discarded and a new one obtained). The other procedure uses the function `random_poset` in package `mccbn` (https://github.com/cbg-ethz/MC-CBN); this function is undocumented, but it returns the transitive reduction of a randomly-filled adjacency matrix for a DAG where the initial number of non-zero connections is equal to the number of possible connections times a constant; we used the default value of that constant (0.15).

*, http://ligarto.org/rdiaz

#### 2. Rough Mount Fuji

In the Rough Mount Fuji fitness landscapes the reference genotype (i.e., the genotype with maximum fitness) was randomly chosen (setting `reference = 'random'` in the `rfitness` function in OncoSimulR). The standard deviation, *sd*, of the random normal variate was set to 0.2 and the decrease in fitness (strictly, birth rate) of a genotype per each unit increase in Hamming distance from the reference genotype, $c$, was chosen from a uniform $U(0, 0.2)$ distribution. This gives a wide variety of fitness landscapes that encompass from close to additive (large values of $c$) to House of Cards ($c$ close to 0), with maximum fitness (birth rate) comparable to those of the representable and local maxima fitness landscapes.

The generated RMF fitness landscape was checked to ensure that all 7 or 10 genes were present in at least one accessible genotype; if they were not, a new fitness landscape was generated (with, possibly, different values of $c$ and reference genotype). Function `rfitness` from the OncoSimulR package [2] was used.

#### 3. Local maxima without reciprocal sign epistasis?

As explained in the paper, creating fitness landscapes with local maxima generally results in creating reciprocal sign epistasis and the number of local fitness maxima is associated with reciprocal sign epistasis —see Figures in section S2 Figure. There were, however, seven cases (out of 420) where introduction of local fitness maxima did not lead to introduction of reciprocal sign epistasis. Is this a contradiction of the statement that reciprocal sign epistasis is a necessary condition for local fitness maxima [4]? No: the key feature of all the models we consider is that back mutation is not allowed, and this explains why scenarios that have local fitness maxima (when no back mutation is allowed) might not have reciprocal sign epistasis.

We will look at one example in detail. The seven cases (which can be seen in S1 Figure) are landscapes with IDs "7E10pIyu7UguIUl8I", "8QlFQCUlUVfC10PZr", " bCsk2Qo5VMVm55fM", "GedZaWDeb1029Mf88", " hw8kQ4g44p4XAkDa", "WpF105HbEDoECa8vs", "t1yUXsv5fVuo10GRi". We will examine in detail fitness landscape "GedZaWDeb1029Mf88" (it is the smallest one — see page 299 in S1 Figure). The fitness of four of the relevant genotypes are

```
ABCDFG : 2.007
ABCDEFG: 1.8
ABCDF: 1.749
ABCDEF: 1.712
```

so there is no reciprocal sign epistasis (use, for example, the graphical criterion in [1] or [3]) and "ABCDFG" is a global maximum.

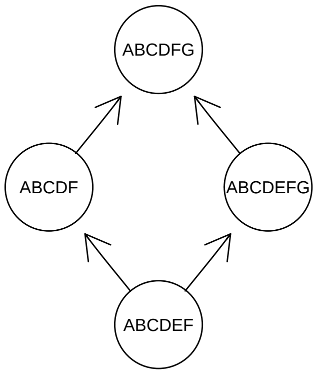


Of course, under an evolutionary model that assumes no back mutations (as is the case of all models considered here), two of those transitions, those that involve loosing "E" ( `ABCDEF -> ABCDF` and `ABCDEFG -> ABCDFG` ) are not allowed, leading to two local fitness maxima (`ABCDFG, ABCDEFG`). To put it differently, under back mutations this fitness landscape would have only a single global maximum (ABCDFG); there is no reciprocal sign epistasis and there would be no local maxima. But once back mutation is not allowed, a local fitness maximum appears as some transitions are not allowed.

Note also that here, for that set of four genotypes, mutating gene E decreases fitness. But mutating E increases fitness in genotypes “ABC” or “ABCD”. Thus, this fitness landscape does not fulfill either the assumption that a mutation never decreases the probability of acquiring other mutations (even if the fraction of pairs of genotypes with reciprocal sign epistasis is 0). Regardless, one can also simply focus on the fact that this fitness landscape contains local maxima (and is missing paths relative to the corresponding fitness graph from the DAG of restrictions).

#### 4. References


[1] Crona, K., Greene, D., Barlow, M., 2013. The peaks and geometry of fitness landscapes. *Journal of Theoretical Biology*, **317**:1–10. doi:10.1016/j.jtbi.2012.09.028. URL http://www.sciencedirect.com/science/article/pii/S0022519312005061.

[2] Diaz-Uriarte, R., 2017. OncoSimulR: Genetic simulation with arbitrary epistasis and mutator genes in asexual populations. *Bioinformatics*, **33(12)**:1898–1899. doi:10.1093/bioinformatics/btx077. URL https://academic.oup.com/bioinformatics/article/33/12/1898/2982052/OncoSimulR-genetic-simulation-with-arbitrary.

[3] Ferretti, L., Schmiegelt, B., Weinreich, D., Yamauchi, A., Kobayashi, Y., Tajima, F., Achaz, G., 2016. Measuring epistasis in fitness landscapes: The correlation of fitness effects of mutations. *Journal of Theoretical Biology*, **396**:132–143. doi:10.1016/j.jtbi.2016.01.037. URL http://www.sciencedirect.com/science/article/pii/S0022519316000771.

[4] Poelwijk, F. J., Tănase-Nicola, S., Kiviet, D. J., Tans, S. J., 2011. Reciprocal sign epistasis is a necessary condition for multi-peaked fitness landscapes. *Journal of Theoretical Biology*, **272(1)**:141–144. doi:10.1016/j.jtbi.2010.12.015. URL http://www.sciencedirect.com/science/article/pii/S0022519310006703.
