## Supplementary material for "Every which way? On predicting tumor evolution using cancer progression models": S3_Text

### Supporting Information for “Every which way? On predicting tumor evolution using cancer progression models”

2019-04-17

#### S3 Text. Evolutionary simulations.

---

#### Contents

**1 Runs until fixation** **1**
1.1 All genes part of lines of descent with frequency $> 0.001$ . . . . . . . . . . . . . . 2

**2 Detection regimes and sampling** **2**

**3 Other parameters of the simulations** **2**

**4 Number of genes used** **2**

**5 LODs through non-accessible genotypes, LODs that go beyond a local maximum, and moving through fitness valleys** **4**

**6 References** **6**

---

#### 1. Runs until fixation

Simulations of evolutionary processes were run until fixation of a genotype, where the genotype was one of the genotypes among the local maxima (or the single global maximum). We used OncoSimulR, with the `fixation` option (introduced in version 2.9.8 of the program). A genotype was considered to have been fixated if it maintained a proportion $\geq$ 0.98 during 15000 consecutive sampling periods (this means that if after reaching a minimum frequency $\geq$ 0.98, at any time the proportion became smaller than 0.98 the counter of successive periods was reset to 0).

Why not require a proportion of 1.0 as evidence of fixation? Because for local maxima, if mutation rate is larger than 0 and neighboring genotypes have non-zero birth rate, the fixated genotype can occasionally generate descending genotypes that exist, with small frequencies, for short periods of time. Using much shorter number of consecutive sampling periods such as

*, http://ligarto.org/rdiaz

1000 or 5000 did not produce different results over using 15000 in trial runs; however, to err on the safe side and make sure fixation had been established, we used that overly long period.

We excluded these 15000 periods from the computation of clonal interference statistics.

##### 1.1. All genes part of lines of descent with frequency $> 0.001$

When the 20000 simulations were completed, we verified that the frequency of all genes in the last genotypes (i.e., the fixated genotypes or the final genotypes of the LODs) were at least 0.001. If they were not, a new fitness landscape was generated and the processes started again. In other words, we avoided fitness landscapes that have a nominal number of, say, 10 genes, but where a smaller number of genes were effectively ever part of the paths of tumor progression (this issue can affect the local maxima and RMF landscapes). In a sample of 4000 individuals, the probability that a gene with a true frequency of 0.001 is never part of a LOD is about 0.018 (= $(1 - 0.001)^{4000}$), so less than 2%. Of course, at the smallest value of the threshold, some data sets of 4000 might have at least one gene missing, and that probability is larger for data sets of 50 and 200.

#### 2. Detection regimes and sampling

For each detection regime, we generated 20000 random deviates (called $r$, below) from the specified beta distribution ($\mathrm{B}(1, 1)$, $\mathrm{B}(5, 3)$, and $\mathrm{B}(3, 5)$ (for uniform, large, and small, respectively).

Using those random deviates, we defined the target size of each sample as $t = \exp(r\ (\ln(M) - \ln(m)) + \ln(m))$, where $M$ and $m$ are the largest and smallest values, respectively, of population sizes (number of cells) ever attained in any of the 20000 simulations. Thus, we obtain target sizes that are uniform or biased towards large sizes or biased towards small sizes in the log scale. In the model of [10], tumor population size increases logarithmically with number of driver mutations. Therefore, uniform, small, and large biases would correspond to approximately uniform, small, or large in terms of number of driver mutations.

For each of the 20000 simulations, the actual sample was the one corresponding to the first observation period at which the total tumor size achieved a value equal to, or larger than, $t$. If all values of tumor population size were $> t$, we returned the sample with the largest population size, and if all values were $< t$ the sample with smallest size.

This procedure determines at which of the sampling times we make the observation. The actual genotype returned is the single genotype with the largest frequency. Thus, we are not emulating taking a biopsy of the entire tumor or bulk sequencing but, rather, single-cell sampling, and sampling the single most common genotype. No observational error was added to the data.

We carried the above steps using OncoSimulR's function `samplePop`, with the values of $t$ (thresholded as explained for $> t$ and $< t$) as arguments to `popSizeSample` and using `typeSample = 'single'`.

#### 3. Other parameters of the simulations

Simulations used the implementation of the McFarland model in the OncoSimulR package [4]. In addition to the parameters specified in the main text, other parameters for the simulations on the fitness landscapes were (see specific meaning in documentation of OncoSimulR [4]): $finalTime = 10000$, $keepEvery = 1$, $sampleEvery = 0.03$, $max.wall.time = 20$, $max.num.tries = 500$.

#### 4. Number of genes used

We have used seven and ten genes for the simulations. Why? Briefly, with the intention of covering a range that both represents a realistic number of genes users would analyze and is computationally feasible.

Seven and ten are values that cover what many previous studies that combine methodological research on CPM and empirical data have used: [2]: 7; [8]: 7 (modules); [13]: 8; [9]: 9; [7]: 7 to 11 (including gene and core pathways); [6]: 10, 11, 12; [11]: 11 and 12; [12]: 6 and 13; we have also previously used those ranges of genes: [3]: 7, 9, 11; [5]: 7.

The execution time of H-CBN increases steeply with number of genes [3] (see Additional File 2, Section 7), with median execution times of 2000 seconds for 11 genes and sample size of 200 subjects. Our results with the simulations on 7 and 10 genes, and the results for the 22 cancer data sets shows that increasing the number of features leads to a decrease in performance; using more than 10 genes for the simulations would not have lead to more optimistic results. In fact, 10 genes, while being computationally feasible, provides a stark contrast with 7 genes: an apparently small increase in number of genes leads, even in the best of circumstances (e.g., Figure 3A) to major decreases in performance —of course, largely the result of the combinatorial increase in the number of paths.

On the other hand, even if the true number of major drivers in cancer is not over 5 or 6, the presence of mini-drivers will easily place the number of drivers analyzed around 7 or above.

### 5. LODs through non-accessible genotypes, LODs that go beyond a local maximum, and moving through fitness valleys

The following example is from the RMF fitness landscape "16H2tTfJLrFoSFFU" (p. 426 in fl-fg-7.pdf), which was run with variable mutation rate and initial population size of 50000.

The figures below reproduce the fitness landscape and the fitness graph.

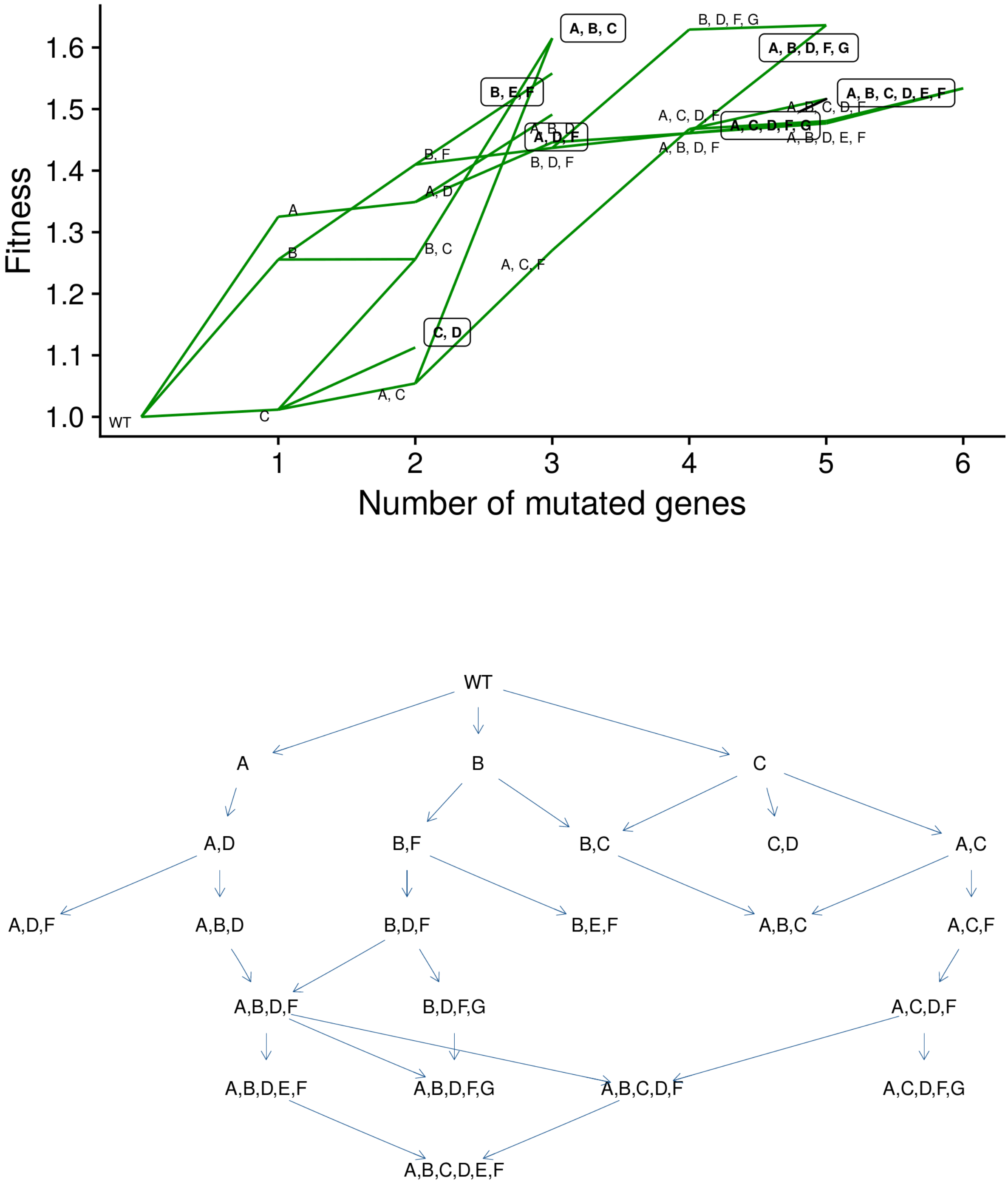


Figure 1: Fitness landscape and fitness graph for fitness landscape "16H2tTfJLrFoSFFU".

As an example, this table shows the LODs (Lines of Descent) that go through genotype ADF. Genotype ADF is a local maximum.

| LOD | $p_i$ |
|---|---|
| WT → A → AD → ADF | 0.22410 |
| WT → A → AF → ADF | 0.00285 |
| WT → D → AD → ADF | 0.00030 |
| WT → A → AD → ADF → ABDF → ABDFG | 0.10935 |
| WT → A → AF → ADF → ABDF → ABDFG | 0.00160 |
| WT → D → AD → ADF → ABDF → ABDFG | 0.00015 |
| WT → A → AD → ADF → ACDF → ACDFG | 0.00085 |
| WT → A → AD → ADF → ADFG → ABDFG | 0.01785 |
| WT → A → AD → ADF → ADFG → ACDFG | 0.00030 |
| WT → A → AF → ADF → ADFG → ABDFG | 0.00040 |
| WT → D → AD → ADF → ADFG → ABDFG | 0.00005 |
| WT → A → AD → ADF → ABDF → ABDEF → ABCDEF | 0.00005 |

Table 1: Proportion of LODs (Lines of Descent) that go through genotype ADF, which is a local maximum. Note that some continue to other local maxima, and some go through non-accessible genotypes. We use "WT" (for "wild type") for the genotype without any mutation in the genes considered; this is the "0000" genotype in Figure 1 in the manuscript.

There are some LODs where ADF is present but for which those LODs then go on to reach other local maxima. Some of those paths actually go through descendant genotypes of lower fitness (e.g., ABDF) and some even go through non-accessible genotypes (e.g., ADFG). This is the birth rate of genotypes with genes A, D, and F mutated:

| Genotype | Birth rate |
|---|---|
| ADF | 1.4912 |
| ABDF | 1.4609 |
| ACDF | 1.4675 |
| ADEF | 1.2575 |
| ADFG | 1.3281 |
| ABCDF | 1.4805 |
| ABDEF | 1.4767 |
| ABDFG | 1.6363 |
| ACDEF | 0.8905 |
| ACDFG | 1.5164 |
| ADEFG | 1.4771 |
| ABCDEF | 1.5337 |
| ABCDFG | 1.3810 |
| ABDEFG | 1.0986 |
| ACDEFG | 0.8672 |
| ABCDEFG | 1.0435 |

Table 2: Birth rate of some of the genotypes depicted in Figure 1 and Table 1.

In a path like "WT → A → AD → ADF → ABDF → ABDFG", a cell with genotype ADF acquires a mutation in gene B; this decreases its fitness slightly relative to its parent (ADF), but ADF is still not fixated, and this descendant eventually acquires another mutation (G), becoming ABDFG, which has higher fitness than ADF, and eventually becomes fixated.

The above example has all genotypes in the LOD as accessible genotypes (it just happened that an intermediate genotype, ABDF, had lower fitness than its parent).

But a path like

"WT → A → AD → ADF → ADFG → ABDFG" goes through ADFG, which is not even an accessible genotype (notice it is not shown in Figure 1). As in the example above, it is possible to go through fitness valleys, specially if the ancestor (here ADF) does not have a very large frequency in the population, in so far as the fitness of the descendant (ADFG) is not too low relative to the average population, and another mutation appears that increases its fitness (here B, so that we end in ABDFG, a local maximum). Note that, in contrast to representable (and local maxima) fitness landscapes, non-accessible genotypes in the RMF landscapes do not necessarily have 0 or tiny birth rates.

As explained in the paper, these types of paths cannot occur under the representable fitness landscapes, as in those fitness landscapes the non-accessible genotypes have a birth rate of 0. In the local maxima fitness landscapes, it is possible to move through a fitness valley if the genotype in the valley is accessible from some other genotype. In the local maxima fitness landscapes, and similar to the representable fitness landscapes, it is not possible to go through non-accessible genotypes: remember that local maxima fitness landscapes are derived from representable ones so the non-accessible genotypes have a birth rate of 0. (Recall that a genotype might be in a valley when coming from some ancestor but in a hill when coming from another ancestor. In Figure 1 in the paper, genotype "1110" is in a valley in the path "1100 → 1110 → 1111", but not in the path "0110 → 1110 → 1111".)

Moving through fitness valleys has been discussed before (see review and references in, for example, 1); how frequent it is depends on populations size (it is generally more common with larger population sizes), mutation rates, and of course how less fit the valleys are compared to the surrounding genotypes.

#### 6. References


[1] de Visser, J. A. G. M., Krug, J., 2014. Empirical fitness landscapes and the predictability of evolution. *Nat Rev Genet*, **15(7)**:480–490. doi:10.1038/nrg3744. URL http://www.nature.com/nrg/journal/v15/n7/full/nrg3744.html.

[2] Desper, R., Jiang, F., Kallioniemi, O. P., Moch, H., Papadimitriou, C. H., Schäffer, A. A., 1999. Inferring tree models for oncogenesis from comparative genome hybridization data. *J Comput Biol*, **6(1)**:37–51. URL http://view.ncbi.nlm.nih.gov/pubmed/10223663.

[3] Diaz-Uriarte, R., 2015. Identifying restrictions in the order of accumulation of mutations during tumor progression: Effects of passengers, evolutionary models, and sampling. *BMC Bioinformatics*, **16(41)**. doi:doi:10.1186/s12859-015-0466-7. URL http://www.biomedcentral.com/1471-2105/16/41/abstract.

[4] Diaz-Uriarte, R., 2017. OncoSimulR: Genetic simulation with arbitrary epistasis and mutator genes in asexual populations. *Bioinformatics*, **33(12)**:1898–1899. doi:10.1093/bioinformatics/btx077. URL https://academic.oup.com/bioinformatics/article/33/12/1898/2982052/OncoSimulR-genetic-simulation-with-arbitrary.

[5] Diaz-Uriarte, R., 2018. Cancer progression models and fitness landscapes: A many-to-many relationship. *Bioinformatics*, **34(5)**:836–844. doi:10.1093/bioinformatics/btx663. URL https://academic.oup.com/bioinformatics/article/34/5/836/4557185.

[6] Gerstung, M., Baudis, M., Moch, H., Beerenwinkel, N., 2009. Quantifying cancer progression with conjunctive Bayesian networks. *Bioinformatics (Oxford, England)*, **25(21)**:2809–2815. doi:10.1093/bioinformatics/btp505. URL http://dx.doi.org/10.1093/bioinformatics/btp505http://www.bsse.ethz.ch/cbg/software/ct-cbn.

[7] Gerstung, M., Eriksson, N., Lin, J., Vogelstein, B., Beerenwinkel, N., 2011. The Temporal Order of Genetic and Pathway Alterations in Tumorigenesis. *PLoS ONE*, **6(11)**:e27136. doi:10.1371/journal.pone.0027136. URL http://dx.plos.org/10.1371/journal.pone.0027136http://www.bsse.ethz.ch/cbg/software/ct-cbn.

[8] Hjelm, M., Höglund, M., Lagergren, J., 2006. New probabilistic network models and algorithms for oncogenesis. *J Comput Biol*, **13(4)**:853–865. doi:10.1089/cmb.2006.13.853. URL http://dx.doi.org/10.1089/cmb.2006.13.853.

[9] Jiang, H. Y., Huang, Z. X., Zhang, X. F., Desper, R., Zhao, T., 2007. Construction and analysis of tree models for chromosomal classification of diffuse large B-cell lymphomas. *World J Gastroenterol*, **13(11)**:1737–1742. URL http://view.ncbi.nlm.nih.gov/pubmed/17461480.

[10] McFarland, C. D., Korolev, K. S., Kryukov, G. V., Sunyaev, S. R., Mirny, L. A., 2013. Impact of deleterious passenger mutations on cancer progression. *Proceedings of the National Academy of Sciences of the United States of America*, **110(8)**:2910–5. doi:10.1073/pnas.1213968110. URL http://www.ncbi.nlm.nih.gov/pubmed/23388632.

[11] Sakoparnig, T., Beerenwinkel, N., 2012. Efficient sampling for Bayesian inference of conjunctive Bayesian networks. *Bioinformatics (Oxford, England)*, **28(18)**:2318–24. doi:10.1093/bioinformatics/bts433. URL http://www.ncbi.nlm.nih.gov/pubmed/22782551http://www.bsse.ethz.ch/cbg/software/bayes-cbn.

[12] Simon, R., Desper, R., Papadimitriou, C. H., Peng, A., Alberts, D. S., Taetle, R., Trent, J. M., Schäffer, A. A., 2000. Chromosome abnormalities in ovarian adenocarcinoma: III. Using breakpoint data to infer and test mathematical models for oncogenesis. *Genes, Chromosomes and Cancer*, **28(1)**:106–120. doi:10.1002/(SICI)1098-2264(200005)28:1\%3C106::AID-GCC13\%3E3.0.CO;2-S. URL http://dx.doi.org/10.1002/(SICI)1098-2264(200005)28:1<106::AID-GCC13>3.0.CO;2-S.

[13] Sweeney, C., Boucher, K. M., Samowitz, W. S., Wolff, R. K., Albertsen, H., Curtin, K., Caan, B. J., Slattery, M. L., 2009. Oncogenetic tree model of somatic mutations and DNA methylation in colon tumors. *Genes, chromosomes & cancer*, **48(1)**:1–9. doi:10.1002/gcc.20614. URL http://www.ncbi.nlm.nih.gov/pubmed/18767147.
