## Supplementary material for "Every which way? On predicting tumor evolution using cancer progression models": S4_Text

### Supporting Information for "Every which way? On predicting tumor evolution using cancer progression models"

2019-04-17

### S4 Text. CPMs: software, probabilities of paths, statistics of performance, linear models.

---

6 Coefficients of linear models 11

7 References 11

---

#### 1. CPM software

Other cancer progression models have been described but either their implementations are too slow for routine use such as [10], or have dependencies on external libraries that are not open source such as DiP [6], or have no software available (e.g., [1, 3]). See further details in [4].

For H-CBN, we used version 0.1.04b from March 2016, and still current as of April 2018, downloaded from https://www.bsse.ethz.ch/cbg/software/ct-cbn.html. We wrote a wrapper to call H-CBN from R, and we used the default settings for temp ($-T = 1$) and steps

*, http://ligarto.org/rdiaz

($-N =$ number of nodes$^2$) —tough we ensure a minimum of 25 steps are used, even if number of nodes is less than five; the simulated annealing search started for the best poset from an initial poset built using OT [11], as preliminary runs suggested this initial poset is as good as, or better than, the default linear poset in [7]. The parameters for the (exponentially distributed) waiting time until the occurrence of a mutation given its restrictions are satisfied ($\lambda$s) were obtained doing an additional run on the fitted model from the previous step, as in [7].

We include the code we used (file ct-cbn-0.1.04b-with-rdu-bug-fix-write-lambda.tar.gz), which we have modified from the original. These are the two changes compared to the file you can download (current as of 2019-03-29) from https://www.bsse.ethz.ch/cbg/software/ct-cbn.html

1. A bug fix for an unjustified stopping when (loglik_new ¡ loglik). These changes are around lines 1434 of ct-cbn.h

2. Force ct-cbn to output a lambda_i, for a lambda from final iteration. The additions are around line 735 of ct-cbn.h

MCCBN was run using version 1.1.9 of the `mccbn` package, downloaded from github (`https://github.com/cbg-ethz/MC-CBN` ).

OT was run using version 0.3.3 of the Oncotree package [11].

For CAPRI and CAPRESE we used version 2.11.0 of the TRONCO BioConductor package, current as of April 2018, downloaded from the official BioConductor site. All options were left at the recommended defaults (e.g., 100 bootstrap samples for the estimation of the selective advantage scores with p-value of 0.05, and heuristic search using Hill Climbing).

We wrote wrapper code for all CPMs to obtain the fitness graphs and, for OT, H-CBN, and MCCBN, the weighted predicted paths. For OT, we use `ot.fit$parent$est.weight` to obtain the probabilities of transition to each descendant genotype; if the OT fit, however, cannot return an error estimate, that operation fails and thus we use the `ot.fit$parent$obs.weight` component.

#### 2. Preprocessing of data for CPMs

Before analyzing data with CPMs, data were preprocessed as follows:

- All columns that had all 0s (i.e., genes that were absent in all samples) were removed. Since these are never present in the data given to the CPMs, no inference can be made about the removed genes, and this necessarily decreases the dimension of the fitness landscape implied by the CPM and the length of the paths to the maximum.

- If two or more columns (genes) were identical over all individuals (i.e., were indistinguishable), all the identical replicate columns, except one, were removed from the analyses[1]. This, of course, will preclude the matching of some (or all) of the true paths since we are constructing the CPM from a data set of smaller dimensionality and the CPM's paths are shorter than they ought to be (see also details in section 5).

  Indistinguishable events will unavoidably create problems. Alternative ways of handling them are not better. If the indistinguishable columns are left in the data, some CPMs (e.g., CAPRI and CAPRESE) complaint about it, whereas others (H-CBN, MCCBN) make the indistinguishable events depend on one another, with a very large $\lambda$, with the order in the DAG depending on the order on the column of data (leftmost events placed as ancestors). OT also places them as independent events (as CAPRI and CAPRESE do).

[1]In more detail, the identical columns were flagged as such by "fusing" the names of the genes, so as to be able to identify them. Then, the paths were post-processed before the analysis to remove the combined name, leaving one of the two, or more, identical names. For example, if we have fused B and C, a path could appear as $WT \rightarrow A \rightarrow A, B\_C \rightarrow A, B\_C, D$. We would then write as $WT \rightarrow A \rightarrow A, B \rightarrow A, B, D$. Here "C" has been dropped (it was indistinguishable from "B").

Note that we use "WT" for the "0000" genotype in Figure 1 in the manuscript.

The consequence of leaving the events as in H-CBN would be similar to expanding the paths of progression, post-analysis, and placing the indistinguishable events one right after the other in the path. The order would, of course, have to be arbitrary and, in most cases, this would actually make matching any true LOD harder. If only one of the replicates is left in the path, the LOD needs to match, by chance, one particular order. If two or more are placed, the probability of matching decreases.

The proportion of data sets with one or more identical columns is about .16, .16, and .11 for the representable, local maxima, and RMF landscapes. They decrease with sample sizes, generally being under 0.05 for sample size 4000.

- Whenever one of more genes were present in all samples, to prevent the removal of these genes present in all cases, we added one case (one "pseudosample") with no mutations to the data set (this is not unlike [4], but we add only one sample, not a fixed percentage, to minimize altering any estimates of probabilities of paths). This allows us to use exactly the same data for all CPMs (CAPRI cannot deal with data where one or more columns are present in all subjects, OT removes them, whereas H-CBN can use this data).

  Even more importantly, this procedure does not decrease the dimensionality of the data set and, thus, does not decrease the length of the CPM's paths to the maximum

  The event that has a frequency of 1 is placed at the top of the DAG of restrictions (it is the first mutation after WT, or "0000" genotype, in all the paths to the maximum). The (very minor) inconvenience is that it has a minor effect on H-CBN's $\lambda$s estimates, but that should be inconsequential for practical purposes.

  The proportion of data sets with pseudosamples added was .30, .21, and .01, for representable, local maxima, and RMF fitness landscapes.

After the data preprocessing, a total of 105 data sets, out of the original 56700, led to data sets in which all CPMs failed to fit a model (e.g., because the final data set contained only a single column). MCCBN, in addition, failed to fit a model to one additional case (to which all other CPMs were able to fit a model).

### 3. Computing probabilities of paths

#### 3.1. CBN (and OT)

The procedure we used is as follows (it might be simpler to understand it by referring to p. i729 of Montazeri et al., 2016 [9]):

1. Obtain the set of genotypes that can exist under the poset (as we will use it in step 3 below).

2. Obtain the set of paths that can exist under the poset (to be used in step 5, below). This itself is obtained from 1.

3. Obtain the transition rate matrix between genotypes from the $\lambda$s (e.g., what is shown in matrix S in Montazeri et al.). As explained in Montazeri et al., "the non-zero off-diagonal elements of the transition matrix are the transition rates from each genotype to its successive genotypes in the genotype lattice, also shown in Figure 1(b)." See also legend of Figure 1: "(b). Directed transition rates among neighboring genotypes are shown on the edges of the lattice".

4. Set the diagonal of the previous matrix to 0[2] and for each row of the transition rate matrix, divide by $\sum \lambda$. Now the entries are probabilities of transition to each descendant genotype given a transition.

[2] In fact, the diagonal entries are never computed explicitly and are always 0.

5. Go over the list of paths (from step 2) and for each path, obtain its probability by multiplying the probabilities of the transitions (from step 4) between the genotypes in a path.

6. (Check: verify that the sum of probabilities of all paths equals 1, within numerical margin of error of machine.)

In the code, steps 3 and 4 above were carried out by creating, from the set of paths to the maximum and the output from CBN (H-CBN or MCCBN), what we called a weighted fitness graph: the fitness graph of paths to the maximum with the $\lambda$s on the edges (or the weighted adjacency matrix corresponding to paths to the maximum where weights are $\lambda$s). This would be Figure 1b in [9]. Dividing by the row sum gives us the transition matrix between genotypes in step 4.

As an example, suppose we obtain from CBN (H-CBN or MCCBN) the following DAG of restrictions and estimated lambdas:

| From | To | $\lambda$ |
|---|---|---|
| Root | A | 2 |
| Root | B | 3 |
| A | C | 4 |
| C | D | 5 |

The paths to the maximum, with their probabilities, are[3]:

| path | probability |
|---|---|
| WT → A → A, B → A, B, C → A, B, C, D | 2/5 * 3/7 * 1 * 1 |
| WT → A → A, C → A, B, C → A, B, C, D | 2/5 * 4/7 * 3/8 * 1 |
| WT → A → A, C → A, C, D → A, B, C, D | 2/5 * 4/7 * 5/8 * 1 |
| WT → B → A, B → A, B, C → A, B, C, D | 3/5 * 1 * 1 * 1 |

For example, from WT with a probability of 2/5 we take the path to A and with 3/5 to B. Once in genotype A, we can either add a B mutation (probability = 3/7) or a C mutation (probability = 4/7). If we add a B mutation, from genotype AB we can only move to ABC (probability 1). Etc. This procedure is equivalent to the one used by Hosseini [8].

An analogous procedure was used with OT.

[3]Recall we indistinctly use "0000", as in Figure 1 in the manuscript, or WT, for wild type, though this is a slight abuse of the concept of wildtype, to denote the genotype without any of the mutations considered.

##### 3.2. CAPRI, CAPRESE, and probabilities of paths of tumor progression

With both OT and CBN if we see a DAG such as

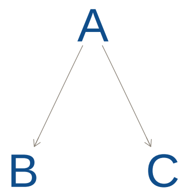

this is saying that, except for errors (errors in the model and observational errors) the genotypes that can exist under the model are only (with our usual notation of using a capital letter to denote that the gene is mutated, and no letter to denote absence of mutation, and "WT" as a shorthand for the genotype without any of the mutations considered):

```
WT
A
AB
AC
ABC
```

and, consequently, there are only two paths to the maximum:

```
WT -> A -> AB -> ABC
WT -> A -> AC -> ABC
```

or

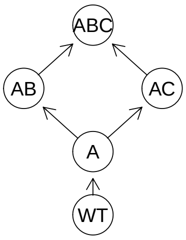

Which means that, for example, under OT and CBN the following paths to the maximum are not allowed under the model:

```
WT -> B -> BC -> ABC
WT -> B -> AB -> ABC
```

This interpretation, however, does not necessarily follow with CAPRI (or CAPRESE). CAPRI returns, as output, a DAG and a set of conditional probability tables (CPTs) associated to each node. What CAPRI seems to say, as can be checked from non-zero entries in the CPTs for B without A or C without A in a DAG as above, is that the DAG says that the most likely mutational paths are

```
WT -> A -> AB -> ABC
WT -> A -> AC -> ABC
```

but other mutational paths could also take place under the model. (This follows directly from seeing CAPRI return a CPT where, say, $P(B|\neg A) = 0.4$, i.e., mutating B without A is certainly not a rare event, even when an arrow exists from A to B). In fact, any path might happen (with some conditional probability tables), and one would seem to need to look at the CPTs to understand which ones can or cannot happen (but see below). And we are given no clear definition of what most likely paths really mean (e.g., "trends of selective advantage among genomic alterations" or "most common evolutionary trajectories" in the wording of 2) in terms of what probabilities are considered or not.

We see a similar issue with the following two DAGs:

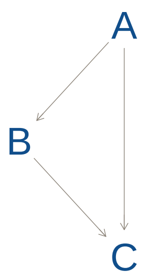

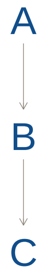

because, of course, the transitive reduction of the first DAG is the second DAG. So from the point of view of paths from the non-mutated to the fully mutated genotype, both DAGs have the same meaning, as both imply that the same order of events needs to happen (or the same restrictions in the accumulation of mutations hold). Yet CAPRI seems to make a distinction between them. A distinction that could only be disentangled, presumably, by looking at the CPTs.

However, no information is available on how to go from CPTs to the conditional probabilities of genotypes implied by the model. In fact, directly using the CPT information seems discouraged and not something that users are supposed to do: access to the CPT has to be done via `TRONCO:::as.bnlearn.network(some_tronco_model)`, i.e., using the `:::` which denotes that we are accessing a non-exported function from the software. (This was the case with v. 2.11.0 and is the case as of July-2018 with version v. 2.13.0).

This contrasts with, say, H-CBN (and MCCBN) and OT which provide, respectively, estimated $\lambda$s and `edge weights` (`object$parent$est.weight`). These conditional probabilities are what we use to obtain the probability-weighted paths implied by the model.

This is a key difference between OT and H-CBN (and MCCBN) on the one hand and CAPRI (and CAPRESE) on the other: OT and H-CBN (and MCCBN) incorporate errors in their models, but they return the estimates of the parameters of their models, i.e., the $\lambda$s or the `est.weights`, that map directly into what can happen under the model, what genotypes can arise and from which other genotypes. (This is analogous to a simple linear regression: there is an error component in the model $y = \alpha + \beta x + \epsilon$, the $\epsilon$, often assumed to be independent and identically distributed from a normal distribution with mean 0 and variance $\sigma^2$, etc, but the model, and the software, return the $\alpha$ and $\beta$ which allow us to predict the expected value of $y$ given $x$, under the model).

In contrast, with CAPRI we can (again, using a non-exported function) obtain the CPTs that correspond to the DAG returned. Remember that essentially what CAPRI is doing is fitting a Bayesian Network to the observational data, with the DAG built so that arrows respect the temporal priority and probability raising restrictions. But the CPTs themselves are not estimated parameters of a model that could be mapped into probabilities of paths. The CPTs of CAPRI seem to be the conditional probabilities of observing what we observe under the DAG and they incorporate errors (model errors and noise in the data), and can contain non-zero entries for child nodes when their parents are absent. Obtaining probabilities of paths from these CPTs is, thus, impossible.

An additional issue that has been noted with CAPRI in the paper is its behavior with increasing sample sizes. That was also noted in [5] and seems related to the penalization in the fits: CAPRI does not seem suited to deal with large data sets as it tends to allow only one, or a few, paths to the maximum when N is 4000. (The transitive reduction of) the DAGs returned from CAPRI is often a linear sequence. This behavior did not change considerably whether we used AIC or BIC.

CAPRESE does not return DAGs, but trees, and in that sense it is simpler to deal with. But still, as with CAPRI, there is no information available on how to go from CPTs to the conditional probabilities of genotypes implied by the model (and accessing the CPT does not seem to be encouraged) and, as for CAPRI, the CPTs seem to mix the underlying model with the error model, and obtaining probabilities of paths from these CPTs is, thus, impossible.

#### 4. Example where perfect recall and precision do not guarantee Jensen-Shannon divergence of 0

Suppose the following set of two paths to the maximum, with predicted and observed probabilities as shown:

| Path | CPM predicted probability | True LOD probability |
|---|---|---|
| WT → A → AB | 0.99 | 0.01 |
| WT → B → AB | 0.01 | 0.99 |

The JS divergence (on a scale 0 to 1) is 0.9192 (remember 1 is the maximum value of divergence), even when 1-recall and 1-precision are both 0 (none of the CPM's paths are missing from the LODs, and none of the LODs are missing from the CPM's paths).

#### 5. Measuring predictability: comparing paths from CPMs and LODs of different lengths

We explain here in detail, and provide a commented example, of the procedure used to compare paths from CPMs and LODs when they might be of different length (this includes the case of LODs that include paths no in the fitness graph).

Let $i$ and $j$ denote two paths, one from the LOD and the other from the CPM, with corresponding probabilities $p_i$ and $q_j$. Here, the $i$ index refers to paths from the LOD, and the $j$ to paths from the CPM (this is in contrast to the paper, where we did not make this specification, to keep the description general).

Let $K_i, K_j$ denote the length of paths $i$ and $j$, respectively. Note that all $K_j$ are equal (as all go up to the genotype with all genes mutated), and we will refer to that unique value of $K_j$ as $K^C$. When there is ambiguity about which $K$ we are referring to, we will use $K_j^C$ for the $K_j$ from the CPM (again, $K_1^C = K_2^C = \ldots = K_C$) and $K_i^L$ for the $K_i$ from the LOD.

The vectors $P$, $Q$, for the computation of JS will have the following types of matching pairs:

1. $p_i, q_j$ when $K_i = K_j$
2. $p_i \frac{K_j}{K_i}, q_j$, when $i$ is partially included in $j$ ($K_i > K_j$), and this accounts for the part of $i$ included in $j$,
3. $p_i, q_j \frac{K_i}{K_j}$ when $j$ is partially included in $i$ ($K_j > K_i$), and this accounts for the part of $j$ included in $i$,

4. $\sum p_i \frac{K_i - K_j}{K_i}, 0$ when $i$ is partially included in $j$ ($K_i > K_j$), and this accounts for the part of $i$ not included in $j$,

5. $0, \sum q_j \frac{K_j - K_i}{K_j}$ when $j$ with is partially included in $i$ ($K_j > K_i$), and this accounts for the part of $j$ not included in $i$,

6. $\sum p_u, 0$, for all paths $u$ among the paths from the LOD that do not match any $j$ (any path from the CPM),

7. $0, \sum q_v$, for all paths $v$ among the paths from the CPM that do not match any $i$ (any path from the LOD).

(Where some notation above, again, can be simplified by noting that all $K_j = K_C$).

We can sum, as appropriate, the relevant entries to simplify computations as the JS is the same for the pair of vectors $P = [p_1, p_2, 0, 0, p_5, p_6], Q = [q_1, q_2, q_3, q_4, 0, 0]$ and the pair of vectors
$P' = [p_1, p_2, 0, p_5 + p_6], Q' = [q_1, q_2, q_3 + q_4, 0]$ (this follows from the definition of JS).

In other words, we have the 0 entry in $Q$ correspond to $\sum p_i \frac{K_i - K_j}{K_i} + \sum p_u$, where the $i$ are the paths in the LOD with some partial match among the paths in the CPM, and the $u$ denote those paths in the LOD that do not match any path in the CPM.

The relevant entries above can be used to compute 1-recall and 1-precision. For example, for 1-recall, $P(\neg DAG|LOD)$, the sum of the probabilities of the paths in the LODs that are not among the paths allowed by the CPMs, we will use $\sum p_i \frac{K_i - K_j}{K_i} + \sum p_u$.

The above expressions are computed for all distinct $k$, the number of mutations of the final genotypes of the true LODs. The final JS (1-recall and 1-precision) are the weighted sum of each of those terms, weighted by $w_k$, the frequency of all paths from the LOD that end at $k$ mutations. But see example below (as we can group together several $k$ for faster computation).

#### 5.1. Commented example for paths of unequal length

To give a specific example, suppose the following paths from a LOD and a CPM. In this example, the CPM has been constructed from a data set that had only mutations A and B (C and D were missing). And we have the three possible cases: some paths from the LOD are shorter than the paths from the CPM, some paths from the LOD are the same length as those from the CPM, and some paths from the LOD are larger than those from the CPM. (Also note that, as commented in section S3 Text , it is possible that some LODs go beyond a local maximum, here

AB, to reach another maximum.)

| Path (i) | LOD | $p_i$ | $K_i$ |
|---|---|---|---|
| 1 | WT → A | 0.1 | 1 |
| 2 | WT → B → AB | 0.3 | 2 |
| 3 | WT → B → BC → BCD | 0.2 | 3 |
| 4 | WT → B → AB → ABC → ABCD | 0.4 | 4 |

| Path (j) | CPM predicted path | $q_j$ | $K_j (= K^C)$ |
|---|---|---|---|
| 1 | WT → A → AB | 0.6 | 2 |
| 2 | WT → B → AB | 0.4 | 2 |

Where $i = 1, 2, 3, 4, 5$ are the four LODs and $j = 1, 2$ the two paths to the maximum from the CPM. $K_j = K^C = 2$. To compute JS, 1-recall and 1-precision, it is much simpler to use an algorithm that splits the cases to be considered into three:

1. $K_i < K_j$ (i.e., $K_i < K^C$), those paths from the LOD that are shorter than the paths from the CPM;
2. $K_i = K_j$ (i.e., $K_i = K^C$), those paths from the LOD that have the same length as the paths from the CPM;
3. $K_i > K_j$ (i.e., $K_i > K_C$), those paths from the LOD that are larger than the paths from the CPM;

We can iterate over all distinct $k$ for $K_i < K_j$, and weight the output by $w_k$, the frequency of all paths from the LOD that end at $k$ mutations; computations for $K_i = K_j$ can be subsumed into those for $K_i < K_j$. Thus, one part of the algorithm iterates over all $k \leq K_j = K_C$ (remember all $K_j$ are identical and equal to $K_C$, the single, unique $K$ at which the paths from the CPM stop). Computations for $K_i > K_j$ can be done in one iteration.

It might also be helpful to think about a cut operation on a path. For instance, for each $k$ where $K_i < K_j$, we can cut the paths from the CPM at $k$ mutations, leaving only the paths from WT up to $k$ mutations (and collapsing, as appropriate, any collection of now indistinguishable subsets of paths, summing their probabilities).

In the exposition below, some computations could be further simplified; they are left as they are for clarity (e.g., we multiply by total frequencies of mutations for the weights when we have previously scaled the total probability by it, so it is 1 in each $k$, etc).

---

**JS**

1. LOD $i = 1$ finishes at one mutated gene. Cutting the CPM path at $k = 1$, the JS for $k = 1$ is obtained from the vectors of probabilities $P = [1, 0, 0]$ (from the LOD) and $Q = [0.6\ 0.5, 0.4\ 0.5, 0.5]$ from the CPM. The last entry in $Q$ is the sum of all the flow through the paths of the CPM that cannot be matched because the length of the CPM paths is $K_C = 2$. And the 1 in $P$ comes from $p_1 / \sum p_i$ for all $i$ that end in $k = 1$ which is only $p_1$.

   The first and second entries are $q_1^1$ and $q_2^1$ multiplied by $k/K_C$, i.e., the probabilities of the fractions of paths from the CPM cut at $k = 1$ mutations.

   The weight, $w_k$, for this value is 0.1 (the frequency of LODs that finish at $k = 1$).

2. LOD $i = 2$ finishes at $k = 2$. Here the comparison is the immediate one for equal length paths and the vectors used for JS are: $P = [1, 0]$ and $Q = [0.4, 0.6]$. $w_2 = 0.3$.

3. Paths $i = 3, 4$ are longer than the CPM paths. The flow $BC \rightarrow BCD$ for $i = 3$ and the flow $AB \rightarrow ABC \rightarrow ABCD$ for $i = 4$, cannot be matched by the CPM.

   Here the total amount of evolutionary flow through the LOD that cannot be captured by the CPM, because the CPM ends prematurely, is $\sum_i p_i(K_i - K^C)/K_i = (0.2 * (1/3) + 0.4 * (1/2)) * (1/0.6)$, for $i = 3, 4$, where the $1/0.6$ scales relative to the total probability in paths $i = 3, 4$.

   Then, to compute JS, the two vectors of probabilities would be
   $P = [0, (1/2)\ 1(0.4/0.6), (1/2)\ 1(0.4/0.6), 1(0.2/0.6)]$, from the LOD, and
   $Q = [0.6, 0.4, 0, 0]$, for the CPM.

   But that is equivalent to using the two vectors
   $P = [0, (1/2)\ (0.4/0.6), (1/2)\ (0.4/0.6)\ +\ 1\ (0.2/0.6)]$, $Q = [0.6, 0.4, 0]$.

   The last entry in $P$ might be easier to see from adding LODs $i = 3$ and the unmatched portion of $i = 4$. Here $w_i = 0.6$

4. We can now add all the JS with their corresponding weights.

---

**1-recall** For 1-recall, the sum of the probabilities of the paths in the LODs that are not among the paths allowed by the CPMs, we have:
$p_3 + p_4 \frac{(K_4 - K_C)}{K_4} = 0.2 + 0.4\ (1/2)\ 0.1 = 0.4$,
where $K_C = 2$ (all CPM paths end at $k = 2$). Note that these same values can be obtained by iterating over $k$ as above, and doing a weighted sum, but in this example it is much simpler to use the computation directly.

Had we used weighted sums, we would have got: $0\ w_1 + 0\ w_2 + (p_3 + p_4 \frac{K_4 - K_C}{K_4}) \frac{1}{0.6} w_{\geq 3} = (0.2 + 0.4 * (1/2)) * (1/0.6) * 0.6 = 0.4$ (where the $\frac{1}{0.6}$ scales so that the probabilities considered when $k \geq 3$ add to 1 —and, sure, we are dividing by 0.6 only to multiply by it because the scaling factor is the weight of the $k_{\geq 3}$ stratum).

---

**1-precision** For 1-precision, the sum of the probabilities of the paths in the CPM's paths that are not among the LOD paths (the paths followed by evolution), it is simpler to use a weighted sum:
$((q_1 \frac{K^C - K_1^L}{K^C}) + q_2)\ w_1 + q_1\ w_2 + q_1\ w_{\geq 3}$.
When $k = 1$, $i = 1$ is $WT \rightarrow A$, and thus all of $q_2$ is not captured, and half of $q_1$ is captured; at $k = 2$, all of $q_2$ but none of $q_1$ is captured; for $k \geq 3$ again path $j = 1$, with $q_1 = 0.6$ is not captured. Thus, we have $(0.6 * (1/2) + 0.4) * 0.1 + 0.6 * 0.3 + 0.6 * 0.6$.

Note that for 1-precision we do not need to re-scale so that probabilities always add up to 1 because they already do (we consider all the $j$). This was not the case for 1-recall (where only some of the $i$ might be considered in turn when we iterate over $k$).

---

To recapitulate, as stated in the paper, we want to compute JS, 1-recall, and 1-precision taking into account that:

1. Any LOD that finishes at $k$ mutations and matches a CPM path up to $k$ mutations is a perfect match, up to $k$; this is the reason we match each LOD with the fitness graph implied by the CPMs, where this fitness graph implied by the CPM has been cut at the number of mutations of the final genotype of the LOD.

2. Any set of LODs that finishes at $k$ mutations when the CPM goes to $K$ with $K > k$ necessarily misses $(K-k)/K$ of the total evolutionary flow, all that which goes from $k+1$ to $K$. This is why we use a category unmatchable by construction.

   Without this, it would be possible to obtain perfect JS from LODs that missed most of the evolutionary flow, for instance very short LODs that finished at one mutation.

   This part of the procedure, thus, accounts for that part of the evolutionary process that the CPM predicts and is not matched by stopping evolution at a local fitness maximum; remember, again, that by construction the CPMs predict that the evolutionary process should go all the way to a global maximum with all genes mutated.

3. A similar reasoning applies when the paths from the CPM are shorter than the paths from the LOD.

#### 6. Coefficients of linear models

Coefficients from the generalized linear mixed-effects models shown in the manuscript (Figure 4) are from overparameterized models, and those are not the models fitted. What we have done is fit the models several times, always with sum-to-zero contrasts, but changing the level of the factor set to $-\Sigma$ (rest of levels) so as to explicitly obtain the coefficients and standard errors for all levels of all terms (e.g., the coefficients that correspond to "Detection, Uniform", "Detection, Small" and "Detection, arge").

#### 7. References

[1] Attolini, C., Cheng, Y., Beroukhim, R., Getz, G., Abdel-Wahab, O., Levine, R. L., Mellinghoff, I. K., Michor, F., 2010. A mathematical framework to determine the temporal sequence of somatic genetic events in cancer. *Proceedings of the National Academy of Sciences*, **107(41)**:17604–17609. doi:10.1073/pnas.1009117107/-/DCSupplemental.www.pnas.org/cgi/doi/10.1073/pnas.1009117107. URL http://www.pnas.org/content/107/41/17604.short.

[2] Caravagna, G., Graudenzi, A., Ramazzotti, D., Sanz-Pamplona, R., Sano, L. D., Mauri, G., Moreno, V., Antoniotti, M., Mishra, B., 2016. Algorithmic methods to infer the evolutionary trajectories in cancer progression. *PNAS*, **113(28)**:E4025–E4034. doi:10.1073/pnas.1520213113. URL http://www.pnas.org/content/113/28/E4025.

[3] Cheng, Y.-K., Beroukhim, R., Levine, R. L., Mellinghoff, I. K., Holland, E. C., Michor, F., 2012. A mathematical methodology for determining the temporal order of pathway alterations arising during gliomagenesis. *PLoS computational biology*, **8(1)**:e1002337. doi:10.1371/journal.pcbi.1002337. URL http://www.pubmedcentral.nih.gov/articlerender.fcgi?artid=3252265&tool=pmcentrez&rendertype=abstract.

[4] Diaz-Uriarte, R., 2015. Identifying restrictions in the order of accumulation of mutations during tumor progression: Effects of passengers, evolutionary models, and sampling. *BMC Bioinformatics*, **16(41)**. doi:doi:10.1186/s12859-015-0466-7. URL http://www.biomedcentral.com/1471-2105/16/41/abstract.

[5] Diaz-Uriarte, R., 2018. Cancer progression models and fitness landscapes: A many-to-many relationship. *Bioinformatics*, **34(5)**:836–844. doi:10.1093/bioinformatics/btx663. URL https://academic.oup.com/bioinformatics/article/34/5/836/4557185.

[6] Farahani, H. S., Lagergren, J., 2013. Learning oncogenetic networks by reducing to mixed integer linear programming. *PloS ONE*, **8(6)**:e65773. doi:10.1371/journal.pone.0065773. URL http://www.pubmedcentral.nih.gov/articlerender.fcgi?artid=3683041&tool=pmcentrez&rendertype=abstract.

[7] Gerstung, M., Eriksson, N., Lin, J., Vogelstein, B., Beerenwinkel, N., 2011. The Temporal Order of Genetic and Pathway Alterations in Tumorigenesis. *PLoS ONE*, **6(11)**:e27136. doi:10.1371/journal.pone.0027136. URL http://dx.plos.org/10.1371/journal.pone.0027136http://www.bsse.ethz.ch/cbg/software/ct-cbn.

[8] Hosseini, S.-R., 2018. Quantifying the predictability of cancer progression using Conjunctive Bayesian Networks. M.Sc. Thesis, Swiss Federal Institute of Technology, Zürich.

[9] Montazeri, H., Kuipers, J., Kouyos, R., Böni, J., Yerly, S., Klimkait, T., Aubert, V., Günthard, H. F., Beerenwinkel, N., Study, T. S. H. C., 2016. Large-scale inference of conjunctive Bayesian networks. *Bioinformatics*, **32(17)**:i727–i735. doi:10.1093/bioinformatics/btw459. URL http://bioinformatics.oxfordjournals.org/content/32/17/i727.

[10] Sakoparnig, T., Beerenwinkel, N., 2012. Efficient sampling for Bayesian inference of conjunctive Bayesian networks. *Bioinformatics (Oxford, England)*, **28(18)**:2318–24. doi:10.1093/bioinformatics/bts433. URL http://www.ncbi.nlm.nih.gov/pubmed/22782551http://www.bsse.ethz.ch/cbg/software/bayes-cbn.

[11] Szabo, A., Pappas, L., 2013. Oncotree: Estimating oncogenetic trees. R package version 0.3.3. URL http://cran.r-project.org/package=Oncotree.
