## Supplementary material for "Every which way? On predicting tumor evolution using cancer progression models": S5_Text

### Supporting Information for "Every which way? On predicting tumor evolution using cancer progression models"

2019-04-17

### S5 Text. Cancer data sets: sources, characteristics, additional results.

---

**3 References** **13**

*, http://ligarto.org/rdiaz

#### List of Figures

1 Results from the cancer data sets analyzed with H-CBN. Data sets have been ordered by increasing sample size, and the x-axis labels provide the acronym (shown in full in the inset legend). Below the data set acronym are the number of subjects and the total number of features, respectively. Analysis were run three times, limiting the number of features analyzed to the 7, 10, and 12 most common ones; the boxplots for each data set are shown in increasing order of number of features. For data sets such as, say, Pancreas genes (Pan_Ge), with 7 features, using 7, 10, or 12 maximum features makes no difference in the number of features analyzed; the three replicate runs show run-to-run variability. A) $JS_{o,b}$: JS statistic for the comparison of the distribution of paths from running H-CBN on the original data set against the distribution of paths from running H-CBN on each one of the bootstrap runs. B) Diamonds show the $S_c$ from the full data, and boxplots the $S_c$ from the bootstrap runs. Right axis labeled by number of equiprobable paths equivalent to the $S_c$. . . . . . . . . . . . . . . . . . . . . . . . . . . 9

2 Cancer data sets: Histograms of number of mutations per subject in the data sets. 10

3 Cancer data sets: Histograms of proportion of individuals in which each mutation is present. For example, in the PP data set, there are four mutations that are present in 80% to 90% of the individuals in the data set, 1 mutation present in 90% to 100% of the individuals, 1 mutation in between 0 and 10% of the individuals, and 1 in between 10% and 15%. . . . . . . . . . . . . . . . . . . . . . . . . . . 11

4 Cancer data sets: scatterplots of the relationship between $JS_{o,b}$, $S_c$, and number of paths to the maximum, using the data labels, using the statistics from analyses with 12 features. . . . . . . . . . . . . . . . . . . . . . . . . . . . . . . . . . . . 12

---

#### 1. Cancer data sets

The cancer data sets used here are a representative example of data sets to which researchers have applied CPMs or data sets to which researchers might want to apply CPMs. All of these data sets (in at least one of their variants) have been used previously in studies with CPMs except for the BRCA data sets, that were obtained *de novo* for this paper.

These data sets vary in:

- sample size (27 to 594 samples);
- data types (nonsynonymous somatic mutations, and copy number aberrations, or both);
- levels of analysis: altered/non-altered pathways —e.g., Pan_pa, GBM_pa, Col_pa, all _pa—, functional modules —GBM_mo–, exclusivity groups [10] —Col_msi_co, Col_mss_co, ACML_co—, genes —e.g., BRCA_ba_s, BRCA_he_s, Pan_ge, GBM_ge, Col_ge, Ov, Lu—, and different types of gene-level events as insertion/deletions, missense point mutations, nonsense point mutations —ACML and ACML_co.
- different procedures for driver selection, from simple frequency-based selection of features (e.g., GBM_ge, Pan_ge, Col_ge) to state-of-the-art methods for the identification of significantly altered genes [31] (e.g., BRCA_he_ s, BRCA_ba_s, Col_msi, Col_mss, GBM_CNA);
- restriction of patient subtypes (with the purpose of achieving sample homogeneity —e.g., BRCA_ba_s, BRCA_he_s, Col_msi, Col_mss);

Thus, in several cases the same source data set has been processed in different ways to produce two different versions. For three of the data sets, two versions, one coded in terms of mutations of genes and one in terms of pathway alterations, were available (Col_ge and

Col_pa, GBM_ge and GBM_pa, Pan_ge and Pan_pa). For three other data sets, we have analyzed both the original data (Col_msi, Col_mss, ACML), and the same data after accounting for so-called "exclusivity relations" (see 10; Col_msi_co, Col_mss_co, ACML_co). Another data set, GBM_CNA, was also analyzed in terms of "functional modules" (GBM_mo).

Other data sets have been obtained from a single source and split to increase subject homogeneity (e.g., BRCA_ba_s and BRCA_he_s; Col_msi, Col_mss).

##### 1.1. Cancer data sets: sources and characteristics

| Name | Source | Original source | Number of features | Number of subjects | Type of event | Abbreviation |
|---|---|---|---|---|---|---|
| All Pathways | [18] | (From sources for colon, glioblastoma, and pancreas genes data sets) | 12 | 268 | candidate mut | all_pa |
| Colon Genes | [18] | [32] | 8 | 95 | candidate mut | Col_ge |
| Colon Pathways | [18] | [32] | 10 | 95 | candidate mut | Col_pa |
| Glioblastoma Genes | [18] | [26] | 8 | 78 | candidate mut | GBM_ge |
| Glioblastoma Pathways | [18] | [26] | 10 | 78 | candidate mut | GBM_pa |
| Pancreas Genes | [18] | [20] | 7 | 90 | candidate mut | Pan_ge |
| Pancreas Pathways | [18] | [20] | 7 | 90 | candidate mut | Pan_pa |
| Lung | [24] | [16] | 51 | 161 | recurrent mut | Lu |
| Ovarian | [24] | [7] | 192 | 326 | recurrent mut | Ov |
| Ovarian driver | [24] | [7] | 9 | 326 | significant mut | Ov_drv |
| Colon MSI | [10] | [8] | 30 | 27 | significant mut and CNA | Col_msi |
| Colon MSS | [10] | [8] | 34 | 152 | significant mut and CNA | Col_mss |
| Colon MSI mutual exclusivity groups collapsed | [10] | [8] | 20 | 27 | significant mut and CNA | Col_msi_co |
| Colon MSS mutual exclusivity groups collapsed | [10] | [8] | 13 | 152 | significant mut and CNA | Col_mss_co |
| ACML | [14, 28] | [27] | 16 | 64 | recurrent mut | ACML |
| ACML mutual exclusivity groups collapsed | [14, 28] | [27] | 11 | 64 | recurrent mut | ACML_co |
| GBM CNA | [12, 17] | [5] | 48 | 563 | significant CNA | GBM_CNA |
| GBM CNA modules | [12, 17] | [5] | 9 | 563 | significant CNA | GBM_mo |
| GBM co-occurrent | [3] | [4, 6] | 3 | 594 | significant co-occurrent CNA | GBM_coo |
| Ovarian CNV | [30] | [21] | 7 | 87 | recurrent arm-level CNA | Ov_CNV |
| BRCA HER2, subtypes | [12, 17] | [9] | 4 | 57 | significant mut | BRCA_he_s |
| BRCA basal-like, subtypes | [12, 17] | [9] | 6 | 81 | significant mut | BRCA_ba_s |

Table 1: Cancer data sets used. Source refers to where the data have been obtained from, generally also the first reference where data set has been used with CPMs. Data sets BRCA_he_s and BRCA_ba_s have been obtained from original sources for this paper. A data set very similar to GBM_CNA was used in [13], but we obtained it from [12, 17], as explained in the text. Type of event: mut: nonsynonymous somatic mutations; CNA: copy number alterations; candidate mut: nonsynonymous mutations on candidate genes [29, 31, 32]; significant mut: nonsynonymous mutations on significant mutated genes, as defined by state-of-the-art algorithms [31] MuSiC [15] or MutSigCV [22]; significant CNA: significant copy number alterations, as defined by GISTIC2.0 [23].

These are further details about how the data were obtained and the rationale for the data processing:

**All Pathways and Colon, Glioblastoma, Pancreas pathways** Data sets Colon Genes, Glioblastoma genes, and Pancreas genes are from [18], with original sources [32], [26], and [20], respectively.

For the corresponding data sets in terms of pathways, the mapping from genes to pathways was done by [18], from the original papers with data sets. Our scripts to reproduce the analysis are provided with the code. Note that for Pancreas pathways we eliminate the four pathways that were present in all subjects (see also [18] and notes in the code for details). For Glioblastoma pathways, two pathways had identical patterns (Apoptosis and Small GTPase-dependent signaling (other than KRAS)) and only one was used.

What we call "All Pathways" here, for brevity, is called "All cancer types" in [18].

**Lung** Original data from [16]. They were obtained from text file `Lung_SM4` from the supplementary material of [24] (file "BMLv1.tar.gz").

**Ovarian** Original data from [7]. They were obtained from text file `OV_SM5` from the supplementary material of [24] (file "BMLv1.tar.gz").

**Ovarian driver** Data come from data set Ovarian, restricting events to 9 significantly mutated genes described in Table 2 of source paper [7].

**Colon MSI** Colorectal cancer, microsatellite unstable tumors. The original data (as well as Colon MSS) come from COADREAD [8]; we obtained them from [10], where the original data were split by tumor subtype into MSI and MSS (see also comments about patient stratification under "BRCA basal-like, subtypes, BRCA HER2, subtypes").

We used GIMP to open the PDF file page (Figure 3 on page 6 of [10]) where the figure was and cropped the grid of the figure and exported it as JPEG with high resolution. Then we imported it in ImageJ(Fiji) (`https://fiji.sc/`), converted it to 8-bit, applied threshold option and set it to B/W, then exported it as text image (matrix as txt). Then we imported the text image in R and used the code in `fig_to_matrix_capri_pnas.R` to convert the text image into a matrix of genotypes. The data were checked against the original figures.

**Colon MSS** Colorectal cancer, microsatellite stable tumors. The original data come from COADREAD [8] and we obtained them from [10], where the original data were split by tumor subtype into MSI and MSS (see above).

From [10]. Same process as for Colon MSI; the figure is Figure S5 from page 16 of the supplementary material to [10]. The authors explain that "Events selected for reconstruction are those involving genes altered in at least 5% of the cases, or part of group of alterations showing an exclusivity trend (see Figure S4)."

**Colon MSI mutual exclusivity groups collapsed, Colon MSS mutual exclusivity groups collapsed** Data sets were obtained from data sets **Colon MSI** and **Colon MSS**, respectively, processed so events showing mutual exclusivity patterns described in [10] were collapsed in a single event representing an exclusivity group.

Mutual exclusivity patterns, as explained in [10], could decrease the performance of CPMs (as CPMs assume no events show exclusivity or otherwise reduce the probability of another event occurring [24]). How to deal with these exclusivity patterns with CPMs such as CBN, OT, CAPRESE, or CAPRI without additional formulas, is not clear, however. For the Colon MSI and Colon MSS data sets, the exclusivity groups identified in Caravagna *et al.* (2016), [10], are supposed to represent fitness-equivalent exclusive sets of alterations. Some of these exclusivity groups share events, some represent "hard" exclusivity relations whereas others represent "soft" exclusivity relations [10]. What we have done is consider each one of the exclusivity groups as analogous to a pathway in Gerstung *et al.* (2011) [18] or a module in Cheng *et al.* (2012) [13] (where the same gene can be part of different pathways/modules or, in this case, different exclusivity groups). This amounts to considering each exclusivity group as a "fitness equivalent" (*sensu* [10]) set of alterations for some "phenotype" shared by the exclusivity group, similar to what [10] did, and should not introduce any additional difficulties for the inference of downstream dependencies.

We removed from the data set any alteration that was a member of one or more exclusivity groups as an individual alteration. Thus, the data sets with exclusivity groups differ from the original ones by adding exclusivity groups and removing alterations that belong to those exclusivity groups. We used the Table S3 of the Supplementary Material of [10] as the canonical source of exclusivity groups. However, notice that there must be the following mistakes in Table S3: in row 6 it says ACVR1B:a, but there is no amplification event that affects ACVR1B in data set **Colon MSI**, according to Figure 3 in the paper (and Figure 5 and Figure S11), but mutation and deletion; in row 7 it says NRAS:a but there is no amplification event that affects NRAS in data set **Colon MSI**, according to Figure 3 in the paper (and Figure 5 and Figure S11), but mutation and deletion; in row 15 it says NRAS:d but there is no deletion event that affects NRAS in data set **Colon MSS**, according to Figure S5 (and Figure S4 and Figure S10), but mutation and amplification. So we assumed it should say ACVR1B:d in row 6, NRAS:d in row 7 and NRAS:a in row 15. Additionally, when a mutual exclusivity group in Table S3 was contained in another (for instance, group in row 5 is part of group of row 1) we used only the bigger one. This procedure results, for **Colon MSI**, in a new data set of 27 subjects and 20 columns (five from the exclusivity groups, and 15 from the 30 alterations in the original data set minus the 15 removed as they are in one or more exclusivity groups). For **Colon MSS**, this results in a new data set of 152 subjects and 13 columns (11 from the exclusivity groups, and 2 from the 34 alterations in the original data set minus the 32 removed as they are in one or more exclusivity groups).

**ACML** Data are originally from [27], and were obtained from the `aCML` data set in R package "TRONCO" [14] and processed to keep the 16 events used in [28]. The data includes alterations with a frequency above 5 % in original data set from [27] and additional selected alterations hypothesized to be part of a functional ACML progression path in the literature and are shown in Figure 5 of [28]. As explained in [28], events are categorized as insertion/deletions, missense point mutations, and nonsense point mutations. This data set shows mutual exclusivity patterns described in Section 4.2 of [28].

**ACML mutual exclusivity groups collapsed** Data come from data set **ACML**, processed so events showing mutual exclusivity patterns described in [28] were collapsed in a single event. As with data sets **Colon MSI** and **Colon MSS**, here we dealt with mutual exclusivity patterns in the data by collapsing individual events in exclusivity groups considered as "fitness equivalent" groups. The two exclusivity groups were obtained from Section 4.2 of [28]: one involves all types of alterations of genes ASXL1 and SF3B1 (ASXL1 nonsense point, ASXL1 ins/del, SF3B1 missense point) and the other involves all types of alterations of genes TET2 and IDH2 (TET2 nonsense point, TET2 missense point, TET2 ins/del, IDH2 missense point); thus, there are 11 columns, 2 from the exclusivity groups, and 9 from the 16 alterations minus the 7 removed as they belong into exclusivity groups.

**GBM CNA** Data come from TCGA GBM PUB CNA data set [5] and were obtained from cBioPortal [12, 17] using the R package "cgdsr" [19], selecting only CNA data from 51 driver genes used by Cheng *et al.* (2012) and detailed in Table 1 of [13], with a GISTIC score of 2 or -2. By doing so, we intended to follow the author's indications to obtain the same data set Cheng *et al.* (2012), [13], used to infer a cancer progression model. Although [13] cite [6] as the source of their data, we understand that the original data set used in [13] should have been the "Provisional" TCGA data set at that time since they got more patients (462) than the total number of patients in the only published TCGA glioblastoma study at the date [6] (206). So, we used the data from the TCGA glioblastoma study published in 2013 [5] on the belief that this is the closest we can get to reproduce the data set used in [13] (the study of 2013, [5], contains 198 patients in common with the study of 2008, [6]). Note that 3 of the 51 genes were not altered in any subject and then were removed from the data set. Also note that, contrary to [13], to avoid mutual exclusivity patterns we only analyze one level of gain/loss per gene; so, here we used high-level amplification (GISTIC score of 2) and homozygous deletion (GISTIC score of - 2).

**GBM CNA modules** Data come from **GBM CNA**, processed so individual alterations were grouped in modules at phenotype level of cancer-related pathways.

We reproduced the procedure used in Cheng *et al.* (2012) [13] to map alterations to functional modules of positive or negative effects within different cancer-related pathways as originally described in [11] and as detailed in Table 1 of [13].

**GBM co-occurrent** Data come originally from [6] and [4] and were obtained from Supplementary Material file `ST01.xls` (GBM_copy_number tab) of Attollini *et al.* (2010) and processed to keep only three highly correlated events described in [3], namely PTEN homozygous deletion, P16 homozygous deletion and EFGR low-level amplification.

**Ovarian CNV** Obtained from data set `ov.cgh` in the R package "Oncotree" [30]. Data are originally from [21]; `ov.cgh` from the "Oncotree" R package has also been used in [25].

**BRCA basal-like, subtypes, BRCA HER2, subtypes** Original data come from TCGA BRCA PUB mutation data set [9] and were obtained from cBioPortal [12, 17] using the R package "cgdsr" [19], restricting the data to subtype-specific significantly mutated genes within patients subtypes.

The data were then split in two, restricting the subjects to those classified as basal-like and HER2-enriched subtypes, respectively. To split the data set according to cancer subtypes we used the patient's classification by the gene expression-based PAM50 technique as detailed in Supplementary Table 1 of [9]. Then, for each subtype, we restricted the features to subtype-specific significantly mutated genes identified by MuSiC algorithm [15] and detailed in Supplementary Table 2 of [9] (Supplementary Tables 1-4.xls file in supplementary file nature11412-s2.zip).

The reason for splitting the data into two subsets is that cancer subtypes are believed to follow distinct evolutionary trajectories, and hence rely on at least some different drivers and/or pathways, and show differences in the chronology of accumulation of alterations [1, 2, 13]. Thus, sample heterogeneity in terms of different cancer subtypes (intertumor heterogeneity) can be confounding and hamper the identification of existing relations in the data. Sample stratification can alleviate this to some extent and should allow to focus on relevant events for specific subsets of subjects [10].

#### 1.2. Bootstrapping on the cancer data sets

If the bootstrapping process resulted in a feature becoming absent from the data, or two or more features having identical patterns (i.e., one feature being identical to another) we discarded the bootstrap sample and obtained a new one; this is done to ensure that all bootstrapped data sets have paths of identical length (see also section "Preprocessing of data for CPMs" in S4 Text). This, therefore, leads to JS values that are more optimistic (smaller).

#### 2. Cancer data sets: additional results, figures

#### 2.1. Cancer data sets: $JS_{ob}$ and unpredictability for the bootstrap runs

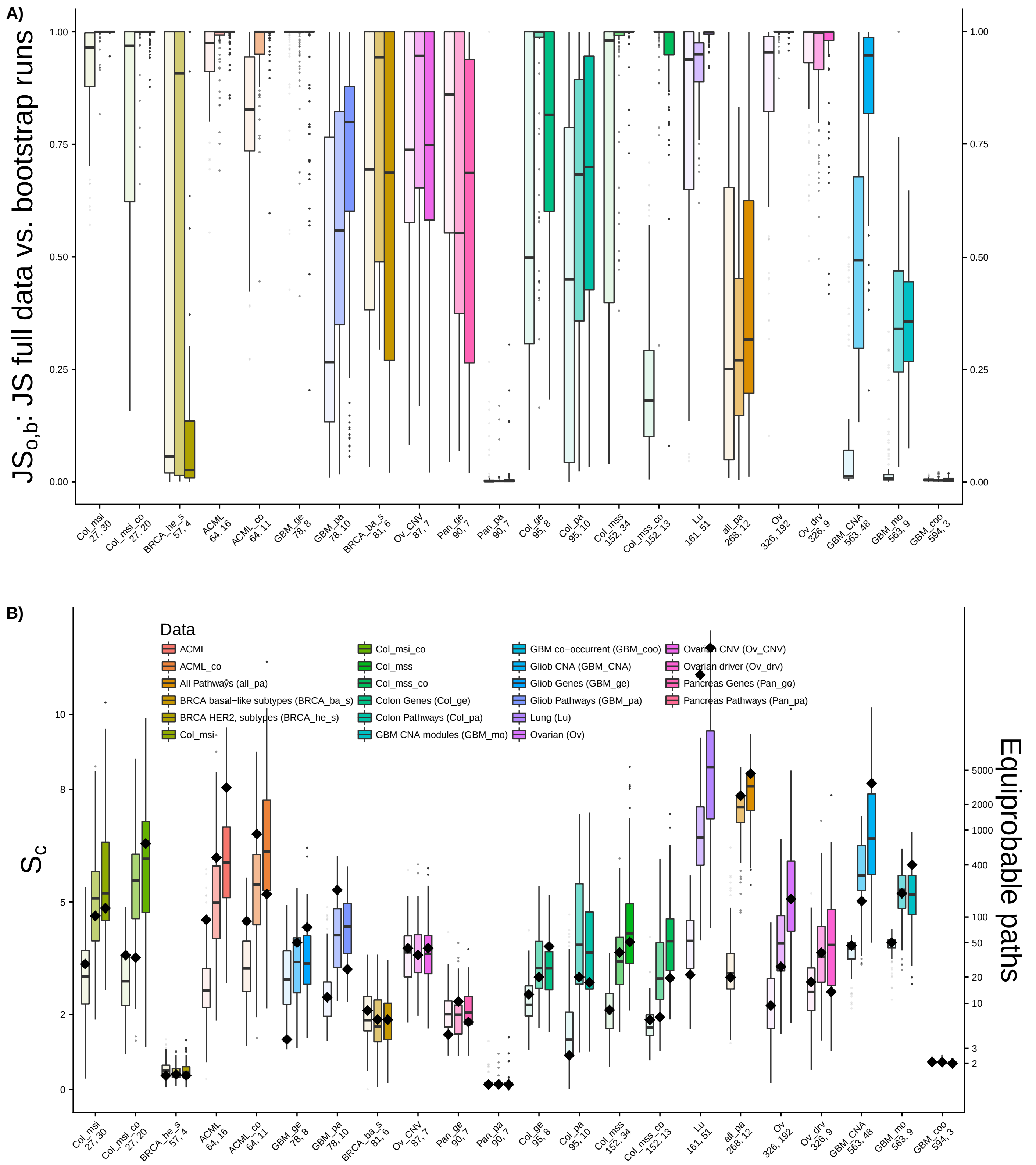

Figure 1: Results from the cancer data sets analyzed with H-CBN. Data sets have been ordered by increasing sample size, and the x-axis labels provide the acronym (shown in full in the inset legend). Below the data set acronym are the number of subjects and the total number of features, respectively. Analysis were run three times, limiting the number of features analyzed to the 7, 10, and 12 most common ones; the boxplots for each data set are shown in increasing order of number of features. For data sets such as, say, Pancreas genes (Pan_Ge), with 7 features, using 7, 10, or 12 maximum features makes no difference in the number of features analyzed; the three replicate runs show run-to-run variability. A) $JS_{o,b}$: JS statistic for the comparison of the distribution of paths from running H-CBN on the original data set against the distribution of paths from running H-CBN on each one of the bootstrap runs. B) Diamonds show the $S_c$ from the full data, and boxplots the $S_c$ from the bootstrap runs. Right axis labeled by number of equiprobable paths equivalent to the $S_c$.

##### 2.1.1. Cancer data sets: distribution of number of mutations per subject

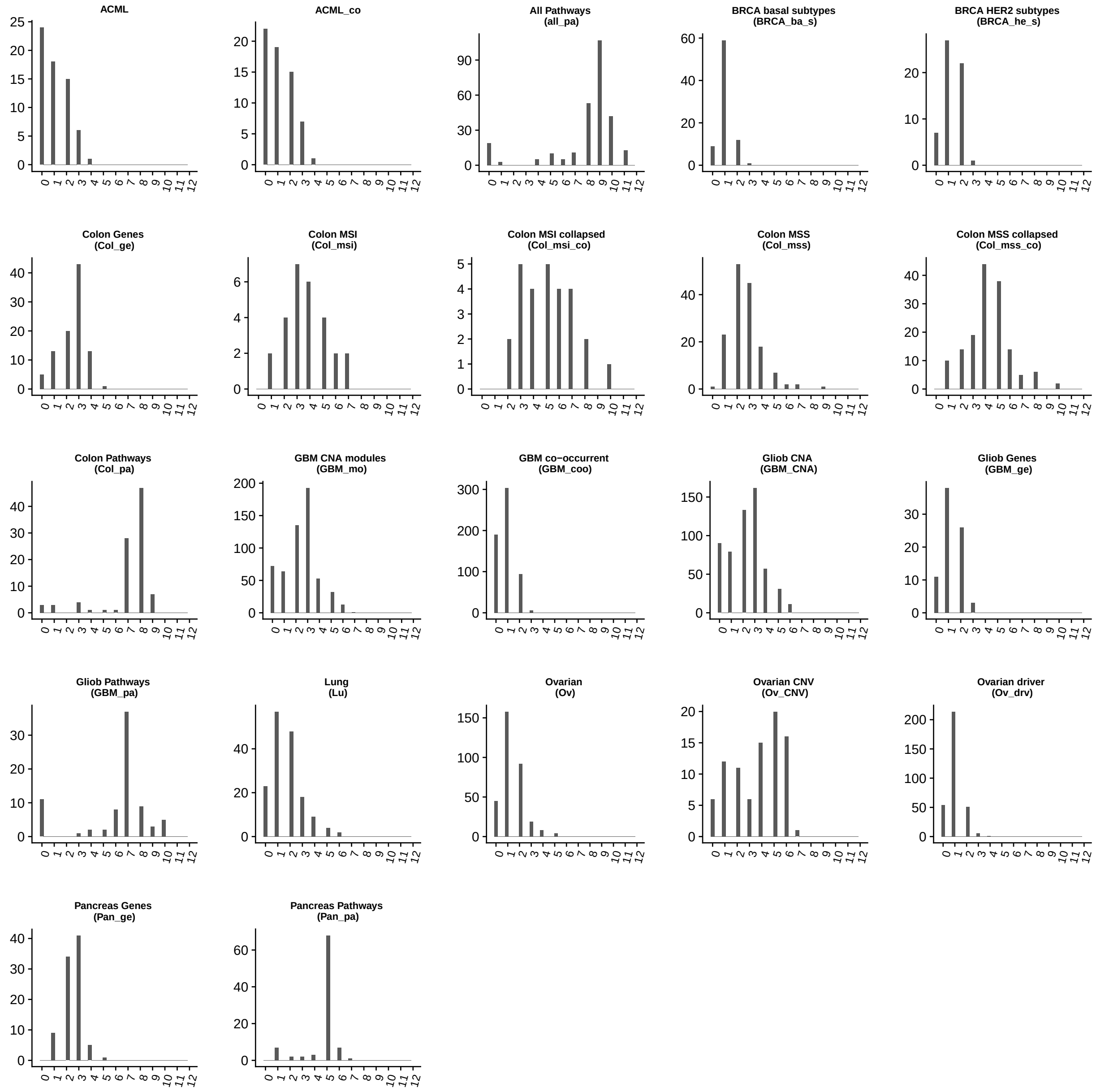

Figure 2: Cancer data sets: Histograms of number of mutations per subject in the data sets.

##### 2.1.2. Cancer data sets: proportion of individuals in which a mutation is present

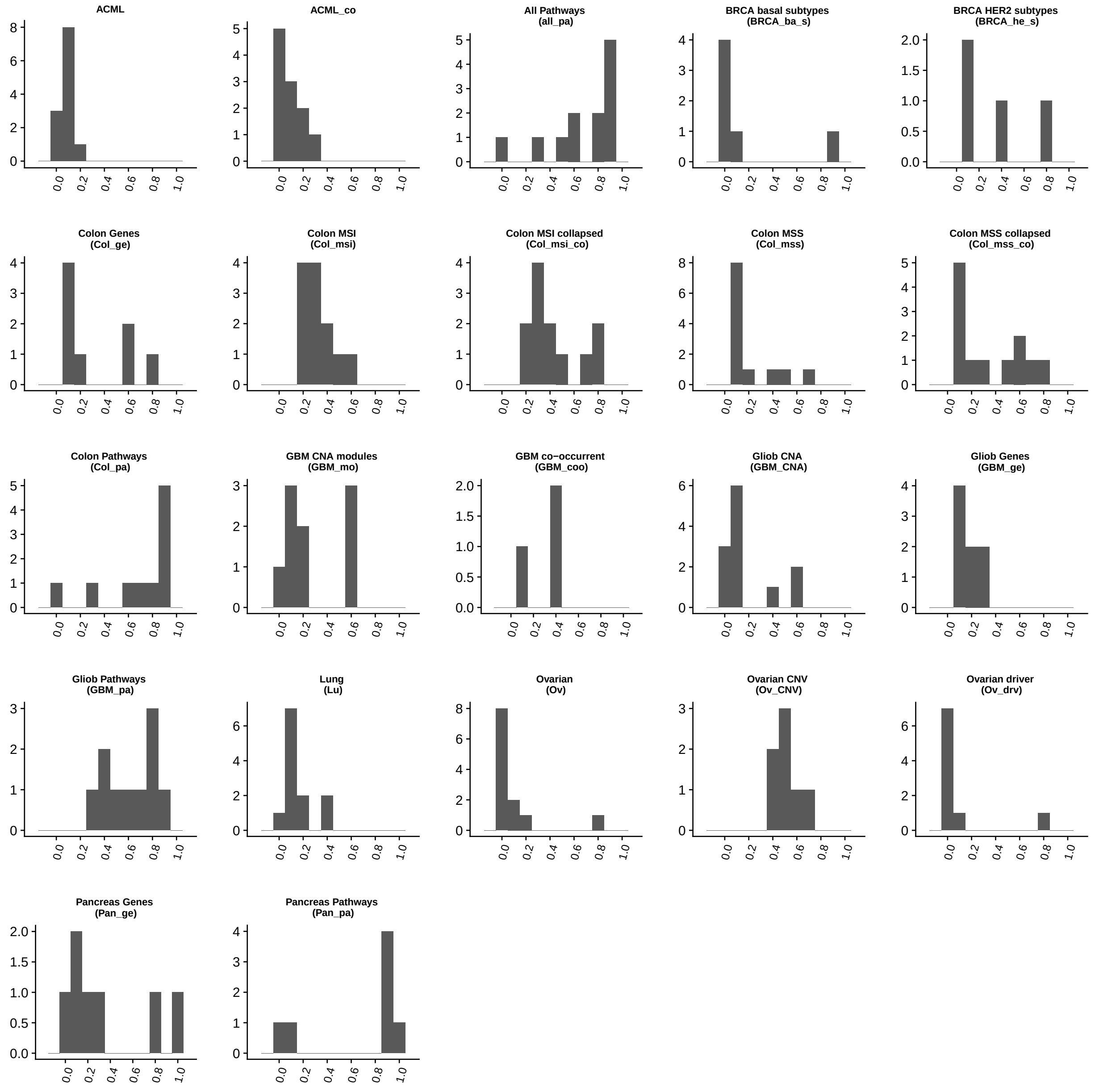

Figure 3: Cancer data sets: Histograms of proportion of individuals in which each mutation is present. For example, in the PP data set, there are four mutations that are present in 80% to 90% of the individuals in the data set, 1 mutation present in 90% to 100% of the individuals, 1 mutation in between 0 and 10% of the individuals, and 1 in between 10% and 15%.

##### 2.1.3. Cancer data sets: scatterplots of $JS_{o,b}$, $S_c$, and number of paths to the maximum

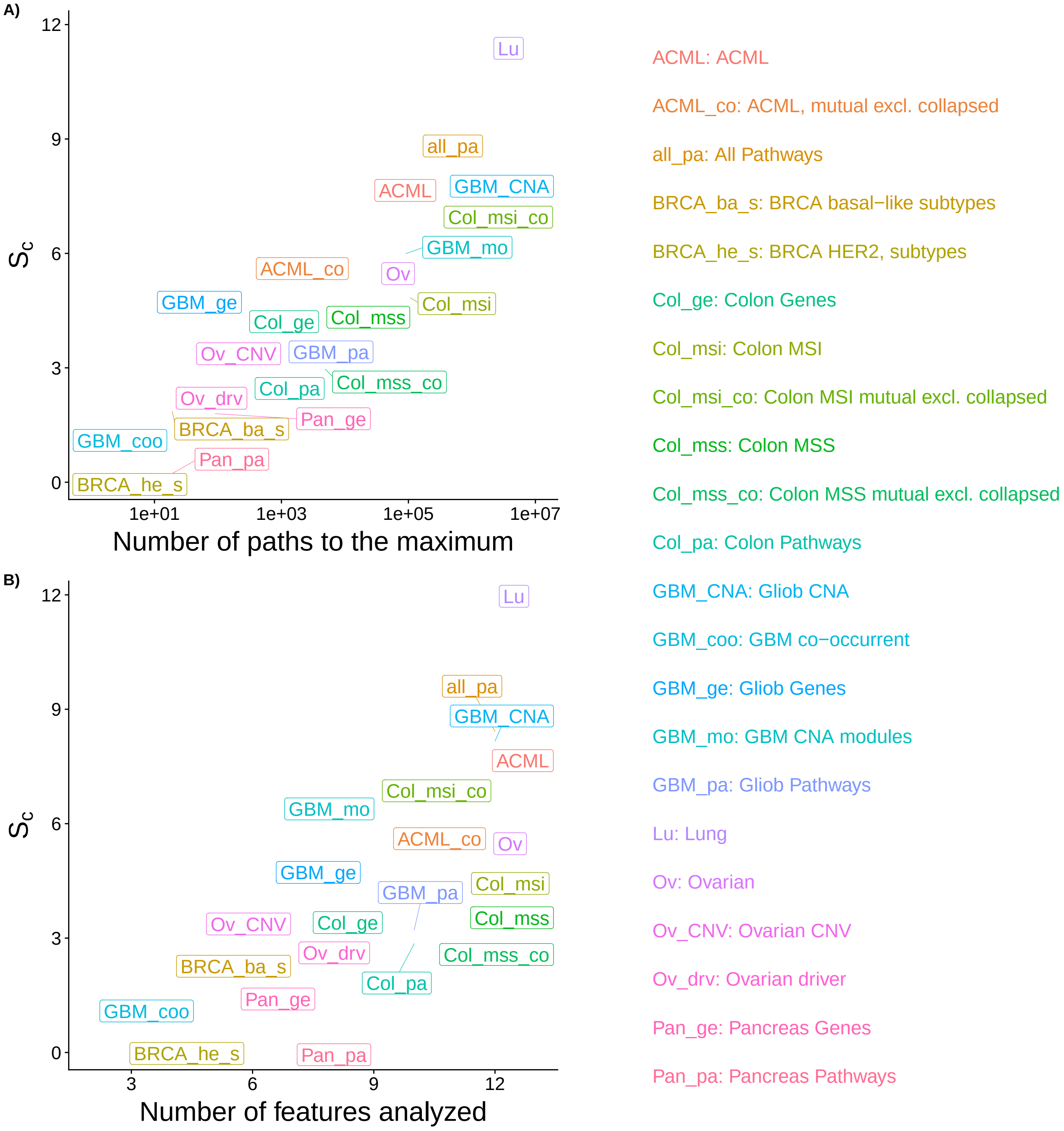

Figure 4: Cancer data sets: scatterplots of the relationship between $JS_{o,b}$, $S_c$, and number of paths to the maximum, using the data labels, using the statistics from analyses with 12 features.

#### 3. References

[1] a. Burrell, R., McGranahan, N., Bartek, J., Swanton, C., 2013. The causes and consequences of genetic heterogeneity in cancer evolution. *Nature*, **501(7467)**:338–345. doi:10.1038/nature12625. URL http://www.nature.com/doifinder/10.1038/nature12625.

[2] Anderson, W. F., Rosenberg, P. S., Prat, A., Perou, C. M., Sherman, M. E., 2014. How many etiological Subtypes of Breast Cancer:two,three, Four, or more? *JNCI: Journal of the National Cancer Institute*, **106(8)**:1–11.

[3] Attolini, C., Cheng, Y., Beroukhim, R., Getz, G., Abdel-Wahab, O., Levine, R. L., Mellinghoff, I. K., Michor, F., 2010. A mathematical framework to determine the temporal sequence of somatic genetic events in cancer. *Proceedings of the National Academy of Sciences*, **107(41)**:17604–17609. doi:10.1073/pnas.1009117107/-/DCSupplemental.www.pnas.org/cgi/doi/10.1073/pnas.1009117107. URL http://www.pnas.org/content/107/41/17604.short.

[4] Bamford, S., Dawson, E., Forbes, S., Clements, J., Pettett, R., Dogan, A., Flanagan, A., Teague, J., Futreal, P. A., Stratton, M. R., Wooster, R., 2004. The COSMIC (Catalogue of Somatic Mutations in Cancer) database and website. *Br J Cancer*, **91(2)**:355–358. doi:10.1038/sj.bjc.6601894. URL https://www.ncbi.nlm.nih.gov/pmc/articles/PMC2409828/.

[5] Brennan, C. W., Verhaak, R. G. W., McKenna, A., Campos, B., Noushmehr, H., Salama, S. R., Zheng, S., Chakravarty, D., Sanborn, J. Z., Berman, S. H., Beroukhim, R., Bernard, B., Wu, C.-J., Genovese, G., Shmulevich, I., Barnholtz-Sloan, J., Zou, L., Vegesna, R., Shukla, S. A., Ciriello, G., Yung, W. K., Zhang, W., Sougnez, C., Mikkelsen, T., Aldape, K., Bigner, D. D., Van Meir, E. G., Prados, M., Sloan, A., Black, K. L., Eschbacher, J., Finocchiaro, G., Friedman, W., Andrews, D. W., Guha, A., Iacocca, M., OtextquoterightNeill, B. P., Foltz, G., Myers, J., Weisenberger, D. J., Penny, R., Kucherlapati, R., Perou, C. M., Hayes, D. N., Gibbs, R., Marra, M., Mills, G. B., Lander, E., Spellman, P., Wilson, R., Sander, C., Weinstein, J., Meyerson, M., Gabriel, S., Laird, P. W., Haussler, D., Getz, G., Chin, L., Benz, C., Barrett, W., Ostrom, Q., Wolinsky, Y., Bose, B., Boulos, P. T., Boulos, M., Brown, J., Czerinski, C., Eppley, M., Kempista, T., Kitko, T., Koyfman, Y., Rabeno, B., Rastogi, P., Sugarman, M., Swanson, P., Yalamanchii, K., Otey, I. P., Liu, Y. S., Xiao, Y., Auman, J. T., Chen, P.-C., Hadjipanayis, A., Lee, E., Lee, S., Park, P. J., Seidman, J., Yang, L., Kalkanis, S., Poisson, L. M., Raghunathan, A., Scarpace, L., Bressler, R., Eakin, A., Iype, L., Kreisberg, R. B., Leinonen, K., Reynolds, S., Rovira, H., Thorsson, V., Annala, M. J., Paulauskis, J., Curley, E., Hatfield, M., Mallery, D., Morris, S., Shelton, T., Shelton, C., Sherman, M., Yena, P., Cuppini, L., DiMeco, F., Eoli, M., Maderna, E., Pollo, B., Saini, M., Balu, S., Hoadley, K. A., Li, L., Miller, C. R., Shi, Y., Topal, M. D., Wu, J., Dunn, G., Giannini, C., Aksoy, B. A., Antipin, Y., Borsu, L., Cerami, E., Gao, J., Gross, B., Jacobsen, A., Ladanyi, M., Lash, A., Liang, Y., Reva, B., Schultz, N., Shen, R., Socci, N. D., Viale, A., Ferguson, M. L., Chen, Q.-R., A, D., Dillon, L. A. L., Shaw, K. R. M., Sheth, M., Tarnuzzer, R., Wang, Z., Yang, L., Davidsen, T., Guyer, M. S., Ozenberger, B. A., Sofia, H. J., Bergsten, J., Eckman, J., Harr, J., Smith, C., Tucker, K., Winemiller, C., Zach, L. A., Ljubimova, J. Y., Eley, G., Ayala, B., Jensen, M. A., Kahn, A., Pihl, T. D., Pot, D. A., Wan, Y., Hansen, N., Hothi, P., Lin, B., Shah, N., Yoon, J.-g., Lau, C., Berens, M., Ardlie, K., Carter, S. L., Cherniack, A. D., Noble, M., Cho, J., Cibulskis, K., DiCara, D., Frazer, S., Gabriel, S. B., Gehlenborg, N., Gentry, J., Heiman, D., Kim, J., Jing, R., Lander, E. S., Lawrence, M., Lin, P., Mallard, W., Onofrio, R. C., Saksena, G., Schumacher, S., Stojanov, P., Tabak, B., Voet, D., Zhang, H., Dees, N. N., Ding, L., Fulton, L. L., Fulton, R. S., Kanchi, K.-L., Mardis, E. R., Wilson, R. K., Baylin, S. B., Harshyne, L., Cohen, M. L., Devine, K., Sloan, A. E., VandenBerg, S. R., Berger, M. S., Carlin, D., Craft, B., Ellrott, K., Goldman, M., Goldstein, T., Grifford, M., Ma, S., Ng, S., Stuart, J., Swatloski, T., Waltman, P., Zhu, J., Foss, R., Frentzen, B., McTiernan, R., Yachnis, A., Mao, Y., Akbani, R., Bogler, O., Fuller, G. N., Liu, W., Liu, Y., Lu, Y., Mills,

G., Protopopov, A., Ren, X., Sun, Y., Yung, W. K. A., Zhang, J., Chen, K., Weinstein, J. N., Bootwalla, ., 2013. The Somatic Genomic Landscape of Glioblastoma. *Cell*, **155(2)**:462–477.

[6] Cancer Genome Atlas Research Network, 2008. Comprehensive genomic characterization defines human glioblastoma genes and core pathways. *Nature*, **455(7216)**:1061–1068.

[7] Cancer Genome Atlas Research Network, 2011. Integrated genomic analyses of ovarian carcinoma. *Nature*, **474(7353)**:609–615. doi:10.1038/nature10166.

[8] Cancer Genome Atlas Research Network, 2012. Comprehensive molecular characterization of human colon and rectal cancer. *Nature*, **487(7407)**:330–337. doi:10.1038/nature11252. URL https://www.nature.com/articles/nature11252.

[9] Cancer Genome Atlas Research Network, 2012. Comprehensive molecular portraits of human breast tumours. *Nature*, **490(7418)**:61–70.

[10] Caravagna, G., Graudenzi, A., Ramazzotti, D., Sanz-Pamplona, R., Sano, L. D., Mauri, G., Moreno, V., Antoniotti, M., Mishra, B., 2016. Algorithmic methods to infer the evolutionary trajectories in cancer progression. *PNAS*, **113(28)**:E4025–E4034. doi:10.1073/pnas.1520213113. URL http://www.pnas.org/content/113/28/E4025.

[11] Cerami, E., Demir, E., Schultz, N., Taylor, B. S., Sander, C., 2010. Automated Network Analysis Identifies Core Pathways in Glioblastoma. *PLoS ONE*, **5(2)**:e8918.

[12] Cerami, E., Gao, J., Dogrusoz, U., Gross, B. E., Sumer, S. O., Aksoy, B. A., Jacobsen, A., Byrne, C. J., Heuer, M. L., Larsson, E., Antipin, Y., Reva, B., Goldberg, A. P., Sander, C., Schultz, N., 2012. The cBio Cancer Genomics Portal: An Open Platform for Exploring Multidimensional Cancer Genomics Data: Figure 1. *Cancer Discovery*, **2(5)**:401–404.

[13] Cheng, Y.-K., Beroukhim, R., Levine, R. L., Mellinghoff, I. K., Holland, E. C., Michor, F., 2012. A mathematical methodology for determining the temporal order of pathway alterations arising during gliomagenesis. *PLoS computational biology*, **8(1)**:e1002337. doi:10.1371/journal.pcbi.1002337. URL http://www.pubmedcentral.nih.gov/articlerender.fcgi?artid=3252265&tool=pmcentrez&rendertype=abstract.

[14] De Sano, L., Caravagna, G., Ramazzotti, D., Graudenzi, A., Mauri, G., Mishra, B., Antoniotti, M., 2016. TRONCO: An R package for the inference of cancer progression models from heterogeneous genomic data. *Bioinformatics*, **32(12)**:1911–1913. doi:10.1093/bioinformatics/btw035.

[15] Dees, N. D., Zhang, Q., Kandoth, C., Wendl, M. C., Schierding, W., Koboldt, D. C., Mooney, T. B., Callaway, M. B., Dooling, D., Mardis, E. R., Wilson, R. K., Ding, L., 2012. MuSiC: Identifying mutational significance in cancer genomes. *Genome Research*, **22(8)**:1589–1598.

[16] Ding, L., Getz, G., Wheeler, D. A., Mardis, E. R., McLellan, M. D., Cibulskis, K., Sougnez, C., Greulich, H., Muzny, D. M., Morgan, M. B., Fulton, L., Fulton, R. S., Zhang, Q., Wendl, M. C., Lawrence, M. S., Larson, D. E., Chen, K., Dooling, D. J., Sabo, A., Hawes, A. C., Shen, H., Jhangiani, S. N., Lewis, L. R., Hall, O., Zhu, Y., Mathew, T., Ren, Y., Yao, J., Scherer, S. E., Clerc, K., Metcalf, G. A., Ng, B., Milosavljevic, A., Gonzalez-Garay, M. L., Osborne, J. R., Meyer, R., Shi, X., Tang, Y., Koboldt, D. C., Lin, L., Abbott, R., Miner, T. L., Pohl, C., Fewell, G., Haipek, C., Schmidt, H., Dunford-Shore, B. H., Kraja, A., Crosby, S. D., Sawyer, C. S., Vickery, T., Sander, S., Robinson, J., Winckler, W., Baldwin, J., Chirieac, L. R., Dutt, A., Fennell, T., Hanna, M., Johnson, B. E., Onofrio, R. C., Thomas, R. K., Tonon, G., Weir, B. A., Zhao, X., Ziaugra, L., Zody, M. C., Giordano, T., Orringer, M. B., Roth, J. A., Spitz, M. R., Wistuba, I. I., Ozenberger, B., Good, P. J., Chang, A. C., Beer, D. G., Watson, M. A., Ladanyi, M., Broderick, S., Yoshizawa, A., Travis, W. D., Pao, W., Province, M. A., Weinstock, G. M., Varmus, H. E., Gabriel, S. B., Lander, E. S., Gibbs, R. A., Meyerson,

M., Wilson, R. K., 2008. Somatic mutations affect key pathways in lung adenocarcinoma. *Nature*, **455(7216)**:1069–1075. doi:10.1038/nature07423.

[17] Gao, J., Aksoy, B. A., Dogrusoz, U., Dresdner, G., Gross, B., Sumer, S. O., Sun, Y., Jacobsen, A., Sinha, R., Larsson, E., Cerami, E., Sander, C., Schultz, N., 2013. Integrative analysis of complex cancer genomics and clinical profiles using the cBioPortal. *Science Signaling*, **6(269)**:pl1. doi:10.1126/scisignal.2004088.

[18] Gerstung, M., Eriksson, N., Lin, J., Vogelstein, B., Beerenwinkel, N., 2011. The Temporal Order of Genetic and Pathway Alterations in Tumorigenesis. *PLoS ONE*, **6(11)**:e27136. doi:10.1371/journal.pone.0027136. URL http://dx.plos.org/10.1371/journal.pone.0027136http://www.bsse.ethz.ch/cbg/software/ct-cbn.

[19] Jacobsen, A., Questions, c., 2018. Cgdsr: R-Based API for Accessing the MSKCC Cancer Genomics Data Server (CGDS). URL https://CRAN.R-project.org/package=cgdsr.

[20] Jones, S., Zhang, X., Parsons, D. W., Lin, J. C.-H., Leary, R. J., Angenendt, P., Mankoo, P., Carter, H., Kamiyama, H., Jimeno, A., Hong, S.-M., Fu, B., Lin, M.-T., Calhoun, E. S., Kamiyama, M., Walter, K., Nikolskaya, T., Nikolsky, Y., Hartigan, J., Smith, D. R., Hidalgo, M., Leach, S. D., Klein, A. P., Jaffee, E. M., Goggins, M., Maitra, A., Iacobuzio-Donahue, C., Eshleman, J. R., Kern, S. E., Hruban, R. H., Karchin, R., Papadopoulos, N., Parmigiani, G., Vogelstein, B., Velculescu, V. E., Kinzler, K. W., 2008. Core signaling pathways in human pancreatic cancers revealed by global genomic analyses. *Science (New York, N.Y.)*, **321(5897)**:1801–6. doi:10.1126/science.1164368. URL http://www.ncbi.nlm.nih.gov/pubmed/18772397.

[21] Knutsen, T., Gobu, V., Knaus, R., Padilla-Nash, H., Augustud, M., Strausberg, R. L., Kirsch, I. R., Sirotkin, K., Ried, T., 2005. The Interactive Online SKY/M-FISH & CGH Database and the Entrez Cancer Chromosomes Search Database: Linkage of Chromosomal Aberrations with the Genome Sequence. *Genes, Chromosomes and Cancer*, **44(1)**:52–64.

[22] Lawrence, M., Huber, W., Pagès, H., Aboyoun, P., Carlson, M., Gentleman, R., Morgan, M. T., Carey, V. J., 2013. Software for computing and annotating genomic ranges. *PLoS computational biology*, **9(8)**:e1003118. doi:10.1371/journal.pcbi.1003118. URL http://www.pubmedcentral.nih.gov/articlerender.fcgi?artid=3738458&tool=pmcentrez&rendertype=abstract.

[23] Mermel, C. H., Schumacher, S. E., Hill, B., Meyerson, M. L., Beroukhim, R., Getz, G., 2011. GISTIC2.0 facilitates sensitive and confident localization of the targets of focal somatic copy-number alteration in human cancers. *Genome Biology*, **12(4)**:R41.

[24] Misra, N., Szczurek, E., Vingron, M., 2014. Inferring the paths of somatic evolution in cancer. *Bioinformatics (Oxford, England)*, **30(17)**:2456–2463. doi:10.1093/bioinformatics/btu319. URL http://www.ncbi.nlm.nih.gov/pubmed/24812340.

[25] Olde Loohuis, L., Caravagna, G., Graudenzi, A., Ramazzotti, D., Mauri, G., Antoniotti, M., Mishra, B., 2014. Inferring Tree Causal Models of Cancer Progression with Probability Raising. *PLOS ONE*, **9(10)**:e108358. doi:10.1371/journal.pone.0108358. URL https://journals.plos.org/plosone/article?id=10.1371/journal.pone.0108358.

[26] Parsons, D. W., Jones, S., Zhang, X., Lin, J. C.-H., Leary, R. J., Angenendt, P., Mankoo, P., Carter, H., Siu, I.-M., Gallia, G. L., Olivi, A., McLendon, R., Rasheed, B. A., Keir, S., Nikolskaya, T., Nikolsky, Y., Busam, D. A., Tekleab, H., Diaz, L. A., Hartigan, J., Smith, D. R., Strausberg, R. L., Marie, S. K. N., Shinjo, S. M. O., Yan, H., Riggins, G. J., Bigner, D. D., Karchin, R., Papadopoulos, N., Parmigiani, G., Vogelstein, B., Velculescu, V. E., Kinzler, K. W., 2008. An Integrated Genomic Analysis of Human Glioblastoma Multiforme. *Science*, **321(5897)**:1807–1812. doi:10.1126/science.1164382. URL http://science.sciencemag.org/content/321/5897/1807.

[27] Piazza, R., Valletta, S., Winkelmann, N., Redaelli, S., Spinelli, R., Pirola, A., Antolini, L., Mologni, L., Donadoni, C., Papaemmanuil, E., Schnittger, S., Kim, D.-W., Boultwood, J., Rossi, F., Gaipa, G., De Martini, G. P., di Celle, P. F., Jang, H. G., Fantin, V., Bignell, G. R., Magistroni, V., Haferlach, T., Pogliani, E. M., Campbell, P. J., Chase, A. J., Tapper, W. J., Cross, N. C. P., Gambacorti-Passerini, C., 2013. Recurrent SETBP1 mutations in atypical chronic myeloid leukemia. *Nature Genetics*, **45(1)**:18–24.

[28] Ramazzotti, D., Caravagna, G., Olde Loohuis, L., Graudenzi, A., Korsunsky, I., Mauri, G., Antoniotti, M., Mishra, B., 2015. CAPRI: Efficient inference of cancer progression models from cross-sectional data. *Bioinformatics*, **31(18)**:3016–3026. doi:10.1093/bioinformatics/btv296. URL https://academic.oup.com/bioinformatics/article-lookup/doi/10.1093/bioinformatics/btv296.

[29] Sjoblom, T., Jones, S., Wood, L. D., Parsons, D. W., Lin, J., Barber, T. D., Mandelker, D., Leary, R. J., Ptak, J., Silliman, N., Szabo, S., Buckhaults, P., Farrell, C., Meeh, P., Markowitz, S. D., Willis, J., Dawson, D., Willson, J. K. V., Gazdar, A. F., Hartigan, J., Wu, L., Liu, C., Parmigiani, G., Park, B. H., Bachman, K. E., Papadopoulos, N., Vogelstein, B., Kinzler, K. W., Velculescu, V. E., 2006. The Consensus Coding Sequences of Human Breast and Colorectal Cancers. *Science*, **314(5797)**:268–274. doi:10.1126/science.1133427. URL http://dx.doi.org/10.1126/science.1133427.

[30] Szabo, A., Pappas, L., 2013. Oncotree: Estimating oncogenetic trees. R package version 0.3.3. URL http://cran.r-project.org/package=Oncotree.

[31] Tokheim, C. J., Papadopoulos, N., Kinzler, K. W., Vogelstein, B., Karchin, R., 2016. Evaluating the evaluation of cancer driver genes. *PNAS*, **113(50)**:14330–14335. doi:10.1073/pnas.1616440113. URL http://www.pnas.org/content/113/50/14330.

[32] Wood, L. D., Parsons, D. W., Jones, S., Lin, J., Sjoblom, T., Leary, R. J., Shen, D., Boca, S. M., Barber, T., Ptak, J., Silliman, N., Szabo, S., Dezso, Z., Ustyanksky, V., Nikolskaya, T., Nikolsky, Y., Karchin, R., Wilson, P. A., Kaminker, J. S., Zhang, Z., Croshaw, R., Willis, J., Dawson, D., Shipitsin, M., Willson, J. K. V., Sukumar, S., Polyak, K., Park, B. H., Pethiyagoda, C. L., Pant, P. V. K., Ballinger, D. G., Sparks, A. B., Hartigan, J., Smith, D. R., Suh, E., Papadopoulos, N., Buckhaults, P., Markowitz, S. D., Parmigiani, G., Kinzler, K. W., Velculescu, V. E., Vogelstein, B., 2007. The Genomic Landscapes of Human Breast and Colorectal Cancers. *Science*, **318(5853)**:1108–1113. doi:10.1126/science.1145720. URL http://dx.doi.org/10.1126/science.1145720.
