## Supplementary material for "Every which way? On predicting tumor evolution using cancer progression models": S6_Text

### Supporting Information for "Every which way? On predicting tumor evolution using cancer progression models"

2019-04-17

### S6 Text. Additional results.

---

#### Contents

**List of Figures** 2

**1 Overall patterns for the six CPM procedures** 3

**2 Probability of recovering the most common LOD** 4

**3 OT and H-CBN, JS, weighted vs. unweighted** 6

**4 CAPRI and H-CBN, 1-precision, unweighted** 7

**5 CAPRESE and OT, 1-precision, unweighted** 8

**6 Coefficient of variation of JS** 9

**7 Number of paths inferred** 10

**8 Slopes of regressions of 1-recall and 1-precision on LOD diversity, $S_p$** 11

**9 Coefficient of variation of $S_c$** 12

**10 Estimated $S_c$ by H-CBN** 13

**11 Analysis of deviance tables for fitted models** 14

*, http://ligarto.org/rdiaz

12 Number of local maxima, mutations of local maxima, reciprocal sign epistasis and performance 29

13 LOD and CPM diversity: ratios and slopes 30

14 References 30

#### List of Figures

1 Summary performance measures for all six CPM procedures for all combinations of sample size, type of landscape, detection regime, and number of genes. For all measures, smaller is better. For OT, H-CBN, and MCCBN, Jensen-Shannon (JS) entropy and 1-precision use probability-weighted predicted paths (see text). Each point represented is the average of 210 points (35 replicates of each one of the six combinations of 3 initial size by 2 mutation rate regimes; we are thus marginalizing over initial size by mutation rate; each one of the 210 points is, itself, the average of five runs on different partitions of the simulated data. . . . . 3

2 Probability of recovering the most common LOD: probability that the most common observed path to the maximum is among the paths allowed by the CPMs. . 4

3 Probability of recovering the most common LOD and 1-recall: relationship. . . . 5

4 Comparison of the performance of OT and H-CBN using weighted and unweighted probabilities of paths to the maximum. . . . . . . . . . . . . . . . . . . . . . 6

5 Comparison of the performance of CAPRI with H-CBN using weighted and unweighted probabilities of paths to the maximum. . . . . . . . . . . . . . . . . . . . . 7

6 Comparison of the performance of CAPRESE with OT using weighted and unweighted probabilities of paths to the maximum. . . . . . . . . . . . . . . . . . . . . 8

7 Coefficient of variation (standard deviation/mean) of JS for each combination of model and type of fitness landscape. The bottom figure represents the same information as the top one but only for the three models shown in the paper. The coefficient of variation has been computed from the five runs for each landscapes on each combination of sample size and detection regime. . . . . . . . . . . . . . 9

8 Number of paths to the maximum according to the CPMs. . . . . . . . . . . . . . 10

9 Slopes of regressions of 1-recall and 1-precision on LOD diversity, $S_p$ . . . . . . . 11

10 Coefficient of variation (standard deviation/mean) of $S_c$ for each combination of model and type of fitness landscape. The coefficient of variation has been computed from the five runs for each landscapes on each combination of sample size and detection regime. For OT and H-CBN, it is computed using the probability-weighted predicted paths (see text). Each point plotted is the average of 210 points. . . . . . . . . . . . . . . . . . . . . . . . . . . . . . . . . . . . . . . 12

11 Estimated $S_c$ by H-CBN for all combinations of sample size by type of landscape by detection regime by number of genes. Each box plot shows 1050 points. . . . . 13

12 Slopes of regression of JS and 1-recall on mean number of mutations of local maxima. . . . . . . . . . . . . . . . . . . . . . . . . . . . . . . . . . . . . . . 29

---

### 1. Overall patterns for the six CPM procedures

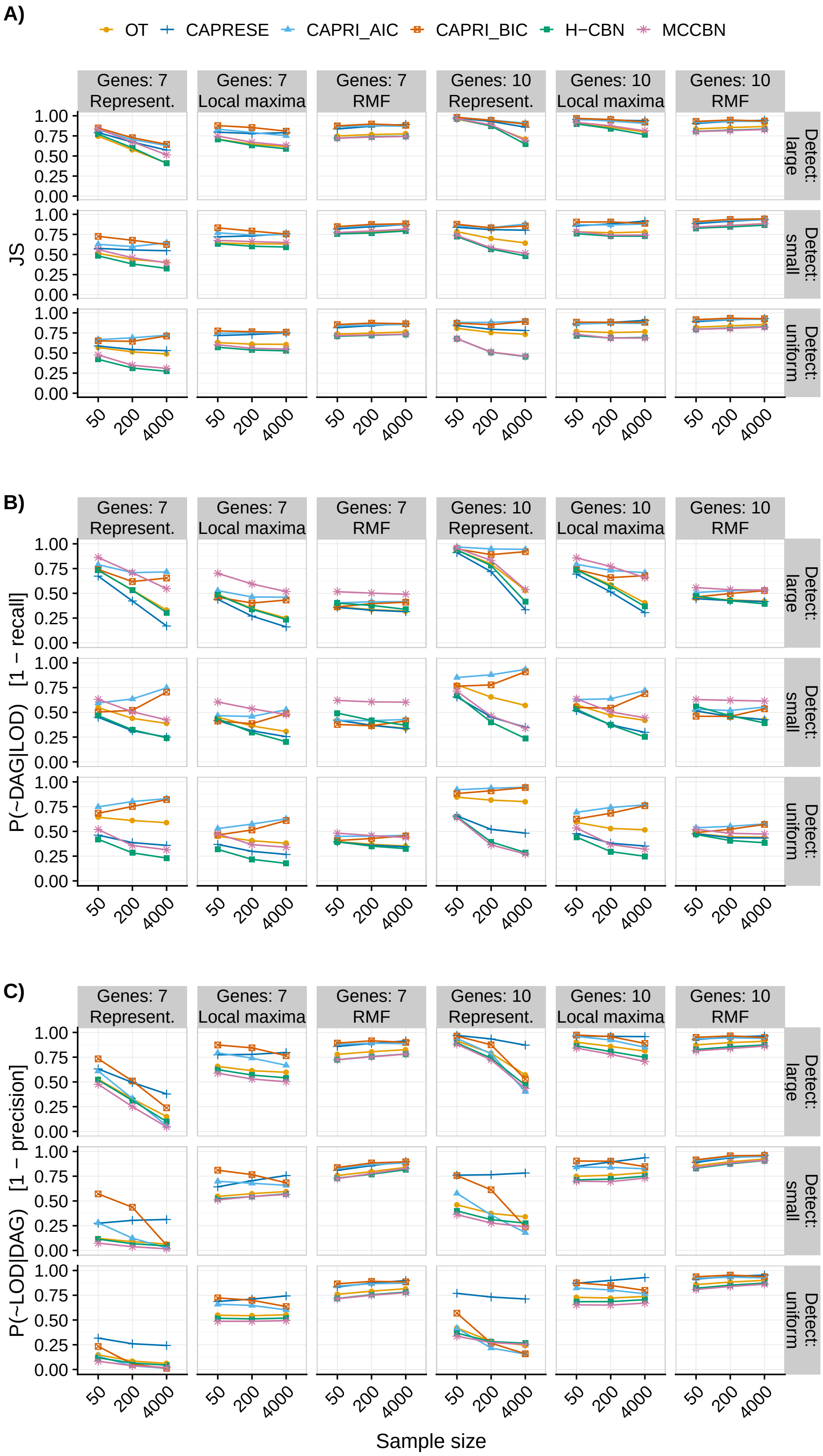

Figure 1: Summary performance measures for all six CPM procedures for all combinations of sample size, type of landscape, detection regime, and number of genes. For all measures, smaller is better. For OT, H-CBN, and MCCBN, Jensen-Shannon (JS) entropy and 1-precision use probability-weighted predicted paths (see text). Each point represented is the average of 210 points (35 replicates of each one of the six combinations of 3 initial size by 2 mutation rate regimes; we are thus marginalizing over initial size by mutation rate; each one of the 210 points is, itself, the average of five runs on different partitions of the simulated data.

#### 2. Probability of recovering the most common LOD

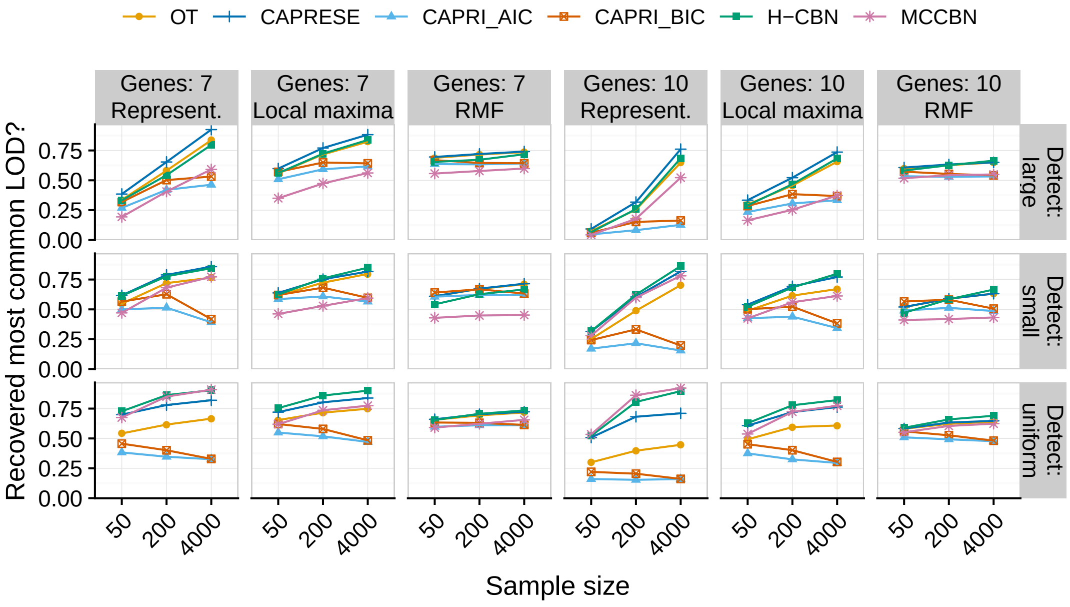

Figure 2: Probability of recovering the most common LOD: probability that the most common observed path to the maximum is among the paths allowed by the CPMs.

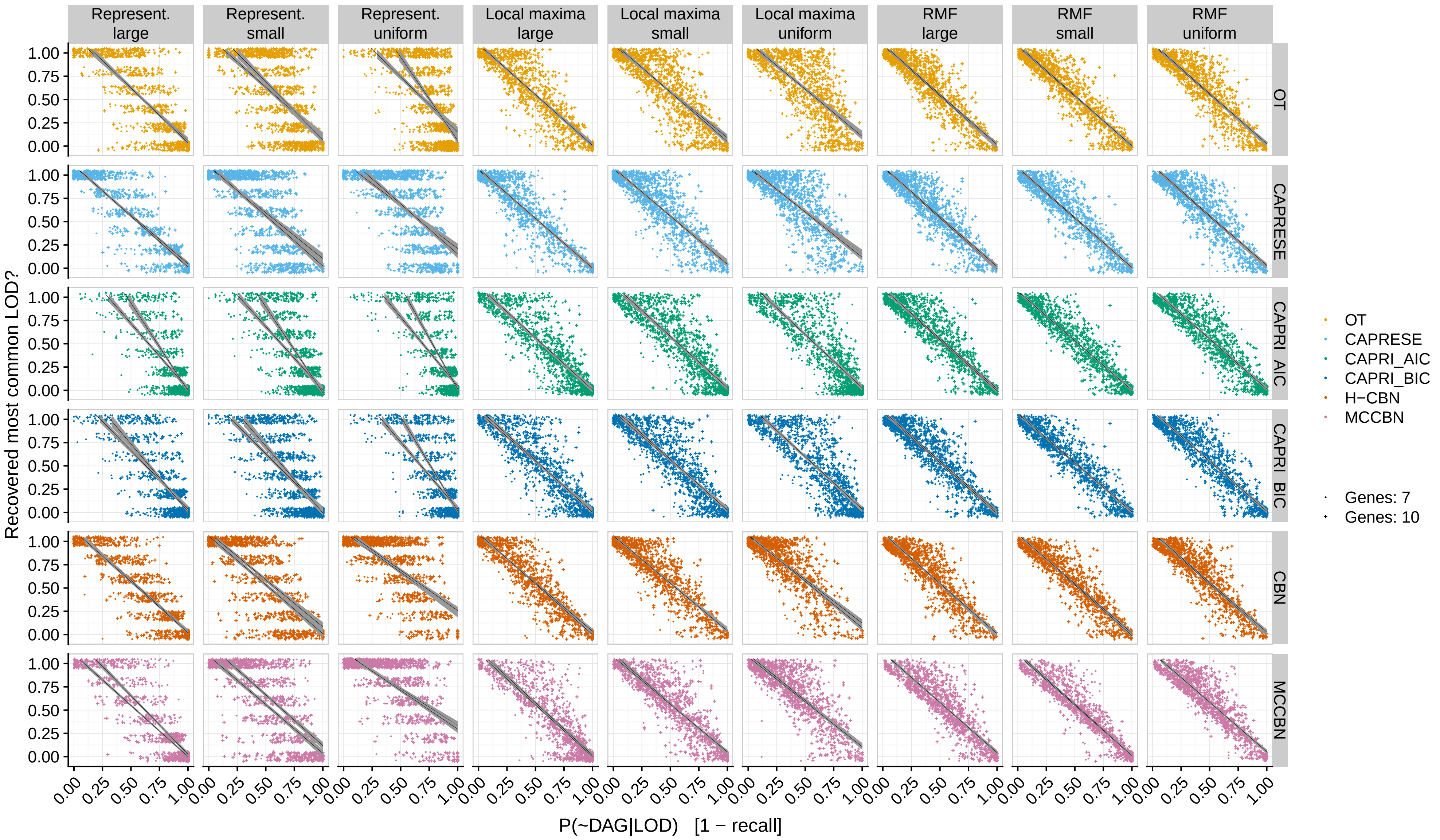

Figure 3: Probability of recovering the most common LOD and 1-recall: relationship.

##### 3. OT and H-CBN, JS, weighted vs. unweighted

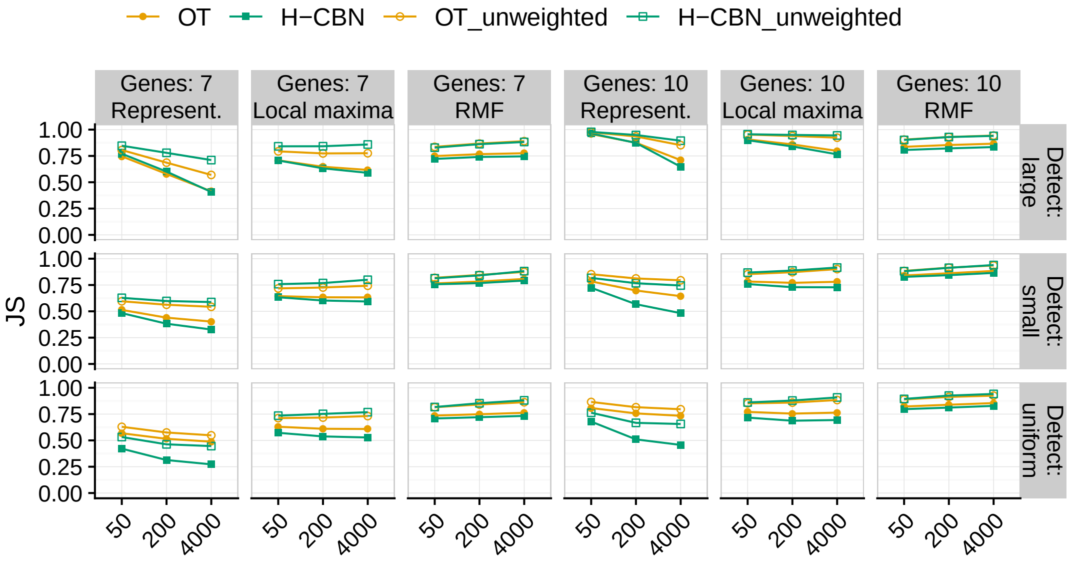

Figure 4: Comparison of the performance of OT and H-CBN using weighted and unweighted probabilities of paths to the maximum.

#### 4. CAPRI and H-CBN, 1-precision, unweighted

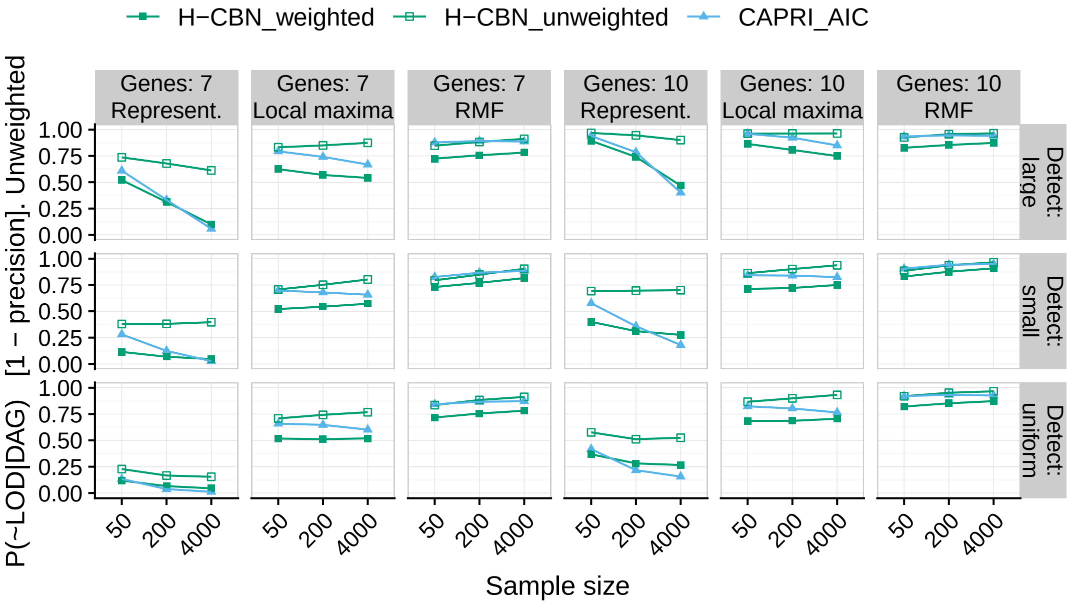

Figure 5: Comparison of the performance of CAPRI with H-CBN using weighted and unweighted probabilities of paths to the maximum.

#### 5. CAPRESE and OT, 1-precision, unweighted

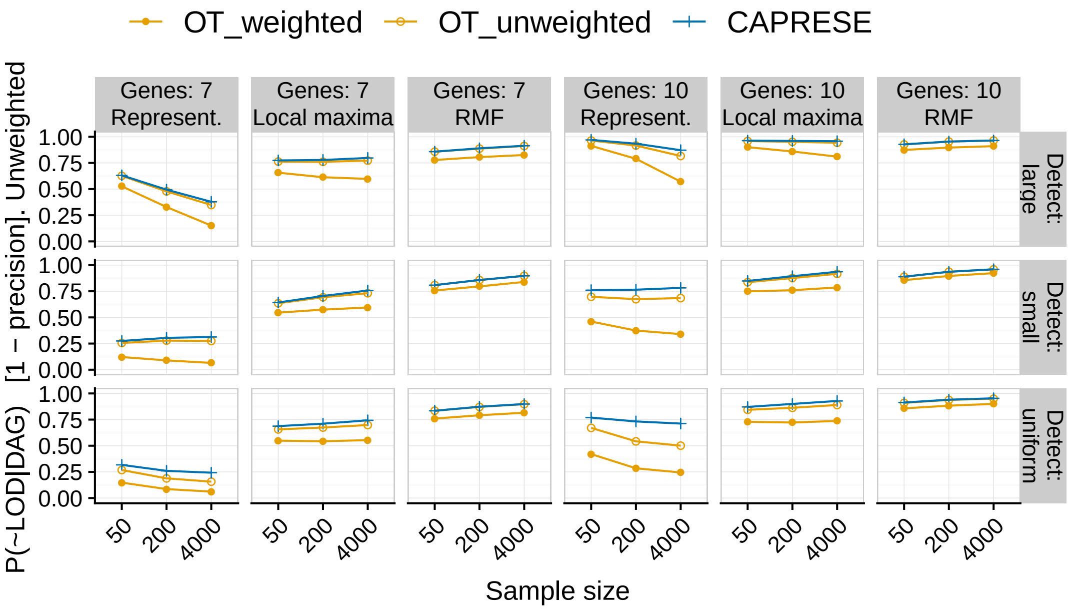

Figure 6: Comparison of the performance of CAPRESE with OT using weighted and unweighted probabilities of paths to the maximum.

The results of OT here are remarkable because OT can only build trees, and therefore cannot reflect the dependency of a mutation on two or more upstream mutations so it is prone to allow more paths to the maximum. The results of OT contrasts with those of CAPRESE, the other model that only builds trees. CAPRESE is building DAGs of restrictions that have too few restrictions and, therefore, allow for too many paths to the maximum. One notable difference between the two models is that with OT it is relatively simple to use a measure of 1-precision that weights by the probability of each path. The performance of OT, even if we use unweighted probabilities of paths, is much better than that of CAPRESE but improves even further when using weighted paths, again highlighting the usefulness of weighting paths to obtain more accurate predictions.

#### 6. Coefficient of variation of JS

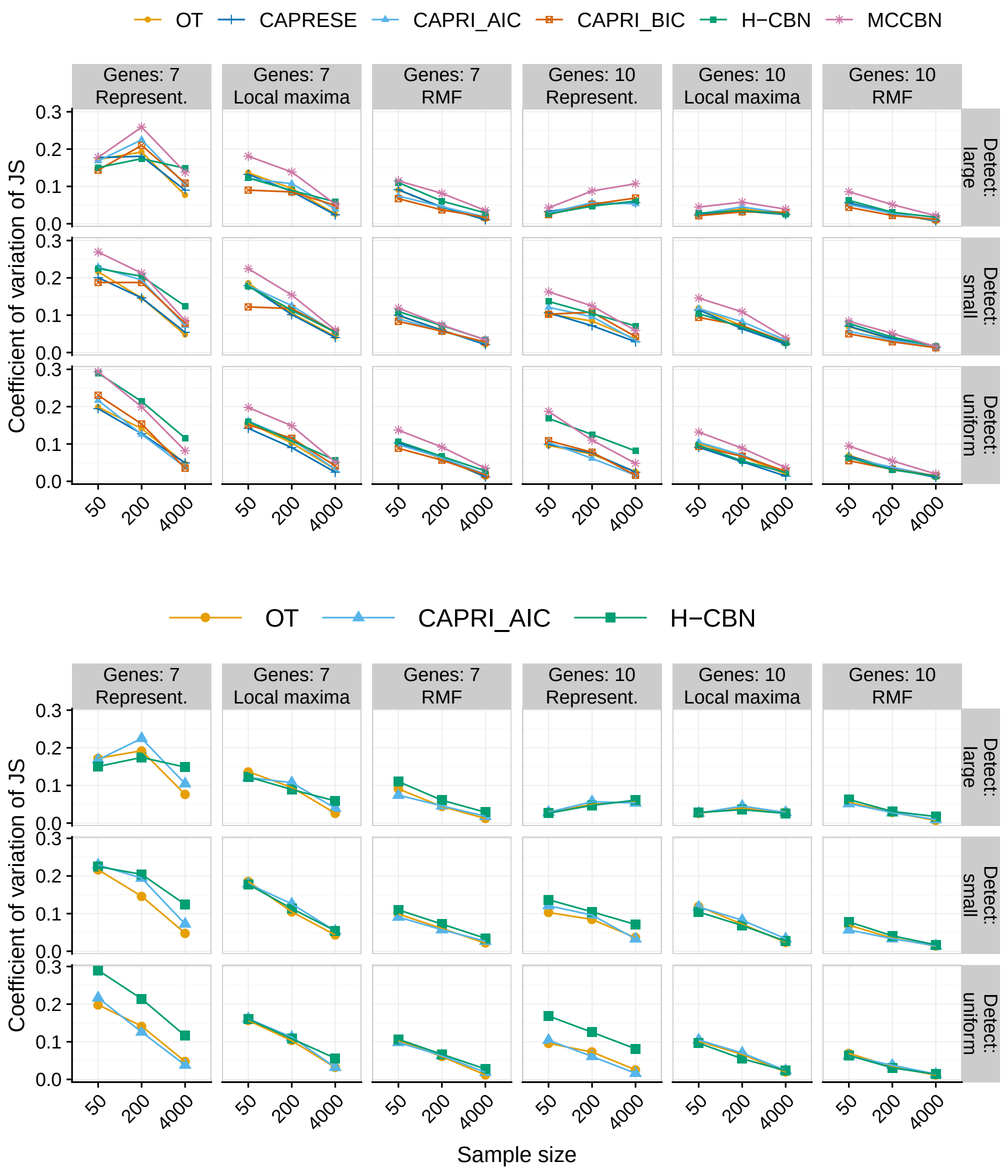

Figure 7: Coefficient of variation (standard deviation/mean) of JS for each combination of model and type of fitness landscape. The bottom figure represents the same information as the top one but only for the three models shown in the paper. The coefficient of variation has been computed from the five runs for each landscapes on each combination of sample size and detection regime.

### 7. Number of paths inferred

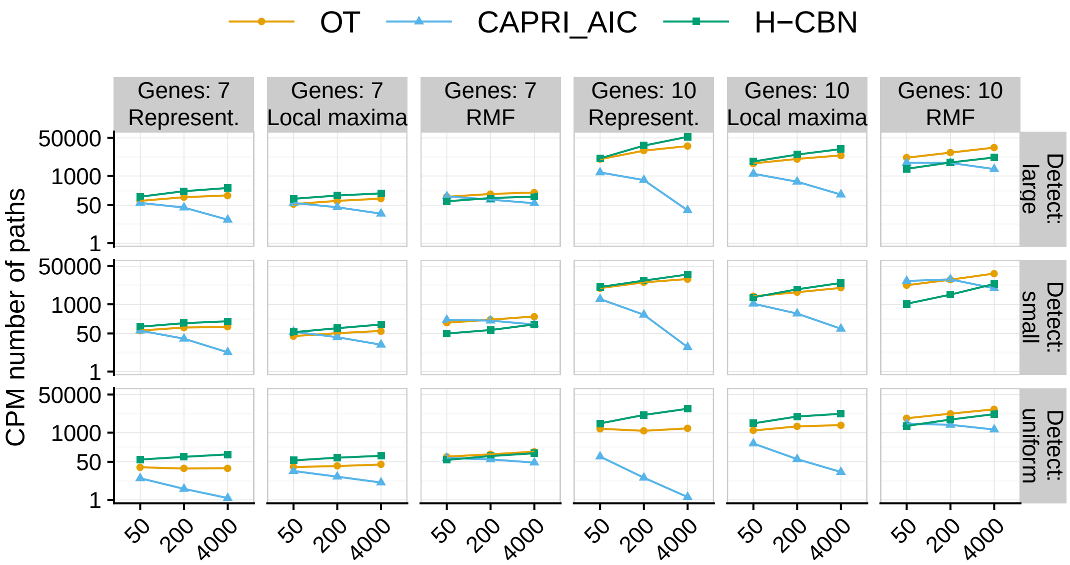

Figure 8: Number of paths to the maximum according to the CPMs.

#### 8. Slopes of regressions of 1-recall and 1-precision on LOD diversity, $S_p$

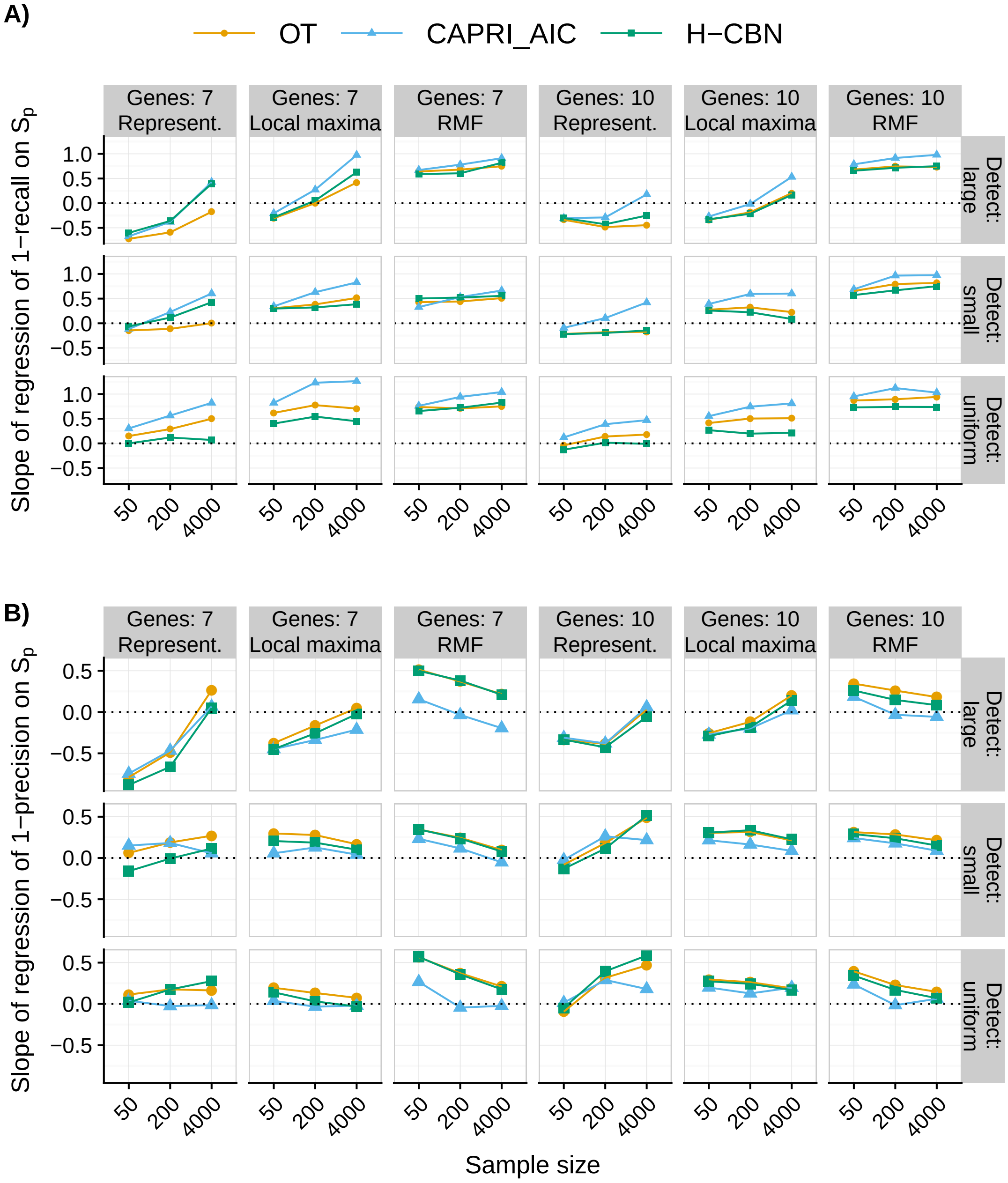

Figure 9: Slopes of regressions of 1-recall and 1-precision on LOD diversity, $S_p$

#### 9. Coefficient of variation of $S_c$

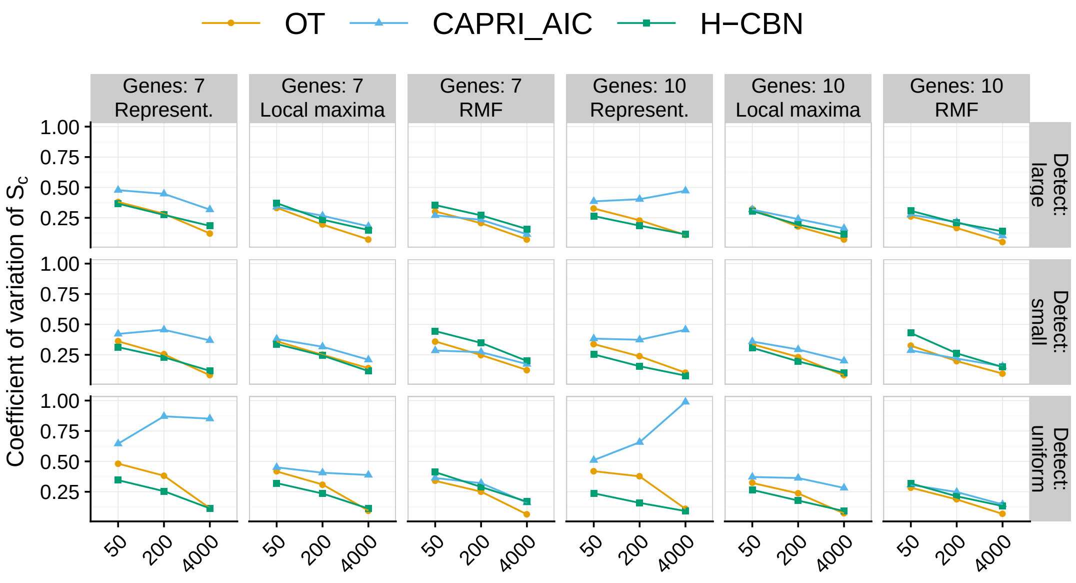

Figure 10: Coefficient of variation (standard deviation/mean) of $S_c$ for each combination of model and type of fitness landscape. The coefficient of variation has been computed from the five runs for each landscapes on each combination of sample size and detection regime. For OT and H-CBN, it is computed using the probability-weighted predicted paths (see text). Each point plotted is the average of 210 points.

#### 10. Estimated $S_c$ by H-CBN

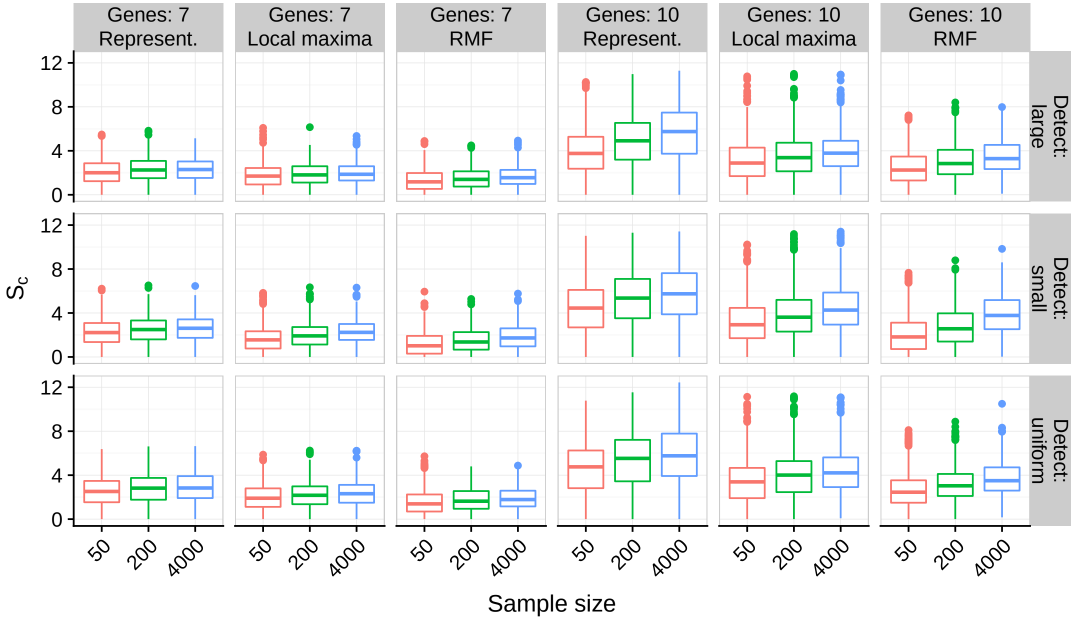

Figure 11: Estimated $S_c$ by H-CBN for all combinations of sample size by type of landscape by detection regime by number of genes. Each box plot shows 1050 points.

### 11. Analysis of deviance tables for fitted models

The tables below show the analysis of deviance tables for the generalized (beta regressions) linear mixed effects models. Models where fitted using the R package glmmTMB [1]. Analysis of deviance tables are from package car [2]. All analysis of deviance tables use Type II Wald chi-square tests.

Analysis have been run on the complete data set (section 11.1), and after splitting for the different combinations of CPM and fitness landscape type (section 11.2).

##### 11.1. Models fitted to the complete data set

(Notice the strong evidence we see for three and four and even five way interactions.)

###### 11.1.1. Two-way interactions

| | Chisq | Df | Pr(>Chisq) |
|---|---|---|---|
| Num. Genes | 223.34 | 1.00 | $< .0001$ |
| CPM | 9459.99 | 2.00 | $< .0001$ |
| Landscape | 322.37 | 2.00 | $< .0001$ |
| Detection | 4768.56 | 2.00 | $< .0001$ |
| Sample Size | 1599.17 | 2.00 | $< .0001$ |
| LOD diversity | 139.98 | 1.00 | $< .0001$ |
| Num. Genes:CPM | 22.98 | 2.00 | $< .0001$ |
| Num. Genes:Landscape | 91.35 | 2.00 | $< .0001$ |
| Num. Genes:Detection | 2318.60 | 2.00 | $< .0001$ |
| Num. Genes:Sample Size | 381.80 | 2.00 | $< .0001$ |
| Num. Genes:LOD diversity | 8.72 | 1.00 | 0.0032 |
| CPM:Landscape | 192.88 | 4.00 | $< .0001$ |
| CPM:Detection | 678.41 | 4.00 | $< .0001$ |
| CPM:Sample Size | 818.96 | 4.00 | $< .0001$ |
| CPM:LOD diversity | 170.20 | 2.00 | $< .0001$ |
| Landscape:Detection | 6534.14 | 4.00 | $< .0001$ |
| Landscape:Sample Size | 3110.94 | 4.00 | $< .0001$ |
| Landscape:LOD diversity | 78.21 | 2.00 | $< .0001$ |
| Detection:Sample Size | 2420.56 | 4.00 | $< .0001$ |
| Detection:LOD diversity | 3303.42 | 2.00 | $< .0001$ |
| Sample Size:LOD diversity | 939.31 | 2.00 | $< .0001$ |

Table 1: Full statistical model (all CPMs), 2-way interactions

##### 11.1.2. Three-way interactions

| | Chisq | Df | Pr(>Chisq) |
|---|---|---|---|
| Num. Genes | 197.19 | 1.00 | $< .0001$ |
| CPM | 11331.58 | 2.00 | $< .0001$ |
| Landscape | 411.00 | 2.00 | $< .0001$ |
| Detection | 4217.33 | 2.00 | $< .0001$ |
| Sample Size | 1536.56 | 2.00 | $< .0001$ |
| LOD diversity | 146.46 | 1.00 | $< .0001$ |
| Num. Genes:CPM | 52.74 | 2.00 | $< .0001$ |
| Num. Genes:Landscape | 80.98 | 2.00 | $< .0001$ |
| Num. Genes:Detection | 2394.91 | 2.00 | $< .0001$ |
| Num. Genes:Sample Size | 354.72 | 2.00 | $< .0001$ |
| Num. Genes:LOD diversity | 8.15 | 1.00 | 0.0043 |
| CPM:Landscape | 285.47 | 4.00 | $< .0001$ |
| CPM:Detection | 707.50 | 4.00 | $< .0001$ |
| CPM:Sample Size | 842.82 | 4.00 | $< .0001$ |
| CPM:LOD diversity | 117.61 | 2.00 | $< .0001$ |
| Landscape:Detection | 6924.74 | 4.00 | $< .0001$ |
| Landscape:Sample Size | 3407.98 | 4.00 | $< .0001$ |
| Landscape:LOD diversity | 77.13 | 2.00 | $< .0001$ |
| Detection:Sample Size | 2507.80 | 4.00 | $< .0001$ |
| Detection:LOD diversity | 4229.09 | 2.00 | $< .0001$ |
| Sample Size:LOD diversity | 1120.46 | 2.00 | $< .0001$ |
| Num. Genes:CPM:Landscape | 90.47 | 4.00 | $< .0001$ |
| Num. Genes:CPM:Detection | 36.64 | 4.00 | $< .0001$ |
| Num. Genes:CPM:Sample Size | 10.43 | 4.00 | 0.0338 |
| Num. Genes:CPM:LOD diversity | 41.35 | 2.00 | $< .0001$ |
| Num. Genes:Landscape:Detection | 1302.06 | 4.00 | $< .0001$ |
| Num. Genes:Landscape:Sample Size | 347.91 | 4.00 | $< .0001$ |
| Num. Genes:Landscape:LOD diversity | 3.70 | 2.00 | 0.1572 |
| Num. Genes:Detection:Sample Size | 553.21 | 4.00 | $< .0001$ |
| Num. Genes:Detection:LOD diversity | 3.91 | 2.00 | 0.1417 |
| Num. Genes:Sample Size:LOD diversity | 56.88 | 2.00 | $< .0001$ |
| CPM:Landscape:Detection | 250.27 | 8.00 | $< .0001$ |
| CPM:Landscape:Sample Size | 192.75 | 8.00 | $< .0001$ |
| CPM:Landscape:LOD diversity | 605.91 | 4.00 | $< .0001$ |
| CPM:Detection:Sample Size | 19.44 | 8.00 | 0.0127 |
| CPM:Detection:LOD diversity | 126.85 | 4.00 | $< .0001$ |
| CPM:Sample Size:LOD diversity | 117.71 | 4.00 | $< .0001$ |
| Landscape:Detection:Sample Size | 2084.94 | 8.00 | $< .0001$ |
| Landscape:Detection:LOD diversity | 867.54 | 4.00 | $< .0001$ |
| Landscape:Sample Size:LOD diversity | 1163.94 | 4.00 | $< .0001$ |
| Detection:Sample Size:LOD diversity | 736.44 | 4.00 | $< .0001$ |

Table 2: Full statistical model (all CPMs), 3-way interactions

##### 11.1.3. Four-way interactions

| | Chisq | Df | Pr(>Chisq) |
|---|---|---|---|
| Num. Genes | 193.68 | 1.00 | < .0001 |
| CPM | 11619.48 | 2.00 | < .0001 |
| Landscape | 426.49 | 2.00 | < .0001 |
| Detection | 4059.78 | 2.00 | < .0001 |
| Sample Size | 1500.59 | 2.00 | < .0001 |
| LOD diversity | 150.27 | 1.00 | < .0001 |
| Num. Genes:CPM | 54.25 | 2.00 | < .0001 |
| Num. Genes:Landscape | 77.19 | 2.00 | < .0001 |
| Num. Genes:Detection | 2370.70 | 2.00 | < .0001 |
| Num. Genes:Sample Size | 357.14 | 2.00 | < .0001 |
| Num. Genes:LOD diversity | 8.47 | 1.00 | 0.0036 |
| CPM:Landscape | 298.90 | 4.00 | < .0001 |
| CPM:Detection | 716.32 | 4.00 | < .0001 |
| CPM:Sample Size | 841.73 | 4.00 | < .0001 |
| CPM:LOD diversity | 119.30 | 2.00 | < .0001 |
| Landscape:Detection | 6697.76 | 4.00 | < .0001 |
| Landscape:Sample Size | 3459.44 | 4.00 | < .0001 |
| Landscape:LOD diversity | 77.85 | 2.00 | < .0001 |
| Detection:Sample Size | 2466.93 | 4.00 | < .0001 |
| Detection:LOD diversity | 4132.54 | 2.00 | < .0001 |
| Sample Size:LOD diversity | 1103.02 | 2.00 | < .0001 |
| Num. Genes:CPM:Landscape | 101.44 | 4.00 | < .0001 |
| Num. Genes:CPM:Detection | 35.22 | 4.00 | < .0001 |
| Num. Genes:CPM:Sample Size | 11.99 | 4.00 | 0.0174 |
| Num. Genes:CPM:LOD diversity | 43.71 | 2.00 | < .0001 |
| Num. Genes:Landscape:Detection | 1262.35 | 4.00 | < .0001 |
| Num. Genes:Landscape:Sample Size | 350.64 | 4.00 | < .0001 |
| Num. Genes:Landscape:LOD diversity | 3.55 | 2.00 | 0.1699 |
| Num. Genes:Detection:Sample Size | 532.95 | 4.00 | < .0001 |
| Num. Genes:Detection:LOD diversity | 2.93 | 2.00 | 0.231 |
| Num. Genes:Sample Size:LOD diversity | 50.25 | 2.00 | < .0001 |
| CPM:Landscape:Detection | 229.42 | 8.00 | < .0001 |
| CPM:Landscape:Sample Size | 215.46 | 8.00 | < .0001 |
| CPM:Landscape:LOD diversity | 627.29 | 4.00 | < .0001 |
| CPM:Detection:Sample Size | 21.27 | 8.00 | 0.0065 |
| CPM:Detection:LOD diversity | 131.95 | 4.00 | < .0001 |
| CPM:Sample Size:LOD diversity | 88.97 | 4.00 | < .0001 |
| Landscape:Detection:Sample Size | 2216.86 | 8.00 | < .0001 |
| Landscape:Detection:LOD diversity | 867.13 | 4.00 | < .0001 |
| Landscape:Sample Size:LOD diversity | 1175.97 | 4.00 | < .0001 |
| Detection:Sample Size:LOD diversity | 819.51 | 4.00 | < .0001 |
| Num. Genes:CPM:Landscape:Detection | 12.73 | 8.00 | 0.1214 |
| Num. Genes:CPM:Landscape:Sample Size | 7.44 | 8.00 | 0.4898 |
| Num. Genes:CPM:Landscape:LOD diversity | 14.84 | 4.00 | 0.0051 |
| Num. Genes:CPM:Detection:Sample Size | 25.87 | 8.00 | 0.0011 |
| Num. Genes:CPM:Detection:LOD diversity | 53.99 | 4.00 | < .0001 |
| Num. Genes:CPM:Sample Size:LOD diversity | 2.51 | 4.00 | 0.6425 |
| Num. Genes:Landscape:Detection:Sample Size | 321.87 | 8.00 | < .0001 |
| Num. Genes:Landscape:Detection:LOD diversity | 74.72 | 4.00 | < .0001 |
| Num. Genes:Landscape:Sample Size:LOD diversity | 101.05 | 4.00 | < .0001 |

| | | | |
|---|---|---|---|
| Num. Genes:Detection:Sample Size:LOD diversity | 115.78 | 4.00 | $< .0001$ |
| CPM:Landscape:Detection:Sample Size | 24.77 | 16.00 | 0.0739 |
| CPM:Landscape:Detection:LOD diversity | 19.36 | 8.00 | 0.013 |
| CPM:Landscape:Sample Size:LOD diversity | 23.93 | 8.00 | 0.0024 |
| CPM:Detection:Sample Size:LOD diversity | 10.02 | 8.00 | 0.2634 |
| Landscape:Detection:Sample Size:LOD diversity | 237.96 | 8.00 | $< .0001$ |

Table 3: Full statistical model (all CPMs), 4-way interactions

##### 11.2. Models fitted to each combination of fitness landscape by CPM

Remember each model uses 3780 observations: 35 replicates, 3 mutation rates, 2 variance settings, 2 number of genes, 3 detection regimes, 3 sample sizes. These correspond to 420 different fitness landscapes: 35 by 3 by 2 by 2. Each observation is itself the average of five different splits of the set of 20000 simulations.

###### 11.2.1. Main effects

| | Chisq | Df | Pr(>Chisq) |
|---|---|---|---|
| Num. Genes | 325.32 | 1.00 | $< .0001$ |
| Sample Size | 797.56 | 2.00 | $< .0001$ |
| Detection | 1101.14 | 2.00 | $< .0001$ |
| LOD diversity | 2.41 | 1.00 | 0.1206 |

Table 4: Represent..OT

| | Chisq | Df | Pr(>Chisq) |
|---|---|---|---|
| Num. Genes | 84.38 | 1.00 | $< .0001$ |
| Sample Size | 179.66 | 2.00 | $< .0001$ |
| Detection | 470.52 | 2.00 | $< .0001$ |
| LOD diversity | 63.71 | 1.00 | $< .0001$ |

Table 5: Local Peaks.OT

| | Chisq | Df | Pr(>Chisq) |
|---|---|---|---|
| Num. Genes | 9.65 | 1.00 | 0.0019 |
| Sample Size | 199.97 | 2.00 | $< .0001$ |
| Detection | 206.52 | 2.00 | $< .0001$ |
| LOD diversity | 162.60 | 1.00 | $< .0001$ |

Table 6: RMF.OT

| | Chisq | Df | Pr(>Chisq) |
|---|---|---|---|
| Num. Genes | 206.70 | 1.00 | $< .0001$ |
| Sample Size | 62.82 | 2.00 | $< .0001$ |
| Detection | 692.92 | 2.00 | $< .0001$ |
| LOD diversity | 79.61 | 1.00 | $< .0001$ |

Table 7: Represent..CAPRI_AIC

| | Chisq | Df | Pr(>Chisq) |
|---|---|---|---|
| Num. Genes | 110.80 | 1.00 | $< .0001$ |
| Sample Size | 48.42 | 2.00 | $< .0001$ |
| Detection | 383.72 | 2.00 | $< .0001$ |
| LOD diversity | 81.86 | 1.00 | $< .0001$ |

Table 8: Local Peaks.CAPRI_AIC

| | Chisq | Df | Pr(>Chisq) |
|---|---|---|---|
| Num. Genes | 35.67 | 1.00 | $< .0001$ |
| Sample Size | 144.12 | 2.00 | $< .0001$ |
| Detection | 48.52 | 2.00 | $< .0001$ |
| LOD diversity | 70.86 | 1.00 | $< .0001$ |

Table 9: RMF.CAPRI_AIC

| | Chisq | Df | Pr(>Chisq) |
|---|---|---|---|
| Num. Genes | 157.75 | 1.00 | $< .0001$ |
| Sample Size | 1331.50 | 2.00 | $< .0001$ |
| Detection | 2493.82 | 2.00 | $< .0001$ |
| LOD diversity | 0.35 | 1.00 | 0.5538 |

Table 10: Represent..H-CBN

| | Chisq | Df | Pr(>Chisq) |
|---|---|---|---|
| Num. Genes | 71.83 | 1.00 | $< .0001$ |
| Sample Size | 315.26 | 2.00 | $< .0001$ |
| Detection | 823.90 | 2.00 | $< .0001$ |
| LOD diversity | 64.31 | 1.00 | $< .0001$ |

Table 11: Local Peaks.H-CBN

| | Chisq | Df | Pr(>Chisq) |
|---|---|---|---|
| Num. Genes | 3.82 | 1.00 | 0.0507 |
| Sample Size | 100.38 | 2.00 | $< .0001$ |
| Detection | 320.43 | 2.00 | $< .0001$ |
| LOD diversity | 151.14 | 1.00 | $< .0001$ |

Table 12: RMF.H-CBN

##### 11.2.2. Two-way interactions

| | Chisq | Df | Pr(>Chisq) |
|---|---|---|---|
| Num. Genes | 229.87 | 1.00 | < .0001 |
| Sample Size | 1084.79 | 2.00 | < .0001 |
| Detection | 1145.03 | 2.00 | < .0001 |
| LOD diversity | 0.02 | 1.00 | 0.8958 |
| Num. Genes:Sample Size | 168.67 | 2.00 | < .0001 |
| Num. Genes:Detection | 705.00 | 2.00 | < .0001 |
| Num. Genes:LOD diversity | 4.46 | 1.00 | 0.0347 |
| Sample Size:Detection | 726.35 | 4.00 | < .0001 |
| Sample Size:LOD diversity | 308.31 | 2.00 | < .0001 |
| Detection:LOD diversity | 1019.12 | 2.00 | < .0001 |

Table 13: Represent..OT

| | Chisq | Df | Pr(>Chisq) |
|---|---|---|---|
| Num. Genes | 66.47 | 1.00 | < .0001 |
| Sample Size | 163.41 | 2.00 | < .0001 |
| Detection | 449.46 | 2.00 | < .0001 |
| LOD diversity | 74.54 | 1.00 | < .0001 |
| Num. Genes:Sample Size | 34.28 | 2.00 | < .0001 |
| Num. Genes:Detection | 443.26 | 2.00 | < .0001 |
| Num. Genes:LOD diversity | 0.02 | 1.00 | 0.895 |
| Sample Size:Detection | 307.15 | 4.00 | < .0001 |
| Sample Size:LOD diversity | 21.38 | 2.00 | < .0001 |
| Detection:LOD diversity | 403.27 | 2.00 | < .0001 |

Table 14: Local Peaks.OT

| | Chisq | Df | Pr(>Chisq) |
|---|---|---|---|
| Num. Genes | 9.65 | 1.00 | 0.0019 |
| Sample Size | 213.84 | 2.00 | < .0001 |
| Detection | 215.42 | 2.00 | < .0001 |
| LOD diversity | 155.94 | 1.00 | < .0001 |
| Num. Genes:Sample Size | 31.75 | 2.00 | < .0001 |
| Num. Genes:Detection | 5.39 | 2.00 | 0.0676 |
| Num. Genes:LOD diversity | 0.05 | 1.00 | 0.8191 |
| Sample Size:Detection | 7.68 | 4.00 | 0.1041 |
| Sample Size:LOD diversity | 190.09 | 2.00 | < .0001 |
| Detection:LOD diversity | 9.29 | 2.00 | 0.0096 |

Table 15: RMF.OT

| | Chisq | Df | Pr(>Chisq) |
|---|---|---|---|
| Num. Genes | 144.41 | 1.00 | < .0001 |
| Sample Size | 70.06 | 2.00 | < .0001 |
| Detection | 720.04 | 2.00 | < .0001 |
| LOD diversity | 89.03 | 1.00 | < .0001 |
| Num. Genes:Sample Size | 154.58 | 2.00 | < .0001 |
| Num. Genes:Detection | 574.17 | 2.00 | < .0001 |
| Num. Genes:LOD diversity | 5.16 | 1.00 | 0.0231 |
| Sample Size:Detection | 691.60 | 4.00 | < .0001 |
| Sample Size:LOD diversity | 734.28 | 2.00 | < .0001 |
| Detection:LOD diversity | 958.49 | 2.00 | < .0001 |

Table 16: Represent..CAPRI_AIC

| | Chisq | Df | Pr(>Chisq) |
|---|---|---|---|
| Num. Genes | 90.86 | 1.00 | < .0001 |
| Sample Size | 36.29 | 2.00 | < .0001 |
| Detection | 351.07 | 2.00 | < .0001 |
| LOD diversity | 89.21 | 1.00 | < .0001 |
| Num. Genes:Sample Size | 8.98 | 2.00 | 0.0112 |
| Num. Genes:Detection | 309.60 | 2.00 | < .0001 |
| Num. Genes:LOD diversity | 0.01 | 1.00 | 0.91 |
| Sample Size:Detection | 262.51 | 4.00 | < .0001 |
| Sample Size:LOD diversity | 80.39 | 2.00 | < .0001 |
| Detection:LOD diversity | 429.26 | 2.00 | < .0001 |

Table 17: Local Peaks.CAPRI_AIC

| | Chisq | Df | Pr(>Chisq) |
|---|---|---|---|
| Num. Genes | 33.69 | 1.00 | < .0001 |
| Sample Size | 164.54 | 2.00 | < .0001 |
| Detection | 50.28 | 2.00 | < .0001 |
| LOD diversity | 65.36 | 1.00 | < .0001 |
| Num. Genes:Sample Size | 34.37 | 2.00 | < .0001 |
| Num. Genes:Detection | 2.23 | 2.00 | 0.3273 |
| Num. Genes:LOD diversity | 1.24 | 1.00 | 0.2663 |
| Sample Size:Detection | 15.51 | 4.00 | 0.0037 |
| Sample Size:LOD diversity | 154.86 | 2.00 | < .0001 |
| Detection:LOD diversity | 40.59 | 2.00 | < .0001 |

Table 18: RMF.CAPRI_AIC

| | Chisq | Df | Pr(>Chisq) |
|---|---|---|---|
| Num. Genes | 111.35 | 1.00 | < .0001 |
| Sample Size | 1727.19 | 2.00 | < .0001 |
| Detection | 2829.04 | 2.00 | < .0001 |
| LOD diversity | 2.47 | 1.00 | 0.116 |
| Num. Genes:Sample Size | 252.55 | 2.00 | < .0001 |
| Num. Genes:Detection | 583.85 | 2.00 | < .0001 |
| Num. Genes:LOD diversity | 11.81 | 1.00 | 6e-04 |
| Sample Size:Detection | 583.96 | 4.00 | < .0001 |
| Sample Size:LOD diversity | 367.88 | 2.00 | < .0001 |
| Detection:LOD diversity | 634.67 | 2.00 | < .0001 |

Table 19: Represent..H-CBN

| | Chisq | Df | Pr(>Chisq) |
|---|---|---|---|
| Num. Genes | 56.06 | 1.00 | $< .0001$ |
| Sample Size | 313.03 | 2.00 | $< .0001$ |
| Detection | 836.24 | 2.00 | $< .0001$ |
| LOD diversity | 76.98 | 1.00 | $< .0001$ |
| Num. Genes:Sample Size | 35.78 | 2.00 | $< .0001$ |
| Num. Genes:Detection | 413.32 | 2.00 | $< .0001$ |
| Num. Genes:LOD diversity | 0.30 | 1.00 | 0.5868 |
| Sample Size:Detection | 296.90 | 4.00 | $< .0001$ |
| Sample Size:LOD diversity | 19.84 | 2.00 | $< .0001$ |
| Detection:LOD diversity | 292.03 | 2.00 | $< .0001$ |

Table 20: Local Peaks.H-CBN

| | Chisq | Df | Pr(>Chisq) |
|---|---|---|---|
| Num. Genes | 3.82 | 1.00 | 0.0506 |
| Sample Size | 107.26 | 2.00 | $< .0001$ |
| Detection | 331.99 | 2.00 | $< .0001$ |
| LOD diversity | 144.95 | 1.00 | $< .0001$ |
| Num. Genes:Sample Size | 21.97 | 2.00 | $< .0001$ |
| Num. Genes:Detection | 4.24 | 2.00 | 0.12 |
| Num. Genes:LOD diversity | 1.16 | 1.00 | 0.2823 |
| Sample Size:Detection | 4.15 | 4.00 | 0.3856 |
| Sample Size:LOD diversity | 181.70 | 2.00 | $< .0001$ |
| Detection:LOD diversity | 1.36 | 2.00 | 0.5071 |

Table 21: RMF.H-CBN

##### 11.2.3. Four-way interactions

| | Chisq | Df | Pr(>Chisq) |
|---|---|---|---|
| Num. Genes | 236.38 | 1.00 | $< .0001$ |
| Sample Size | 1105.42 | 2.00 | $< .0001$ |
| Detection | 1046.08 | 2.00 | $< .0001$ |
| LOD diversity | 0.84 | 1.00 | 0.3593 |
| Num. Genes:Sample Size | 193.98 | 2.00 | $< .0001$ |
| Num. Genes:Detection | 778.62 | 2.00 | $< .0001$ |
| Num. Genes:LOD diversity | 6.38 | 1.00 | 0.0116 |
| Sample Size:Detection | 722.33 | 4.00 | $< .0001$ |
| Sample Size:LOD diversity | 322.04 | 2.00 | $< .0001$ |
| Detection:LOD diversity | 1027.32 | 2.00 | $< .0001$ |
| Num. Genes:Sample Size:Detection | 96.81 | 4.00 | $< .0001$ |
| Num. Genes:Sample Size:LOD diversity | 28.18 | 2.00 | $< .0001$ |
| Num. Genes:Detection:LOD diversity | 26.39 | 2.00 | $< .0001$ |
| Sample Size:Detection:LOD diversity | 116.00 | 4.00 | $< .0001$ |
| Num. Genes:Sample Size:Detection:LOD diversity | 76.54 | 4.00 | $< .0001$ |

Table 22: Represent..OT

| | Chisq | Df | Pr(>Chisq) |
|---|---|---|---|
| Num. Genes | 57.63 | 1.00 | $< .0001$ |
| Sample Size | 166.26 | 2.00 | $< .0001$ |
| Detection | 380.88 | 2.00 | $< .0001$ |
| LOD diversity | 72.98 | 1.00 | $< .0001$ |
| Num. Genes:Sample Size | 35.33 | 2.00 | $< .0001$ |
| Num. Genes:Detection | 409.75 | 2.00 | $< .0001$ |
| Num. Genes:LOD diversity | 0.04 | 1.00 | 0.8394 |
| Sample Size:Detection | 326.02 | 4.00 | $< .0001$ |
| Sample Size:LOD diversity | 24.07 | 2.00 | $< .0001$ |
| Detection:LOD diversity | 412.53 | 2.00 | $< .0001$ |
| Num. Genes:Sample Size:Detection | 198.32 | 4.00 | $< .0001$ |
| Num. Genes:Sample Size:LOD diversity | 1.85 | 2.00 | 0.3967 |
| Num. Genes:Detection:LOD diversity | 5.28 | 2.00 | 0.0715 |
| Sample Size:Detection:LOD diversity | 219.97 | 4.00 | $< .0001$ |
| Num. Genes:Sample Size:Detection:LOD diversity | 8.77 | 4.00 | 0.0672 |

Table 23: Local Peaks.OT

| | Chisq | Df | Pr(>Chisq) |
|---:|---:|---:|:---|
| Num. Genes | 9.69 | 1.00 | 0.0019 |
| Sample Size | 215.06 | 2.00 | < .0001 |
| Detection | 215.20 | 2.00 | < .0001 |
| LOD diversity | 155.03 | 1.00 | < .0001 |
| Num. Genes:Sample Size | 32.40 | 2.00 | < .0001 |
| Num. Genes:Detection | 5.40 | 2.00 | 0.0673 |
| Num. Genes:LOD diversity | 0.05 | 1.00 | 0.8191 |
| Sample Size:Detection | 7.55 | 4.00 | 0.1096 |
| Sample Size:LOD diversity | 191.25 | 2.00 | < .0001 |
| Detection:LOD diversity | 9.34 | 2.00 | 0.0094 |
| Num. Genes:Sample Size:Detection | 0.26 | 4.00 | 0.9925 |
| Num. Genes:Sample Size:LOD diversity | 1.73 | 2.00 | 0.4219 |
| Num. Genes:Detection:LOD diversity | 32.05 | 2.00 | < .0001 |
| Sample Size:Detection:LOD diversity | 1.65 | 4.00 | 0.799 |
| Num. Genes:Sample Size:Detection:LOD diversity | 1.63 | 4.00 | 0.8035 |

Table 24: RMF.OT

| | Chisq | Df | Pr(>Chisq) |
|---:|---:|---:|:---|
| Num. Genes | 163.81 | 1.00 | < .0001 |
| Sample Size | 53.20 | 2.00 | < .0001 |
| Detection | 661.33 | 2.00 | < .0001 |
| LOD diversity | 100.78 | 1.00 | < .0001 |
| Num. Genes:Sample Size | 170.47 | 2.00 | < .0001 |
| Num. Genes:Detection | 654.22 | 2.00 | < .0001 |
| Num. Genes:LOD diversity | 5.37 | 1.00 | 0.0205 |
| Sample Size:Detection | 691.21 | 4.00 | < .0001 |
| Sample Size:LOD diversity | 750.44 | 2.00 | < .0001 |
| Detection:LOD diversity | 972.35 | 2.00 | < .0001 |
| Num. Genes:Sample Size:Detection | 48.16 | 4.00 | < .0001 |
| Num. Genes:Sample Size:LOD diversity | 24.57 | 2.00 | < .0001 |
| Num. Genes:Detection:LOD diversity | 42.41 | 2.00 | < .0001 |
| Sample Size:Detection:LOD diversity | 116.19 | 4.00 | < .0001 |
| Num. Genes:Sample Size:Detection:LOD diversity | 53.68 | 4.00 | < .0001 |

Table 25: Represent..CAPRI_AIC

| | Chisq | Df | Pr(>Chisq) |
|---|---|---|---|
| Num. Genes | 87.57 | 1.00 | $< .0001$ |
| Sample Size | 35.06 | 2.00 | $< .0001$ |
| Detection | 329.14 | 2.00 | $< .0001$ |
| LOD diversity | 88.97 | 1.00 | $< .0001$ |
| Num. Genes:Sample Size | 9.80 | 2.00 | 0.0074 |
| Num. Genes:Detection | 309.31 | 2.00 | $< .0001$ |
| Num. Genes:LOD diversity | 0.03 | 1.00 | 0.8725 |
| Sample Size:Detection | 272.21 | 4.00 | $< .0001$ |
| Sample Size:LOD diversity | 87.34 | 2.00 | $< .0001$ |
| Detection:LOD diversity | 437.56 | 2.00 | $< .0001$ |
| Num. Genes:Sample Size:Detection | 63.36 | 4.00 | $< .0001$ |
| Num. Genes:Sample Size:LOD diversity | 1.44 | 2.00 | 0.4879 |
| Num. Genes:Detection:LOD diversity | 4.16 | 2.00 | 0.1248 |
| Sample Size:Detection:LOD diversity | 73.43 | 4.00 | $< .0001$ |
| Num. Genes:Sample Size:Detection:LOD diversity | 2.63 | 4.00 | 0.6213 |

Table 26: Local Peaks.CAPRI_AIC

| | Chisq | Df | Pr(>Chisq) |
|---|---|---|---|
| Num. Genes | 33.91 | 1.00 | $< .0001$ |
| Sample Size | 165.45 | 2.00 | $< .0001$ |
| Detection | 50.85 | 2.00 | $< .0001$ |
| LOD diversity | 65.50 | 1.00 | $< .0001$ |
| Num. Genes:Sample Size | 34.42 | 2.00 | $< .0001$ |
| Num. Genes:Detection | 2.30 | 2.00 | 0.3163 |
| Num. Genes:LOD diversity | 1.25 | 1.00 | 0.2641 |
| Sample Size:Detection | 15.66 | 4.00 | 0.0035 |
| Sample Size:LOD diversity | 155.70 | 2.00 | $< .0001$ |
| Detection:LOD diversity | 41.21 | 2.00 | $< .0001$ |
| Num. Genes:Sample Size:Detection | 1.76 | 4.00 | 0.7805 |
| Num. Genes:Sample Size:LOD diversity | 9.35 | 2.00 | 0.0093 |
| Num. Genes:Detection:LOD diversity | 3.06 | 2.00 | 0.2169 |
| Sample Size:Detection:LOD diversity | 2.52 | 4.00 | 0.6414 |
| Num. Genes:Sample Size:Detection:LOD diversity | 0.83 | 4.00 | 0.9342 |

Table 27: RMF.CAPRI_AIC

| | Chisq | Df | Pr(>Chisq) |
|---|---|---|---|
| Num. Genes | 109.99 | 1.00 | $<.0001$ |
| Sample Size | 1798.63 | 2.00 | $<.0001$ |
| Detection | 2781.42 | 2.00 | $<.0001$ |
| LOD diversity | 4.29 | 1.00 | 0.0384 |
| Num. Genes:Sample Size | 292.35 | 2.00 | $<.0001$ |
| Num. Genes:Detection | 633.77 | 2.00 | $<.0001$ |
| Num. Genes:LOD diversity | 13.30 | 1.00 | 3e-04 |
| Sample Size:Detection | 577.47 | 4.00 | $<.0001$ |
| Sample Size:LOD diversity | 383.39 | 2.00 | $<.0001$ |
| Detection:LOD diversity | 647.75 | 2.00 | $<.0001$ |
| Num. Genes:Sample Size:Detection | 91.32 | 4.00 | $<.0001$ |
| Num. Genes:Sample Size:LOD diversity | 48.42 | 2.00 | $<.0001$ |
| Num. Genes:Detection:LOD diversity | 16.31 | 2.00 | 3e-04 |
| Sample Size:Detection:LOD diversity | 97.29 | 4.00 | $<.0001$ |
| Num. Genes:Sample Size:Detection:LOD diversity | 98.90 | 4.00 | $<.0001$ |

Table 28: Represent..H-CBN

| | Chisq | Df | Pr(>Chisq) |
|---|---|---|---|
| Num. Genes | 46.47 | 1.00 | $<.0001$ |
| Sample Size | 326.11 | 2.00 | $<.0001$ |
| Detection | 762.69 | 2.00 | $<.0001$ |
| LOD diversity | 74.96 | 1.00 | $<.0001$ |
| Num. Genes:Sample Size | 35.59 | 2.00 | $<.0001$ |
| Num. Genes:Detection | 377.15 | 2.00 | $<.0001$ |
| Num. Genes:LOD diversity | 0.43 | 1.00 | 0.5135 |
| Sample Size:Detection | 315.03 | 4.00 | $<.0001$ |
| Sample Size:LOD diversity | 21.87 | 2.00 | $<.0001$ |
| Detection:LOD diversity | 296.12 | 2.00 | $<.0001$ |
| Num. Genes:Sample Size:Detection | 213.53 | 4.00 | $<.0001$ |
| Num. Genes:Sample Size:LOD diversity | 2.25 | 2.00 | 0.3248 |
| Num. Genes:Detection:LOD diversity | 20.79 | 2.00 | $<.0001$ |
| Sample Size:Detection:LOD diversity | 233.91 | 4.00 | $<.0001$ |
| Num. Genes:Sample Size:Detection:LOD diversity | 4.47 | 4.00 | 0.3461 |

Table 29: Local Peaks.H-CBN

| | Chisq | Df | Pr(>Chisq) |
|---|---|---|---|
| Num. Genes | 3.85 | 1.00 | 0.0499 |
| Sample Size | 107.17 | 2.00 | $< .0001$ |
| Detection | 332.57 | 2.00 | $< .0001$ |
| LOD diversity | 143.99 | 1.00 | $< .0001$ |
| Num. Genes:Sample Size | 22.27 | 2.00 | $< .0001$ |
| Num. Genes:Detection | 4.25 | 2.00 | 0.1195 |
| Num. Genes:LOD diversity | 1.16 | 1.00 | 0.2811 |
| Sample Size:Detection | 4.19 | 4.00 | 0.3803 |
| Sample Size:LOD diversity | 182.19 | 2.00 | $< .0001$ |
| Detection:LOD diversity | 1.29 | 2.00 | 0.5253 |
| Num. Genes:Sample Size:Detection | 0.42 | 4.00 | 0.9807 |
| Num. Genes:Sample Size:LOD diversity | 0.44 | 2.00 | 0.8014 |
| Num. Genes:Detection:LOD diversity | 24.00 | 2.00 | $< .0001$ |
| Sample Size:Detection:LOD diversity | 8.52 | 4.00 | 0.0744 |
| Num. Genes:Sample Size:Detection:LOD diversity | 1.49 | 4.00 | 0.8284 |

Table 30: RMF.H-CBN

#### 12. Number of local maxima, mutations of local maxima, reciprocal sign epistasis and performance

(When interpreting these figures, you should focus on sign of slope and comparisons between models within panel. Comparisons between panels in terms of magnitudes of slopes are complicated because the range of the independent variable can vary a lot between panels —e.g., the range of mean number of mutations in local fitness maxima in the local maxima vs. RMF fitness landscapes.)

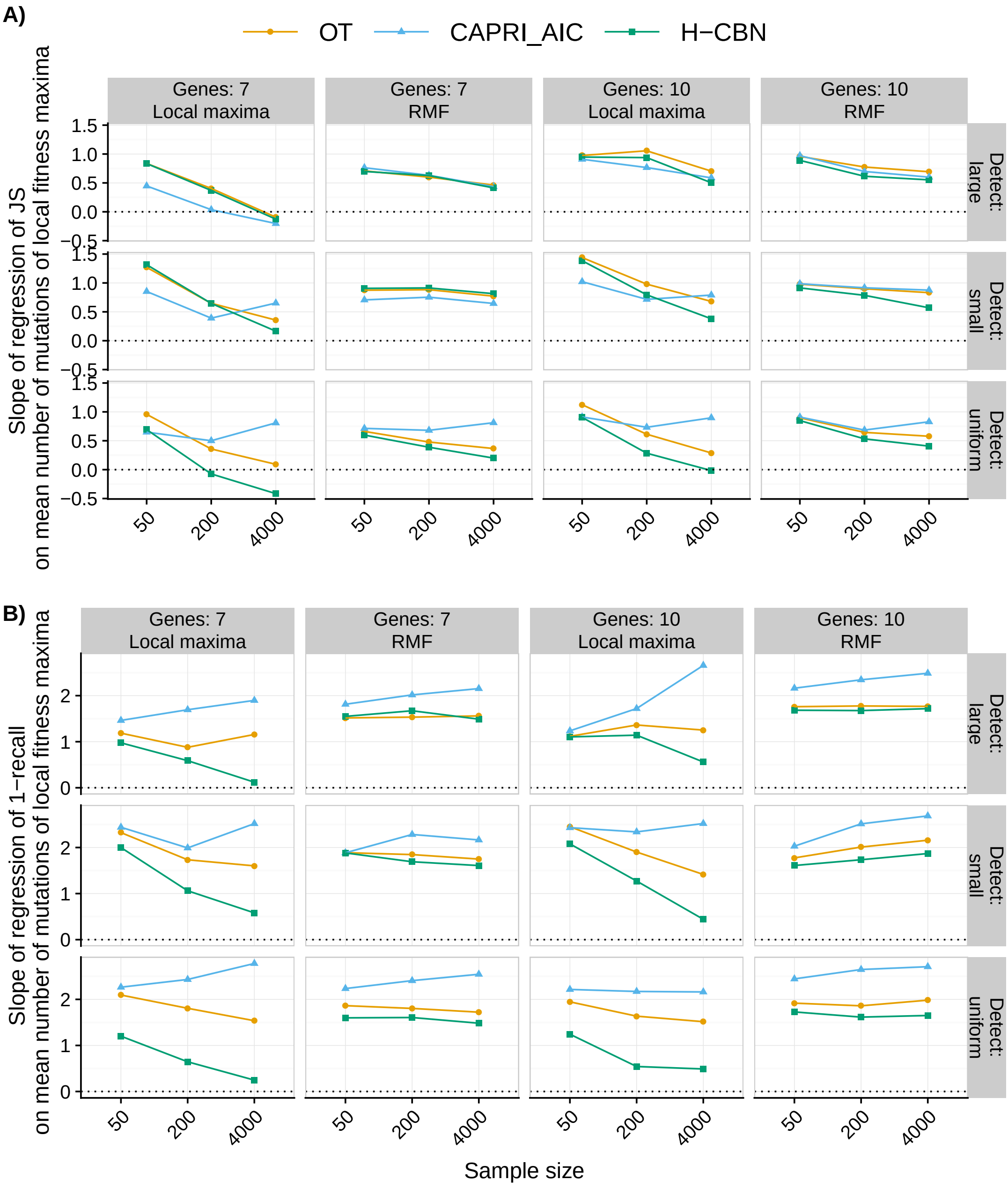

Figure 12: Slopes of regression of JS and 1-recall on mean number of mutations of local maxima.

#### 13. LOD and CPM diversity: ratios and slopes

The following R code will, via a simple example, show that it is easy to have data where the average of the ratios is larger than one whereas the slope of the regression is negative:

```
a <- 10
n <- 100
sd <- 0.5
x <- runif(n, min = 1, max = 5)
y <- -1 * x + a + rnorm(n, mean = 0, sd = sd)
plot(y ~ x)
summary(lm(y ~ x))
mean(y/x)
```

#### 14. References

[1] Brooks, M. E., Kristensen, K., van Benthem, K. J., Magnusson, A., Berg, C. W., Nielsen, A., Skaug, H. J., Maechler, M., Bolker, B. M., 2017. glmmTMB Balances Speed and Flexibility Among Packages for Zero-inflated Generalized Linear Mixed Modeling. *The R Journal*, **9(2)**:378–400. URL https://journal.r-project.org/archive/2017/RJ-2017-066/index.html.

[2] Fox, J., Weisberg, S., 2011. *An R Companion to Applied Regression, 2nd Ed*. Sage, Thousand Oaks, CA.
