## Supplementary material for "Every which way? On predicting tumor evolution using cancer progression models": S7_Text

### Supporting Information for "Every which way? On predicting tumor evolution using cancer progression models"

2019-04-17

#### S7 Text. Data and code availability

All data for this article, along with source code, is available from the compressed zip files. Because of size limits during upload, this compressed file has been split into two, `S1_Dataset.zip` and `S1_Dataset.z01`. Because of naming conventions for file names, `S1_Dataset.z01` has had to be renamed to `S2_Dataset.z01`.

Thus, the very first thing you need to do is **rename** `S2_Dataset.z01` **to** `S1_Dataset.z01`, so that decompression programs know they are dealing with the split files from a single original zip archive.

Once you have renamed `S2_Dataset.z01` to `S1_Dataset.z01`:

- Under **Windows**:

  Open `S1_Dataset.zip` with 7-Zip (https://www.7-zip.org/) (or WinRAR or WinZIP). Alternatively, click on the file, and specify it to be extracted with 7-zip/WinRAR/WinZIP.

- Under **GNU/Linux** or **OS X**:

  You first have to combine the split archive into a single archive, and then unzip (you will need the `zip` program to be installed):

```
zip -s 0 S1_Dataset.zip --out unsplit-S1_Dataset.zip
unzip unsplit-S1_Dataset.zip
```

- Under **GNU/Linux**, you can also use `7z` directly (if you have it installed) (in Debian and Ubuntu systems, you will need to install `p7zip-full`):

```
7z x S1_Dataset.zip
```

  (see further details in, for example, https://superuser.com/a/517758 and https://unix.stackexchange.com/q/40480).

*, http://ligarto.org/rdiaz
