## Supplementary material for "Every which way? On predicting tumor evolution using cancer progression models": S1_Figure

### Supporting Information for “Every which way? On predicting tumor evolution using cancer progression models”

2019-04-17

#### S1 Figure. Plots of simulated fitness landscapes and fitness graphs.

---

#### Contents

1 Fitness landscapes: explanation of plots 1

2 Seven genes 3

3 Ten genes 634

---

#### 1. Fitness landscapes: explanation of plots

Here we show the 1260 fitness landscapes used. The first section, “Seven genes”, section 2, shows those for 7 genes, the second section, “Ten genes’, ’ section 3, those for 10 genes. In each section, the first 210 fitness landscapes are fully representable fitness landscapes, the next 210 are the “Local maxima” fitness landscapes. The next 210 correspond to Rough Mount Fuji (RMF) fitness landscapes. Landscapes are not ordered according to any criterion within type of fitness landscape.

In each page, the following figures are shown:

**DAG of restrictions (top left)** The true DAG of restrictions. This only applies properly to the representable fitness landscapes. In the local maxima fitness landscapes, not all paths between genotypes are available. (See below). For the RMF landscapes this, of course, is not available, since there is no underlying DAG of restrictions.

The true DAG of restrictions could be a tree. Among the representable fitness landscapes, 30% and 2%, of the landscapes with 7 and 10 genes, respectively, were trees (i.e., can be represented by OT and CAPRESE).

**Fitness landscape (bottom left)** As it says, the fitness landscape. Boxes surround the fitness maxima in the 7-genes landscapes. To minimize clutter, genotype labels are not shown in the 10-genes landscapes.

*, http://ligarto.org/rdiaz

**ID and landscape characteristics (center)** The ID of the fitness landscape (a random string that matches the value of ID in the data tables), the number of accessible genotypes, the number of fitness maxima or peaks. Values for "removed edges" and "prop rem edges" denote, respectively, the number and proportion of edges in the fitness graph that were removed; this only applies to the "Local maxima" landscapes; thus, these values are 0 for "Representable" fitness landscapes, and NA for RMF fitness landscapes.

**Fitness graph from DAG of restrictions (top right)** The fitness graph implied by the DAG of restrictions. Not available for RMF (since there is no DAG of restrictions).

**True fitness graph (bottom right)** The actual fitness graph that corresponds to the fitness landscape. For "Representable" fitness landscapes this would be the same as the Fitness graph from DAG of restrictions, so we do not show it (to make the size of the files smaller).

Thus, a simple way to tell what kind of landscape you are looking at in any random page is this:

- If there is a fitness graph on the top right and no fitness graph in the bottom right corner: a representable fitness landscape.
- If there is a fitness graph on the top right and another fitness graph in the bottom right corner (with, necessarily, fewer paths than the graph in the upper right corner and at least two local maxima): a local maxima fitness landscape.
- If there is only a fitness graph on the bottom right: a RMF fitness landscape.

#### 2. Seven genes

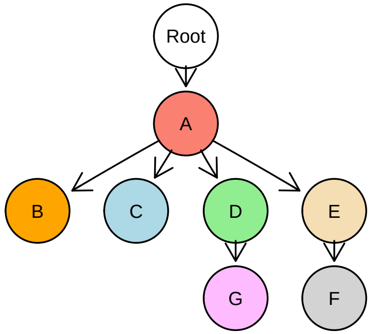


| | |
|---|---|
| *ID* | 1010kGD9gL10Sq9tJsP |
| *acc genotypes* | 37 |
| *peaks no back* | 1 |
| *removed edges* | 0 |
| *prop rem edges* | 0 |

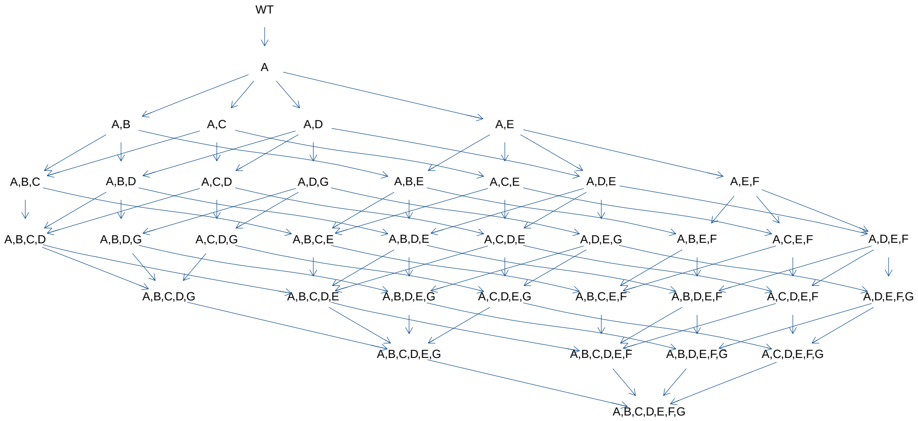


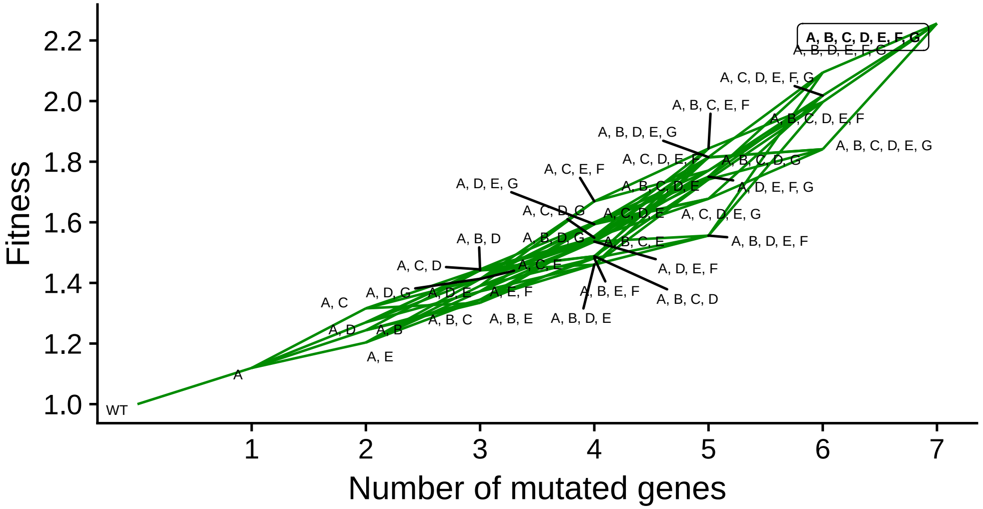

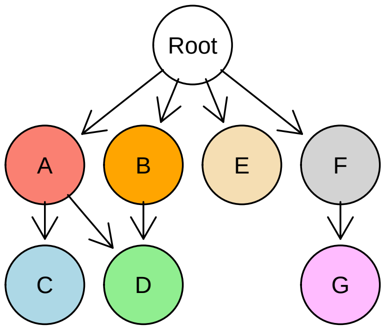


| | |
|---|---|
| *ID* | 10Dgx1l4YRhmToSuW |
| *acc genotypes* | 48 |
| *peaks no back* | 1 |
| *removed edges* | 0 |
| *prop rem edges* | 0 |

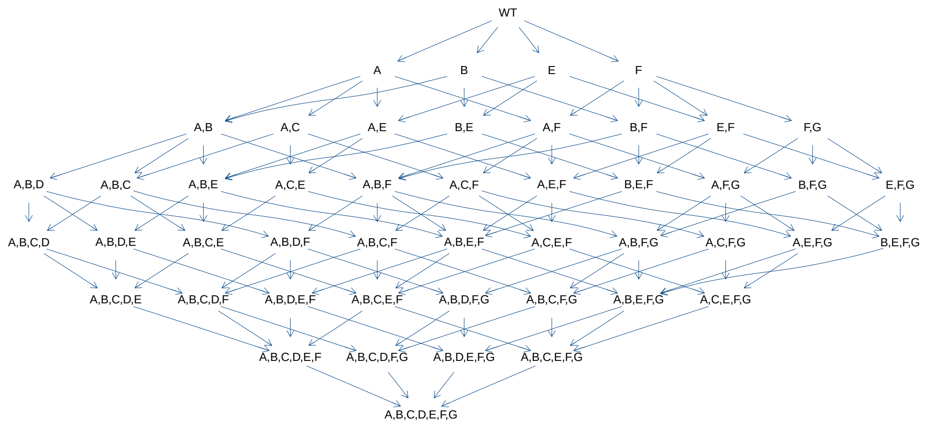


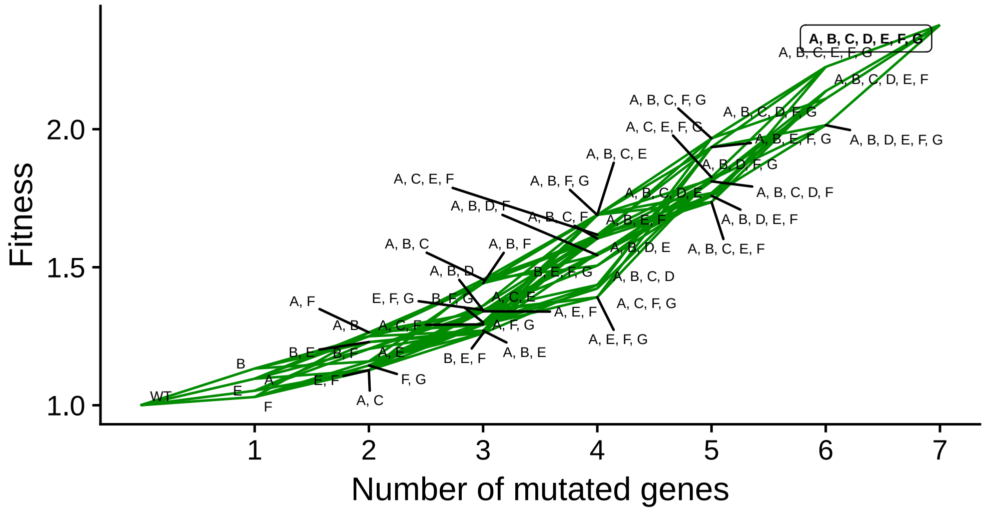

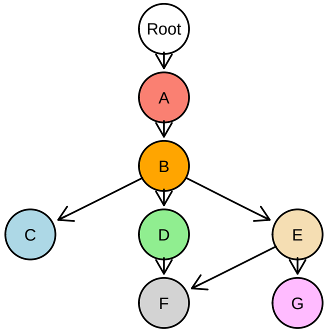


| | |
|---|---|
| *ID* | 1g3AE5zjGYmnuTbC |
| *acc genotypes* | 18 |
| *peaks no back* | 1 |
| *removed edges* | 0 |
| *prop rem edges* | 0 |

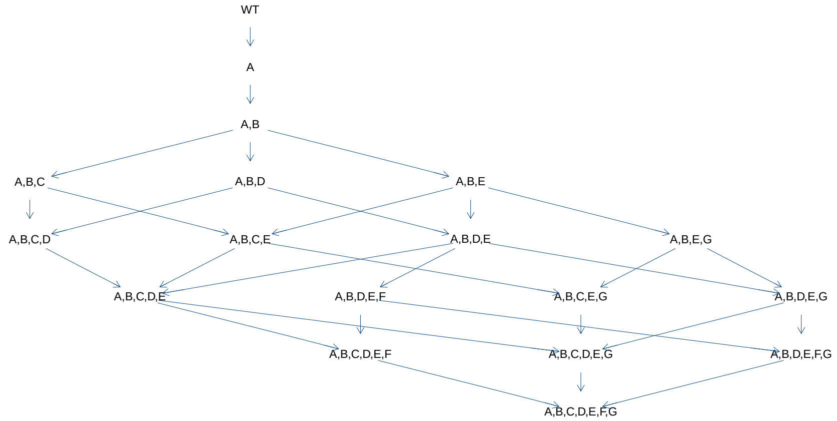


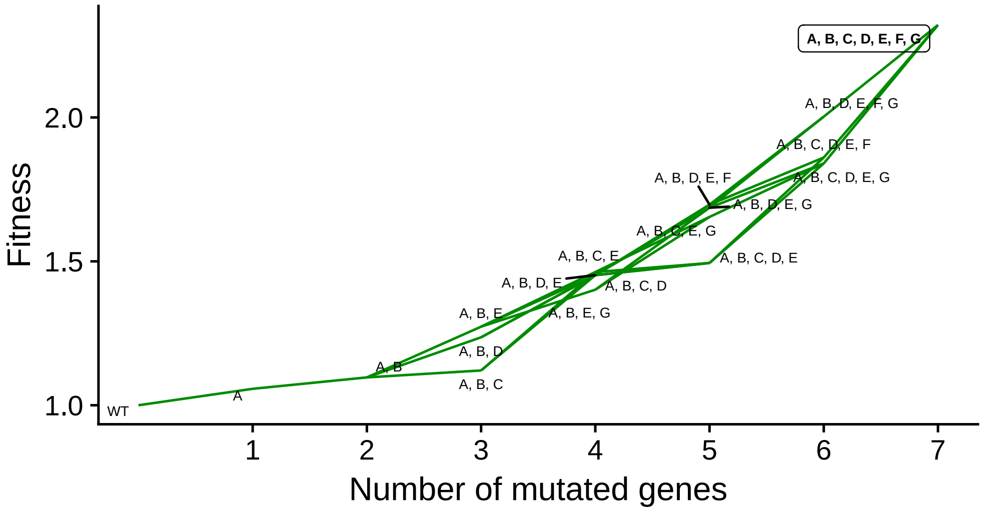

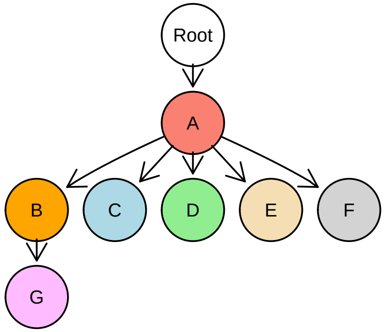


| | |
|---|---|
| *ID* | 1HutaalyylPzhYlD |
| *acc genotypes* | 49 |
| *peaks no back* | 1 |
| *removed edges* | 0 |
| *prop rem edges* | 0 |

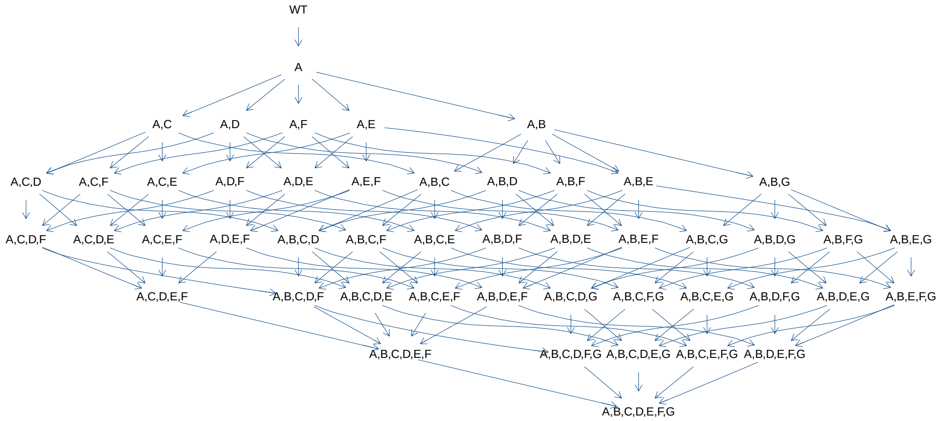


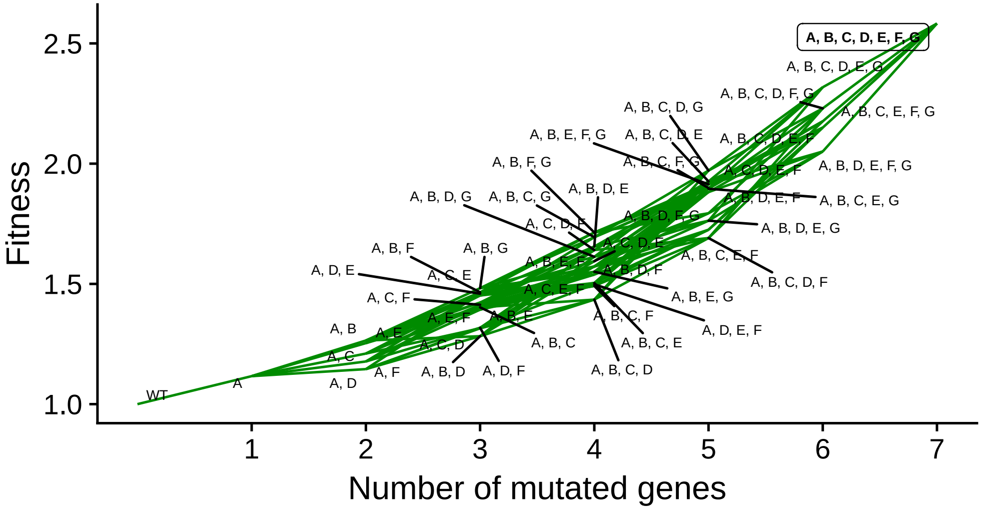

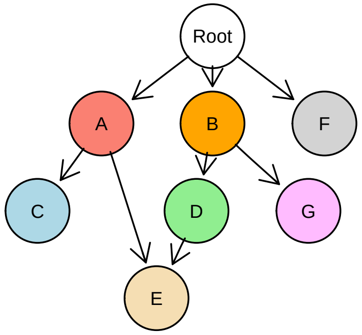


| | |
|---|---|
| *ID* | 1Mj1yzbmqyN510otm |
| *acc genotypes* | 38 |
| *peaks no back* | 1 |
| *removed edges* | 0 |
| *prop rem edges* | 0 |

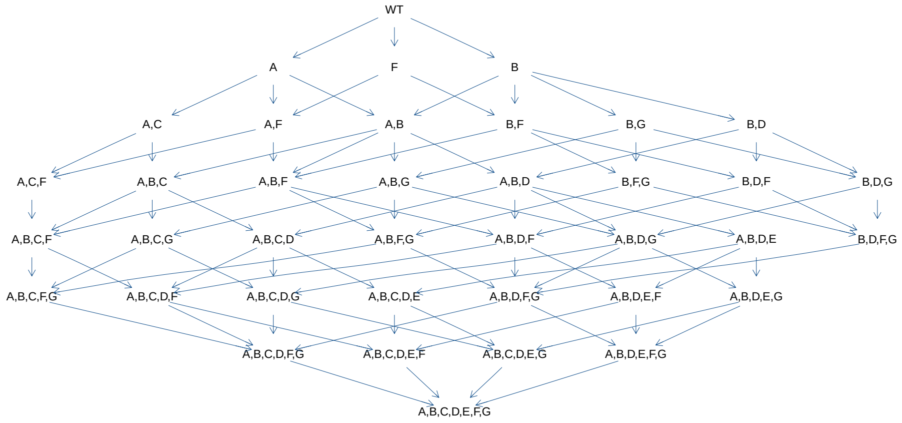


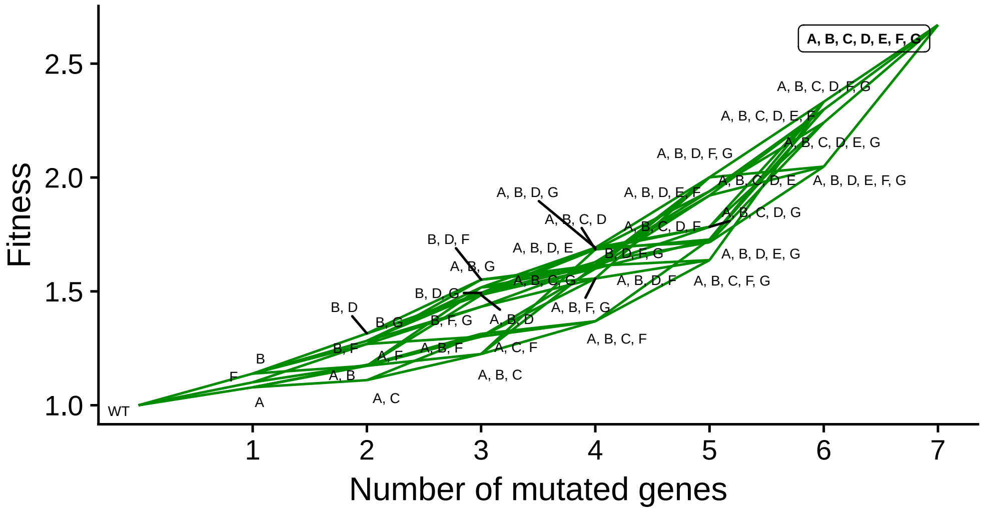

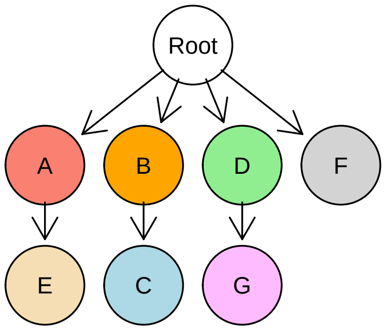


| | |
|---|---|
| *ID* | 1OIh71InKRui4v2e |
| *acc genotypes* | 54 |
| *peaks no back* | 1 |
| *removed edges* | 0 |
| *prop rem edges* | 0 |

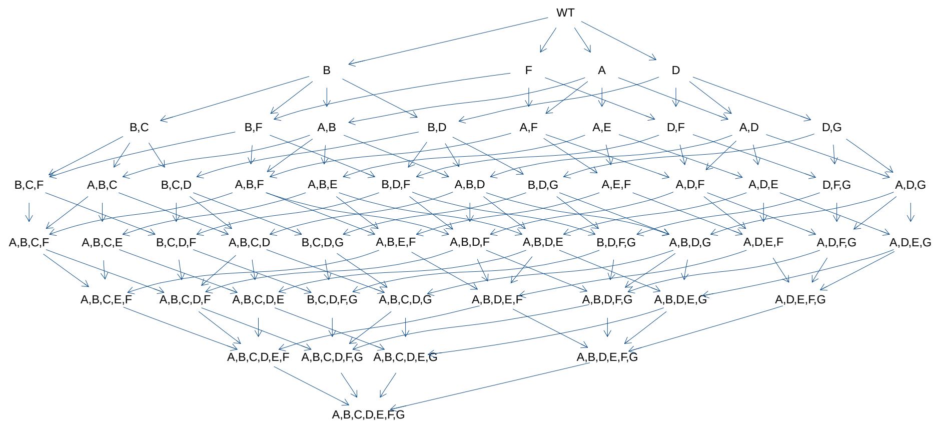


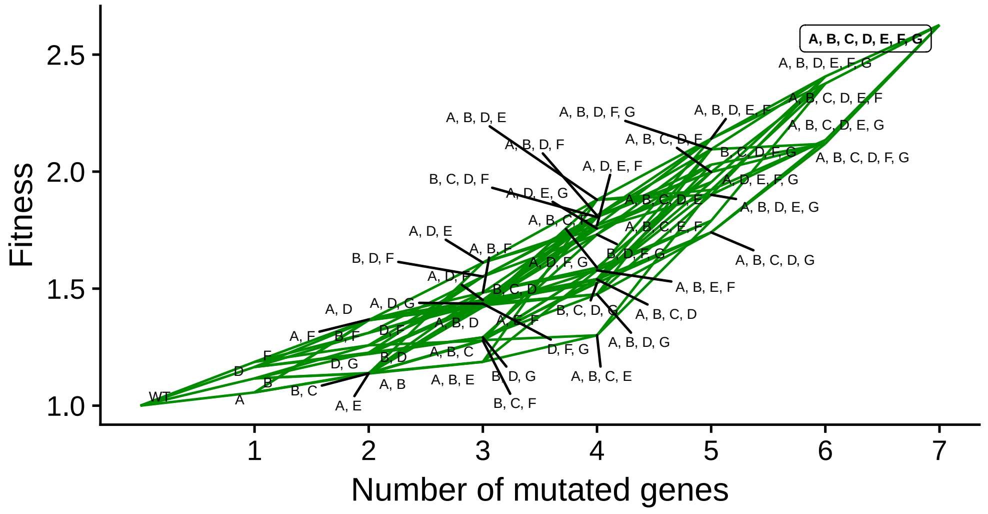

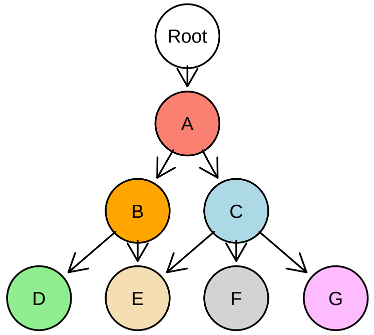


| | |
|---|---|
| *ID* | 1uosnPLLoevoyoqh |
| *acc genotypes* | 24 |
| *peaks no back* | 1 |
| *removed edges* | 0 |
| *prop rem edges* | 0 |

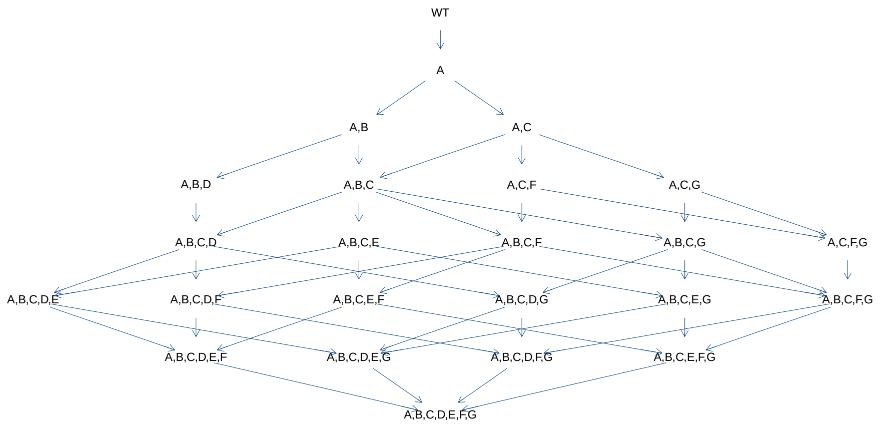


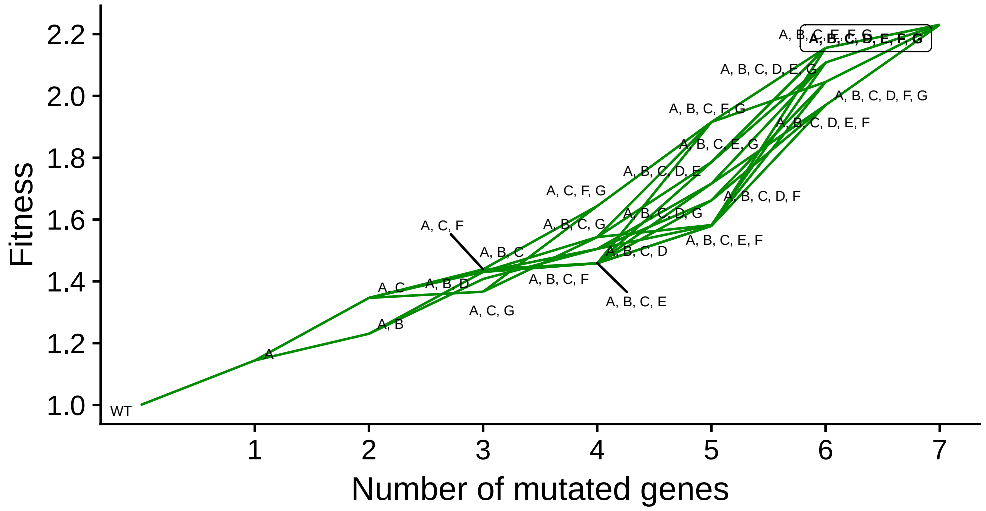

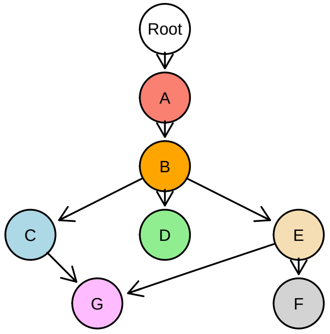


*ID* 29lH82hKYXw3zuT10
*acc genotypes* 18
*peaks no back* 1
*removed edges* 0
*prop rem edges* 0

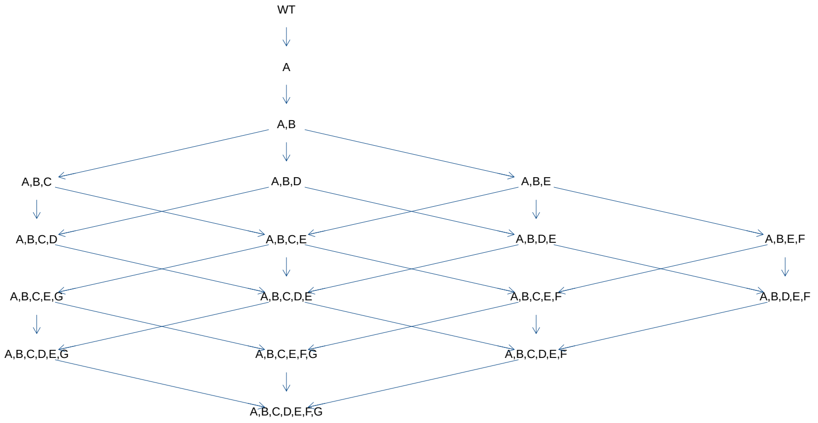


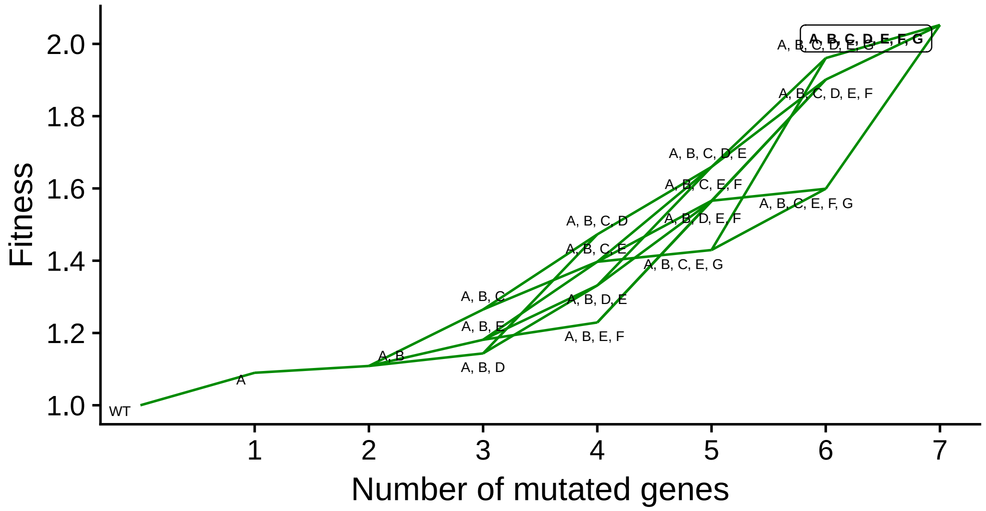

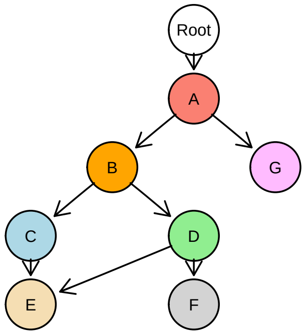


| | |
|---|---|
| *ID* | 2FyhSdCFo4Nq5dT10 |
| *acc genotypes* | 19 |
| *peaks no back* | 1 |
| *removed edges* | 0 |
| *prop rem edges* | 0 |

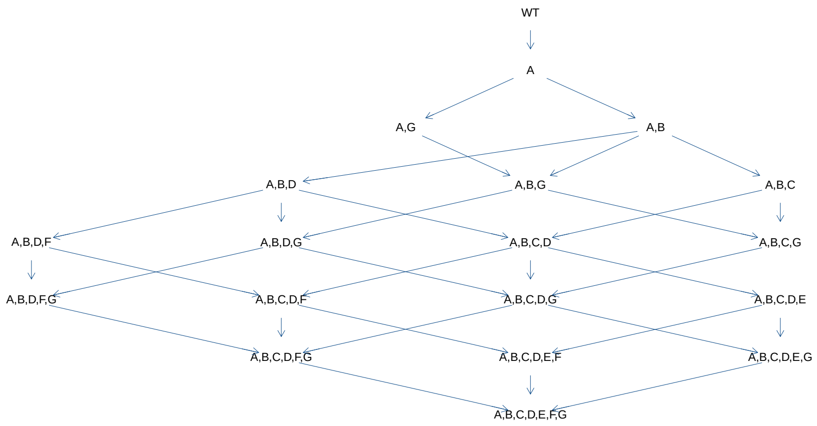


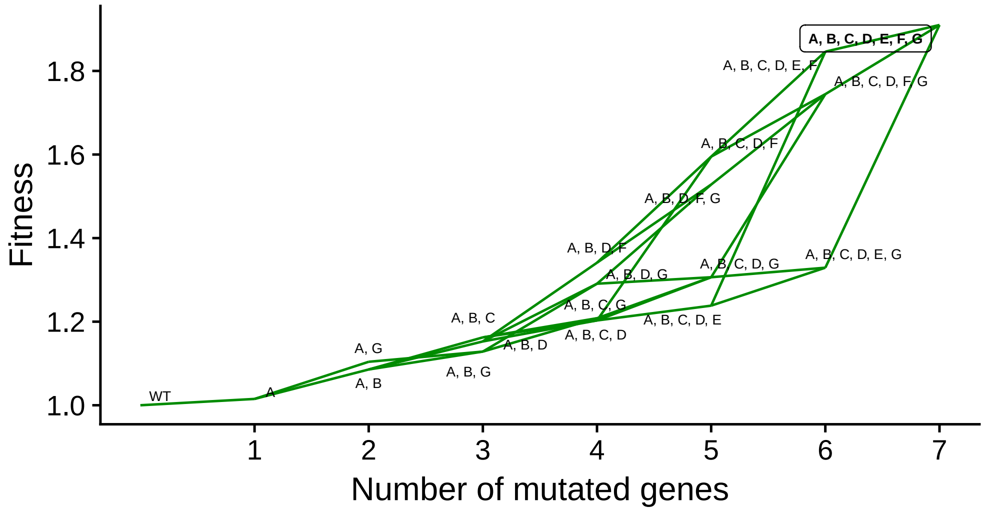

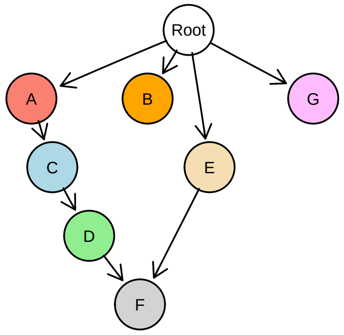


| | |
|---|---|
| *ID* | 2qPuA9kEjZvpHzWR |
| *acc genotypes* | 36 |
| *peaks no back* | 1 |
| *removed edges* | 0 |
| *prop rem edges* | 0 |

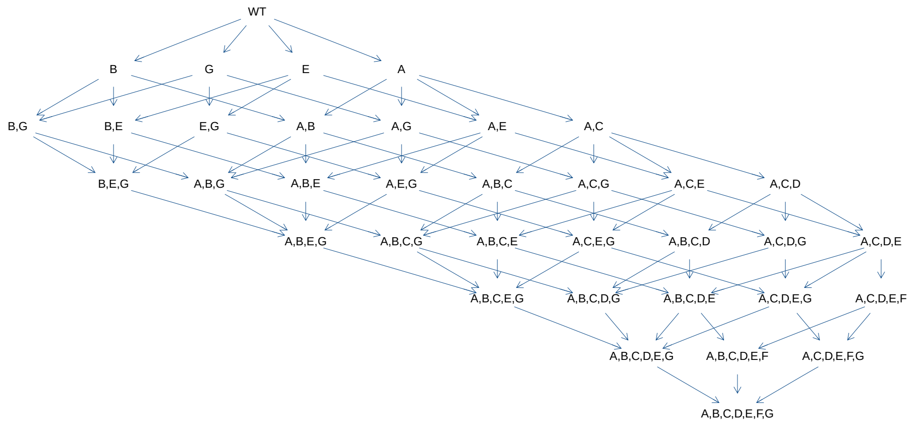


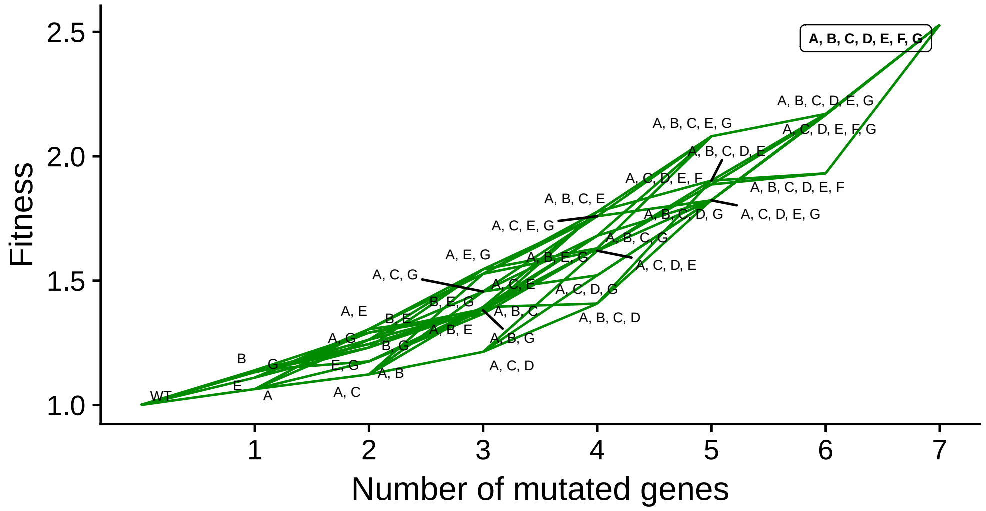

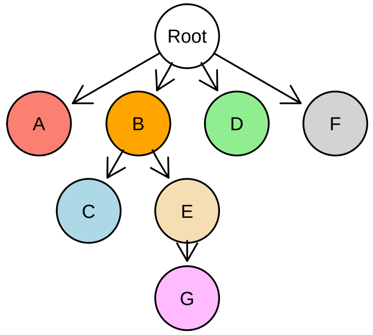


*ID* 2yfABs8hSv10jftX2
*acc genotypes* 56
*peaks no back* 1
*removed edges* 0
*prop rem edges* 0

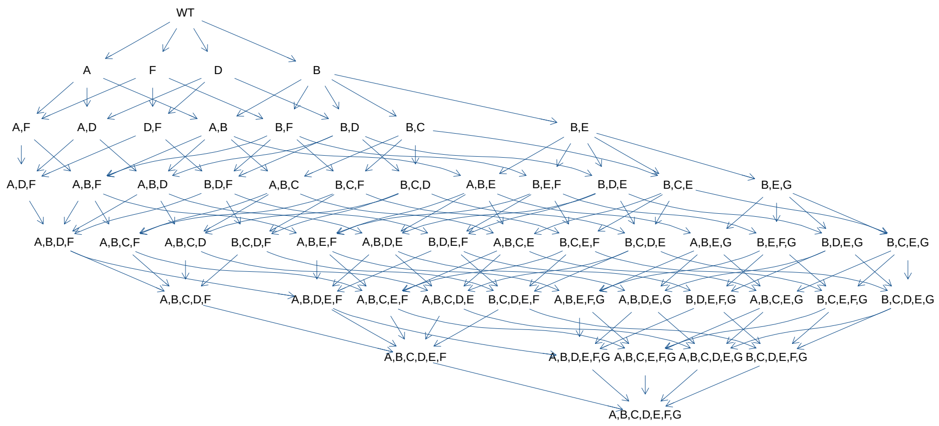


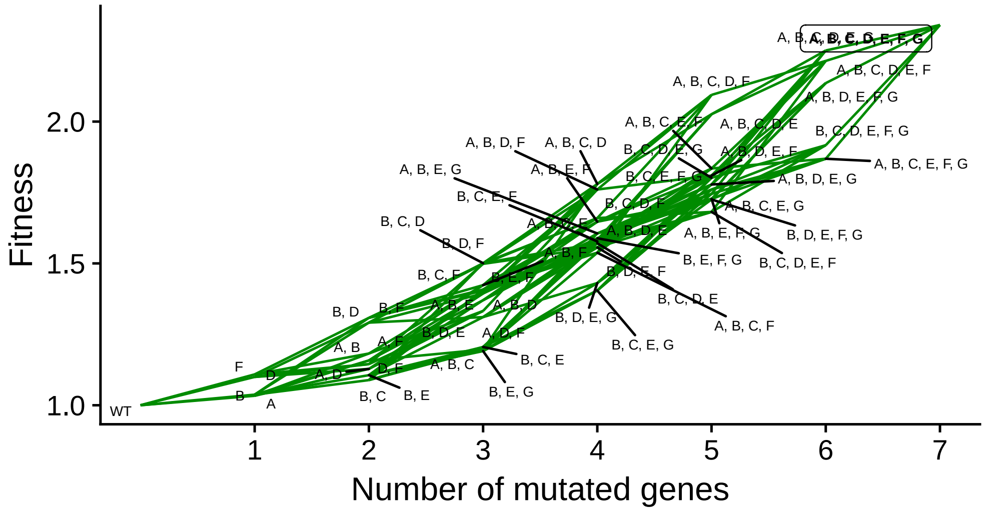

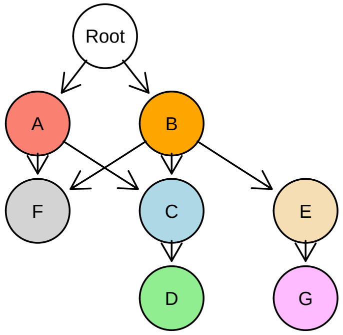


| | |
|---|---|
| *ID* | 3PMBKfaRwTHJEyd8 |
| *acc genotypes* | 23 |
| *peaks no back* | 1 |
| *removed edges* | 0 |
| *prop rem edges* | 0 |

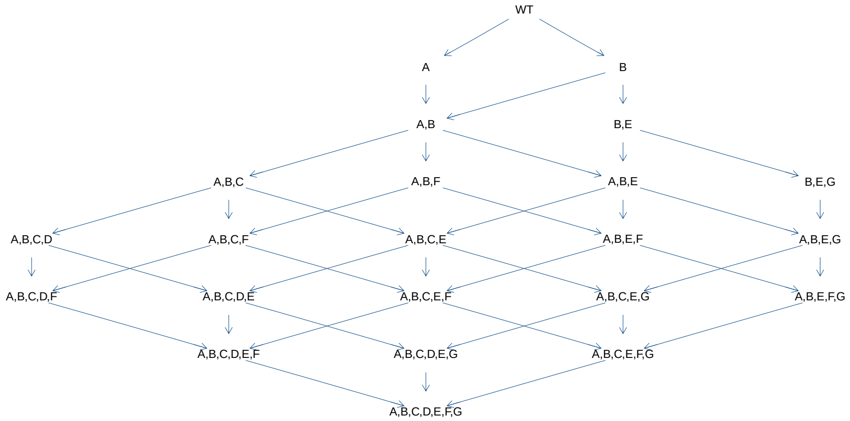


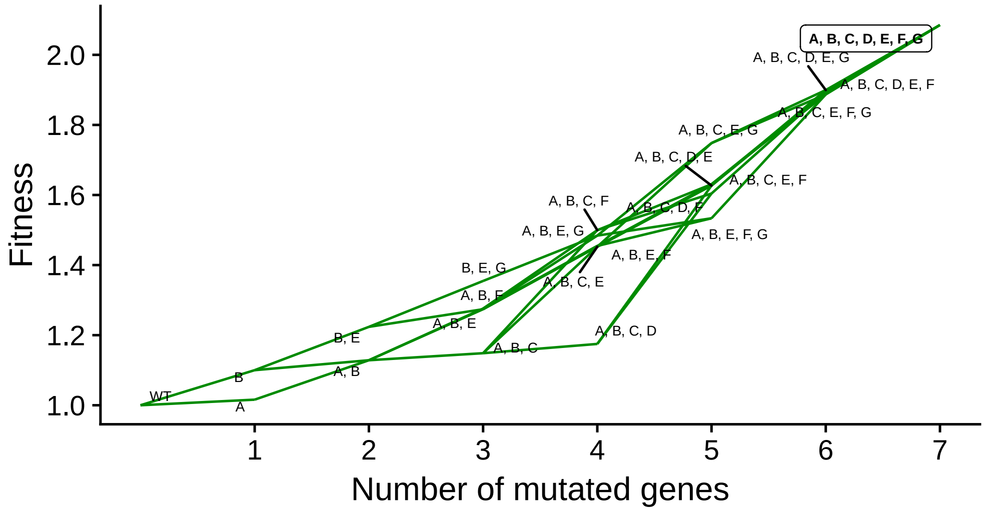

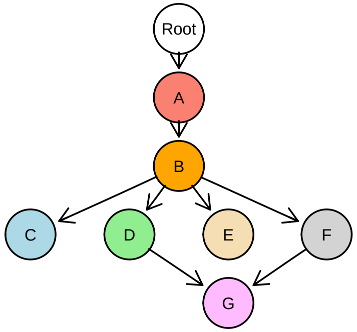


| | |
|---|---|
| *ID* | 3qyRJOswye1eG84H |
| *acc genotypes* | 22 |
| *peaks no back* | 1 |
| *removed edges* | 0 |
| *prop rem edges* | 0 |

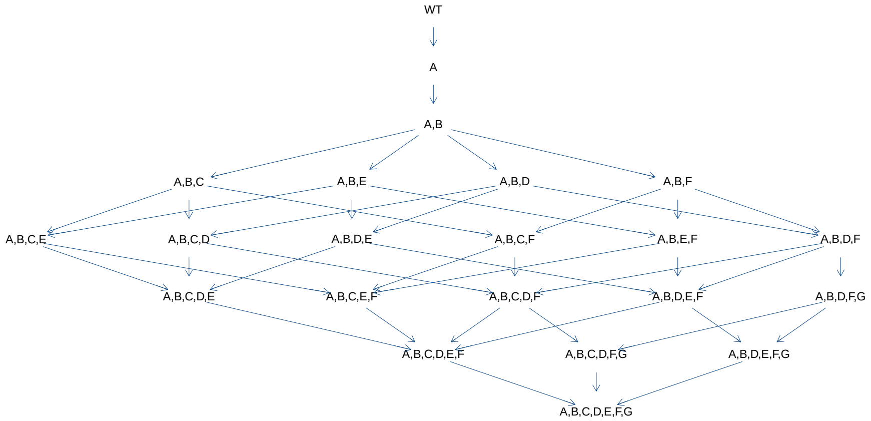


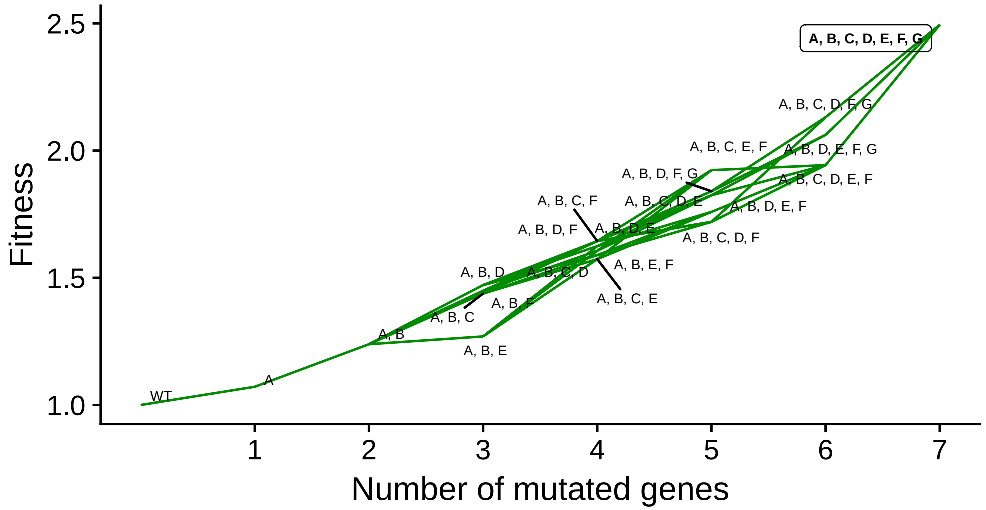

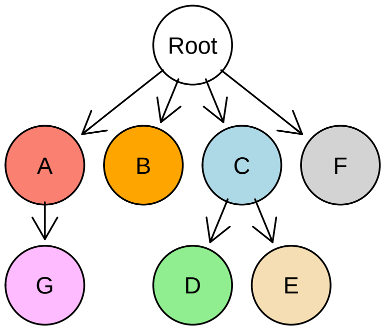


| | |
|---|---|
| *ID* | 3YPBDYDpA3ERbdMp |
| *acc genotypes* | 60 |
| *peaks no back* | 1 |
| *removed edges* | 0 |
| *prop rem edges* | 0 |

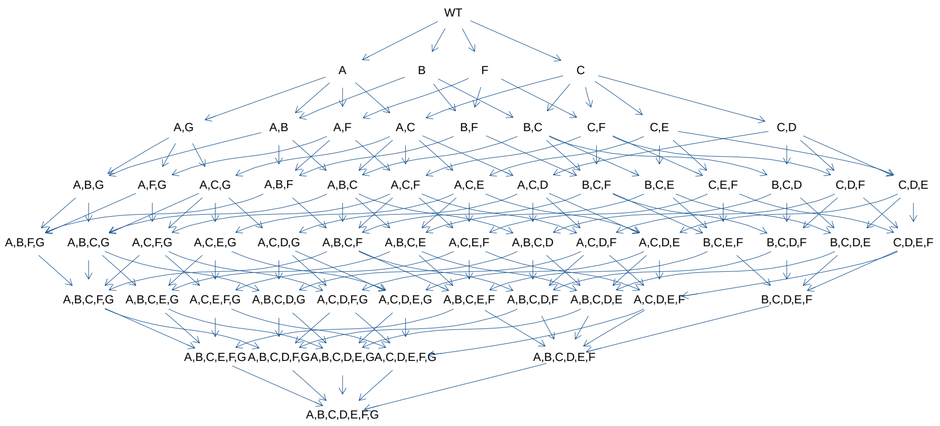


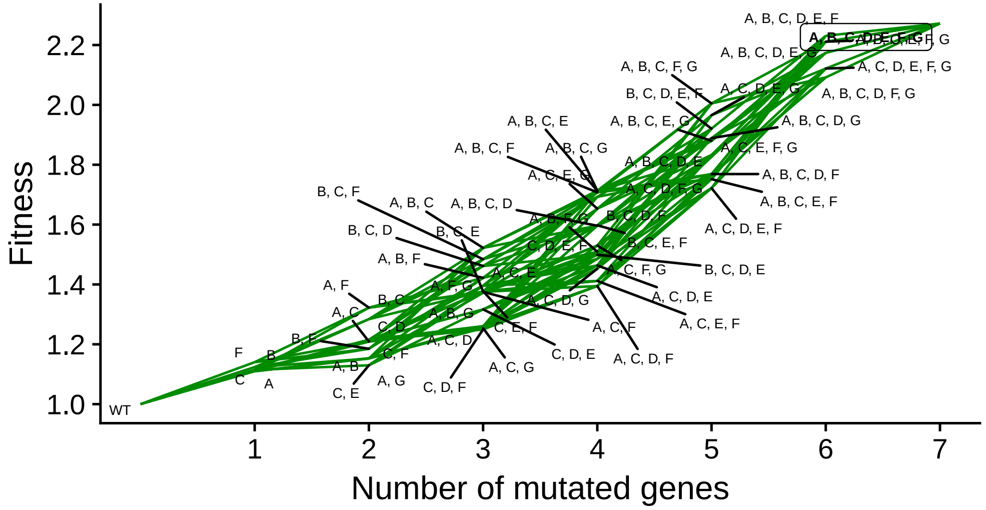

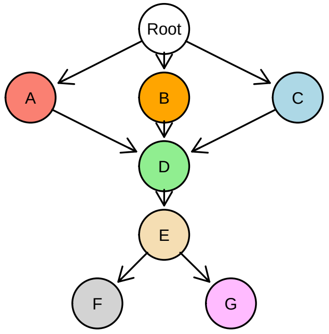


| | |
|---|---|
| *ID* | 417teASz4uBqi4qQ |
| *acc genotypes* | 13 |
| *peaks no back* | 1 |
| *removed edges* | 0 |
| *prop rem edges* | 0 |

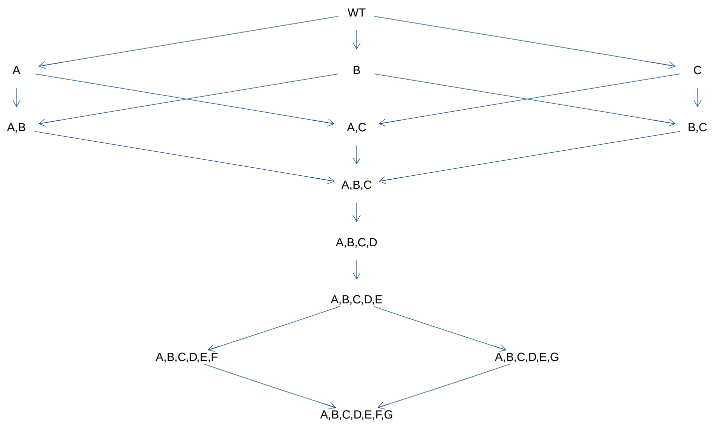


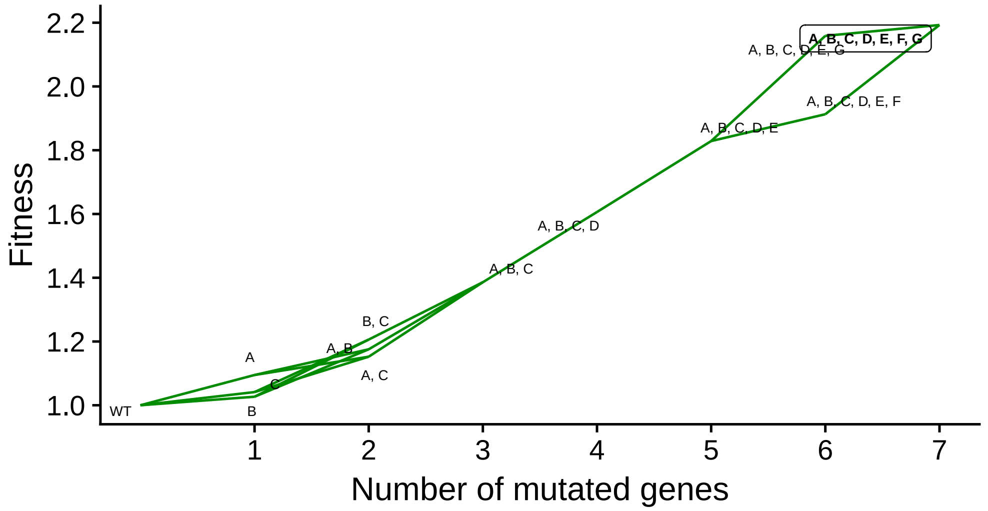

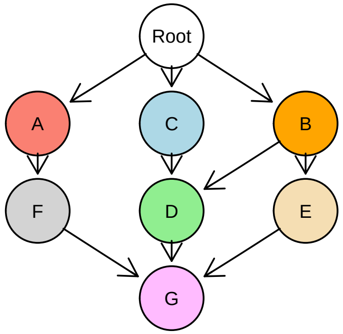


| | |
|---|---|
| *ID* | 46rFIBbA10BhomWGN |
| *acc genotypes* | 25 |
| *peaks no back* | 1 |
| *removed edges* | 0 |
| *prop rem edges* | 0 |

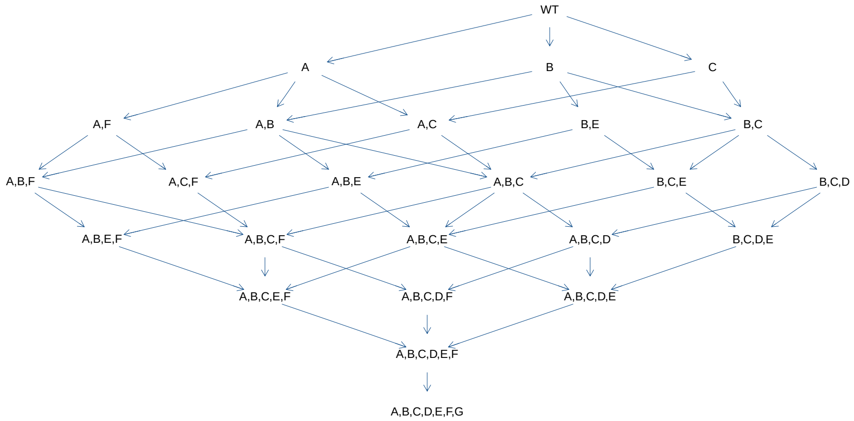


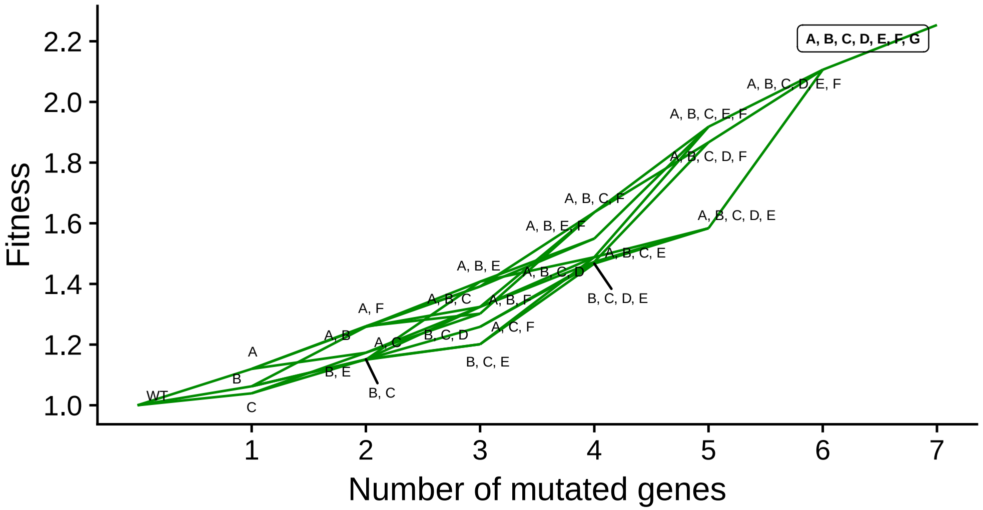

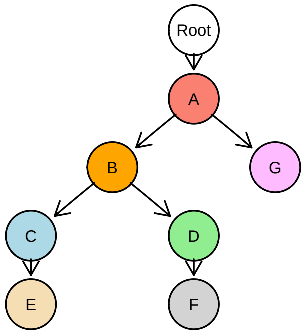


| | |
|---|---|
| *ID* | 4mouiGQselPBUfsX |
| *acc genotypes* | 21 |
| *peaks no back* | 1 |
| *removed edges* | 0 |
| *prop rem edges* | 0 |

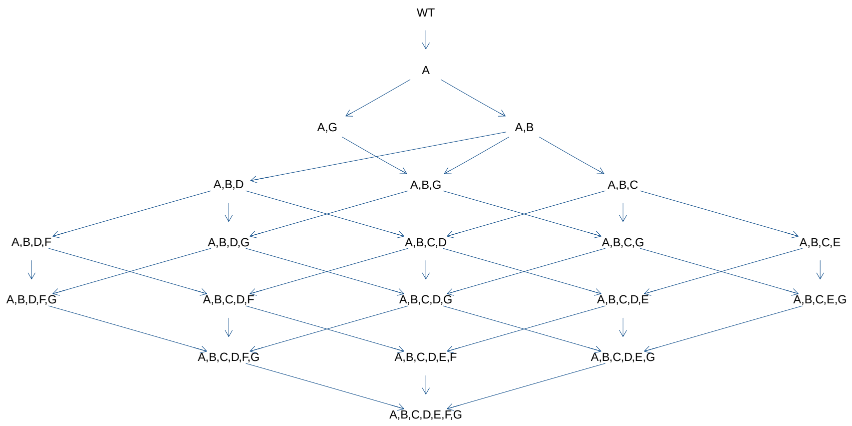


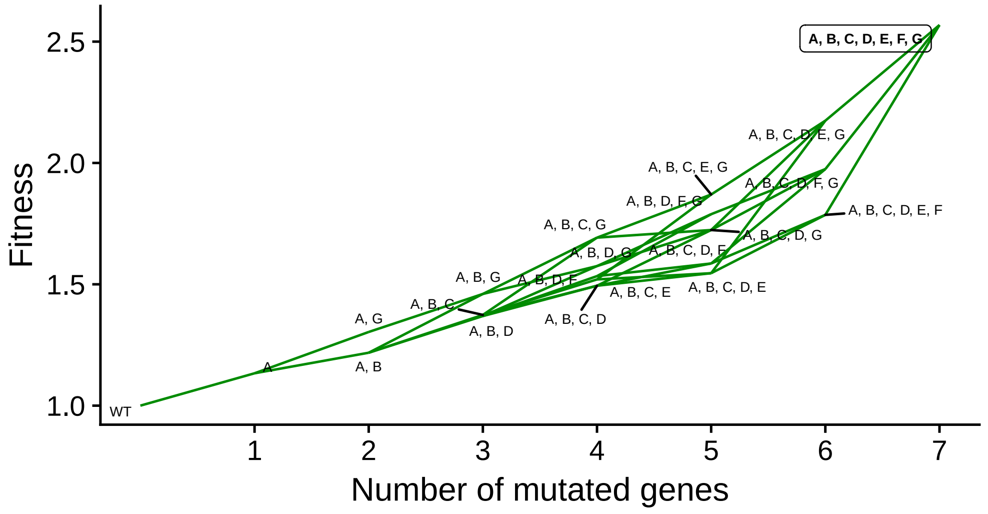

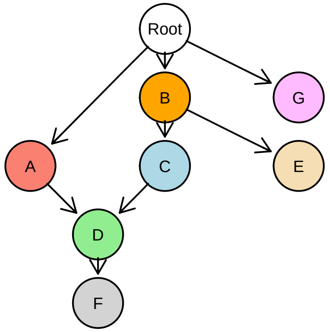


| | |
|---|---|
| *ID* | 4RoAEsprOB10Yv3U6 |
| *acc genotypes* | 28 |
| *peaks no back* | 1 |
| *removed edges* | 0 |
| *prop rem edges* | 0 |

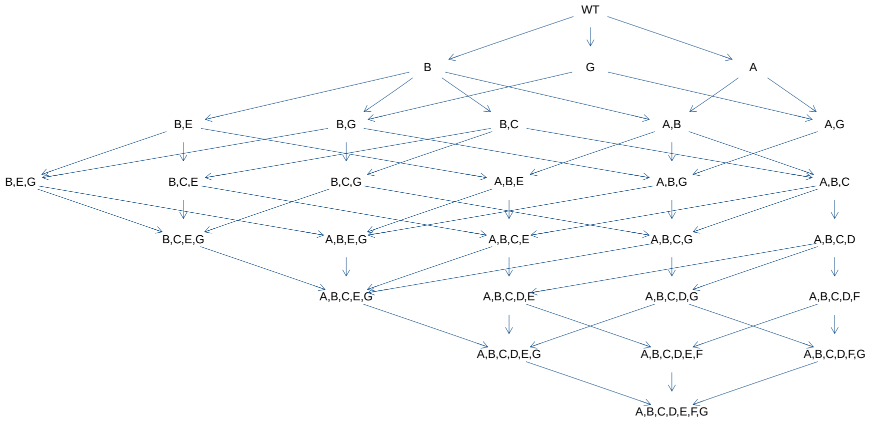


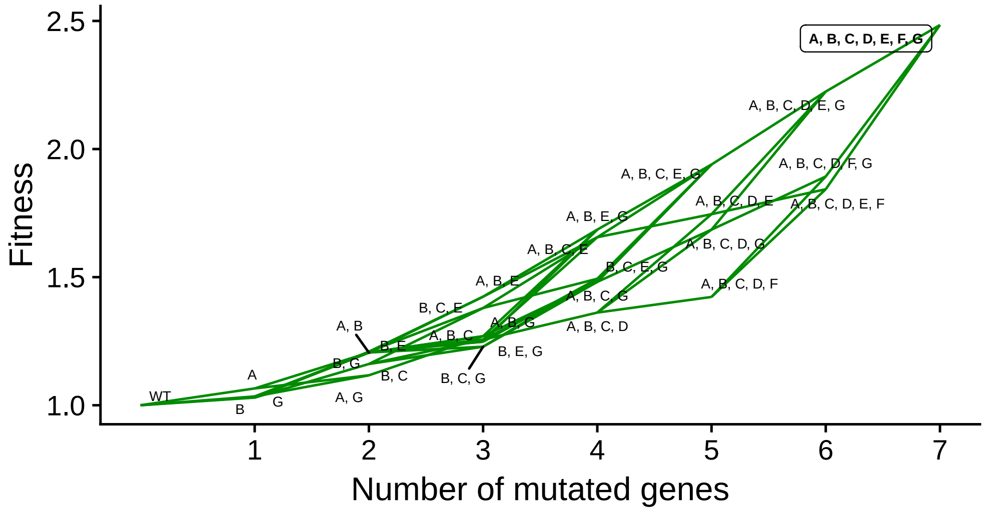

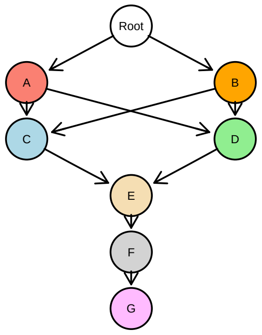


| | |
|---|---|
| *ID* | 4yUgjvZI1UgonSqo |
| *acc genotypes* | 10 |
| *peaks no back* | 1 |
| *removed edges* | 0 |
| *prop rem edges* | 0 |

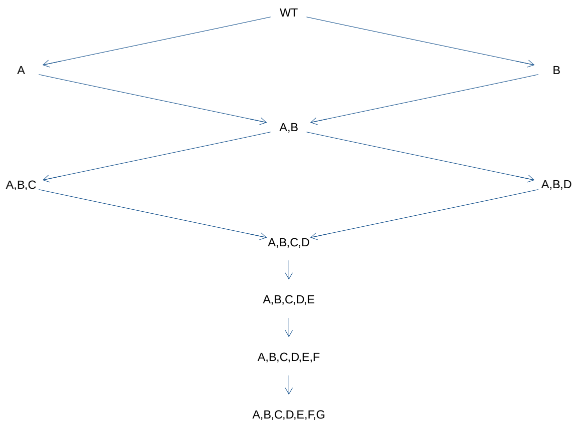


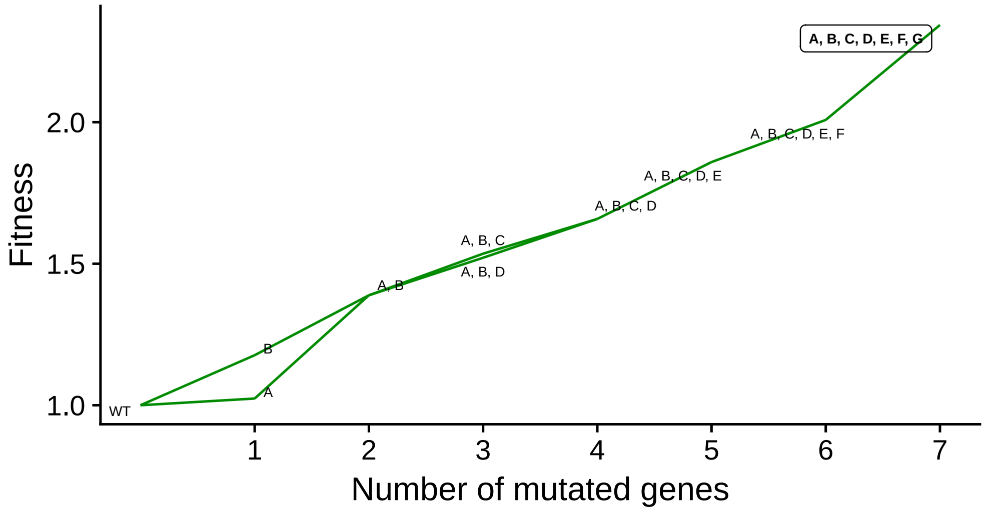

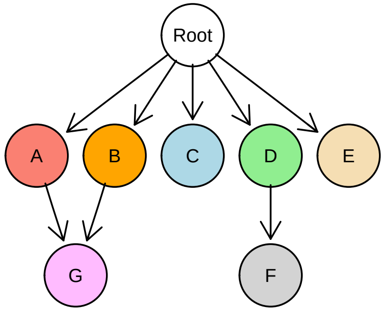


| | |
|---|---|
| *ID* | 5fz101Z59qIyO10Rxn |
| *acc genotypes* | 60 |
| *peaks no back* | 1 |
| *removed edges* | 0 |
| *prop rem edges* | 0 |

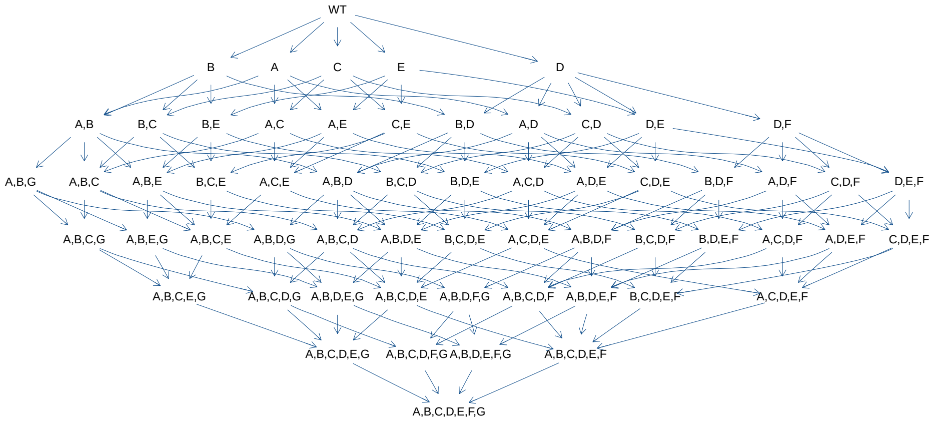


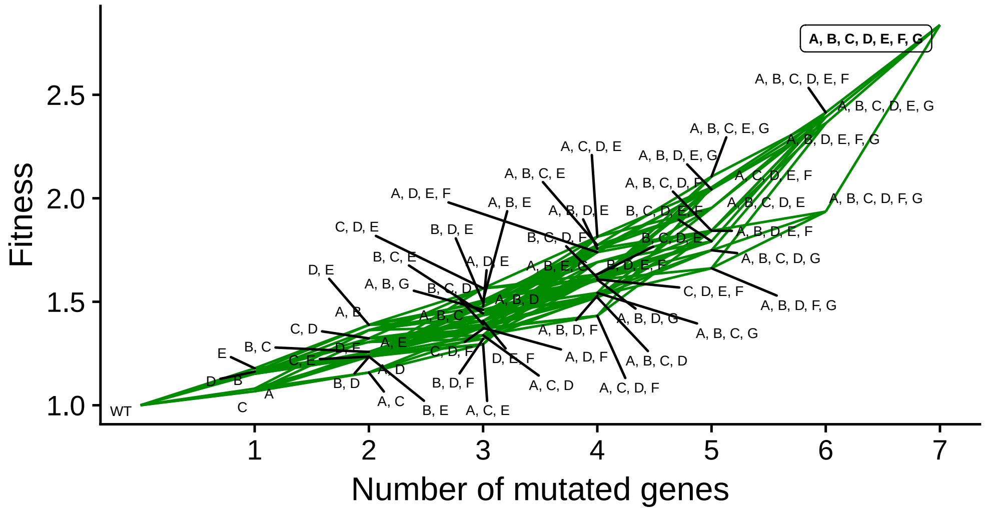

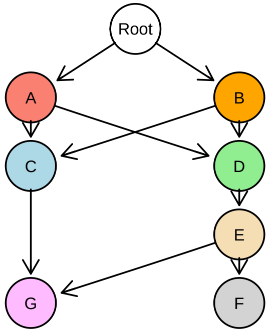


| | |
|---|---|
| *ID* | 6pjAHfBKJ10Ocigjg |
| *acc genotypes* | 13 |
| *peaks no back* | 1 |
| *removed edges* | 0 |
| *prop rem edges* | 0 |

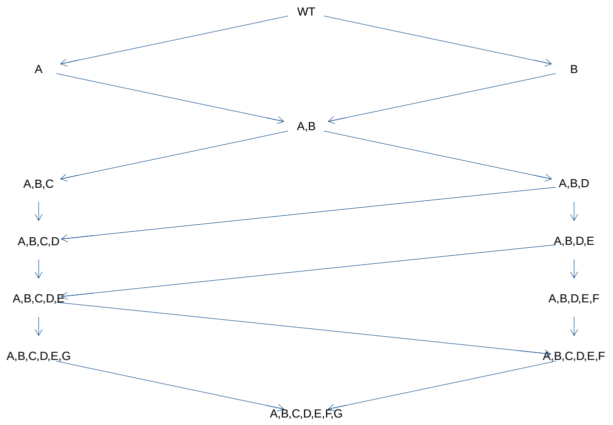


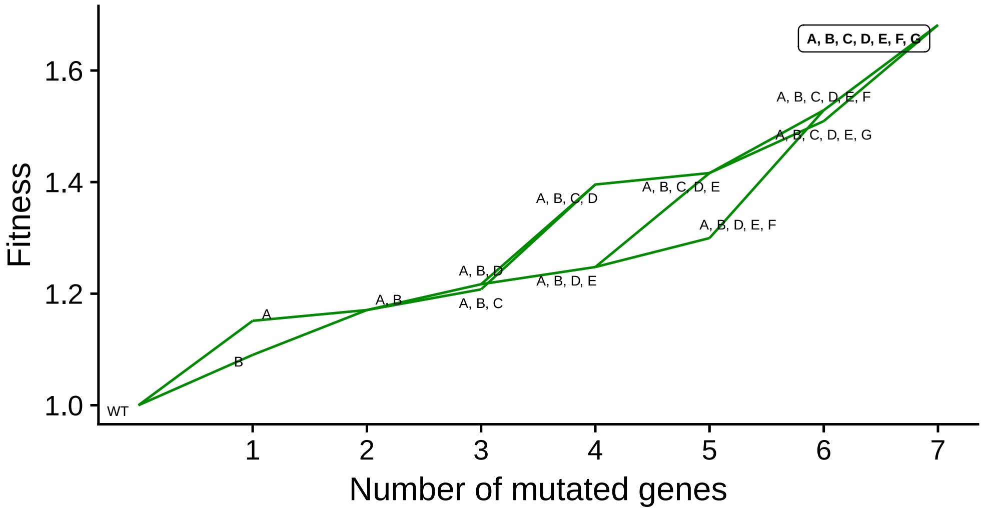

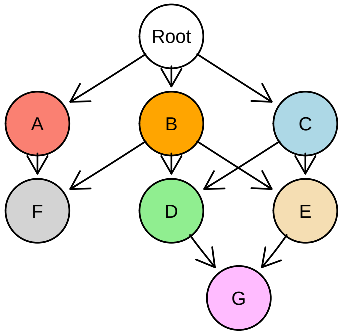


| | |
|---|---|
| *ID* | 718HviJL2vTZgZyJ |
| *acc genotypes* | 22 |
| *peaks no back* | 1 |
| *removed edges* | 0 |
| *prop rem edges* | 0 |

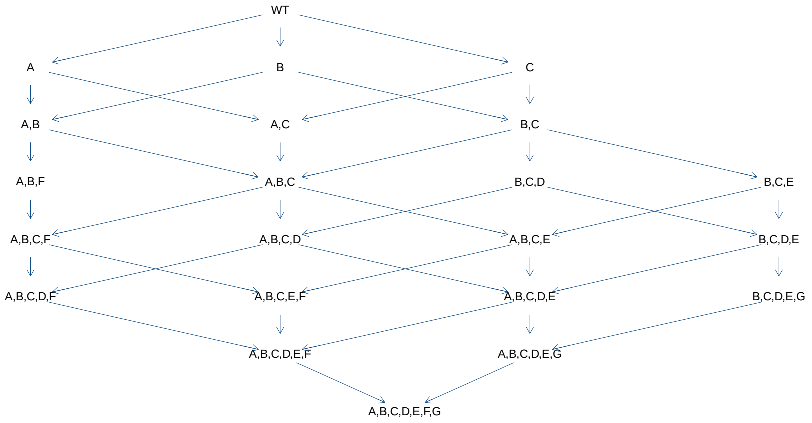


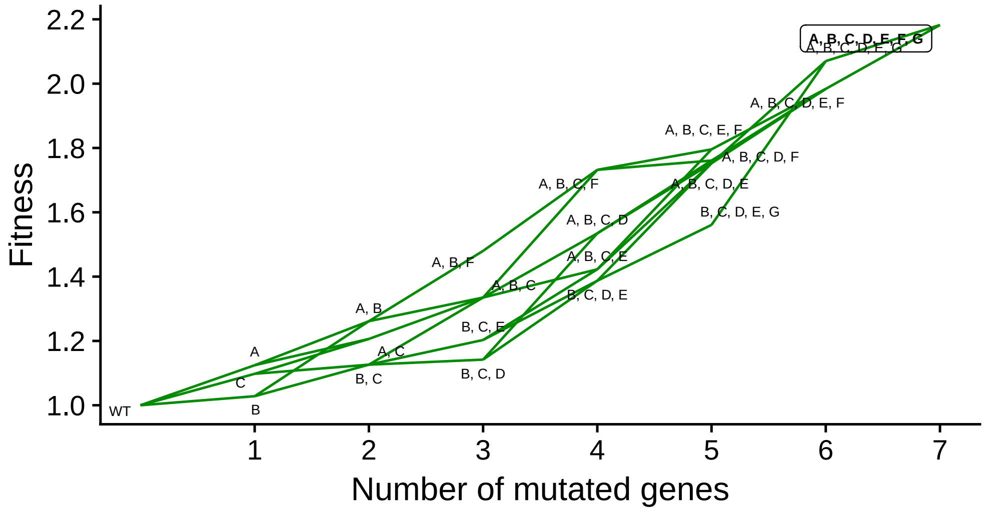

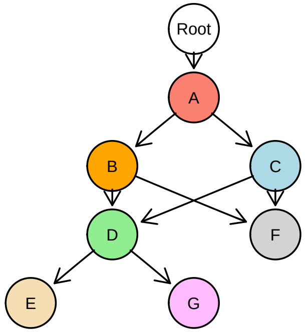


| | |
|---|---|
| *ID* | 7GFrRtiQ5Cyt3w8k |
| *acc genotypes* | 14 |
| *peaks no back* | 1 |
| *removed edges* | 0 |
| *prop rem edges* | 0 |

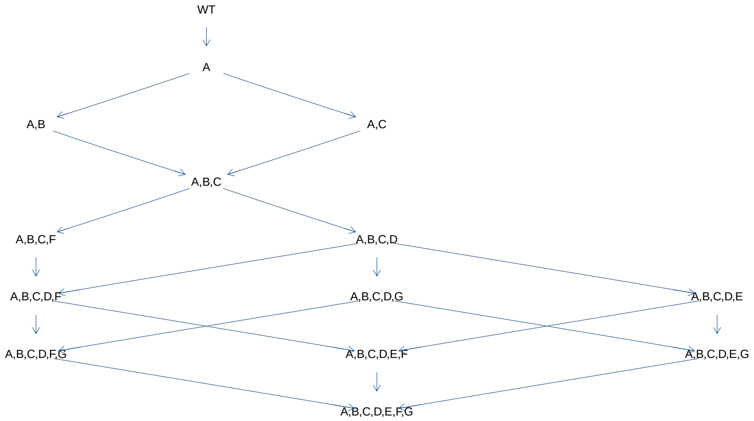


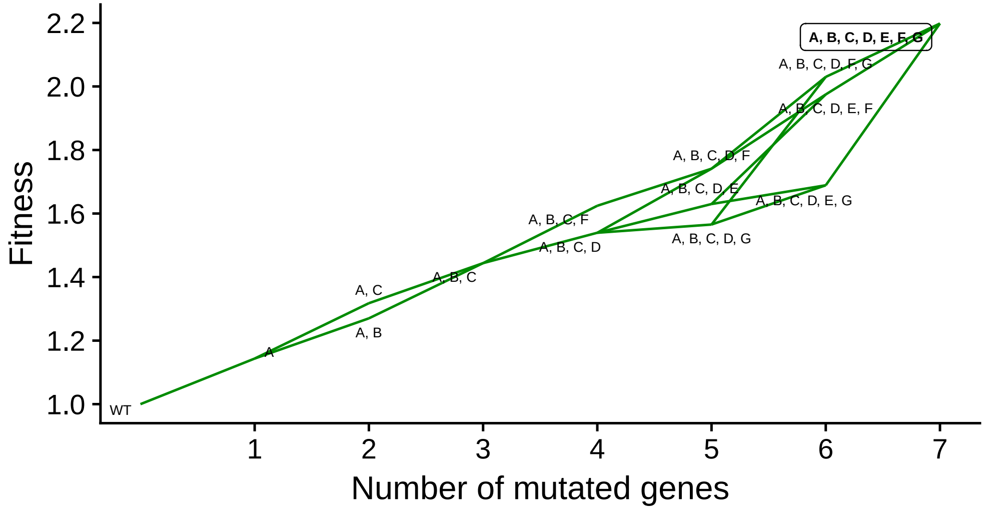

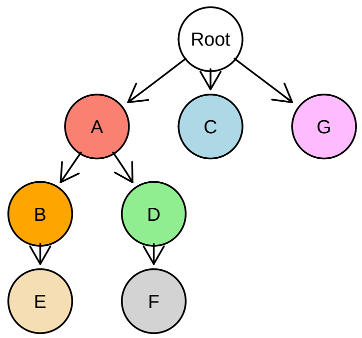


| | |
|---|---|
| *ID* | 7O6M3PSFd6vyICJi |
| *acc genotypes* | 40 |
| *peaks no back* | 1 |
| *removed edges* | 0 |
| *prop rem edges* | 0 |

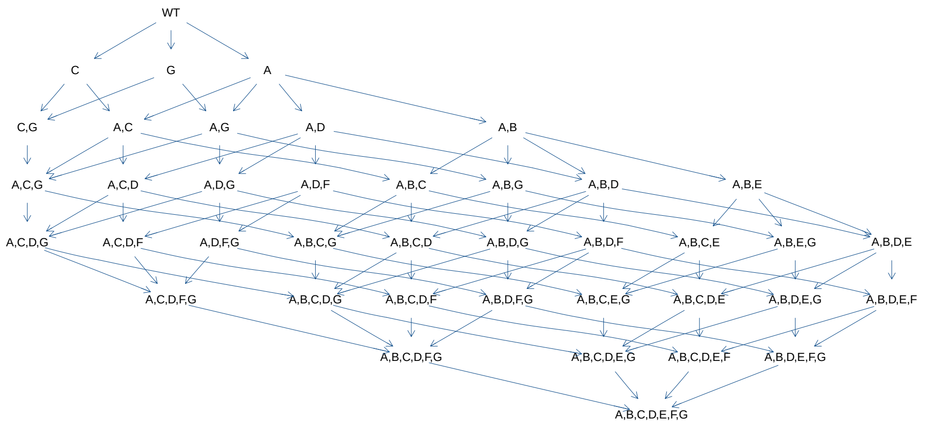


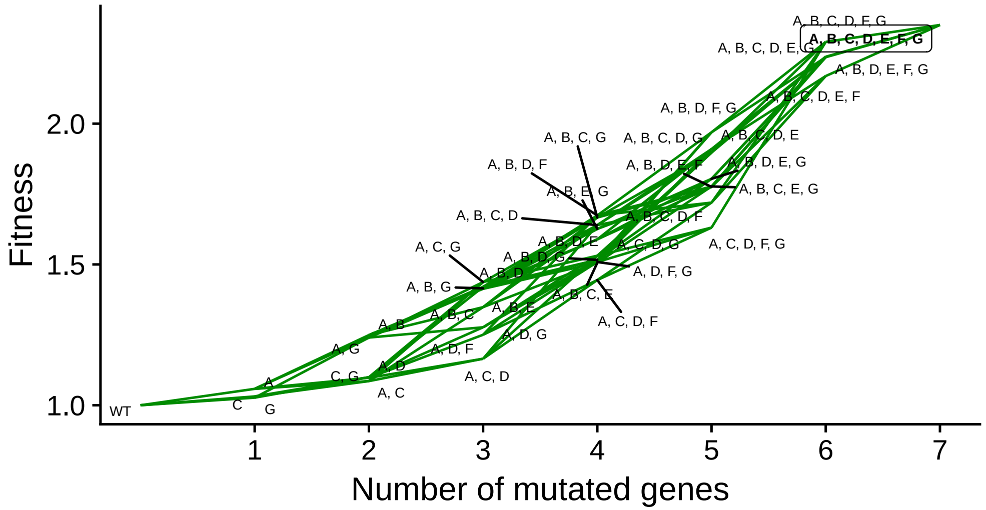

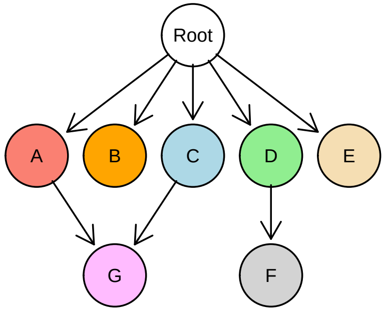


*ID* 8GEzhn1FLKorAfUm
*acc genotypes* 60
*peaks no back* 1
*removed edges* 0
*prop rem edges* 0

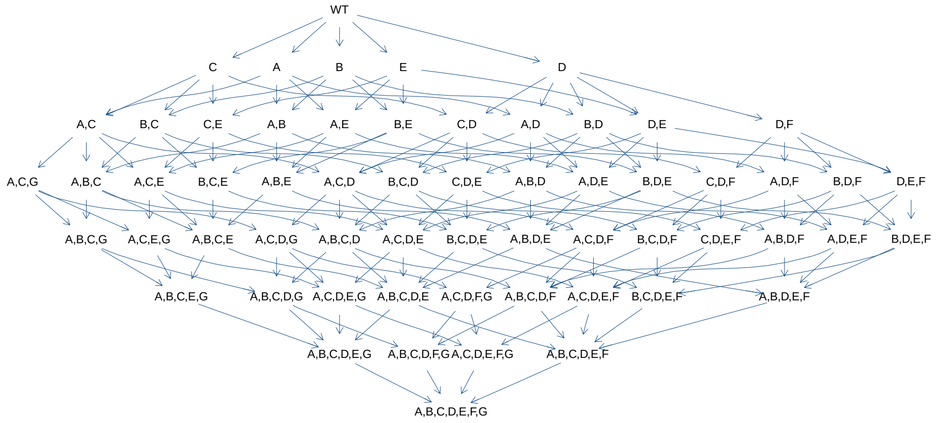


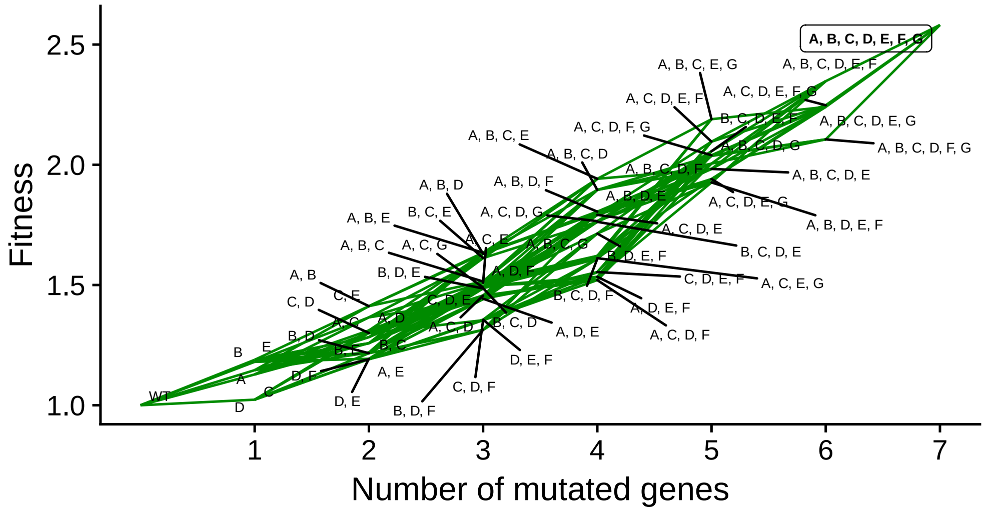

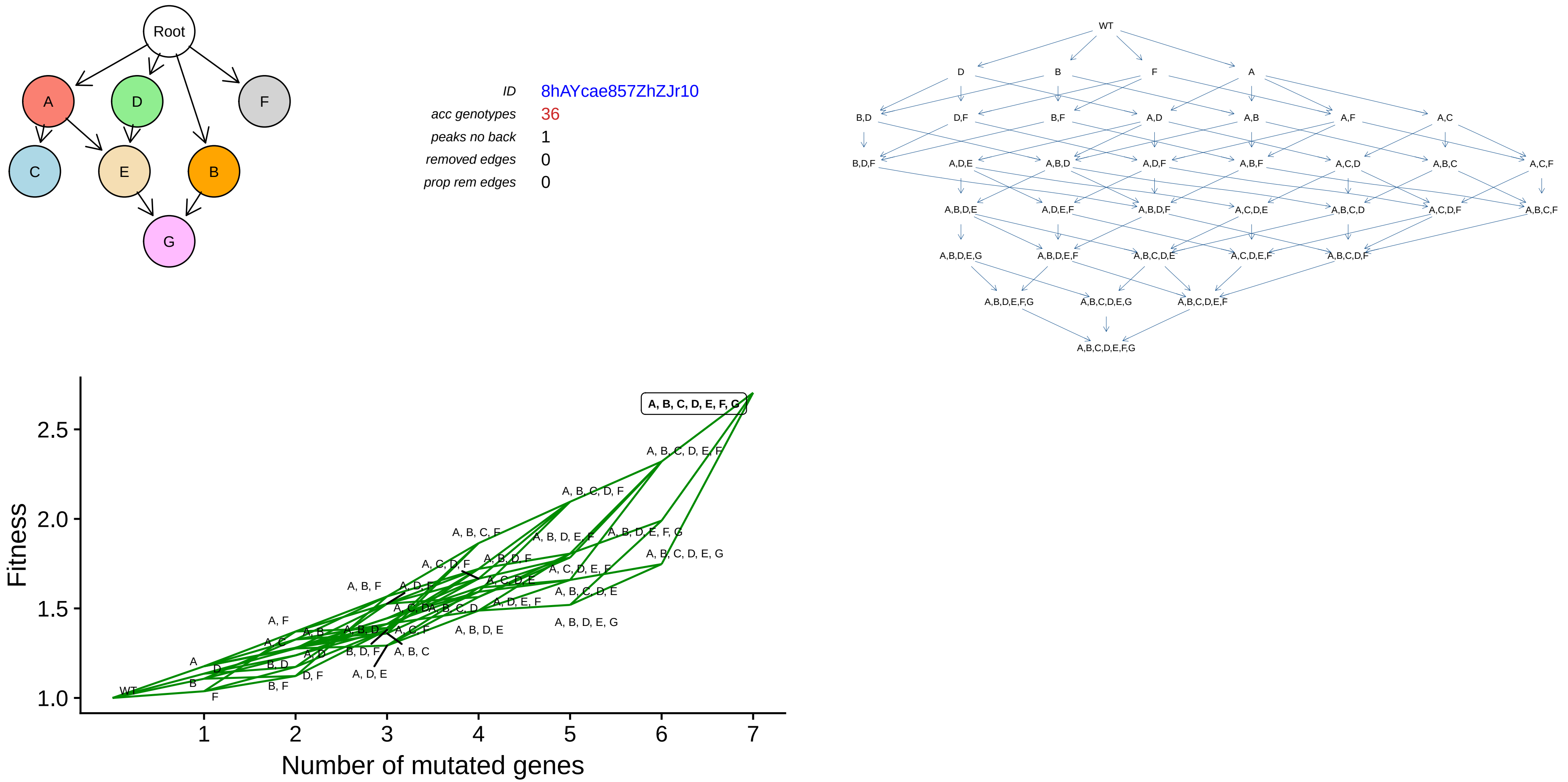
Root
A
D
F
C
E
B
G
ID 8hAYcae857ZhZJr10
acc genotypes 36
peaks no back 1
removed edges 0
prop rem edges 0
A, B, C, D, E, F, G
Fitness
Number of mutated genes

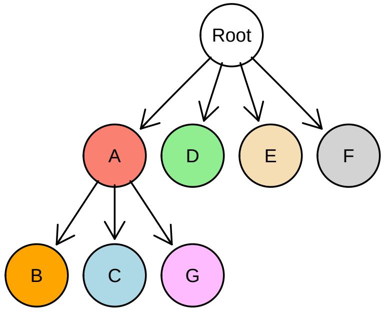


| | |
|---|---|
| *ID* | 8Zr6qLC9yn9cyhAY |
| *acc genotypes* | 72 |
| *peaks no back* | 1 |
| *removed edges* | 0 |
| *prop rem edges* | 0 |

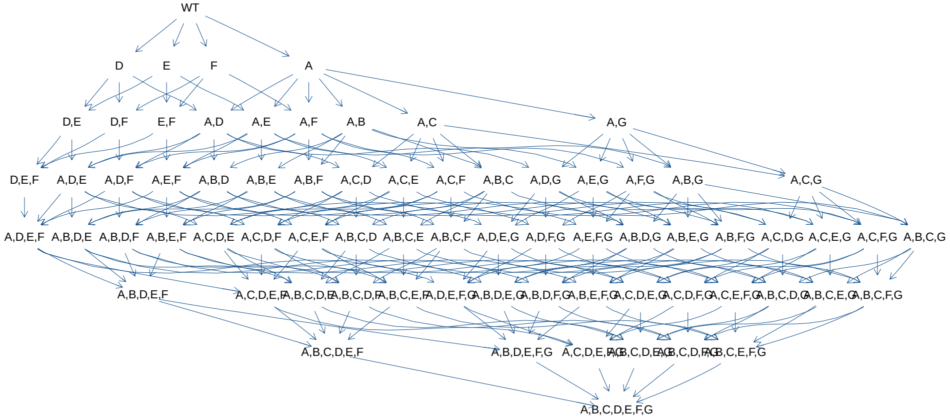


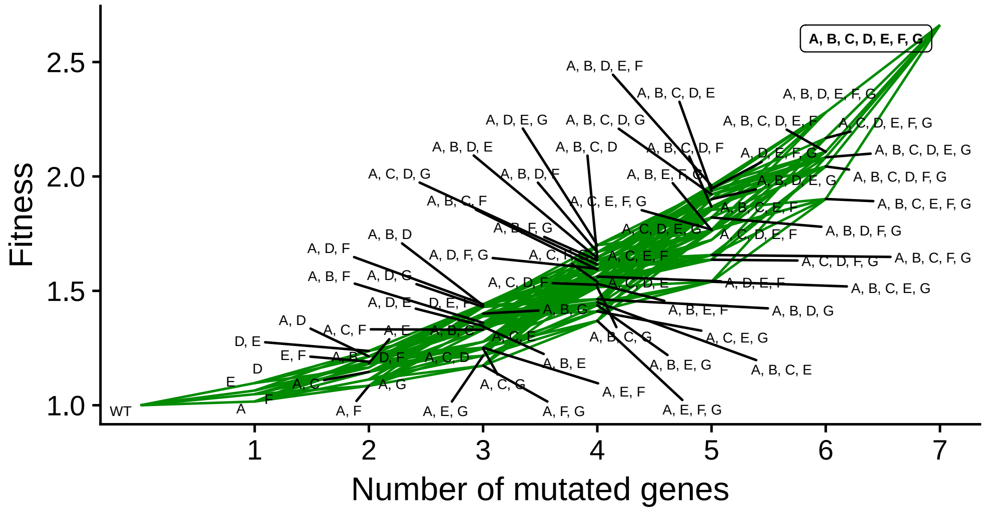

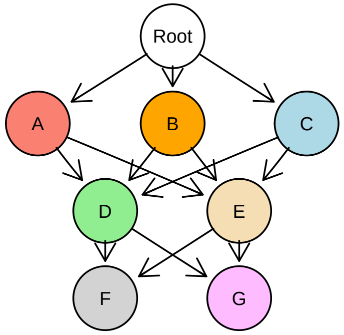


*ID* 9o10QUimqcWrPHbGH
*acc genotypes* 14
*peaks no back* 1
*removed edges* 0
*prop rem edges* 0

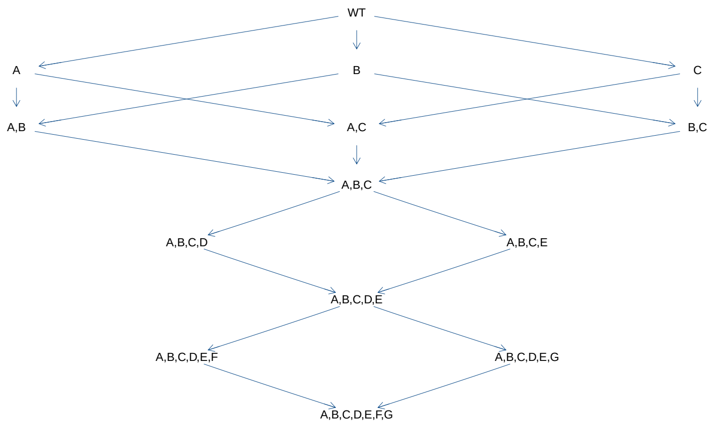


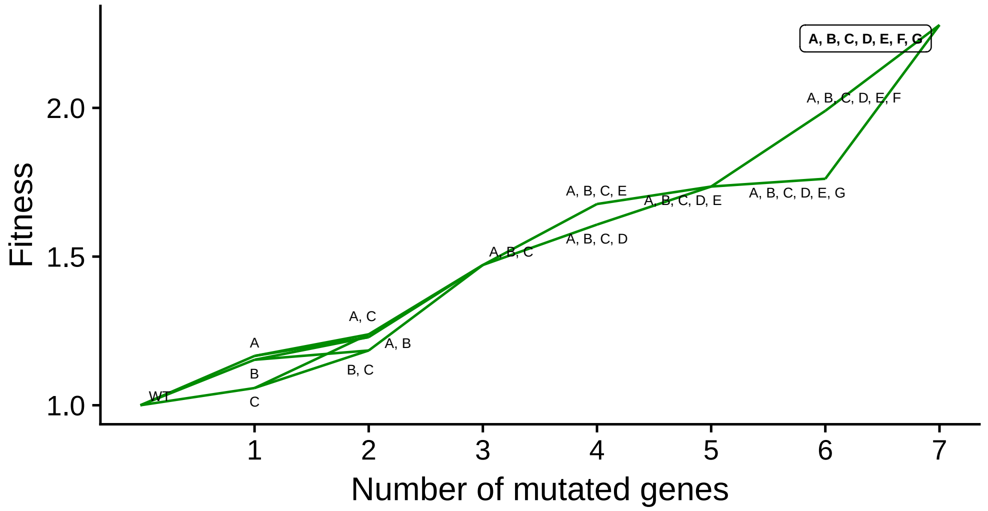

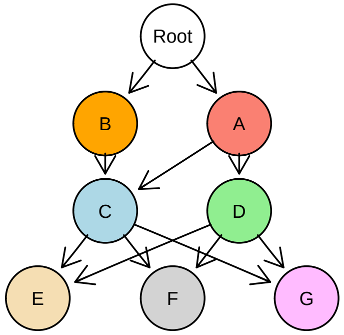


| | |
|---|---|
| *ID* | a2tStINqtyWzD10eQ |
| *acc genotypes* | 15 |
| *peaks no back* | 1 |
| *removed edges* | 0 |
| *prop rem edges* | 0 |

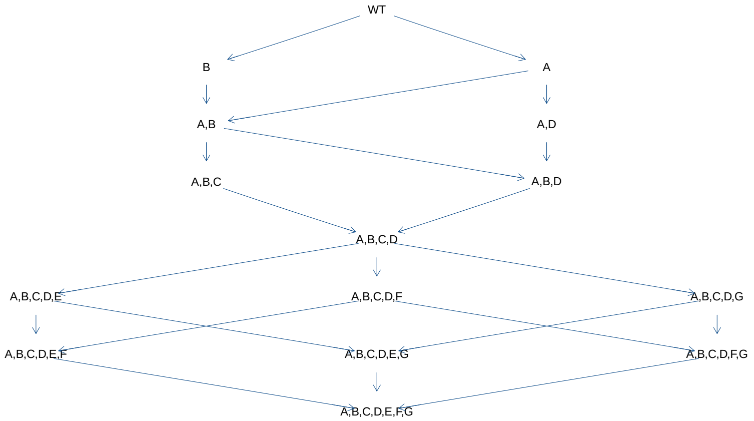


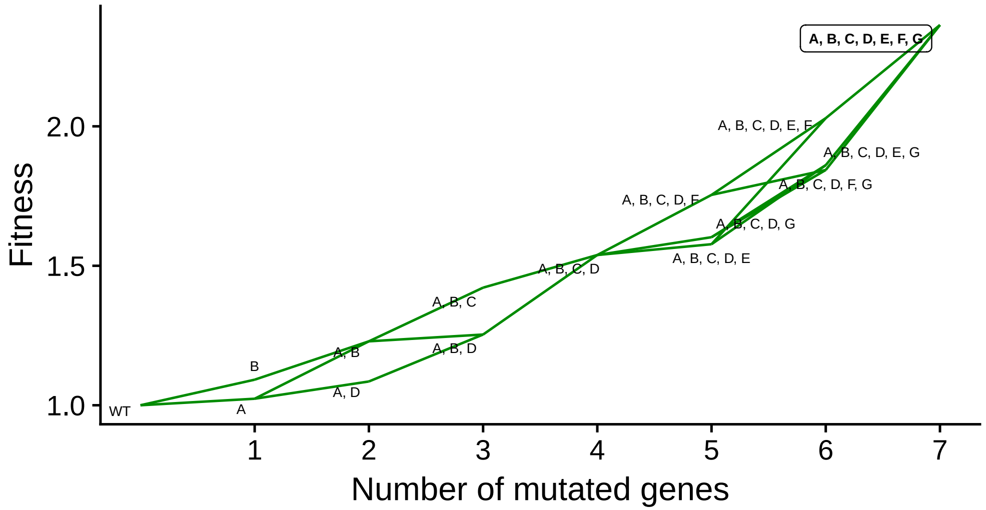

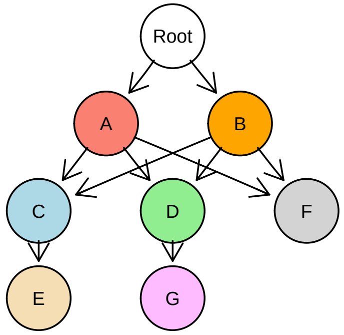


| | |
|---|---|
| *ID* | a7XZXptDjjWuSFhl |
| *acc genotypes* | 21 |
| *peaks no back* | 1 |
| *removed edges* | 0 |
| *prop rem edges* | 0 |

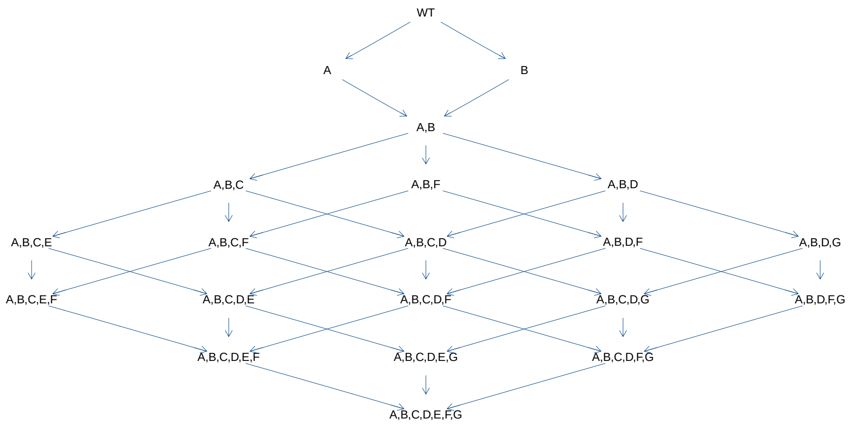


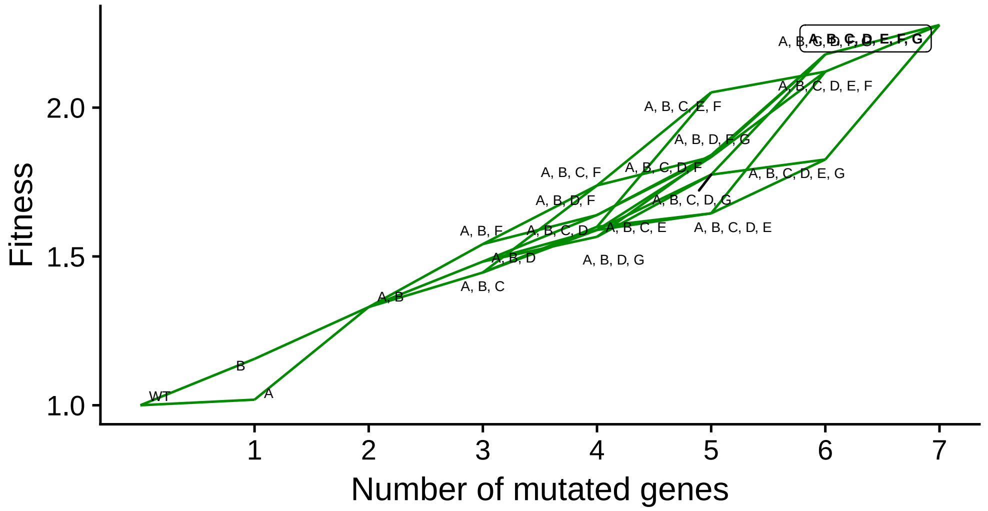

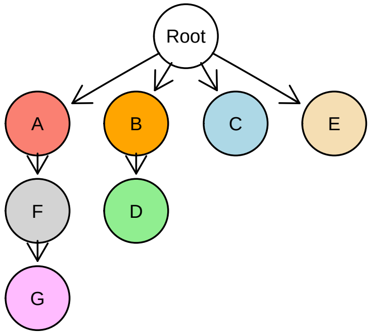


| | |
|---|---|
| *ID* | aGBVZklQ4oOGxWTb |
| *acc genotypes* | 48 |
| *peaks no back* | 1 |
| *removed edges* | 0 |
| *prop rem edges* | 0 |

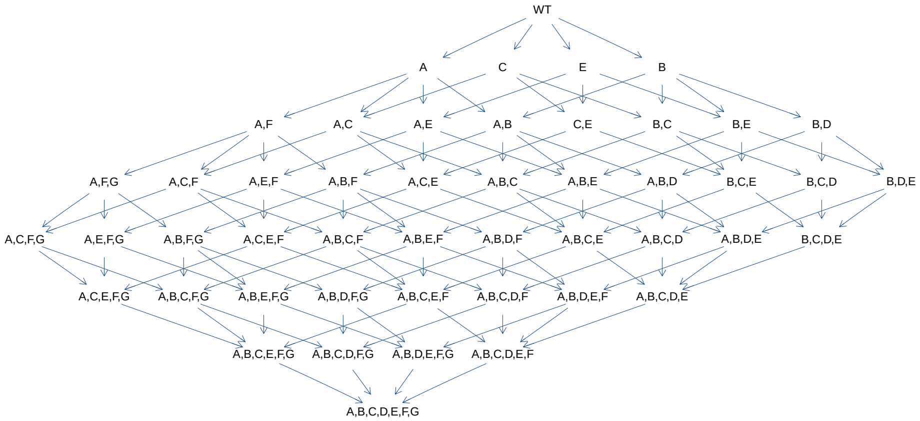


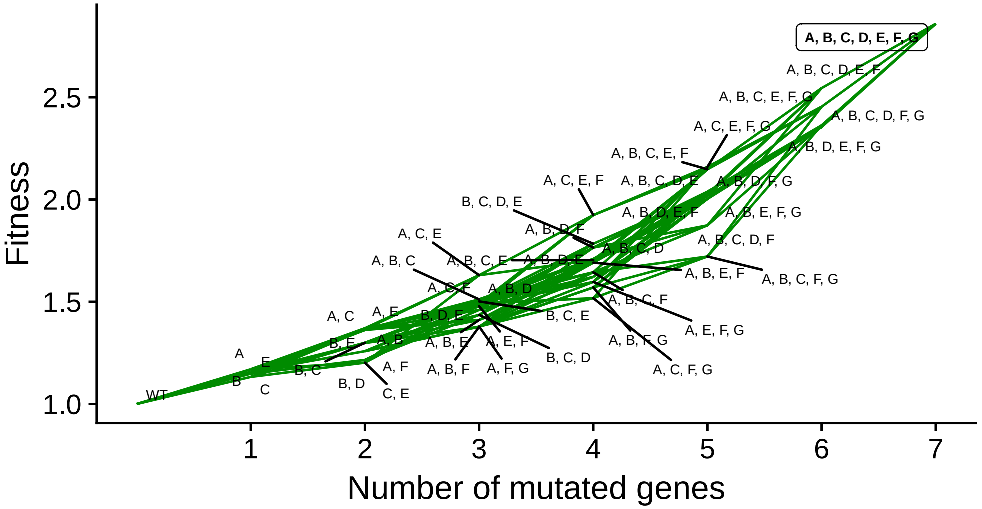

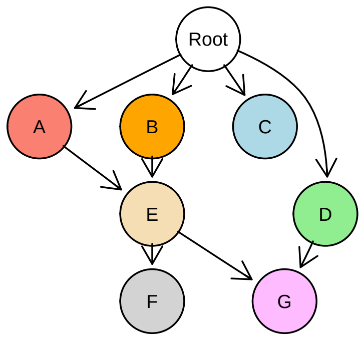


| | |
|---|---|
| *ID* | AlBDGRuJ29dXk10wK |
| *acc genotypes* | 28 |
| *peaks no back* | 1 |
| *removed edges* | 0 |
| *prop rem edges* | 0 |

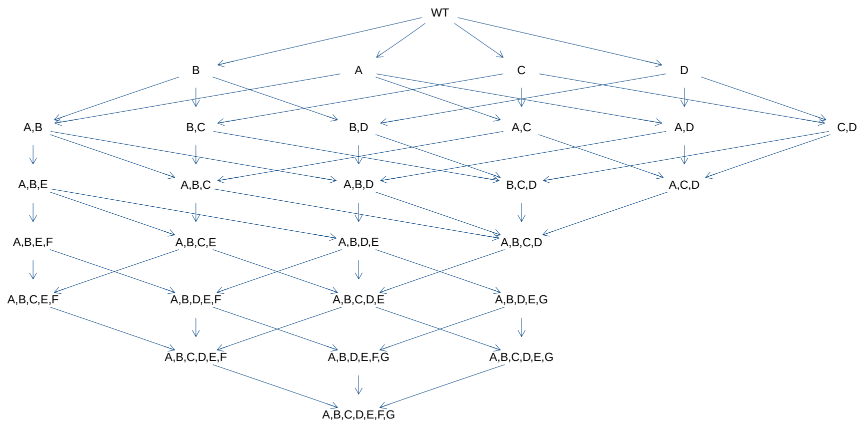


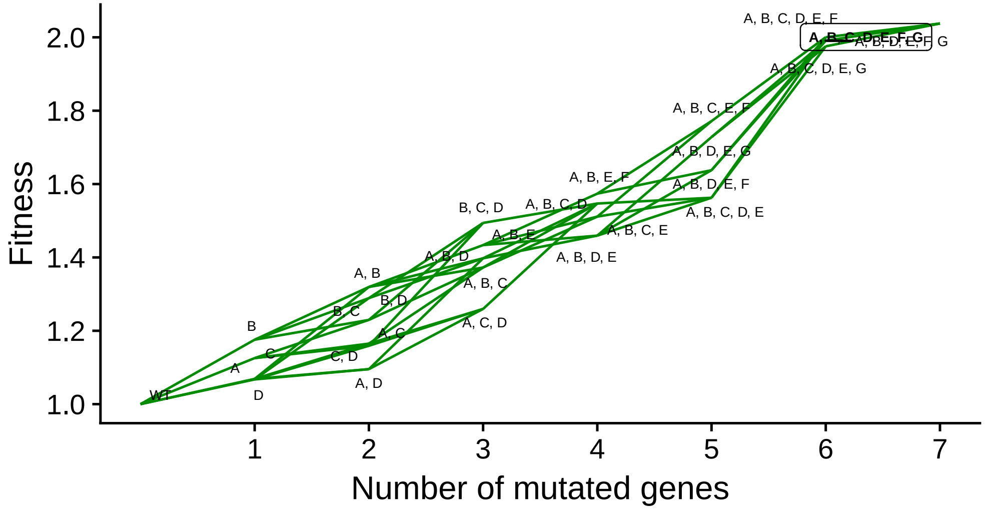

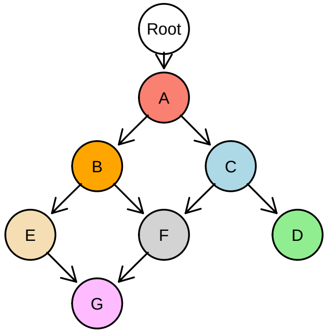


*ID* ALtLgLoV6JFZgc9s
*acc genotypes* 16
*peaks no back* 1
*removed edges* 0
*prop rem edges* 0

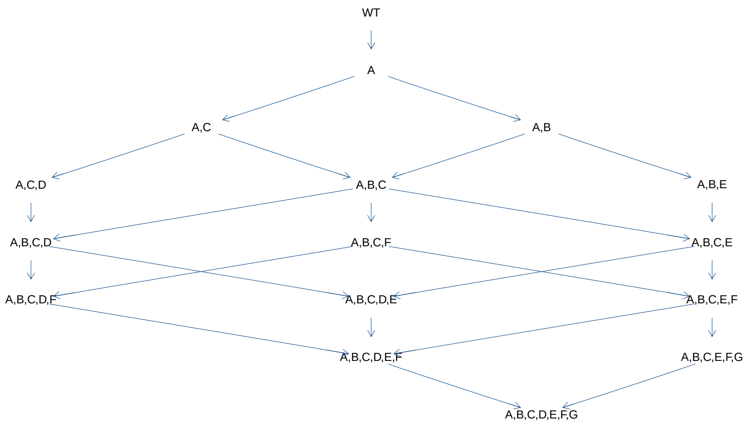


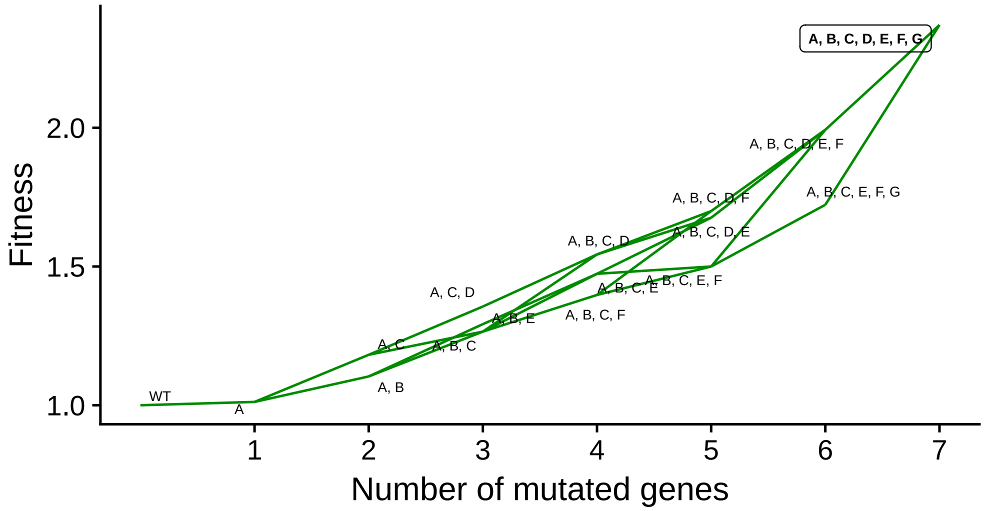

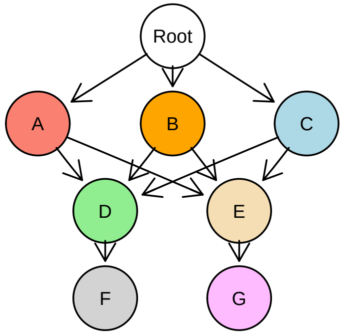


| | |
|---|---|
| *ID* | AomkJD2bjXlfY6bL |
| *acc genotypes* | 16 |
| *peaks no back* | 1 |
| *removed edges* | 0 |
| *prop rem edges* | 0 |

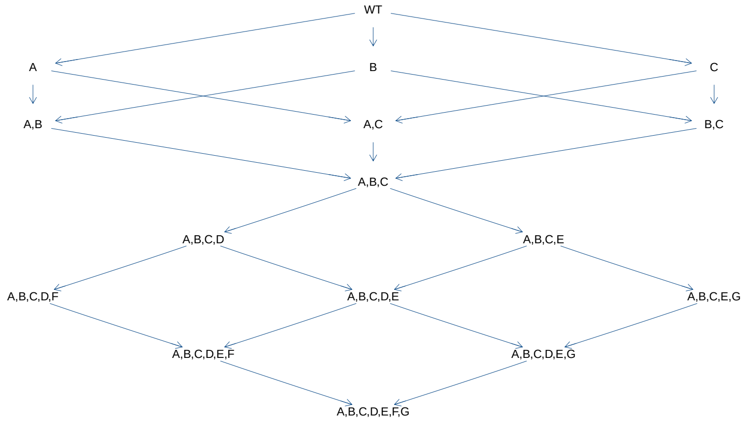


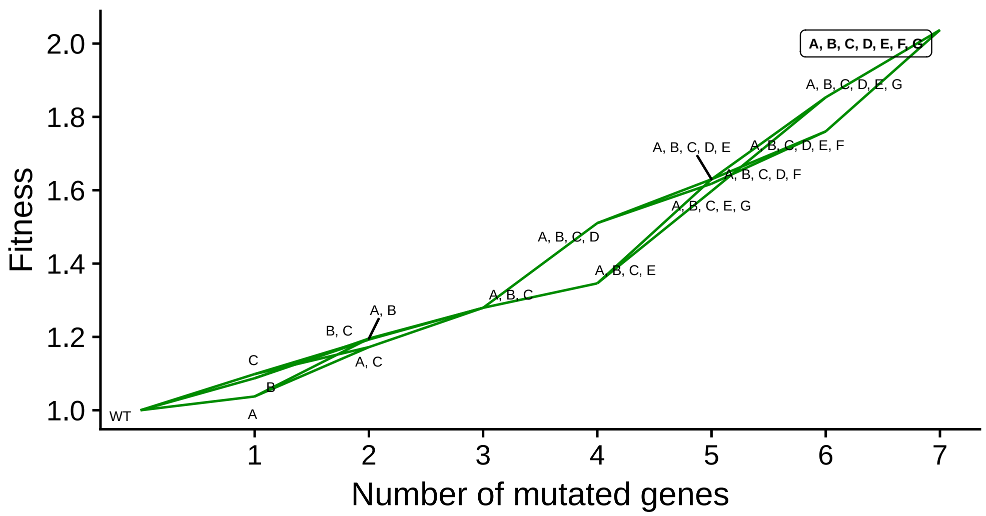

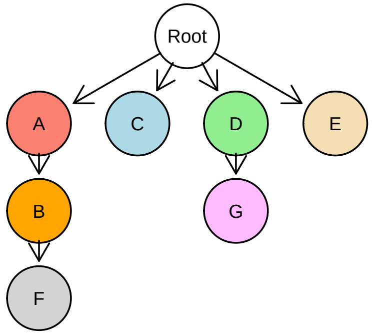


| | |
|---|---|
| *ID* | AQcGRsclzSVWoU7Z |
| *acc genotypes* | 48 |
| *peaks no back* | 1 |
| *removed edges* | 0 |
| *prop rem edges* | 0 |

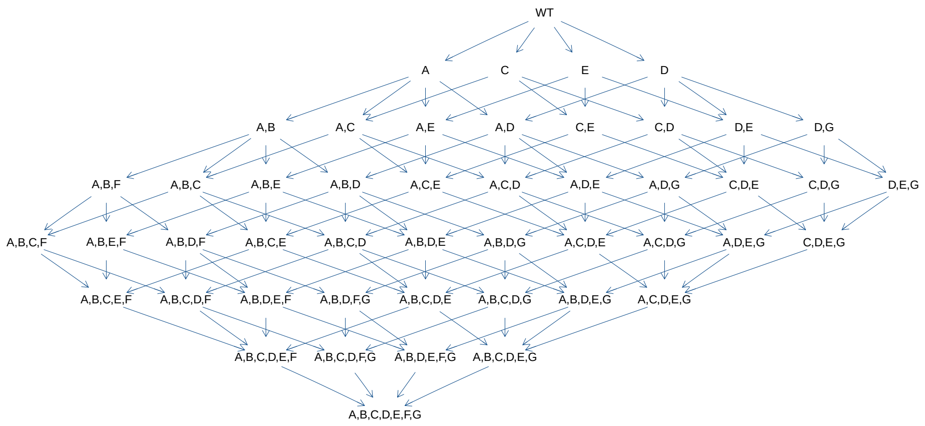


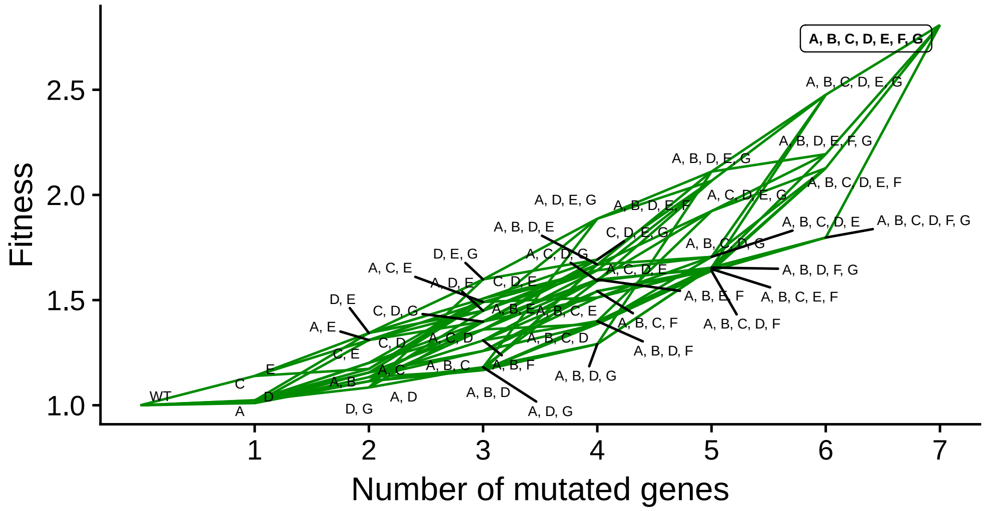

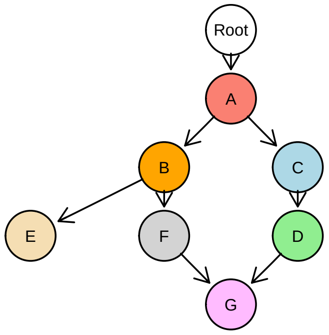


| | |
|---|---|
| *ID* | Azj8j6q13euWbW53 |
| *acc genotypes* | 18 |
| *peaks no back* | 1 |
| *removed edges* | 0 |
| *prop rem edges* | 0 |

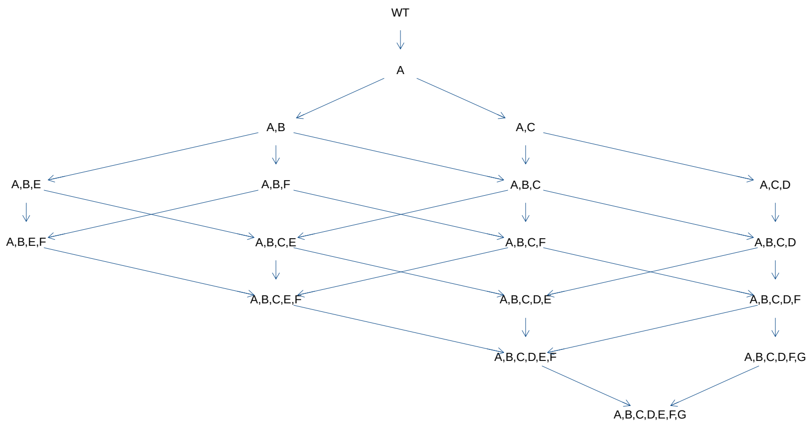


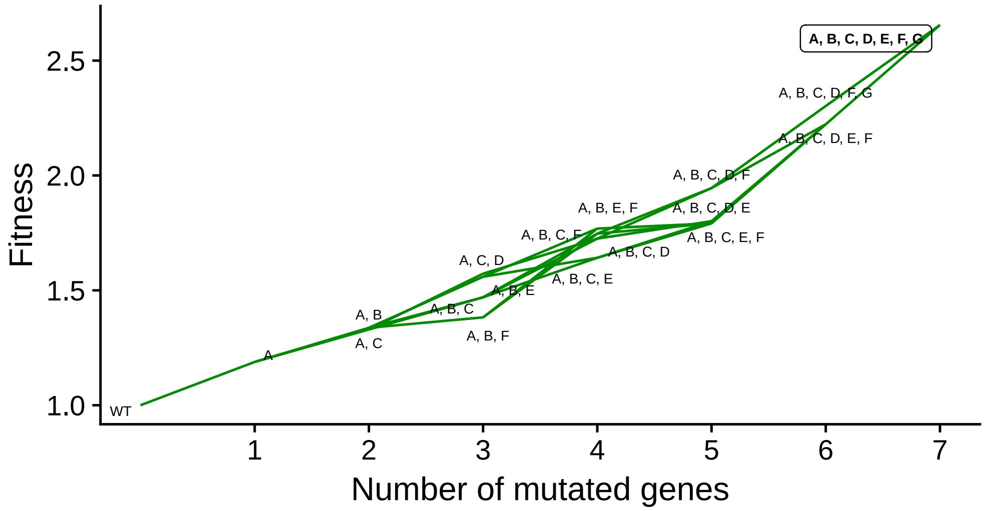

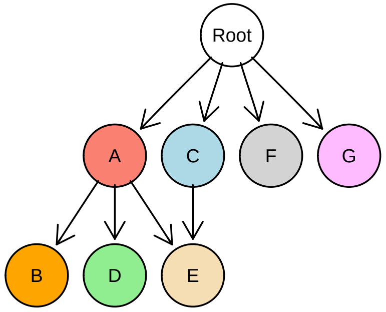


| | |
|---|---|
| *ID* | B3I8D10NQwXKC9sLs |
| *acc genotypes* | 56 |
| *peaks no back* | 1 |
| *removed edges* | 0 |
| *prop rem edges* | 0 |

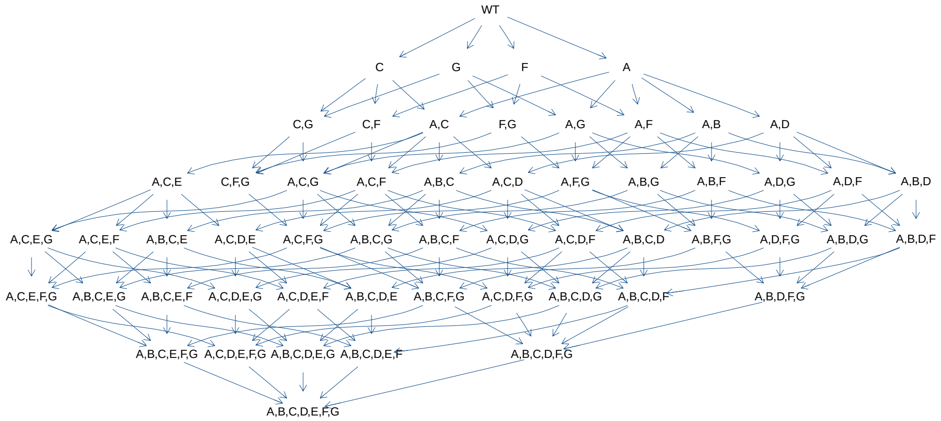


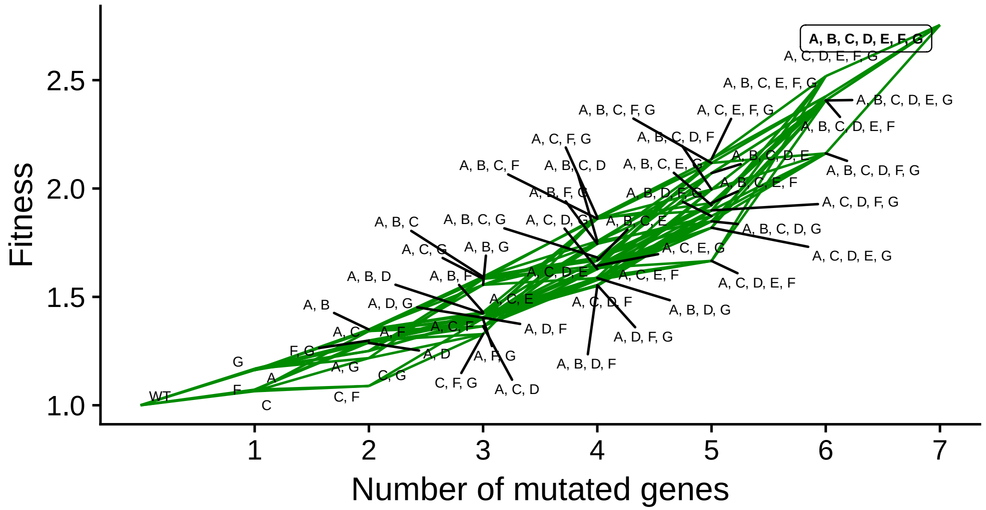

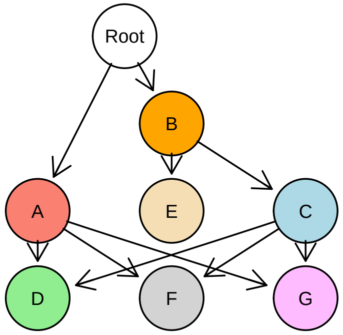


| | |
|---|---|
| *ID* | BawvXOS7sG8kOijC |
| *acc genotypes* | 24 |
| *peaks no back* | 1 |
| *removed edges* | 0 |
| *prop rem edges* | 0 |

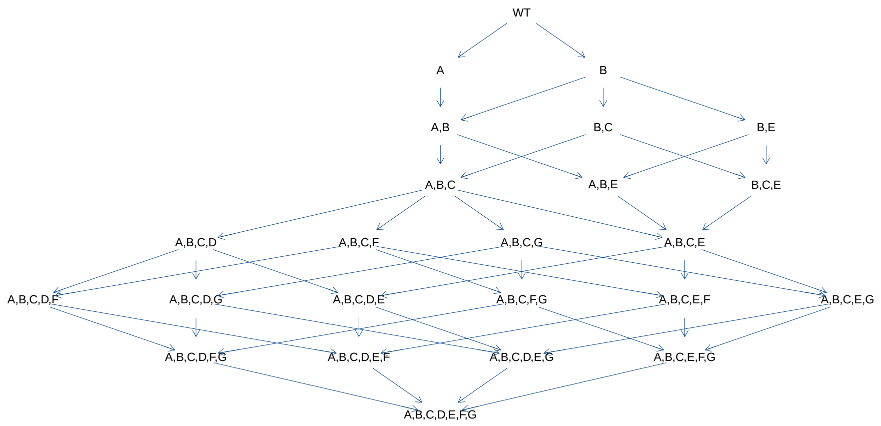


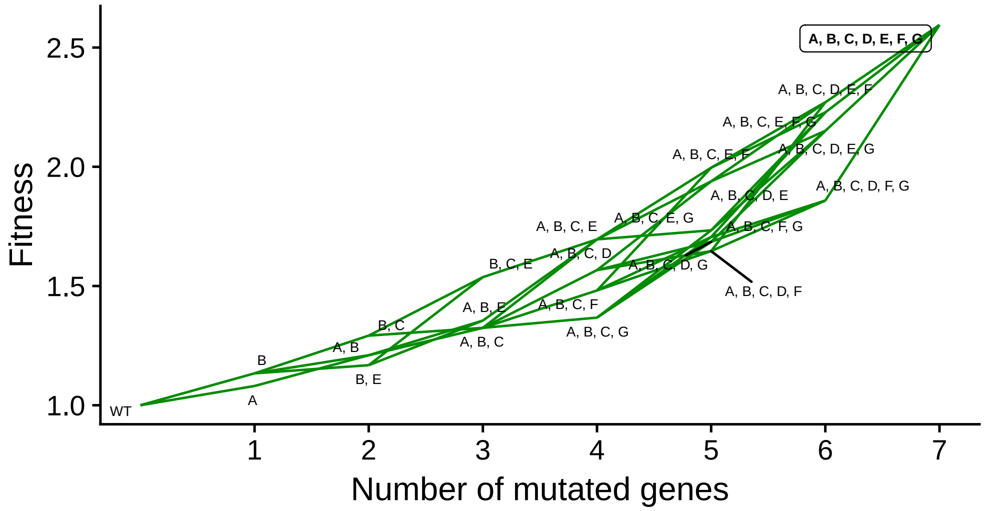

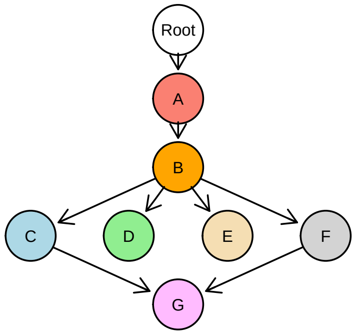


| | |
|---|---|
| *ID* | BFElrFle1os3fm18 |
| *acc genotypes* | 22 |
| *peaks no back* | 1 |
| *removed edges* | 0 |
| *prop rem edges* | 0 |

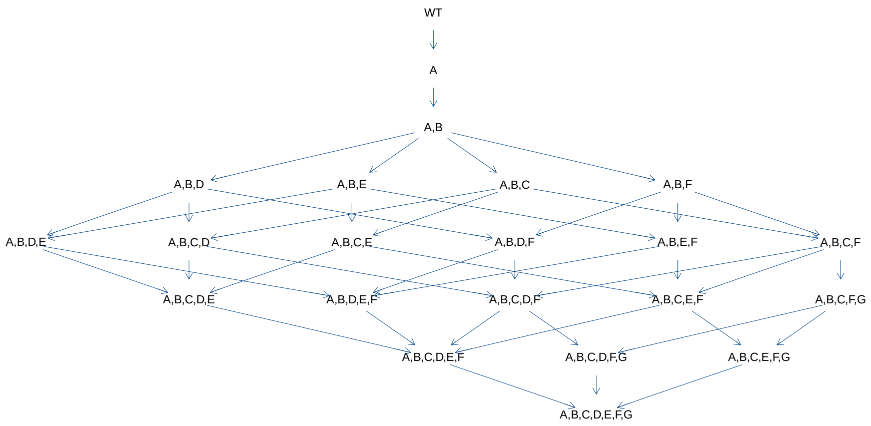


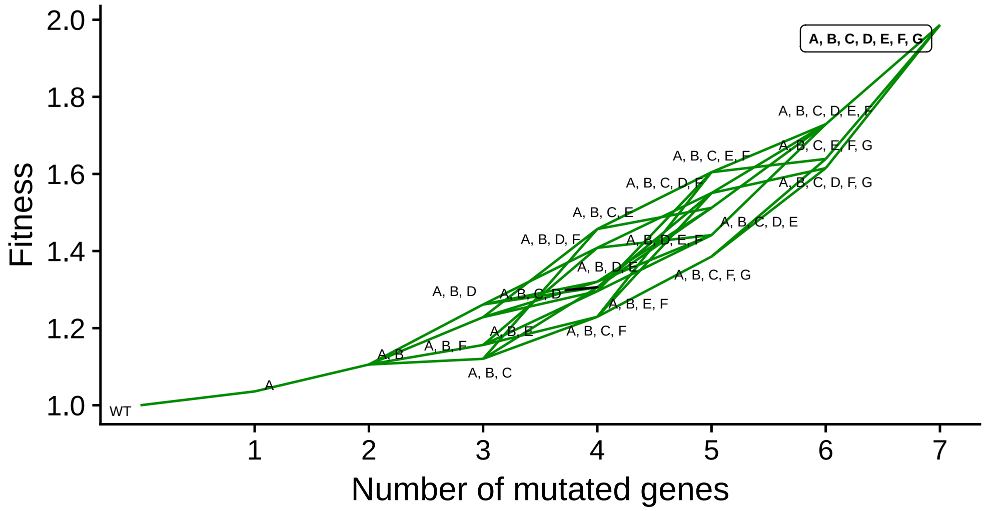

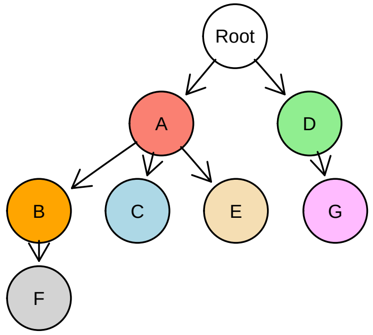


| | |
|---|---|
| *ID* | bFmYC2KrBktypLw3 |
| *acc genotypes* | 39 |
| *peaks no back* | 1 |
| *removed edges* | 0 |
| *prop rem edges* | 0 |

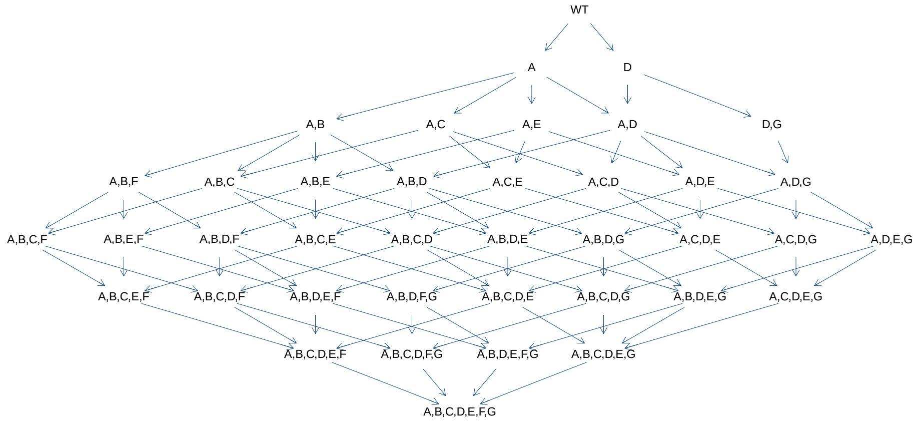


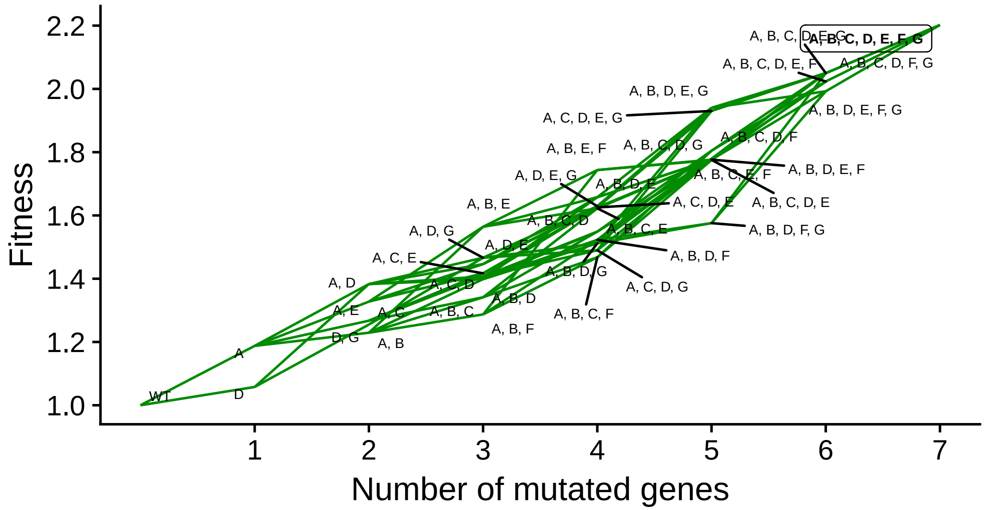

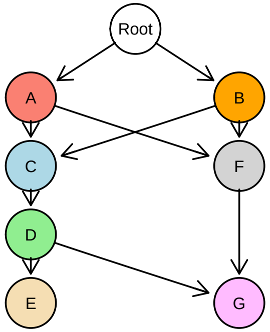


| | |
|---|---|
| *ID* | BfsxwvogFr6VDw4S |
| *acc genotypes* | 13 |
| *peaks no back* | 1 |
| *removed edges* | 0 |
| *prop rem edges* | 0 |

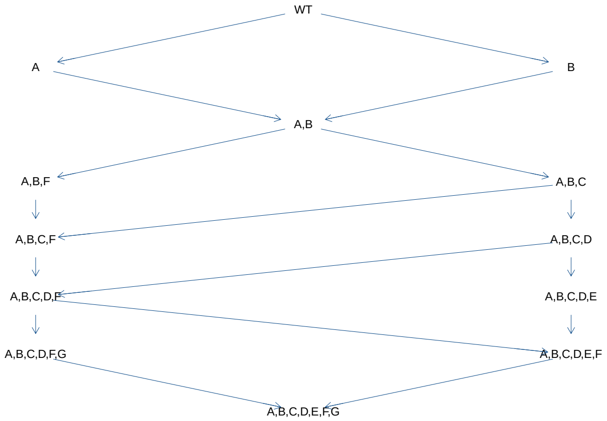


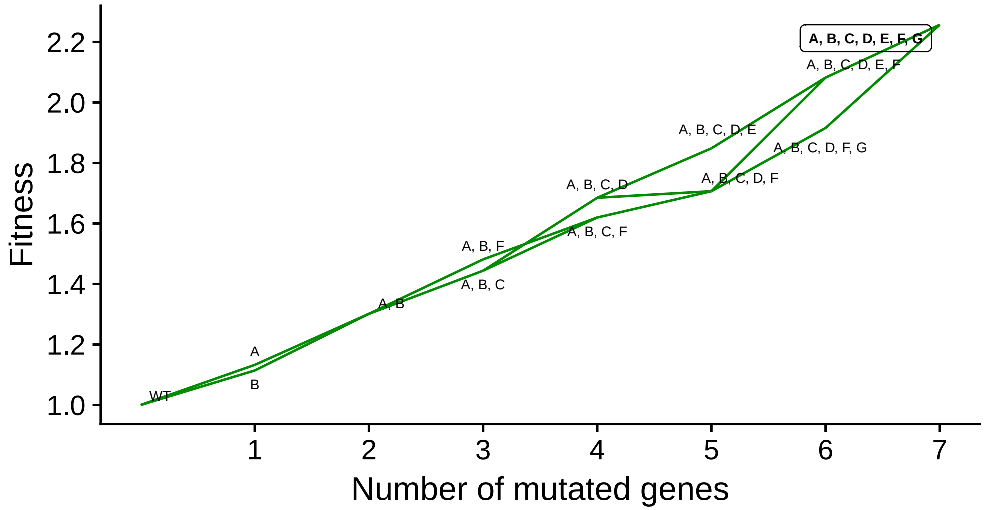

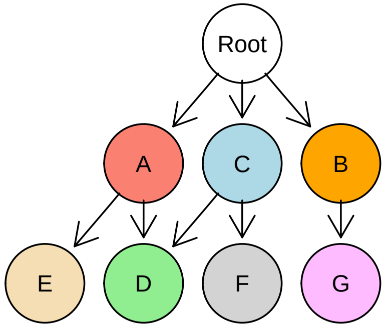


| | |
|---|---|
| *ID* | bJlgffVcPbF6GUJl |
| *acc genotypes* | 39 |
| *peaks no back* | 1 |
| *removed edges* | 0 |
| *prop rem edges* | 0 |

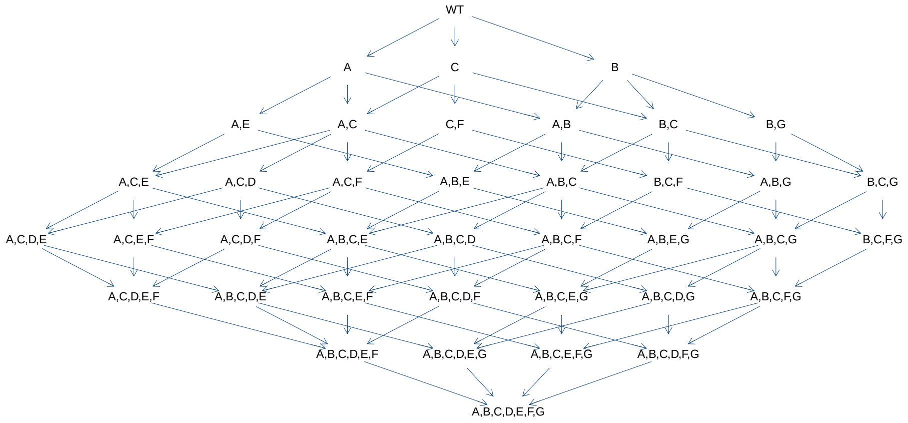


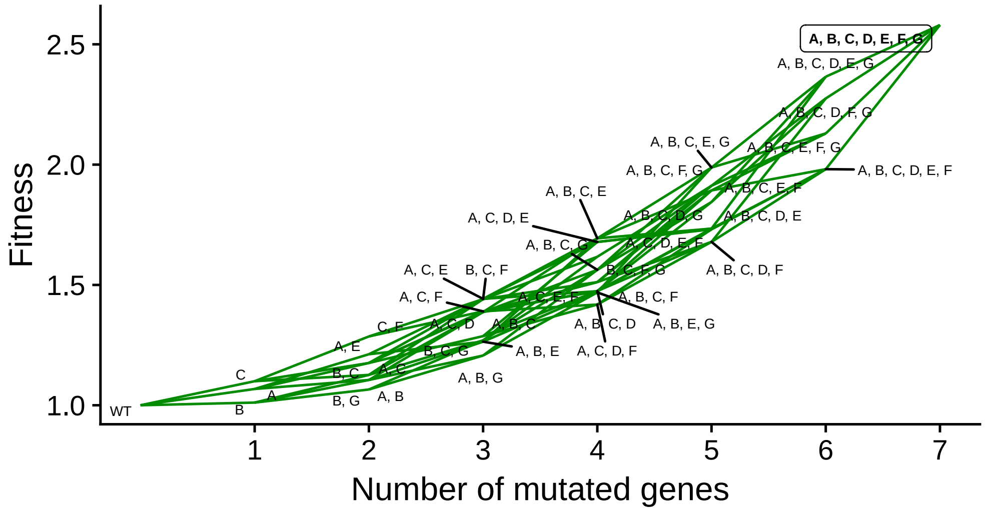

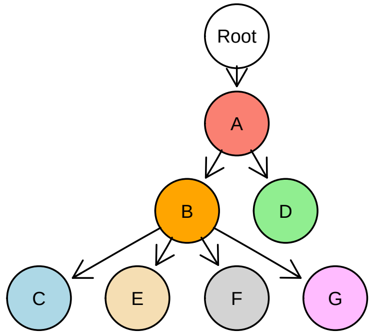


| | |
|---|---|
| *ID* | bqjCnNng38uQA1OY |
| *acc genotypes* | 35 |
| *peaks no back* | 1 |
| *removed edges* | 0 |
| *prop rem edges* | 0 |

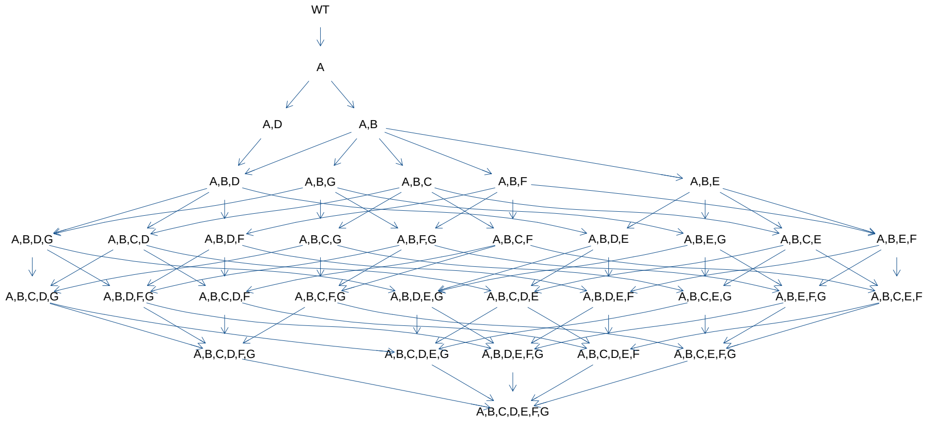


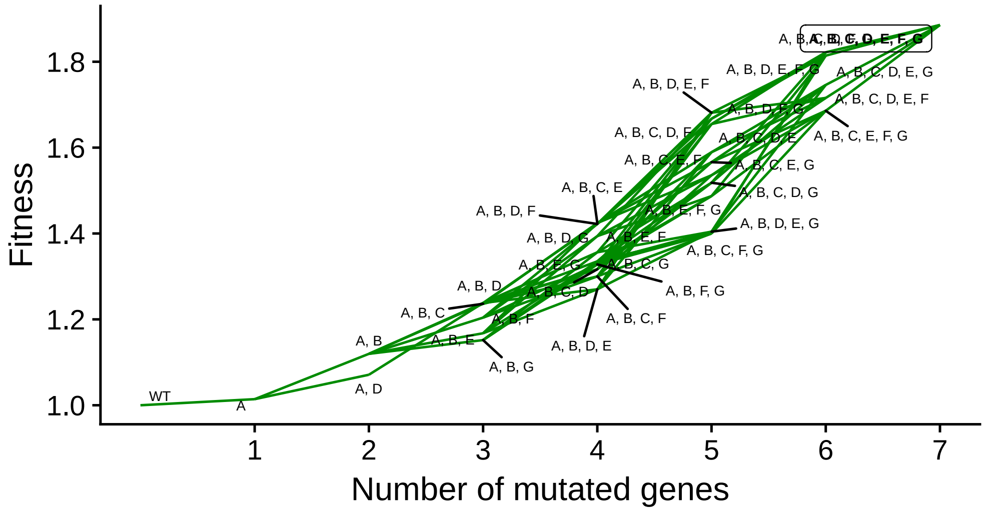

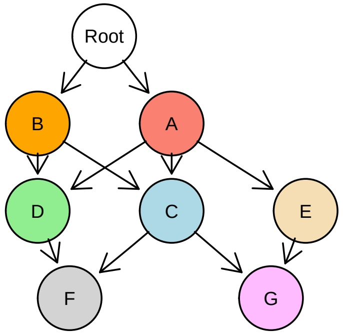


| | |
|---|---|
| *ID* | btsq9bPlvHef8E4b |
| *acc genotypes* | 17 |
| *peaks no back* | 1 |
| *removed edges* | 0 |
| *prop rem edges* | 0 |

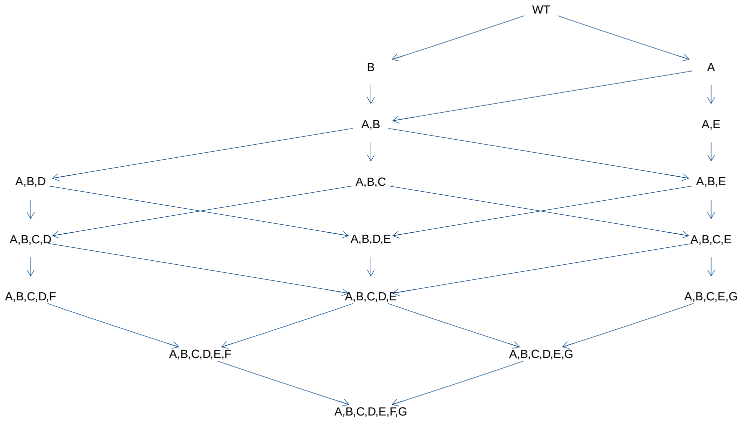


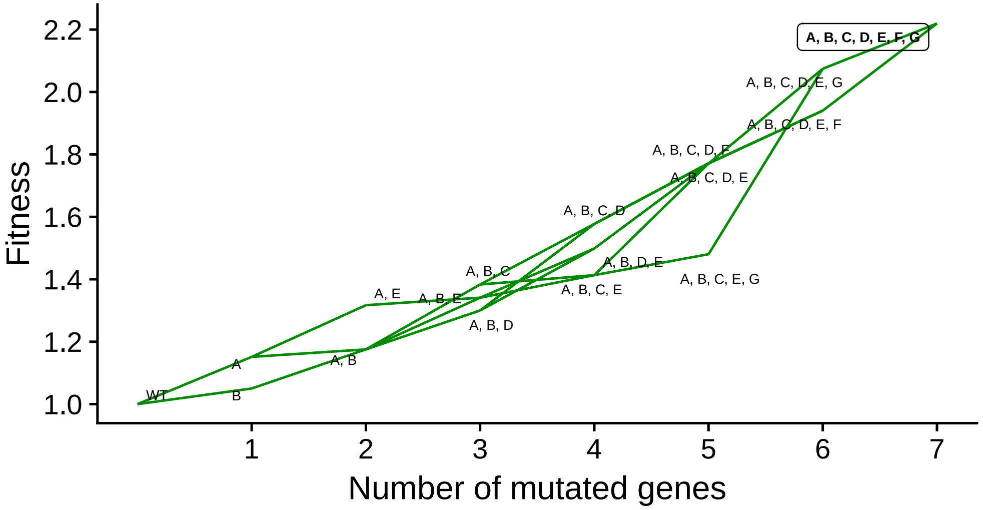

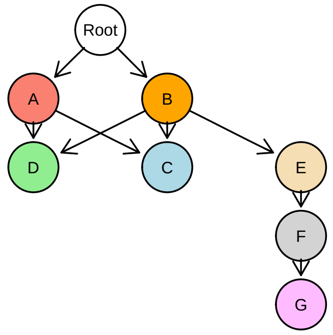


| | |
|---|---|
| *ID* | BuXQklRXr18pe1ec |
| *acc genotypes* | 22 |
| *peaks no back* | 1 |
| *removed edges* | 0 |
| *prop rem edges* | 0 |

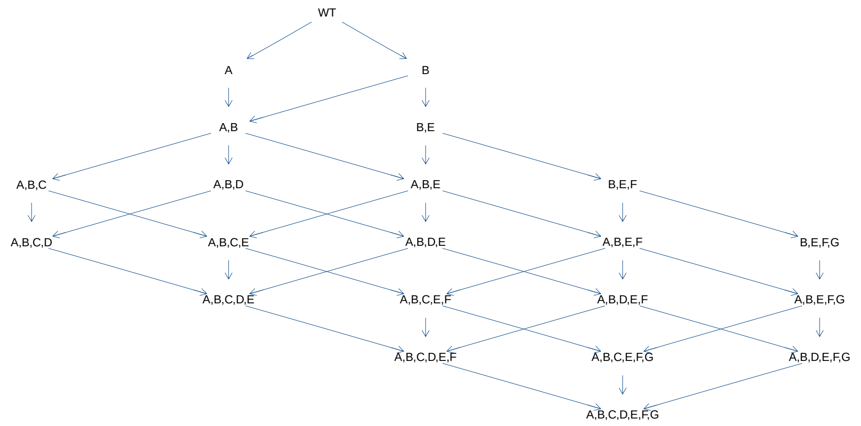


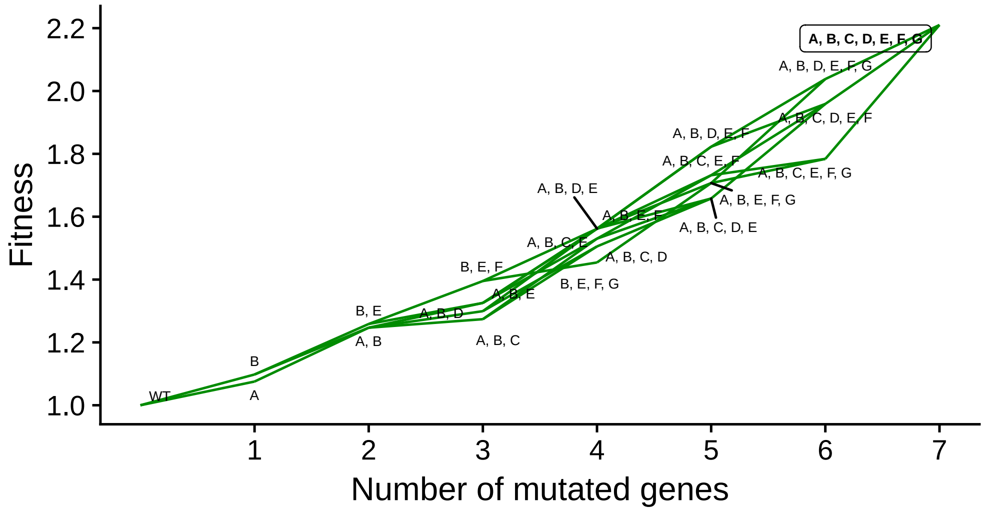

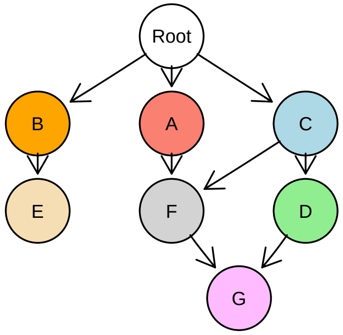


| | |
|---|---|
| *ID* | cPY2aEYEnXRtcOVx |
| *acc genotypes* | 27 |
| *peaks no back* | 1 |
| *removed edges* | 0 |
| *prop rem edges* | 0 |

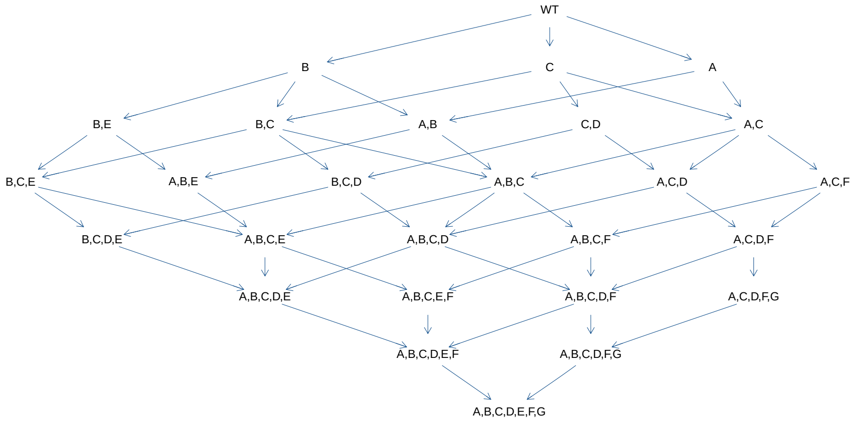


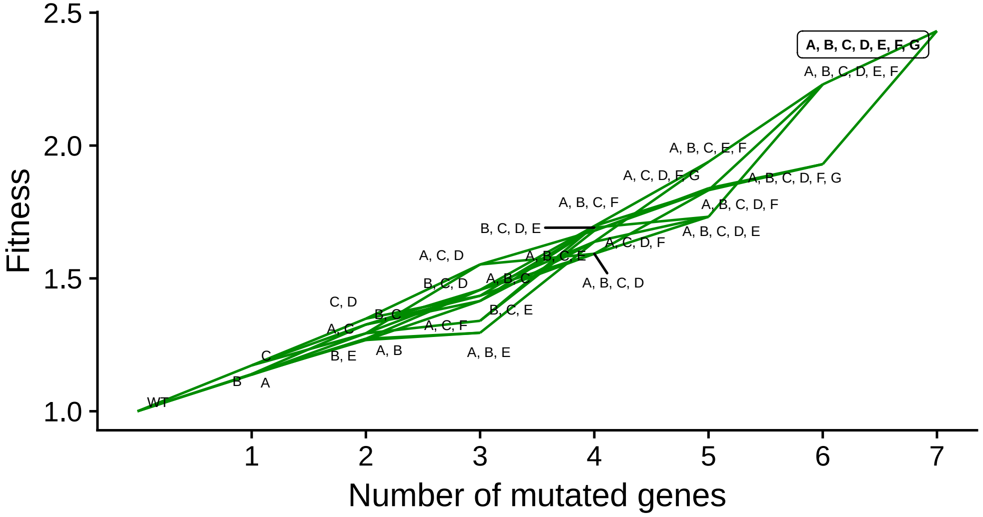

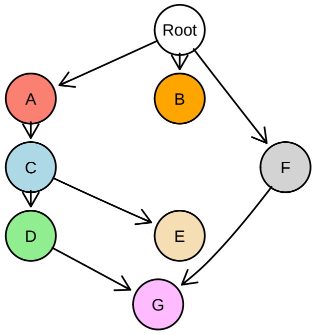


*ID* cTShsHTaSM2pFU3Z
*acc genotypes* 28
*peaks no back* 1
*removed edges* 0
*prop rem edges* 0

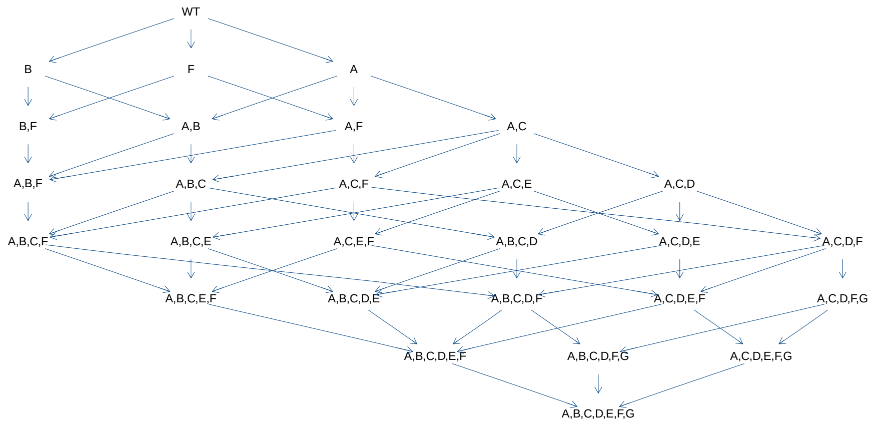


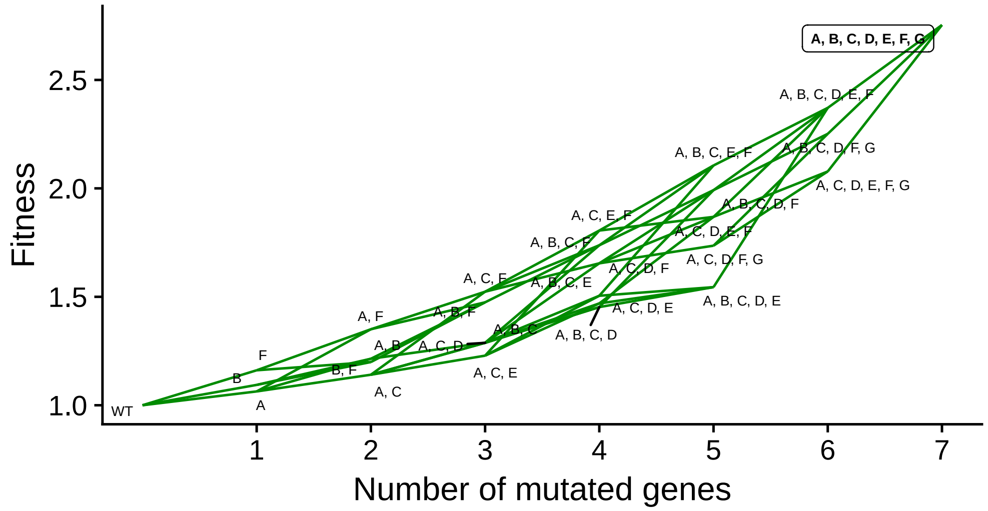

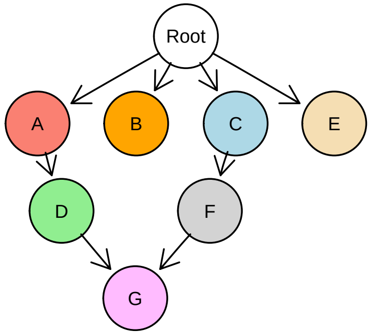


| | |
|---|---|
| *ID* | cZ5tgt8lsWr92I8O |
| *acc genotypes* | 40 |
| *peaks no back* | 1 |
| *removed edges* | 0 |
| *prop rem edges* | 0 |

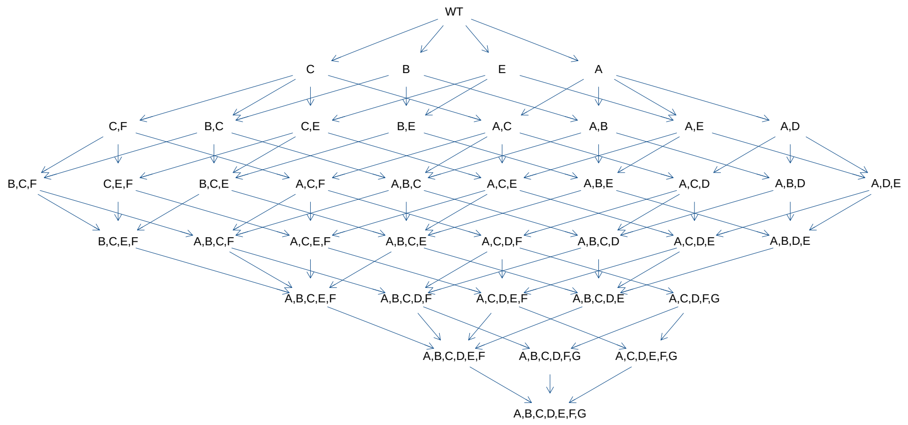


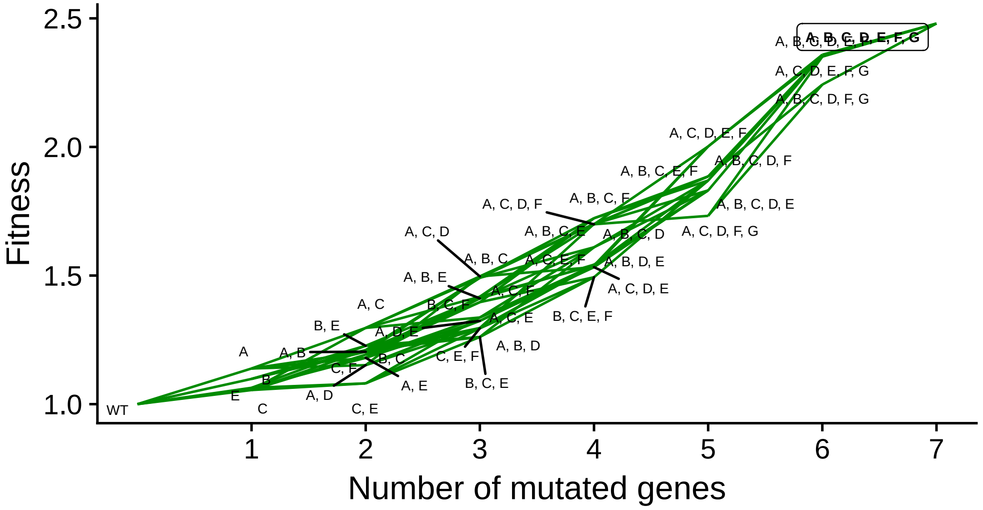

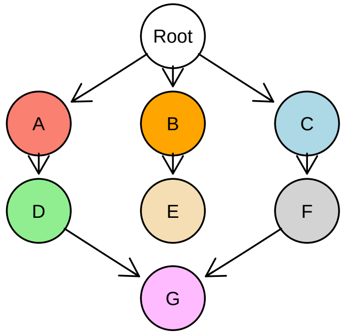


| | |
|---|---|
| *ID* | D10upTTNDLyeed8b9 |
| *acc genotypes* | 30 |
| *peaks no back* | 1 |
| *removed edges* | 0 |
| *prop rem edges* | 0 |

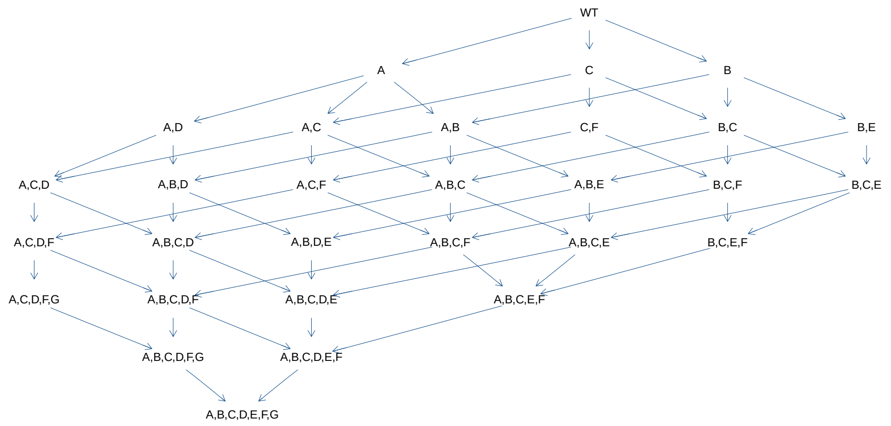


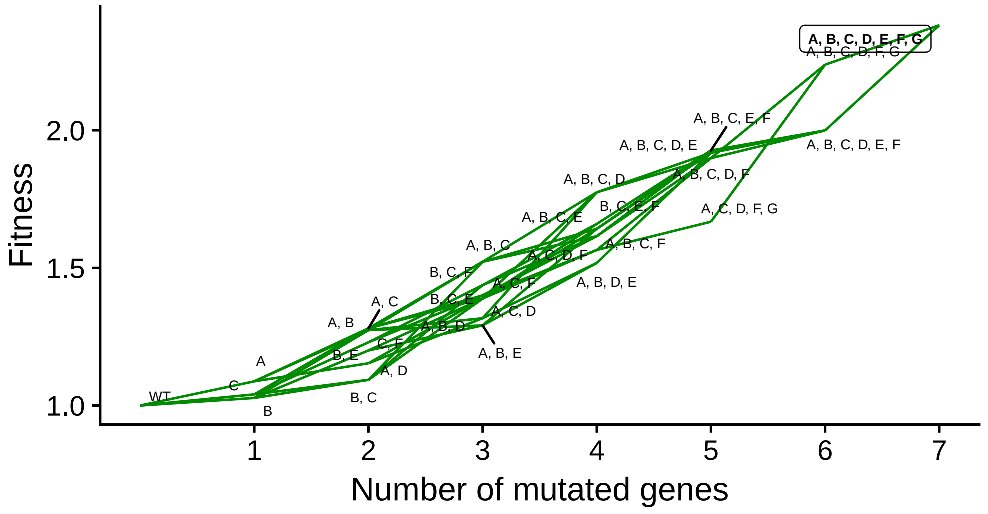

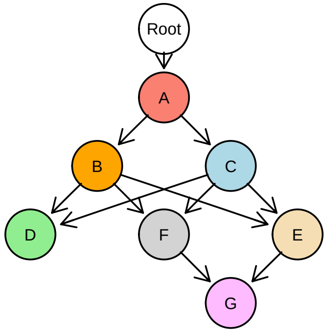


| | |
|---|---|
| *ID* | DaQ1lMqJ1ktP10uHB |
| *acc genotypes* | 14 |
| *peaks no back* | 1 |
| *removed edges* | 0 |
| *prop rem edges* | 0 |

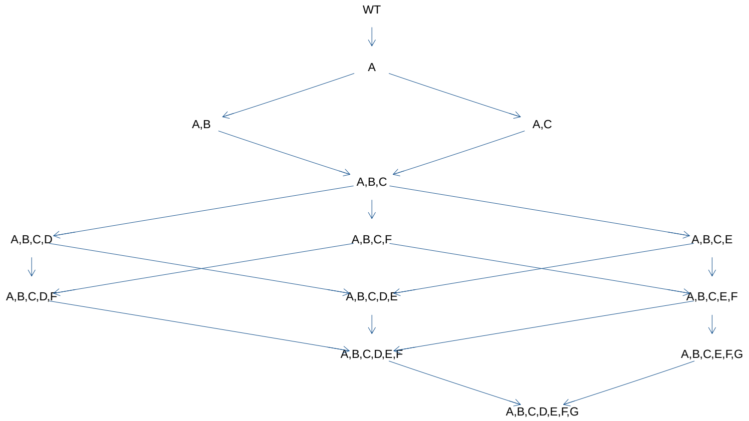


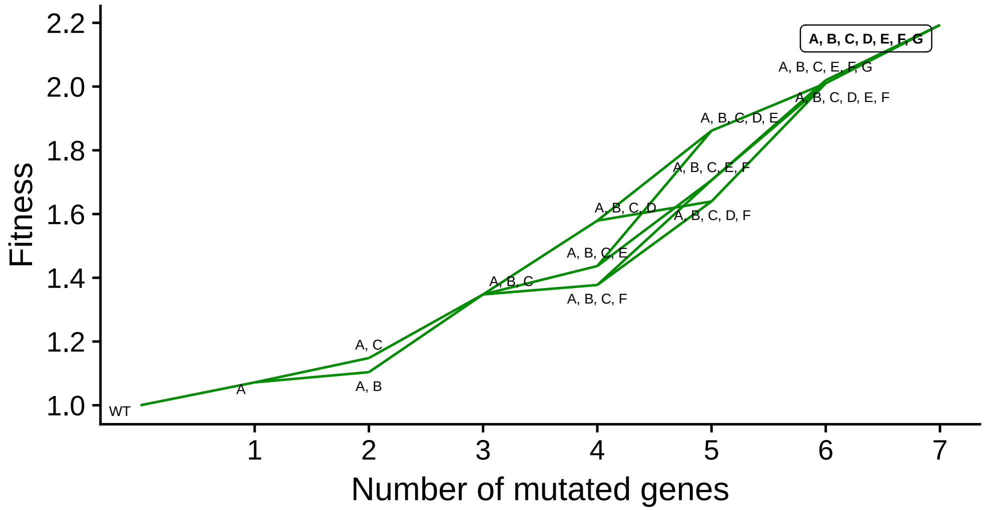

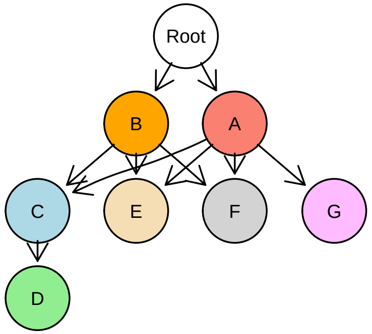


| | |
|---|---|
| *ID* | dG5EKqdkRReFl4LP |
| *acc genotypes* | 28 |
| *peaks no back* | 1 |
| *removed edges* | 0 |
| *prop rem edges* | 0 |

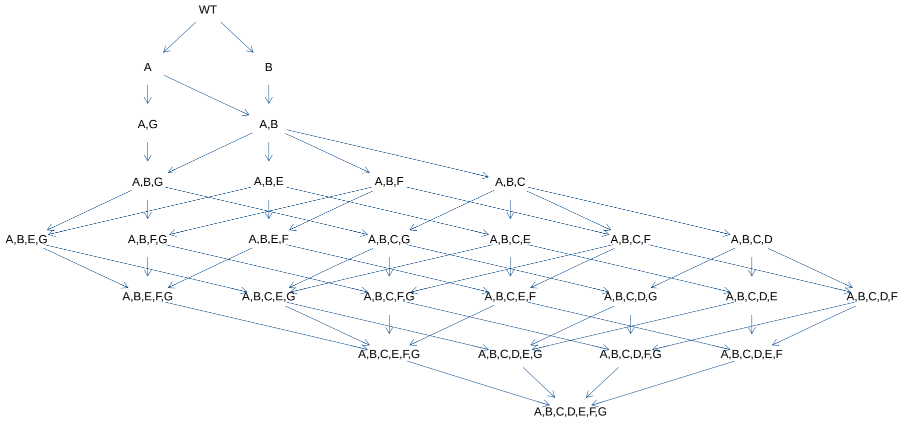


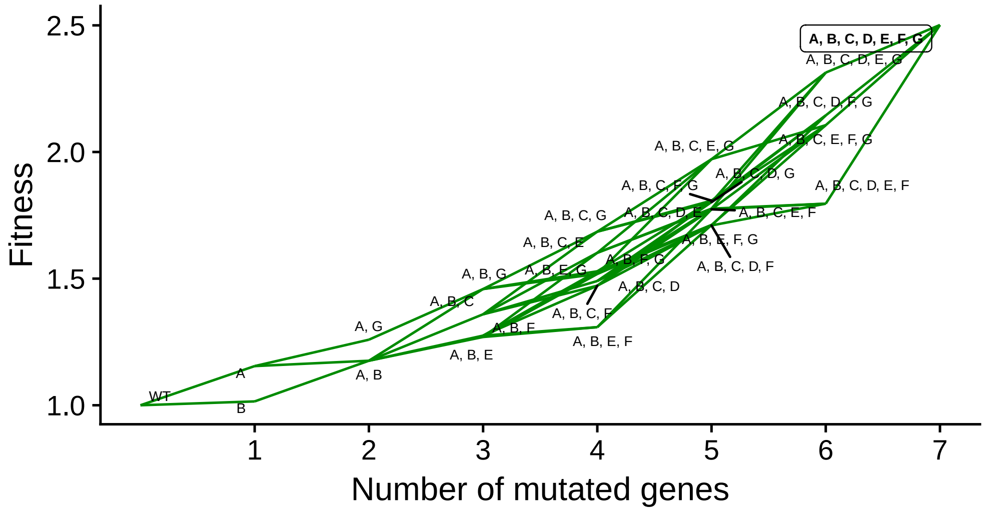

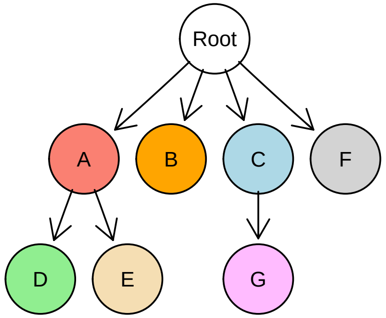


| | |
|---|---|
| *ID* | DMErt7pbZBAcNnQg |
| *acc genotypes* | 60 |
| *peaks no back* | 1 |
| *removed edges* | 0 |
| *prop rem edges* | 0 |

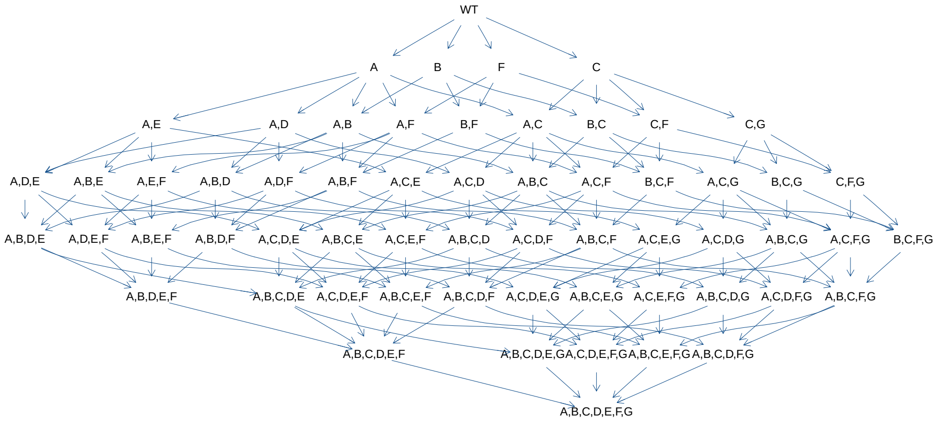

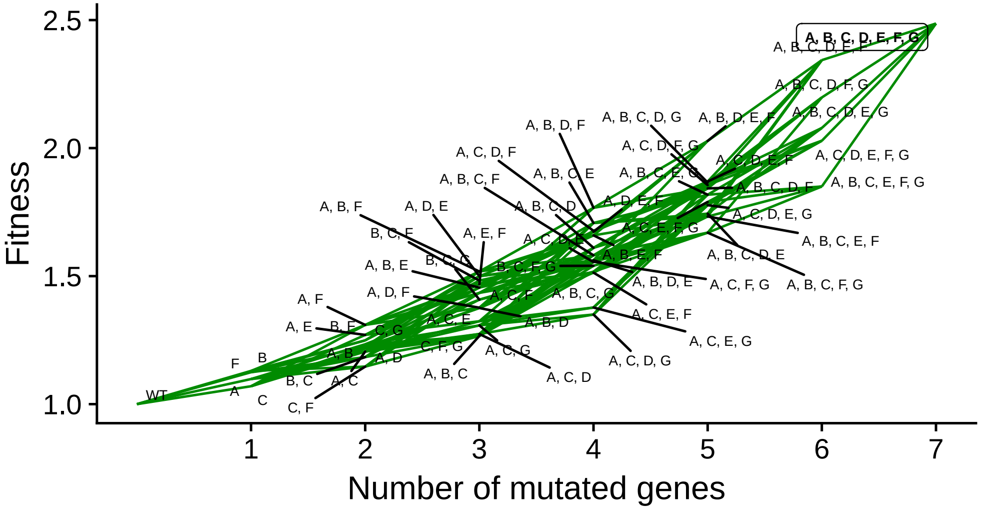

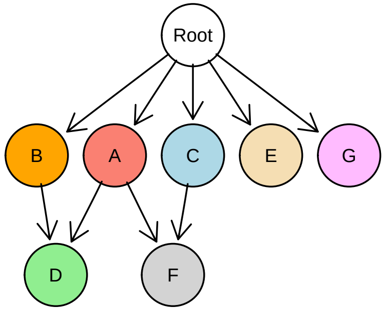


| | |
|---|---|
| *ID* | dmZHV2wAZ3zFiHOq |
| *acc genotypes* | 52 |
| *peaks no back* | 1 |
| *removed edges* | 0 |
| *prop rem edges* | 0 |

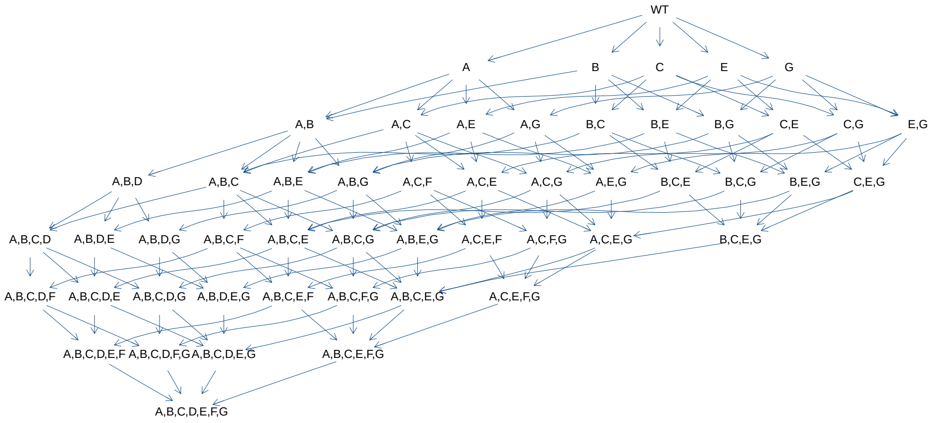


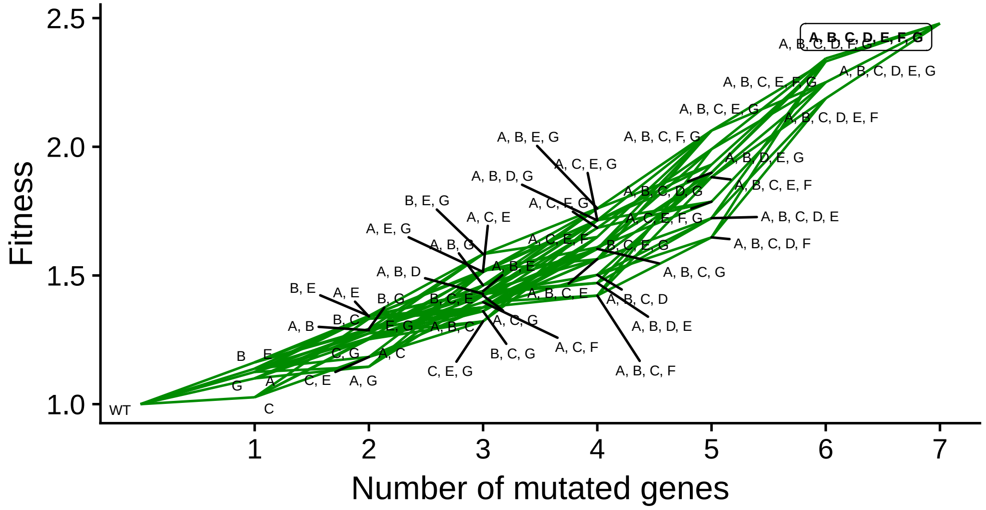

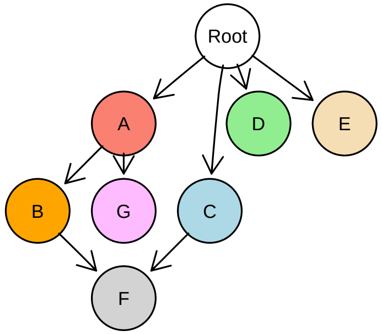


| | |
|---|---|
| *ID* | DNdxF6nsuNX9Tqlx |
| *acc genotypes* | 48 |
| *peaks no back* | 1 |
| *removed edges* | 0 |
| *prop rem edges* | 0 |

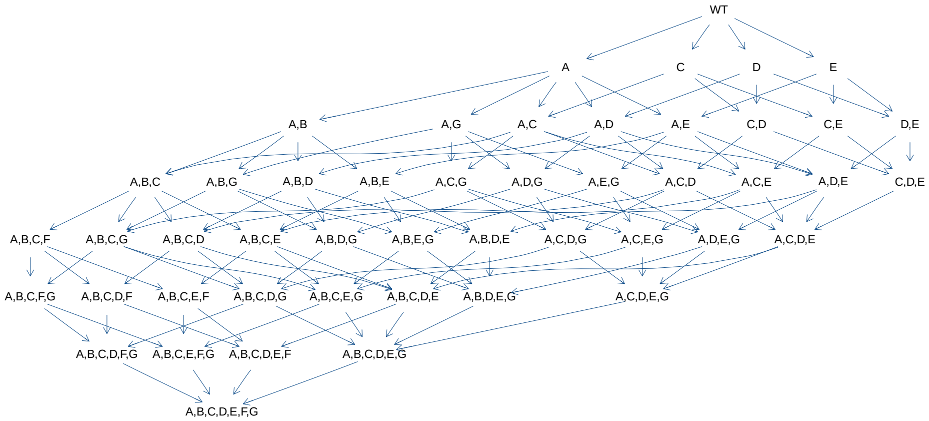


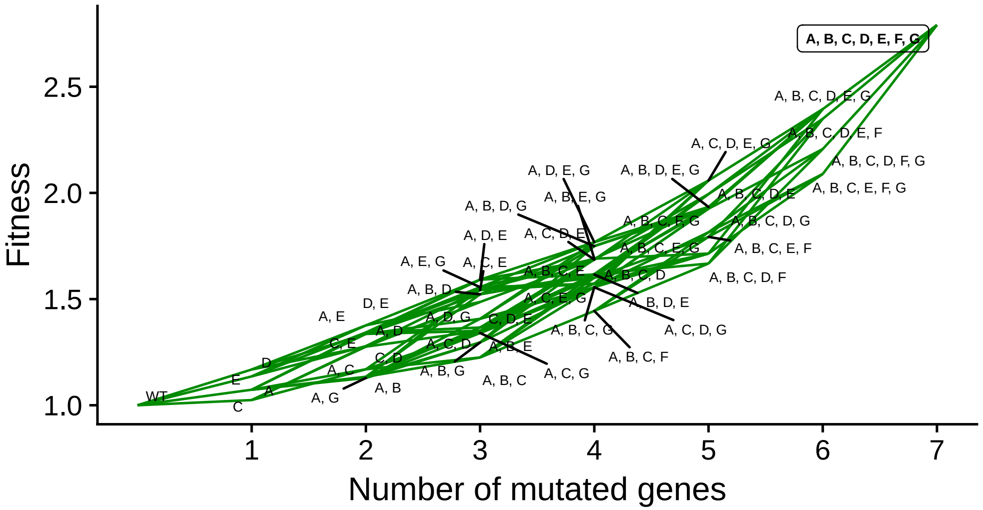

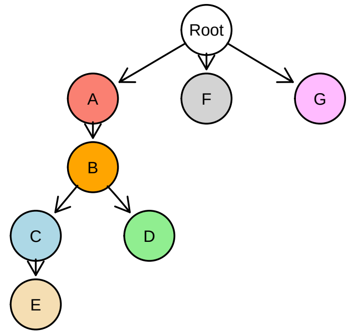


*ID* Dv8B7RHWDguRzH1W
*acc genotypes* 32
*peaks no back* 1
*removed edges* 0
*prop rem edges* 0

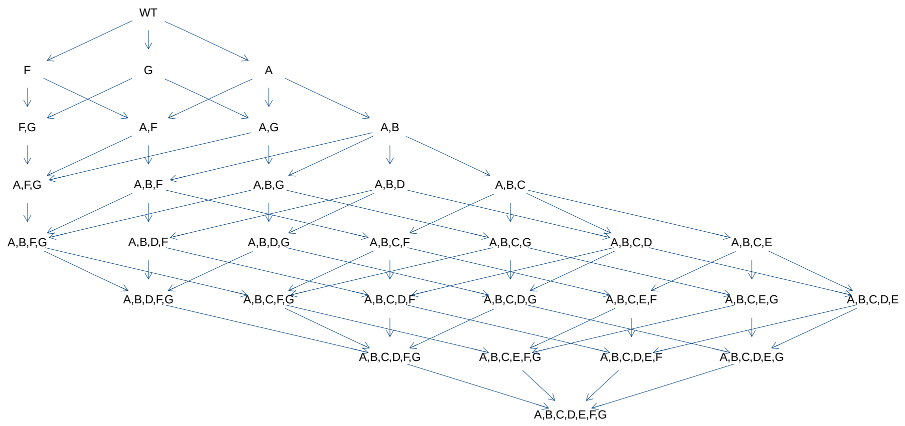


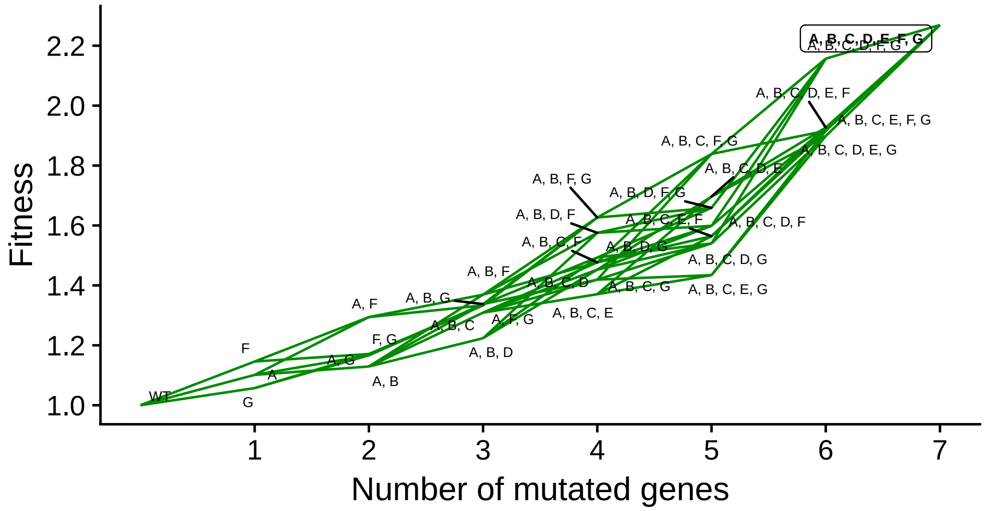

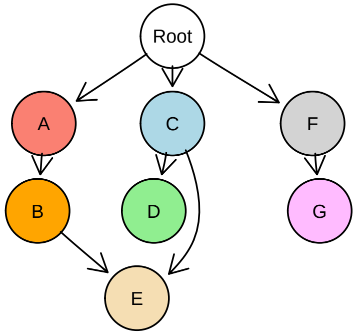


| | |
|---|---|
| *ID* | e6d6sLGltLVqnRO4 |
| *acc genotypes* | 33 |
| *peaks no back* | 1 |
| *removed edges* | 0 |
| *prop rem edges* | 0 |

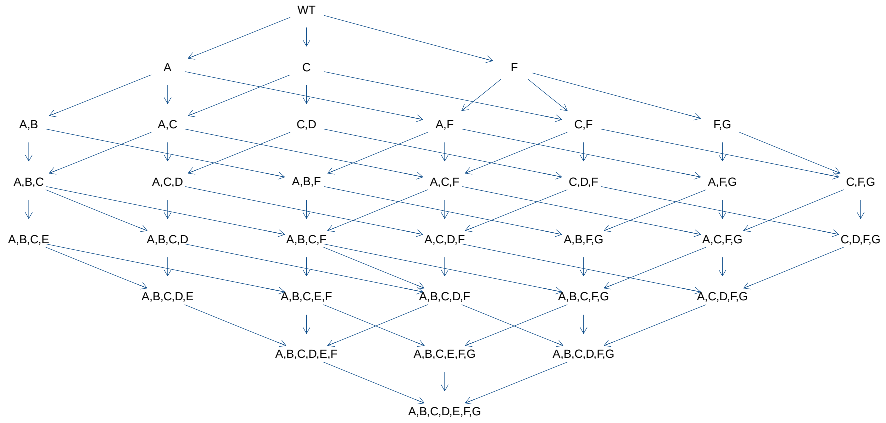


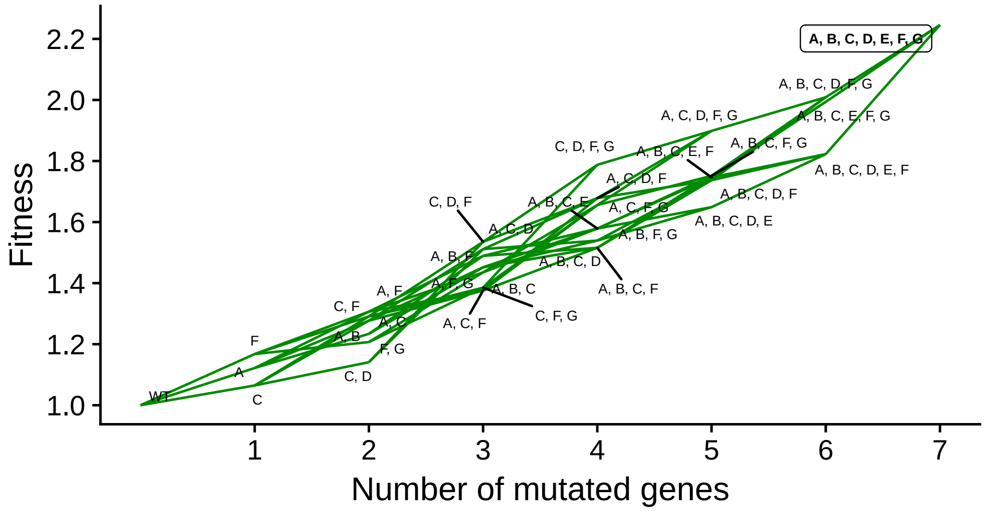

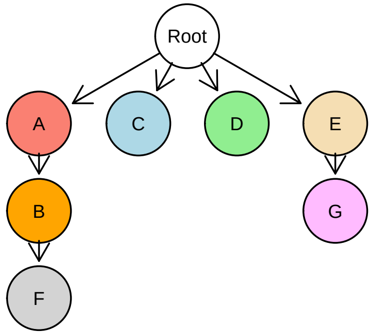


| | |
|---|---|
| *ID* | E7PUOGD47fKBfgGx |
| *acc genotypes* | 48 |
| *peaks no back* | 1 |
| *removed edges* | 0 |
| *prop rem edges* | 0 |

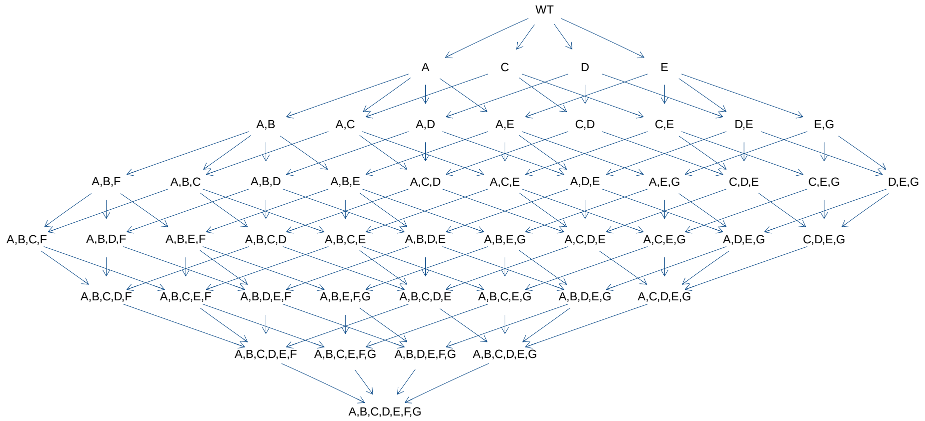


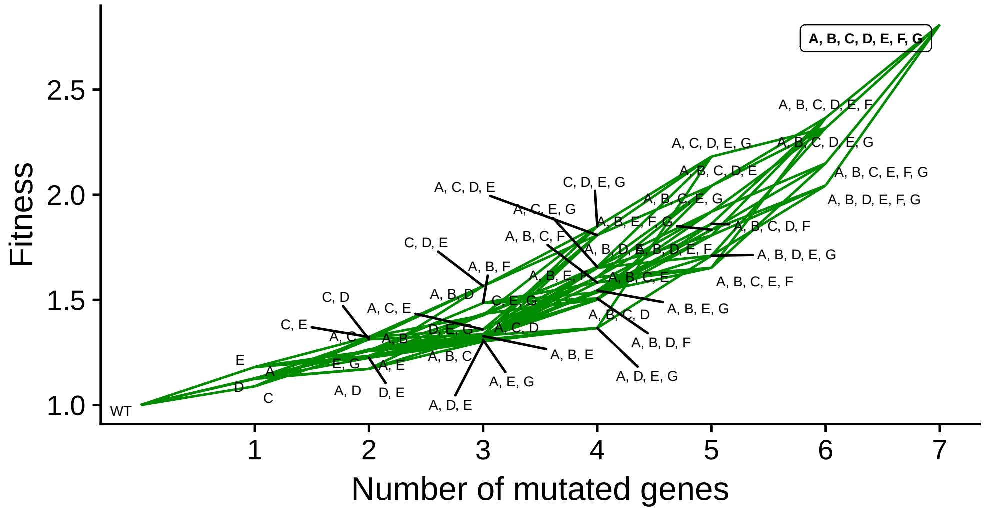

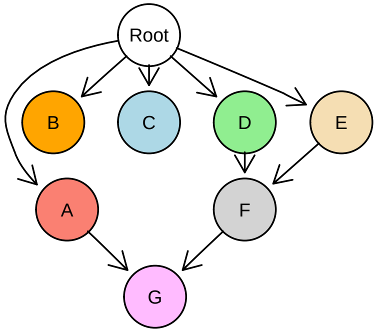


| | |
|---|---|
| *ID* | eEiiU6IXTAitvbXD |
| *acc genotypes* | 44 |
| *peaks no back* | 1 |
| *removed edges* | 0 |
| *prop rem edges* | 0 |

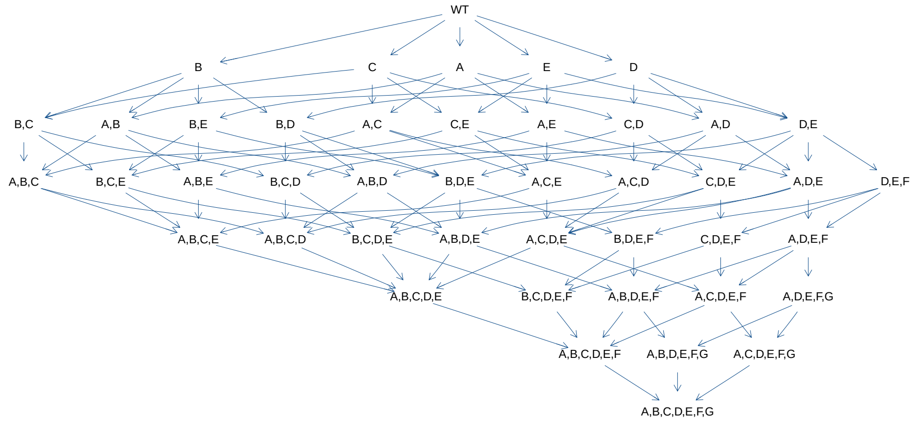


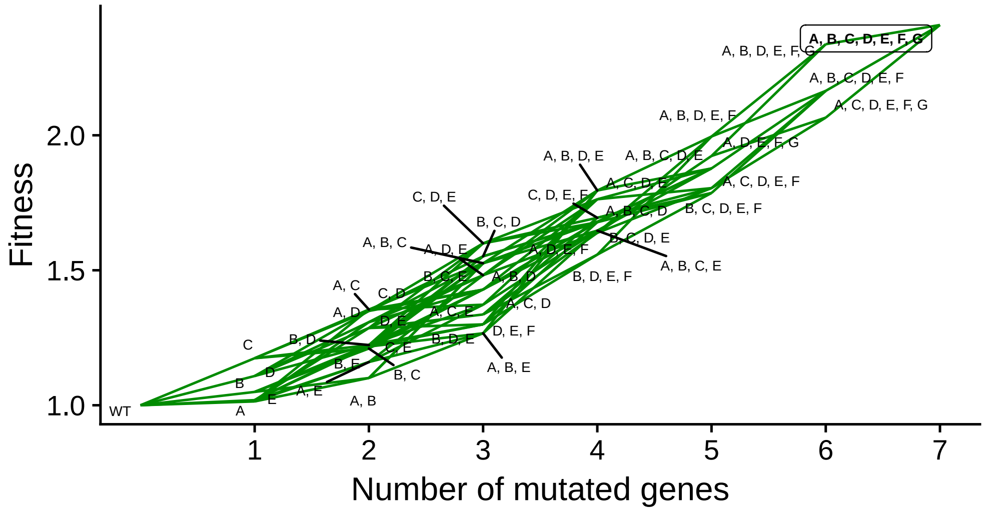

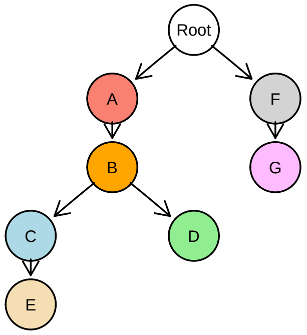


*ID* EF3dPOLYSD4SFyjp
*acc genotypes* 24
*peaks no back* 1
*removed edges* 0
*prop rem edges* 0

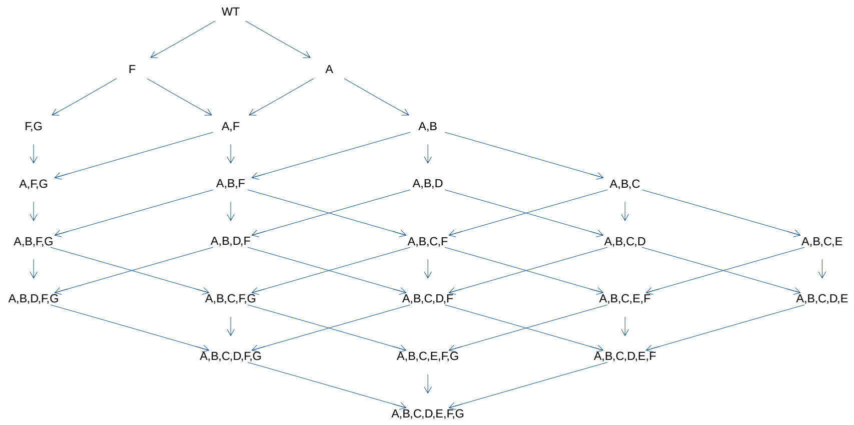


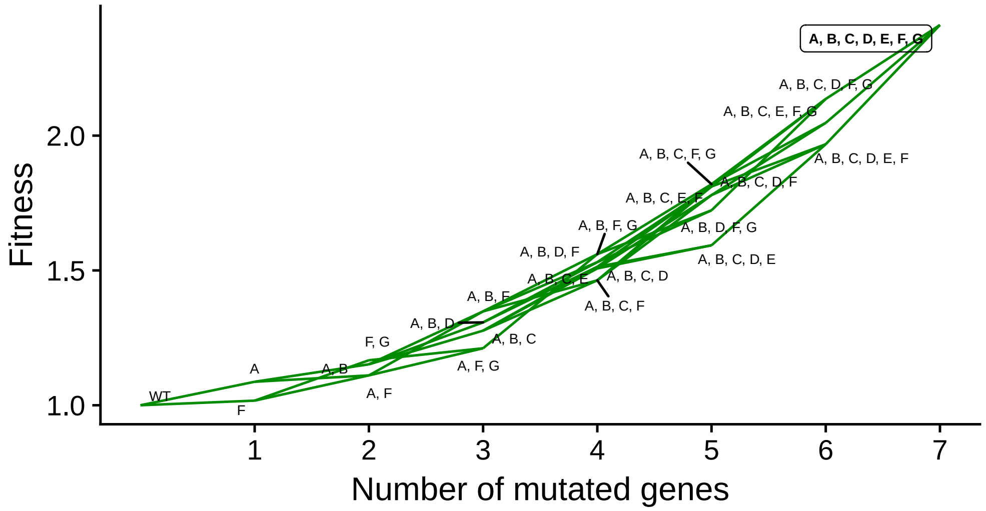

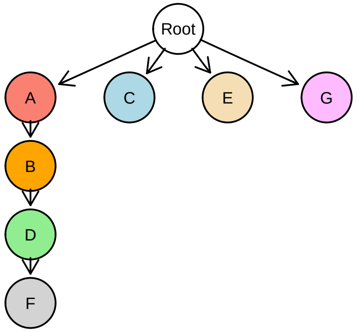


| | |
|---|---|
| *ID* | EikpyAkVdLnWHzZF |
| *acc genotypes* | 40 |
| *peaks no back* | 1 |
| *removed edges* | 0 |
| *prop rem edges* | 0 |

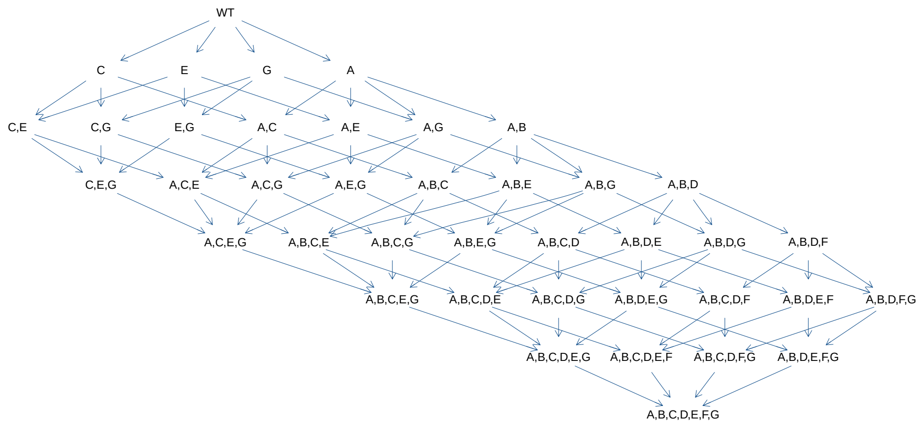


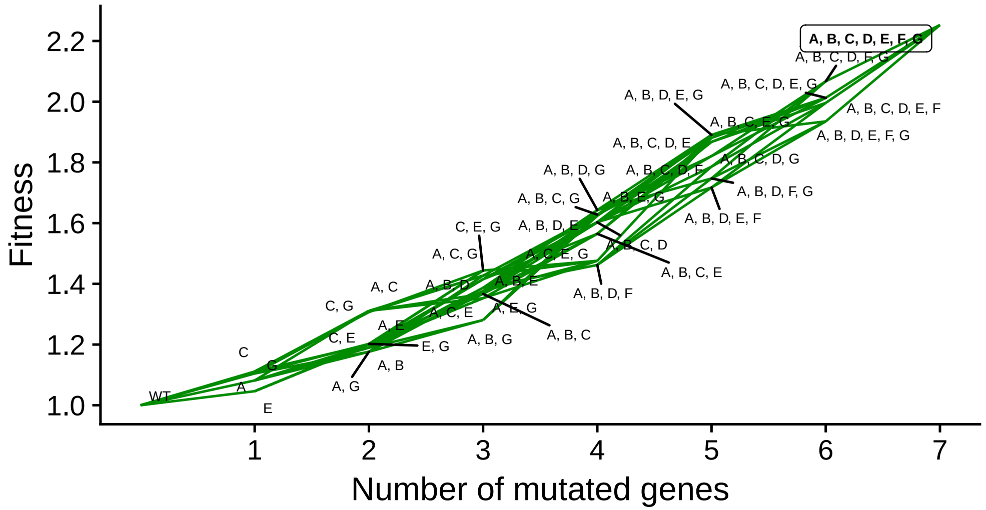

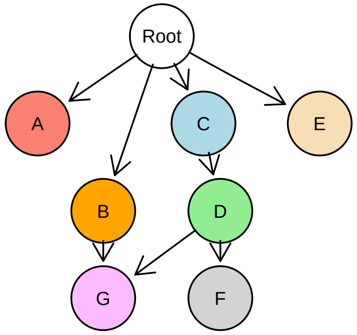


| | |
|---|---|
| *ID* | EL1Qu5ixDsldHIqT |
| *acc genotypes* | 40 |
| *peaks no back* | 1 |
| *removed edges* | 0 |
| *prop rem edges* | 0 |

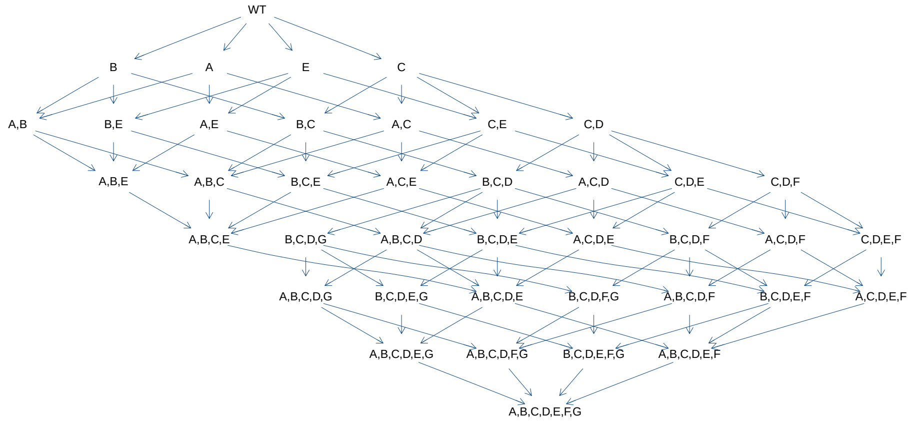


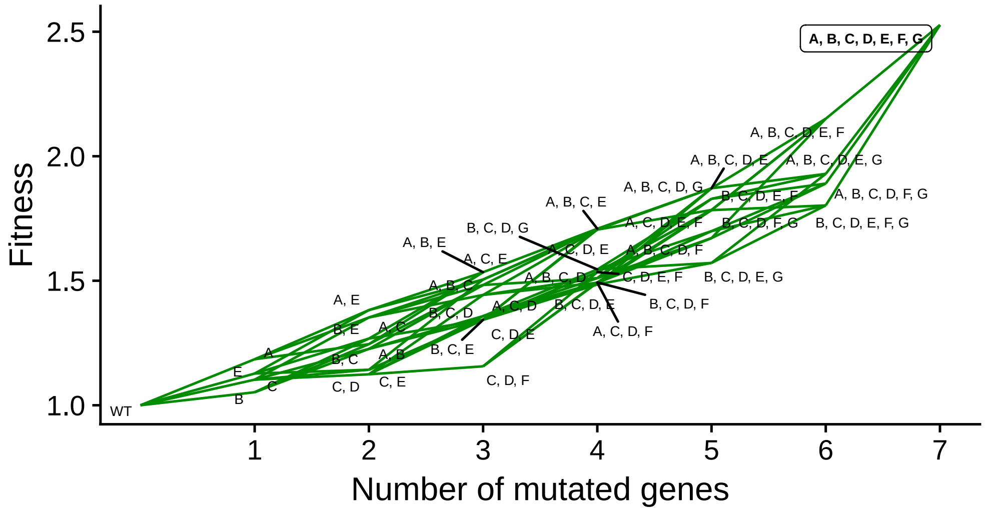

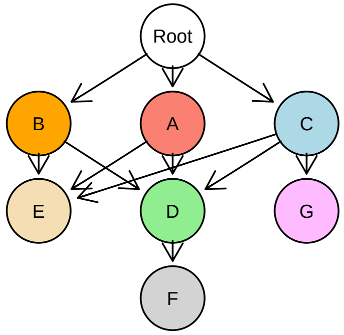


| | |
|---|---|
| *ID* | eyXg2MFYlT9dyyLY |
| *acc genotypes* | 22 |
| *peaks no back* | 1 |
| *removed edges* | 0 |
| *prop rem edges* | 0 |

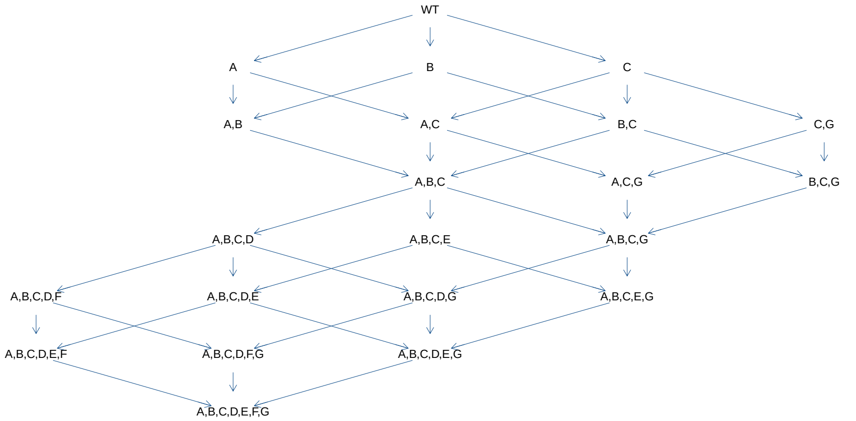


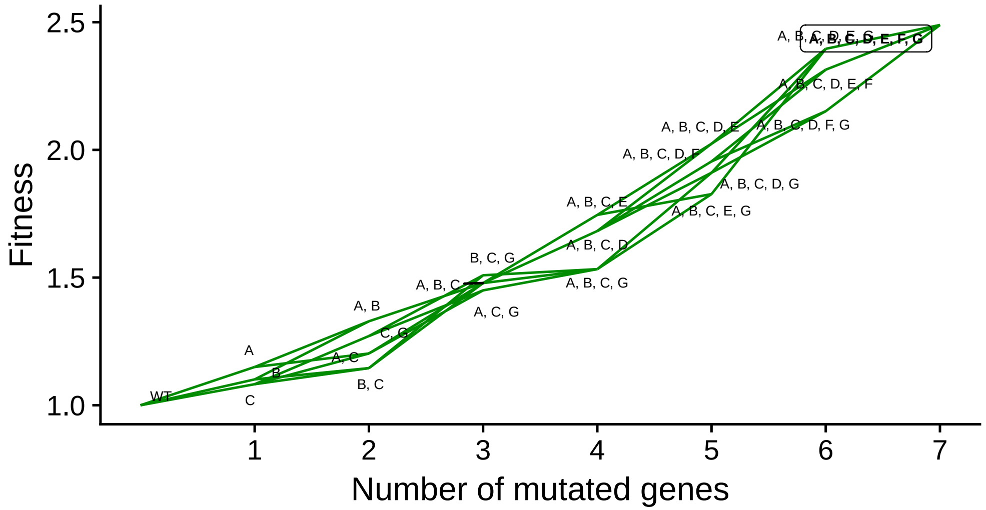

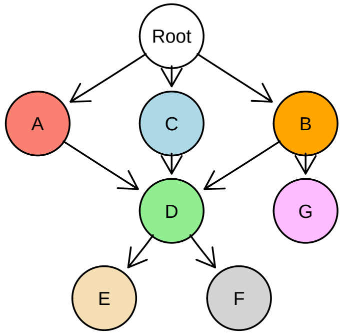


| | |
|---|---|
| *ID* | FcKyPNlgfuFUAoV10 |
| *acc genotypes* | 20 |
| *peaks no back* | 1 |
| *removed edges* | 0 |
| *prop rem edges* | 0 |

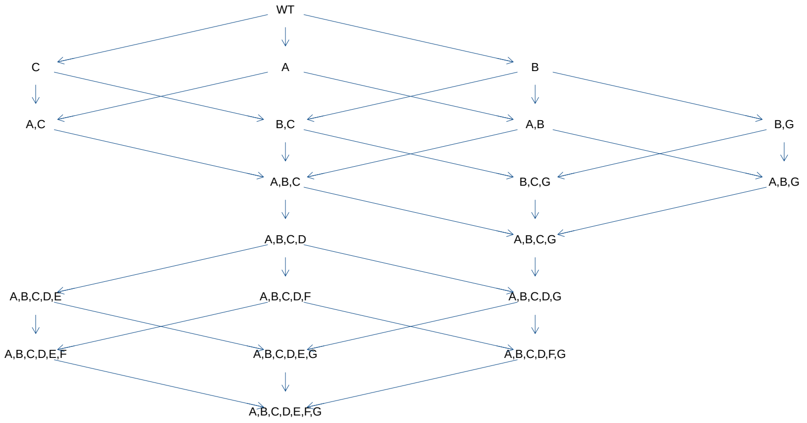


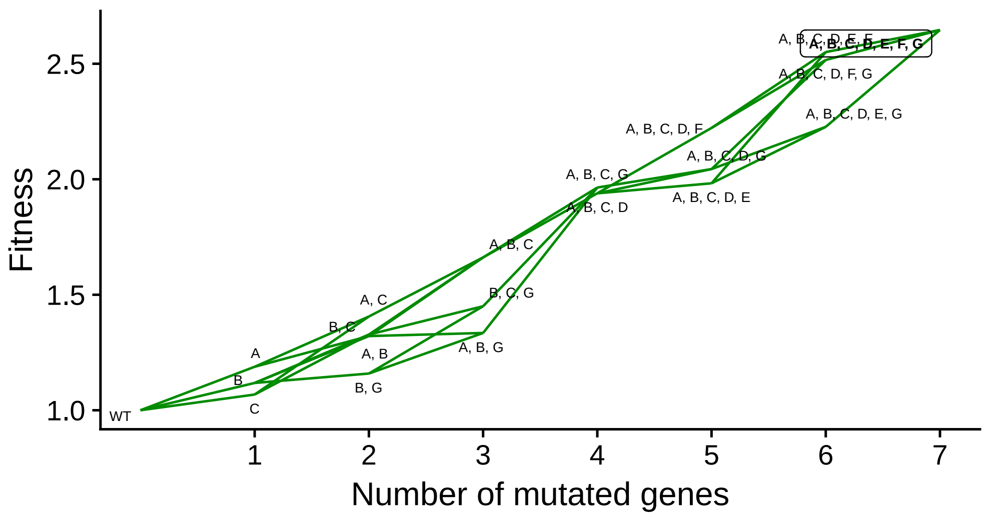

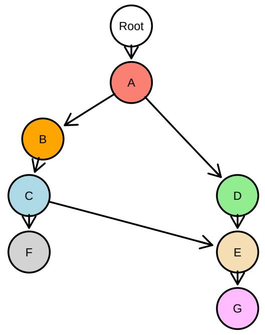


| | |
|---|---|
| *ID* | FL34ukOM10yOTdr2W |
| *acc genotypes* | 13 |
| *peaks no back* | 1 |
| *removed edges* | 0 |
| *prop rem edges* | 0 |

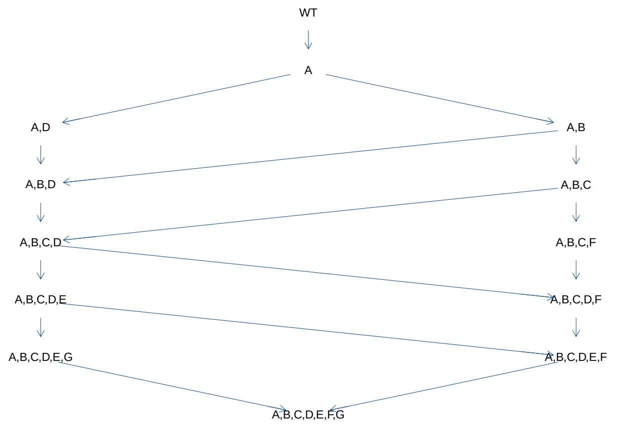


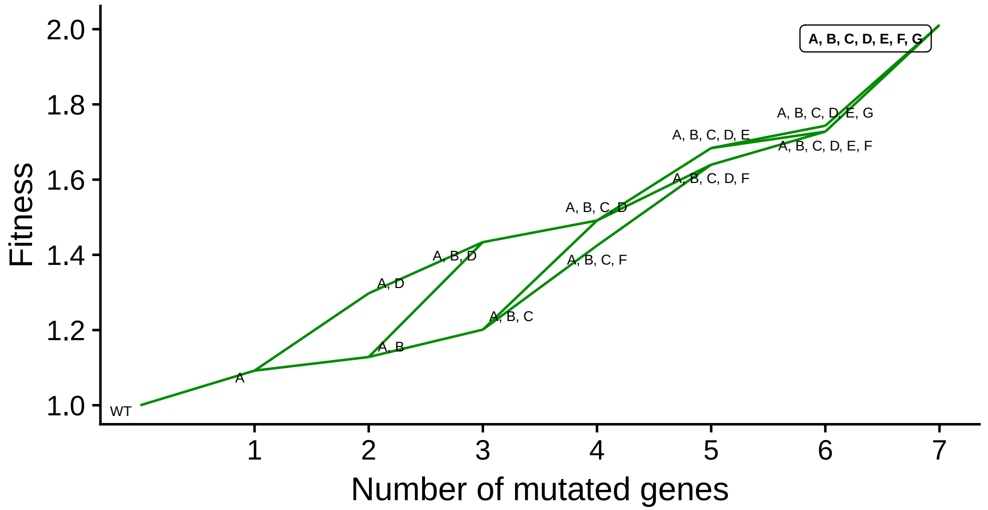

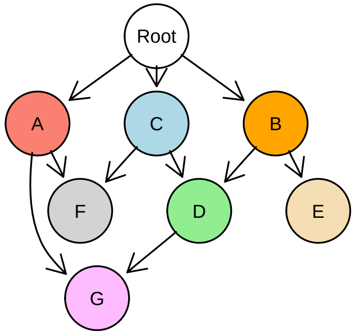


| | |
|---|---|
| *ID* | fMD9ePnbEbDWW2mR |
| *acc genotypes* | 25 |
| *peaks no back* | 1 |
| *removed edges* | 0 |
| *prop rem edges* | 0 |

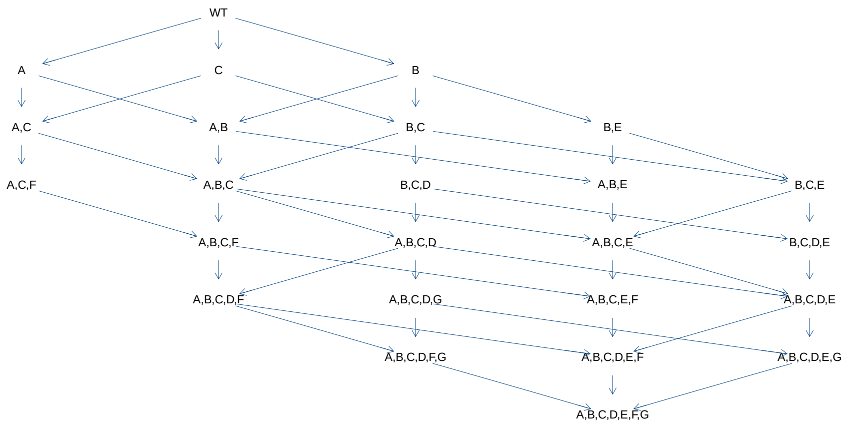


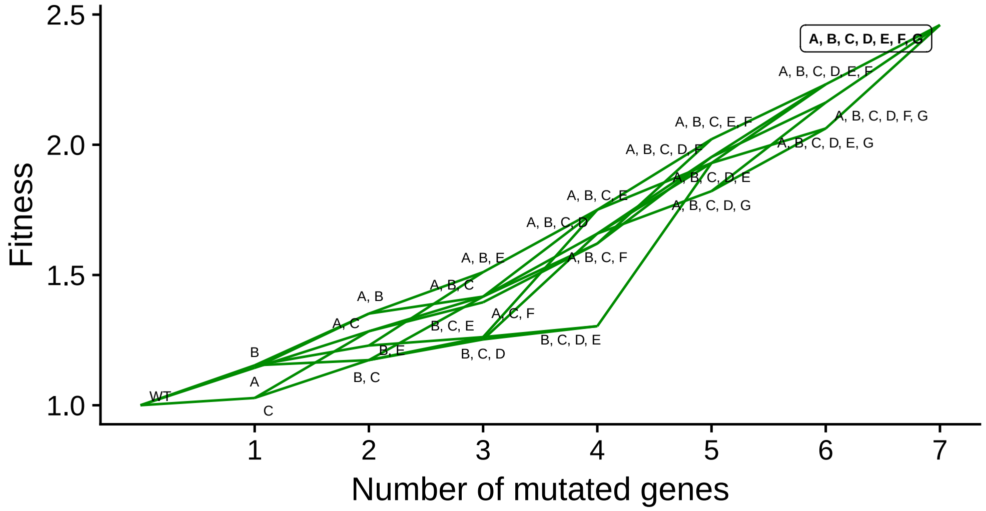

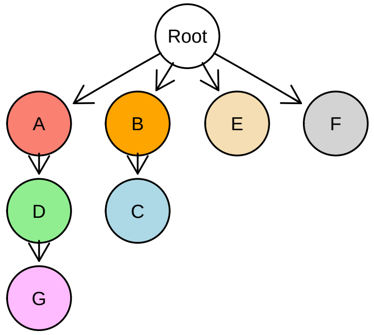


| | |
|---|---|
| *ID* | fO3JEhzhobZEAWxH |
| *acc genotypes* | 48 |
| *peaks no back* | 1 |
| *removed edges* | 0 |
| *prop rem edges* | 0 |

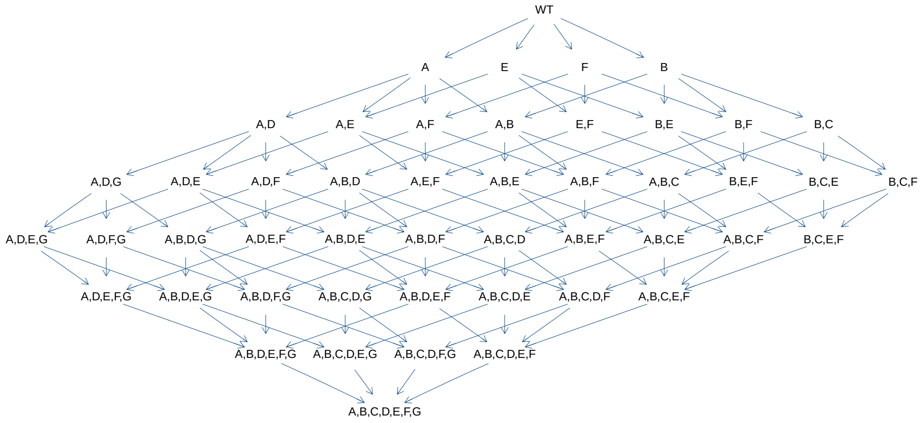


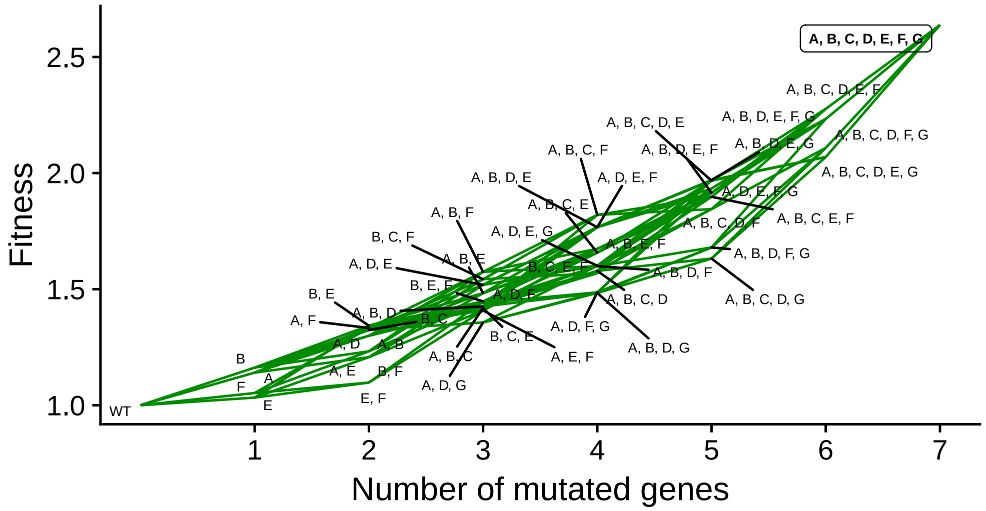

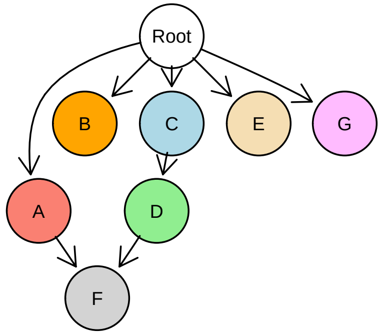


| | |
|---|---|
| *ID* | FoHYHTCWCEdt7JW8 |
| *acc genotypes* | 56 |
| *peaks no back* | 1 |
| *removed edges* | 0 |
| *prop rem edges* | 0 |

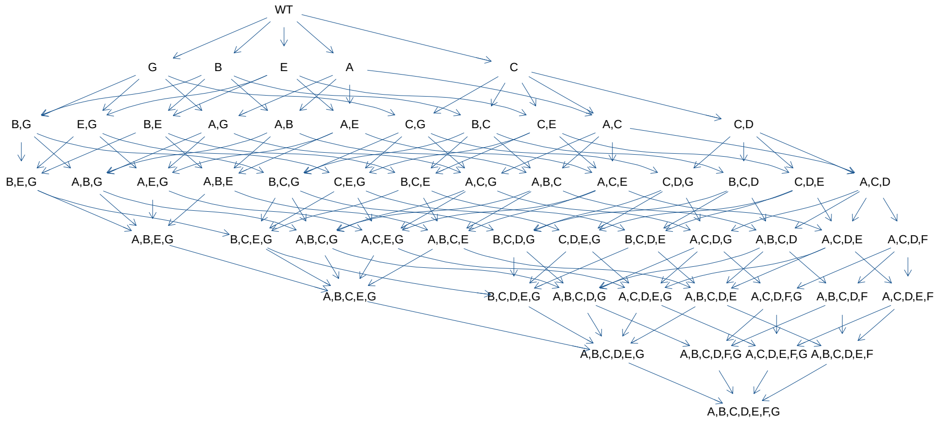


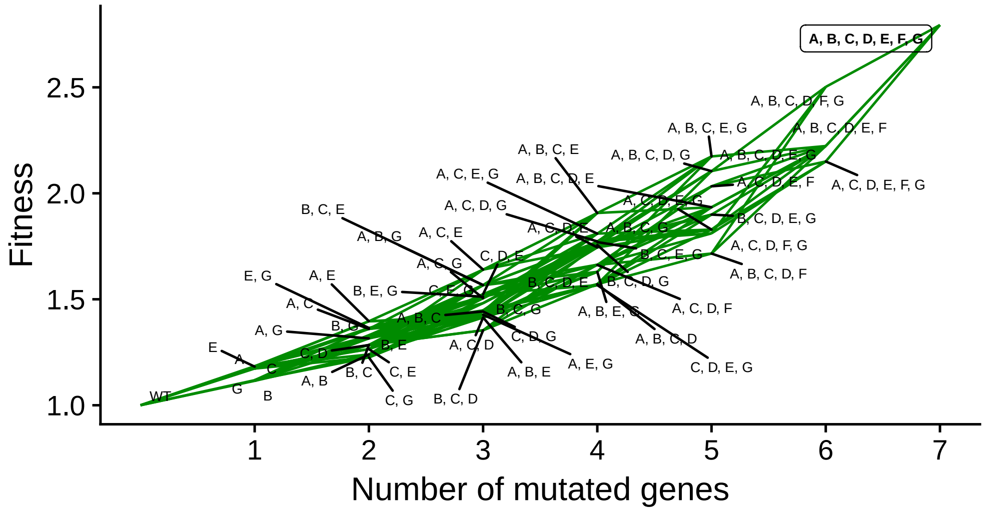

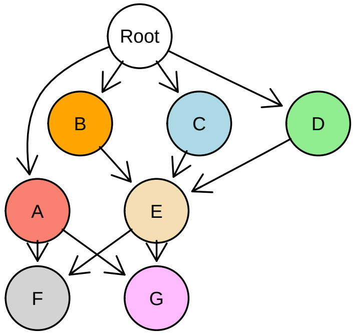


| | |
|---|---|
| *ID* | FoocHDFetllLLSqj |
| *acc genotypes* | 21 |
| *peaks no back* | 1 |
| *removed edges* | 0 |
| *prop rem edges* | 0 |

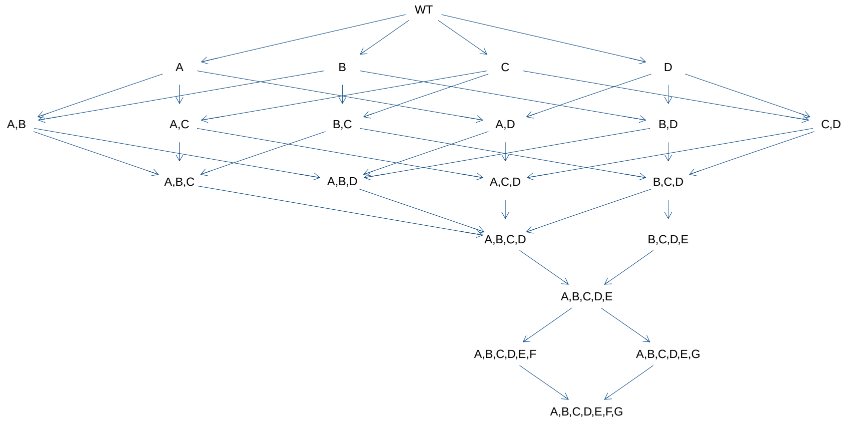


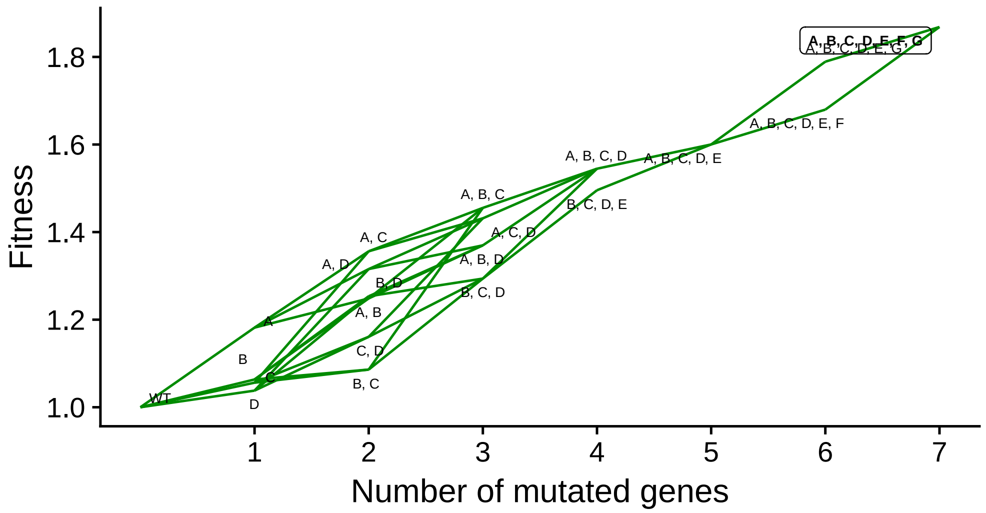

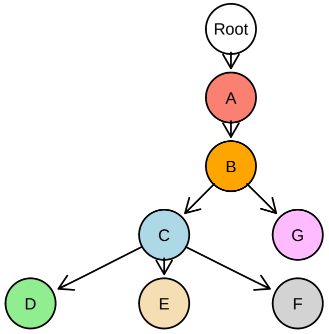


| | |
|---|---|
| *ID* | Gb23niBOgzn9YAzw |
| *acc genotypes* | 20 |
| *peaks no back* | 1 |
| *removed edges* | 0 |
| *prop rem edges* | 0 |

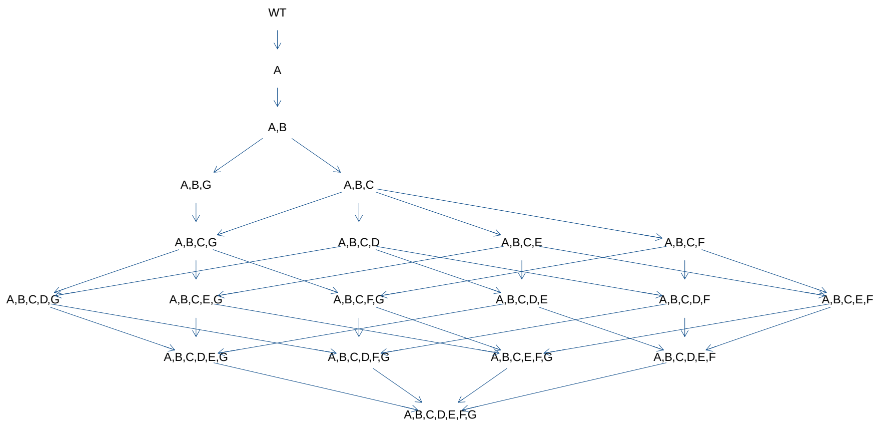


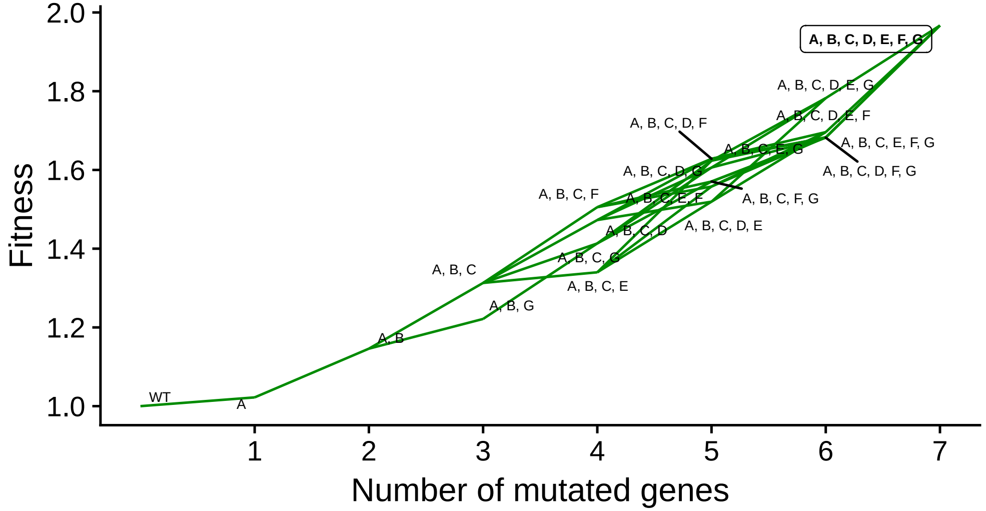

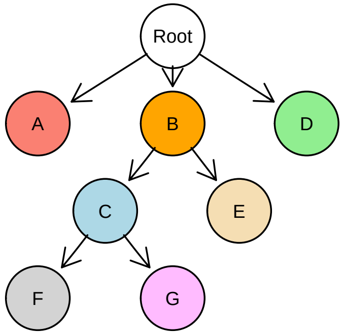


| | |
|---|---|
| *ID* | GIWxSFRzKt1hZIMN |
| *acc genotypes* | 44 |
| *peaks no back* | 1 |
| *removed edges* | 0 |
| *prop rem edges* | 0 |

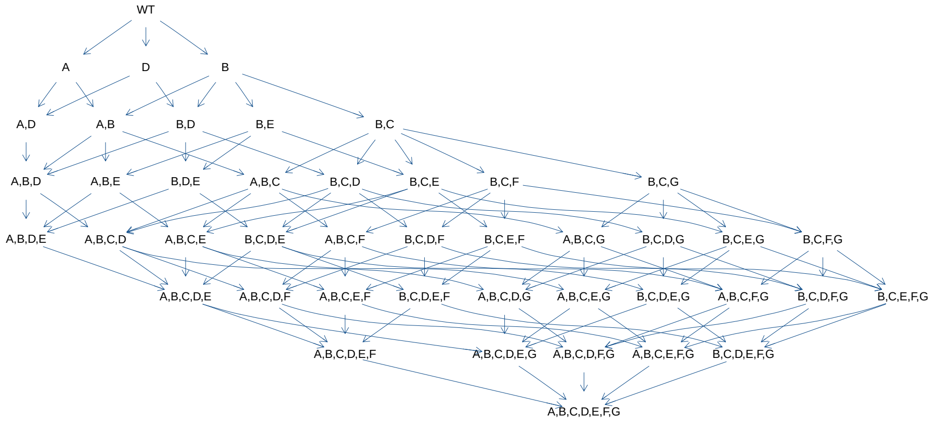


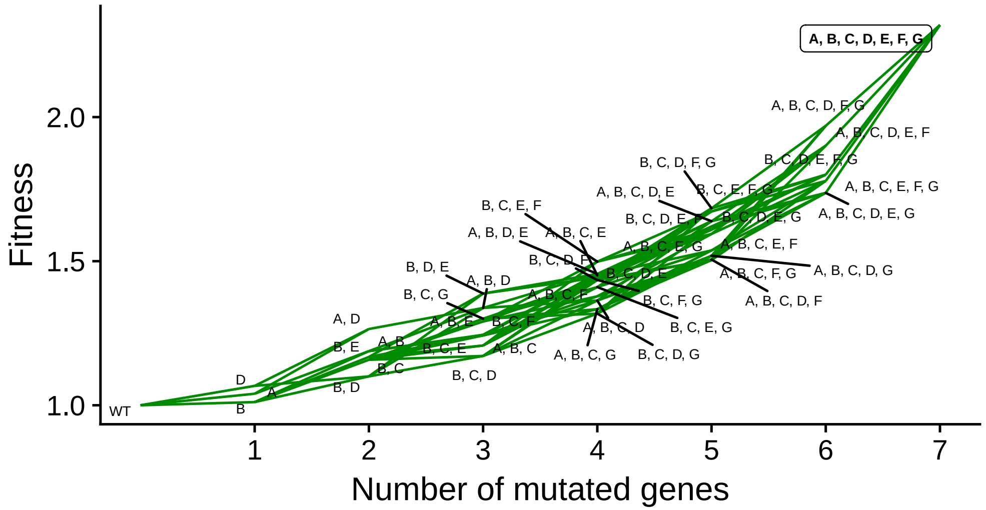

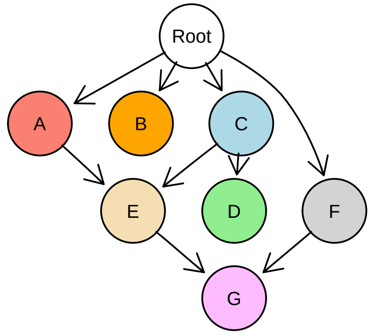


| | |
|---|---|
| *ID* | gLwa1VlNXzVMwvea |
| *acc genotypes* | 36 |
| *peaks no back* | 1 |
| *removed edges* | 0 |
| *prop rem edges* | 0 |

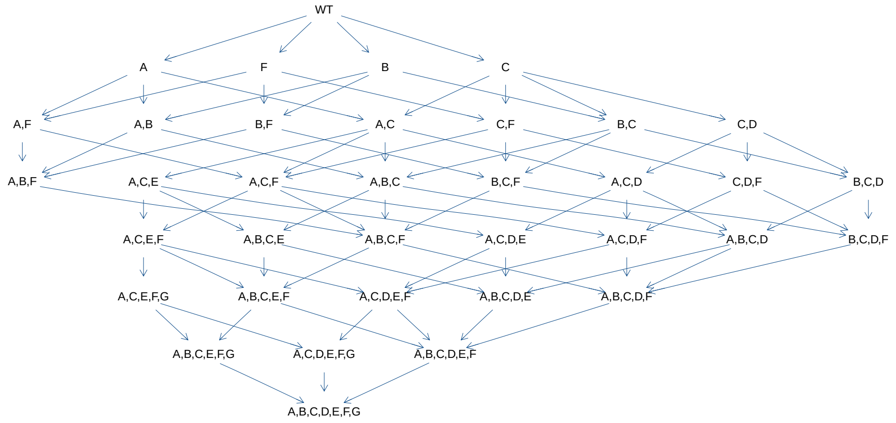


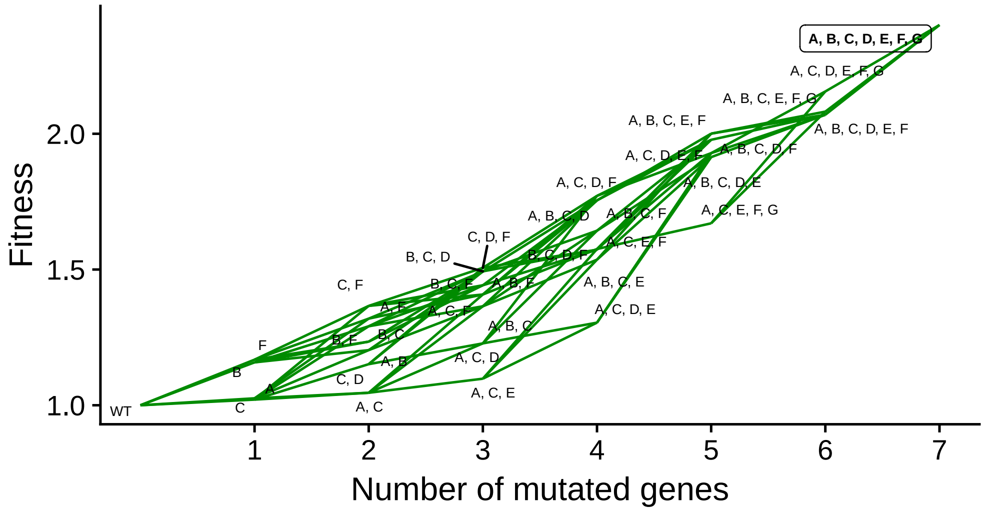

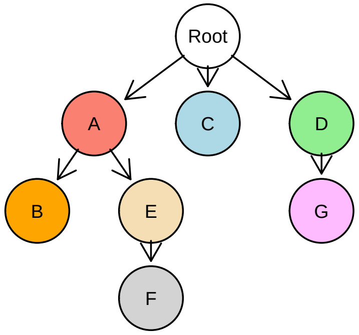


| | |
|---|---|
| *ID* | Gn2HkZc73V1xJWfm |
| *acc genotypes* | 42 |
| *peaks no back* | 1 |
| *removed edges* | 0 |
| *prop rem edges* | 0 |

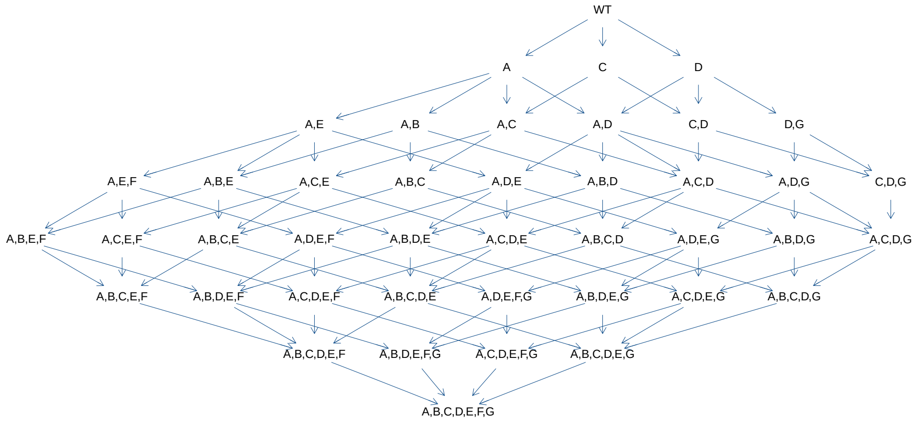


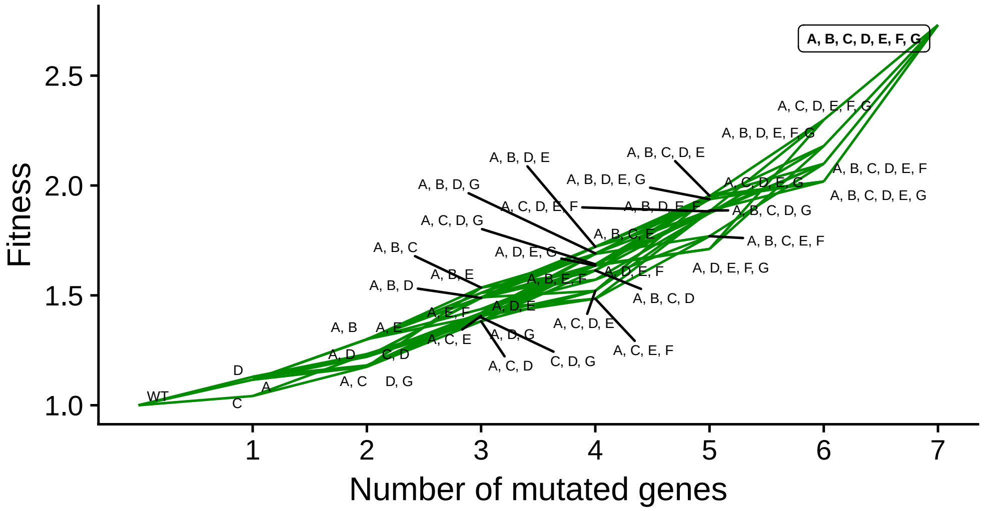

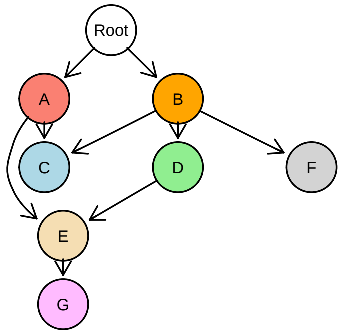


| | |
|---|---|
| *ID* | GNq25Sjl71oPTH7O |
| *acc genotypes* | 22 |
| *peaks no back* | 1 |
| *removed edges* | 0 |
| *prop rem edges* | 0 |

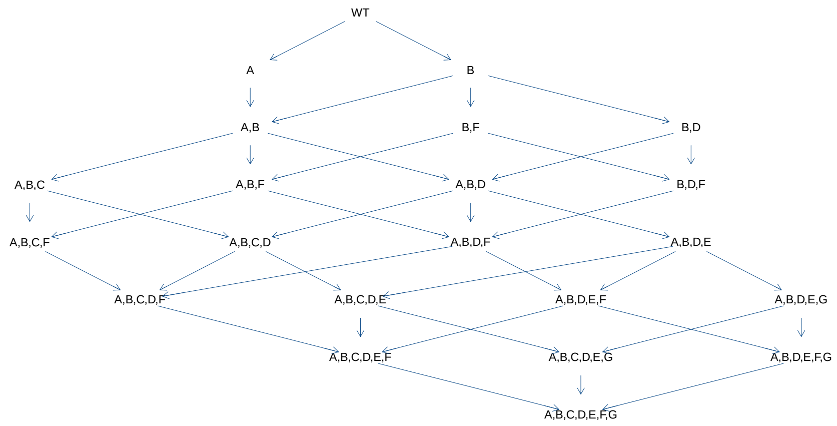


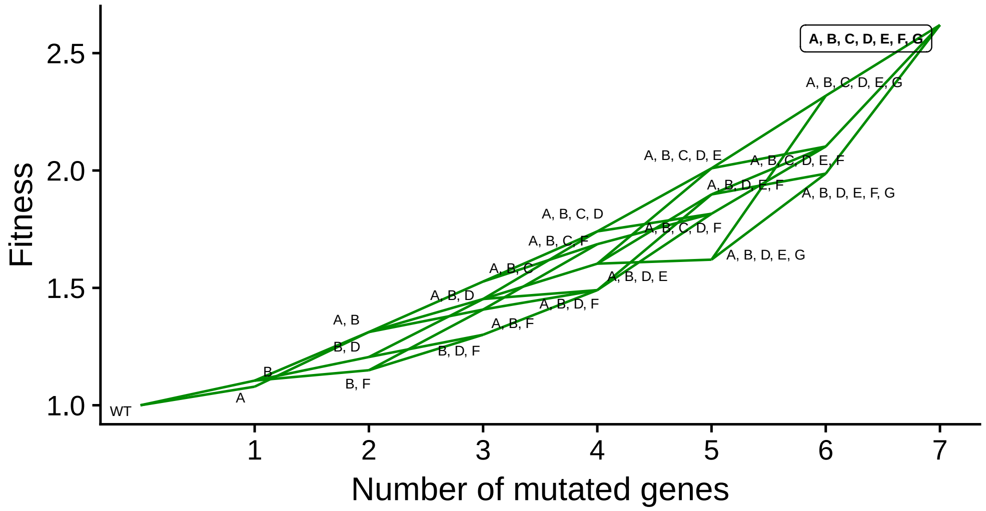

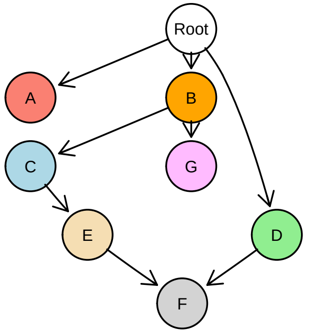


*ID* gNsBkqVxNevMiAxL
*acc genotypes* 32
*peaks no back* 1
*removed edges* 0
*prop rem edges* 0

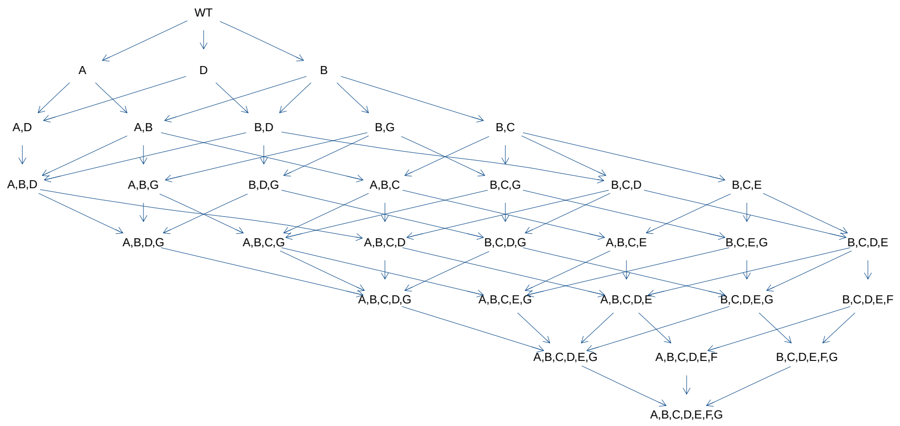


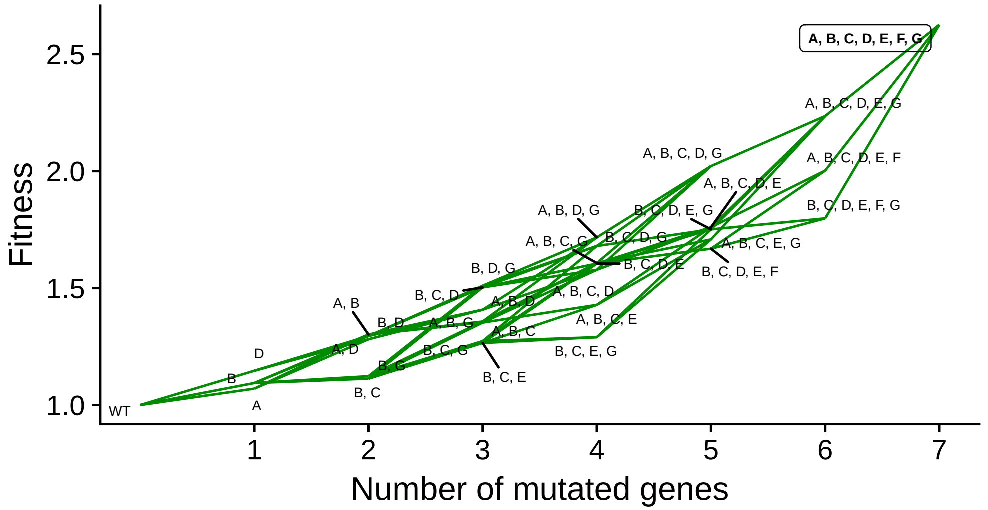

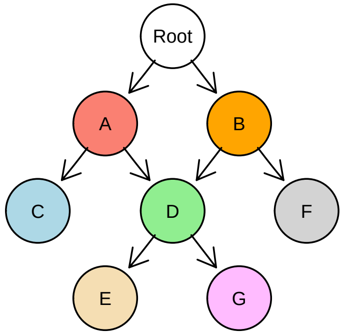


| | |
|---|---|
| *ID* | Gp543eGHWygA1P8H |
| *acc genotypes* | 25 |
| *peaks no back* | 1 |
| *removed edges* | 0 |
| *prop rem edges* | 0 |

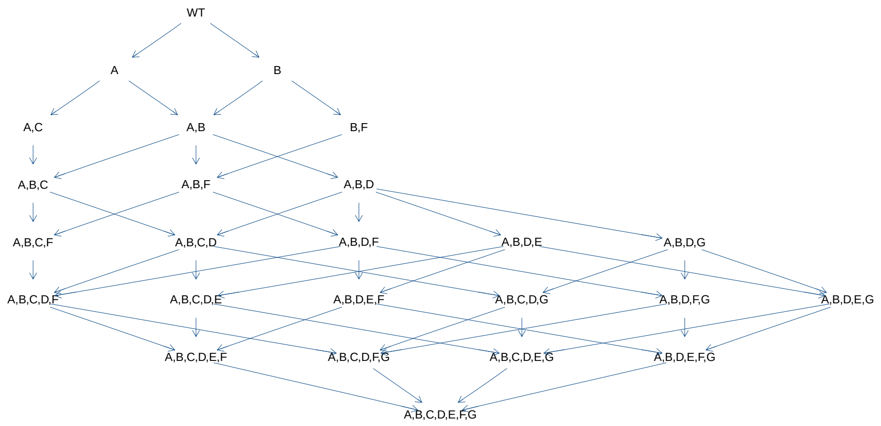


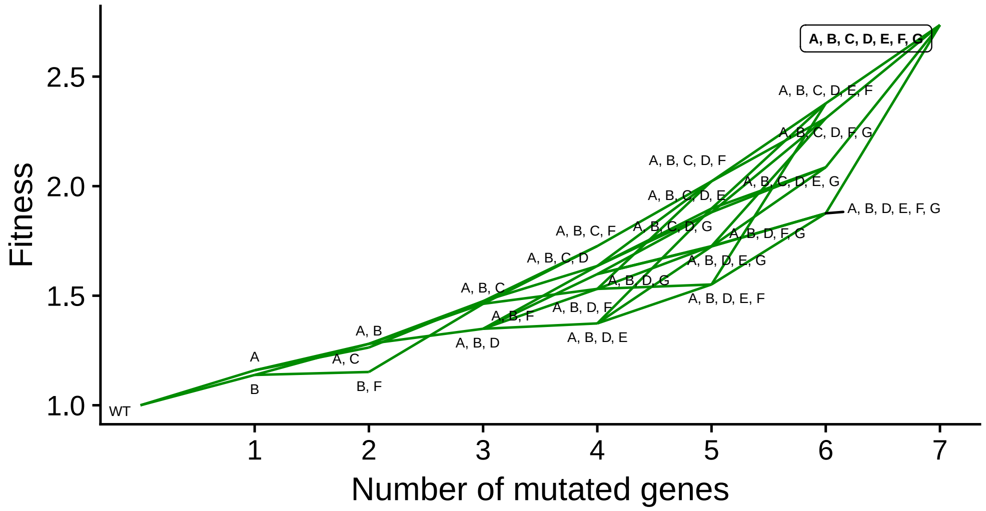

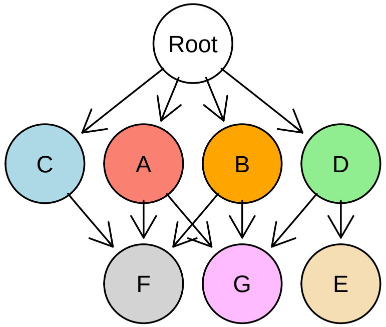


| | |
|---|---|
| *ID* | gtuJ3lZP3MzeoW31 |
| *acc genotypes* | 33 |
| *peaks no back* | 1 |
| *removed edges* | 0 |
| *prop rem edges* | 0 |

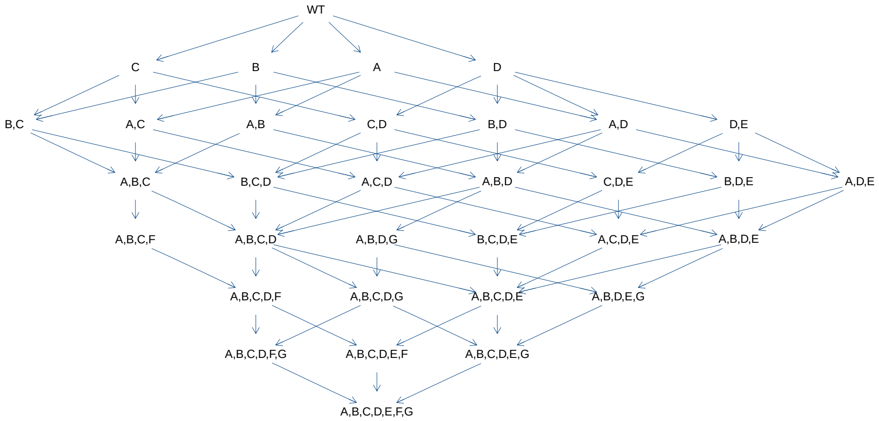


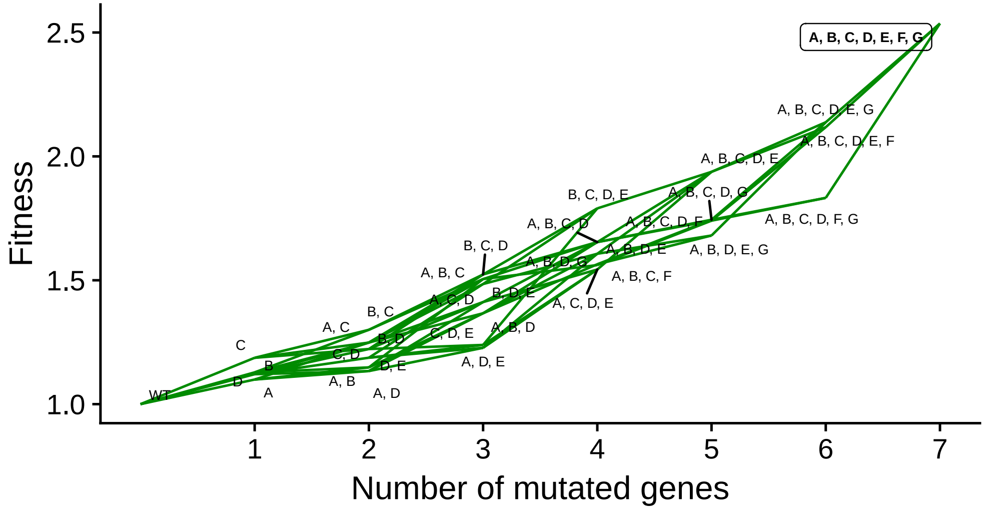

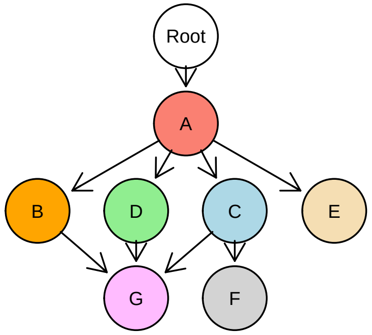


| | |
|---|---|
| *ID* | GyBZiyDvewoQU6ri |
| *acc genotypes* | 29 |
| *peaks no back* | 1 |
| *removed edges* | 0 |
| *prop rem edges* | 0 |

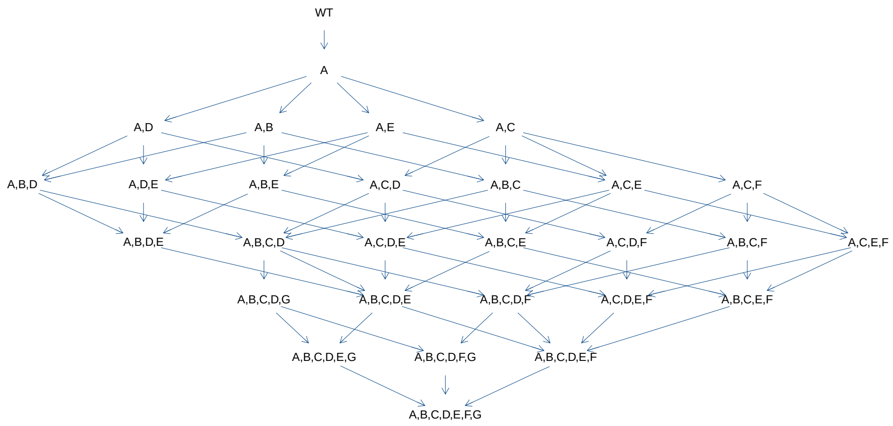


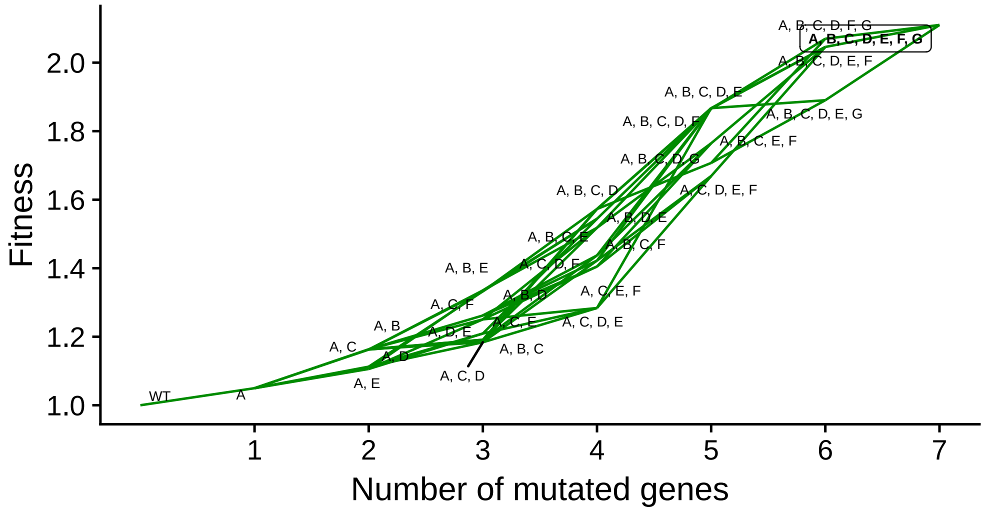

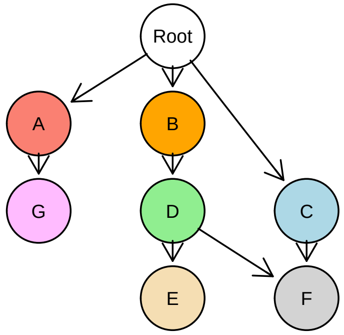


*ID* h5XuDSnadWt672QY
*acc genotypes* 30
*peaks no back* 1
*removed edges* 0
*prop rem edges* 0

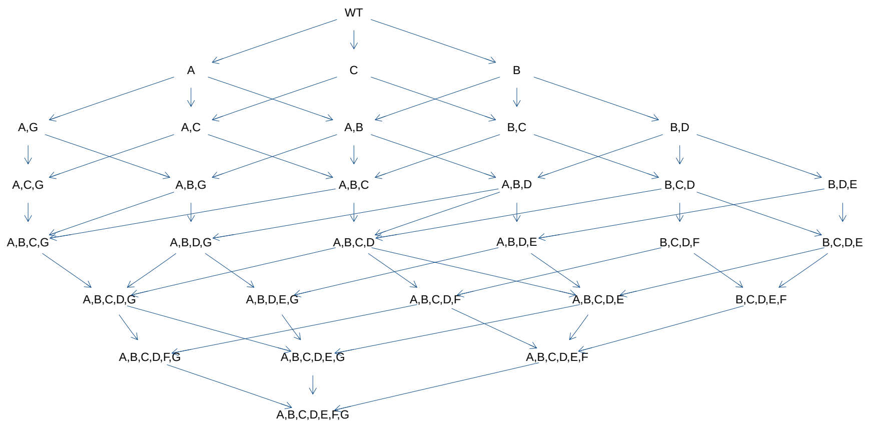


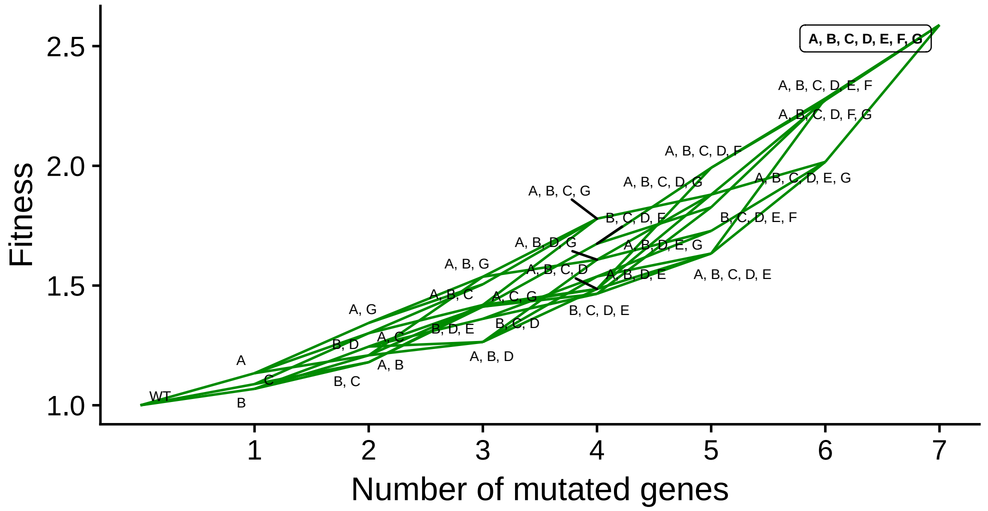

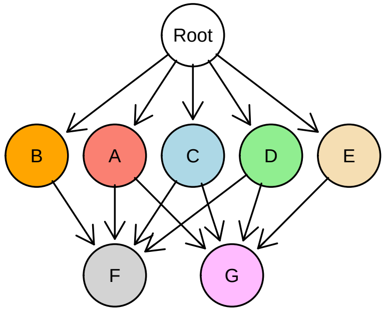


| | |
|---|---|
| *ID* | h88O5DNbc34DtJRm |
| *acc genotypes* | 37 |
| *peaks no back* | 1 |
| *removed edges* | 0 |
| *prop rem edges* | 0 |

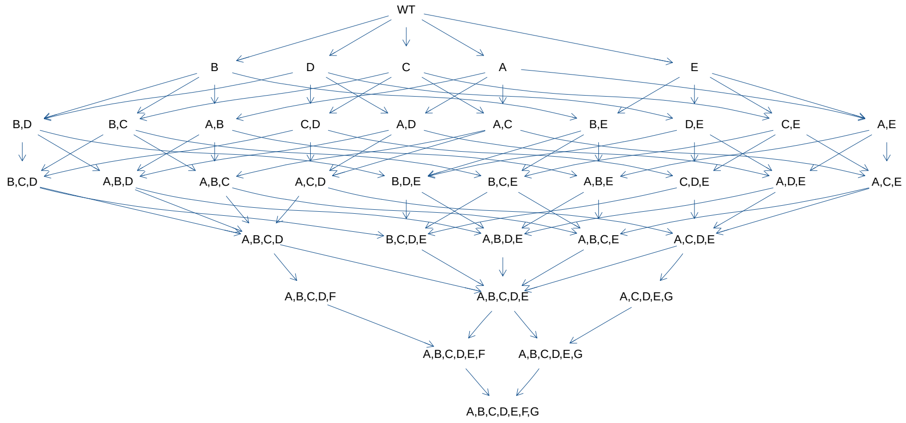


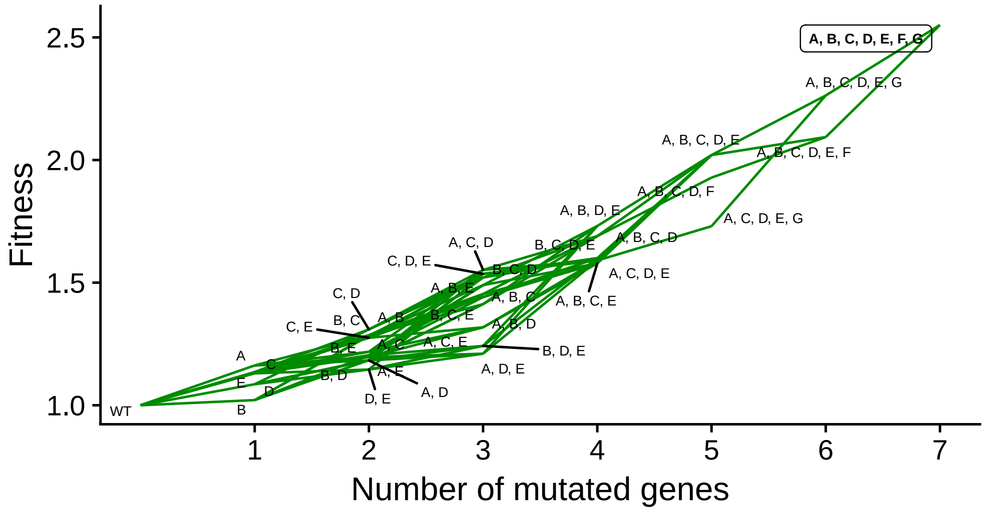

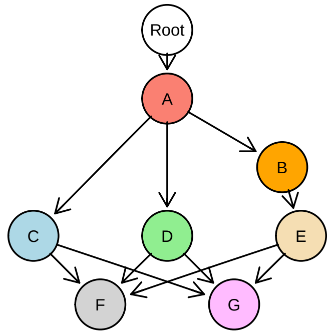


| | |
|---|---|
| *ID* | Hbk7A4ZWDH110SIas |
| *acc genotypes* | 16 |
| *peaks no back* | 1 |
| *removed edges* | 0 |
| *prop rem edges* | 0 |

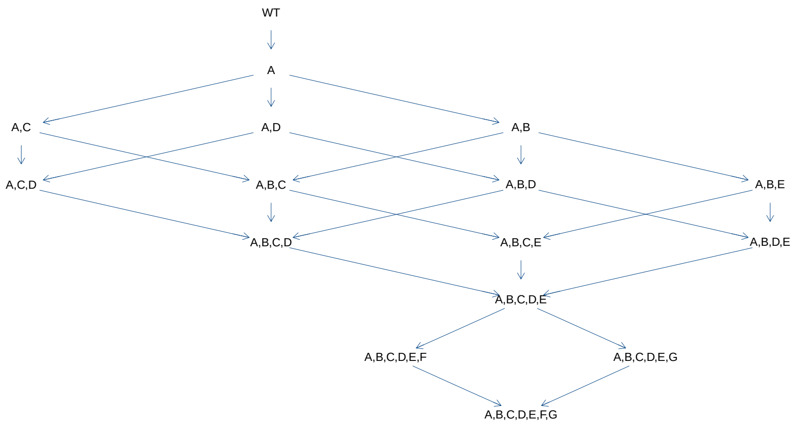


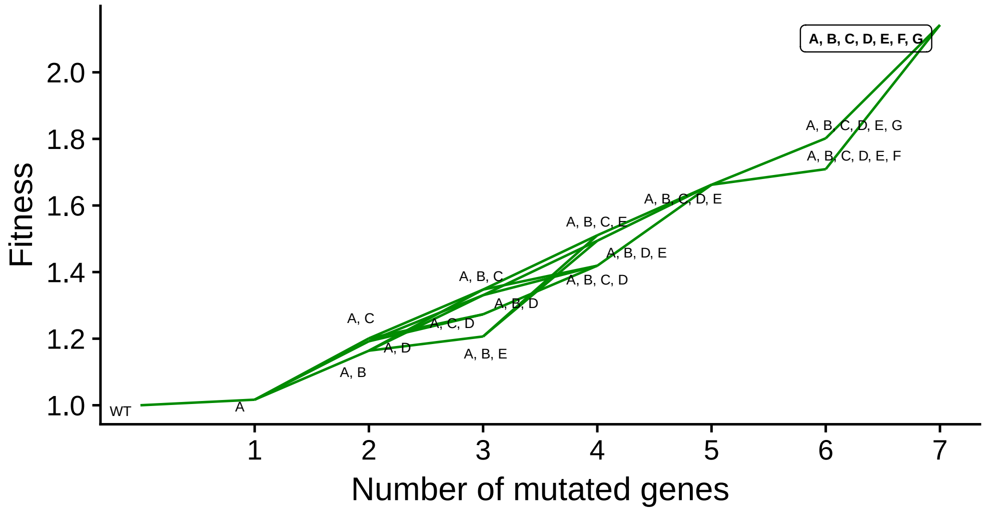

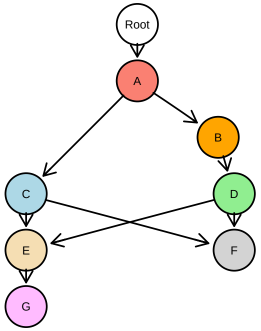


| | |
|---|---|
| *ID* | HCfuosyRn2Mnwy2Y |
| *acc genotypes* | 12 |
| *peaks no back* | 1 |
| *removed edges* | 0 |
| *prop rem edges* | 0 |

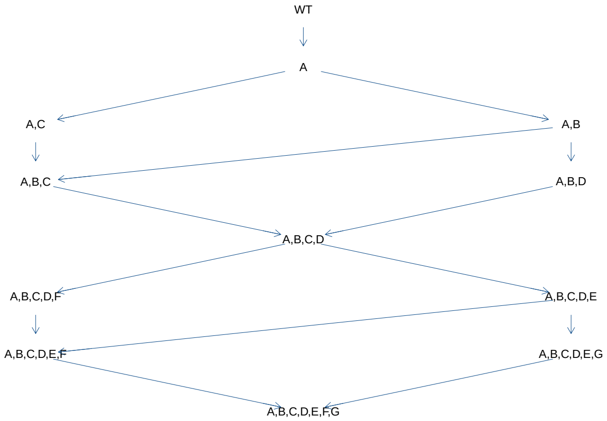


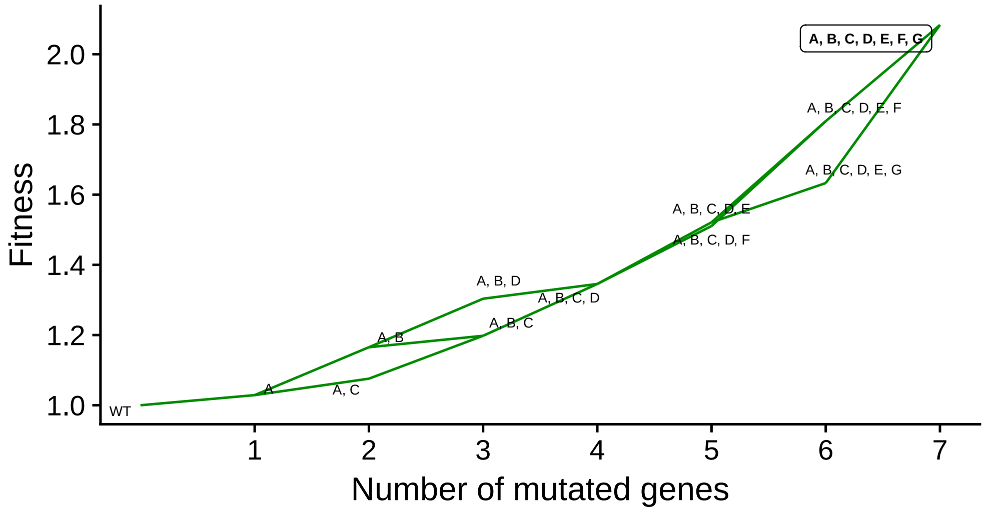

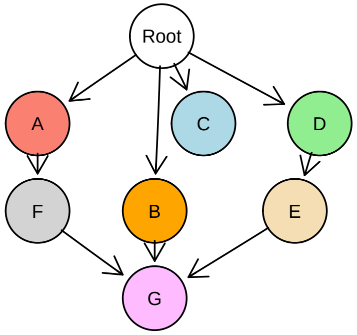


*ID* hEfBL8MoVyBlRoMA
*acc genotypes* 38
*peaks no back* 1
*removed edges* 0
*prop rem edges* 0

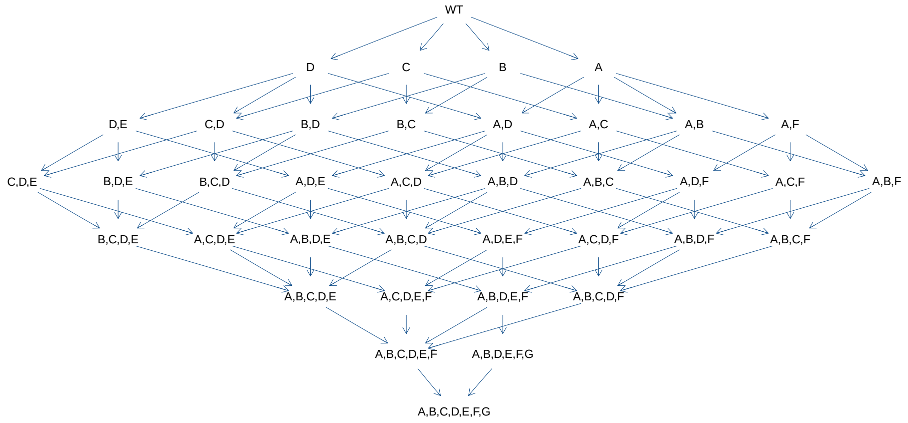


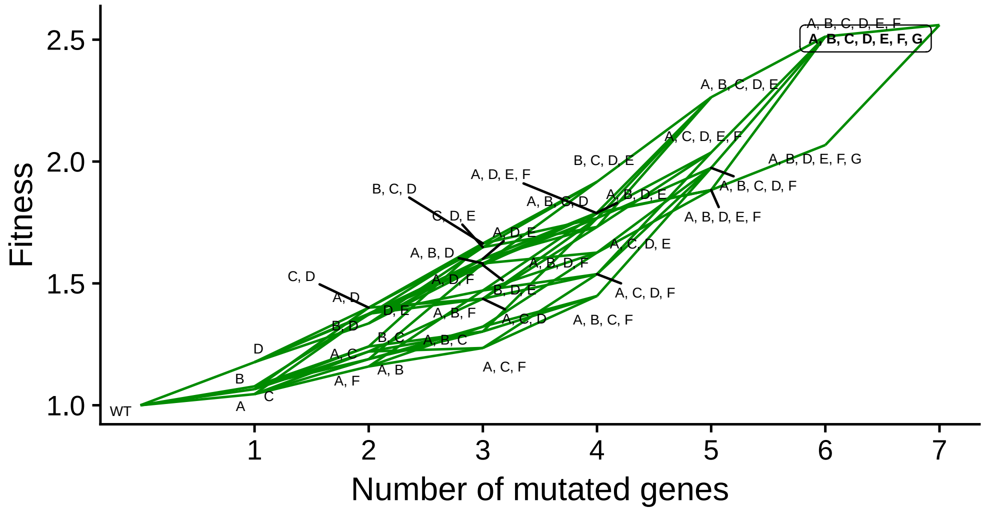

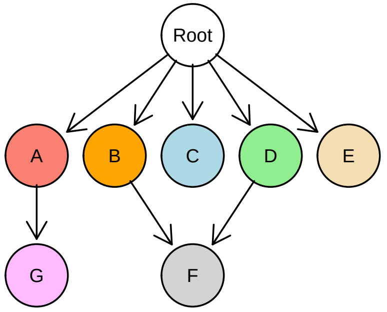


| | |
|---|---|
| *ID* | HF510lGzHhUvaK1Wc |
| *acc genotypes* | 60 |
| *peaks no back* | 1 |
| *removed edges* | 0 |
| *prop rem edges* | 0 |

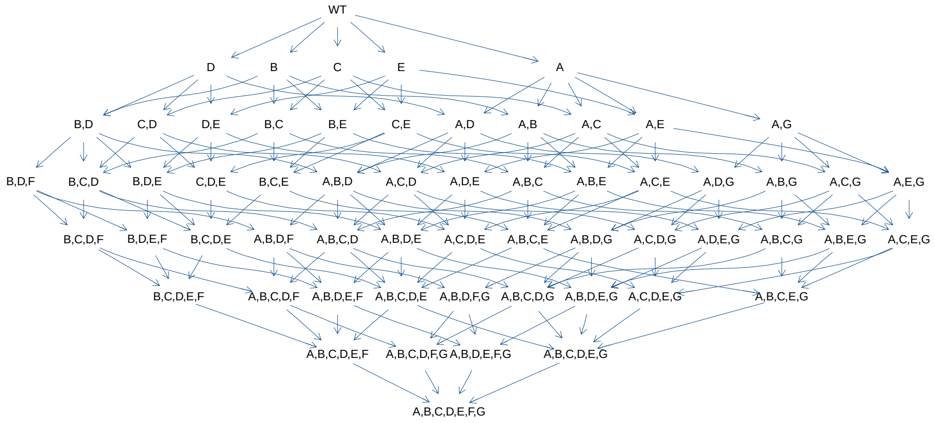


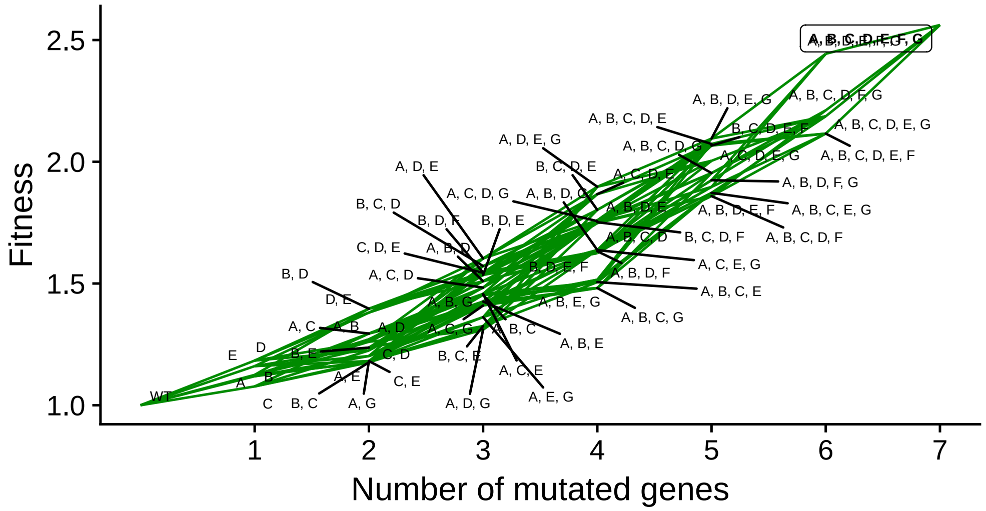

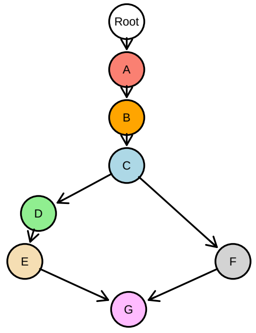


| | |
|---|---|
| *ID* | HfNvtnFRbnBZJbsa |
| *acc genotypes* | 10 |
| *peaks no back* | 1 |
| *removed edges* | 0 |
| *prop rem edges* | 0 |

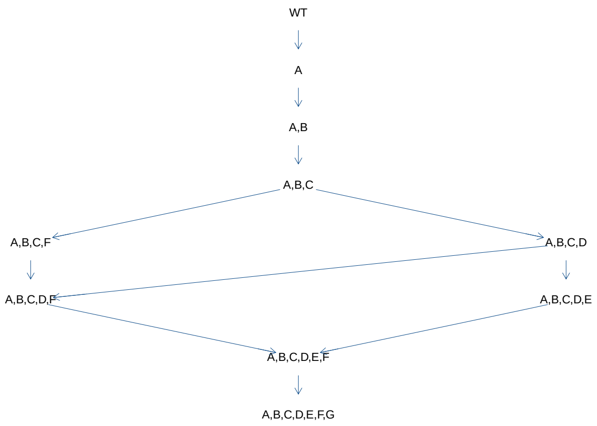


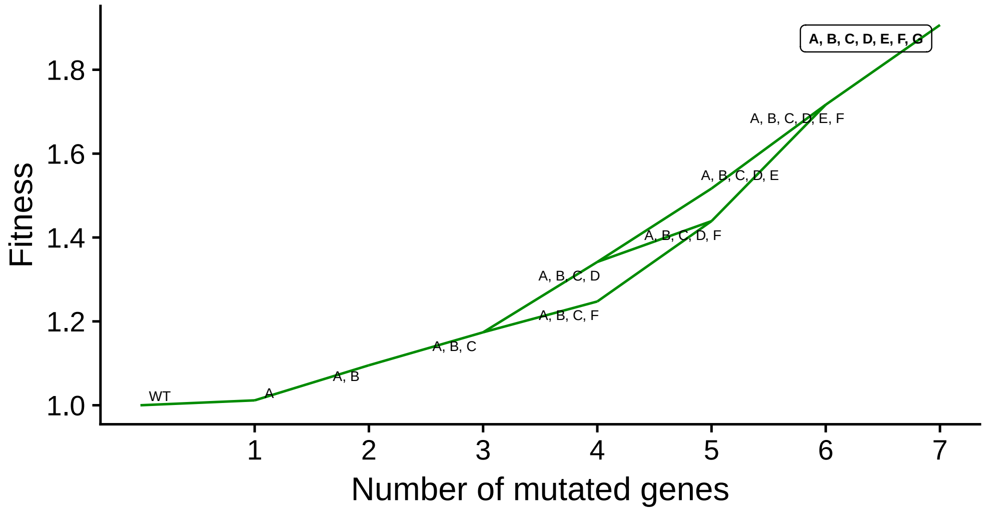

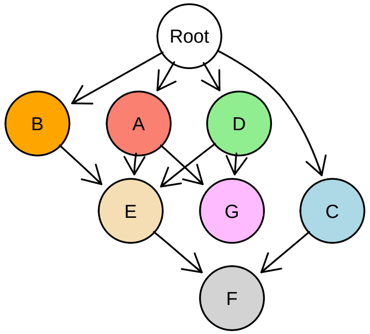


| | |
|---|---|
| *ID* | HGZXE3cU1toI8NwB |
| *acc genotypes* | 26 |
| *peaks no back* | 1 |
| *removed edges* | 0 |
| *prop rem edges* | 0 |

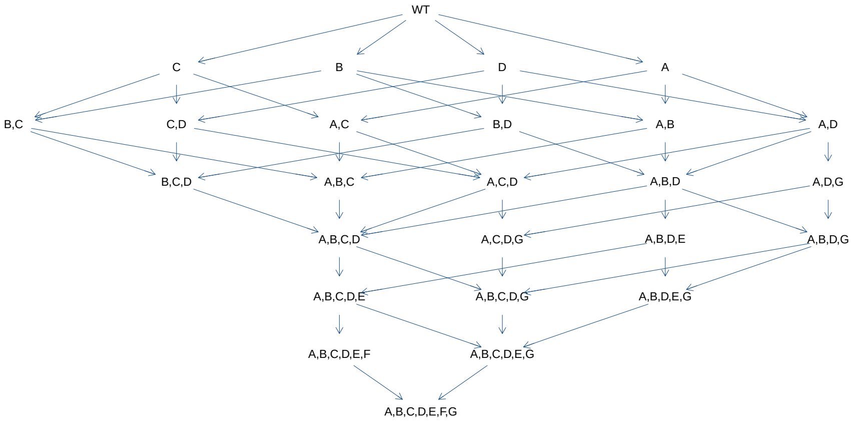


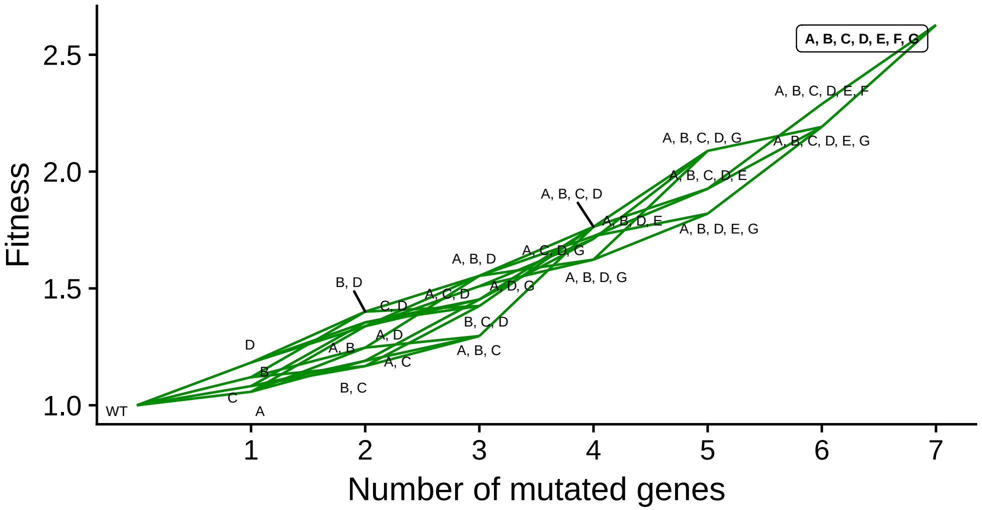

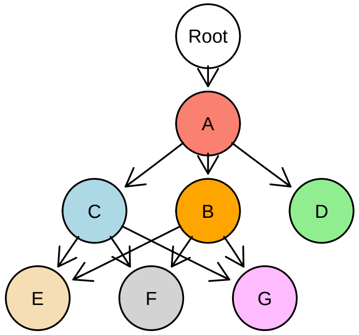


| | |
|---|---|
| *ID* | HhTnaJyFFi4y9Ptx |
| *acc genotypes* | 23 |
| *peaks no back* | 1 |
| *removed edges* | 0 |
| *prop rem edges* | 0 |

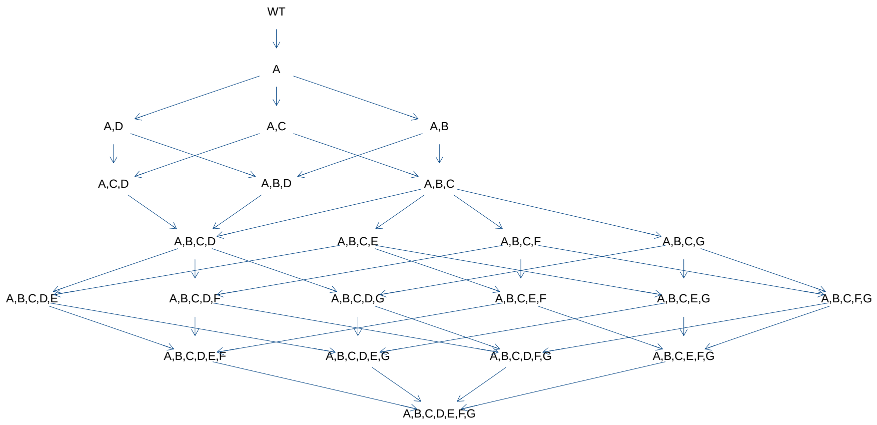


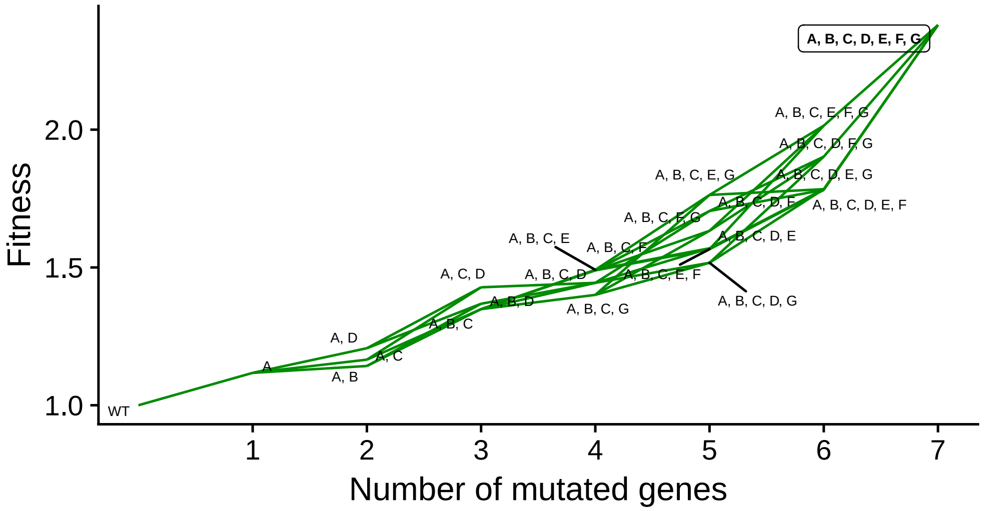

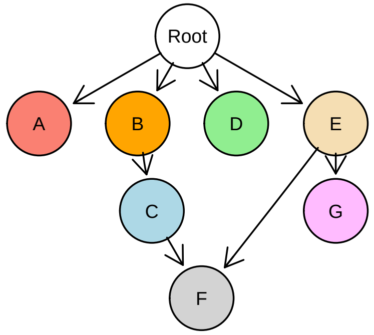


| | |
|---|---|
| *ID* | hiqO72oQ9p210Vxcp |
| *acc genotypes* | 44 |
| *peaks no back* | 1 |
| *removed edges* | 0 |
| *prop rem edges* | 0 |

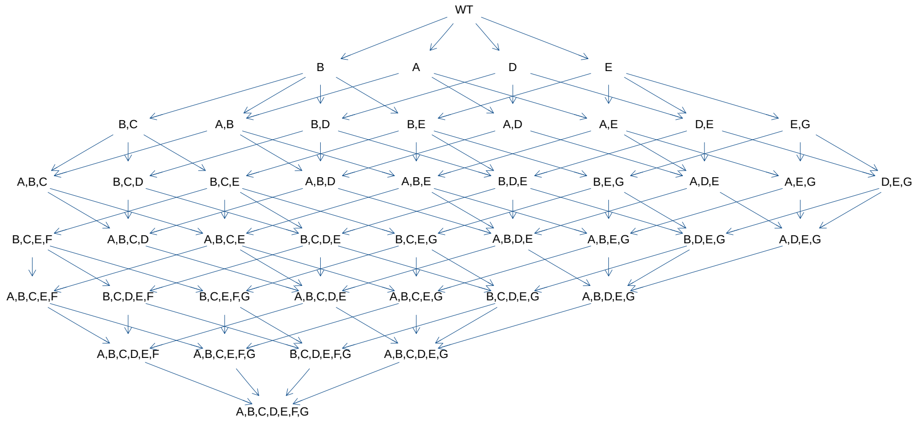


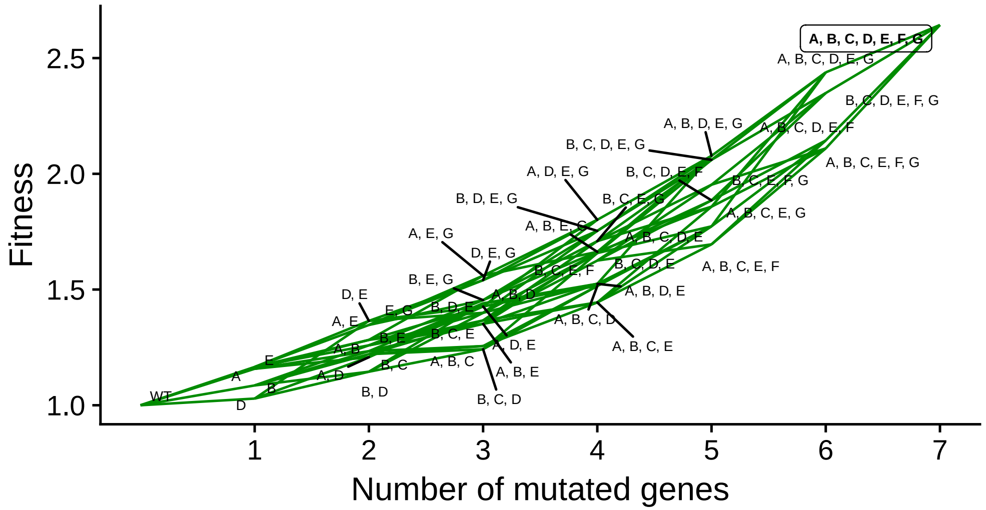

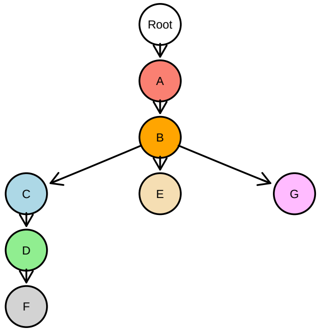


| | |
|---|---|
| *ID* | hTuov2KGeLAIrrac |
| *acc genotypes* | 18 |
| *peaks no back* | 1 |
| *removed edges* | 0 |
| *prop rem edges* | 0 |

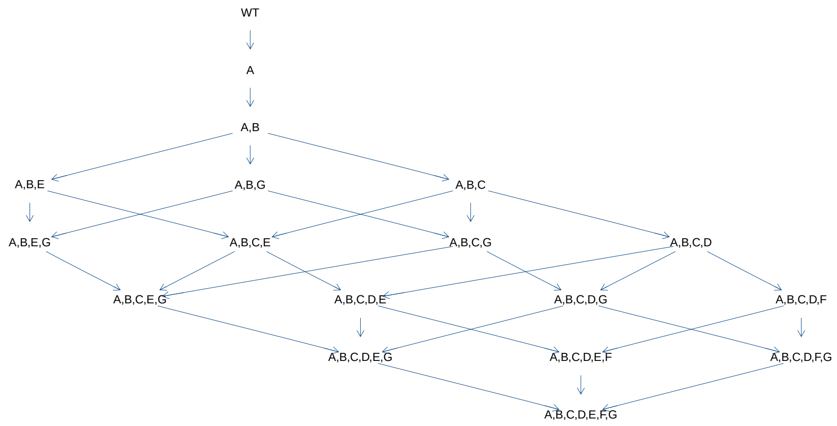


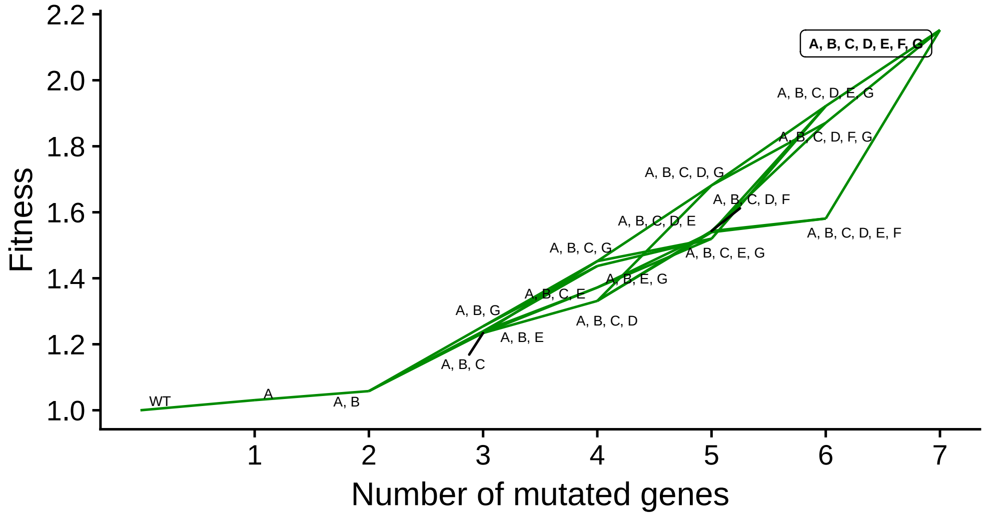

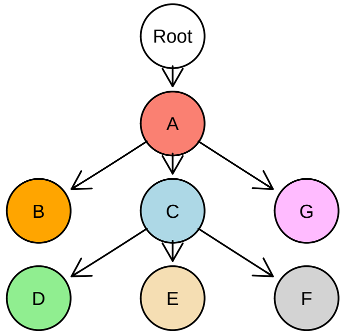


| | |
|---|---|
| *ID* | hyGvt7bXxXmlBy4c |
| *acc genotypes* | 37 |
| *peaks no back* | 1 |
| *removed edges* | 0 |
| *prop rem edges* | 0 |

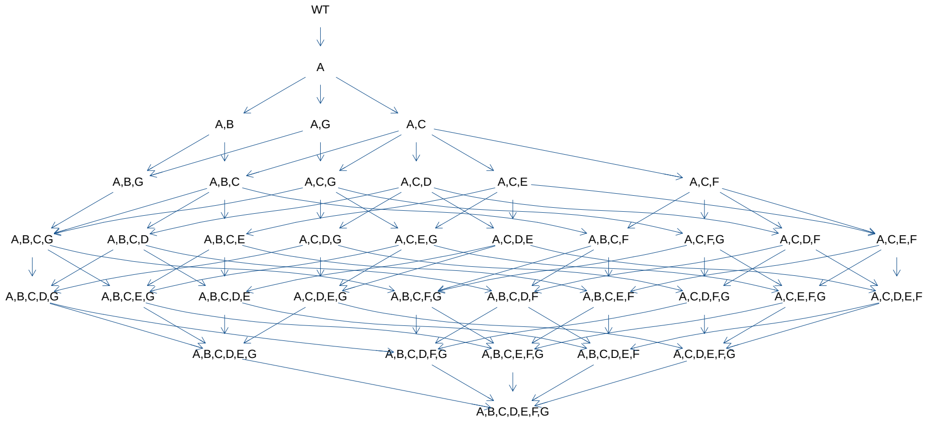


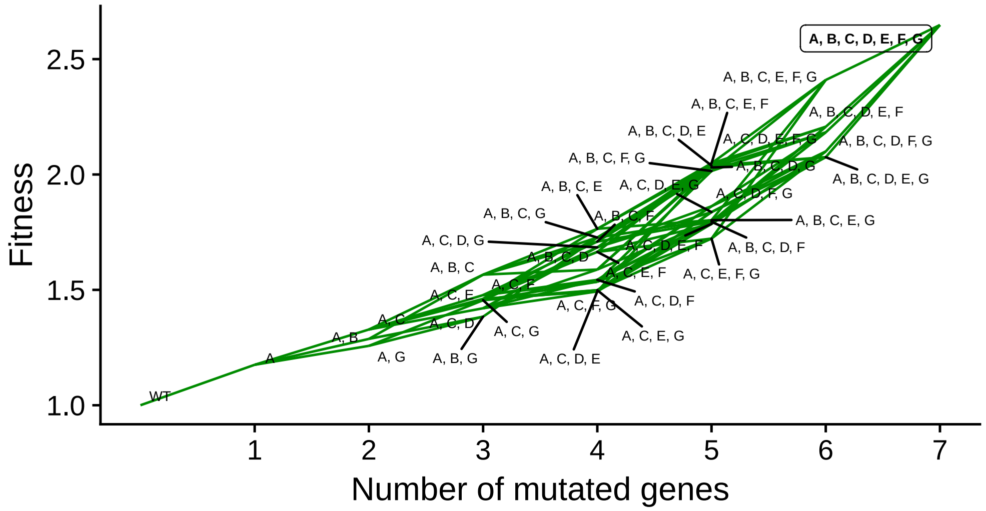

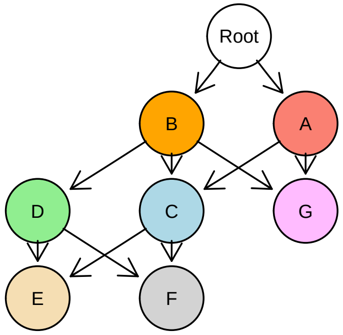


*ID* HYyO10PwiQqDuWK2r
*acc genotypes* 18
*peaks no back* 1
*removed edges* 0
*prop rem edges* 0

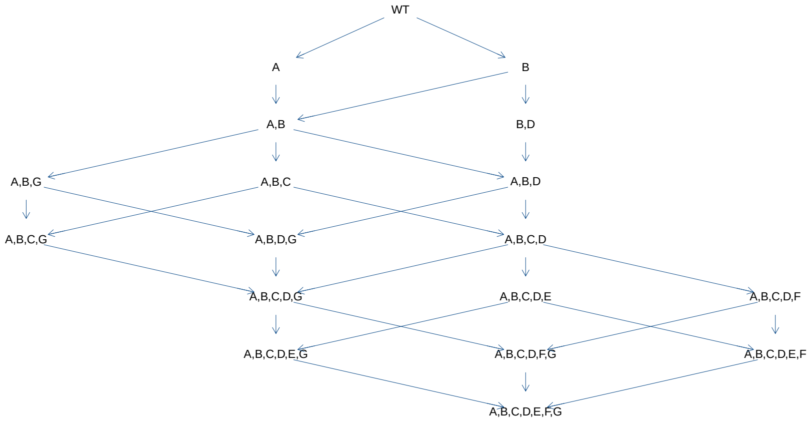


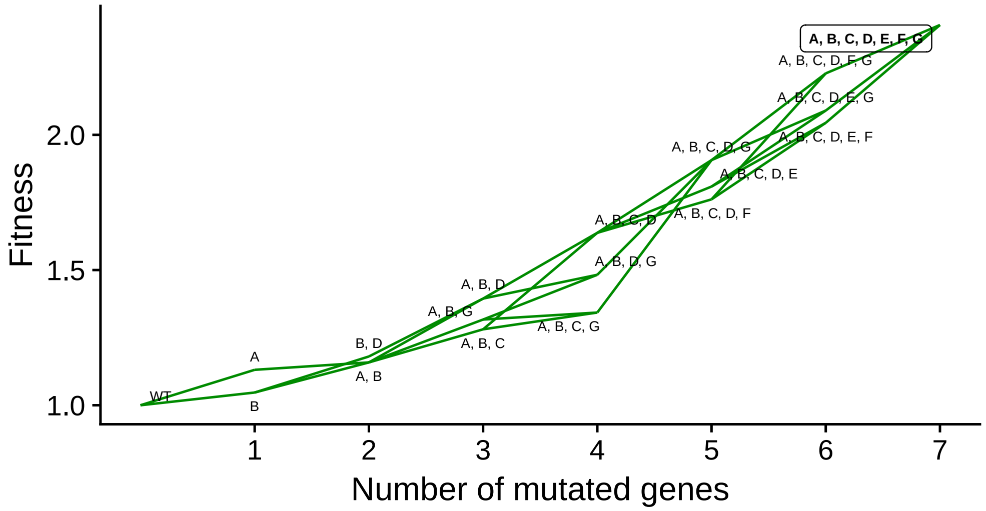

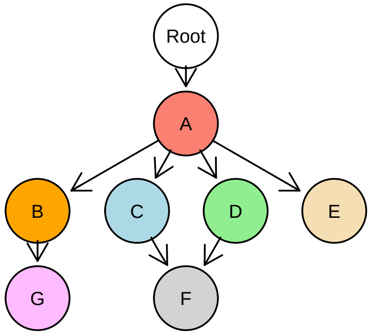


| | |
|---|---|
| *ID* | IFOFBmVbzrLBfqR1 |
| *acc genotypes* | 31 |
| *peaks no back* | 1 |
| *removed edges* | 0 |
| *prop rem edges* | 0 |

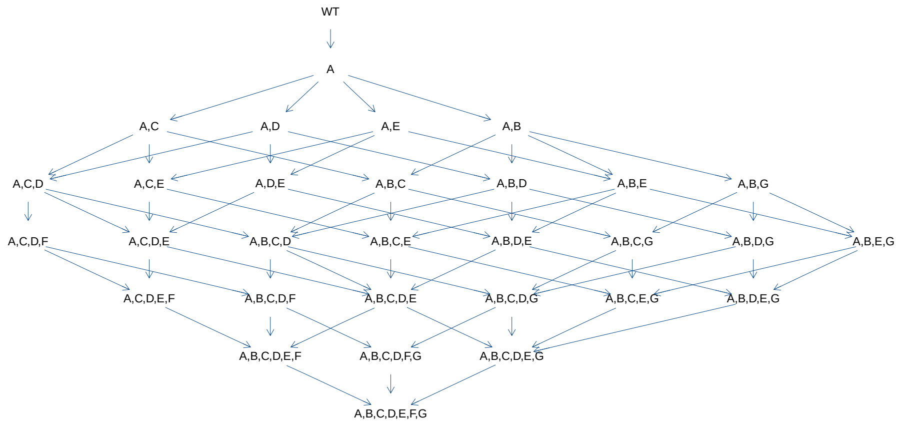


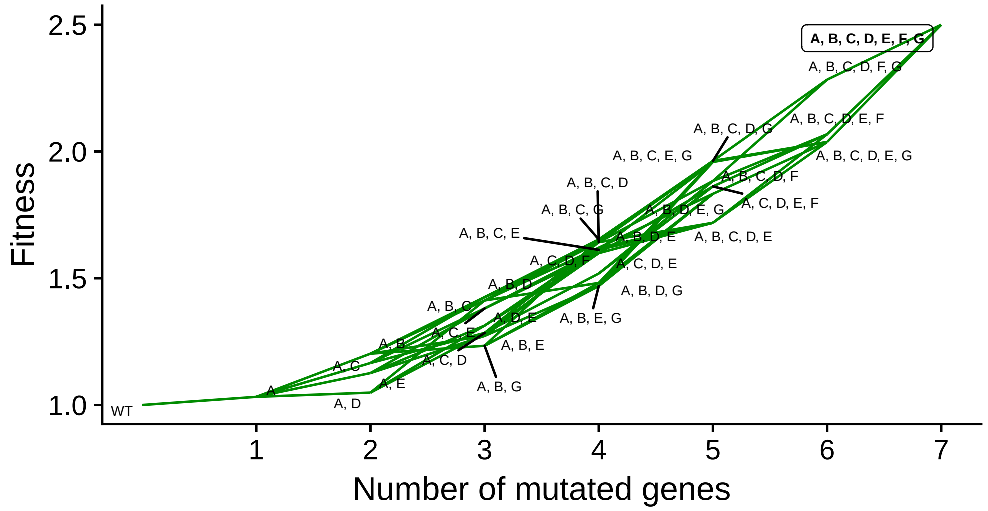

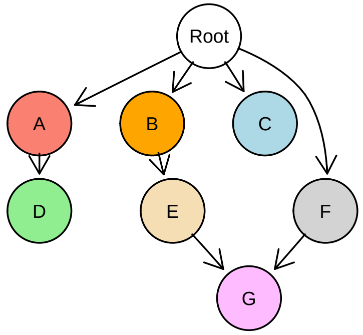


| | |
|---|---|
| *ID* | IJ15PYr1XTbhonDL |
| *acc genotypes* | 42 |
| *peaks no back* | 1 |
| *removed edges* | 0 |
| *prop rem edges* | 0 |

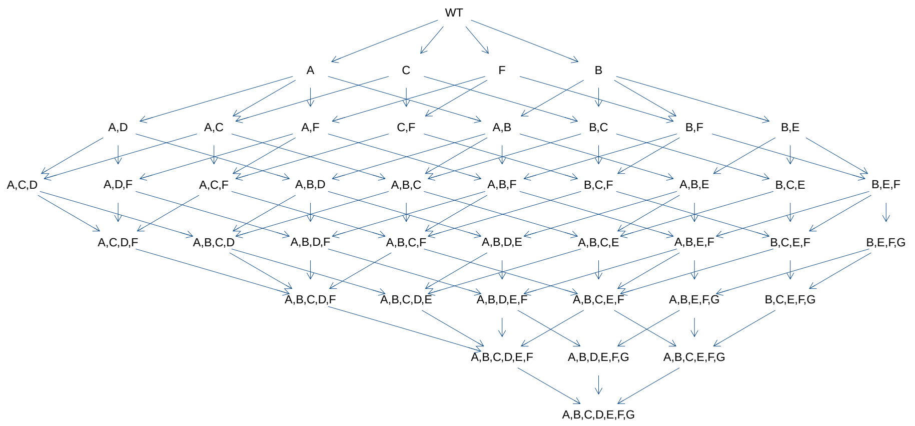


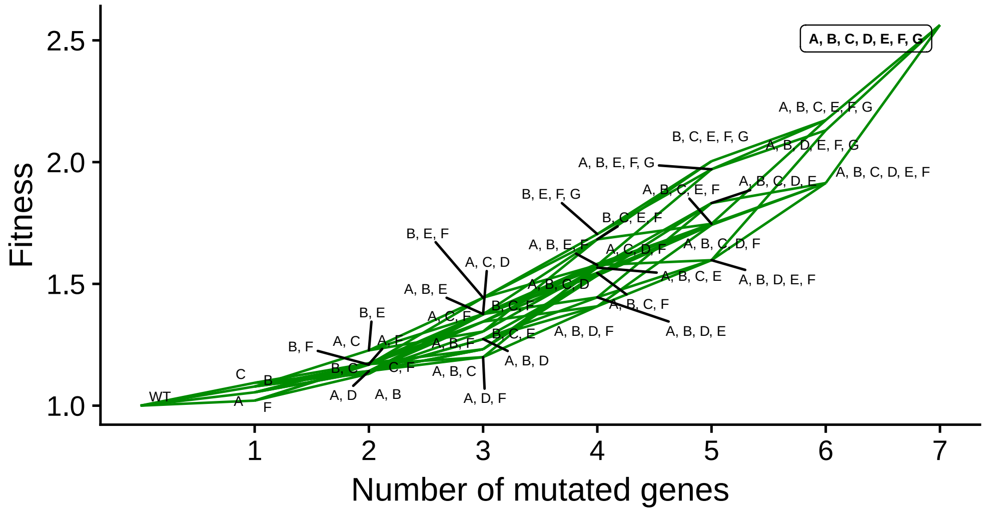

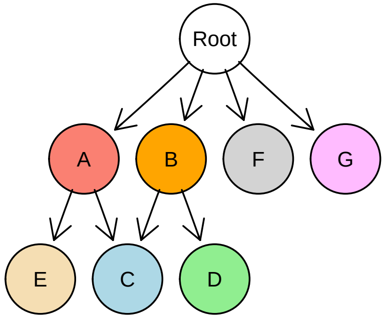


| | |
|---|---|
| *ID* | iKHz2mPyuFYjaiPo |
| *acc genotypes* | 52 |
| *peaks no back* | 1 |
| *removed edges* | 0 |
| *prop rem edges* | 0 |

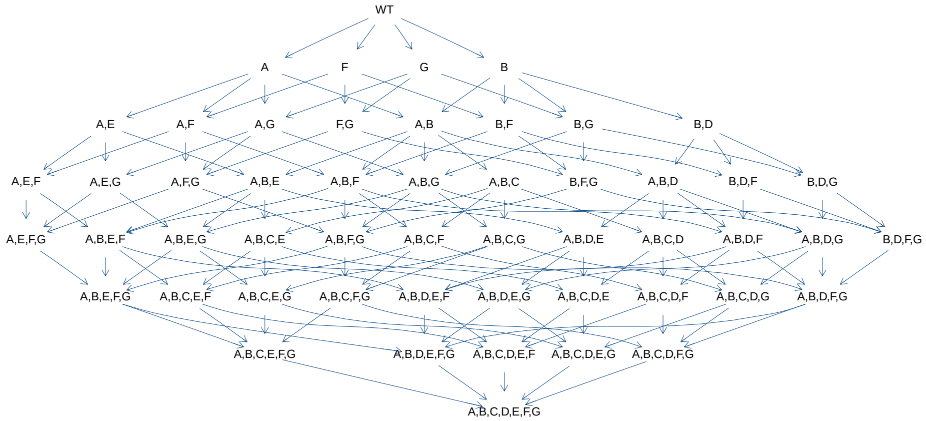


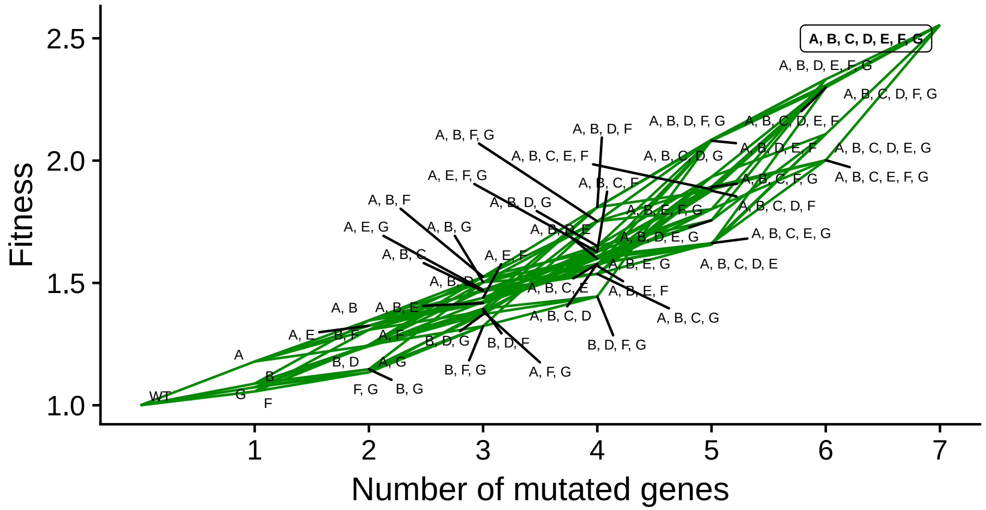

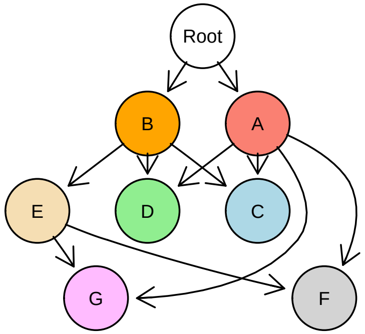


| | |
|---|---|
| *ID* | imGyIoiov7jJGQPM |
| *acc genotypes* | 24 |
| *peaks no back* | 1 |
| *removed edges* | 0 |
| *prop rem edges* | 0 |

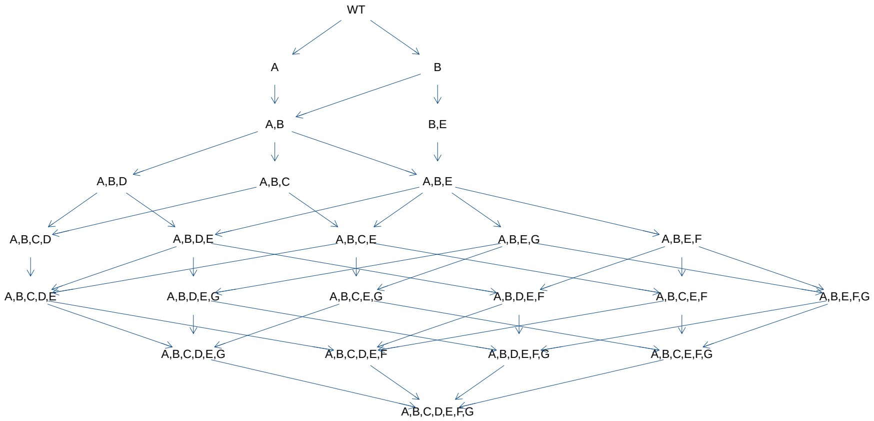


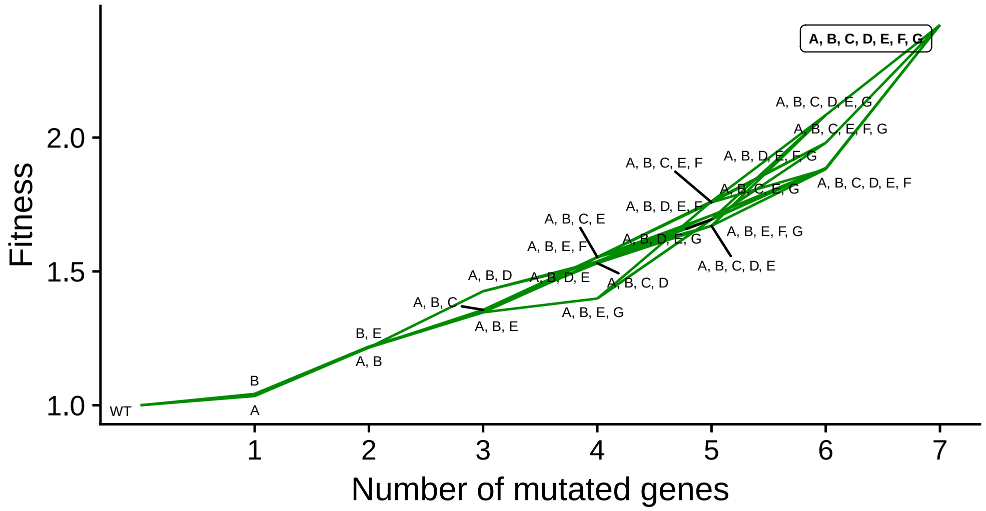

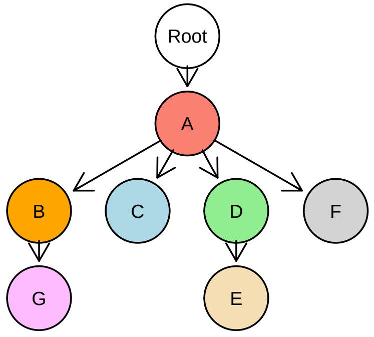


| | |
|---|---|
| *ID* | Iq2ILDZ3MG3ndder |
| *acc genotypes* | 37 |
| *peaks no back* | 1 |
| *removed edges* | 0 |
| *prop rem edges* | 0 |

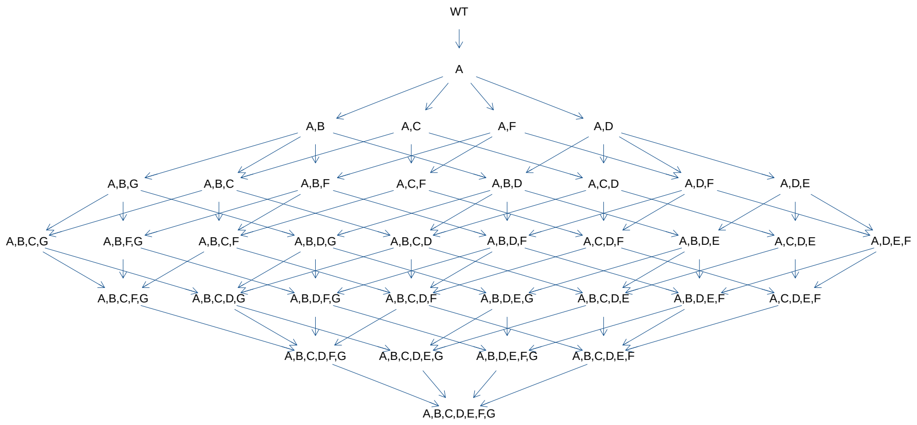


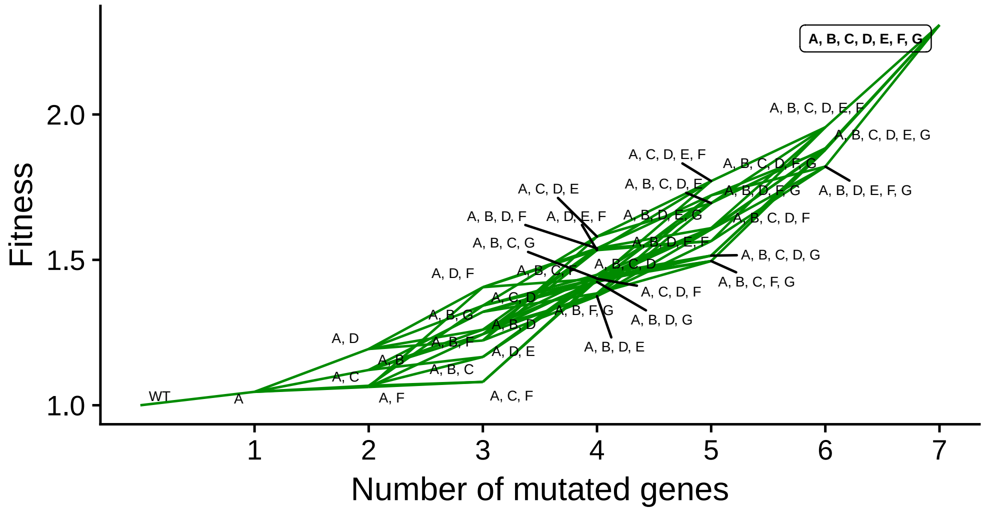

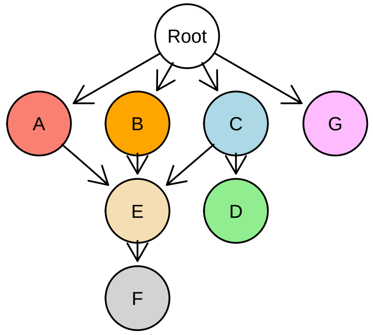


| | |
|---|---|
| *ID* | IRpnRnTLcDvmGnyE |
| *acc genotypes* | 32 |
| *peaks no back* | 1 |
| *removed edges* | 0 |
| *prop rem edges* | 0 |

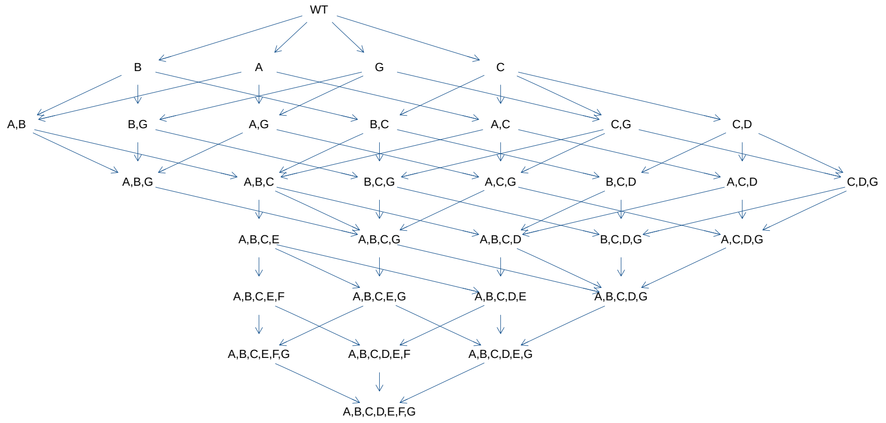


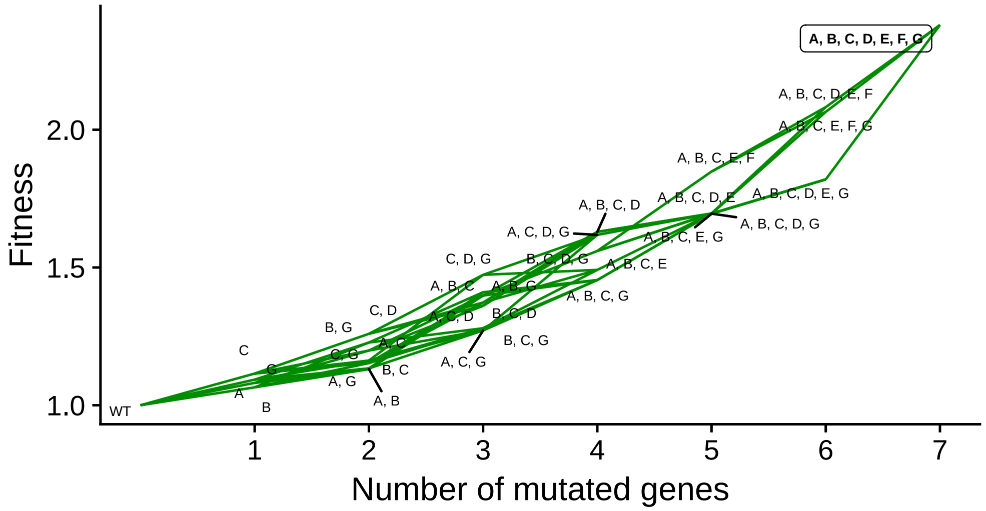

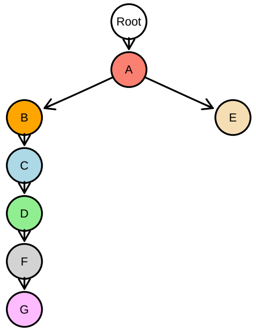


| | |
|---|---|
| *ID* | iU2gBfqvhEyguOWr |
| *acc genotypes* | 13 |
| *peaks no back* | 1 |
| *removed edges* | 0 |
| *prop rem edges* | 0 |

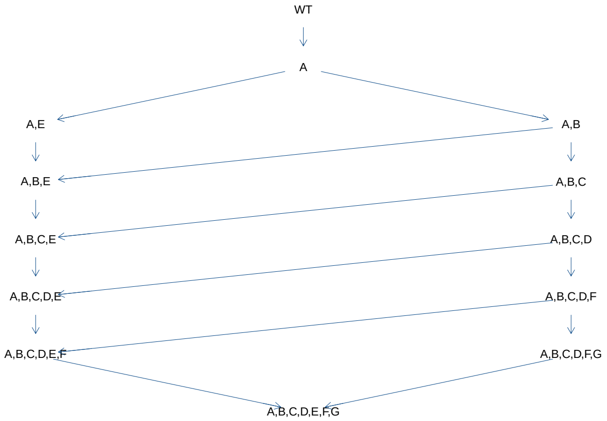


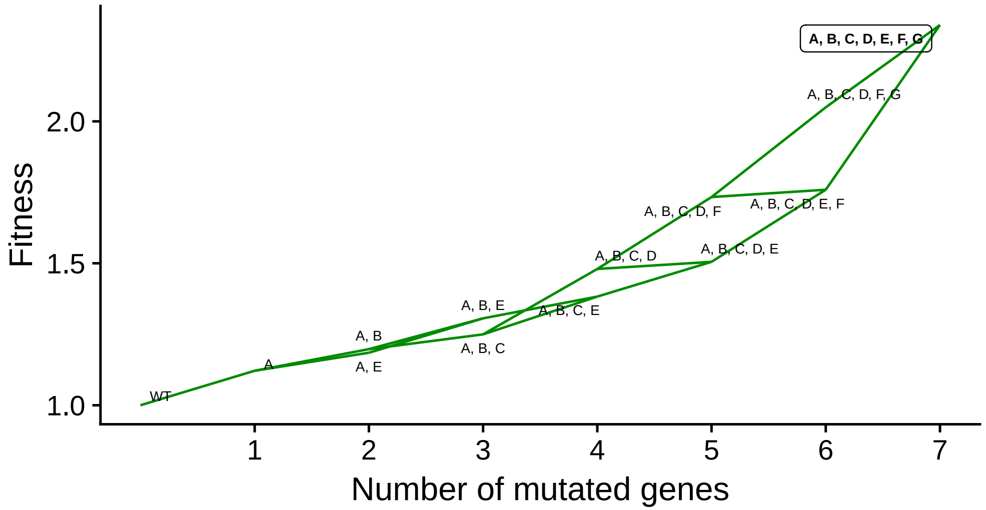

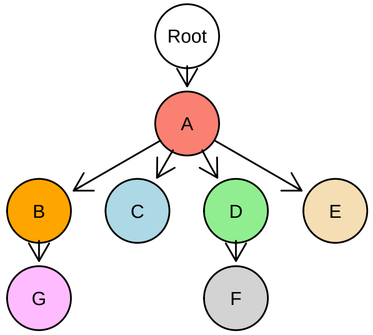


| | |
|---|---|
| *ID* | IySPgbGJgYzQfV5k |
| *acc genotypes* | 37 |
| *peaks no back* | 1 |
| *removed edges* | 0 |
| *prop rem edges* | 0 |

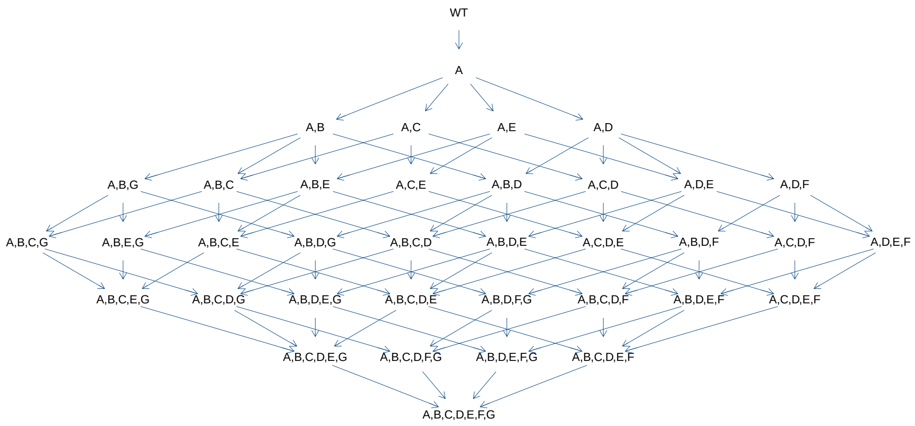


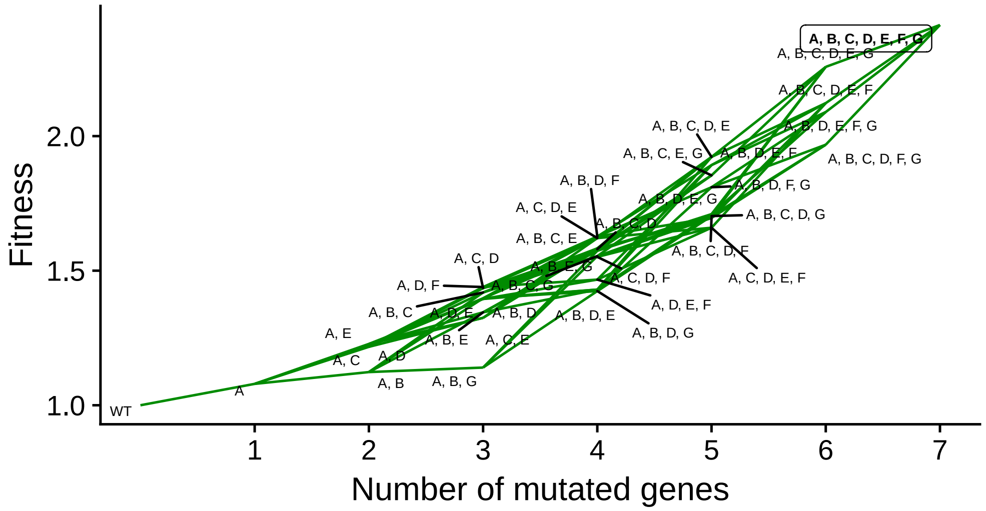

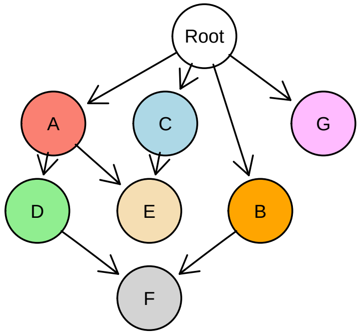


| | |
|---|---|
| *ID* | J5q7C41n6m4aEUp10 |
| *acc genotypes* | 38 |
| *peaks no back* | 1 |
| *removed edges* | 0 |
| *prop rem edges* | 0 |

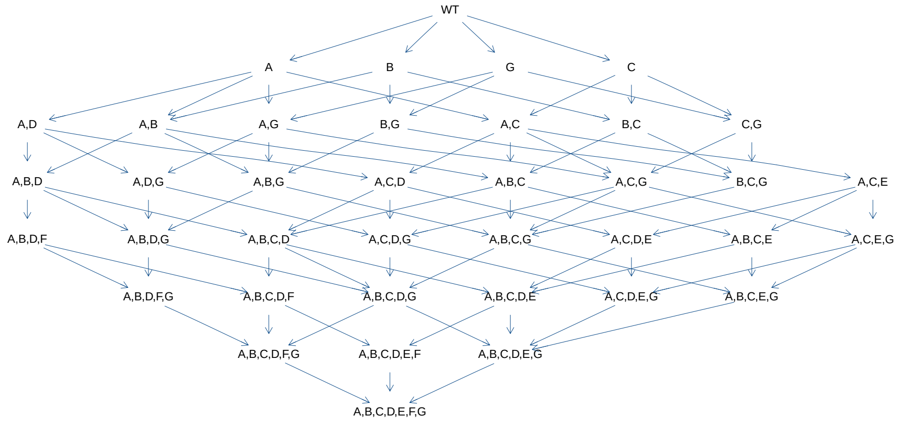


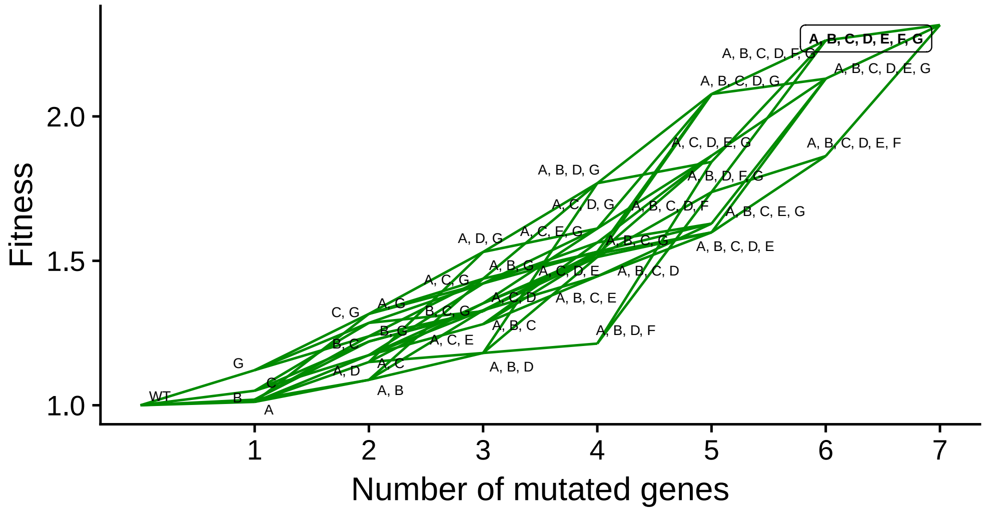

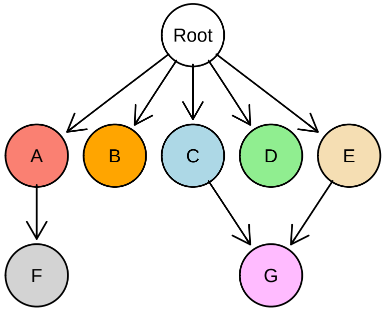


*ID* j5YLGuLuDa6cvNb2
*acc genotypes* 60
*peaks no back* 1
*removed edges* 0
*prop rem edges* 0

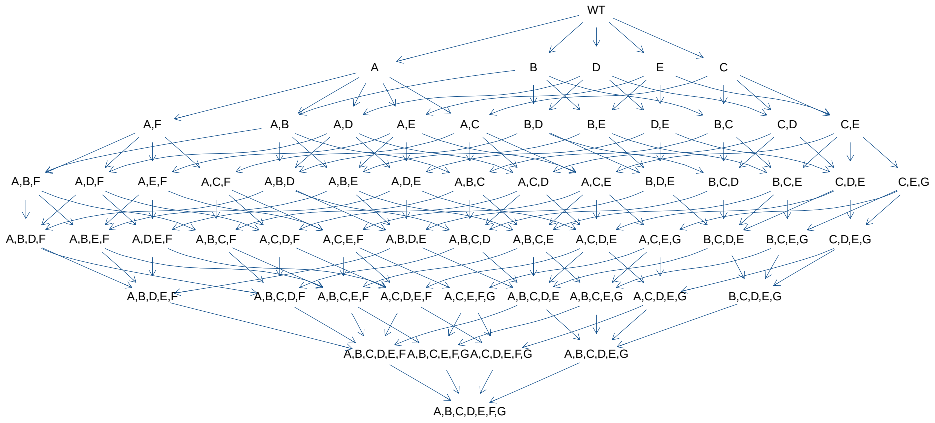


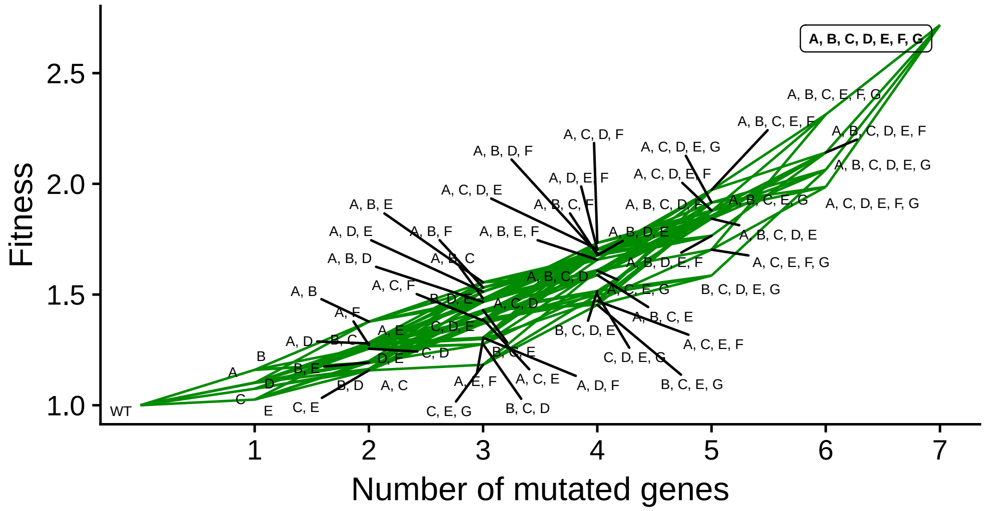

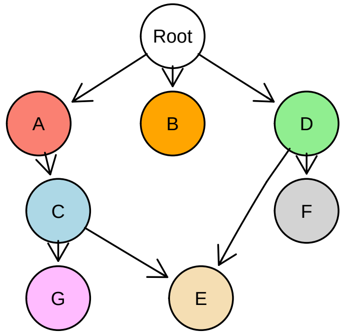


*ID* J6Itoaqms6HIPc7B
*acc genotypes* 32
*peaks no back* 1
*removed edges* 0
*prop rem edges* 0

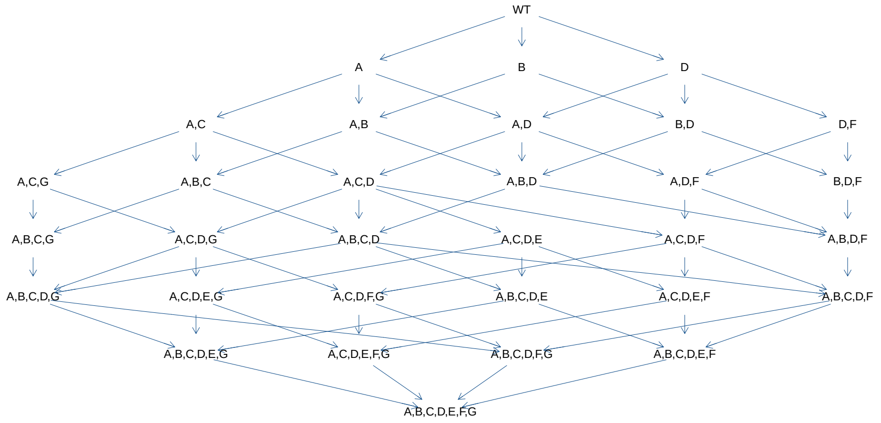


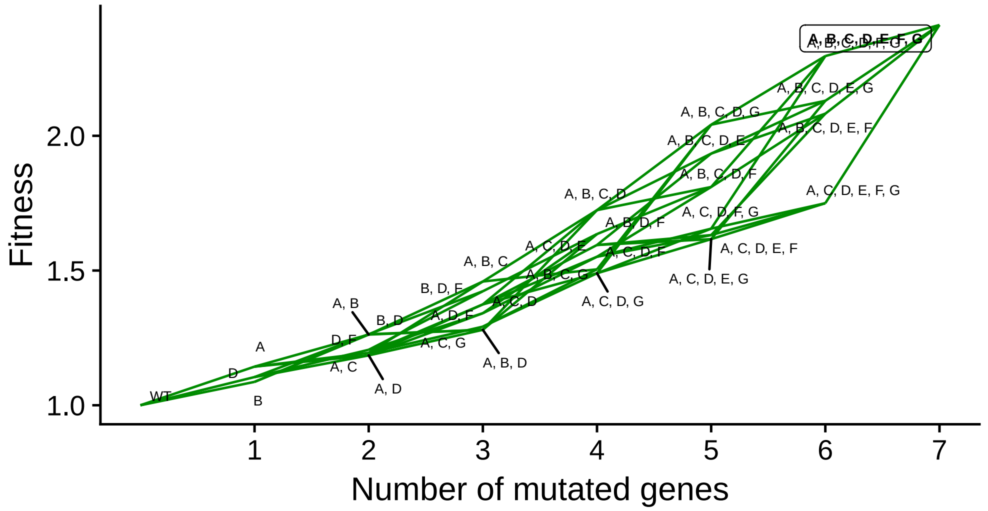

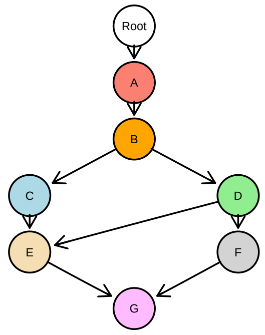


| | |
|---|---|
| *ID* | j71sbZK1fQ6ihd8O |
| *acc genotypes* | 11 |
| *peaks no back* | 1 |
| *removed edges* | 0 |
| *prop rem edges* | 0 |

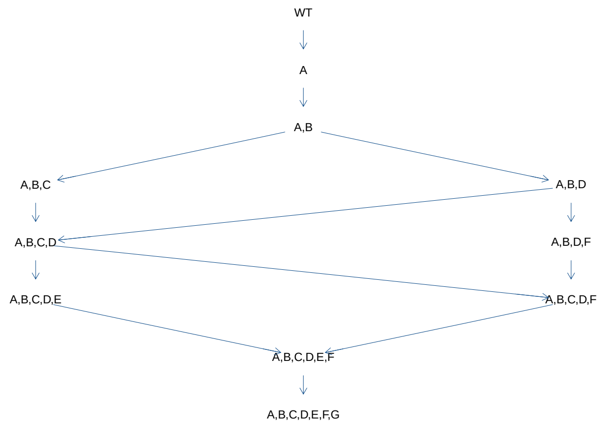


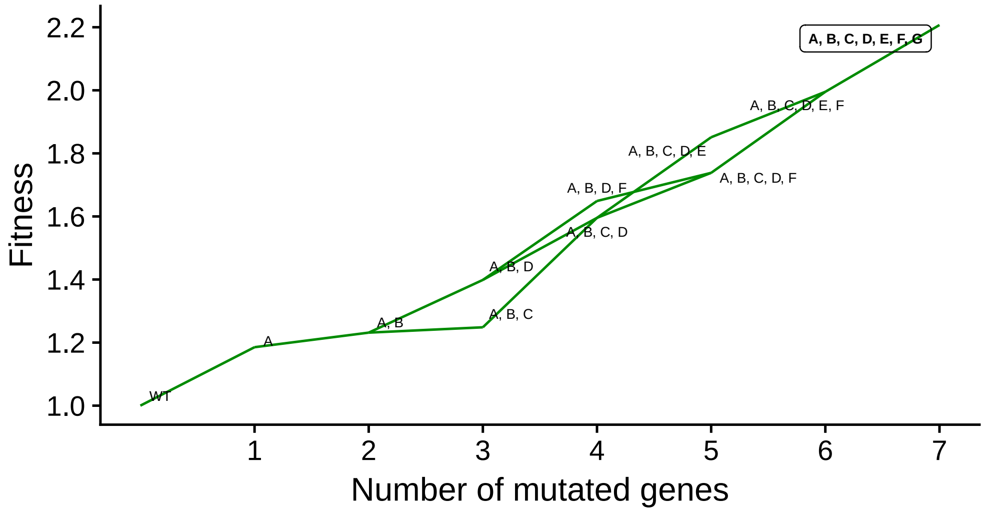

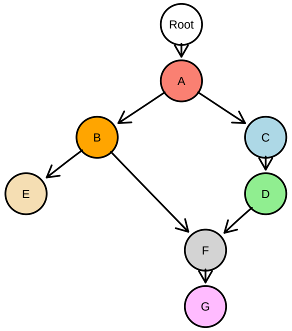


| | |
|---|---|
| *ID* | JLERmKvl2w2lquvc |
| *acc genotypes* | 14 |
| *peaks no back* | 1 |
| *removed edges* | 0 |
| *prop rem edges* | 0 |

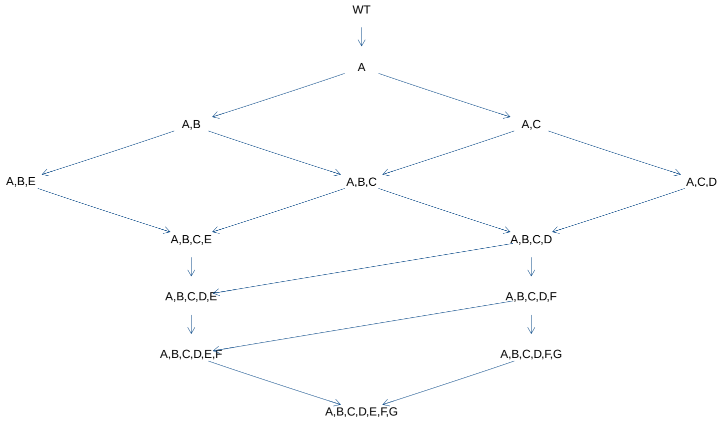


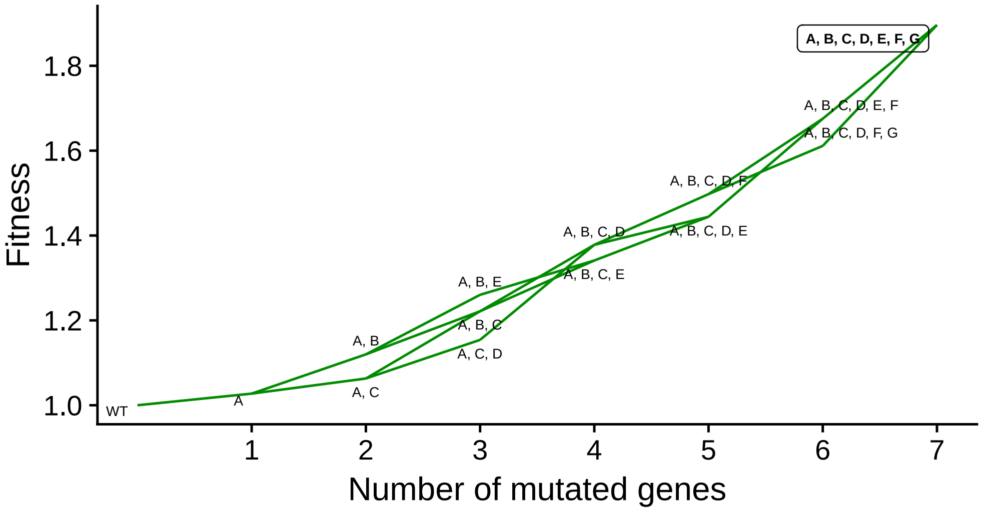

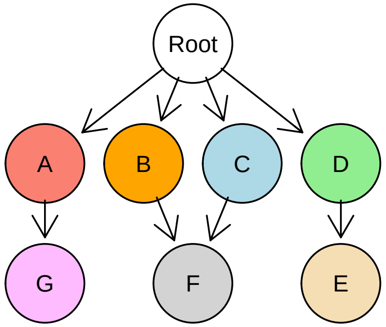


| | |
|---|---|
| *ID* | JoTSpt4LngDDMASj |
| *acc genotypes* | 45 |
| *peaks no back* | 1 |
| *removed edges* | 0 |
| *prop rem edges* | 0 |

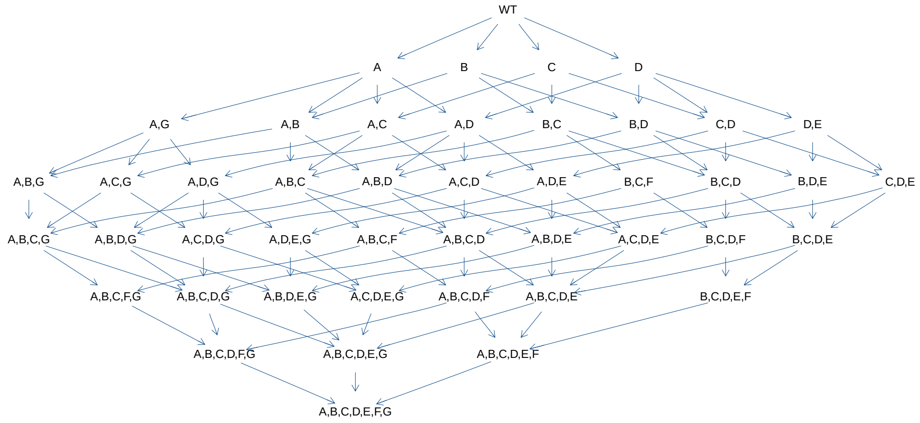


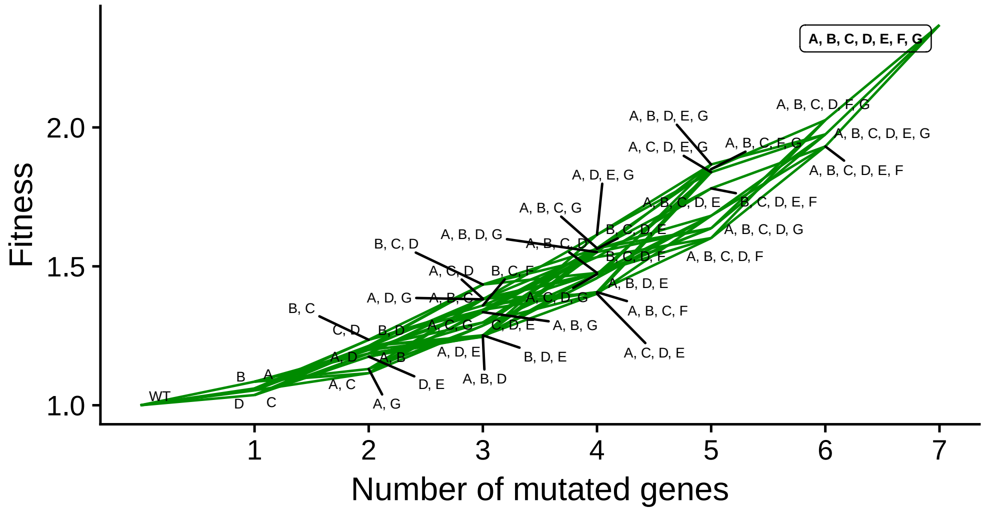

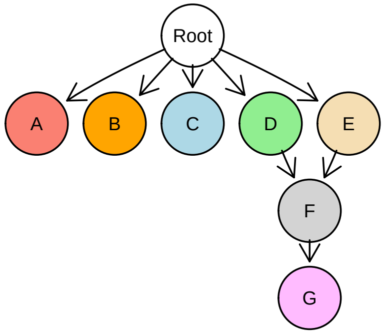


| | |
|---|---|
| *ID* | jq7mqYXKFpls2uXm |
| *acc genotypes* | 48 |
| *peaks no back* | 1 |
| *removed edges* | 0 |
| *prop rem edges* | 0 |

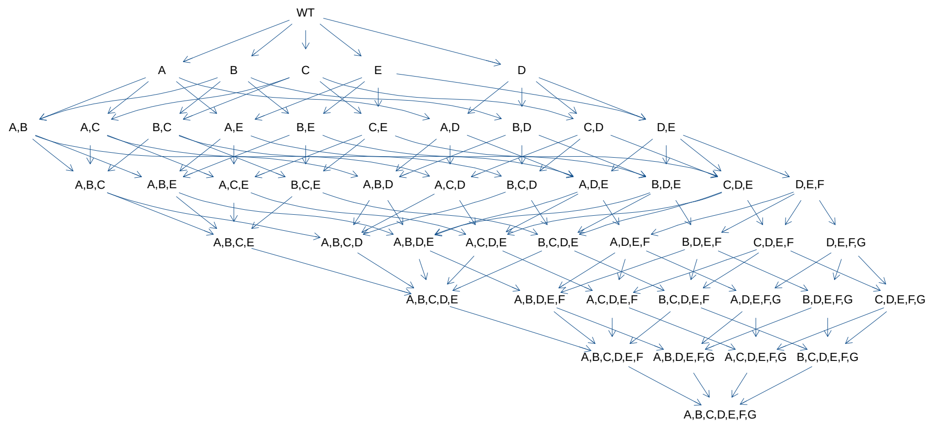


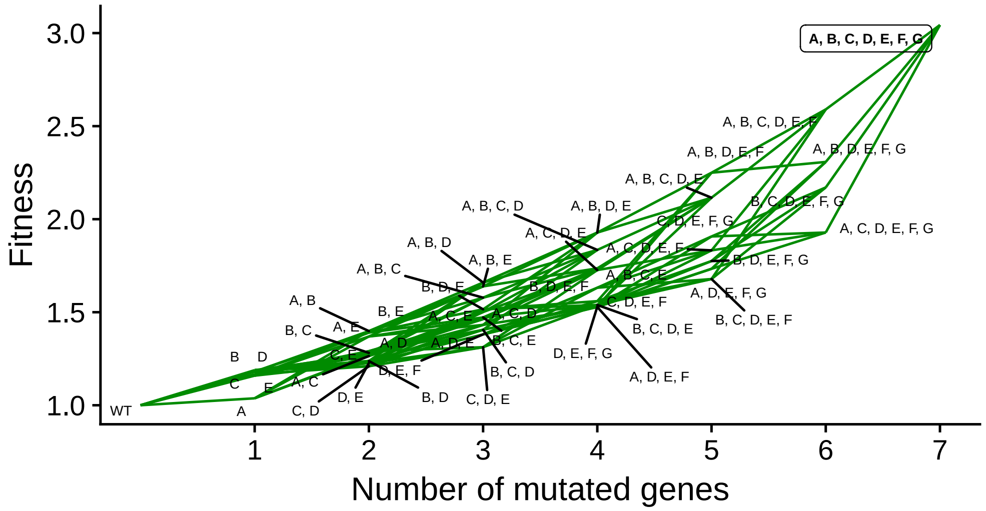

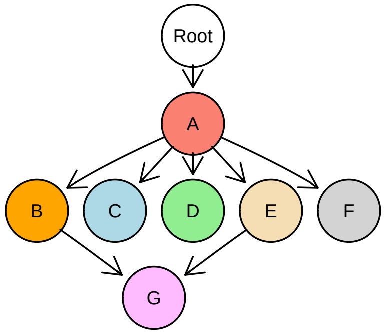


*ID* JVEFSk5ORIOz9YE7
*acc genotypes* 41
*peaks no back* 1
*removed edges* 0
*prop rem edges* 0

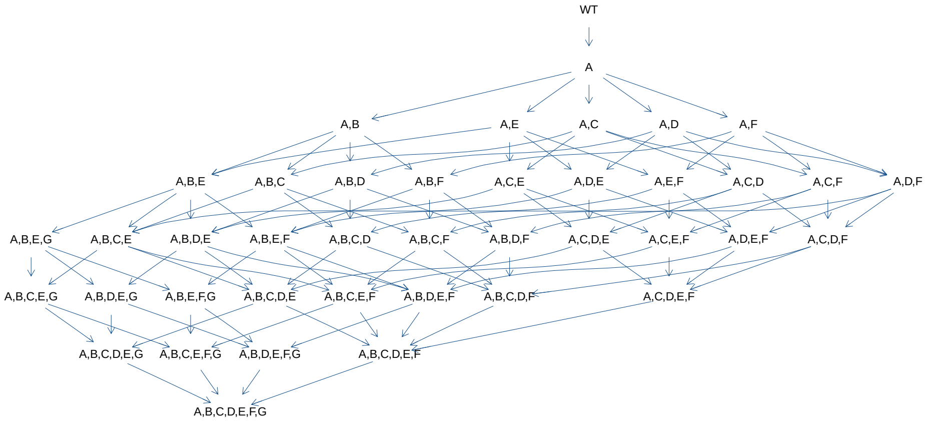


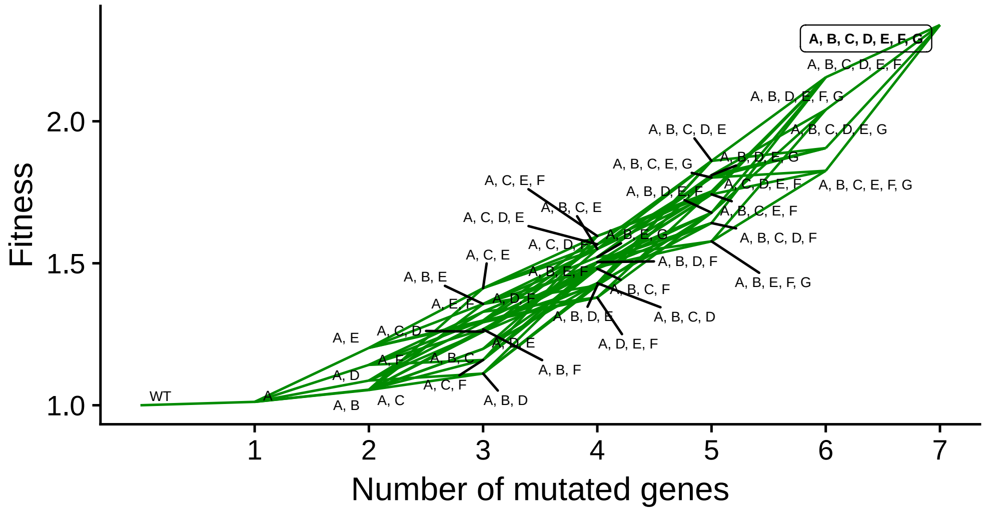

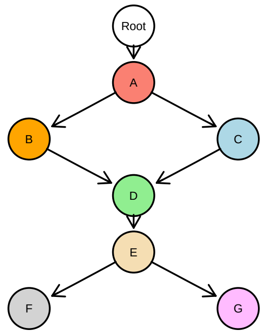


| | |
|---|---|
| *ID* | K9x9g2eq1VT8fICv |
| *acc genotypes* | 10 |
| *peaks no back* | 1 |
| *removed edges* | 0 |
| *prop rem edges* | 0 |

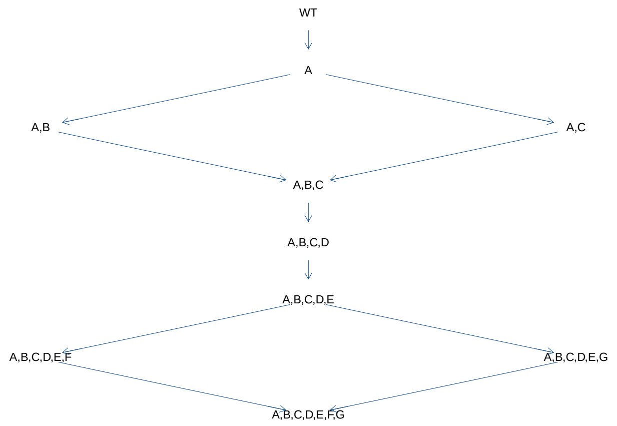


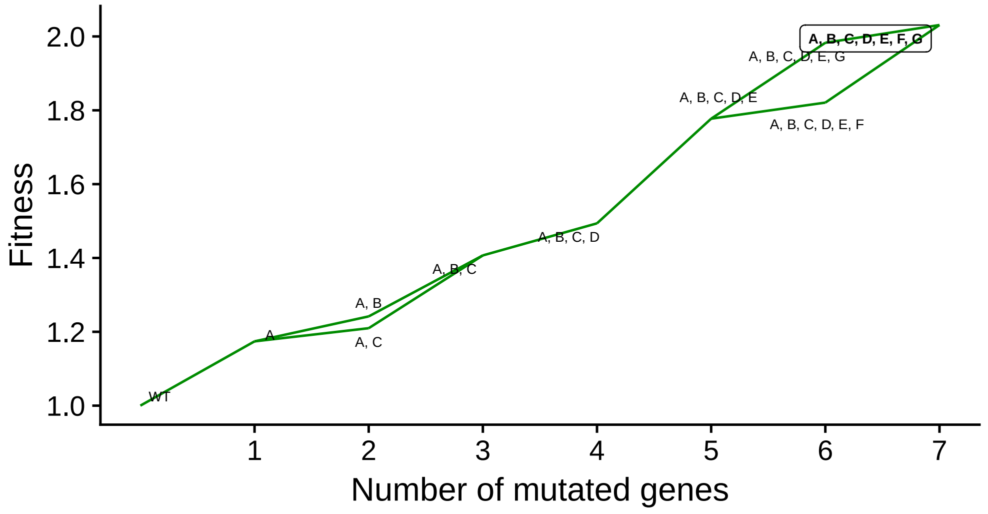

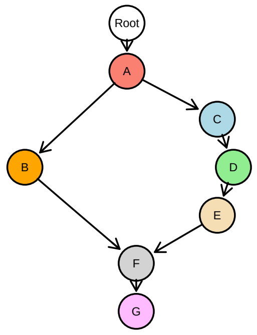


| | |
|---|---|
| *ID* | kBK5zr8jiuD8Etoo |
| *acc genotypes* | 11 |
| *peaks no back* | 1 |
| *removed edges* | 0 |
| *prop rem edges* | 0 |

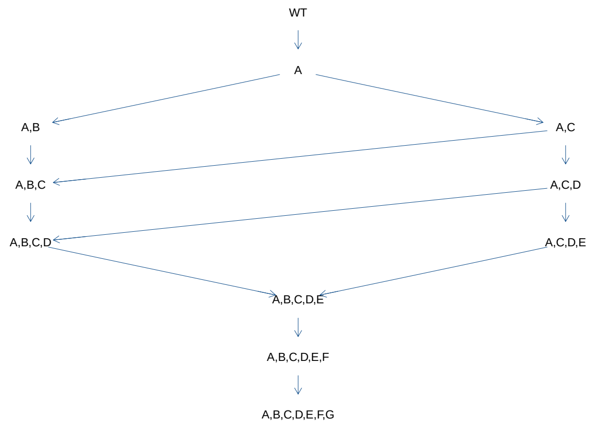


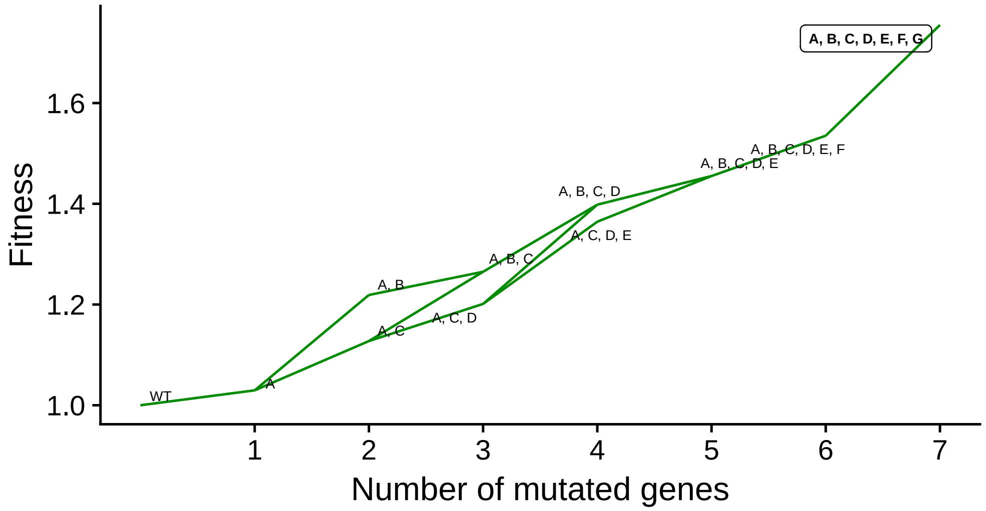

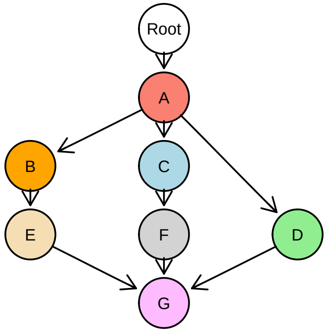


| | |
|---|---|
| *ID* | KctNTtEFQr1oMvyT |
| *acc genotypes* | 20 |
| *peaks no back* | 1 |
| *removed edges* | 0 |
| *prop rem edges* | 0 |

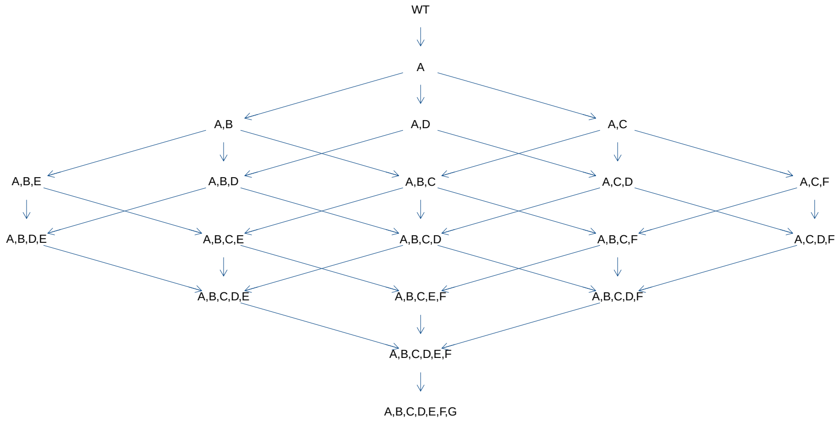


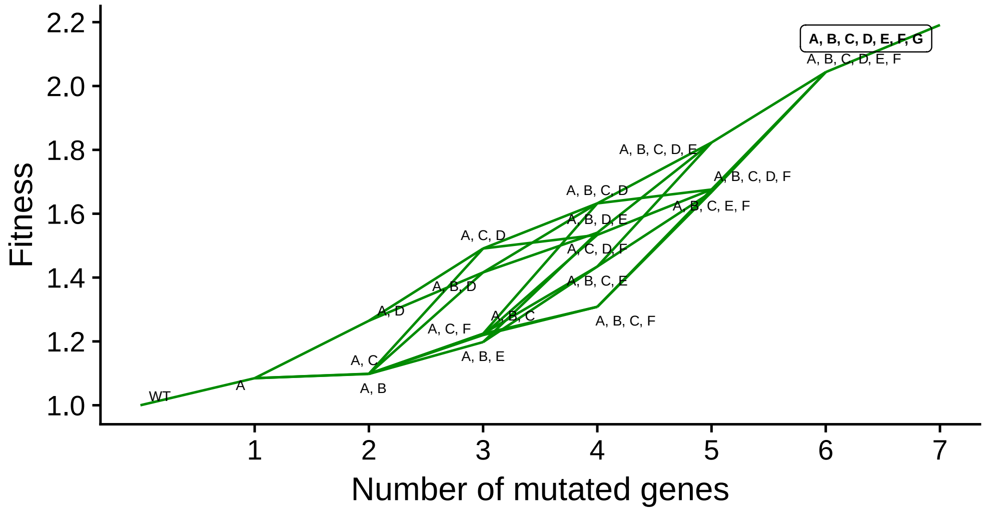

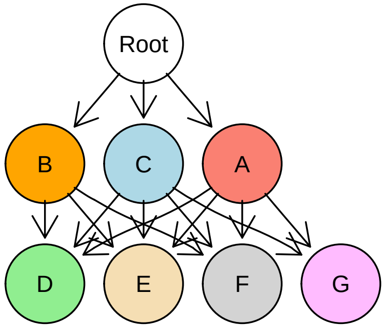


| | |
|---|---|
| *ID* | kf9mgLp5ZKwjcVhz |
| *acc genotypes* | 24 |
| *peaks no back* | 1 |
| *removed edges* | 0 |
| *prop rem edges* | 0 |

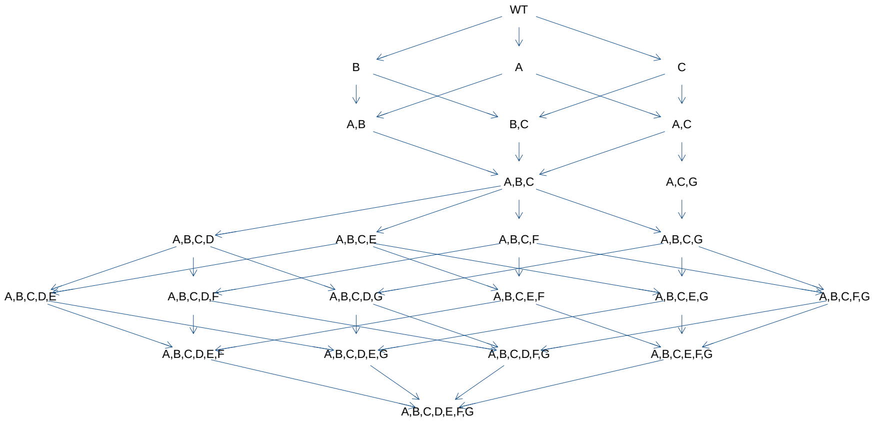


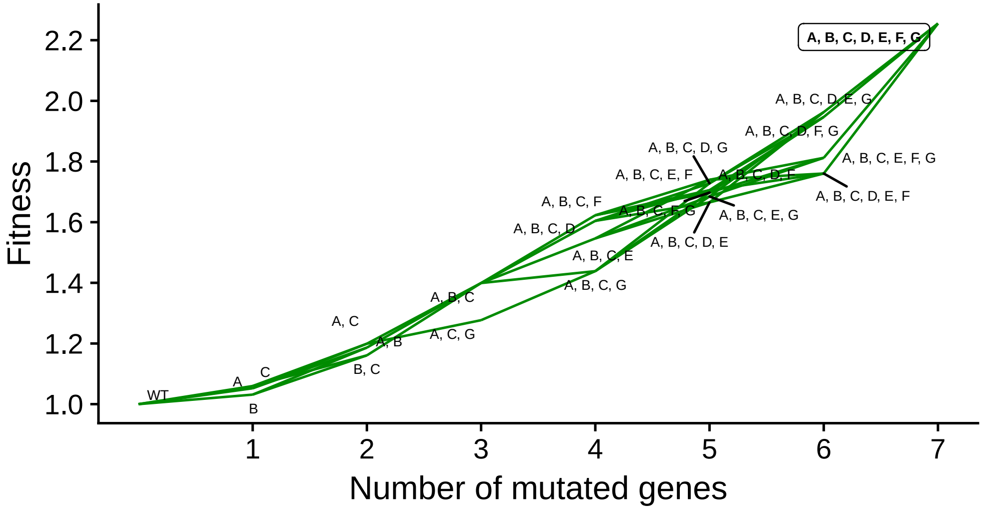

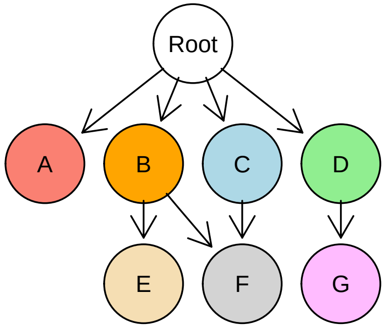


| | |
|---|---|
| *ID* | KtmVfIOzxH44NUrR |
| *acc genotypes* | 48 |
| *peaks no back* | 1 |
| *removed edges* | 0 |
| *prop rem edges* | 0 |

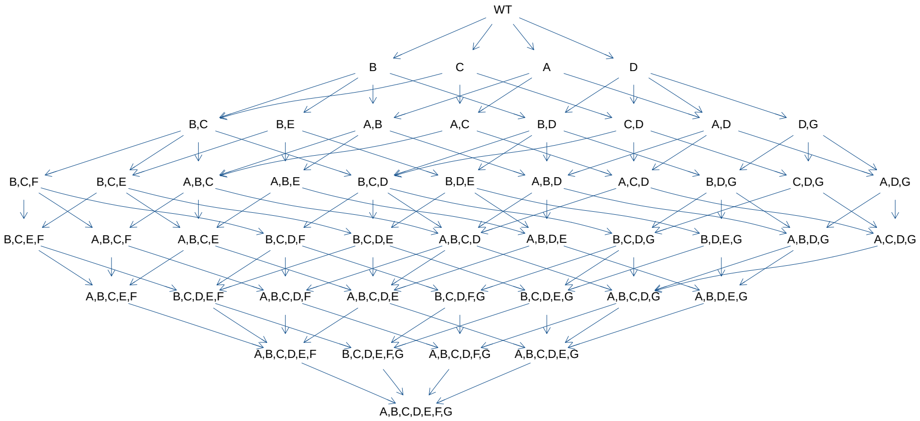

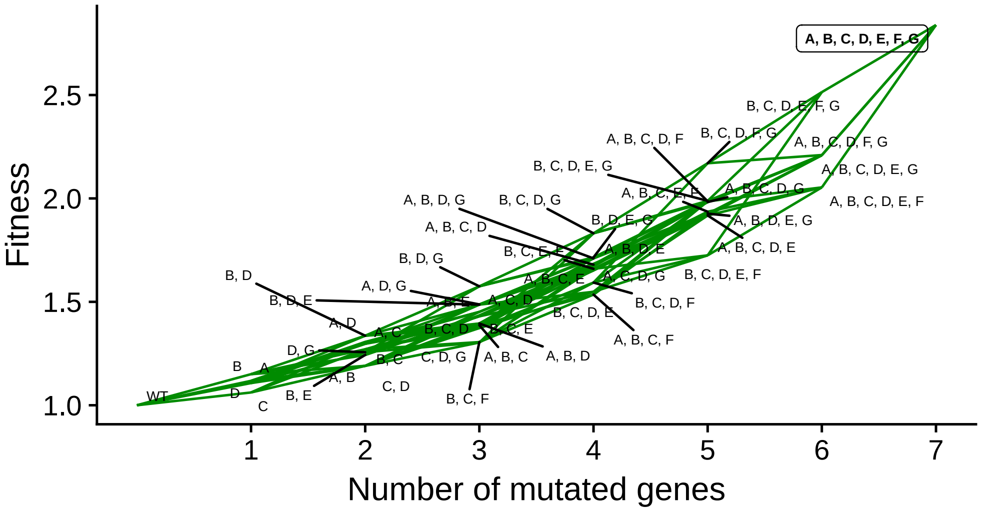

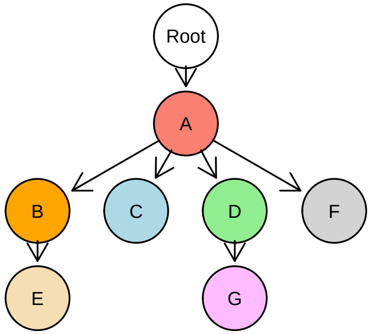


| | |
|---|---|
| *ID* | KZkcV1r2fzi9JPSr |
| *acc genotypes* | 37 |
| *peaks no back* | 1 |
| *removed edges* | 0 |
| *prop rem edges* | 0 |

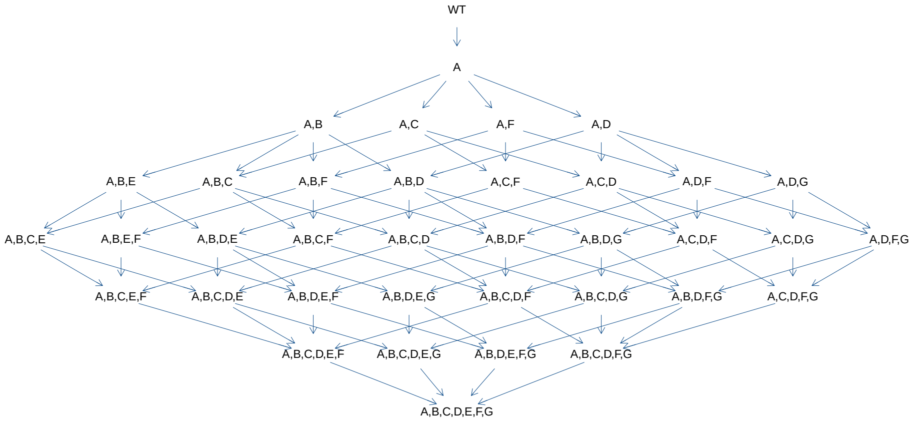


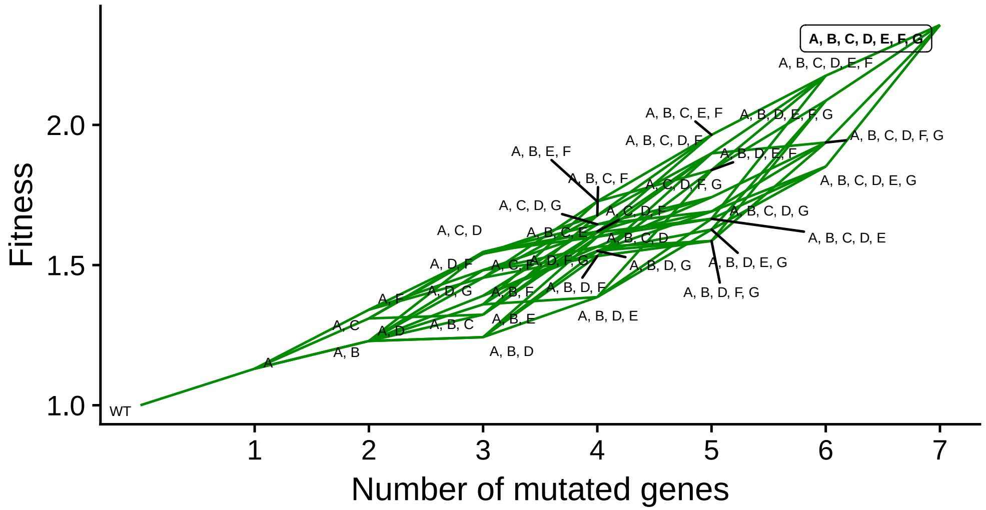

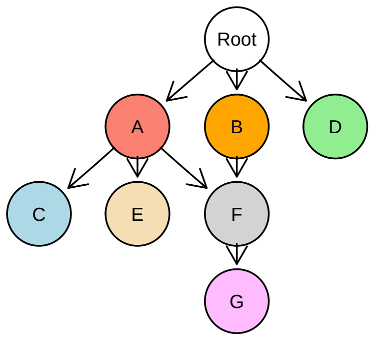


| | |
|---|---|
| *ID* | Lke10loDf84oef4BC |
| *acc genotypes* | 36 |
| *peaks no back* | 1 |
| *removed edges* | 0 |
| *prop rem edges* | 0 |

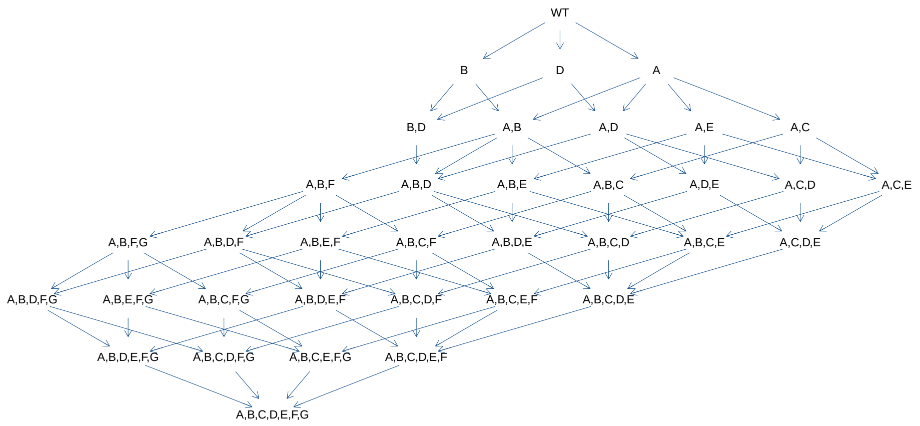


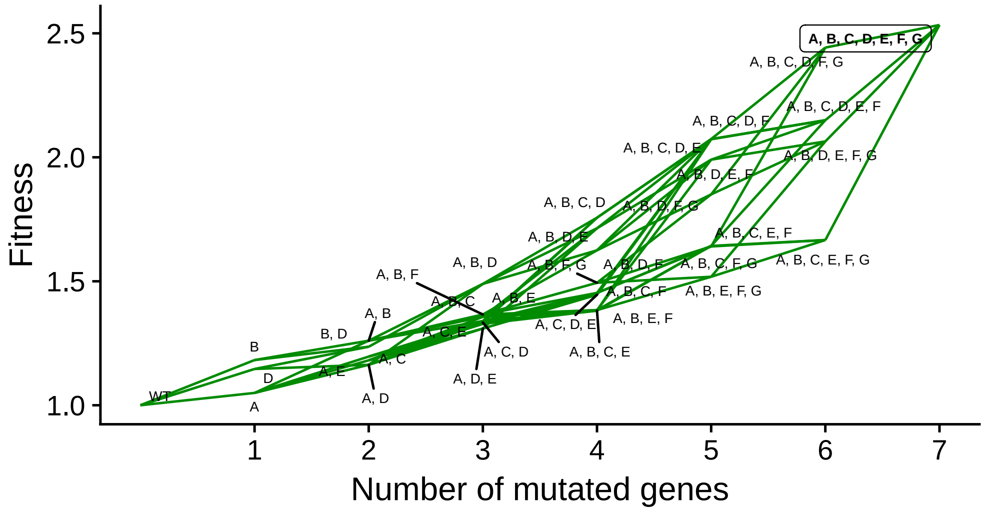

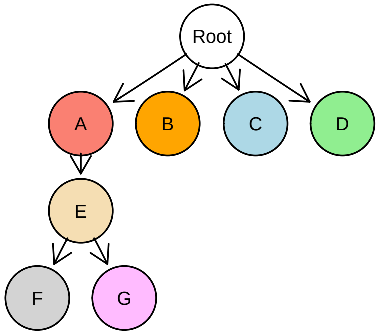


| | |
|---|---|
| *ID* | LIVHZ106EHMqYUsuF |
| *acc genotypes* | 48 |
| *peaks no back* | 1 |
| *removed edges* | 0 |
| *prop rem edges* | 0 |

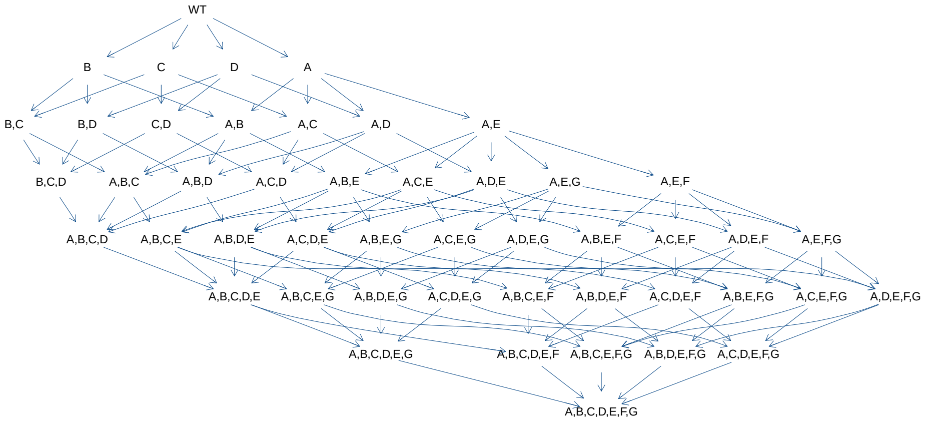


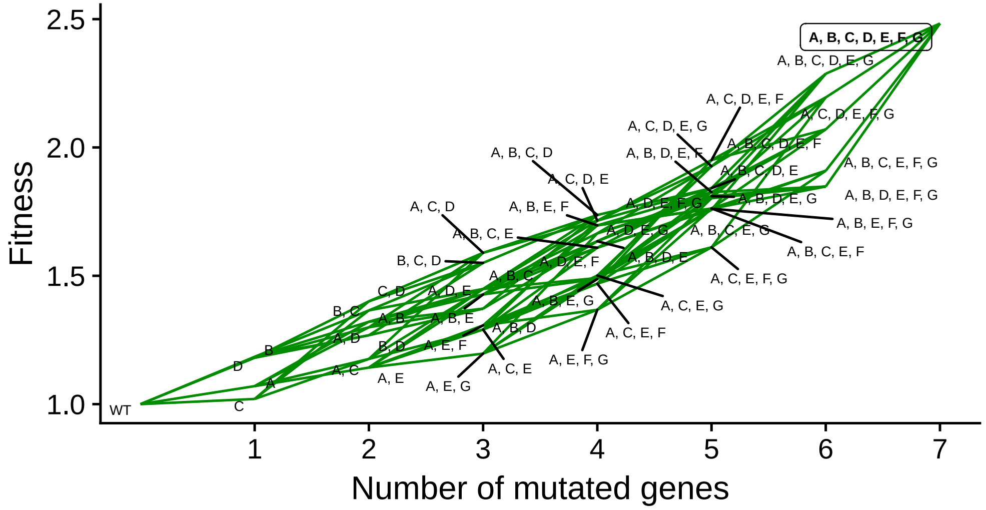

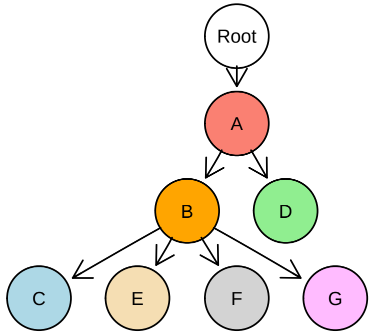


| | |
|---|---|
| *ID* | LpANxOHMhj1cYLtN |
| *acc genotypes* | 35 |
| *peaks no back* | 1 |
| *removed edges* | 0 |
| *prop rem edges* | 0 |

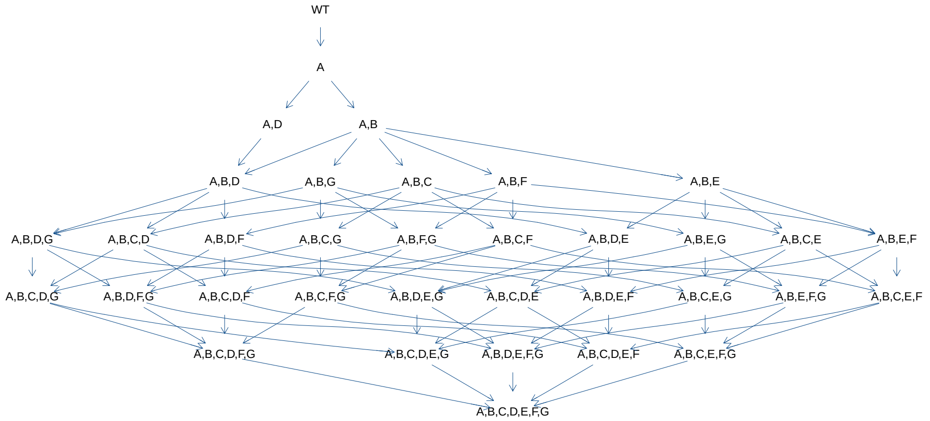


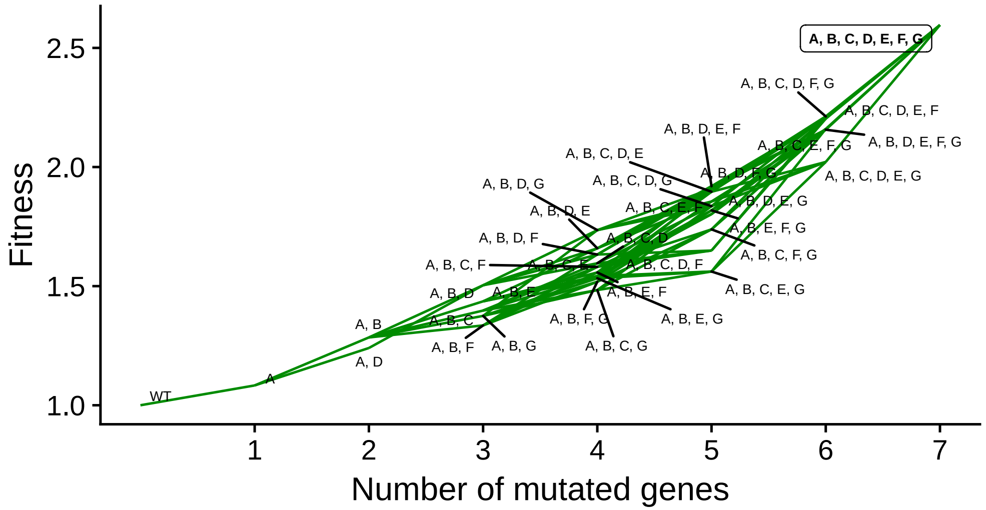

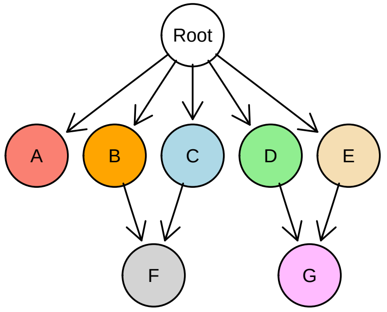


| | |
|---|---|
| *ID* | LTgRqknZ6BY1cfLB |
| *acc genotypes* | 50 |
| *peaks no back* | 1 |
| *removed edges* | 0 |
| *prop rem edges* | 0 |

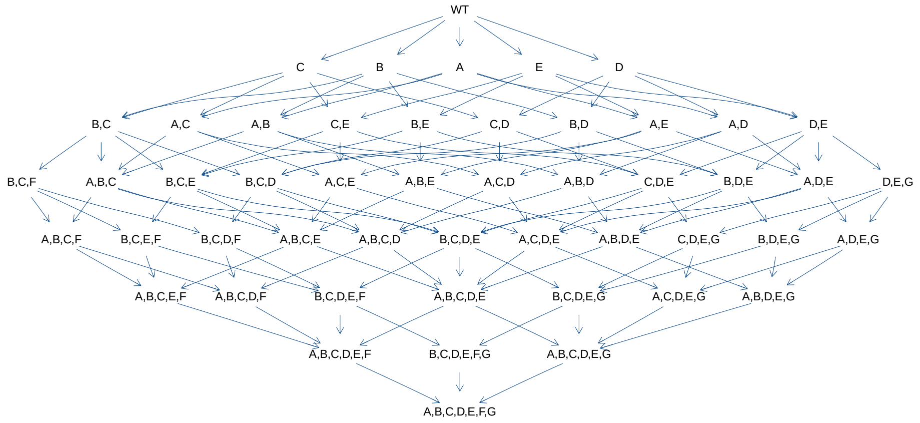


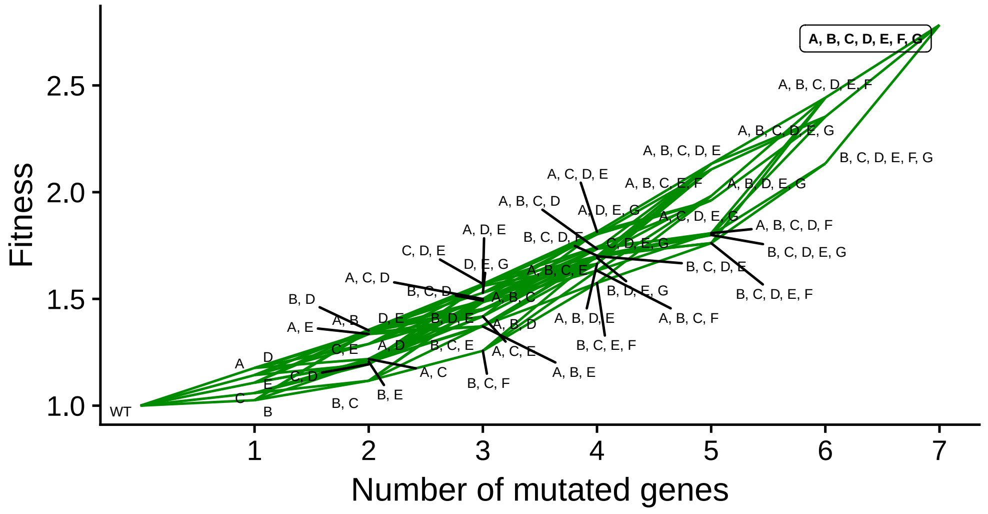

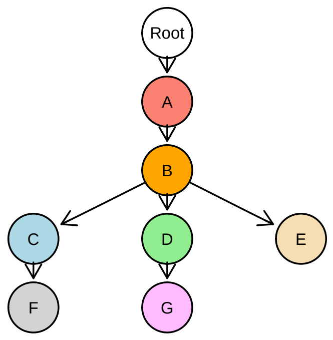


| | |
|---|---|
| *ID* | Mdi5kYcCCepKSprh |
| *acc genotypes* | 20 |
| *peaks no back* | 1 |
| *removed edges* | 0 |
| *prop rem edges* | 0 |

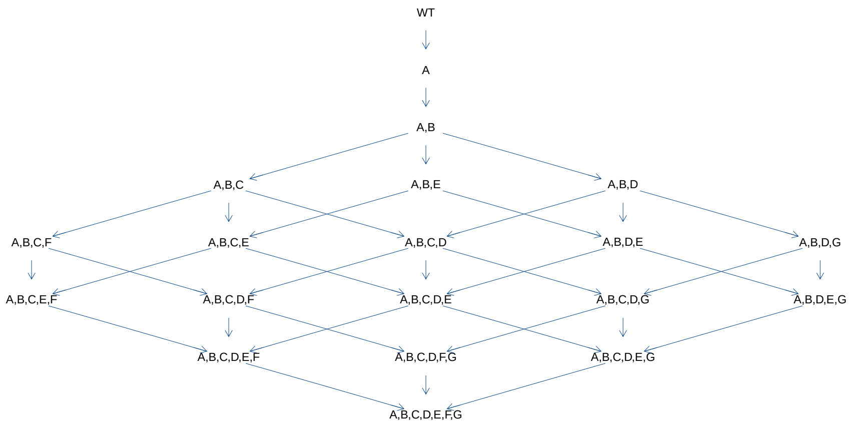


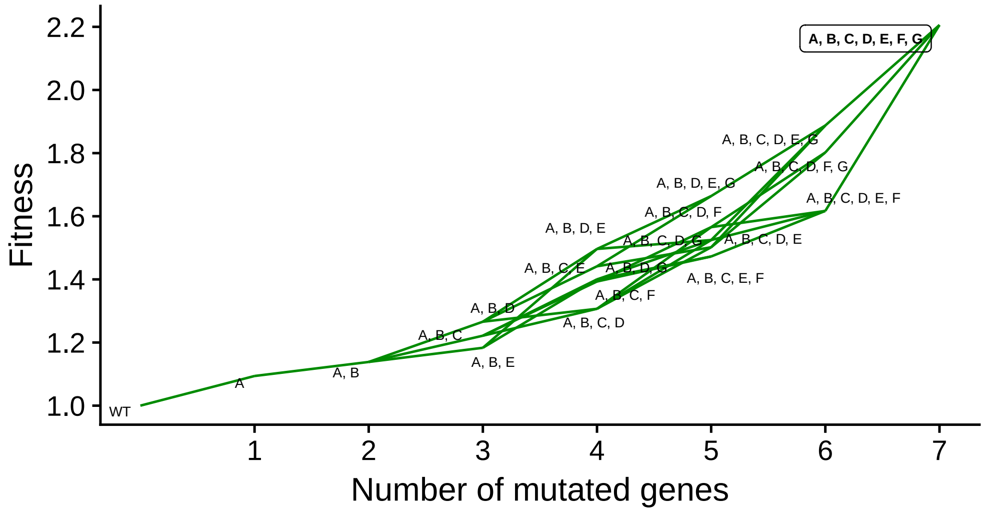

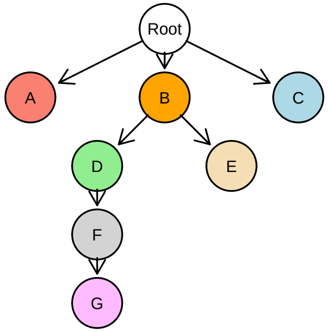


*ID* mEX5RjVtCmtkvI6C
*acc genotypes* 36
*peaks no back* 1
*removed edges* 0
*prop rem edges* 0

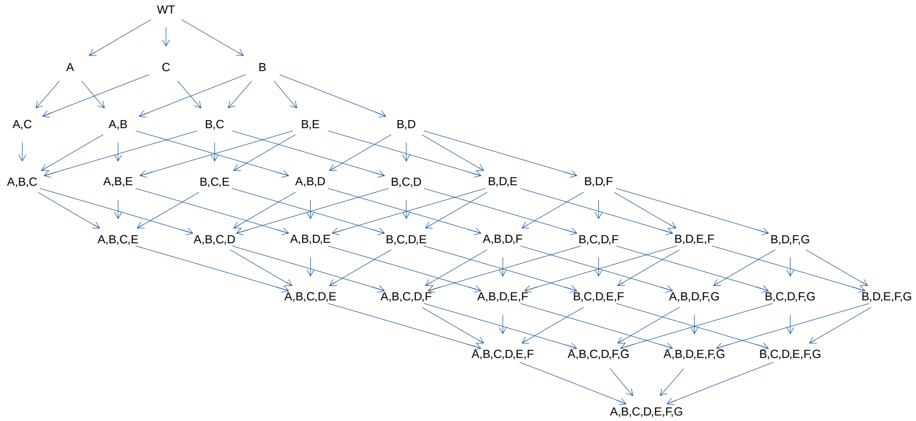


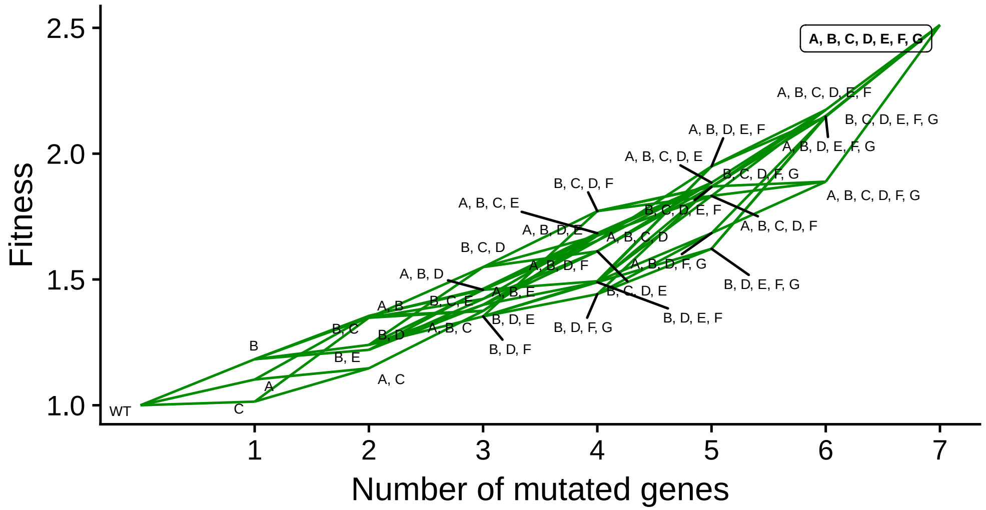

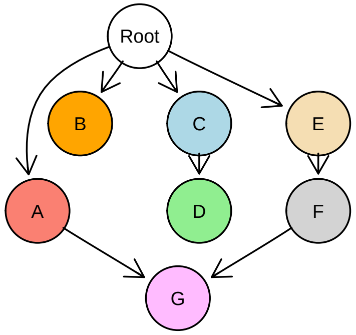


*ID* MHHdzPsMBi26wD8j
*acc genotypes* 42
*peaks no back* 1
*removed edges* 0
*prop rem edges* 0

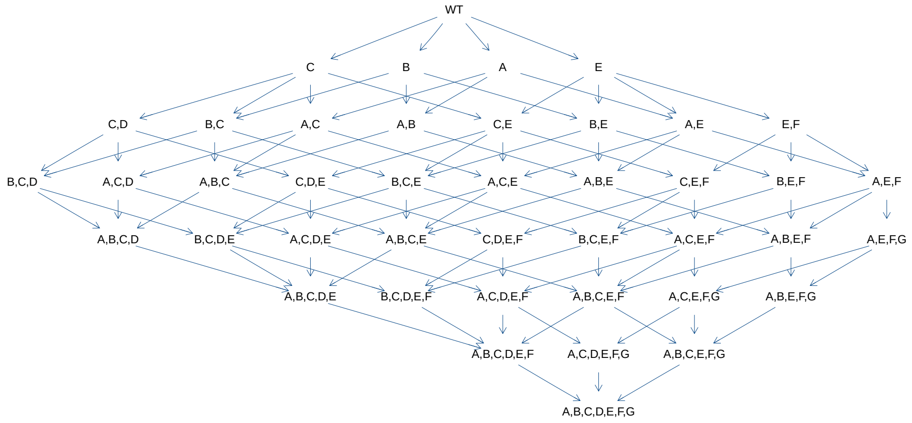


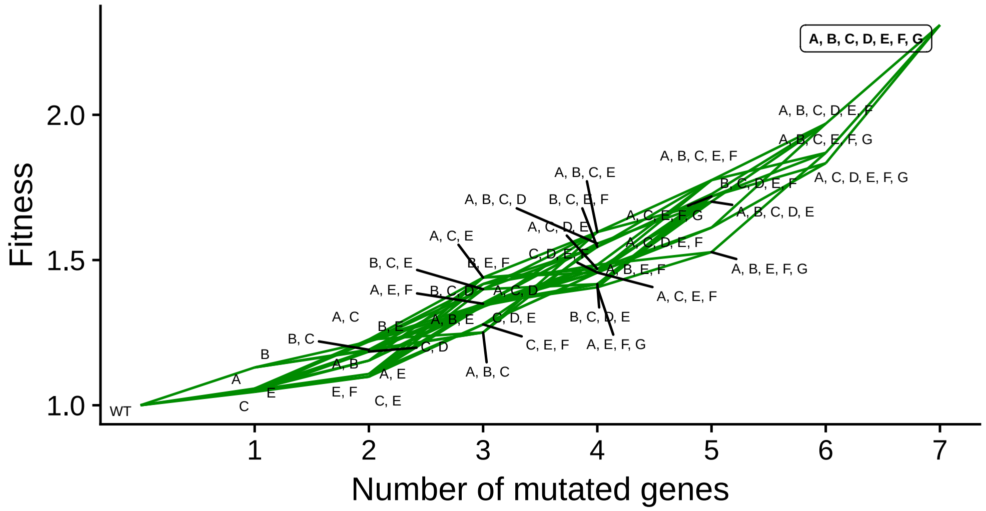

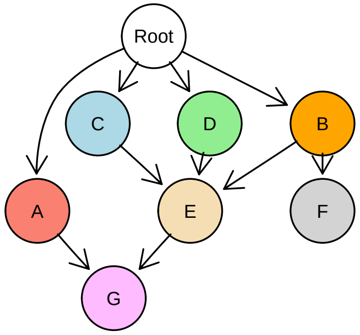


| | |
|---|---|
| *ID* | MiiXBqQH10YjcXVMI |
| *acc genotypes* | 30 |
| *peaks no back* | 1 |
| *removed edges* | 0 |
| *prop rem edges* | 0 |

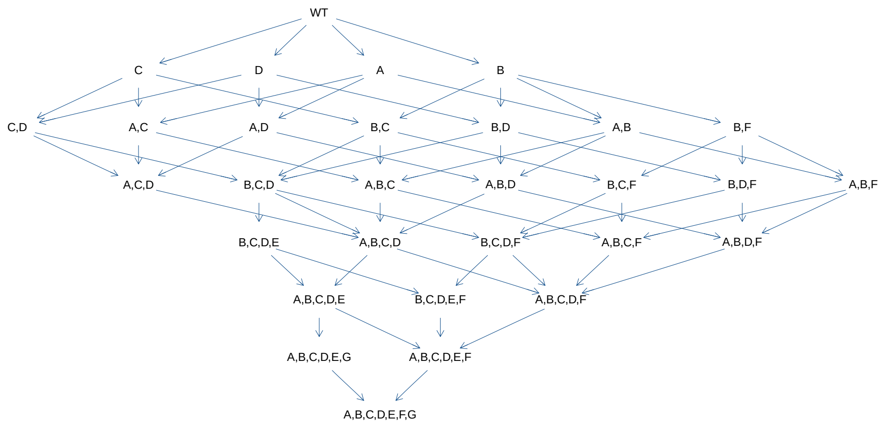


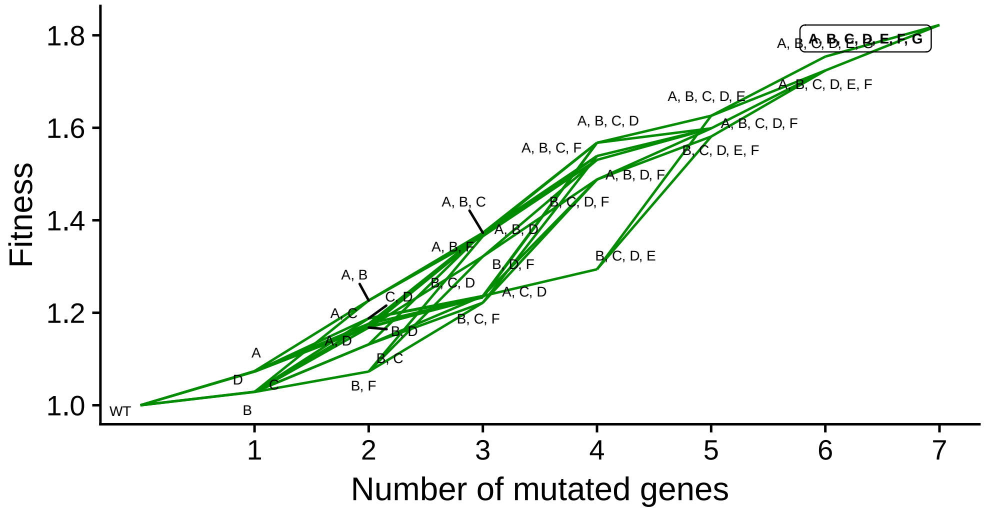

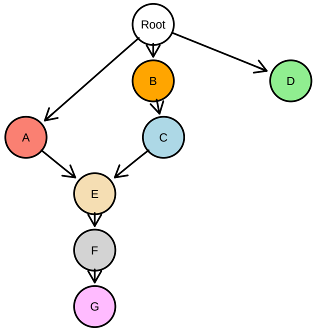


| | |
|---|---|
| *ID* | MJEstF1cdVazEIfA |
| *acc genotypes* | 18 |
| *peaks no back* | 1 |
| *removed edges* | 0 |
| *prop rem edges* | 0 |

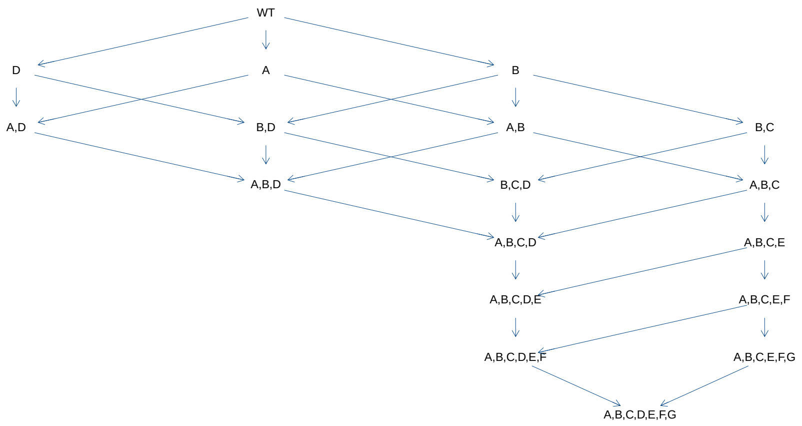


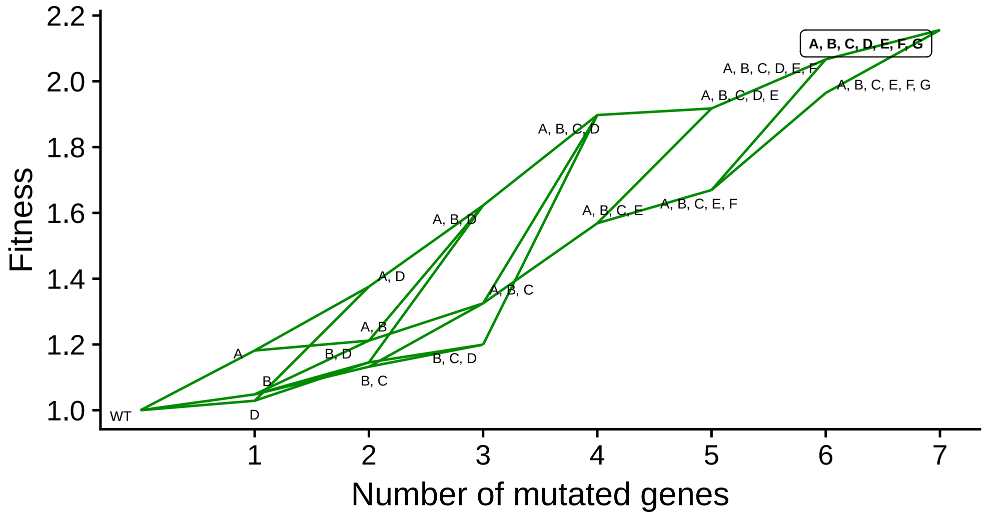

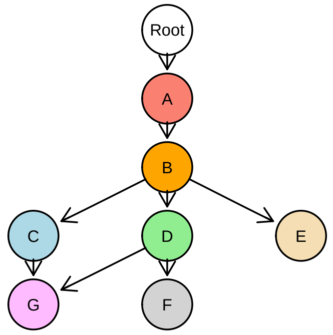


| | |
|---|---|
| *ID* | MJMydL6WRmXU5Lio |
| *acc genotypes* | 18 |
| *peaks no back* | 1 |
| *removed edges* | 0 |
| *prop rem edges* | 0 |

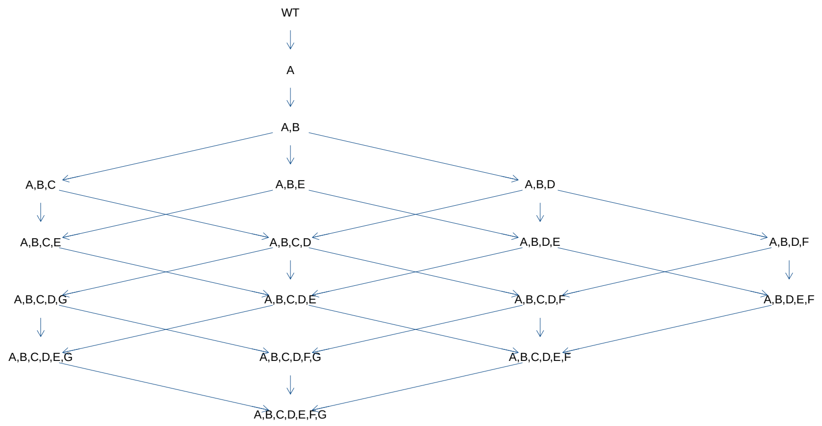


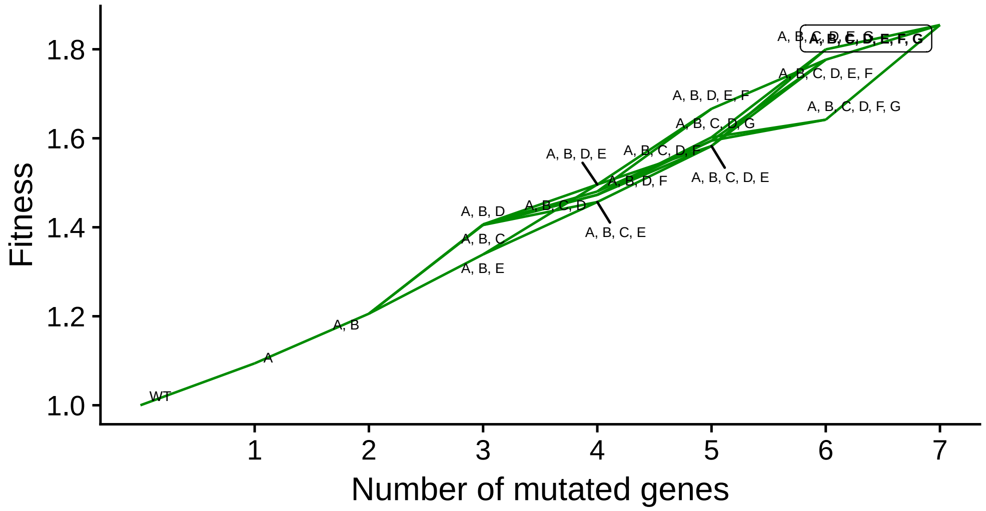

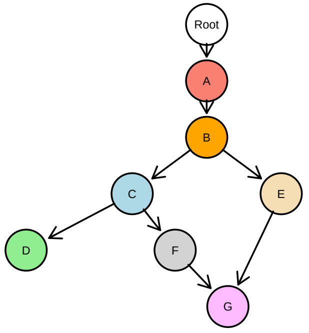


*ID* mJyMPts2CCp9bFCw
*acc genotypes* 14
*peaks no back* 1
*removed edges* 0
*prop rem edges* 0

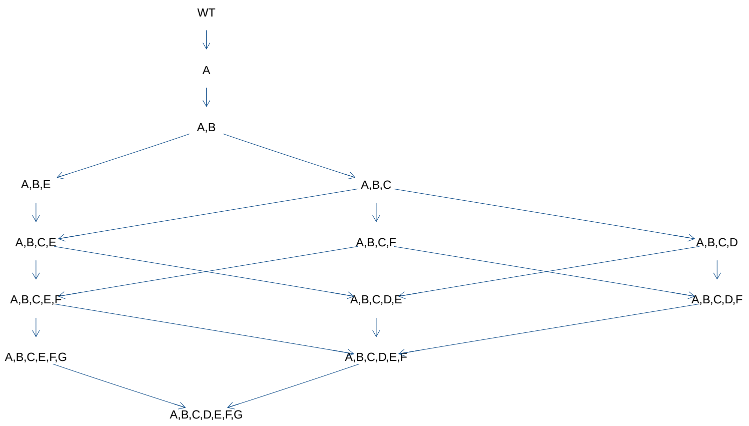


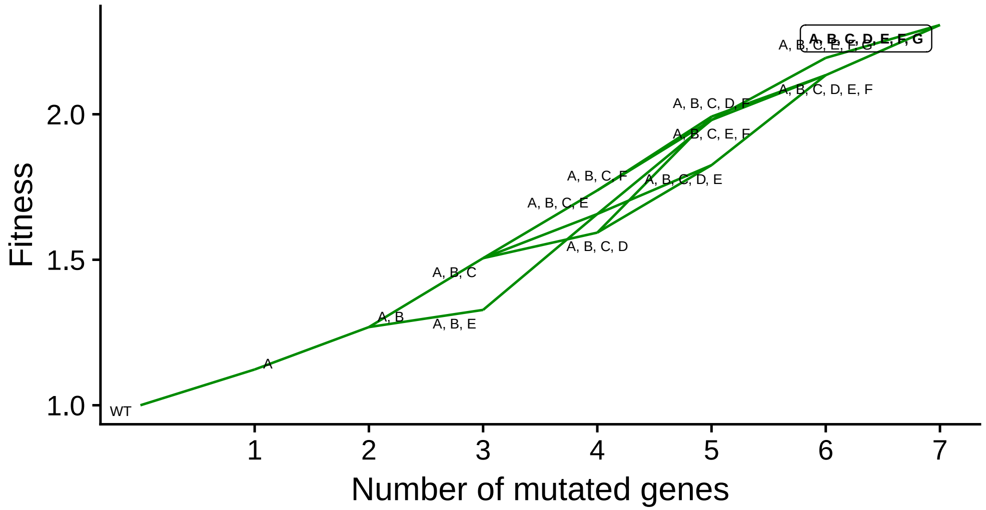

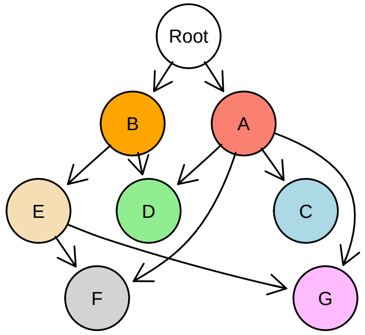


| | |
|---|---|
| *ID* | MmQTibreqvyLDuMK |
| *acc genotypes* | 25 |
| *peaks no back* | 1 |
| *removed edges* | 0 |
| *prop rem edges* | 0 |

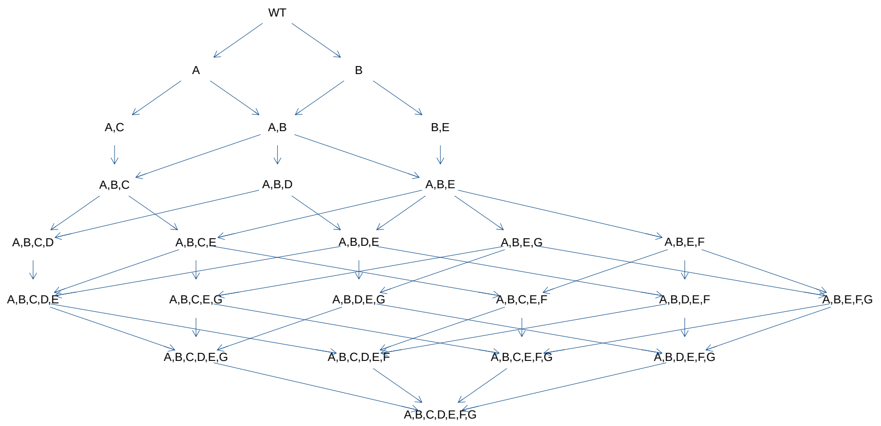


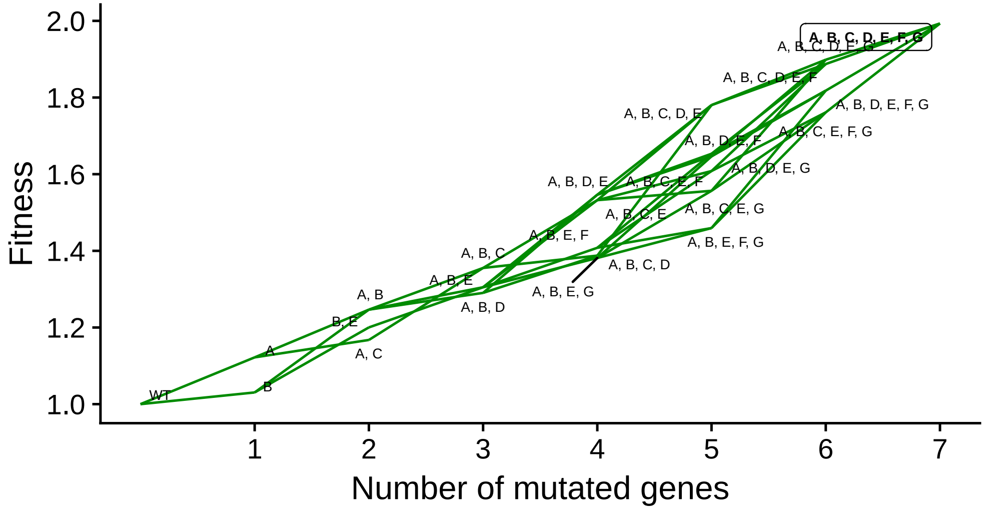

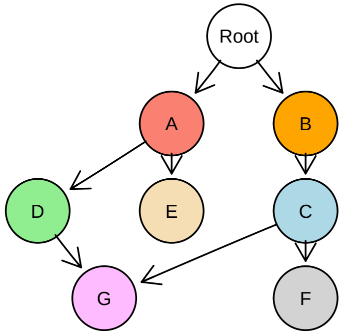


| | |
|---|---|
| *ID* | msMIql4eo3nmvydj |
| *acc genotypes* | 24 |
| *peaks no back* | 1 |
| *removed edges* | 0 |
| *prop rem edges* | 0 |

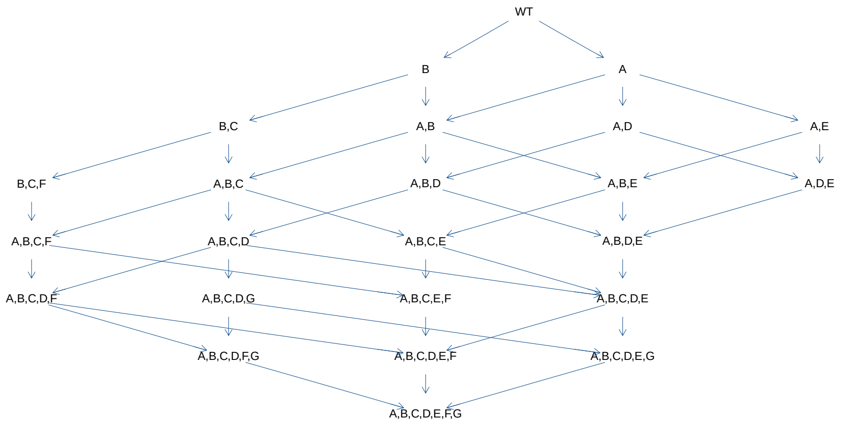


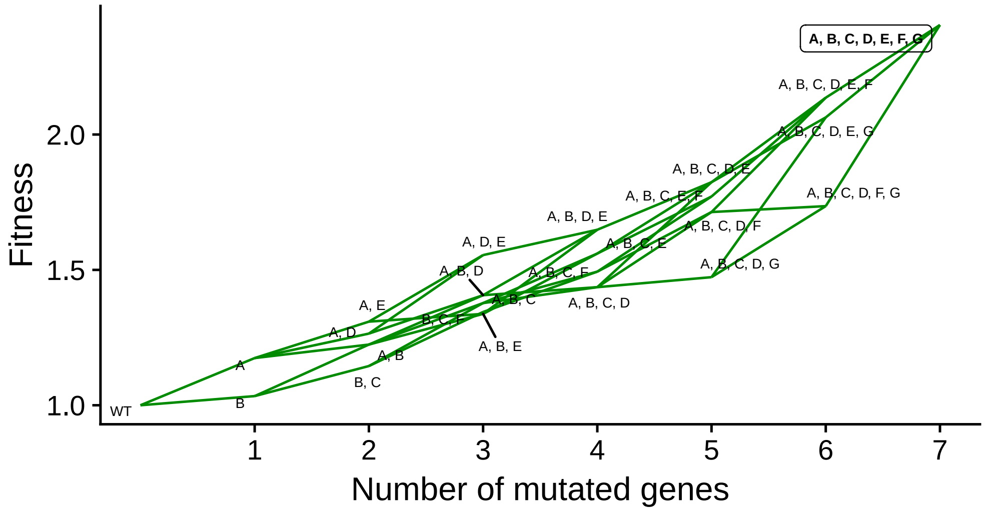

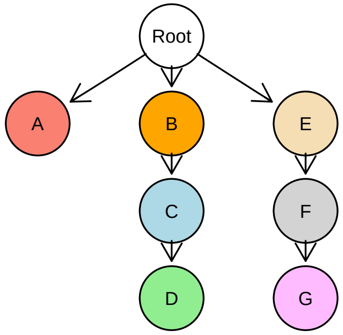


| | |
|---|---|
| *ID* | mzgsSEJyju3Unz2A |
| *acc genotypes* | 32 |
| *peaks no back* | 1 |
| *removed edges* | 0 |
| *prop rem edges* | 0 |

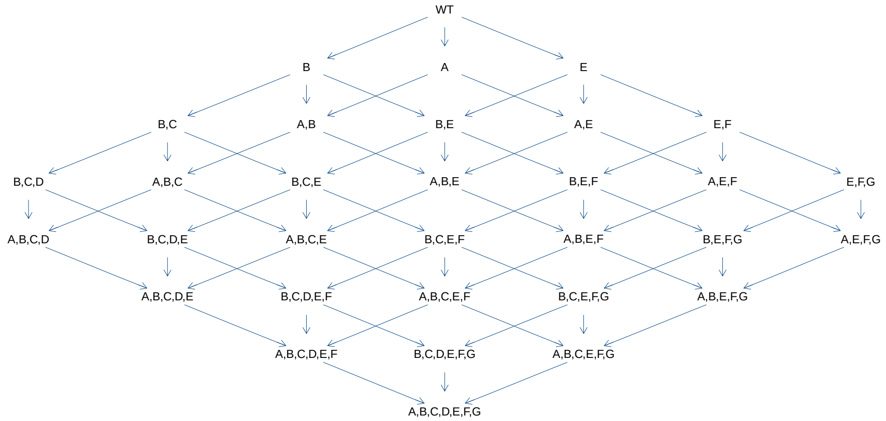


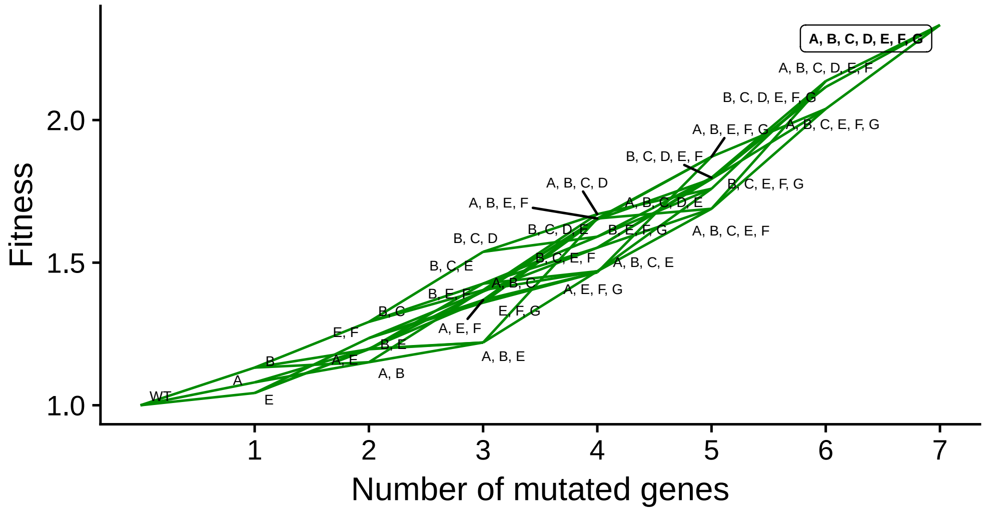

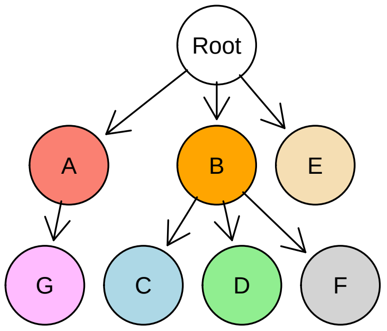


| | |
|---|---|
| *ID* | N4iitsjj5WGvCNiO |
| *acc genotypes* | 54 |
| *peaks no back* | 1 |
| *removed edges* | 0 |
| *prop rem edges* | 0 |

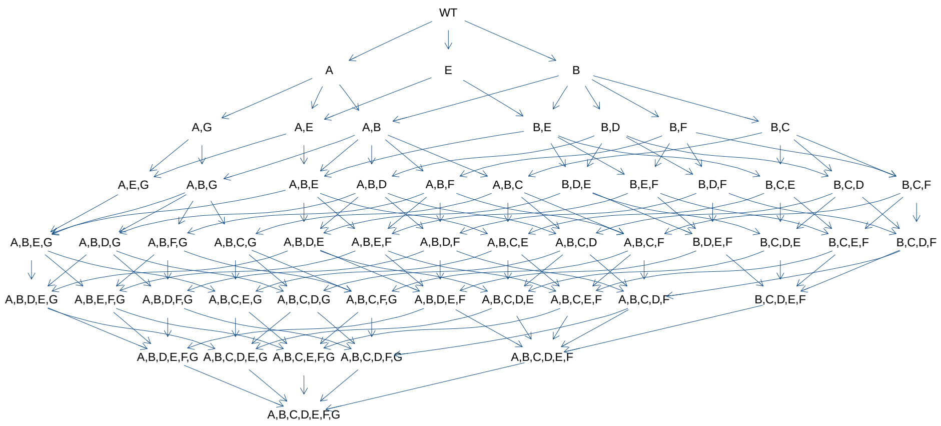


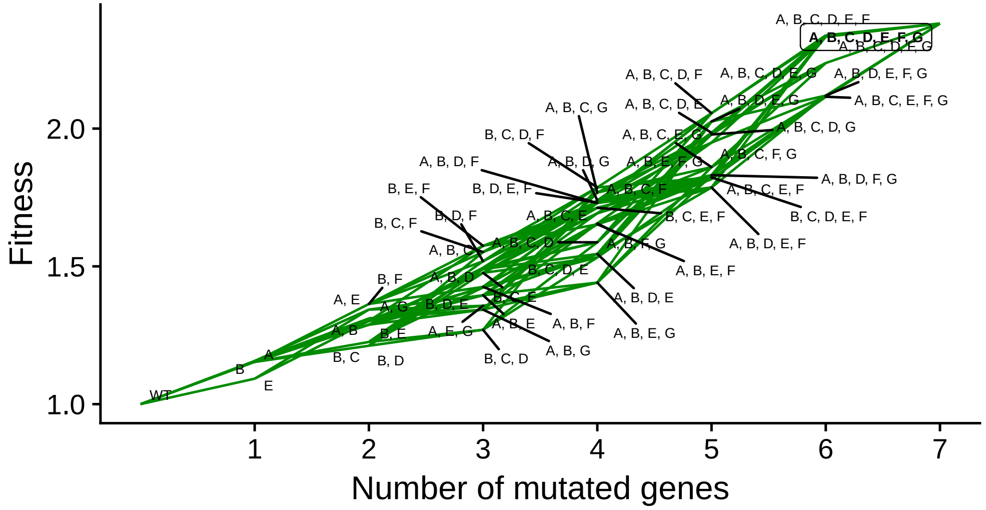

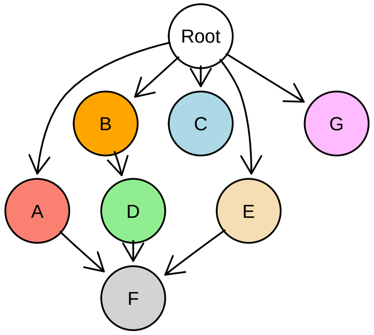


| | |
|---|---|
| *ID* | N9wjEenhGOlChUML |
| *acc genotypes* | 52 |
| *peaks no back* | 1 |
| *removed edges* | 0 |
| *prop rem edges* | 0 |

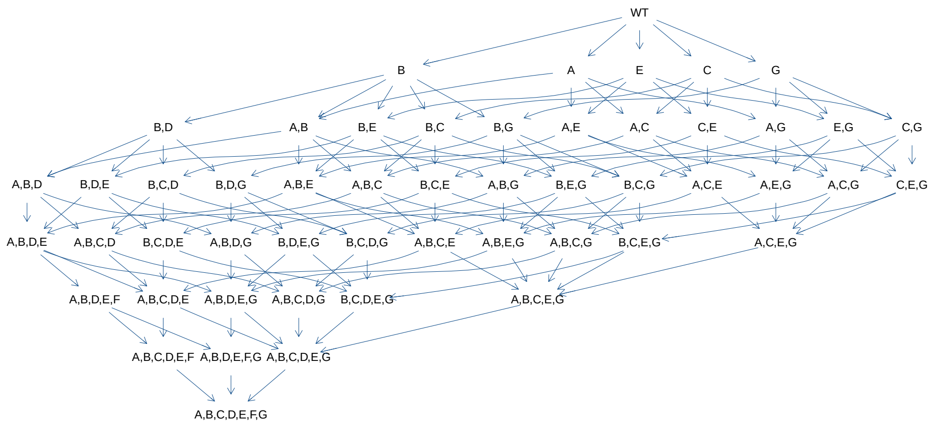


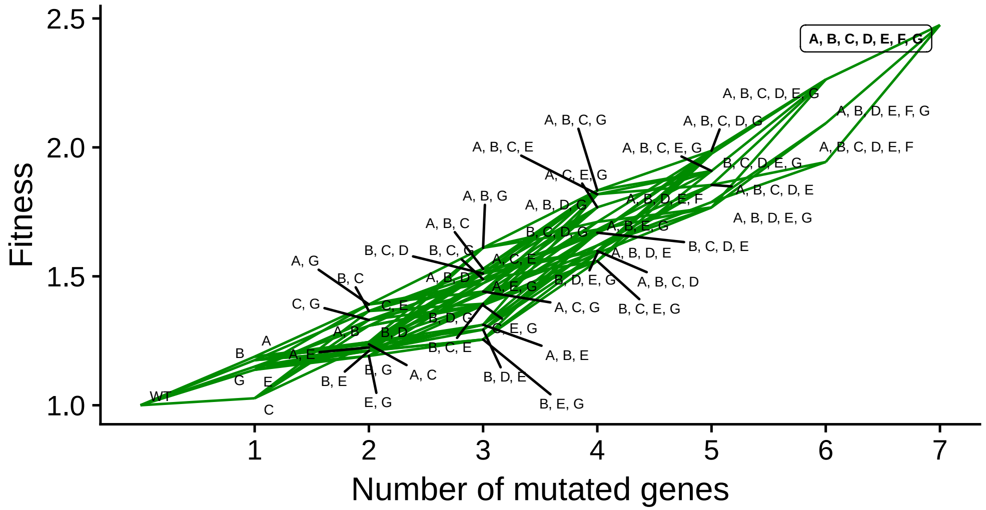

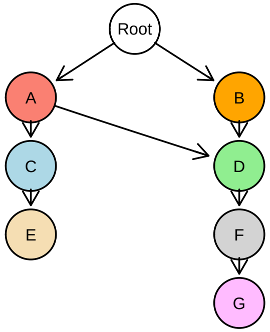


| | |
|---|---|
| *ID* | nbY47YBy9tCkfebb |
| *acc genotypes* | 17 |
| *peaks no back* | 1 |
| *removed edges* | 0 |
| *prop rem edges* | 0 |

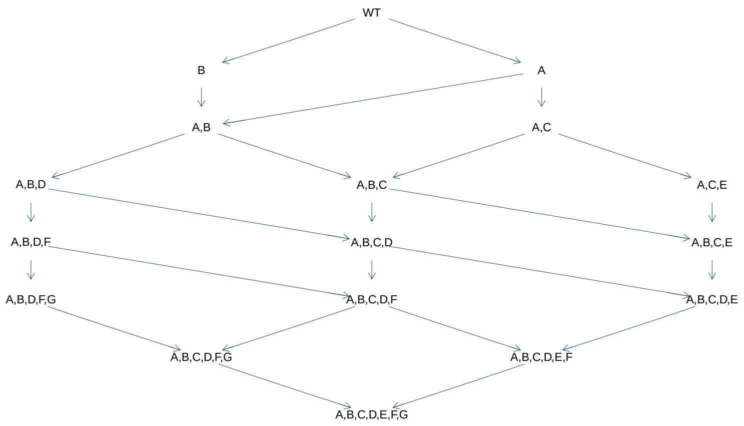


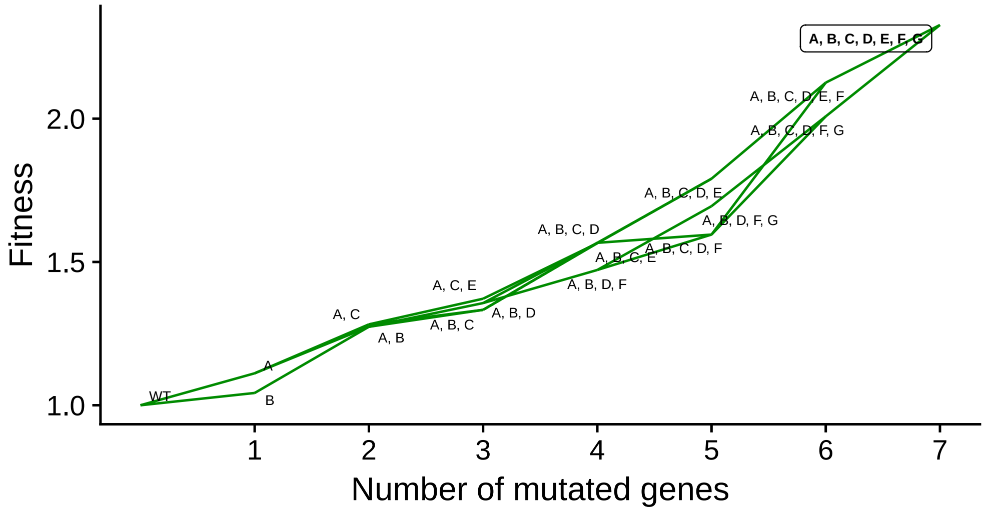

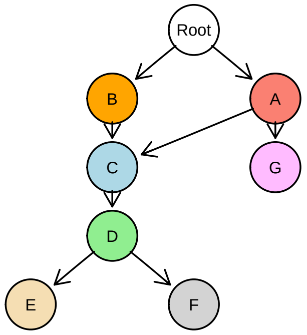


| | |
|---|---|
| *ID* | NEtTcxoCuFV4dbsx |
| *acc genotypes* | 16 |
| *peaks no back* | 1 |
| *removed edges* | 0 |
| *prop rem edges* | 0 |

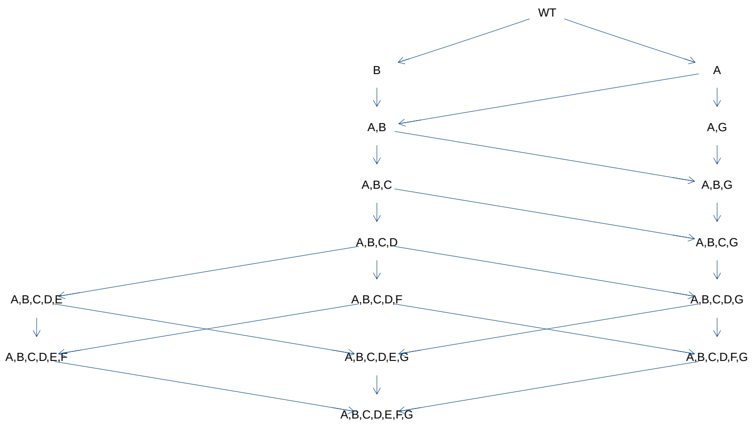


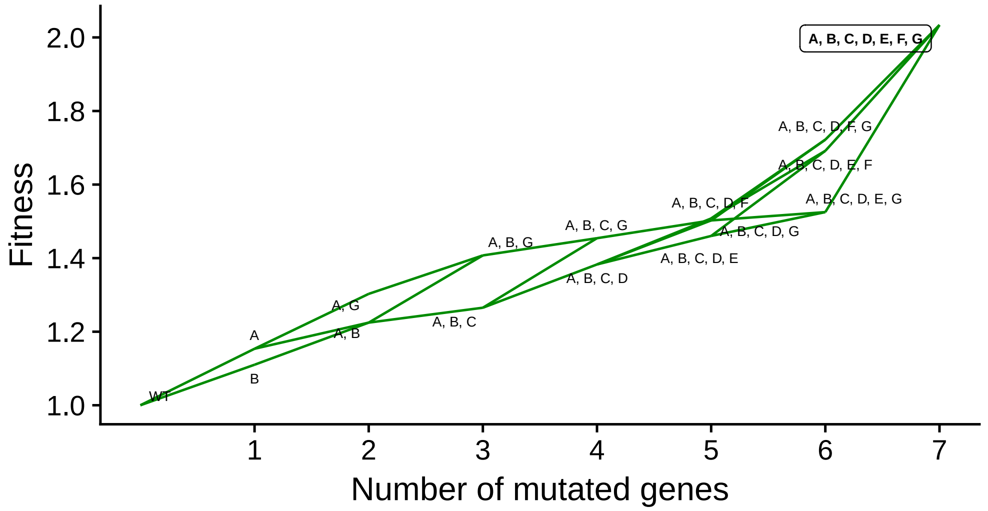

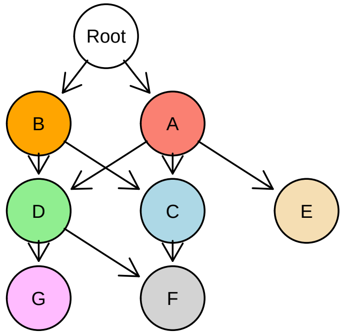


*ID* NL2plam10LkyAR93o
*acc genotypes* 20
*peaks no back* 1
*removed edges* 0
*prop rem edges* 0

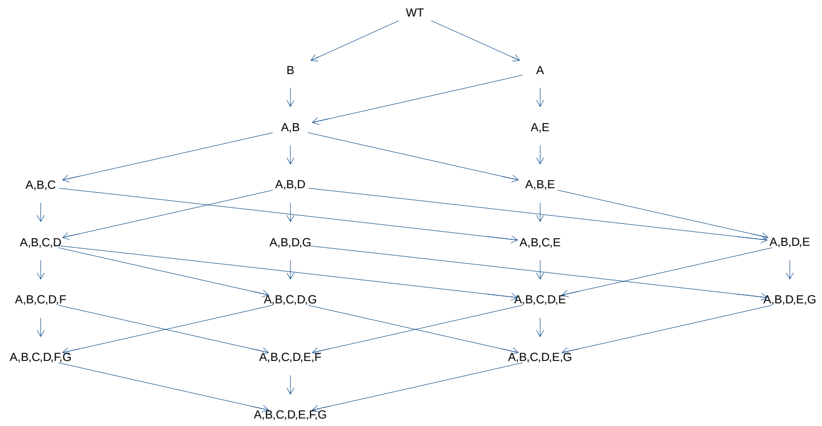


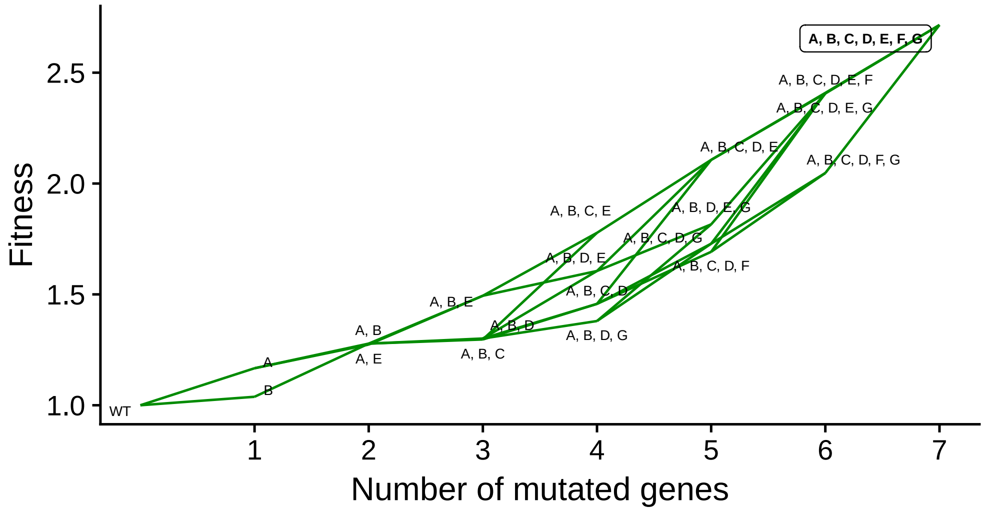

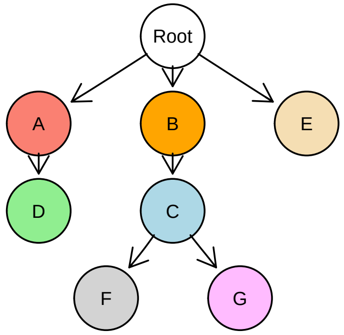


| | |
|---|---|
| *ID* | nNXWsbAmror4SleI |
| *acc genotypes* | 36 |
| *peaks no back* | 1 |
| *removed edges* | 0 |
| *prop rem edges* | 0 |

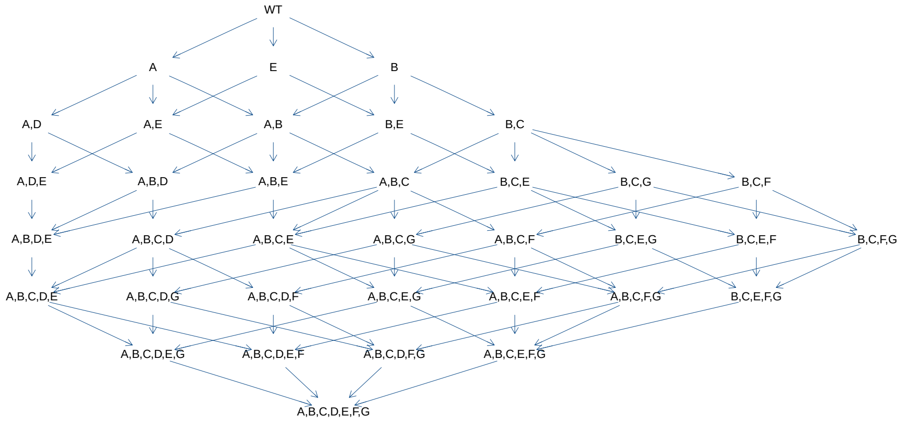


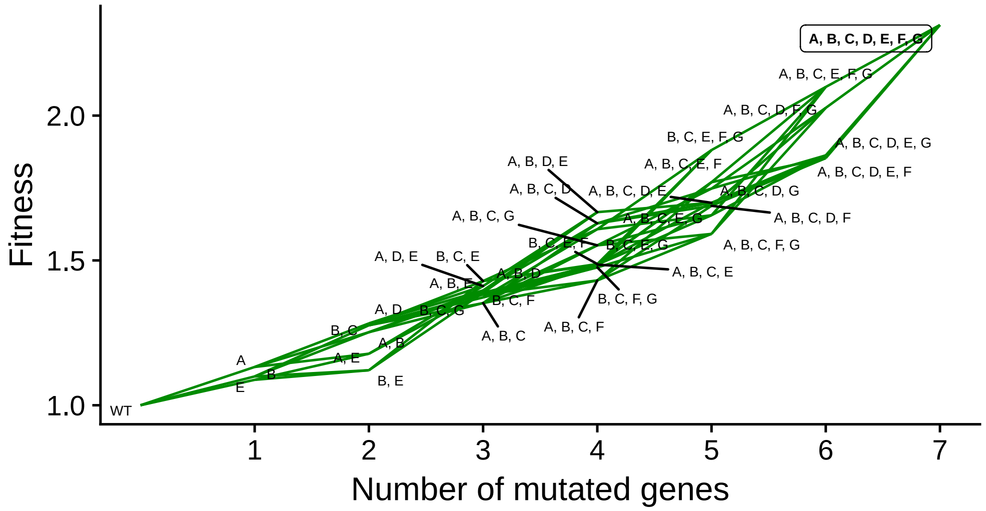

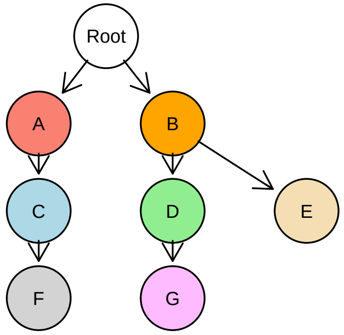


| | |
|---|---|
| *ID* | NP6G6AnYJEoAMZzC |
| *acc genotypes* | 28 |
| *peaks no back* | 1 |
| *removed edges* | 0 |
| *prop rem edges* | 0 |

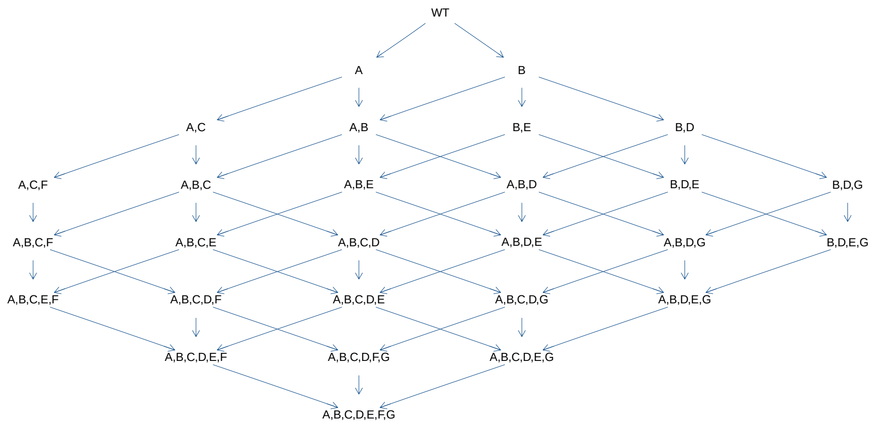


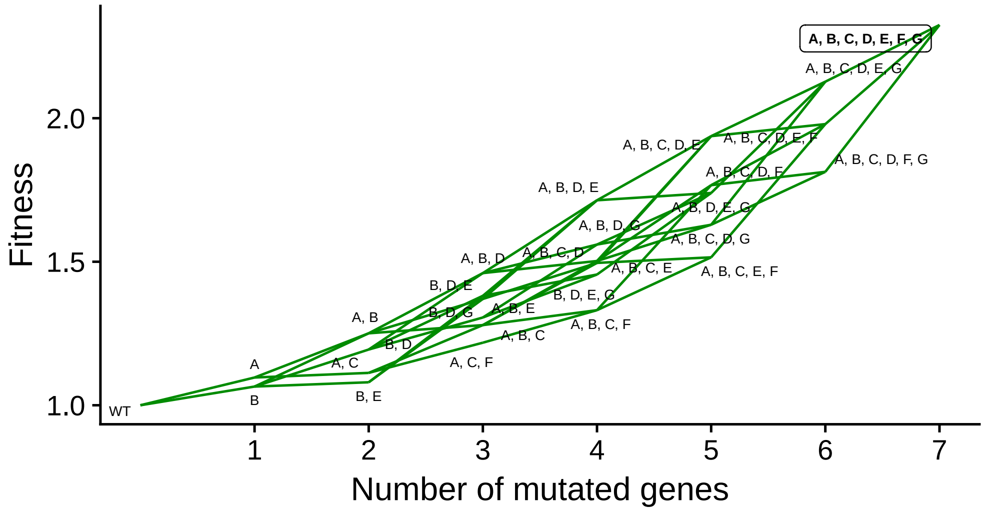

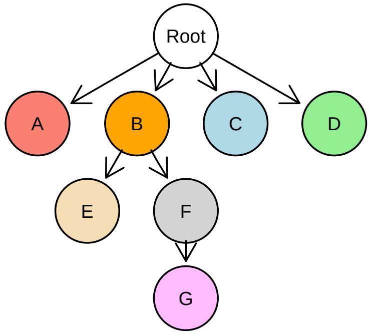


| | |
|---|---|
| *ID* | NQkZsXiFwR5kJZQb |
| *acc genotypes* | 56 |
| *peaks no back* | 1 |
| *removed edges* | 0 |
| *prop rem edges* | 0 |

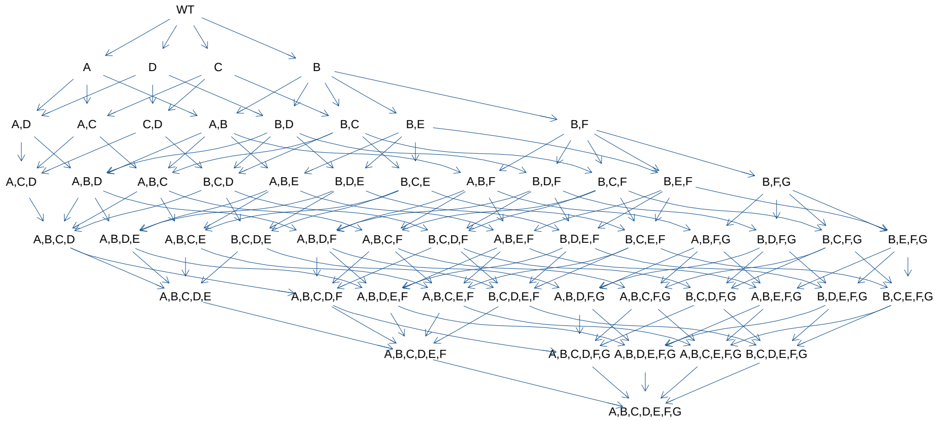


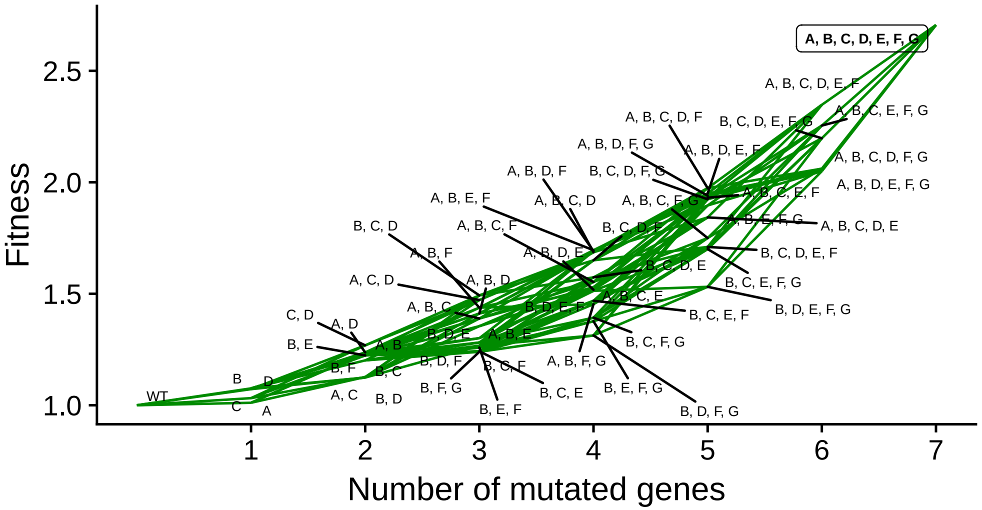

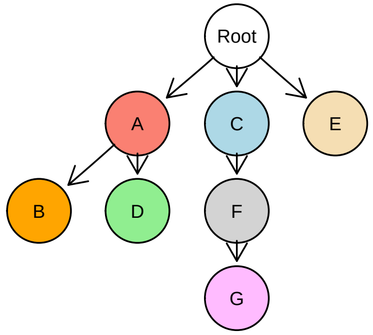


| | |
|---|---|
| *ID* | O9dkG7dXRCSpbF7s |
| *acc genotypes* | 40 |
| *peaks no back* | 1 |
| *removed edges* | 0 |
| *prop rem edges* | 0 |

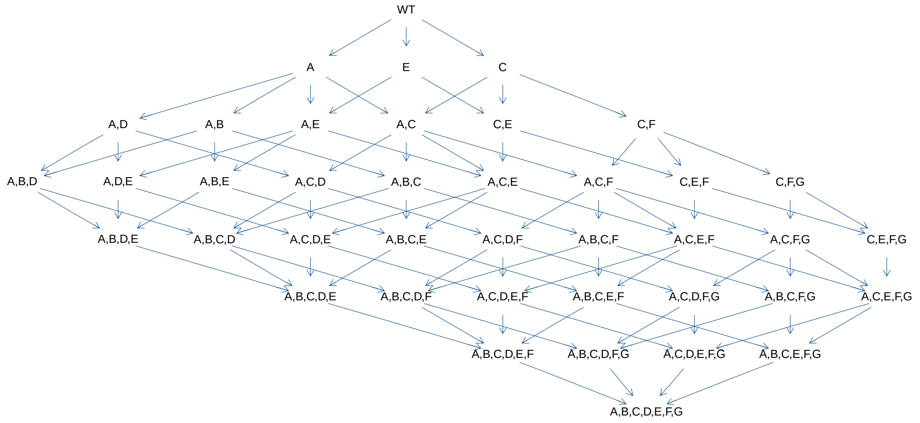


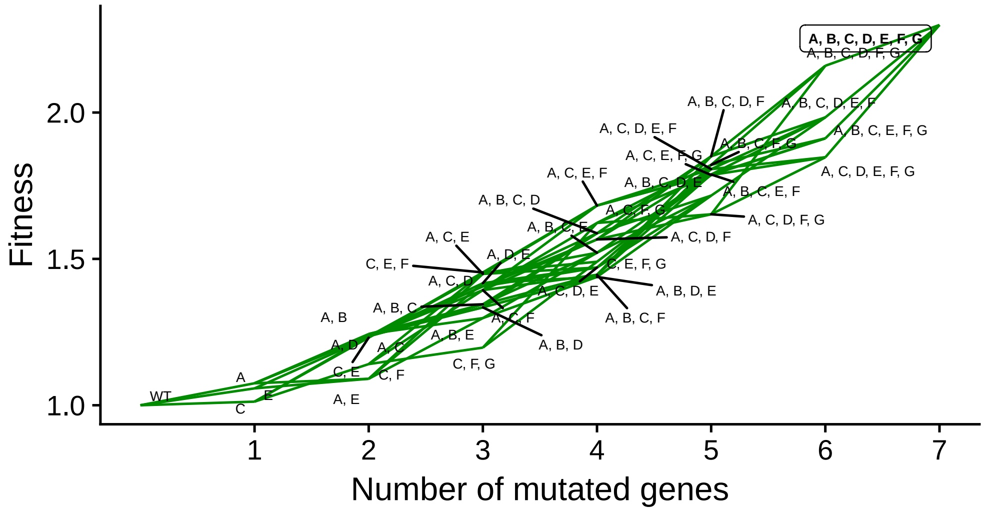

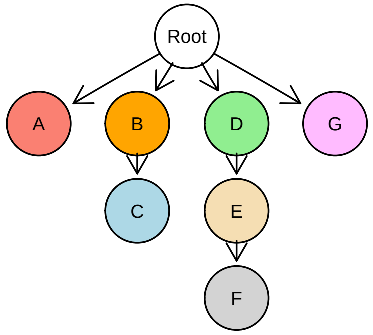


| | |
|---|---|
| *ID* | o9zATJwfW4X1dLDs |
| *acc genotypes* | 48 |
| *peaks no back* | 1 |
| *removed edges* | 0 |
| *prop rem edges* | 0 |

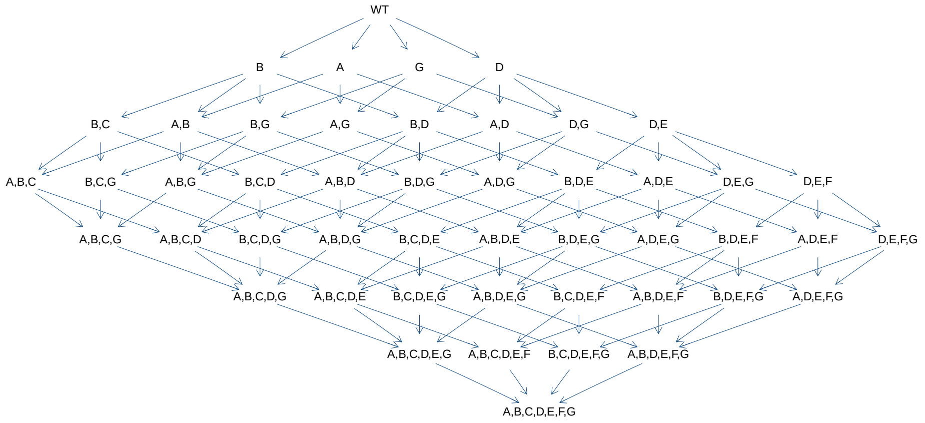


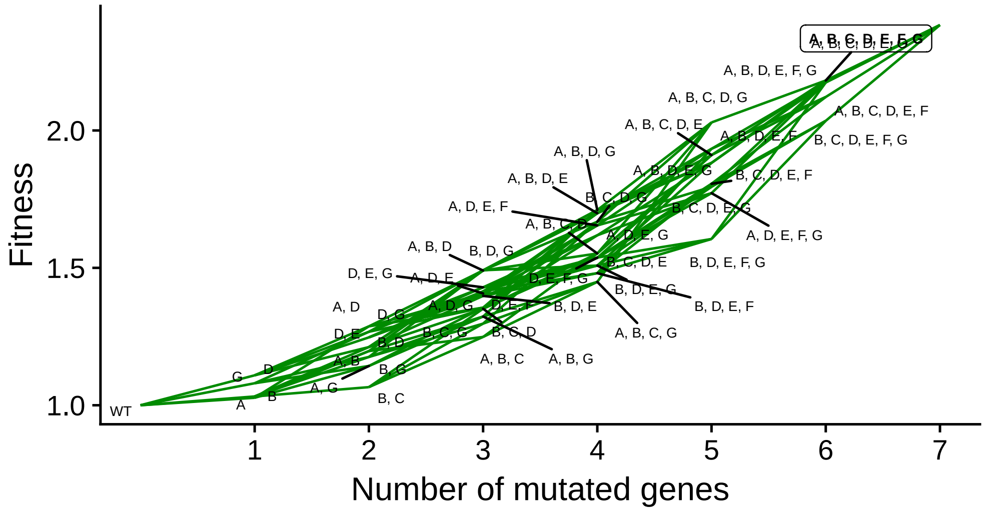

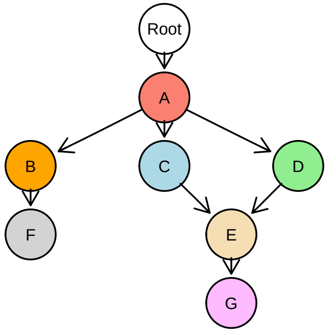


| | |
|---|---|
| *ID* | oZm9q2fpYypg1gEQ |
| *acc genotypes* | 19 |
| *peaks no back* | 1 |
| *removed edges* | 0 |
| *prop rem edges* | 0 |

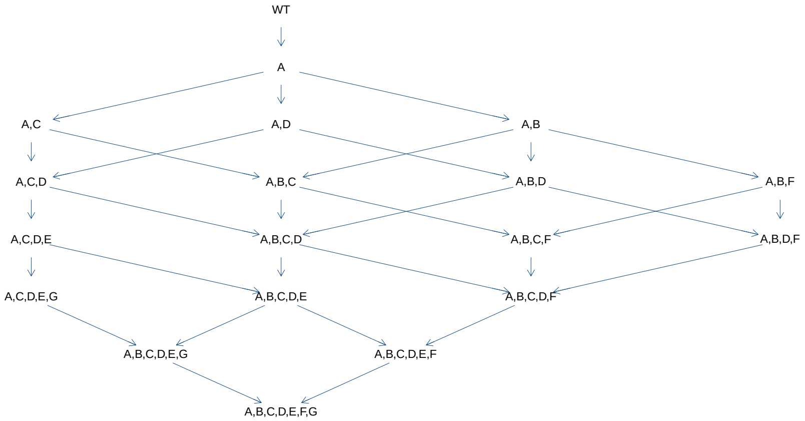


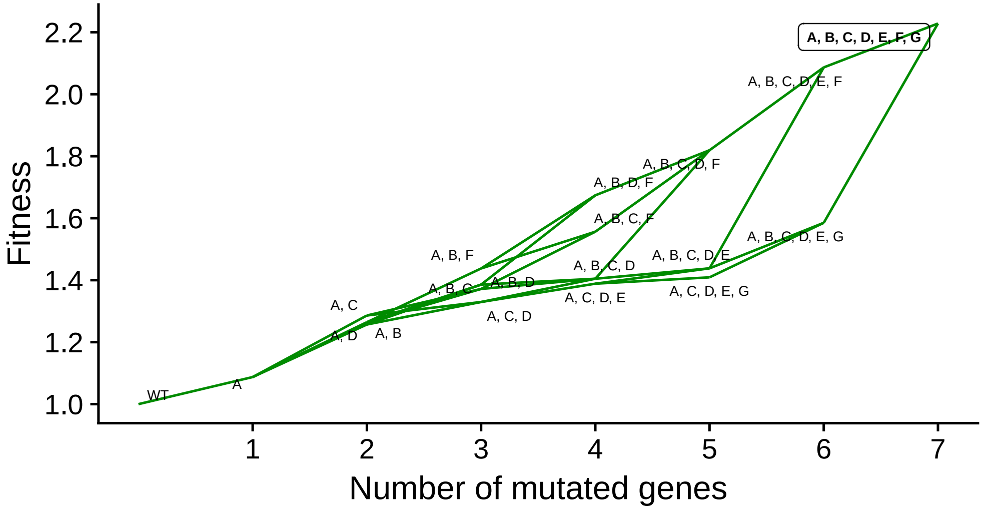

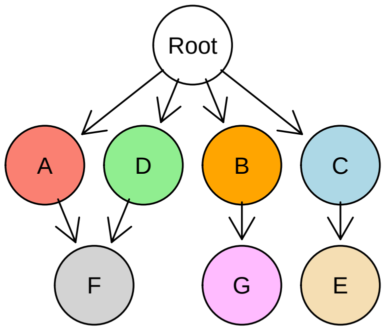


*ID* P6fkPLOb9jfitZfv
*acc genotypes* 45
*peaks no back* 1
*removed edges* 0
*prop rem edges* 0

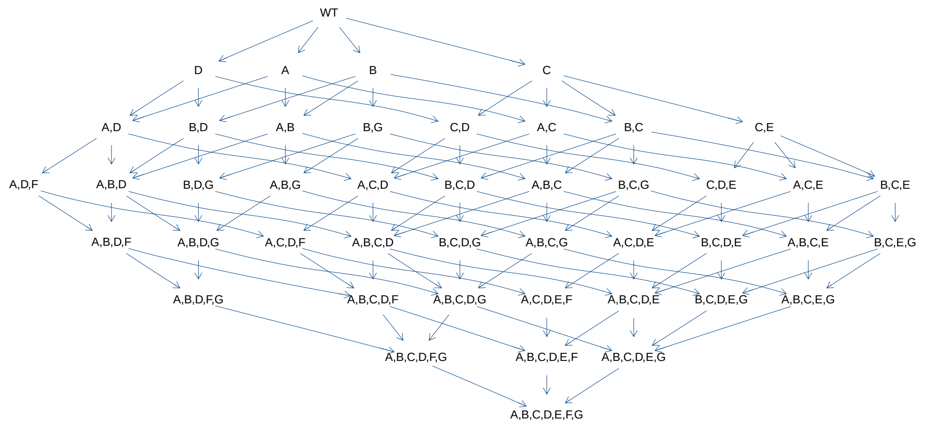


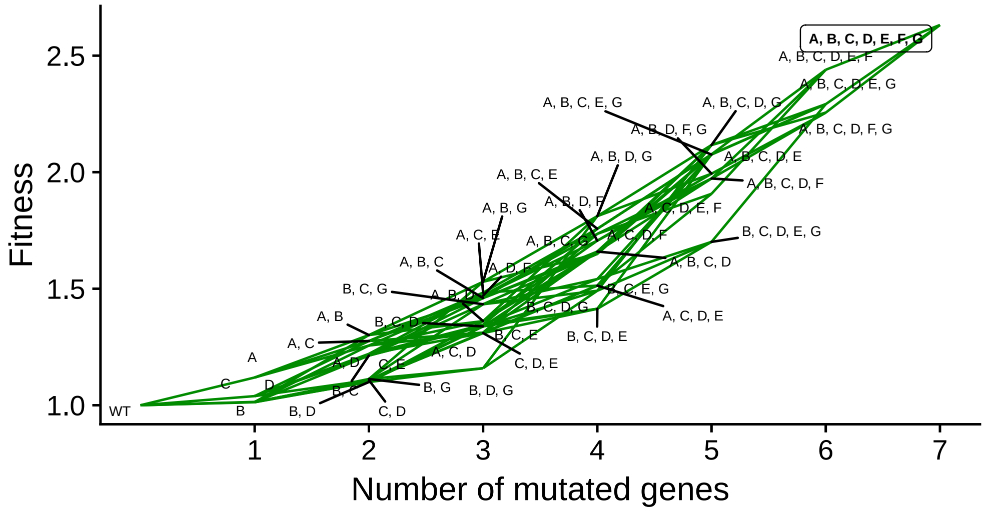

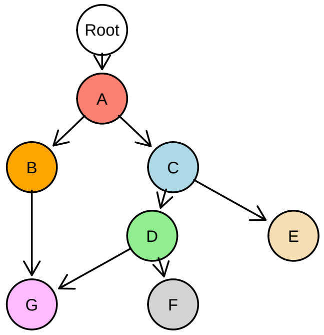


| | |
|---|---|
| *ID* | pDRZFntEs1roodY8 |
| *acc genotypes* | 19 |
| *peaks no back* | 1 |
| *removed edges* | 0 |
| *prop rem edges* | 0 |

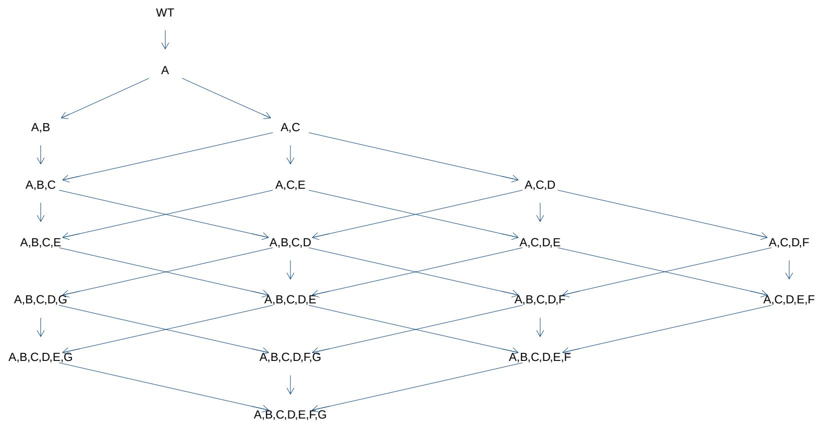


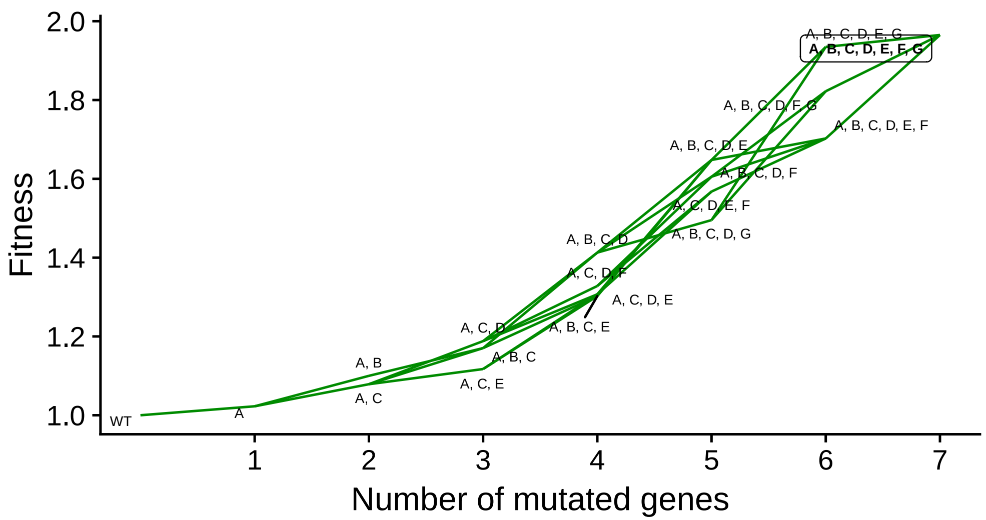

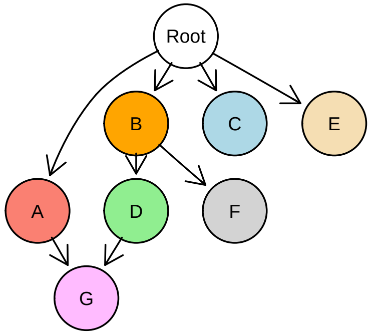


| | |
|---|---|
| *ID* | pJAvDquRk5XGCjrp |
| *acc genotypes* | 48 |
| *peaks no back* | 1 |
| *removed edges* | 0 |
| *prop rem edges* | 0 |

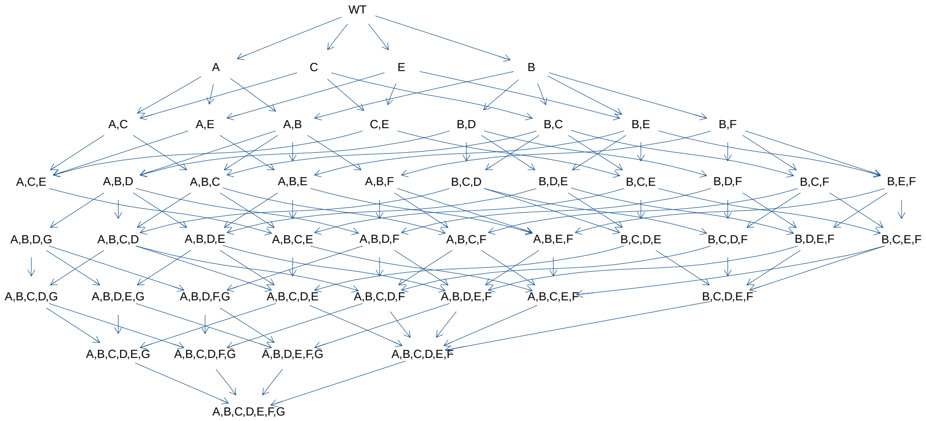


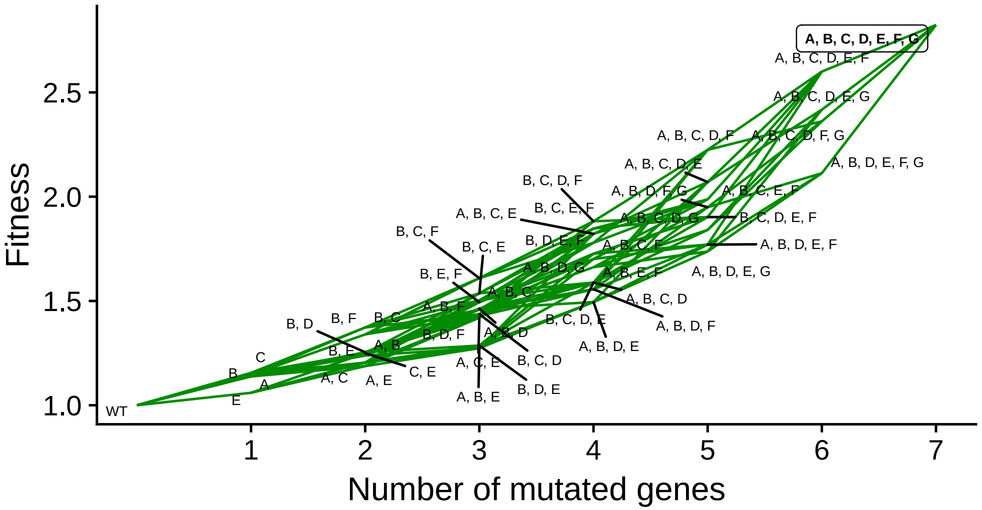

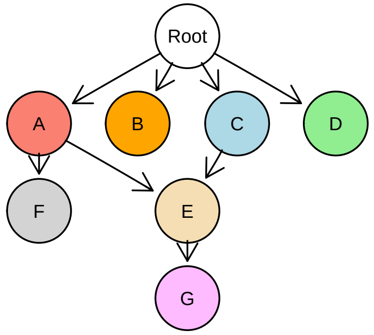


| | |
|---|---|
| *ID* | PQUizfzfBmdE7gFY |
| *acc genotypes* | 40 |
| *peaks no back* | 1 |
| *removed edges* | 0 |
| *prop rem edges* | 0 |

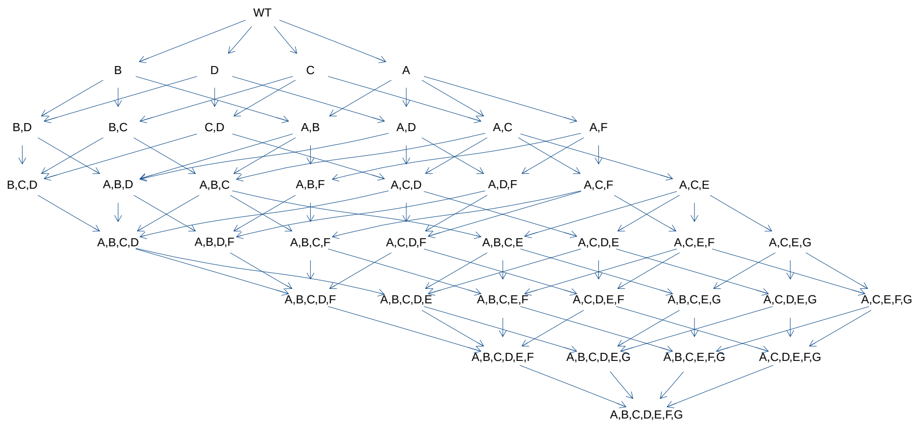


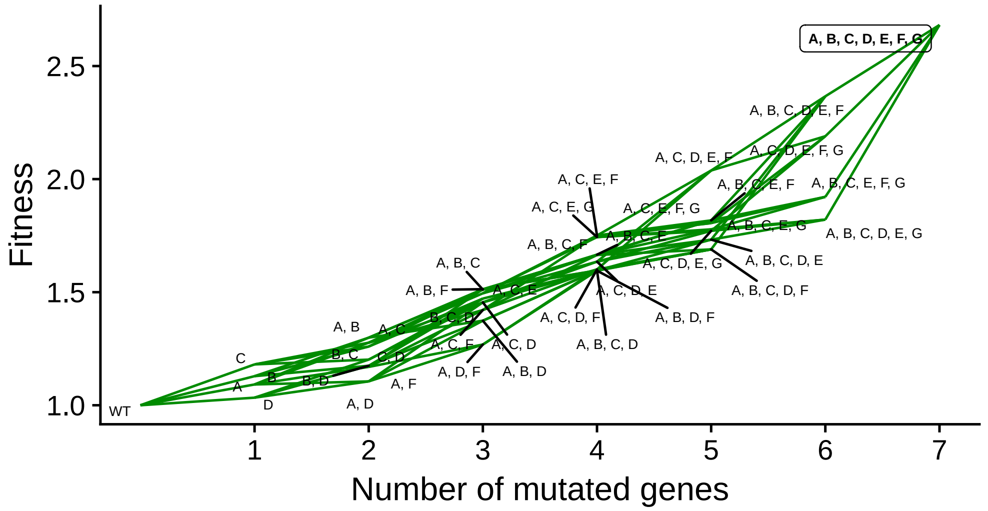

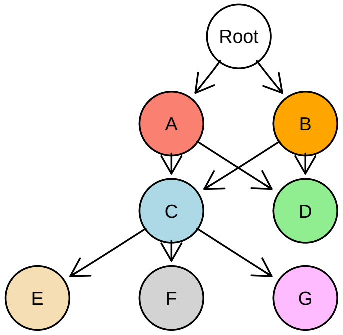


| | |
|---|---|
| *ID* | q10sOtkvPmaBzPDOV |
| *acc genotypes* | 21 |
| *peaks no back* | 1 |
| *removed edges* | 0 |
| *prop rem edges* | 0 |

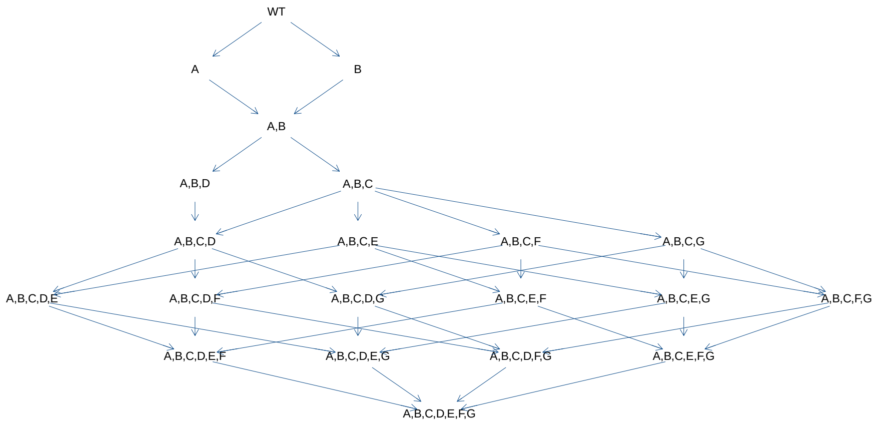


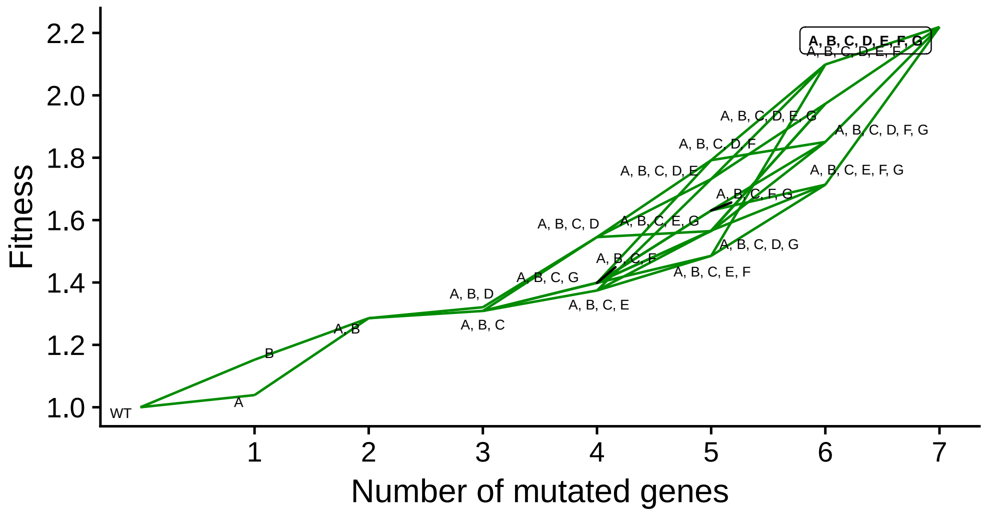

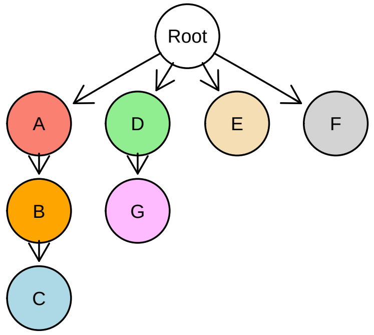


| | |
|---|---|
| *ID* | q10upXvME3iKLaWcZ |
| *acc genotypes* | 48 |
| *peaks no back* | 1 |
| *removed edges* | 0 |
| *prop rem edges* | 0 |

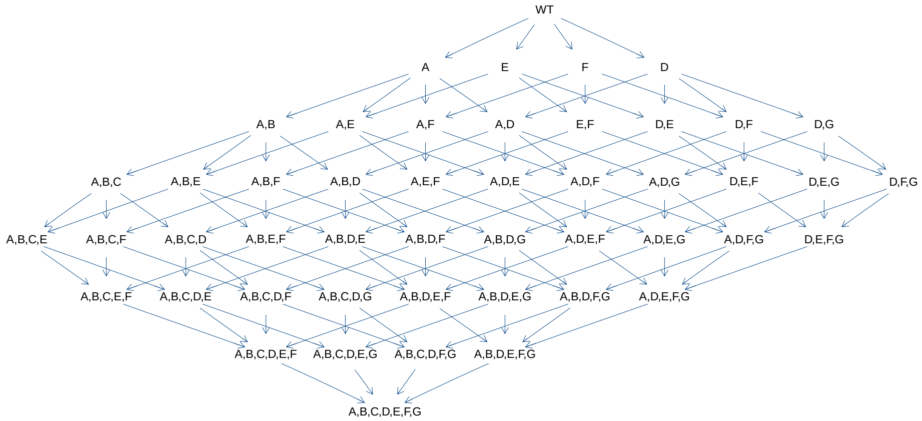


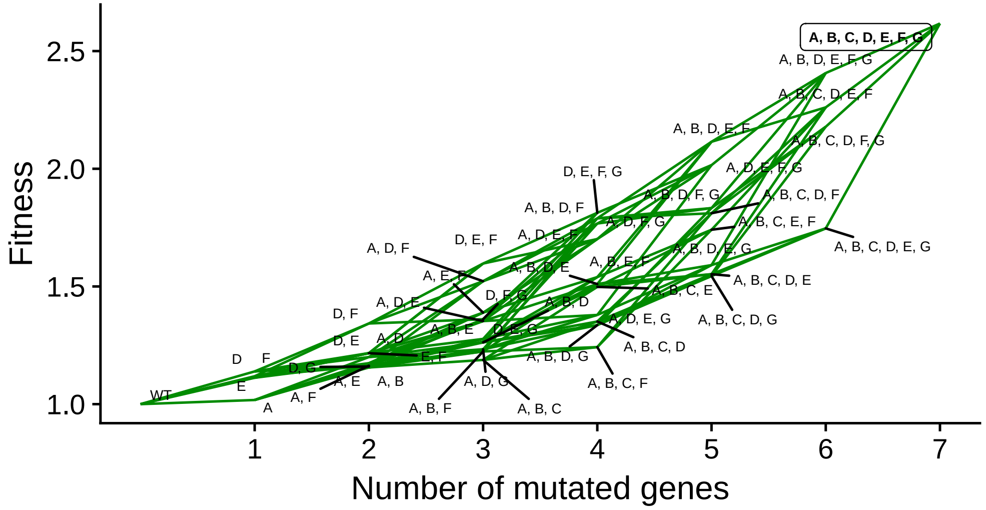

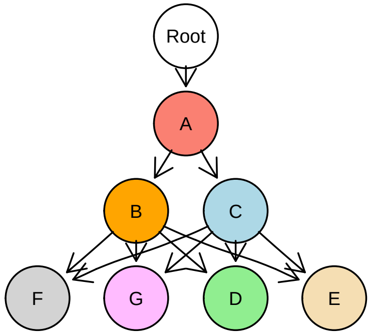


| | |
|---|---|
| *ID* | q3bB6O4aHcWy2GNL |
| *acc genotypes* | 20 |
| *peaks no back* | 1 |
| *removed edges* | 0 |
| *prop rem edges* | 0 |

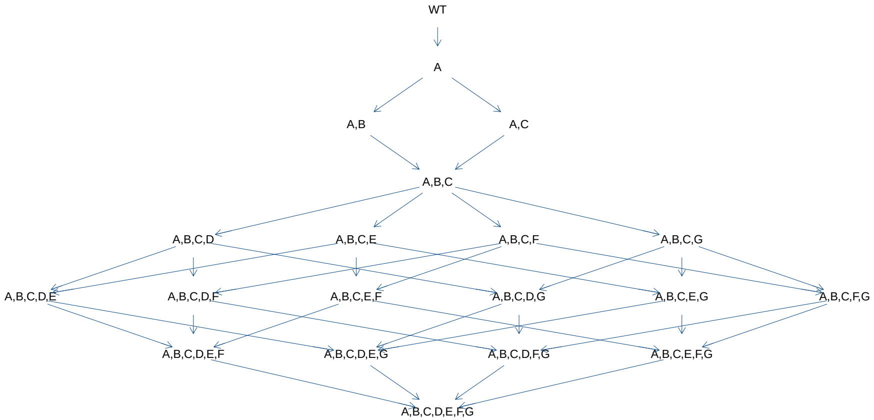


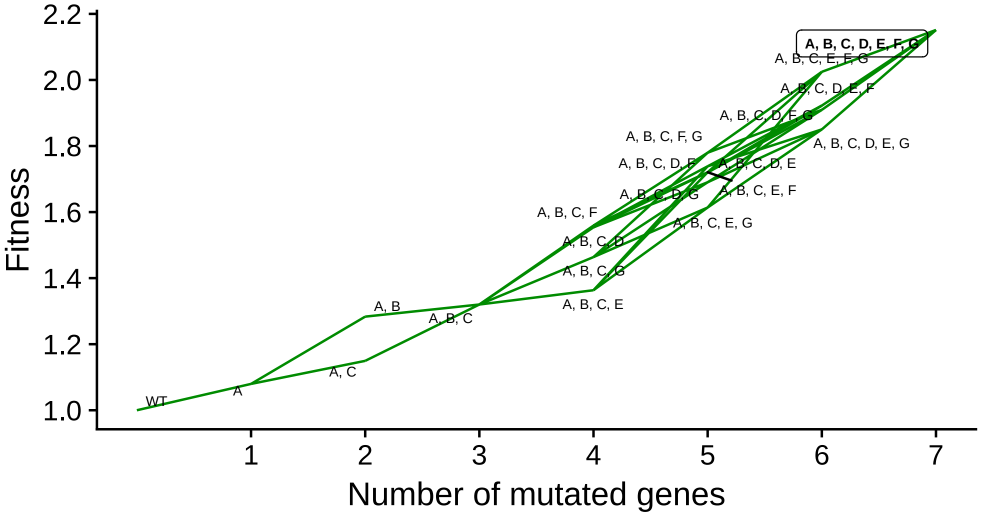

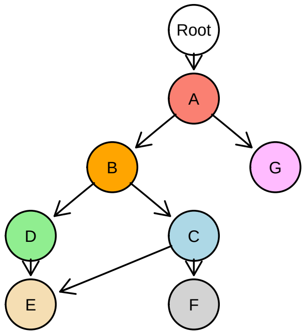


| | |
|---|---|
| *ID* | qDTXWJ10Vx4dad6eV |
| *acc genotypes* | 19 |
| *peaks no back* | 1 |
| *removed edges* | 0 |
| *prop rem edges* | 0 |

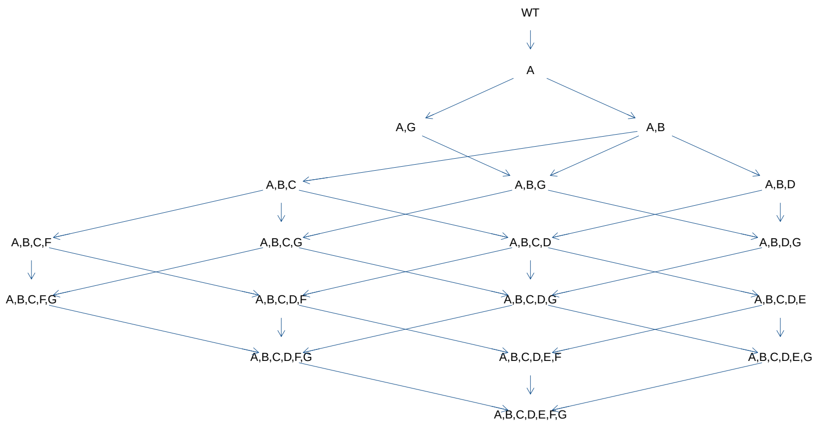


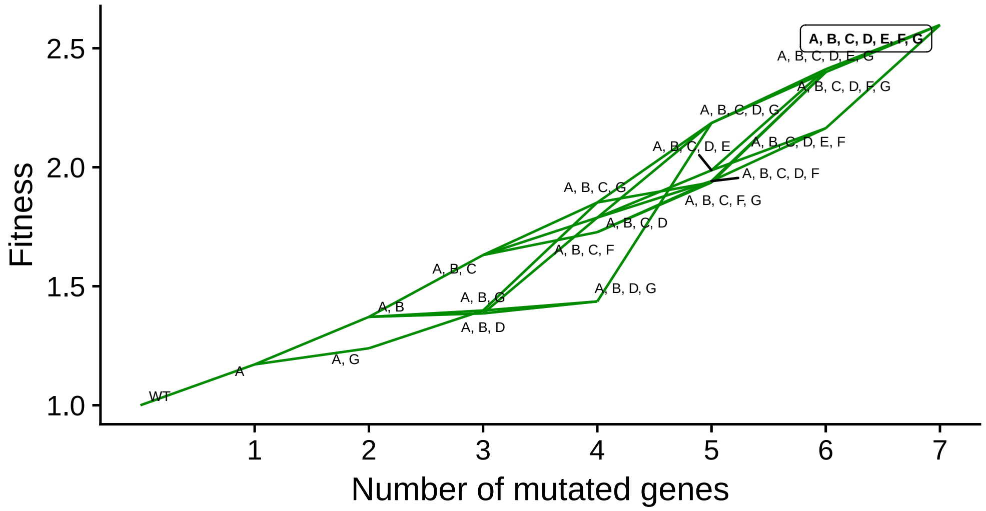

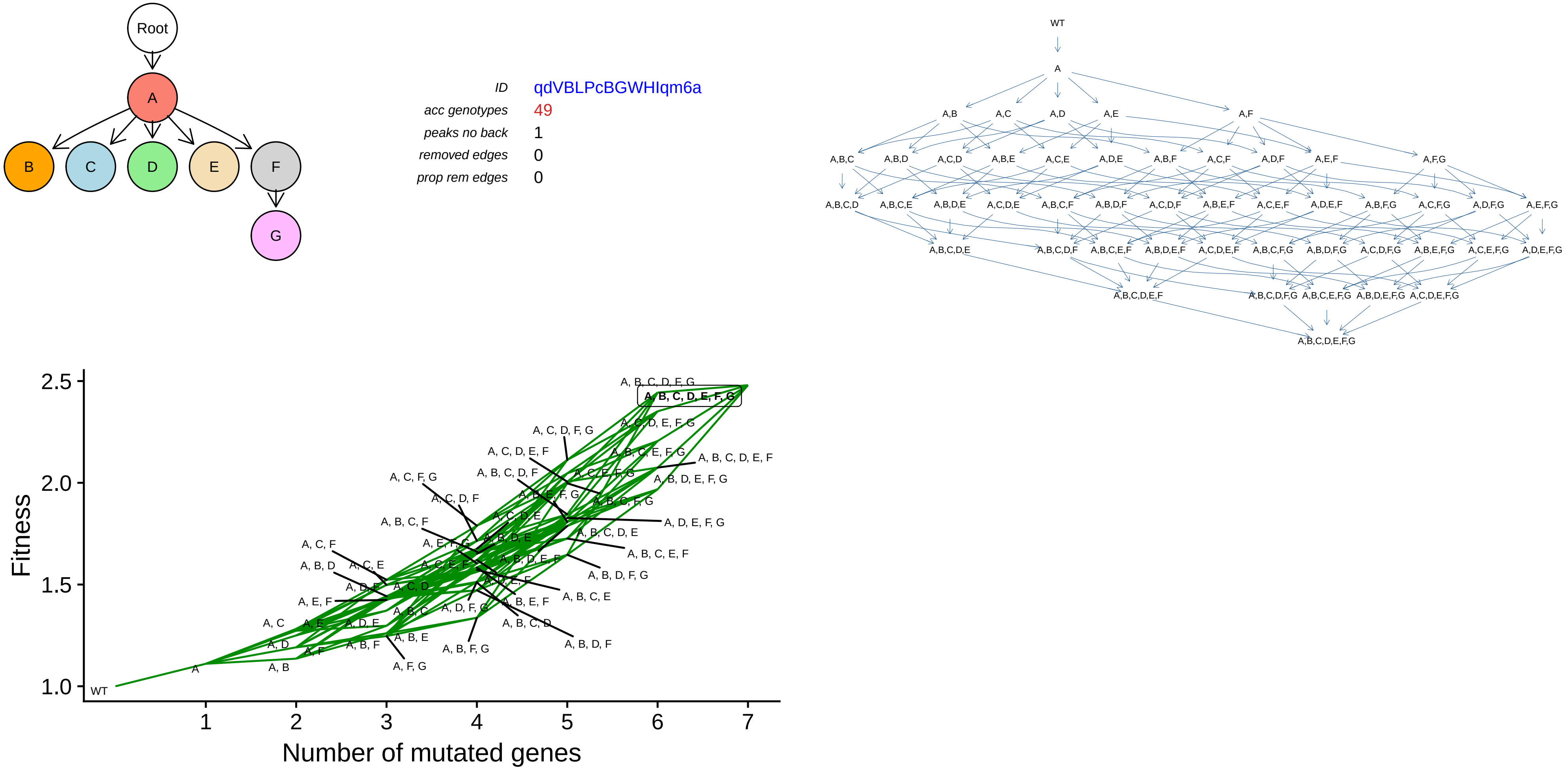

Root
A
B
C
D
E
F
G
ID qdVBLPcBGWHIqm6a
acc genotypes 49
peaks no back 1
removed edges 0
prop rem edges 0
WT
Fitness
Number of mutated genes
A, B, C, D, E, F, G

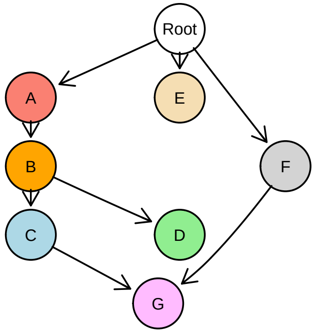


*ID* QfKhJqFLfL10D8u110
*acc genotypes* 28
*peaks no back* 1
*removed edges* 0
*prop rem edges* 0

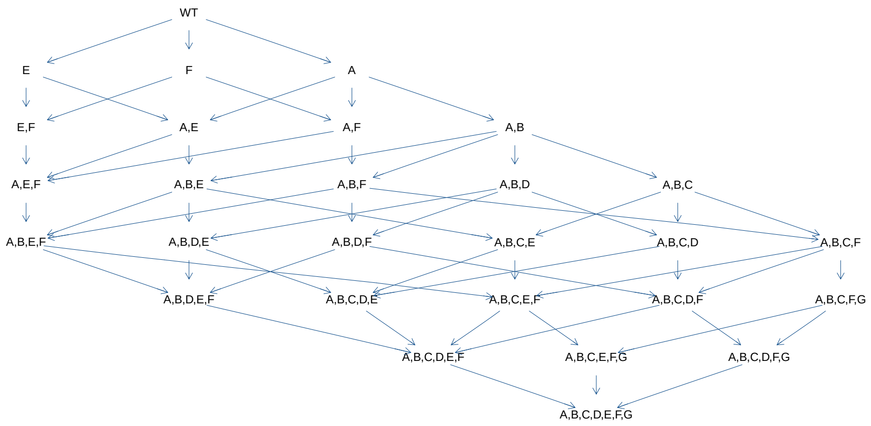


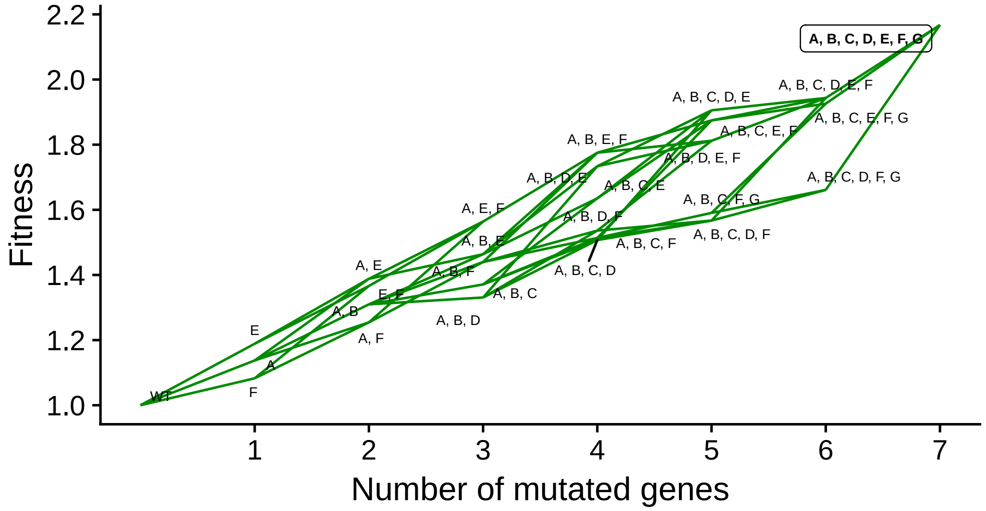

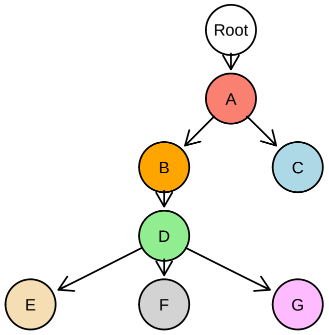


| | |
|---|---|
| *ID* | qhzHLgirL8adNfAJ |
| *acc genotypes* | 21 |
| *peaks no back* | 1 |
| *removed edges* | 0 |
| *prop rem edges* | 0 |

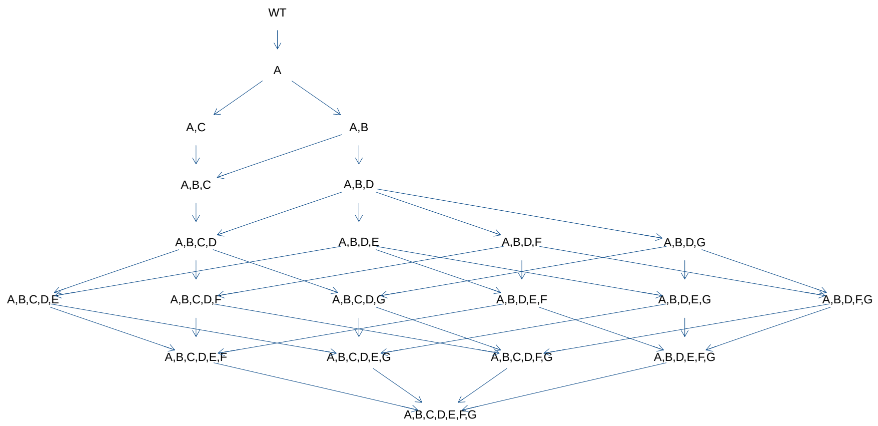


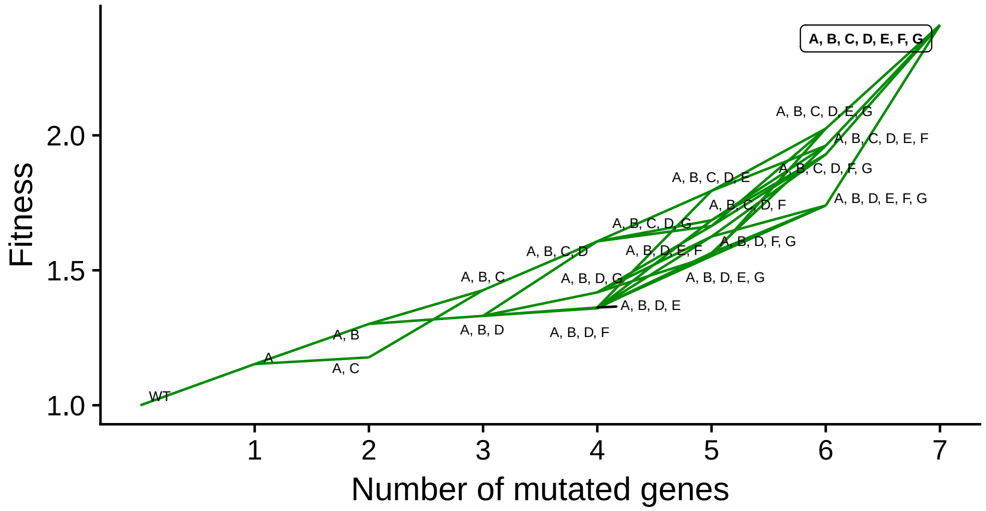

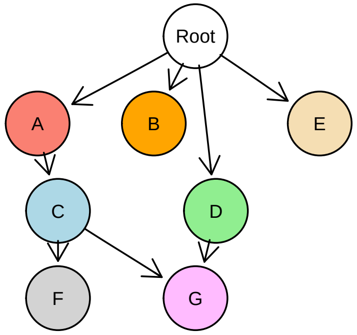


| | |
|---|---|
| *ID* | qIEabZH111rt10AQ2 |
| *acc genotypes* | 40 |
| *peaks no back* | 1 |
| *removed edges* | 0 |
| *prop rem edges* | 0 |

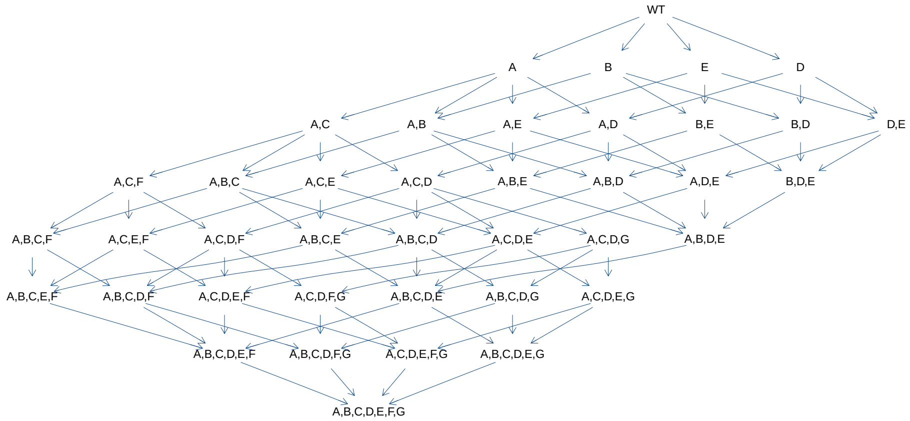


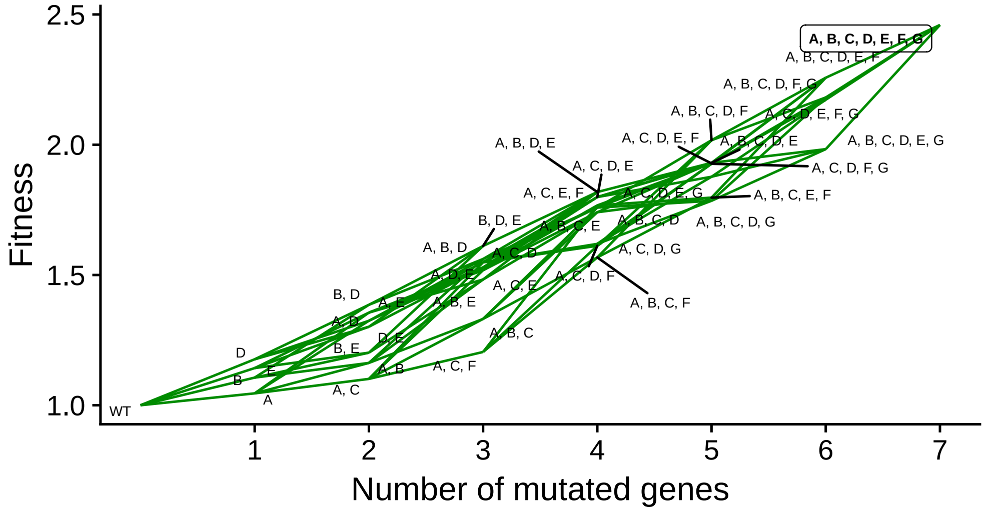

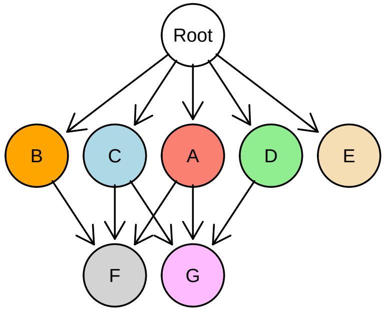


| | |
|---|---|
| *ID* | qsBbMjGmsMVjLZVW |
| *acc genotypes* | 42 |
| *peaks no back* | 1 |
| *removed edges* | 0 |
| *prop rem edges* | 0 |

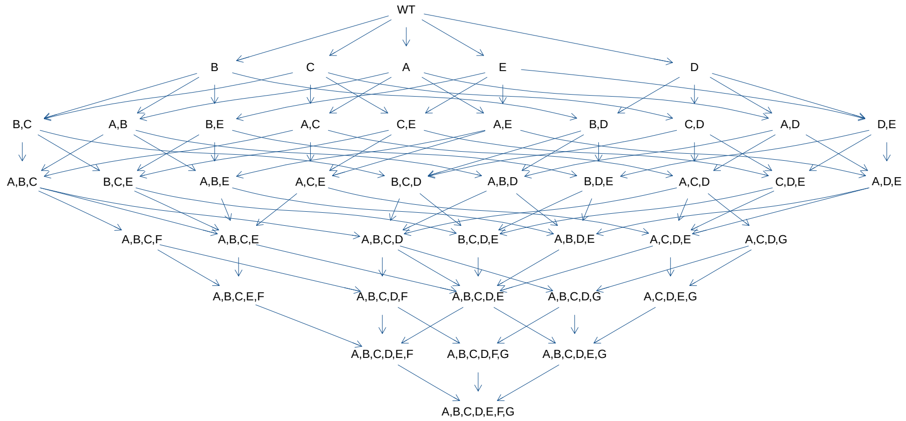


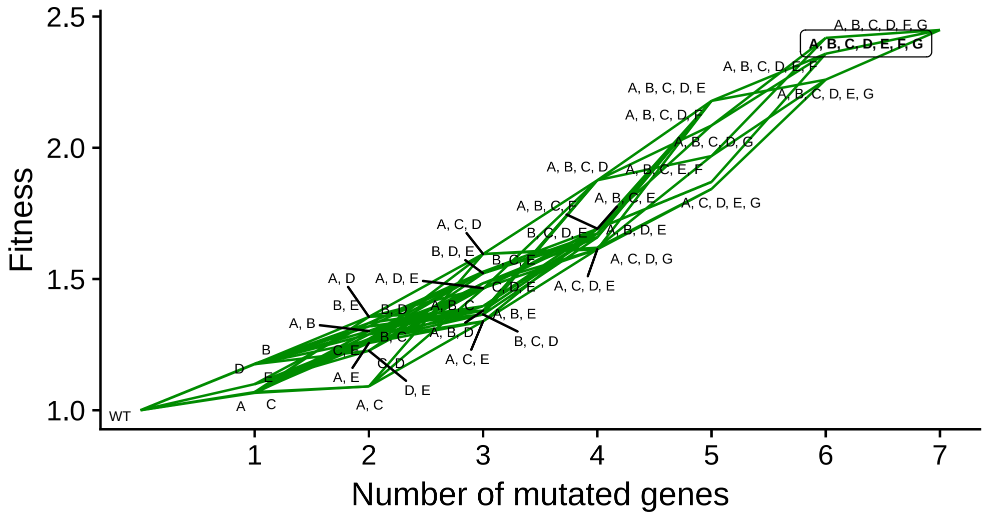

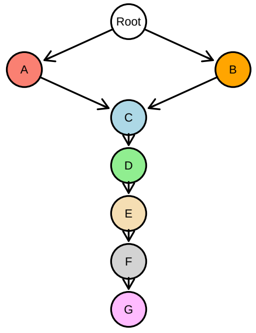


| | |
|---|---|
| *ID* | QTVh104soQUgOBfKi |
| *acc genotypes* | 9 |
| *peaks no back* | 1 |
| *removed edges* | 0 |
| *prop rem edges* | 0 |

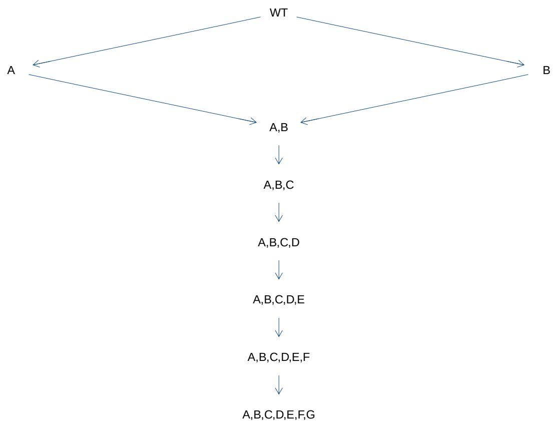


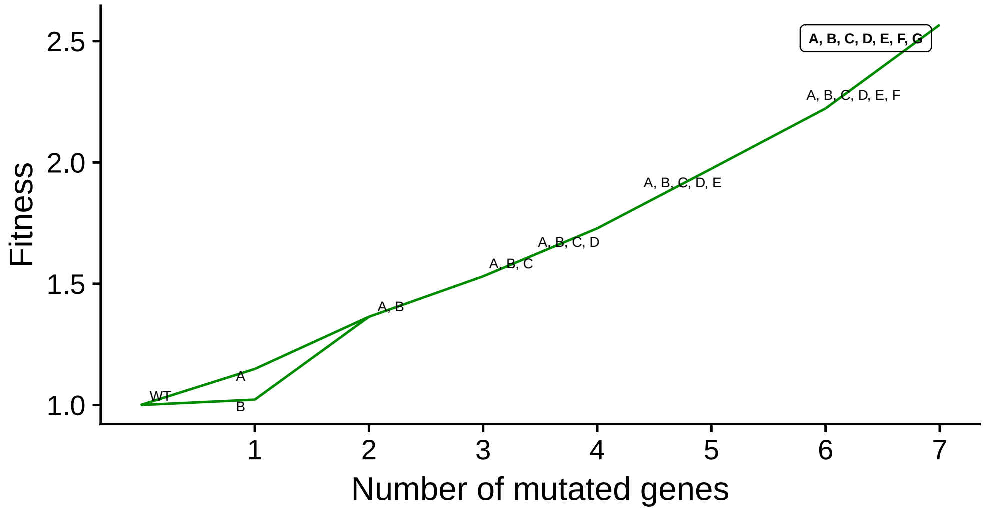

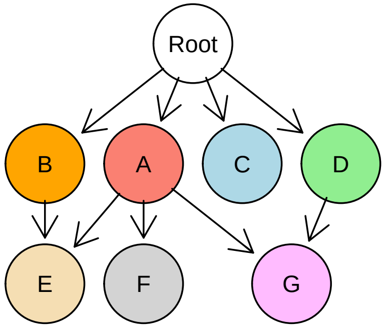


| | |
|---|---|
| *ID* | qYgVxb5ntdQ1oibs |
| *acc genotypes* | 44 |
| *peaks no back* | 1 |
| *removed edges* | 0 |
| *prop rem edges* | 0 |

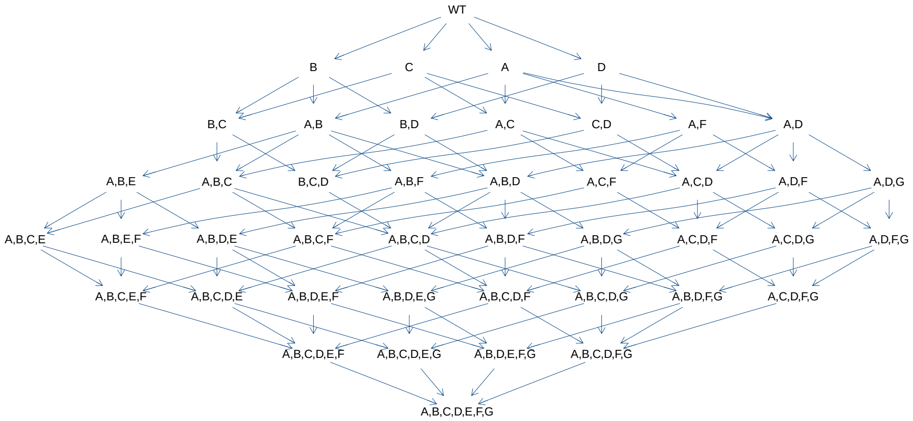


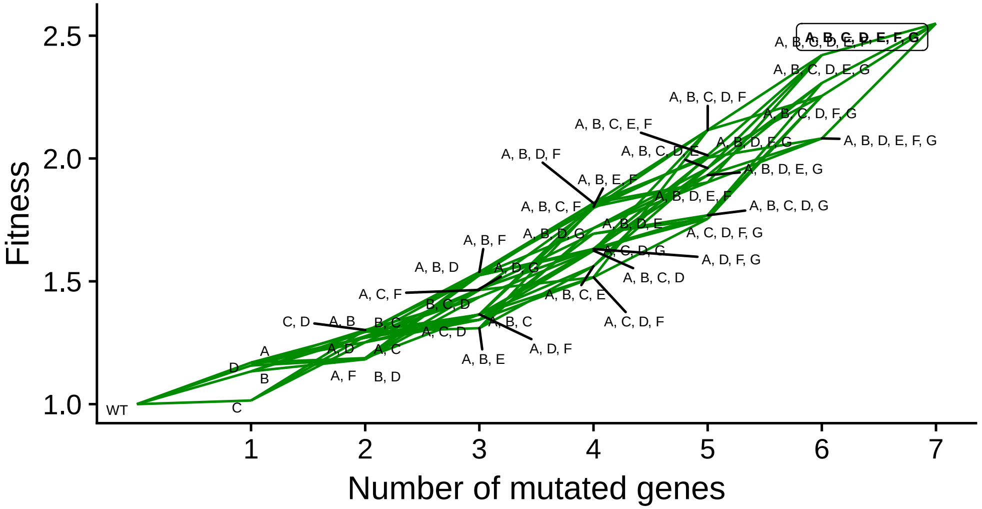

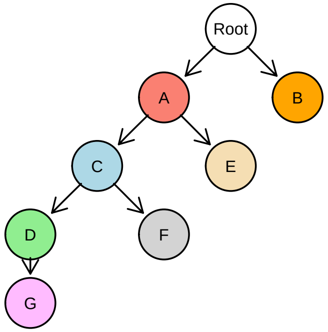


| | |
|---|---|
| *ID* | r6gSASnFWWRSp10wF |
| *acc genotypes* | 30 |
| *peaks no back* | 1 |
| *removed edges* | 0 |
| *prop rem edges* | 0 |

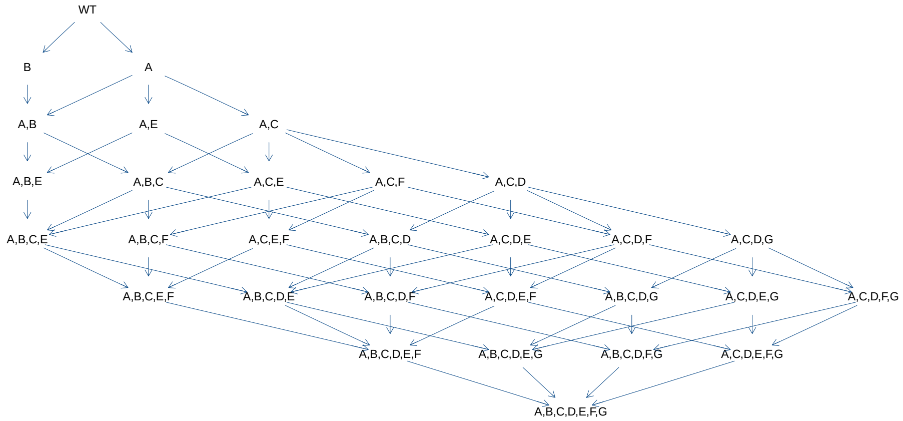


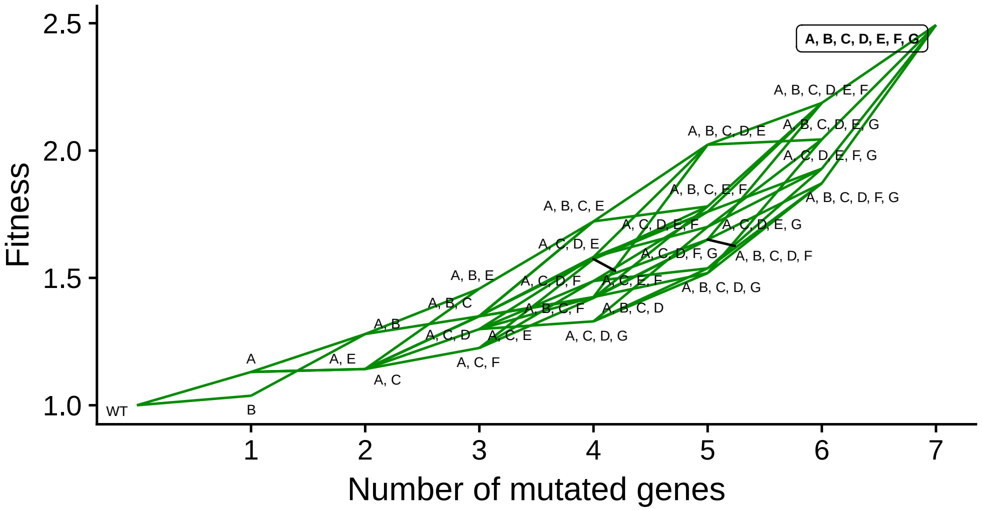

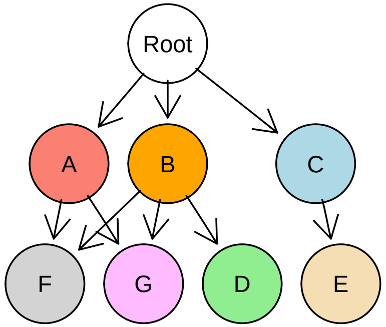


| | |
|---|---|
| *ID* | rcySfhqbP6f5Ys45 |
| *acc genotypes* | 36 |
| *peaks no back* | 1 |
| *removed edges* | 0 |
| *prop rem edges* | 0 |

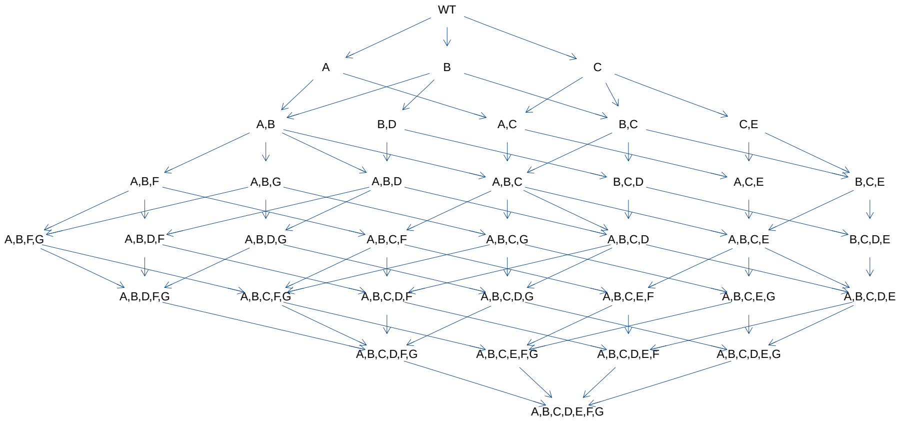


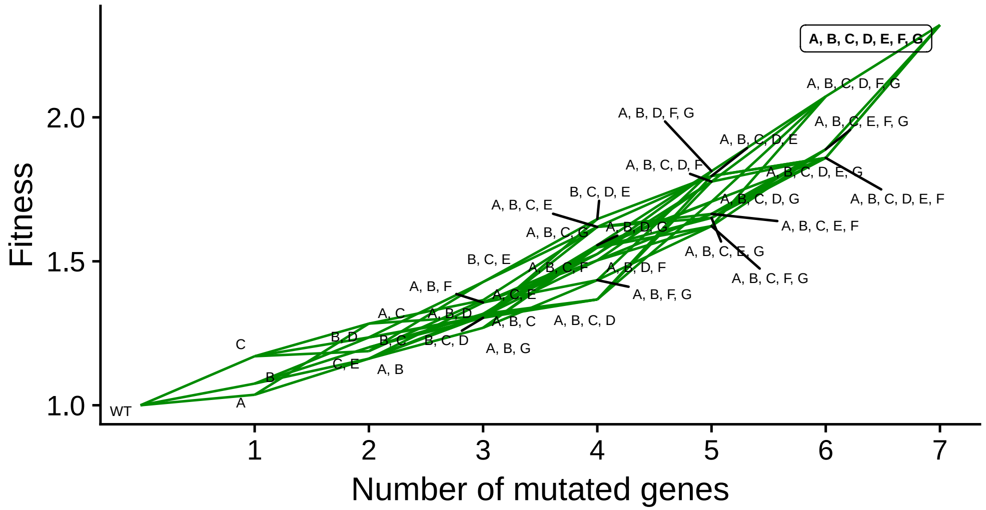

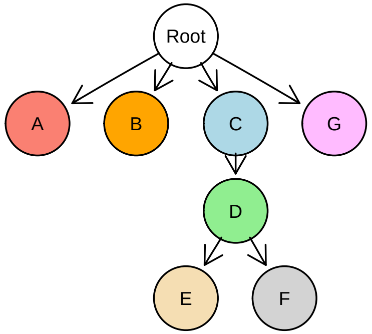


| | |
|---|---|
| *ID* | RgRwHiCY10NBQ5ya4 |
| *acc genotypes* | 48 |
| *peaks no back* | 1 |
| *removed edges* | 0 |
| *prop rem edges* | 0 |

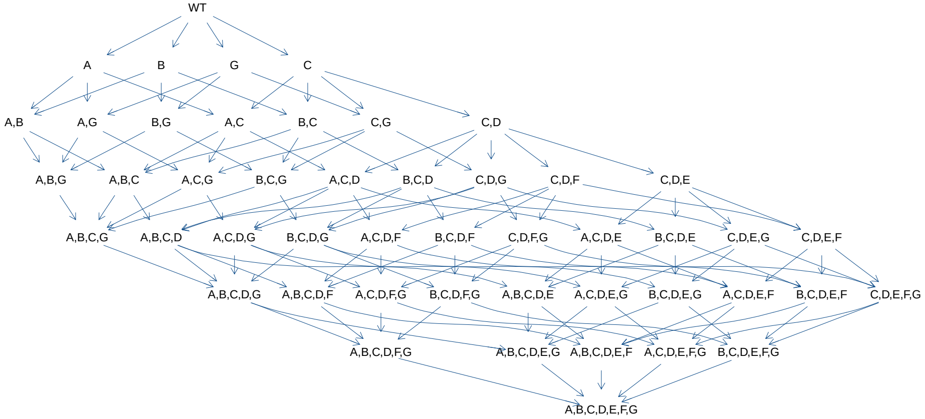


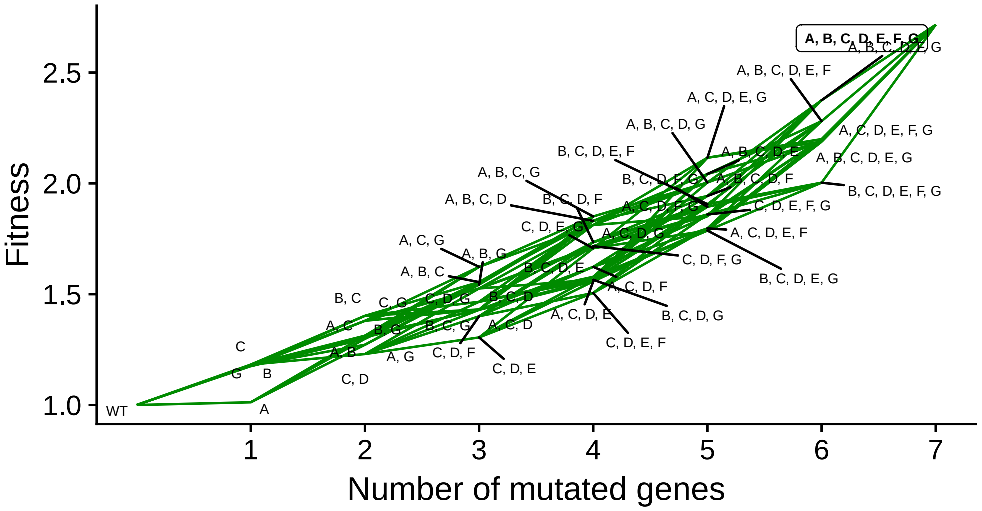

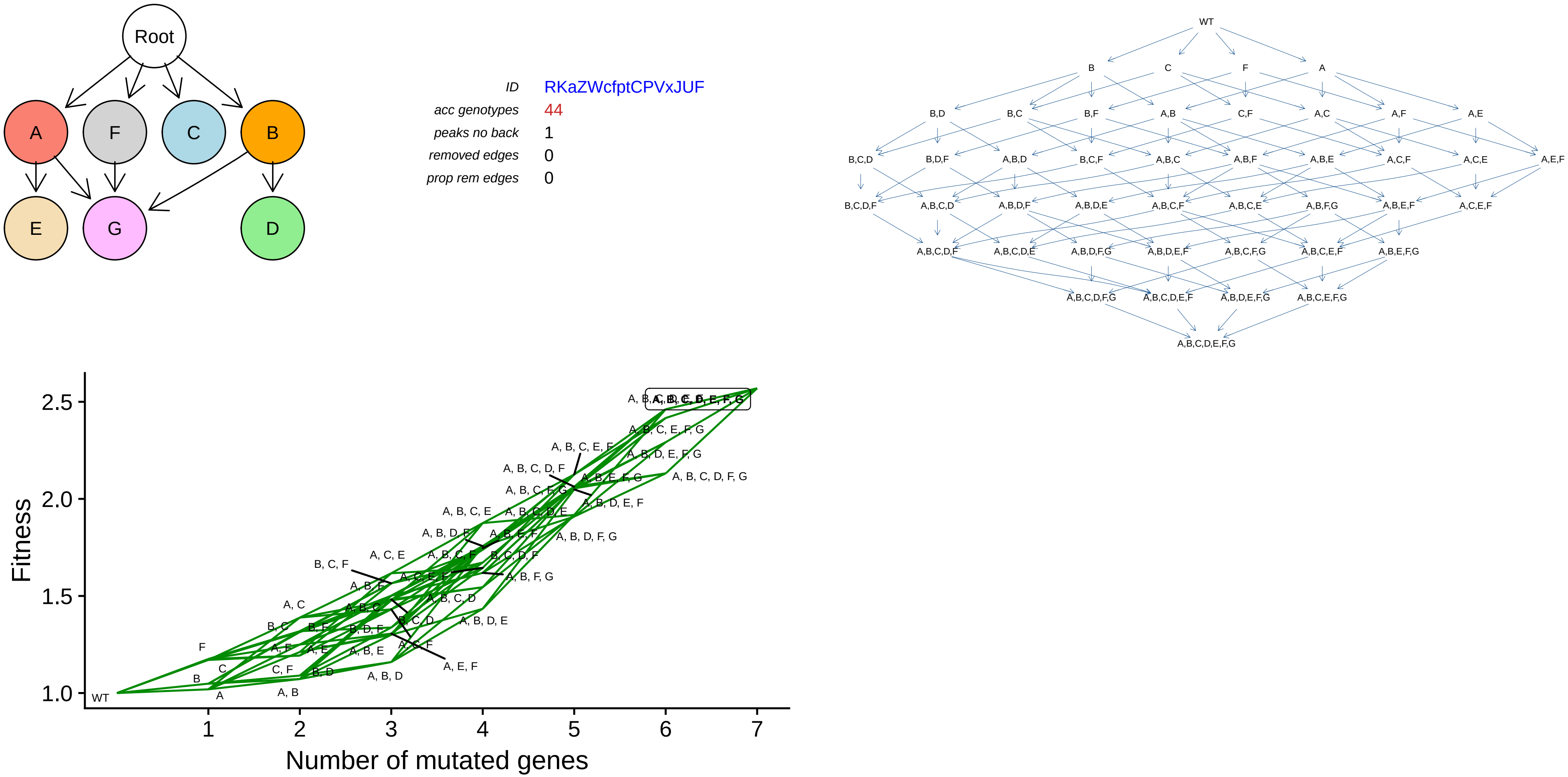

Root
A
F
C
B
E
G
D
ID RKaZWcfptCPVxJUF
acc genotypes 44
peaks no back 1
removed edges 0
prop rem edges 0
Fitness
Number of mutated genes
WT
A, B, C, D, F, G
A, B, C, E, F, G
A, B, D, E, F, G
A, B, C, D, E, F
A, B, C, D, F
A, B, C, F, G
A, B, E, F, G
A, B, D, E, F
A, B, C, E
A, B, C, D, E
A, B, D, F
A, B, E, F
A, B, D, F, G
A, B, C, F
B, C, D, F
A, B, F, G
A, C, E
B, C, F
A, C, E, F
A, B, F
A, B, C, D
A, C
A, B, C
B, C, D
A, B, D, E
B, C
B, F
B, D, F
A, F
A, E
A, B, E
A, C, F
A, E, F
C, F
B, D
A, B, D
A, B

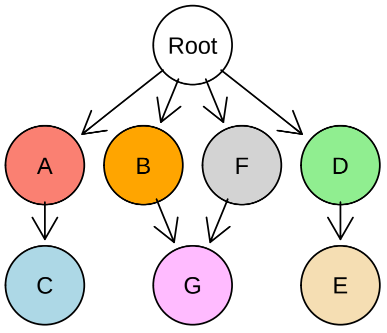


| | |
|---|---|
| *ID* | rokOVrDugL5VXneF |
| *acc genotypes* | 45 |
| *peaks no back* | 1 |
| *removed edges* | 0 |
| *prop rem edges* | 0 |

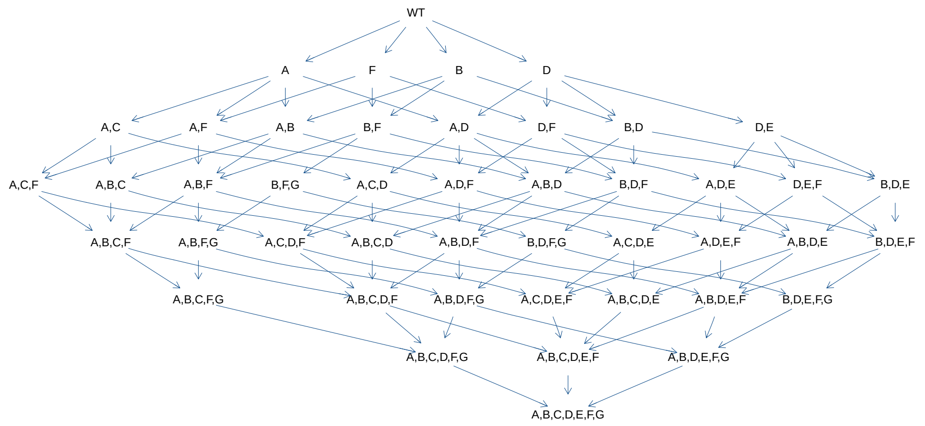


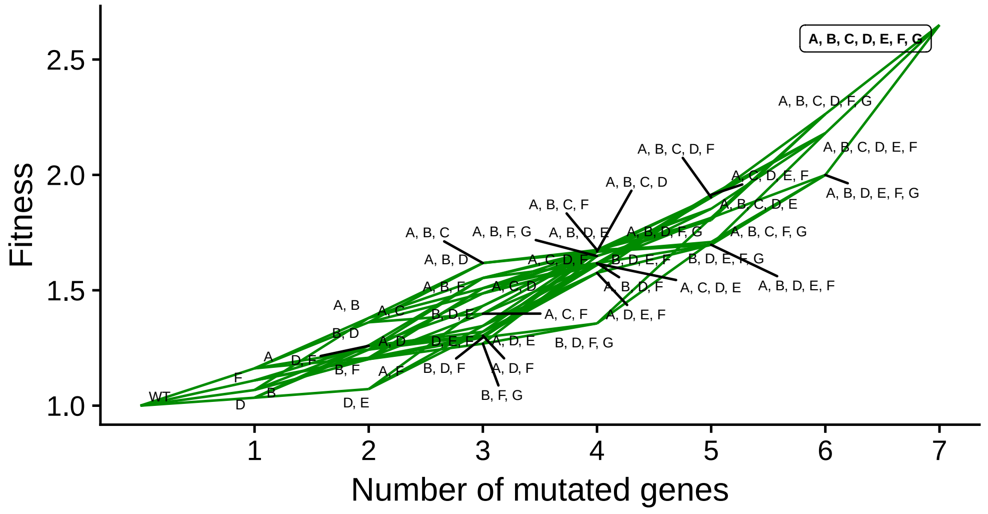

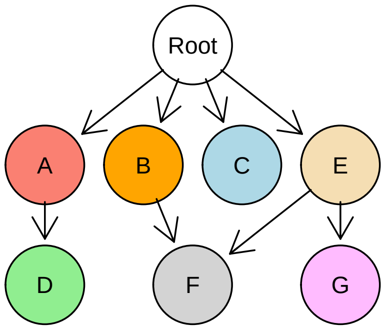


| | |
|---|---|
| *ID* | ryICF10Z10RhI71ONm |
| *acc genotypes* | 48 |
| *peaks no back* | 1 |
| *removed edges* | 0 |
| *prop rem edges* | 0 |

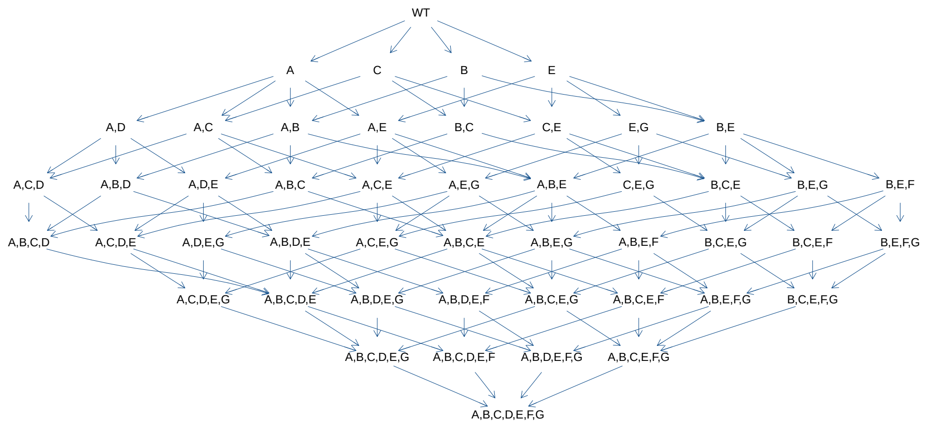


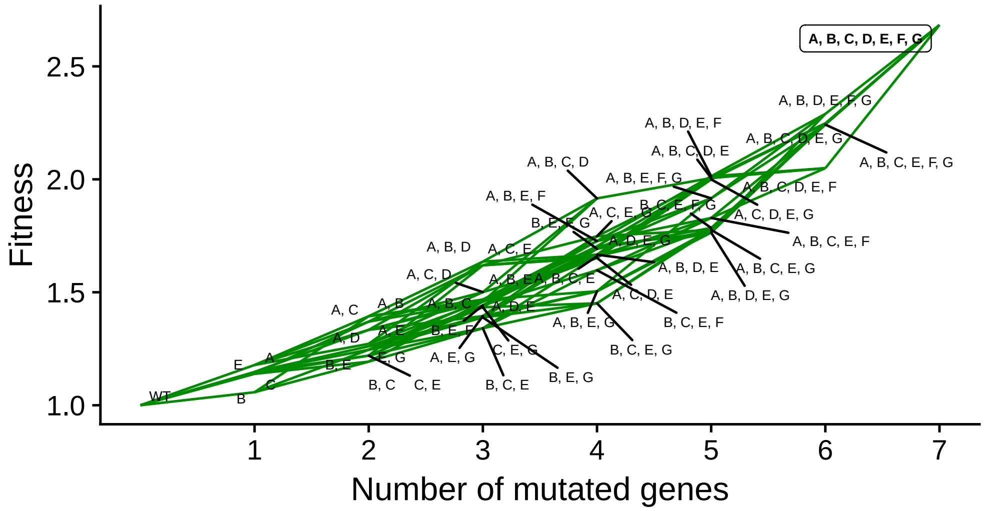

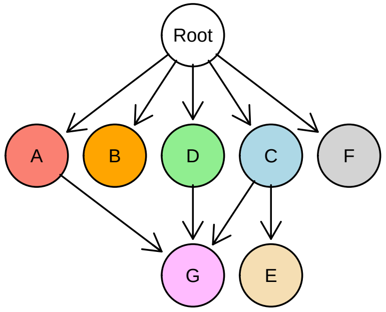


| | |
|---|---|
| *ID* | s6QENCe4O6bJtquJ |
| *acc genotypes* | 56 |
| *peaks no back* | 1 |
| *removed edges* | 0 |
| *prop rem edges* | 0 |

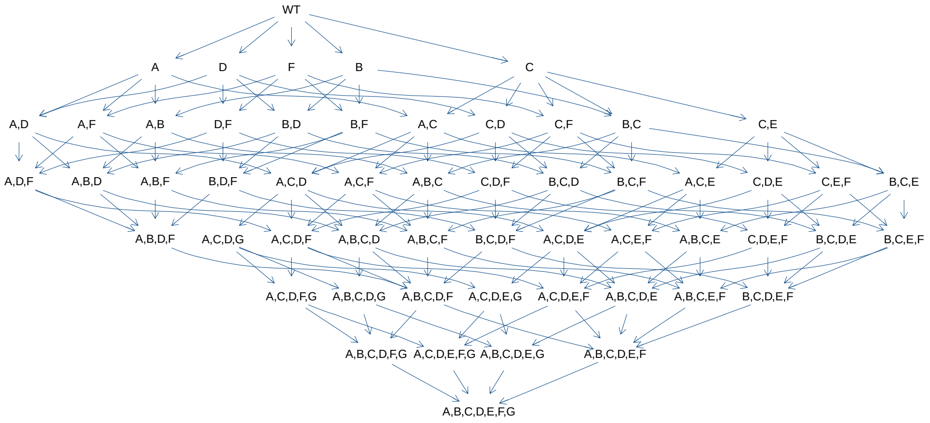


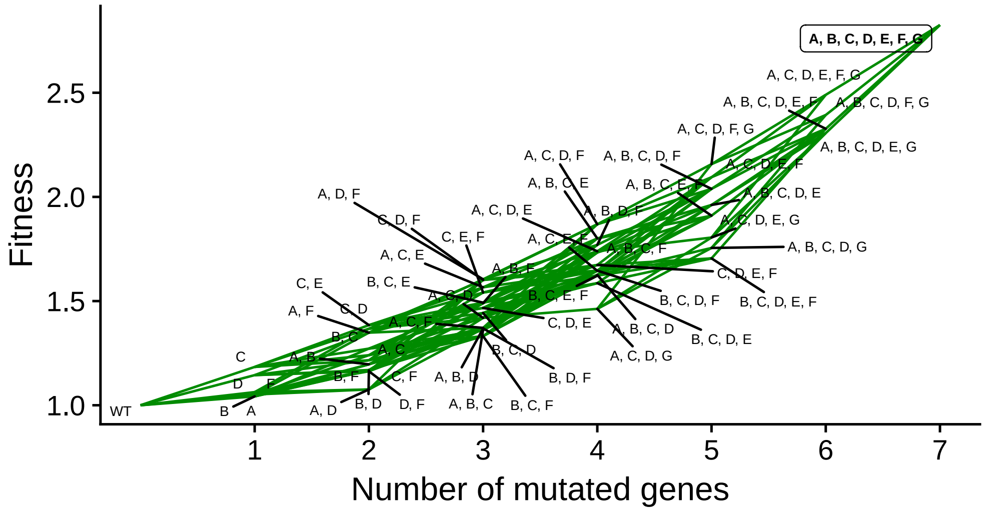

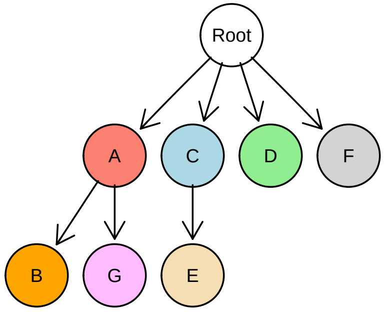


| | |
|---|---|
| *ID* | SeULzsEELispb2JY |
| *acc genotypes* | 60 |
| *peaks no back* | 1 |
| *removed edges* | 0 |
| *prop rem edges* | 0 |

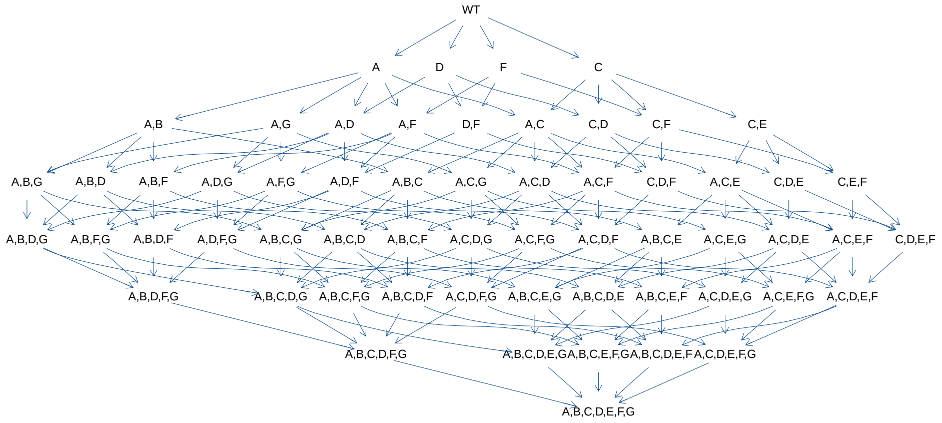


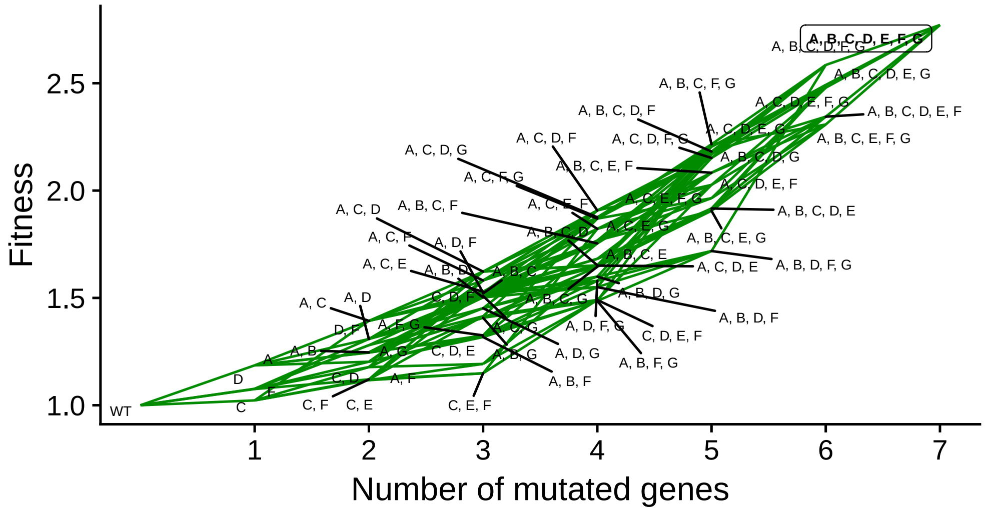

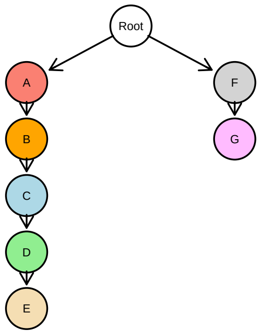


*ID* sJxeTmNhXKbZaZnQ
*acc genotypes* 18
*peaks no back* 1
*removed edges* 0
*prop rem edges* 0

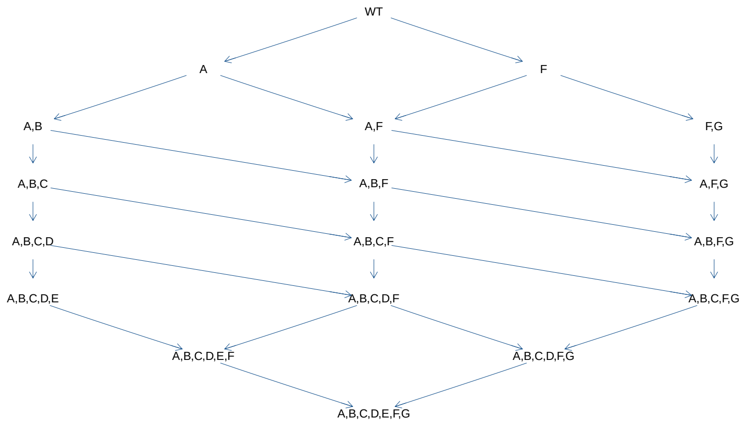


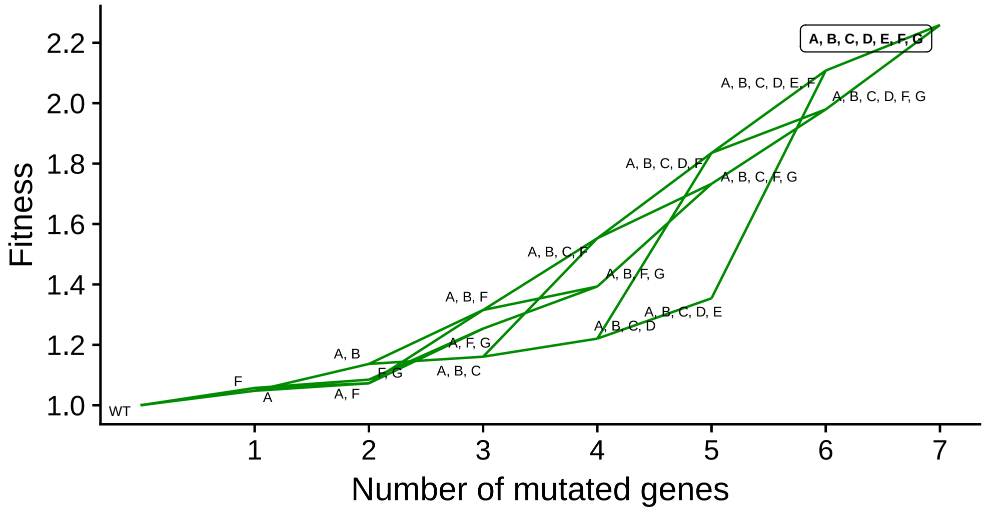

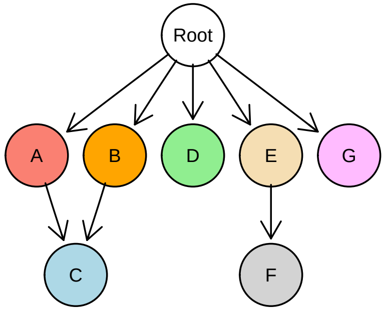


| | |
|---|---|
| *ID* | SUTWyy1ERdbEqE8j |
| *acc genotypes* | 60 |
| *peaks no back* | 1 |
| *removed edges* | 0 |
| *prop rem edges* | 0 |

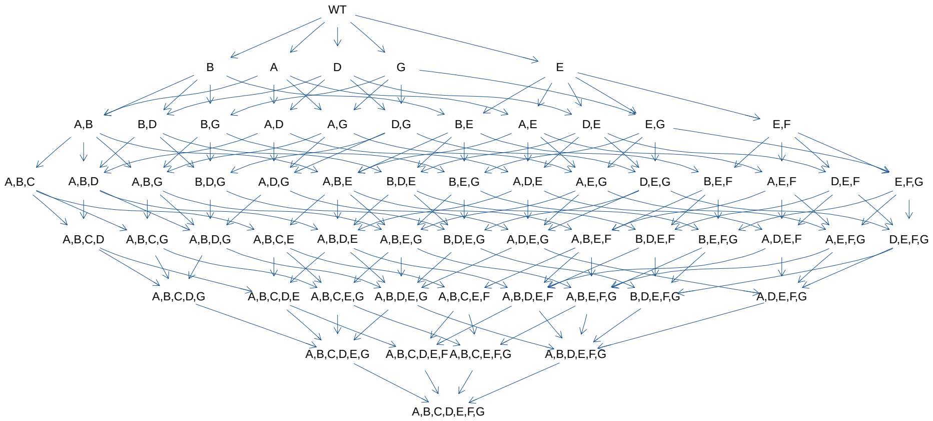


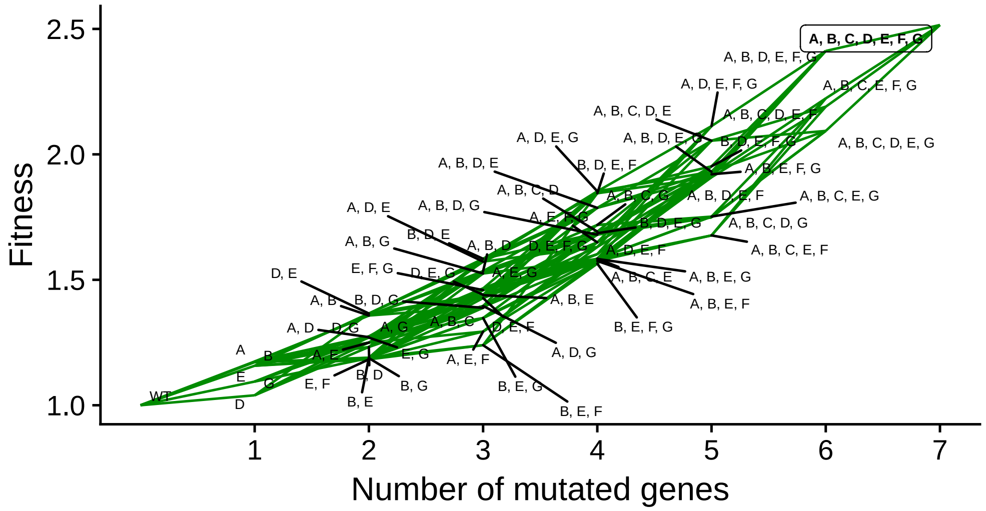

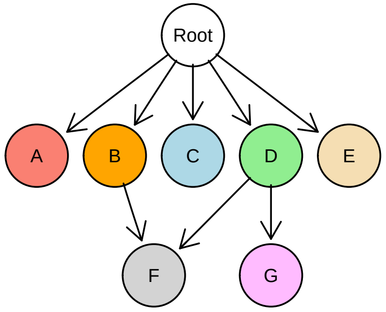


| | |
|---|---|
| *ID* | TEOeR5kKW73StsGe |
| *acc genotypes* | 64 |
| *peaks no back* | 1 |
| *removed edges* | 0 |
| *prop rem edges* | 0 |

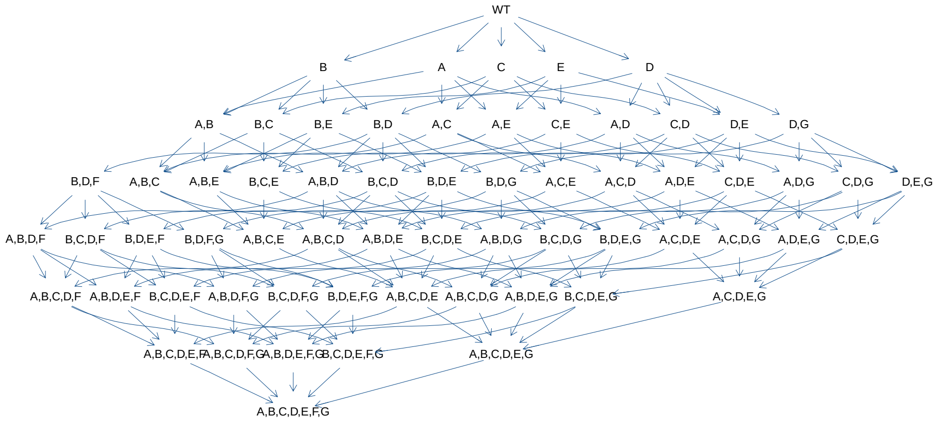


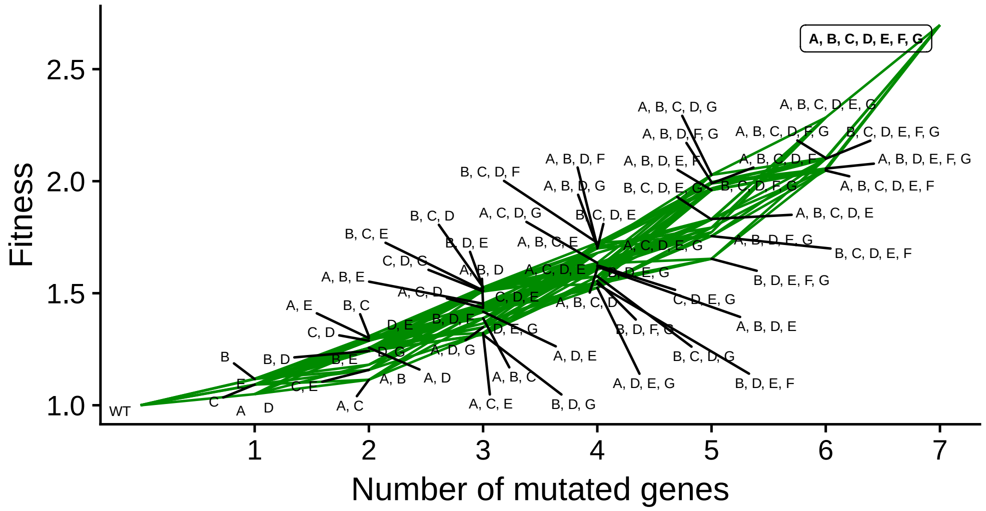

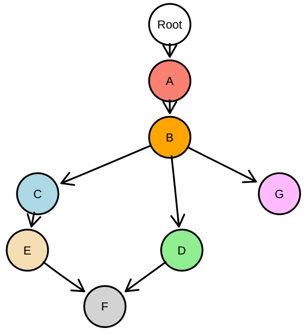


| | |
|---|---|
| *ID* | TqrLOOQde8L7lZjK |
| *acc genotypes* | 16 |
| *peaks no back* | 1 |
| *removed edges* | 0 |
| *prop rem edges* | 0 |

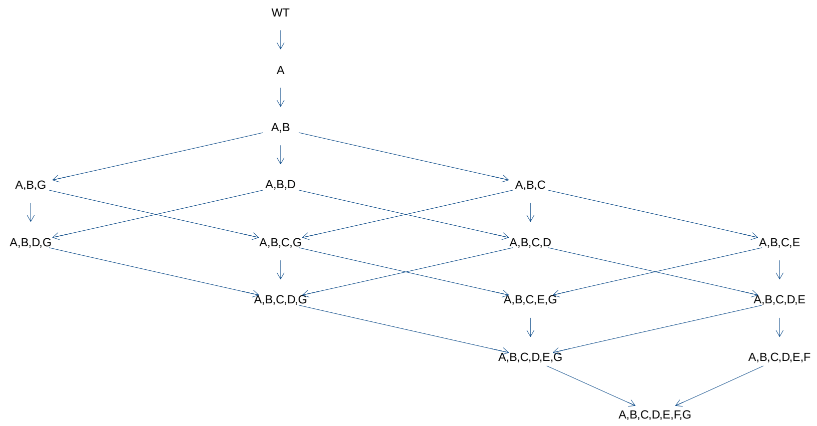


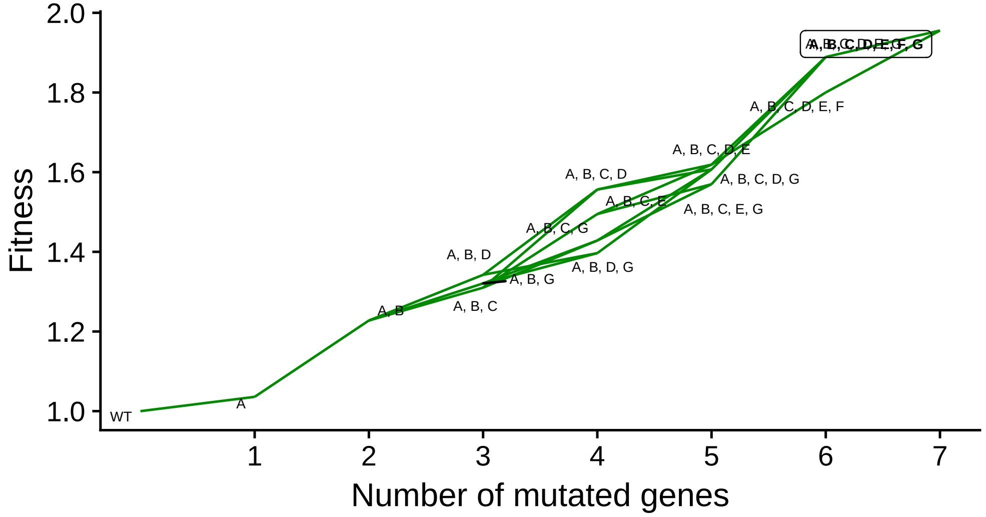

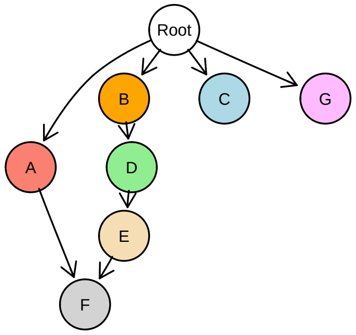


| | |
|---|---|
| *ID* | tRehYFGQCIcbWhJZ |
| *acc genotypes* | 36 |
| *peaks no back* | 1 |
| *removed edges* | 0 |
| *prop rem edges* | 0 |

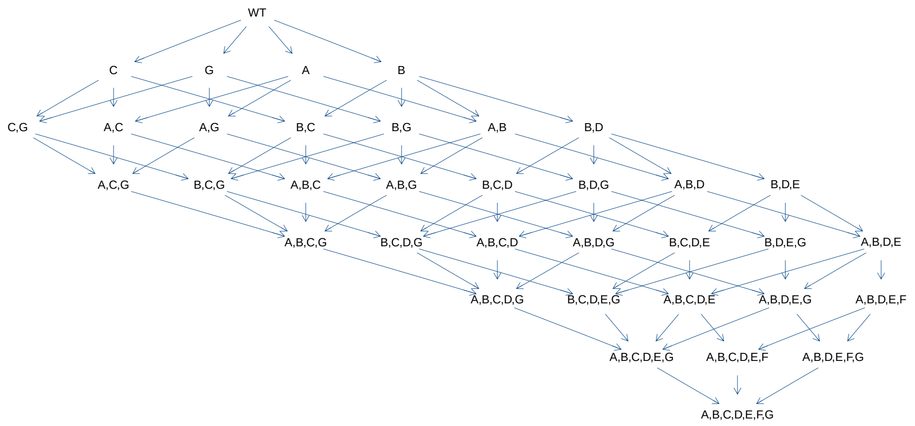


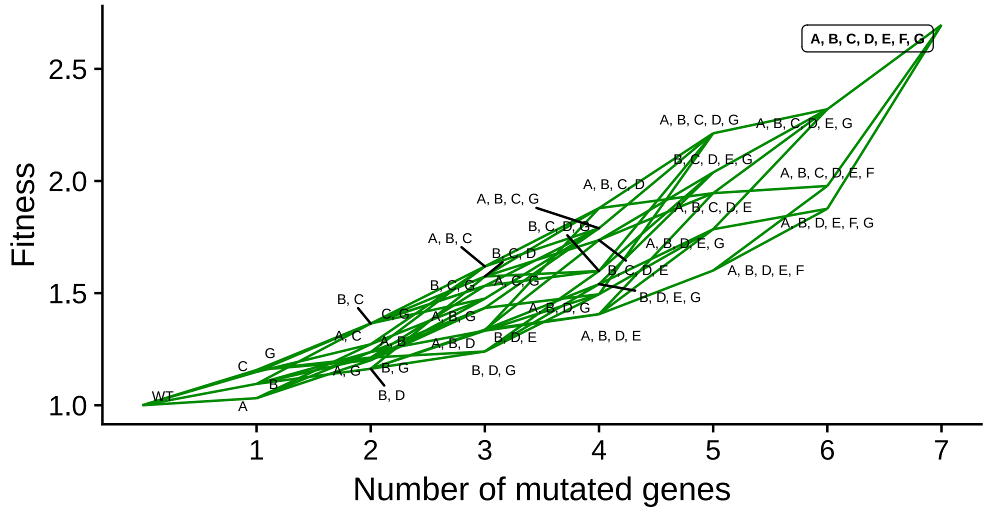

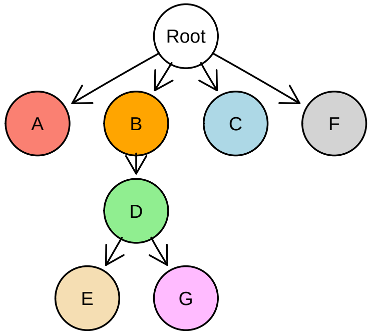


| | |
|---|---|
| *ID* | TX10VAp1TpO185dpx |
| *acc genotypes* | 48 |
| *peaks no back* | 1 |
| *removed edges* | 0 |
| *prop rem edges* | 0 |

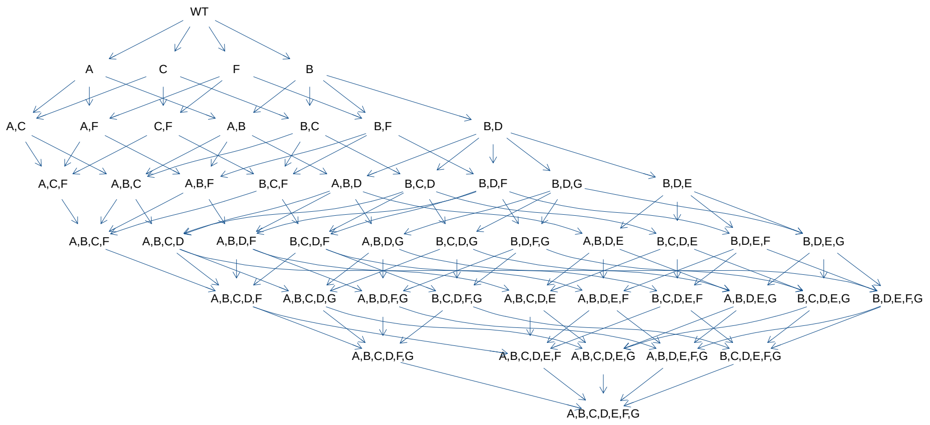


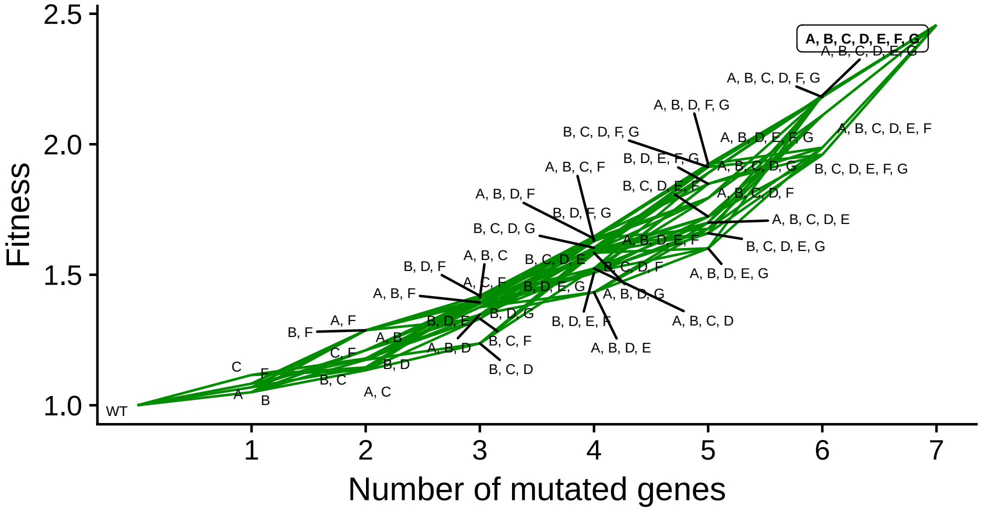

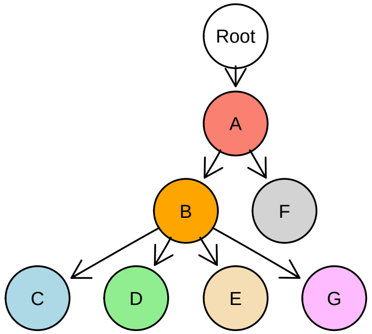


| | |
|---|---|
| *ID* | tZE7JJadDSWCdZhf |
| *acc genotypes* | 35 |
| *peaks no back* | 1 |
| *removed edges* | 0 |
| *prop rem edges* | 0 |

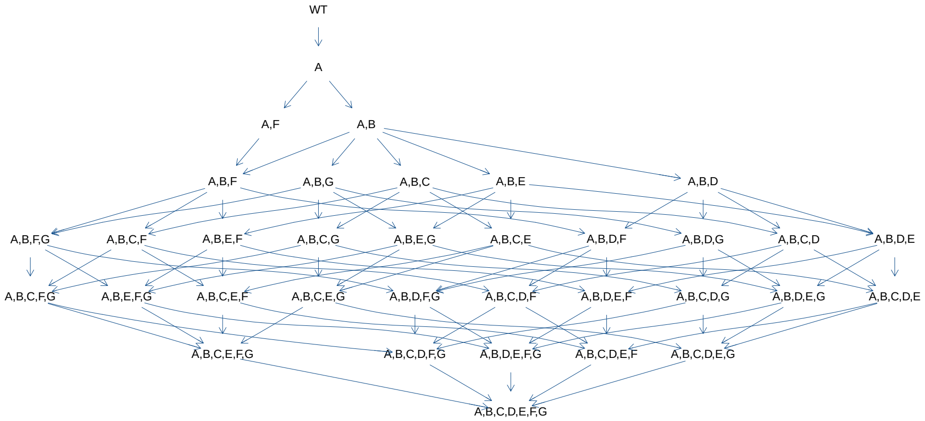


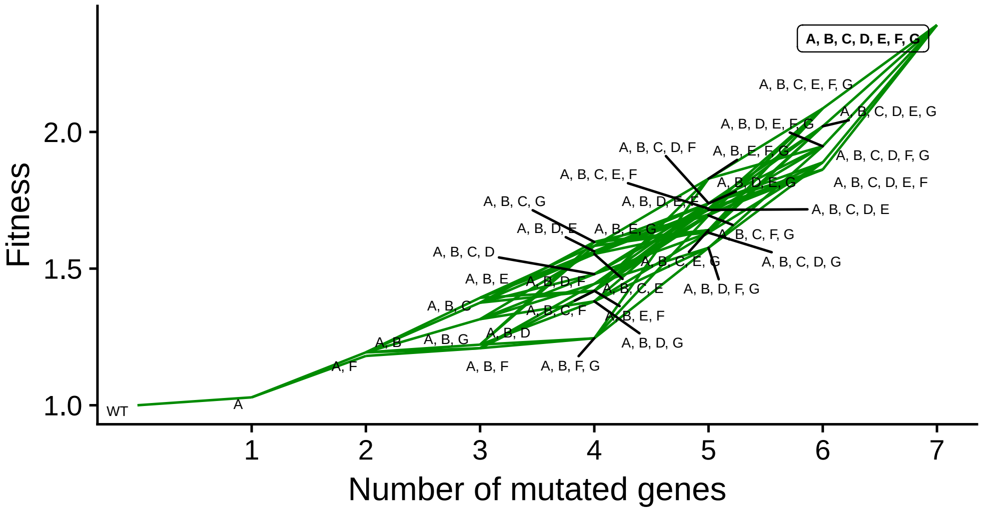

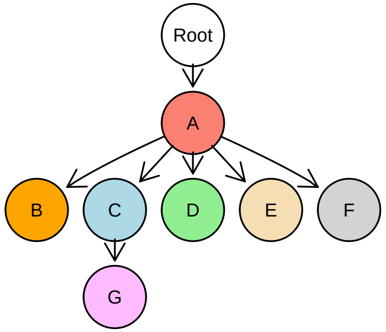


| | |
|---|---|
| *ID* | TZvpRpDnbfQvlt4G |
| *acc genotypes* | 49 |
| *peaks no back* | 1 |
| *removed edges* | 0 |
| *prop rem edges* | 0 |

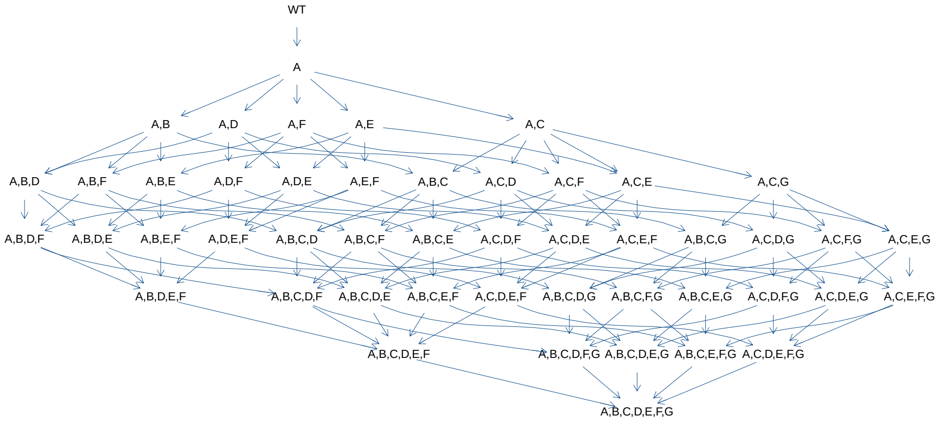


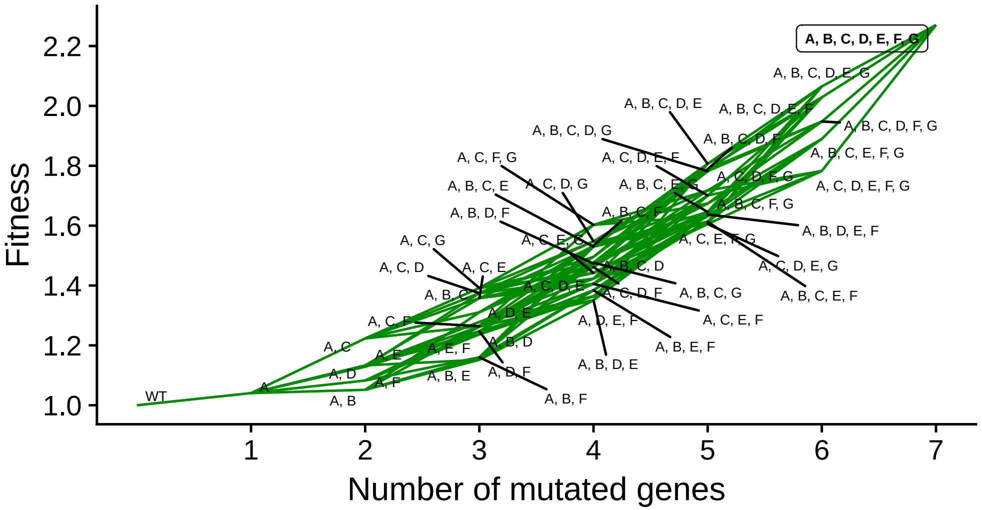

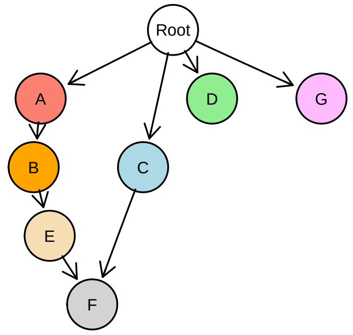


| | |
|---|---|
| *ID* | U9FVyqSn82tFhAtl |
| *acc genotypes* | 36 |
| *peaks no back* | 1 |
| *removed edges* | 0 |
| *prop rem edges* | 0 |

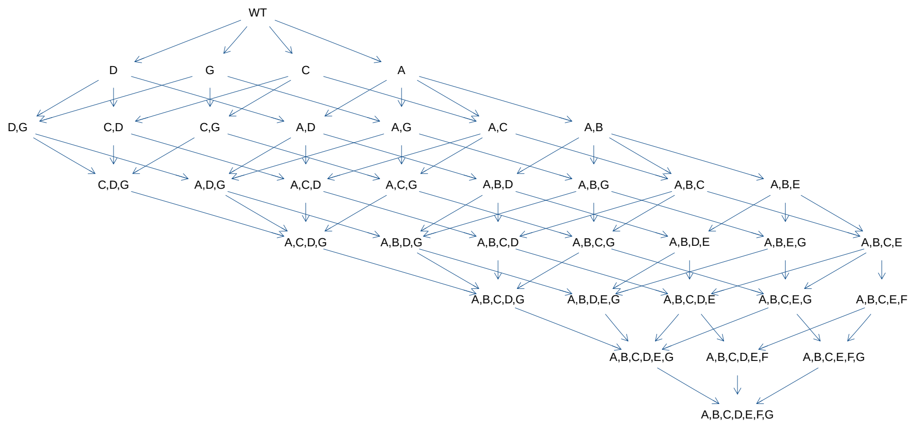


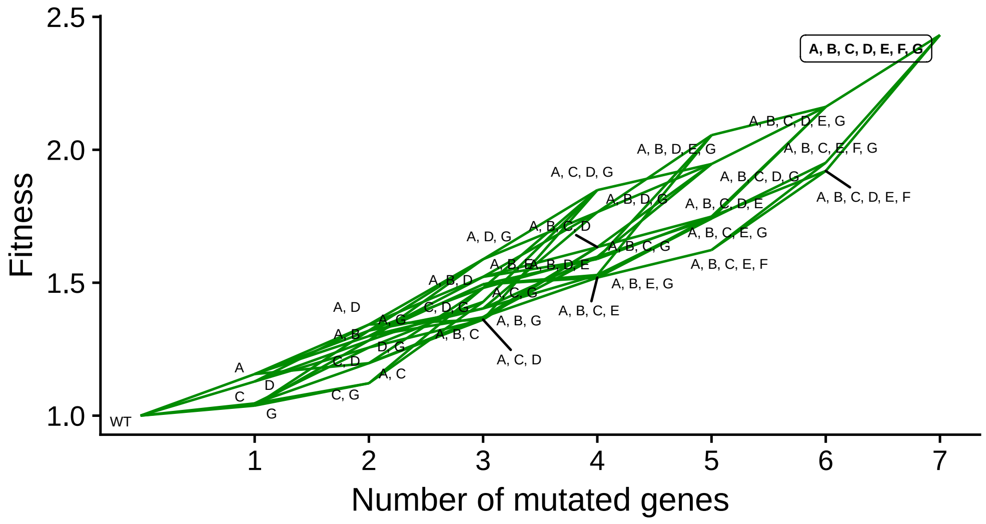

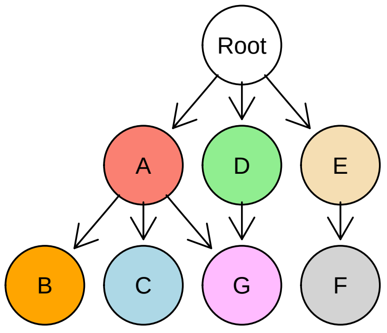


| | |
|---|---|
| *ID* | U9tqKXHuH1LkO5mF |
| *acc genotypes* | 42 |
| *peaks no back* | 1 |
| *removed edges* | 0 |
| *prop rem edges* | 0 |

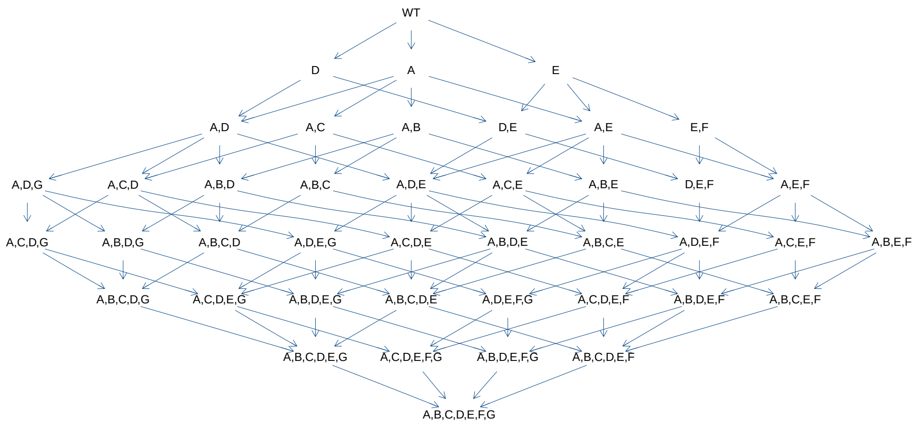


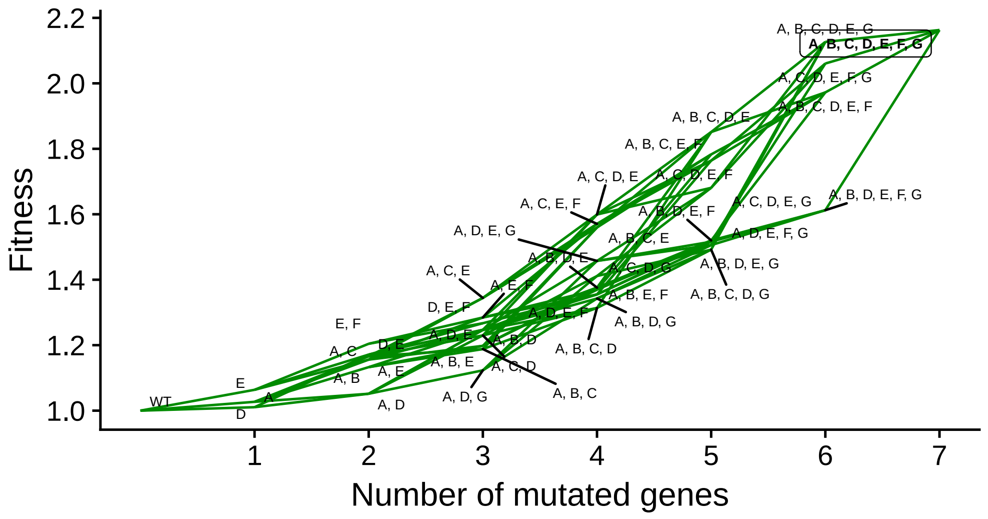

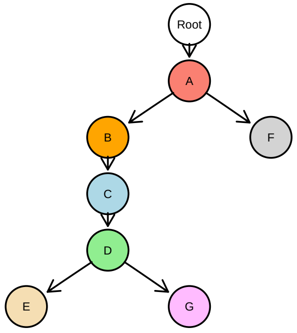


| | |
|---|---|
| *ID* | Ua2QSlSPNSRgJaJY |
| *acc genotypes* | 15 |
| *peaks no back* | 1 |
| *removed edges* | 0 |
| *prop rem edges* | 0 |

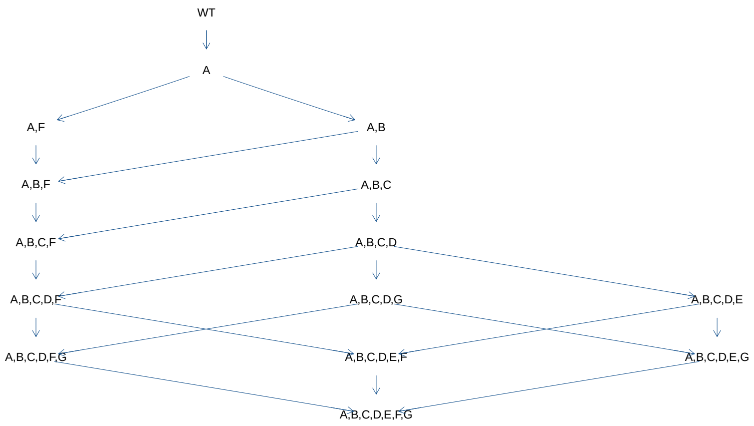


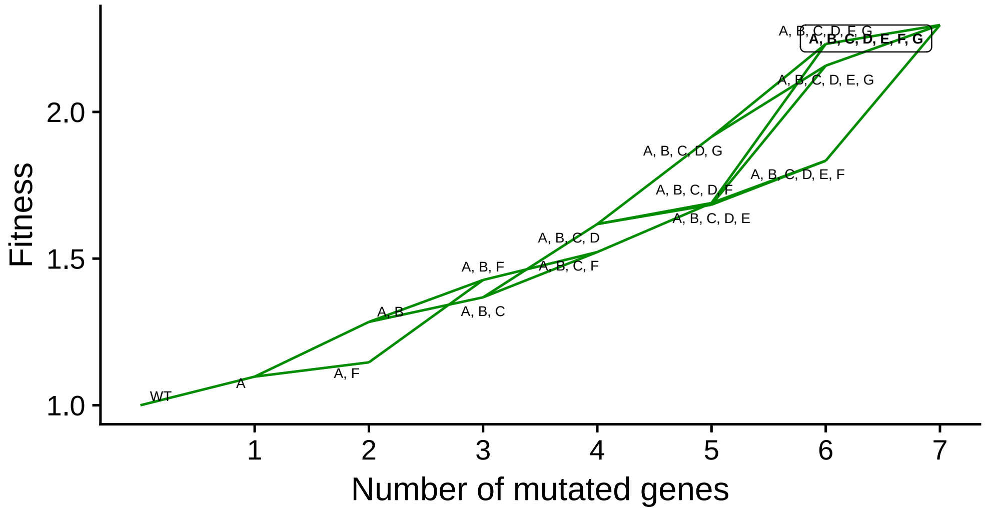

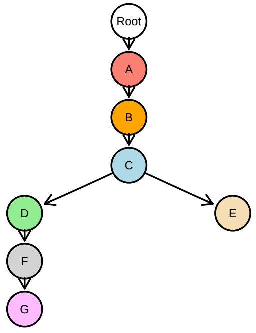


| | |
|---|---|
| *ID* | UDkNV5Dxa4w10lh6h |
| *acc genotypes* | 11 |
| *peaks no back* | 1 |
| *removed edges* | 0 |
| *prop rem edges* | 0 |

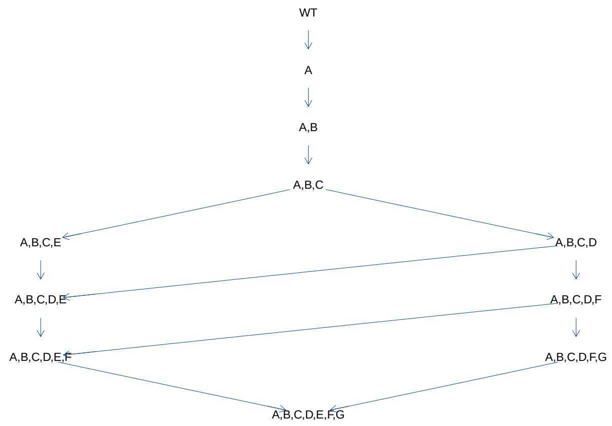


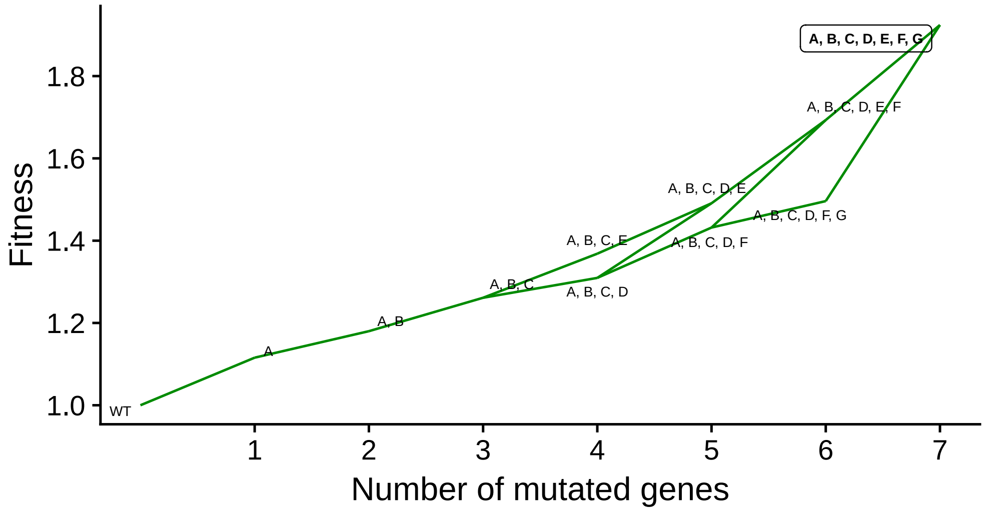

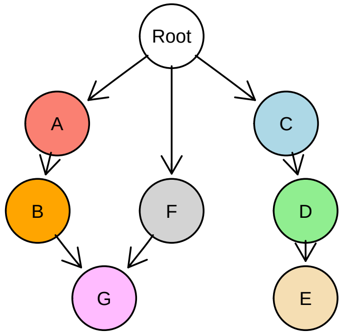


| | |
|---|---|
| *ID* | uggvrwXSAwaMkYSi |
| *acc genotypes* | 28 |
| *peaks no back* | 1 |
| *removed edges* | 0 |
| *prop rem edges* | 0 |

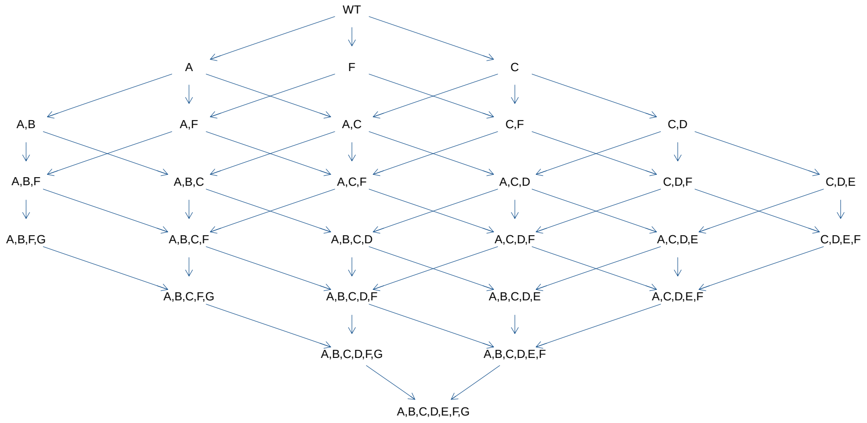


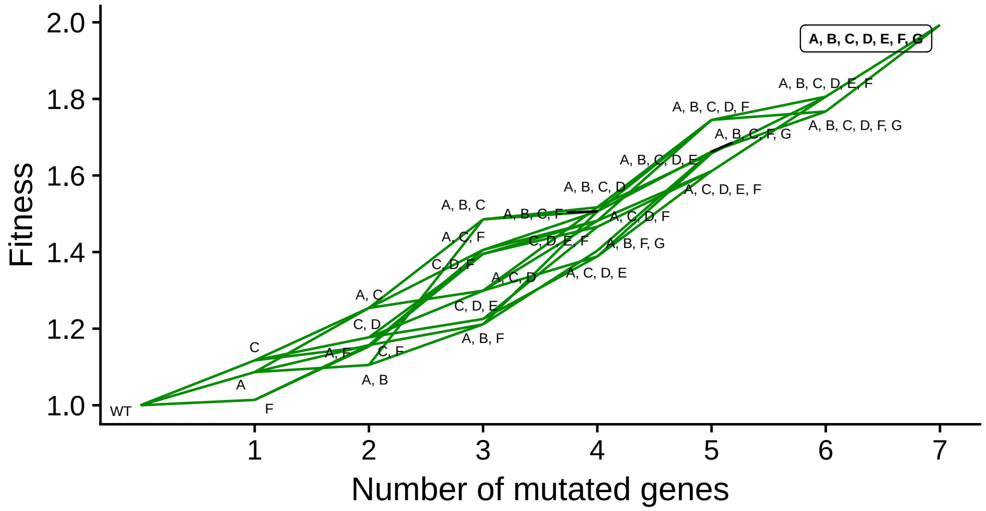

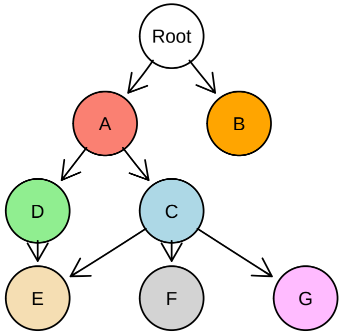


| | |
|---|---|
| *ID* | UqGRanXLIlQe1z2o |
| *acc genotypes* | 30 |
| *peaks no back* | 1 |
| *removed edges* | 0 |
| *prop rem edges* | 0 |

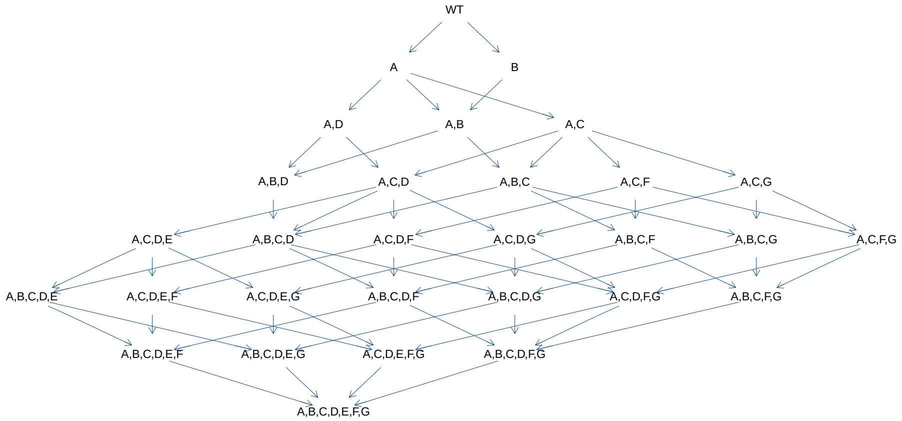


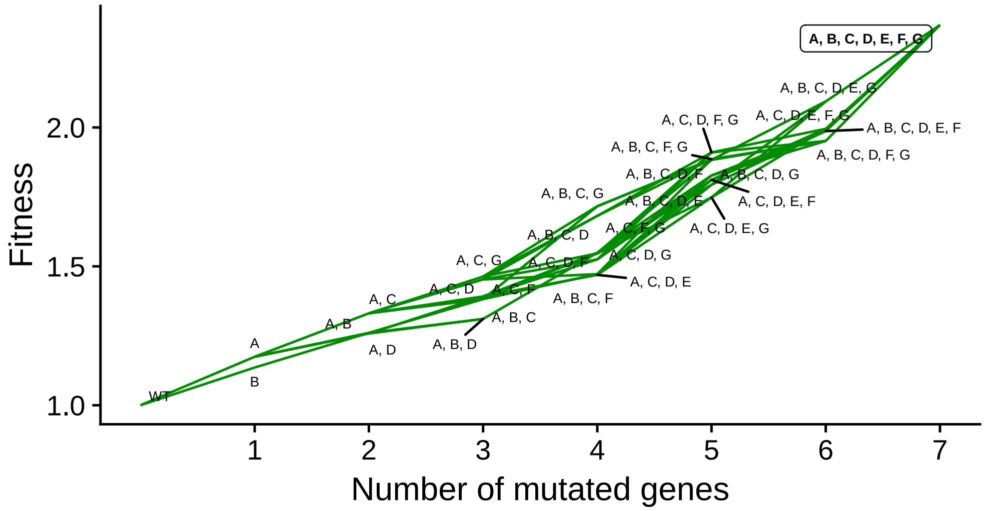

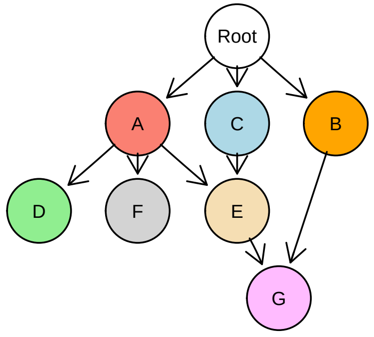


| | |
|---|---|
| *ID* | uvOx4rGT8Tr6iHvm |
| *acc genotypes* | 32 |
| *peaks no back* | 1 |
| *removed edges* | 0 |
| *prop rem edges* | 0 |

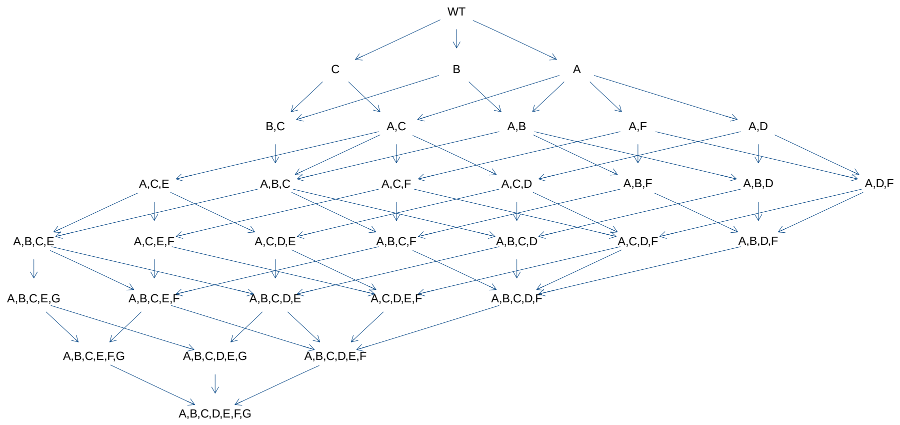


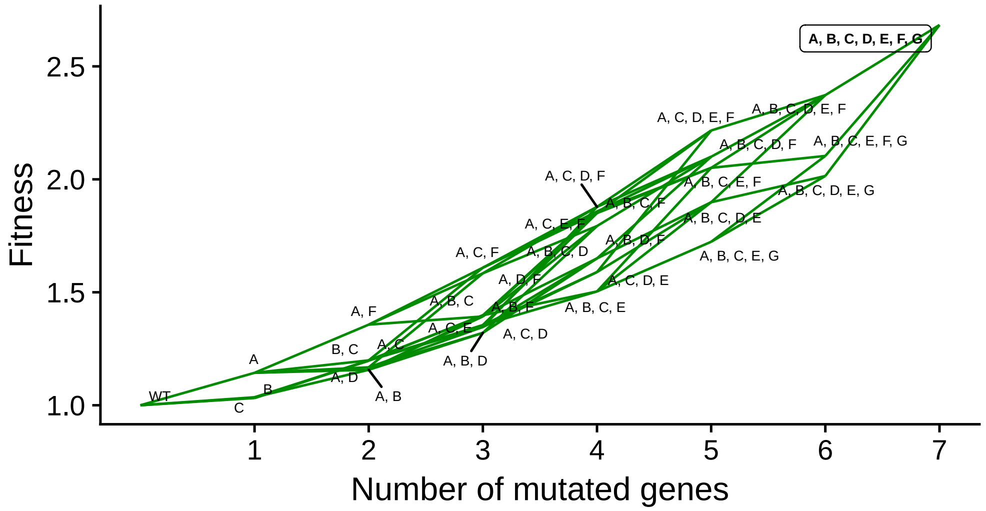

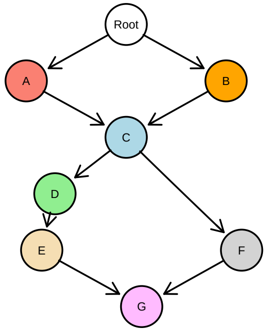


| | |
|---|---|
| *ID* | uW1bRozHBgg4IHvp |
| *acc genotypes* | 11 |
| *peaks no back* | 1 |
| *removed edges* | 0 |
| *prop rem edges* | 0 |

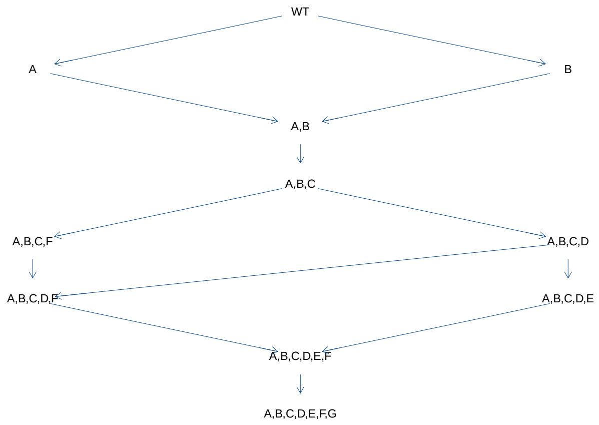


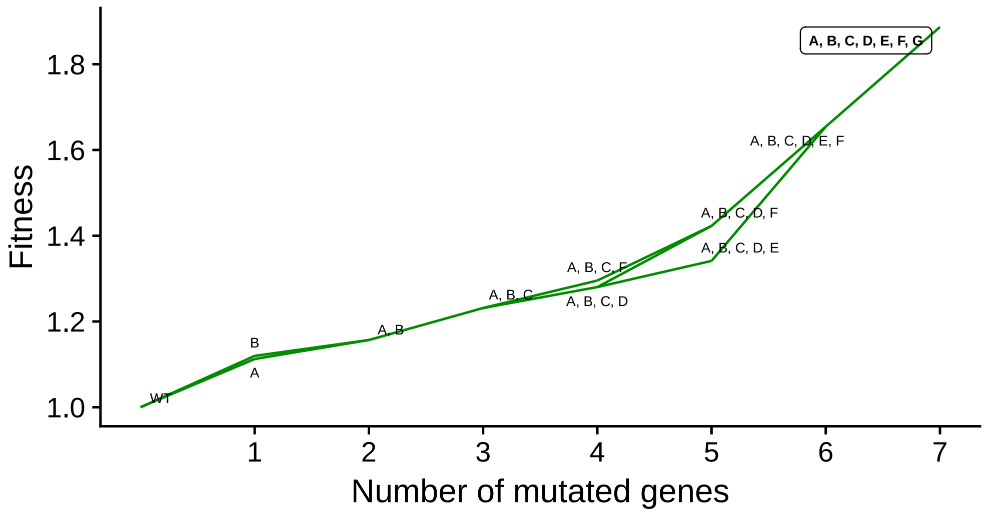

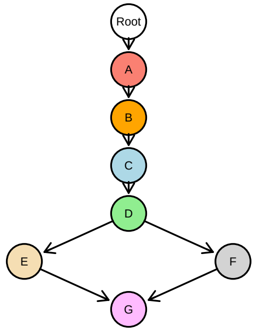


| | |
|---|---|
| *ID* | uXxo7pBZAOc10w5w3 |
| *acc genotypes* | 9 |
| *peaks no back* | 1 |
| *removed edges* | 0 |
| *prop rem edges* | 0 |

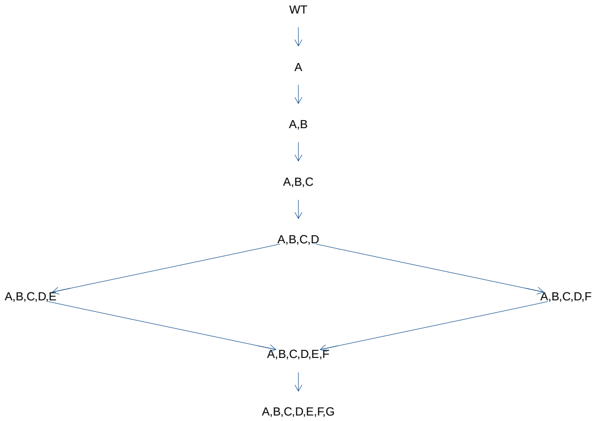


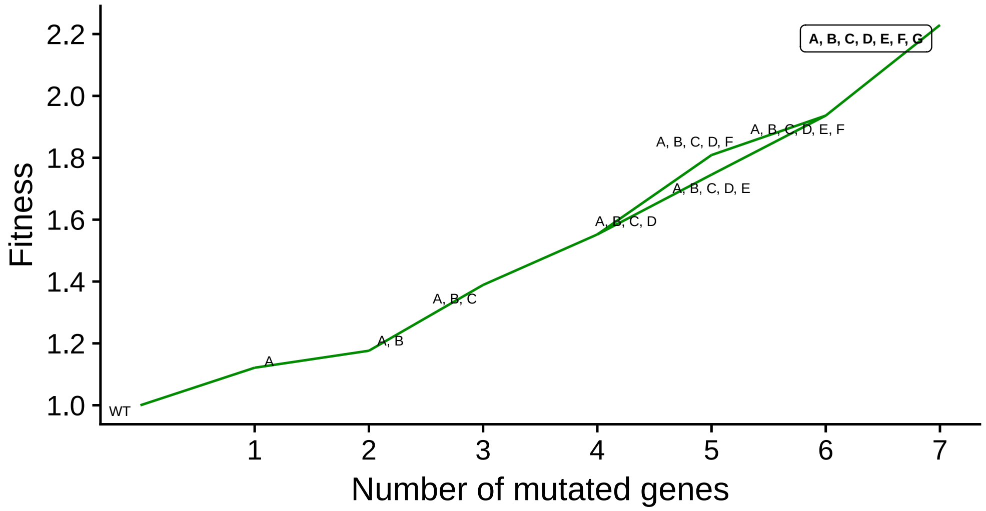

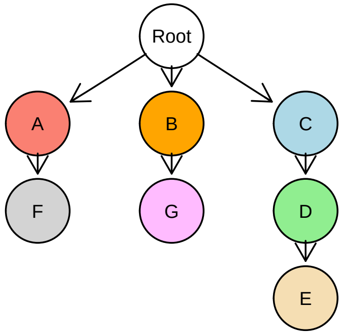


| | |
|---|---|
| *ID* | vG10cQRZVOxdasUPn |
| *acc genotypes* | 36 |
| *peaks no back* | 1 |
| *removed edges* | 0 |
| *prop rem edges* | 0 |

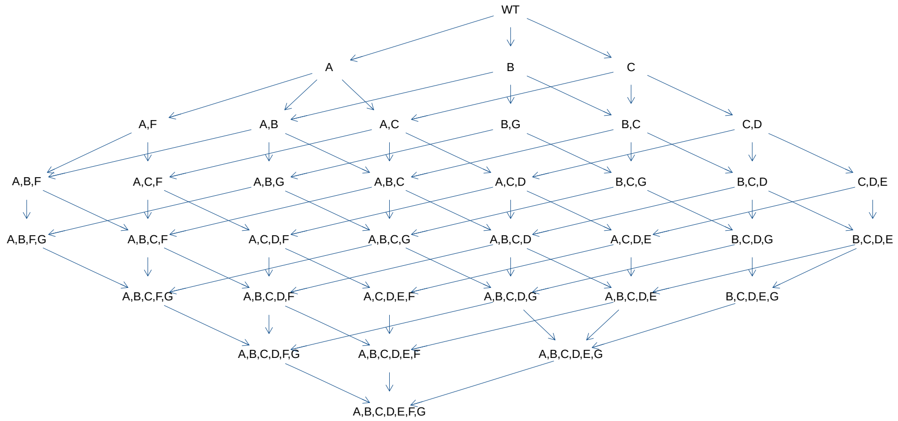


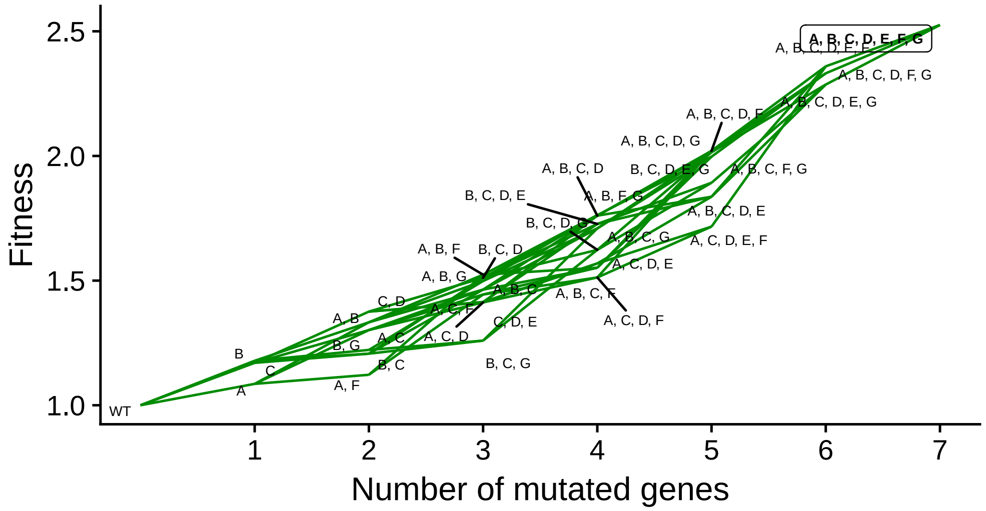

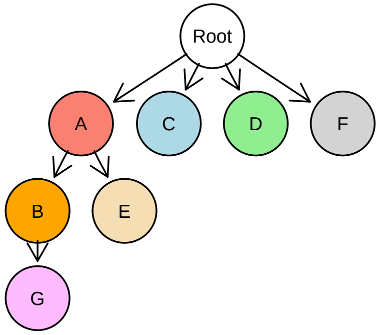


| | |
|---|---|
| *ID* | vpP1t7lvrEU3dpj1 |
| *acc genotypes* | 56 |
| *peaks no back* | 1 |
| *removed edges* | 0 |
| *prop rem edges* | 0 |

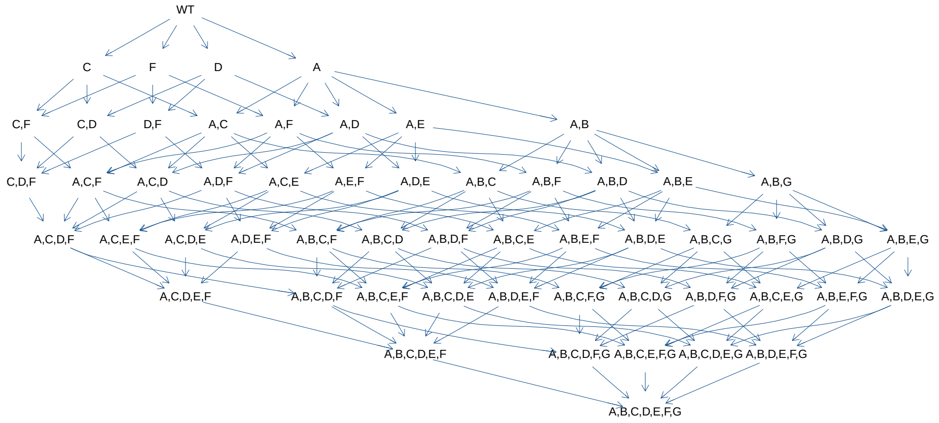


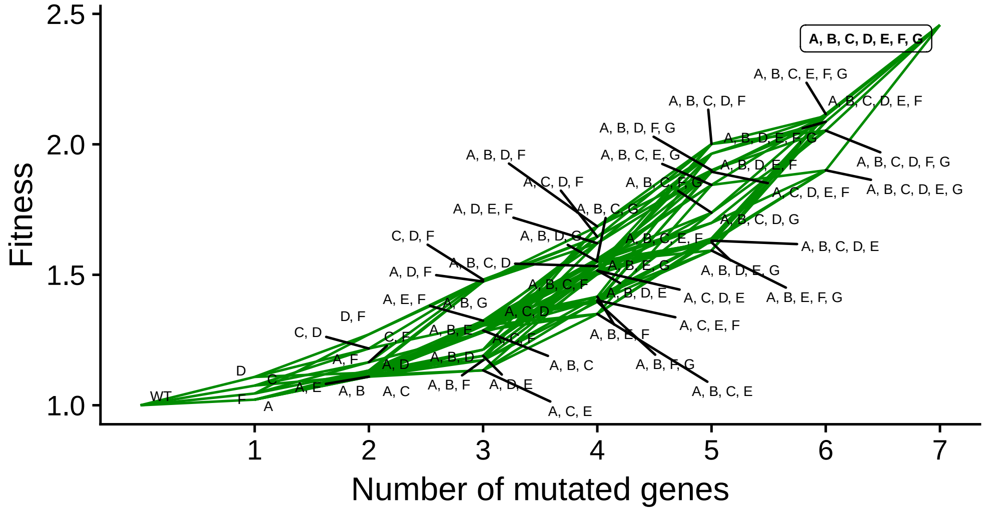

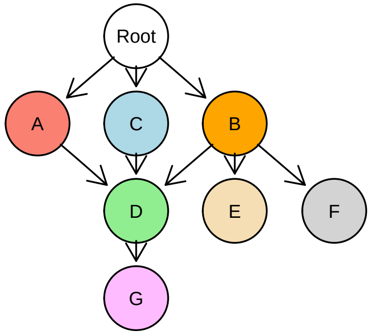


| | |
|---|---|
| *ID* | vq10rX5oSabdGhDLE |
| *acc genotypes* | 28 |
| *peaks no back* | 1 |
| *removed edges* | 0 |
| *prop rem edges* | 0 |

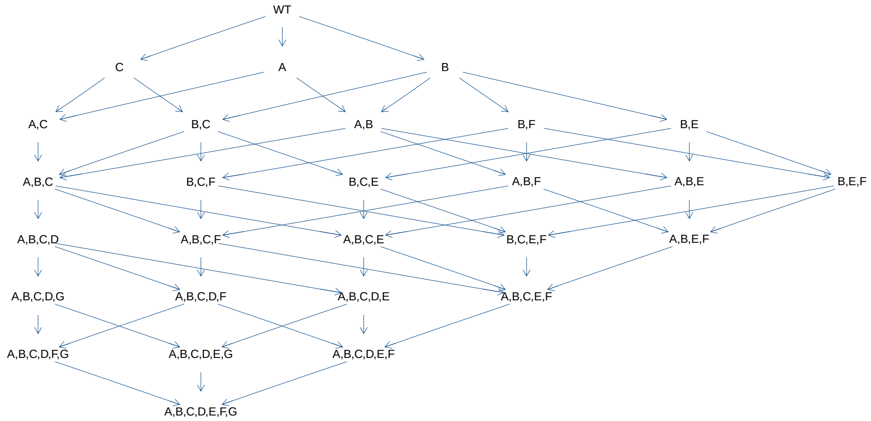


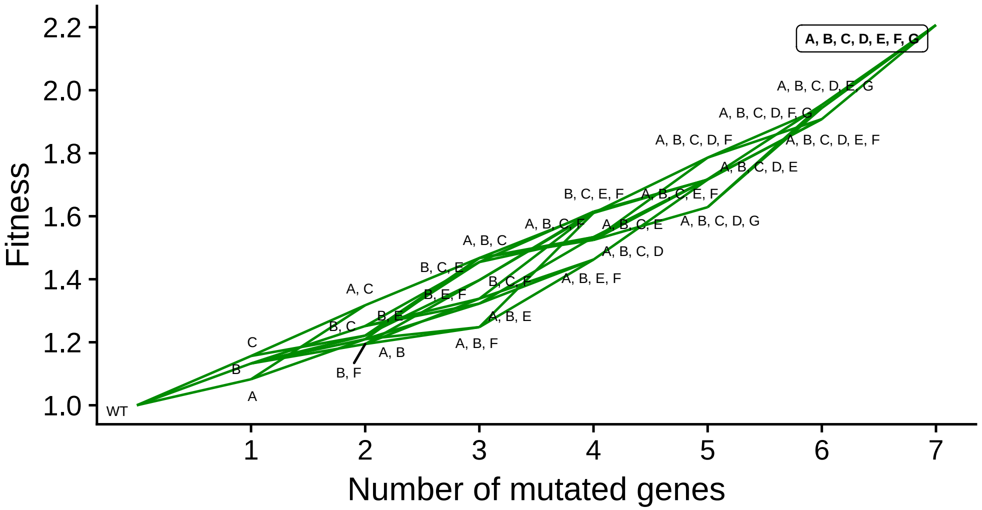

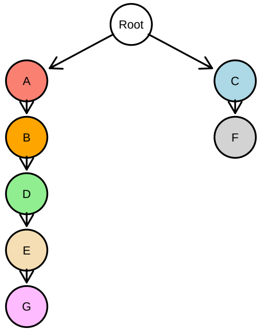


| | |
|---|---|
| *ID* | vrlakeKZSaVC1Lzx |
| *acc genotypes* | 18 |
| *peaks no back* | 1 |
| *removed edges* | 0 |
| *prop rem edges* | 0 |

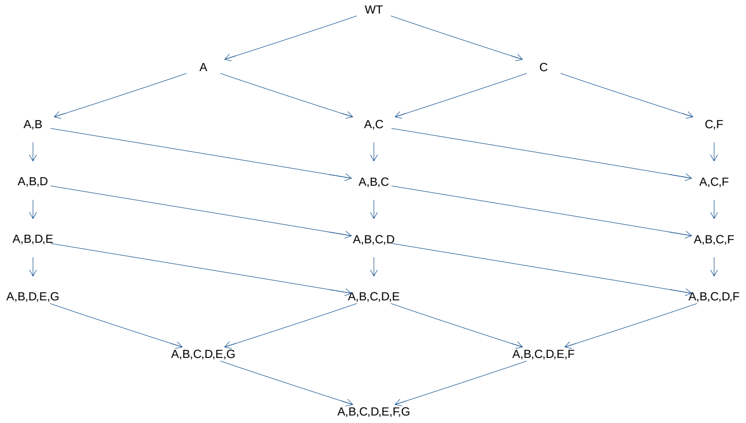


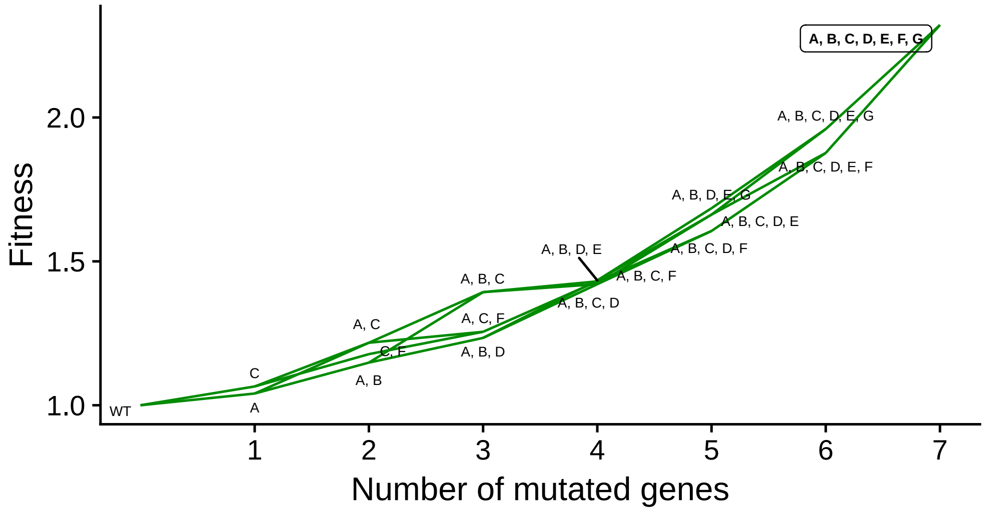

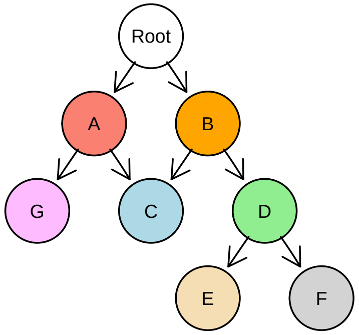


*ID* VYbhtDvalyAvnLsQ

*acc genotypes* 28

*peaks no back* 1

*removed edges* 0

*prop rem edges* 0

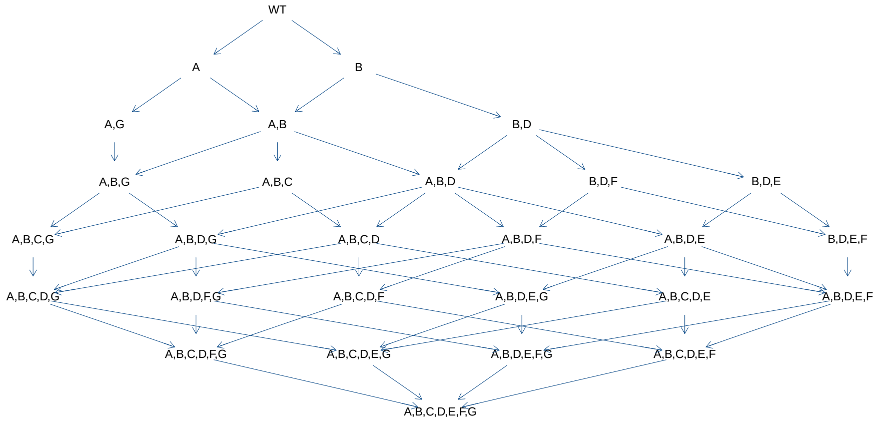


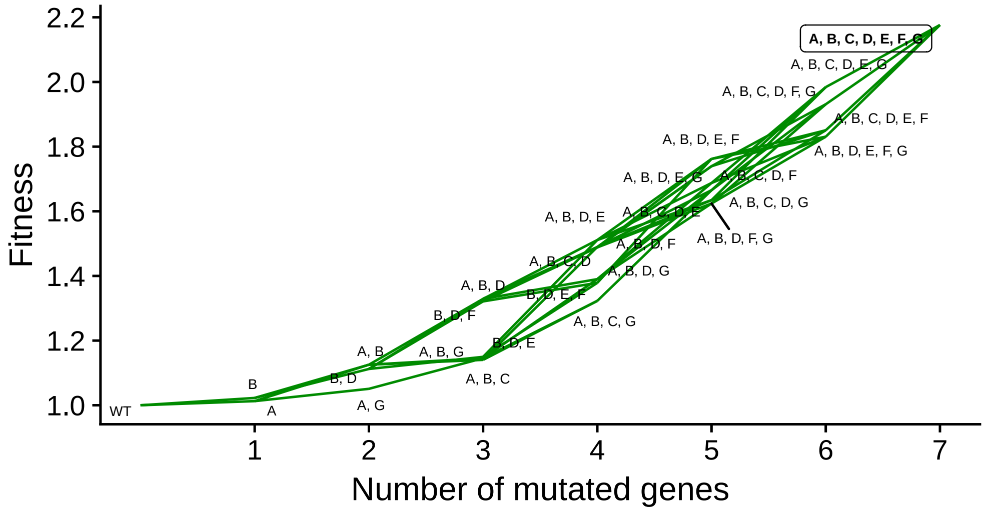

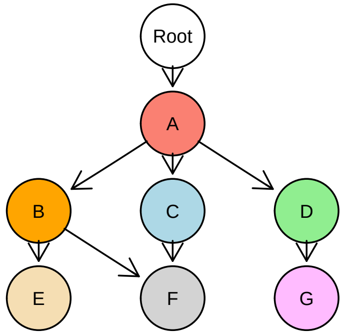


| | |
|---|---|
| *ID* | Wc5FDMqc5Bb6oeDH |
| *acc genotypes* | 25 |
| *peaks no back* | 1 |
| *removed edges* | 0 |
| *prop rem edges* | 0 |

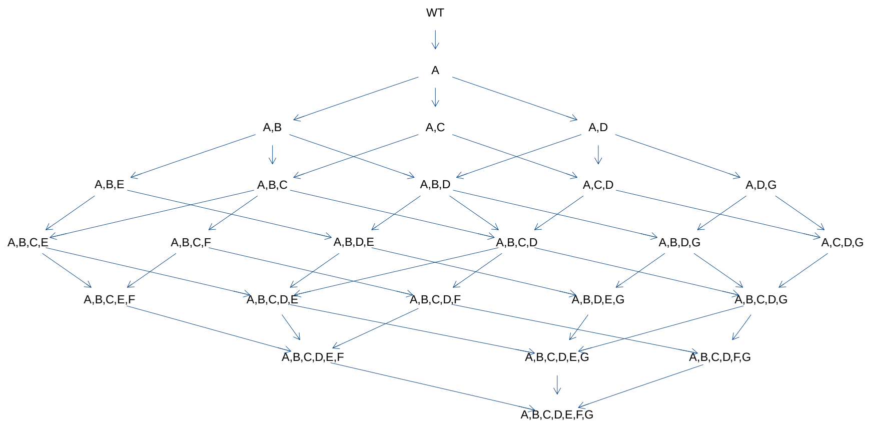


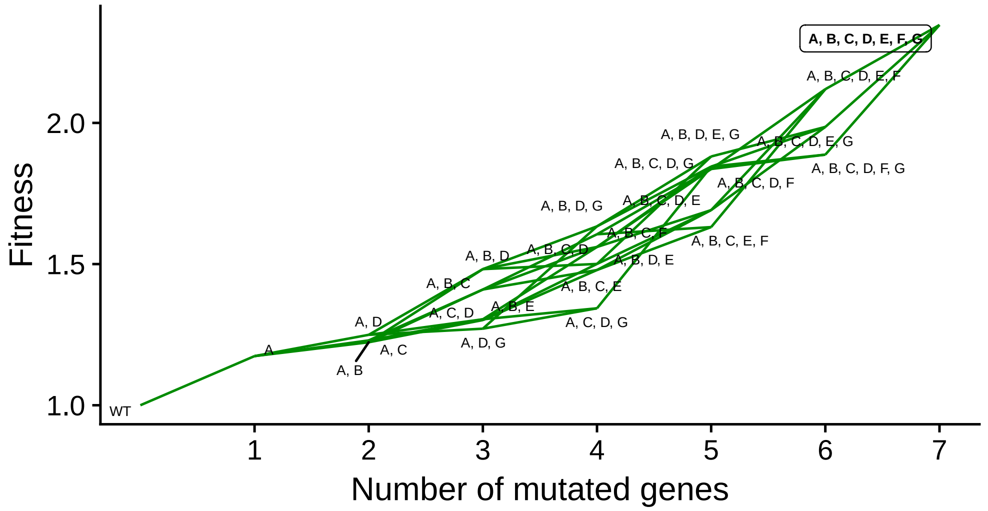

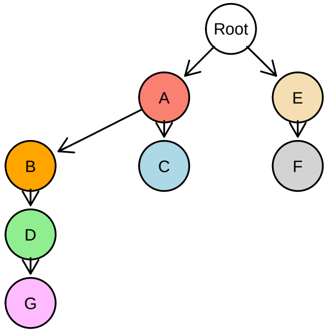


| | |
|---|---|
| *ID* | Wh8iukI5tW3rKboV |
| *acc genotypes* | 27 |
| *peaks no back* | 1 |
| *removed edges* | 0 |
| *prop rem edges* | 0 |

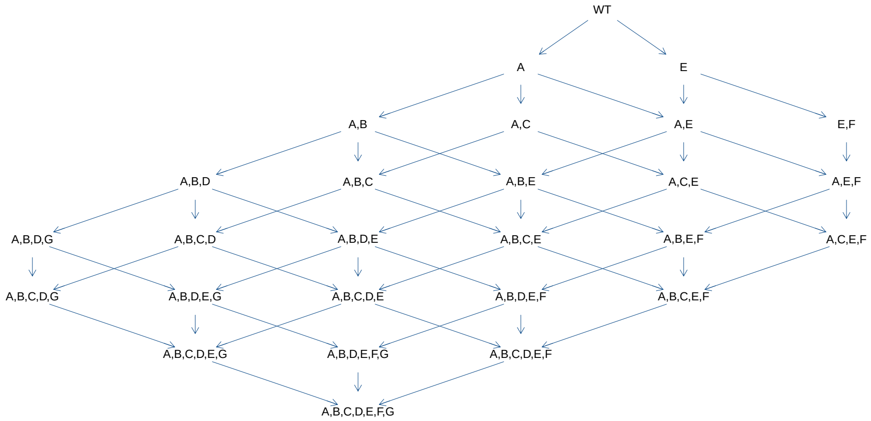


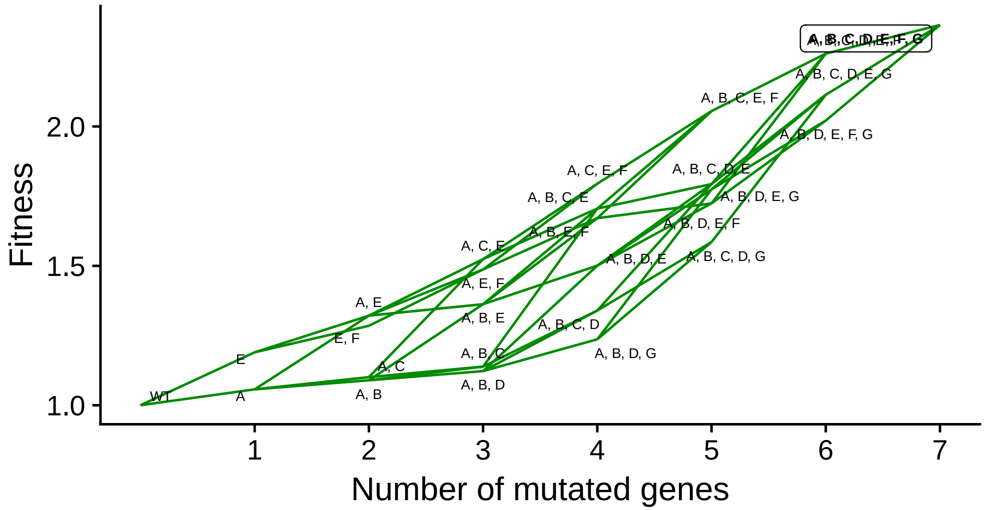

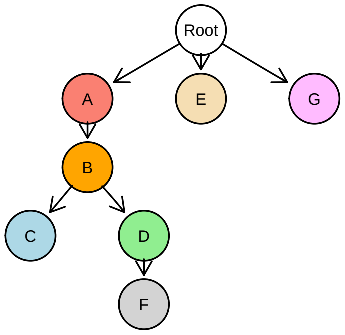


*ID* wKElTkPsWorfdIcX
*acc genotypes* 32
*peaks no back* 1
*removed edges* 0
*prop rem edges* 0

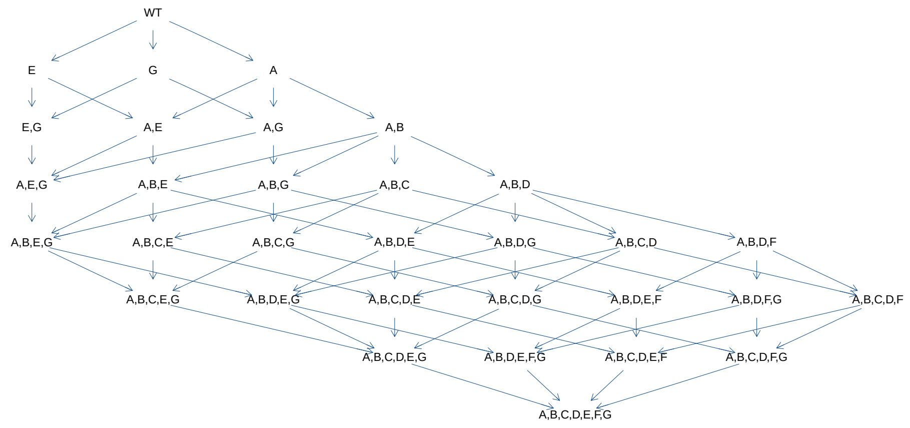


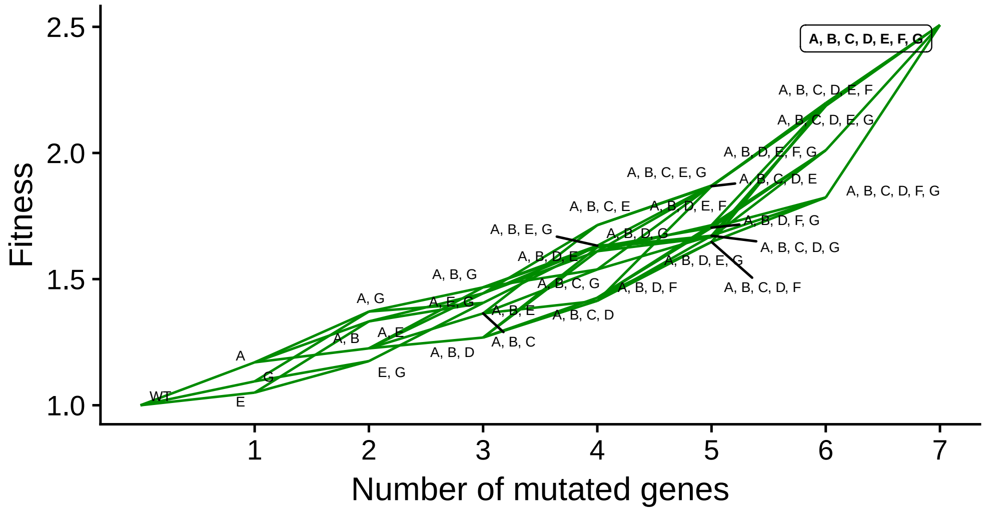

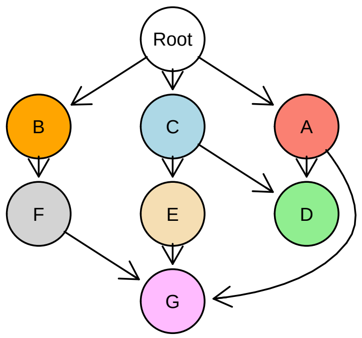


| | |
|---|---|
| *ID* | wLlgb17DLATLHcdE |
| *acc genotypes* | 26 |
| *peaks no back* | 1 |
| *removed edges* | 0 |
| *prop rem edges* | 0 |

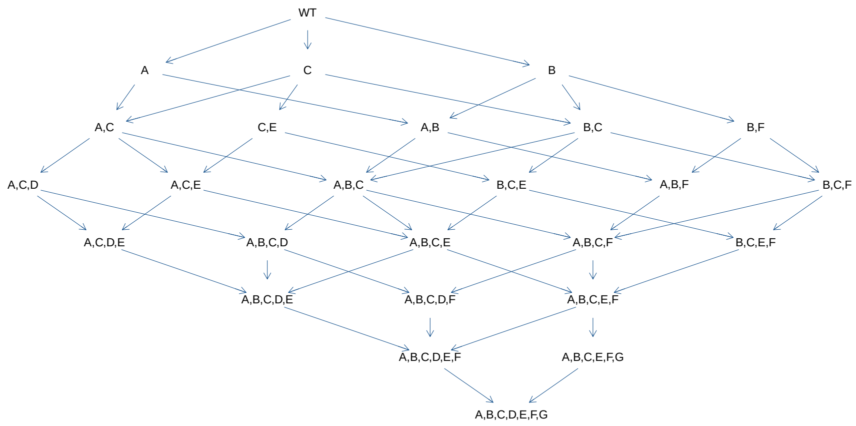


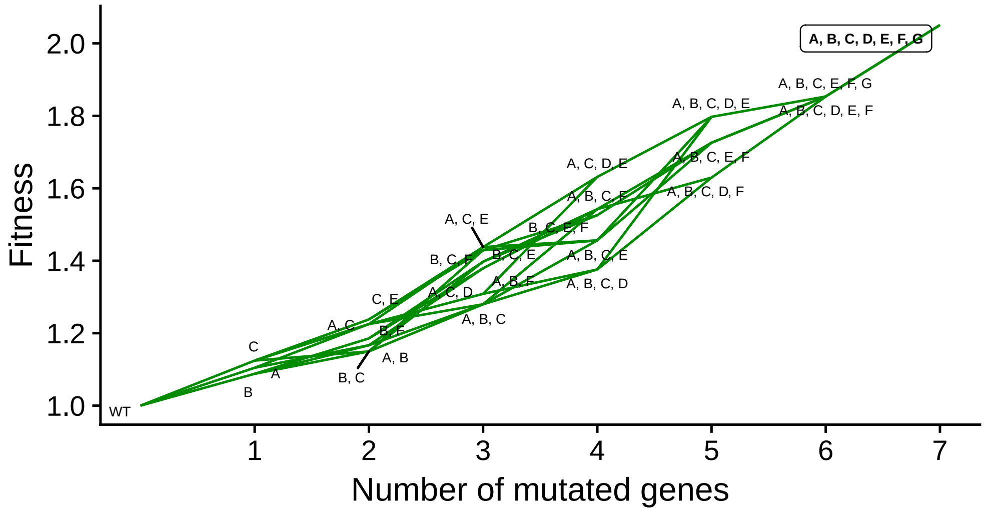

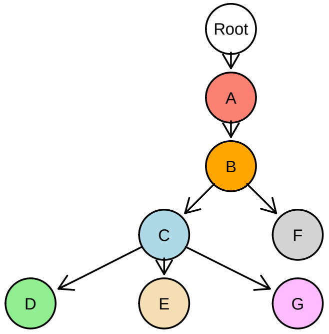


| | |
|---|---|
| *ID* | wNAatqlHhqMbIvMb |
| *acc genotypes* | 20 |
| *peaks no back* | 1 |
| *removed edges* | 0 |
| *prop rem edges* | 0 |

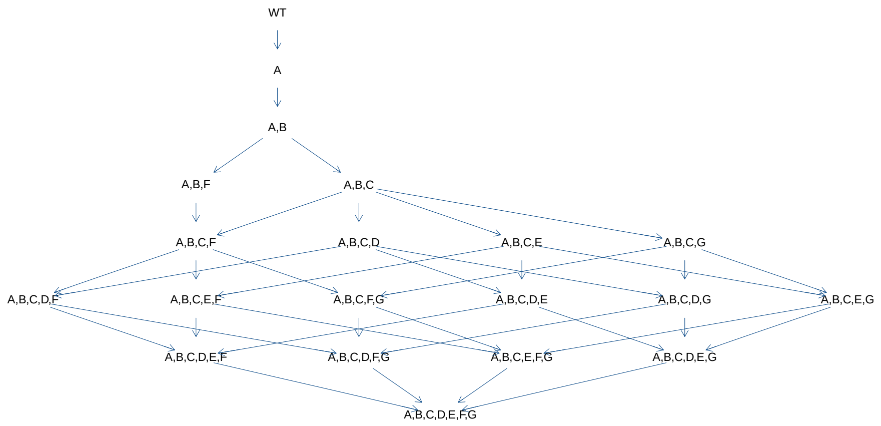


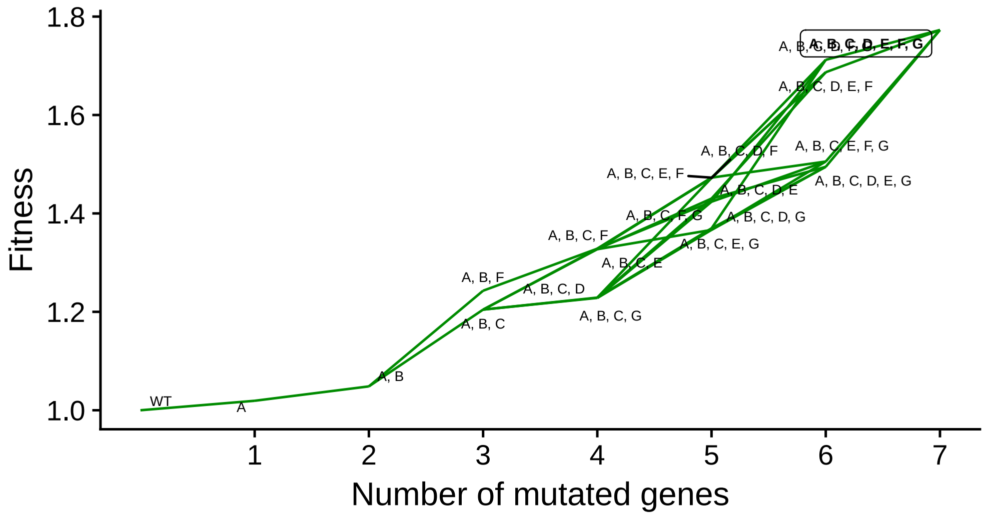

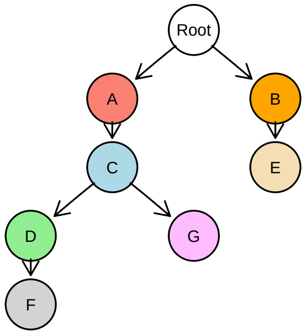


| | |
|---|---|
| *ID* | Wo7CkQUAYwfIkfX4 |
| *acc genotypes* | 24 |
| *peaks no back* | 1 |
| *removed edges* | 0 |
| *prop rem edges* | 0 |

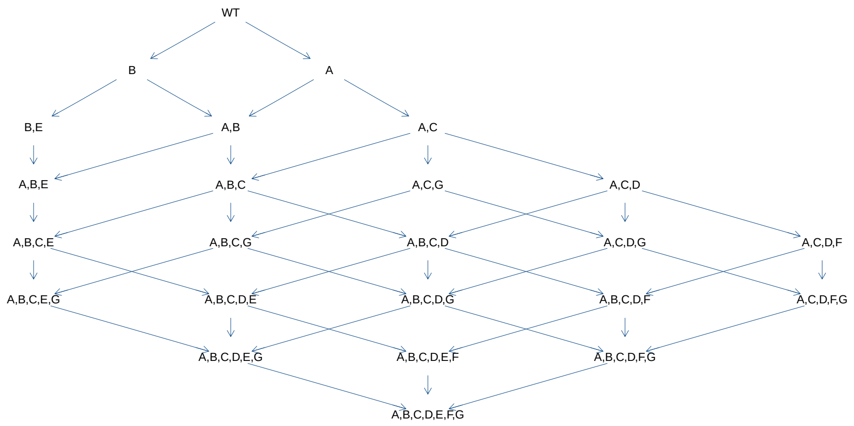


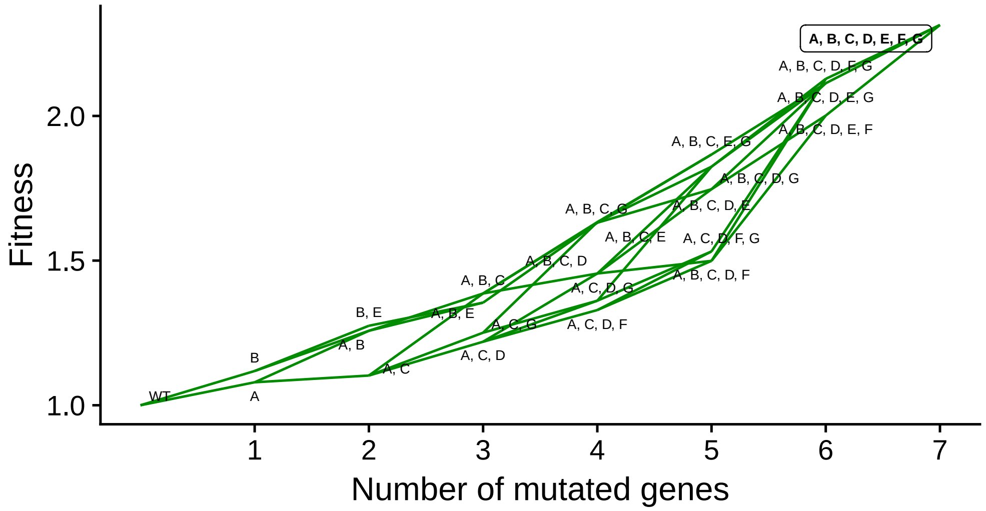

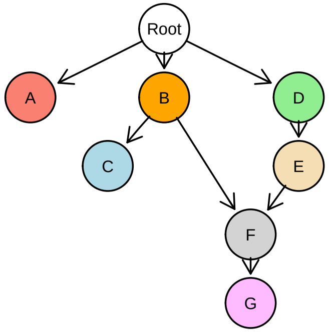


| | |
|---|---|
| *ID* | WoqvN8oF47jbnoBK |
| *acc genotypes* | 26 |
| *peaks no back* | 1 |
| *removed edges* | 0 |
| *prop rem edges* | 0 |

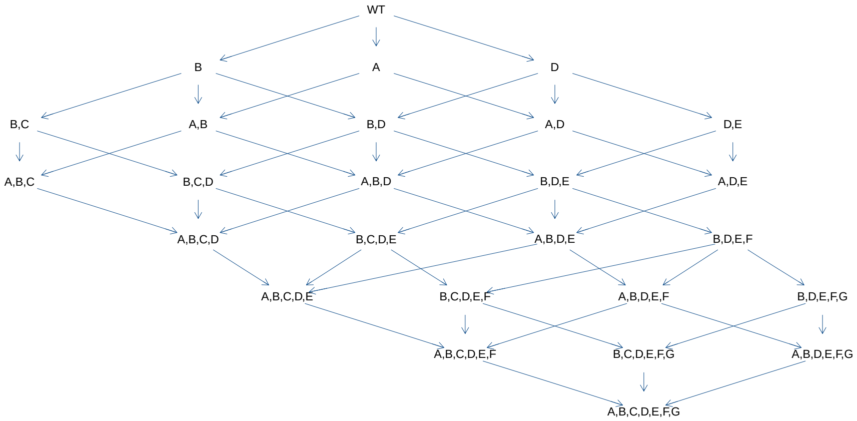


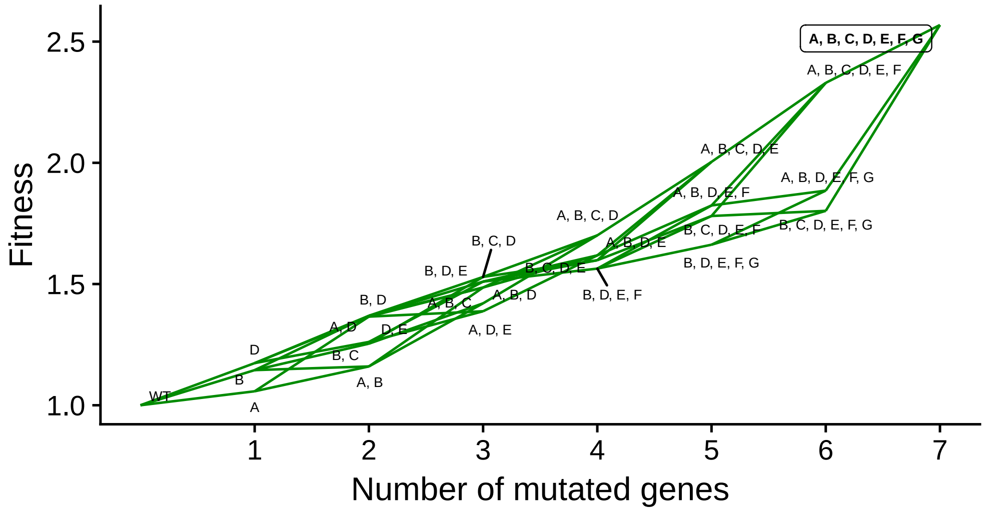

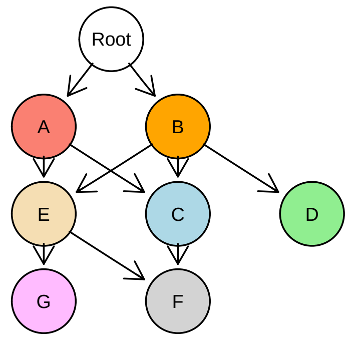


| | |
|---|---|
| *ID* | WQ5ixaPdtGNgofTu |
| *acc genotypes* | 20 |
| *peaks no back* | 1 |
| *removed edges* | 0 |
| *prop rem edges* | 0 |

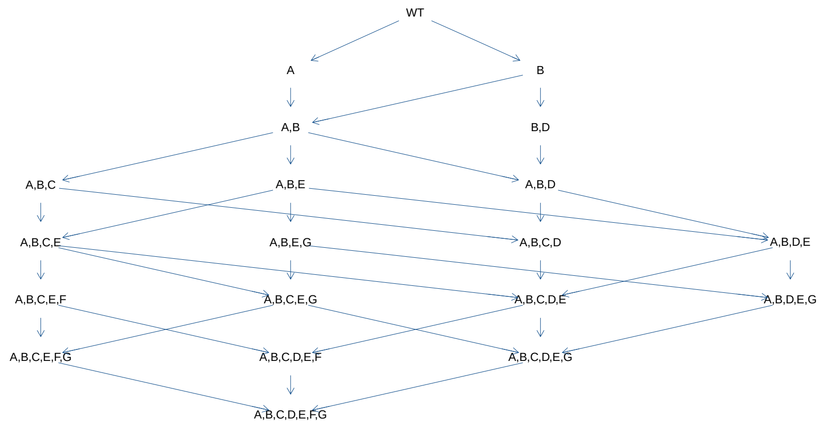


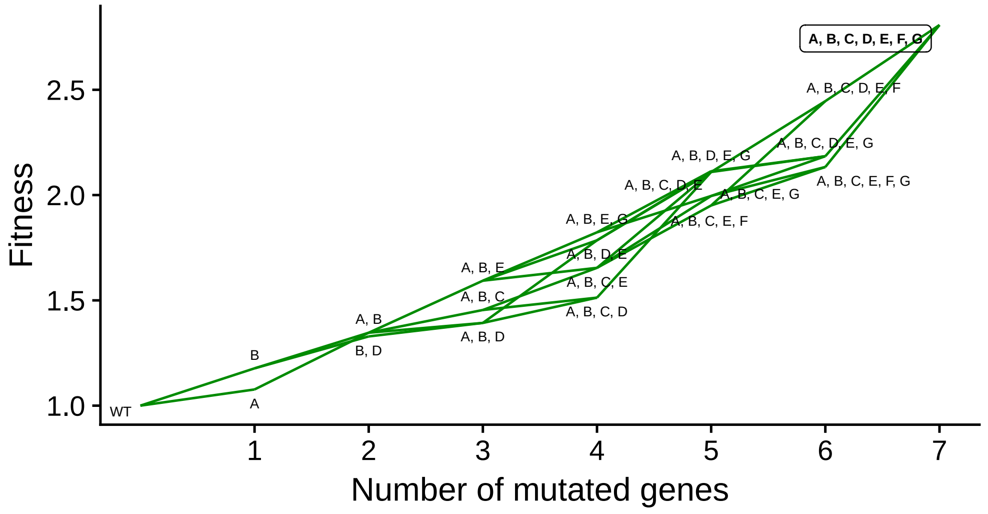

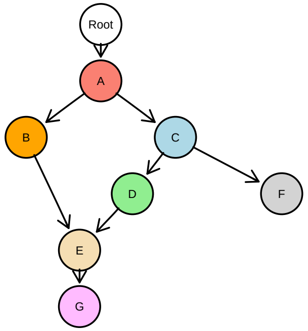


| | |
|---|---|
| *ID* | wrcr1dFF7SQNSwts |
| *acc genotypes* | 15 |
| *peaks no back* | 1 |
| *removed edges* | 0 |
| *prop rem edges* | 0 |

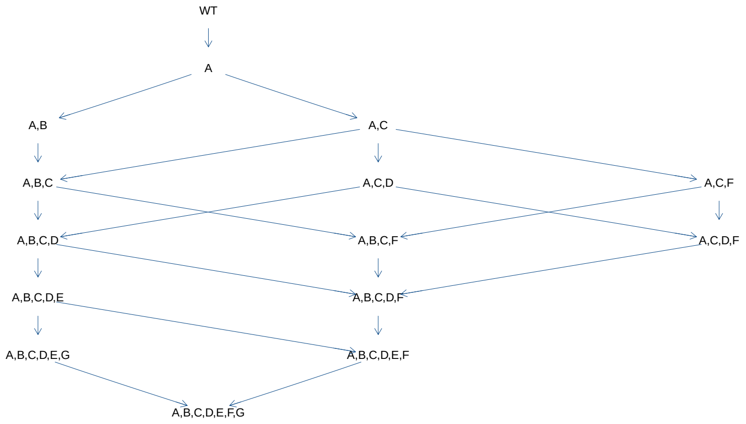


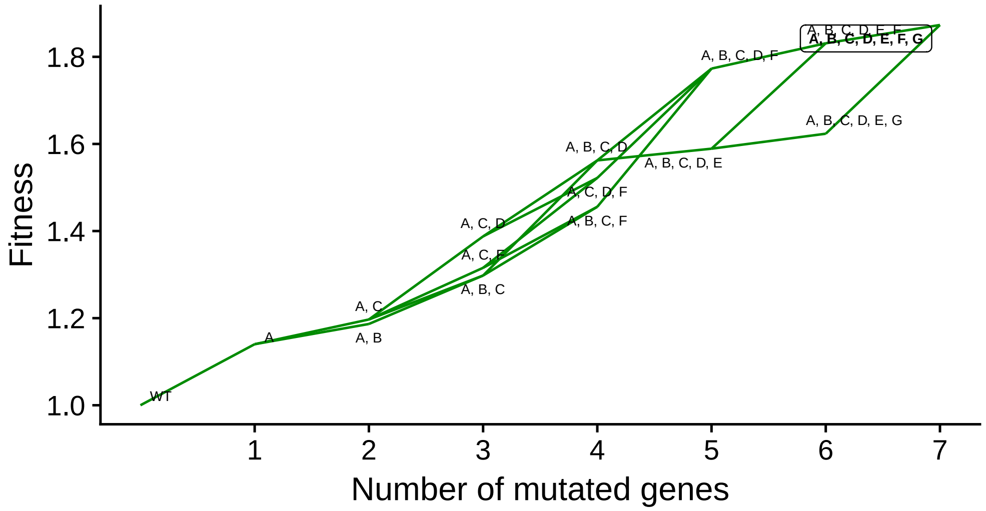

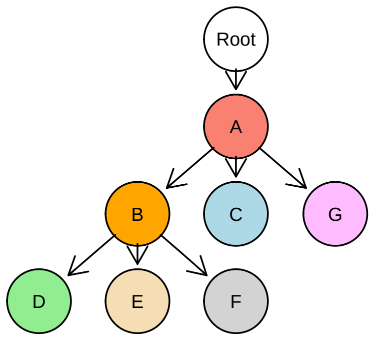


| | |
|---|---|
| *ID* | wrRTzMwK5mQOC85K |
| *acc genotypes* | 37 |
| *peaks no back* | 1 |
| *removed edges* | 0 |
| *prop rem edges* | 0 |

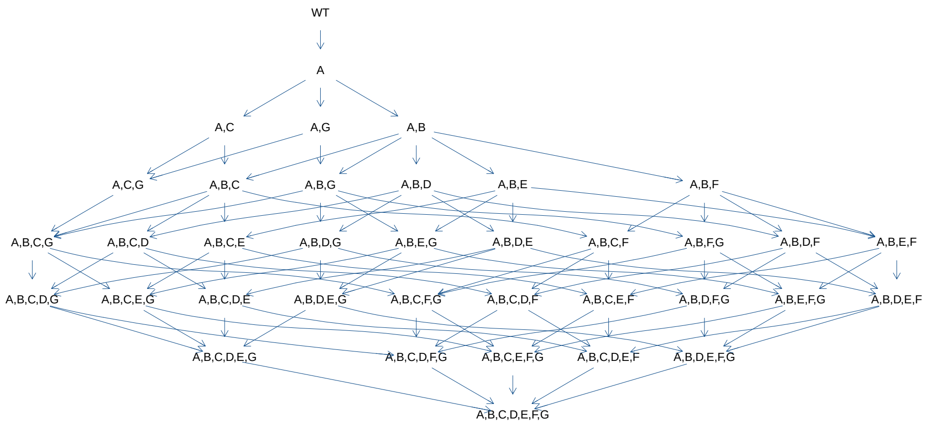


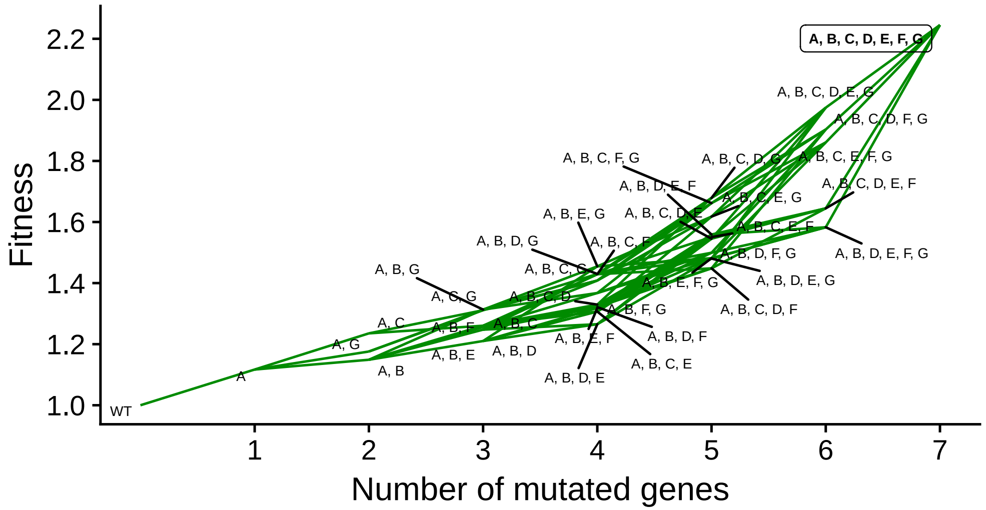

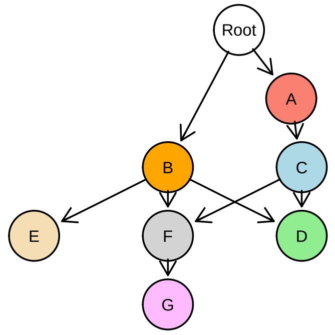


*ID* WTwHv3VrTVwsxPNF
*acc genotypes* 19
*peaks no back* 1
*removed edges* 0
*prop rem edges* 0

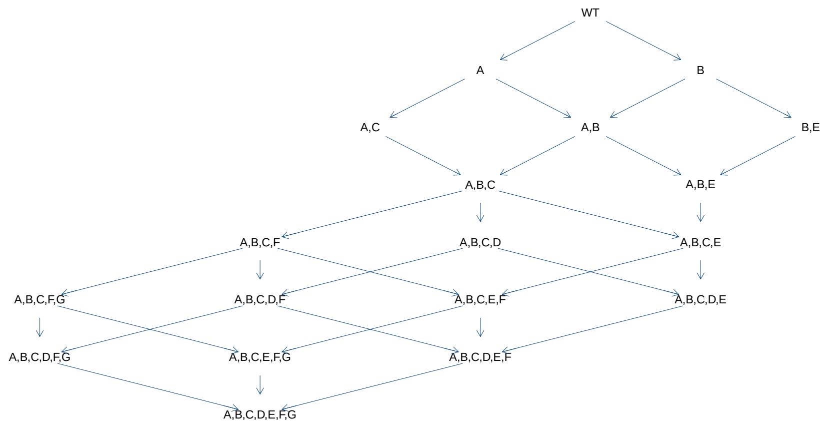


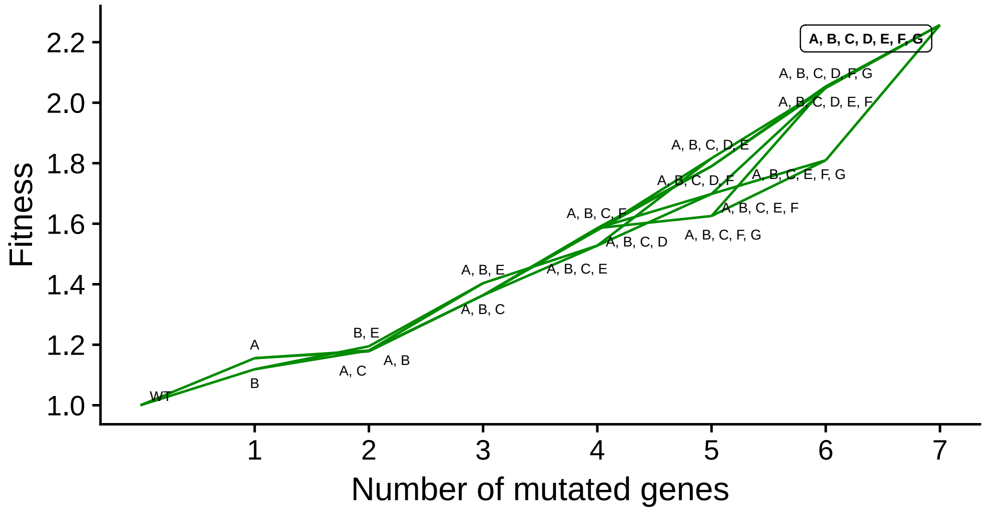

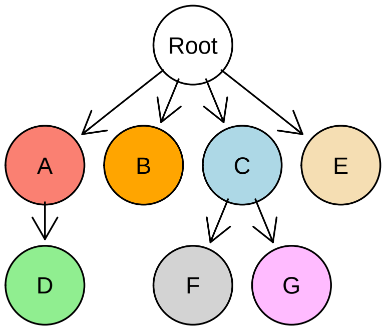


| | |
|---|---|
| *ID* | wu54FEWu694xA1Ko |
| *acc genotypes* | 60 |
| *peaks no back* | 1 |
| *removed edges* | 0 |
| *prop rem edges* | 0 |

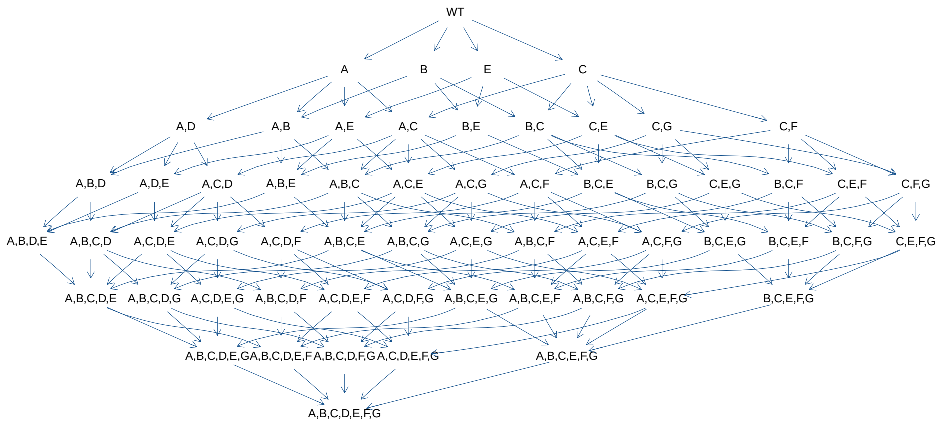


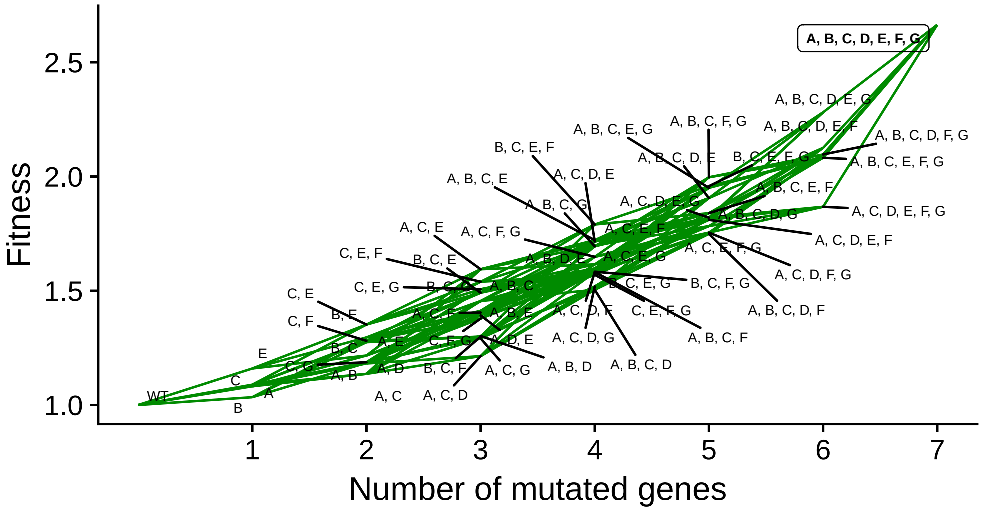

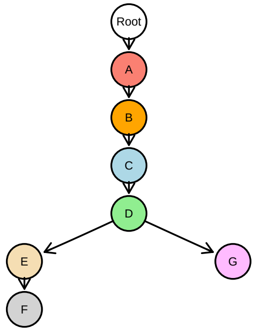


| | |
|---|---|
| *ID* | wXqQ210QBkvlJzh1K |
| *acc genotypes* | 10 |
| *peaks no back* | 1 |
| *removed edges* | 0 |
| *prop rem edges* | 0 |

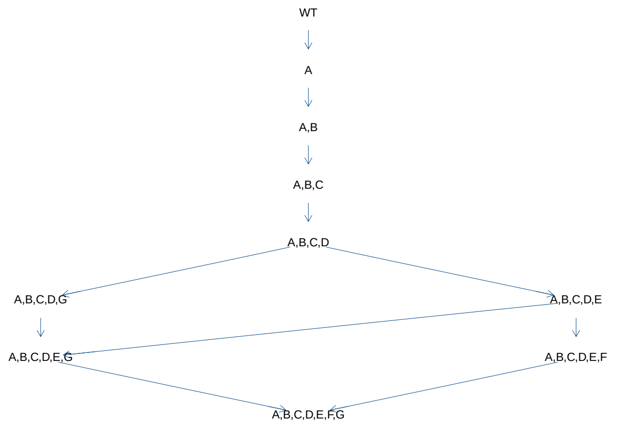


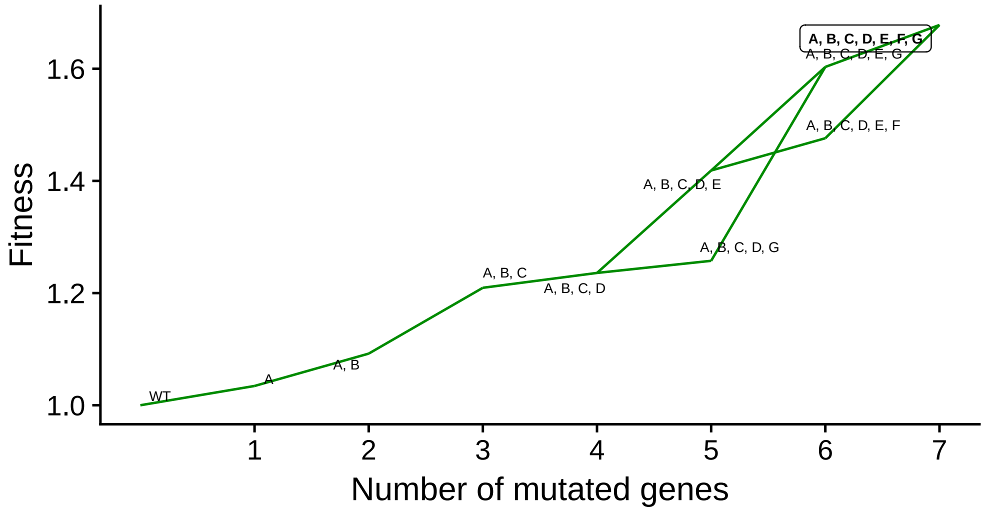

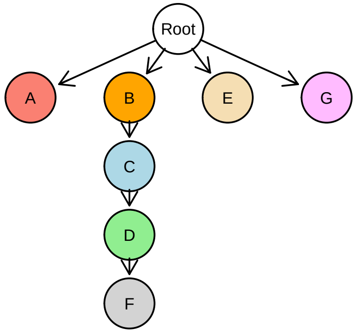


| | |
|---|---|
| *ID* | WzFbSkNFewLZM2Lc |
| *acc genotypes* | 40 |
| *peaks no back* | 1 |
| *removed edges* | 0 |
| *prop rem edges* | 0 |

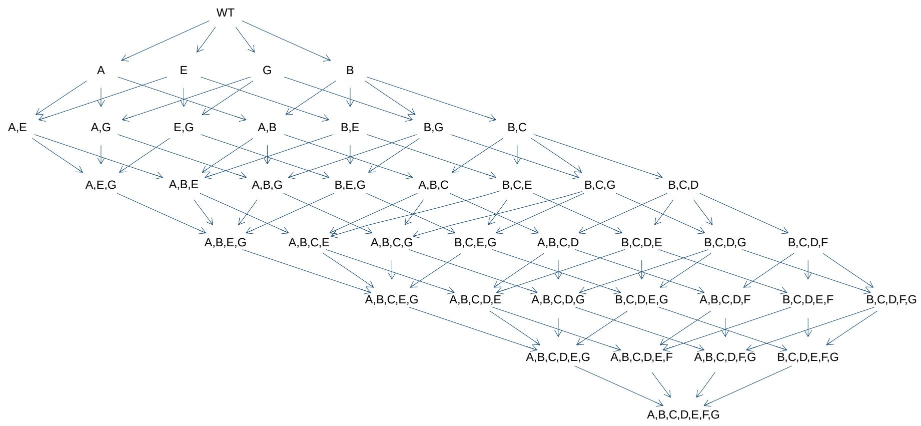


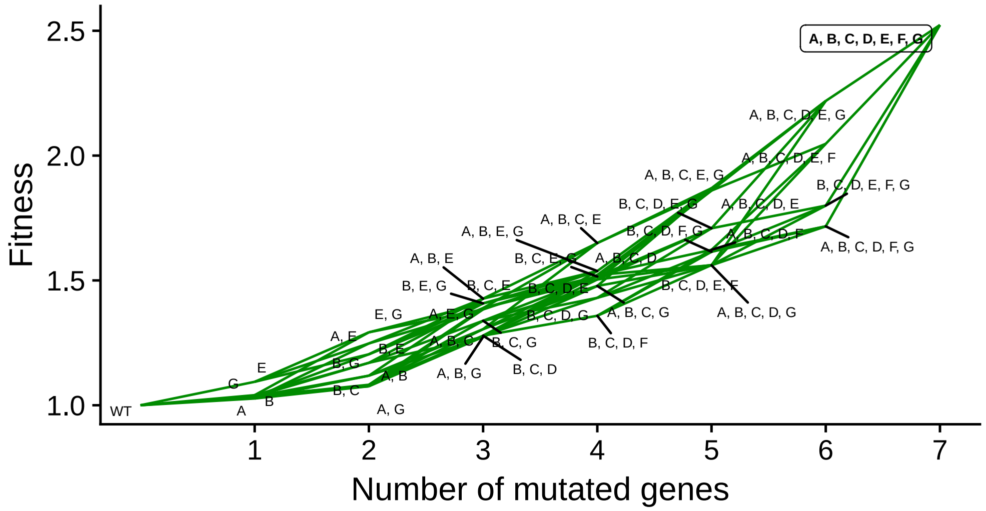

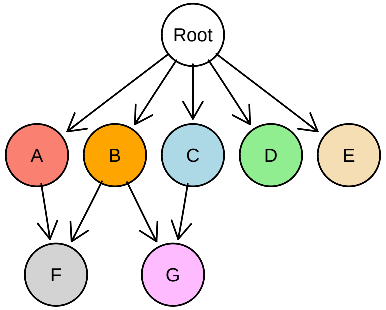


*ID* wZHRlWE1jjVBM1mQ

*acc genotypes* 52

*peaks no back* 1

*removed edges* 0

*prop rem edges* 0

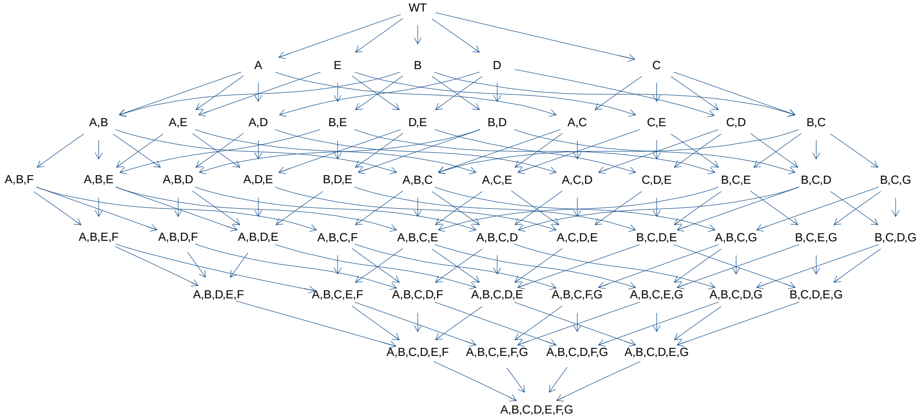


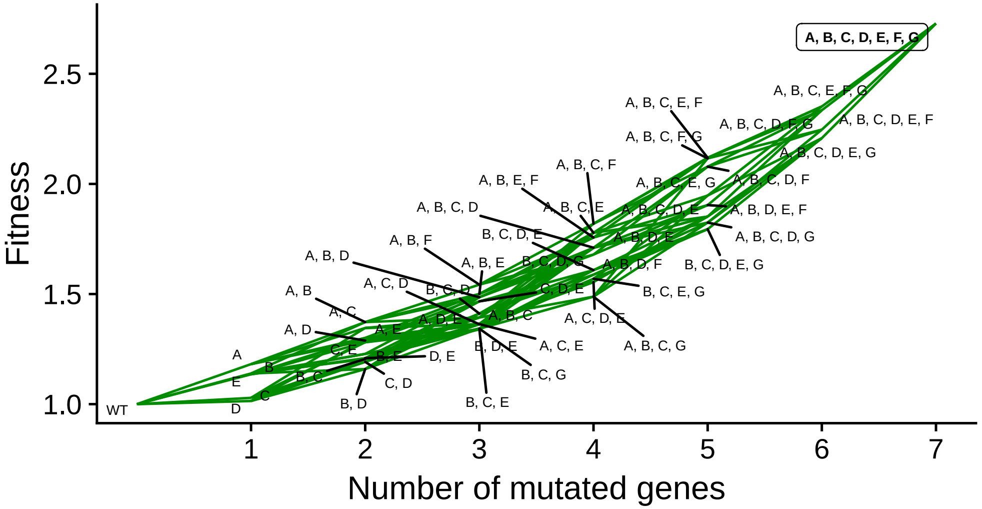

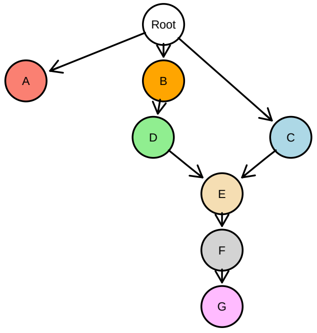


| | |
|---|---|
| *ID* | xBMt9dOS3zBvChSy |
| *acc genotypes* | 18 |
| *peaks no back* | 1 |
| *removed edges* | 0 |
| *prop rem edges* | 0 |

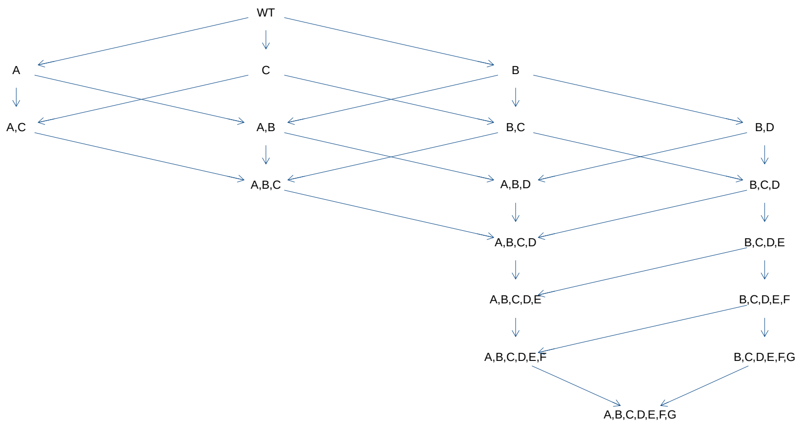


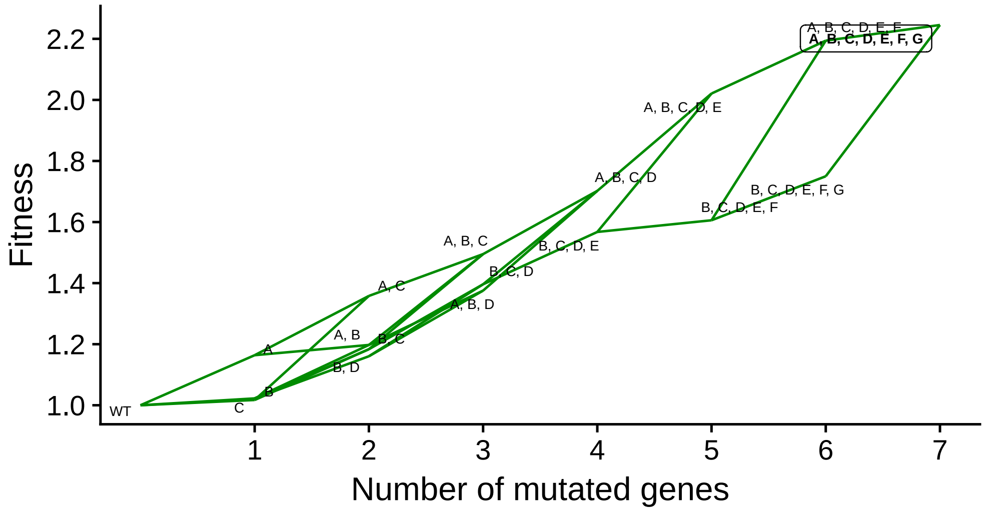

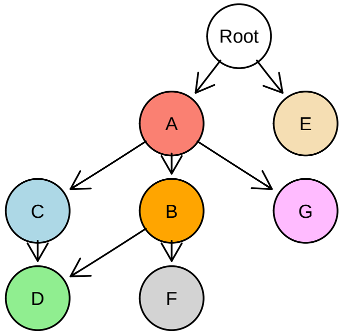


| | |
|---|---|
| *ID* | xkgudeGhJGOlP7X7 |
| *acc genotypes* | 34 |
| *peaks no back* | 1 |
| *removed edges* | 0 |
| *prop rem edges* | 0 |

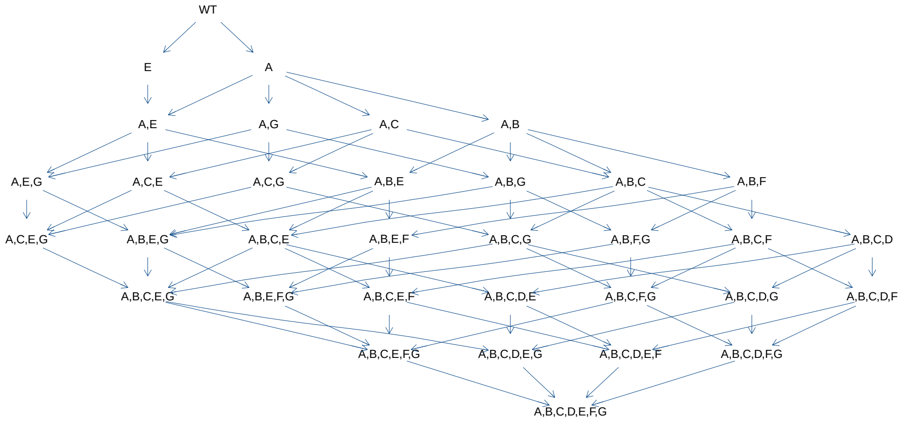


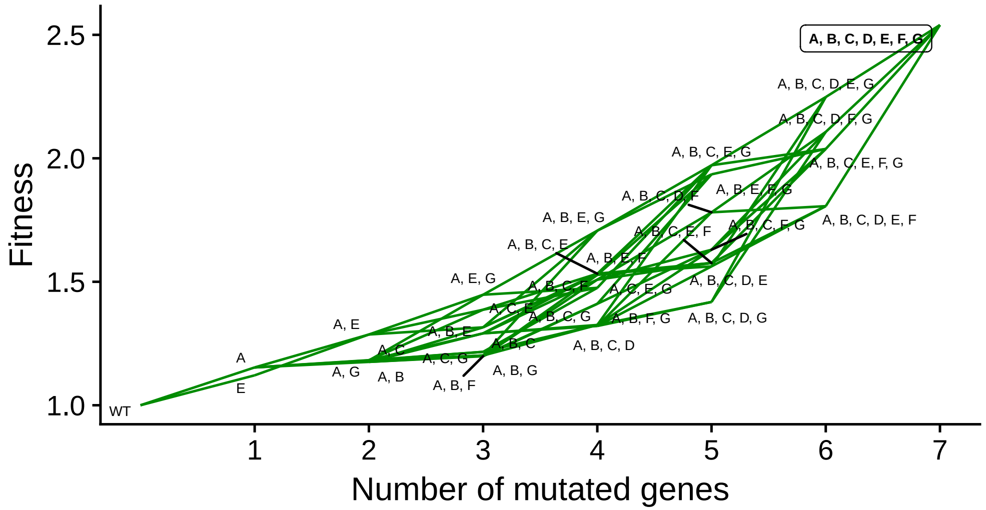

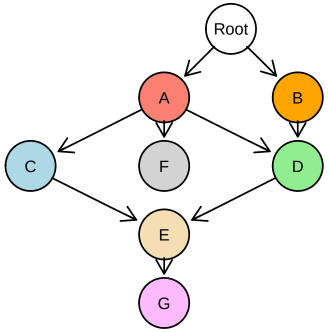


| | |
|---|---|
| *ID* | XolBQBDPoWH3DwDQ |
| *acc genotypes* | 18 |
| *peaks no back* | 1 |
| *removed edges* | 0 |
| *prop rem edges* | 0 |

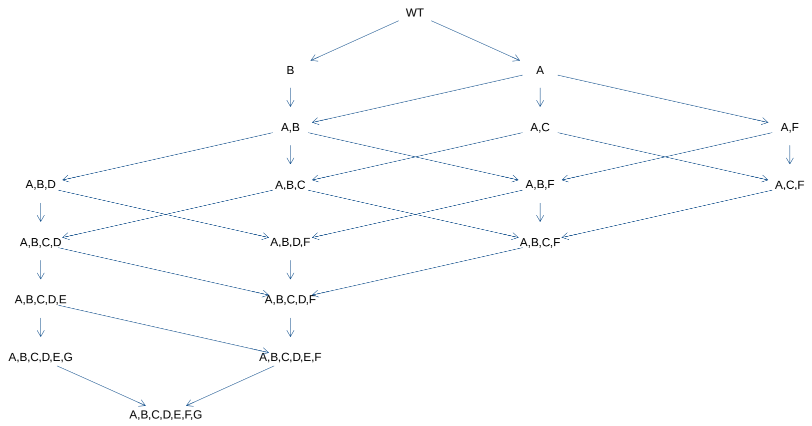


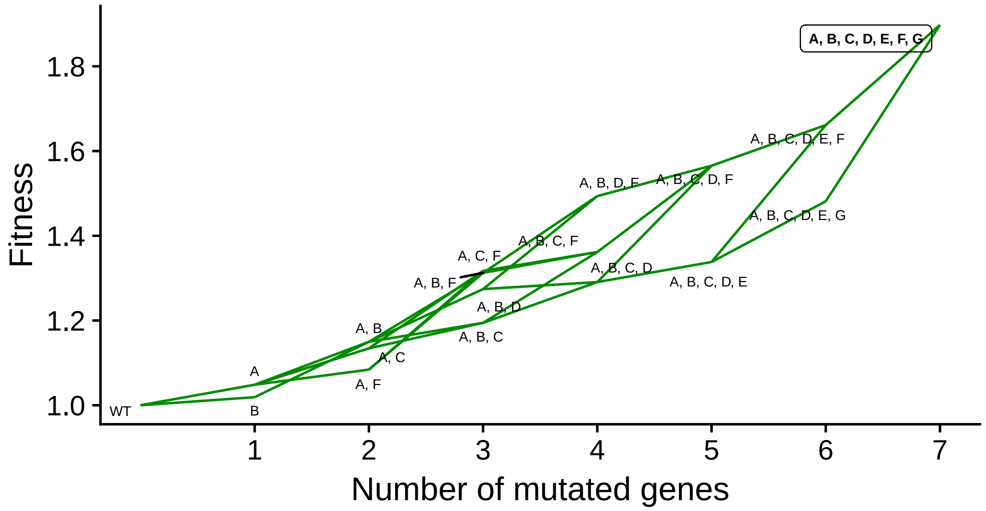

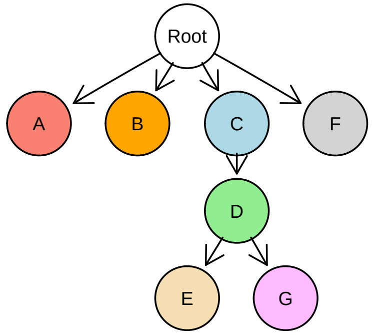


| | |
|---|---|
| *ID* | XRovuAJWlNYiVJHn |
| *acc genotypes* | 48 |
| *peaks no back* | 1 |
| *removed edges* | 0 |
| *prop rem edges* | 0 |

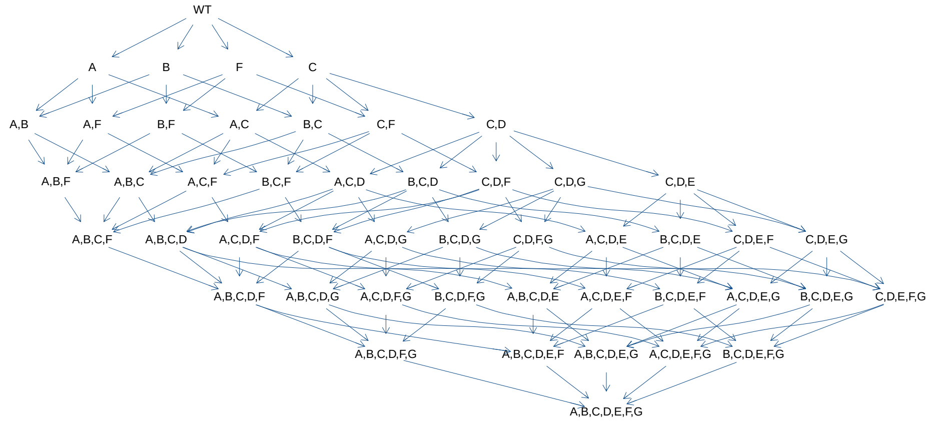


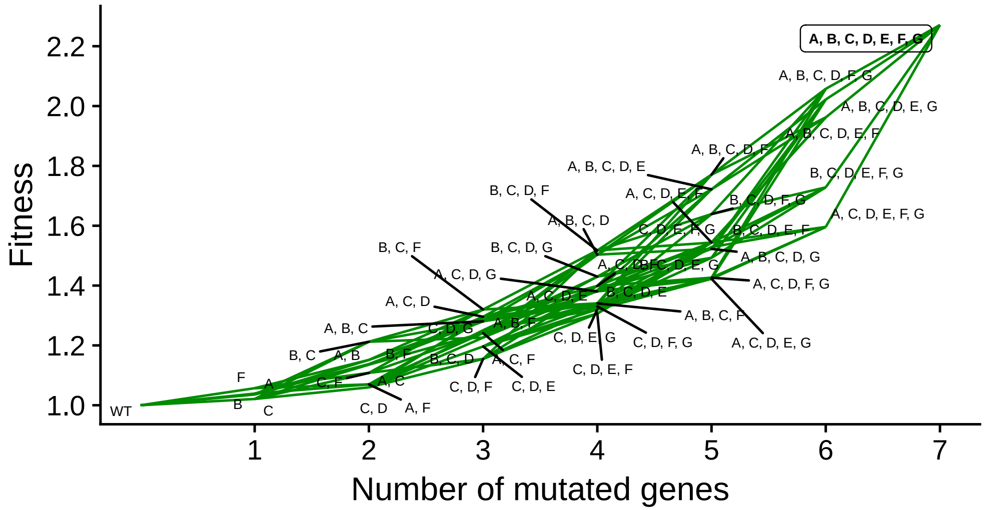

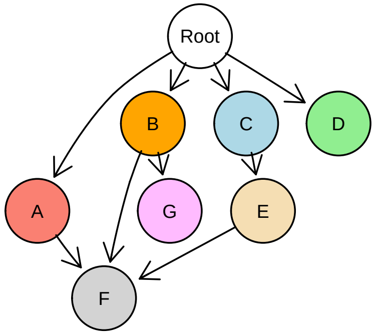


| | |
|---|---|
| *ID* | xVDfqEWHyqTK7PeA |
| *acc genotypes* | 40 |
| *peaks no back* | 1 |
| *removed edges* | 0 |
| *prop rem edges* | 0 |

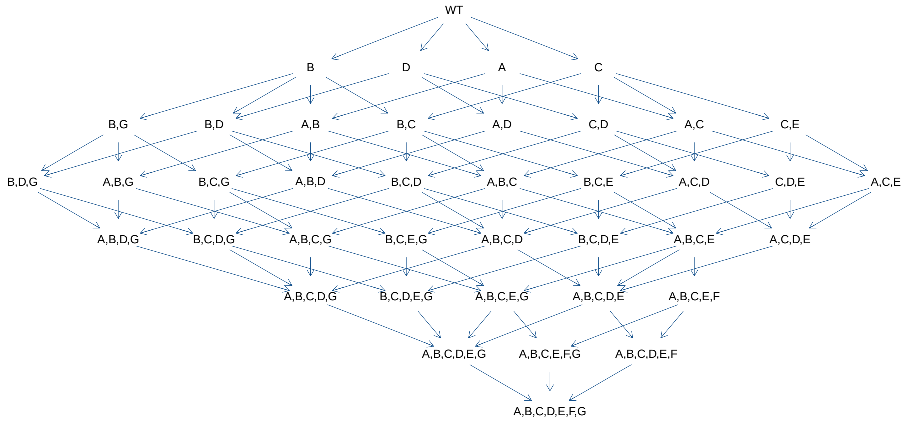


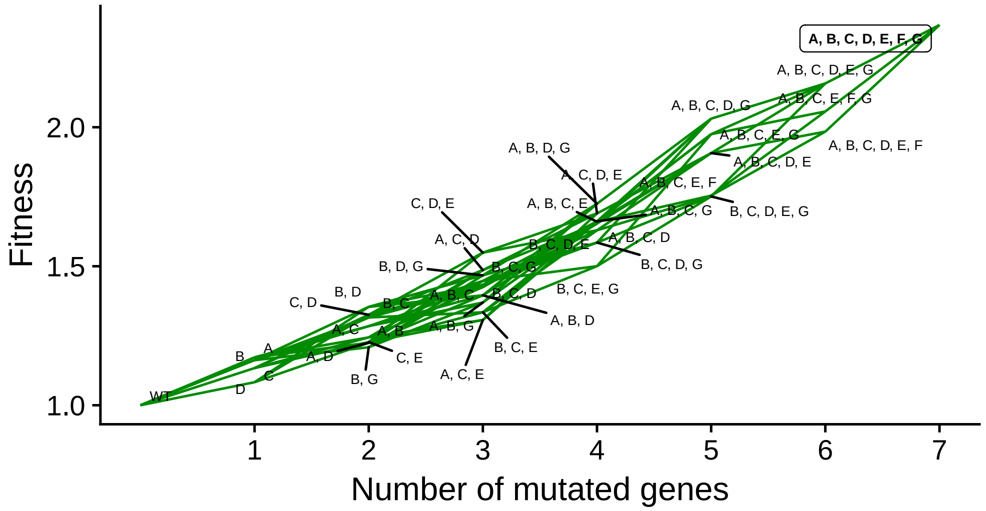

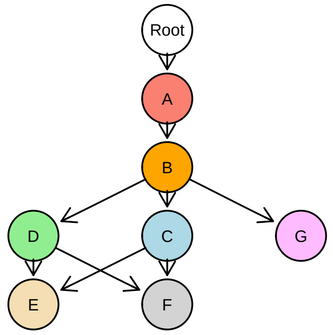


| | |
|---|---|
| *ID* | y46oBk106Lnsh10yaQ |
| *acc genotypes* | 16 |
| *peaks no back* | 1 |
| *removed edges* | 0 |
| *prop rem edges* | 0 |

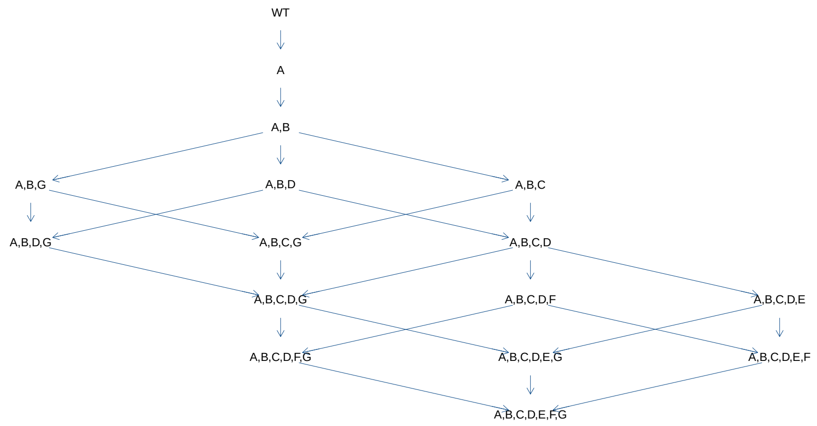


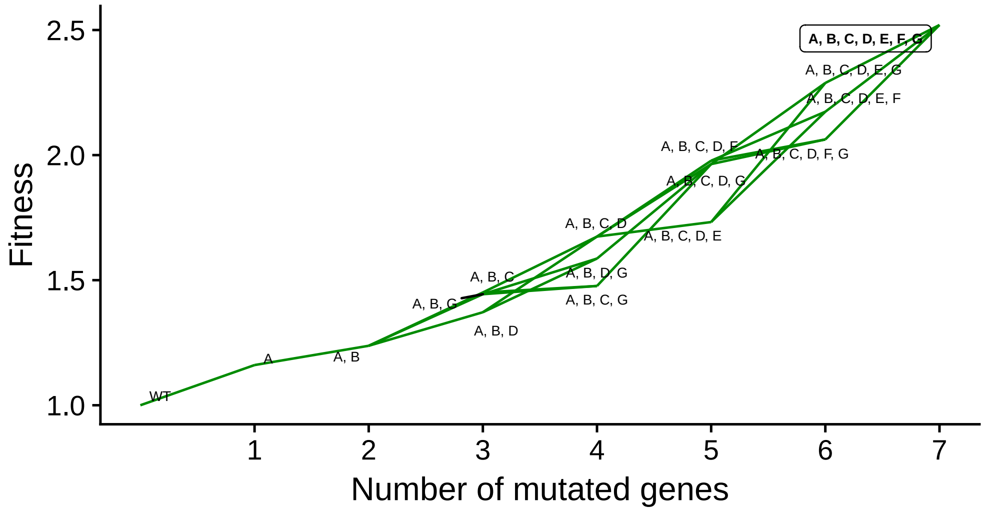

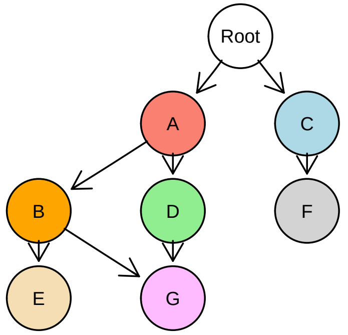


| | |
|---|---|
| *ID* | YbCf9soBzZRhuiw10 |
| *acc genotypes* | 27 |
| *peaks no back* | 1 |
| *removed edges* | 0 |
| *prop rem edges* | 0 |

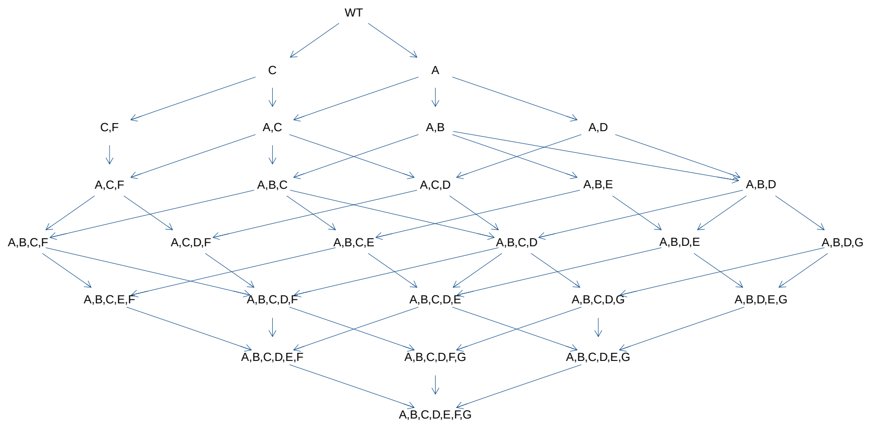


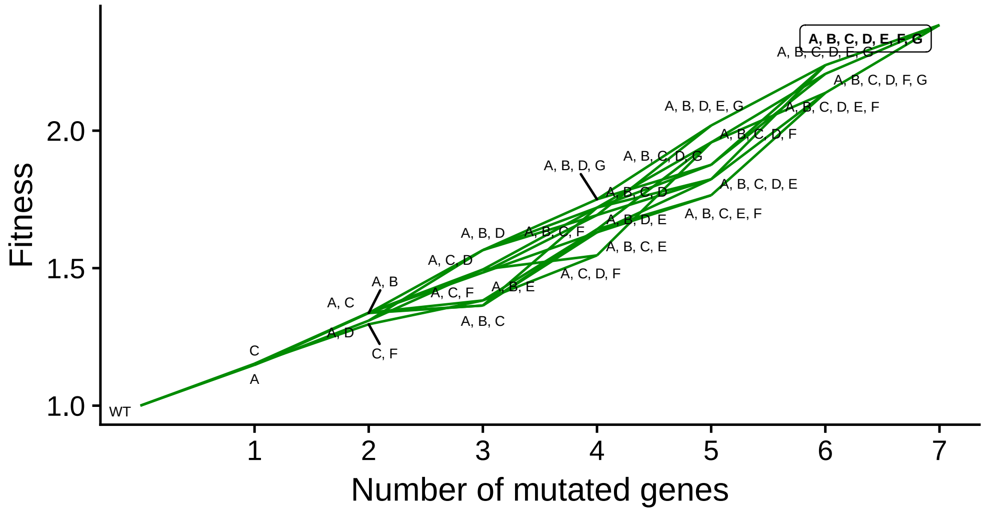

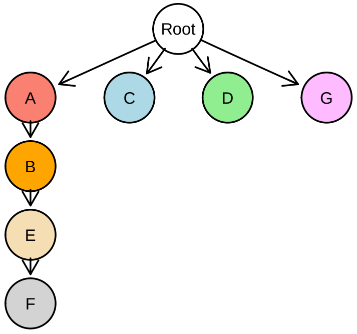


| | |
|---|---|
| *ID* | YFmY1csTgroJgPCG |
| *acc genotypes* | 40 |
| *peaks no back* | 1 |
| *removed edges* | 0 |
| *prop rem edges* | 0 |

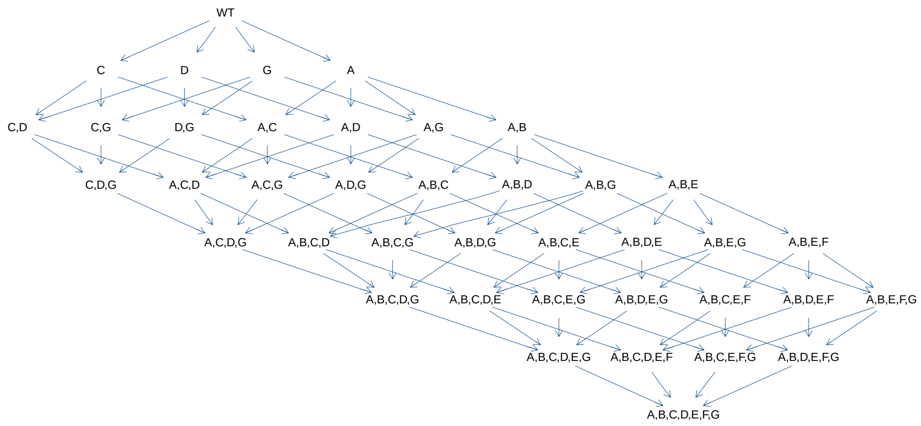


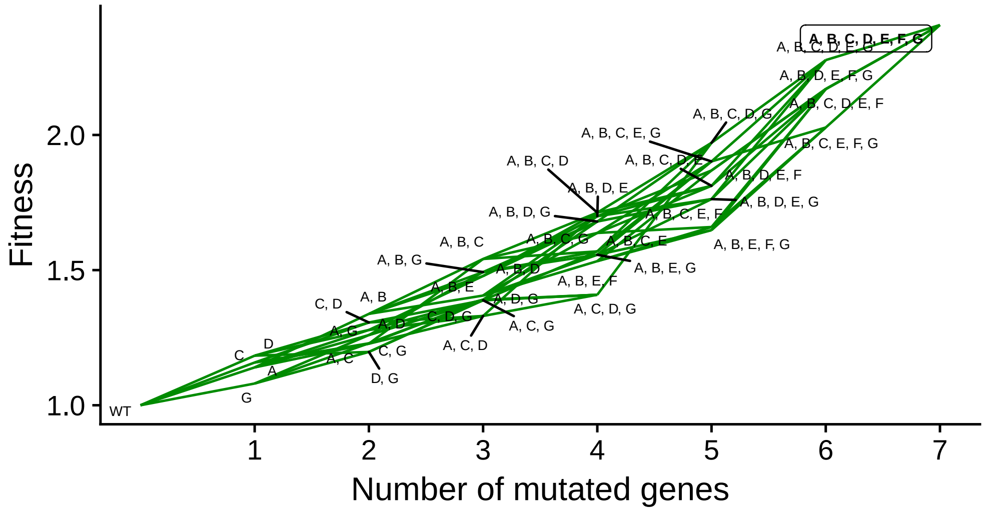

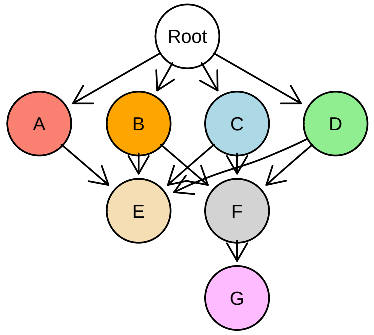


| | |
|---|---|
| *ID* | yjwFeMSZVW68gVPI |
| *acc genotypes* | 23 |
| *peaks no back* | 1 |
| *removed edges* | 0 |
| *prop rem edges* | 0 |

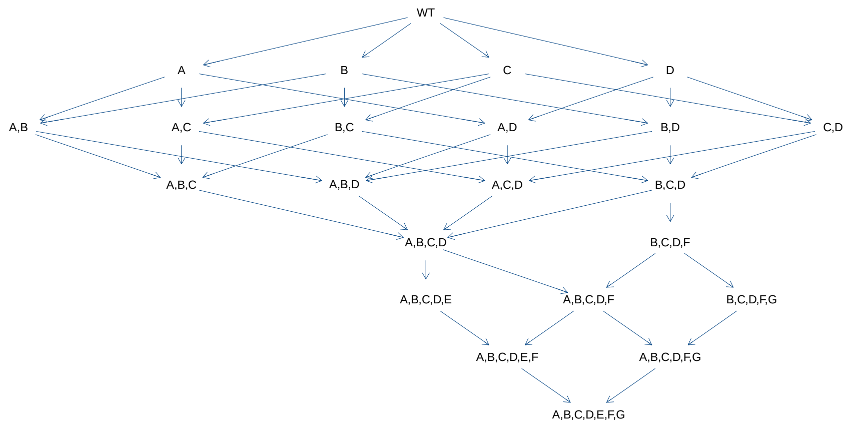


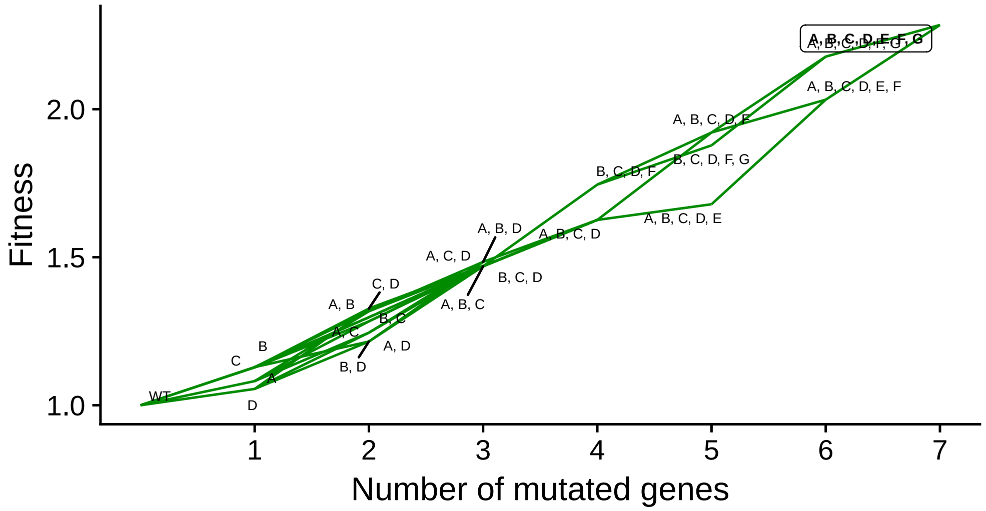

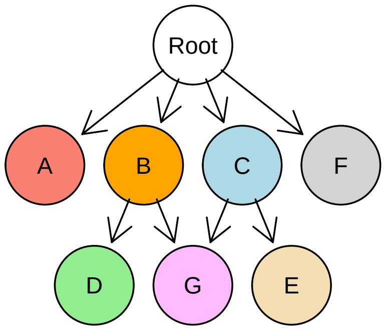


| | |
|---|---|
| *ID* | yVlAr6nny7pIzy1a |
| *acc genotypes* | 52 |
| *peaks no back* | 1 |
| *removed edges* | 0 |
| *prop rem edges* | 0 |

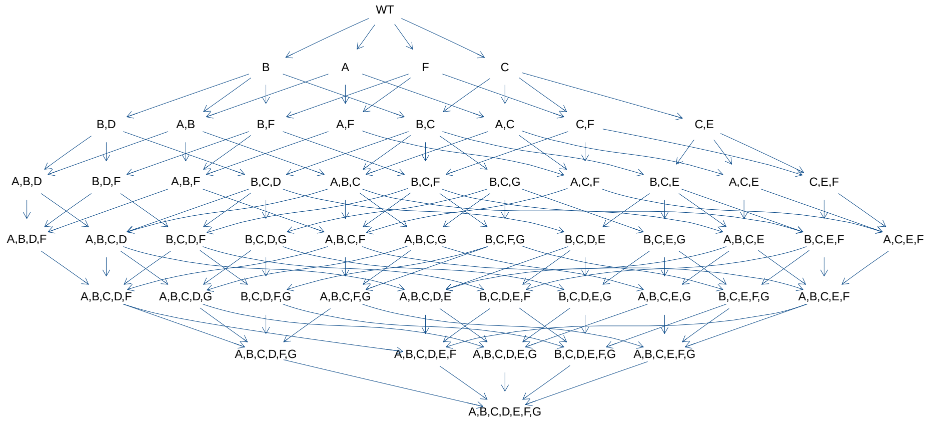


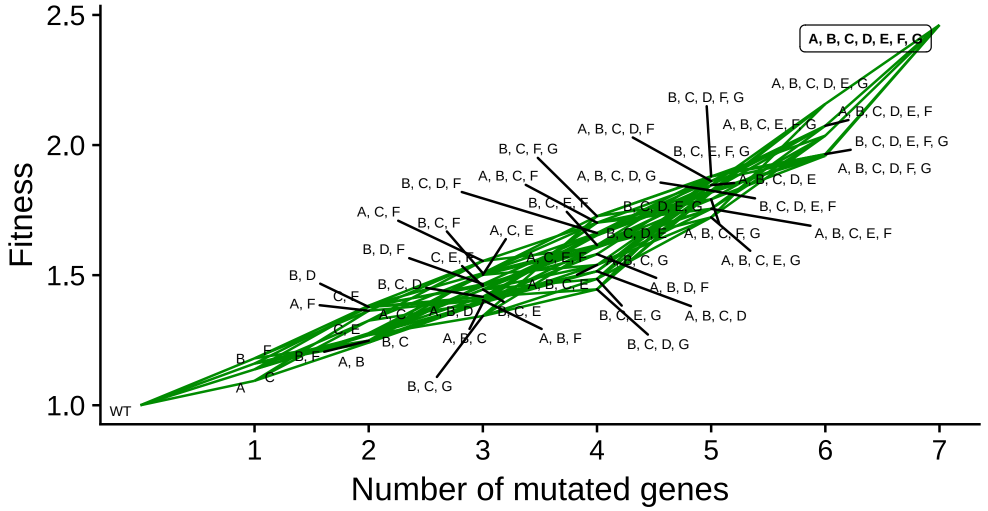

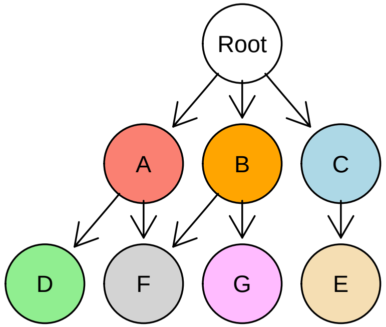


*ID* YZtw9jhxSkZdMuV2
*acc genotypes* 39
*peaks no back* 1
*removed edges* 0
*prop rem edges* 0

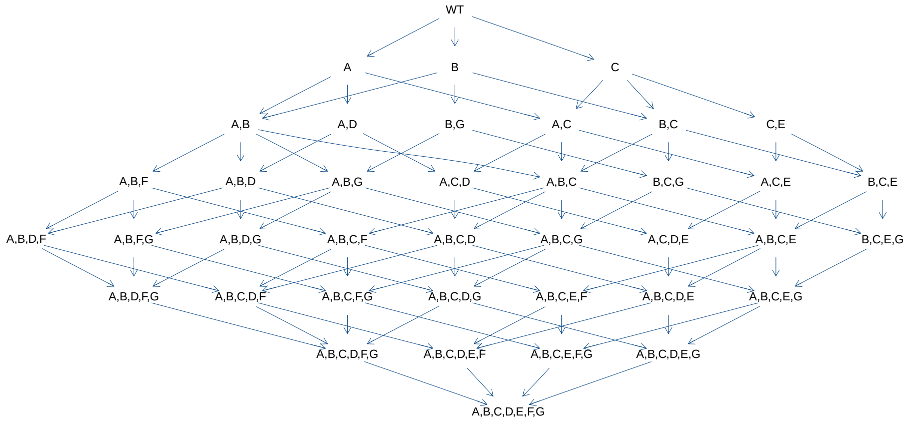


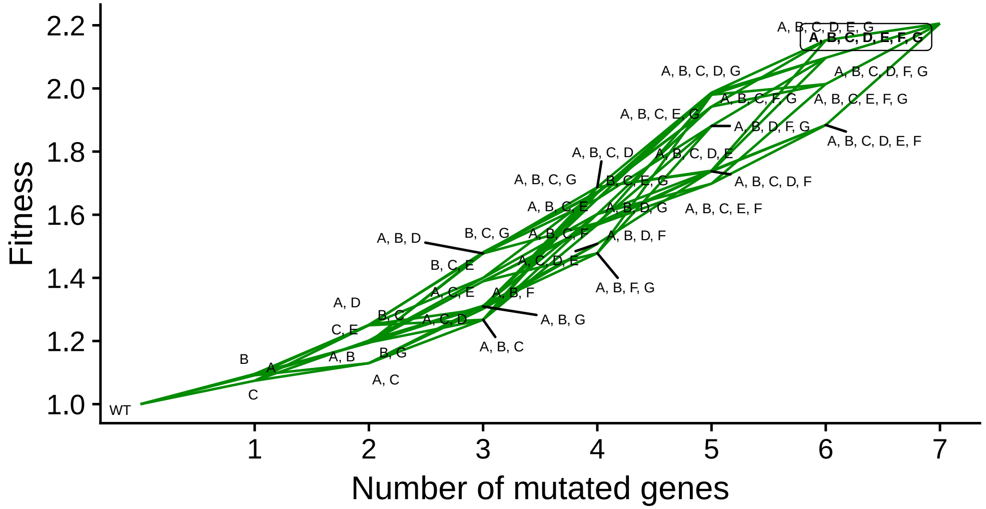

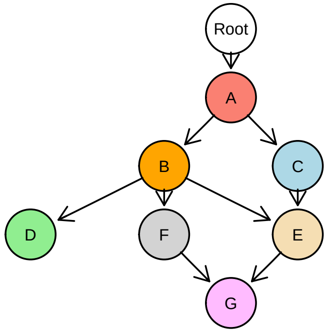


*ID* Z5HeEO7UheuYVWra
*acc genotypes* 17
*peaks no back* 1
*removed edges* 0
*prop rem edges* 0

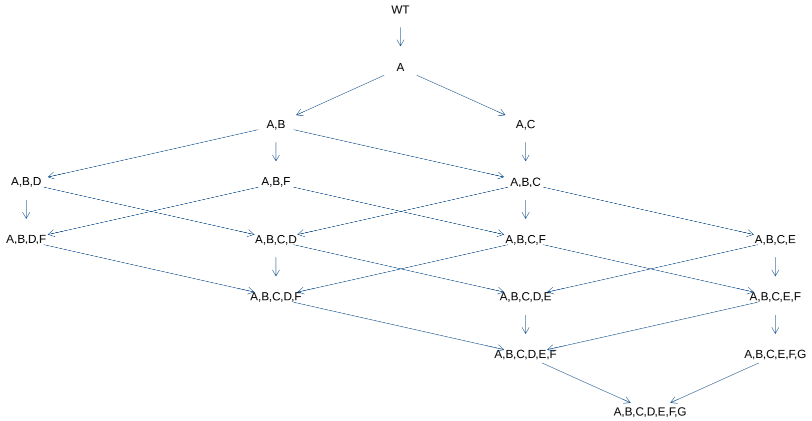


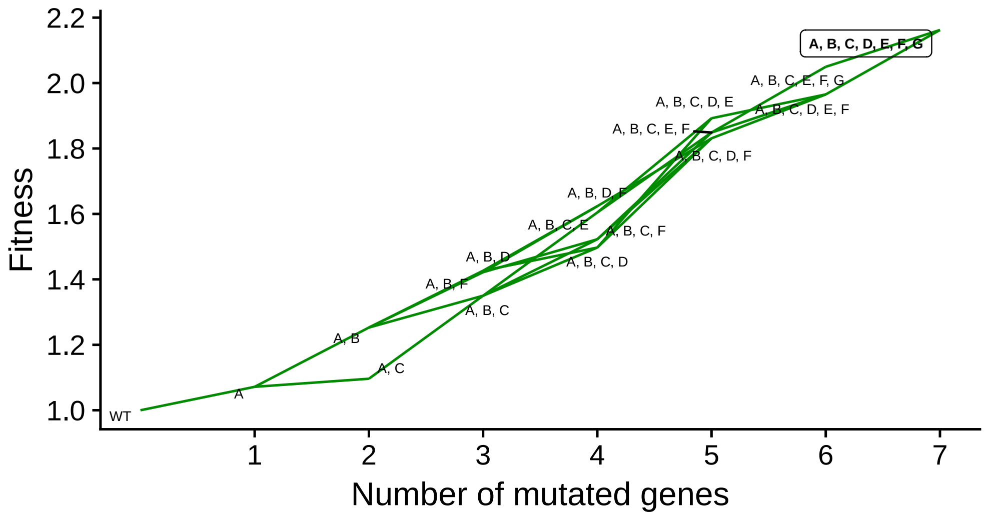

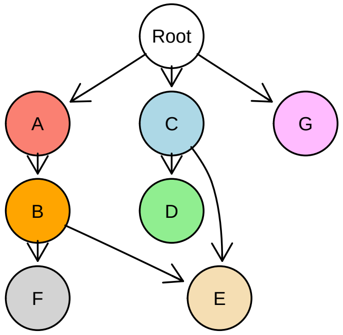


| | |
|---|---|
| *ID* | zy1cmG2IWriY25OZ |
| *acc genotypes* | 32 |
| *peaks no back* | 1 |
| *removed edges* | 0 |
| *prop rem edges* | 0 |

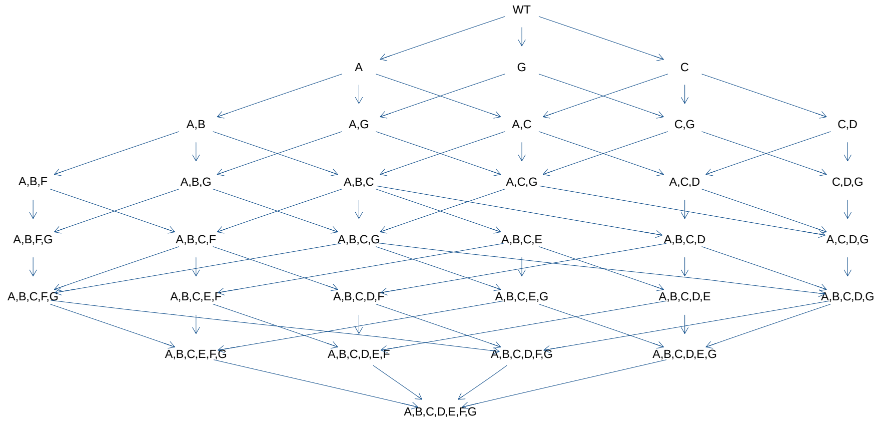


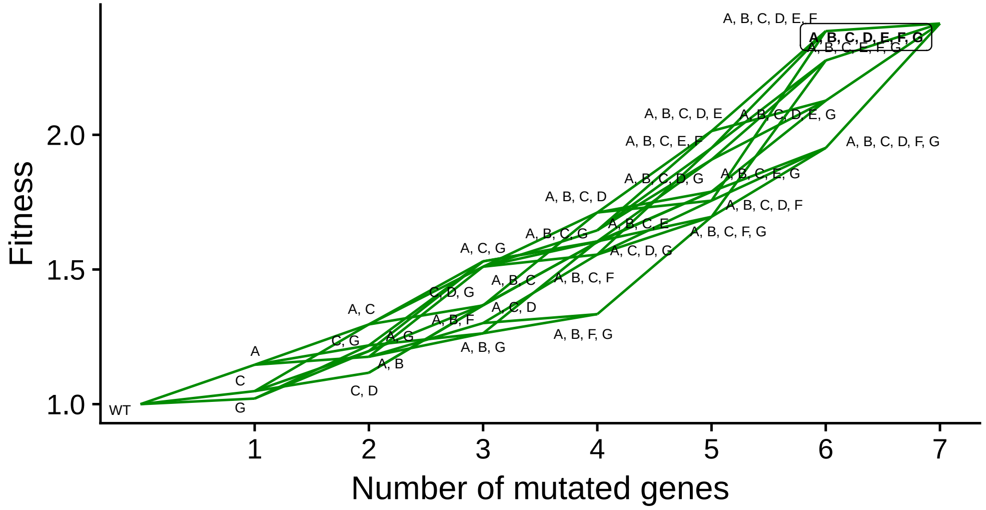

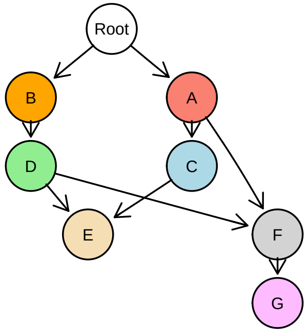


| | |
|---|---|
| *ID* | 10cxjmXXiW5qEyv73 |
| *acc genotypes* | 16 |
| *peaks no back* | 3 |
| *removed edges* | 2 |
| *prop rem edges* | 0.087 |

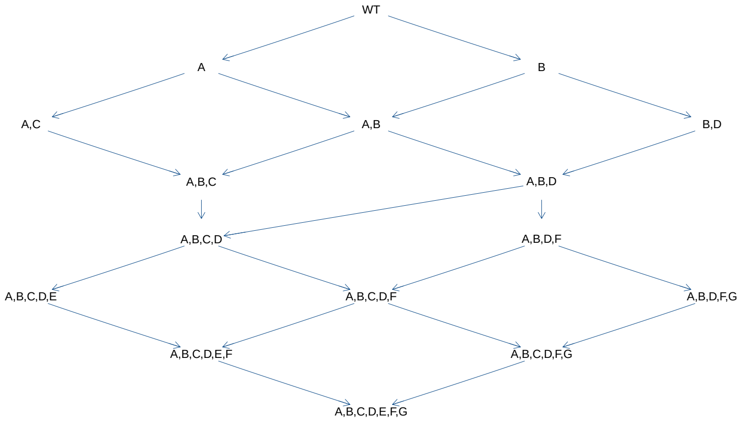


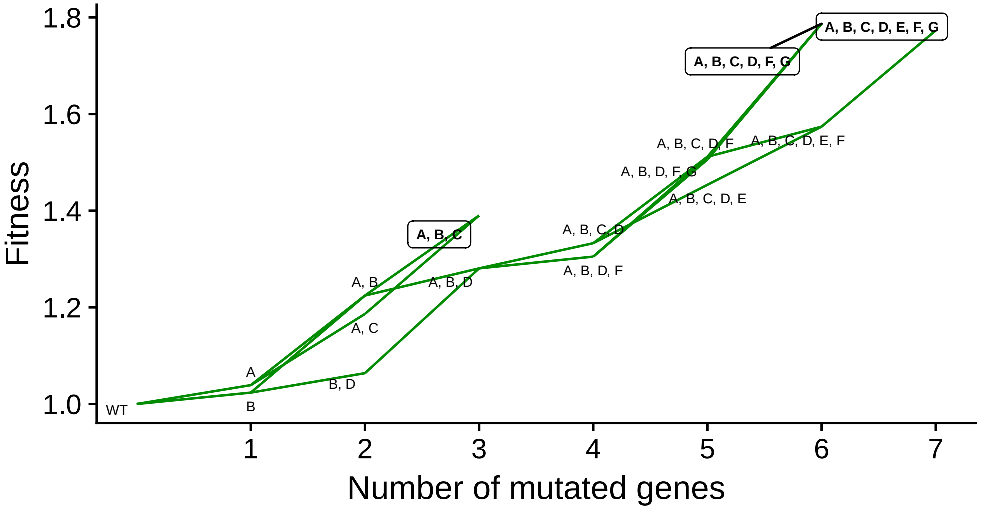


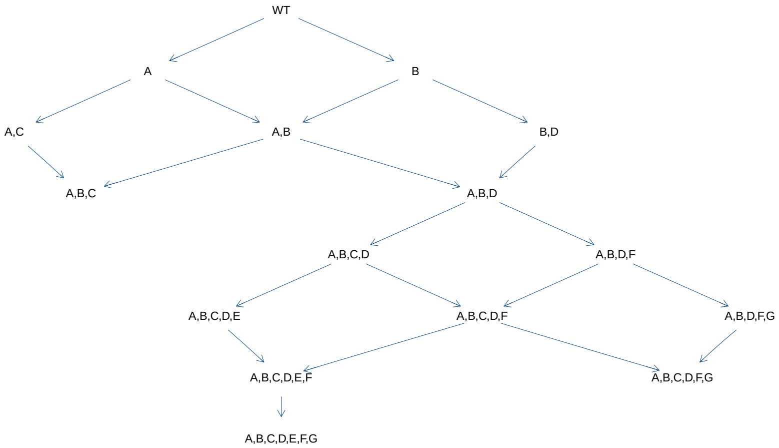

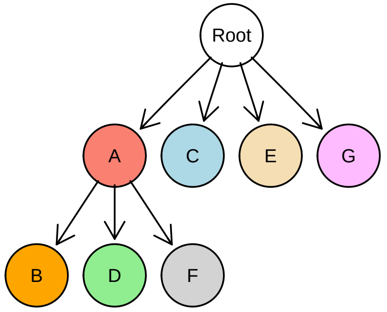


| | |
|---|---|
| *ID* | 1QsHDEe4ASsHeCxX |
| *acc genotypes* | 72 |
| *peaks no back* | 9 |
| *removed edges* | 27 |
| *prop rem edges* | 0.127 |

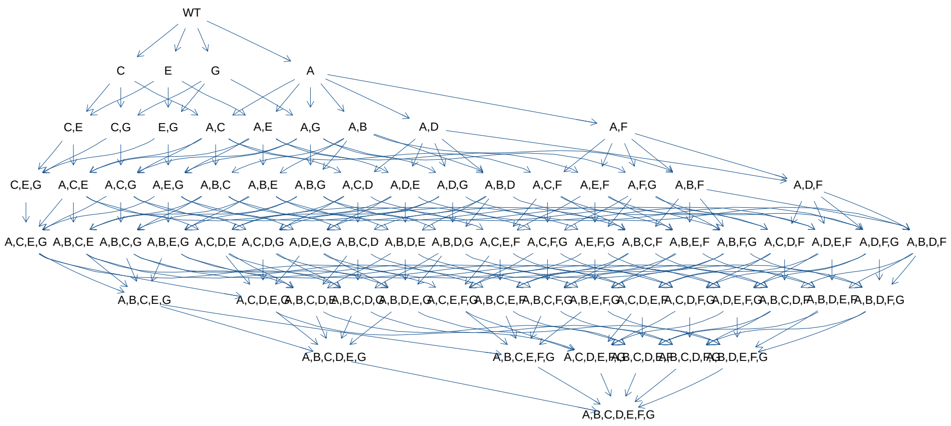


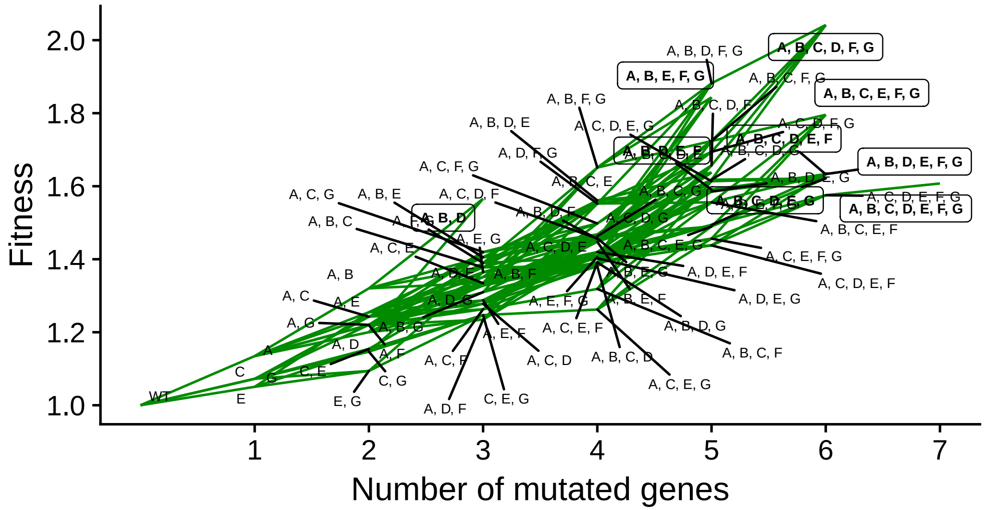


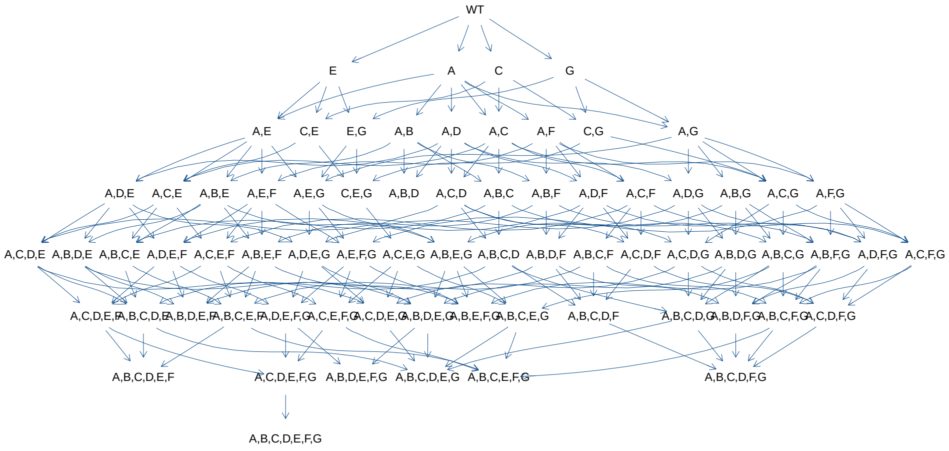

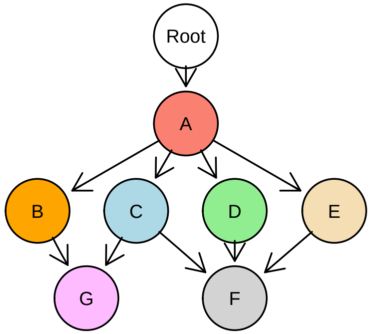


| | |
|---|---|
| *ID* | 2hKTyl9dwrqbrkDP |
| *acc genotypes* | 24 |
| *peaks no back* | 4 |
| *removed edges* | 8 |
| *prop rem edges* | 0.174 |

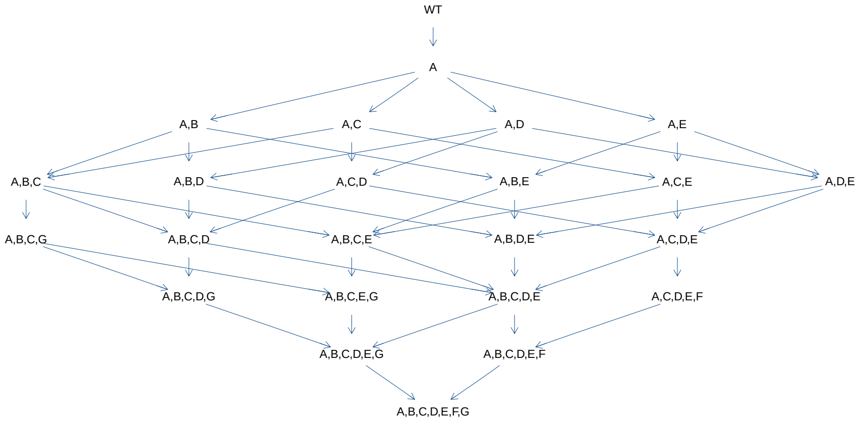


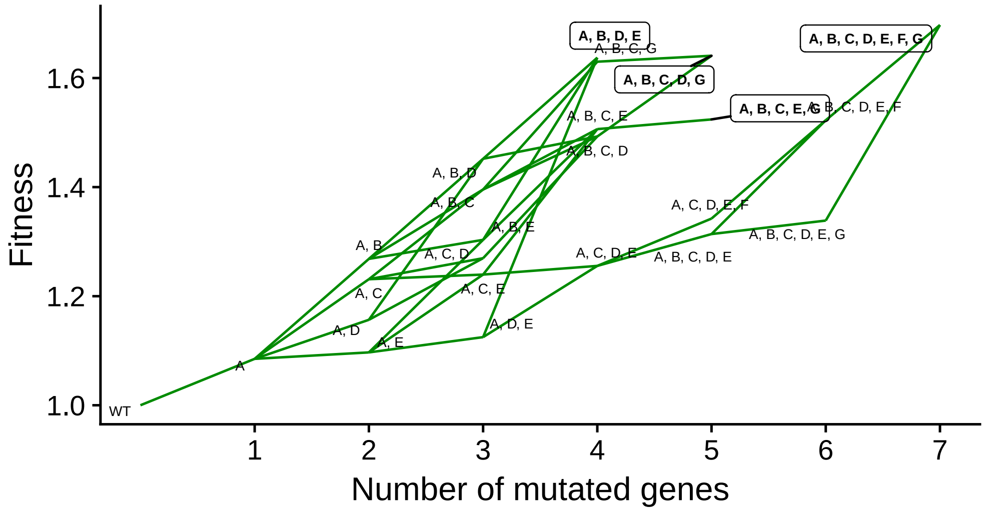


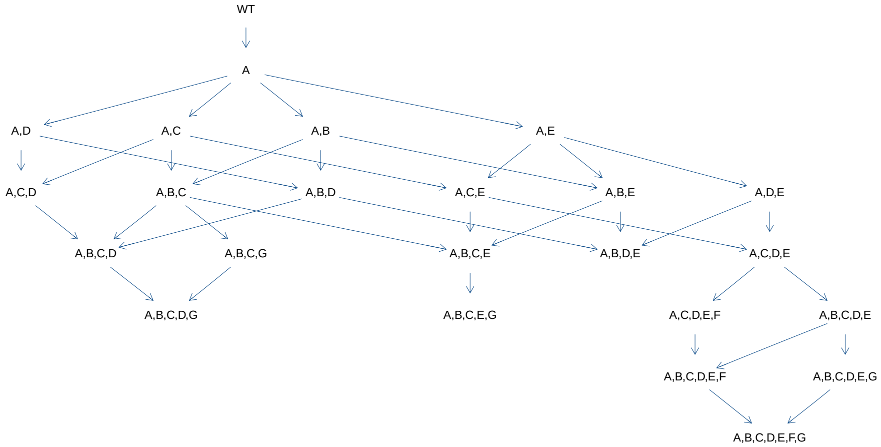

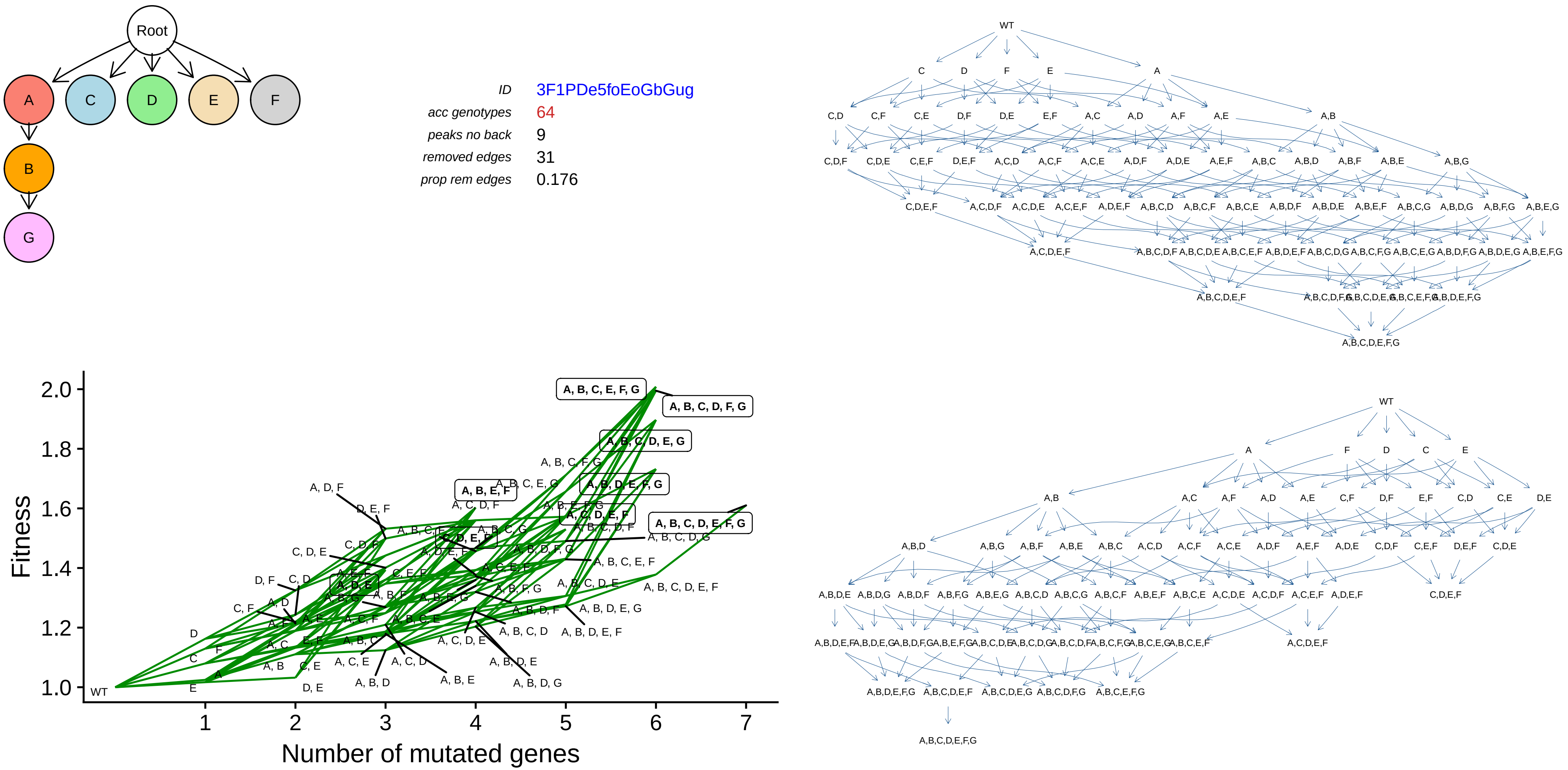
Root
A
C
D
E
F
B
G
ID 3F1PDe5foEoGbGug
acc genotypes 64
peaks no back 9
removed edges 31
prop rem edges 0.176
Fitness
Number of mutated genes
A, B, C, E, F, G
A, B, C, D, F, G
A, B, C, D, E, G
A, B, D, E, F, G
A, B, C, D, E, F, G
A, B, E, F
A, C, D, E, F
A, D, E, F
A, D, E
WT

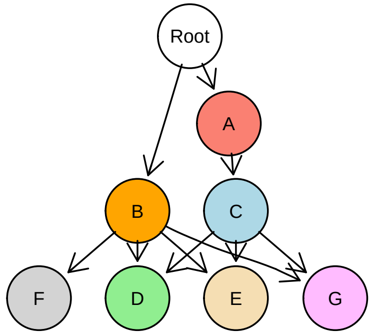


| | |
|---|---|
| *ID* | 3HeBeujvSFoM1FDd |
| *acc genotypes* | 23 |
| *peaks no back* | 5 |
| *removed edges* | 7 |
| *prop rem edges* | 0.163 |

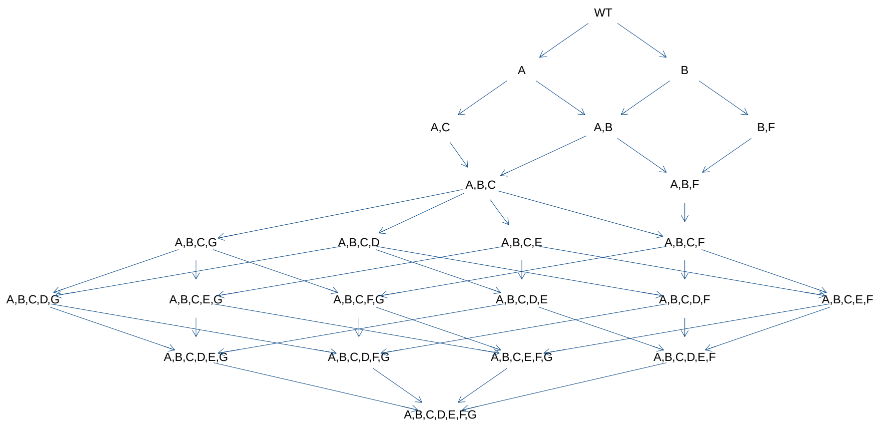


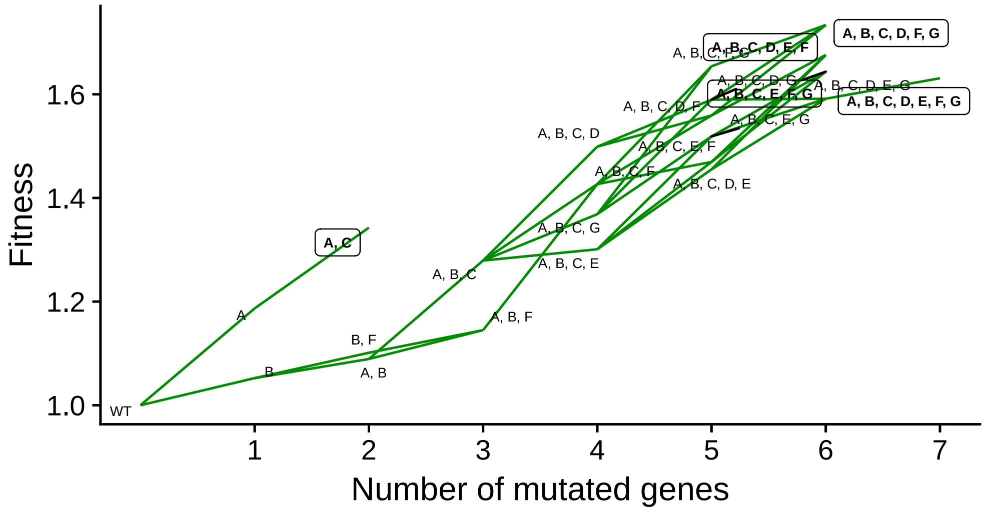


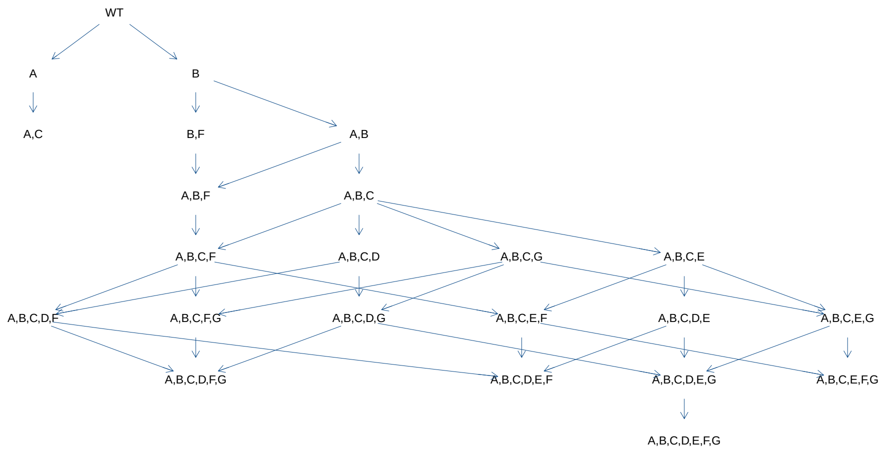

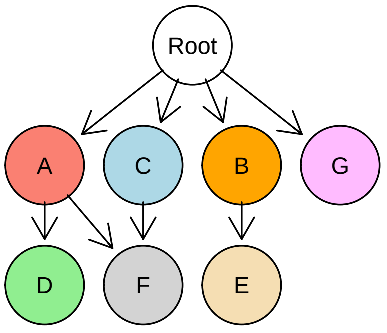


| | |
|---|---|
| *ID* | 3zLLGRM6bZ1renKV |
| *acc genotypes* | 48 |
| *peaks no back* | 7 |
| *removed edges* | 25 |
| *prop rem edges* | 0.216 |

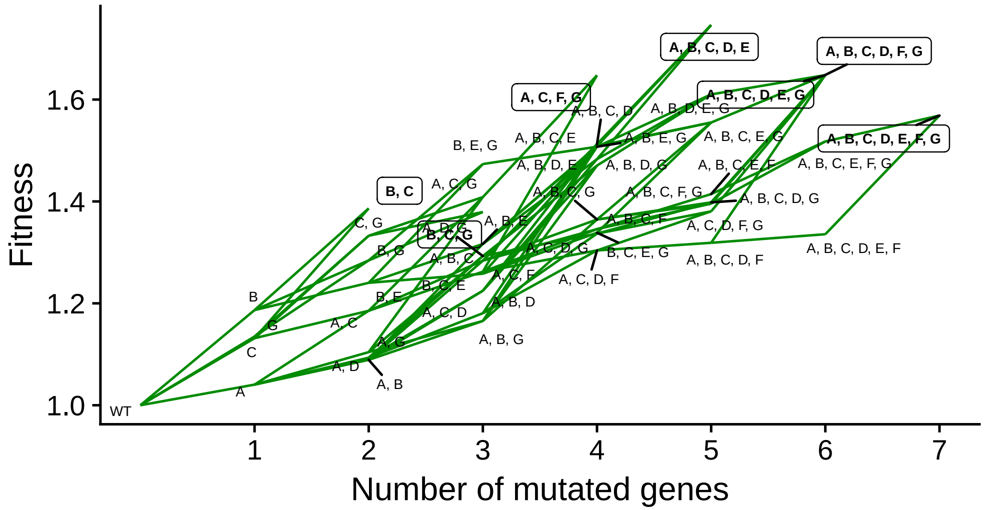


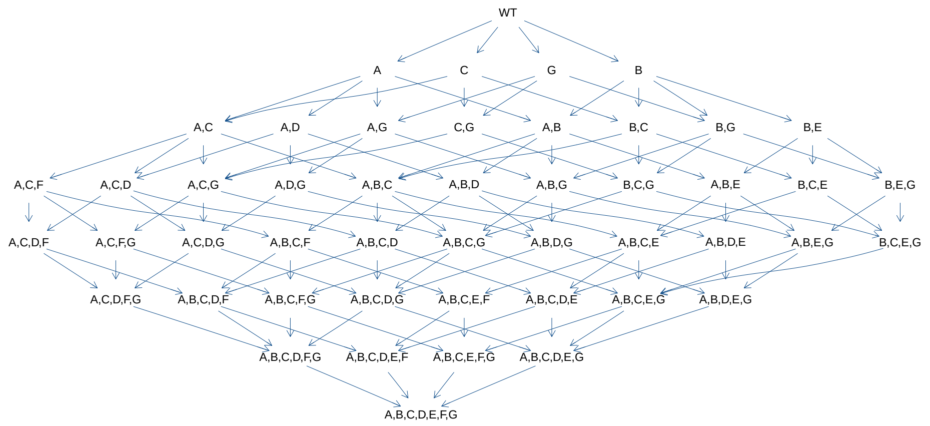


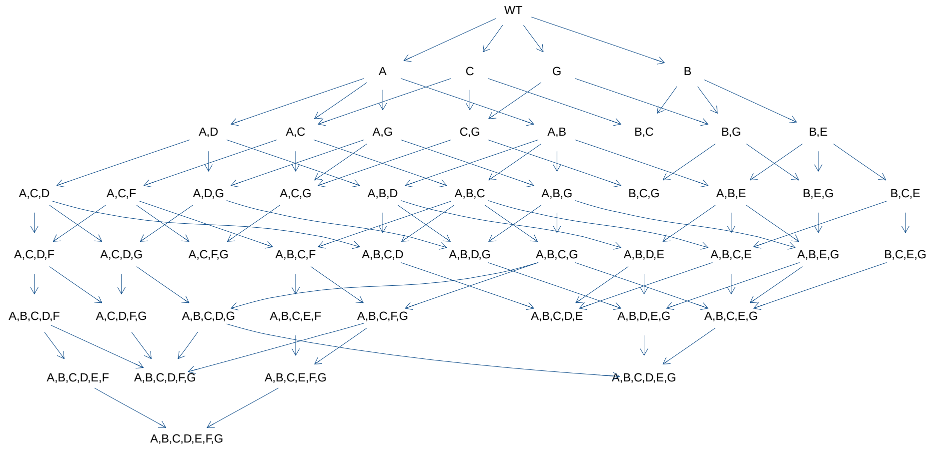

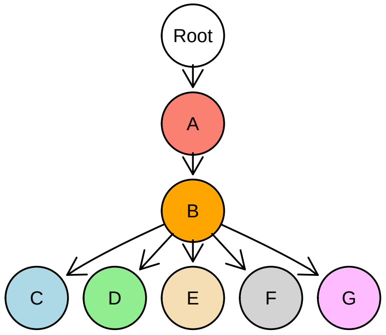


*ID* 410cr63nve1pNTOmC
*acc genotypes* 34
*peaks no back* 5
*removed edges* 5
*prop rem edges* 0.061

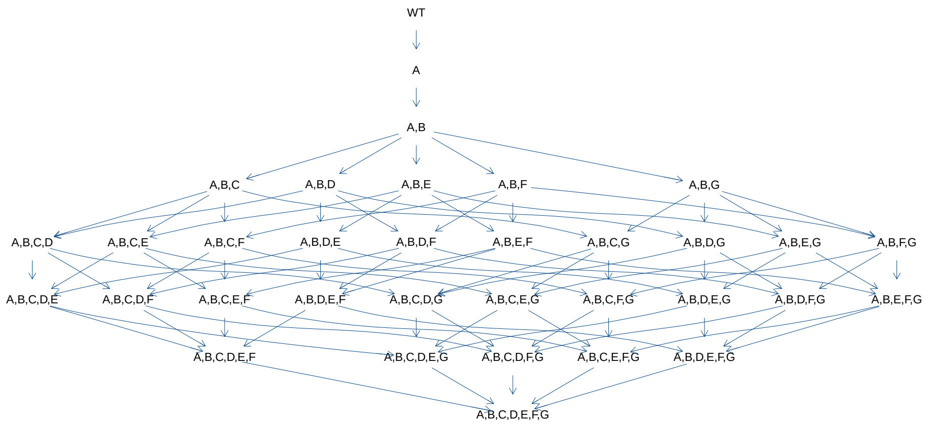


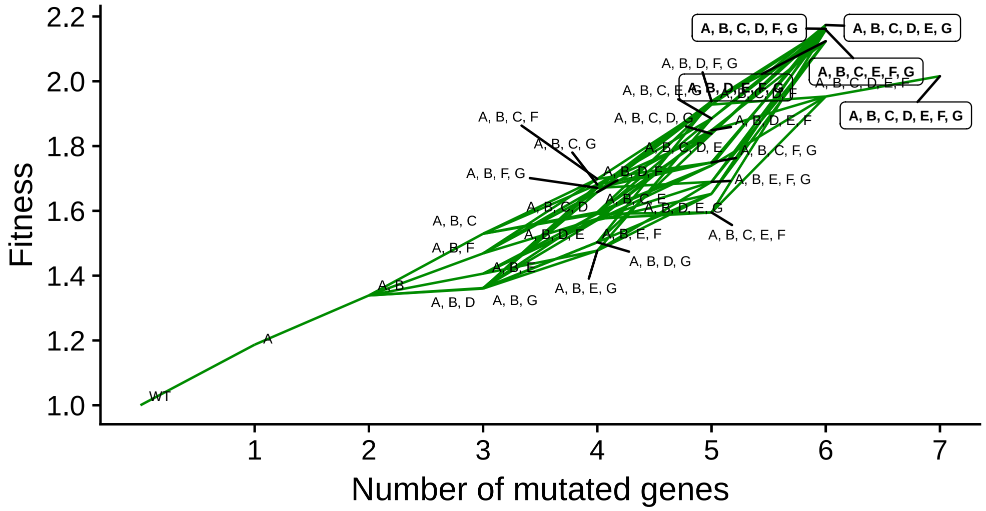


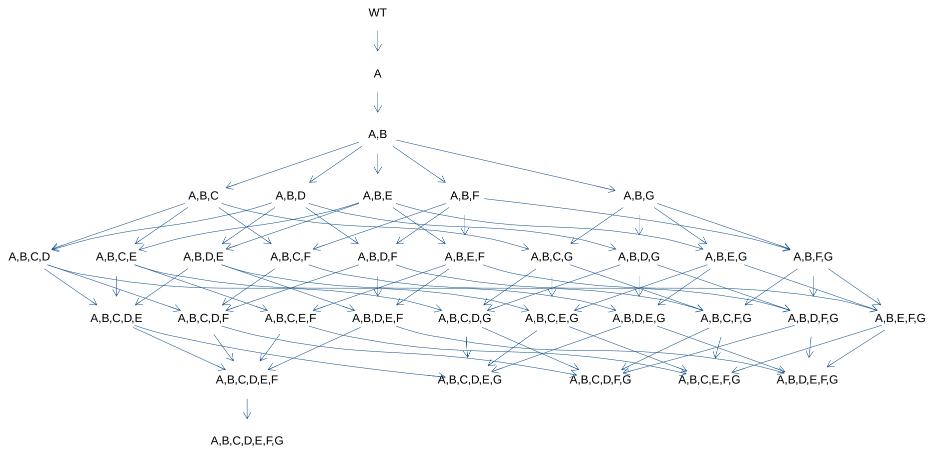

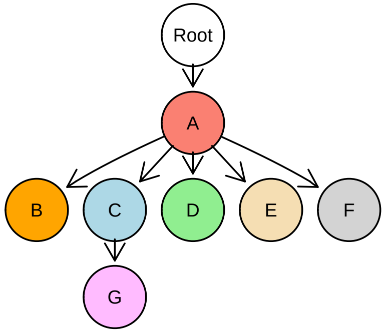


*ID* 4K18VlZE2xqKOhrH
*acc genotypes* 49
*peaks no back* 8
*removed edges* 22
*prop rem edges* 0.171

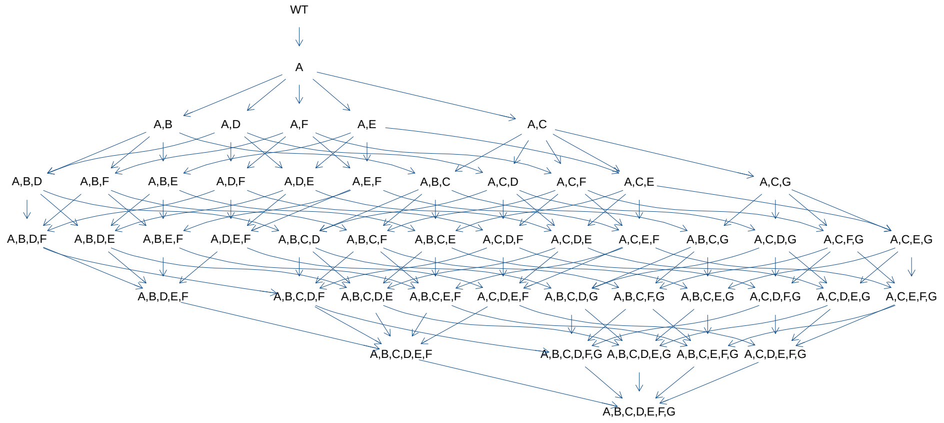

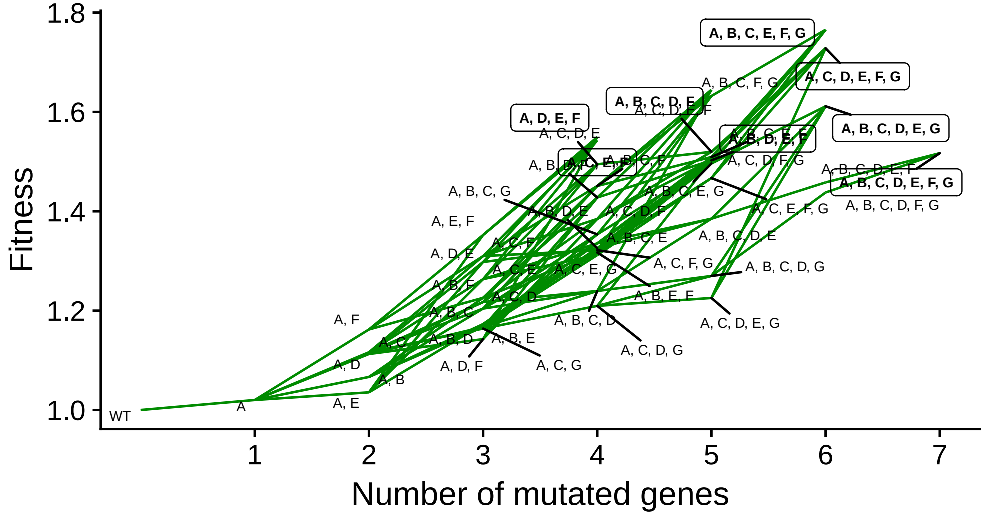


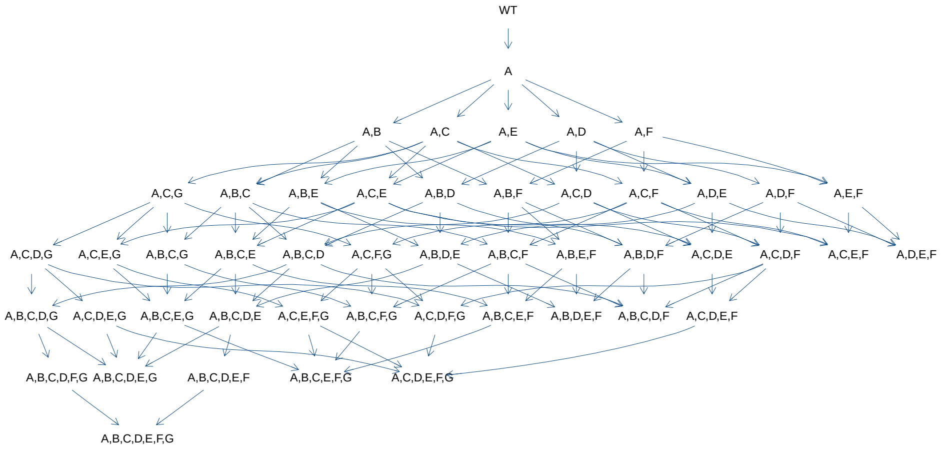

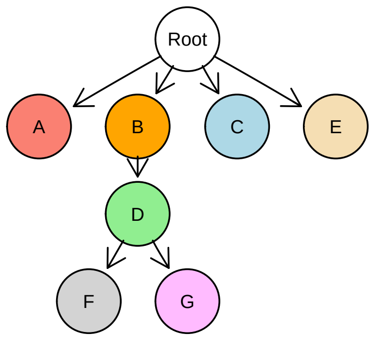


| | |
|---|---|
| *ID* | 4KzuP1MMEwOUSymI |
| *acc genotypes* | 48 |
| *peaks no back* | 10 |
| *removed edges* | 26 |
| *prop rem edges* | 0.217 |

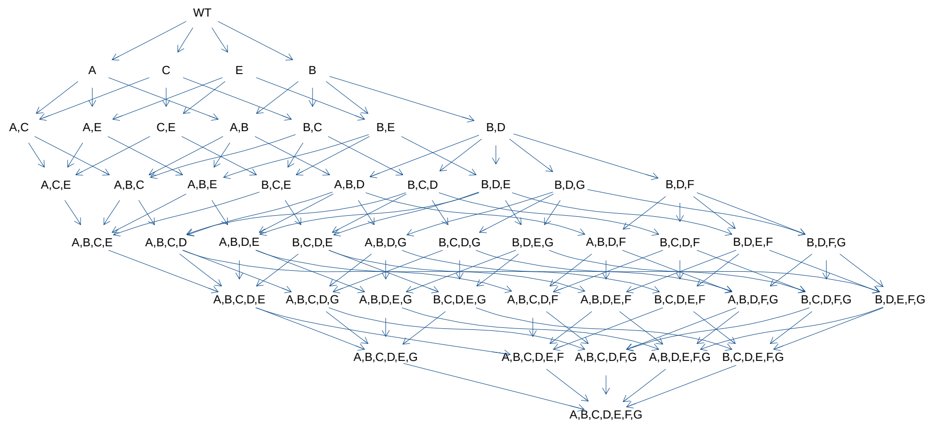


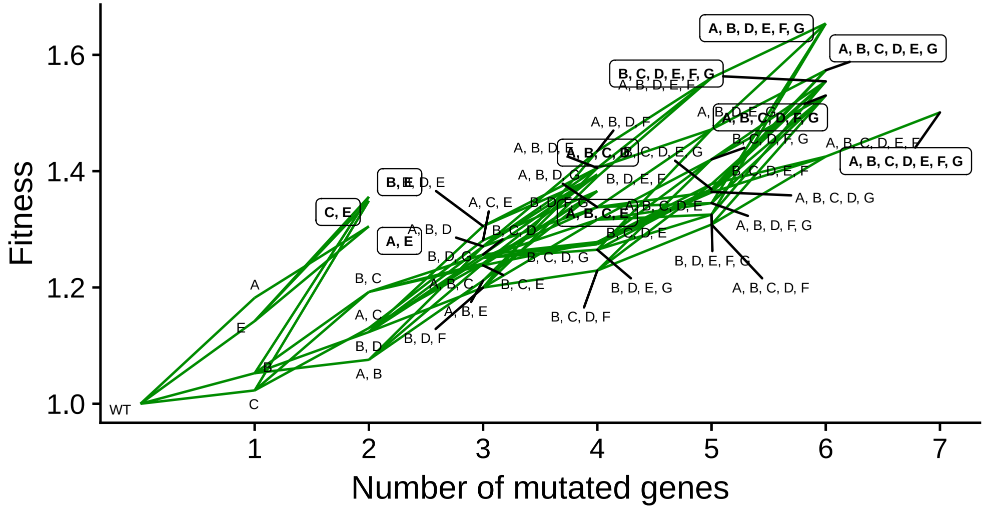


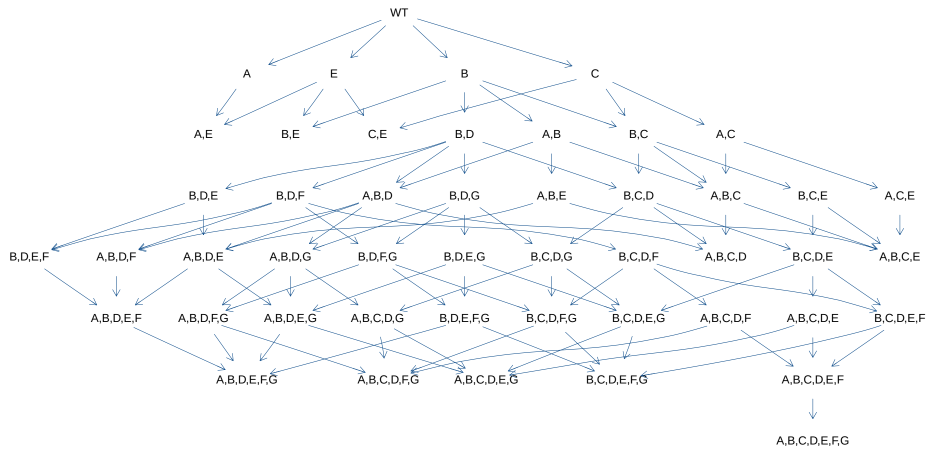

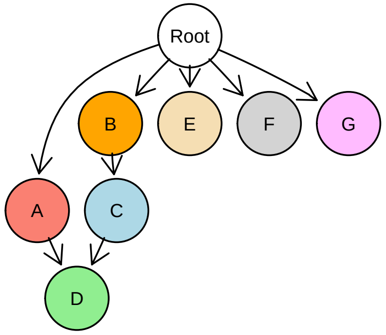


| | |
|---|---|
| *ID* | 4uRd77KBIQ1QZeh9 |
| *acc genotypes* | 56 |
| *peaks no back* | 9 |
| *removed edges* | 36 |
| *prop rem edges* | 0.243 |

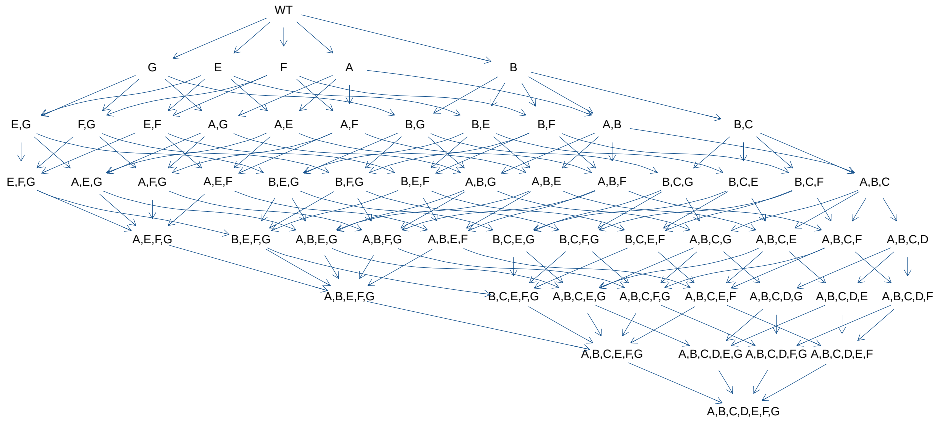


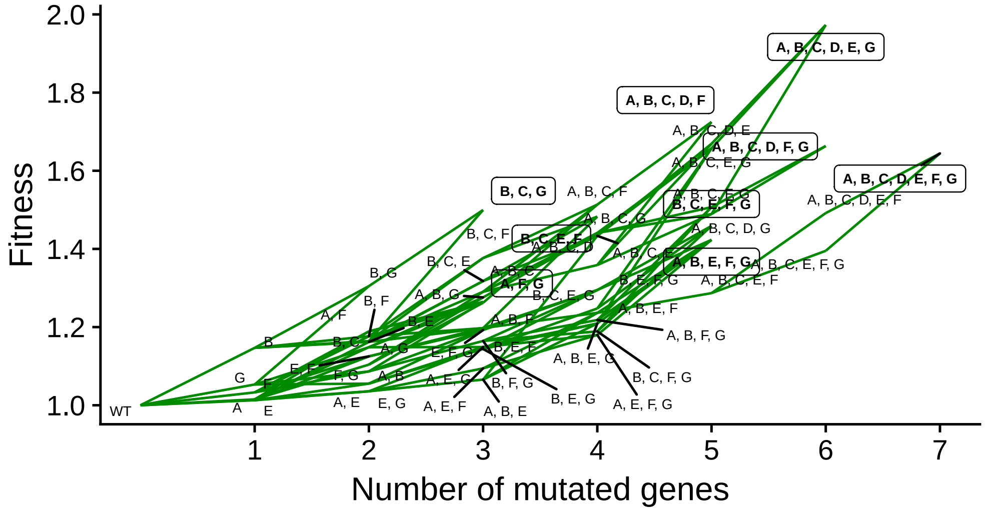


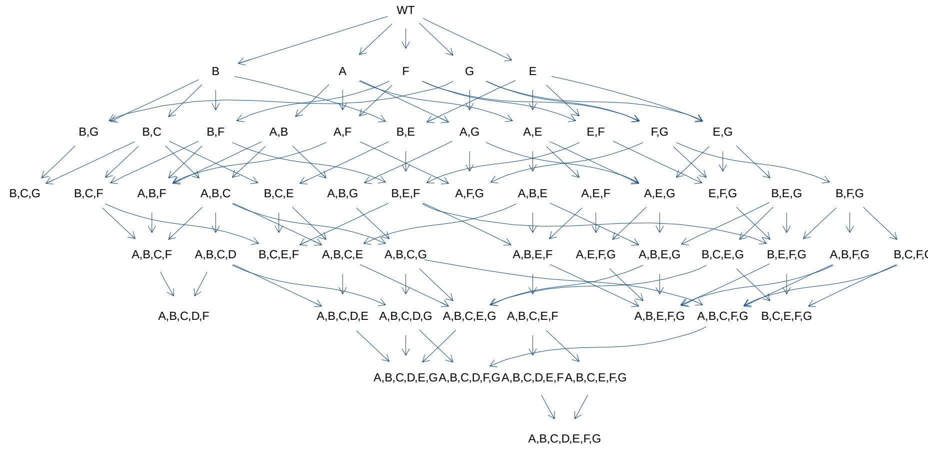

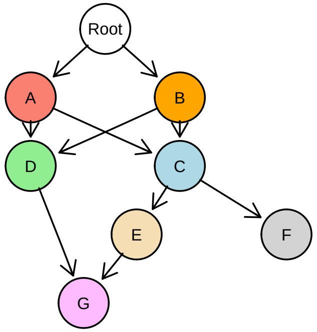


| | |
|---|---|
| *ID* | 4xM3Wofst2xw10jJ6 |
| *acc genotypes* | 15 |
| *peaks no back* | 2 |
| *removed edges* | 2 |
| *prop rem edges* | 0.091 |

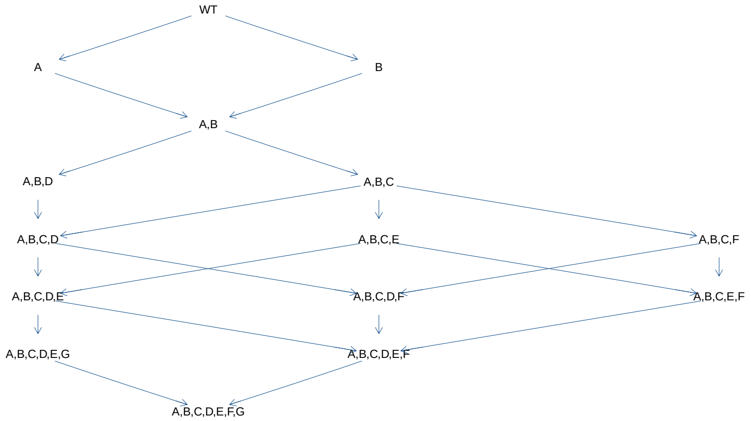


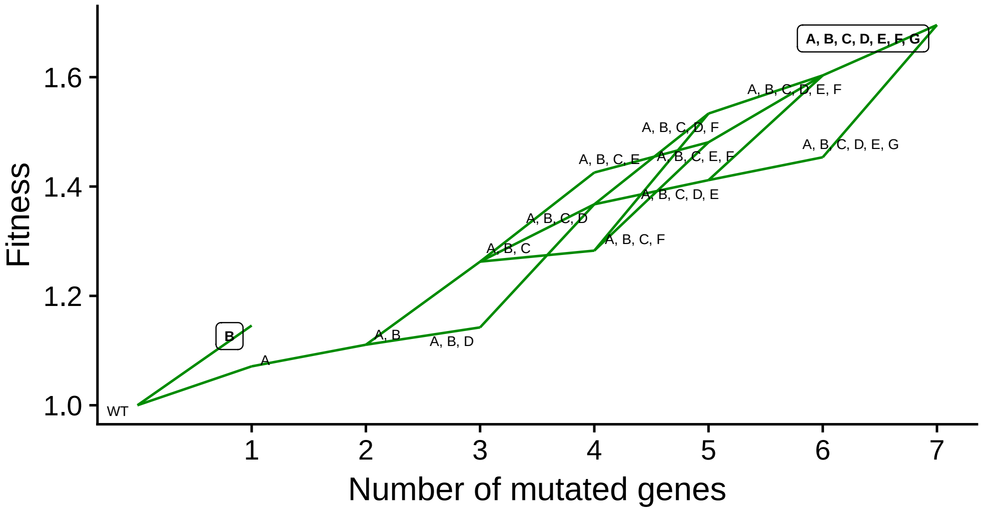


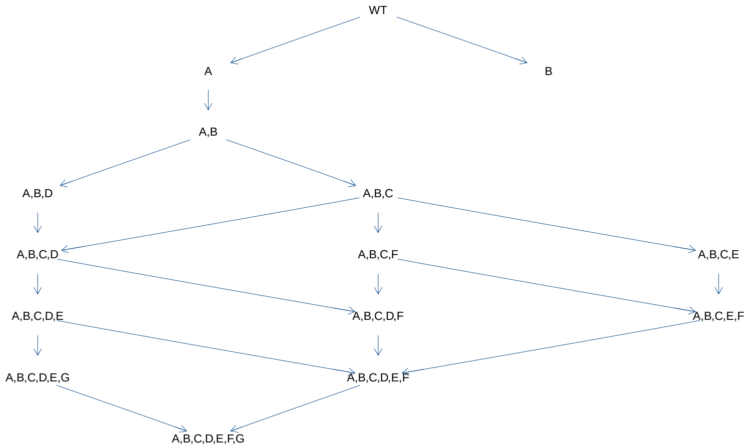

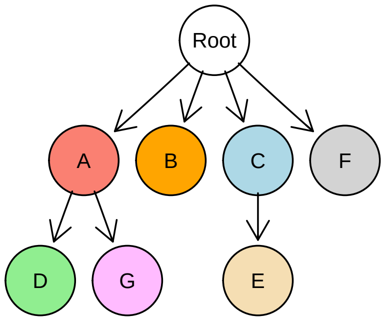


*ID* 4YkXkc14paEyTomW
*acc genotypes* 60
*peaks no back* 8
*removed edges* 28
*prop rem edges* 0.175

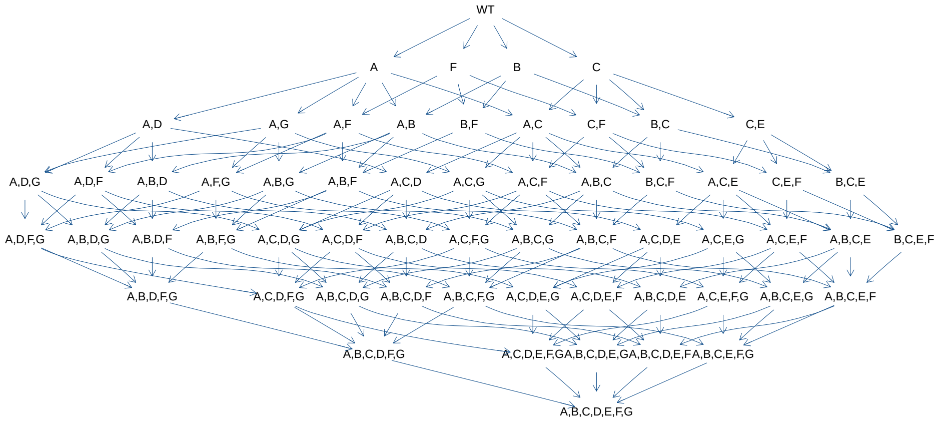


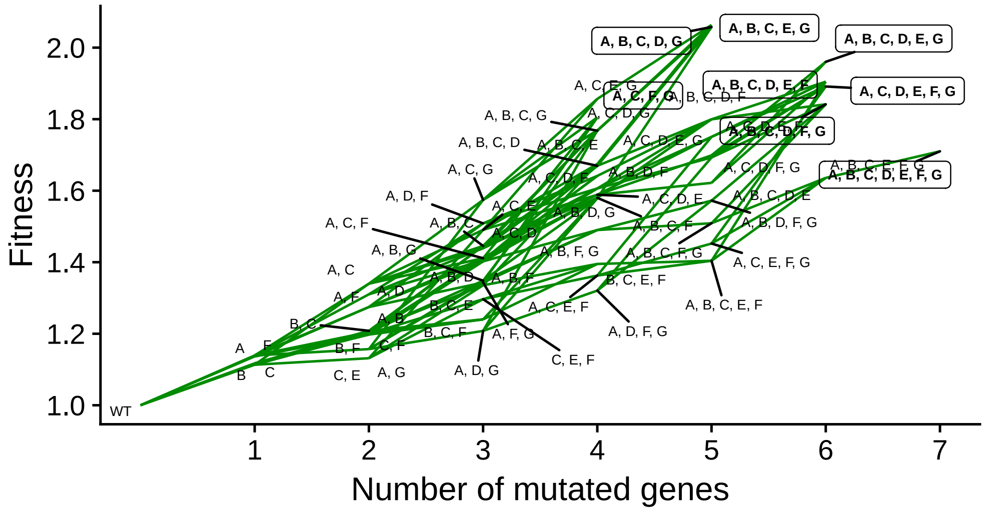


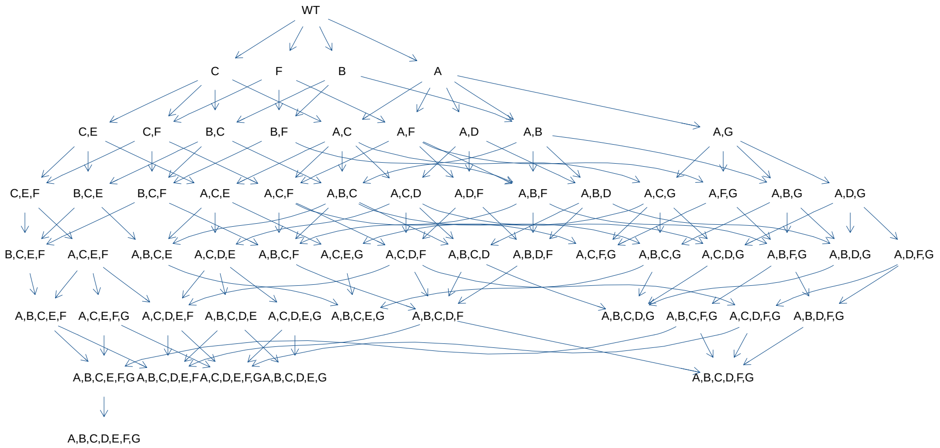

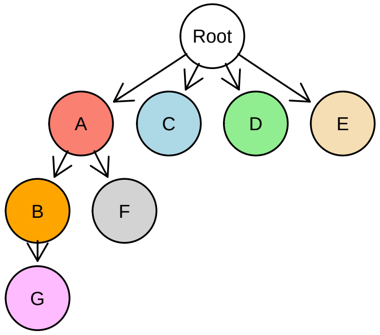


| | |
|---|---|
| *ID* | 5cLLbdWPenssPhaD |
| *acc genotypes* | 56 |
| *peaks no back* | 7 |
| *removed edges* | 20 |
| *prop rem edges* | 0.135 |

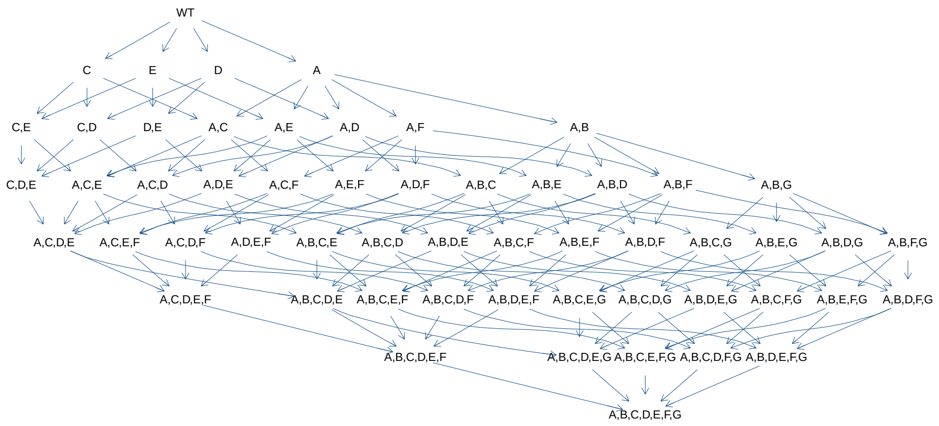


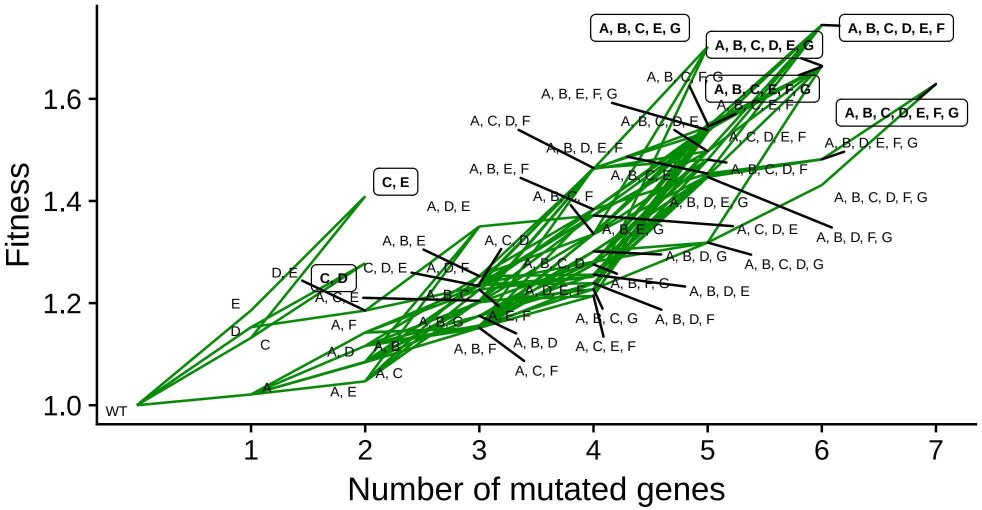


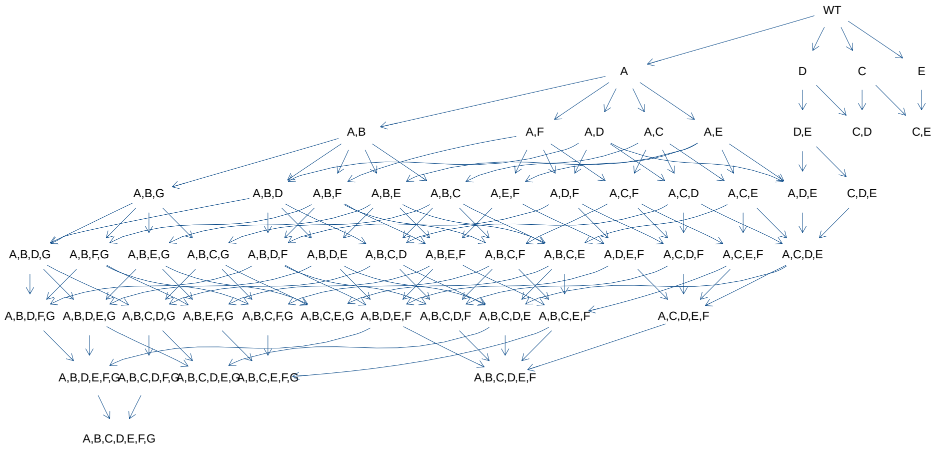

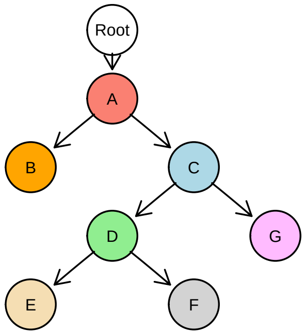


*ID* 5EI6S7TMXAPy65pq
*acc genotypes* 23
*peaks no back* 2
*removed edges* 3
*prop rem edges* 0.068

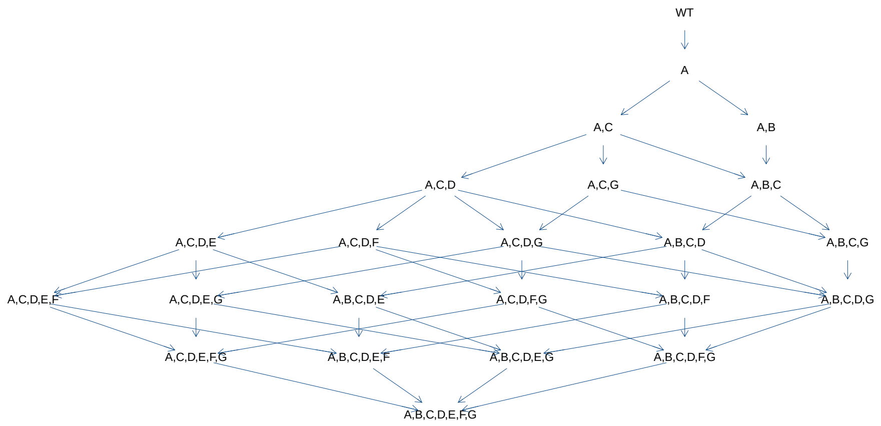


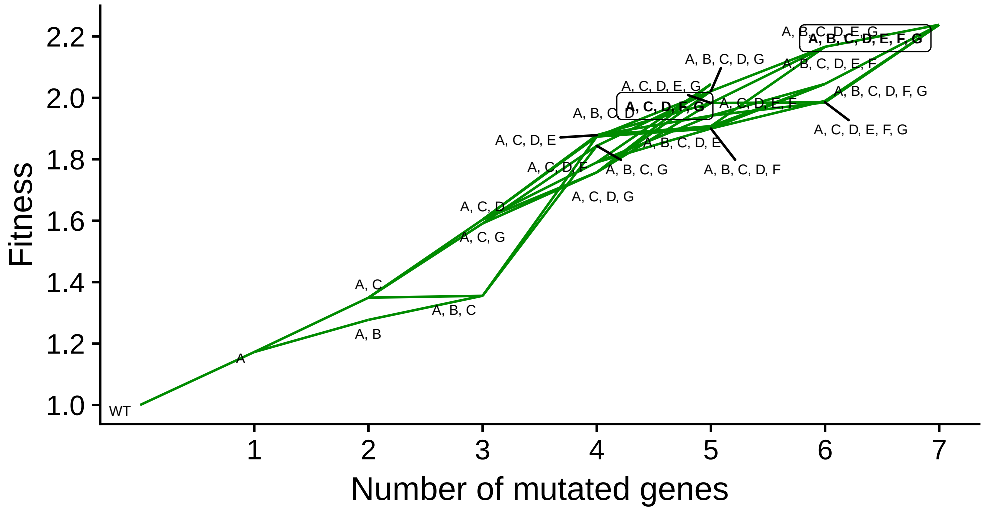


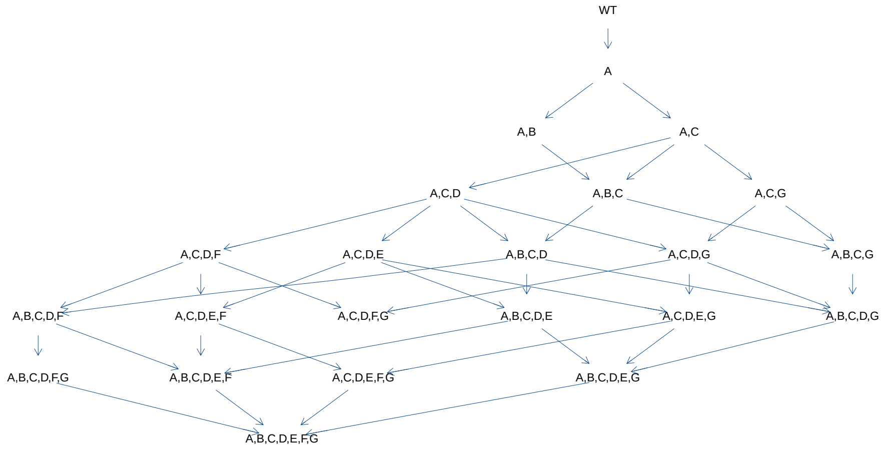

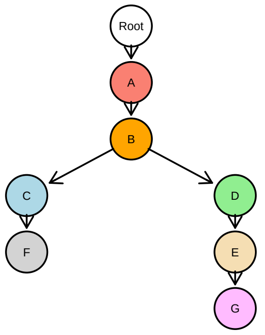


| | |
|---|---|
| *ID* | 5gEdOQwKmmjGCuwA |
| *acc genotypes* | 14 |
| *peaks no back* | 3 |
| *removed edges* | 4 |
| *prop rem edges* | 0.211 |

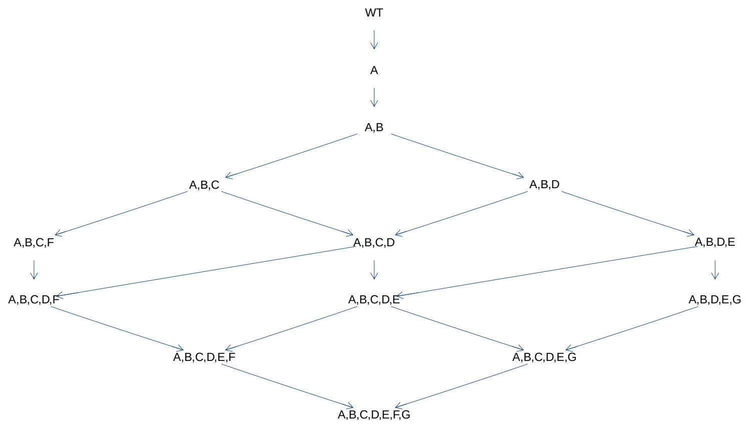


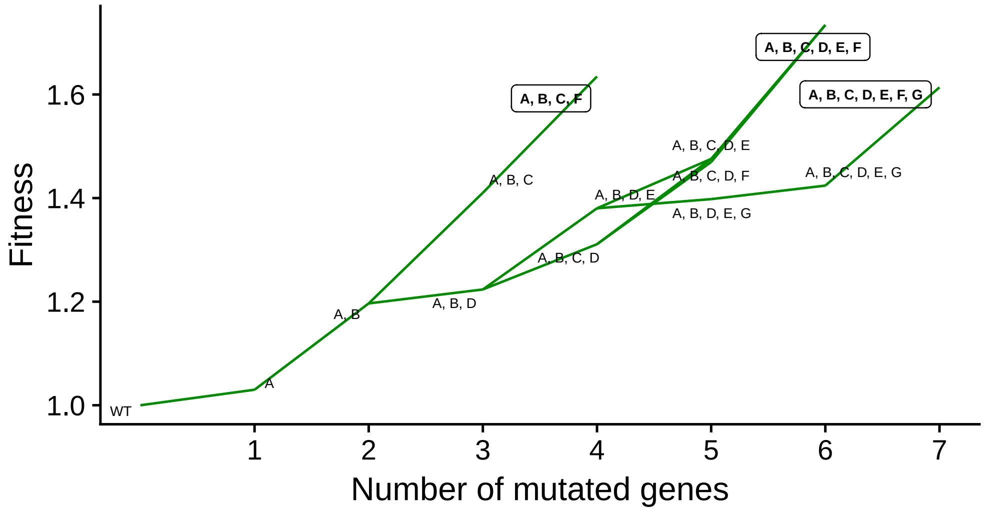


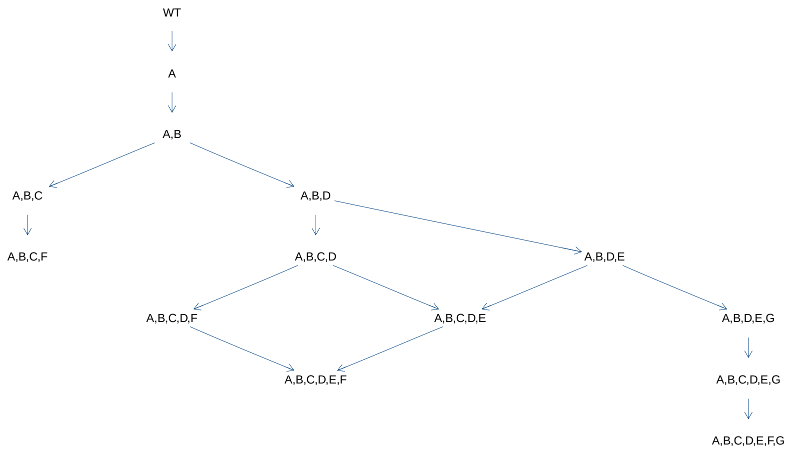

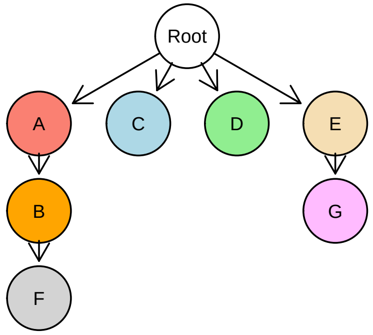


*ID* 5kZqKEOHzl10BWV78
*acc genotypes* 48
*peaks no back* 7
*removed edges* 16
*prop rem edges* 0.138

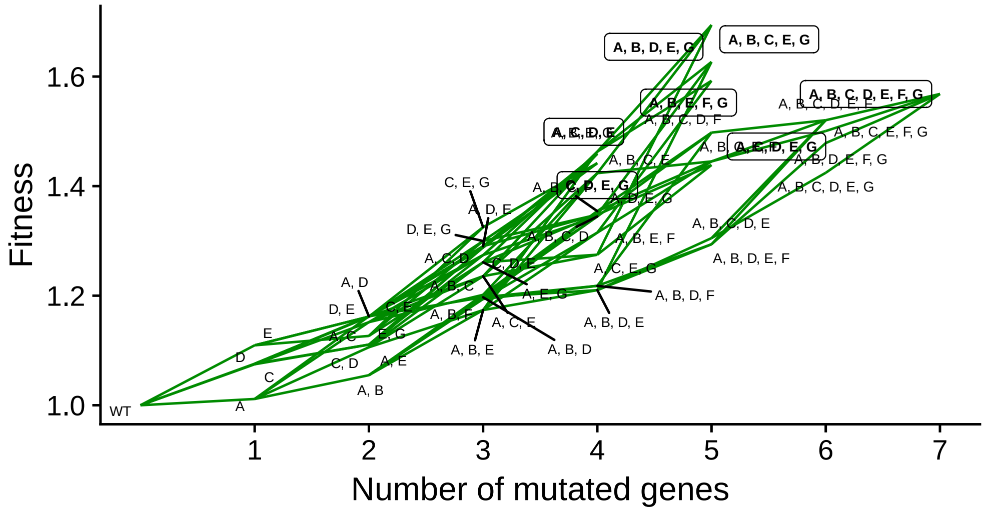


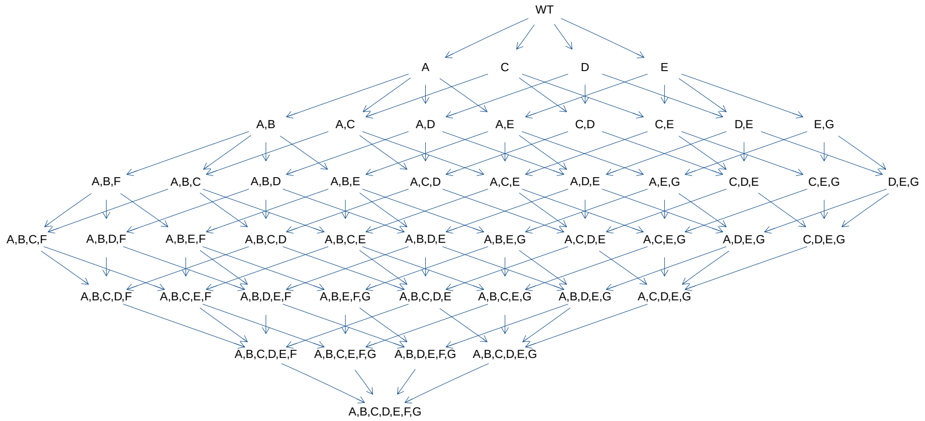


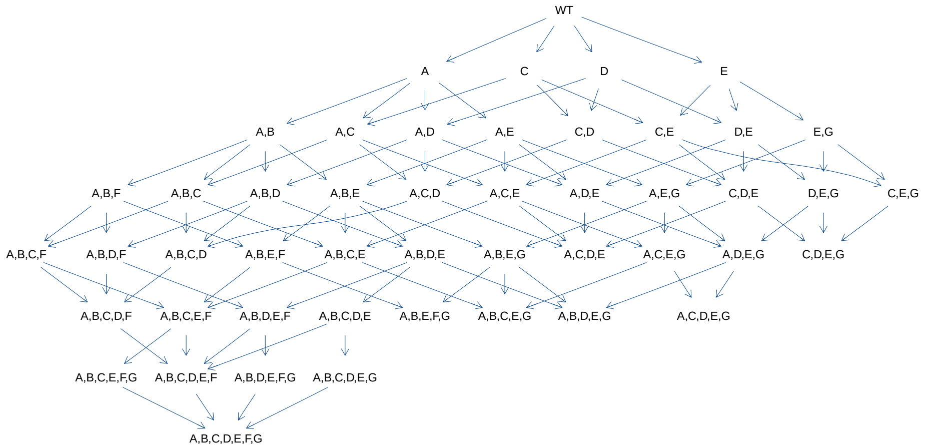

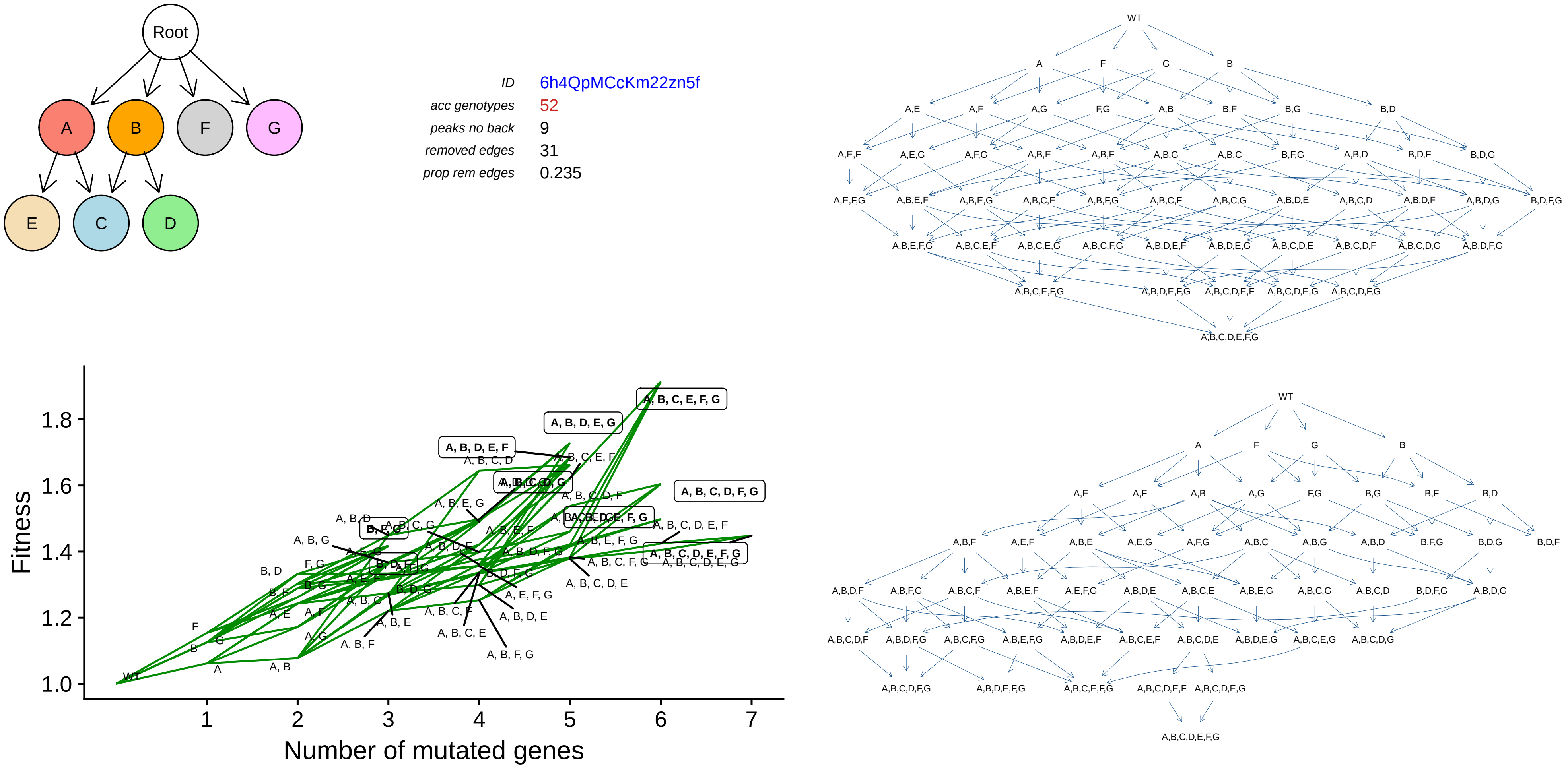
Root
A
B
F
G
E
C
D
ID
6h4QpMCcKm22zn5f
acc genotypes
52
peaks no back
9
removed edges
31
prop rem edges
0.235
Fitness
Number of mutated genes
A, B, C, E, F, G
A, B, D, E, G
A, B, D, E, F
A, B, C, D, F, G
A, B, C, D, E, F, G
WT
A,B,C,D,E,F,G

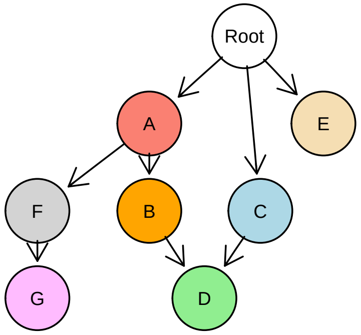


| | |
|---|---|
| *ID* | 6Hn6i2YqHUFLoALB |
| *acc genotypes* | 34 |
| *peaks no back* | 5 |
| *removed edges* | 15 |
| *prop rem edges* | 0.205 |

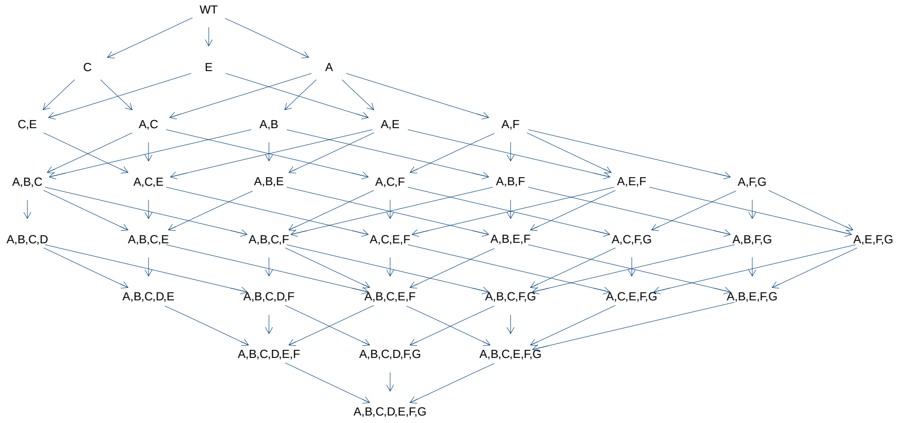


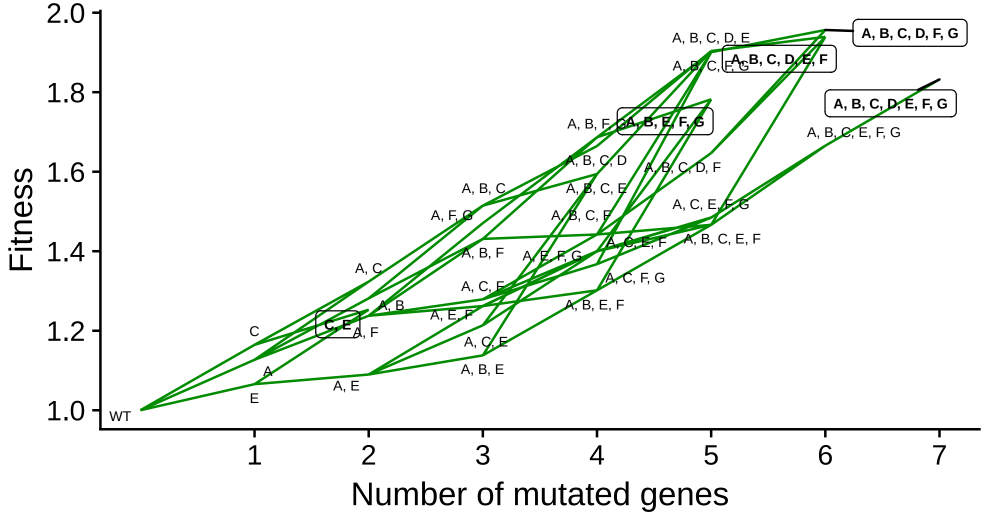


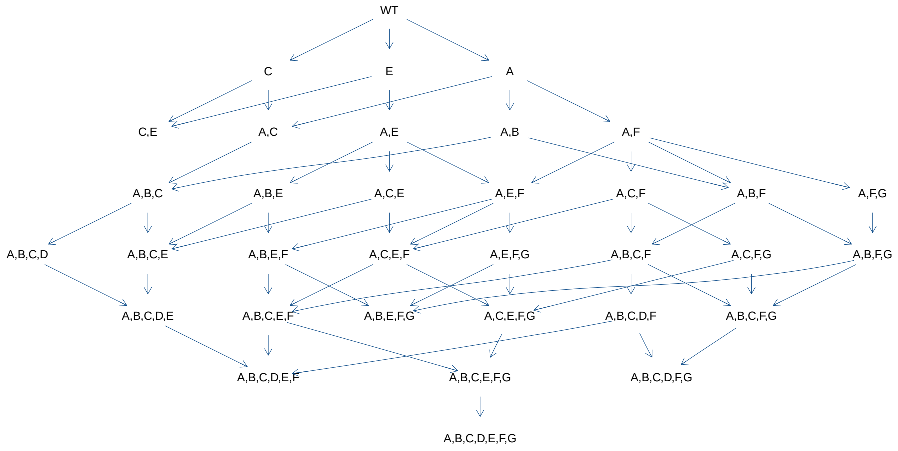

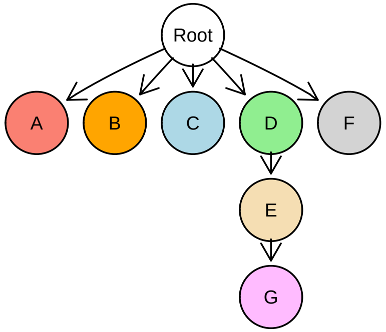


| | |
|---|---|
| *ID* | 6JONUdvaeCCGnS7Y |
| *acc genotypes* | 64 |
| *peaks no back* | 8 |
| *removed edges* | 34 |
| *prop rem edges* | 0.193 |

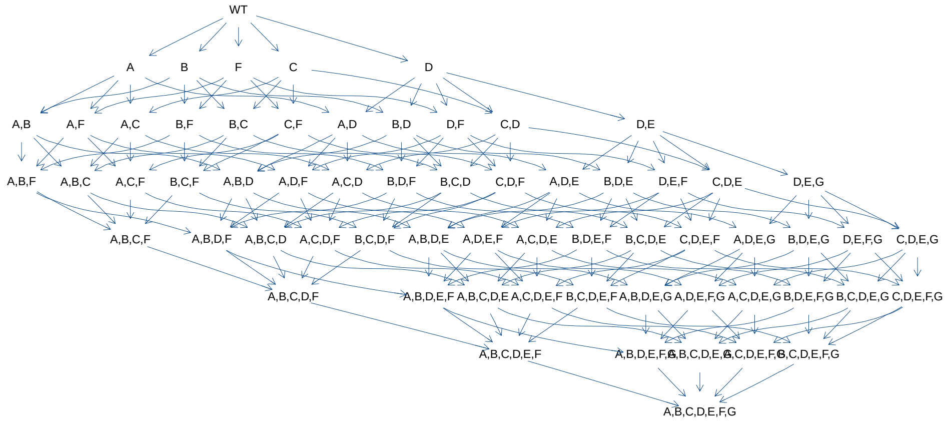


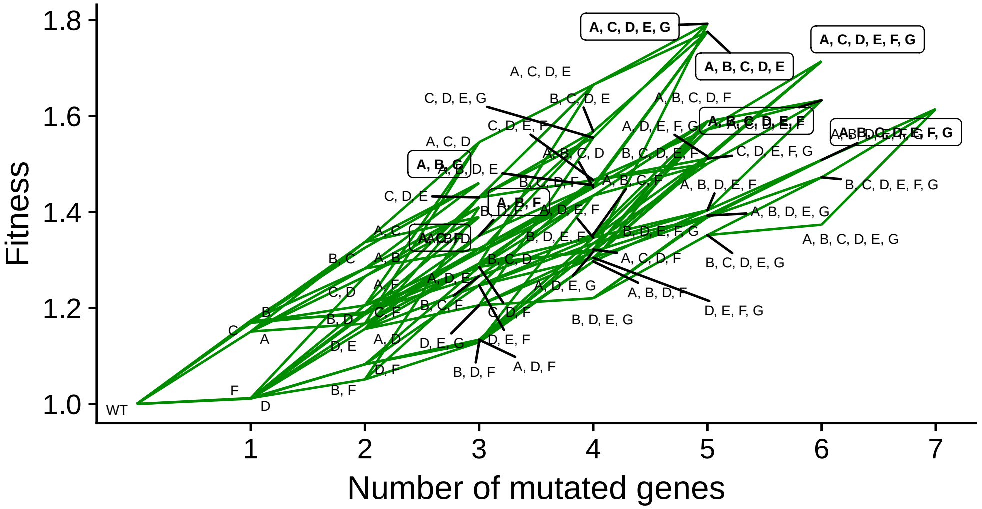


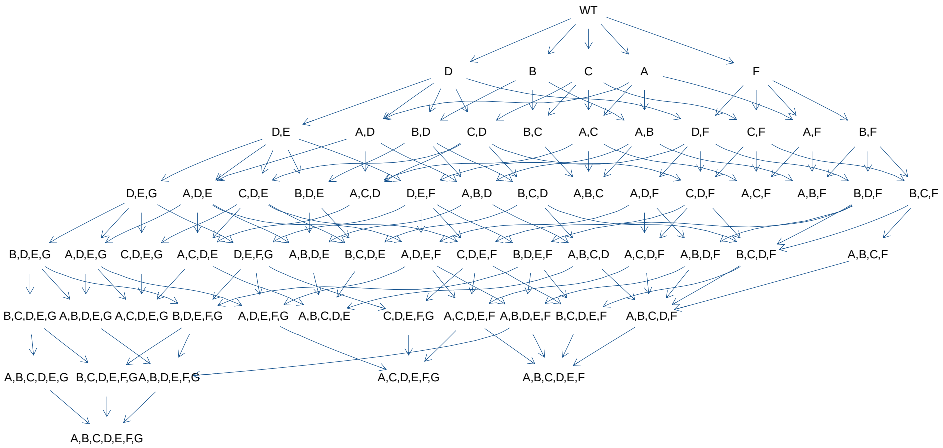

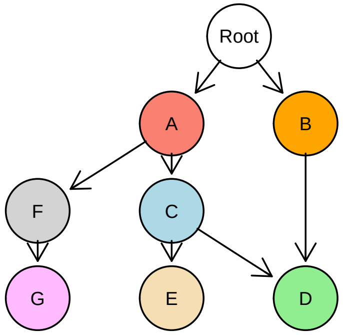


| | |
|---|---|
| *ID* | 6V3IXXX77zzMpI58 |
| *acc genotypes* | 26 |
| *peaks no back* | 2 |
| *removed edges* | 9 |
| *prop rem edges* | 0.184 |

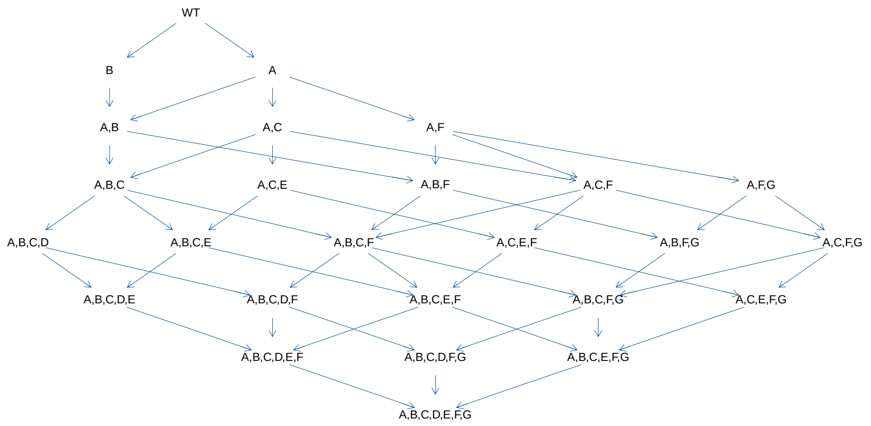

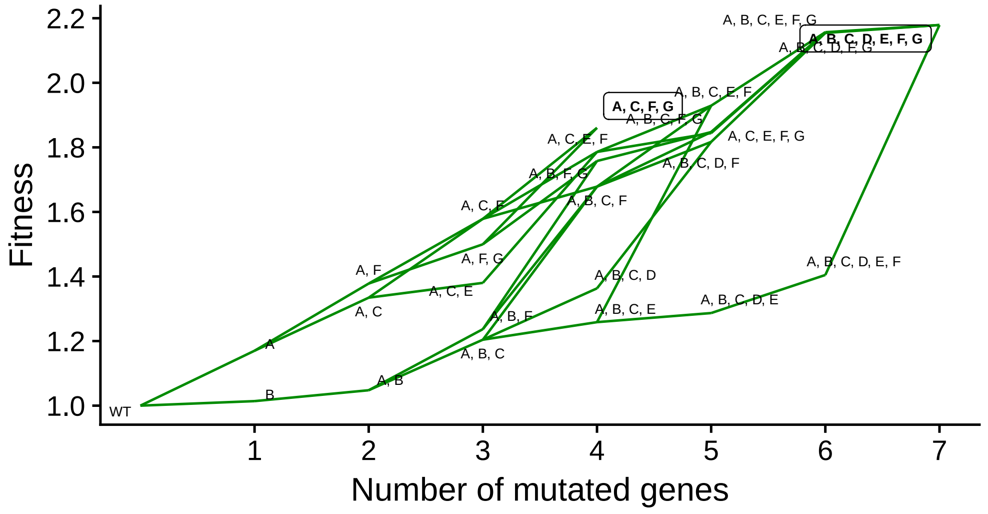


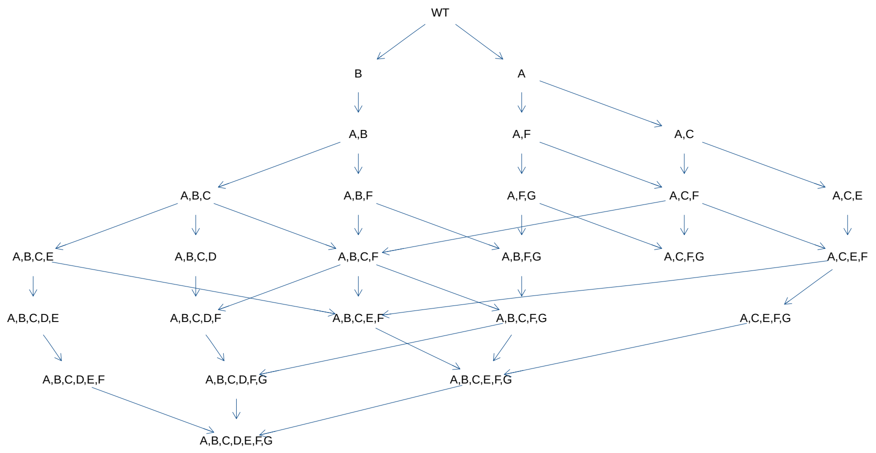

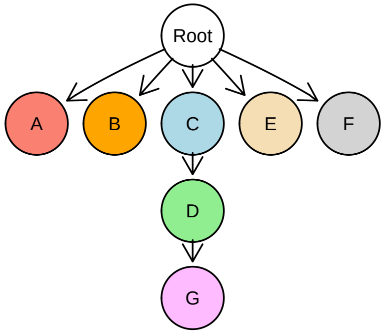


| | |
|---|---|
| *ID* | 738zqMpeBSiZXfez |
| *acc genotypes* | 64 |
| *peaks no back* | 7 |
| *removed edges* | 26 |
| *prop rem edges* | 0.148 |

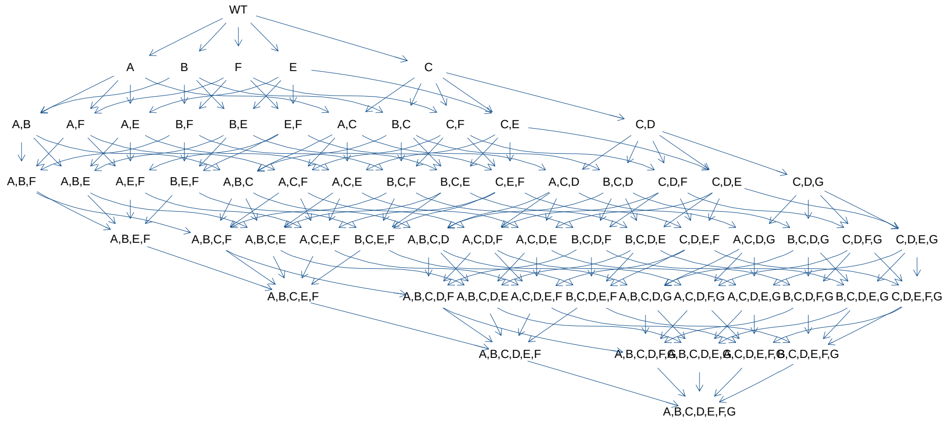


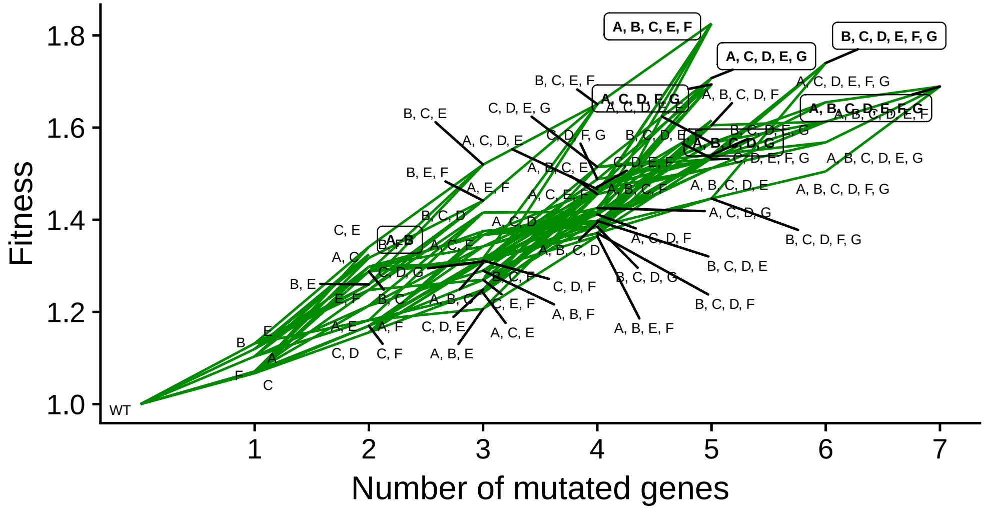


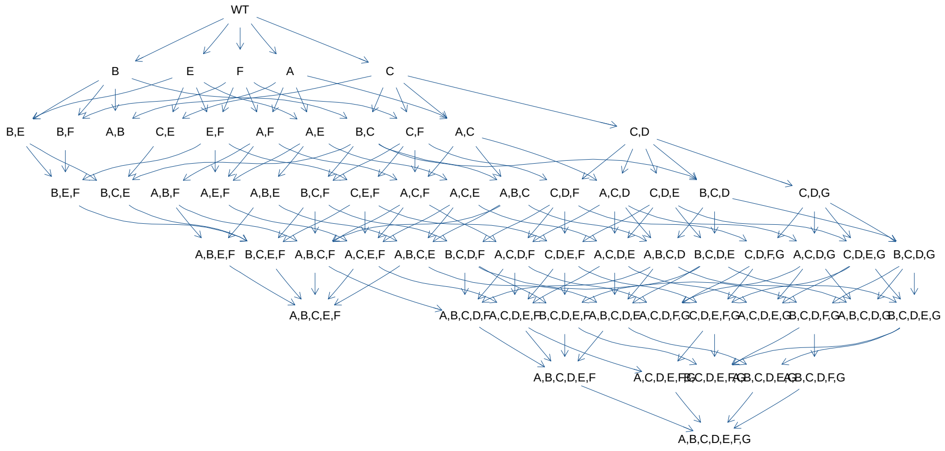

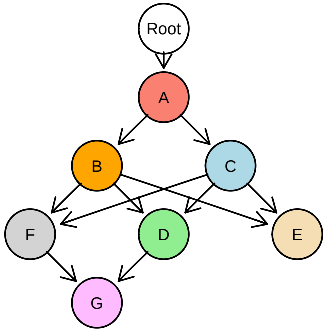


| | |
|---|---|
| *ID* | 7E10pIyu7UguIUl8I |
| *acc genotypes* | 14 |
| *peaks no back* | 2 |
| *removed edges* | 1 |
| *prop rem edges* | 0.05 |

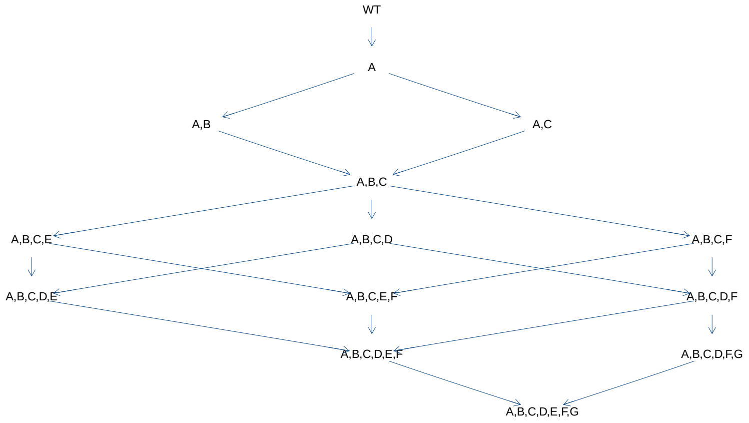


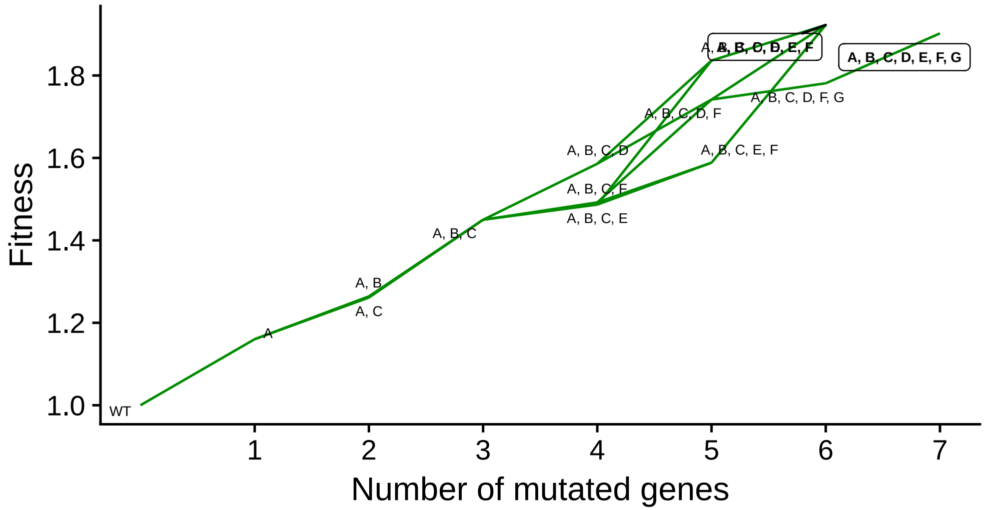


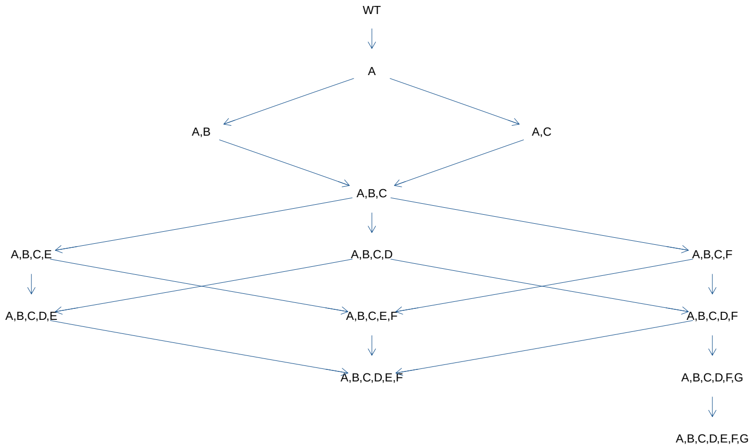

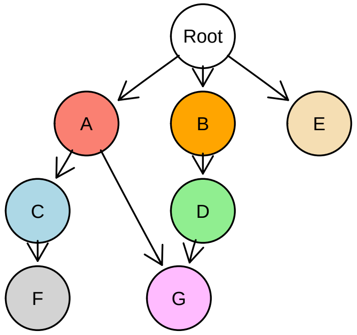


*ID* 7L5tLW7iVltNxTJ6
*acc genotypes* 30
*peaks no back* 6
*removed edges* 14
*prop rem edges* 0.237

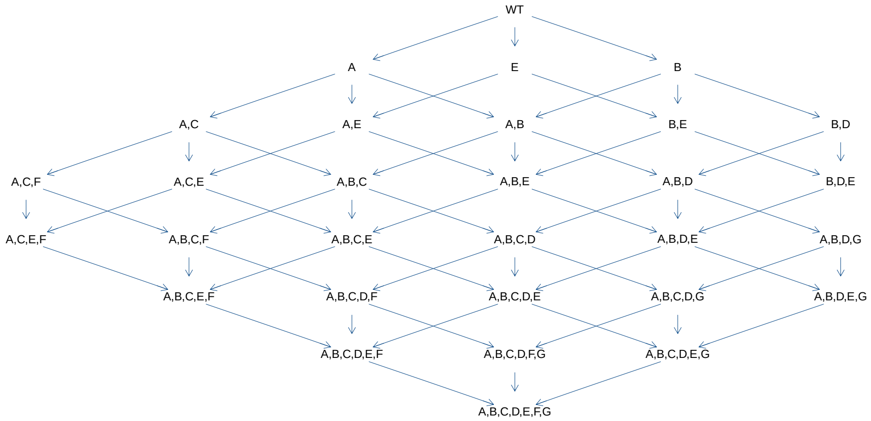


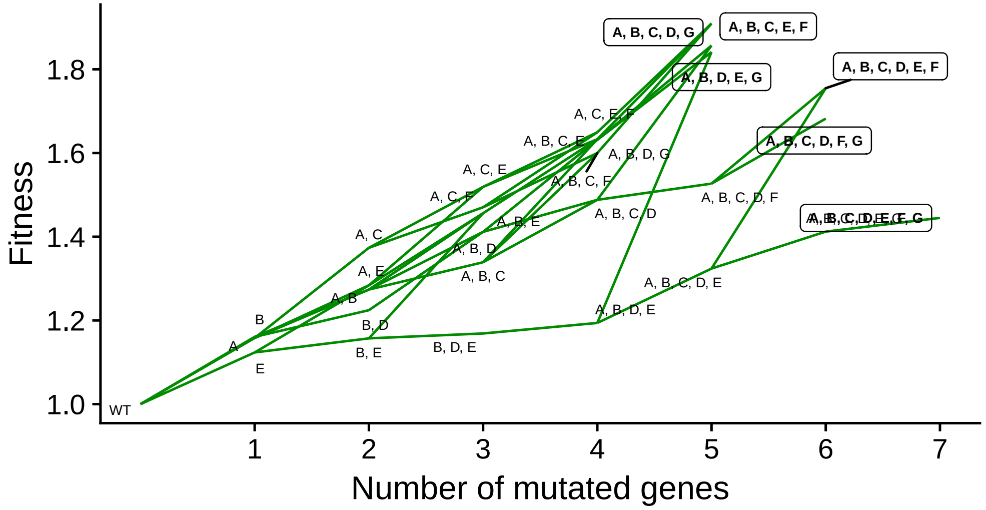


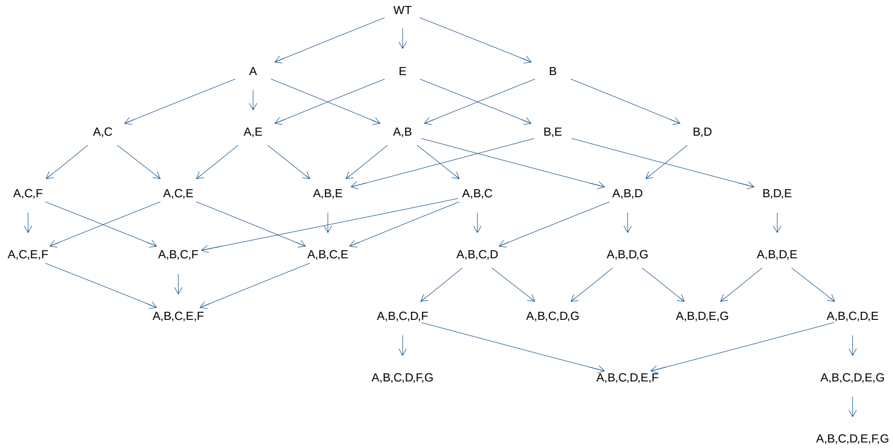

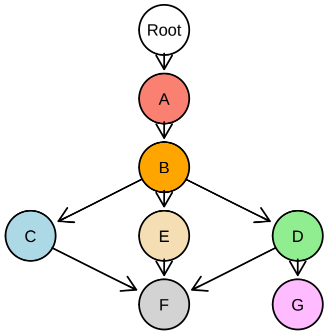


| | |
|---|---|
| *ID* | 7N2SgGN48X5QWxk9 |
| *acc genotypes* | 16 |
| *peaks no back* | 2 |
| *removed edges* | 5 |
| *prop rem edges* | 0.2 |

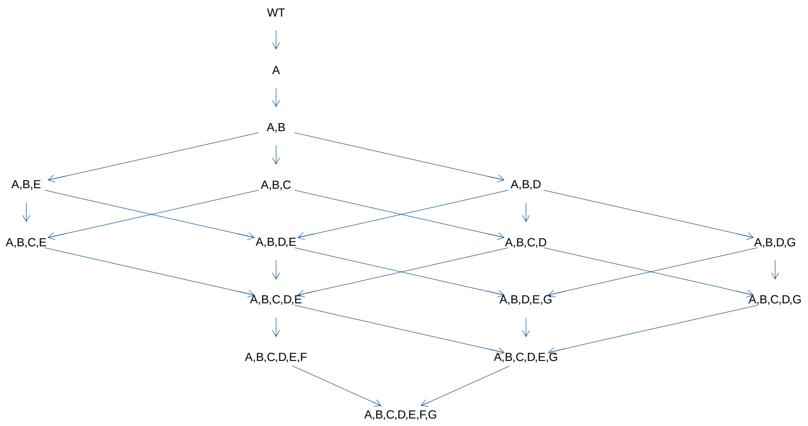


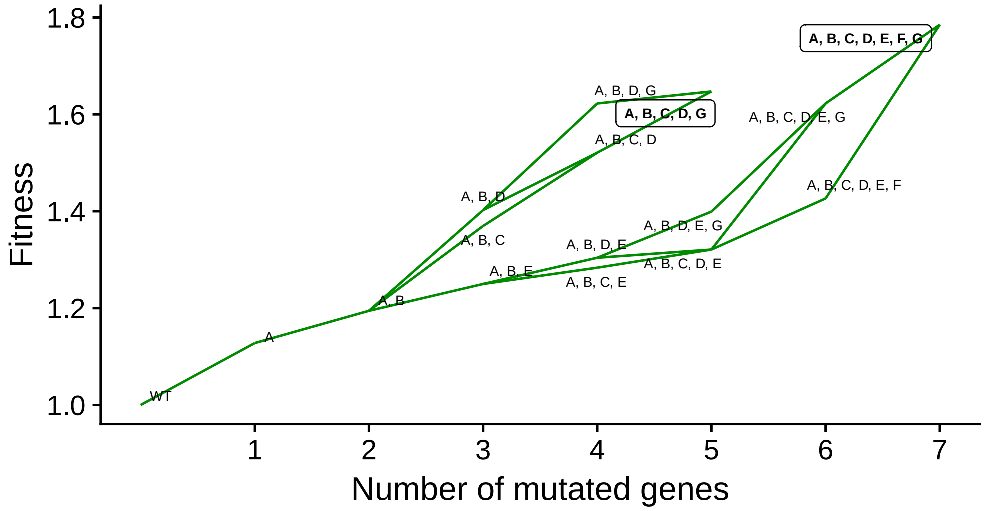


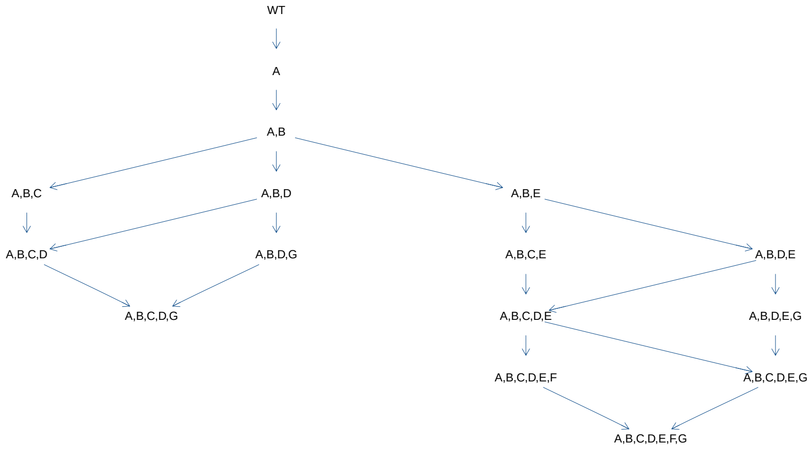

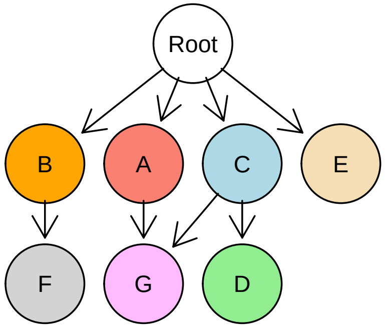


| | |
|---|---|
| *ID* | 7N9Bf18TzdbZ2zab |
| *acc genotypes* | 48 |
| *peaks no back* | 6 |
| *removed edges* | 24 |
| *prop rem edges* | 0.207 |

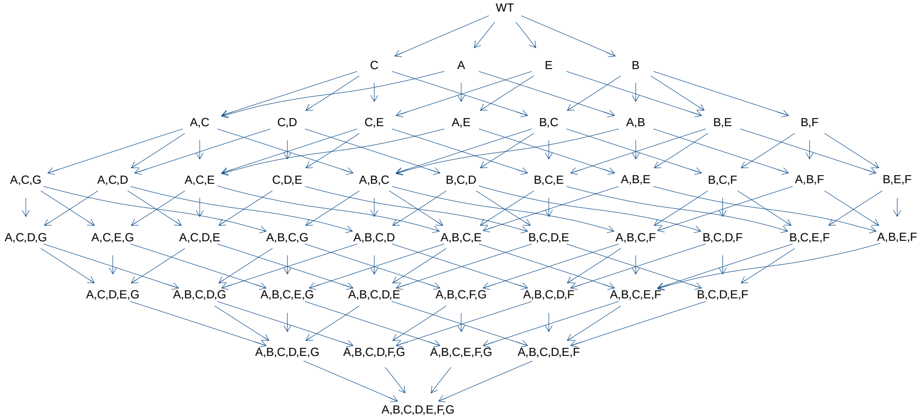

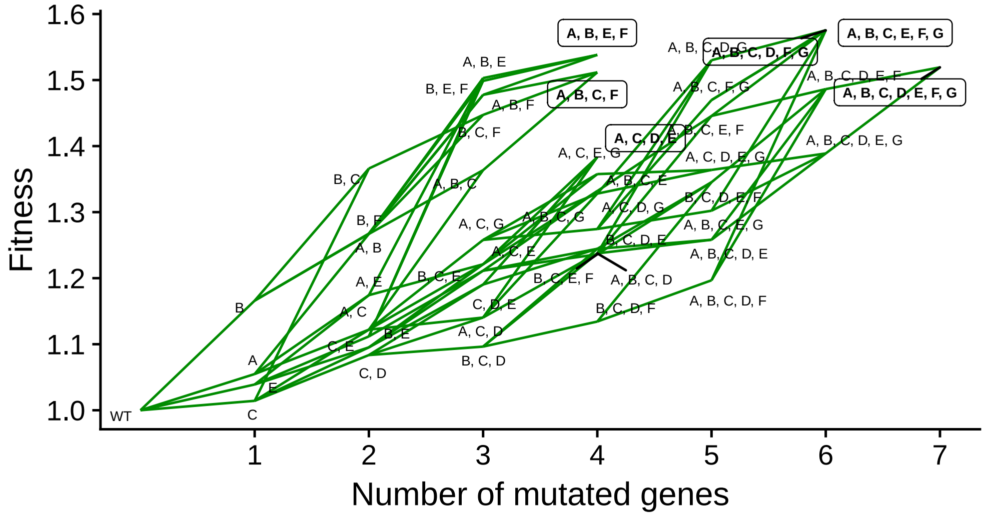


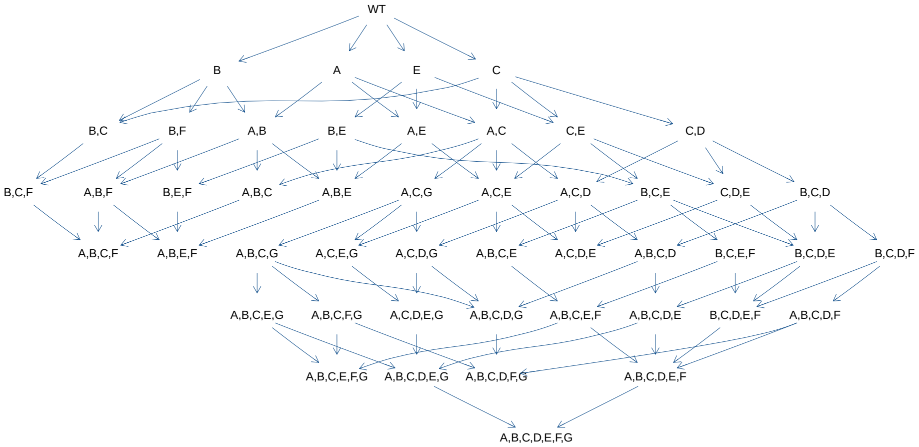

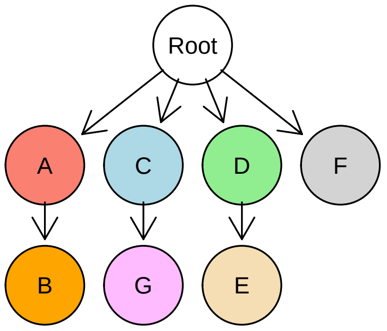


| | |
|---|---|
| *ID* | 7wKXobmcjD38LD6b |
| *acc genotypes* | 54 |
| *peaks no back* | 8 |
| *removed edges* | 29 |
| *prop rem edges* | 0.215 |

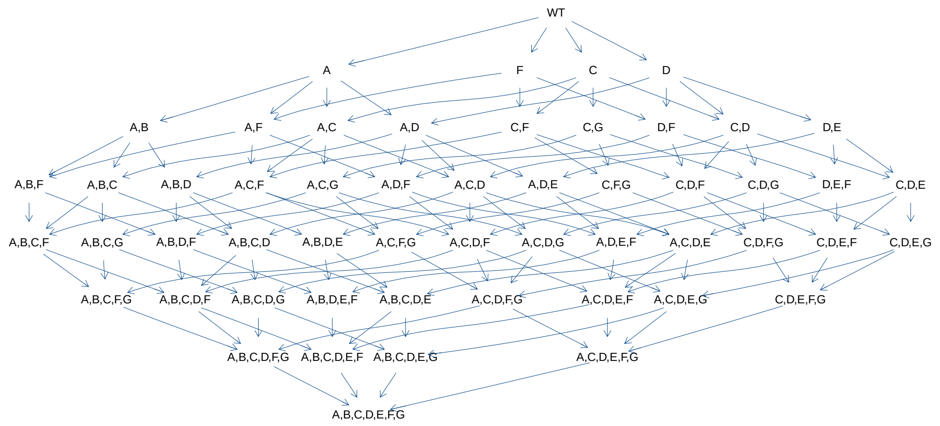


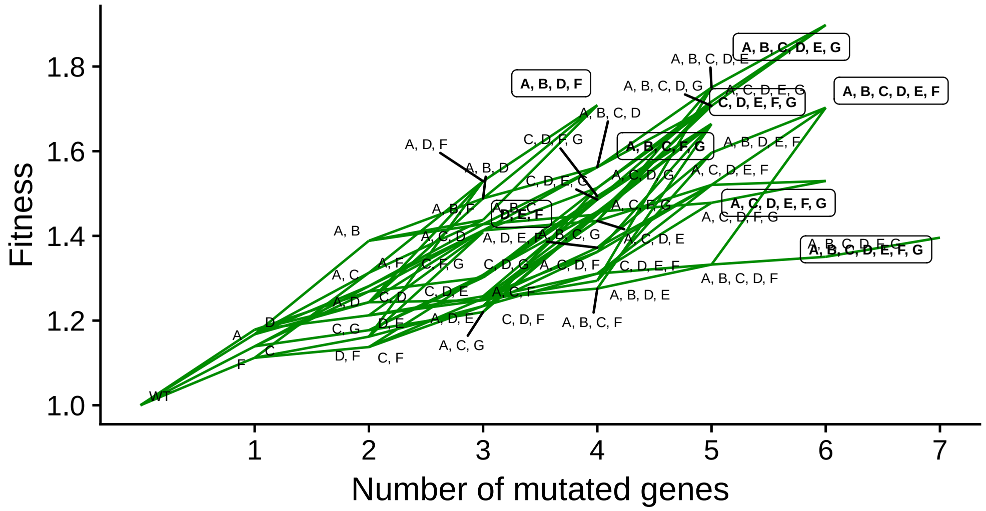


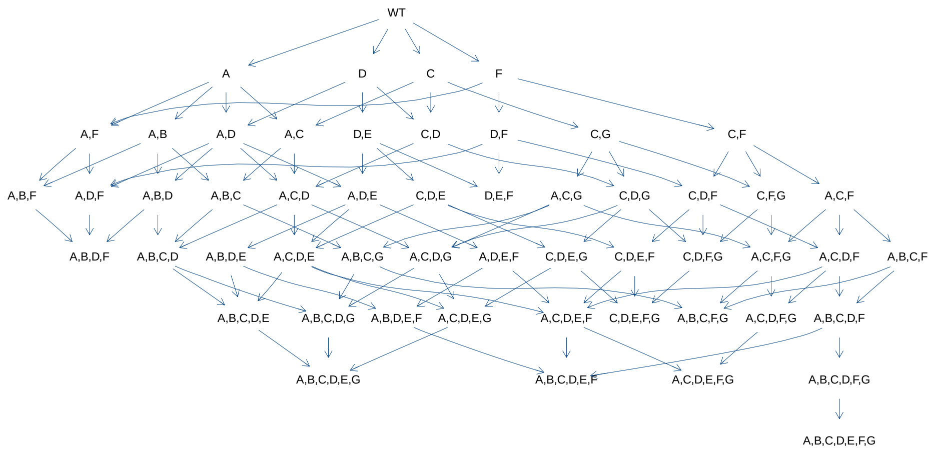

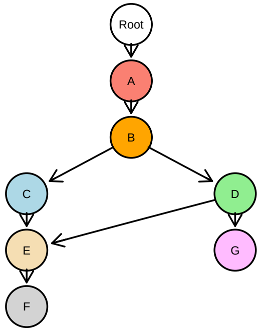


| | |
|---|---|
| *ID* | 8jUIKcUqRbPGoMif |
| *acc genotypes* | 12 |
| *peaks no back* | 3 |
| *removed edges* | 2 |
| *prop rem edges* | 0.133 |

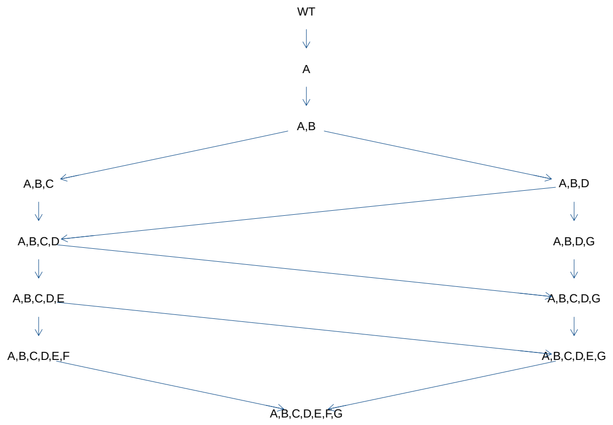


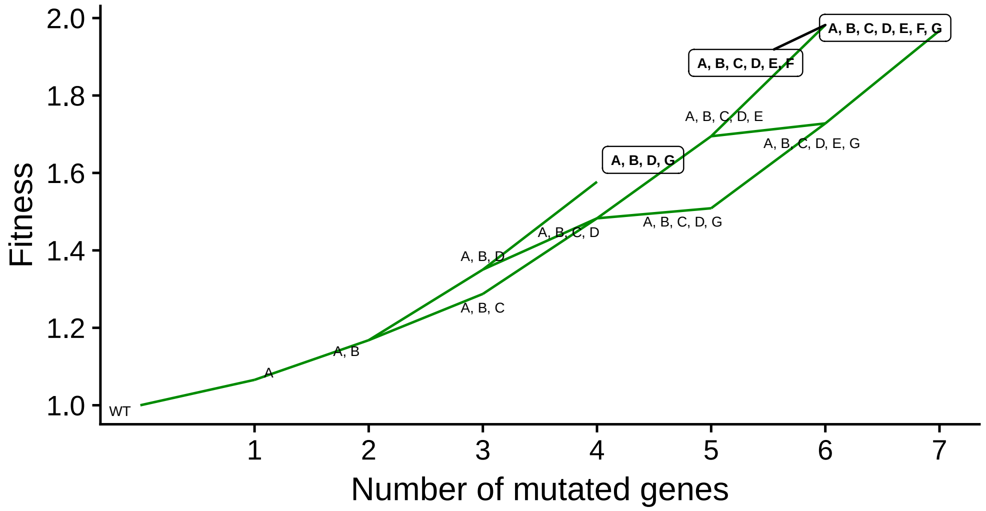


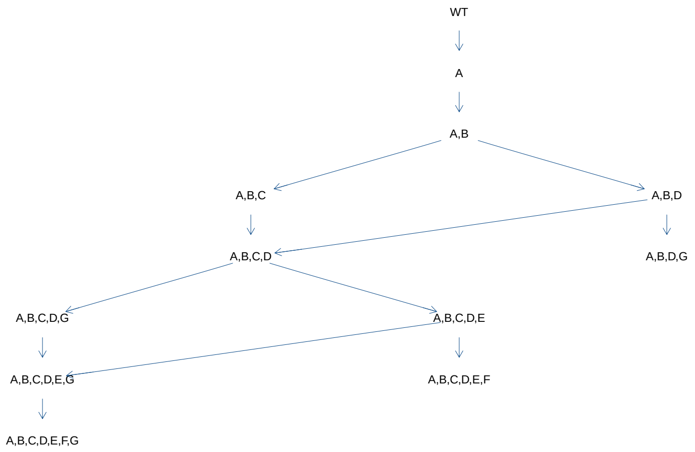

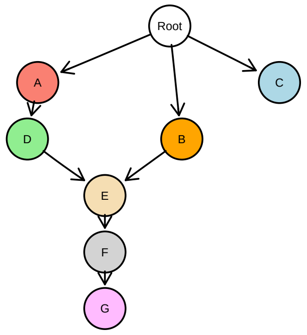


| | |
|---|---|
| *ID* | 8mSMlsAi8LuLuzZC |
| *acc genotypes* | 18 |
| *peaks no back* | 4 |
| *removed edges* | 4 |
| *prop rem edges* | 0.138 |

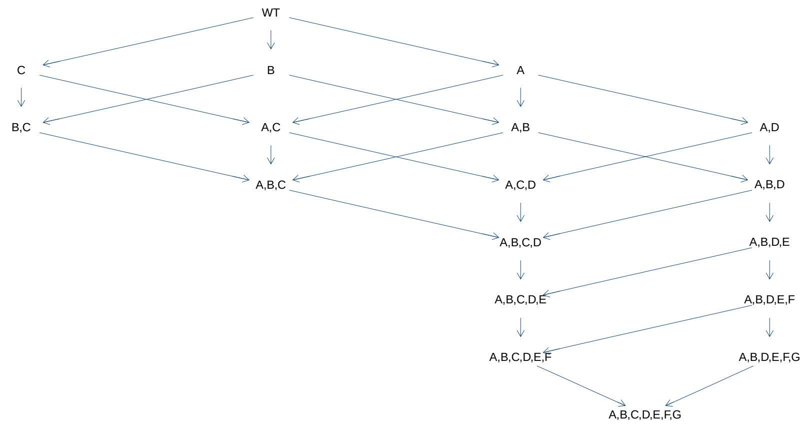


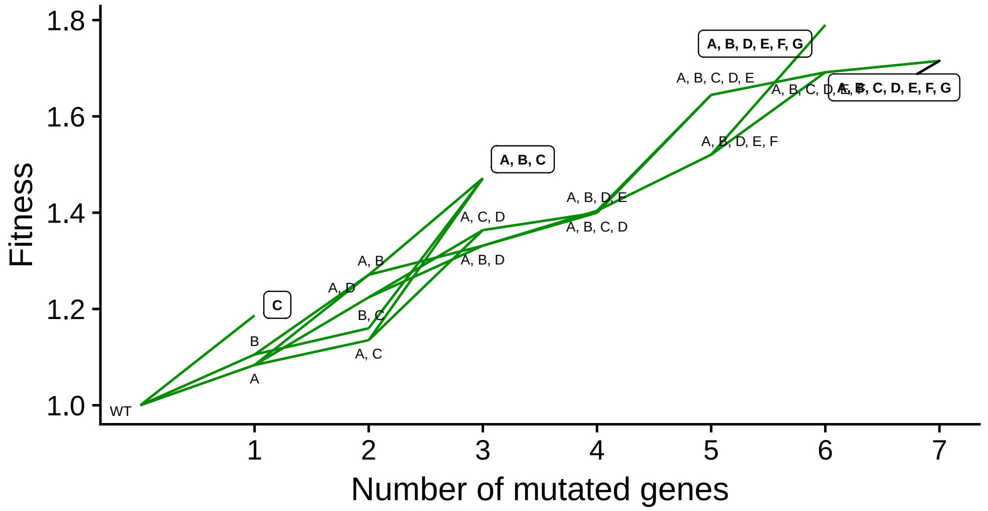


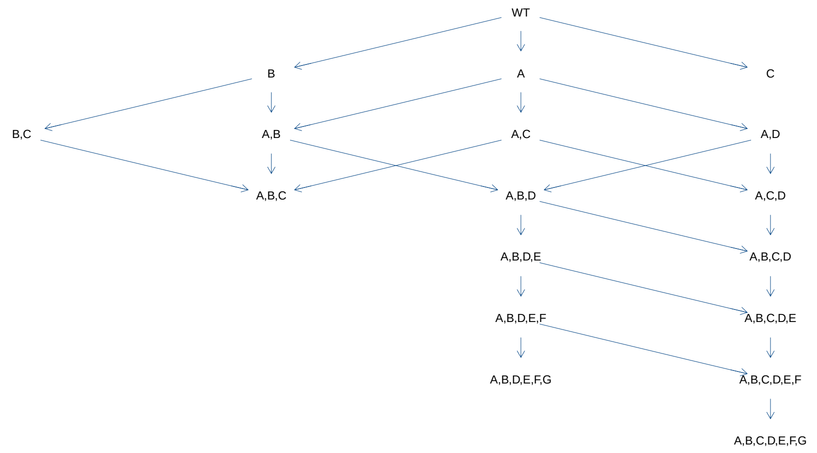

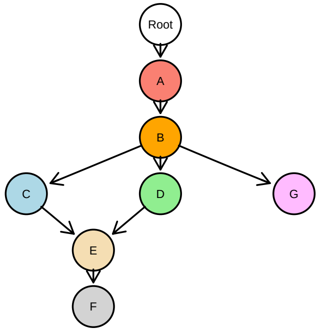


*ID* 8QIFQCUlUVfC10PZr
*acc genotypes* 14
*peaks no back* 2
*removed edges* 1
*prop rem edges* 0.05

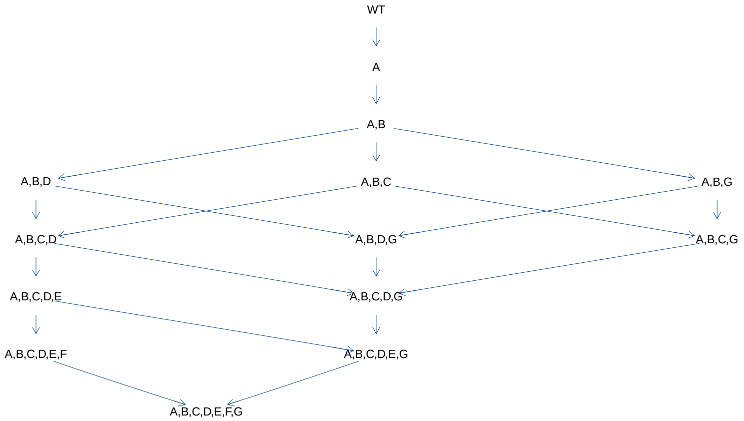


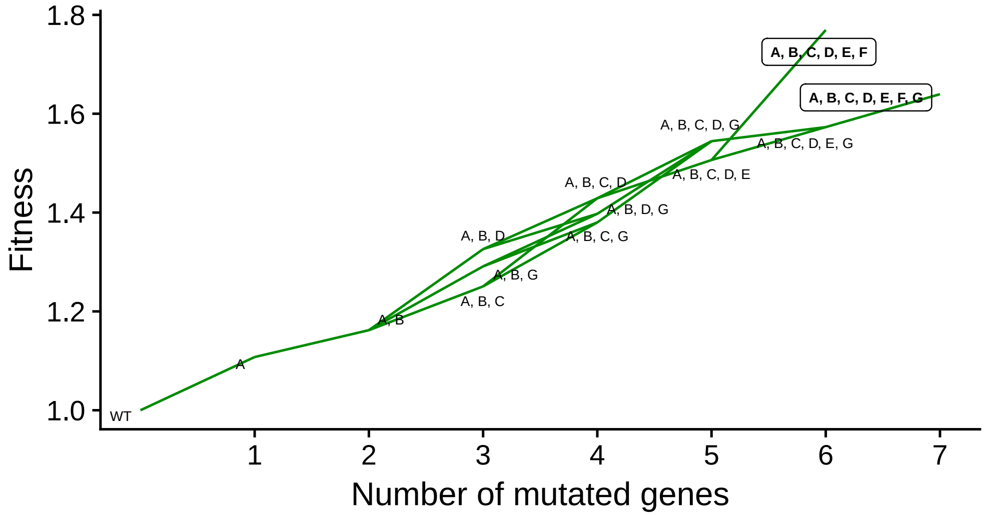


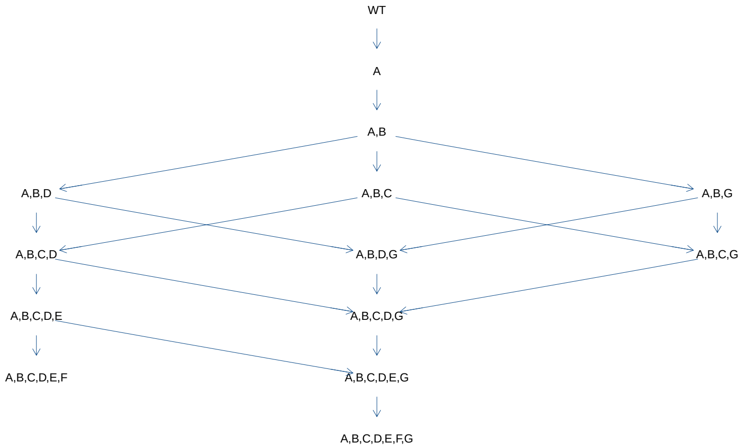

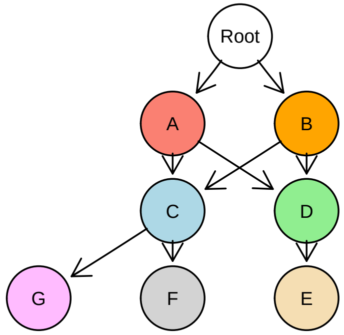


| | |
|---|---|
| *ID* | 8u1FzrFscU1z8xqB |
| *acc genotypes* | 18 |
| *peaks no back* | 4 |
| *removed edges* | 3 |
| *prop rem edges* | 0.103 |

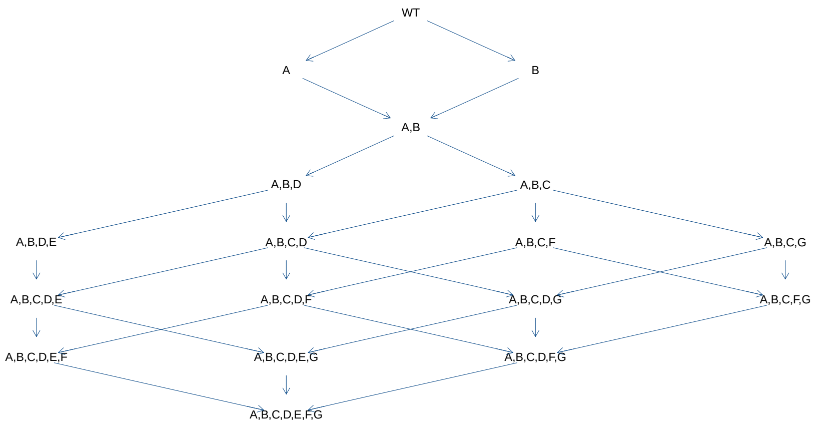


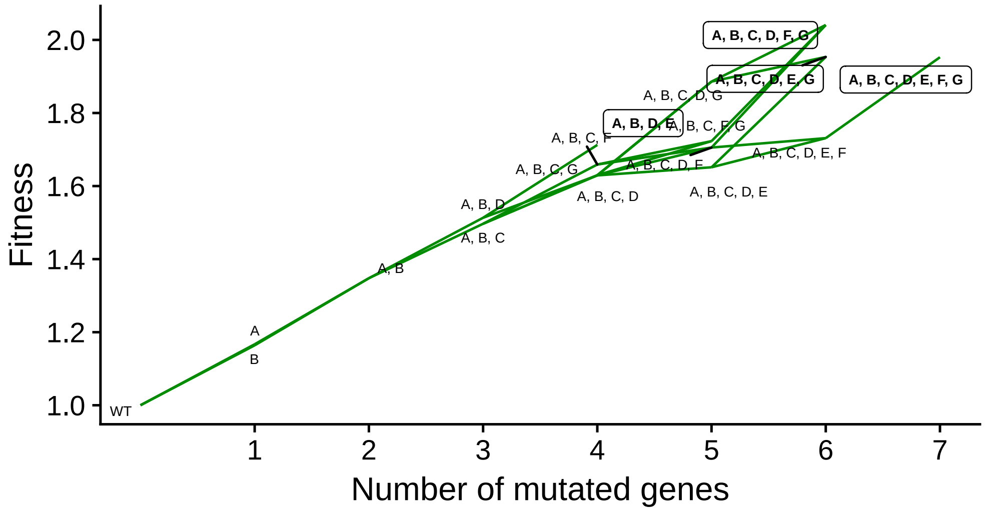


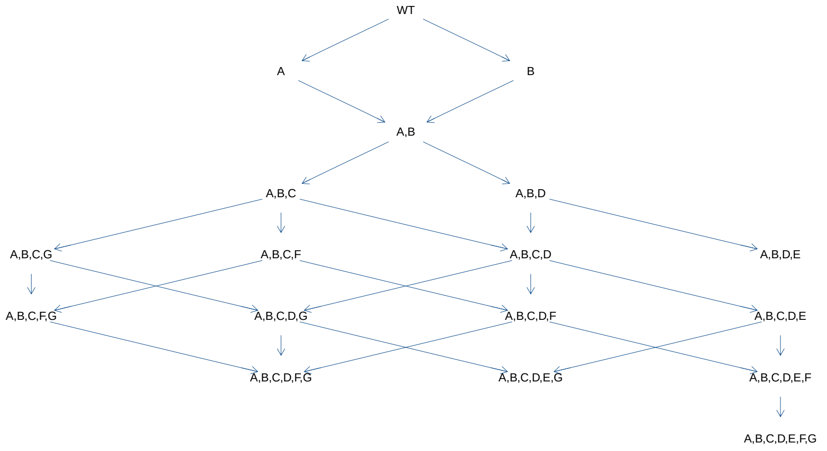

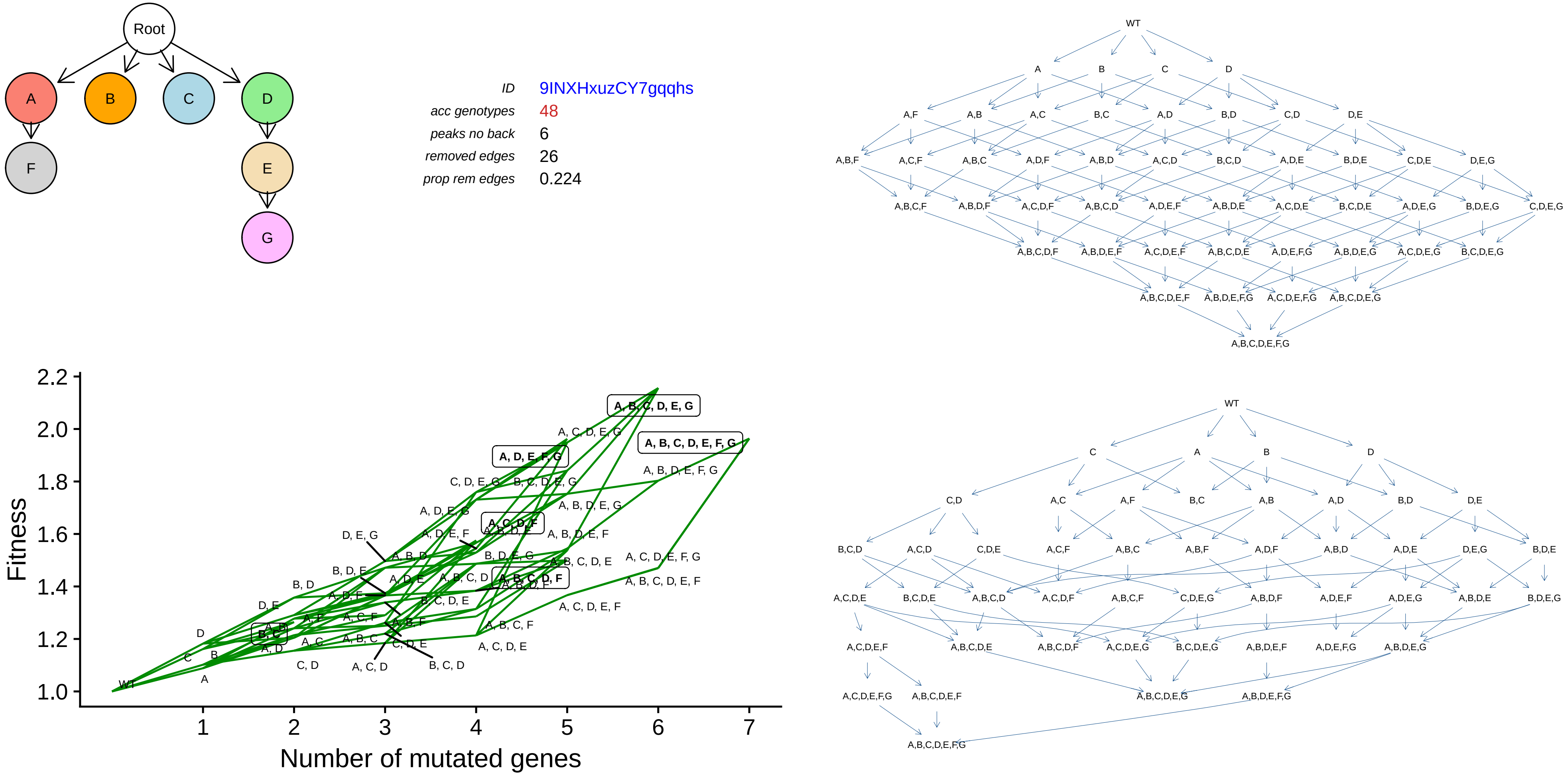
Root
A
B
C
D
F
E
G
ID 9INXHxuzCY7gqqhs
acc genotypes 48
peaks no back 6
removed edges 26
prop rem edges 0.224
Fitness
Number of mutated genes
A, B, C, D, E, G
A, B, C, D, E, F, G
A, D, E, F, G
A, C, D, F
A, B, C, D, F
B, C
WT

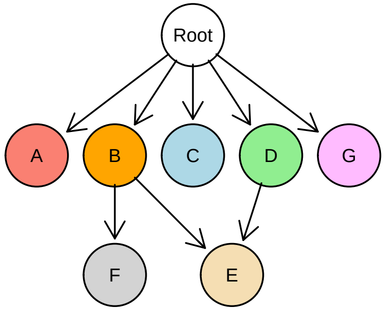


| | |
|---|---|
| *ID* | 9p9QxtT6PnU5iRpy |
| *acc genotypes* | 64 |
| *peaks no back* | 9 |
| *removed edges* | 37 |
| *prop rem edges* | 0.21 |

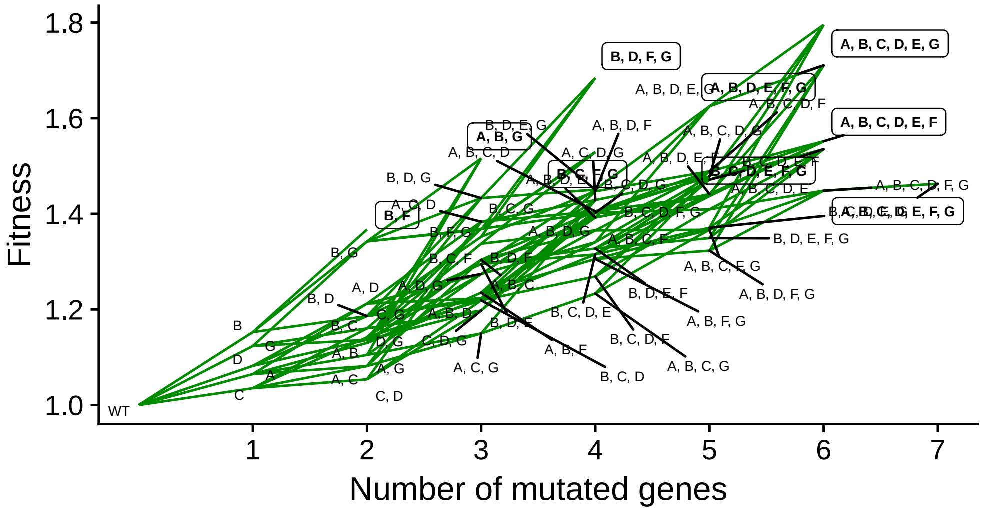


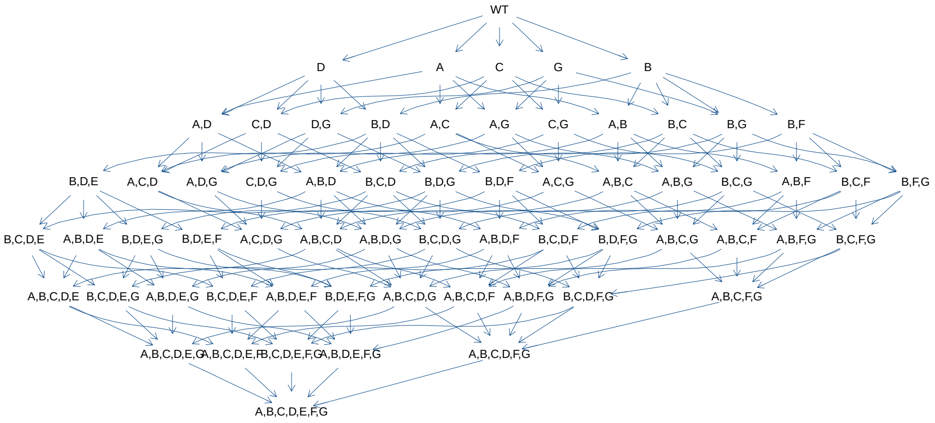

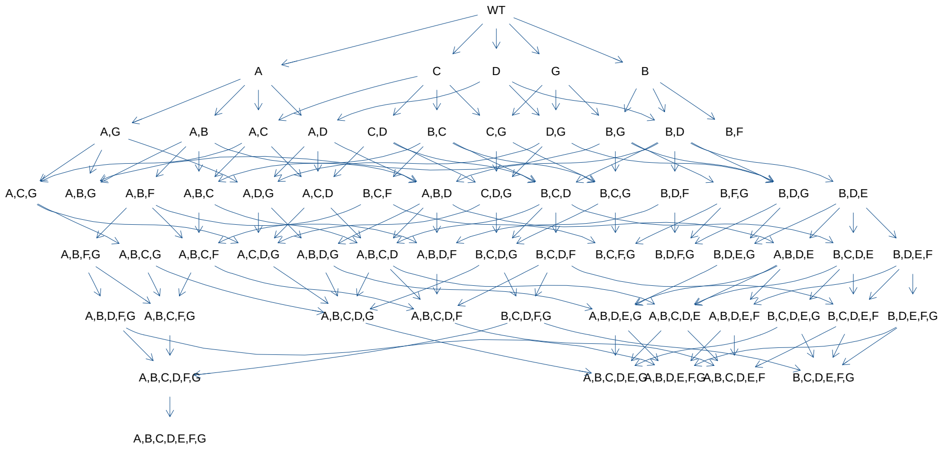

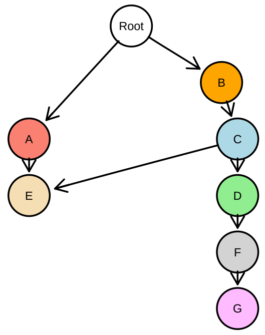


*ID* 9xXGITgcqHNk6QAO
*acc genotypes* 16
*peaks no back* 4
*removed edges* 4
*prop rem edges* 0.174

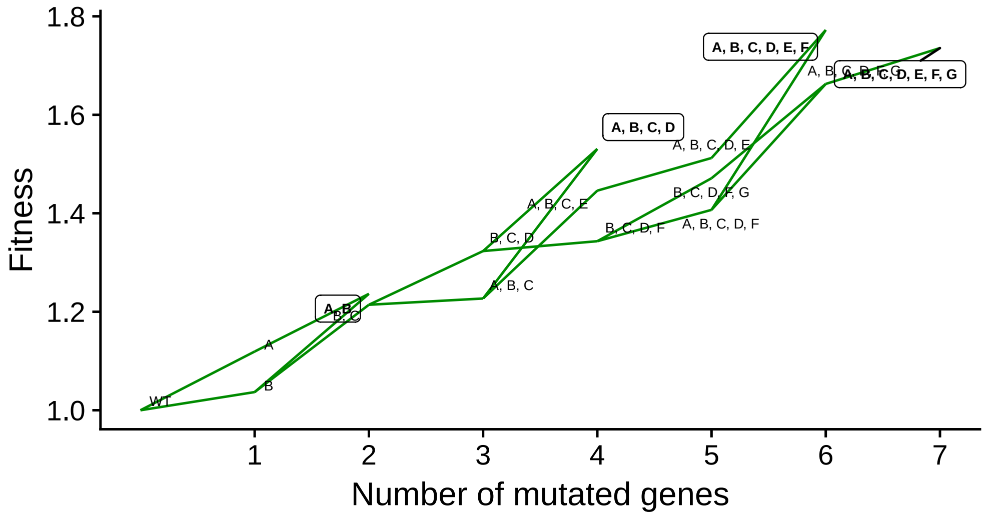


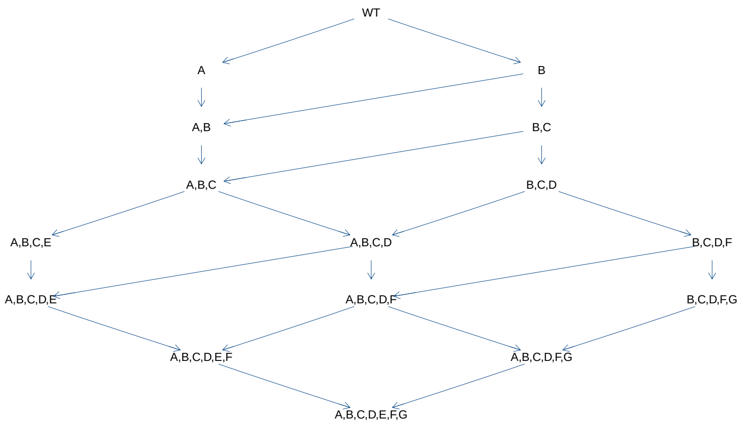


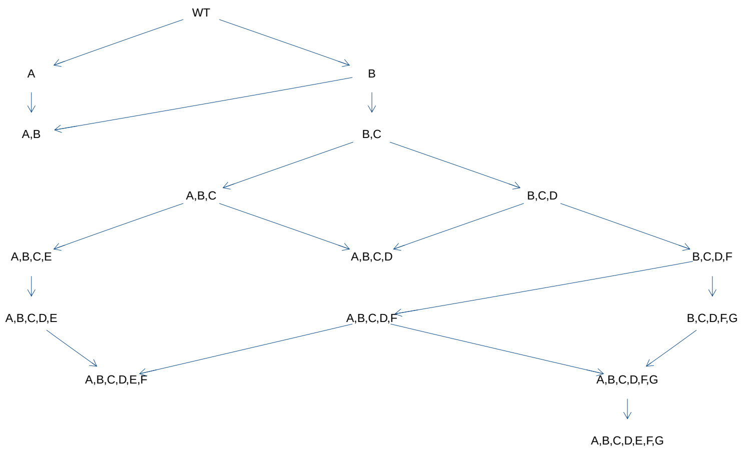

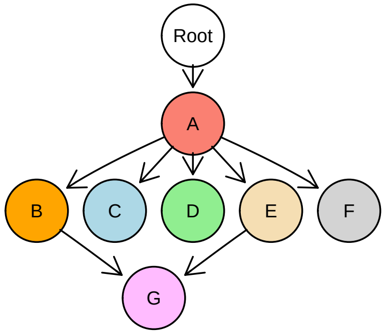


| | |
|---|---|
| *ID* | 9yJVB9KVObV7syFm |
| *acc genotypes* | 41 |
| *peaks no back* | 6 |
| *removed edges* | 16 |
| *prop rem edges* | 0.158 |

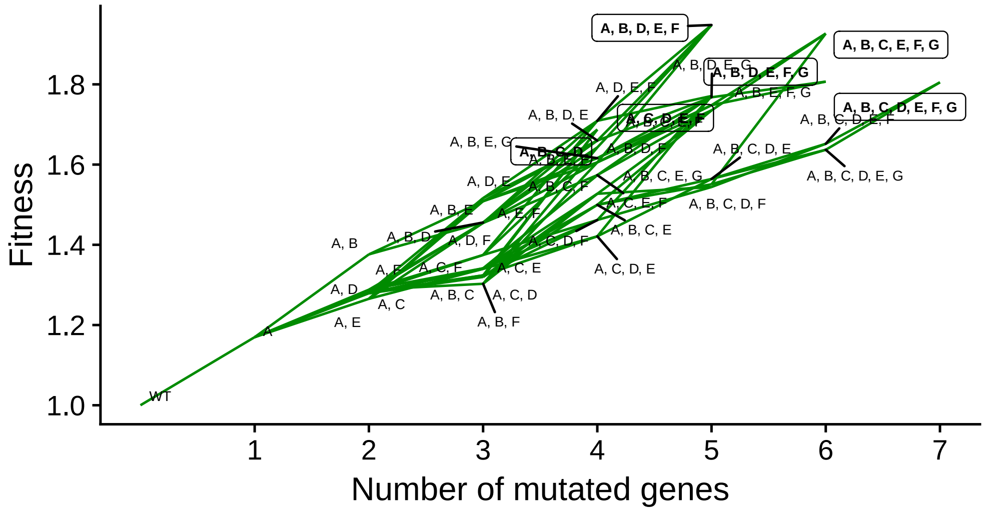


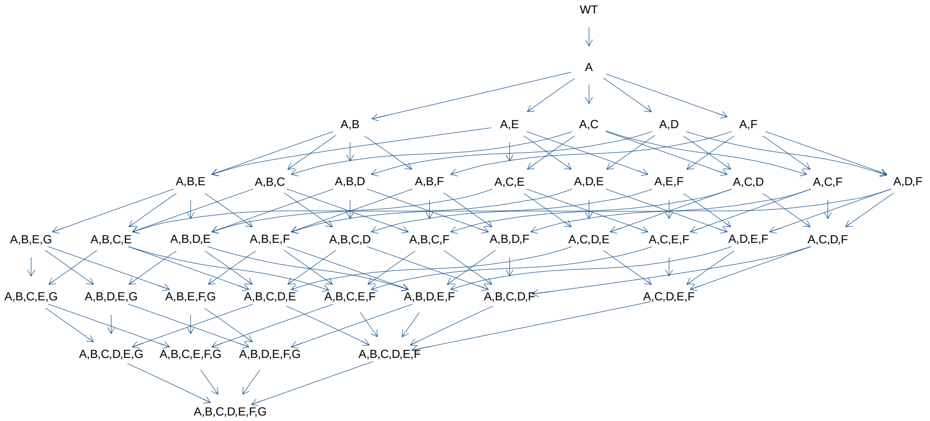


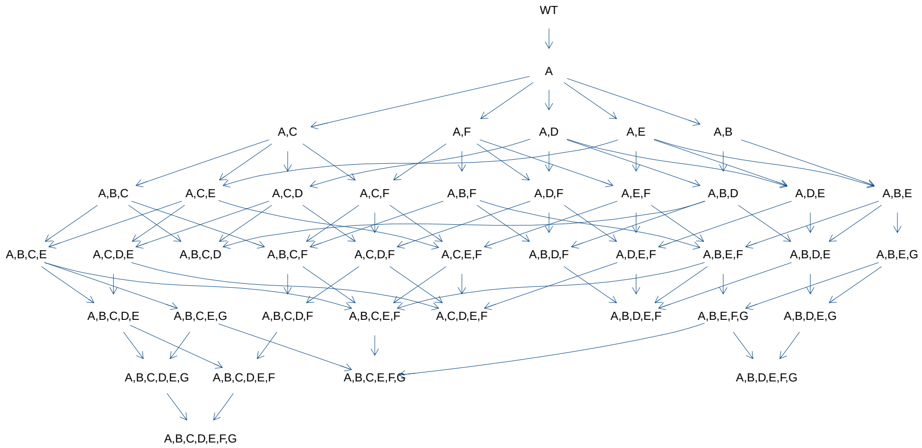

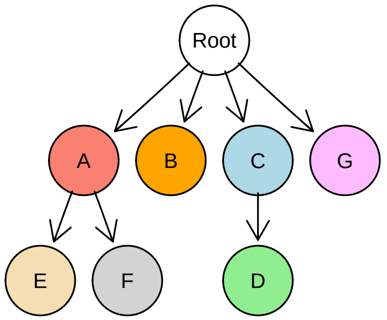


| | |
|---|---|
| *ID* | a71ZlUjjTtaY98zg |
| *acc genotypes* | 60 |
| *peaks no back* | 7 |
| *removed edges* | 21 |
| *prop rem edges* | 0.131 |

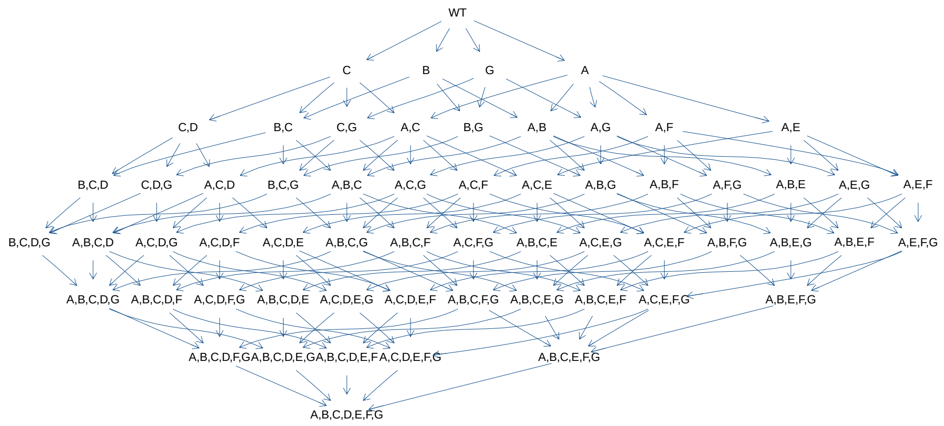


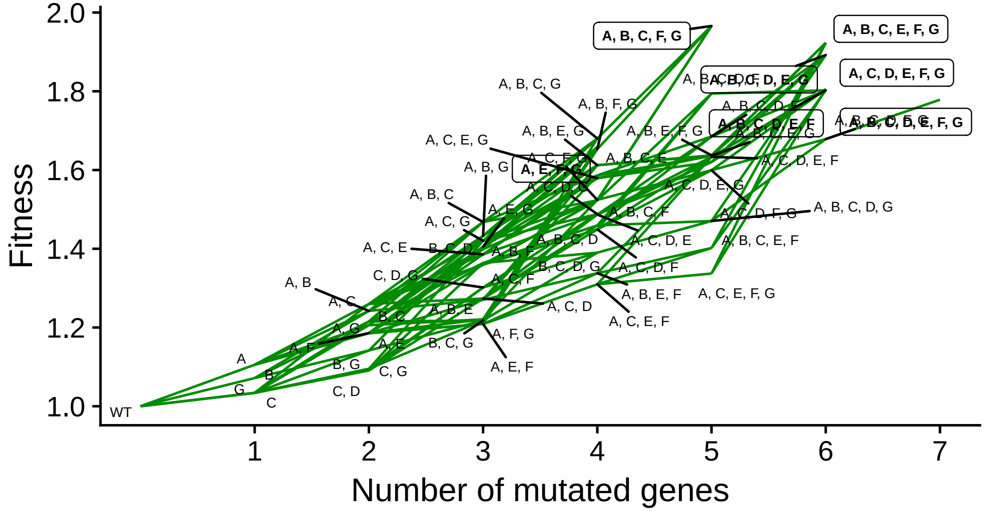


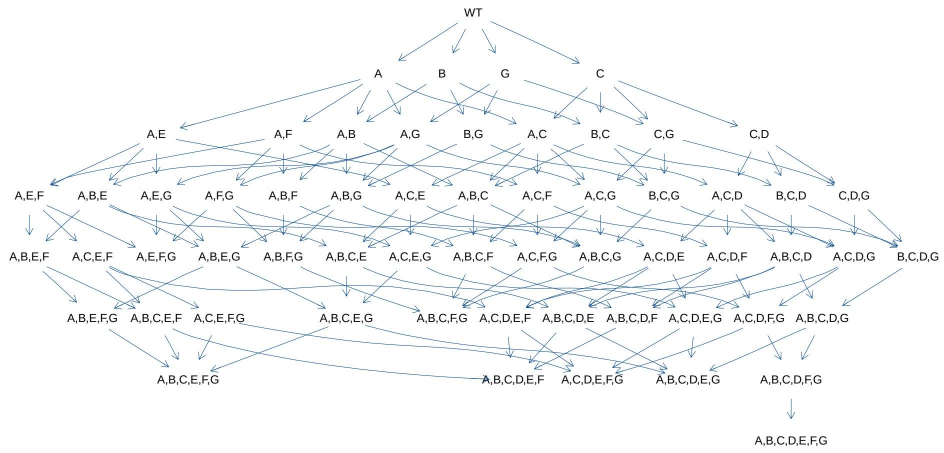

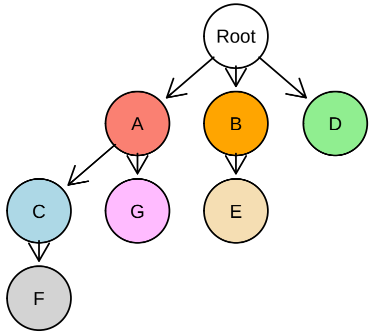


| | |
|---|---|
| *ID* | aABp8Hfk3fLTX4co |
| *acc genotypes* | 42 |
| *peaks no back* | 7 |
| *removed edges* | 17 |
| *prop rem edges* | 0.175 |

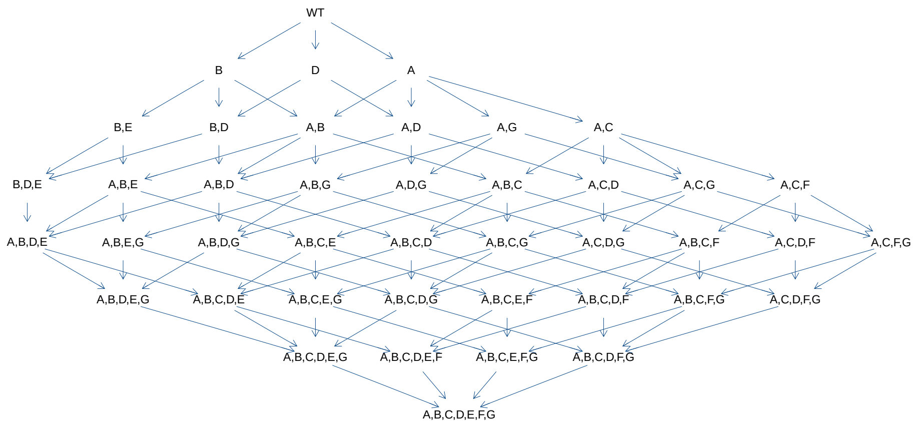


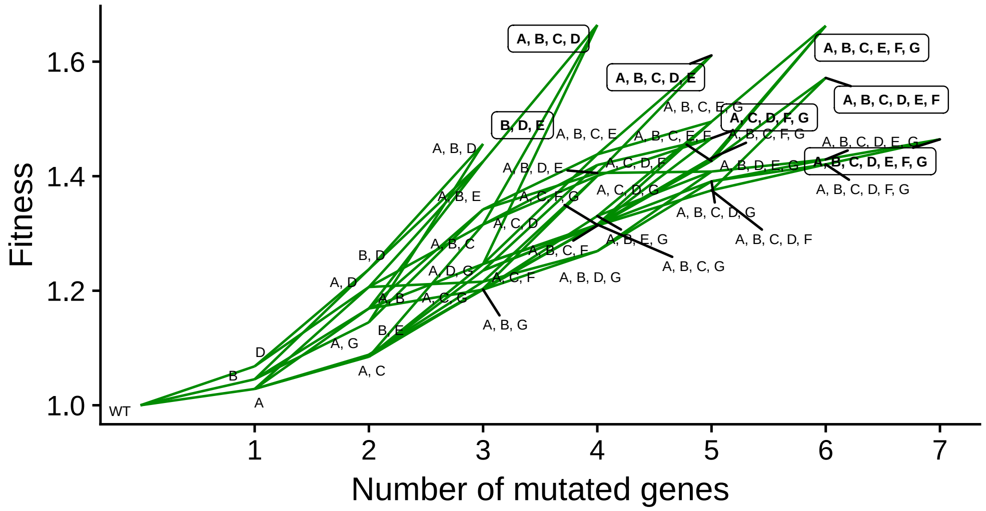


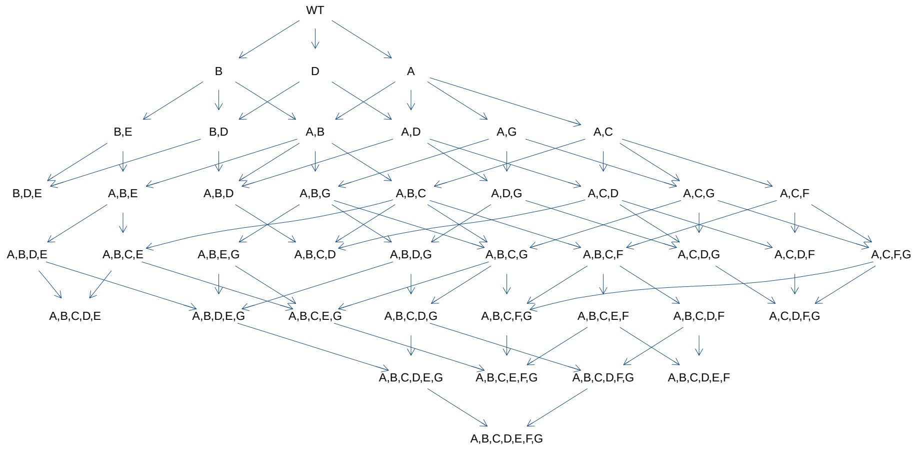

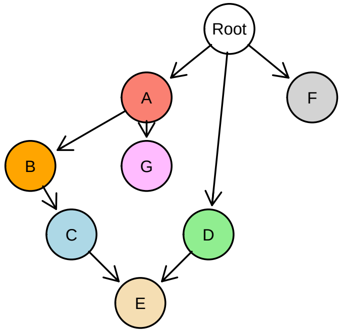


*ID* AeBhYdSJ32SIaHgW
*acc genotypes* 32
*peaks no back* 3
*removed edges* 8
*prop rem edges* 0.118

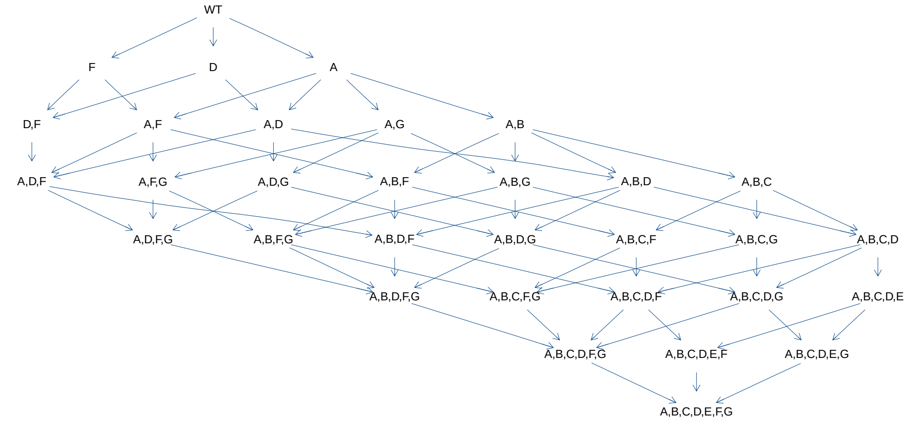


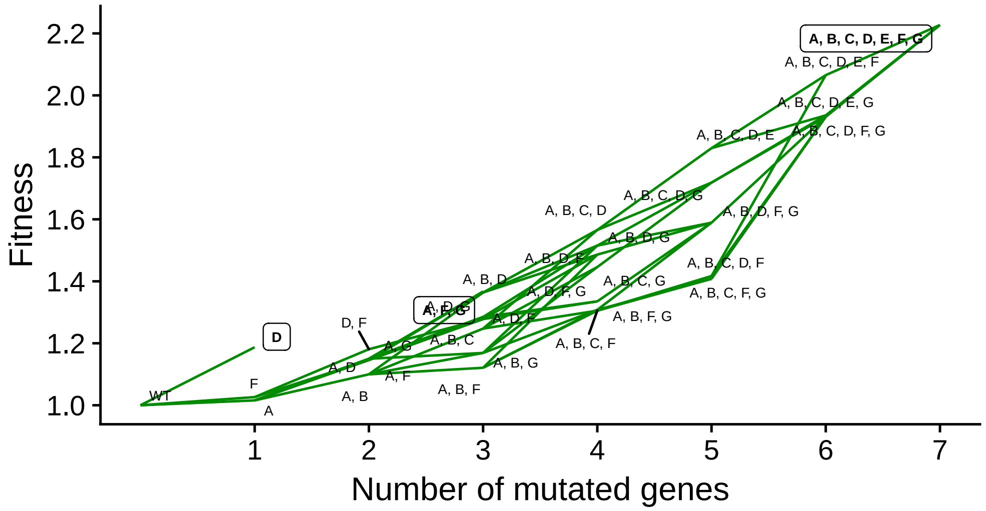


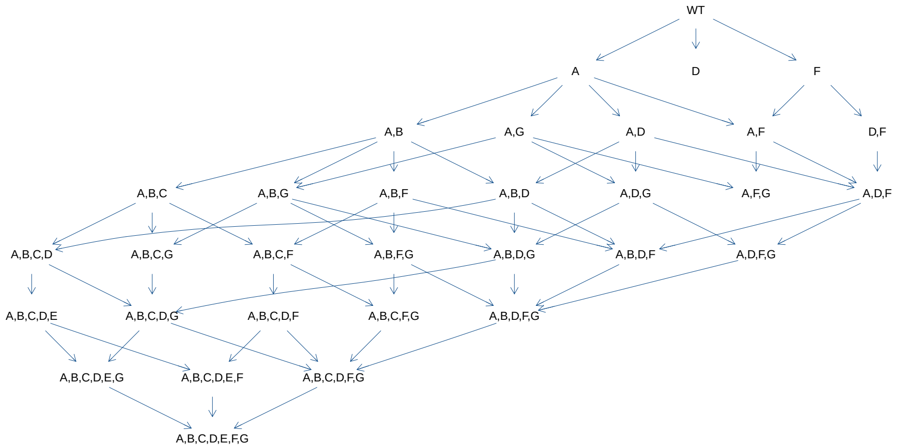

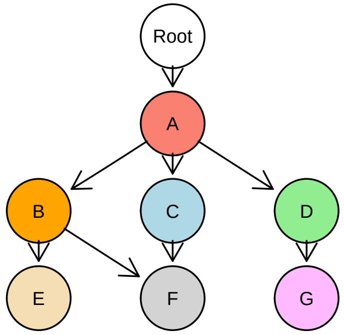


| | |
|---|---|
| *ID* | ANTtoA35KKWUX10uQ |
| *acc genotypes* | 25 |
| *peaks no back* | 3 |
| *removed edges* | 5 |
| *prop rem edges* | 0.106 |

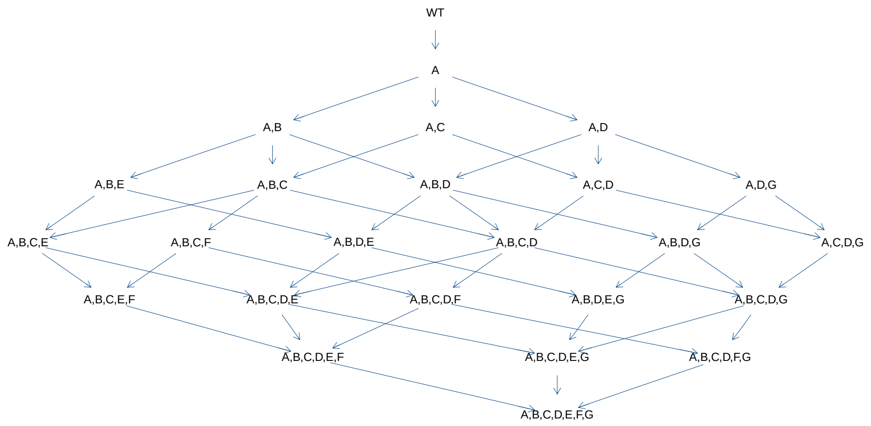


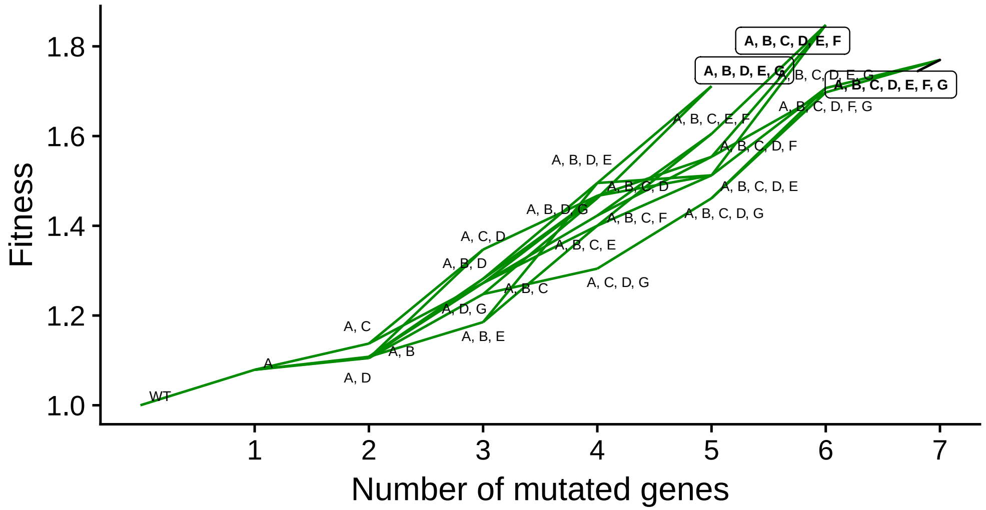


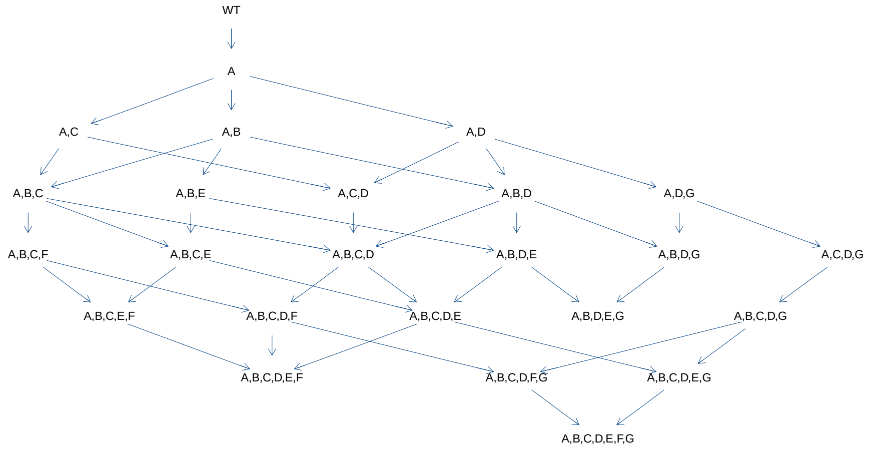

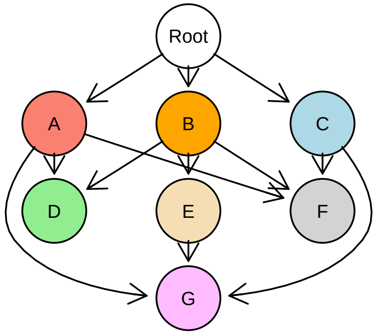


| | |
|---|---|
| *ID* | Asdds4Nn2MDRrNBH |
| *acc genotypes* | 24 |
| *peaks no back* | 5 |
| *removed edges* | 6 |
| *prop rem edges* | 0.136 |

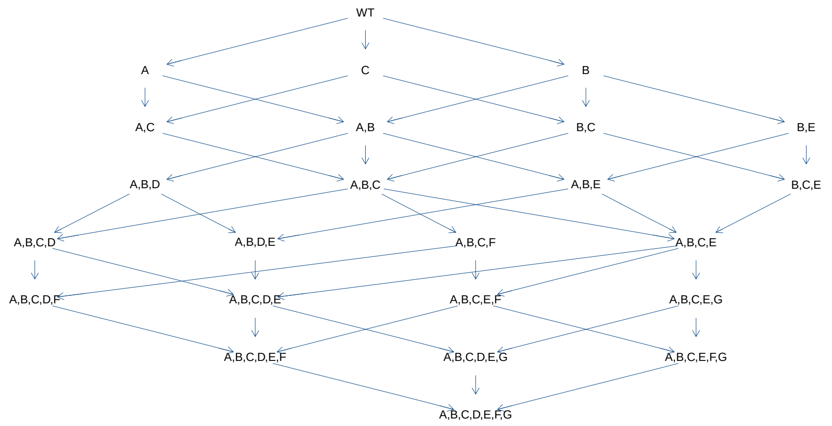


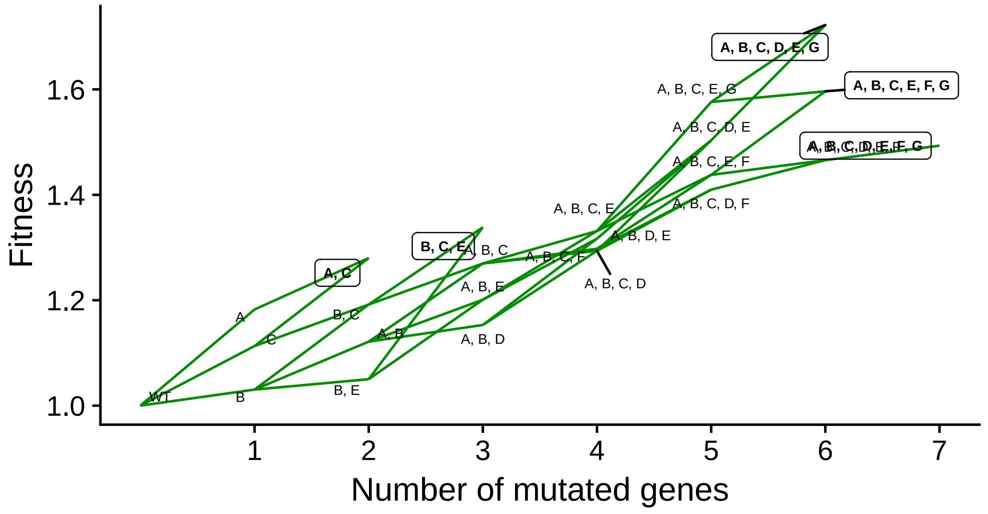


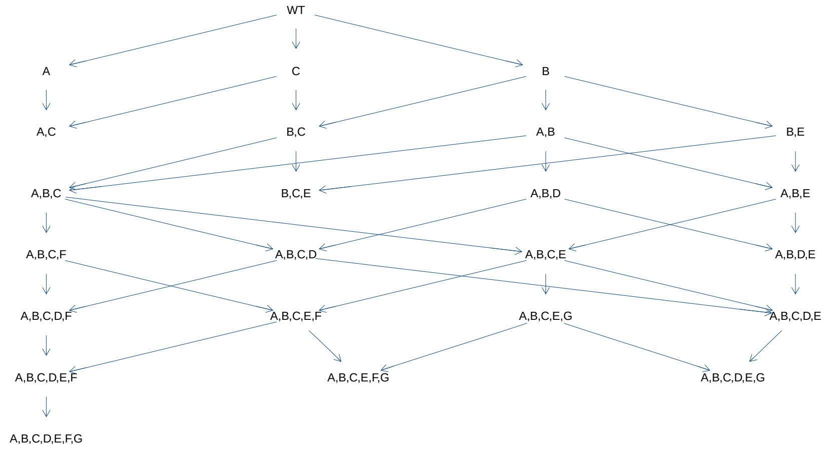

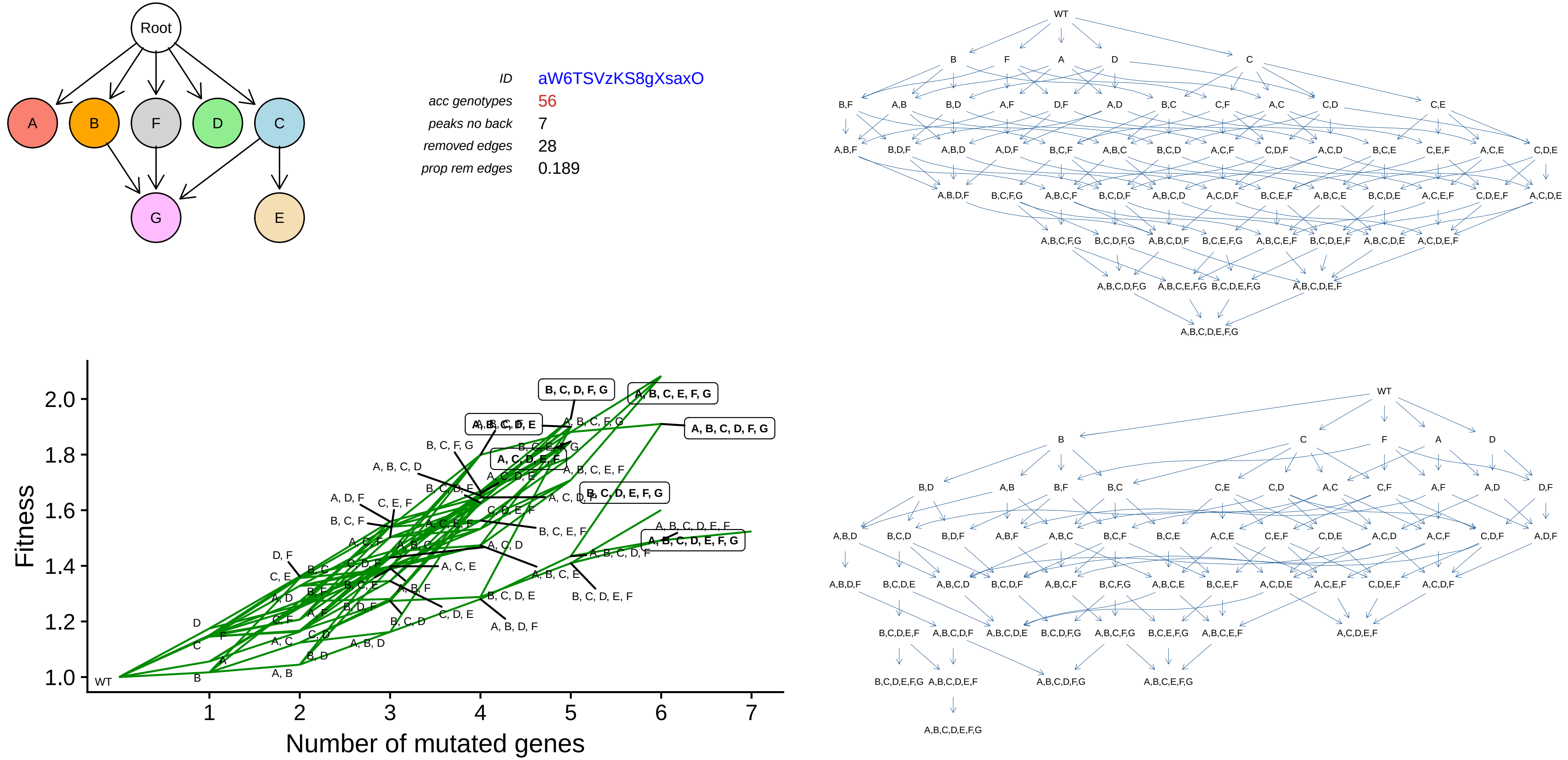
Root
A
B
F
D
C
G
E
ID aW6TSVzKS8gXsaxO
acc genotypes 56
peaks no back 7
removed edges 28
prop rem edges 0.189
Fitness
Number of mutated genes
B, C, D, F, G
A, B, C, E, F, G
A, B, C, D, F, G
A, C, D, E, F
B, C, D, E, F, G
A, B, C, D, E, F, G
WT
A,B,C,D,E,F,G

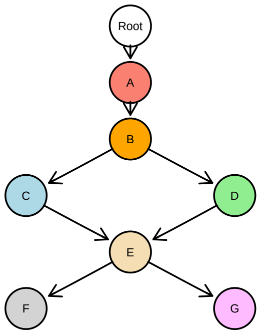


| | |
|---|---|
| *ID* | AXnGmWZxzz5kfly4 |
| *acc genotypes* | 10 |
| *peaks no back* | 2 |
| *removed edges* | 1 |
| *prop rem edges* | 0.091 |

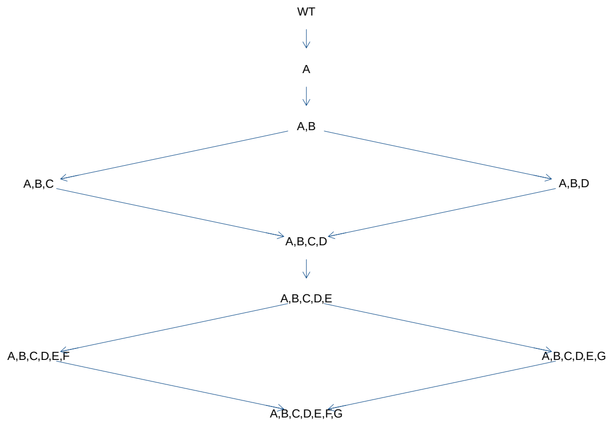


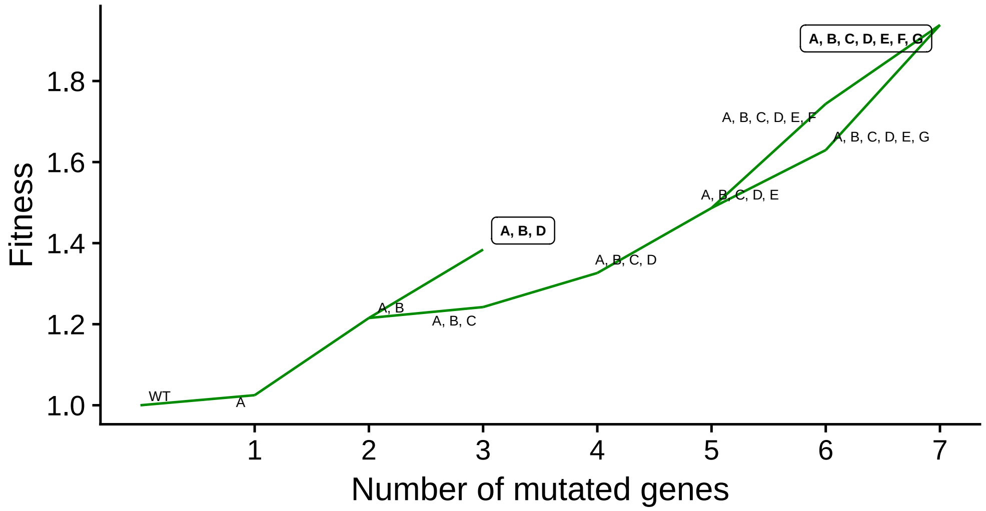


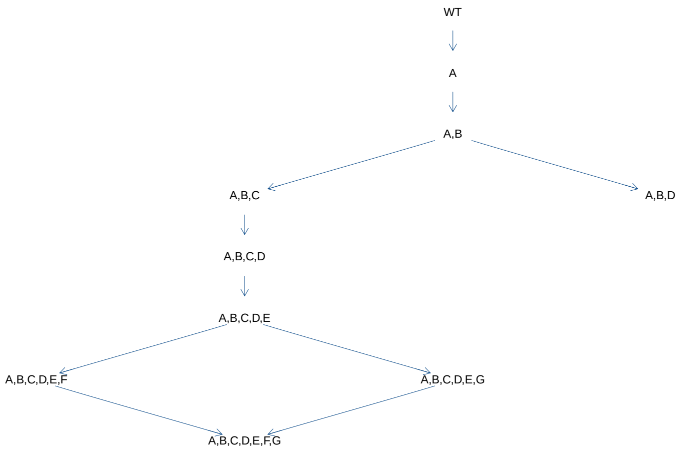

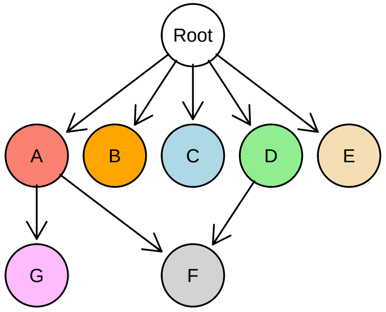


| | |
|---|---|
| *ID* | AYfLkIxUEKLpymWH |
| *acc genotypes* | 64 |
| *peaks no back* | 9 |
| *removed edges* | 33 |
| *prop rem edges* | 0.188 |

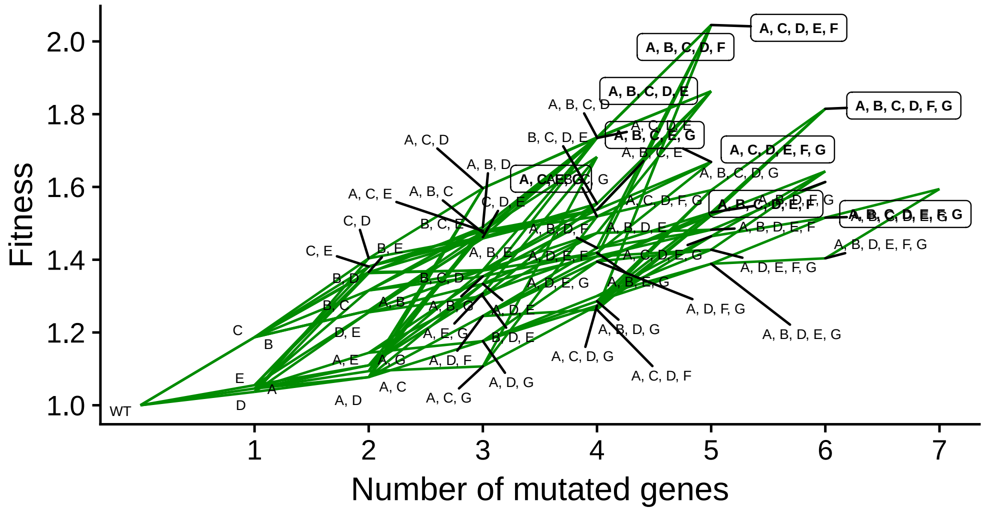


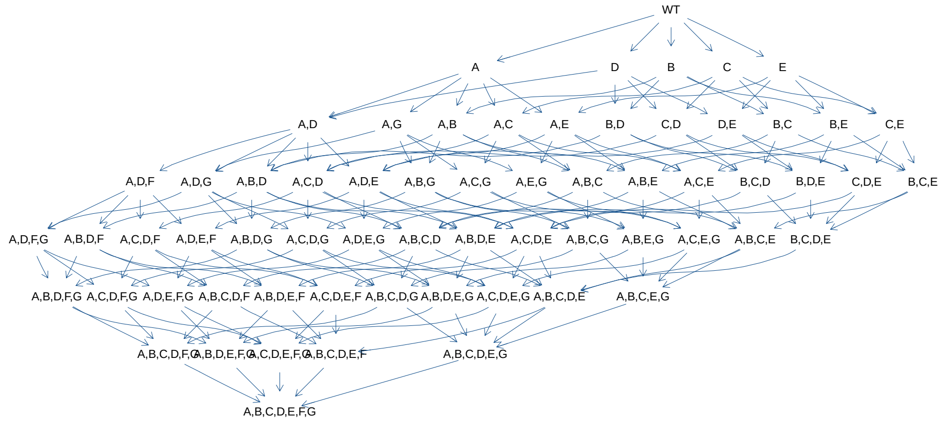


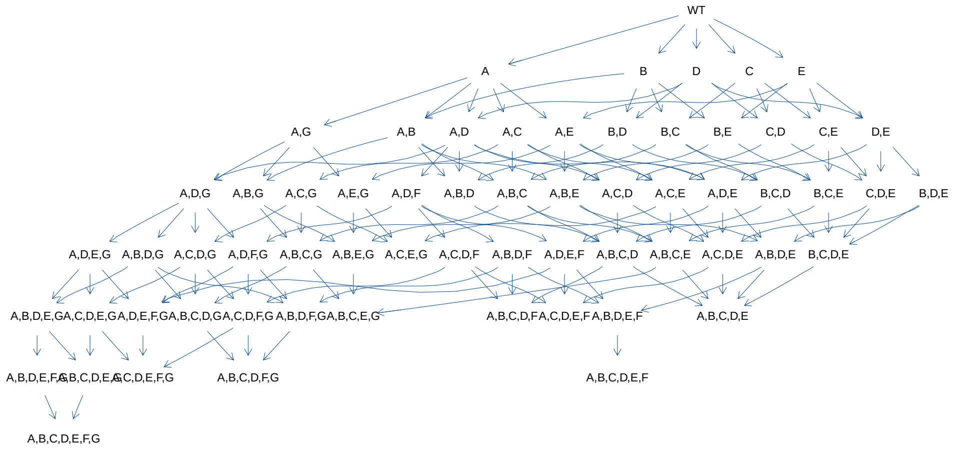

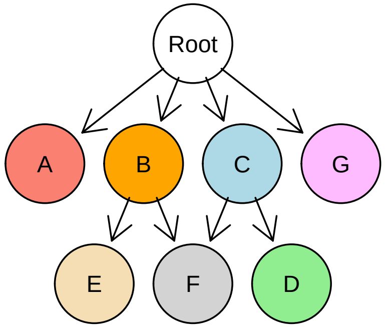


| | |
|---|---|
| *ID* | b8dcfCJtzgN6gupz |
| *acc genotypes* | 52 |
| *peaks no back* | 7 |
| *removed edges* | 20 |
| *prop rem edges* | 0.152 |

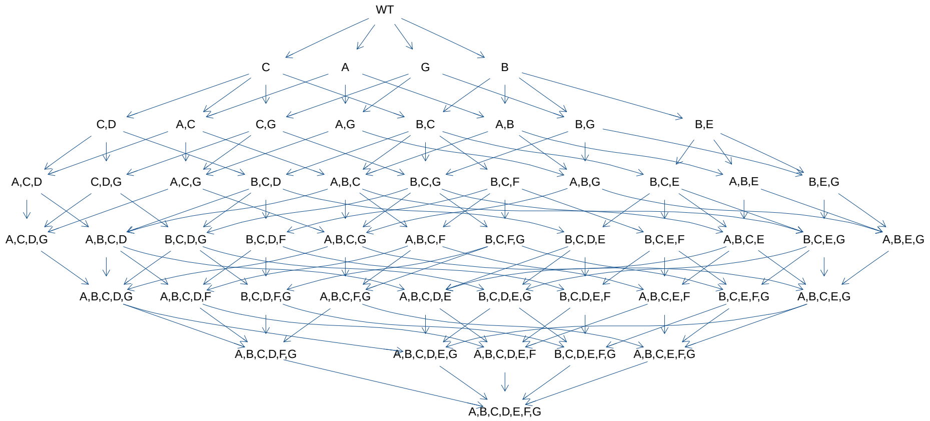

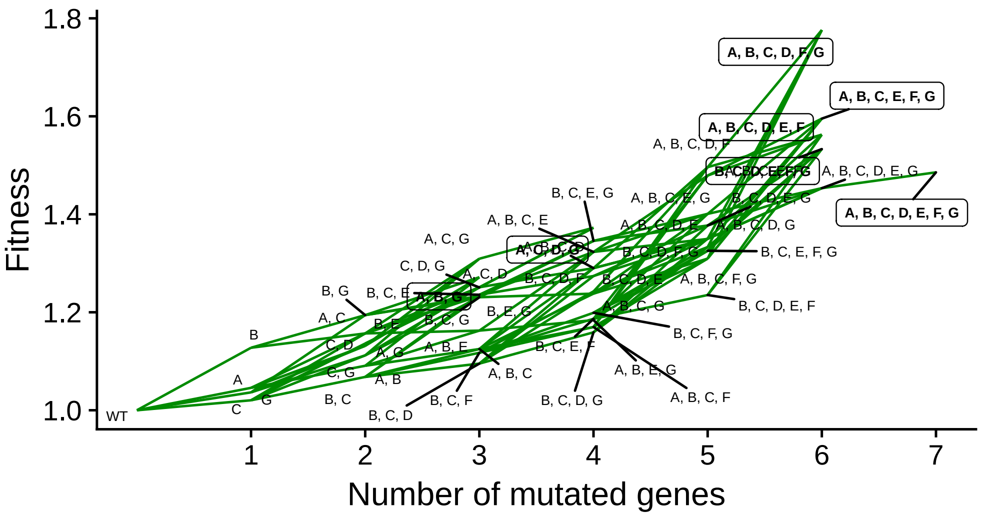


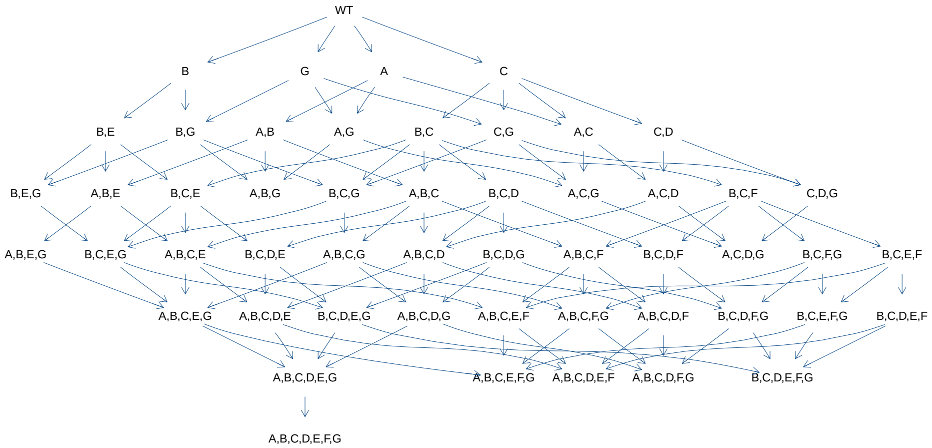

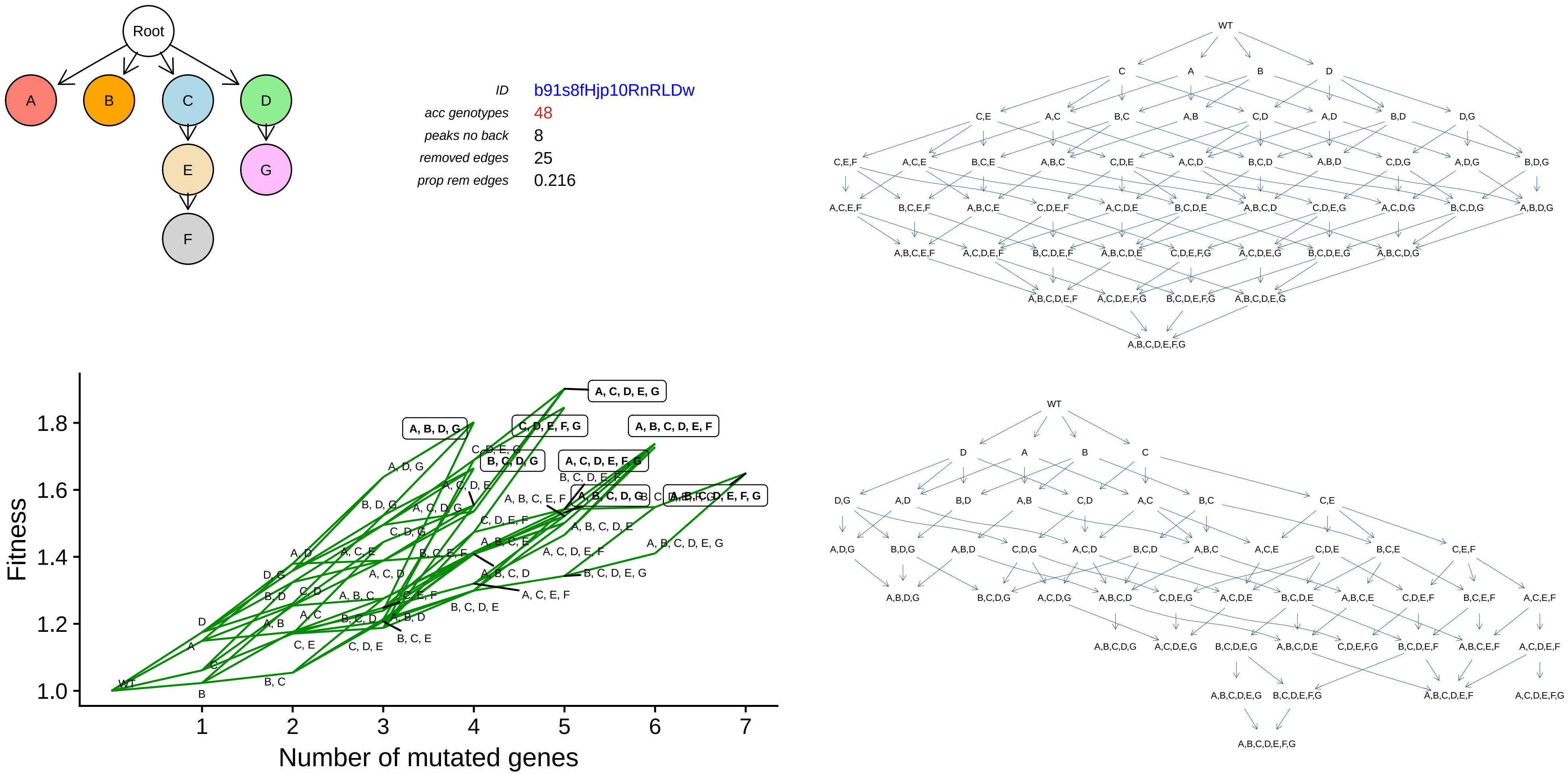
Root
A
B
C
D
E
G
F
ID b91s8fHjp10RnRLDw
acc genotypes 48
peaks no back 8
removed edges 25
prop rem edges 0.216
Fitness
Number of mutated genes
A, C, D, E, G
A, B, D, G
C, D, E, F, G
A, B, C, D, E, F
B, C, D, G
A, C, D, E, F, G
A, B, C, D, G
A, B, C, D, E, F, G
WT
A,B,C,D,E,F,G

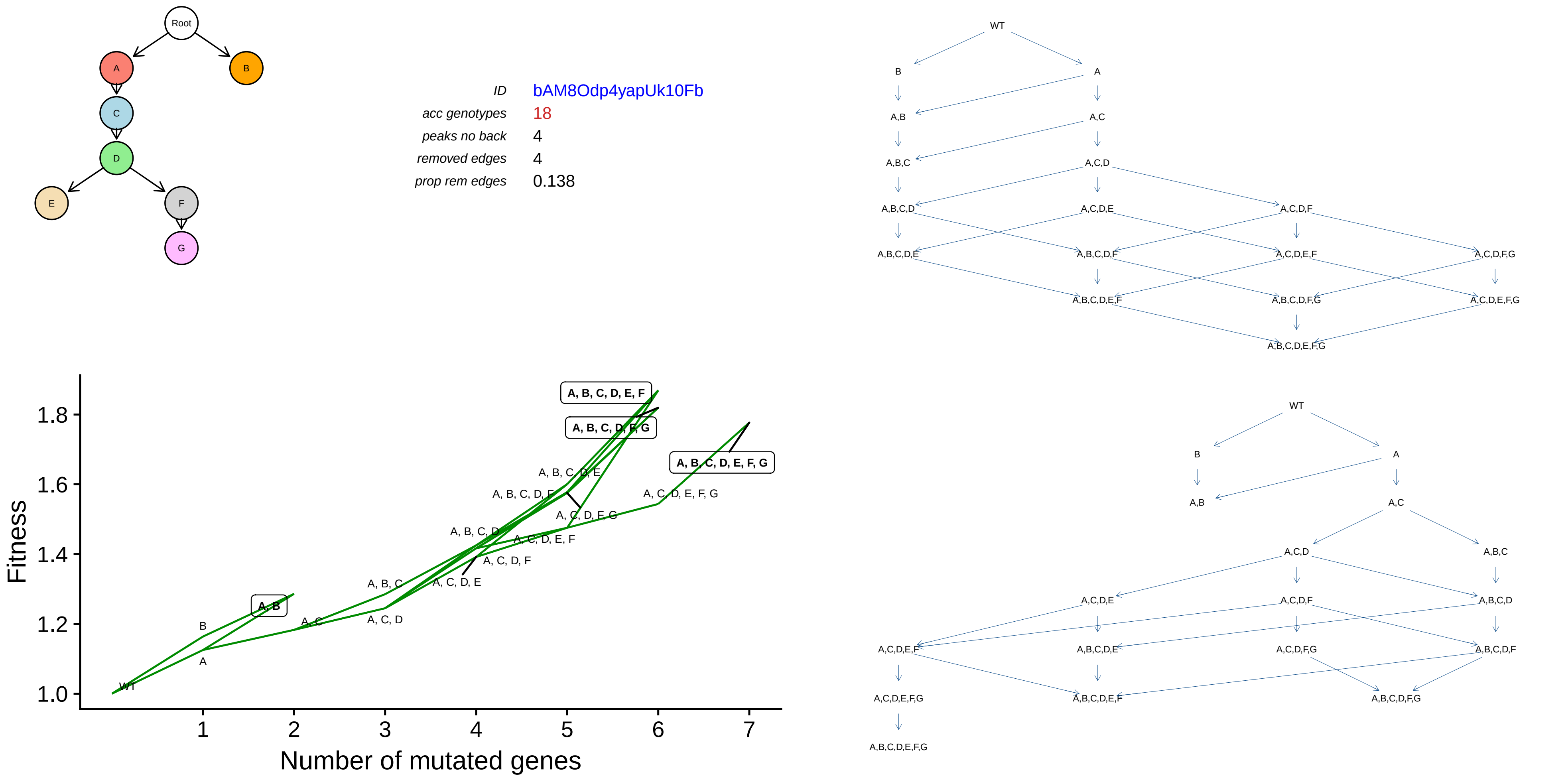

Root
A
B
C
D
E
F
G
ID bAM8Odp4yapUk10Fb
acc genotypes 18
peaks no back 4
removed edges 4
prop rem edges 0.138
Fitness
Number of mutated genes
A, B
A, B, C, D, E, F
A, B, C, D, F, G
A, B, C, D, E, F, G
WT
A,B,C,D,E,F,G

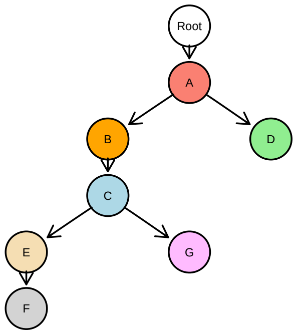


| | |
|---|---|
| *ID* | bCsk2Qo5VMVm55fM |
| *acc genotypes* | 17 |
| *peaks no back* | 2 |
| *removed edges* | 3 |
| *prop rem edges* | 0.111 |

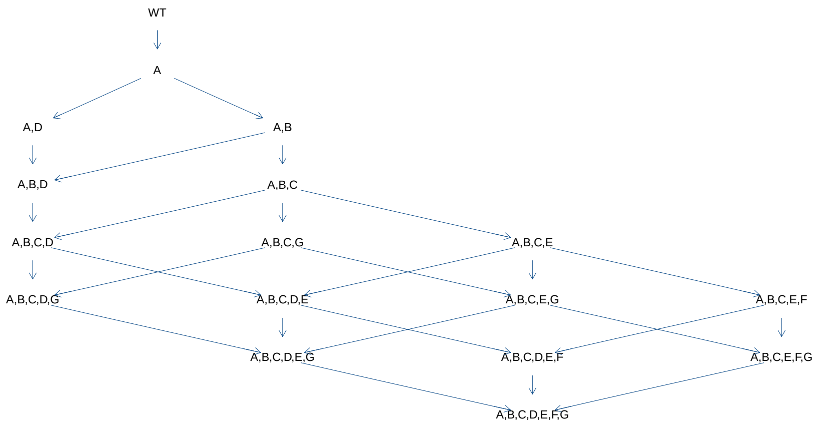


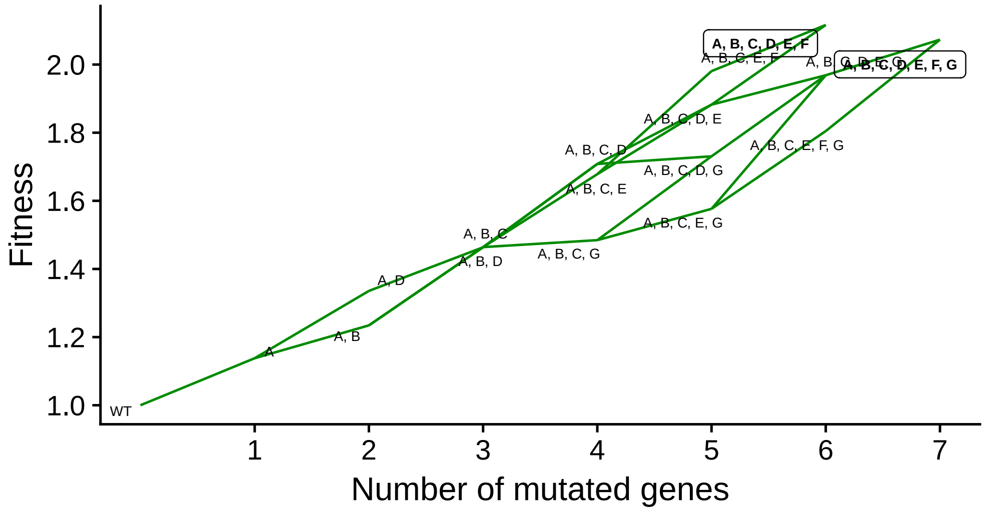


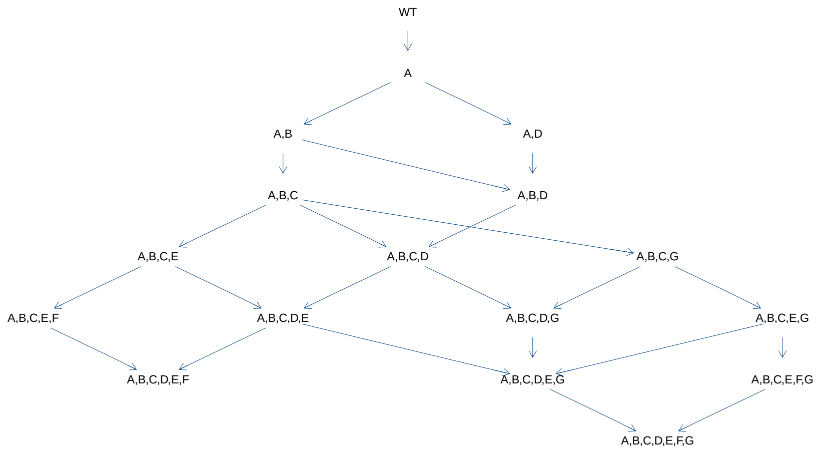

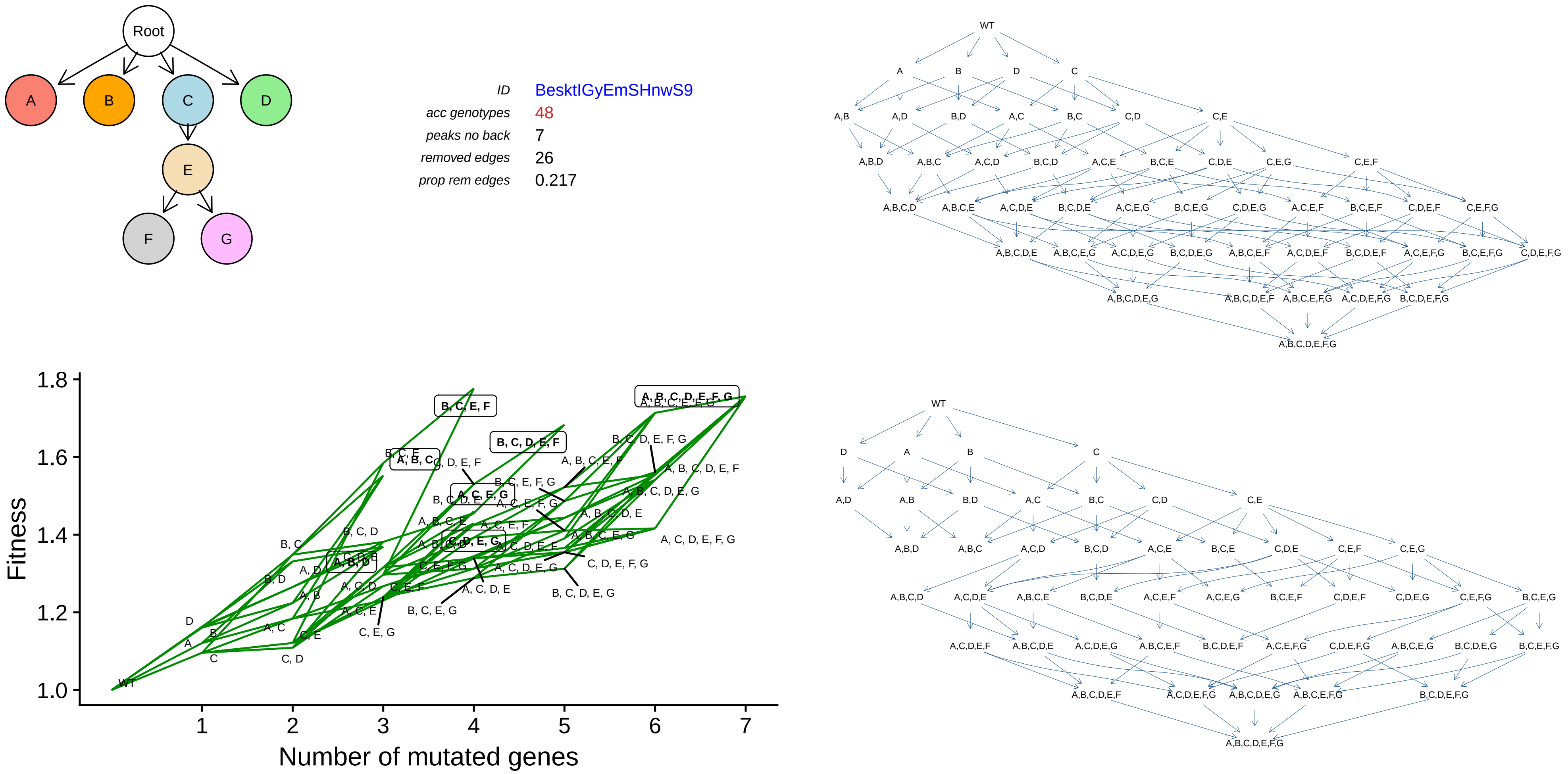

Root
A
B
C
D
E
F
G
ID BesktIGyEmSHnwS9
acc genotypes 48
peaks no back 7
removed edges 26
prop rem edges 0.217
Fitness
Number of mutated genes
1.0
1.2
1.4
1.6
1.8
1
2
3
4
5
6
7
A, B, C, D, E, F, G
B, C, E, F
B, C, D, E, F
WT
A,B,C,D,E,F,G

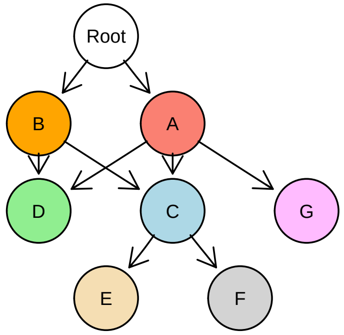


| | |
|---|---|
| *ID* | Bm107M6KzXSHrdvsu |
| *acc genotypes* | 24 |
| *peaks no back* | 2 |
| *removed edges* | 3 |
| *prop rem edges* | 0.065 |

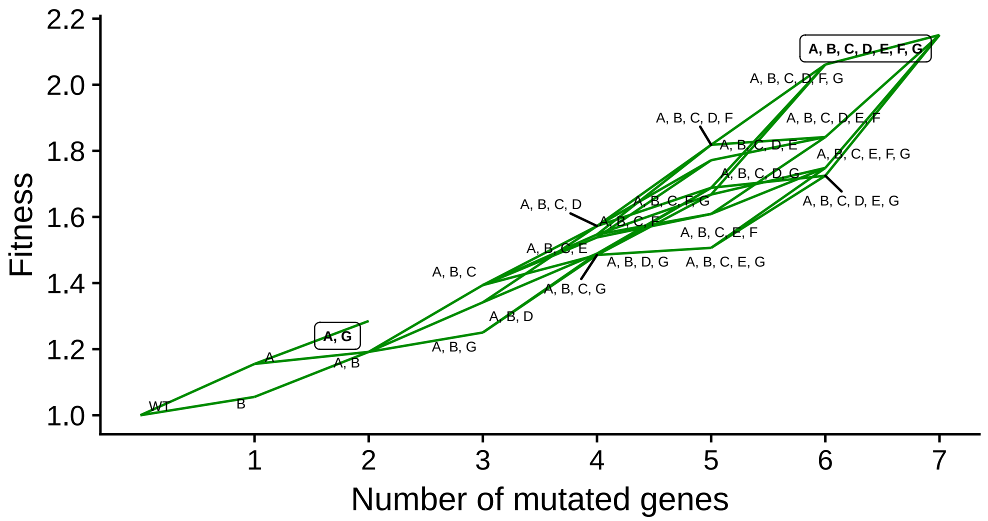


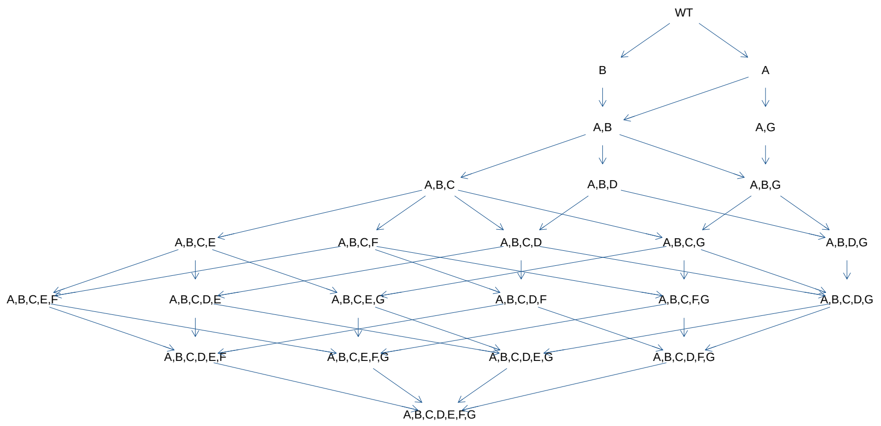


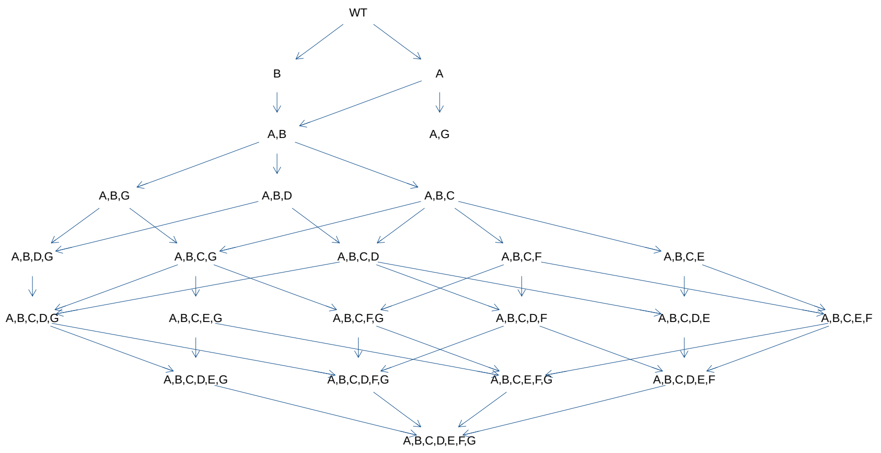

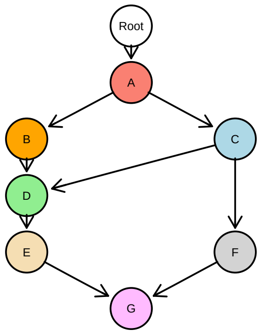


| | |
|---|---|
| *ID* | Bm9gSkay1l4nALeA |
| *acc genotypes* | 12 |
| *peaks no back* | 2 |
| *removed edges* | 1 |
| *prop rem edges* | 0.067 |

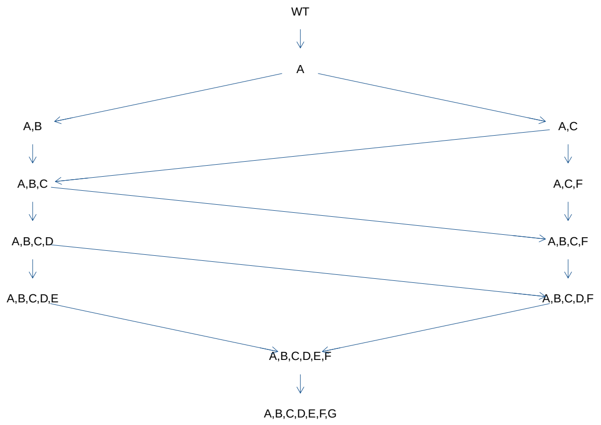


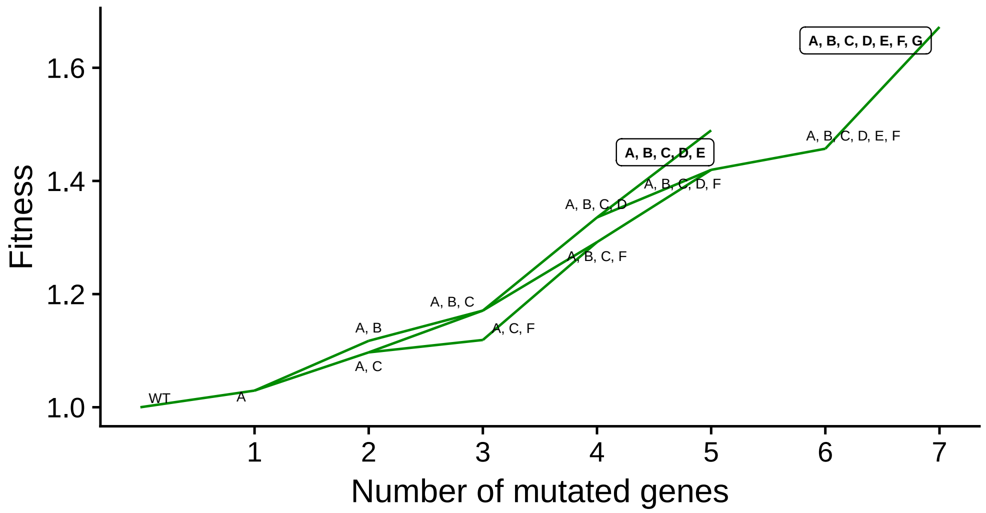


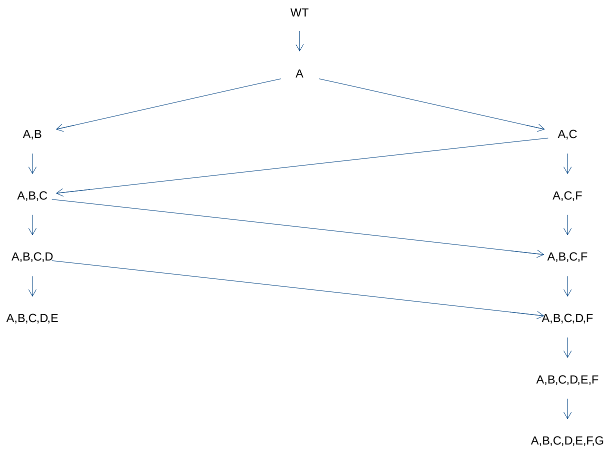

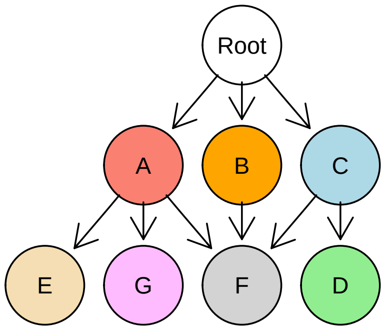


| | |
|---|---|
| *ID* | bpHispMQClsXVnvs |
| *acc genotypes* | 38 |
| *peaks no back* | 6 |
| *removed edges* | 9 |
| *prop rem edges* | 0.106 |

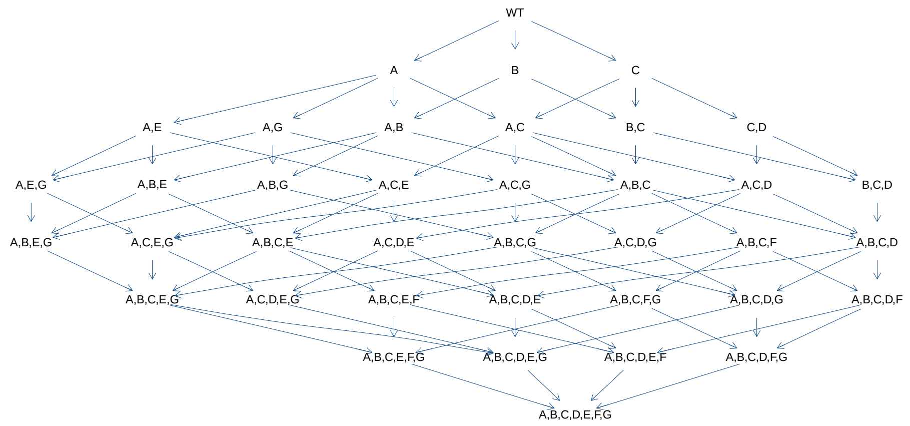


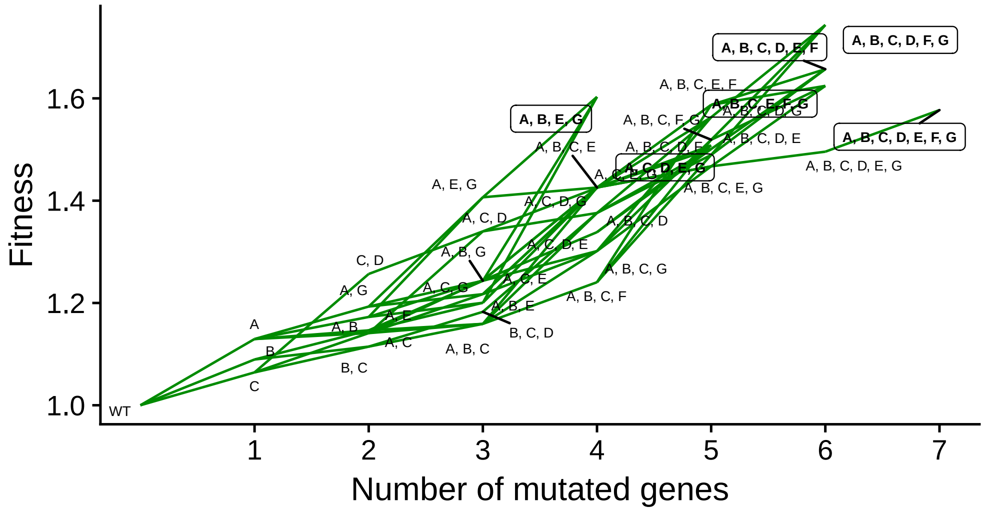


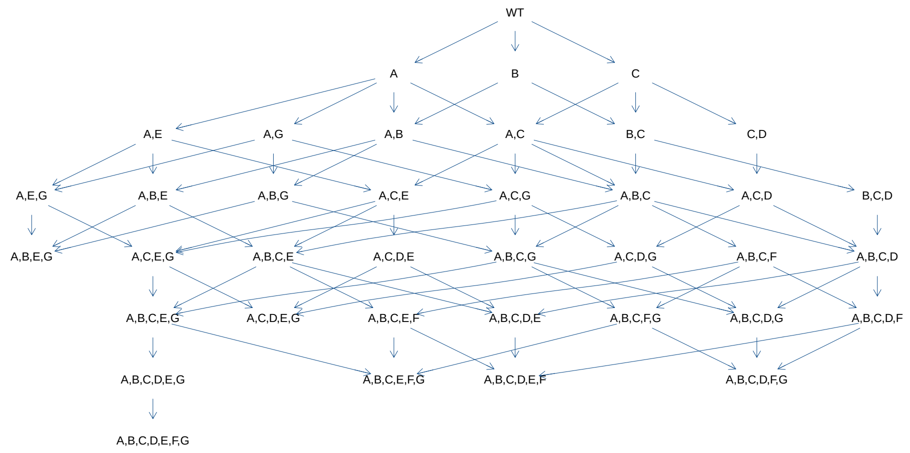

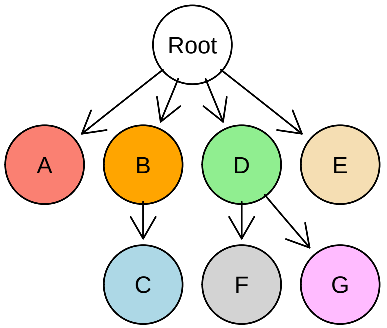


| | |
|---|---|
| *ID* | BQW2TKpC1B6DJ7jN |
| *acc genotypes* | 60 |
| *peaks no back* | 4 |
| *removed edges* | 27 |
| *prop rem edges* | 0.169 |

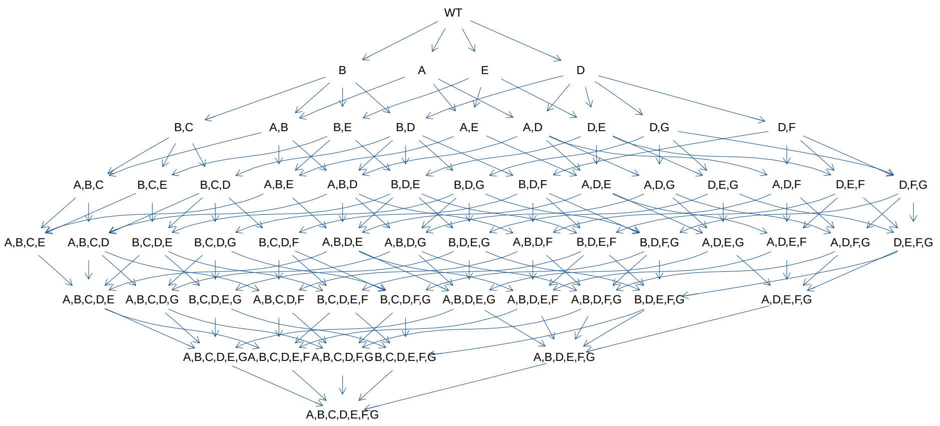

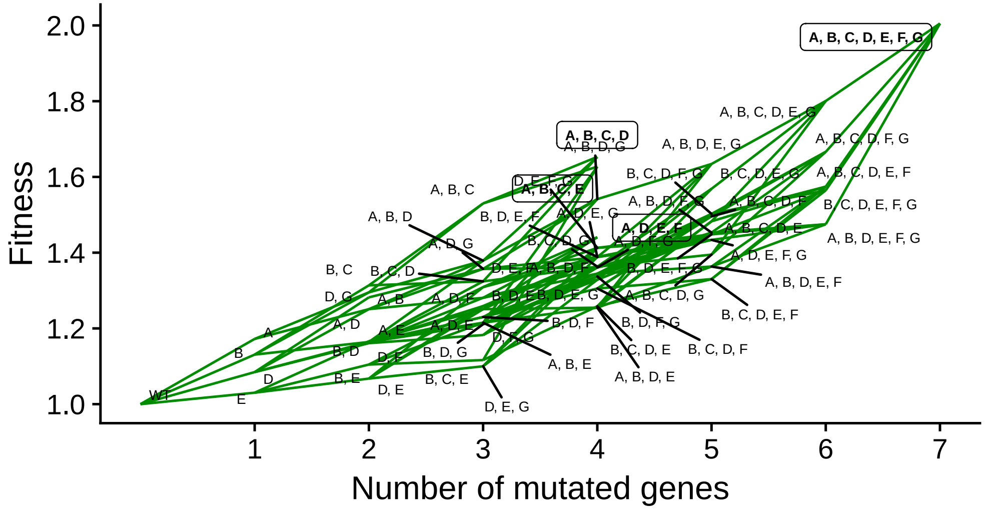


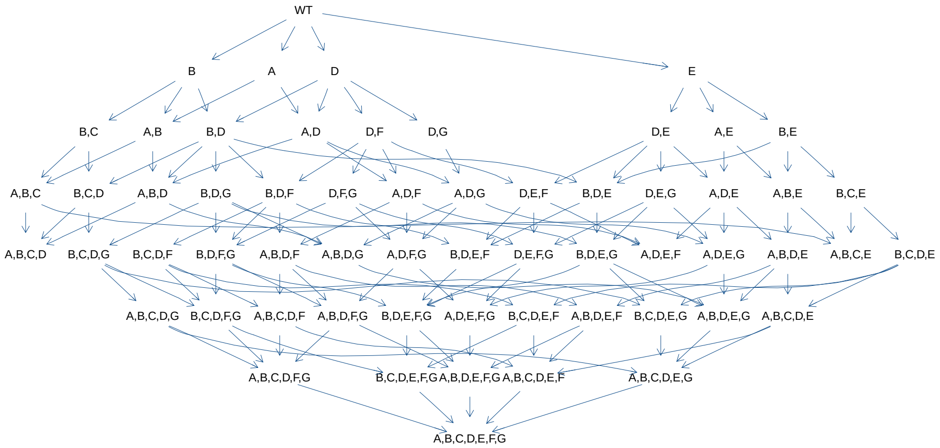

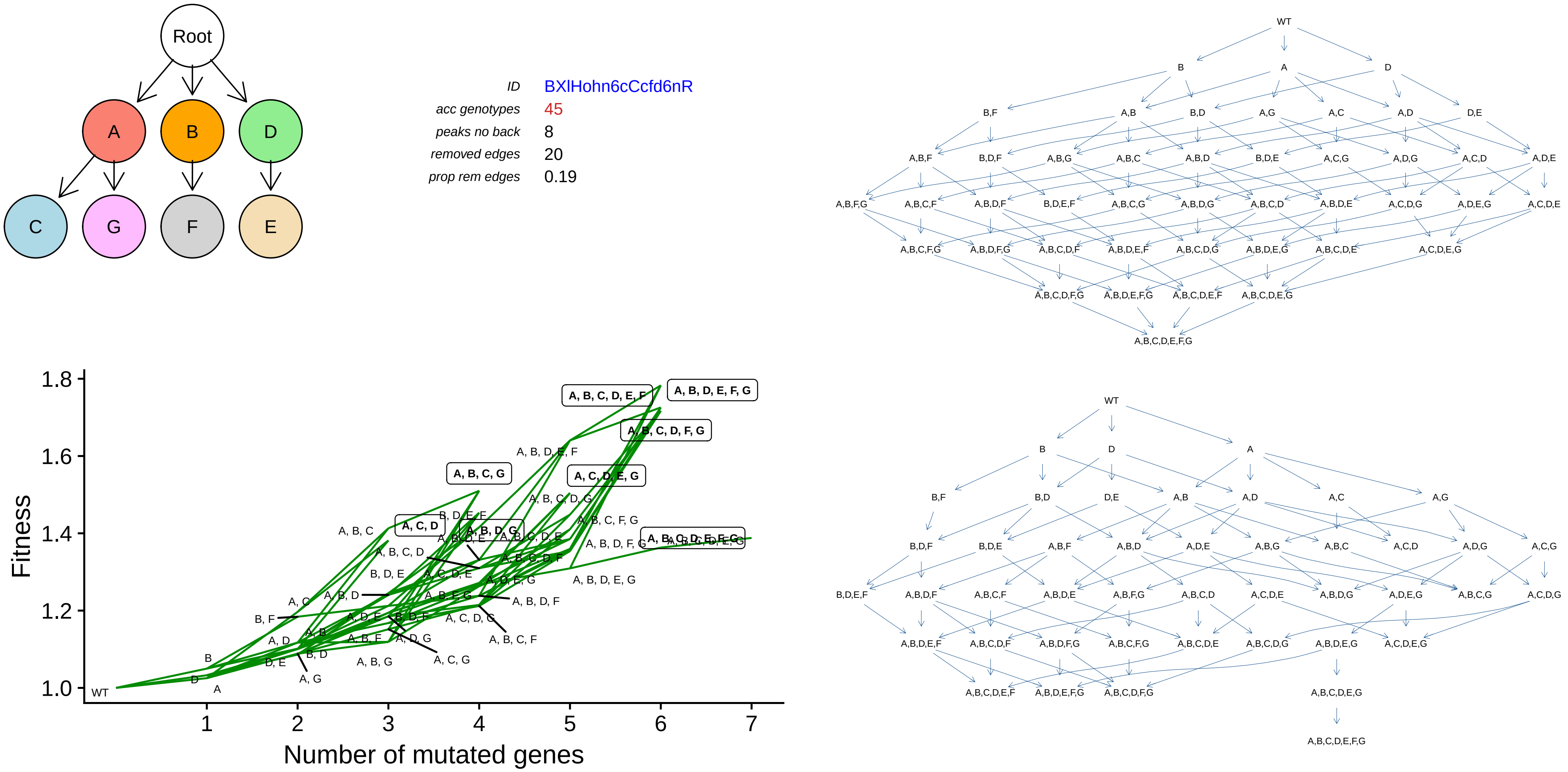

Root
A
B
D
C
G
F
E
ID BXlHohn6cCcfd6nR
acc genotypes 45
peaks no back 8
removed edges 20
prop rem edges 0.19
Fitness
Number of mutated genes
A, B, C, D, E, F
A, B, D, E, F, G
A, B, C, D, F, G
A, B, C, G
A, C, D, E, G
A, C, D
A, B, D, G
A, B, C, D, E, F, G

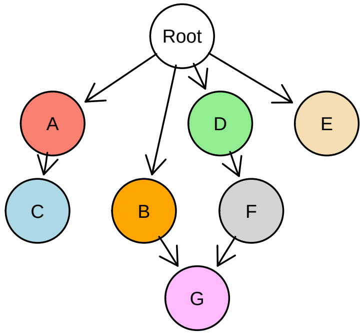


| | |
|---|---|
| *ID* | BzCzics10JlcdE102O |
| *acc genotypes* | 42 |
| *peaks no back* | 6 |
| *removed edges* | 16 |
| *prop rem edges* | 0.165 |

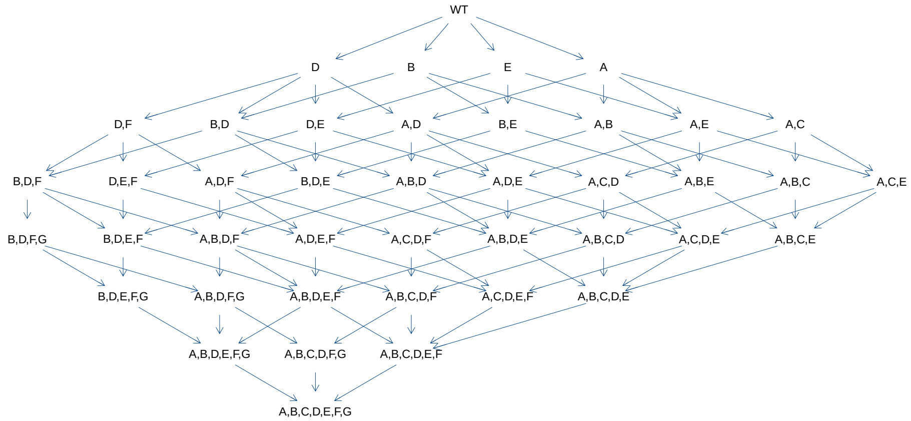


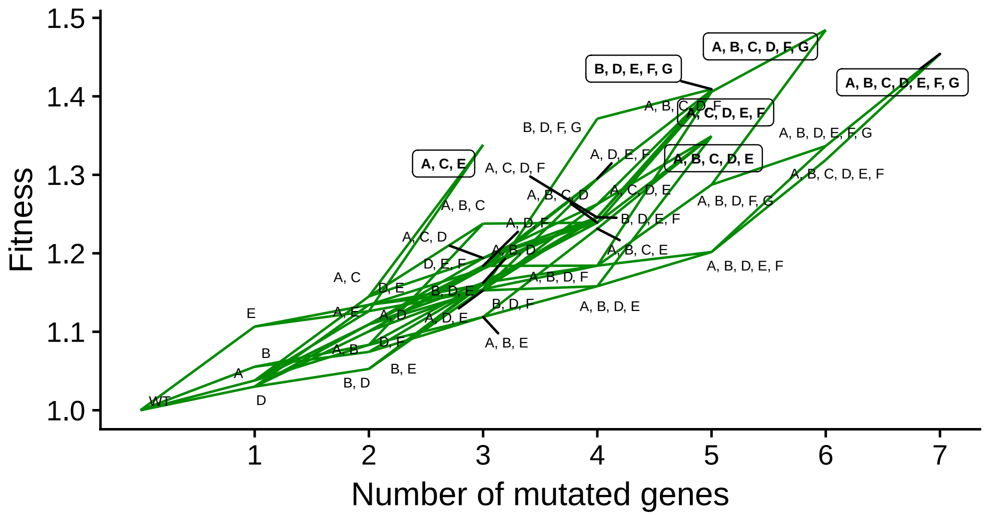


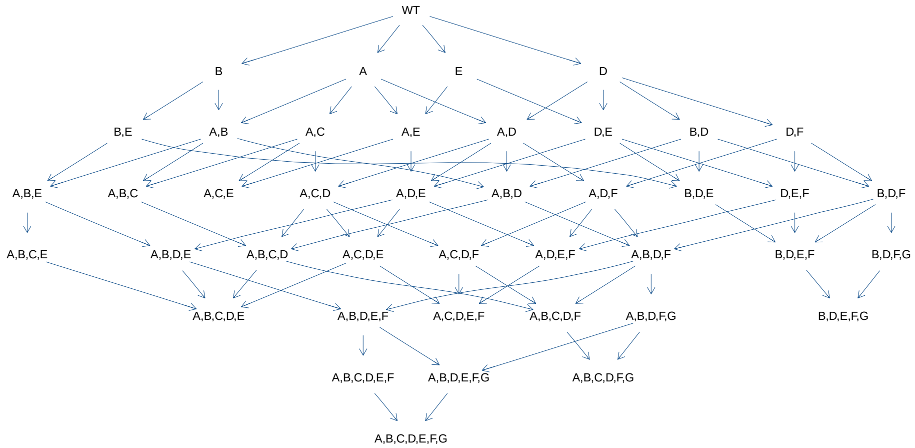

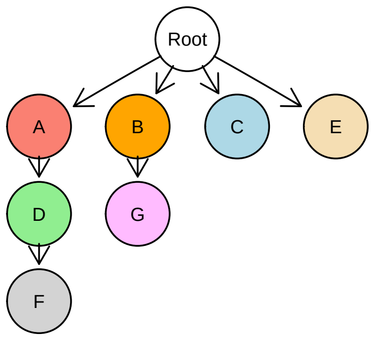


*ID* c9GVdRVUtkHv3yR7
*acc genotypes* 48
*peaks no back* 4
*removed edges* 23
*prop rem edges* 0.198

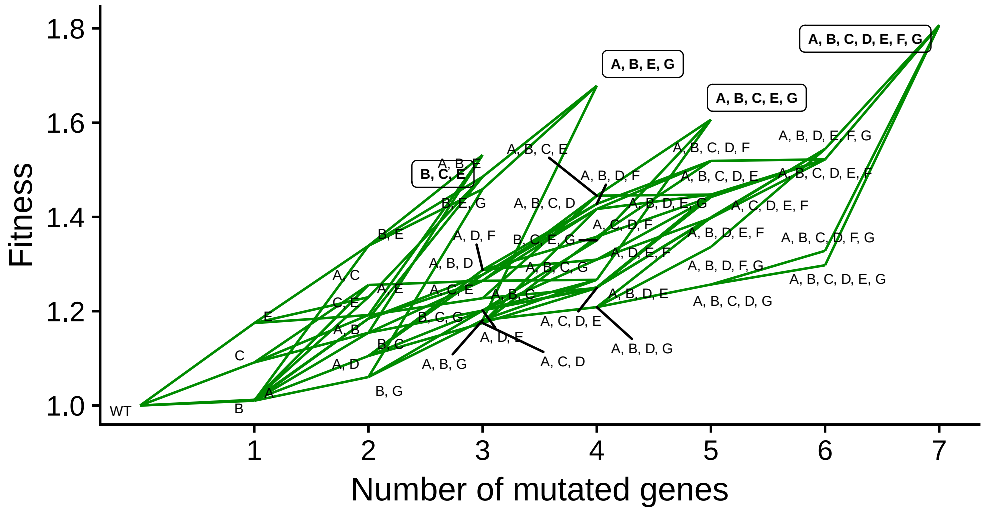


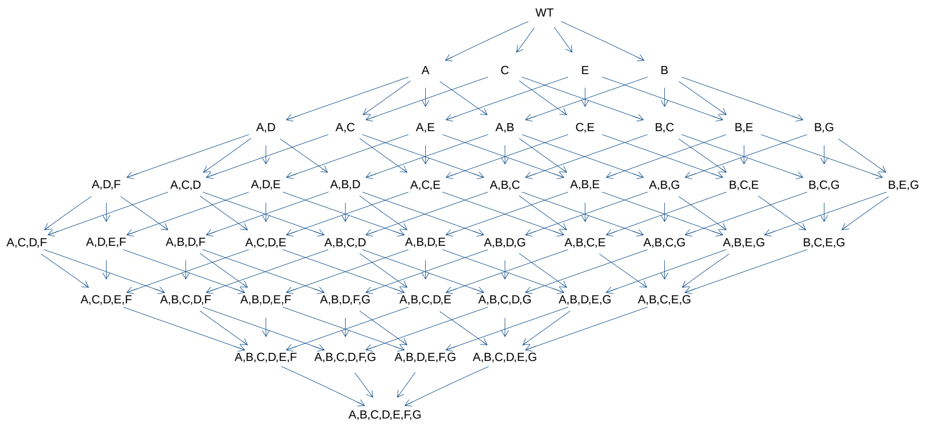


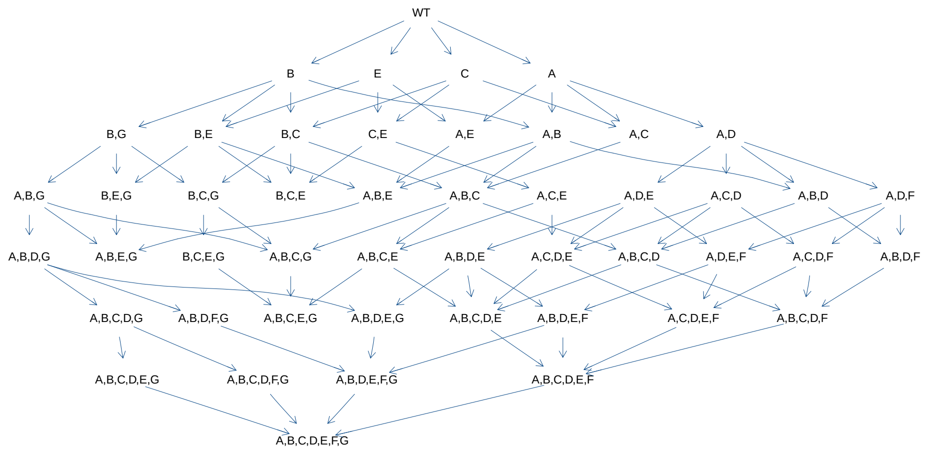

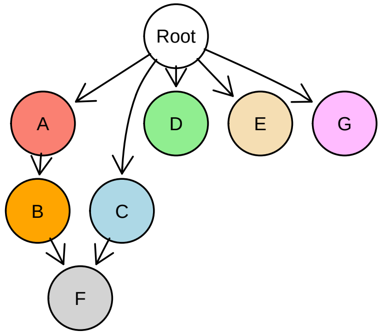


| | |
|---|---|
| *ID* | C9MQvf10YWmVwuUpW |
| *acc genotypes* | 56 |
| *peaks no back* | 9 |
| *removed edges* | 32 |
| *prop rem edges* | 0.216 |

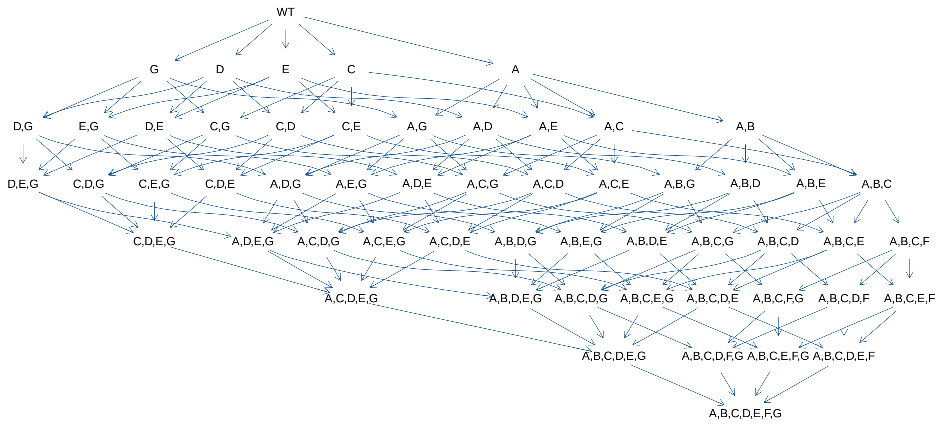

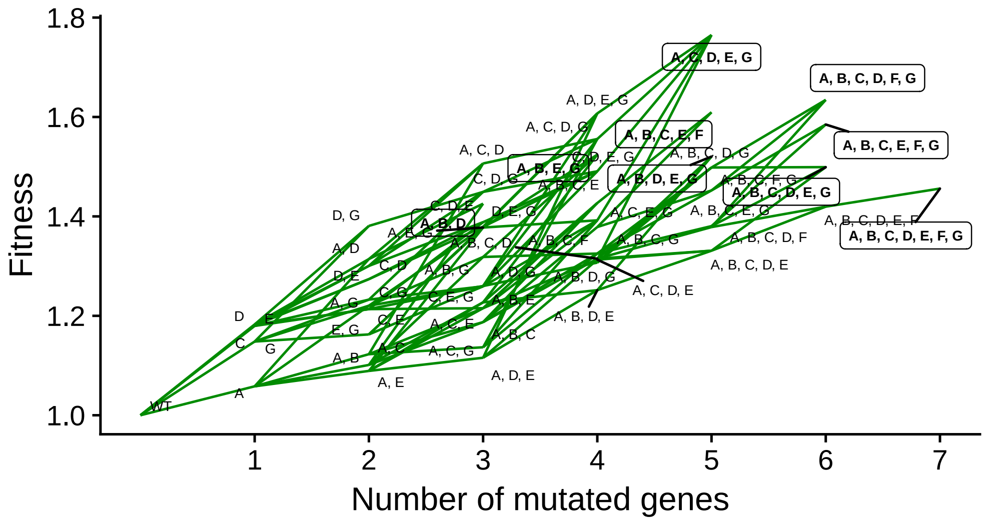


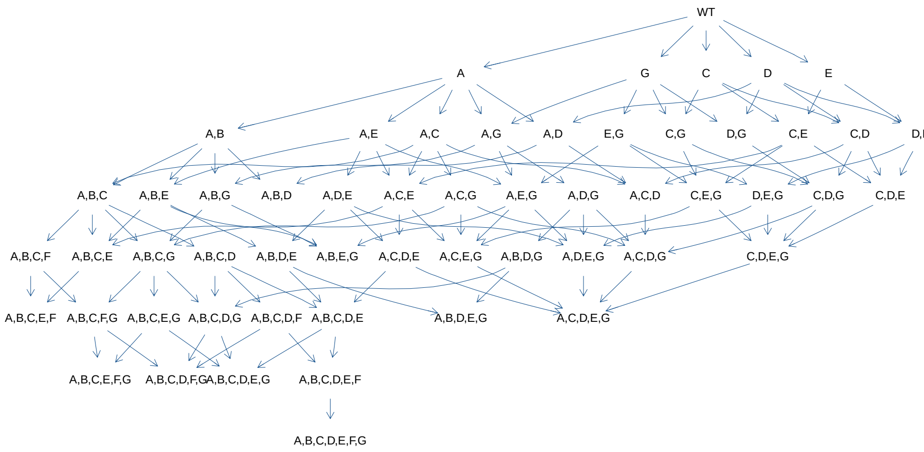

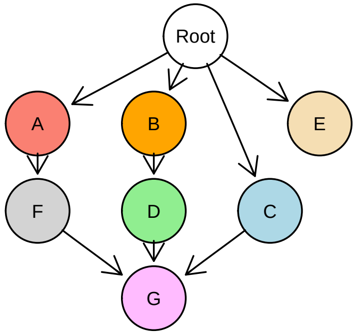


| | |
|---|---|
| *ID* | ce4Pp9mhcgxMJhs9 |
| *acc genotypes* | 38 |
| *peaks no back* | 3 |
| *removed edges* | 9 |
| *prop rem edges* | 0.103 |

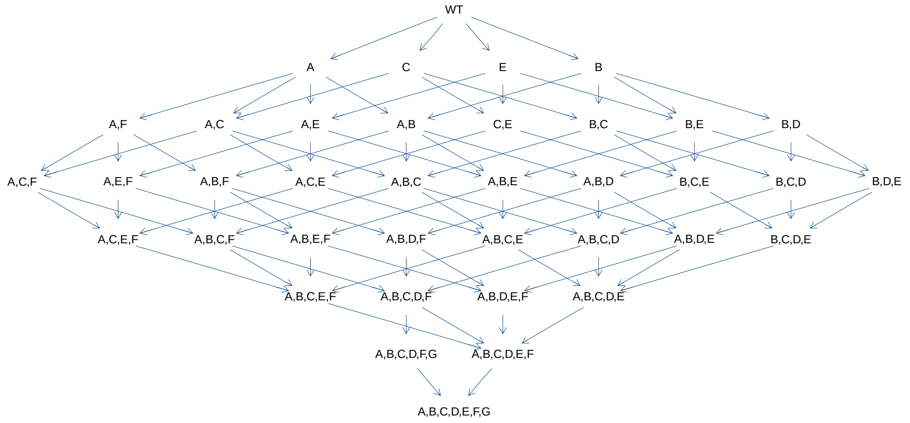


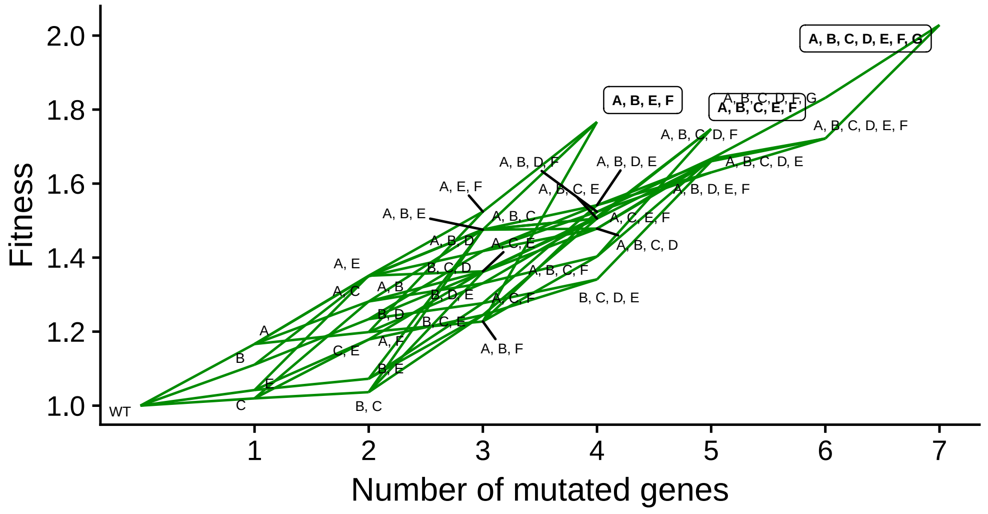


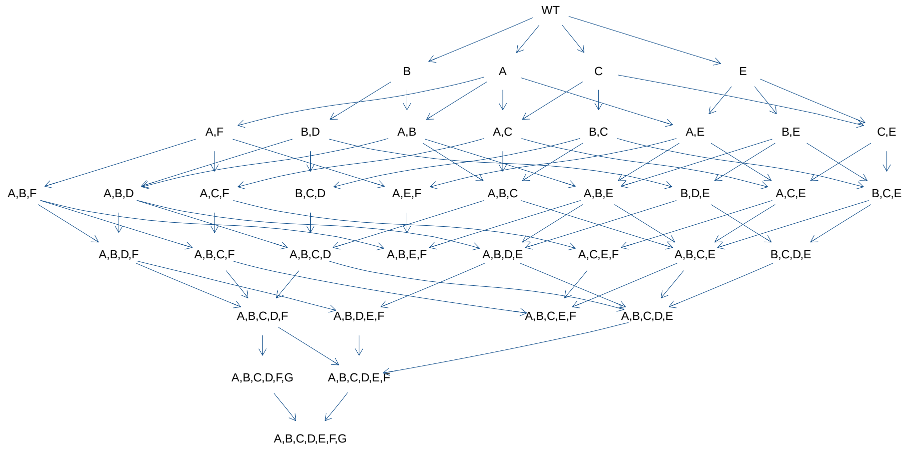

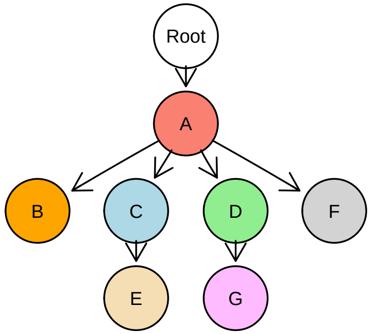


| | |
|---|---|
| *ID* | cfODc4wi4uNS9zkM |
| *acc genotypes* | 37 |
| *peaks no back* | 6 |
| *removed edges* | 16 |
| *prop rem edges* | 0.188 |

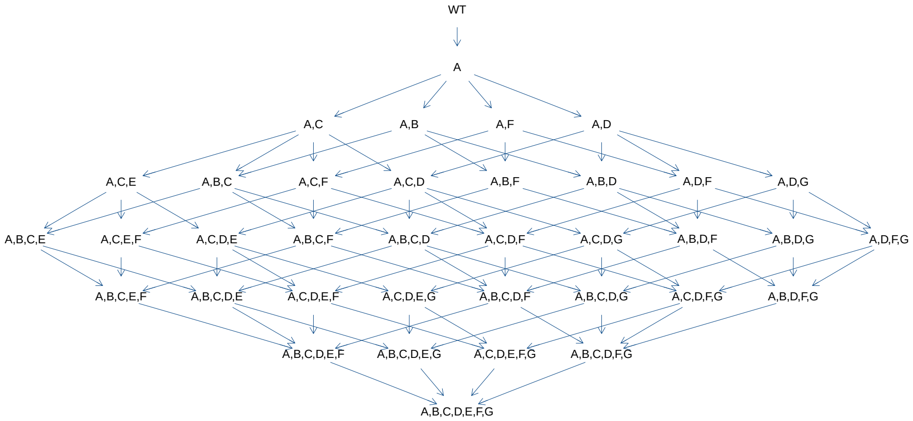


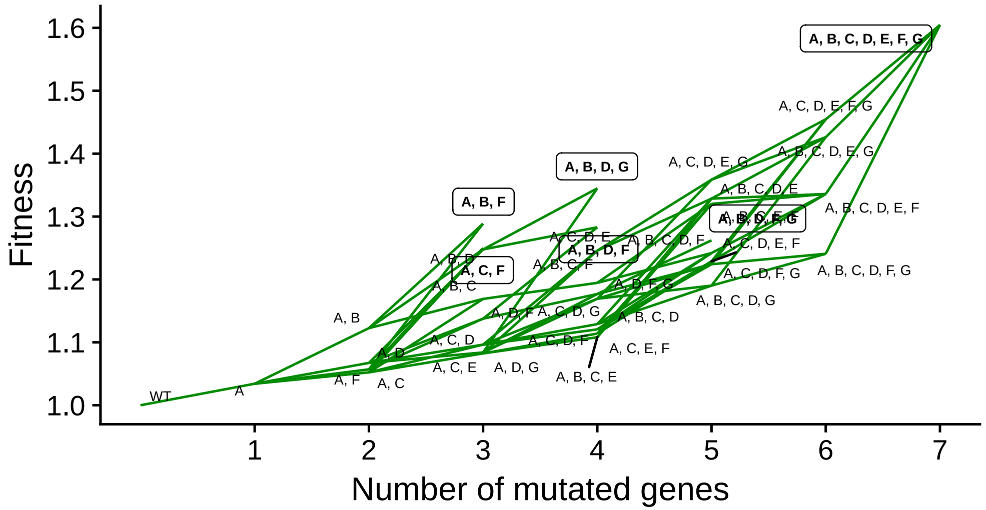


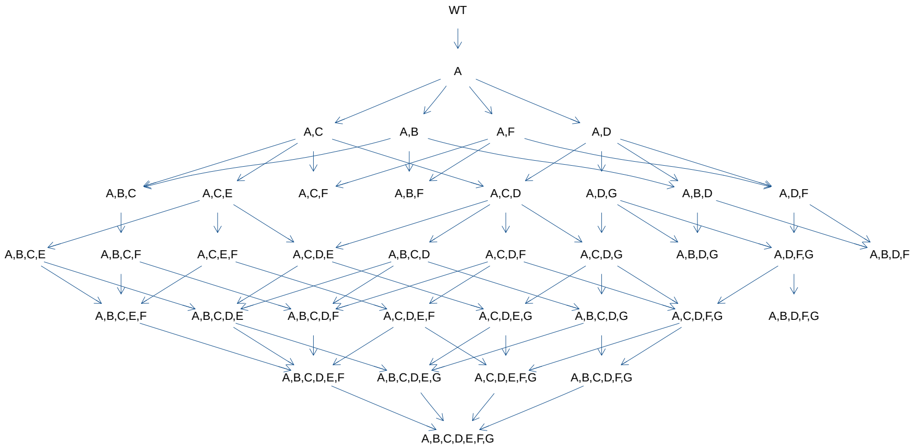

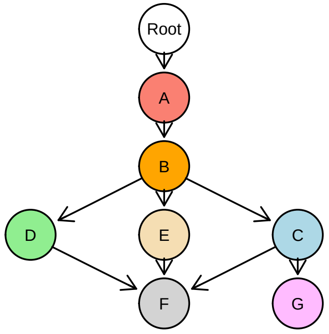


| | |
|---|---|
| *ID* | chTnx8SjPrqKECWb |
| *acc genotypes* | 16 |
| *peaks no back* | 3 |
| *removed edges* | 5 |
| *prop rem edges* | 0.2 |

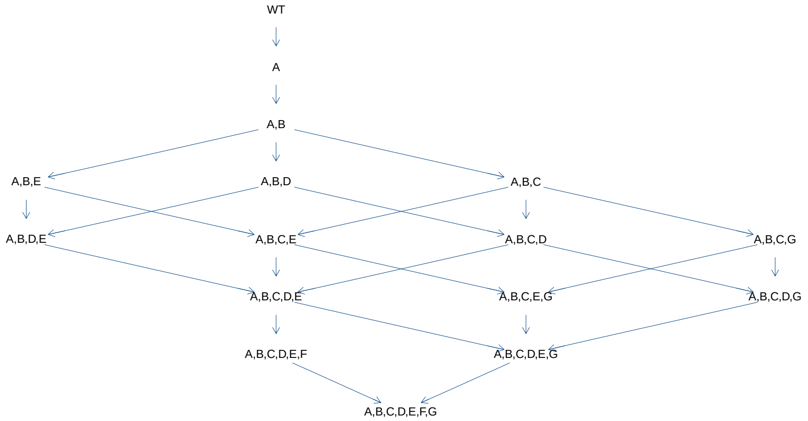


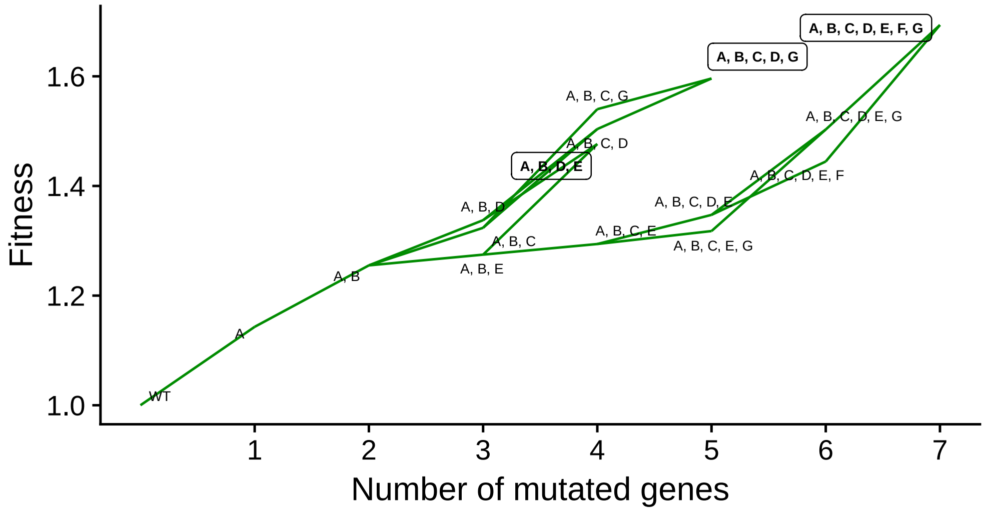


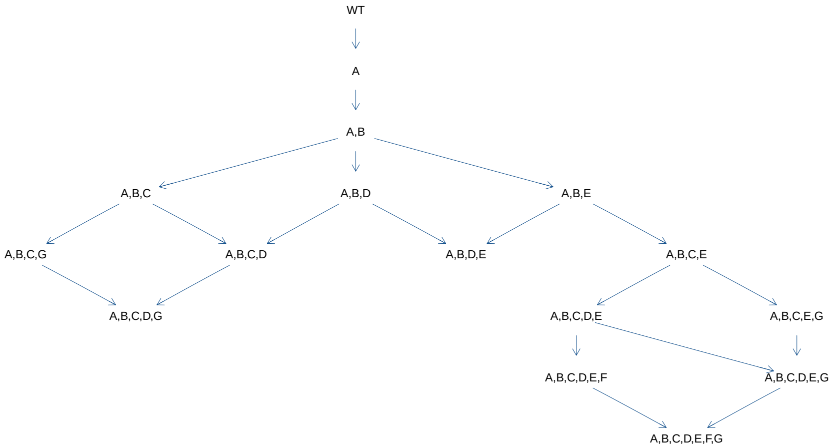

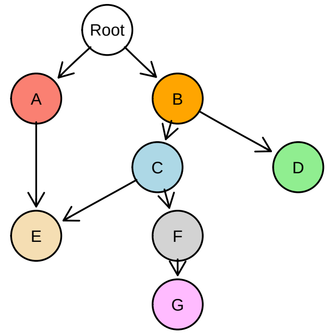


| | |
|---|---|
| *ID* | ClxqJzXIXlPDwCzC |
| *acc genotypes* | 24 |
| *peaks no back* | 5 |
| *removed edges* | 7 |
| *prop rem edges* | 0.159 |

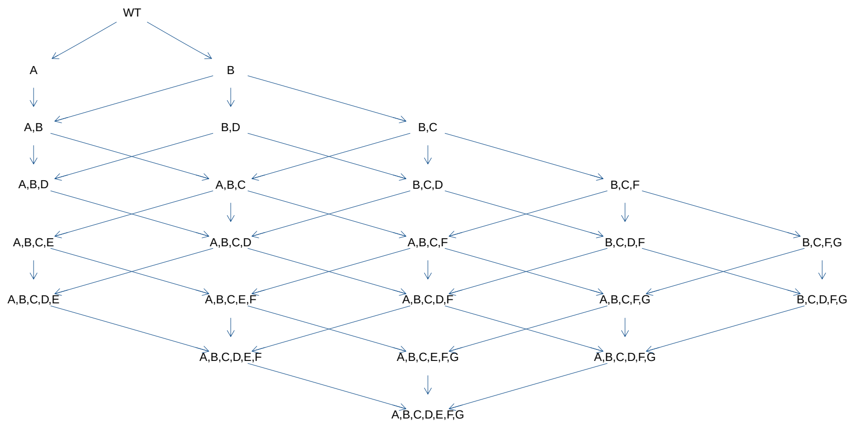


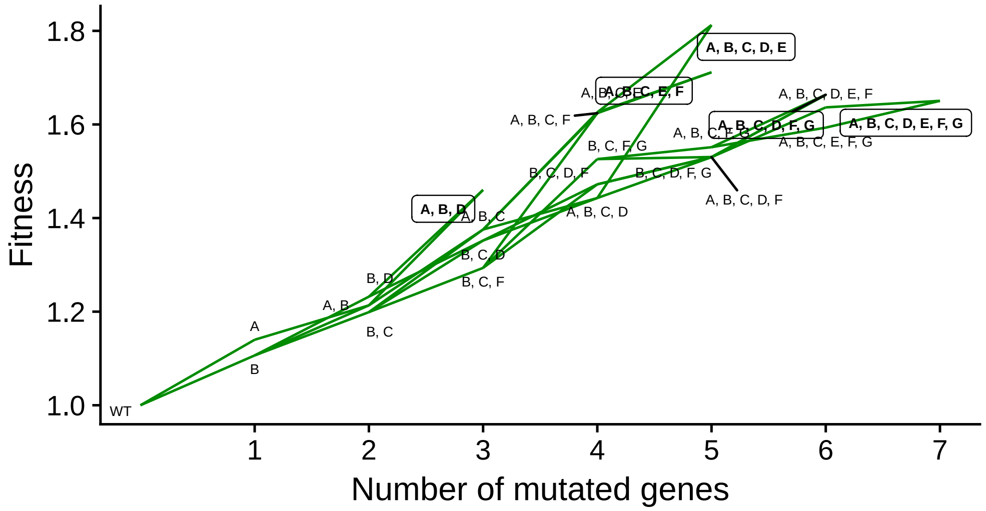


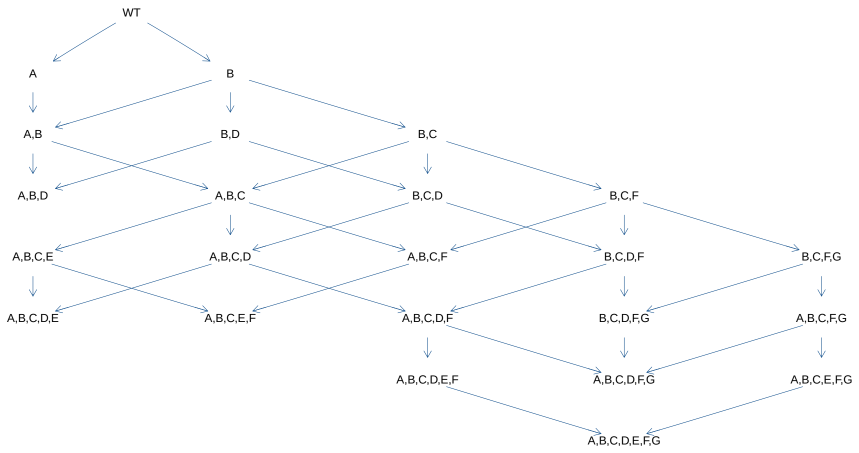

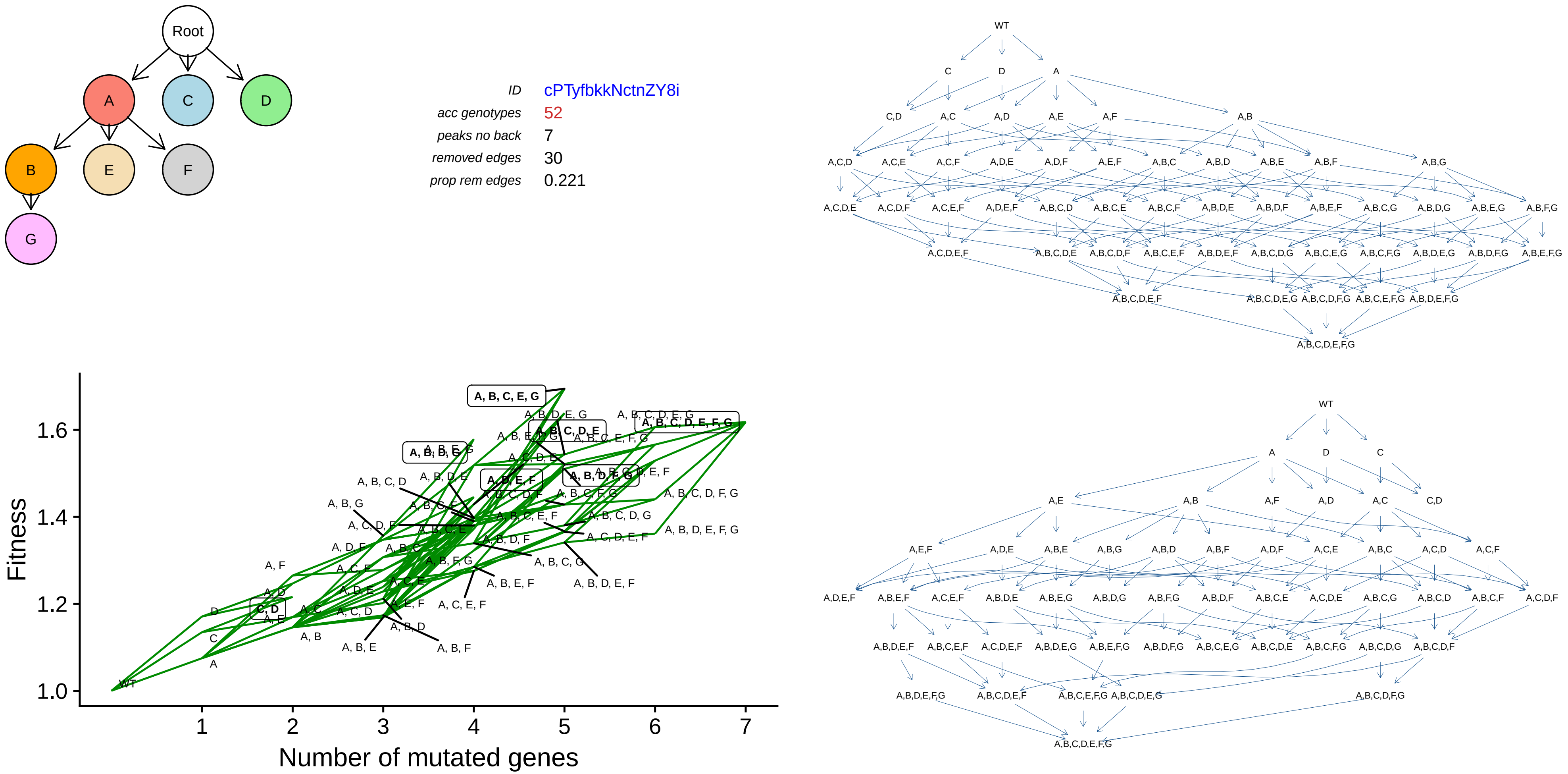

Root
A
C
D
B
E
F
G
ID
cPTyfbkkNctnZY8i
acc genotypes
52
peaks no back
7
removed edges
30
prop rem edges
0.221
Fitness
Number of mutated genes
A, B, C, E, G
A, B, C, D, E, F, G
WT

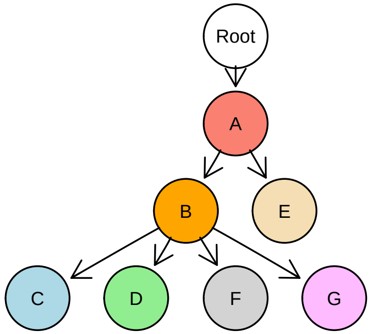


| | |
|---|---|
| *ID* | cPvlM3BD3obpDIZu |
| *acc genotypes* | 35 |
| *peaks no back* | 4 |
| *removed edges* | 6 |
| *prop rem edges* | 0.071 |

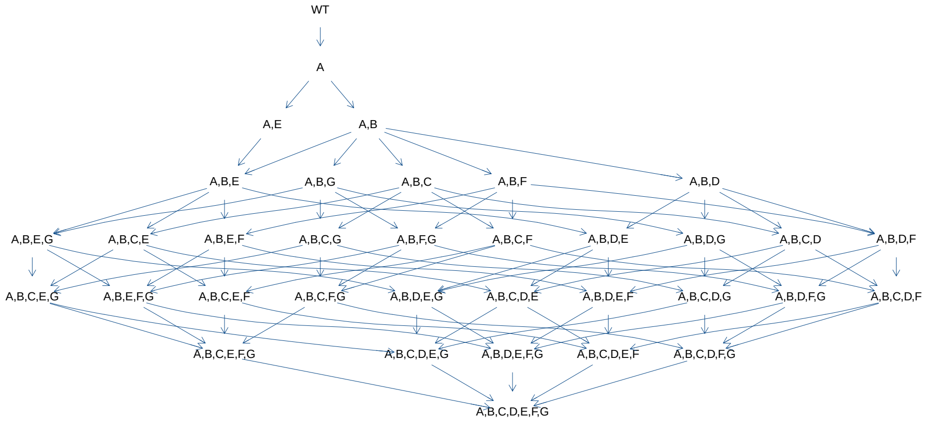


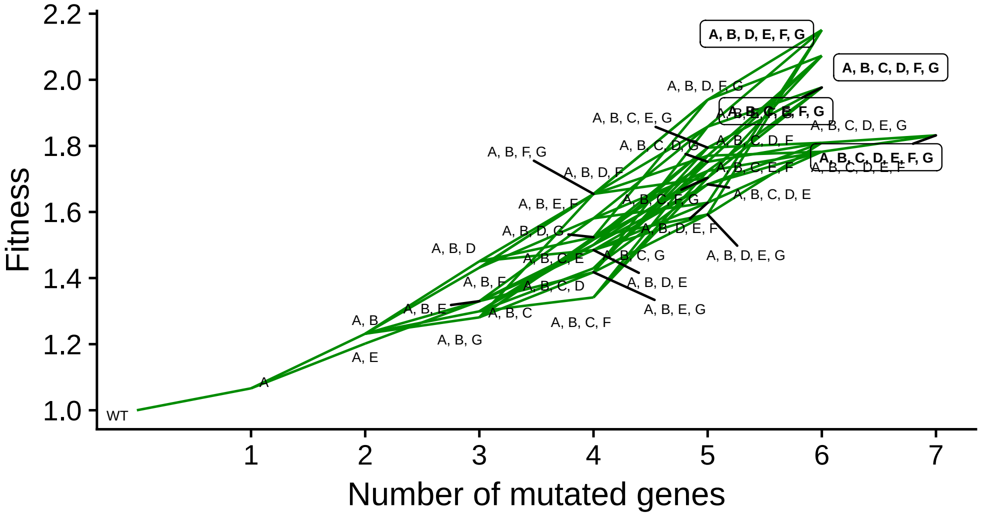


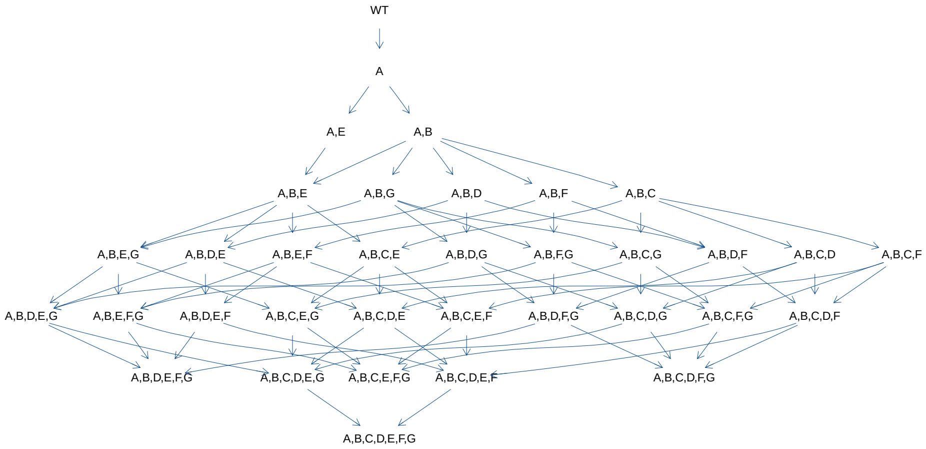

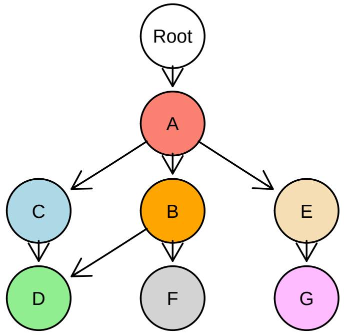


| | |
|---|---|
| *ID* | CQaJm9lTvQh26obo |
| *acc genotypes* | 25 |
| *peaks no back* | 4 |
| *removed edges* | 11 |
| *prop rem edges* | 0.234 |

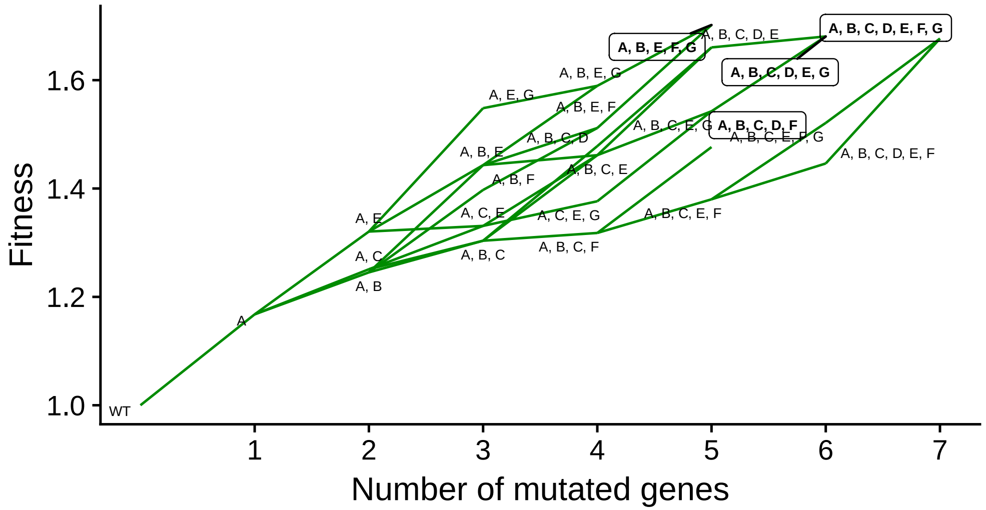


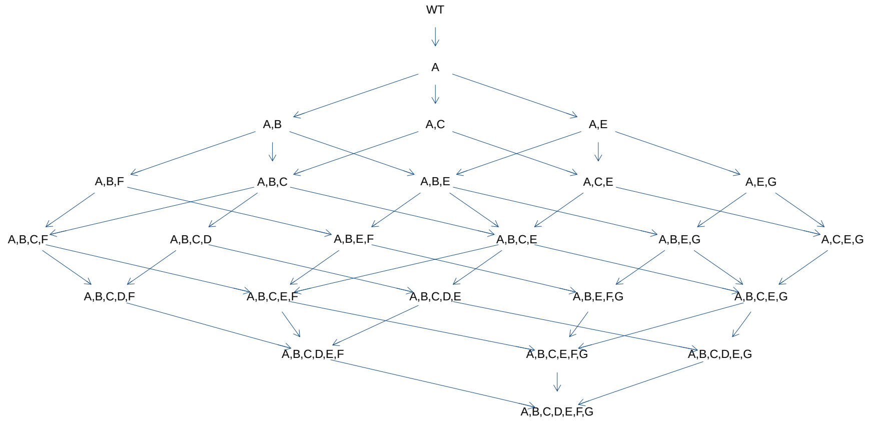


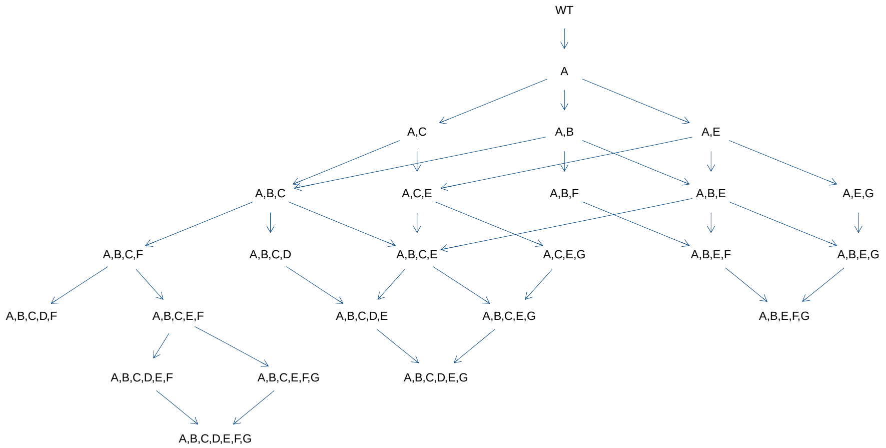

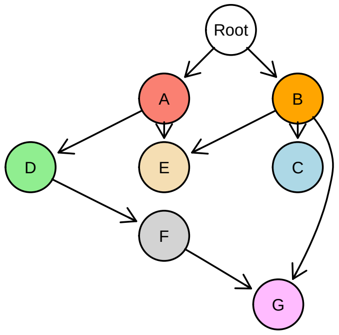


*ID* cQdnO9XMvDNKOW39
*acc genotypes* 22
*peaks no back* 4
*removed edges* 5
*prop rem edges* 0.132

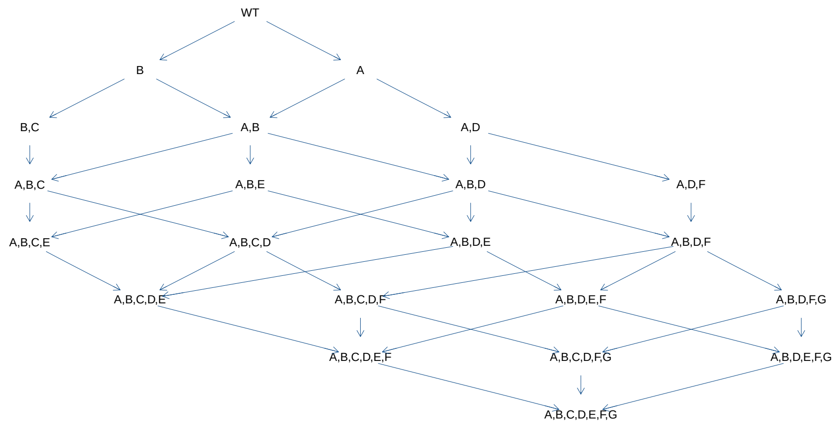


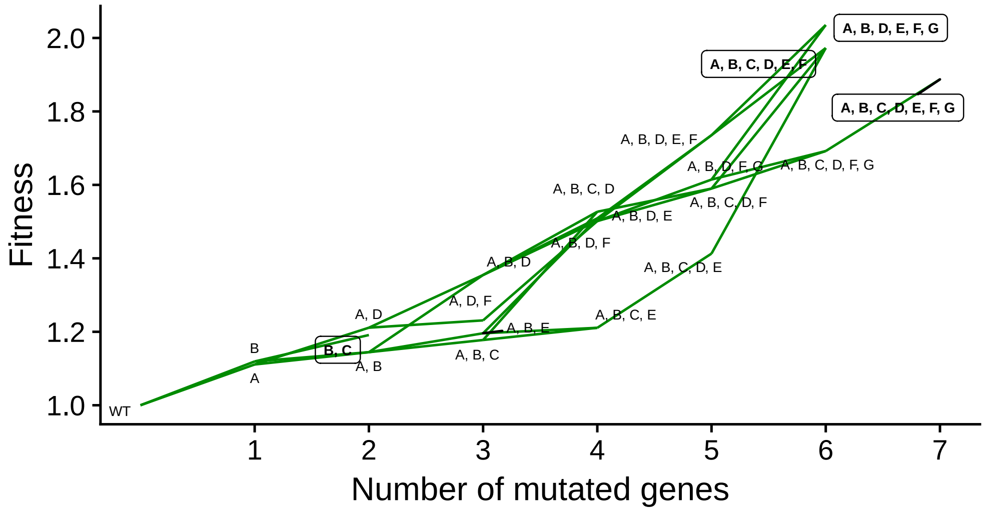


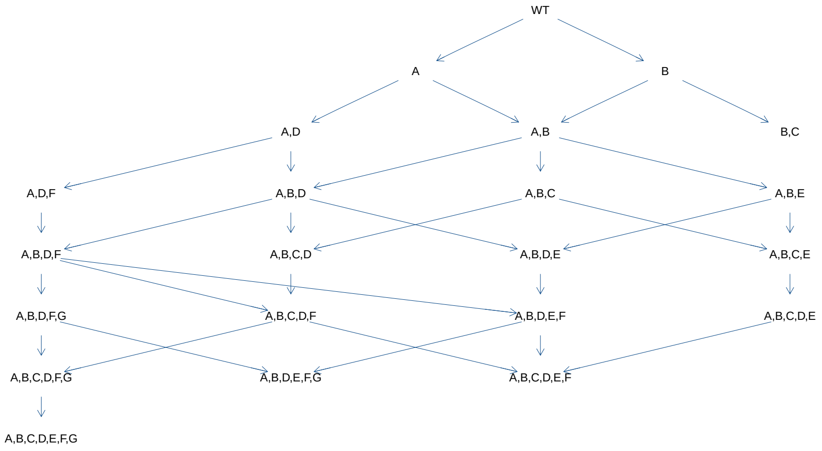

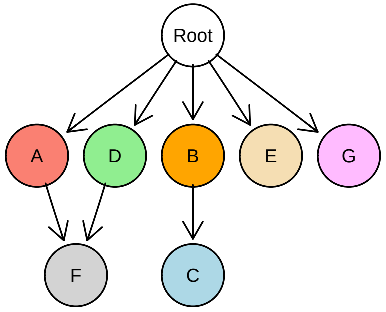


| | |
|---|---|
| *ID* | cwFWkTsrBGFO2dfv |
| *acc genotypes* | 60 |
| *peaks no back* | 7 |
| *removed edges* | 29 |
| *prop rem edges* | 0.181 |

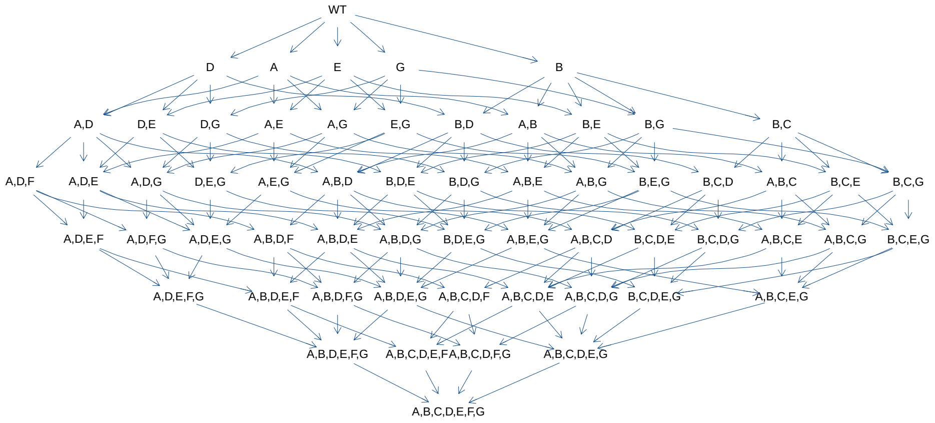

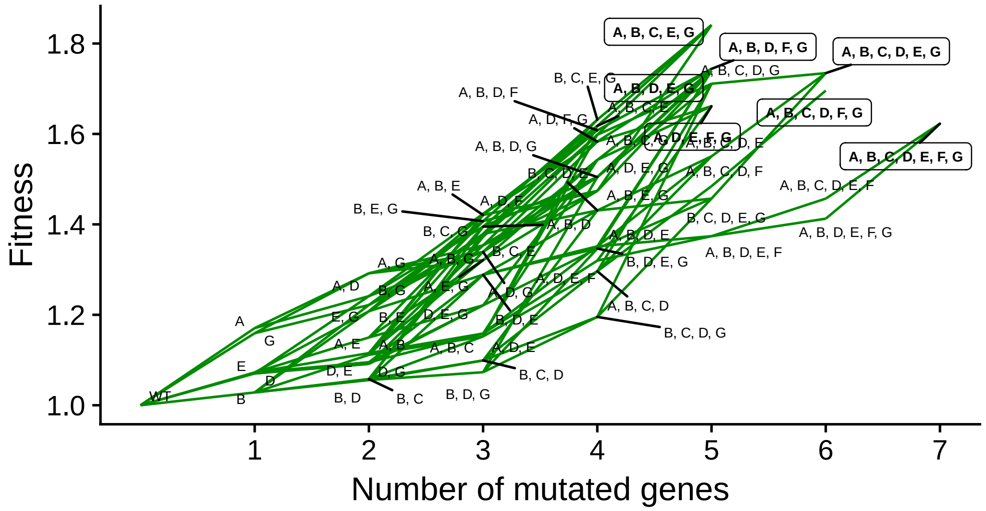


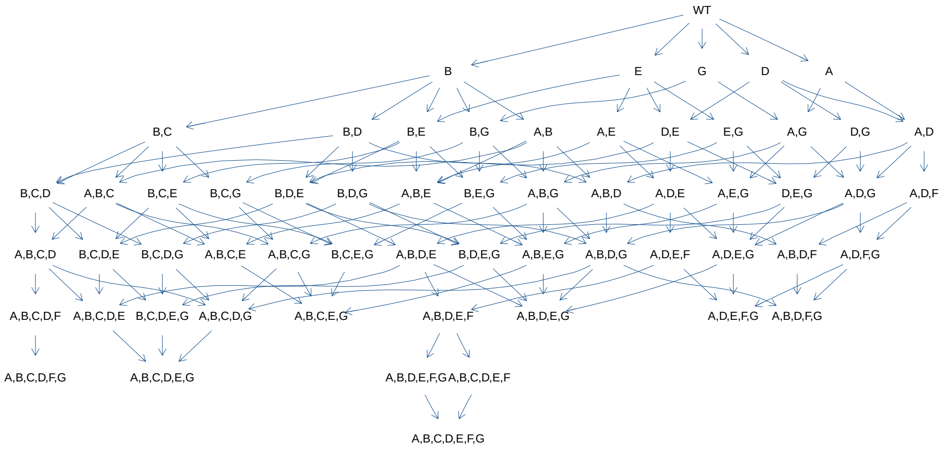

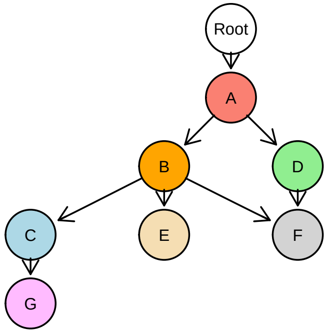


| | |
|---|---|
| *ID* | D1wA9X4HhUWYpfHR |
| *acc genotypes* | 21 |
| *peaks no back* | 5 |
| *removed edges* | 8 |
| *prop rem edges* | 0.216 |

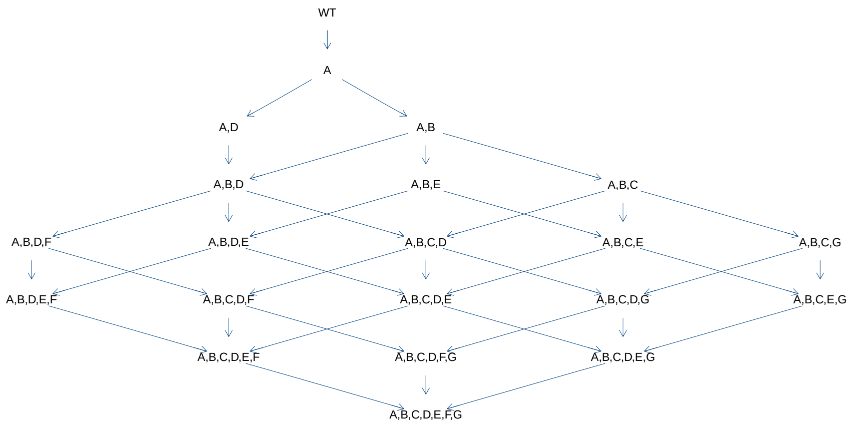


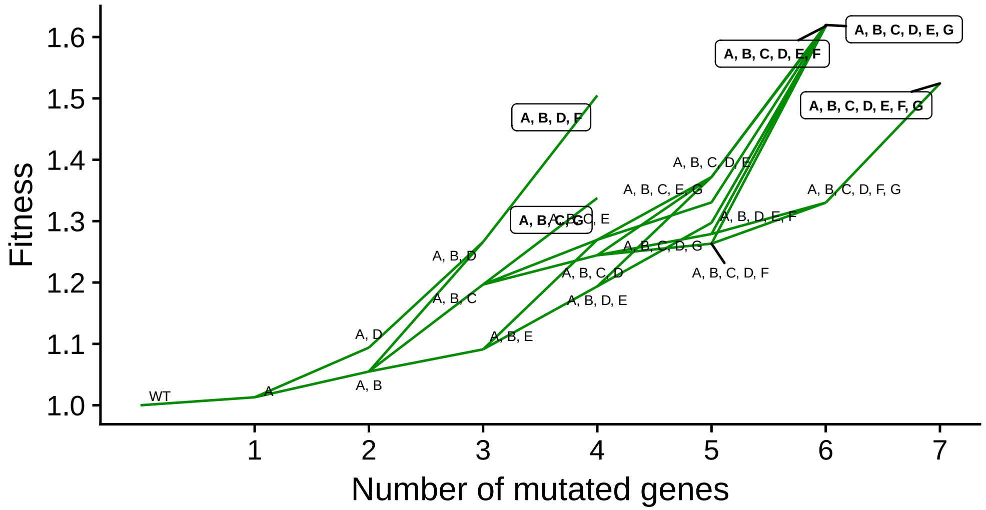


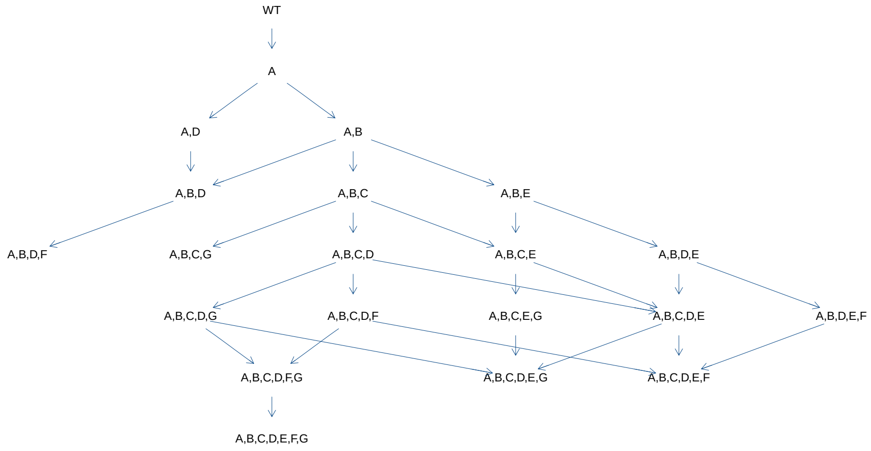

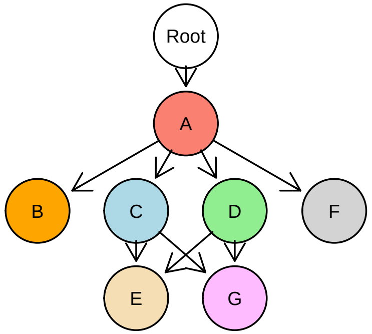


| | |
|---|---|
| *ID* | DBuyPIOfmeKuioDc |
| *acc genotypes* | 29 |
| *peaks no back* | 5 |
| *removed edges* | 9 |
| *prop rem edges* | 0.148 |

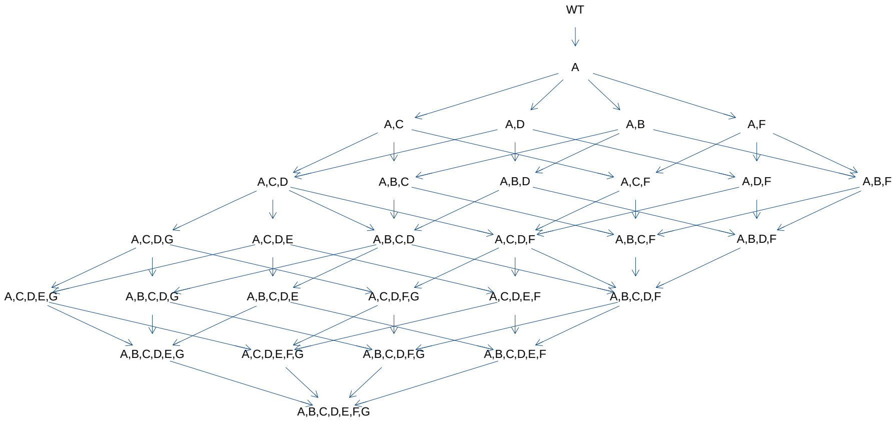


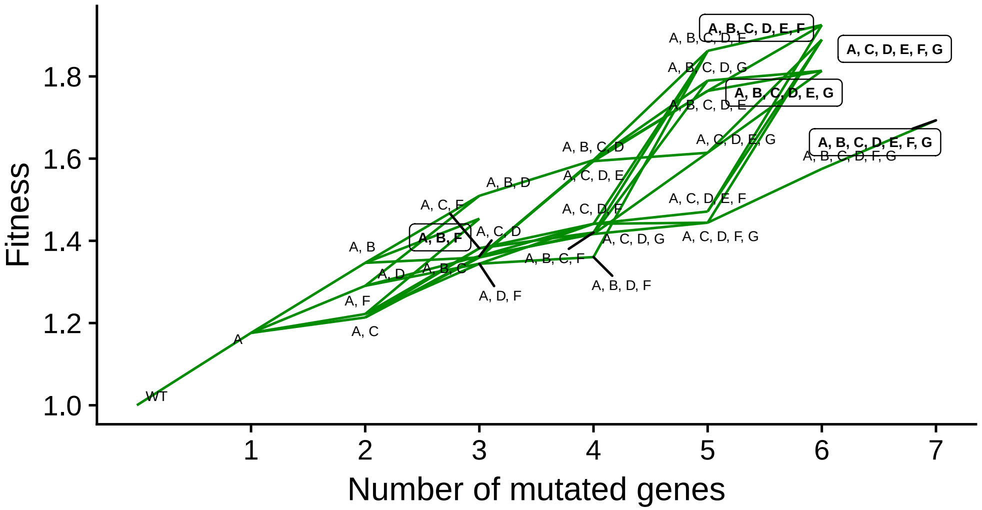


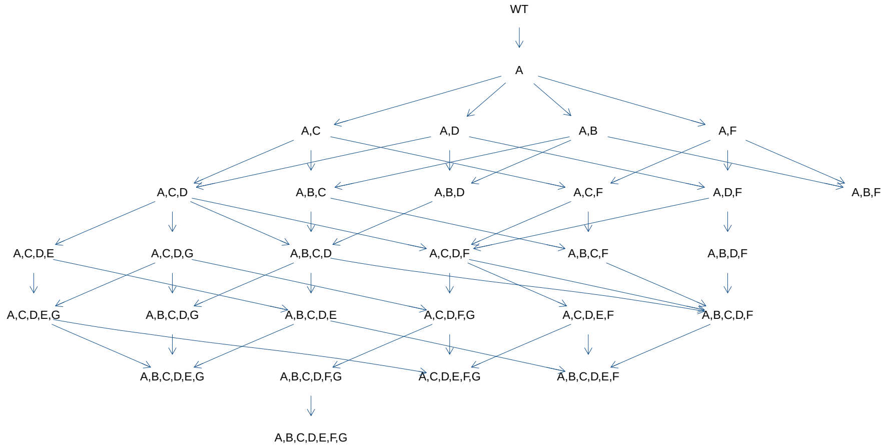

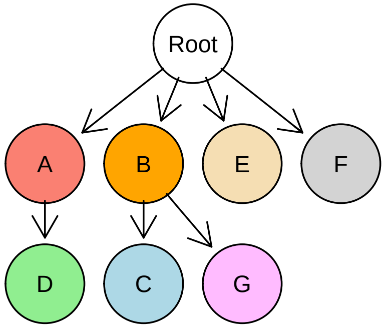


| | |
|---|---|
| *ID* | DDAFd9YlDa3wcgyX |
| *acc genotypes* | 60 |
| *peaks no back* | 8 |
| *removed edges* | 32 |
| *prop rem edges* | 0.2 |

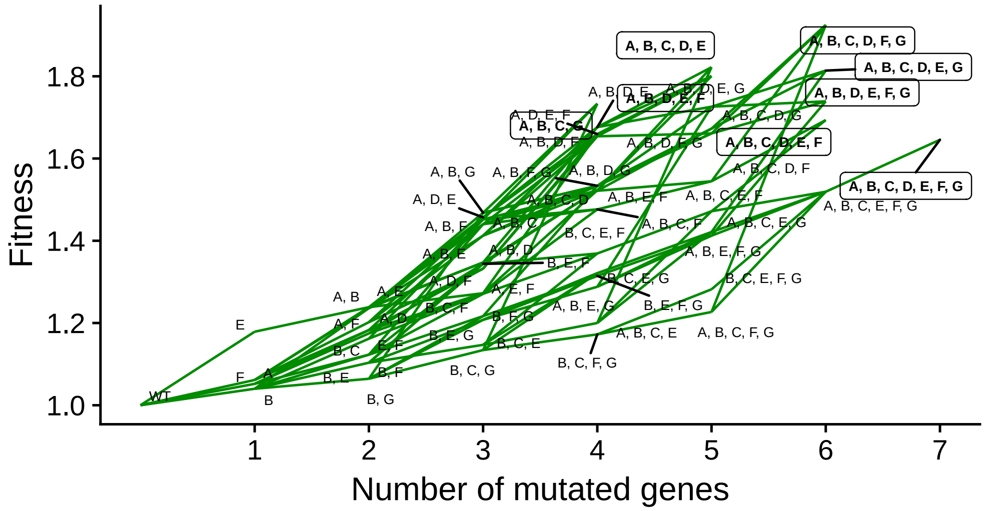


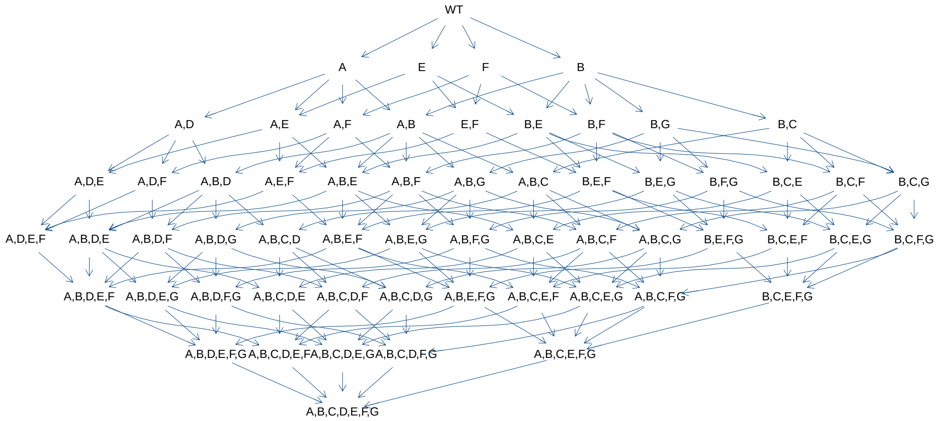


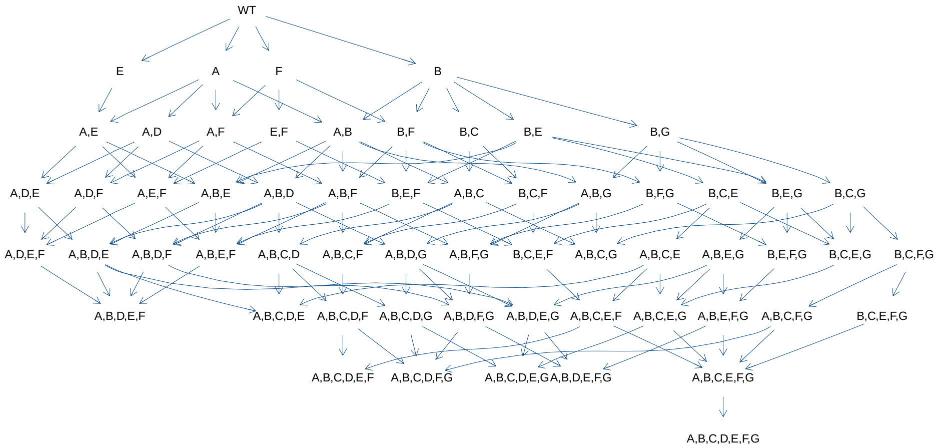

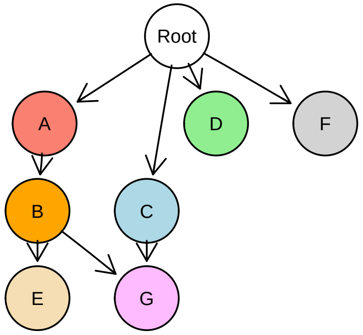


| | |
|---|---|
| *ID* | DE10Mak4NoN6QJYUY |
| *acc genotypes* | 40 |
| *peaks no back* | 4 |
| *removed edges* | 13 |
| *prop rem edges* | 0.141 |

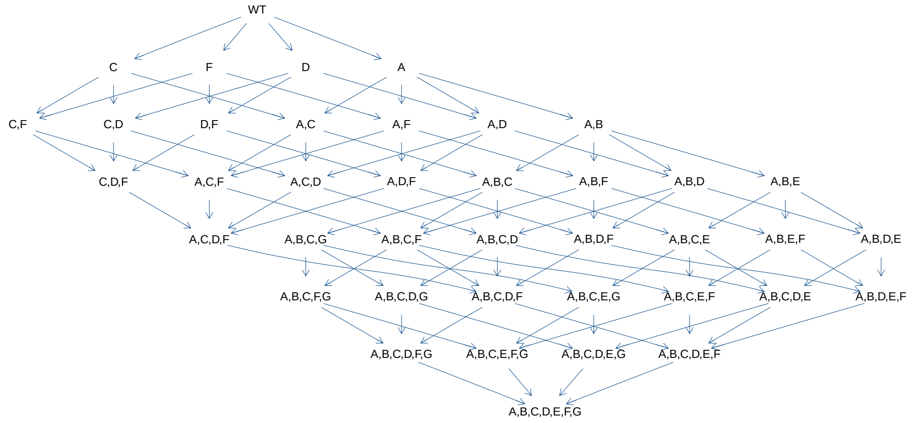


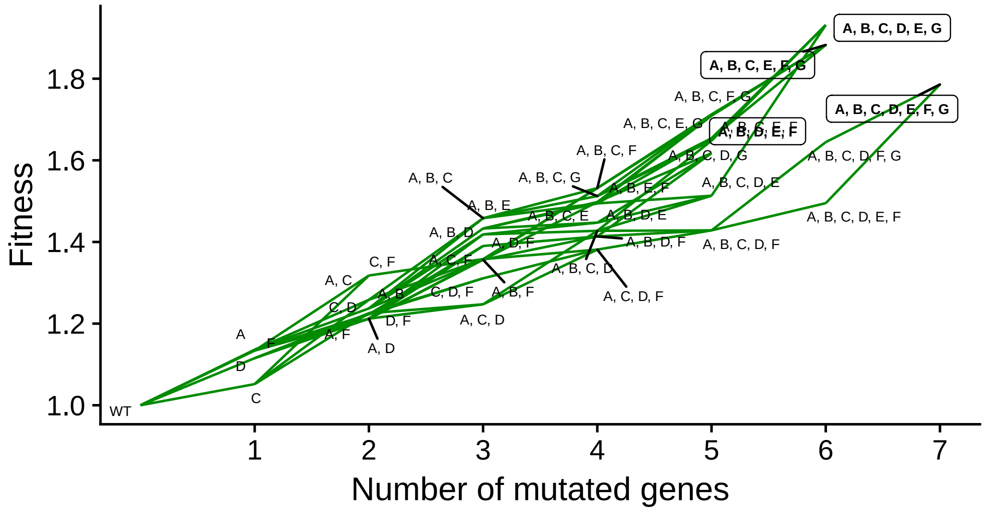


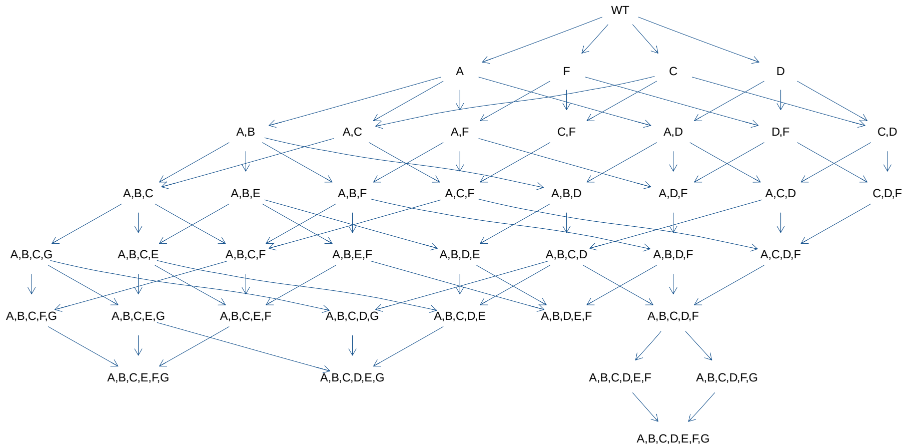

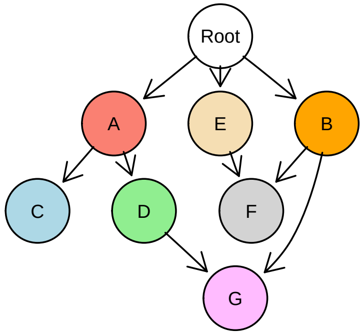


*ID* dJOynYb7GGPbx910F
*acc genotypes* 31
*peaks no back* 5
*removed edges* 12
*prop rem edges* 0.19

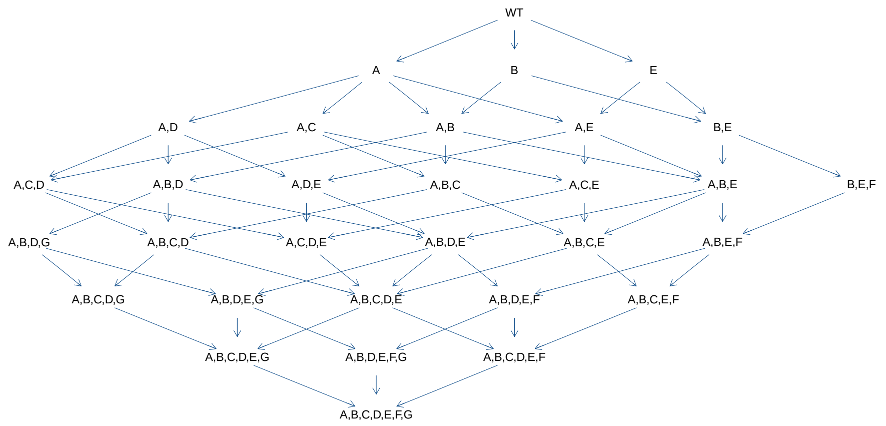


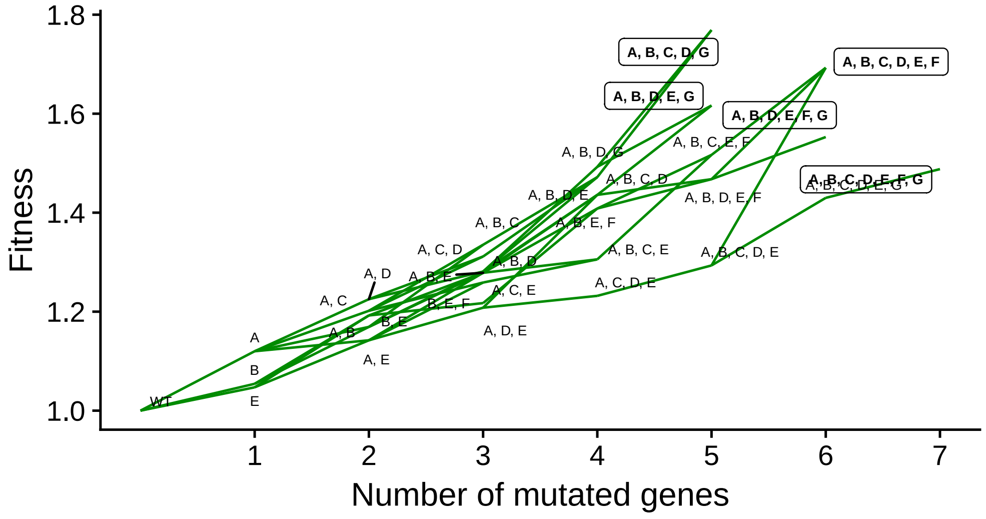


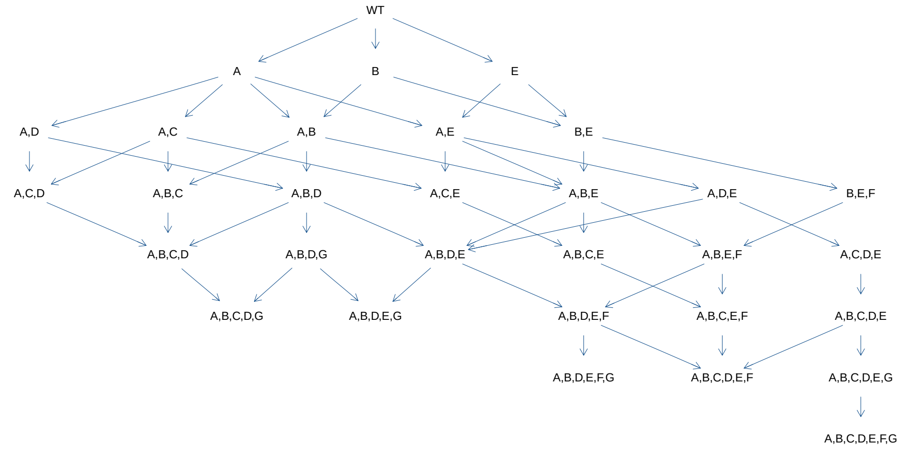

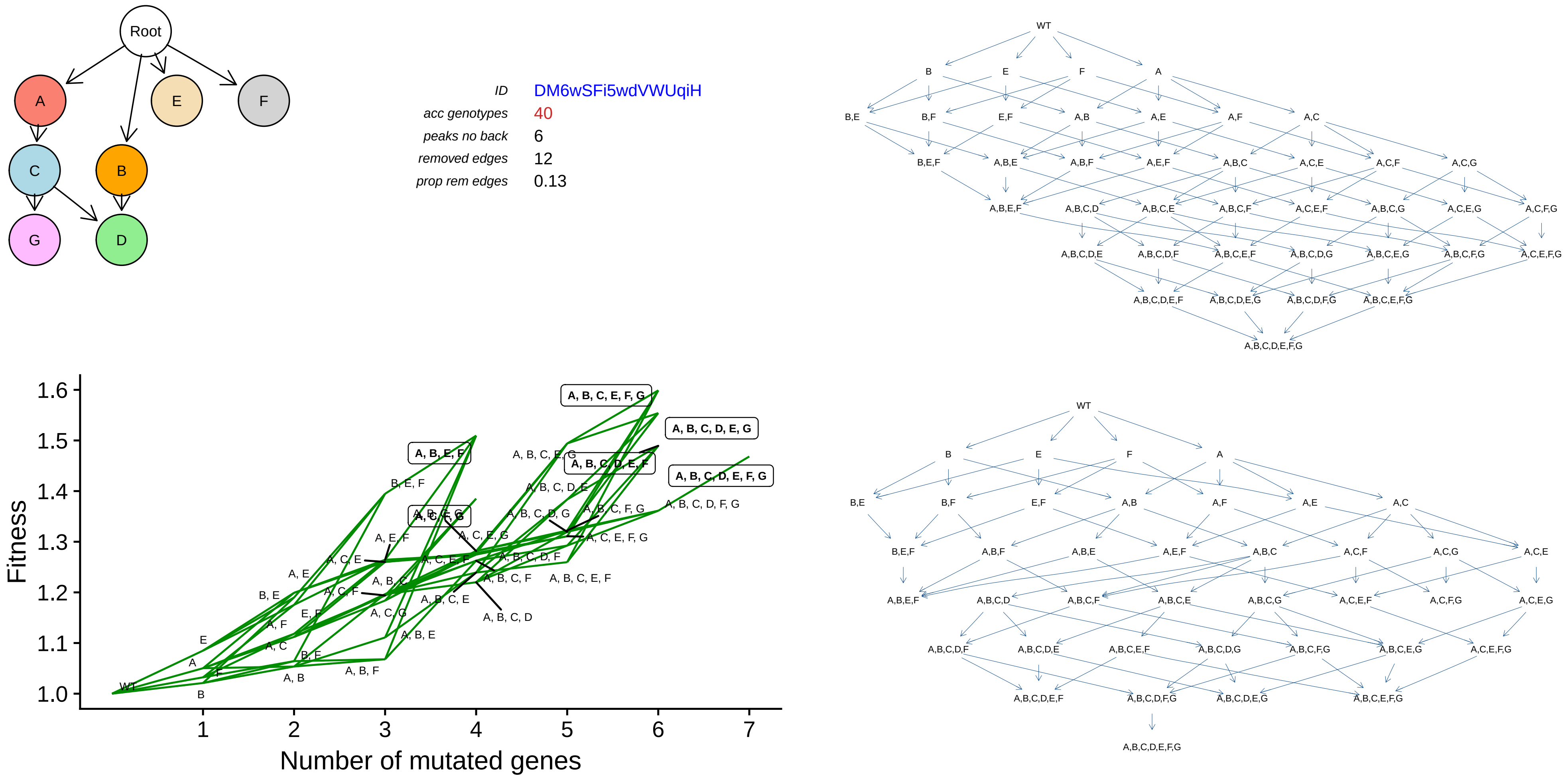

Root
A
E
F
C
B
G
D
ID DM6wSFi5wdVWUqiH
acc genotypes 40
peaks no back 6
removed edges 12
prop rem edges 0.13
Fitness
Number of mutated genes
A, B, C, E, F, G
A, B, C, D, E, G
A, B, C, D, E, F
A, B, C, D, E, F, G
A, B, E, F
A, C, F, G
WT

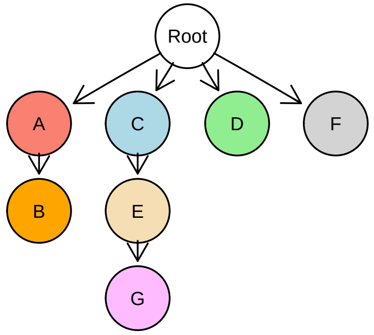


| | |
|---|---|
| *ID* | dO8v6amgefufggoE |
| *acc genotypes* | 48 |
| *peaks no back* | 7 |
| *removed edges* | 24 |
| *prop rem edges* | 0.207 |

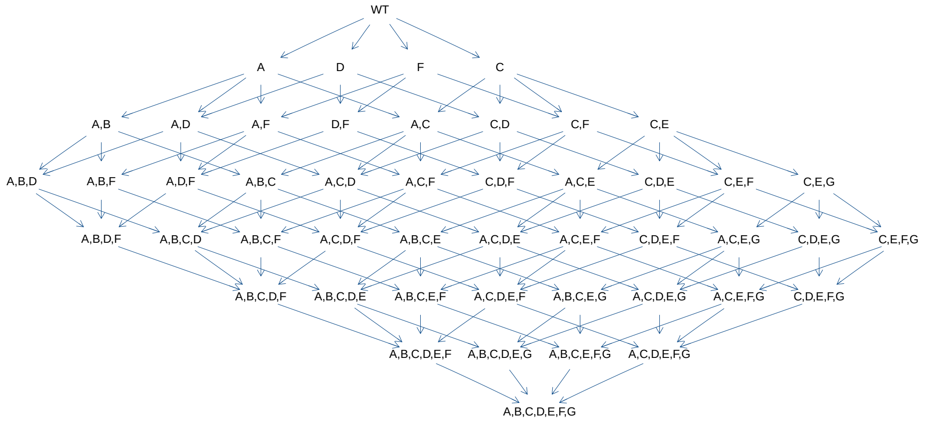


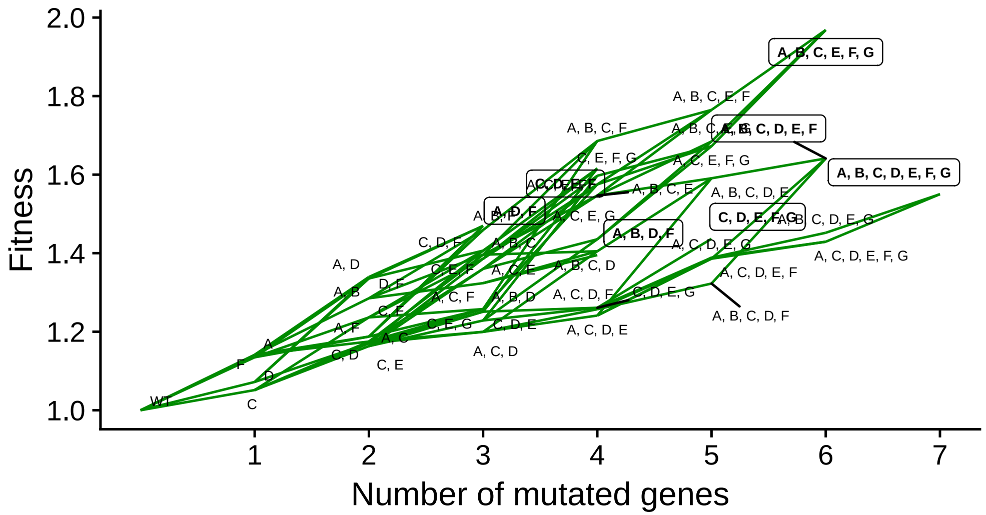


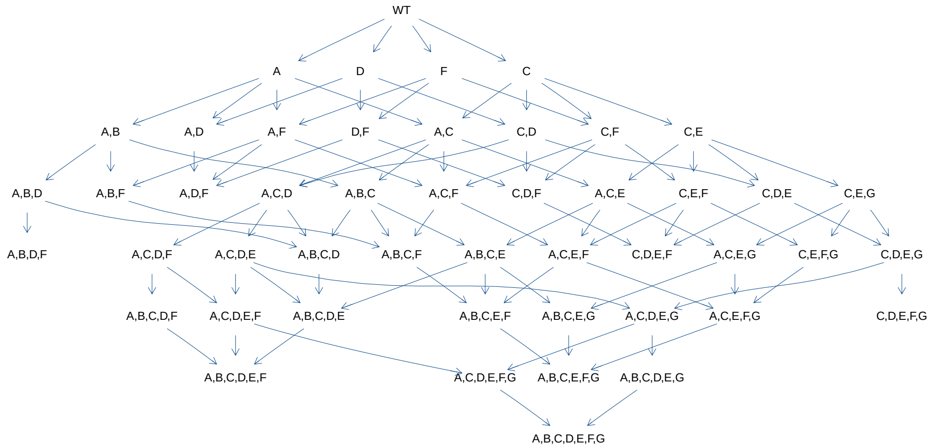

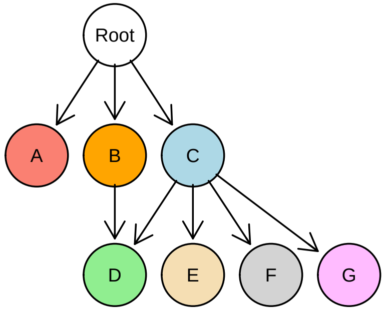


| | |
|---|---|
| *ID* | dPDOh3Us3MOolJnT |
| *acc genotypes* | 52 |
| *peaks no back* | 9 |
| *removed edges* | 26 |
| *prop rem edges* | 0.191 |

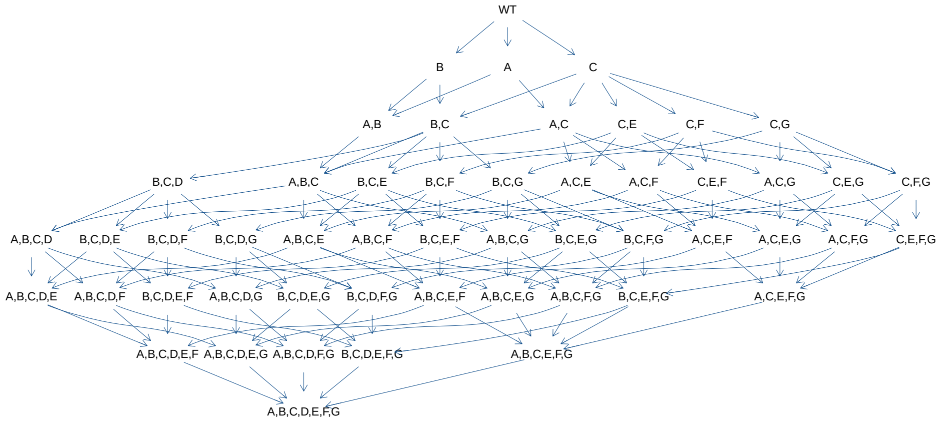

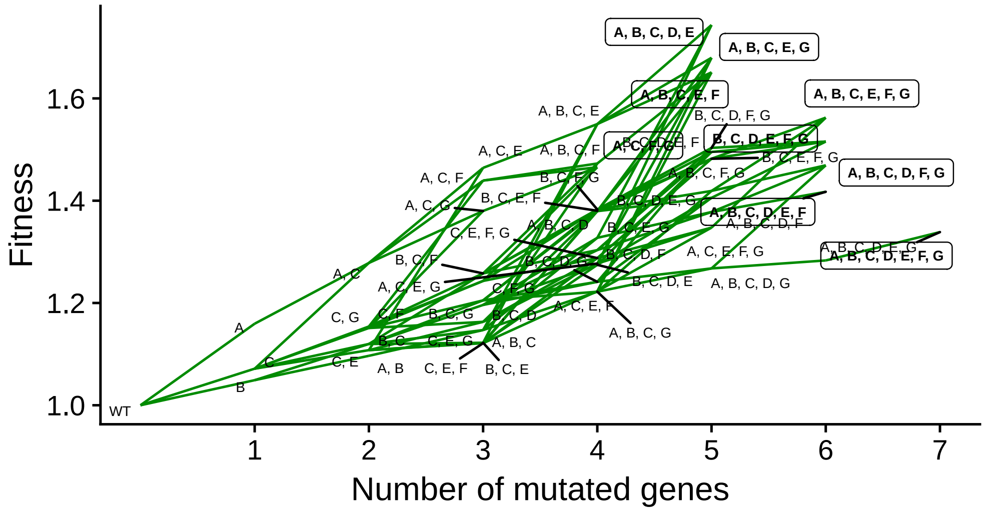


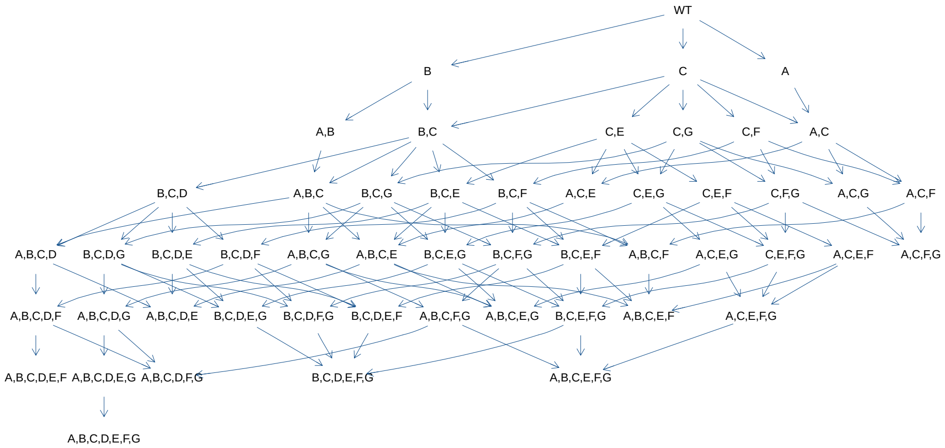

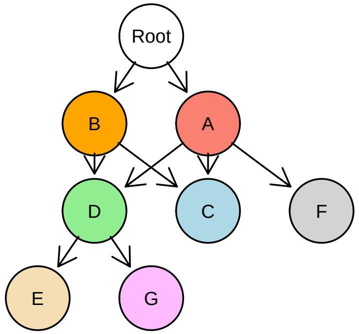


| | |
|---|---|
| *ID* | E4Ngwmi2x89iIda9 |
| *acc genotypes* | 24 |
| *peaks no back* | 4 |
| *removed edges* | 4 |
| *prop rem edges* | 0.087 |

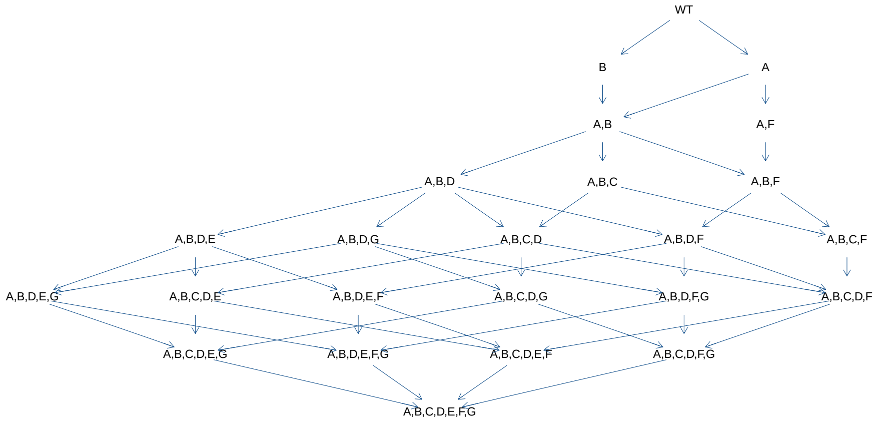


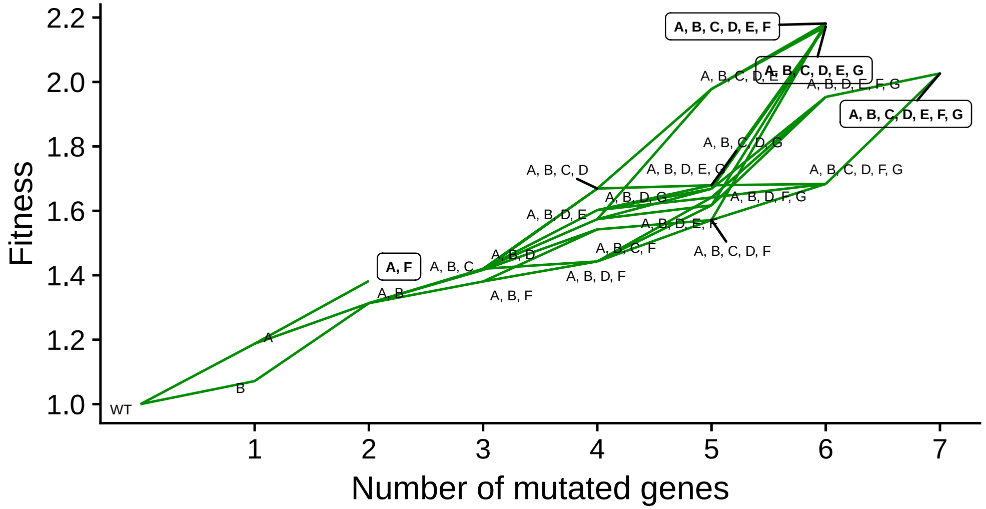


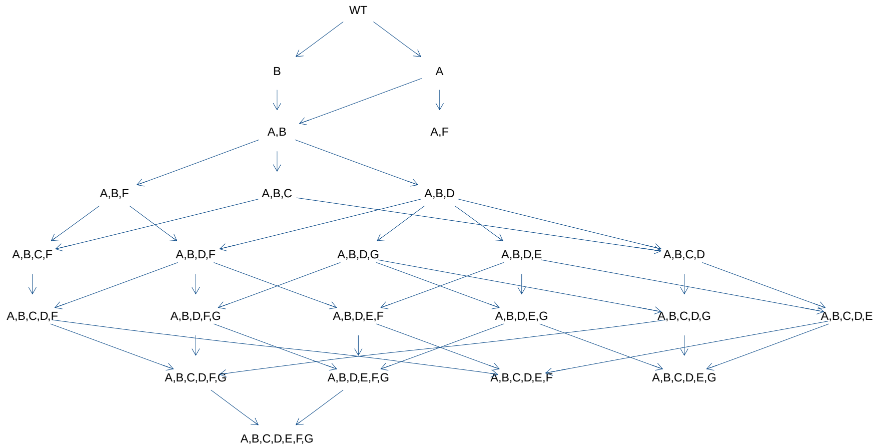

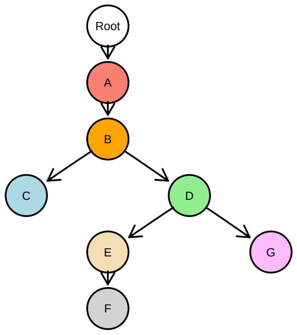


| | |
|---|---|
| *ID* | ejQdvM3QTzcrMUgC |
| *acc genotypes* | 16 |
| *peaks no back* | 4 |
| *removed edges* | 4 |
| *prop rem edges* | 0.16 |

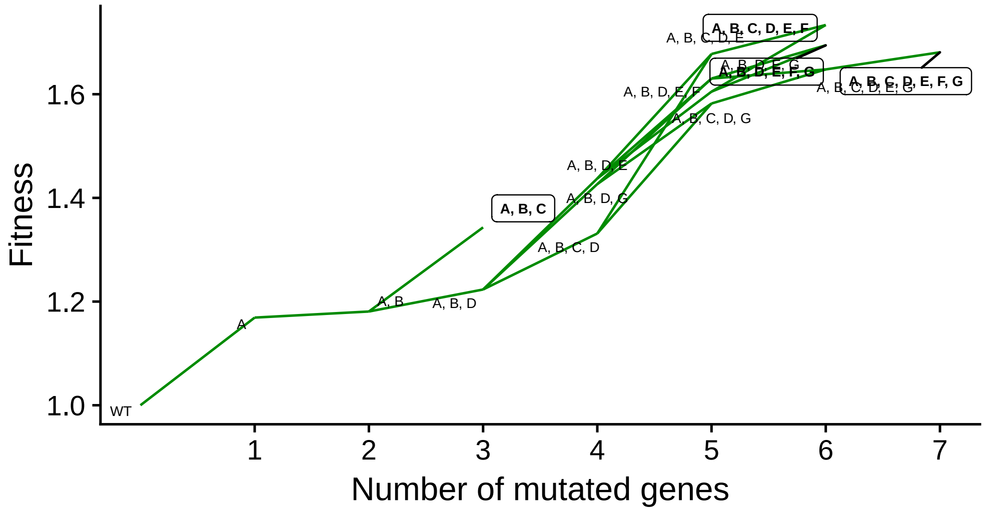


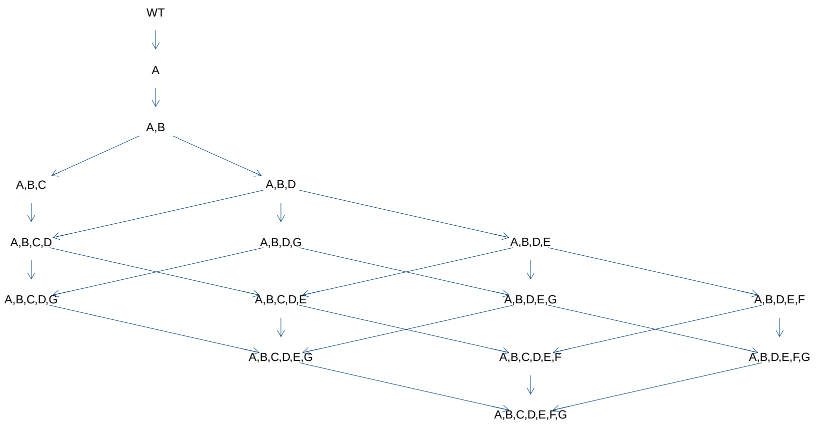


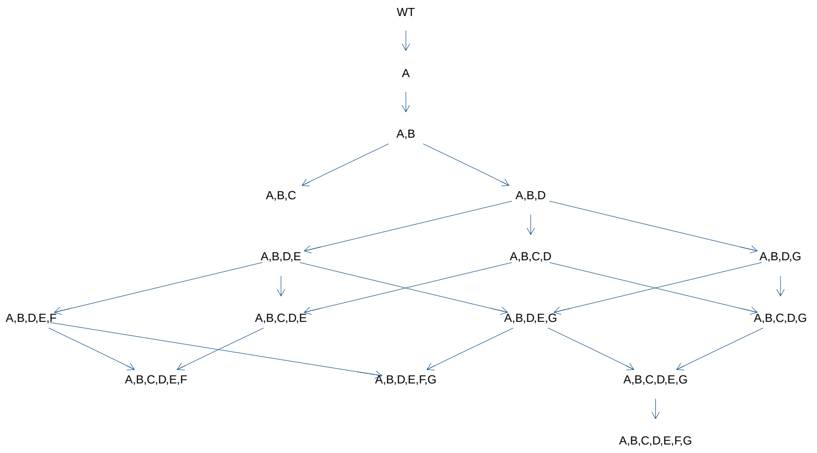

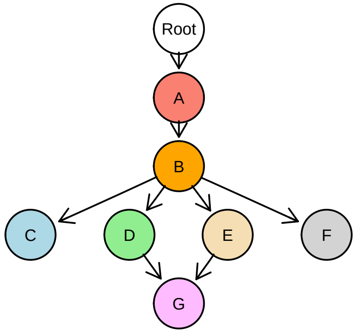


| | |
|---|---|
| *ID* | eNAunG4FXTpwgDbz |
| *acc genotypes* | 22 |
| *peaks no back* | 4 |
| *removed edges* | 9 |
| *prop rem edges* | 0.214 |

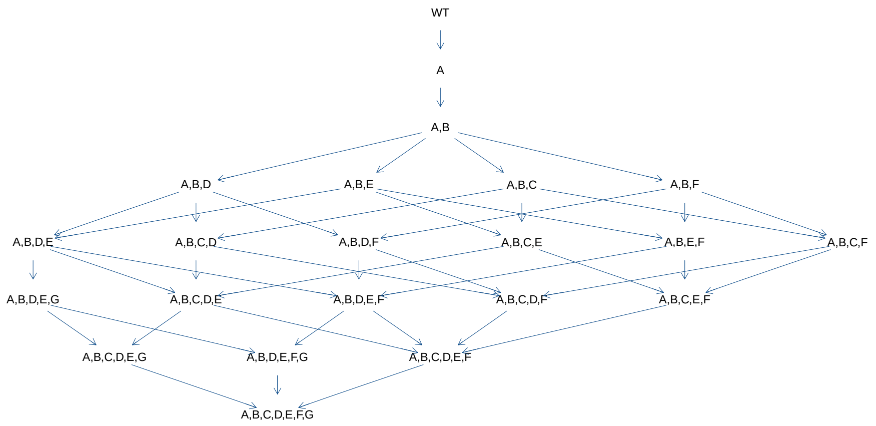


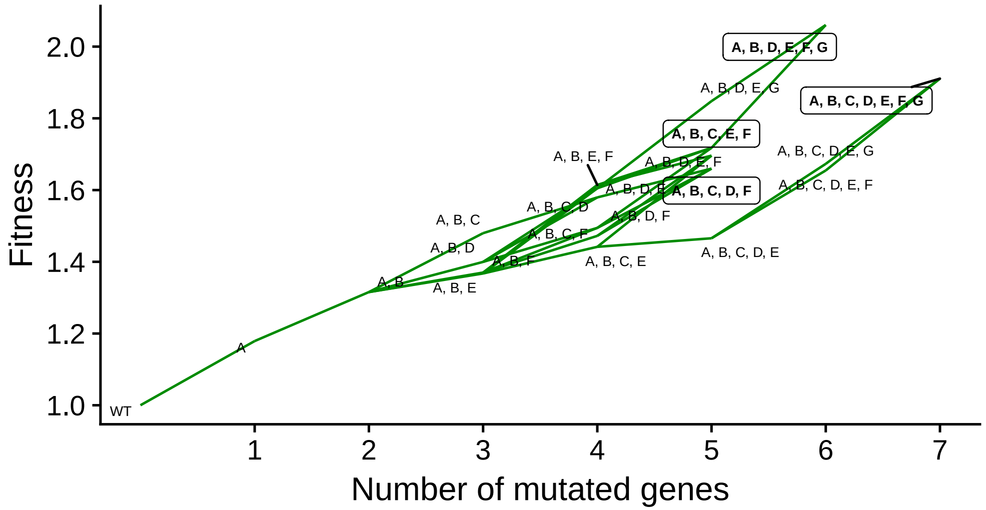


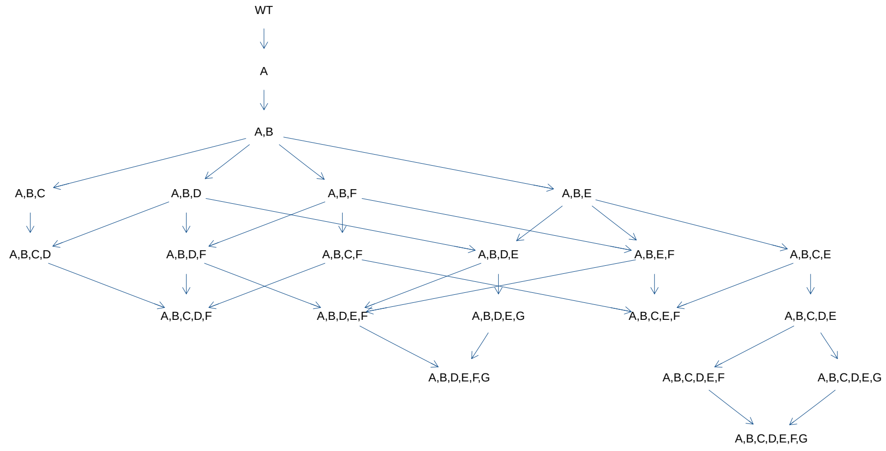

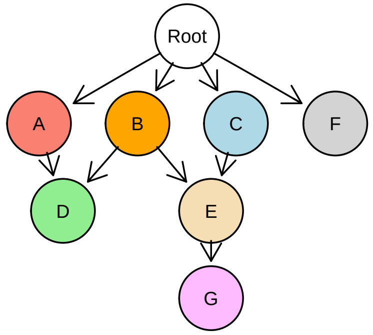


*ID* eO96nNtKZRtCYFxB
*acc genotypes* 32
*peaks no back* 6
*removed edges* 10
*prop rem edges* 0.152

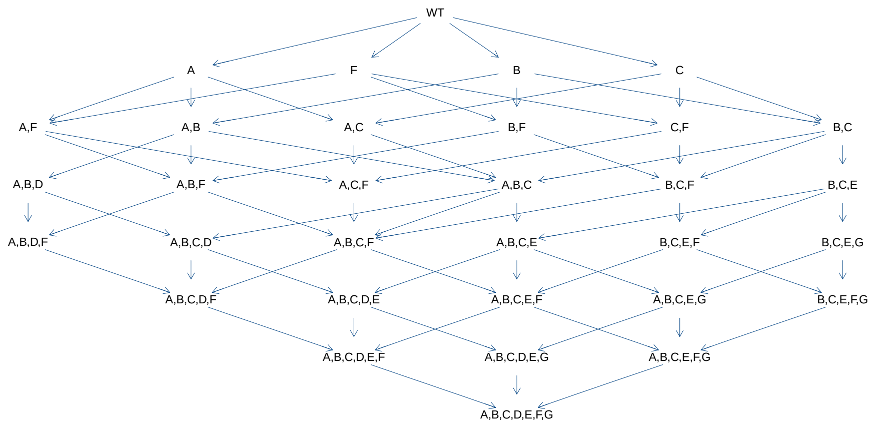


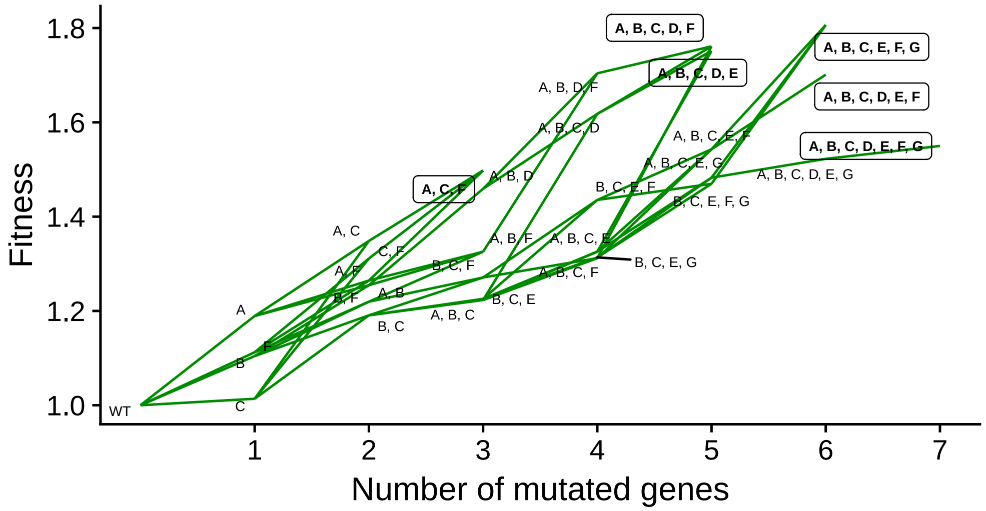


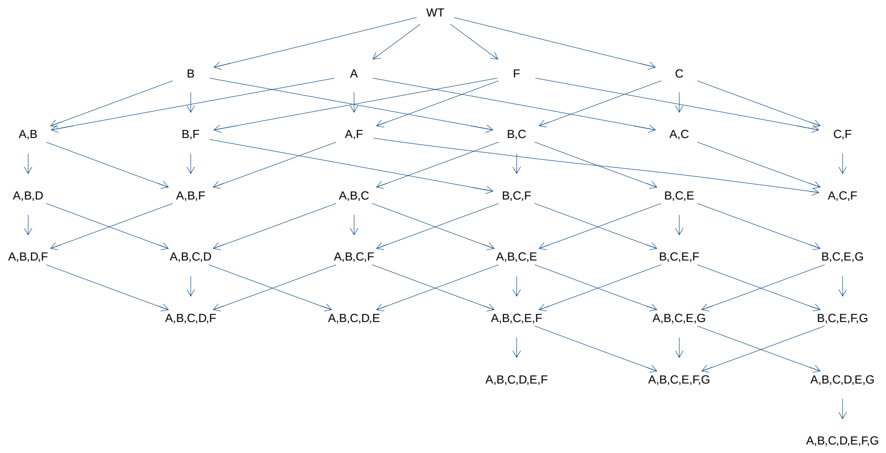

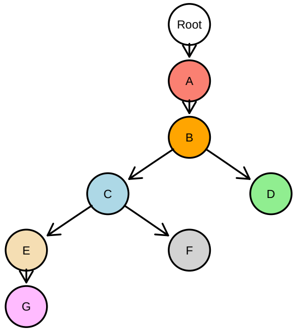


| | |
|---|---|
| *ID* | EtcHaQHBlLbIjaZQ |
| *acc genotypes* | 16 |
| *peaks no back* | 2 |
| *removed edges* | 2 |
| *prop rem edges* | 0.08 |

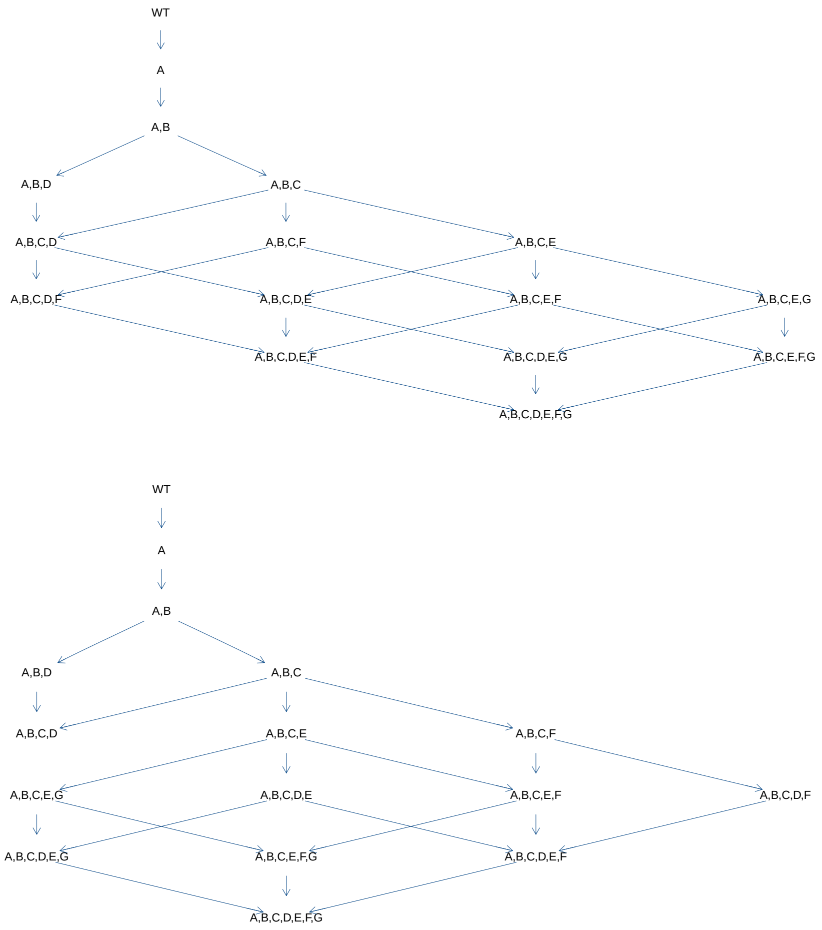


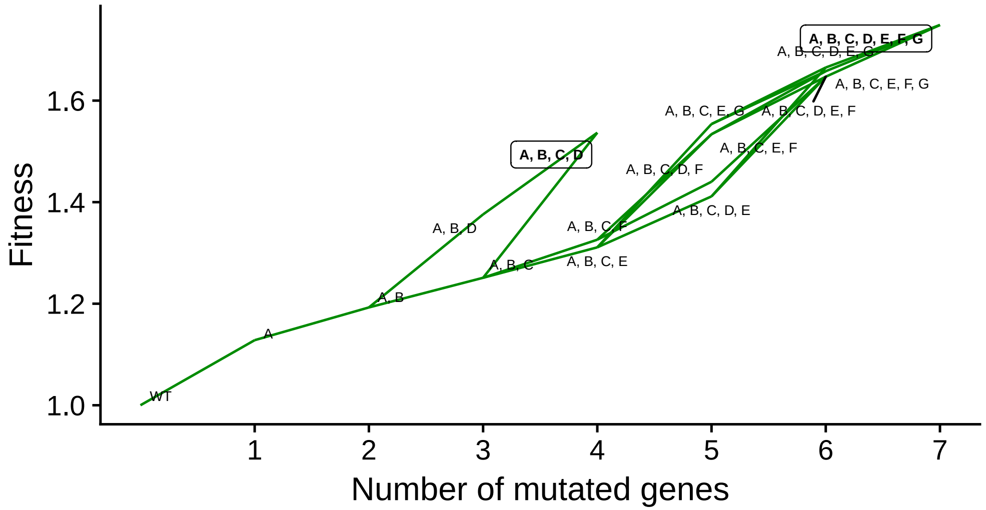

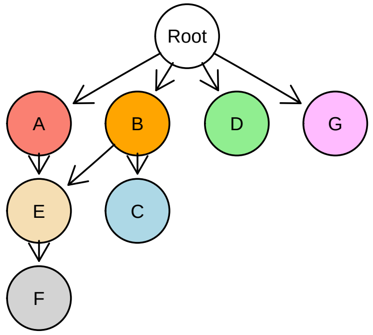


*ID* f103lT84L6WhInZWQ
*acc genotypes* 40
*peaks no back* 5
*removed edges* 13
*prop rem edges* 0.141

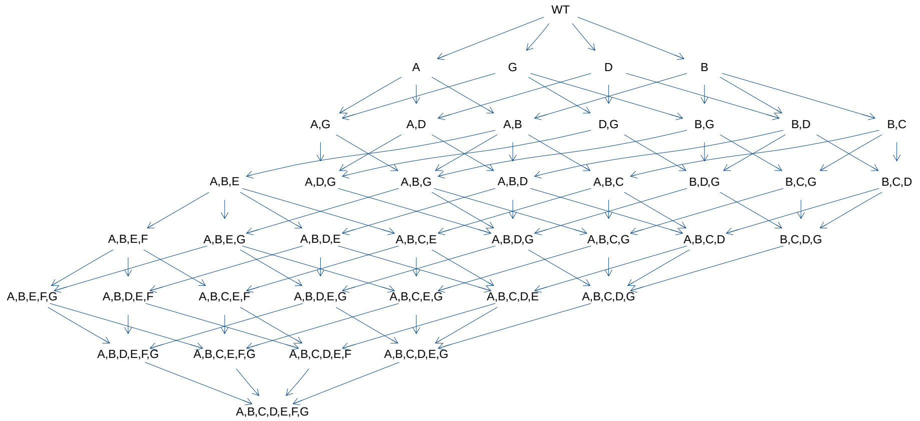


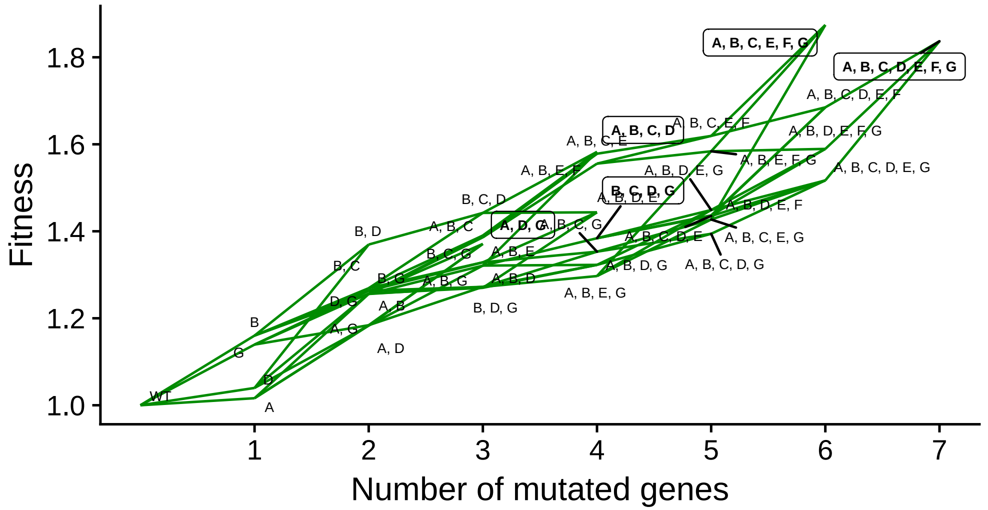


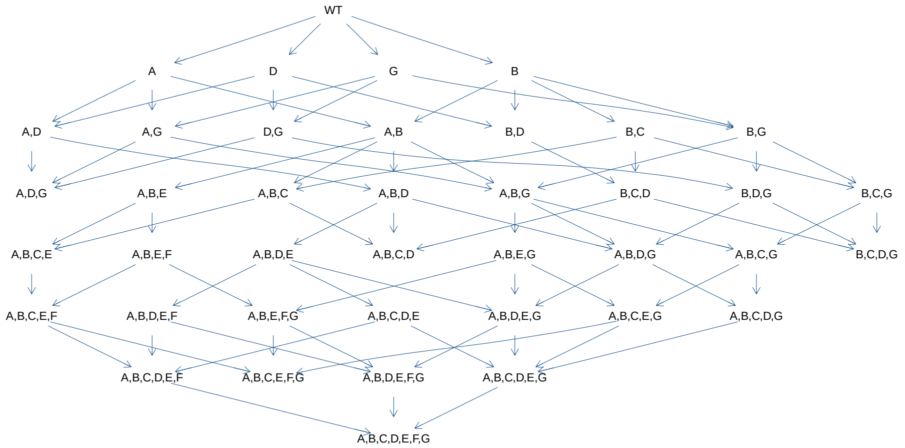

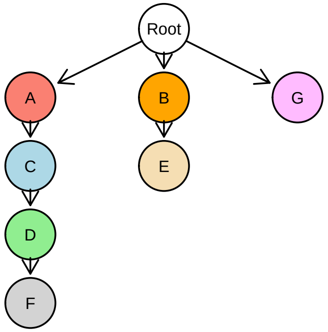


| | |
|---|---|
| *ID* | Fhypo3U5P7710LvXb |
| *acc genotypes* | 30 |
| *peaks no back* | 6 |
| *removed edges* | 8 |
| *prop rem edges* | 0.136 |

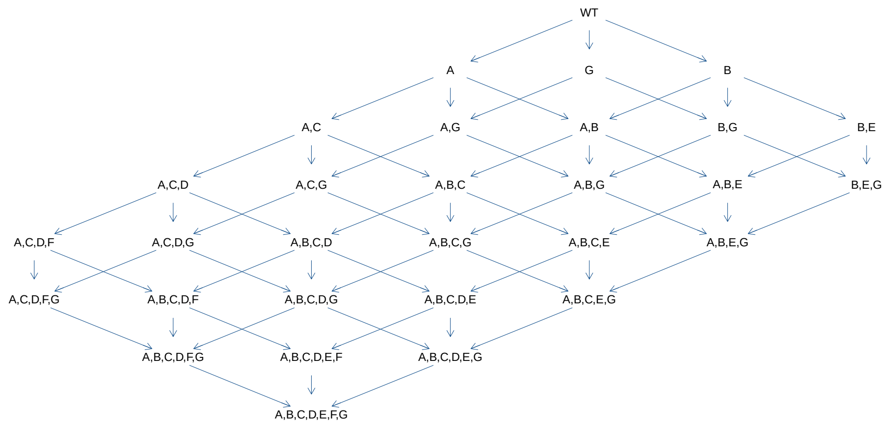


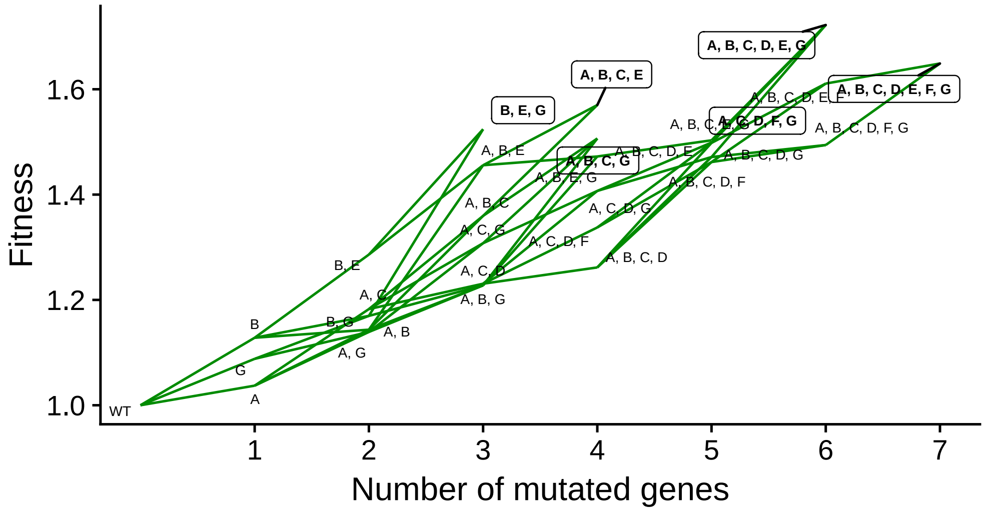


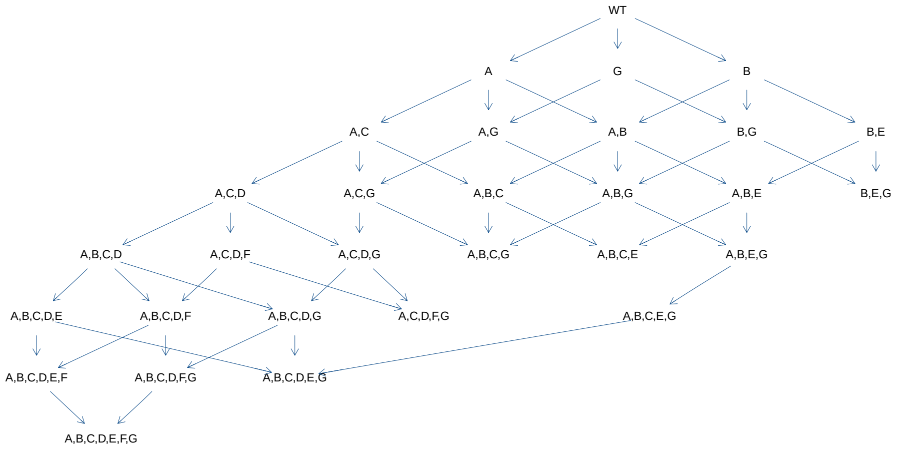

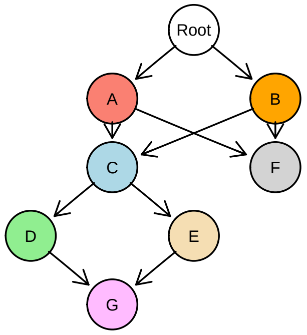


| | |
|---|---|
| *ID* | FJcdcp4ZT1UXlBCC |
| *acc genotypes* | 15 |
| *peaks no back* | 3 |
| *removed edges* | 3 |
| *prop rem edges* | 0.136 |

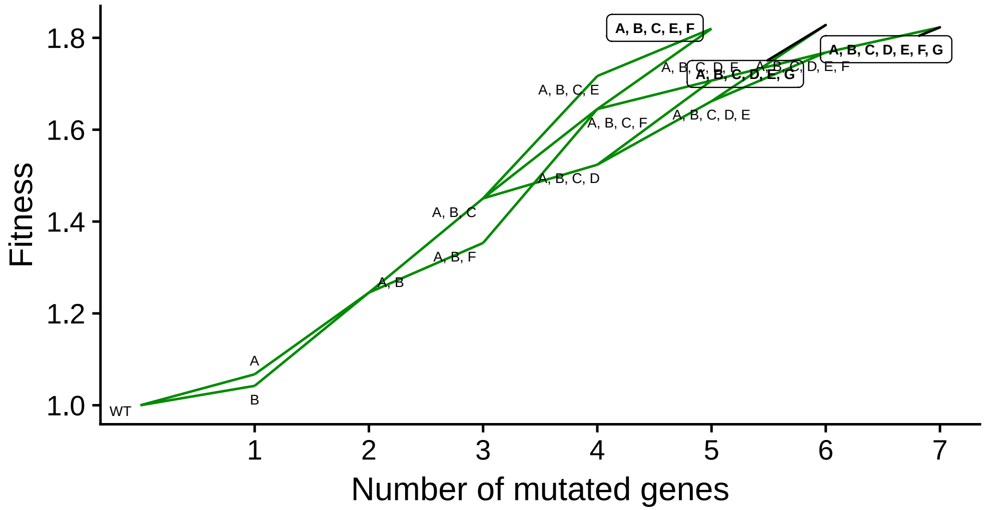


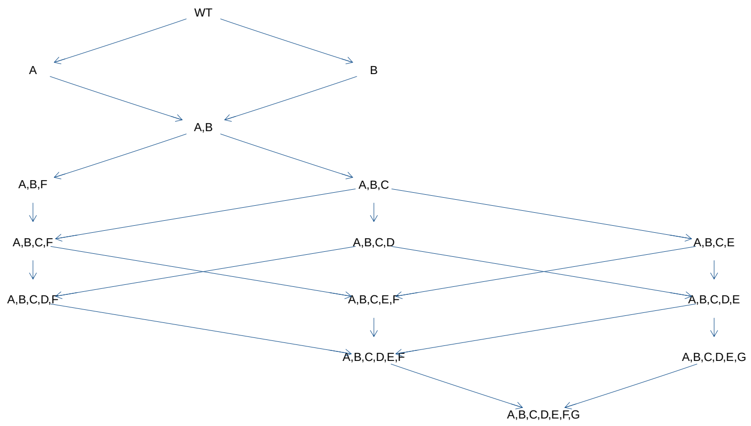


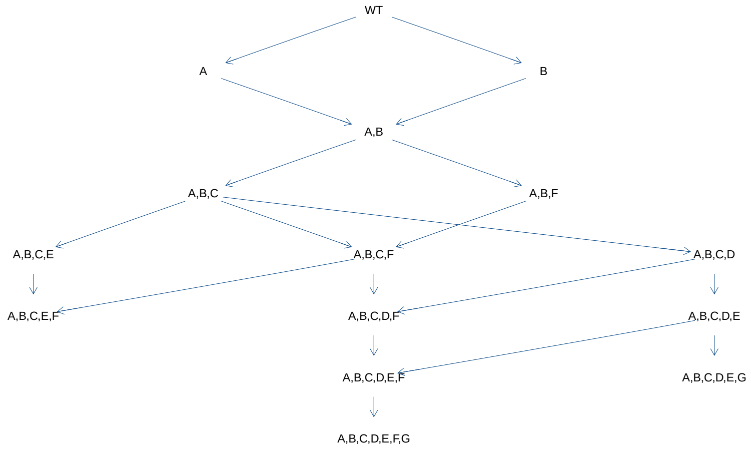

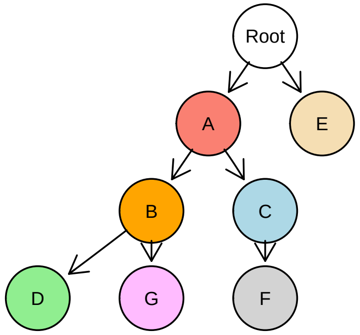


| | |
|---|---|
| *ID* | FMNyhwmR9SOSPPYf |
| *acc genotypes* | 32 |
| *peaks no back* | 6 |
| *removed edges* | 15 |
| *prop rem edges* | 0.221 |

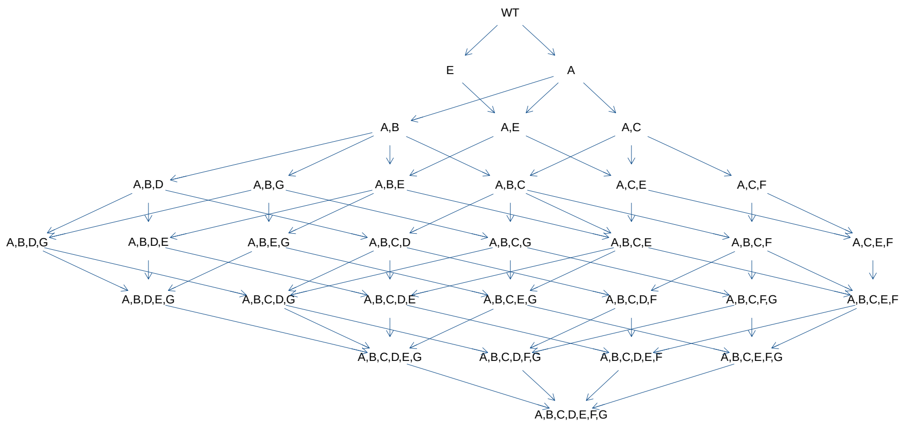


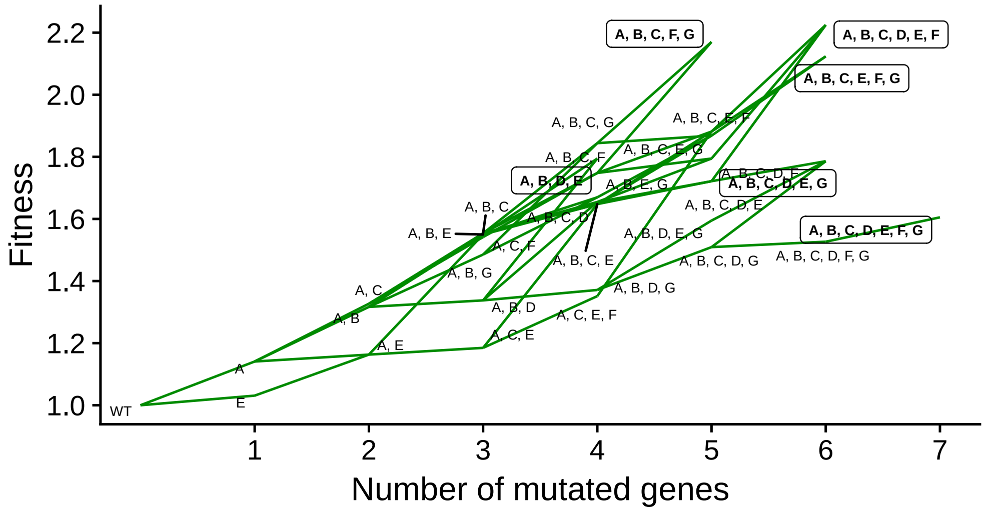


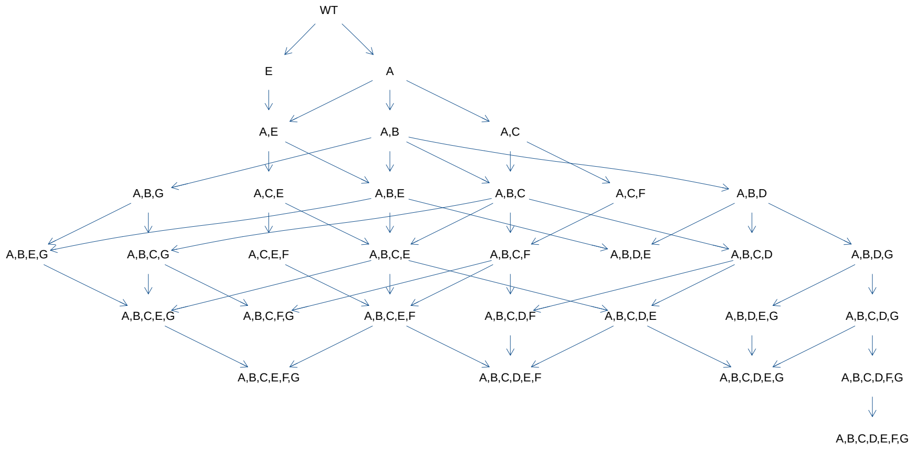

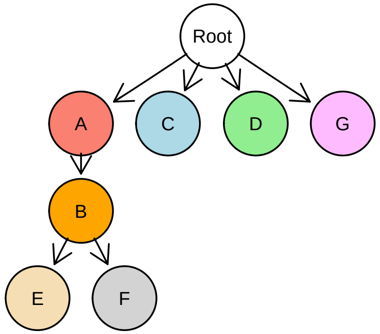


| | |
|---|---|
| *ID* | FTNjYXrURUcCxqz9 |
| *acc genotypes* | 48 |
| *peaks no back* | 9 |
| *removed edges* | 27 |
| *prop rem edges* | 0.225 |

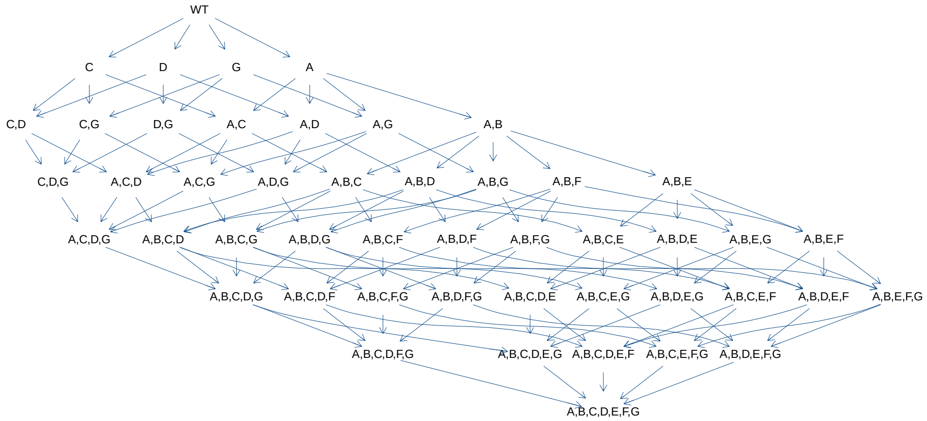

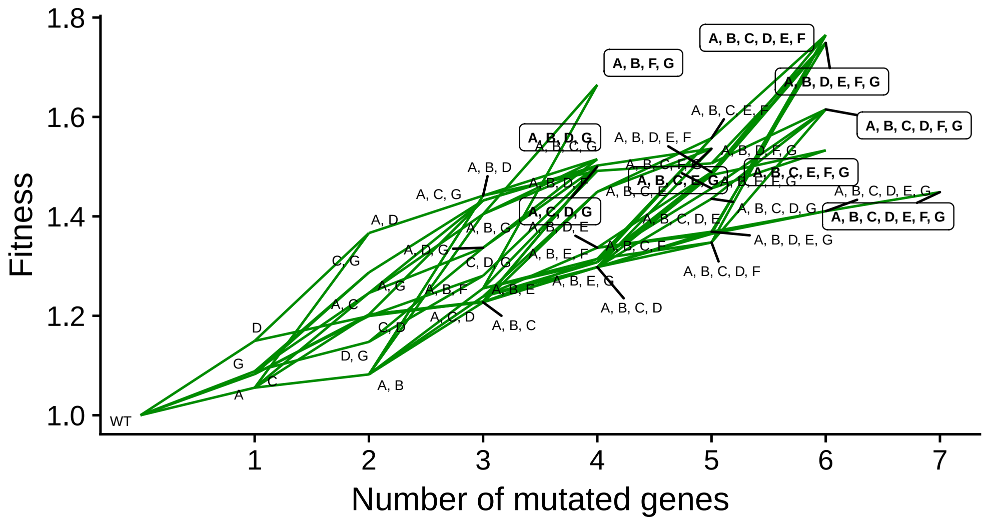


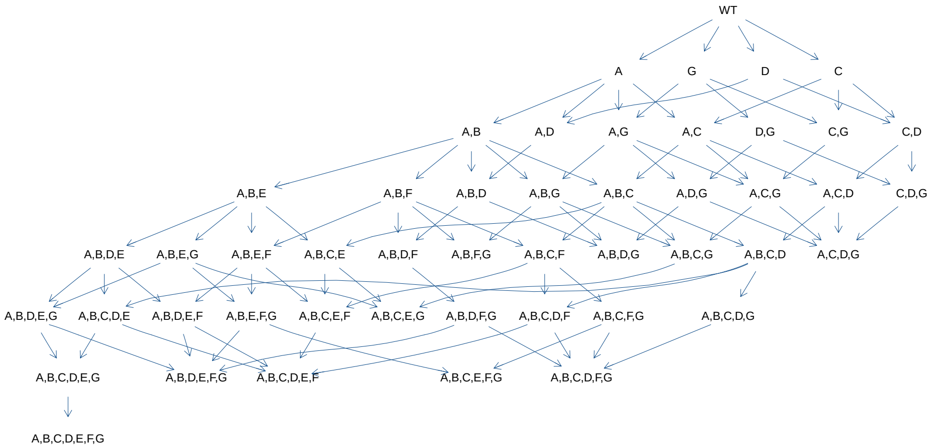

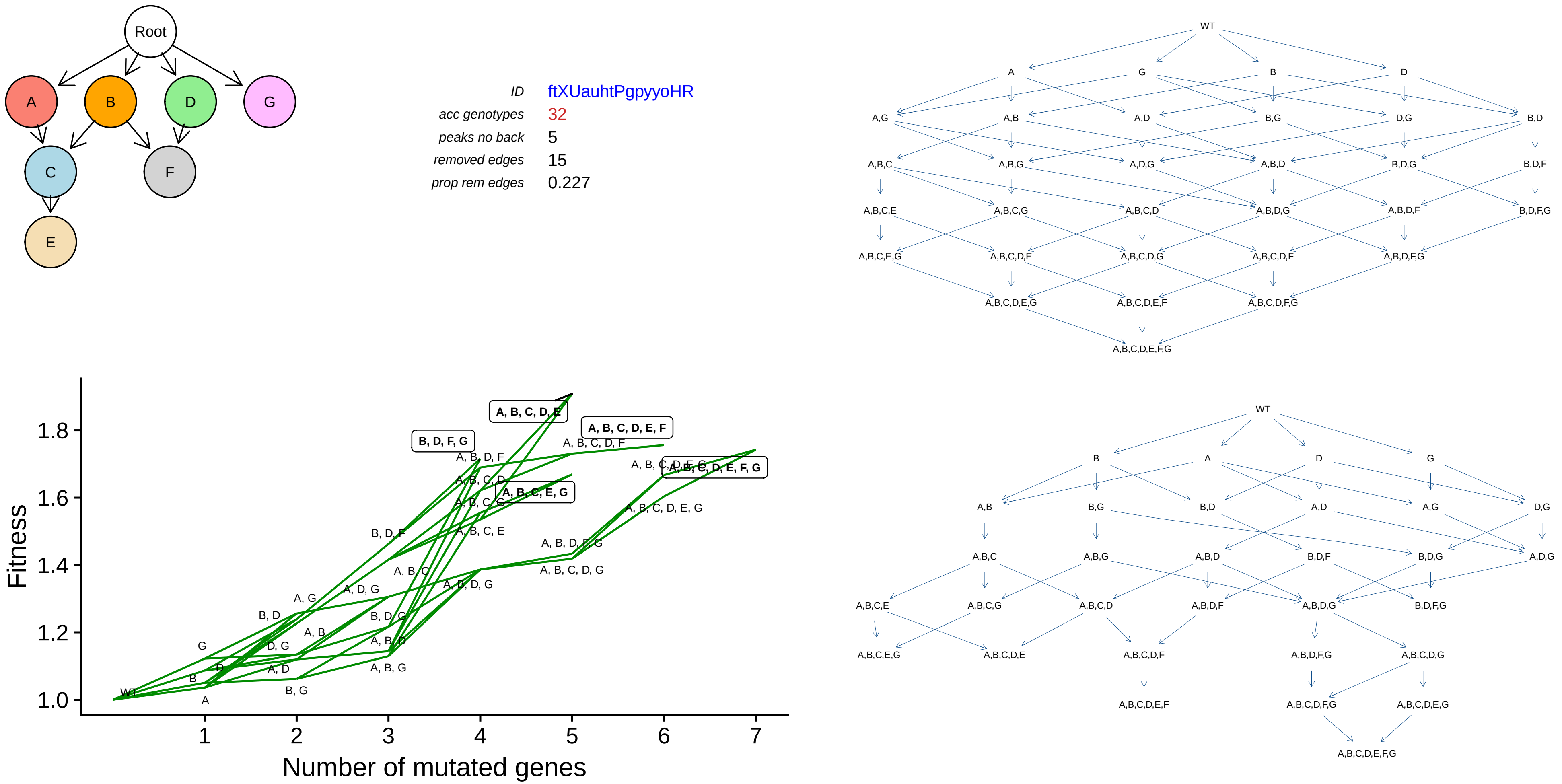

Root
A
B
D
G
C
F
E
ID ftXUauhtPgpyyoHR
acc genotypes 32
peaks no back 5
removed edges 15
prop rem edges 0.227
Fitness
Number of mutated genes
A, B, C, D, E
B, D, F, G
A, B, C, D, E, F
A, B, C, E, G
A, B, C, D, E, F, G
WT

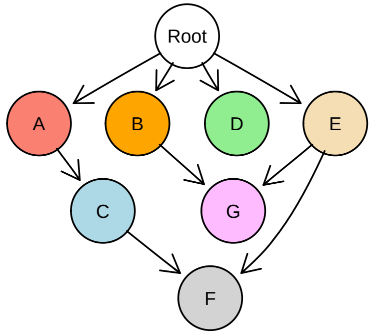


| | |
|---|---|
| *ID* | fW757Yty4nJQzn3B |
| *acc genotypes* | 36 |
| *peaks no back* | 5 |
| *removed edges* | 10 |
| *prop rem edges* | 0.128 |

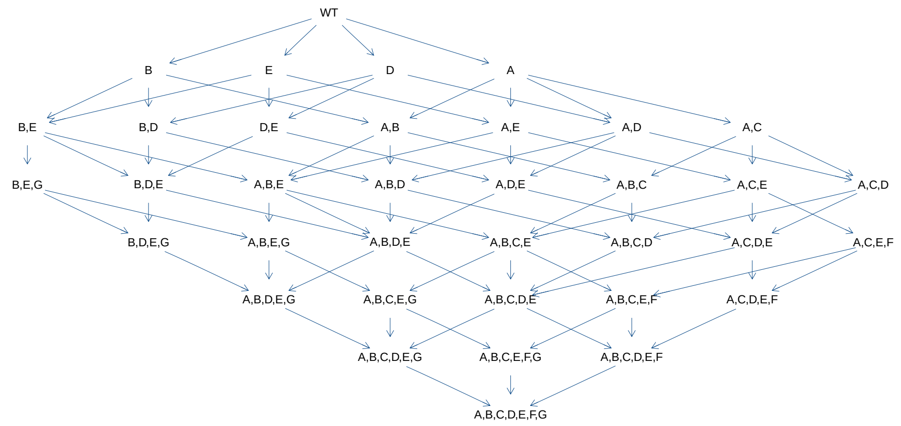


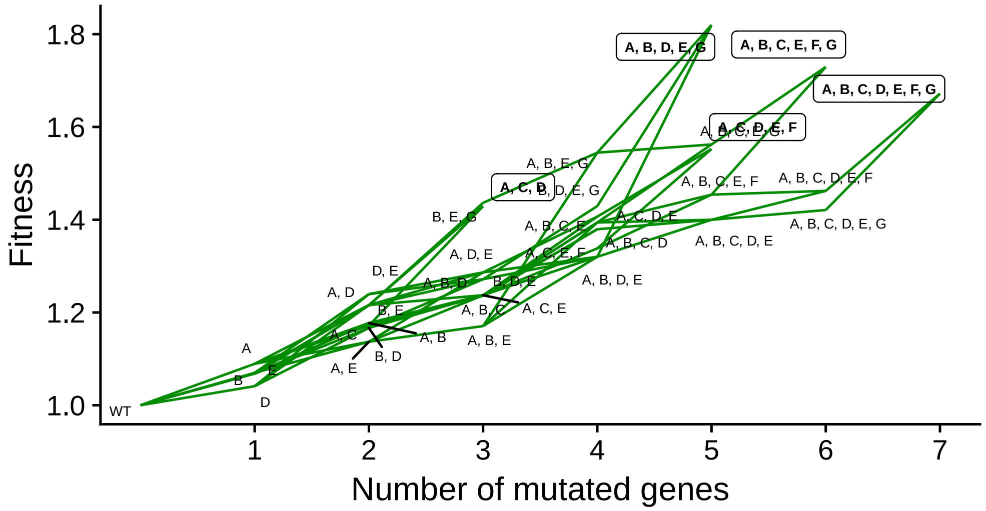


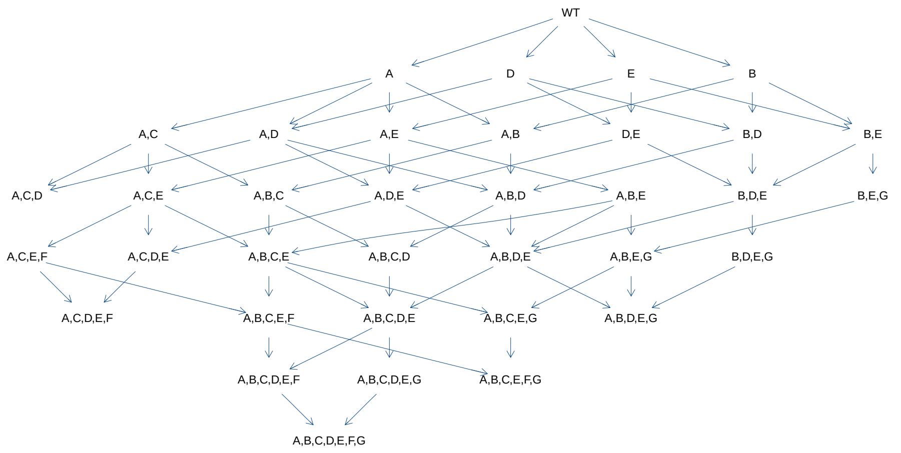

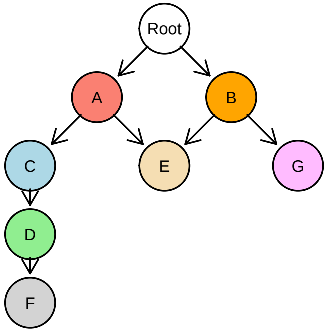


| | |
|---|---|
| *ID* | g38nzkCLq72TPfvC |
| *acc genotypes* | 23 |
| *peaks no back* | 5 |
| *removed edges* | 8 |
| *prop rem edges* | 0.2 |

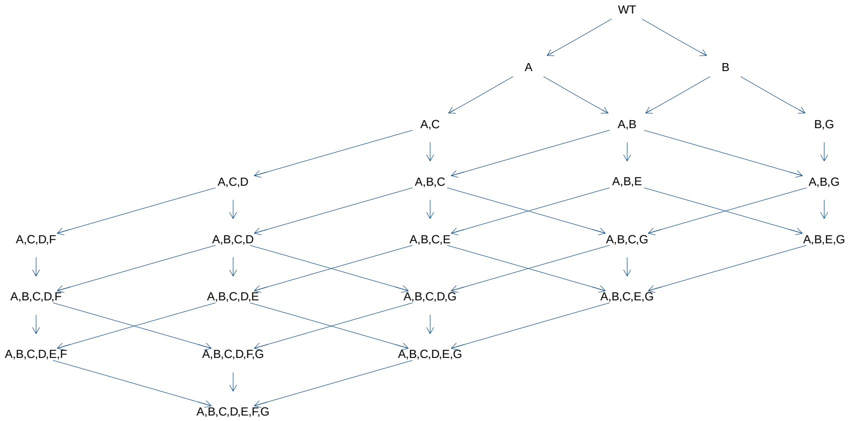


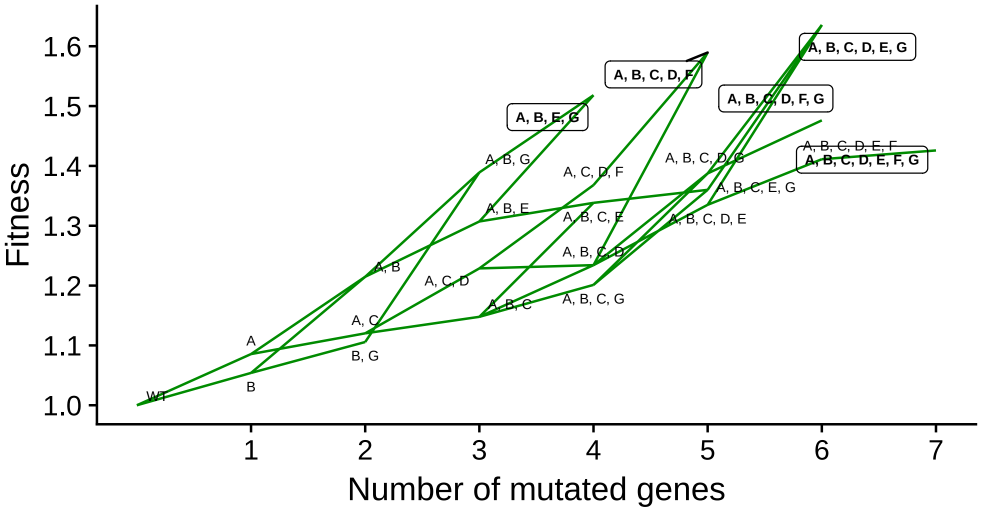


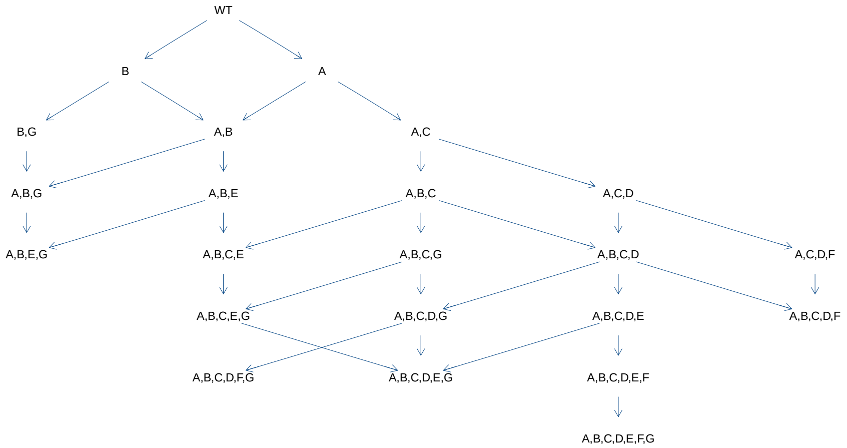

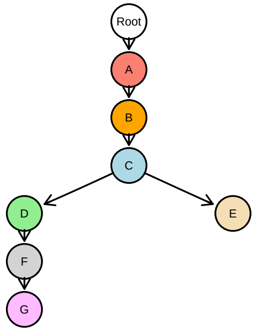


| | |
|---|---|
| *ID* | GedZaWDeb1029Mf88 |
| *acc genotypes* | 11 |
| *peaks no back* | 2 |
| *removed edges* | 2 |
| *prop rem edges* | 0.154 |

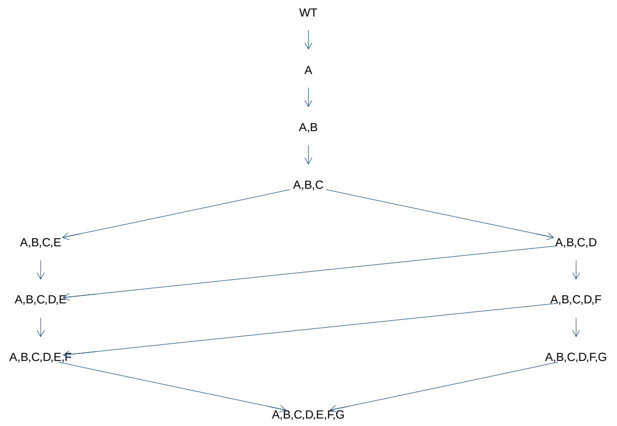


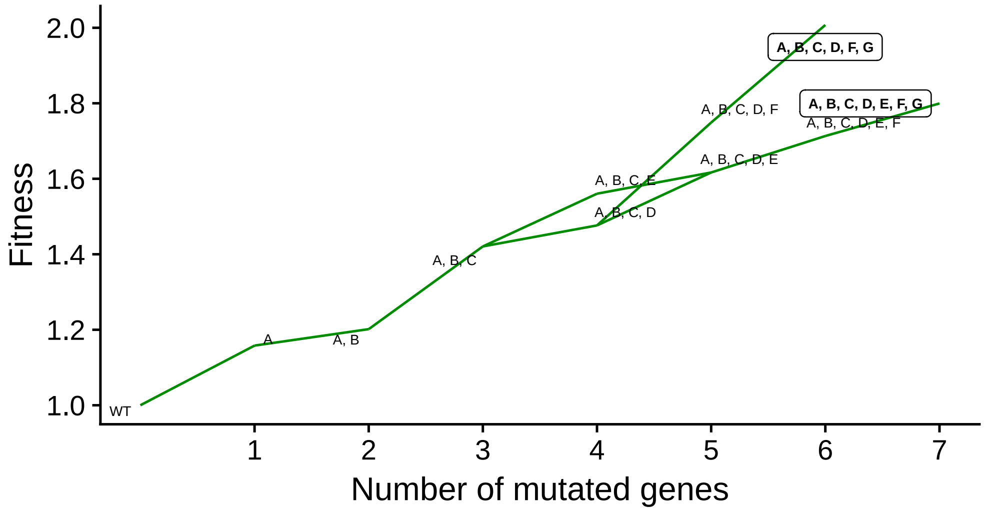


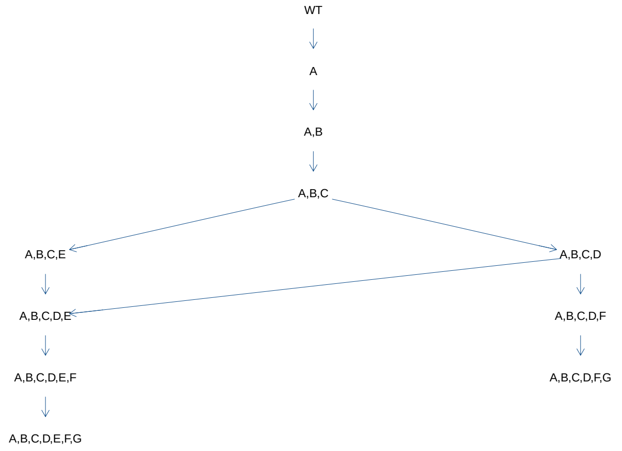

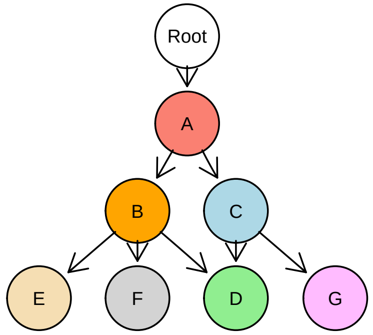


| | |
|---|---|
| *ID* | gfkgOLnyr7gjNQkC |
| *acc genotypes* | 24 |
| *peaks no back* | 5 |
| *removed edges* | 10 |
| *prop rem edges* | 0.217 |

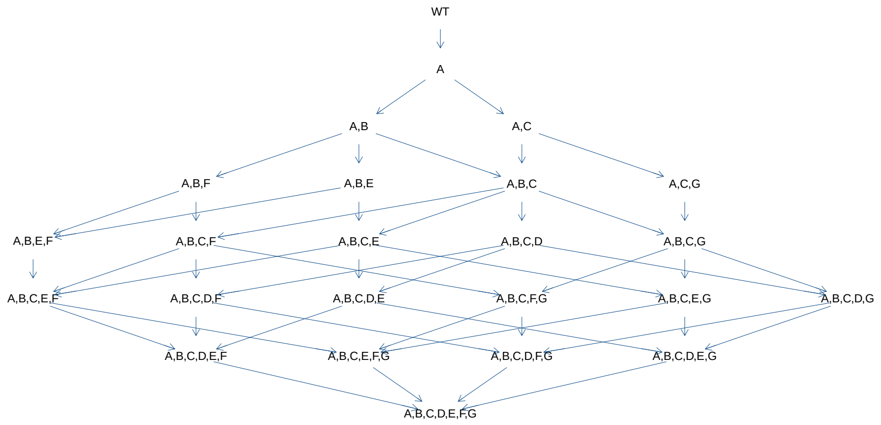


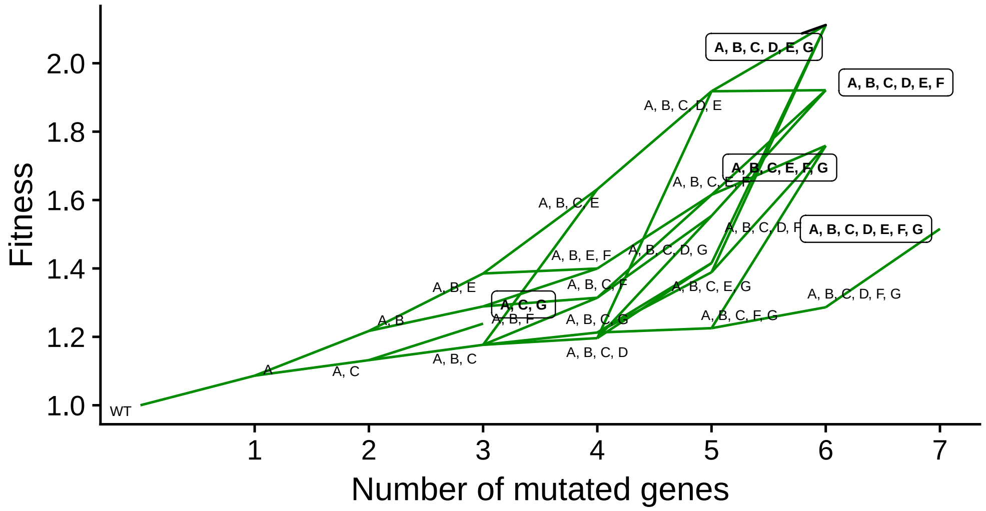


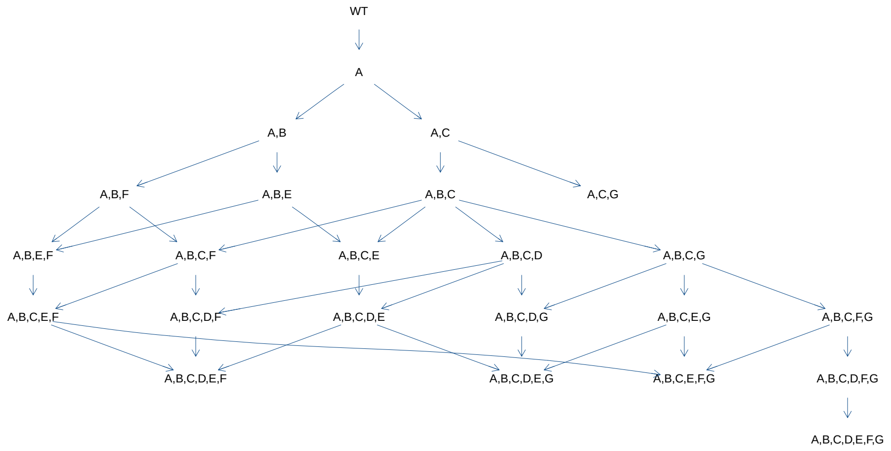

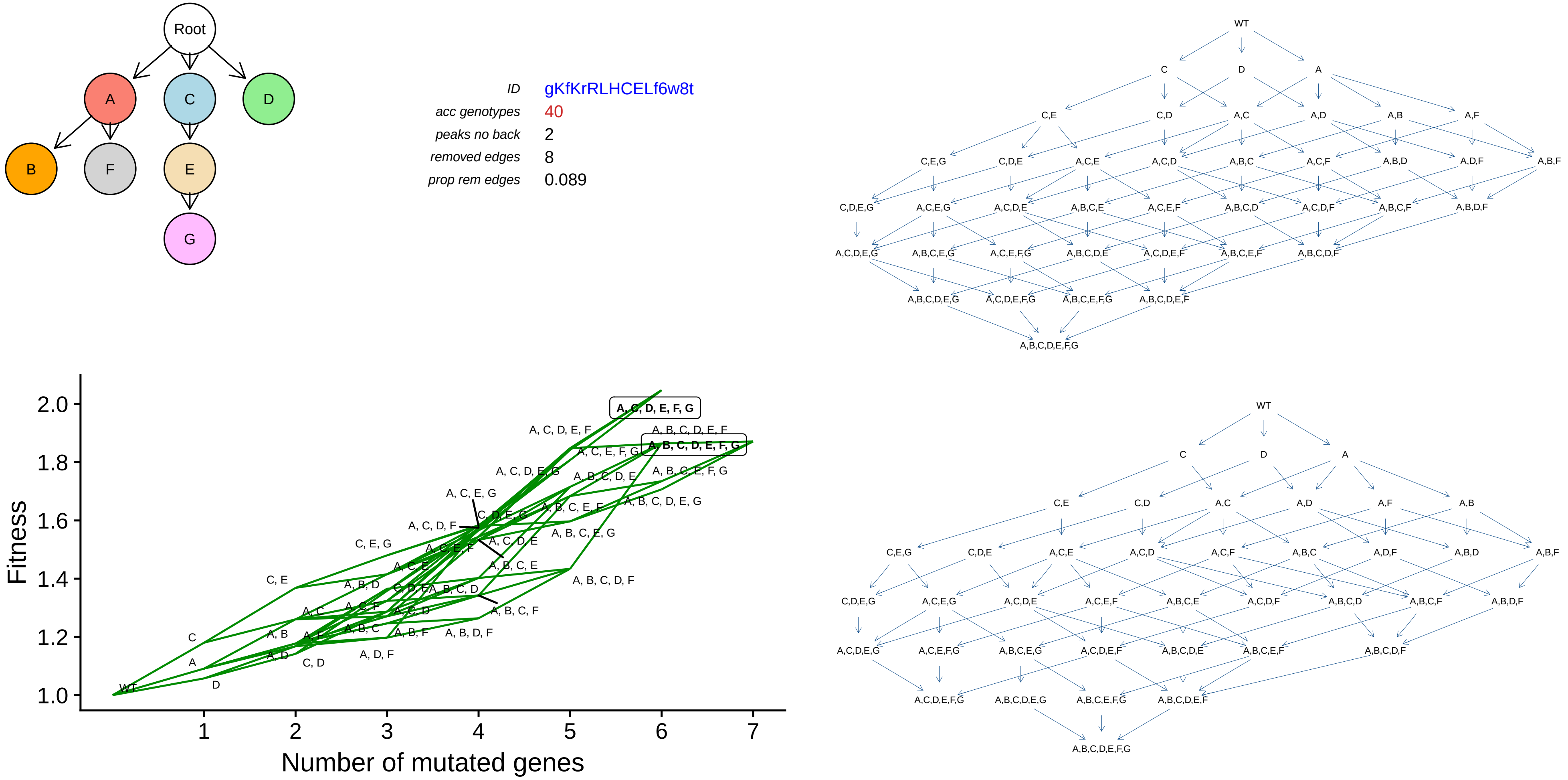
Root
A
C
D
B
F
E
G
ID gKfKrRLHCELf6w8t
acc genotypes 40
peaks no back 2
removed edges 8
prop rem edges 0.089
A, C, D, E, F, G
A, B, C, D, E, F, G
Fitness
Number of mutated genes
1.0
1.2
1.4
1.6
1.8
2.0
1
2
3
4
5
6
7
WT
A,B,C,D,E,F,G

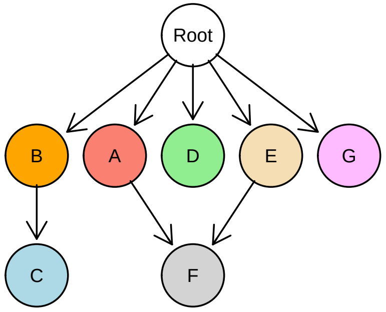


| | |
|---|---|
| *ID* | GM3TL5IyxTmLPLFB |
| *acc genotypes* | 60 |
| *peaks no back* | 9 |
| *removed edges* | 25 |
| *prop rem edges* | 0.156 |

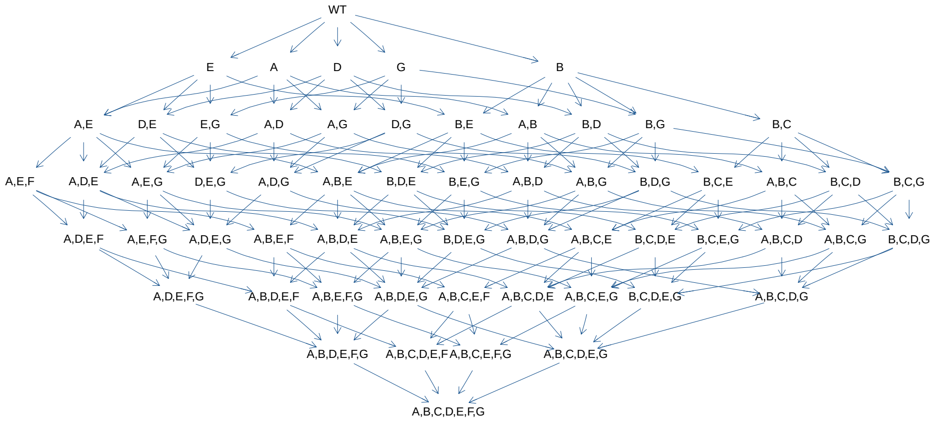

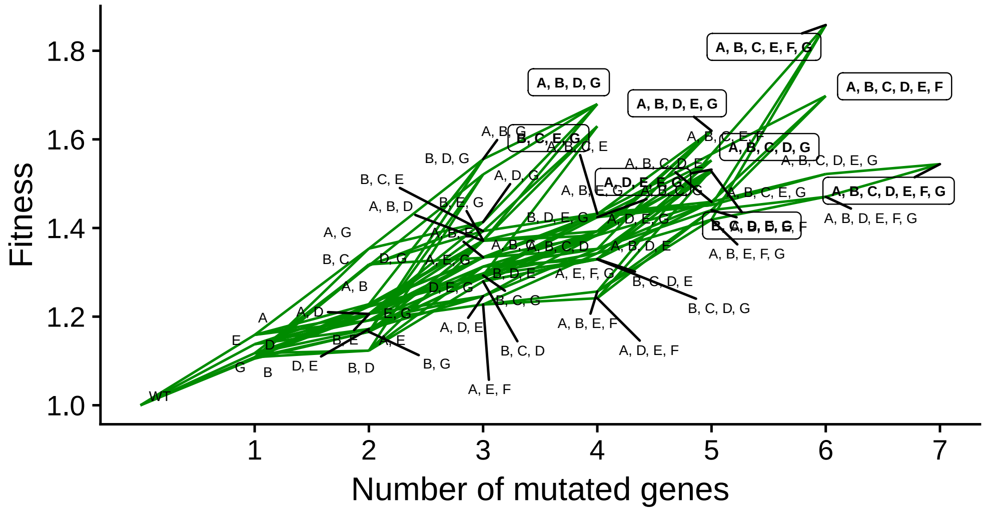


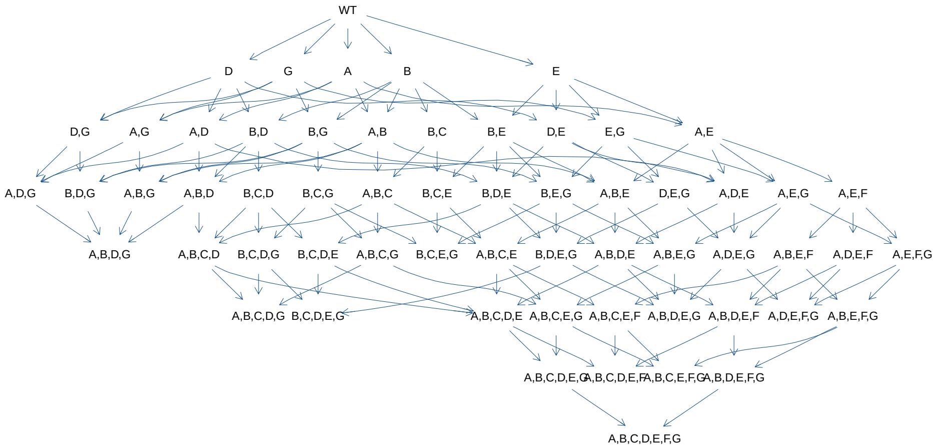

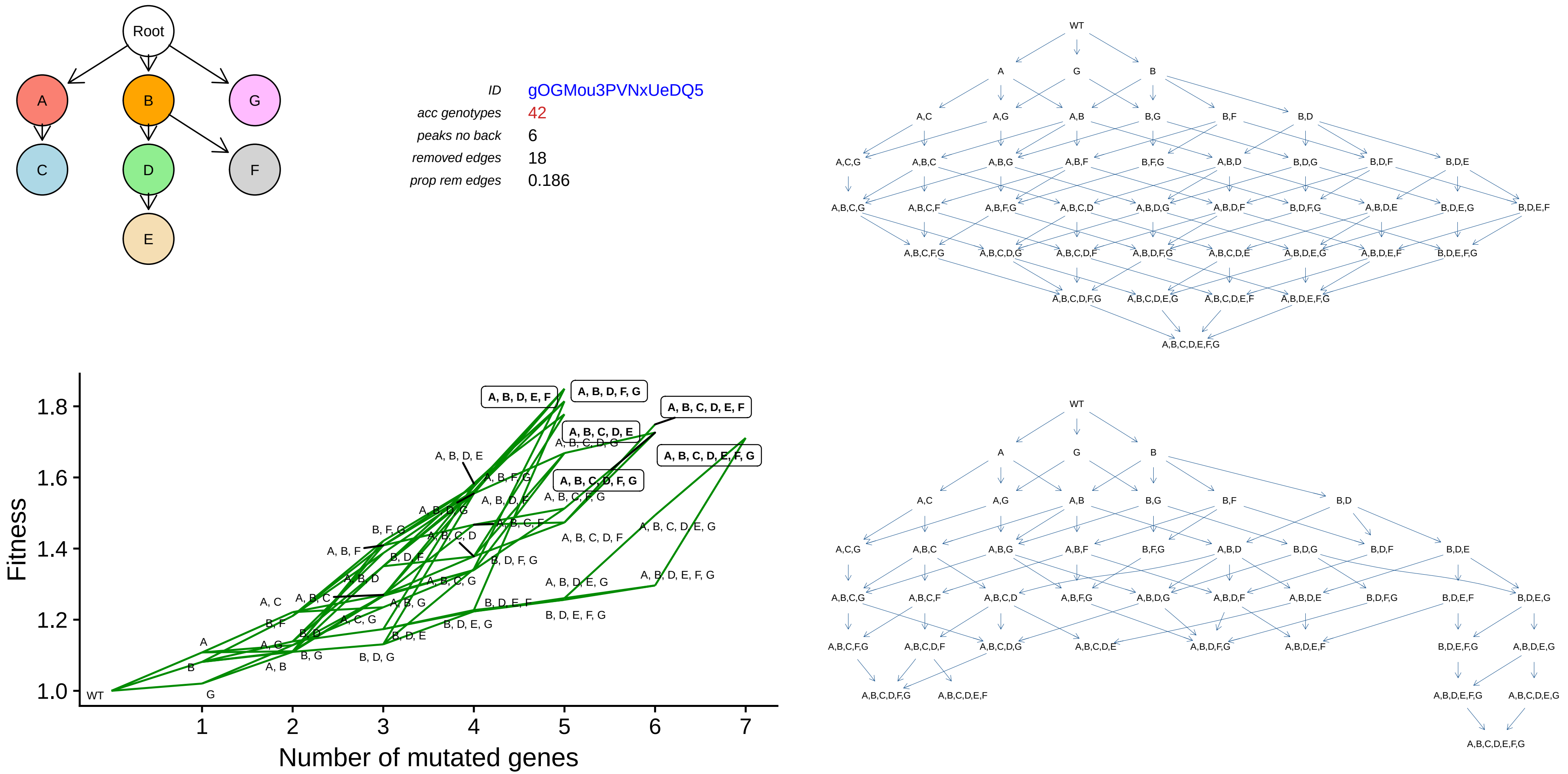

Root
A
B
G
C
D
F
E
ID gOGMou3PVNxUeDQ5
acc genotypes 42
peaks no back 6
removed edges 18
prop rem edges 0.186
Fitness
Number of mutated genes
A, B, D, E, F
A, B, D, F, G
A, B, C, D, E, F
A, B, C, D, E
A, B, C, D, E, F, G
A, B, C, D, F, G
WT

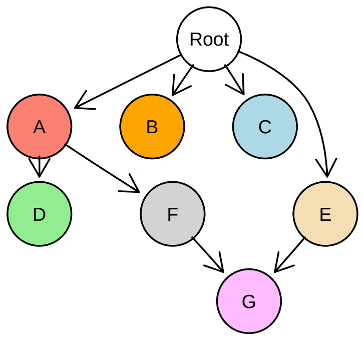


*ID* GwFwG10TqvAnPNr82
*acc genotypes* 48
*peaks no back* 7
*removed edges* 33
*prop rem edges* 0.275

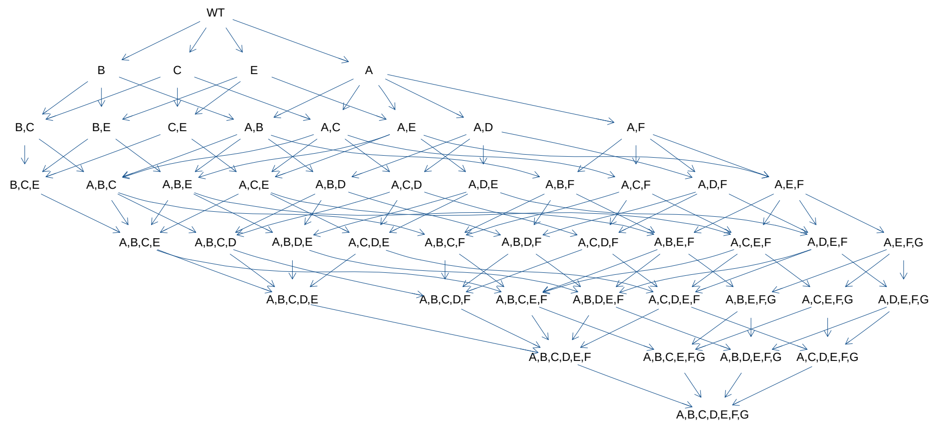


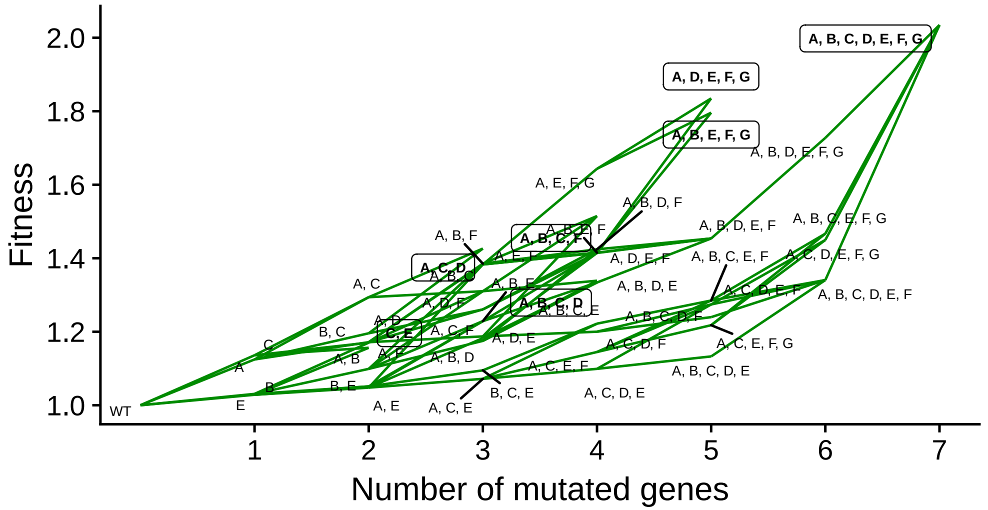


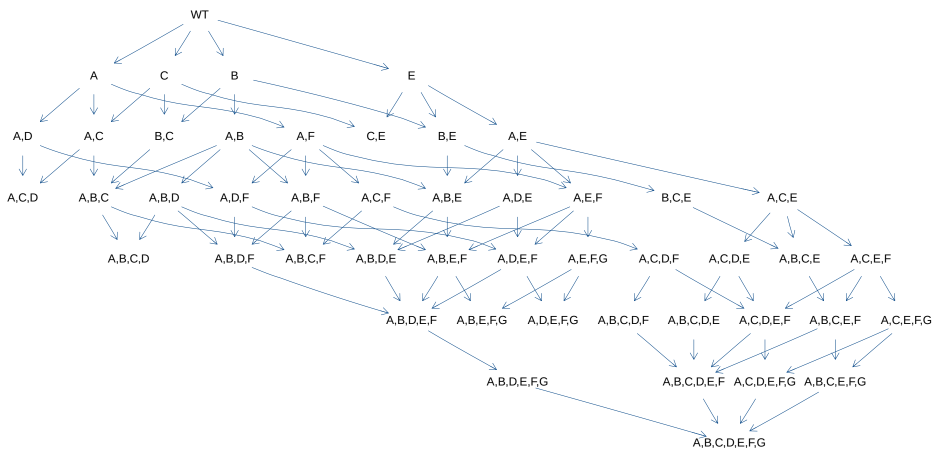

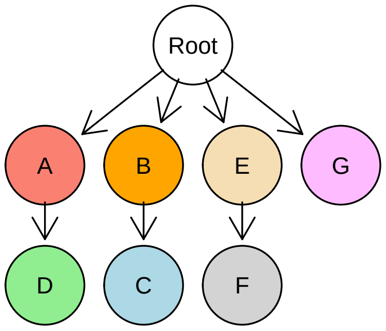


*ID* haC6AnZXOHVkOVUX
*acc genotypes* 54
*peaks no back* 6
*removed edges* 21
*prop rem edges* 0.156

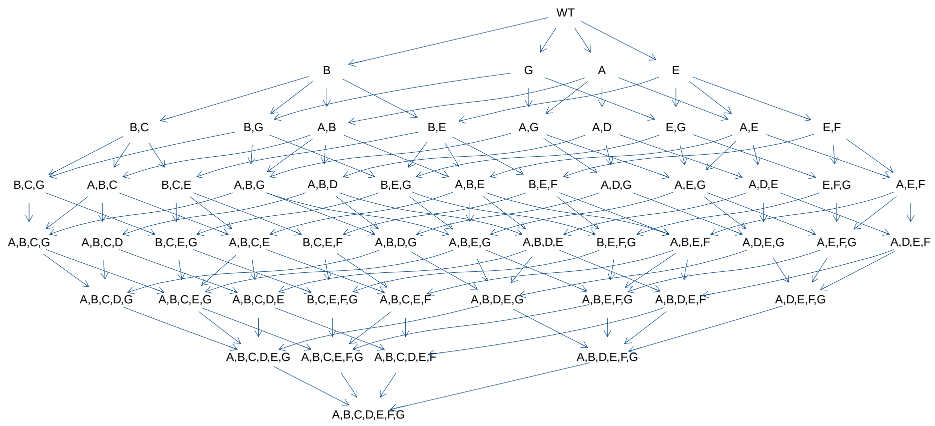


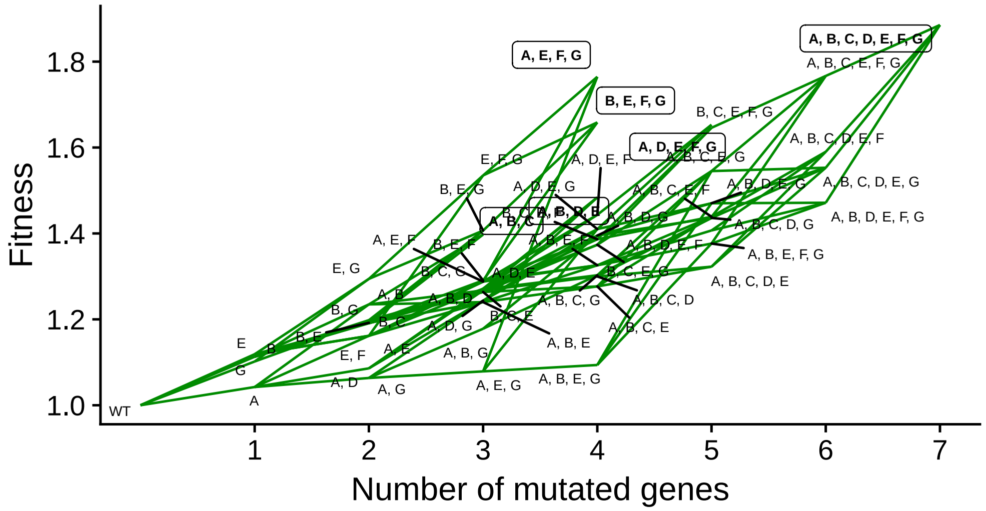


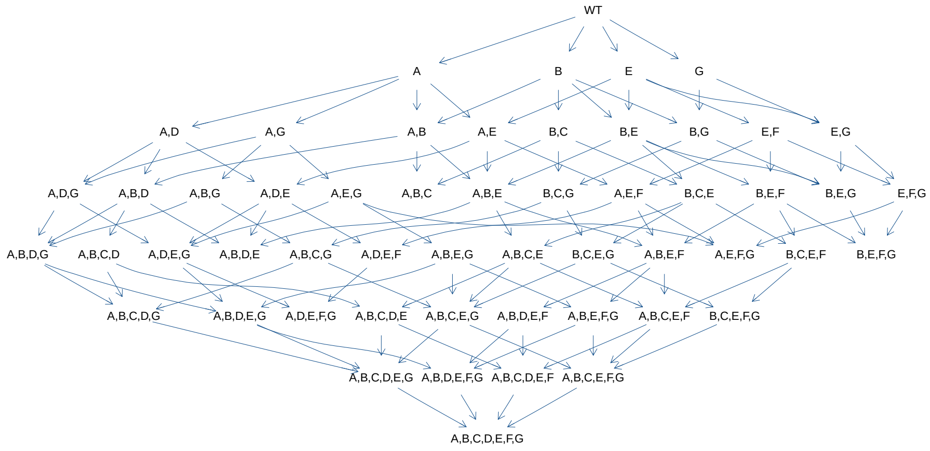

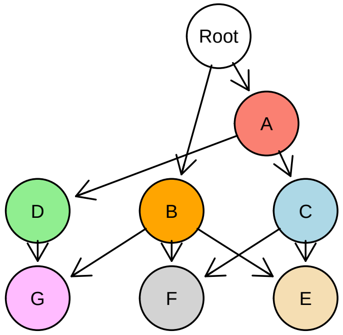


| | |
|---|---|
| *ID* | Hc4dDsccE1OXyhxX |
| *acc genotypes* | 21 |
| *peaks no back* | 5 |
| *removed edges* | 7 |
| *prop rem edges* | 0.194 |

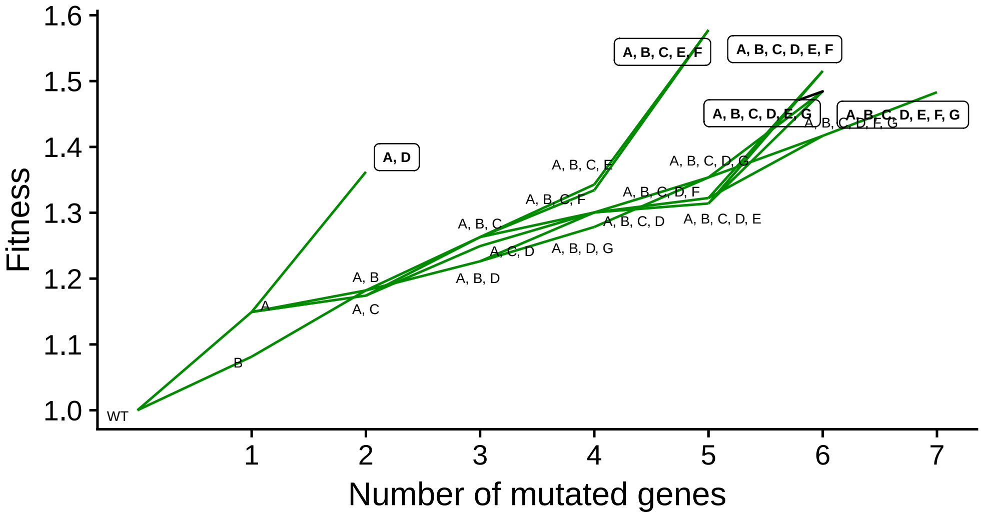


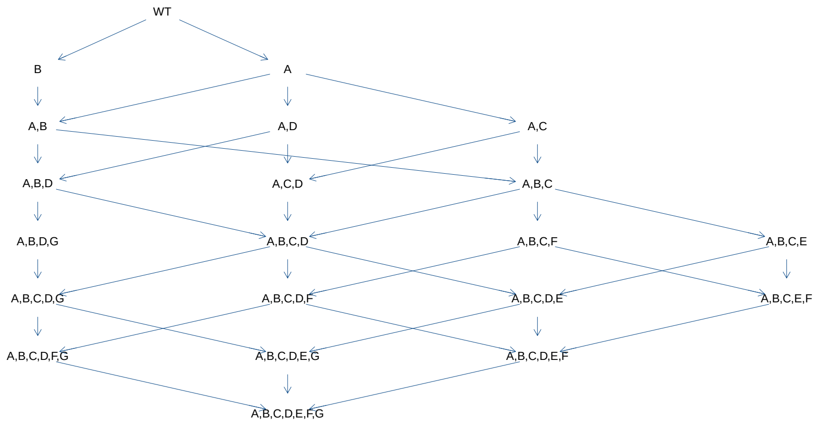


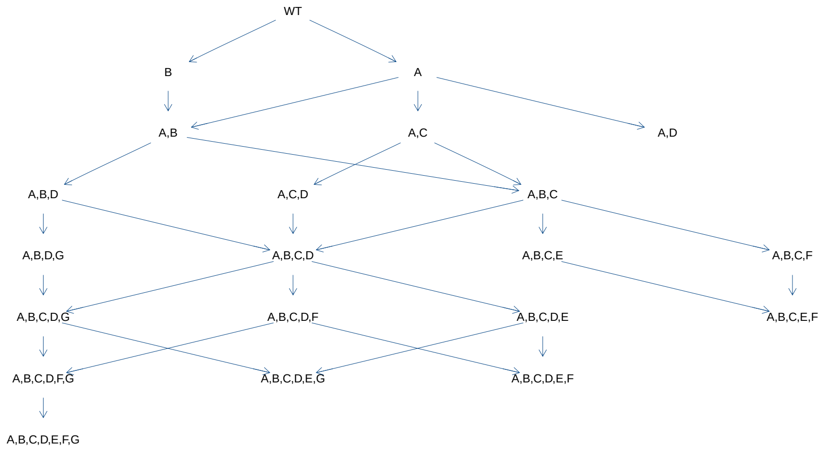

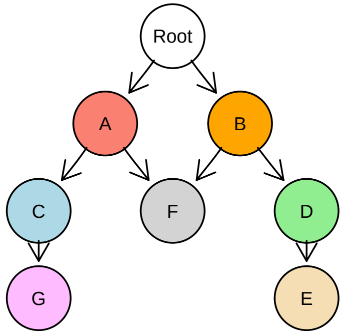


| | |
|---|---|
| *ID* | HFe7qJ1sPtvbaUtG |
| *acc genotypes* | 25 |
| *peaks no back* | 4 |
| *removed edges* | 5 |
| *prop rem edges* | 0.111 |

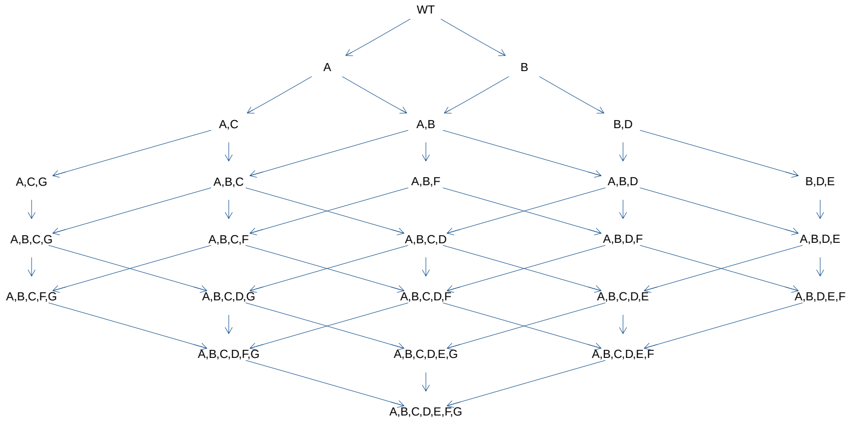


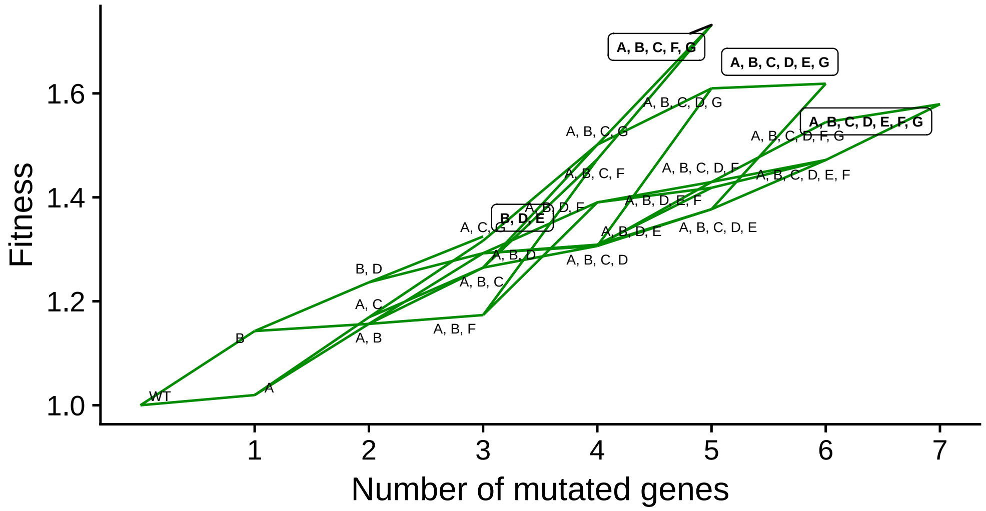


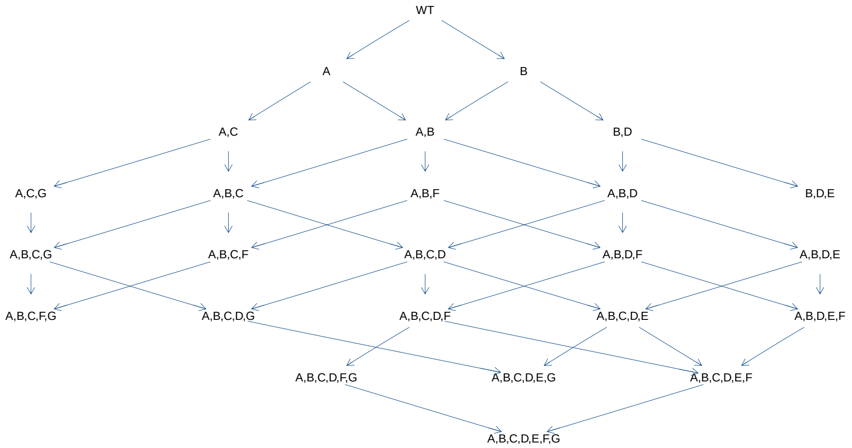

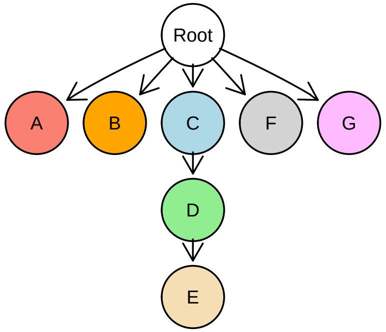


| | |
|---|---|
| *ID* | hloXRsRm4d7aXmN4 |
| *acc genotypes* | 64 |
| *peaks no back* | 2 |
| *removed edges* | 22 |
| *prop rem edges* | 0.125 |

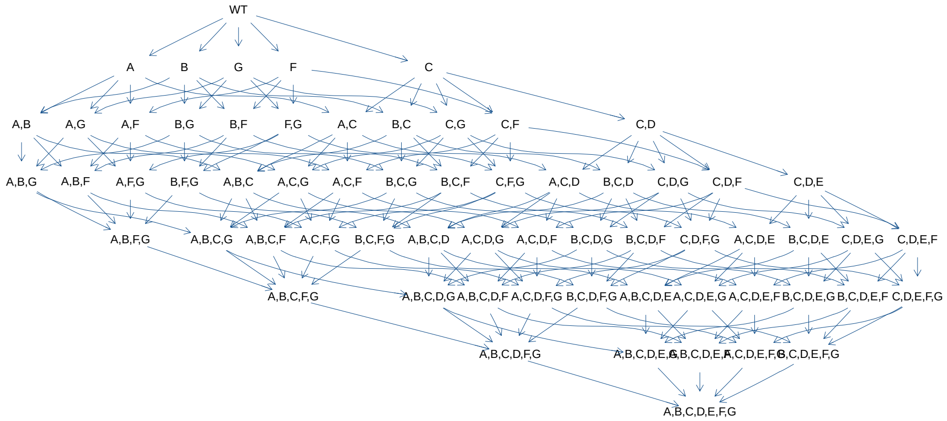

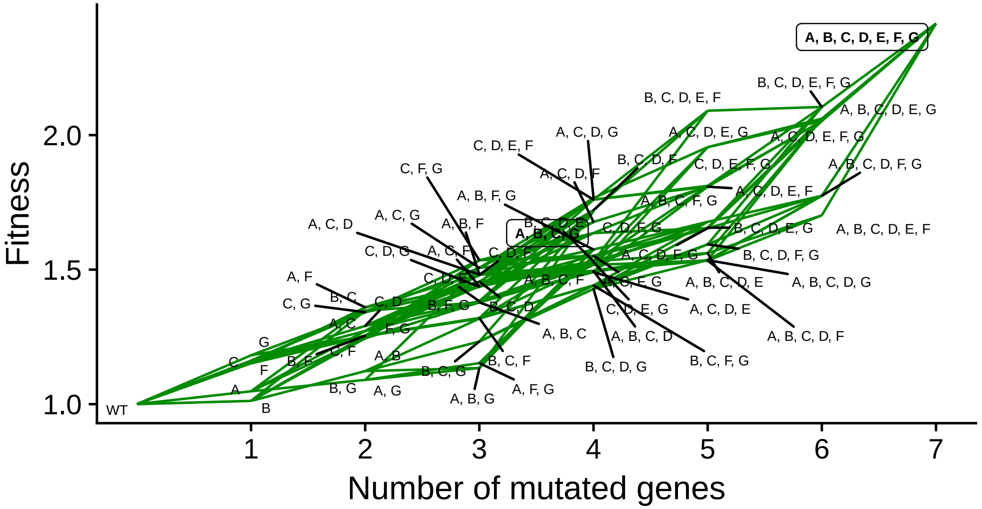


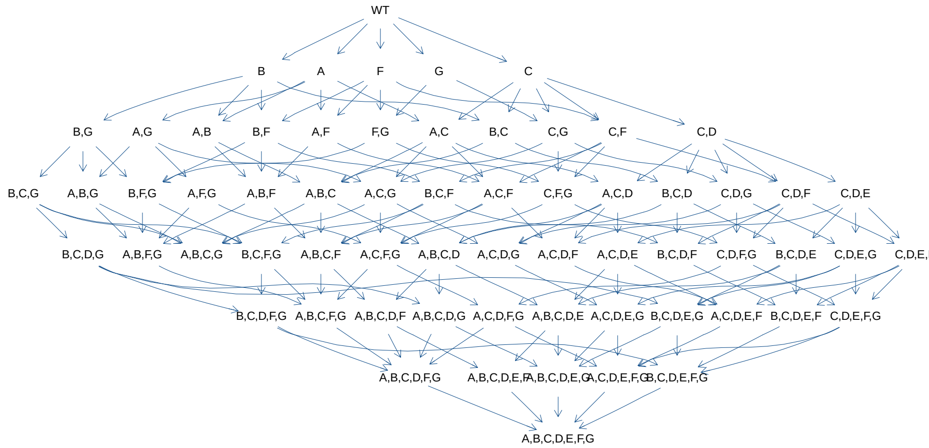

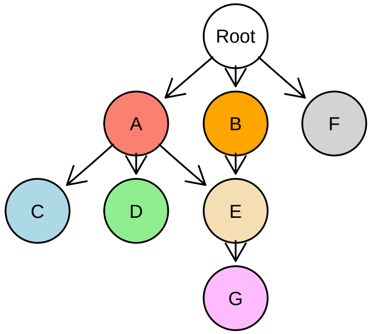


*ID* HizemNX910W6f2YLA
*acc genotypes* 36
*peaks no back* 5
*removed edges* 12
*prop rem edges* 0.15

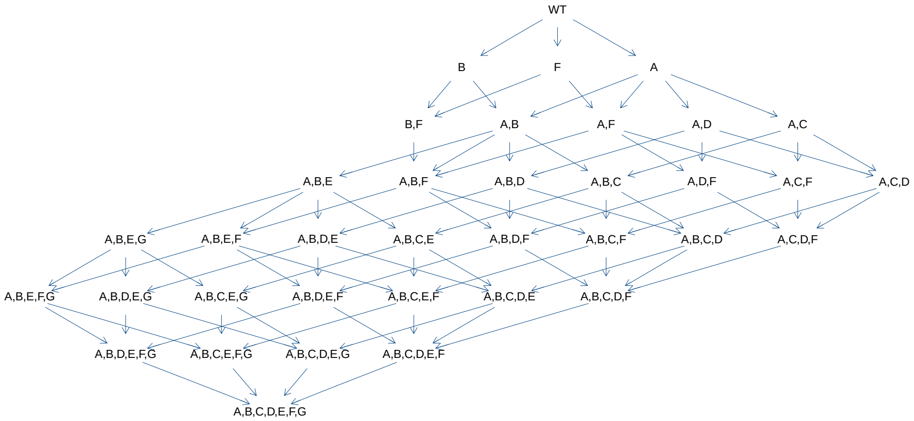


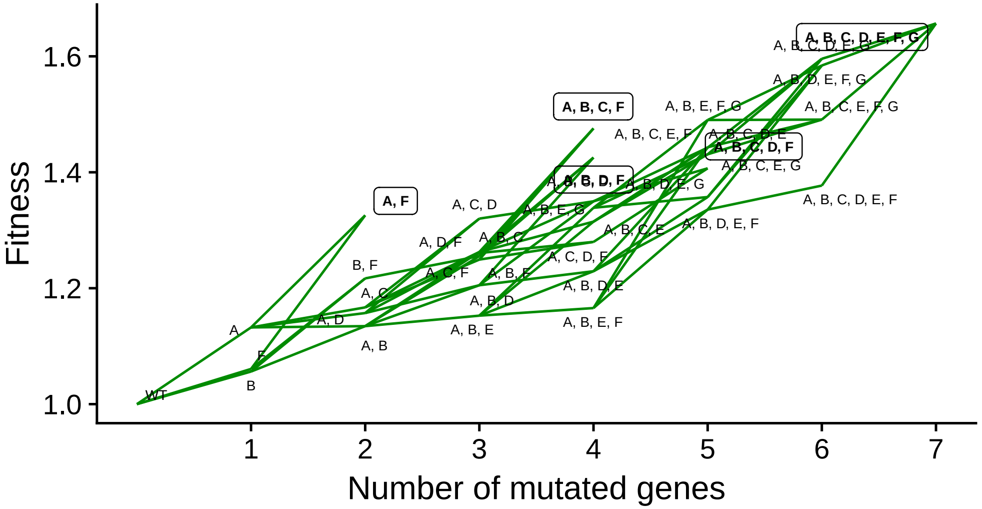


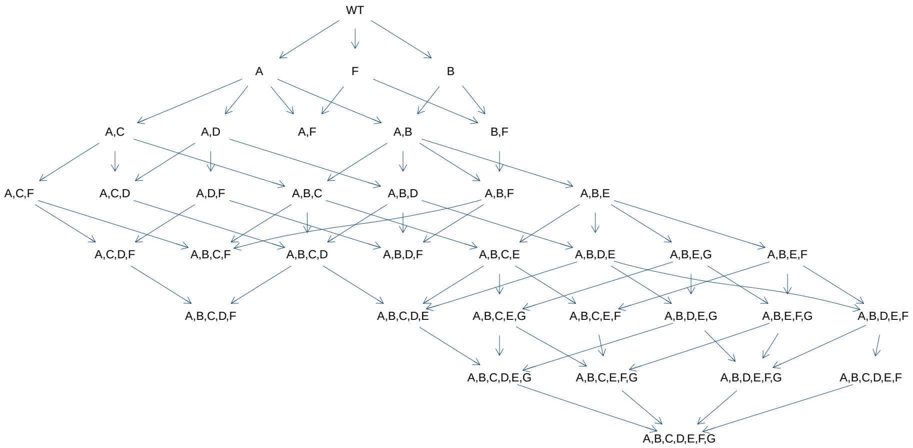

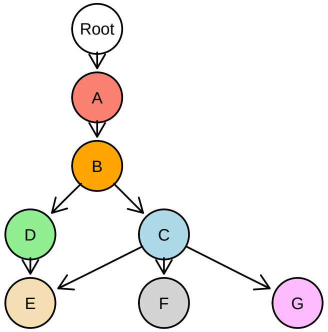


*ID* hw8kQ4g44p4XAkDa
*acc genotypes* 16
*peaks no back* 2
*removed edges* 3
*prop rem edges* 0.12

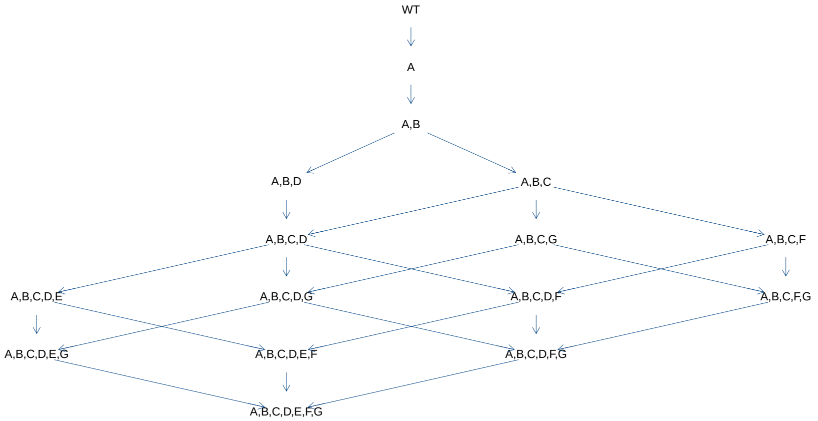


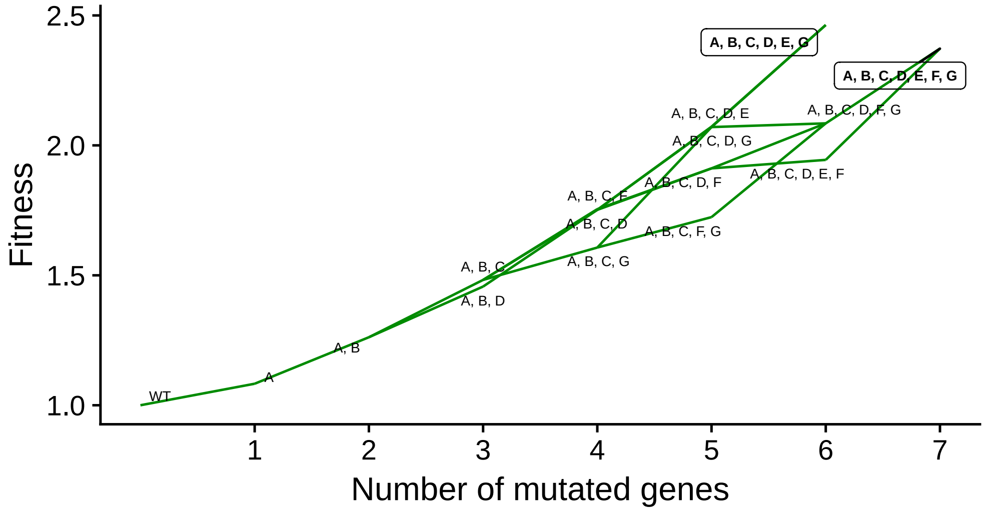


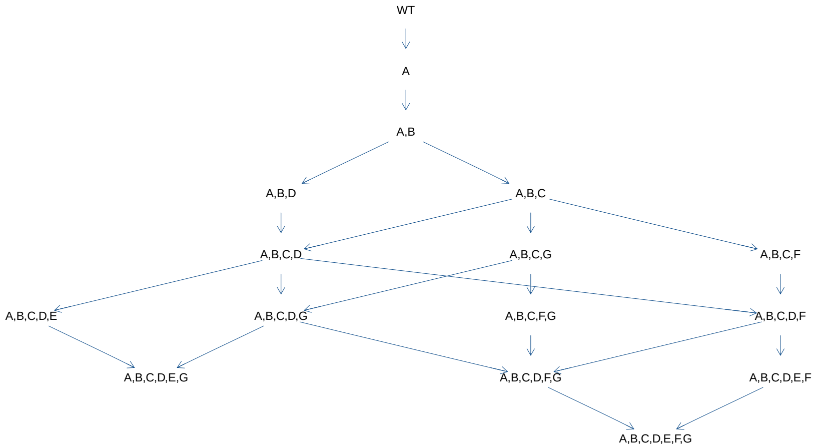

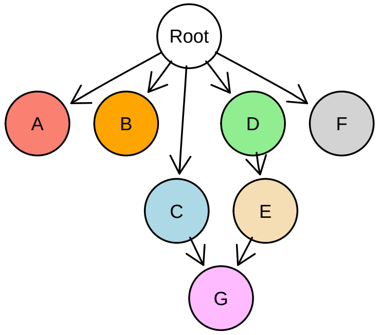


*ID* Hy5Py5ryXo10HeUYG
*acc genotypes* 56
*peaks no back* 8
*removed edges* 35
*prop rem edges* 0.236

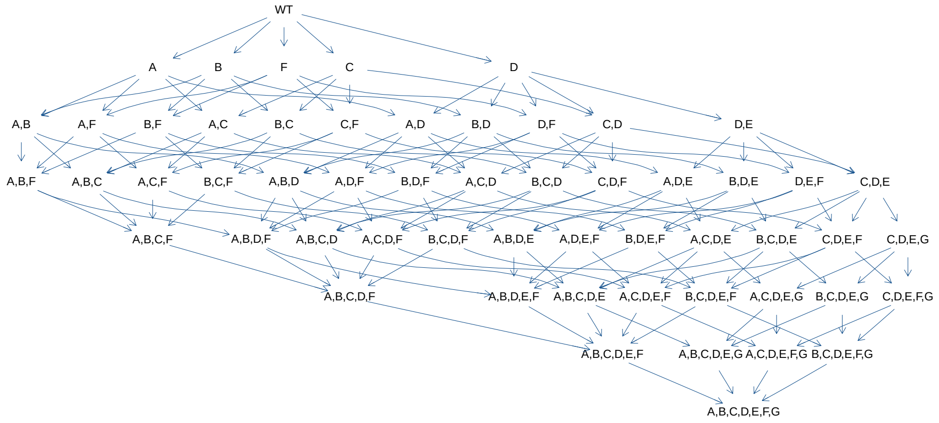


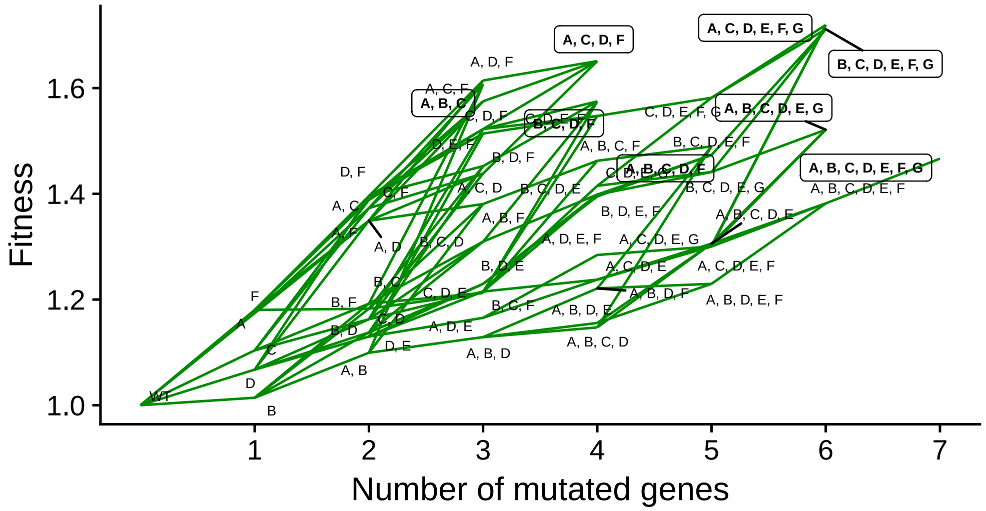


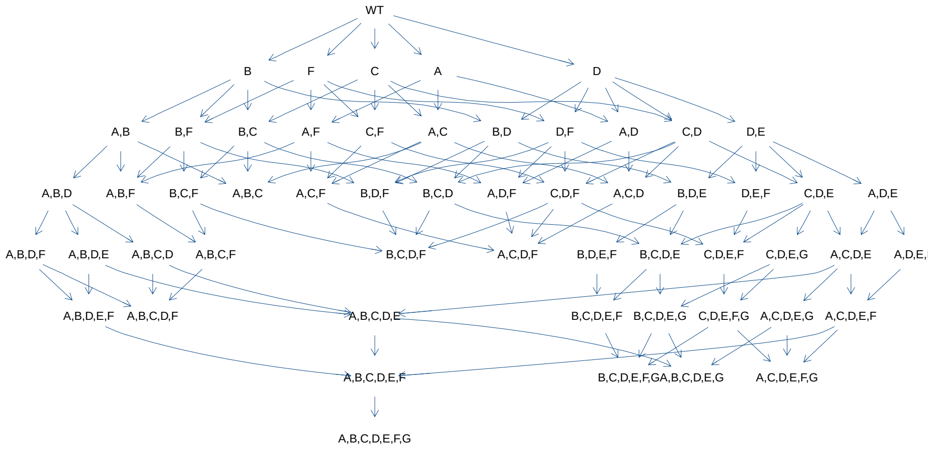

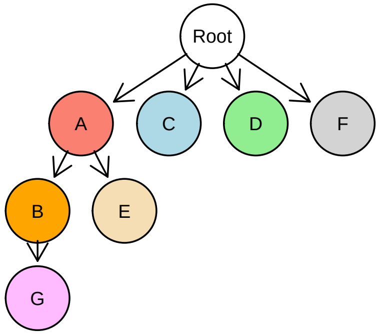


| | |
|---|---|
| *ID* | Id85CzgL2g6AV3510 |
| *acc genotypes* | 56 |
| *peaks no back* | 9 |
| *removed edges* | 38 |
| *prop rem edges* | 0.257 |

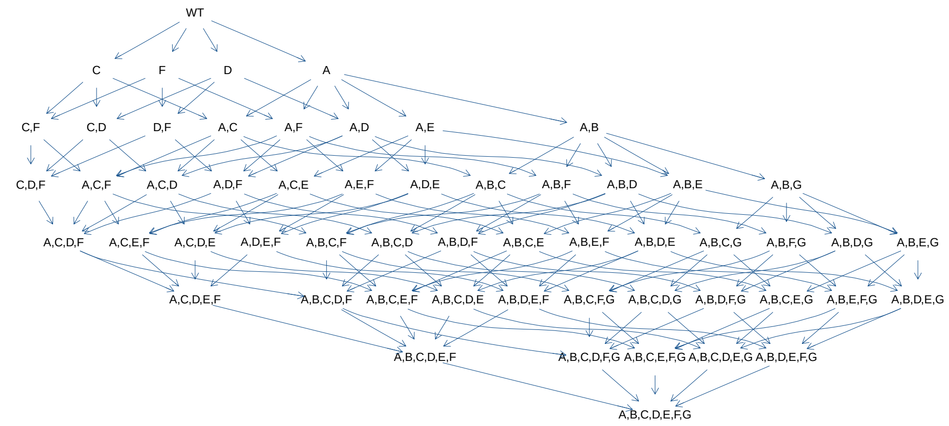


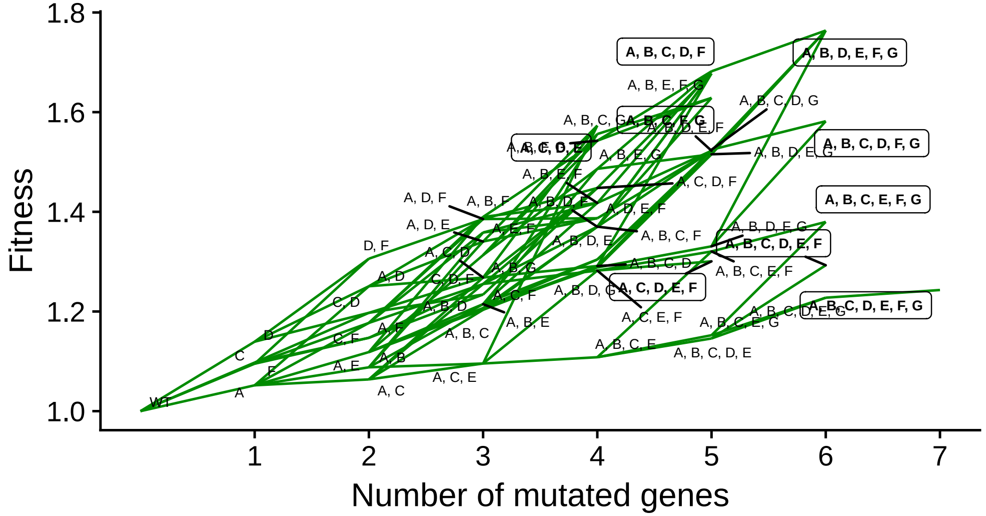


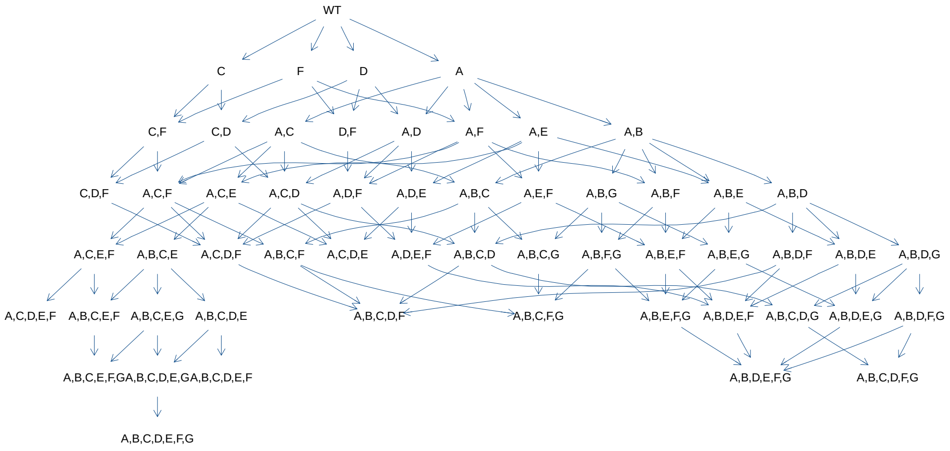

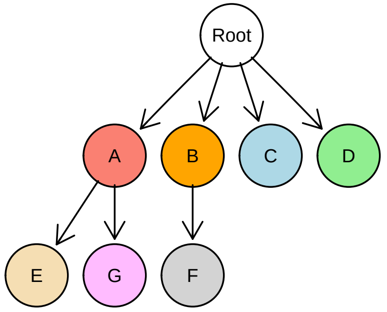


| | |
|---|---|
| *ID* | ILWtHZyJ9o3F1Qgp |
| *acc genotypes* | 60 |
| *peaks no back* | 3 |
| *removed edges* | 20 |
| *prop rem edges* | 0.125 |

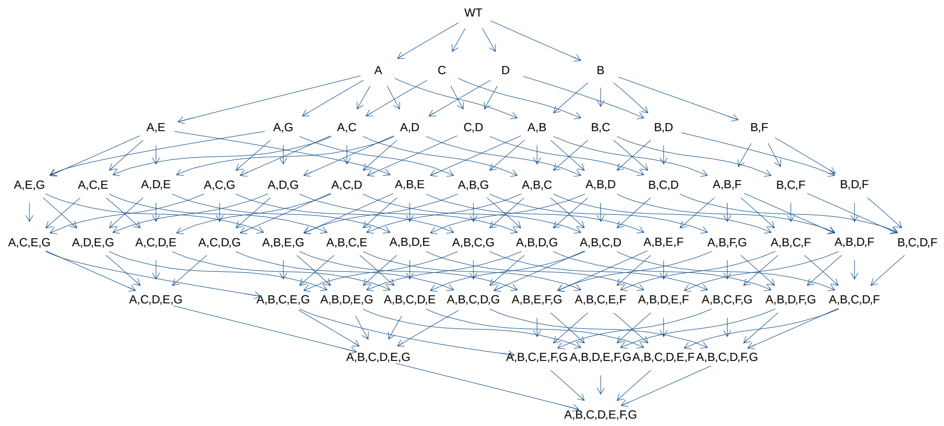

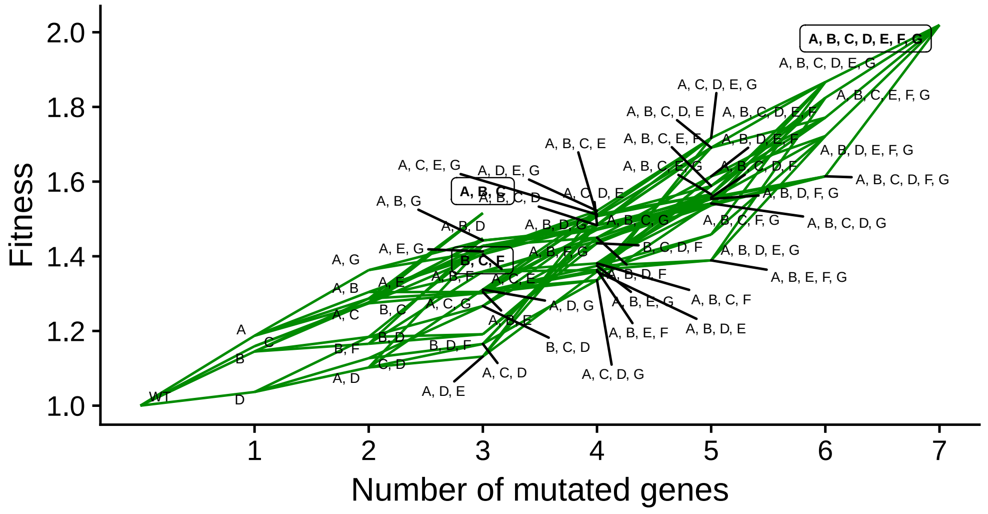


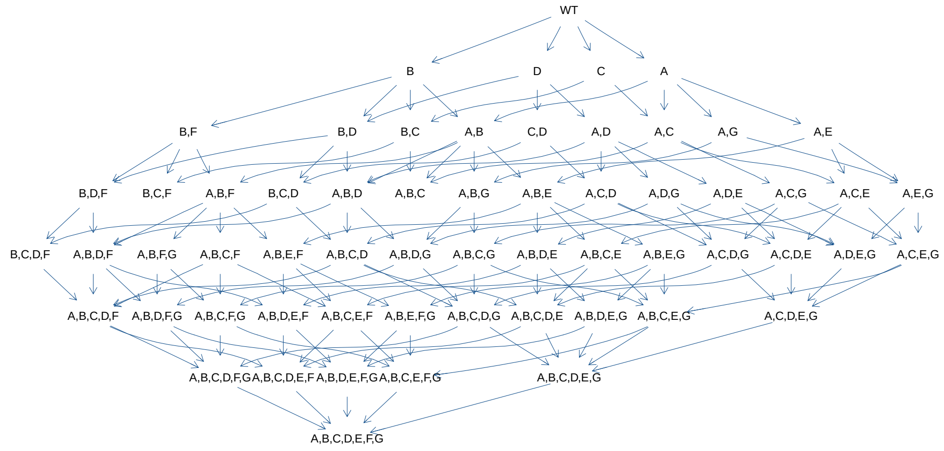

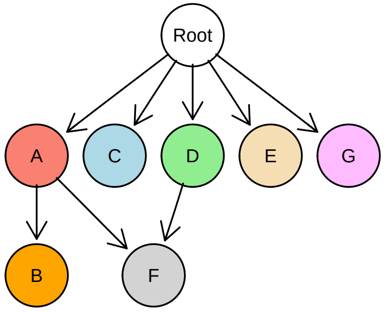


| | |
|---|---|
| *ID* | JABP9YQJqeNrAKVt |
| *acc genotypes* | 64 |
| *peaks no back* | 8 |
| *removed edges* | 35 |
| *prop rem edges* | 0.199 |

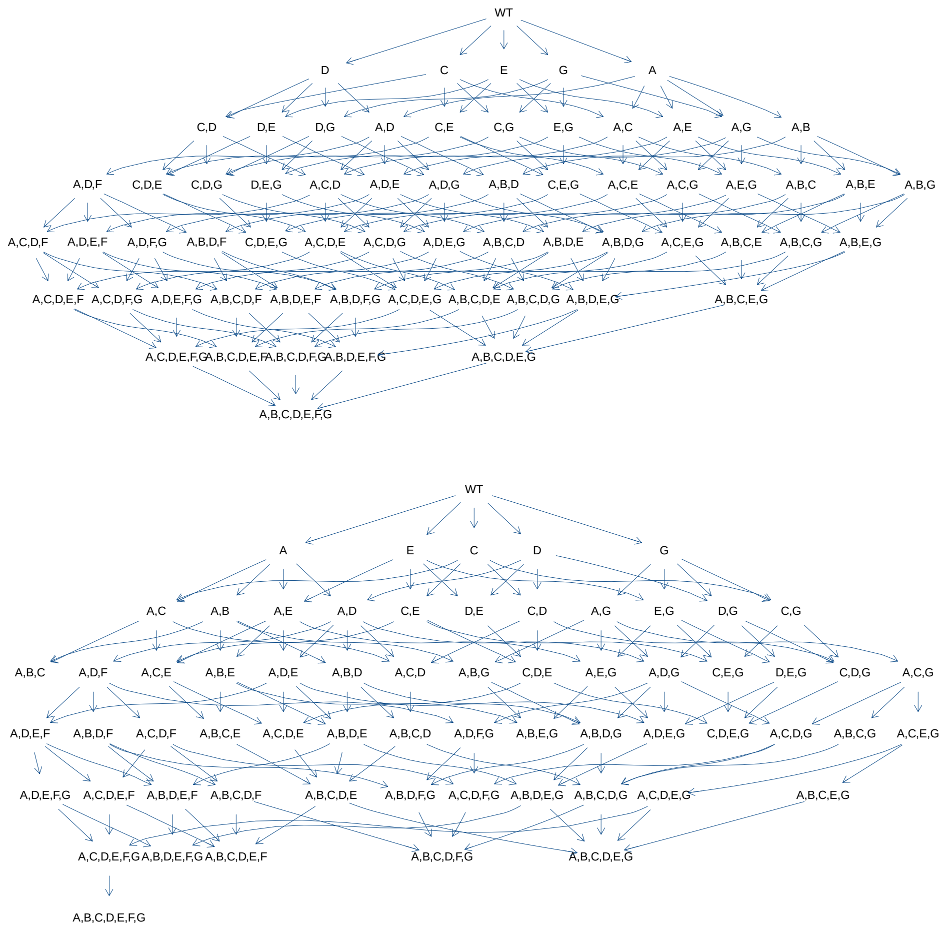

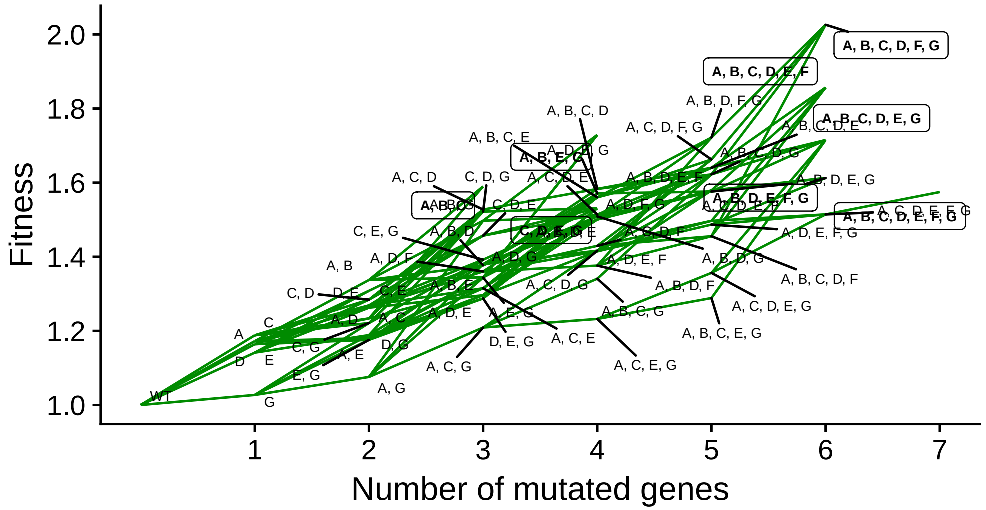

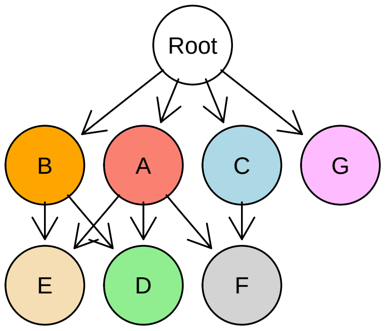


| | |
|---|---|
| *ID* | jFHaUyfw10mPN2R1P |
| *acc genotypes* | 38 |
| *peaks no back* | 4 |
| *removed edges* | 15 |
| *prop rem edges* | 0.176 |

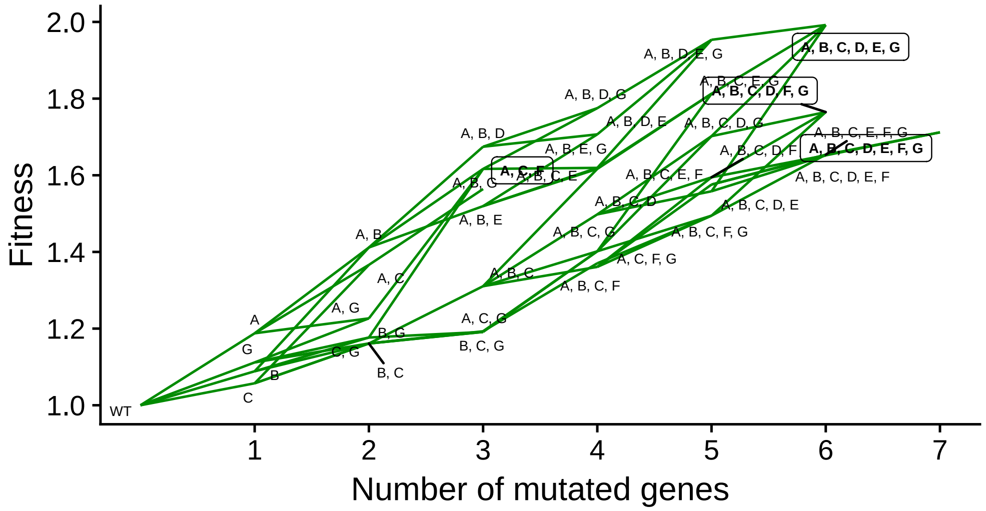


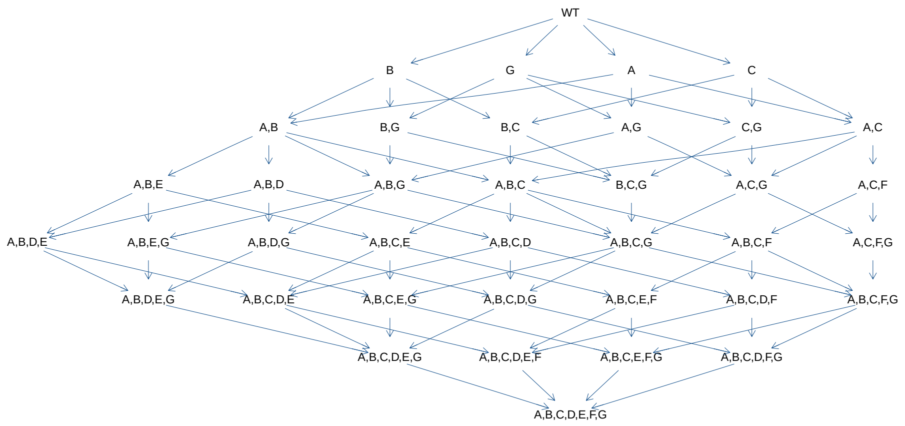


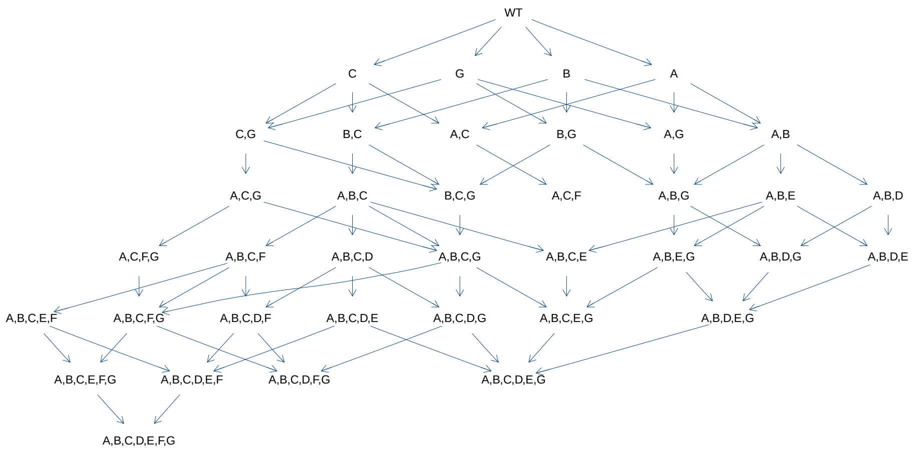

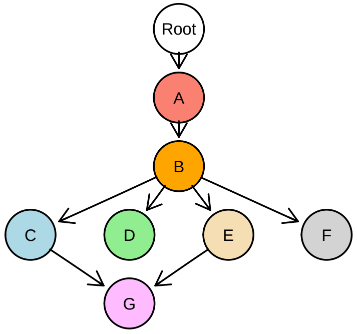


| | |
|---|---|
| *ID* | jj397ue810e35pKQe |
| *acc genotypes* | 22 |
| *peaks no back* | 3 |
| *removed edges* | 5 |
| *prop rem edges* | 0.119 |

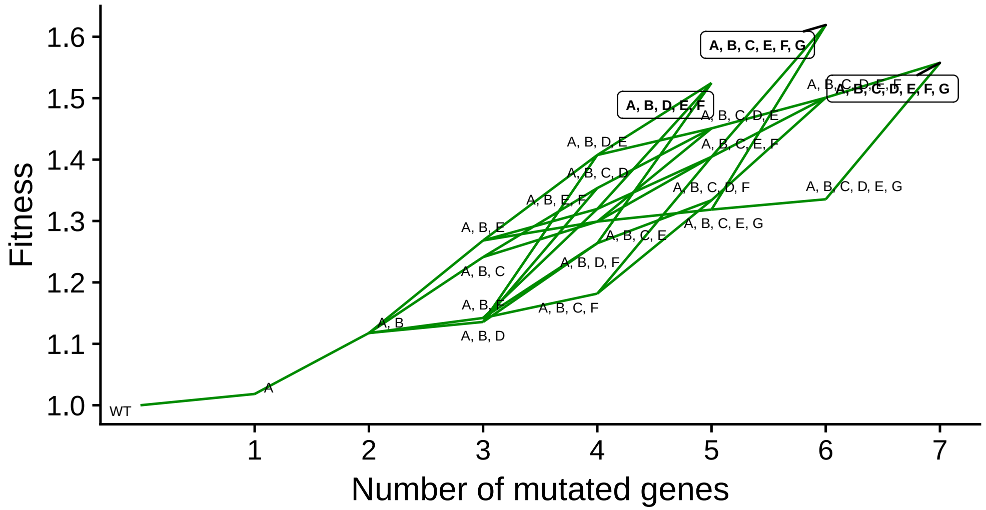


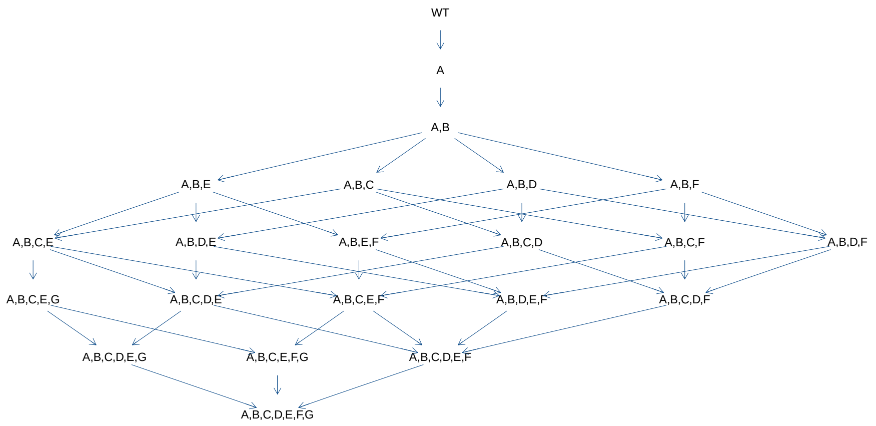


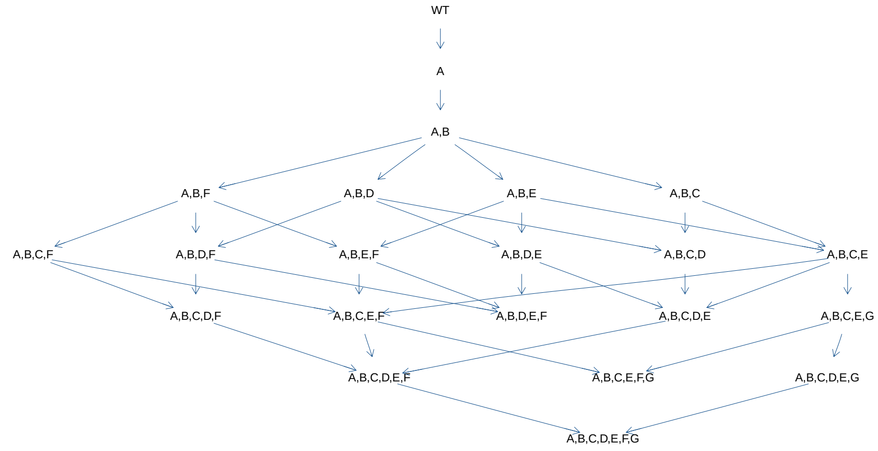

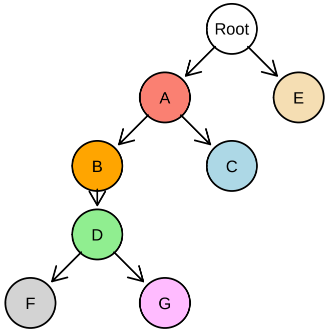


| | |
|---|---|
| *ID* | jle1t255I110i164X |
| *acc genotypes* | 26 |
| *peaks no back* | 3 |
| *removed edges* | 4 |
| *prop rem edges* | 0.078 |

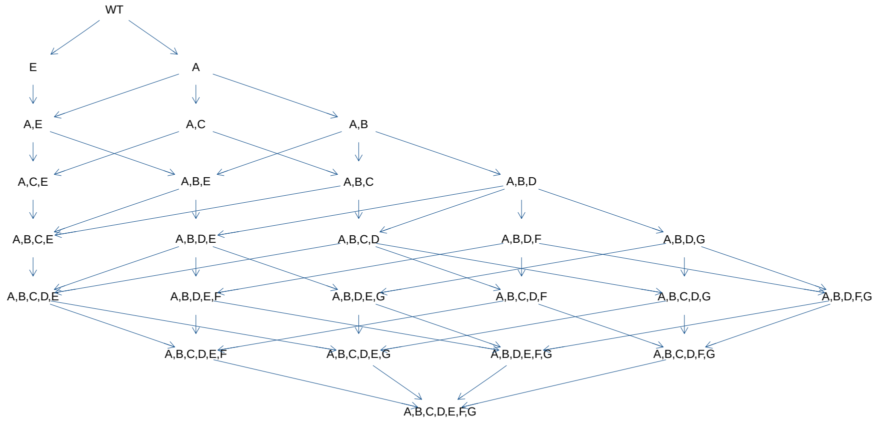


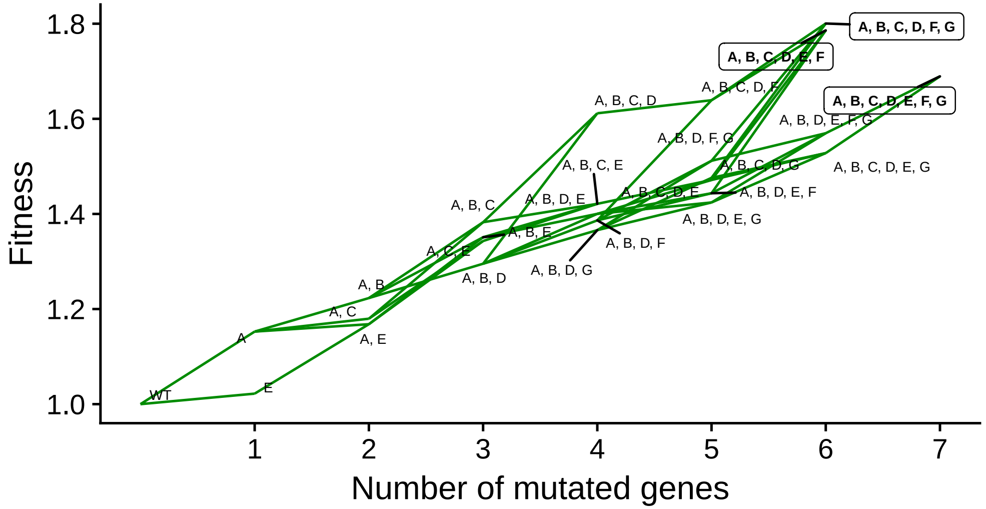


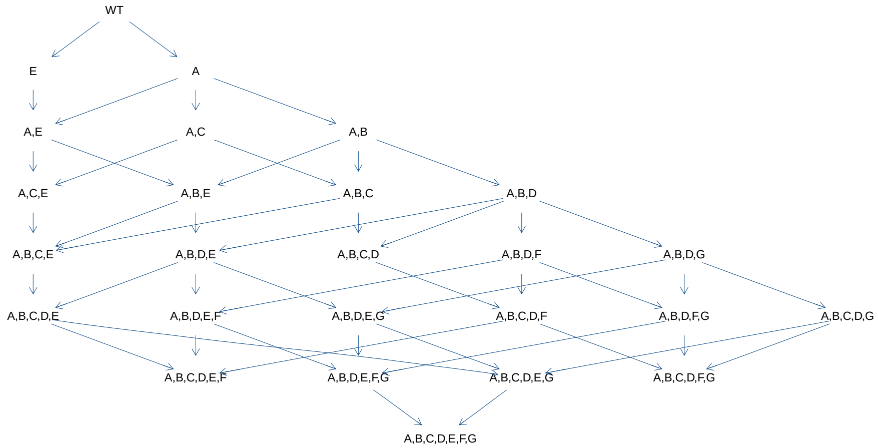

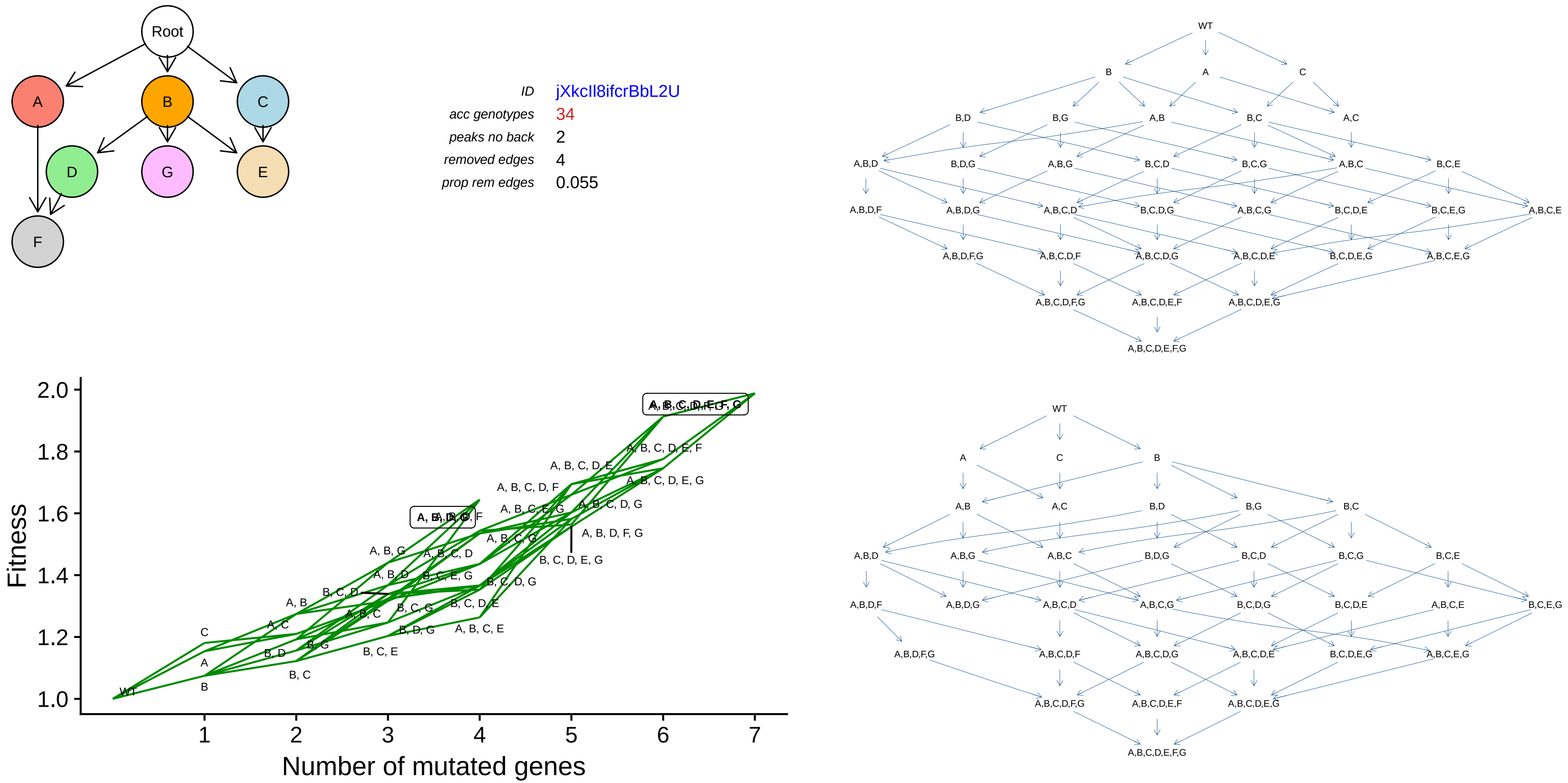

Root
A
B
C
D
G
E
F
ID jXkcIl8ifcrBbL2U
acc genotypes 34
peaks no back 2
removed edges 4
prop rem edges 0.055
Fitness
Number of mutated genes

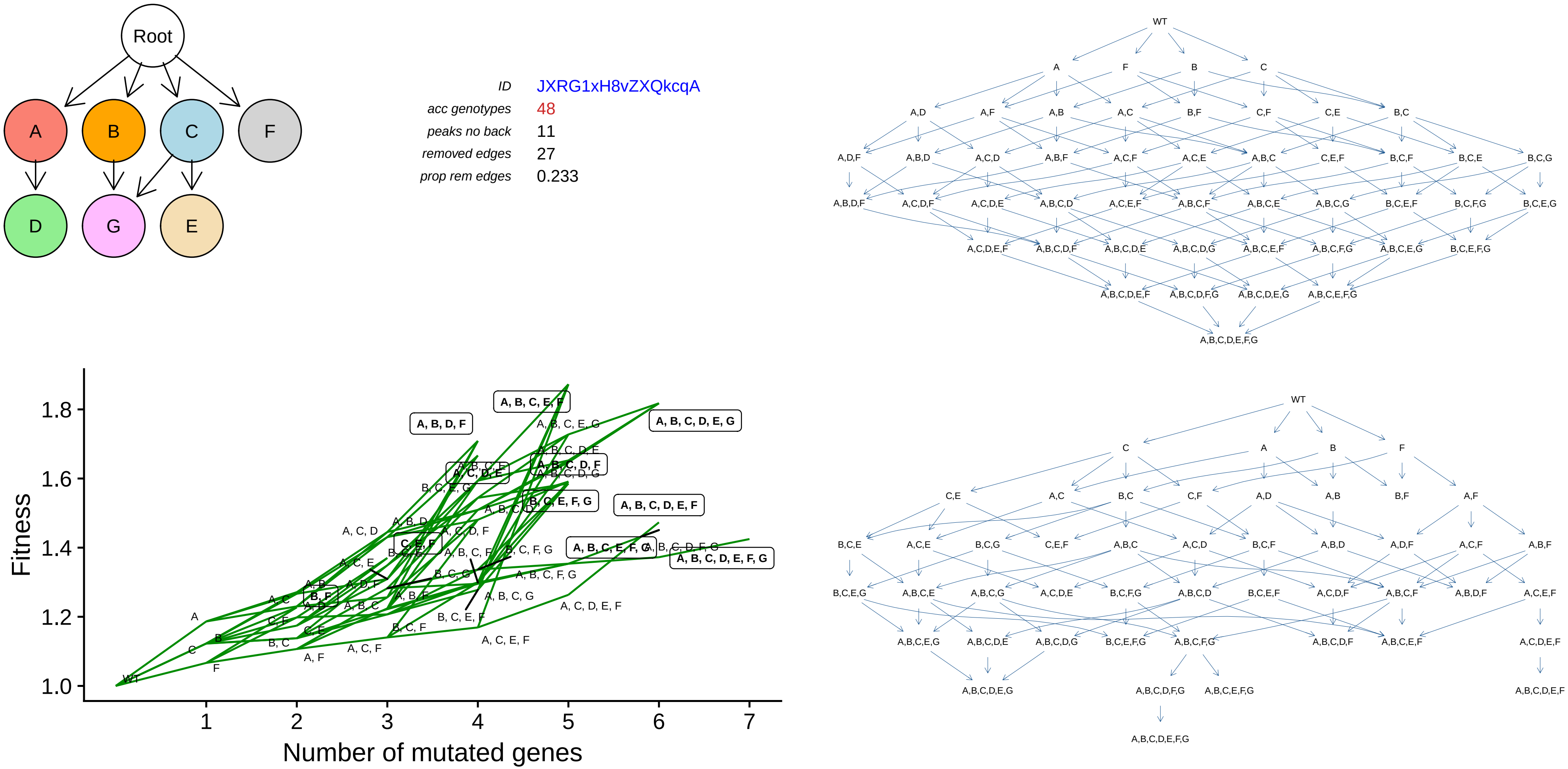

Root
A
B
C
F
D
G
E
ID JXRG1xH8vZXQkcqA
acc genotypes 48
peaks no back 11
removed edges 27
prop rem edges 0.233
Fitness
Number of mutated genes
A, B, D, F
A, B, C, E, F
A, B, C, D, E, G
A, B, C, D, F
A, C, D, E
B, C, E, F, G
A, B, C, D, E, F
C, E, F
A, B, C, E, F, G
A, B, C, D, E, F, G
B, F

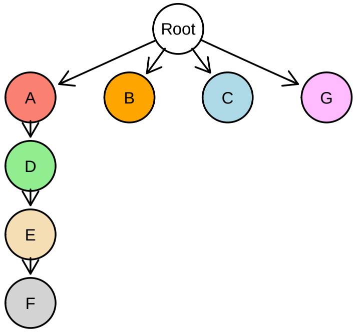


*ID* k58HtWC5MFWCpF19
*acc genotypes* 40
*peaks no back* 7
*removed edges* 12
*prop rem edges* 0.13

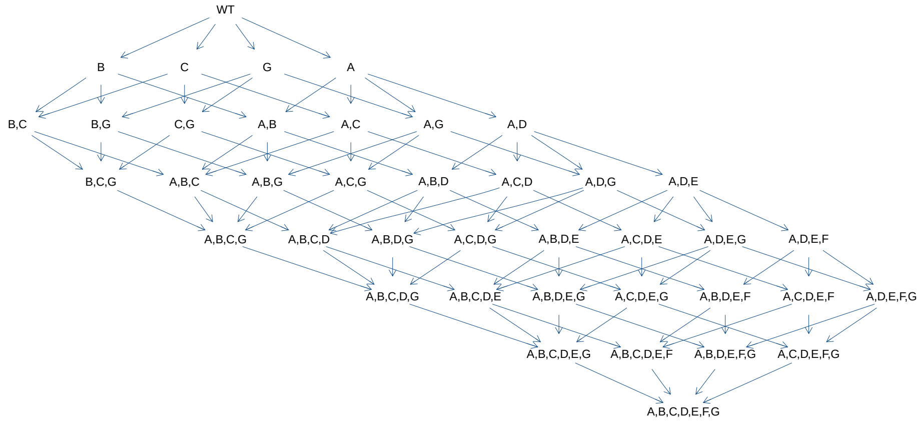


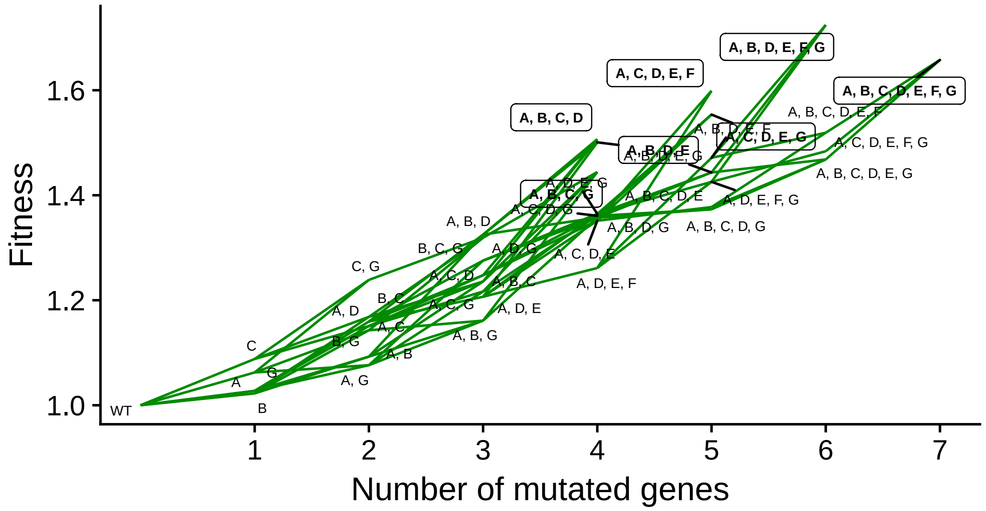


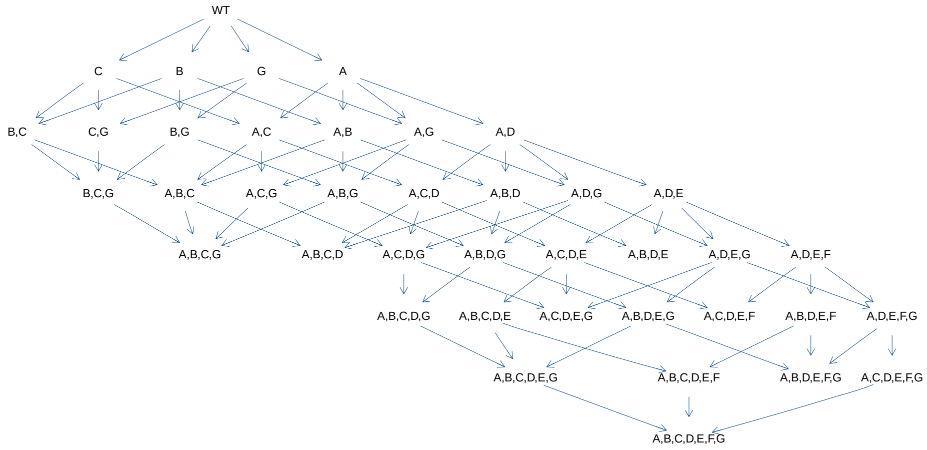

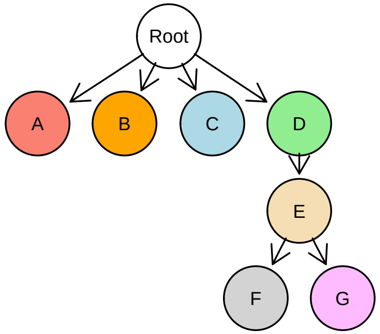


| | |
|---|---|
| *ID* | kHBNd5rumamnFMto |
| *acc genotypes* | 48 |
| *peaks no back* | 7 |
| *removed edges* | 25 |
| *prop rem edges* | 0.208 |

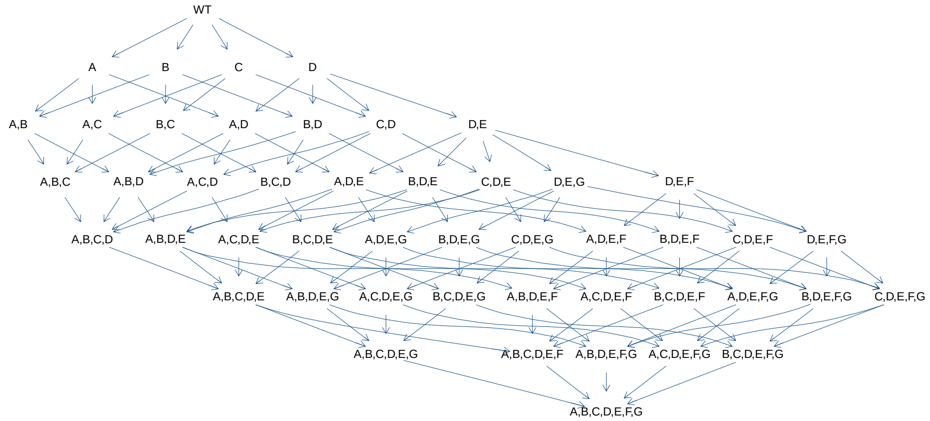

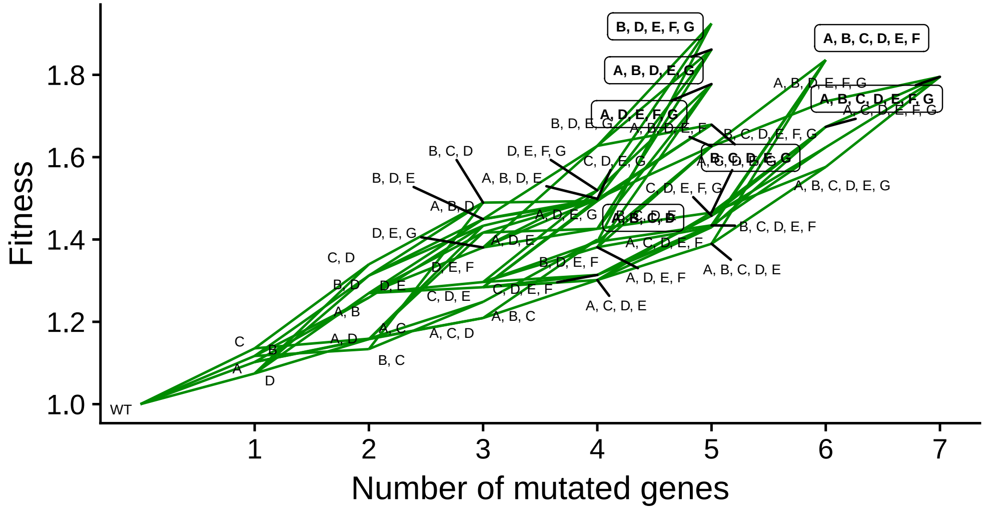


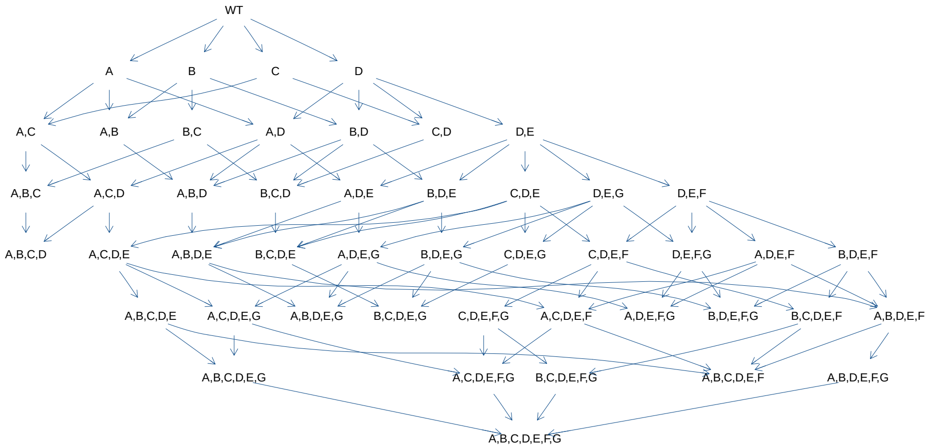

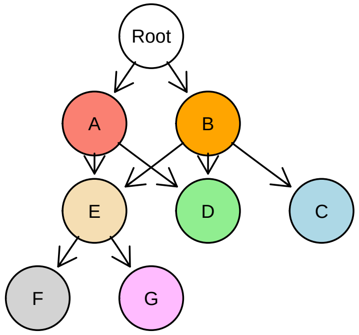


| | |
|---|---|
| *ID* | kj6bITZoZLt3OYX2 |
| *acc genotypes* | 24 |
| *peaks no back* | 5 |
| *removed edges* | 8 |
| *prop rem edges* | 0.174 |

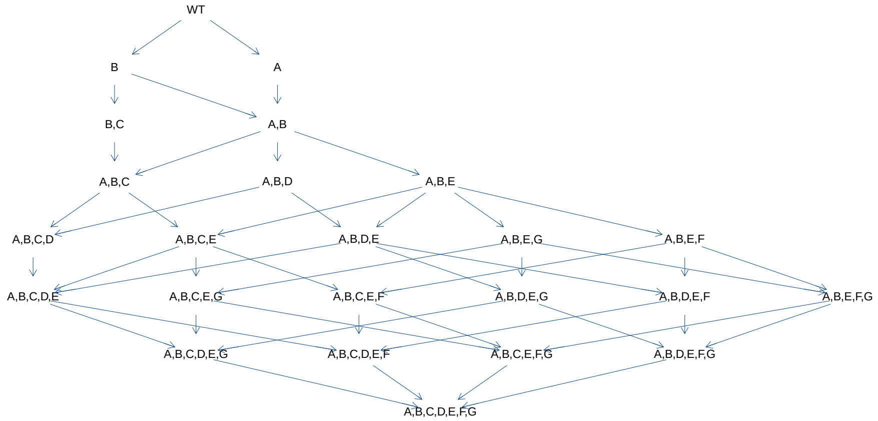


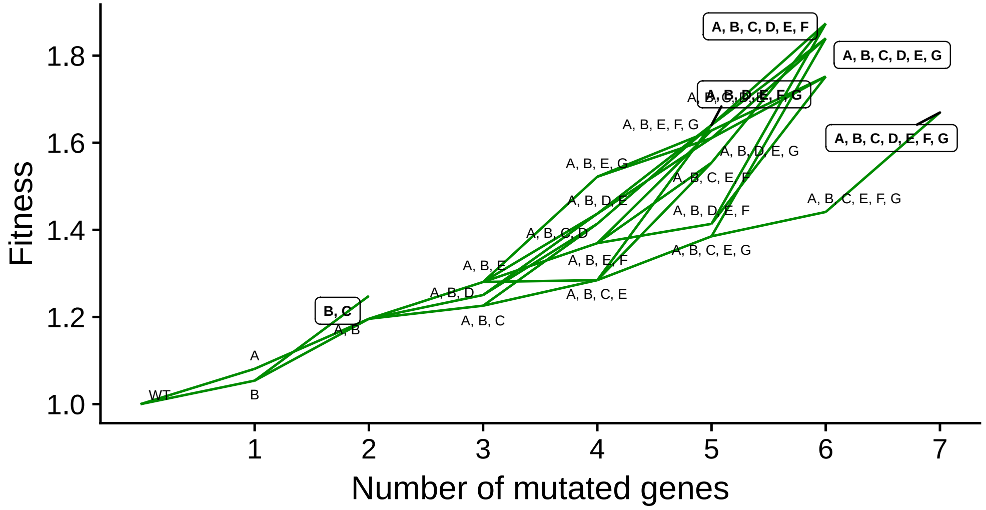


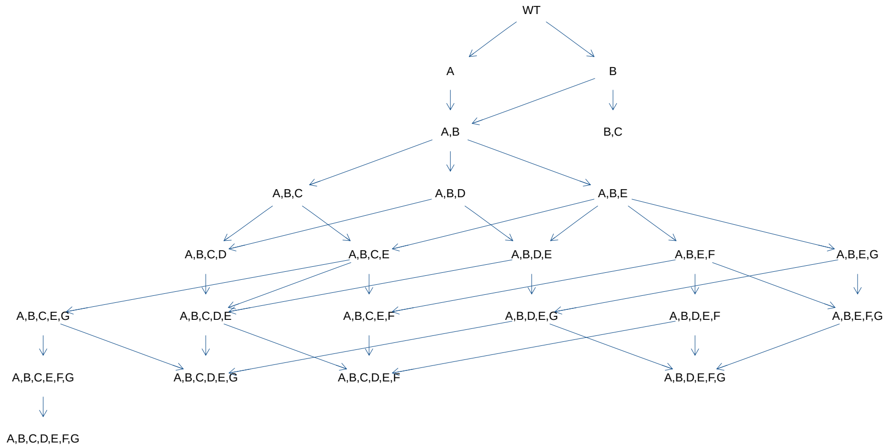

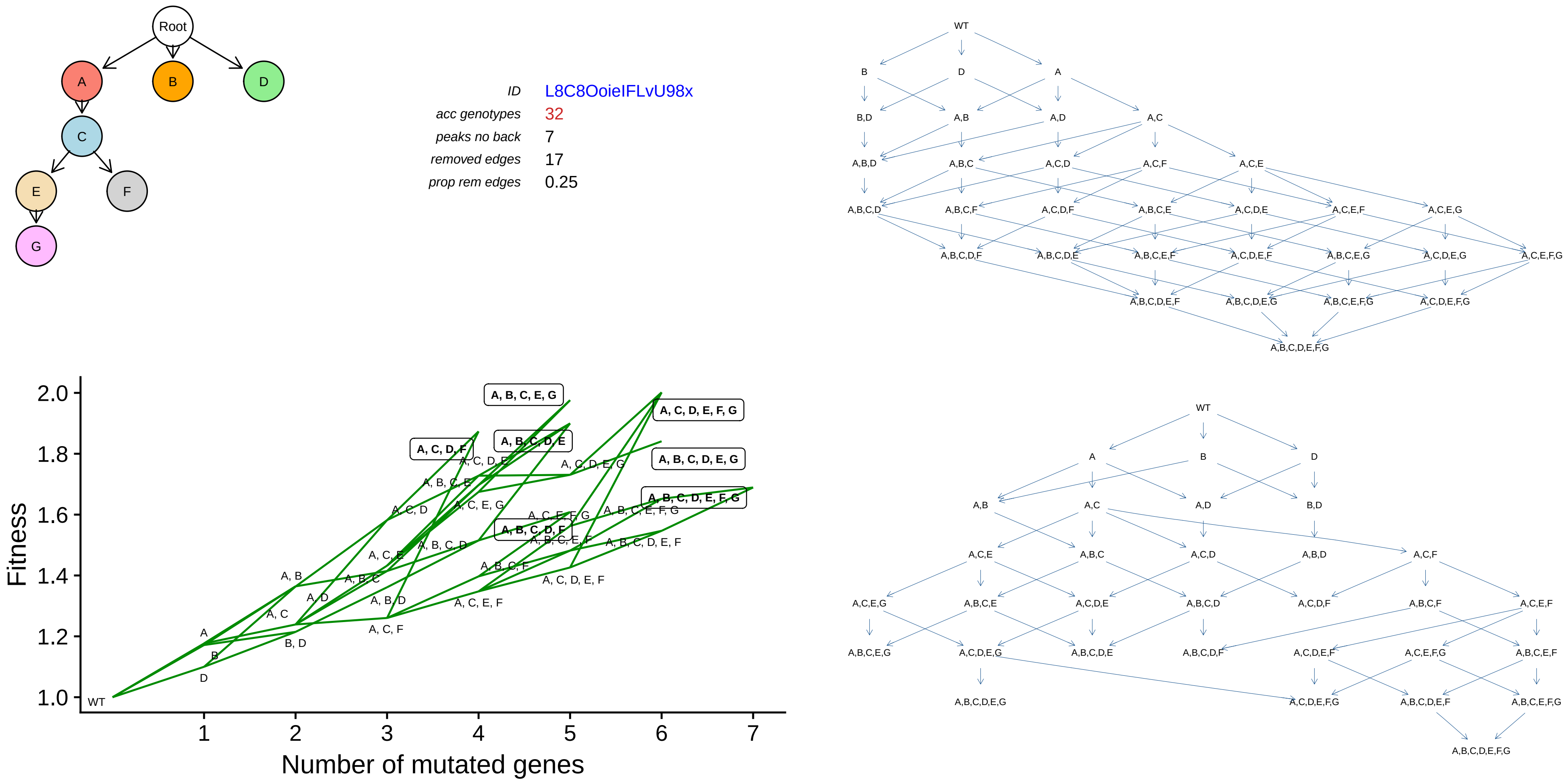

Root
A
B
D
C
E
F
G
ID L8C8OoieIFLvU98x
acc genotypes 32
peaks no back 7
removed edges 17
prop rem edges 0.25
Fitness
Number of mutated genes
A, B, C, E, G
A, C, D, F
A, B, C, D, E
A, C, D, E, F, G
A, B, C, D, E, G
A, B, C, D, E, F, G
A, B, C, D, F
WT

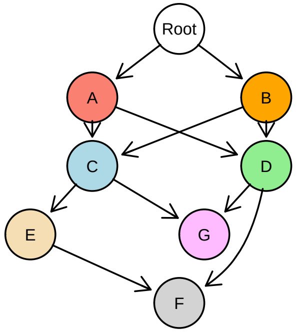


| | |
|---|---|
| *ID* | L9U6EXaxa4AgbWdQ |
| *acc genotypes* | 13 |
| *peaks no back* | 3 |
| *removed edges* | 2 |
| *prop rem edges* | 0.118 |

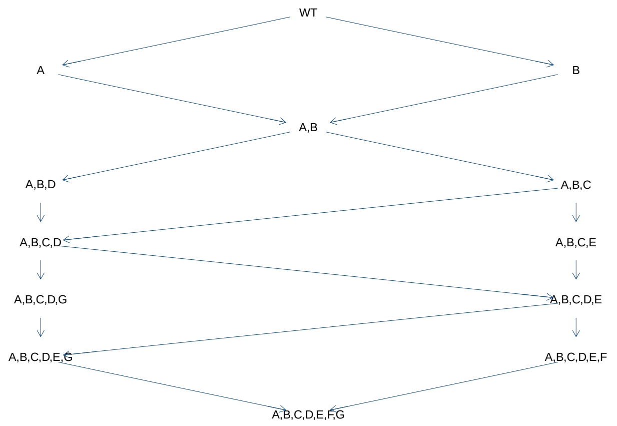


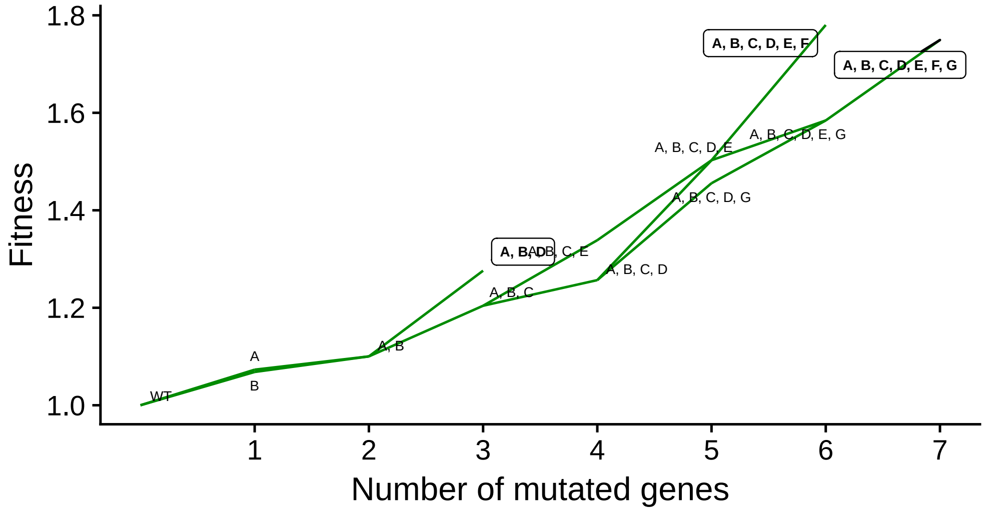


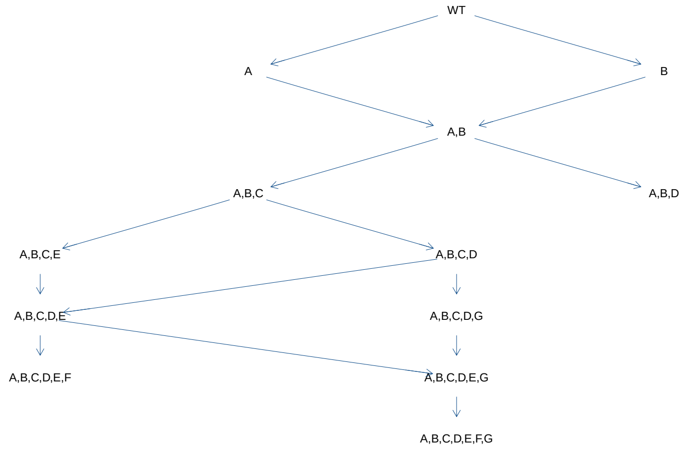

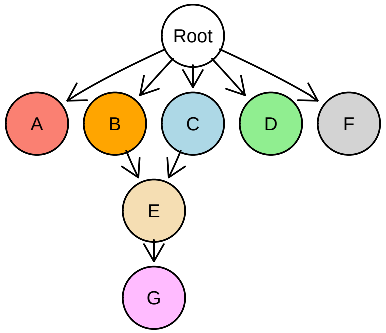


| | |
|---|---|
| *ID* | IDsDxkW7aWMDoTYv |
| *acc genotypes* | 48 |
| *peaks no back* | 8 |
| *removed edges* | 27 |
| *prop rem edges* | 0.225 |

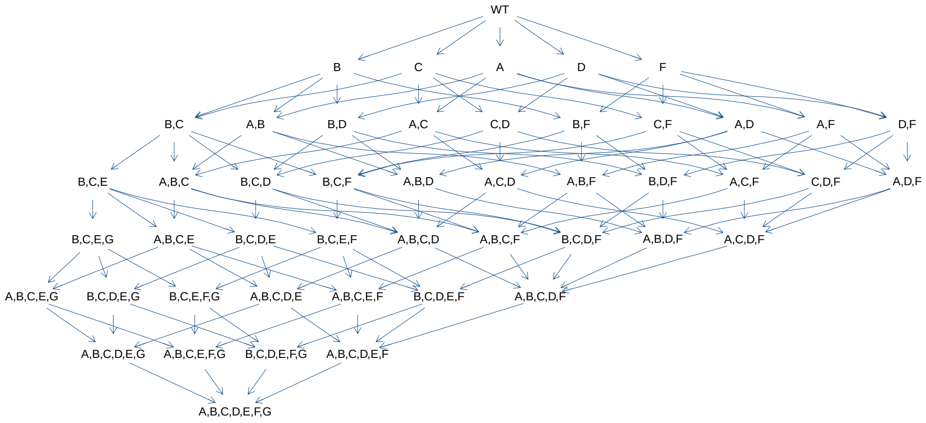


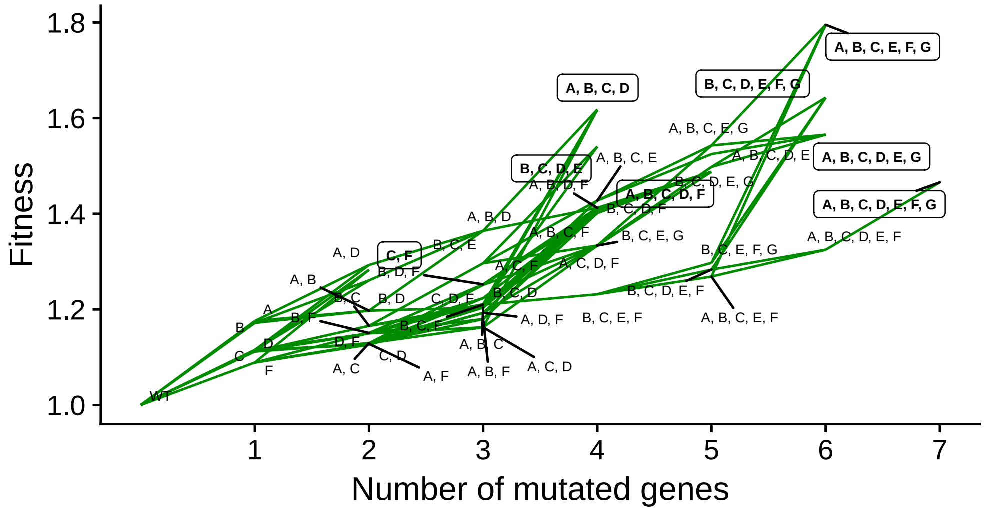


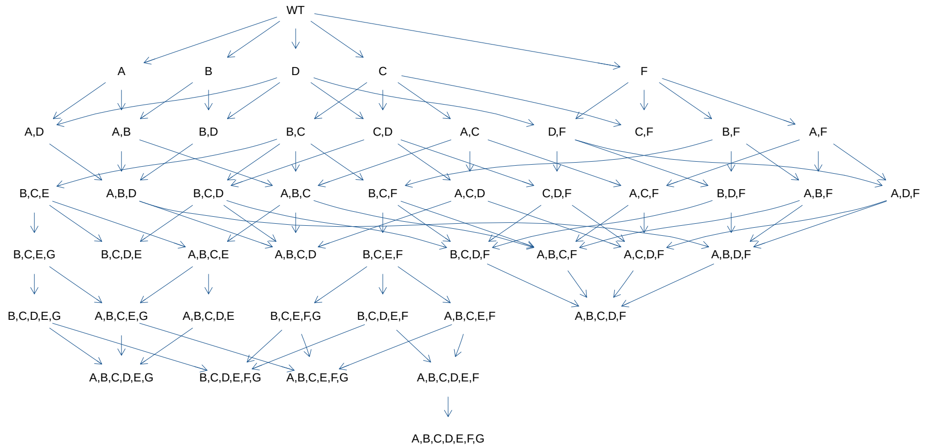

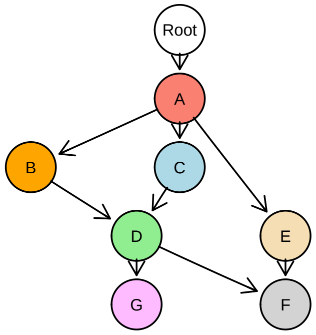


| | |
|---|---|
| *ID* | Lh10CtFMPrzzV6ihq |
| *acc genotypes* | 15 |
| *peaks no back* | 2 |
| *removed edges* | 1 |
| *prop rem edges* | 0.045 |

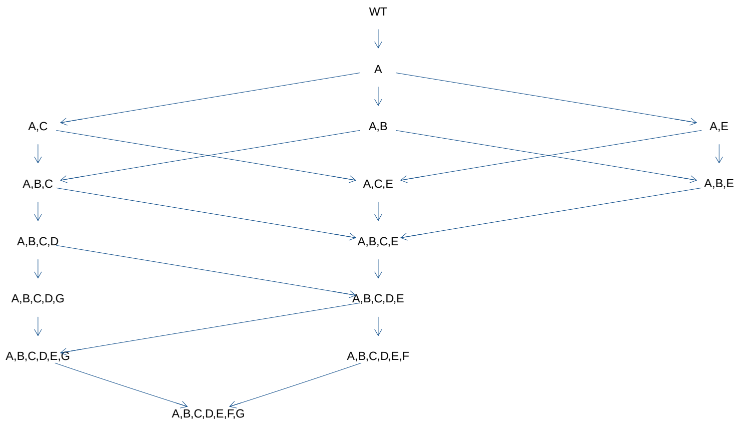


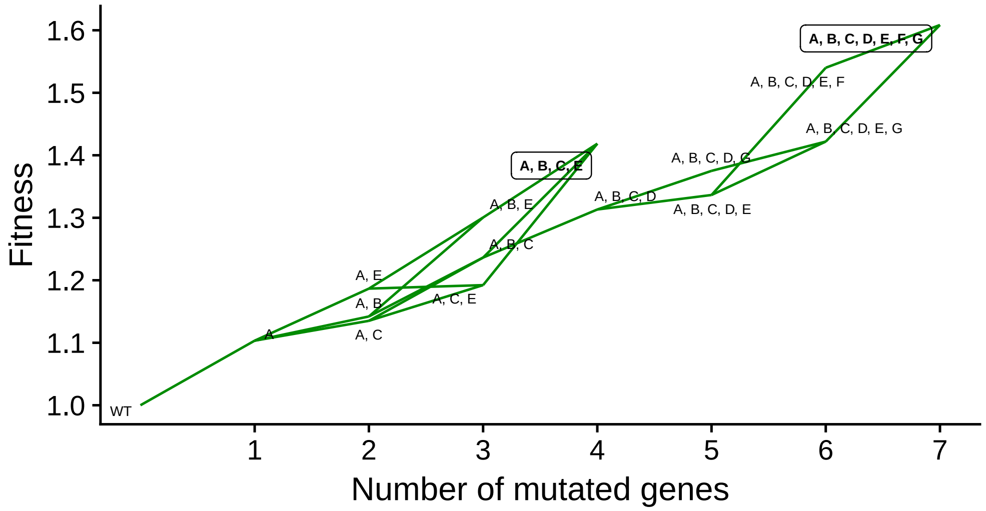


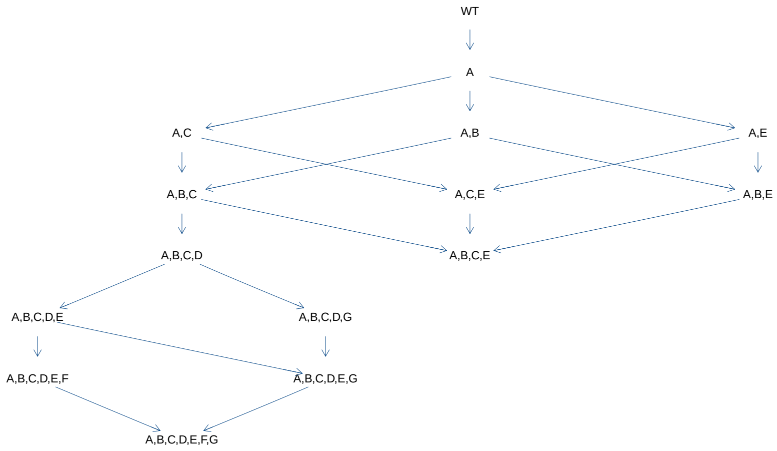

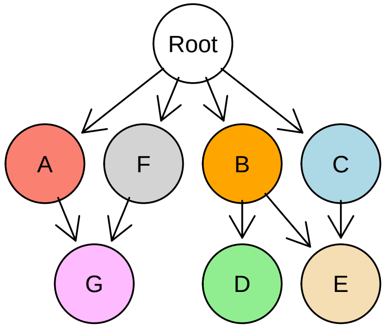


| | |
|---|---|
| *ID* | LqgKBG79CNzADijg |
| *acc genotypes* | 40 |
| *peaks no back* | 6 |
| *removed edges* | 14 |
| *prop rem edges* | 0.156 |

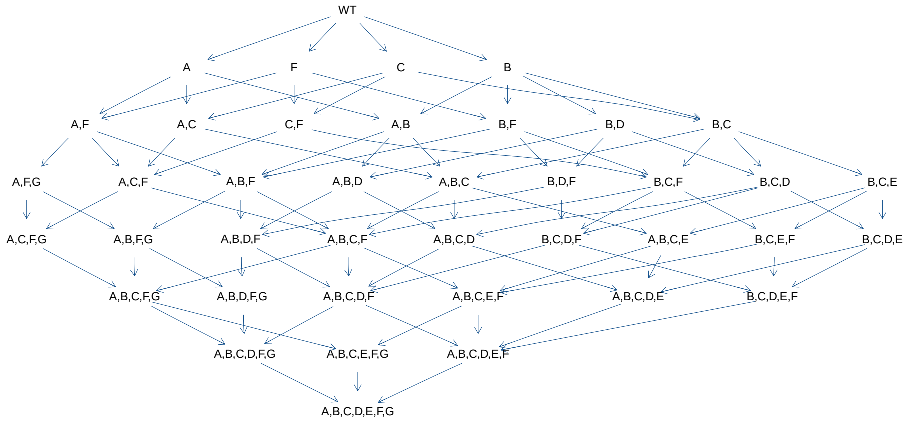


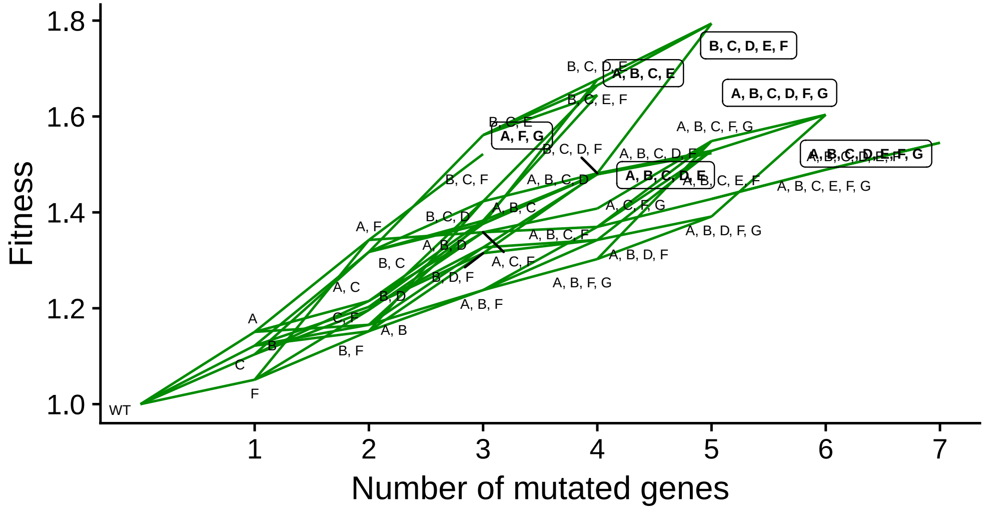


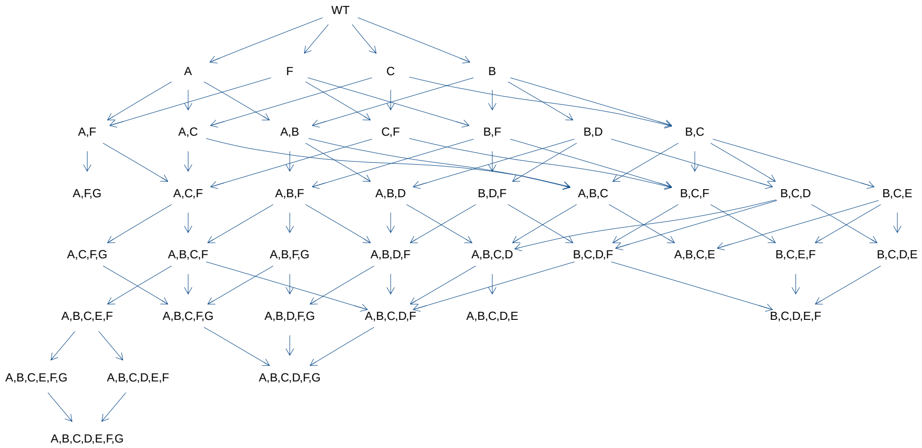

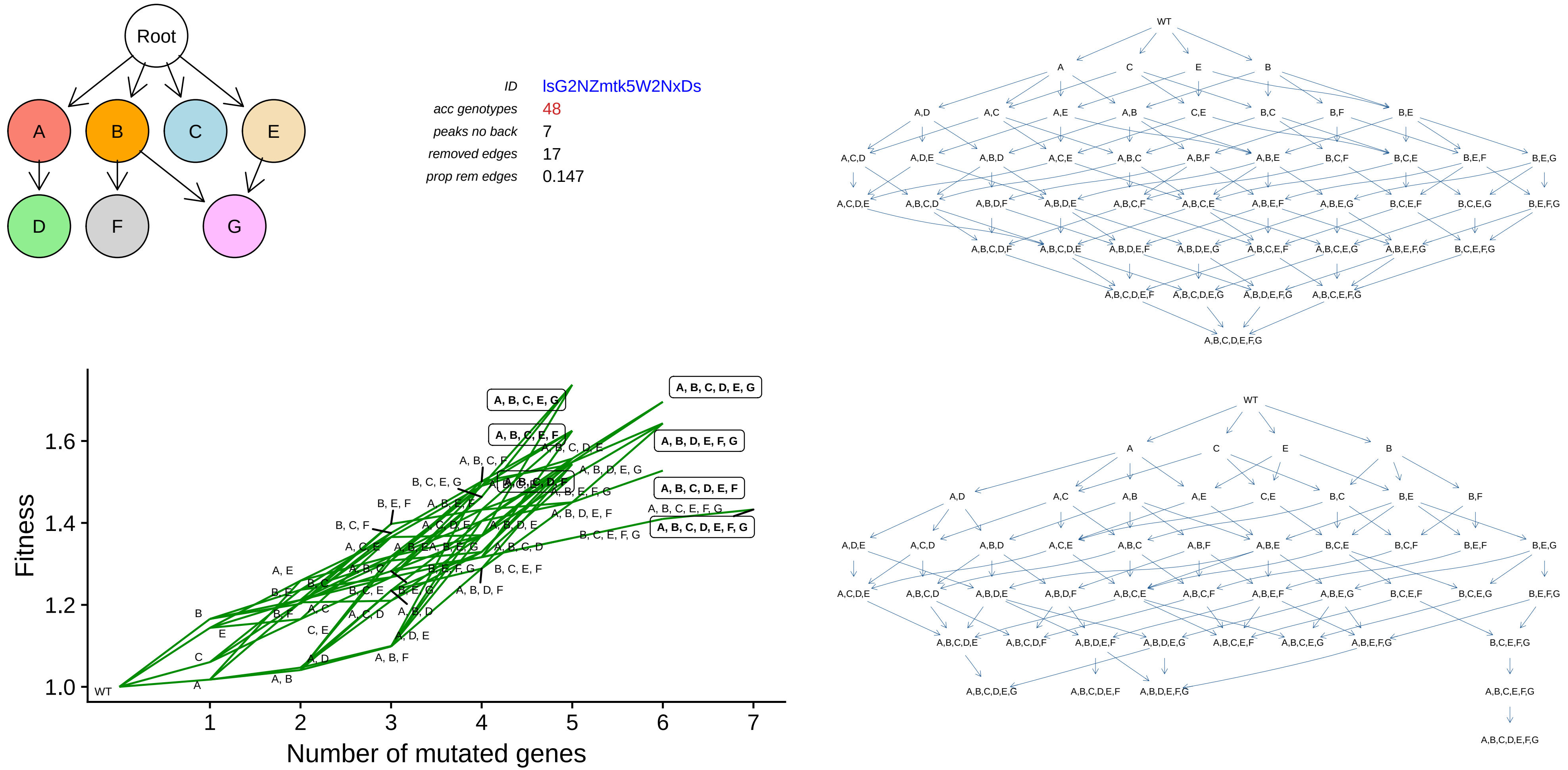
Root
A
B
C
E
D
F
G
ID lsG2NZmtk5W2NxDs
acc genotypes 48
peaks no back 7
removed edges 17
prop rem edges 0.147
A, B, C, E, G
A, B, C, D, E, G
A, B, C, E, F
A, B, D, E, F, G
A, B, C, D, E, F
A, B, C, D, E, F, G
Fitness
Number of mutated genes
WT
A,B,C,D,E,F,G

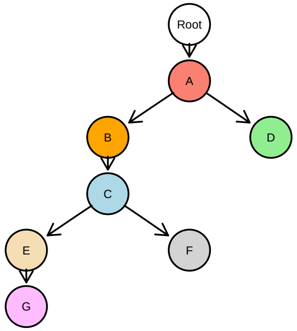


*ID* lsjclB1mAC8mHR9Q
*acc genotypes* 17
*peaks no back* 3
*removed edges* 3
*prop rem edges* 0.111

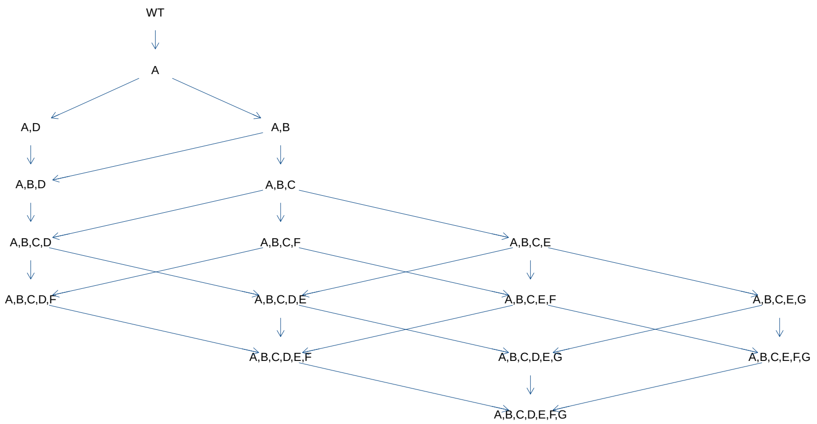


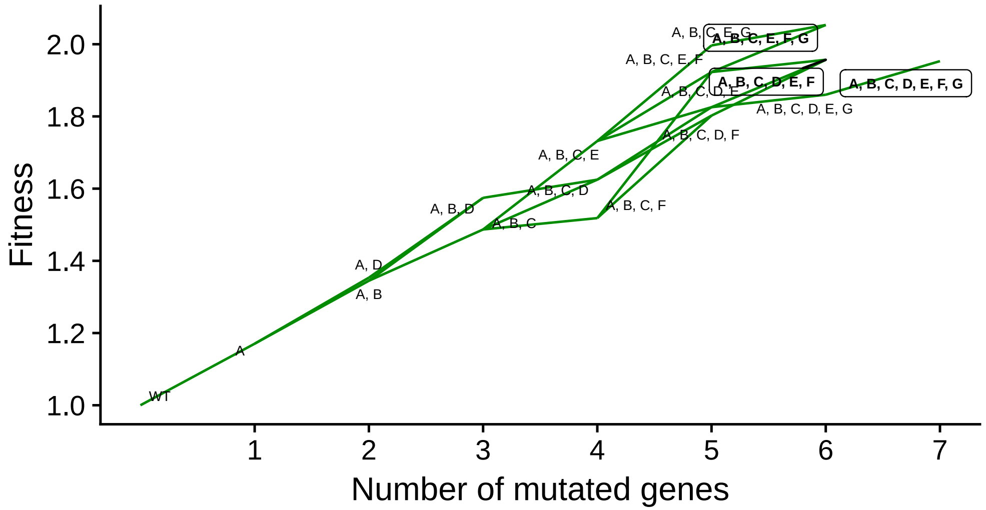


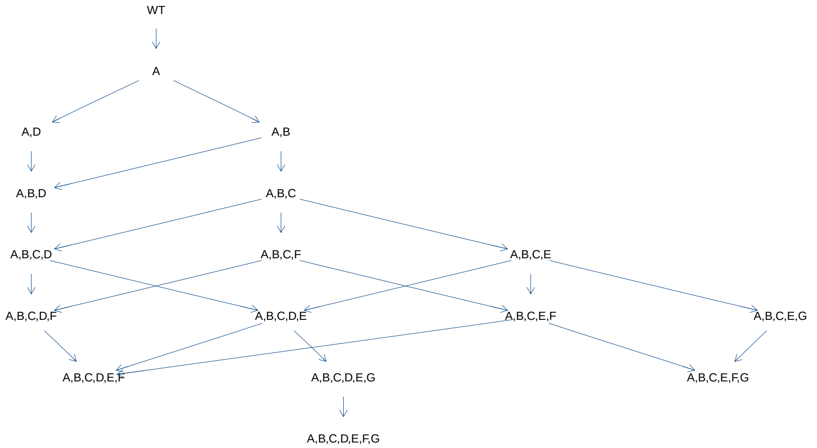

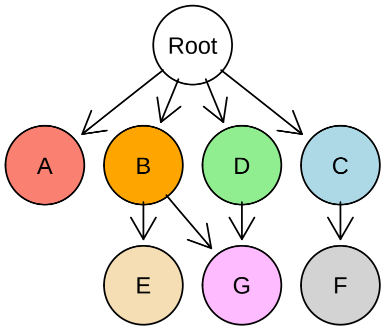


*ID* LSnOGiz5m5AoyDGr
*acc genotypes* 48
*peaks no back* 7
*removed edges* 17
*prop rem edges* 0.147

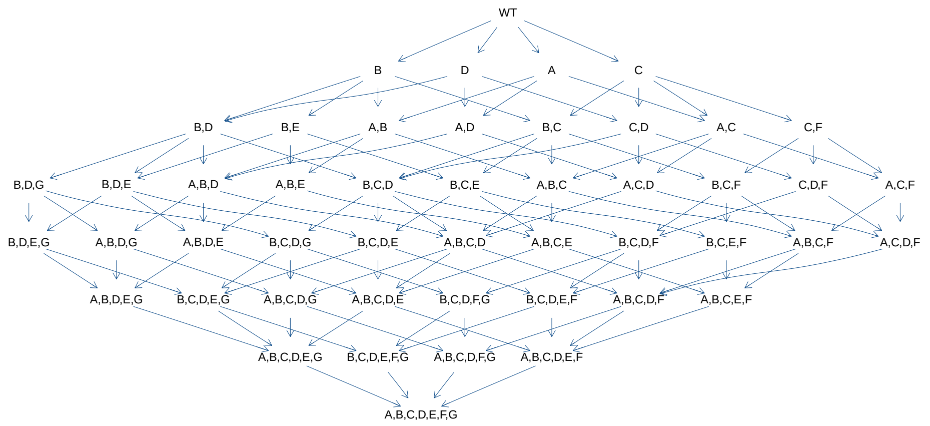


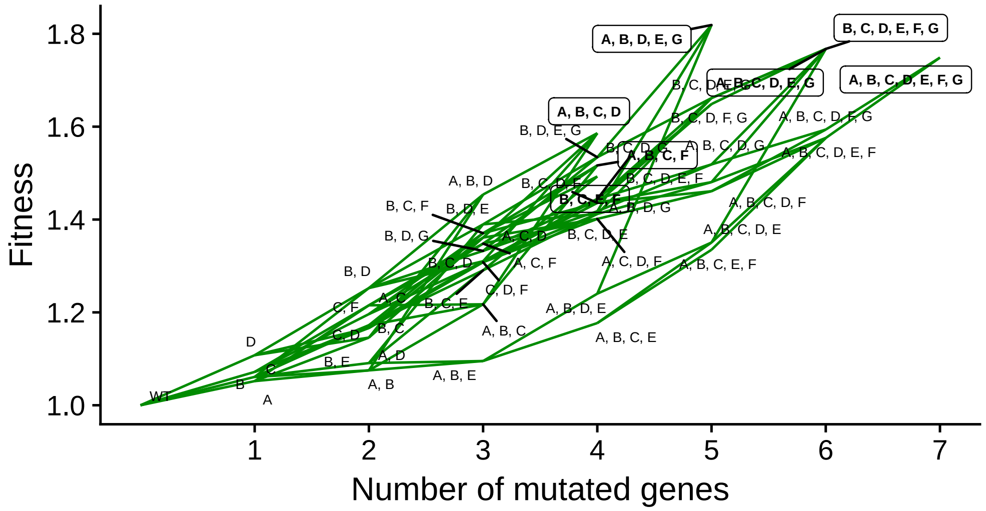


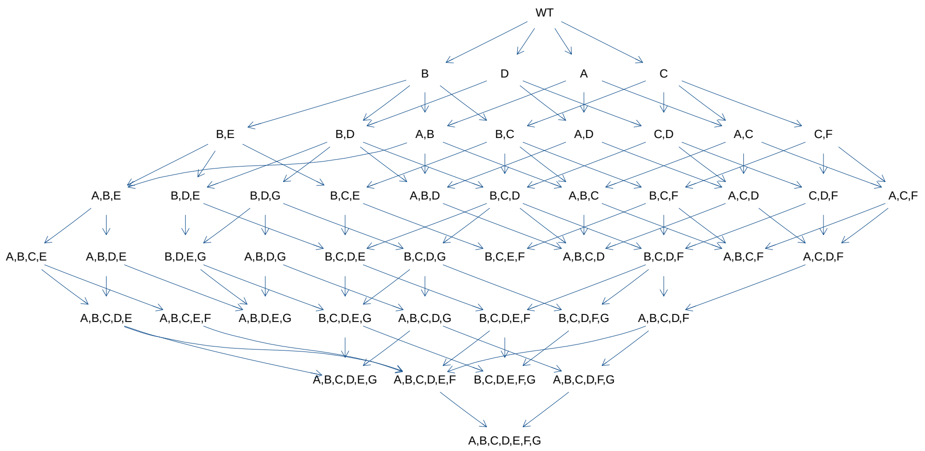

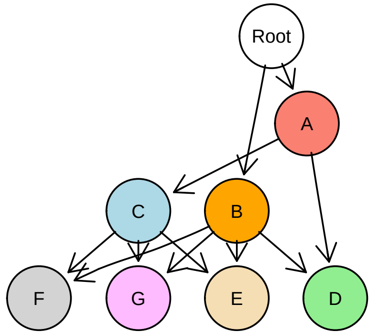


| | |
|---|---|
| *ID* | LSXwTnPZnOfXSdzA |
| *acc genotypes* | 22 |
| *peaks no back* | 2 |
| *removed edges* | 2 |
| *prop rem edges* | 0.049 |

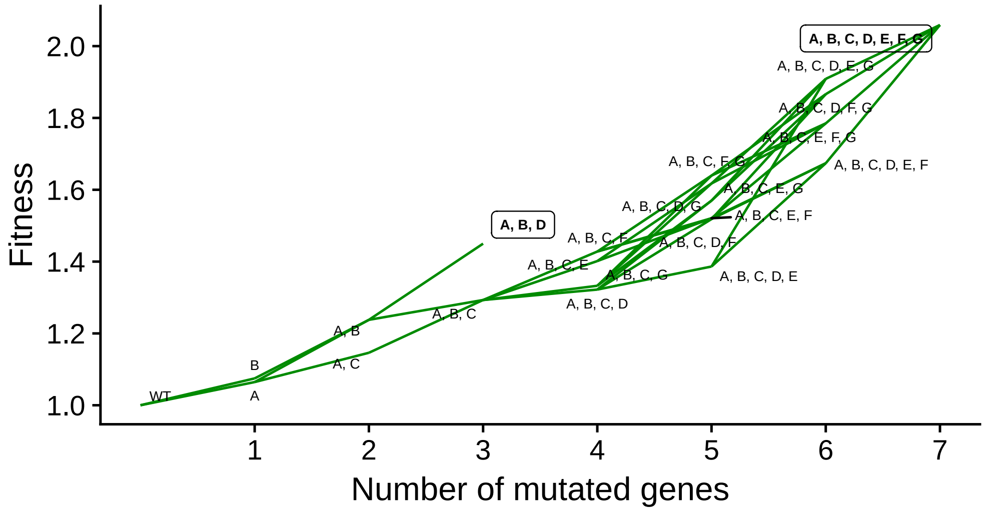


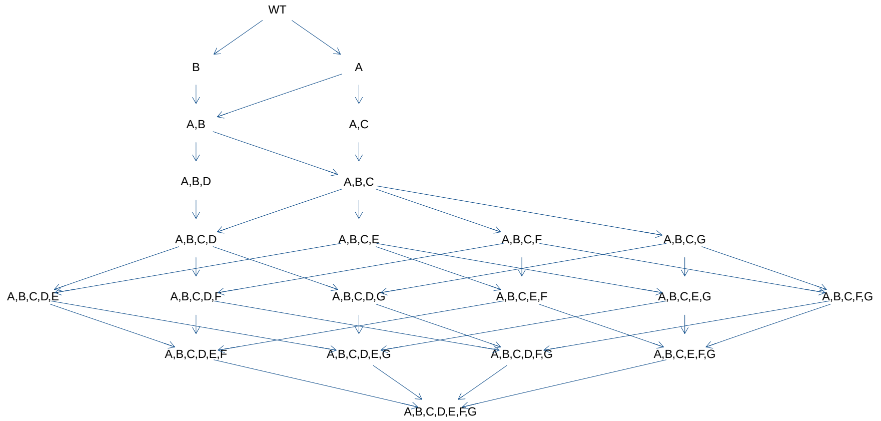


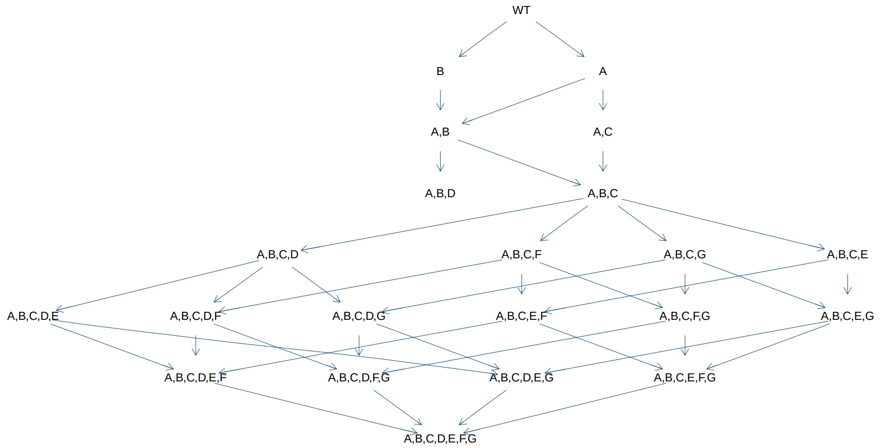

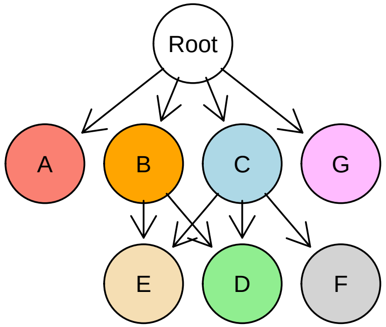


| | |
|---|---|
| *ID* | lsZUuCCknMZno10B8 |
| *acc genotypes* | 48 |
| *peaks no back* | 8 |
| *removed edges* | 21 |
| *prop rem edges* | 0.175 |

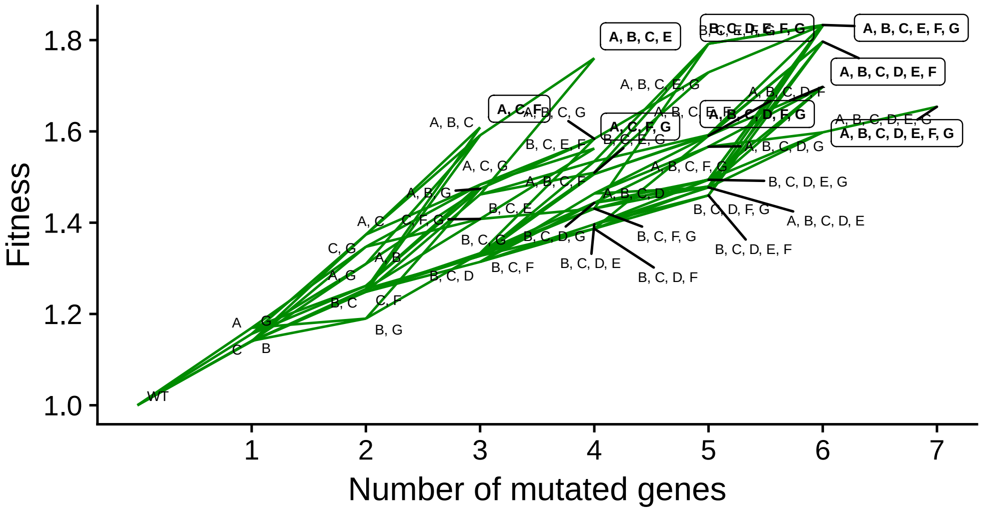


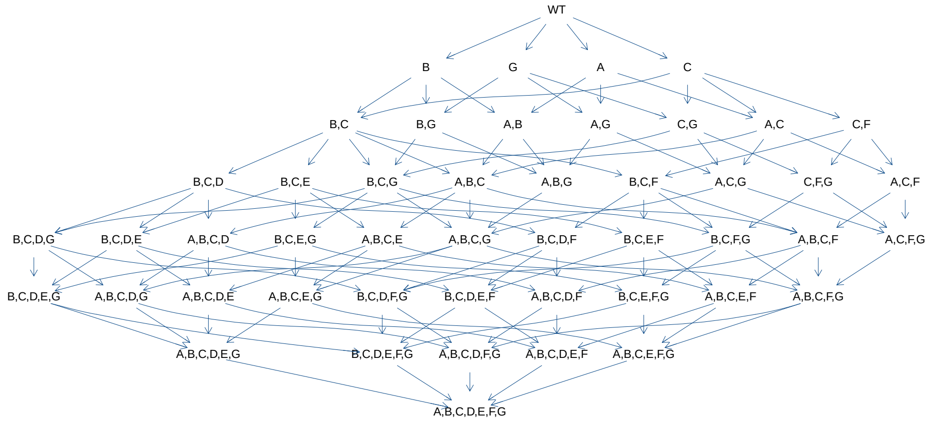


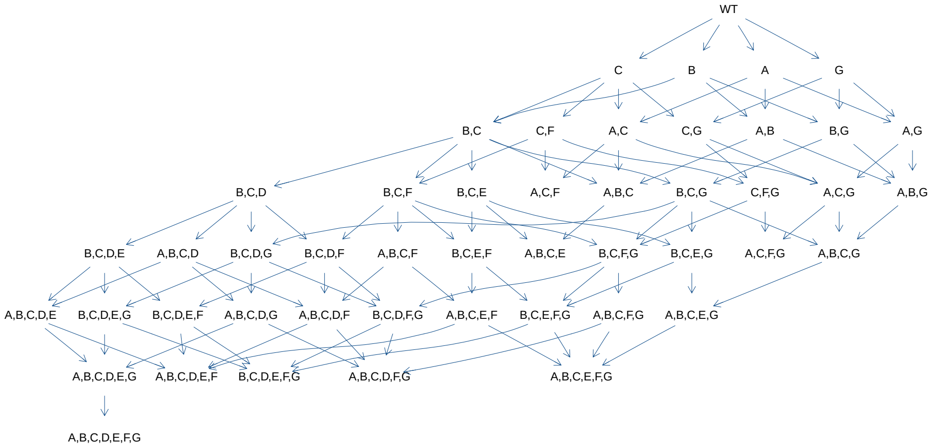

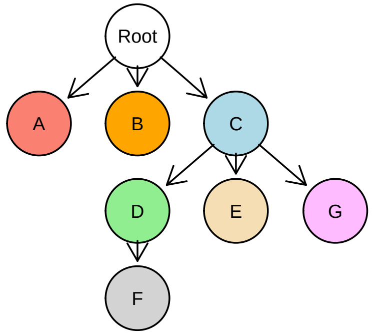


| | |
|---|---|
| *ID* | m3T5PDQjmlID9CNk |
| *acc genotypes* | 52 |
| *peaks no back* | 3 |
| *removed edges* | 17 |
| *prop rem edges* | 0.125 |

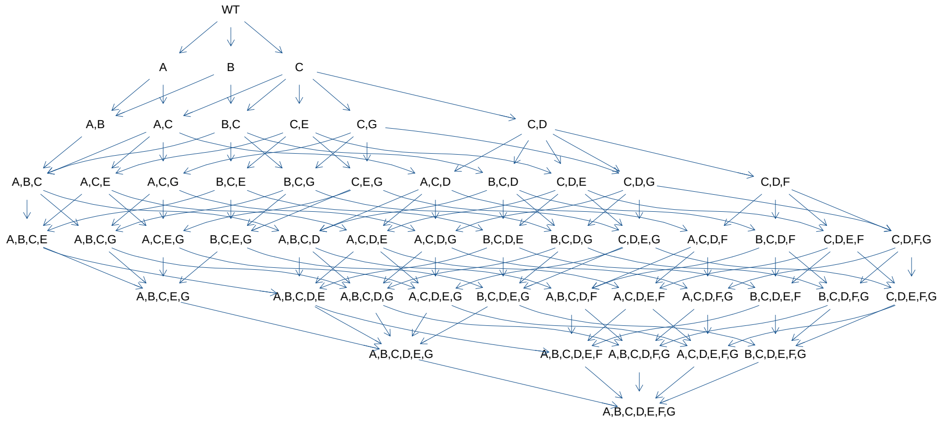

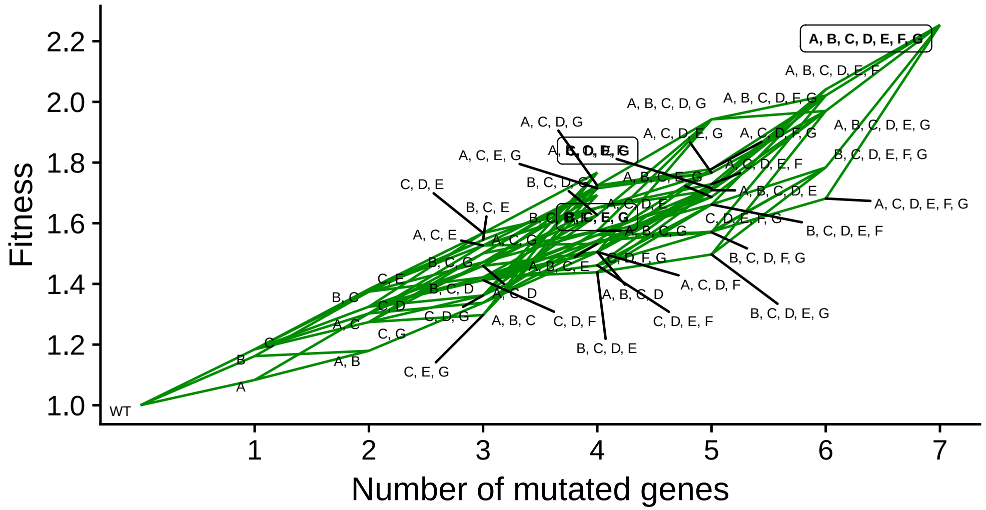


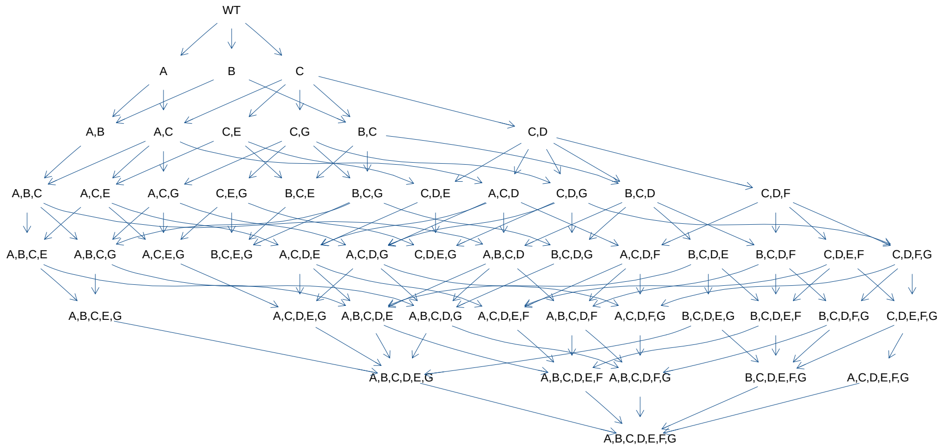

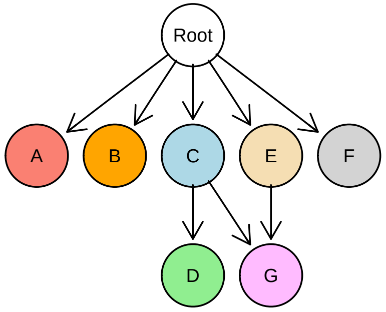


*ID* m8kM7ayaSoitwqmo
*acc genotypes* 64
*peaks no back* 11
*removed edges* 36
*prop rem edges* 0.205

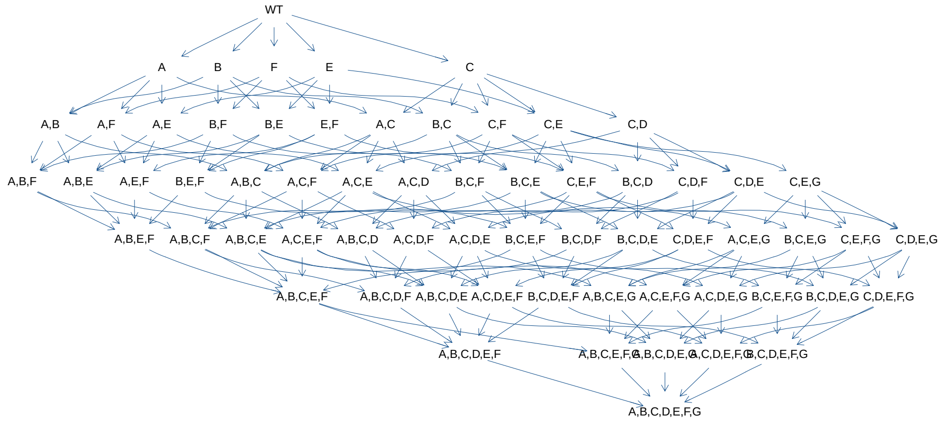

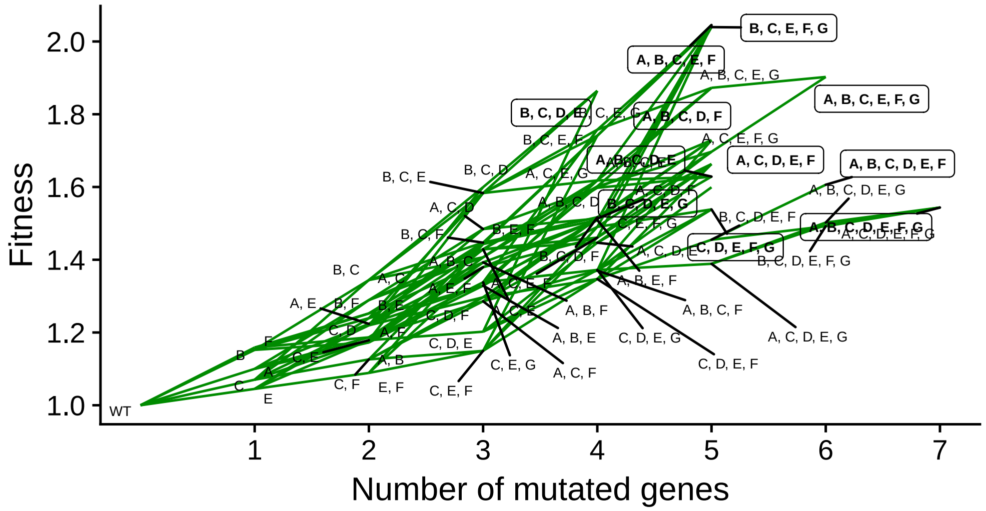


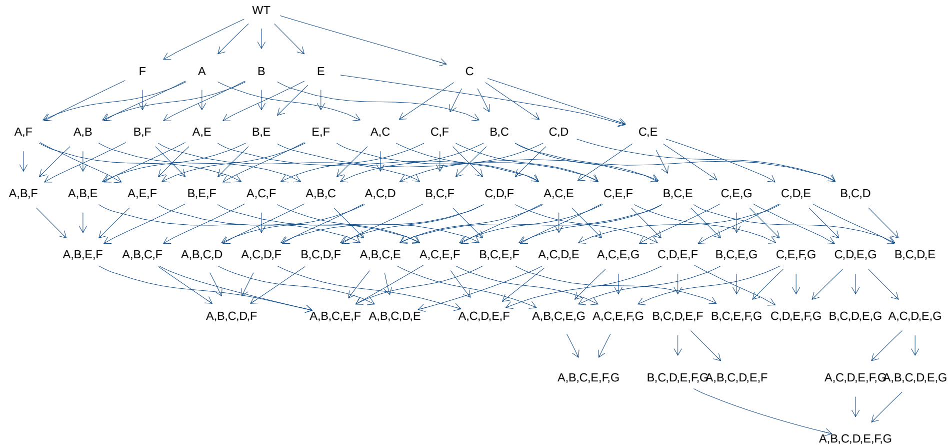

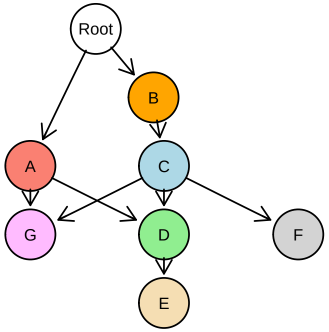


| | |
|---|---|
| *ID* | MHkrL757u9iPj10kv |
| *acc genotypes* | 18 |
| *peaks no back* | 4 |
| *removed edges* | 4 |
| *prop rem edges* | 0.138 |

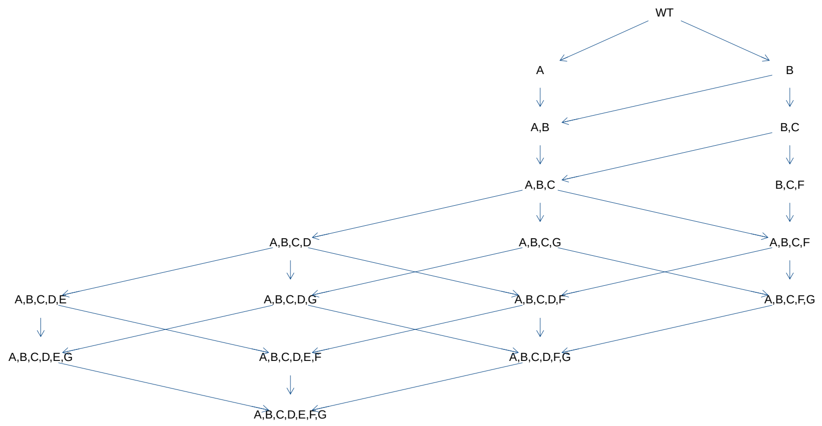


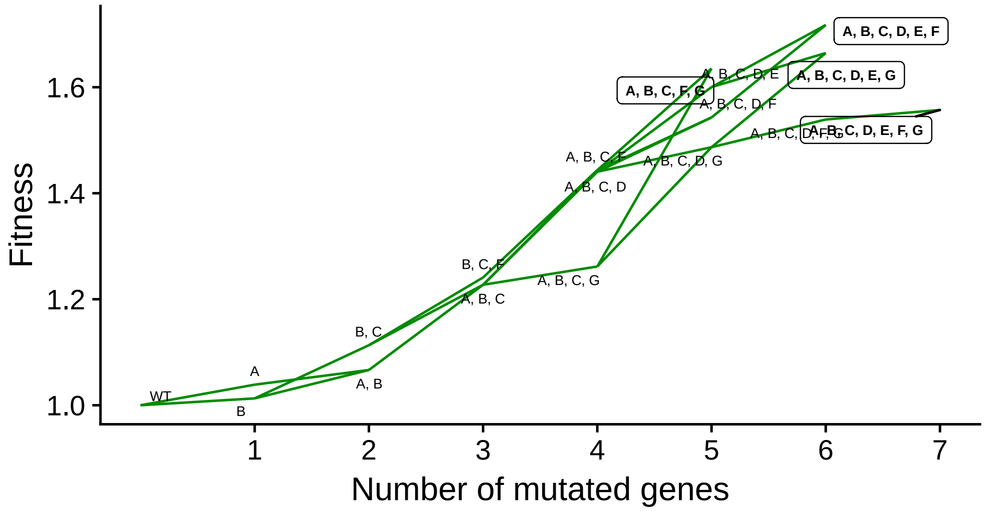


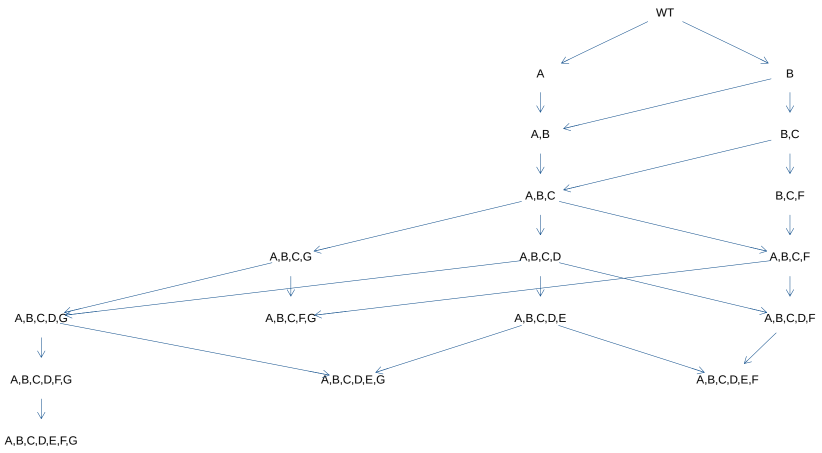

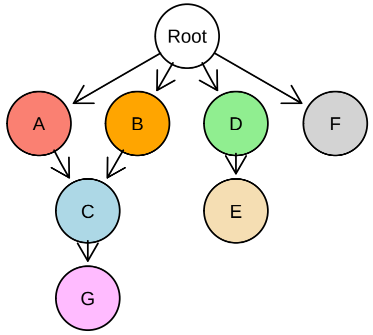


| | |
|---|---|
| *ID* | MifrHKJzEqocbcMH |
| *acc genotypes* | 36 |
| *peaks no back* | 3 |
| *removed edges* | 9 |
| *prop rem edges* | 0.115 |

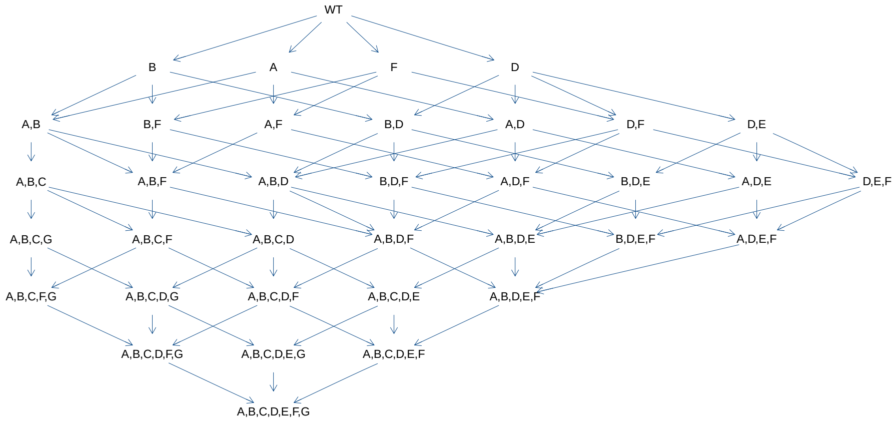


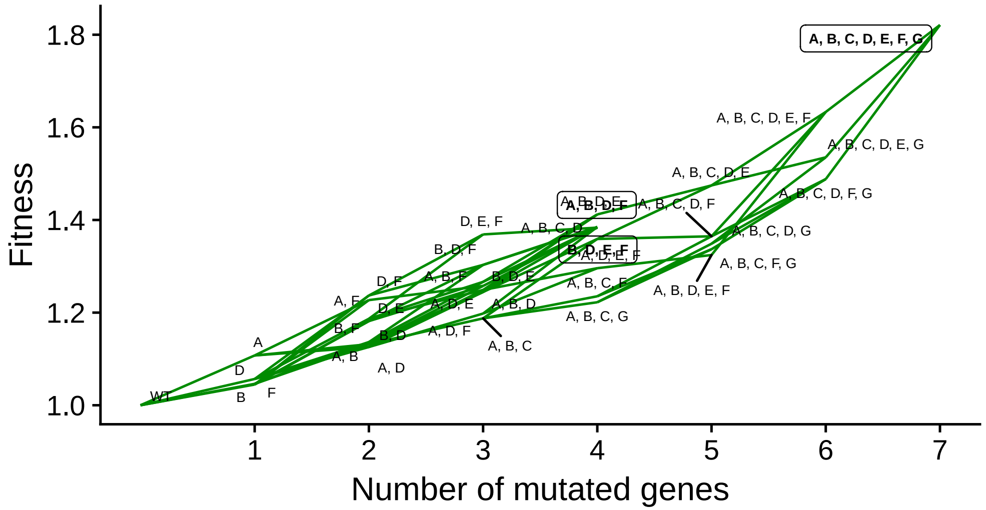


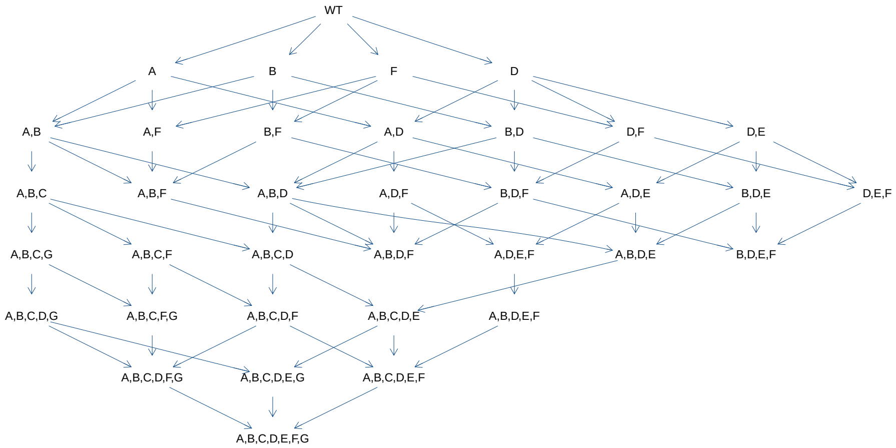

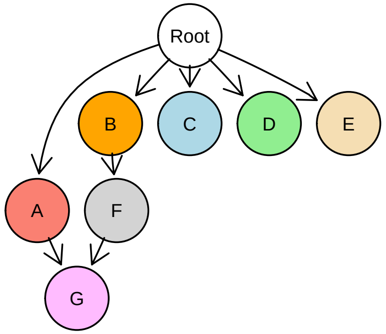


| | |
|---|---|
| *ID* | mtZbHQ97NJy8xIsf |
| *acc genotypes* | 56 |
| *peaks no back* | 10 |
| *removed edges* | 26 |
| *prop rem edges* | 0.176 |

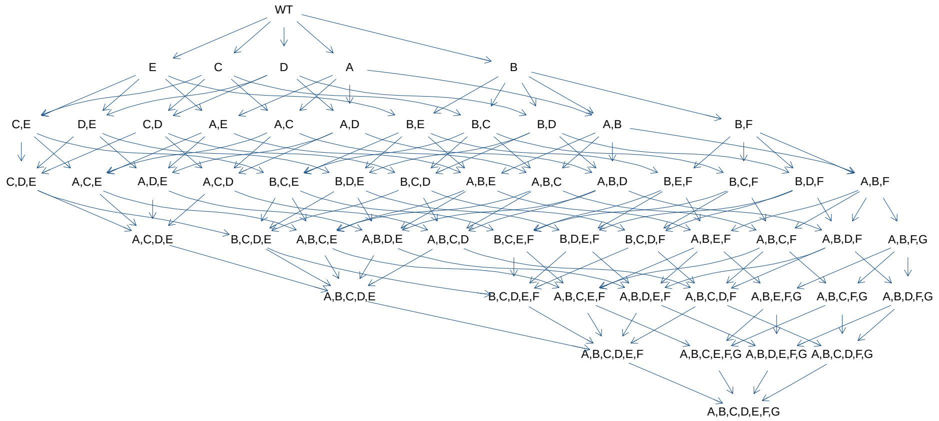

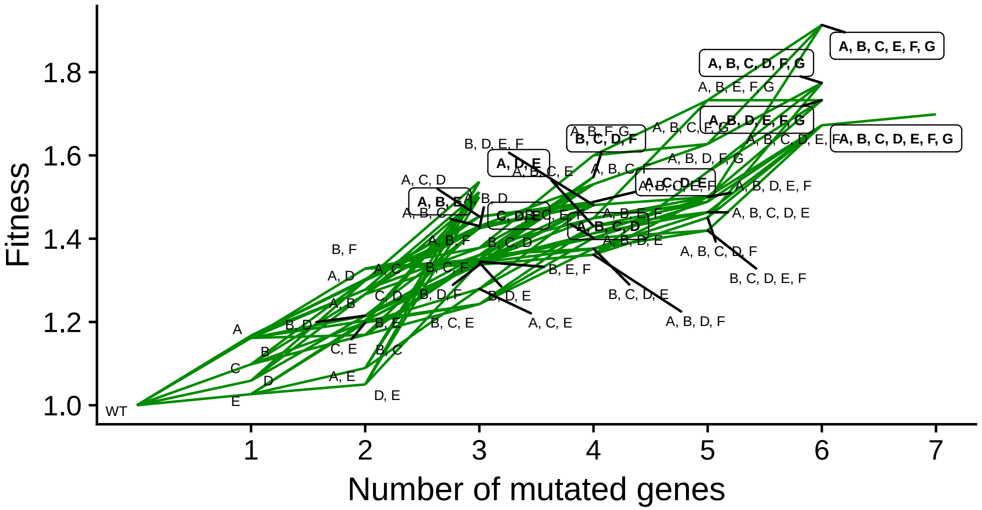


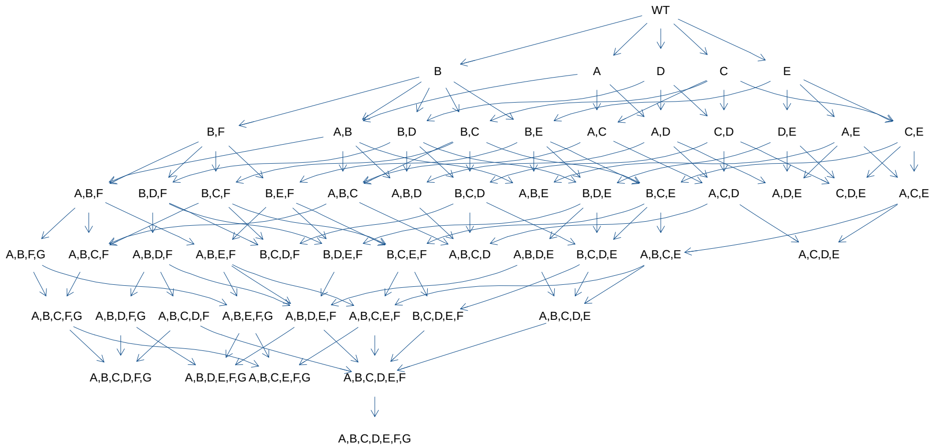

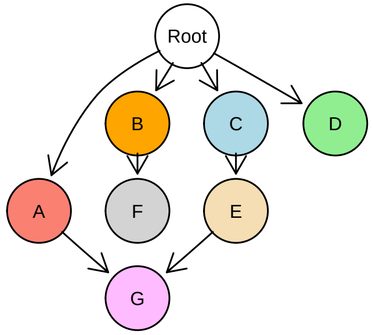


*ID* MuAVAUt10hjGNt5AD
*acc genotypes* 42
*peaks no back* 7
*removed edges* 15
*prop rem edges* 0.155

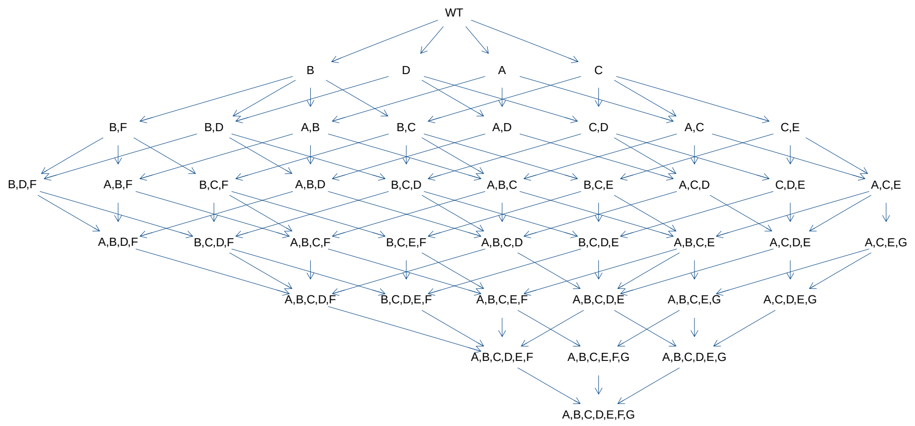


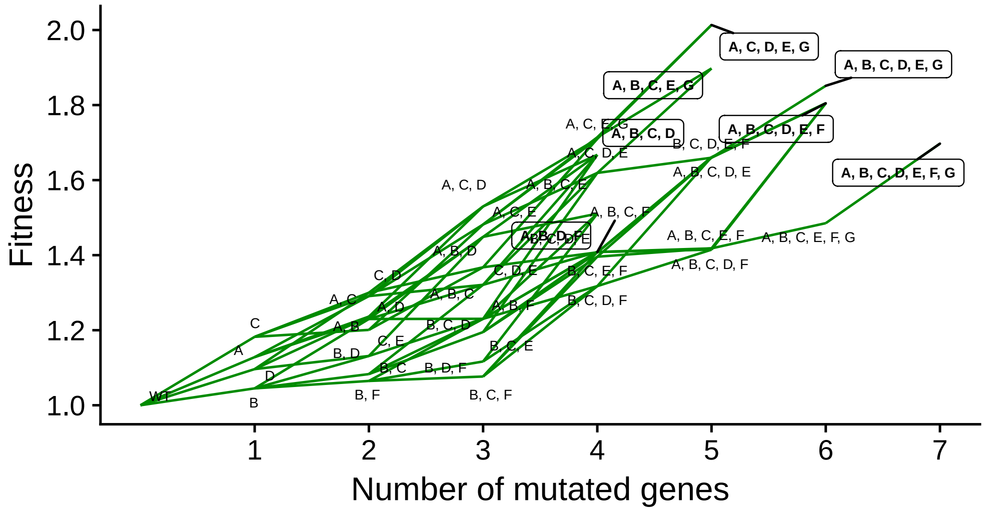


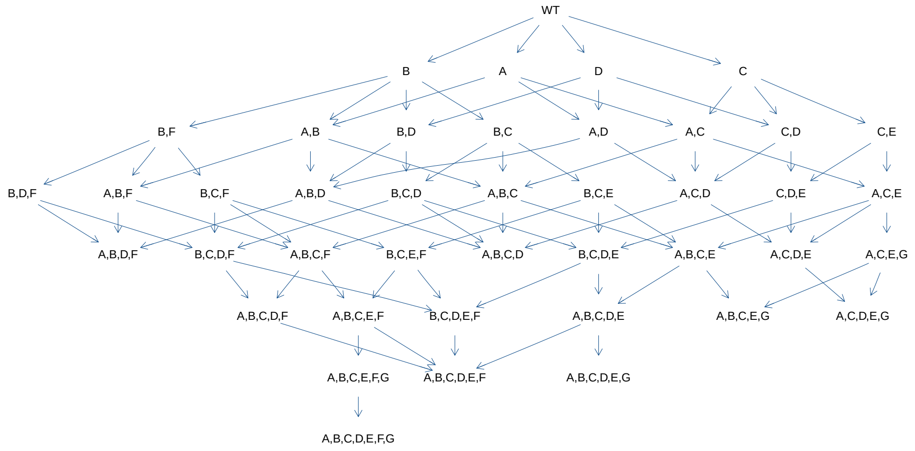

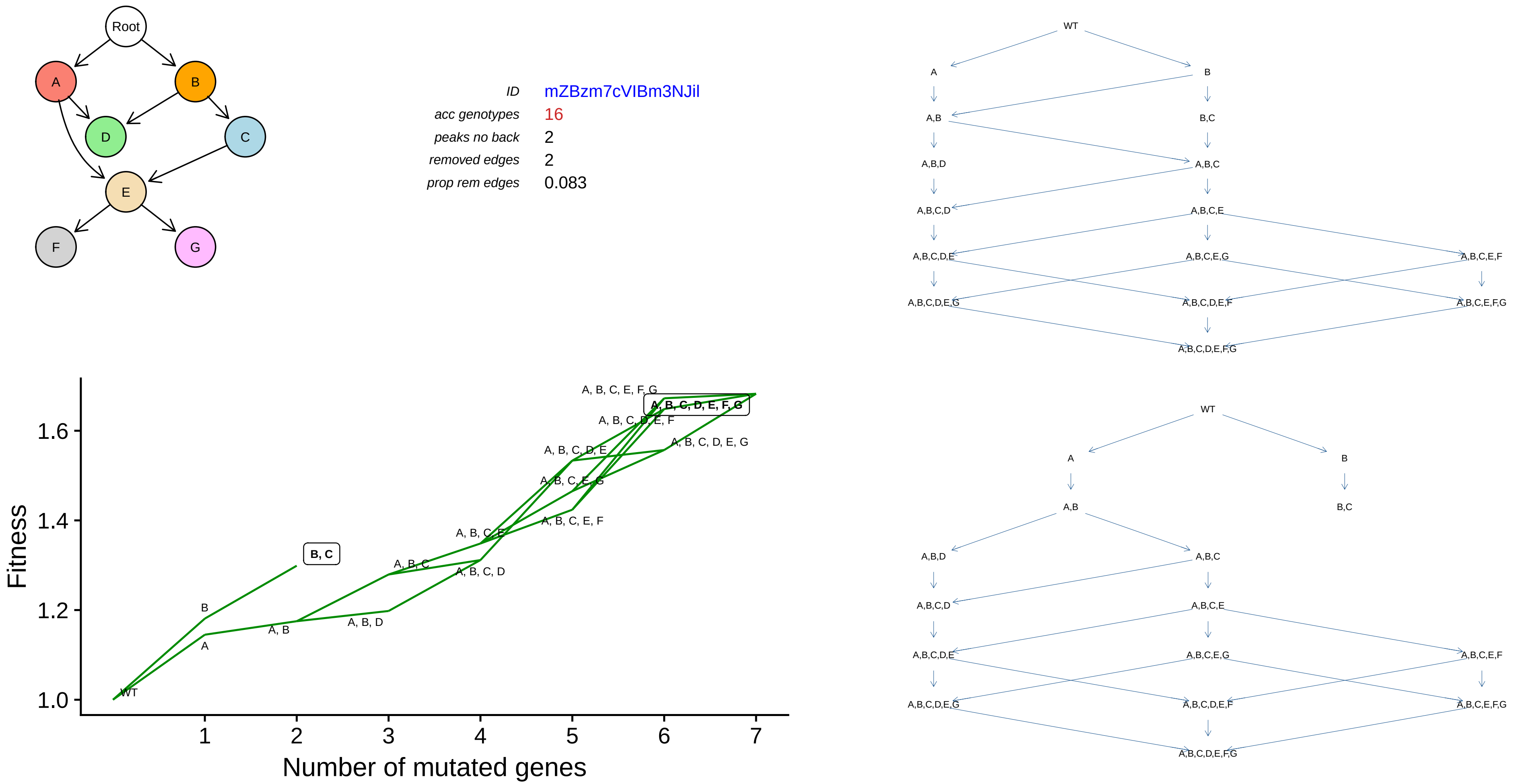
Root
A
B
C
D
E
F
G
ID mZBzm7cVIBm3NJil
acc genotypes 16
peaks no back 2
removed edges 2
prop rem edges 0.083
Fitness
Number of mutated genes
1.0
1.2
1.4
1.6
1
2
3
4
5
6
7
WT
A
B
A, B
B, C
A, B, C
A, B, D
A, B, C, D
A, B, C, E
A, B, C, D, E
A, B, C, E, G
A, B, C, E, F
A, B, C, D, E, F
A, B, C, D, E, G
A, B, C, E, F, G
A, B, C, D, E, F, G
WT
A
B
A,B
B,C
A,B,D
A,B,C
A,B,C,D
A,B,C,E
A,B,C,D,E
A,B,C,E,G
A,B,C,E,F
A,B,C,D,E,G
A,B,C,D,E,F
A,B,C,E,F,G
A,B,C,D,E,F,G

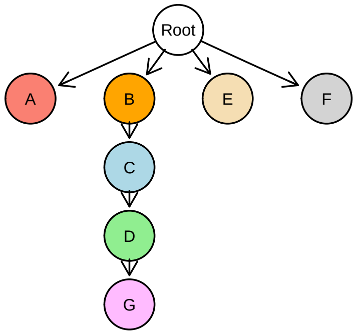


| | |
|---|---|
| *ID* | N7LwhGSvS3wDt8P2 |
| *acc genotypes* | 40 |
| *peaks no back* | 7 |
| *removed edges* | 13 |
| *prop rem edges* | 0.141 |

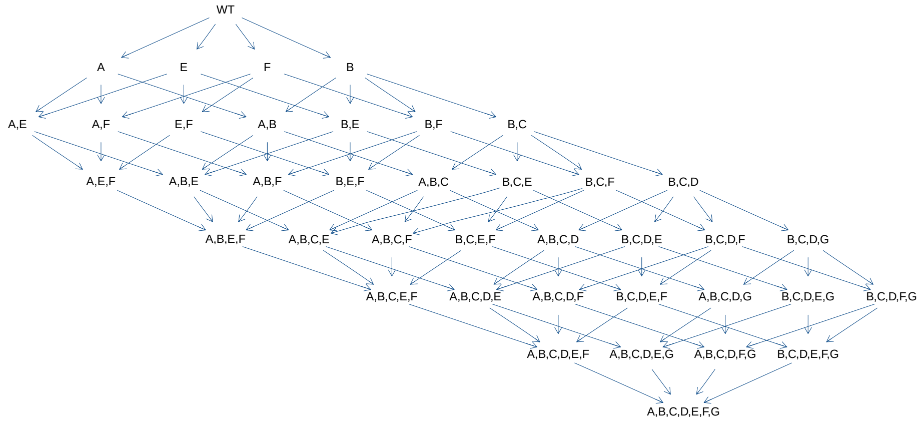


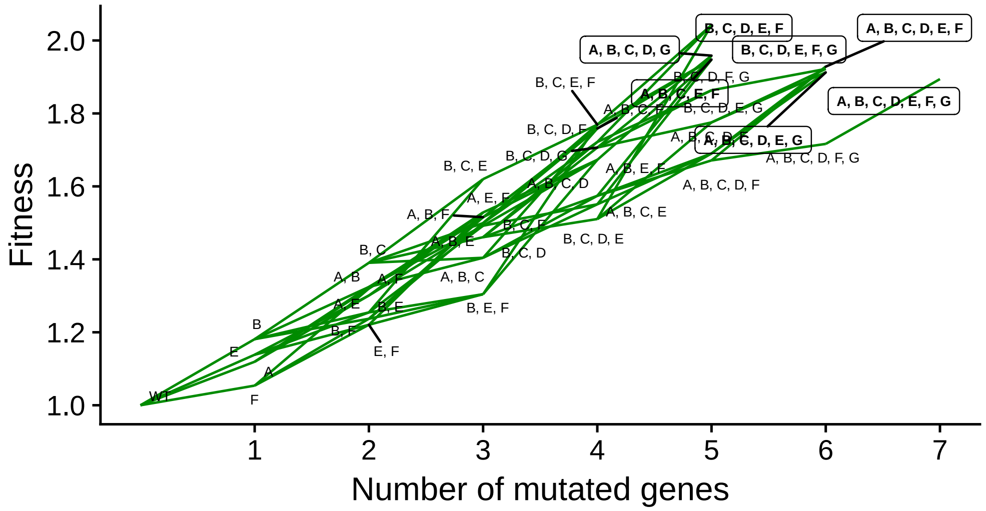


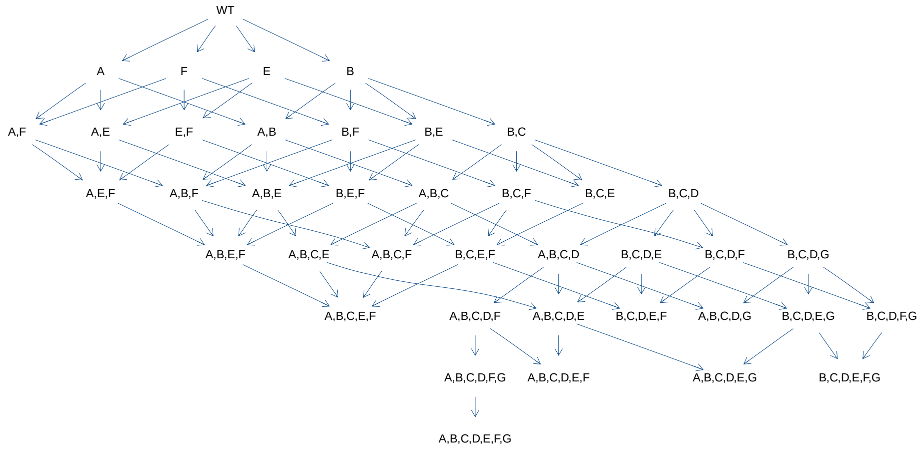

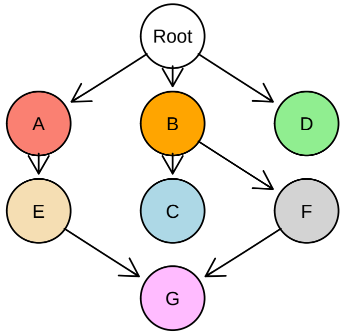


| | |
|---|---|
| *ID* | nAB3TW1qrCNzLA92 |
| *acc genotypes* | 34 |
| *peaks no back* | 5 |
| *removed edges* | 10 |
| *prop rem edges* | 0.137 |

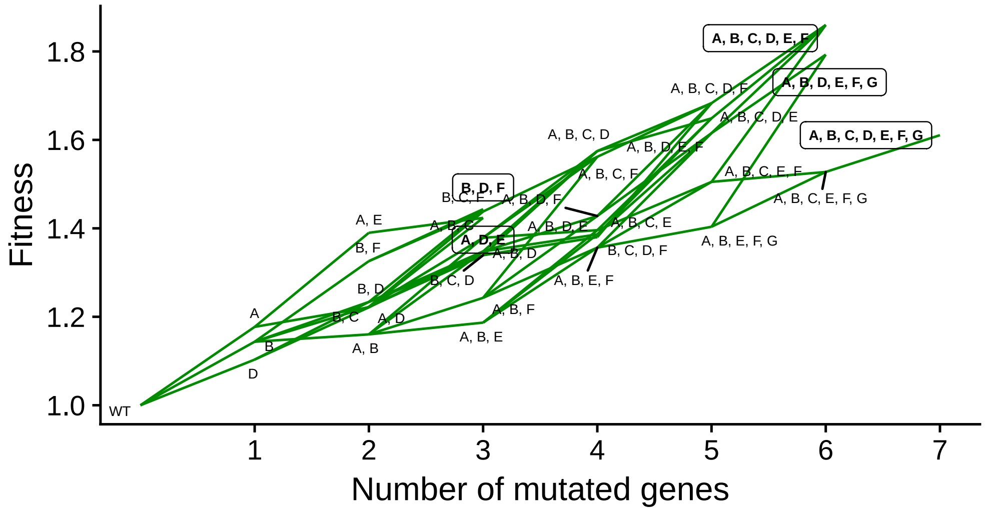


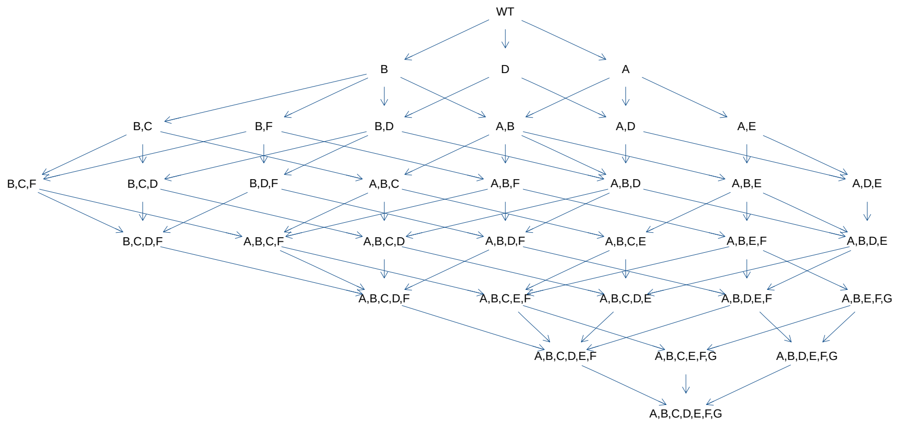

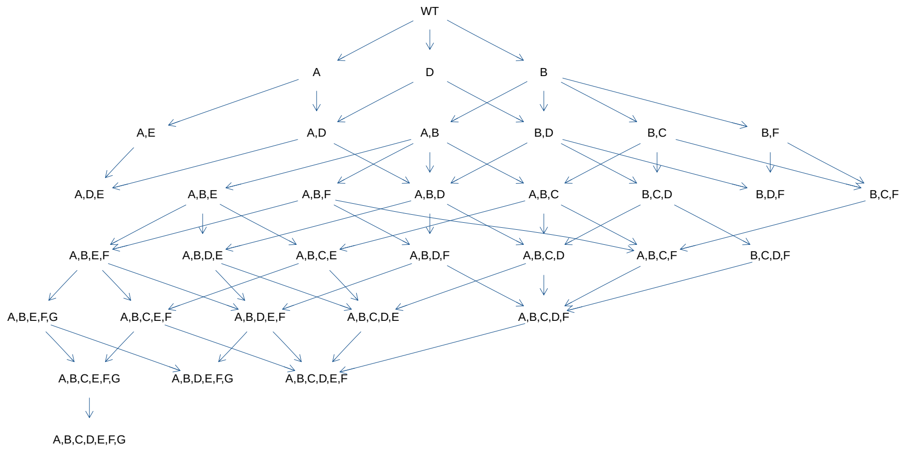

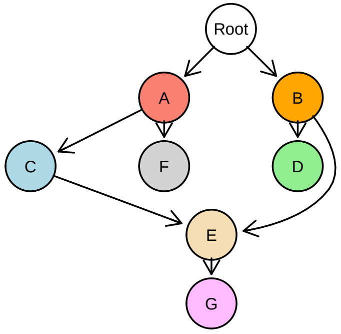


| | |
|---|---|
| *ID* | Nc8hEZJTul6Jz10xm |
| *acc genotypes* | 23 |
| *peaks no back* | 5 |
| *removed edges* | 9 |
| *prop rem edges* | 0.22 |

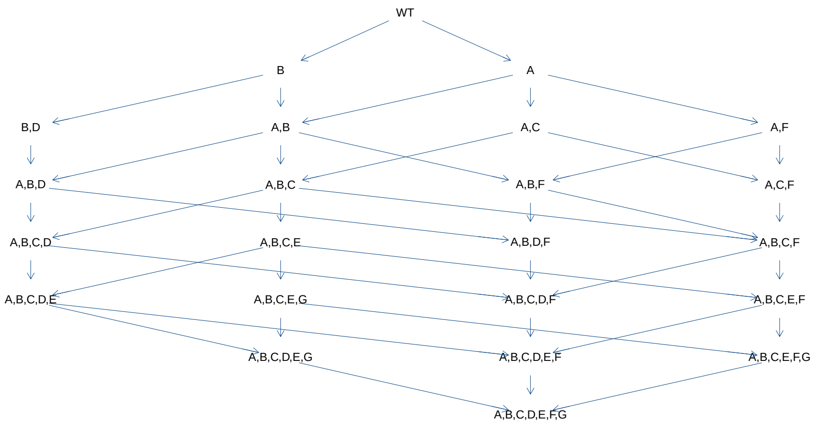


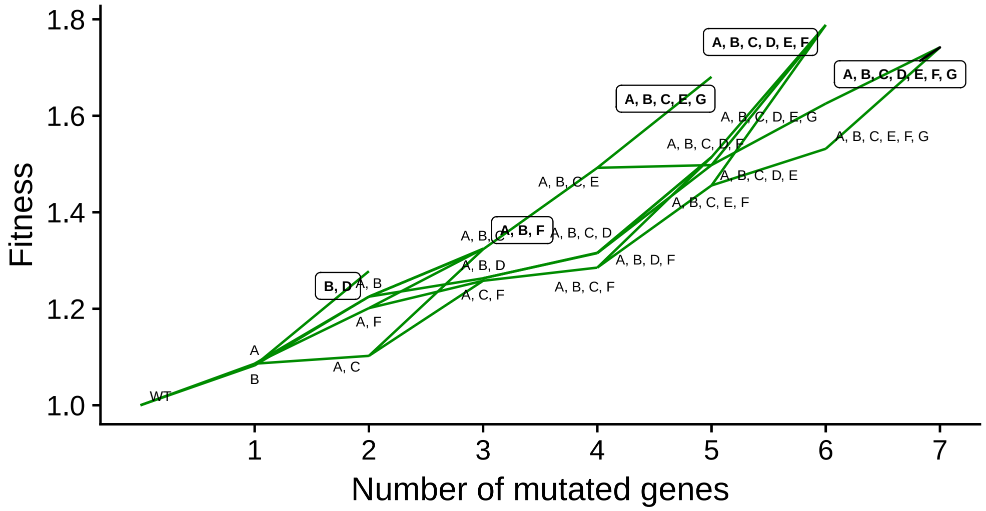


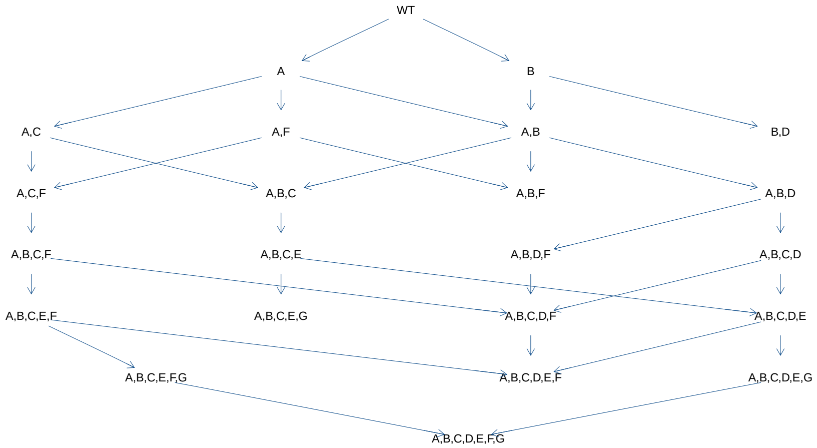

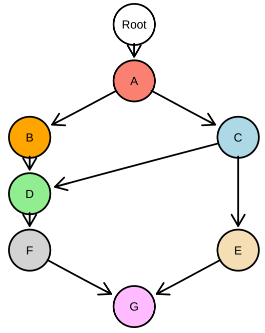


| | |
|---|---|
| *ID* | nCI8mAj78ZzZKlBn |
| *acc genotypes* | 12 |
| *peaks no back* | 2 |
| *removed edges* | 1 |
| *prop rem edges* | 0.067 |

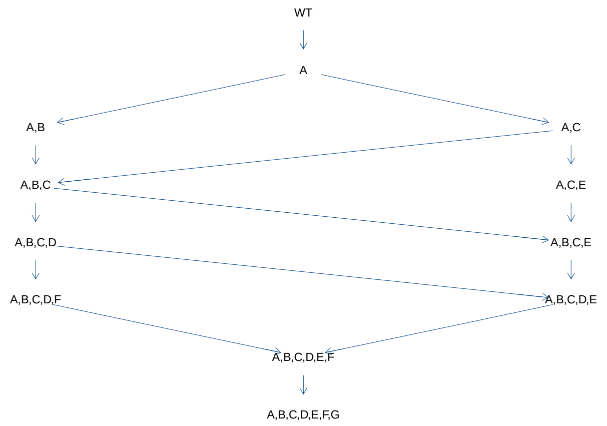


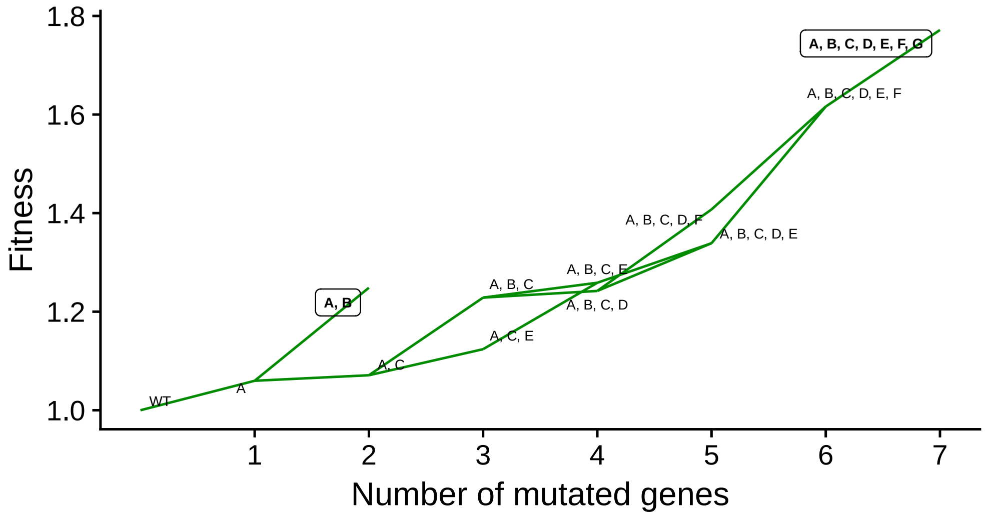


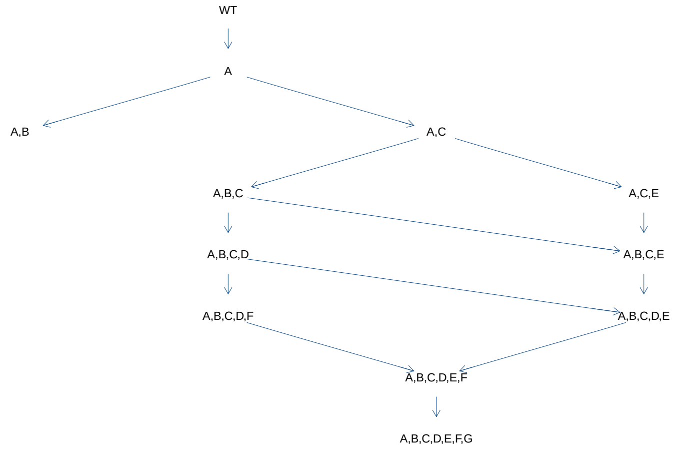

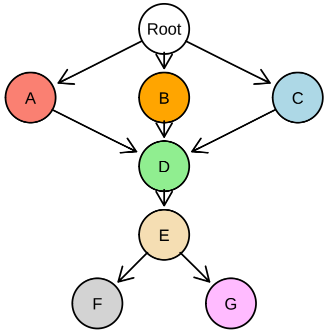


| | |
|---|---|
| *ID* | nKSE4LBW1z9FriV10 |
| *acc genotypes* | 13 |
| *peaks no back* | 2 |
| *removed edges* | 2 |
| *prop rem edges* | 0.111 |

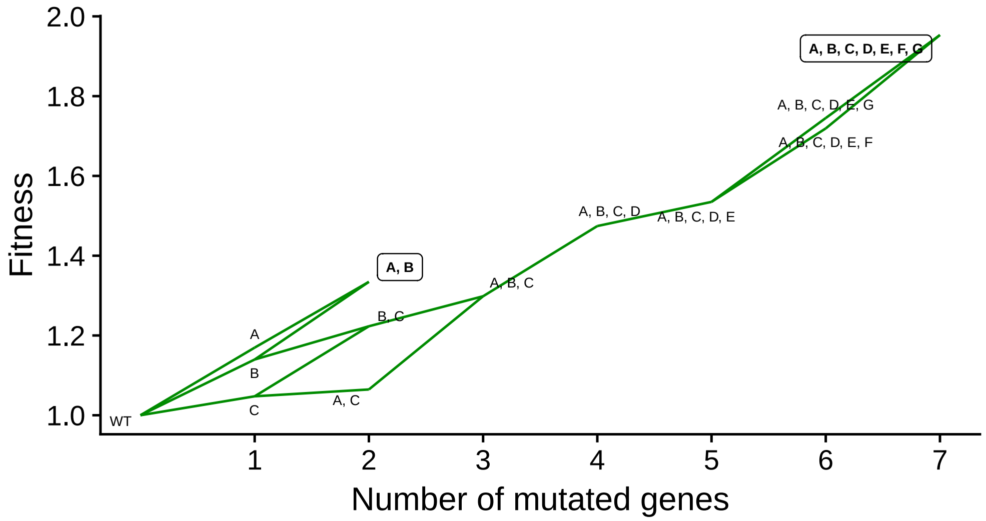


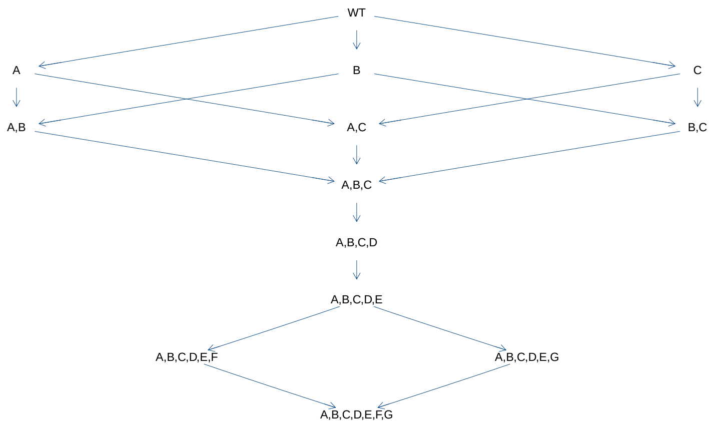


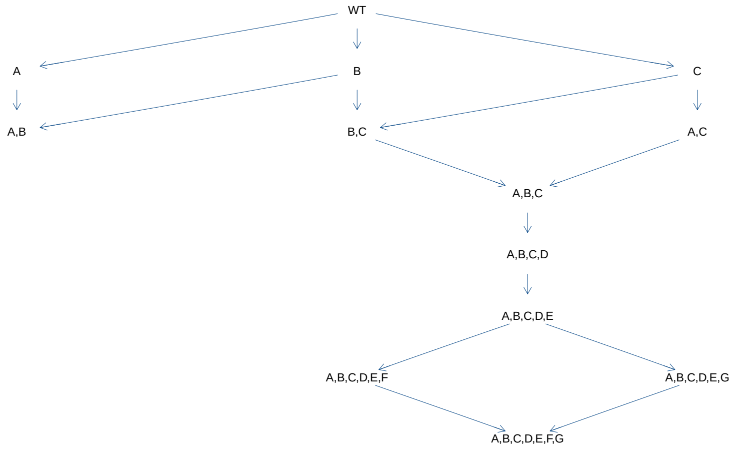

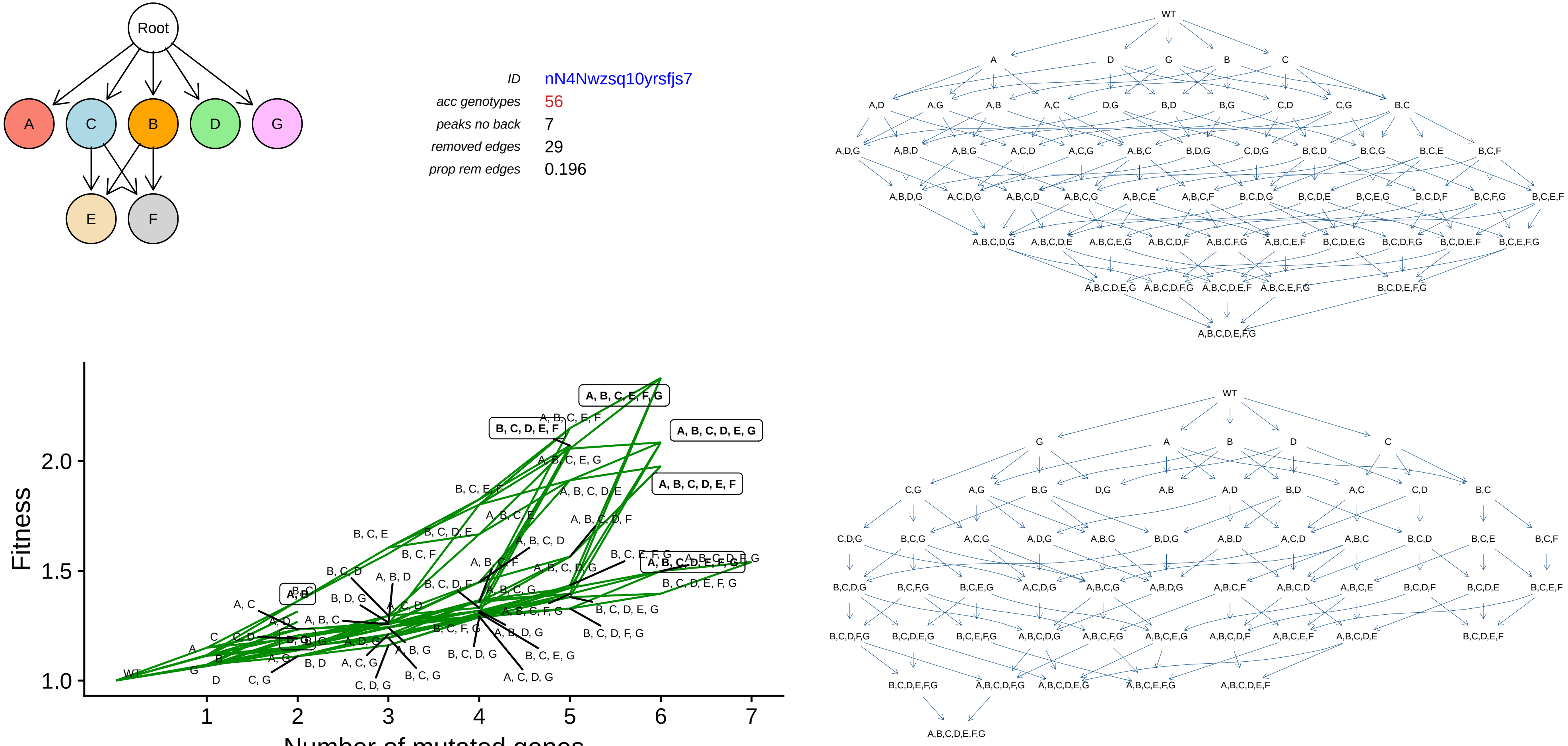

Root
A
C
B
D
G
E
F
ID nN4Nwzsq10yrsfjs7
acc genotypes 56
peaks no back 7
removed edges 29
prop rem edges 0.196
Fitness
Number of mutated genes
A, B, C, E, F, G
B, C, D, E, F
A, B, C, D, E, G
A, B, C, D, E, F
WT

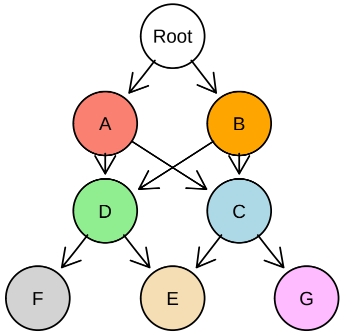


| | |
|---|---|
| *ID* | NnQqInZk8XUxrSDx |
| *acc genotypes* | 16 |
| *peaks no back* | 5 |
| *removed edges* | 5 |
| *prop rem edges* | 0.208 |

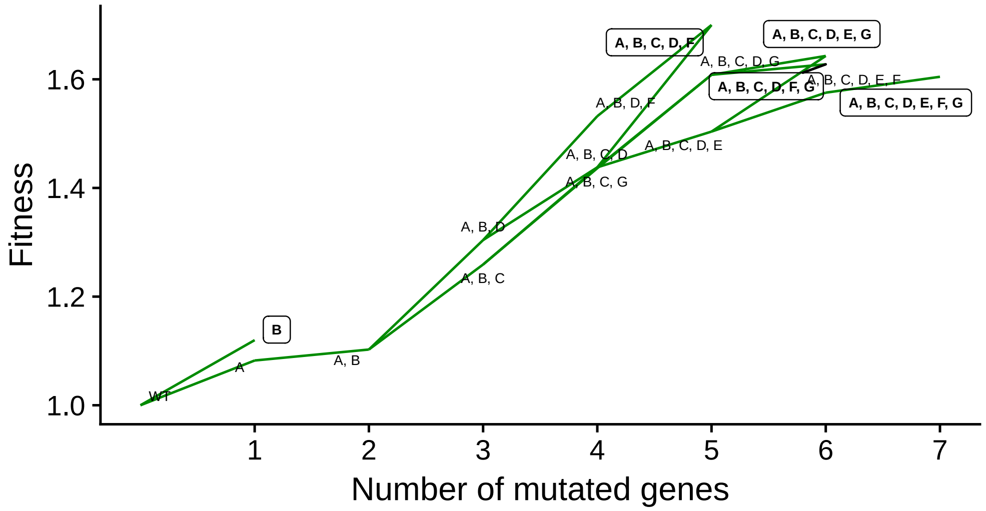


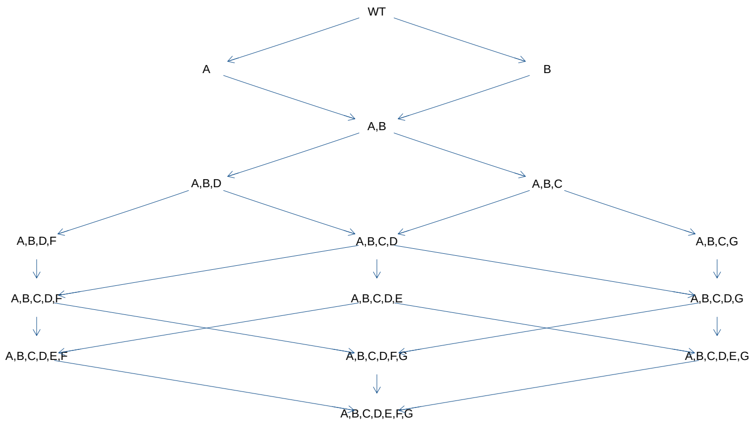


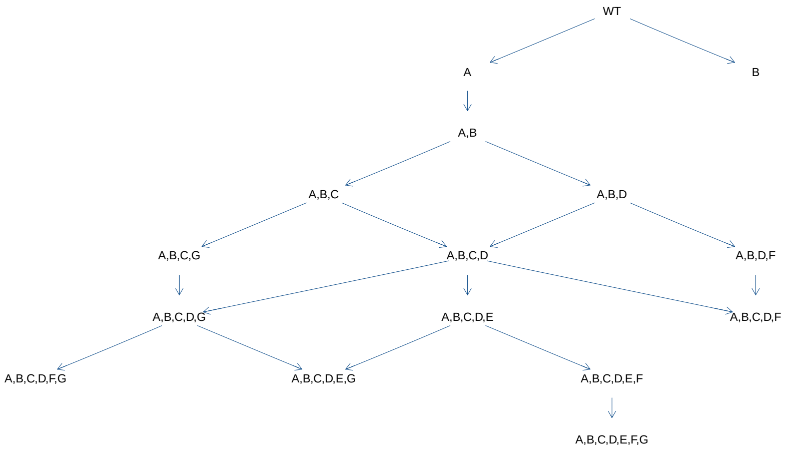

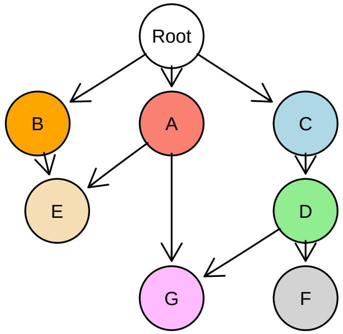


| | |
|---|---|
| *ID* | nX7dlkO9zvyl4JCM |
| *acc genotypes* | 26 |
| *peaks no back* | 2 |
| *removed edges* | 5 |
| *prop rem edges* | 0.104 |

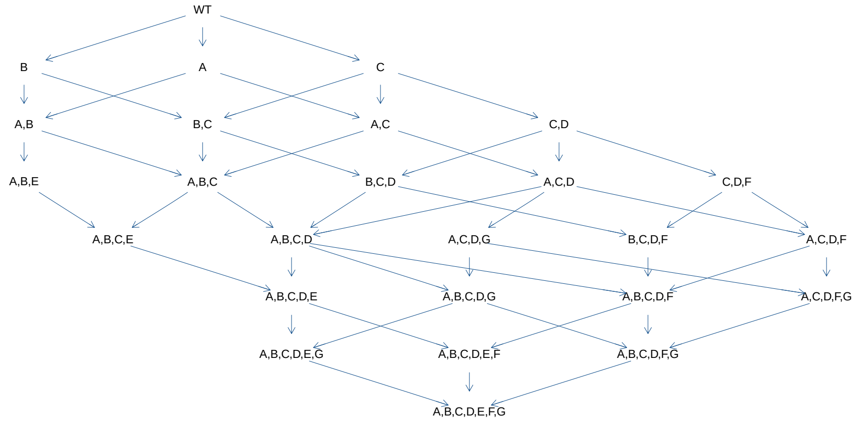


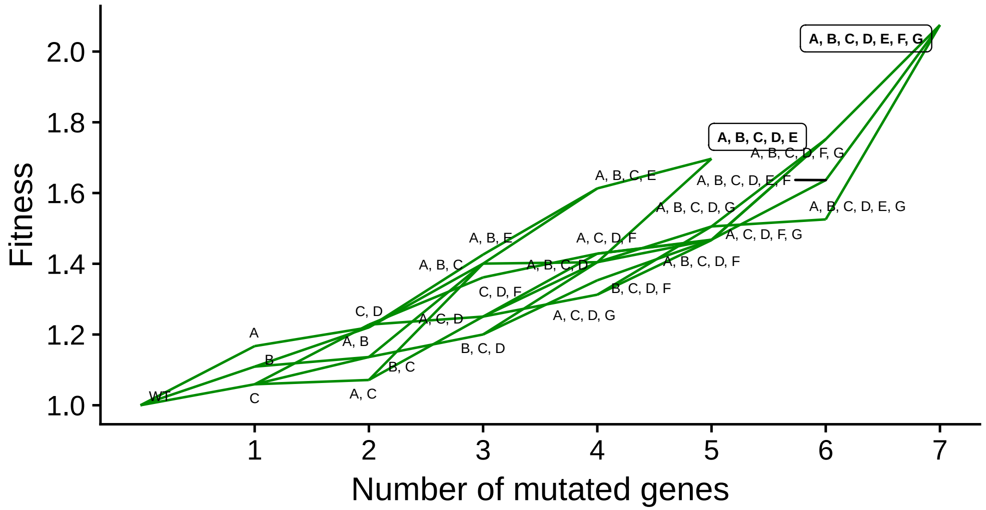


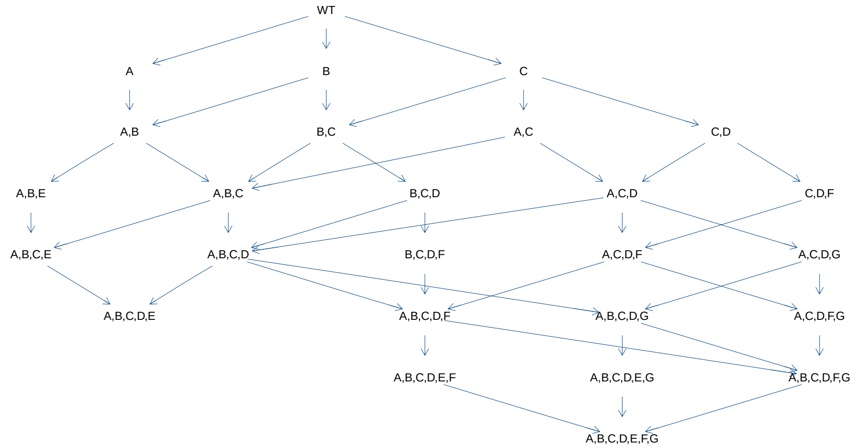

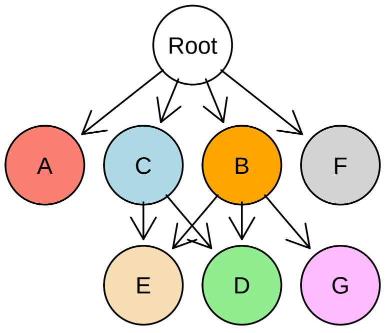


| | |
|---|---|
| *ID* | nYEr3NE5NcmF25mT |
| *acc genotypes* | 48 |
| *peaks no back* | 9 |
| *removed edges* | 17 |
| *prop rem edges* | 0.142 |

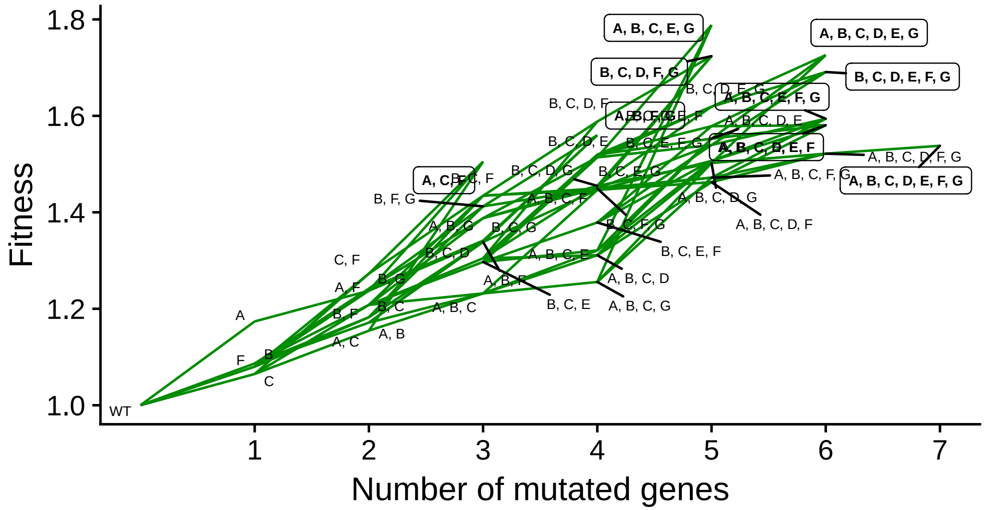


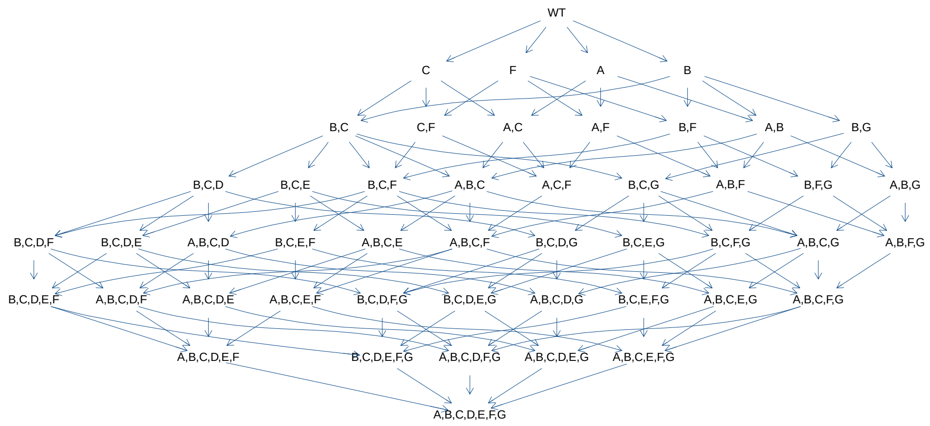

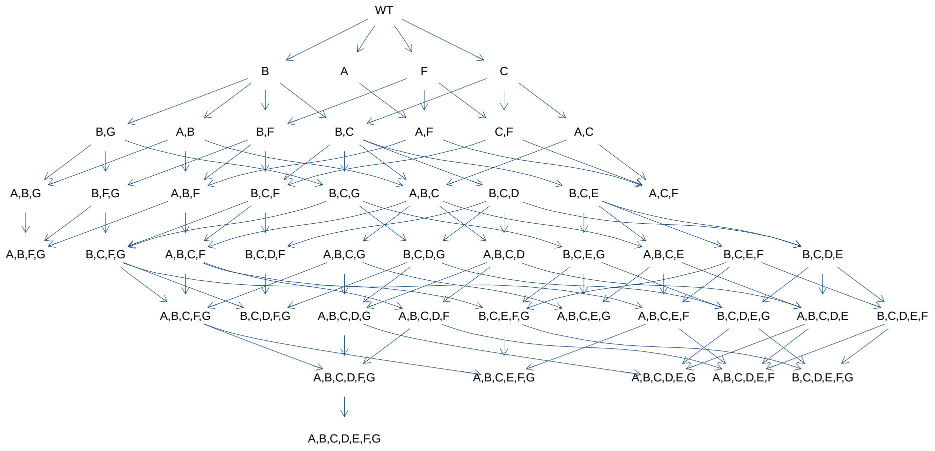

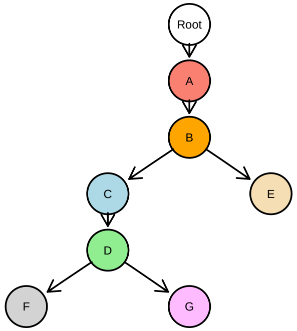


| | |
|---|---|
| *ID* | o4Y3hPxIjQmaWRx1 |
| *acc genotypes* | 14 |
| *peaks no back* | 2 |
| *removed edges* | 1 |
| *prop rem edges* | 0.05 |

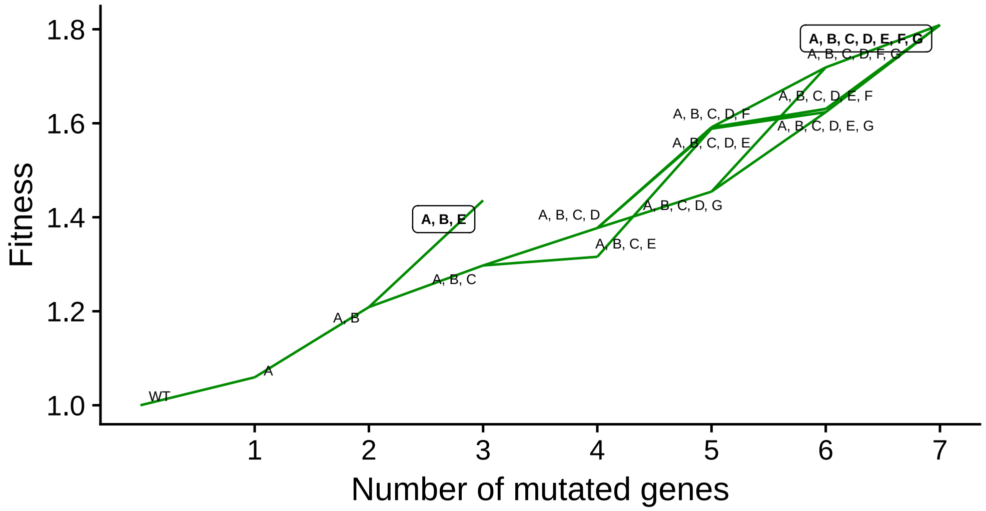


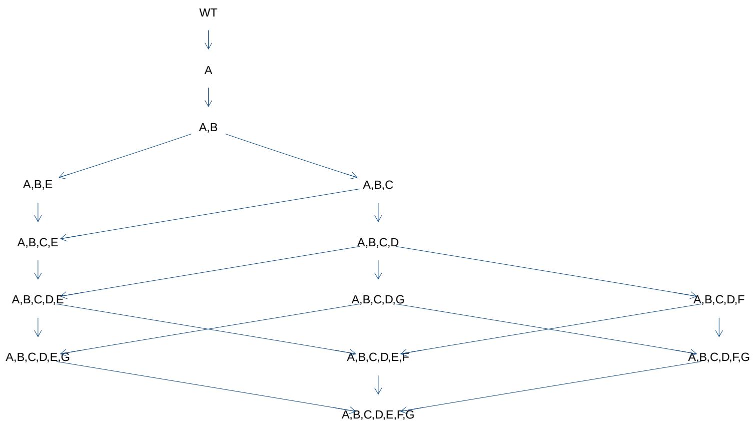


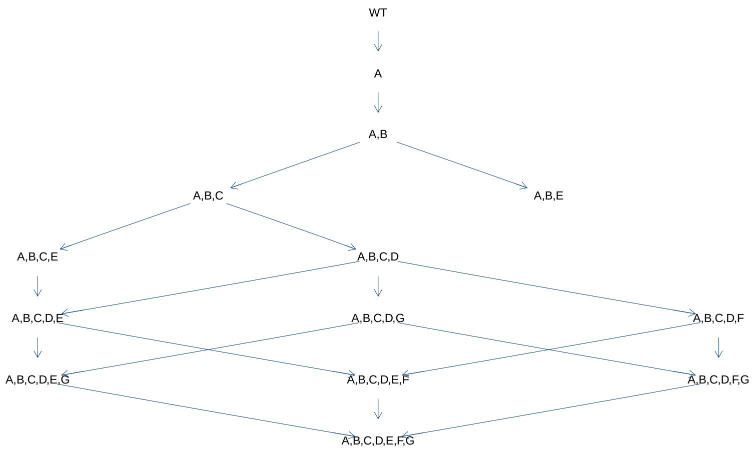

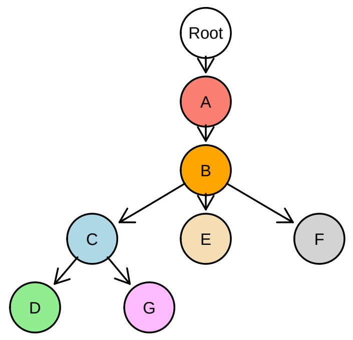


| | |
|---|---|
| *ID* | OaMMWW2waVOxtGK3 |
| *acc genotypes* | 22 |
| *peaks no back* | 5 |
| *removed edges* | 5 |
| *prop rem edges* | 0.119 |

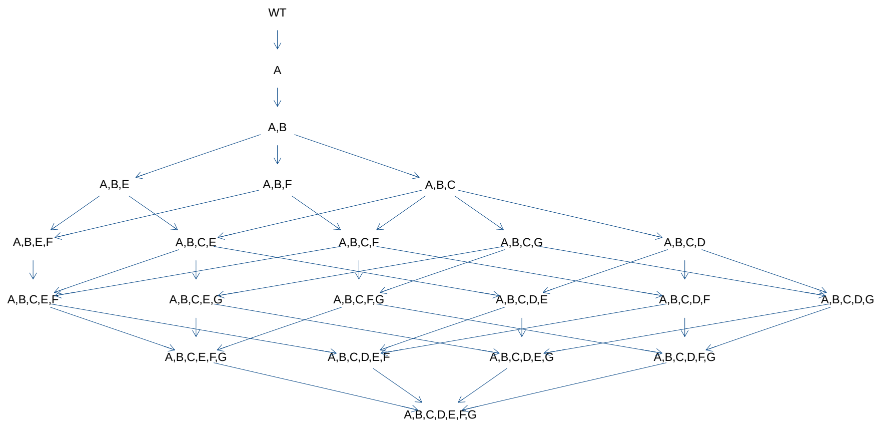


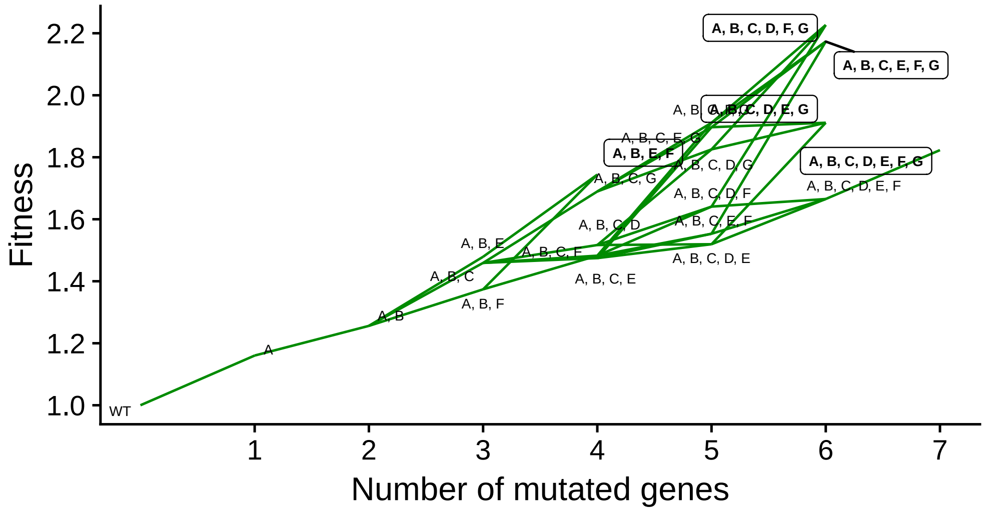


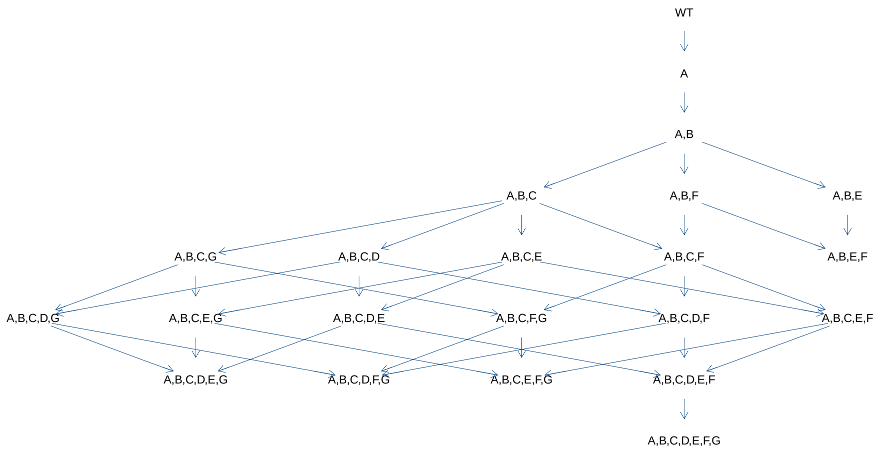

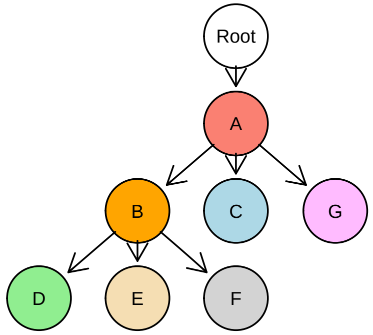


*ID* ODbe43XDGuHvXueq
*acc genotypes* 37
*peaks no back* 3
*removed edges* 6
*prop rem edges* 0.067

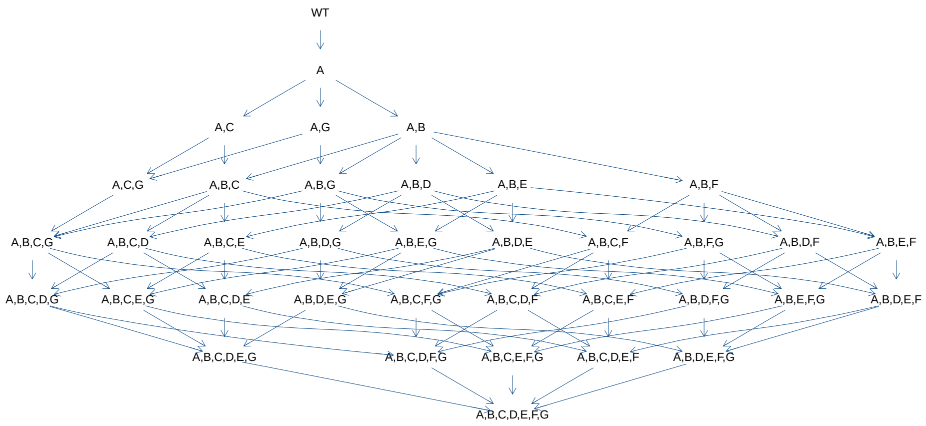


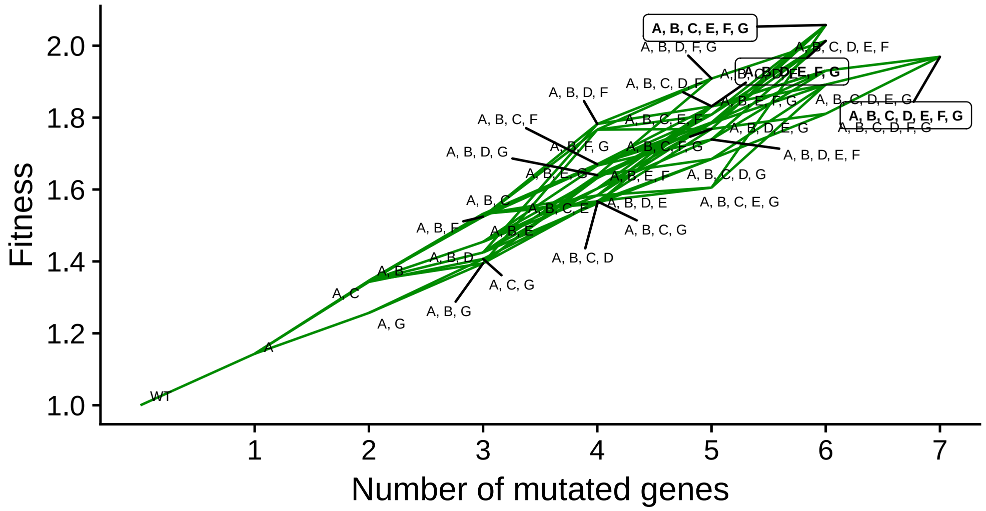


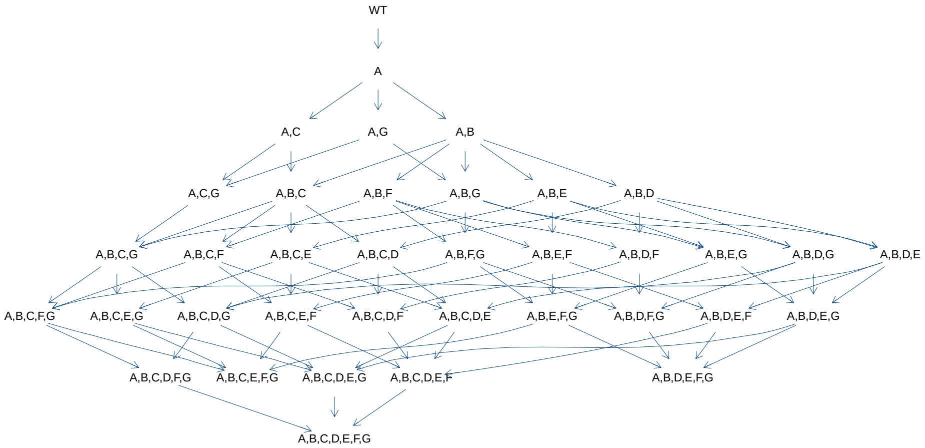

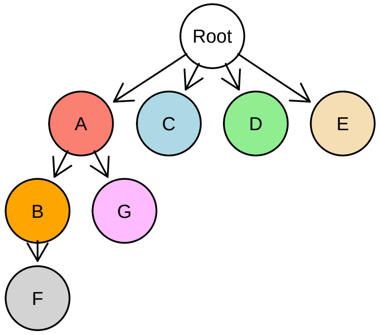


*ID* OflKufVm9LrZy9VH

*acc genotypes* 56

*peaks no back* 8

*removed edges* 27

*prop rem edges* 0.182

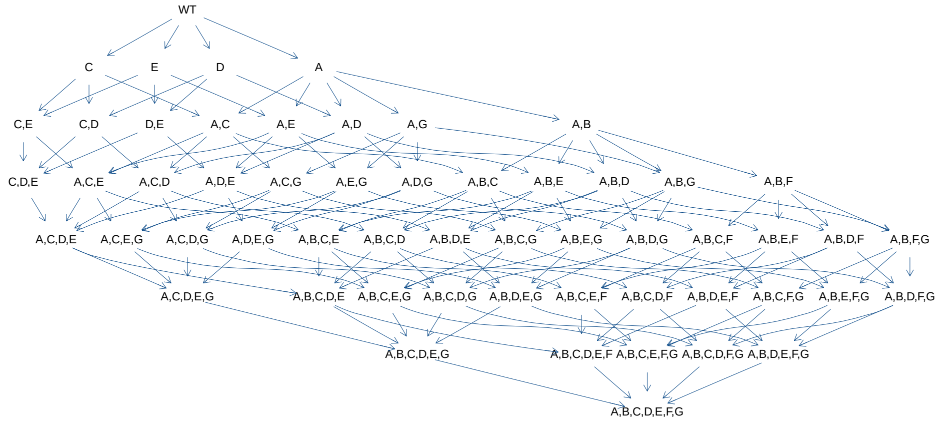


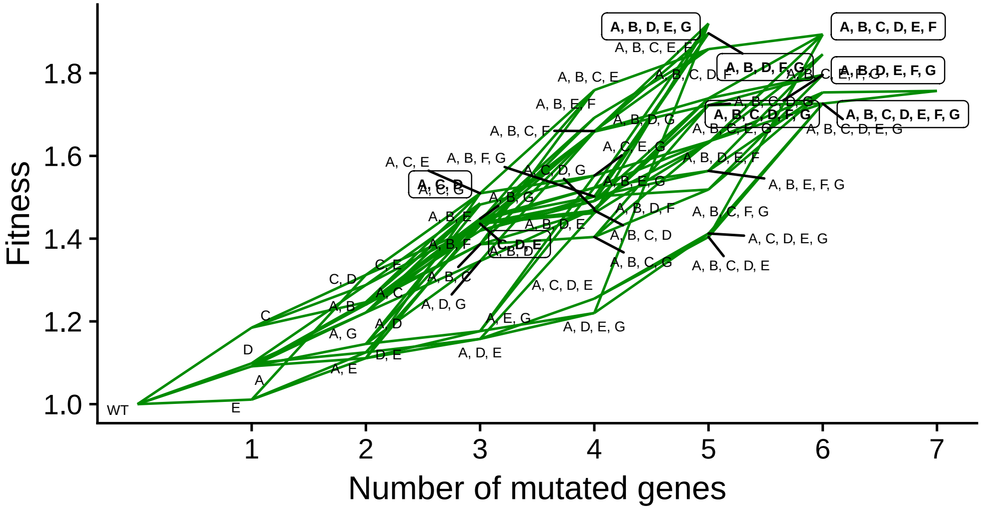


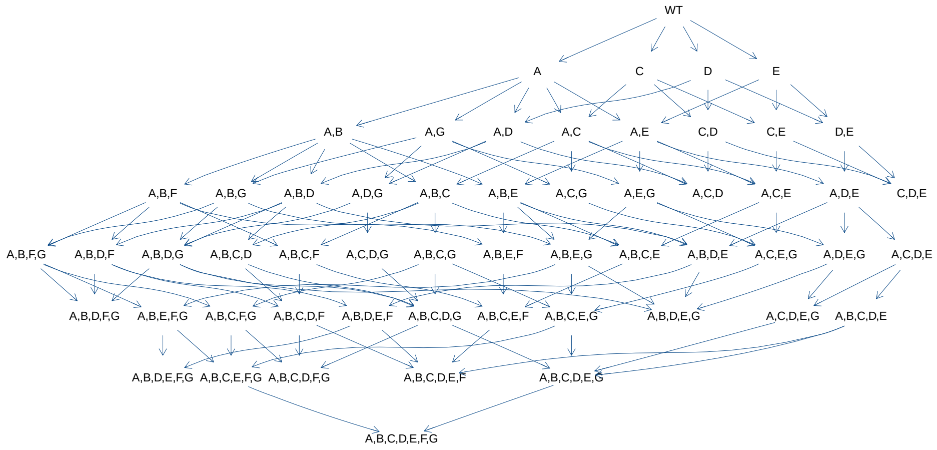

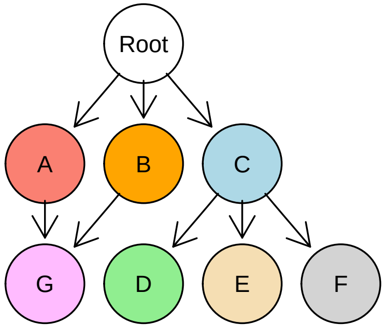


| | |
|---|---|
| *ID* | OMDWvgSAoX1WSQy1 |
| *acc genotypes* | 45 |
| *peaks no back* | 3 |
| *removed edges* | 13 |
| *prop rem edges* | 0.118 |

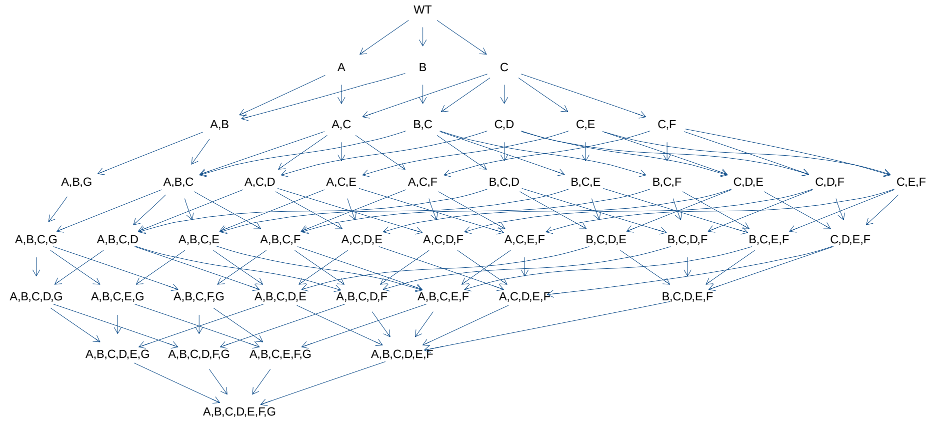


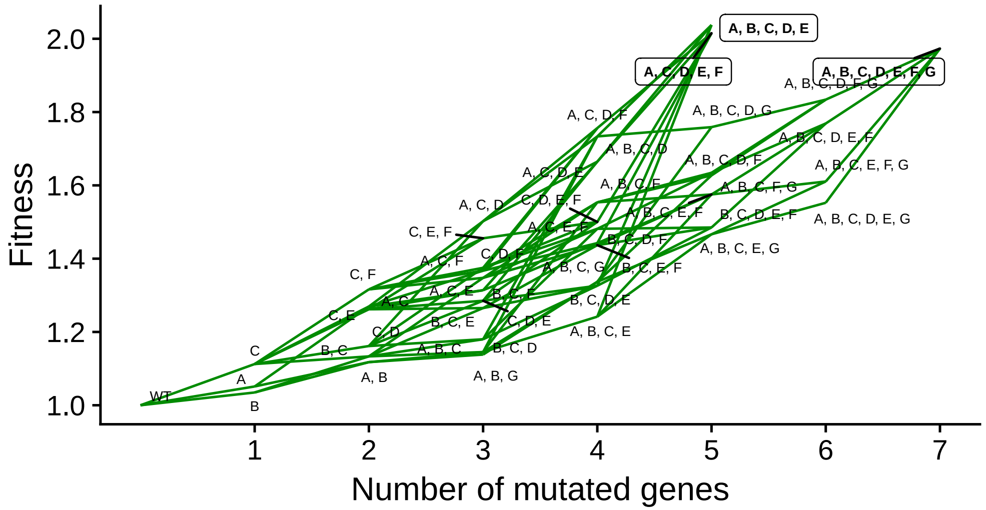


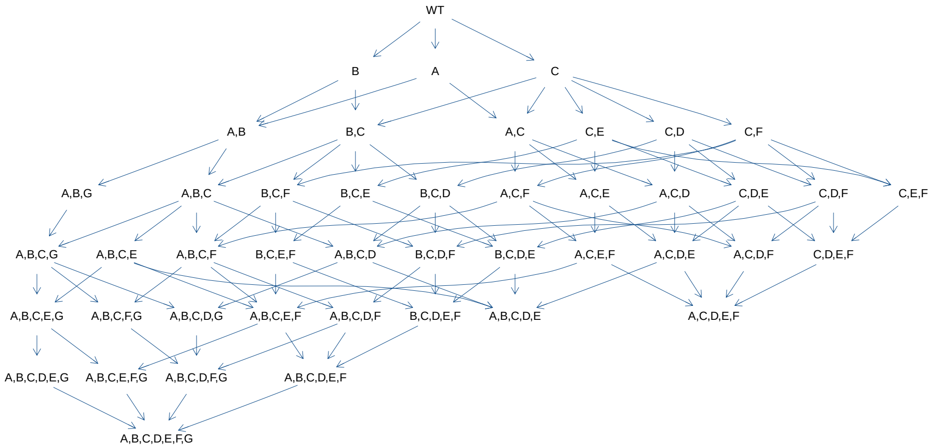

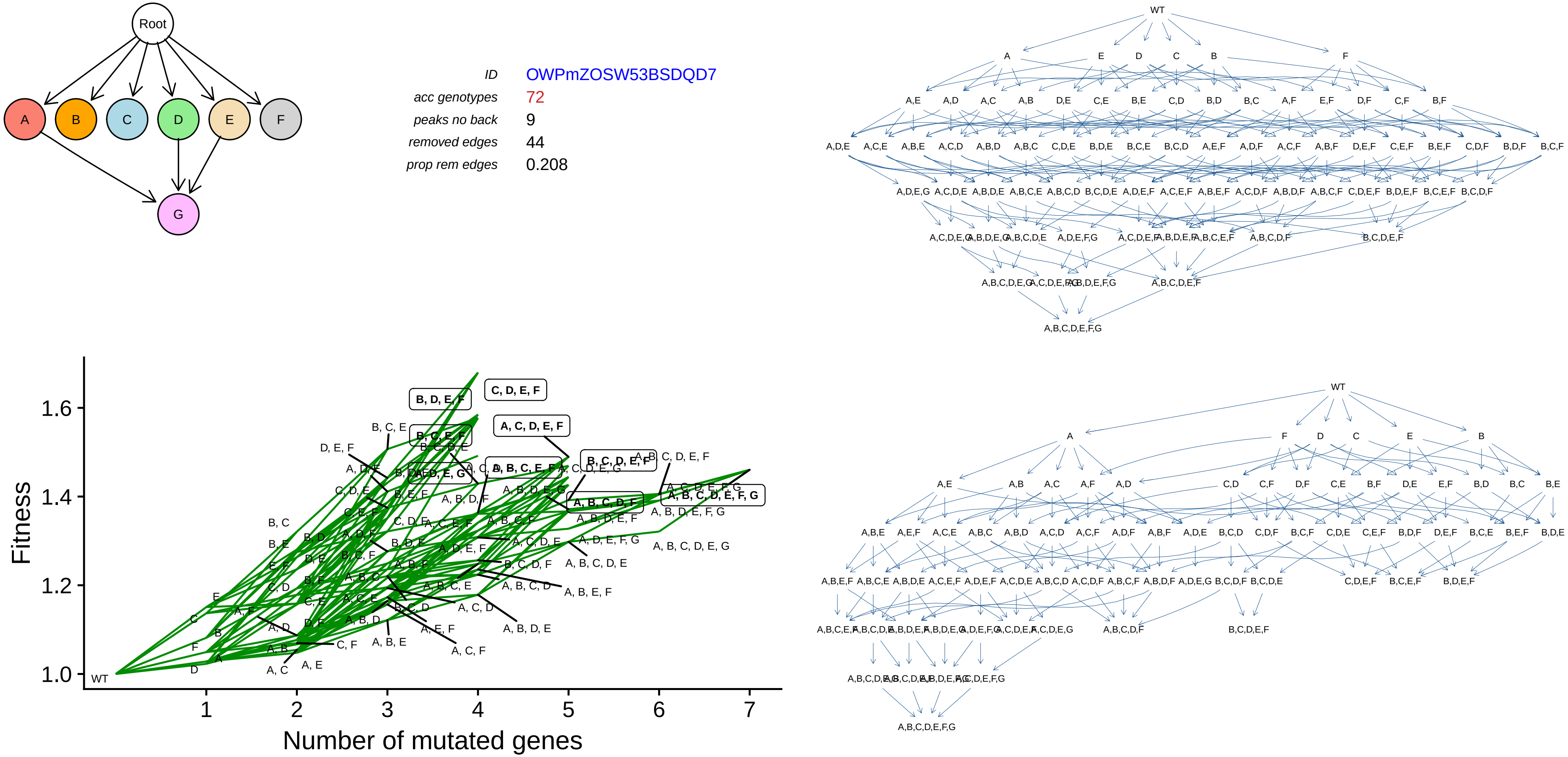

Root
A
B
C
D
E
F
G
ID OWPmZOSW53BSDQD7
acc genotypes 72
peaks no back 9
removed edges 44
prop rem edges 0.208
Fitness
Number of mutated genes
WT
A,B,C,D,E,F,G

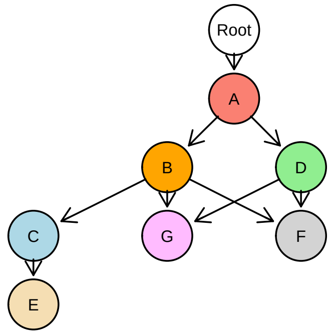


| | |
|---|---|
| *ID* | OyMB3UqWrH3e9QQY |
| *acc genotypes* | 18 |
| *peaks no back* | 4 |
| *removed edges* | 3 |
| *prop rem edges* | 0.103 |

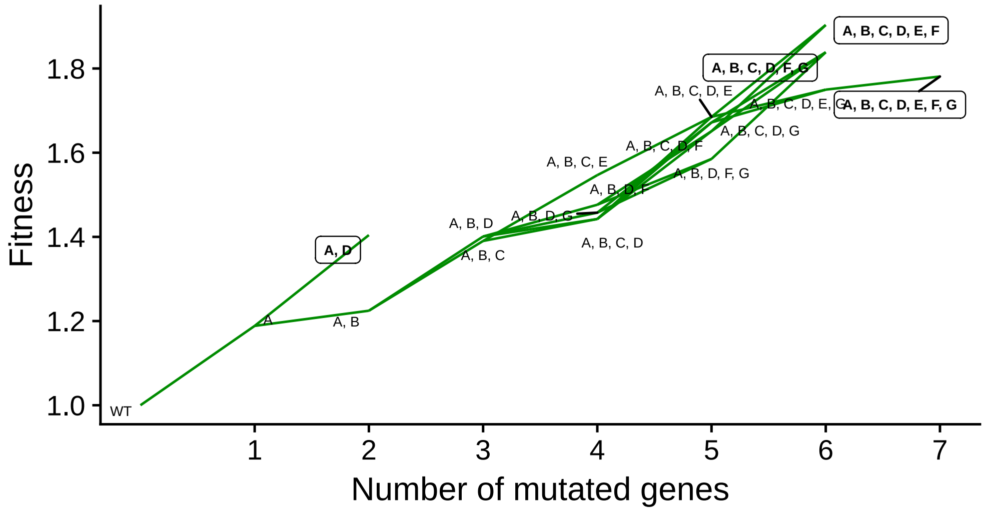


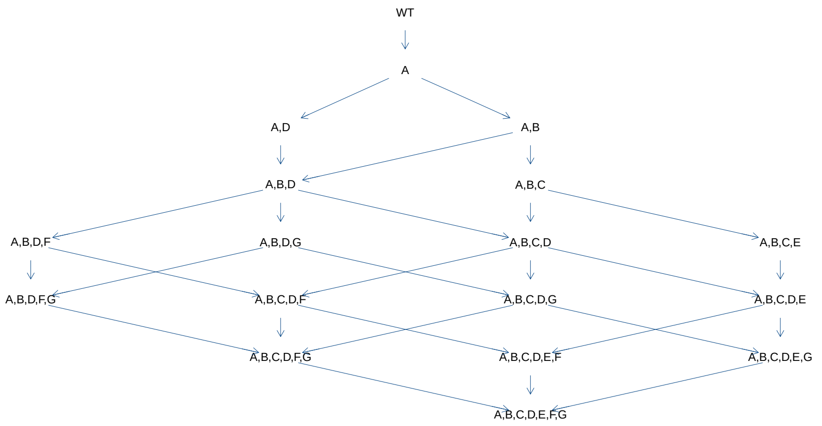


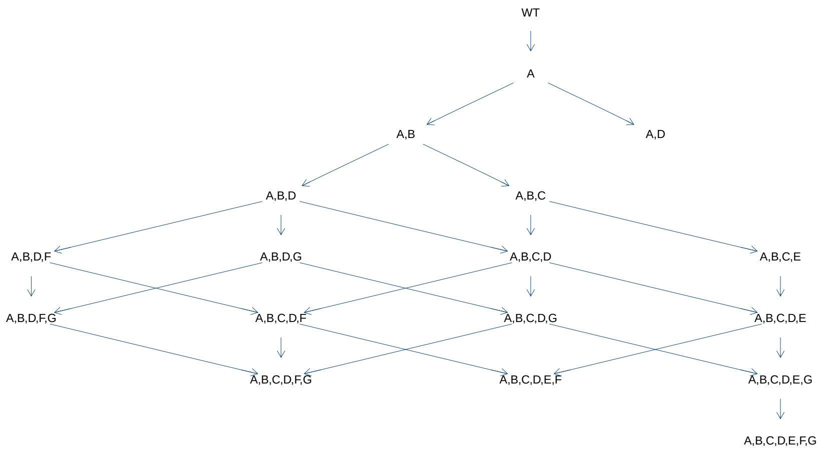

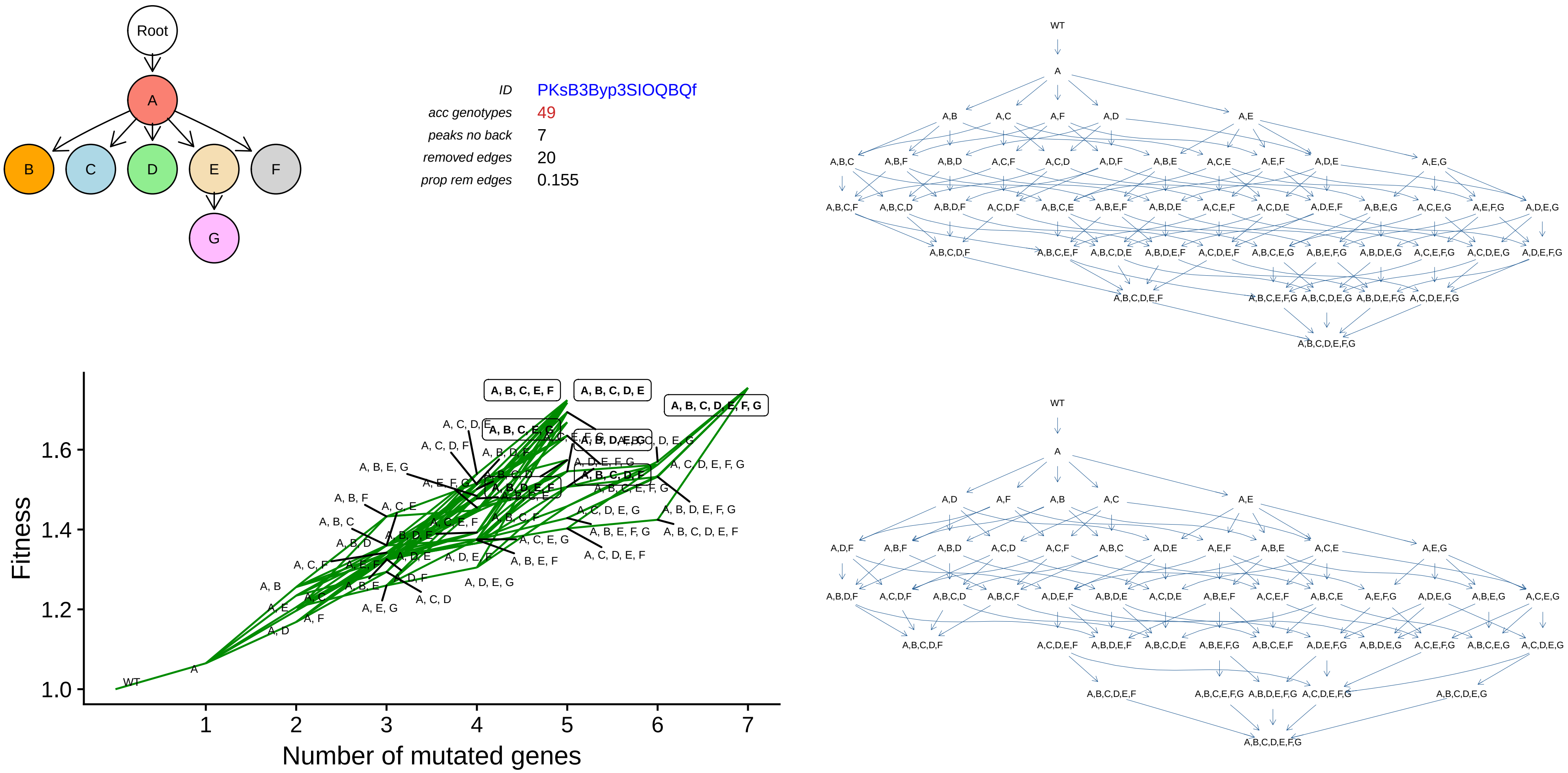

Root
A
B
C
D
E
F
G
ID PKsB3Byp3SIOQBQf
acc genotypes 49
peaks no back 7
removed edges 20
prop rem edges 0.155
Fitness
Number of mutated genes
1.0
1.2
1.4
1.6
1
2
3
4
5
6
7
WT
A
A, B
A, E
A, D
A, F
A, C
A, B, C, E, F
A, B, C, D, E
A, B, C, D, E, F, G
A, B, C, E, G
A, B, D, E, G
A, B, C, D, F
A, B, C, D, E
A, B, C, D, E
A, C, D, E, F, G
A, B, D, E, F, G
A, B, C, D, E, F
A, C, D, E, F
A, B, E, F, G
A, C, D, E, G
A, B, C, E, F, G
A, D, E, F, G
A, C, D, E
A, C, D, F
A, B, E, G
A, B, D, F
A, B, F
A, C, E
A, B, C
A, B, D
A, C, F
A, E, F
A, B, E
A, D, F
A, C, D
A, E, G
A, D, E, G
A, B, E, F
A, C, E, G
A, D, E, F
A, B, C, F
A, C, E, F
A, E, F, G
WT
A
A,B
A,C
A,F
A,D
A,E
A,B,C
A,B,F
A,B,D
A,C,F
A,C,D
A,D,F
A,B,E
A,C,E
A,E,F
A,D,E
A,E,G
A,B,C,F
A,B,C,D
A,B,D,F
A,C,D,F
A,B,C,E
A,B,E,F
A,B,D,E
A,C,E,F
A,C,D,E
A,D,E,F
A,B,E,G
A,C,E,G
A,E,F,G
A,D,E,G
A,B,C,D,F
A,B,C,E,F
A,B,C,D,E
A,B,D,E,F
A,C,D,E,F
A,B,C,E,G
A,B,E,F,G
A,B,D,E,G
A,C,E,F,G
A,C,D,E,G
A,D,E,F,G
A,B,C,D,E,F
A,B,C,E,F,G
A,B,C,D,E,G
A,B,D,E,F,G
A,C,D,E,F,G
A,B,C,D,E,F,G

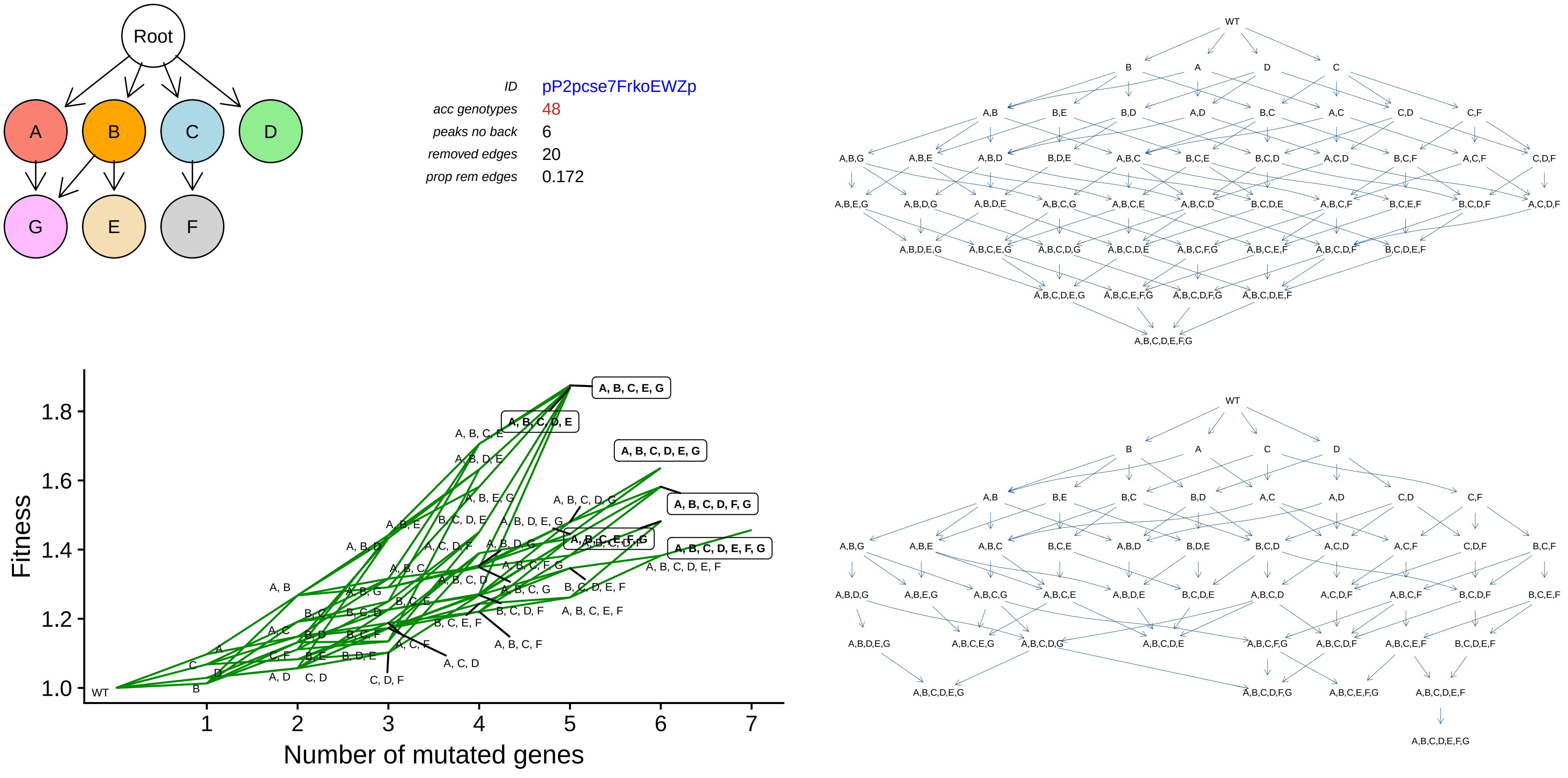
Root
A
B
C
D
G
E
F
ID pP2pcse7FrkoEWZp
acc genotypes 48
peaks no back 6
removed edges 20
prop rem edges 0.172
Fitness
Number of mutated genes
A, B, C, E, G
A, B, C, D, E
A, B, C, D, E, G
A, B, C, D, F, G
A, B, C, E, F, G
A, B, C, D, E, F, G
WT
A,B,C,D,E,F,G

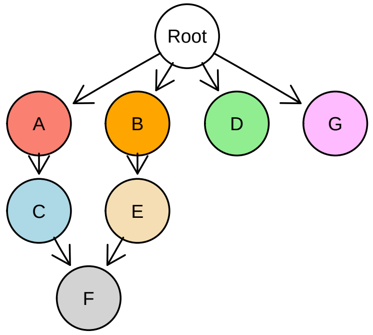


| | |
|---|---|
| *ID* | PPvpOC9kPl3xygTj |
| *acc genotypes* | 40 |
| *peaks no back* | 8 |
| *removed edges* | 19 |
| *prop rem edges* | 0.207 |

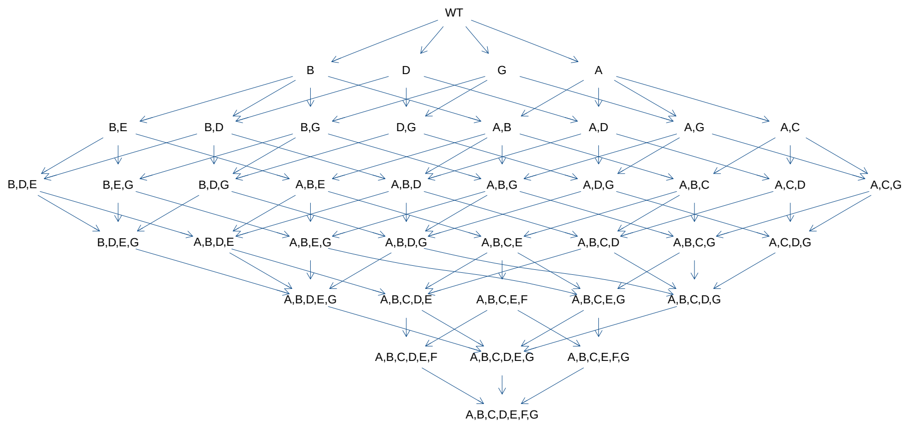


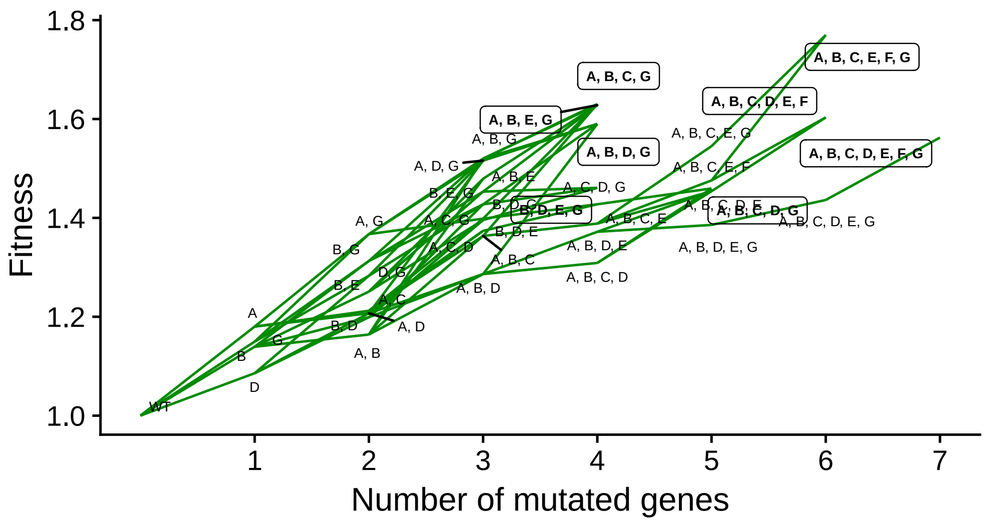


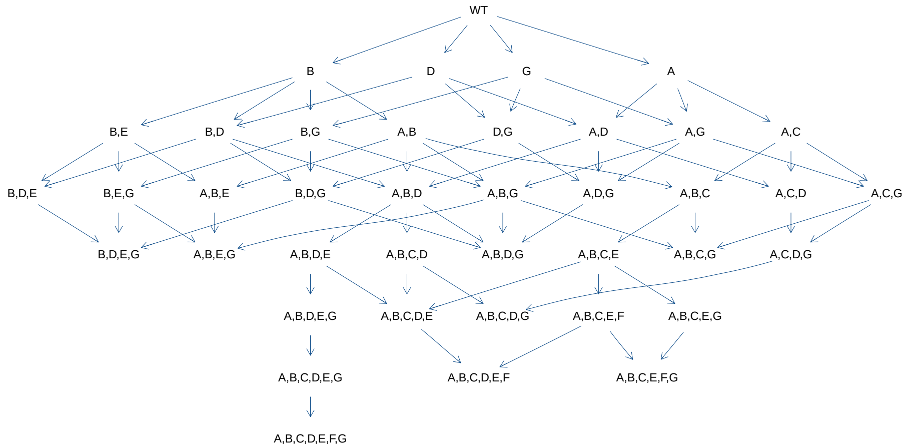

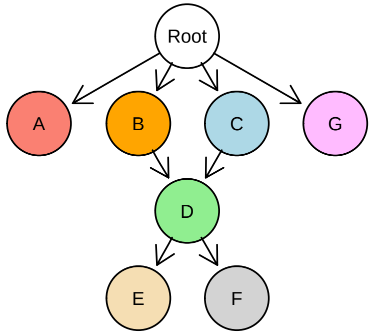


*ID* pSBATEcePfnX7JWh
*acc genotypes* 32
*peaks no back* 2
*removed edges* 7
*prop rem edges* 0.103

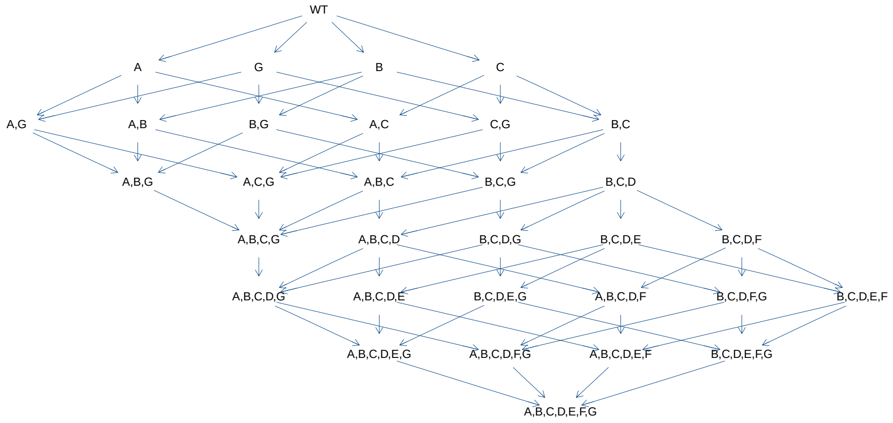


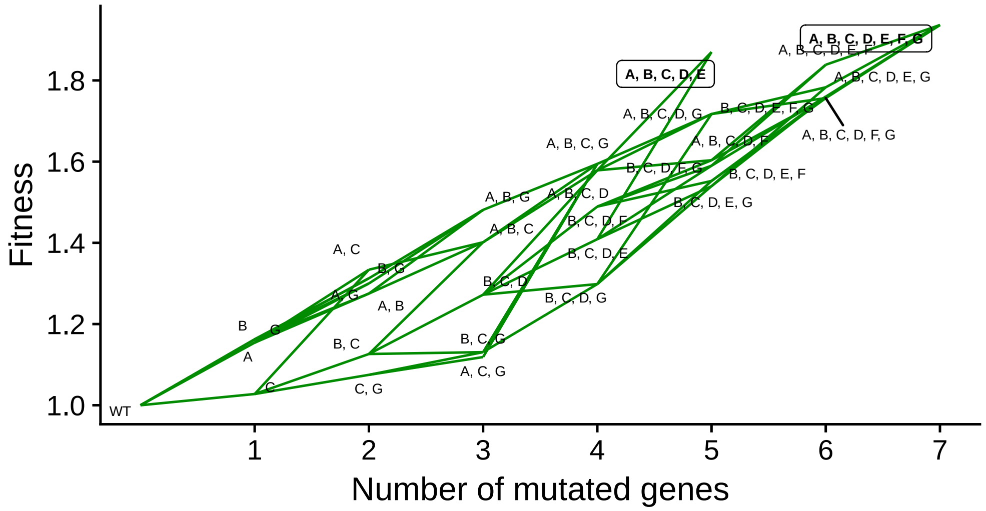


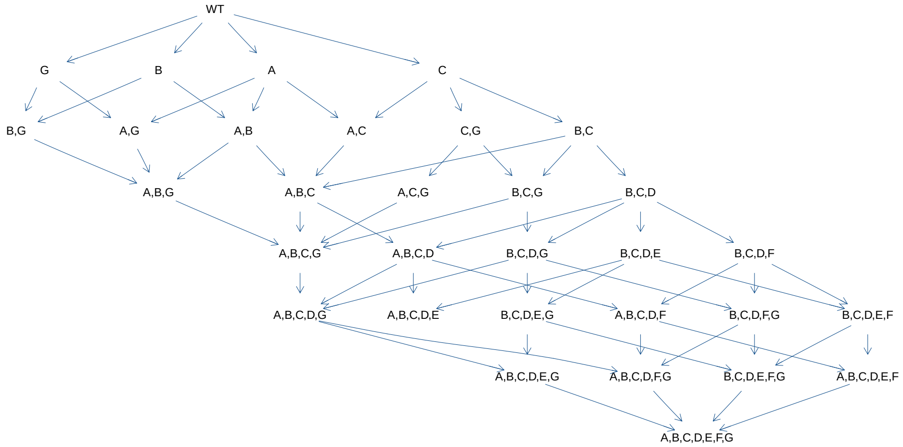

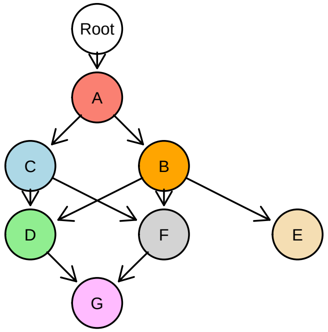


| | |
|---|---|
| *ID* | pui6npZKf4iG9VmR |
| *acc genotypes* | 15 |
| *peaks no back* | 3 |
| *removed edges* | 3 |
| *prop rem edges* | 0.136 |

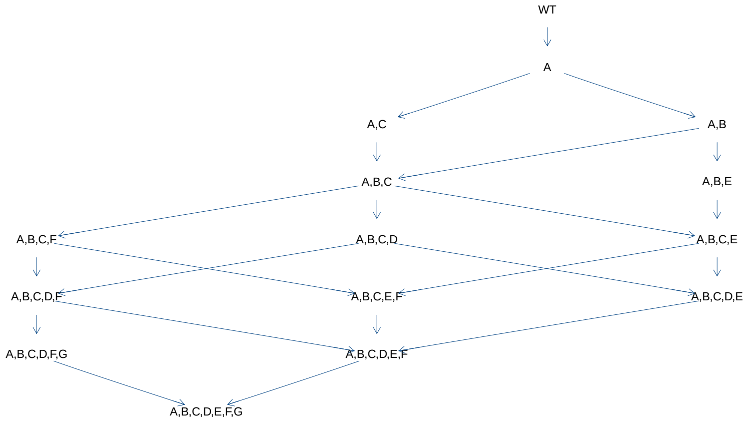


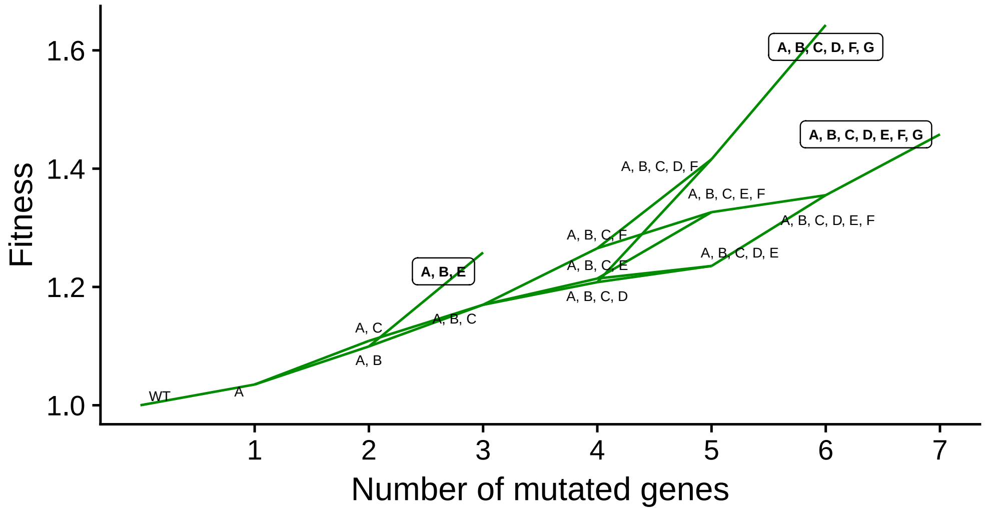


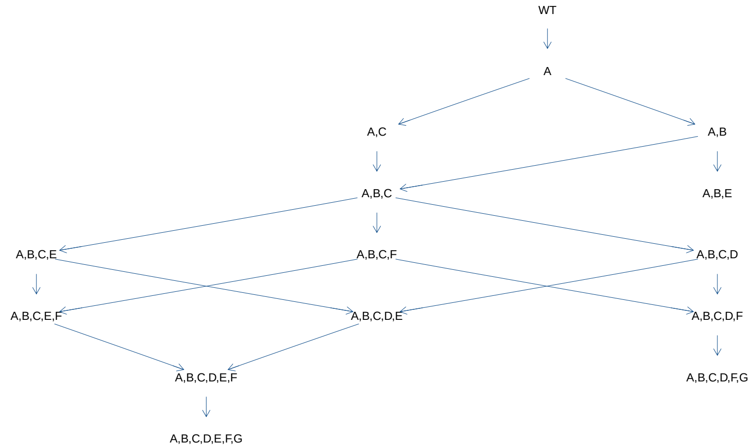

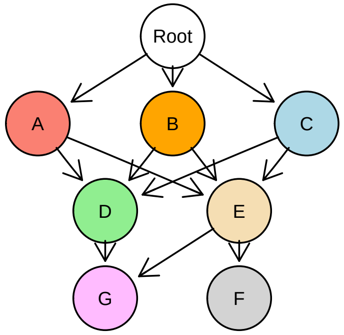


| | |
|---|---|
| *ID* | Q7PLBKdNNwZTd7om |
| *acc genotypes* | 15 |
| *peaks no back* | 2 |
| *removed edges* | 2 |
| *prop rem edges* | 0.091 |

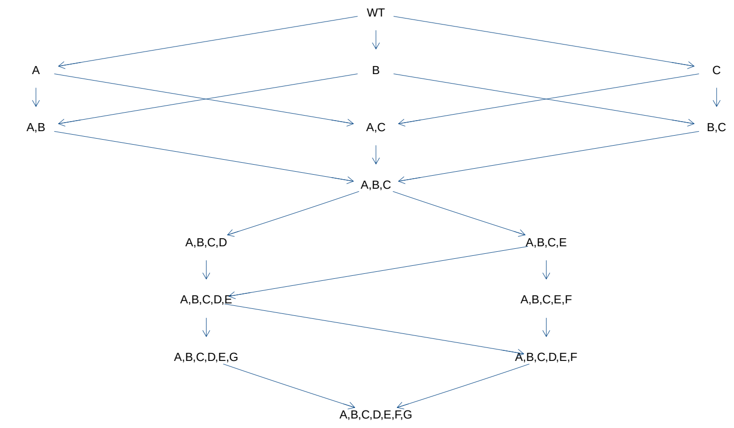


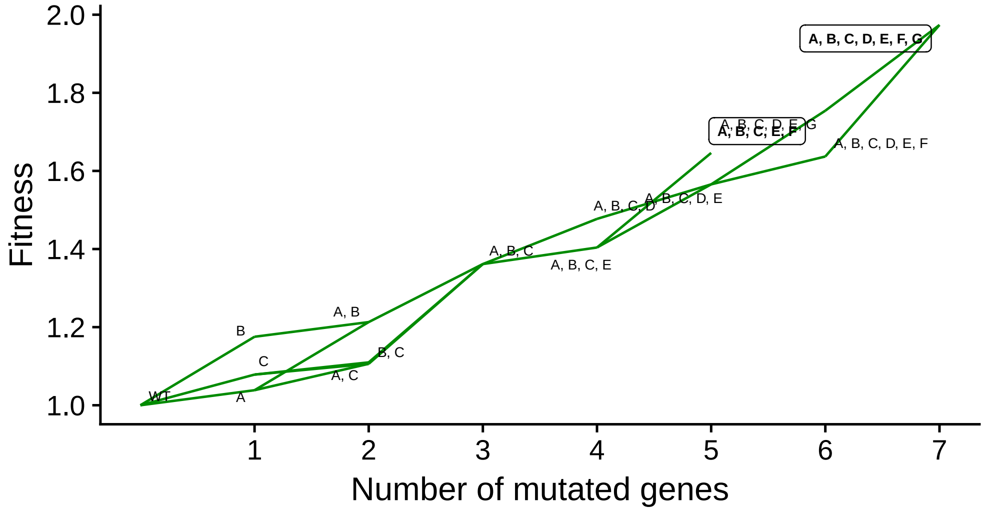


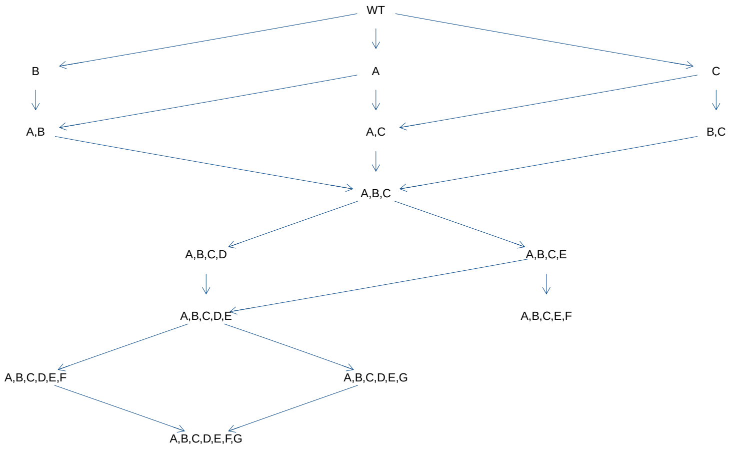

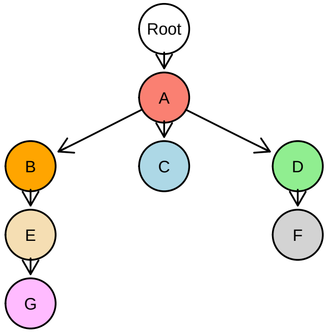


| | |
|---|---|
| *ID* | qEpRgGAzFDFL2rE5 |
| *acc genotypes* | 25 |
| *peaks no back* | 3 |
| *removed edges* | 10 |
| *prop rem edges* | 0.213 |

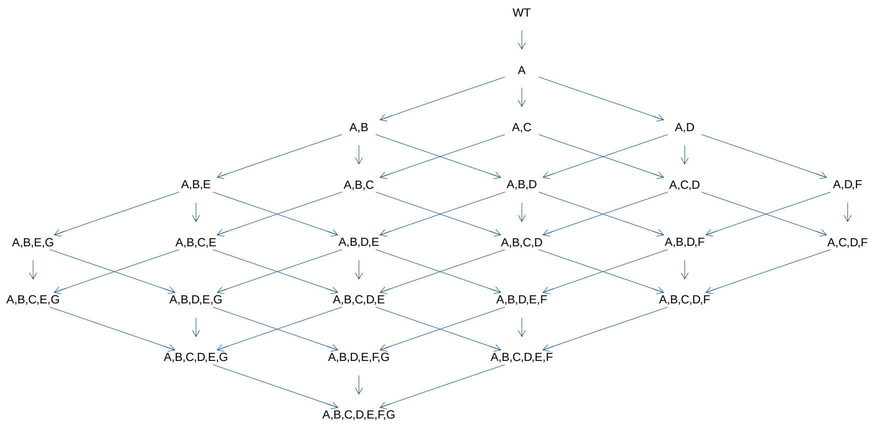


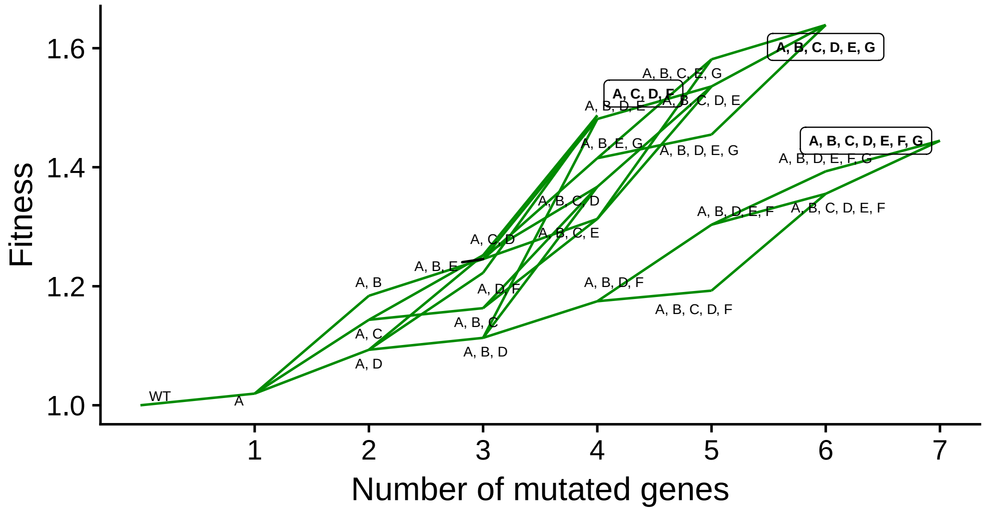


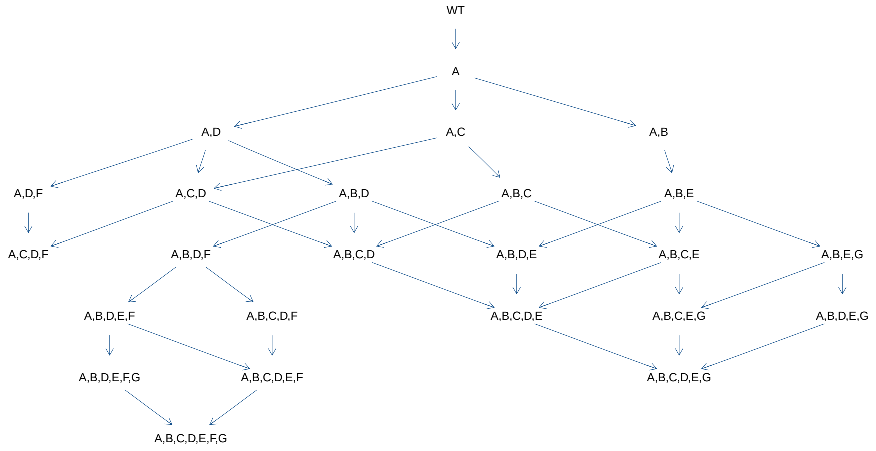

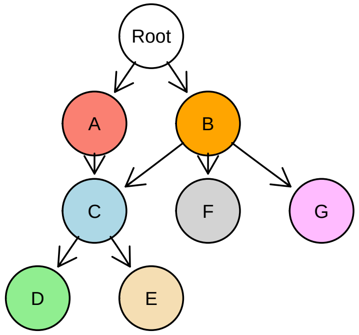


*ID* qEvXADpjSKIVHQcX
*acc genotypes* 26
*peaks no back* 2
*removed edges* 4
*prop rem edges* 0.078

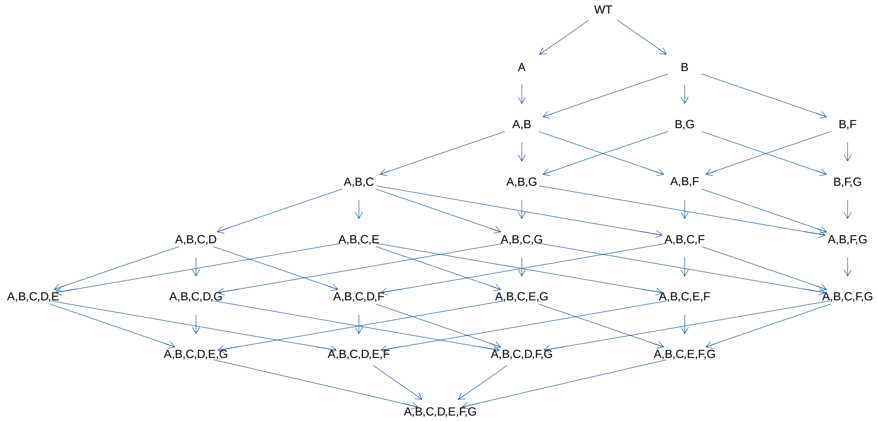


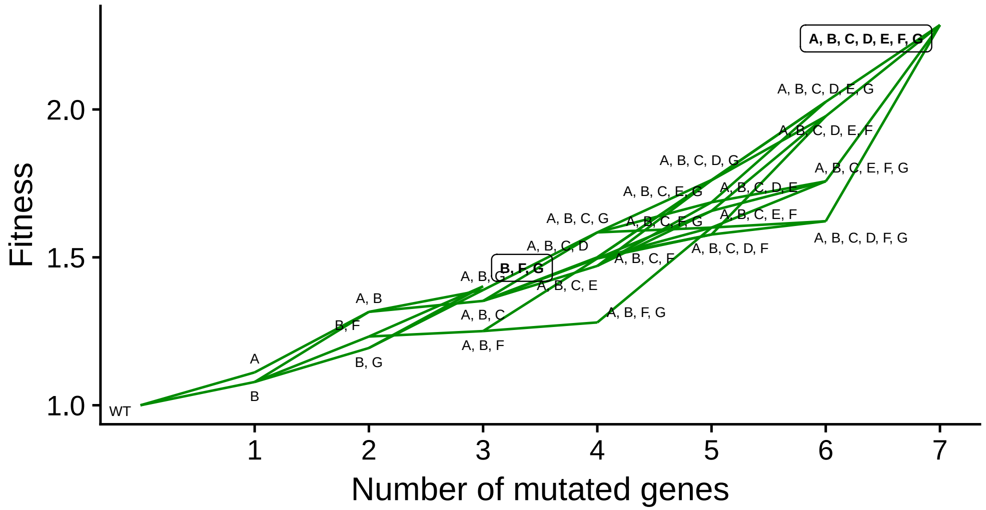


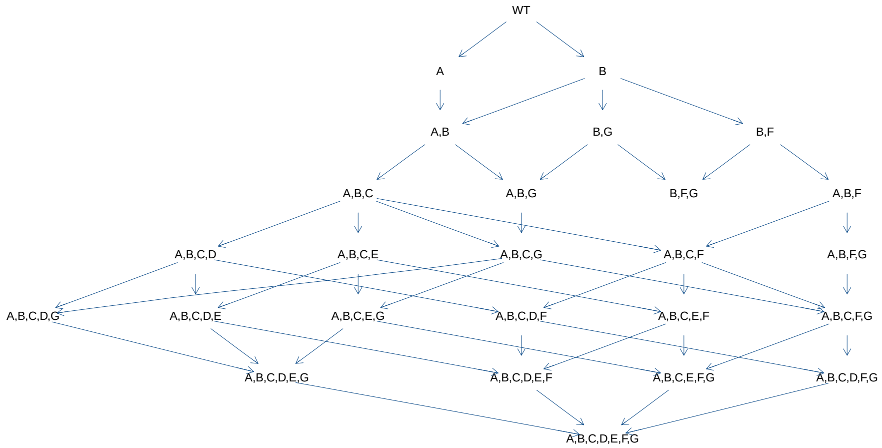

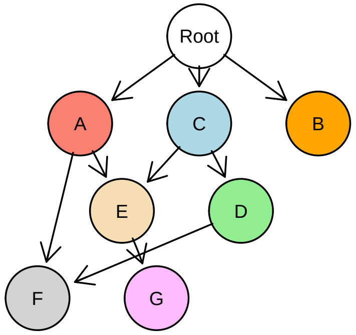


*ID* qp7i3jRVfJ15Ghst
*acc genotypes* 26
*peaks no back* 6
*removed edges* 12
*prop rem edges* 0.245

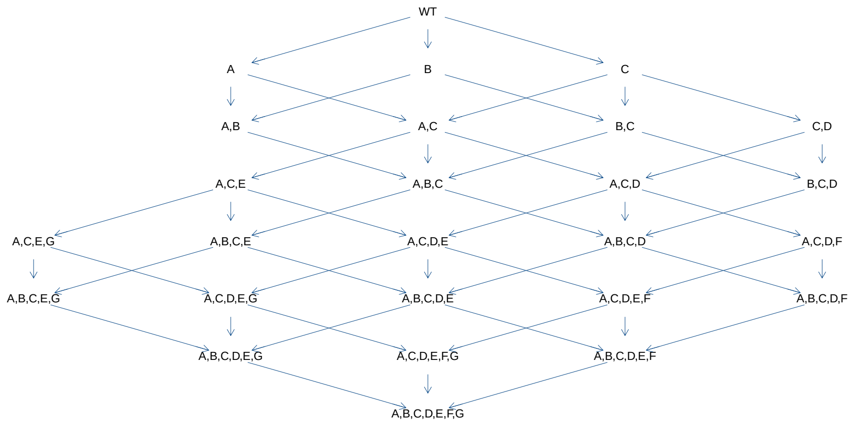


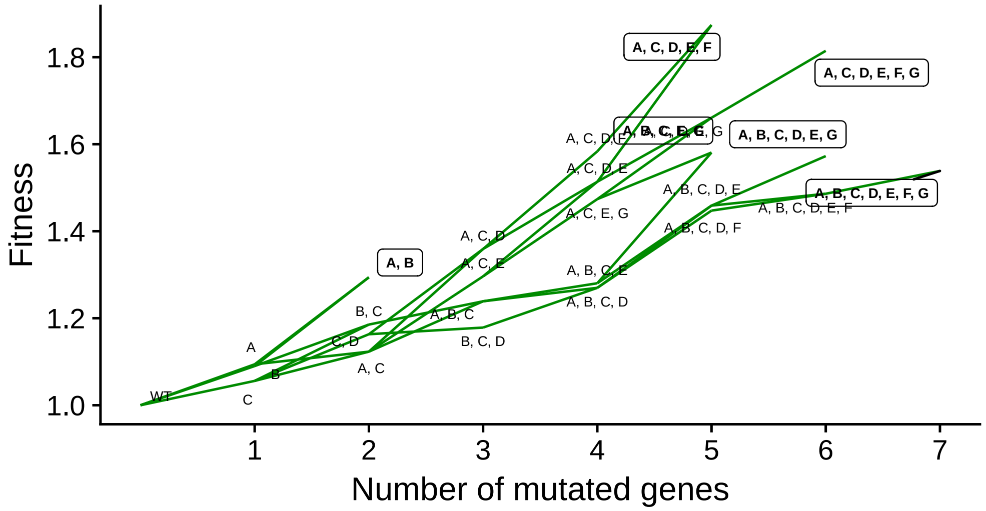


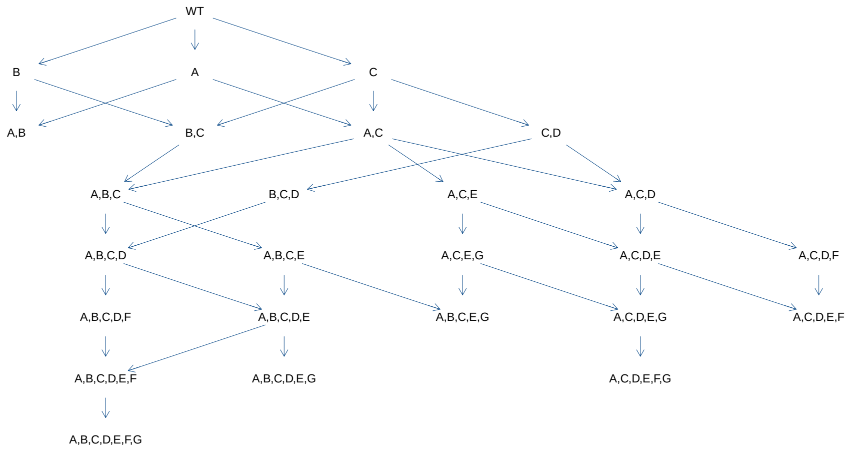

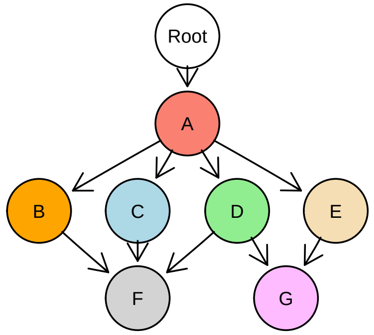


*ID* QqSOOn7ihVdtBpuj
*acc genotypes* 24
*peaks no back* 4
*removed edges* 4
*prop rem edges* 0.087

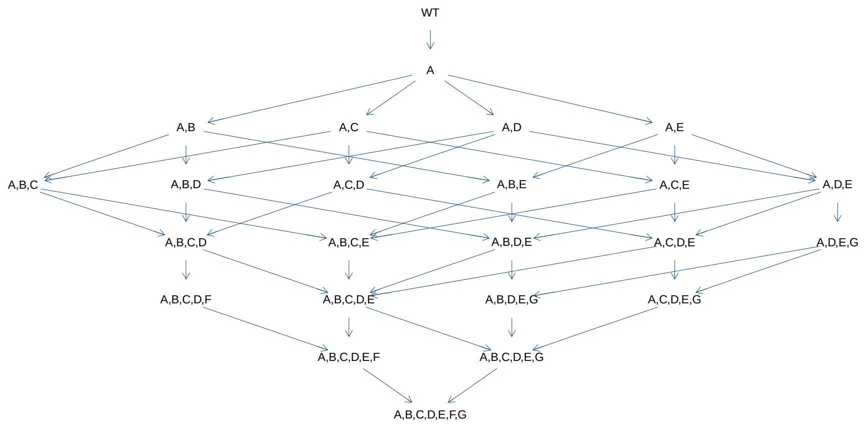


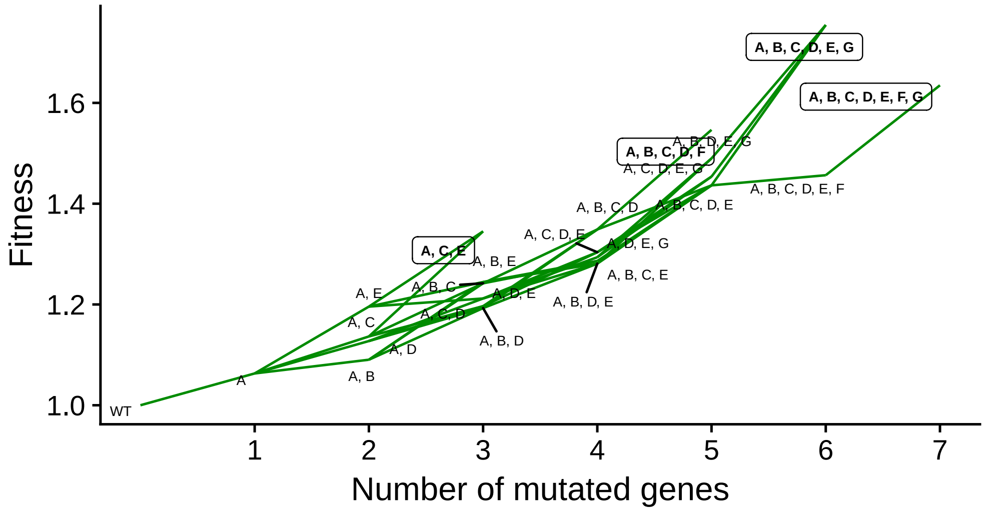


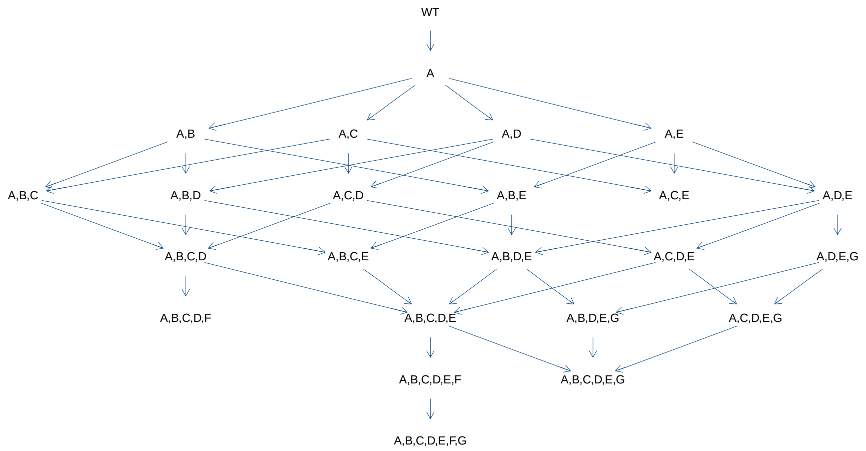

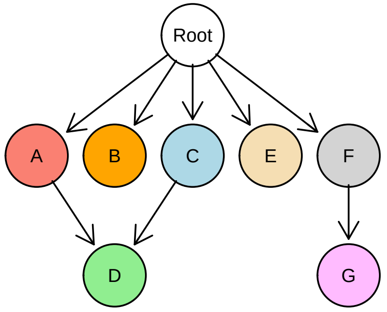


| | |
|---|---|
| *ID* | qUS7LZwsUxmyH3in |
| *acc genotypes* | 60 |
| *peaks no back* | 7 |
| *removed edges* | 32 |
| *prop rem edges* | 0.2 |

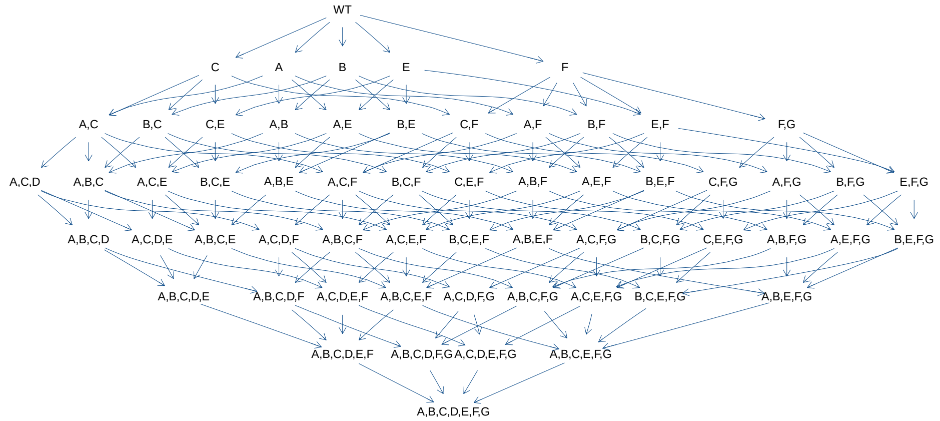


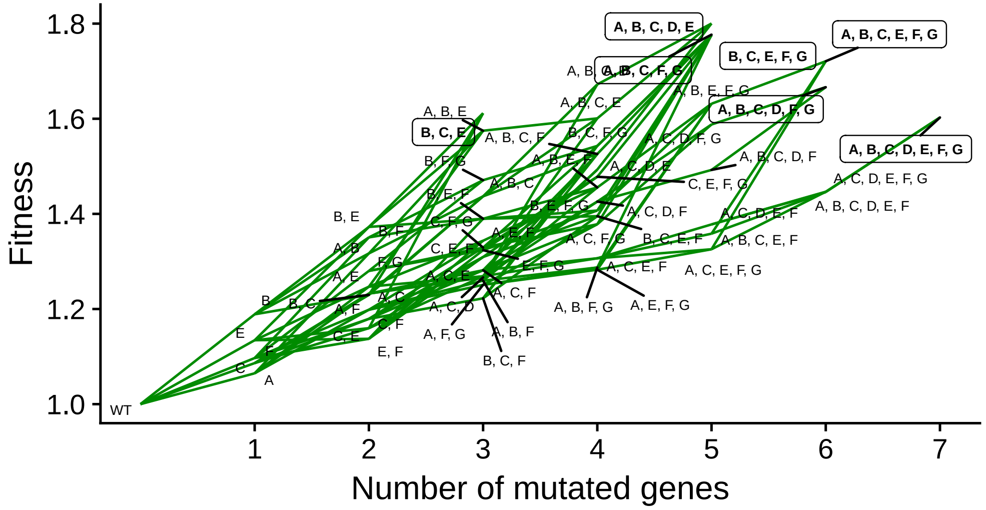


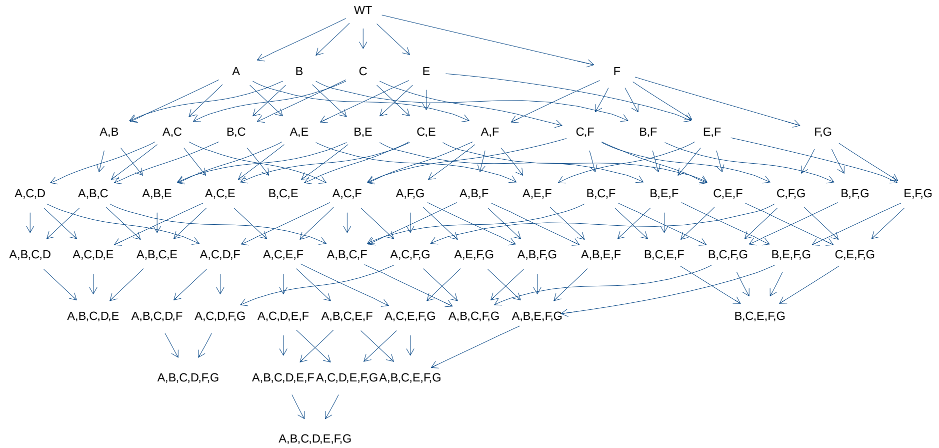

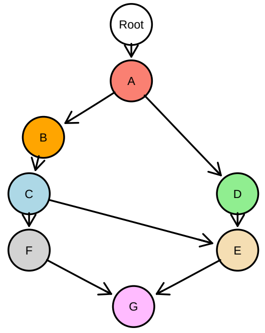


| | |
|---|---|
| *ID* | R102qzAOOheDkzBdz |
| *acc genotypes* | 12 |
| *peaks no back* | 2 |
| *removed edges* | 1 |
| *prop rem edges* | 0.067 |

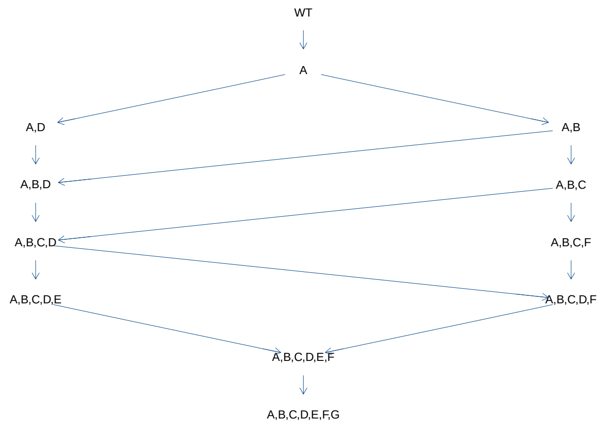


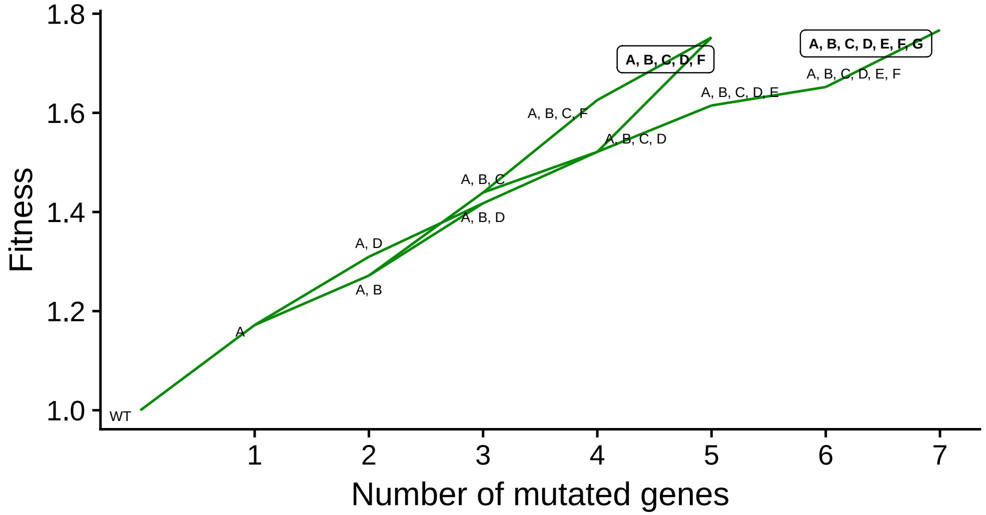


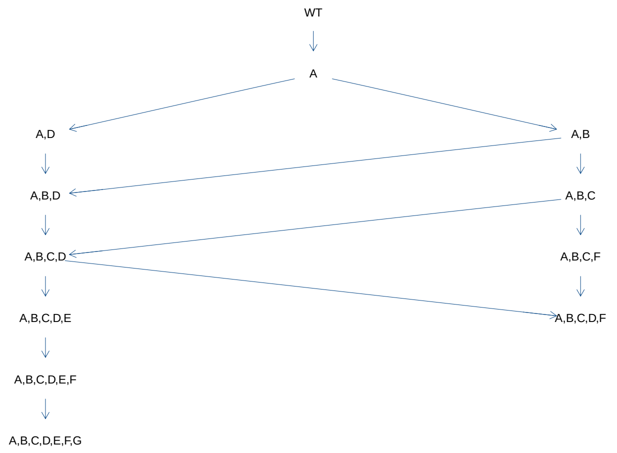

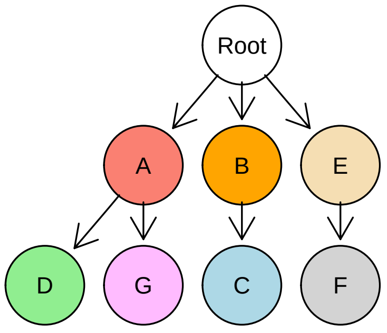


*ID* RXEIHdhCdDnzsWrB
*acc genotypes* 45
*peaks no back* 6
*removed edges* 13
*prop rem edges* 0.124

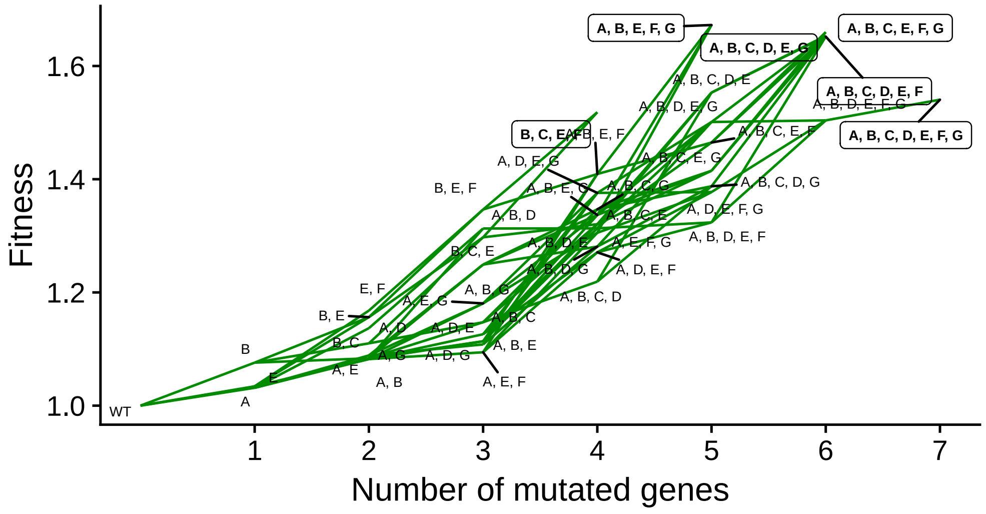


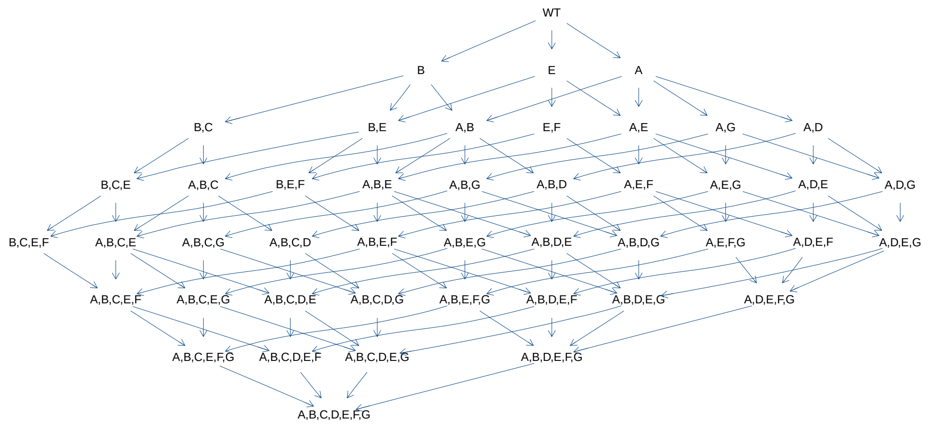


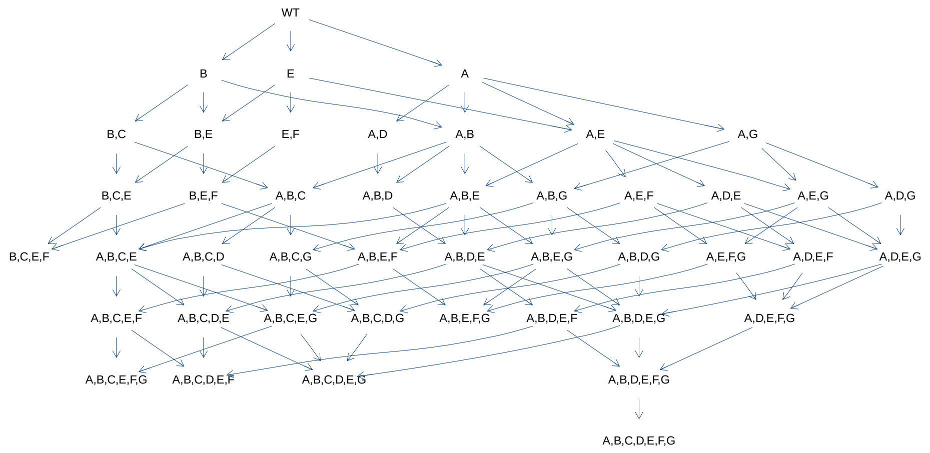

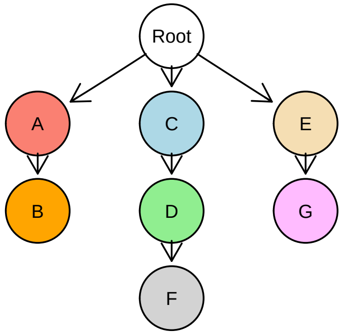


| | |
|---|---|
| *ID* | ScvT9EQPUJ5Ak10em |
| *acc genotypes* | 36 |
| *peaks no back* | 2 |
| *removed edges* | 8 |
| *prop rem edges* | 0.107 |

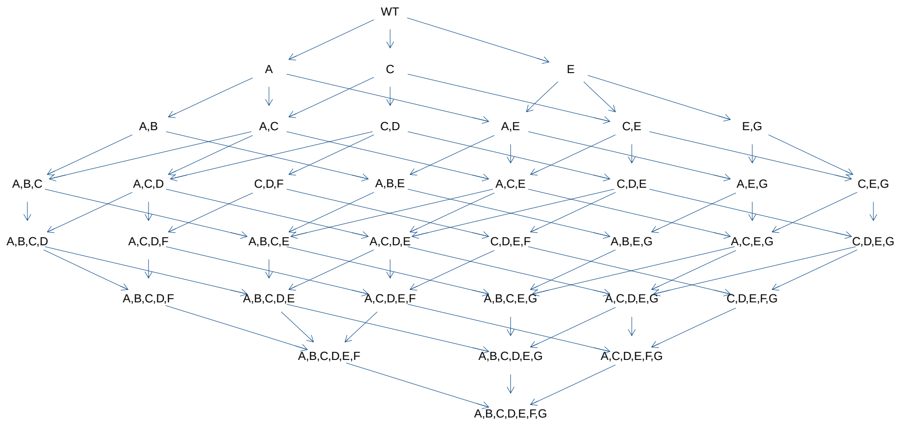


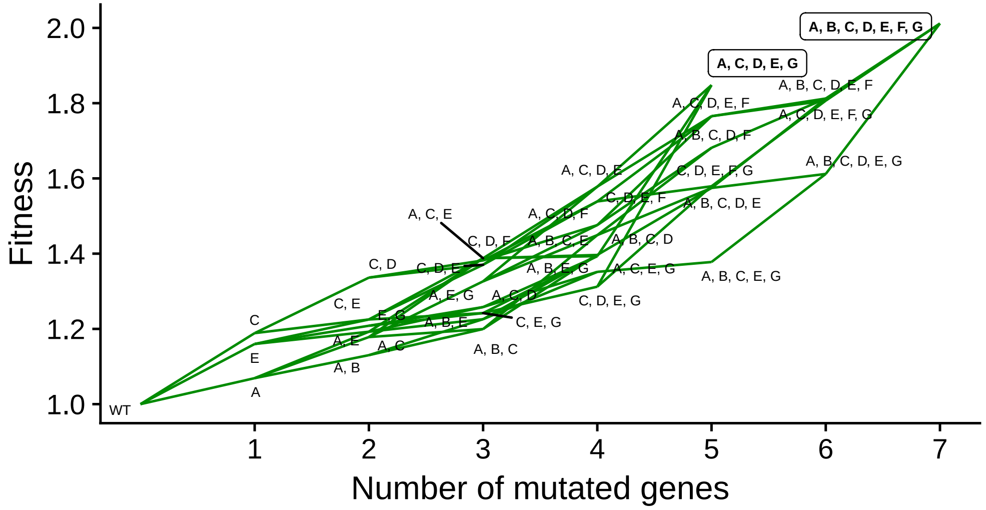


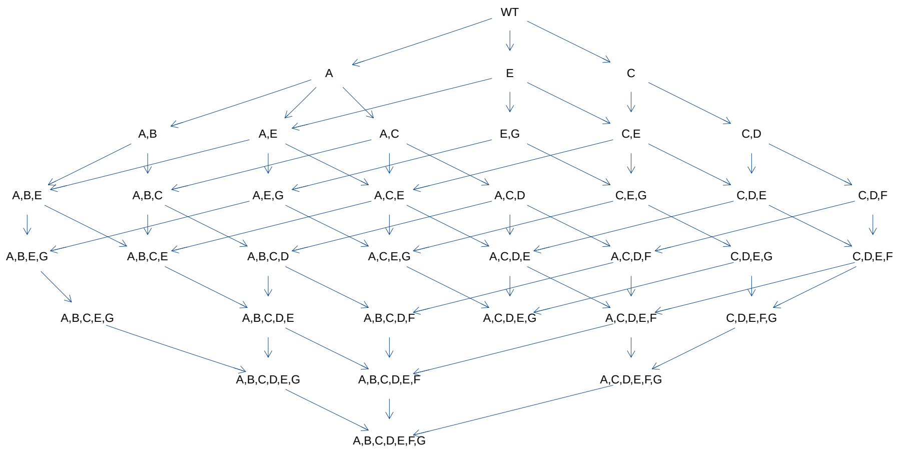

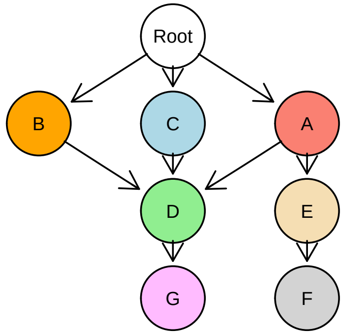


*ID* sLsugLzPftgJ7O1e
*acc genotypes* 22
*peaks no back* 2
*removed edges* 3
*prop rem edges* 0.079

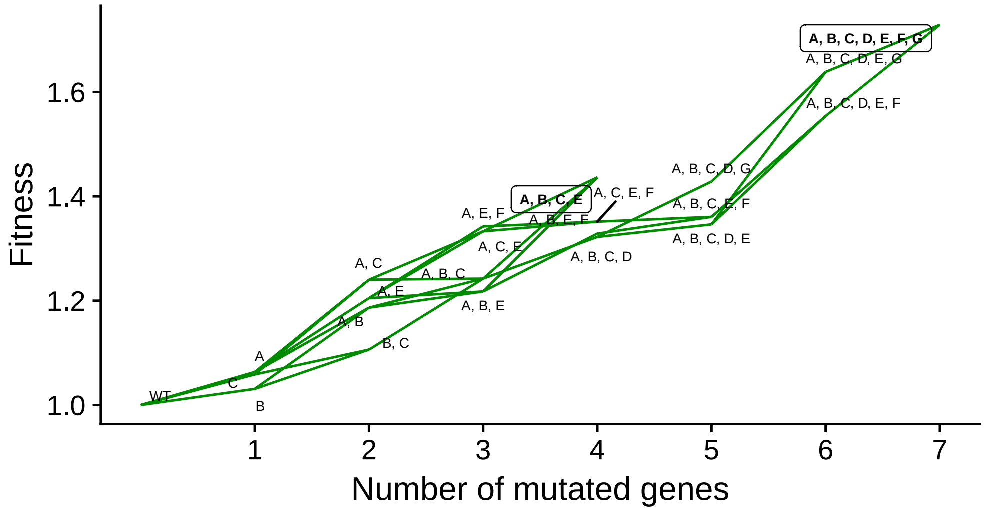


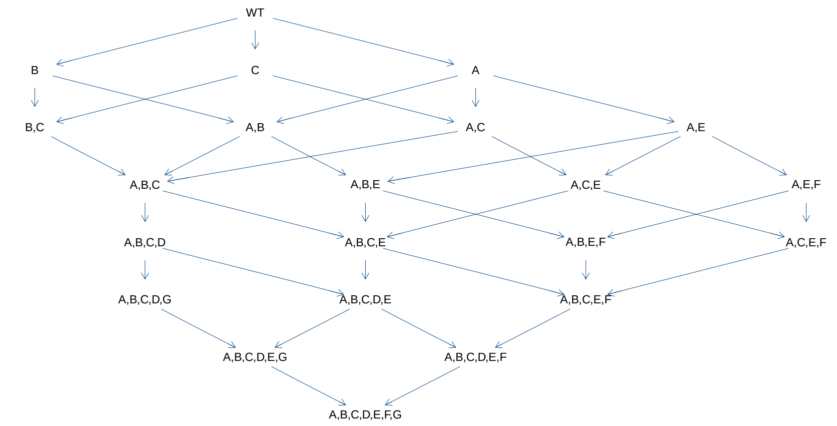


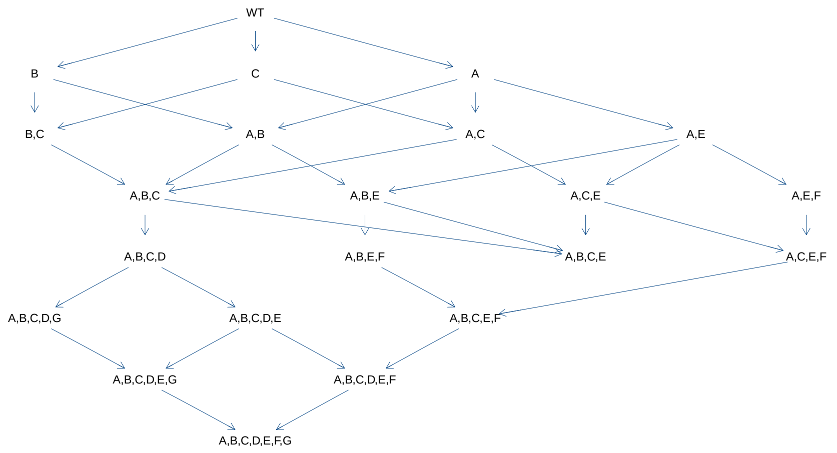

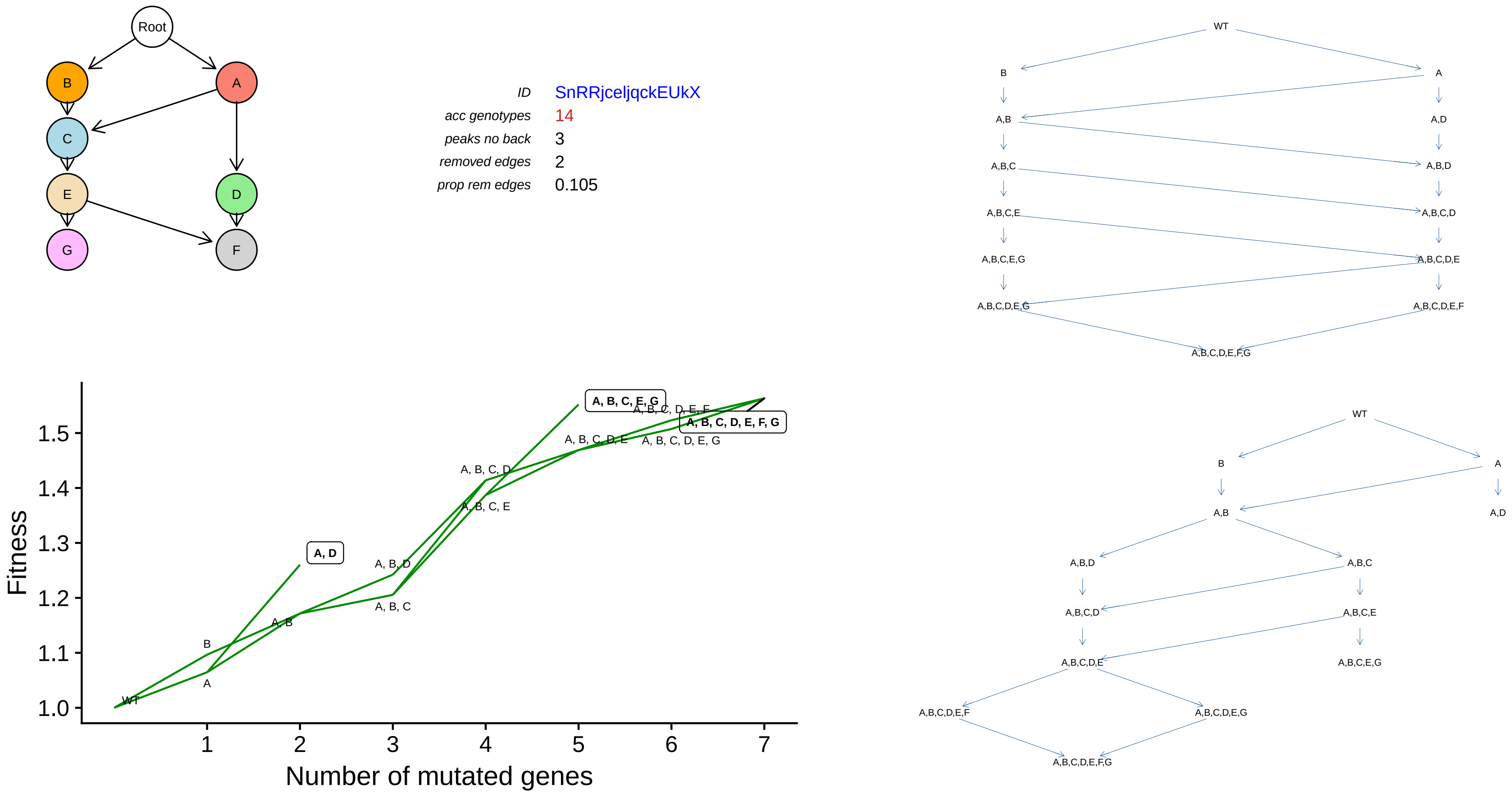

Root
B
A
C
E
D
G
F
ID SnRRjceljqckEUkX
acc genotypes 14
peaks no back 3
removed edges 2
prop rem edges 0.105
Fitness
Number of mutated genes
1.0
1.1
1.2
1.3
1.4
1.5
1
2
3
4
5
6
7
WT
A
B
A, B
A, D
A, B, C
A, B, D
A, B, C, D
A, B, C, E
A, B, C, D, E
A, B, C, E, G
A, B, C, D, E, F
A, B, C, D, E, G
A, B, C, D, E, F, G
WT
B
A
A,B
A,D
A,B,C
A,B,D
A,B,C,E
A,B,C,D
A,B,C,E,G
A,B,C,D,E
A,B,C,D,E,G
A,B,C,D,E,F
A,B,C,D,E,F,G
WT
B
A
A,B
A,D
A,B,D
A,B,C
A,B,C,D
A,B,C,E
A,B,C,D,E
A,B,C,E,G
A,B,C,D,E,F
A,B,C,D,E,G
A,B,C,D,E,F,G

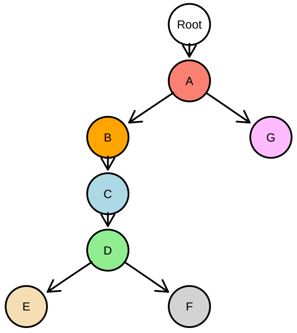


| | |
|---|---|
| *ID* | t2kQYfWf61wq10313 |
| *acc genotypes* | 15 |
| *peaks no back* | 2 |
| *removed edges* | 2 |
| *prop rem edges* | 0.091 |

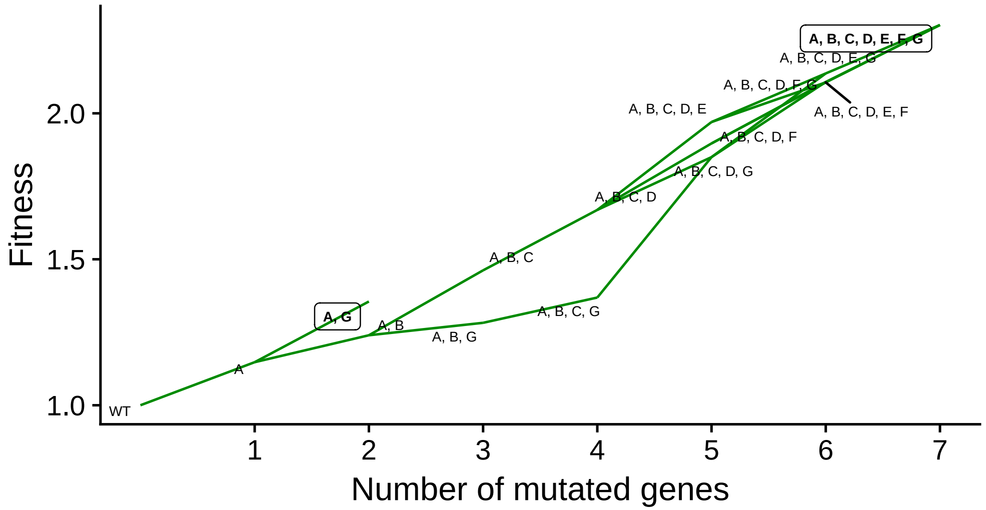


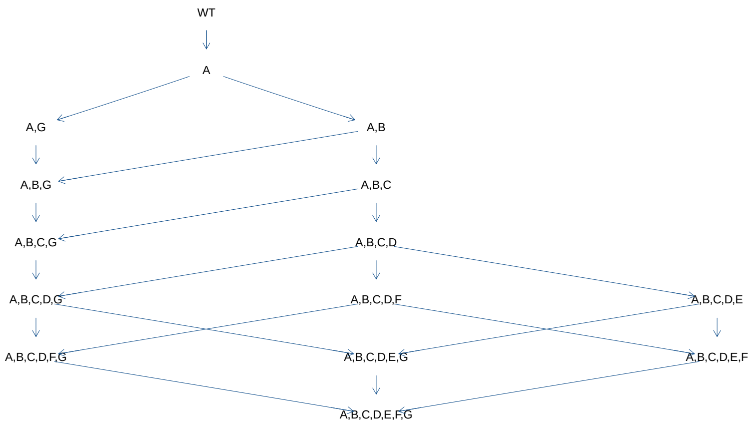


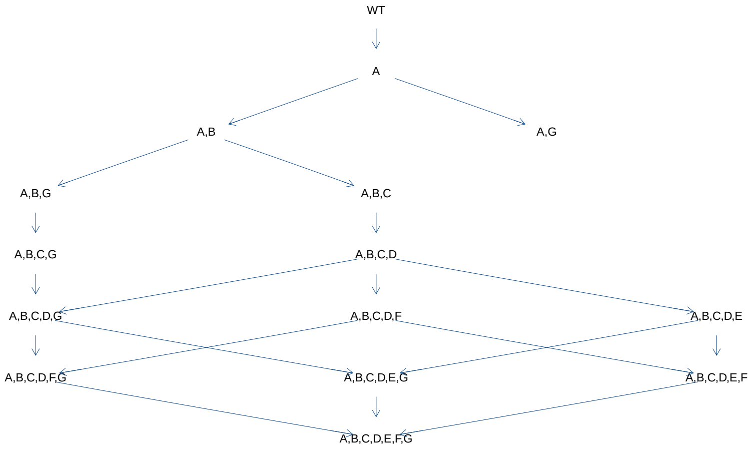

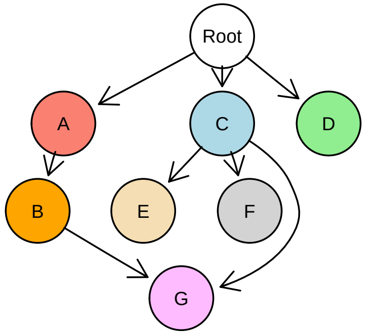


| | |
|---|---|
| *ID* | t4BmCLHxEiv103ES6 |
| *acc genotypes* | 38 |
| *peaks no back* | 8 |
| *removed edges* | 17 |
| *prop rem edges* | 0.2 |

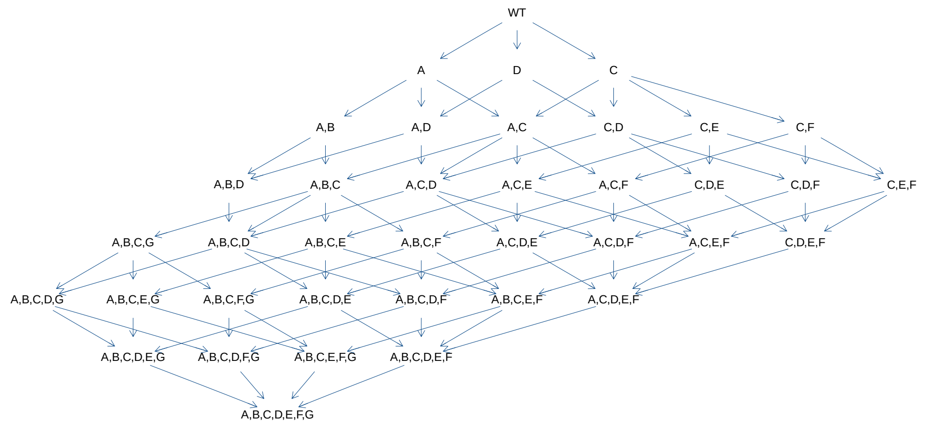


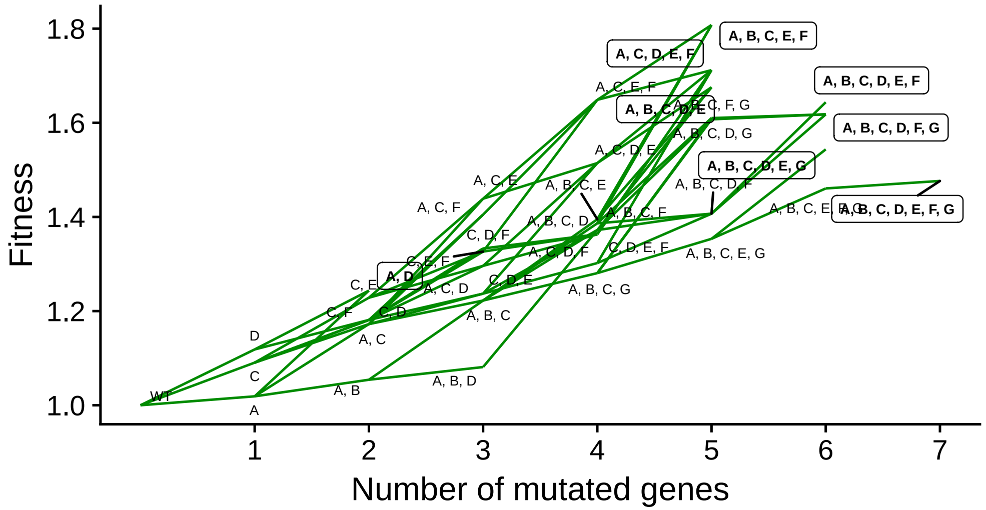


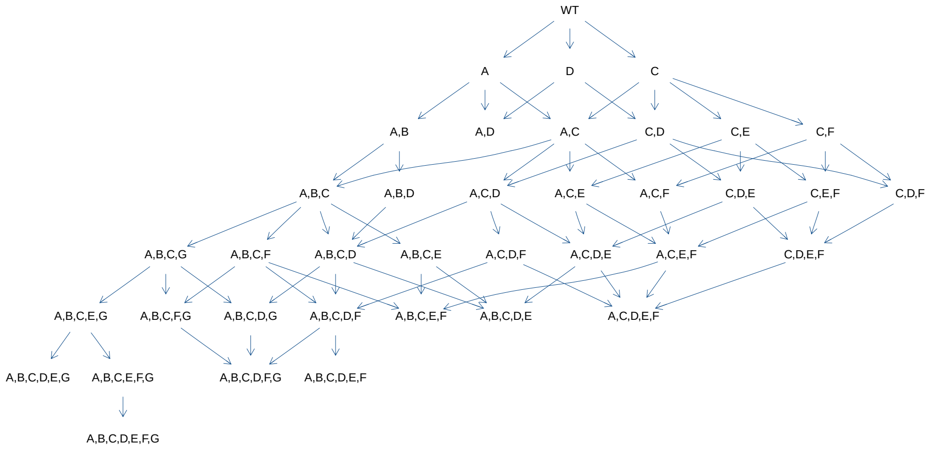

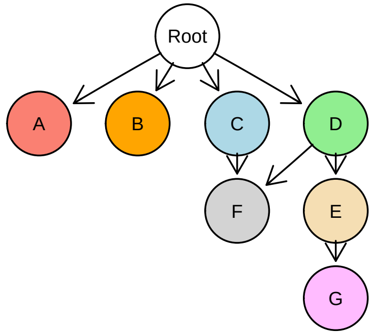


| | |
|---|---|
| *ID* | T9n9yW1qk7hj10GjU |
| *acc genotypes* | 44 |
| *peaks no back* | 7 |
| *removed edges* | 25 |
| *prop rem edges* | 0.24 |

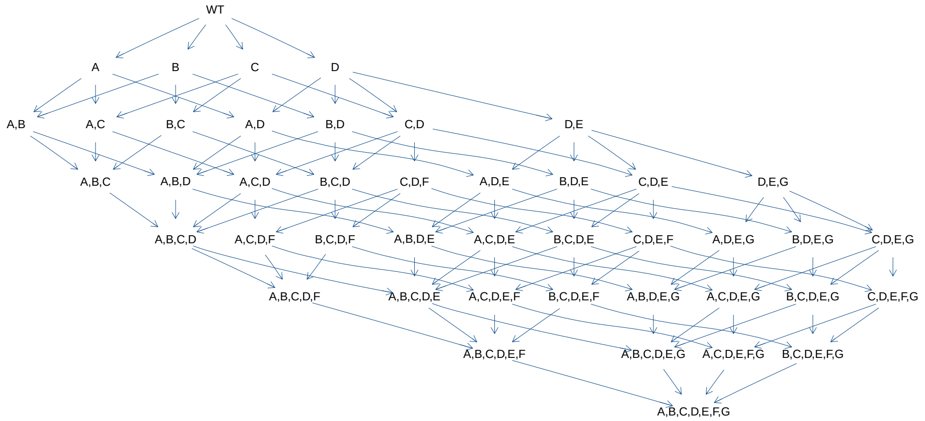


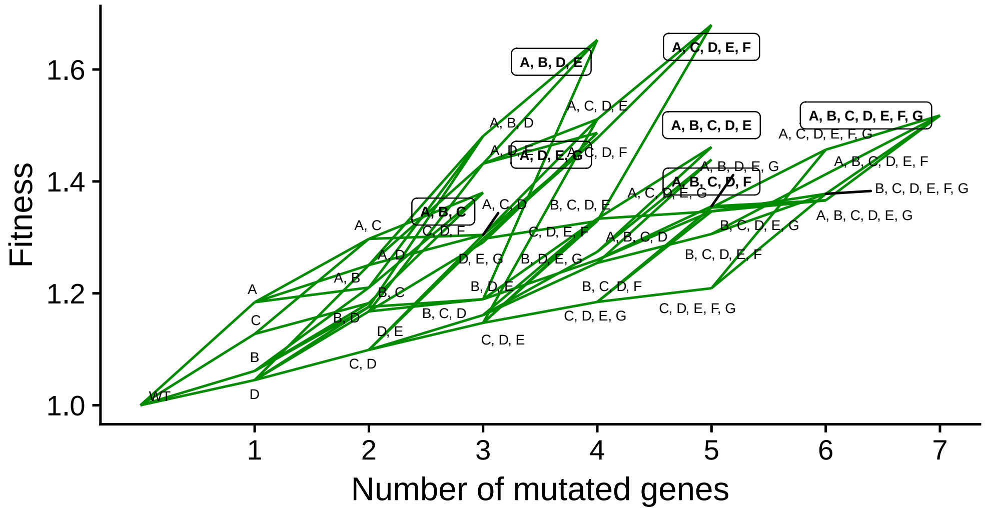


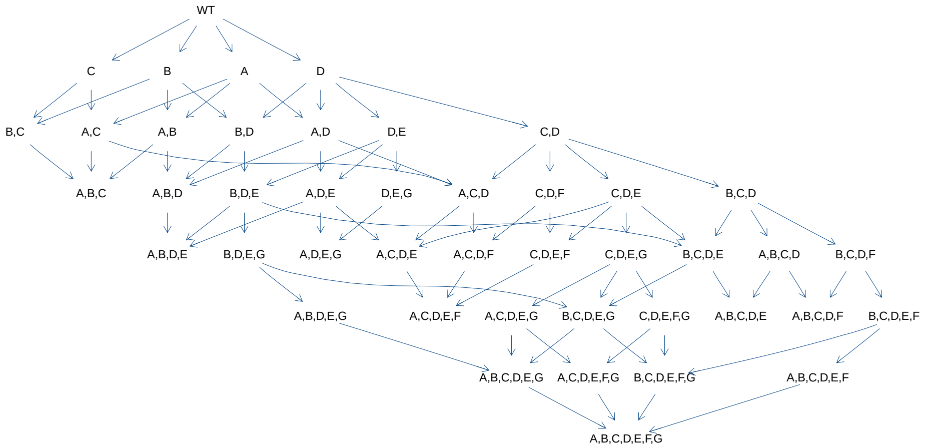

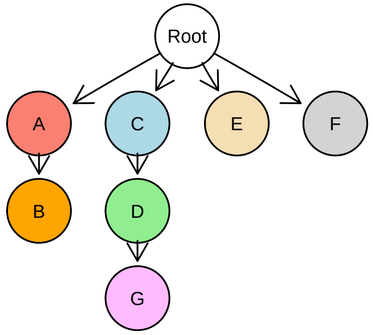


| | |
|---|---|
| *ID* | TcQu1TxfmYIxeOT4 |
| *acc genotypes* | 48 |
| *peaks no back* | 6 |
| *removed edges* | 19 |
| *prop rem edges* | 0.164 |

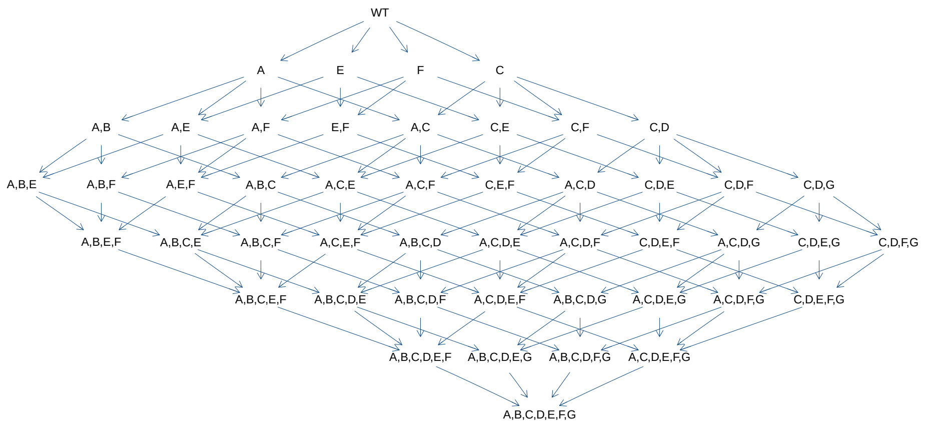

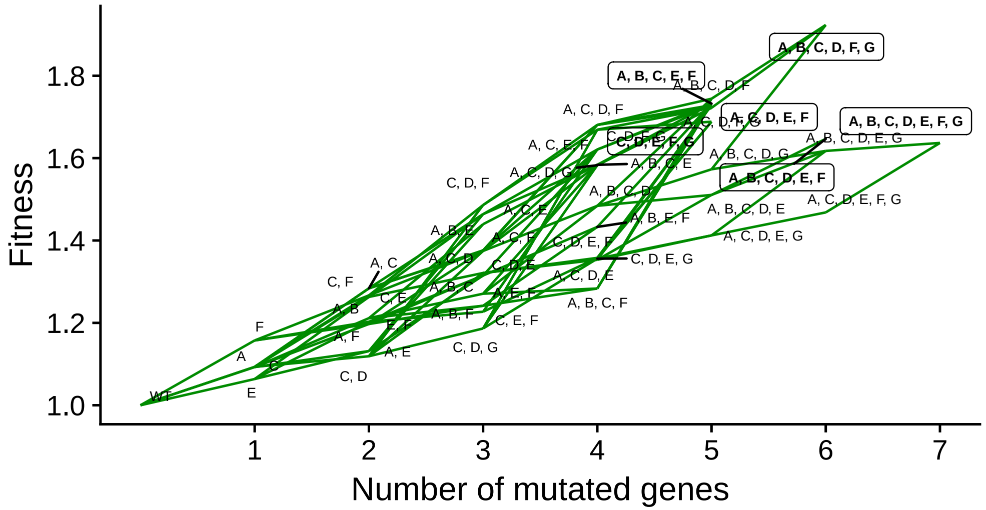


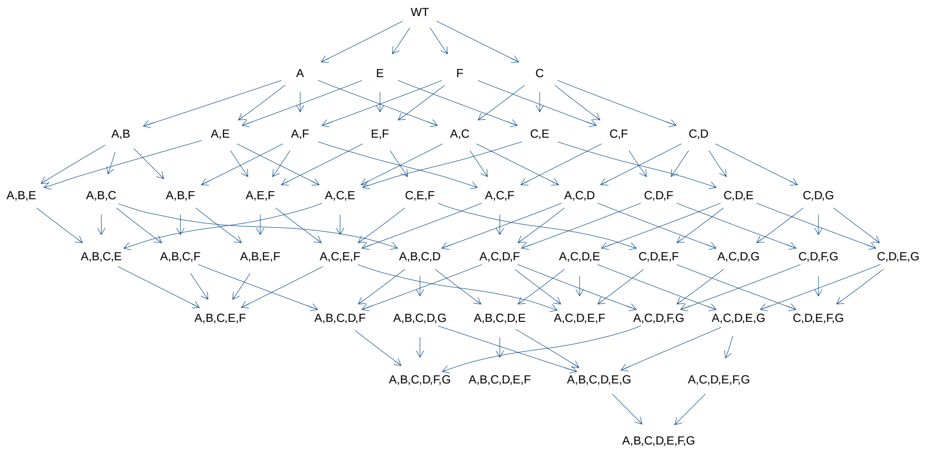

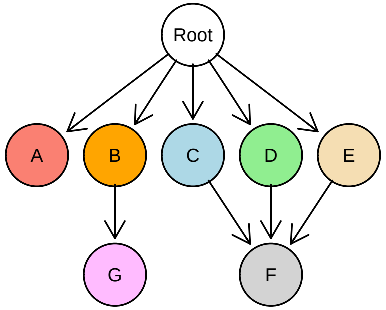


*ID* tdjsoypZIFJzlC5O
*acc genotypes* 54
*peaks no back* 7
*removed edges* 23
*prop rem edges* 0.163

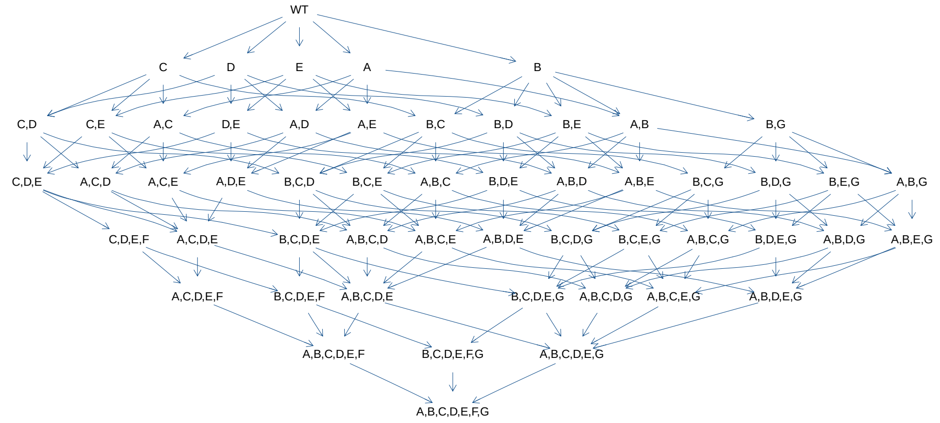


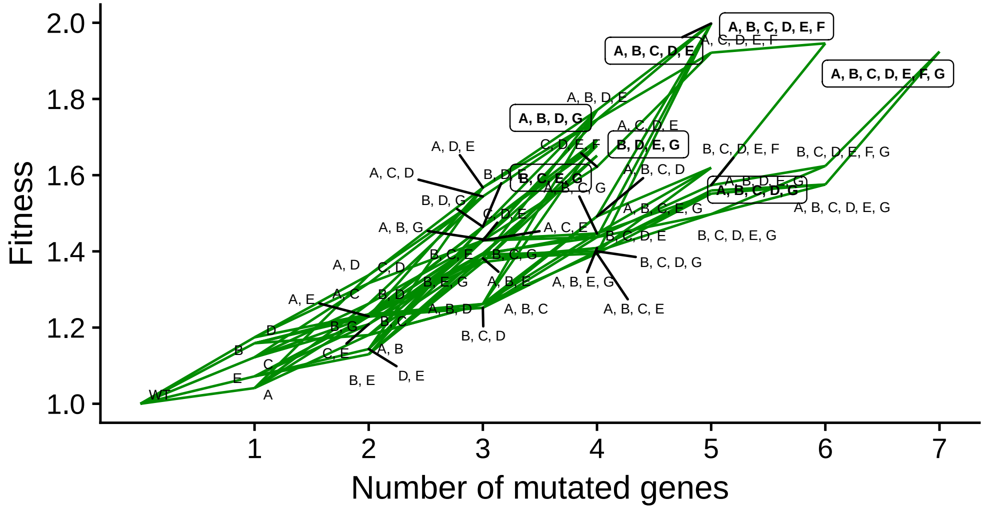


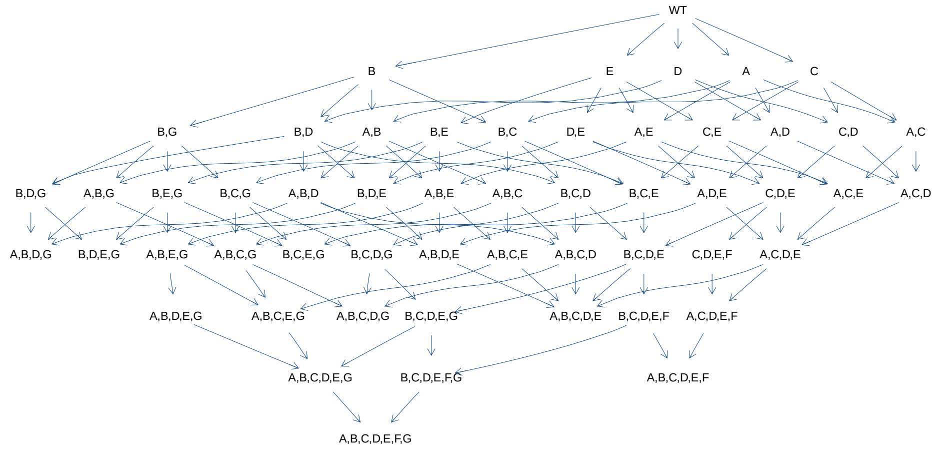

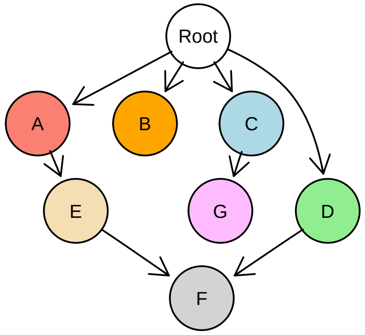


| | |
|---|---|
| *ID* | THM3Re8DzitZ9K5X |
| *acc genotypes* | 42 |
| *peaks no back* | 6 |
| *removed edges* | 12 |
| *prop rem edges* | 0.124 |

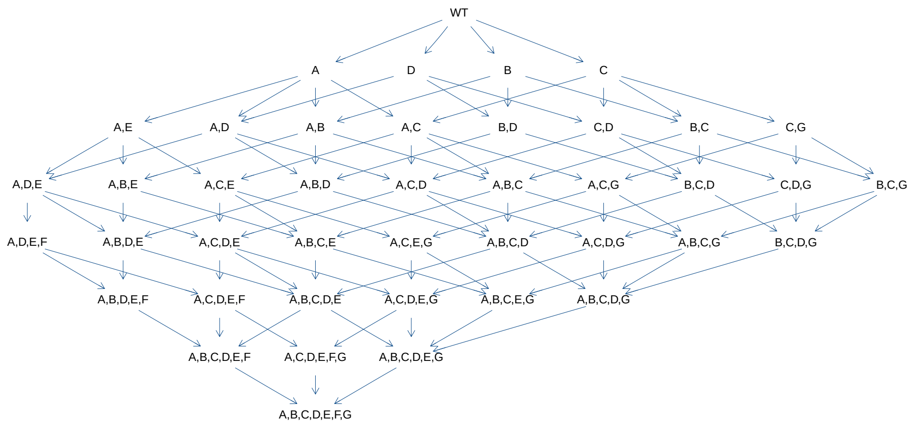


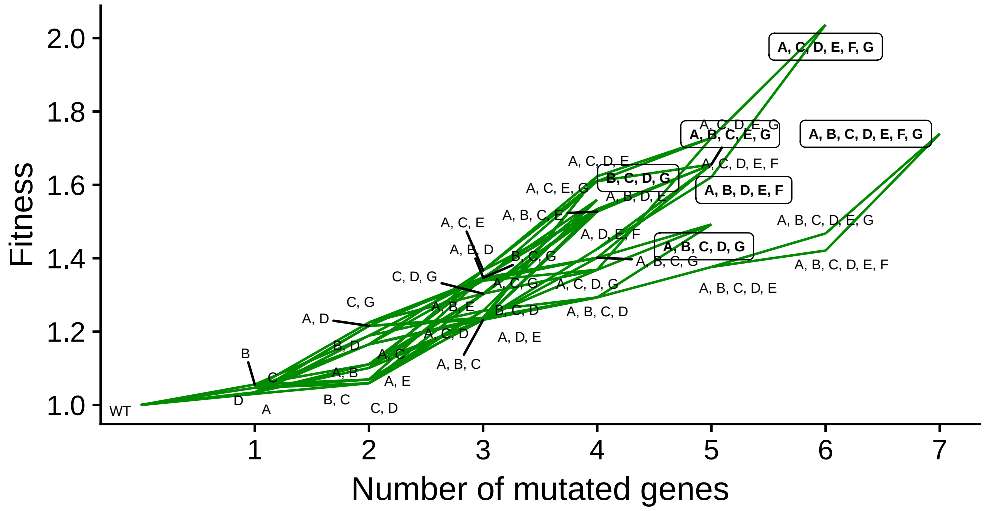


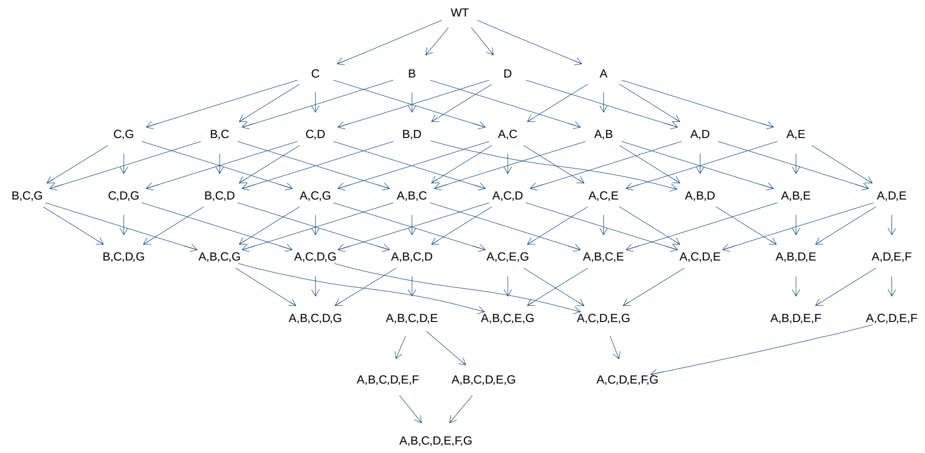

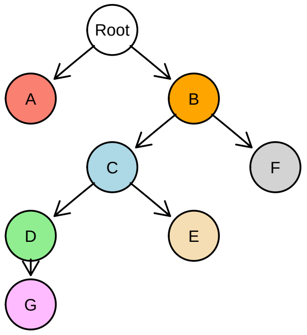


| | |
|---|---|
| *ID* | tPr5Vg10103qrkePxM |
| *acc genotypes* | 30 |
| *peaks no back* | 5 |
| *removed edges* | 10 |
| *prop rem edges* | 0.159 |

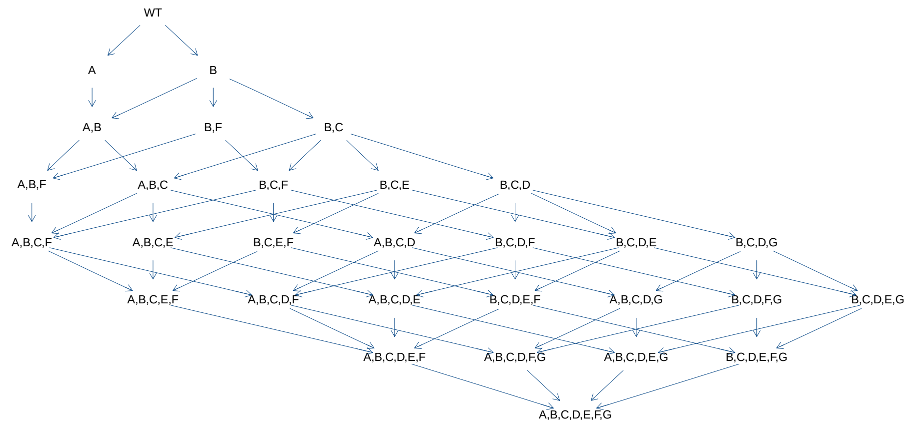


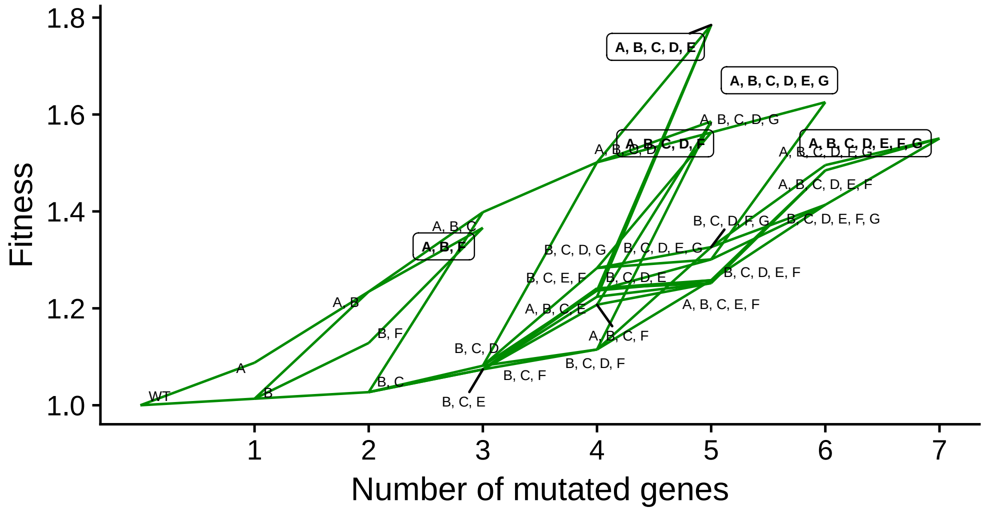


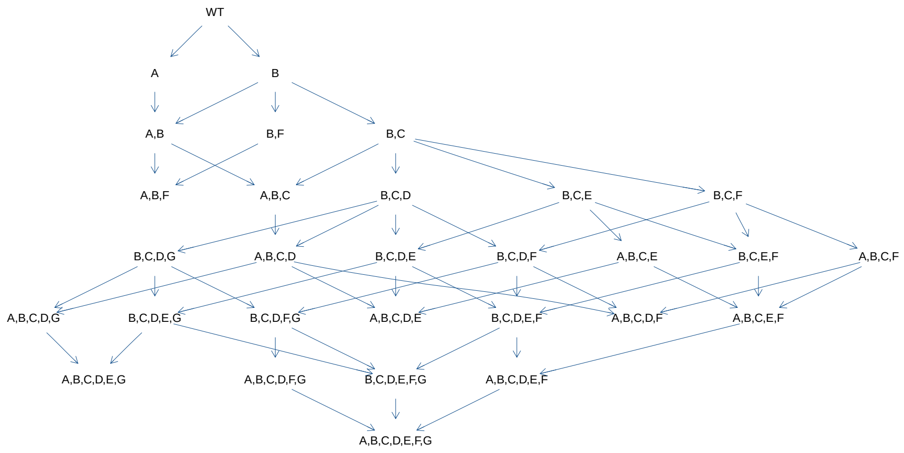

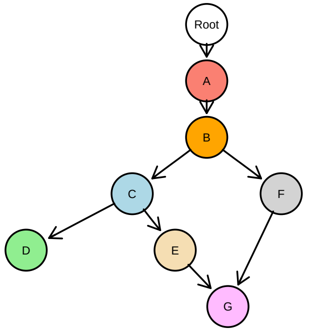


| | |
|---|---|
| *ID* | tZ722GsITVDN5PNF |
| *acc genotypes* | 14 |
| *peaks no back* | 4 |
| *removed edges* | 4 |
| *prop rem edges* | 0.2 |

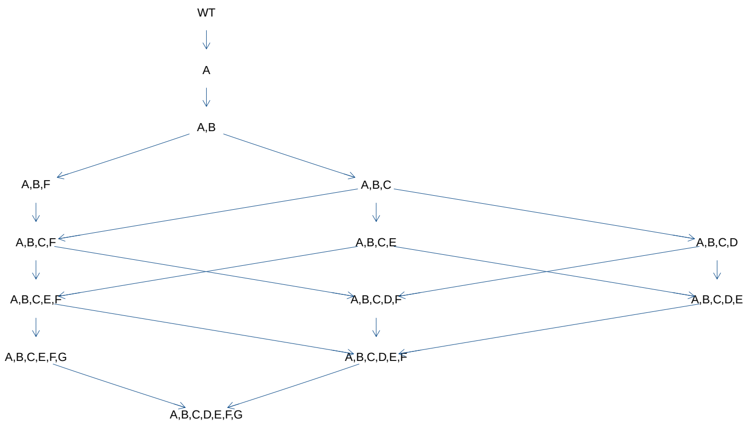


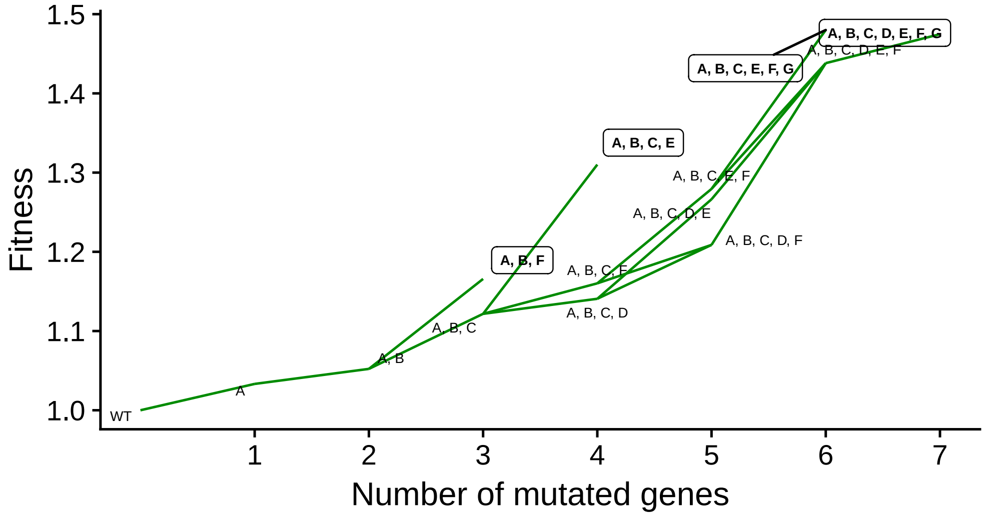


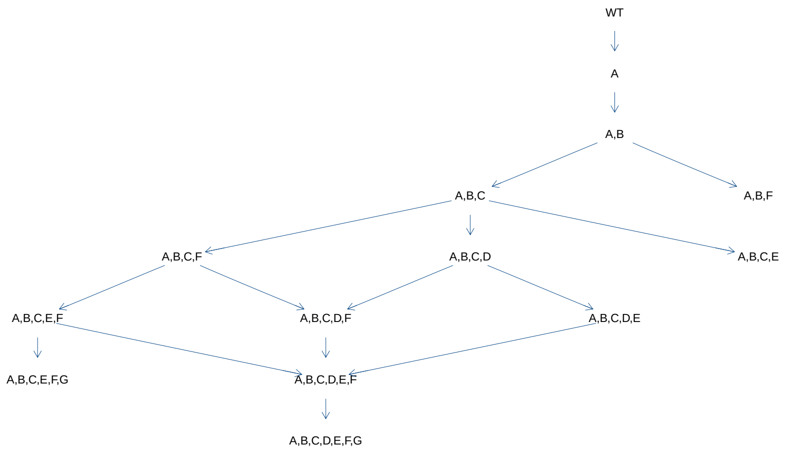

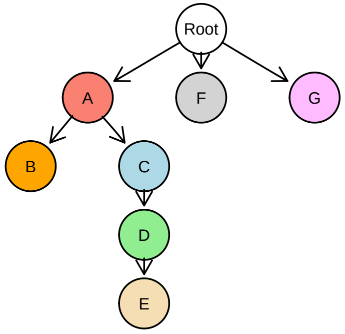


*ID* U3F9k2e17pCATfZA
*acc genotypes* 36
*peaks no back* 6
*removed edges* 9
*prop rem edges* 0.112

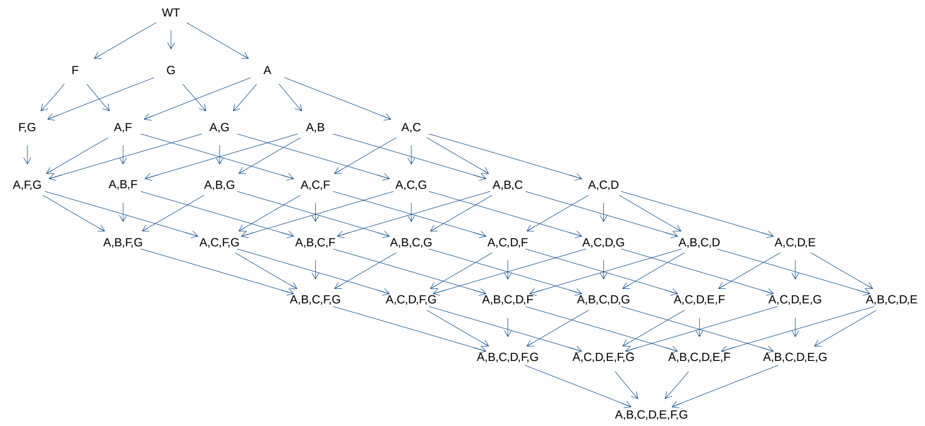


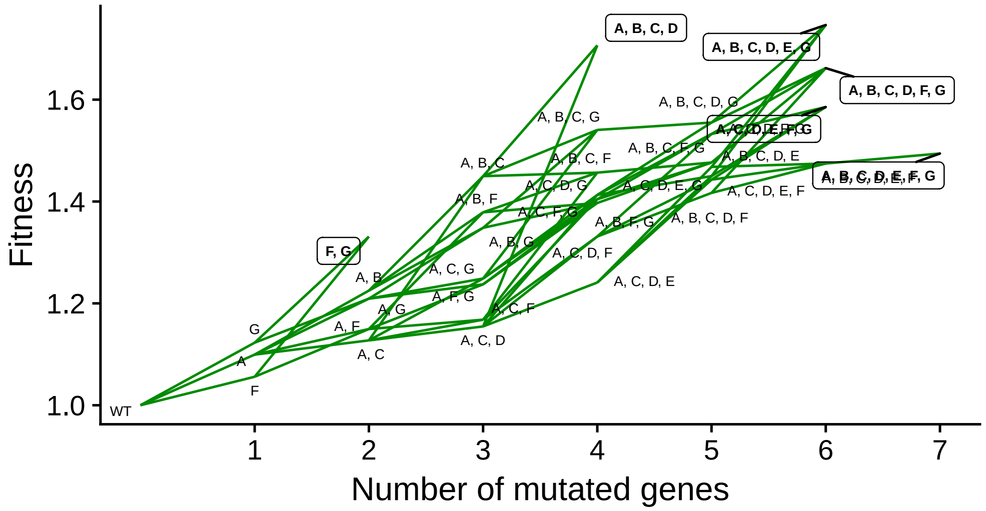


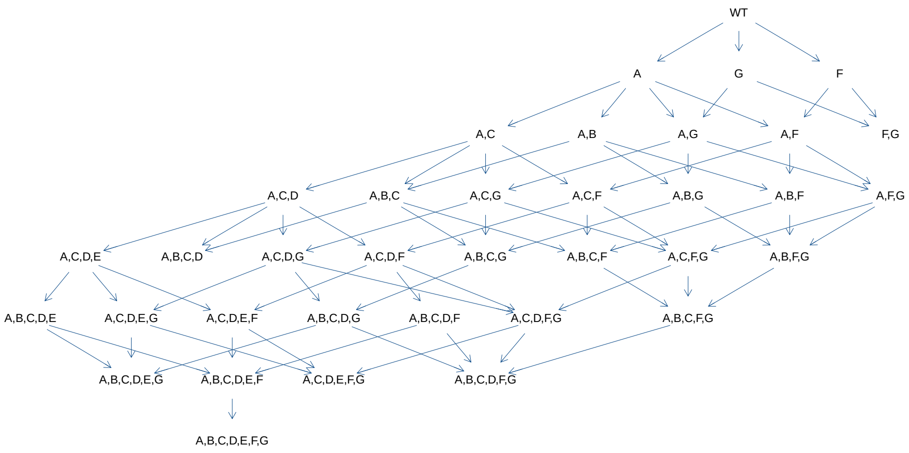

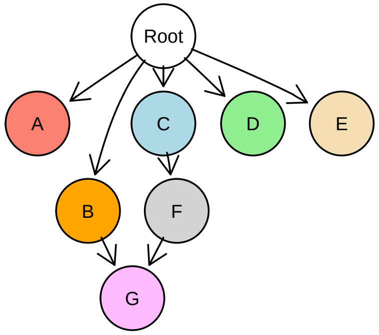


| | |
|---|---|
| *ID* | u5c4a1CX10bL5RtgV |
| *acc genotypes* | 56 |
| *peaks no back* | 7 |
| *removed edges* | 27 |
| *prop rem edges* | 0.182 |

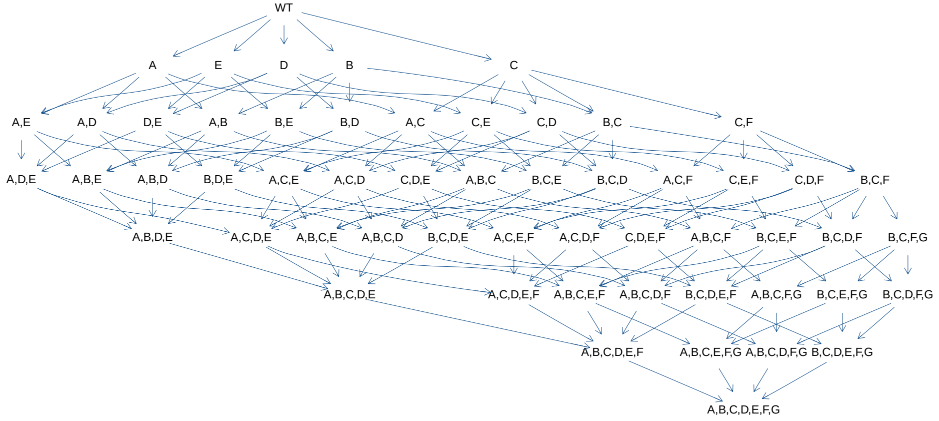


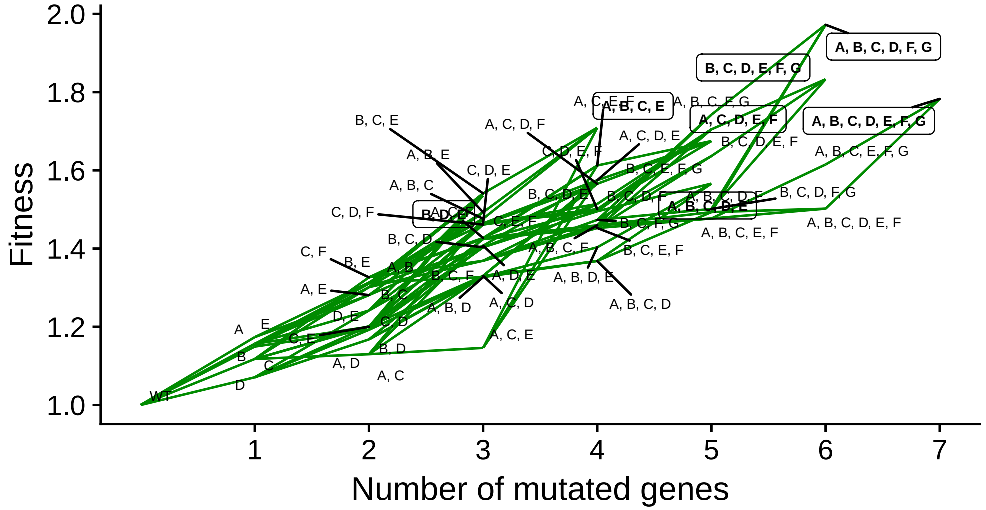


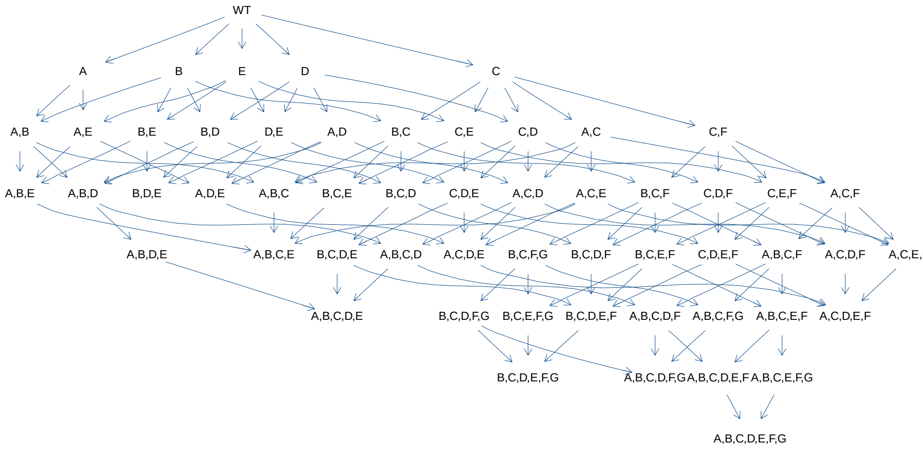

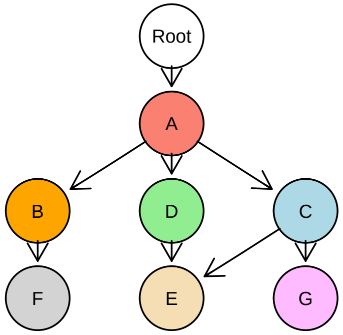


| | |
|---|---|
| *ID* | U7zMFhVmpGvNZWah |
| *acc genotypes* | 25 |
| *peaks no back* | 6 |
| *removed edges* | 9 |
| *prop rem edges* | 0.191 |

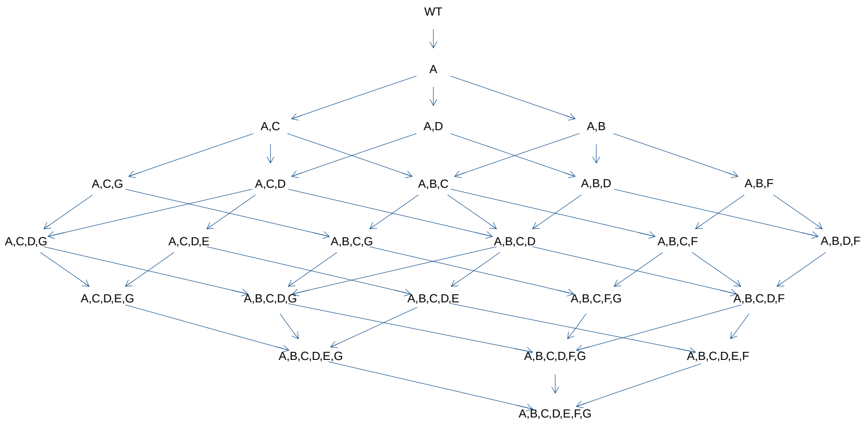


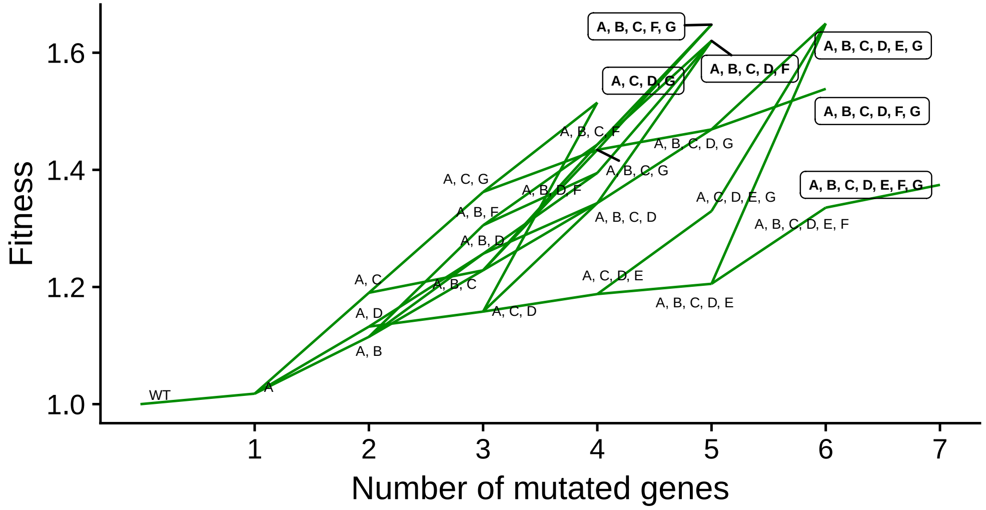


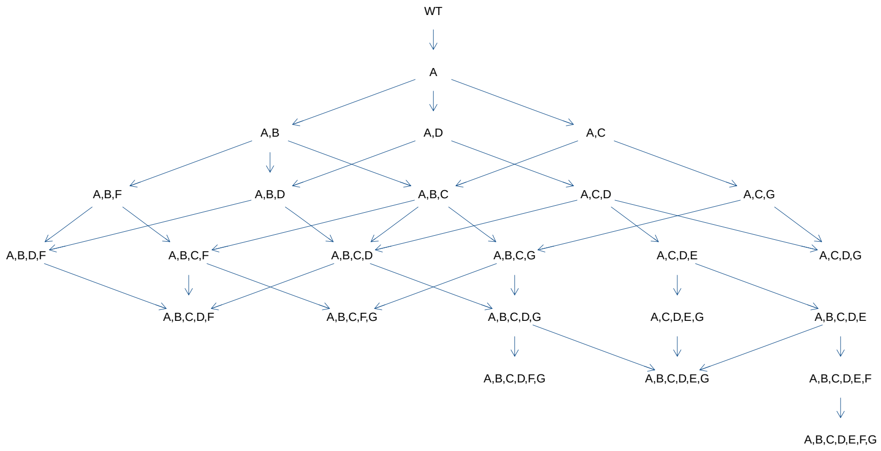

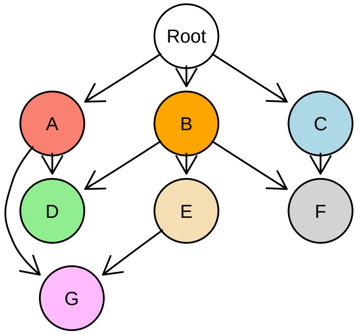


*ID* UlCZ6OQWroBIY8ER
*acc genotypes* 28
*peaks no back* 3
*removed edges* 4
*prop rem edges* 0.074

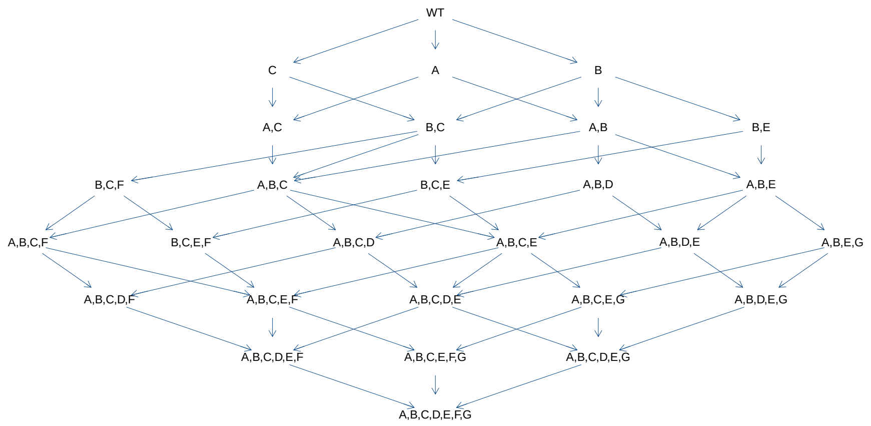


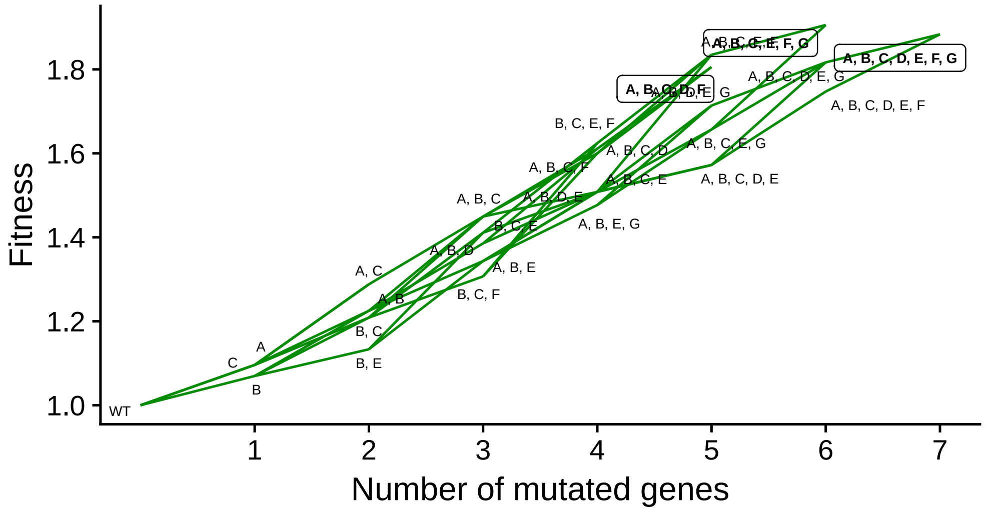


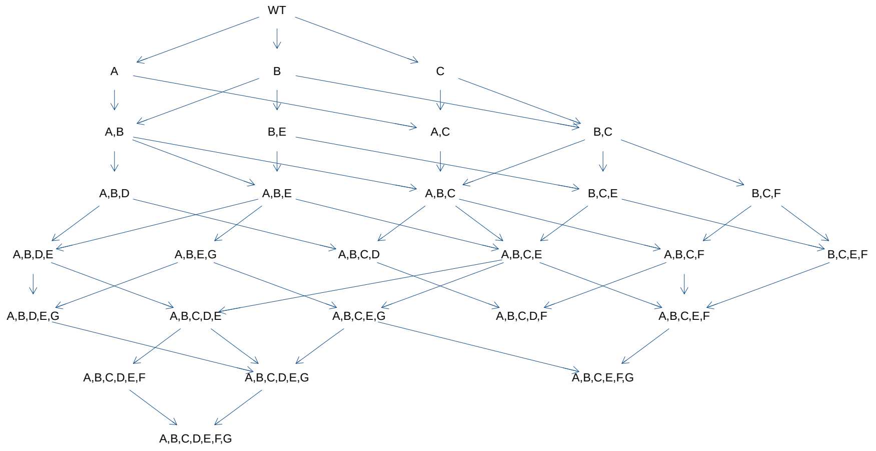

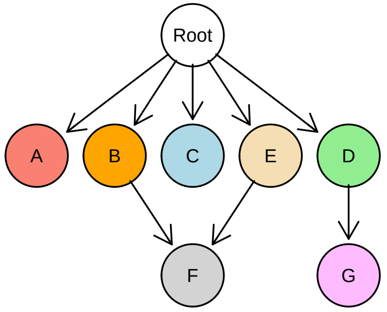


| | |
|---|---|
| *ID* | unh1zVBv7V9gaHC5 |
| *acc genotypes* | 60 |
| *peaks no back* | 7 |
| *removed edges* | 29 |
| *prop rem edges* | 0.181 |

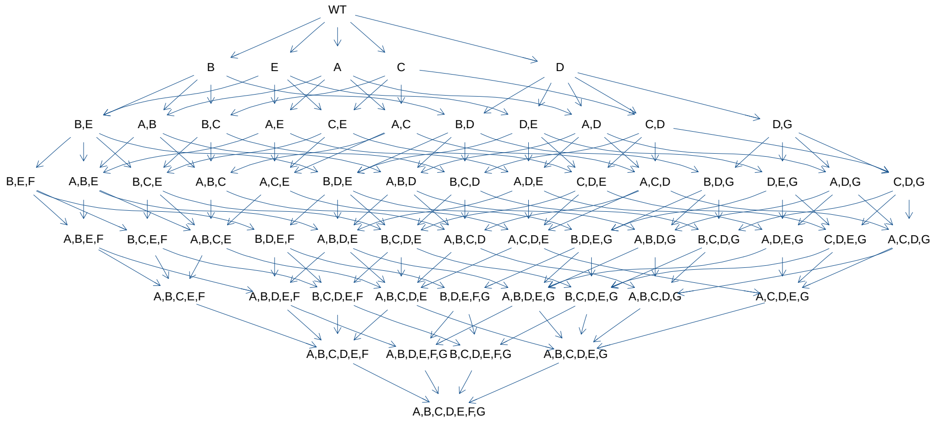


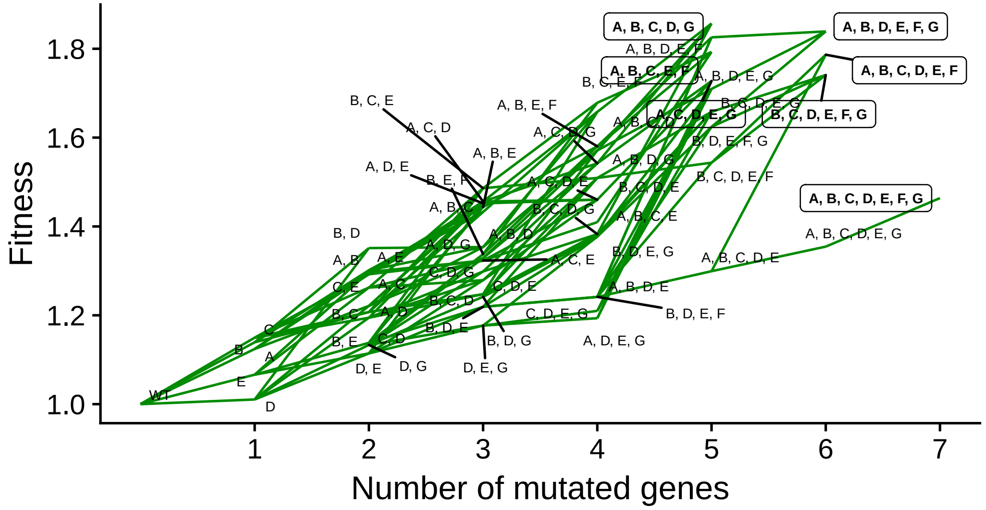


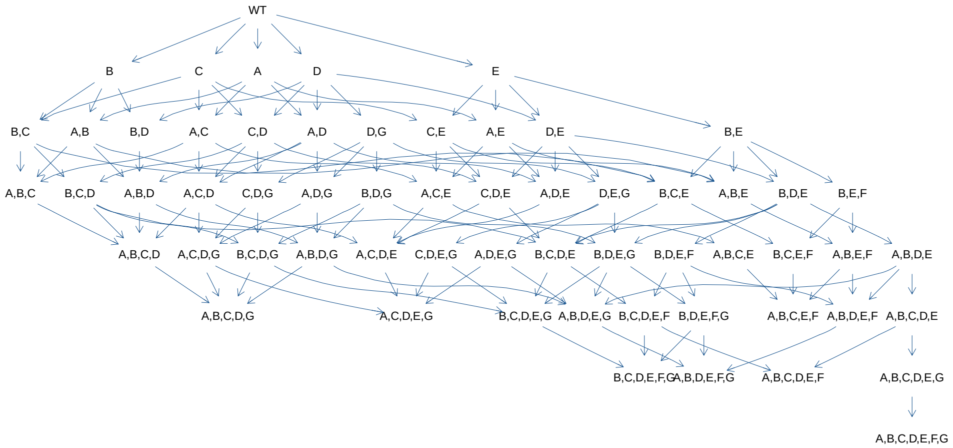

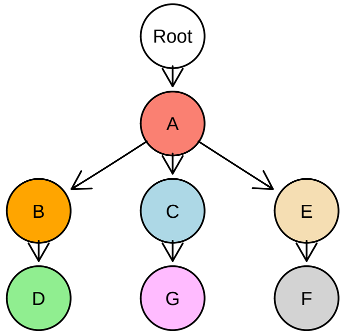


| | |
|---|---|
| *ID* | uOql5ivL2U4HWUKS |
| *acc genotypes* | 28 |
| *peaks no back* | 4 |
| *removed edges* | 4 |
| *prop rem edges* | 0.073 |

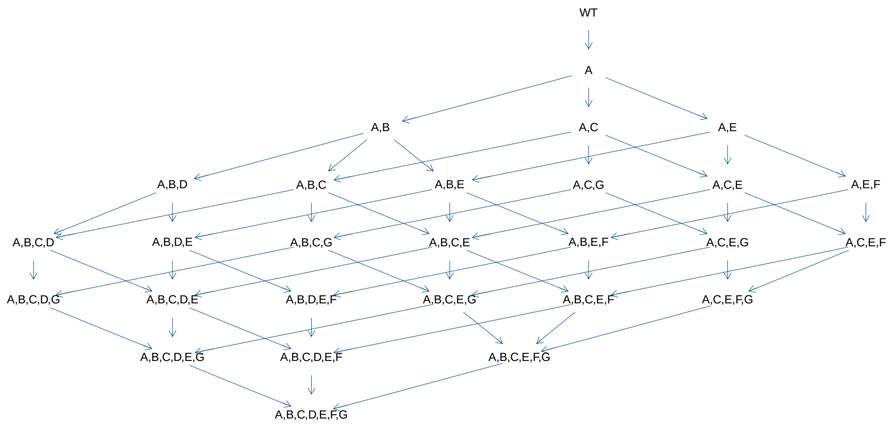


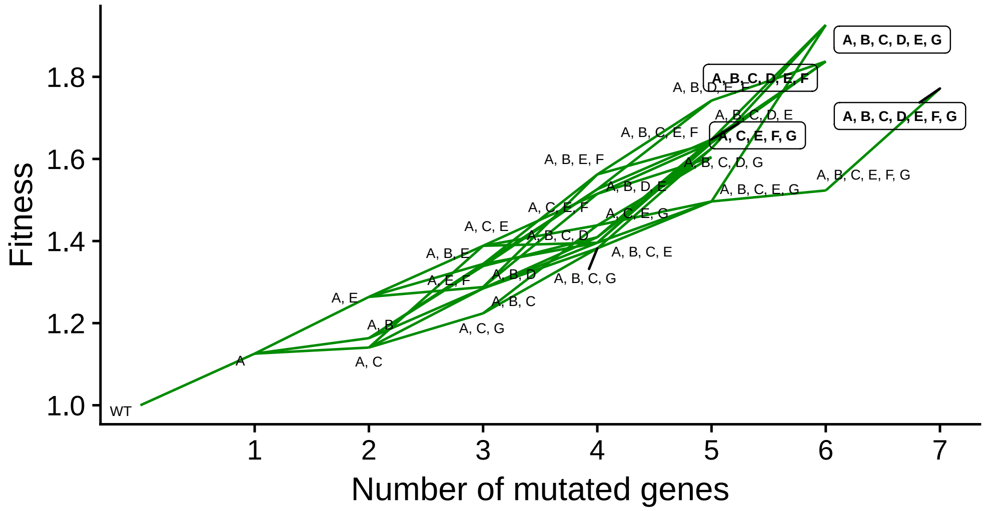


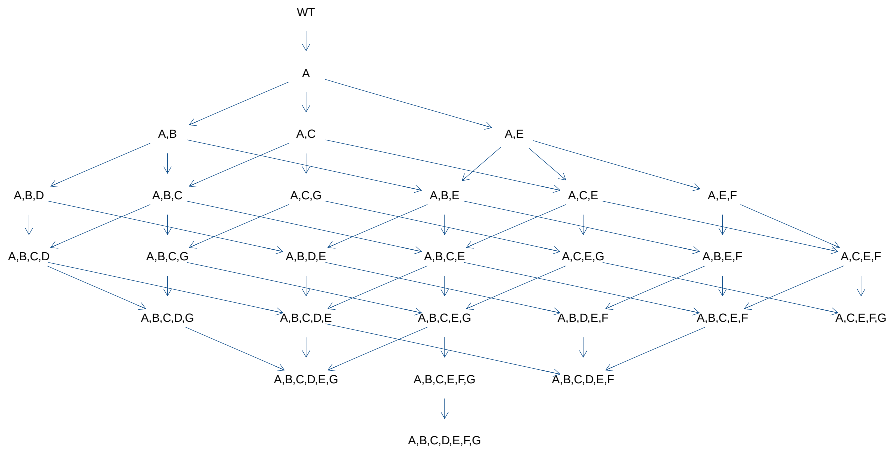

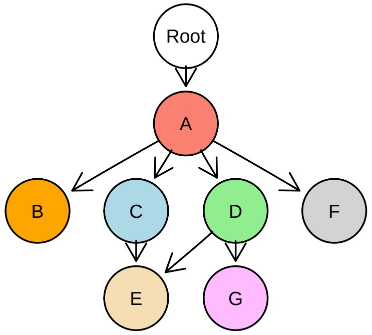


| | |
|---|---|
| *ID* | upEZISo6oFVQcSEK |
| *acc genotypes* | 33 |
| *peaks no back* | 6 |
| *removed edges* | 10 |
| *prop rem edges* | 0.137 |

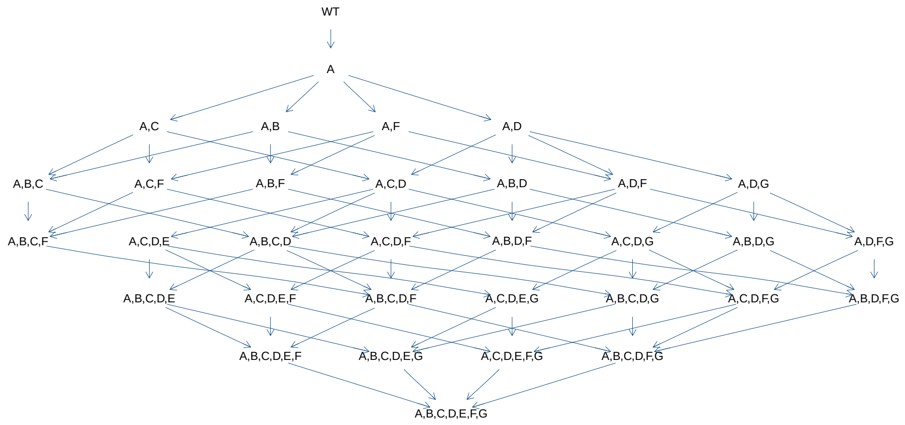


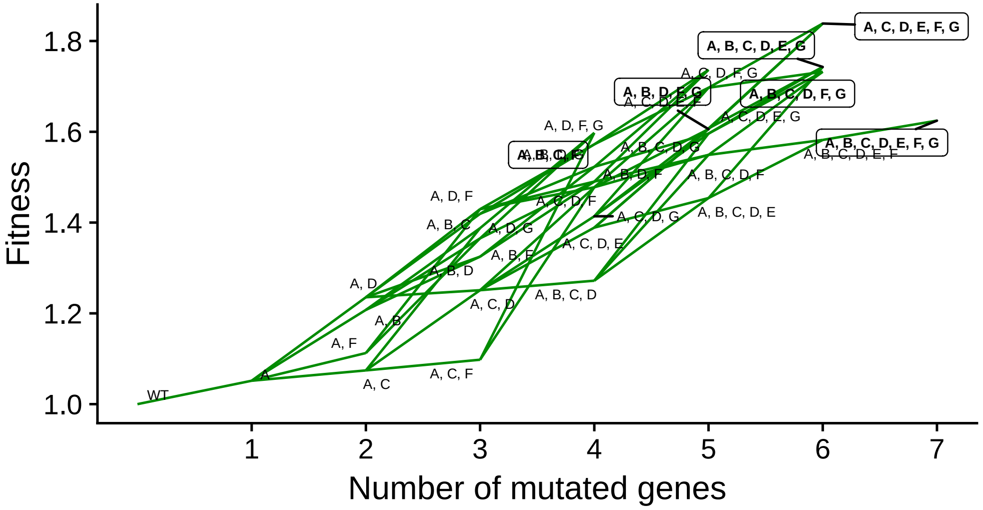


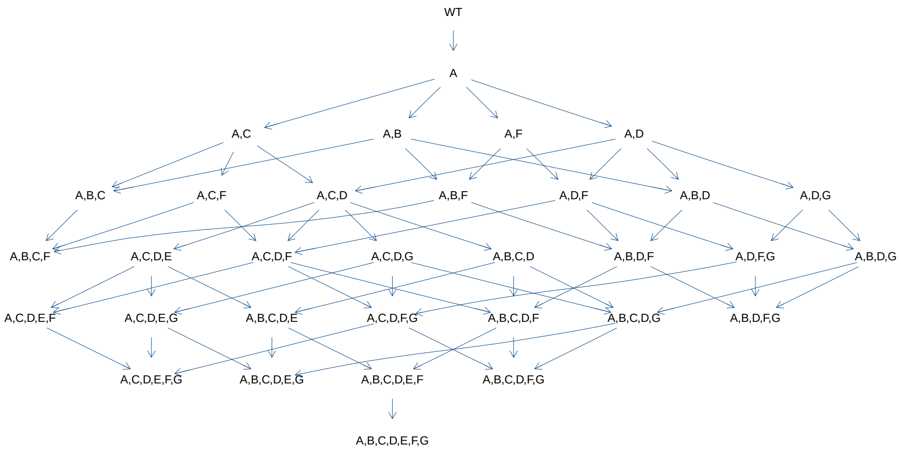

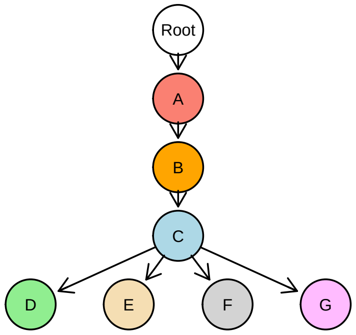


| | |
|---|---|
| *ID* | UQu5n10aPzWHnGf59 |
| *acc genotypes* | 19 |
| *peaks no back* | 5 |
| *removed edges* | 6 |
| *prop rem edges* | 0.171 |

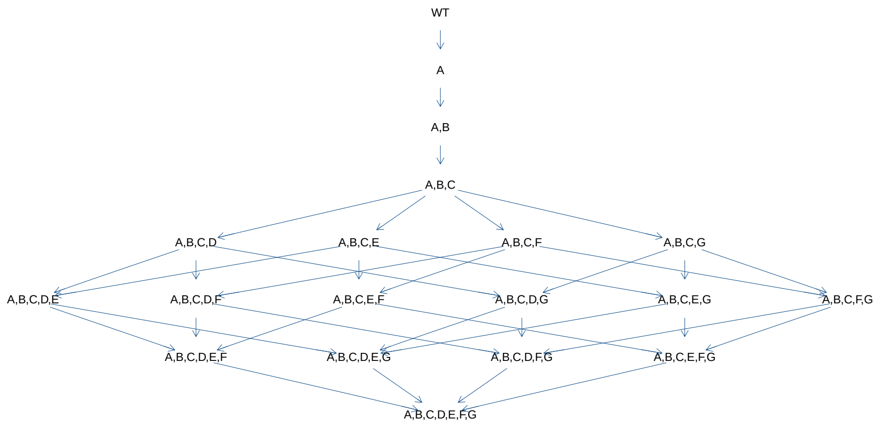


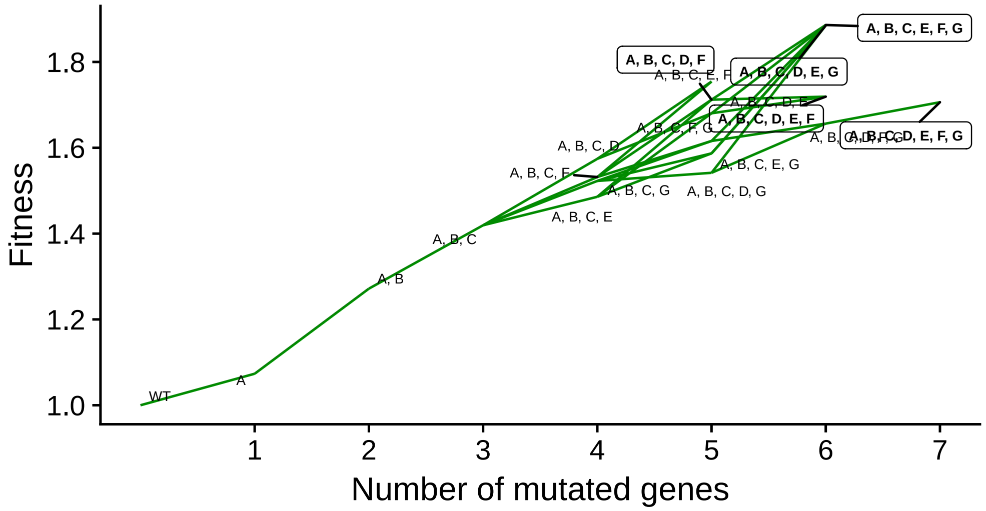


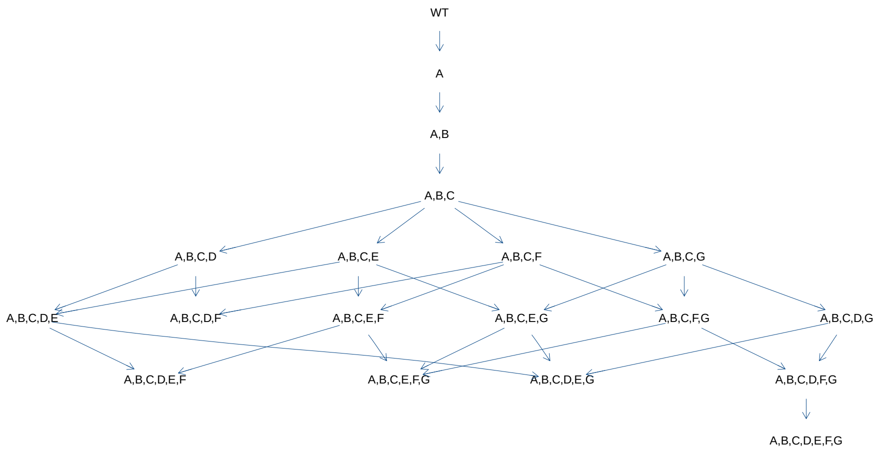

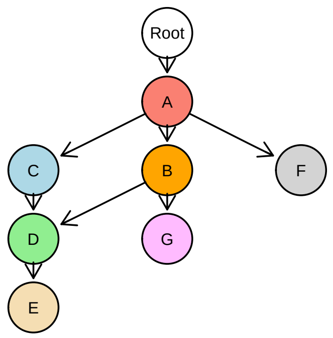


| | |
|---|---|
| *ID* | uw7YHXhohAPISHwZ |
| *acc genotypes* | 21 |
| *peaks no back* | 2 |
| *removed edges* | 8 |
| *prop rem edges* | 0.216 |

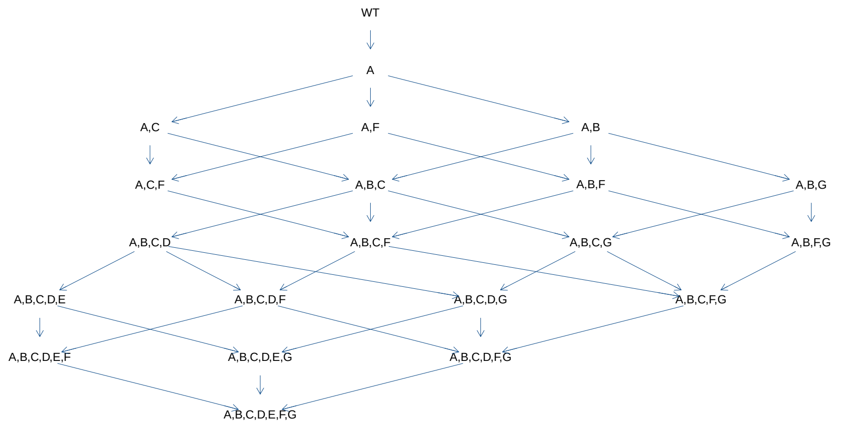


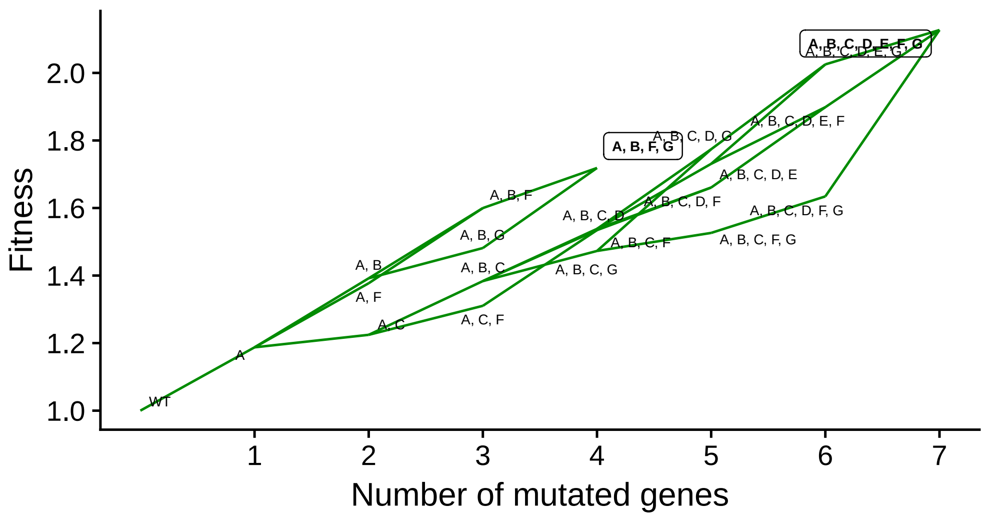


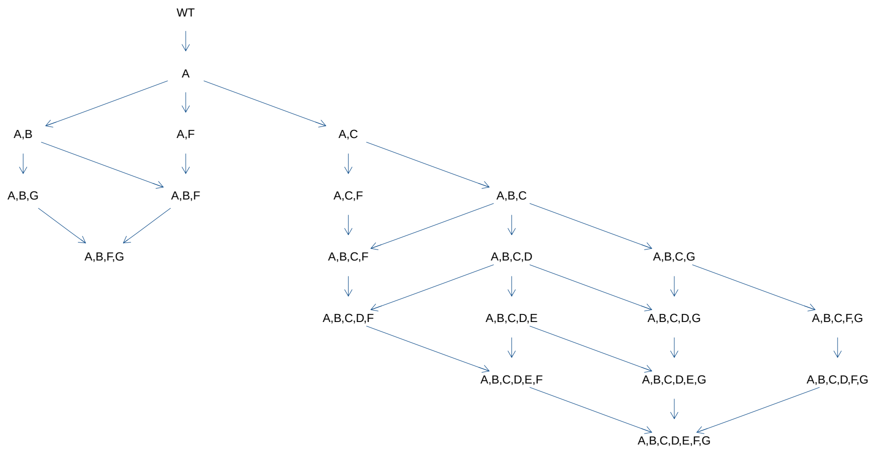

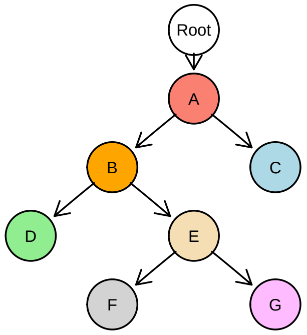


| | |
|---|---|
| *ID* | V4JKndnKWM310nITX |
| *acc genotypes* | 23 |
| *peaks no back* | 4 |
| *removed edges* | 5 |
| *prop rem edges* | 0.114 |

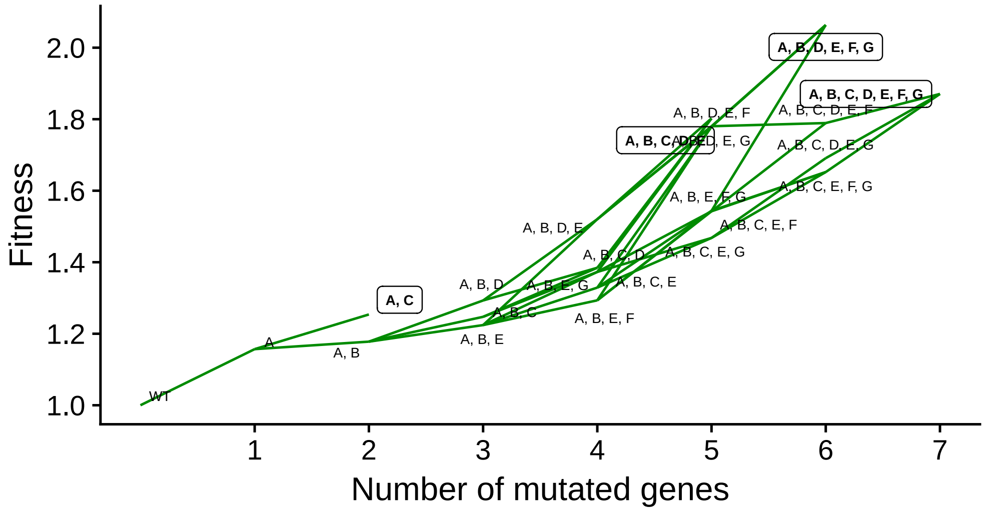


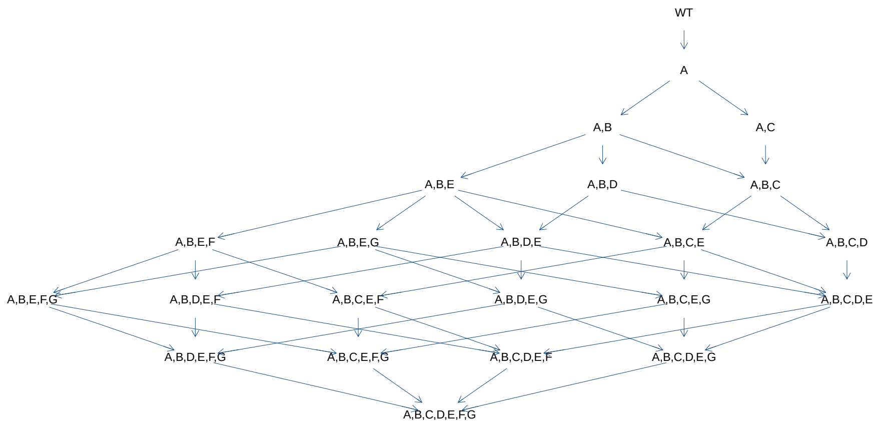


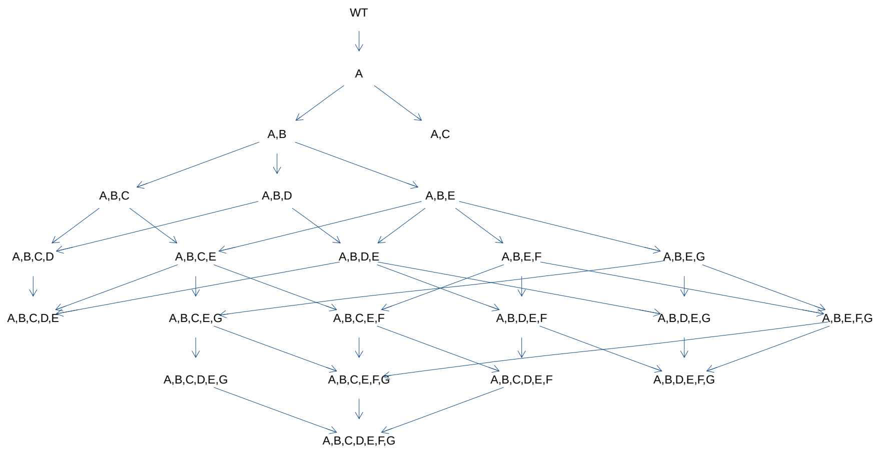

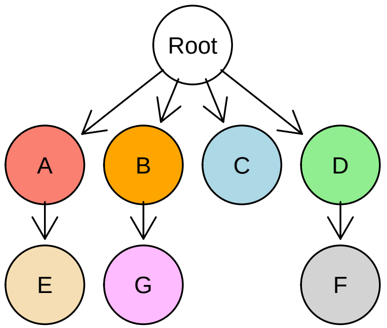


| | |
|---|---|
| *ID* | V4TR5vUVudRPVWmM |
| *acc genotypes* | 54 |
| *peaks no back* | 8 |
| *removed edges* | 24 |
| *prop rem edges* | 0.178 |

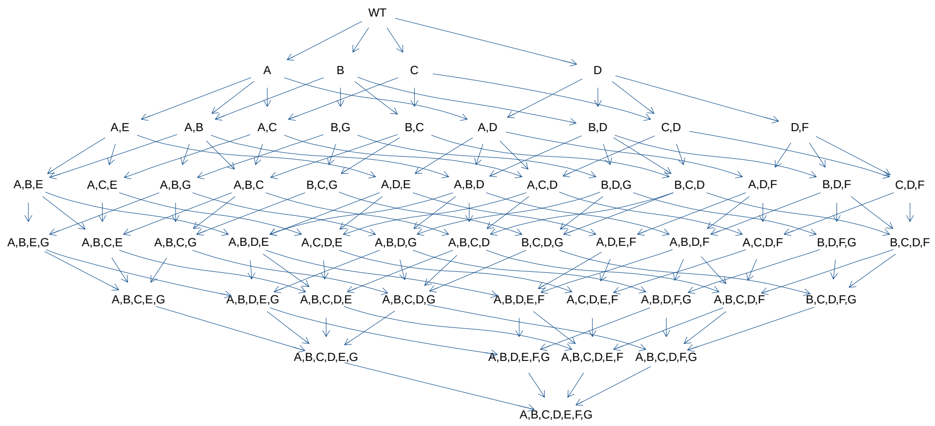

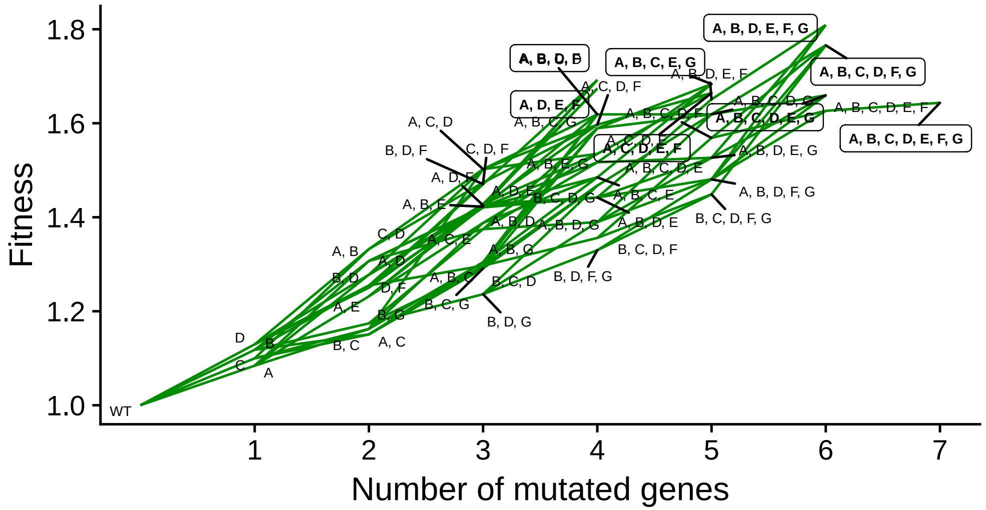


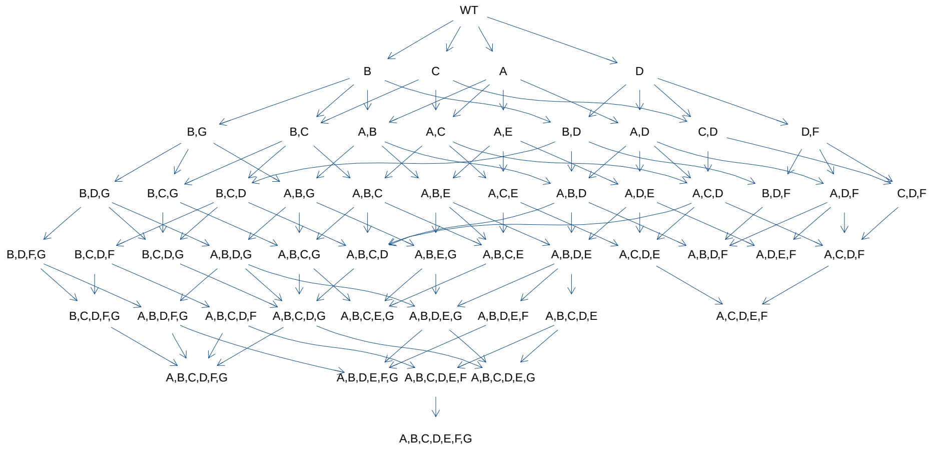

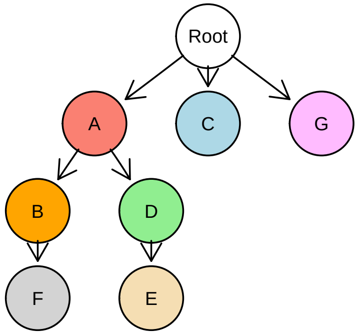


| | |
|---|---|
| *ID* | vDqUv4dtGIoB94NH |
| *acc genotypes* | 40 |
| *peaks no back* | 8 |
| *removed edges* | 13 |
| *prop rem edges* | 0.141 |

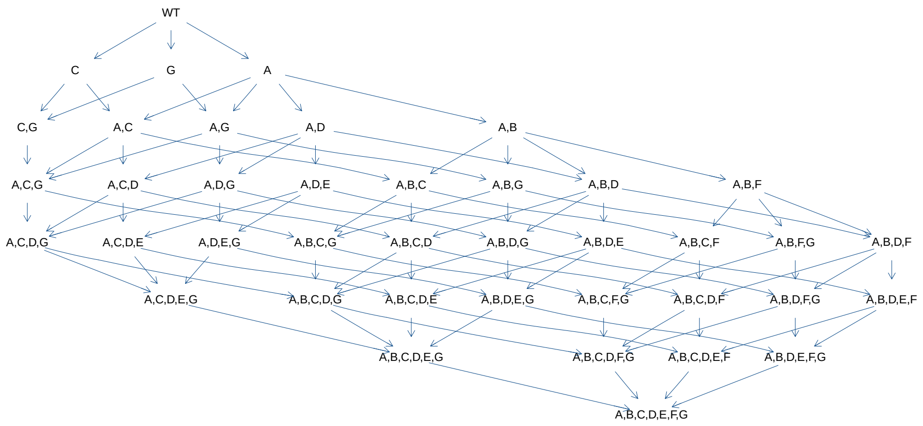


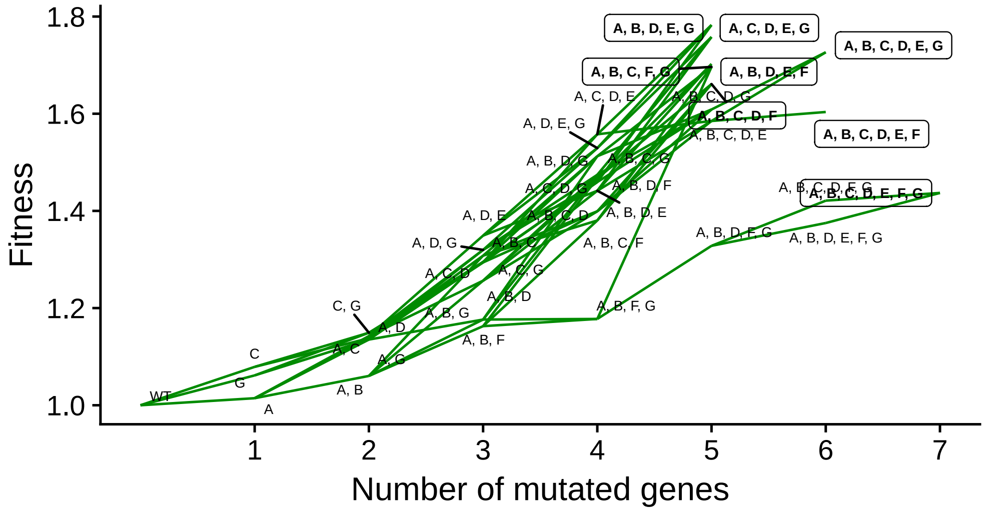


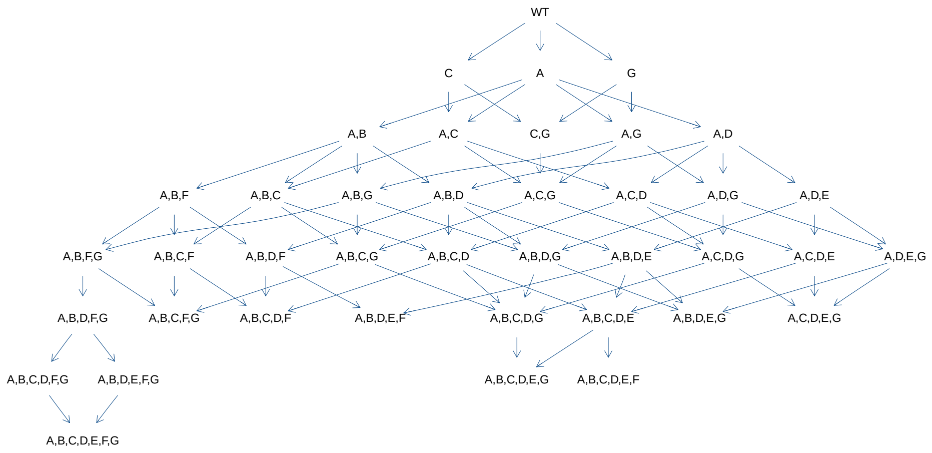

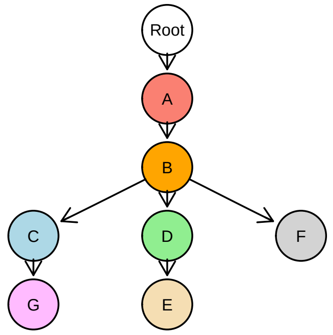


| | |
|---|---|
| *ID* | VIFJpVZjq2HKeP6z |
| *acc genotypes* | 20 |
| *peaks no back* | 5 |
| *removed edges* | 8 |
| *prop rem edges* | 0.229 |

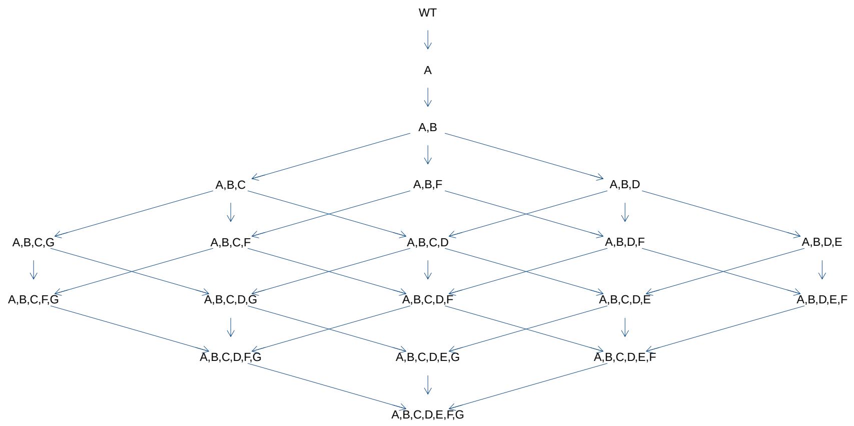


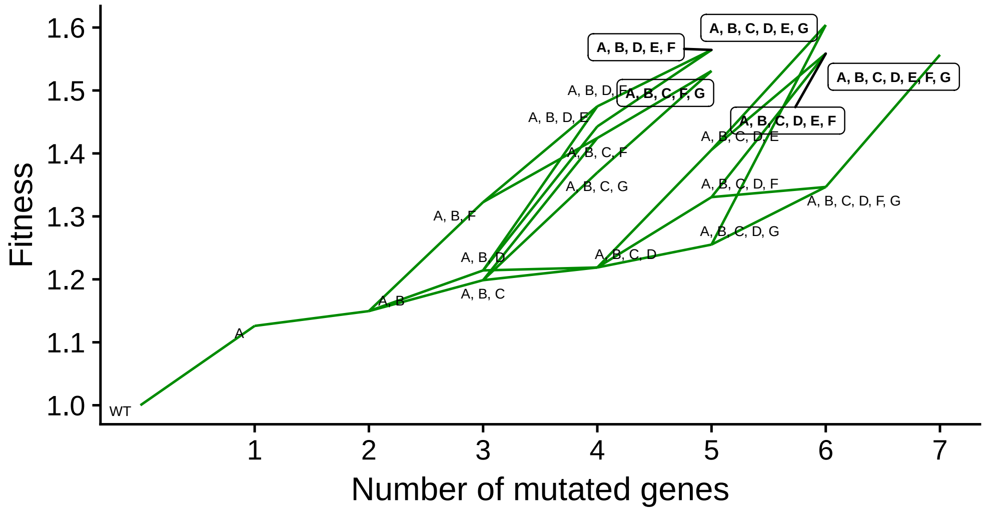


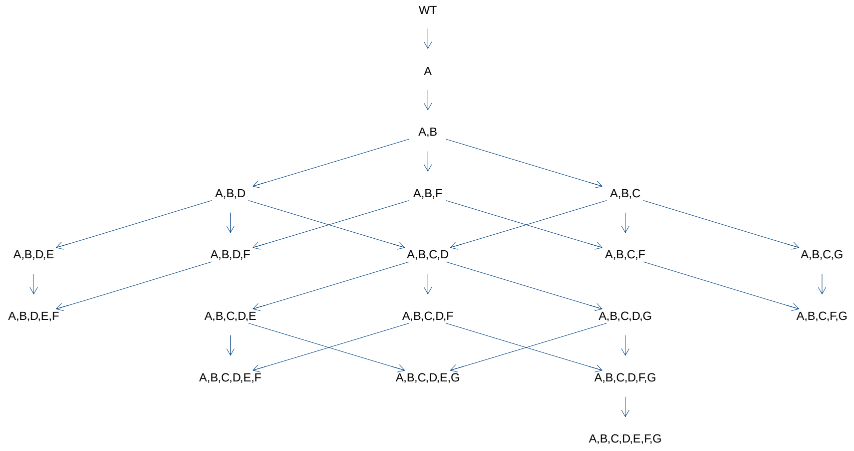

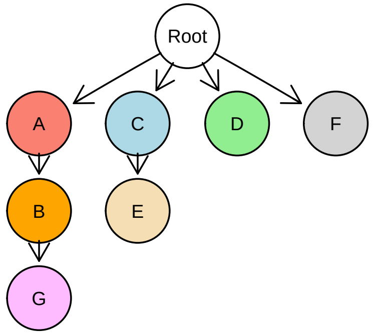


| | |
|---|---|
| *ID* | vK7dEdwIvdYYs5rB |
| *acc genotypes* | 48 |
| *peaks no back* | 10 |
| *removed edges* | 24 |
| *prop rem edges* | 0.207 |

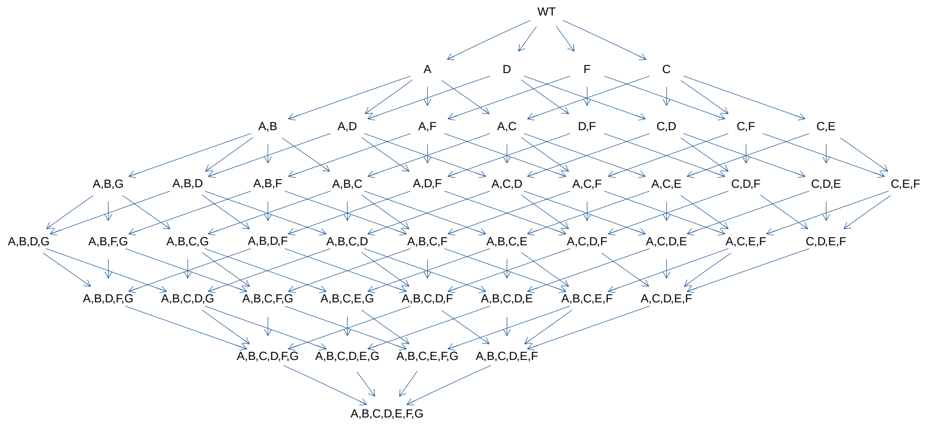


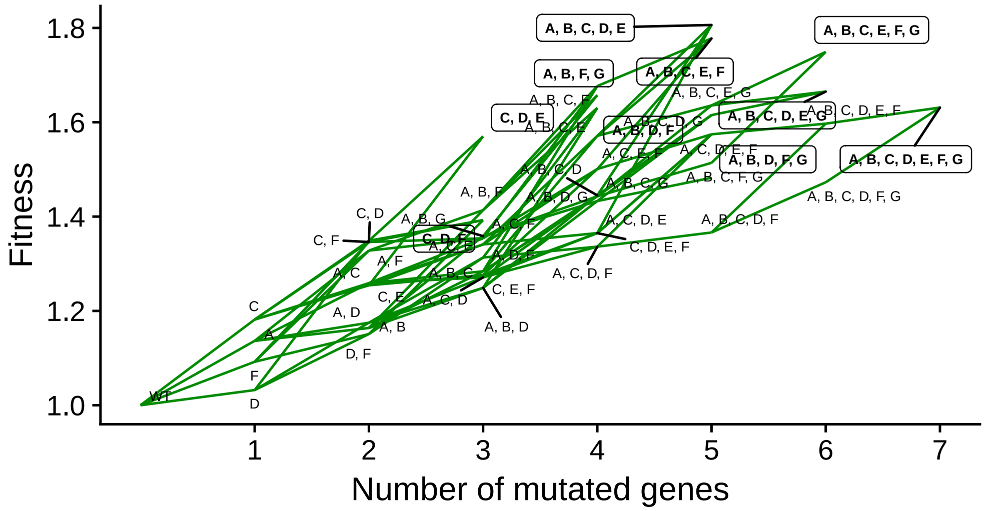


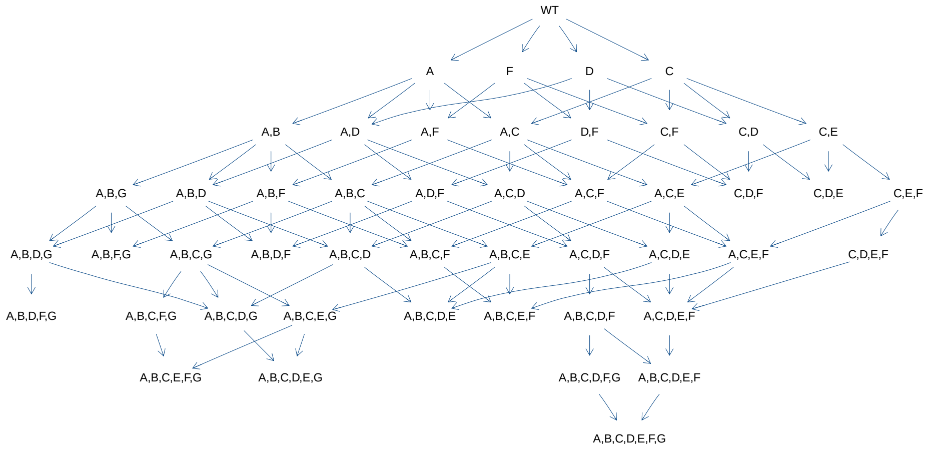

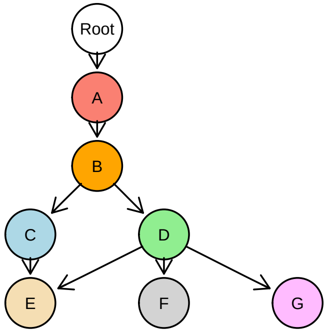


| | |
|---|---|
| *ID* | VqVZDfELXTXwJzx10 |
| *acc genotypes* | 16 |
| *peaks no back* | 3 |
| *removed edges* | 2 |
| *prop rem edges* | 0.08 |

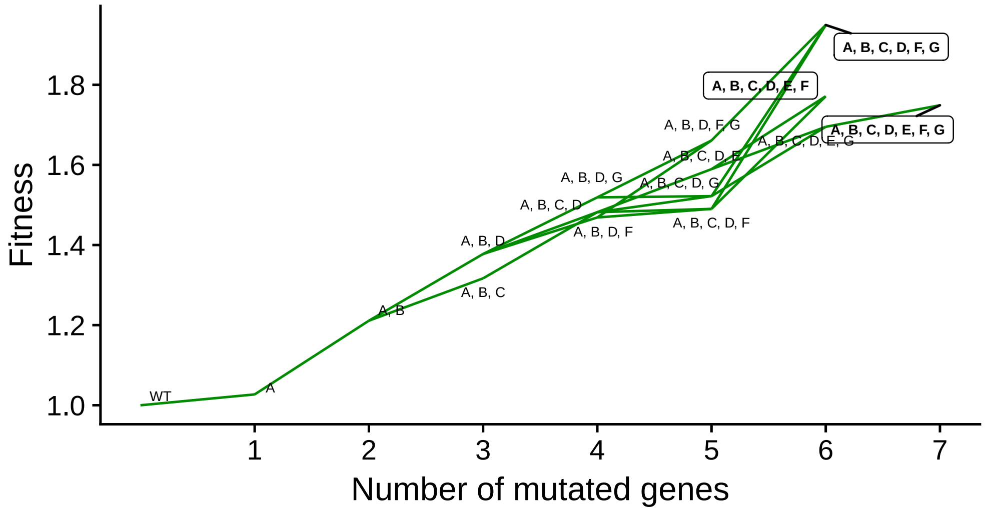


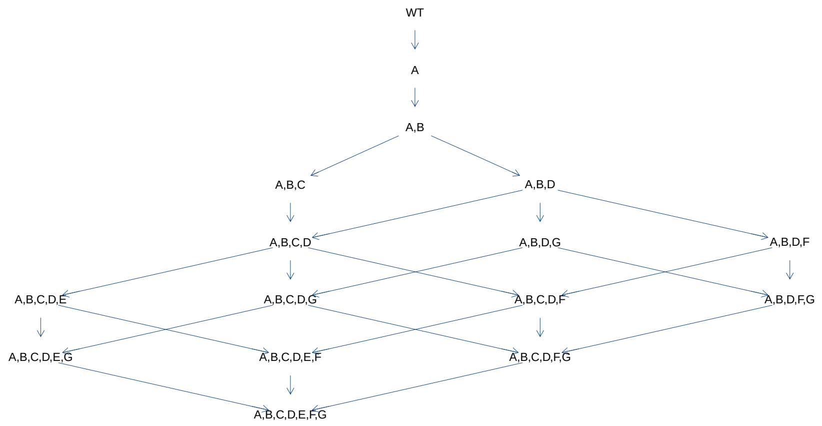


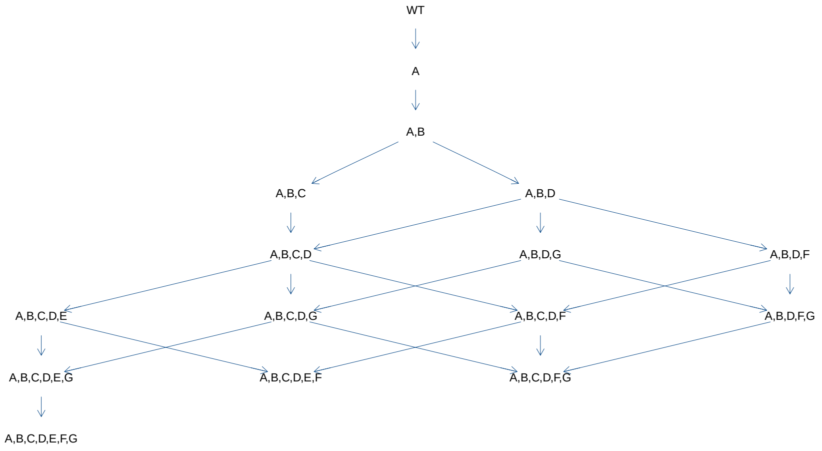

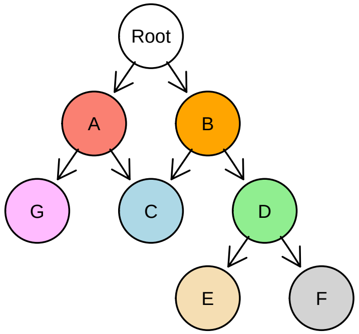


| | |
|---|---|
| *ID* | vviAaqqVbYrKTfxK |
| *acc genotypes* | 28 |
| *peaks no back* | 7 |
| *removed edges* | 12 |
| *prop rem edges* | 0.218 |

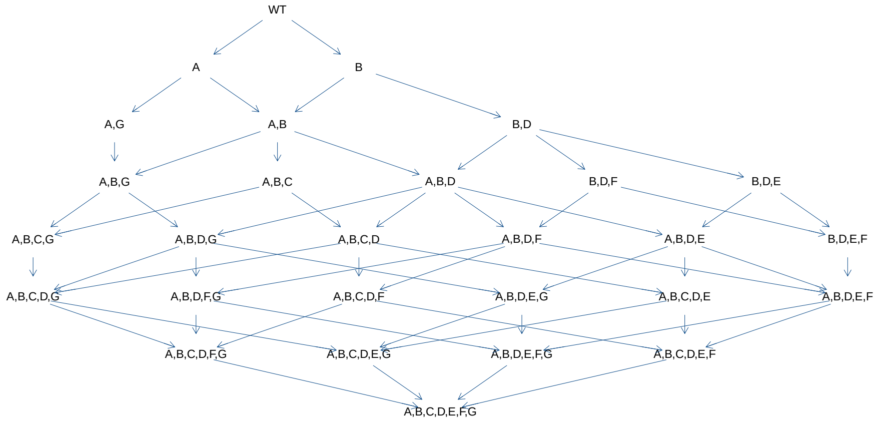


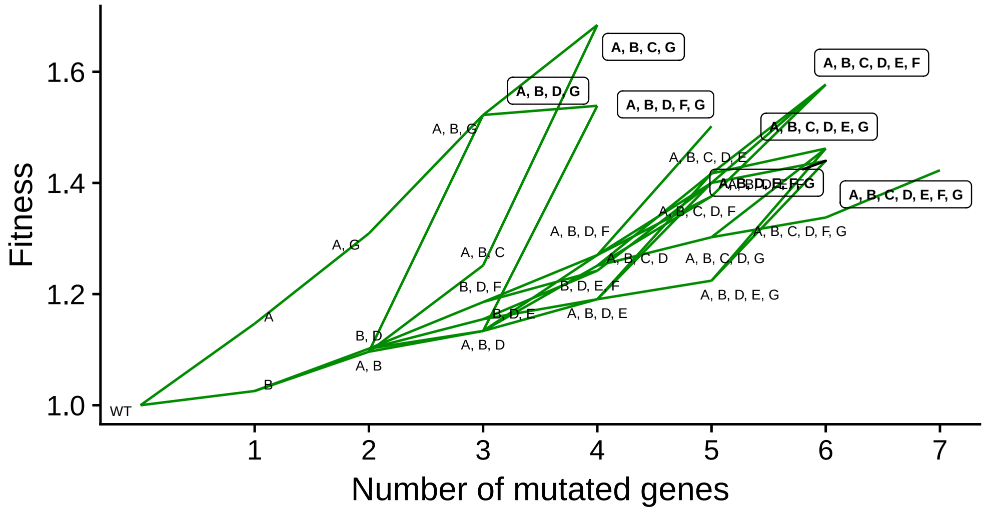


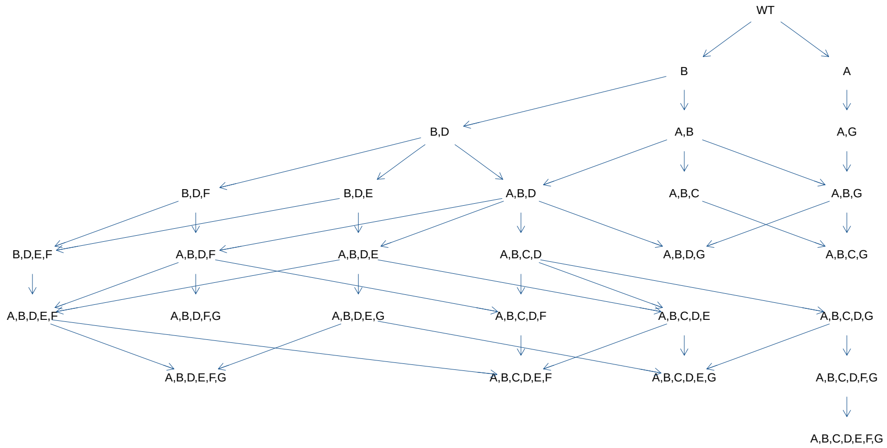

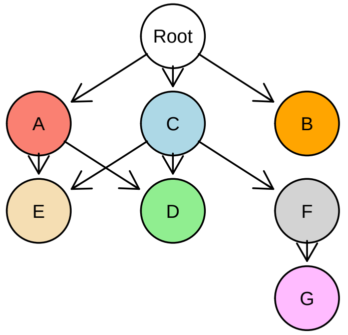


| | |
|---|---|
| *ID* | VYKTwc6PGOfo5RH5 |
| *acc genotypes* | 34 |
| *peaks no back* | 5 |
| *removed edges* | 12 |
| *prop rem edges* | 0.164 |

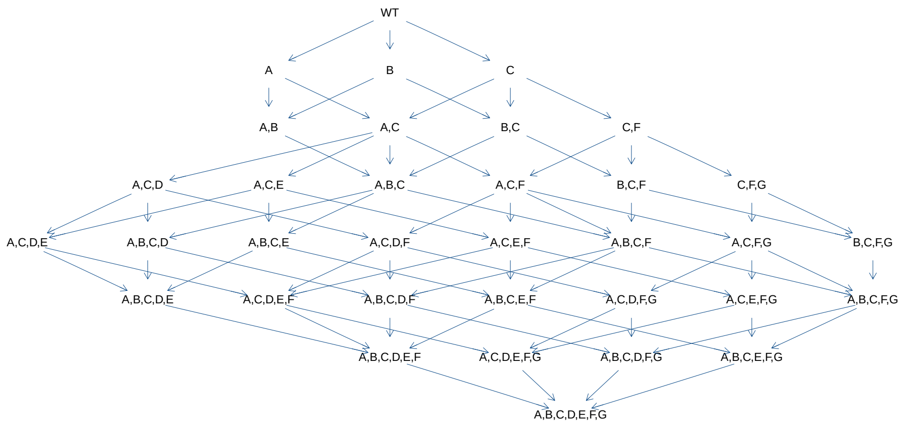


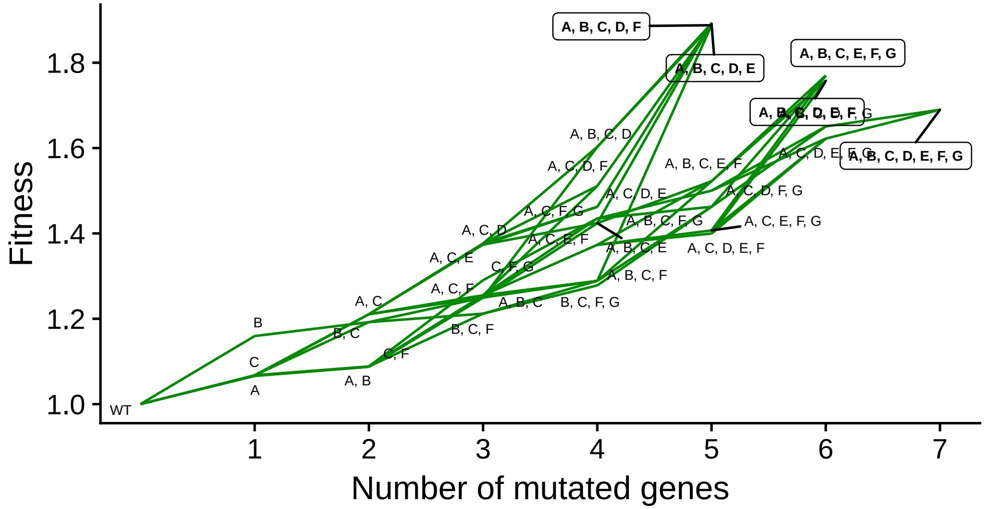


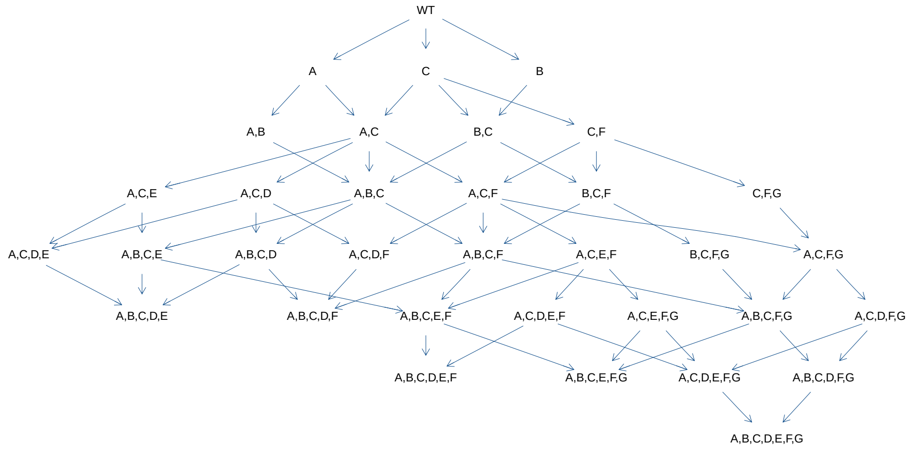

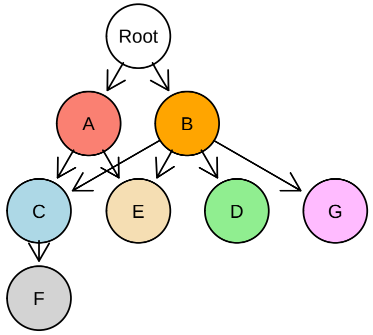


| | |
|---|---|
| *ID* | vz7pdw2T4KJ7Ey10T |
| *acc genotypes* | 30 |
| *peaks no back* | 6 |
| *removed edges* | 14 |
| *prop rem edges* | 0.222 |

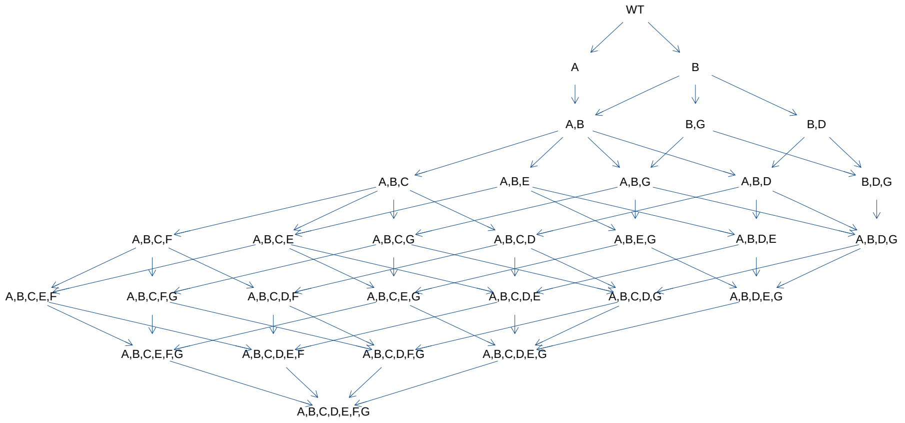


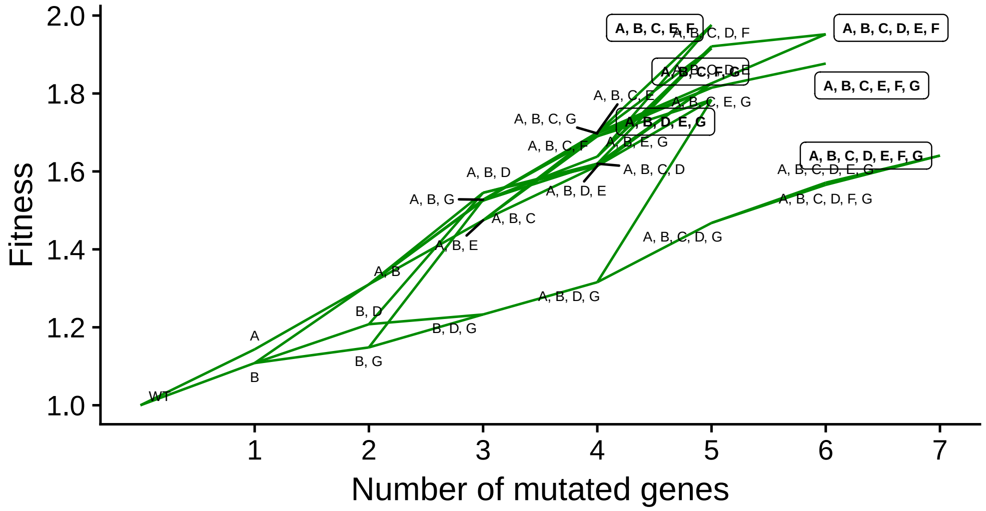


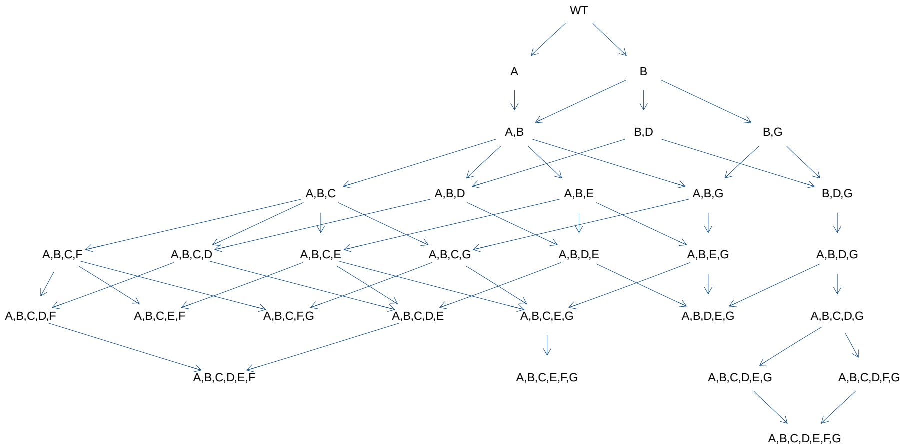

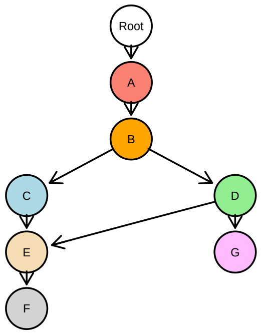


| | |
|---|---|
| *ID* | WCthf7pnLrf65210C |
| *acc genotypes* | 12 |
| *peaks no back* | 2 |
| *removed edges* | 1 |
| *prop rem edges* | 0.067 |

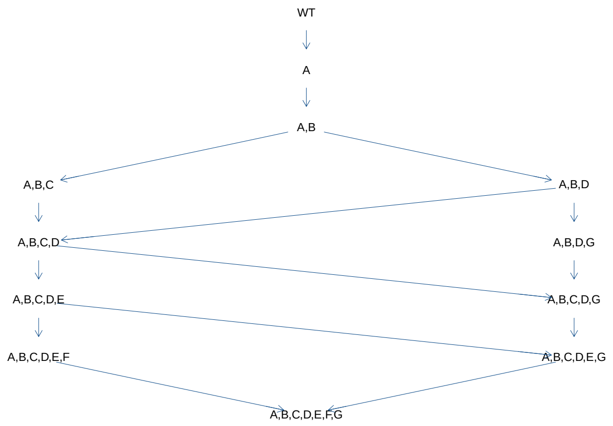


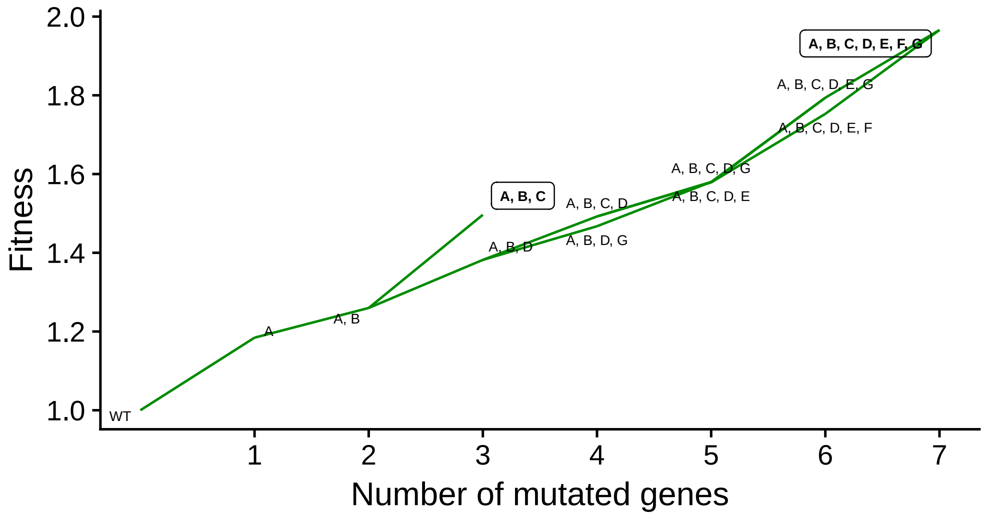


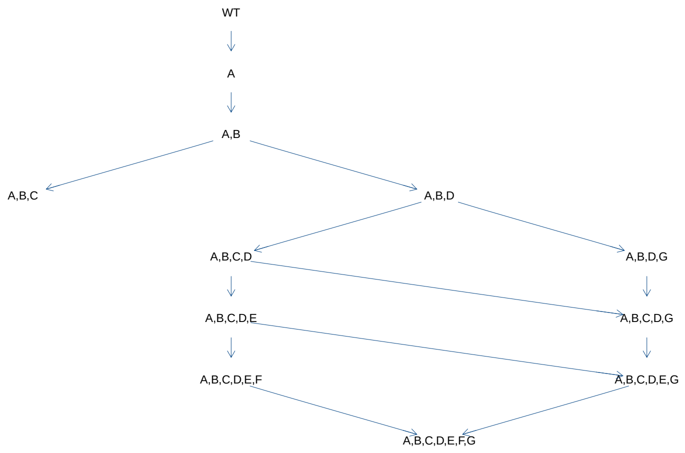

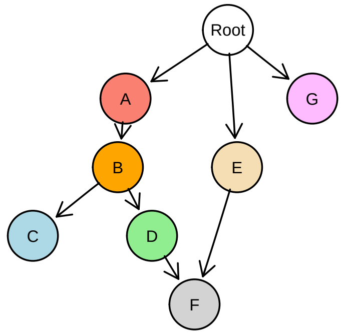


| | |
|---|---|
| *ID* | wGM2GvMSYVAE2KwT |
| *acc genotypes* | 28 |
| *peaks no back* | 6 |
| *removed edges* | 13 |
| *prop rem edges* | 0.232 |

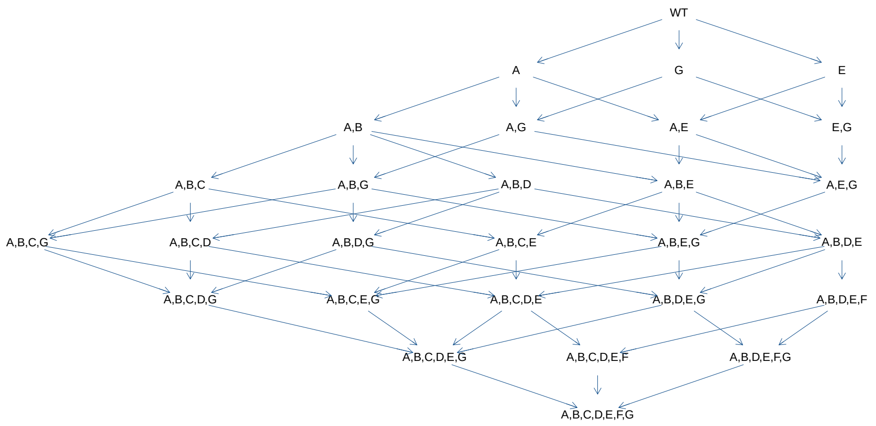


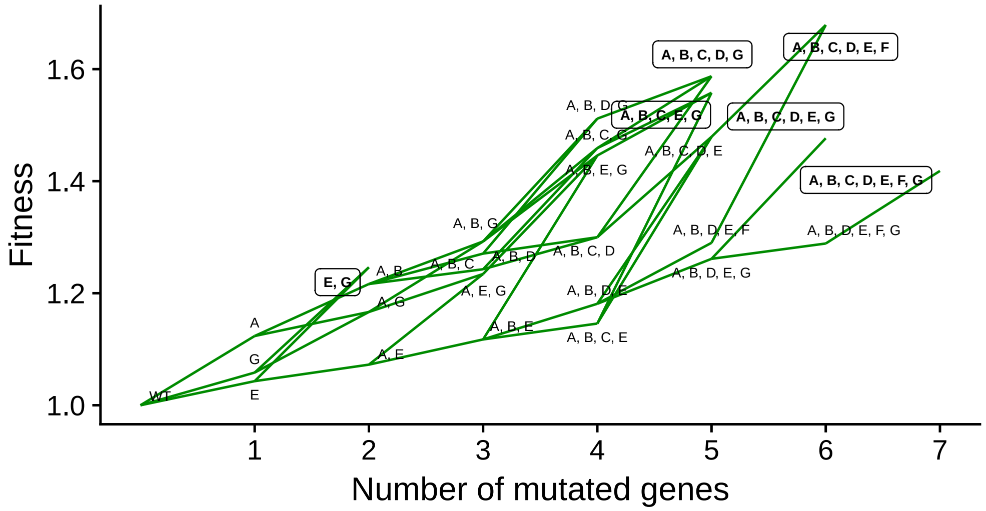


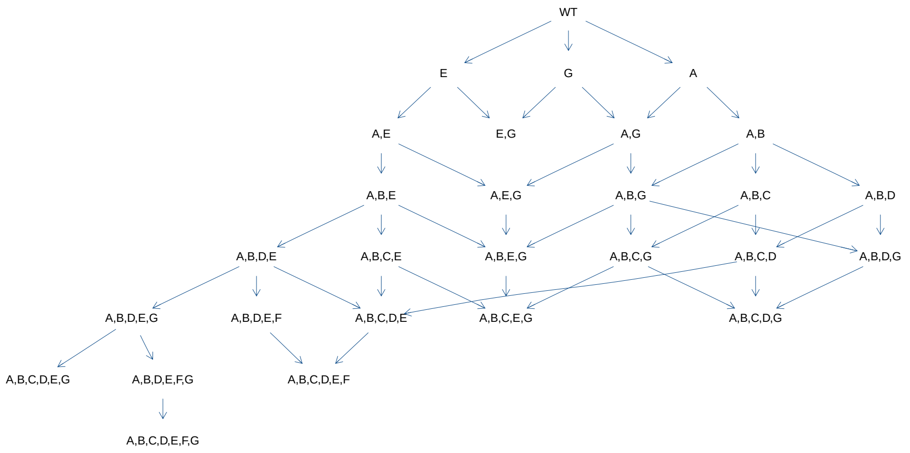

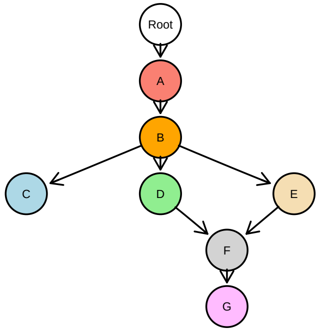


| | |
|---|---|
| *ID* | WpF105HbEDoECa8vs |
| *acc genotypes* | 14 |
| *peaks no back* | 2 |
| *removed edges* | 1 |
| *prop rem edges* | 0.05 |

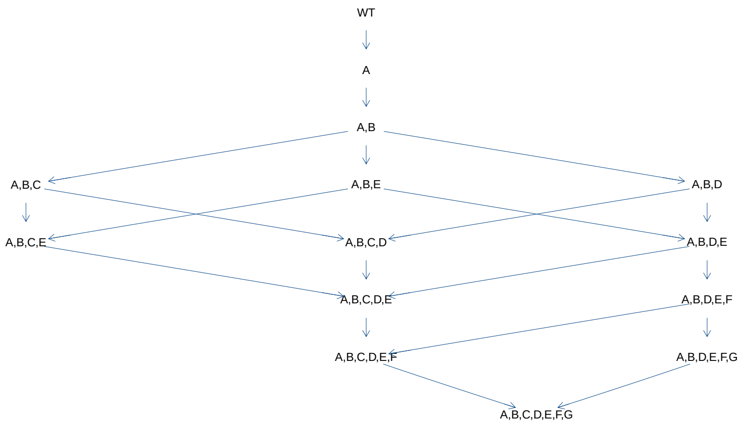


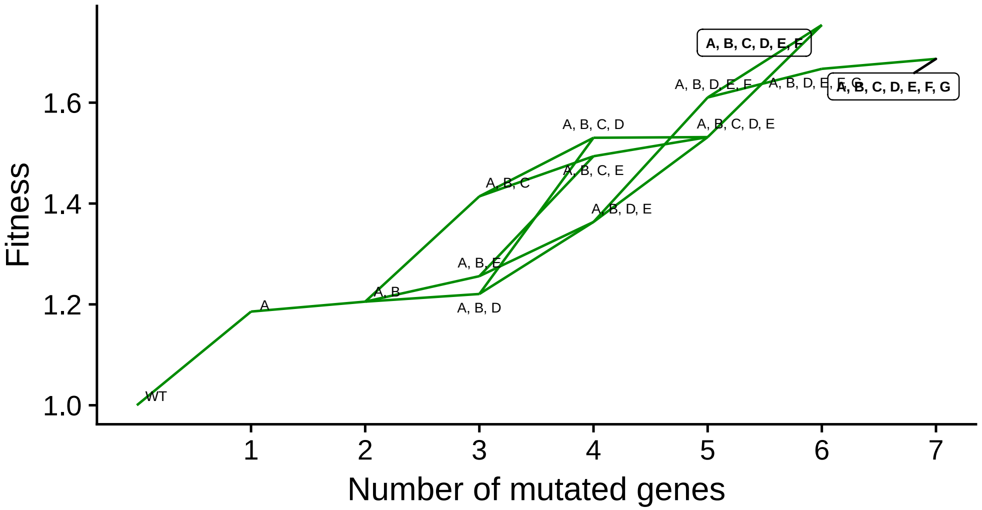


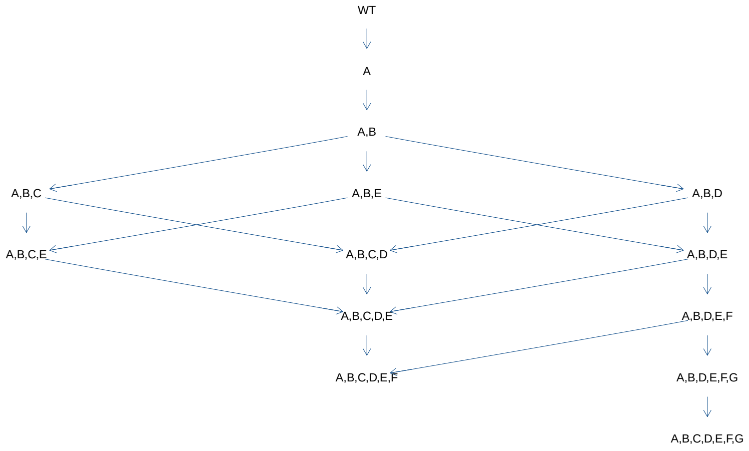

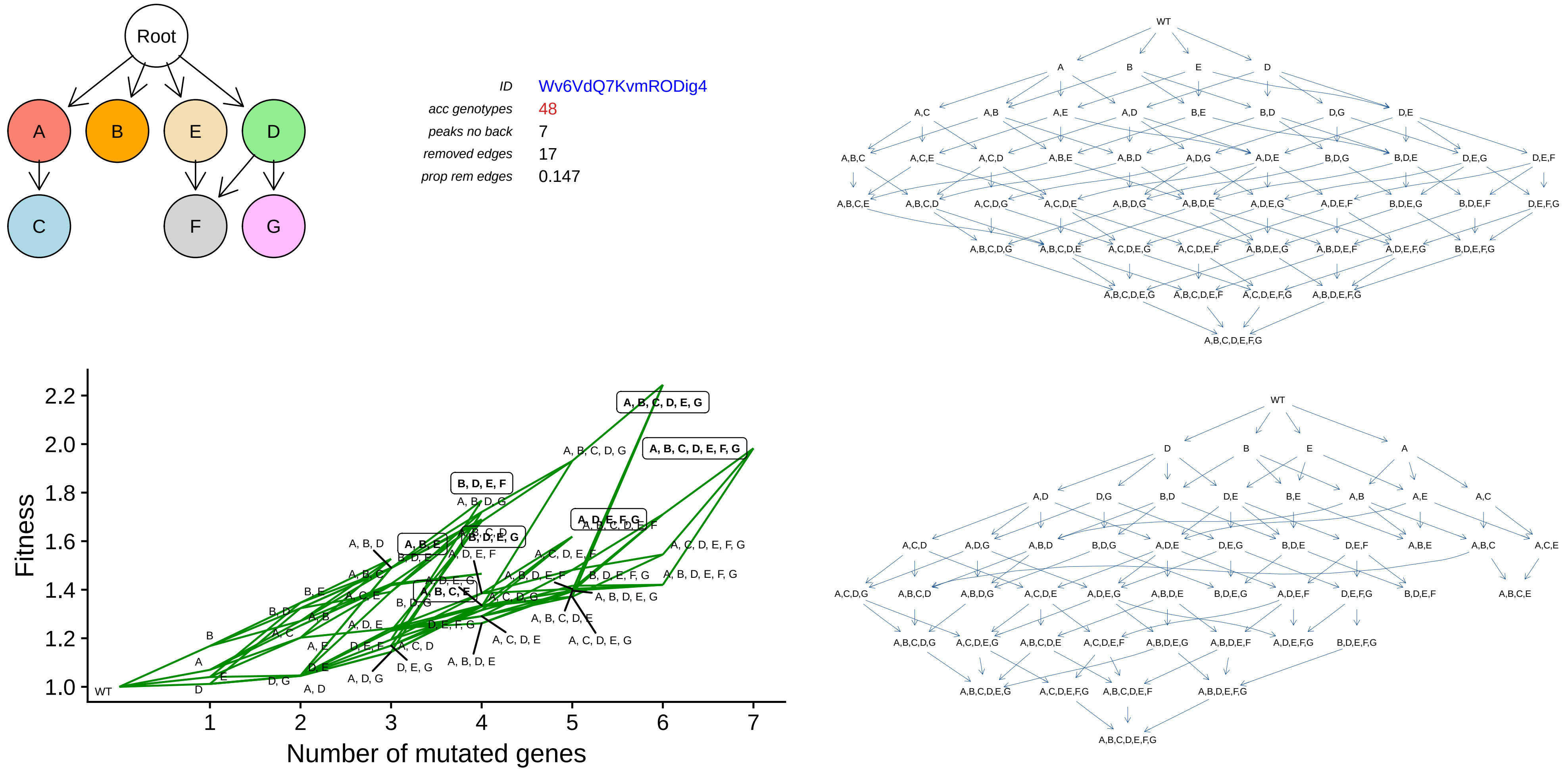
Root
A
B
E
D
C
F
G
ID Wv6VdQ7KvmRODig4
acc genotypes 48
peaks no back 7
removed edges 17
prop rem edges 0.147
Fitness
Number of mutated genes
1.0
1.2
1.4
1.6
1.8
2.0
2.2
1
2
3
4
5
6
7
WT
A, B, C, D, E, G
A, B, C, D, E, F, G
B, D, E, F
A, D, E, F, G
A, B, E
B, D, E, G
A, B, C, E
A,B,C,D,E,F,G

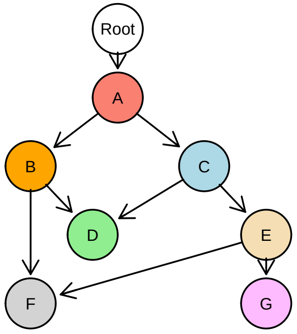


| | |
|---|---|
| *ID* | wXZP1BcX68smcT7W |
| *acc genotypes* | 16 |
| *peaks no back* | 4 |
| *removed edges* | 4 |
| *prop rem edges* | 0.167 |

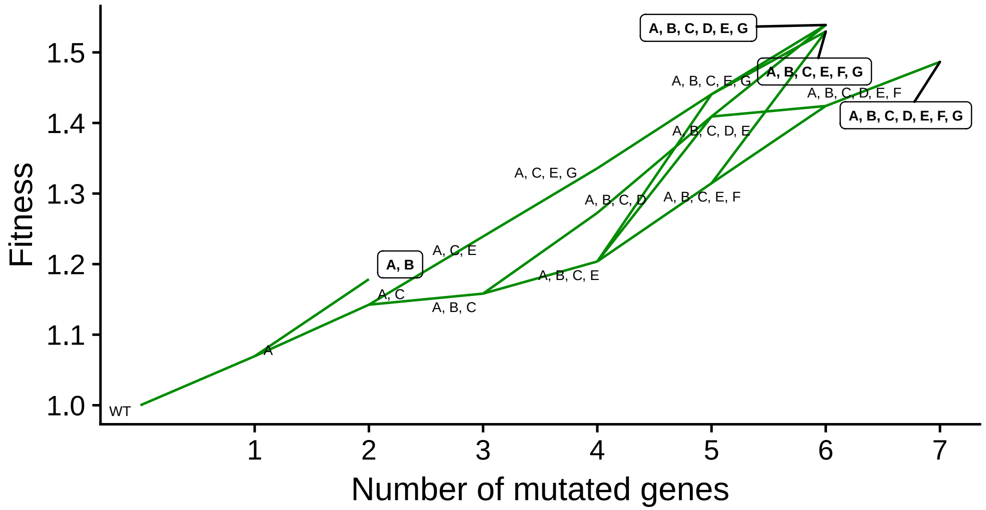


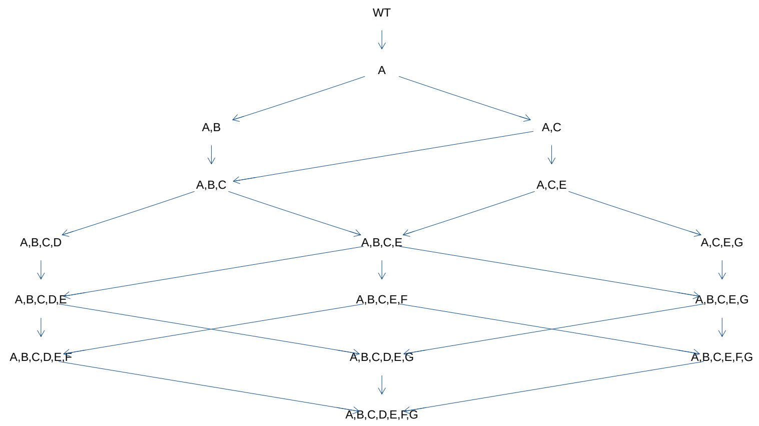


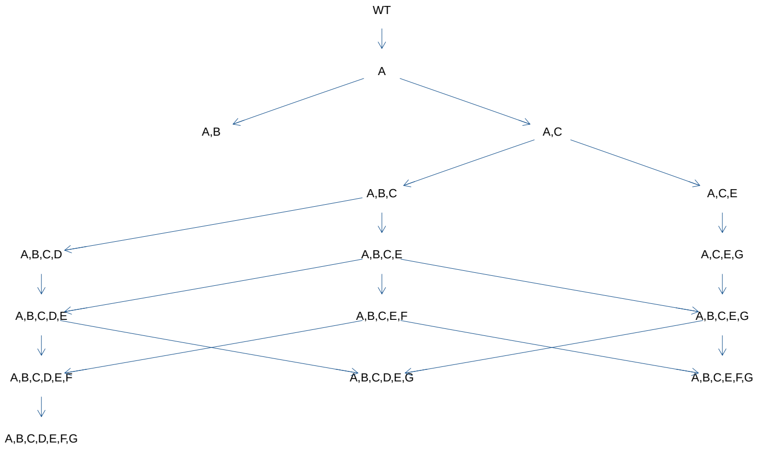

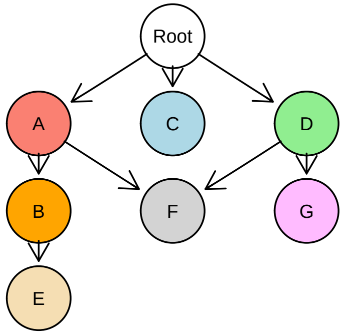


| | |
|---|---|
| *ID* | WzbE1eRFpaclNCxU |
| *acc genotypes* | 36 |
| *peaks no back* | 6 |
| *removed edges* | 14 |
| *prop rem edges* | 0.179 |

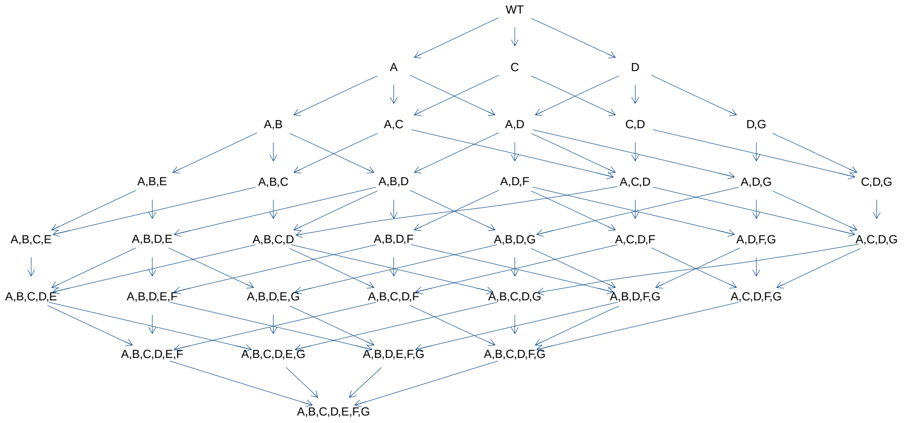


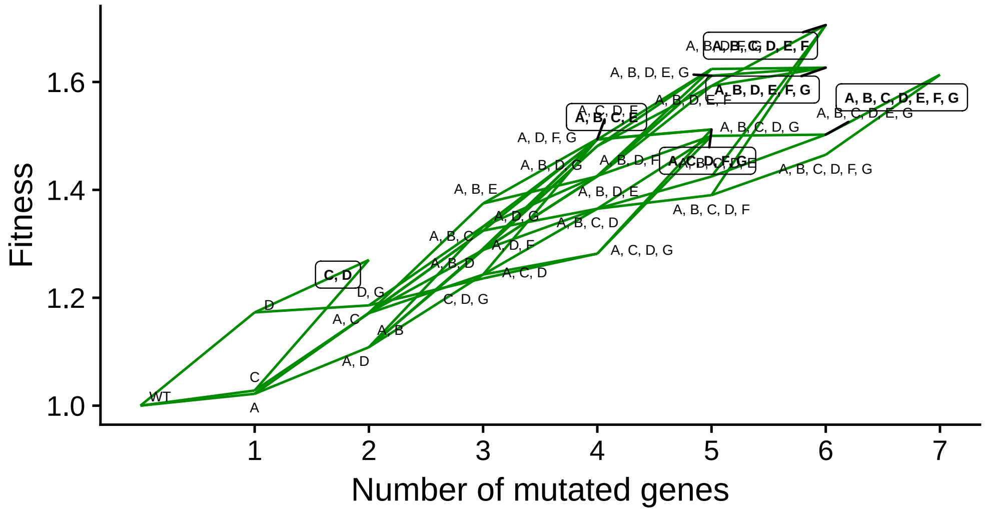


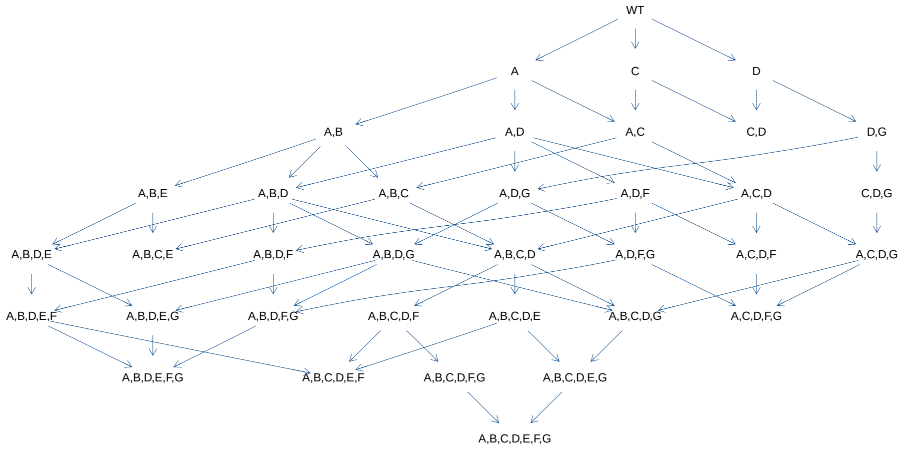

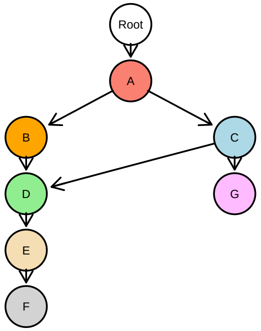


| | |
|---|---|
| *ID* | wzBFbd7oN5KHrhdR |
| *acc genotypes* | 13 |
| *peaks no back* | 2 |
| *removed edges* | 1 |
| *prop rem edges* | 0.059 |

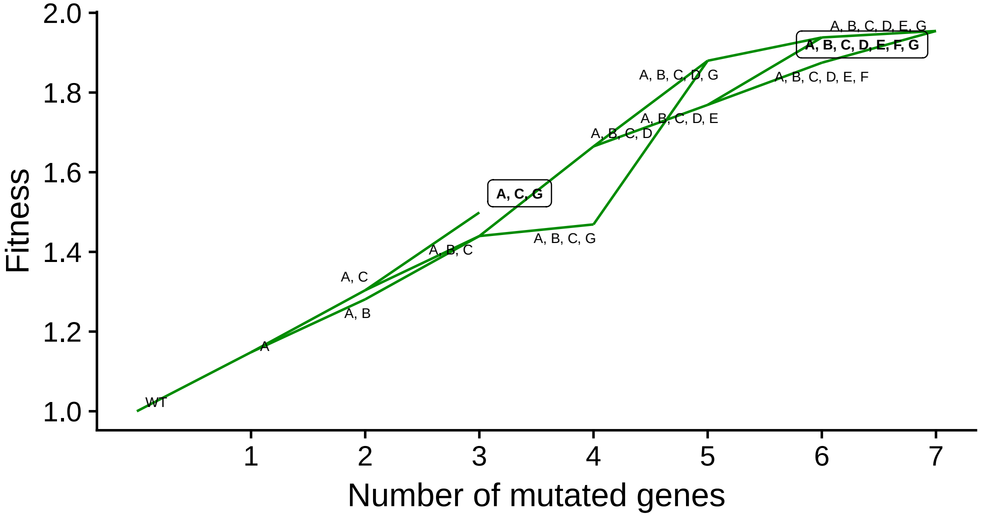


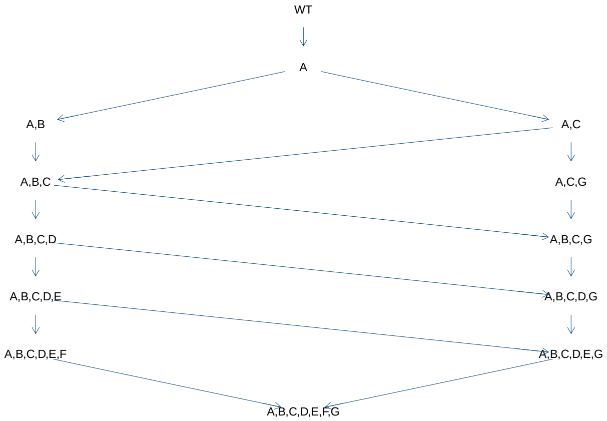


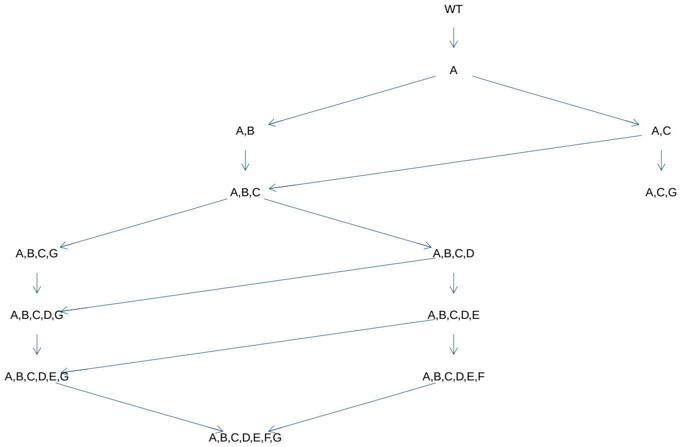

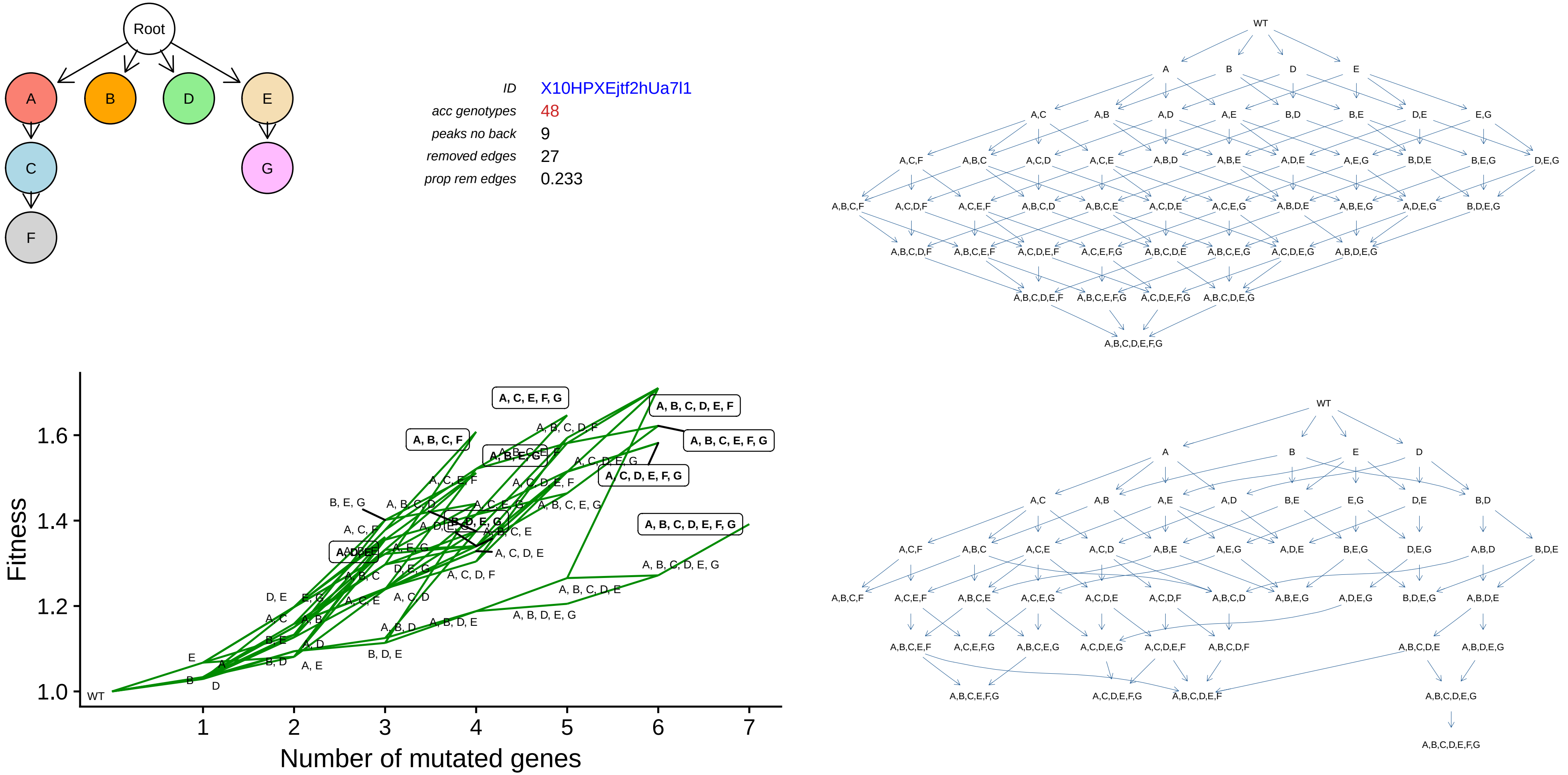
Root
A
B
D
E
C
G
F
ID X10HPXEjtf2hUa7l1
acc genotypes 48
peaks no back 9
removed edges 27
prop rem edges 0.233
Fitness
Number of mutated genes
A, C, E, F, G
A, B, C, D, E, F
A, B, C, F
A, B, C, E, F, G
A, C, D, E, F, G
A, B, C, D, E, F, G
WT
A,B,C,D,E,F,G

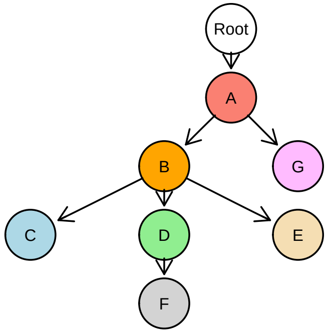


| | |
|---|---|
| *ID* | x2Dh5bR5ksqmvSQp |
| *acc genotypes* | 27 |
| *peaks no back* | 5 |
| *removed edges* | 14 |
| *prop rem edges* | 0.25 |

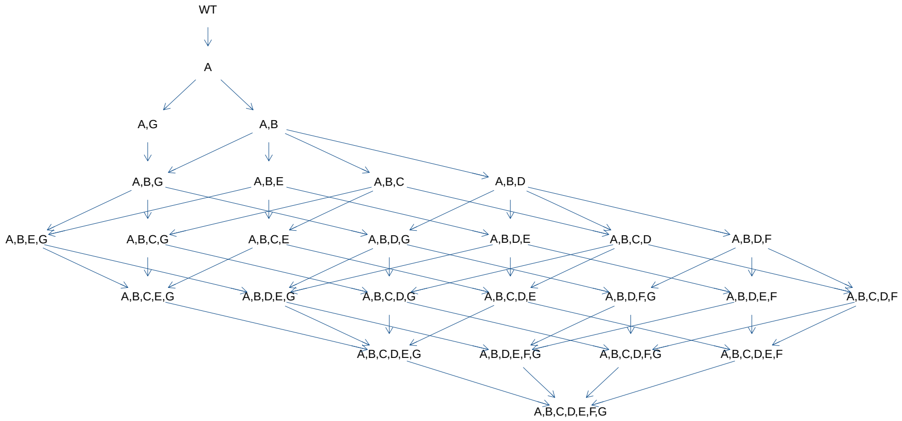


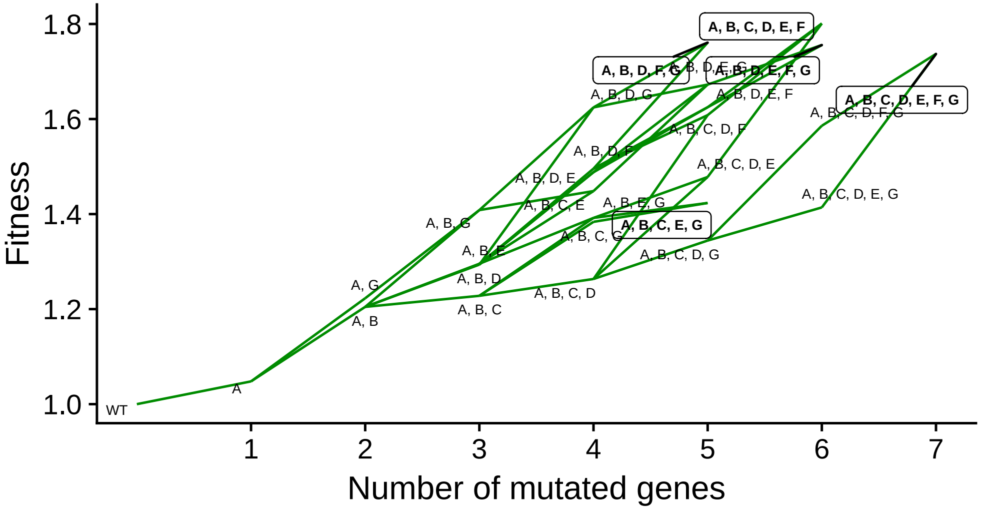


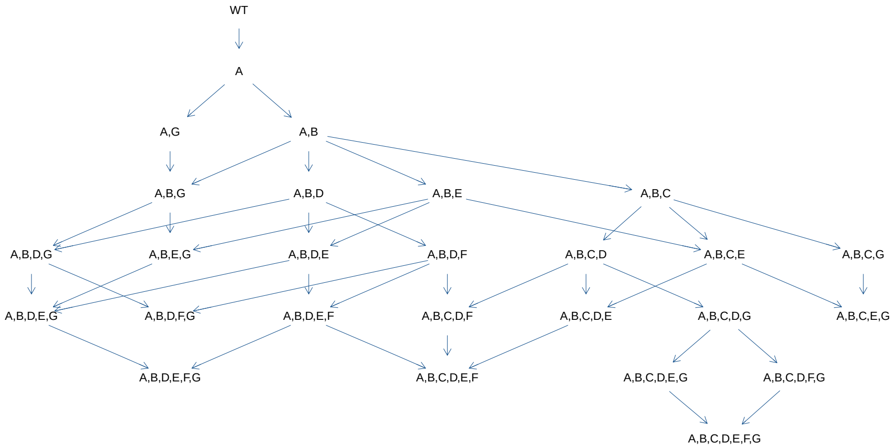

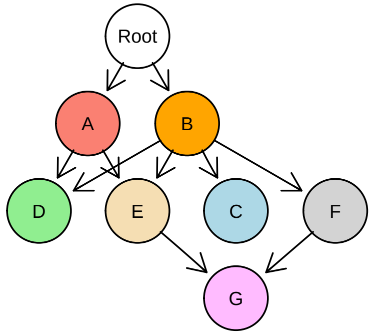


| | |
|---|---|
| *ID* | XkAnRfxPG4FT103ju |
| *acc genotypes* | 26 |
| *peaks no back* | 6 |
| *removed edges* | 12 |
| *prop rem edges* | 0.235 |

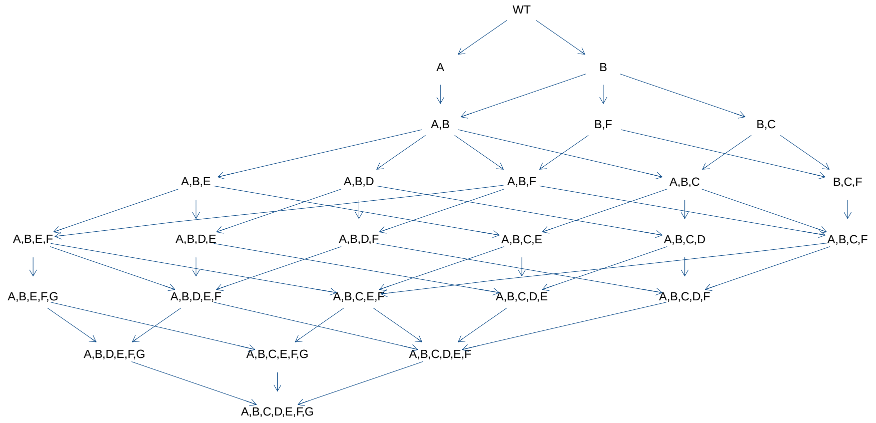


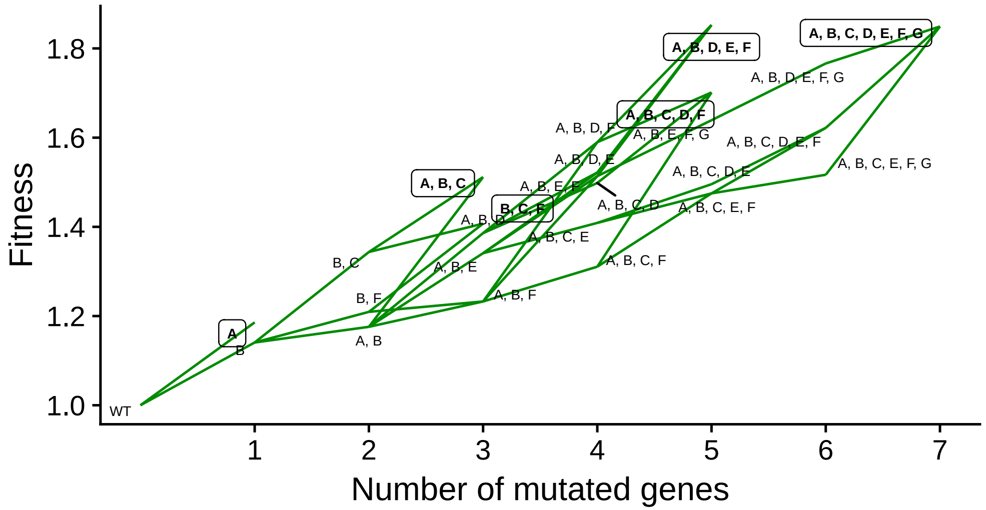


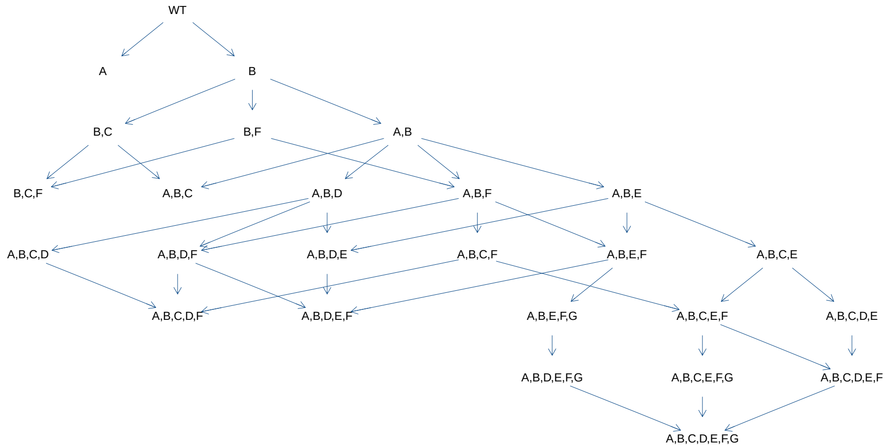

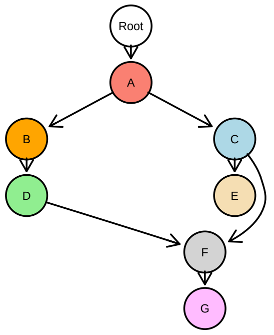


| | |
|---|---|
| *ID* | XMtNwZBJ7EWu8yH7 |
| *acc genotypes* | 14 |
| *peaks no back* | 2 |
| *removed edges* | 2 |
| *prop rem edges* | 0.105 |

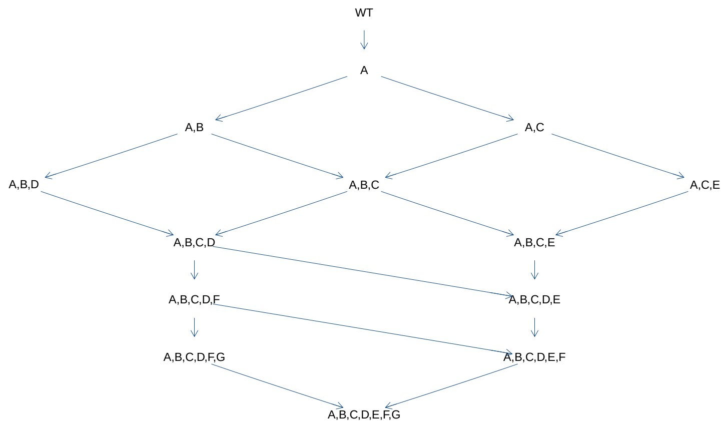


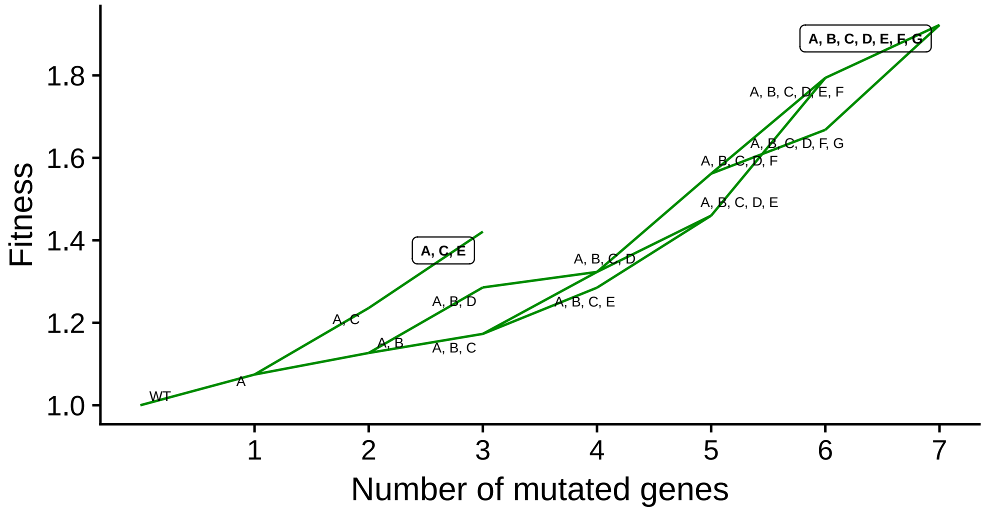


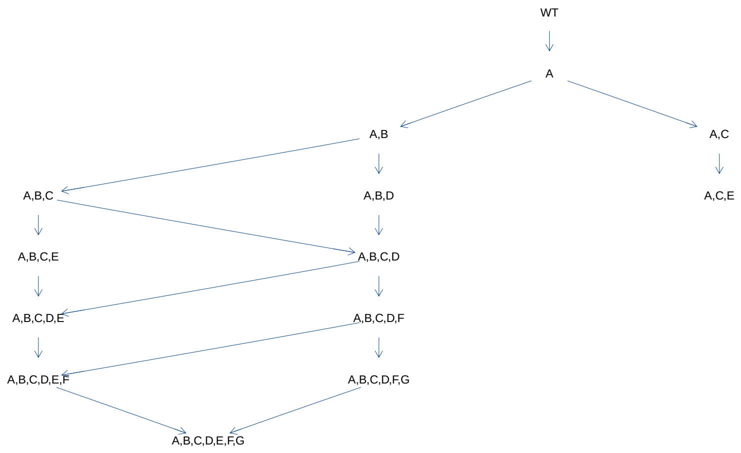

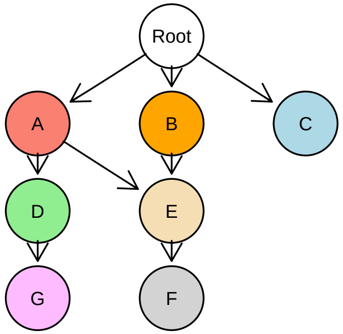


| | |
|---|---|
| *ID* | xN6jmjSdlYDl6ALk |
| *acc genotypes* | 28 |
| *peaks no back* | 3 |
| *removed edges* | 5 |
| *prop rem edges* | 0.093 |

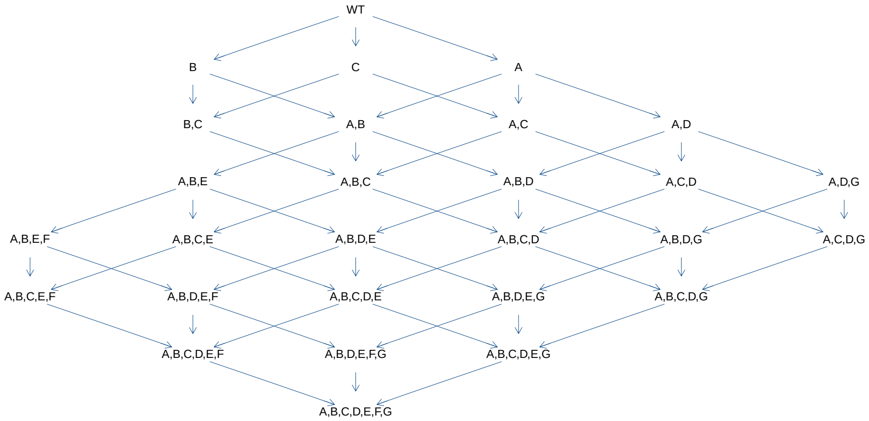


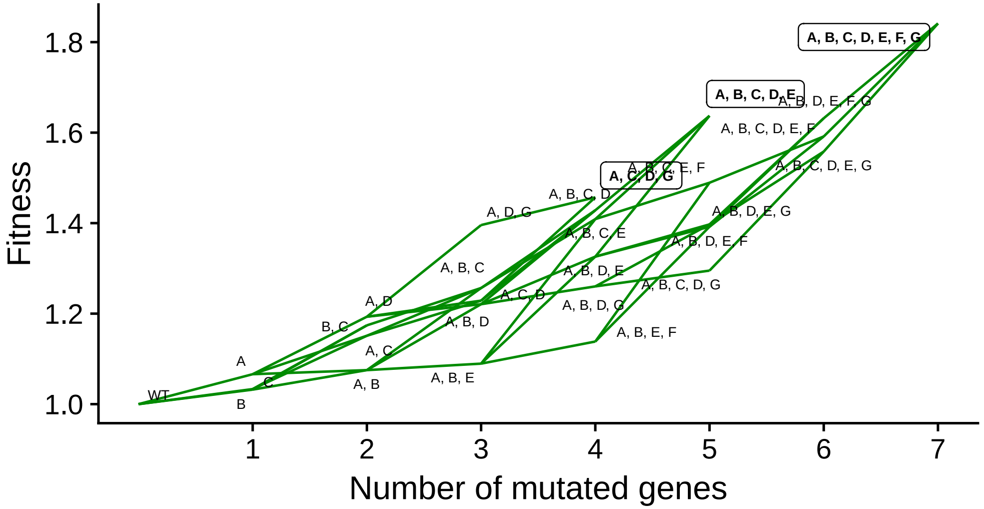


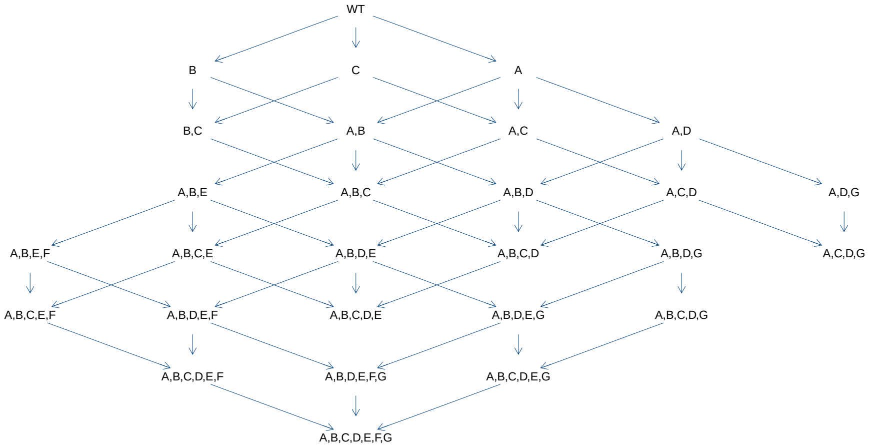

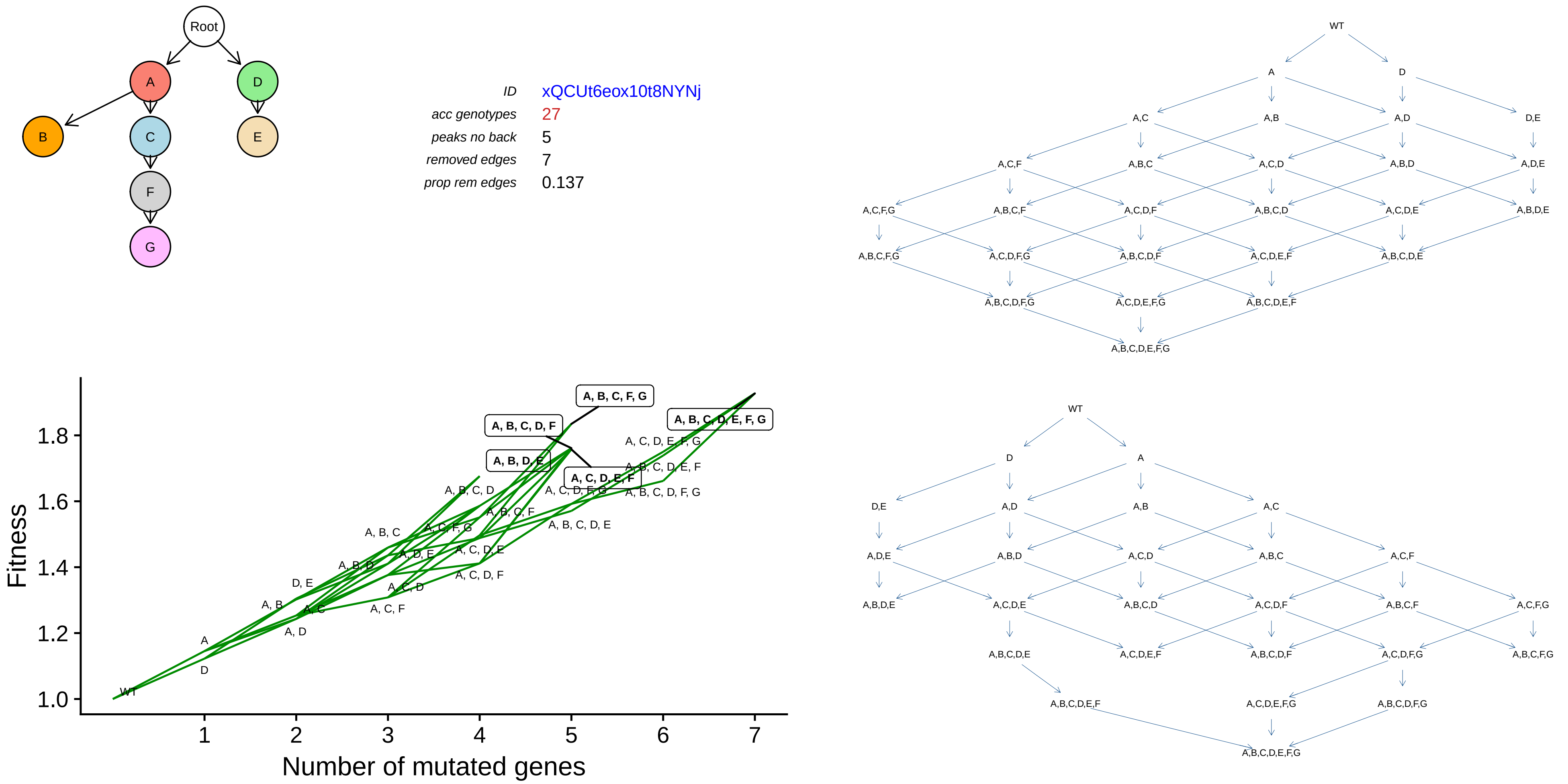

Root
A
B
C
D
E
F
G
ID
xQCUt6eox10t8NYNj
acc genotypes
27
peaks no back
5
removed edges
7
prop rem edges
0.137
Fitness
Number of mutated genes
A, B, C, F, G
A, B, C, D, F
A, B, D, E
A, C, D, E, F
A, B, C, D, E, F, G
WT

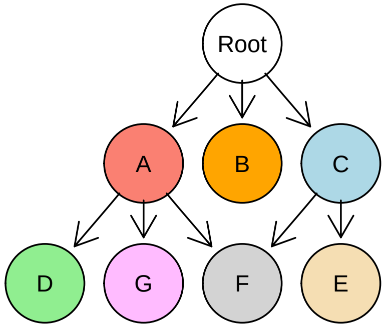


| | |
|---|---|
| *ID* | xsLEpm8mUqbWuEtX |
| *acc genotypes* | 46 |
| *peaks no back* | 6 |
| *removed edges* | 22 |
| *prop rem edges* | 0.195 |

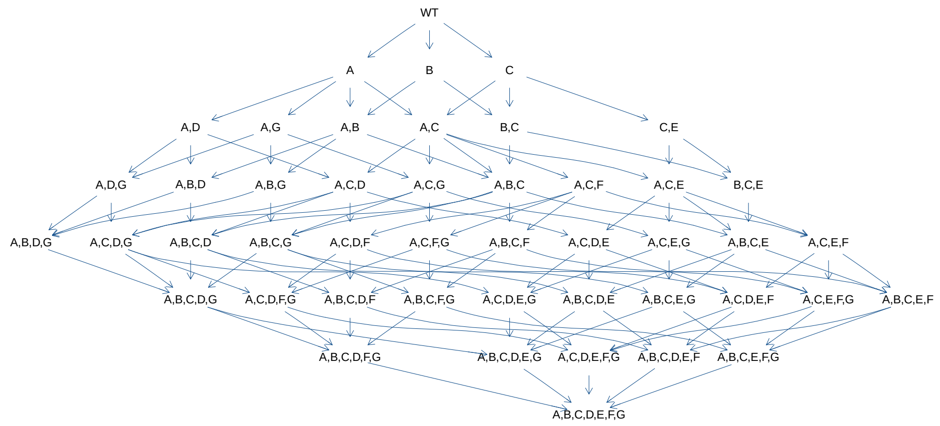


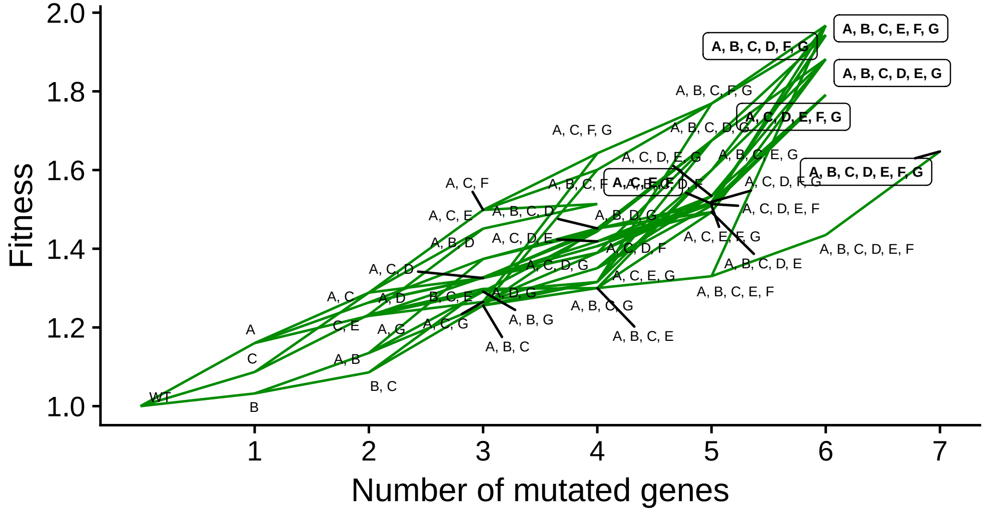


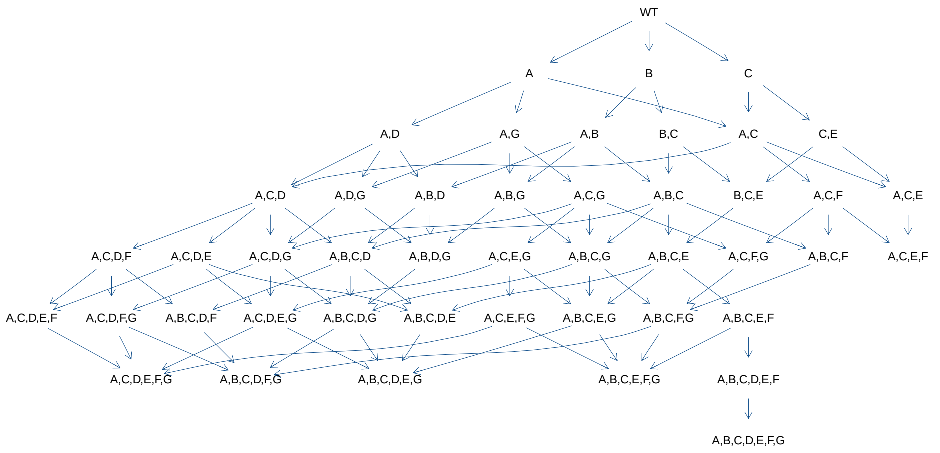

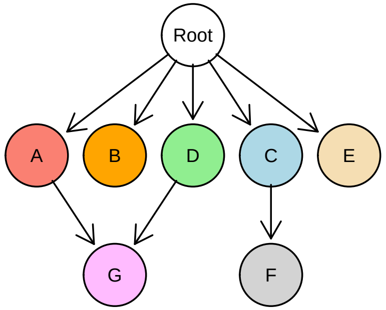


| | |
|---|---|
| *ID* | XUvwmwc1Jrh7g5gR |
| *acc genotypes* | 60 |
| *peaks no back* | 7 |
| *removed edges* | 26 |
| *prop rem edges* | 0.162 |

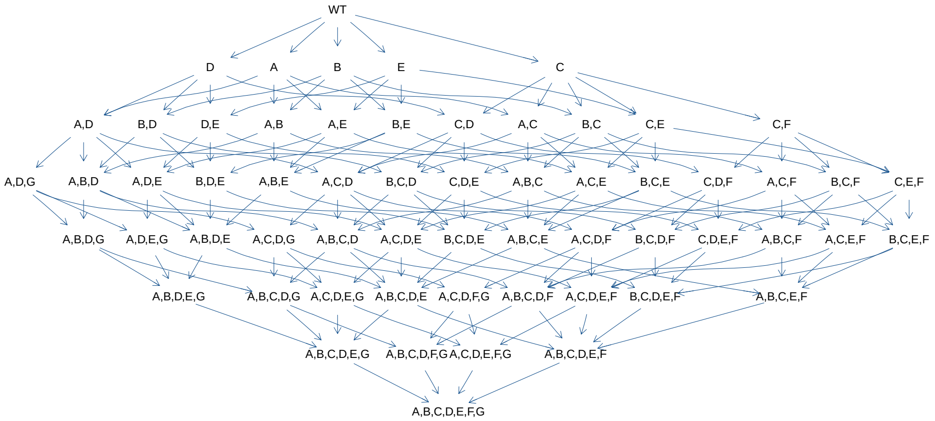


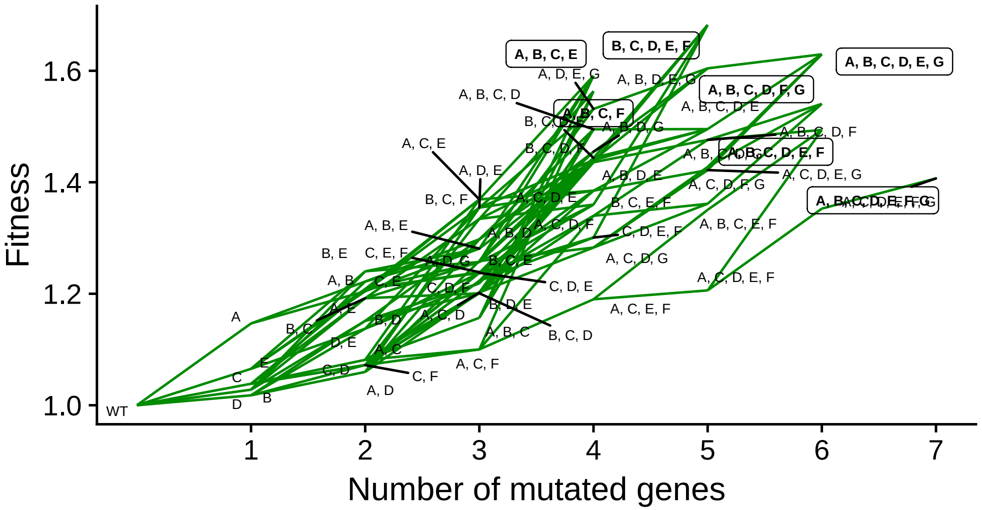


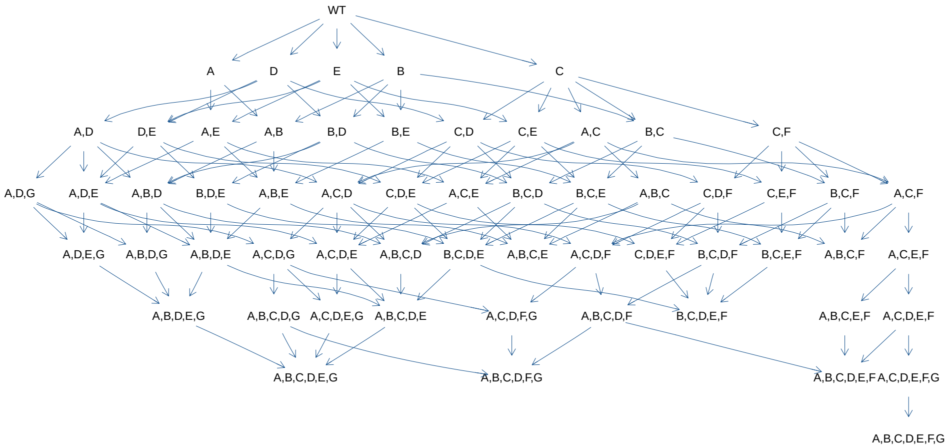

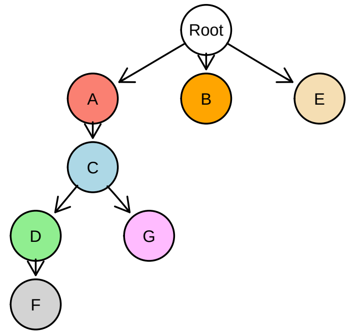


*ID* XvPBLPU6GY62cXv3
*acc genotypes* 32
*peaks no back* 6
*removed edges* 11
*prop rem edges* 0.162

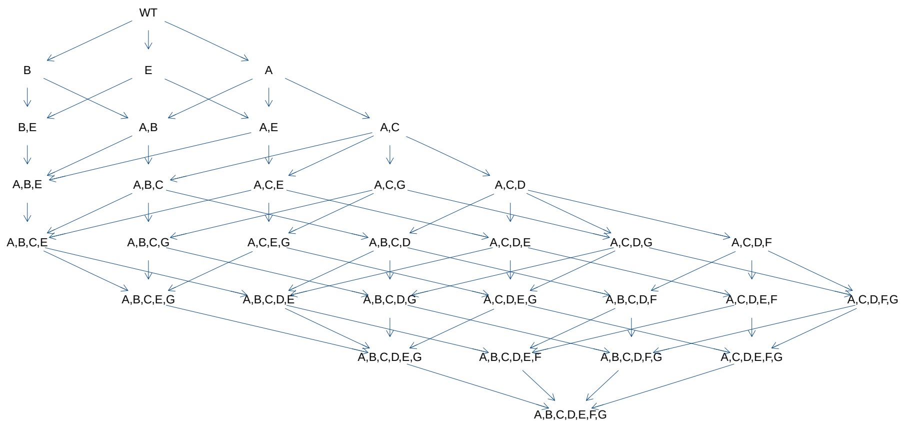


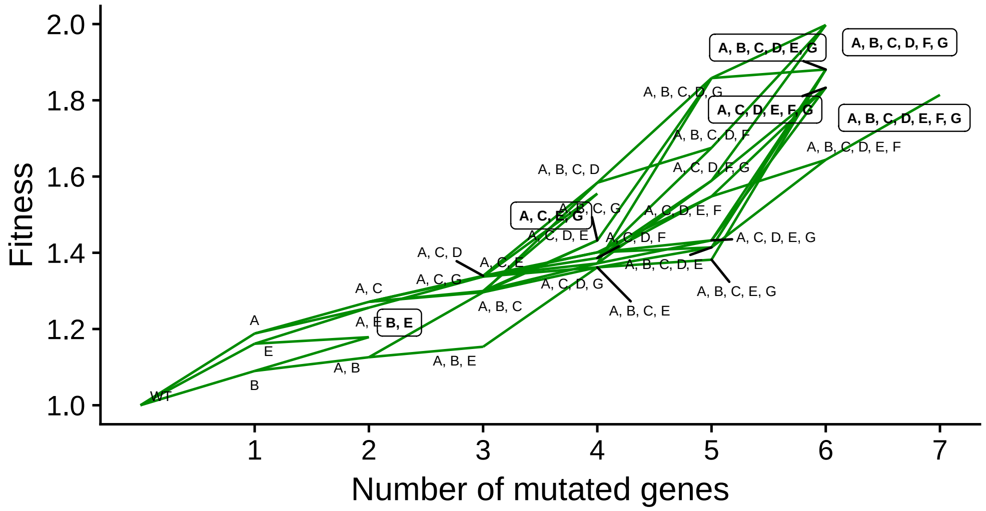


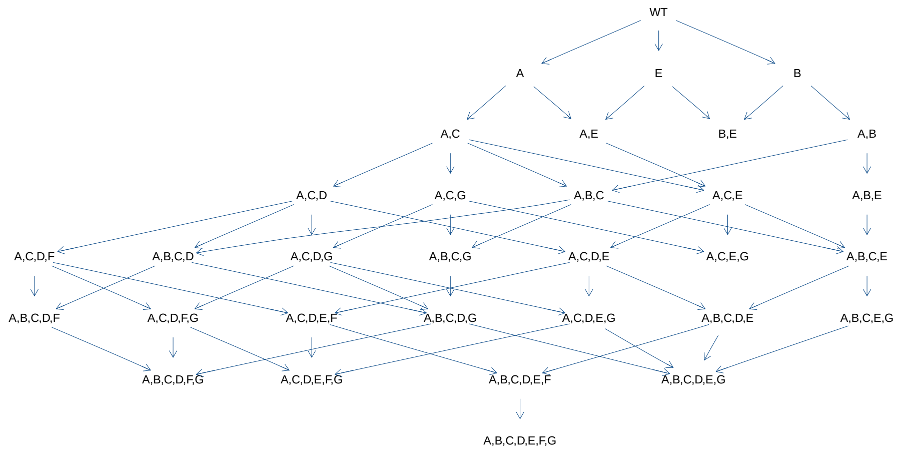

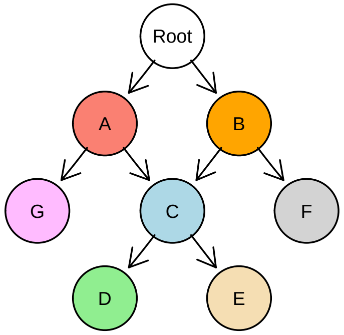


| | |
|---|---|
| *ID* | yBXLyZECBuzOCwPP |
| *acc genotypes* | 25 |
| *peaks no back* | 2 |
| *removed edges* | 4 |
| *prop rem edges* | 0.083 |

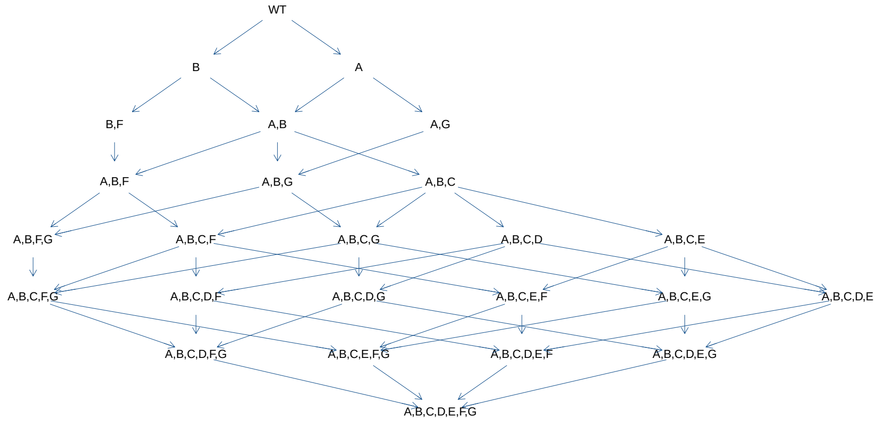


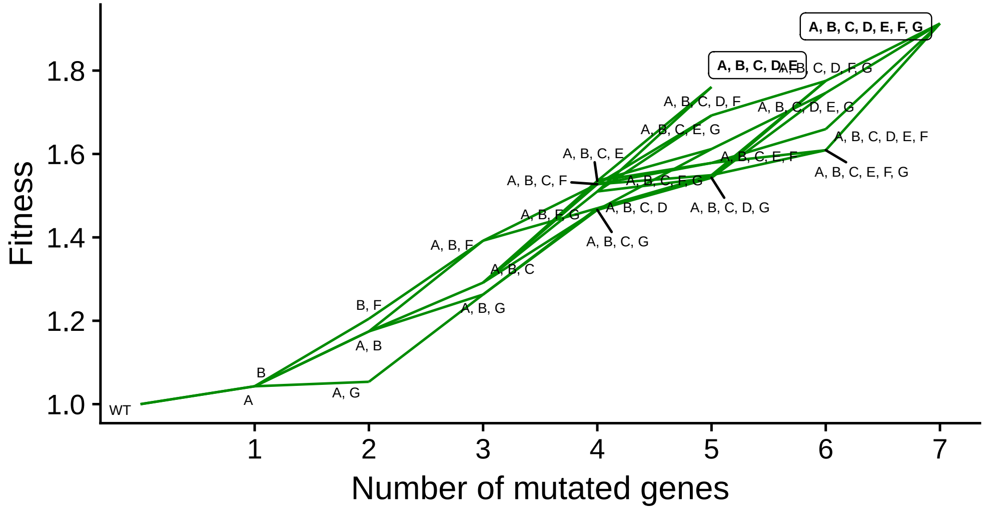


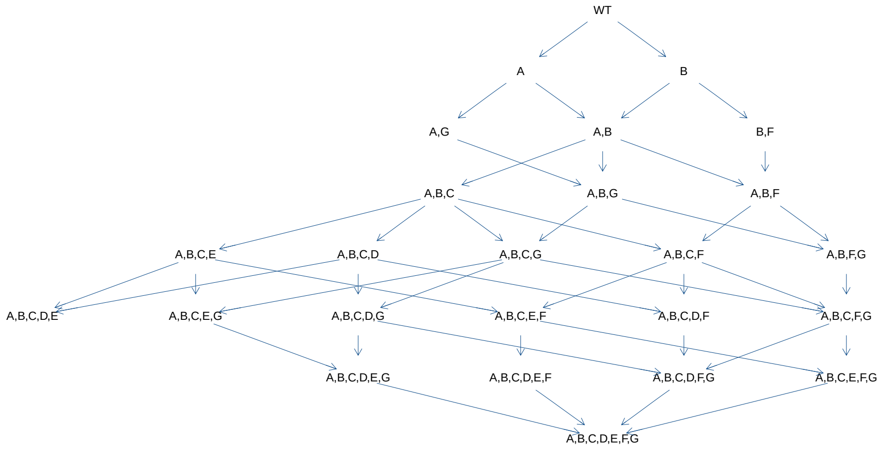

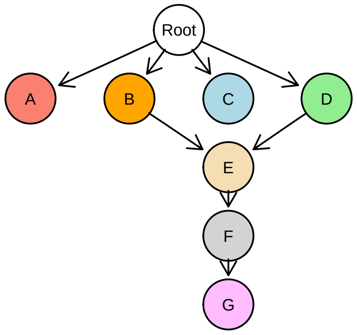


| | |
|---|---|
| *ID* | ycfMUnx16qqaBjIW |
| *acc genotypes* | 28 |
| *peaks no back* | 6 |
| *removed edges* | 11 |
| *prop rem edges* | 0.196 |

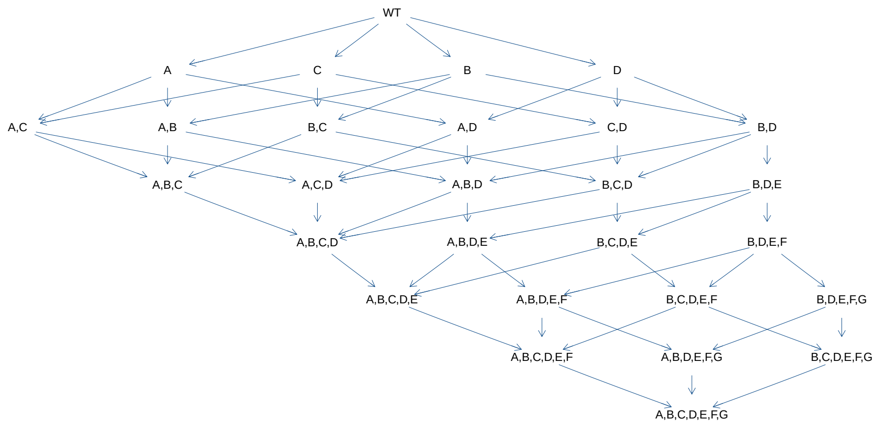


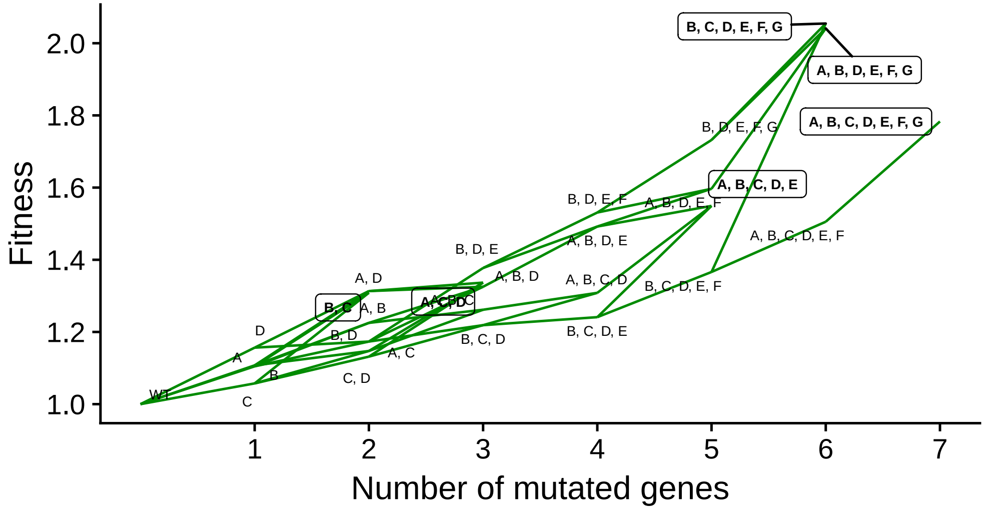


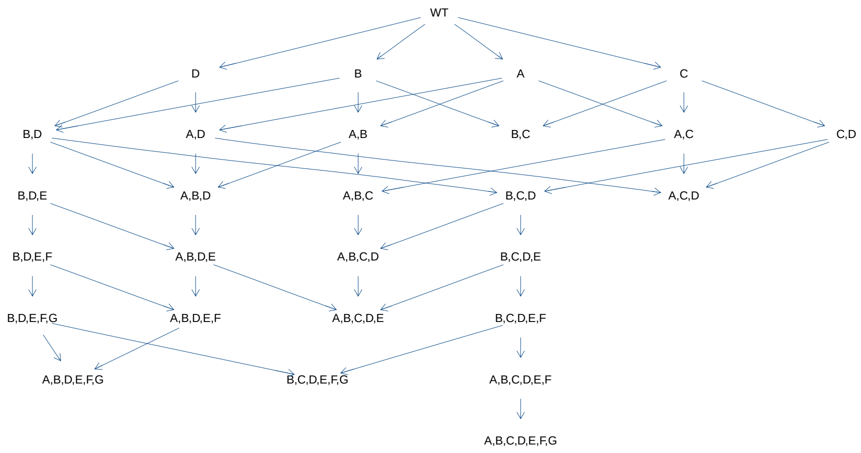

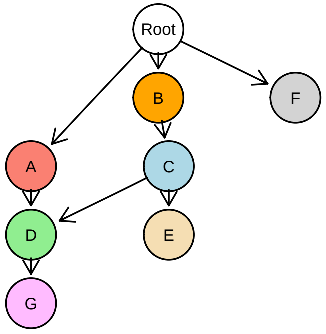


| | |
|---|---|
| *ID* | yCxbC75Zji2W7wiD |
| *acc genotypes* | 24 |
| *peaks no back* | 7 |
| *removed edges* | 11 |
| *prop rem edges* | 0.25 |

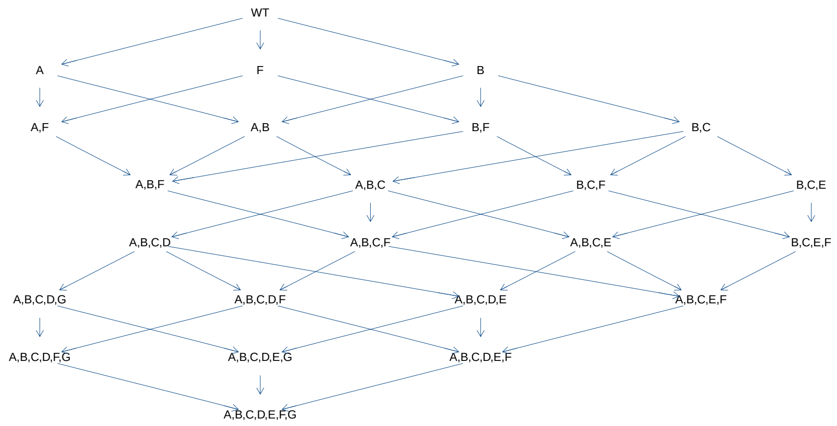


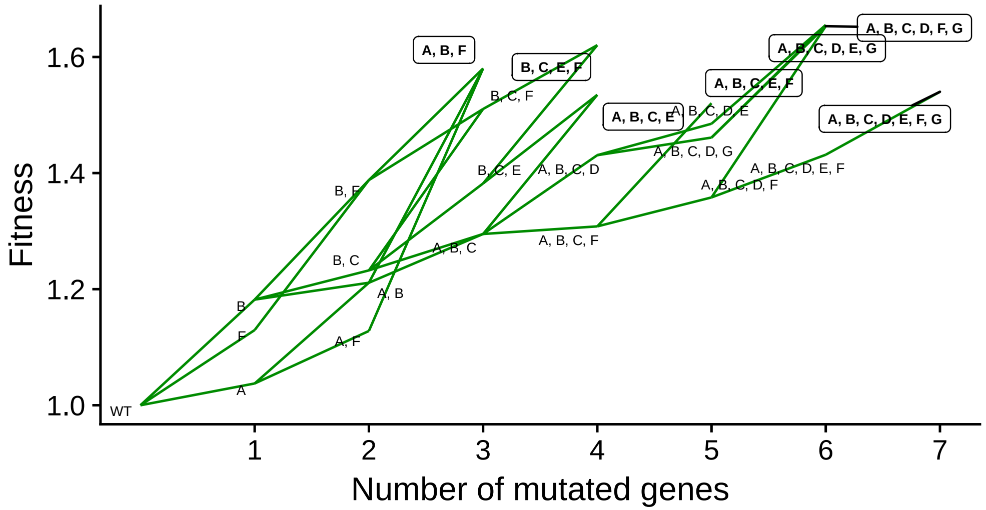


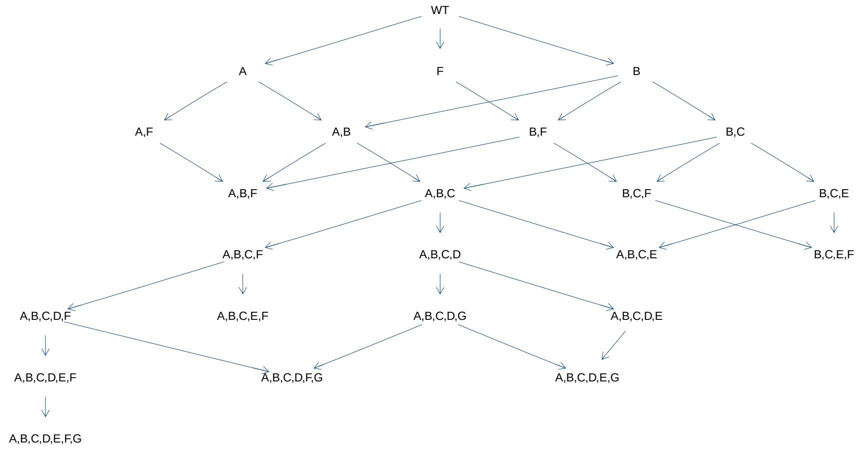

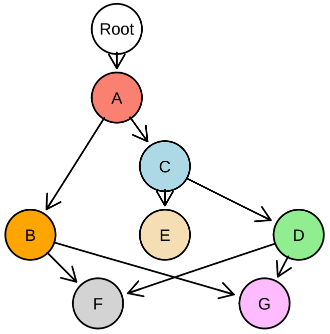


| | |
|---|---|
| *ID* | yoDlXpd2iByH7zZf |
| *acc genotypes* | 17 |
| *peaks no back* | 5 |
| *removed edges* | 6 |
| *prop rem edges* | 0.222 |

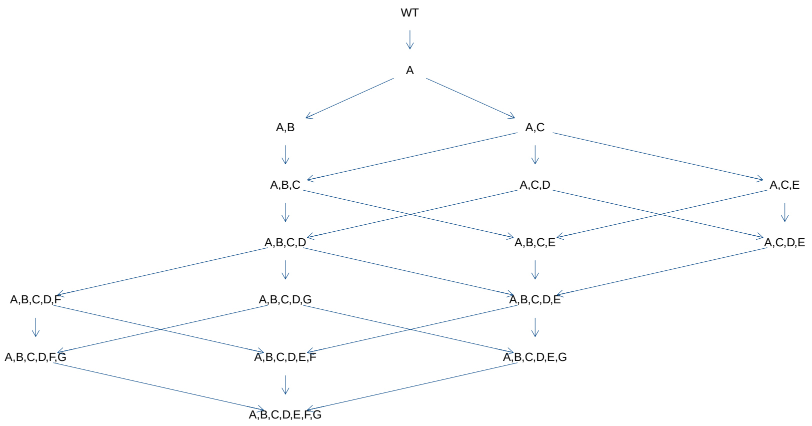


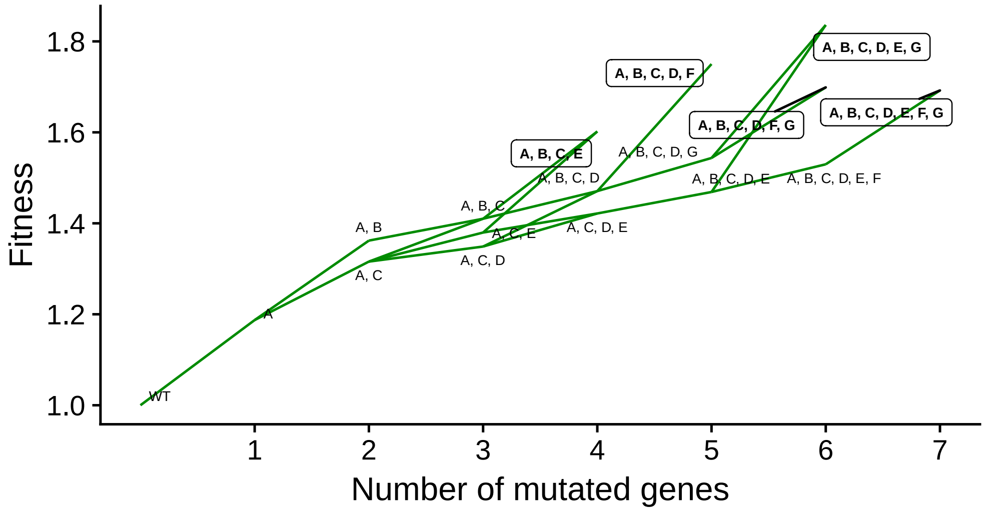


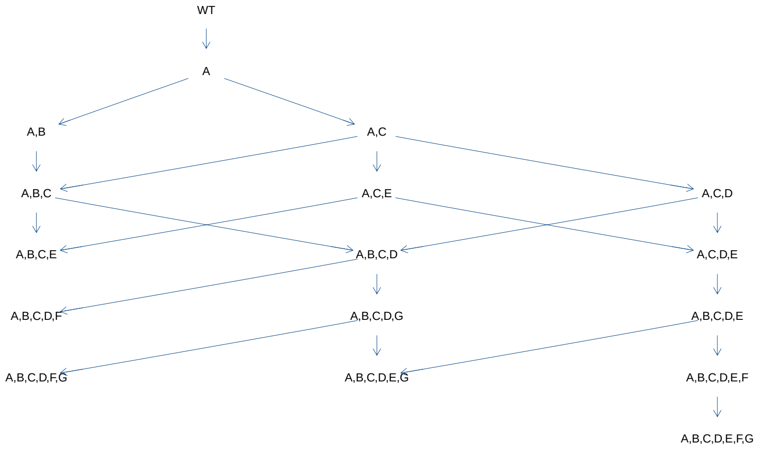

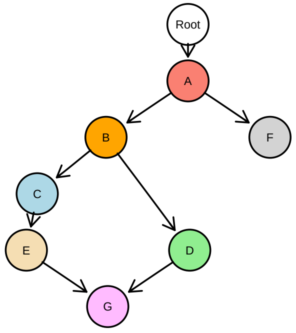


| | |
|---|---|
| *ID* | YqJMzkvAccJ4gHYm |
| *acc genotypes* | 17 |
| *peaks no back* | 3 |
| *removed edges* | 3 |
| *prop rem edges* | 0.111 |

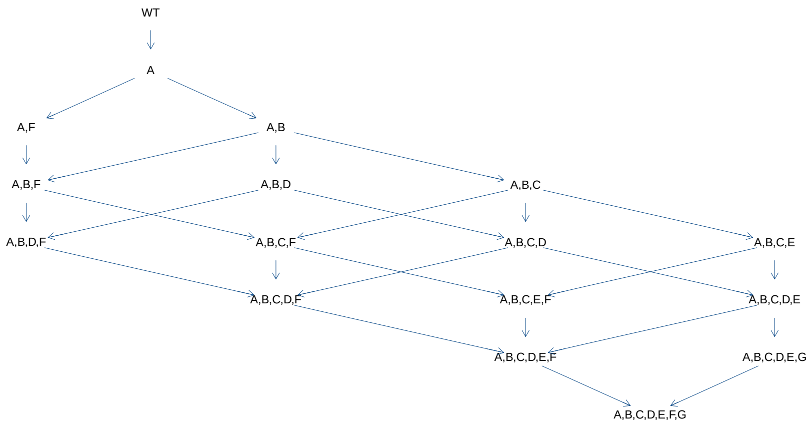


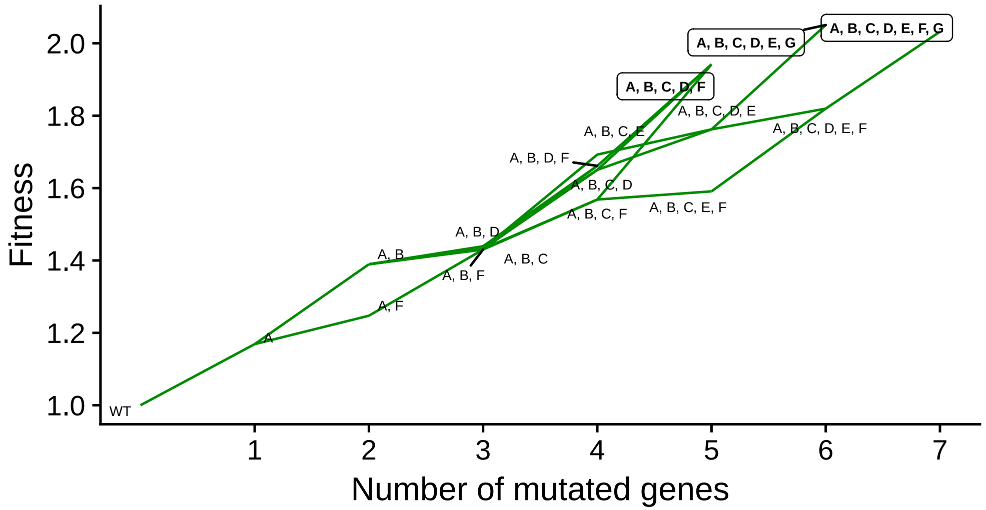


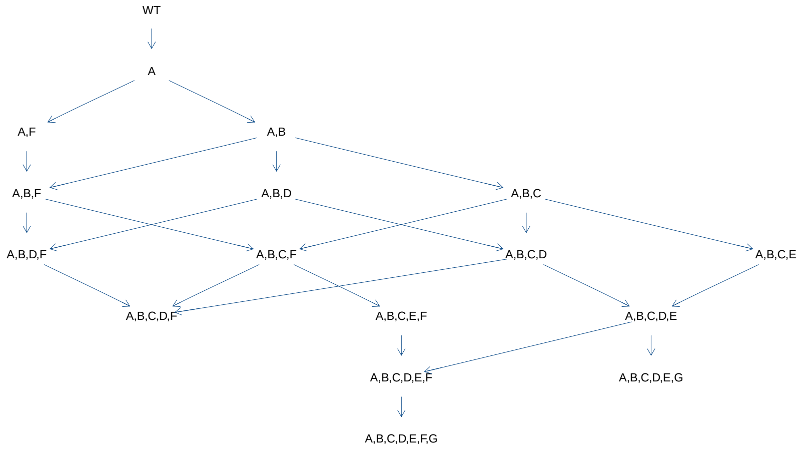

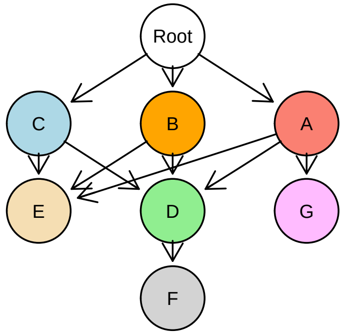


*ID* yX5bopH3PiXCr10r7
*acc genotypes* 22
*peaks no back* 4
*removed edges* 8
*prop rem edges* 0.205

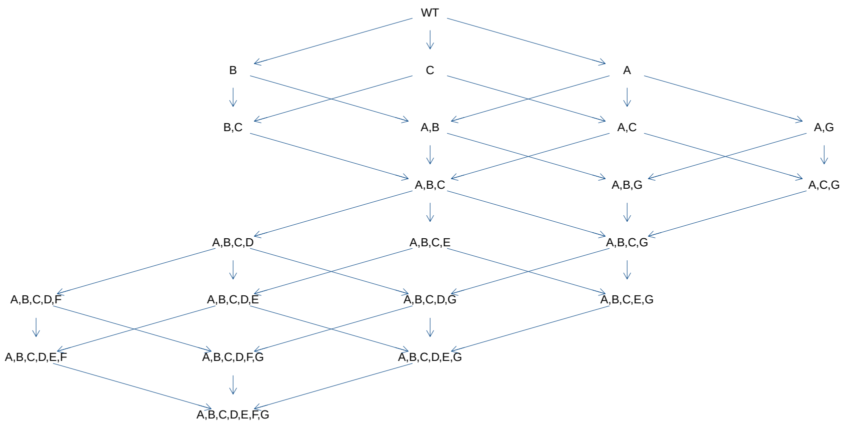


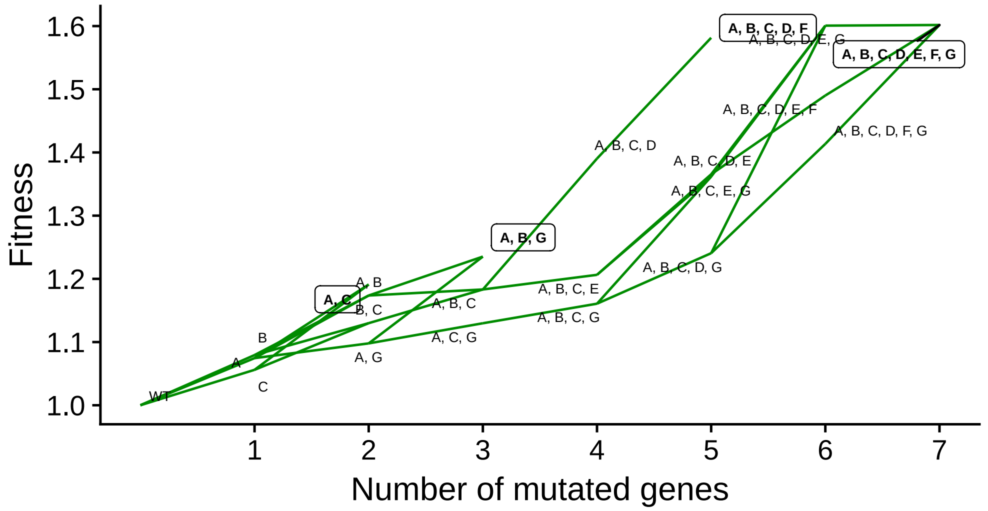


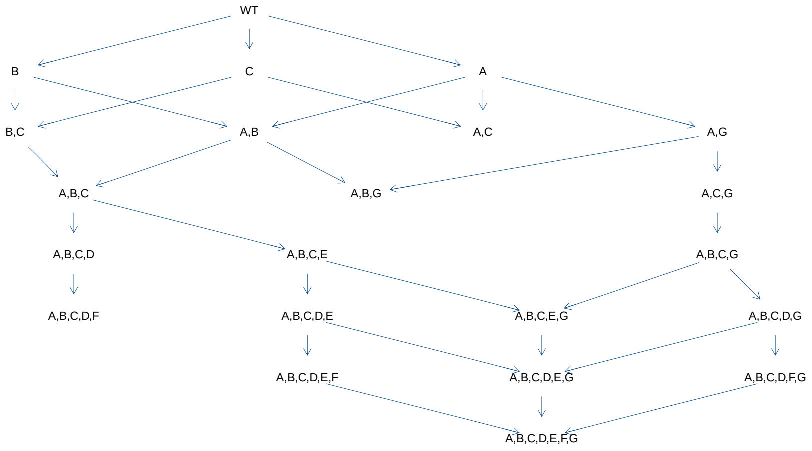

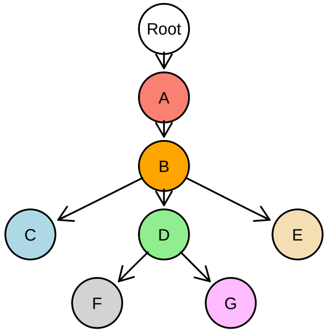


| | |
|---|---|
| *ID* | yy6dYwpU1ktFm49a |
| *acc genotypes* | 22 |
| *peaks no back* | 6 |
| *removed edges* | 10 |
| *prop rem edges* | 0.238 |

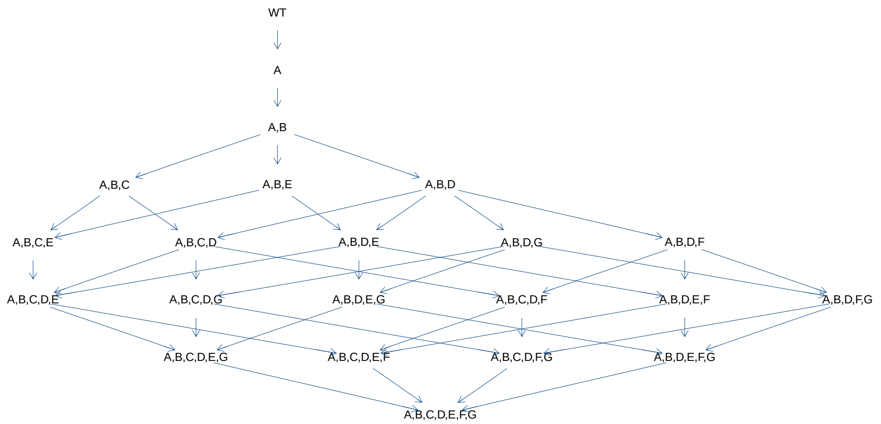


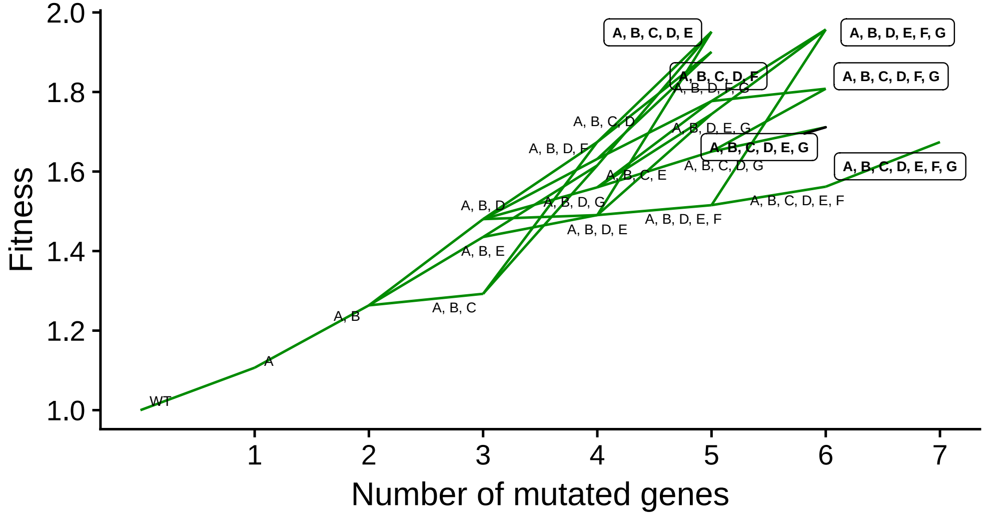


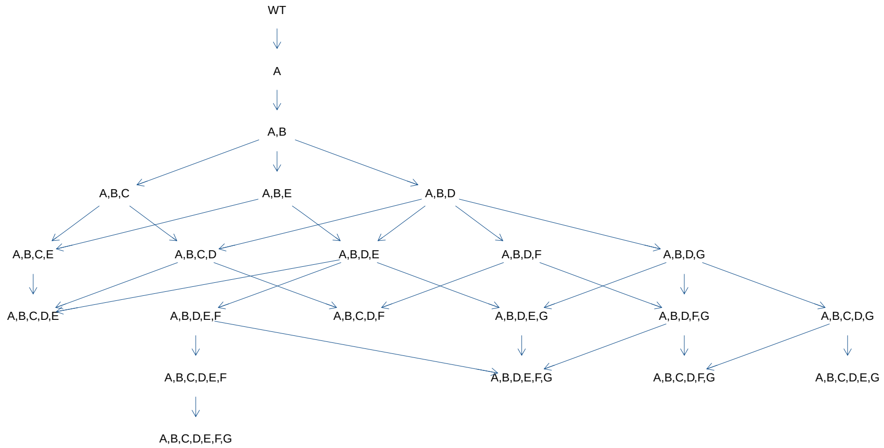

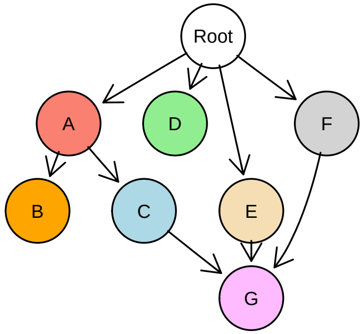


| | |
|---|---|
| *ID* | zE6cfkQmrWiDYn5D |
| *acc genotypes* | 44 |
| *peaks no back* | 5 |
| *removed edges* | 18 |
| *prop rem edges* | 0.167 |

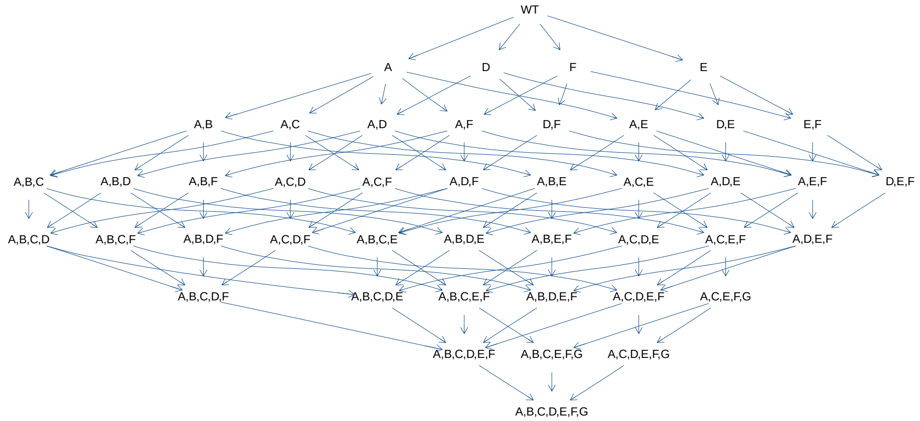


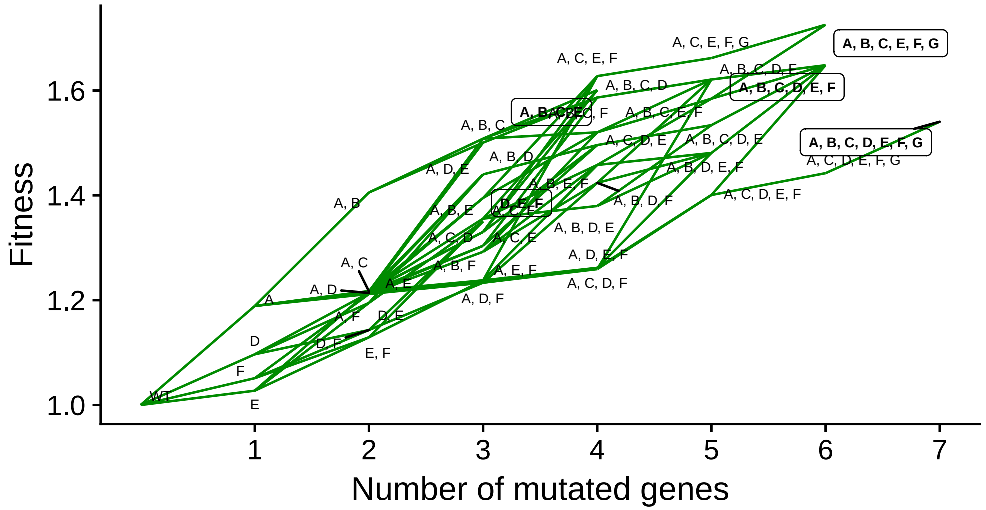


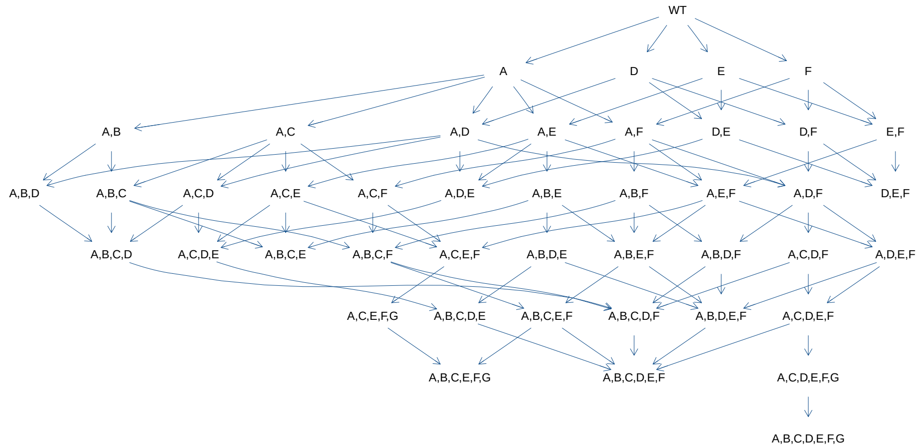

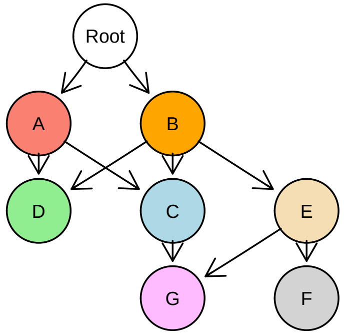


*ID* zHGCza3O5qL97M76
*acc genotypes* 21
*peaks no back* 2
*removed edges* 3
*prop rem edges* 0.083

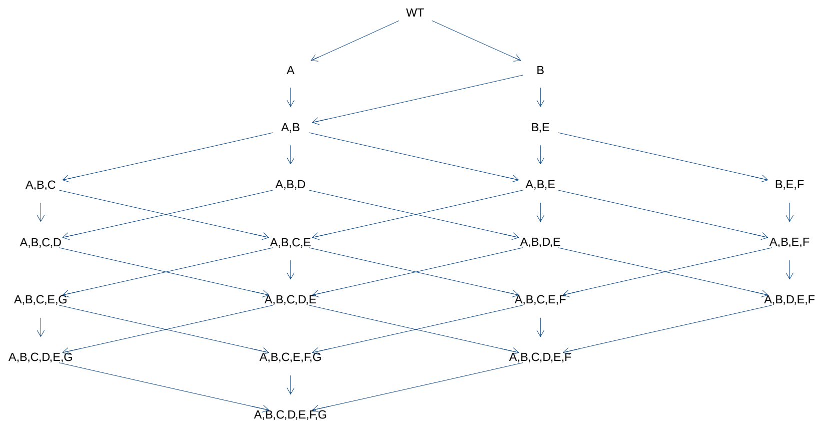


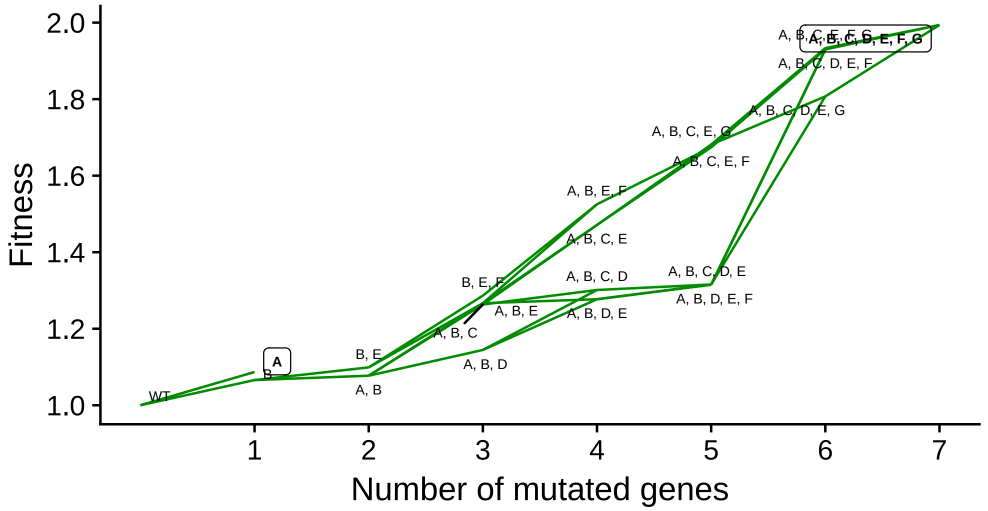


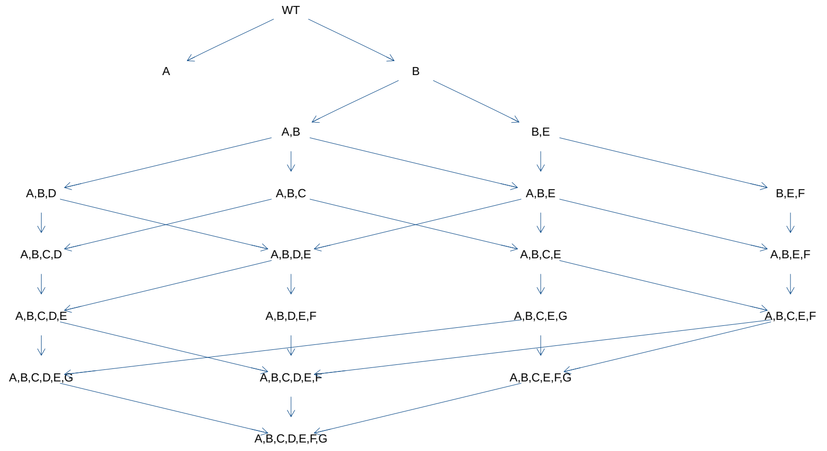

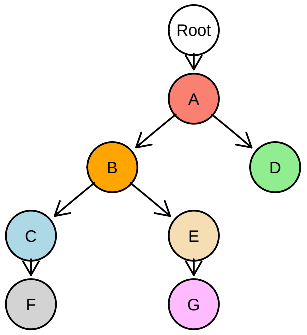


| | |
|---|---|
| *ID* | ZkjYaBc10QMnrUp8W |
| *acc genotypes* | 21 |
| *peaks no back* | 3 |
| *removed edges* | 2 |
| *prop rem edges* | 0.054 |

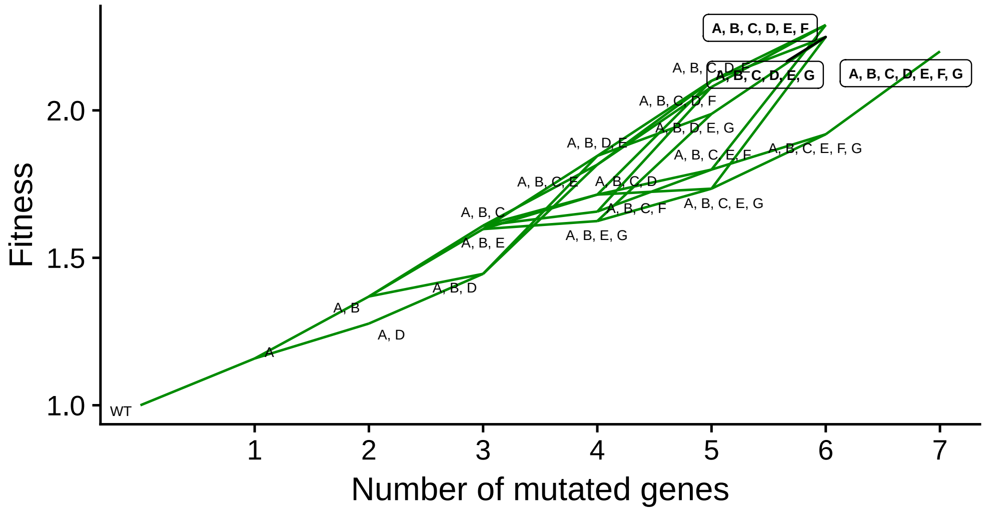


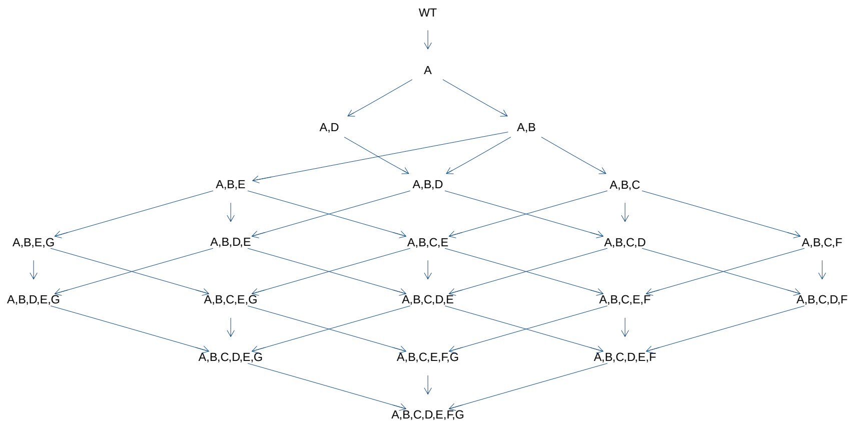


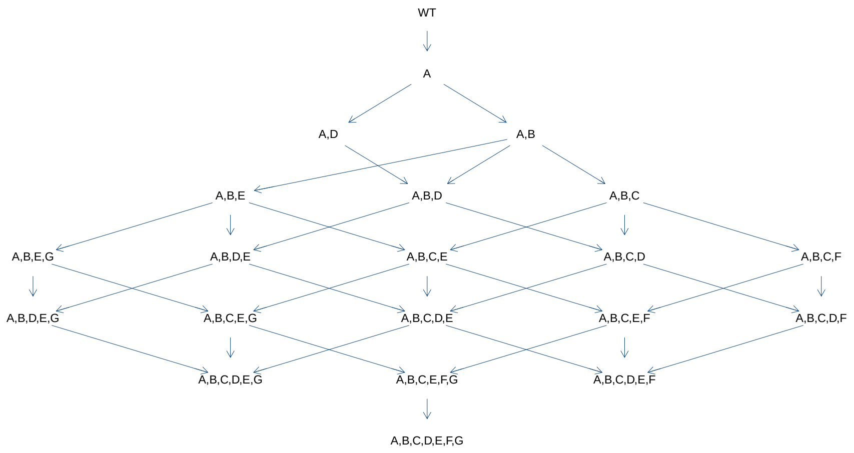

| | |
|---|---|
| *ID* | 10bxK1gc1IxCjpaJM |
| *acc genotypes* | 24 |
| *peaks no back* | 9 |
| *removed edges* | NA |
| *prop rem edges* | NA |

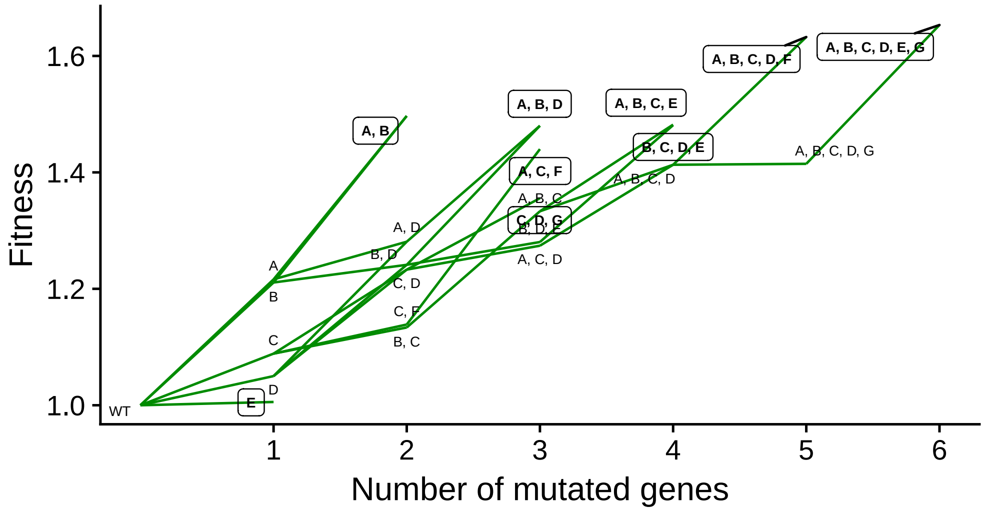


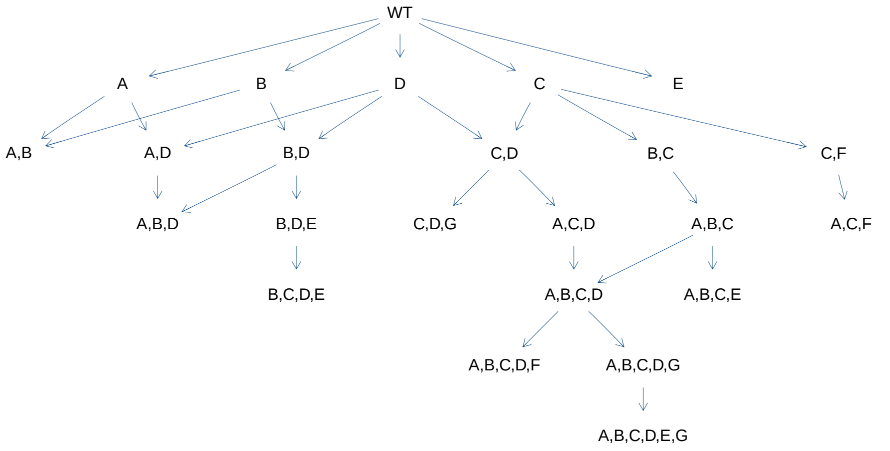

| | |
|---|---|
| *ID* | 10fs6bc54hwzLal8K |
| *acc genotypes* | 99 |
| *peaks no back* | 14 |
| *removed edges* | NA |
| *prop rem edges* | NA |

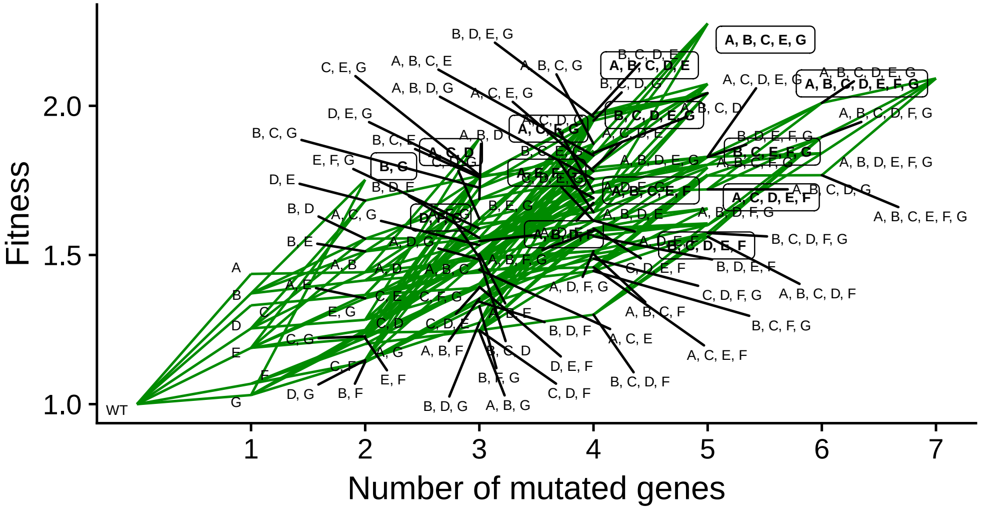


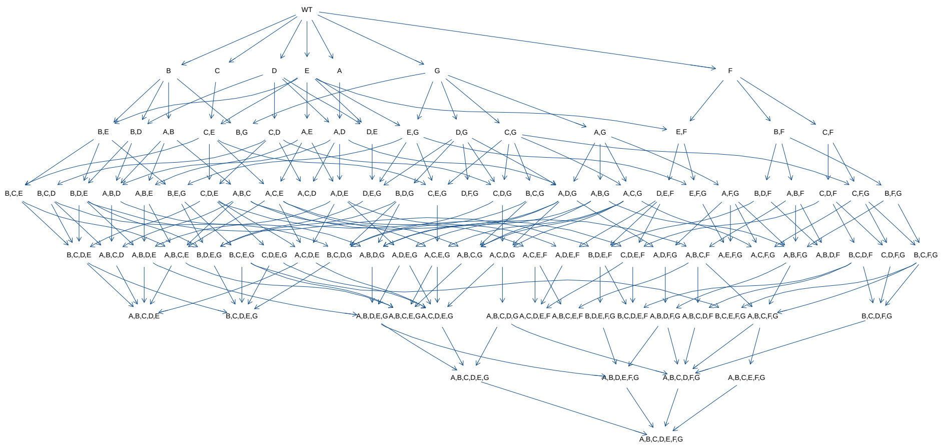

| | |
|---|---|
| *ID* | 10H8qiQAOoVrnY9T3 |
| *acc genotypes* | 42 |
| *peaks no back* | 15 |
| *removed edges* | NA |
| *prop rem edges* | NA |

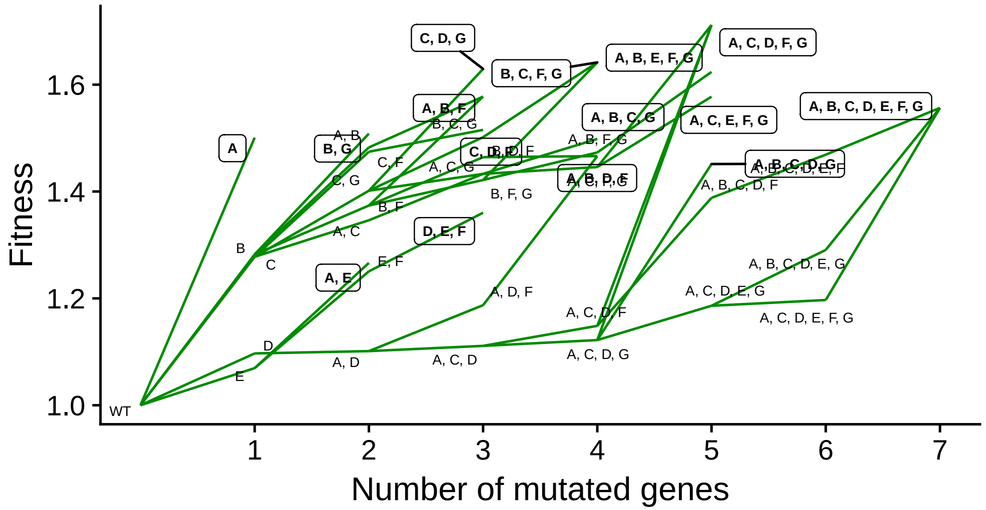


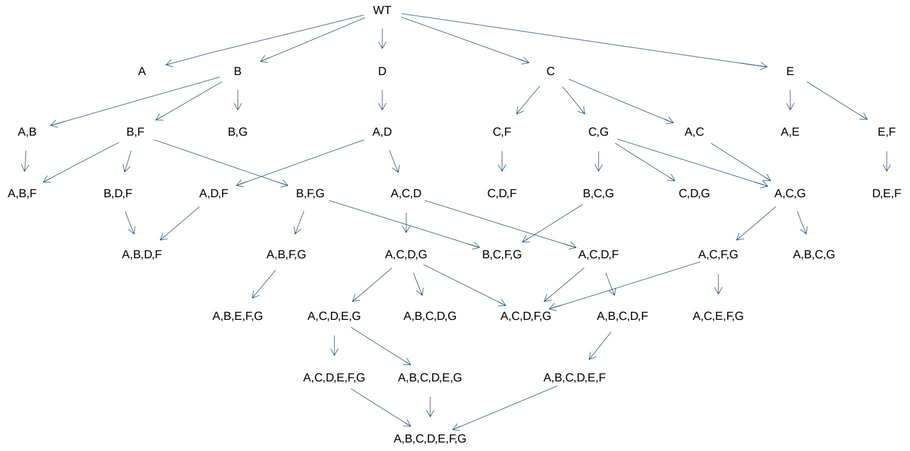

| | |
|---|---|
| *ID* | 10rLLqPV6rJMo5c6I |
| *acc genotypes* | 28 |
| *peaks no back* | 9 |
| *removed edges* | NA |
| *prop rem edges* | NA |

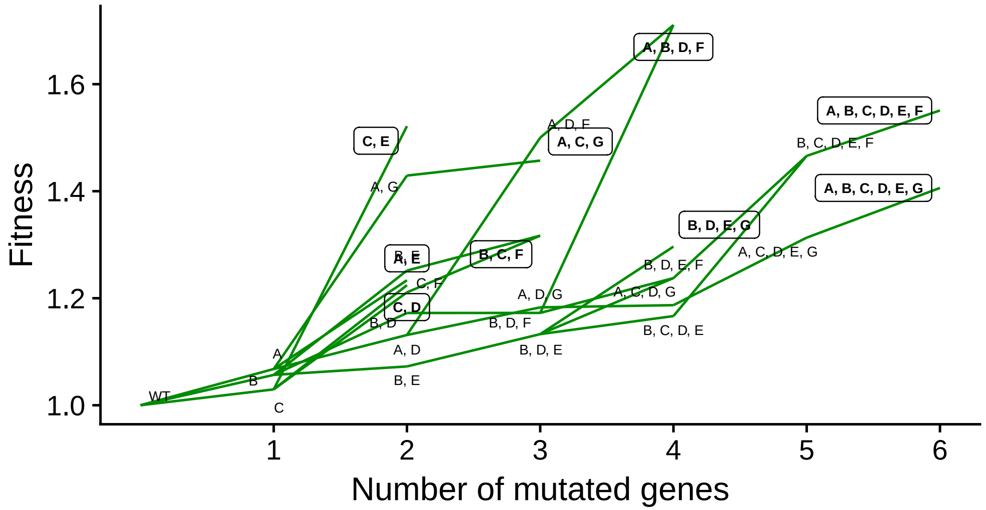


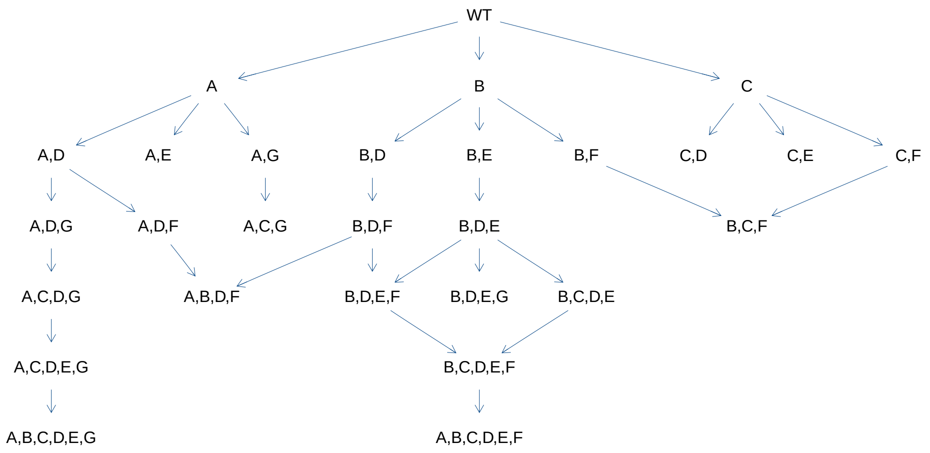

| | |
|---|---|
| *ID* | 10uENjADj4VYkG10Z6 |
| *acc genotypes* | 58 |
| *peaks no back* | 26 |
| *removed edges* | NA |
| *prop rem edges* | NA |

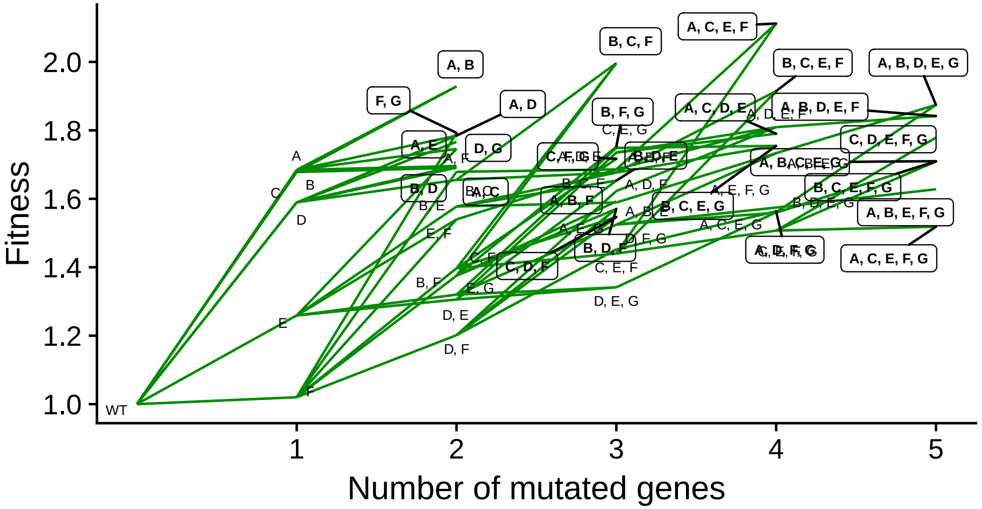


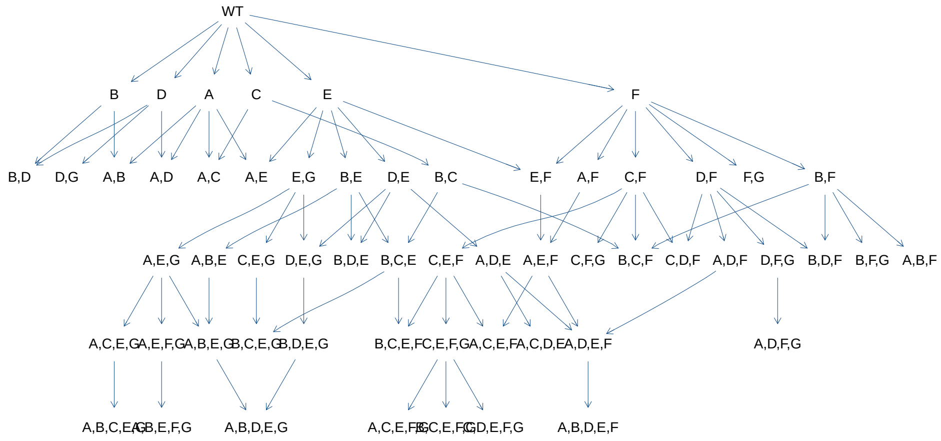

| | |
|---|---|
| *ID* | 16H2tTfJLrFoSFFU |
| *acc genotypes* | 23 |
| *peaks no back* | 7 |
| *removed edges* | NA |
| *prop rem edges* | NA |

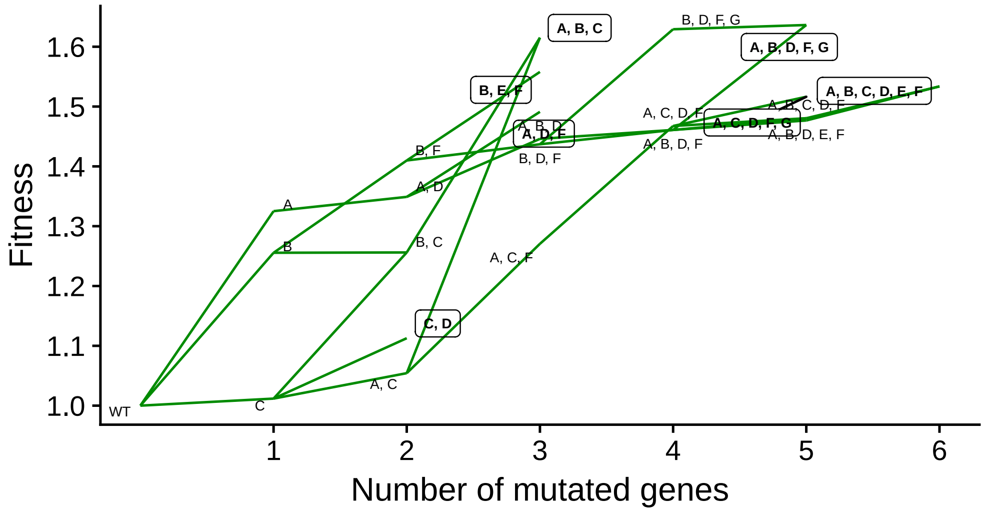


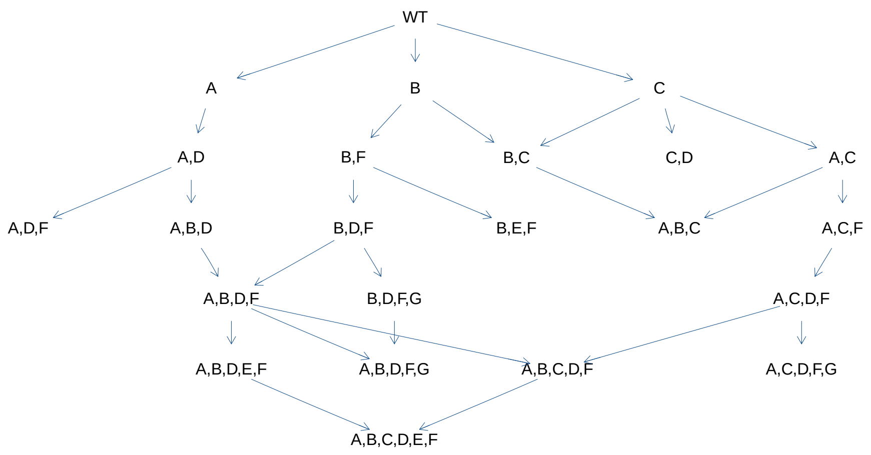

| | |
|---|---|
| *ID* | 18VIJjttIjPc2A62 |
| *acc genotypes* | 24 |
| *peaks no back* | 10 |
| *removed edges* | NA |
| *prop rem edges* | NA |

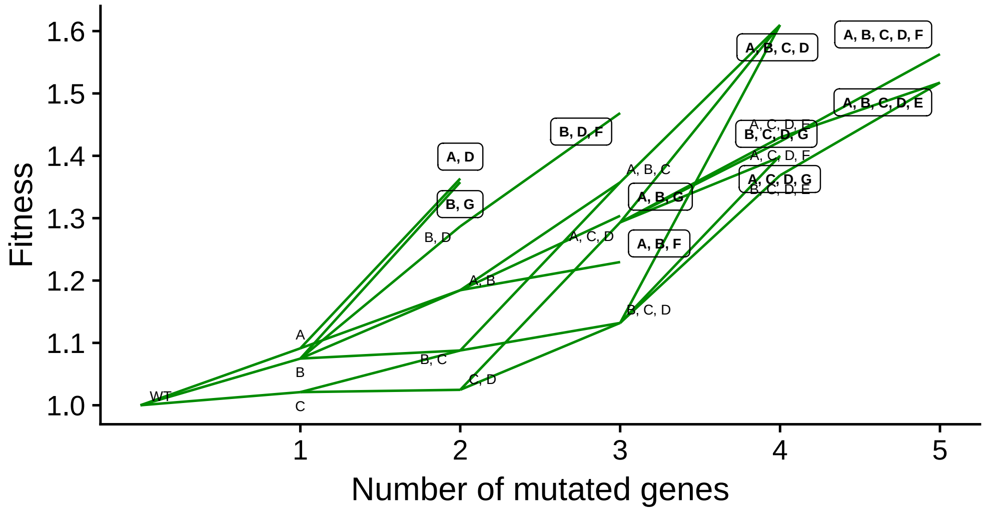


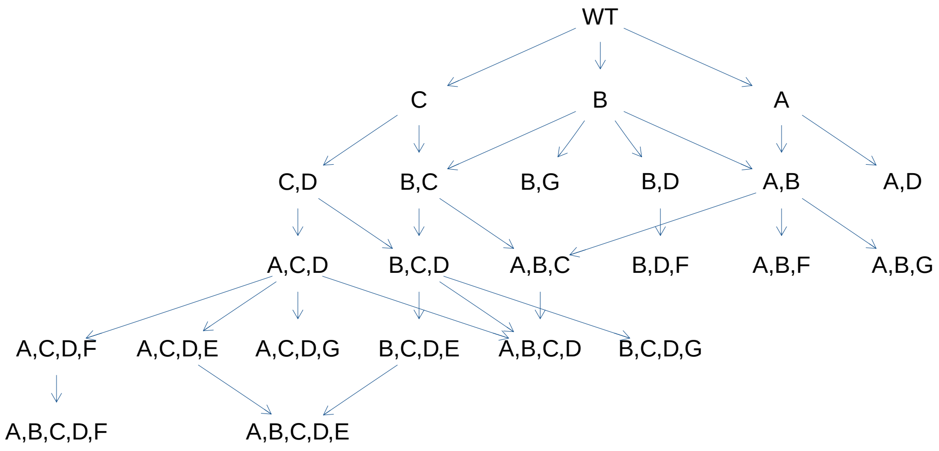

| | |
|---|---|
| *ID* | 1ylOxuiyzdWquD2u |
| *acc genotypes* | 27 |
| *peaks no back* | 12 |
| *removed edges* | NA |
| *prop rem edges* | NA |

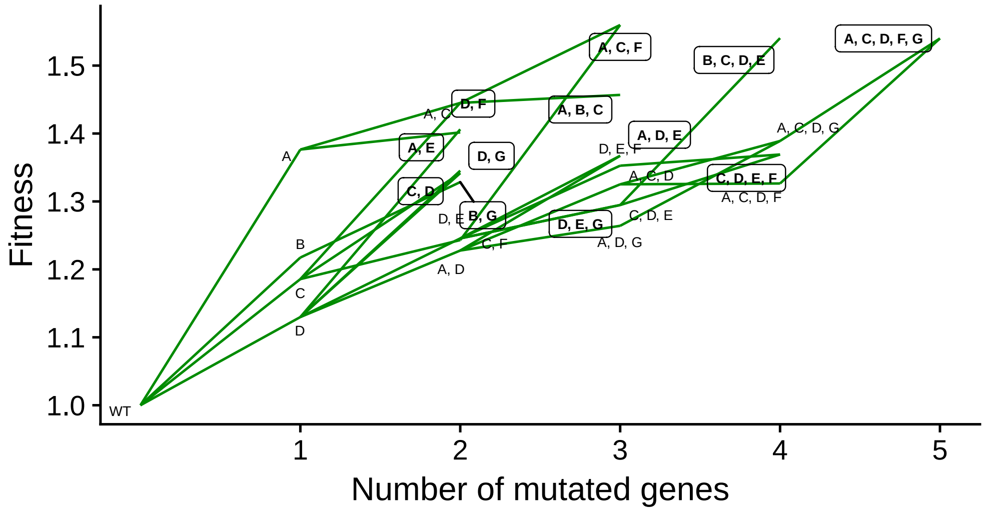


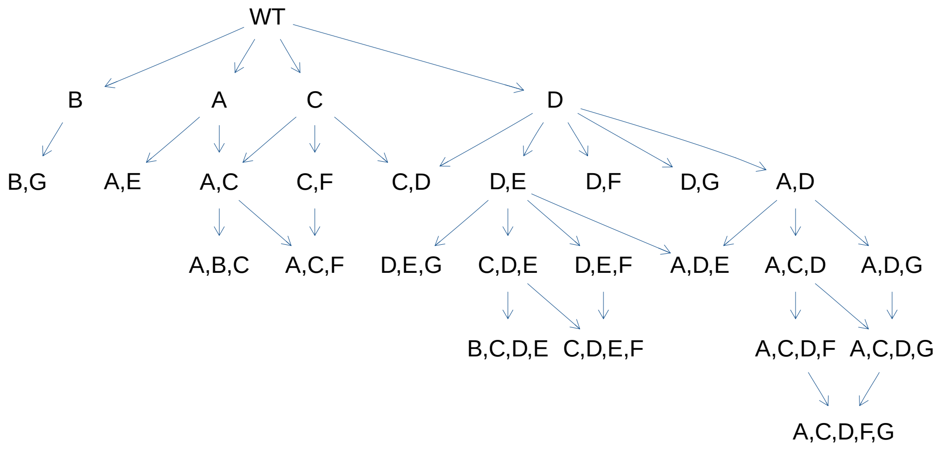

| | |
|---|---|
| *ID* | 28U7vxmKq8g5SSwV |
| *acc genotypes* | 26 |
| *peaks no back* | 11 |
| *removed edges* | NA |
| *prop rem edges* | NA |

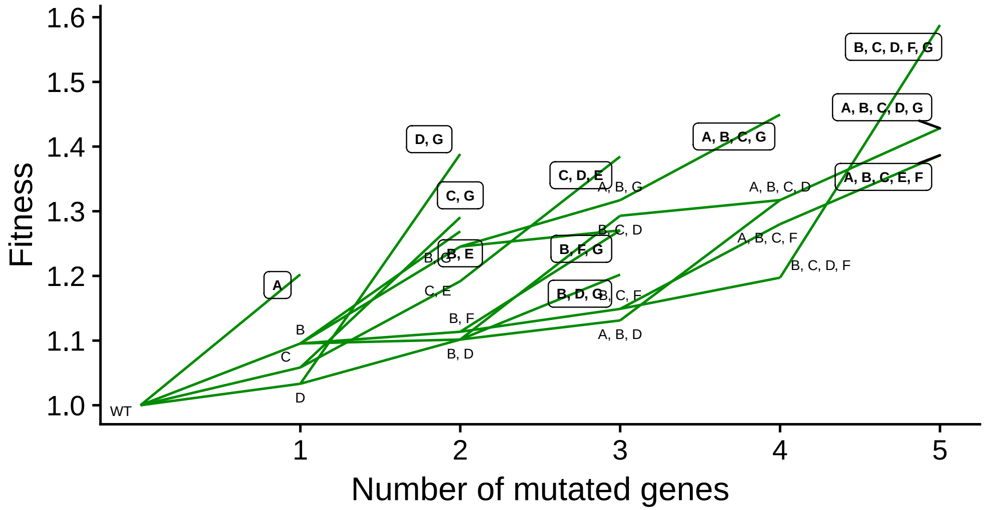


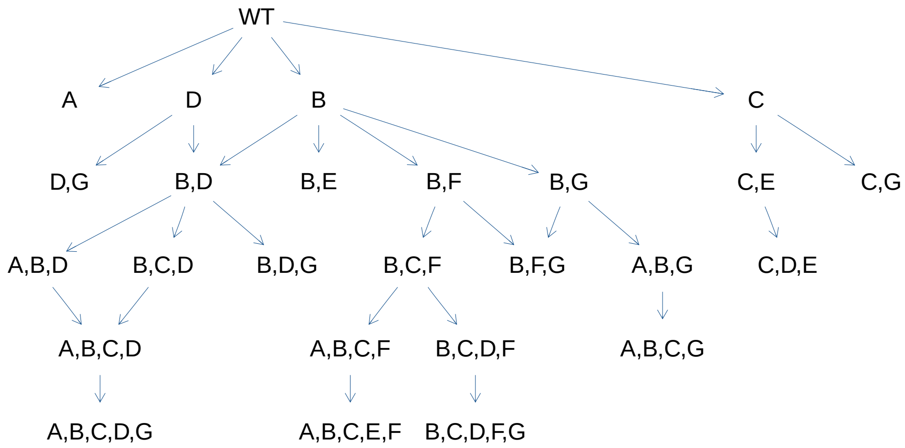

| | |
|---|---|
| *ID* | 2PO6NSrcnBJAxivm |
| *acc genotypes* | 12 |
| *peaks no back* | 5 |
| *removed edges* | NA |
| *prop rem edges* | NA |

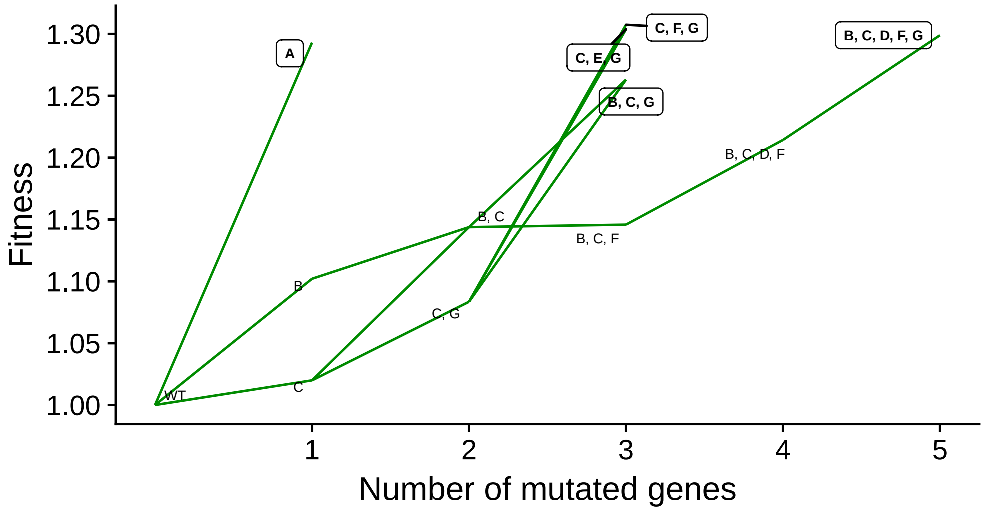


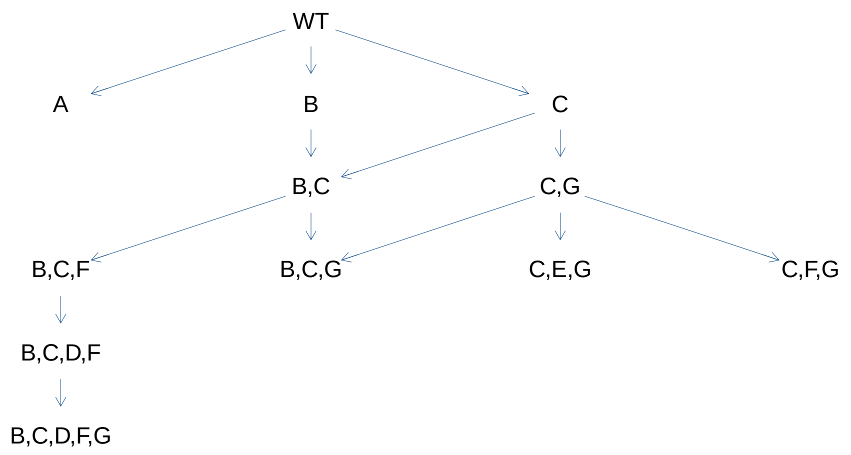

| | |
|---|---|
| *ID* | 344NmuCSF610HIunl |
| *acc genotypes* | 44 |
| *peaks no back* | 17 |
| *removed edges* | NA |
| *prop rem edges* | NA |

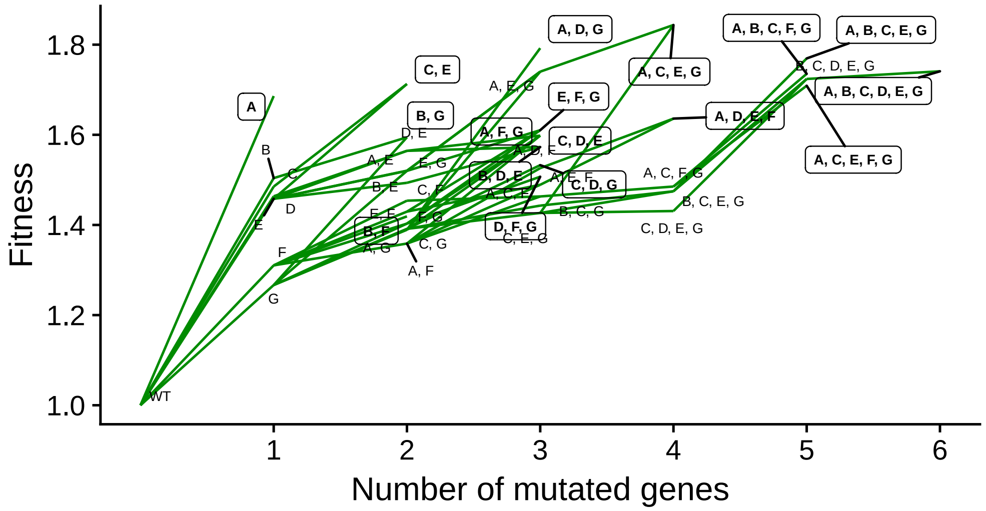


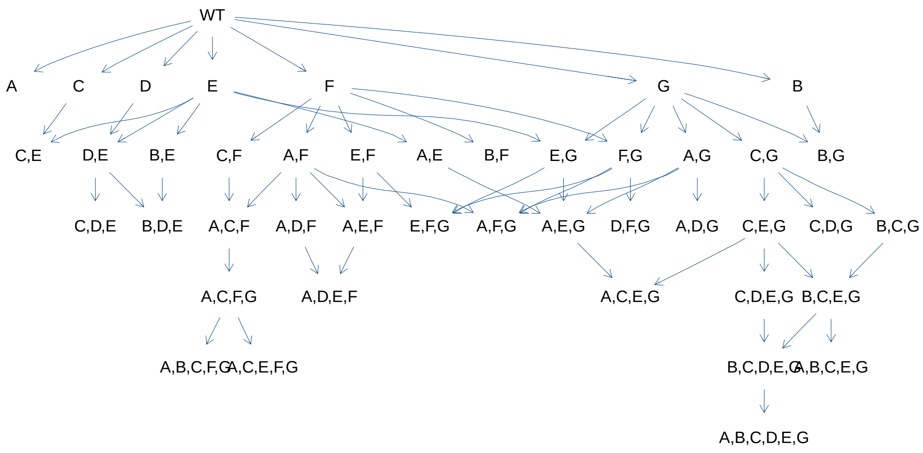

| | |
|---|---|
| *ID* | 3JtCUkHFLmBXxzdb |
| *acc genotypes* | 25 |
| *peaks no back* | 13 |
| *removed edges* | NA |
| *prop rem edges* | NA |

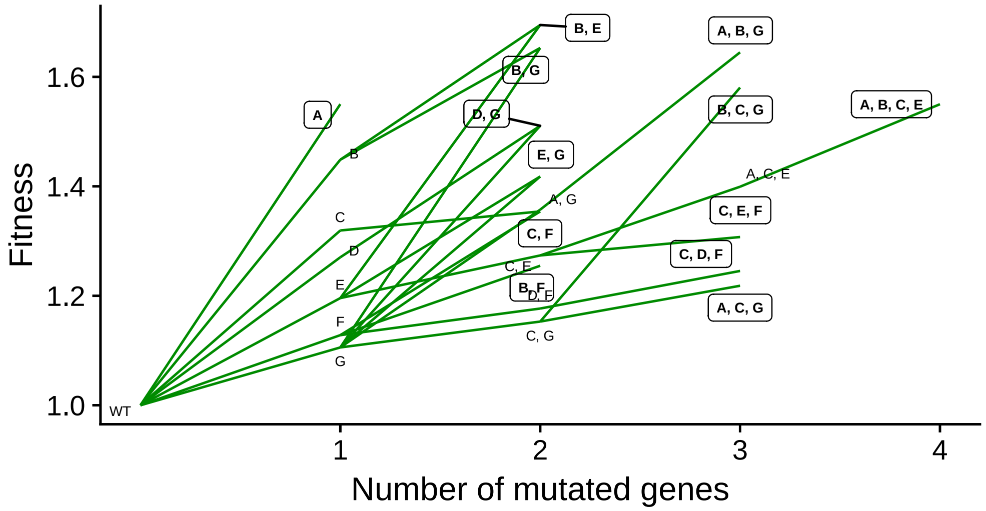


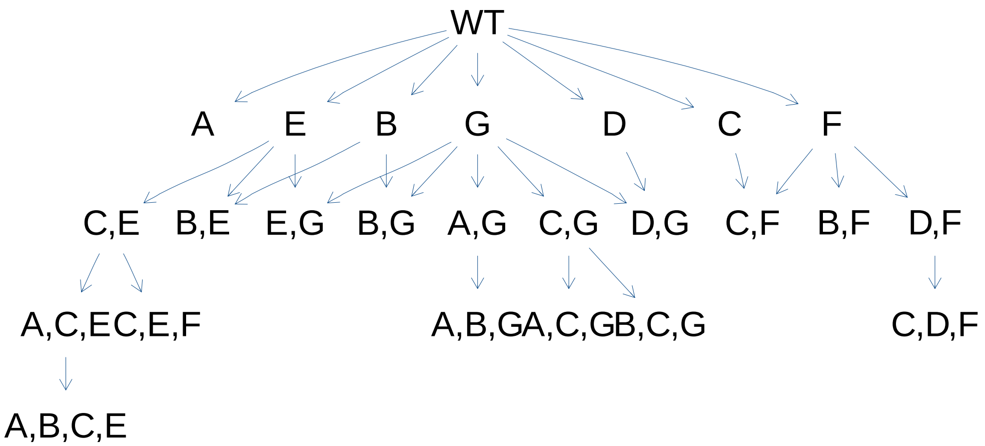

| | |
|---|---|
| *ID* | 3LxjVpequ1QHM105O |
| *acc genotypes* | 26 |
| *peaks no back* | 13 |
| *removed edges* | NA |
| *prop rem edges* | NA |

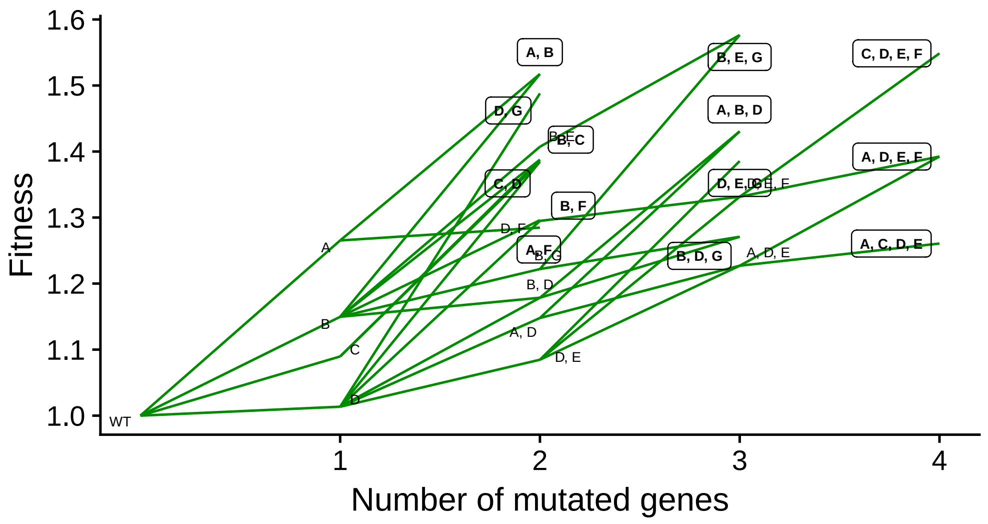


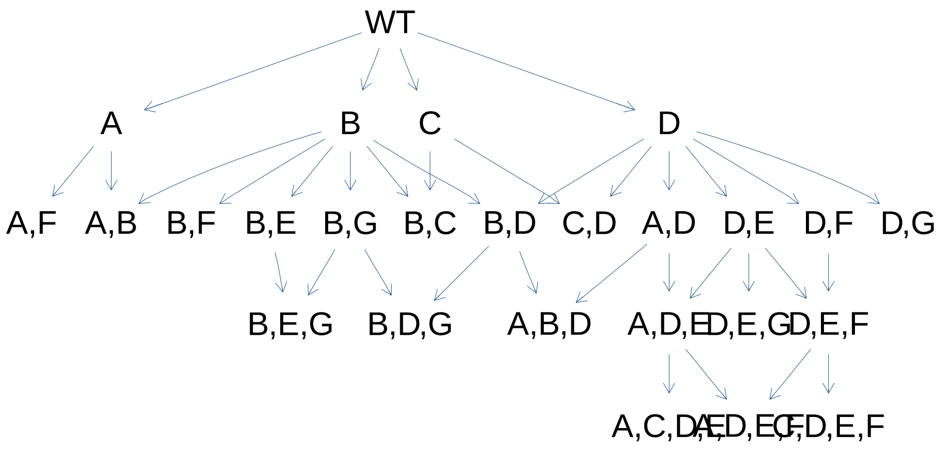

| | |
|---|---|
| *ID* | 3NwMO8WF7CzUnn4q |
| *acc genotypes* | 57 |
| *peaks no back* | 15 |
| *removed edges* | NA |
| *prop rem edges* | NA |

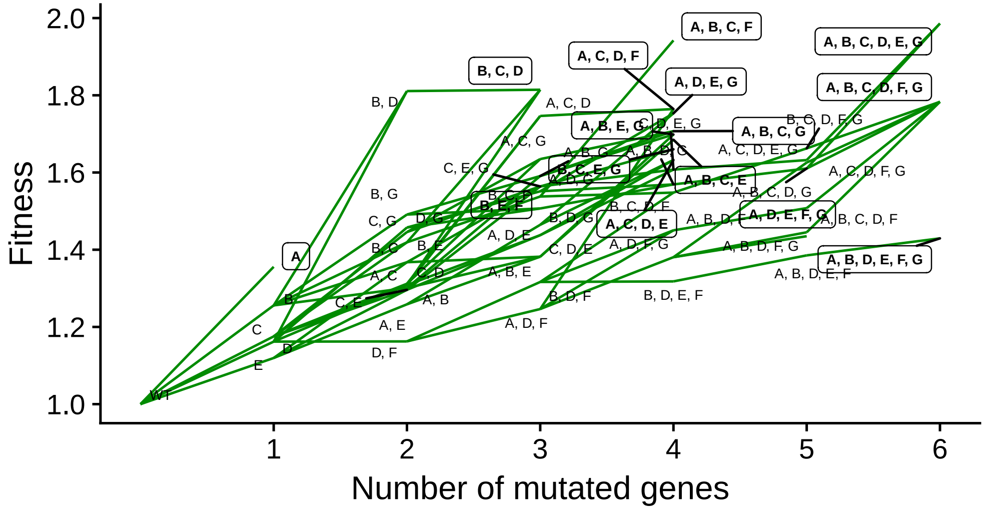


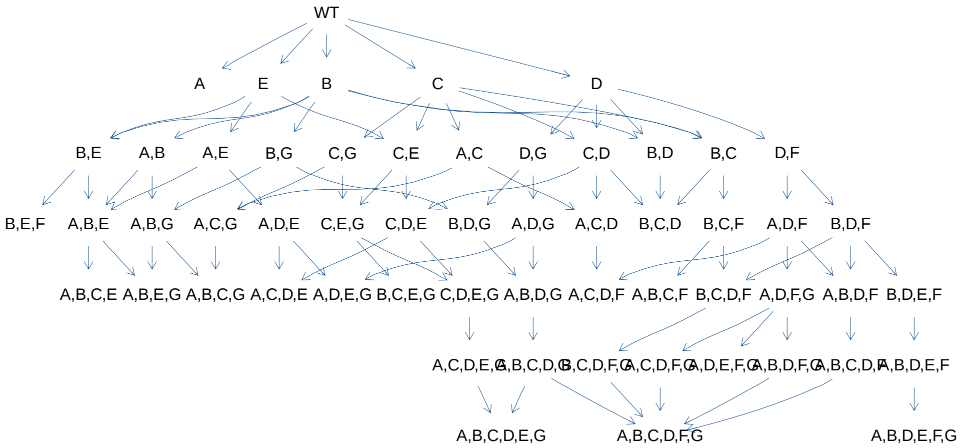

| | |
|---|---|
| *ID* | 4b99AWrdV4410hdsO |
| *acc genotypes* | 78 |
| *peaks no back* | 21 |
| *removed edges* | NA |
| *prop rem edges* | NA |

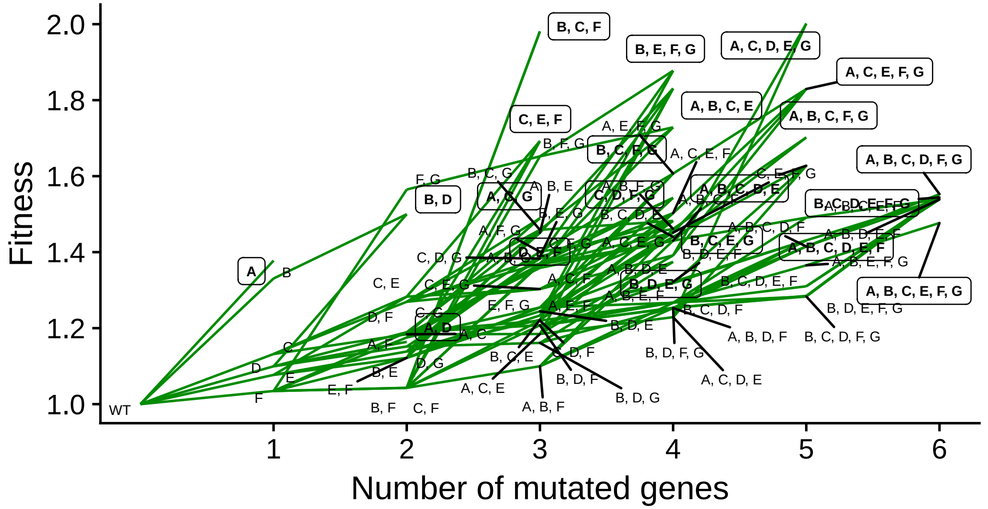


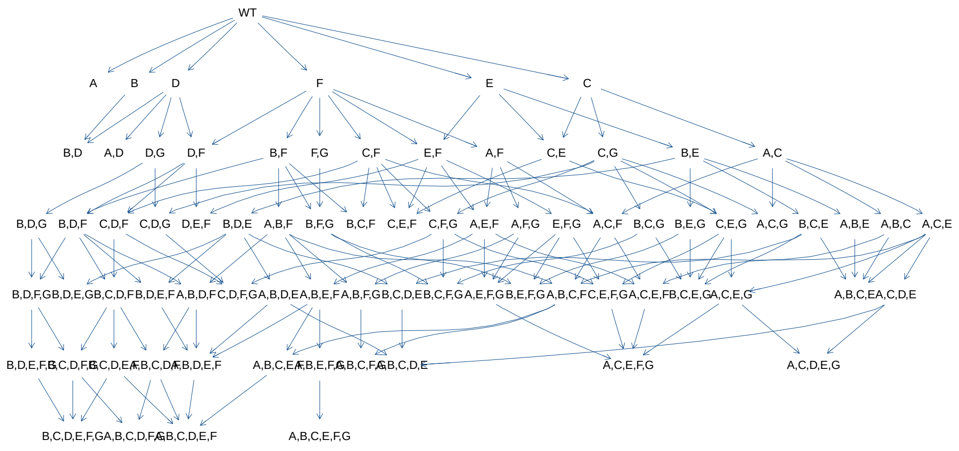

| | |
|---|---|
| *ID* | 4cDYPNJ8oYbtJWIe |
| *acc genotypes* | 50 |
| *peaks no back* | 13 |
| *removed edges* | NA |
| *prop rem edges* | NA |

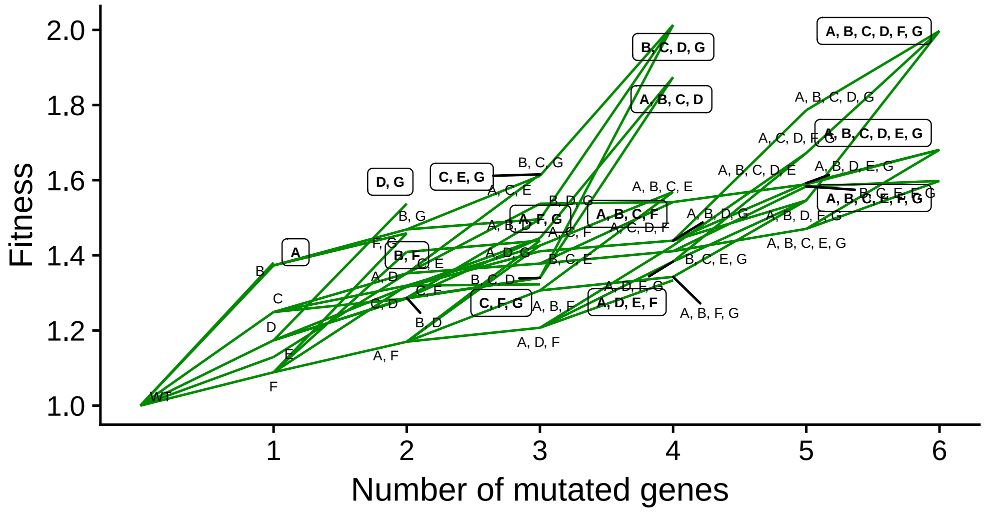


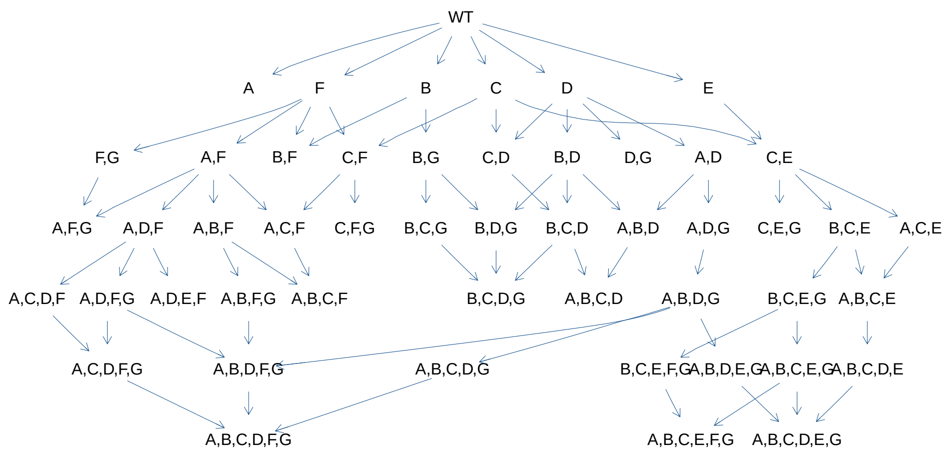

| | |
|---|---|
| *ID* | 4oM8TrevISckCyjy |
| *acc genotypes* | 17 |
| *peaks no back* | 11 |
| *removed edges* | NA |
| *prop rem edges* | NA |

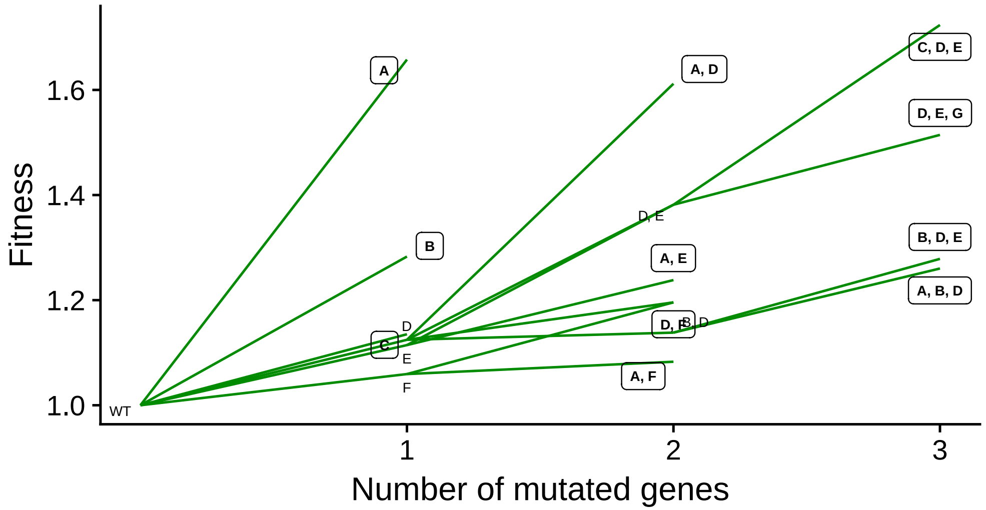


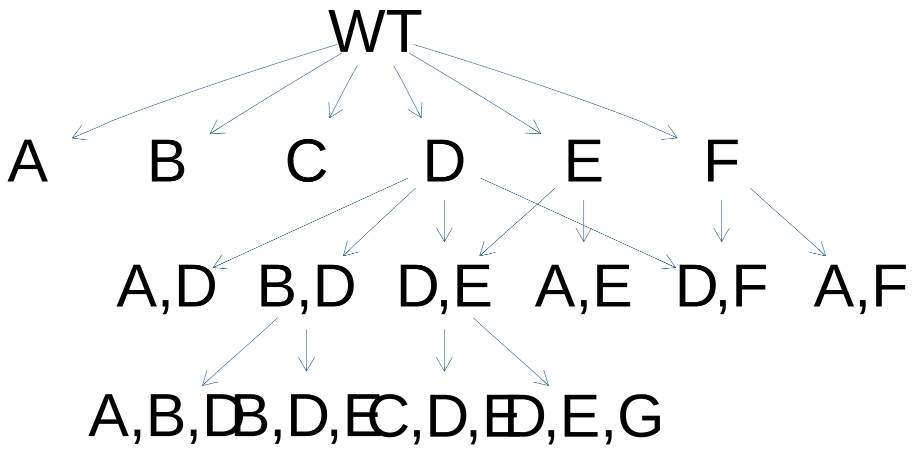

| | |
|---|---|
| *ID* | 4oYbhN10xm4ltLDHm |
| *acc genotypes* | 115 |
| *peaks no back* | 12 |
| *removed edges* | NA |
| *prop rem edges* | NA |

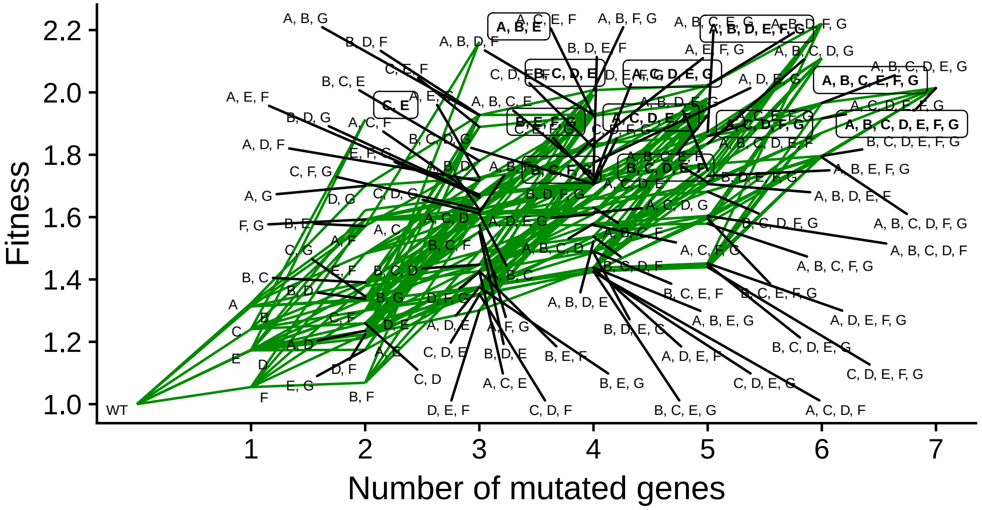


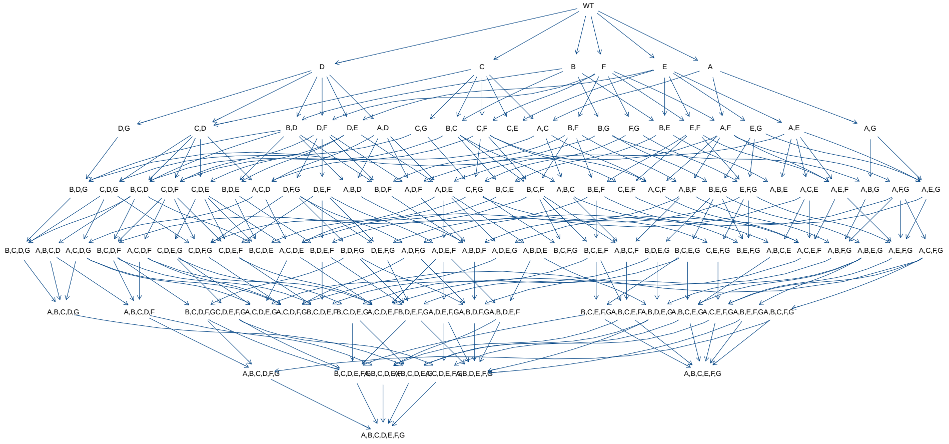

| | |
|---|---|
| *ID* | 4UvCBrp2StvG6sde |
| *acc genotypes* | 68 |
| *peaks no back* | 27 |
| *removed edges* | NA |
| *prop rem edges* | NA |

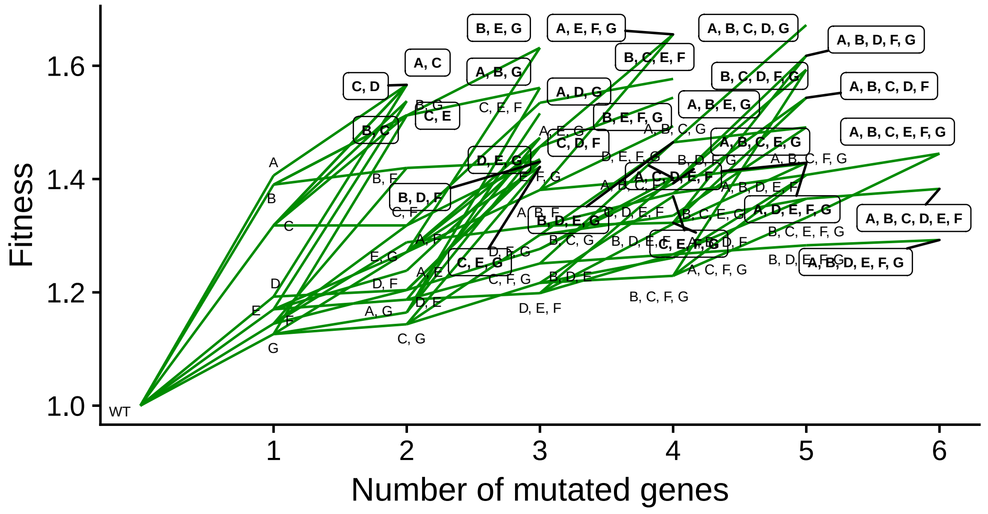


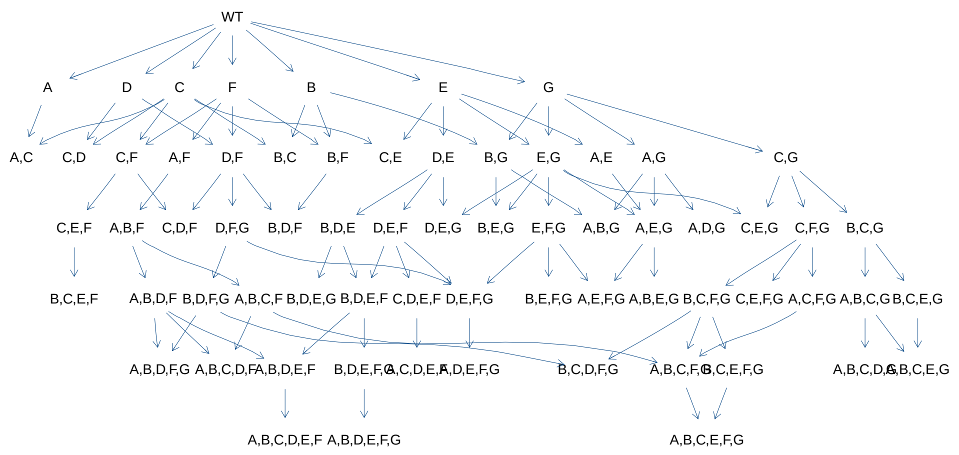

| | |
|---|---|
| *ID* | 4xoT2TA10PlN1Y8f9 |
| *acc genotypes* | 43 |
| *peaks no back* | 17 |
| *removed edges* | NA |
| *prop rem edges* | NA |

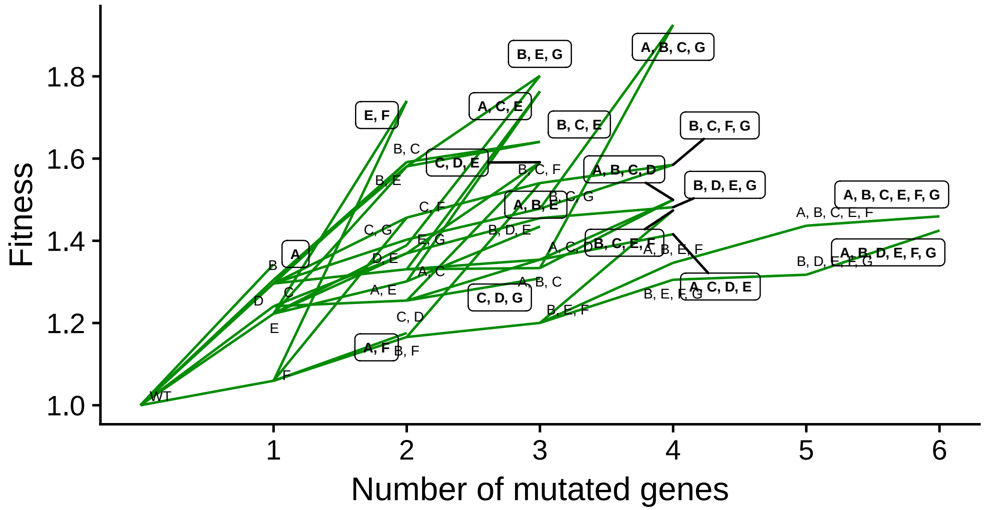


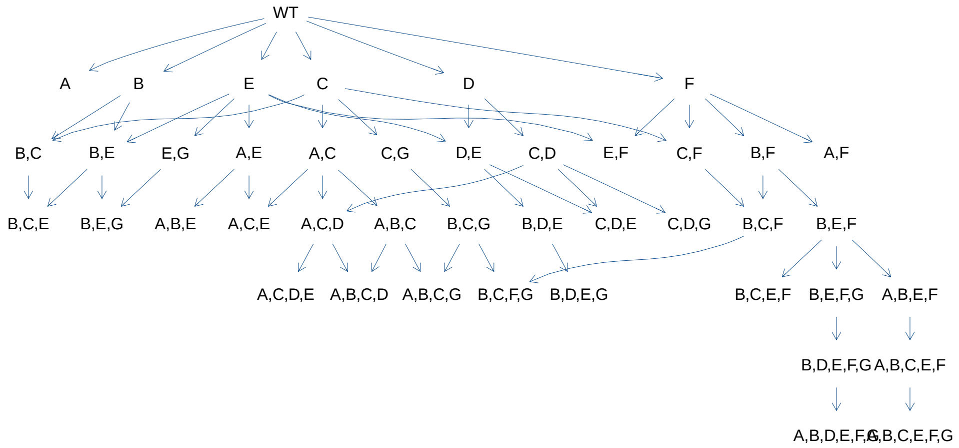

| | |
|---|---|
| *ID* | 5cropZmgpoQ1ehPe |
| *acc genotypes* | 33 |
| *peaks no back* | 13 |
| *removed edges* | NA |
| *prop rem edges* | NA |

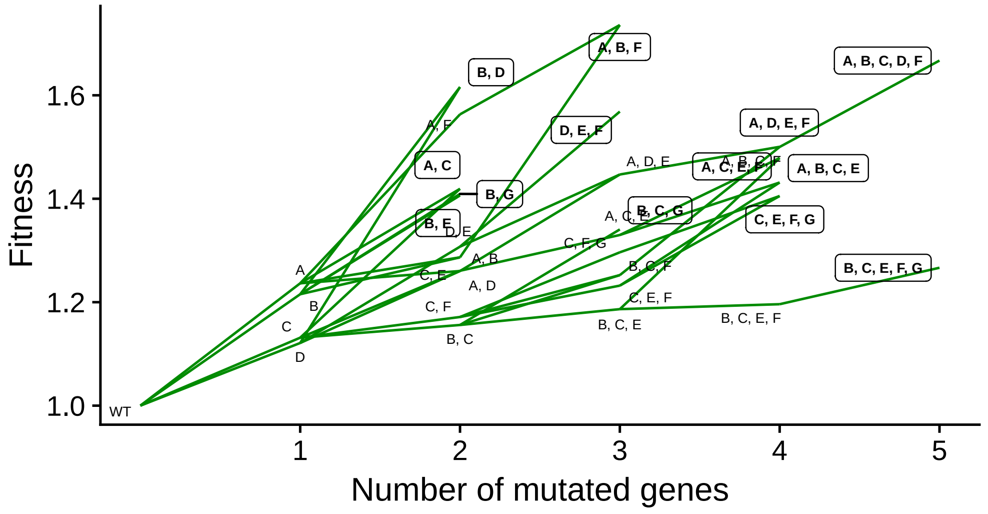


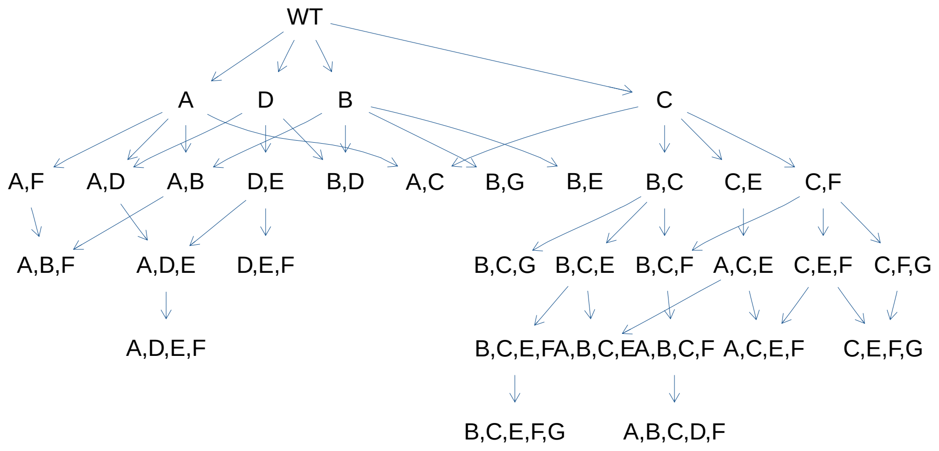

*ID* 5epf10NPUUcYzNrU9
*acc genotypes* 46
*peaks no back* 7
*removed edges* NA
*prop rem edges* NA

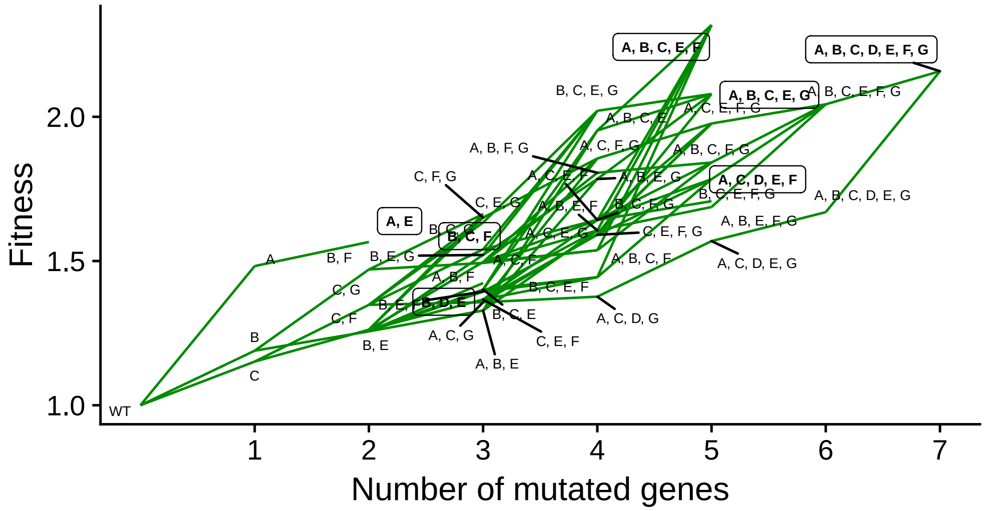


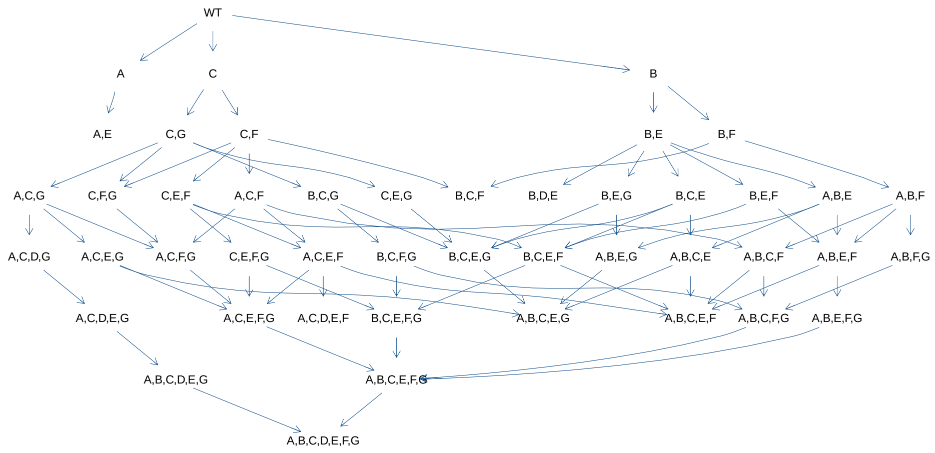

| | |
|---|---|
| *ID* | 5J10fJoXgI3k5FMVe |
| *acc genotypes* | 81 |
| *peaks no back* | 18 |
| *removed edges* | NA |
| *prop rem edges* | NA |

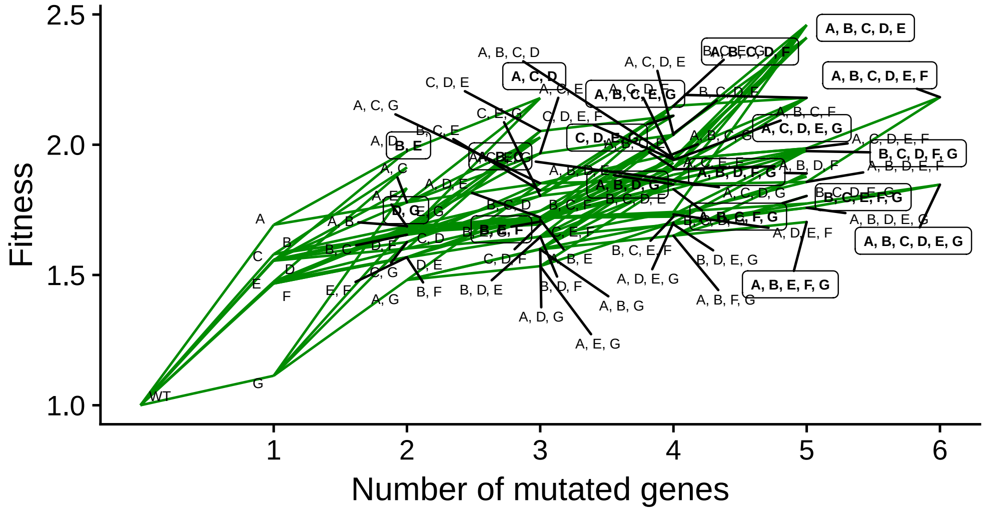


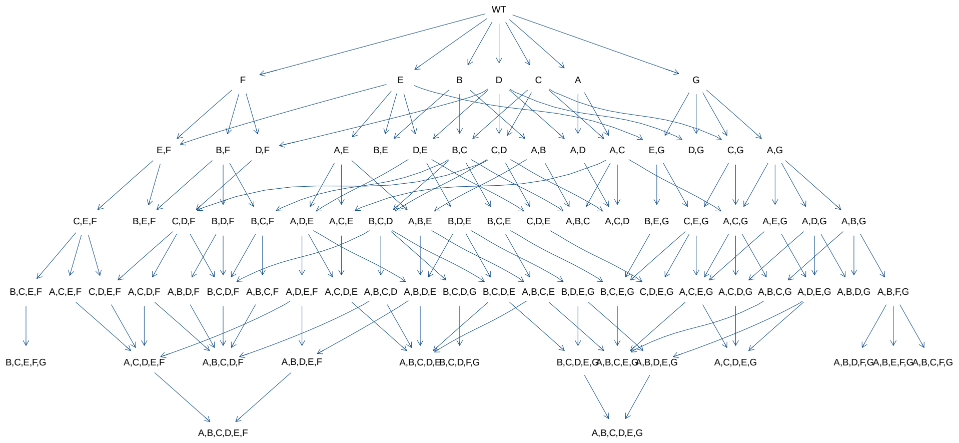

| | |
|---|---|
| *ID* | 5J6VWoMy7ROl8aOu |
| *acc genotypes* | 72 |
| *peaks no back* | 23 |
| *removed edges* | NA |
| *prop rem edges* | NA |

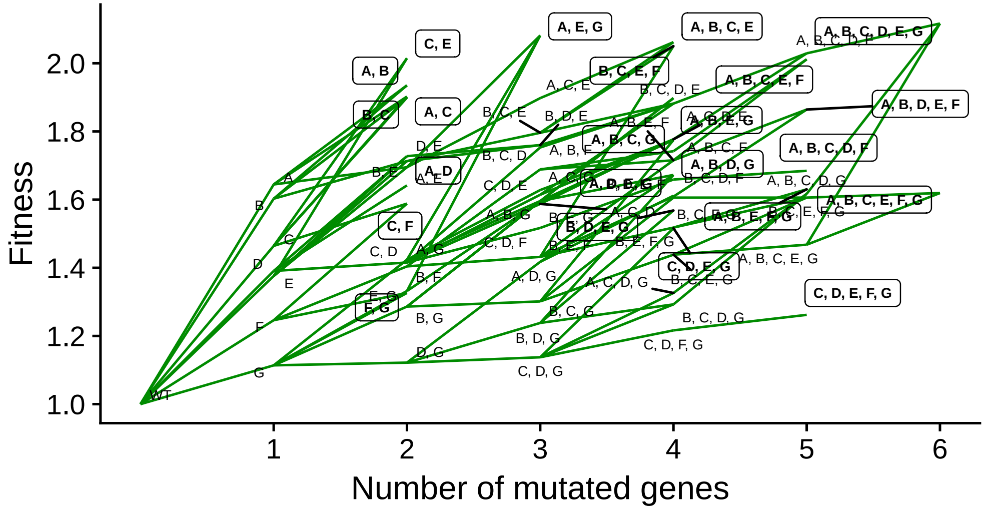


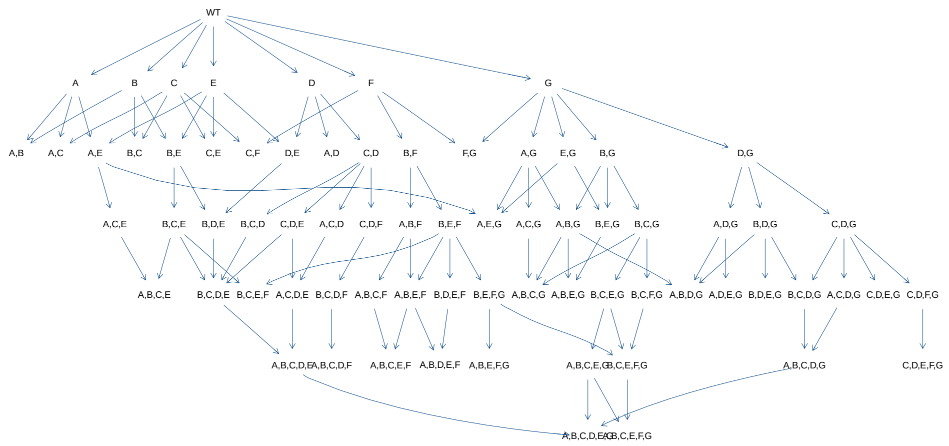

| | |
|---|---|
| *ID* | 5MozgxpY4MNYCtSZ |
| *acc genotypes* | 80 |
| *peaks no back* | 28 |
| *removed edges* | NA |
| *prop rem edges* | NA |

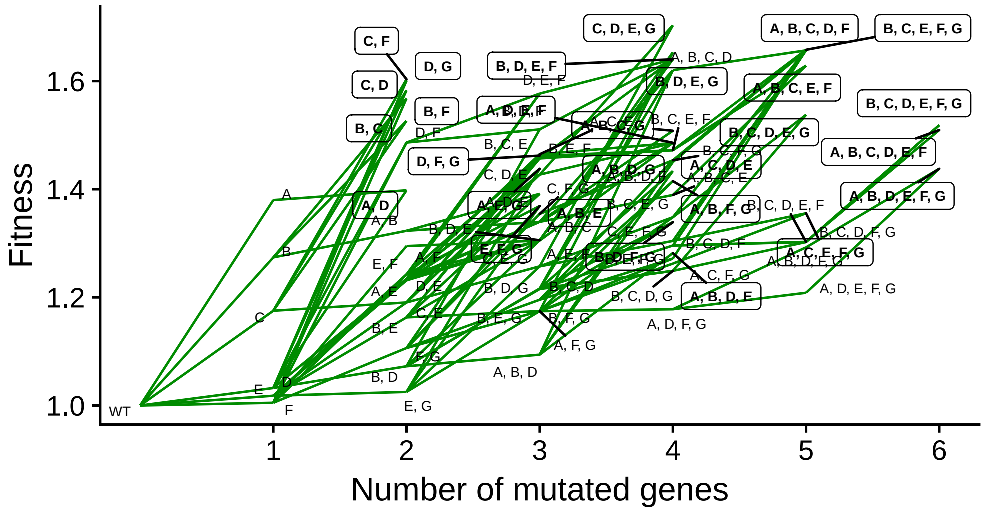


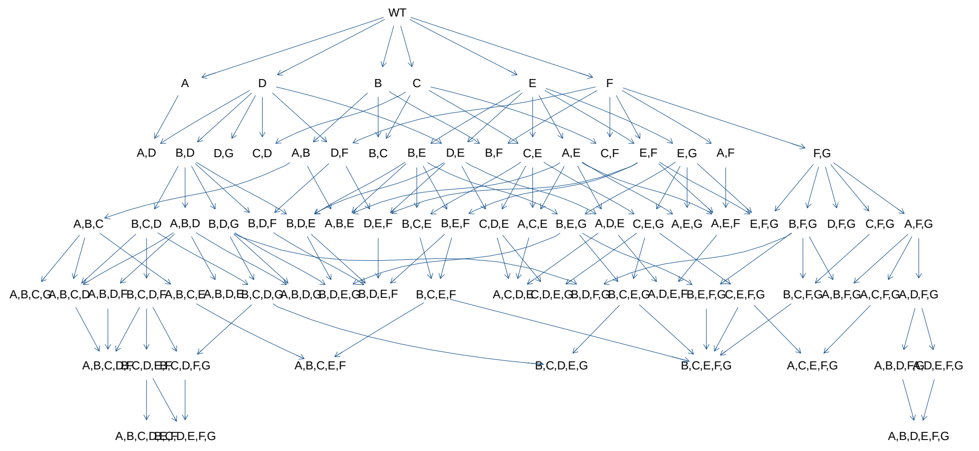

*ID* 6ARgJvilfN23NjA4
*acc genotypes* 81
*peaks no back* 22
*removed edges* NA
*prop rem edges* NA

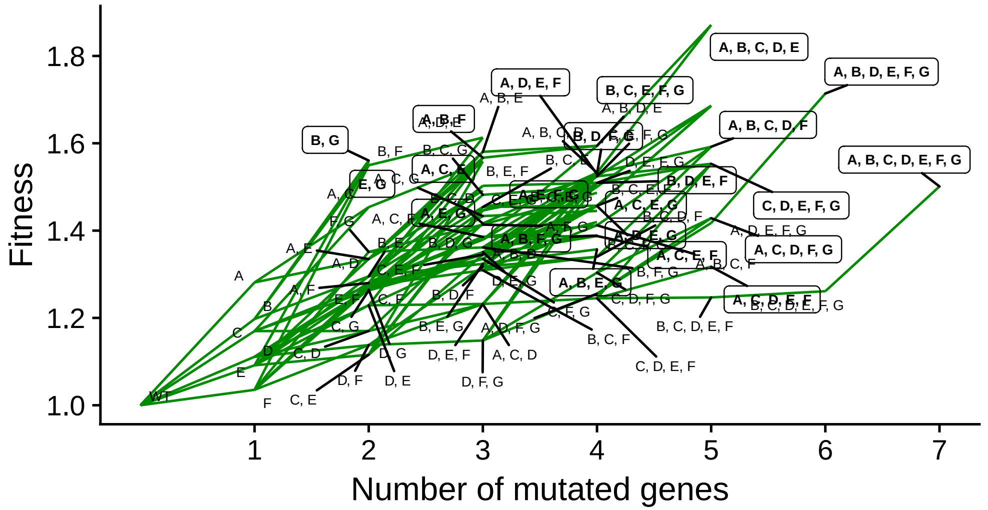


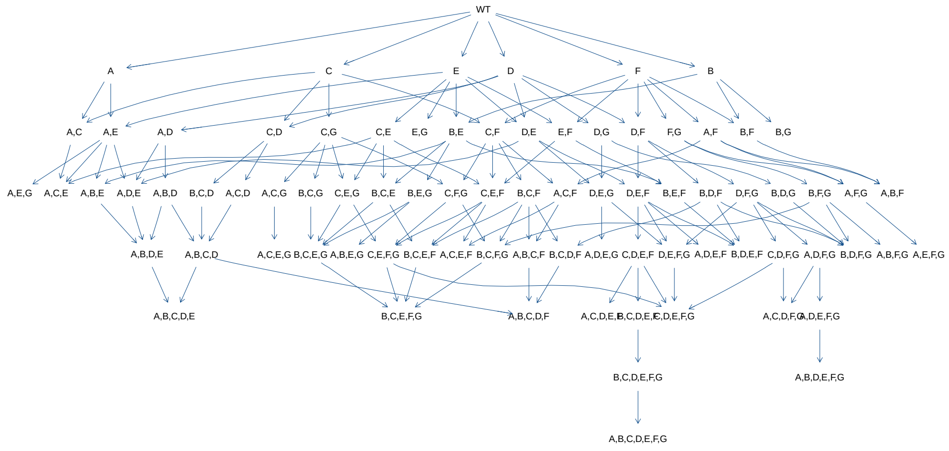

| | |
|---|---|
| *ID* | 6grkdzP10lm6kQ42g |
| *acc genotypes* | 81 |
| *peaks no back* | 12 |
| *removed edges* | NA |
| *prop rem edges* | NA |

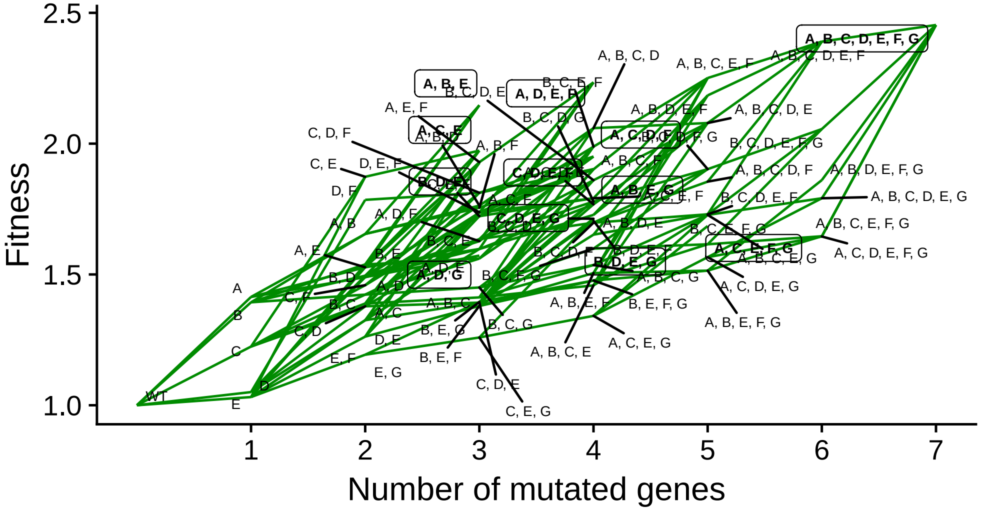


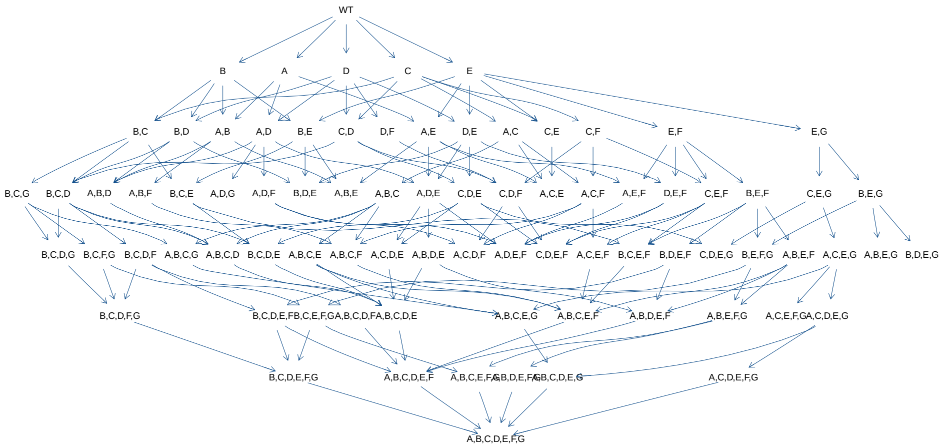

| | |
|---|---|
| *ID* | 75PXrtvV107d4XbmV |
| *acc genotypes* | 70 |
| *peaks no back* | 19 |
| *removed edges* | NA |
| *prop rem edges* | NA |

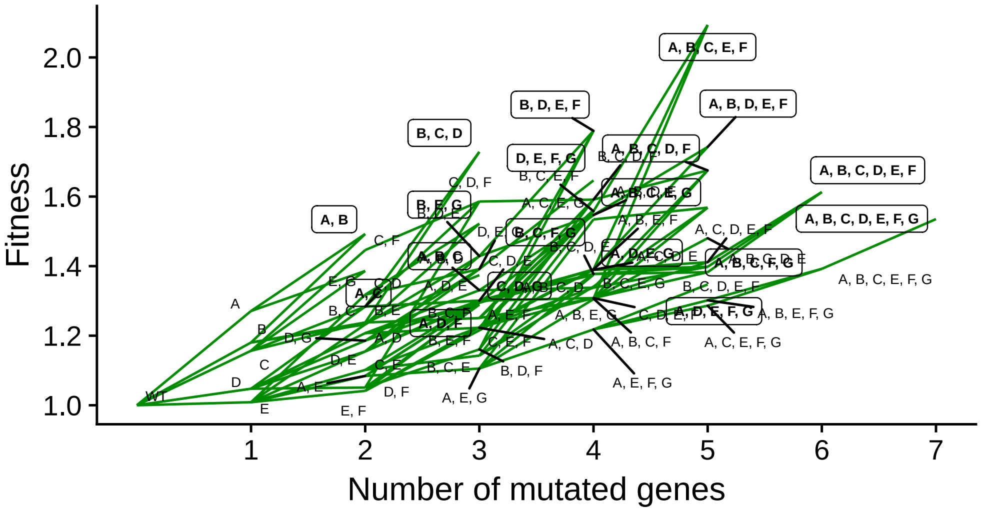


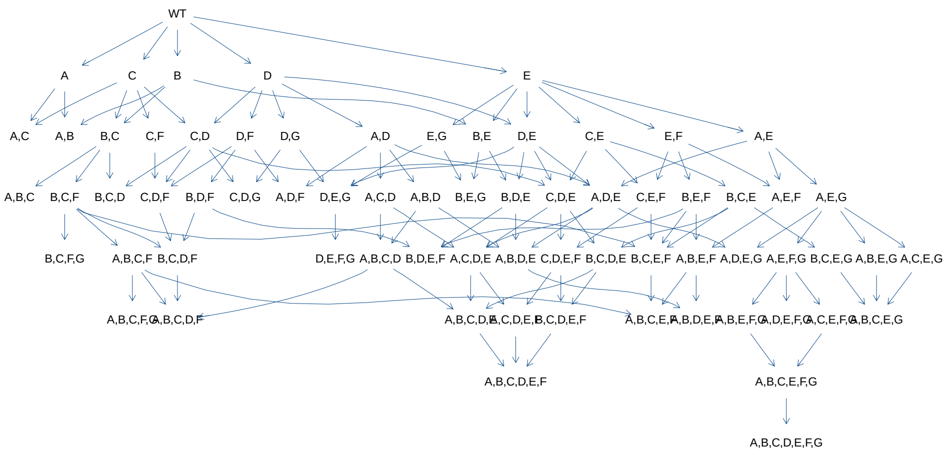

| | |
|---|---|
| *ID* | 84eRNCZIduWDrCW4 |
| *acc genotypes* | 39 |
| *peaks no back* | 15 |
| *removed edges* | NA |
| *prop rem edges* | NA |

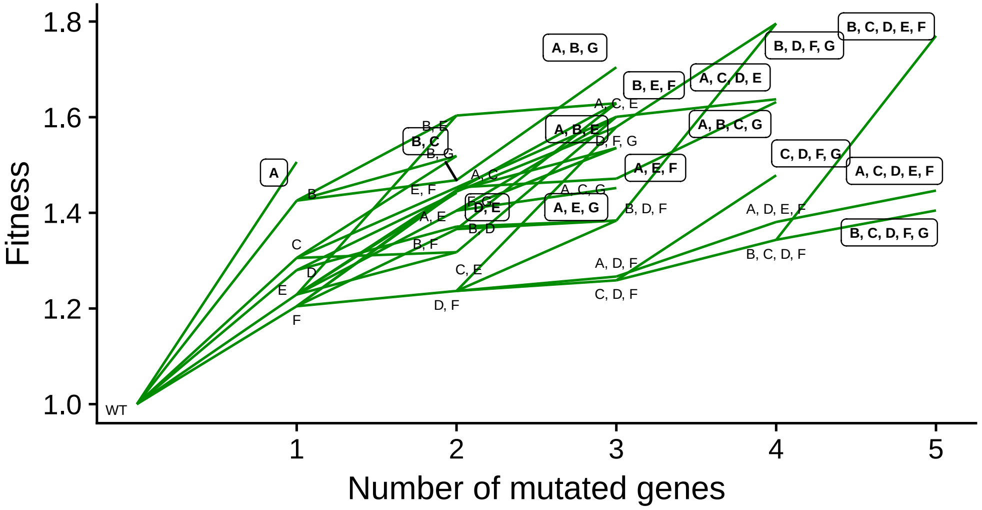


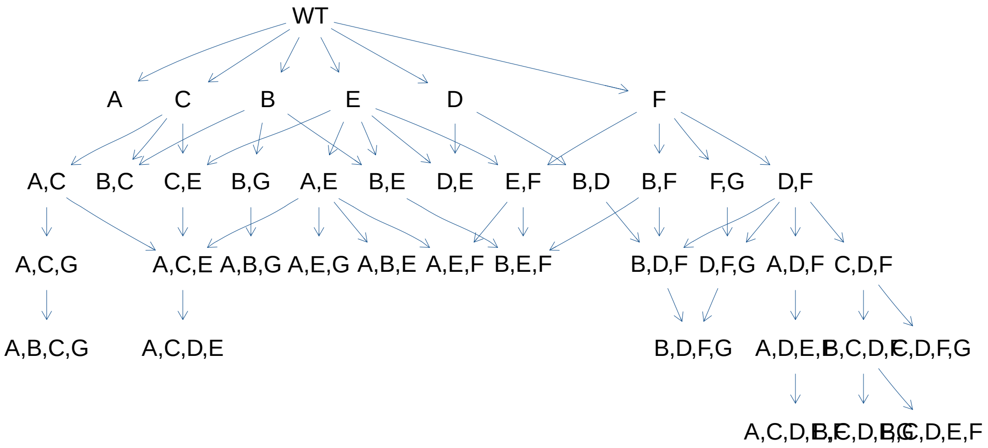

| | |
|---|---|
| *ID* | 84uIeheQaYaVNux8 |
| *acc genotypes* | 18 |
| *peaks no back* | 6 |
| *removed edges* | NA |
| *prop rem edges* | NA |

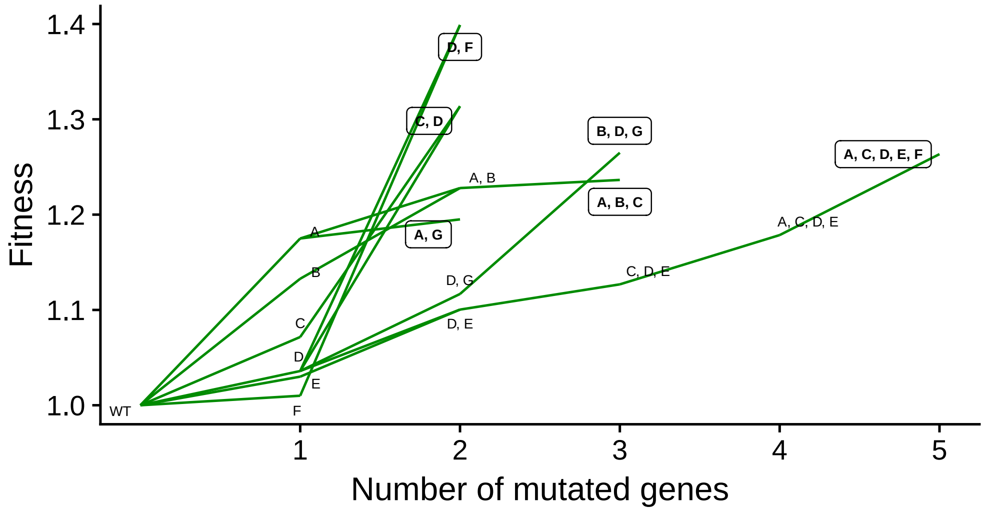


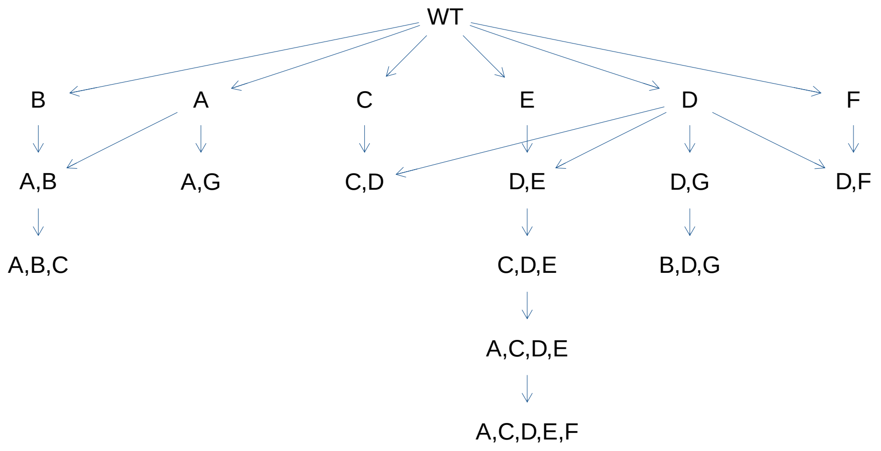

| | |
|---|---|
| *ID* | 8a9vNJyhKyFzgtGl |
| *acc genotypes* | 29 |
| *peaks no back* | 6 |
| *removed edges* | NA |
| *prop rem edges* | NA |

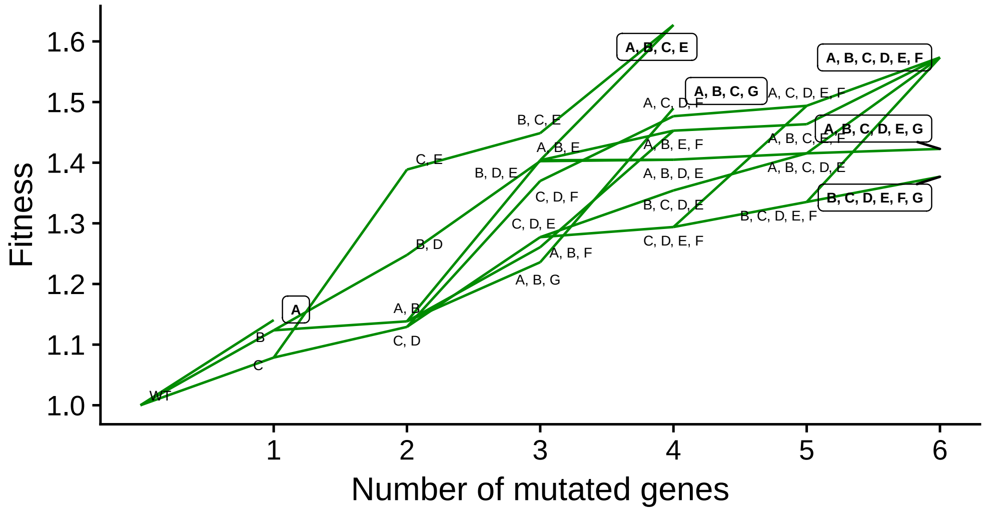


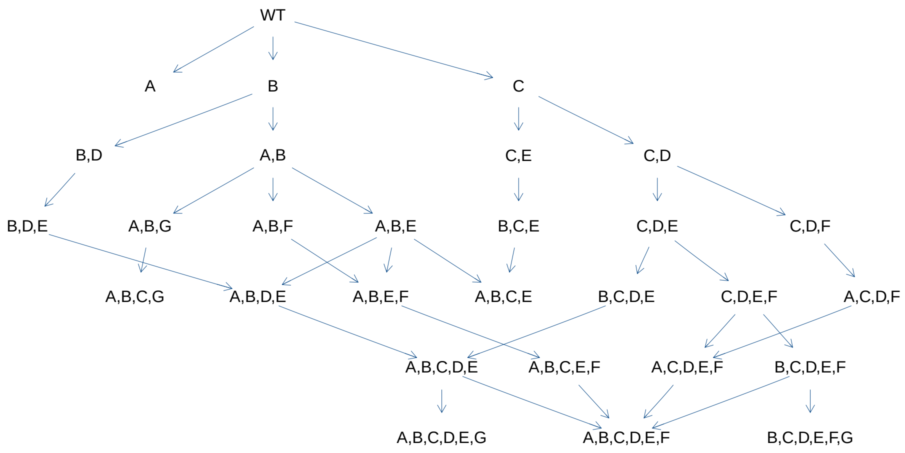

| | |
|---|---|
| *ID* | 8UysAOd1WpNJ6JEA |
| *acc genotypes* | 67 |
| *peaks no back* | 15 |
| *removed edges* | NA |
| *prop rem edges* | NA |

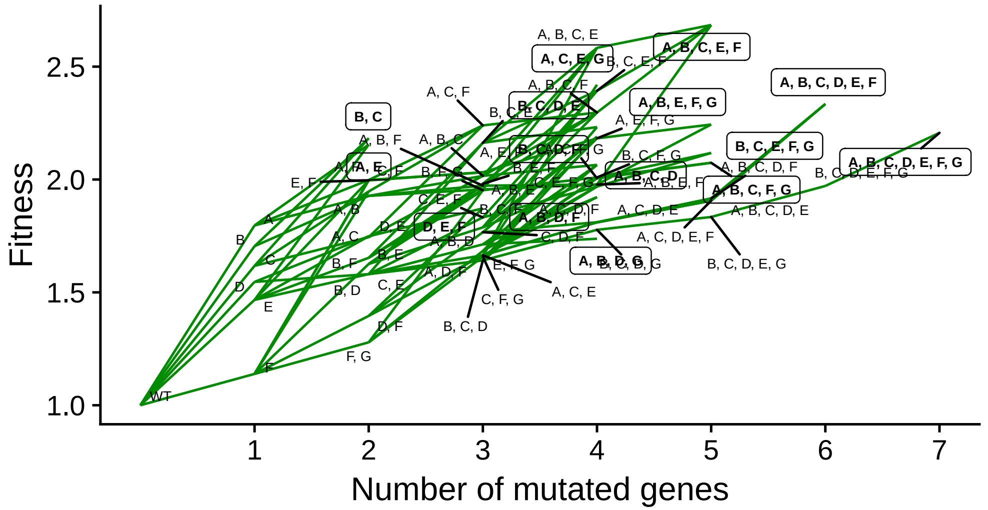


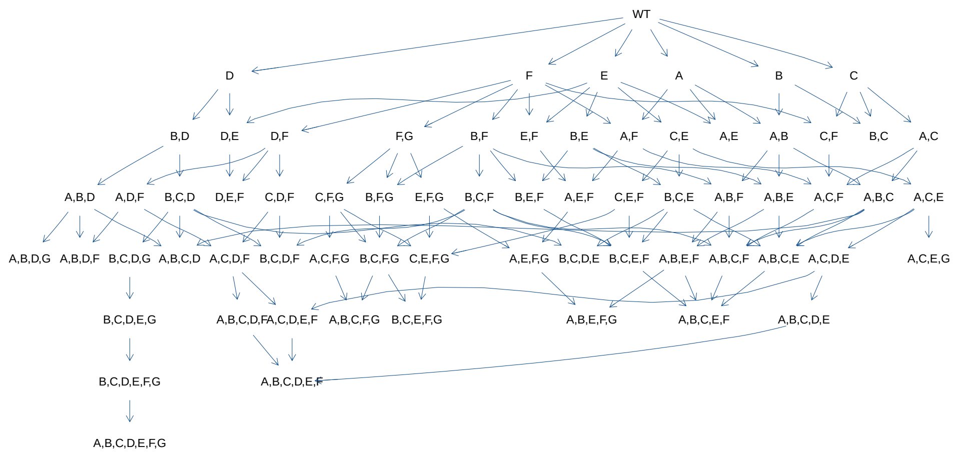

| | |
|---|---|
| *ID* | 8yQFlKWNFgRltIOW |
| *acc genotypes* | 40 |
| *peaks no back* | 18 |
| *removed edges* | NA |
| *prop rem edges* | NA |

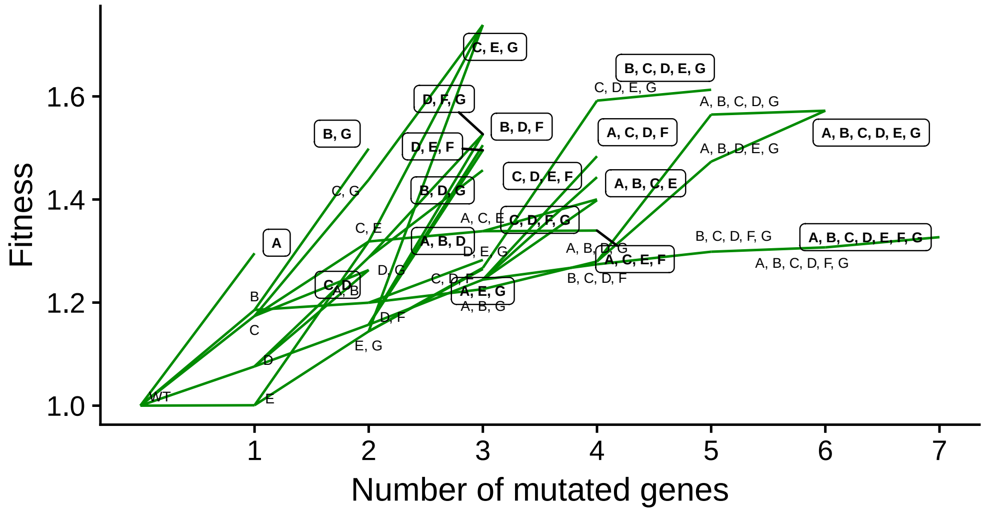


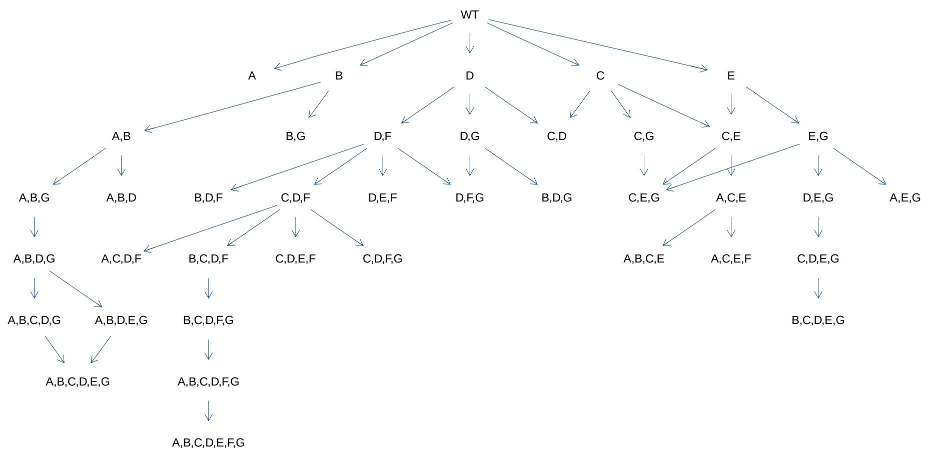

| | |
|---|---|
| *ID* | 98Ltpa3lXTnc4zbC |
| *acc genotypes* | 29 |
| *peaks no back* | 12 |
| *removed edges* | NA |
| *prop rem edges* | NA |

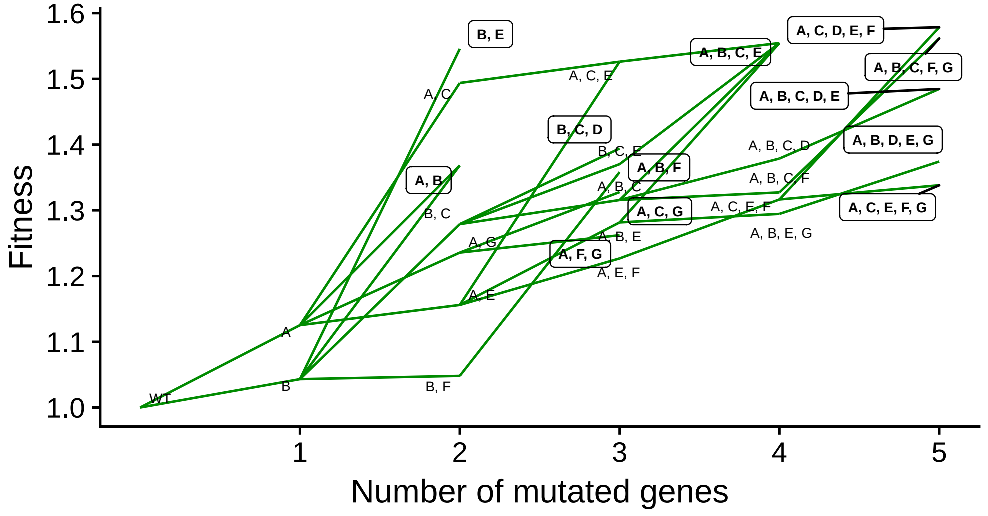


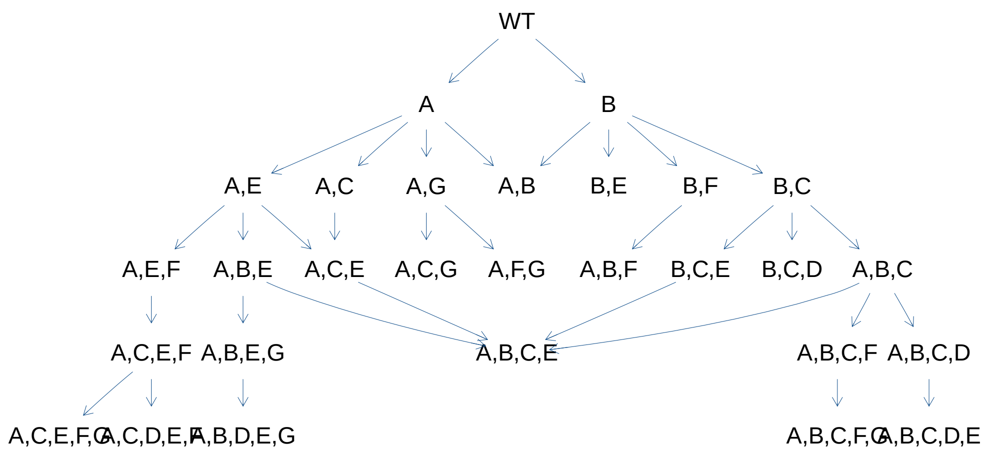

| | |
|---|---|
| *ID* | 9KrYrcUaMBo5BoOd |
| *acc genotypes* | 89 |
| *peaks no back* | 13 |
| *removed edges* | NA |
| *prop rem edges* | NA |

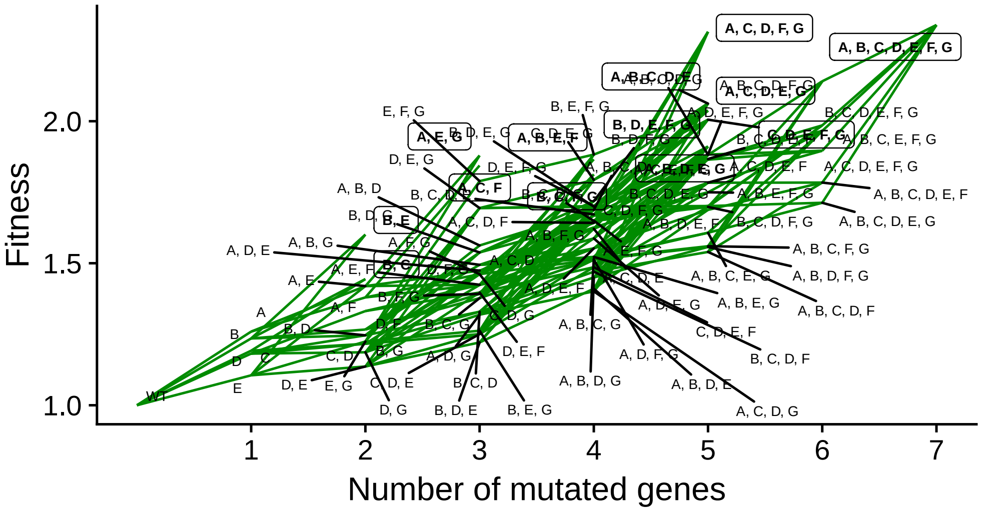


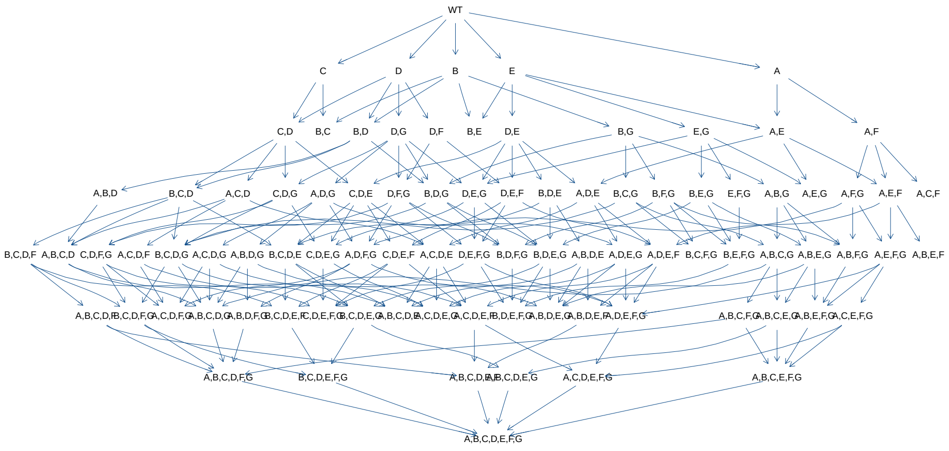

| | |
|---|---|
| *ID* | 9MBO3qg10BvSTzUqM |
| *acc genotypes* | 52 |
| *peaks no back* | 15 |
| *removed edges* | NA |
| *prop rem edges* | NA |

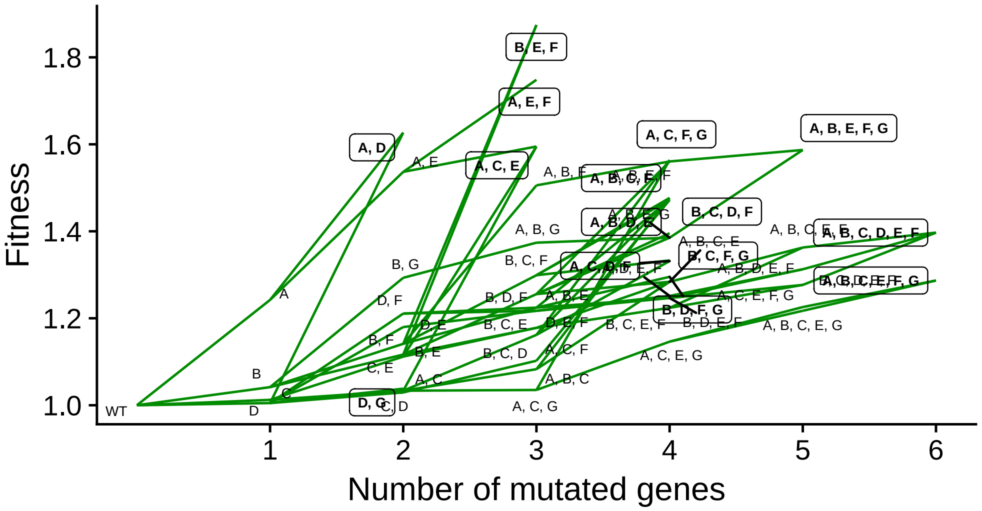


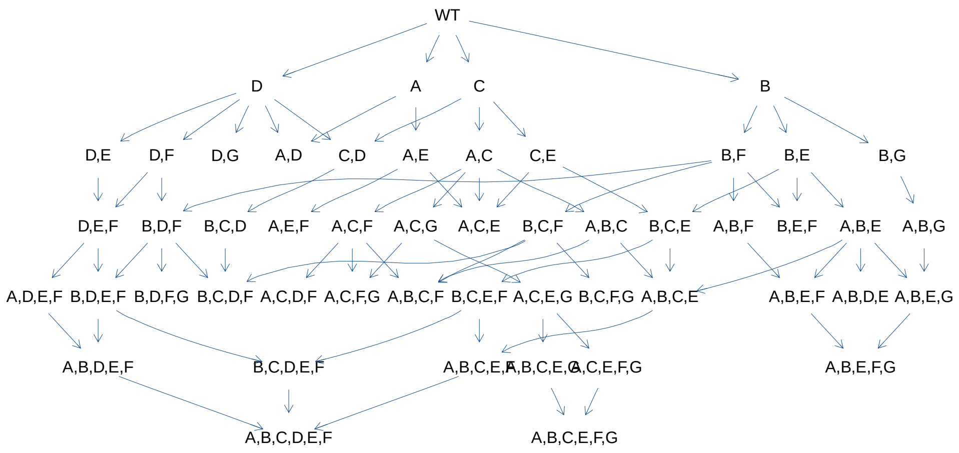

| | |
|---|---|
| *ID* | 9rebvjtcu7f5fc6O |
| *acc genotypes* | 38 |
| *peaks no back* | 12 |
| *removed edges* | NA |
| *prop rem edges* | NA |

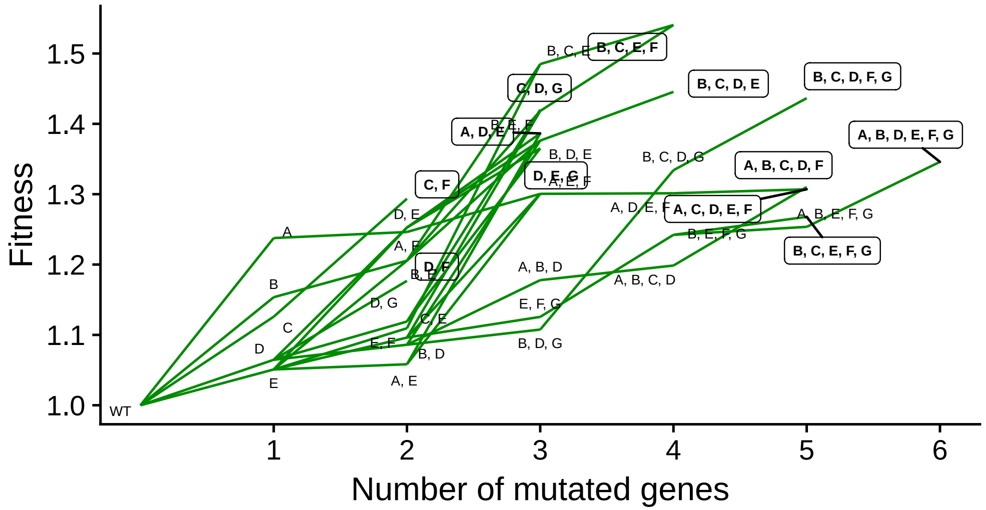


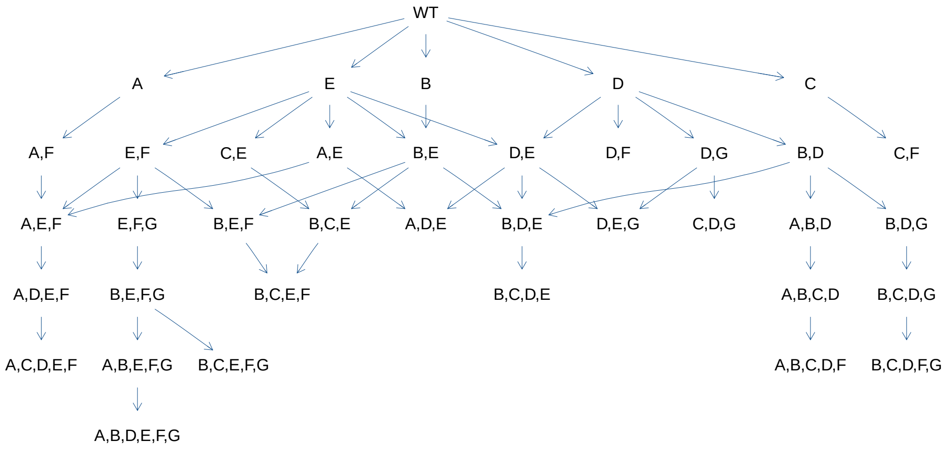

| | |
|---|---|
| *ID* | A3LrldXjUIno4iNV |
| *acc genotypes* | 77 |
| *peaks no back* | 19 |
| *removed edges* | NA |
| *prop rem edges* | NA |

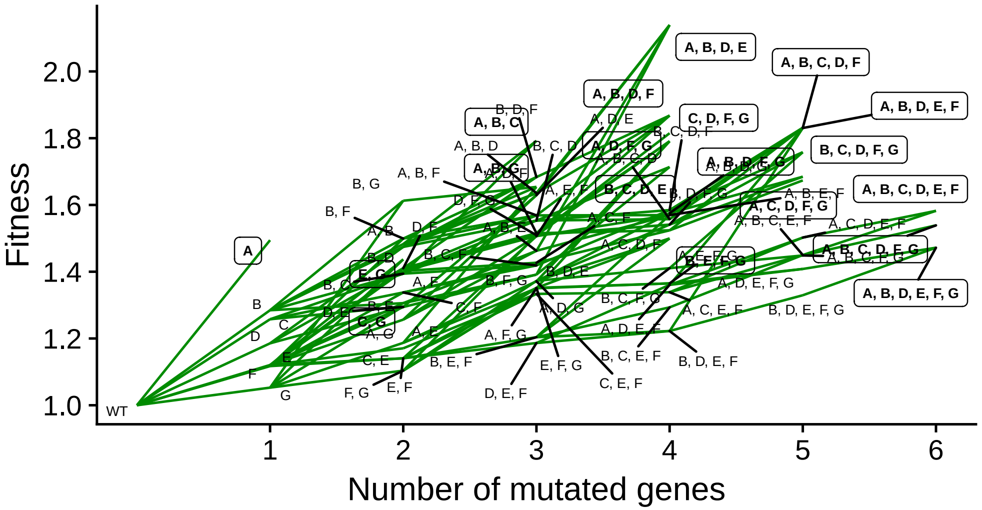


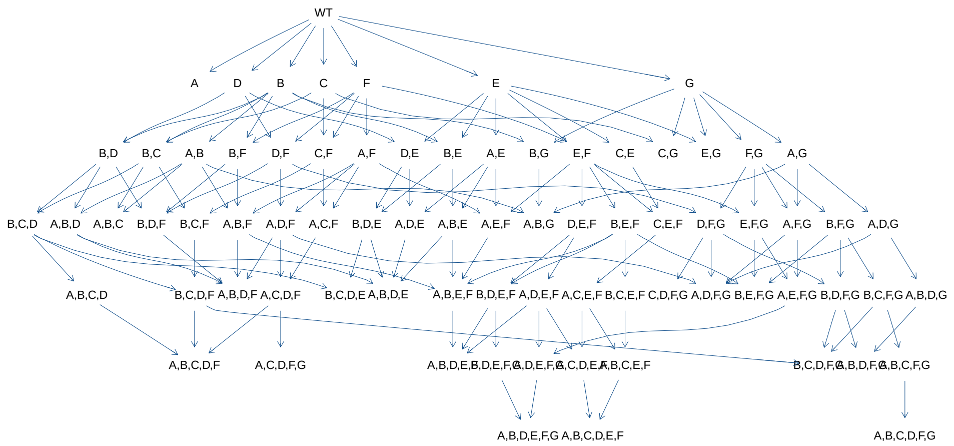

| | |
|---|---|
| *ID* | aA3KFQazfmWZiKVN |
| *acc genotypes* | 54 |
| *peaks no back* | 20 |
| *removed edges* | NA |
| *prop rem edges* | NA |

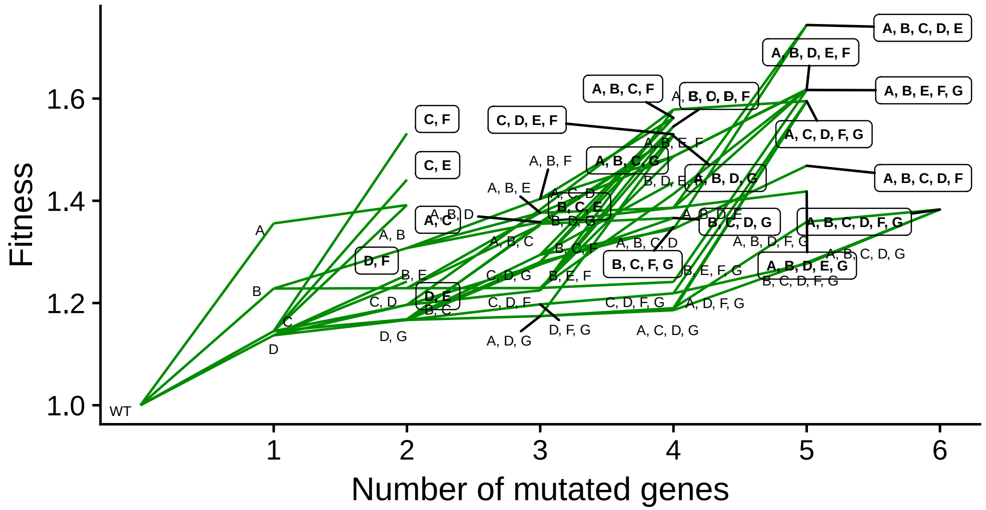


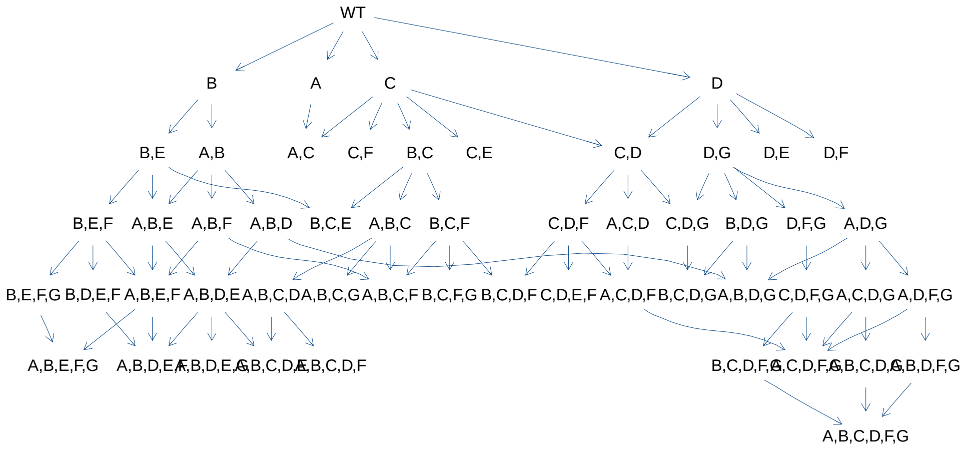

| | |
|---|---|
| *ID* | ACGaLODBO3L7xXPK |
| *acc genotypes* | 70 |
| *peaks no back* | 19 |
| *removed edges* | NA |
| *prop rem edges* | NA |

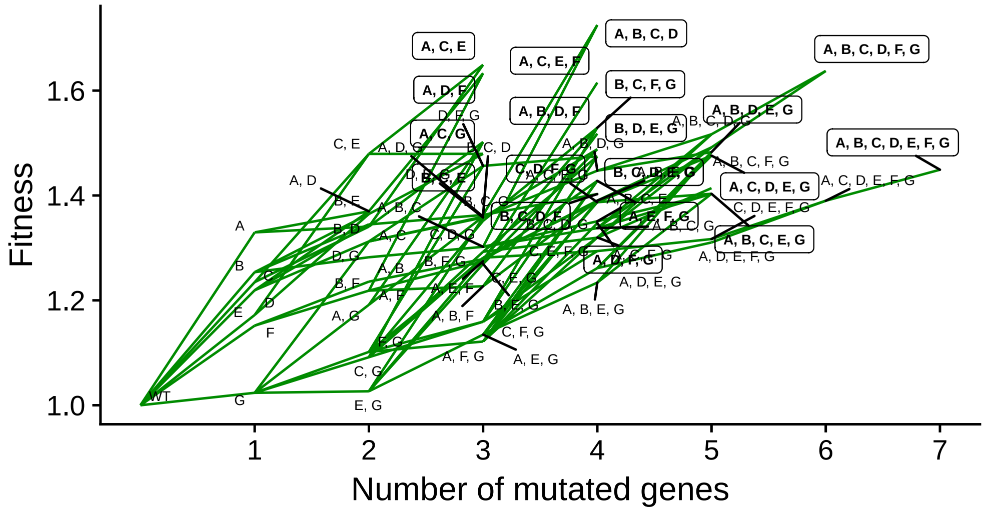


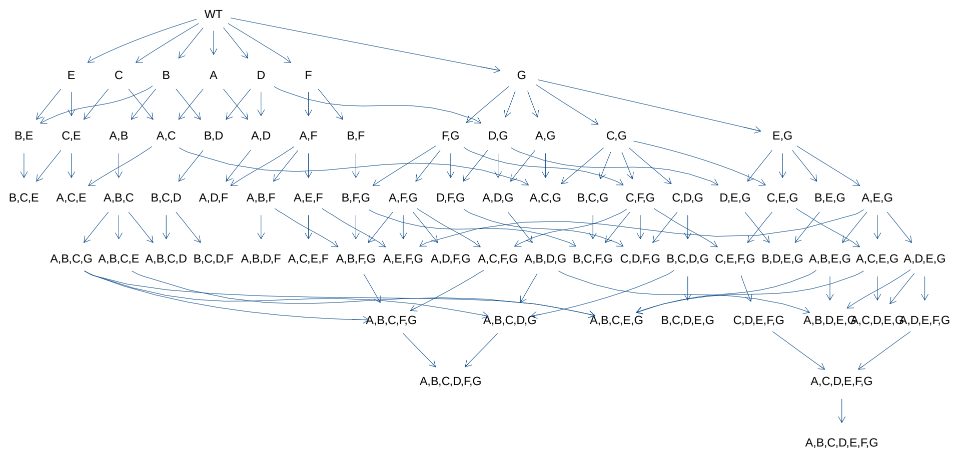

| | |
|---|---|
| *ID* | AdlPqP6mmr9aElyf |
| *acc genotypes* | 19 |
| *peaks no back* | 8 |
| *removed edges* | NA |
| *prop rem edges* | NA |

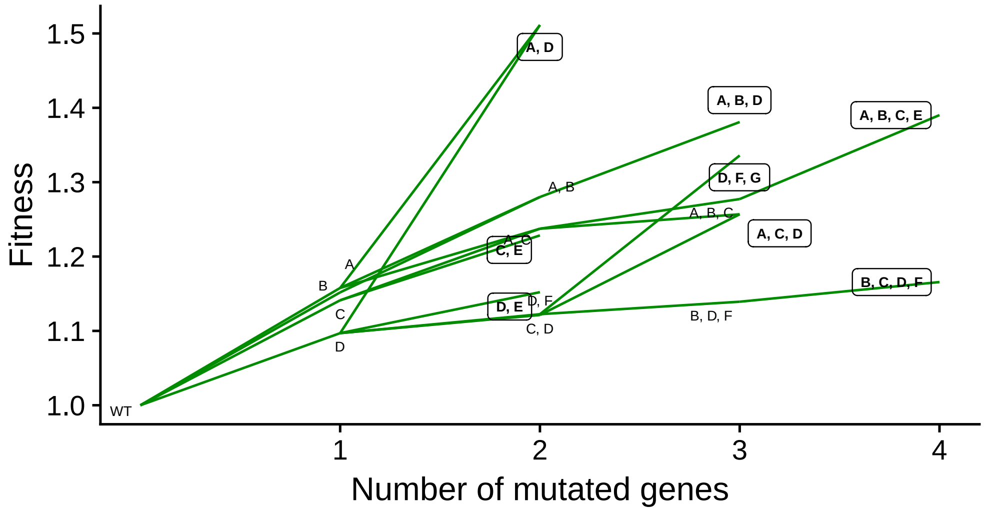


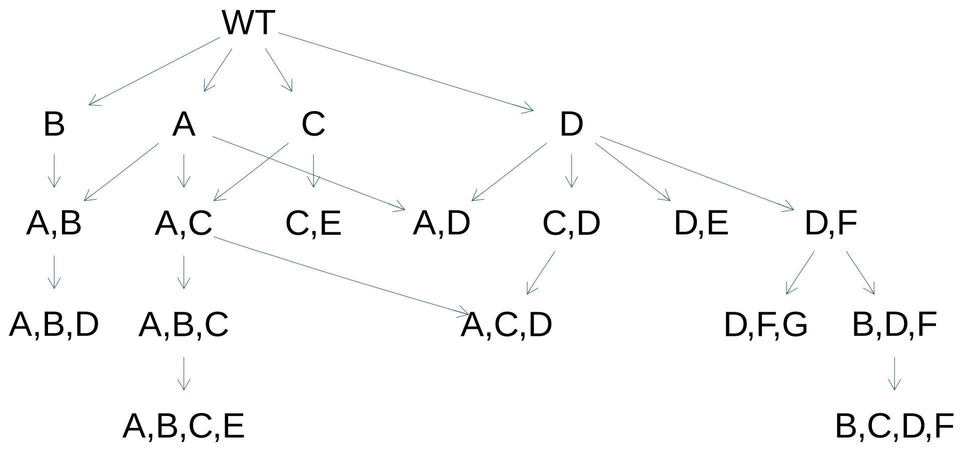

*ID* aeQvEQMAOMlDD5Ee
*acc genotypes* 49
*peaks no back* 20
*removed edges* NA
*prop rem edges* NA

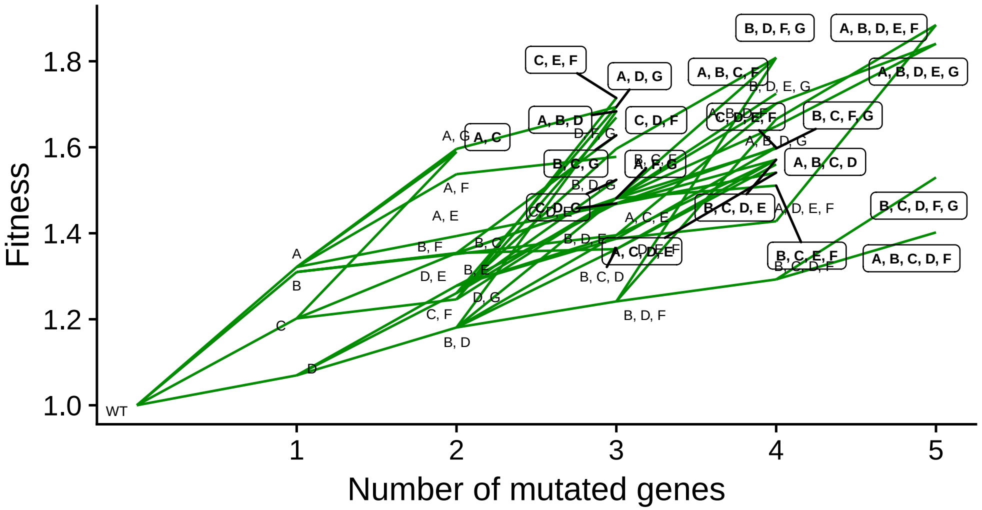


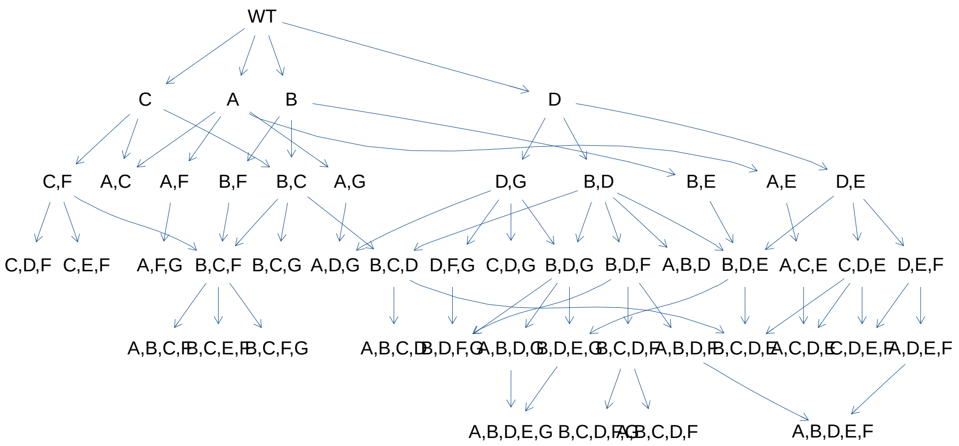

| | |
|---|---|
| *ID* | AqnbiWCAFm6eogfg |
| *acc genotypes* | 17 |
| *peaks no back* | 6 |
| *removed edges* | NA |
| *prop rem edges* | NA |

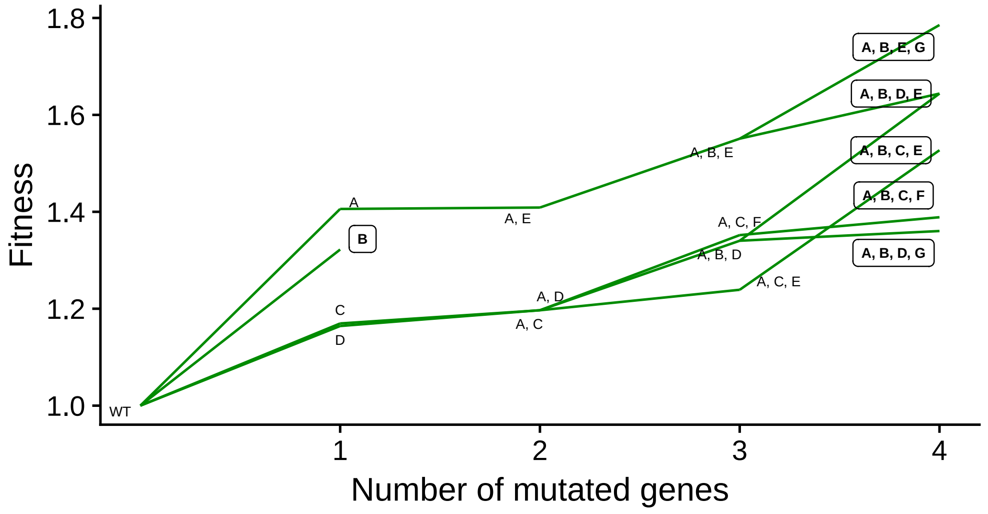


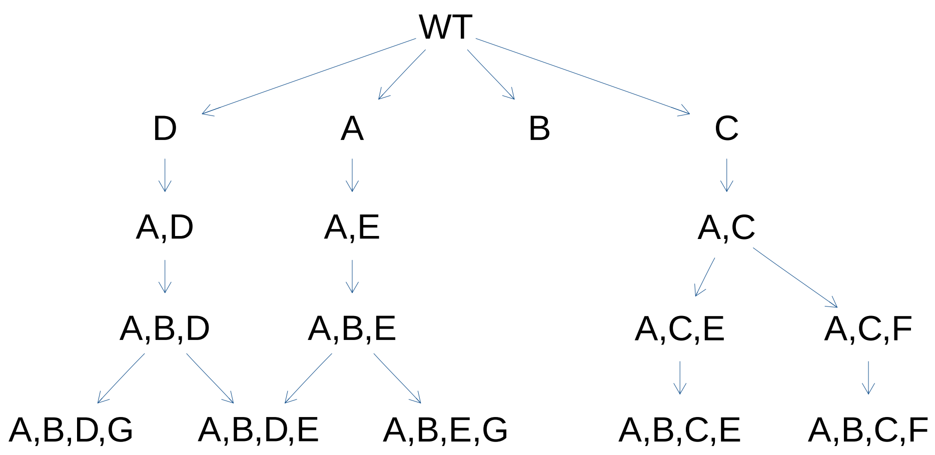

| | |
|---|---|
| *ID* | aVdtnwMxlBa9JGAI |
| *acc genotypes* | 48 |
| *peaks no back* | 14 |
| *removed edges* | NA |
| *prop rem edges* | NA |

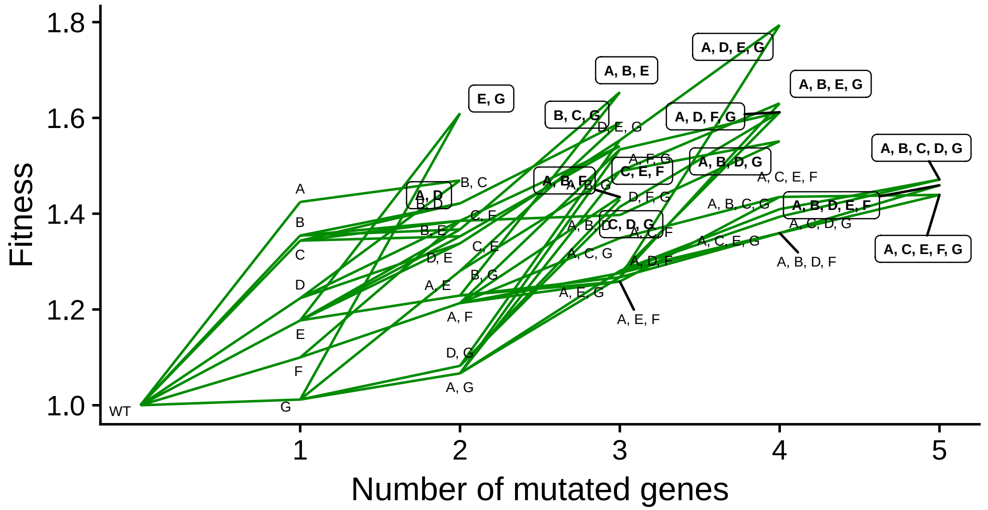


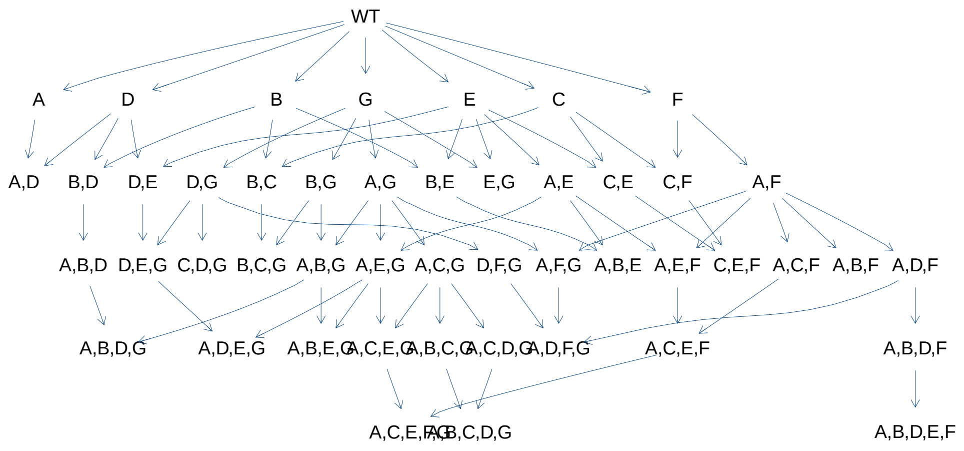

*ID* Avmue778lBrop2vO
*acc genotypes* 44
*peaks no back* 11
*removed edges* NA
*prop rem edges* NA

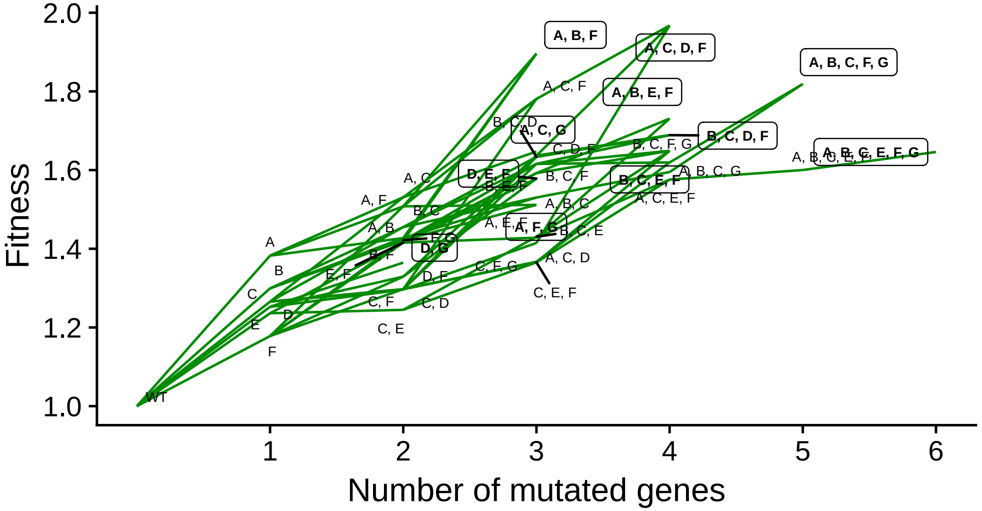


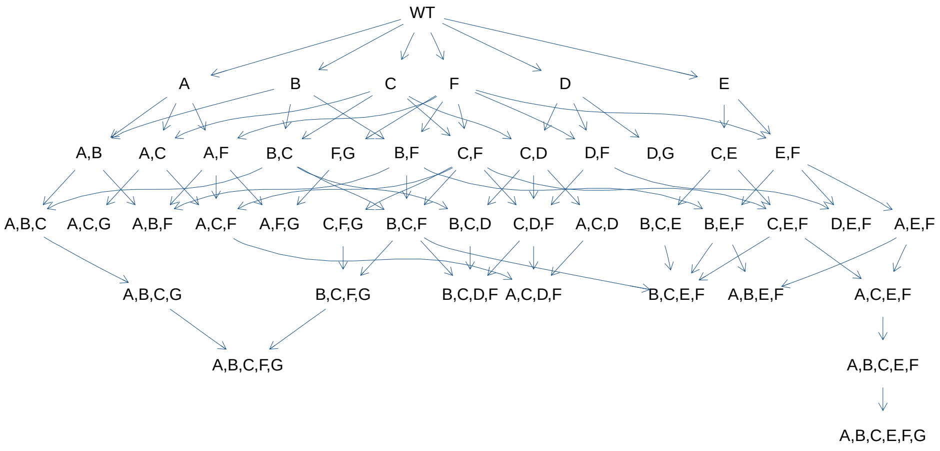

| | |
|---|---|
| *ID* | AyannCS2nwfQ2TDc |
| *acc genotypes* | 25 |
| *peaks no back* | 9 |
| *removed edges* | NA |
| *prop rem edges* | NA |

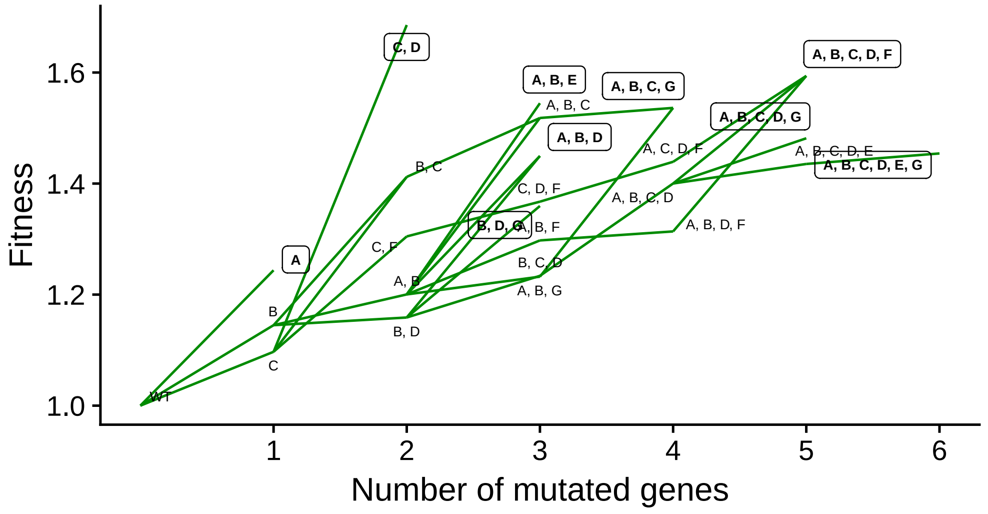


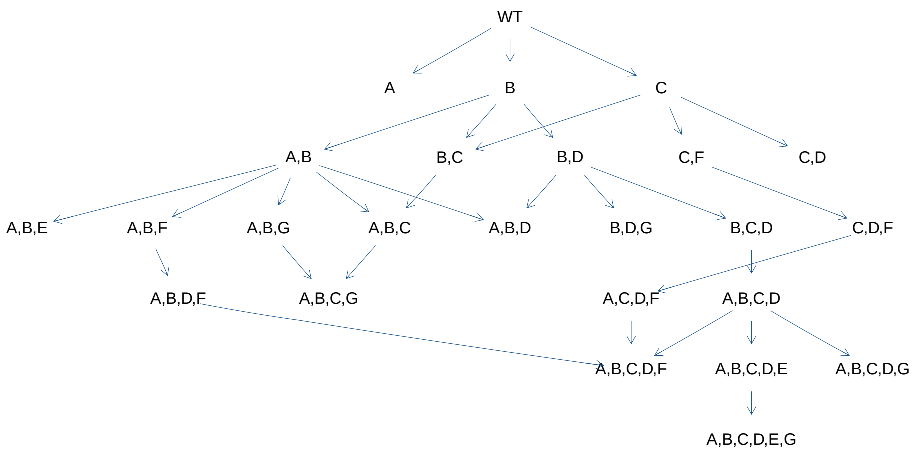

| | |
|---|---|
| *ID* | aZQI57105TWpukXl1 |
| *acc genotypes* | 22 |
| *peaks no back* | 10 |
| *removed edges* | NA |
| *prop rem edges* | NA |

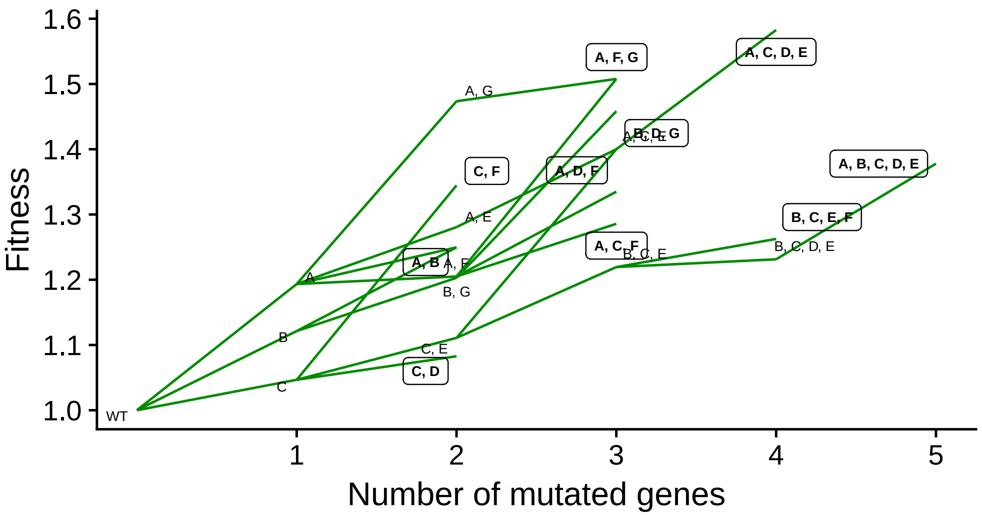


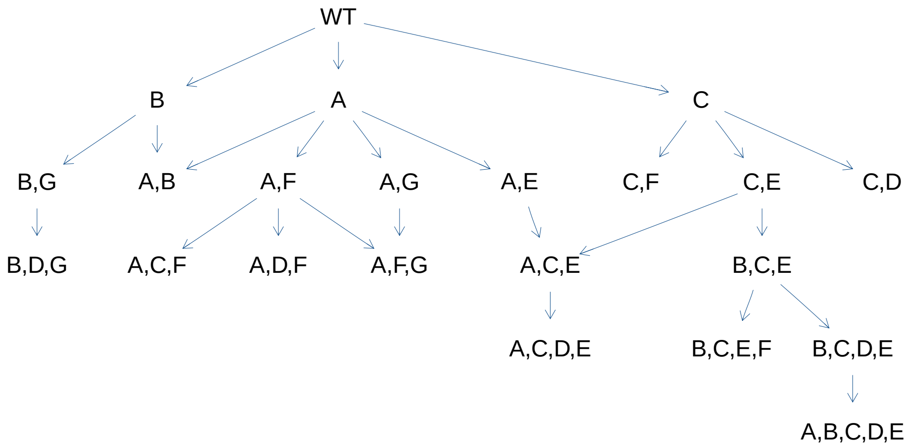

| | |
|---|---|
| *ID* | BD3duuYOmGuZZNVr |
| *acc genotypes* | 14 |
| *peaks no back* | 6 |
| *removed edges* | NA |
| *prop rem edges* | NA |

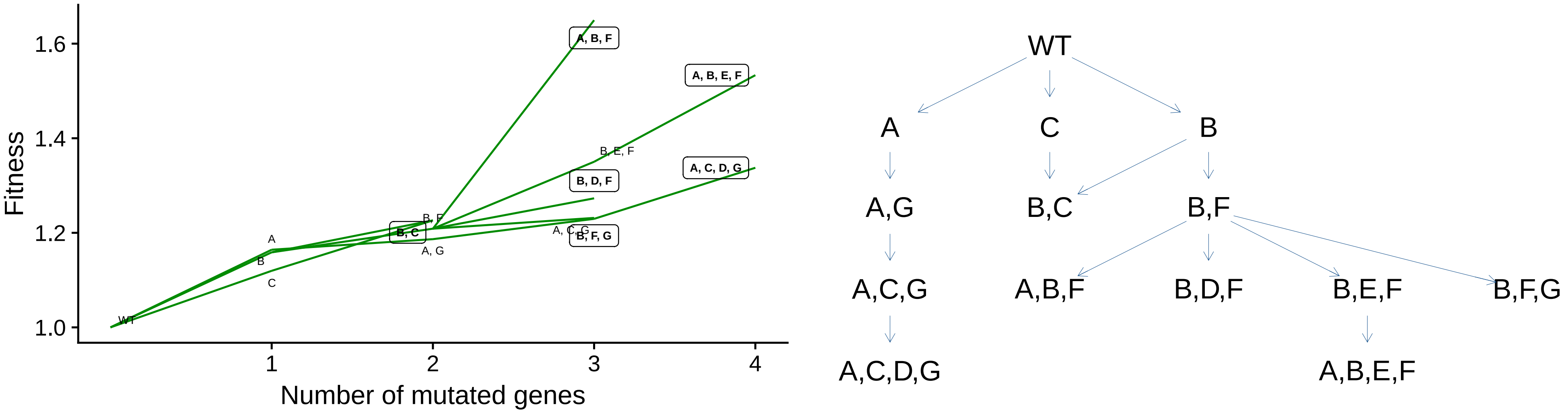

| | |
|---|---|
| *ID* | BfGQkzq8zBGWdITj |
| *acc genotypes* | 18 |
| *peaks no back* | 7 |
| *removed edges* | NA |
| *prop rem edges* | NA |

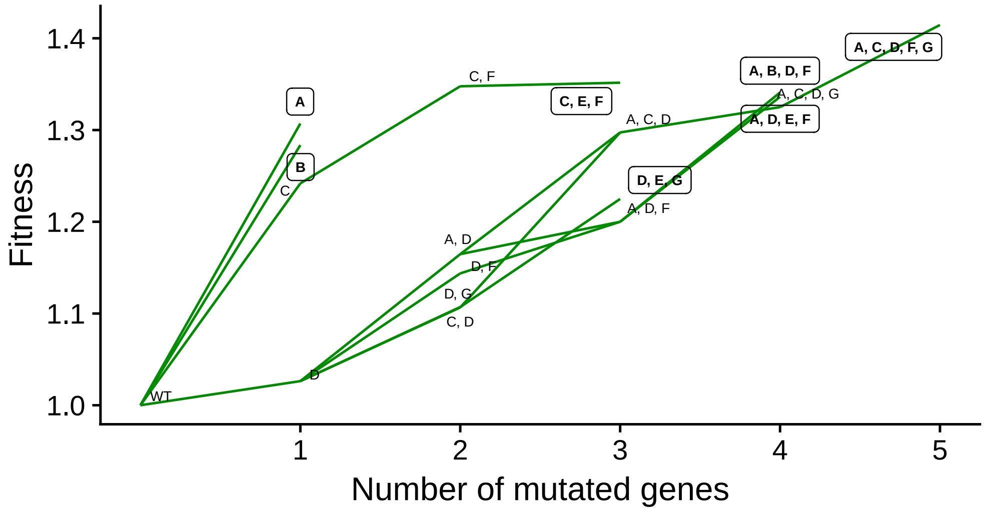


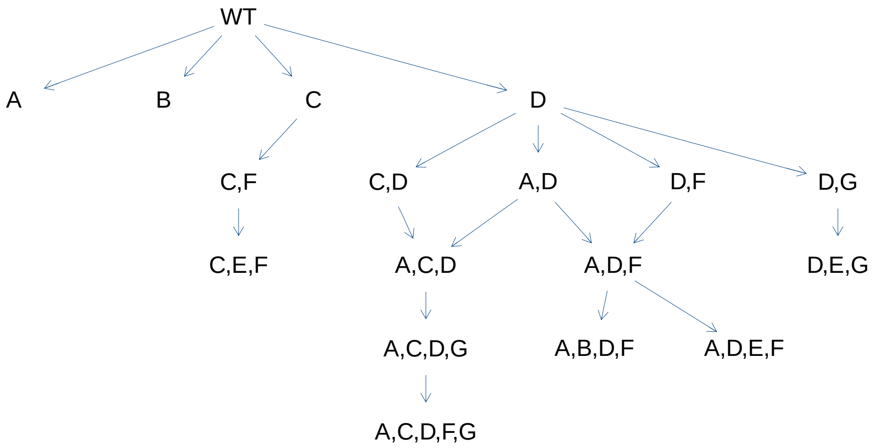

| | |
|---|---|
| *ID* | bIGKwHhjkye10mnvq |
| *acc genotypes* | 85 |
| *peaks no back* | 10 |
| *removed edges* | NA |
| *prop rem edges* | NA |

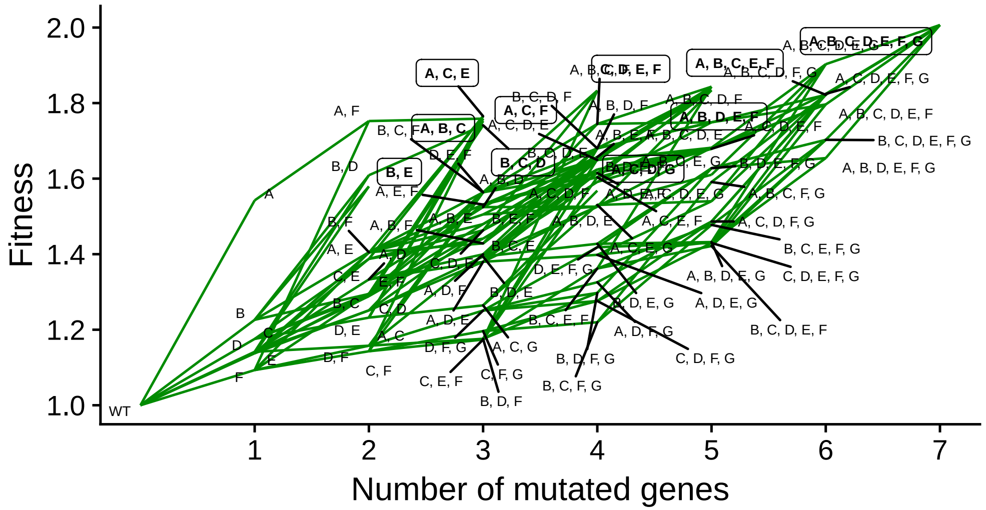


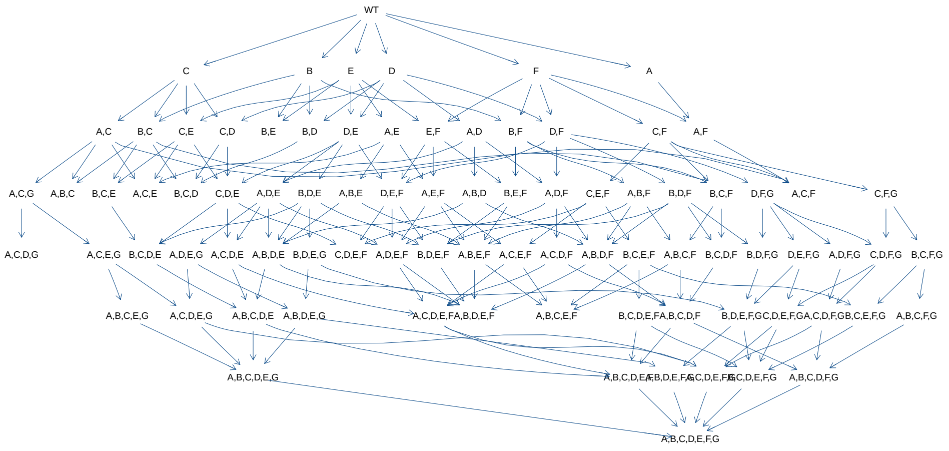

*ID* bpM86wHSqemVZQ1G

*acc genotypes* 57

*peaks no back* 18

*removed edges* NA

*prop rem edges* NA

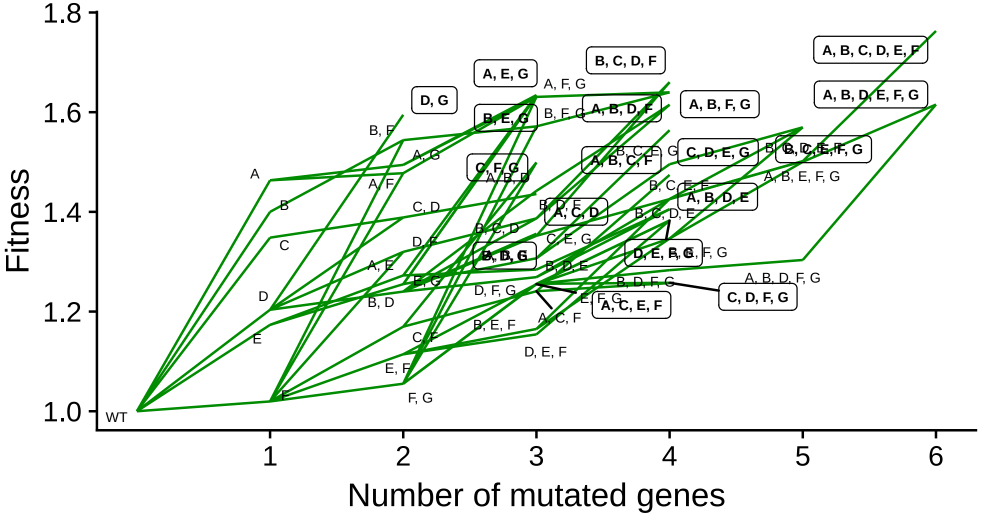


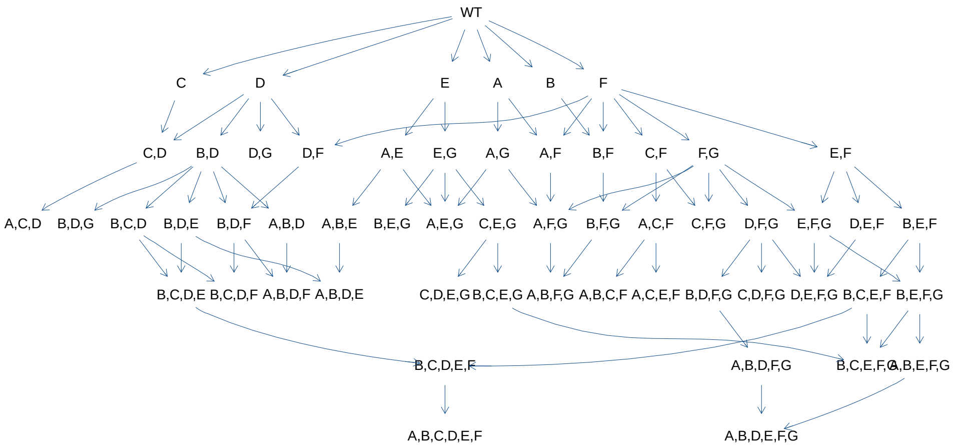

| | |
|---|---|
| *ID* | BTJyOApuNPFWEPsU |
| *acc genotypes* | 17 |
| *peaks no back* | 11 |
| *removed edges* | NA |
| *prop rem edges* | NA |

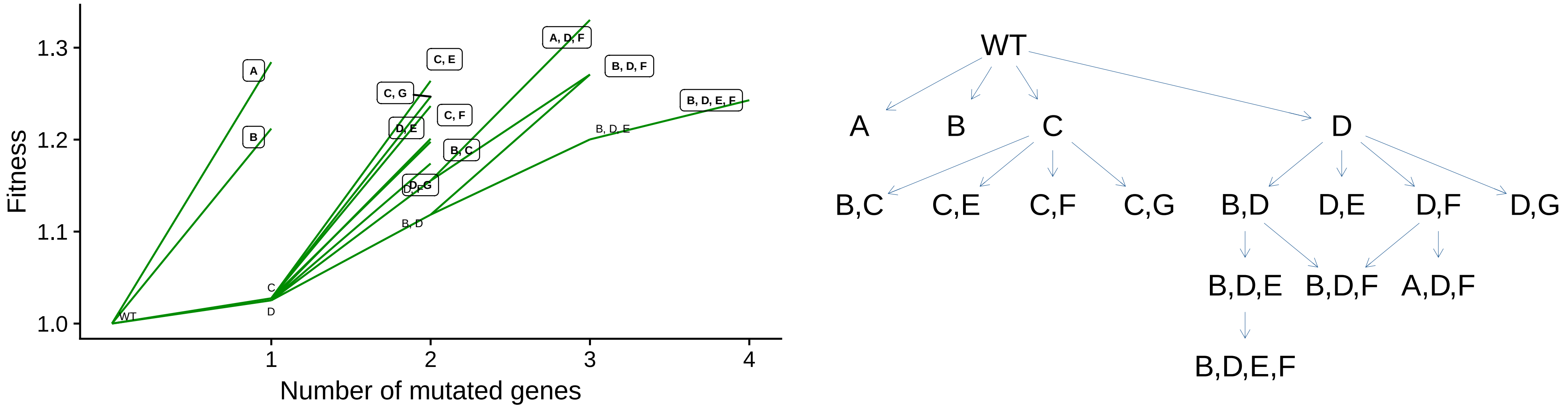

| | |
|---|---|
| *ID* | BxBVJz10wDpOmersD |
| *acc genotypes* | 90 |
| *peaks no back* | 21 |
| *removed edges* | NA |
| *prop rem edges* | NA |

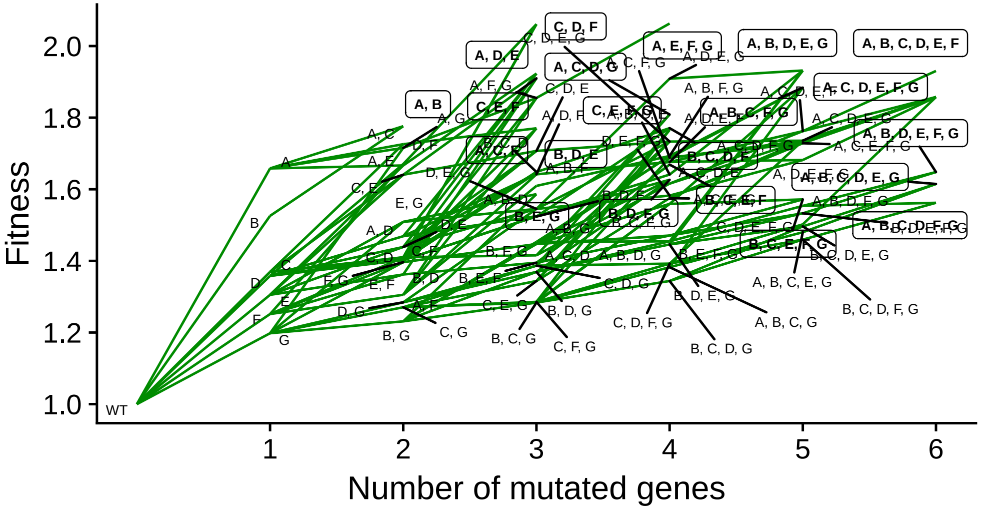


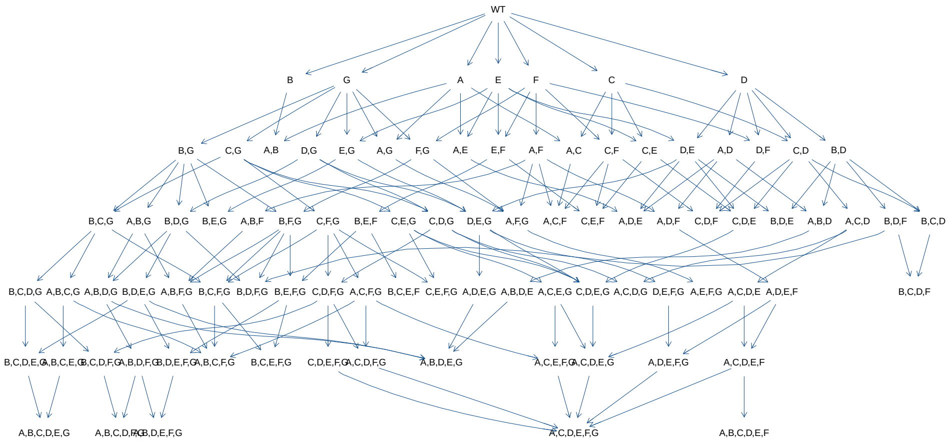

*ID* c42Quu4ADM6baevo
*acc genotypes* 60
*peaks no back* 14
*removed edges* NA
*prop rem edges* NA

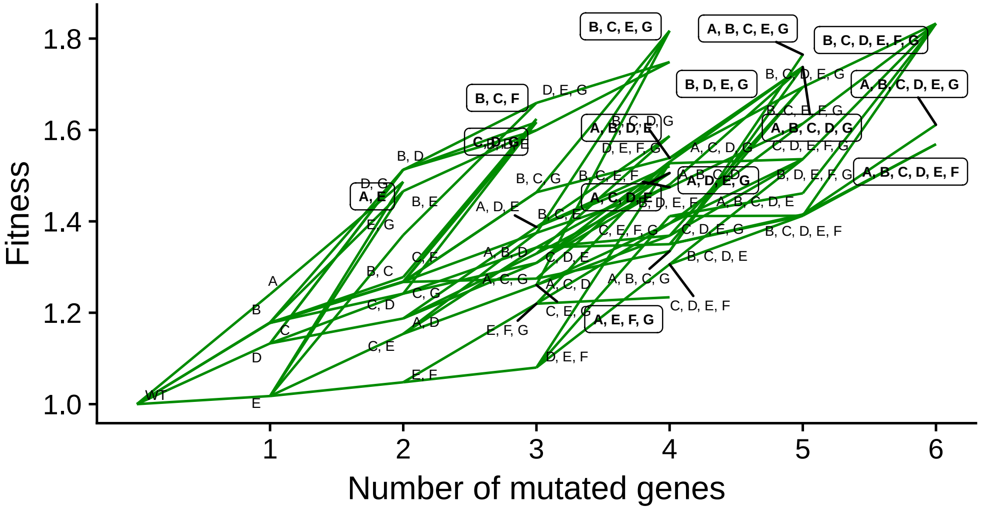


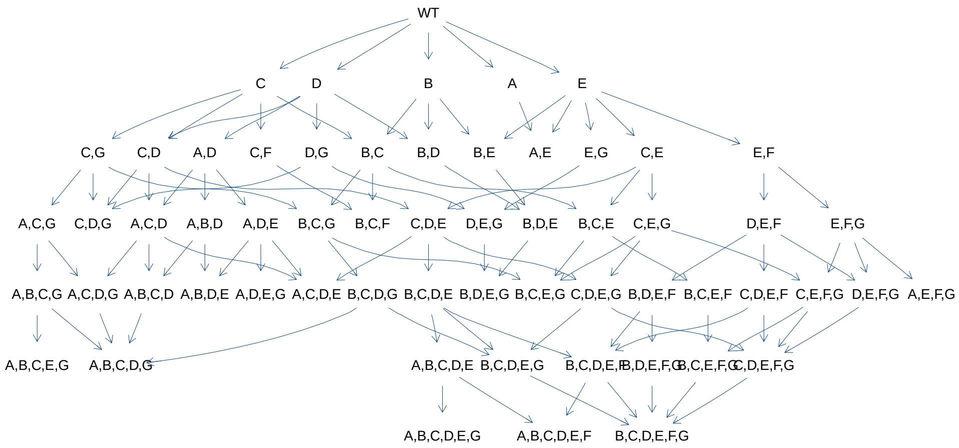

| | |
|---|---|
| *ID* | c534FhXAH8kOwR3J |
| *acc genotypes* | 53 |
| *peaks no back* | 24 |
| *removed edges* | NA |
| *prop rem edges* | NA |

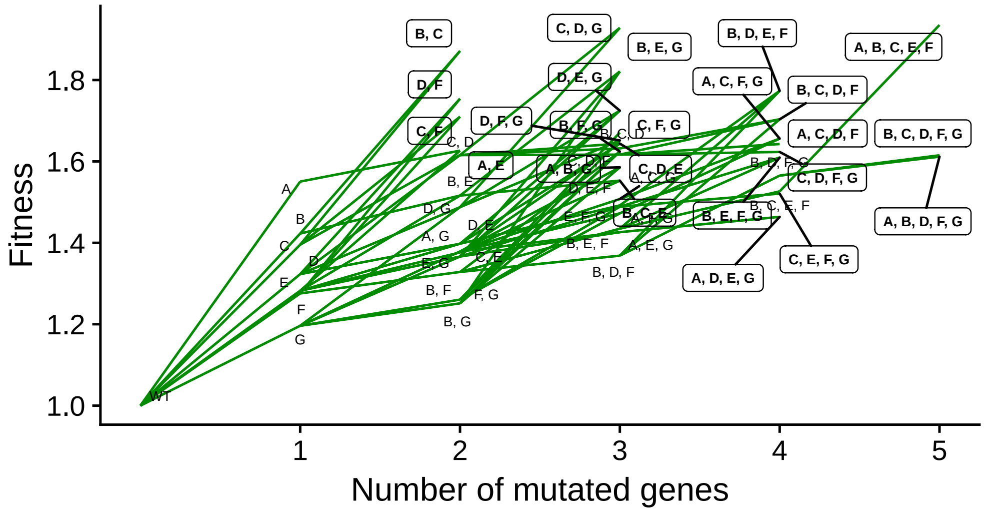


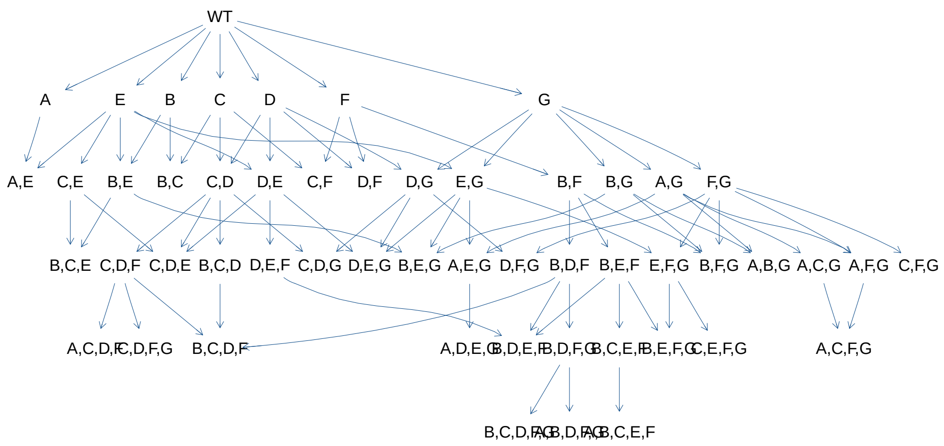

| | |
|---|---|
| *ID* | CfrC3ZAnJWi3s9E3 |
| *acc genotypes* | 38 |
| *peaks no back* | 15 |
| *removed edges* | NA |
| *prop rem edges* | NA |

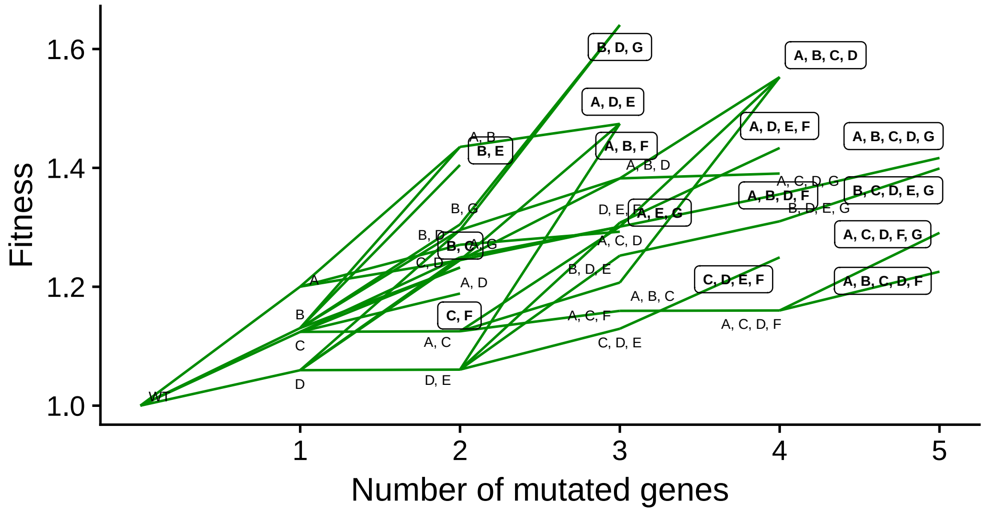


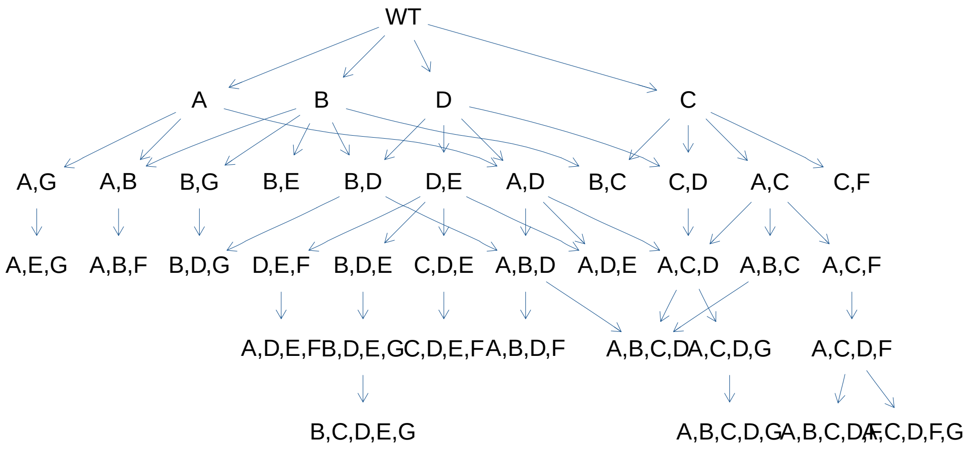

| | |
|---|---|
| *ID* | CHYQvcfXo103swhhd |
| *acc genotypes* | 26 |
| *peaks no back* | 9 |
| *removed edges* | NA |
| *prop rem edges* | NA |

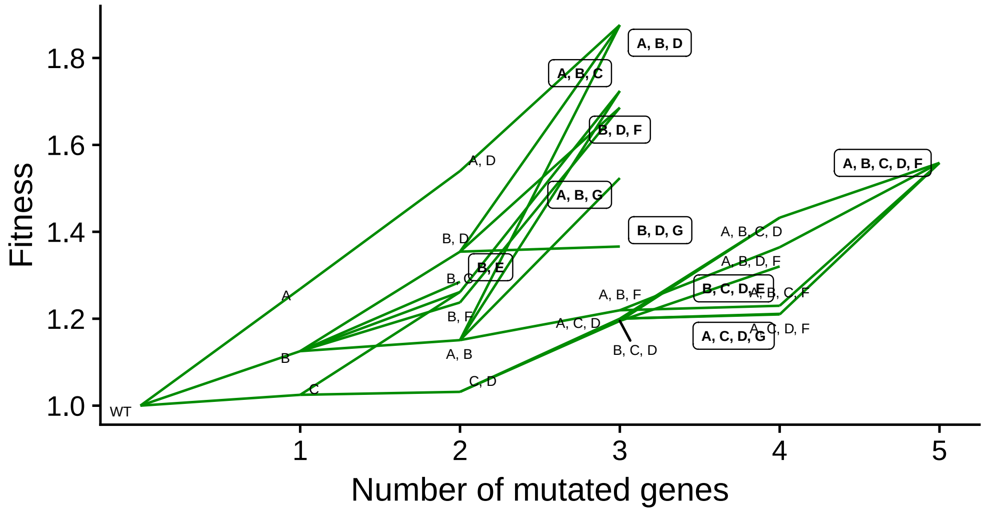


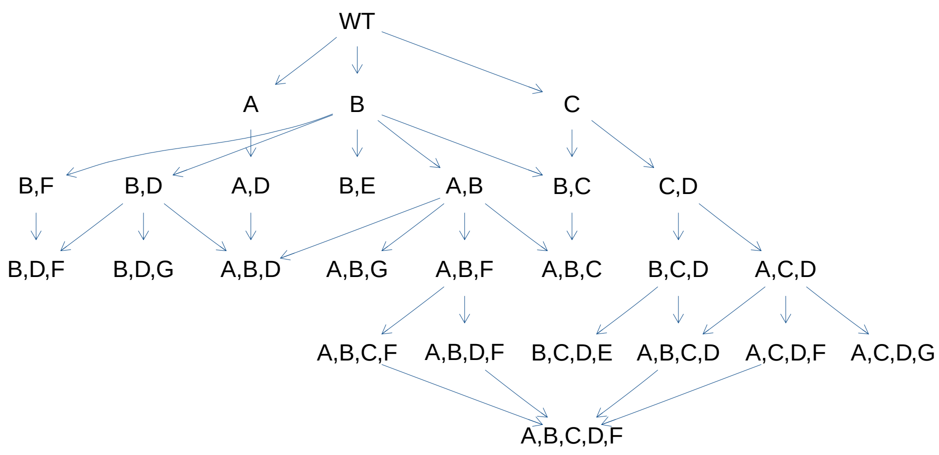

| | |
|---|---|
| *ID* | cI8oWQEqqPY7cX2o |
| *acc genotypes* | 22 |
| *peaks no back* | 13 |
| *removed edges* | NA |
| *prop rem edges* | NA |

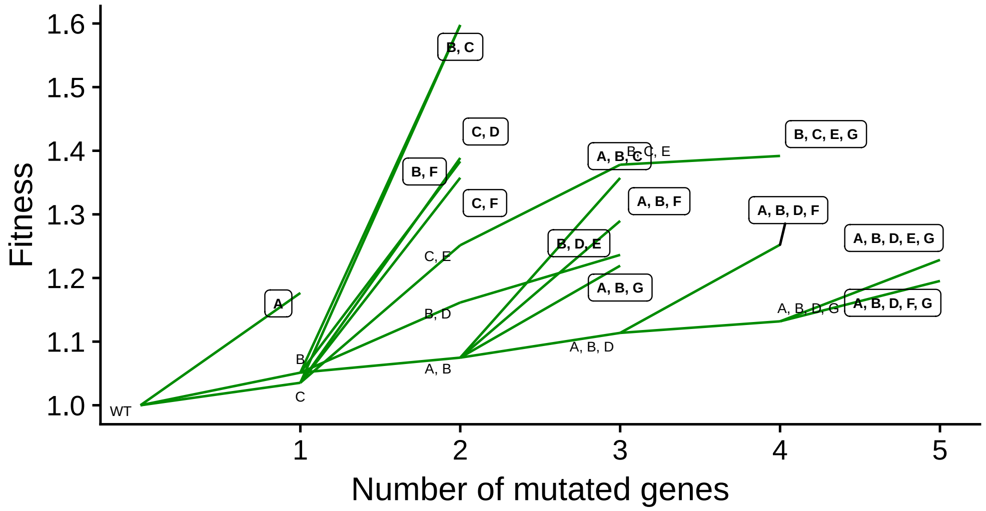


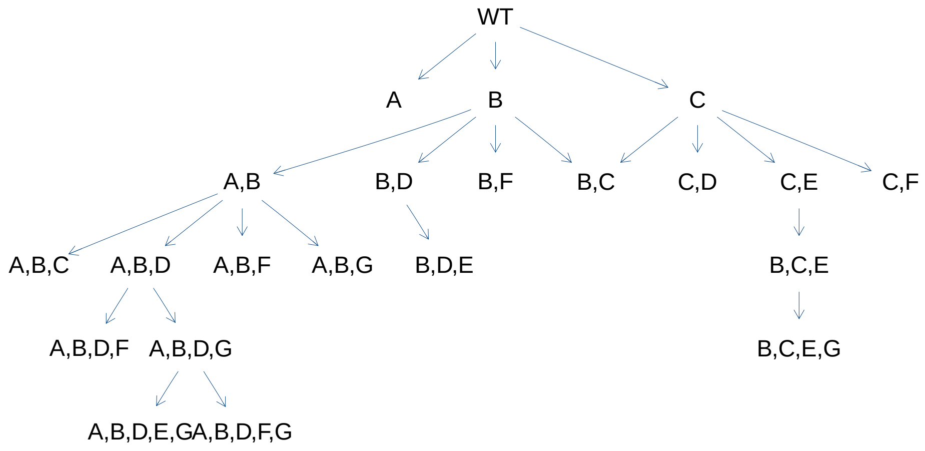

| | |
|---|---|
| *ID* | CjzXJbJTcI5BNmE1 |
| *acc genotypes* | 74 |
| *peaks no back* | 21 |
| *removed edges* | NA |
| *prop rem edges* | NA |

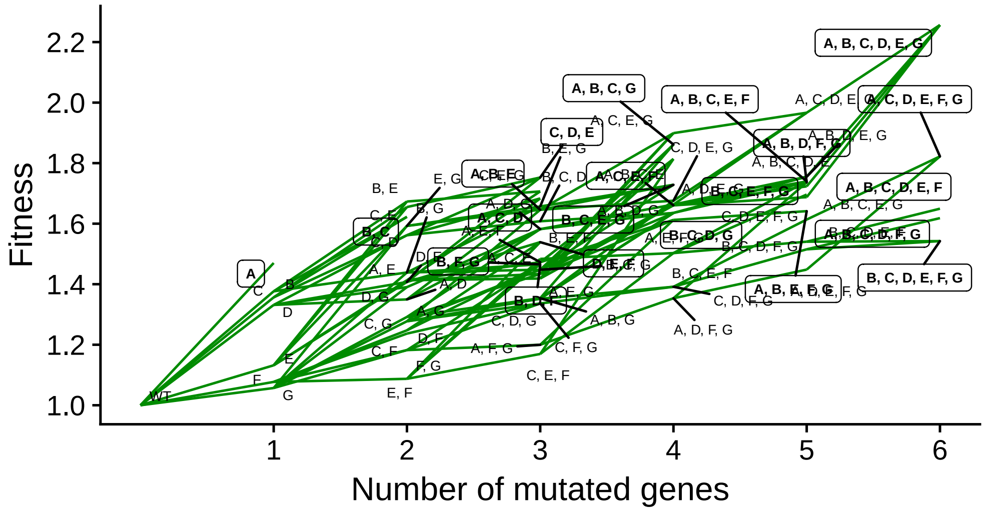


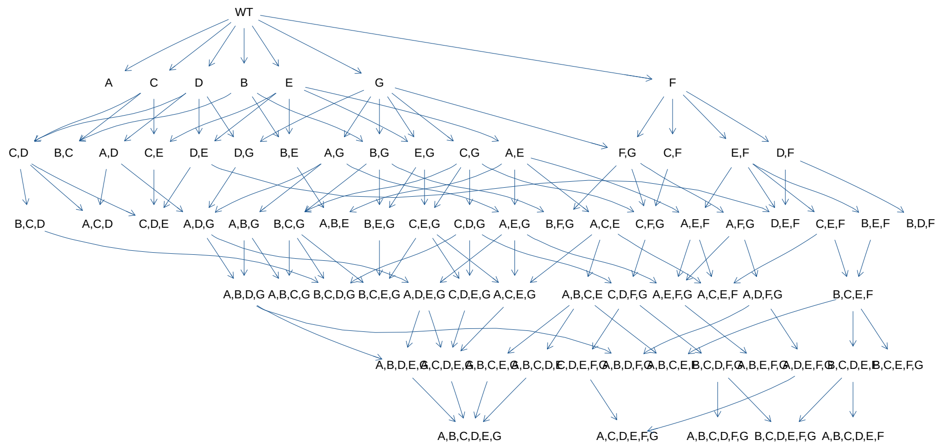

| | |
|---|---|
| *ID* | Ck1zf9wAa7Dm9QHw |
| *acc genotypes* | 24 |
| *peaks no back* | 9 |
| *removed edges* | NA |
| *prop rem edges* | NA |

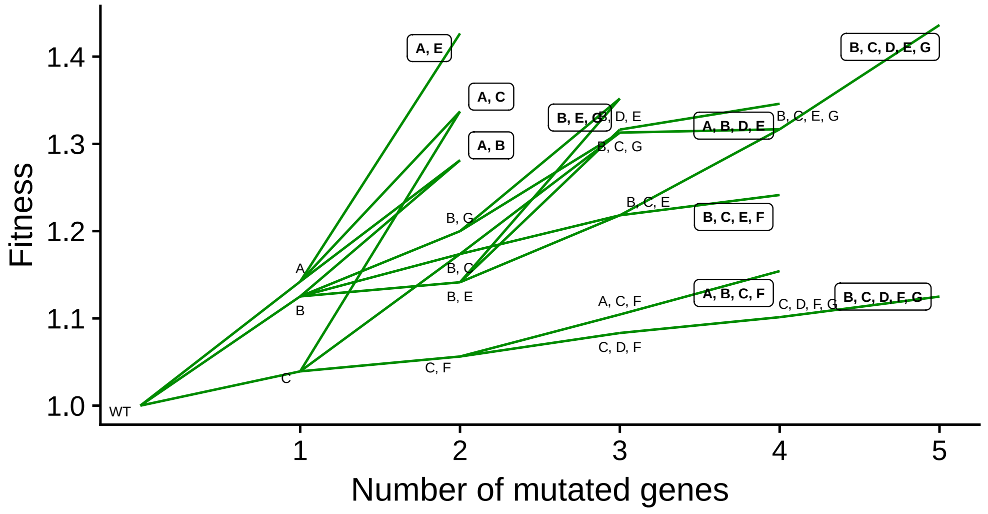


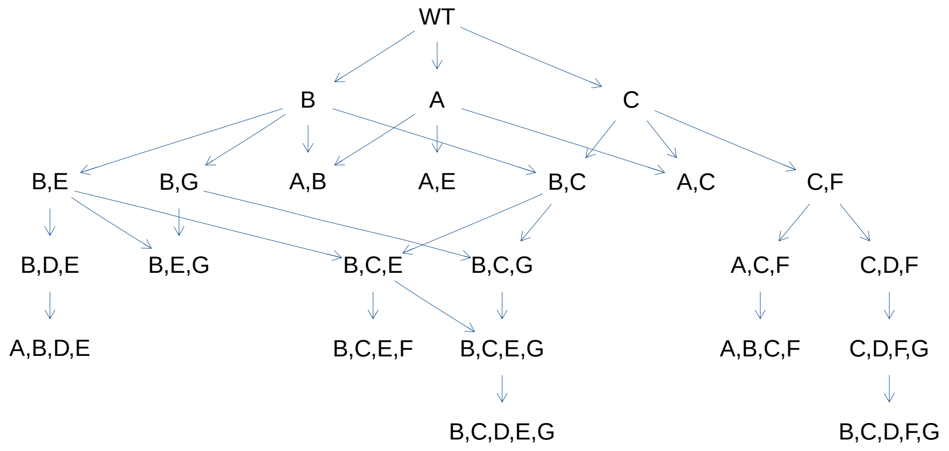

| | |
|---|---|
| *ID* | CsQCk5oitWE3VWiq |
| *acc genotypes* | 90 |
| *peaks no back* | 25 |
| *removed edges* | NA |
| *prop rem edges* | NA |

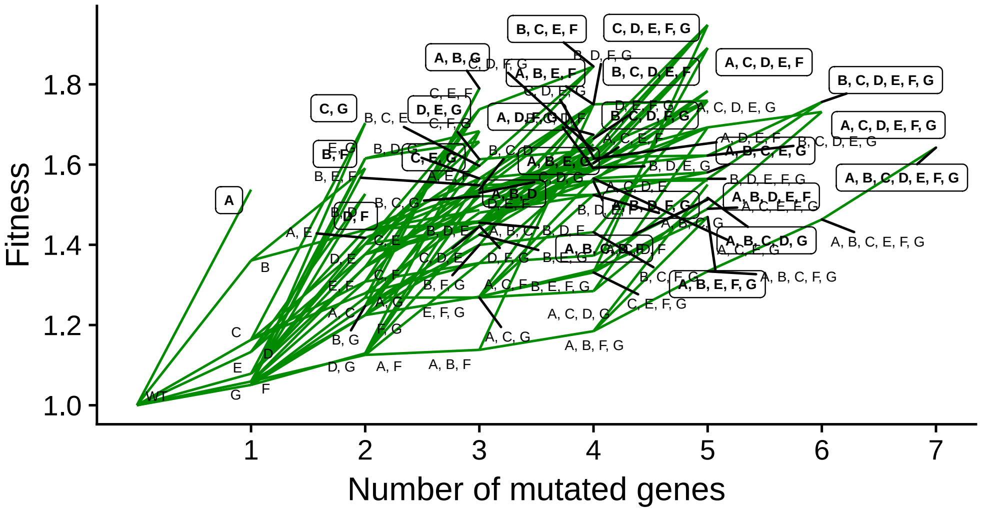


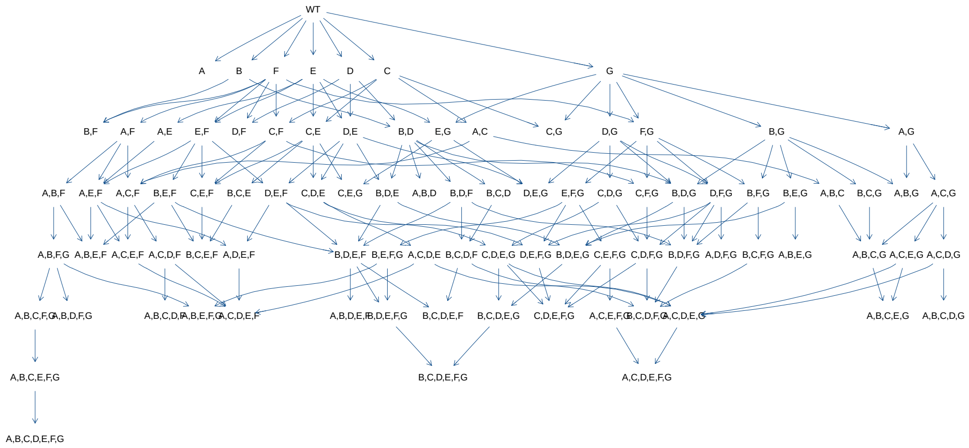

| | |
|---|---|
| *ID* | cueZ258JKmvObVdj |
| *acc genotypes* | 67 |
| *peaks no back* | 15 |
| *removed edges* | NA |
| *prop rem edges* | NA |

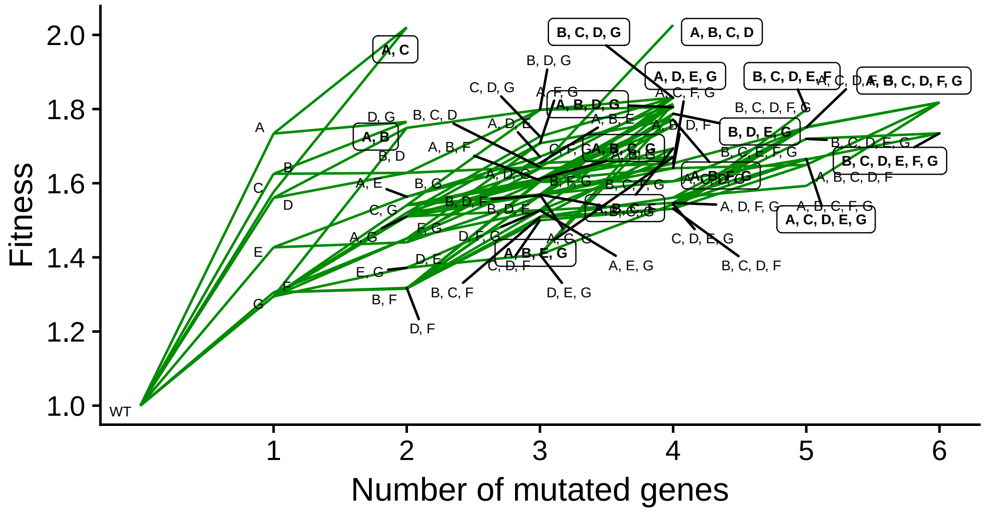


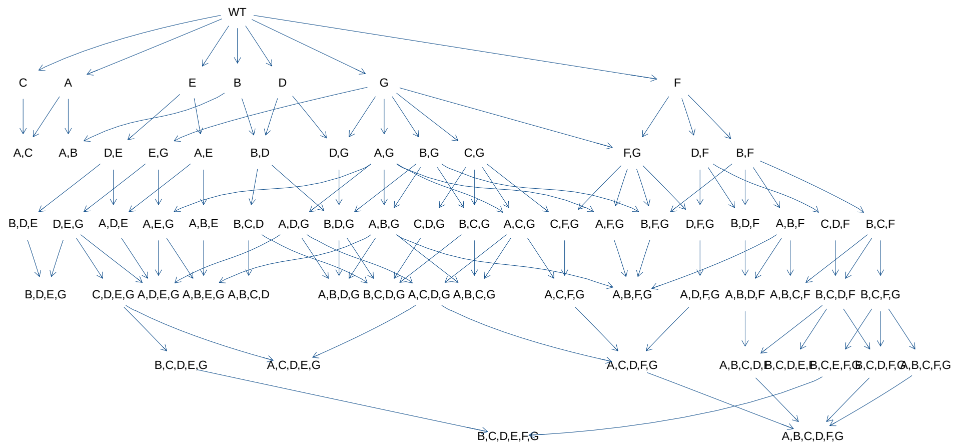

| | |
|---|---|
| *ID* | CvnM3L5EYGnoEGpc |
| *acc genotypes* | 44 |
| *peaks no back* | 19 |
| *removed edges* | NA |
| *prop rem edges* | NA |

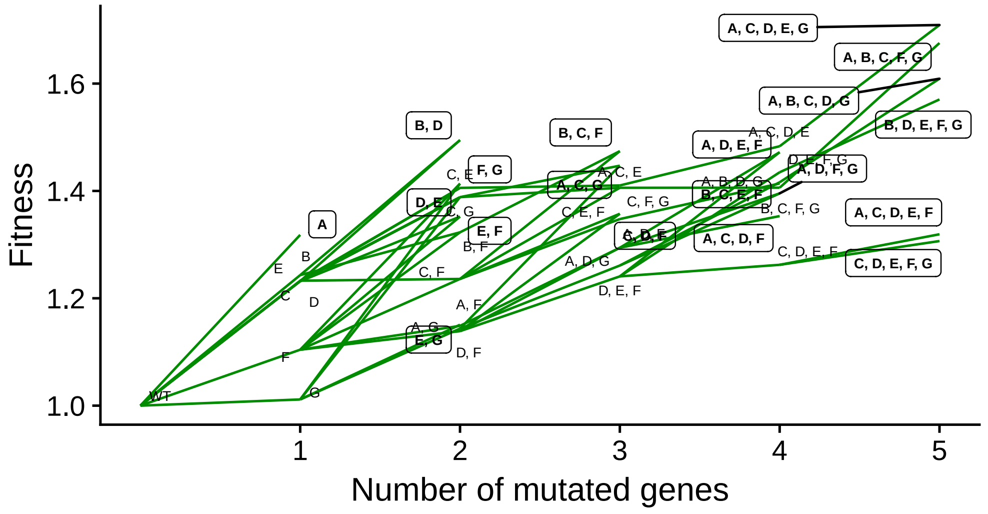


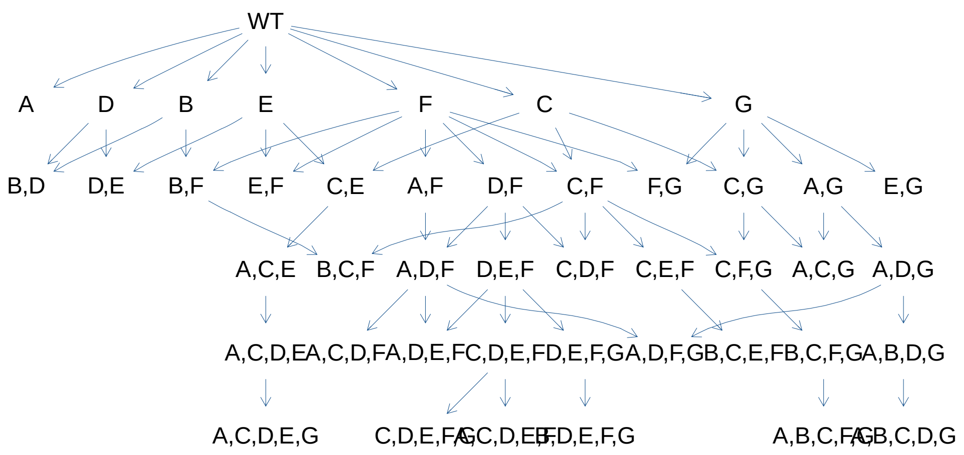

| | |
|---|---|
| *ID* | D8vky9fRuaEmBWIV |
| *acc genotypes* | 45 |
| *peaks no back* | 16 |
| *removed edges* | NA |
| *prop rem edges* | NA |

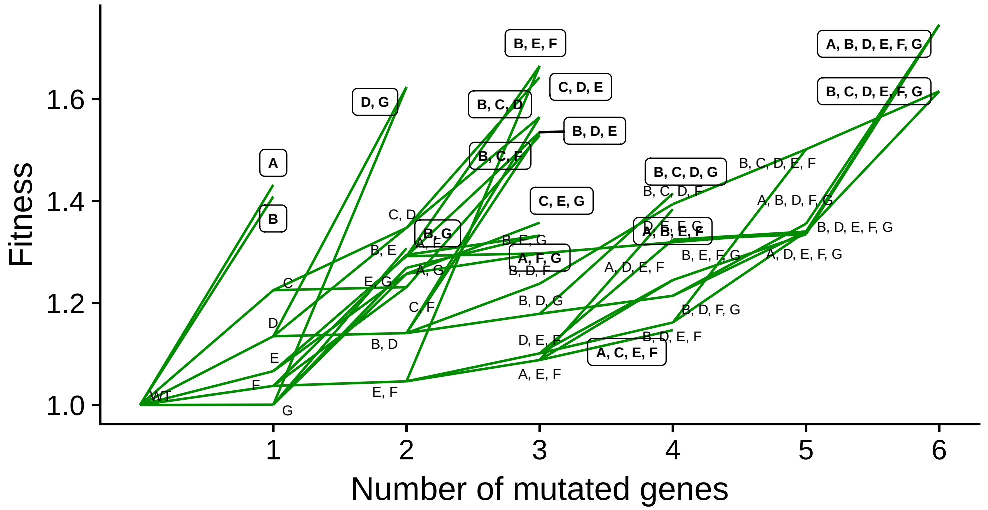

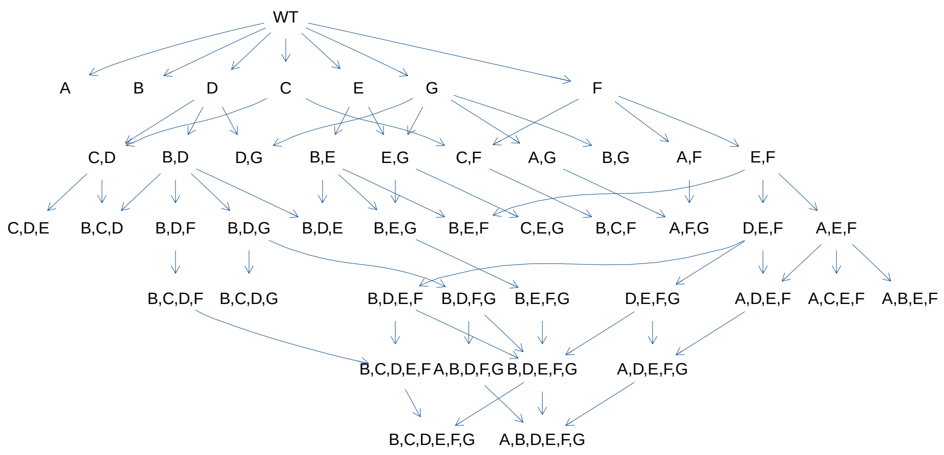

| | |
|---|---|
| *ID* | Daq9vjIG10xrsjP23 |
| *acc genotypes* | 22 |
| *peaks no back* | 13 |
| *removed edges* | NA |
| *prop rem edges* | NA |

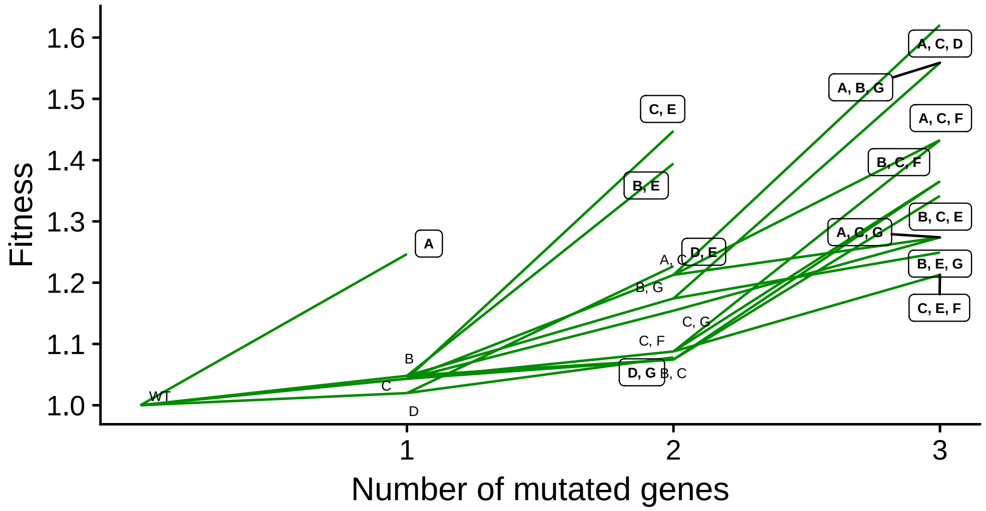


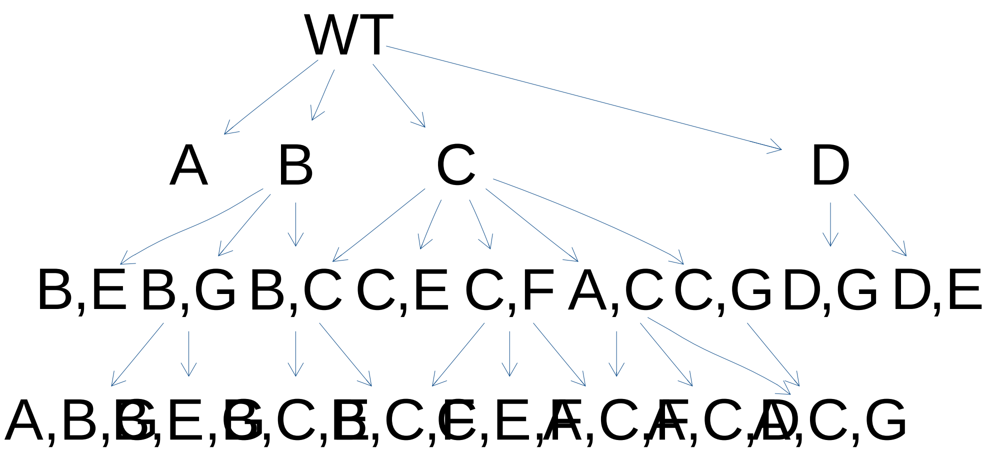

| | |
|---|---|
| *ID* | deoj10TADCLDLOA5o |
| *acc genotypes* | 30 |
| *peaks no back* | 8 |
| *removed edges* | NA |
| *prop rem edges* | NA |

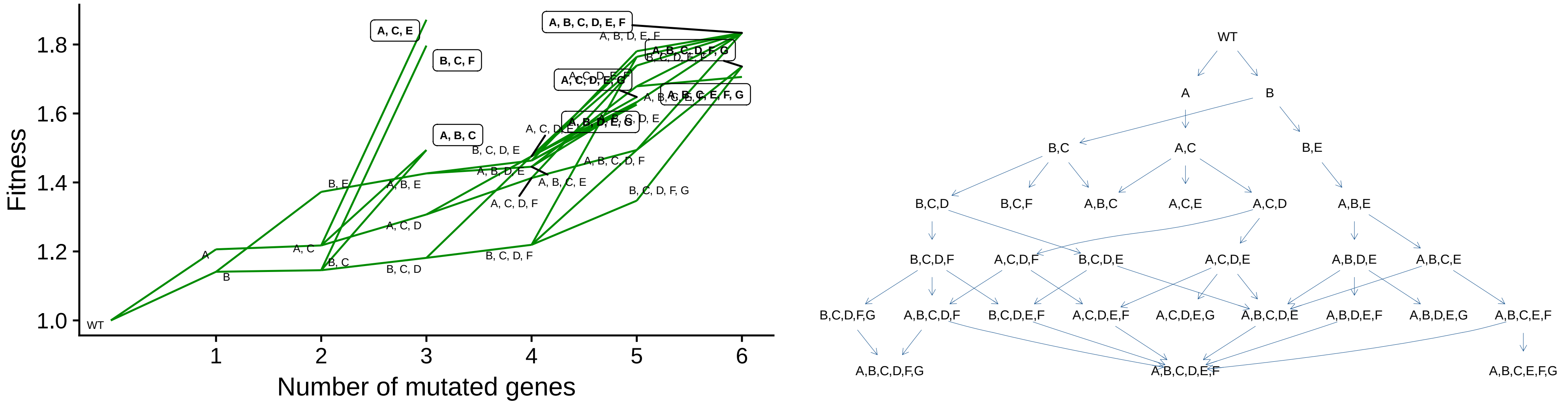

| | |
|---|---|
| *ID* | dTwzjRHQb3XHk1Qo |
| *acc genotypes* | 39 |
| *peaks no back* | 14 |
| *removed edges* | NA |
| *prop rem edges* | NA |

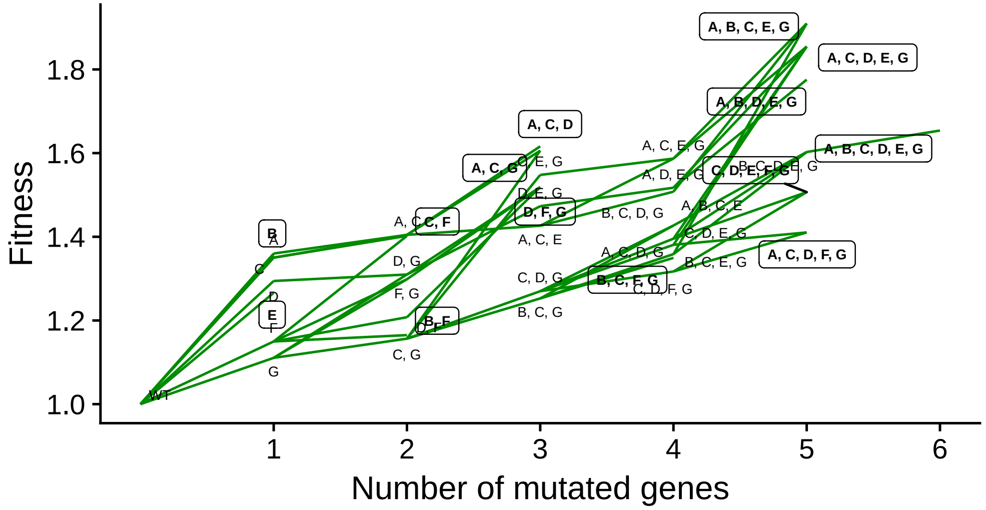


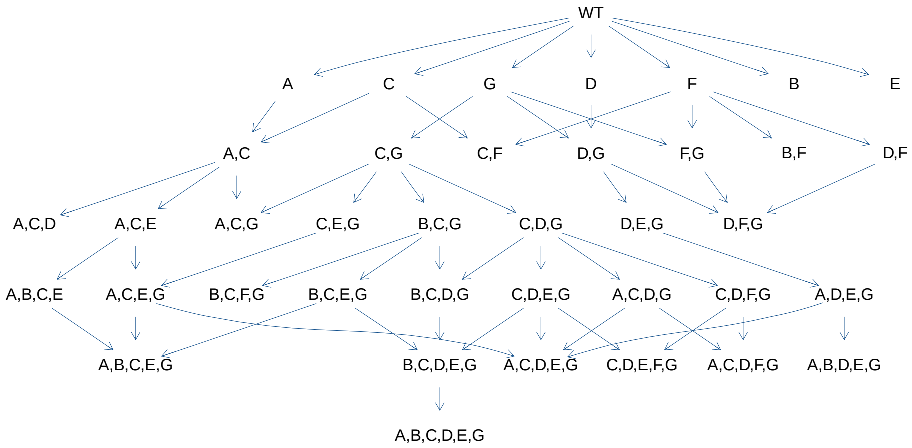

| | |
|---|---|
| *ID* | duoNzOQwAe8B75y5 |
| *acc genotypes* | 42 |
| *peaks no back* | 8 |
| *removed edges* | NA |
| *prop rem edges* | NA |

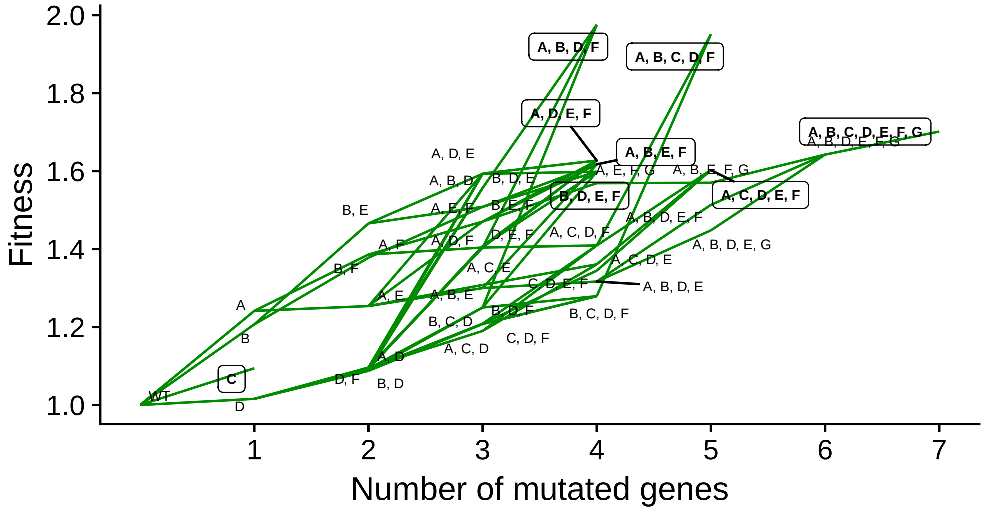


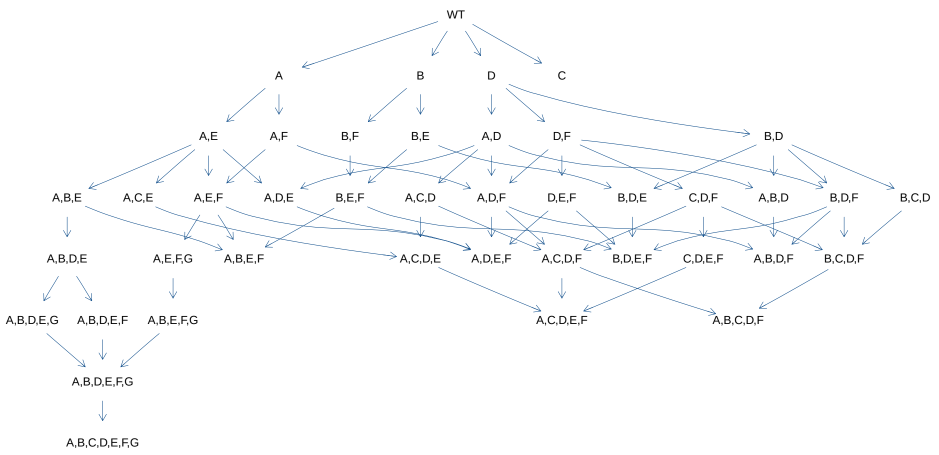

| | |
|---|---|
| *ID* | Dv4AjSOStEZrnaox |
| *acc genotypes* | 62 |
| *peaks no back* | 17 |
| *removed edges* | NA |
| *prop rem edges* | NA |

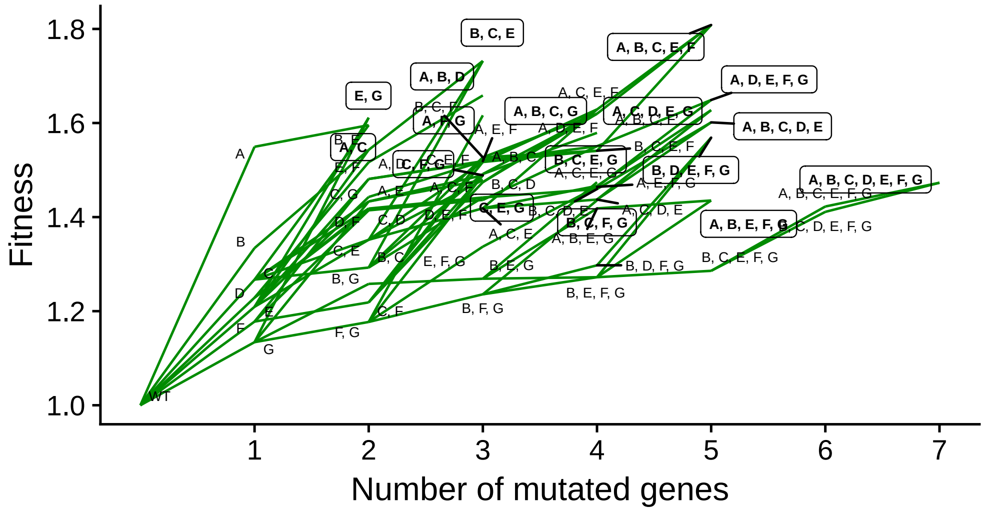


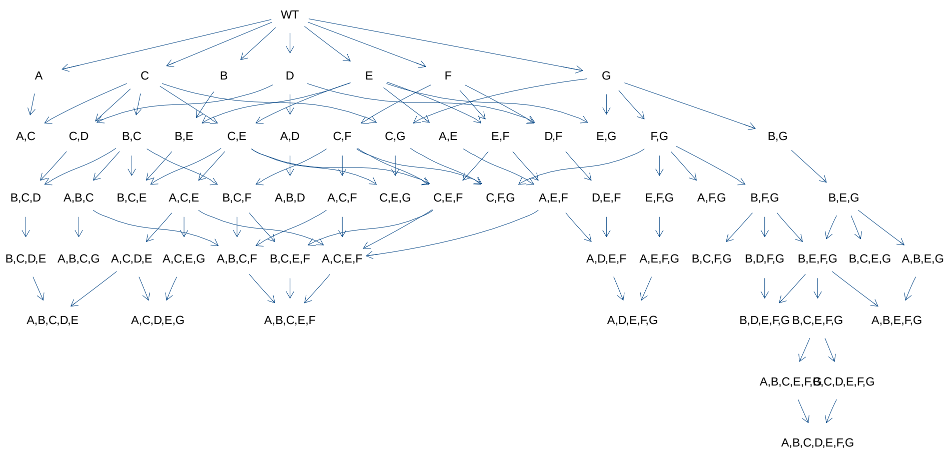

| | |
|---|---|
| *ID* | E1088yM4b7OyoaM3C |
| *acc genotypes* | 96 |
| *peaks no back* | 12 |
| *removed edges* | NA |
| *prop rem edges* | NA |

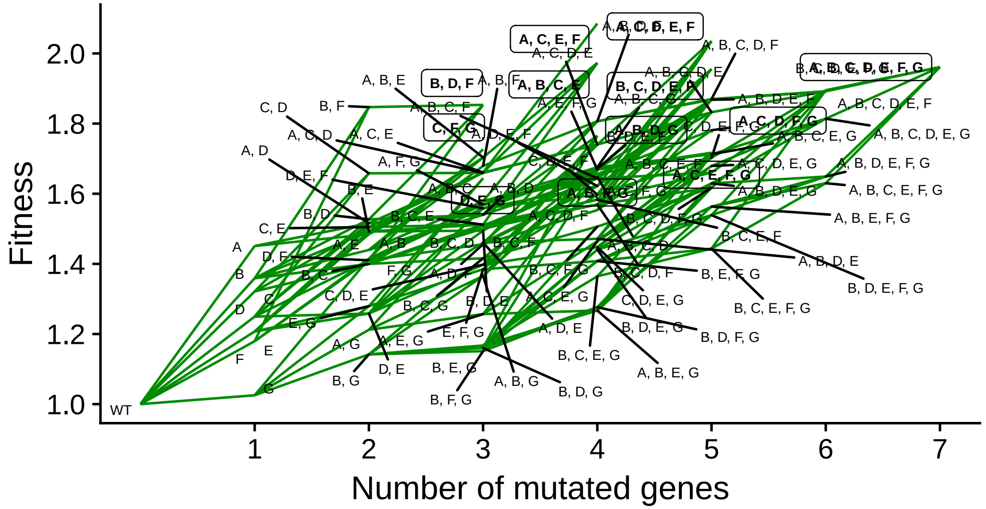


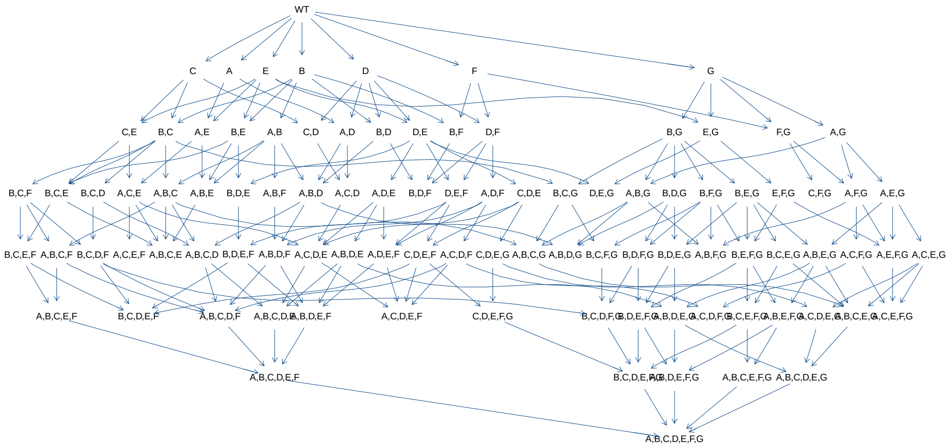

| | |
|---|---|
| *ID* | EAI9G10BQGPjhOaTd |
| *acc genotypes* | 20 |
| *peaks no back* | 9 |
| *removed edges* | NA |
| *prop rem edges* | NA |

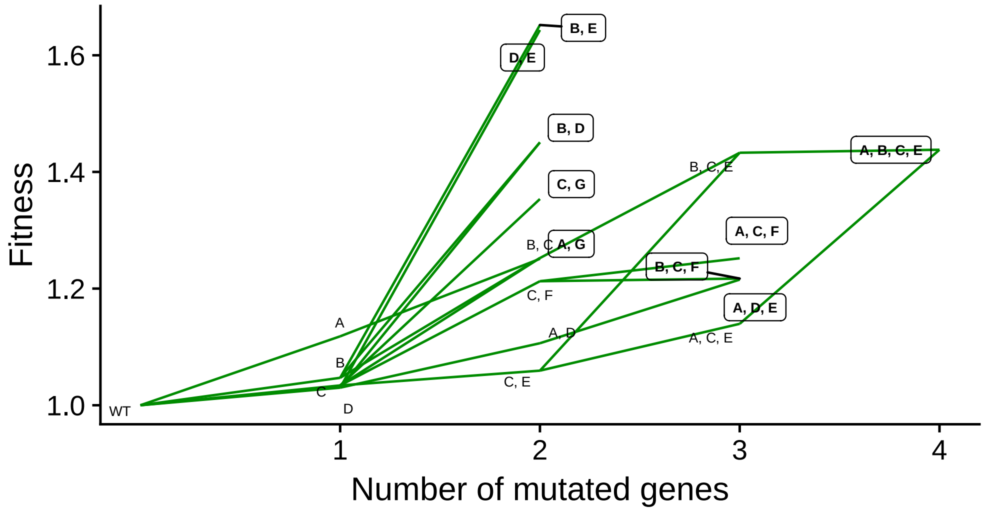


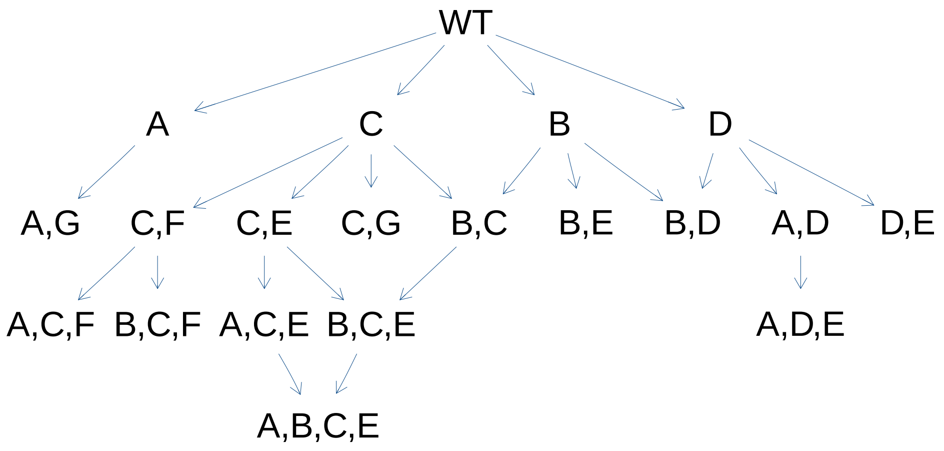

| | |
|---|---|
| *ID* | eb4moeiVT2ClnVLV |
| *acc genotypes* | 74 |
| *peaks no back* | 28 |
| *removed edges* | NA |
| *prop rem edges* | NA |

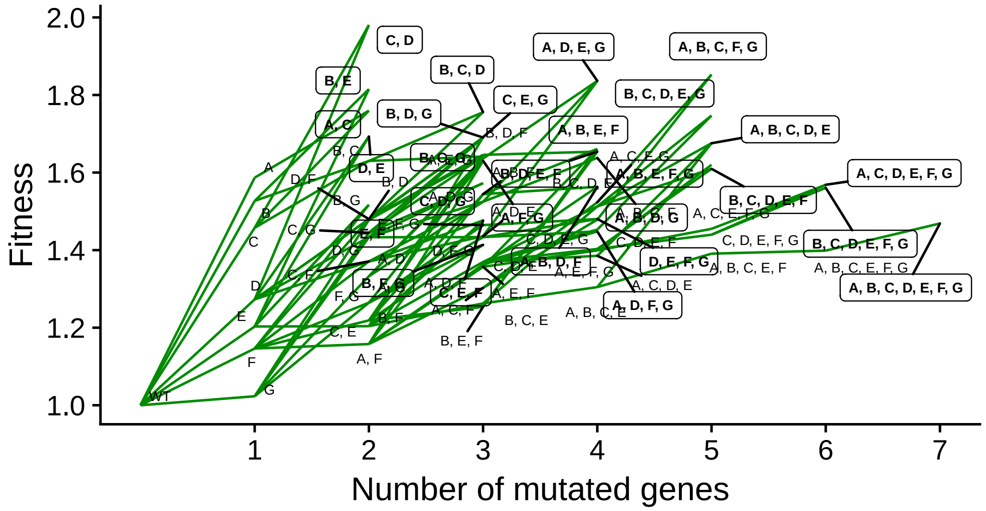


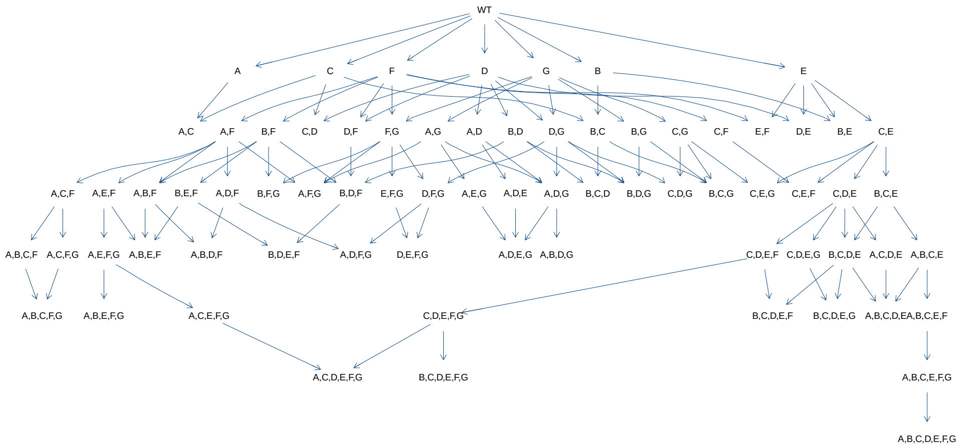

| | |
|---|---|
| *ID* | EdoyPlJWzxCpBrys |
| *acc genotypes* | 31 |
| *peaks no back* | 14 |
| *removed edges* | NA |
| *prop rem edges* | NA |

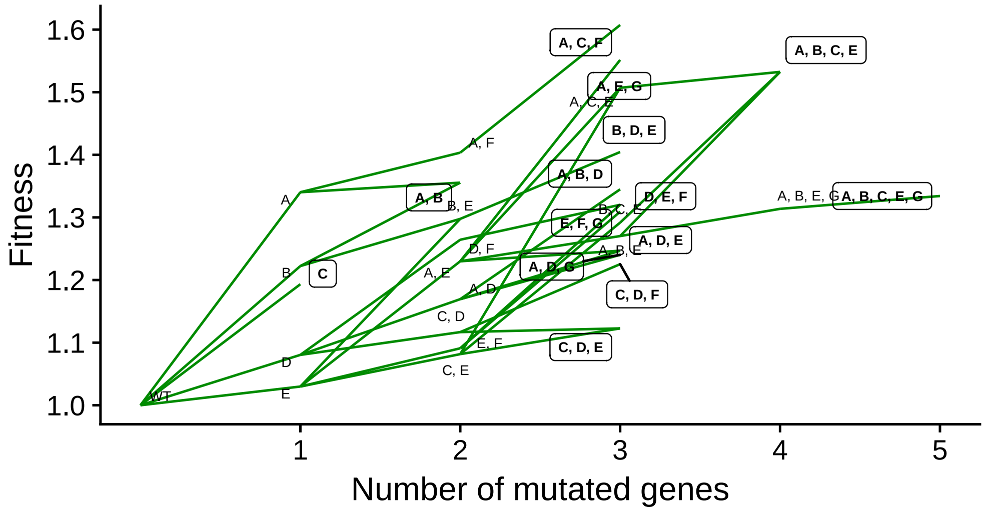


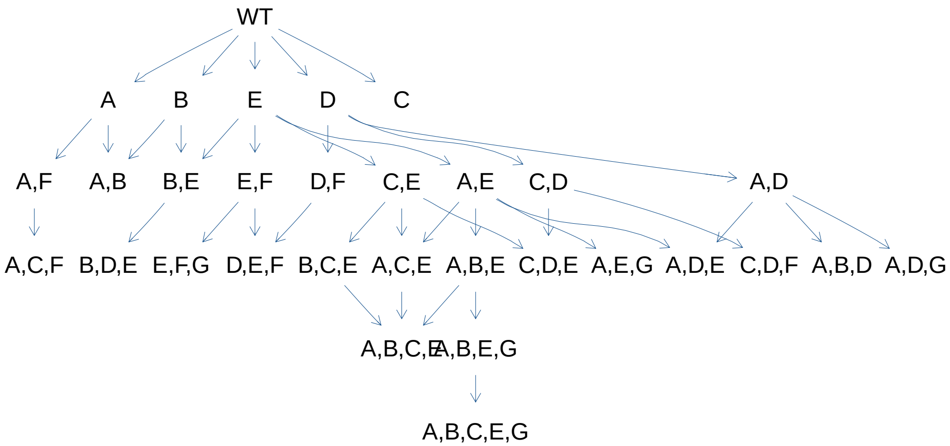

*ID* EKkcYkyuQzPjnUDy
*acc genotypes* 54
*peaks no back* 16
*removed edges* NA
*prop rem edges* NA

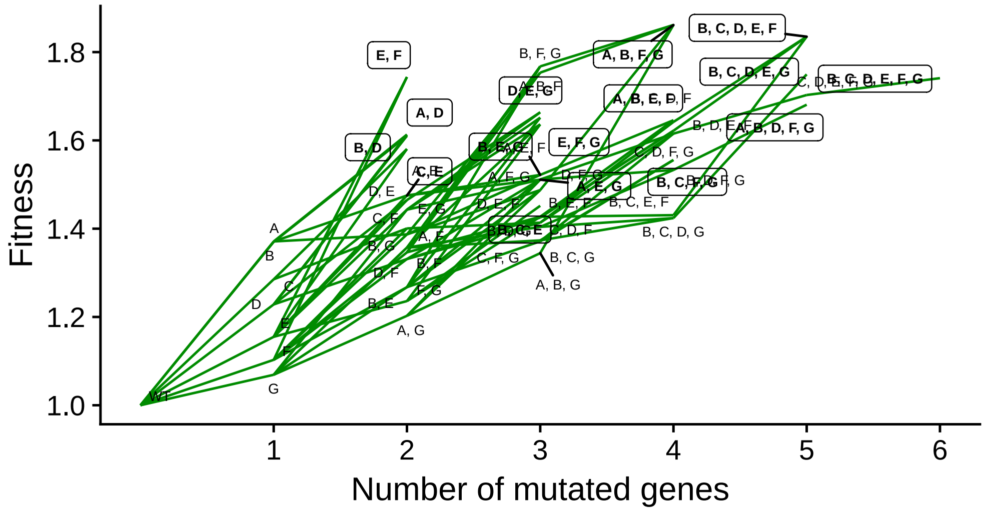


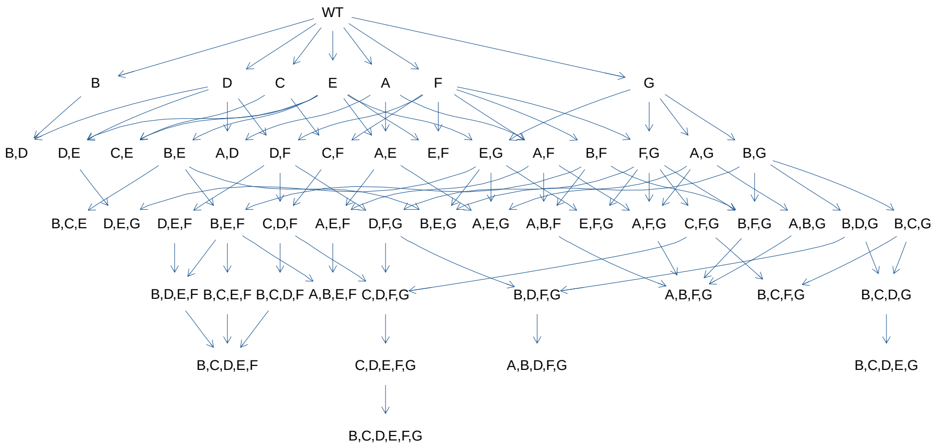

| | |
|---|---|
| *ID* | EM68B9s53CXWI9KL |
| *acc genotypes* | 41 |
| *peaks no back* | 16 |
| *removed edges* | NA |
| *prop rem edges* | NA |

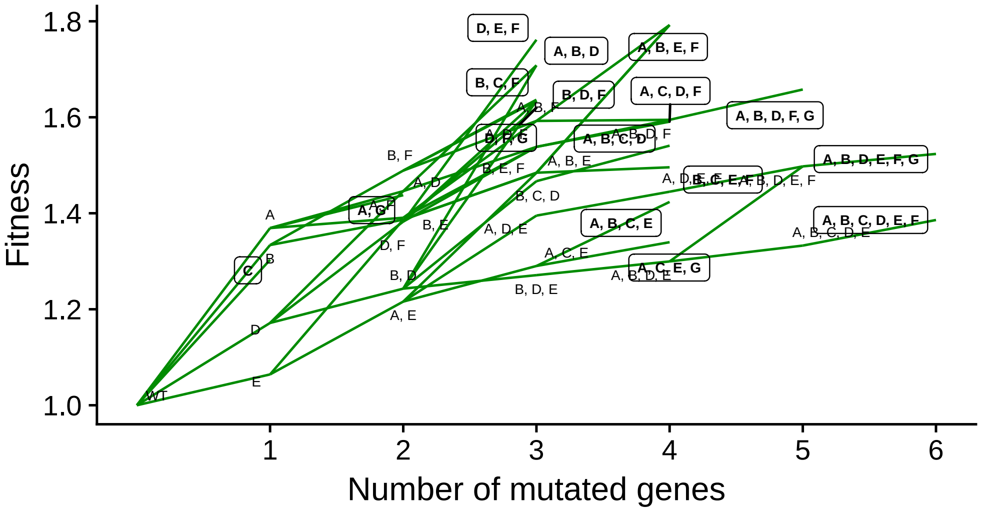


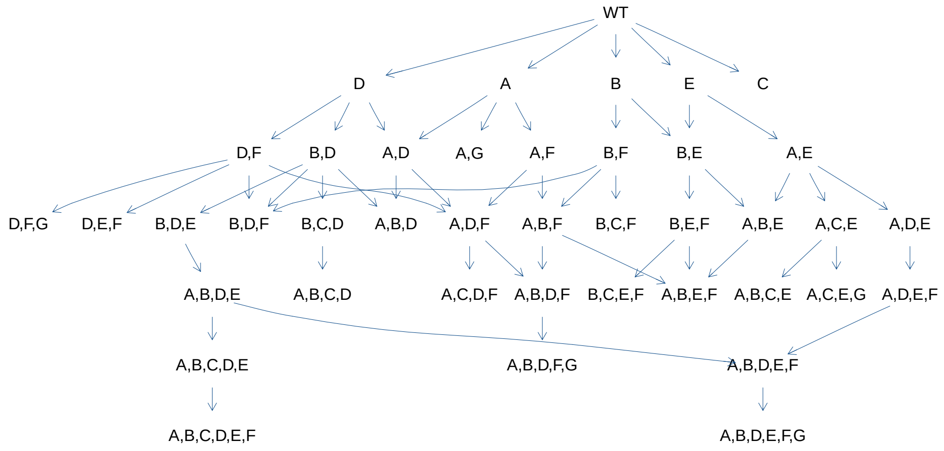

| | |
|---|---|
| *ID* | eQOmKNmXnLoUPYw5 |
| *acc genotypes* | 30 |
| *peaks no back* | 14 |
| *removed edges* | NA |
| *prop rem edges* | NA |

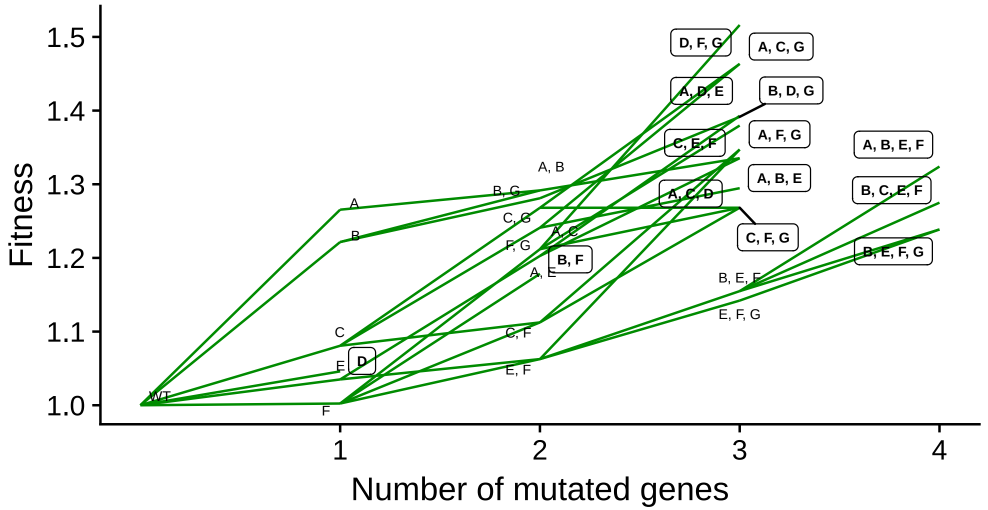


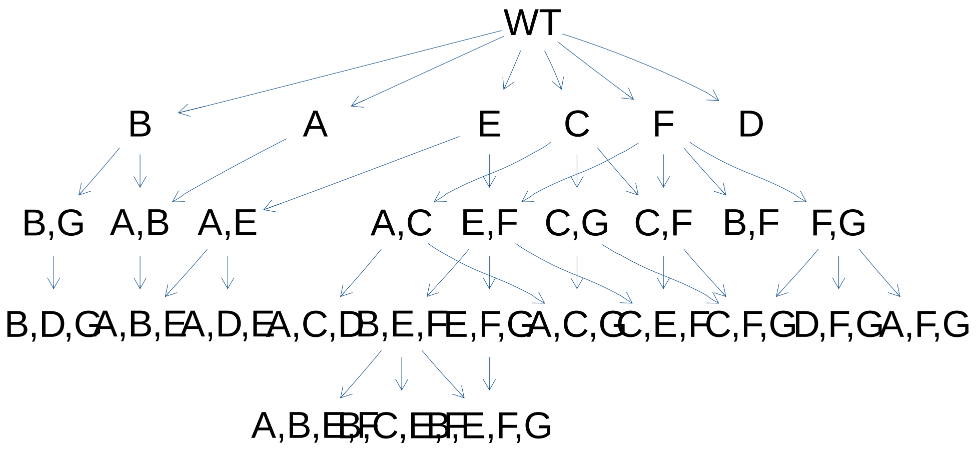

| | |
|---|---|
| *ID* | EsTjWIE7vDhzVqNd |
| *acc genotypes* | 69 |
| *peaks no back* | 20 |
| *removed edges* | NA |
| *prop rem edges* | NA |

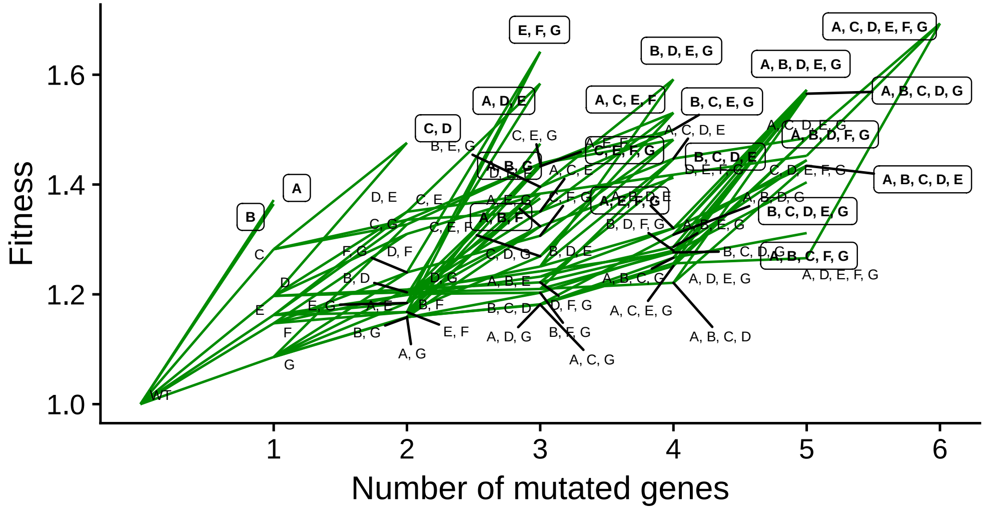


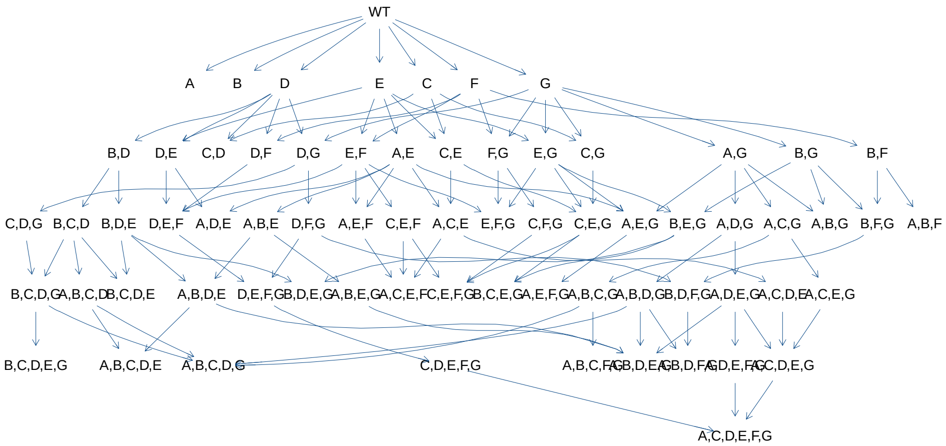

| | |
|---|---|
| *ID* | ETH7102TvoUx6htCa |
| *acc genotypes* | 21 |
| *peaks no back* | 9 |
| *removed edges* | NA |
| *prop rem edges* | NA |

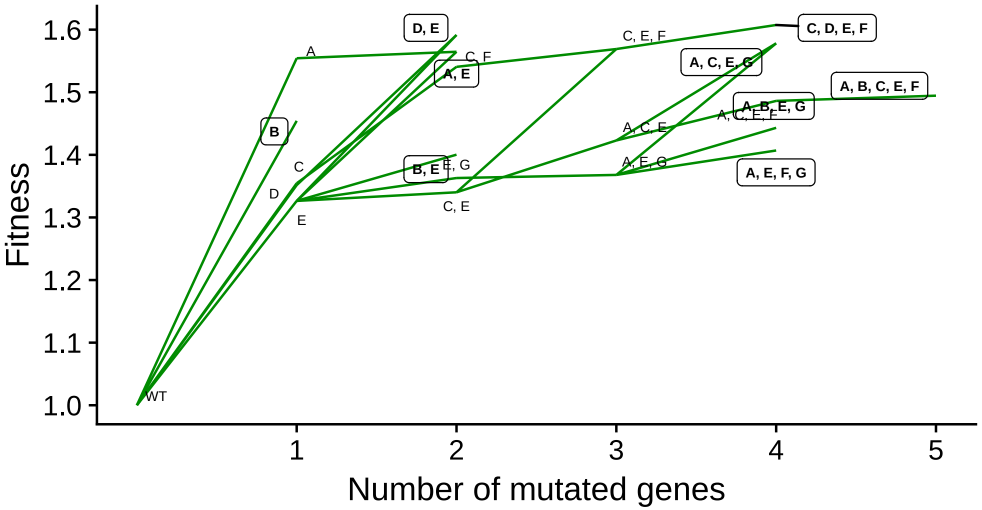


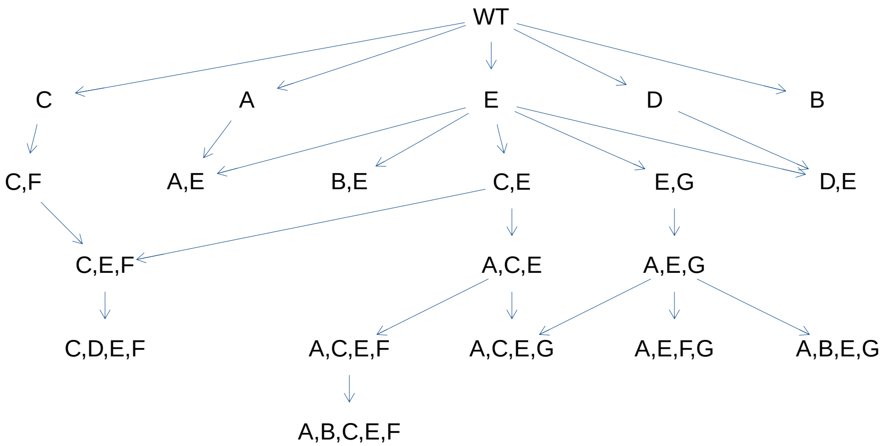

| | |
|---|---|
| *ID* | eUzrnqLvhVknhTTj |
| *acc genotypes* | 40 |
| *peaks no back* | 11 |
| *removed edges* | NA |
| *prop rem edges* | NA |

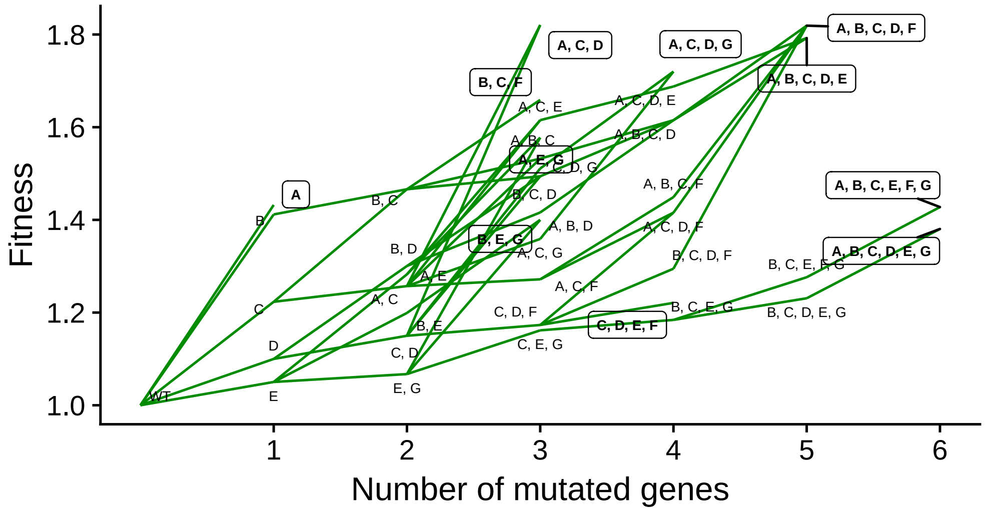


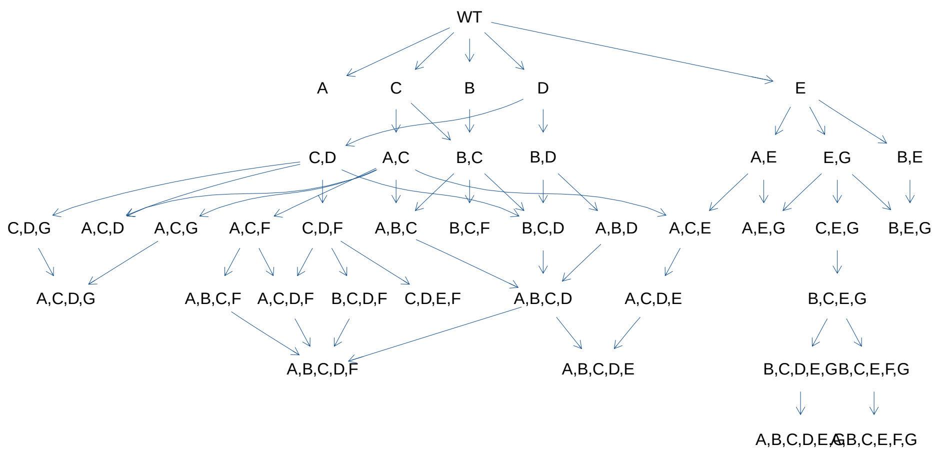

| | |
|---|---|
| *ID* | fdZVeDlLcm54cJxM |
| *acc genotypes* | 41 |
| *peaks no back* | 17 |
| *removed edges* | NA |
| *prop rem edges* | NA |

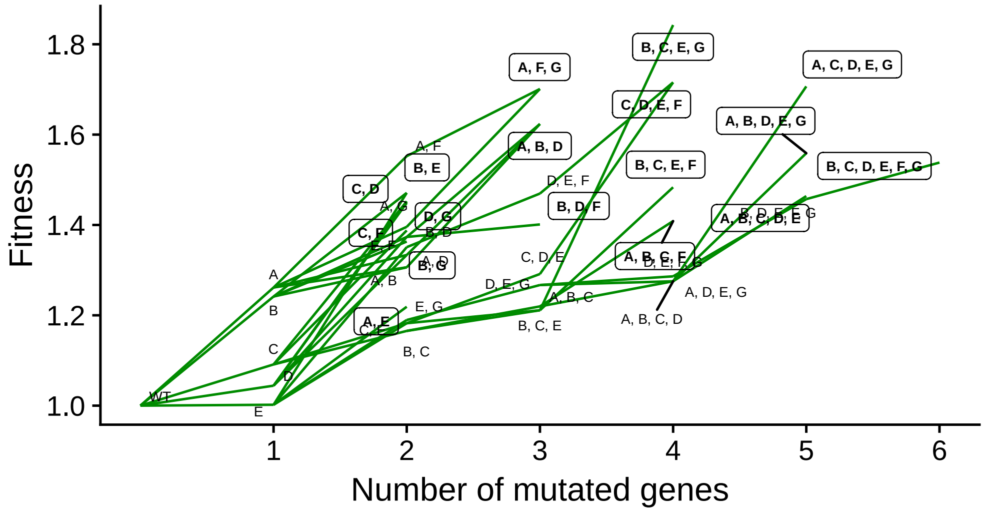


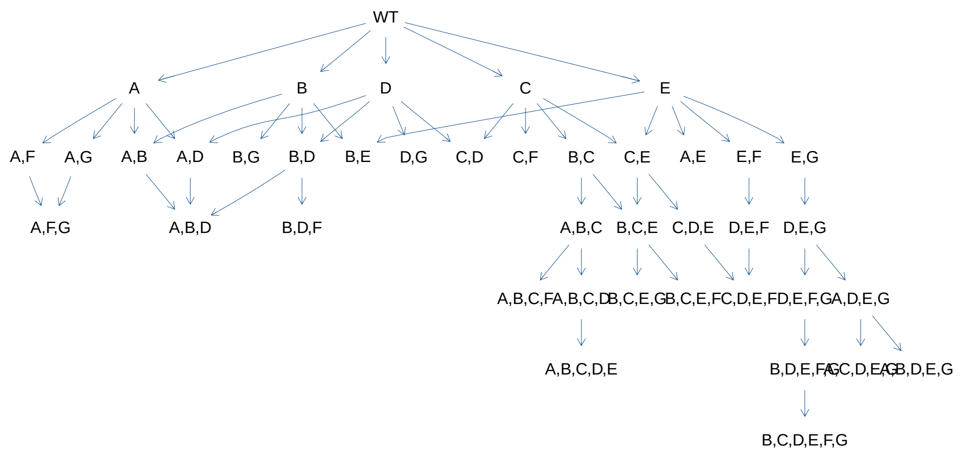

| | |
|---|---|
| *ID* | fE3LkSfz106ZntdBy |
| *acc genotypes* | 48 |
| *peaks no back* | 13 |
| *removed edges* | NA |
| *prop rem edges* | NA |

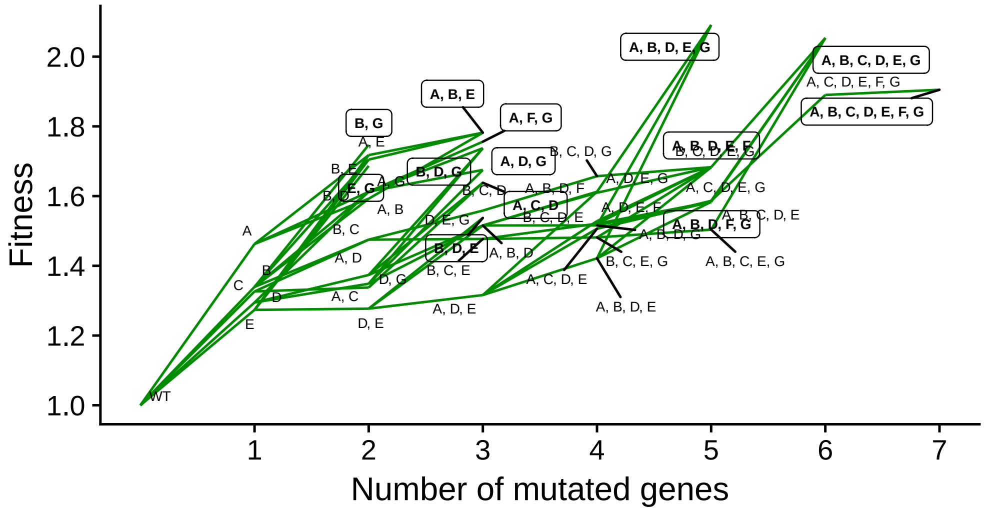


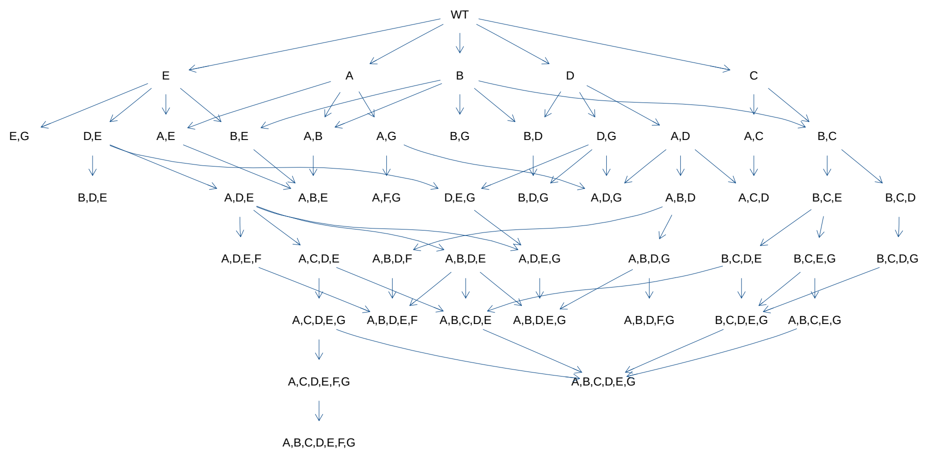

| | |
|---|---|
| *ID* | fEwT1uohy1ShL5nH |
| *acc genotypes* | 38 |
| *peaks no back* | 17 |
| *removed edges* | NA |
| *prop rem edges* | NA |

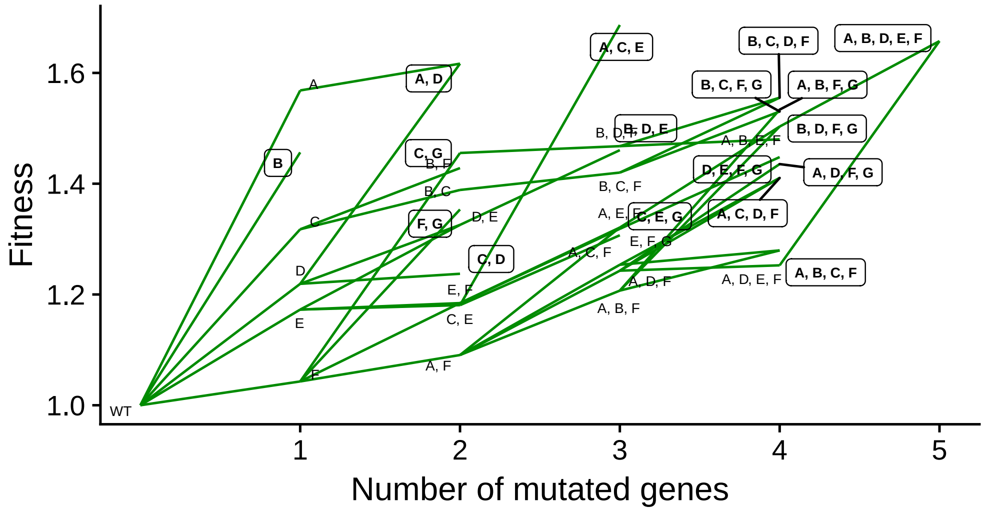


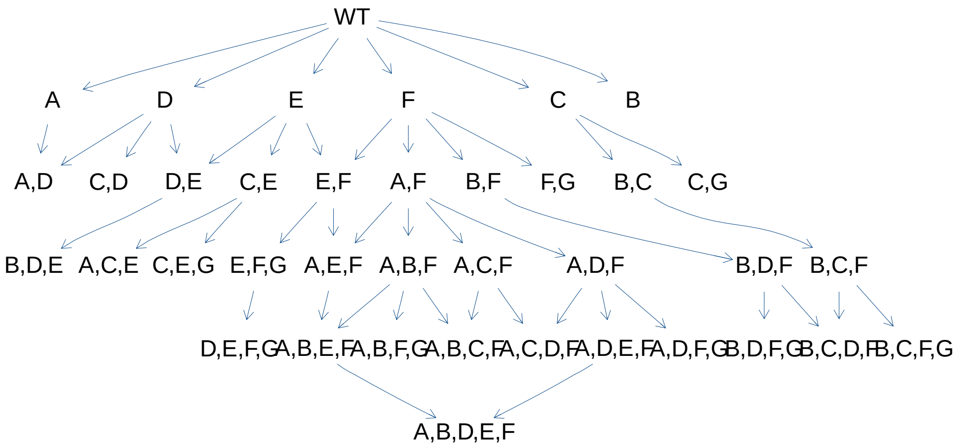

| | |
|---|---|
| *ID* | fS10Ju2gMFkEFOANk |
| *acc genotypes* | 20 |
| *peaks no back* | 8 |
| *removed edges* | NA |
| *prop rem edges* | NA |

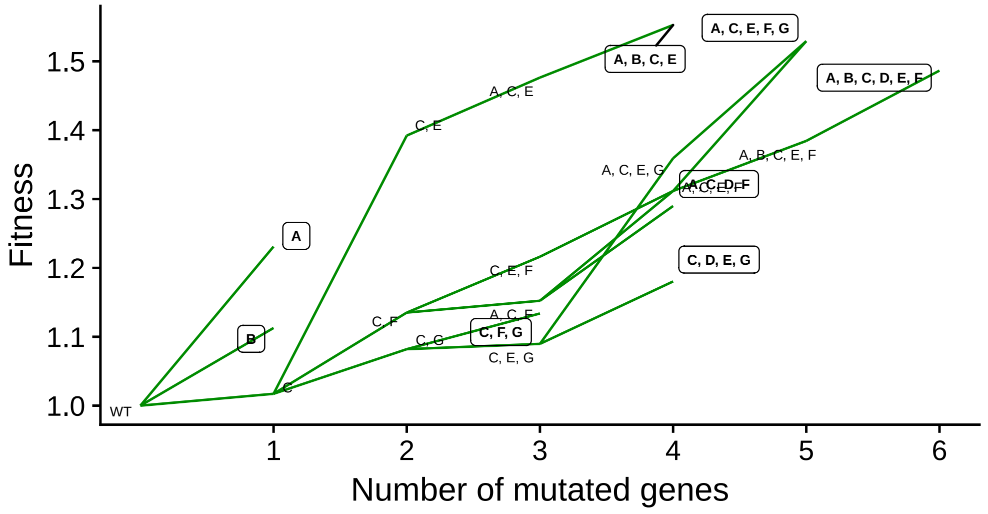


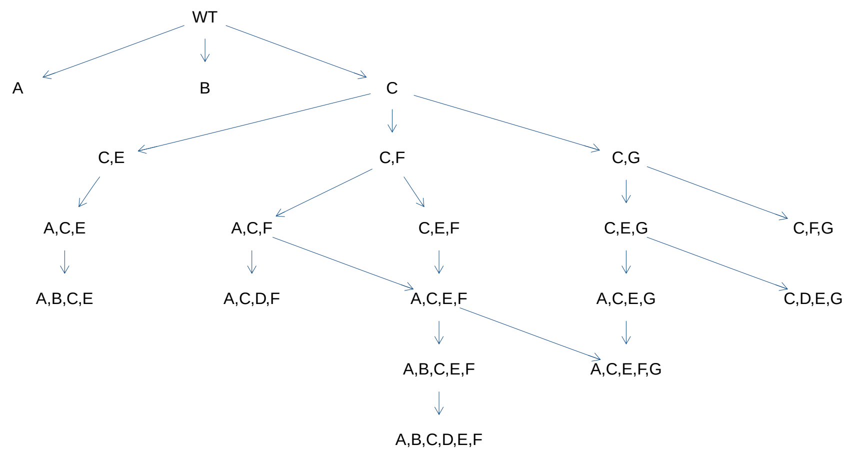

| | |
|---|---|
| *ID* | Fwg3jWZQftuLfTc8 |
| *acc genotypes* | 43 |
| *peaks no back* | 13 |
| *removed edges* | NA |
| *prop rem edges* | NA |

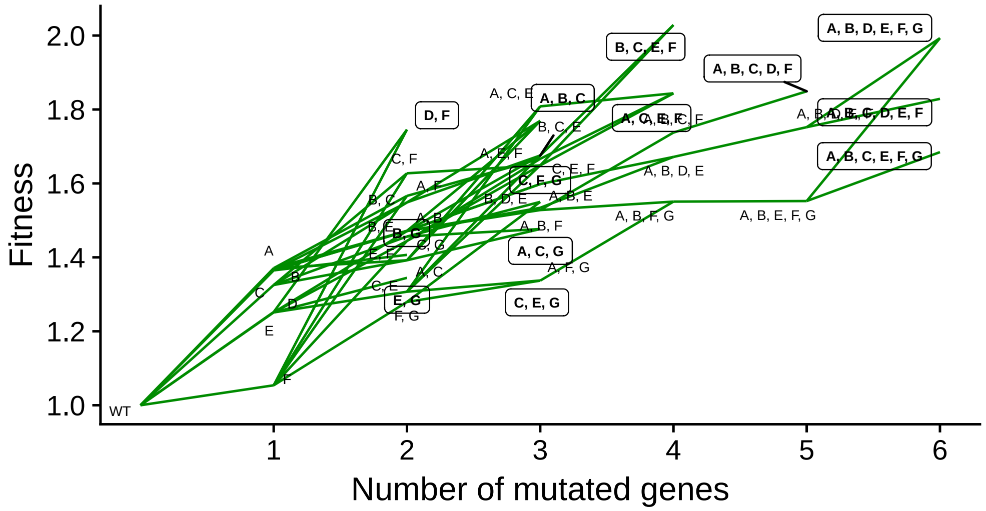


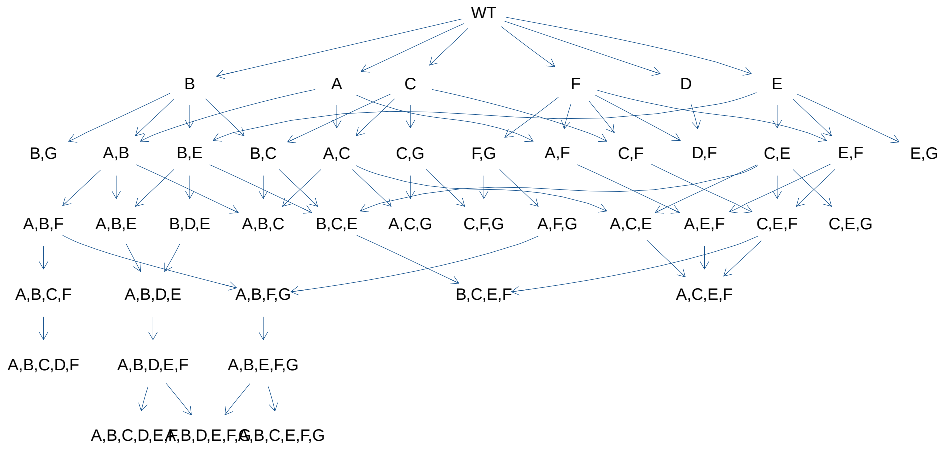

| | |
|---|---|
| *ID* | fzwb5MOAwOWpF4B9 |
| *acc genotypes* | 62 |
| *peaks no back* | 18 |
| *removed edges* | NA |
| *prop rem edges* | NA |

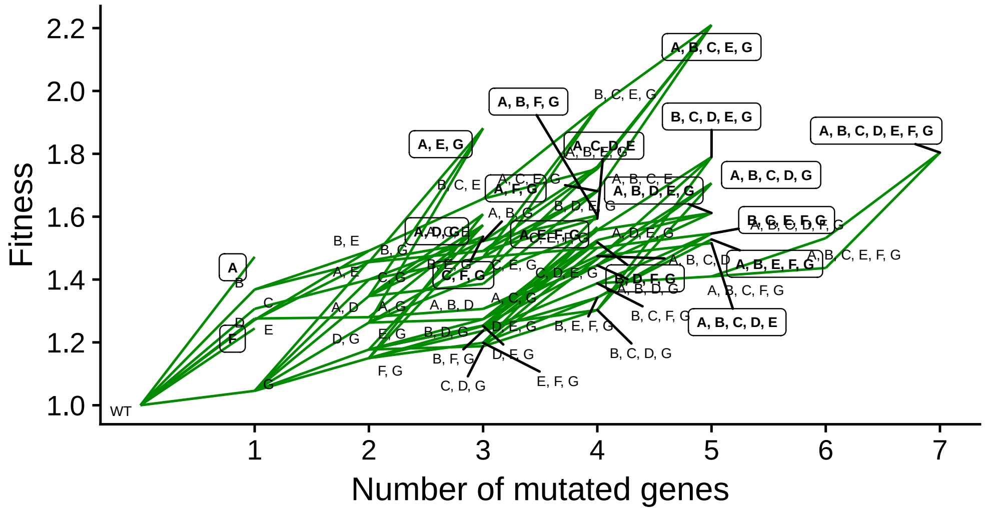


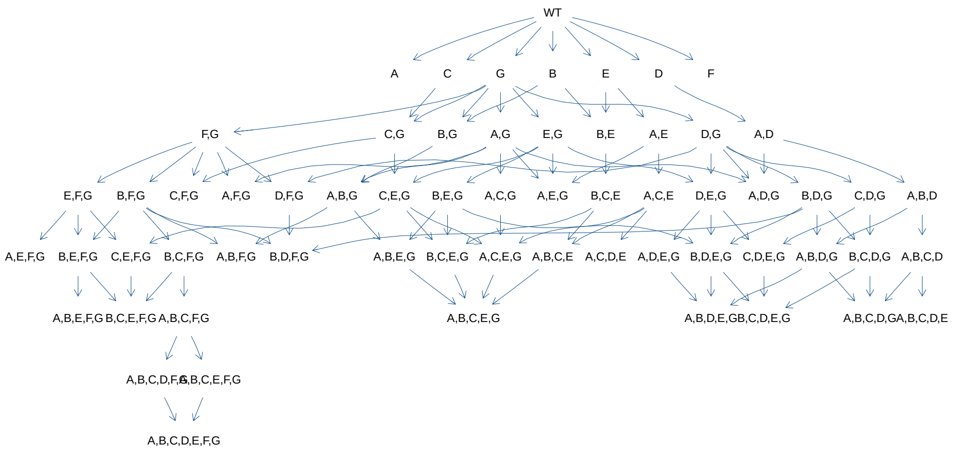

| | |
|---|---|
| *ID* | GDVo4gmjaDr6lo1c |
| *acc genotypes* | 51 |
| *peaks no back* | 16 |
| *removed edges* | NA |
| *prop rem edges* | NA |

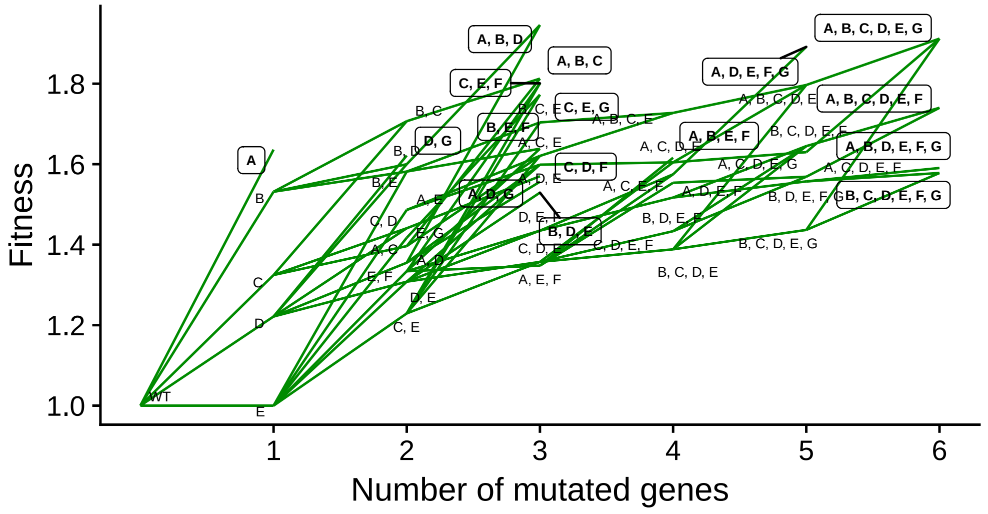

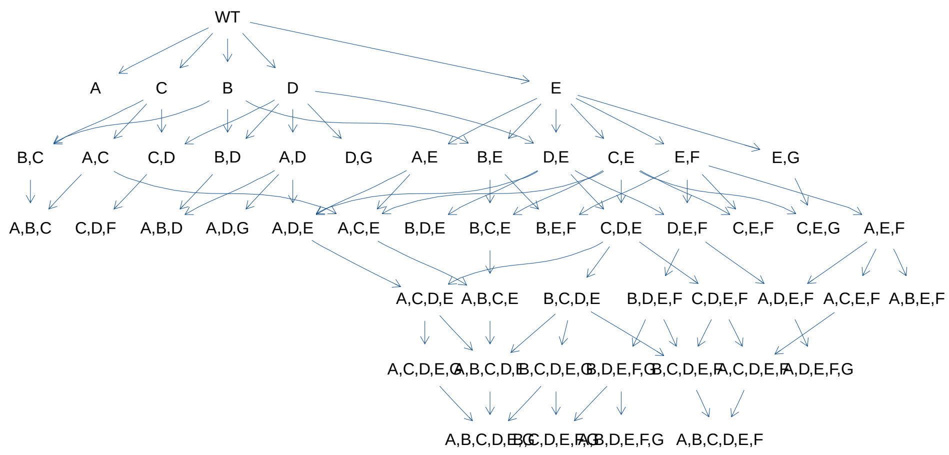

| | |
|---|---|
| *ID* | gHFY6JsOn10PnBYf3 |
| *acc genotypes* | 40 |
| *peaks no back* | 13 |
| *removed edges* | NA |
| *prop rem edges* | NA |

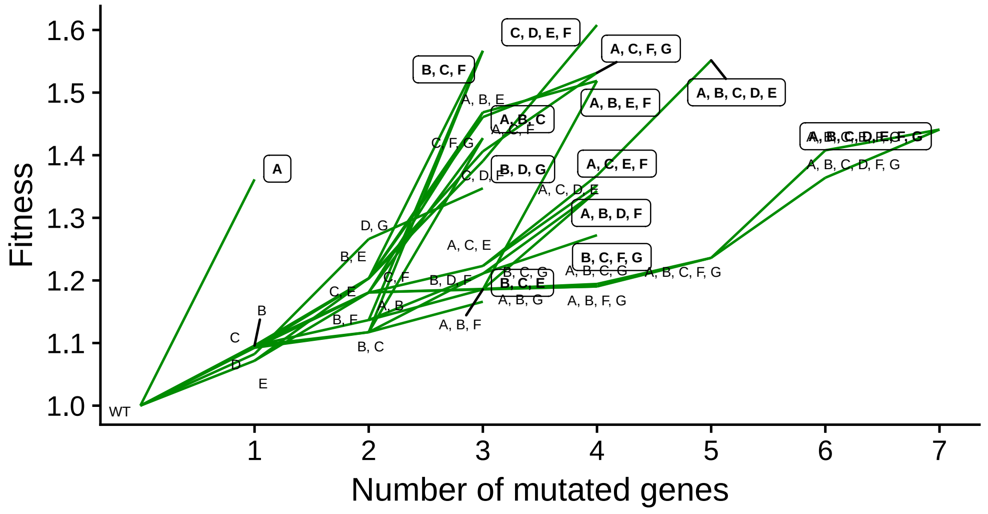


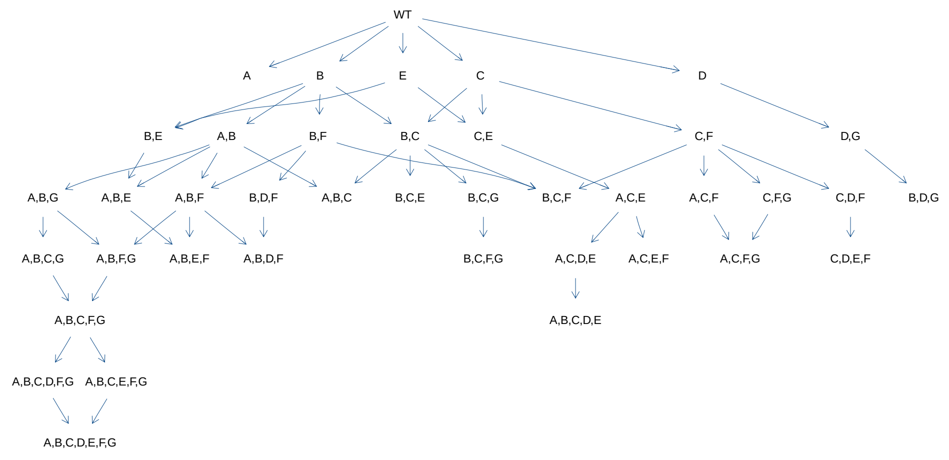

| | |
|---|---|
| *ID* | GJH85dNm2AtrzoUI |
| *acc genotypes* | 30 |
| *peaks no back* | 13 |
| *removed edges* | NA |
| *prop rem edges* | NA |

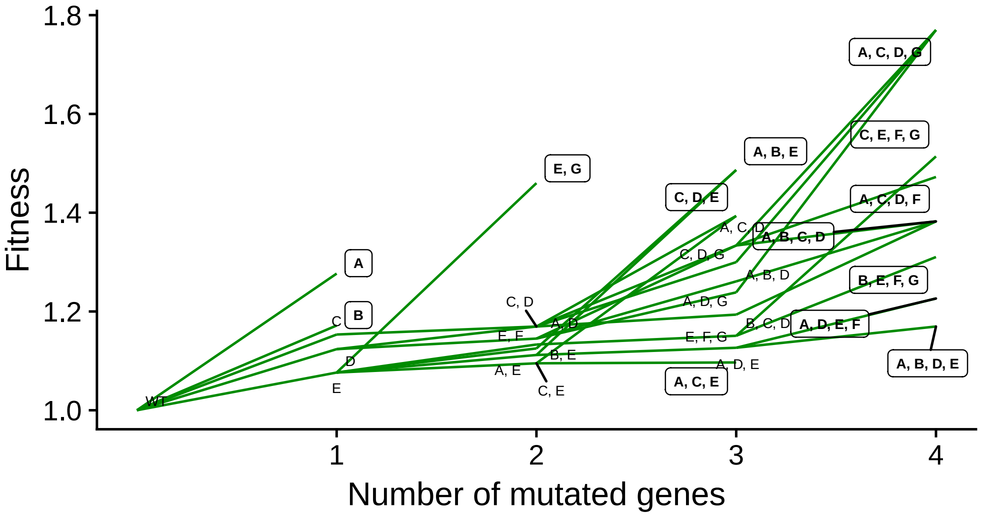


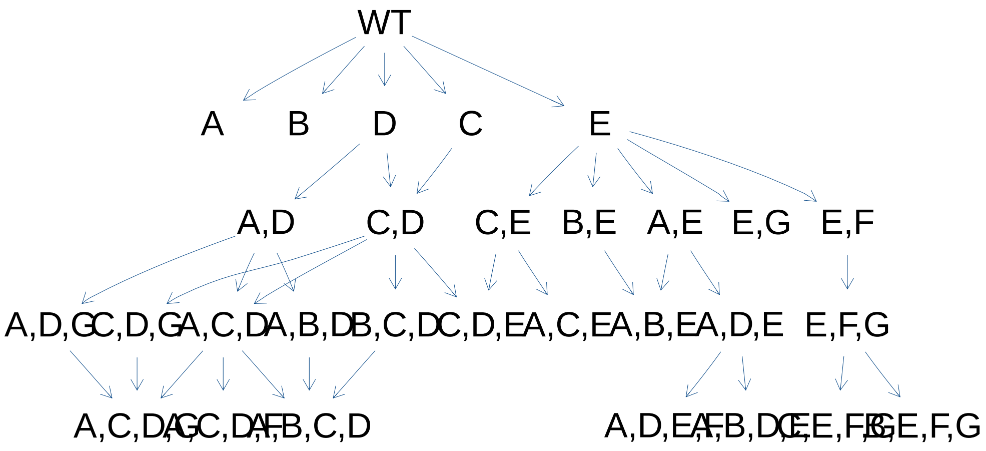

| | |
|---|---|
| *ID* | GrnJzITUvPUix6BN |
| *acc genotypes* | 51 |
| *peaks no back* | 19 |
| *removed edges* | NA |
| *prop rem edges* | NA |

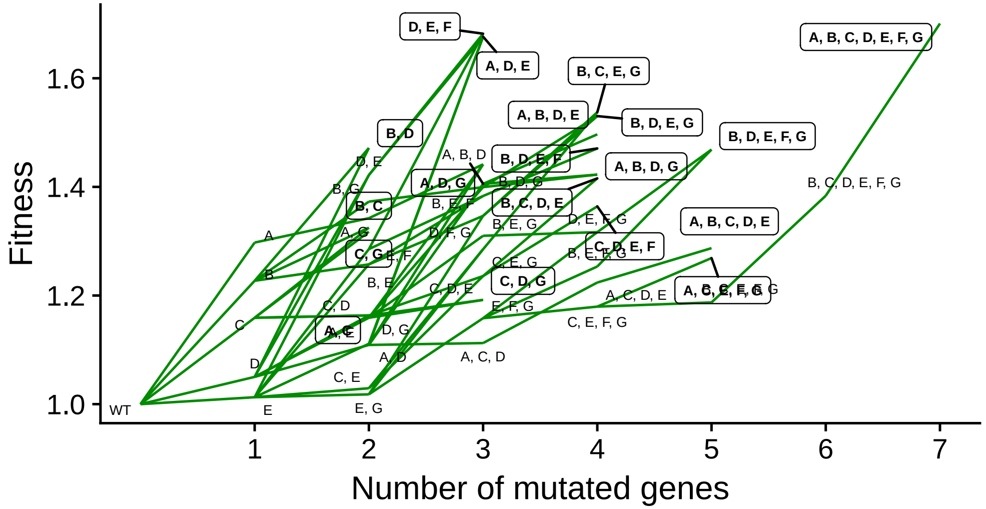


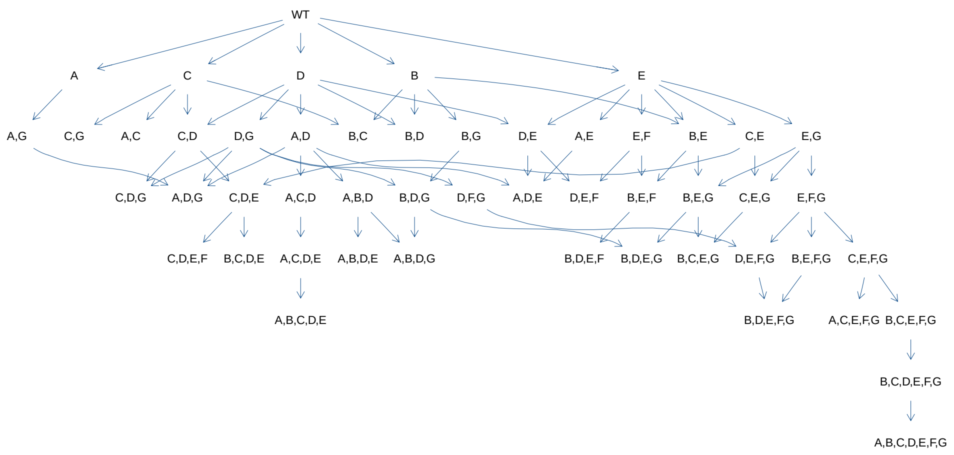

| | |
|---|---|
| *ID* | Hm3E9vk10sP85Ykfb |
| *acc genotypes* | 14 |
| *peaks no back* | 6 |
| *removed edges* | NA |
| *prop rem edges* | NA |

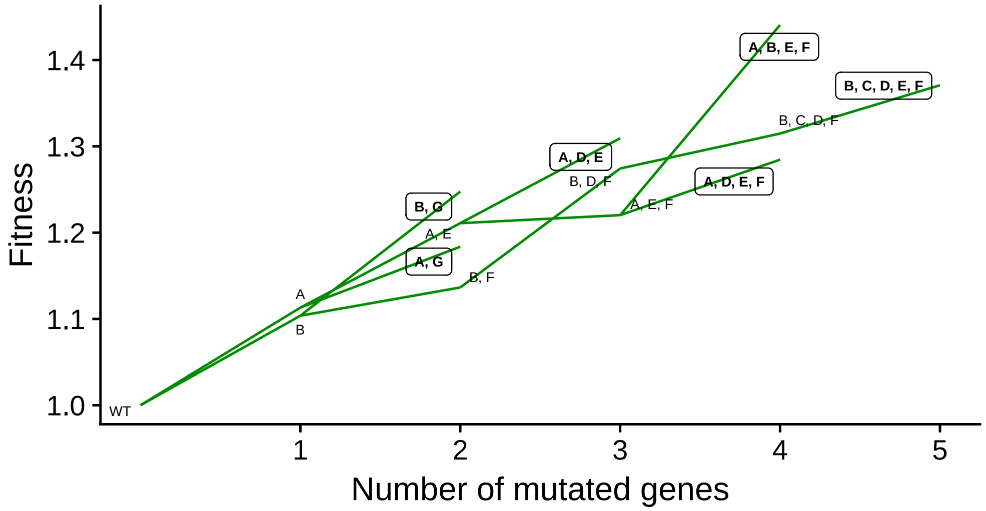


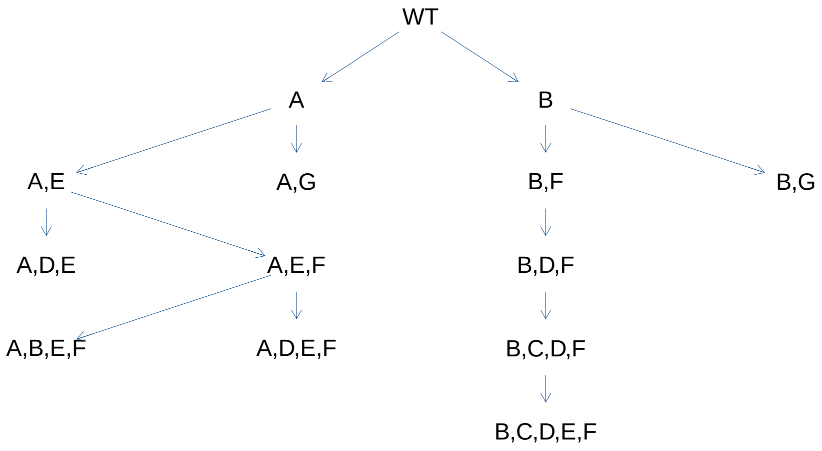

| | |
|---|---|
| *ID* | hnKNIikQ9aKghXCm |
| *acc genotypes* | 36 |
| *peaks no back* | 14 |
| *removed edges* | NA |
| *prop rem edges* | NA |

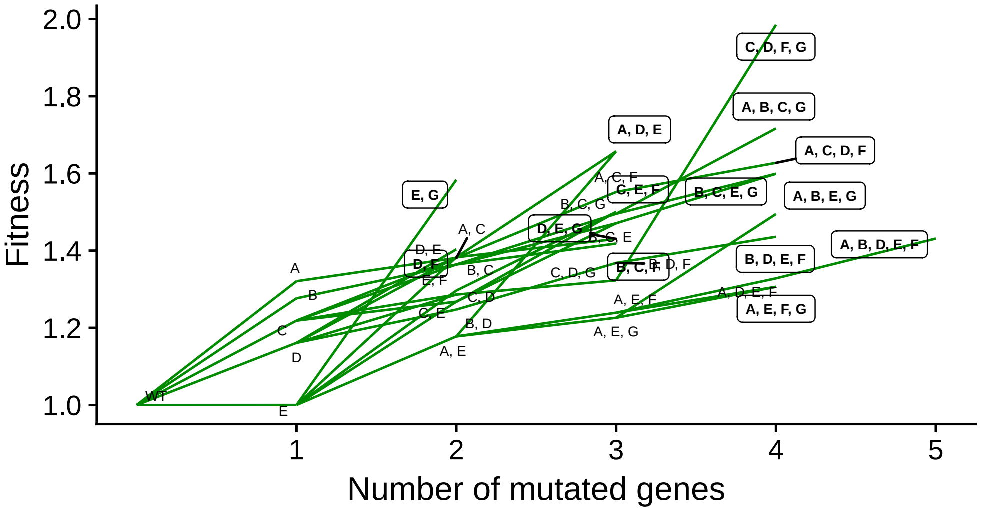


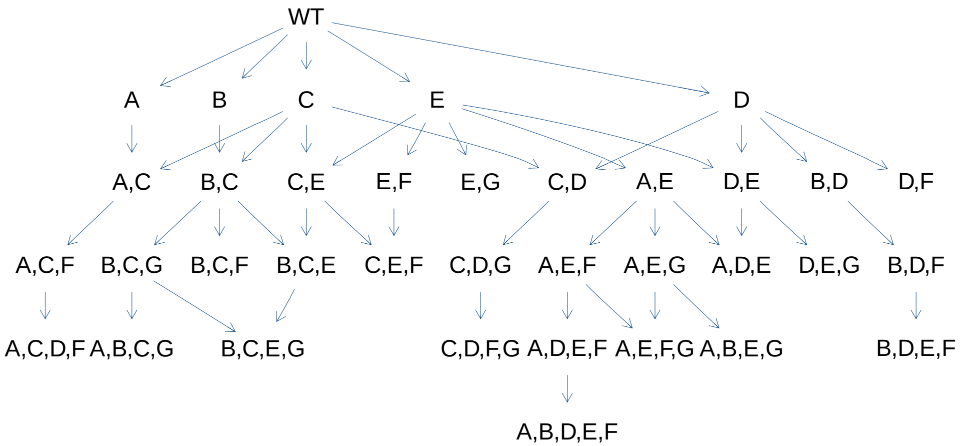

| | |
|---|---|
| *ID* | HUGD4mPfT3Y9PL8i |
| *acc genotypes* | 18 |
| *peaks no back* | 4 |
| *removed edges* | NA |
| *prop rem edges* | NA |

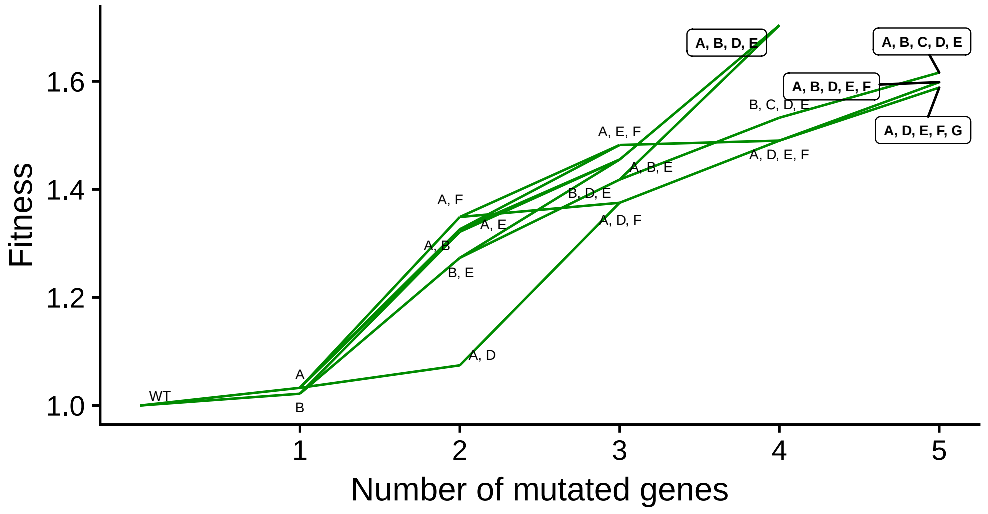


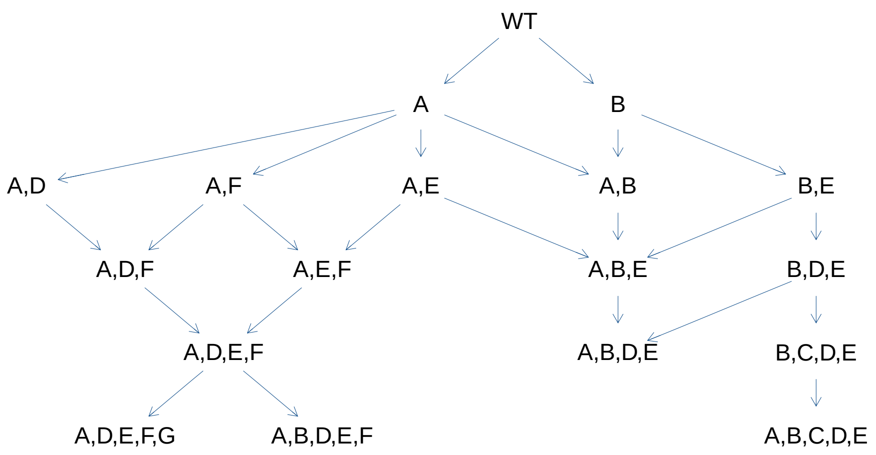

| | |
|---|---|
| *ID* | HVUm1KRpYXeSvmPO |
| *acc genotypes* | 28 |
| *peaks no back* | 13 |
| *removed edges* | NA |
| *prop rem edges* | NA |

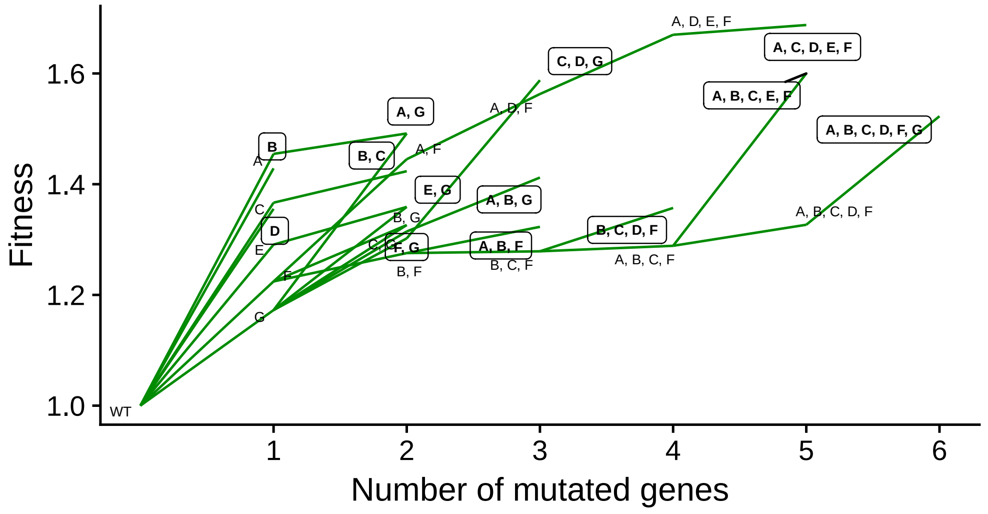


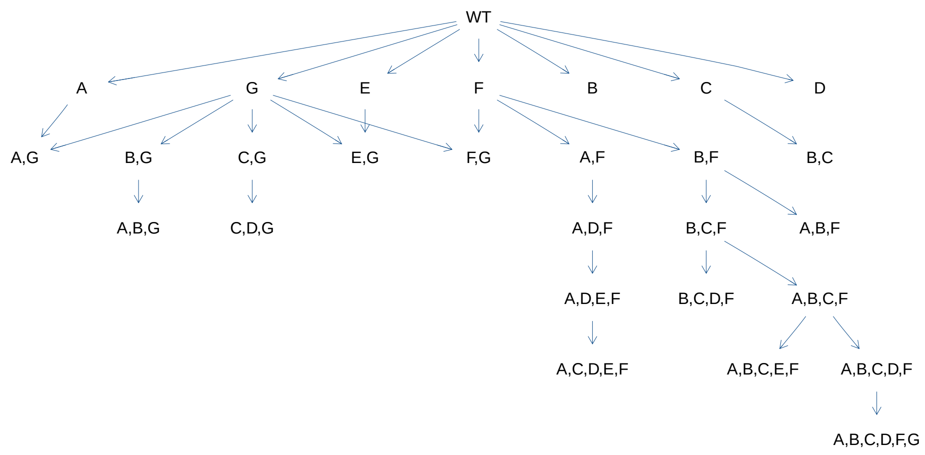

| | |
|---|---|
| *ID* | hwWOucBKvmrZN9ue |
| *acc genotypes* | 55 |
| *peaks no back* | 15 |
| *removed edges* | NA |
| *prop rem edges* | NA |

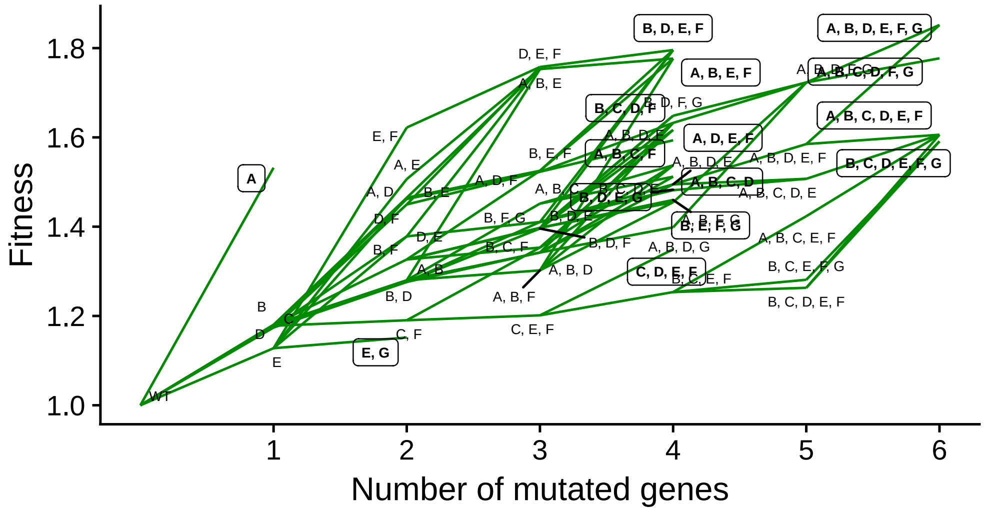


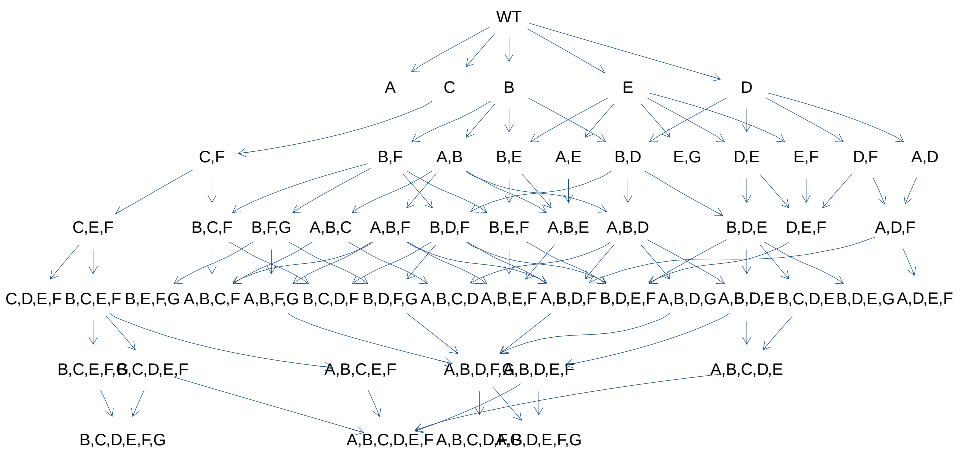

| | |
|---|---|
| *ID* | hXnNgnV34hLXDvEz |
| *acc genotypes* | 49 |
| *peaks no back* | 16 |
| *removed edges* | NA |
| *prop rem edges* | NA |

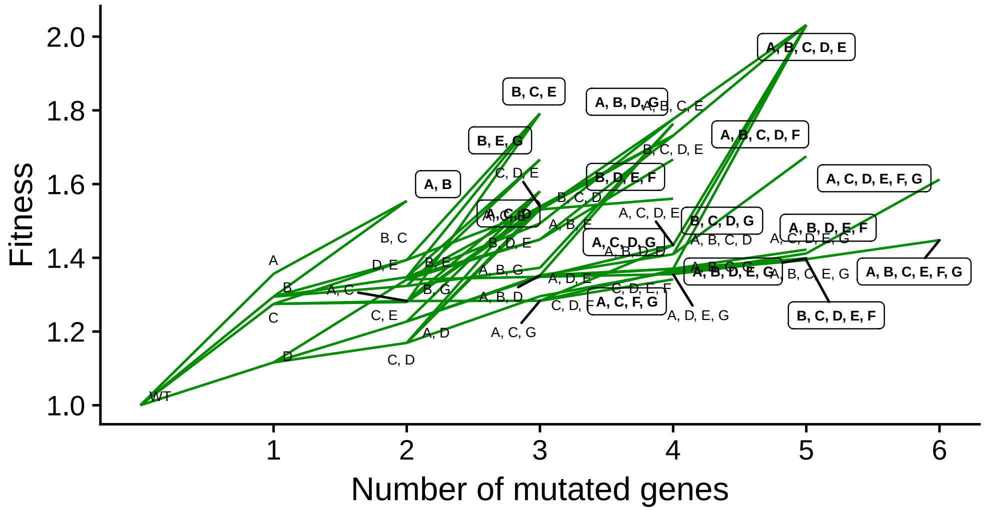

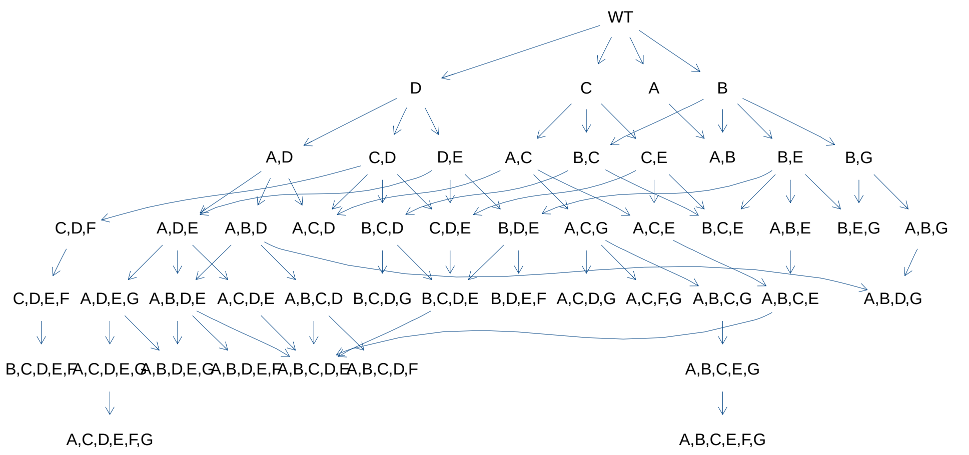

| | |
|---|---|
| *ID* | hz9XzL5TRKBXl8Xk |
| *acc genotypes* | 45 |
| *peaks no back* | 21 |
| *removed edges* | NA |
| *prop rem edges* | NA |

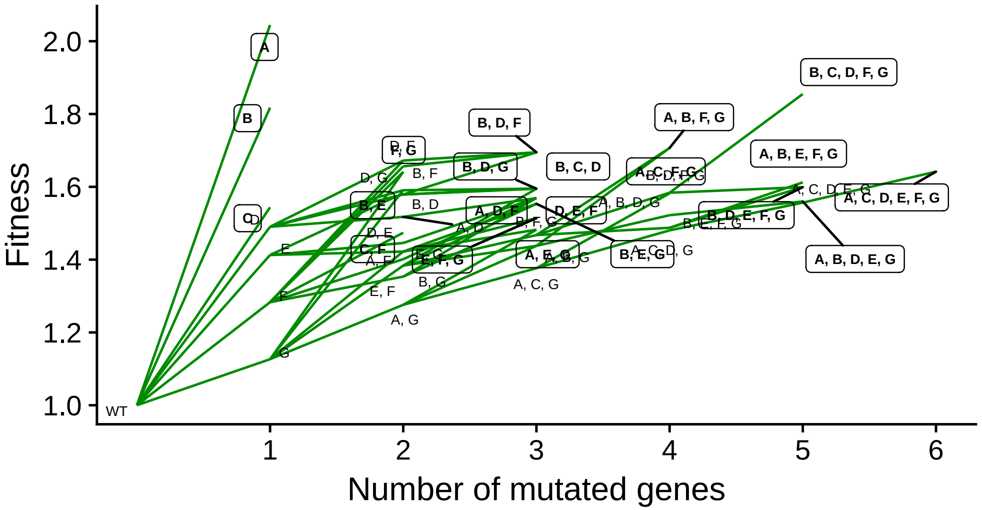


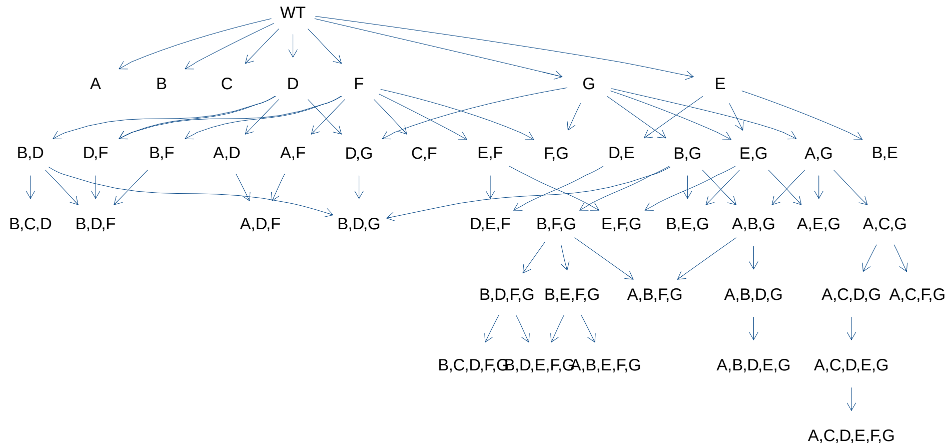

| | |
|---|---|
| *ID* | i102joVz69wXudIlf |
| *acc genotypes* | 26 |
| *peaks no back* | 11 |
| *removed edges* | NA |
| *prop rem edges* | NA |

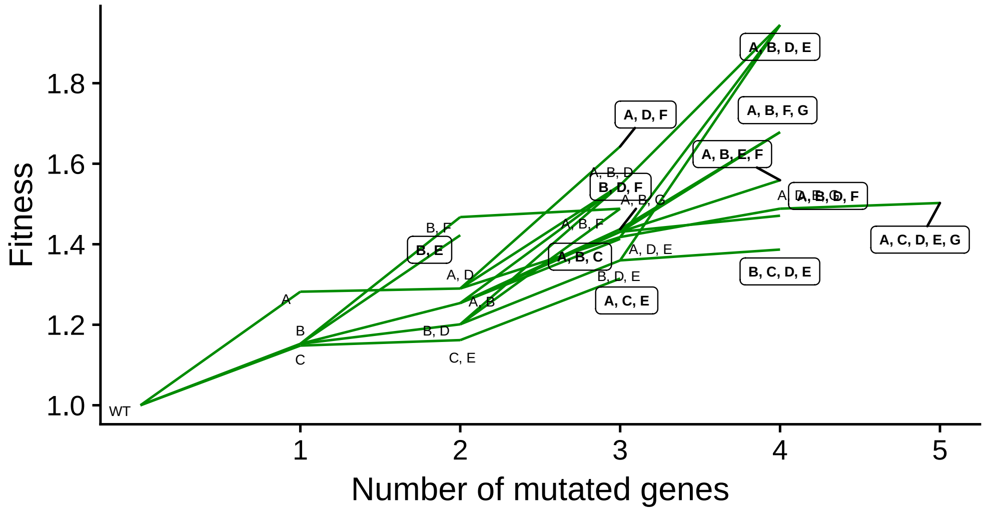


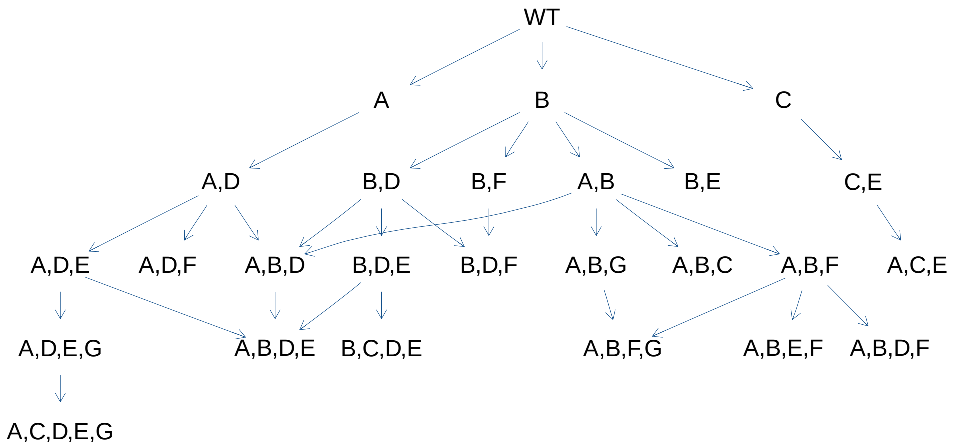

| | |
|---|---|
| *ID* | I4wGMzShjuo8XyuN |
| *acc genotypes* | 43 |
| *peaks no back* | 18 |
| *removed edges* | NA |
| *prop rem edges* | NA |

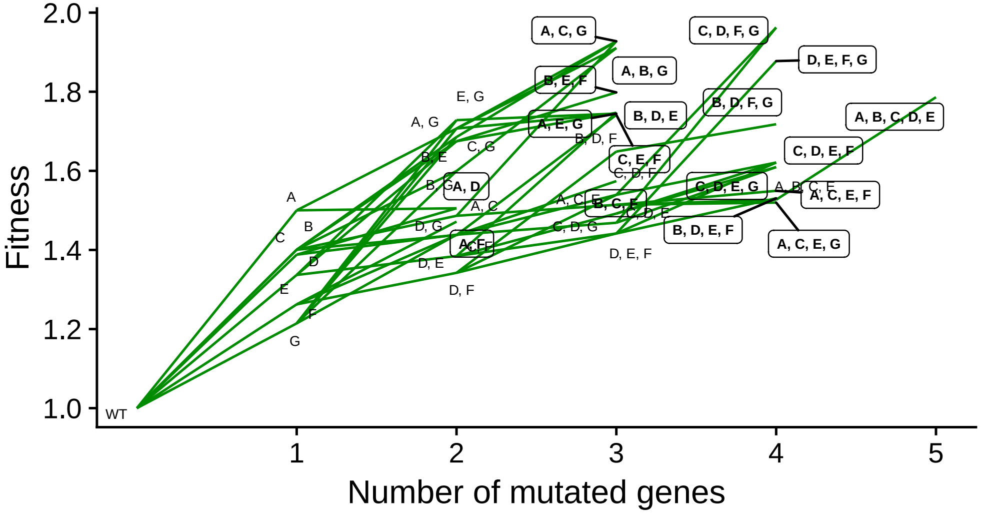


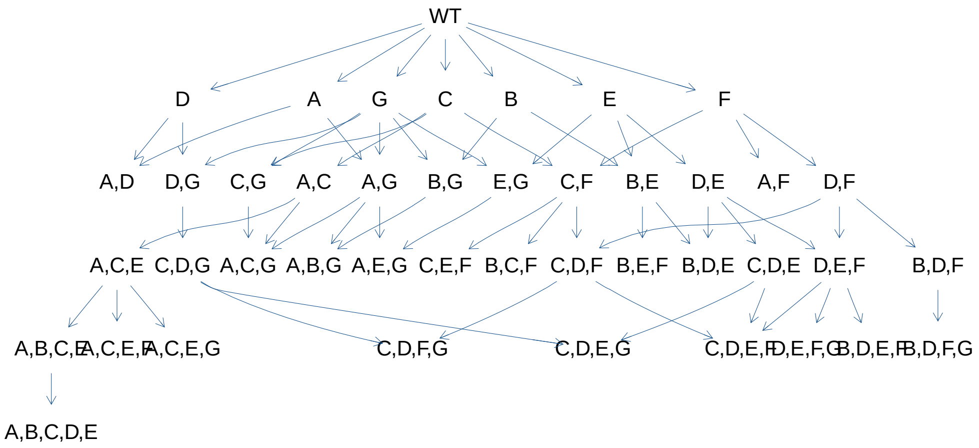

| | |
|---|---|
| *ID* | IDLT5qGFT3uM1rEn |
| *acc genotypes* | 50 |
| *peaks no back* | 19 |
| *removed edges* | NA |
| *prop rem edges* | NA |

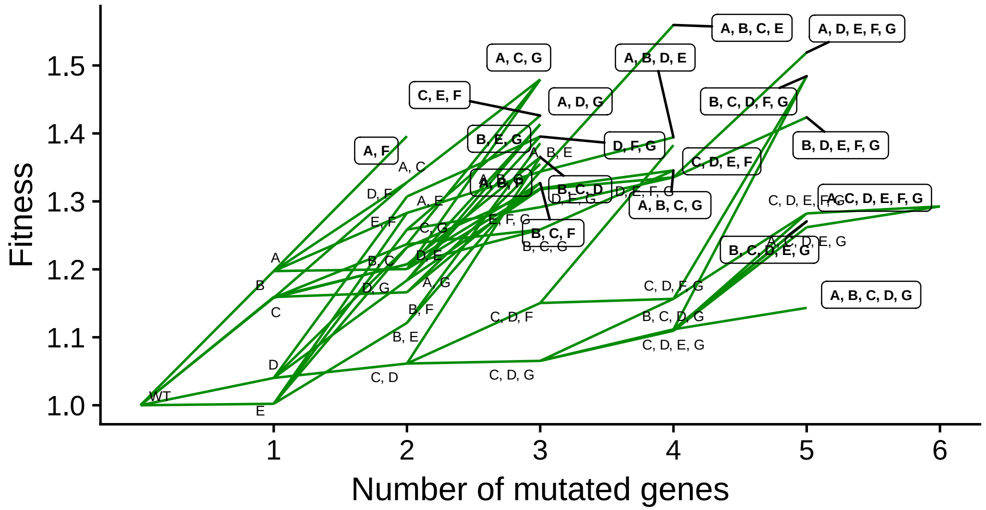


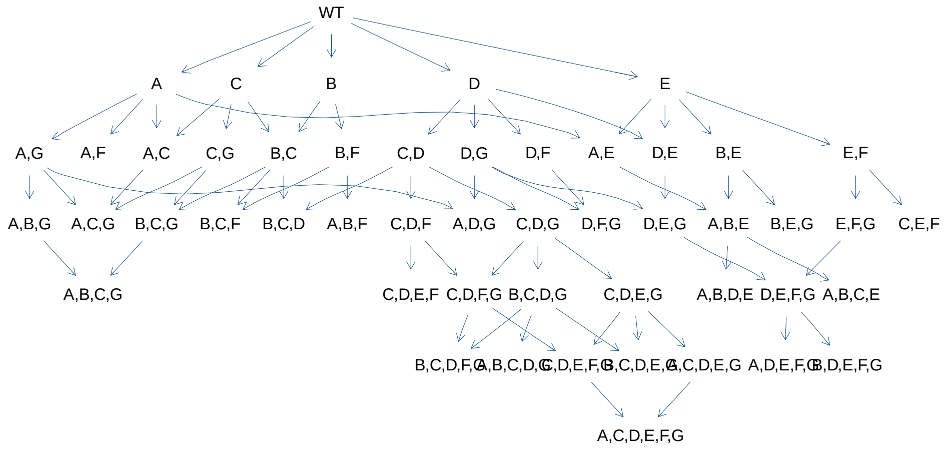

| | |
|---|---|
| *ID* | IJl10hVtdsgYgvoV7 |
| *acc genotypes* | 46 |
| *peaks no back* | 10 |
| *removed edges* | NA |
| *prop rem edges* | NA |

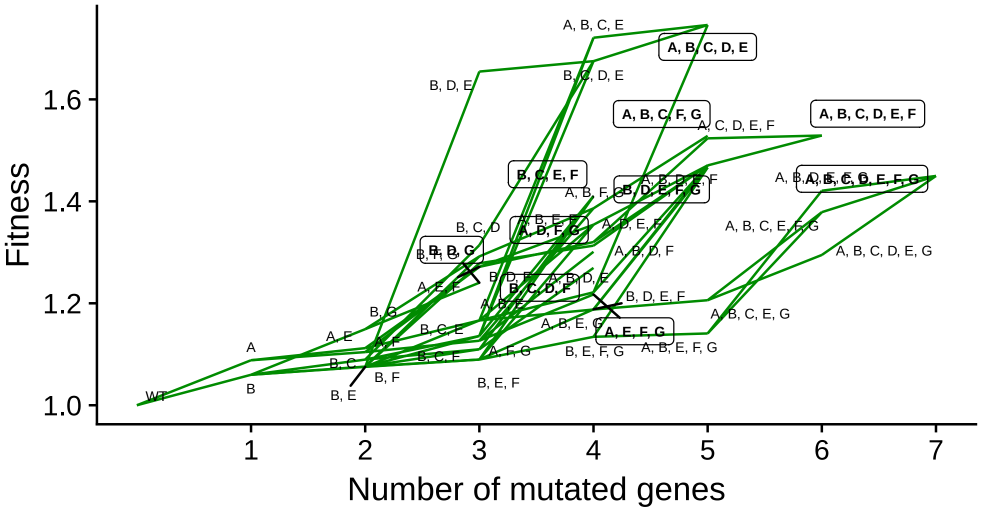


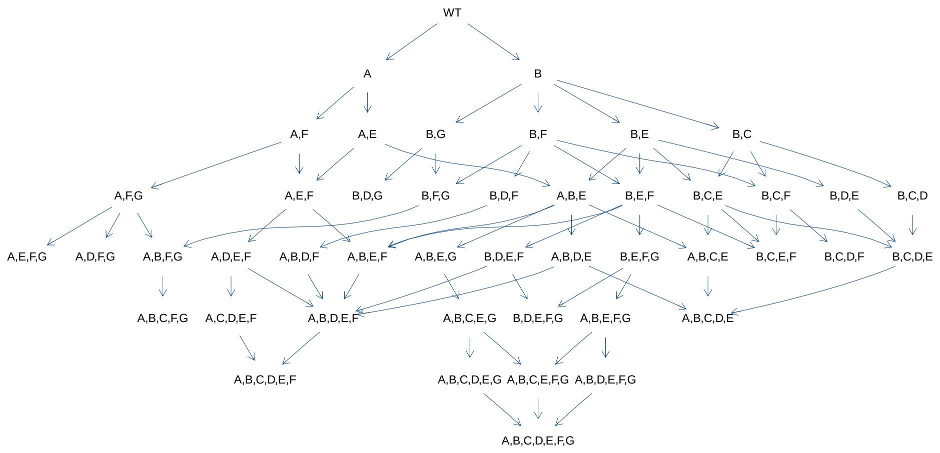

| | |
|---|---|
| *ID* | iuizHj29wSFmqntf |
| *acc genotypes* | 39 |
| *peaks no back* | 17 |
| *removed edges* | NA |
| *prop rem edges* | NA |

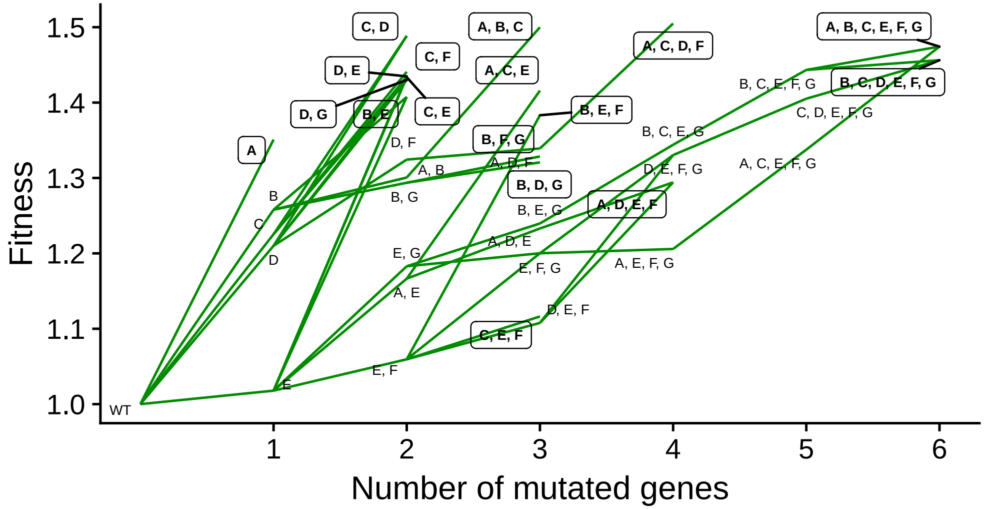


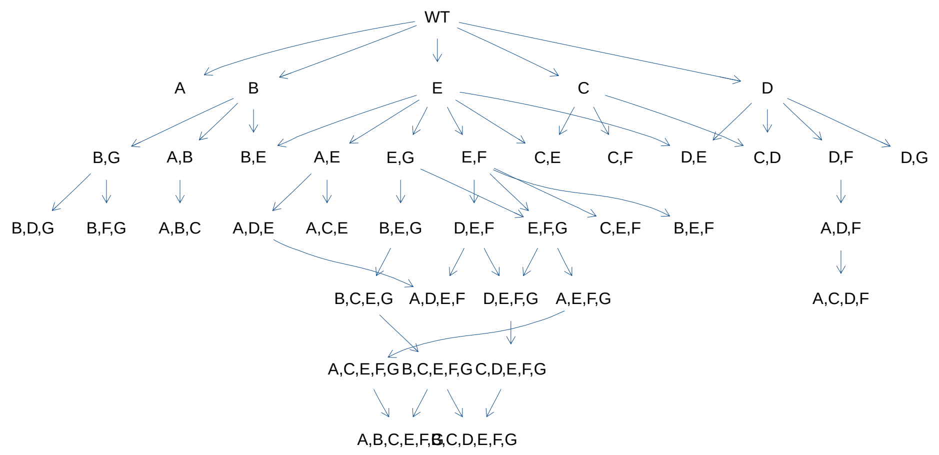

| | |
|---|---|
| *ID* | jARDfKjvS5xgIbjy |
| *acc genotypes* | 32 |
| *peaks no back* | 12 |
| *removed edges* | NA |
| *prop rem edges* | NA |

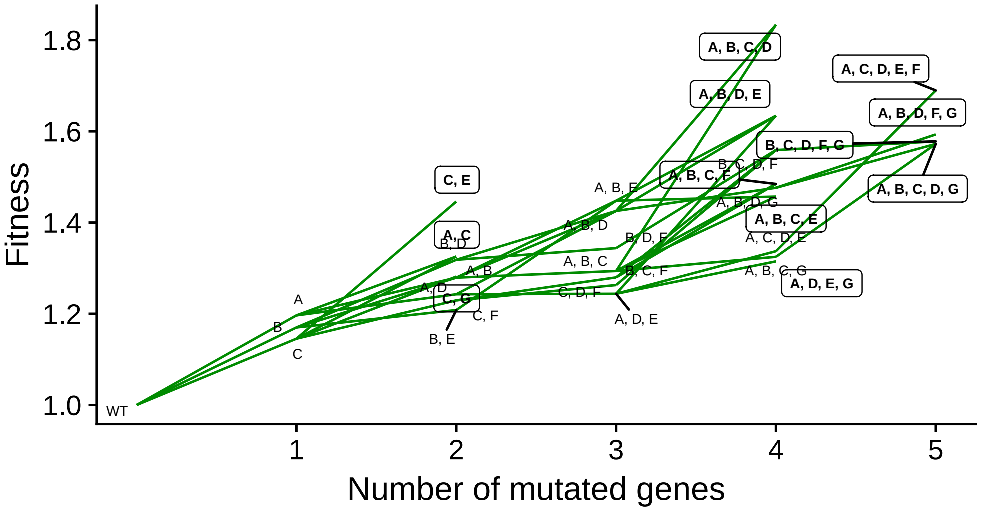


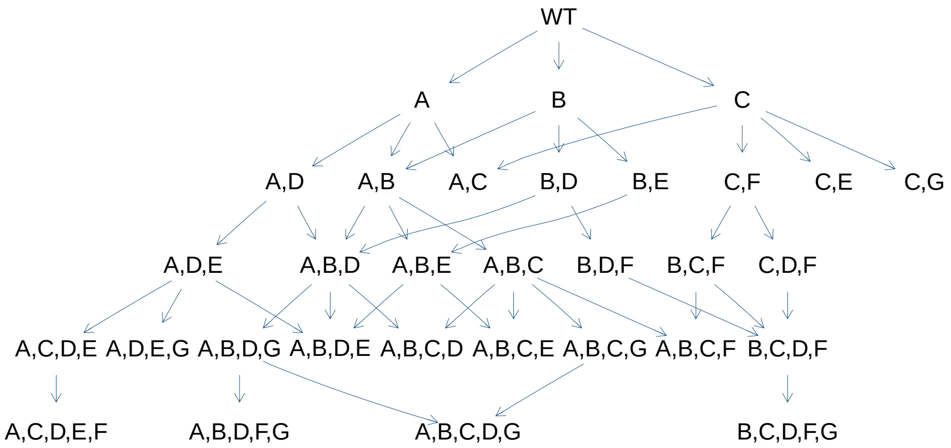

| | |
|---|---|
| *ID* | JC7USXL8yt3B9M10N |
| *acc genotypes* | 36 |
| *peaks no back* | 15 |
| *removed edges* | NA |
| *prop rem edges* | NA |

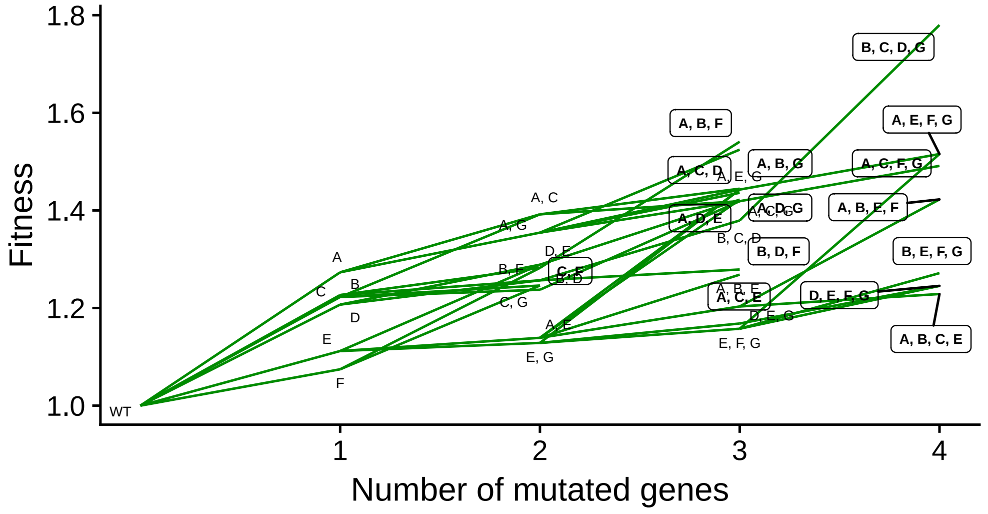


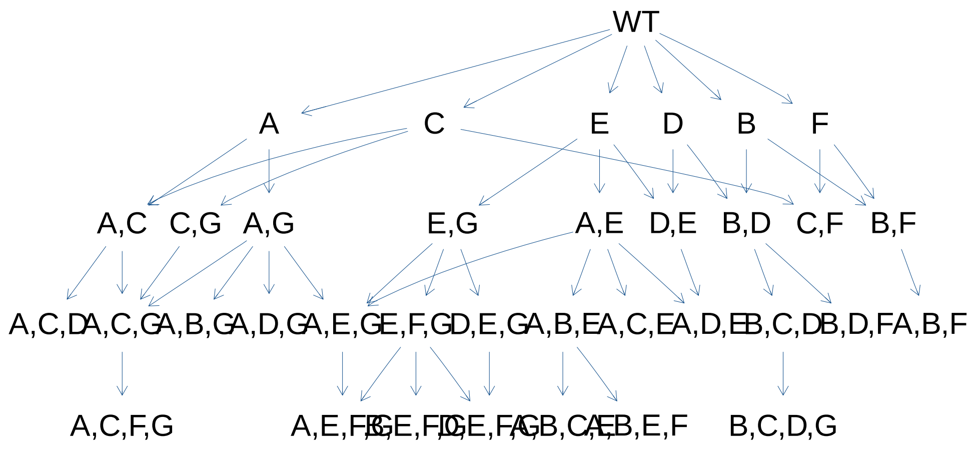

| | |
|---|---|
| *ID* | JdIeKg10Z6a5TlQp5 |
| *acc genotypes* | 21 |
| *peaks no back* | 7 |
| *removed edges* | NA |
| *prop rem edges* | NA |

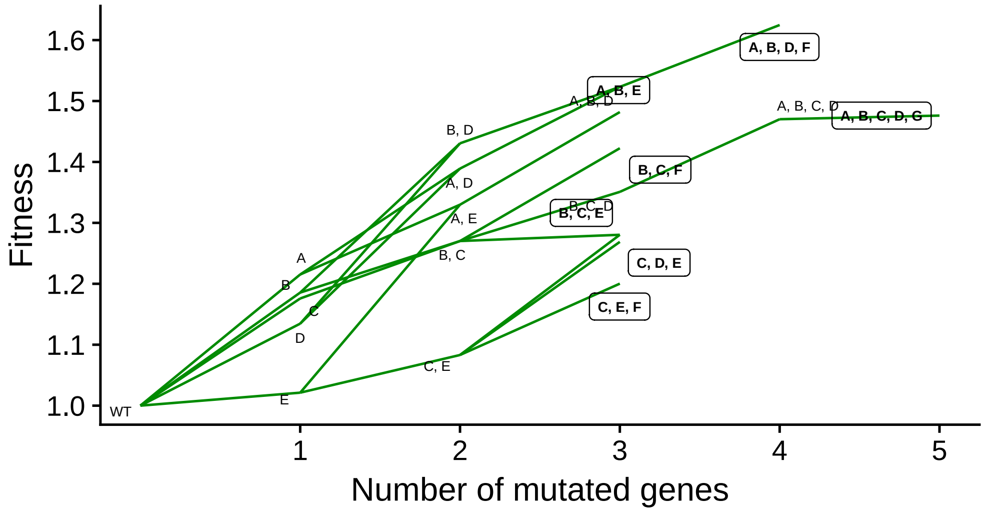


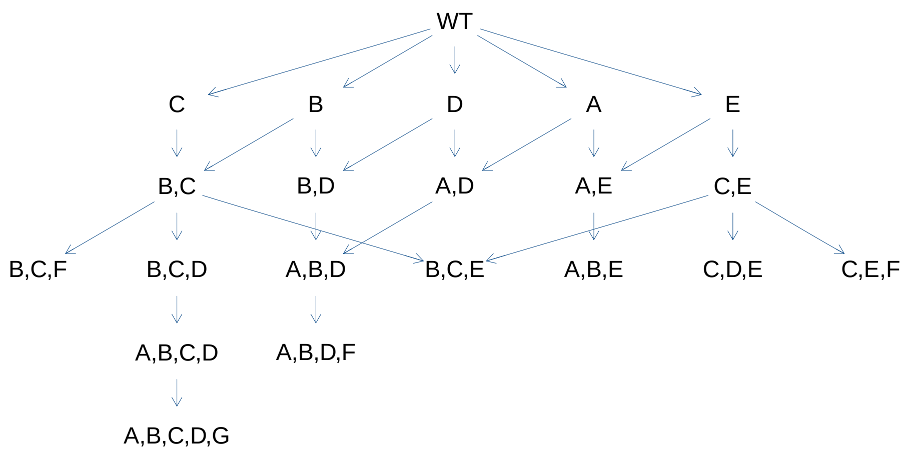

| | |
|---|---|
| *ID* | jJkQq5oUGT4cTflw |
| *acc genotypes* | 11 |
| *peaks no back* | 5 |
| *removed edges* | NA |
| *prop rem edges* | NA |

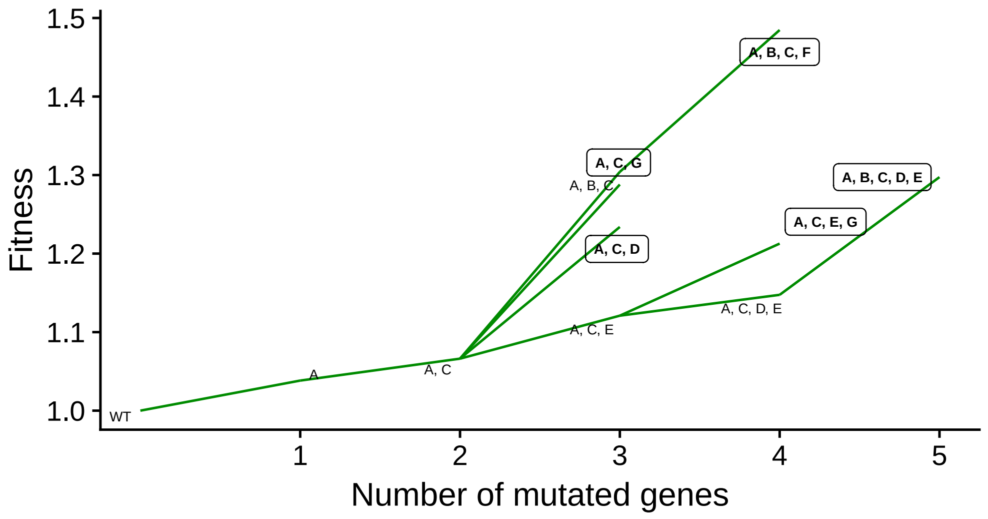


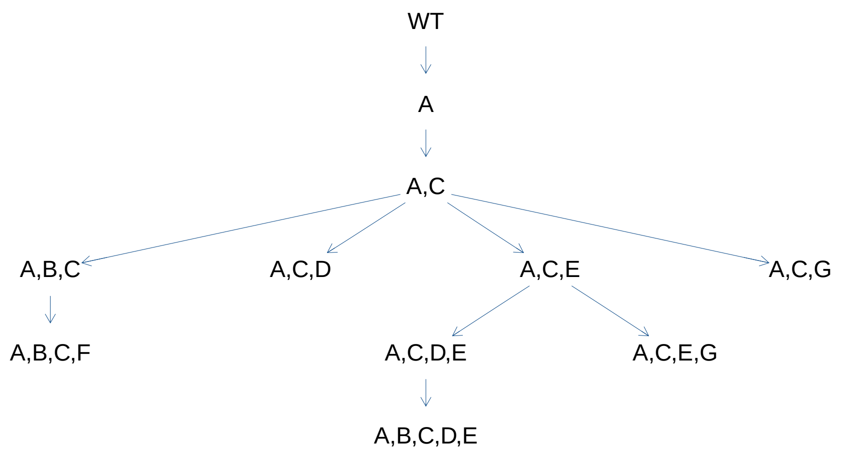

| | |
|---|---|
| *ID* | JqytlTKKThh5QBbf |
| *acc genotypes* | 11 |
| *peaks no back* | 5 |
| *removed edges* | NA |
| *prop rem edges* | NA |

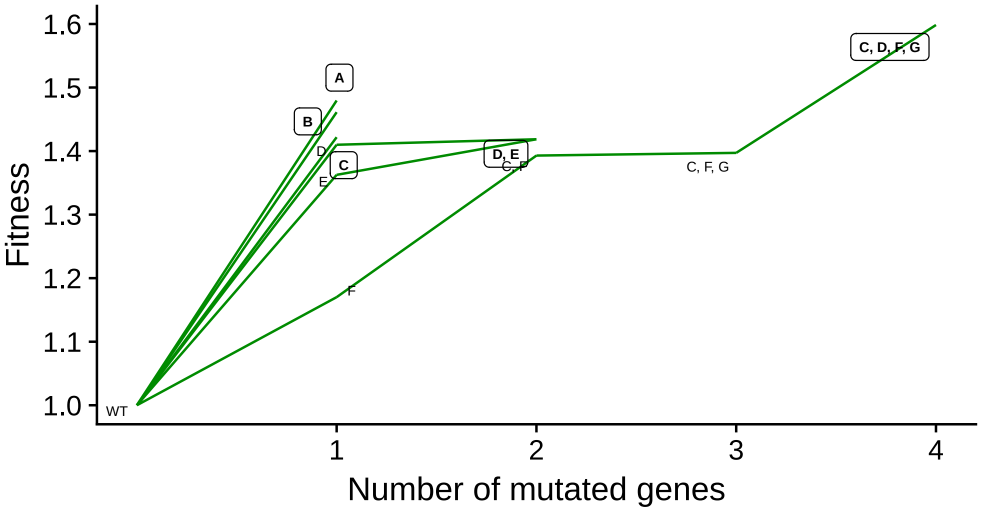


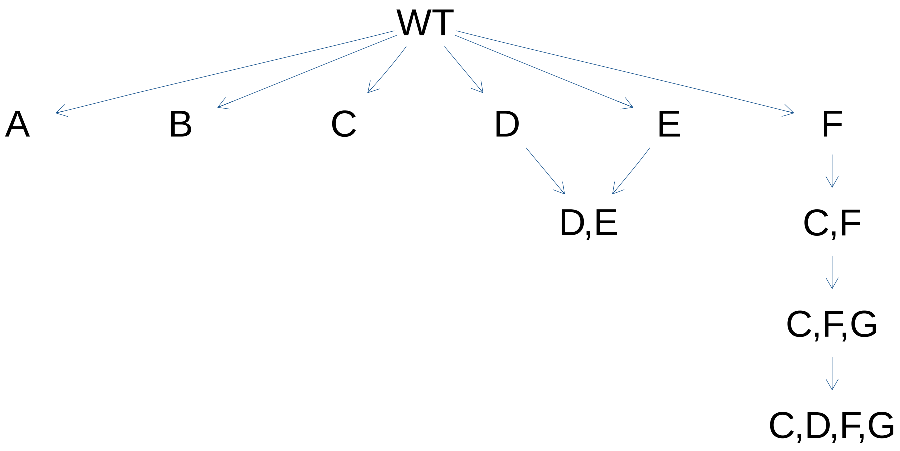

| | |
|---|---|
| *ID* | jvGYmalfWVHjDSsr |
| *acc genotypes* | 39 |
| *peaks no back* | 15 |
| *removed edges* | NA |
| *prop rem edges* | NA |

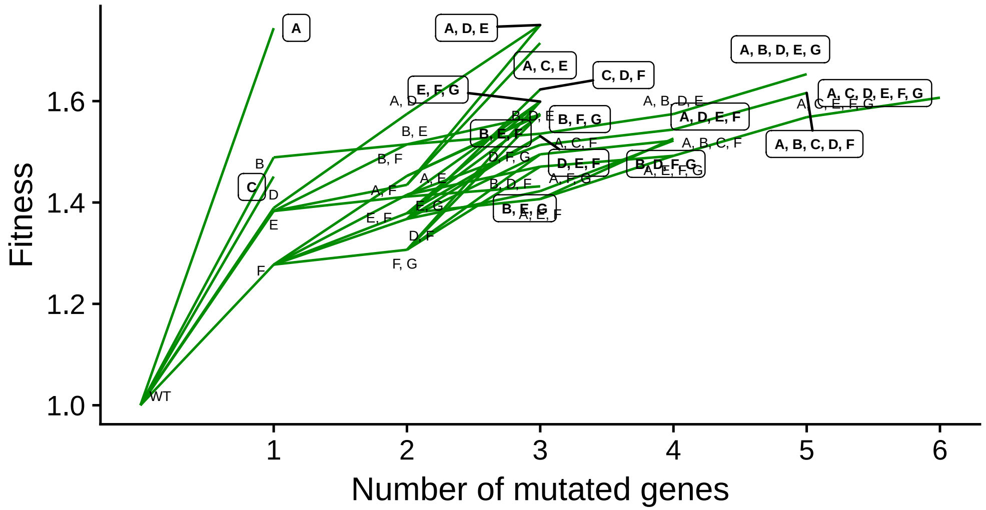


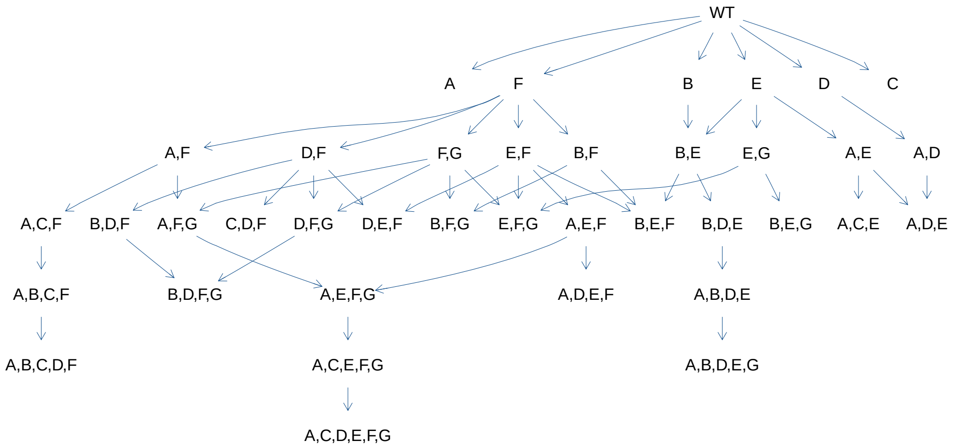

| | |
|---|---|
| *ID* | jvTSHGXJvbnFtVLt |
| *acc genotypes* | 30 |
| *peaks no back* | 9 |
| *removed edges* | NA |
| *prop rem edges* | NA |

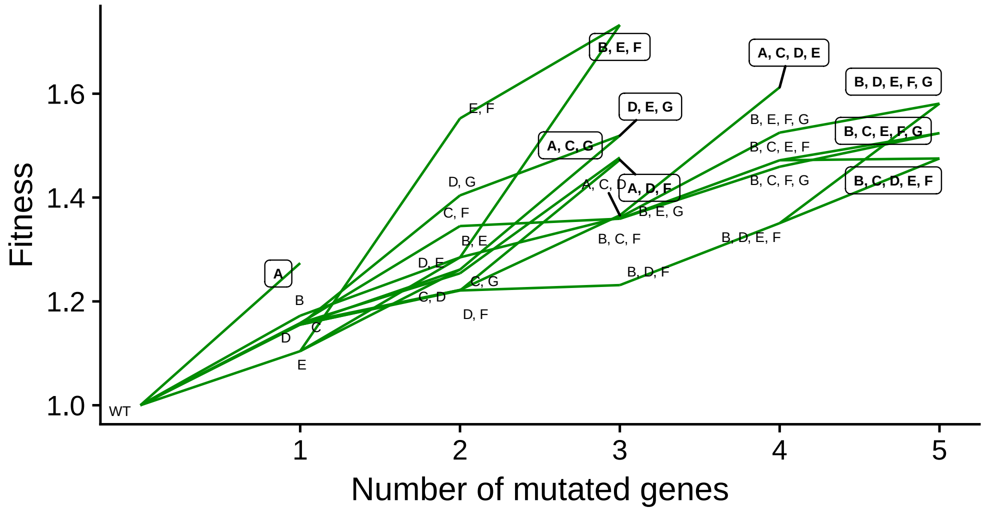


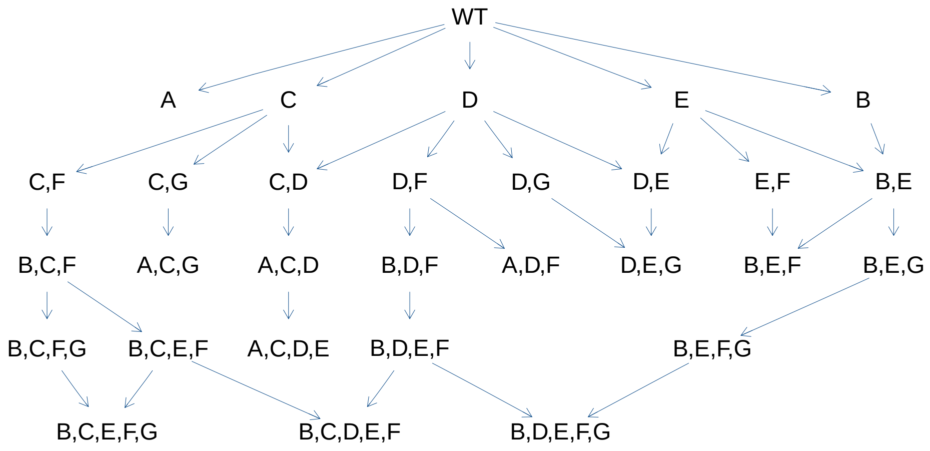

| | |
|---|---|
| *ID* | k4wHgtkweyDrnCn8 |
| *acc genotypes* | 80 |
| *peaks no back* | 17 |
| *removed edges* | NA |
| *prop rem edges* | NA |

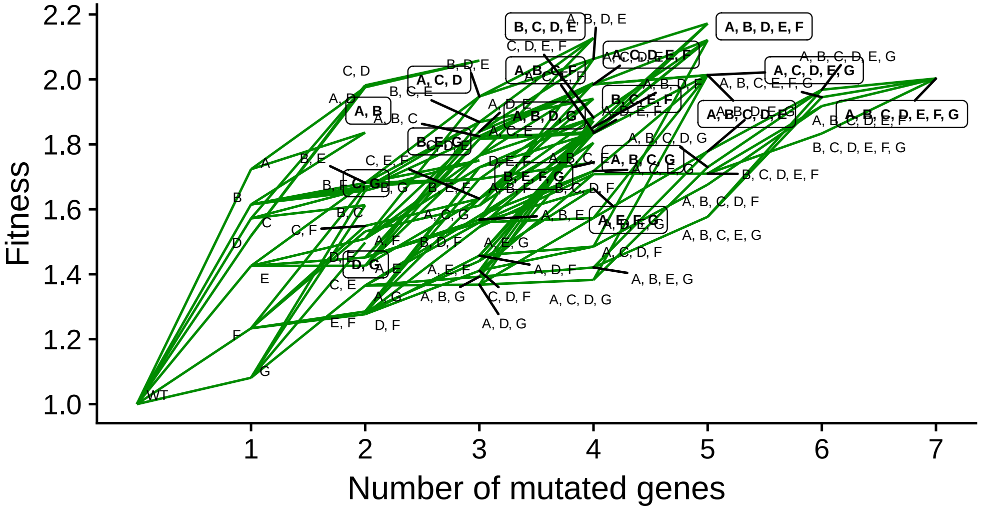


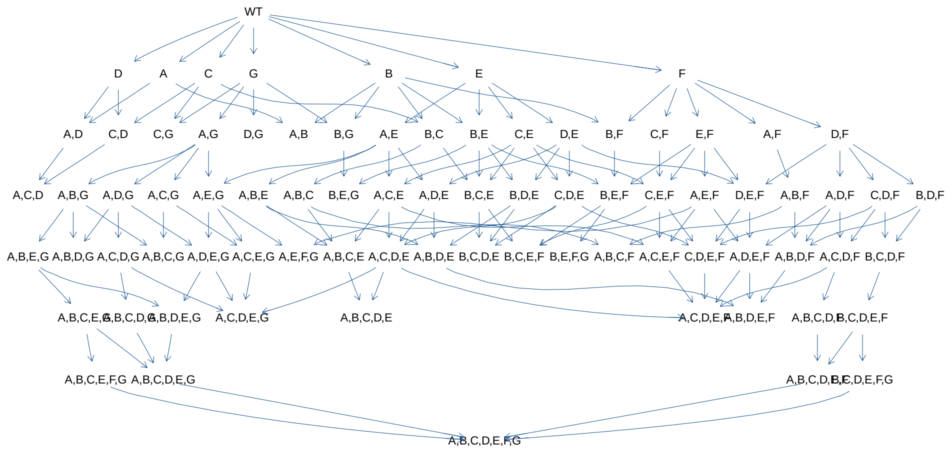

| | |
|---|---|
| *ID* | kd10vK9jGcoOkaAMm |
| *acc genotypes* | 75 |
| *peaks no back* | 7 |
| *removed edges* | NA |
| *prop rem edges* | NA |

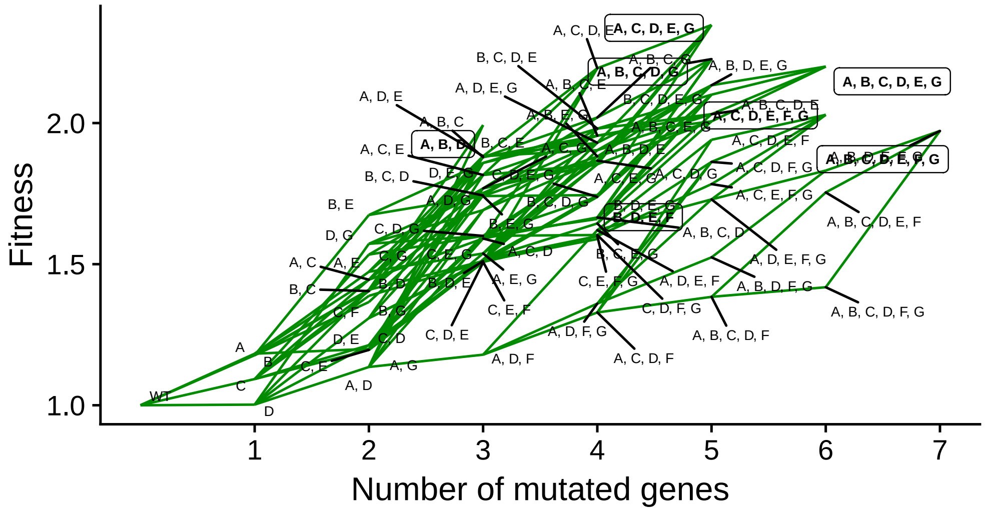


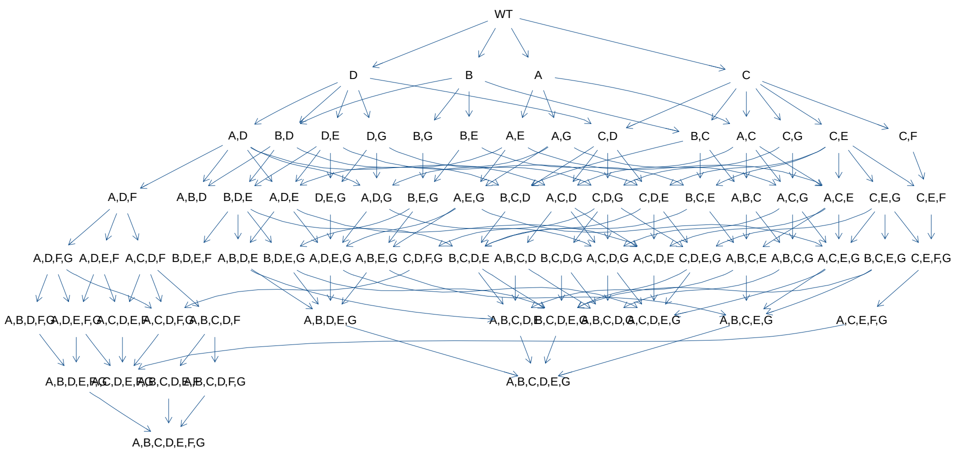

| | |
|---|---|
| *ID* | kfB7NOk1J247FNsy |
| *acc genotypes* | 18 |
| *peaks no back* | 8 |
| *removed edges* | NA |
| *prop rem edges* | NA |

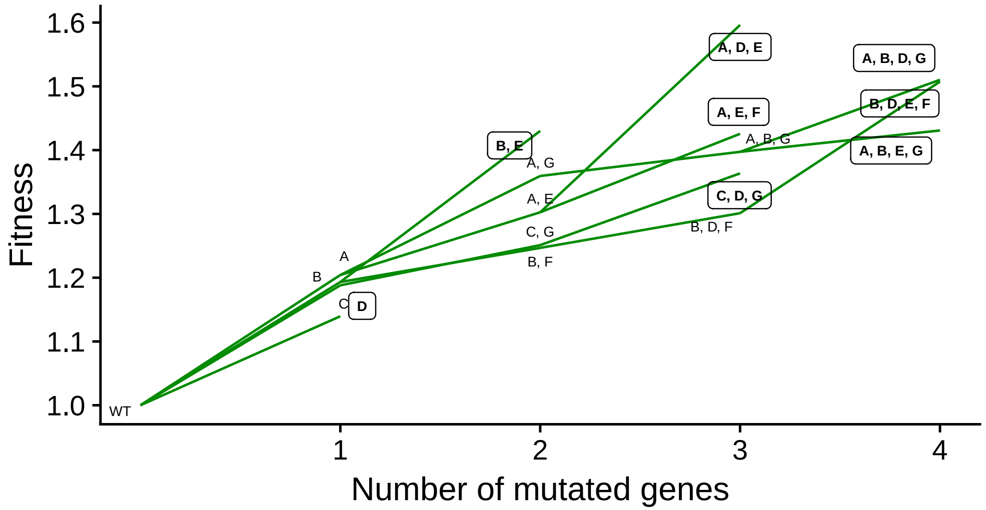


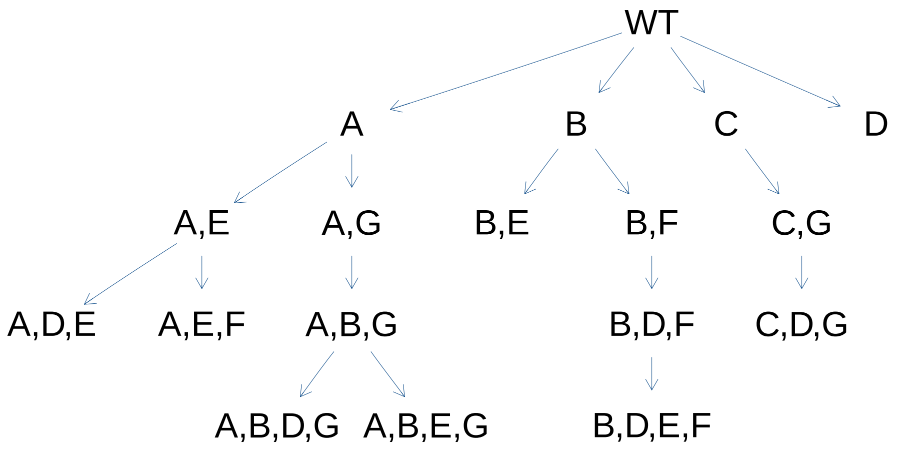

| | |
|---|---|
| *ID* | kfC76EeHnueEMKzE |
| *acc genotypes* | 70 |
| *peaks no back* | 19 |
| *removed edges* | NA |
| *prop rem edges* | NA |

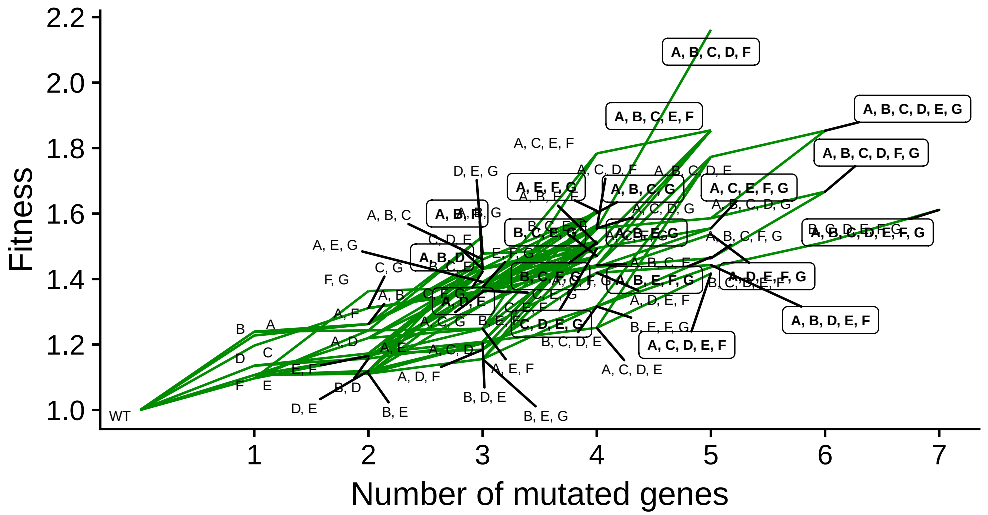

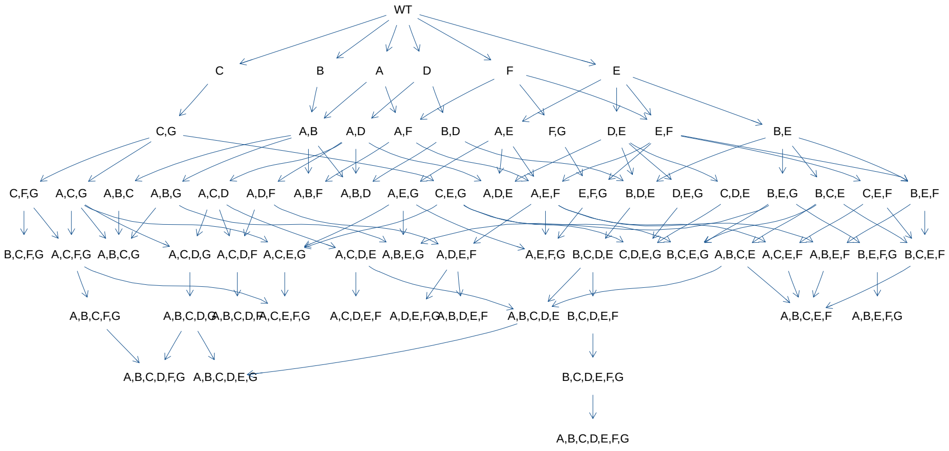

| | |
|---|---|
| *ID* | knzBi4rUYms1ZmCY |
| *acc genotypes* | 57 |
| *peaks no back* | 20 |
| *removed edges* | NA |
| *prop rem edges* | NA |

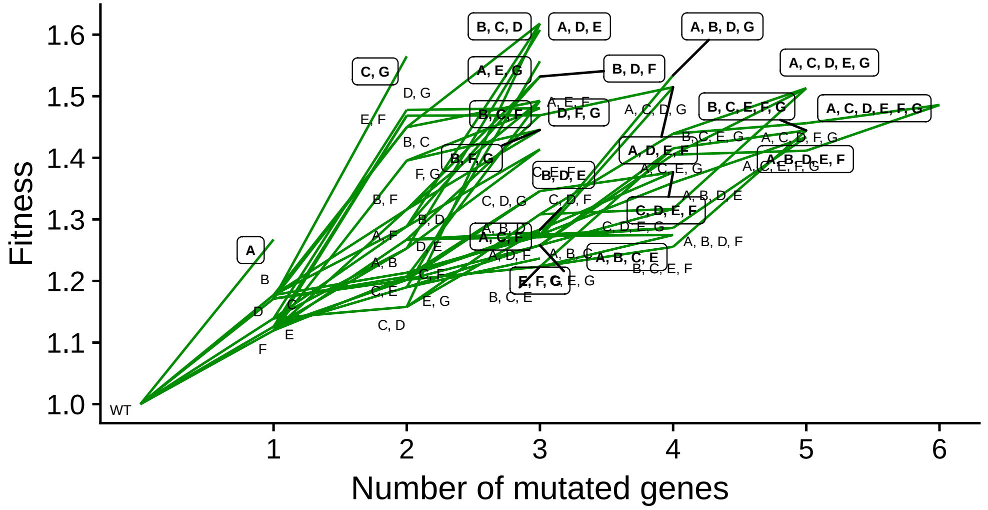


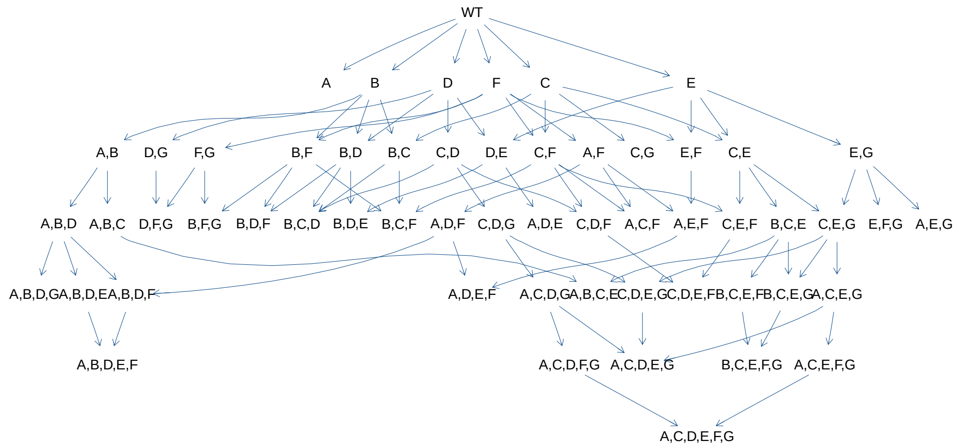

| | |
|---|---|
| *ID* | KRi7SXjpurE910L5q |
| *acc genotypes* | 37 |
| *peaks no back* | 13 |
| *removed edges* | NA |
| *prop rem edges* | NA |

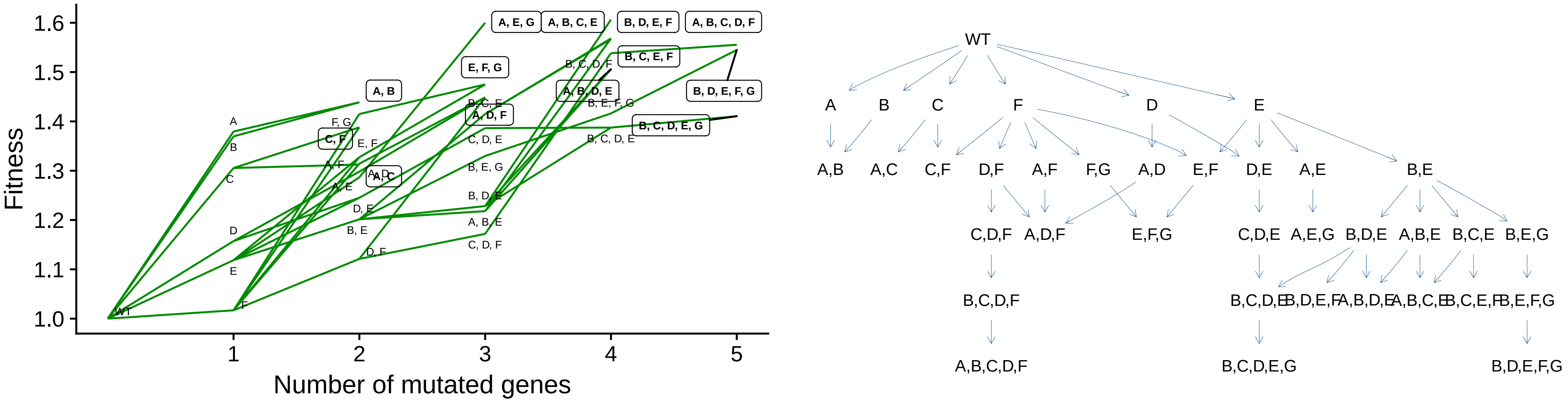

| | |
|---|---|
| *ID* | KwdXNcrapMR7uPi7 |
| *acc genotypes* | 31 |
| *peaks no back* | 10 |
| *removed edges* | NA |
| *prop rem edges* | NA |

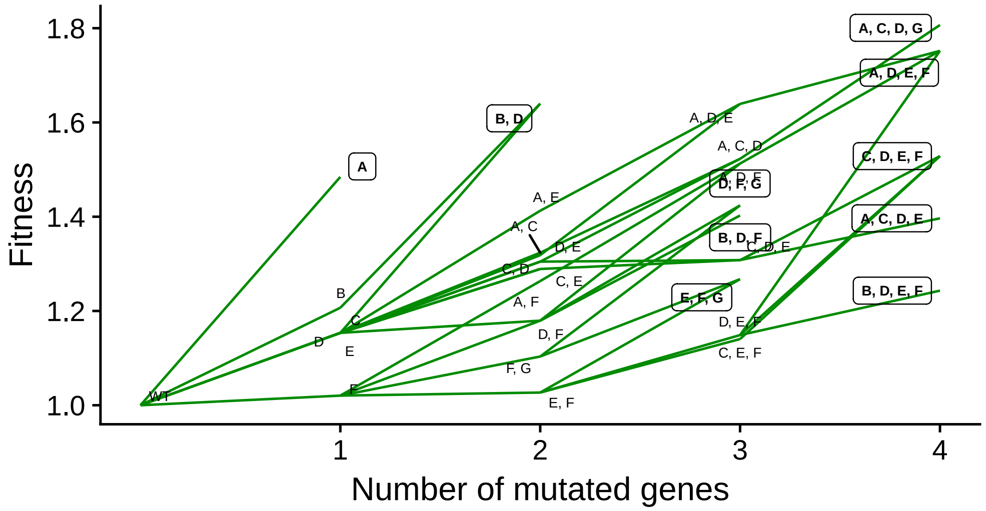


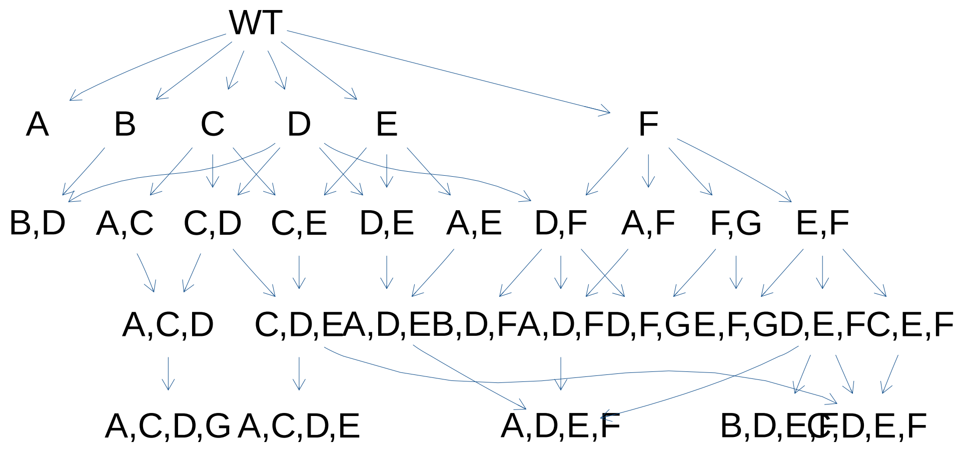

| | |
|---|---|
| *ID* | L53nf1rd3BsJMUlj |
| *acc genotypes* | 53 |
| *peaks no back* | 20 |
| *removed edges* | NA |
| *prop rem edges* | NA |

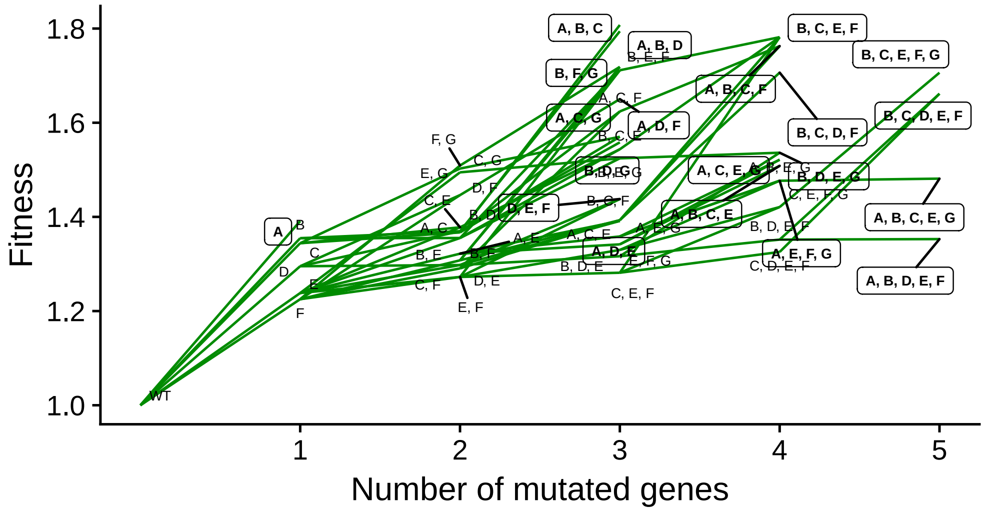


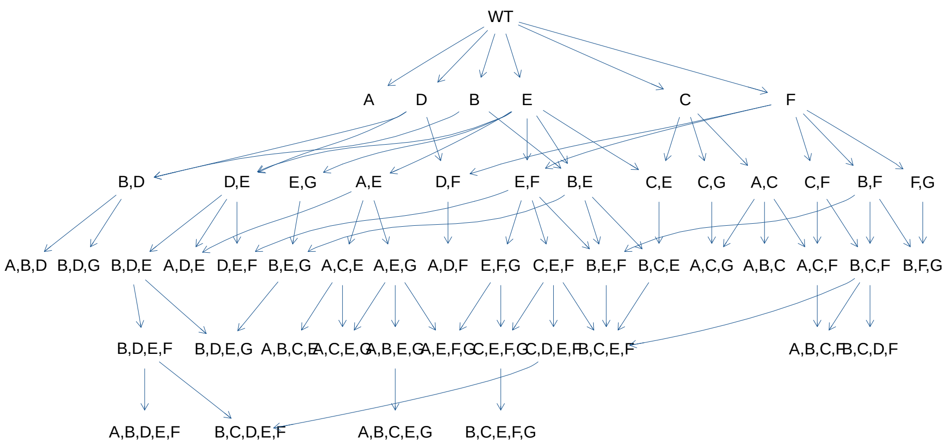

| | |
|---|---|
| *ID* | L96VgnsLKl1UgcXs |
| *acc genotypes* | 29 |
| *peaks no back* | 10 |
| *removed edges* | NA |
| *prop rem edges* | NA |

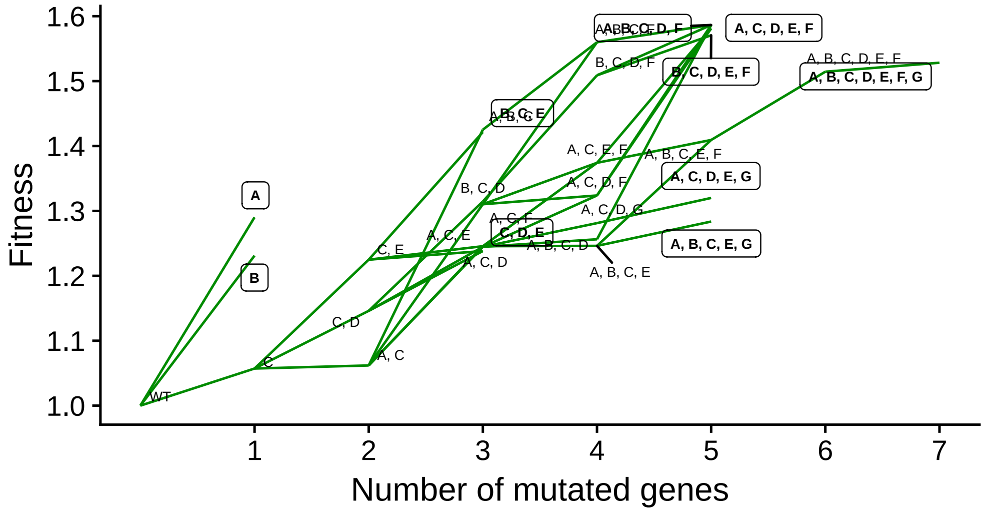


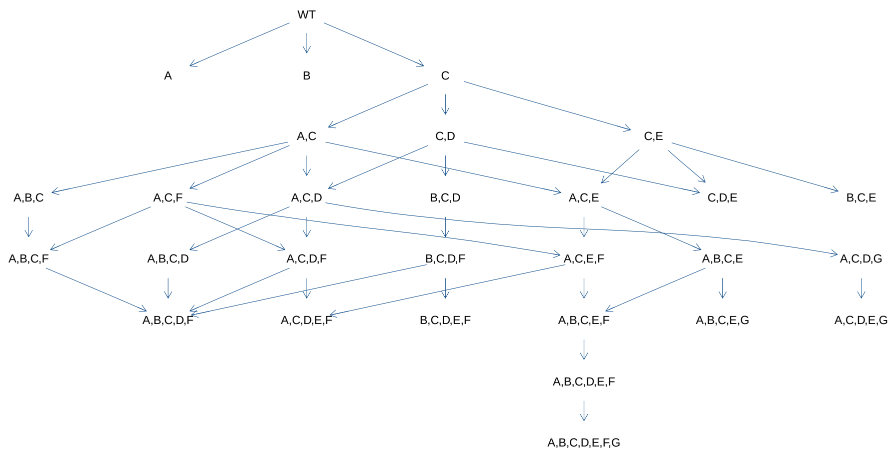

| | |
|---|---|
| *ID* | IG7VsdckSOb10Utw8 |
| *acc genotypes* | 40 |
| *peaks no back* | 13 |
| *removed edges* | NA |
| *prop rem edges* | NA |

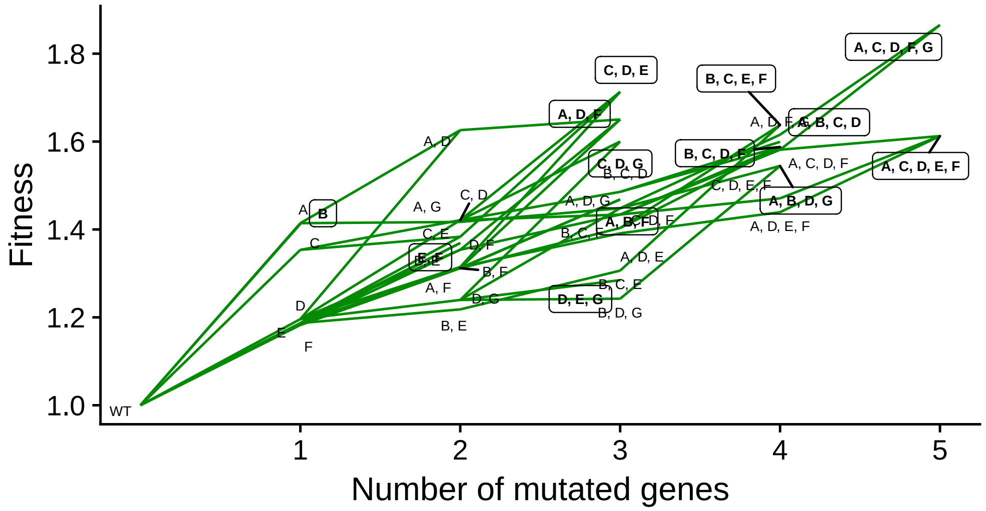


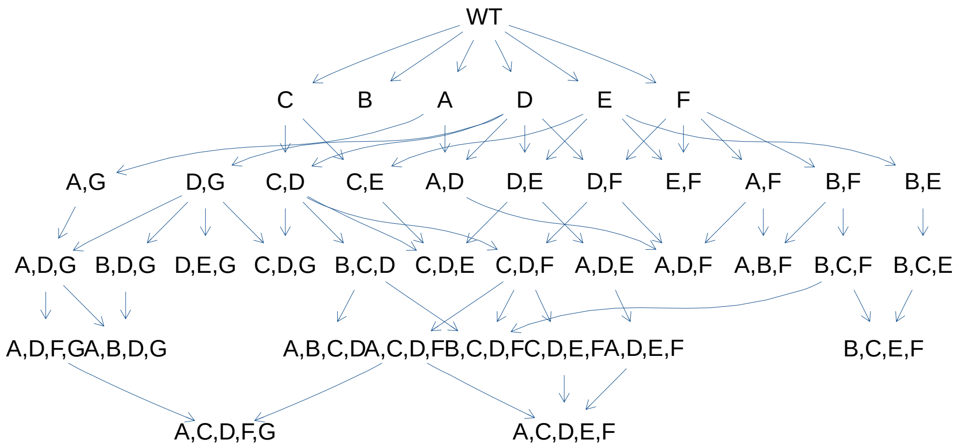

| | |
|---|---|
| *ID* | LJ27jI8rSPxKS2A9 |
| *acc genotypes* | 52 |
| *peaks no back* | 19 |
| *removed edges* | NA |
| *prop rem edges* | NA |

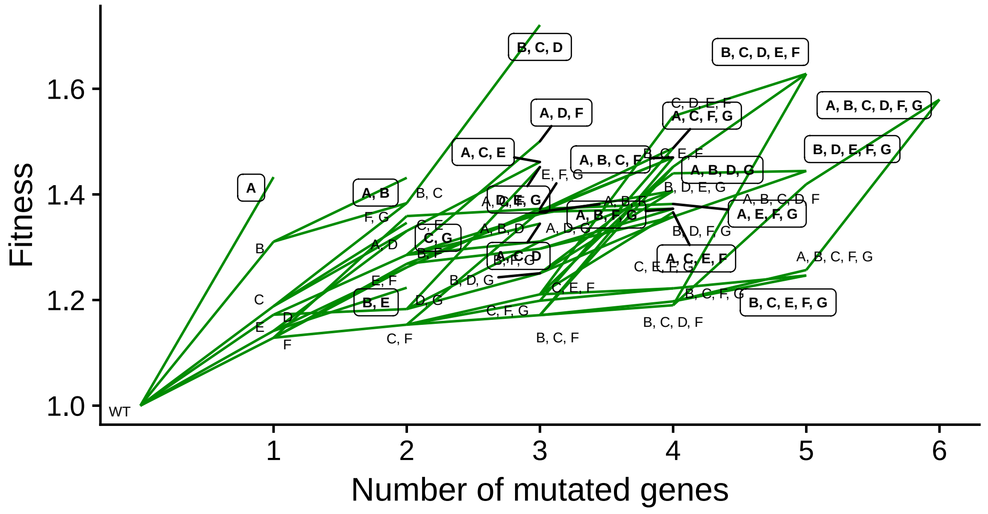


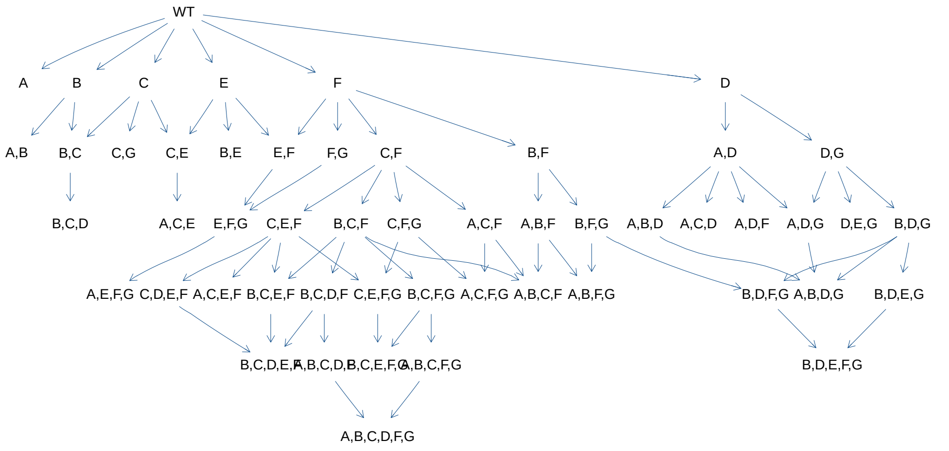

| | |
|---|---|
| *ID* | M19kX109AWS4lET93 |
| *acc genotypes* | 21 |
| *peaks no back* | 10 |
| *removed edges* | NA |
| *prop rem edges* | NA |

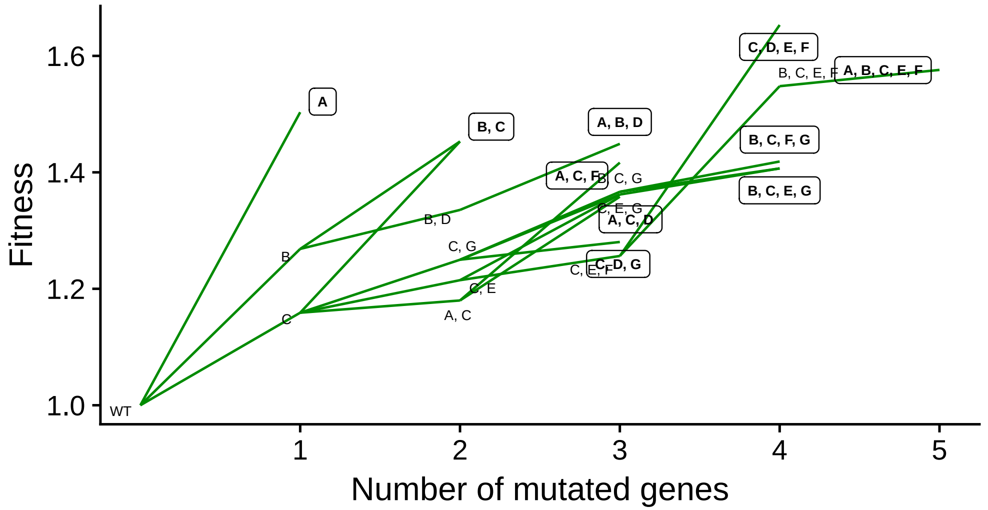


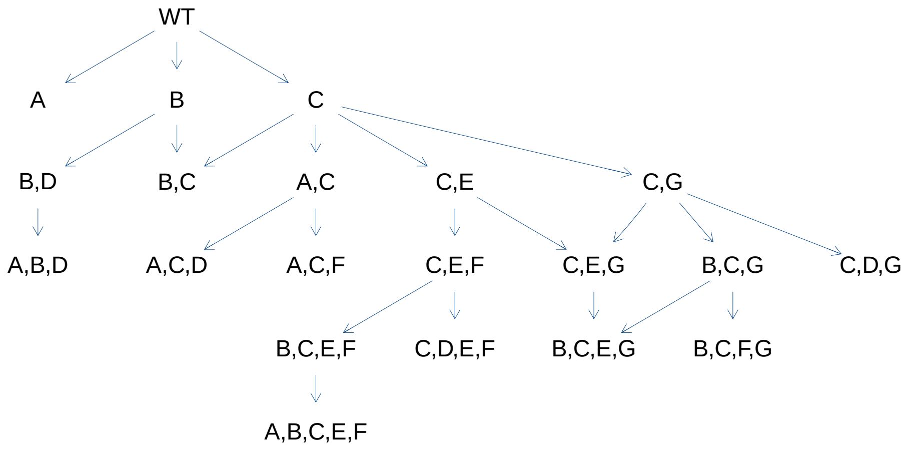

| | |
|---|---|
| *ID* | mBMwivnTLA1F1fOQ |
| *acc genotypes* | 26 |
| *peaks no back* | 11 |
| *removed edges* | NA |
| *prop rem edges* | NA |

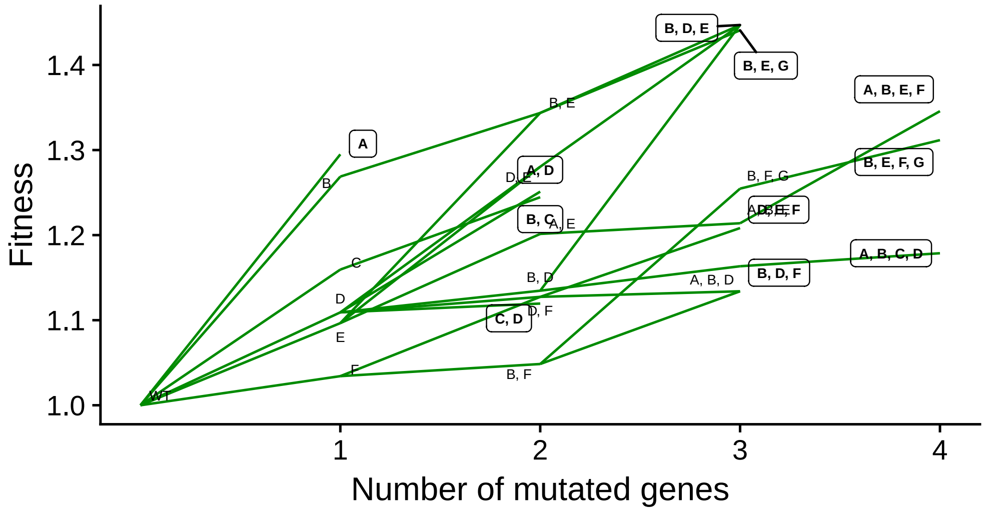


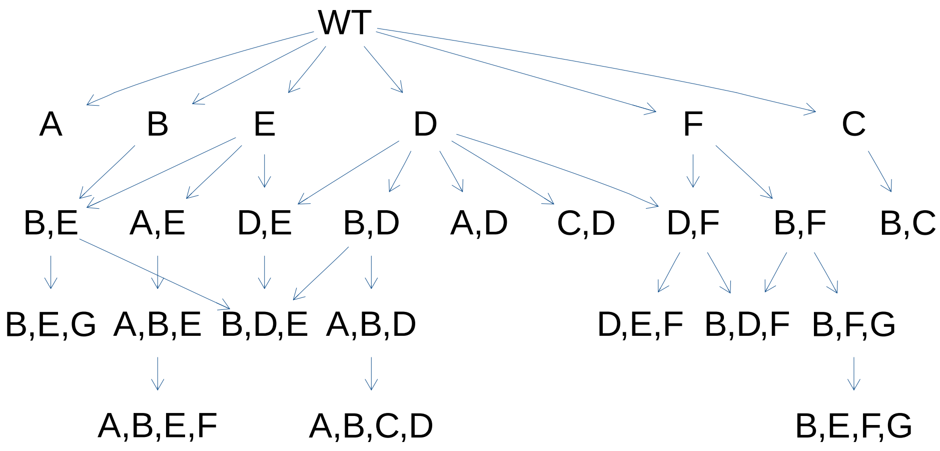

| | |
|---|---|
| *ID* | MPJBgvRZOA1Xp4SQ |
| *acc genotypes* | 53 |
| *peaks no back* | 19 |
| *removed edges* | NA |
| *prop rem edges* | NA |

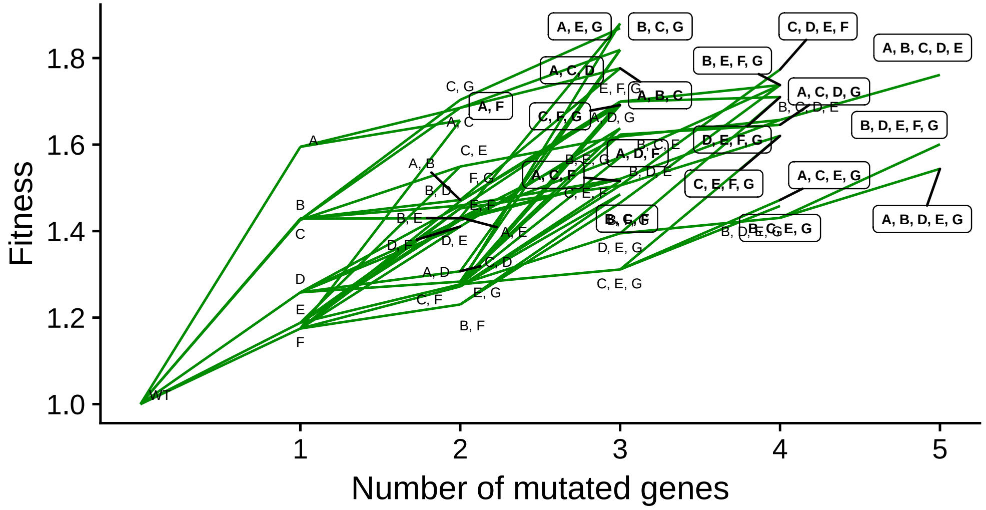


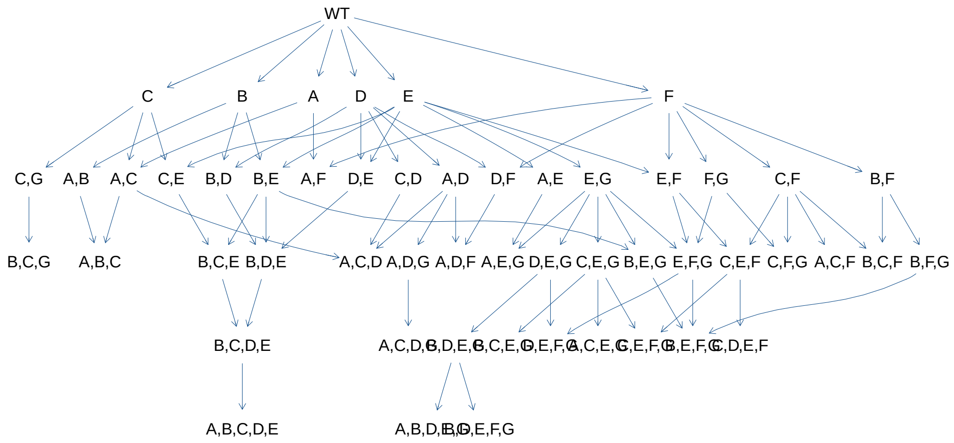

| | |
|---|---|
| *ID* | Mq3VEO6q8h6j5Q2K |
| *acc genotypes* | 43 |
| *peaks no back* | 17 |
| *removed edges* | NA |
| *prop rem edges* | NA |

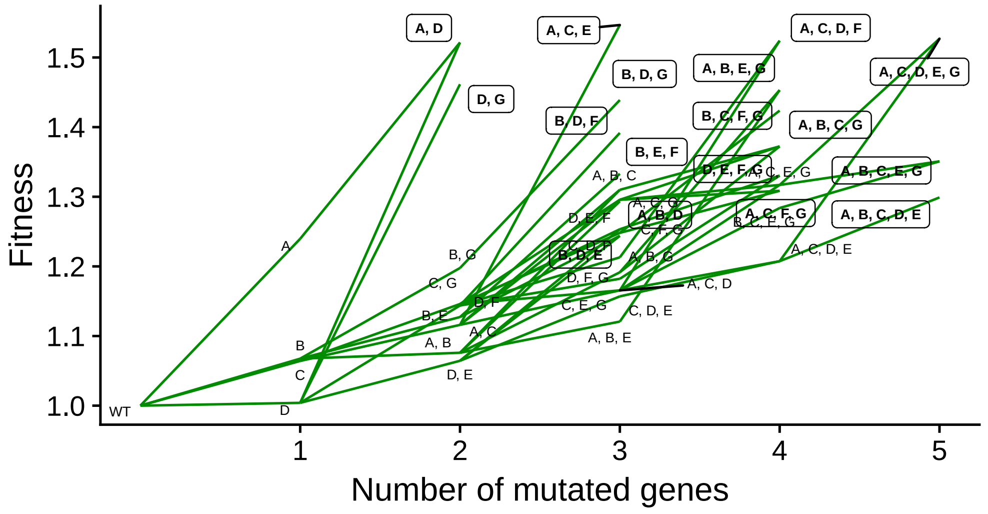


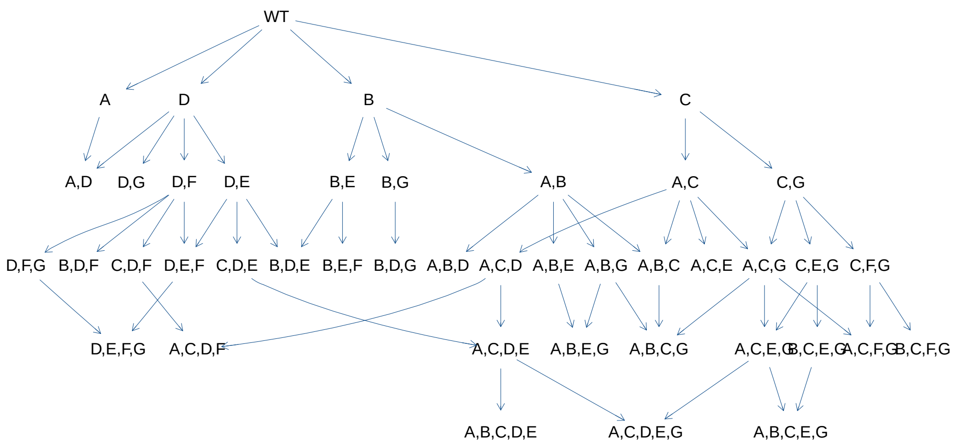

| | |
|---|---|
| *ID* | mrmstEuzwuuKXnIi |
| *acc genotypes* | 64 |
| *peaks no back* | 18 |
| *removed edges* | NA |
| *prop rem edges* | NA |

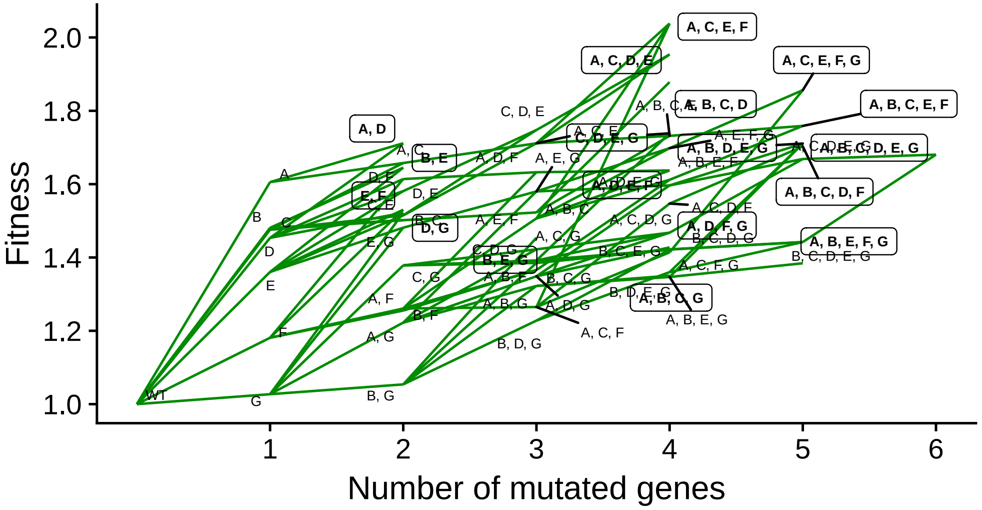


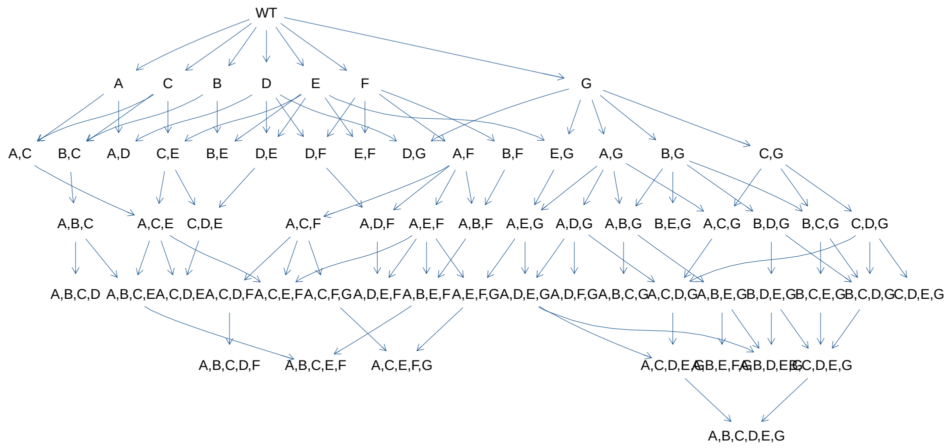

| | |
|---|---|
| *ID* | mTWr1XzBqvHzZVjW |
| *acc genotypes* | 63 |
| *peaks no back* | 22 |
| *removed edges* | NA |
| *prop rem edges* | NA |

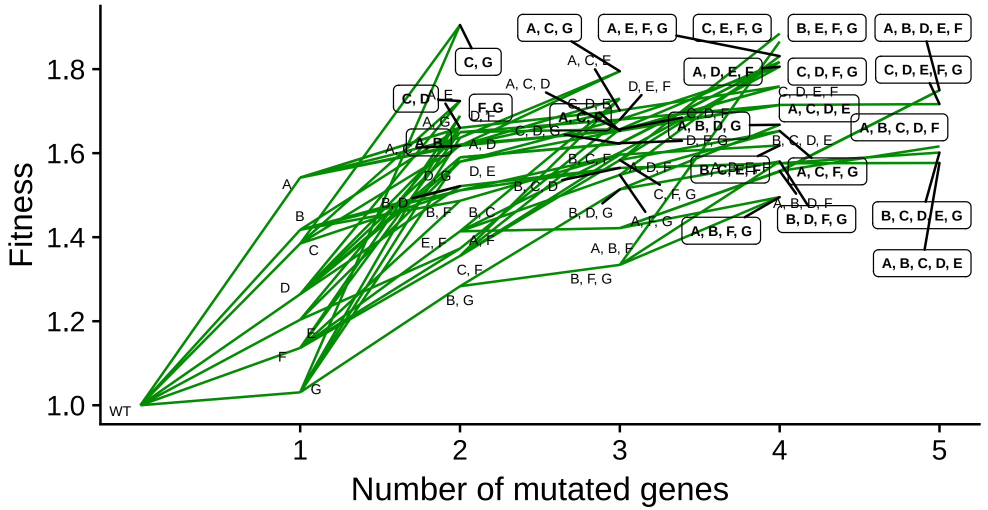


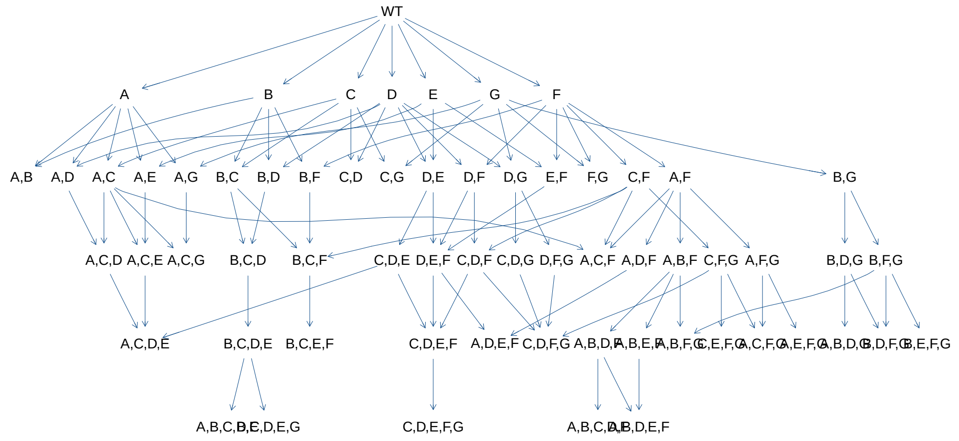

| | |
|---|---|
| *ID* | mwHTaW8c10WKVxeLt |
| *acc genotypes* | 71 |
| *peaks no back* | 22 |
| *removed edges* | NA |
| *prop rem edges* | NA |

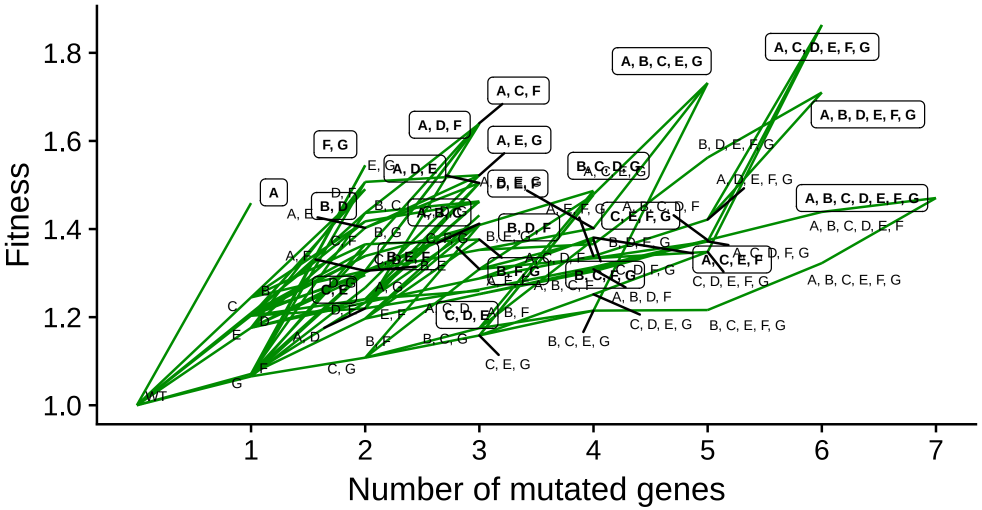


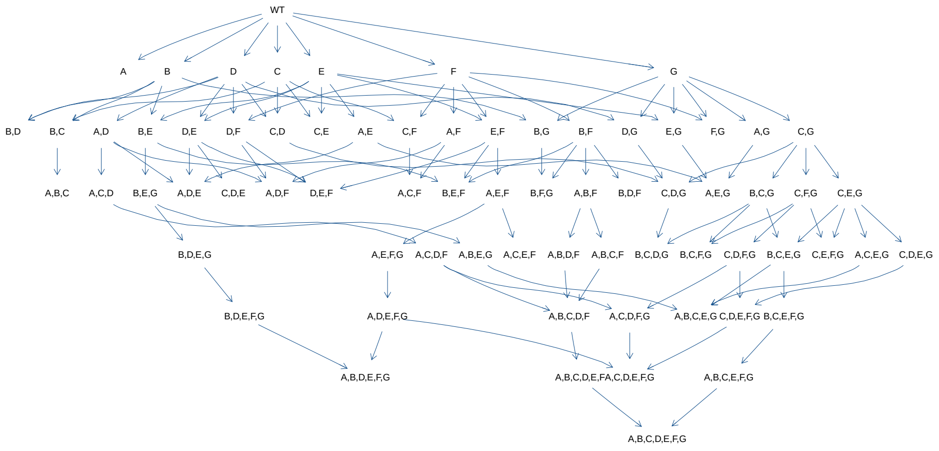

| | |
|---|---|
| *ID* | n2WFR10n5VPYOSSSt |
| *acc genotypes* | 61 |
| *peaks no back* | 17 |
| *removed edges* | NA |
| *prop rem edges* | NA |

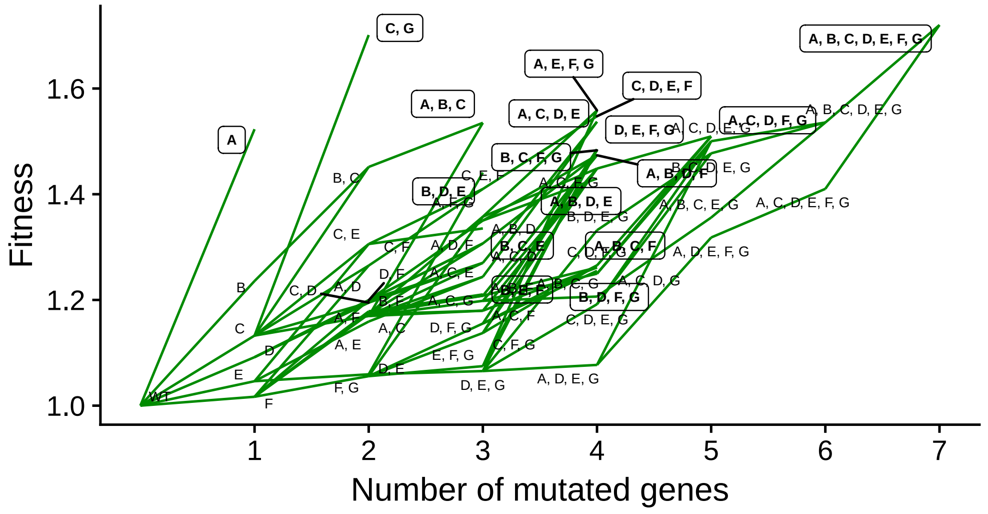


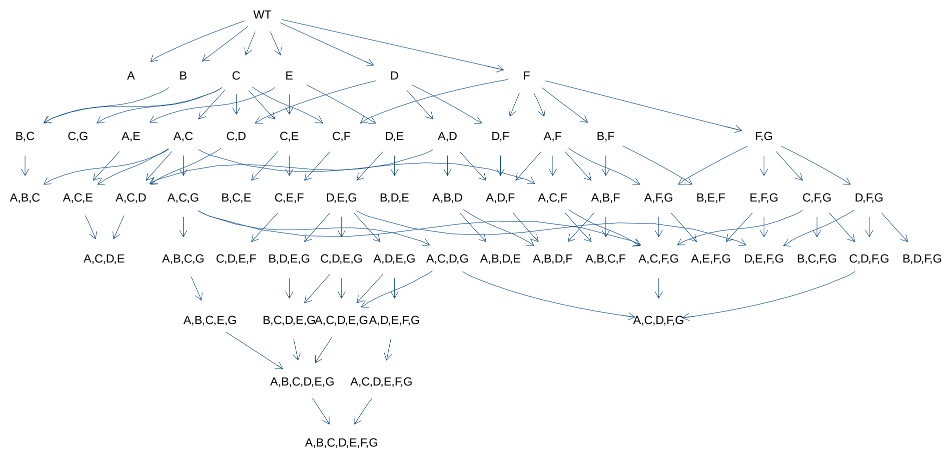

| | |
|---|---|
| *ID* | NejgQ4lbqE6syfEk |
| *acc genotypes* | 60 |
| *peaks no back* | 22 |
| *removed edges* | NA |
| *prop rem edges* | NA |

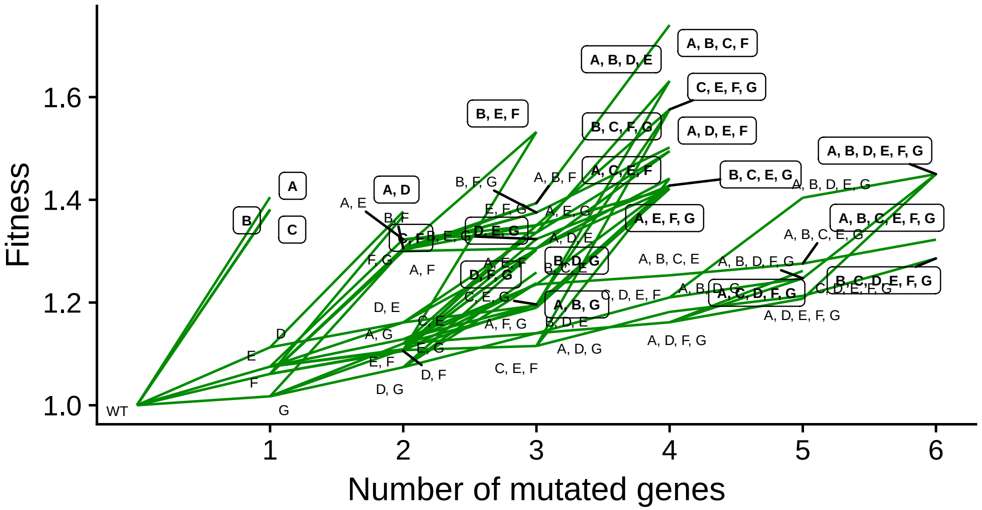


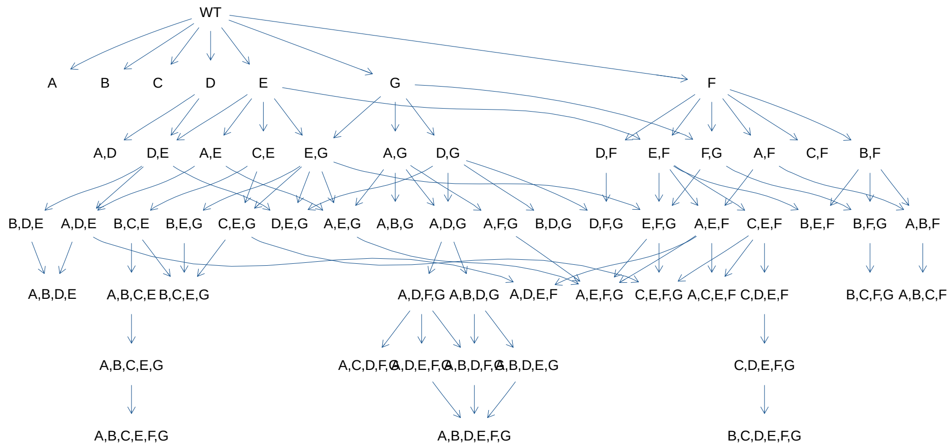

*ID* nIkUt9Az1n9o6UtN
*acc genotypes* 40
*peaks no back* 17
*removed edges* NA
*prop rem edges* NA

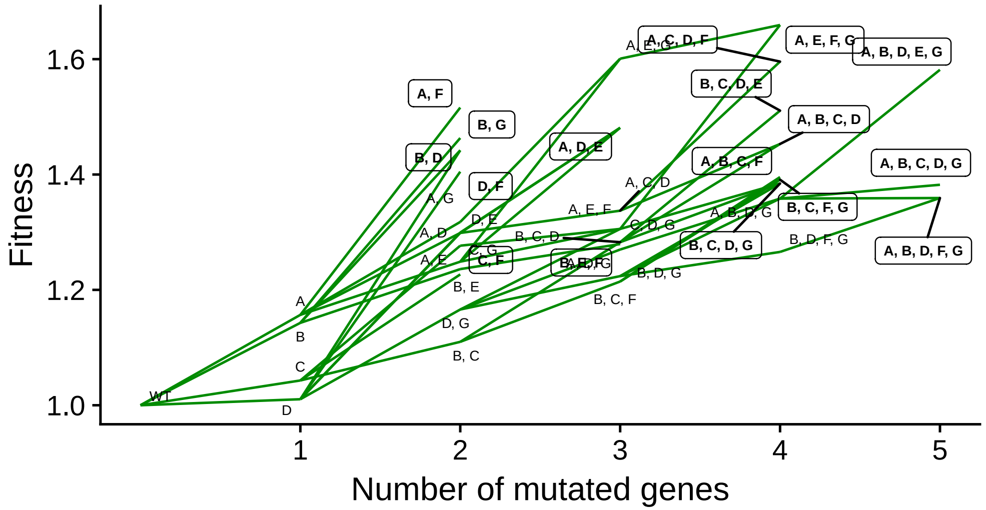


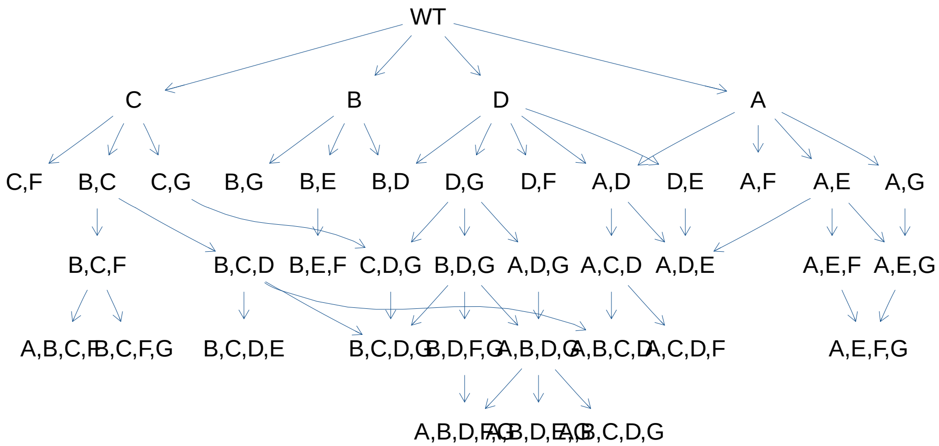

| | |
|---|---|
| *ID* | NLcu10ZEMImemfBBs |
| *acc genotypes* | 32 |
| *peaks no back* | 14 |
| *removed edges* | NA |
| *prop rem edges* | NA |

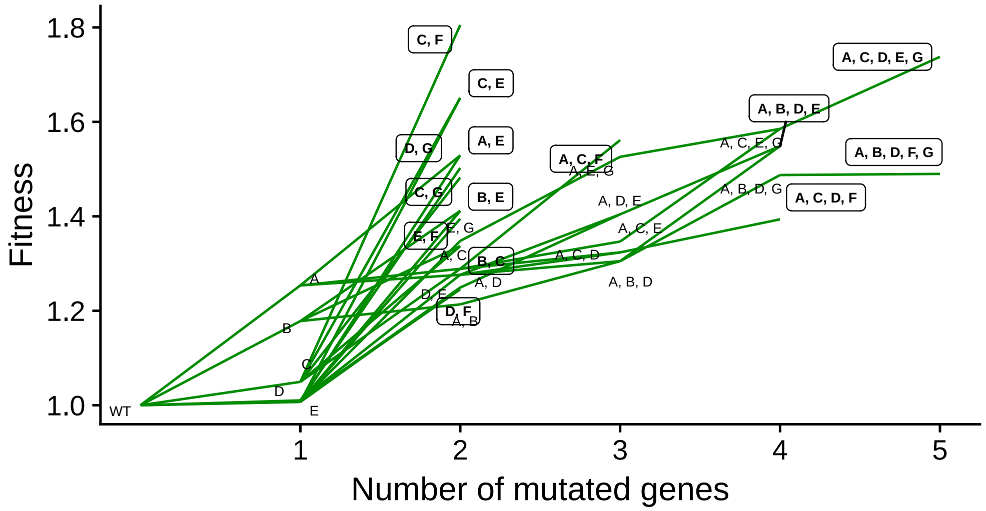


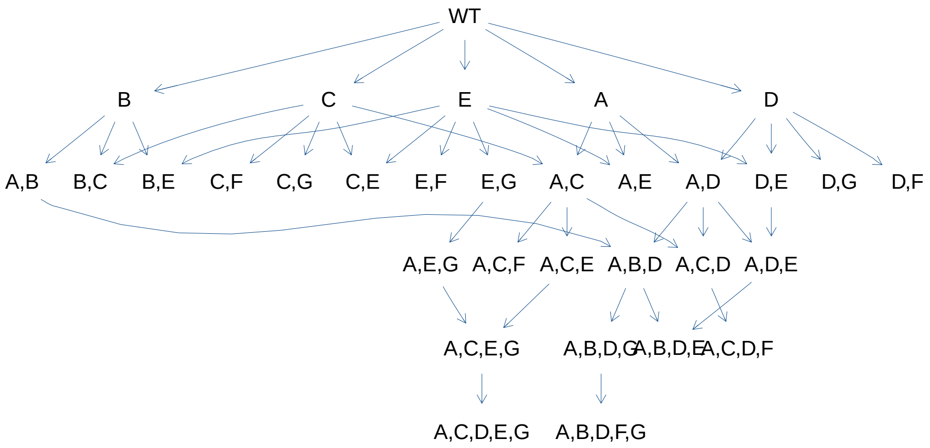

| | |
|---|---|
| *ID* | nlx10k3xX8bswraYG |
| *acc genotypes* | 19 |
| *peaks no back* | 5 |
| *removed edges* | NA |
| *prop rem edges* | NA |

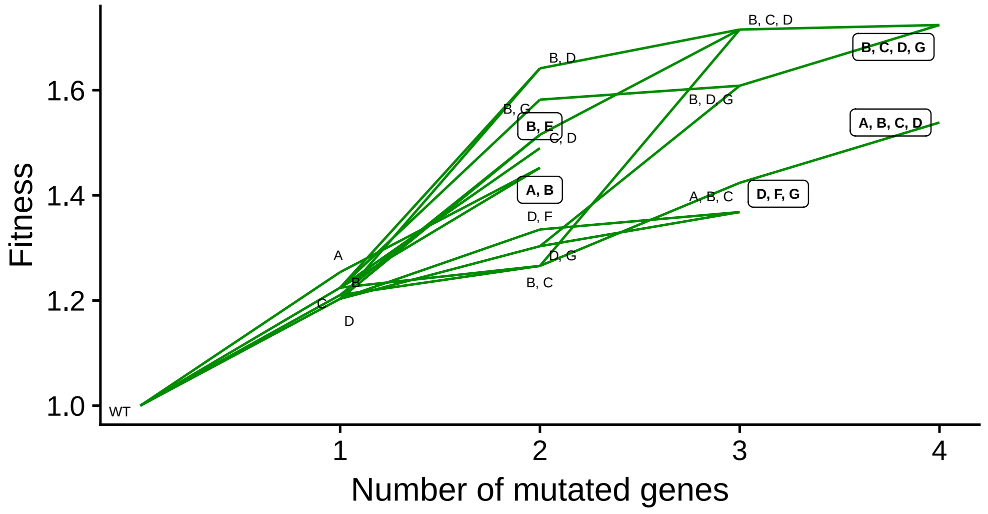


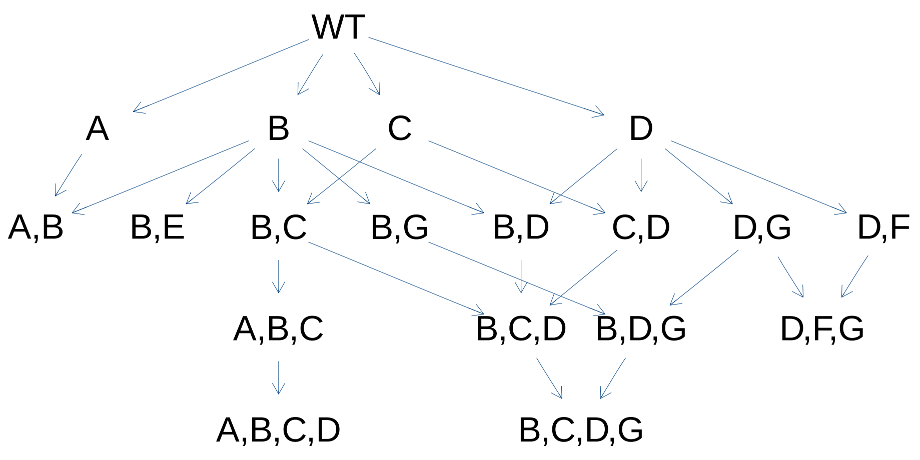

| | |
|---|---|
| *ID* | Nq9C7m8MciTio6I4 |
| *acc genotypes* | 45 |
| *peaks no back* | 13 |
| *removed edges* | NA |
| *prop rem edges* | NA |

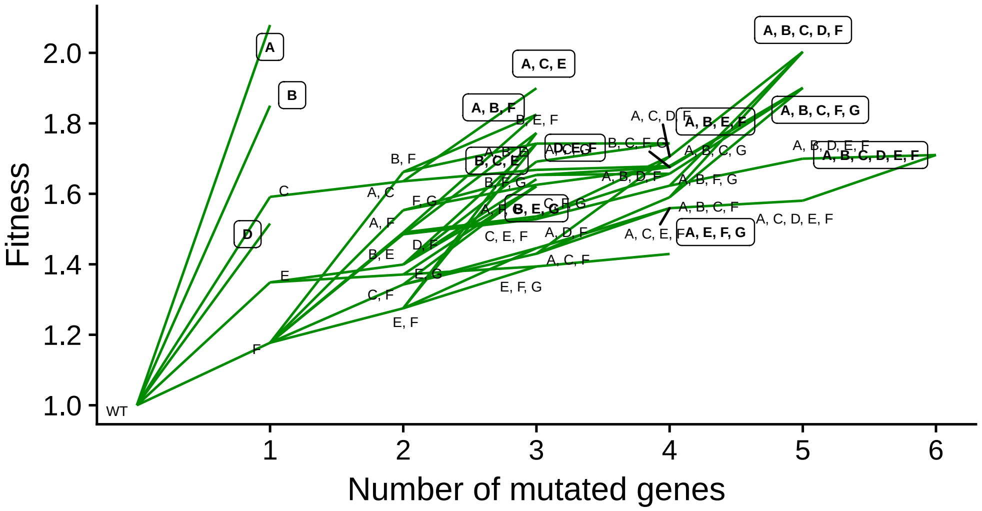


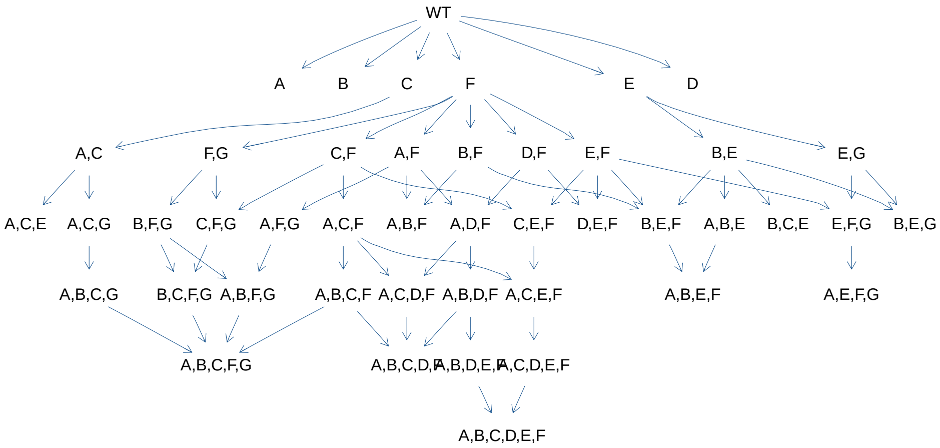

| | |
|---|---|
| *ID* | NuYHUmW2Daw10xoxW |
| *acc genotypes* | 61 |
| *peaks no back* | 15 |
| *removed edges* | NA |
| *prop rem edges* | NA |

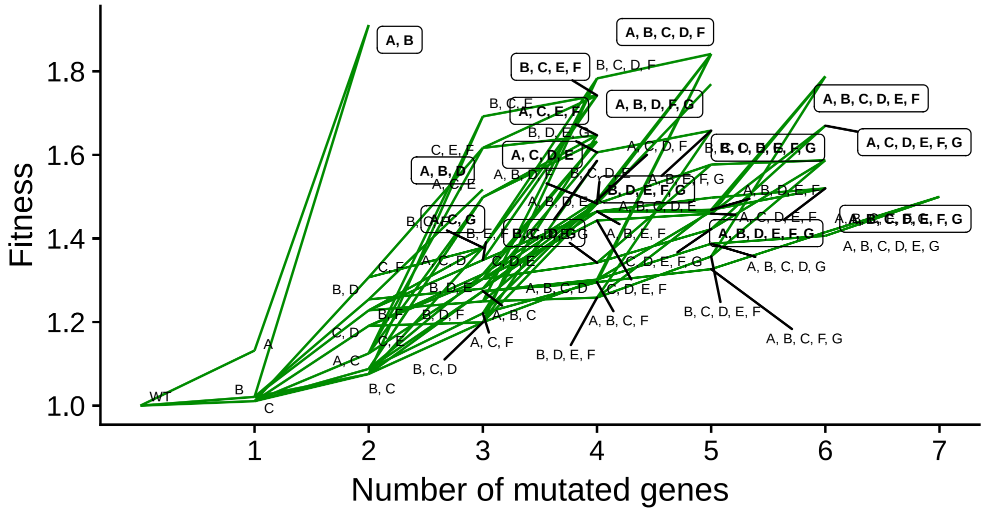


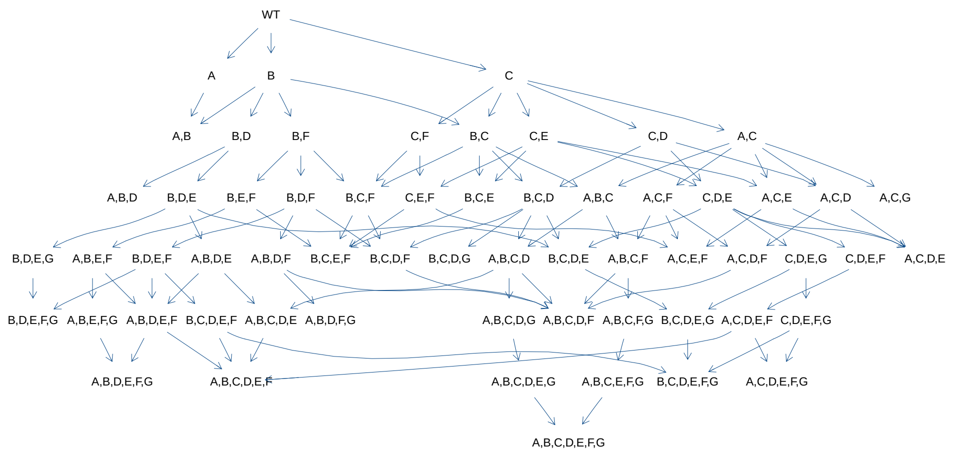

| | |
|---|---|
| *ID* | nwLrcLgueac4e3RH |
| *acc genotypes* | 57 |
| *peaks no back* | 16 |
| *removed edges* | NA |
| *prop rem edges* | NA |

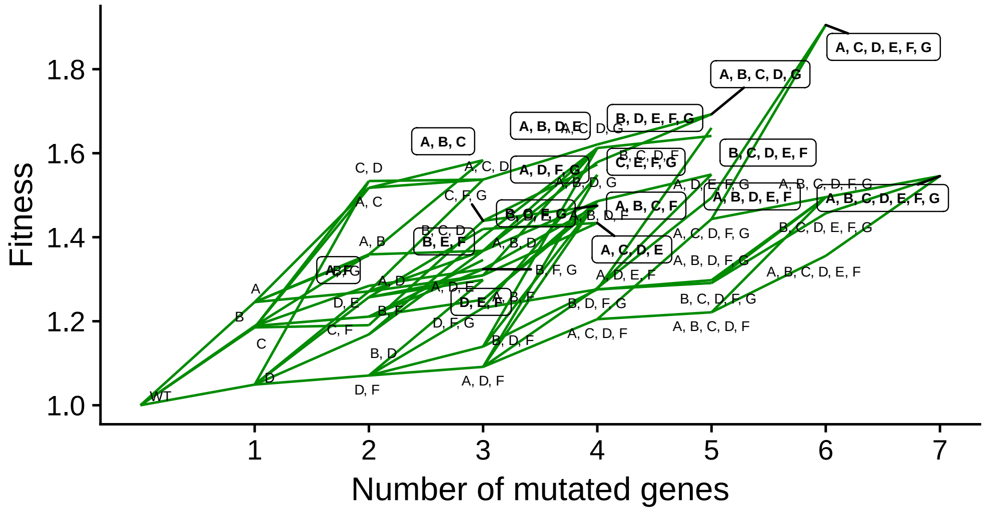


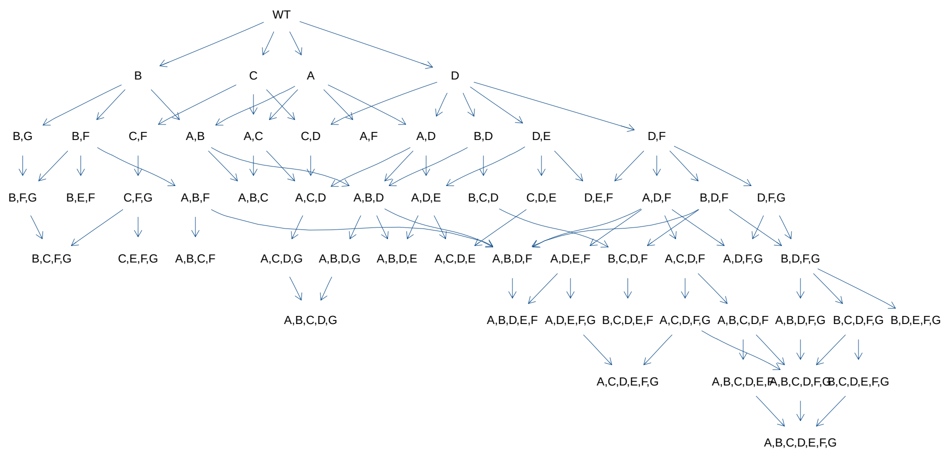

| | |
|---|---|
| *ID* | om3Pj5IphjvaQhAk |
| *acc genotypes* | 82 |
| *peaks no back* | 27 |
| *removed edges* | NA |
| *prop rem edges* | NA |

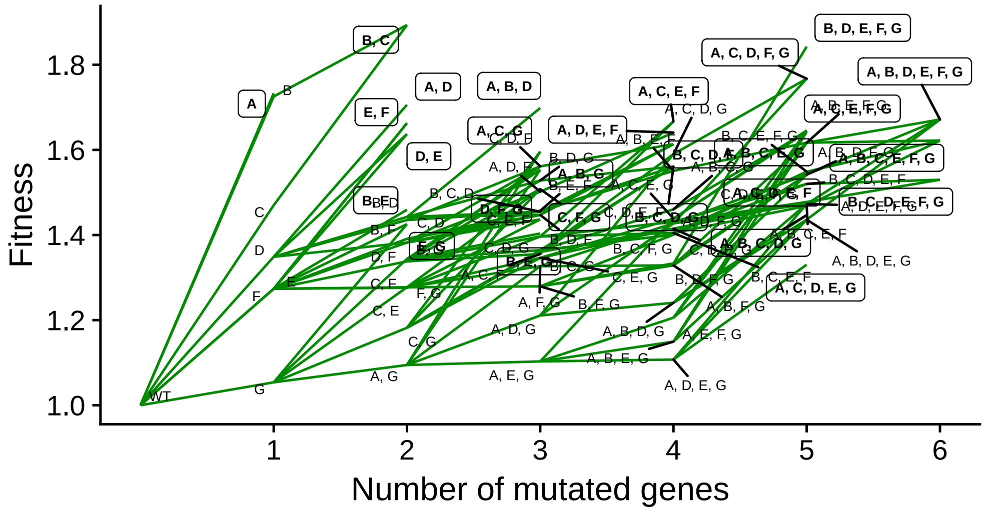


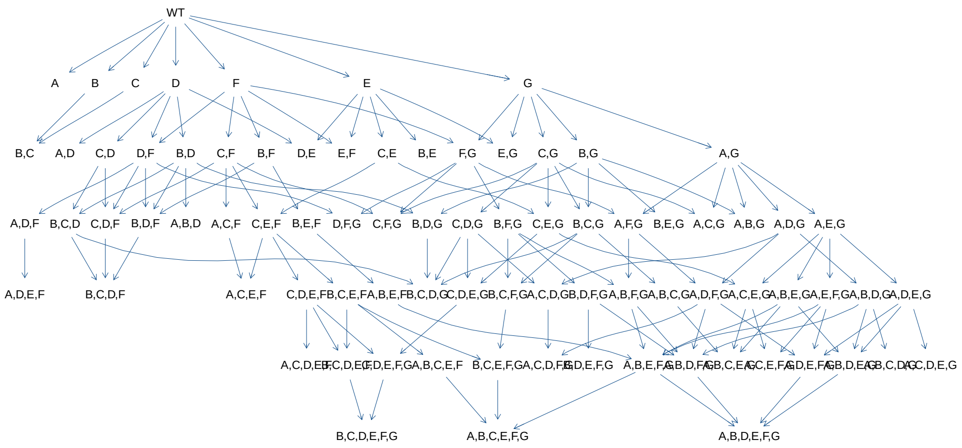

| | |
|---|---|
| *ID* | OsDpSWcfqKrGoOGX |
| *acc genotypes* | 31 |
| *peaks no back* | 10 |
| *removed edges* | NA |
| *prop rem edges* | NA |

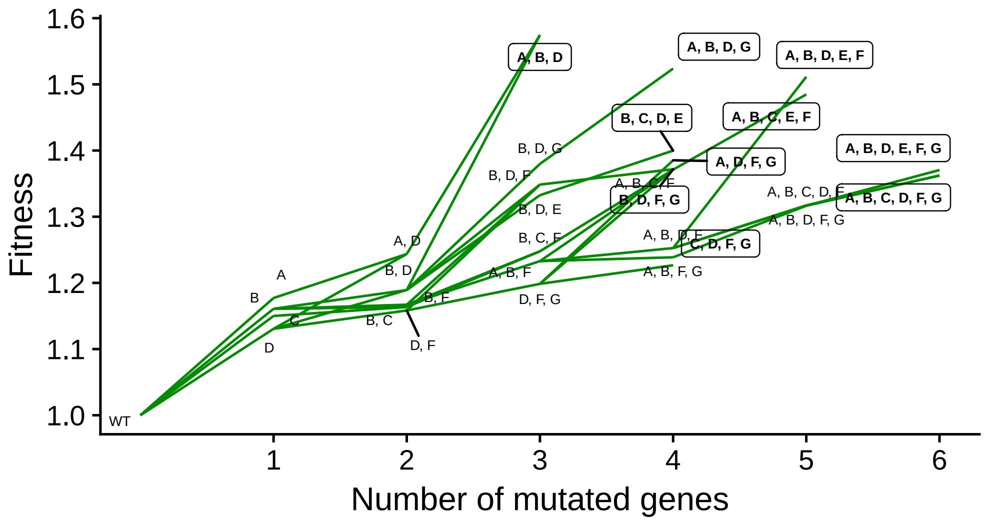


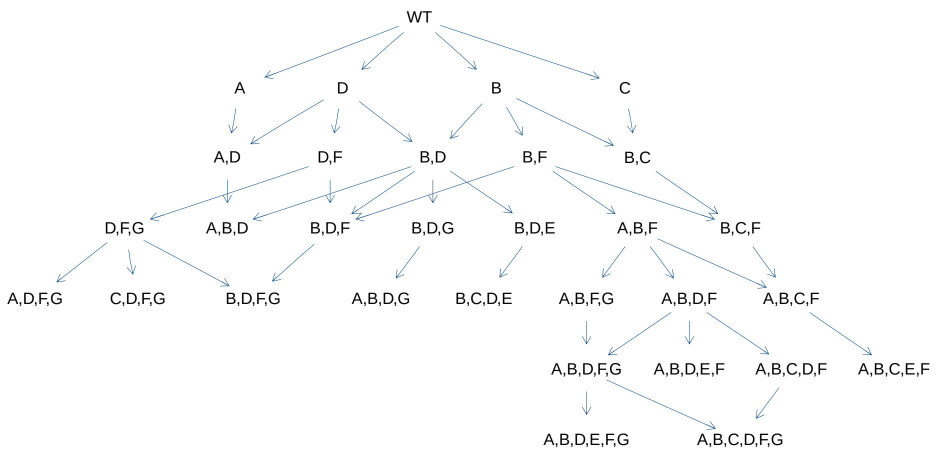

| | |
|---|---|
| *ID* | ozA8bz3JzJBjMoZP |
| *acc genotypes* | 38 |
| *peaks no back* | 11 |
| *removed edges* | NA |
| *prop rem edges* | NA |

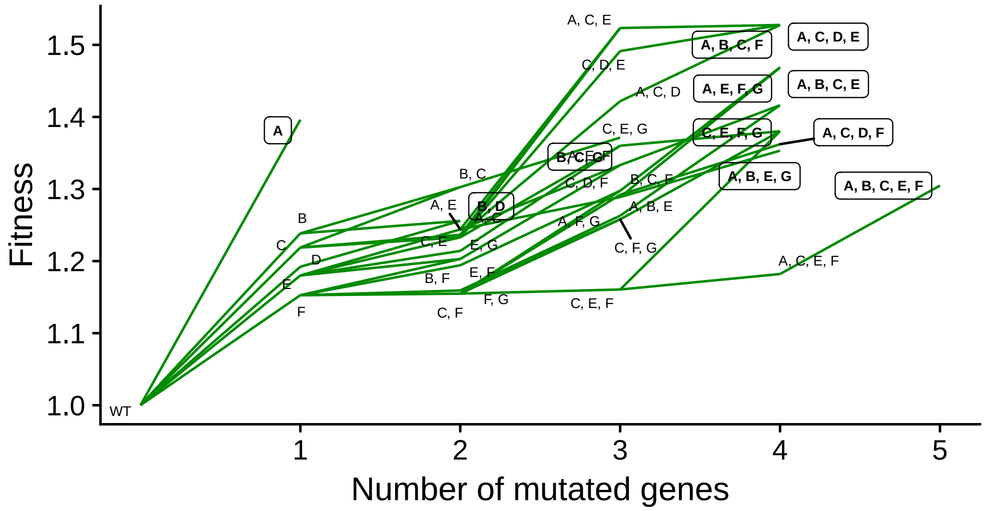


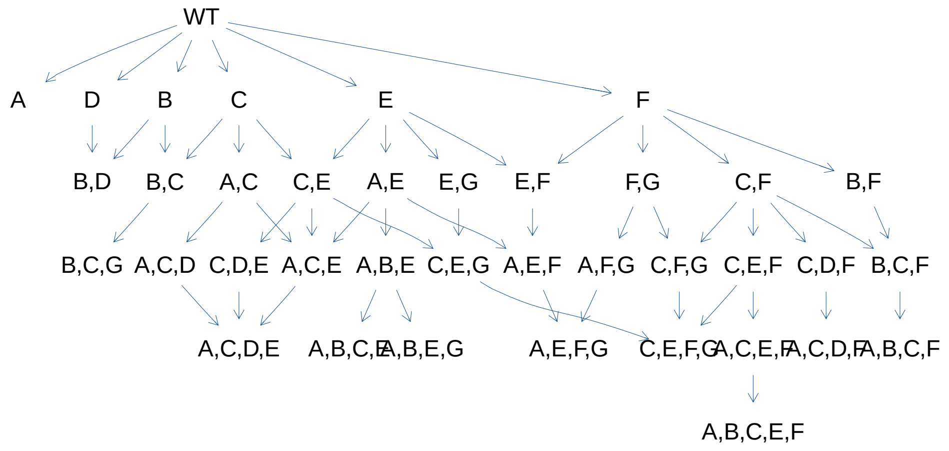

| | |
|---|---|
| *ID* | p2rZTfzQSZaEzk7D |
| *acc genotypes* | 42 |
| *peaks no back* | 11 |
| *removed edges* | NA |
| *prop rem edges* | NA |

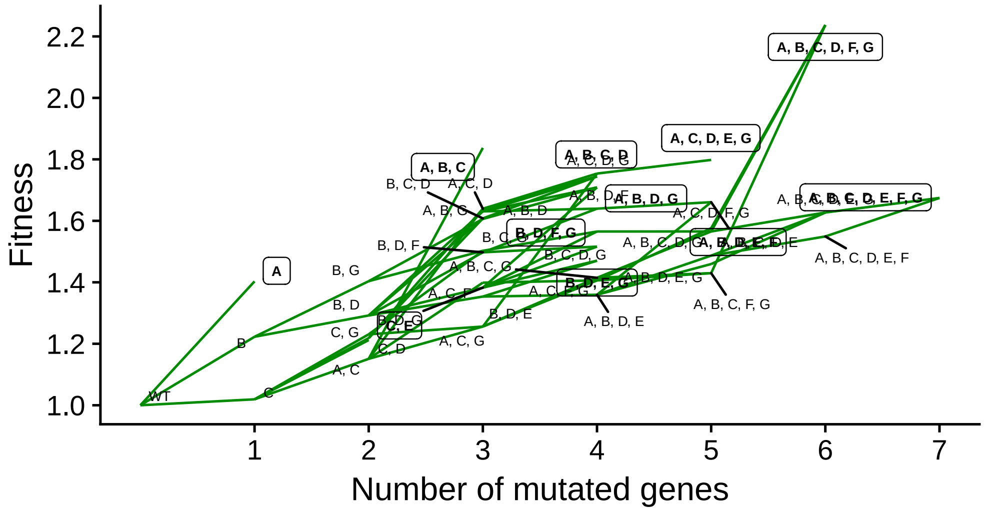


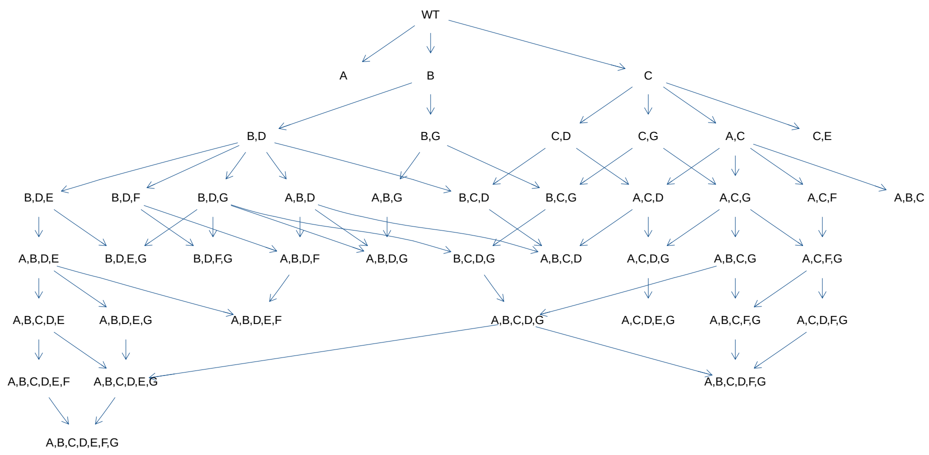

| | |
|---|---|
| *ID* | PdBS51bgDdPXdV8W |
| *acc genotypes* | 33 |
| *peaks no back* | 10 |
| *removed edges* | NA |
| *prop rem edges* | NA |

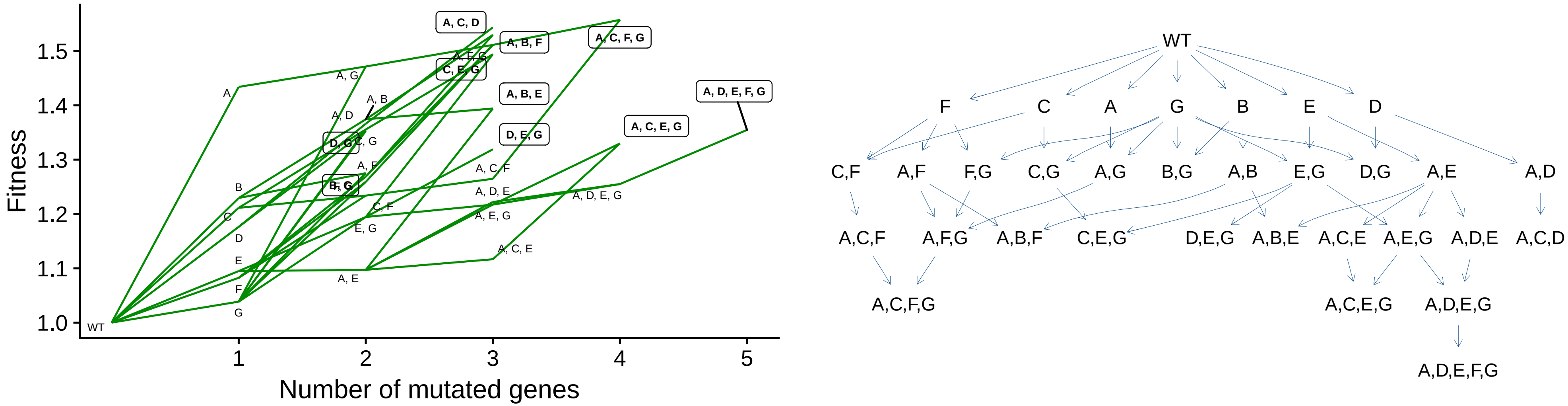

*ID* PdEpnOBgly104Dp4A
*acc genotypes* 50
*peaks no back* 16
*removed edges* NA
*prop rem edges* NA

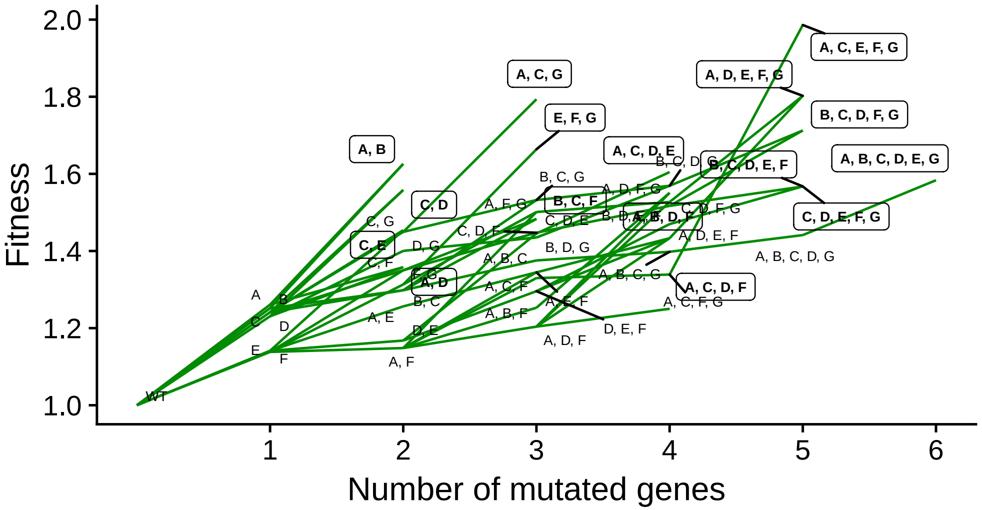


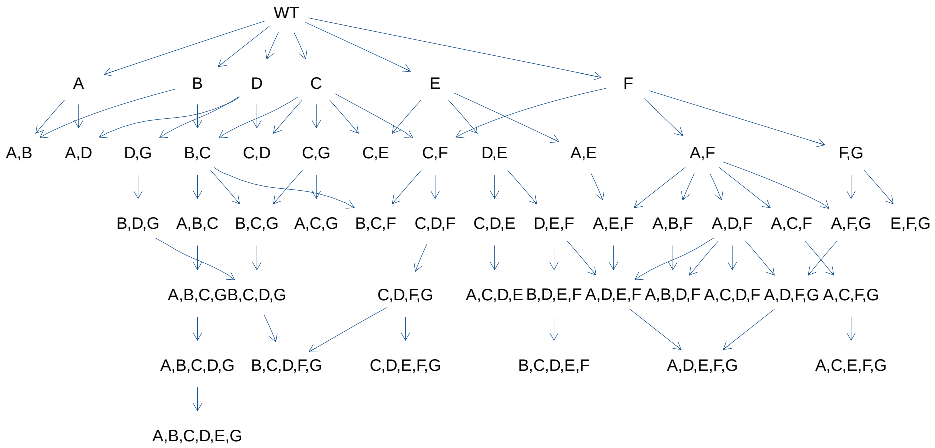

| | |
|---|---|
| *ID* | pdLy5HbuIEcGMFw8 |
| *acc genotypes* | 39 |
| *peaks no back* | 12 |
| *removed edges* | NA |
| *prop rem edges* | NA |

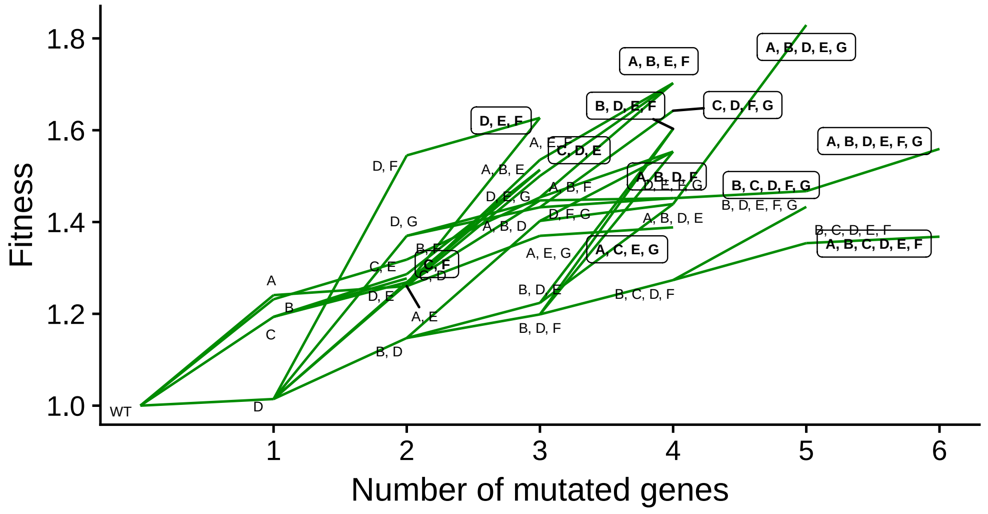


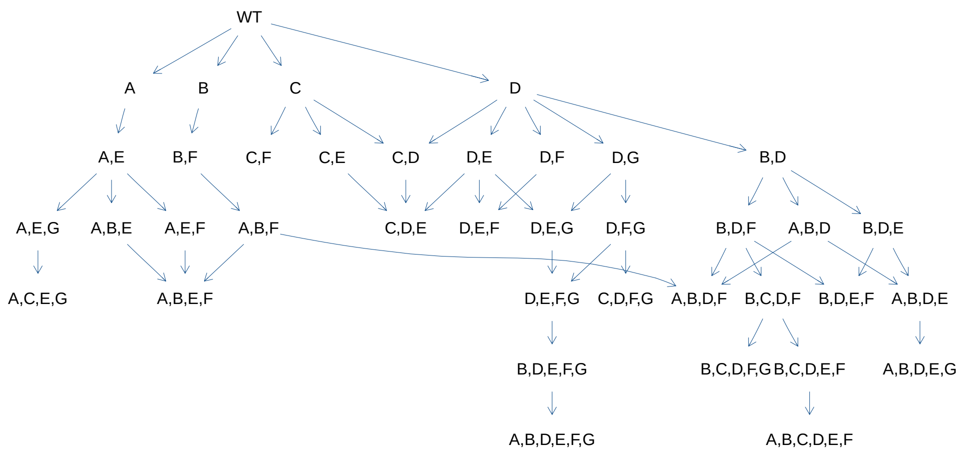

| | |
|---|---|
| *ID* | PMnxoLN1P10vNrmKW |
| *acc genotypes* | 22 |
| *peaks no back* | 8 |
| *removed edges* | NA |
| *prop rem edges* | NA |

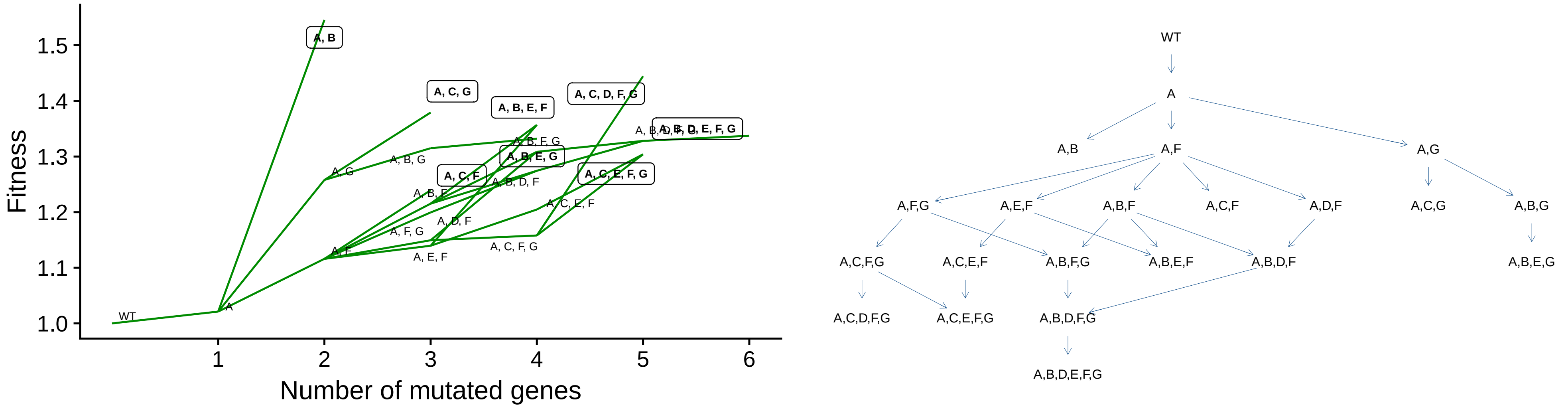

| | |
|---|---|
| *ID* | poffFlI3VRWbB7qo |
| *acc genotypes* | 49 |
| *peaks no back* | 17 |
| *removed edges* | NA |
| *prop rem edges* | NA |

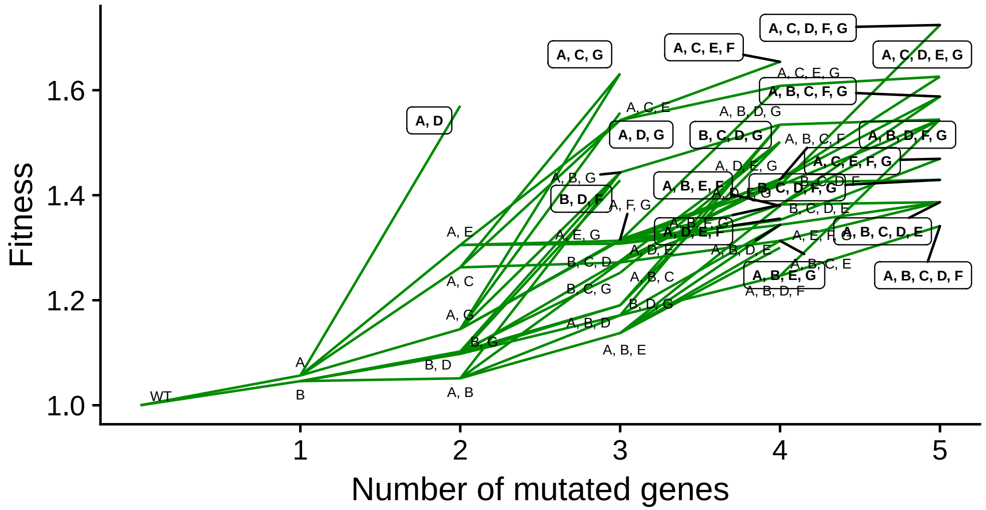


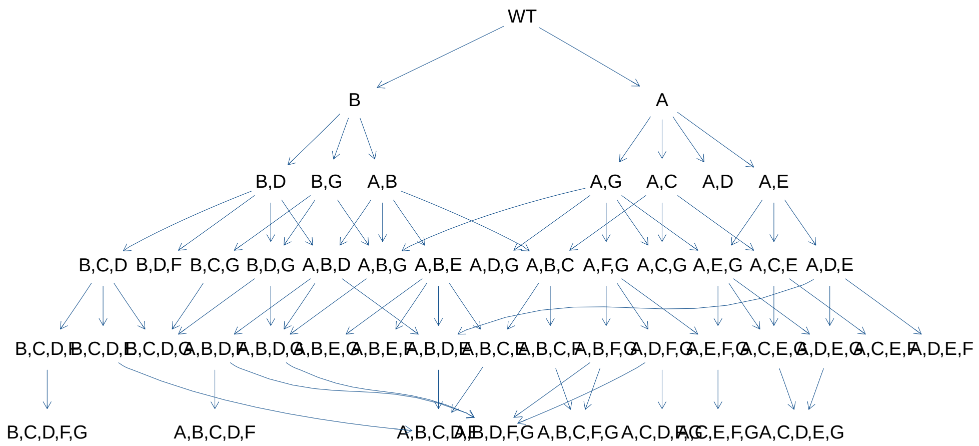

| | |
|---|---|
| *ID* | pZAMLLsskT5d10Q10B |
| *acc genotypes* | 41 |
| *peaks no back* | 16 |
| *removed edges* | NA |
| *prop rem edges* | NA |

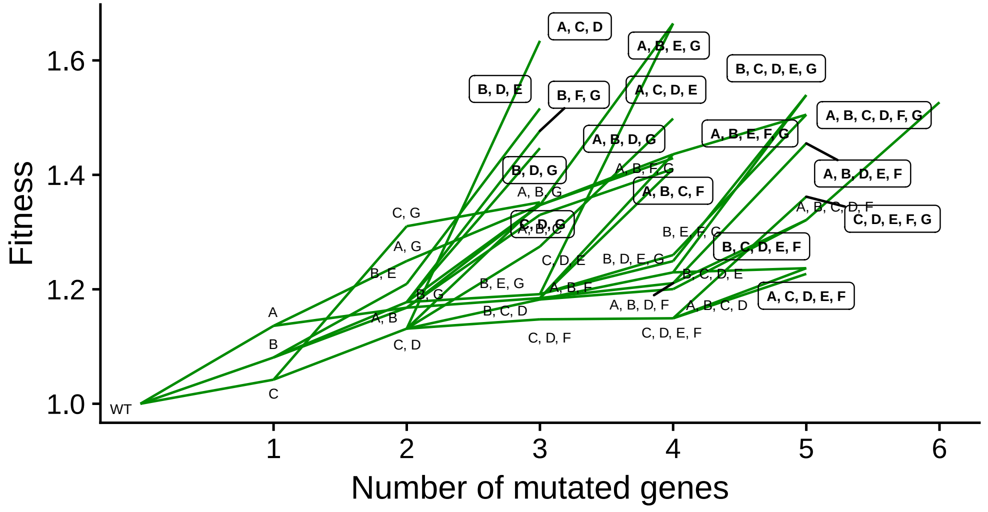


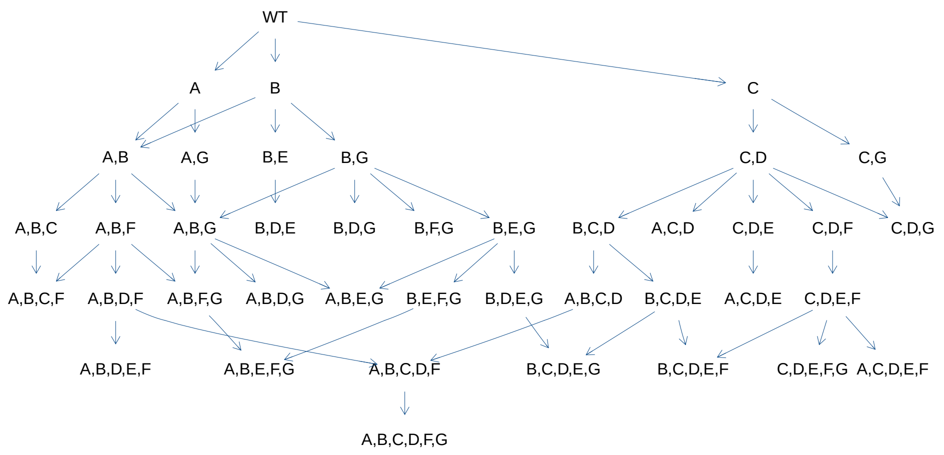

| | |
|---|---|
| *ID* | QdO42eriCLzJlKPT |
| *acc genotypes* | 56 |
| *peaks no back* | 22 |
| *removed edges* | NA |
| *prop rem edges* | NA |

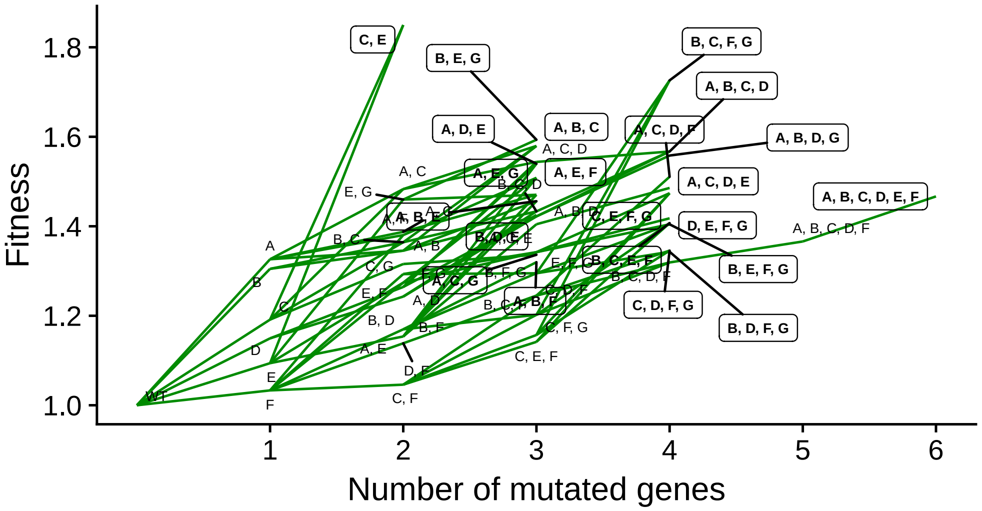


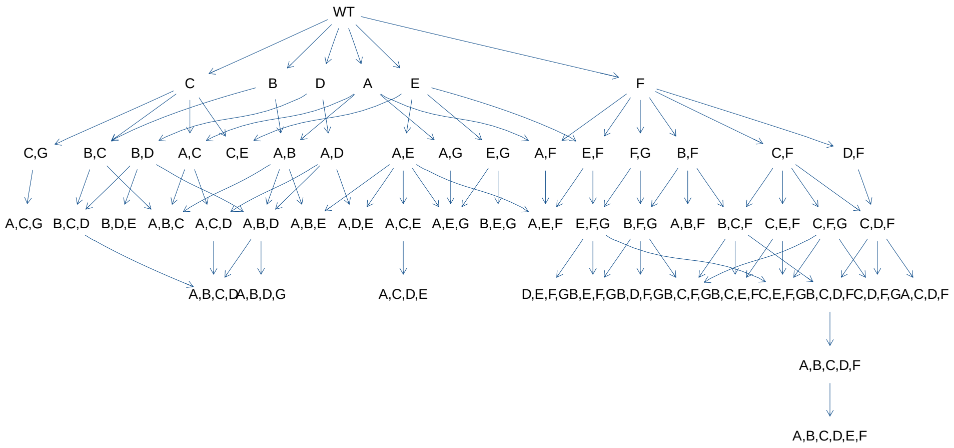

| | |
|---|---|
| *ID* | QDPtbXAp8i6koPnT |
| *acc genotypes* | 18 |
| *peaks no back* | 8 |
| *removed edges* | NA |
| *prop rem edges* | NA |

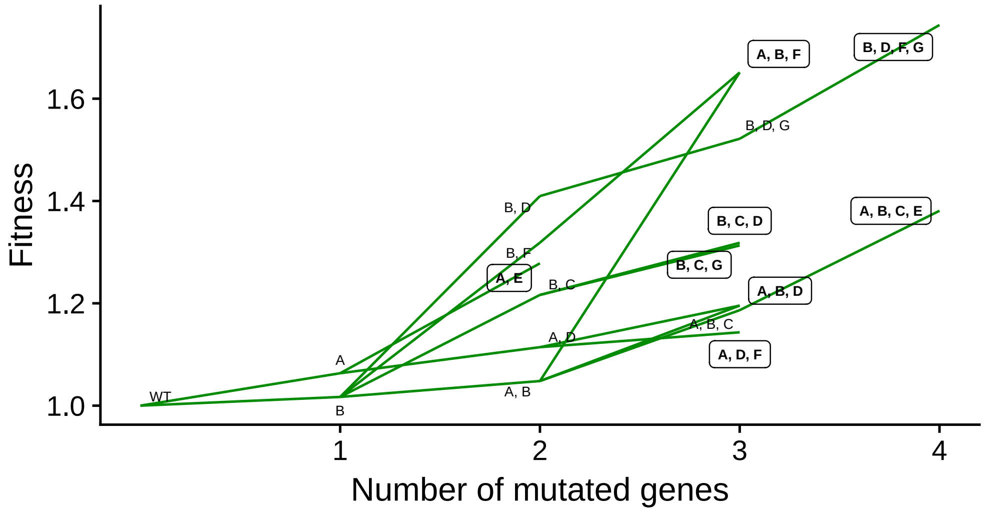


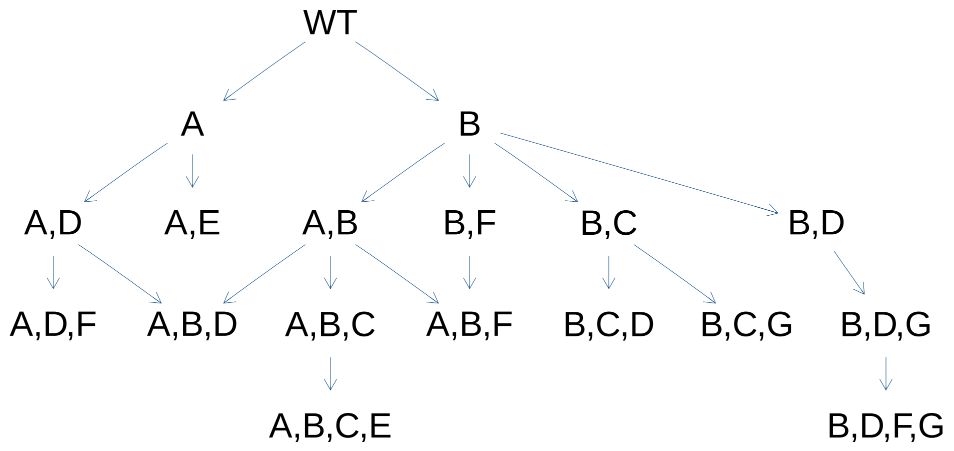

| | |
|---|---|
| *ID* | QIQaxajtj6kjit1V |
| *acc genotypes* | 27 |
| *peaks no back* | 10 |
| *removed edges* | NA |
| *prop rem edges* | NA |

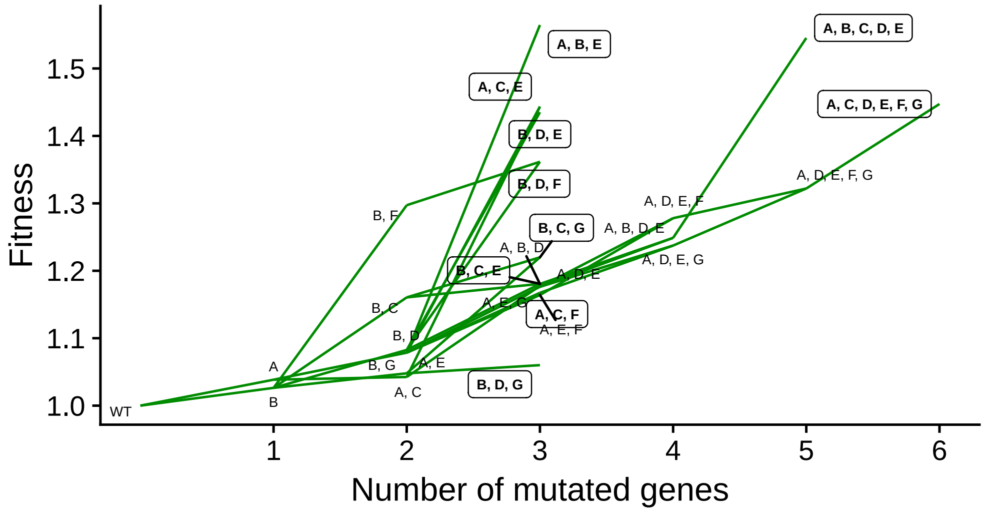


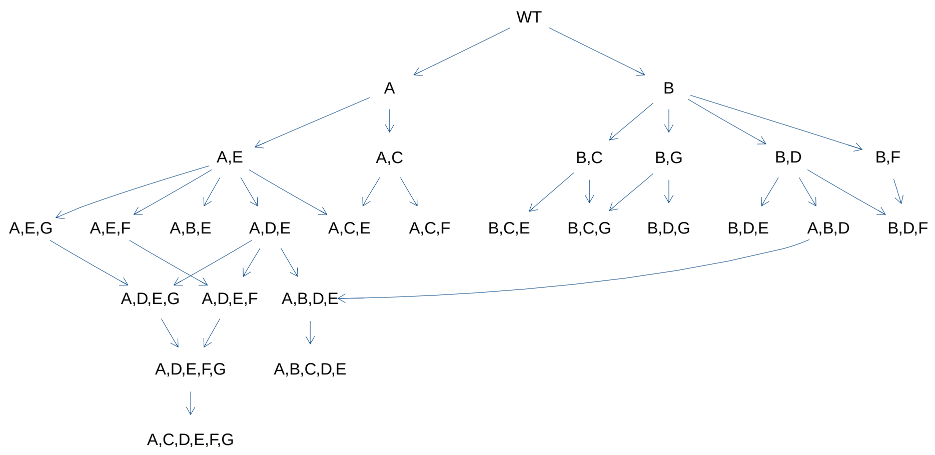

| | |
|---|---|
| *ID* | QTFRtRudotyAGuqB |
| *acc genotypes* | 55 |
| *peaks no back* | 18 |
| *removed edges* | NA |
| *prop rem edges* | NA |

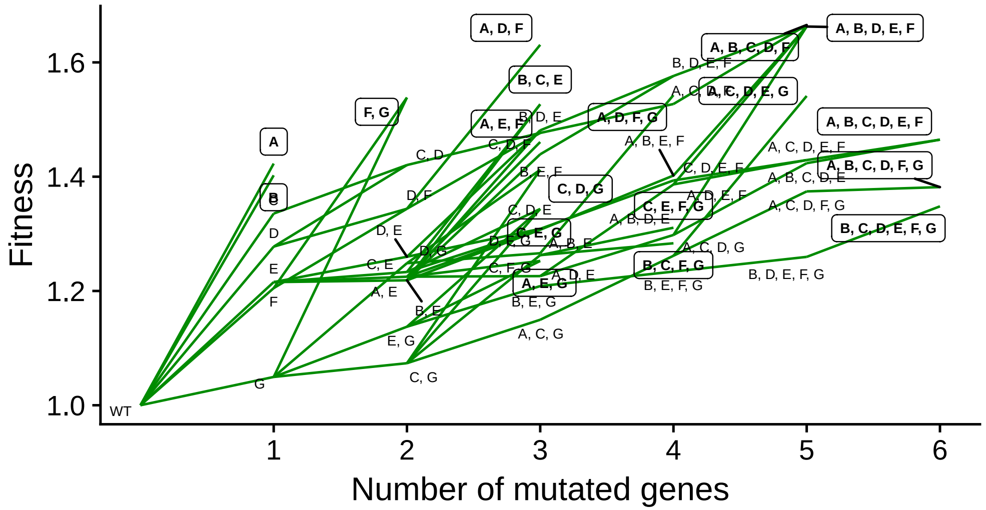


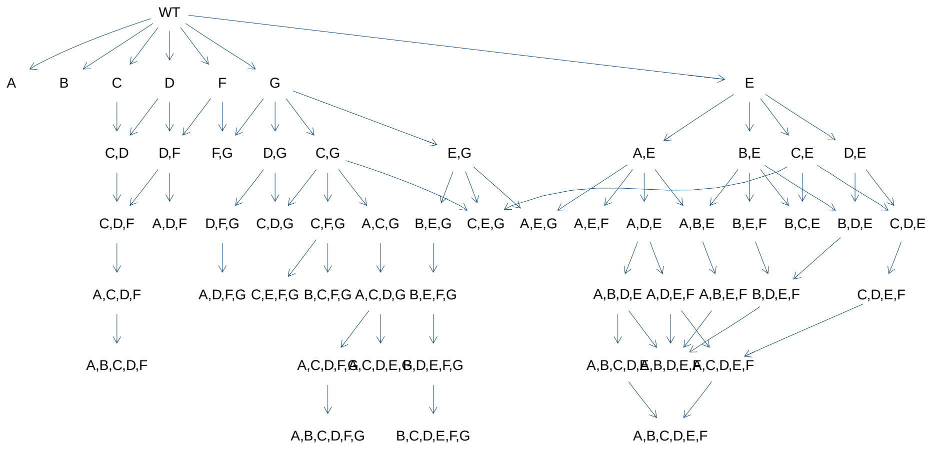

| | |
|---|---|
| *ID* | qUmaRIOlDCO4dtyu |
| *acc genotypes* | 44 |
| *peaks no back* | 17 |
| *removed edges* | NA |
| *prop rem edges* | NA |

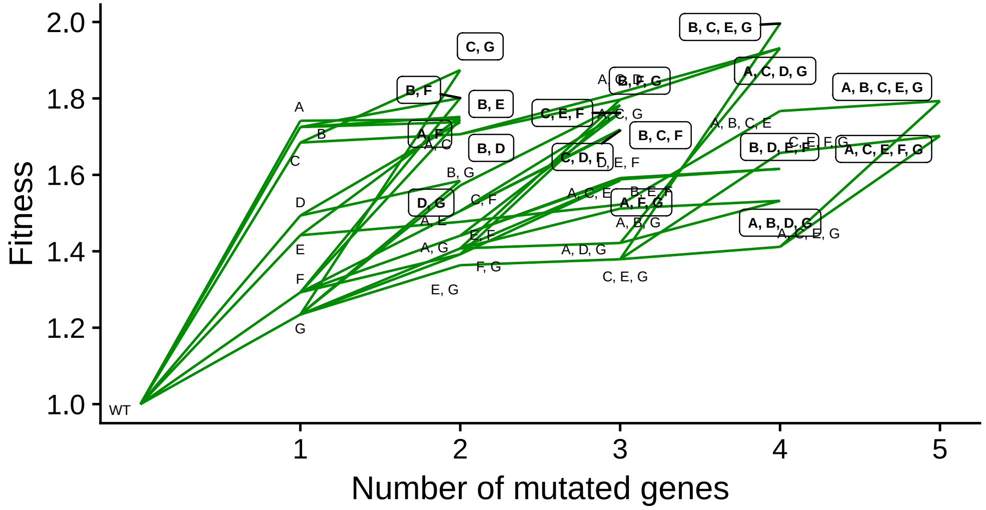


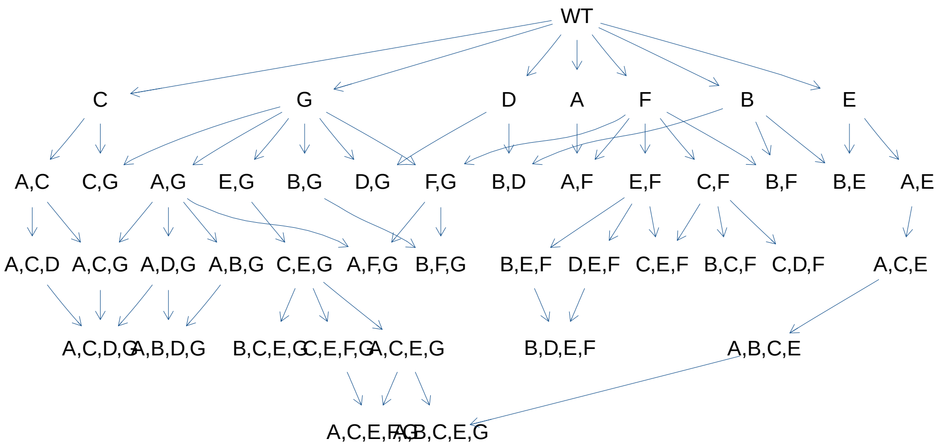

| | |
|---|---|
| *ID* | r4JwqMTj8Y3k7yRZ |
| *acc genotypes* | 52 |
| *peaks no back* | 15 |
| *removed edges* | NA |
| *prop rem edges* | NA |

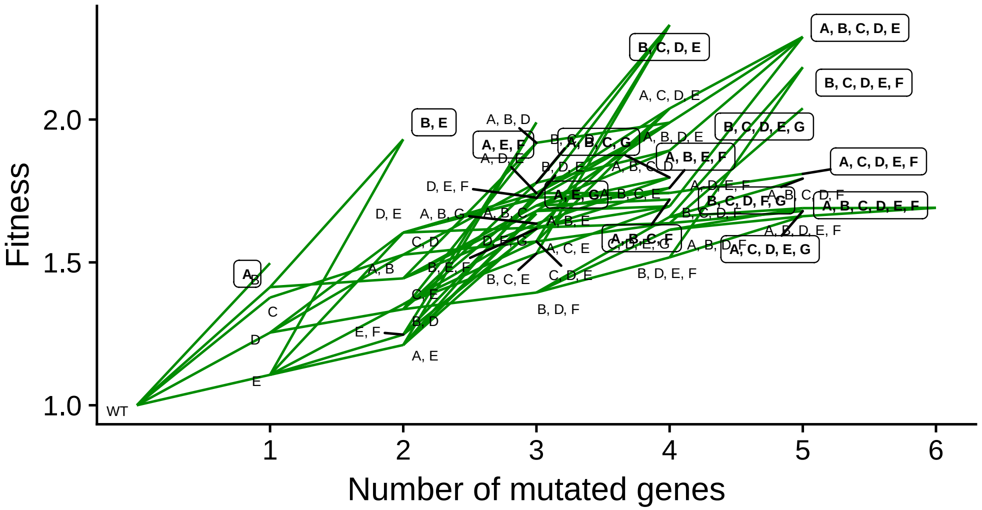


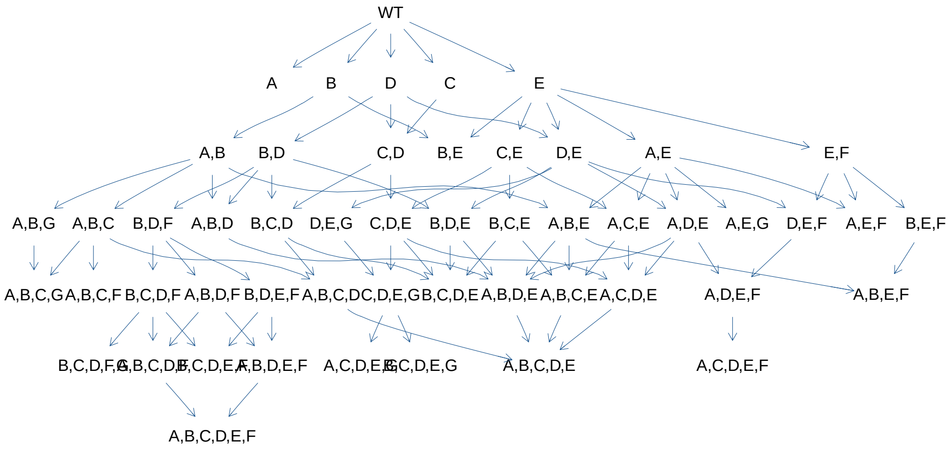

| | |
|---|---|
| *ID* | rB10EATzKOO2QwOIv |
| *acc genotypes* | 23 |
| *peaks no back* | 9 |
| *removed edges* | NA |
| *prop rem edges* | NA |

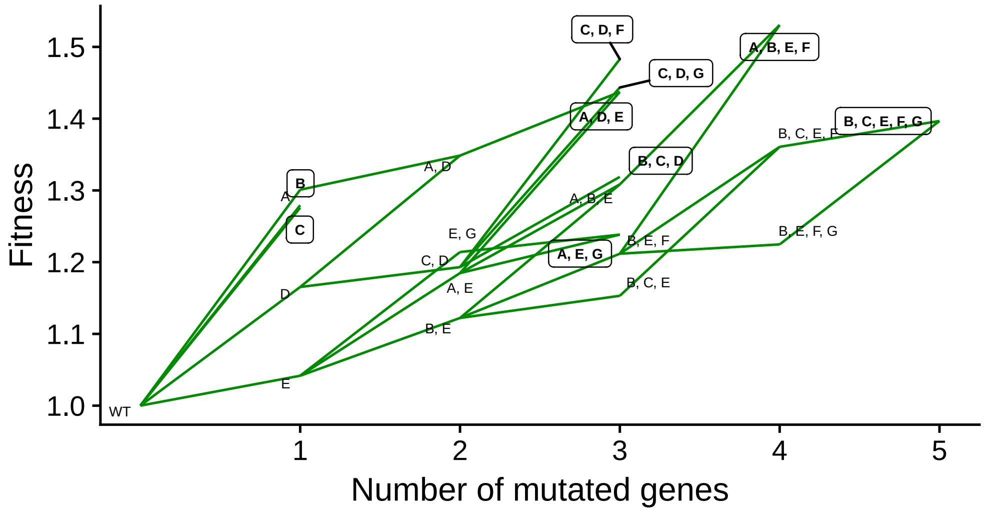


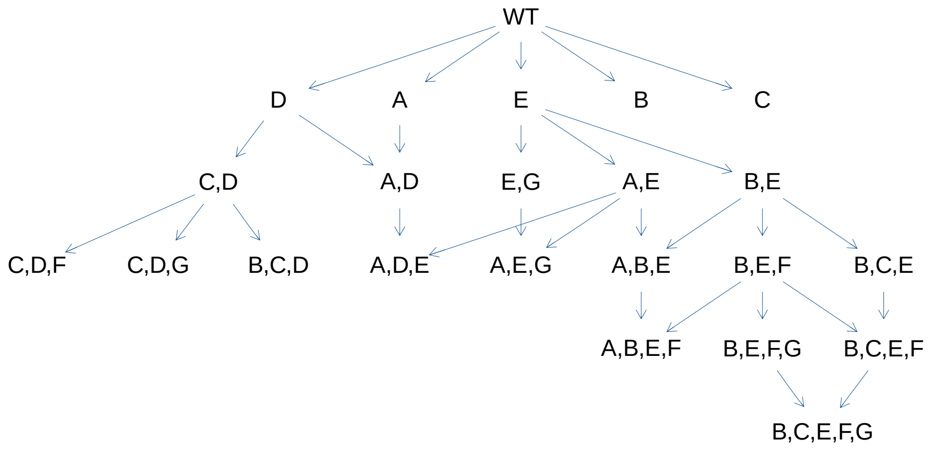

| | |
|---|---|
| *ID* | rCDeBXmZNnImUHl1 |
| *acc genotypes* | 60 |
| *peaks no back* | 24 |
| *removed edges* | NA |
| *prop rem edges* | NA |

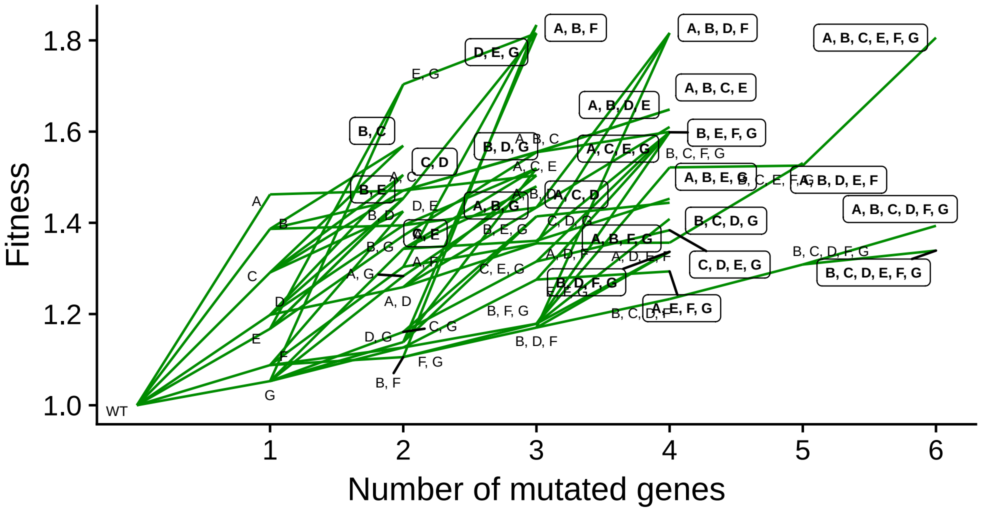


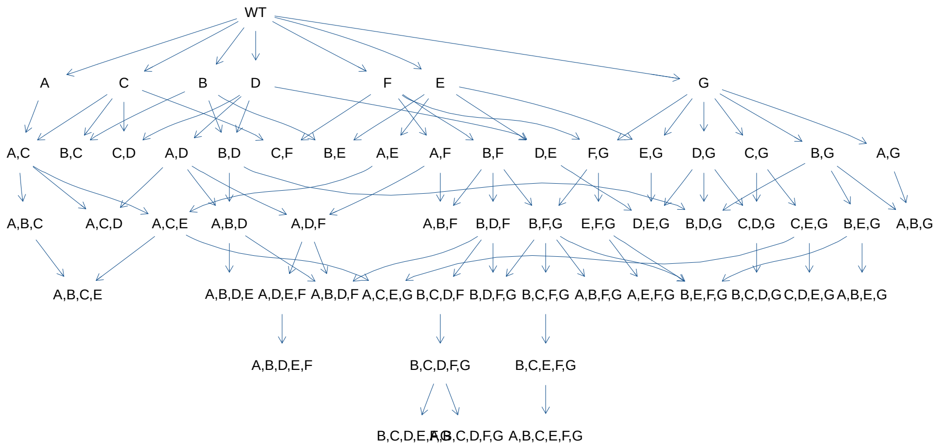

| | |
|---|---|
| *ID* | ReBiTNyW6Sg6eJwN |
| *acc genotypes* | 75 |
| *peaks no back* | 13 |
| *removed edges* | NA |
| *prop rem edges* | NA |

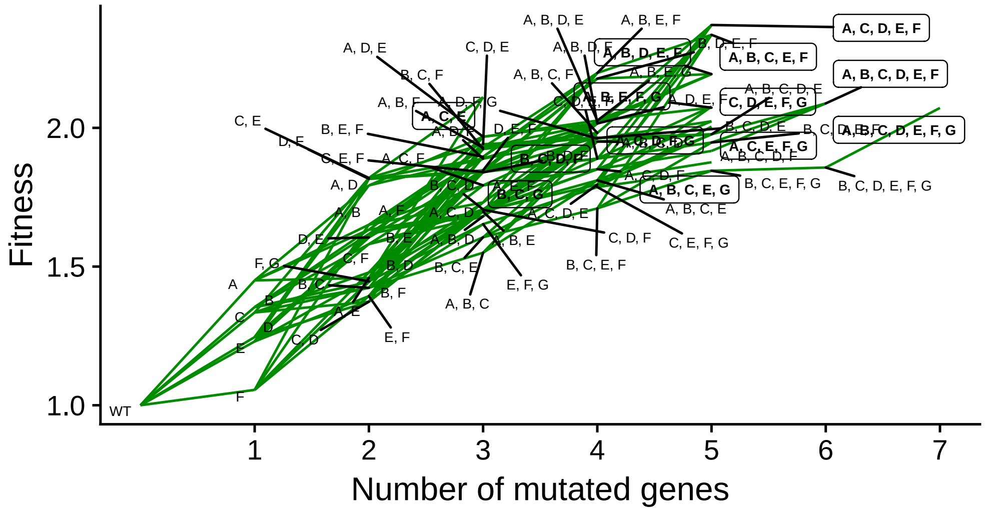


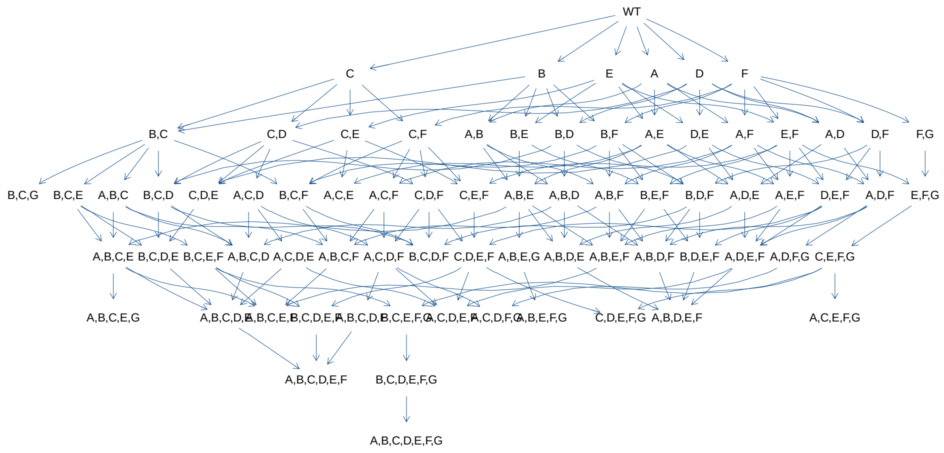

| | |
|---|---|
| *ID* | rhc6UcRhymI47xh4 |
| *acc genotypes* | 12 |
| *peaks no back* | 6 |
| *removed edges* | NA |
| *prop rem edges* | NA |

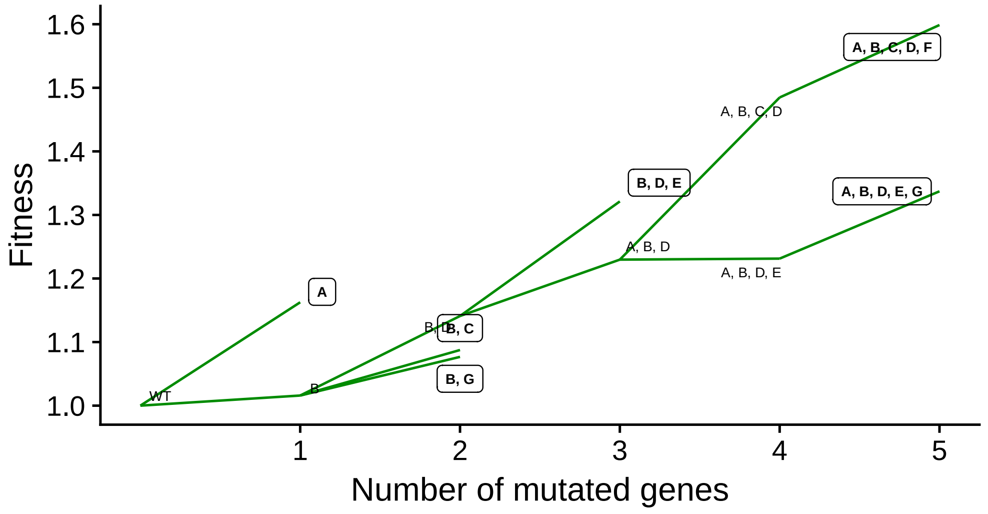


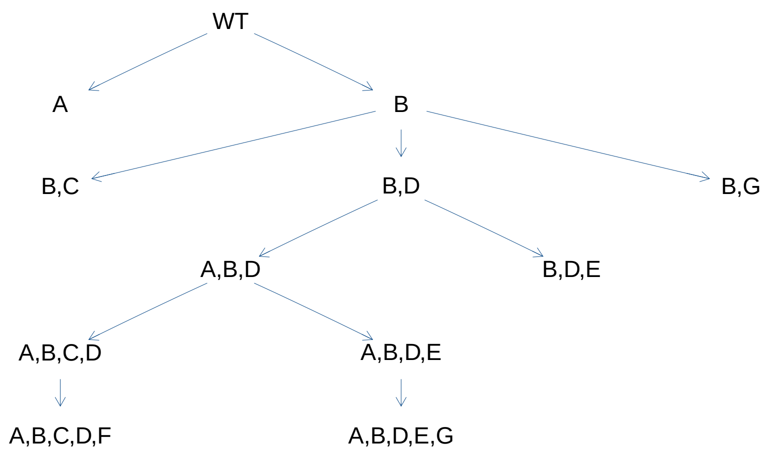

| | |
|---|---|
| *ID* | RmugDTsuCjx1AI9r |
| *acc genotypes* | 14 |
| *peaks no back* | 6 |
| *removed edges* | NA |
| *prop rem edges* | NA |

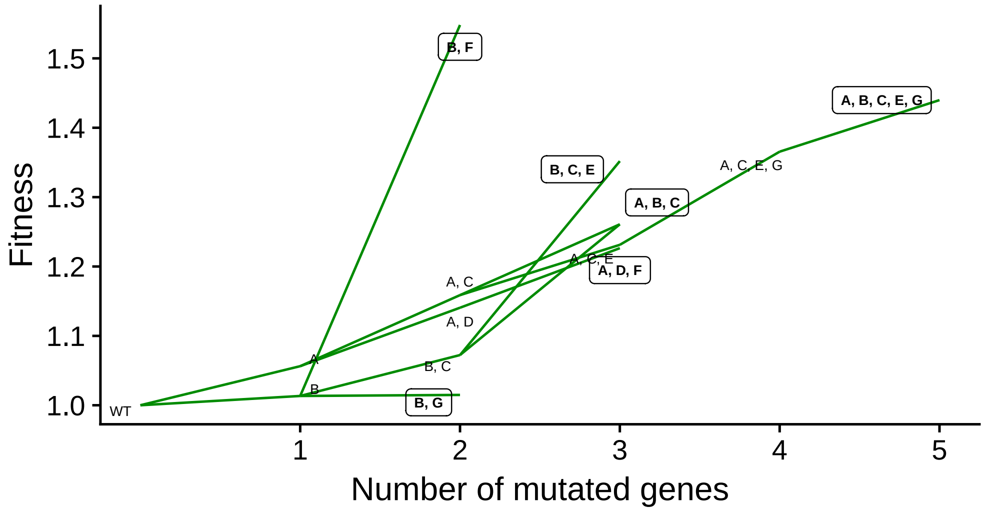


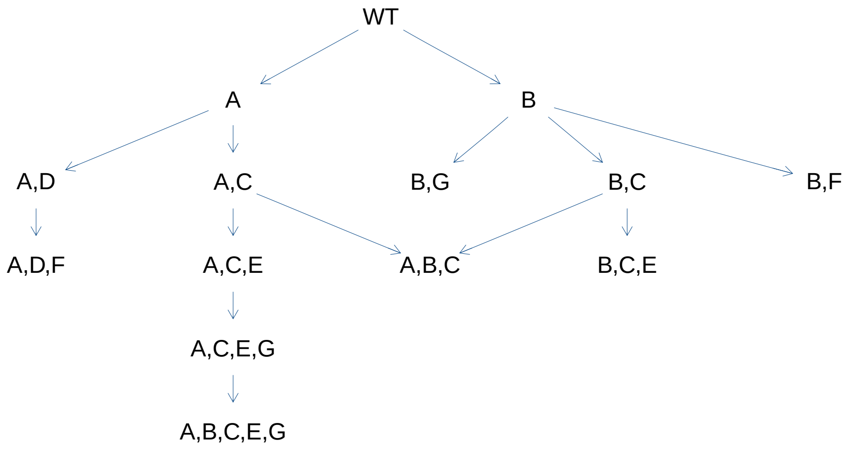

| | |
|---|---|
| *ID* | RnHnxYc271fXVfMs |
| *acc genotypes* | 57 |
| *peaks no back* | 15 |
| *removed edges* | NA |
| *prop rem edges* | NA |

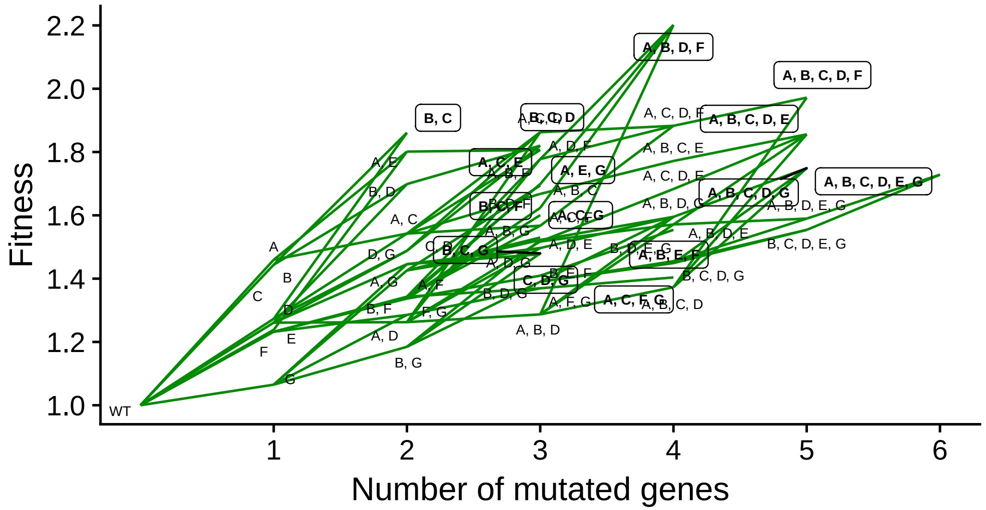


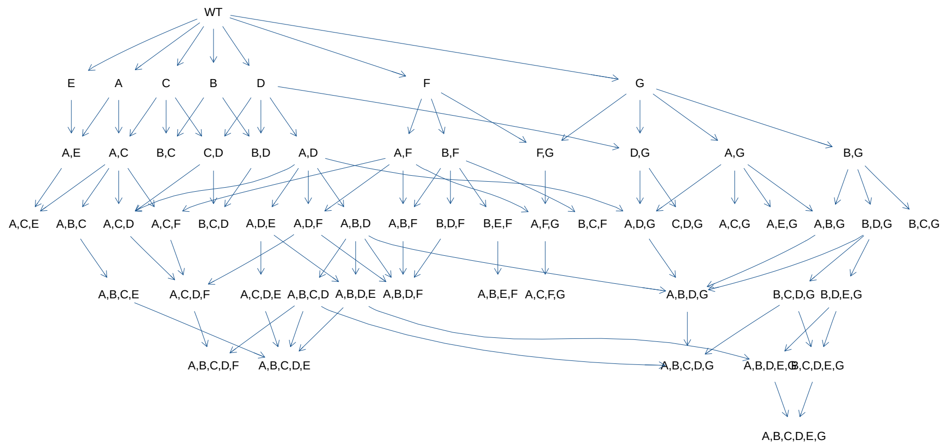

| | |
|---|---|
| *ID* | rWquwMtuP4hFOLRB |
| *acc genotypes* | 66 |
| *peaks no back* | 15 |
| *removed edges* | NA |
| *prop rem edges* | NA |

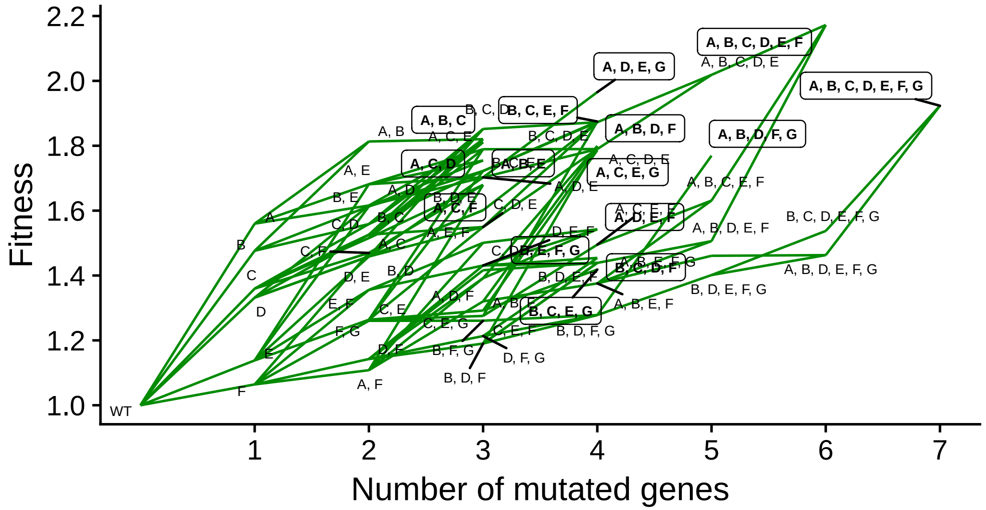


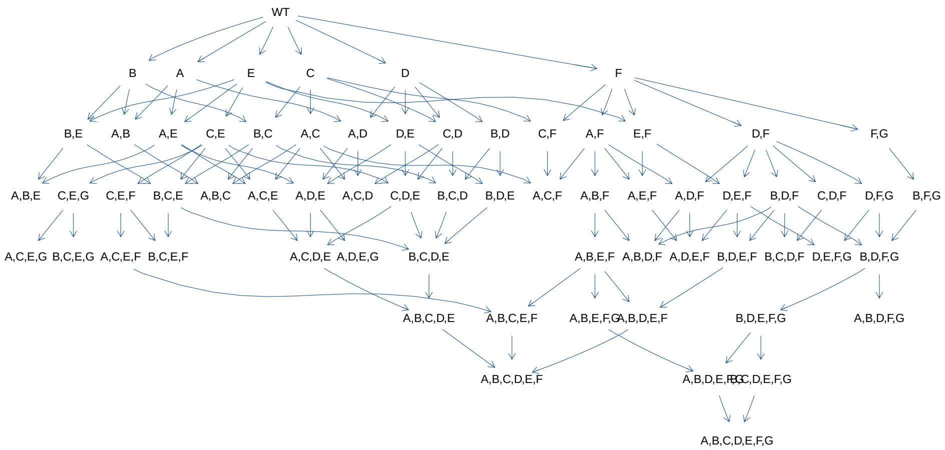

| | |
|---|---|
| *ID* | rX57ibCOZcMSMm4z |
| *acc genotypes* | 30 |
| *peaks no back* | 12 |
| *removed edges* | NA |
| *prop rem edges* | NA |

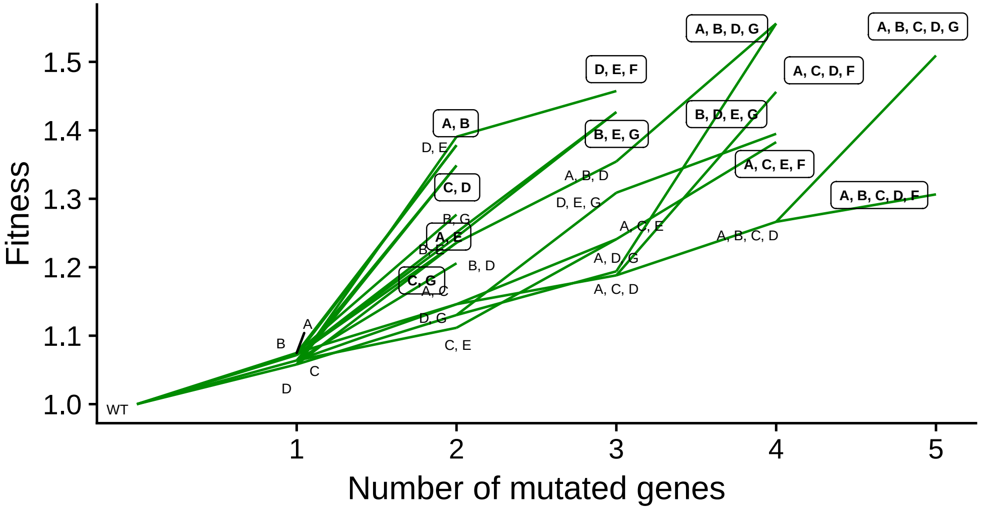


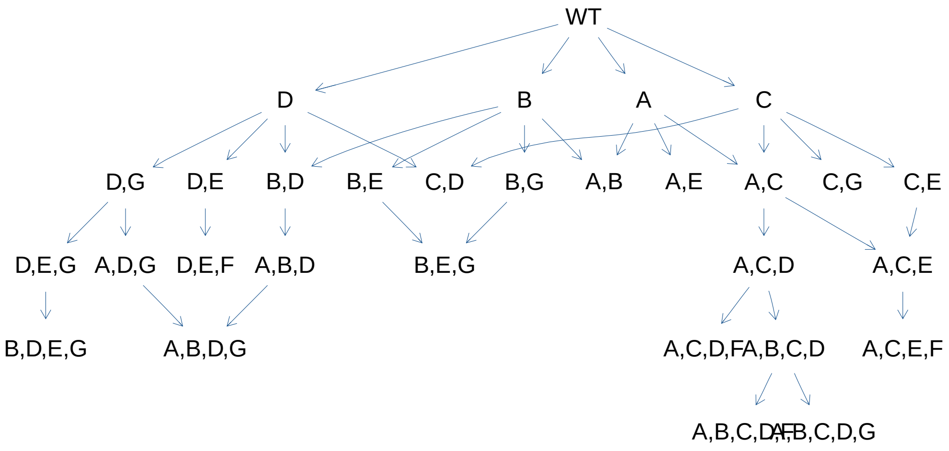

| | |
|---|---|
| *ID* | s10ieWttHAueSTqaL |
| *acc genotypes* | 74 |
| *peaks no back* | 25 |
| *removed edges* | NA |
| *prop rem edges* | NA |

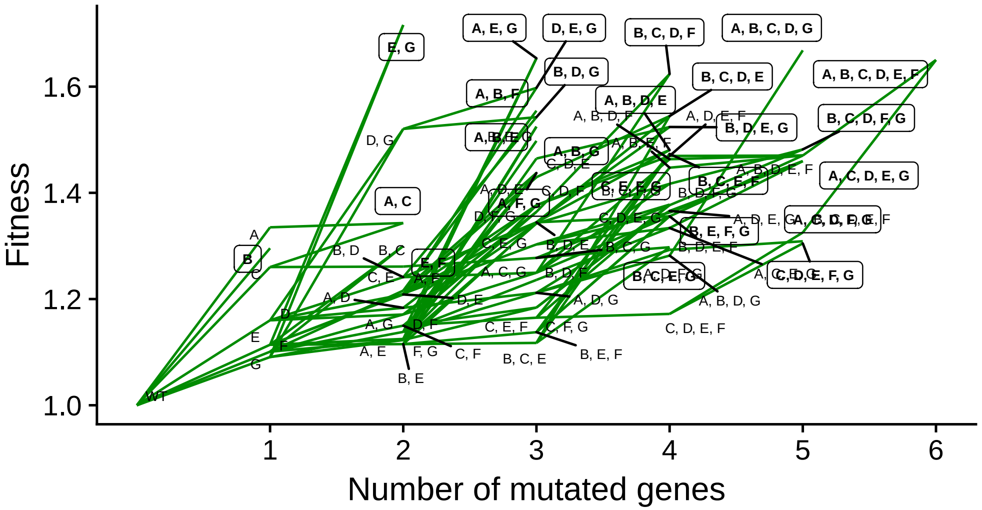


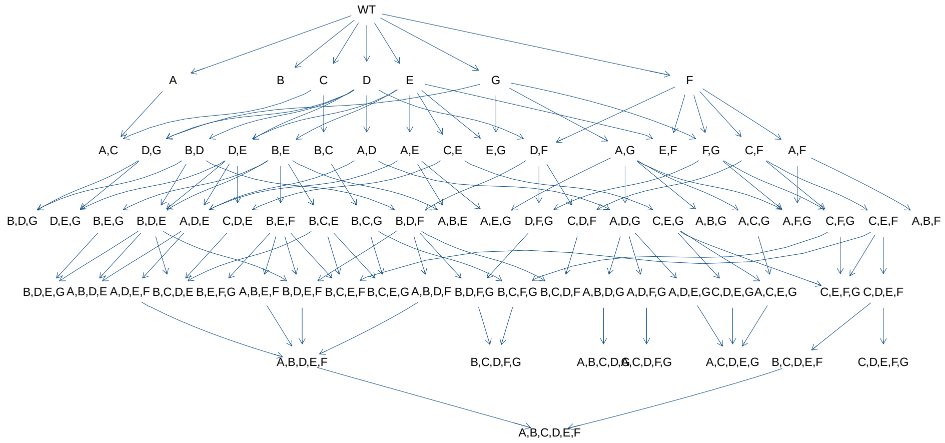

| | |
|---|---|
| *ID* | s5lzMrcmZRB5b6Mo |
| *acc genotypes* | 28 |
| *peaks no back* | 10 |
| *removed edges* | NA |
| *prop rem edges* | NA |

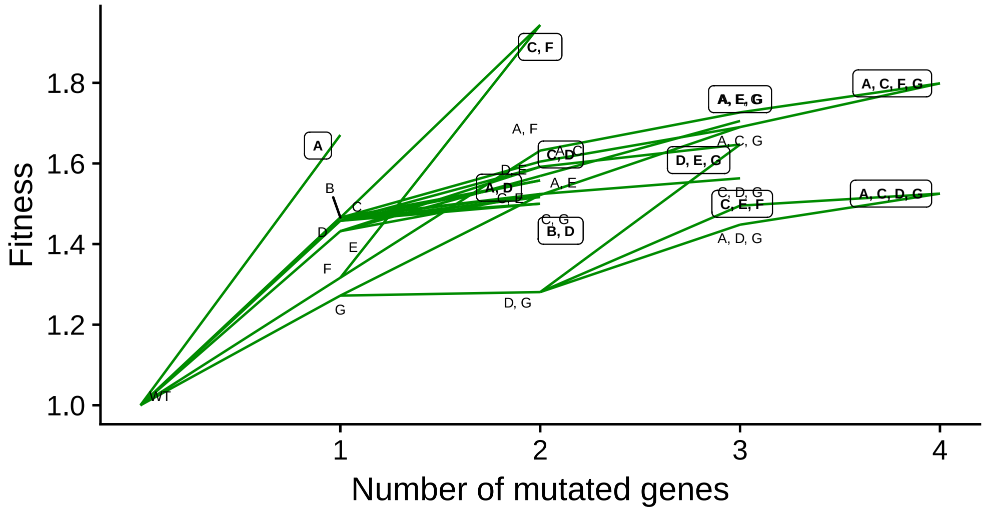


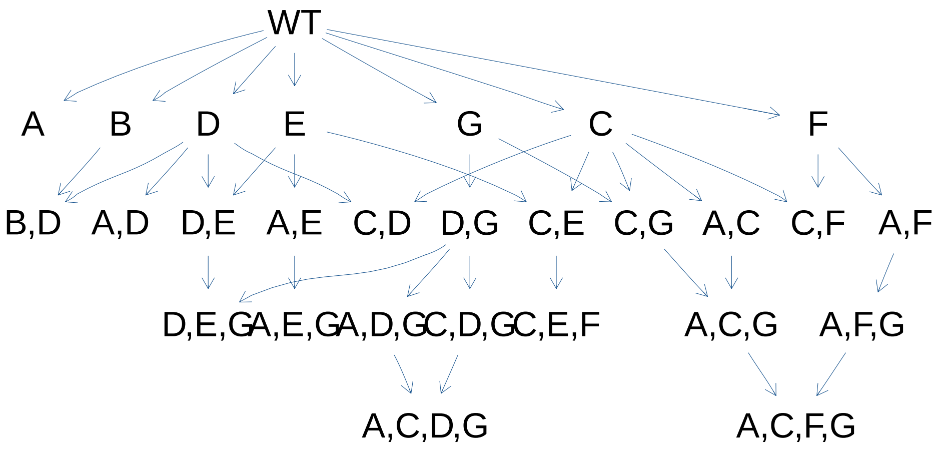

| | |
|---|---|
| *ID* | sgP24AiGrN2flVGV |
| *acc genotypes* | 43 |
| *peaks no back* | 10 |
| *removed edges* | NA |
| *prop rem edges* | NA |

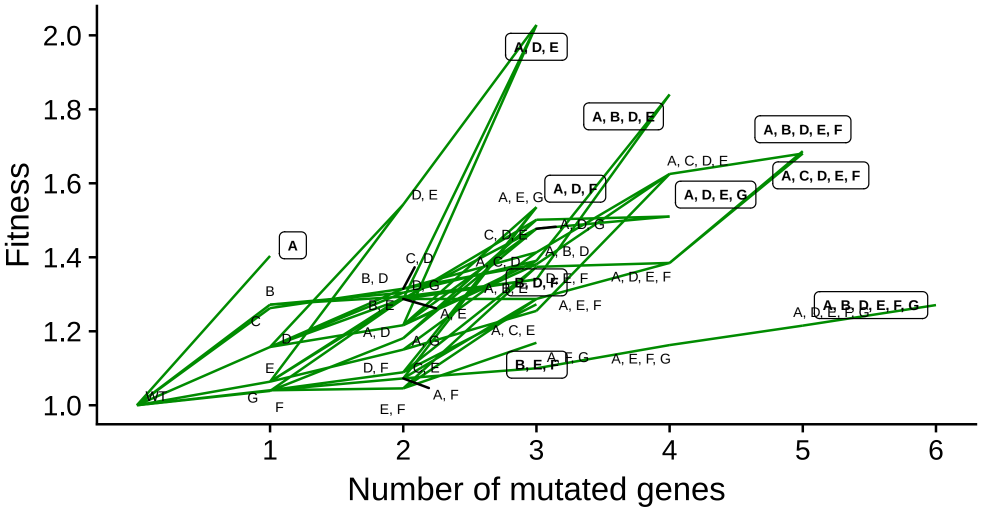


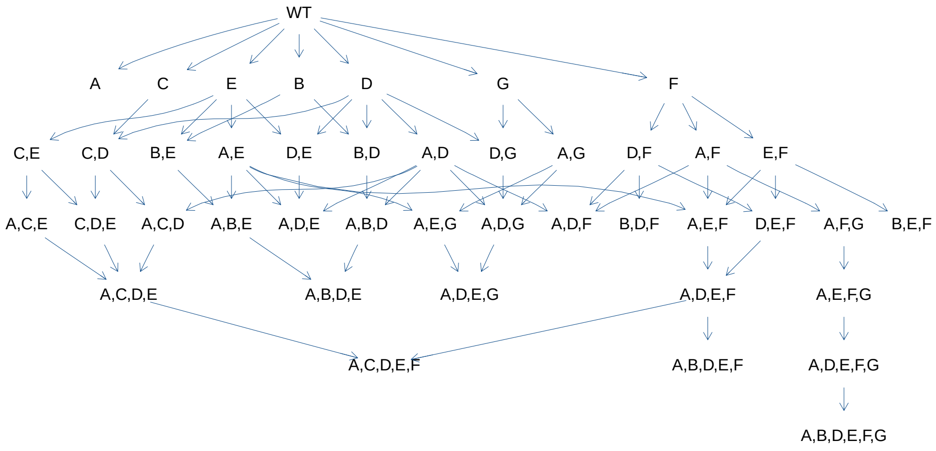

| | |
|---|---|
| *ID* | sJhH8DjY2yo2ZQjo |
| *acc genotypes* | 19 |
| *peaks no back* | 8 |
| *removed edges* | NA |
| *prop rem edges* | NA |

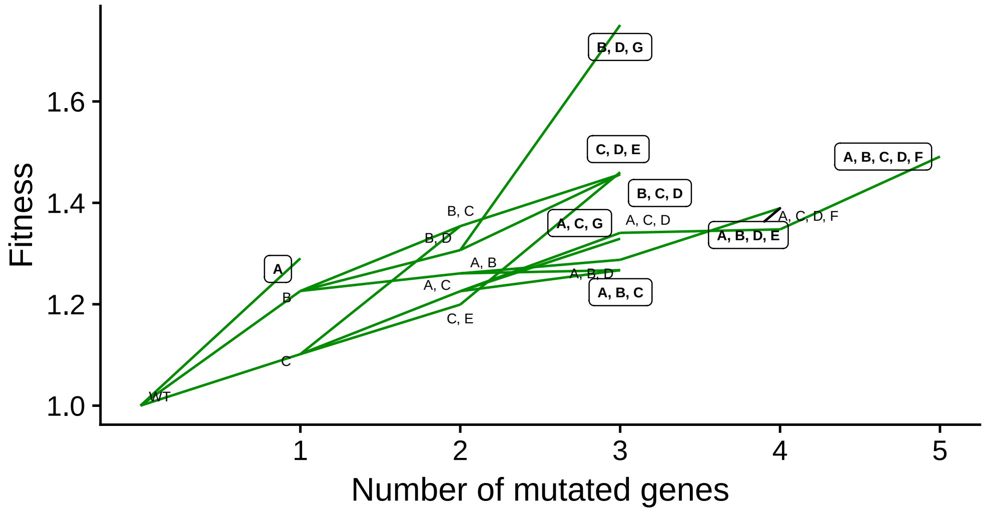


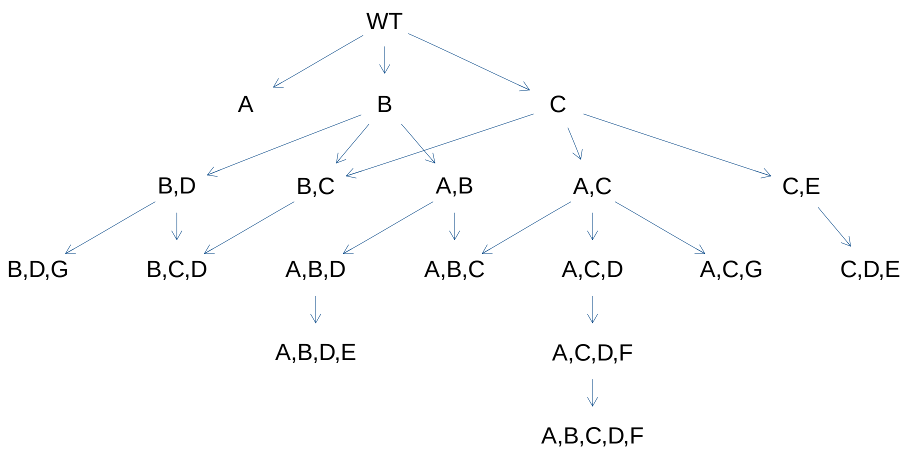

| | |
|---|---|
| *ID* | SpvxDYG1010zub43g5 |
| *acc genotypes* | 25 |
| *peaks no back* | 12 |
| *removed edges* | NA |
| *prop rem edges* | NA |

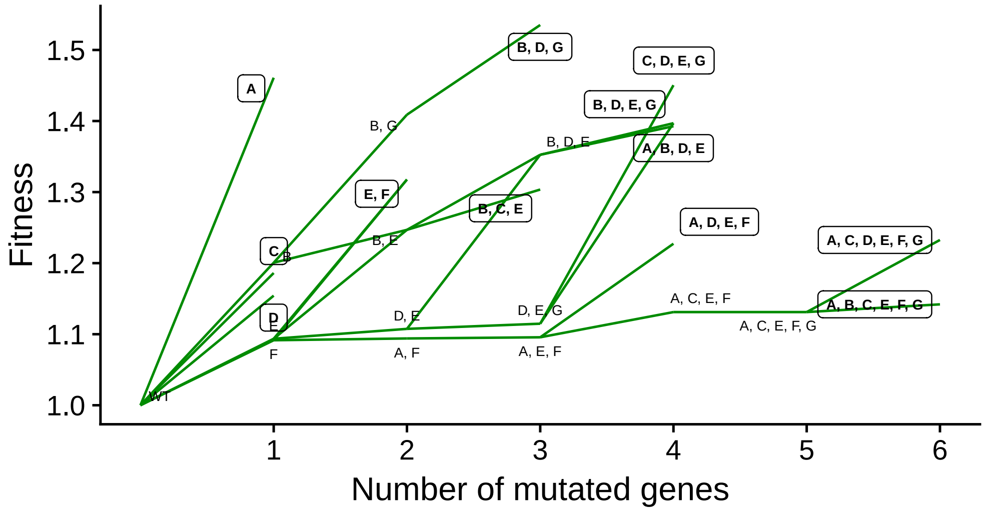


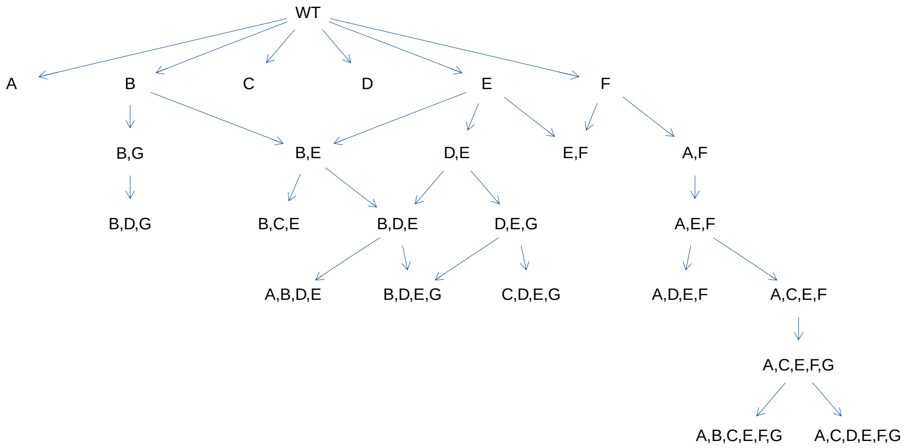

| | |
|---|---|
| *ID* | SRBzzEiA3CEspfsm |
| *acc genotypes* | 51 |
| *peaks no back* | 12 |
| *removed edges* | NA |
| *prop rem edges* | NA |

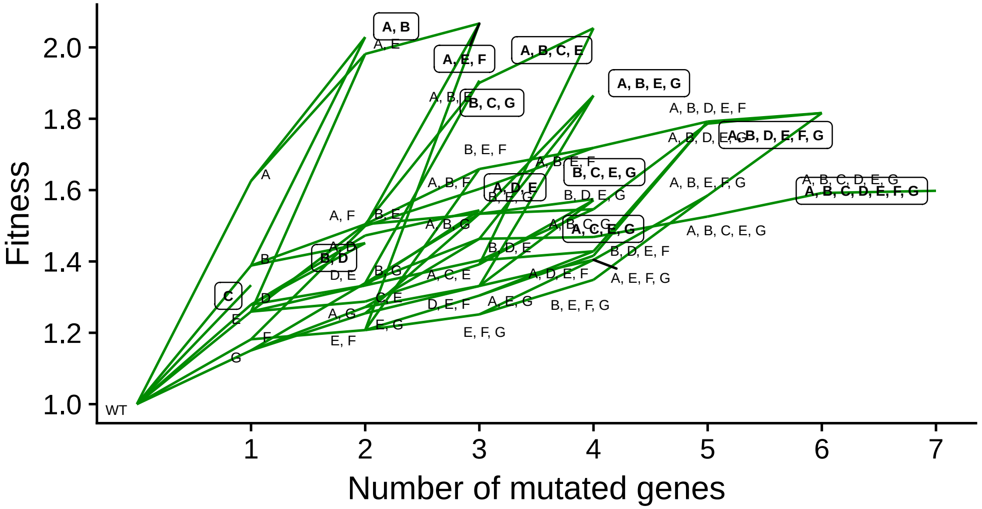


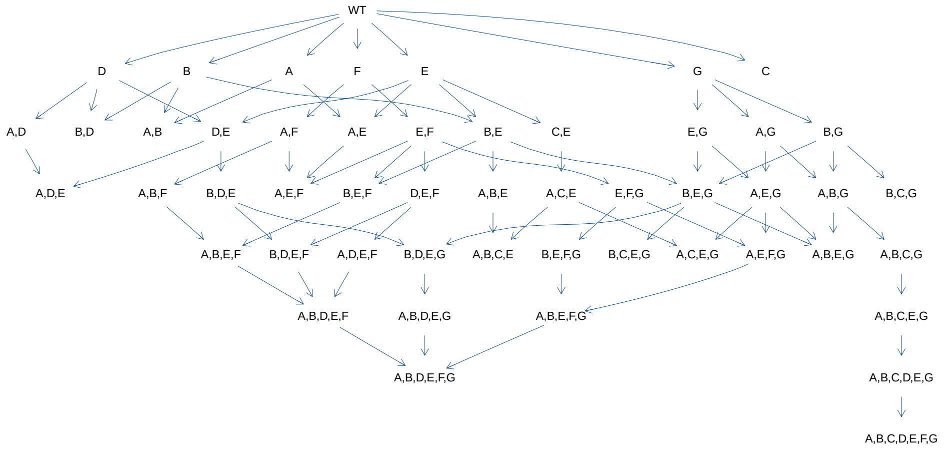

| | |
|---|---|
| *ID* | SWE6nweARORL1yFW |
| *acc genotypes* | 19 |
| *peaks no back* | 6 |
| *removed edges* | NA |
| *prop rem edges* | NA |

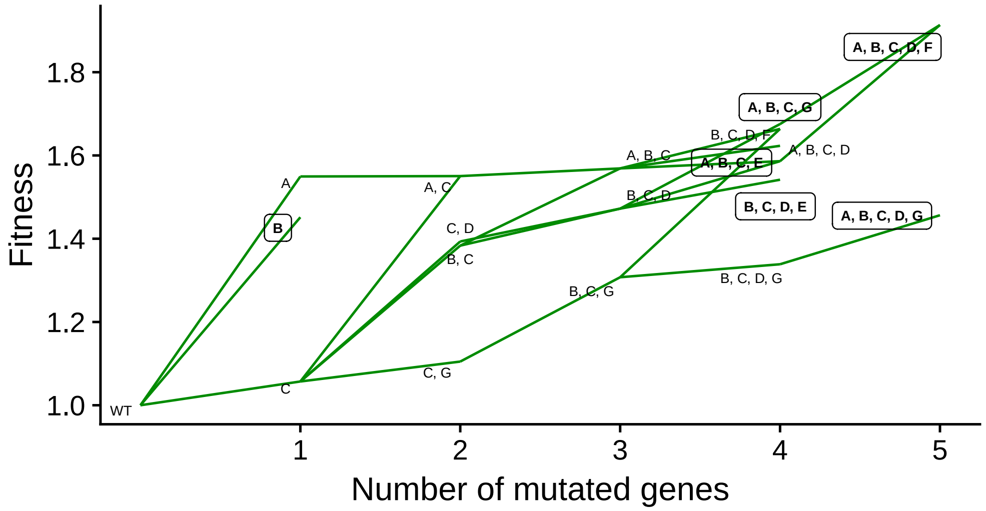


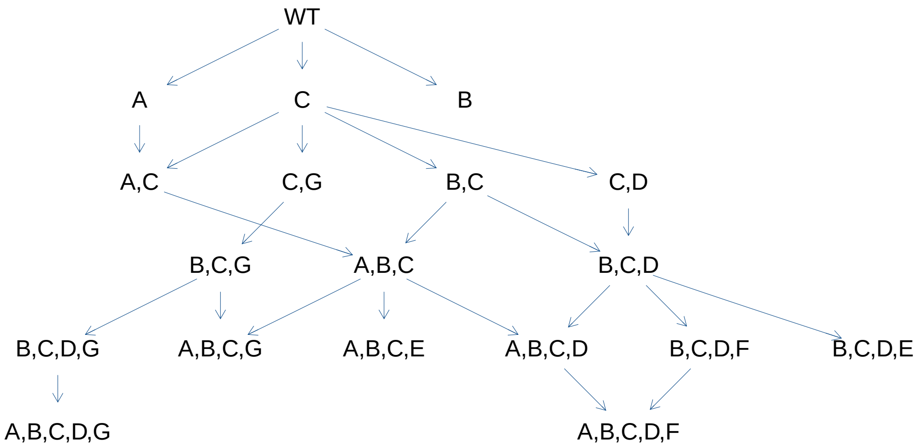

*ID* t10CCmCuCK9esa9Rx
*acc genotypes* 55
*peaks no back* 20
*removed edges* NA
*prop rem edges* NA

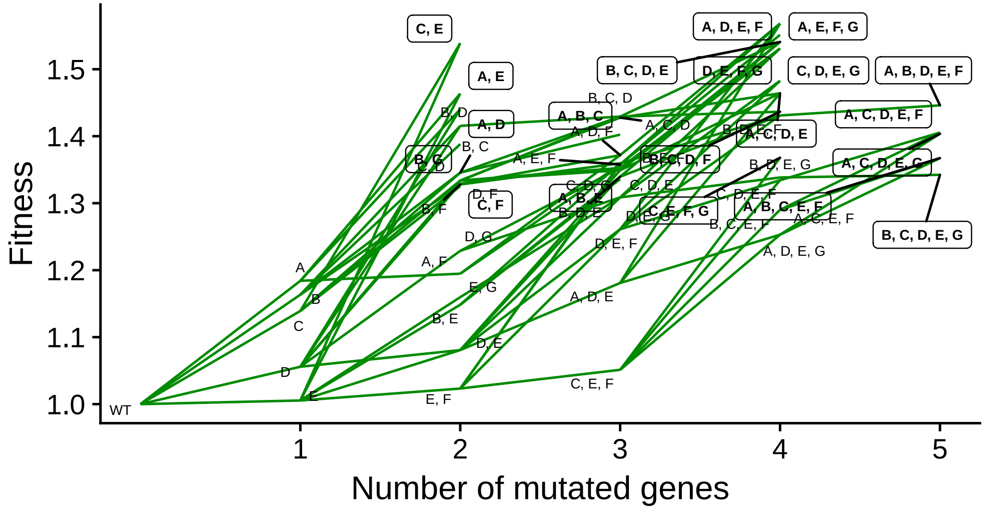


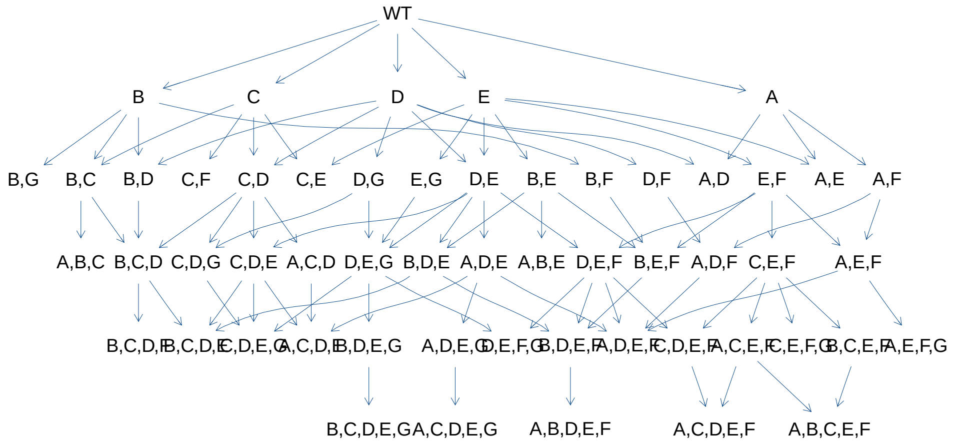

| | |
|---|---|
| *ID* | tgqvbY64a2p366jT |
| *acc genotypes* | 46 |
| *peaks no back* | 14 |
| *removed edges* | NA |
| *prop rem edges* | NA |

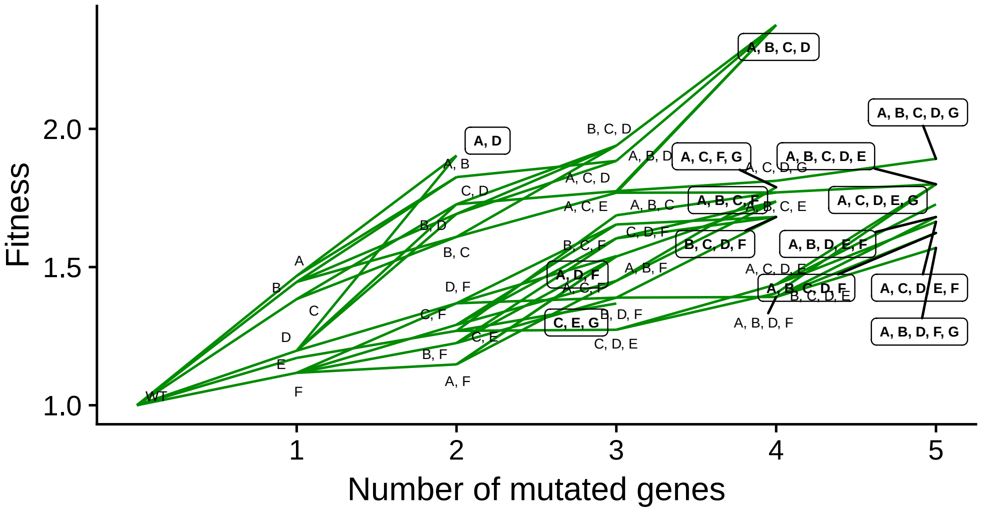


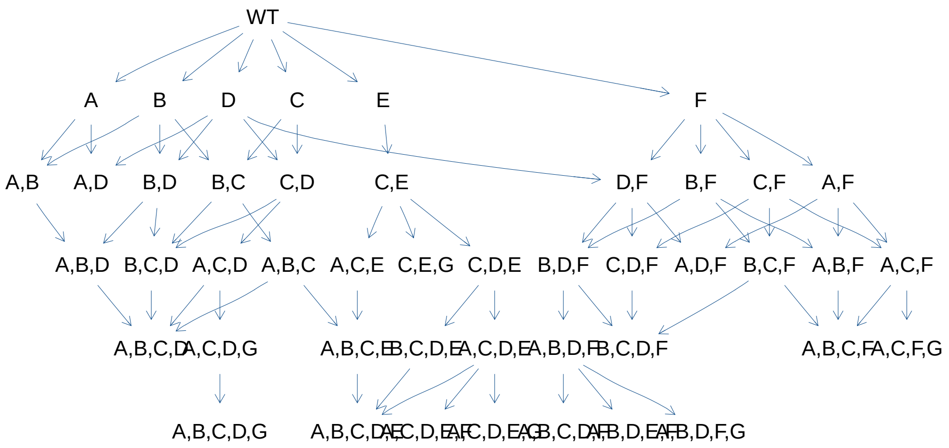

| | |
|---|---|
| *ID* | TgUTHKueceu4QsBH |
| *acc genotypes* | 18 |
| *peaks no back* | 9 |
| *removed edges* | NA |
| *prop rem edges* | NA |

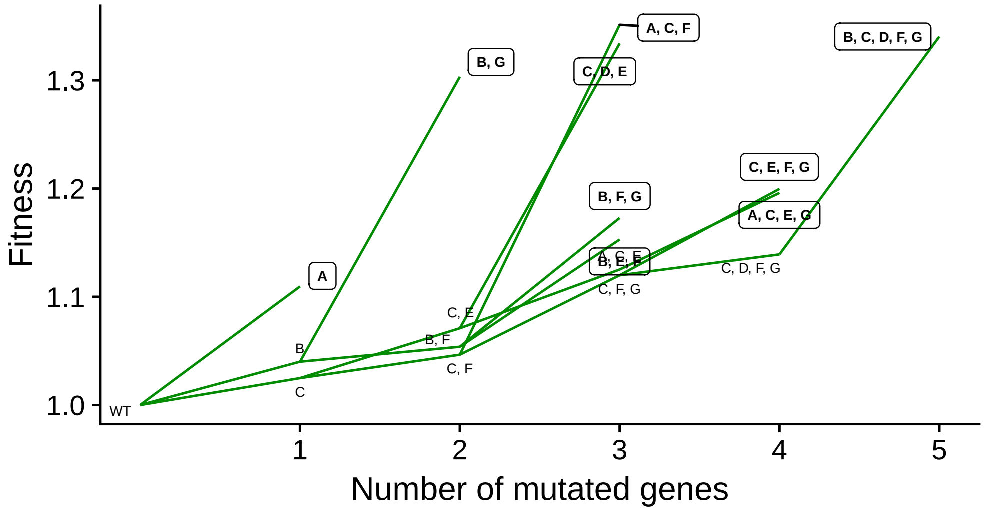


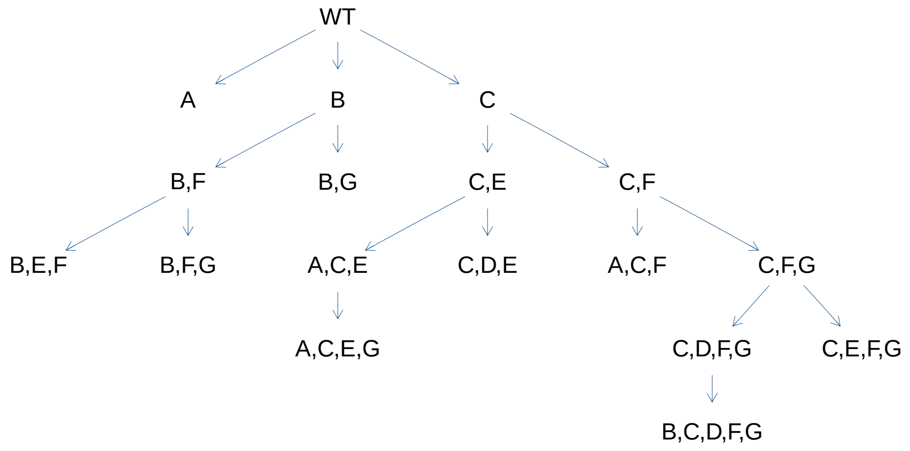

| | |
|---|---|
| *ID* | TgvwxIvA3IzbdsYC |
| *acc genotypes* | 38 |
| *peaks no back* | 14 |
| *removed edges* | NA |
| *prop rem edges* | NA |

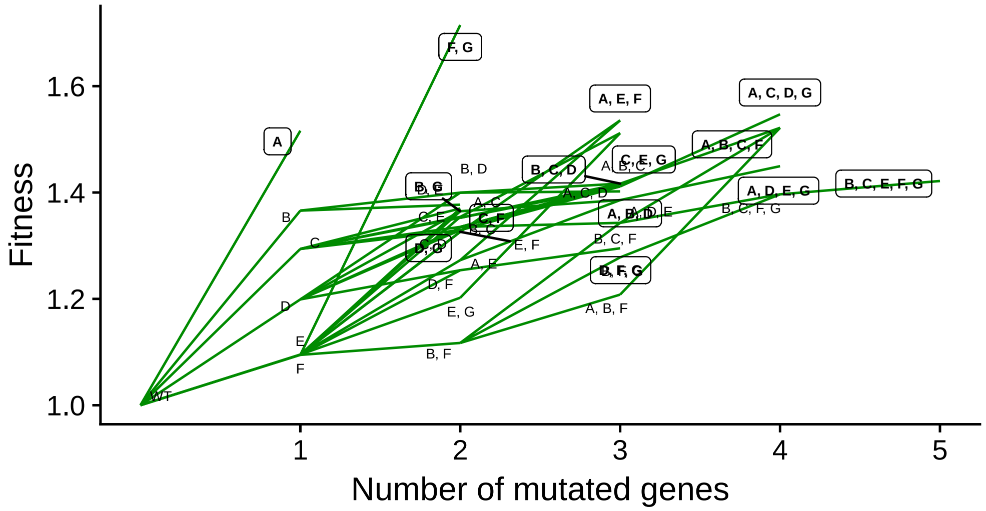


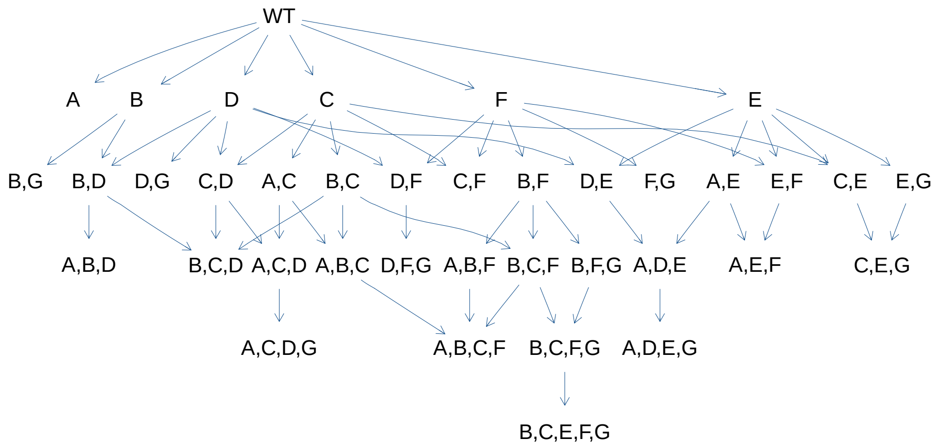

| | |
|---|---|
| *ID* | tHfMAysqIiLPDZQ5 |
| *acc genotypes* | 23 |
| *peaks no back* | 7 |
| *removed edges* | NA |
| *prop rem edges* | NA |

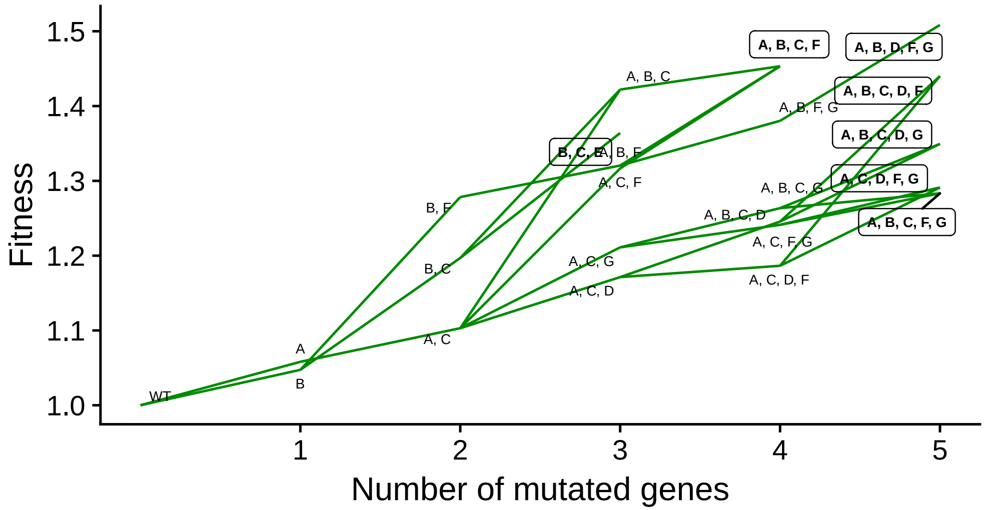


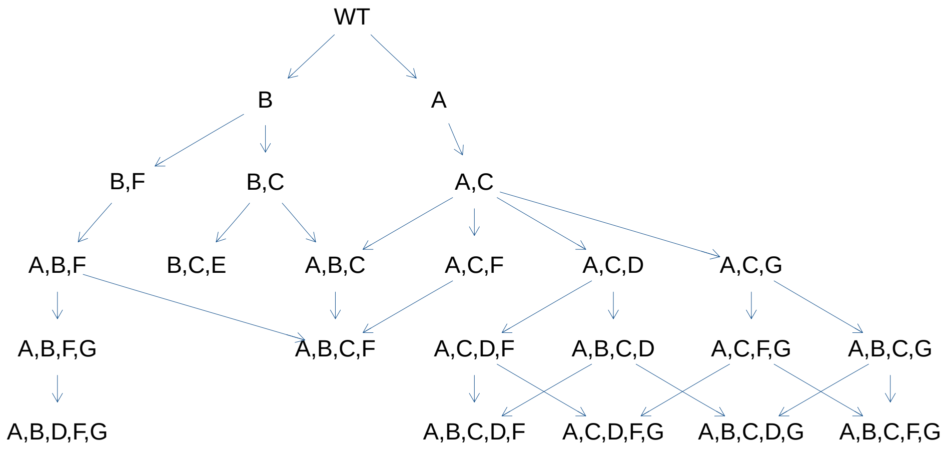

| | |
|---|---|
| *ID* | TJaoNibPlMSgtl1f |
| *acc genotypes* | 24 |
| *peaks no back* | 12 |
| *removed edges* | NA |
| *prop rem edges* | NA |

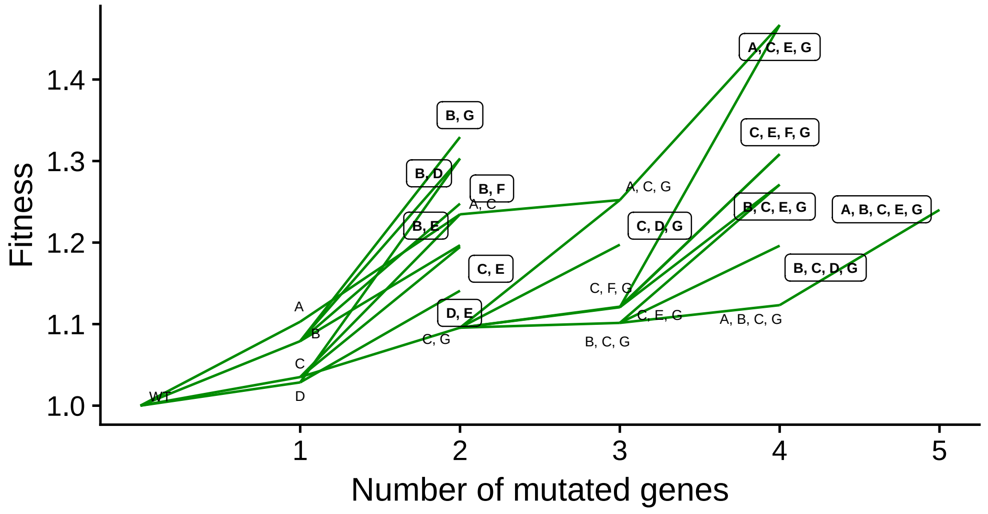


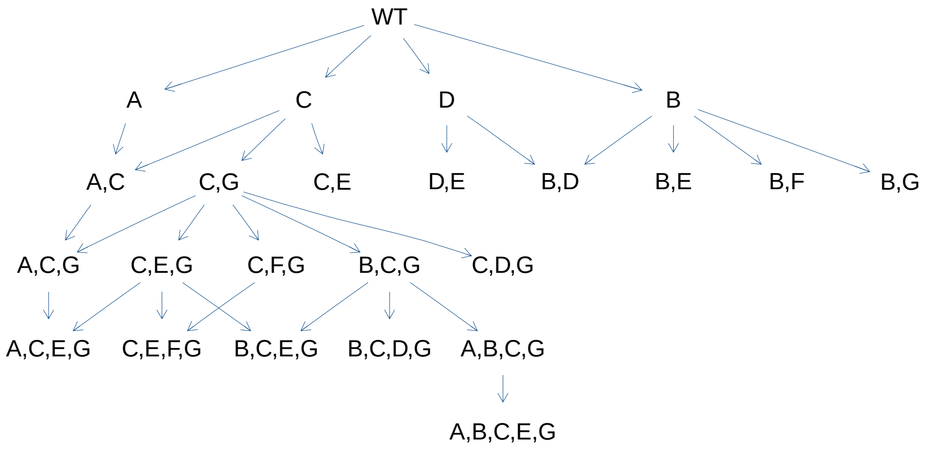

| | |
|---|---|
| *ID* | TI10ATX82Nhqew8T7 |
| *acc genotypes* | 80 |
| *peaks no back* | 15 |
| *removed edges* | NA |
| *prop rem edges* | NA |

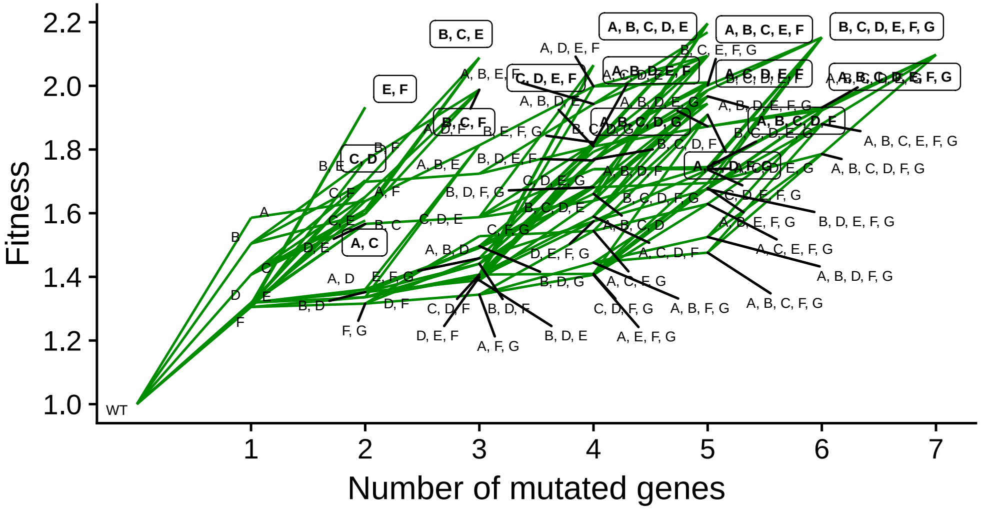


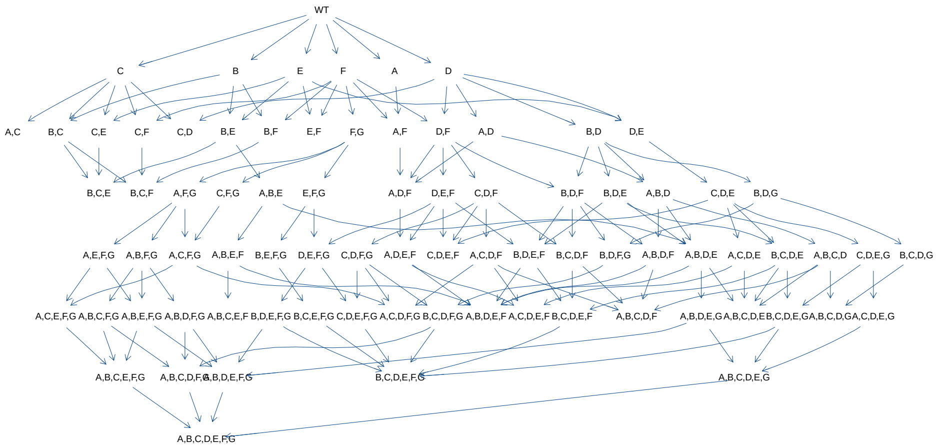

| | |
|---|---|
| *ID* | tljFPA395CywDkbx |
| *acc genotypes* | 17 |
| *peaks no back* | 8 |
| *removed edges* | NA |
| *prop rem edges* | NA |

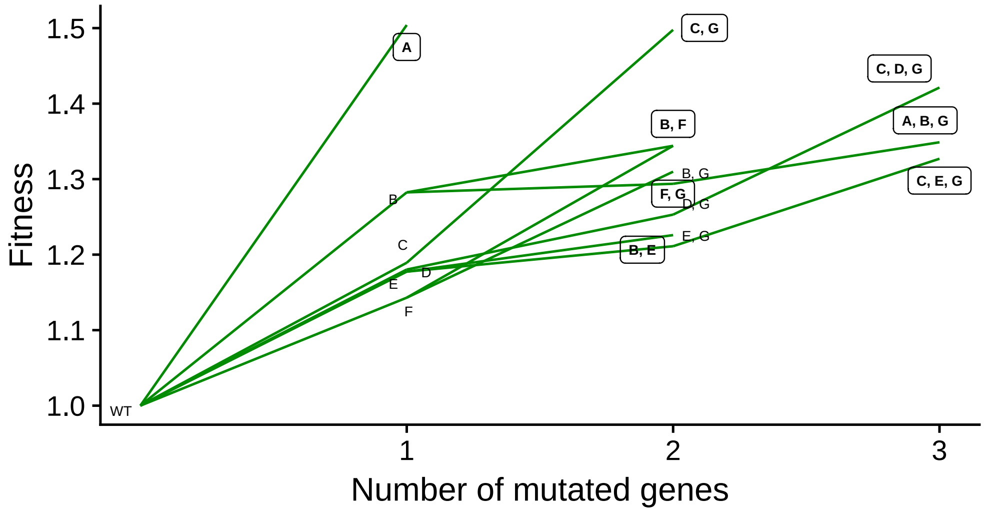


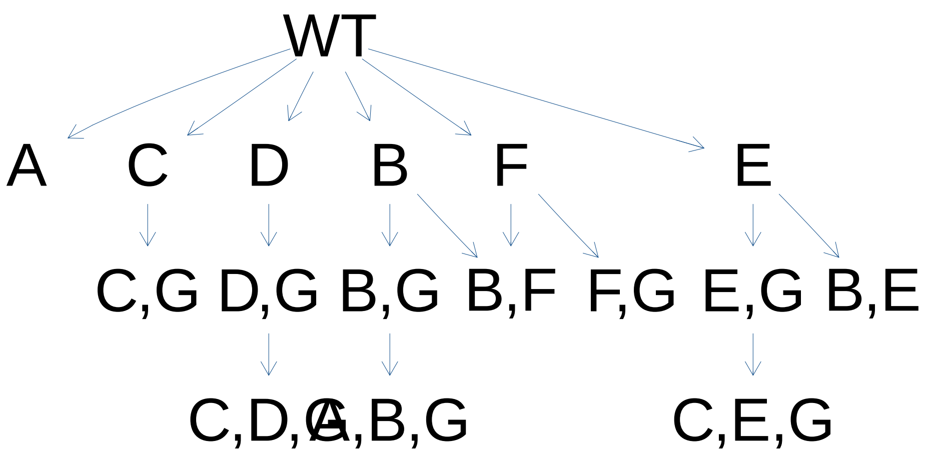

| | |
|---|---|
| *ID* | TonSIXDkiuvbpNr6 |
| *acc genotypes* | 64 |
| *peaks no back* | 16 |
| *removed edges* | NA |
| *prop rem edges* | NA |

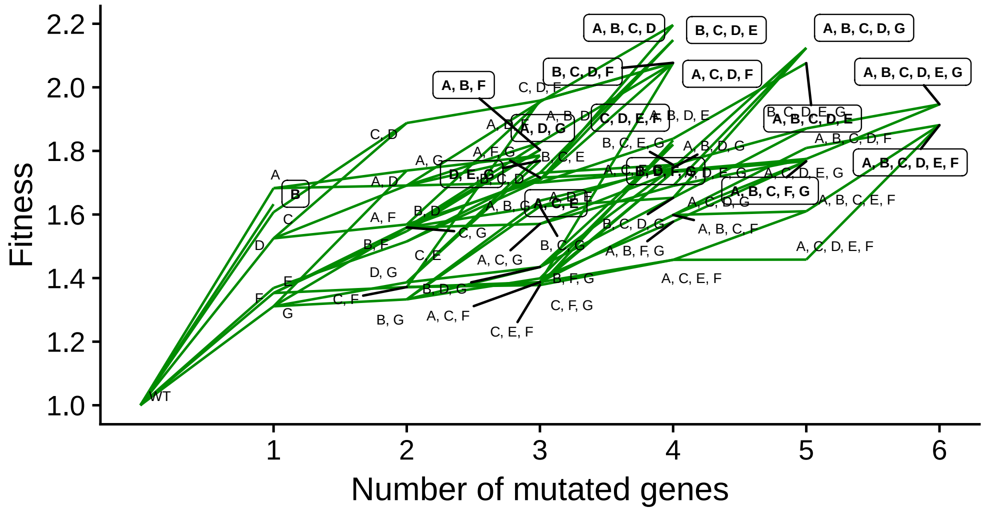


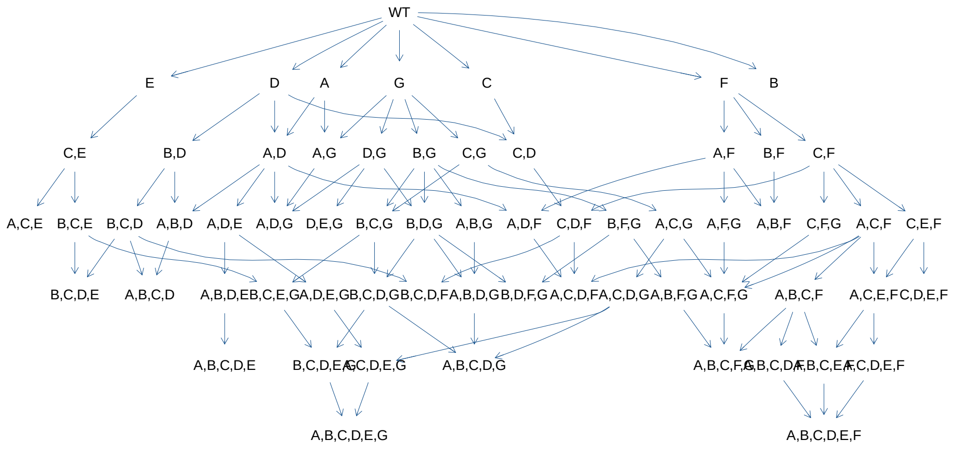

| | |
|---|---|
| *ID* | TuigQ6ZYErsjqG8r |
| *acc genotypes* | 19 |
| *peaks no back* | 9 |
| *removed edges* | NA |
| *prop rem edges* | NA |

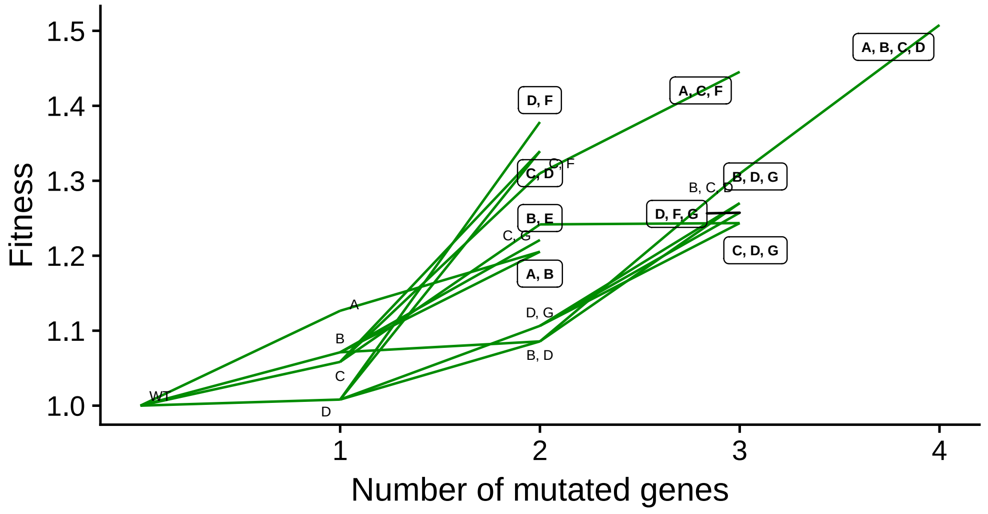


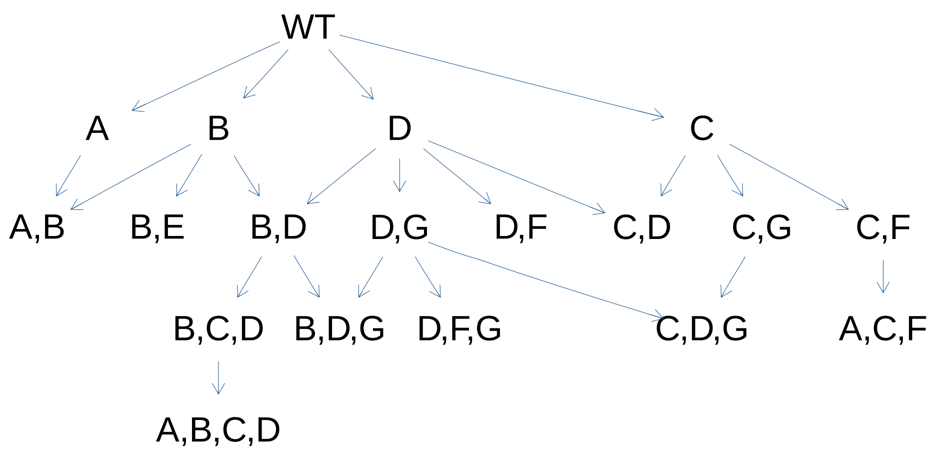

| | |
|---|---|
| *ID* | U2qBdGfdtF10VB9Ww |
| *acc genotypes* | 30 |
| *peaks no back* | 12 |
| *removed edges* | NA |
| *prop rem edges* | NA |

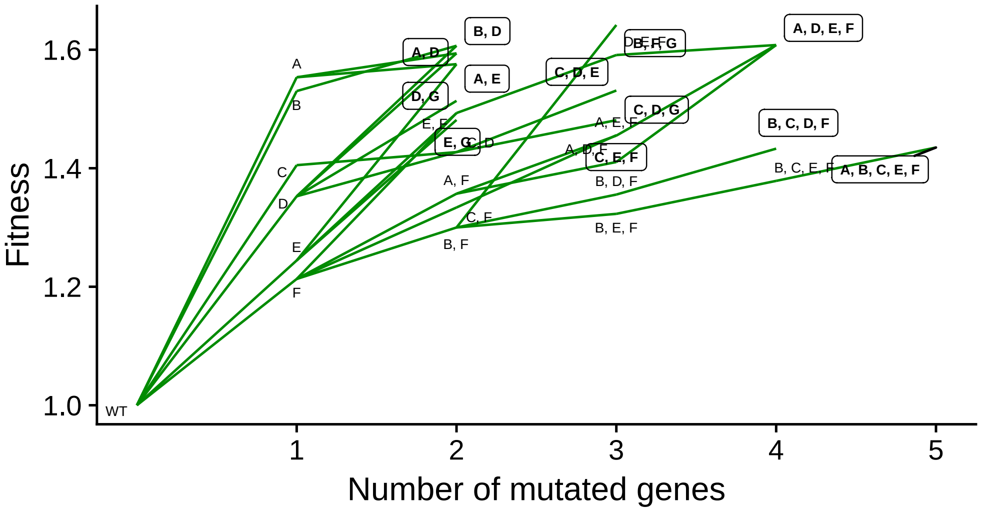


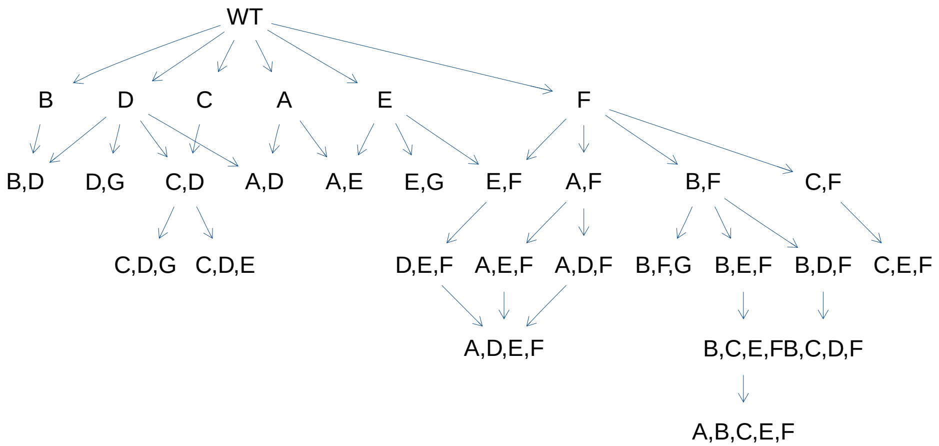

*ID* U7R64nQkJqBxJIju

*acc genotypes* 43

*peaks no back* 14

*removed edges* NA

*prop rem edges* NA

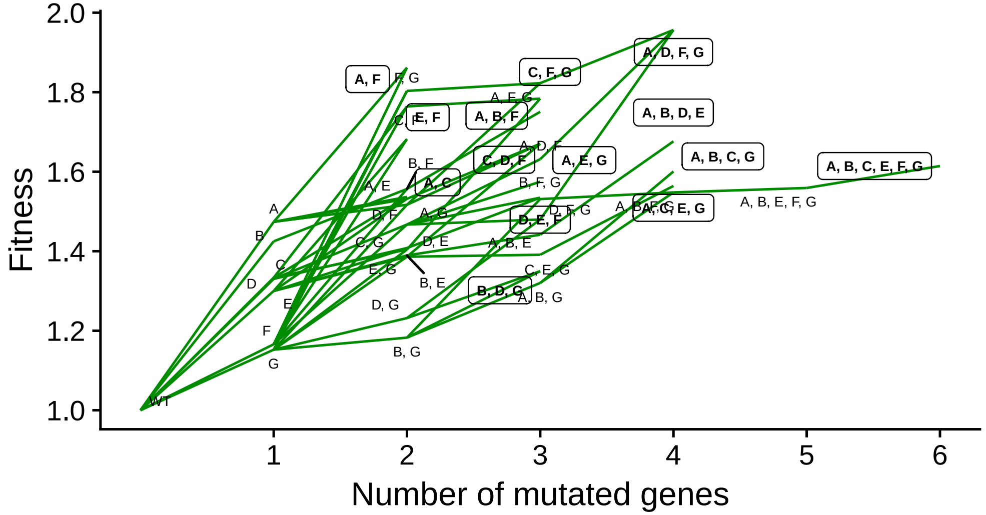


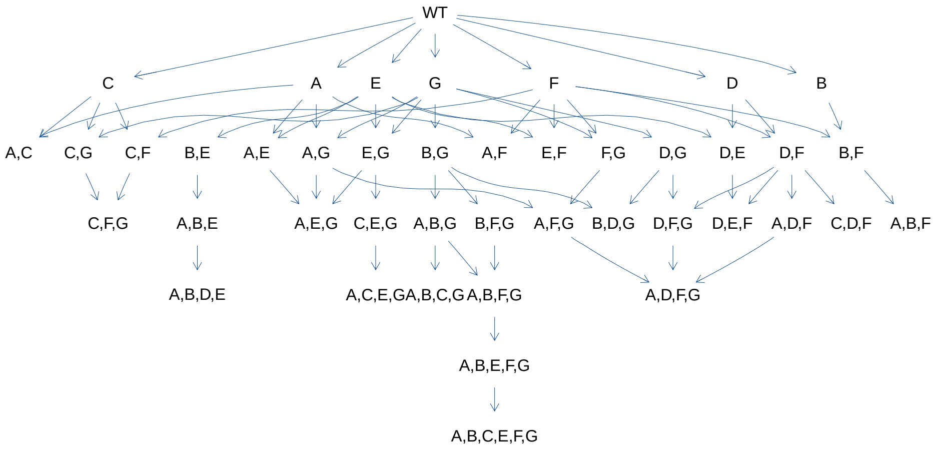

| | |
|---|---|
| *ID* | usBeU10iBdpLaKISX |
| *acc genotypes* | 14 |
| *peaks no back* | 7 |
| *removed edges* | NA |
| *prop rem edges* | NA |

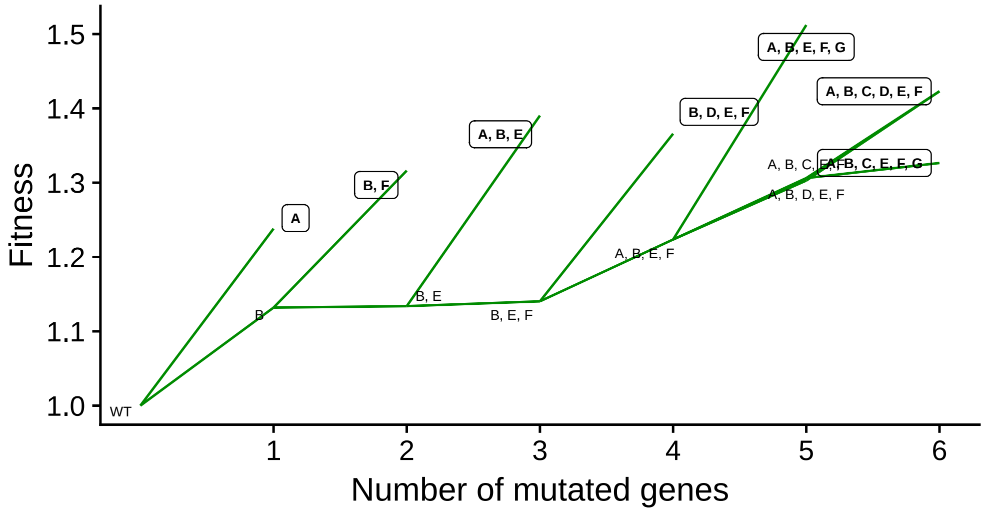


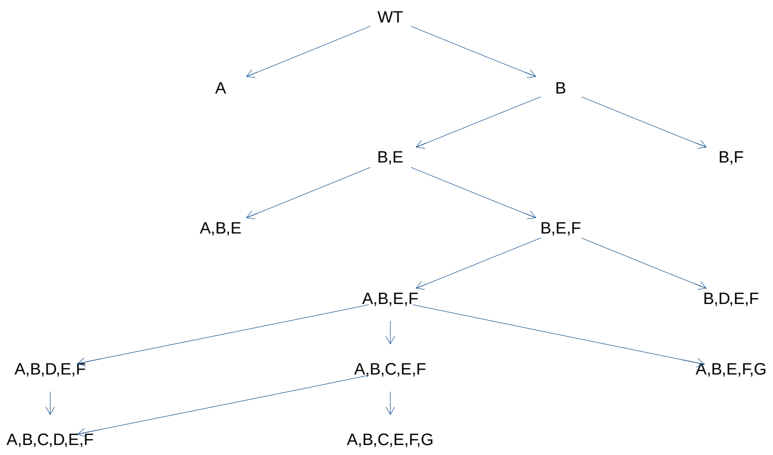

| | |
|---|---|
| *ID* | uTuVU1GMy3Xg8Zye |
| *acc genotypes* | 43 |
| *peaks no back* | 17 |
| *removed edges* | NA |
| *prop rem edges* | NA |

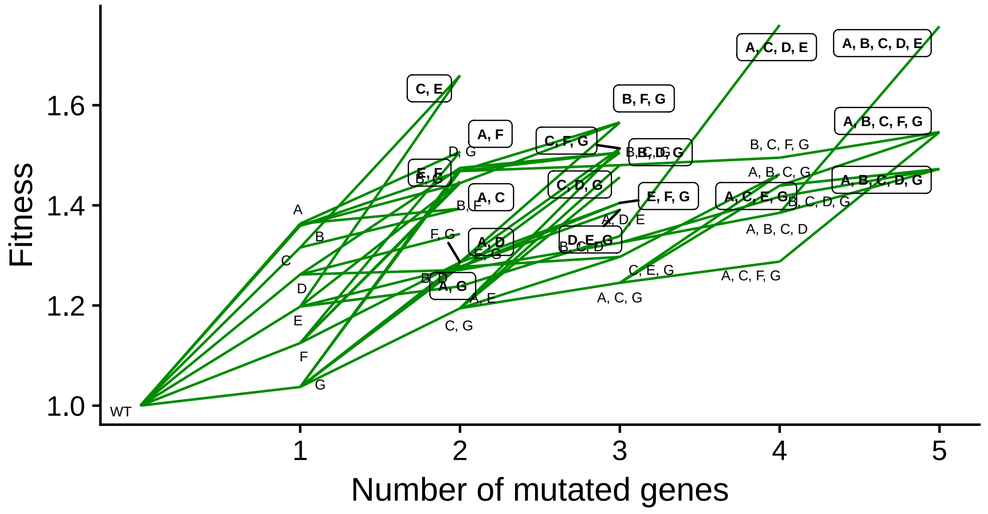


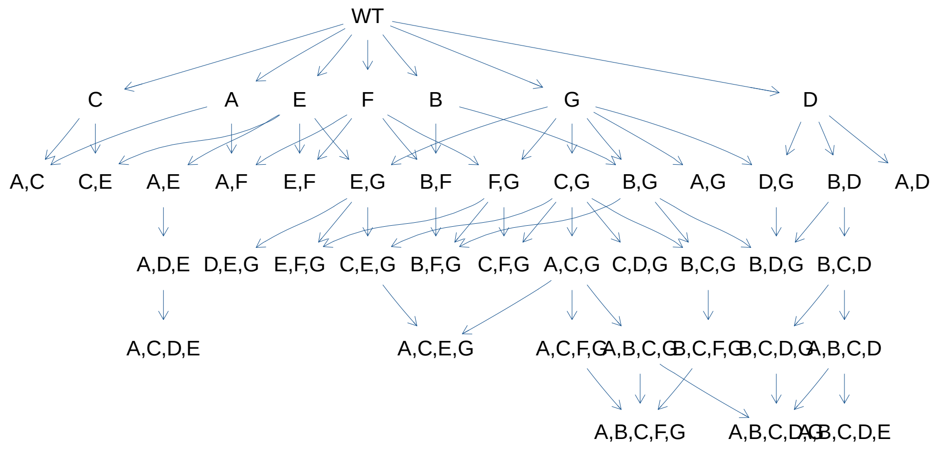

| | |
|---|---|
| *ID* | uxdKQzl2TafUiD7b |
| *acc genotypes* | 42 |
| *peaks no back* | 17 |
| *removed edges* | NA |
| *prop rem edges* | NA |

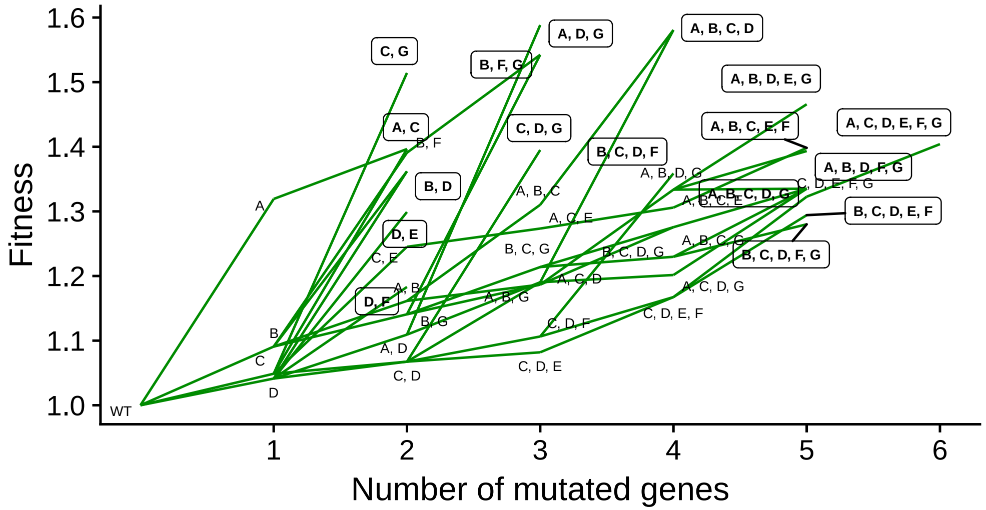


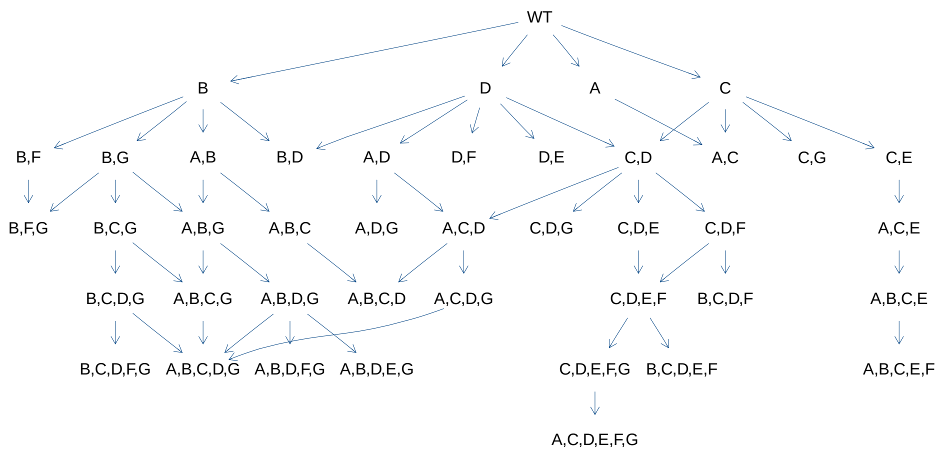

*ID* vFhJDZxTJ3k4epfY
*acc genotypes* 49
*peaks no back* 19
*removed edges* NA
*prop rem edges* NA

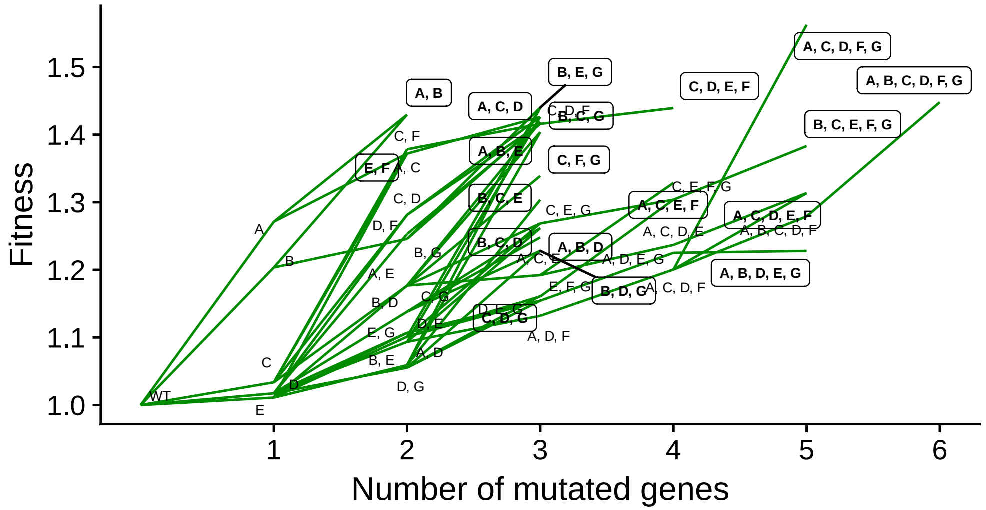


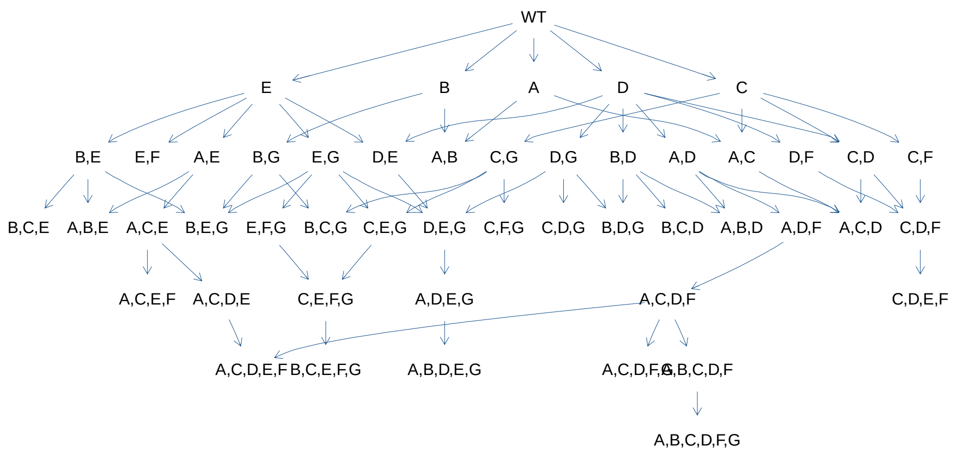

| | |
|---|---|
| *ID* | vmQ3luxZBIZ10PMCg |
| *acc genotypes* | 25 |
| *peaks no back* | 9 |
| *removed edges* | NA |
| *prop rem edges* | NA |

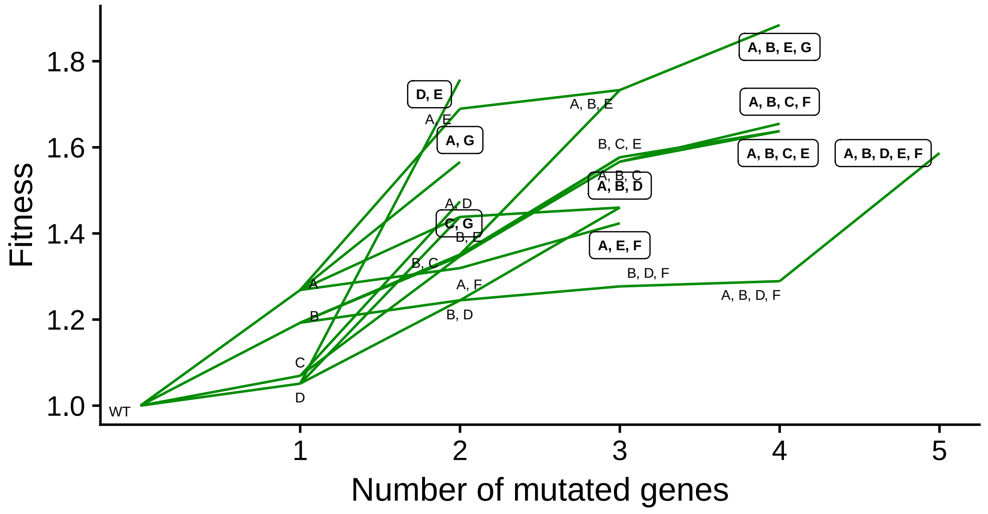


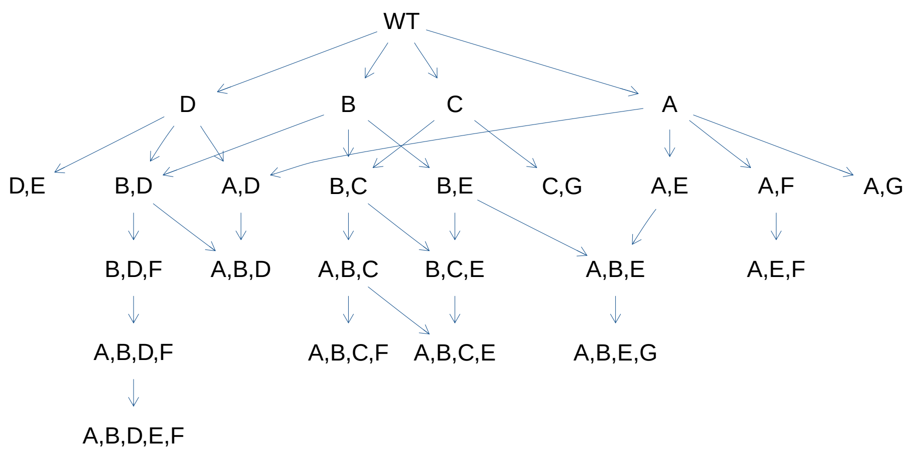

| | |
|---|---|
| *ID* | vt56Tsvc76JVWmej |
| *acc genotypes* | 20 |
| *peaks no back* | 5 |
| *removed edges* | NA |
| *prop rem edges* | NA |

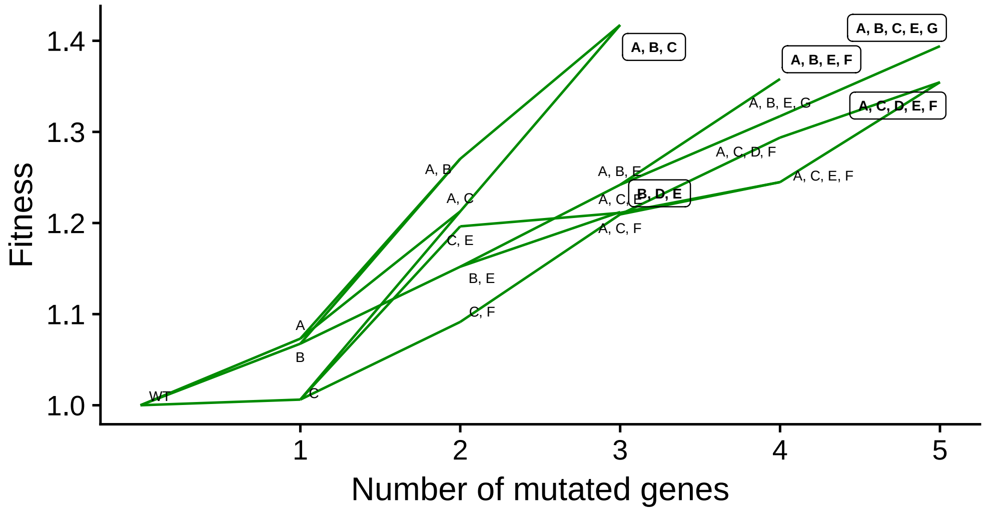


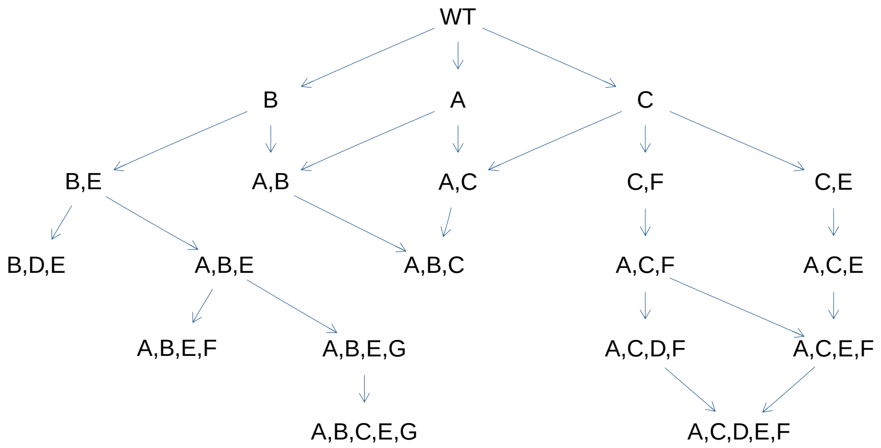

| | |
|---|---|
| *ID* | vweucMX7s10v2ZeDw |
| *acc genotypes* | 57 |
| *peaks no back* | 13 |
| *removed edges* | NA |
| *prop rem edges* | NA |

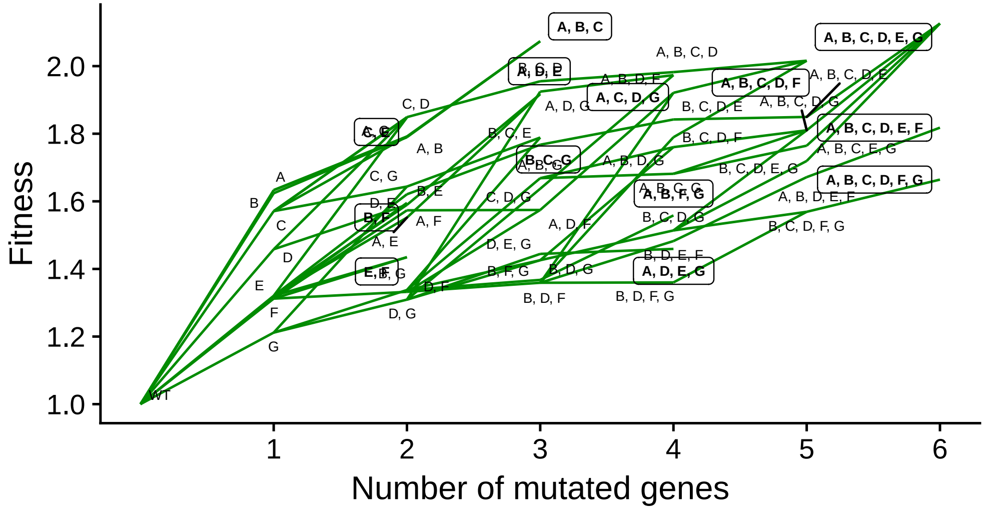


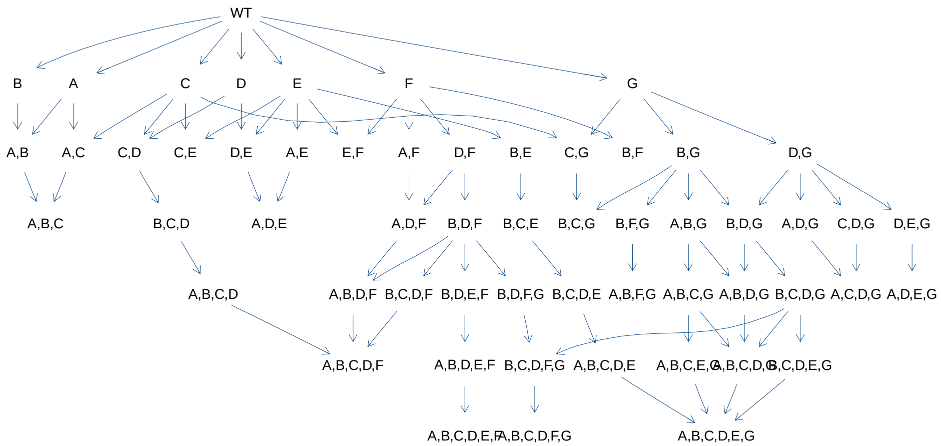

| | |
|---|---|
| *ID* | w3y65IfSHEfh1Qrj |
| *acc genotypes* | 45 |
| *peaks no back* | 15 |
| *removed edges* | NA |
| *prop rem edges* | NA |

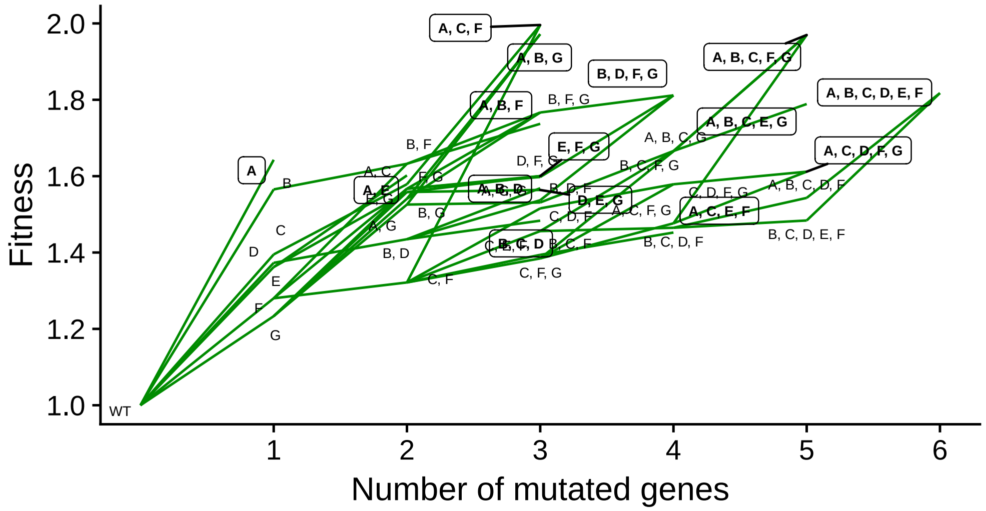


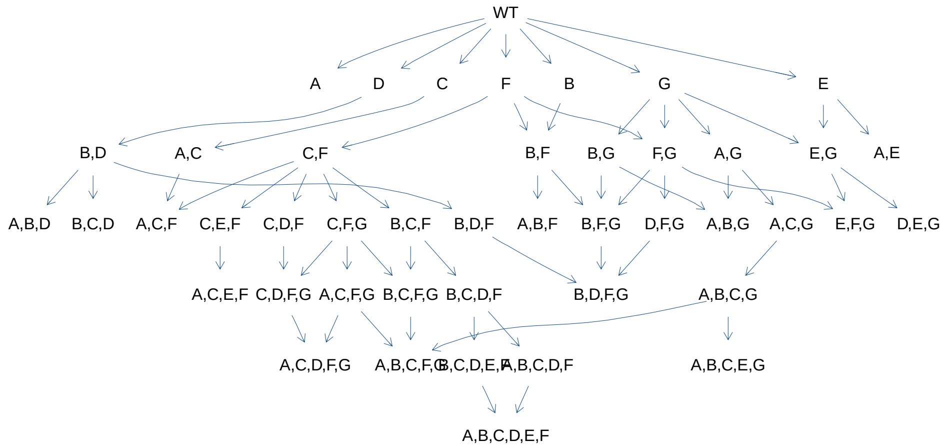

| | |
|---|---|
| *ID* | W7VVpn96d4J5b6gp |
| *acc genotypes* | 43 |
| *peaks no back* | 13 |
| *removed edges* | NA |
| *prop rem edges* | NA |

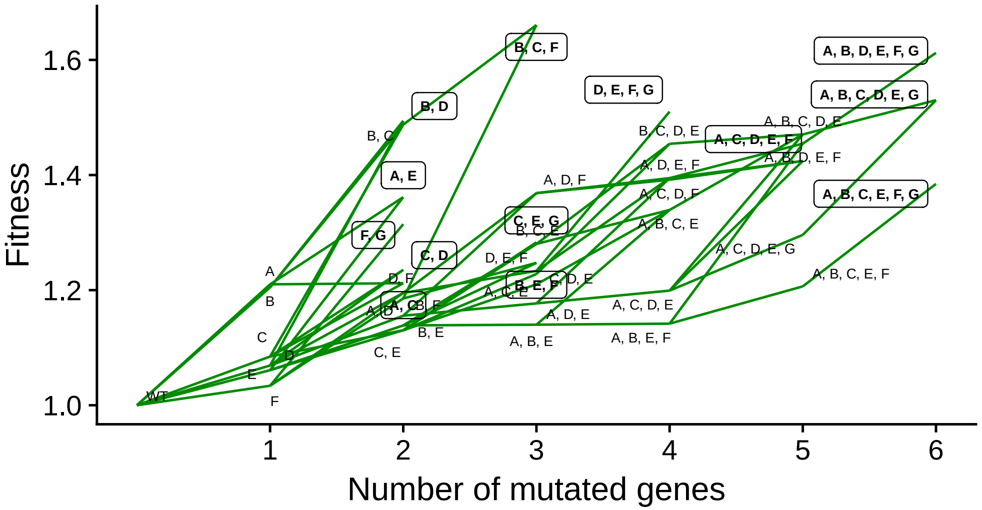


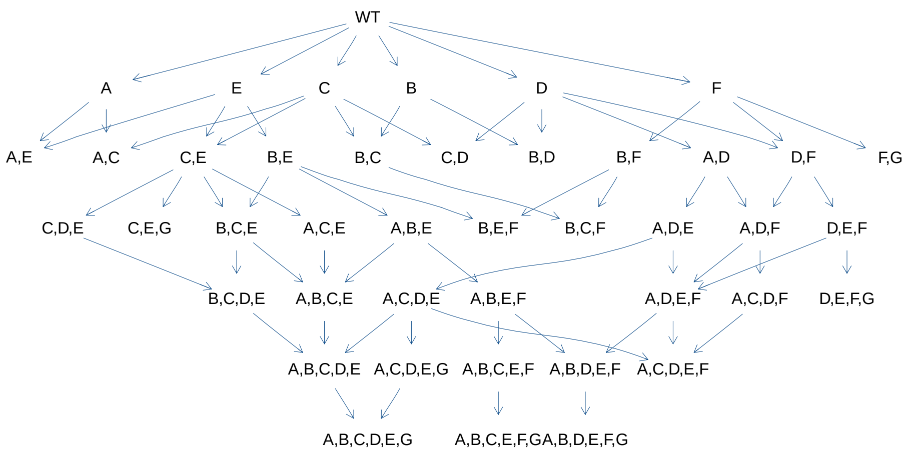

| | |
|---|---|
| *ID* | w8IlHfJBXuIGpKP10 |
| *acc genotypes* | 57 |
| *peaks no back* | 18 |
| *removed edges* | NA |
| *prop rem edges* | NA |

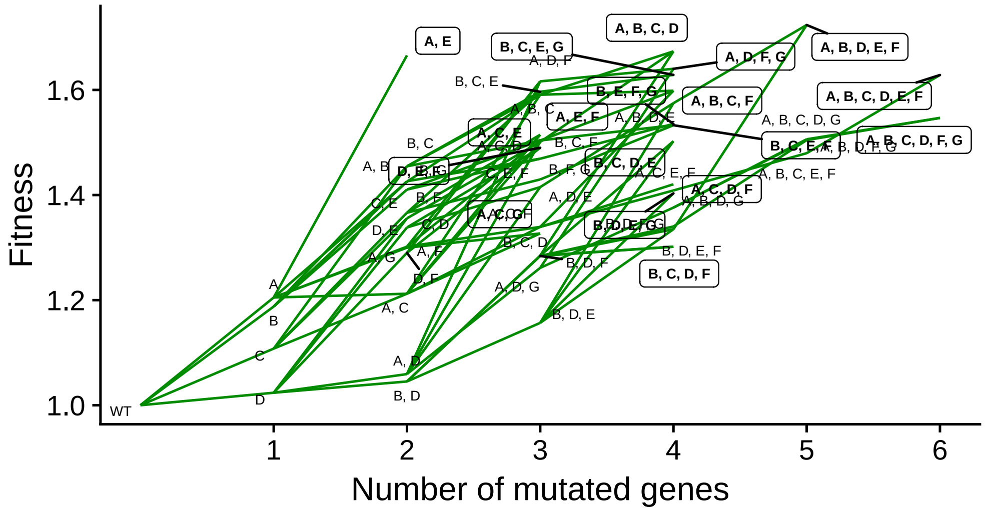


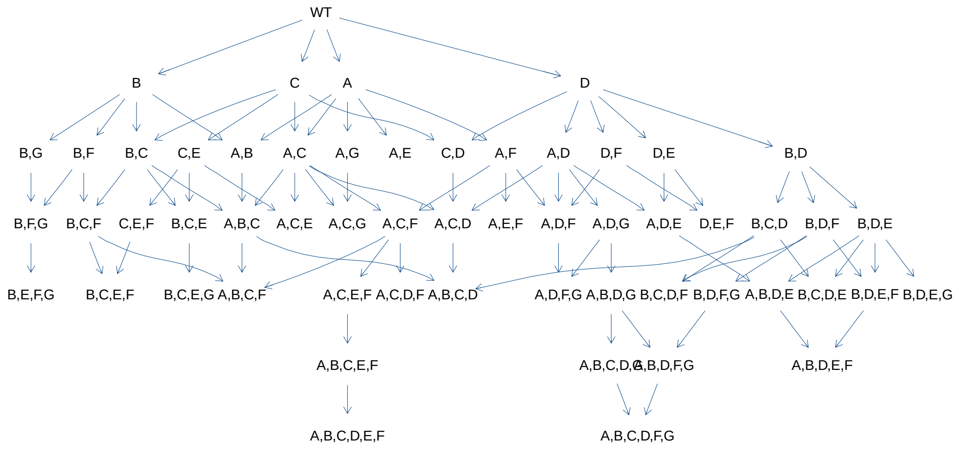

| | |
|---|---|
| *ID* | WTbc1BmerHIsalgq |
| *acc genotypes* | 38 |
| *peaks no back* | 15 |
| *removed edges* | NA |
| *prop rem edges* | NA |

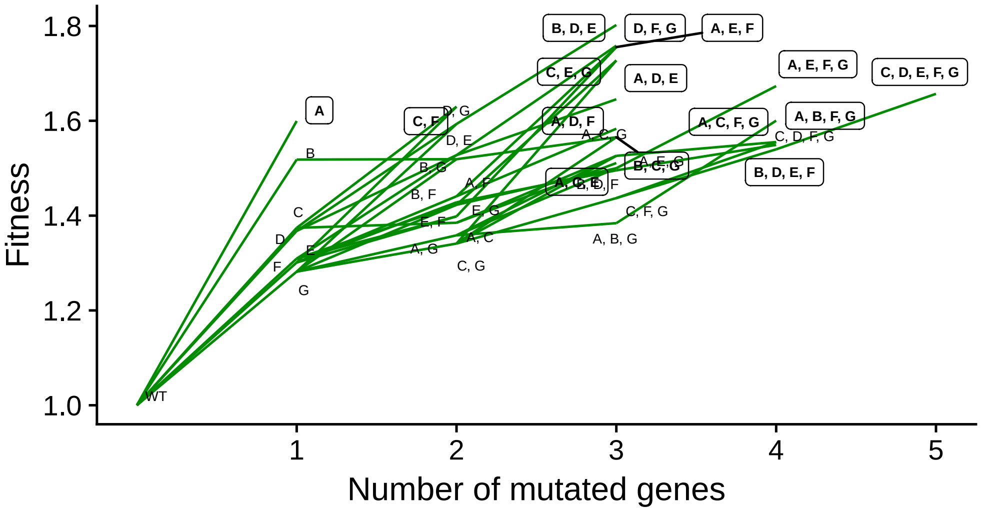


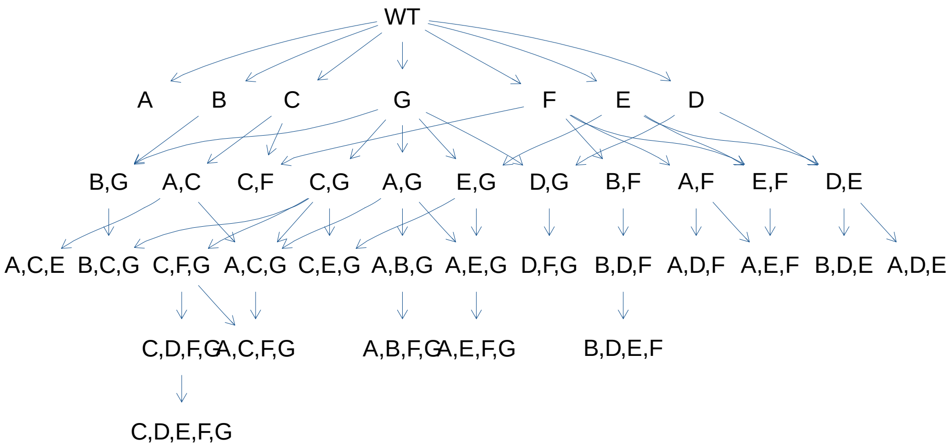

| | |
|---|---|
| *ID* | WuYVFvHnmpxvVWQi |
| *acc genotypes* | 72 |
| *peaks no back* | 20 |
| *removed edges* | NA |
| *prop rem edges* | NA |

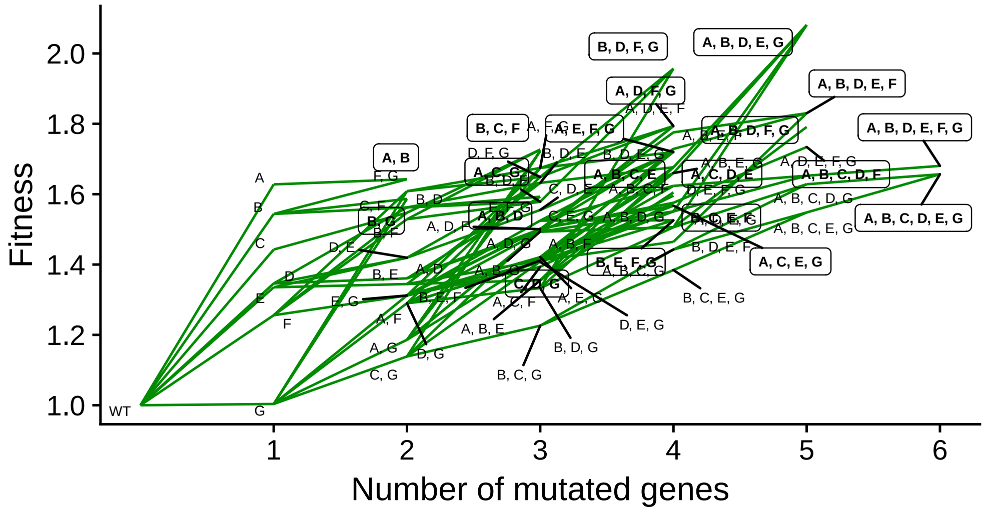


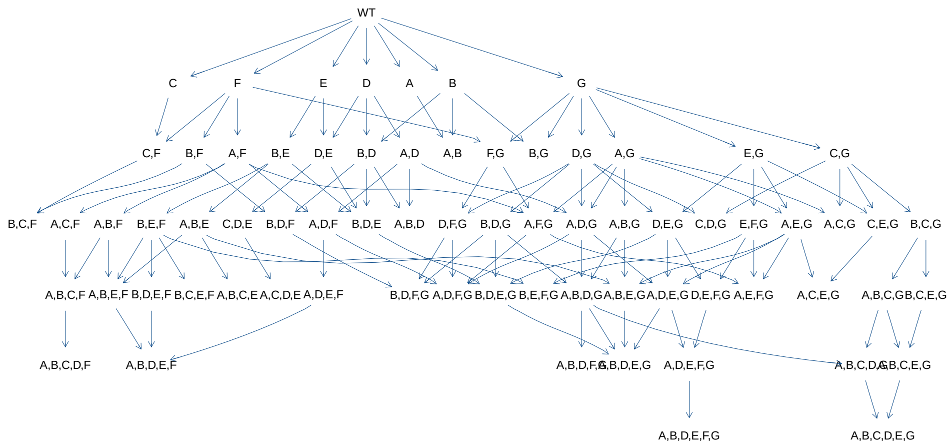

| | |
|---|---|
| *ID* | WXvNSe1lPVS5wf7B |
| *acc genotypes* | 46 |
| *peaks no back* | 14 |
| *removed edges* | NA |
| *prop rem edges* | NA |

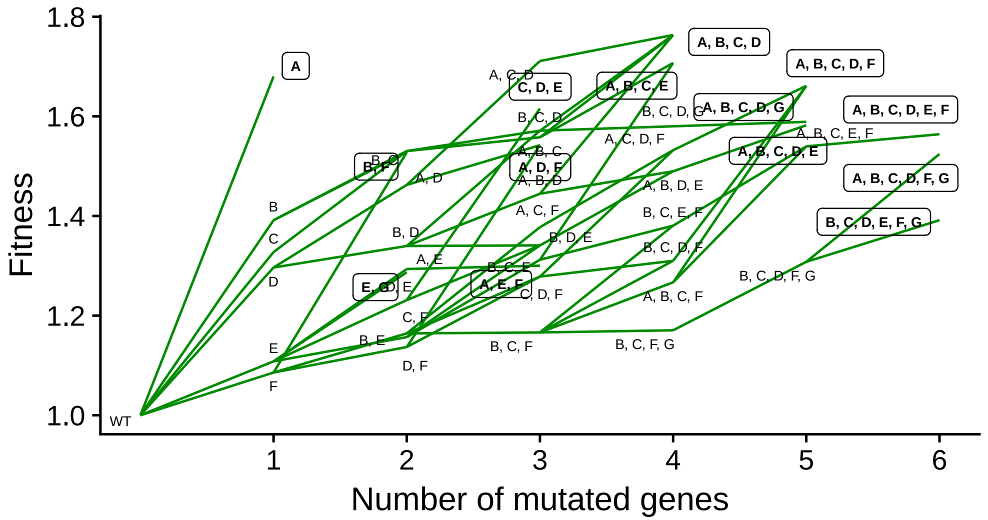


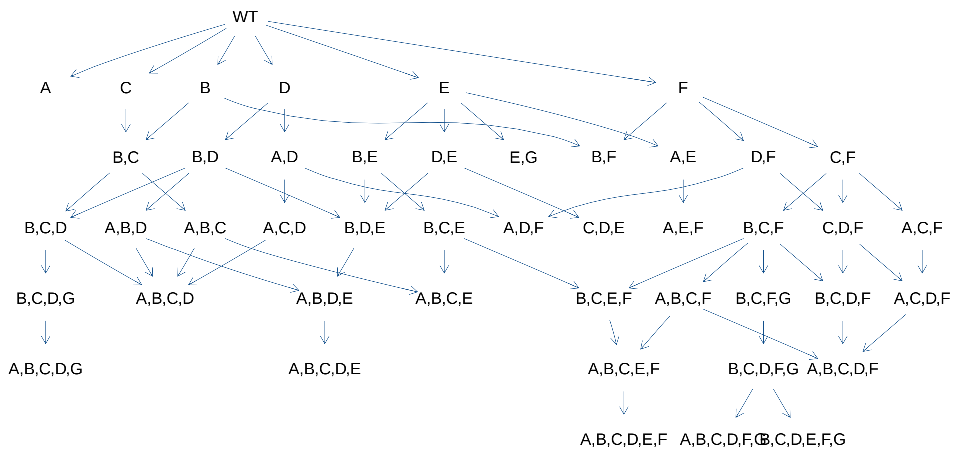

| | |
|---|---|
| *ID* | x4ZFzf8D5q4Z4anR |
| *acc genotypes* | 11 |
| *peaks no back* | 4 |
| *removed edges* | NA |
| *prop rem edges* | NA |

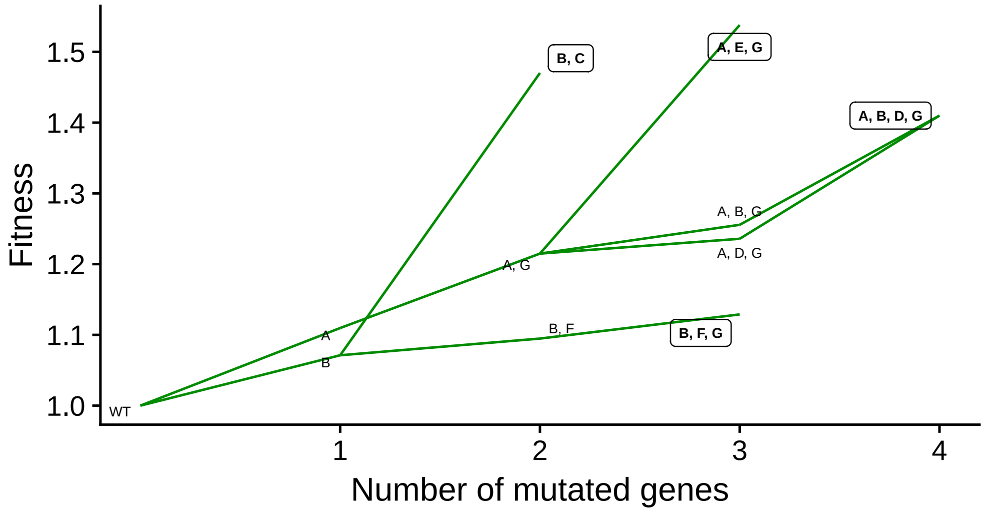


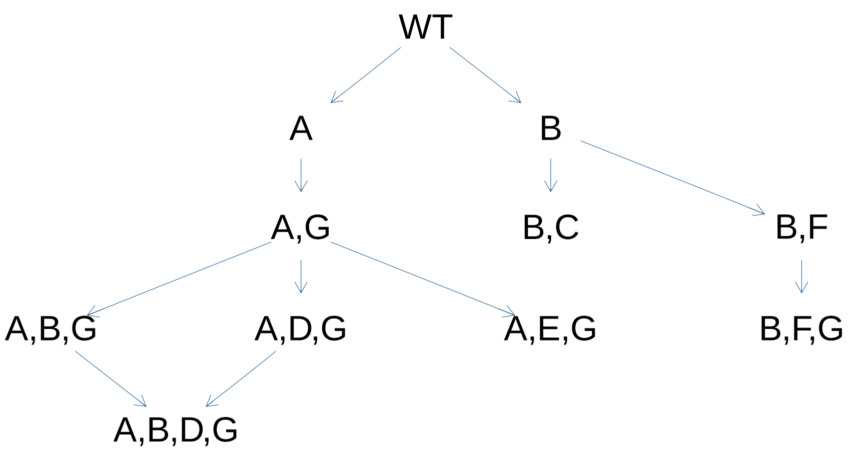

| | |
|---|---|
| *ID* | x53kISylIWMWcjly |
| *acc genotypes* | 22 |
| *peaks no back* | 6 |
| *removed edges* | NA |
| *prop rem edges* | NA |

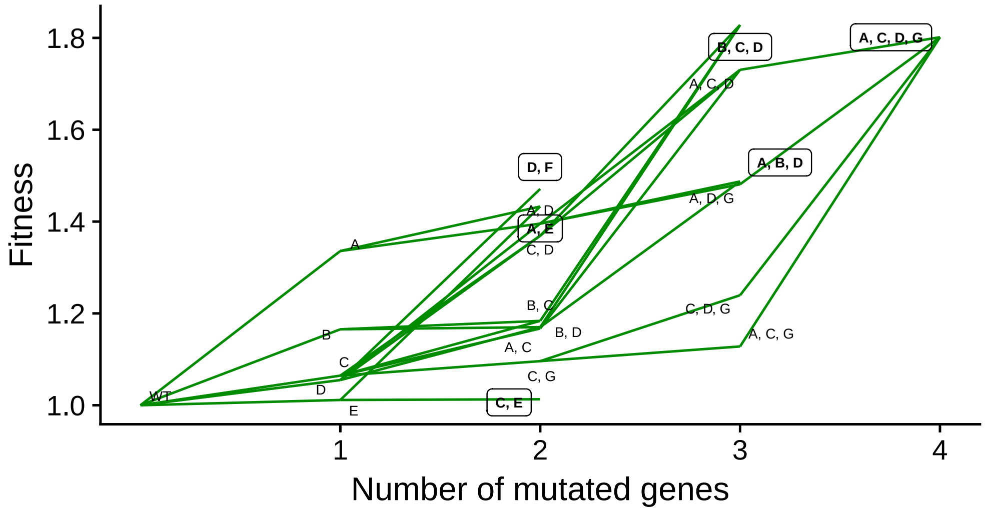


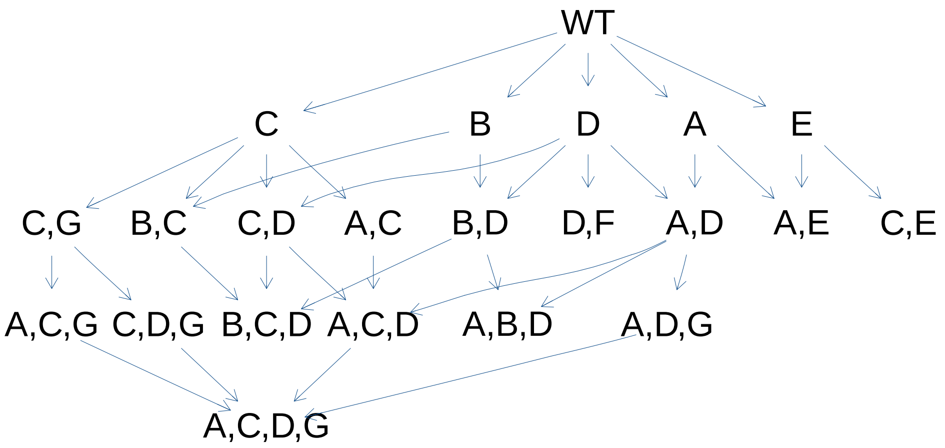

| | |
|---|---|
| *ID* | x63vO10FZvDdTuarD |
| *acc genotypes* | 20 |
| *peaks no back* | 9 |
| *removed edges* | NA |
| *prop rem edges* | NA |

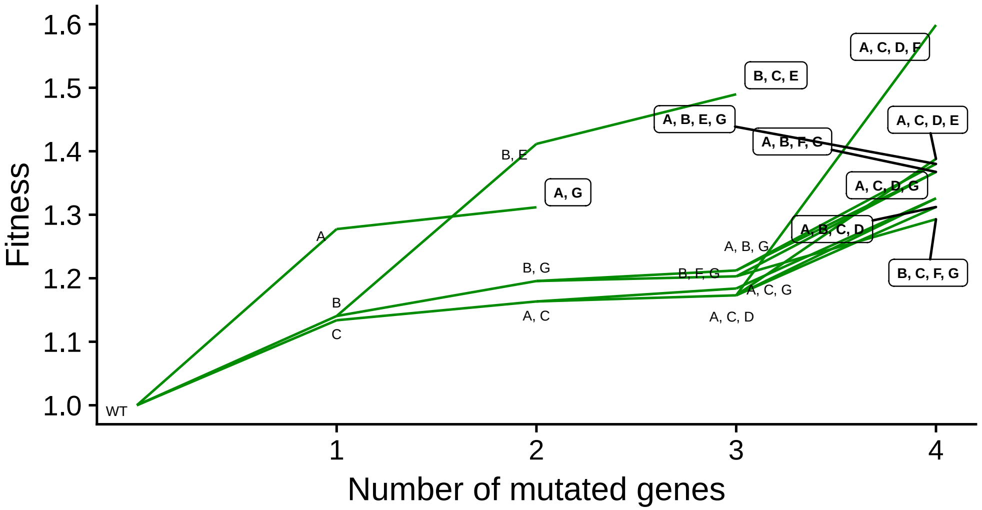


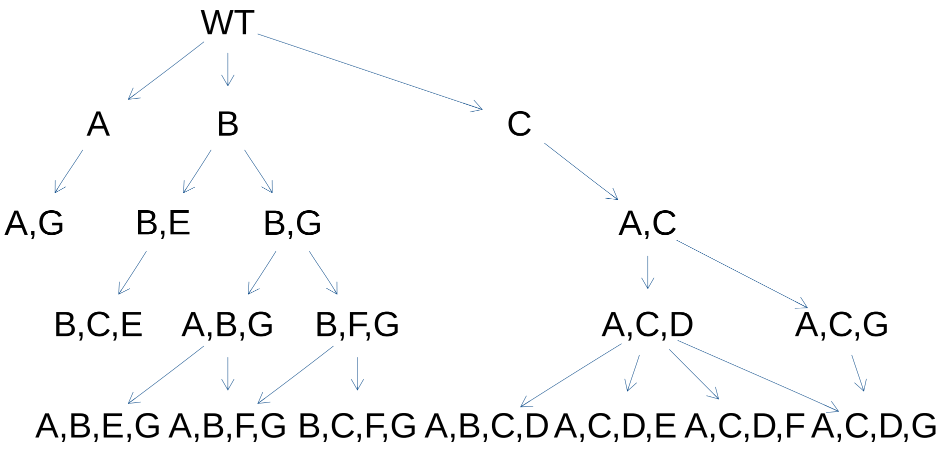

| | |
|---|---|
| *ID* | XhJcAoB3WdJKttP7 |
| *acc genotypes* | 55 |
| *peaks no back* | 21 |
| *removed edges* | NA |
| *prop rem edges* | NA |

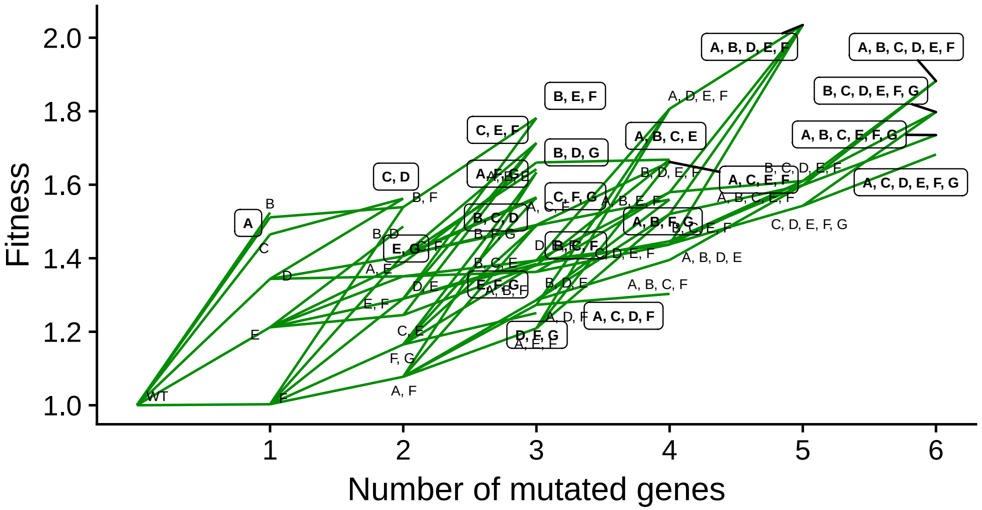


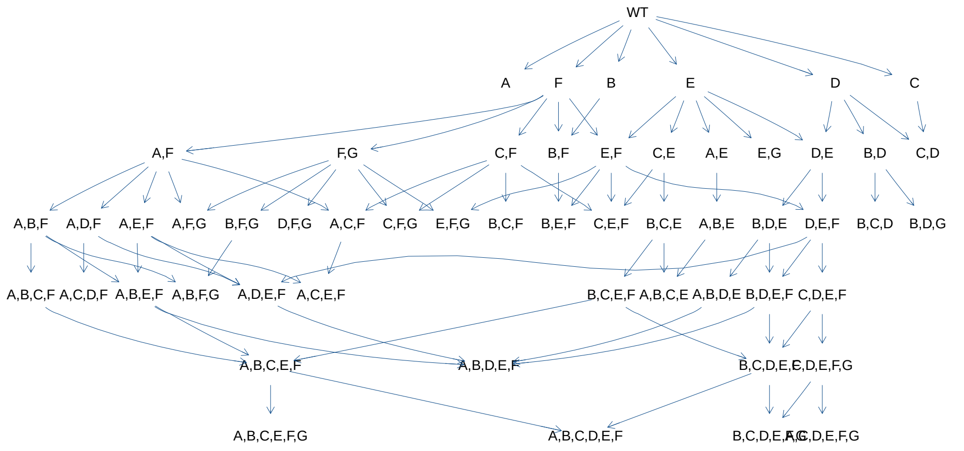

| | |
|---|---|
| *ID* | xp6hoPCA8glPKMzG |
| *acc genotypes* | 62 |
| *peaks no back* | 16 |
| *removed edges* | NA |
| *prop rem edges* | NA |

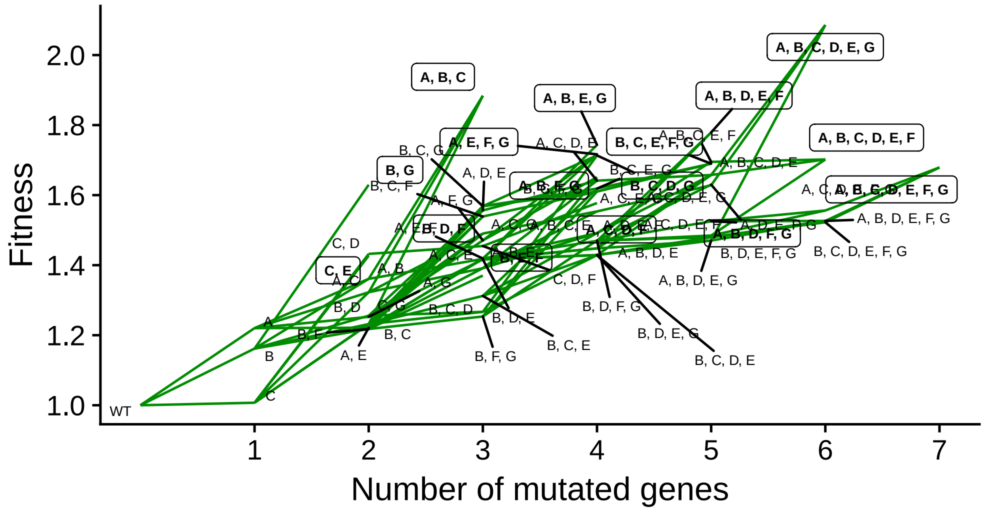


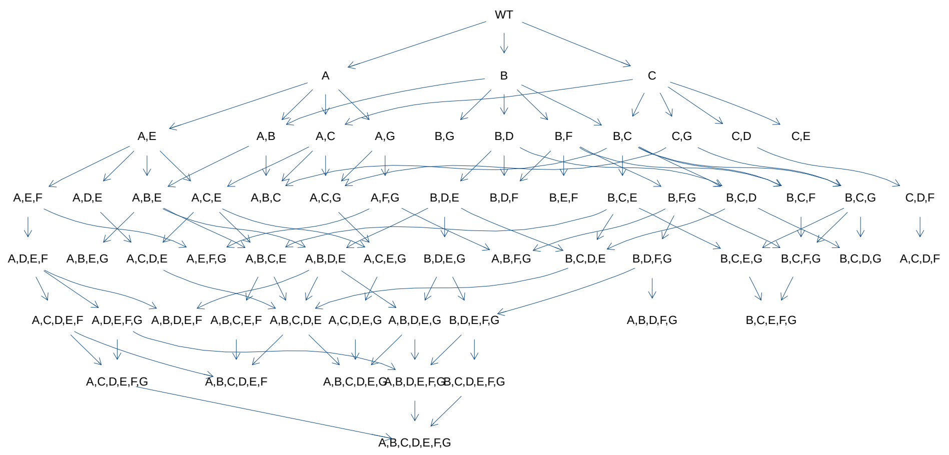

| | |
|---|---|
| *ID* | xU2TICTIsth9hYb3 |
| *acc genotypes* | 23 |
| *peaks no back* | 7 |
| *removed edges* | NA |
| *prop rem edges* | NA |

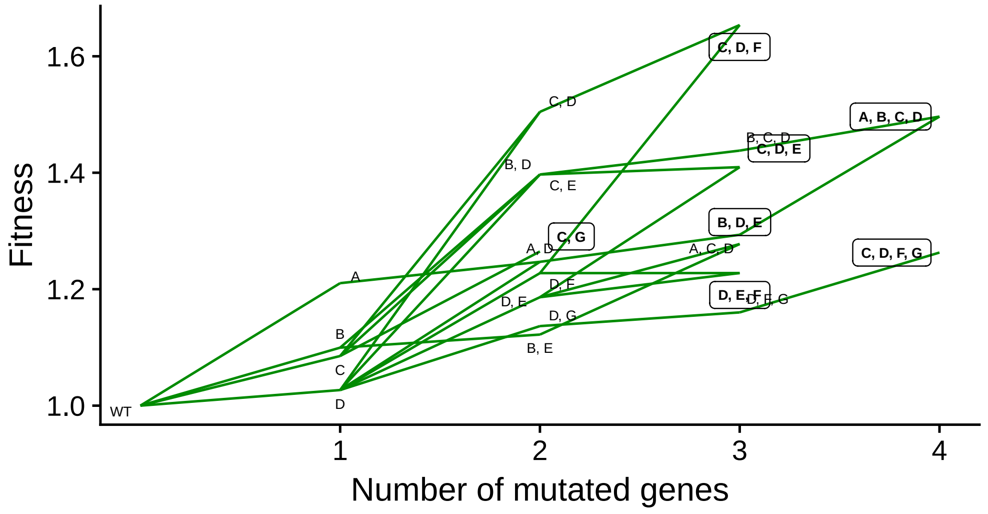


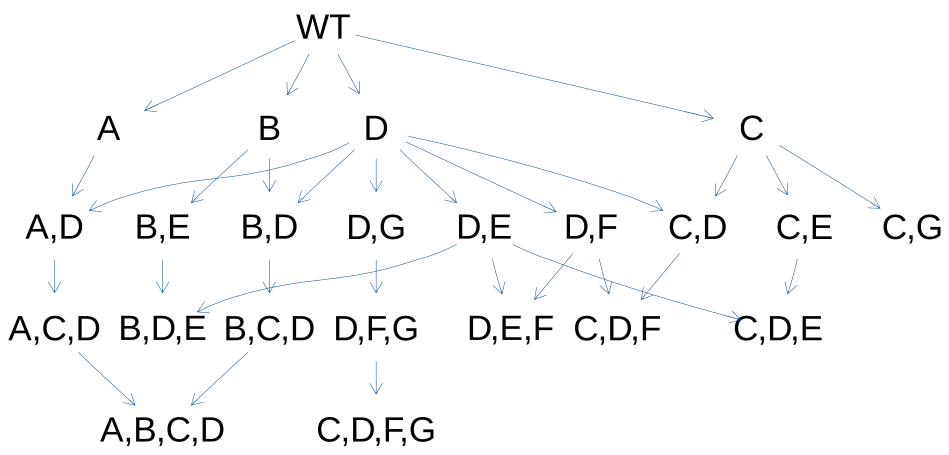

| | |
|---|---|
| *ID* | xZGO107xRn107k9sKc |
| *acc genotypes* | 37 |
| *peaks no back* | 12 |
| *removed edges* | NA |
| *prop rem edges* | NA |

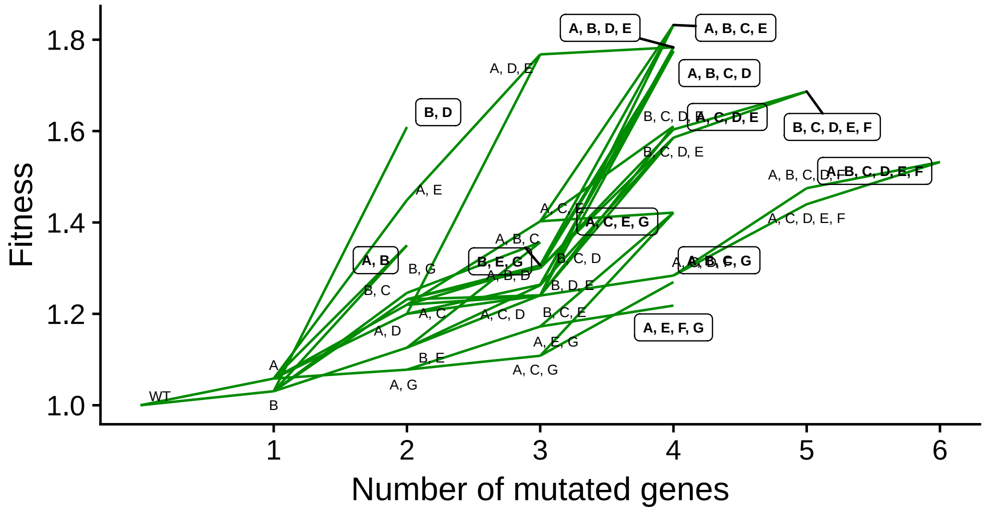


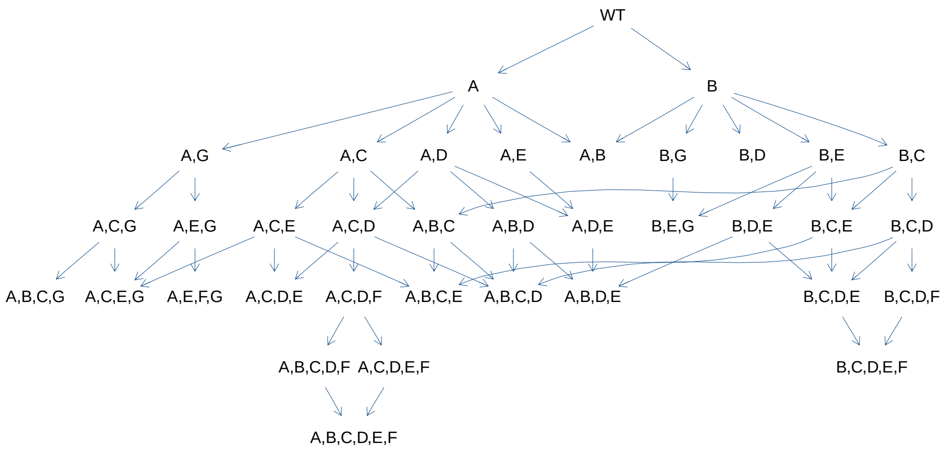

| | |
|---|---|
| *ID* | Y8pEslk1p3jDN3PC |
| *acc genotypes* | 16 |
| *peaks no back* | 7 |
| *removed edges* | NA |
| *prop rem edges* | NA |

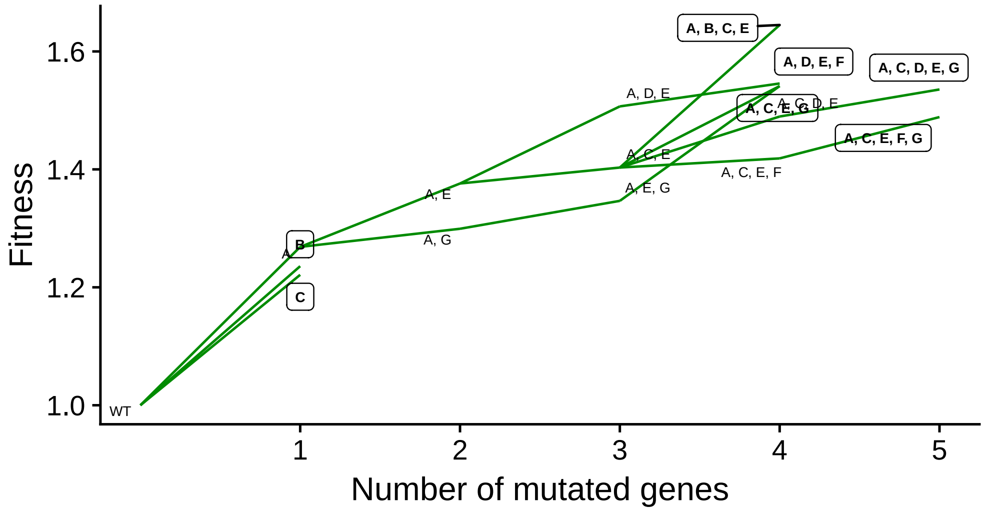


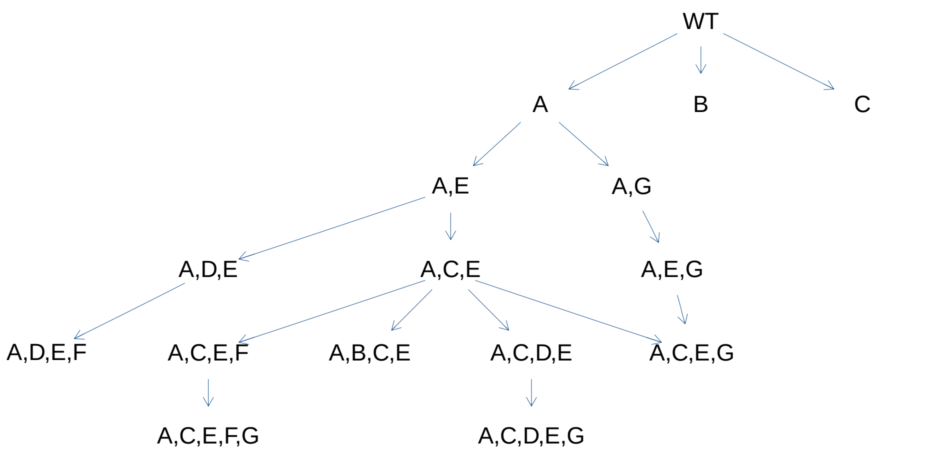

| | |
|---|---|
| *ID* | yc851VadCH5z6cH10 |
| *acc genotypes* | 30 |
| *peaks no back* | 14 |
| *removed edges* | NA |
| *prop rem edges* | NA |

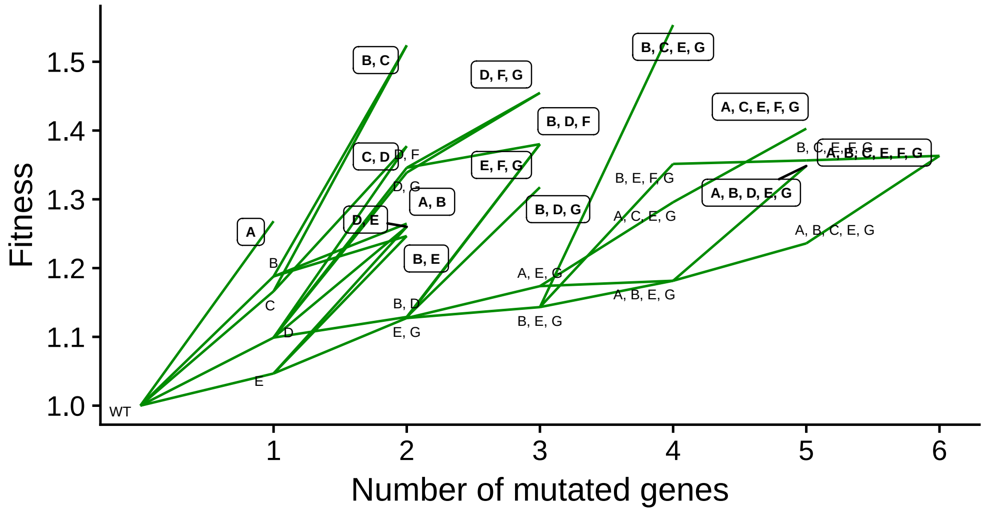


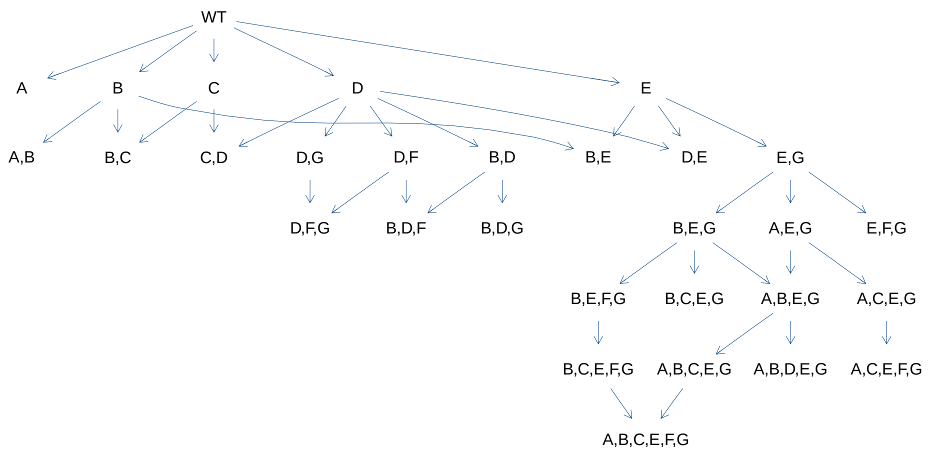

| | |
|---|---|
| *ID* | yeCk19IGNHzq9Zia |
| *acc genotypes* | 44 |
| *peaks no back* | 18 |
| *removed edges* | NA |
| *prop rem edges* | NA |

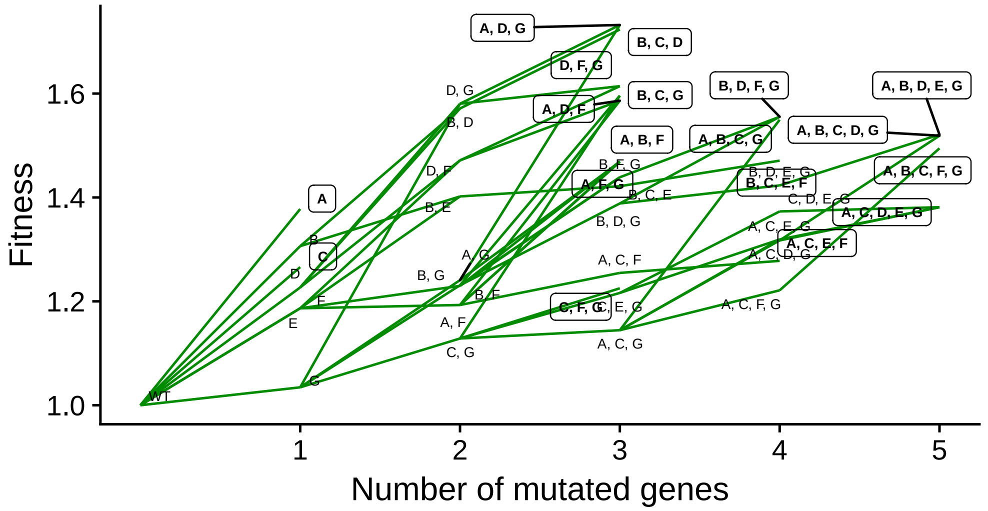


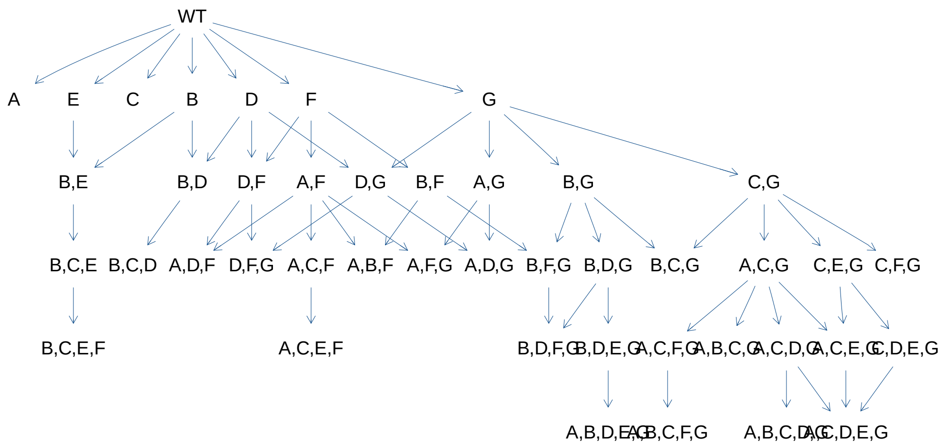

| | |
|---|---|
| *ID* | YpVNspihDcE5kIsm |
| *acc genotypes* | 48 |
| *peaks no back* | 12 |
| *removed edges* | NA |
| *prop rem edges* | NA |

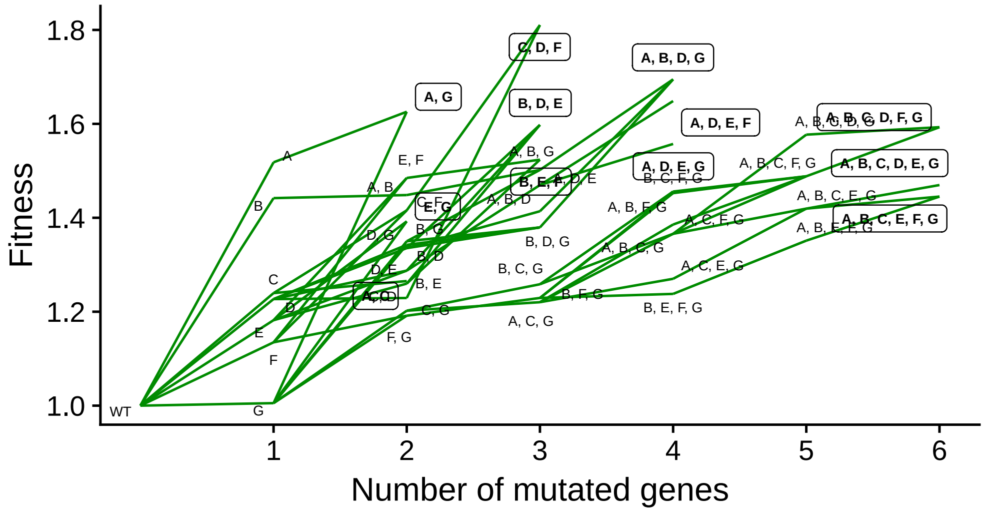


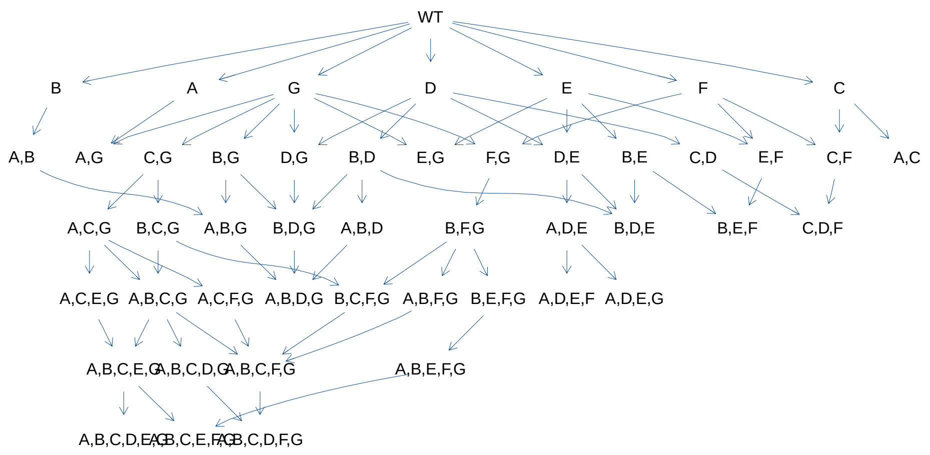

| | |
|---|---|
| *ID* | YuMhOqw10Xkx33JOs |
| *acc genotypes* | 61 |
| *peaks no back* | 17 |
| *removed edges* | NA |
| *prop rem edges* | NA |

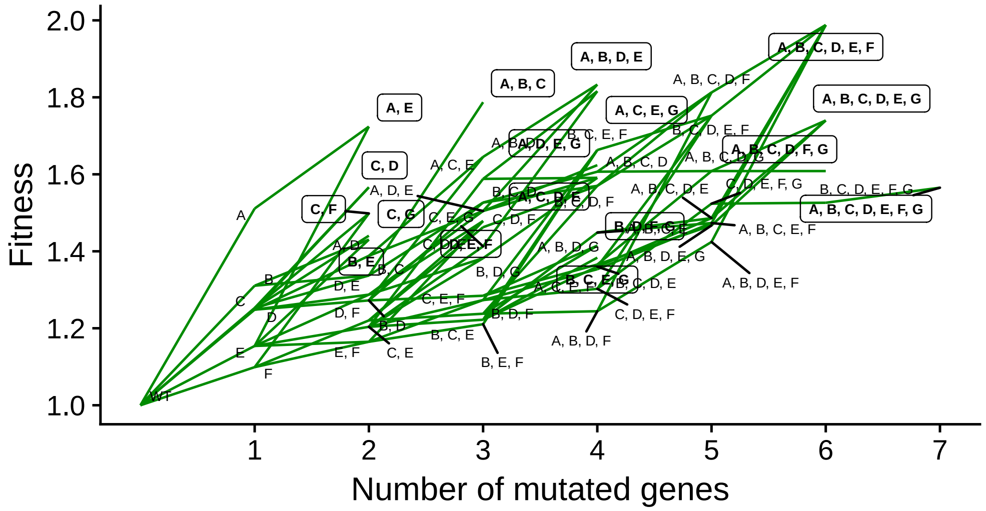


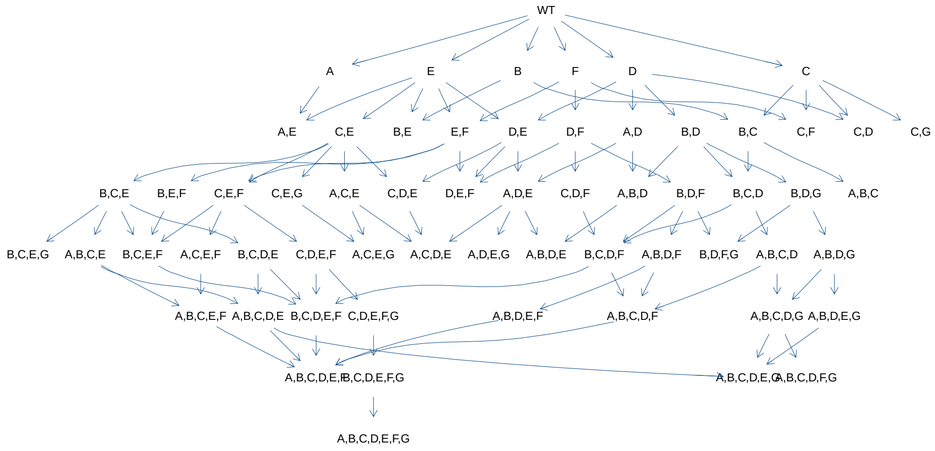

| | |
|---|---|
| *ID* | yYOyPKhl3NOTcbHe |
| *acc genotypes* | 50 |
| *peaks no back* | 18 |
| *removed edges* | NA |
| *prop rem edges* | NA |

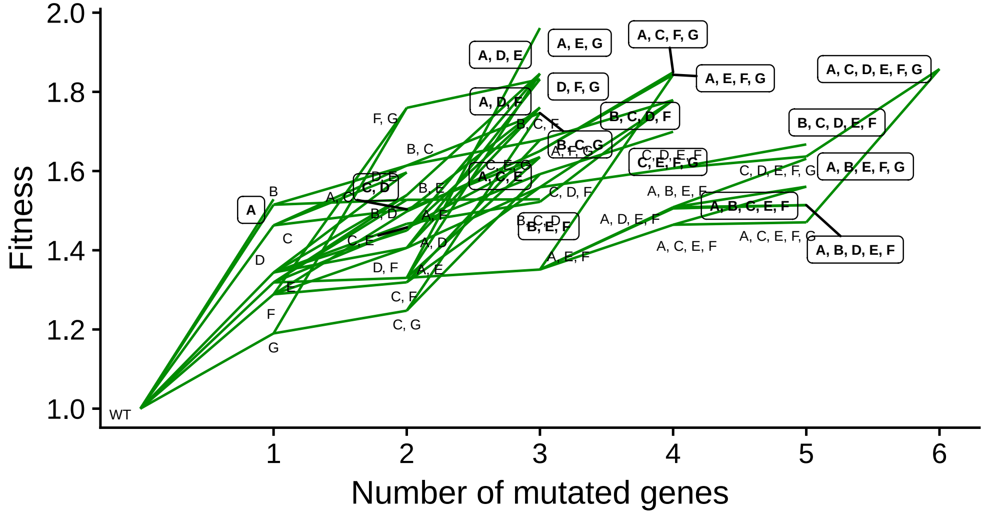


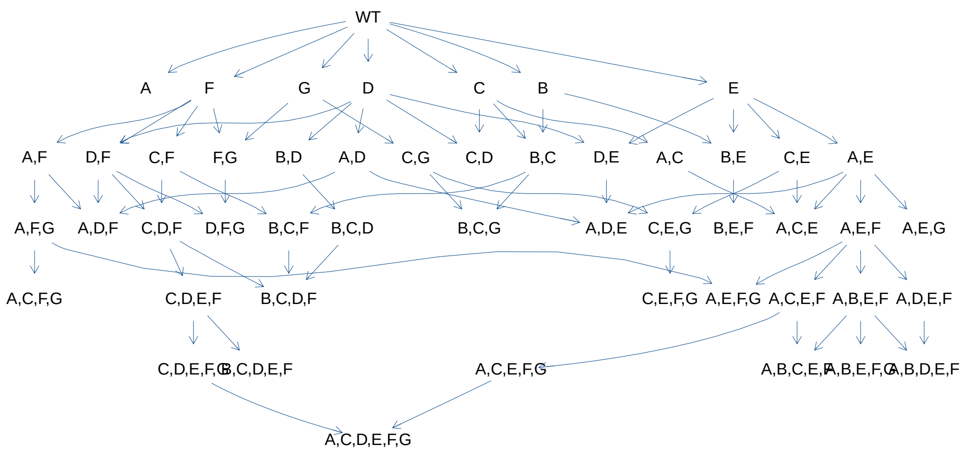

| | |
|---|---|
| *ID* | Z10gb2ZT22WwOQVNk |
| *acc genotypes* | 30 |
| *peaks no back* | 12 |
| *removed edges* | NA |
| *prop rem edges* | NA |

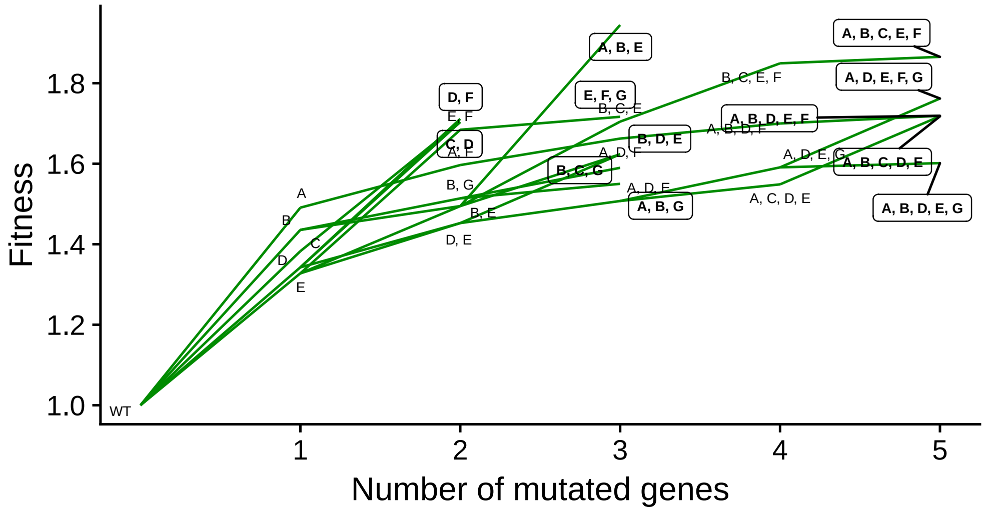


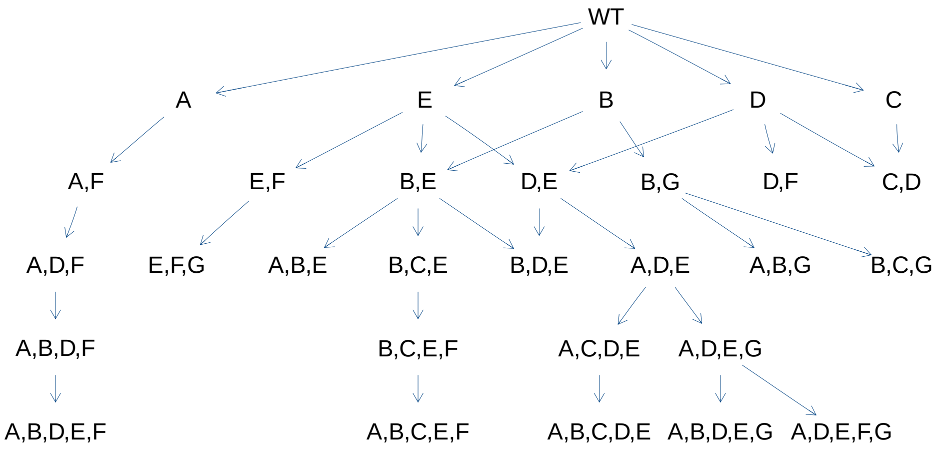

| | |
|---|---|
| *ID* | Z7IpusjT8G9uuSKU |
| *acc genotypes* | 59 |
| *peaks no back* | 18 |
| *removed edges* | NA |
| *prop rem edges* | NA |

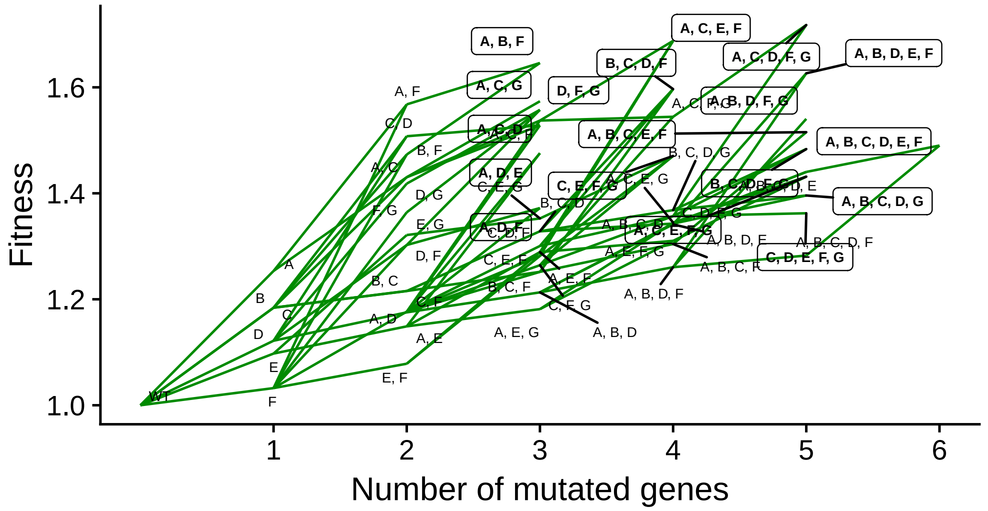


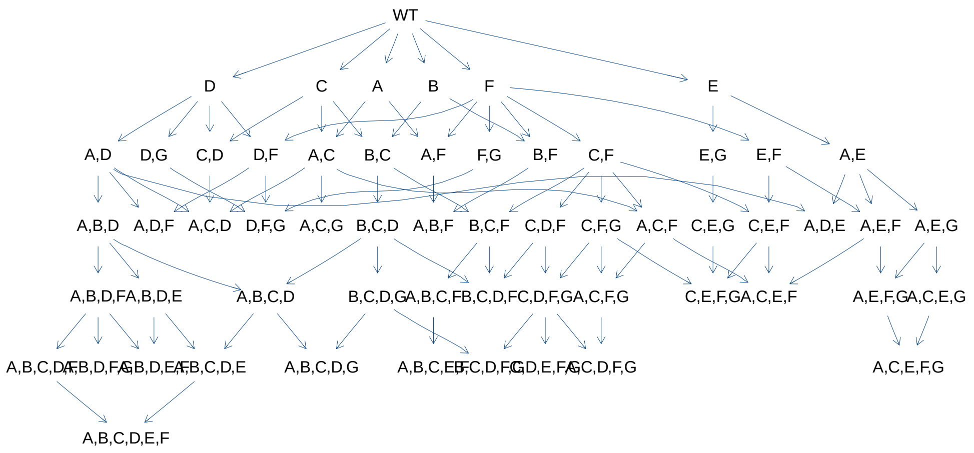

| | |
|---|---|
| *ID* | ZGkYkR2uDqq44mge |
| *acc genotypes* | 34 |
| *peaks no back* | 14 |
| *removed edges* | NA |
| *prop rem edges* | NA |

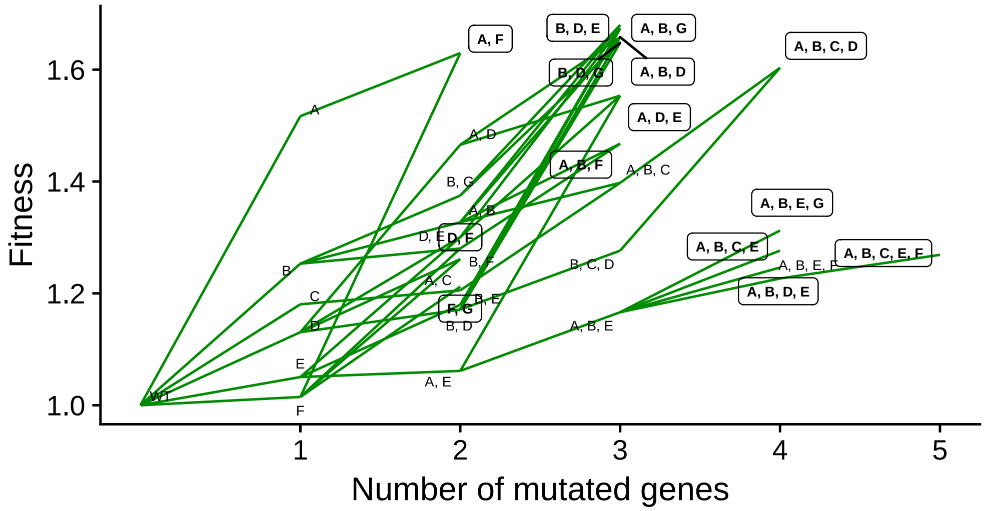


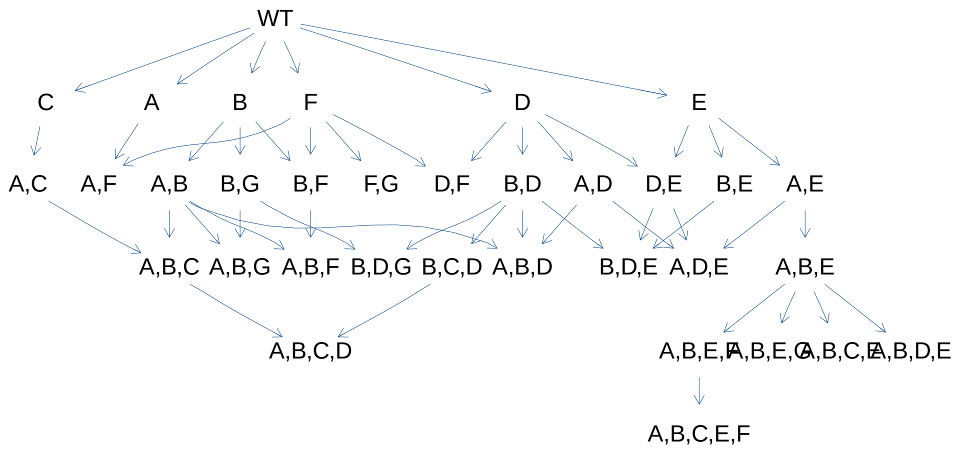

| | |
|---|---|
| *ID* | ZjLMUPFuopk2WZUI |
| *acc genotypes* | 22 |
| *peaks no back* | 7 |
| *removed edges* | NA |
| *prop rem edges* | NA |

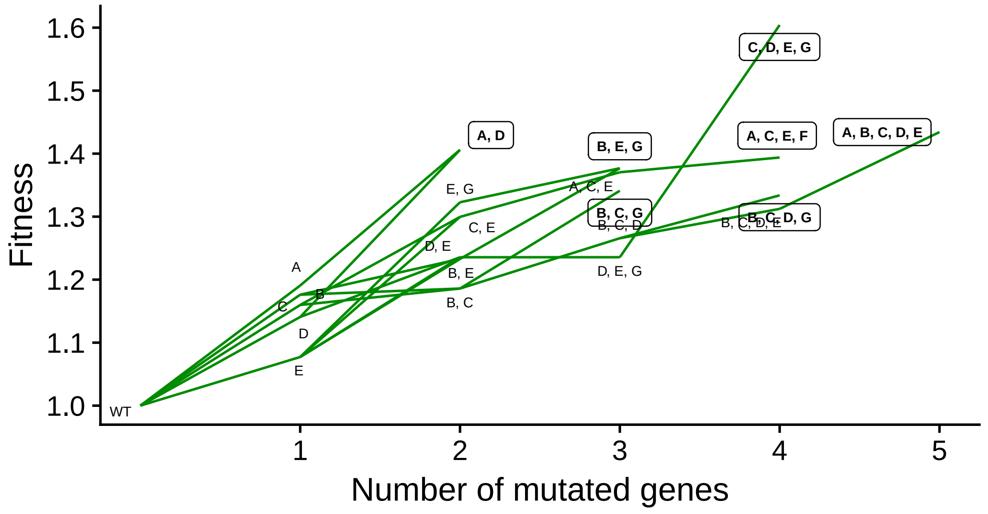


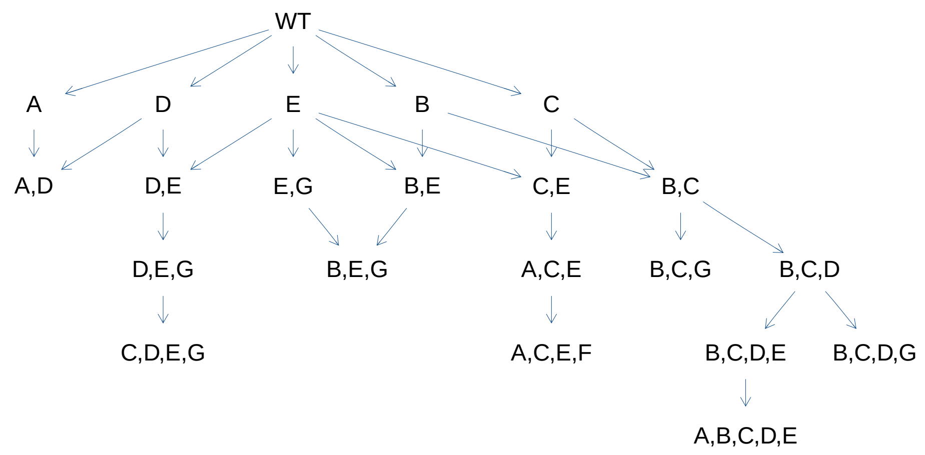

| | |
|---|---|
| *ID* | ZVWECQo9V7rYLRNy |
| *acc genotypes* | 72 |
| *peaks no back* | 10 |
| *removed edges* | NA |
| *prop rem edges* | NA |

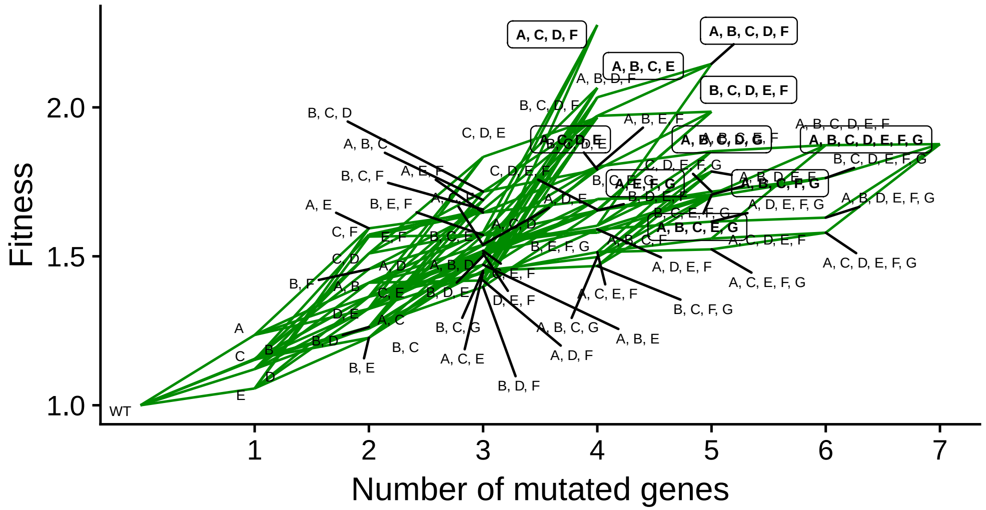


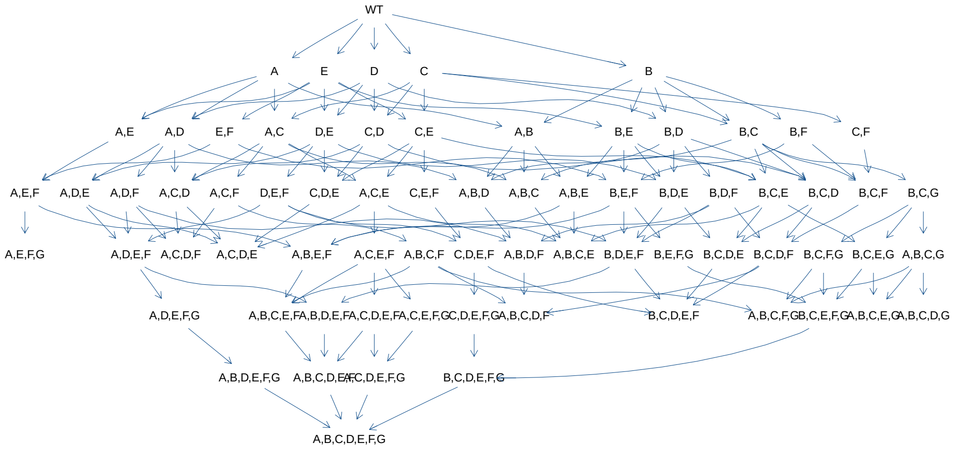

| | |
|---|---|
| *ID* | ZWDP3y7TKpRZUzKJ |
| *acc genotypes* | 46 |
| *peaks no back* | 15 |
| *removed edges* | NA |
| *prop rem edges* | NA |

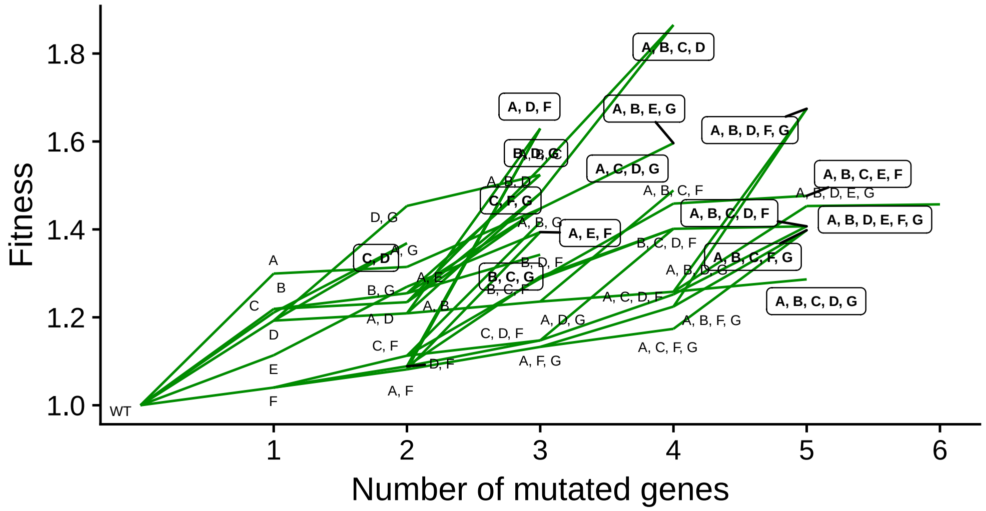


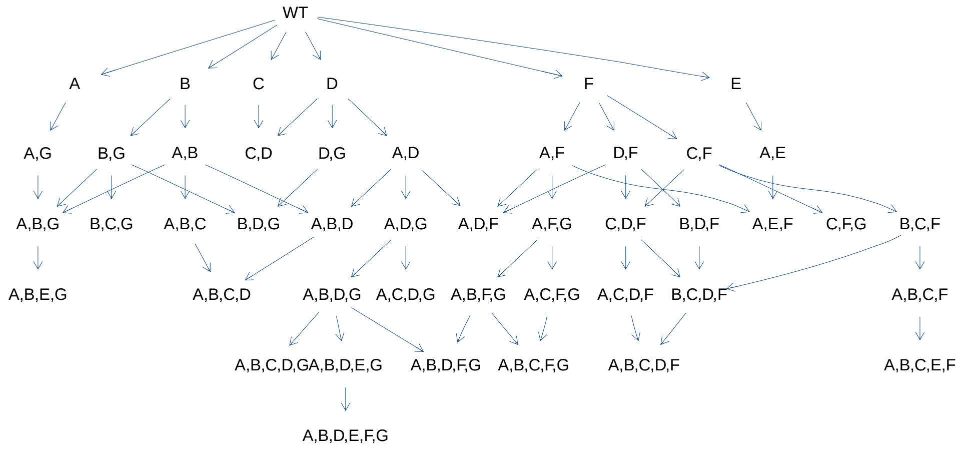

| | |
|---|---|
| *ID* | ZXShHtwLyLLxewaU |
| *acc genotypes* | 51 |
| *peaks no back* | 22 |
| *removed edges* | NA |
| *prop rem edges* | NA |

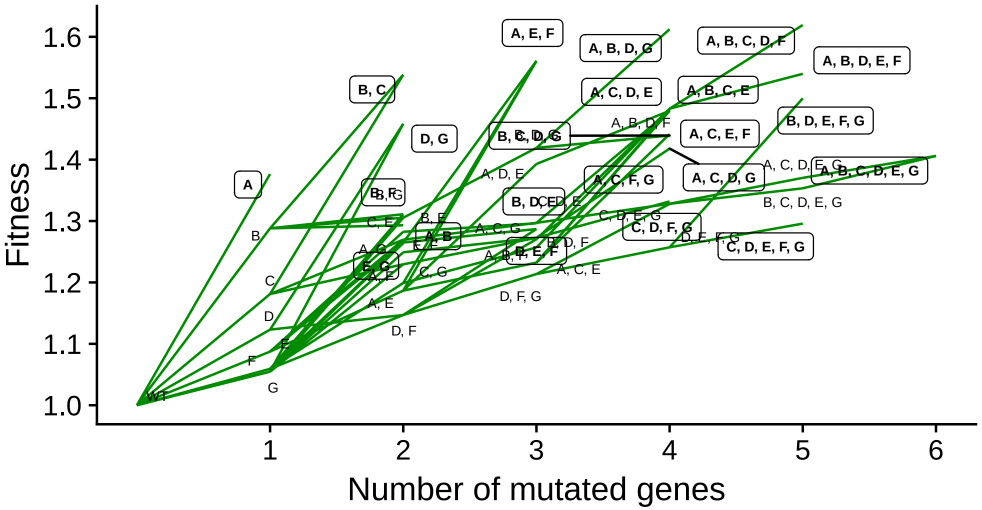


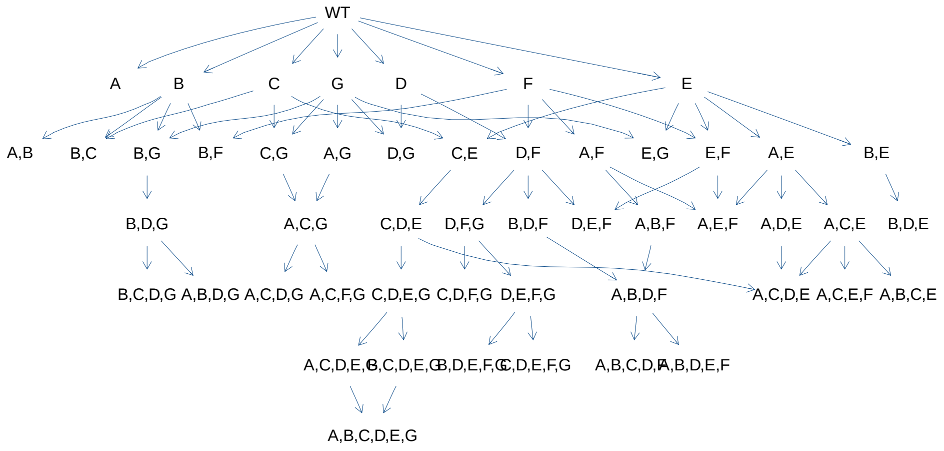

#### 3. Ten genes

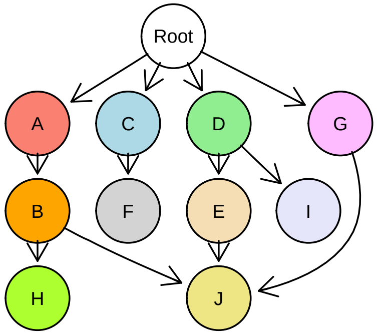


| | |
|---|---|
| *ID* | 10Ic5x6106RnZOmZ6Z |
| *acc genotypes* | 132 |
| *peaks no back* | 1 |
| *removed edges* | 0 |
| *prop rem edges* | 0 |

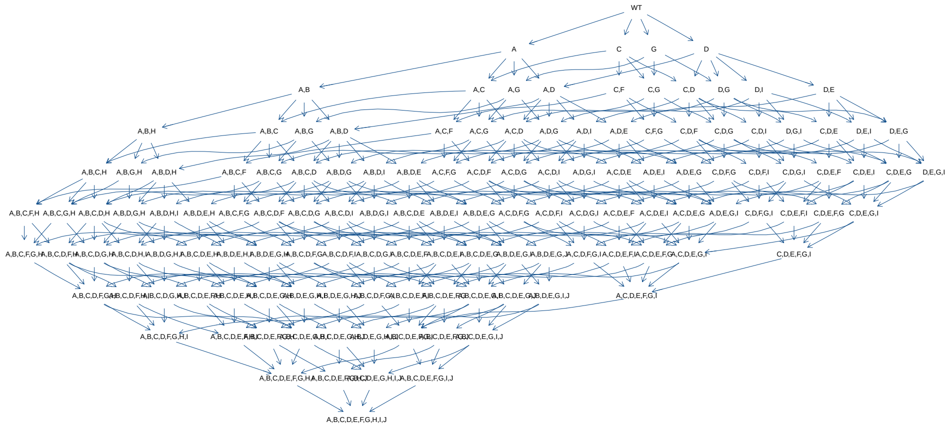

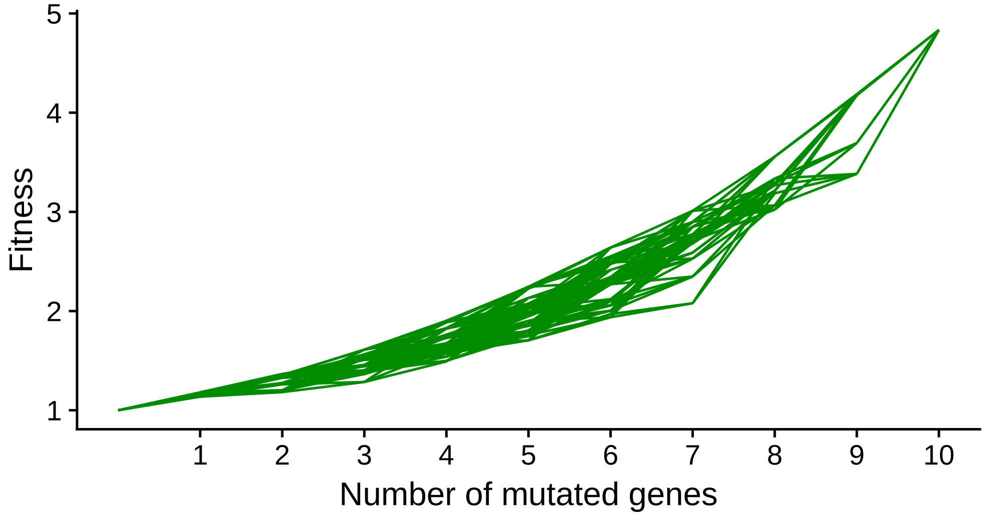

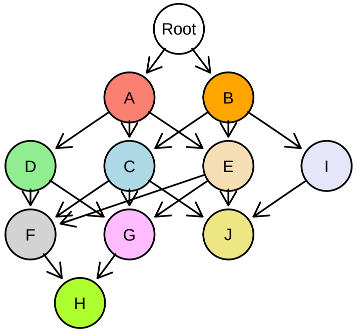


| | |
|---|---|
| *ID* | 10sA3fCTx7METho38 |
| *acc genotypes* | 35 |
| *peaks no back* | 1 |
| *removed edges* | 0 |
| *prop rem edges* | 0 |

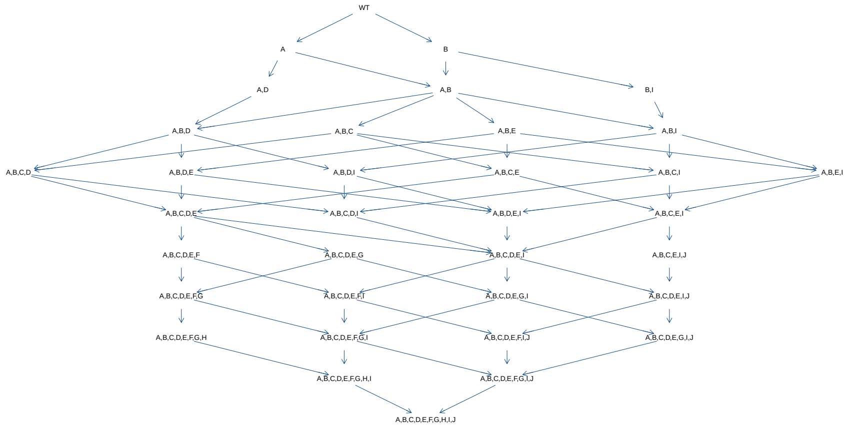

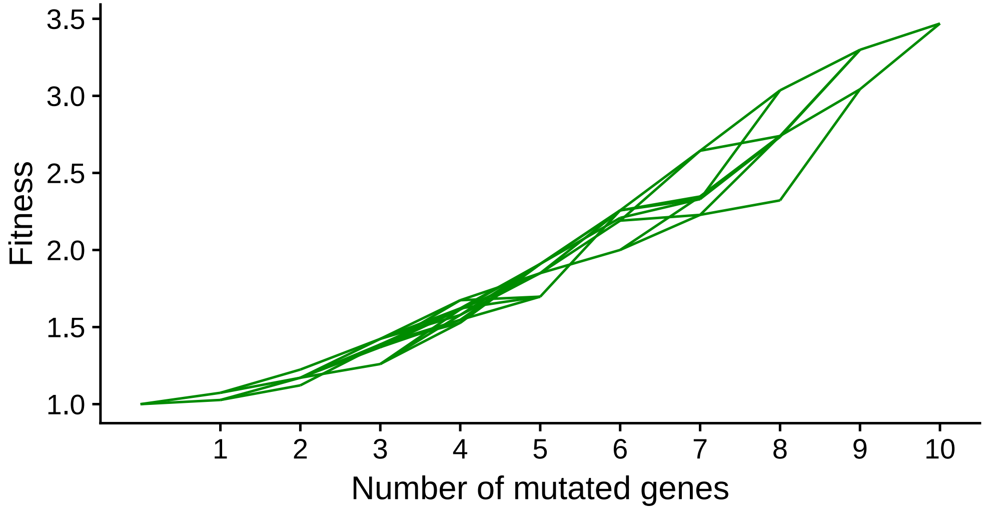

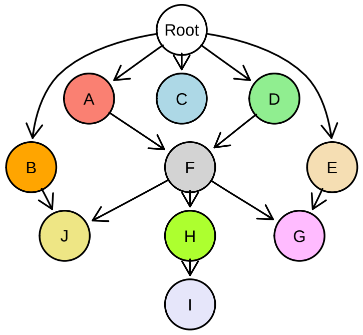


*ID* 1MaScdYnWRHseOZ7

*acc genotypes* 86

*peaks no back* 1

*removed edges* 0

*prop rem edges* 0

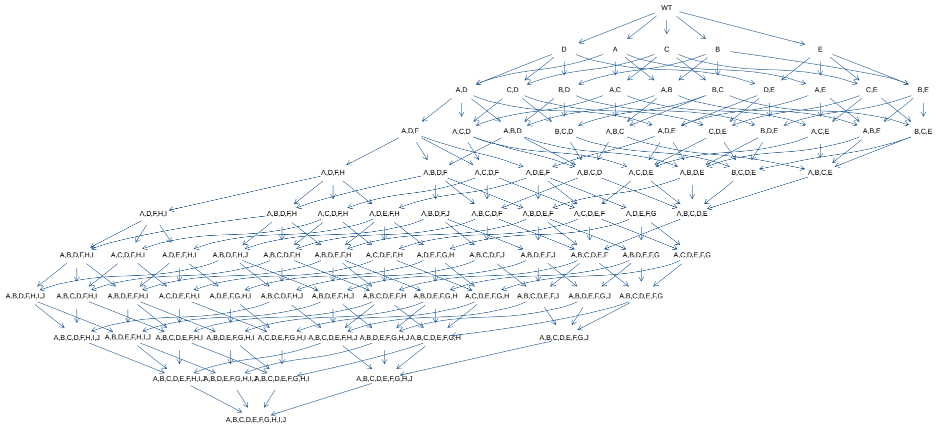

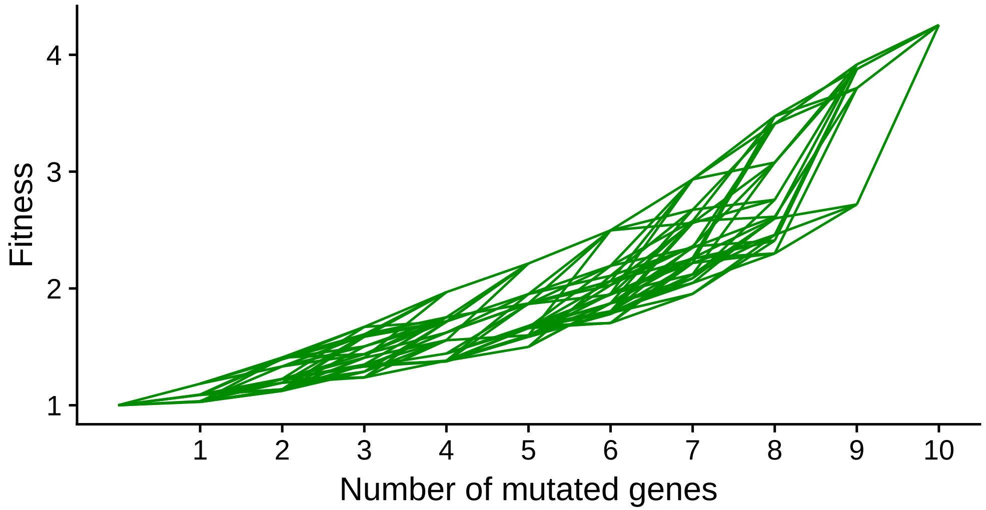

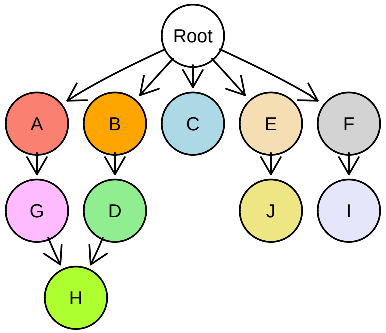


| | |
|---|---|
| *ID* | 1Nl1Lblu3EuEdbw8 |
| *acc genotypes* | 180 |
| *peaks no back* | 1 |
| *removed edges* | 0 |
| *prop rem edges* | 0 |

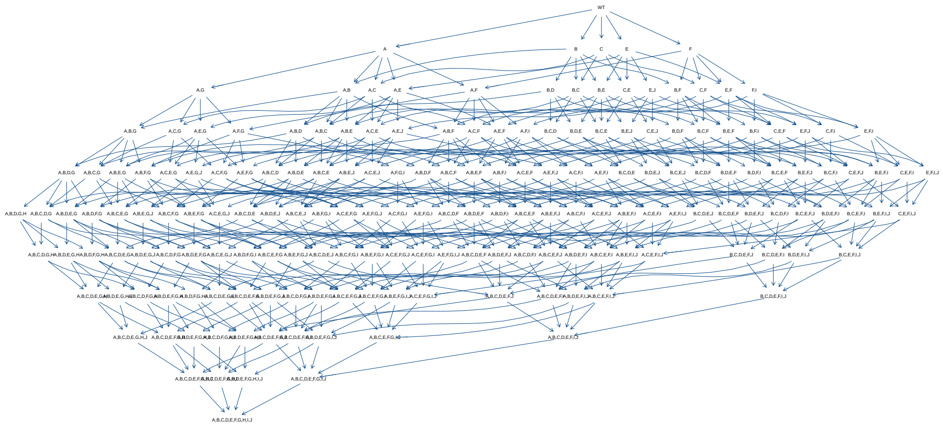

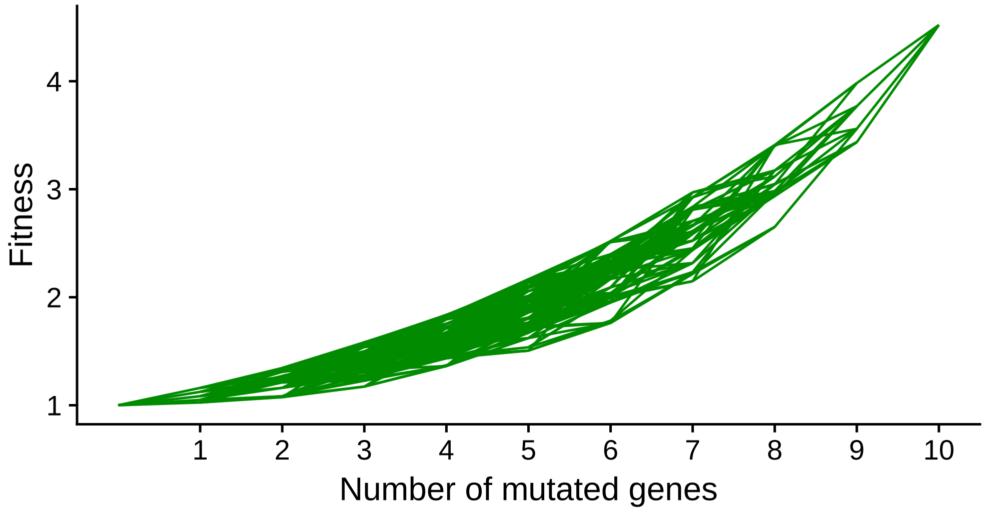

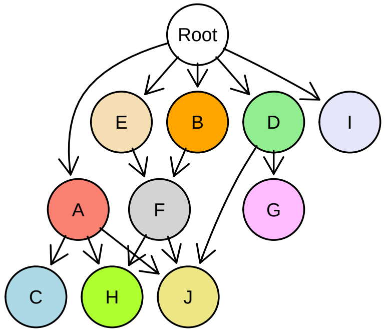


| | |
|---|---|
| *ID* | 2PhVbwXtsibgHoCR |
| *acc genotypes* | 118 |
| *peaks no back* | 1 |
| *removed edges* | 0 |
| *prop rem edges* | 0 |

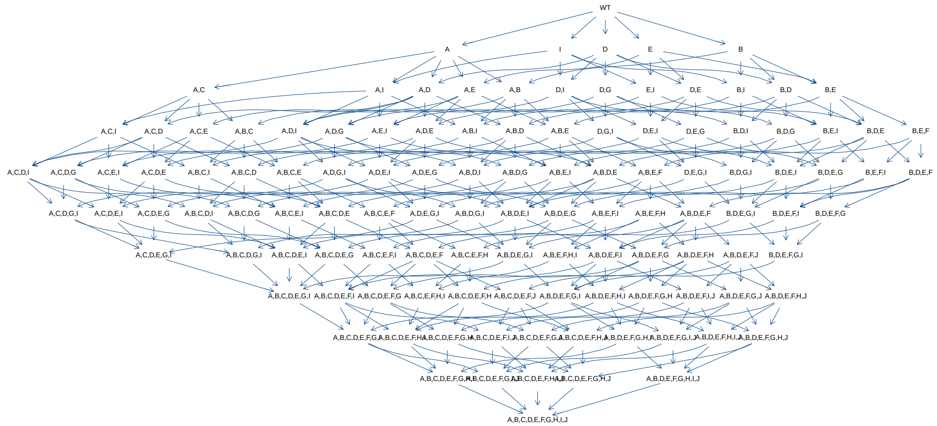

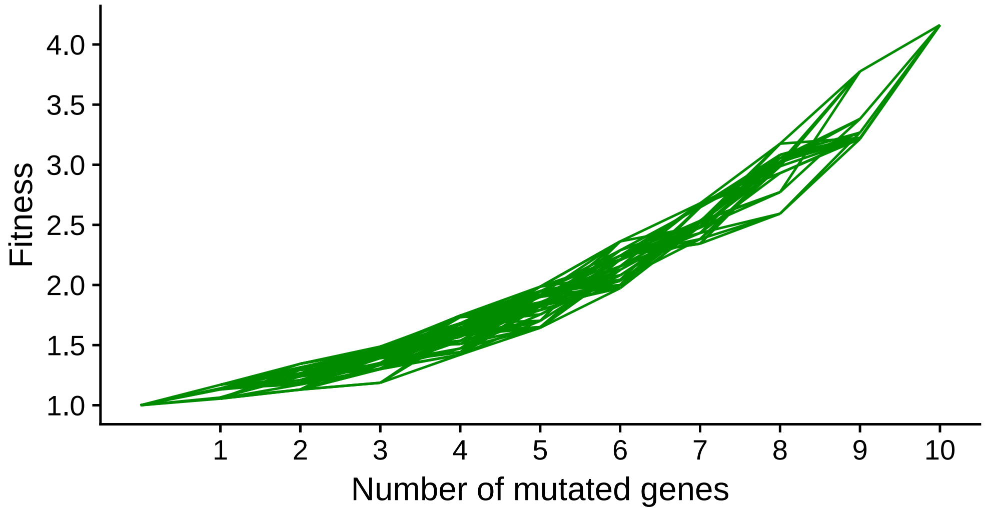

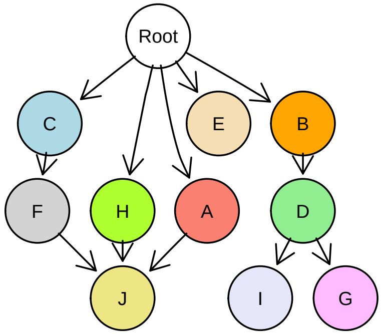


*ID* 2QFqp77NSVxKUuRw
*acc genotypes* 156
*peaks no back* 1
*removed edges* 0
*prop rem edges* 0

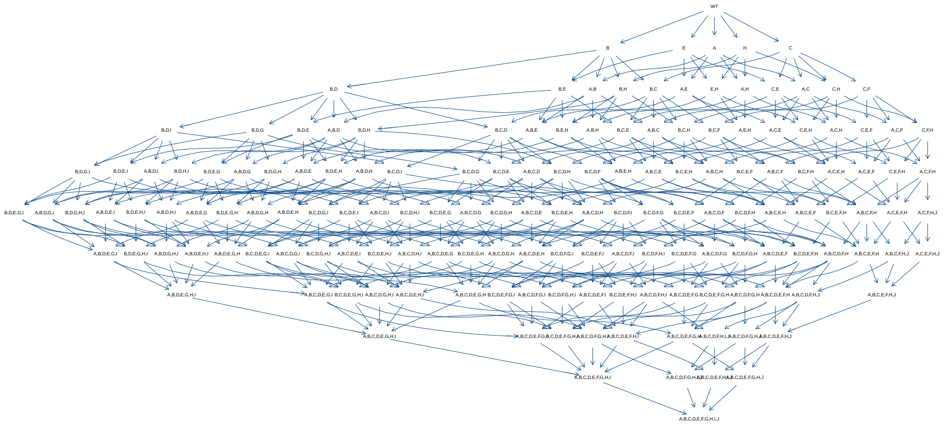

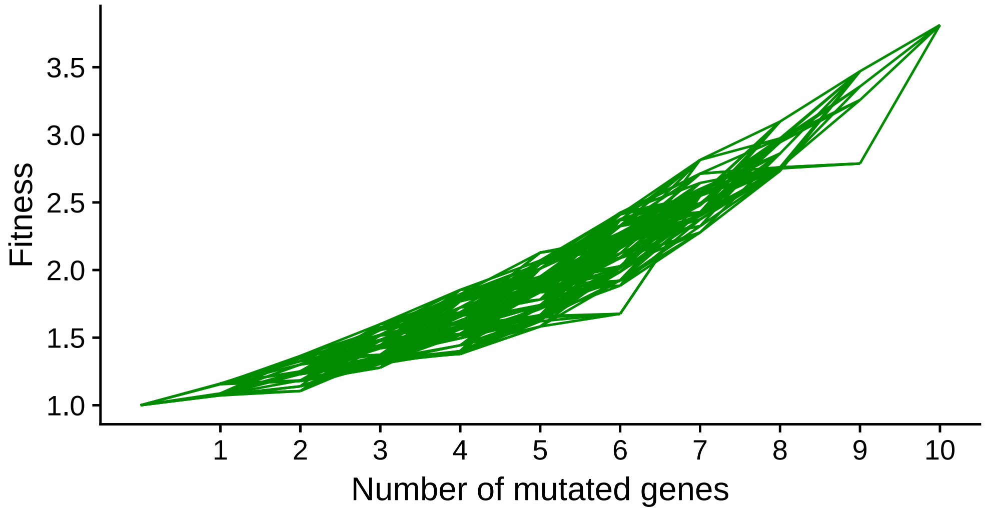

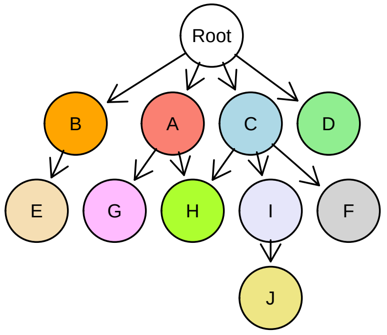


| | |
|---|---|
| *ID* | 3LaNuAy8eWU4GMwm |
| *acc genotypes* | 198 |
| *peaks no back* | 1 |
| *removed edges* | 0 |
| *prop rem edges* | 0 |

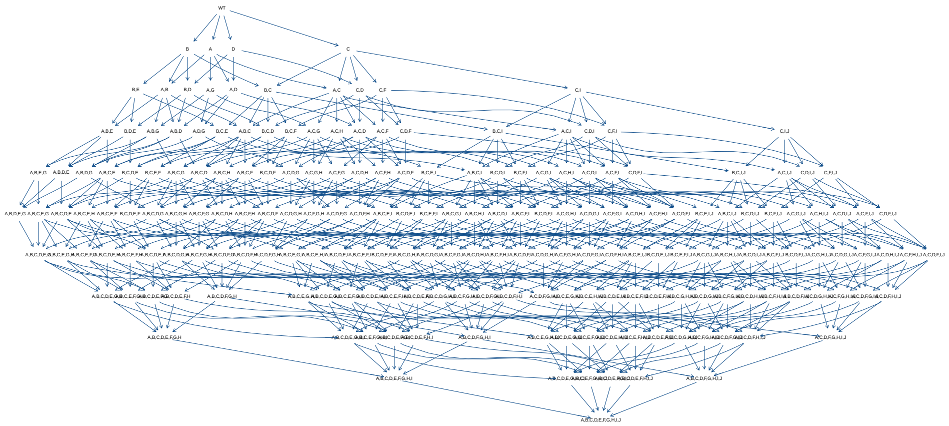

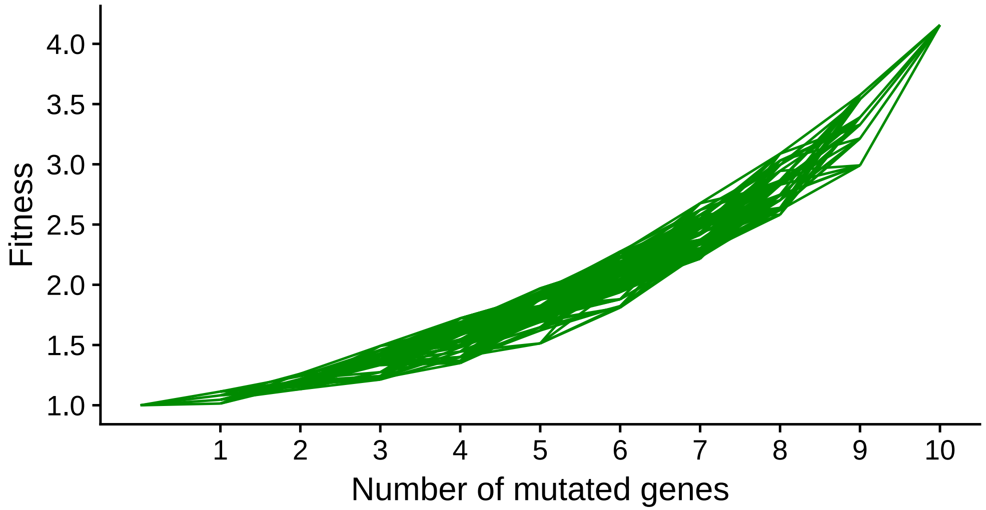

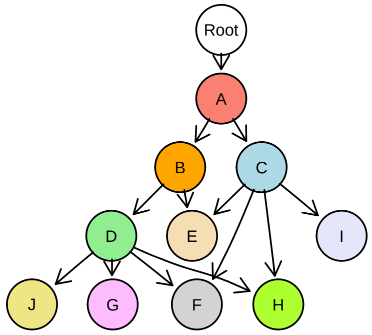


| | |
|---|---|
| *ID* | 3qwClNpvS1TW6yT3 |
| *acc genotypes* | 77 |
| *peaks no back* | 1 |
| *removed edges* | 0 |
| *prop rem edges* | 0 |

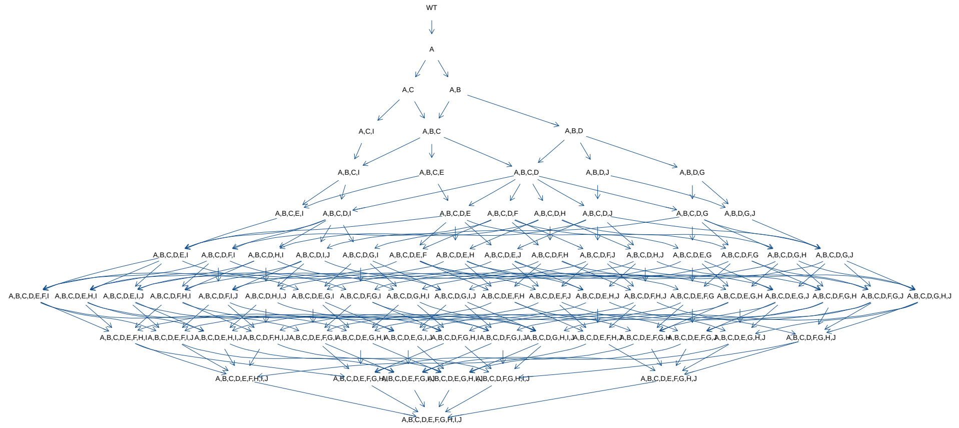

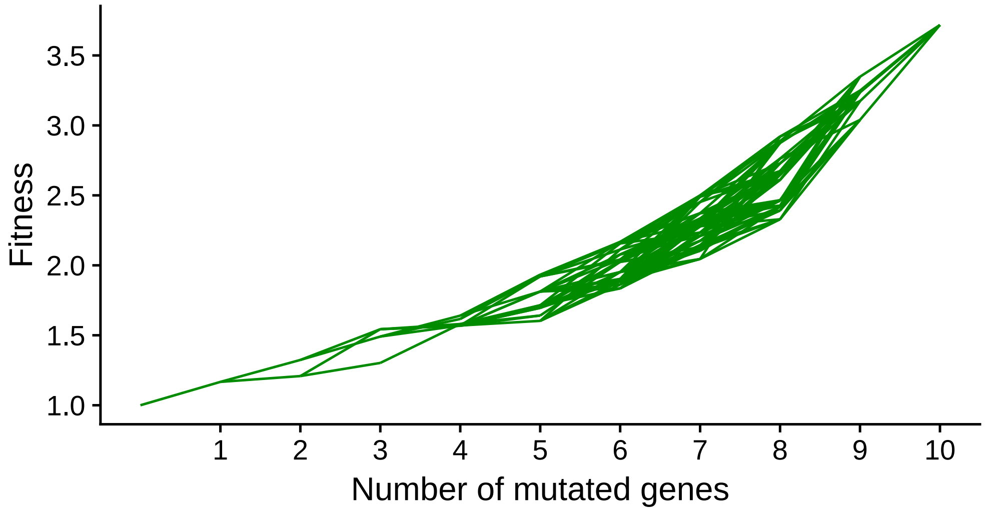

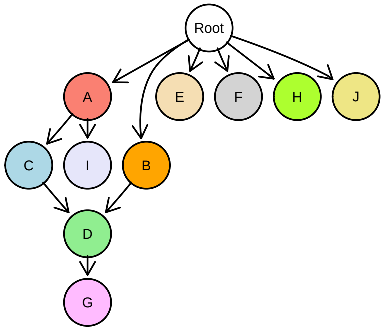


*ID* 3wRHh10MSegWf4SVc
*acc genotypes* 224
*peaks no back* 1
*removed edges* 0
*prop rem edges* 0

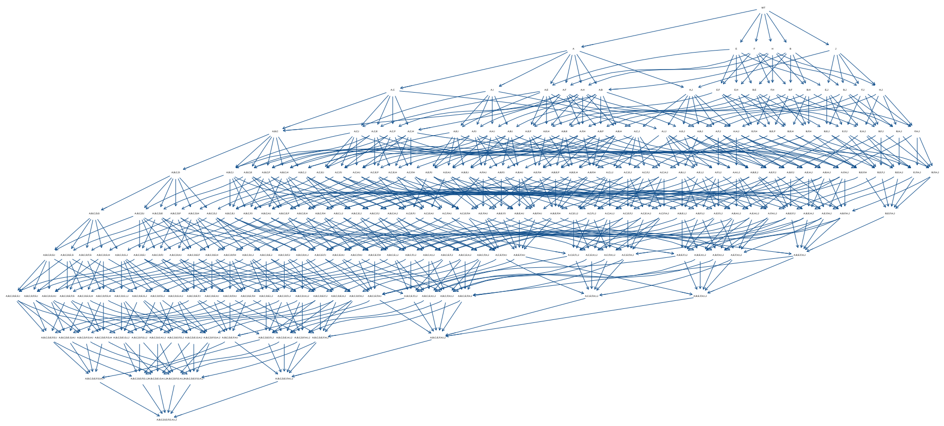

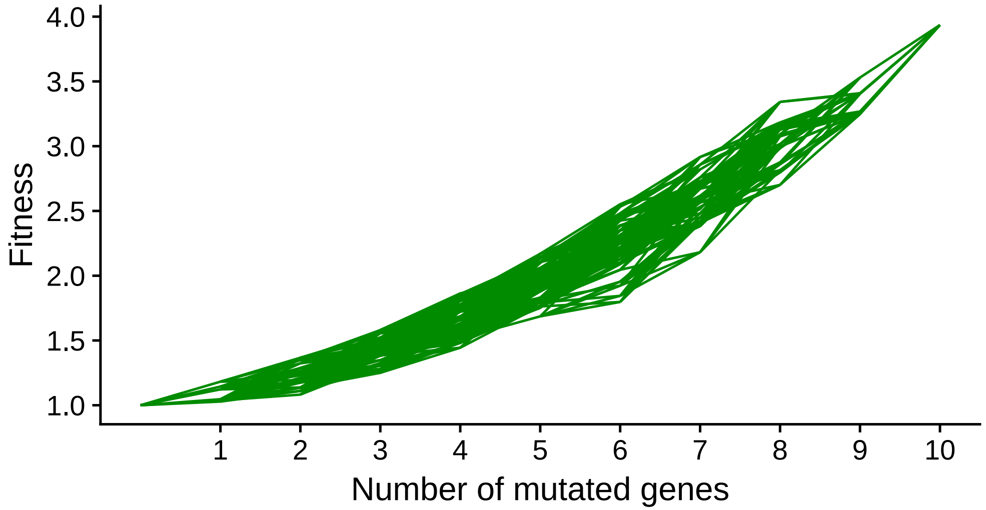

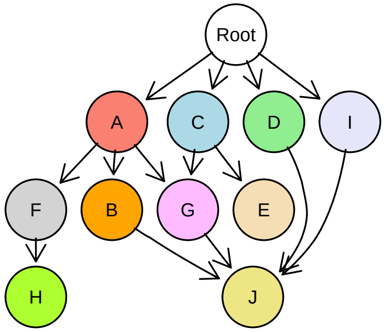


| | |
|---|---|
| *ID* | 3xUzonJpTvNpS9cY |
| *acc genotypes* | 138 |
| *peaks no back* | 1 |
| *removed edges* | 0 |
| *prop rem edges* | 0 |

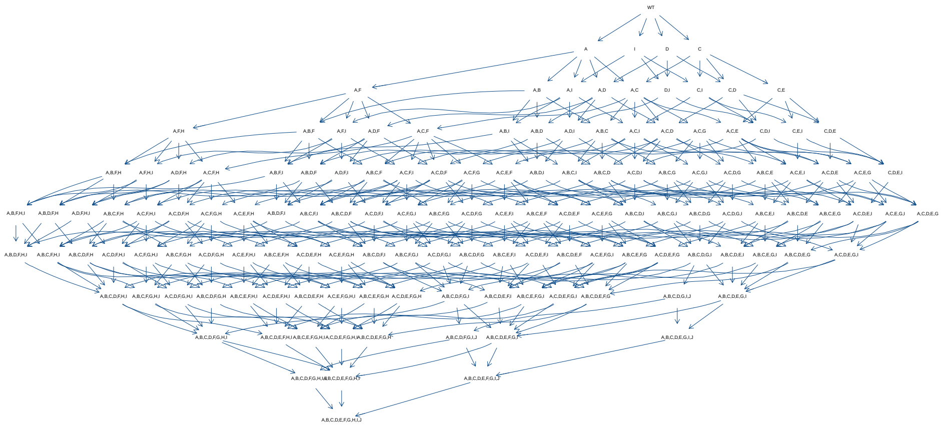

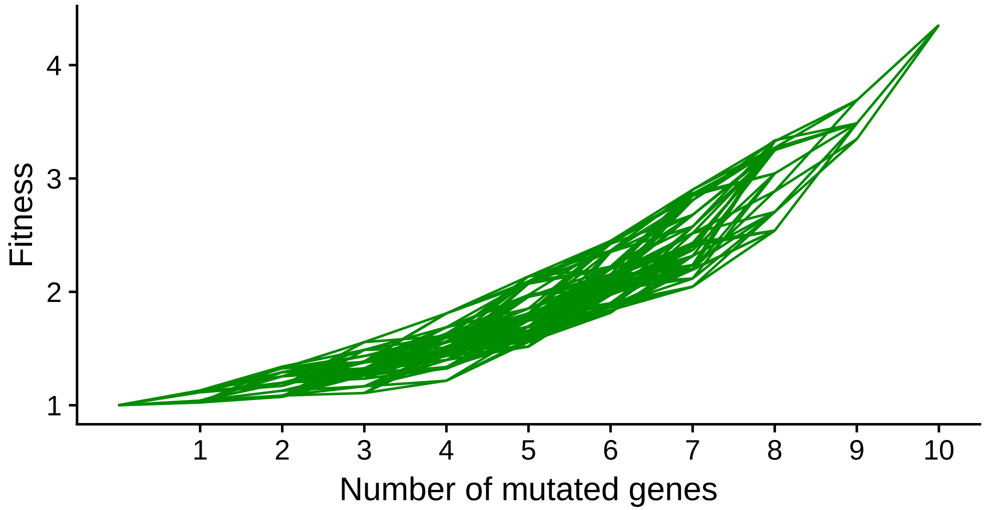

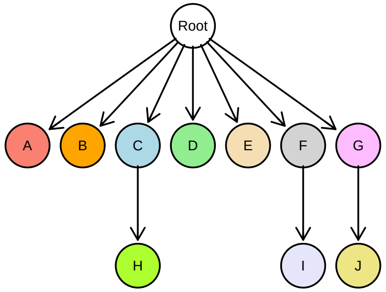


| | |
|---|---|
| *ID* | 41wj4xc10qjq6tByv |
| *acc genotypes* | 432 |
| *peaks no back* | 1 |
| *removed edges* | 0 |
| *prop rem edges* | 0 |

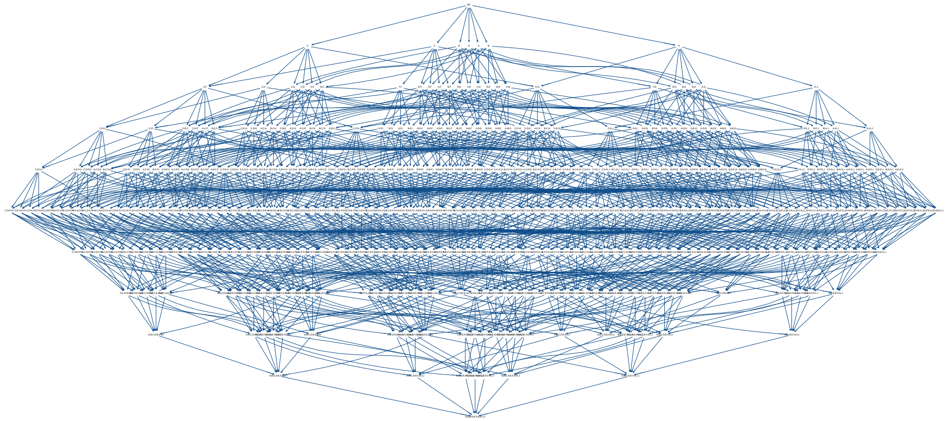

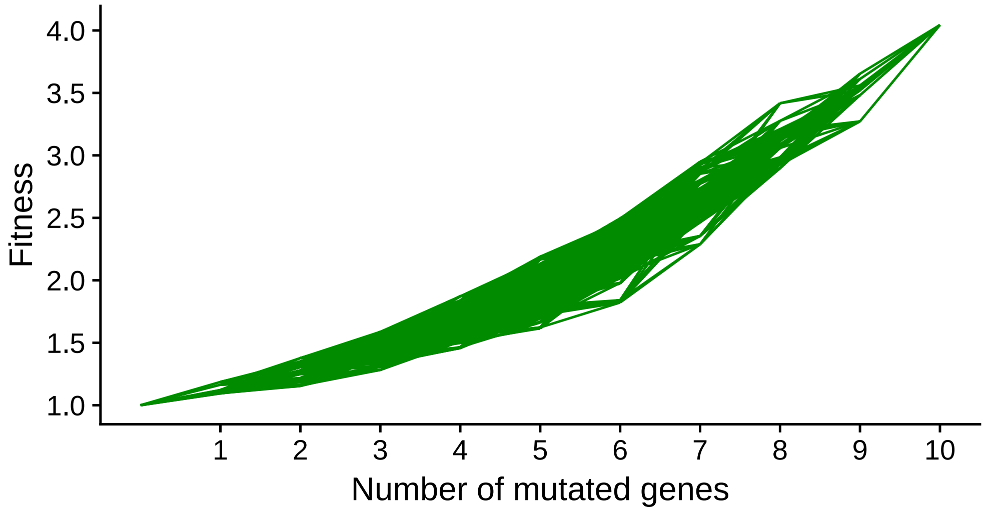

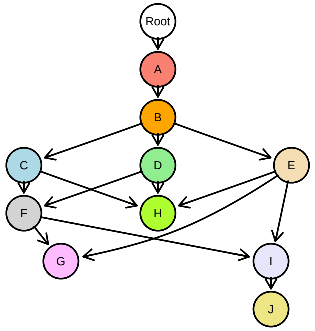


| | |
|---|---|
| *ID* | 43e2wOk4101CSHhe7 |
| *acc genotypes* | 24 |
| *peaks no back* | 1 |
| *removed edges* | 0 |
| *prop rem edges* | 0 |

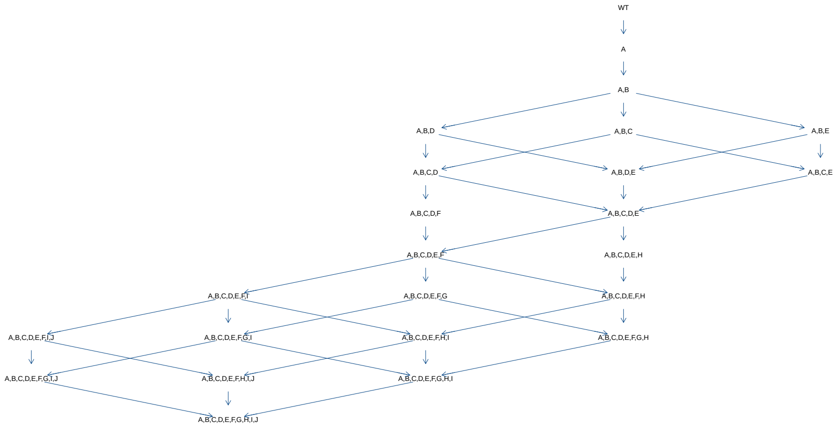

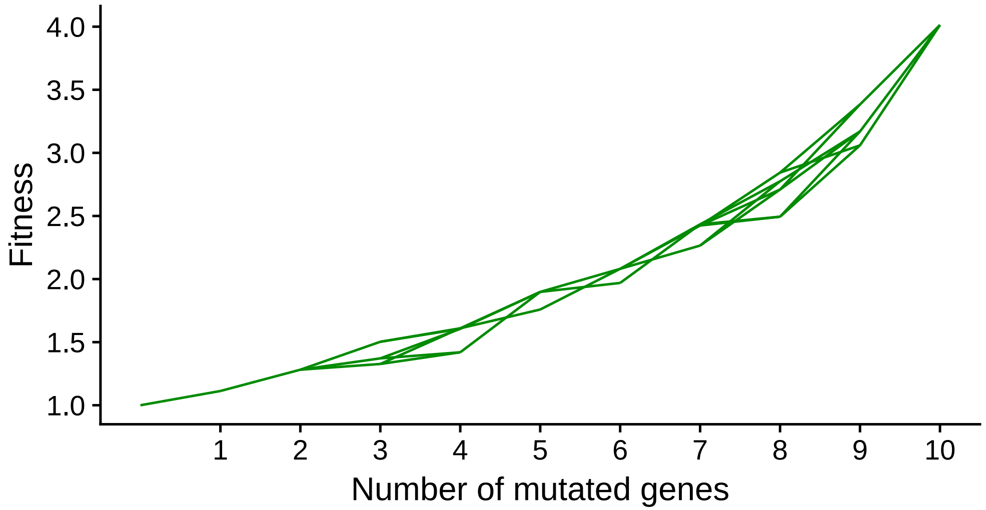

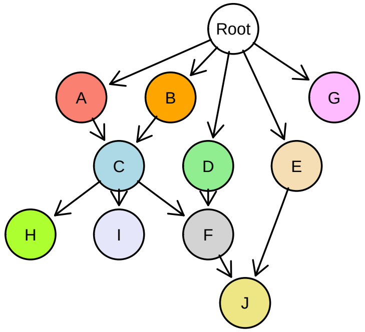


| | |
|---|---|
| *ID* | 45ZtV5kGH2IwAPAq |
| *acc genotypes* | 88 |
| *peaks no back* | 1 |
| *removed edges* | 0 |
| *prop rem edges* | 0 |

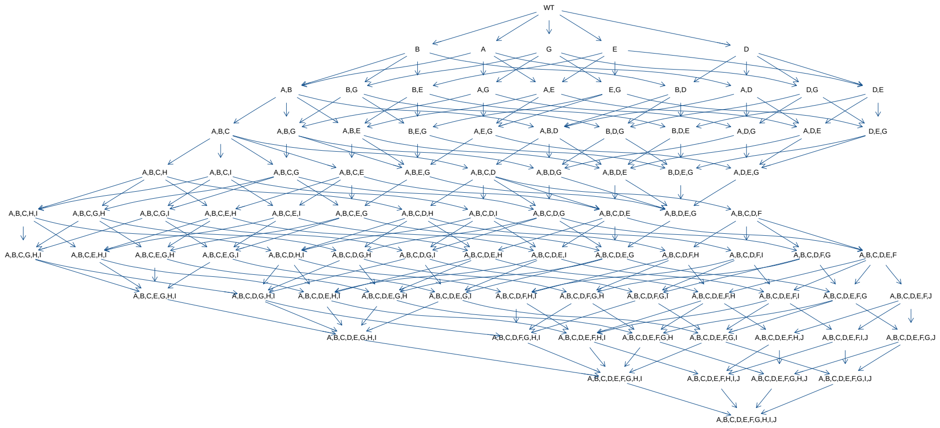

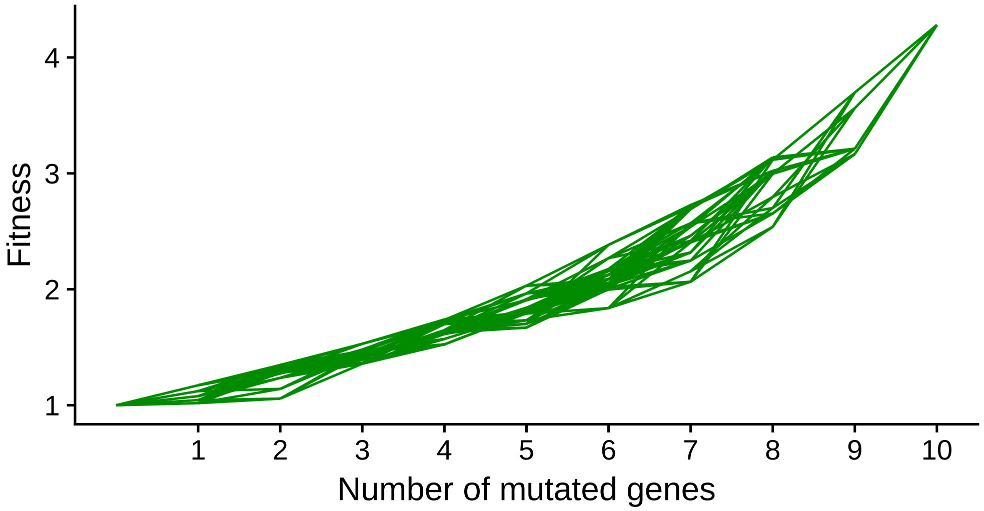

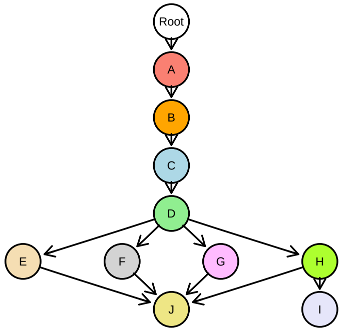


*ID* 510skoIlHNus2jjBB
*acc genotypes* 30
*peaks no back* 1
*removed edges* 0
*prop rem edges* 0

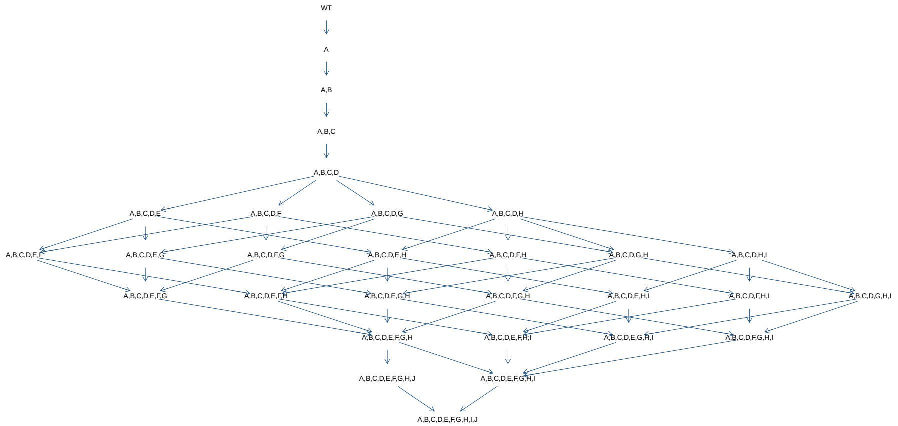


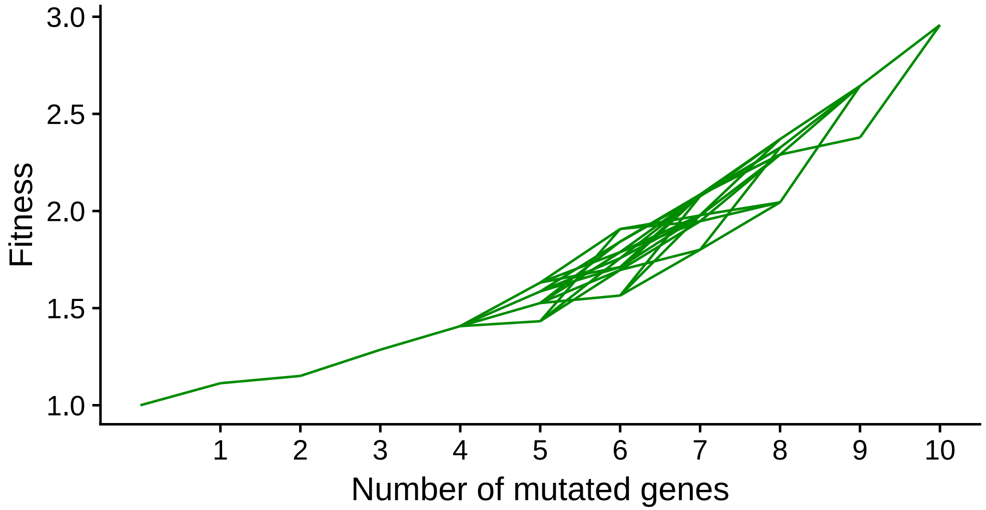

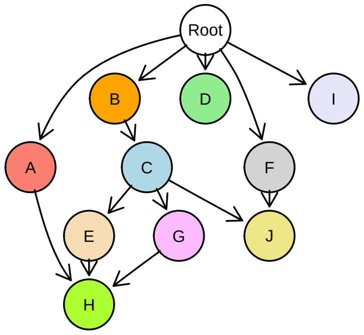


| | |
|---|---|
| *ID* | 56QJuQAd106HujMS5 |
| *acc genotypes* | 140 |
| *peaks no back* | 1 |
| *removed edges* | 0 |
| *prop rem edges* | 0 |

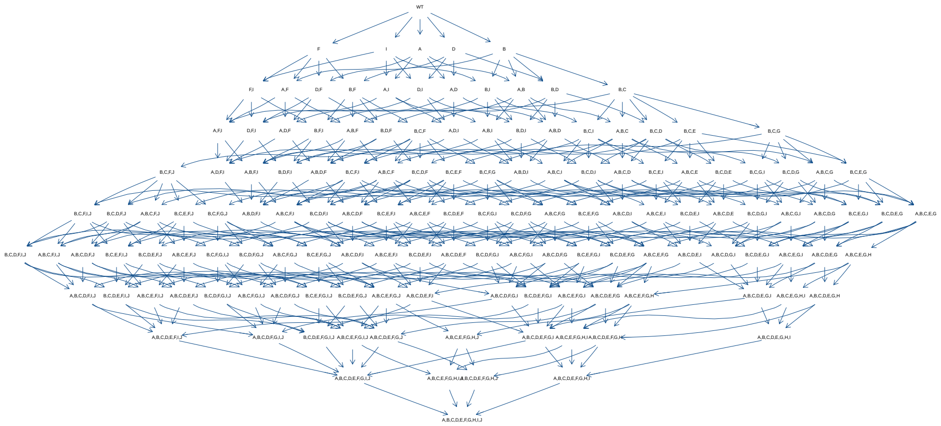

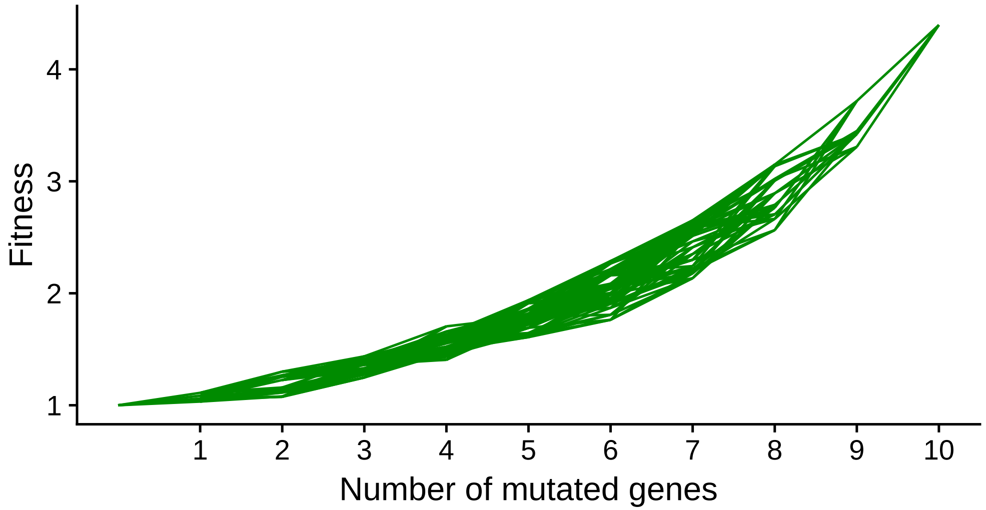

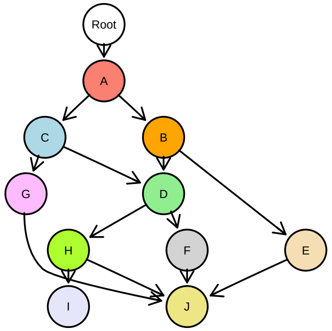


| | |
|---|---|
| *ID* | 59d4W3MnYvIGCNDg |
| *acc genotypes* | 36 |
| *peaks no back* | 1 |
| *removed edges* | 0 |
| *prop rem edges* | 0 |

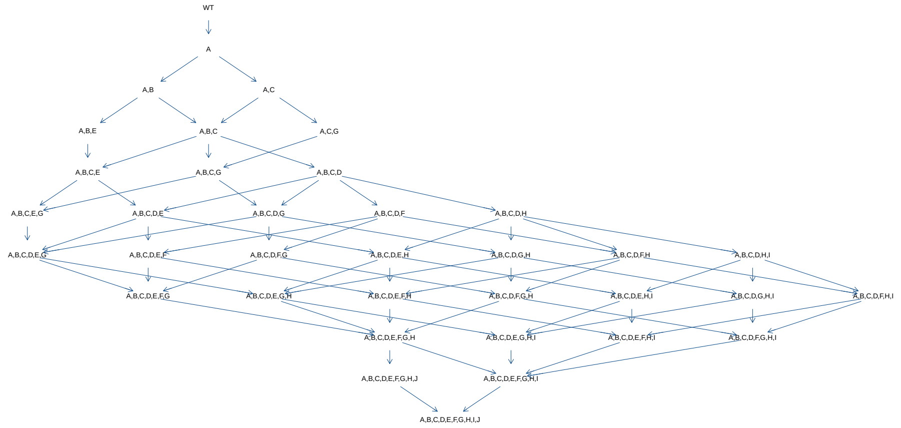

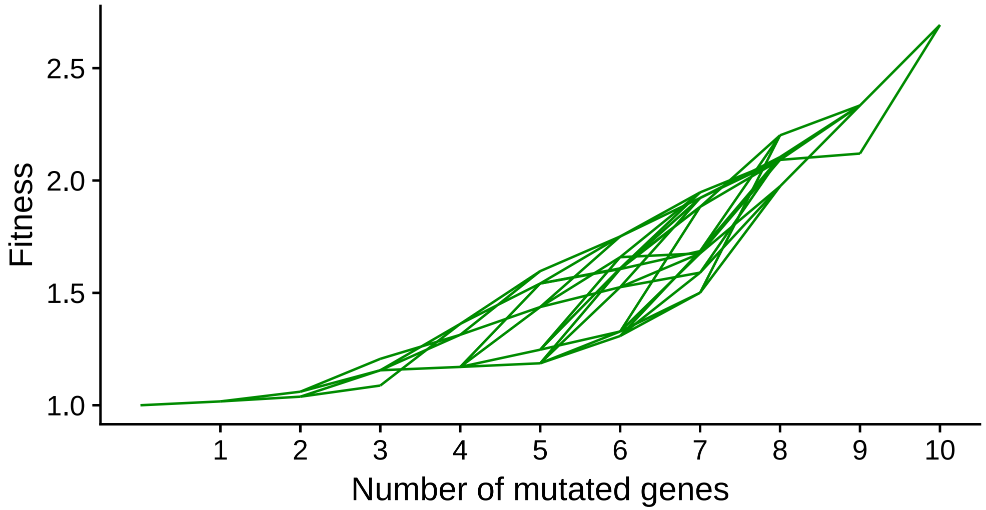

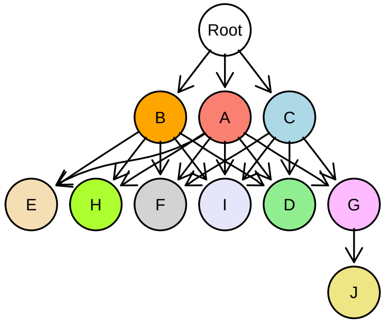


| | |
|---|---|
| *ID* | 5aGJFJcQGLZfPkJM |
| *acc genotypes* | 108 |
| *peaks no back* | 1 |
| *removed edges* | 0 |
| *prop rem edges* | 0 |

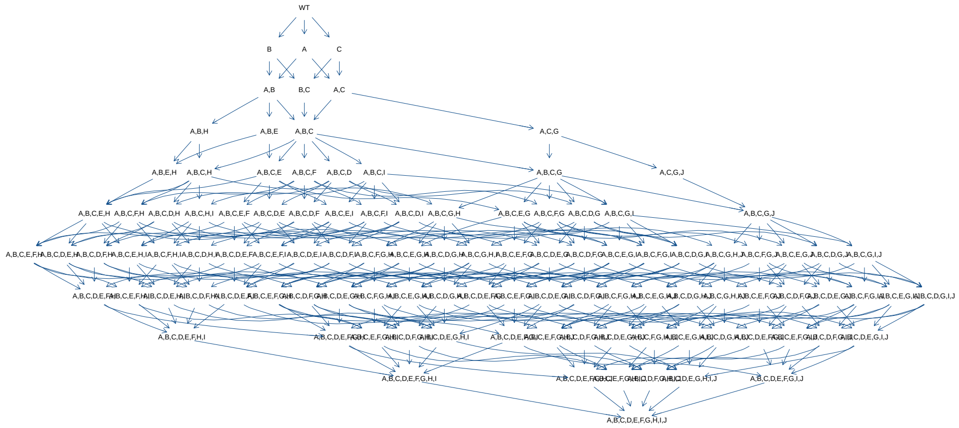

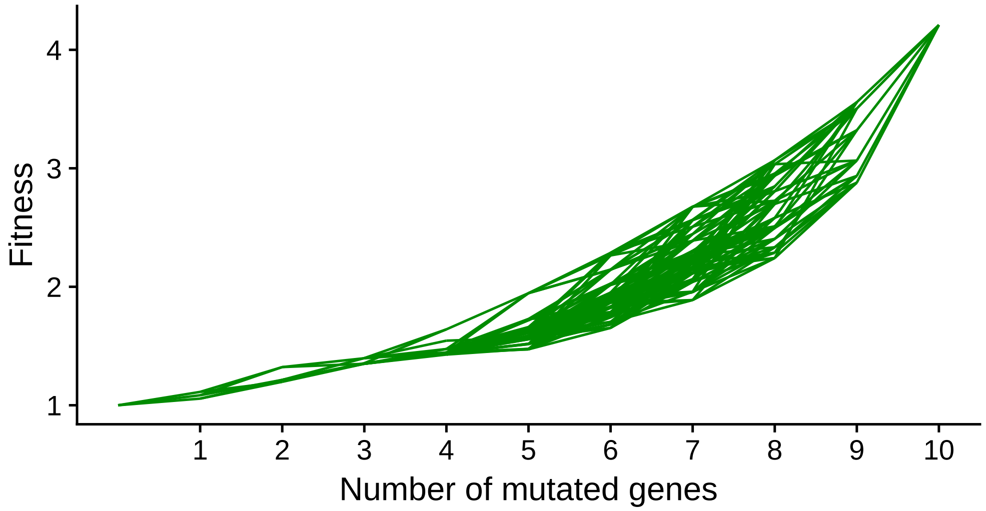

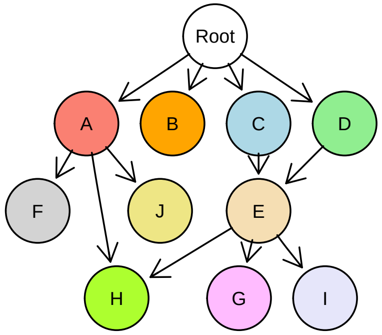


| | |
|---|---|
| *ID* | 5ism4PKhL1Vt6vub |
| *acc genotypes* | 112 |
| *peaks no back* | 1 |
| *removed edges* | 0 |
| *prop rem edges* | 0 |

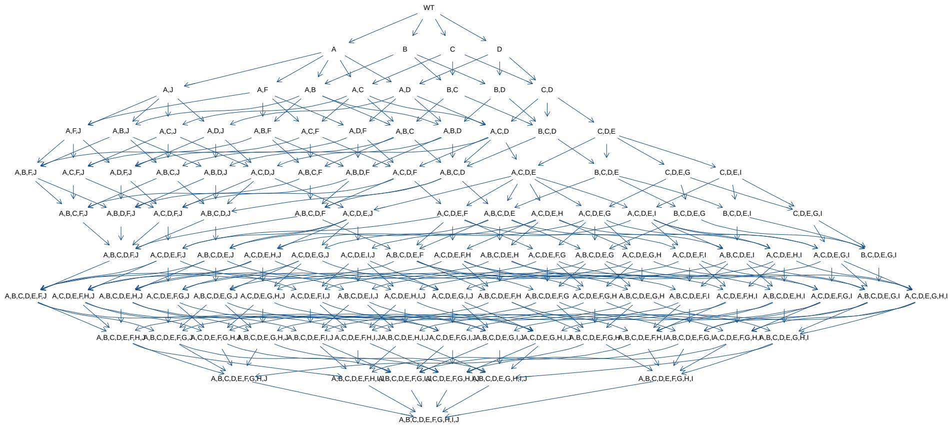

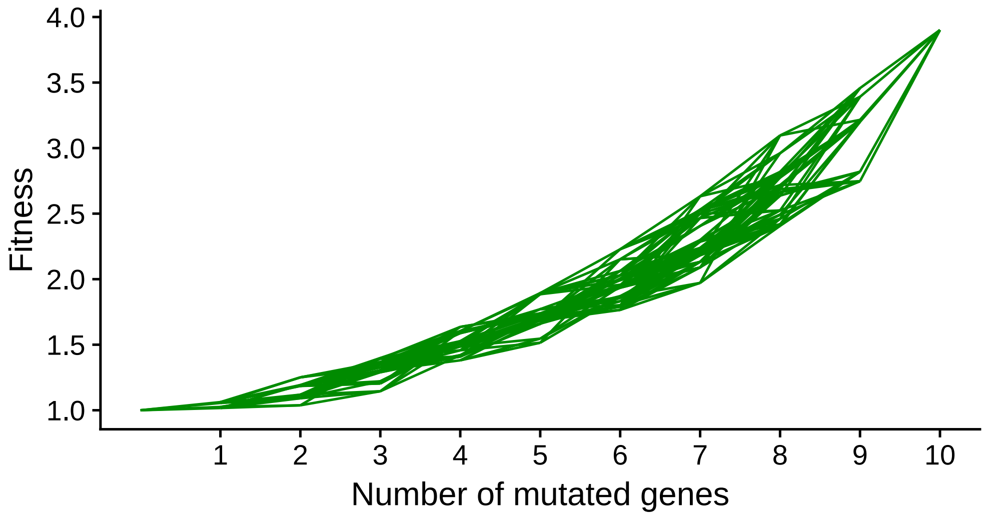

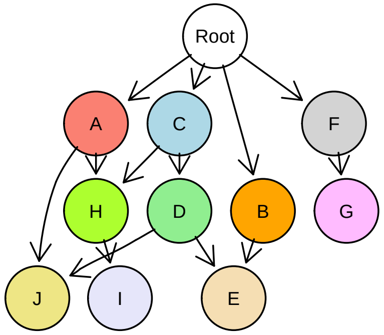


| | |
|---|---|
| *ID* | 5j8b10ekasUqQVF9H |
| *acc genotypes* | 99 |
| *peaks no back* | 1 |
| *removed edges* | 0 |
| *prop rem edges* | 0 |

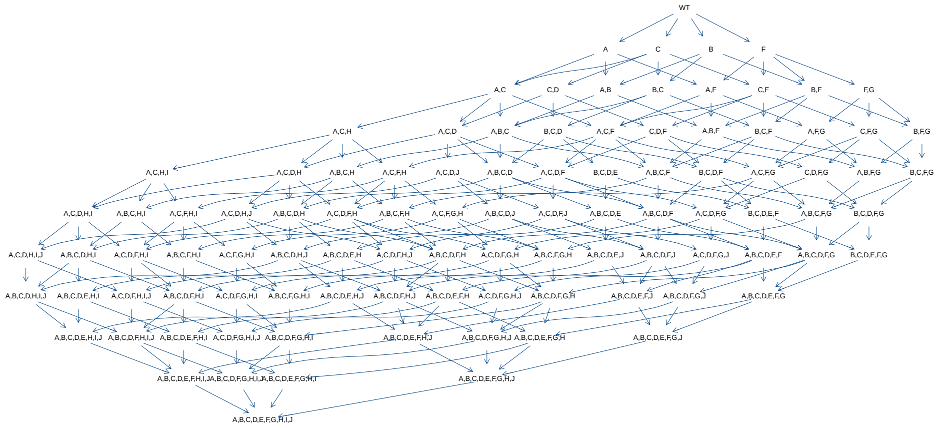

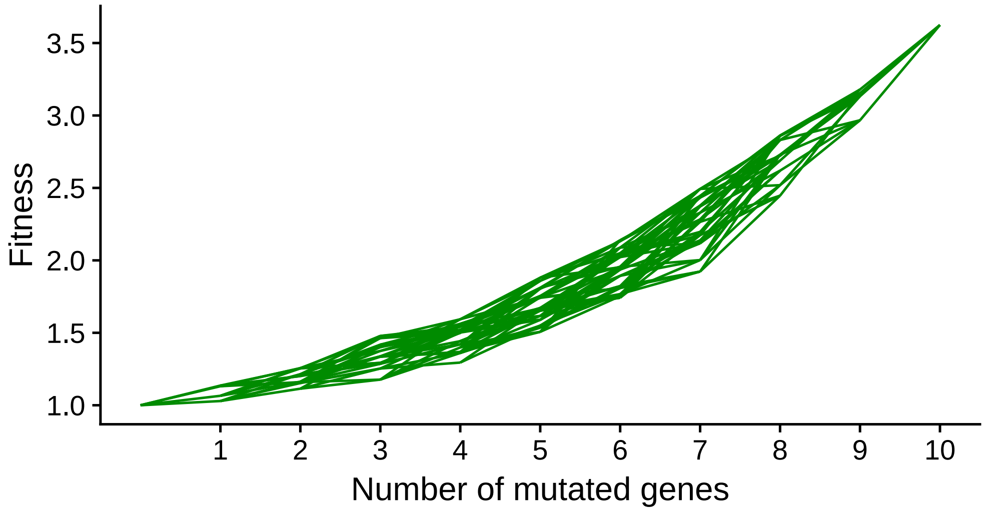

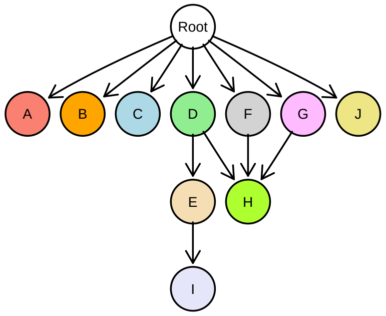


| | |
|---|---|
| *ID* | 5RbSm92BvC5zV10ss |
| *acc genotypes* | 304 |
| *peaks no back* | 1 |
| *removed edges* | 0 |
| *prop rem edges* | 0 |

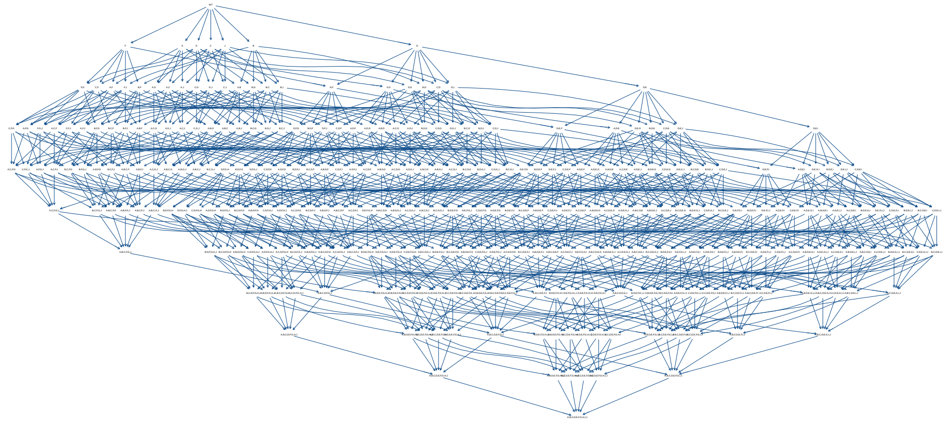

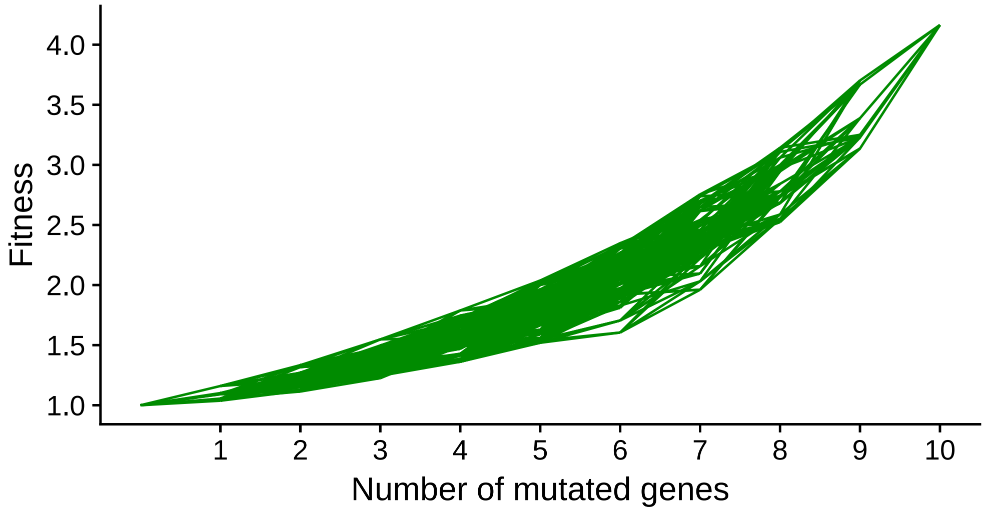

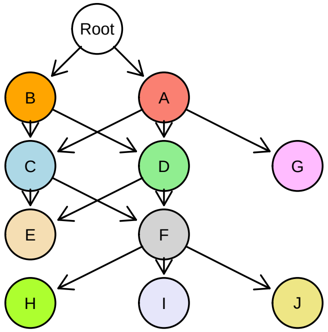


| | |
|---|---|
| *ID* | 6Lxt9hxPDEYs7HrO |
| *acc genotypes* | 46 |
| *peaks no back* | 1 |
| *removed edges* | 0 |
| *prop rem edges* | 0 |

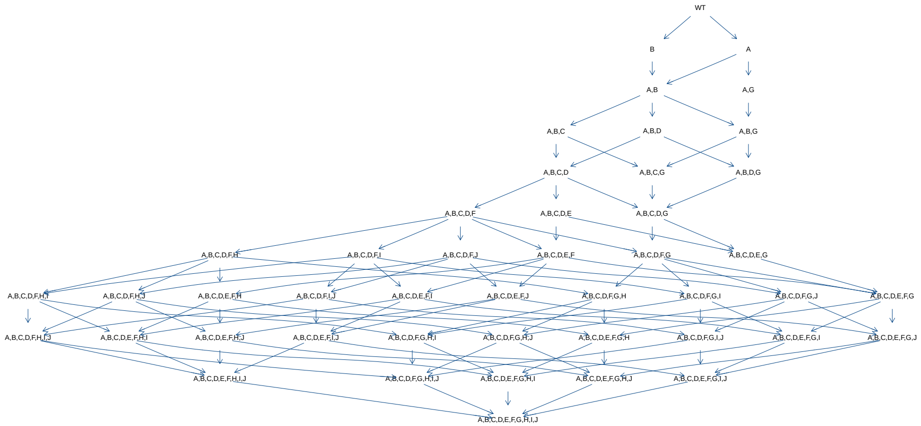

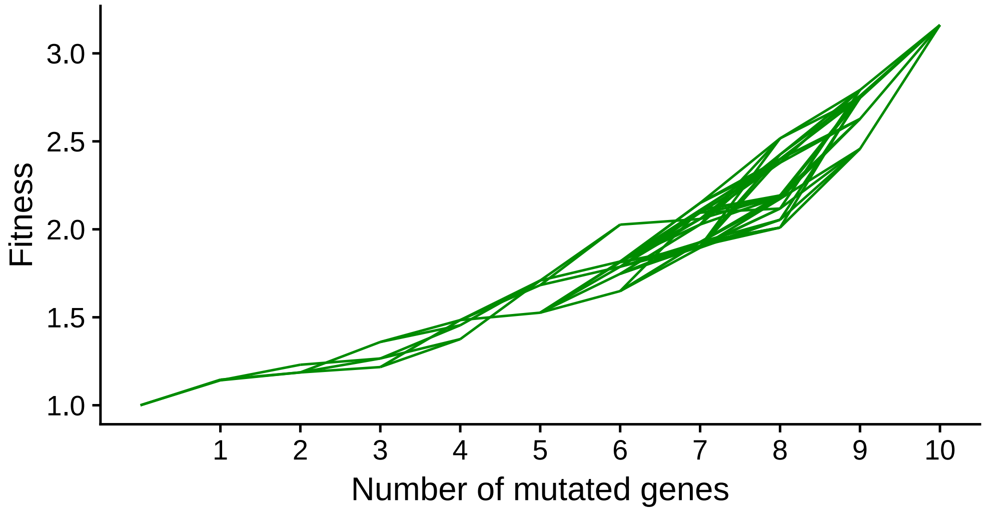

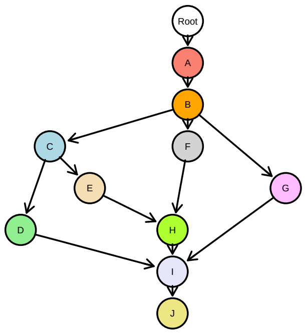


| | |
|---|---|
| *ID* | 7Fj7Va75pN1rV57O |
| *acc genotypes* | 28 |
| *peaks no back* | 1 |
| *removed edges* | 0 |
| *prop rem edges* | 0 |

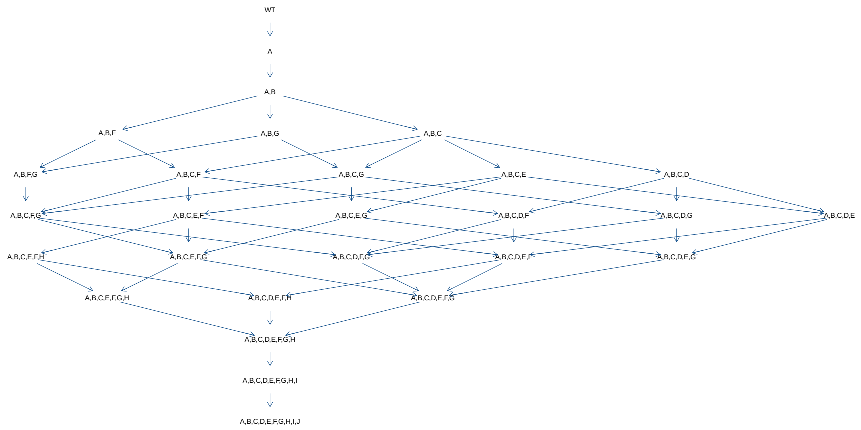

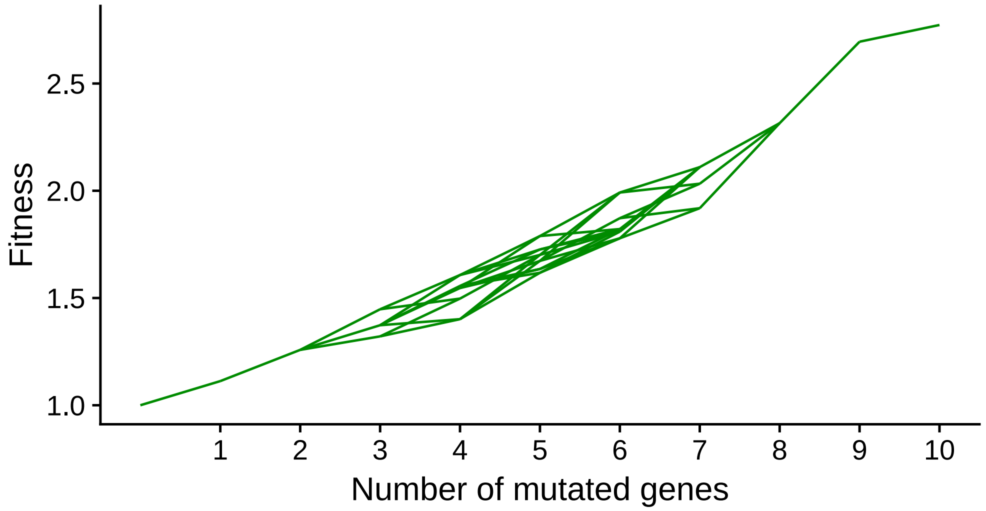

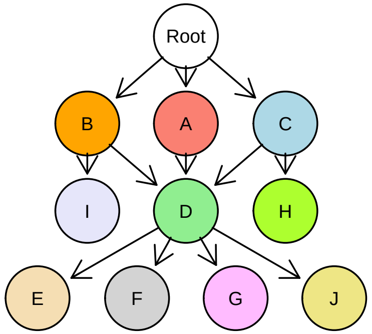


*ID* 7MYmvXyPDesfmMhs
*acc genotypes* 82
*peaks no back* 1
*removed edges* 0
*prop rem edges* 0

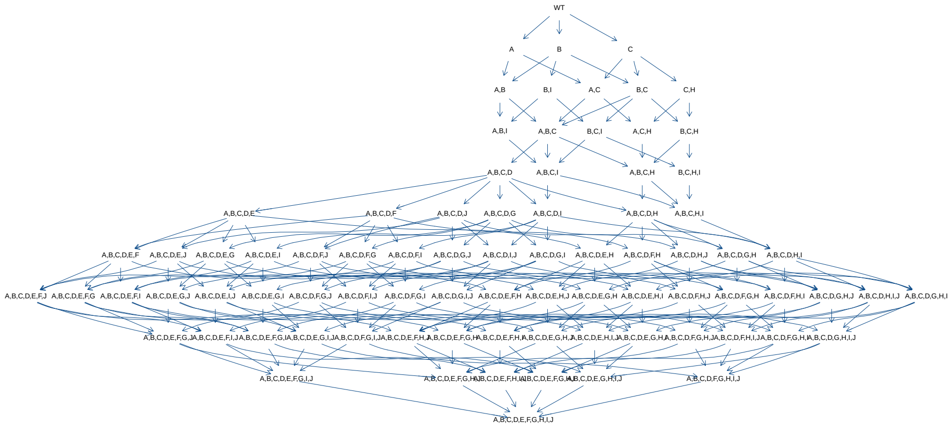

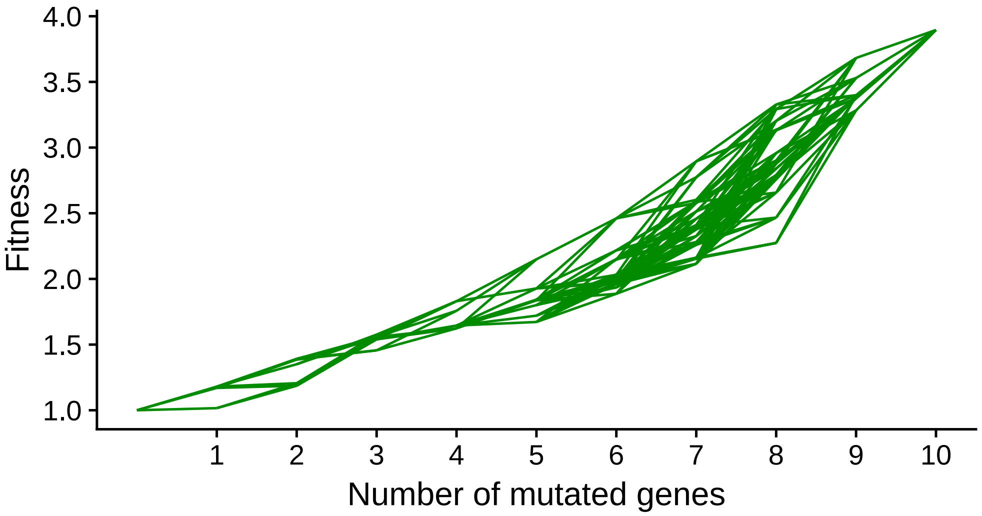

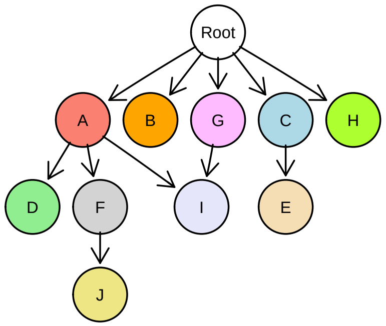


| | |
|---|---|
| *ID* | 7xha8OlAIqIZ4SXr |
| *acc genotypes* | 240 |
| *peaks no back* | 1 |
| *removed edges* | 0 |
| *prop rem edges* | 0 |

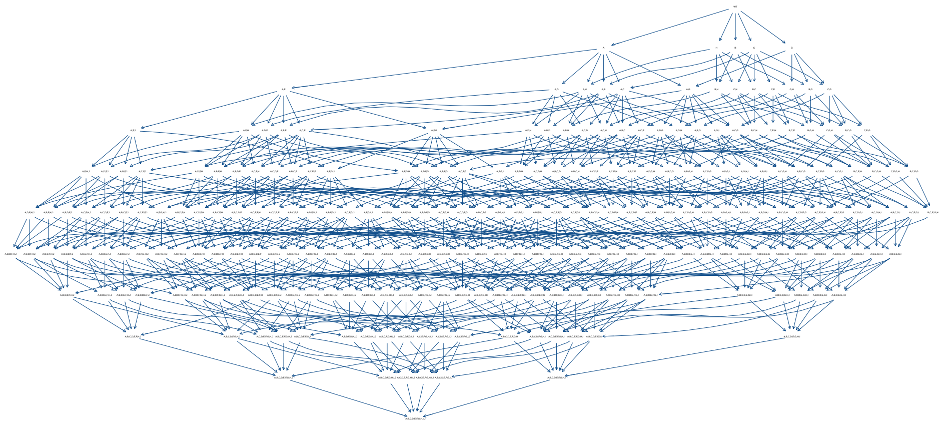

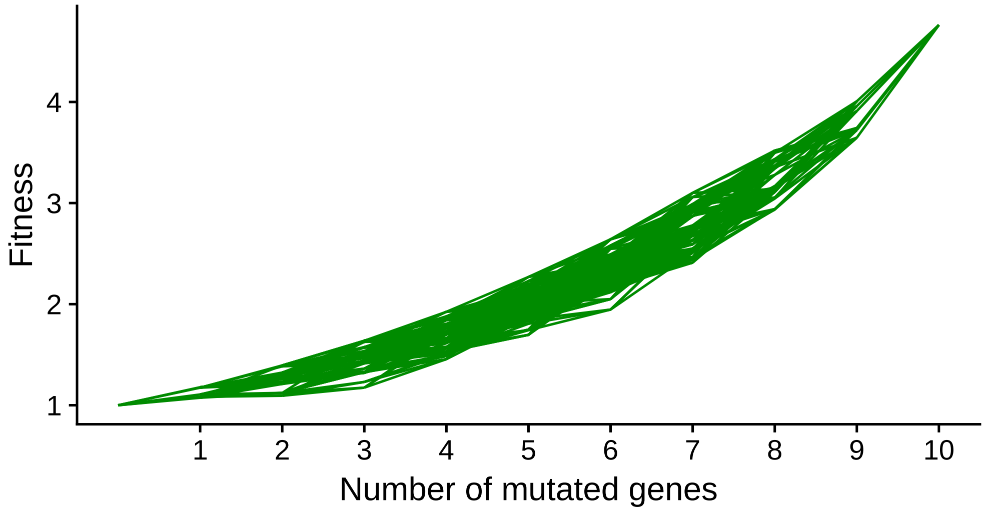

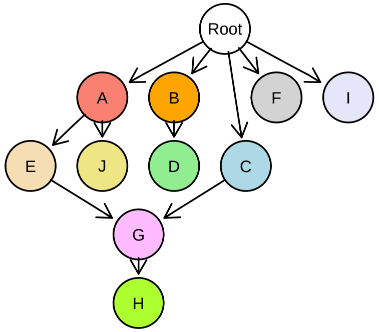


| | |
|---|---|
| *ID* | 84w3Yj1QSkf4zJx4 |
| *acc genotypes* | 168 |
| *peaks no back* | 1 |
| *removed edges* | 0 |
| *prop rem edges* | 0 |

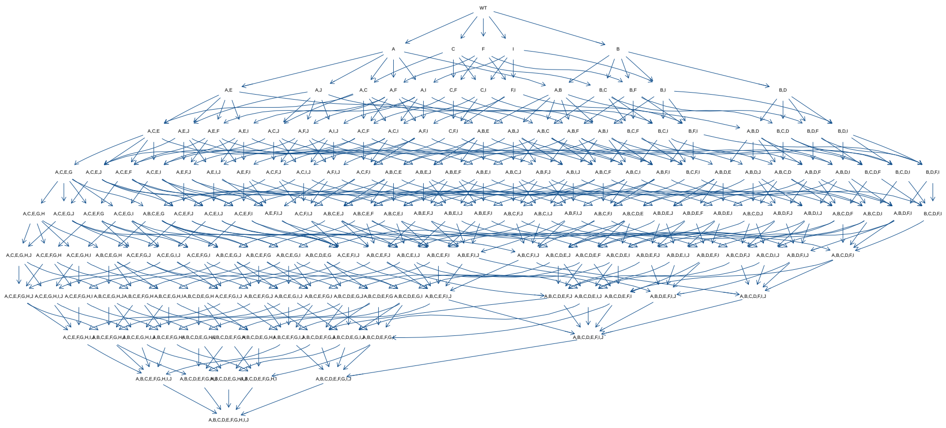

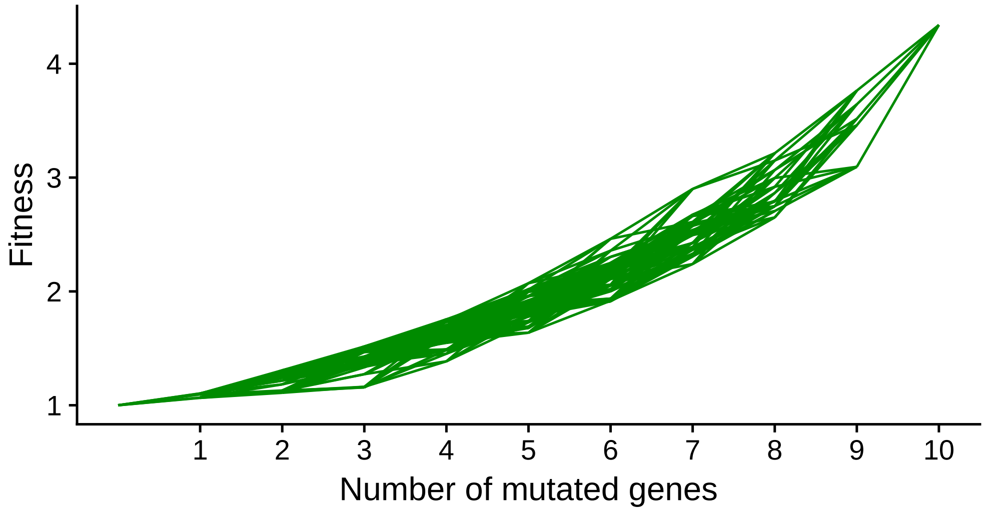

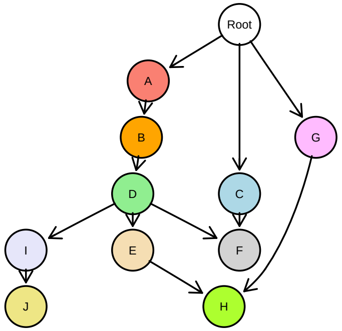


| | |
|---|---|
| *ID* | 8Libw9yeJgfs5Q95 |
| *acc genotypes* | 57 |
| *peaks no back* | 1 |
| *removed edges* | 0 |
| *prop rem edges* | 0 |

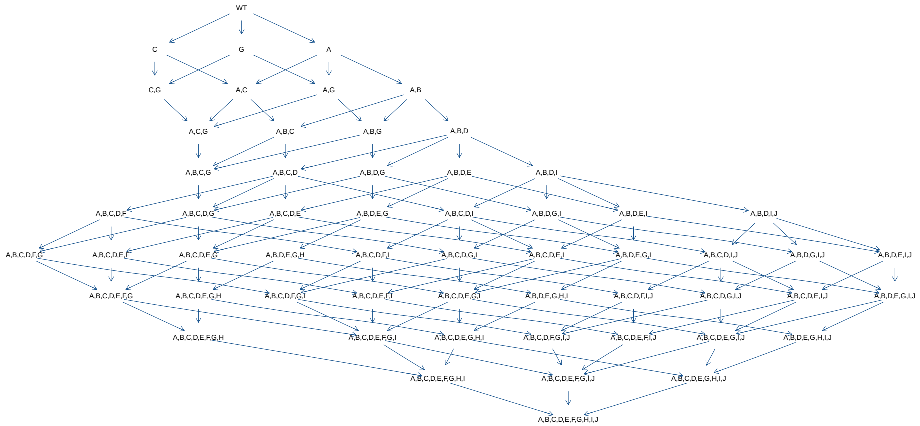

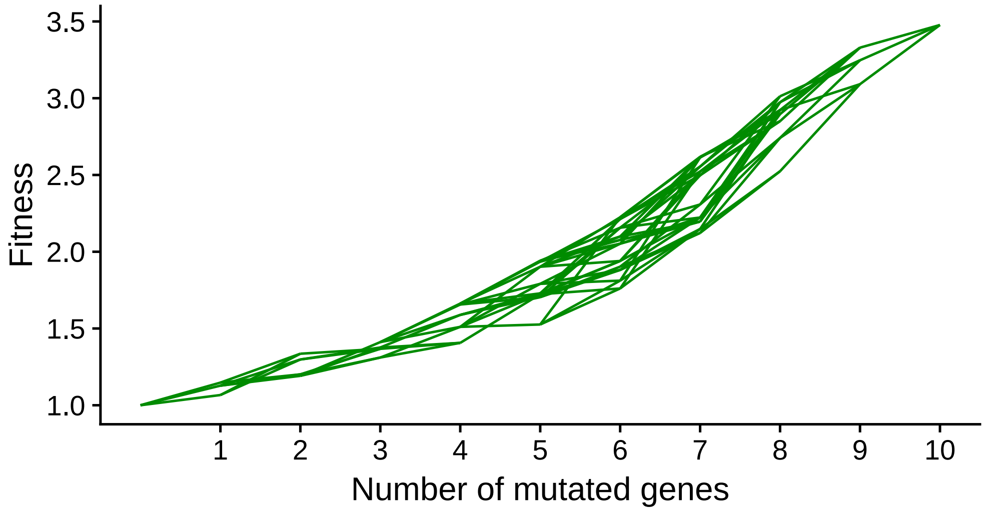

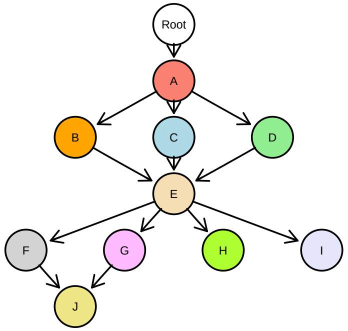


*ID* 9wKr6Qeu5I92VwmA
*acc genotypes* 29
*peaks no back* 1
*removed edges* 0
*prop rem edges* 0

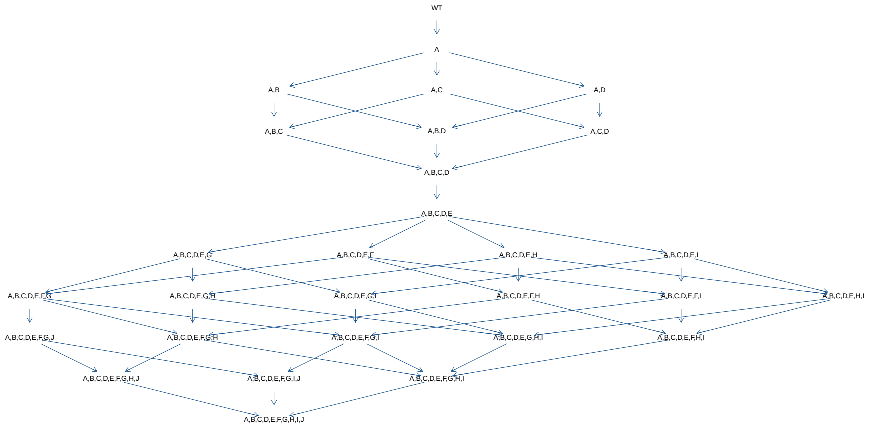

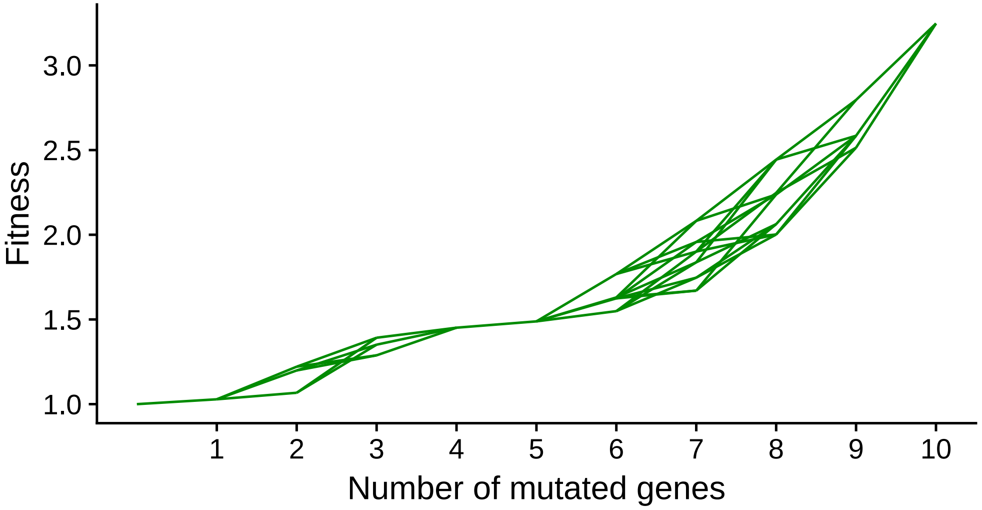

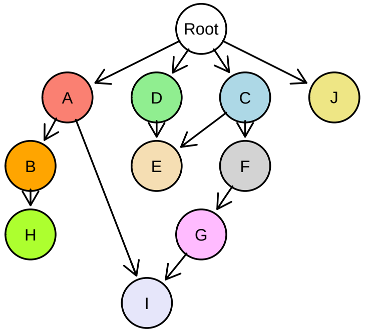


| | |
|---|---|
| *ID* | ad9ndCIhE585lPSN |
| *acc genotypes* | 106 |
| *peaks no back* | 1 |
| *removed edges* | 0 |
| *prop rem edges* | 0 |

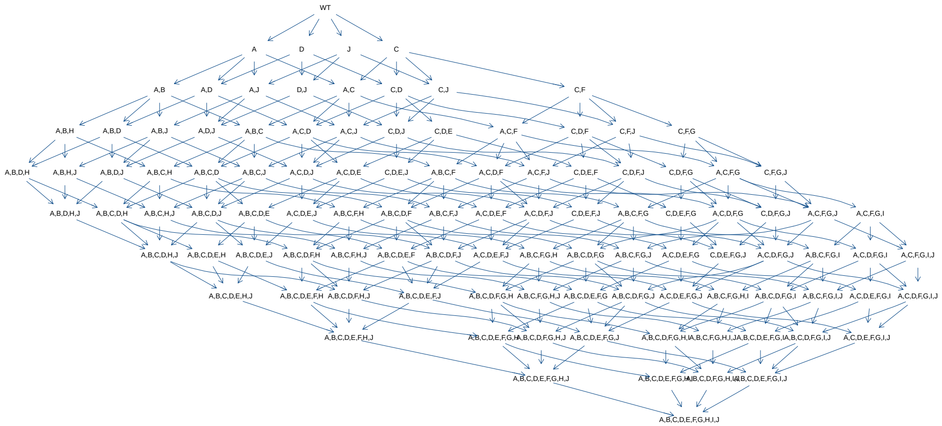

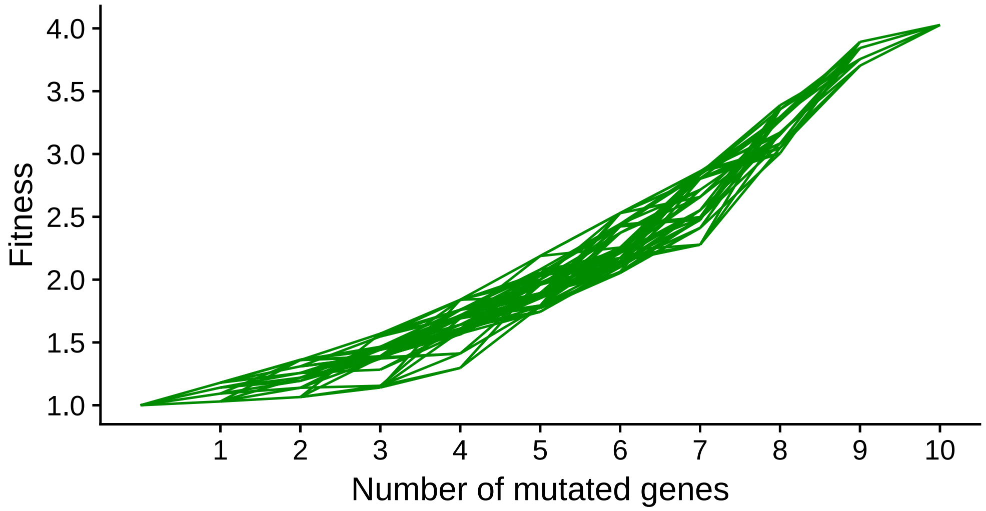

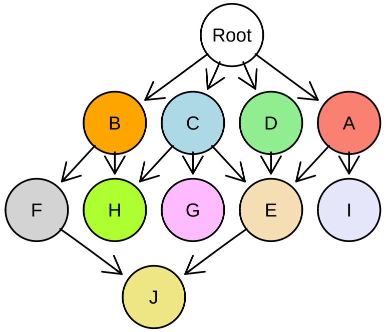


| | |
|---|---|
| *ID* | AFZHDDg9eZ1wABOd |
| *acc genotypes* | 106 |
| *peaks no back* | 1 |
| *removed edges* | 0 |
| *prop rem edges* | 0 |

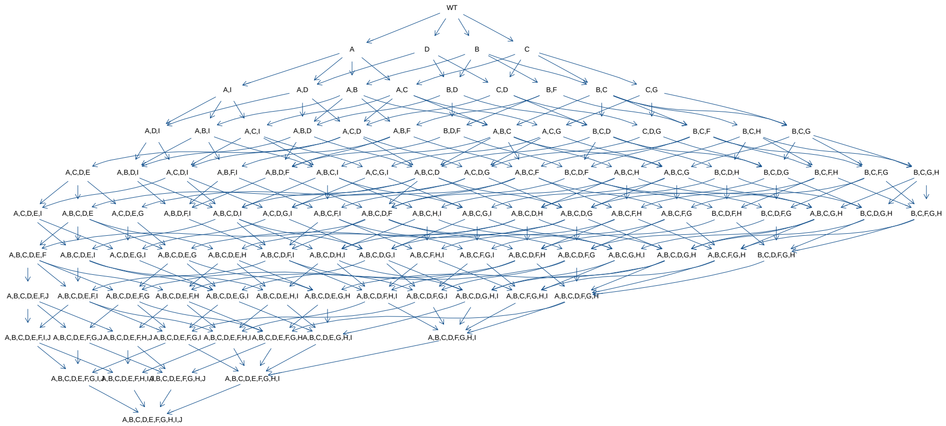

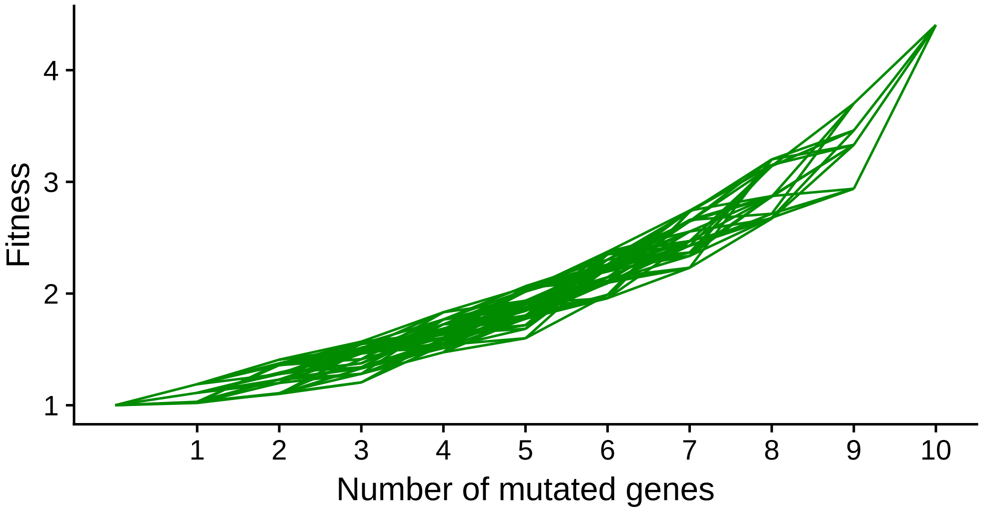

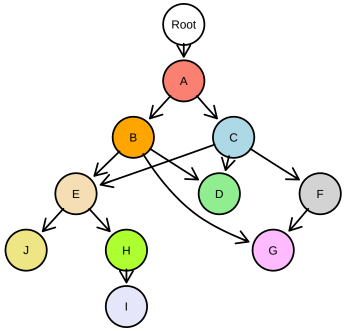


| | |
|---|---|
| *ID* | ALSuQeDPTtbq8lNG |
| *acc genotypes* | 47 |
| *peaks no back* | 1 |
| *removed edges* | 0 |
| *prop rem edges* | 0 |

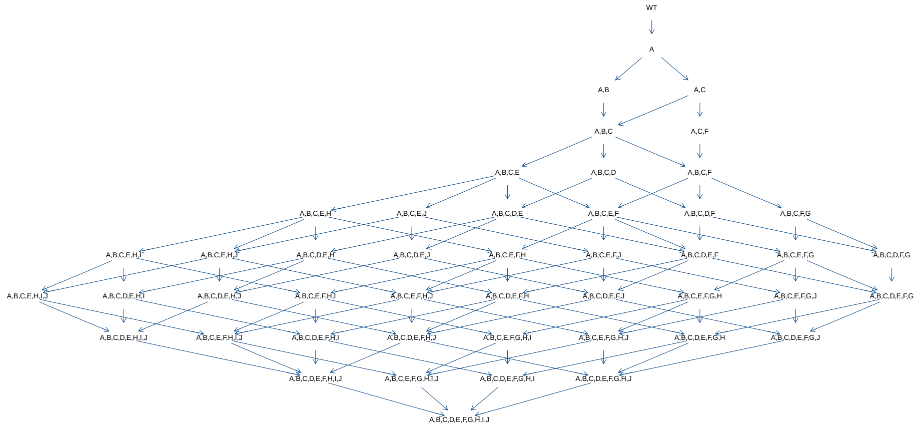

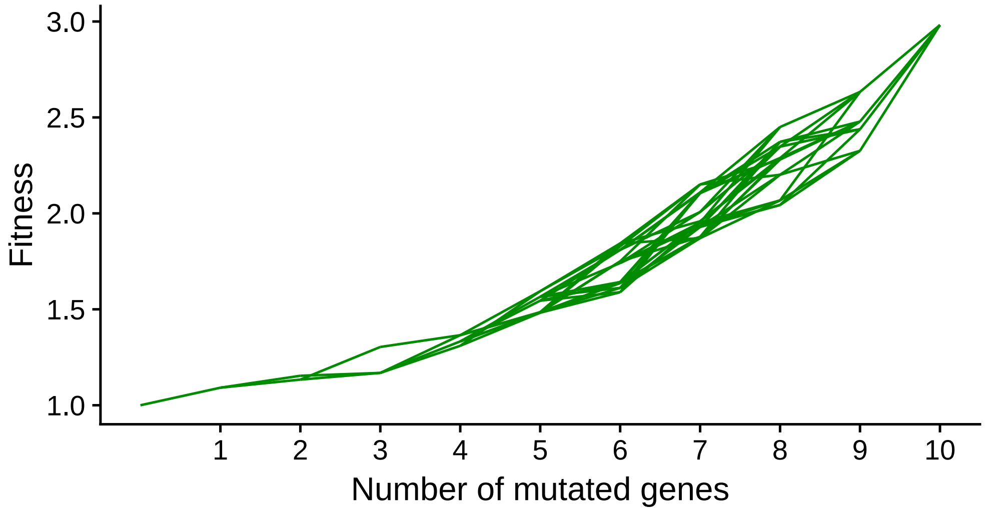

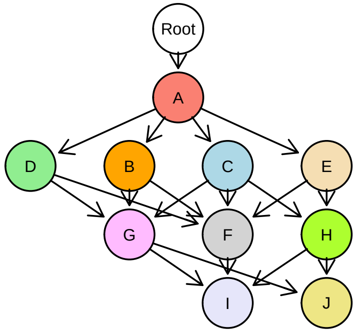


| | |
|---|---|
| *ID* | AV7D5TvJW78FIxWP |
| *acc genotypes* | 32 |
| *peaks no back* | 1 |
| *removed edges* | 0 |
| *prop rem edges* | 0 |

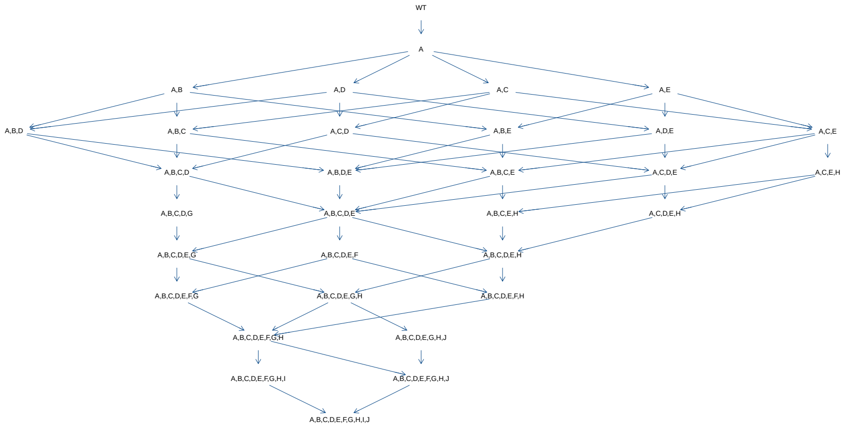

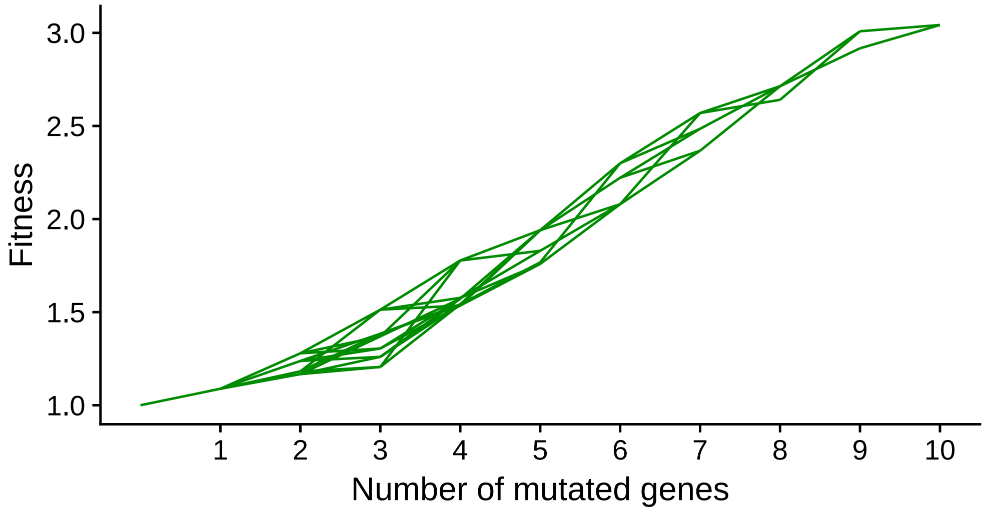

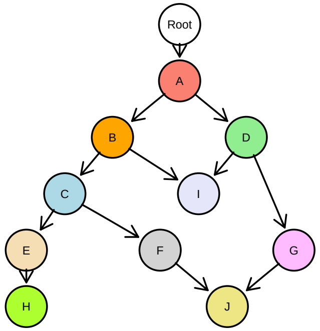


| | |
|---|---|
| *ID* | AzcqFM9IjyJEevU2 |
| *acc genotypes* | 45 |
| *peaks no back* | 1 |
| *removed edges* | 0 |
| *prop rem edges* | 0 |

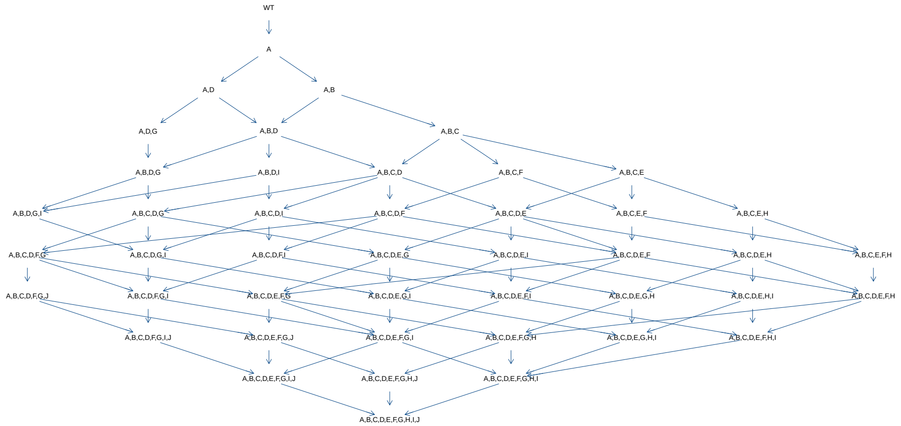

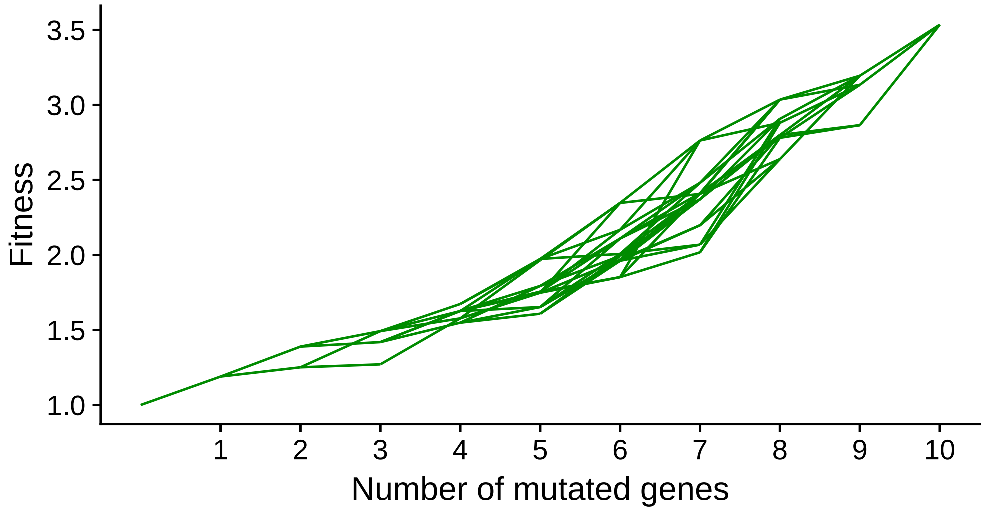

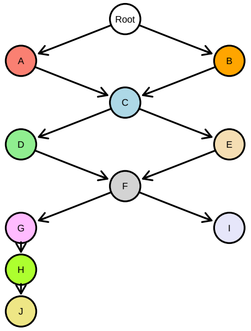


*ID* B2Xyqnt5VTgzAZsS
*acc genotypes* 16
*peaks no back* 1
*removed edges* 0
*prop rem edges* 0

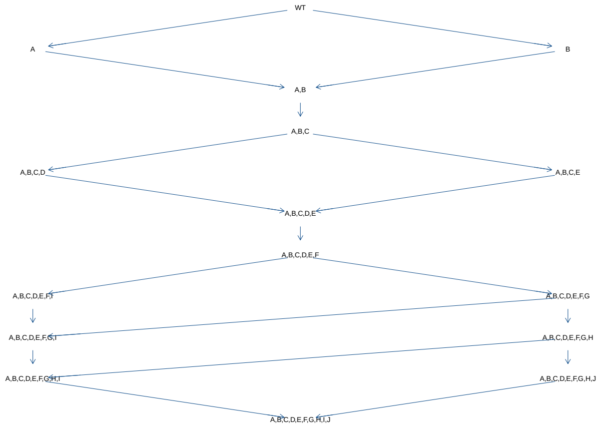


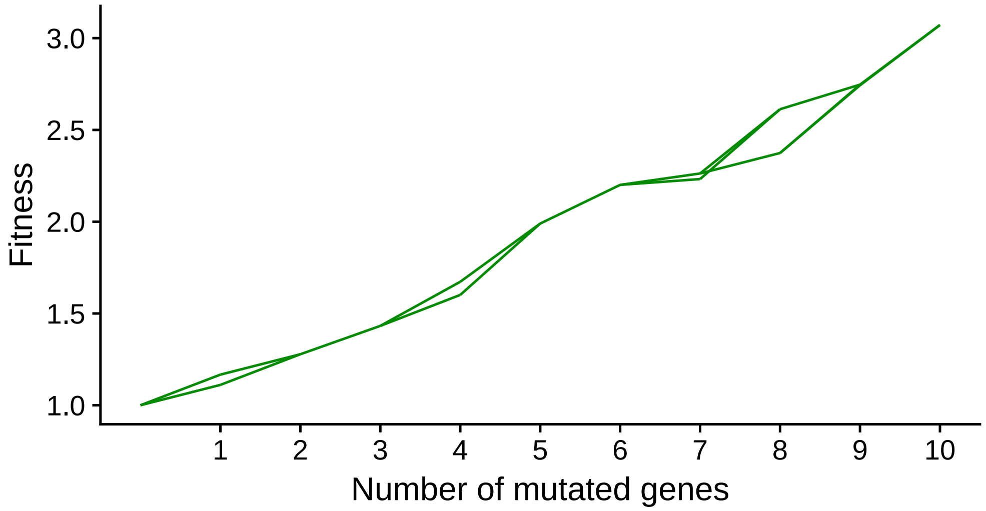

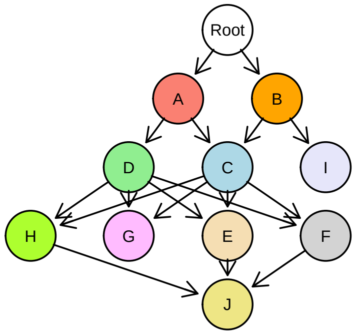


*ID* b6IqOPrfURmxrTRJ
*acc genotypes* 47
*peaks no back* 1
*removed edges* 0
*prop rem edges* 0

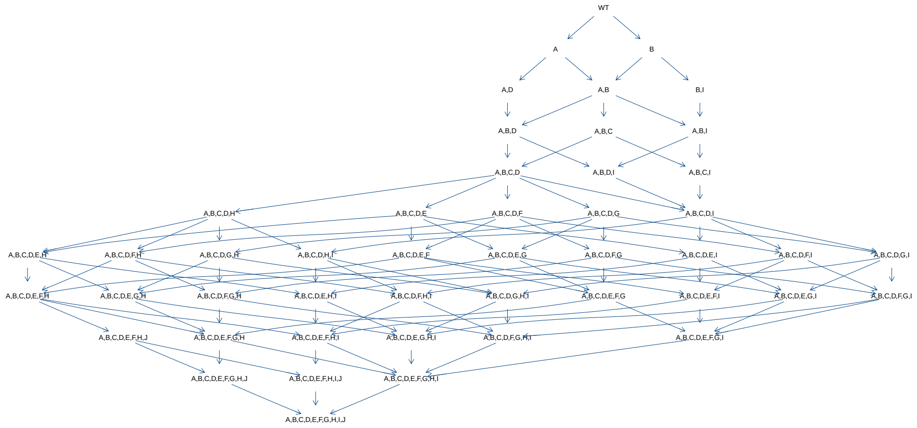

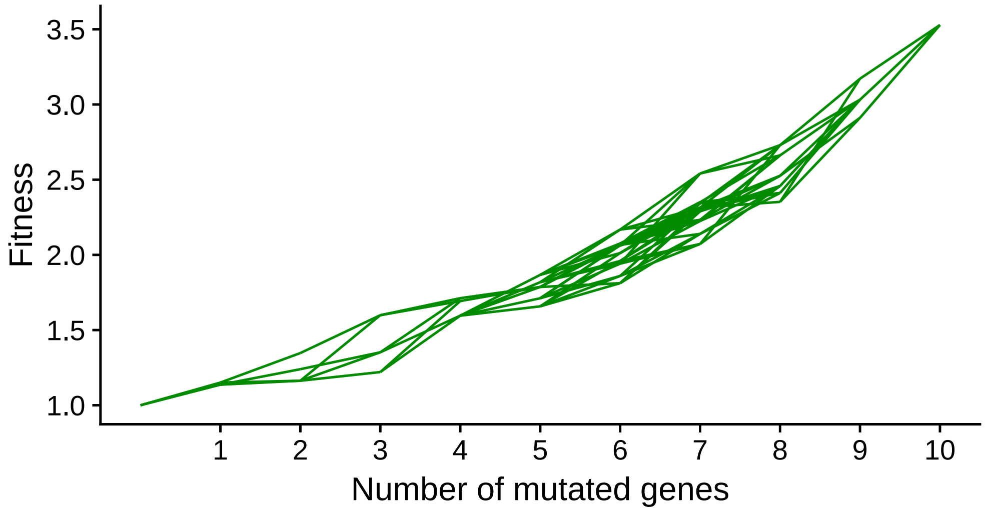

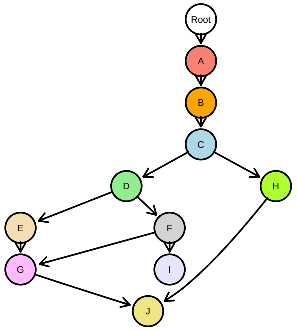


| | |
|---|---|
| *ID* | BkSjd8cAWeN4eVyO |
| *acc genotypes* | 23 |
| *peaks no back* | 1 |
| *removed edges* | 0 |
| *prop rem edges* | 0 |

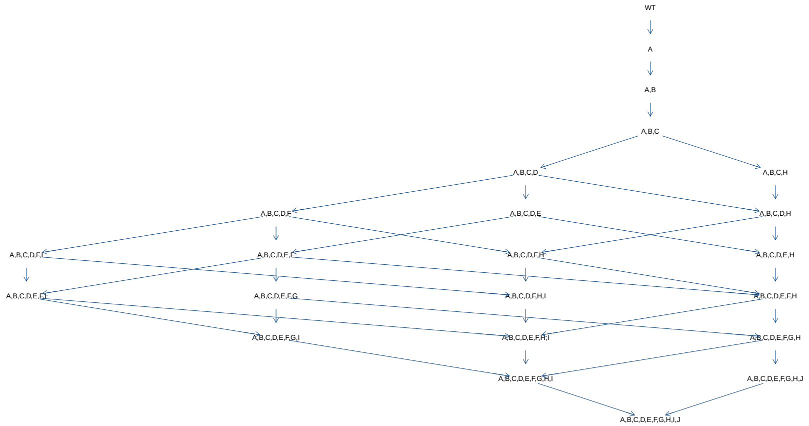


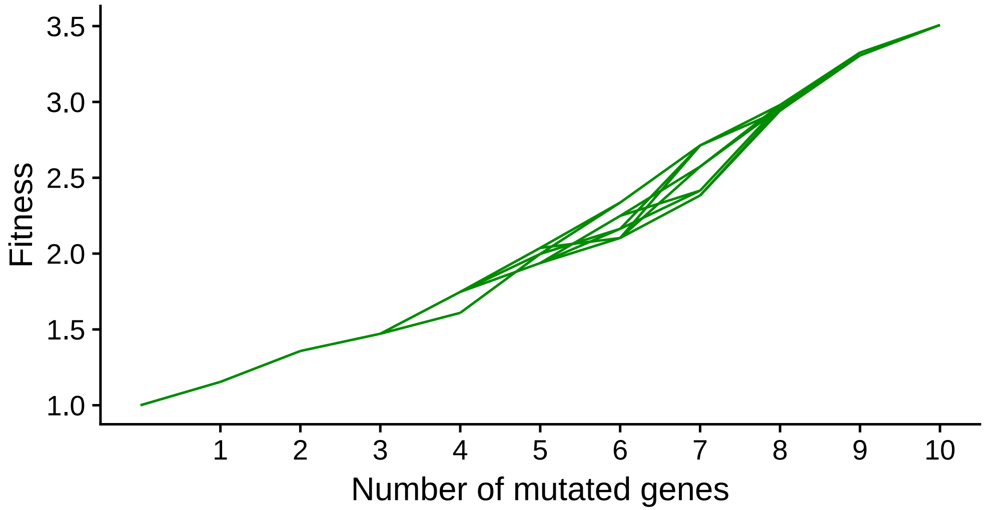

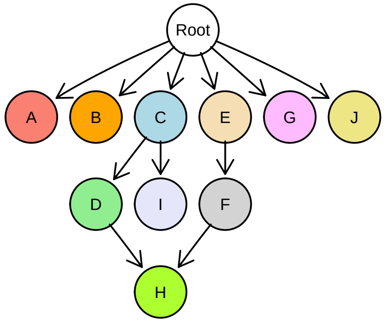


| | |
|---|---|
| *ID* | BqTdapi2oHdGJNeQ |
| *acc genotypes* | 272 |
| *peaks no back* | 1 |
| *removed edges* | 0 |
| *prop rem edges* | 0 |

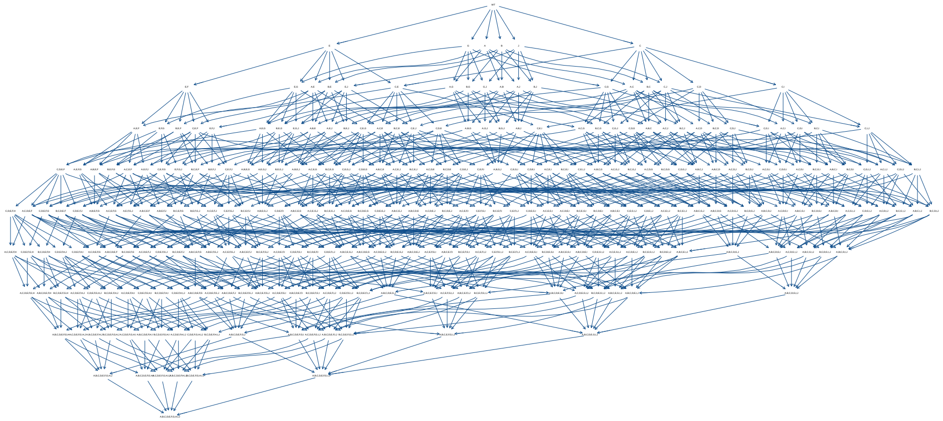

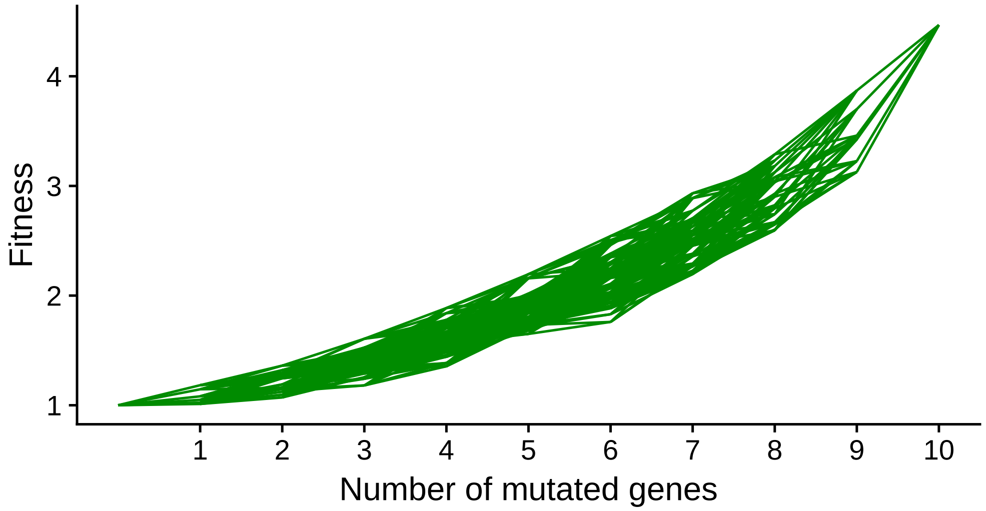

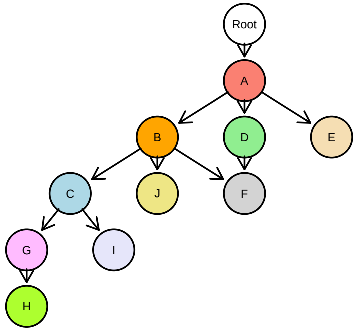


| | |
|---|---|
| *ID* | bTi93VZQ3TjAOVaY |
| *acc genotypes* | 89 |
| *peaks no back* | 1 |
| *removed edges* | 0 |
| *prop rem edges* | 0 |

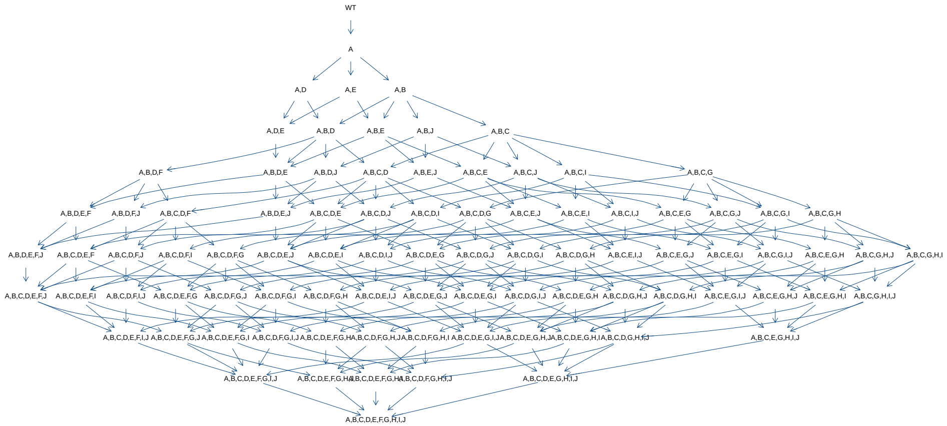

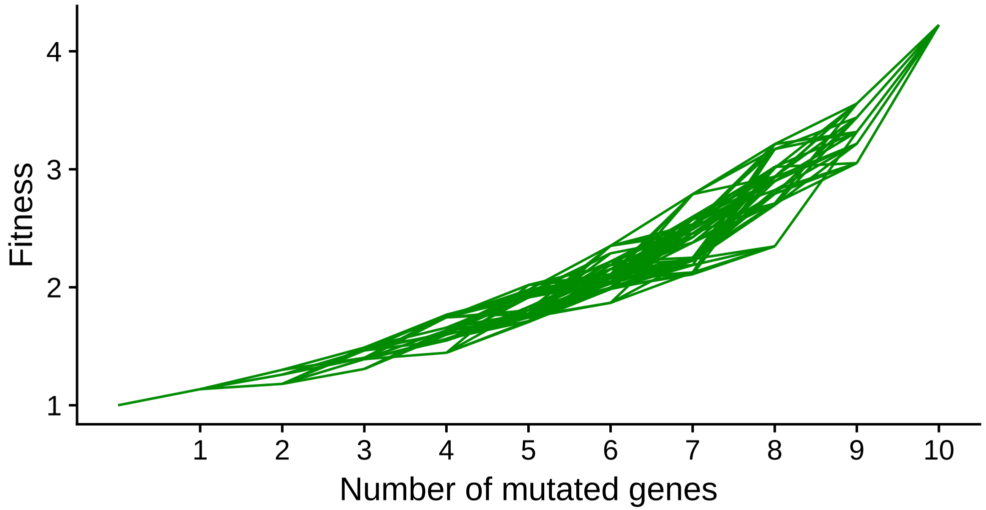

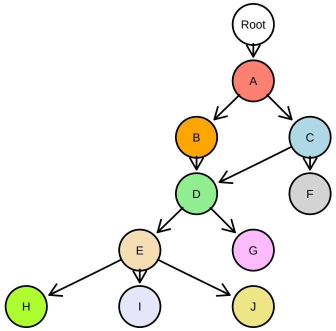


| | |
|---|---|
| *ID* | BU2srhsWr1UElYvS |
| *acc genotypes* | 43 |
| *peaks no back* | 1 |
| *removed edges* | 0 |
| *prop rem edges* | 0 |

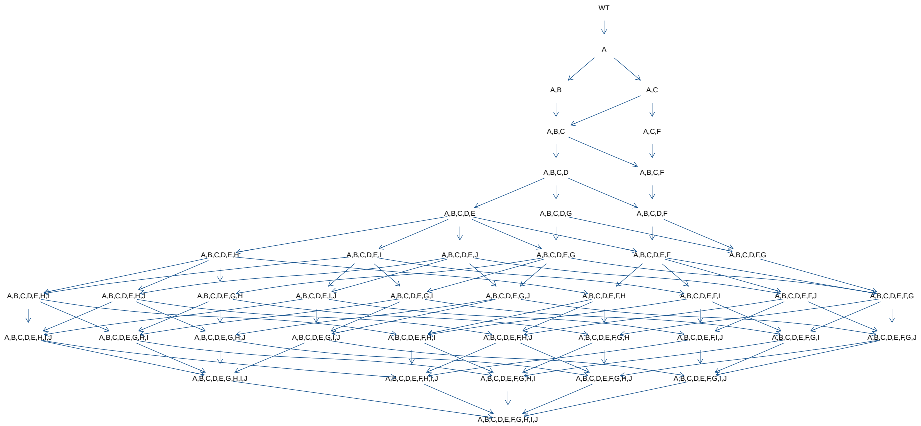

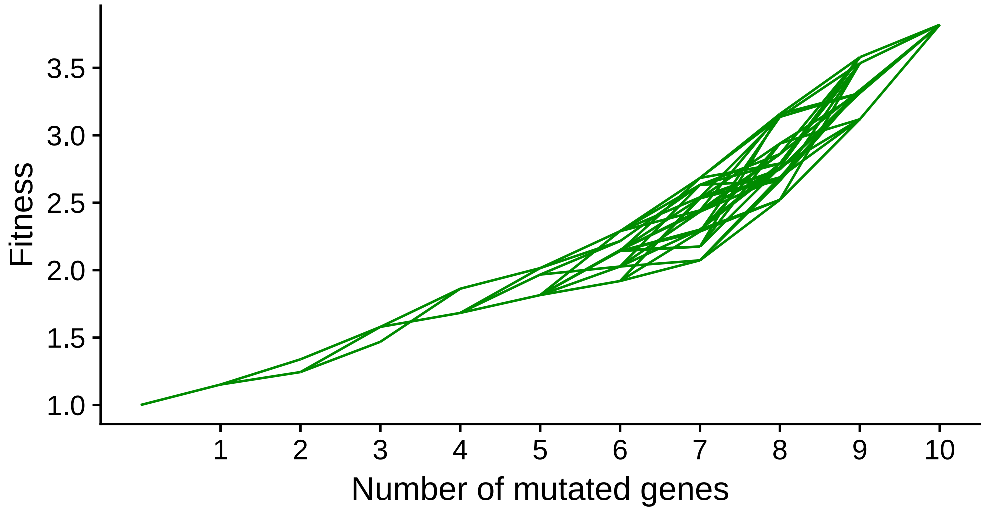

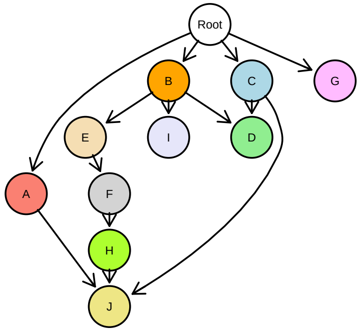


| | |
|---|---|
| *ID* | bv105Mh6lDH10HPdOB |
| *acc genotypes* | 112 |
| *peaks no back* | 1 |
| *removed edges* | 0 |
| *prop rem edges* | 0 |

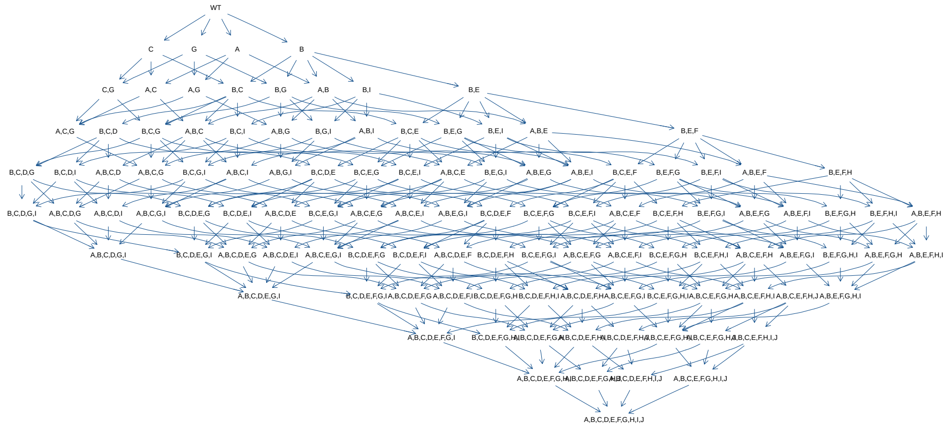

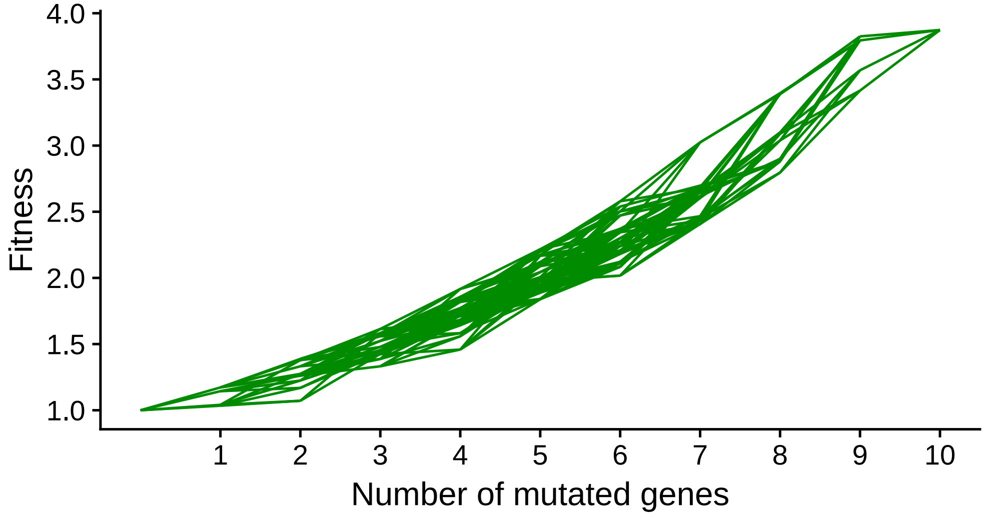

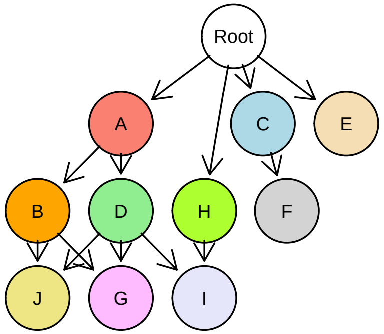


| | |
|---|---|
| *ID* | BWkhGPzysoNbGoq1 |
| *acc genotypes* | 126 |
| *peaks no back* | 1 |
| *removed edges* | 0 |
| *prop rem edges* | 0 |

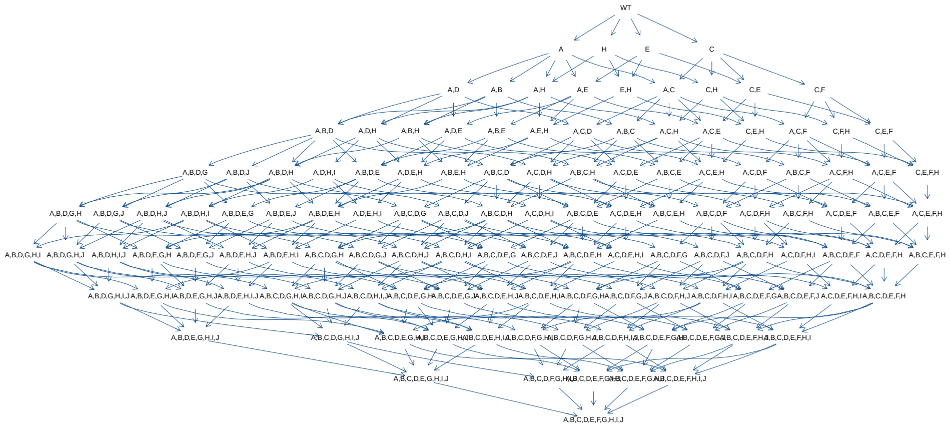

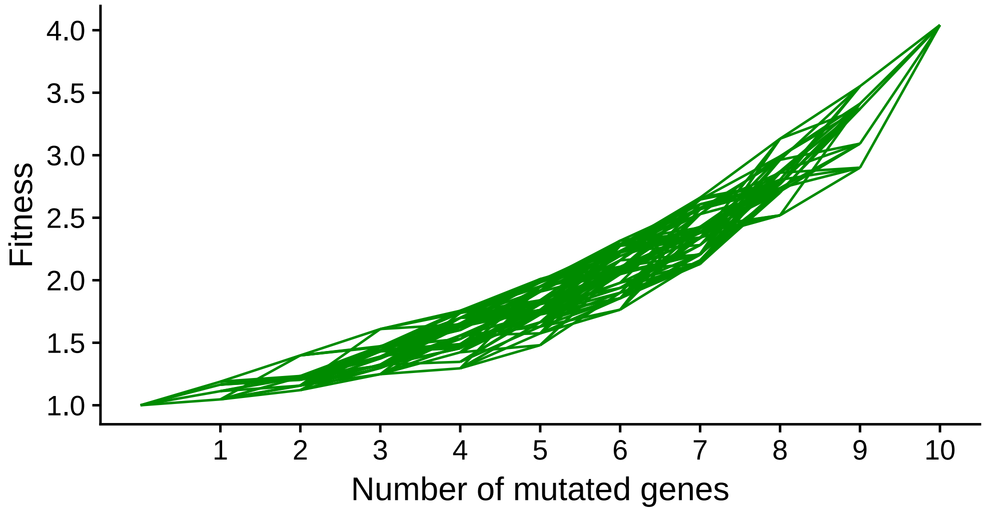

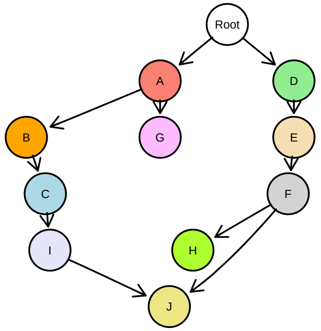


*ID* bXvsBjdeGDgvk10nj
*acc genotypes* 49
*peaks no back* 1
*removed edges* 0
*prop rem edges* 0

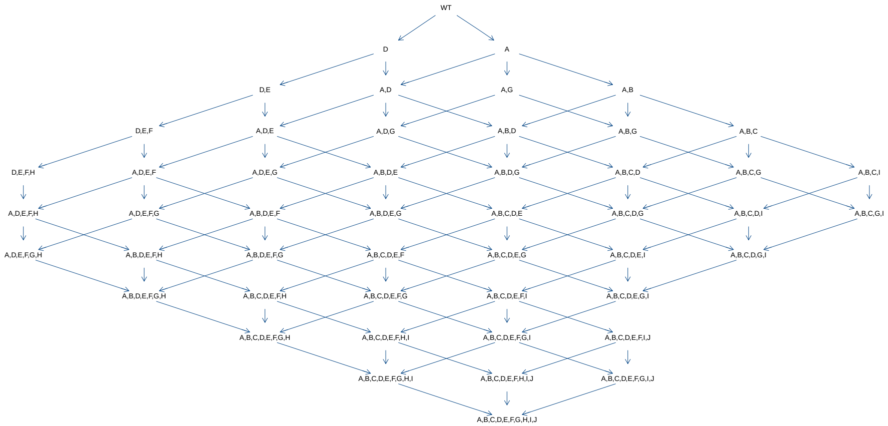

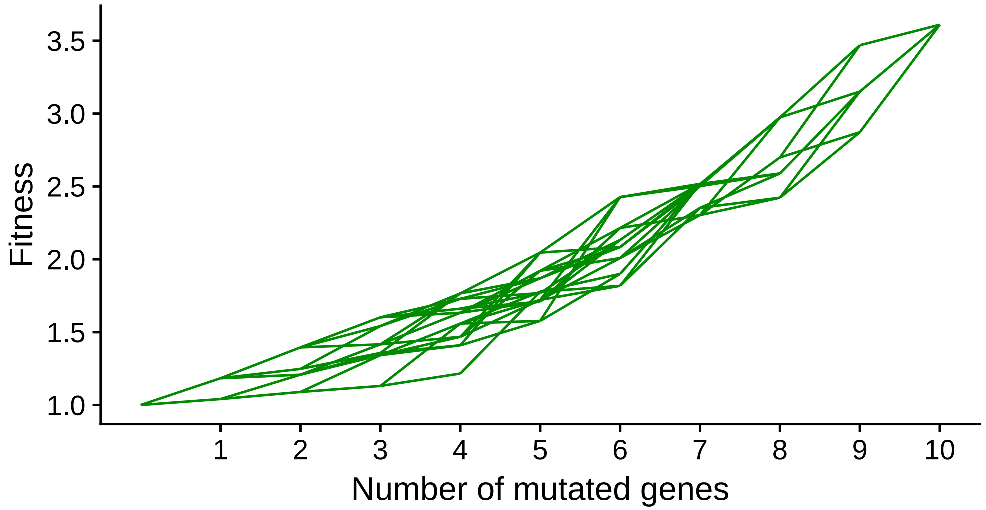

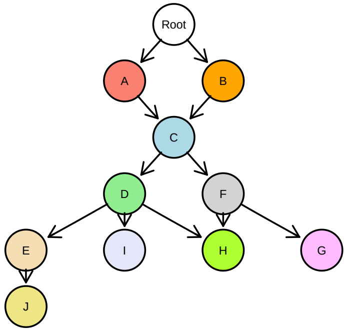


| | |
|---|---|
| *ID* | c4ImcRlDBXQAiAza |
| *acc genotypes* | 37 |
| *peaks no back* | 1 |
| *removed edges* | 0 |
| *prop rem edges* | 0 |

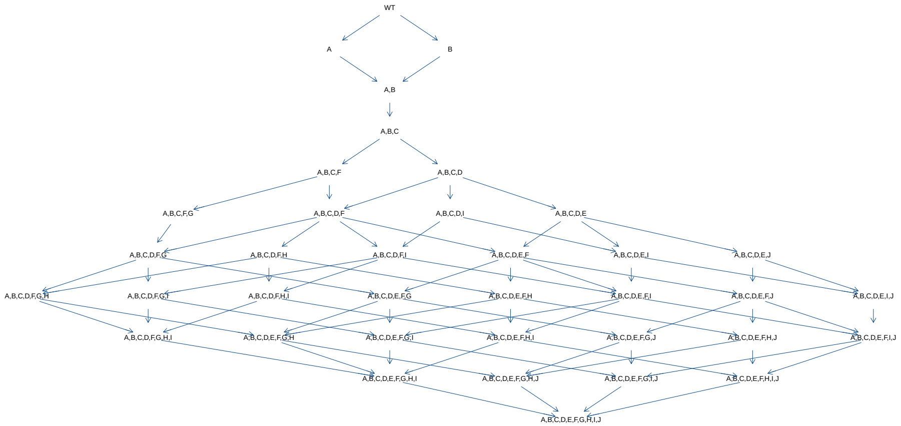

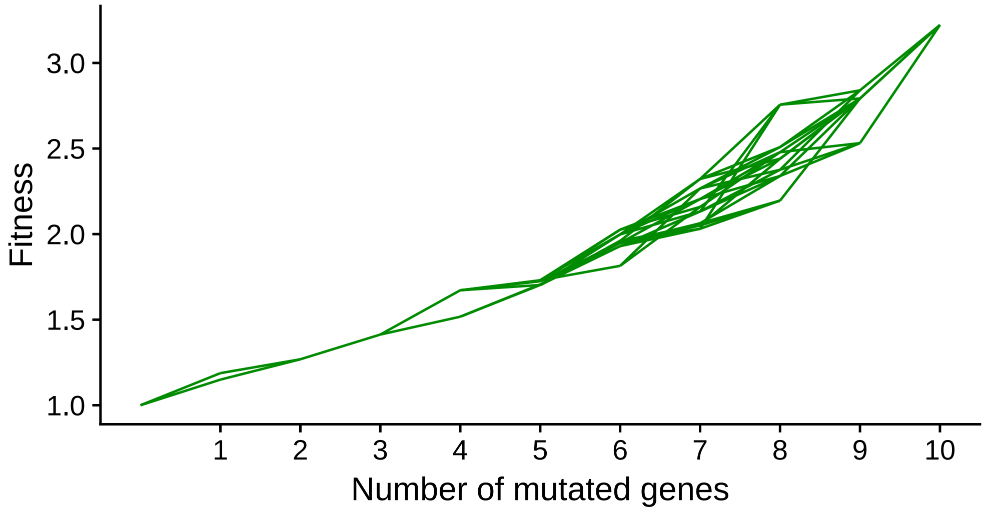

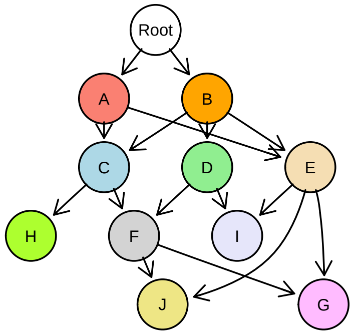


| | |
|---|---|
| *ID* | cczLUqQeSfB6Czbr |
| *acc genotypes* | 37 |
| *peaks no back* | 1 |
| *removed edges* | 0 |
| *prop rem edges* | 0 |

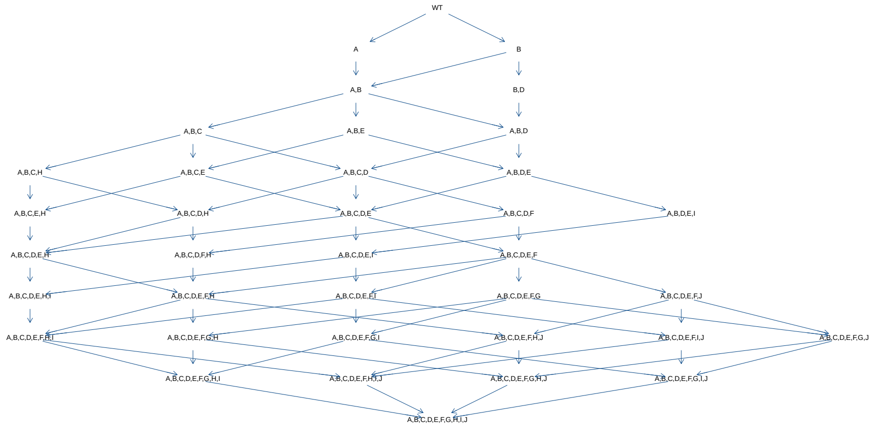

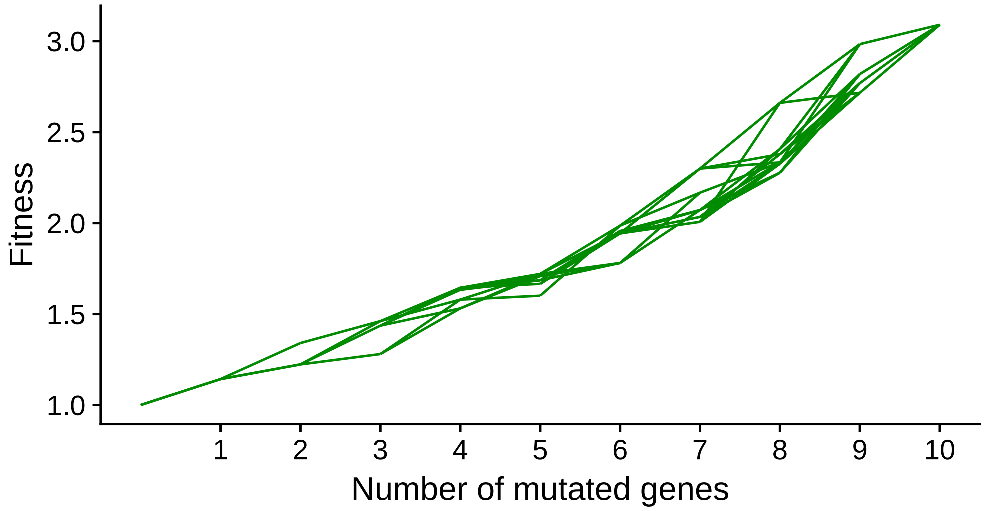

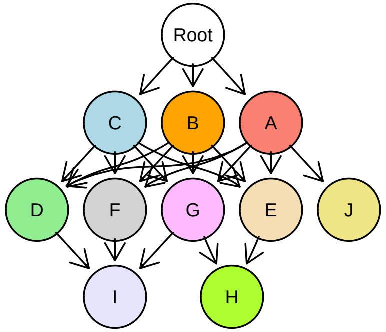


| | |
|---|---|
| *ID* | CEKKz1011cQcW3hT7 |
| *acc genotypes* | 56 |
| *peaks no back* | 1 |
| *removed edges* | 0 |
| *prop rem edges* | 0 |

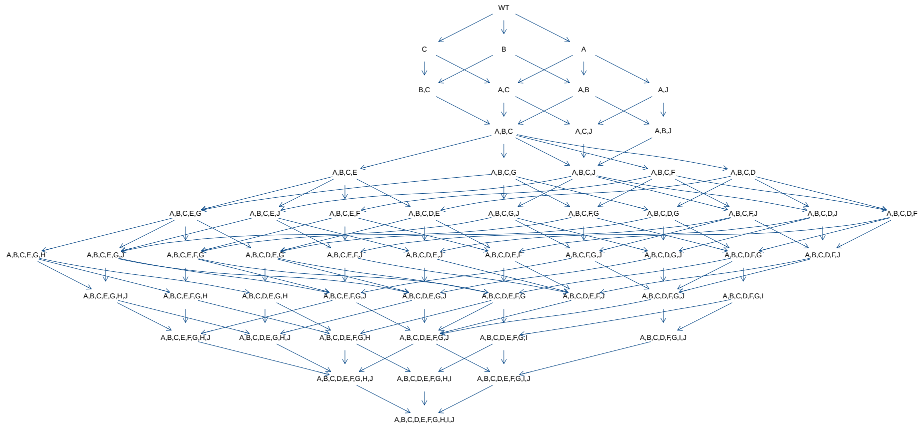

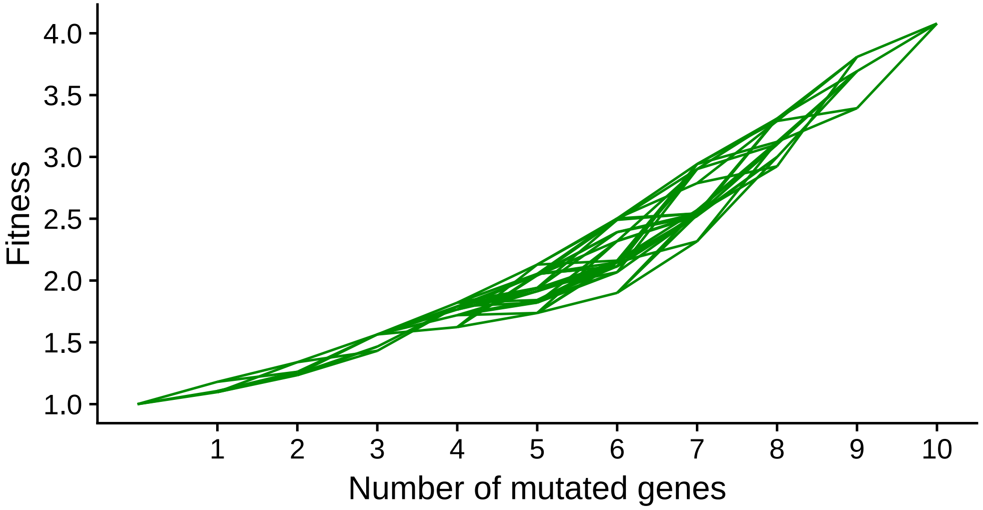

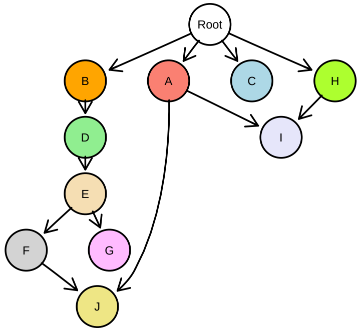


| | |
|---|---|
| *ID* | COhbmk810lA2tC32X |
| *acc genotypes* | 82 |
| *peaks no back* | 1 |
| *removed edges* | 0 |
| *prop rem edges* | 0 |

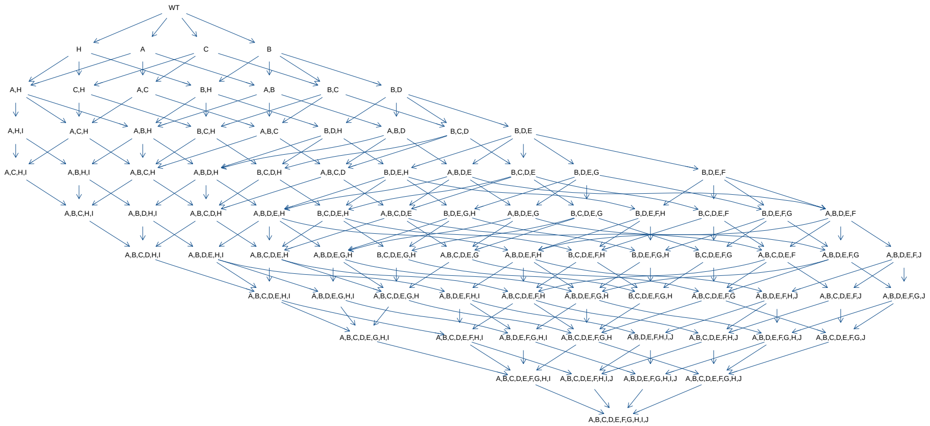

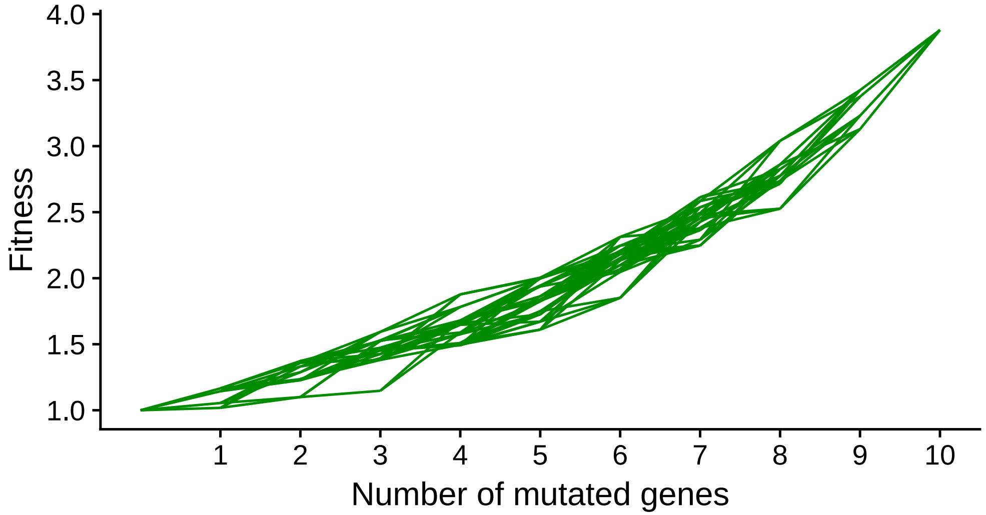

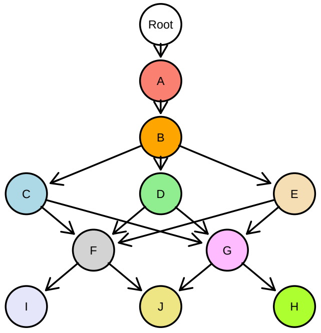


| | |
|---|---|
| *ID* | CRSLV974aXfmgnX10 |
| *acc genotypes* | 22 |
| *peaks no back* | 1 |
| *removed edges* | 0 |
| *prop rem edges* | 0 |

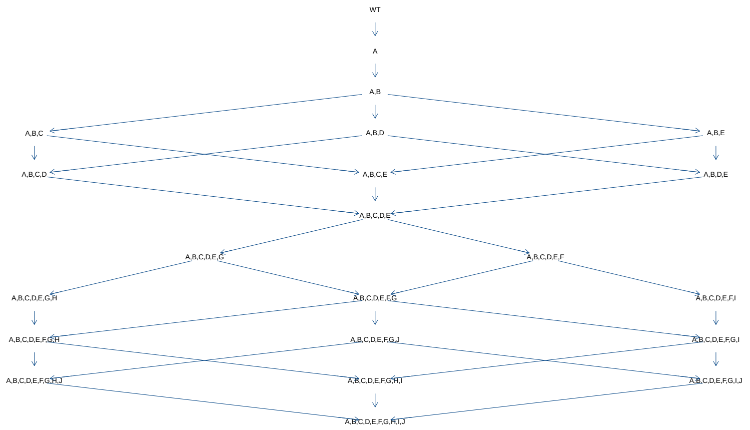


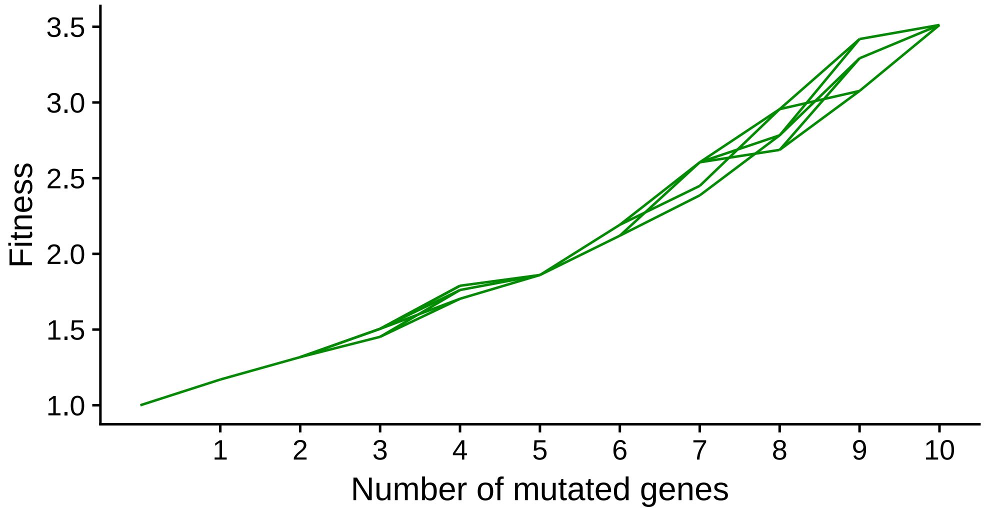

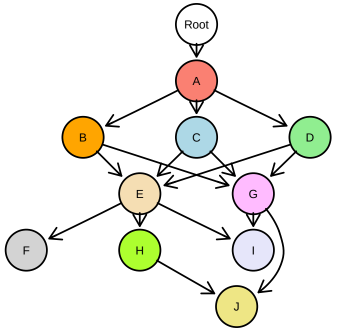


| | |
|---|---|
| *ID* | cWla4xvRzx8wVebz |
| *acc genotypes* | 26 |
| *peaks no back* | 1 |
| *removed edges* | 0 |
| *prop rem edges* | 0 |

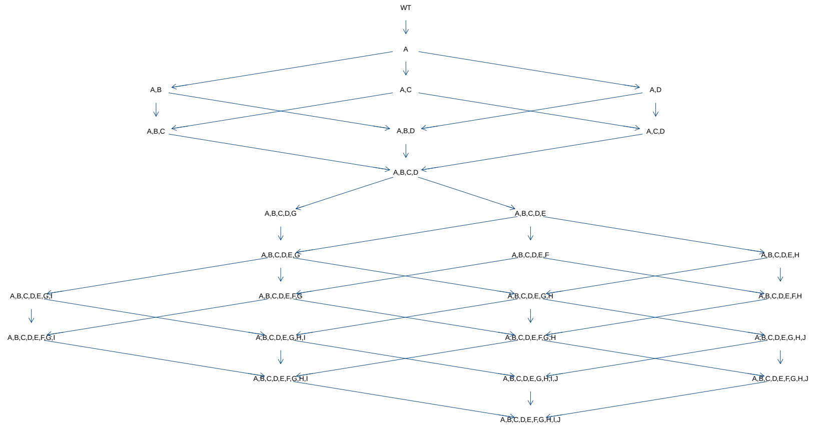


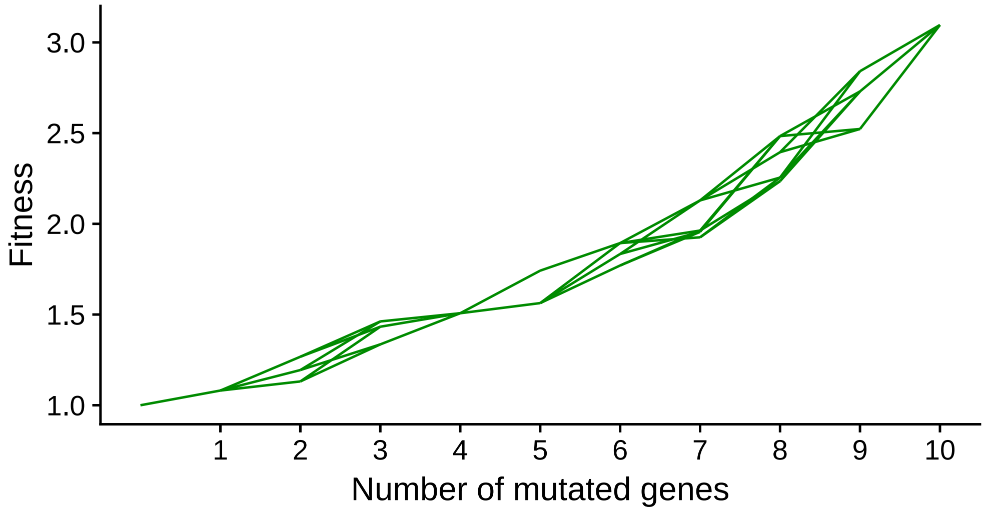

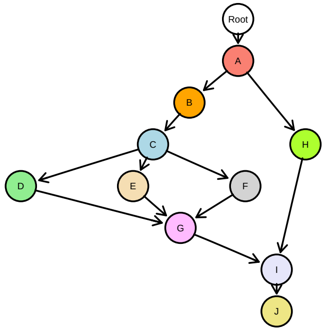


| | |
|---|---|
| *ID* | d1i86X4sy10ttNPpW |
| *acc genotypes* | 25 |
| *peaks no back* | 1 |
| *removed edges* | 0 |
| *prop rem edges* | 0 |

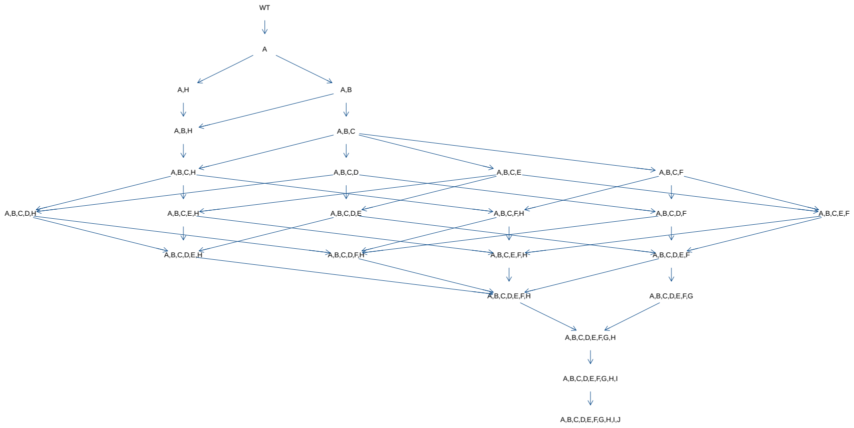


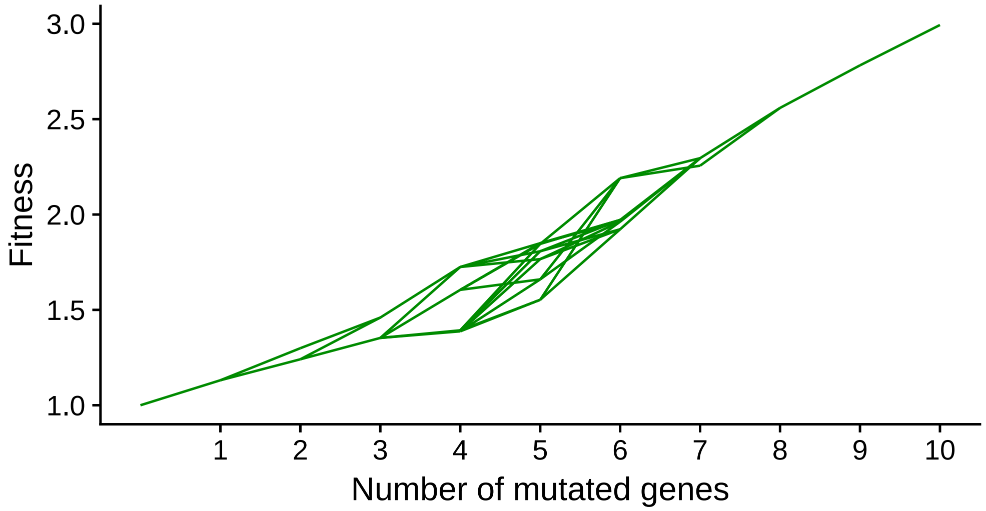

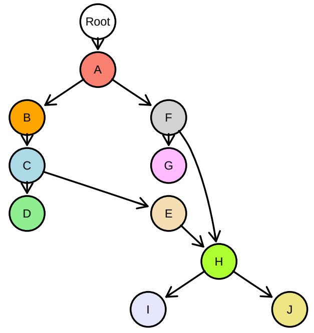


| | |
|---|---|
| *ID* | D3yYb2Y5LZWHMlC9 |
| *acc genotypes* | 35 |
| *peaks no back* | 1 |
| *removed edges* | 0 |
| *prop rem edges* | 0 |

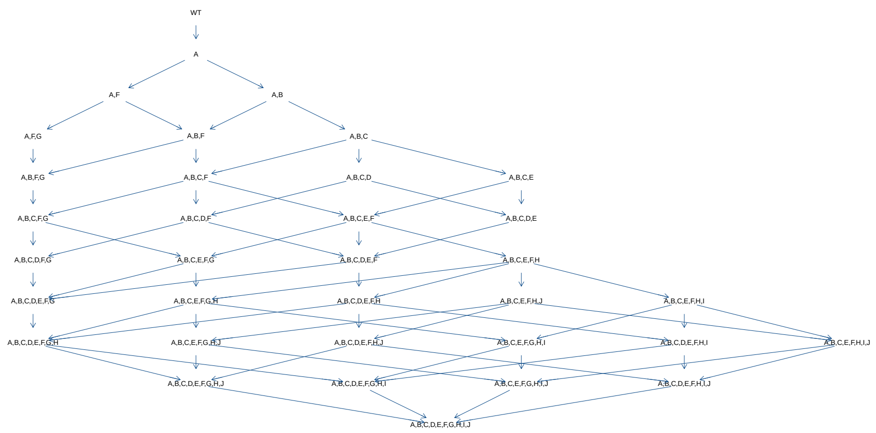

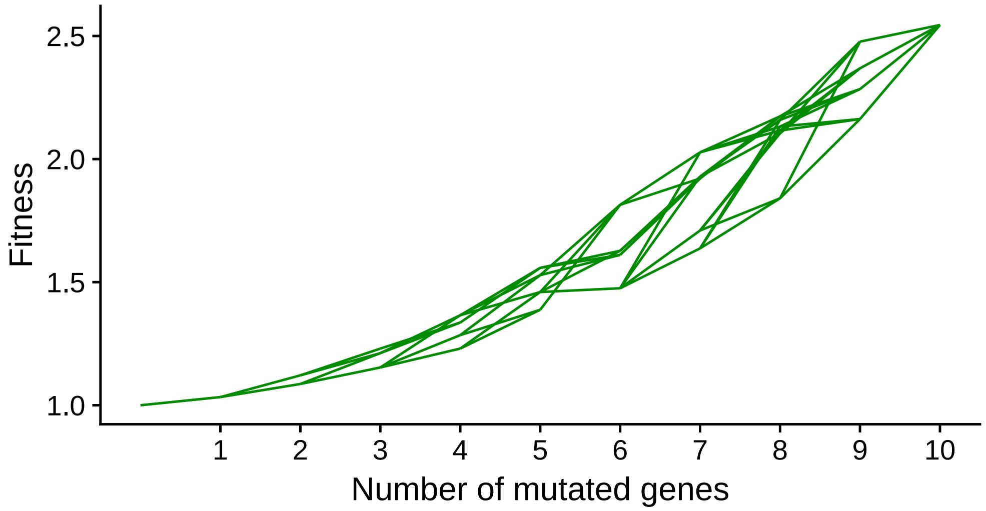

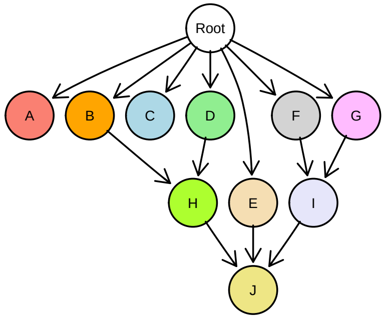


| | |
|---|---|
| *ID* | DishuUtqtuc3gwcU |
| *acc genotypes* | 204 |
| *peaks no back* | 1 |
| *removed edges* | 0 |
| *prop rem edges* | 0 |

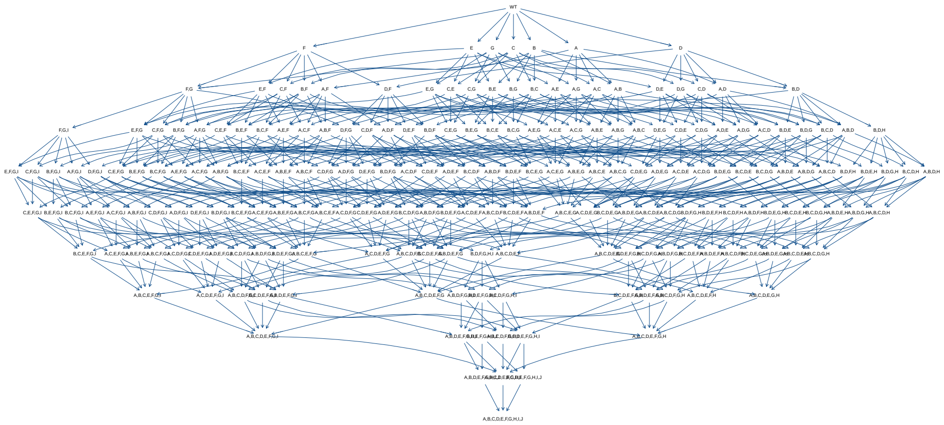

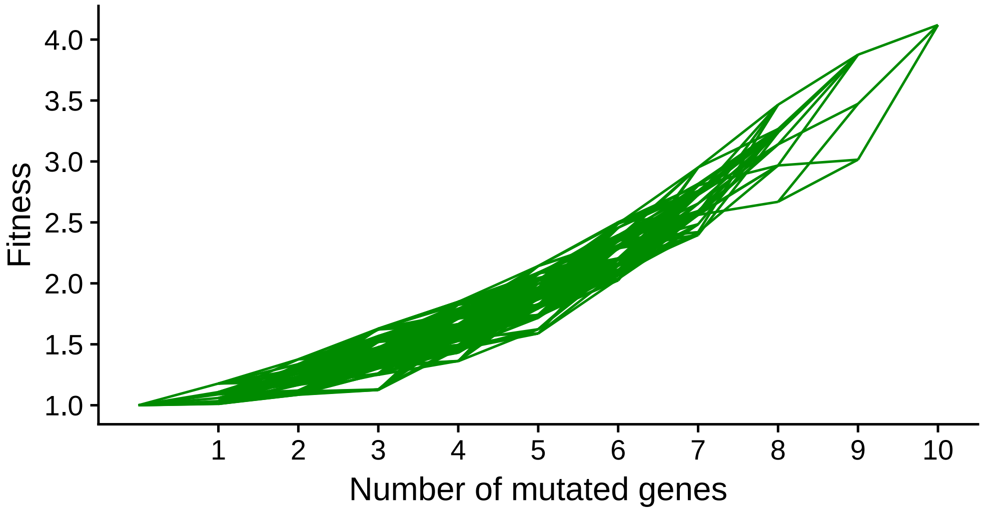

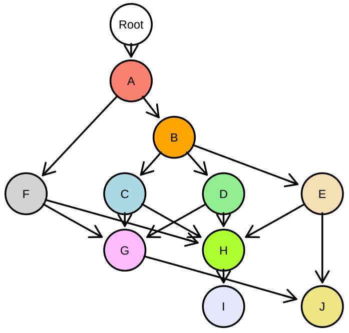


| | |
|---|---|
| *ID* | DLJ64zEF6RPBgmZU |
| *acc genotypes* | 28 |
| *peaks no back* | 1 |
| *removed edges* | 0 |
| *prop rem edges* | 0 |

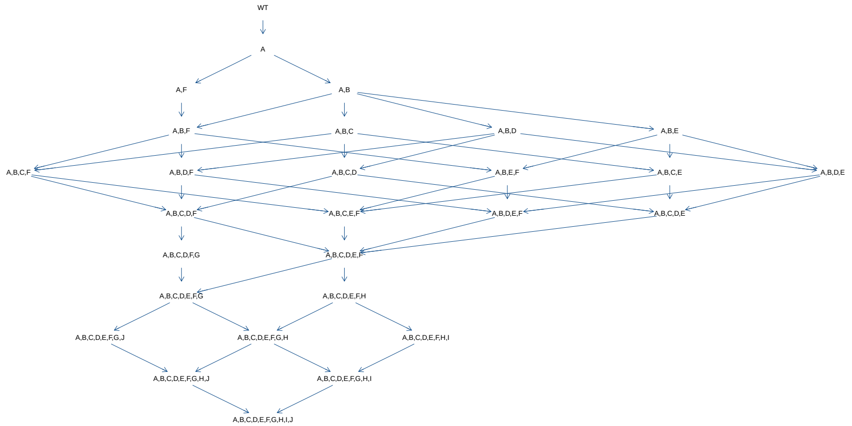


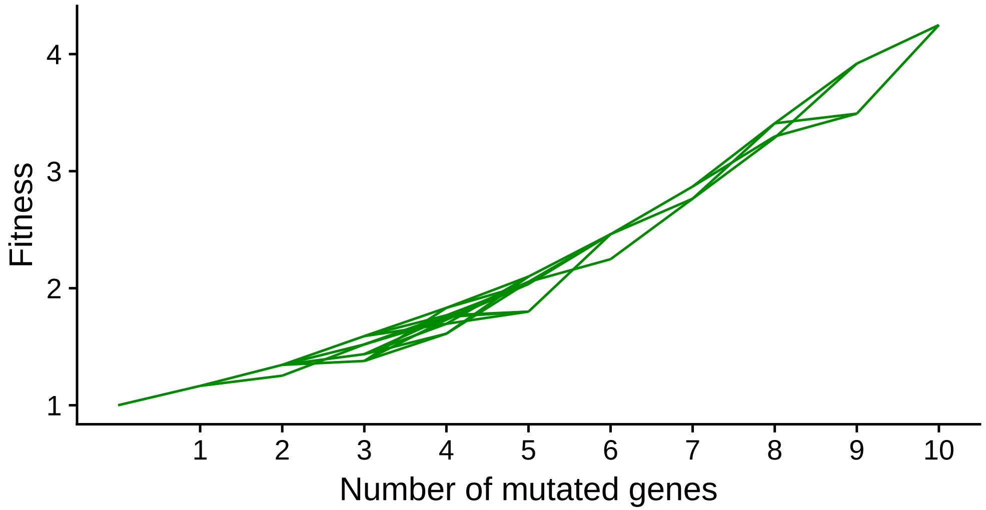

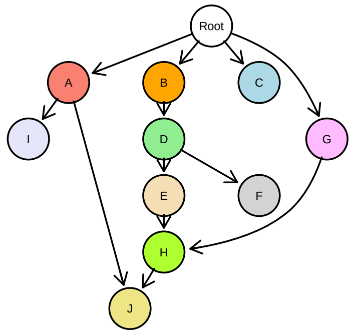


| | |
|---|---|
| *ID* | dvBzM4I8nentNLyl |
| *acc genotypes* | 92 |
| *peaks no back* | 1 |
| *removed edges* | 0 |
| *prop rem edges* | 0 |

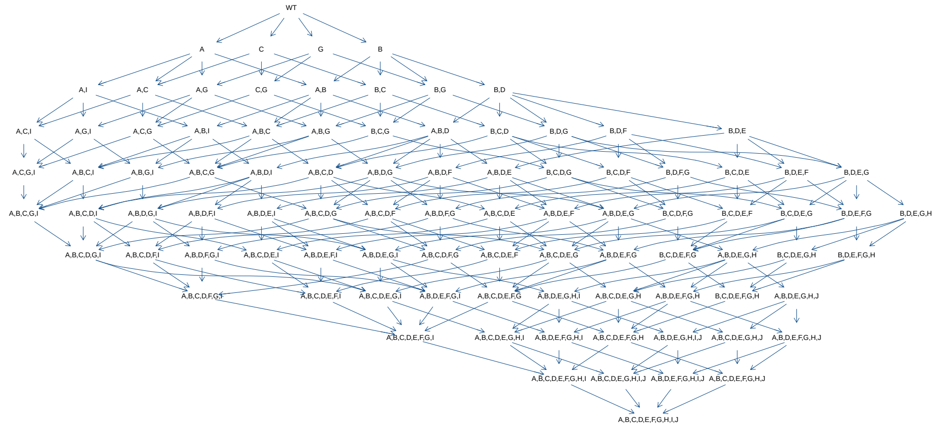

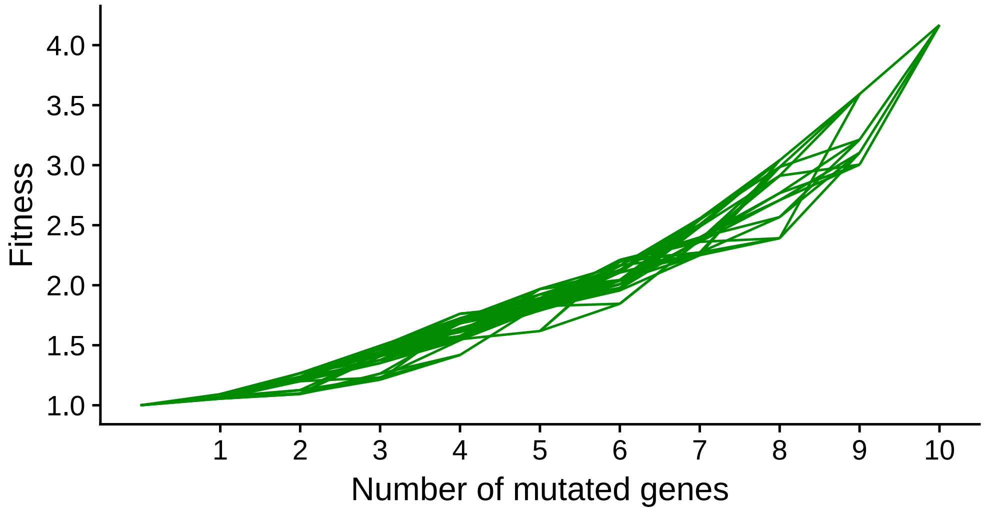

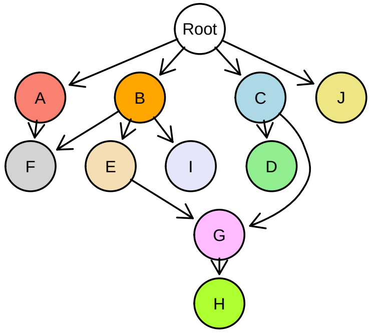


| | |
|---|---|
| *ID* | dvQzUhETlllrU97u |
| *acc genotypes* | 132 |
| *peaks no back* | 1 |
| *removed edges* | 0 |
| *prop rem edges* | 0 |

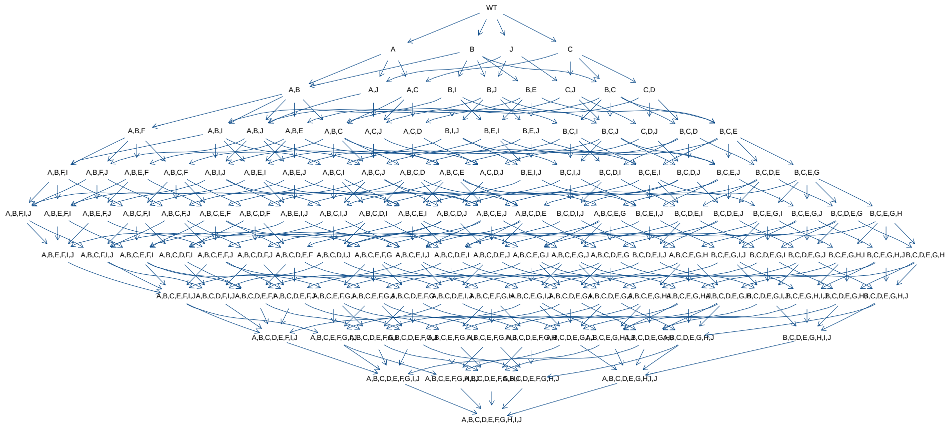

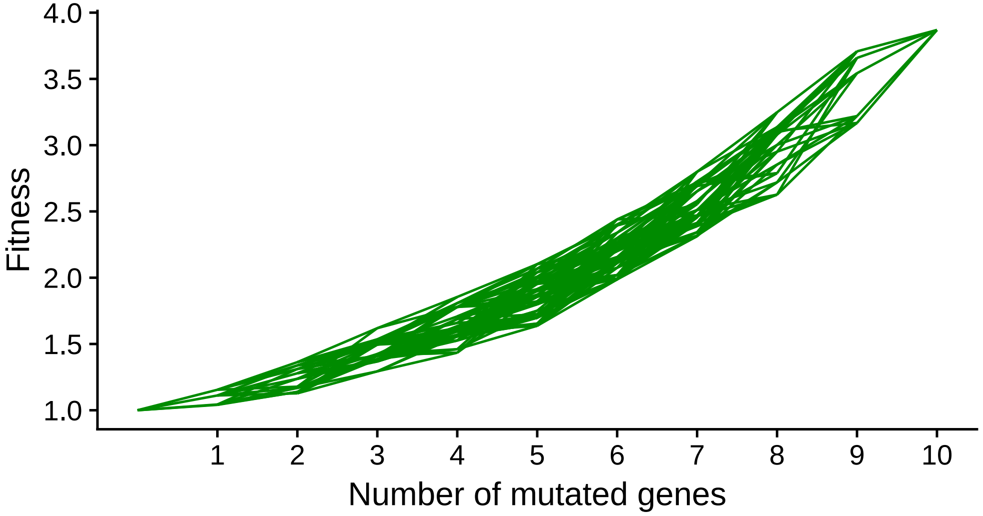

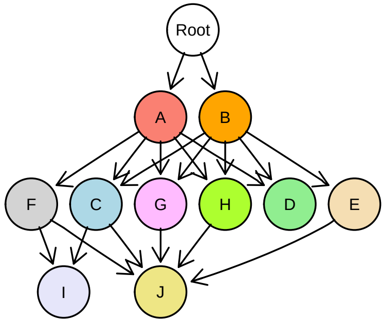


*ID* EBt810tAfY9D105FkY
*acc genotypes* 89
*peaks no back* 1
*removed edges* 0
*prop rem edges* 0

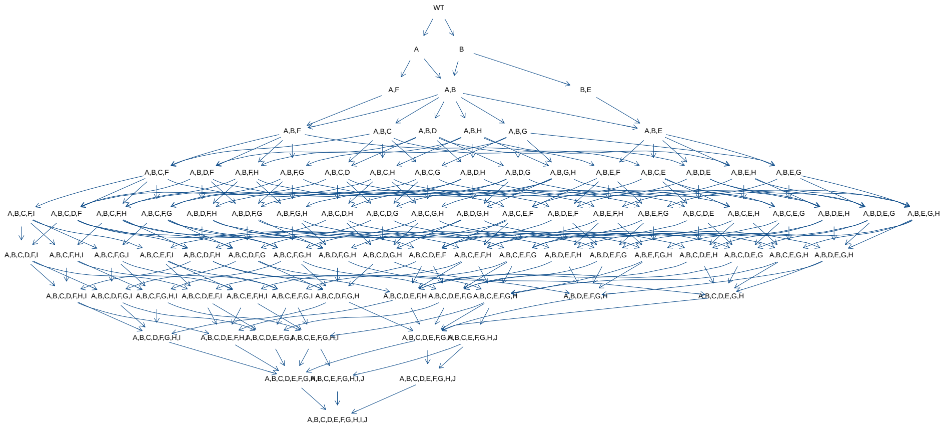

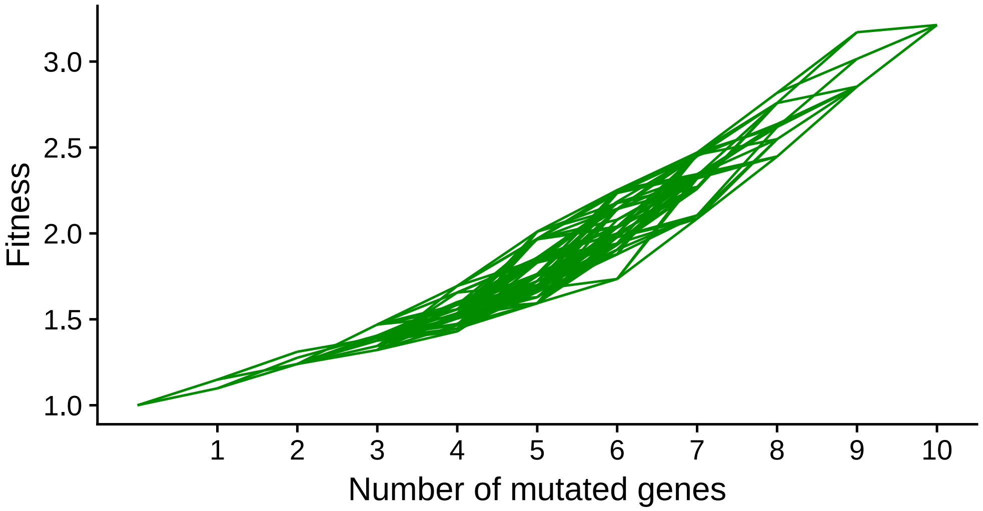

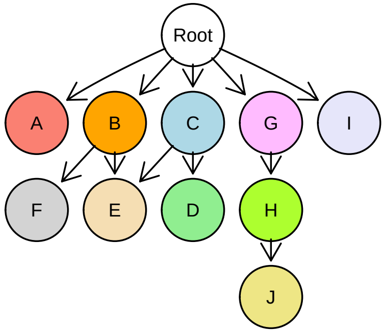


| | |
|---|---|
| *ID* | ekTHm442hqA10lVJm |
| *acc genotypes* | 208 |
| *peaks no back* | 1 |
| *removed edges* | 0 |
| *prop rem edges* | 0 |

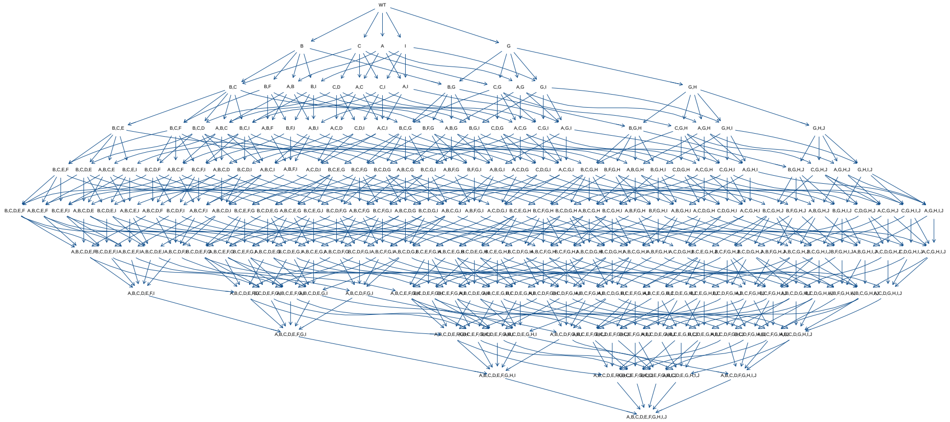

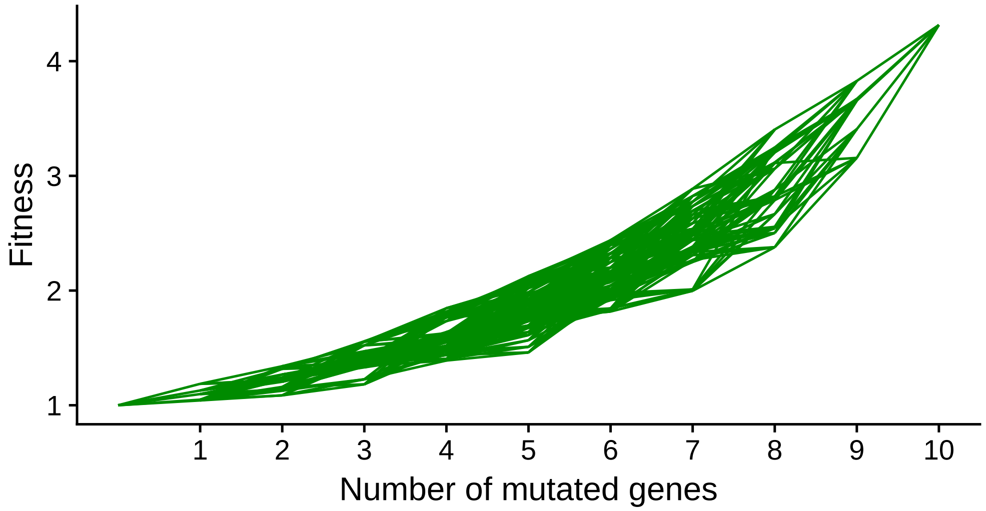

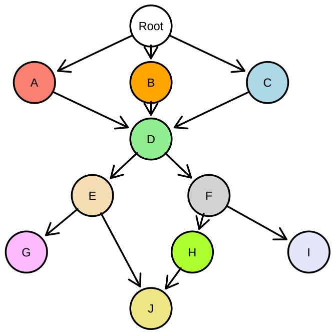


| | |
|---|---|
| *ID* | EkuM7k8AJLiPR6hS |
| *acc genotypes* | 27 |
| *peaks no back* | 1 |
| *removed edges* | 0 |
| *prop rem edges* | 0 |

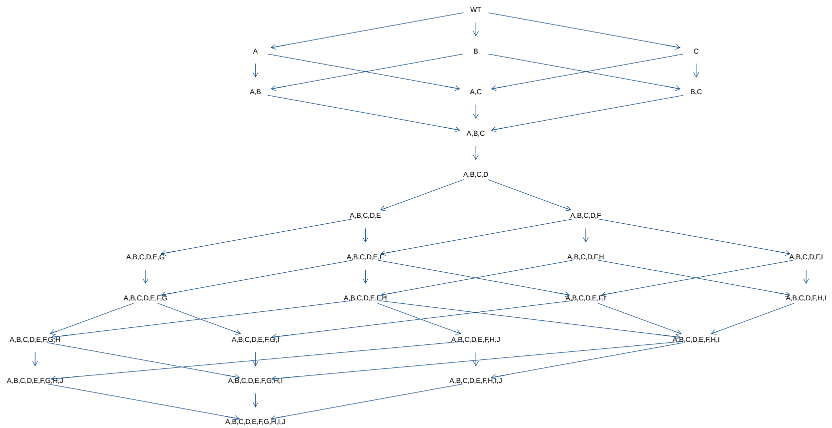


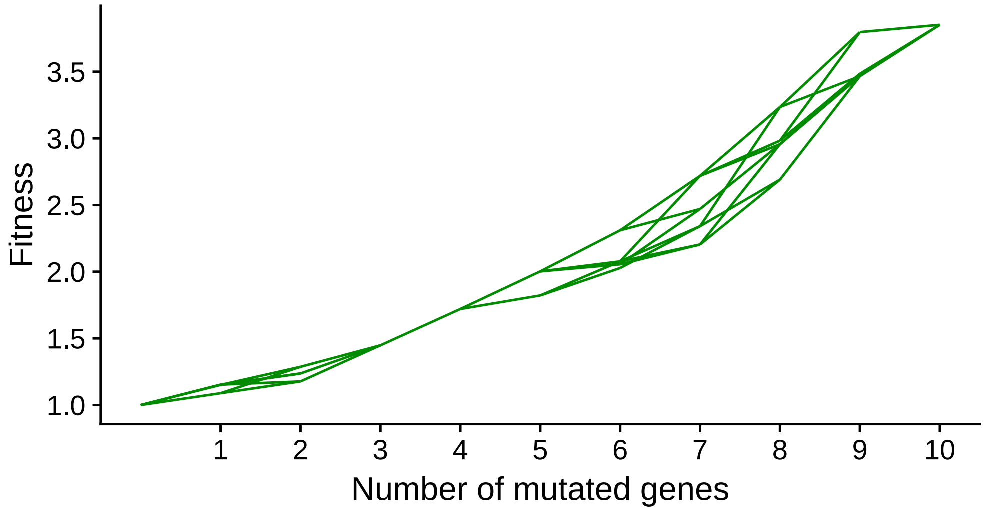

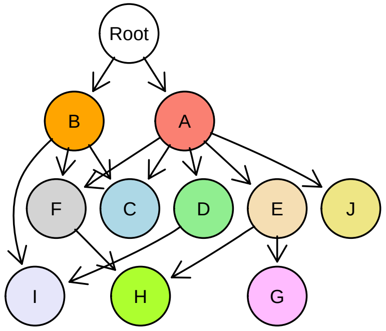


| | |
|---|---|
| *ID* | eNPbGhByo2yD8VoG |
| *acc genotypes* | 110 |
| *peaks no back* | 1 |
| *removed edges* | 0 |
| *prop rem edges* | 0 |

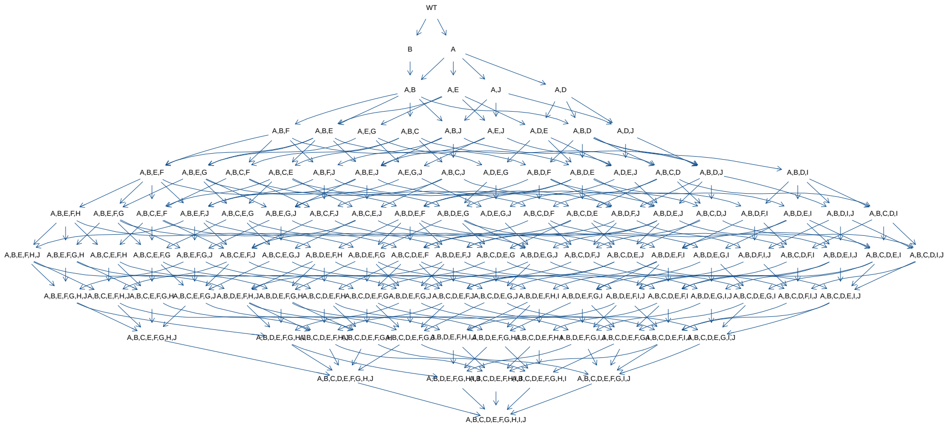


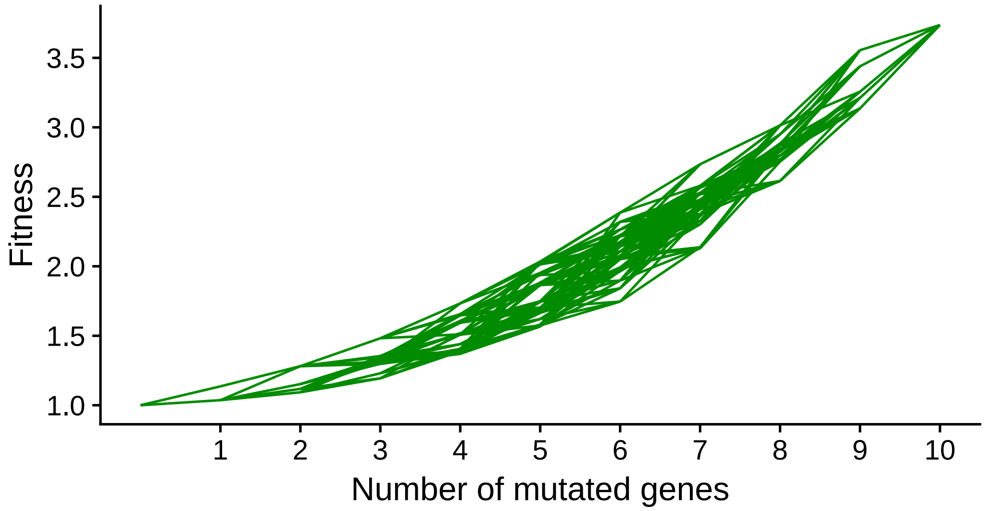

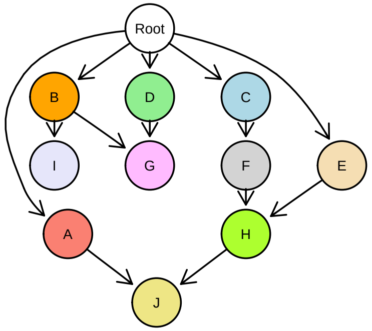


| | |
|---|---|
| *ID* | EoCzYRJurCmFz168 |
| *acc genotypes* | 120 |
| *peaks no back* | 1 |
| *removed edges* | 0 |
| *prop rem edges* | 0 |

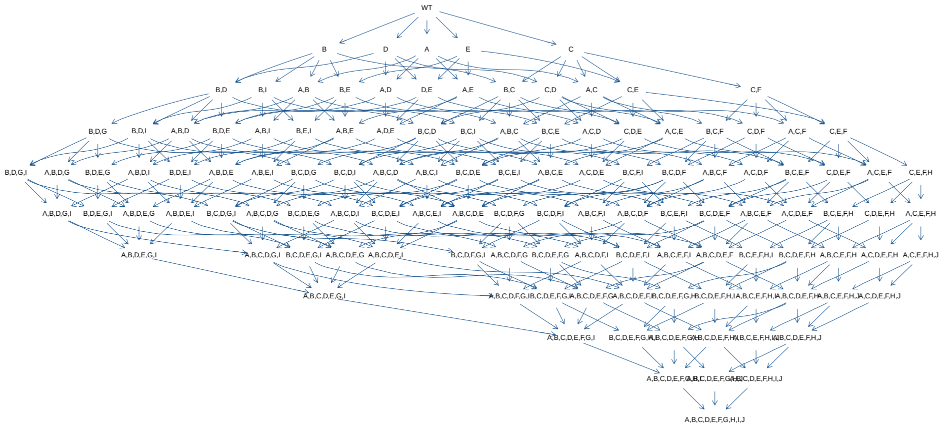

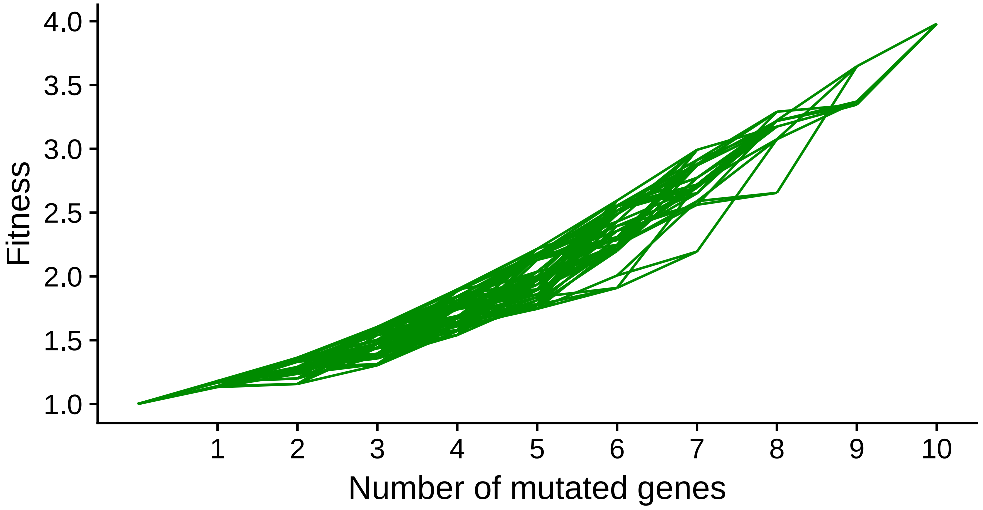

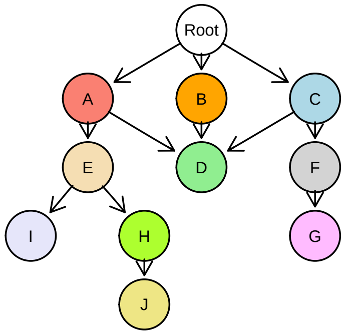


| | |
|---|---|
| *ID* | eOGzHUxNXJg7737x |
| *acc genotypes* | 85 |
| *peaks no back* | 1 |
| *removed edges* | 0 |
| *prop rem edges* | 0 |

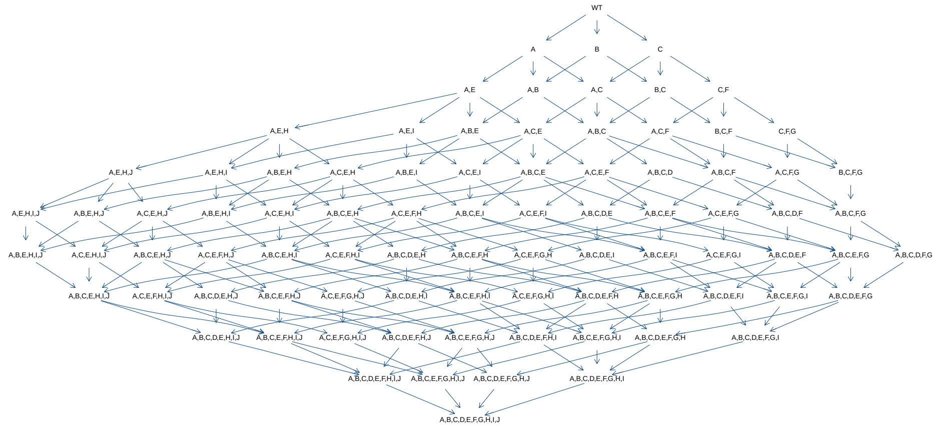

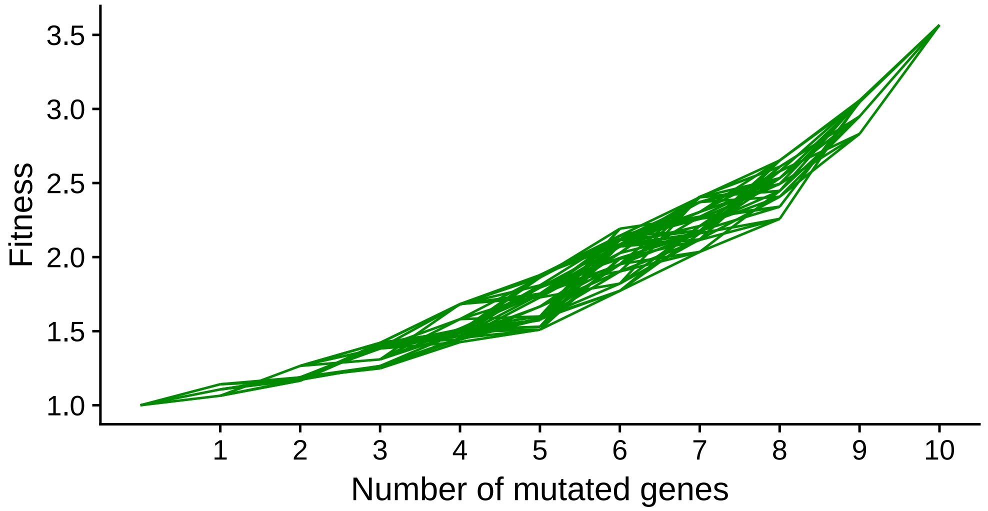

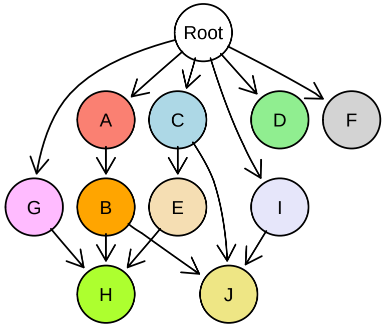


| | |
|---|---|
| *ID* | EScCMp2k63ciS5e1 |
| *acc genotypes* | 172 |
| *peaks no back* | 1 |
| *removed edges* | 0 |
| *prop rem edges* | 0 |

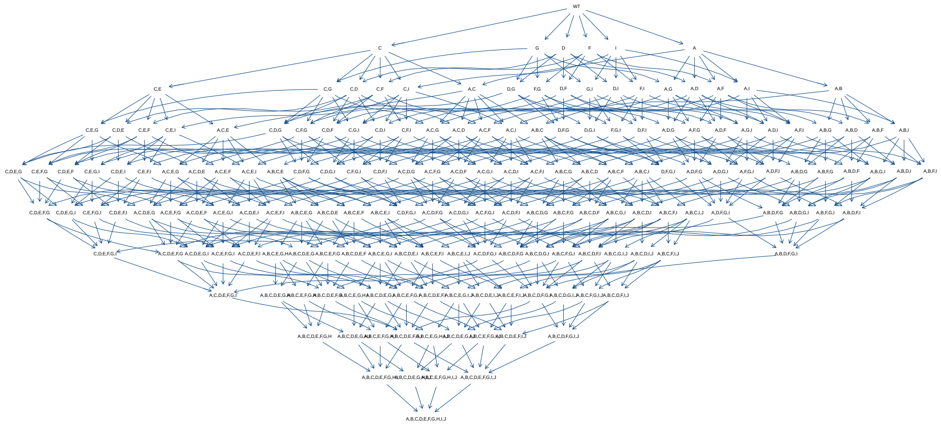

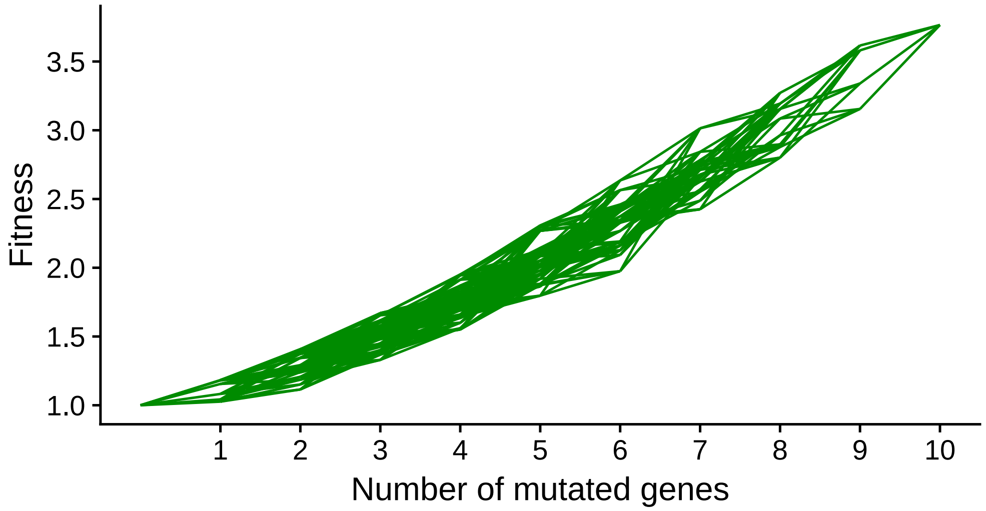

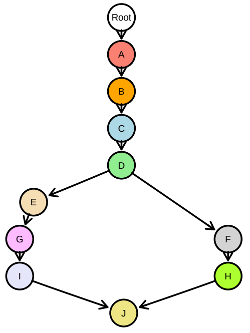


| | |
|---|---|
| *ID* | EUl10dI8bbHyhh4nE |
| *acc genotypes* | 17 |
| *peaks no back* | 1 |
| *removed edges* | 0 |
| *prop rem edges* | 0 |

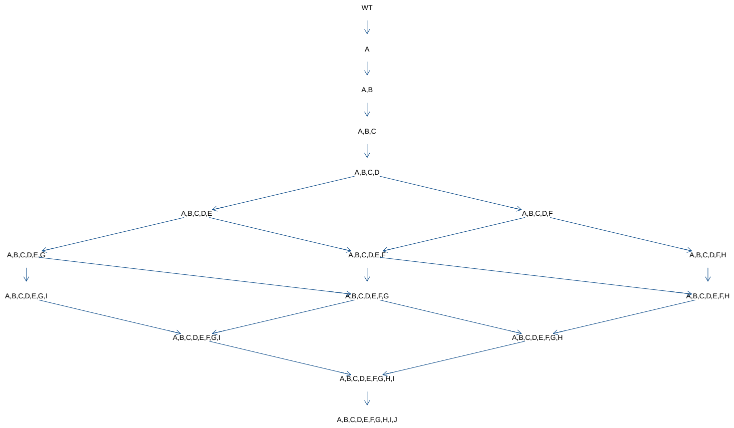


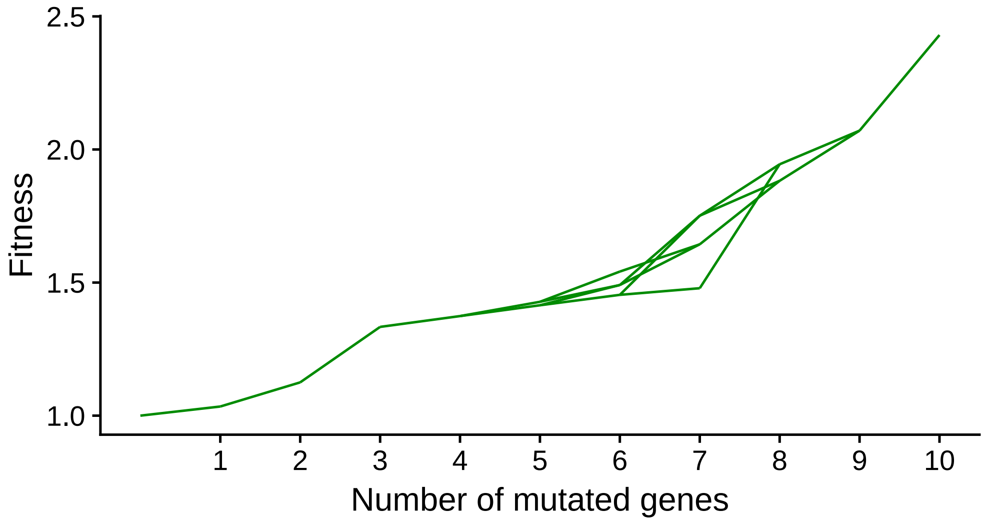

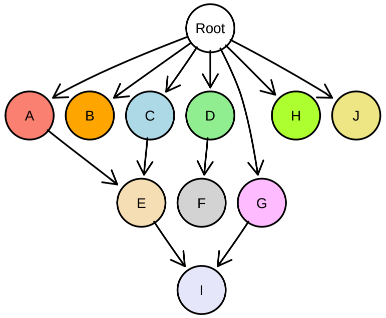


| | |
|---|---|
| *ID* | ExaZNcadWsiZp1os |
| *acc genotypes* | 264 |
| *peaks no back* | 1 |
| *removed edges* | 0 |
| *prop rem edges* | 0 |

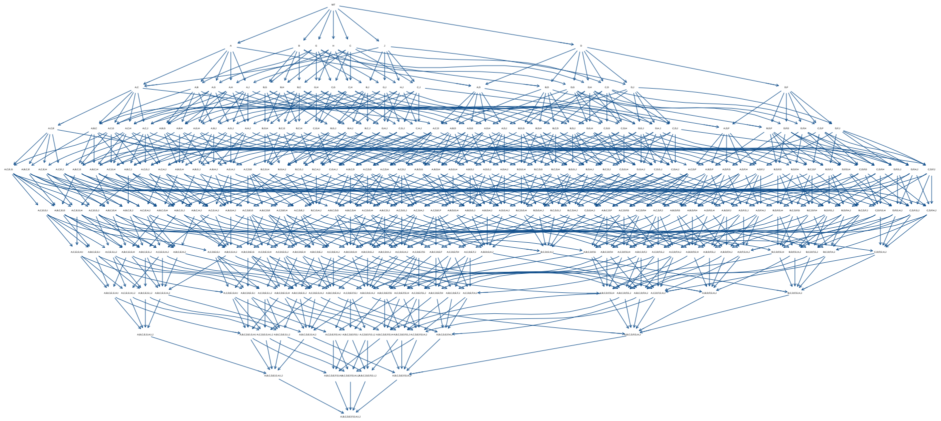

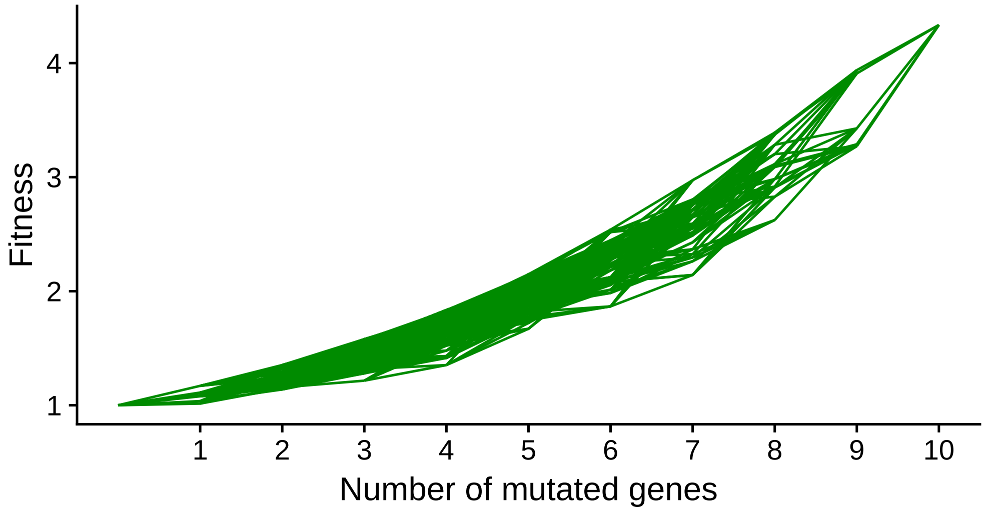

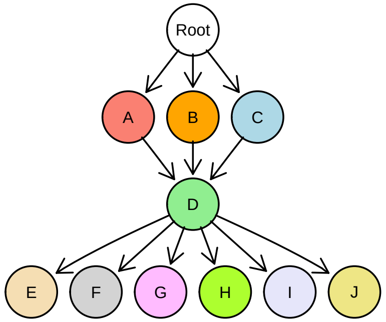


*ID* F74OrUlE2JUNgUl10
*acc genotypes* 72
*peaks no back* 1
*removed edges* 0
*prop rem edges* 0

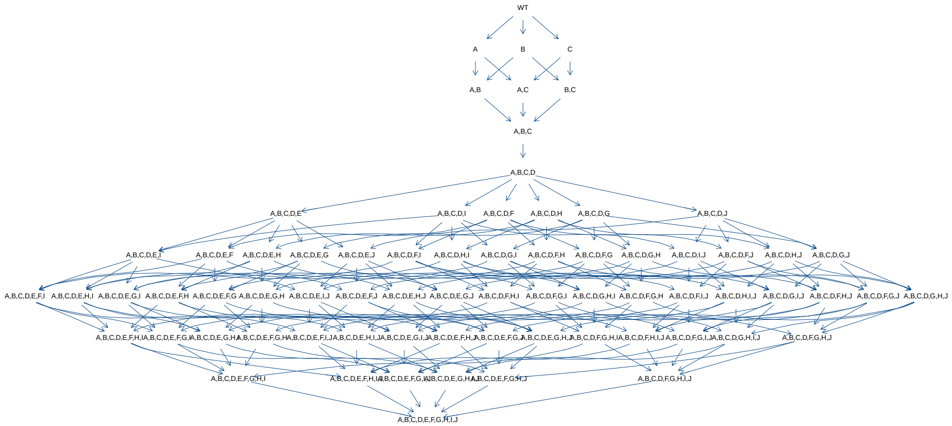

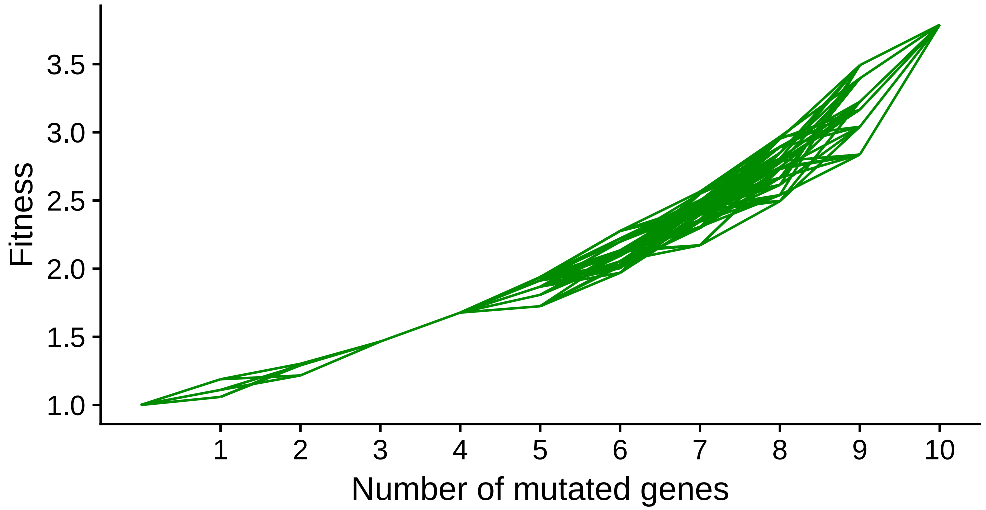

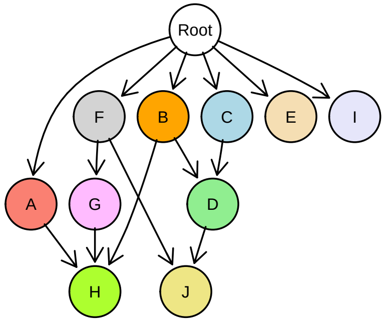


| | |
|---|---|
| *ID* | fDVyfwz5EixVTGkl |
| *acc genotypes* | 152 |
| *peaks no back* | 1 |
| *removed edges* | 0 |
| *prop rem edges* | 0 |

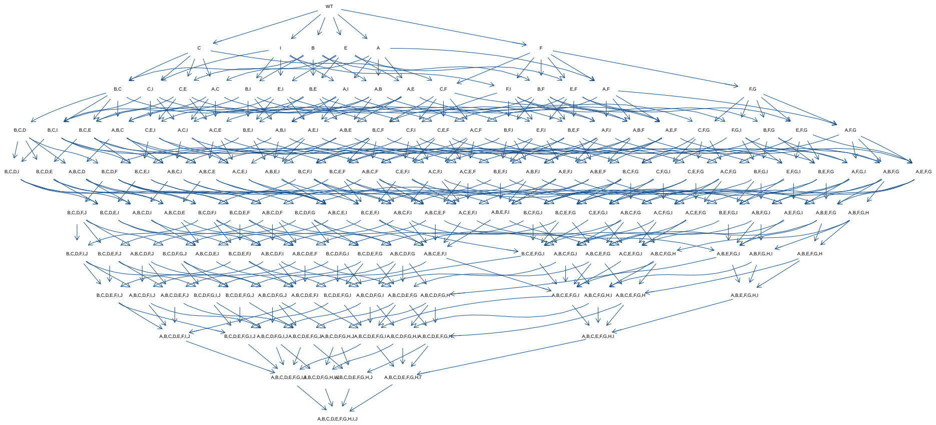

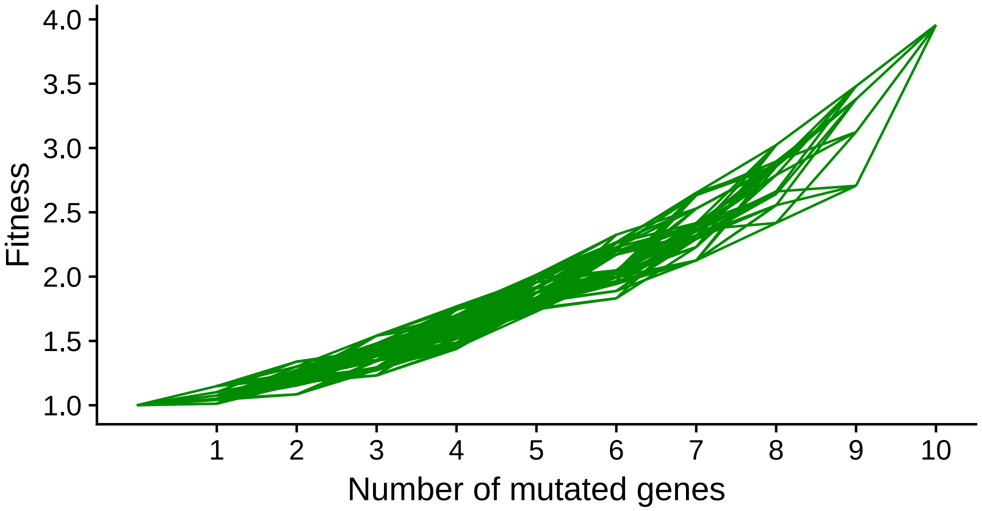

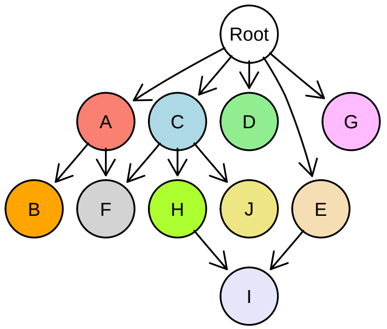


| | |
|---|---|
| *ID* | FMa3on8bA8mNavrZ |
| *acc genotypes* | 224 |
| *peaks no back* | 1 |
| *removed edges* | 0 |
| *prop rem edges* | 0 |

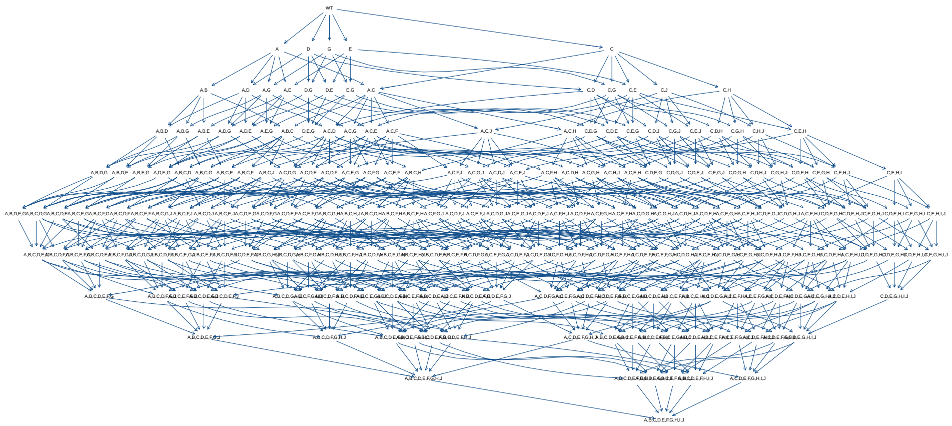

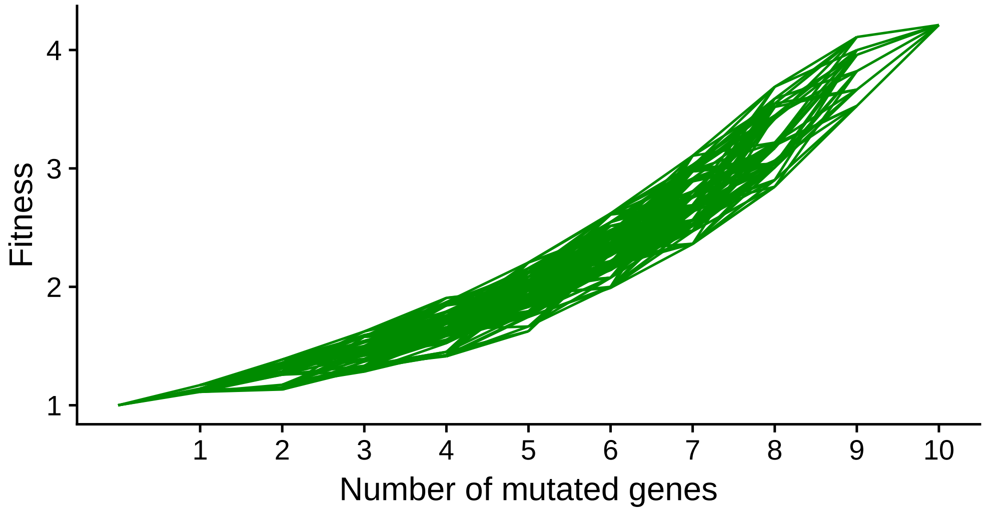

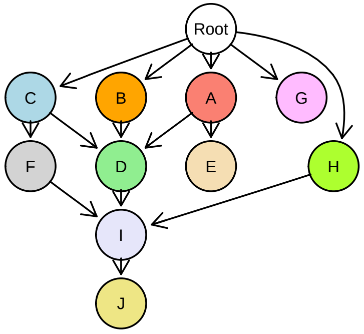


| | |
|---|---|
| *ID* | Fsqw1LKsGEDExuVa |
| *acc genotypes* | 96 |
| *peaks no back* | 1 |
| *removed edges* | 0 |
| *prop rem edges* | 0 |

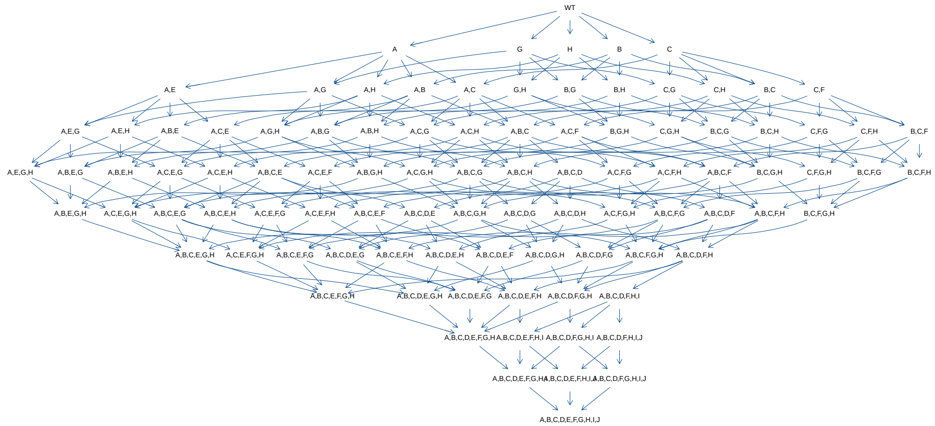


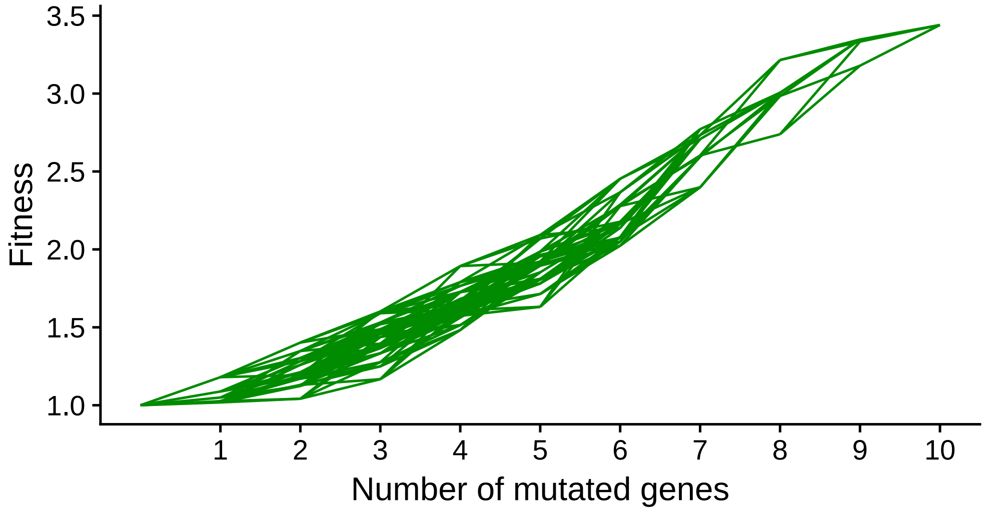

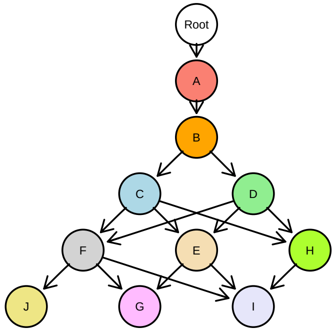


| | |
|---|---|
| *ID* | FYcNi4WyweZ3YLZu |
| *acc genotypes* | 25 |
| *peaks no back* | 1 |
| *removed edges* | 0 |
| *prop rem edges* | 0 |

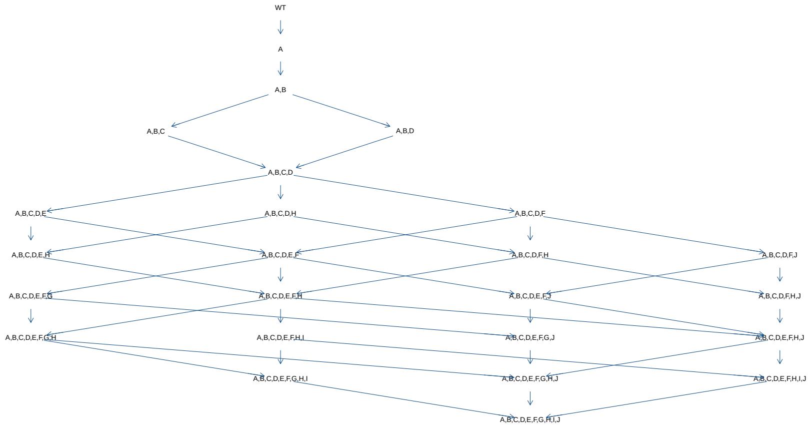


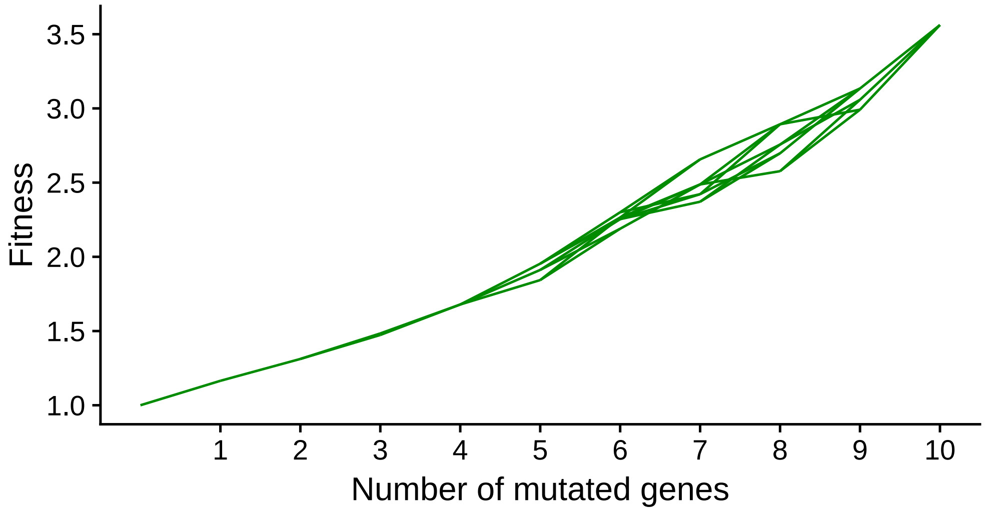

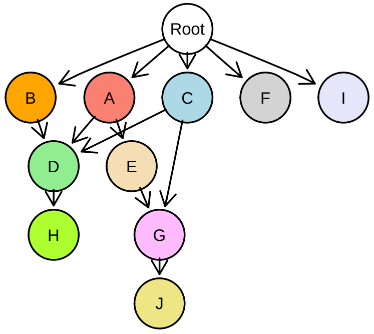


*ID* G3GJI10gesv99QKDi
*acc genotypes* 96
*peaks no back* 1
*removed edges* 0
*prop rem edges* 0

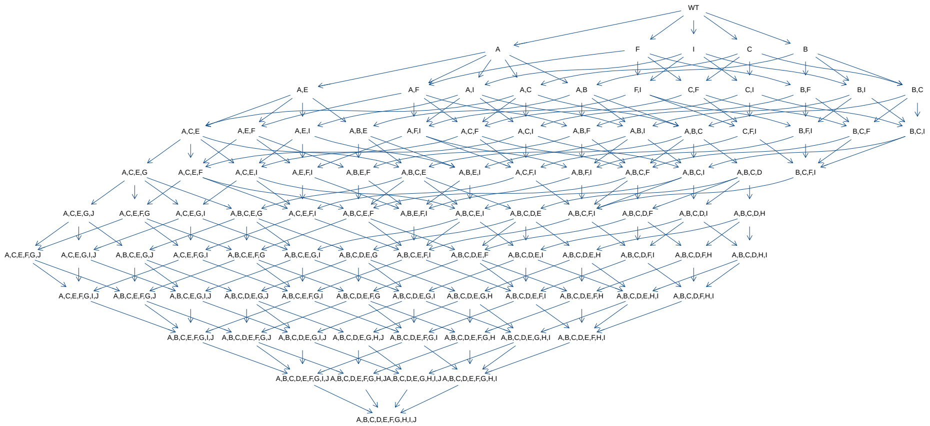

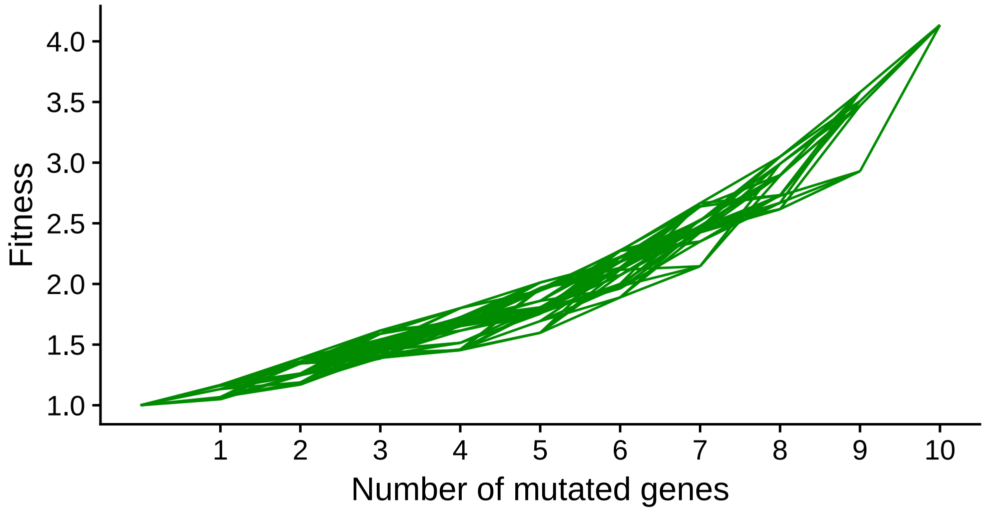

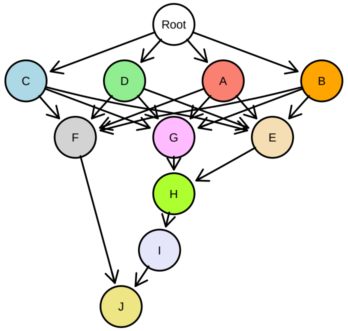


| | |
|---|---|
| *ID* | gcBjSEP8ZTo7NX2w |
| *acc genotypes* | 28 |
| *peaks no back* | 1 |
| *removed edges* | 0 |
| *prop rem edges* | 0 |

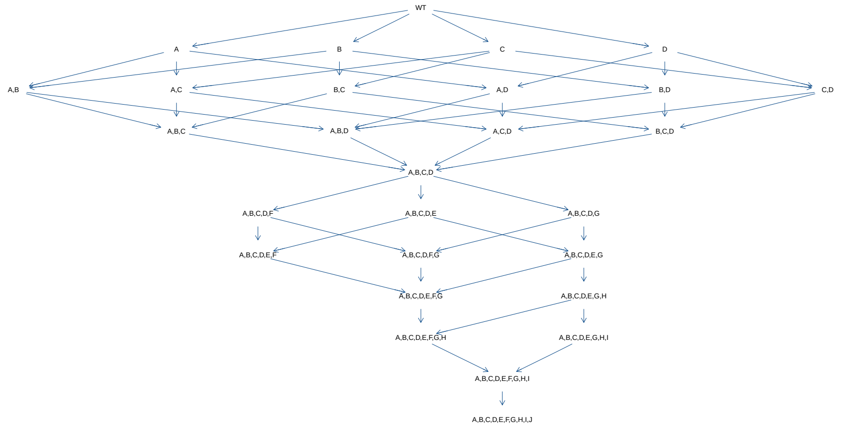


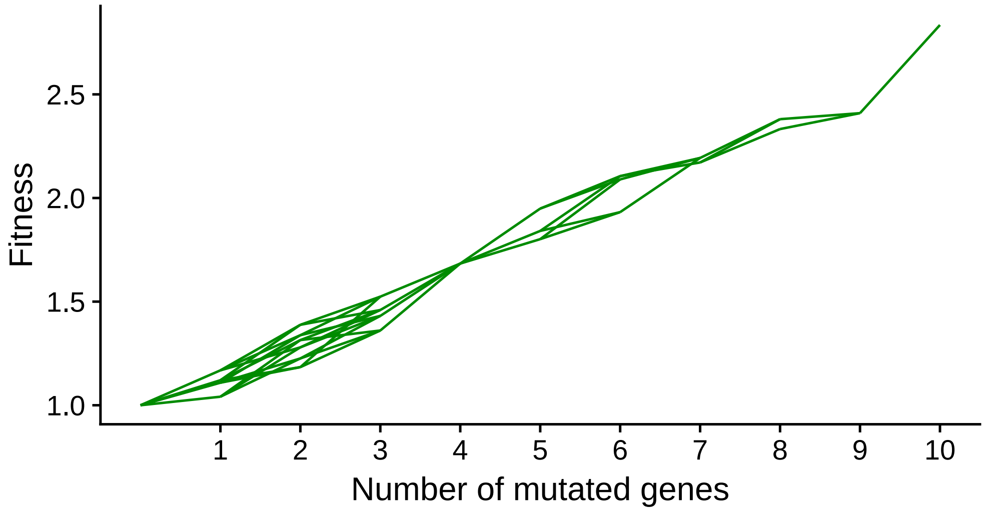

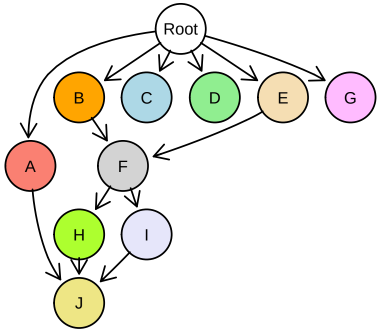


| | |
|---|---|
| *ID* | GdKyxu7pbMBmXsha |
| *acc genotypes* | 136 |
| *peaks no back* | 1 |
| *removed edges* | 0 |
| *prop rem edges* | 0 |

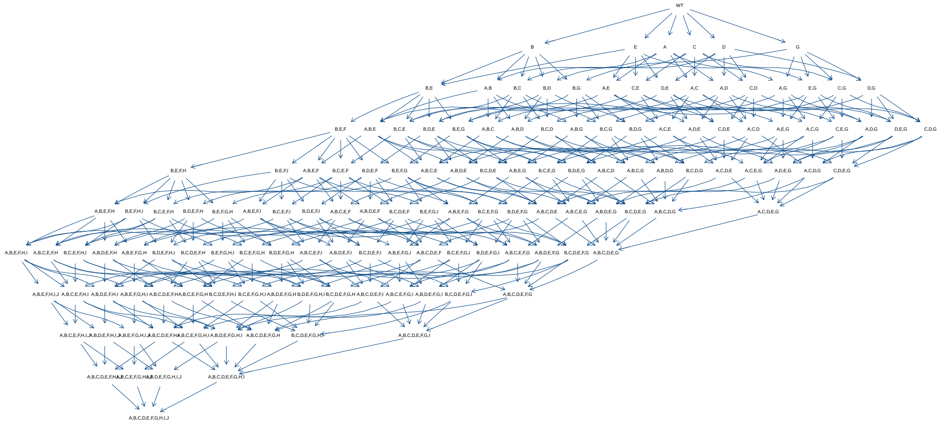

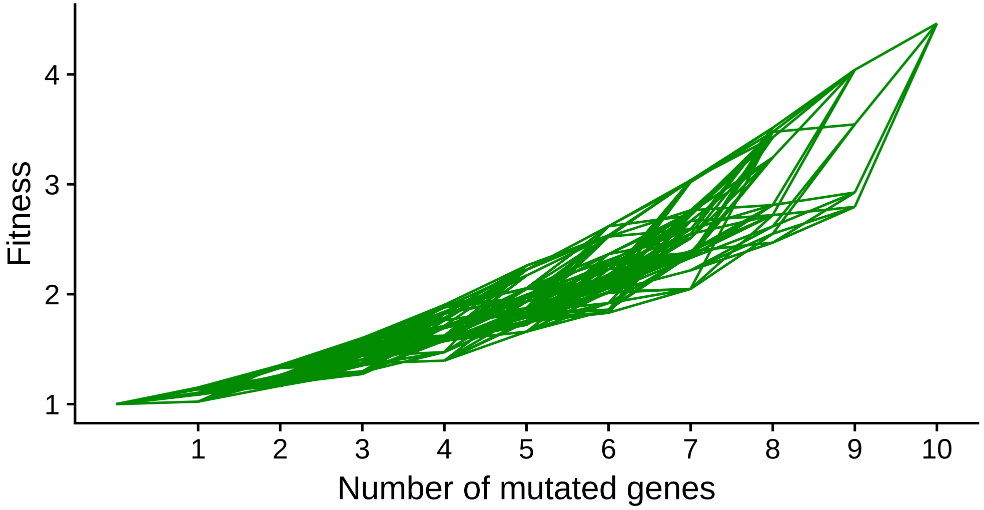

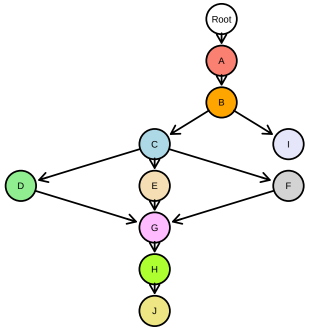


| | |
|---|---|
| *ID* | GDWChzATMq5hpozJ |
| *acc genotypes* | 26 |
| *peaks no back* | 1 |
| *removed edges* | 0 |
| *prop rem edges* | 0 |

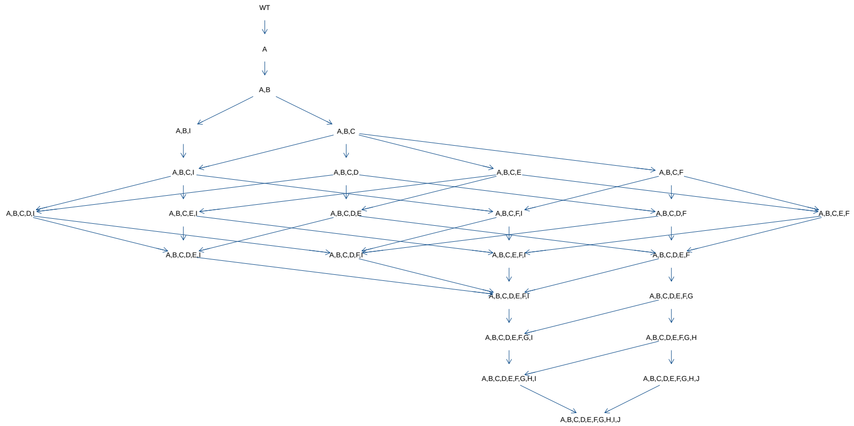

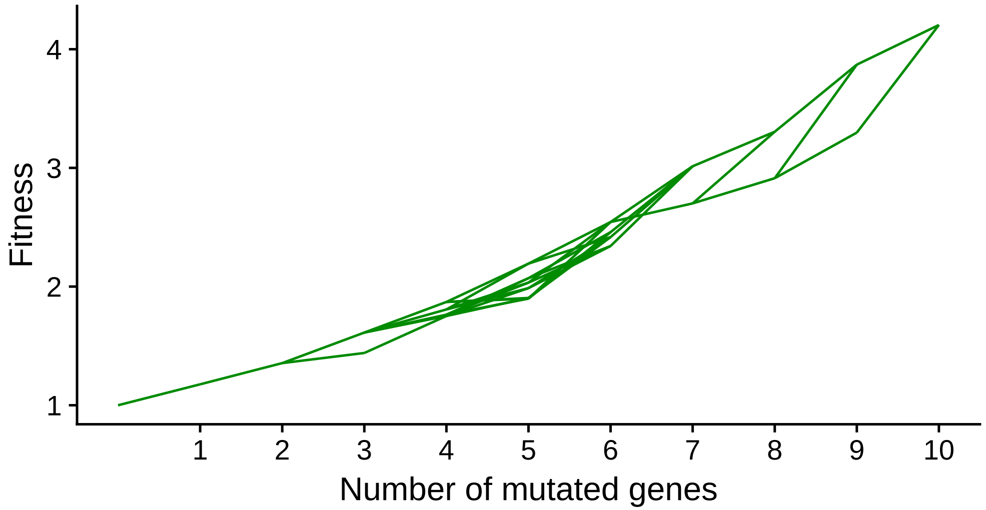

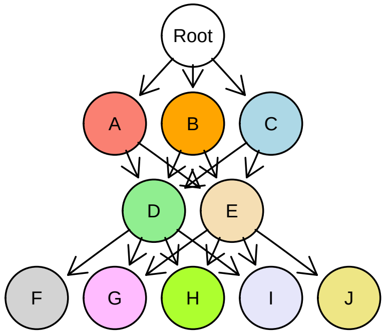


| | |
|---|---|
| *ID* | gNDvvHs3jYcerc7l |
| *acc genotypes* | 44 |
| *peaks no back* | 1 |
| *removed edges* | 0 |
| *prop rem edges* | 0 |

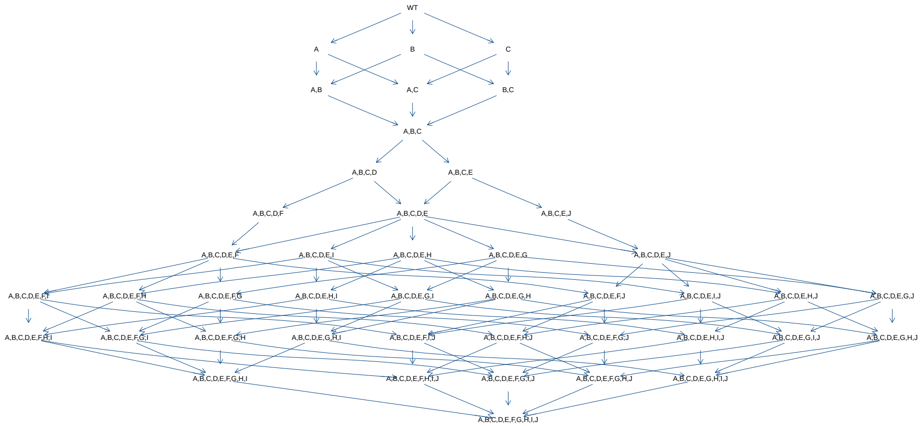

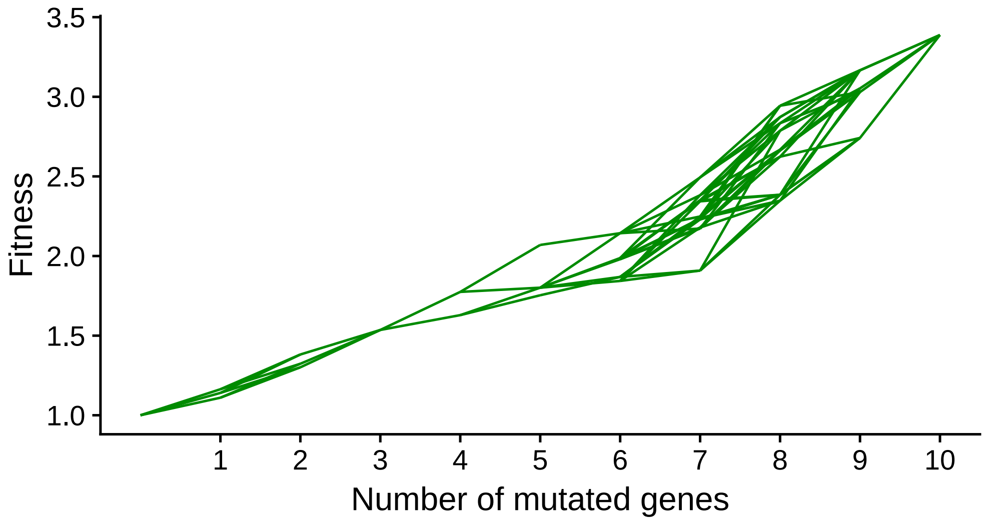

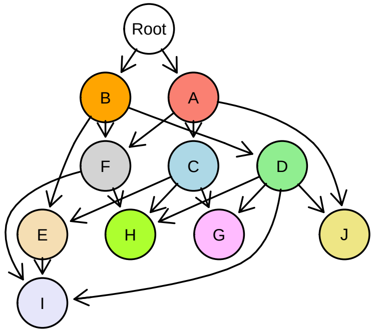


| | |
|---|---|
| *ID* | gqmarqZNpxa6L2Zx |
| *acc genotypes* | 47 |
| *peaks no back* | 1 |
| *removed edges* | 0 |
| *prop rem edges* | 0 |

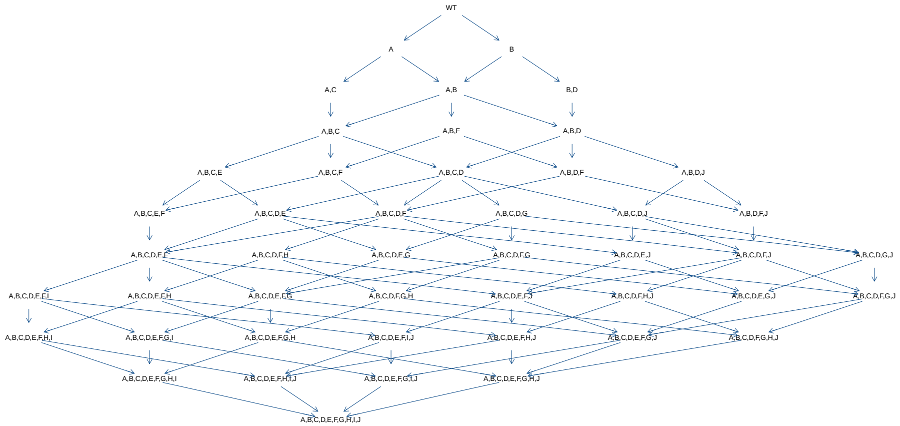

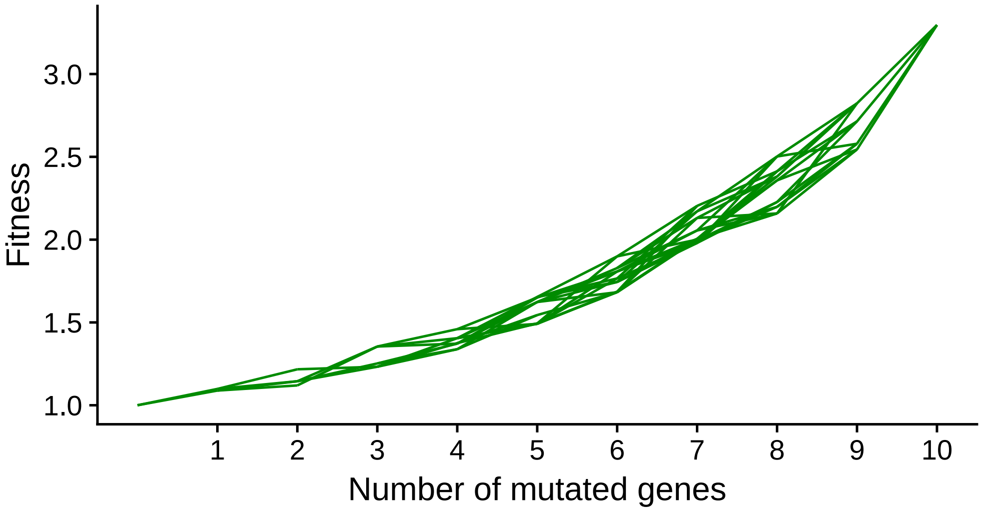

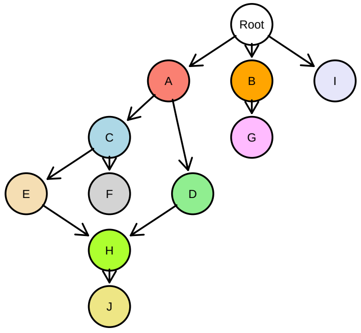


*ID* gYZAxs6hKlMQf3Ca
*acc genotypes* 90
*peaks no back* 1
*removed edges* 0
*prop rem edges* 0

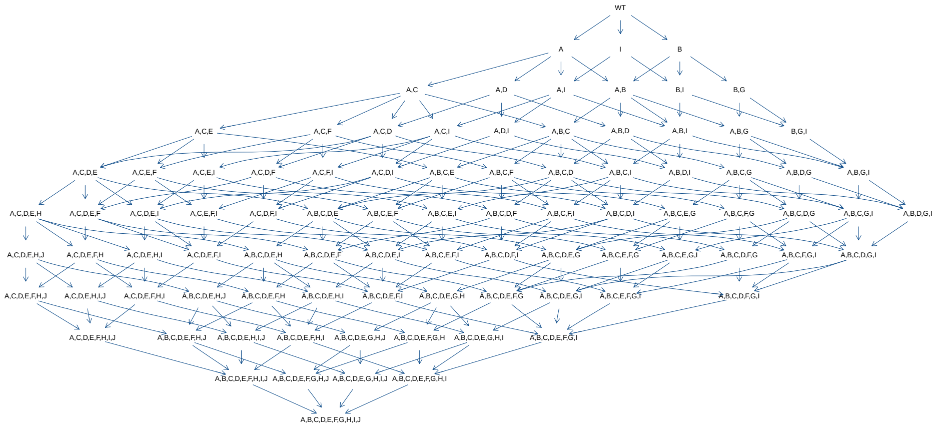

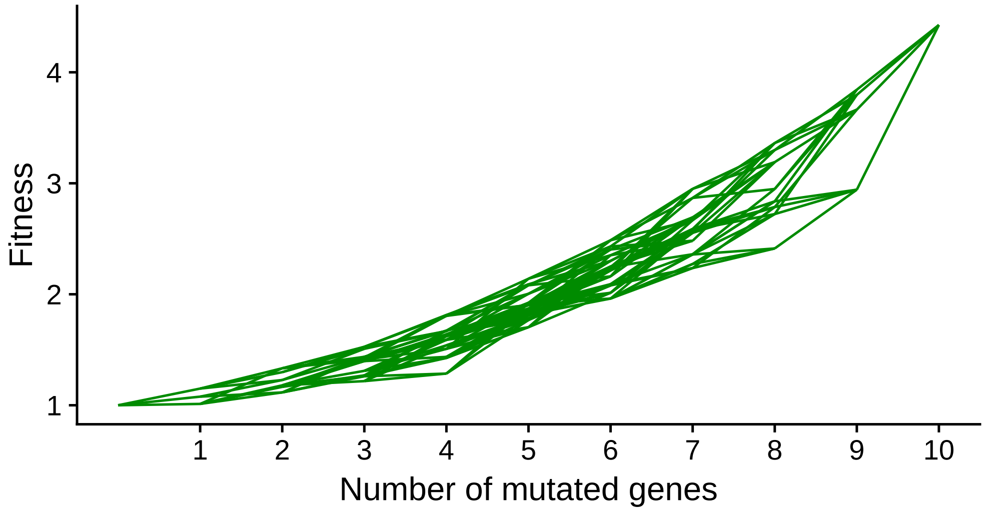

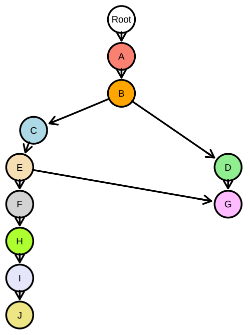


| | |
|---|---|
| *ID* | H10RfX4lukGLEX1kv |
| *acc genotypes* | 21 |
| *peaks no back* | 1 |
| *removed edges* | 0 |
| *prop rem edges* | 0 |

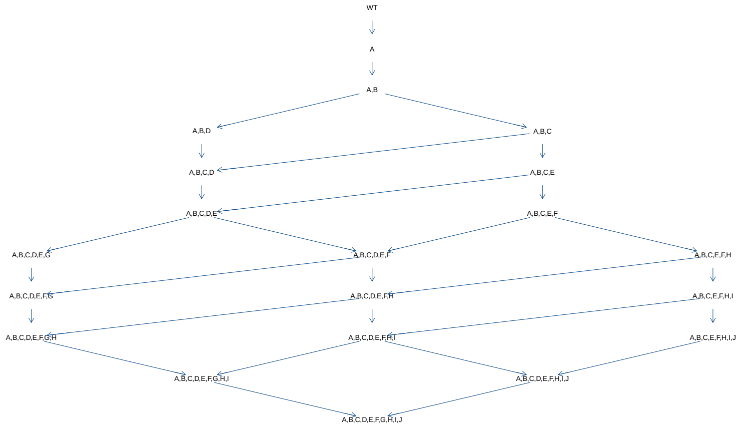


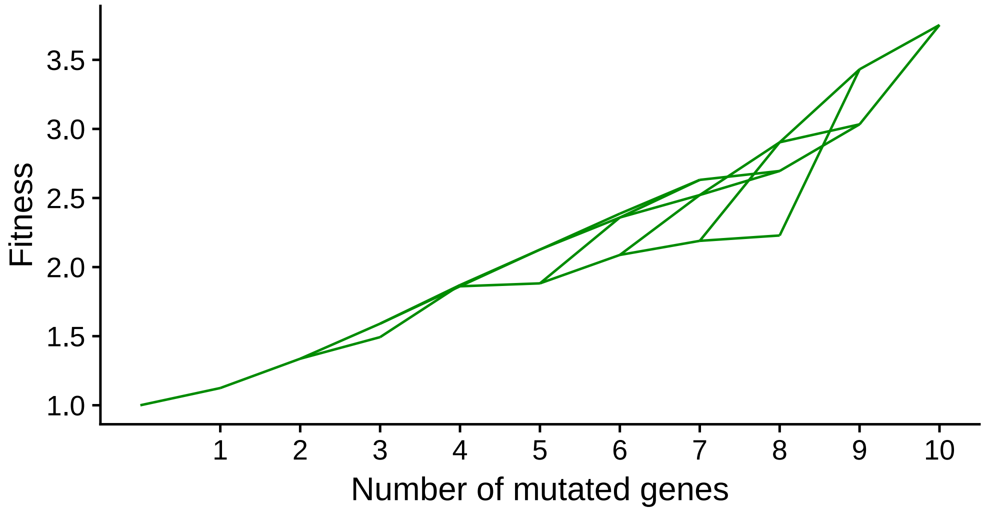

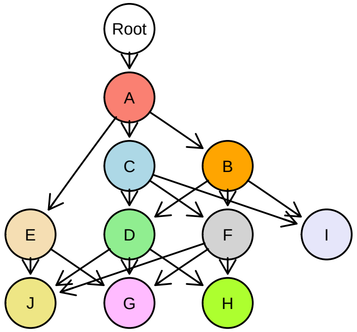


| | |
|---|---|
| *ID* | HaVk9pjpjvDygF8e |
| *acc genotypes* | 39 |
| *peaks no back* | 1 |
| *removed edges* | 0 |
| *prop rem edges* | 0 |

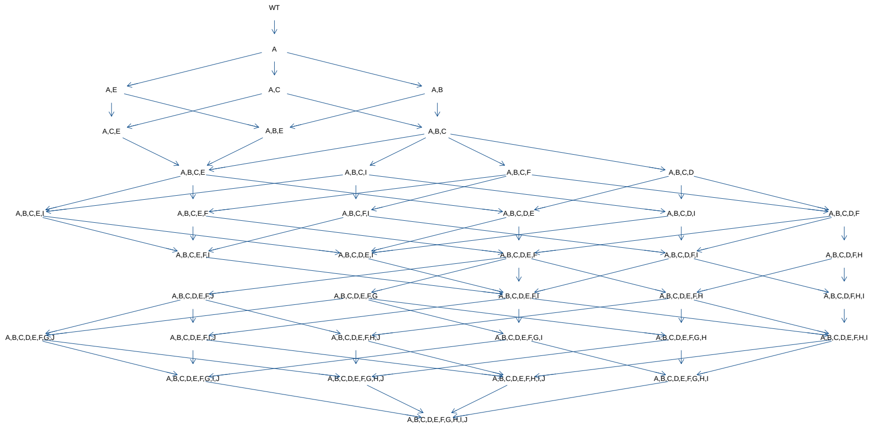

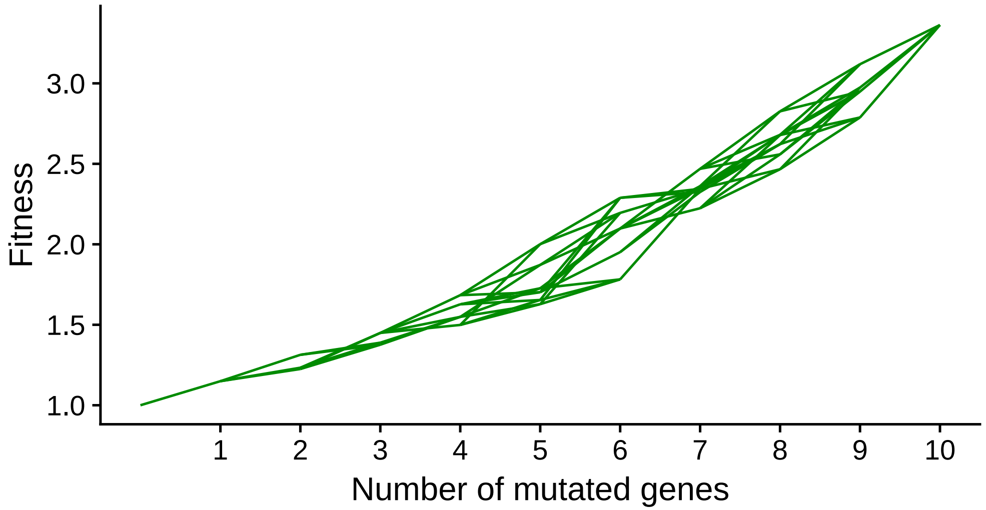

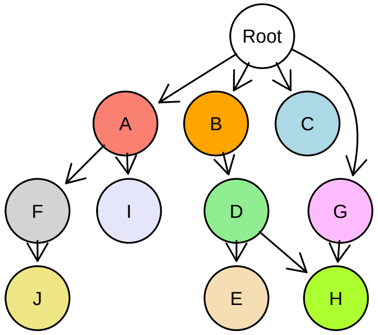


*ID* hB10YoS6mVpFbv6aL
*acc genotypes* 140
*peaks no back* 1
*removed edges* 0
*prop rem edges* 0

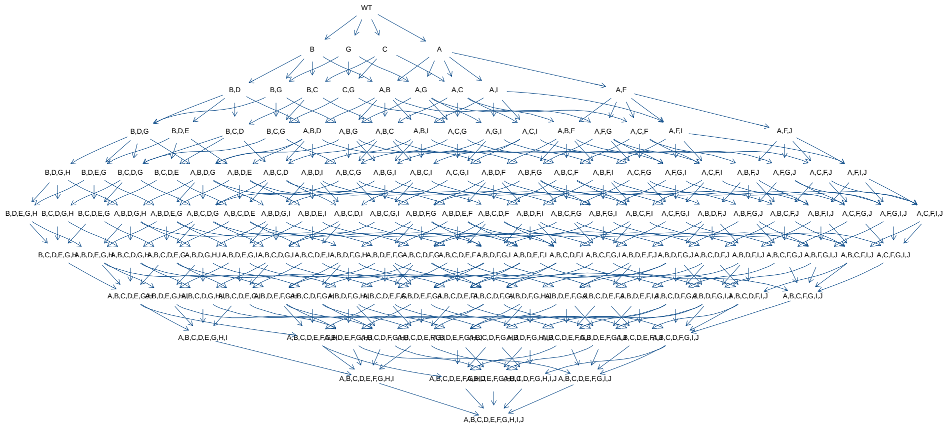

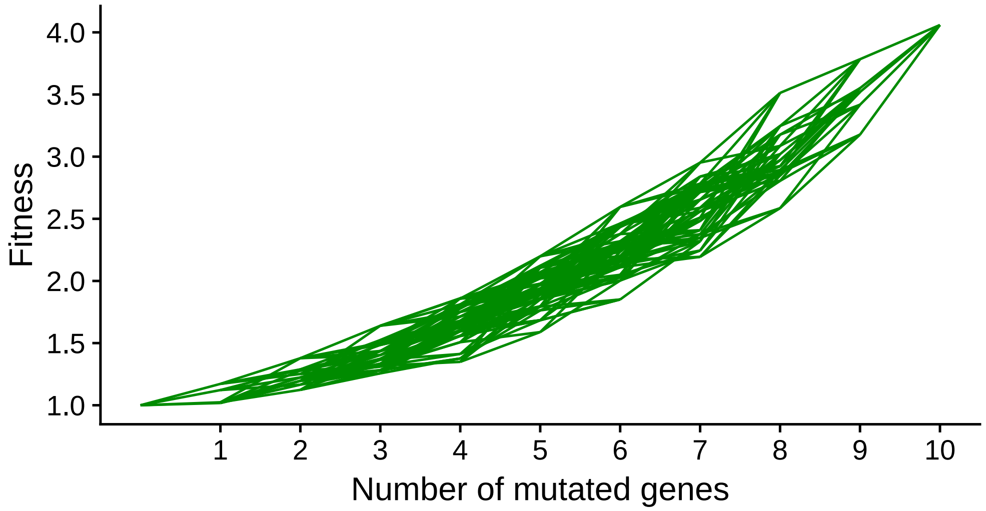

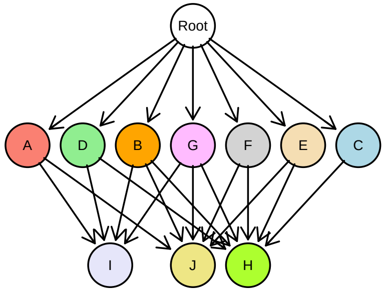


| | |
|---|---|
| *ID* | hiImr3F710LYpQsGL |
| *acc genotypes* | 147 |
| *peaks no back* | 1 |
| *removed edges* | 0 |
| *prop rem edges* | 0 |

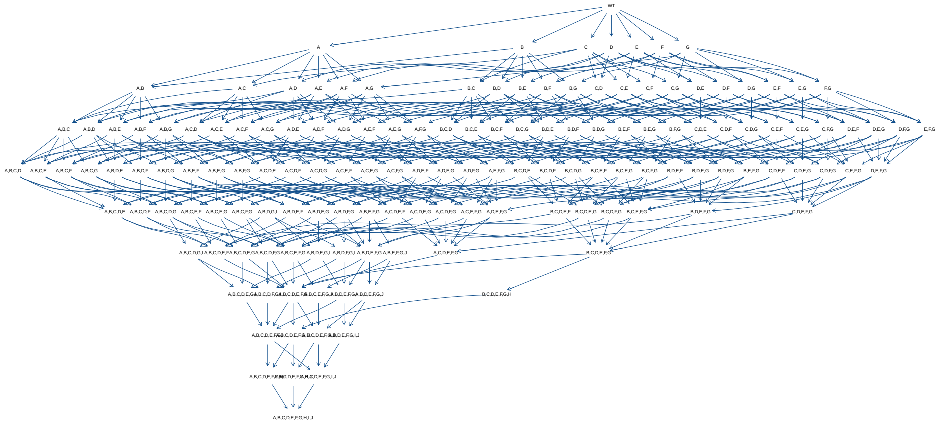

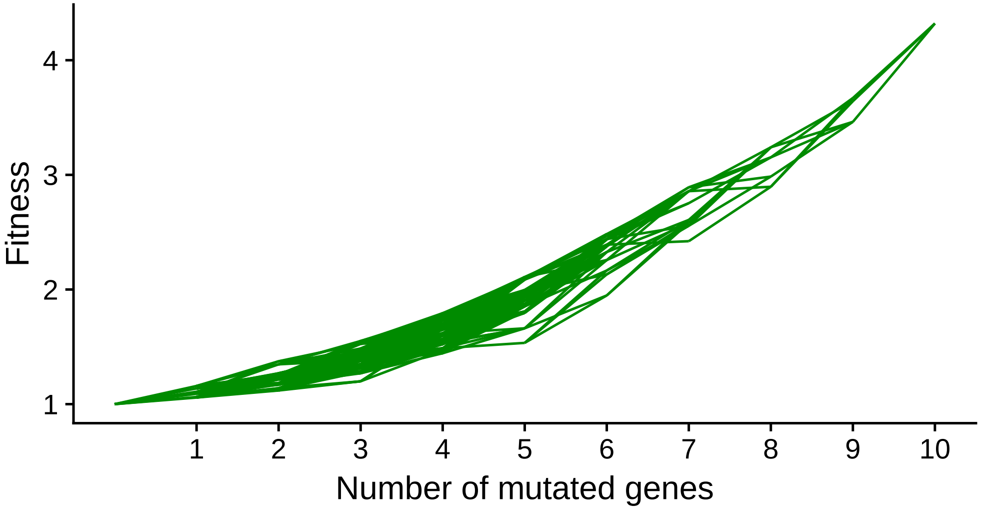

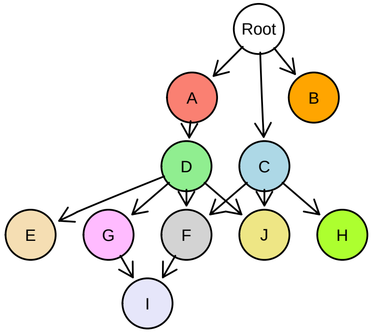


*ID* hNfoY8xLhb3Nf7ZZ
*acc genotypes* 100
*peaks no back* 1
*removed edges* 0
*prop rem edges* 0

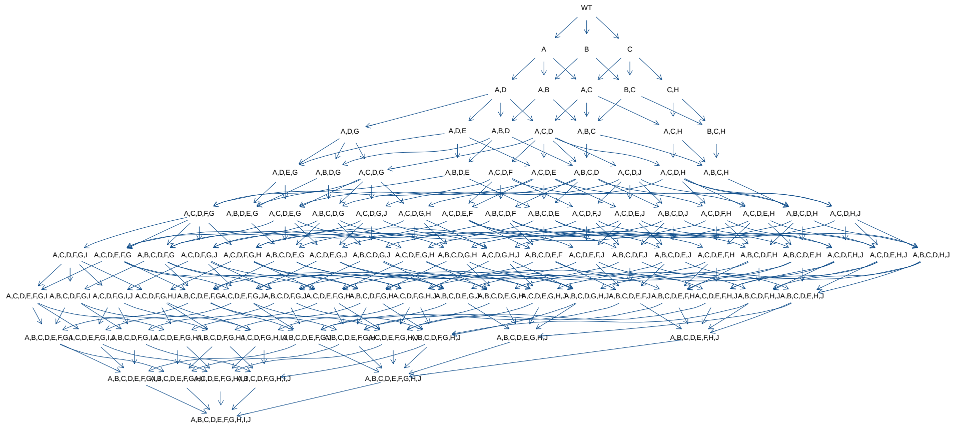

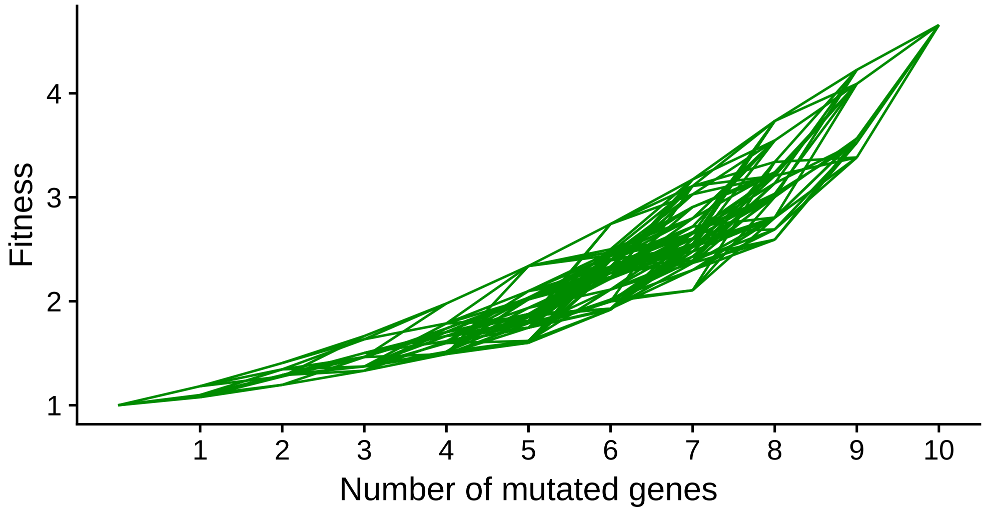

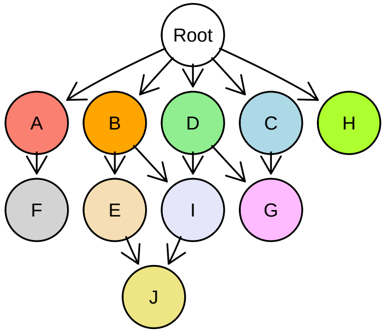


| | |
|---|---|
| *ID* | HnZ10pnWSr3M1MXbv |
| *acc genotypes* | 144 |
| *peaks no back* | 1 |
| *removed edges* | 0 |
| *prop rem edges* | 0 |

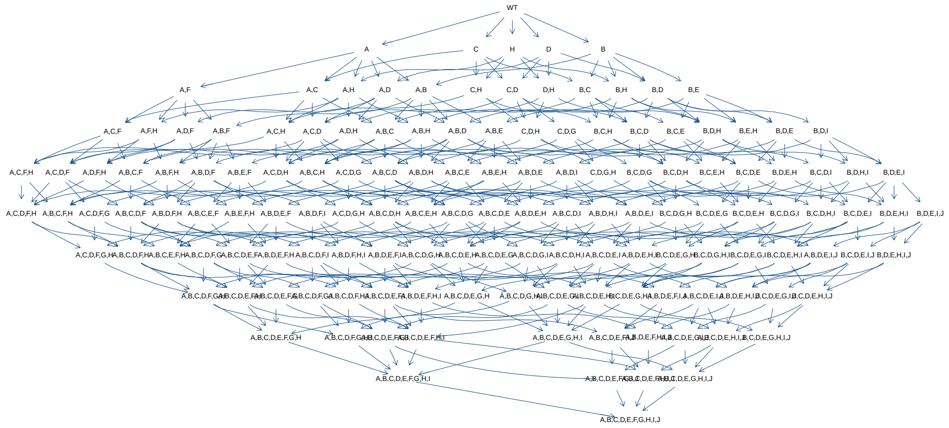


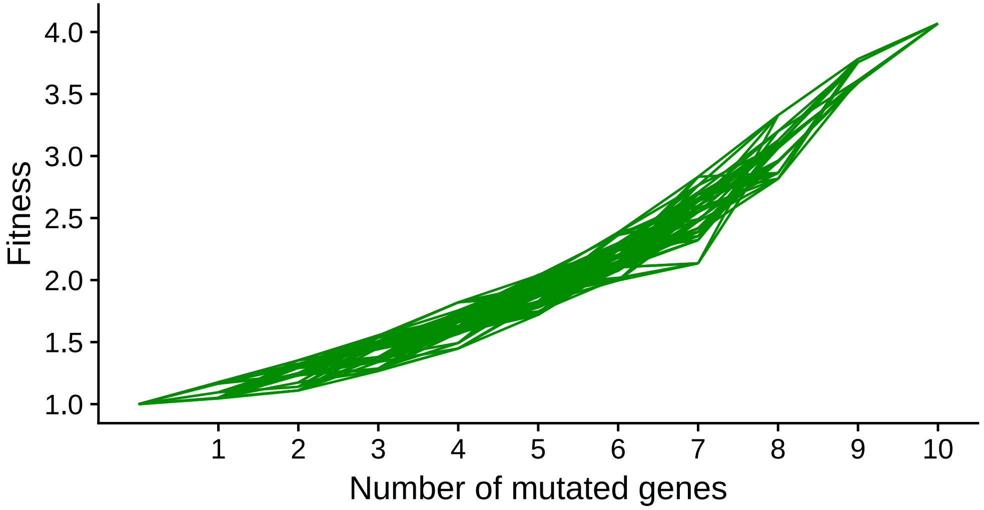

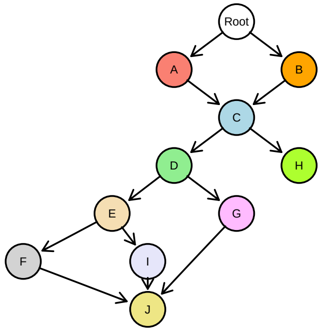


| | |
|---|---|
| *ID* | Hon2Ud6RNbFJ826U |
| *acc genotypes* | 28 |
| *peaks no back* | 1 |
| *removed edges* | 0 |
| *prop rem edges* | 0 |

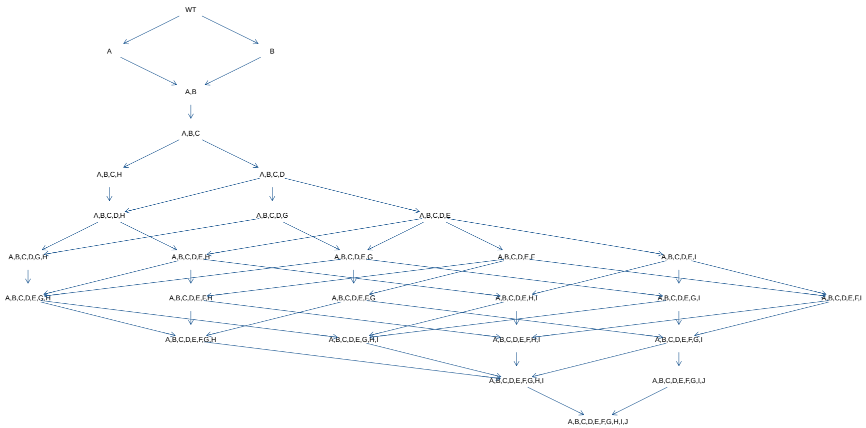


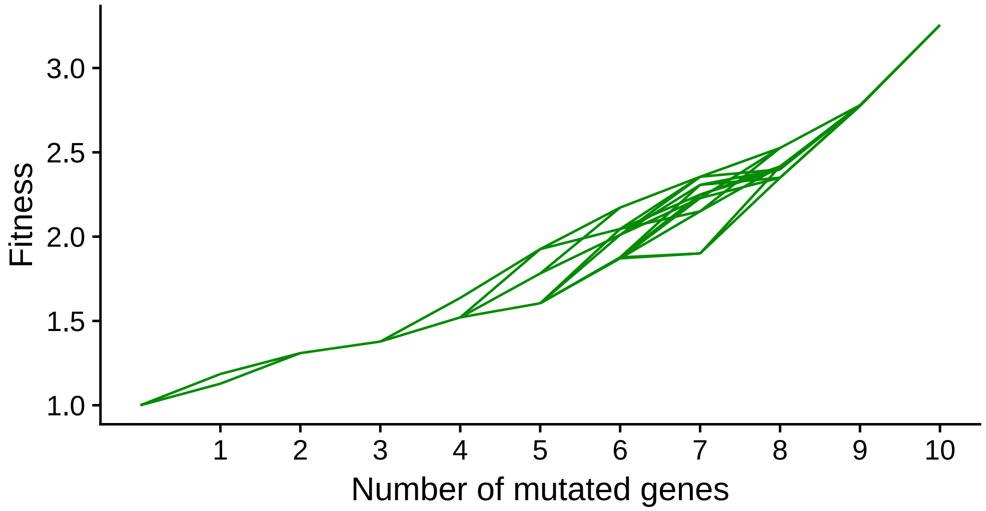

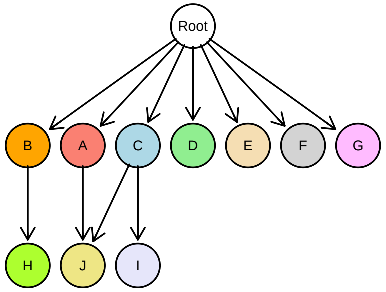


| | |
|---|---|
| *ID* | hqhSp8AzeBqMadTe |
| *acc genotypes* | 384 |
| *peaks no back* | 1 |
| *removed edges* | 0 |
| *prop rem edges* | 0 |

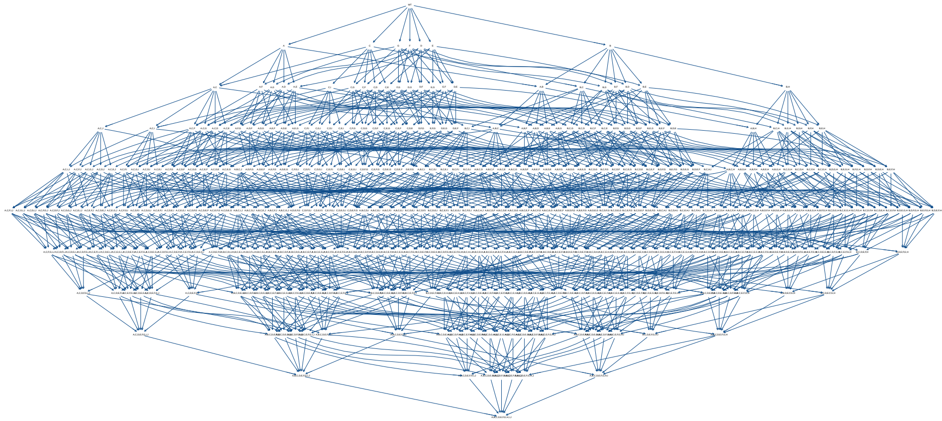

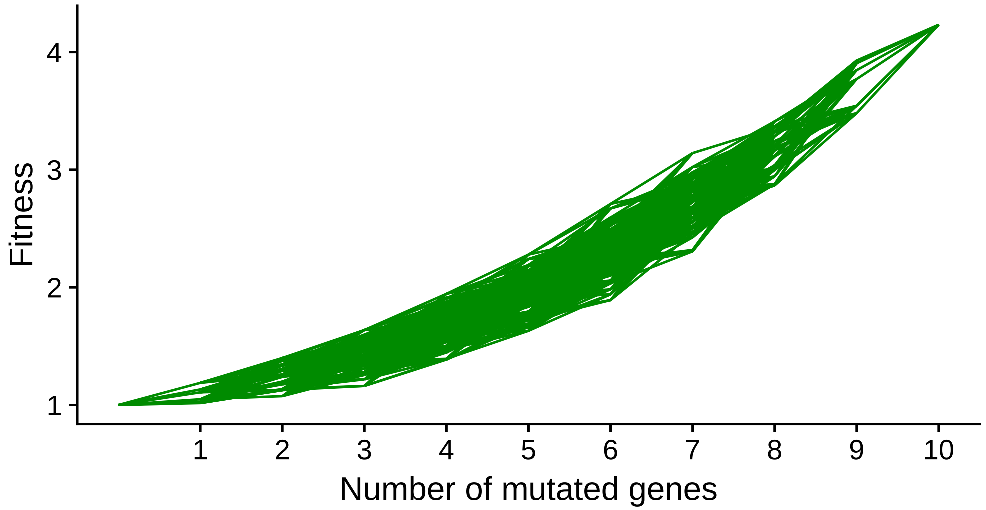

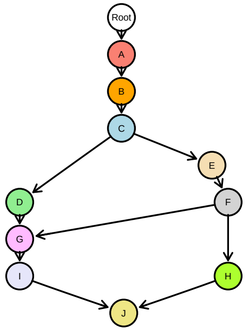


| | |
|---|---|
| *ID* | hshAK10qbNh6UkHAB |
| *acc genotypes* | 16 |
| *peaks no back* | 1 |
| *removed edges* | 0 |
| *prop rem edges* | 0 |

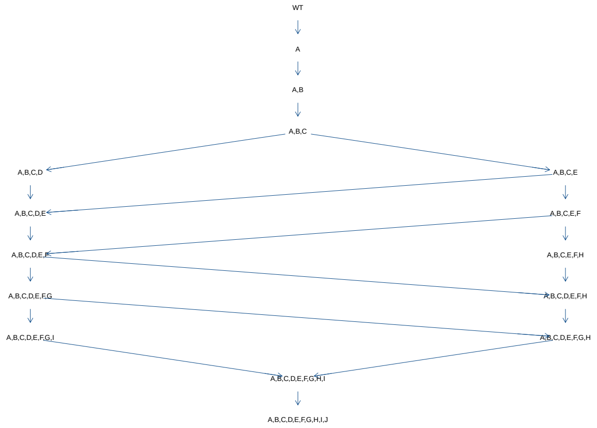


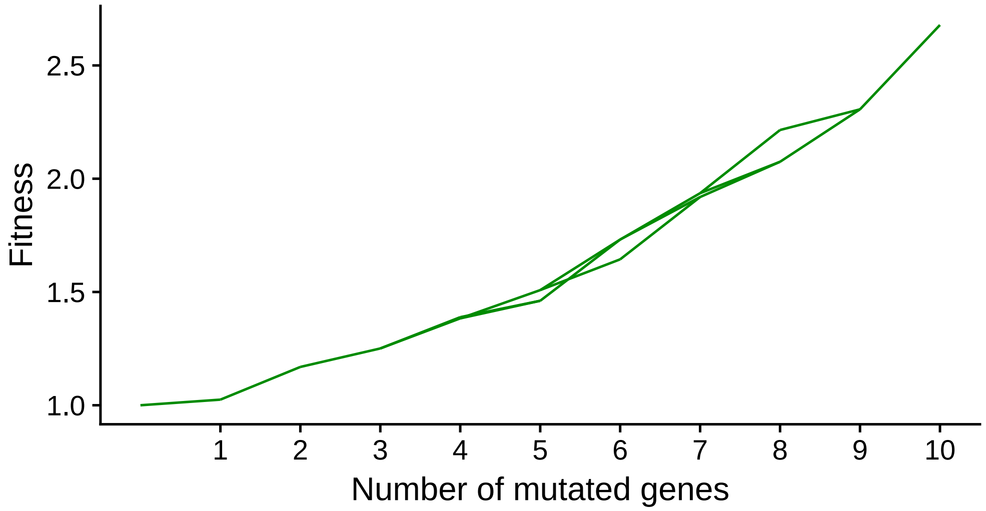

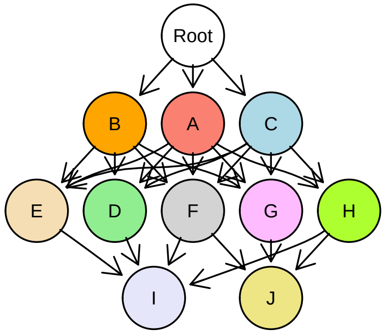


| | |
|---|---|
| *ID* | HXhpKGzcwUflLLeF |
| *acc genotypes* | 47 |
| *peaks no back* | 1 |
| *removed edges* | 0 |
| *prop rem edges* | 0 |

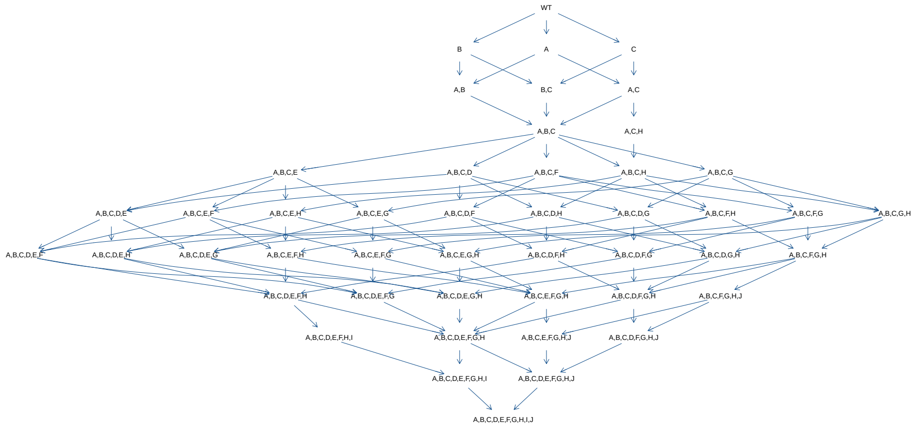

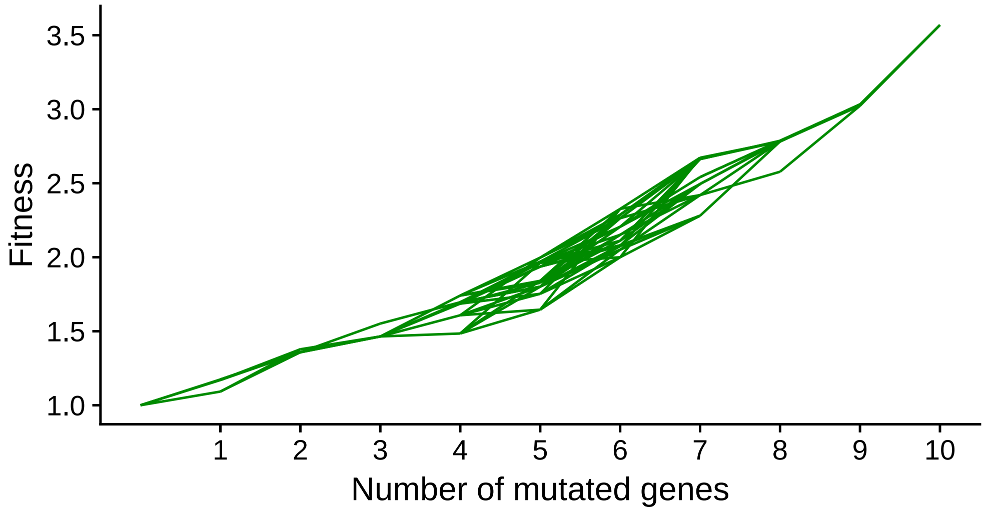

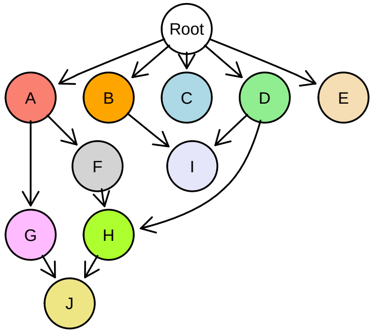


| | |
|---|---|
| *ID* | HXibbT4wtp8elqYm |
| *acc genotypes* | 136 |
| *peaks no back* | 1 |
| *removed edges* | 0 |
| *prop rem edges* | 0 |

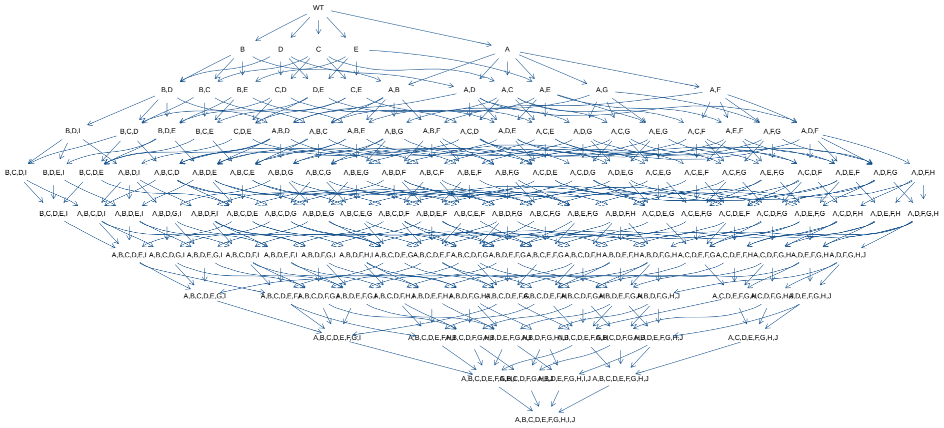

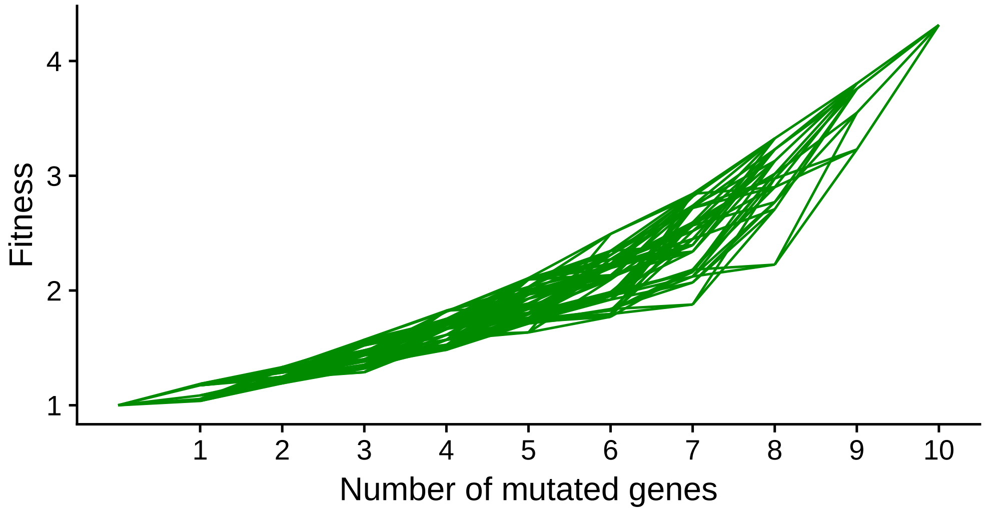

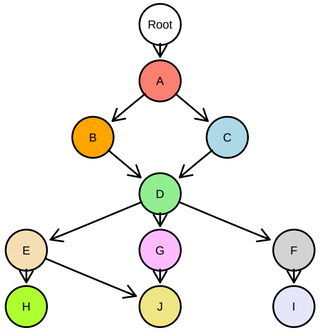


*ID* IBfkuKvC5LnGkILI
*acc genotypes* 29
*peaks no back* 1
*removed edges* 0
*prop rem edges* 0

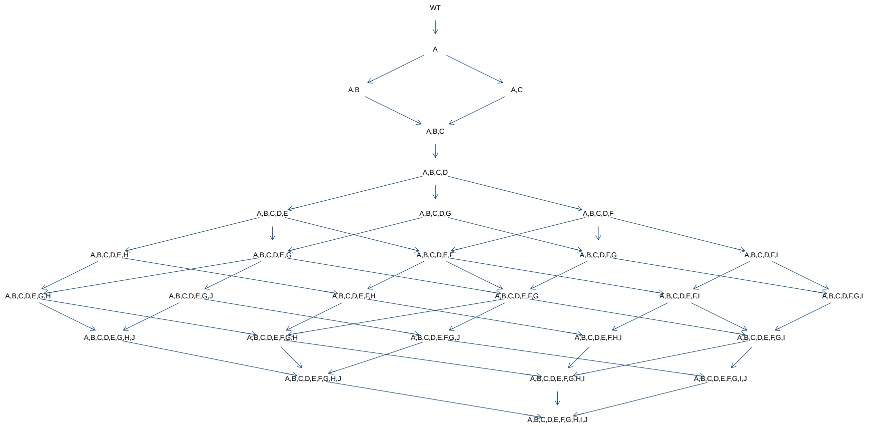

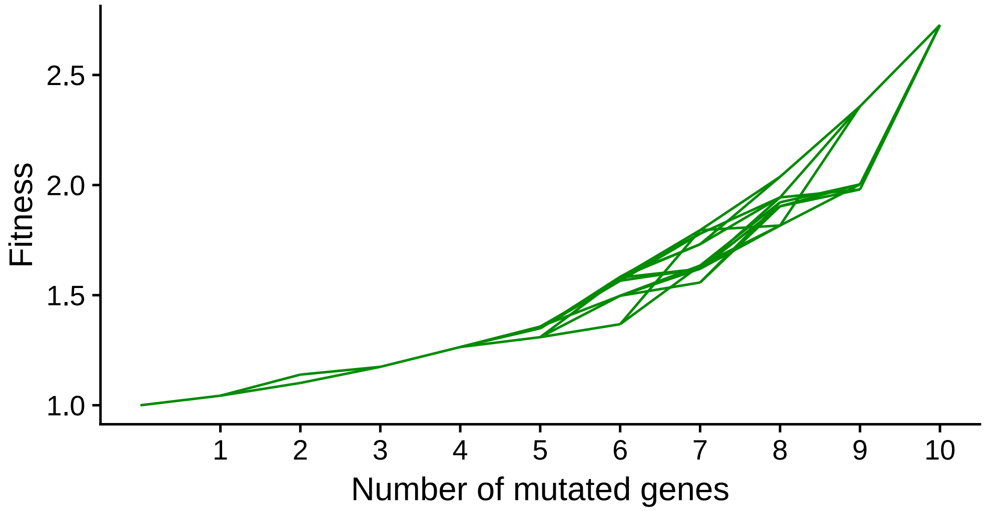

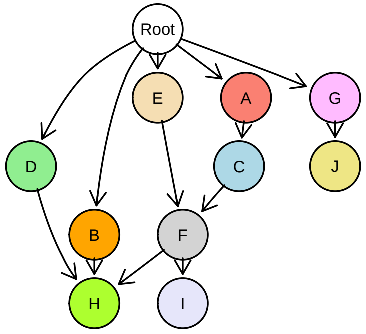


| | |
|---|---|
| *ID* | ibqIgMnpLmhx8hJy |
| *acc genotypes* | 102 |
| *peaks no back* | 1 |
| *removed edges* | 0 |
| *prop rem edges* | 0 |

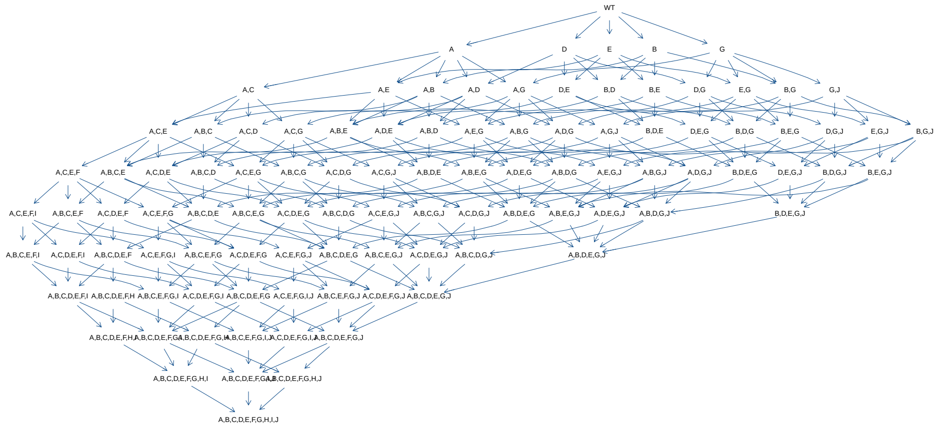

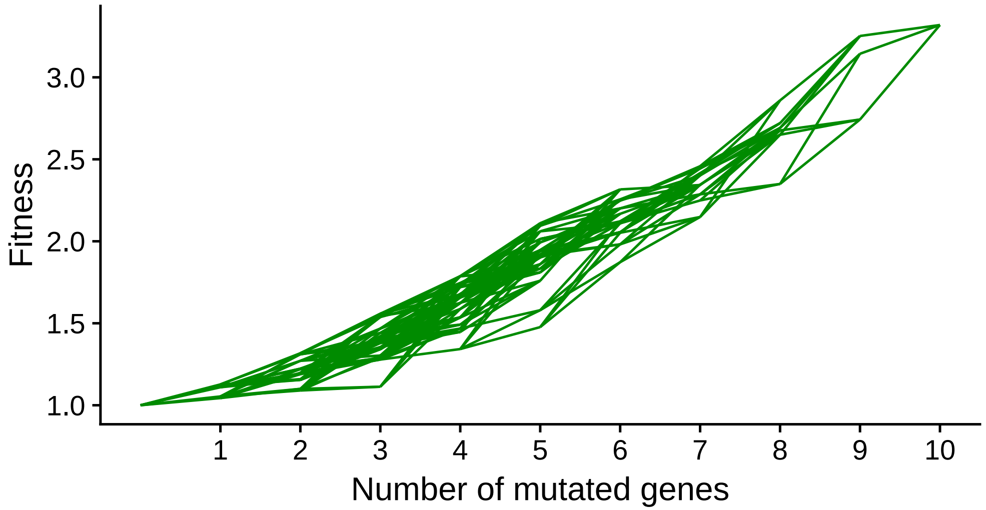

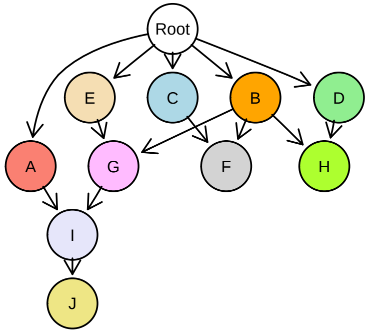


| | |
|---|---|
| *ID* | IFzpx6g9rMSHRpvL |
| *acc genotypes* | 88 |
| *peaks no back* | 1 |
| *removed edges* | 0 |
| *prop rem edges* | 0 |

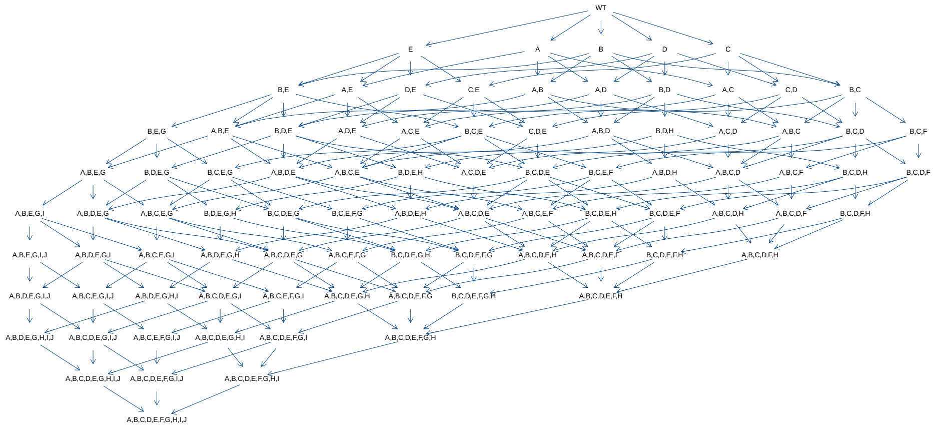

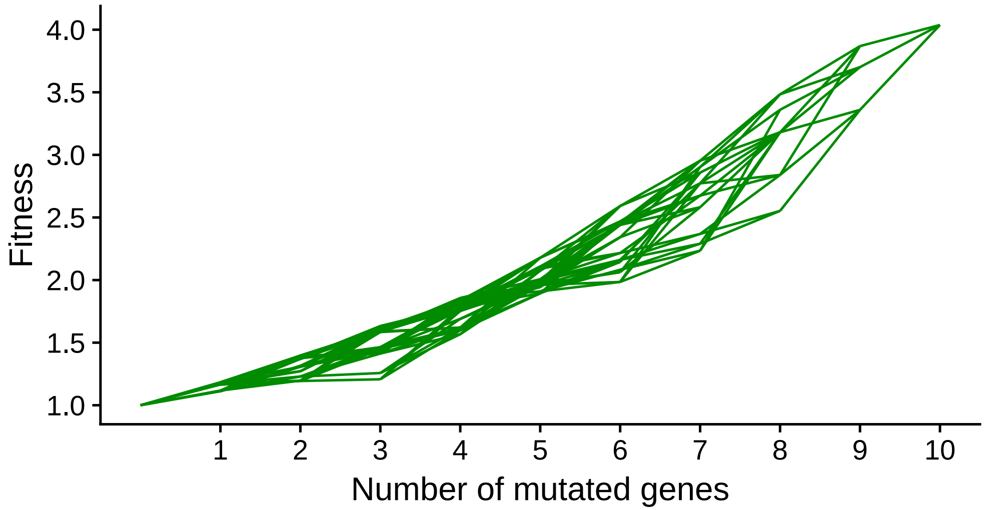

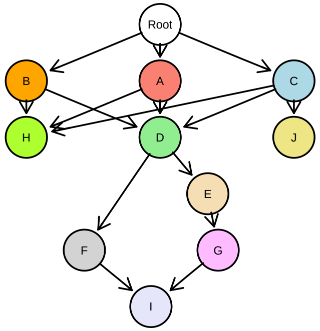


*ID* IGNIamDyyL6pJkAH
*acc genotypes* 42
*peaks no back* 1
*removed edges* 0
*prop rem edges* 0

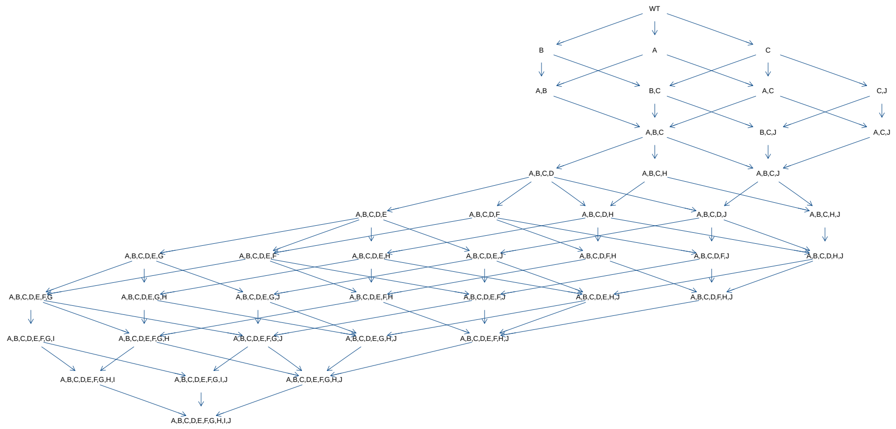


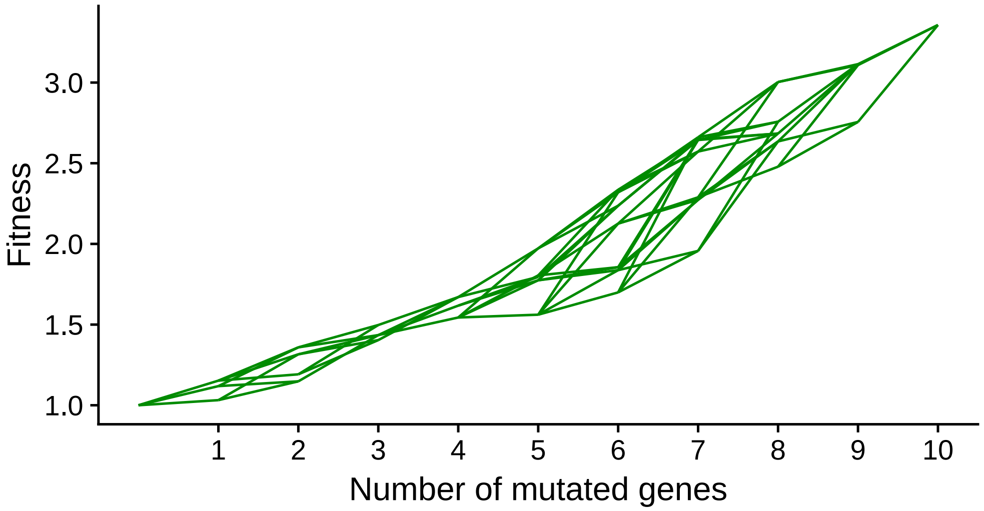

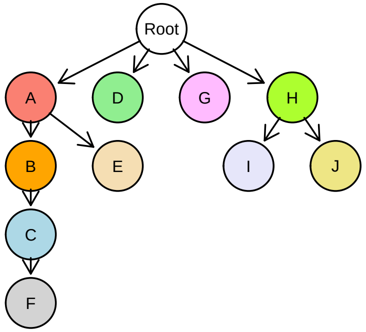


| | |
|---|---|
| *ID* | IIY6ewqbwnTGL76Q |
| *acc genotypes* | 180 |
| *peaks no back* | 1 |
| *removed edges* | 0 |
| *prop rem edges* | 0 |

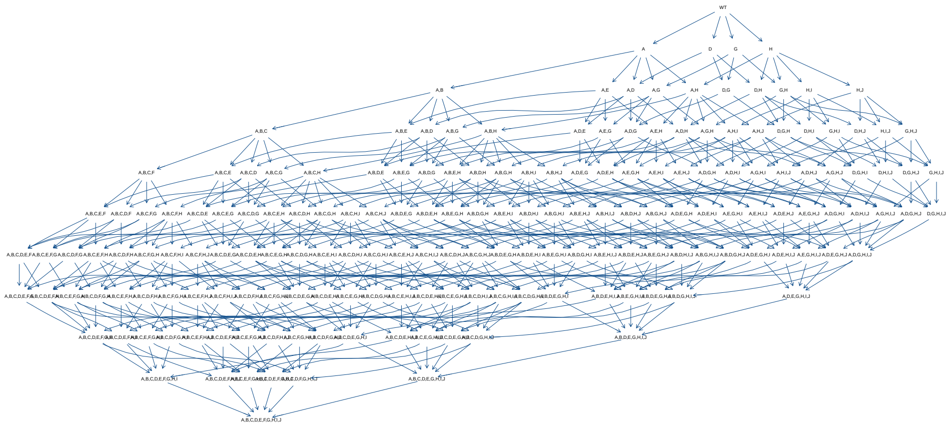

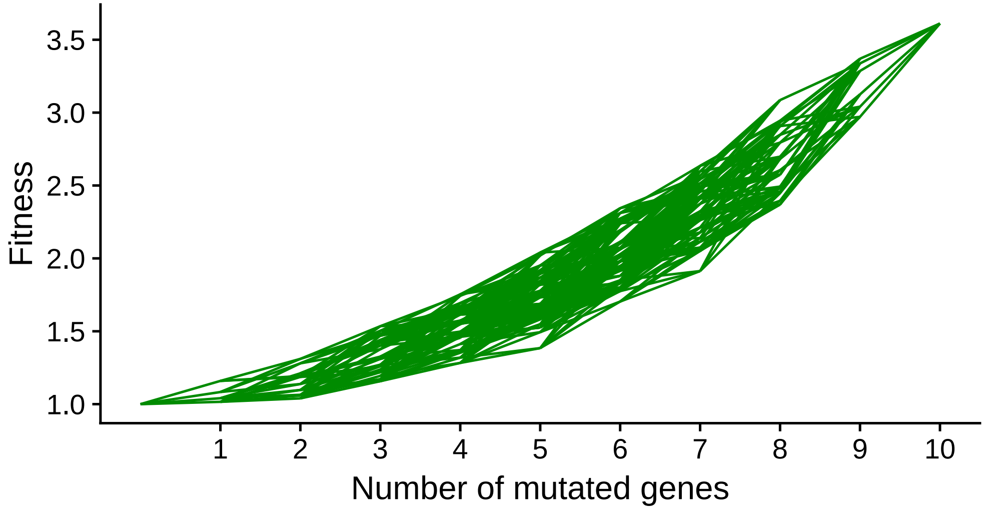

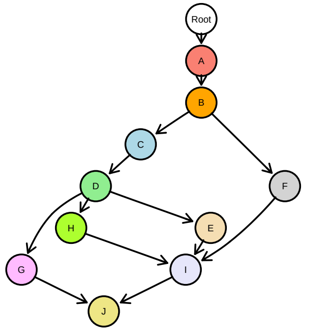


| | |
|---|---|
| *ID* | IJ82SrhcD1ULypf8 |
| *acc genotypes* | 25 |
| *peaks no back* | 1 |
| *removed edges* | 0 |
| *prop rem edges* | 0 |

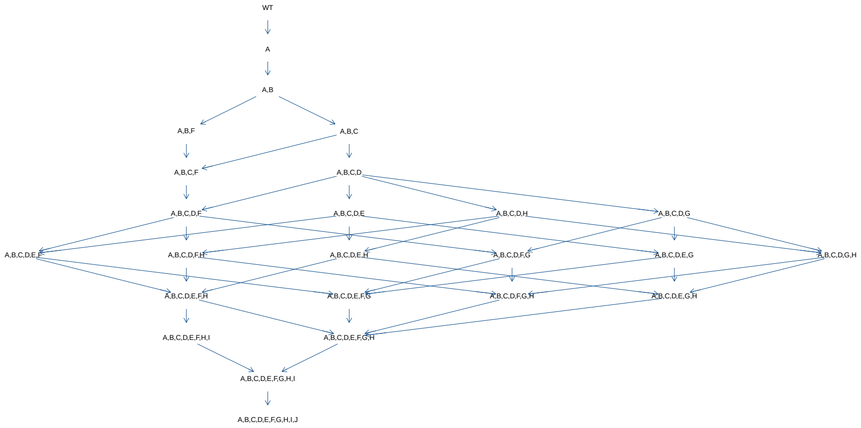

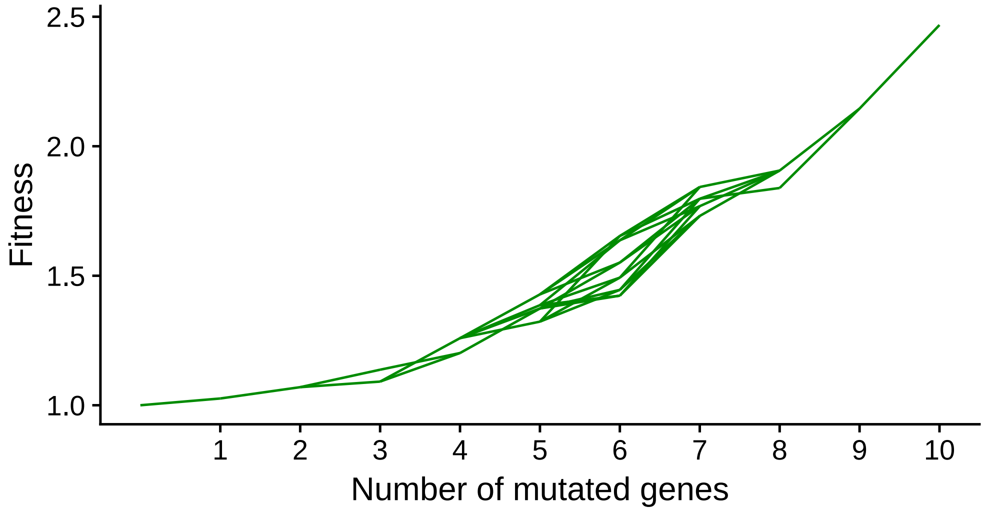

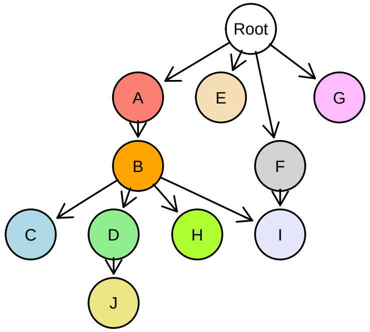


| | |
|---|---|
| *ID* | IqyqZ4siy10VLgJ5s |
| *acc genotypes* | 160 |
| *peaks no back* | 1 |
| *removed edges* | 0 |
| *prop rem edges* | 0 |

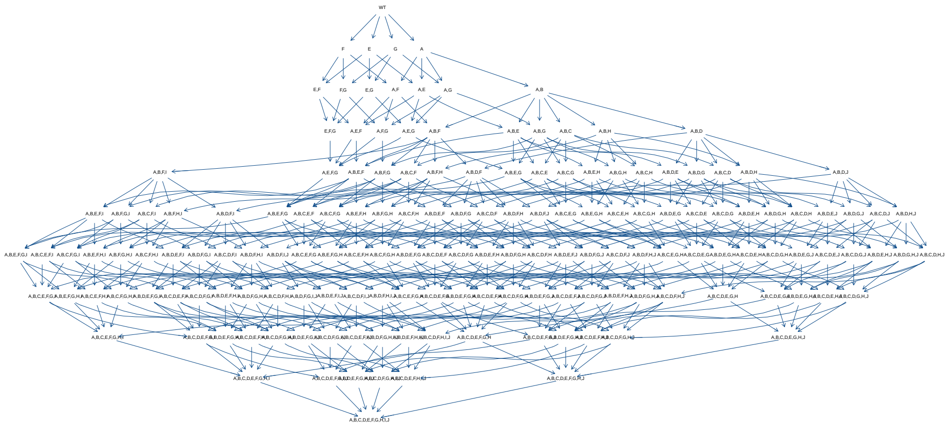

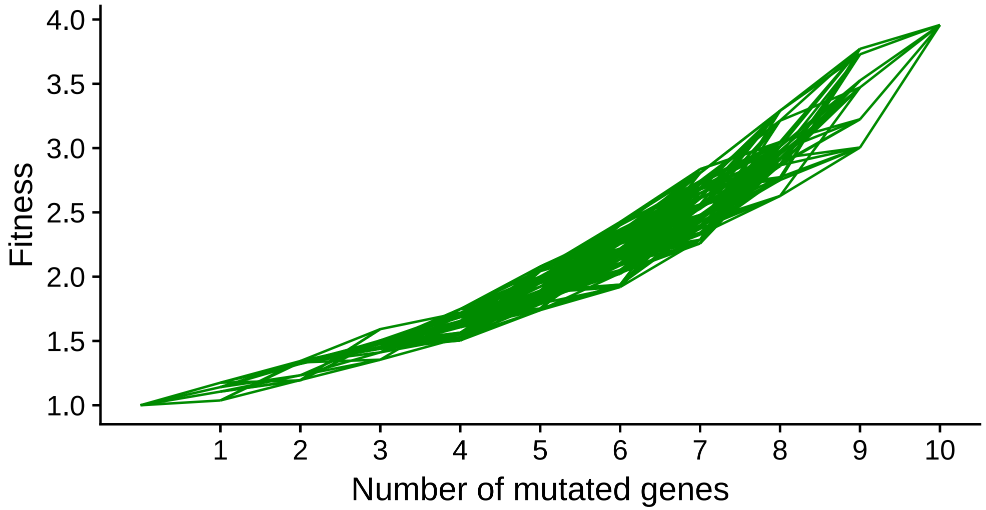

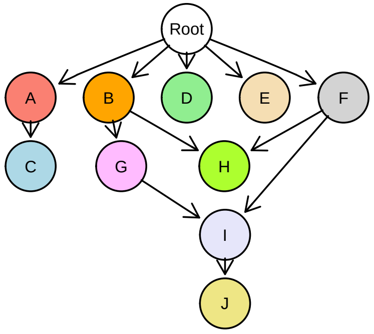


| | |
|---|---|
| *ID* | ITJu10YflRKiTyZLt |
| *acc genotypes* | 144 |
| *peaks no back* | 1 |
| *removed edges* | 0 |
| *prop rem edges* | 0 |

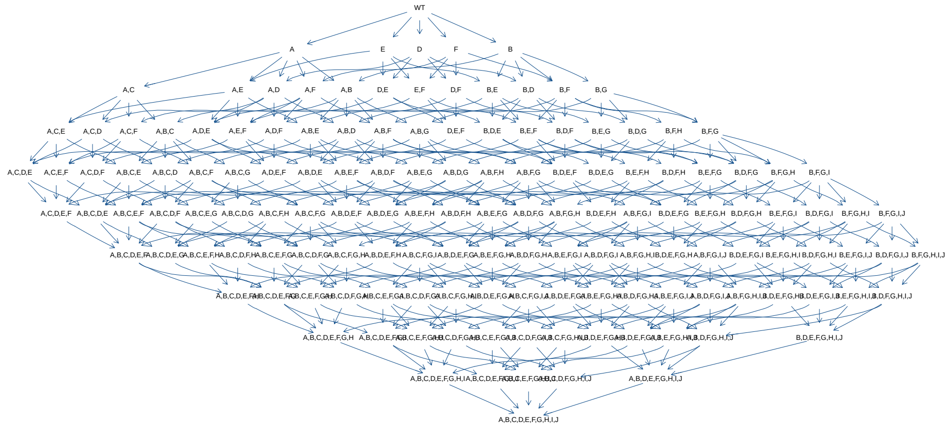

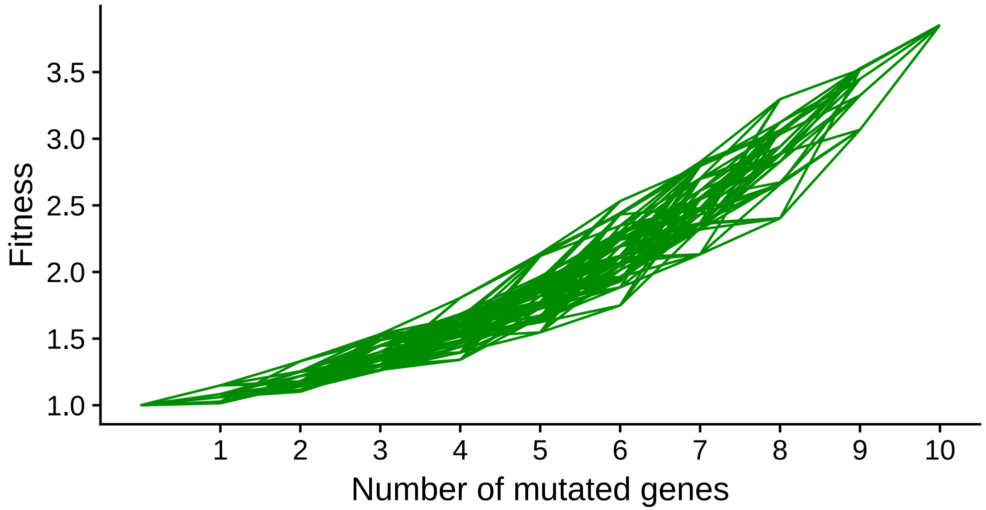

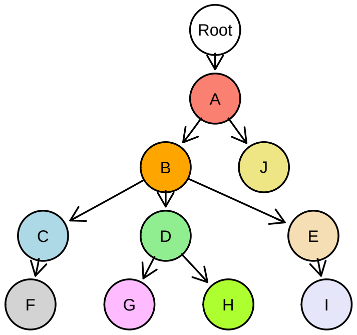


| | |
|---|---|
| *ID* | j5cMndnUwm51TTqp |
| *acc genotypes* | 93 |
| *peaks no back* | 1 |
| *removed edges* | 0 |
| *prop rem edges* | 0 |

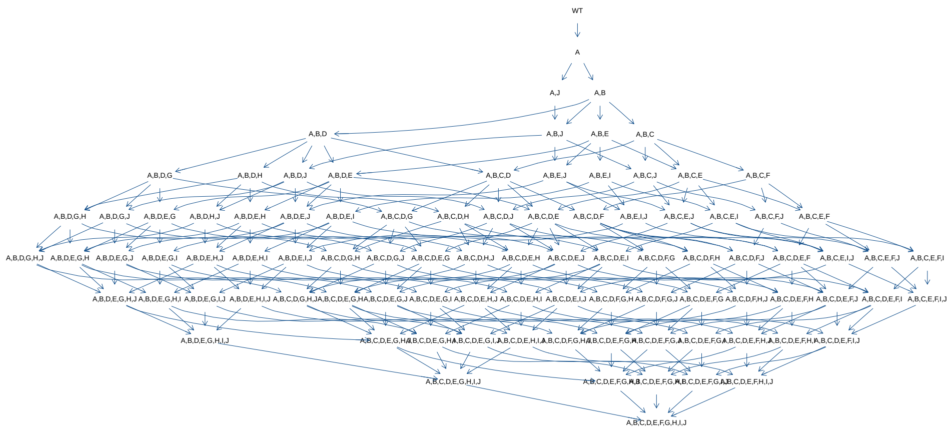


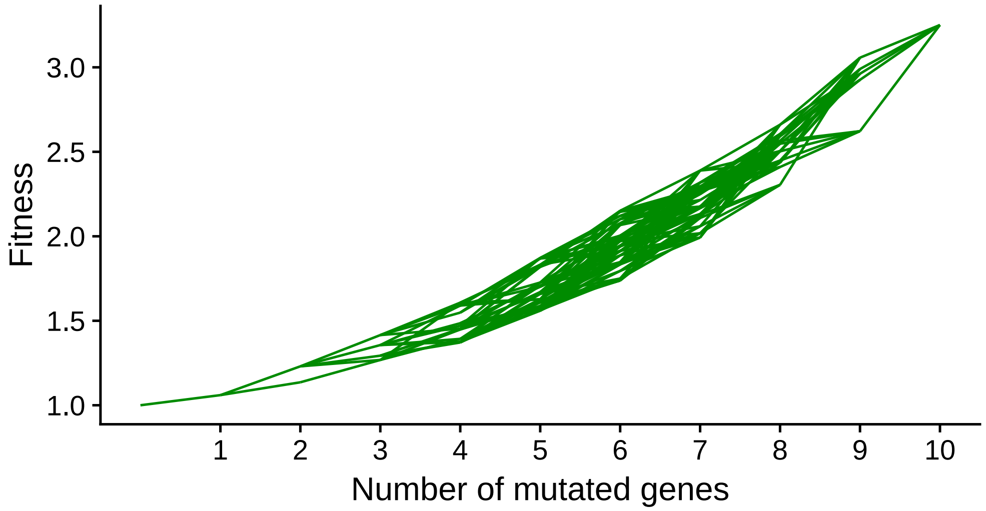

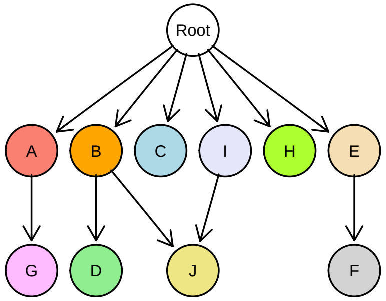


*ID* j6HW2RoAhJ2N68DN
*acc genotypes* 288
*peaks no back* 1
*removed edges* 0
*prop rem edges* 0

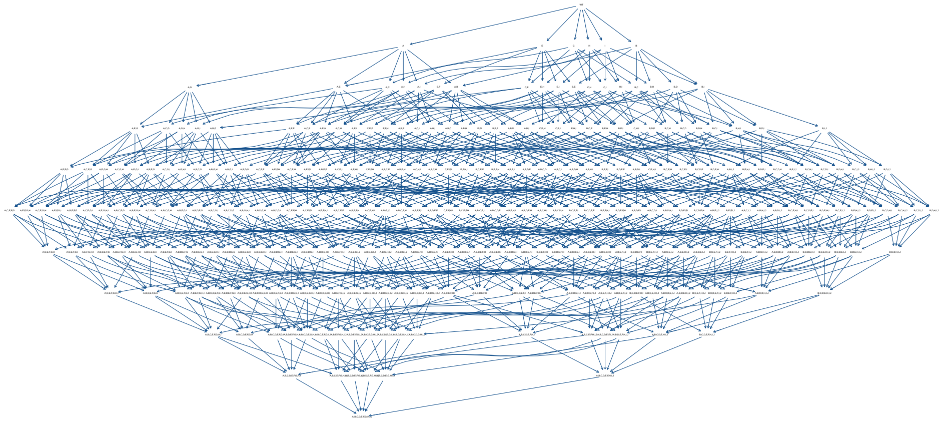

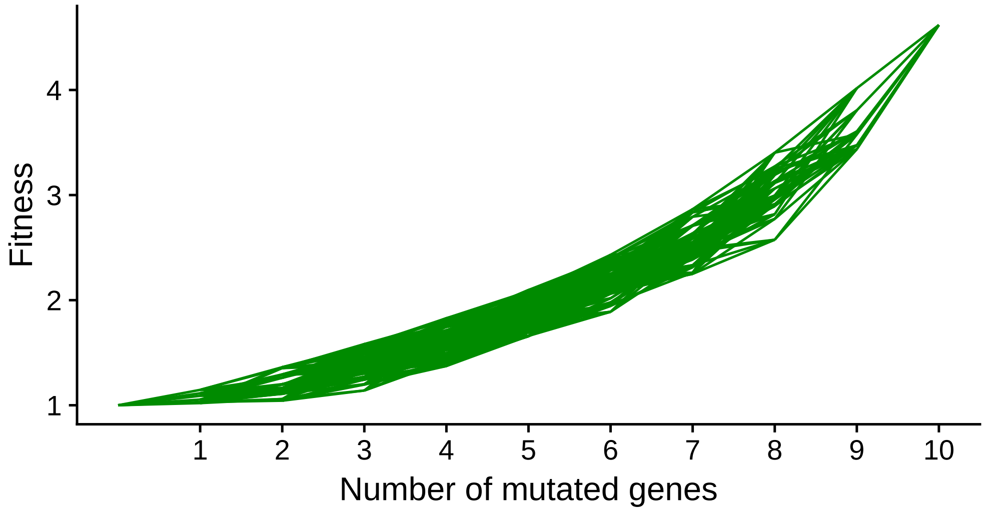

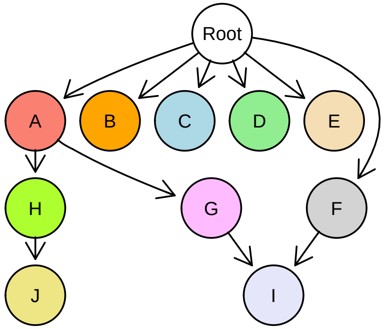


| | |
|---|---|
| *ID* | jAZ9fNKs3jEXePB4 |
| *acc genotypes* | 272 |
| *peaks no back* | 1 |
| *removed edges* | 0 |
| *prop rem edges* | 0 |

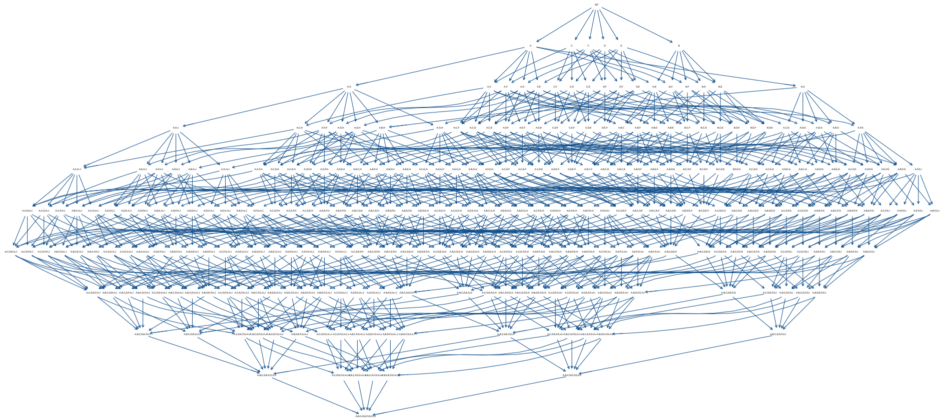

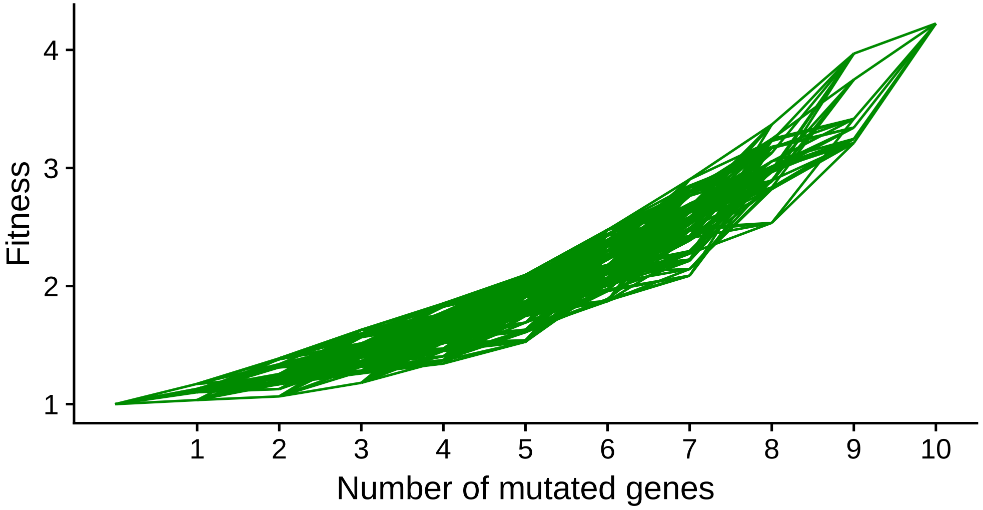

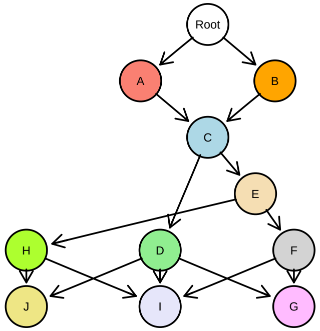


| | |
|---|---|
| *ID* | jD4PcR3DEFwIK6qC |
| *acc genotypes* | 23 |
| *peaks no back* | 1 |
| *removed edges* | 0 |
| *prop rem edges* | 0 |

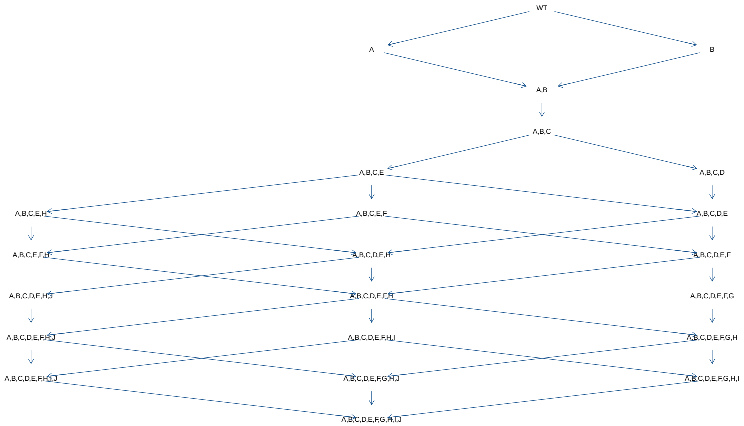


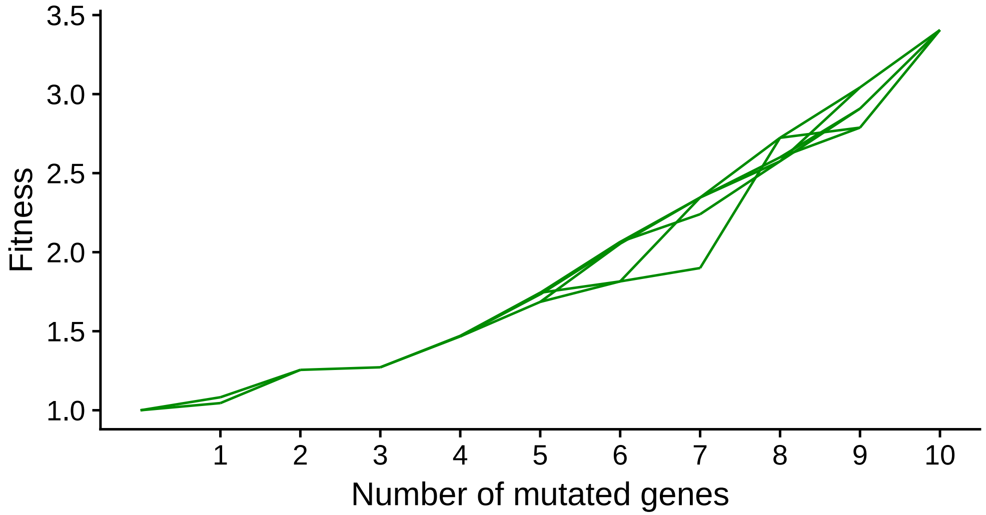

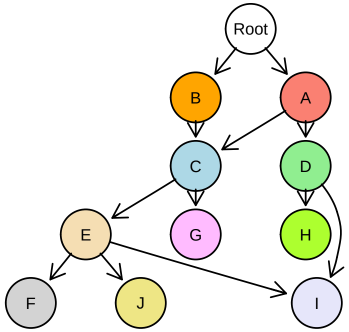


| | |
|---|---|
| *ID* | Jh8glYdX10audNWJF |
| *acc genotypes* | 54 |
| *peaks no back* | 1 |
| *removed edges* | 0 |
| *prop rem edges* | 0 |

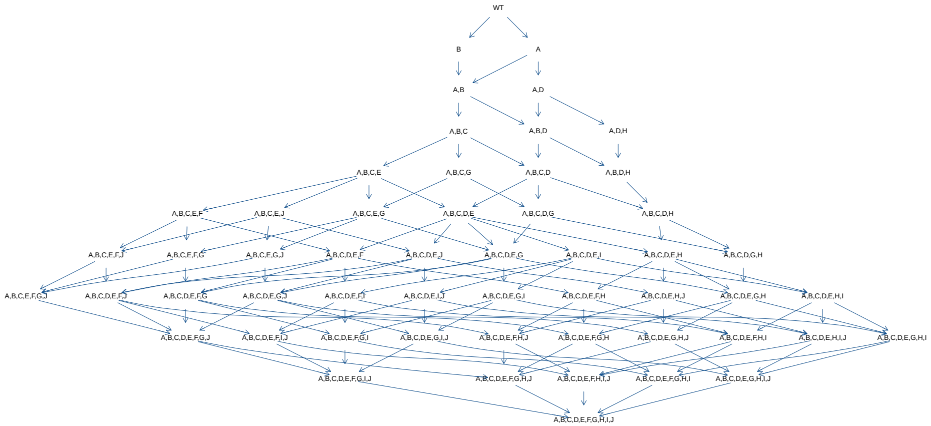

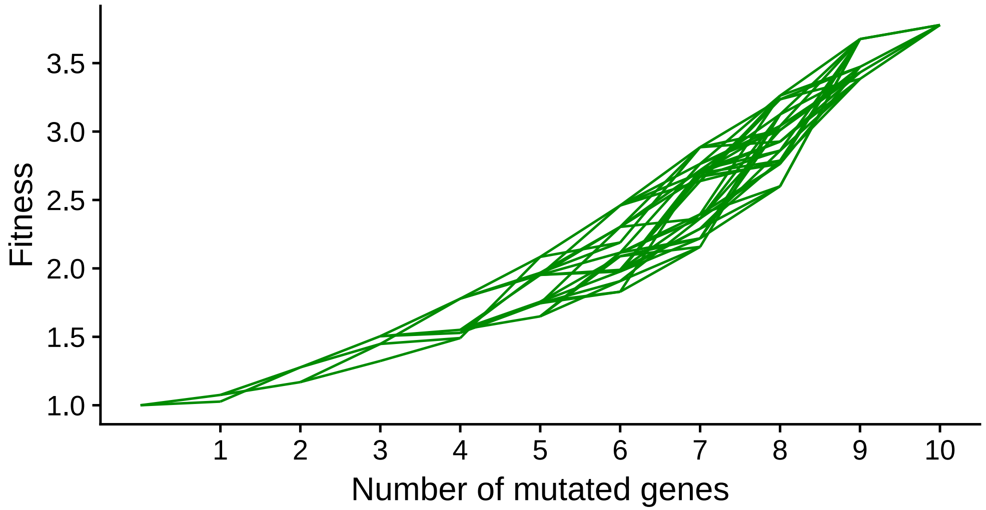

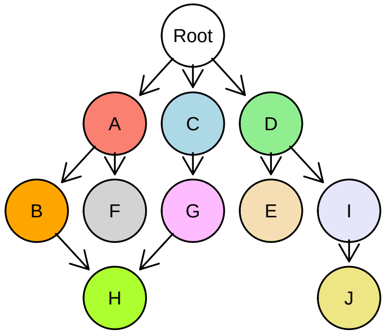


| | |
|---|---|
| *ID* | JhPPnqoKAJCL8pvR |
| *acc genotypes* | 119 |
| *peaks no back* | 1 |
| *removed edges* | 0 |
| *prop rem edges* | 0 |

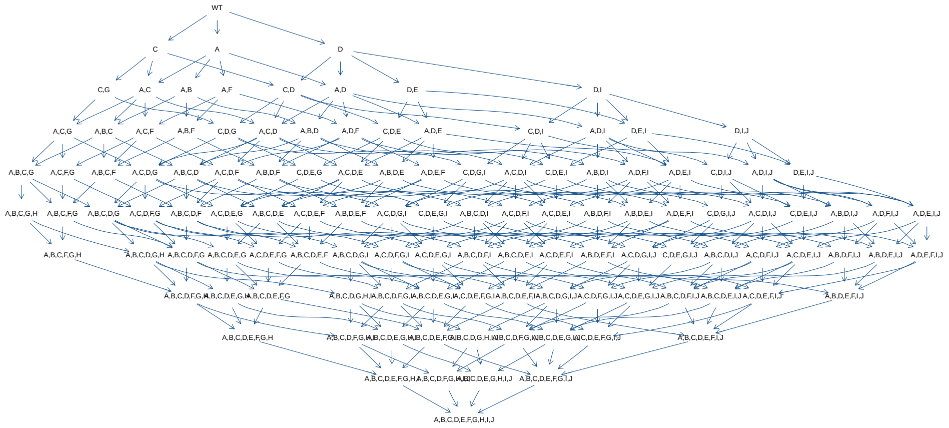

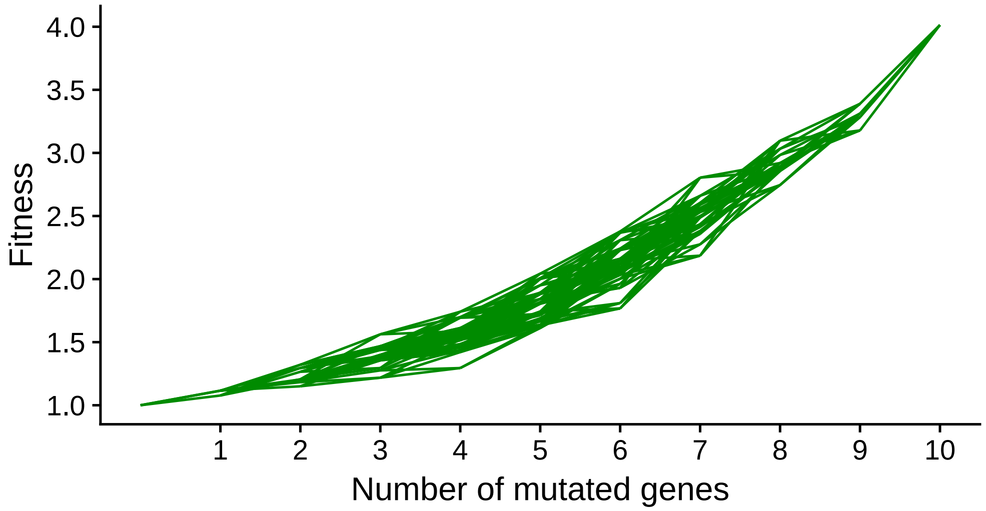

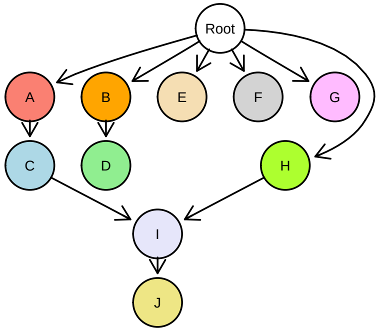


| | |
|---|---|
| *ID* | JIKf7ckdhSo6ArQQ |
| *acc genotypes* | 192 |
| *peaks no back* | 1 |
| *removed edges* | 0 |
| *prop rem edges* | 0 |

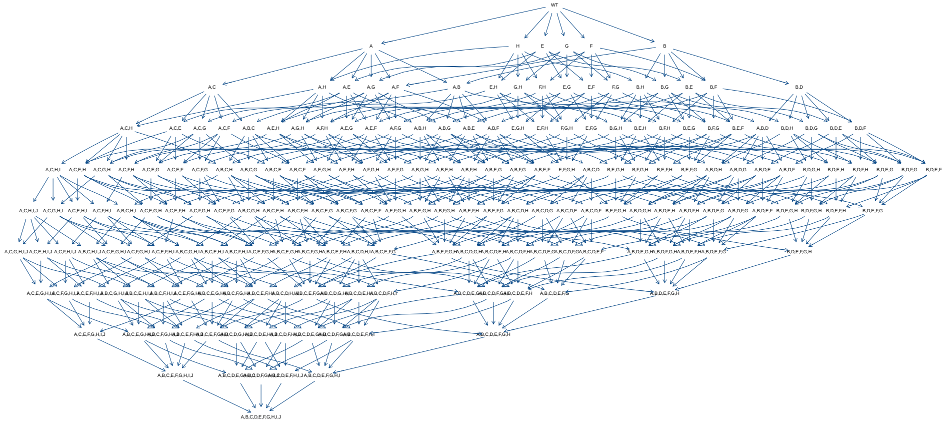

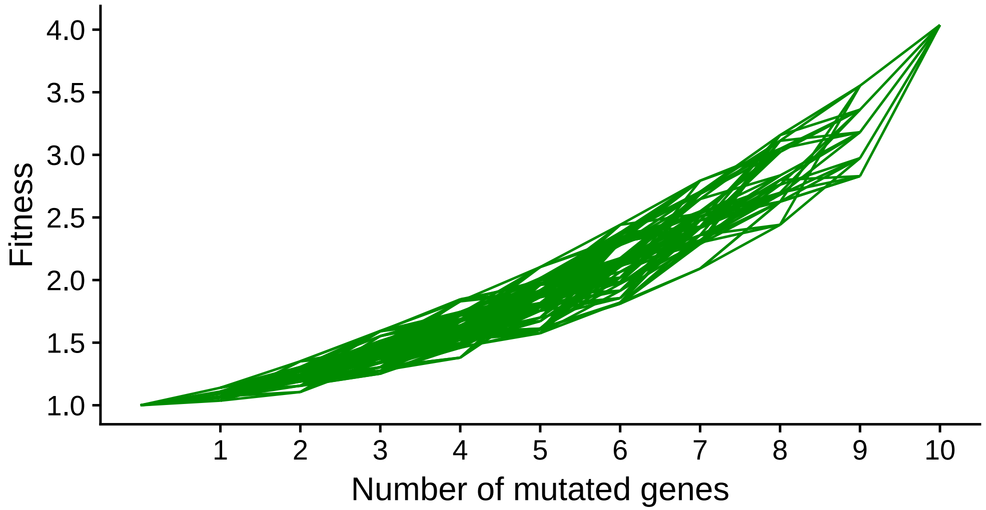

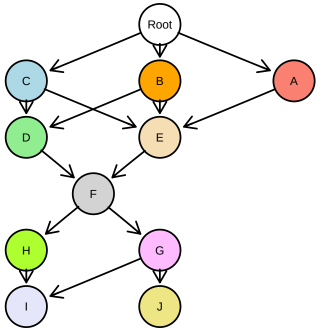


| | |
|---|---|
| *ID* | JoiUrmmVqi67SyTm |
| *acc genotypes* | 20 |
| *peaks no back* | 1 |
| *removed edges* | 0 |
| *prop rem edges* | 0 |

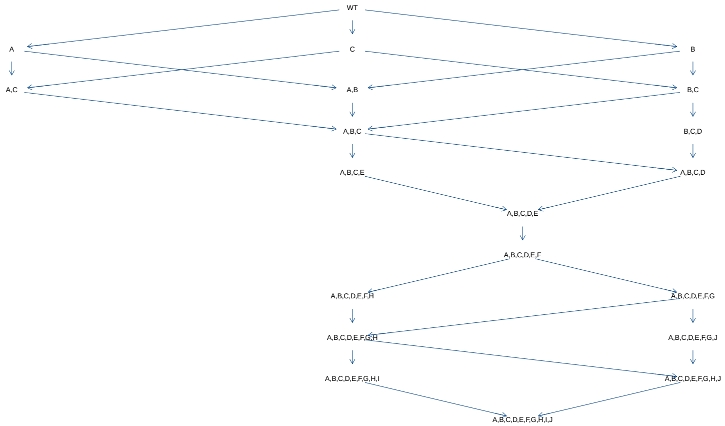


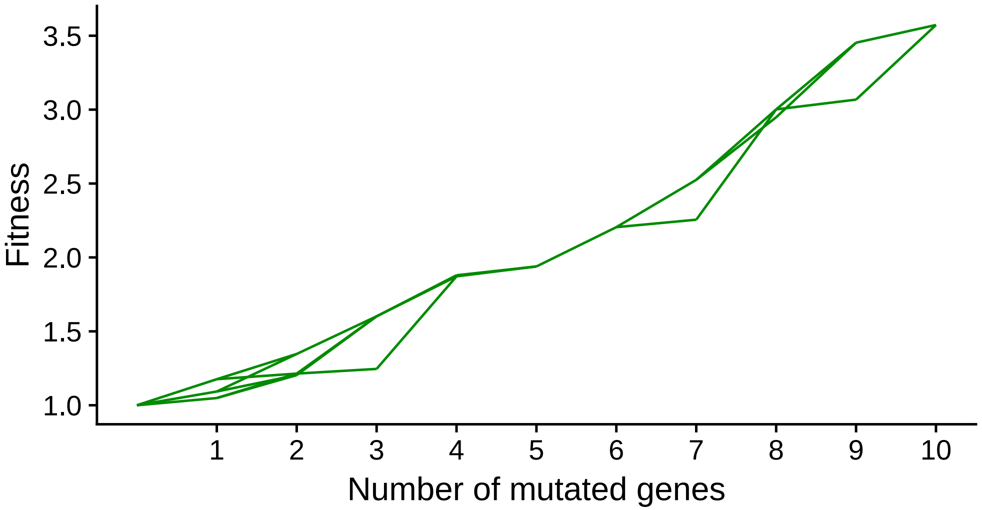

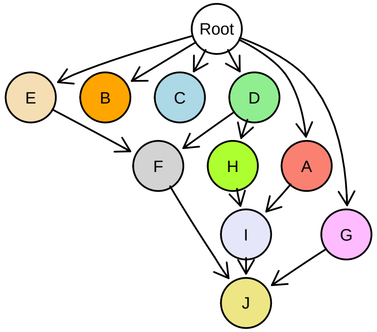


| | |
|---|---|
| *ID* | jSQmuyLWDxqE106yj |
| *acc genotypes* | 156 |
| *peaks no back* | 1 |
| *removed edges* | 0 |
| *prop rem edges* | 0 |

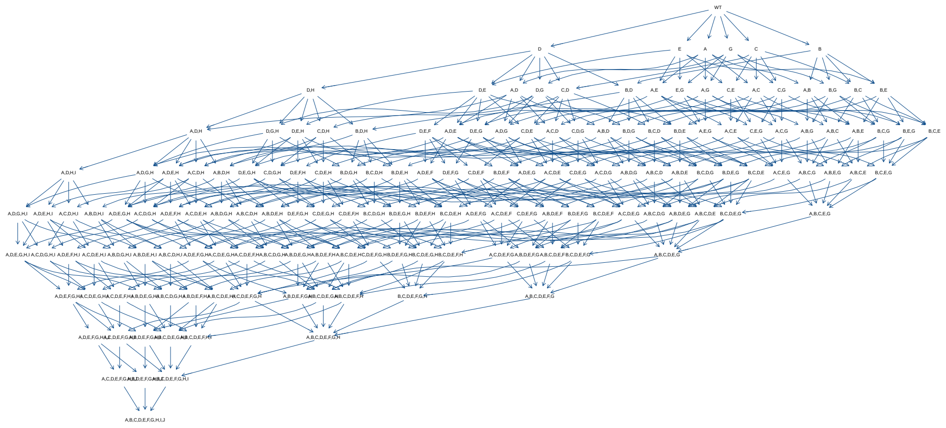

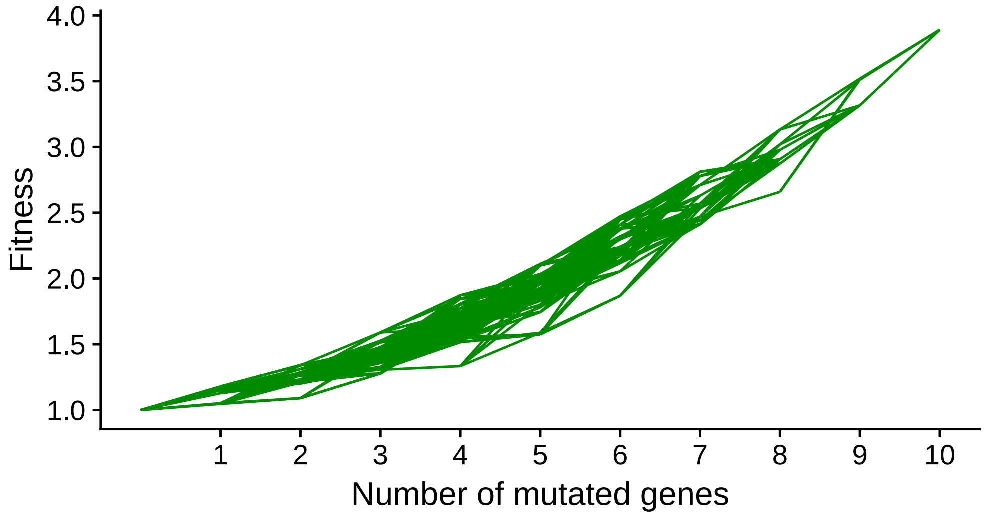

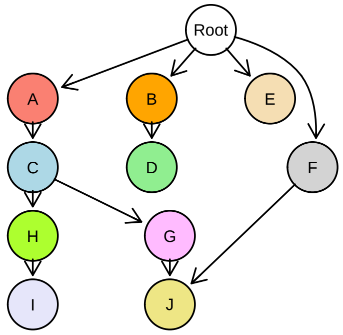


| | |
|---|---|
| *ID* | jXDzA3wcvwQmXJvD |
| *acc genotypes* | 114 |
| *peaks no back* | 1 |
| *removed edges* | 0 |
| *prop rem edges* | 0 |

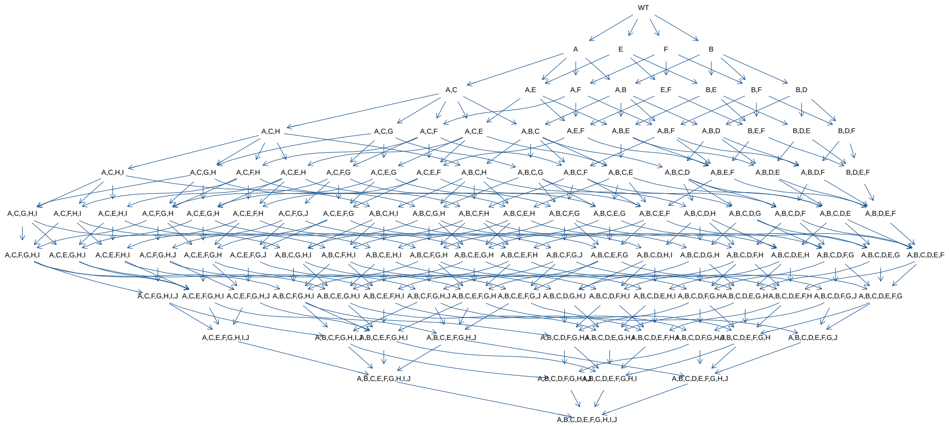

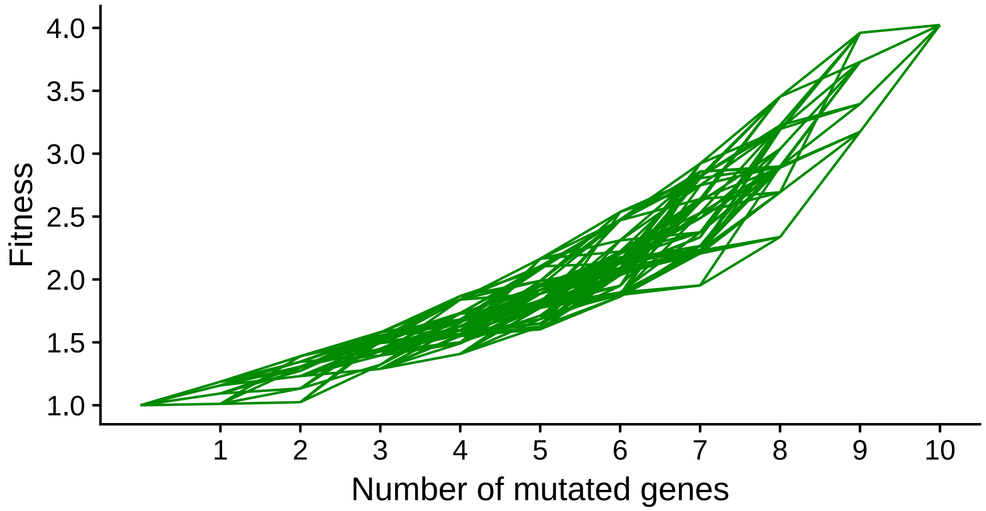

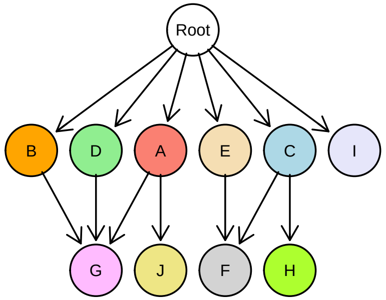


| | |
|---|---|
| *ID* | K1jCHK7t4zCUPD6B |
| *acc genotypes* | 224 |
| *peaks no back* | 1 |
| *removed edges* | 0 |
| *prop rem edges* | 0 |

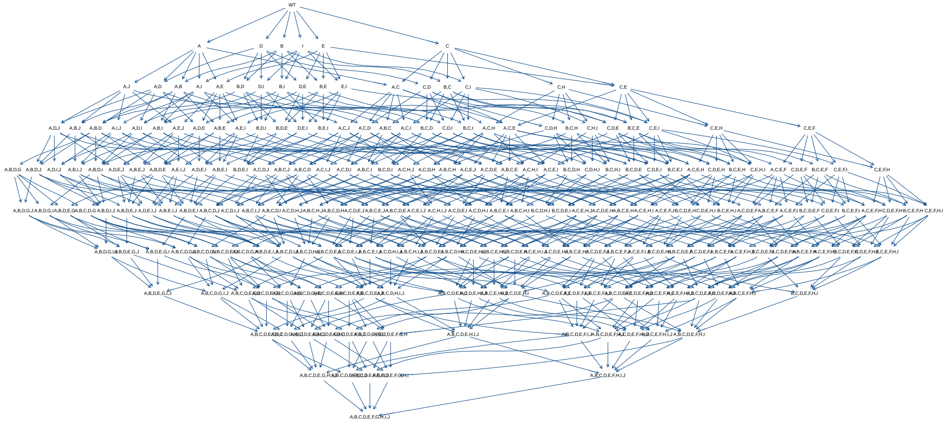

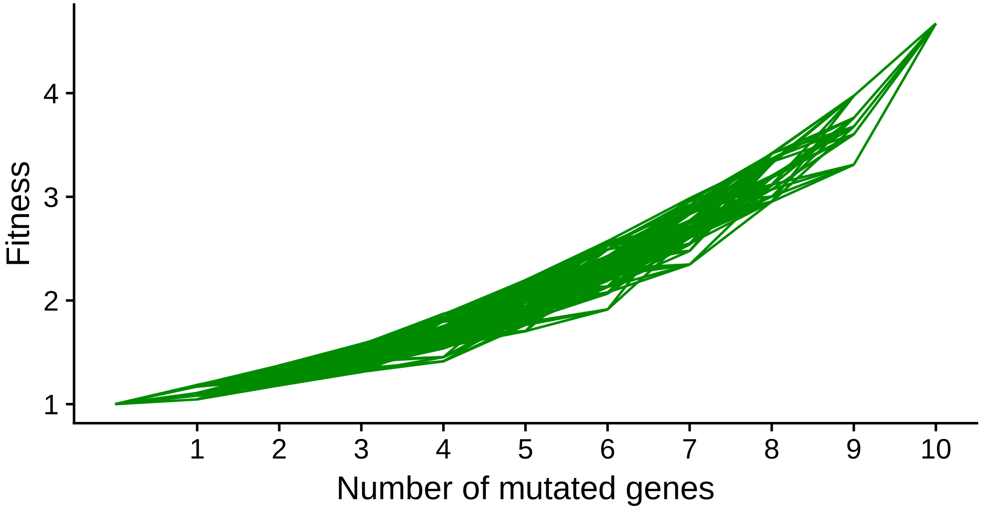

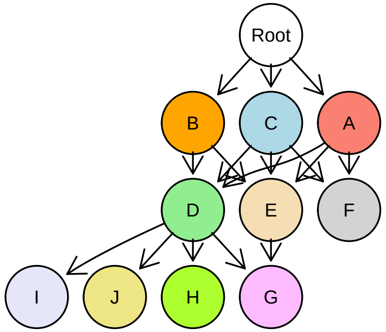


| | |
|---|---|
| *ID* | KkQL102YepOlao9Bb |
| *acc genotypes* | 60 |
| *peaks no back* | 1 |
| *removed edges* | 0 |
| *prop rem edges* | 0 |

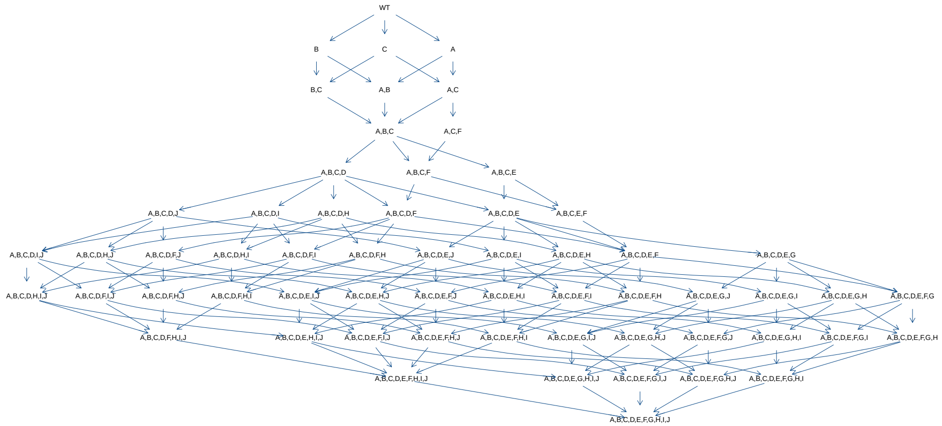

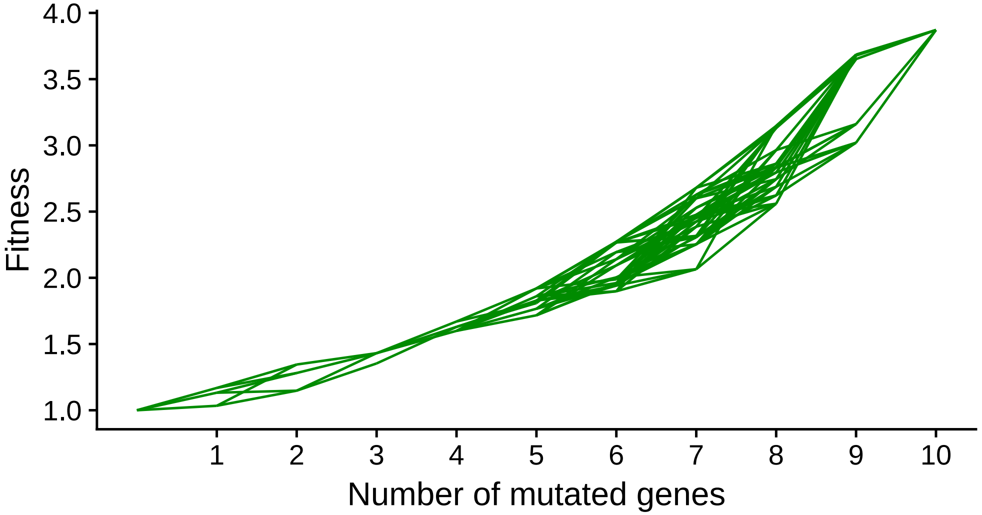

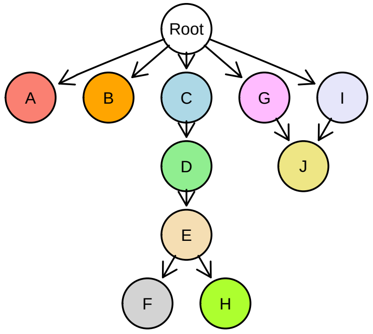


| | |
|---|---|
| *ID* | kpRFNq2Q10Ex8oSMG |
| *acc genotypes* | 140 |
| *peaks no back* | 1 |
| *removed edges* | 0 |
| *prop rem edges* | 0 |

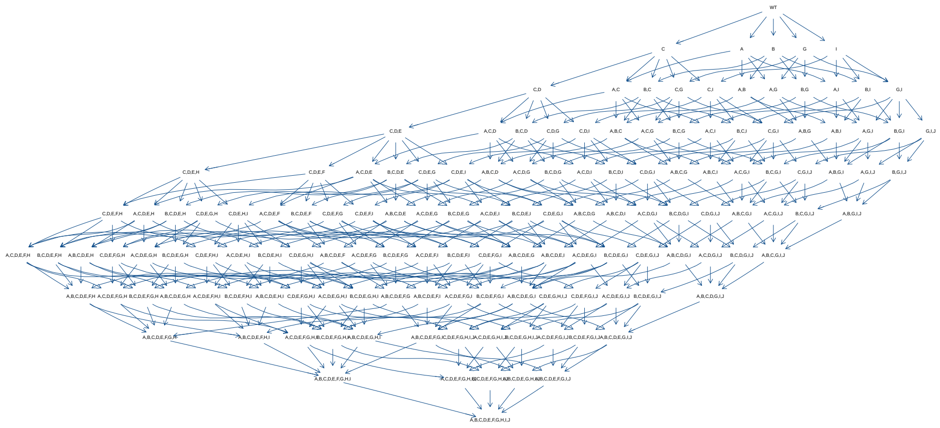

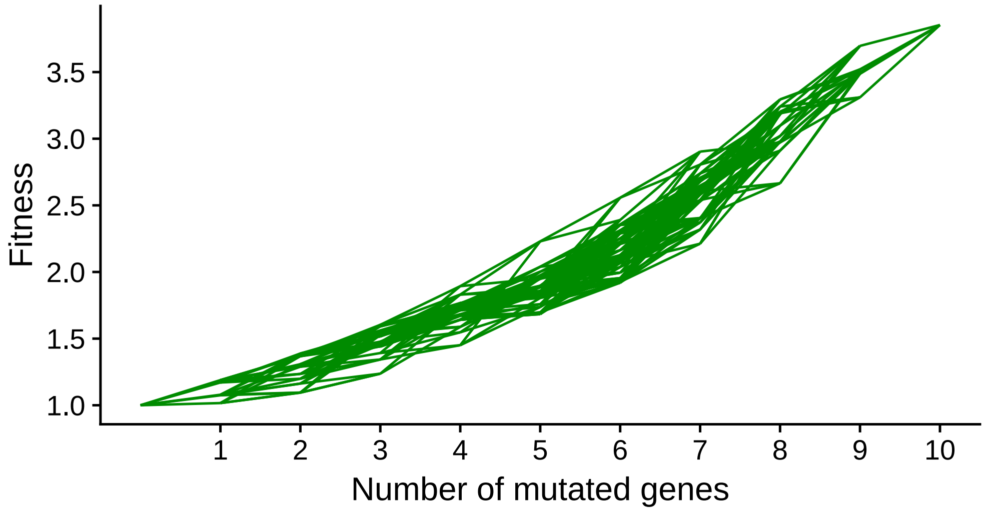

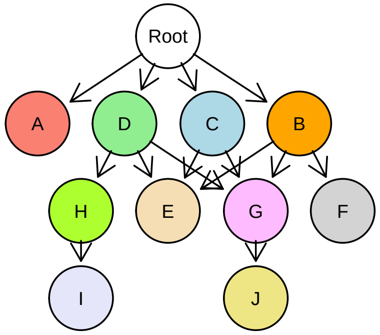


*ID* ktOHeXaq4zp10UQNO
*acc genotypes* 108
*peaks no back* 1
*removed edges* 0
*prop rem edges* 0

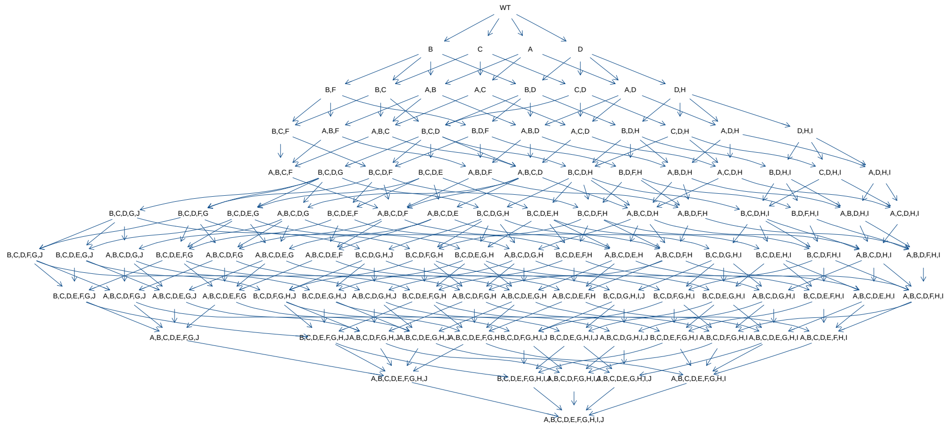

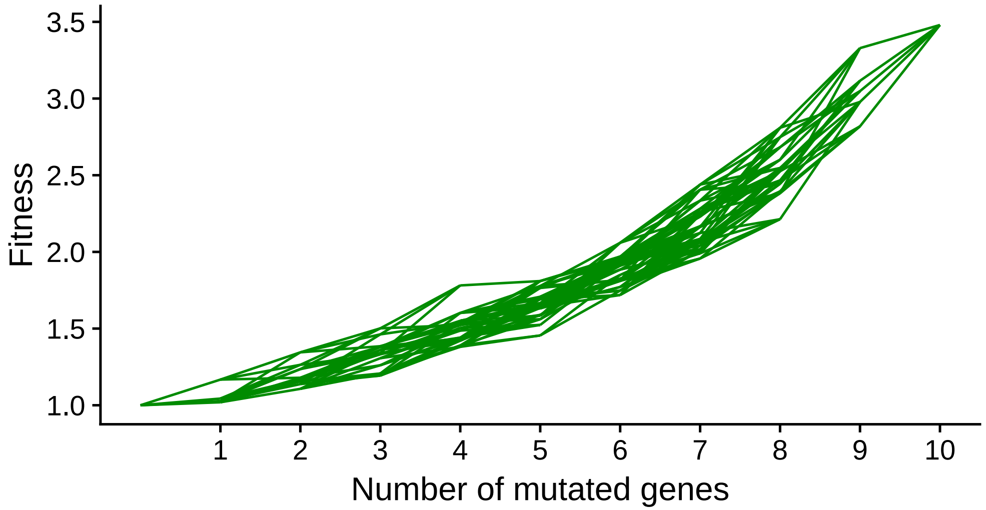

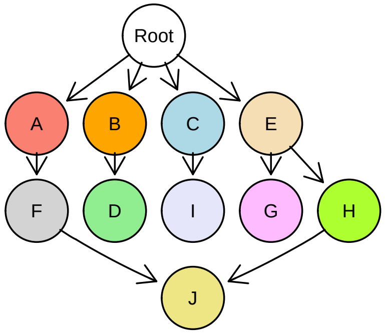


| | |
|---|---|
| *ID* | kvDLqWZKOAz5w43U |
| *acc genotypes* | 153 |
| *peaks no back* | 1 |
| *removed edges* | 0 |
| *prop rem edges* | 0 |

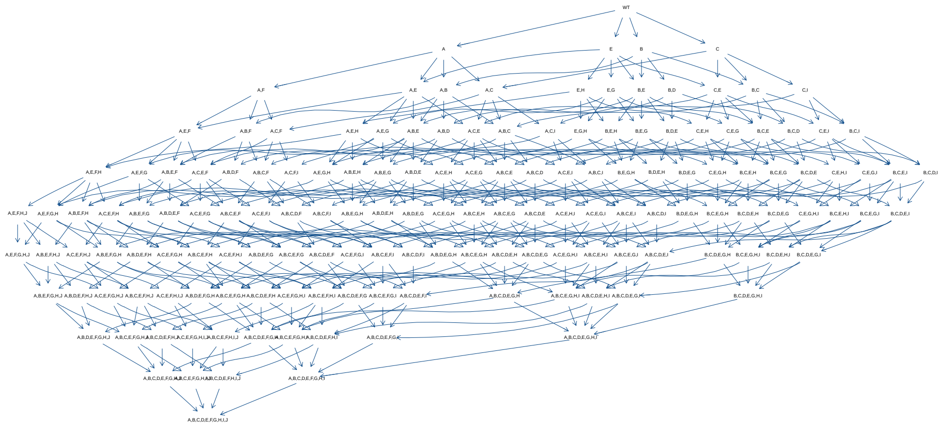

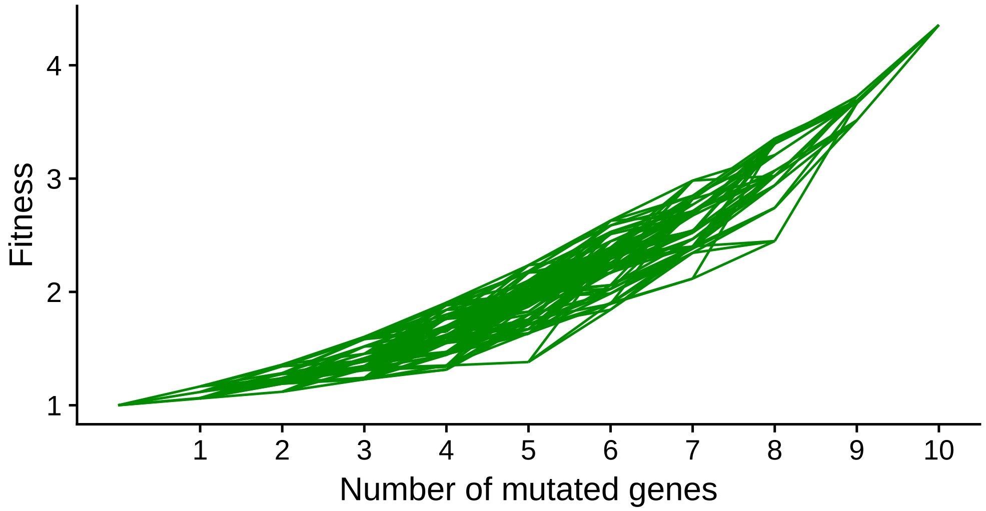

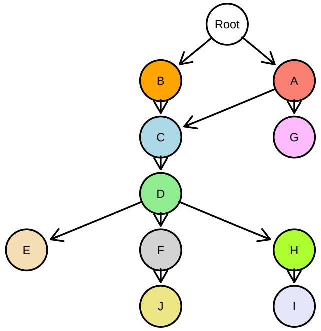


| | |
|---|---|
| *ID* | LAiwaf4n74KiIOZv |
| *acc genotypes* | 44 |
| *peaks no back* | 1 |
| *removed edges* | 0 |
| *prop rem edges* | 0 |

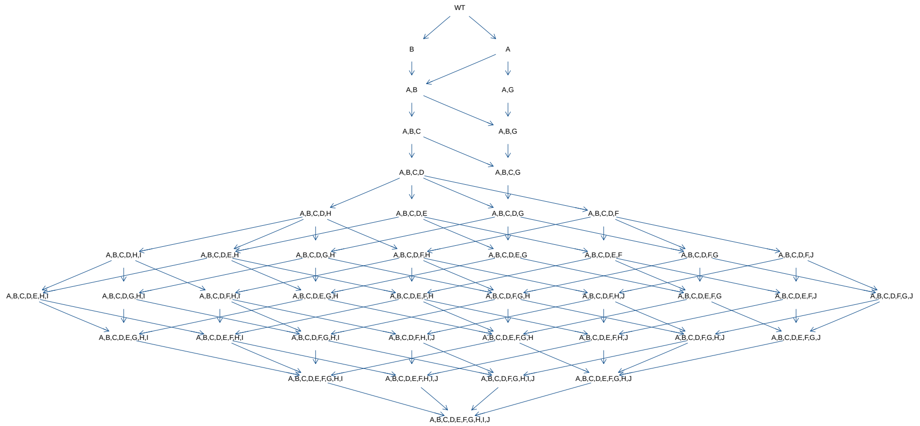

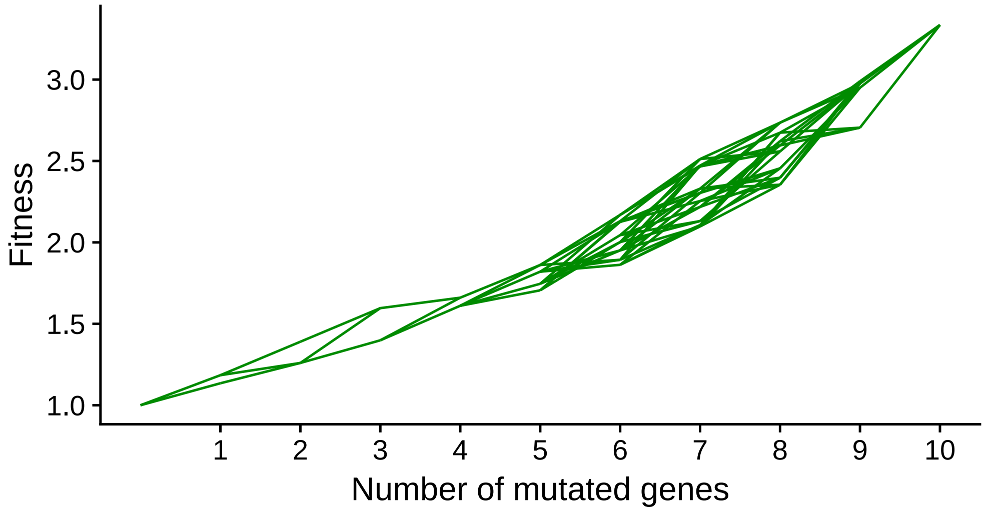

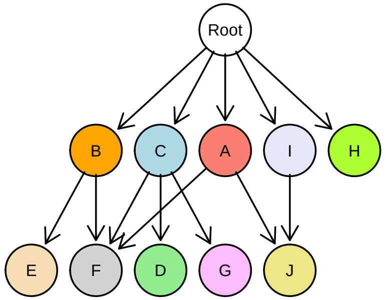


| | |
|---|---|
| *ID* | LgHU63NzyxJ8cVAR |
| *acc genotypes* | 198 |
| *peaks no back* | 1 |
| *removed edges* | 0 |
| *prop rem edges* | 0 |

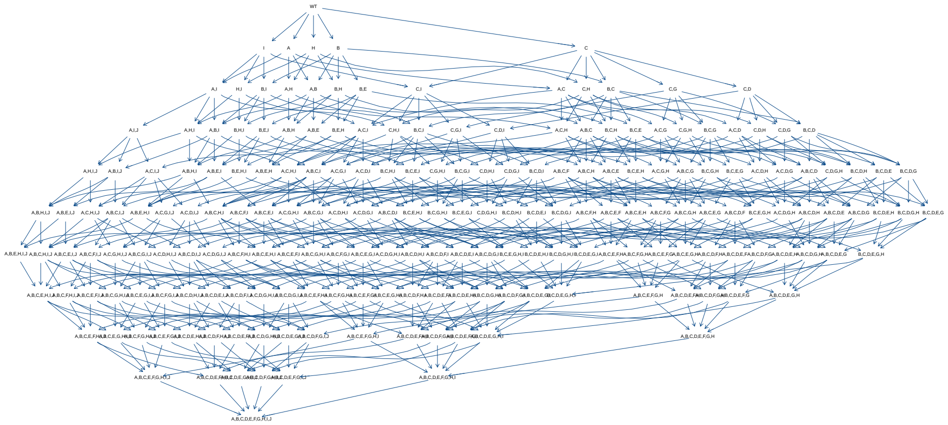

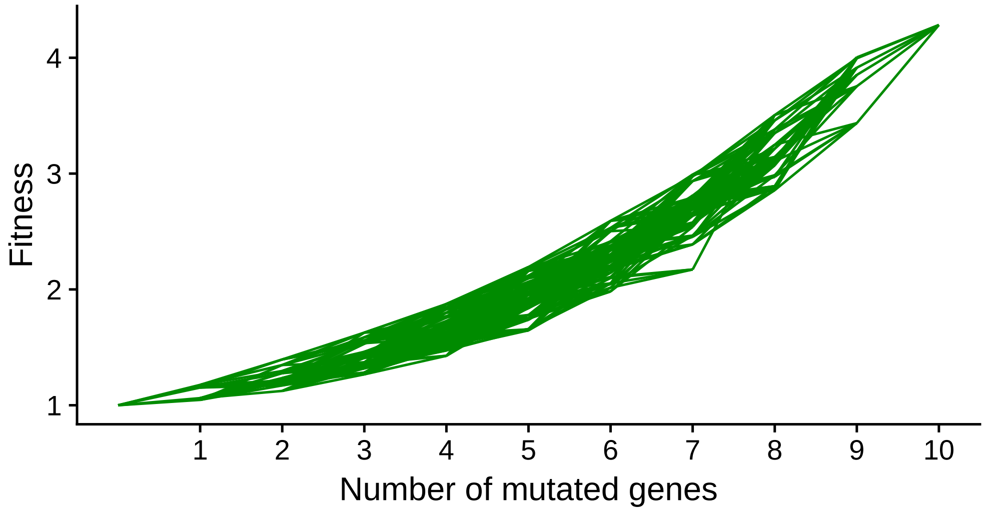

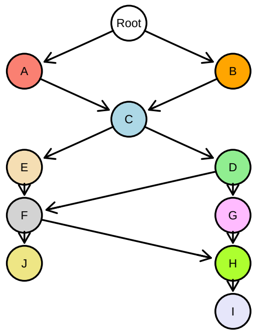


| | |
|---|---|
| *ID* | lIOath10wN5BQcyee |
| *acc genotypes* | 18 |
| *peaks no back* | 1 |
| *removed edges* | 0 |
| *prop rem edges* | 0 |

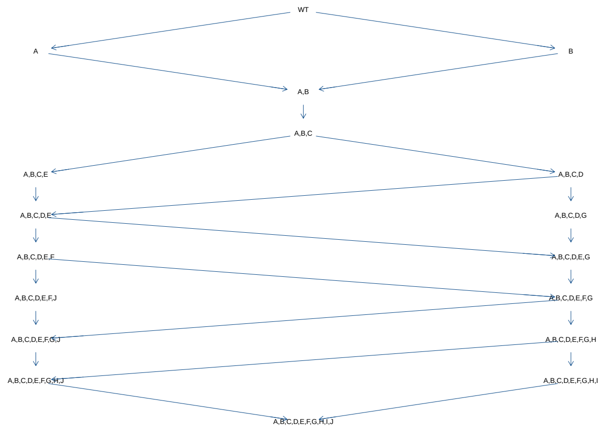


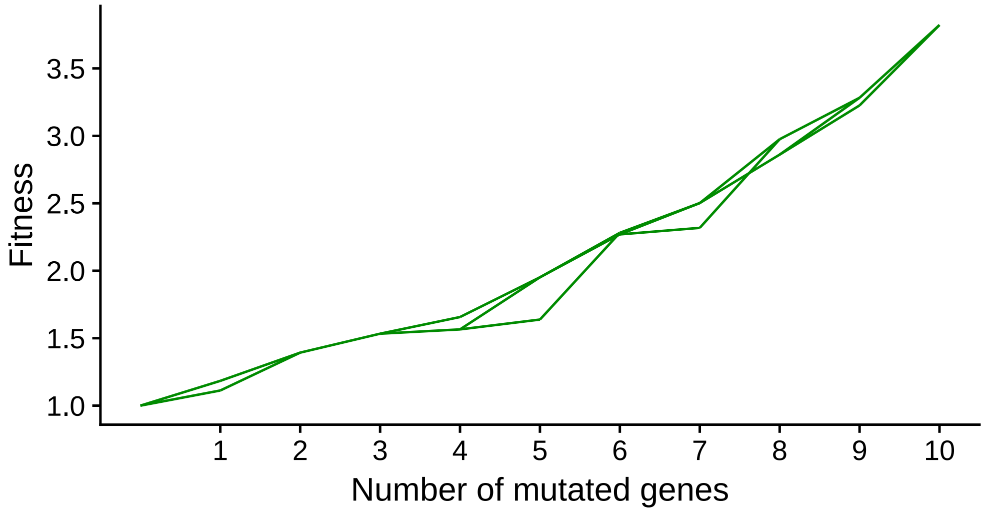

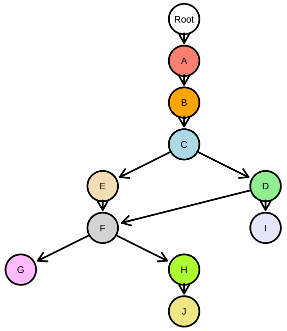


| | |
|---|---|
| *ID* | LiqVyEk5185giJ87 |
| *acc genotypes* | 21 |
| *peaks no back* | 1 |
| *removed edges* | 0 |
| *prop rem edges* | 0 |

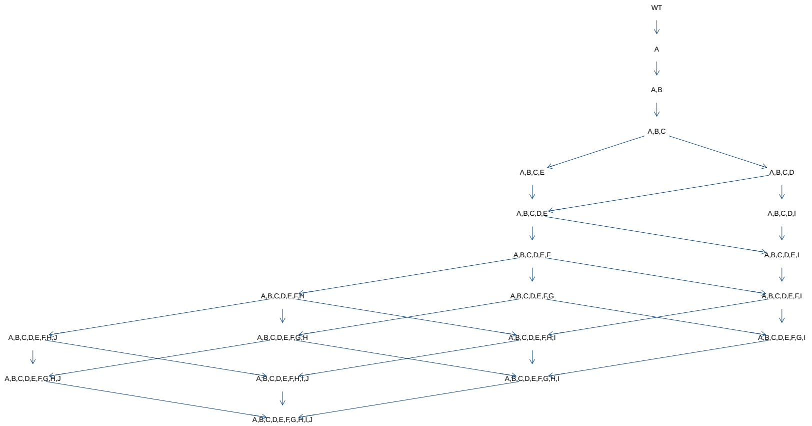

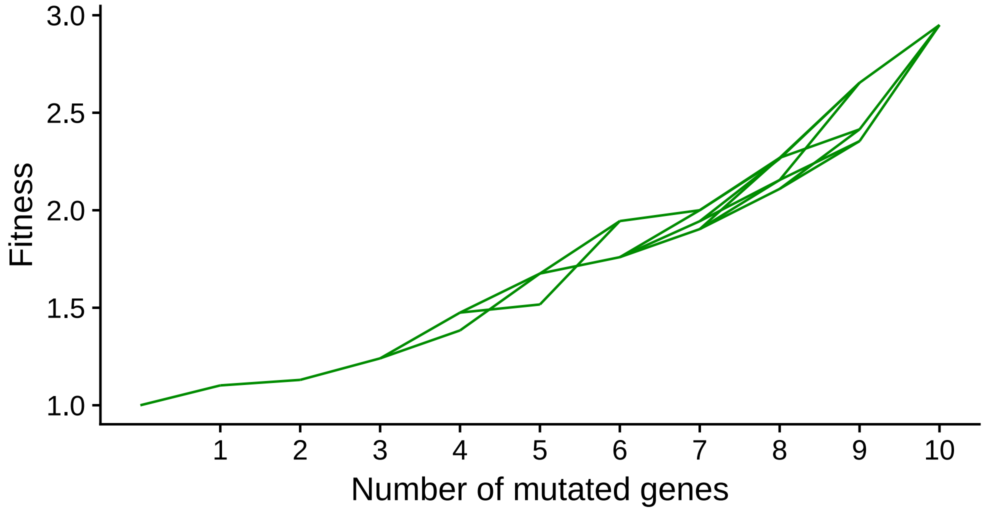

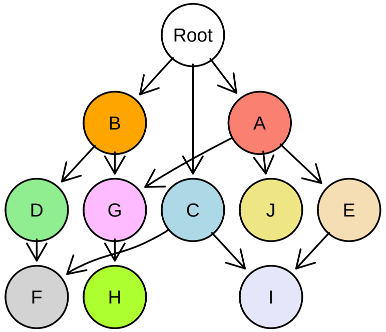


| | |
|---|---|
| *ID* | IJ3M9zaA46iBzYO2 |
| *acc genotypes* | 95 |
| *peaks no back* | 1 |
| *removed edges* | 0 |
| *prop rem edges* | 0 |

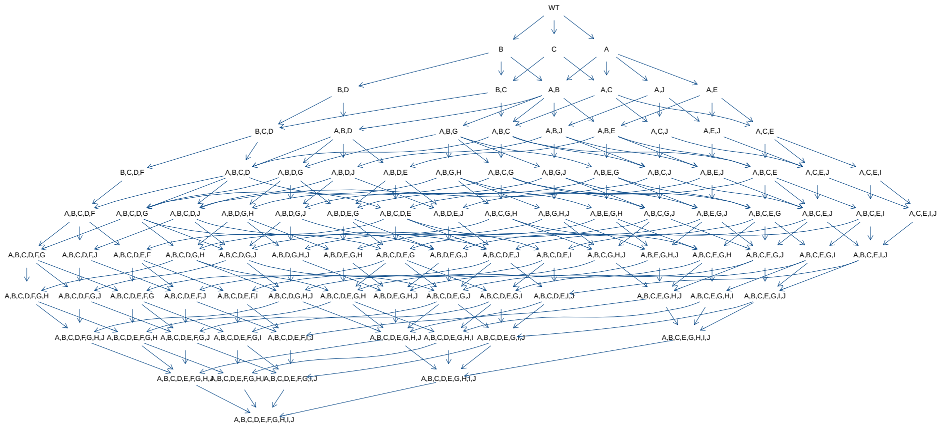

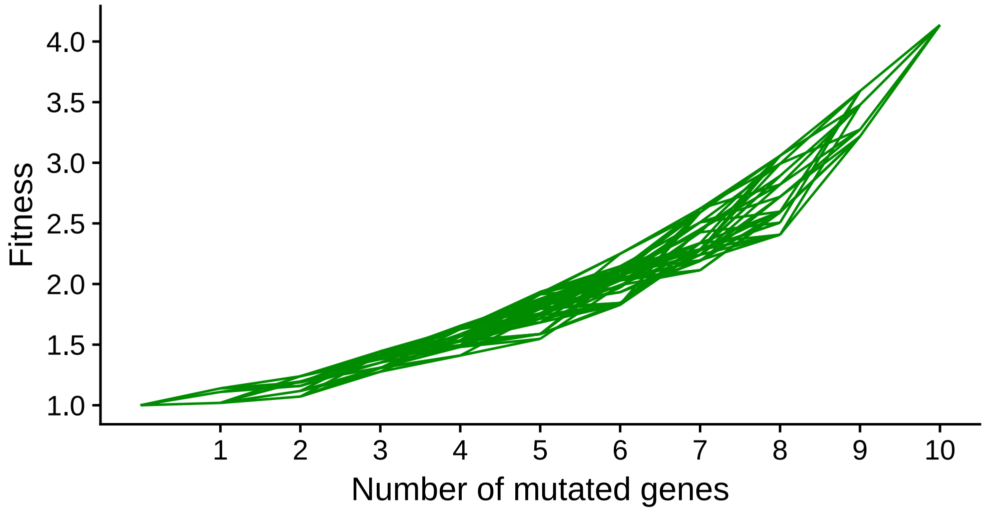

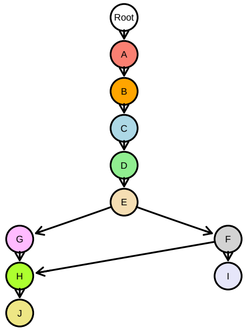


| | |
|---|---|
| *ID* | LKzTjEtauR10FkWvI |
| *acc genotypes* | 15 |
| *peaks no back* | 1 |
| *removed edges* | 0 |
| *prop rem edges* | 0 |

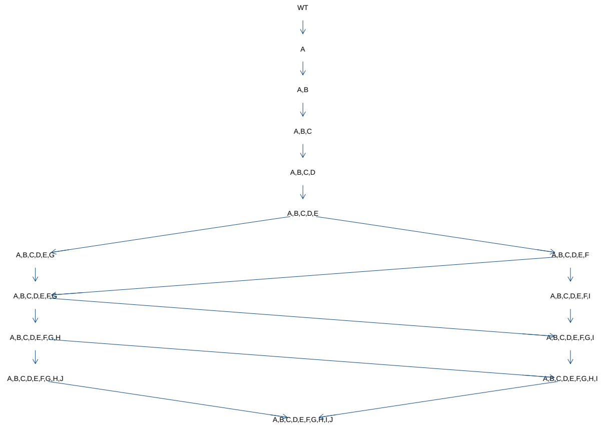


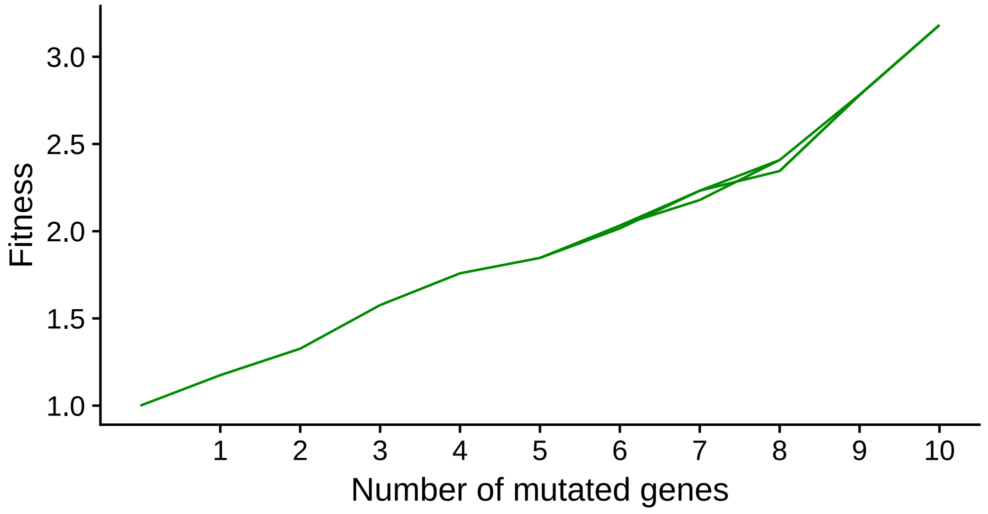

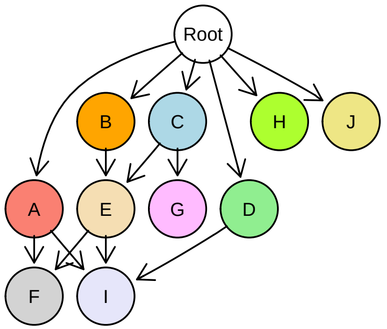


| | |
|---|---|
| *ID* | LLARXmNmlvPm7fLH |
| *acc genotypes* | 160 |
| *peaks no back* | 1 |
| *removed edges* | 0 |
| *prop rem edges* | 0 |

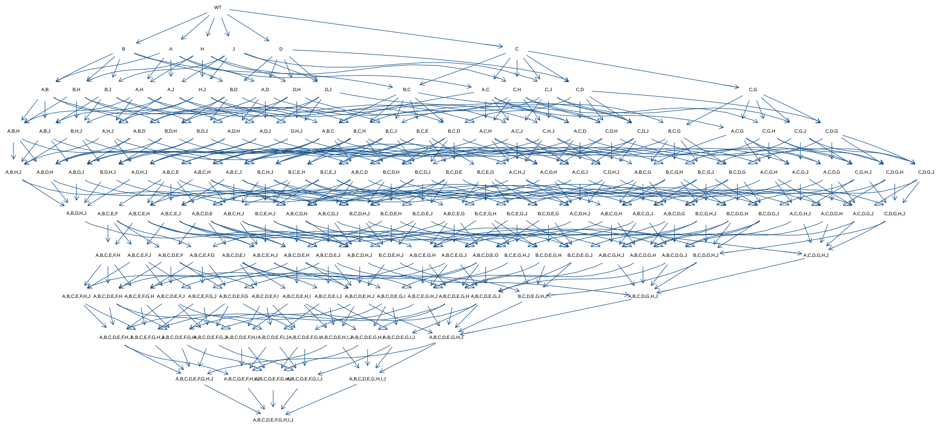

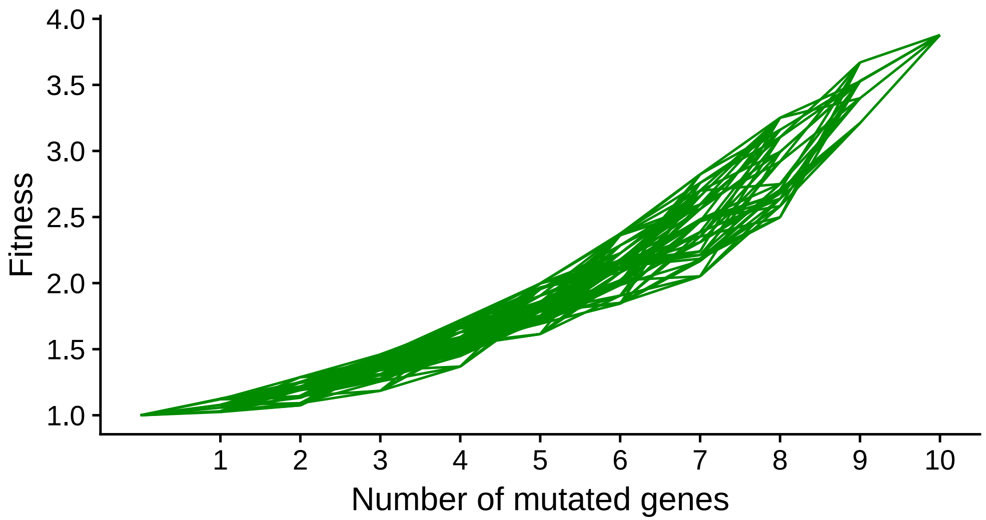

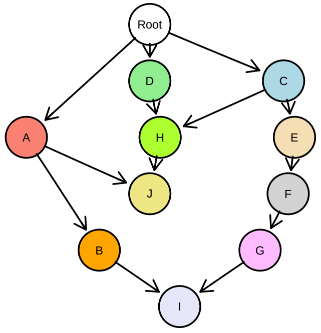


| | |
|---|---|
| *ID* | LoVQjzyAttkUHwB8 |
| *acc genotypes* | 54 |
| *peaks no back* | 1 |
| *removed edges* | 0 |
| *prop rem edges* | 0 |

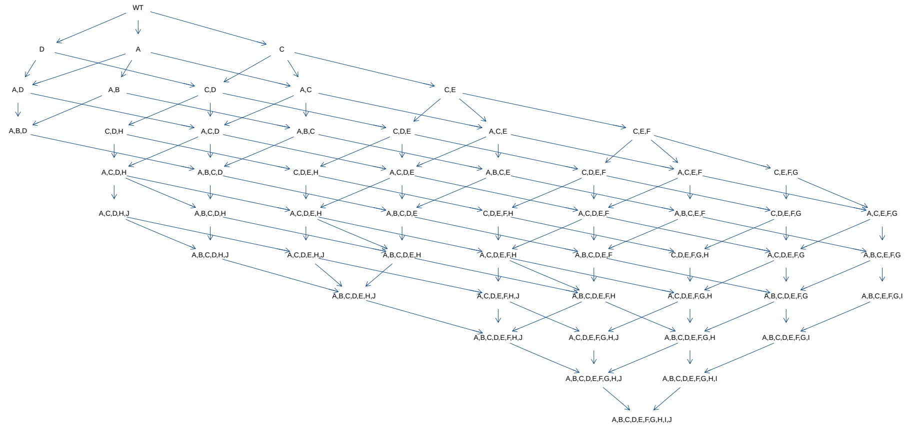

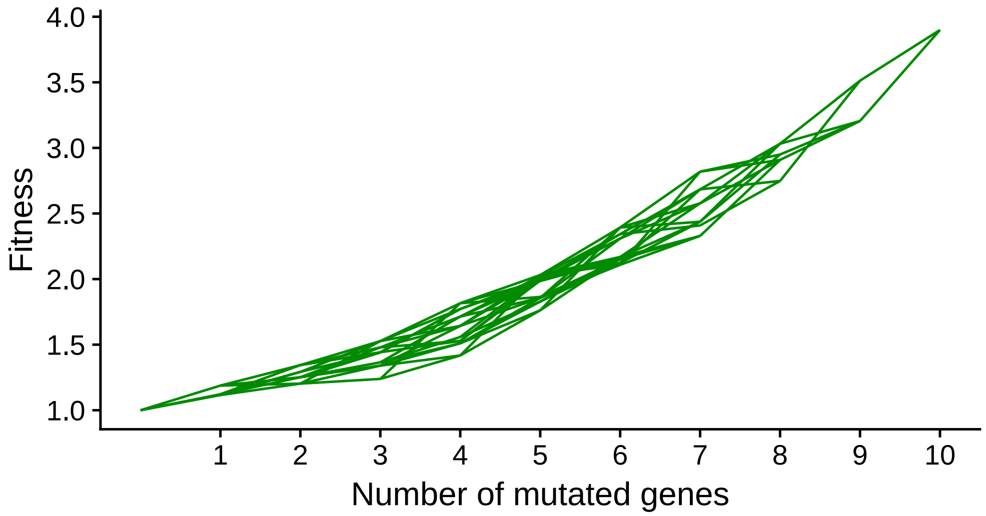

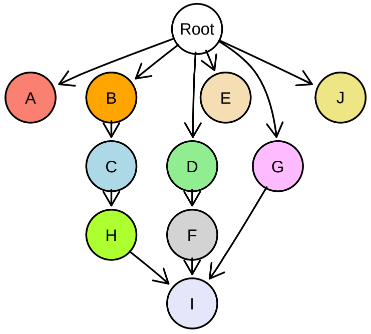


| | |
|---|---|
| *ID* | m2LJE5DdVSyhA9R9 |
| *acc genotypes* | 200 |
| *peaks no back* | 1 |
| *removed edges* | 0 |
| *prop rem edges* | 0 |

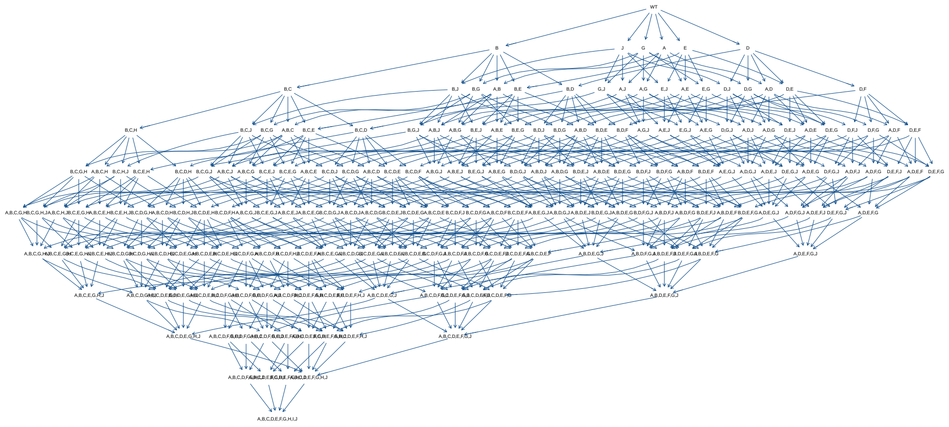

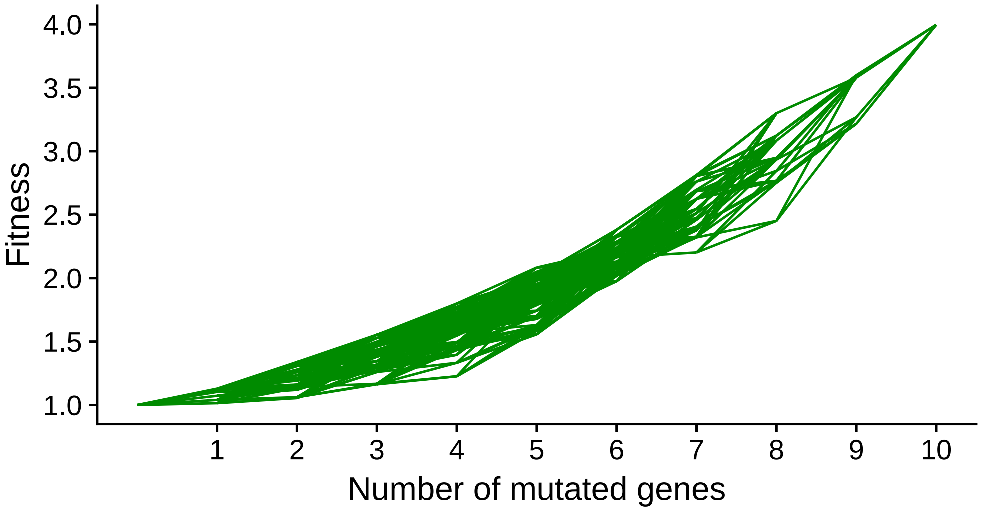

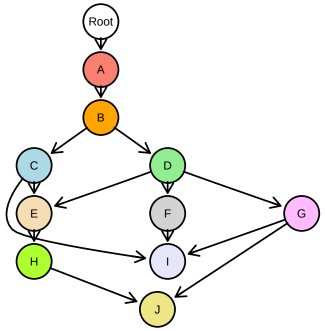


| | |
|---|---|
| *ID* | M5NjxUHUHAkU7nQF |
| *acc genotypes* | 26 |
| *peaks no back* | 1 |
| *removed edges* | 0 |
| *prop rem edges* | 0 |

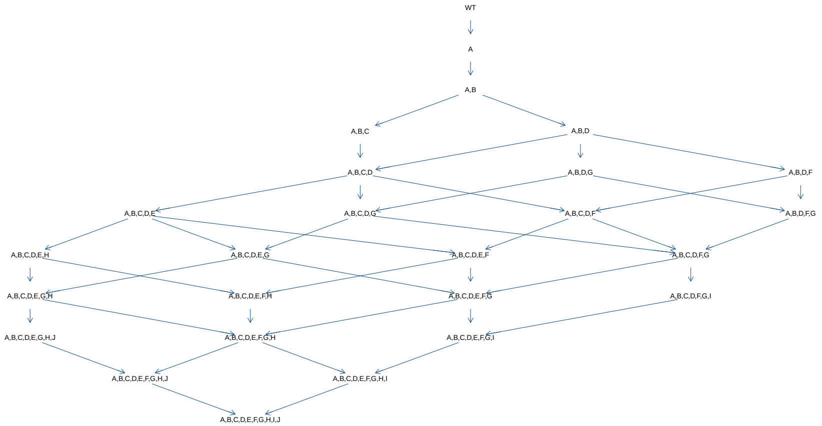


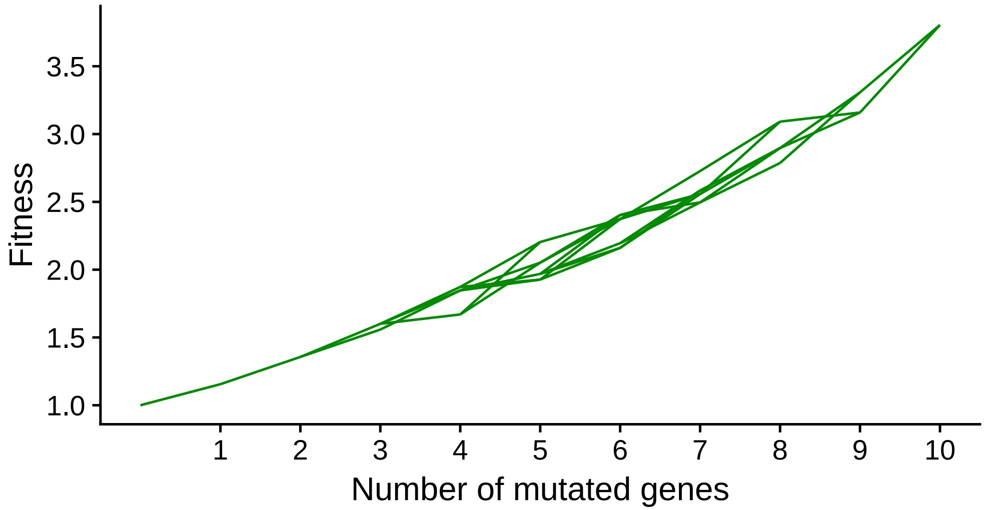

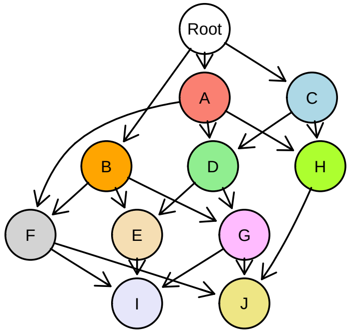


| | |
|---|---|
| *ID* | MJxO4QrzOUXVWzXM |
| *acc genotypes* | 36 |
| *peaks no back* | 1 |
| *removed edges* | 0 |
| *prop rem edges* | 0 |

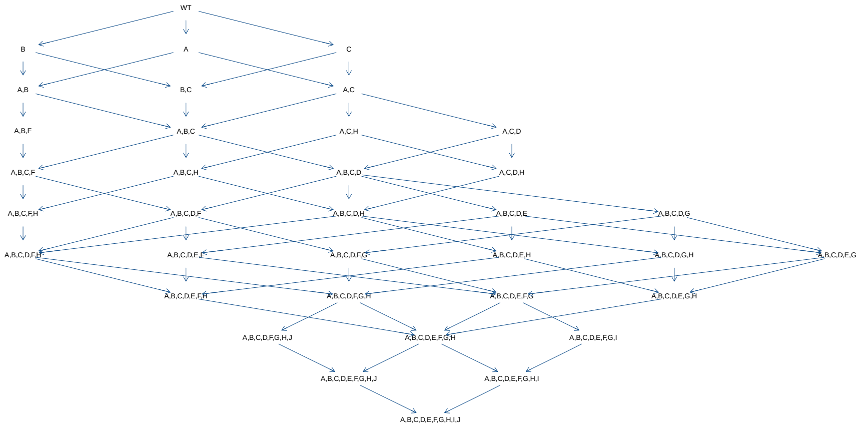

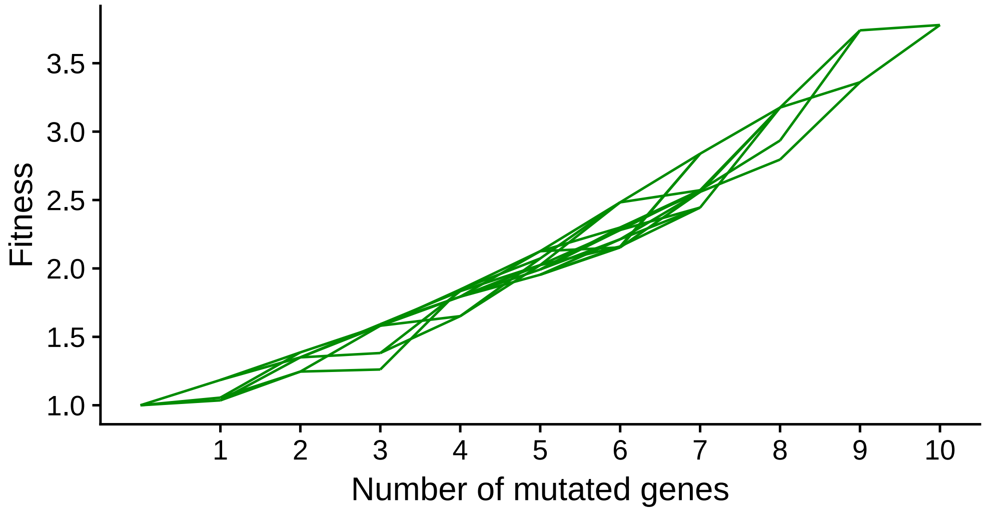

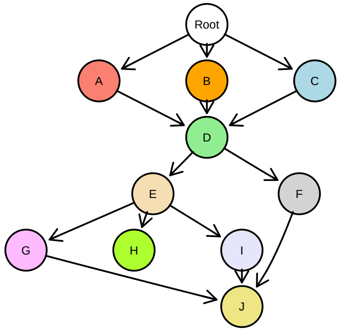


| | |
|---|---|
| *ID* | MkJPmR2HWGDw34xG |
| *acc genotypes* | 28 |
| *peaks no back* | 1 |
| *removed edges* | 0 |
| *prop rem edges* | 0 |

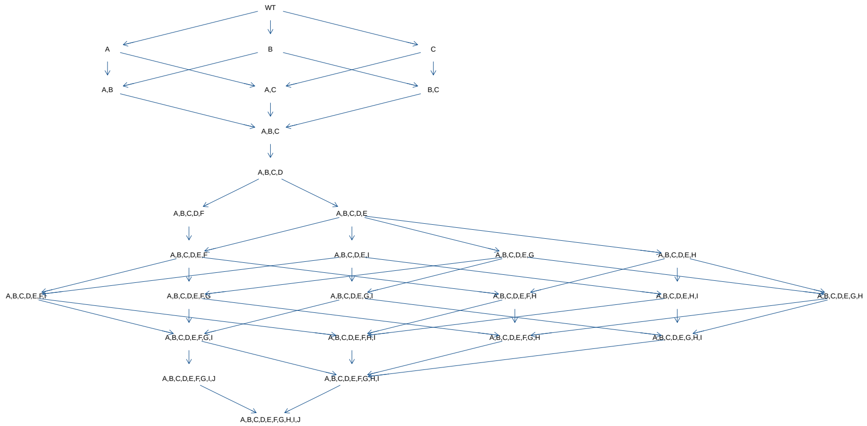


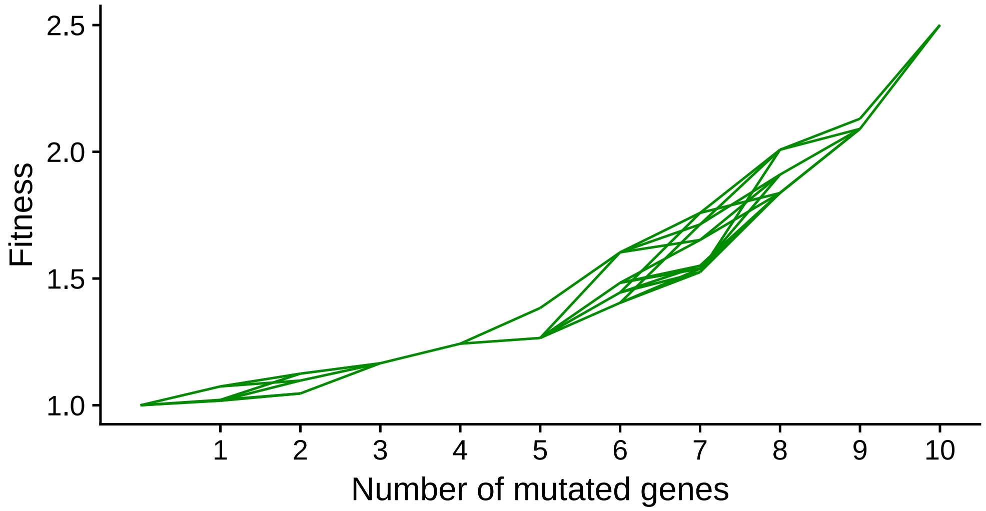

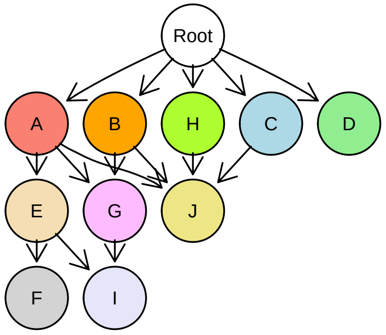


| | |
|---|---|
| *ID* | MocksvQ5er102Pjpk |
| *acc genotypes* | 120 |
| *peaks no back* | 1 |
| *removed edges* | 0 |
| *prop rem edges* | 0 |

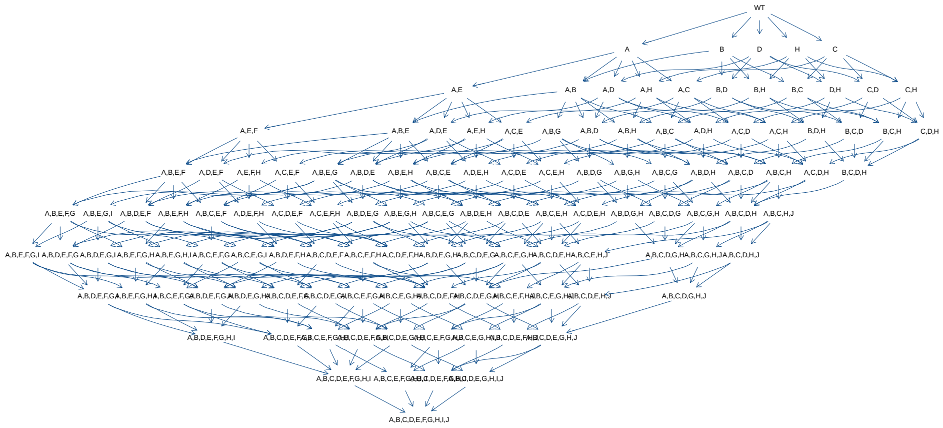

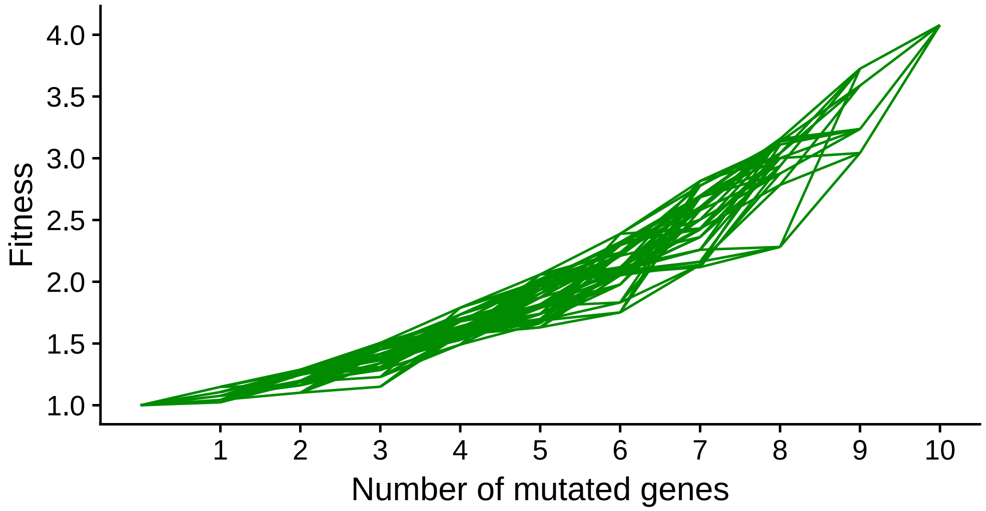

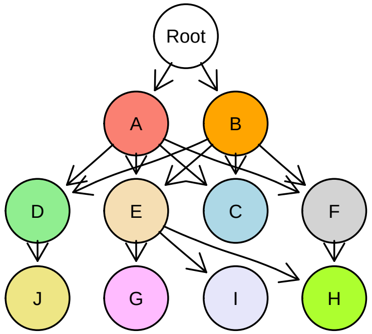


| | |
|---|---|
| *ID* | MPXnkjvgPoPYXZbV |
| *acc genotypes* | 87 |
| *peaks no back* | 1 |
| *removed edges* | 0 |
| *prop rem edges* | 0 |

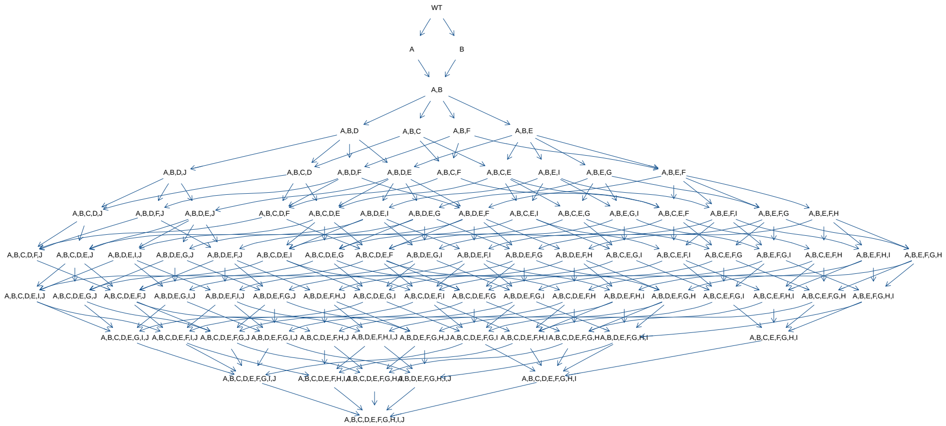

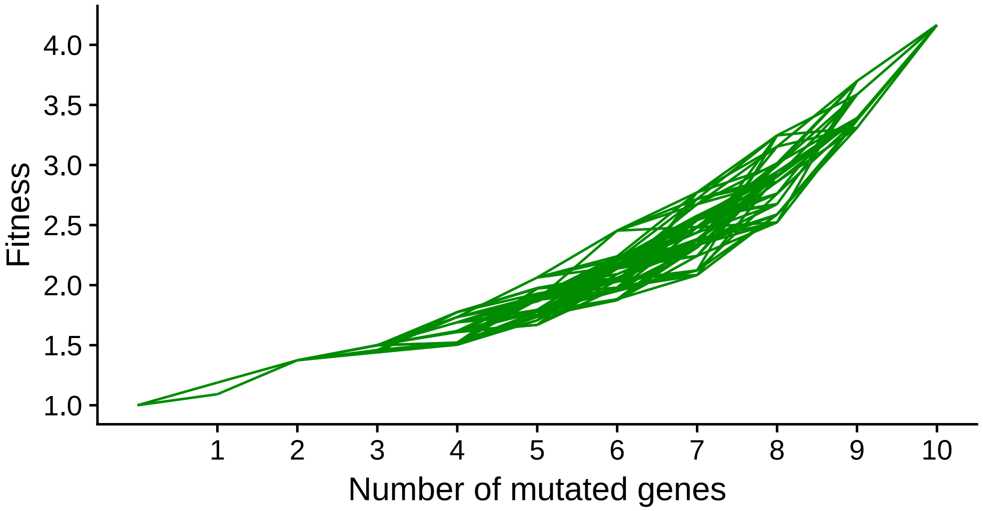

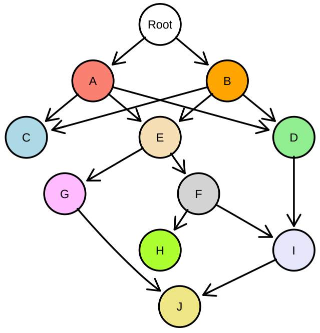


| | |
|---|---|
| *ID* | msWBh10MLRgzEH10BX |
| *acc genotypes* | 43 |
| *peaks no back* | 1 |
| *removed edges* | 0 |
| *prop rem edges* | 0 |

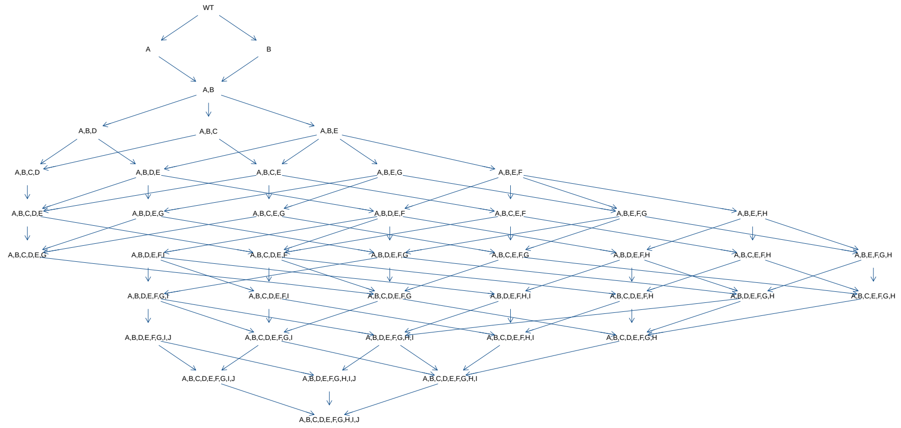

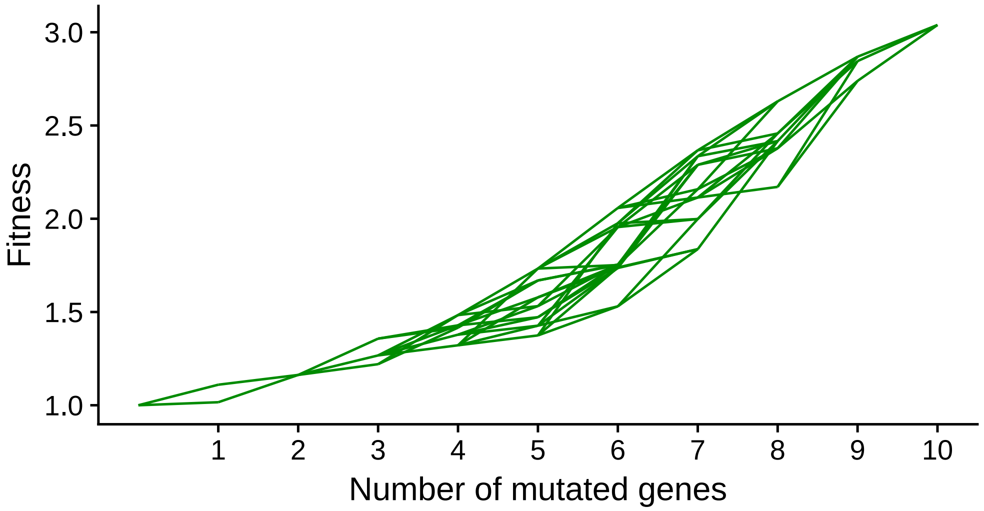

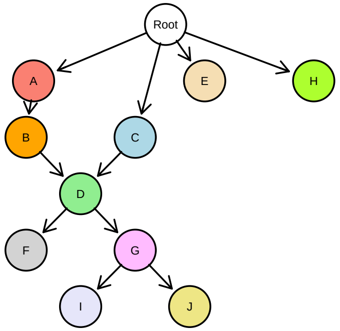


| | |
|---|---|
| *ID* | MT7WzjxZkqx10snFO |
| *acc genotypes* | 64 |
| *peaks no back* | 1 |
| *removed edges* | 0 |
| *prop rem edges* | 0 |

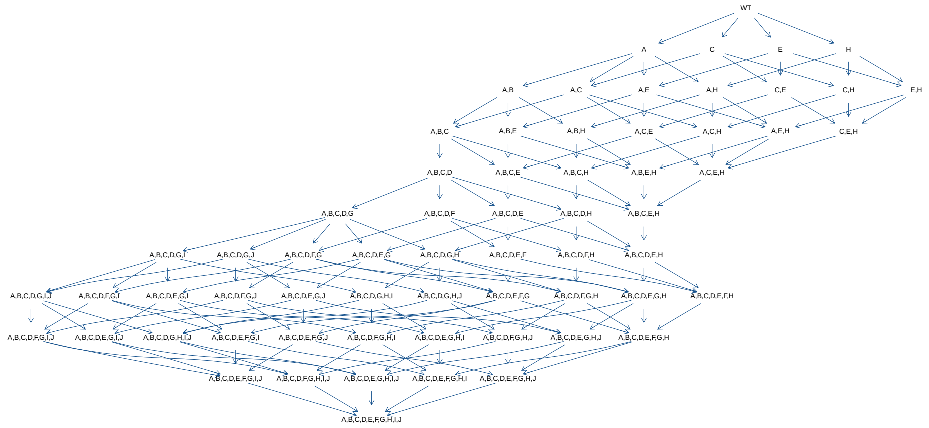

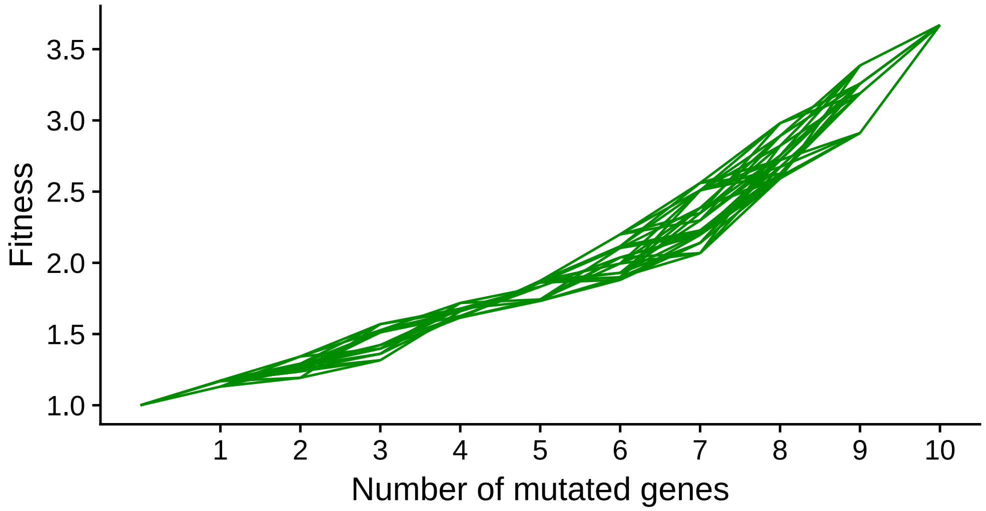

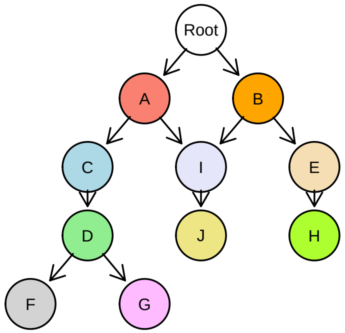


| | |
|---|---|
| *ID* | n210BxEwpJXJV8VL6 |
| *acc genotypes* | 64 |
| *peaks no back* | 1 |
| *removed edges* | 0 |
| *prop rem edges* | 0 |

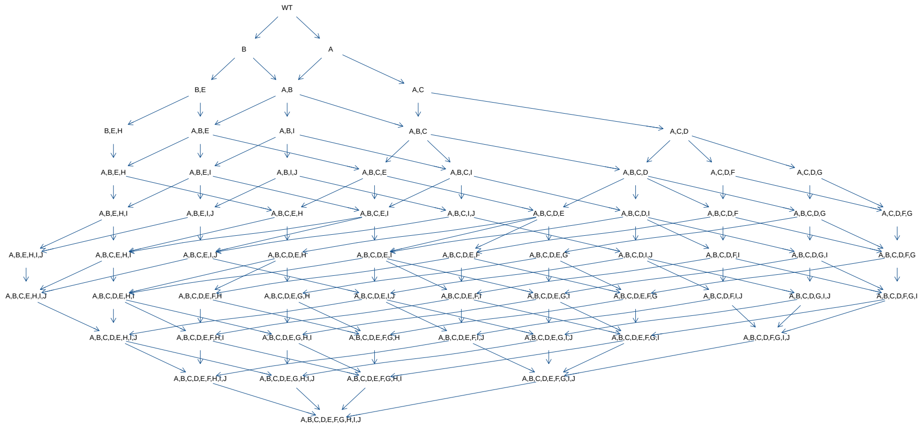

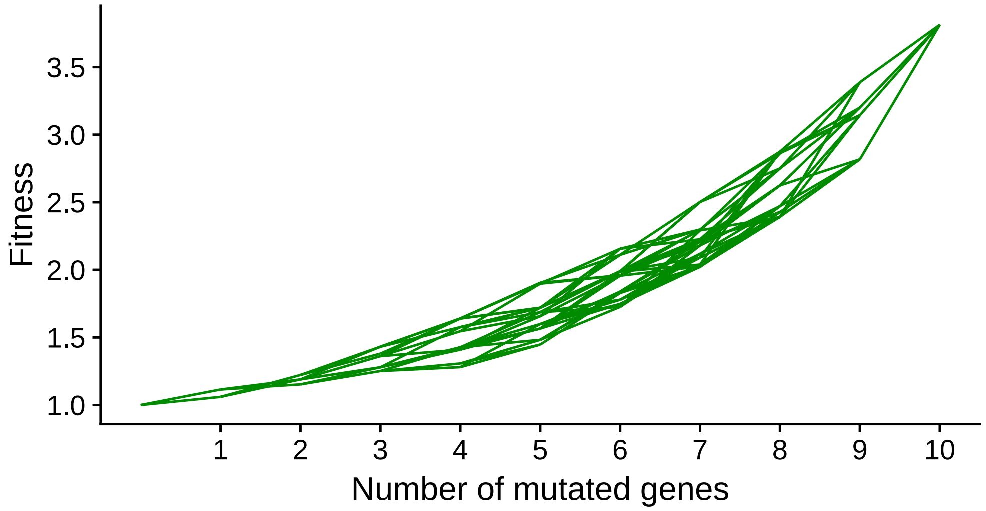

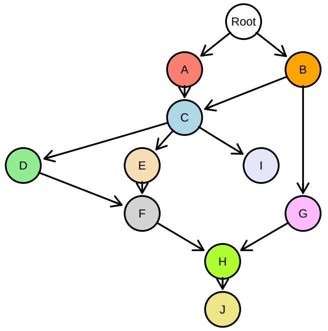


*ID* Nd5tXtGonmFk3JtF
*acc genotypes* 30
*peaks no back* 1
*removed edges* 0
*prop rem edges* 0

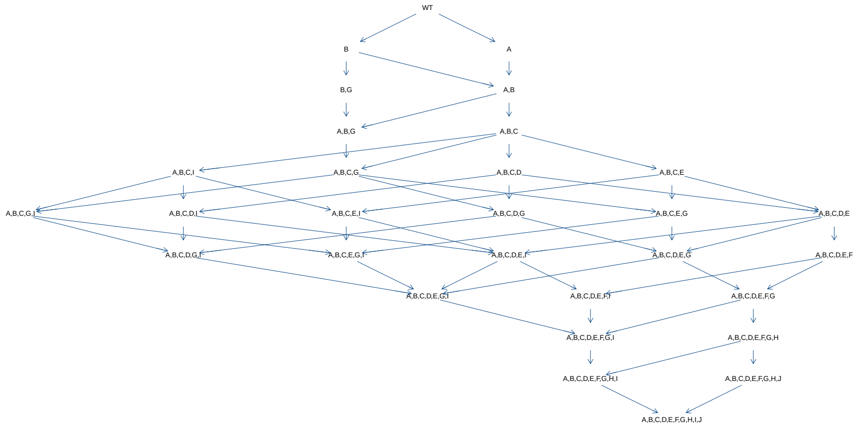


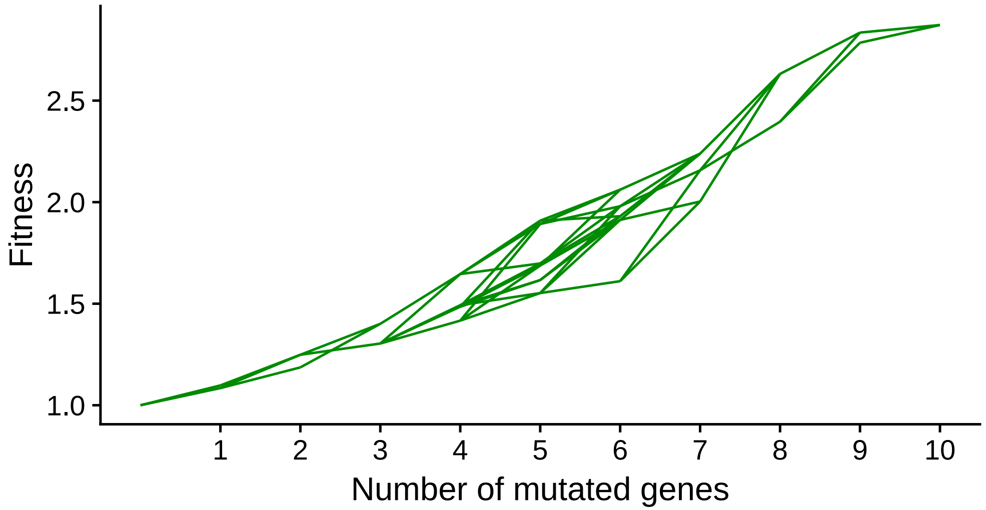

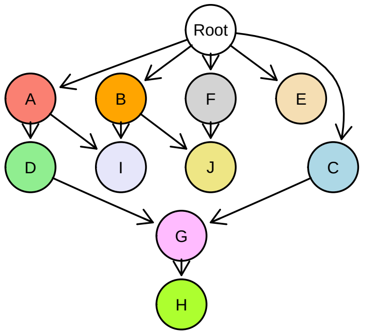


| | |
|---|---|
| *ID* | nfjDxDAtFc4uTrco |
| *acc genotypes* | 116 |
| *peaks no back* | 1 |
| *removed edges* | 0 |
| *prop rem edges* | 0 |

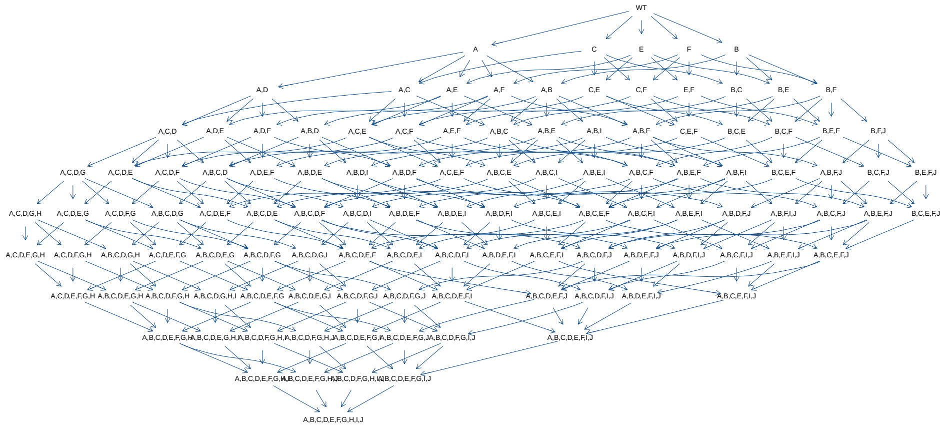

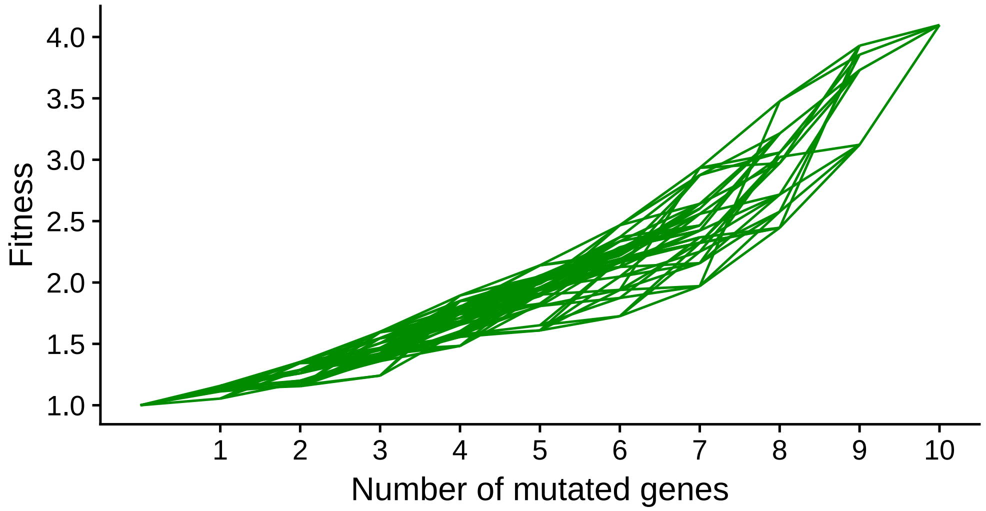

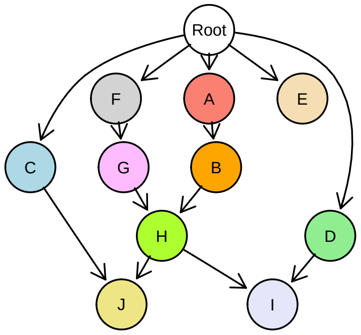


| | |
|---|---|
| *ID* | nO5yu7D15ZhqKnN9 |
| *acc genotypes* | 90 |
| *peaks no back* | 1 |
| *removed edges* | 0 |
| *prop rem edges* | 0 |

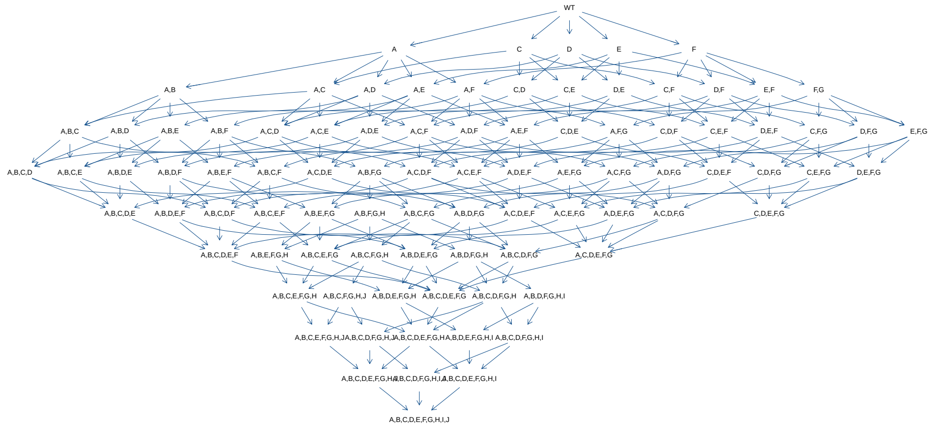


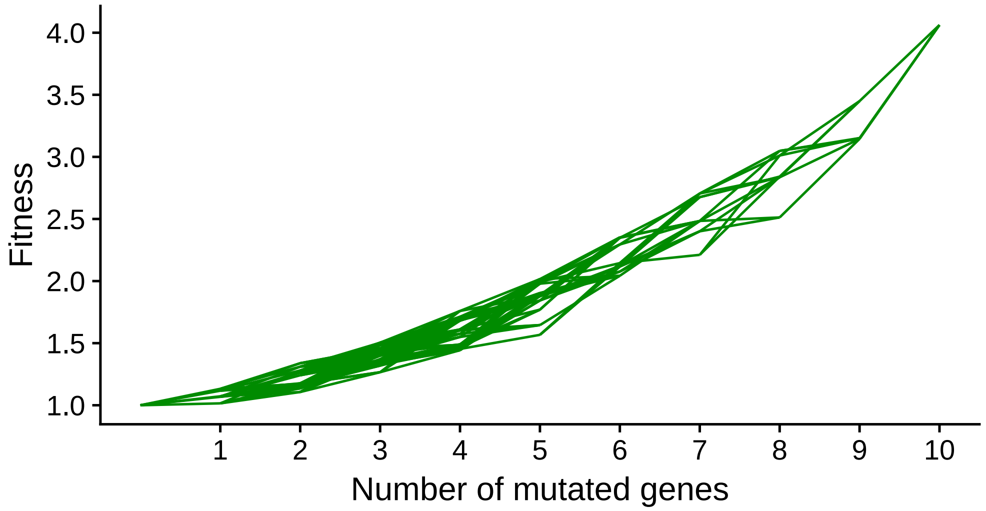

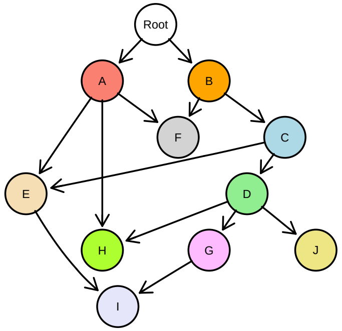


| | |
|---|---|
| *ID* | NP7ssqpSWvIesNBx |
| *acc genotypes* | 54 |
| *peaks no back* | 1 |
| *removed edges* | 0 |
| *prop rem edges* | 0 |

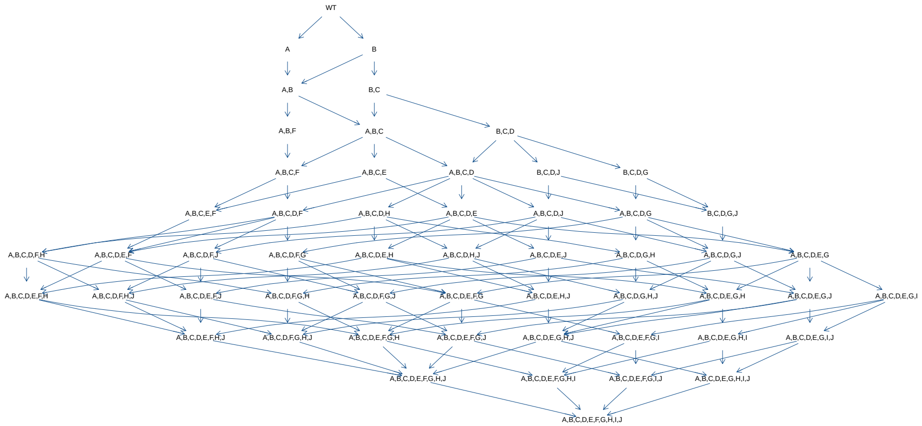

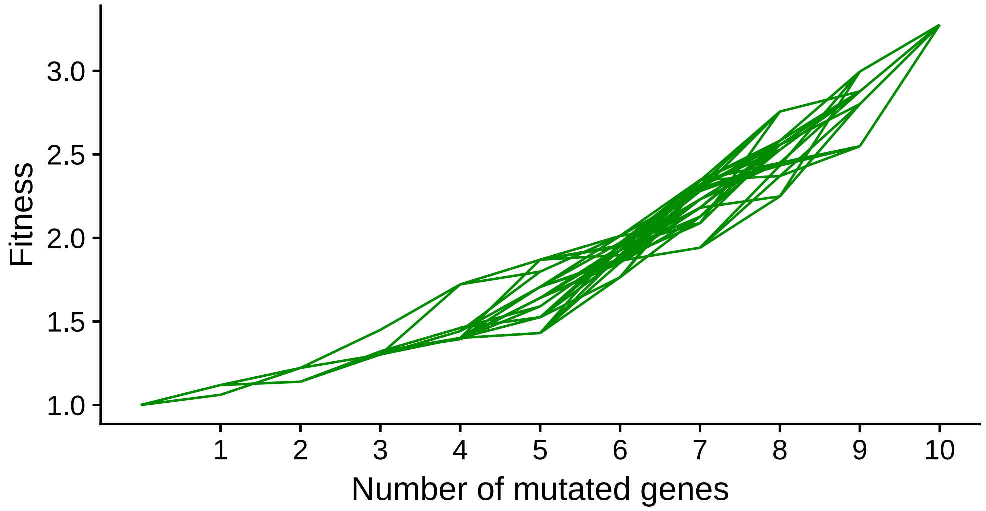

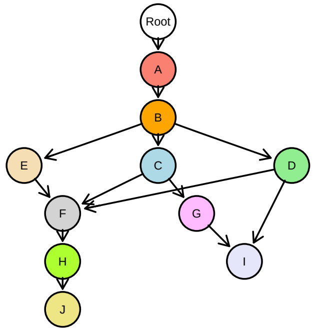


| | |
|---|---|
| *ID* | NyMcE3XWsePNxuEs |
| *acc genotypes* | 25 |
| *peaks no back* | 1 |
| *removed edges* | 0 |
| *prop rem edges* | 0 |

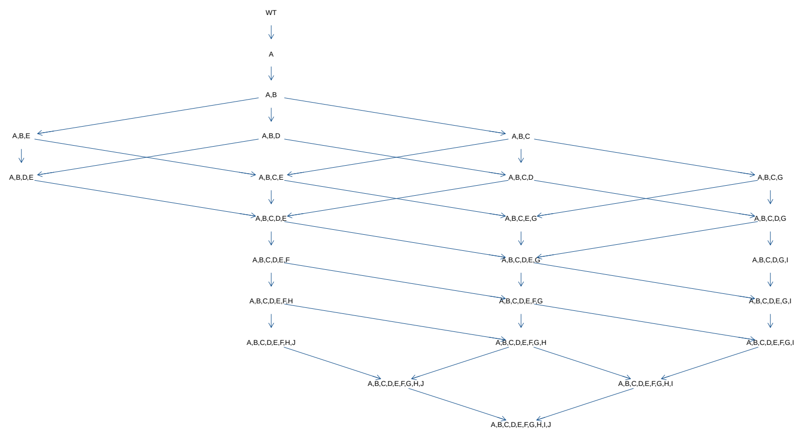


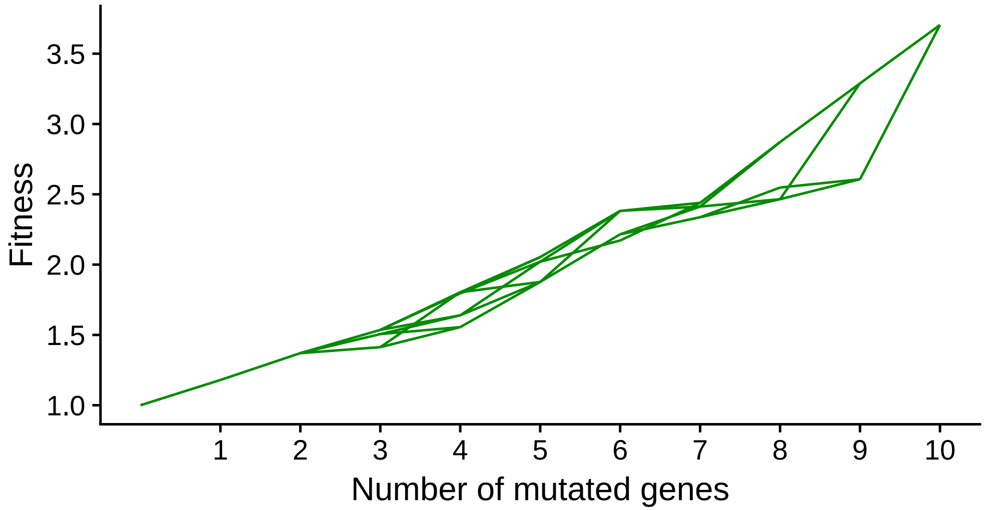

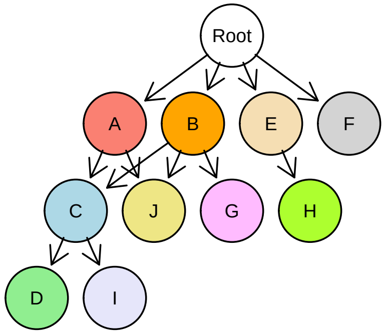


| | |
|---|---|
| *ID* | o10BaslJKzTznq5sp |
| *acc genotypes* | 144 |
| *peaks no back* | 1 |
| *removed edges* | 0 |
| *prop rem edges* | 0 |

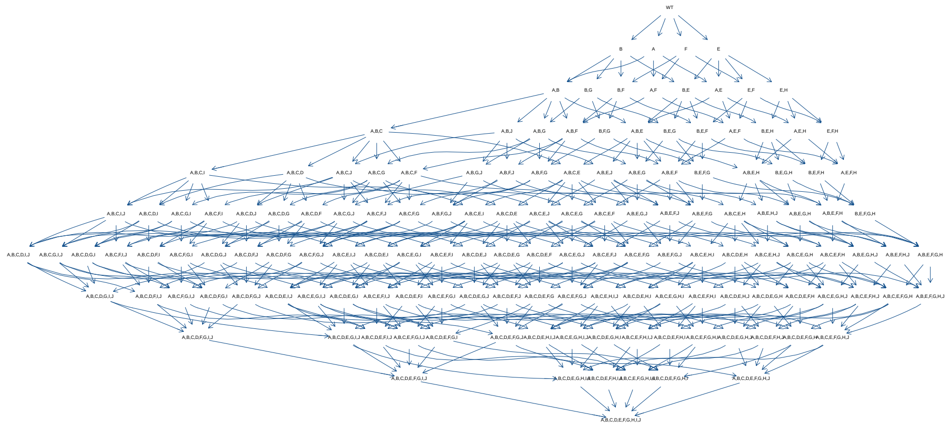

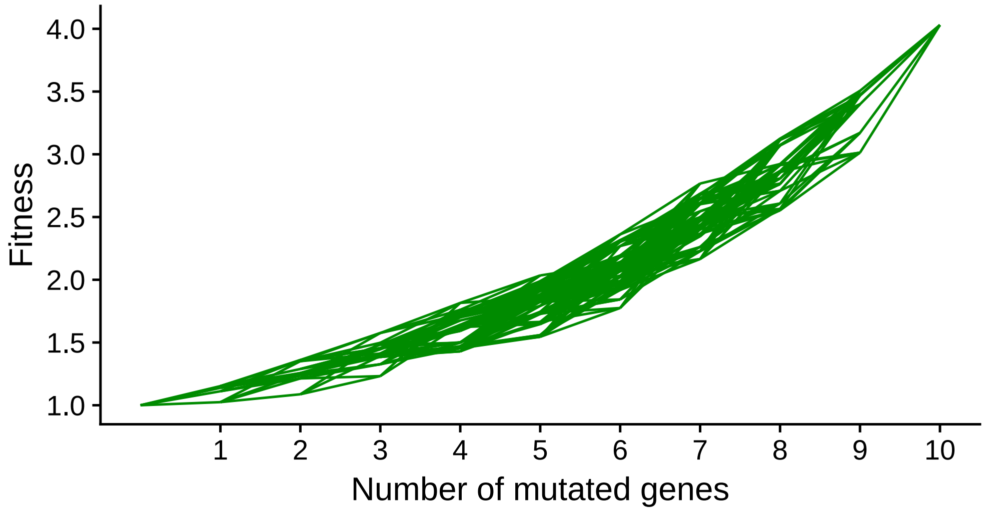

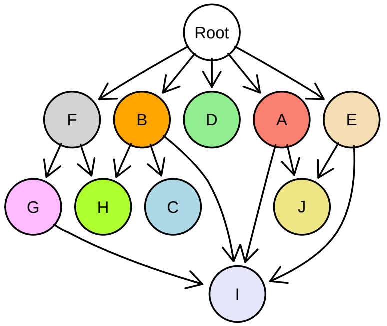


| | |
|---|---|
| *ID* | o89L6ak94kWOZRNA |
| *acc genotypes* | 146 |
| *peaks no back* | 1 |
| *removed edges* | 0 |
| *prop rem edges* | 0 |

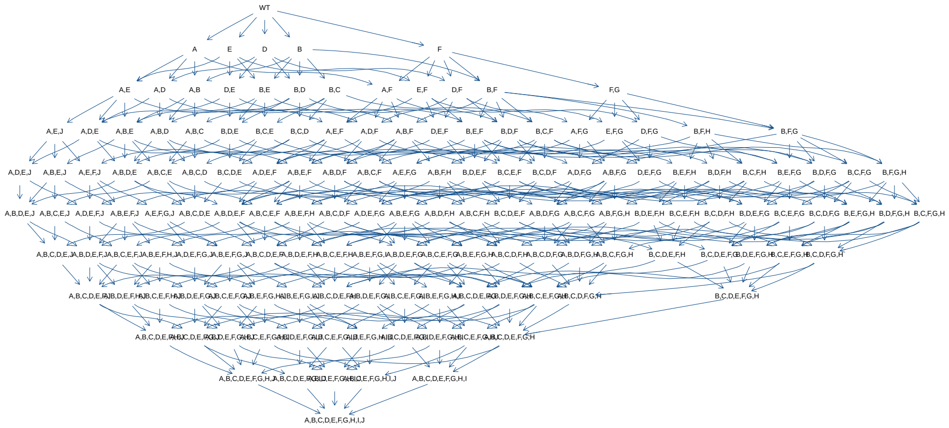

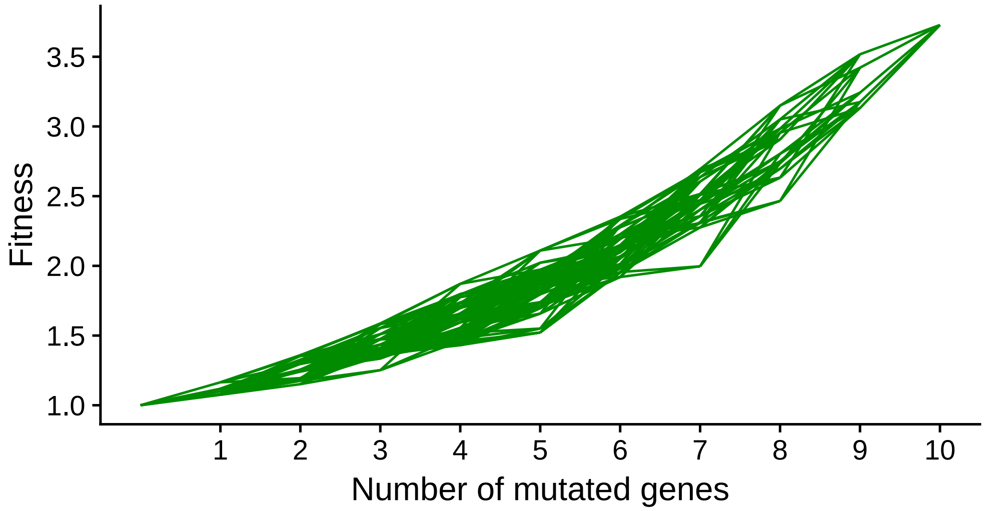

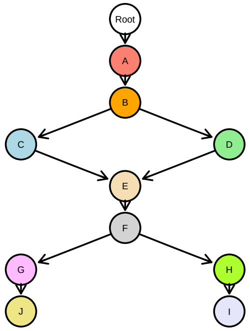


| | |
|---|---|
| *ID* | ocx2Aw4wa94NsR64 |
| *acc genotypes* | 16 |
| *peaks no back* | 1 |
| *removed edges* | 0 |
| *prop rem edges* | 0 |

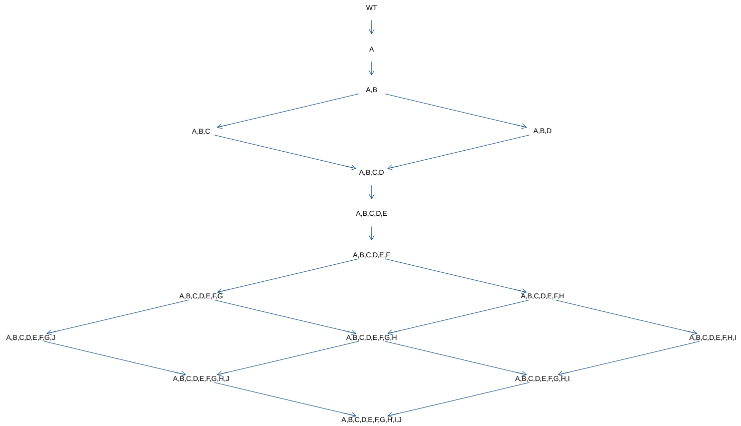


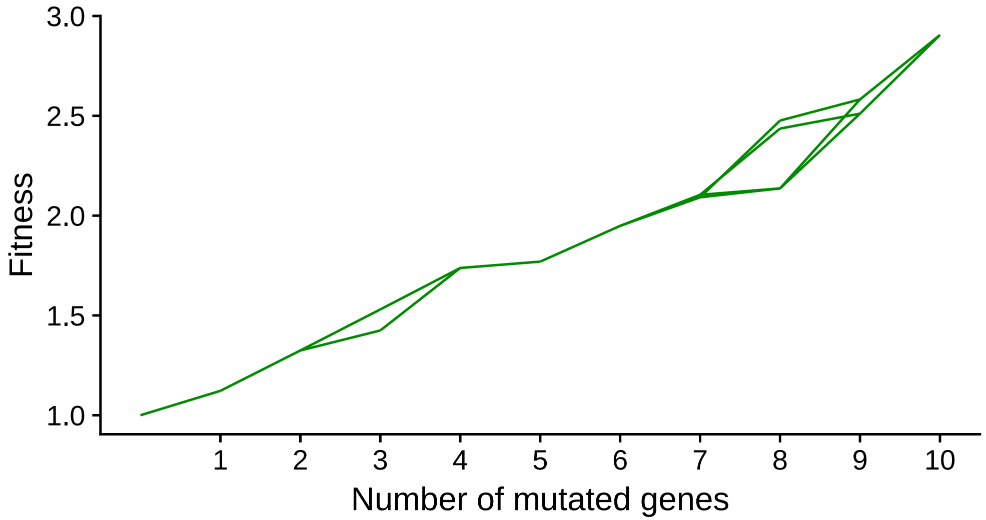

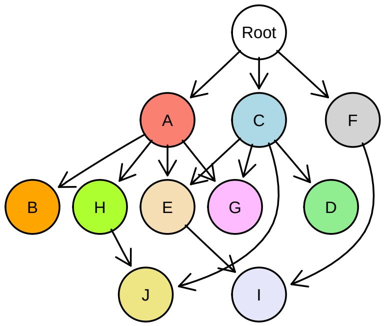


*ID* OGeUcOzxp4J7ocOW
*acc genotypes* 134
*peaks no back* 1
*removed edges* 0
*prop rem edges* 0

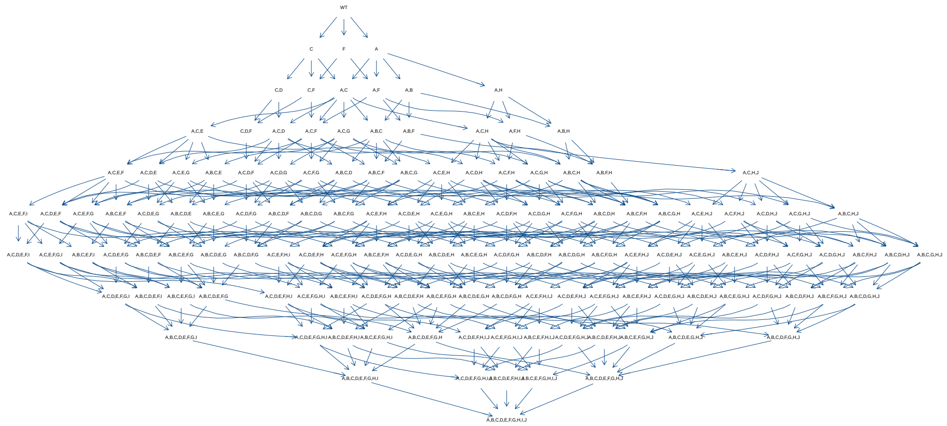


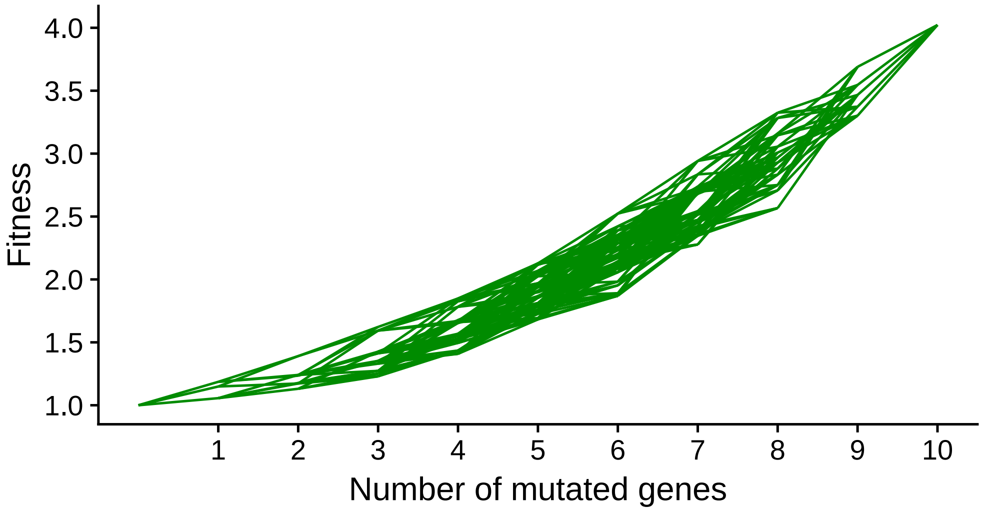

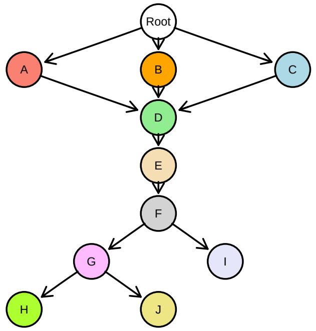


*ID* oGhD99FEVEPwA8X8
*acc genotypes* 20
*peaks no back* 1
*removed edges* 0
*prop rem edges* 0

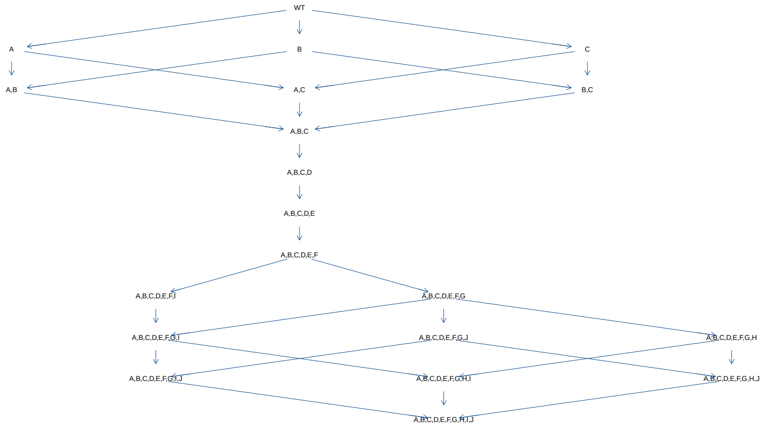


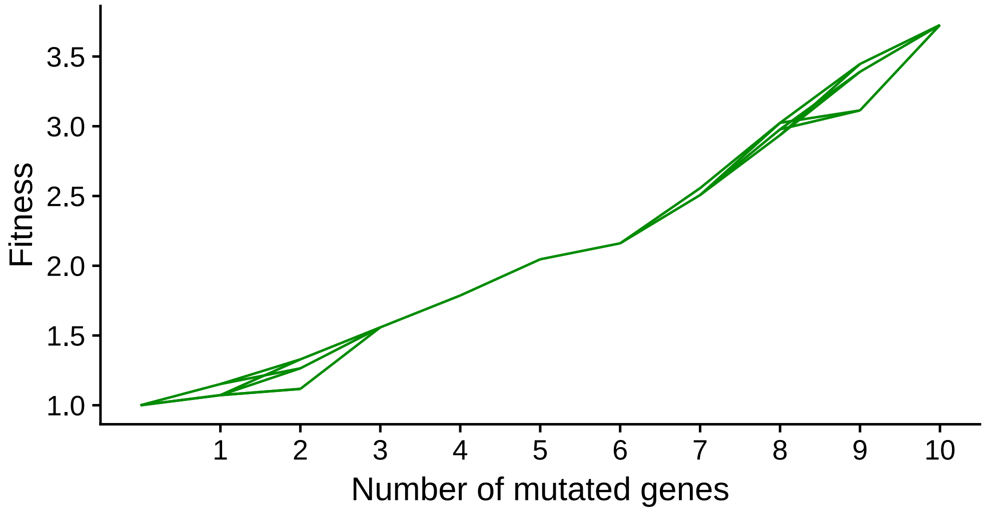

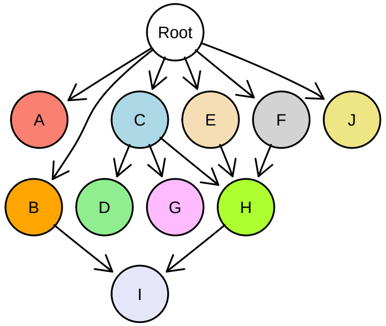


| | |
|---|---|
| *ID* | ohfmWmj2mPmIjDEY |
| *acc genotypes* | 208 |
| *peaks no back* | 1 |
| *removed edges* | 0 |
| *prop rem edges* | 0 |

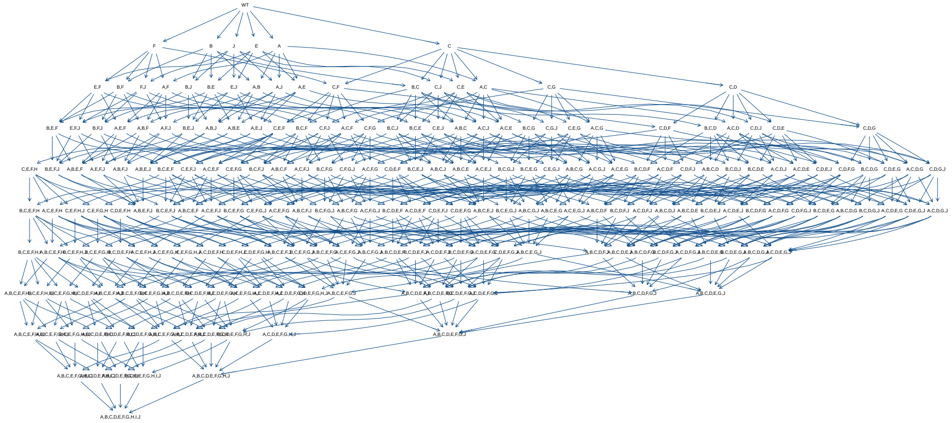

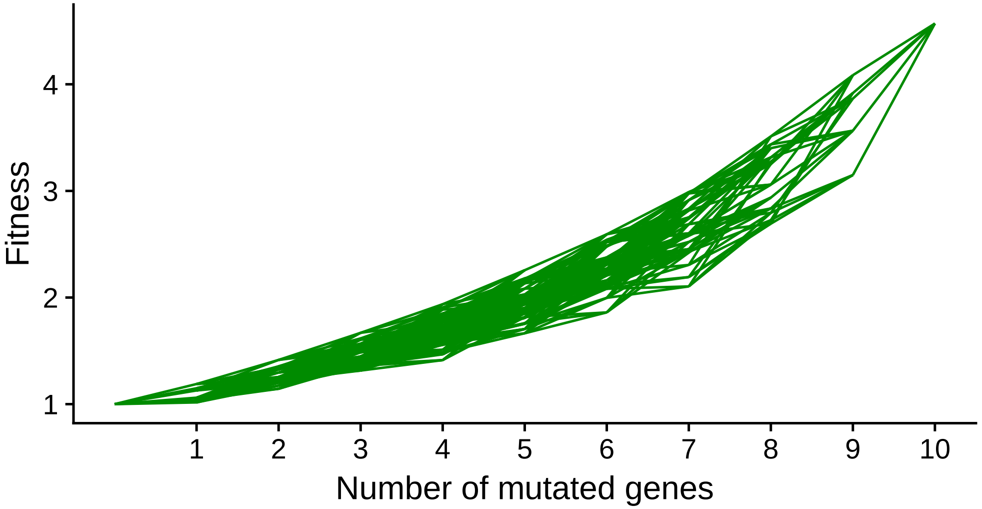

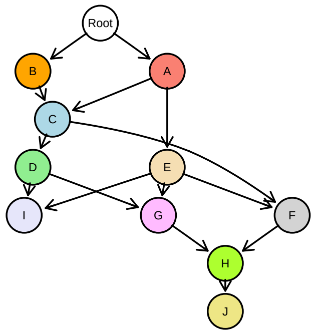


| | |
|---|---|
| *ID* | oRJlh1e10mdF10GFUm |
| *acc genotypes* | 22 |
| *peaks no back* | 1 |
| *removed edges* | 0 |
| *prop rem edges* | 0 |

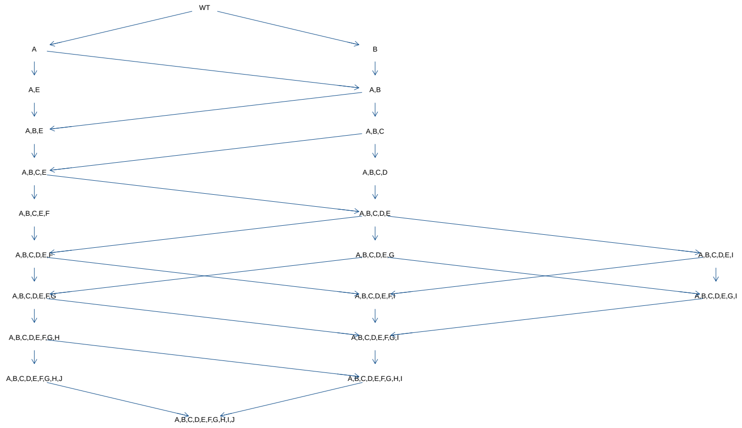


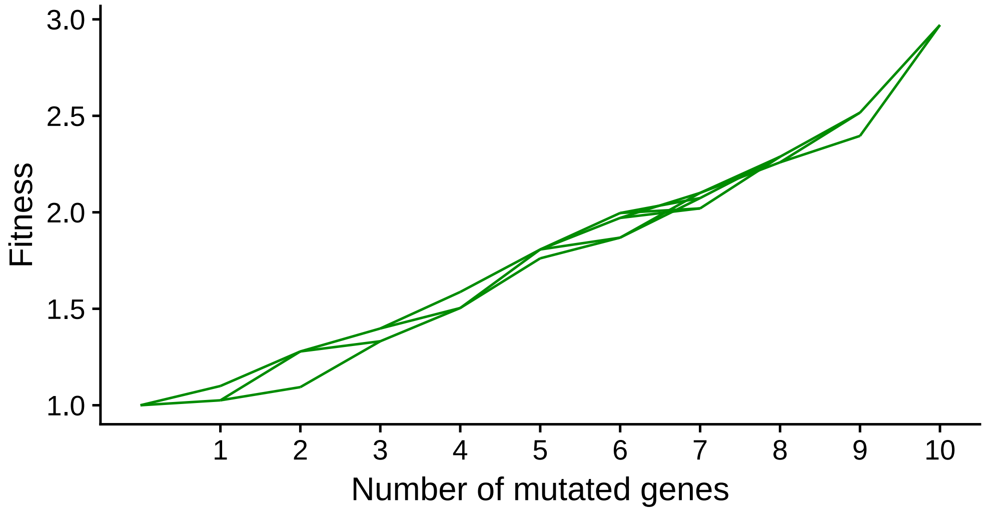

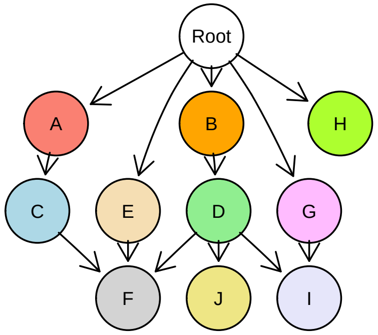


| | |
|---|---|
| *ID* | PEUXKMJo1Ym7Cv4Z |
| *acc genotypes* | 132 |
| *peaks no back* | 1 |
| *removed edges* | 0 |
| *prop rem edges* | 0 |

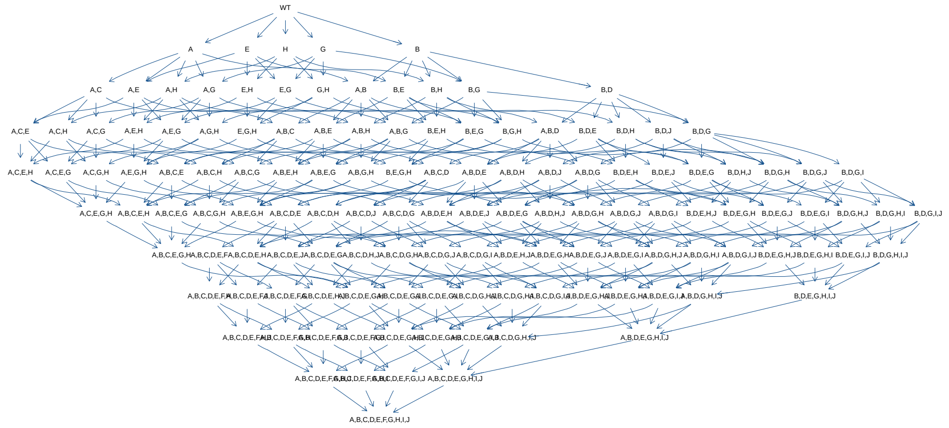

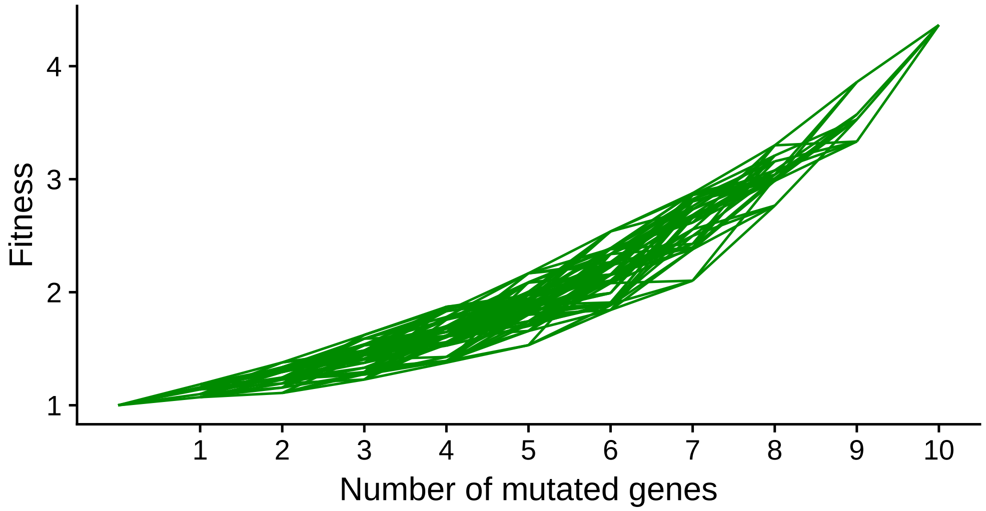

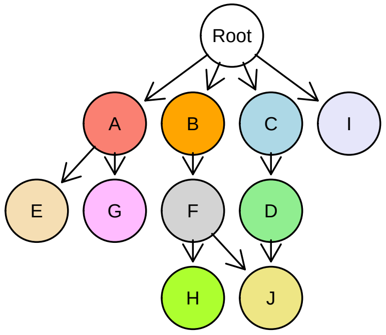


| | |
|---|---|
| *ID* | PhaG1vE1MZA6aPS7 |
| *acc genotypes* | 140 |
| *peaks no back* | 1 |
| *removed edges* | 0 |
| *prop rem edges* | 0 |

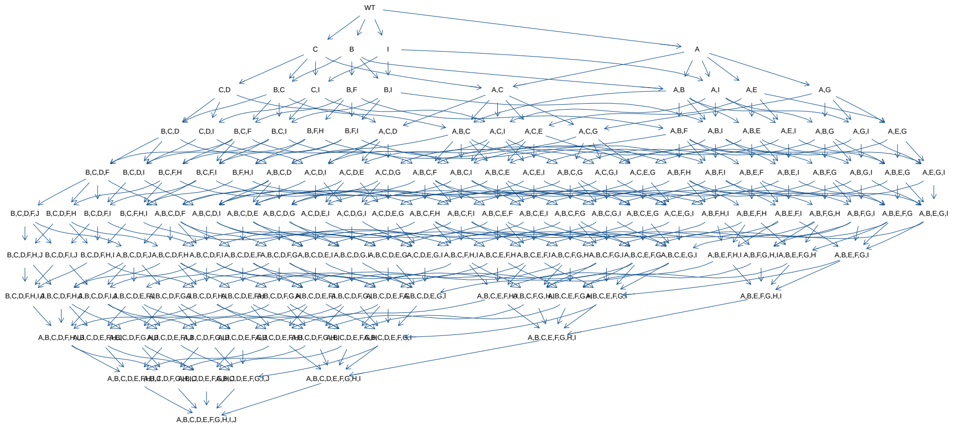

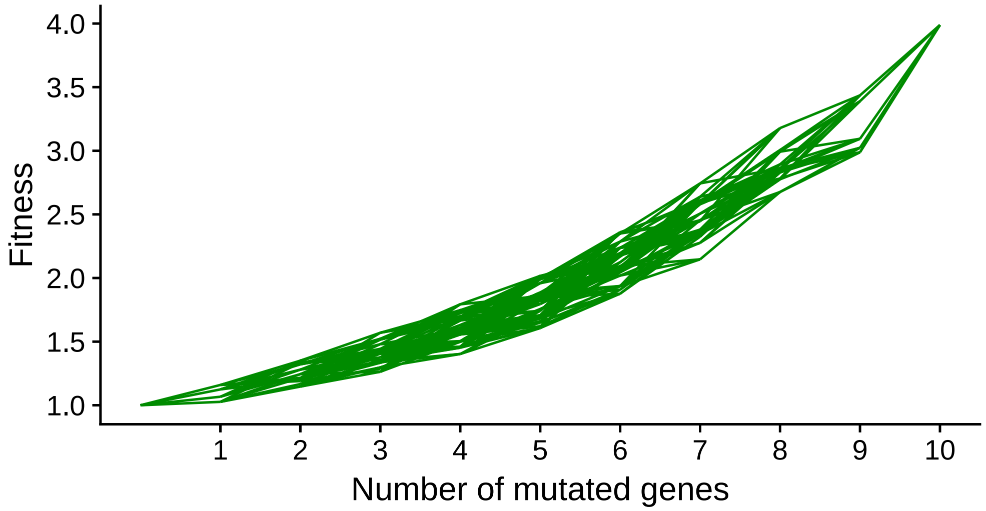

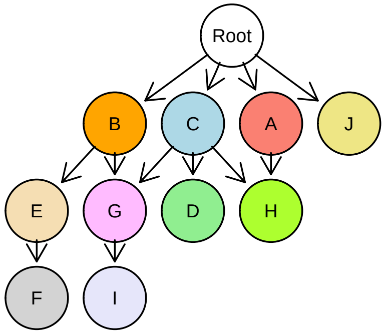


| | |
|---|---|
| *ID* | Pl10ybVHbxKrShrRr |
| *acc genotypes* | 136 |
| *peaks no back* | 1 |
| *removed edges* | 0 |
| *prop rem edges* | 0 |

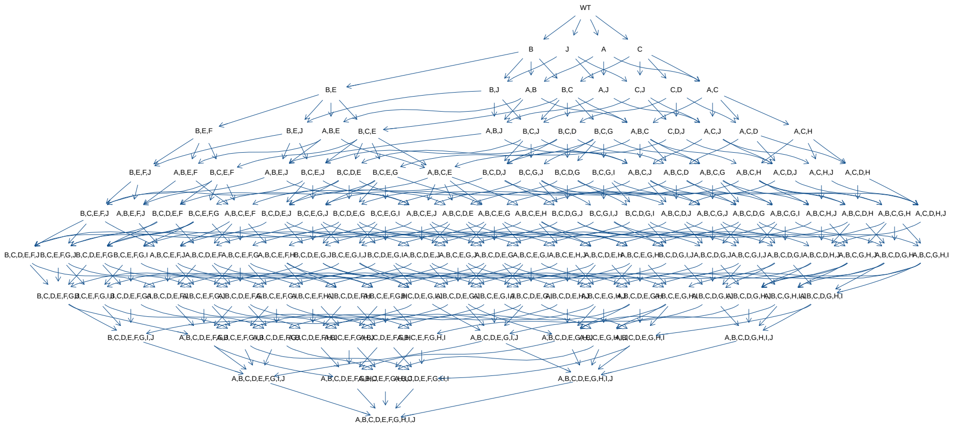

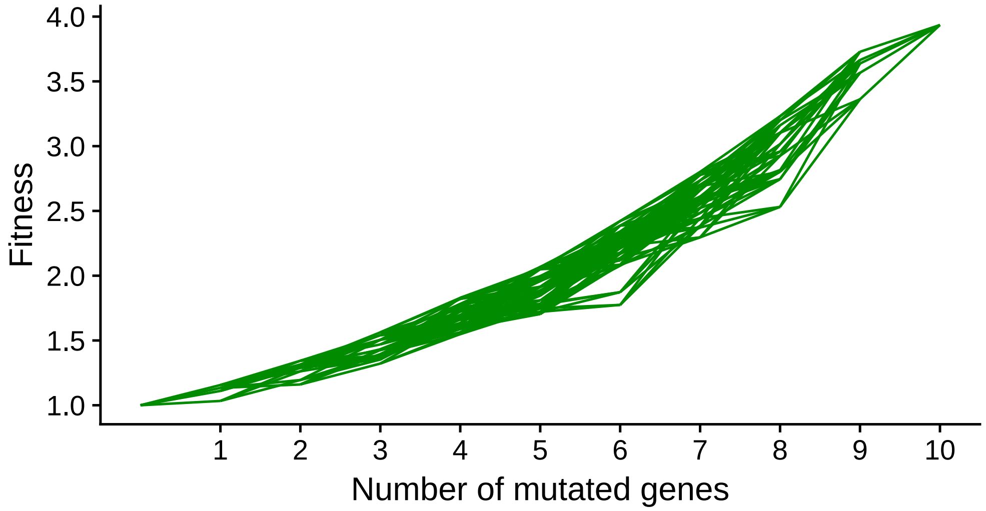

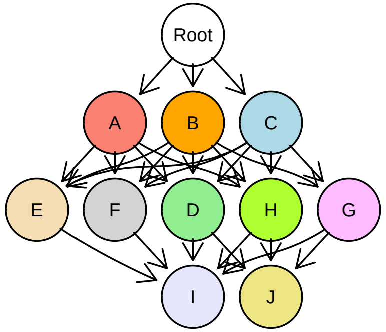


*ID* pTNzMT9igvRrzlPi
*acc genotypes* 46
*peaks no back* 1
*removed edges* 0
*prop rem edges* 0

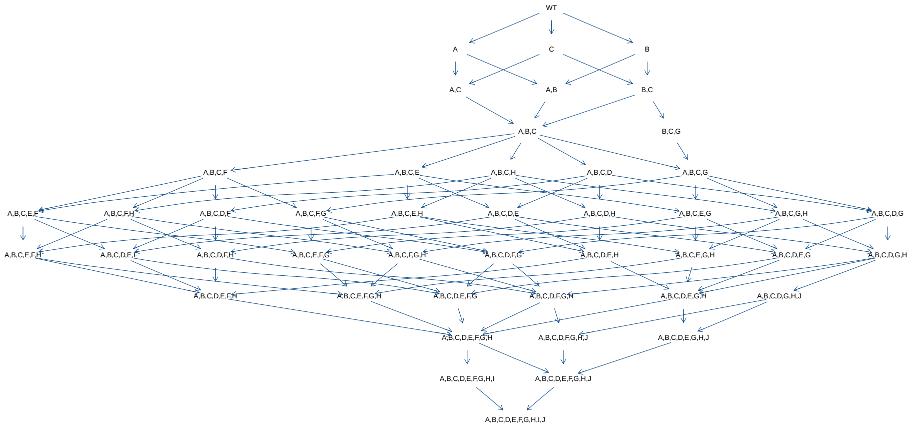

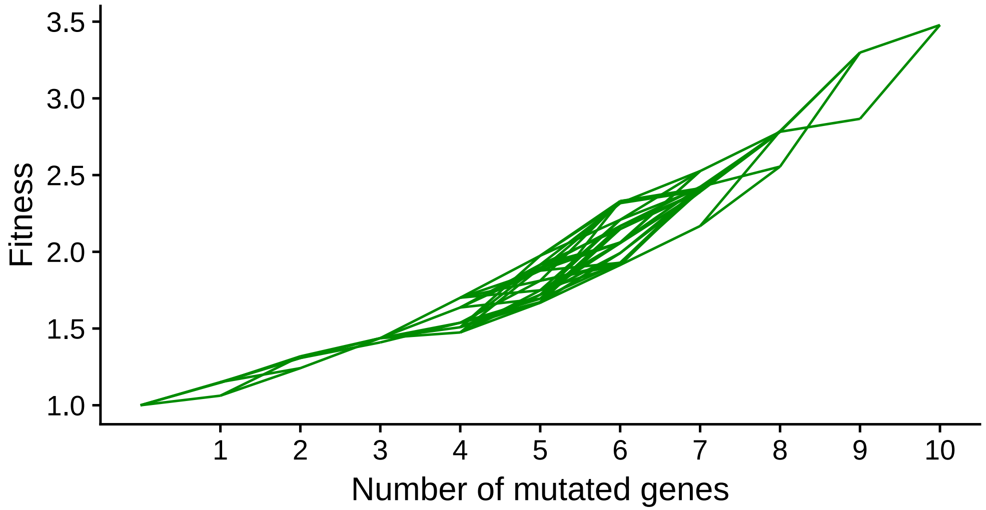

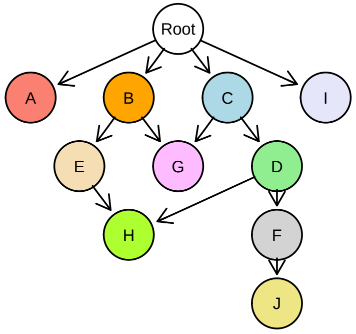


*ID* Pu5M3F61IUoT6zBD
*acc genotypes* 116
*peaks no back* 1
*removed edges* 0
*prop rem edges* 0

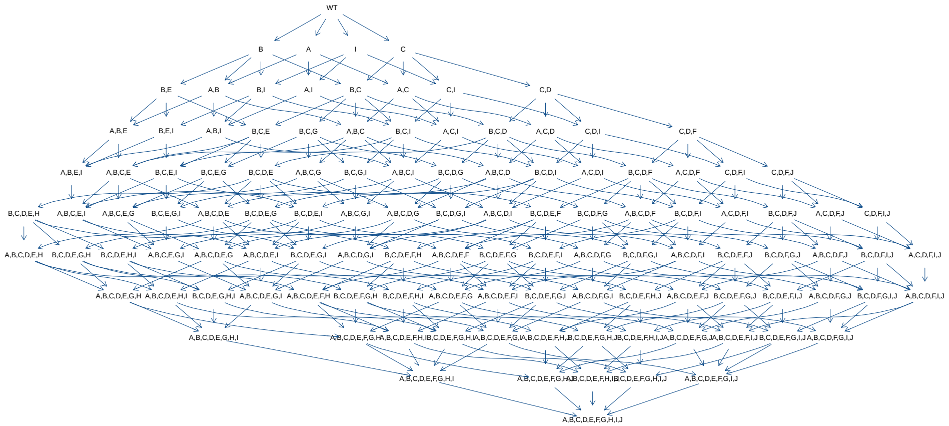

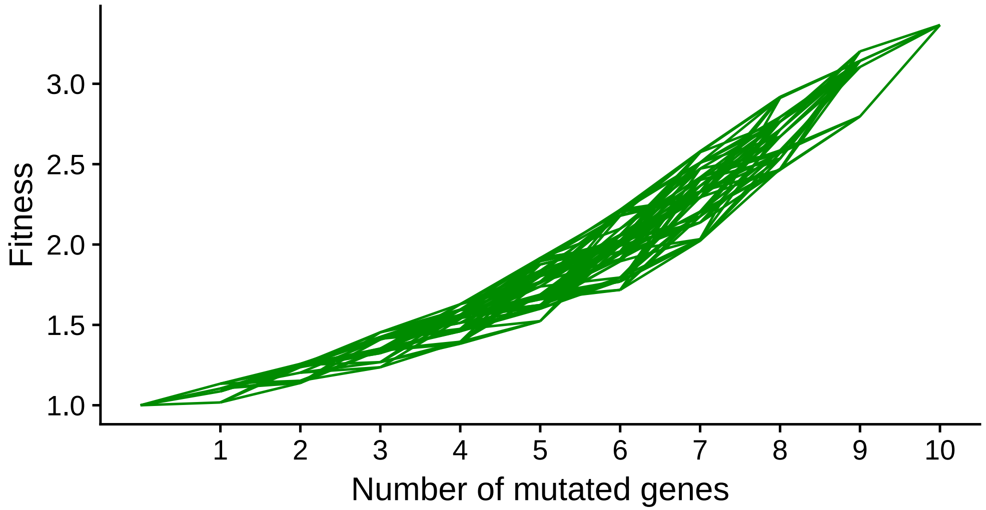

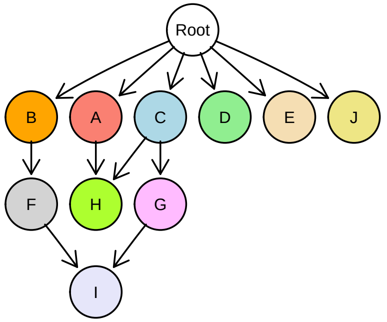


| | |
|---|---|
| *ID* | PVfJTlNJUgHLfOjP |
| *acc genotypes* | 216 |
| *peaks no back* | 1 |
| *removed edges* | 0 |
| *prop rem edges* | 0 |

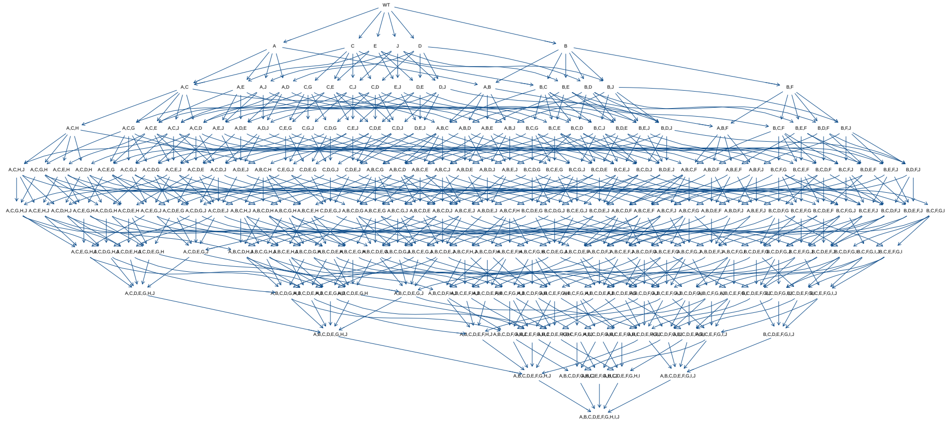

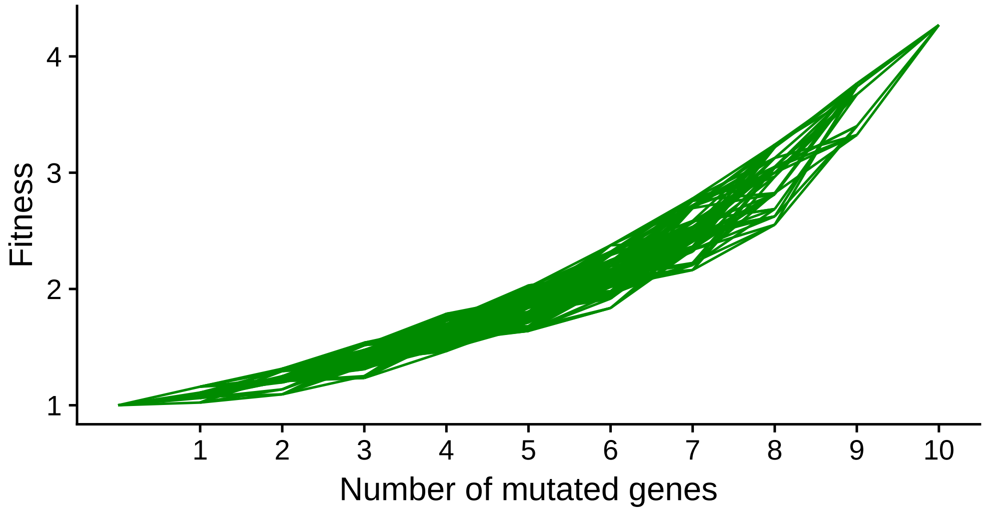

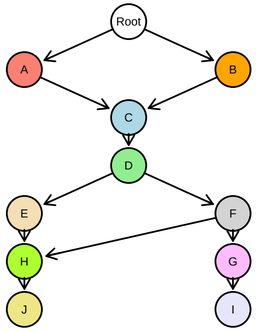


| | |
|---|---|
| *ID* | pwyaau7tK7LZumPR |
| *acc genotypes* | 19 |
| *peaks no back* | 1 |
| *removed edges* | 0 |
| *prop rem edges* | 0 |

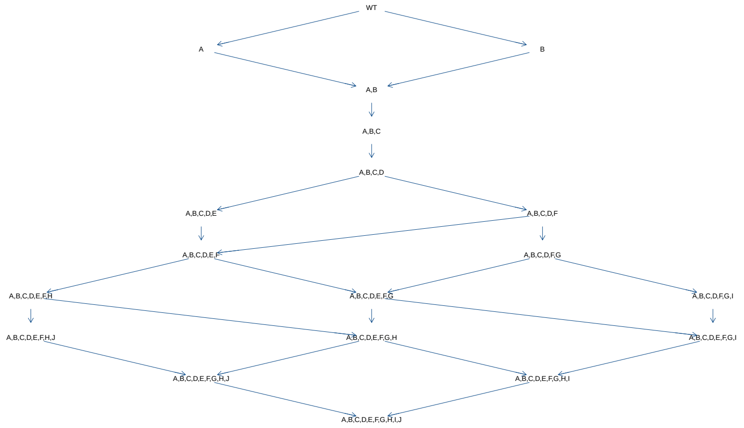


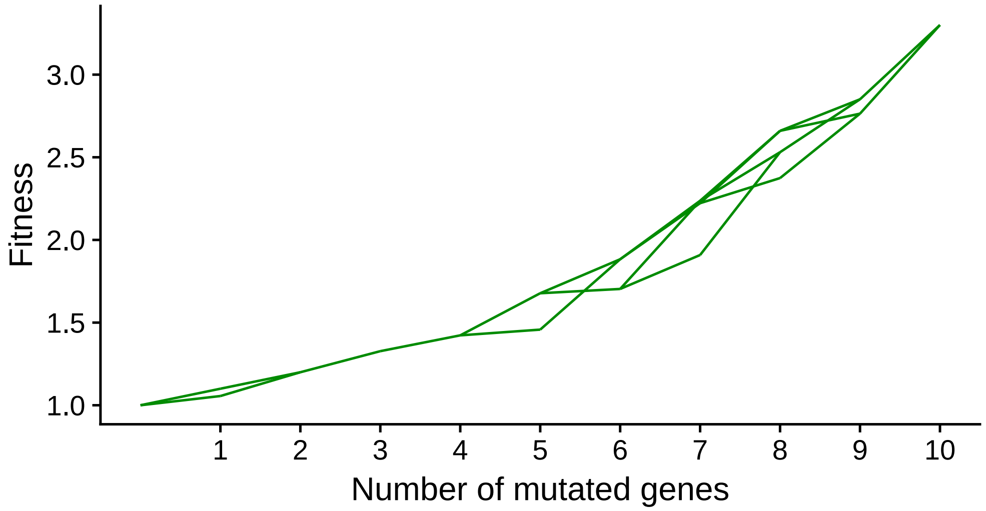

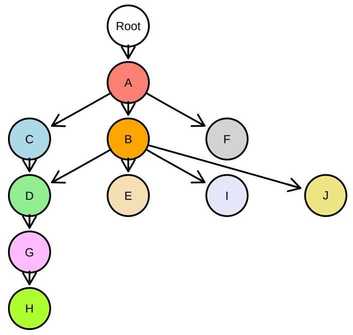


| | |
|---|---|
| *ID* | QePxqp7BQd10uZ5Ko |
| *acc genotypes* | 85 |
| *peaks no back* | 1 |
| *removed edges* | 0 |
| *prop rem edges* | 0 |

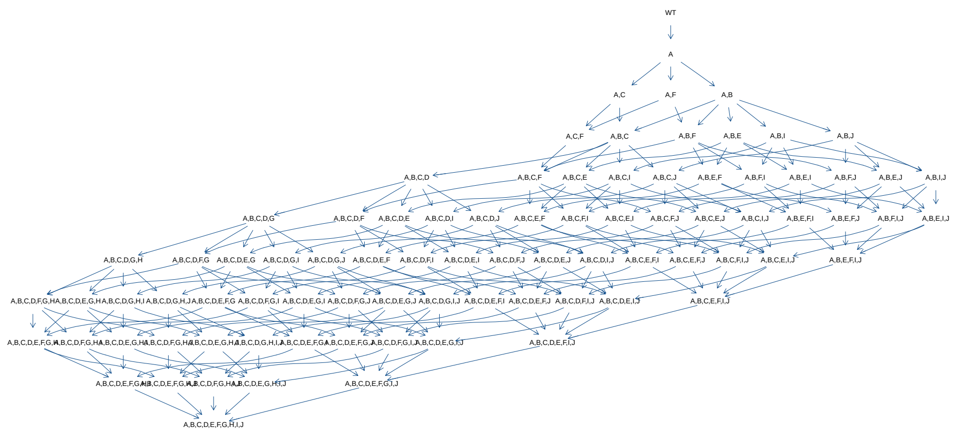

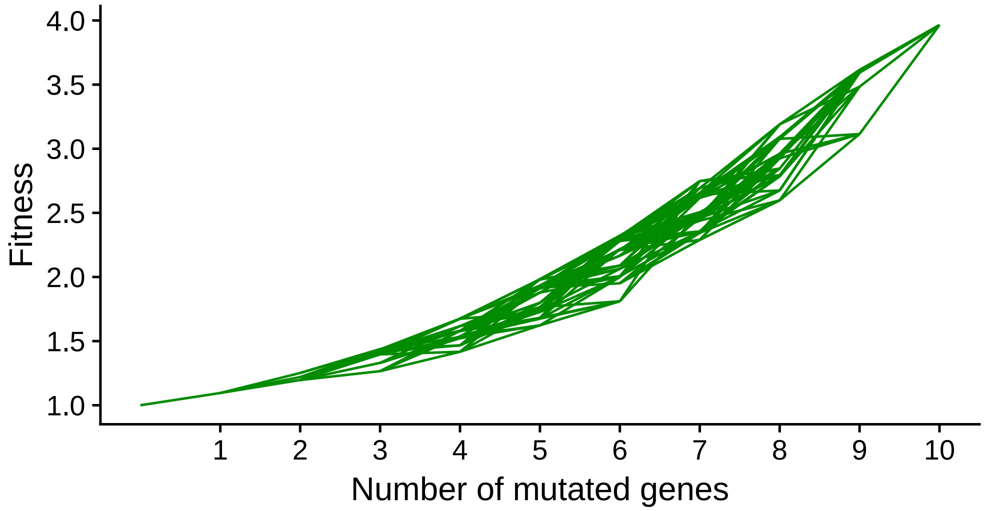

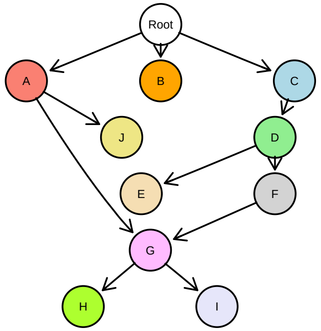


| | |
|---|---|
| *ID* | QGRVlBiM3bkC395C |
| *acc genotypes* | 68 |
| *peaks no back* | 1 |
| *removed edges* | 0 |
| *prop rem edges* | 0 |

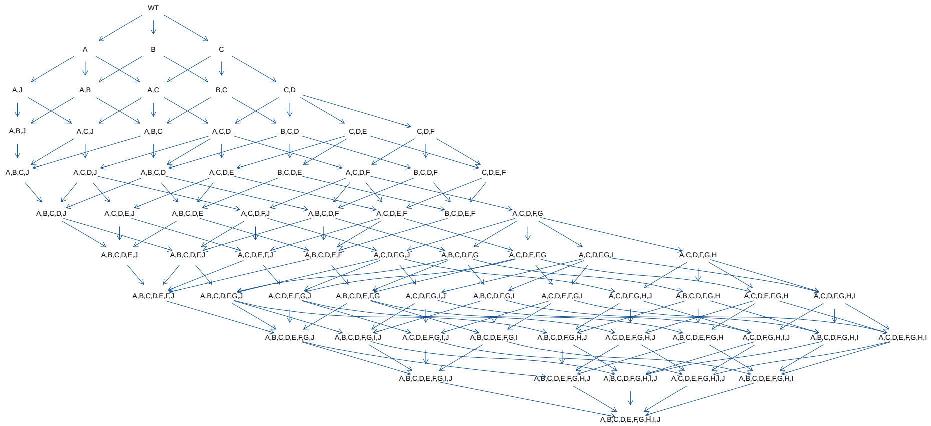

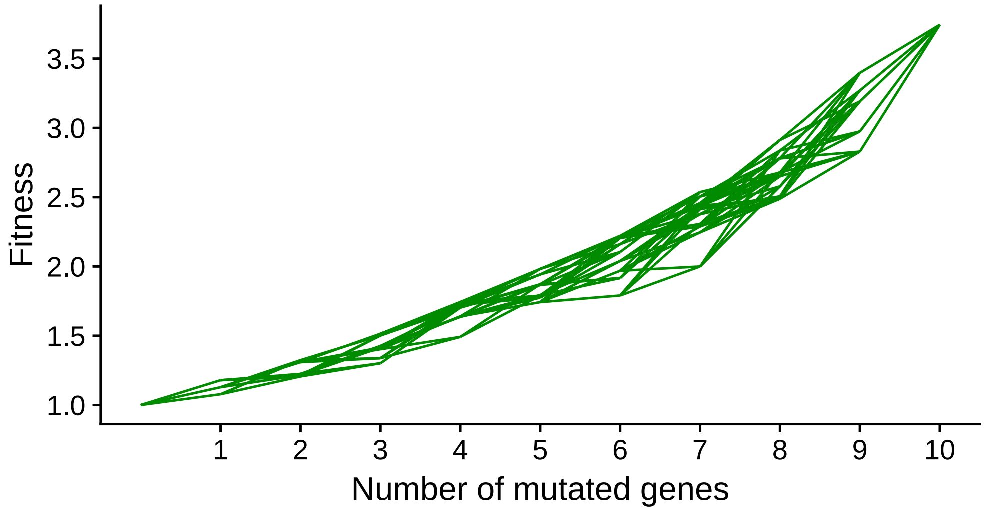

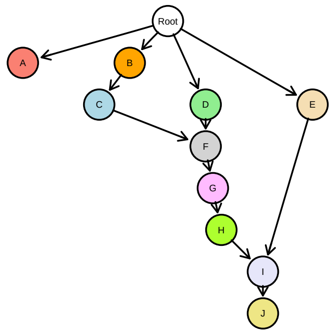


| | |
|---|---|
| *ID* | Qt7h8PhrOcDVjbUJ |
| *acc genotypes* | 40 |
| *peaks no back* | 1 |
| *removed edges* | 0 |
| *prop rem edges* | 0 |

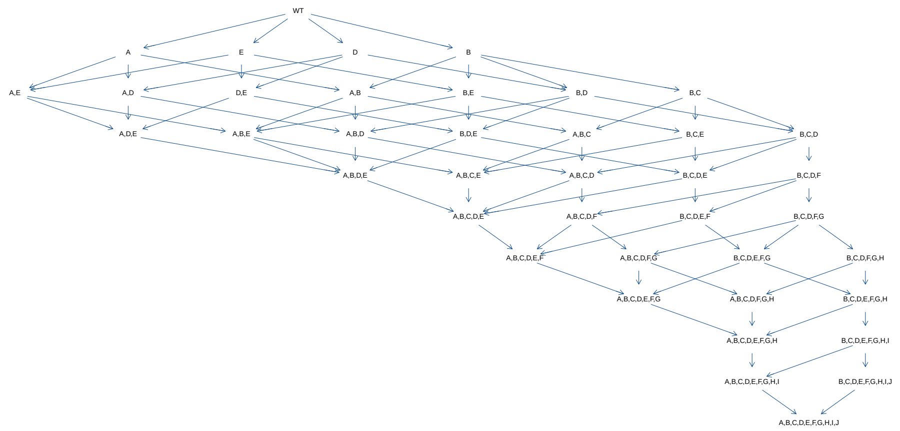

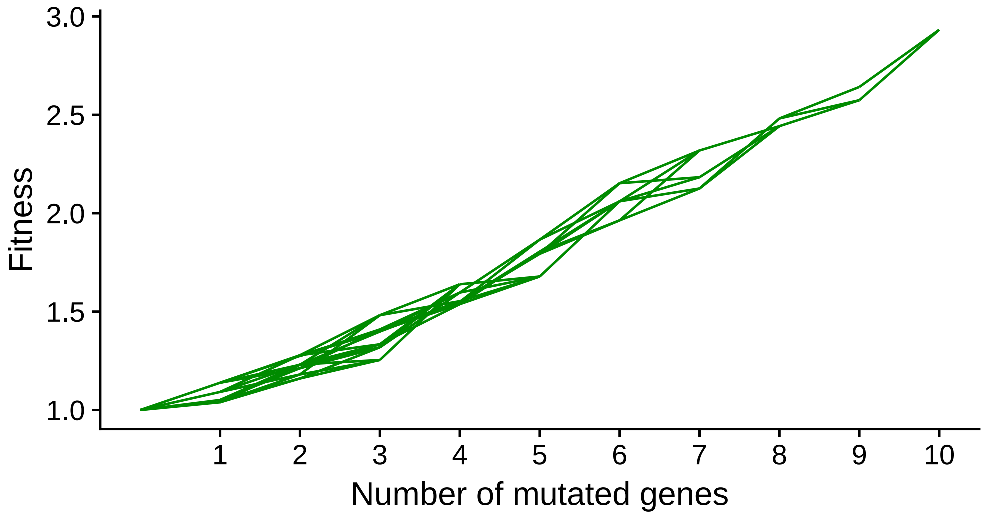

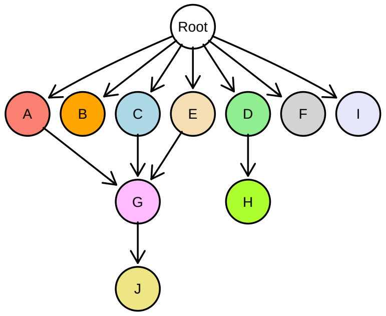


| | |
|---|---|
| *ID* | Rgqi10zX3omumQ9oh |
| *acc genotypes* | 240 |
| *peaks no back* | 1 |
| *removed edges* | 0 |
| *prop rem edges* | 0 |

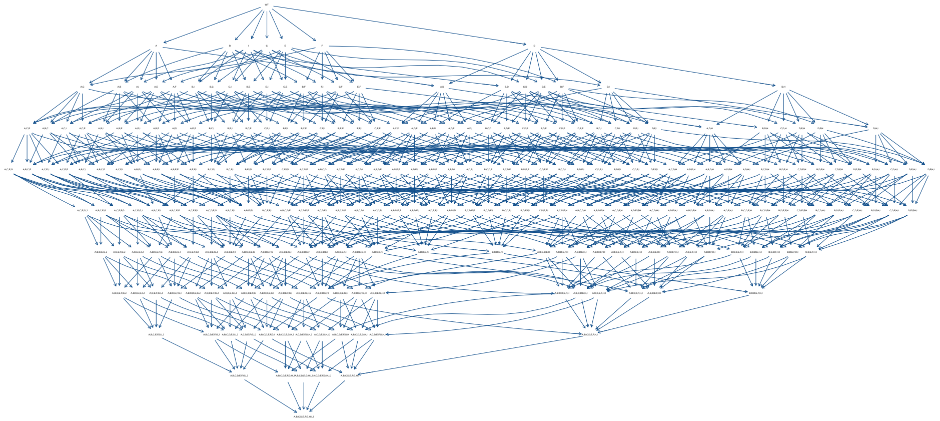

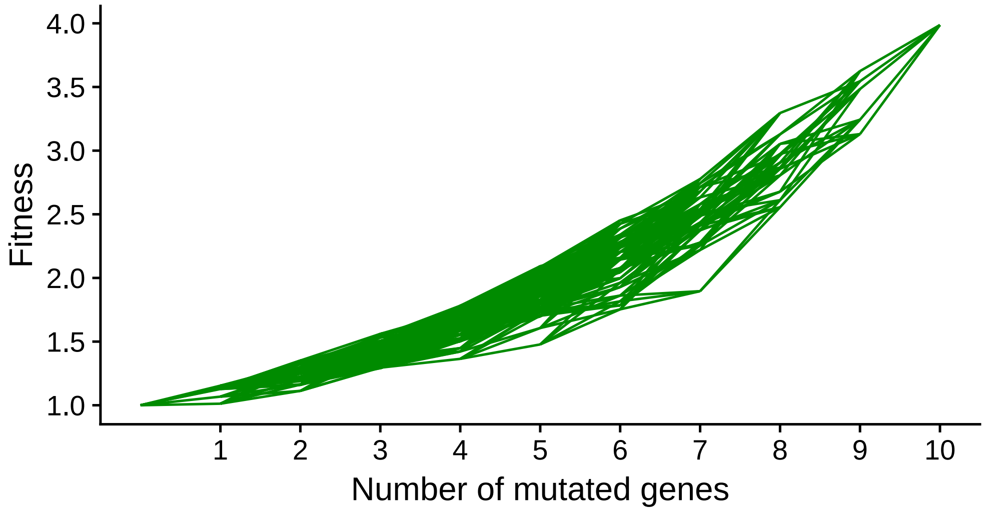

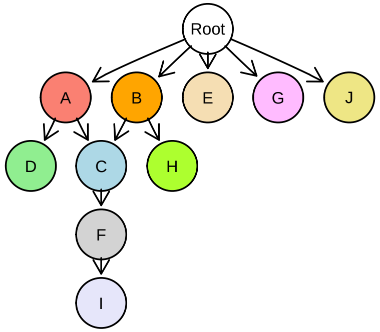


| | |
|---|---|
| *ID* | RHdC3iuVYPR6IBD10 |
| *acc genotypes* | 168 |
| *peaks no back* | 1 |
| *removed edges* | 0 |
| *prop rem edges* | 0 |

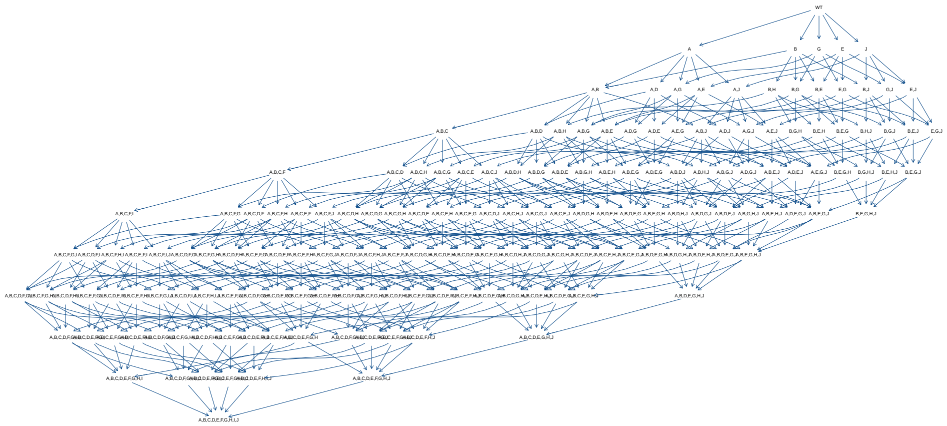

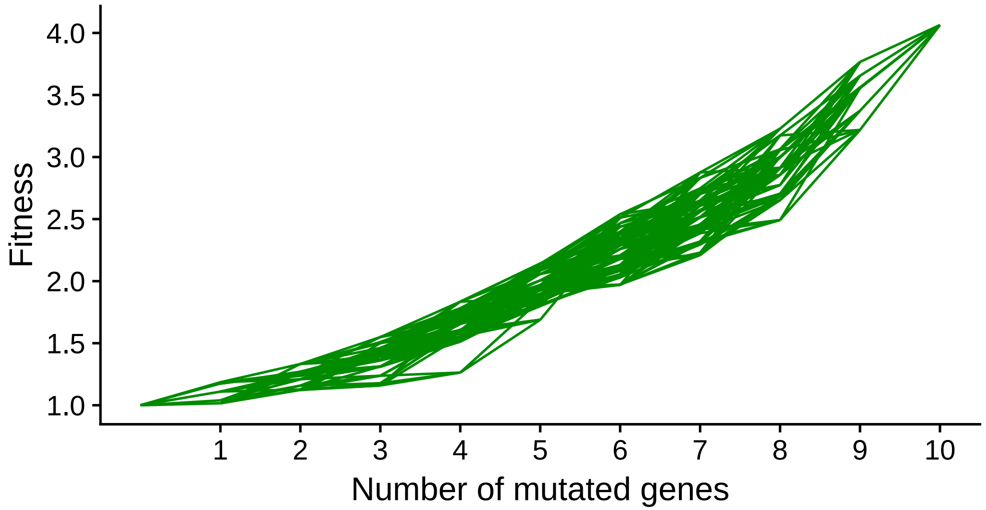

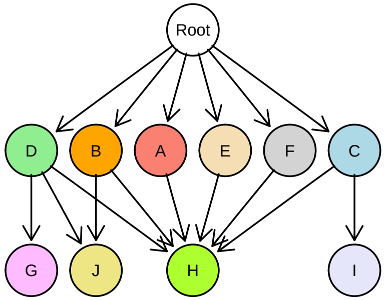


| | |
|---|---|
| *ID* | RM2aZBkyRpQtZvKY |
| *acc genotypes* | 200 |
| *peaks no back* | 1 |
| *removed edges* | 0 |
| *prop rem edges* | 0 |

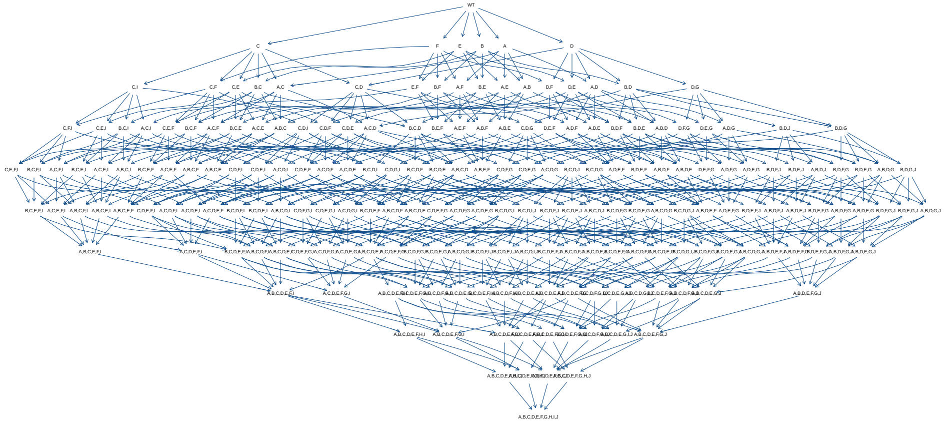

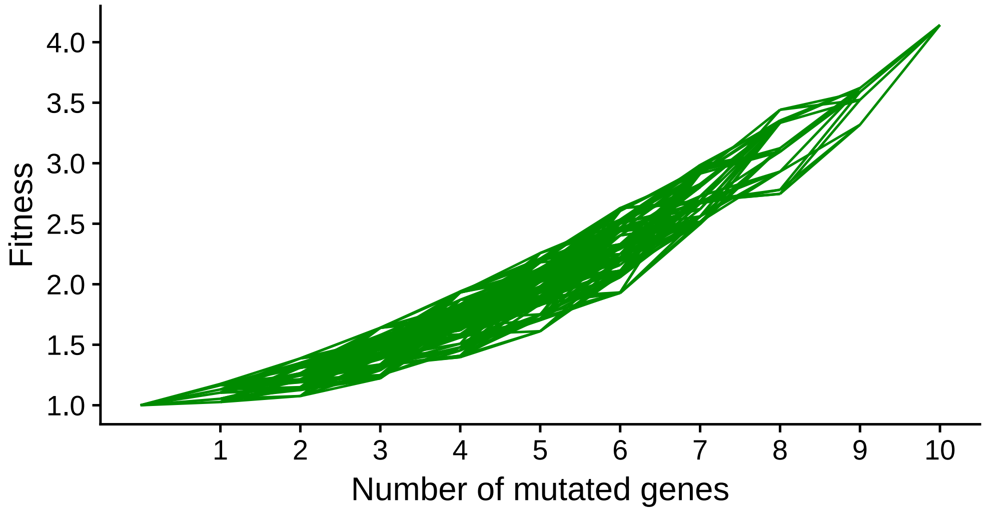

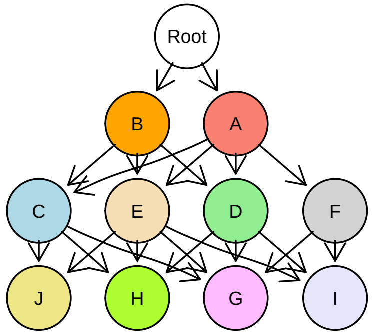


*ID* RNqpQsaM6eeS7RWA
*acc genotypes* 41
*peaks no back* 1
*removed edges* 0
*prop rem edges* 0

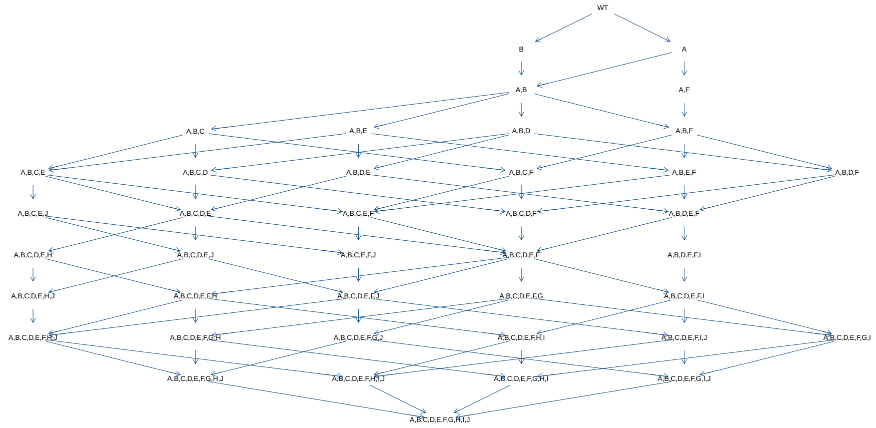

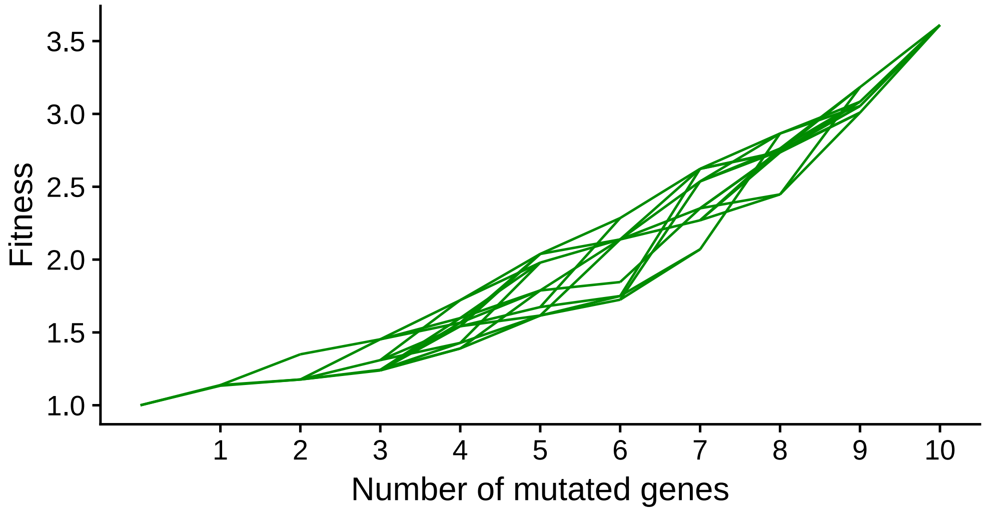

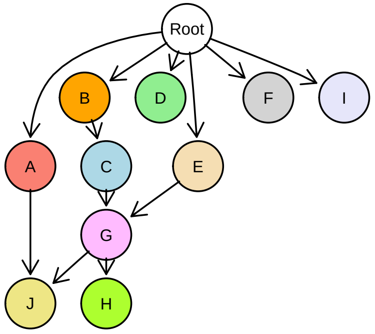


| | |
|---|---|
| *ID* | rRVesv4UlXP10d31k |
| *acc genotypes* | 144 |
| *peaks no back* | 1 |
| *removed edges* | 0 |
| *prop rem edges* | 0 |

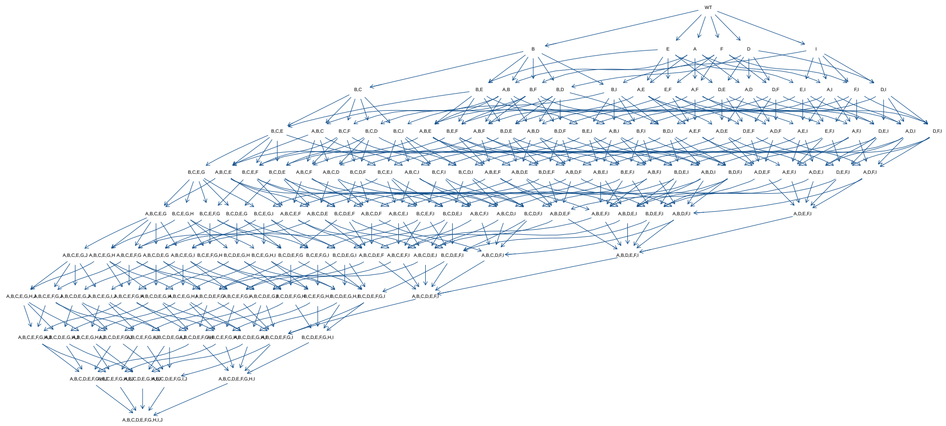

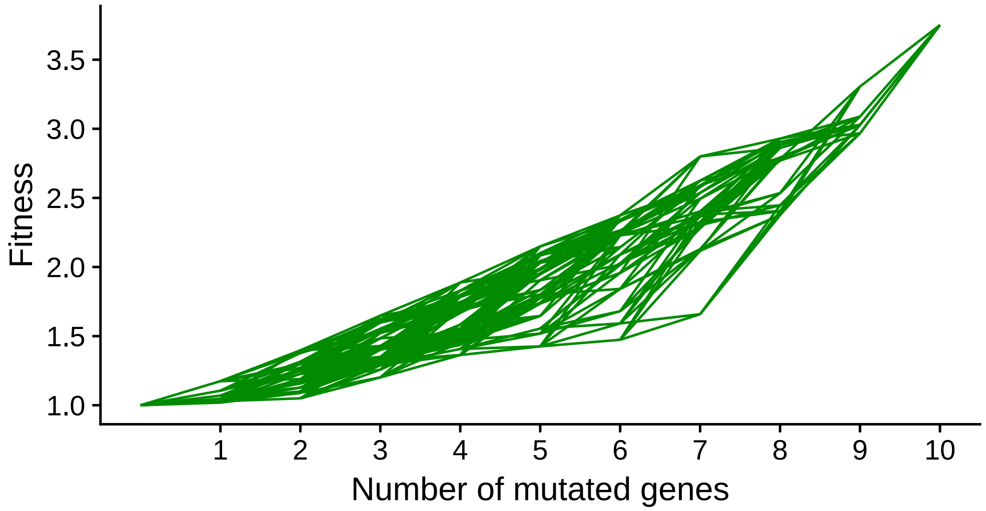

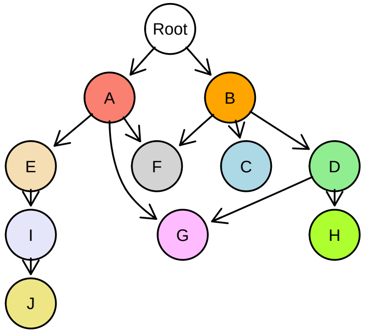


| | |
|---|---|
| *ID* | rsk7tIpjfEVKOLgb |
| *acc genotypes* | 91 |
| *peaks no back* | 1 |
| *removed edges* | 0 |
| *prop rem edges* | 0 |

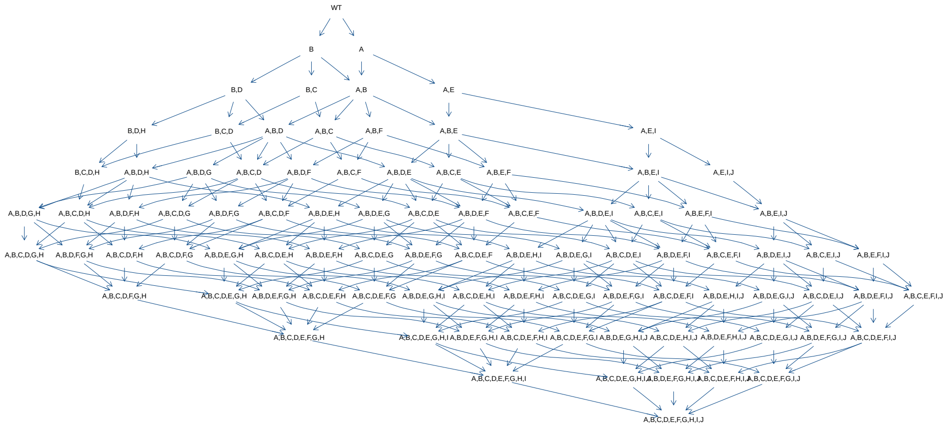


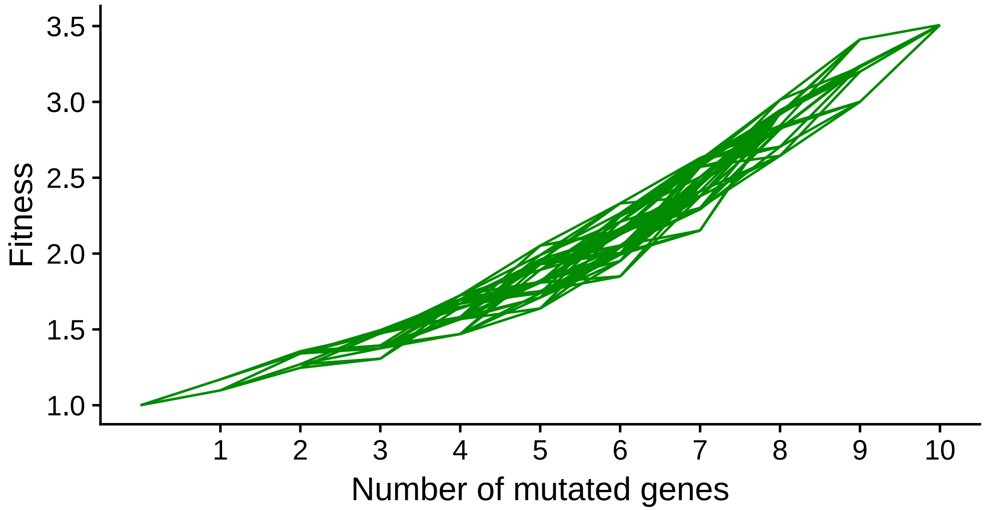

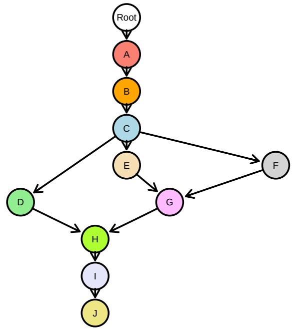


| | |
|---|---|
| *ID* | rXEM4KMif4W386fC |
| *acc genotypes* | 16 |
| *peaks no back* | 1 |
| *removed edges* | 0 |
| *prop rem edges* | 0 |

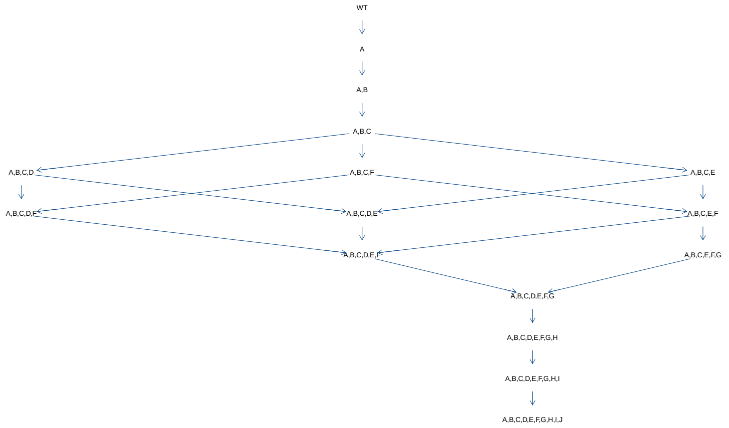


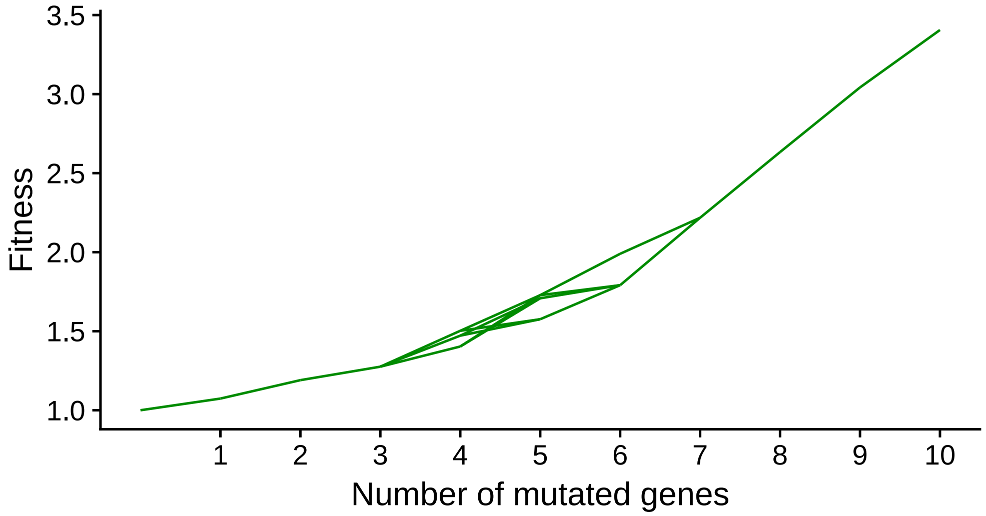

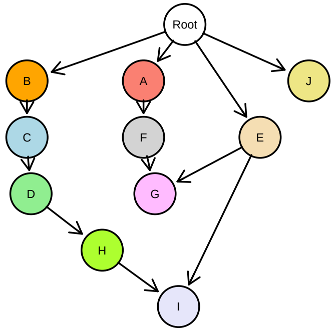


| | |
|---|---|
| *ID* | rZYDGyZC7mmbrvG7 |
| *acc genotypes* | 78 |
| *peaks no back* | 1 |
| *removed edges* | 0 |
| *prop rem edges* | 0 |

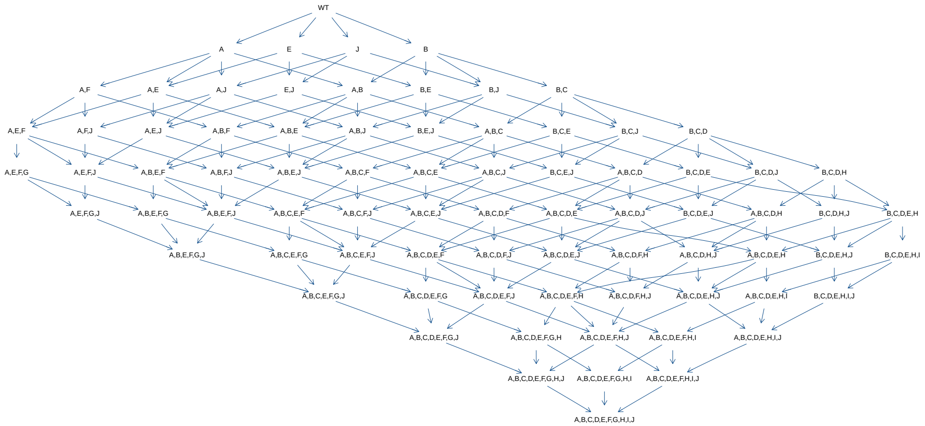

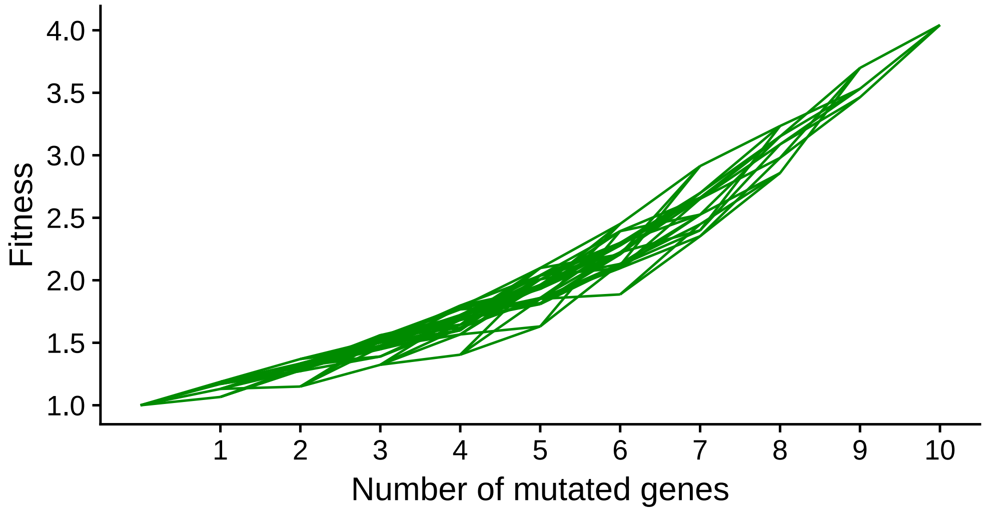

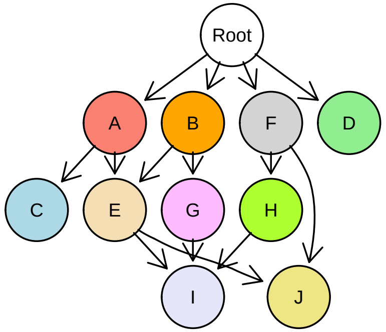


| | |
|---|---|
| *ID* | sH934jpdql10yYeSf |
| *acc genotypes* | 102 |
| *peaks no back* | 1 |
| *removed edges* | 0 |
| *prop rem edges* | 0 |

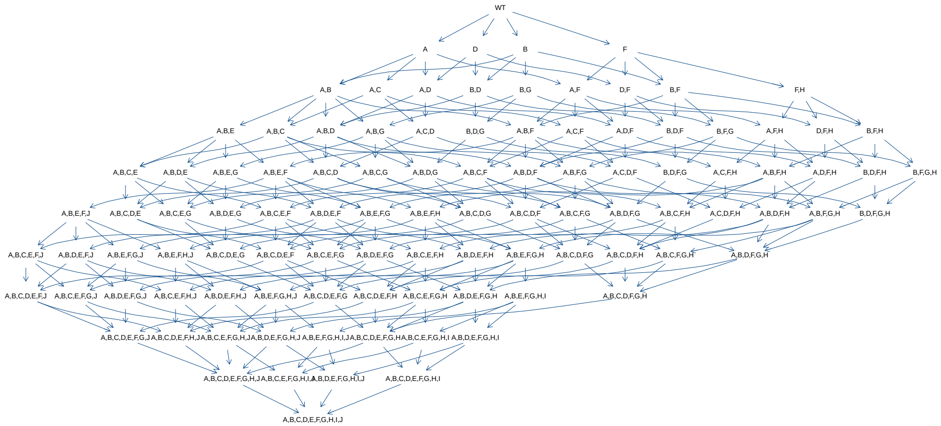

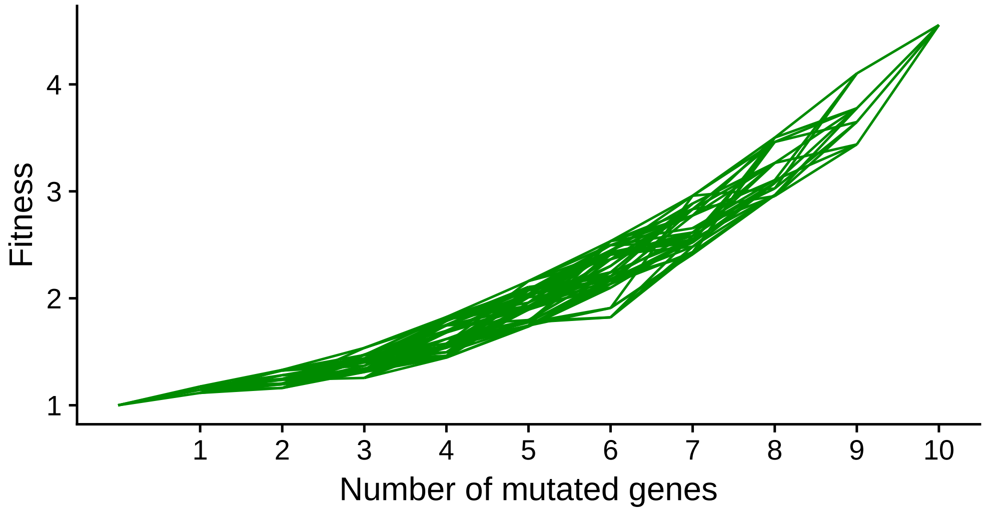

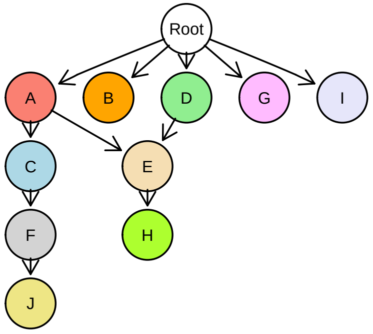


| | |
|---|---|
| *ID* | sisQewhG7QPpthsD |
| *acc genotypes* | 144 |
| *peaks no back* | 1 |
| *removed edges* | 0 |
| *prop rem edges* | 0 |

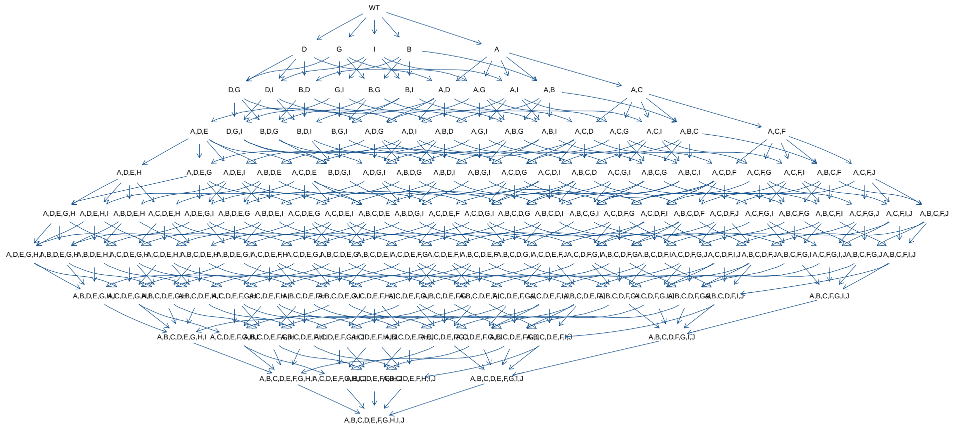

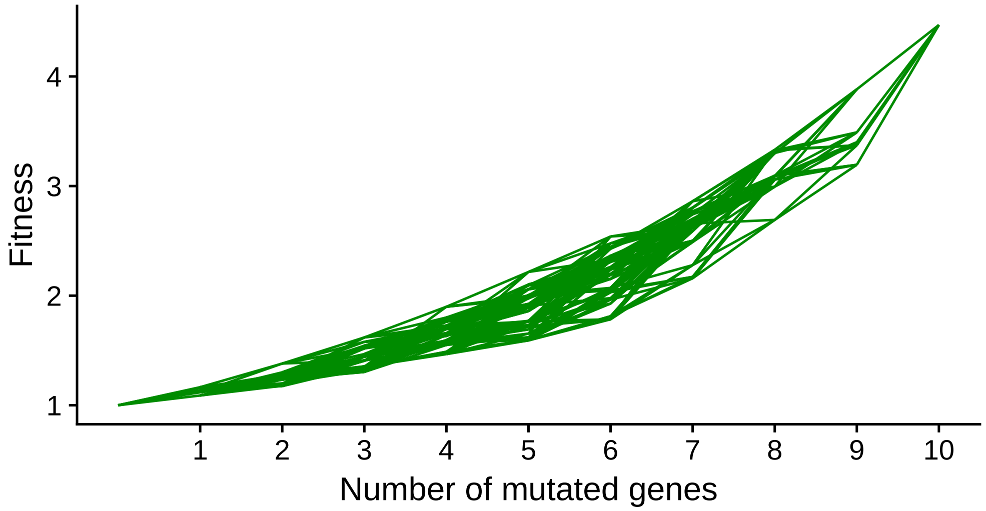

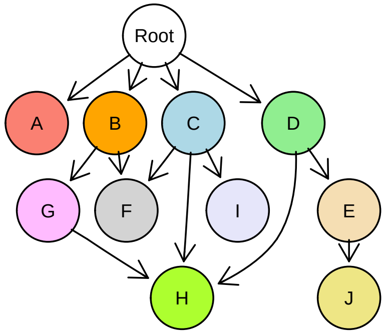


| | |
|---|---|
| *ID* | sNgShm95dlSfotH4 |
| *acc genotypes* | 128 |
| *peaks no back* | 1 |
| *removed edges* | 0 |
| *prop rem edges* | 0 |

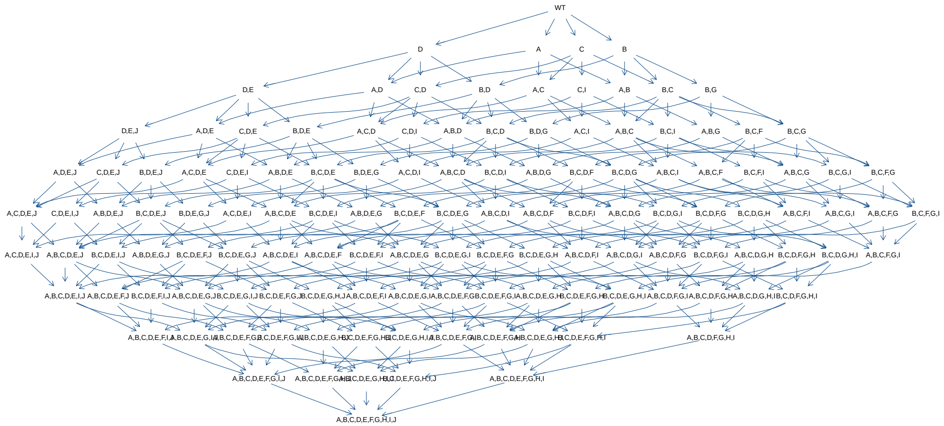

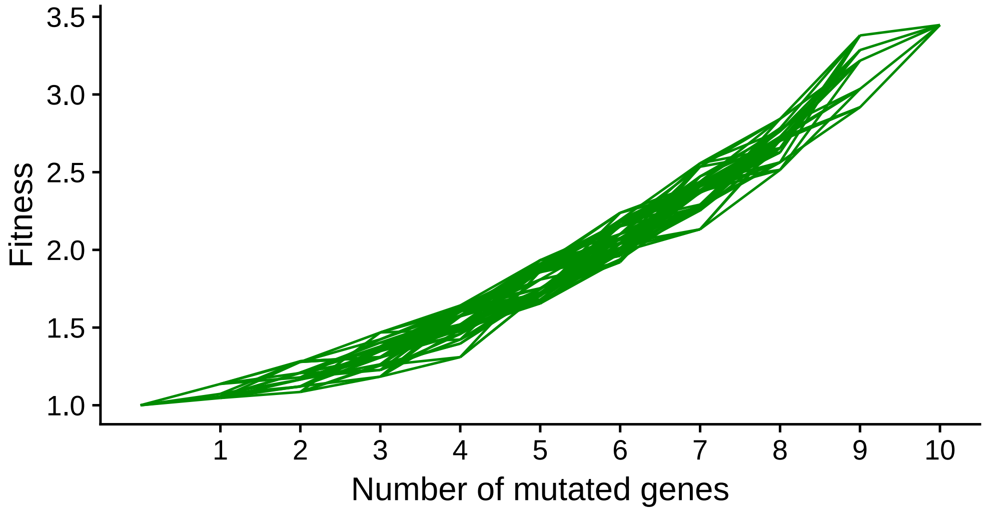

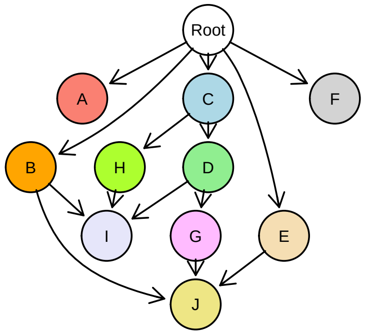


| | |
|---|---|
| *ID* | sQiN7joZyxfMKsXn |
| *acc genotypes* | 140 |
| *peaks no back* | 1 |
| *removed edges* | 0 |
| *prop rem edges* | 0 |

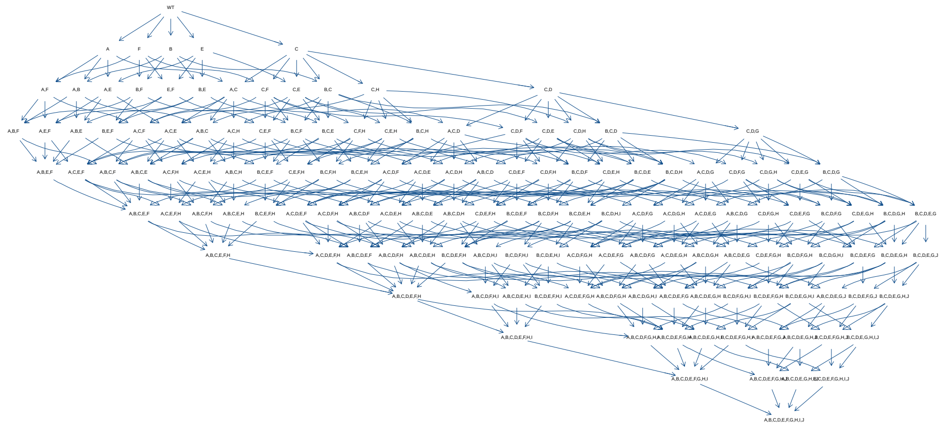

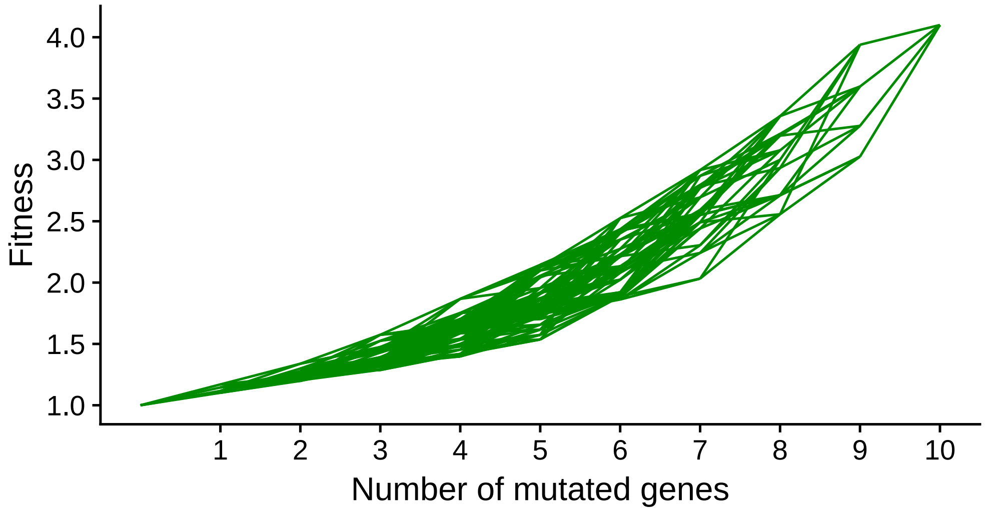

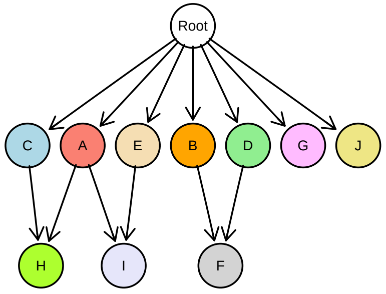


| | |
|---|---|
| *ID* | sUfgZw6OOPt5SCir |
| *acc genotypes* | 260 |
| *peaks no back* | 1 |
| *removed edges* | 0 |
| *prop rem edges* | 0 |

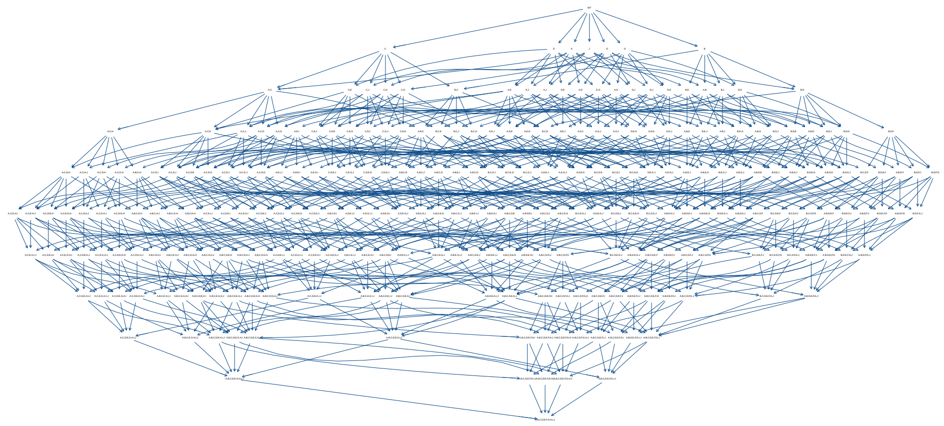

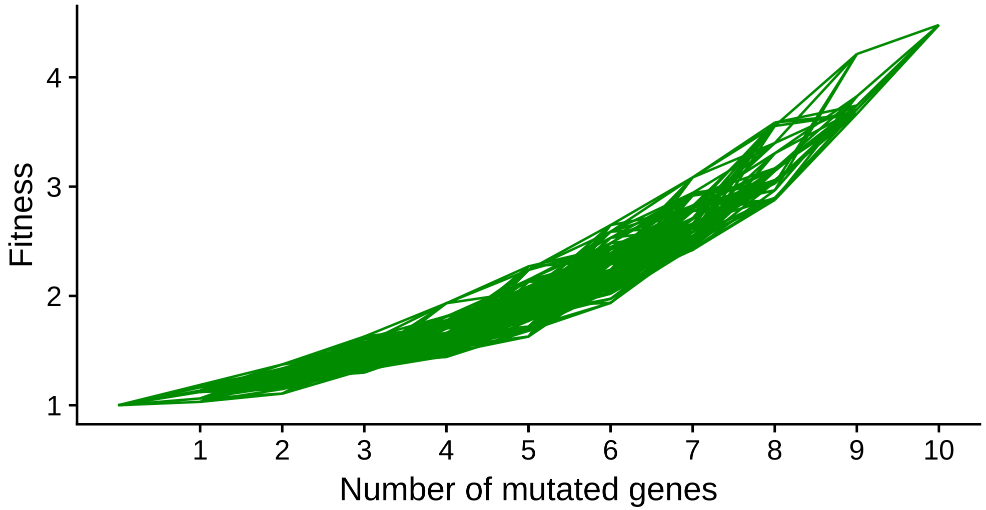

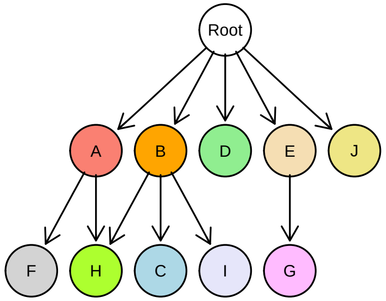


| | |
|---|---|
| *ID* | svUNVWKM4yHG4ZG6 |
| *acc genotypes* | 276 |
| *peaks no back* | 1 |
| *removed edges* | 0 |
| *prop rem edges* | 0 |

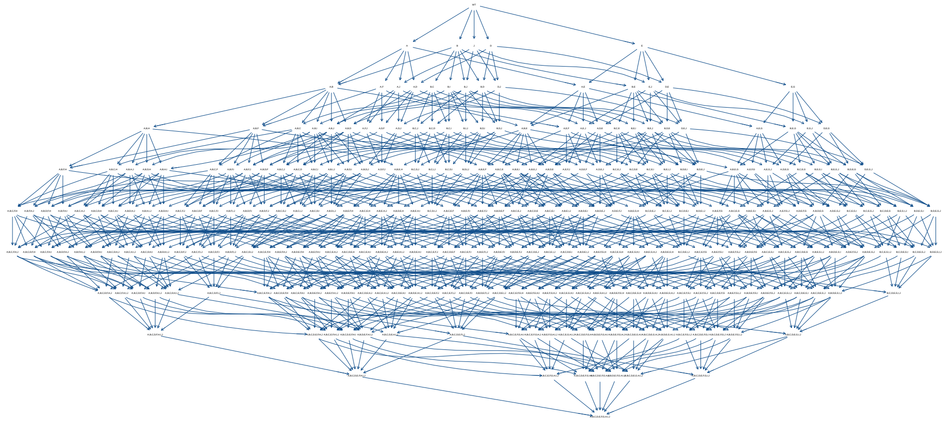

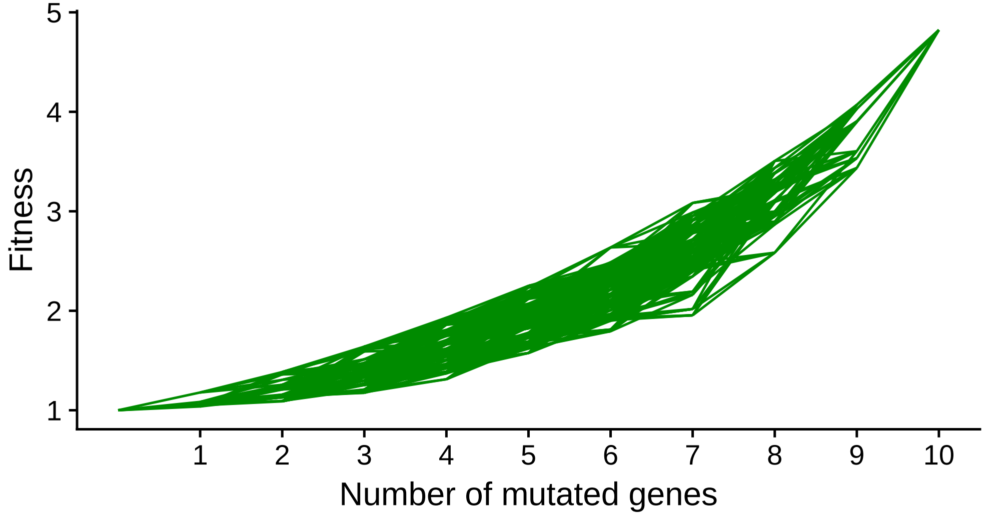

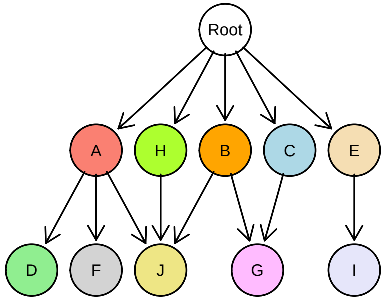


| | |
|---|---|
| *ID* | tbhoAhVYmdTD8tmZ |
| *acc genotypes* | 186 |
| *peaks no back* | 1 |
| *removed edges* | 0 |
| *prop rem edges* | 0 |

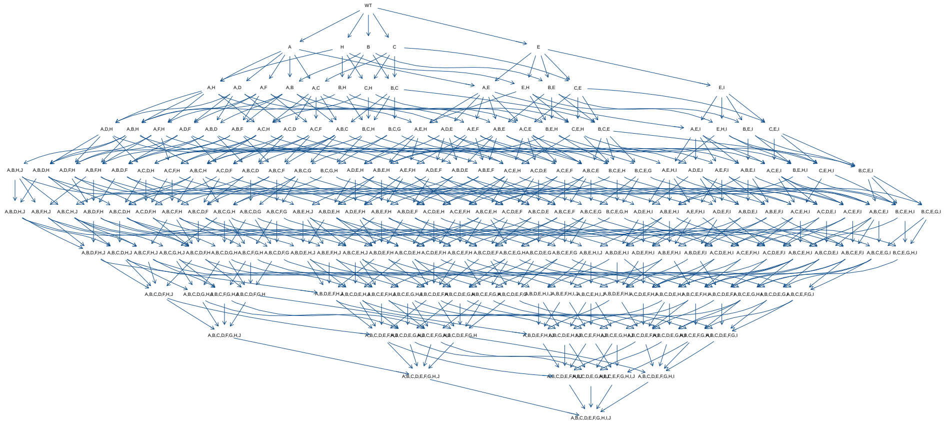

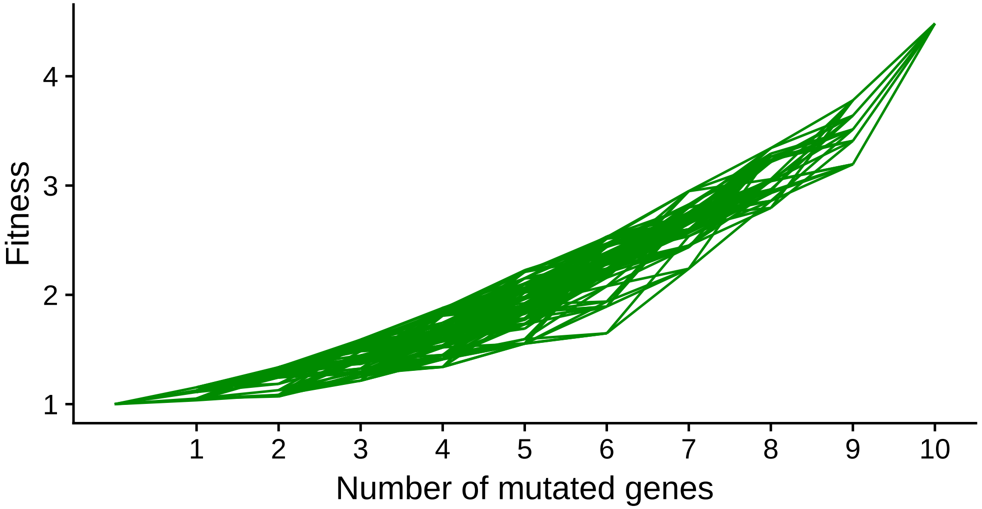

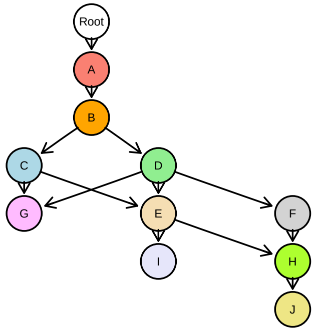


| | |
|---|---|
| *ID* | TFxs2KPm5WO9Abi10 |
| *acc genotypes* | 26 |
| *peaks no back* | 1 |
| *removed edges* | 0 |
| *prop rem edges* | 0 |

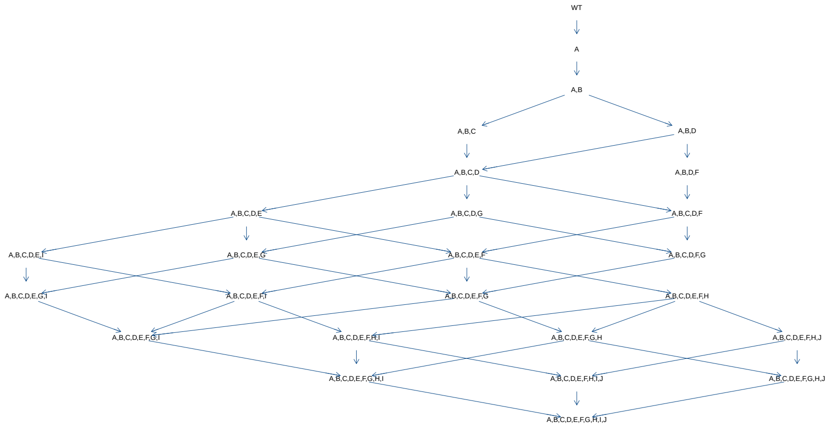

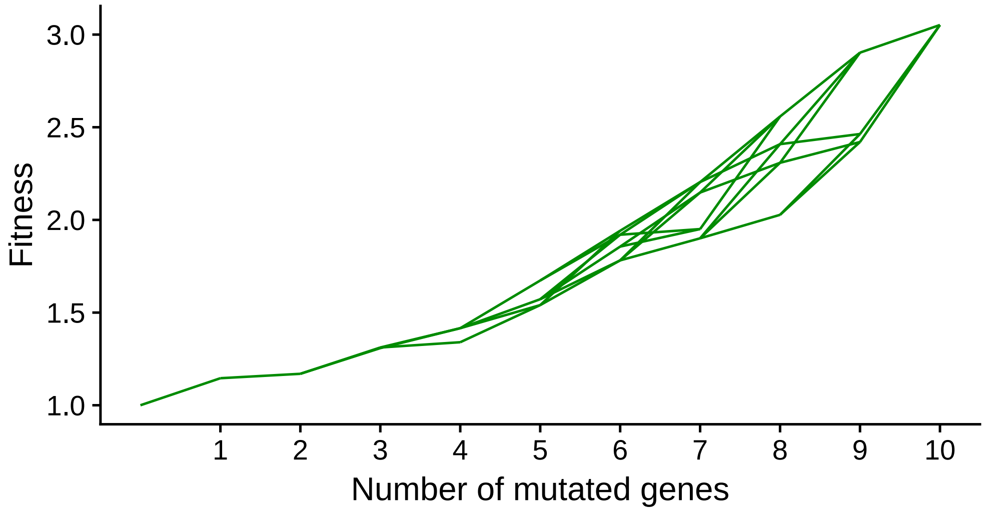

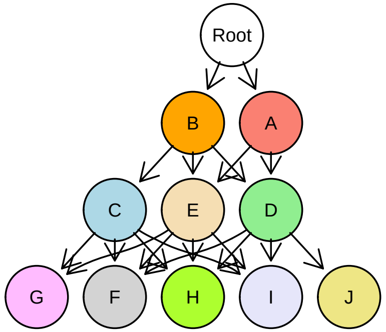


| | |
|---|---|
| *ID* | TH4wayDxtpPQJ3zV |
| *acc genotypes* | 47 |
| *peaks no back* | 1 |
| *removed edges* | 0 |
| *prop rem edges* | 0 |

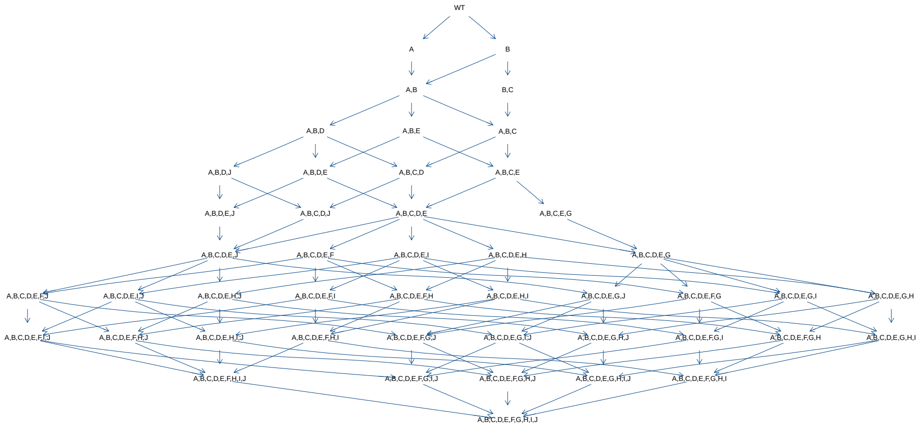

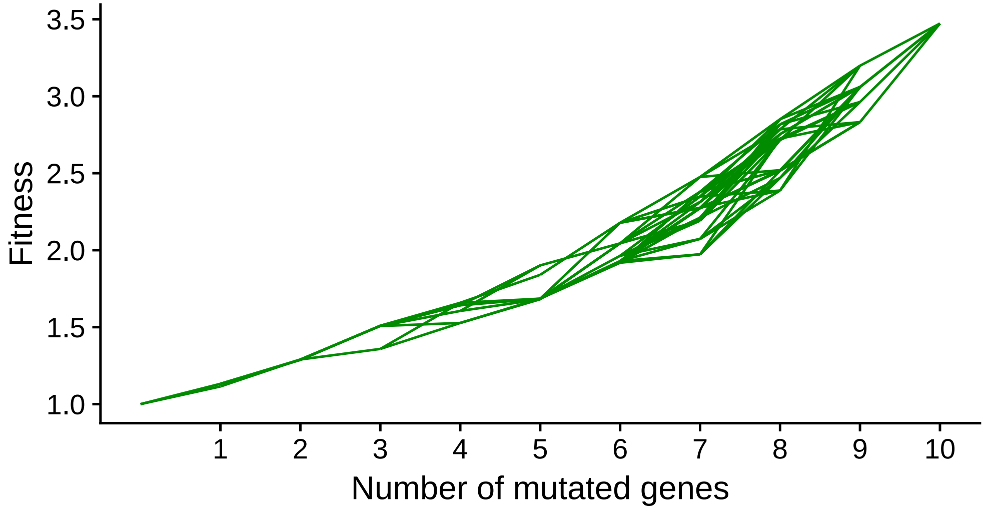

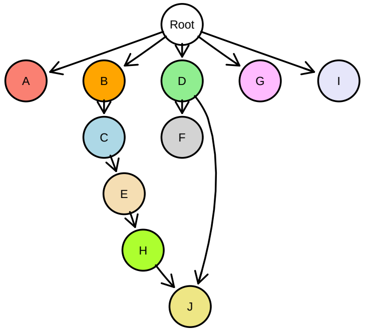


| | |
|---|---|
| *ID* | tl7D6tClLmHkt9fG |
| *acc genotypes* | 136 |
| *peaks no back* | 1 |
| *removed edges* | 0 |
| *prop rem edges* | 0 |

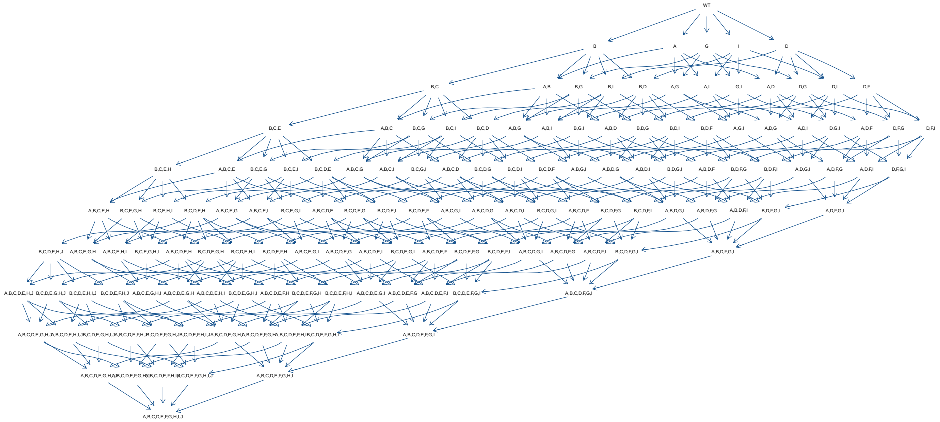

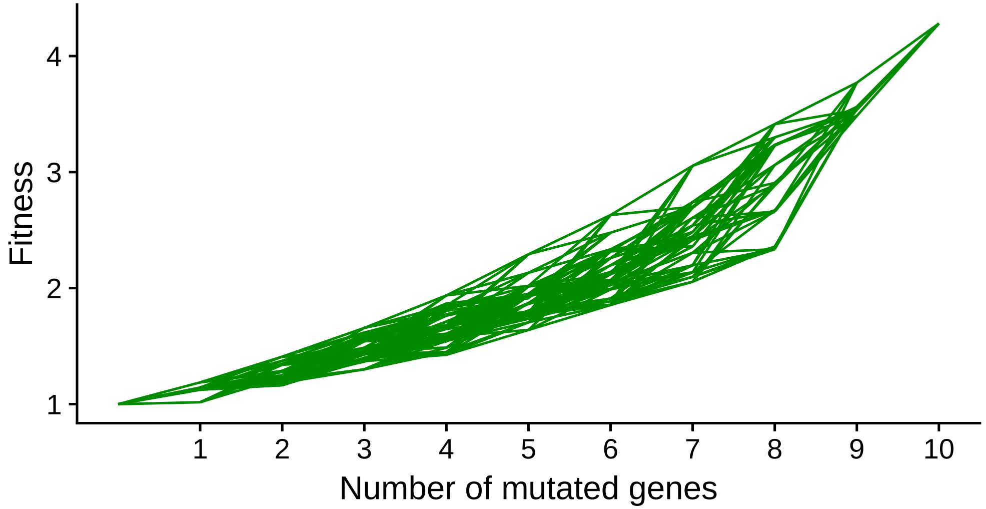

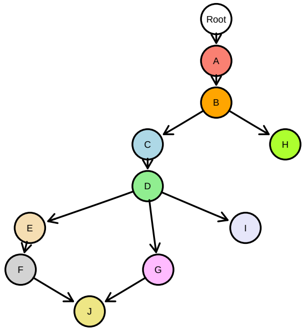


| | |
|---|---|
| *ID* | tq7jgfG17AM4lqGW |
| *acc genotypes* | 34 |
| *peaks no back* | 1 |
| *removed edges* | 0 |
| *prop rem edges* | 0 |

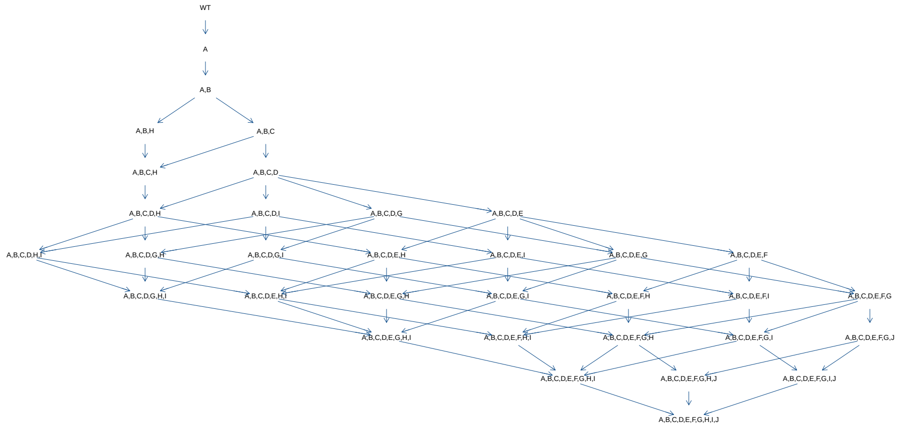

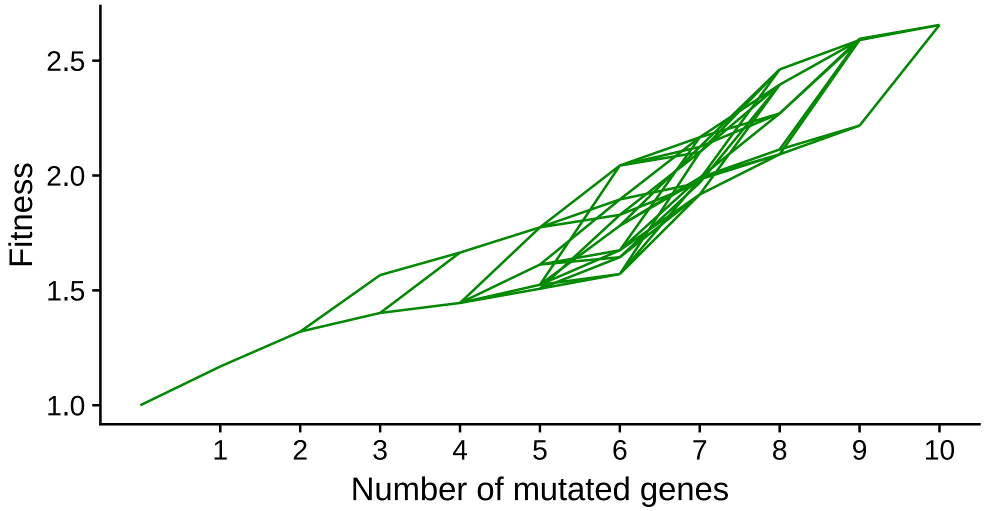

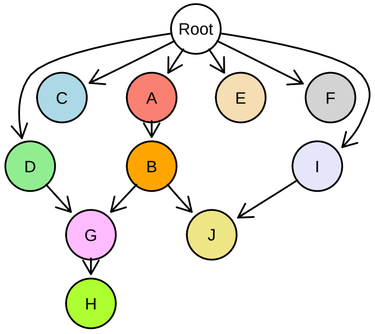


| | |
|---|---|
| *ID* | TVJsMGLUm85XS67g |
| *acc genotypes* | 160 |
| *peaks no back* | 1 |
| *removed edges* | 0 |
| *prop rem edges* | 0 |

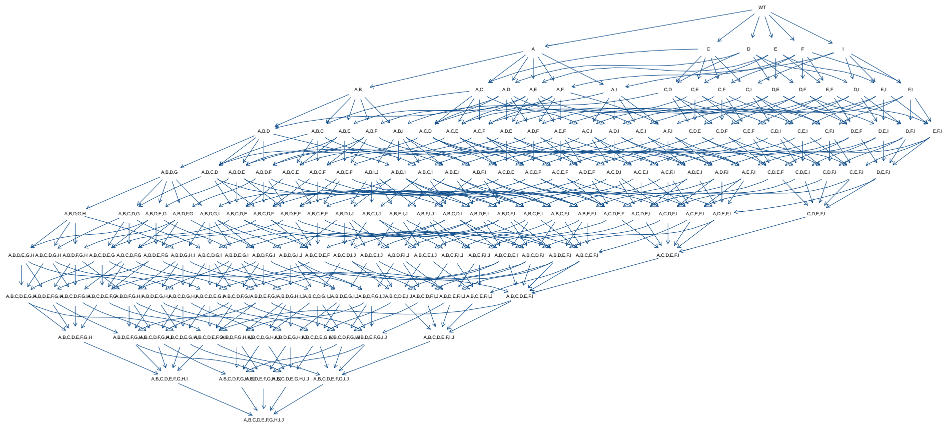

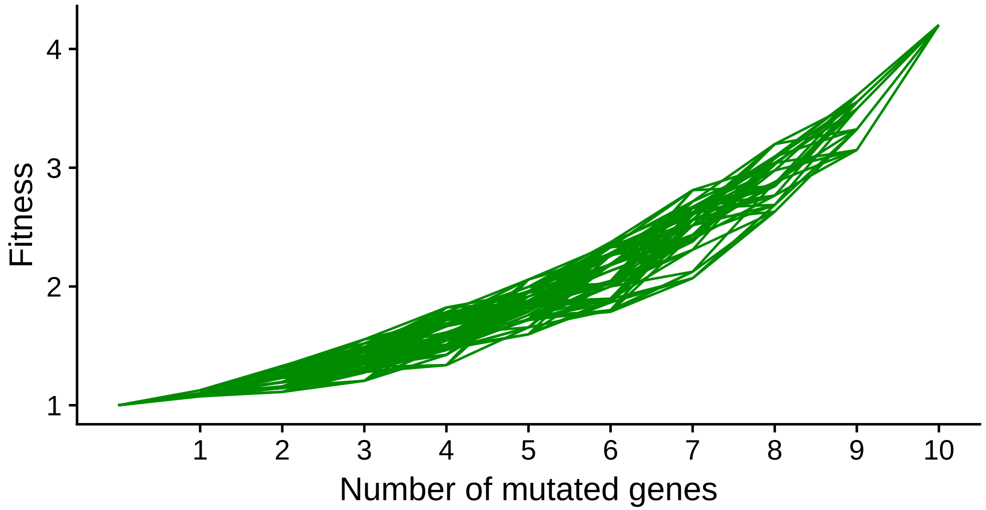

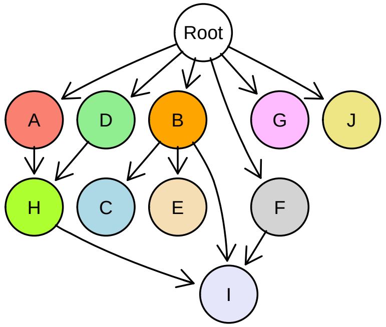


*ID* TXSYGzI7tgHRqH7B
*acc genotypes* 216
*peaks no back* 1
*removed edges* 0
*prop rem edges* 0

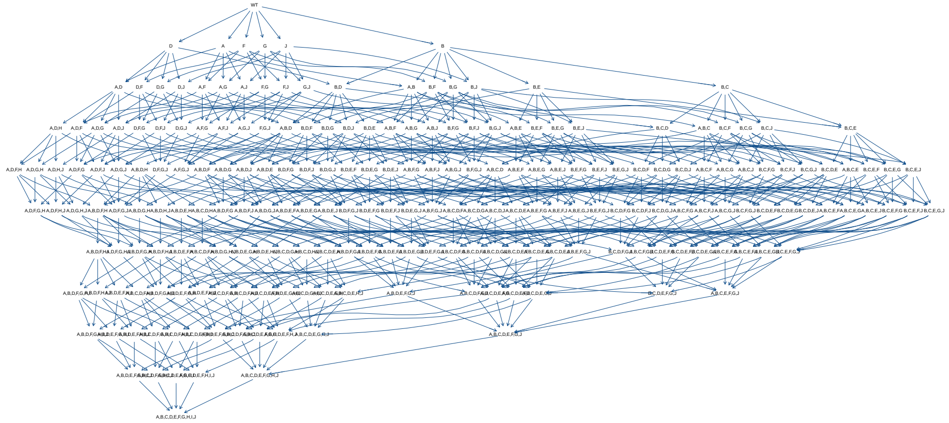


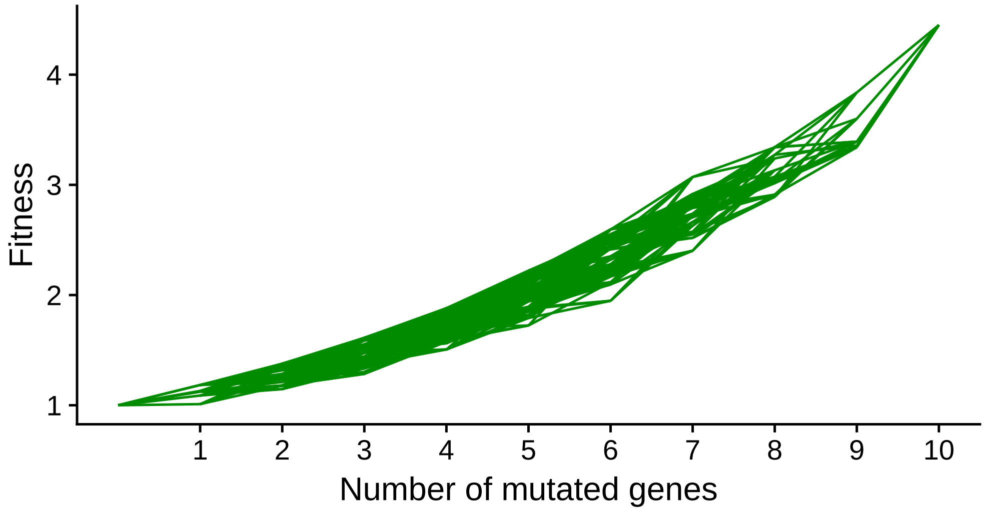

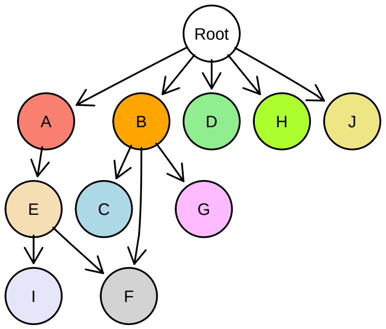


*ID* u1PuIFhBeoVLcUwQ
*acc genotypes* 224
*peaks no back* 1
*removed edges* 0
*prop rem edges* 0

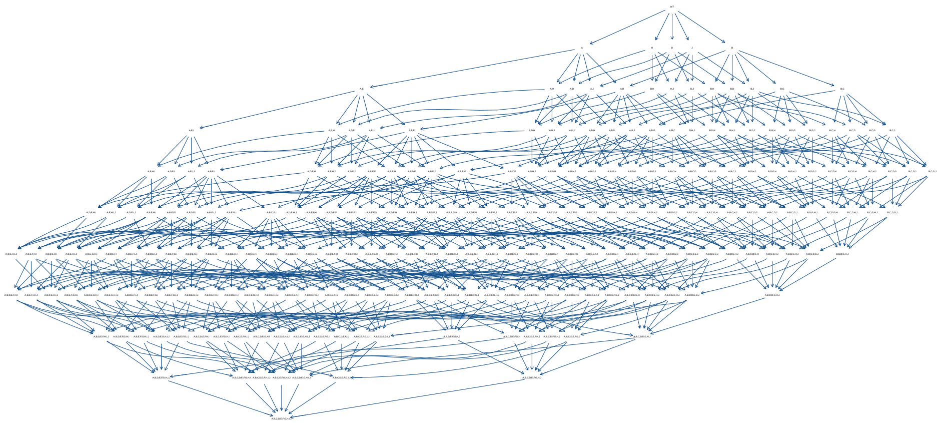

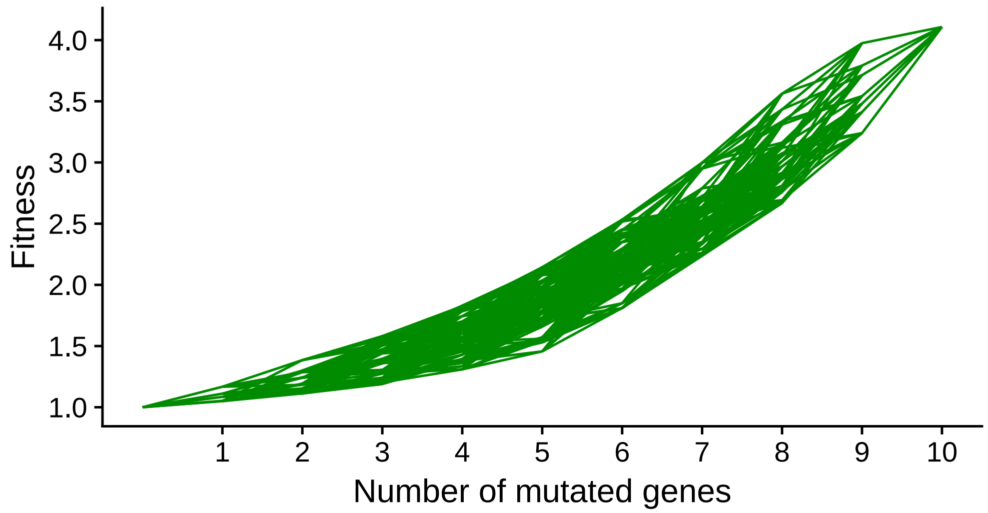

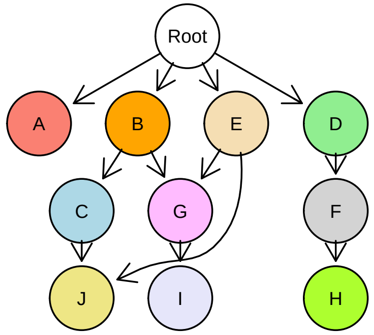


| | |
|---|---|
| *ID* | u7NcTlp7H7vgrd6Z |
| *acc genotypes* | 104 |
| *peaks no back* | 1 |
| *removed edges* | 0 |
| *prop rem edges* | 0 |

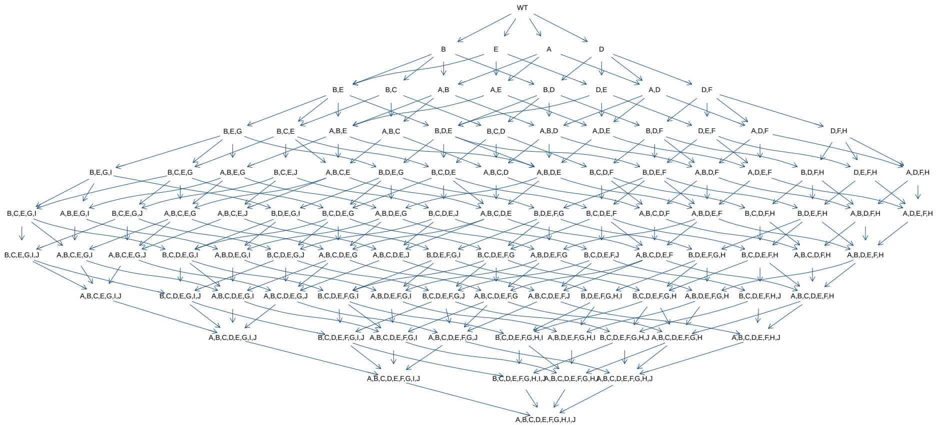

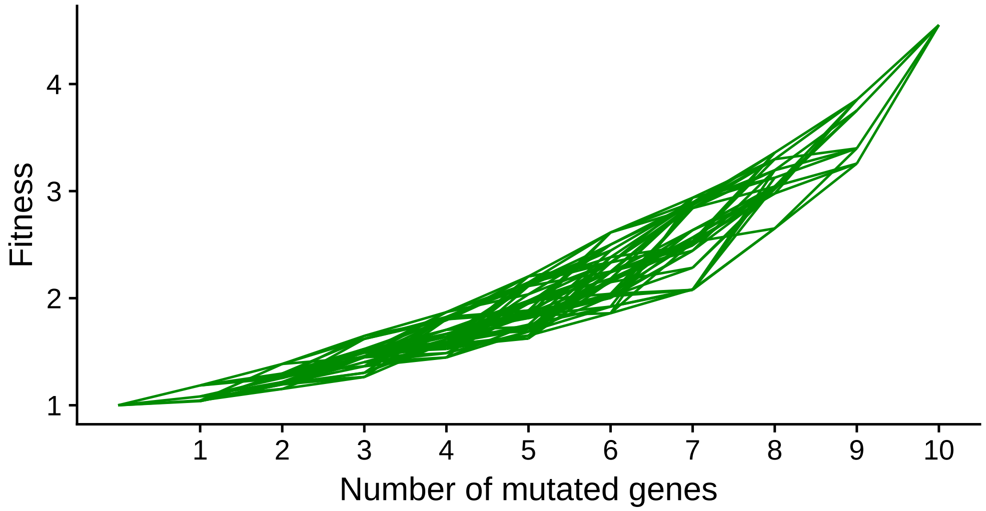

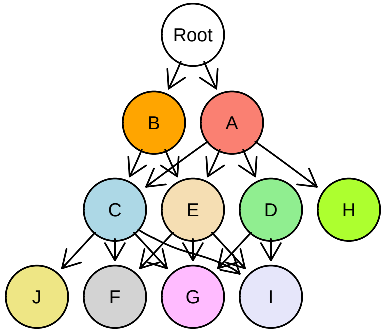


| | |
|---|---|
| *ID* | u8LSkOke6emtVQOi |
| *acc genotypes* | 62 |
| *peaks no back* | 1 |
| *removed edges* | 0 |
| *prop rem edges* | 0 |

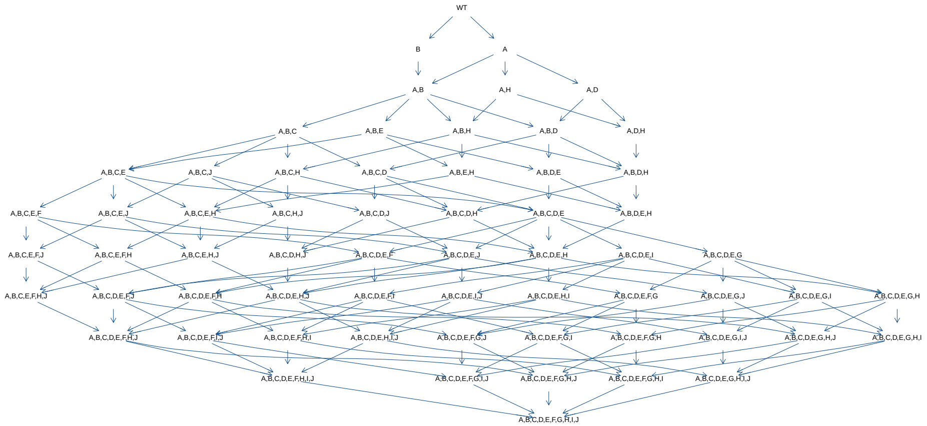

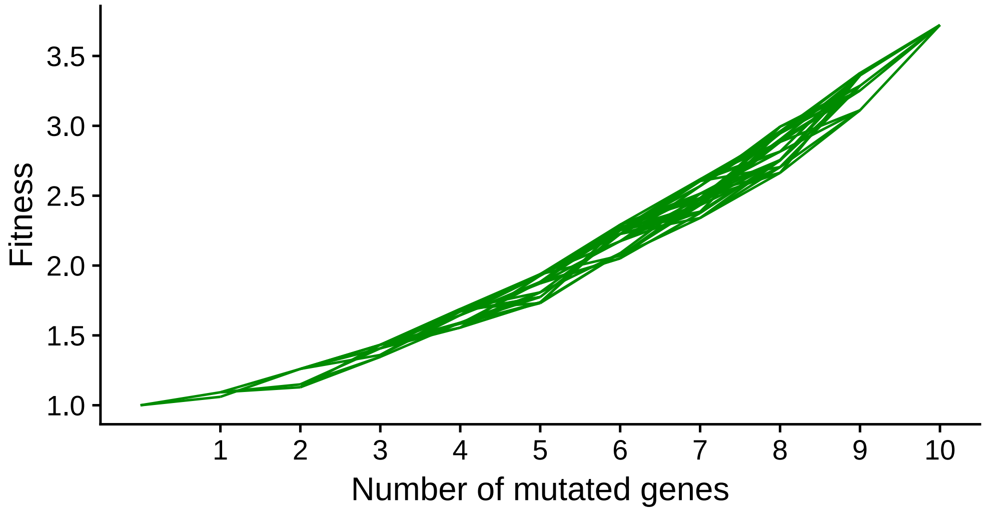

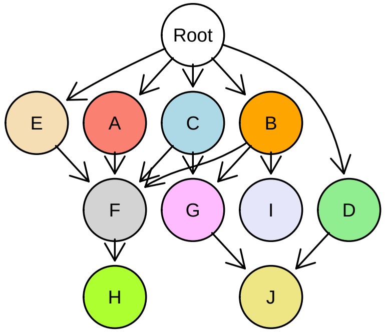


*ID* UjXjH4wCYwemR9Ik
*acc genotypes* 92
*peaks no back* 1
*removed edges* 0
*prop rem edges* 0

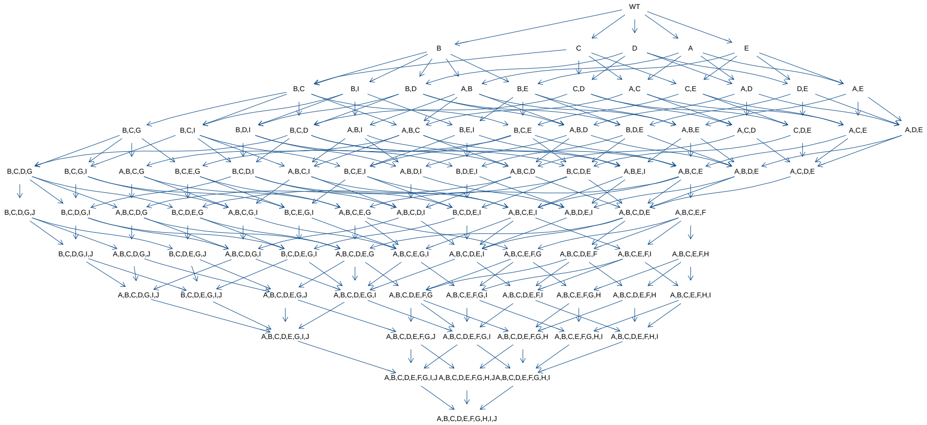


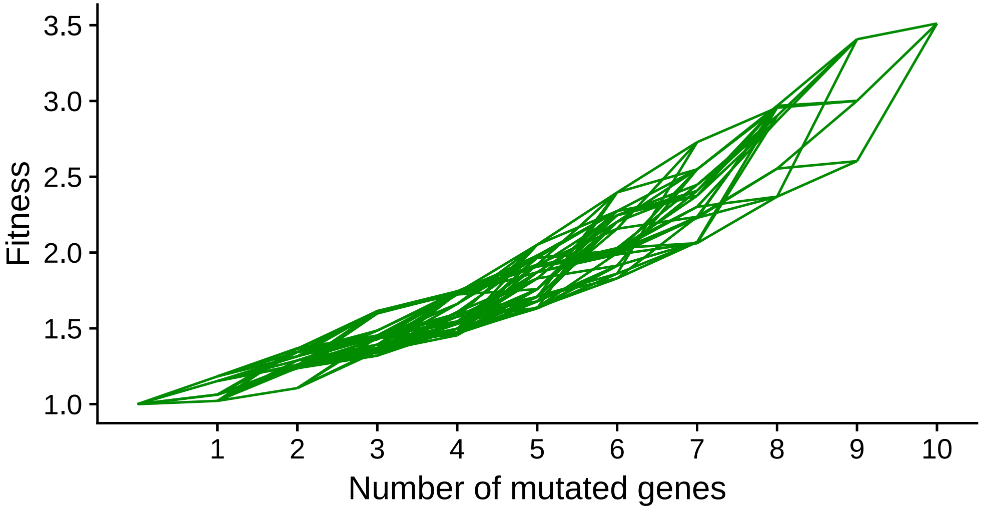

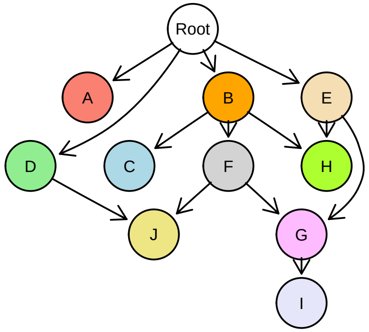


| | |
|---|---|
| *ID* | UPacqPOLUgev2Qbz |
| *acc genotypes* | 116 |
| *peaks no back* | 1 |
| *removed edges* | 0 |
| *prop rem edges* | 0 |

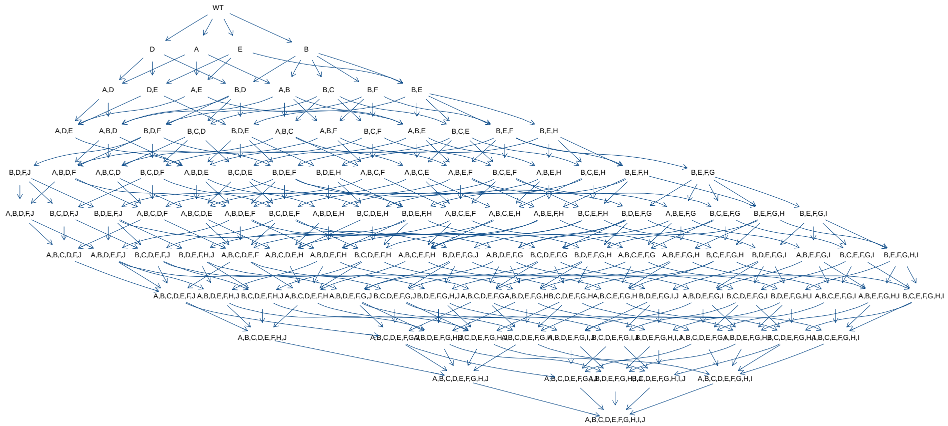

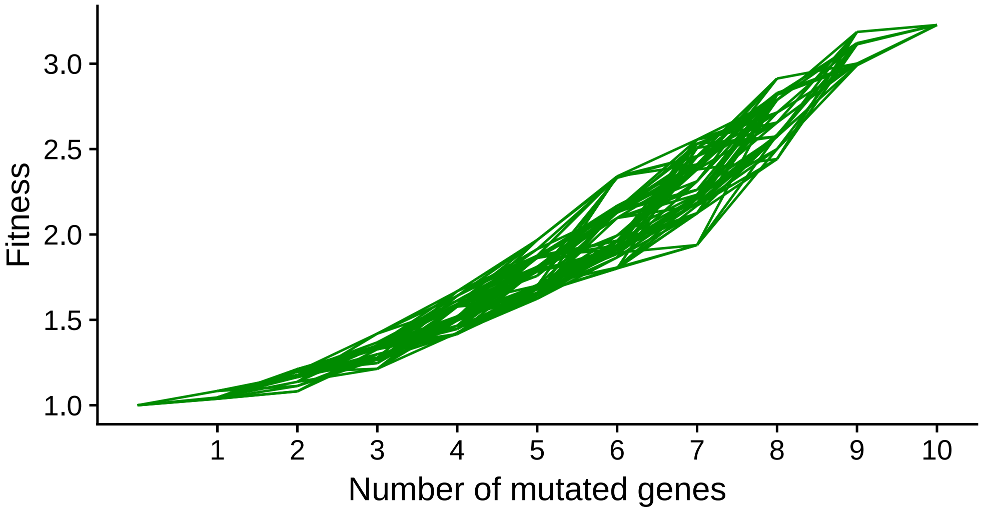

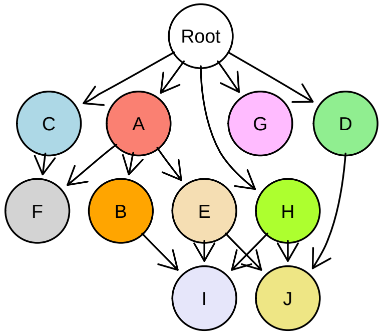


| | |
|---|---|
| *ID* | UtTlZN4ssUPI1042T |
| *acc genotypes* | 142 |
| *peaks no back* | 1 |
| *removed edges* | 0 |
| *prop rem edges* | 0 |

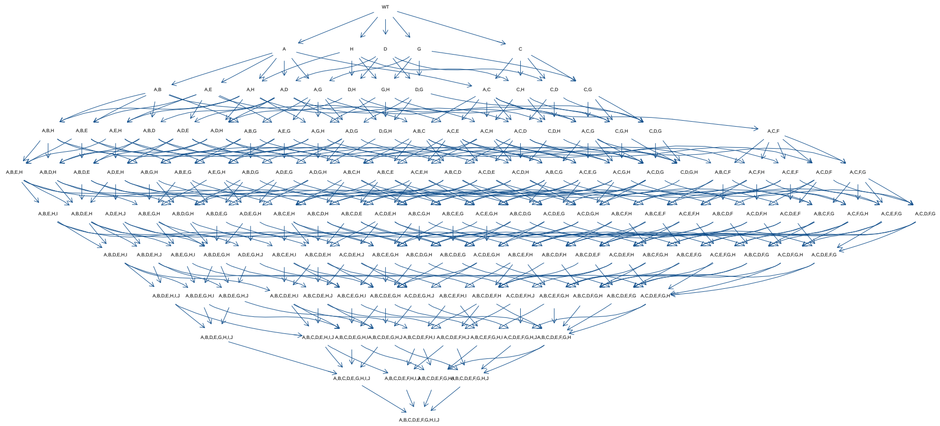

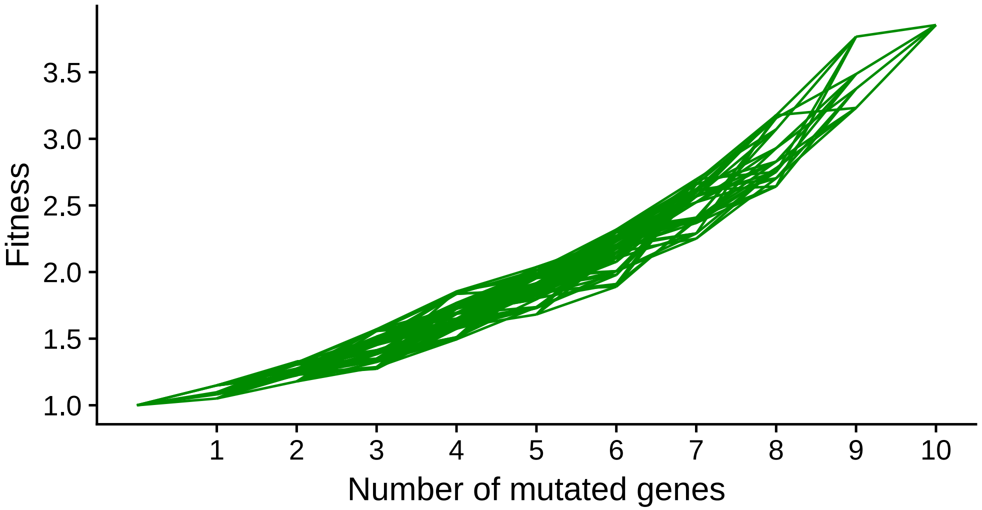

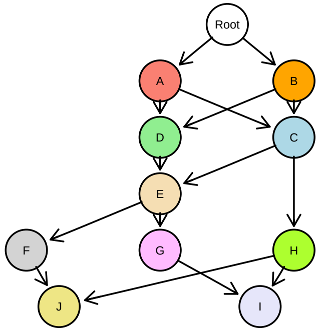


| | |
|---|---|
| *ID* | uU3eWnOFxxJy7Vct |
| *acc genotypes* | 22 |
| *peaks no back* | 1 |
| *removed edges* | 0 |
| *prop rem edges* | 0 |

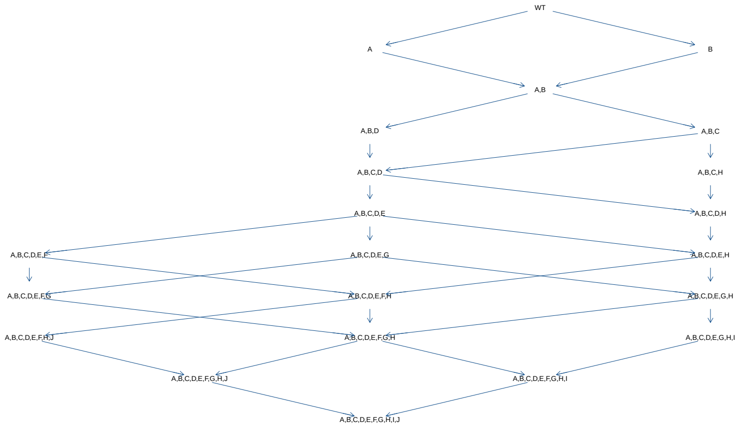


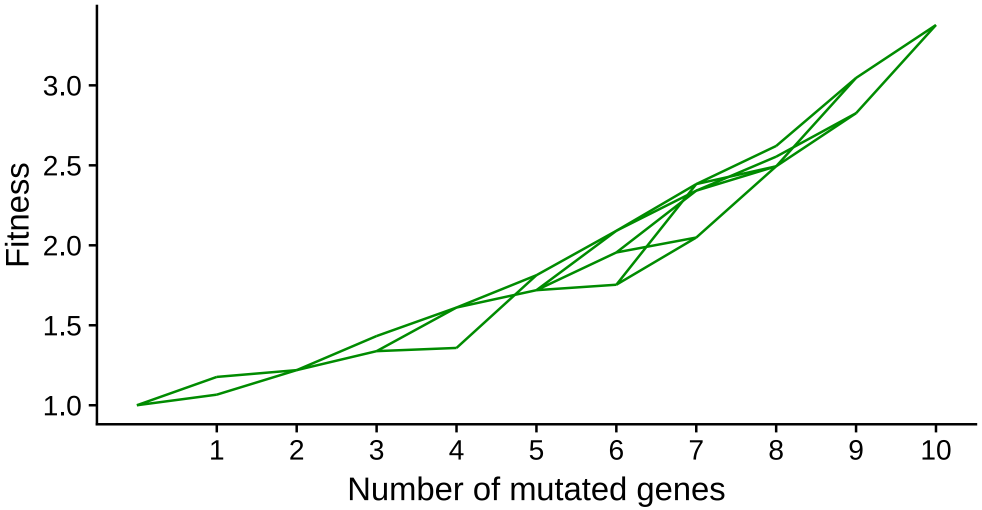

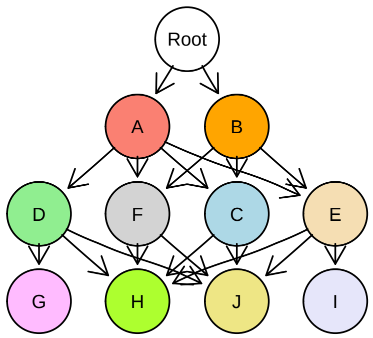


| | |
|---|---|
| *ID* | UvbbY5R10xG3o3Fbv |
| *acc genotypes* | 53 |
| *peaks no back* | 1 |
| *removed edges* | 0 |
| *prop rem edges* | 0 |

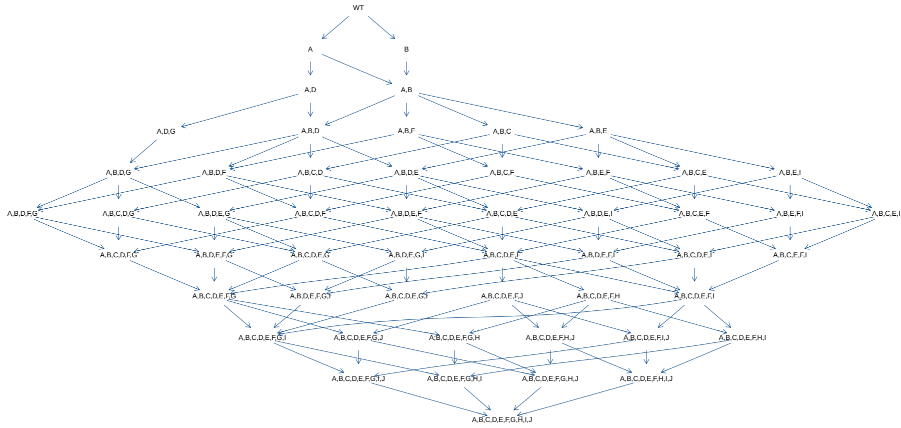

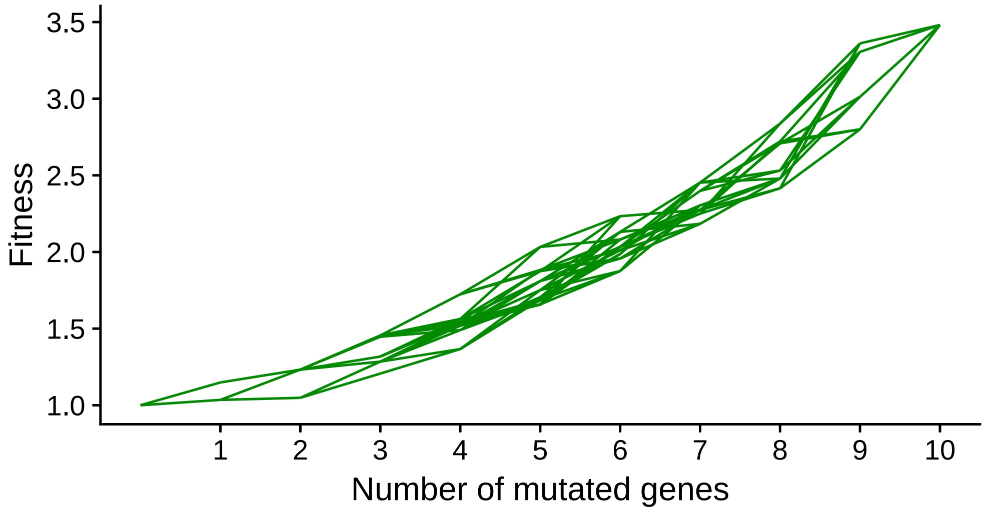

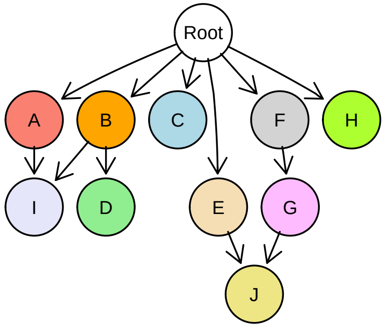


| | |
|---|---|
| *ID* | uVPTmh9a12FoeCLG |
| *acc genotypes* | 224 |
| *peaks no back* | 1 |
| *removed edges* | 0 |
| *prop rem edges* | 0 |

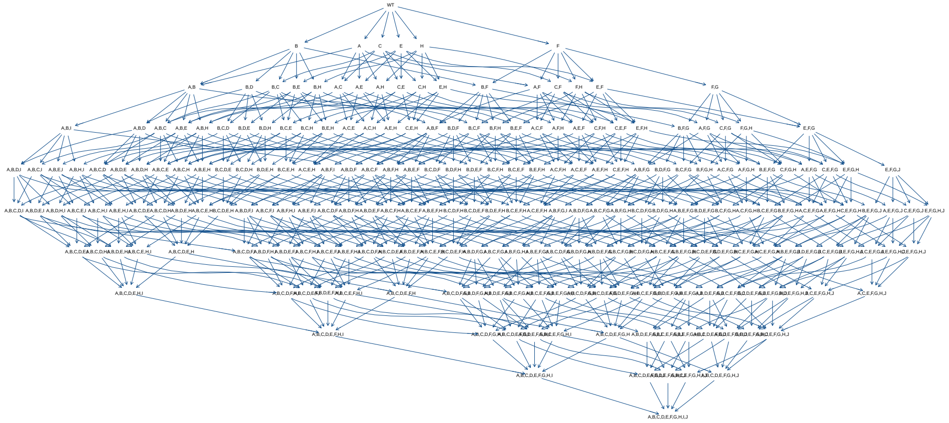

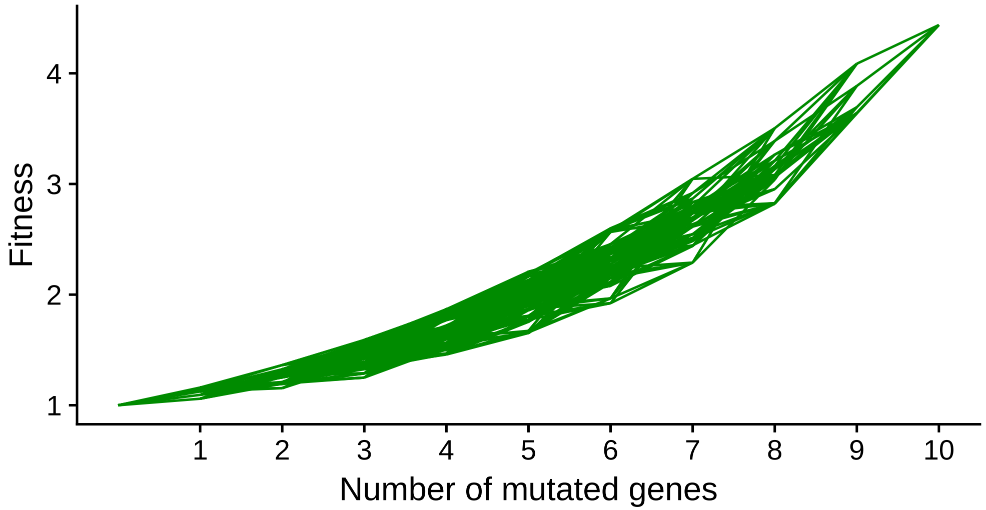

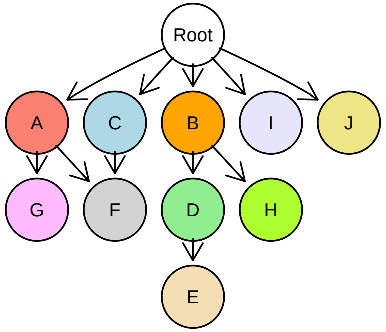


| | |
|---|---|
| *ID* | uWAn38F57JFUrKmr |
| *acc genotypes* | 224 |
| *peaks no back* | 1 |
| *removed edges* | 0 |
| *prop rem edges* | 0 |

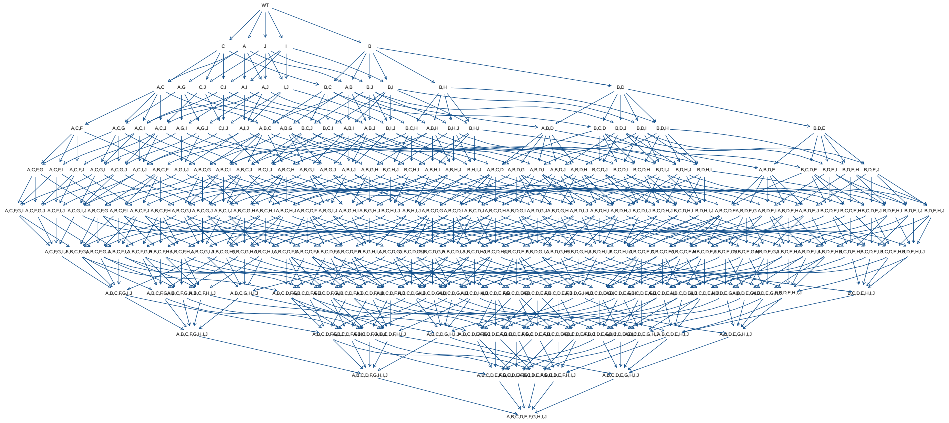

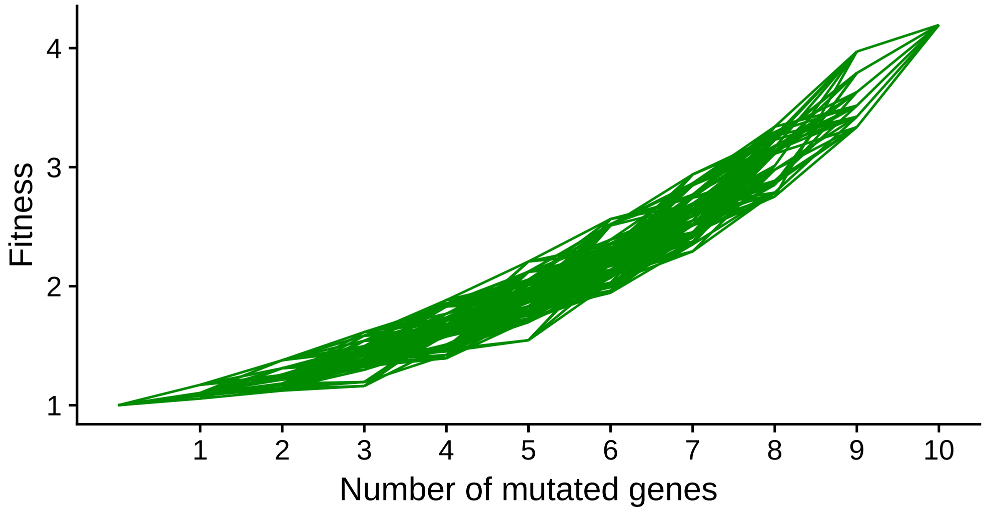

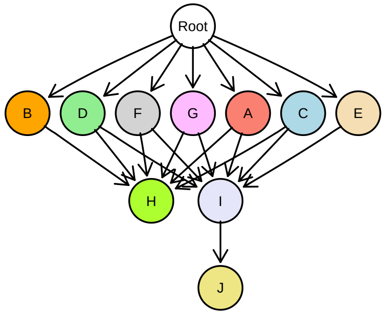


| | |
|---|---|
| *ID* | UXtvbte61fs7Xr8g |
| *acc genotypes* | 136 |
| *peaks no back* | 1 |
| *removed edges* | 0 |
| *prop rem edges* | 0 |

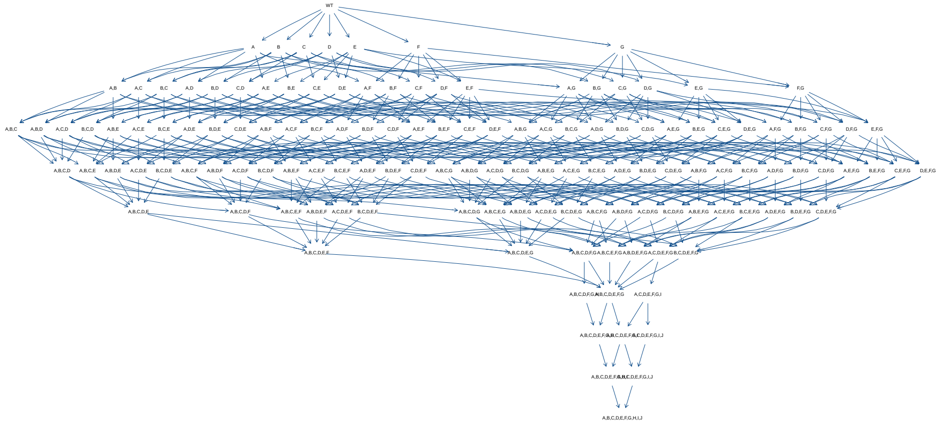

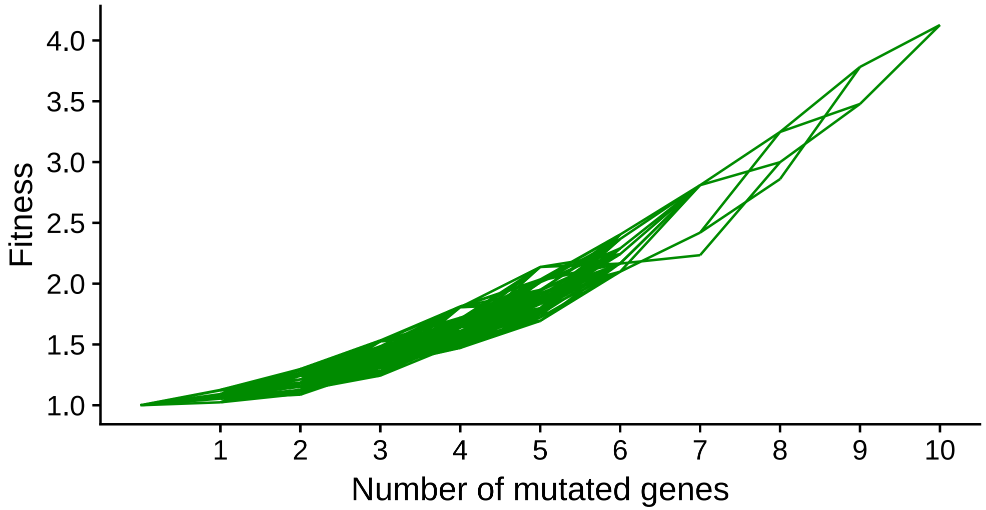

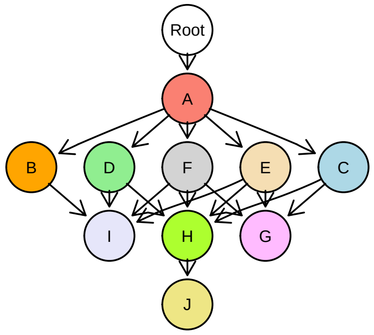


| | |
|---|---|
| *ID* | v7v1qJcKO4kxs78T |
| *acc genotypes* | 52 |
| *peaks no back* | 1 |
| *removed edges* | 0 |
| *prop rem edges* | 0 |

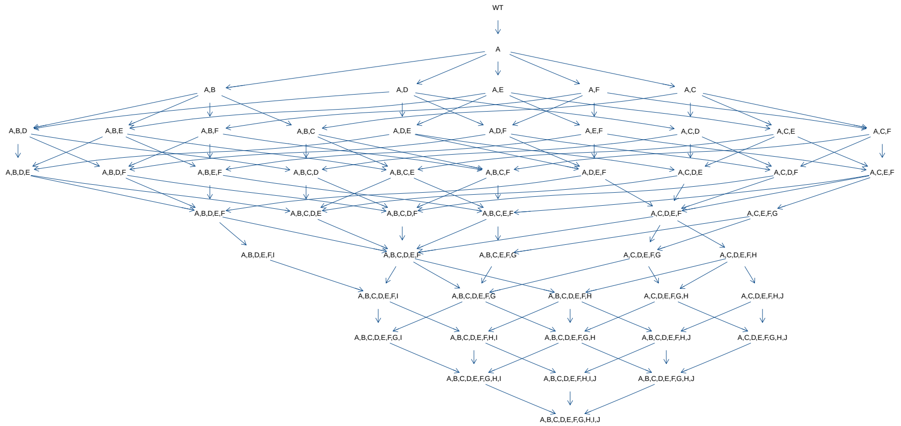

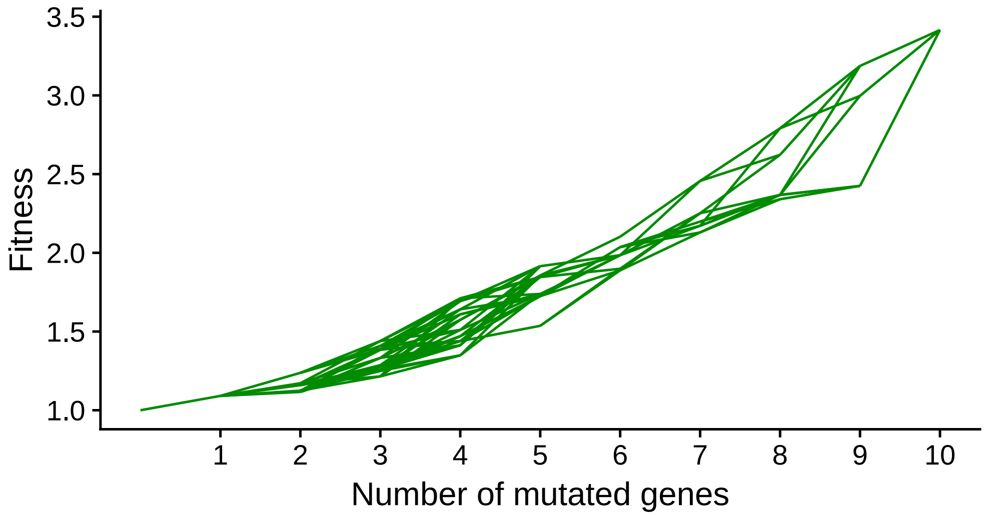

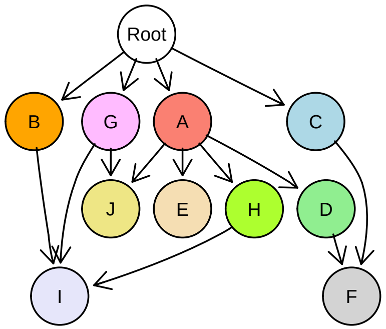


| | |
|---|---|
| *ID* | VdRwaw10j2G10mqCxP |
| *acc genotypes* | 148 |
| *peaks no back* | 1 |
| *removed edges* | 0 |
| *prop rem edges* | 0 |

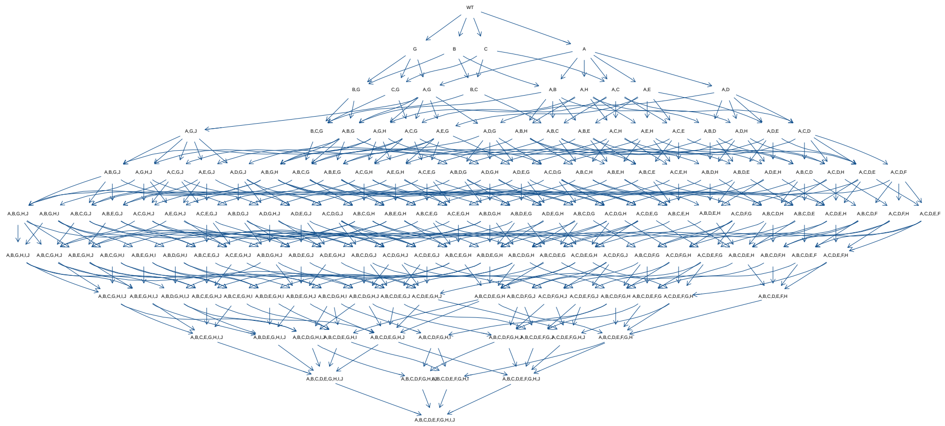

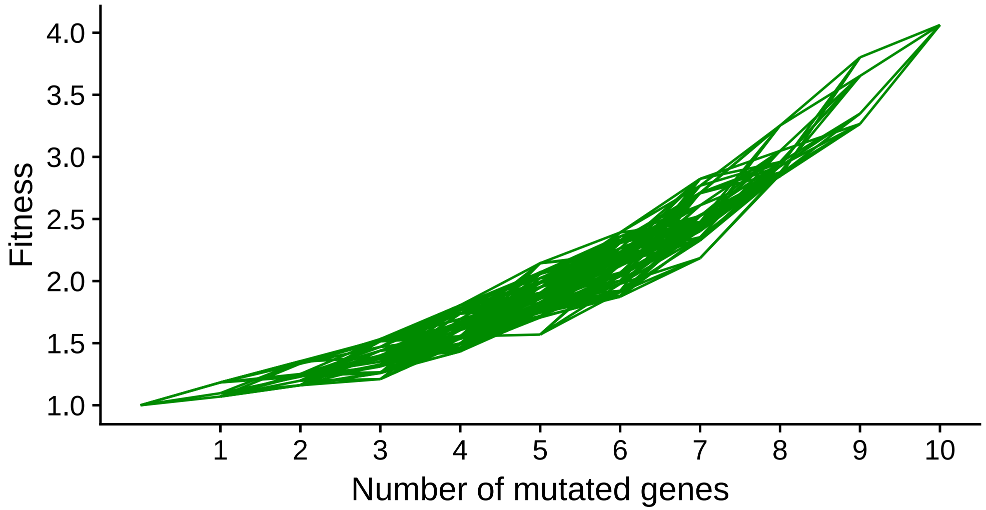

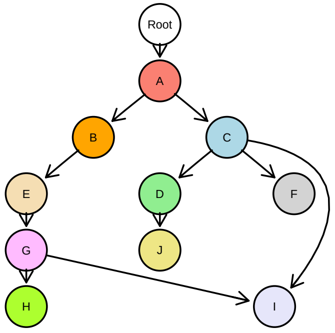


| | |
|---|---|
| *ID* | VIlrIHndJDehvx8p |
| *acc genotypes* | 48 |
| *peaks no back* | 1 |
| *removed edges* | 0 |
| *prop rem edges* | 0 |

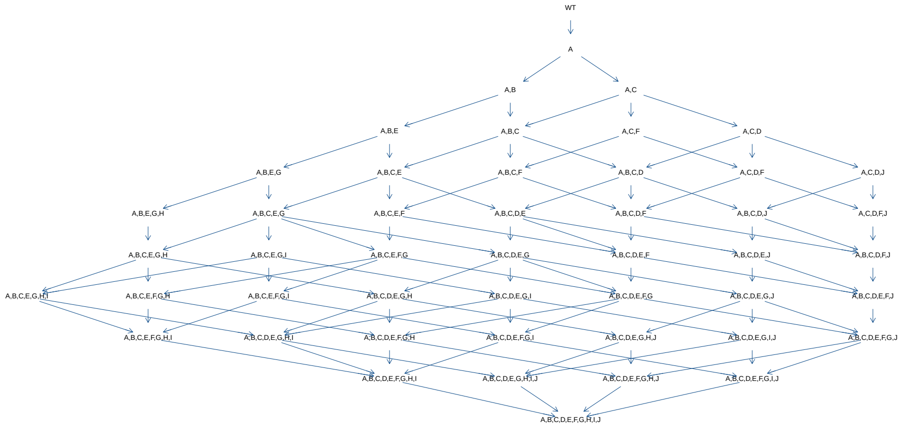


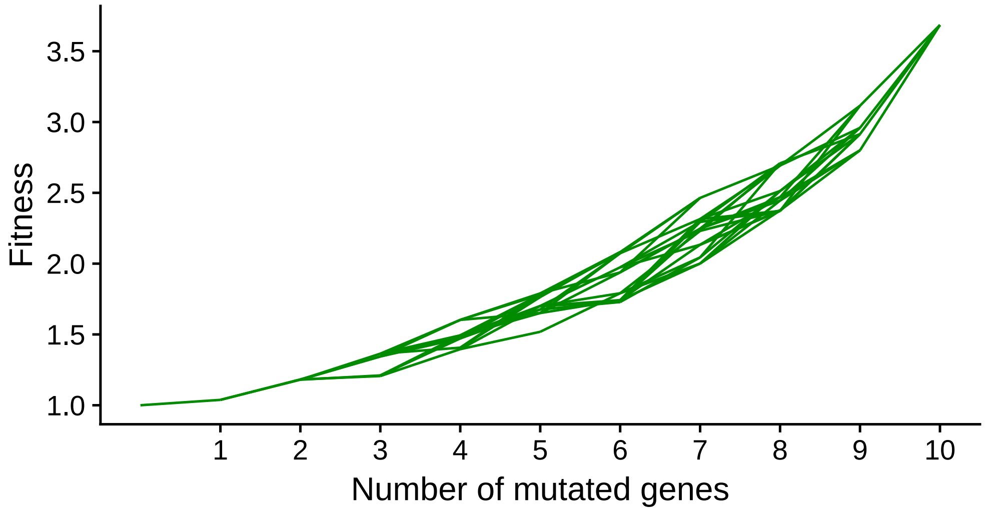

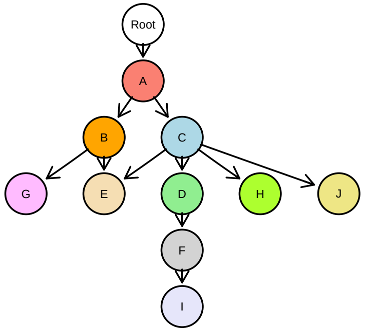


| | |
|---|---|
| *ID* | vKqR1PxaeGDS6o2G |
| *acc genotypes* | 84 |
| *peaks no back* | 1 |
| *removed edges* | 0 |
| *prop rem edges* | 0 |

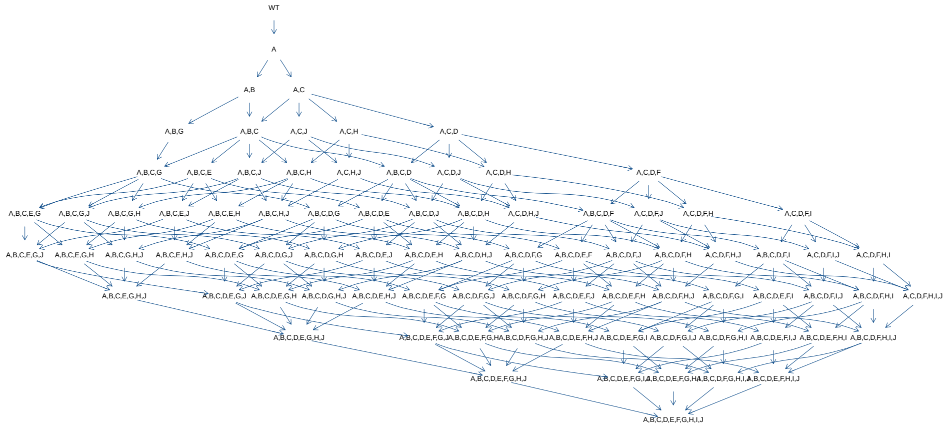

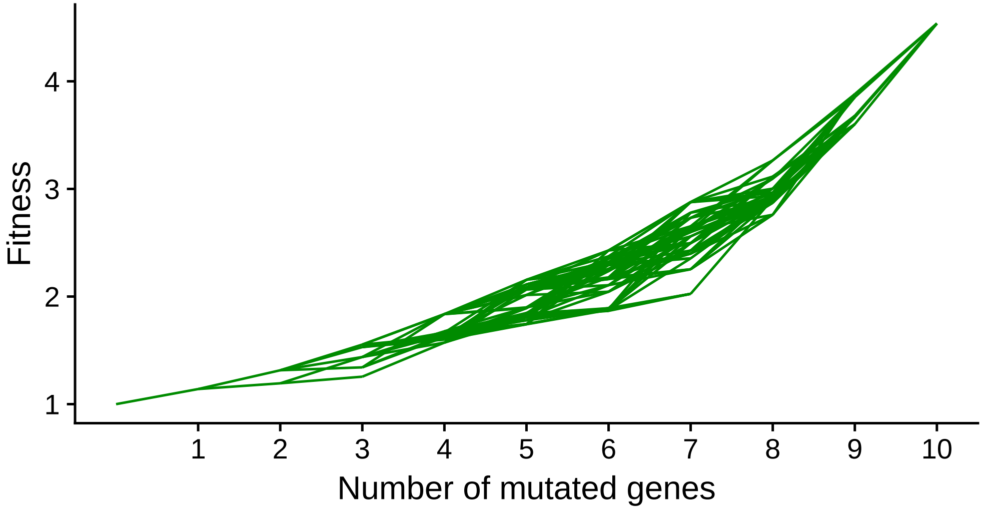

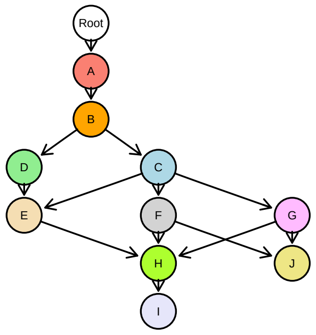


| | |
|---|---|
| *ID* | vWa8dLmVr7VaMVrU |
| *acc genotypes* | 23 |
| *peaks no back* | 1 |
| *removed edges* | 0 |
| *prop rem edges* | 0 |

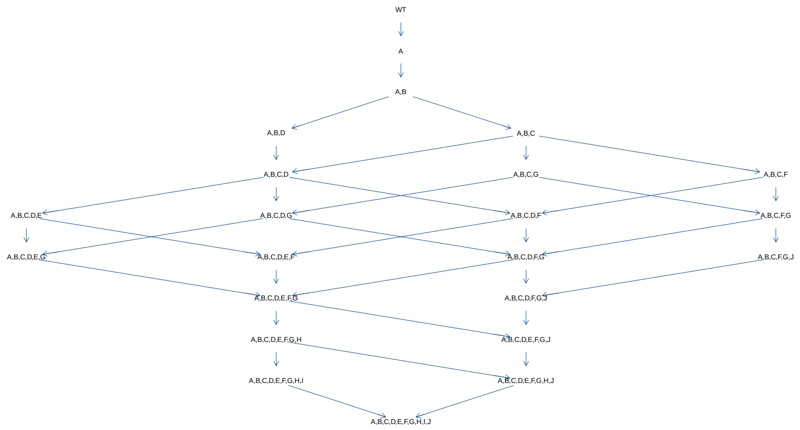


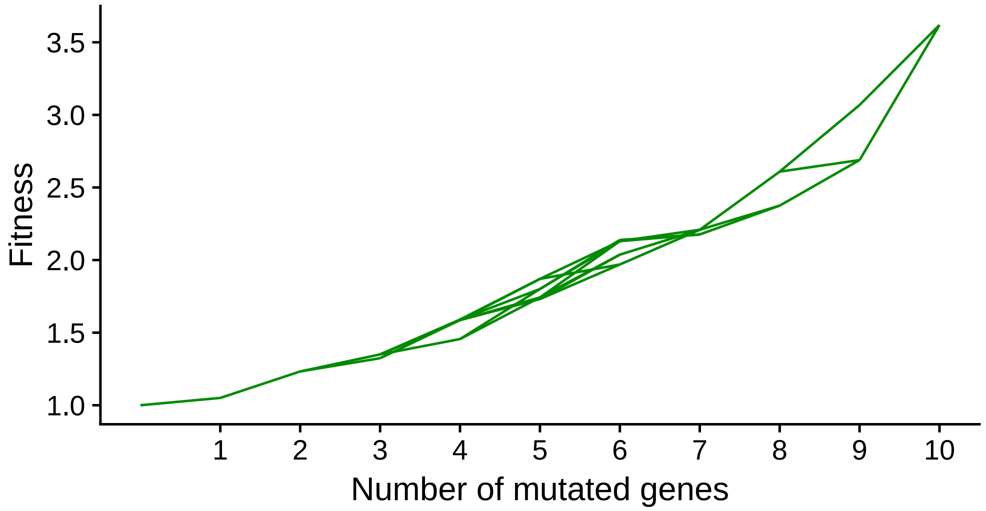

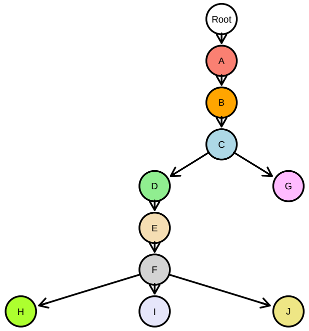


| | |
|---|---|
| *ID* | W10RExqb1hvnBZhkz |
| *acc genotypes* | 25 |
| *peaks no back* | 1 |
| *removed edges* | 0 |
| *prop rem edges* | 0 |

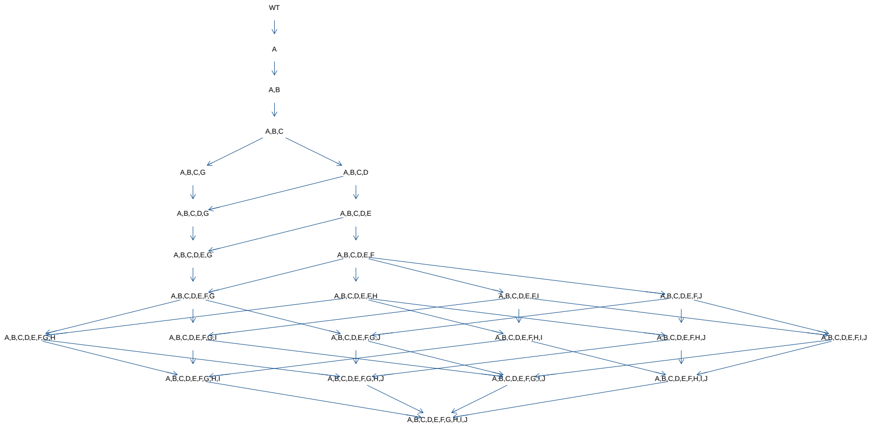

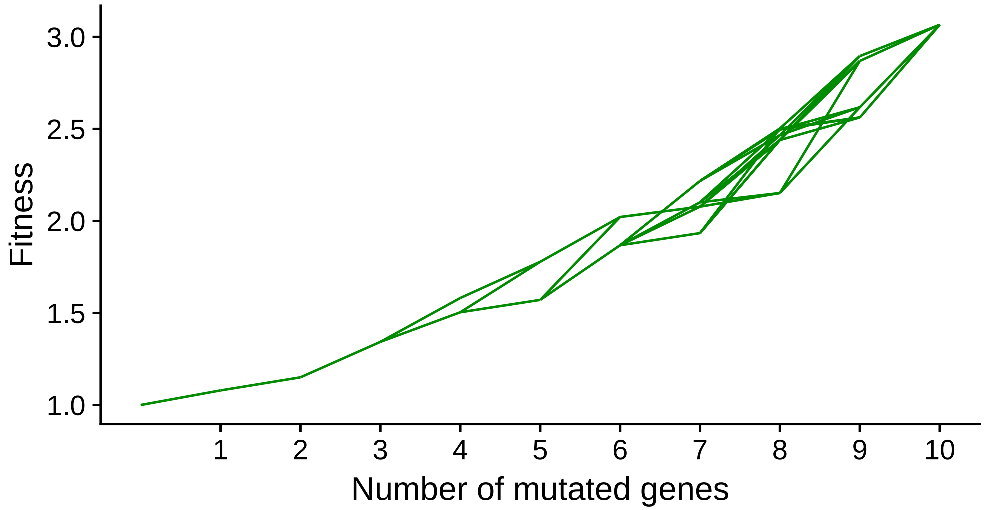

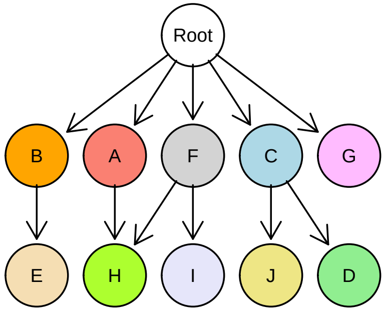


| | |
|---|---|
| *ID* | w2LGyrZeMILULslK |
| *acc genotypes* | 240 |
| *peaks no back* | 1 |
| *removed edges* | 0 |
| *prop rem edges* | 0 |

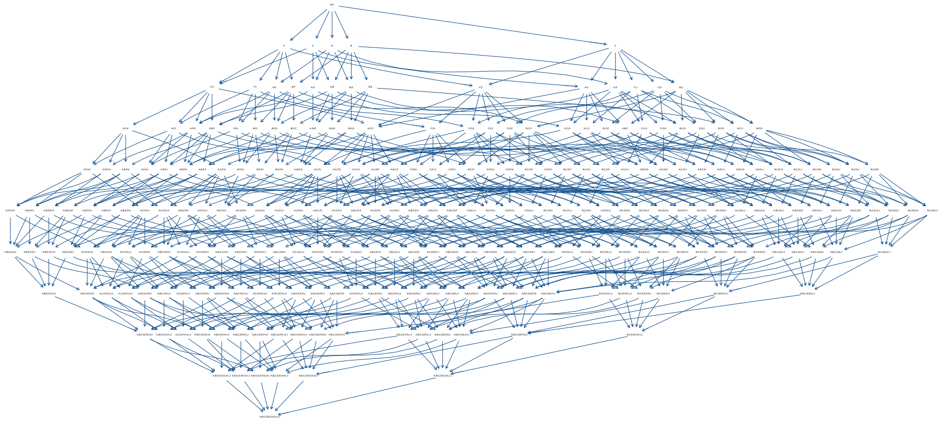

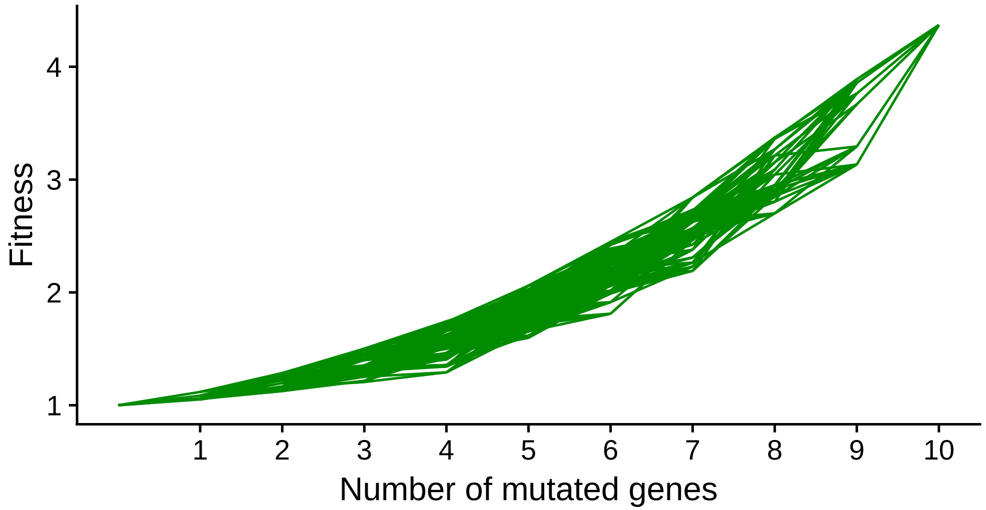

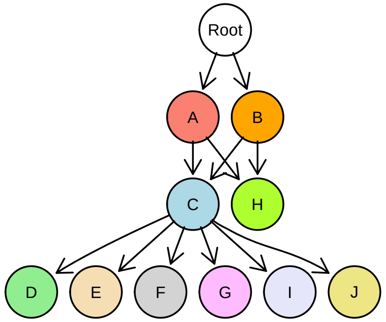


| | |
|---|---|
| *ID* | wOVnm2kwBXXY2Fm4 |
| *acc genotypes* | 133 |
| *peaks no back* | 1 |
| *removed edges* | 0 |
| *prop rem edges* | 0 |

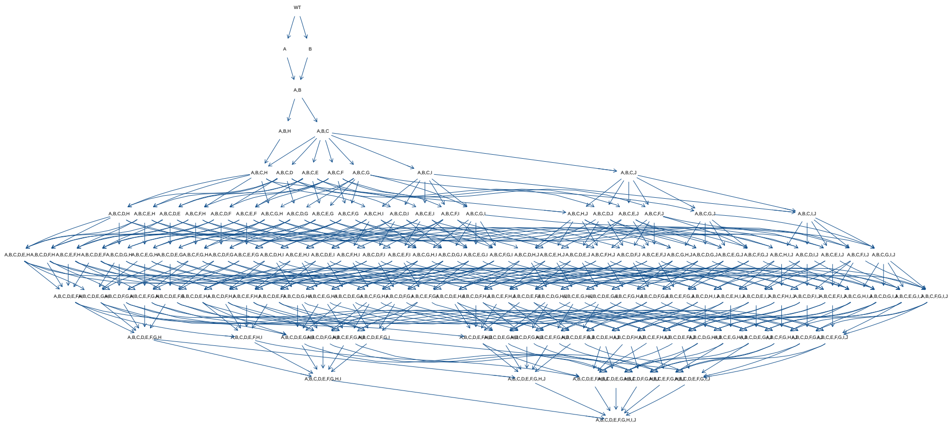

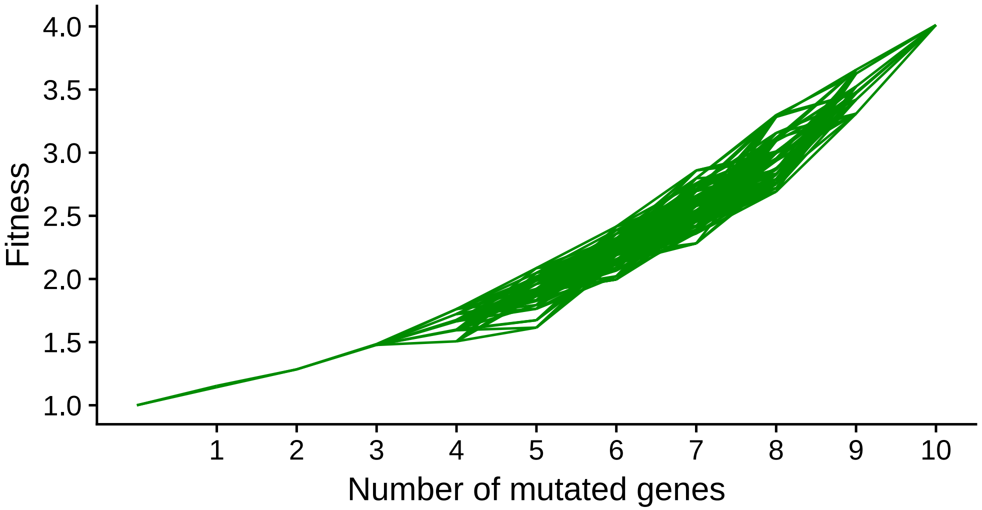

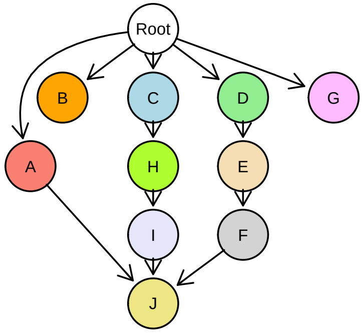


| | |
|---|---|
| *ID* | WX2rq5CKiTz10Vbzs |
| *acc genotypes* | 132 |
| *peaks no back* | 1 |
| *removed edges* | 0 |
| *prop rem edges* | 0 |

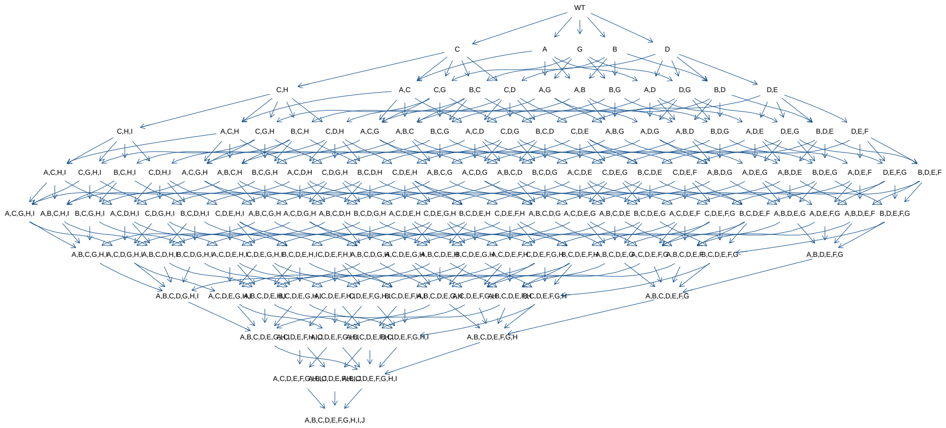


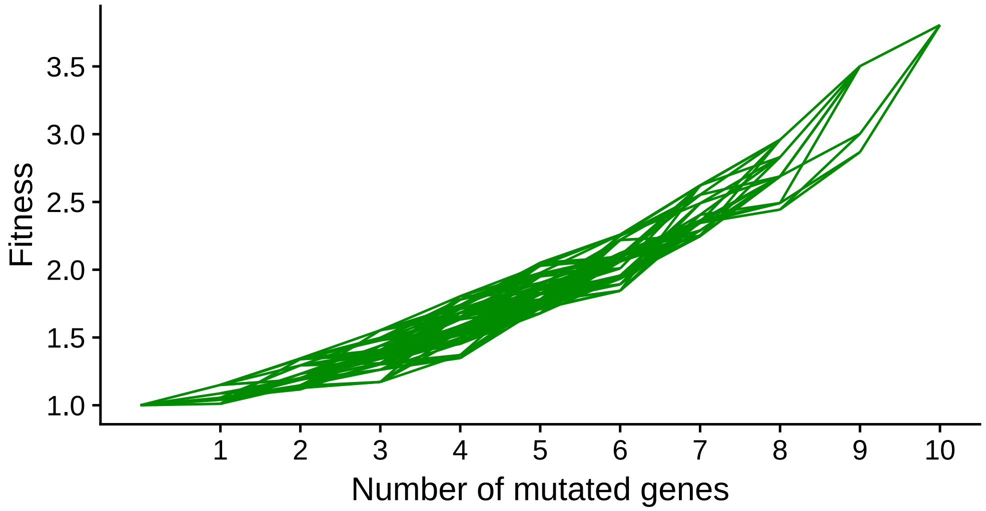

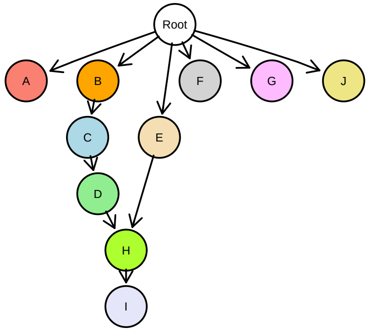


| | |
|---|---|
| *ID* | X73CthbrahOJ65dM |
| *acc genotypes* | 160 |
| *peaks no back* | 1 |
| *removed edges* | 0 |
| *prop rem edges* | 0 |

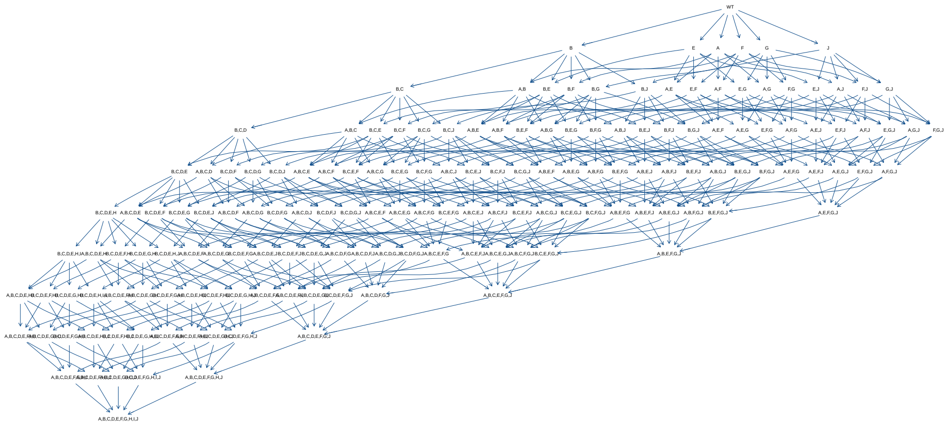

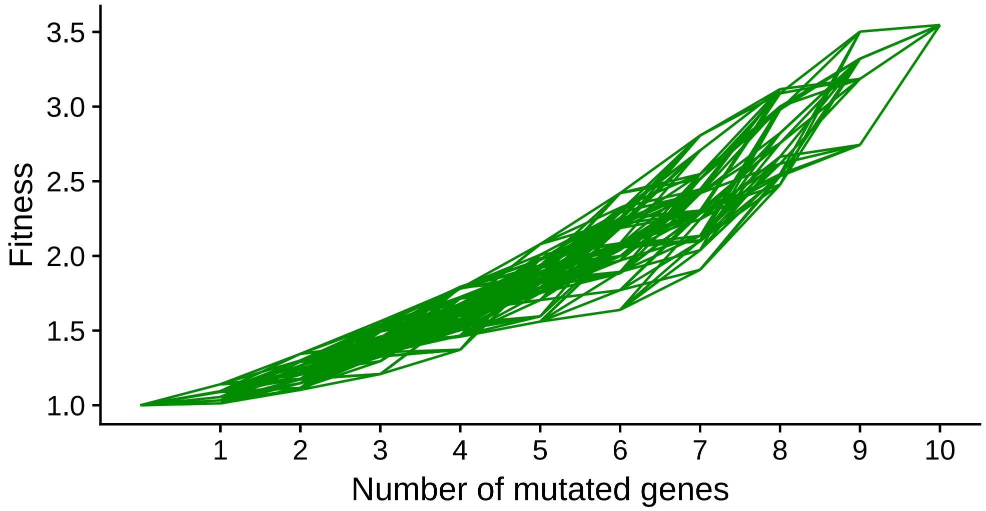

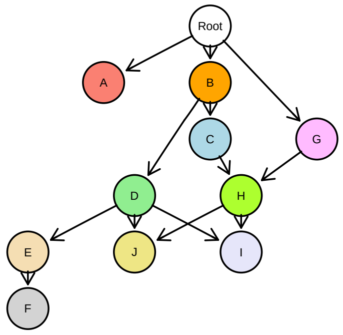


| | |
|---|---|
| *ID* | xghzdPbC2Ste8yA4 |
| *acc genotypes* | 62 |
| *peaks no back* | 1 |
| *removed edges* | 0 |
| *prop rem edges* | 0 |

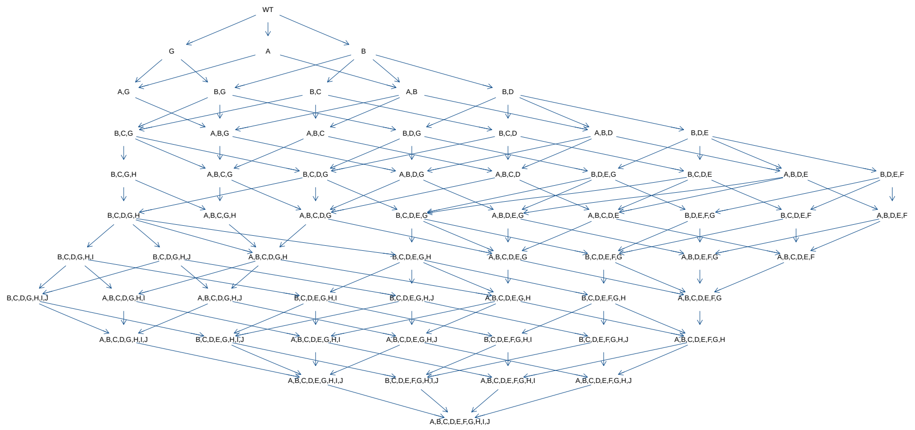

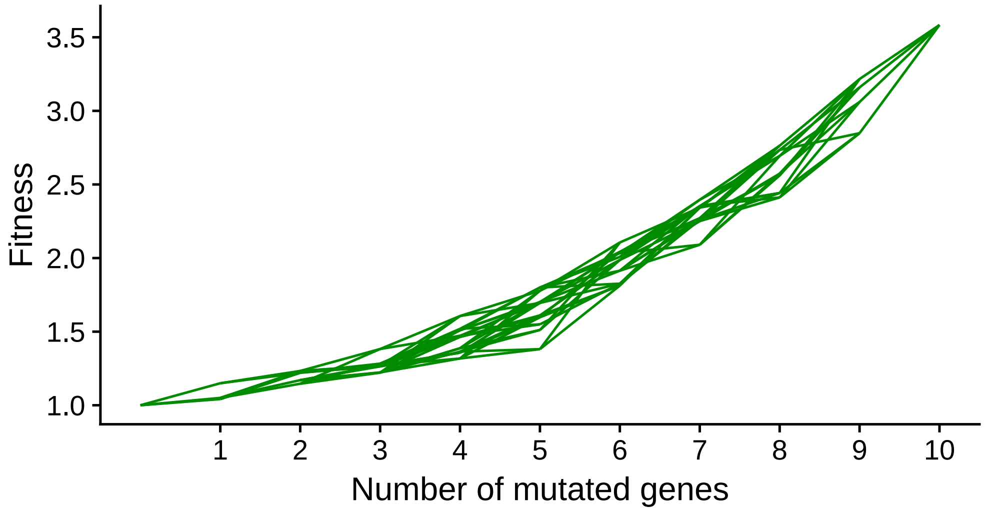

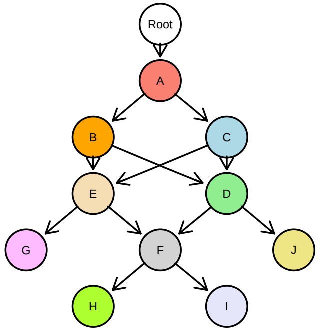


| | |
|---|---|
| *ID* | xsBe1fvNRQrAYKEK |
| *acc genotypes* | 29 |
| *peaks no back* | 1 |
| *removed edges* | 0 |
| *prop rem edges* | 0 |

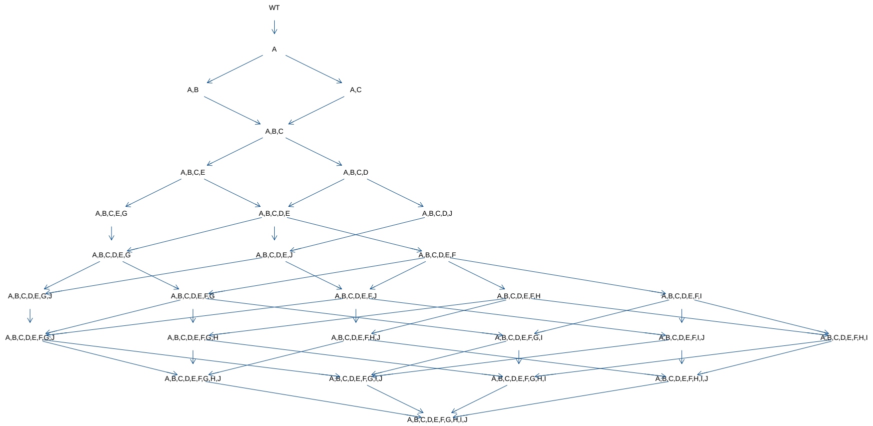


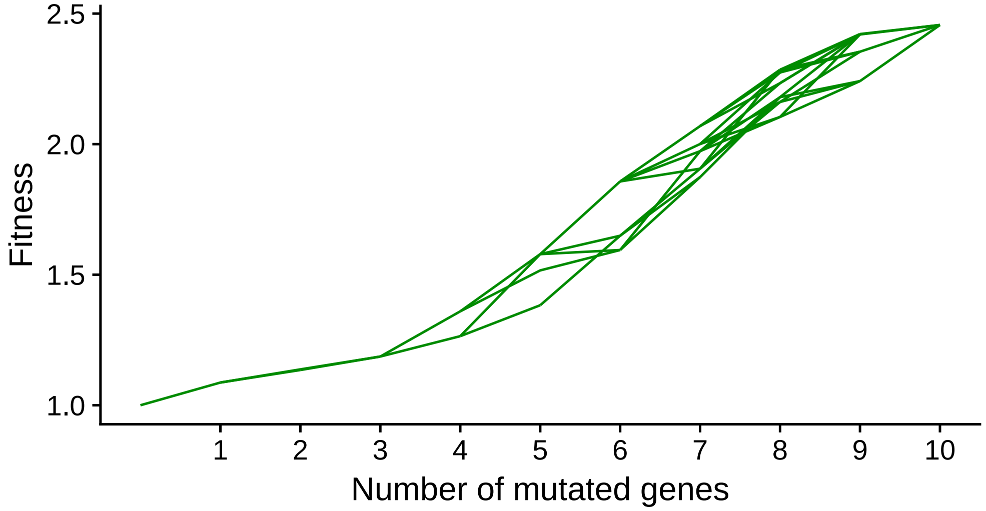

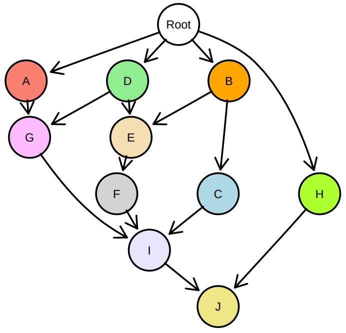


| | |
|---|---|
| *ID* | xTeo6GEUfqWOarjY |
| *acc genotypes* | 57 |
| *peaks no back* | 1 |
| *removed edges* | 0 |
| *prop rem edges* | 0 |

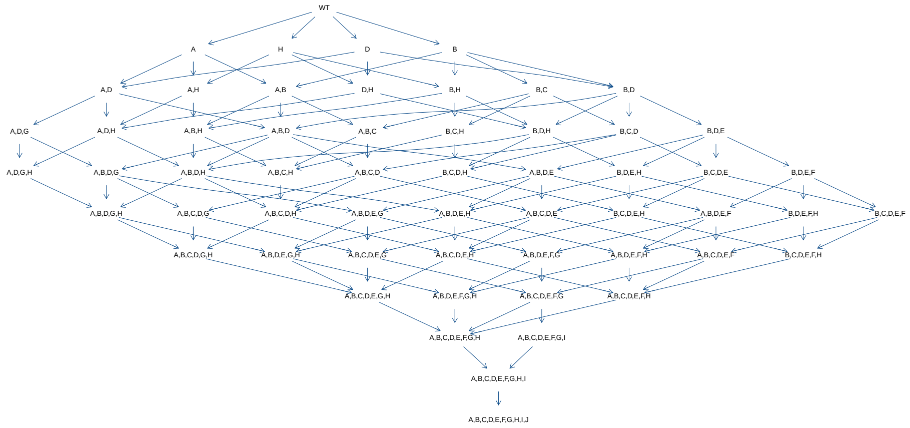

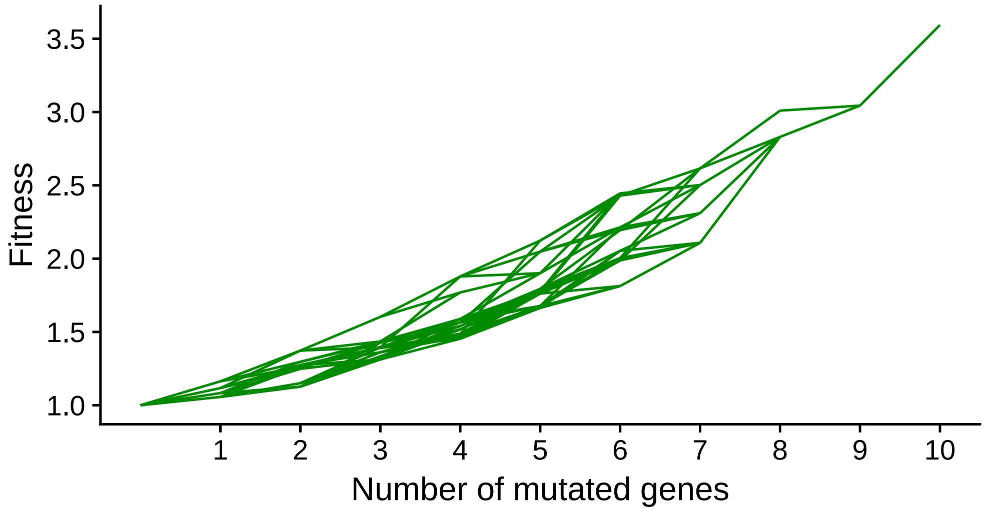

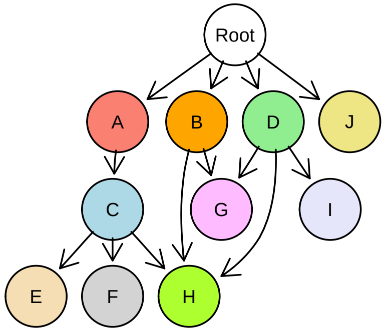


| | |
|---|---|
| *ID* | xUB10YVej3P32RKIi |
| *acc genotypes* | 128 |
| *peaks no back* | 1 |
| *removed edges* | 0 |
| *prop rem edges* | 0 |

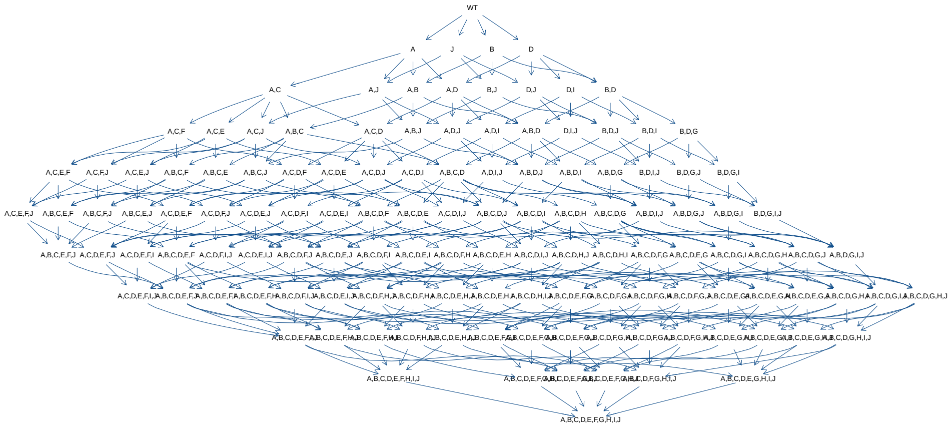

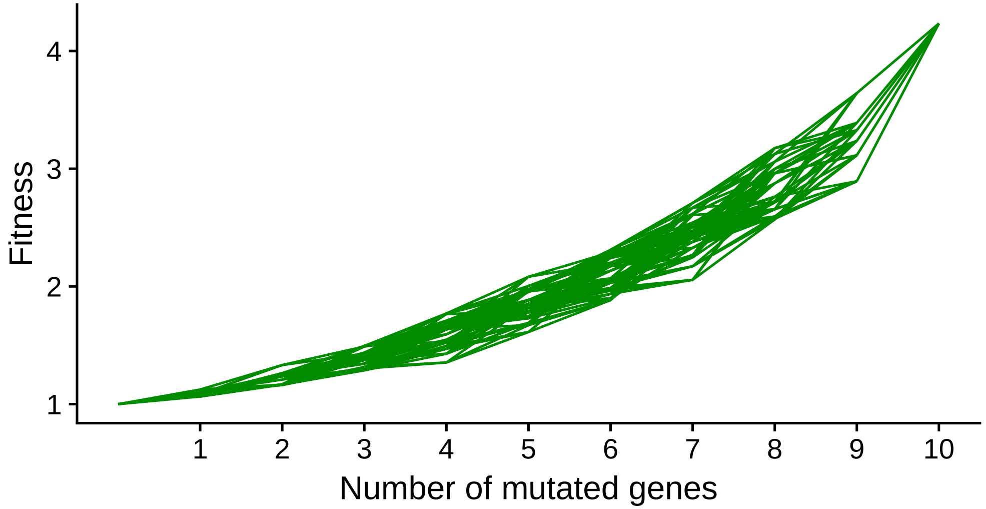

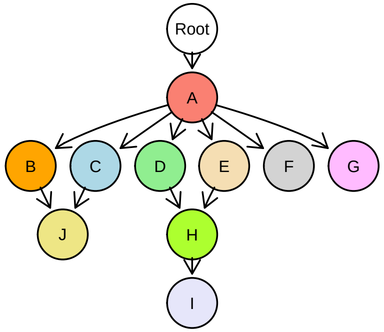


| | |
|---|---|
| *ID* | xVYnsd10xBjzI3Lt2 |
| *acc genotypes* | 121 |
| *peaks no back* | 1 |
| *removed edges* | 0 |
| *prop rem edges* | 0 |

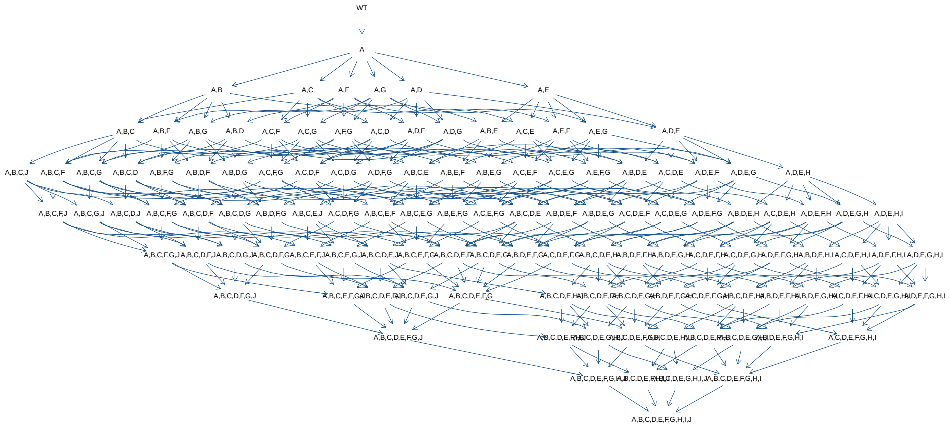

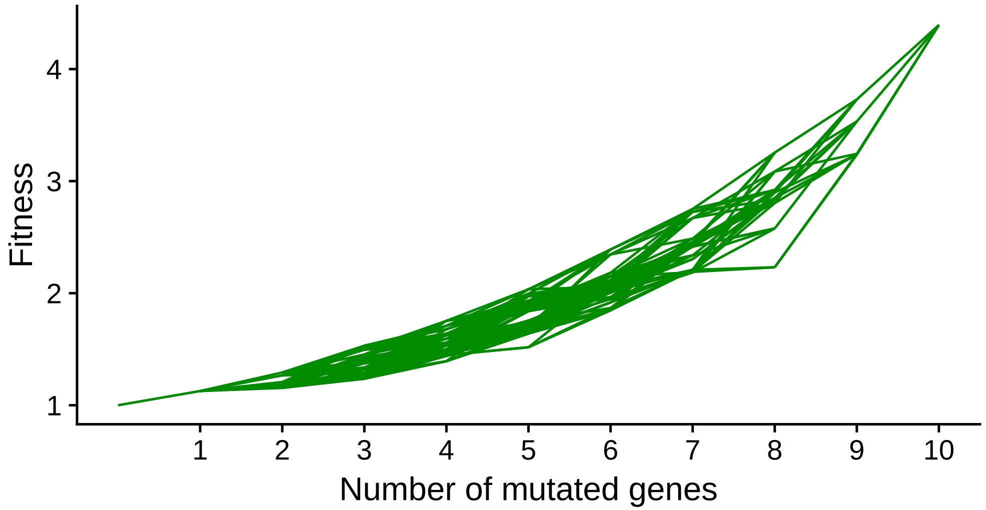

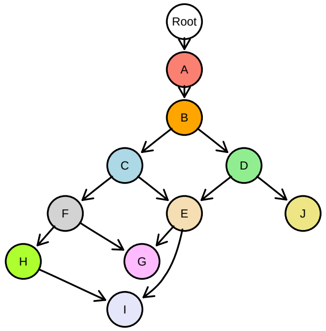


| | |
|---|---|
| *ID* | xWR3MCuUy8RKrLHF |
| *acc genotypes* | 28 |
| *peaks no back* | 1 |
| *removed edges* | 0 |
| *prop rem edges* | 0 |

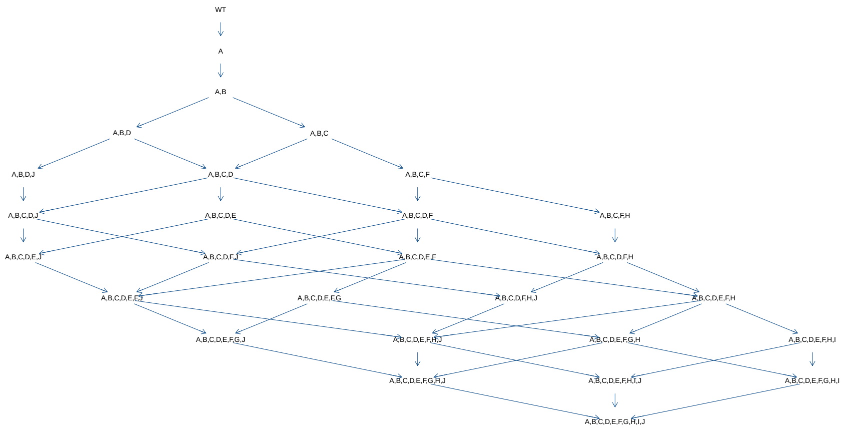


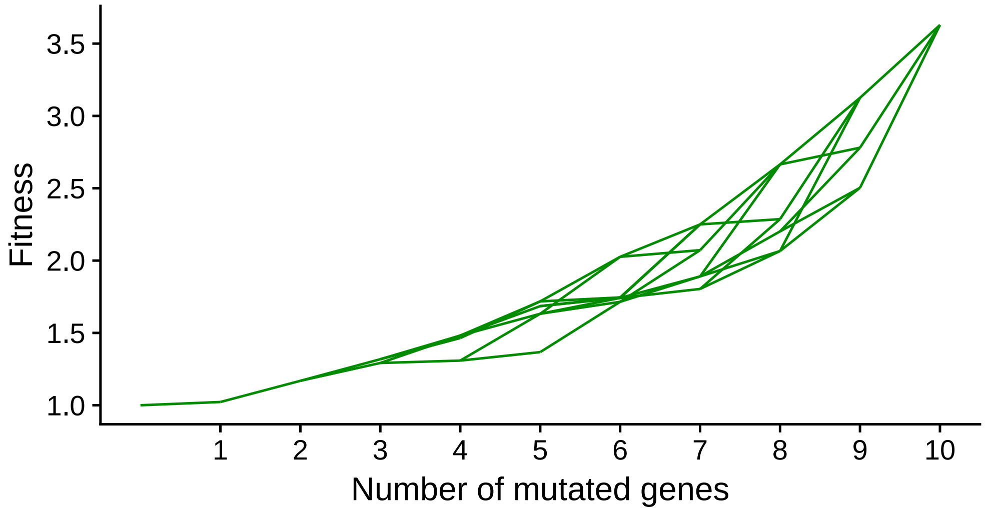

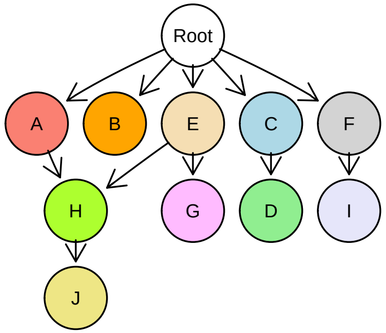


| | |
|---|---|
| *ID* | XYVDPIvrNBJywJzR |
| *acc genotypes* | 180 |
| *peaks no back* | 1 |
| *removed edges* | 0 |
| *prop rem edges* | 0 |

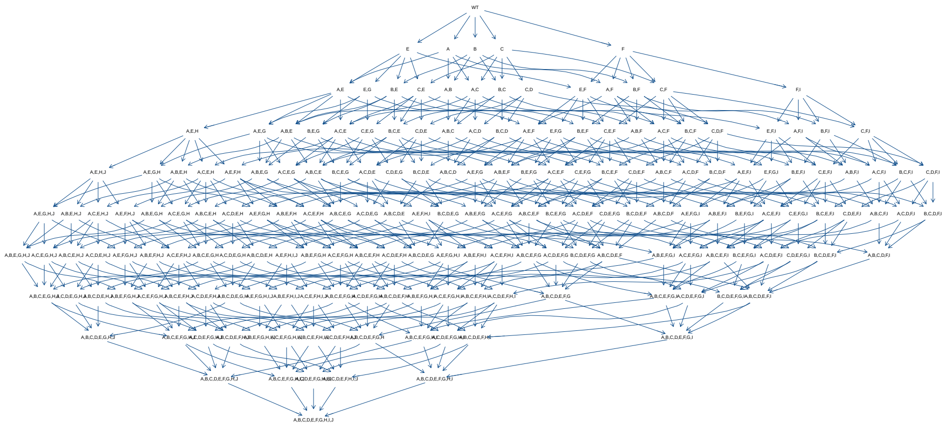

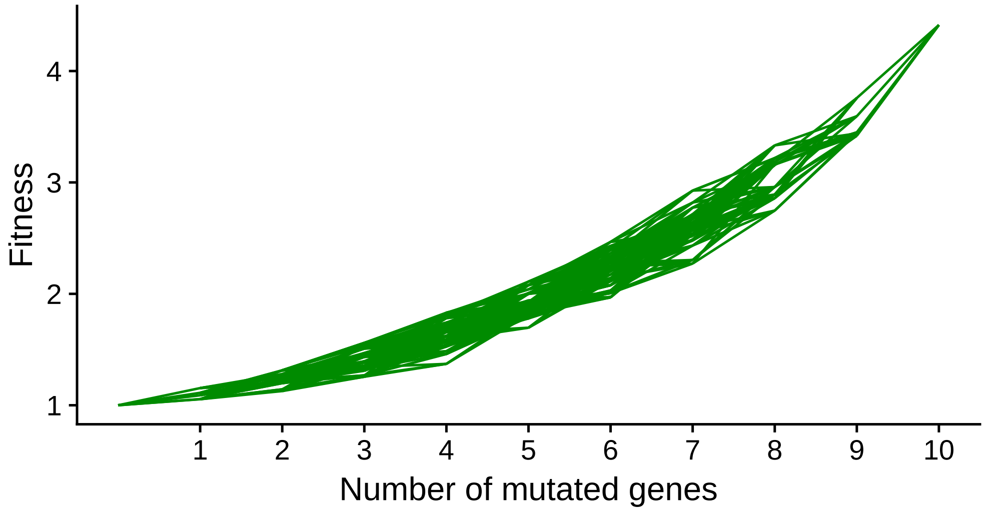

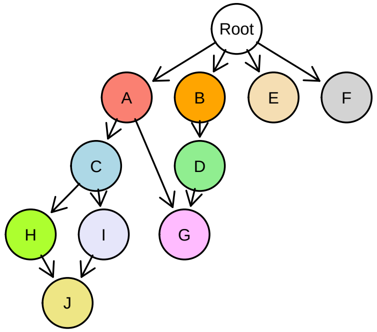


| | |
|---|---|
| *ID* | y1DWAEMgCsc8ELl2 |
| *acc genotypes* | 108 |
| *peaks no back* | 1 |
| *removed edges* | 0 |
| *prop rem edges* | 0 |

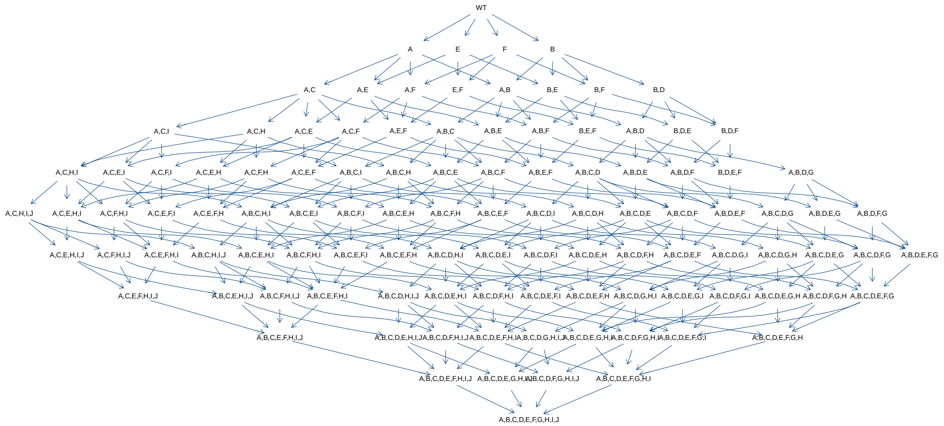

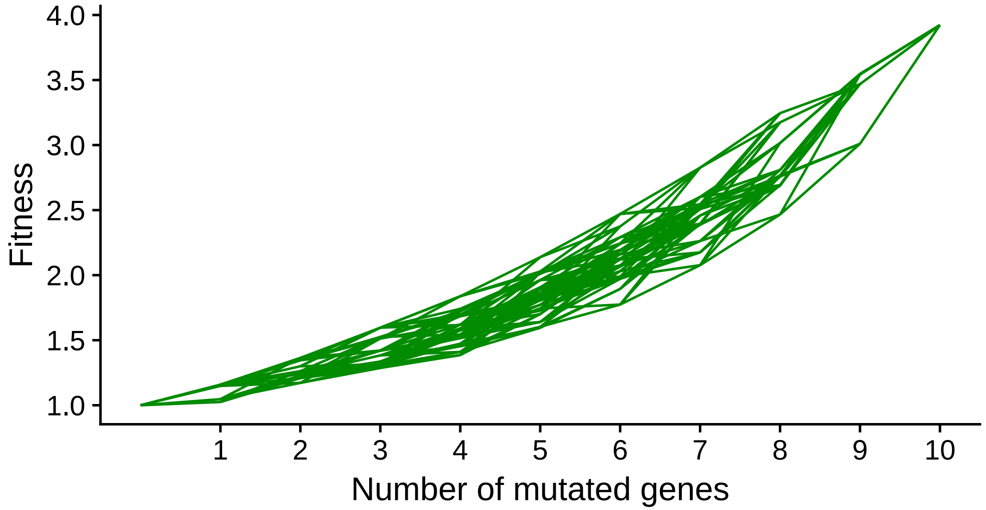

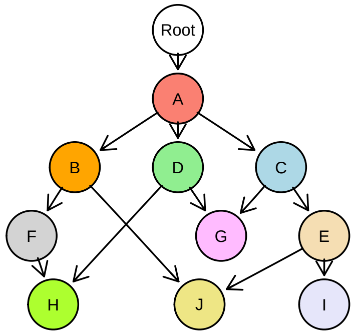


| | |
|---|---|
| *ID* | y910g6HDlnpD8GSSB |
| *acc genotypes* | 57 |
| *peaks no back* | 1 |
| *removed edges* | 0 |
| *prop rem edges* | 0 |

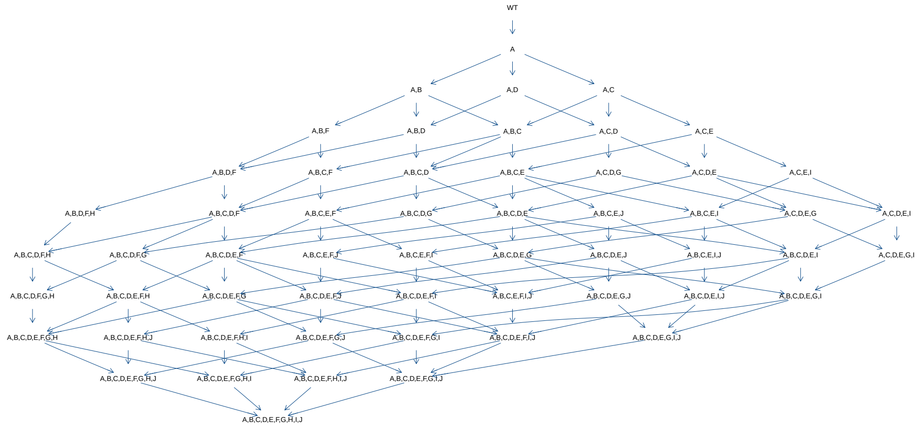

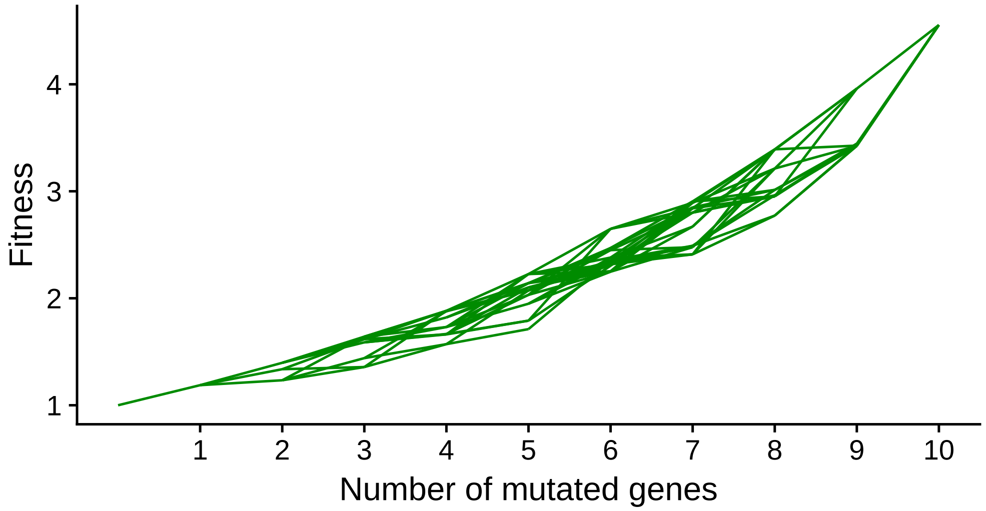

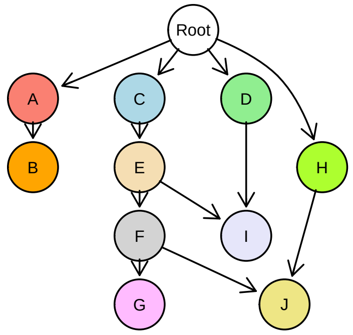


| | |
|---|---|
| *ID* | YbPjwENMtPWo53FW |
| *acc genotypes* | 96 |
| *peaks no back* | 1 |
| *removed edges* | 0 |
| *prop rem edges* | 0 |

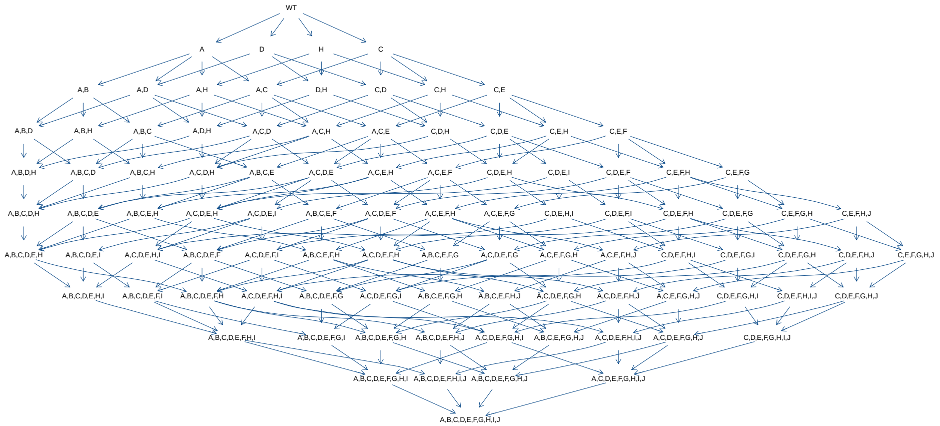

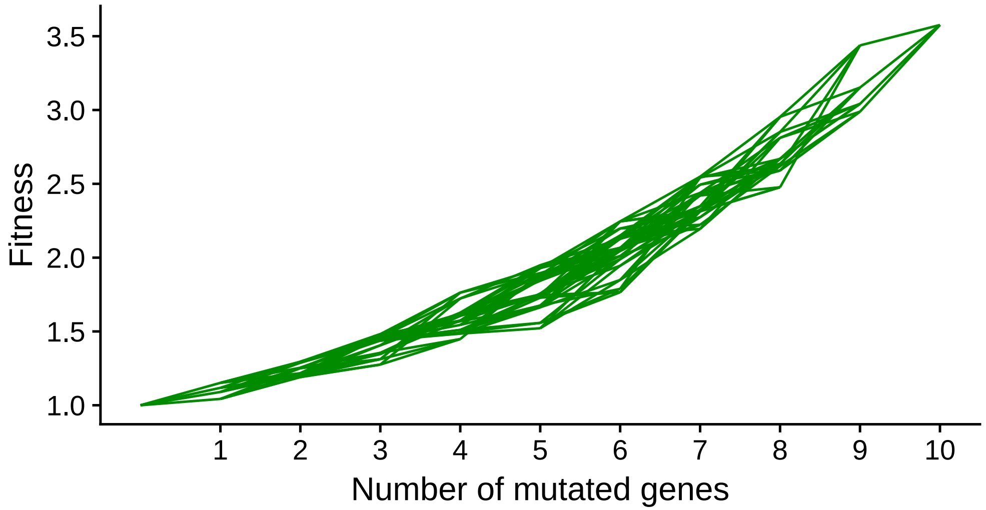

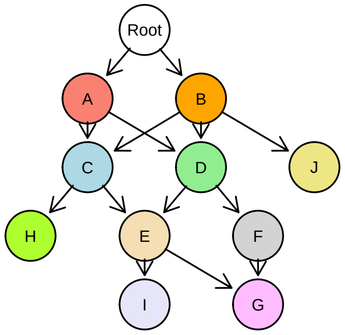


| | |
|---|---|
| *ID* | YDOzLX10m8IHyrpAK |
| *acc genotypes* | 46 |
| *peaks no back* | 1 |
| *removed edges* | 0 |
| *prop rem edges* | 0 |

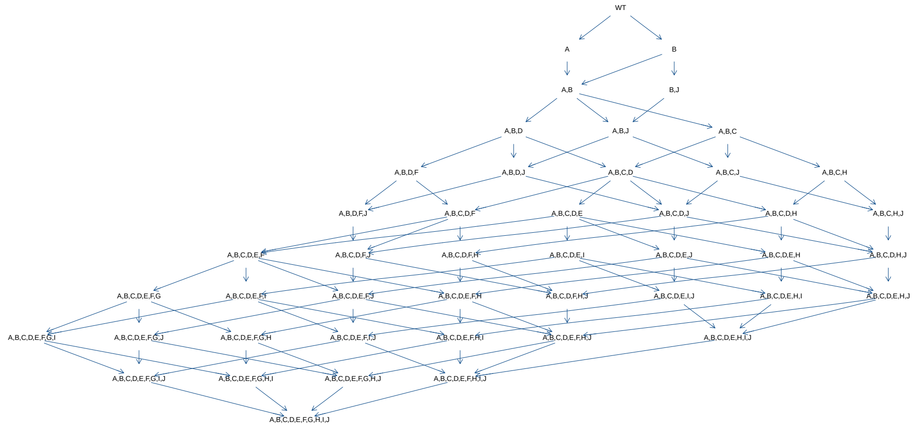

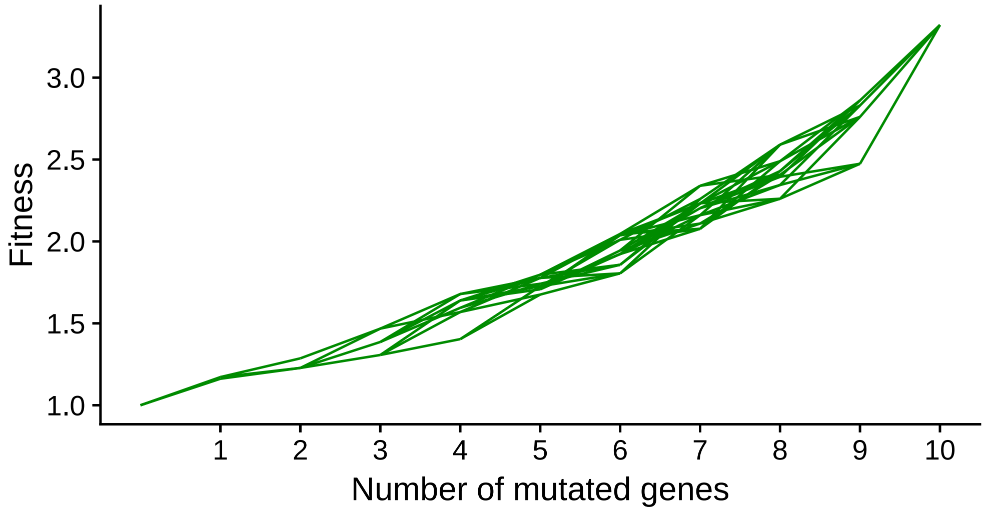

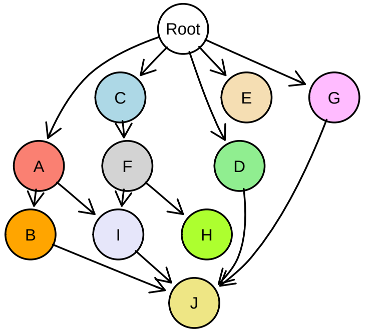


| | |
|---|---|
| *ID* | yKg7RfKP7wBk10kV3 |
| *acc genotypes* | 132 |
| *peaks no back* | 1 |
| *removed edges* | 0 |
| *prop rem edges* | 0 |

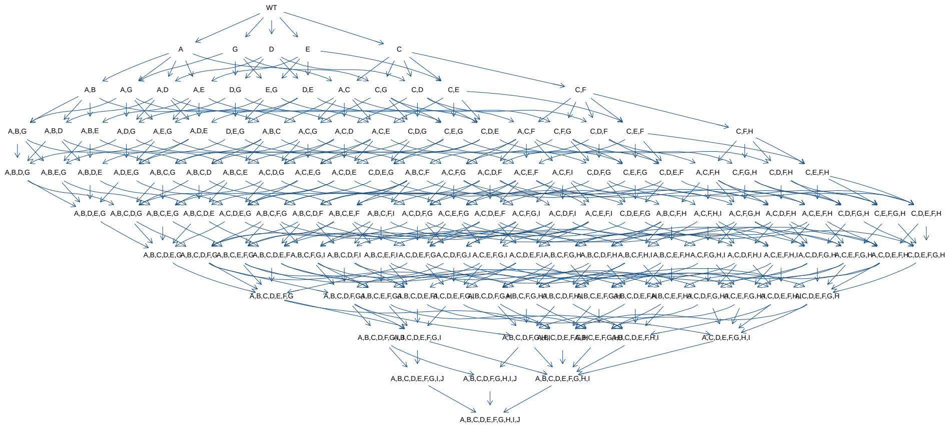

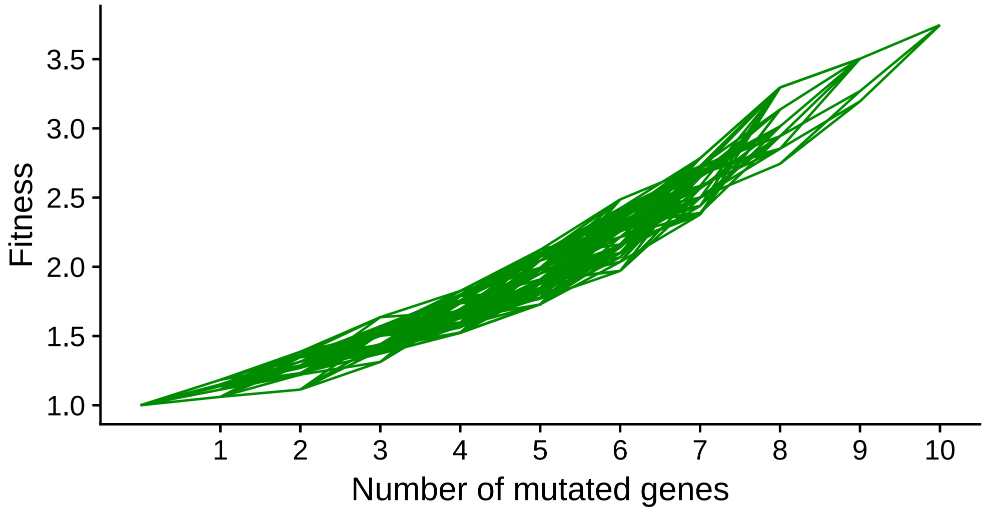

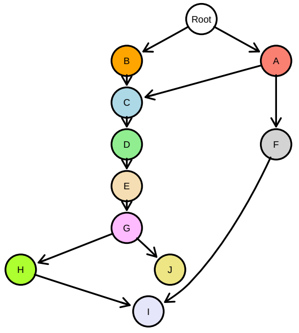


| | |
|---|---|
| *ID* | YlNcYnp363stUXfS |
| *acc genotypes* | 22 |
| *peaks no back* | 1 |
| *removed edges* | 0 |
| *prop rem edges* | 0 |

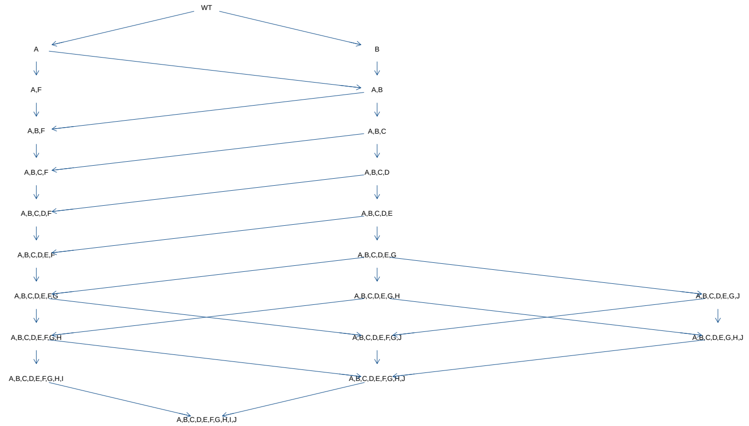

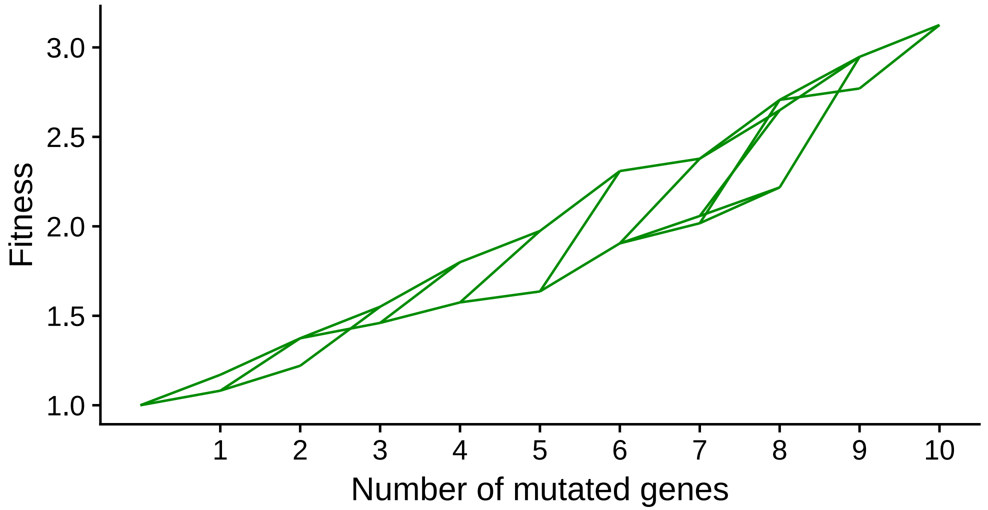

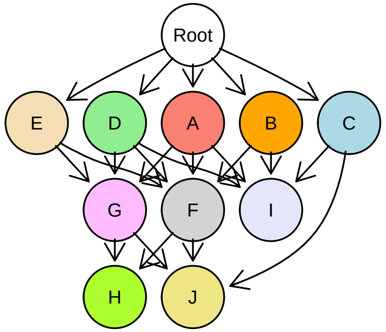


*ID* YlnEDjC9bTAwZFRa
*acc genotypes* 52
*peaks no back* 1
*removed edges* 0
*prop rem edges* 0

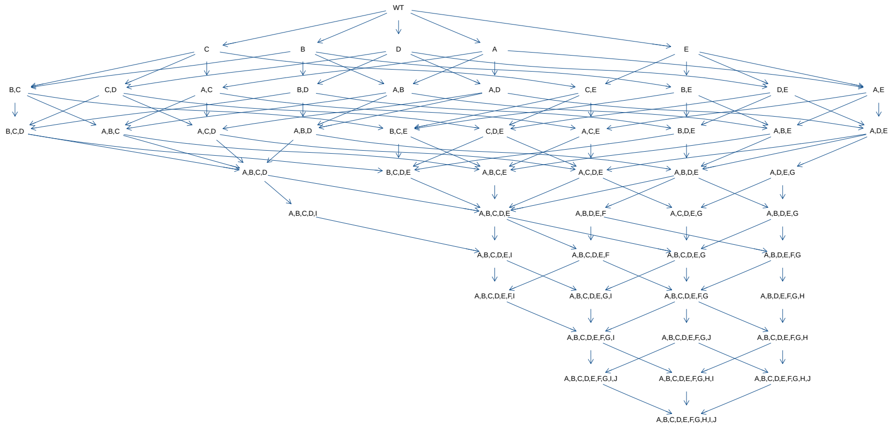

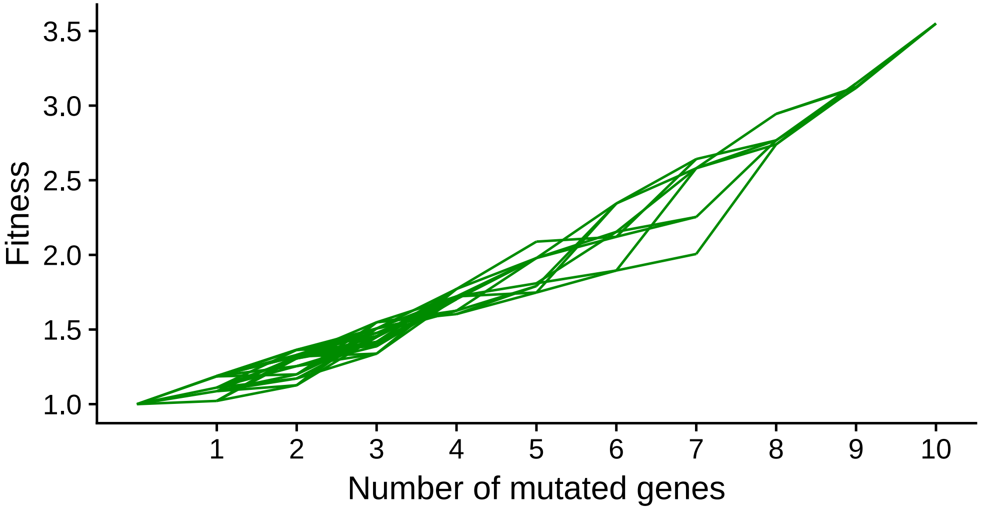

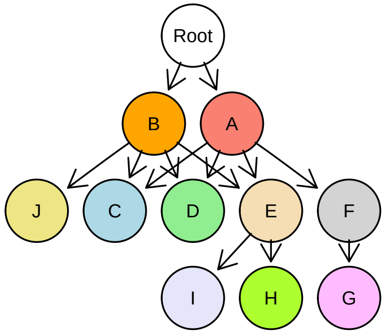


| | |
|---|---|
| *ID* | YsJ3QBe72KOpkYgG |
| *acc genotypes* | 126 |
| *peaks no back* | 1 |
| *removed edges* | 0 |
| *prop rem edges* | 0 |

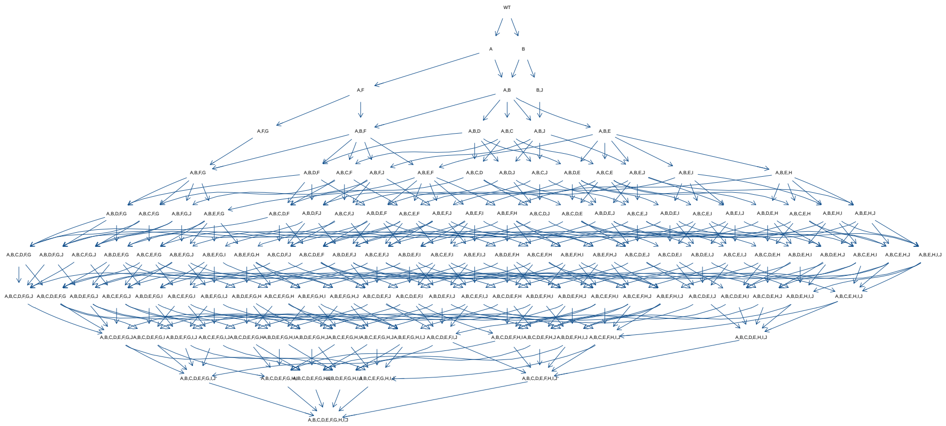

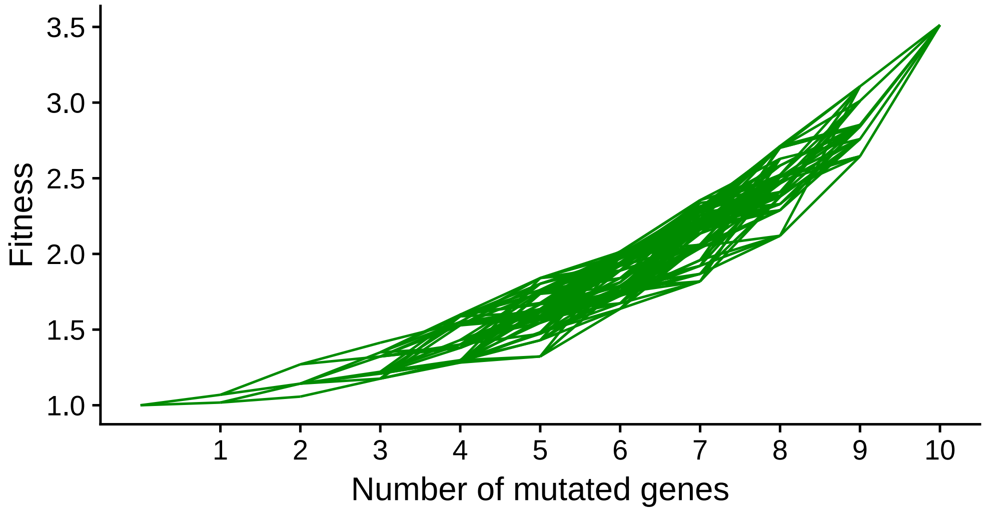

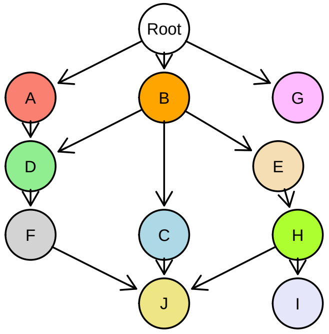


*ID* yUkCSfeekAS3UsYq
*acc genotypes* 72
*peaks no back* 1
*removed edges* 0
*prop rem edges* 0

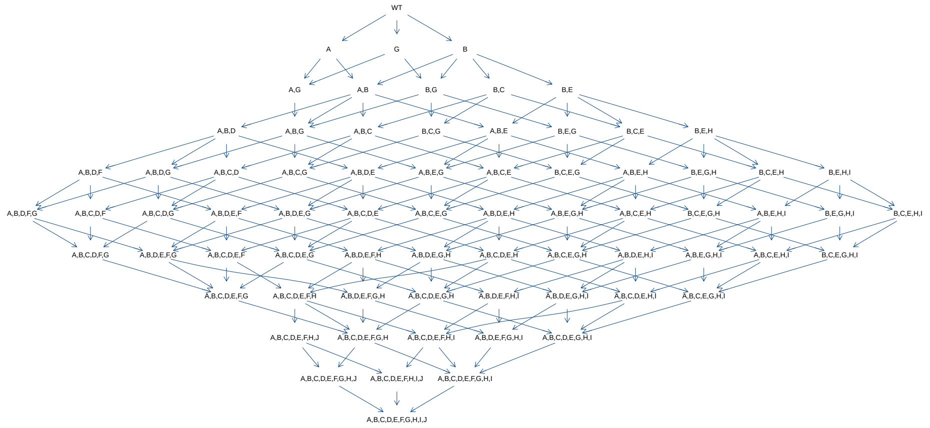

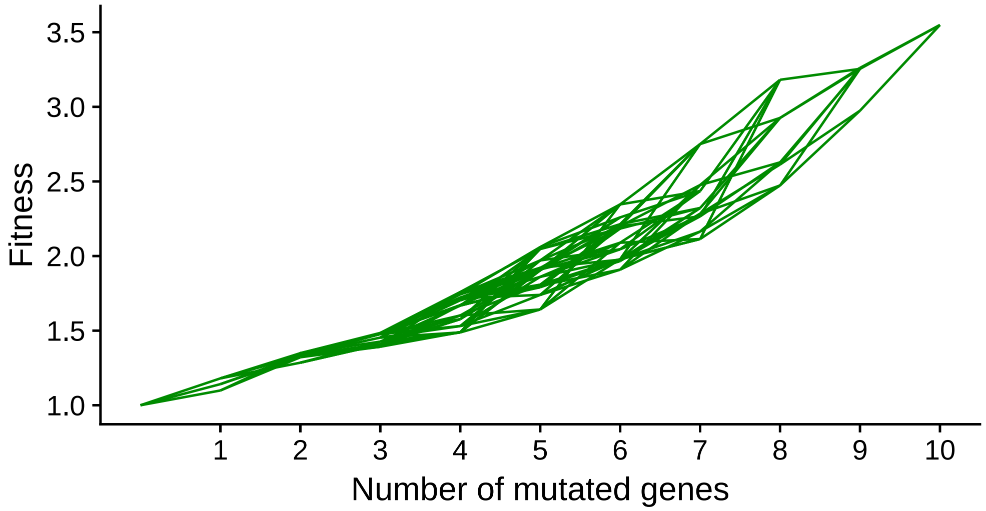

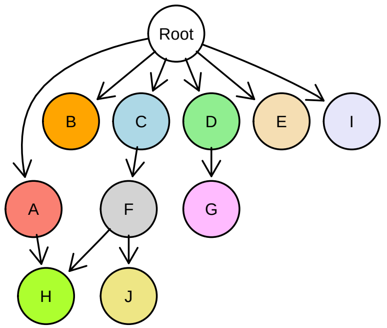


| | |
|---|---|
| *ID* | z1gqmdhtodgm6biP |
| *acc genotypes* | 240 |
| *peaks no back* | 1 |
| *removed edges* | 0 |
| *prop rem edges* | 0 |

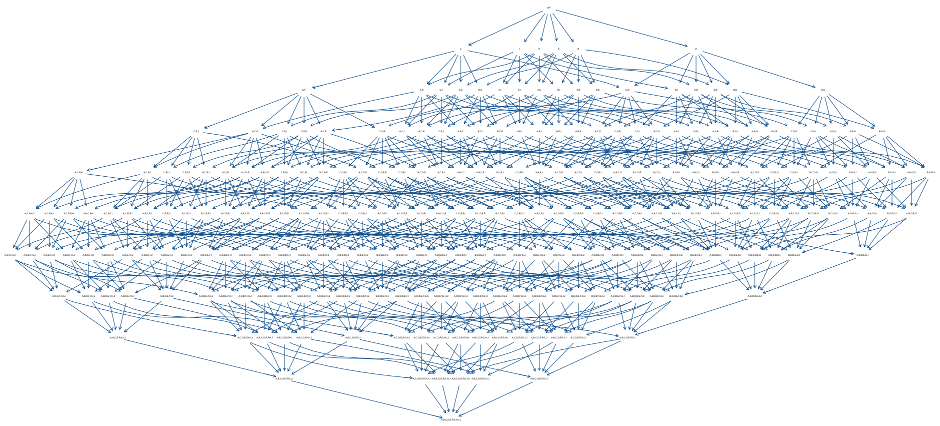

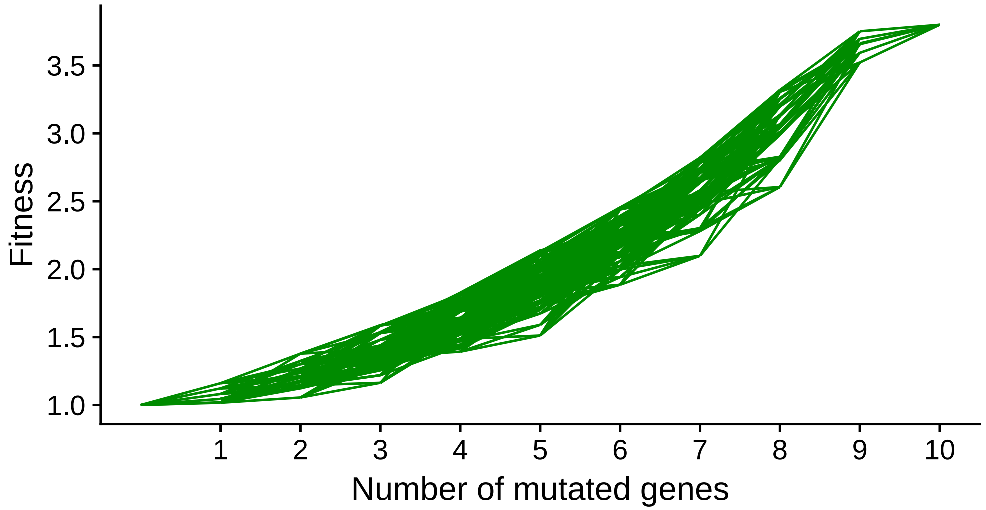

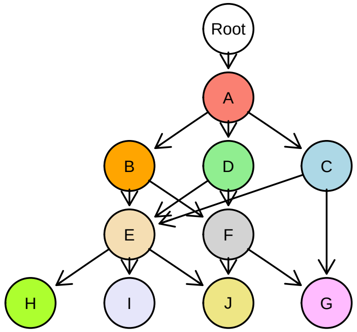


| | |
|---|---|
| *ID* | z55GMs5PYH8kzvrX |
| *acc genotypes* | 32 |
| *peaks no back* | 1 |
| *removed edges* | 0 |
| *prop rem edges* | 0 |

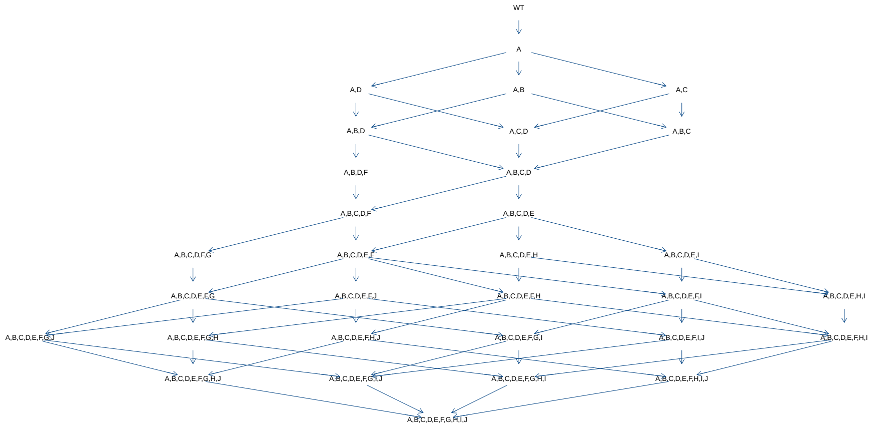

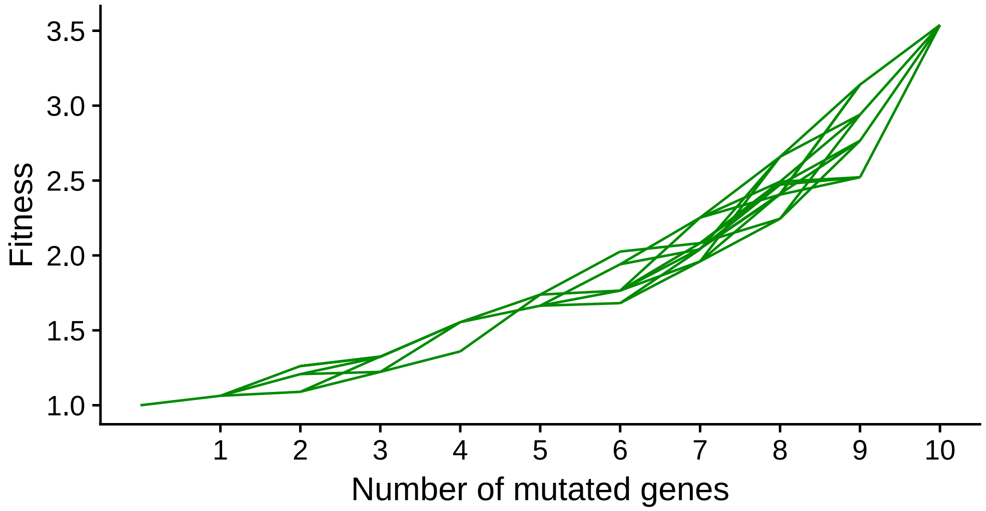

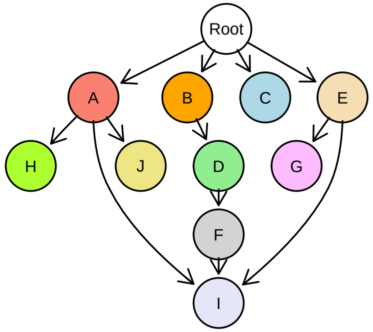


*ID* ZEOZqZ8hAfpl29IT
*acc genotypes* 136
*peaks no back* 1
*removed edges* 0
*prop rem edges* 0

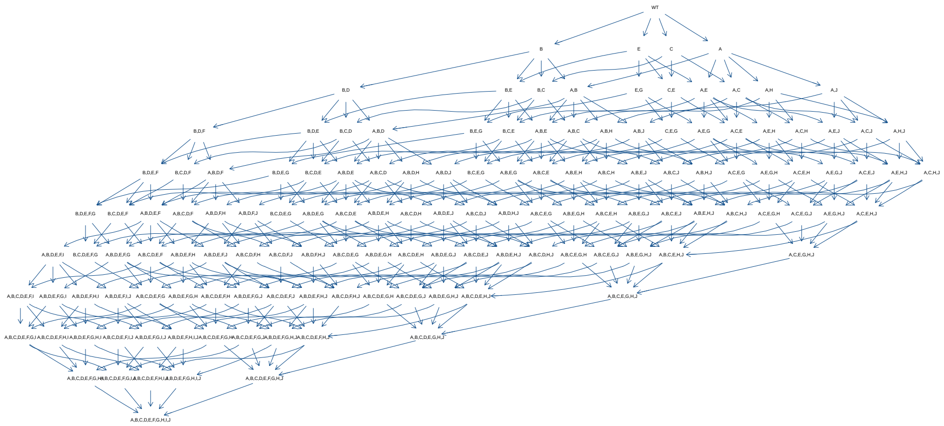

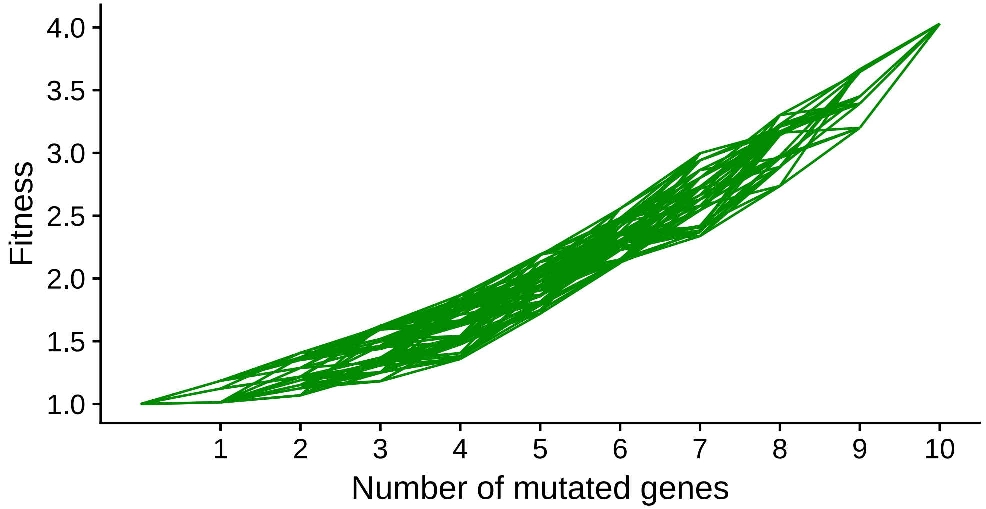

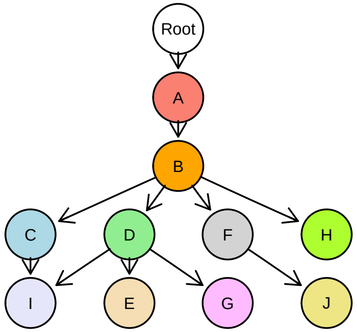


| | |
|---|---|
| *ID* | zibjAEA10MPGwv10fw |
| *acc genotypes* | 86 |
| *peaks no back* | 1 |
| *removed edges* | 0 |
| *prop rem edges* | 0 |

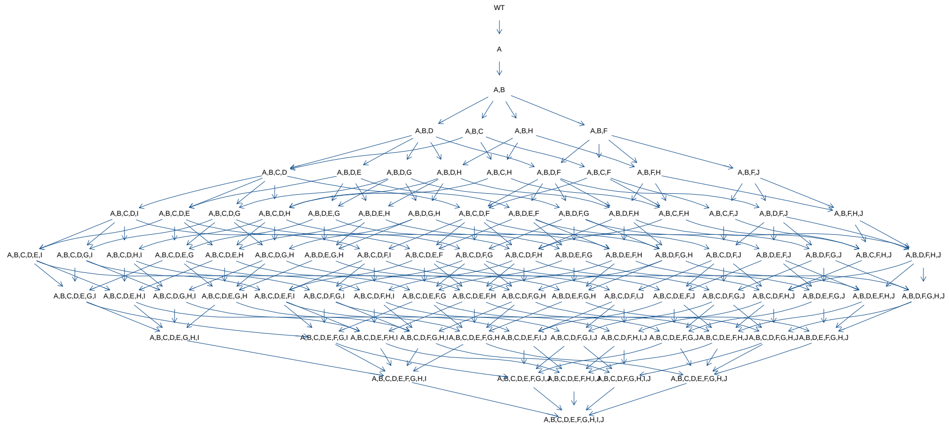


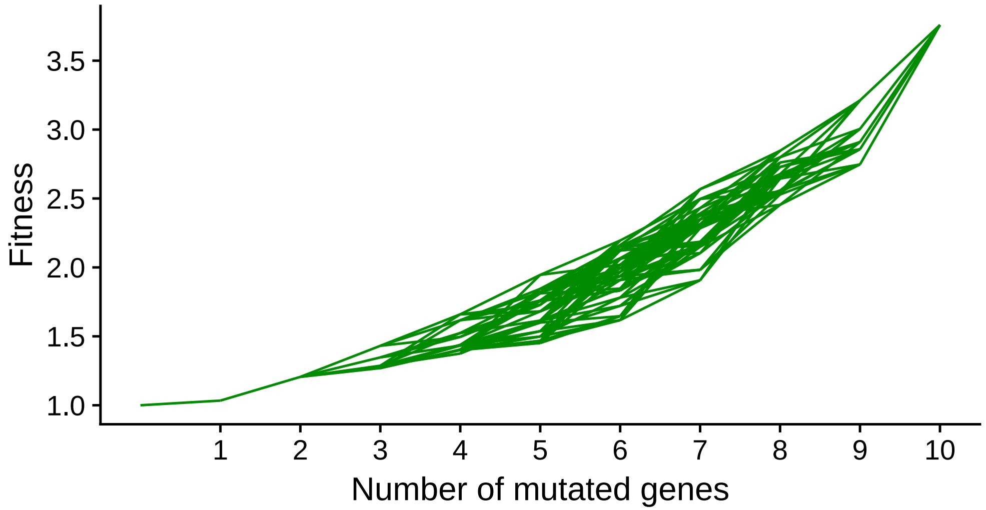

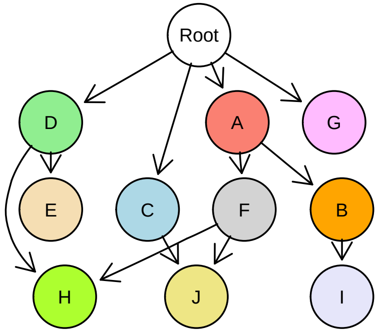


*ID* zwBgTR5JXPWHnI1B
*acc genotypes* 138
*peaks no back* 1
*removed edges* 0
*prop rem edges* 0

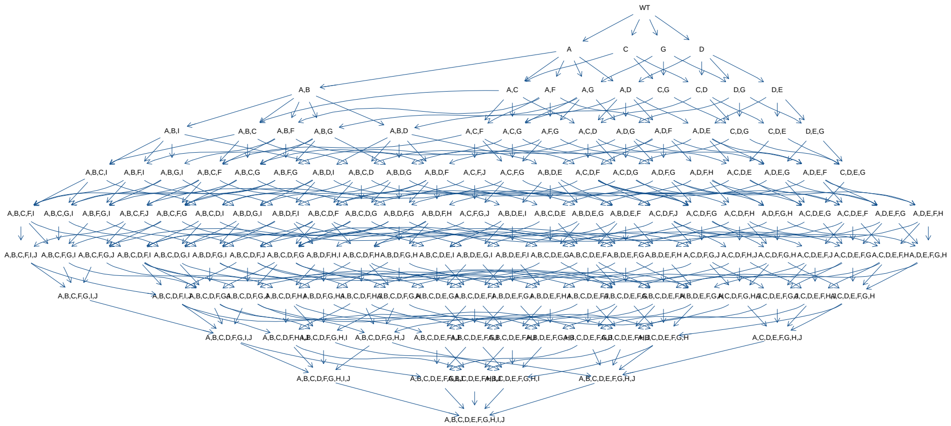

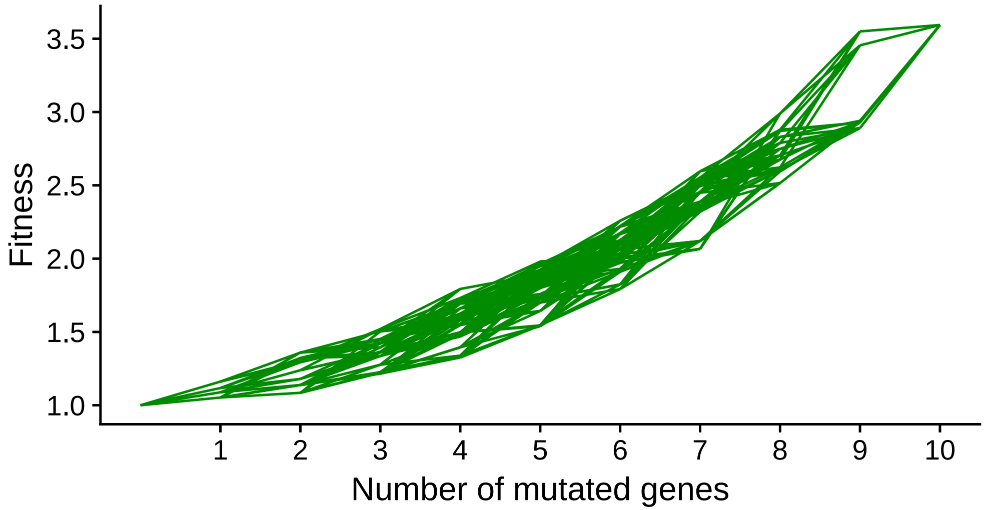

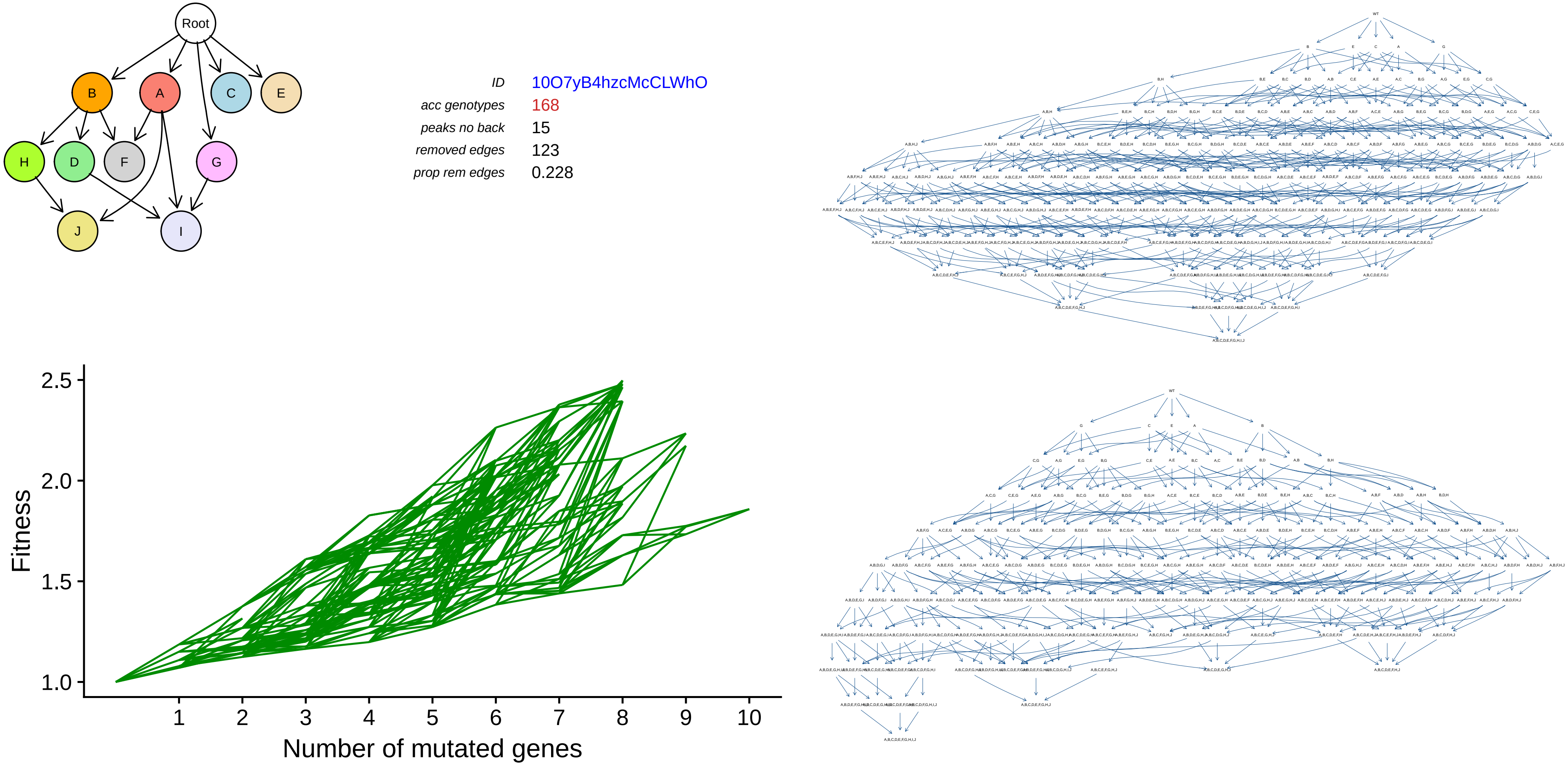

Root
B
A
C
E
H
D
F
G
J
I
ID 10O7yB4hzcMcCLWhO
acc genotypes 168
peaks no back 15
removed edges 123
prop rem edges 0.228
Fitness
2.5
2.0
1.5
1.0
1
2
3
4
5
6
7
8
9
10
Number of mutated genes

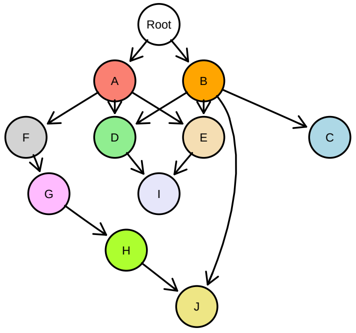


| | |
|---|---|
| *ID* | 1cfjEhrDN713Rg8k |
| *acc genotypes* | 57 |
| *peaks no back* | 7 |
| *removed edges* | 19 |
| *prop rem edges* | 0.15 |

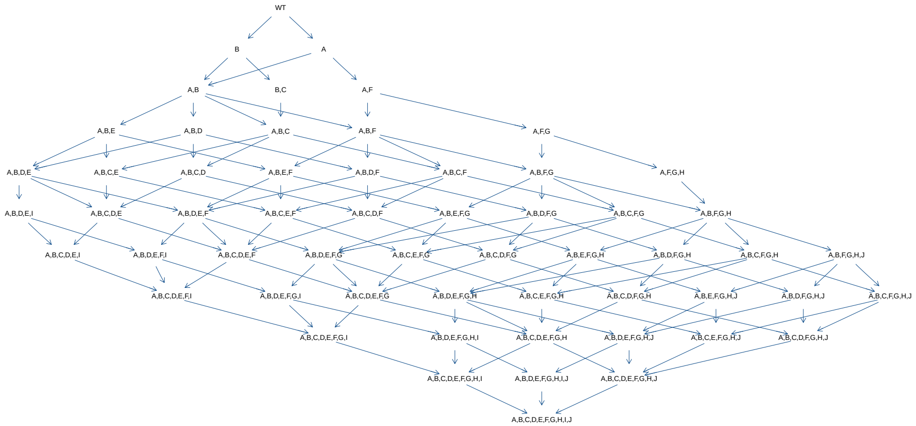

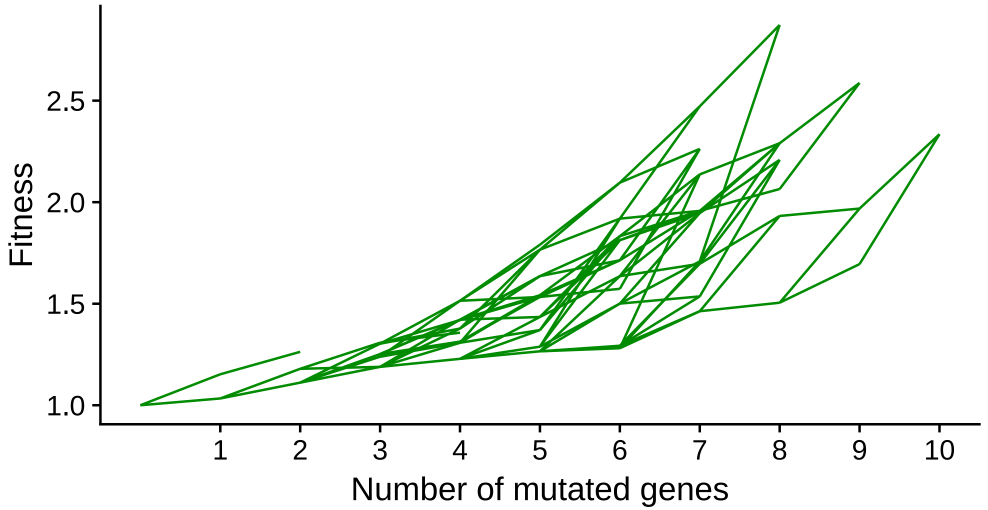


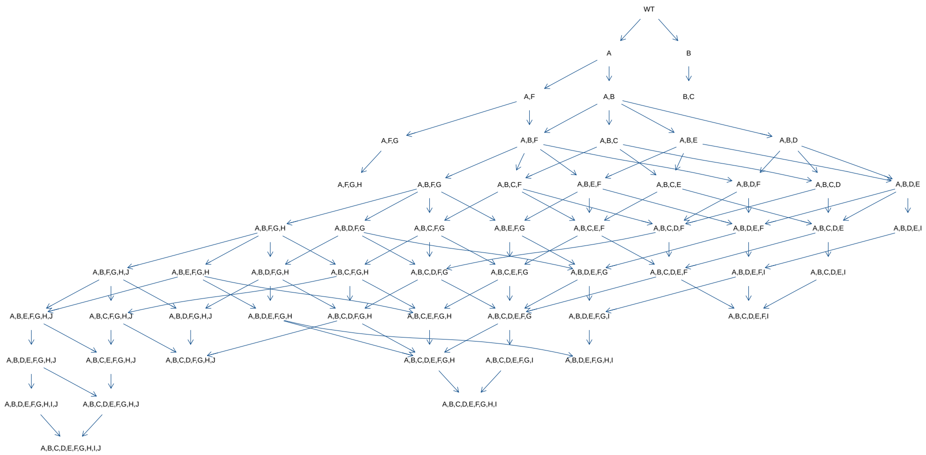

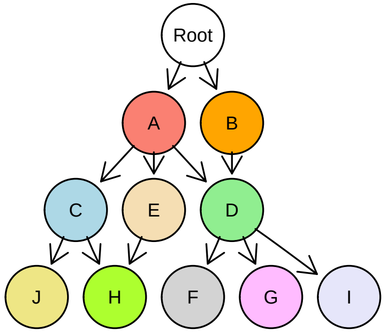


| | |
|---|---|
| *ID* | 1flJ9gvVIxHu10l4I |
| *acc genotypes* | 82 |
| *peaks no back* | 10 |
| *removed edges* | 36 |
| *prop rem edges* | 0.167 |

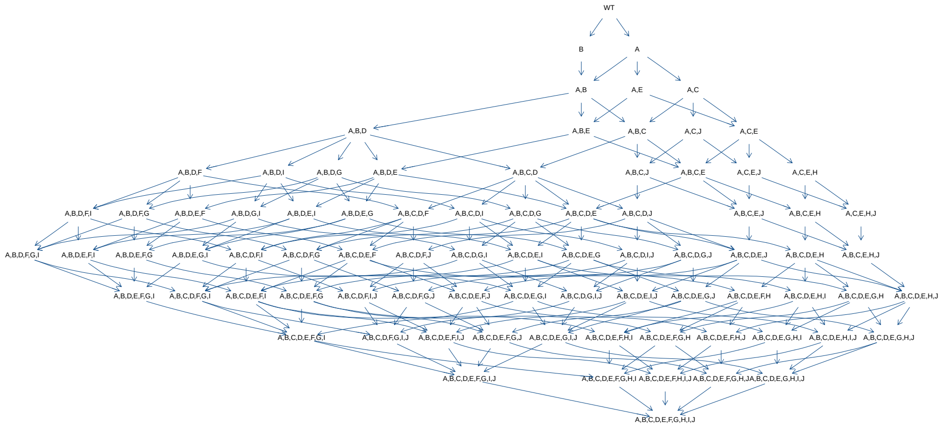

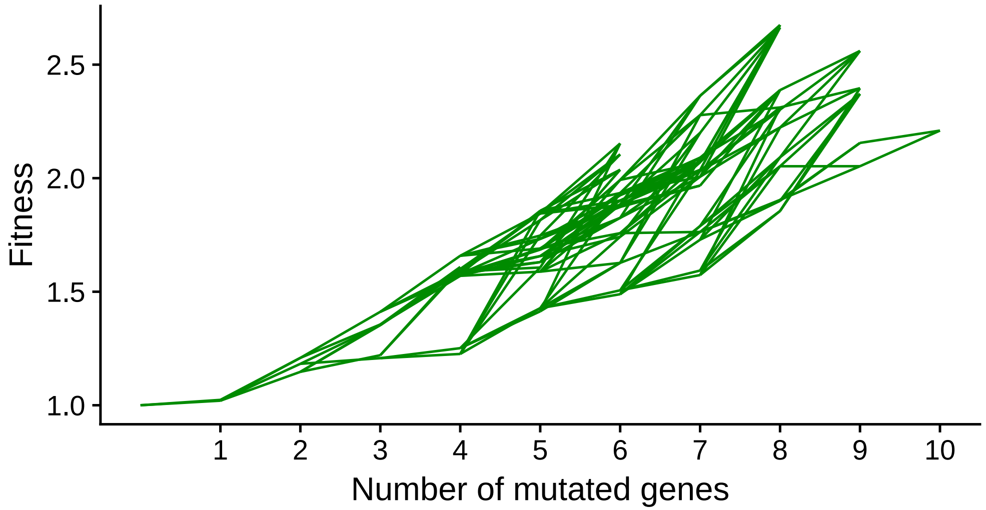


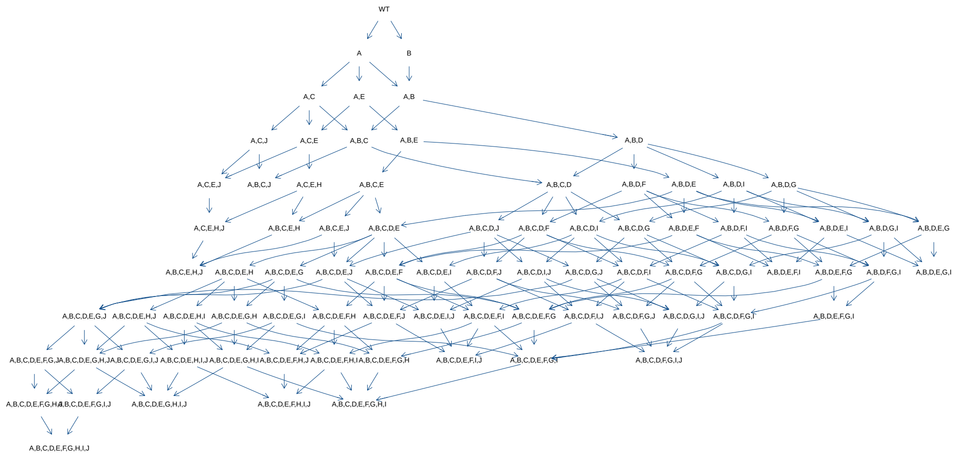

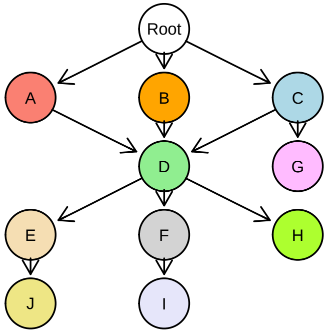


| | |
|---|---|
| *ID* | 1LRNl4YXQOtosQeO |
| *acc genotypes* | 48 |
| *peaks no back* | 7 |
| *removed edges* | 20 |
| *prop rem edges* | 0.189 |

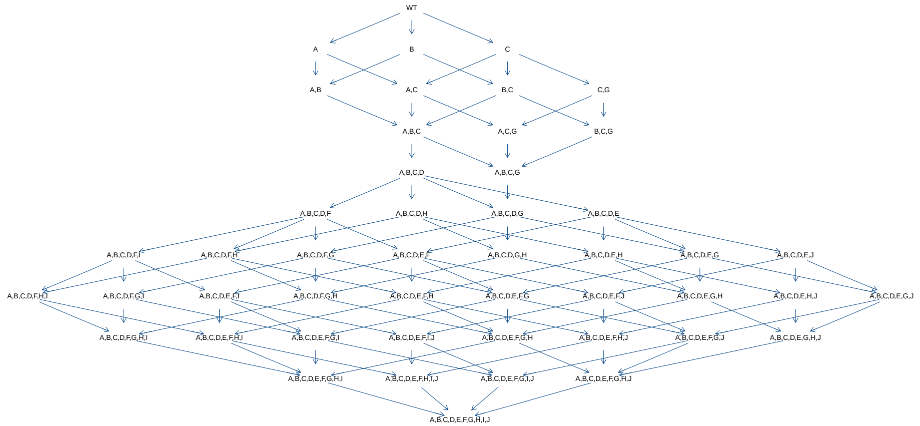

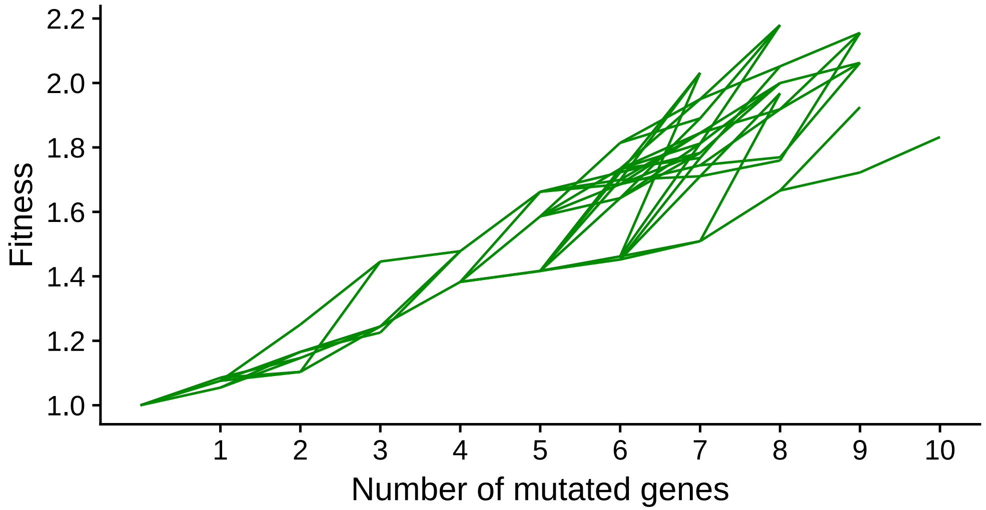


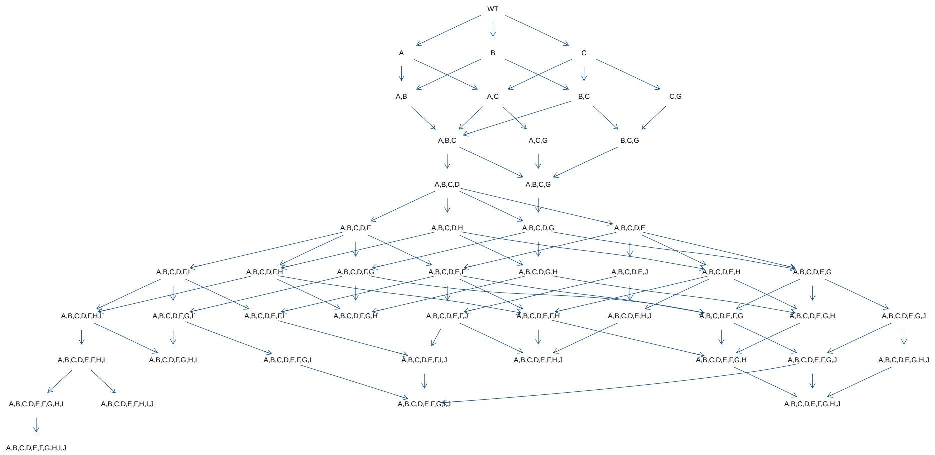

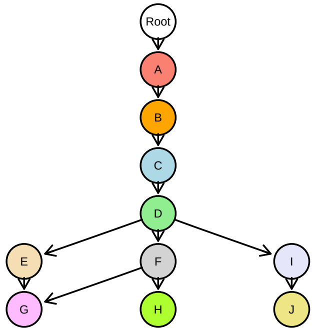


| | |
|---|---|
| *ID* | 24psYb2Z5uoeEIgL |
| *acc genotypes* | 28 |
| *peaks no back* | 5 |
| *removed edges* | 9 |
| *prop rem edges* | 0.18 |

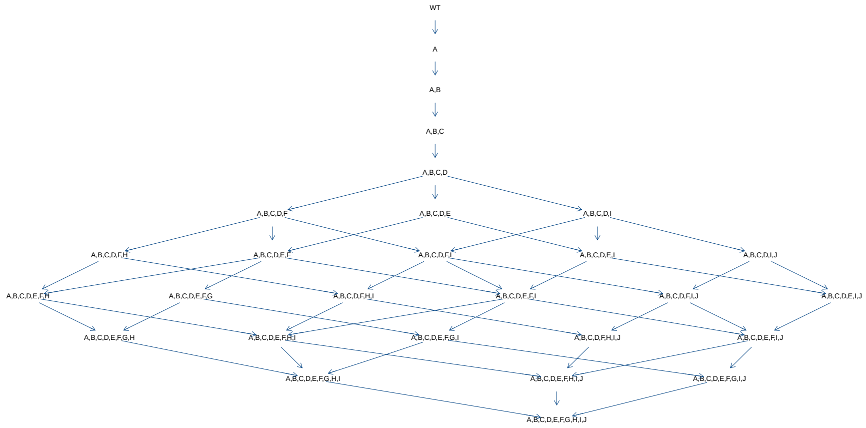

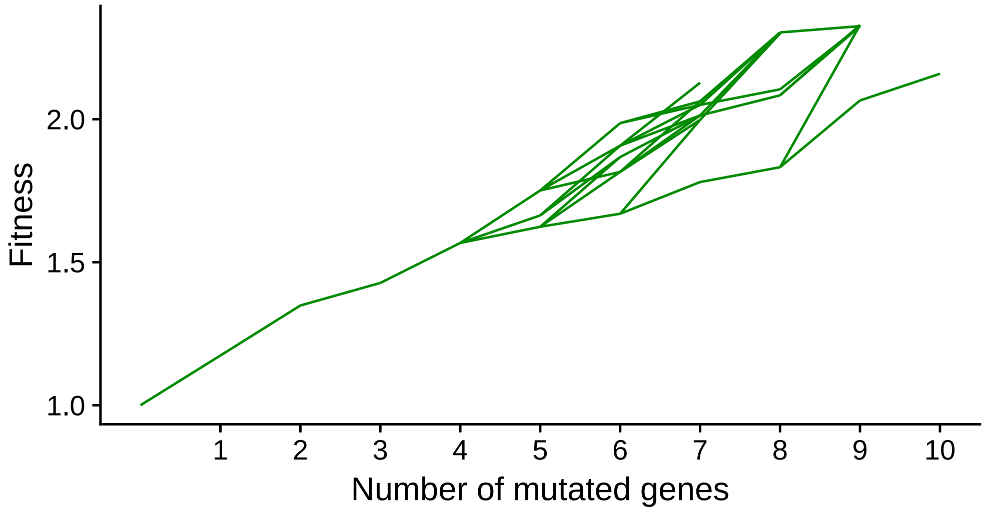


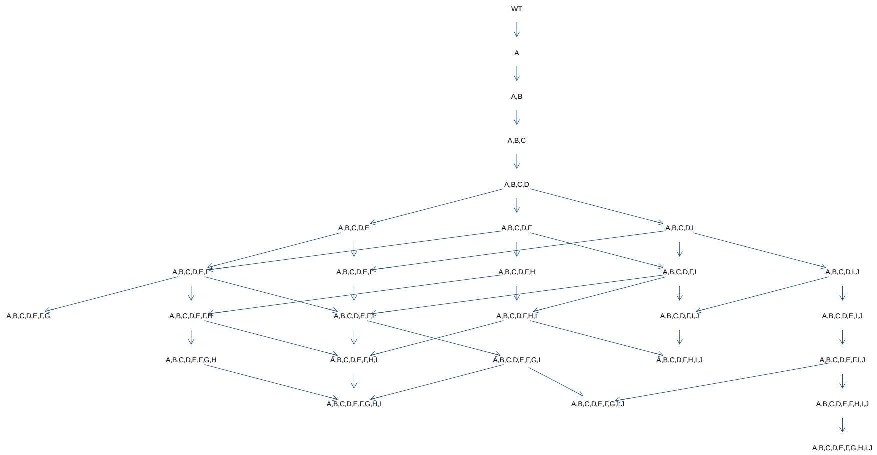

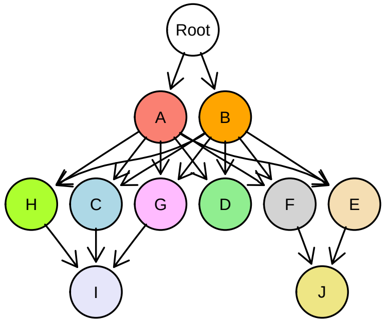


| | |
|---|---|
| *ID* | 26j4DR9Ab9swRweC |
| *acc genotypes* | 93 |
| *peaks no back* | 10 |
| *removed edges* | 40 |
| *prop rem edges* | 0.149 |

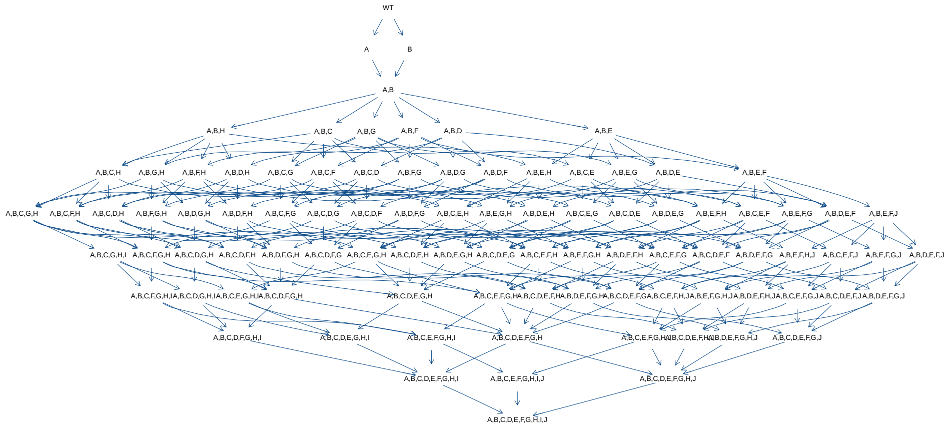

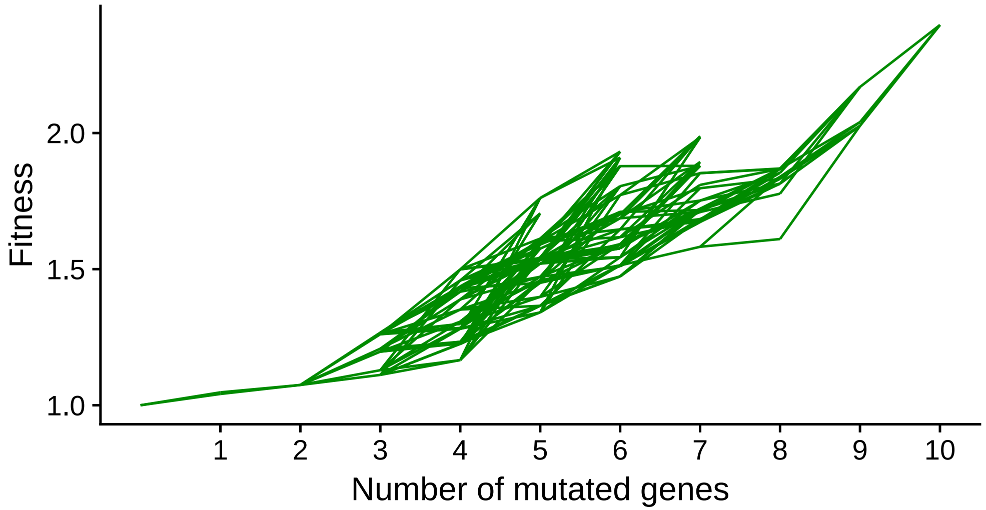


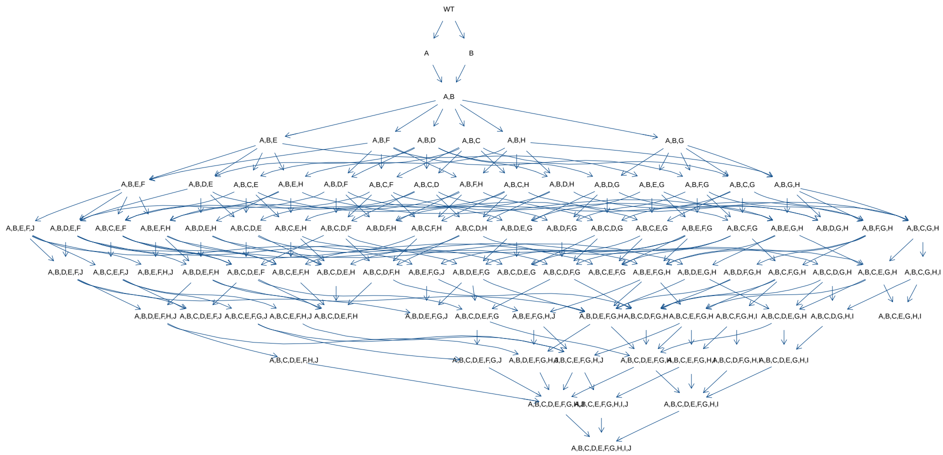

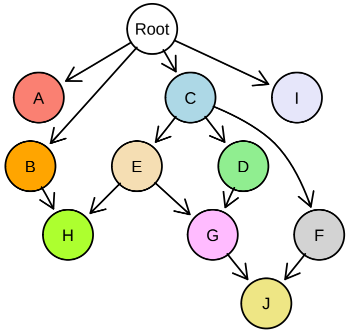


| | |
|---|---|
| *ID* | 29J8aMCesoGoSpvM |
| *acc genotypes* | 124 |
| *peaks no back* | 10 |
| *removed edges* | 55 |
| *prop rem edges* | 0.149 |

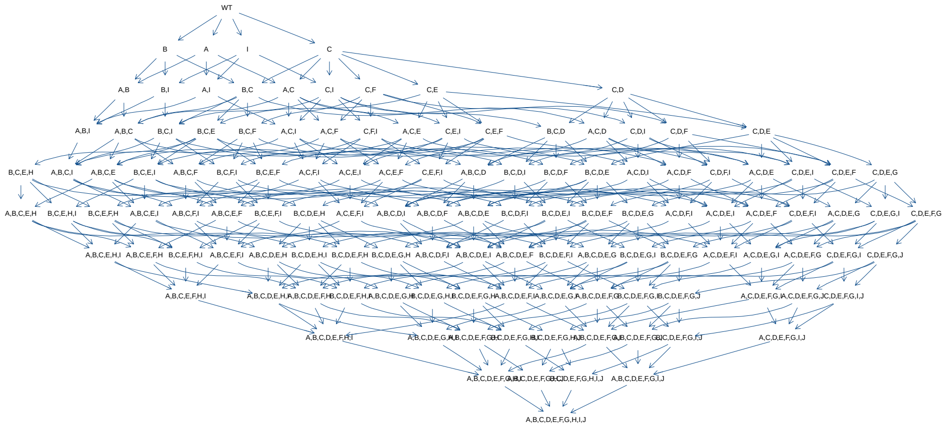

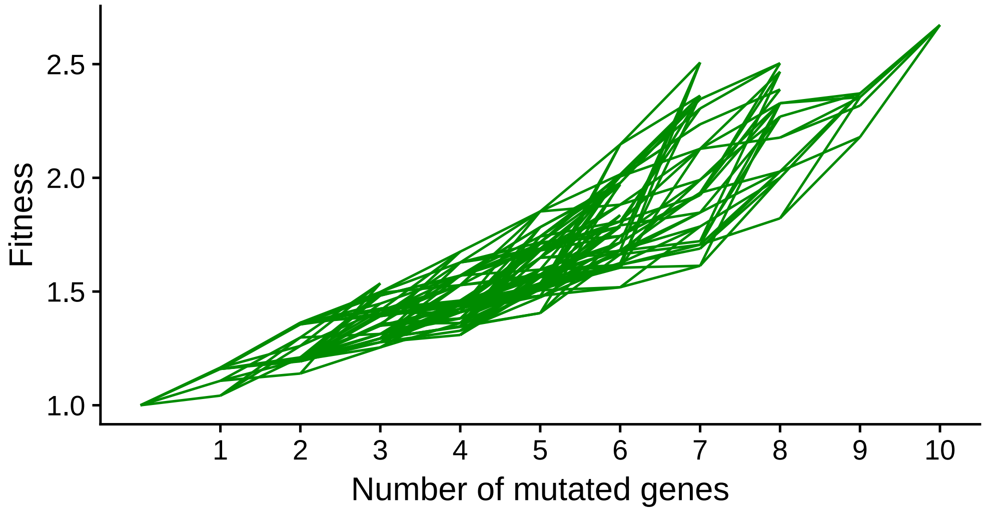


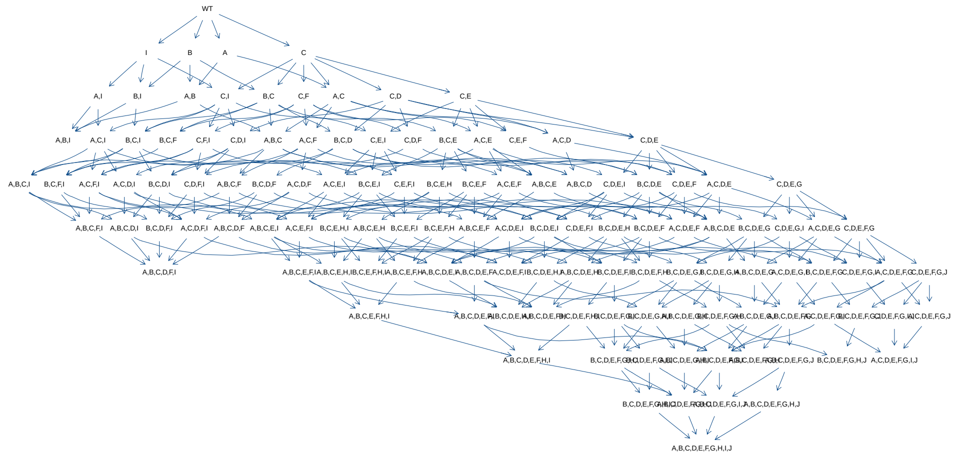

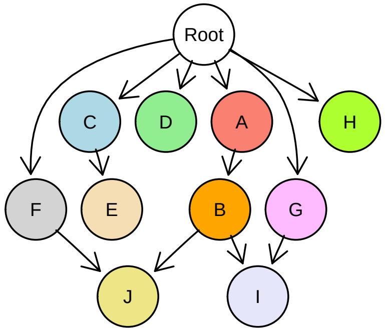


| | |
|---|---|
| *ID* | 2e8NWr3WSIIdSHqm |
| *acc genotypes* | 204 |
| *peaks no back* | 16 |
| *removed edges* | 139 |
| *prop rem edges* | 0.206 |

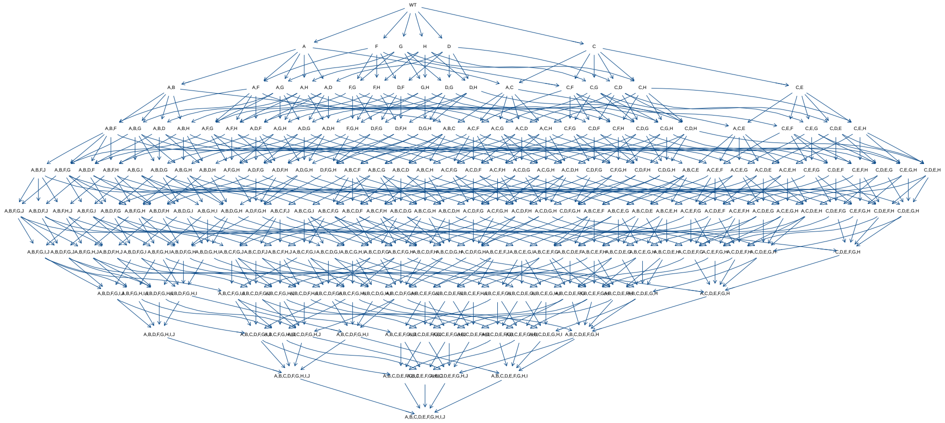

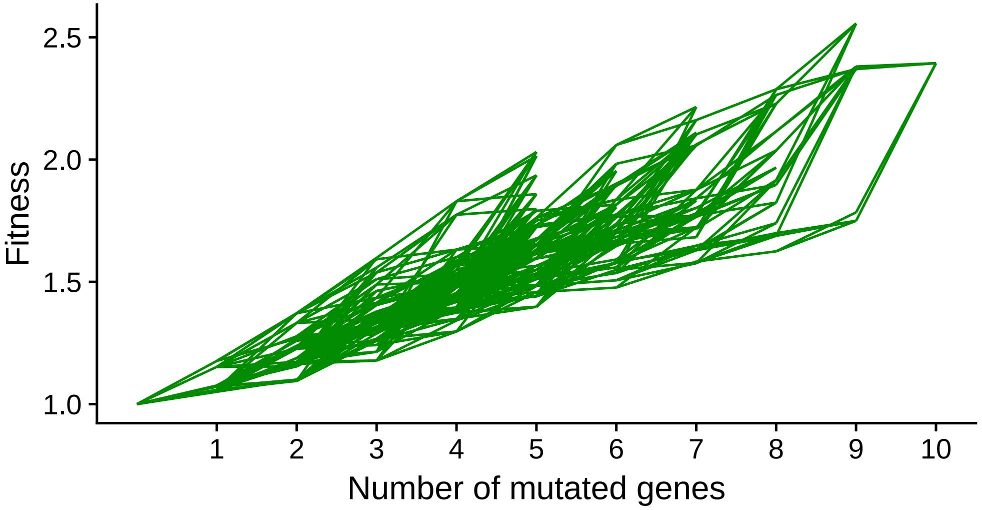


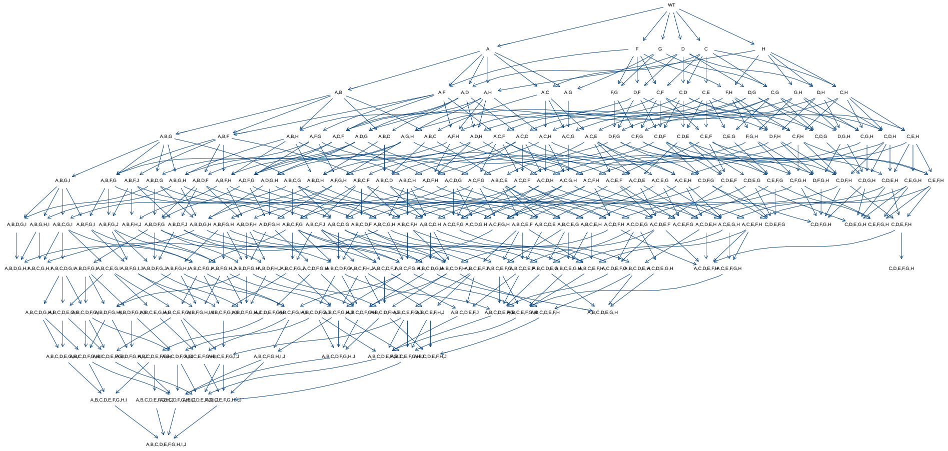

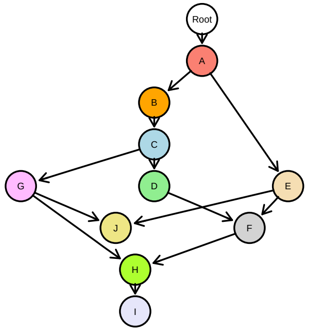


*ID* 2NKDNCMNshVULJ2E
*acc genotypes* 22
*peaks no back* 2
*removed edges* 3
*prop rem edges* 0.091

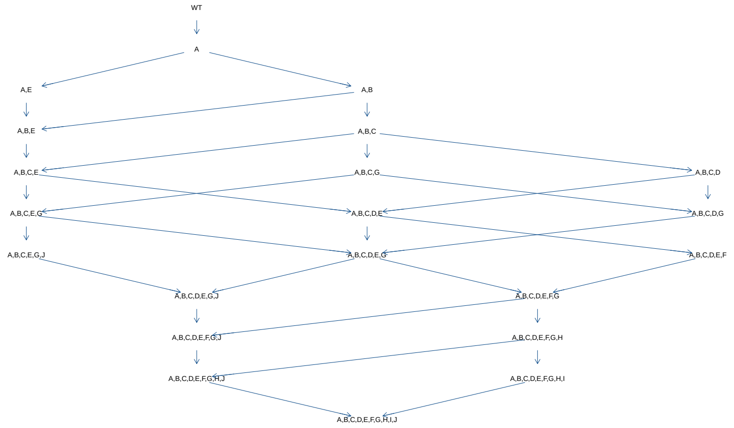

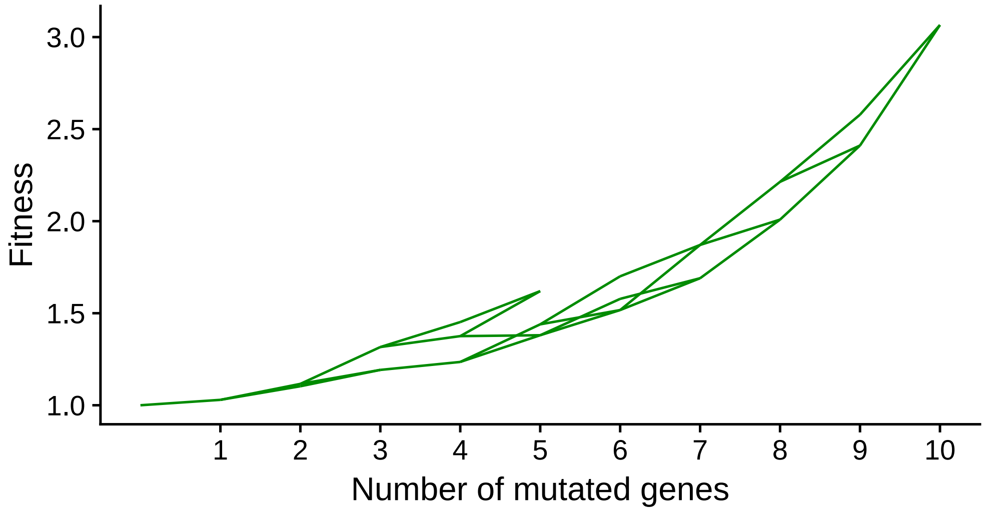


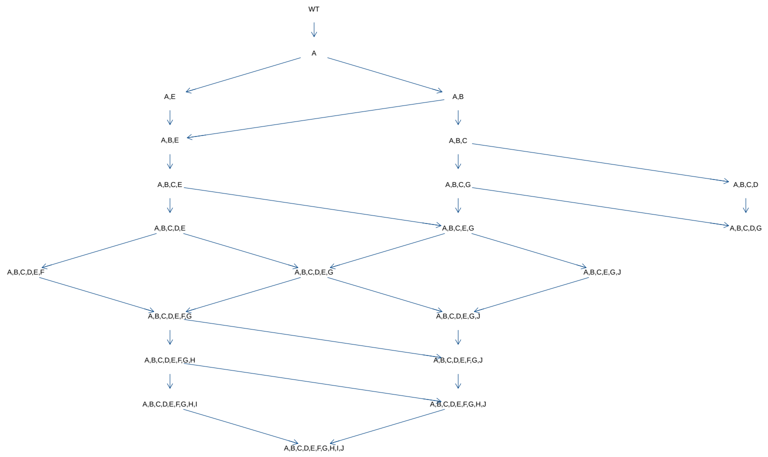

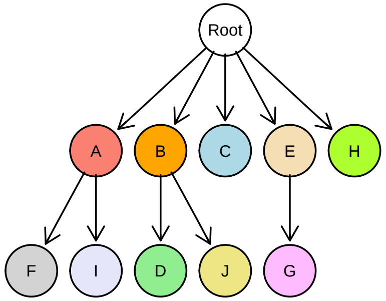


| | |
|---|---|
| *ID* | 2ti105XmgduWlCPHP |
| *acc genotypes* | 300 |
| *peaks no back* | 24 |
| *removed edges* | 245 |
| *prop rem edges* | 0.223 |

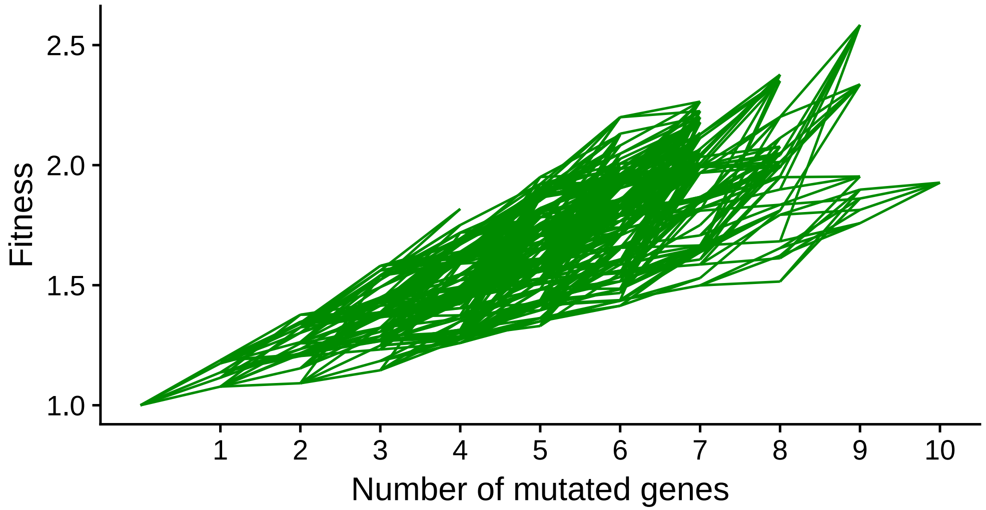


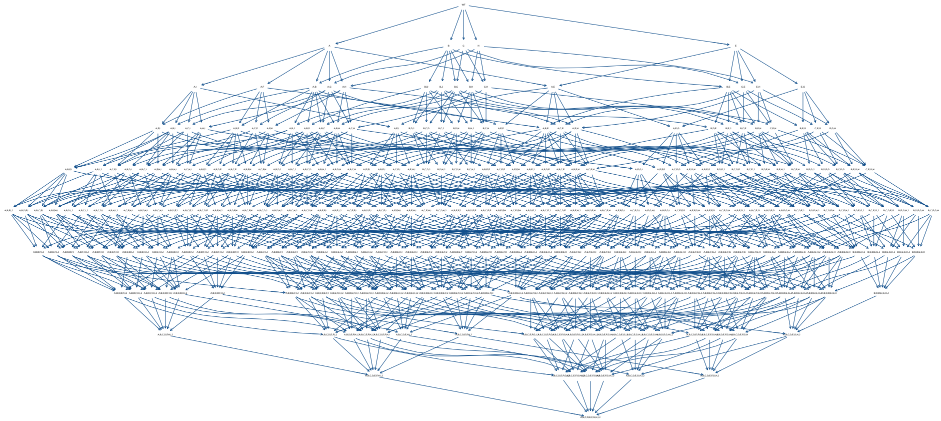

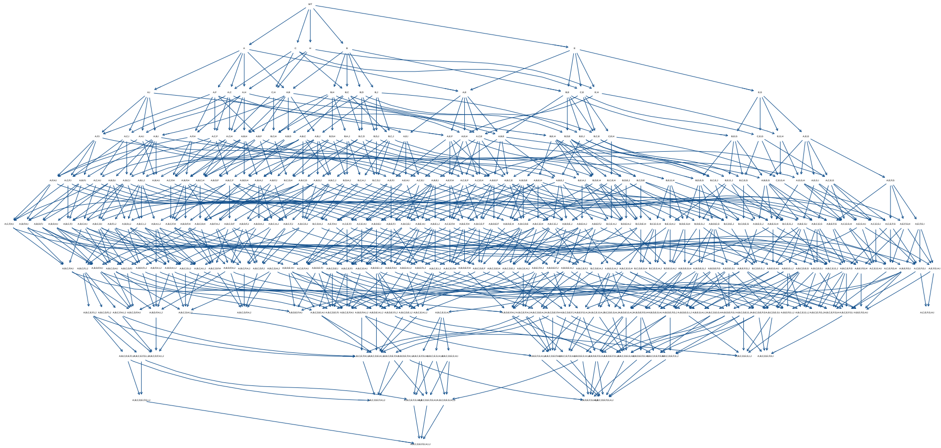

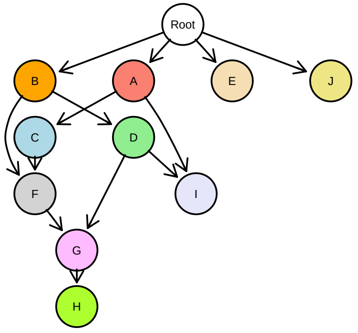


| | |
|---|---|
| *ID* | 3a7jMZPzBxEpCMra |
| *acc genotypes* | 72 |
| *peaks no back* | 11 |
| *removed edges* | 42 |
| *prop rem edges* | 0.239 |

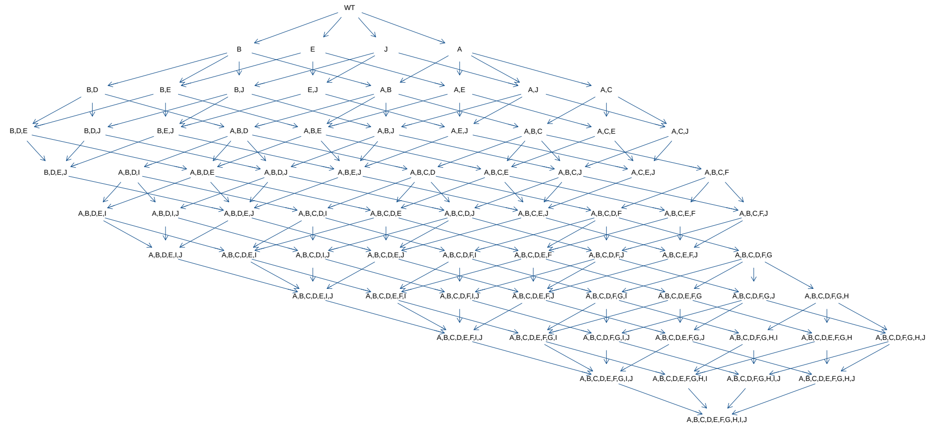

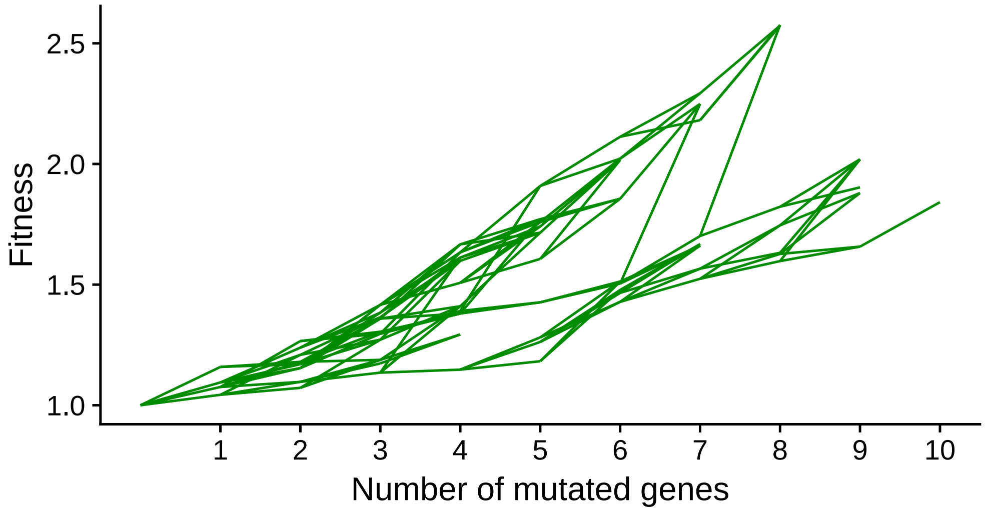


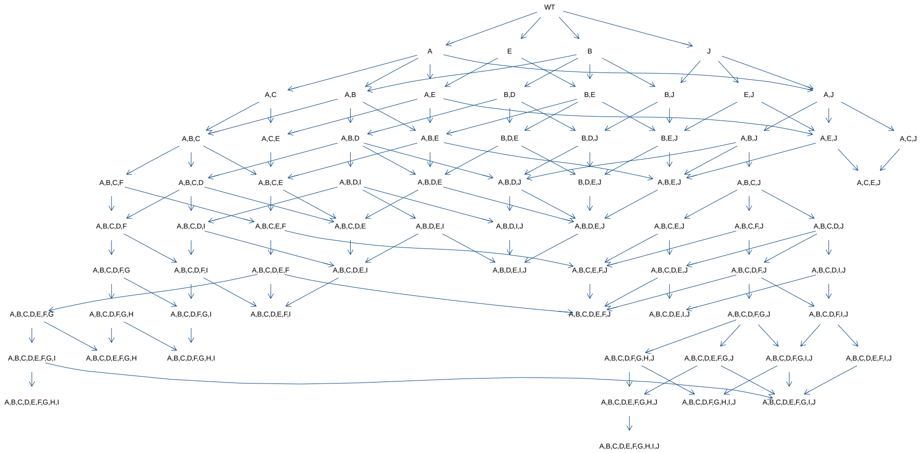

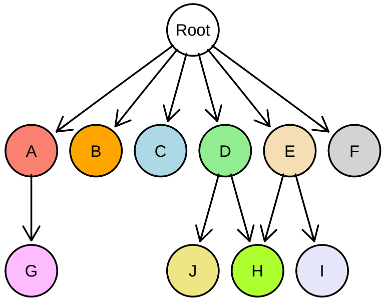


*ID* 46TSoH89nM3Ac4Jd
*acc genotypes* 312
*peaks no back* 23
*removed edges* 210
*prop rem edges* 0.182

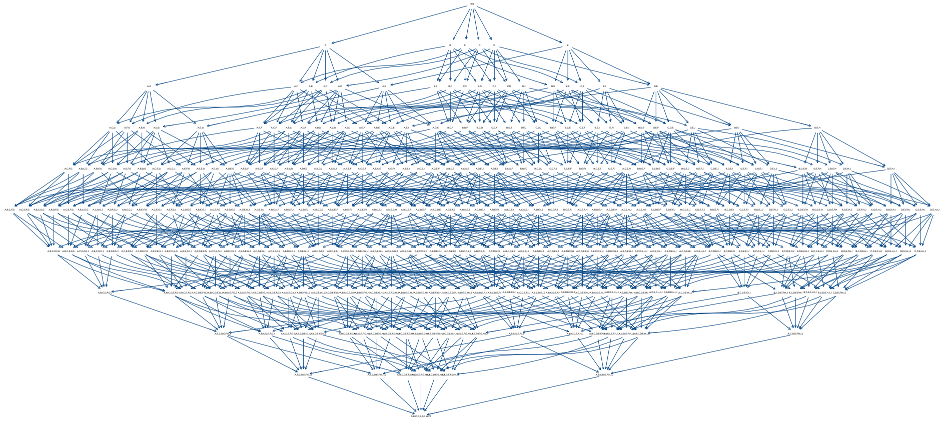

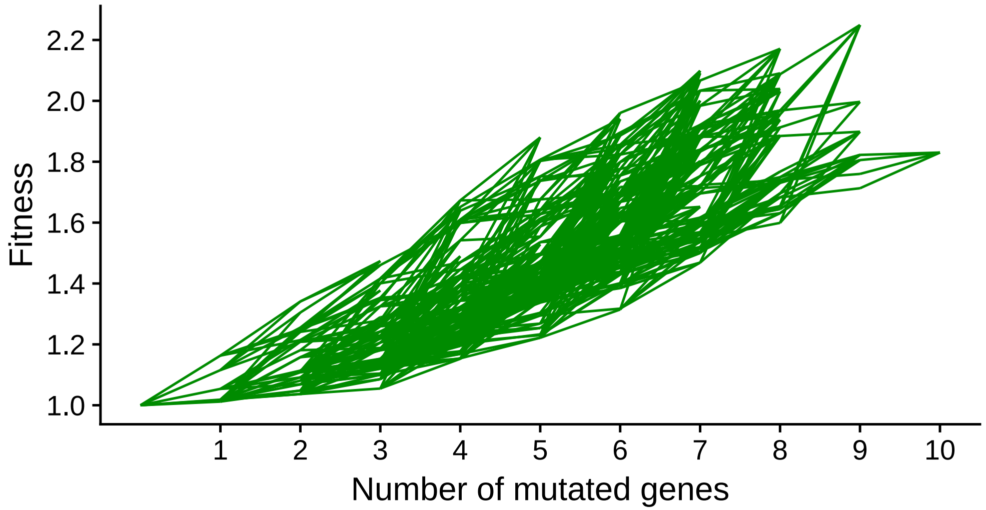


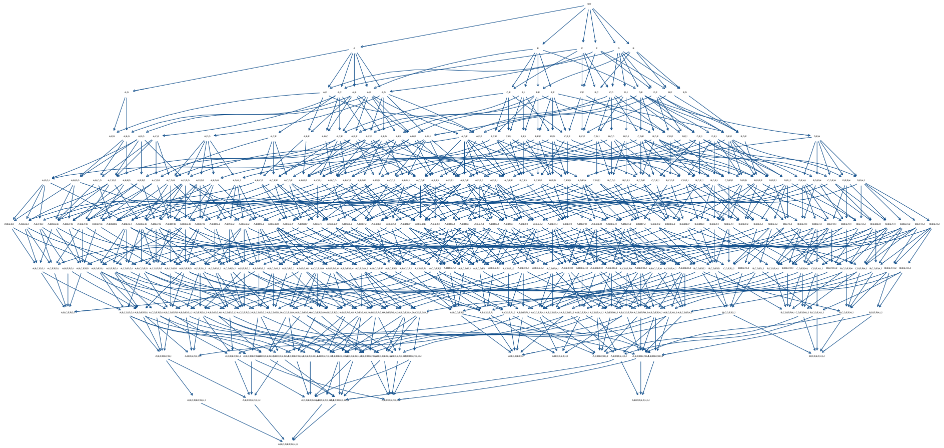

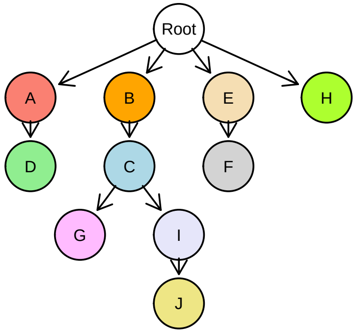


| | |
|---|---|
| *ID* | 4KWL9eg4fUjg1ndt |
| *acc genotypes* | 144 |
| *peaks no back* | 15 |
| *removed edges* | 97 |
| *prop rem edges* | 0.228 |

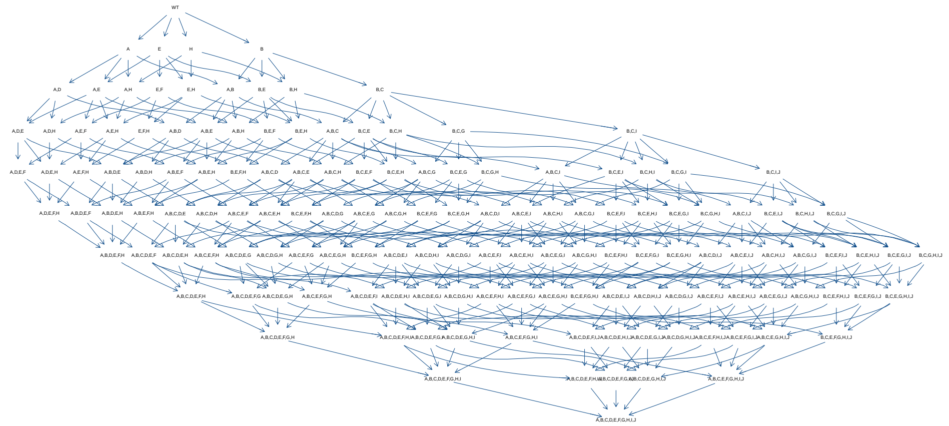

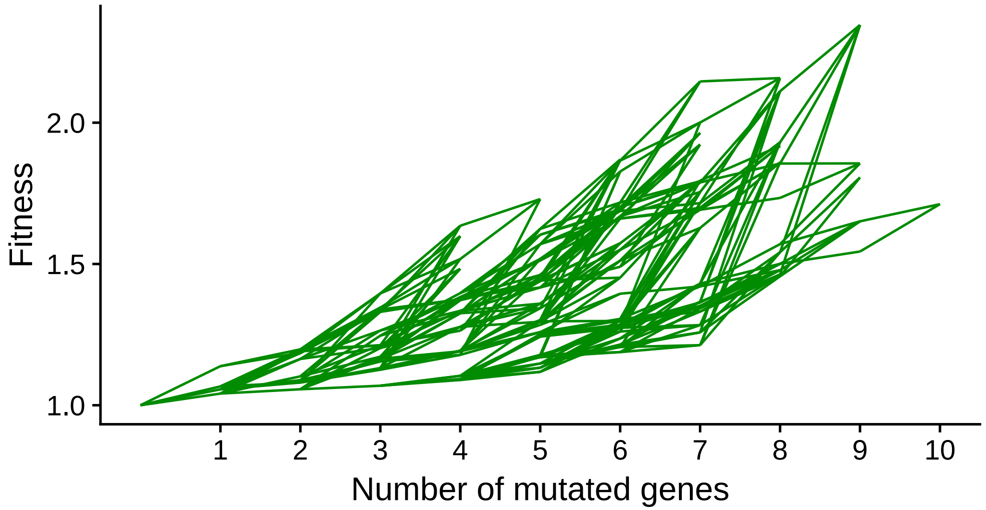


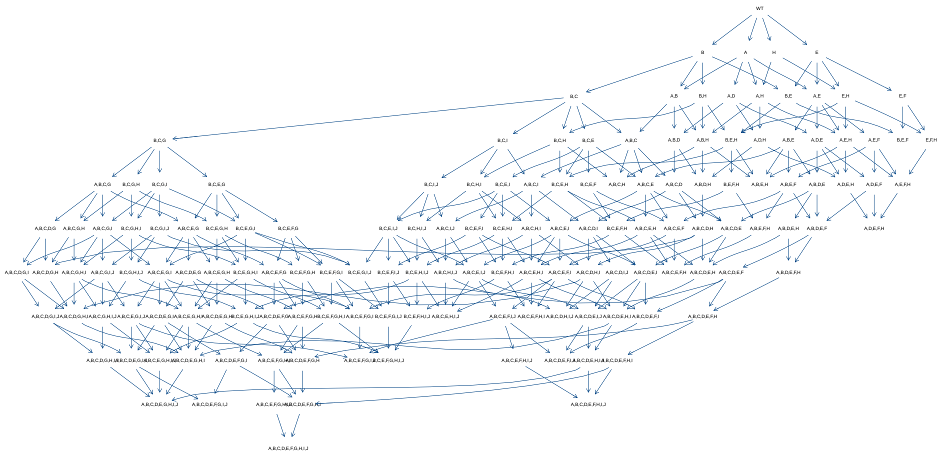

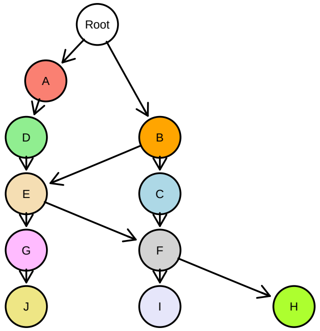


| | |
|---|---|
| *ID* | 4r7diBoGWUyNOWMY |
| *acc genotypes* | 27 |
| *peaks no back* | 5 |
| *removed edges* | 6 |
| *prop rem edges* | 0.136 |

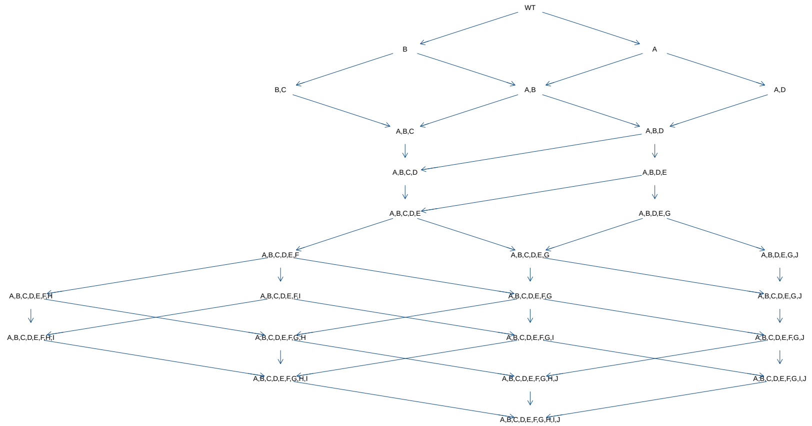

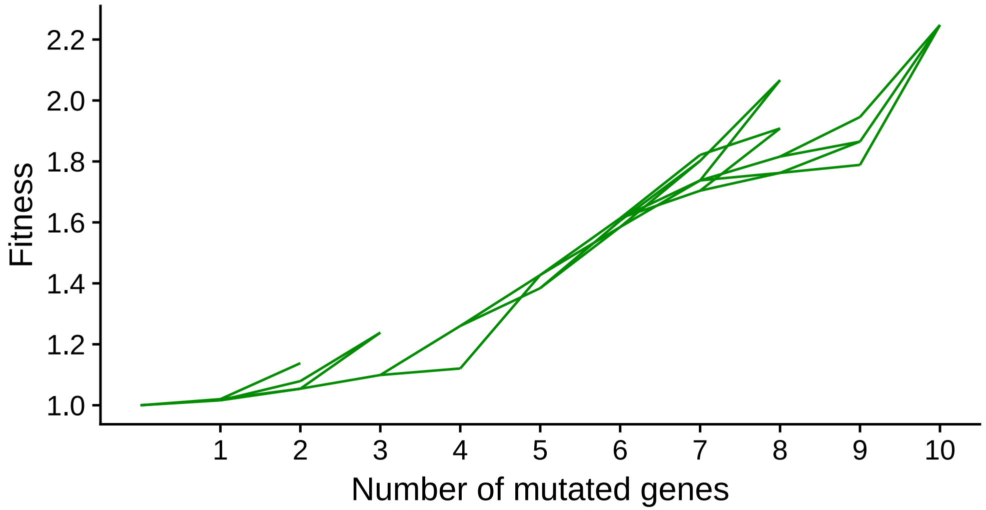


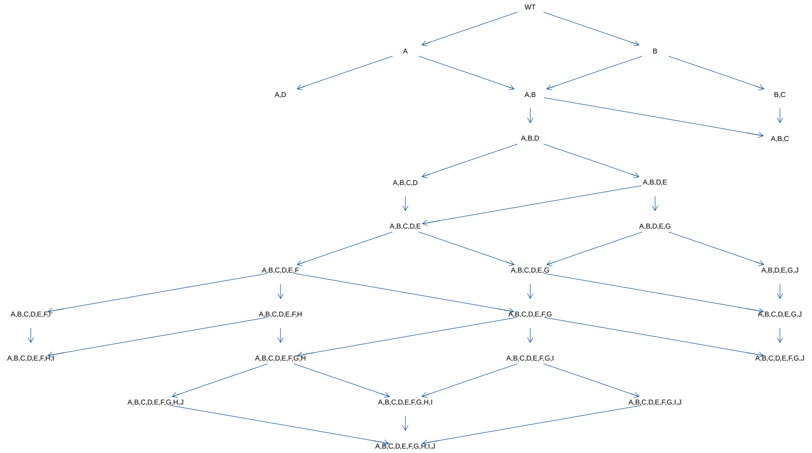

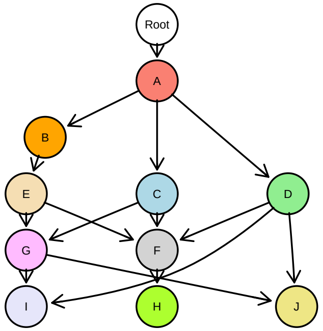


*ID* 56zMgtzUdnUxiwN9
*acc genotypes* 28
*peaks no back* 6
*removed edges* 8
*prop rem edges* 0.167

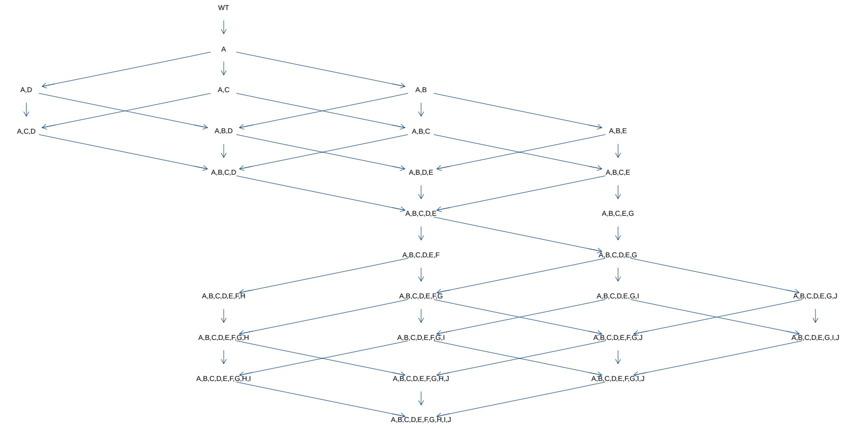

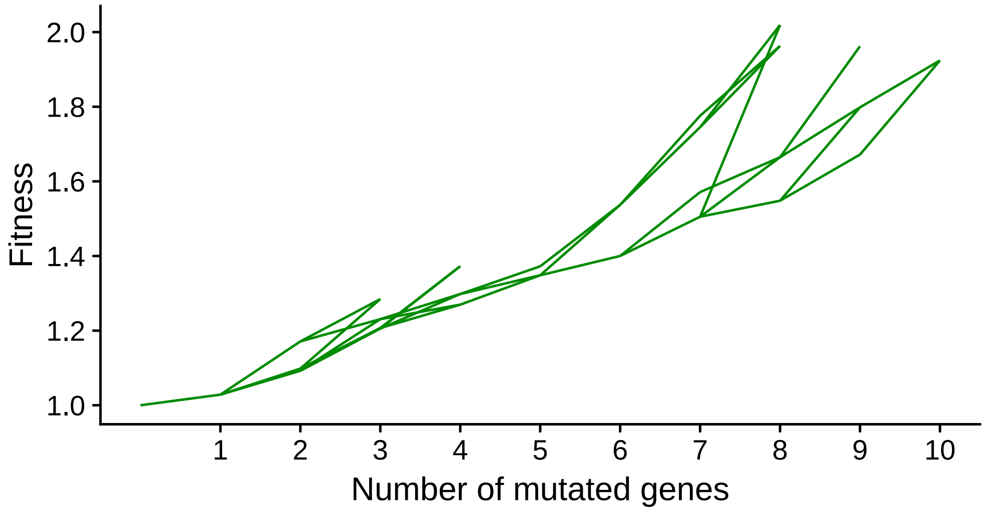


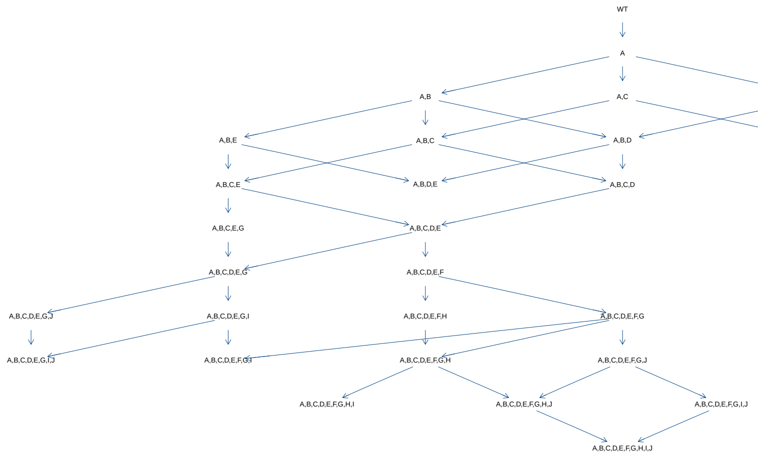

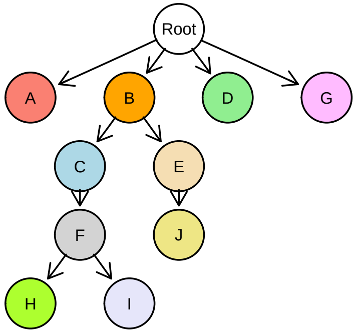


| | |
|---|---|
| *ID* | 5D4EKkcfeVoj4fHS |
| *acc genotypes* | 152 |
| *peaks no back* | 14 |
| *removed edges* | 94 |
| *prop rem edges* | 0.197 |

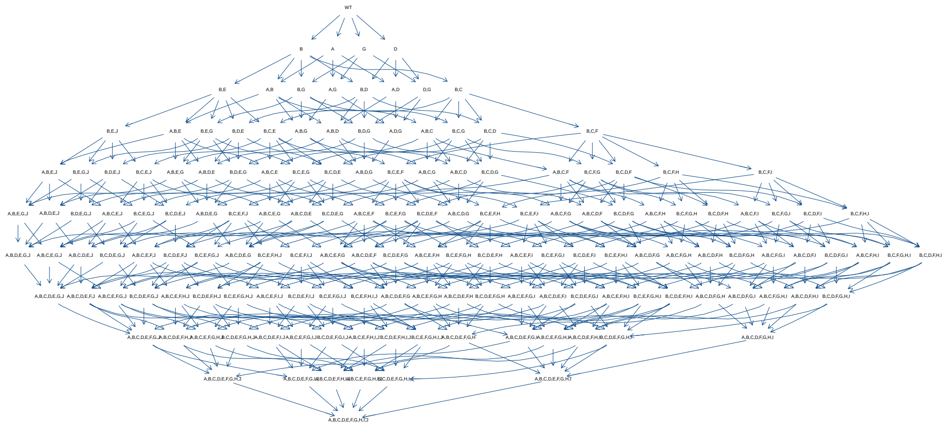


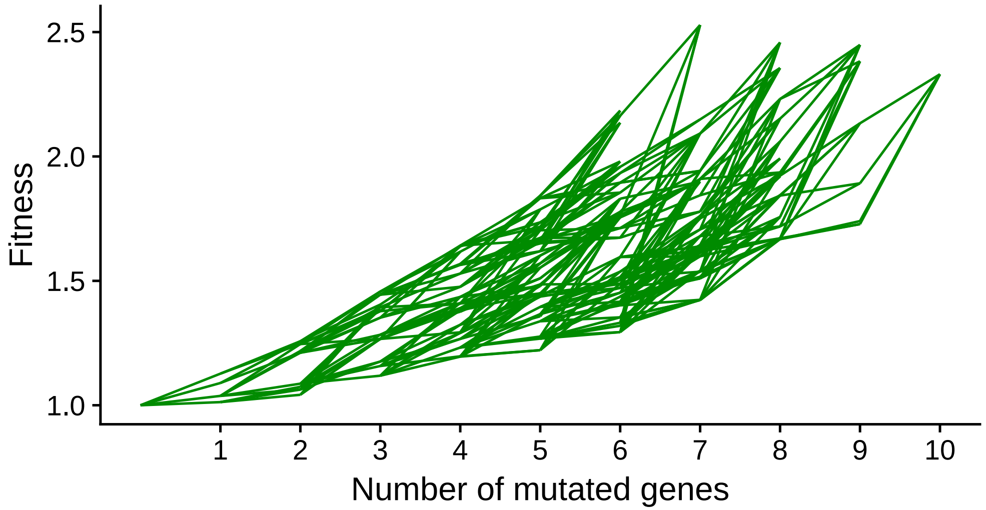


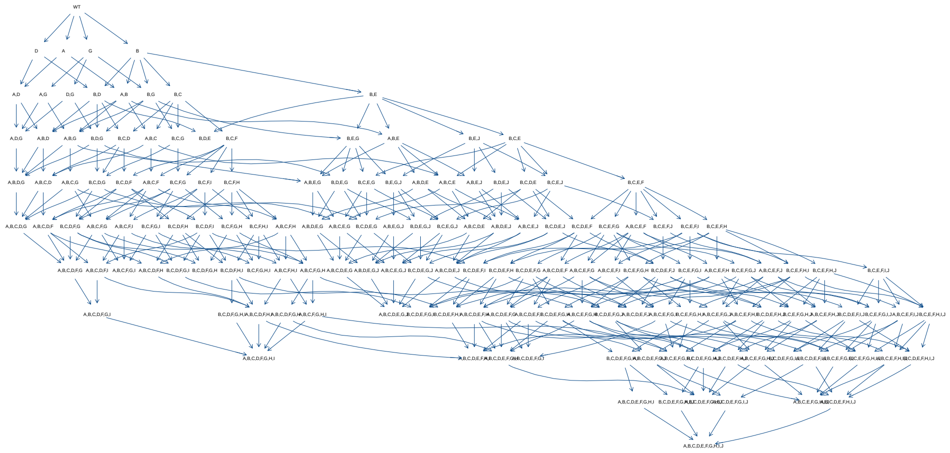

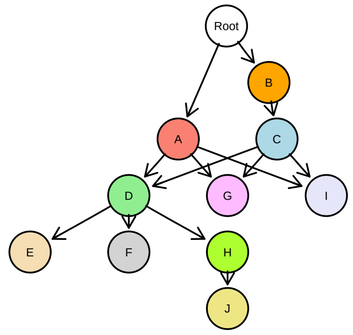


| | |
|---|---|
| *ID* | 5DSlnm8xyZVZ5wT9 |
| *acc genotypes* | 57 |
| *peaks no back* | 8 |
| *removed edges* | 20 |
| *prop rem edges* | 0.14 |

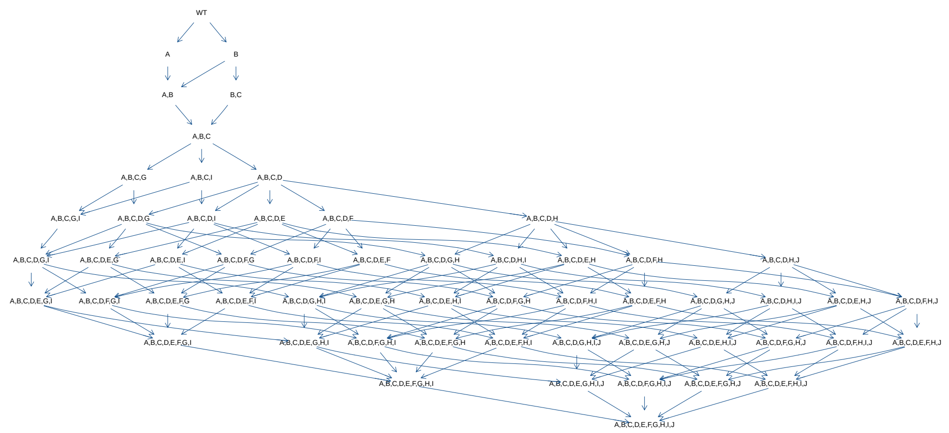

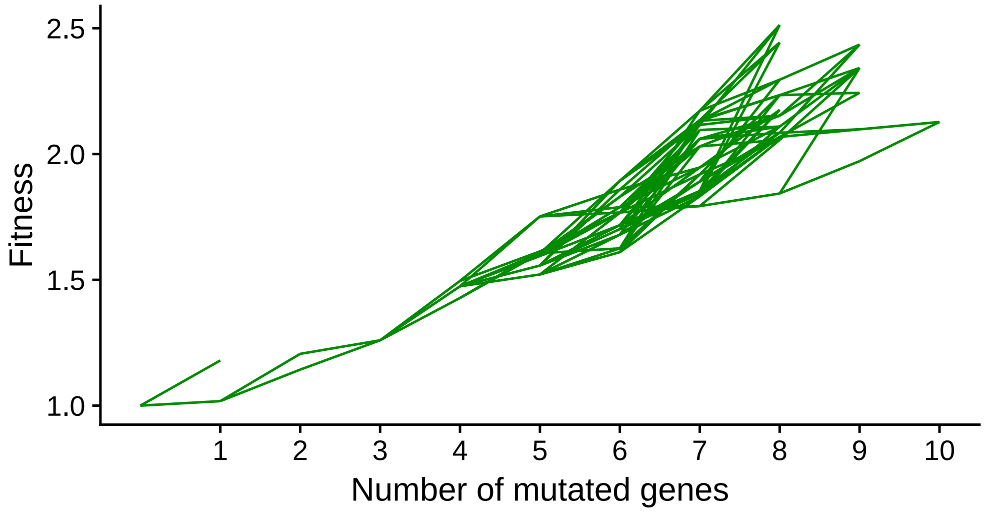


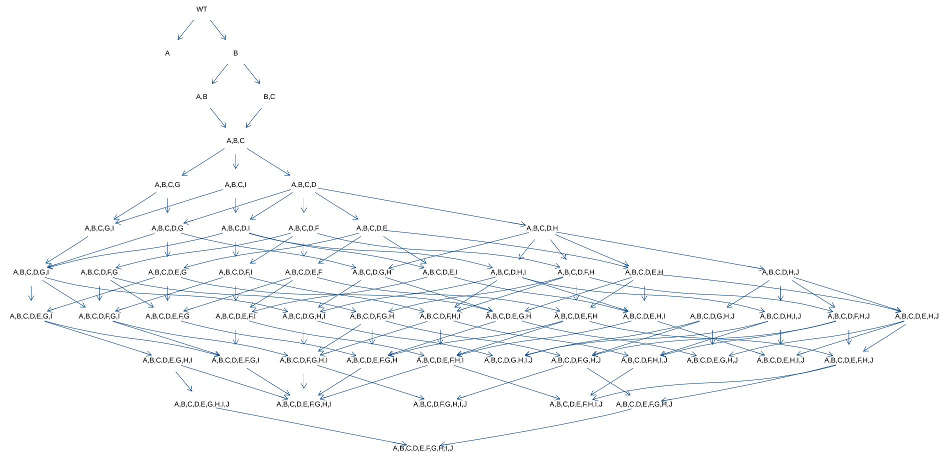

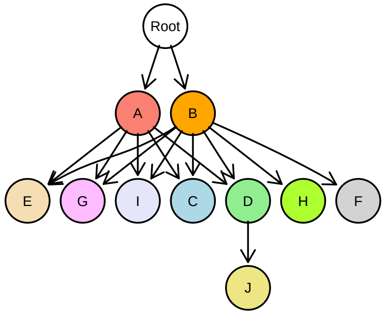


| | |
|---|---|
| *ID* | 5ervwavRLQyVUmM4 |
| *acc genotypes* | 198 |
| *peaks no back* | 19 |
| *removed edges* | 124 |
| *prop rem edges* | 0.173 |

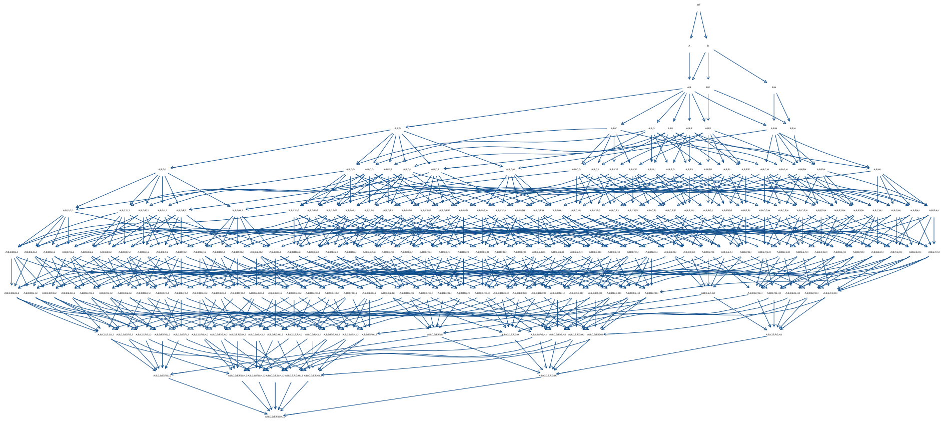

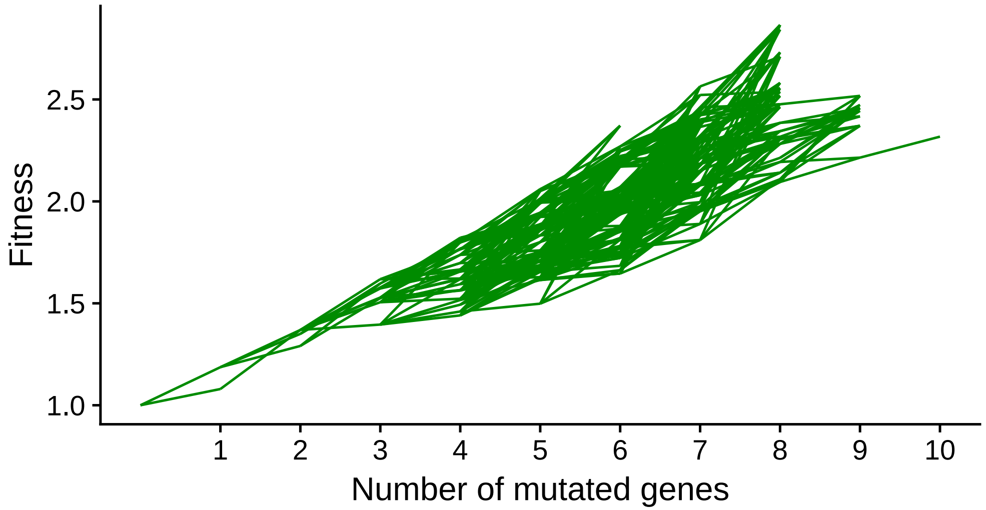


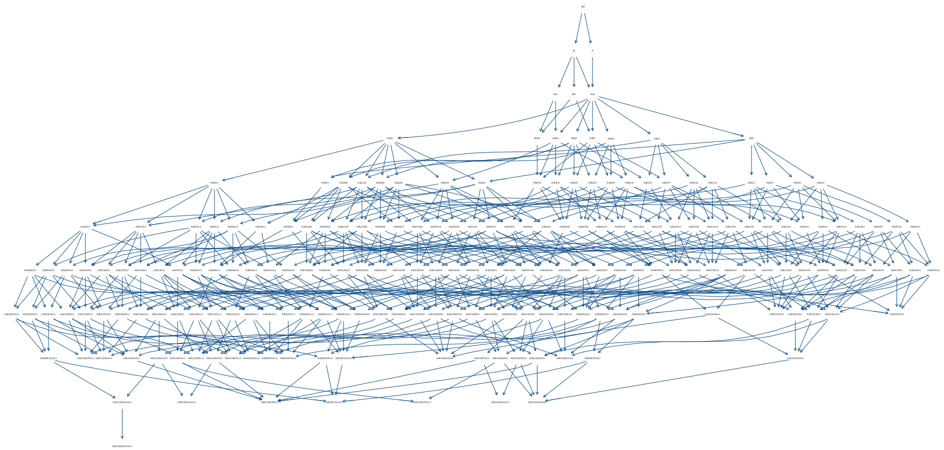

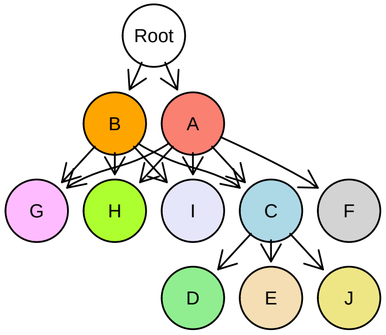


| | |
|---|---|
| *ID* | 5Kk3ENaa6v1KQQvo |
| *acc genotypes* | 148 |
| *peaks no back* | 18 |
| *removed edges* | 100 |
| *prop rem edges* | 0.199 |

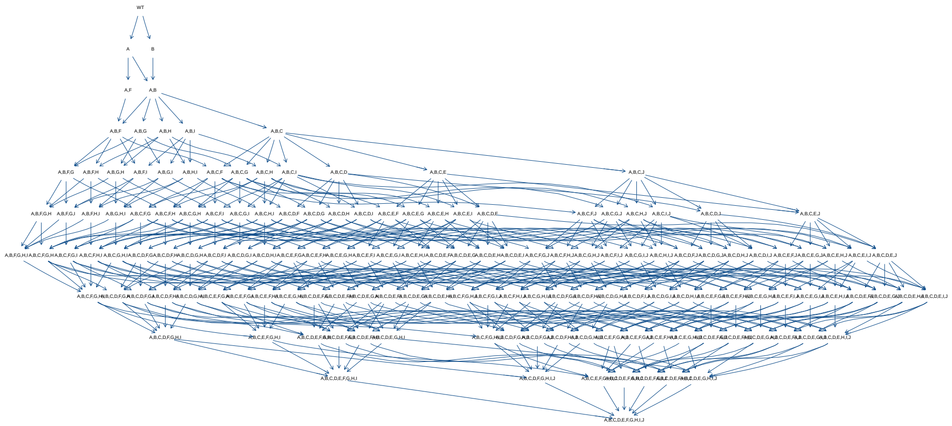

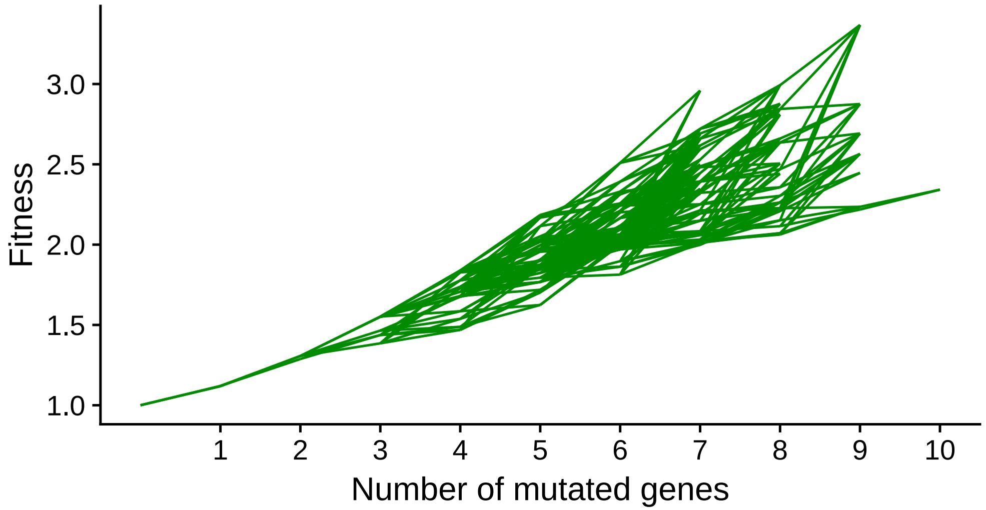


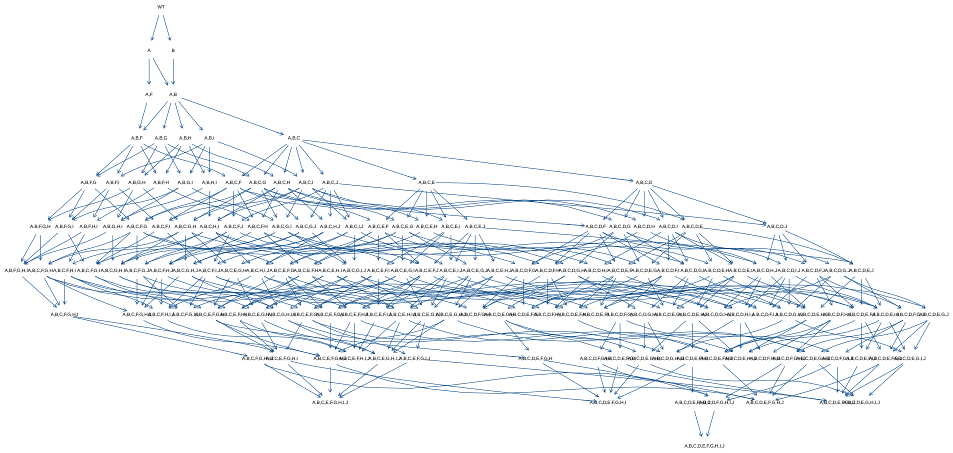

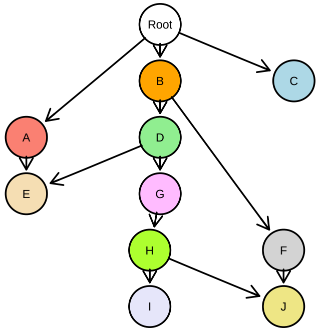


| | |
|---|---|
| *ID* | 5MCNYeerEvdy7pit |
| *acc genotypes* | 72 |
| *peaks no back* | 8 |
| *removed edges* | 33 |
| *prop rem edges* | 0.188 |

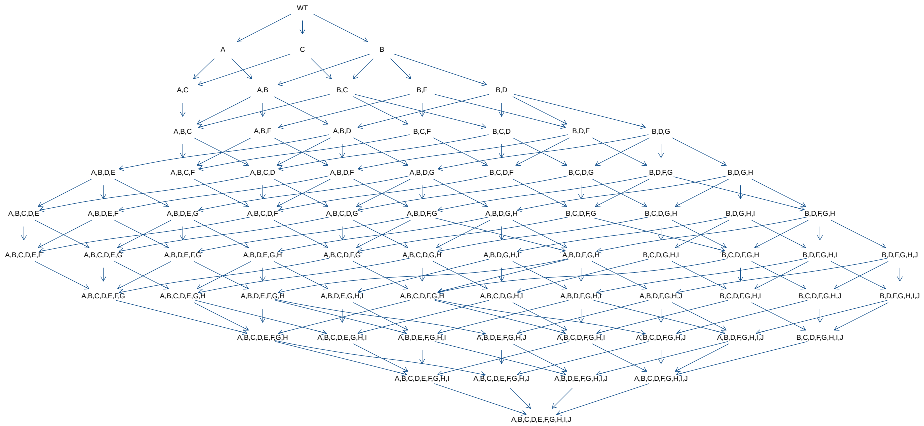

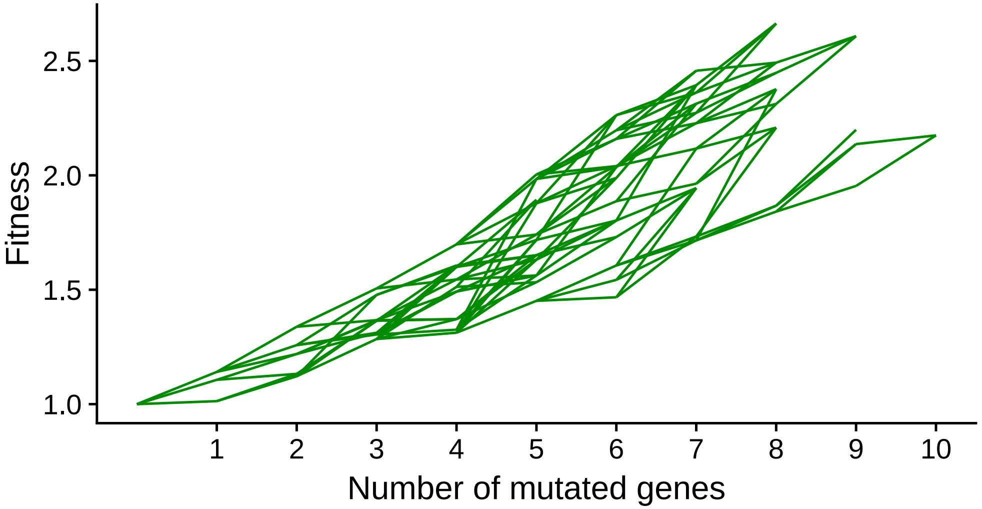


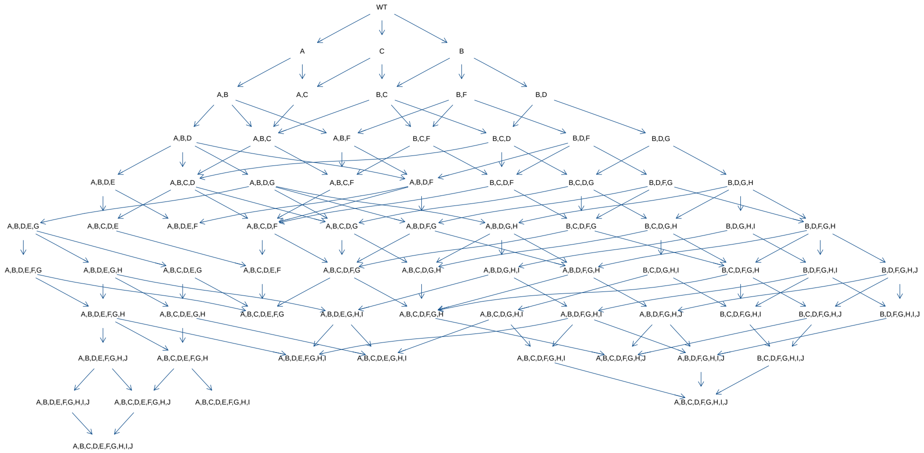

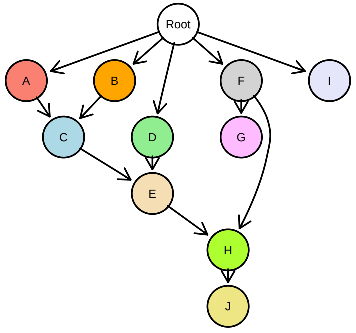


| | |
|---|---|
| *ID* | 5MxLARdmCMPqgUDW |
| *acc genotypes* | 74 |
| *peaks no back* | 9 |
| *removed edges* | 35 |
| *prop rem edges* | 0.185 |

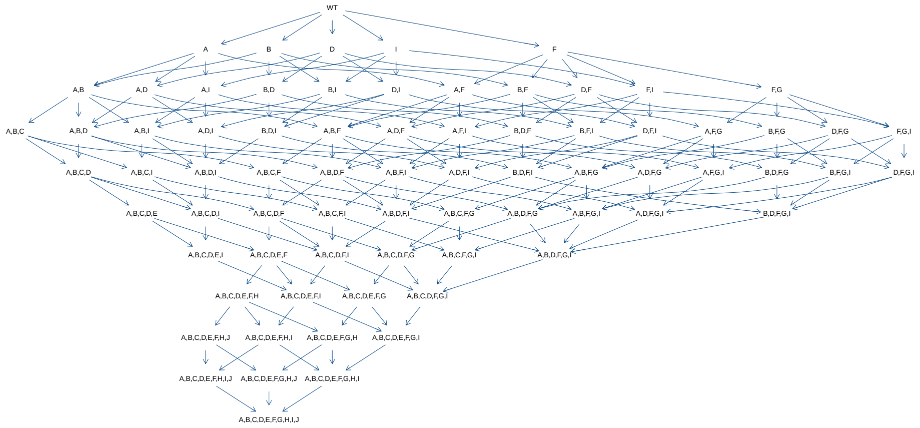

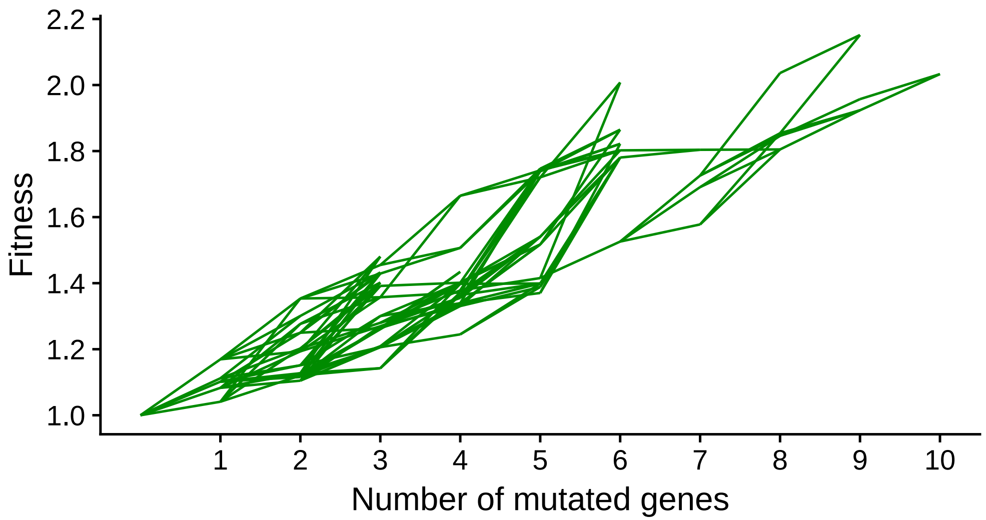


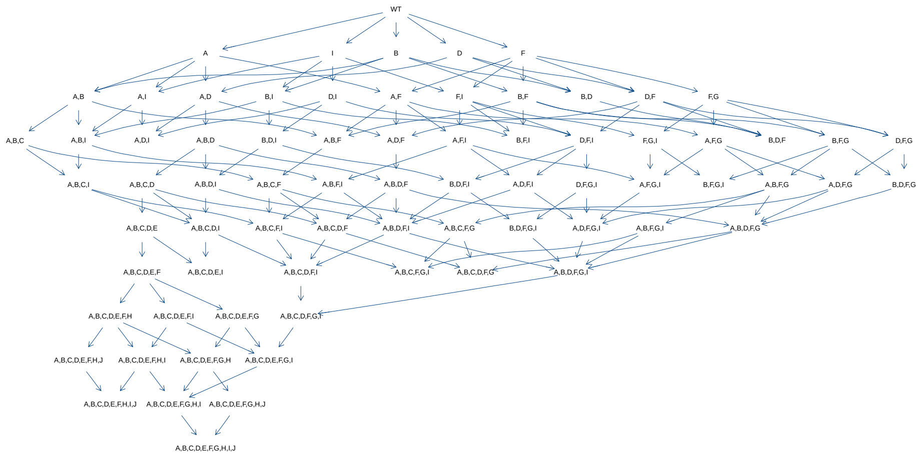

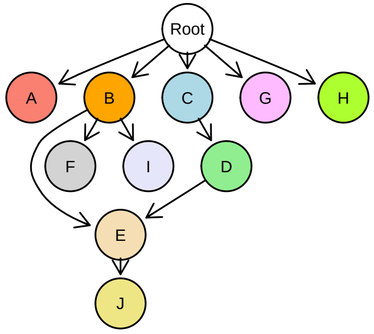


| | |
|---|---|
| *ID* | 5URt4ktIFolpEsp3 |
| *acc genotypes* | 184 |
| *peaks no back* | 19 |
| *removed edges* | 120 |
| *prop rem edges* | 0.199 |

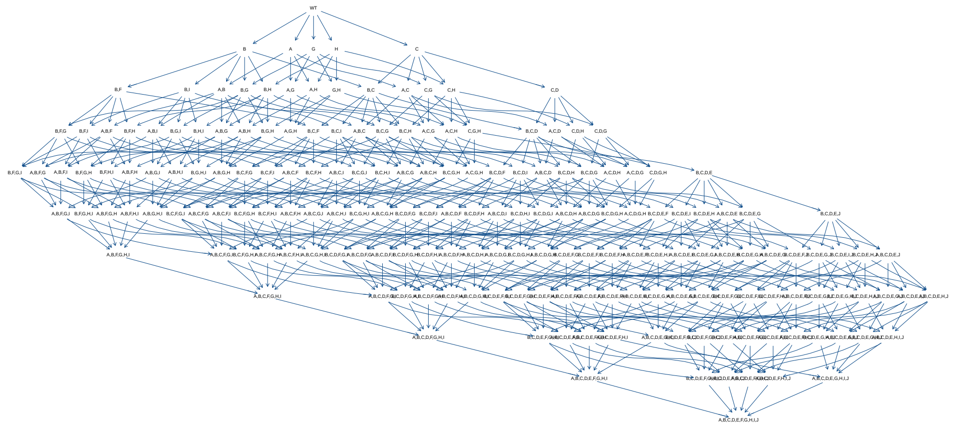

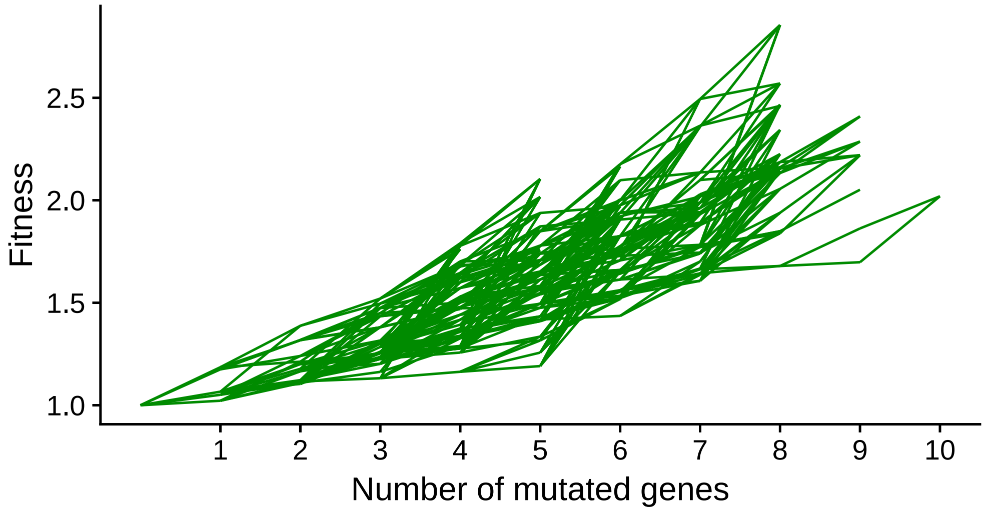


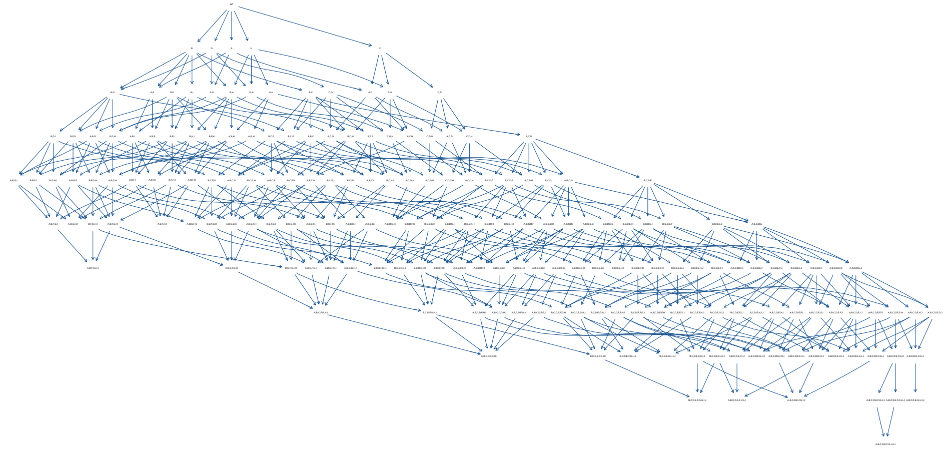

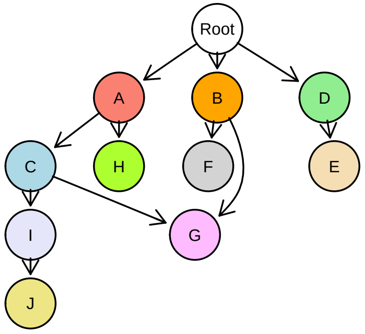


| | |
|---|---|
| *ID* | 64xXOCLCH1gfTB10v |
| *acc genotypes* | 117 |
| *peaks no back* | 12 |
| *removed edges* | 78 |
| *prop rem edges* | 0.239 |

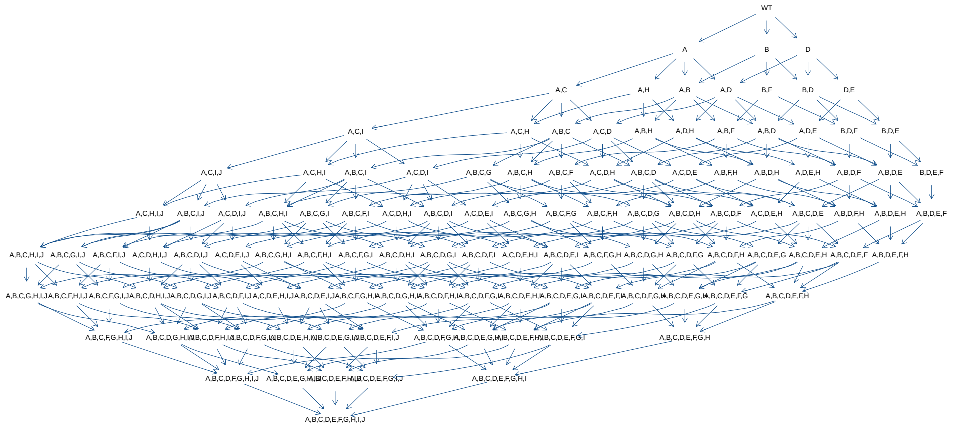

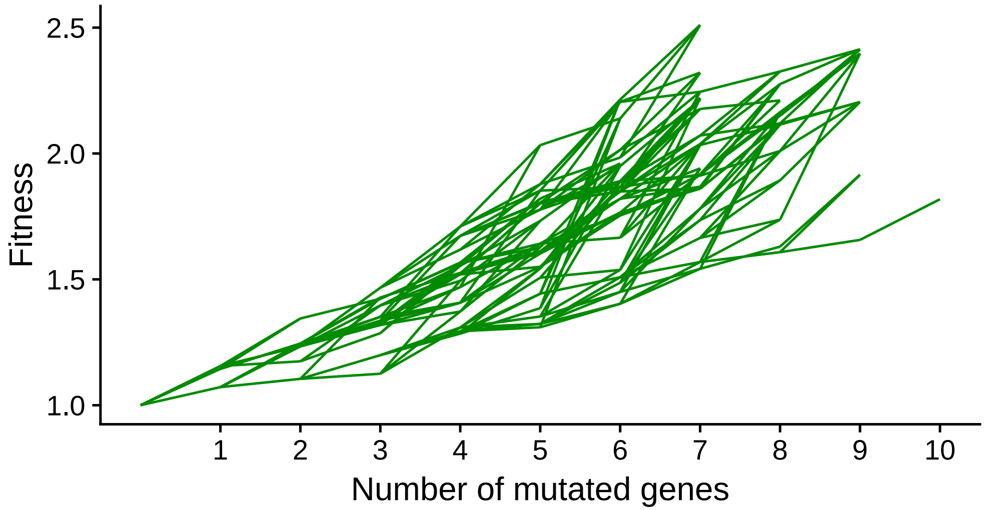


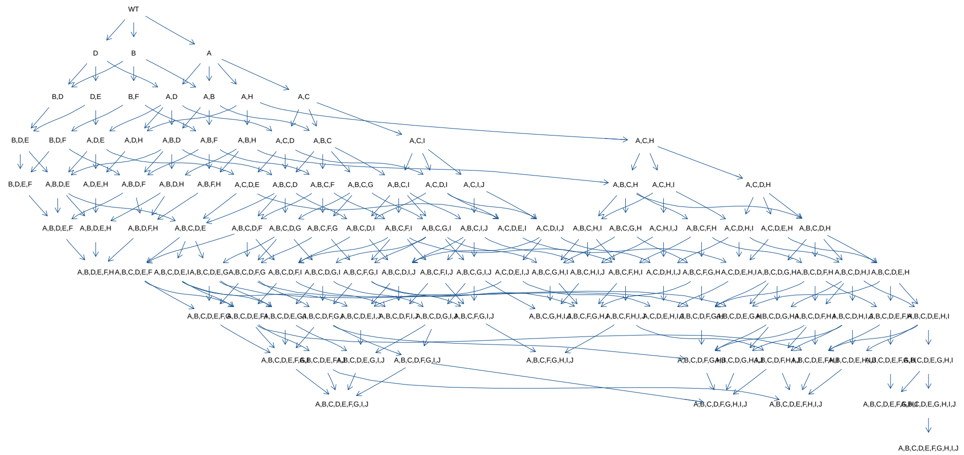

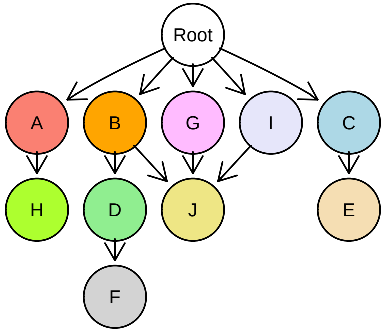


| | |
|---|---|
| *ID* | 69GLOdUeTxOq6MVW |
| *acc genotypes* | 171 |
| *peaks no back* | 13 |
| *removed edges* | 102 |
| *prop rem edges* | 0.194 |

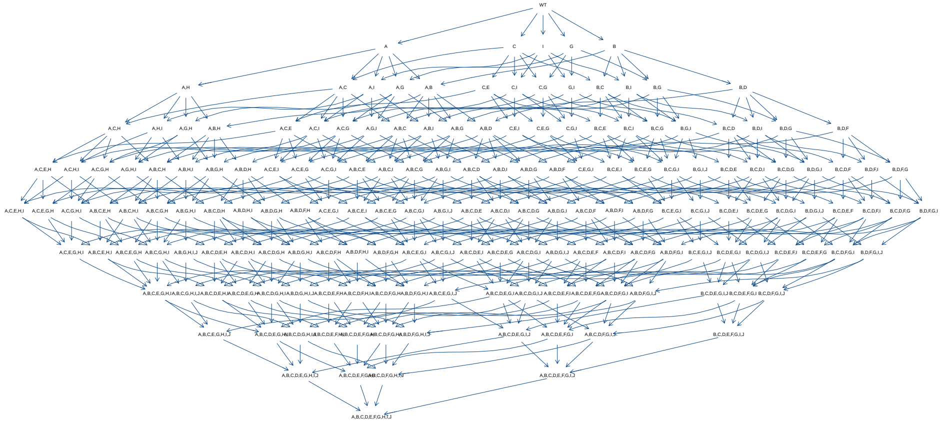

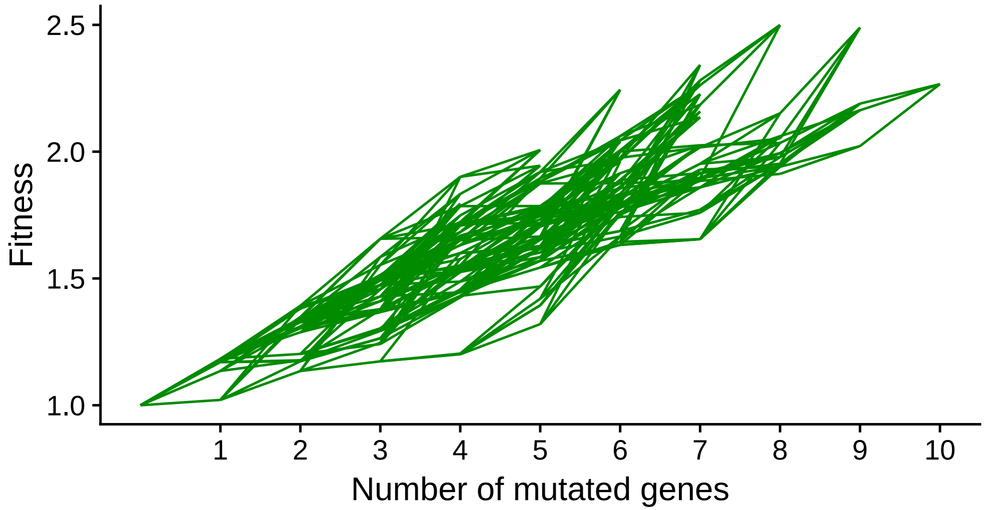


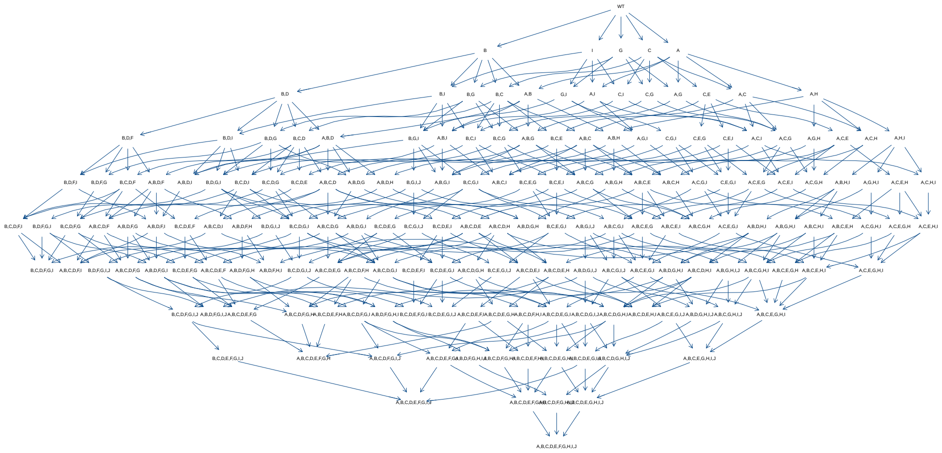

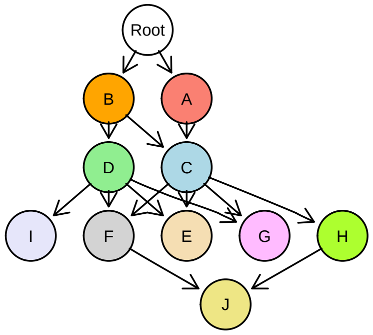


| | |
|---|---|
| *ID* | 6dNxcxBCKj6SRe7c |
| *acc genotypes* | 50 |
| *peaks no back* | 7 |
| *removed edges* | 19 |
| *prop rem edges* | 0.164 |

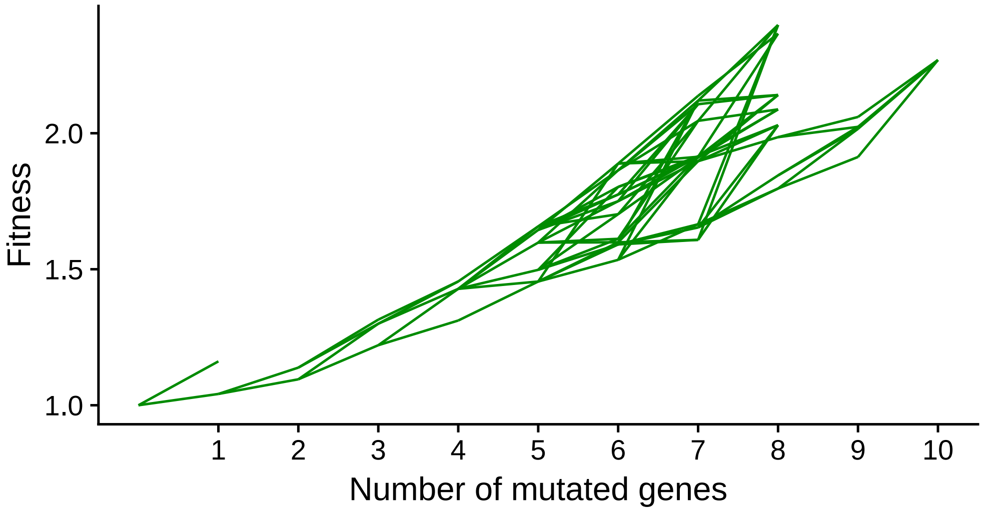


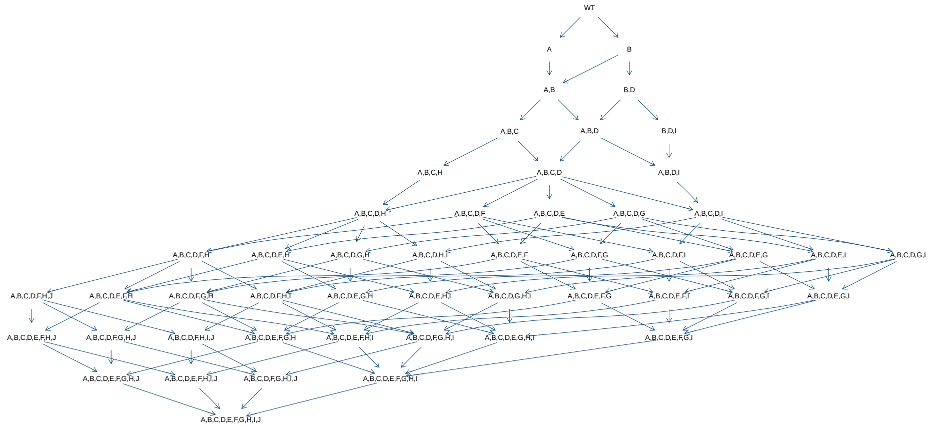

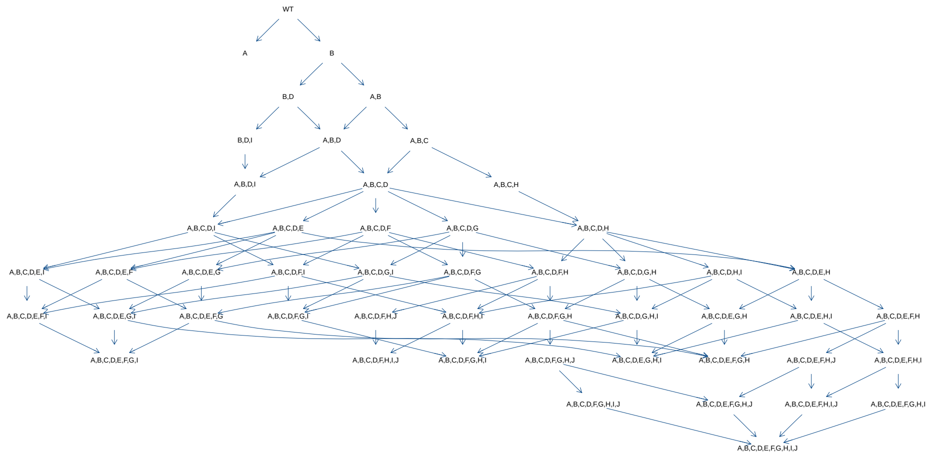

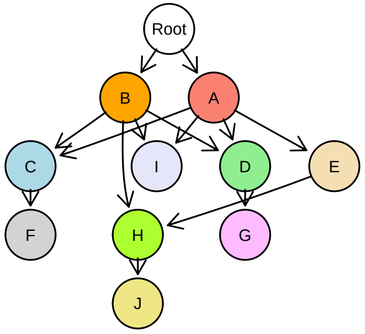


| | |
|---|---|
| *ID* | 6faIe6UPTkVnVhjC |
| *acc genotypes* | 76 |
| *peaks no back* | 7 |
| *removed edges* | 24 |
| *prop rem edges* | 0.125 |

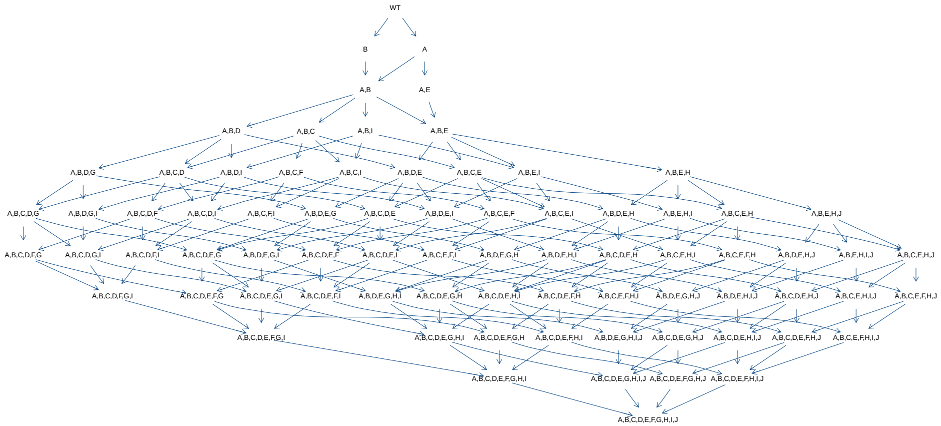

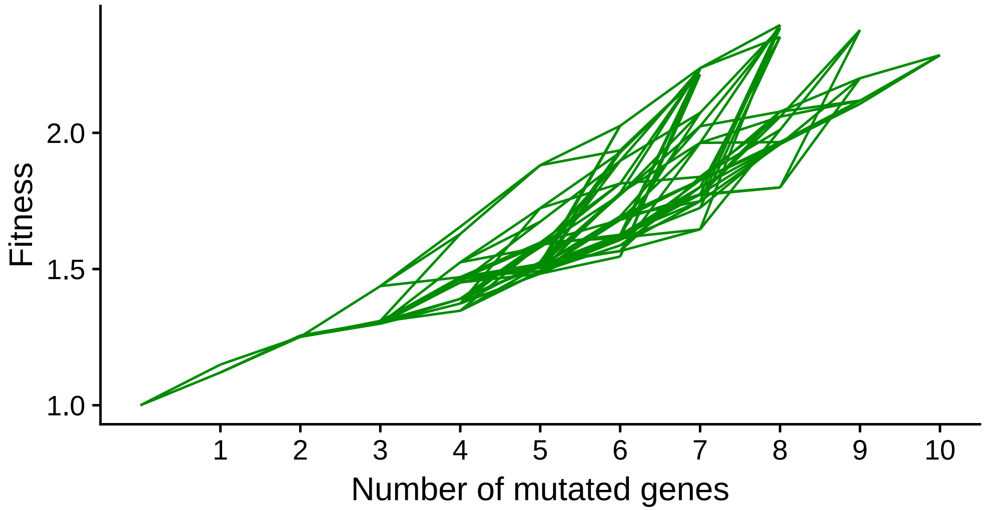


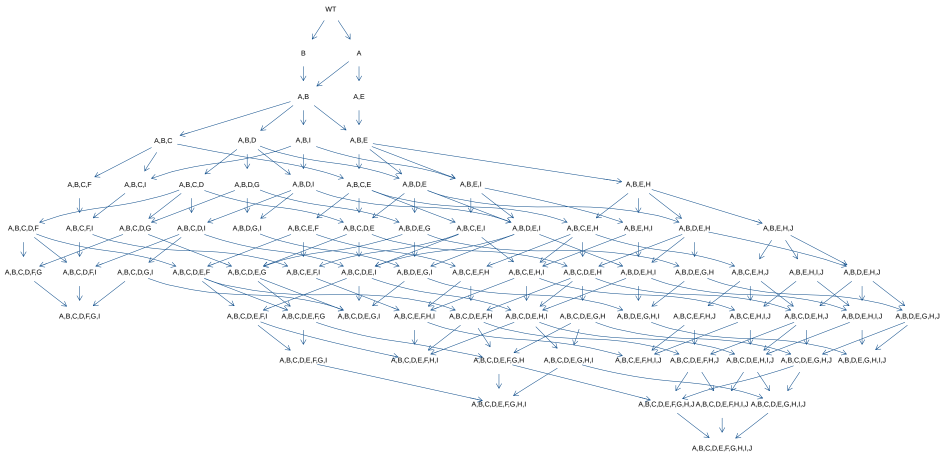

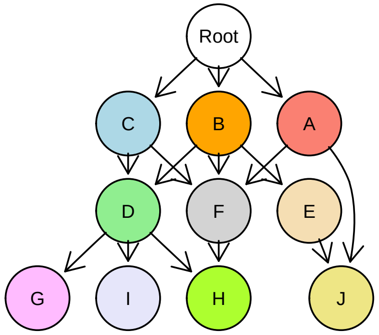


| | |
|---|---|
| *ID* | 6j2DyMAhErFeSJJe |
| *acc genotypes* | 61 |
| *peaks no back* | 9 |
| *removed edges* | 26 |
| *prop rem edges* | 0.186 |

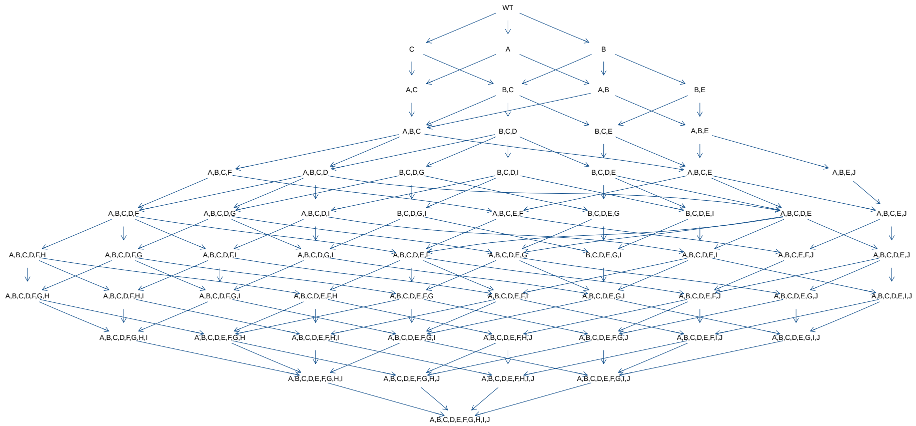

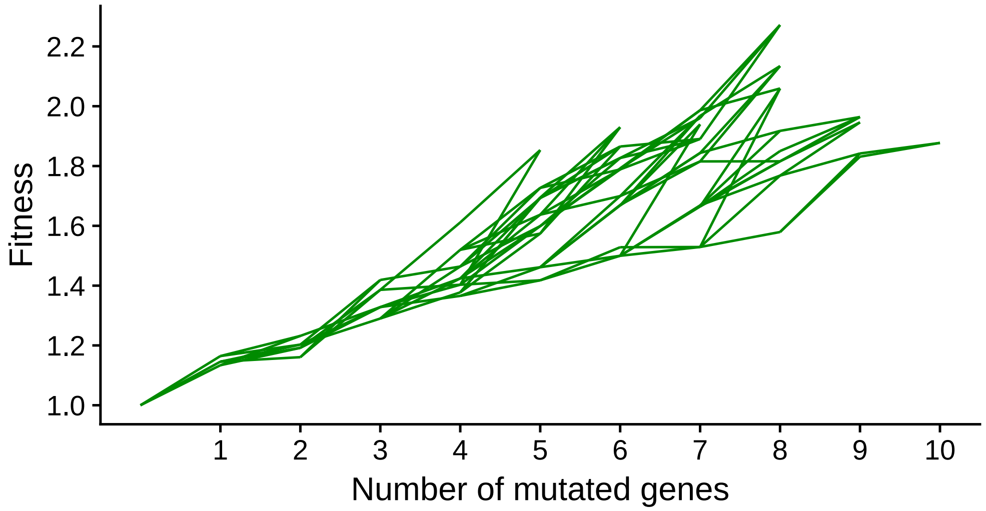


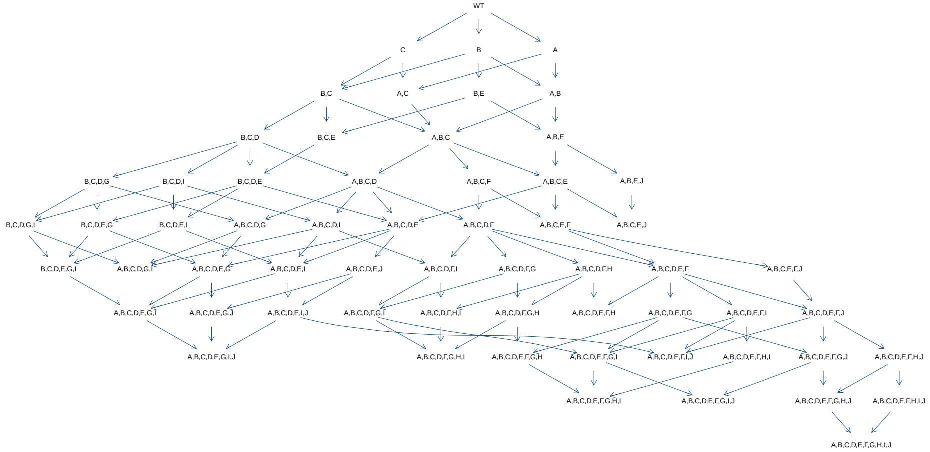

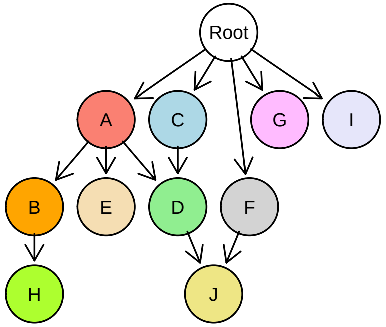


| | |
|---|---|
| *ID* | 7wUPQMRQPYFpPVSq |
| *acc genotypes* | 184 |
| *peaks no back* | 17 |
| *removed edges* | 135 |
| *prop rem edges* | 0.224 |

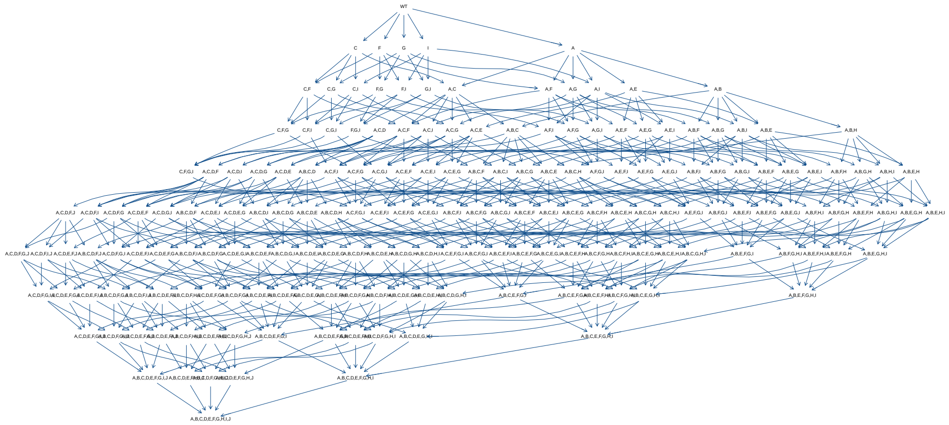

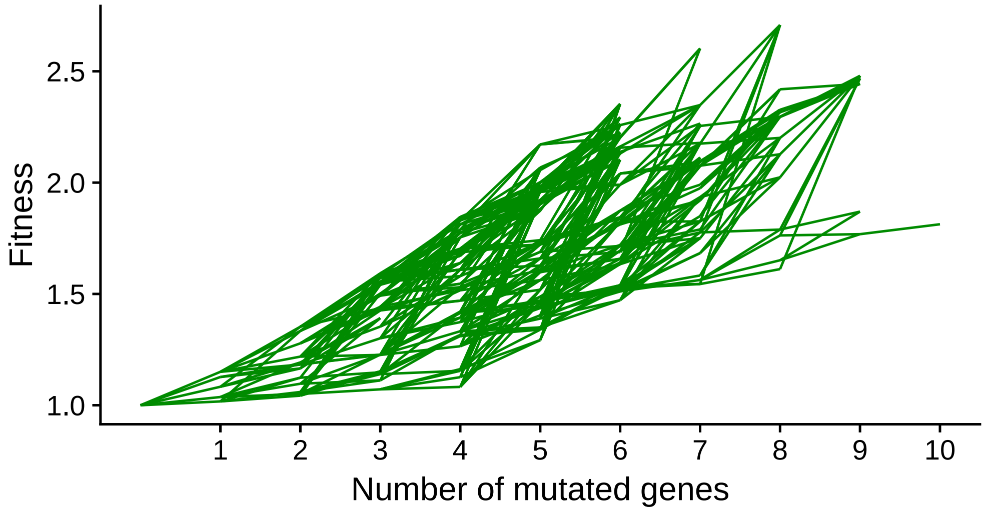


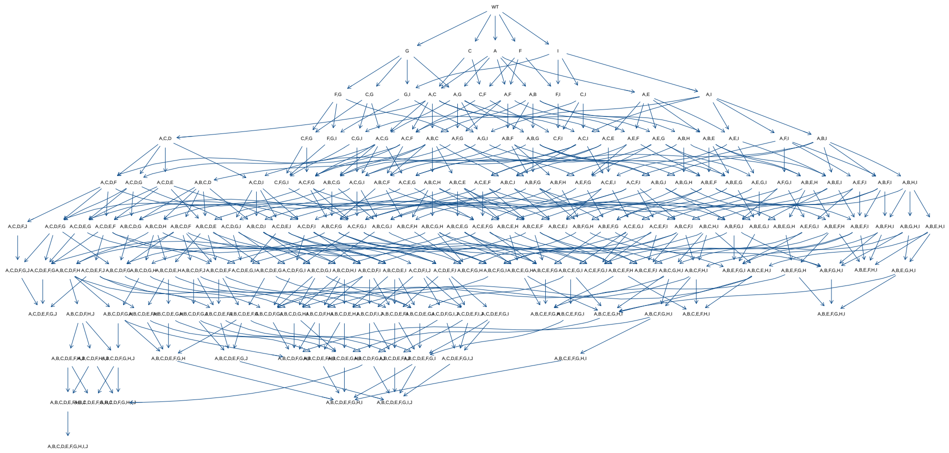

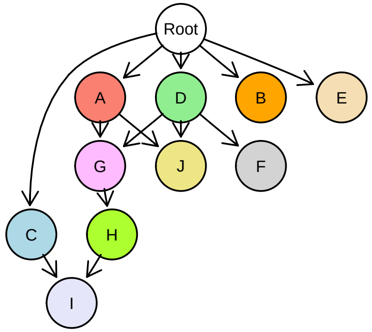


| | |
|---|---|
| *ID* | 87fcnp96NbP7zN7N |
| *acc genotypes* | 144 |
| *peaks no back* | 13 |
| *removed edges* | 63 |
| *prop rem edges* | 0.141 |

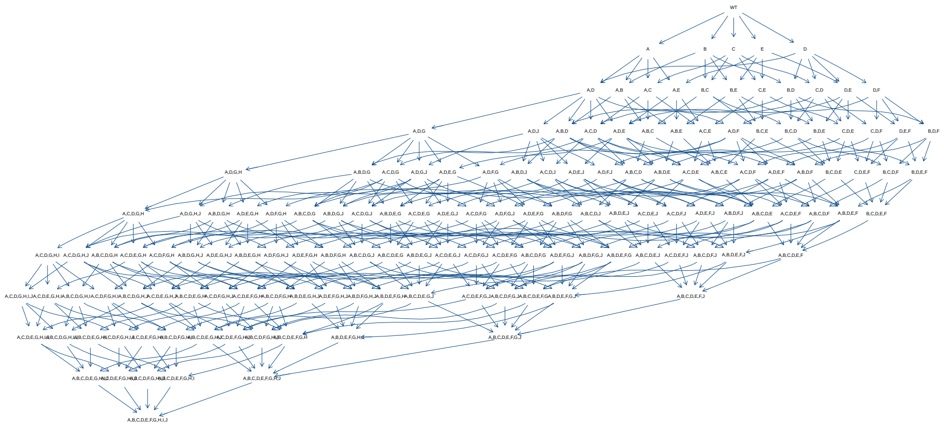

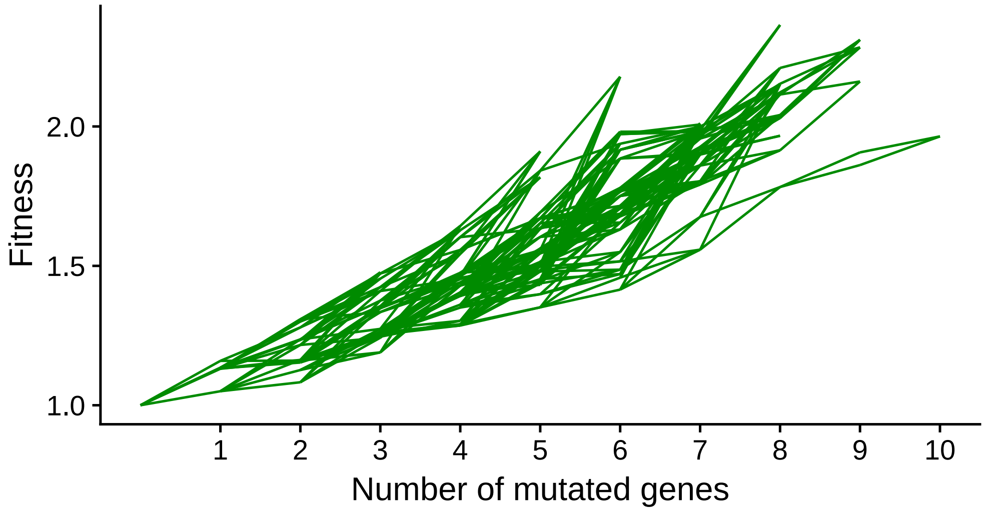


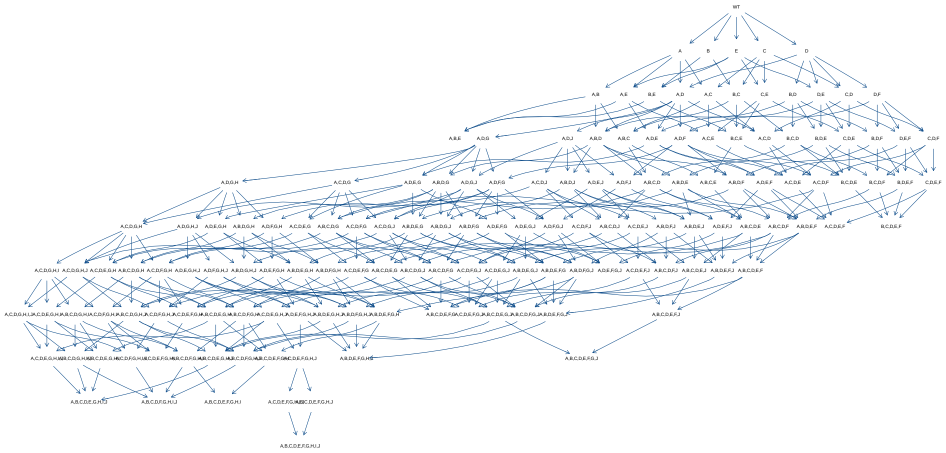

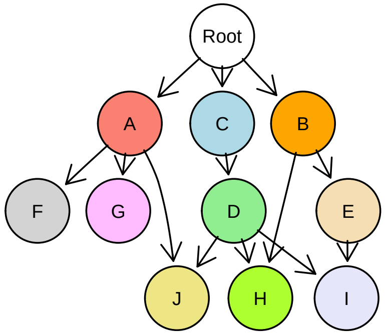


| | |
|---|---|
| *ID* | 8GlpgW8kSqHXEOja |
| *acc genotypes* | 93 |
| *peaks no back* | 9 |
| *removed edges* | 40 |
| *prop rem edges* | 0.165 |

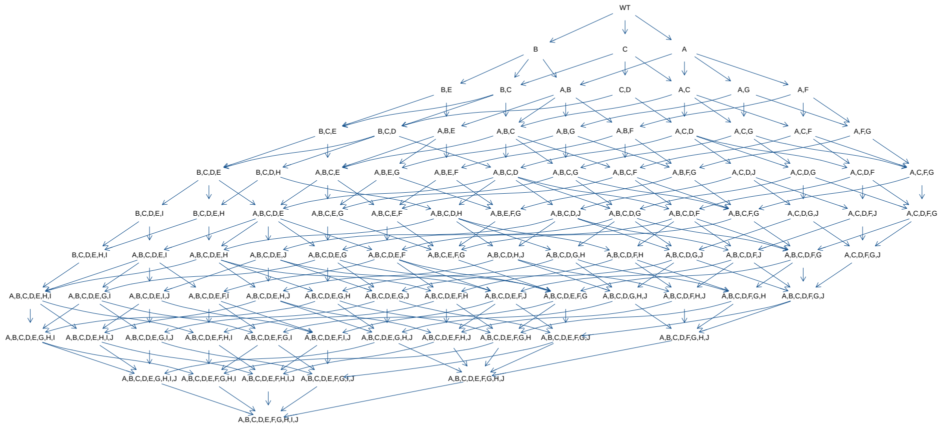

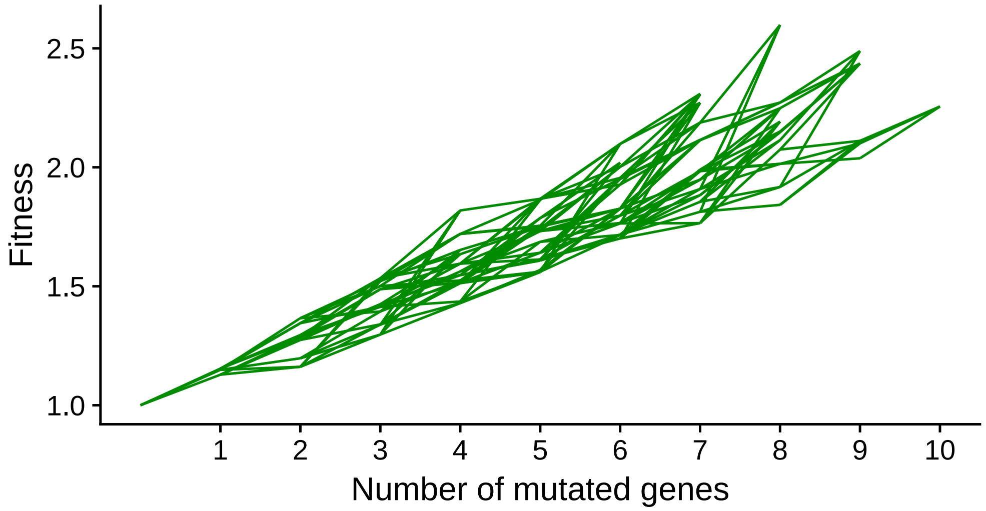


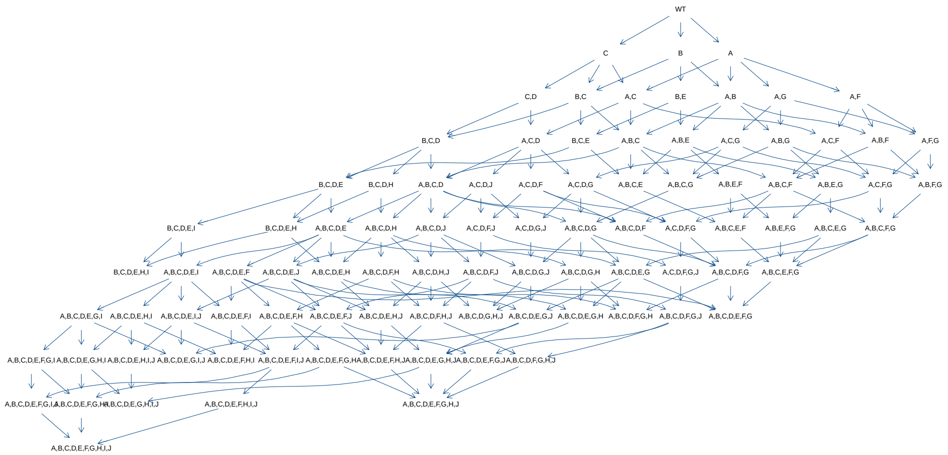

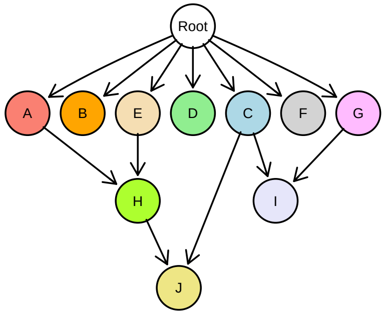


| | |
|---|---|
| *ID* | 8OAYTmnWfxxdHTRV |
| *acc genotypes* | 224 |
| *peaks no back* | 18 |
| *removed edges* | 163 |
| *prop rem edges* | 0.21 |

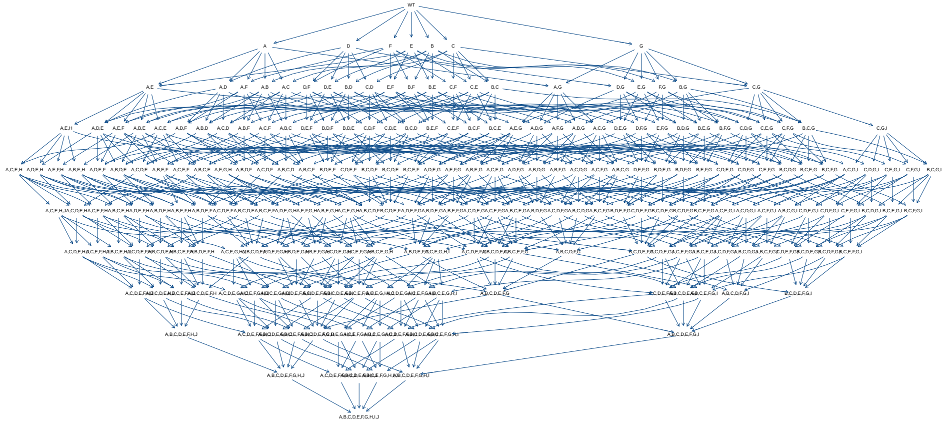

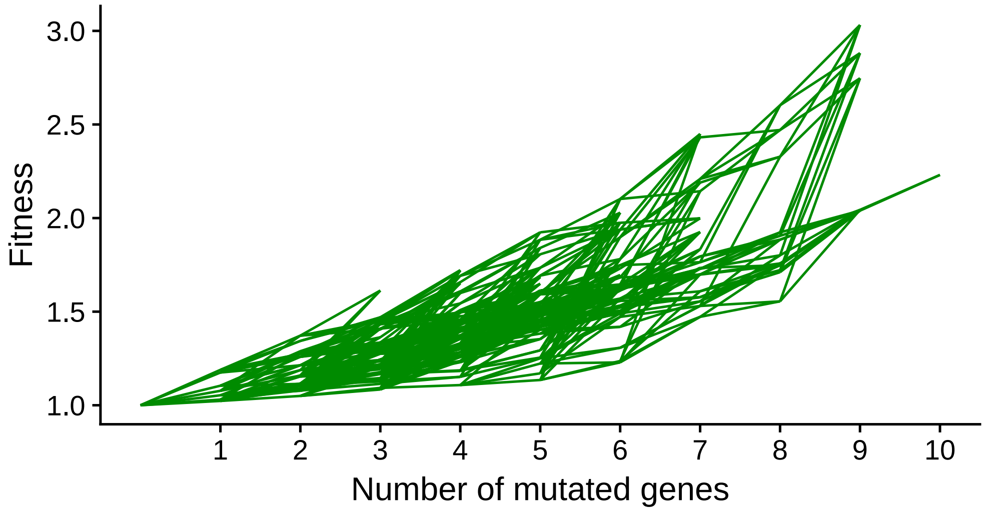


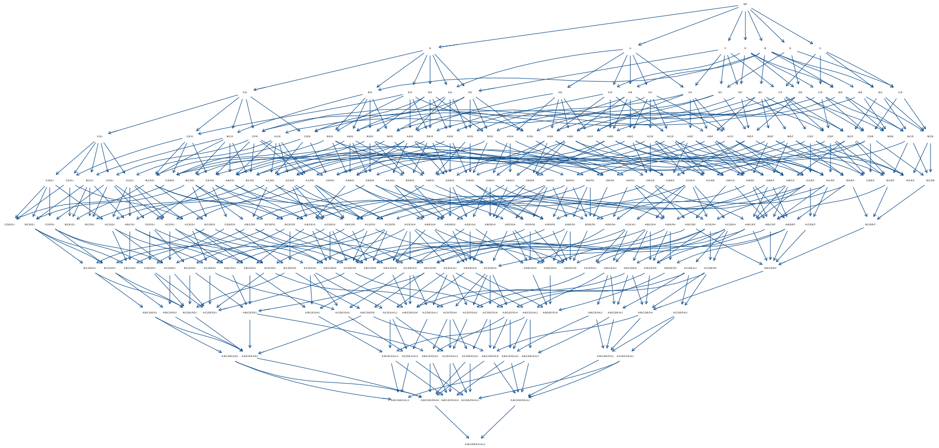

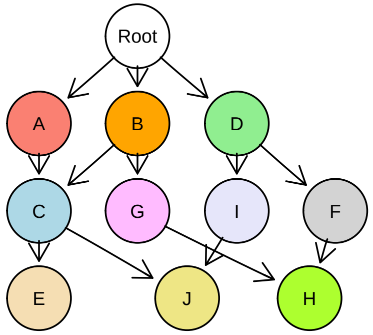


| | |
|---|---|
| *ID* | 8pBC5Dt22yGwrF10I |
| *acc genotypes* | 68 |
| *peaks no back* | 2 |
| *removed edges* | 15 |
| *prop rem edges* | 0.095 |

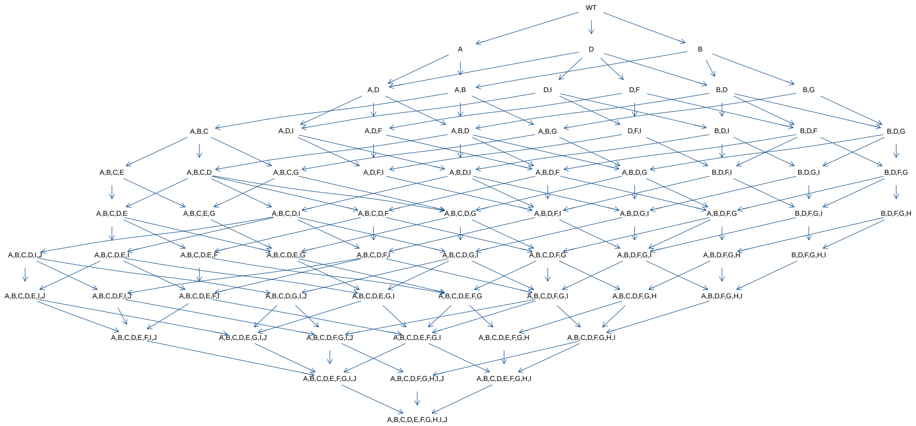

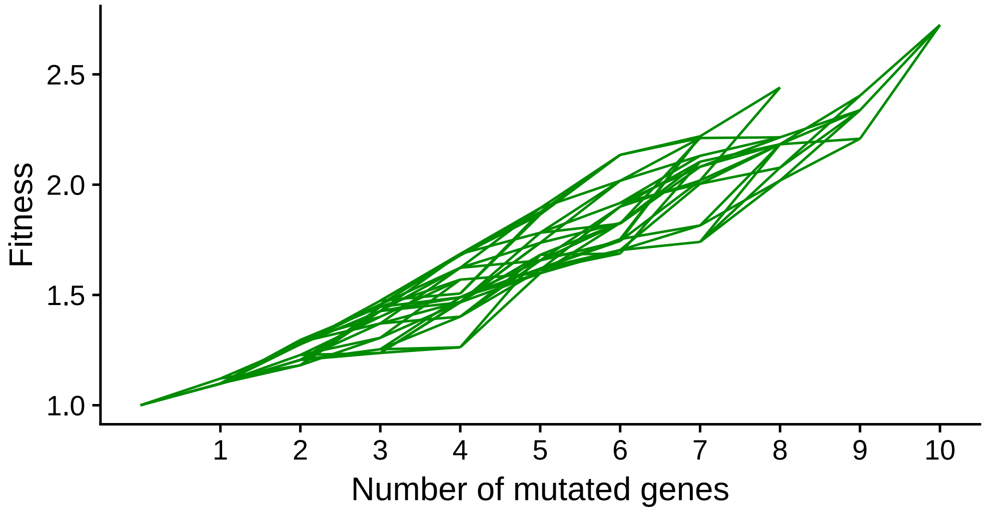


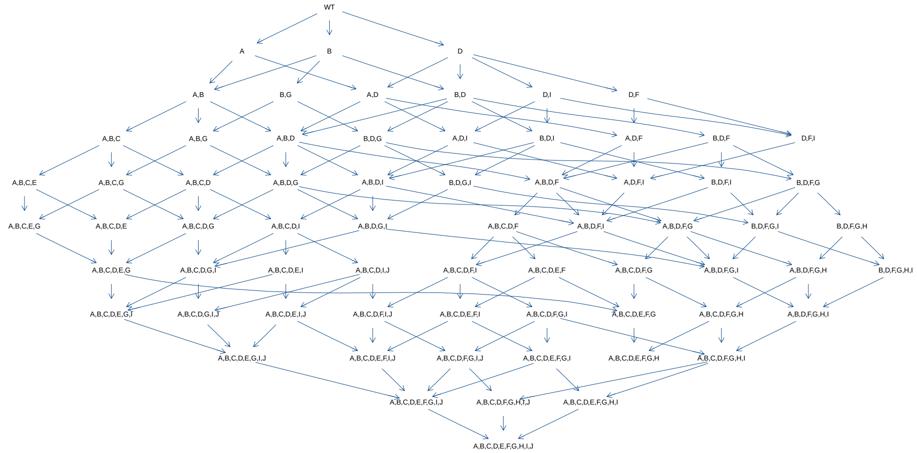

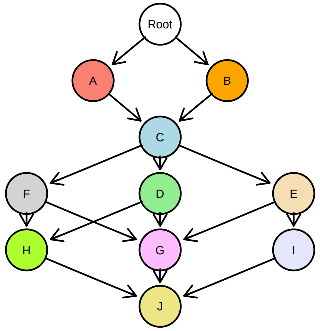


| | |
|---|---|
| *ID* | 8zRUVMJqJUMsBBkl |
| *acc genotypes* | 24 |
| *peaks no back* | 4 |
| *removed edges* | 3 |
| *prop rem edges* | 0.077 |

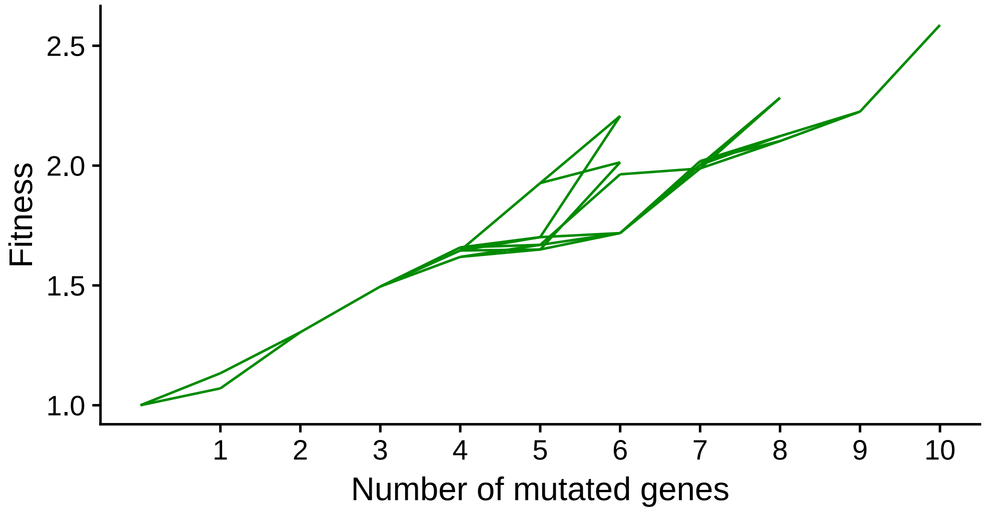


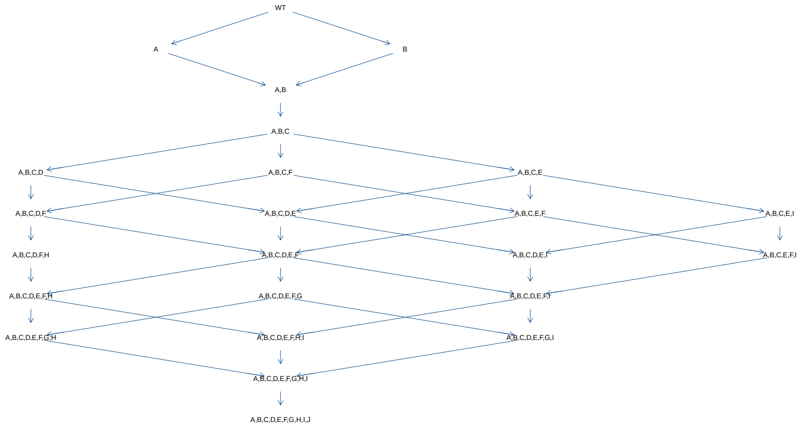

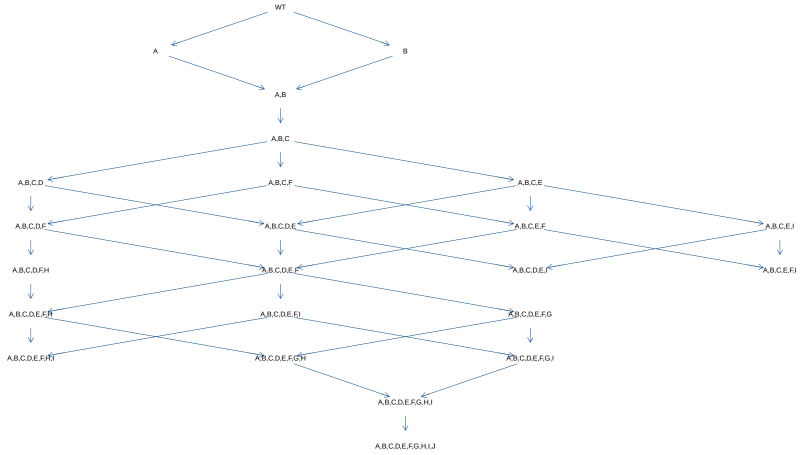

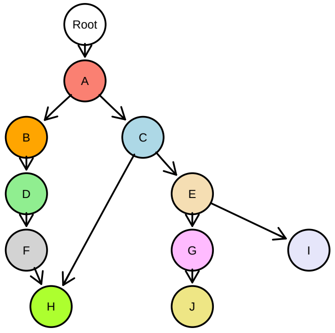


| | |
|---|---|
| *ID* | 8ZUcIGXESgYkqacc |
| *acc genotypes* | 40 |
| *peaks no back* | 6 |
| *removed edges* | 13 |
| *prop rem edges* | 0.171 |

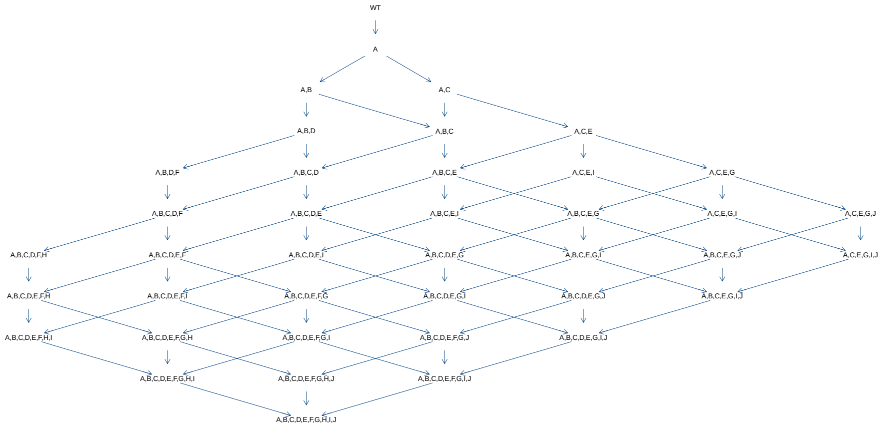

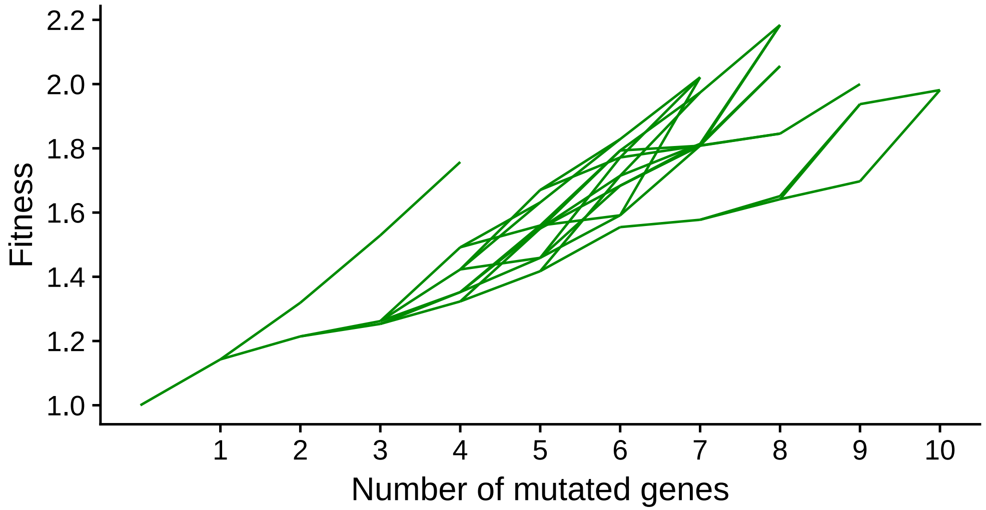


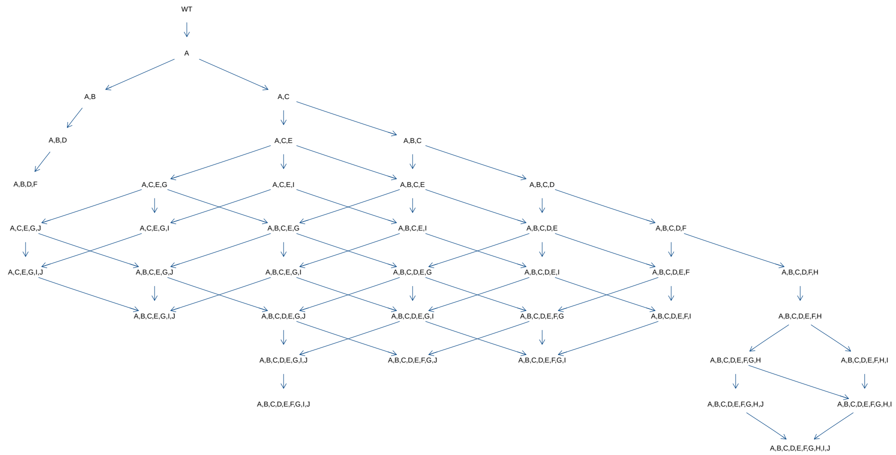

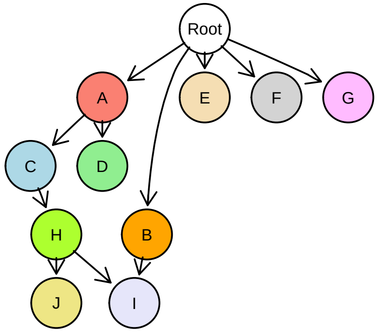


*ID* 92mZfe3axm10pyB10O

*acc genotypes* 176

*peaks no back* 15

*removed edges* 108

*prop rem edges* 0.188

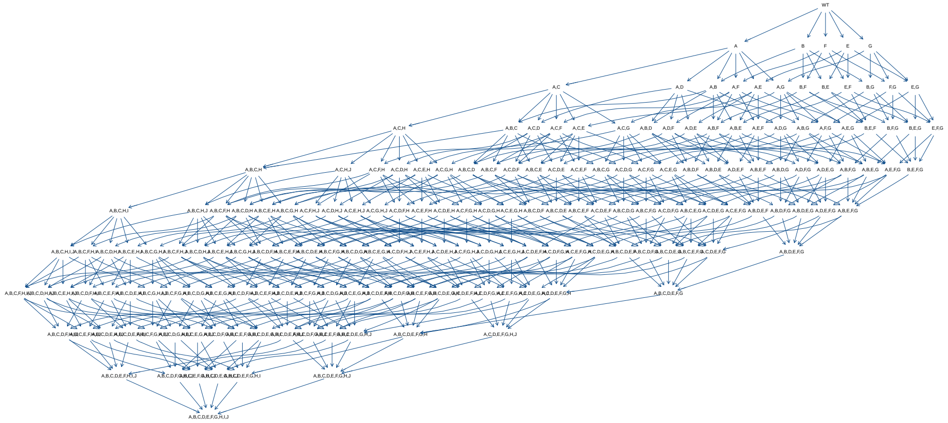

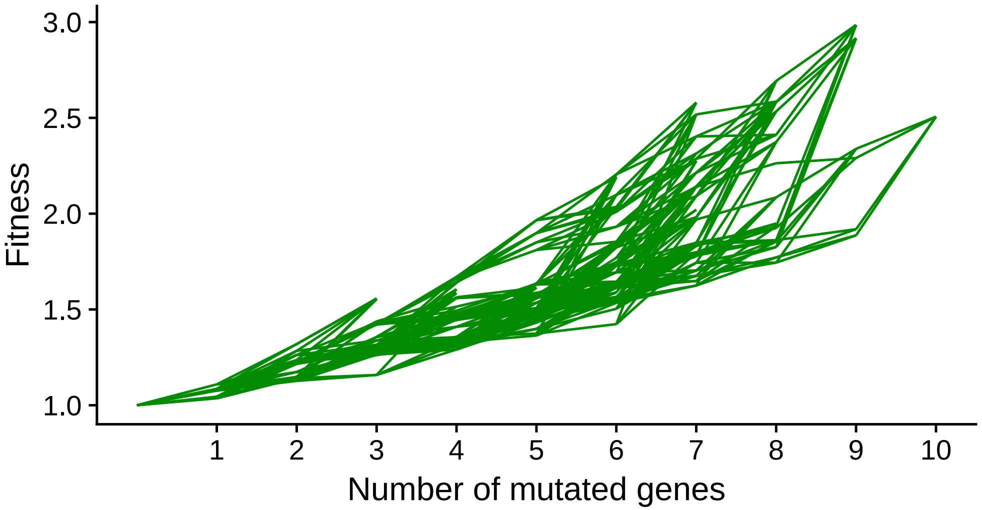


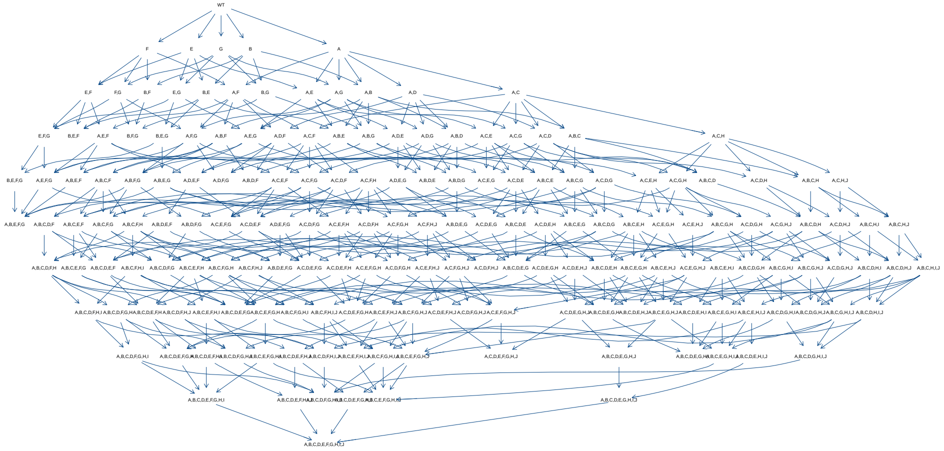

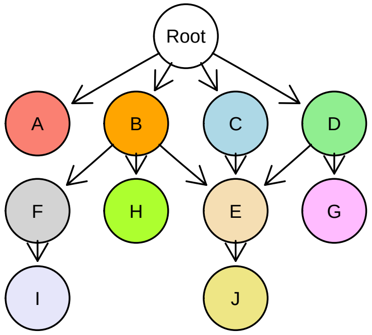


| | |
|---|---|
| *ID* | 9tzPJQwcPFgCvZpS |
| *acc genotypes* | 132 |
| *peaks no back* | 12 |
| *removed edges* | 78 |
| *prop rem edges* | 0.201 |

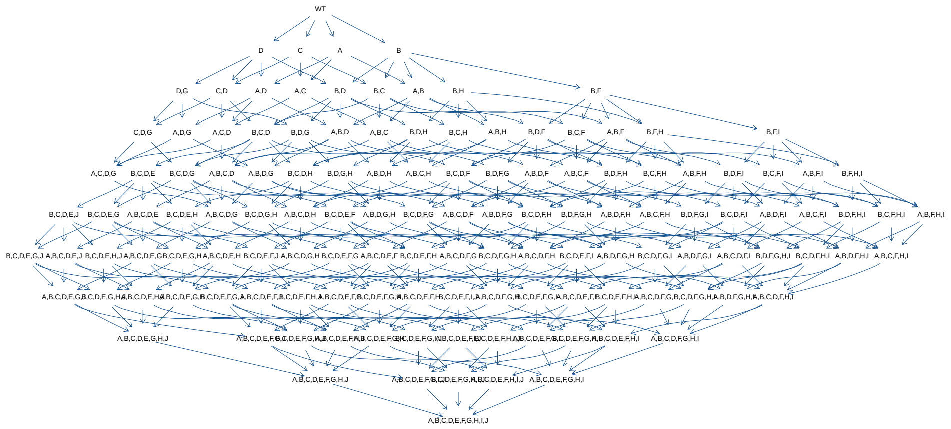

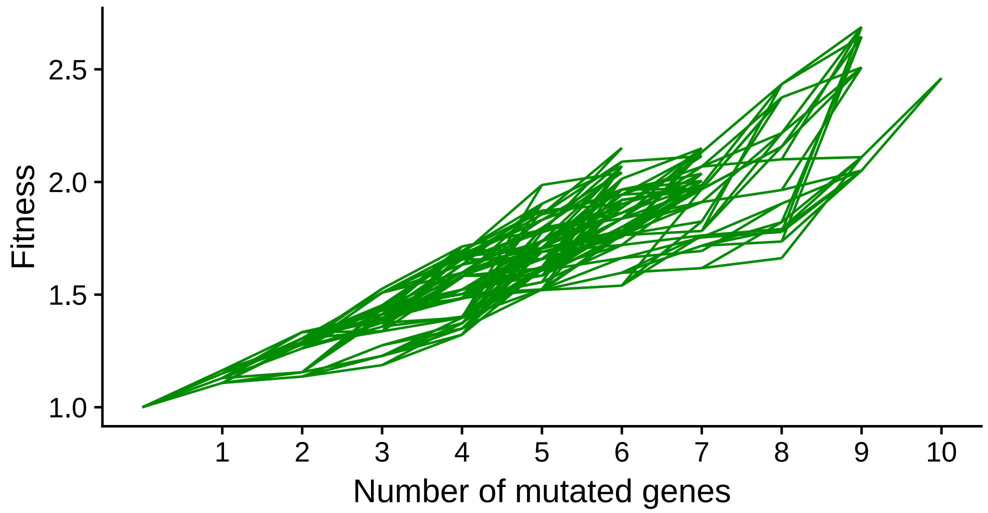


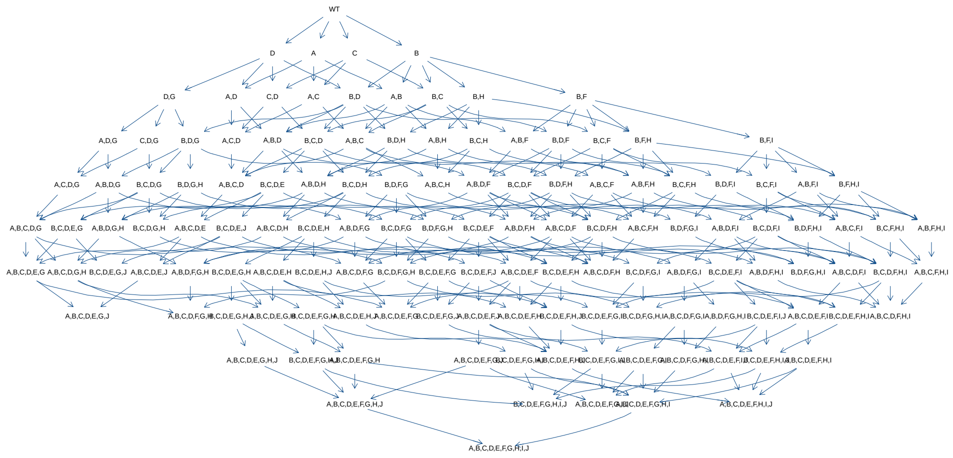

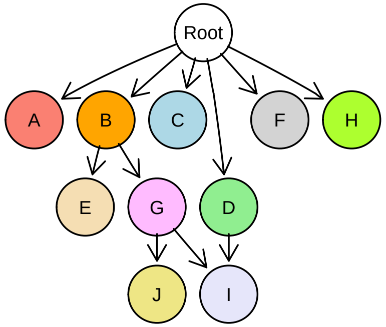


| | |
|---|---|
| *ID* | 9XhdPR4OnXtyF10ls |
| *acc genotypes* | 288 |
| *peaks no back* | 21 |
| *removed edges* | 224 |
| *prop rem edges* | 0.209 |

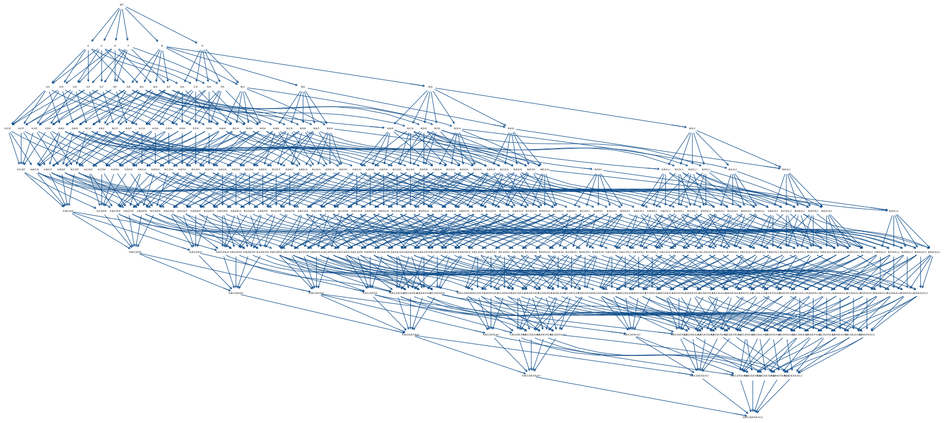

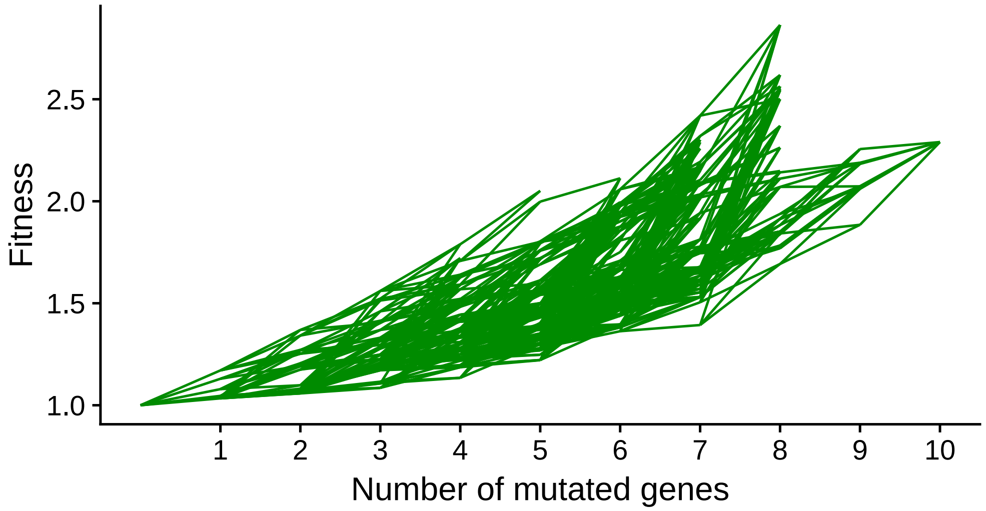


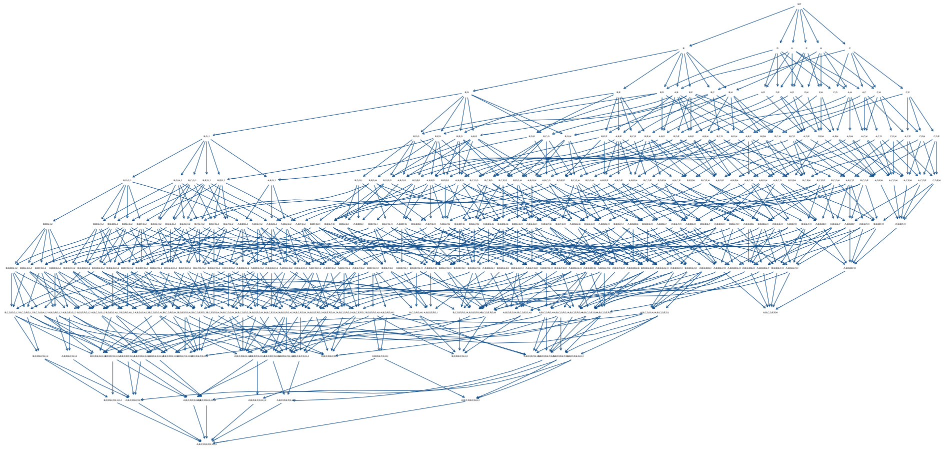

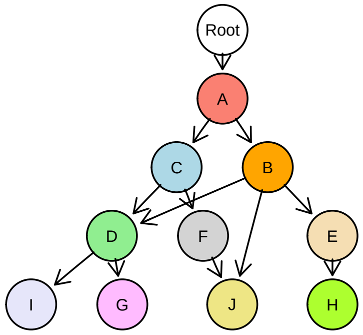


| | |
|---|---|
| *ID* | AAvENuRd7EV7ymhh |
| *acc genotypes* | 52 |
| *peaks no back* | 8 |
| *removed edges* | 21 |
| *prop rem edges* | 0.181 |

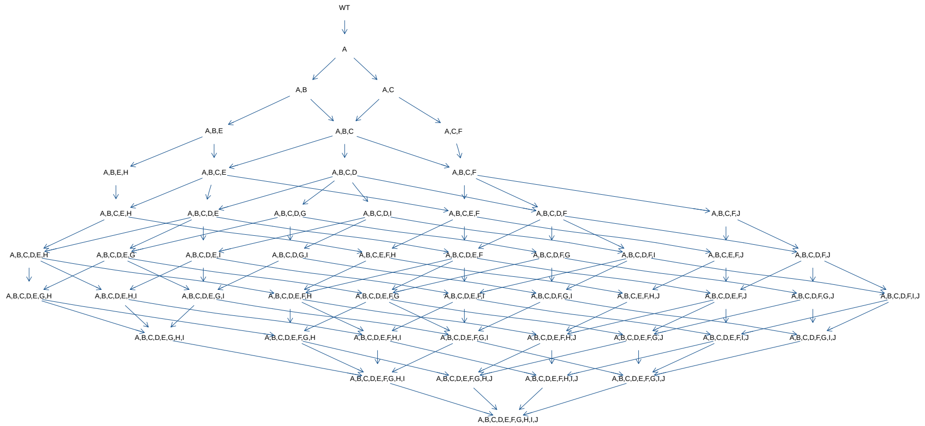

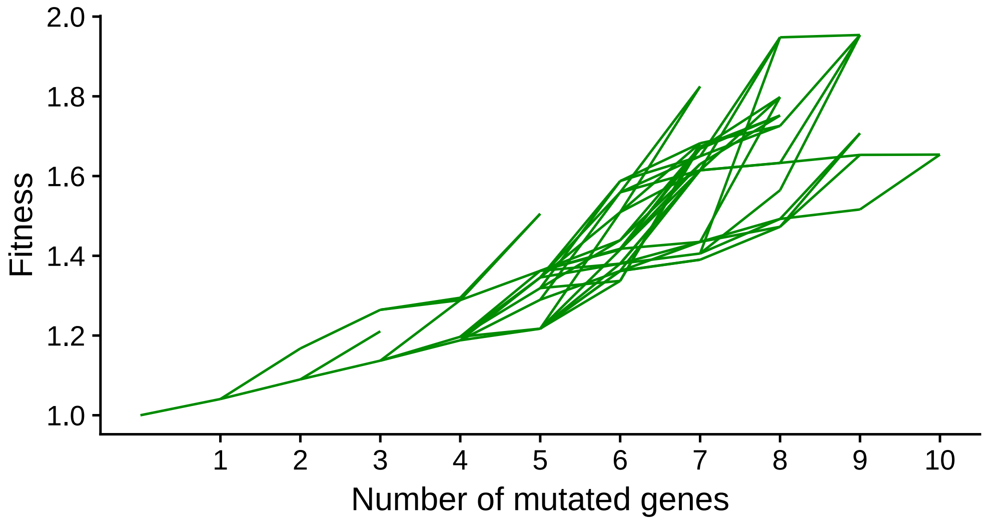


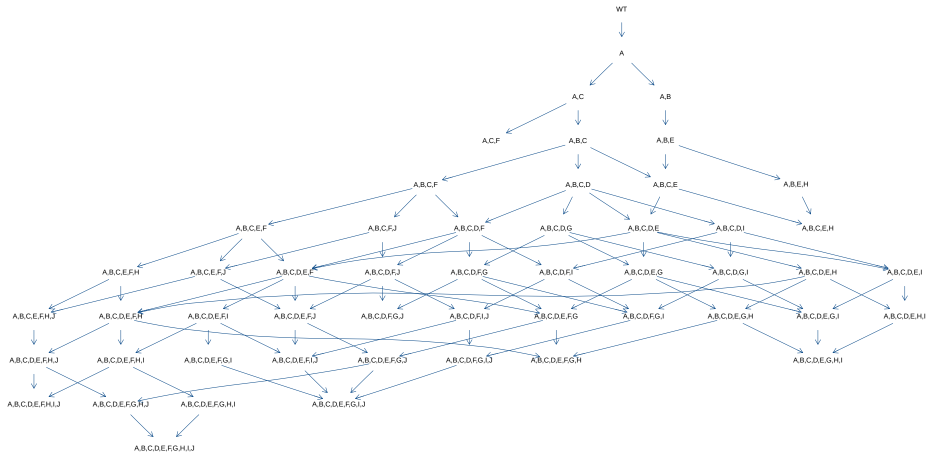

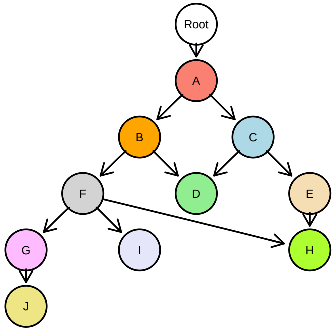


*ID* aeSijOttBLlSw1Z1
*acc genotypes* 51
*peaks no back* 8
*removed edges* 21
*prop rem edges* 0.186

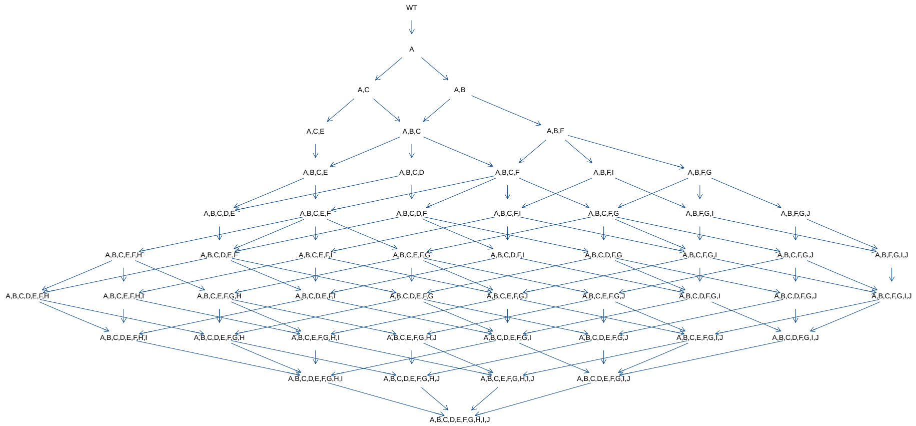

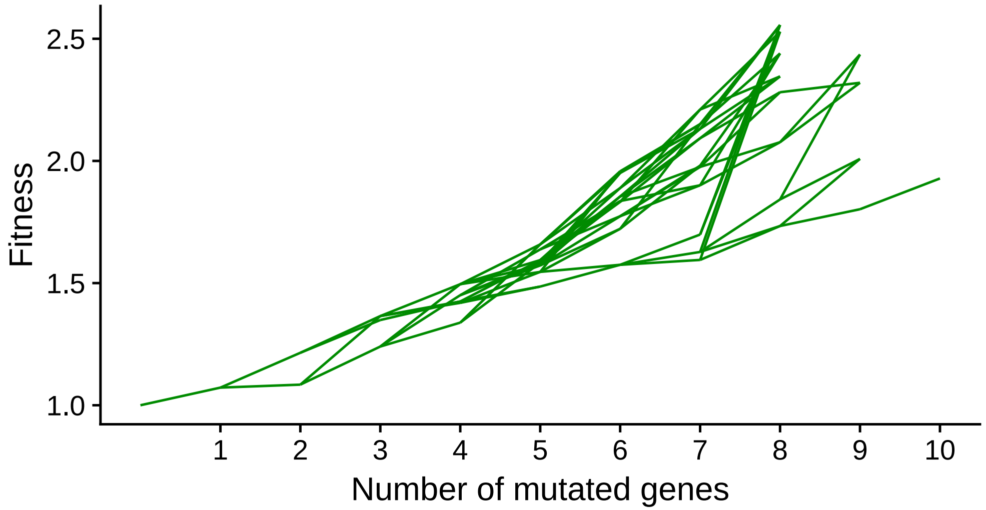


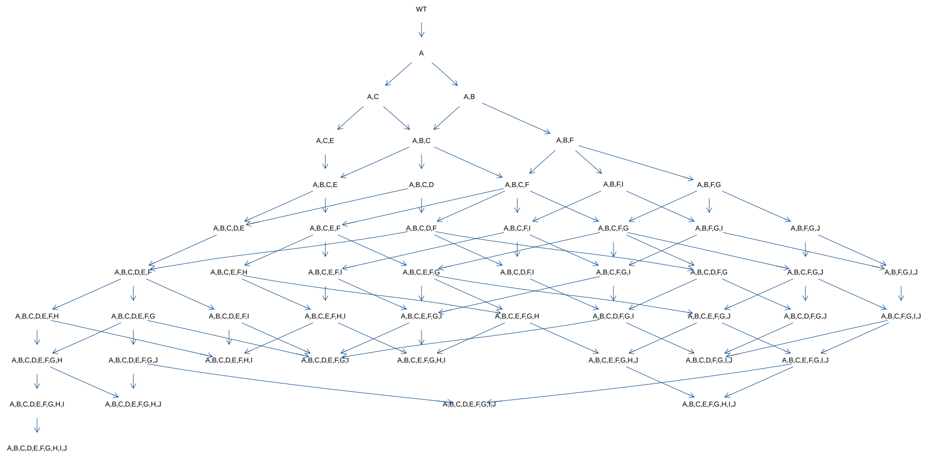

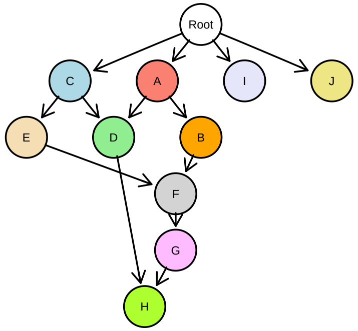


| | |
|---|---|
| *ID* | aFSJF5G7y4DBZWSr |
| *acc genotypes* | 72 |
| *peaks no back* | 8 |
| *removed edges* | 39 |
| *prop rem edges* | 0.217 |

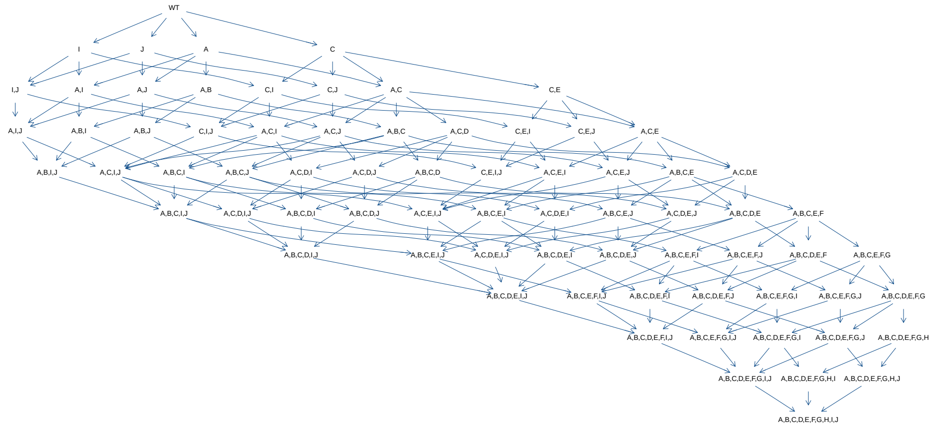

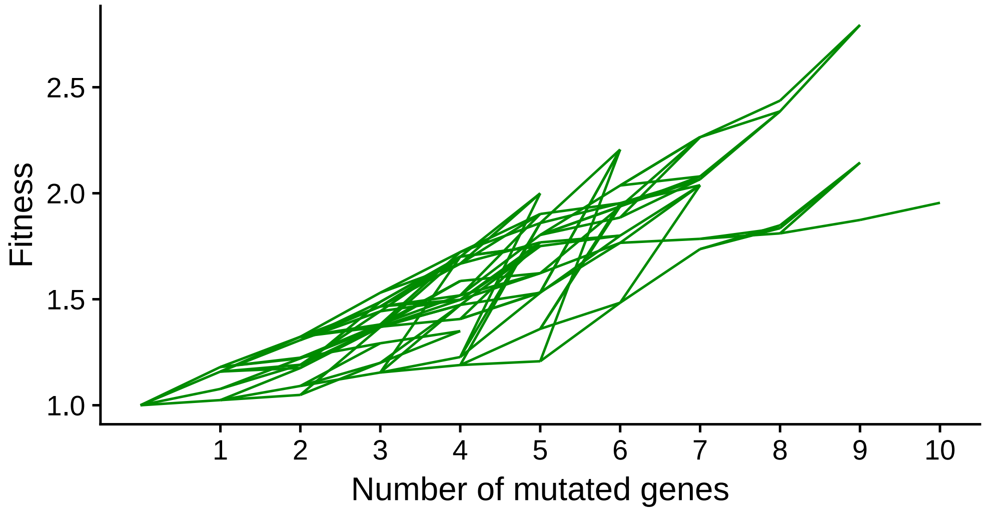


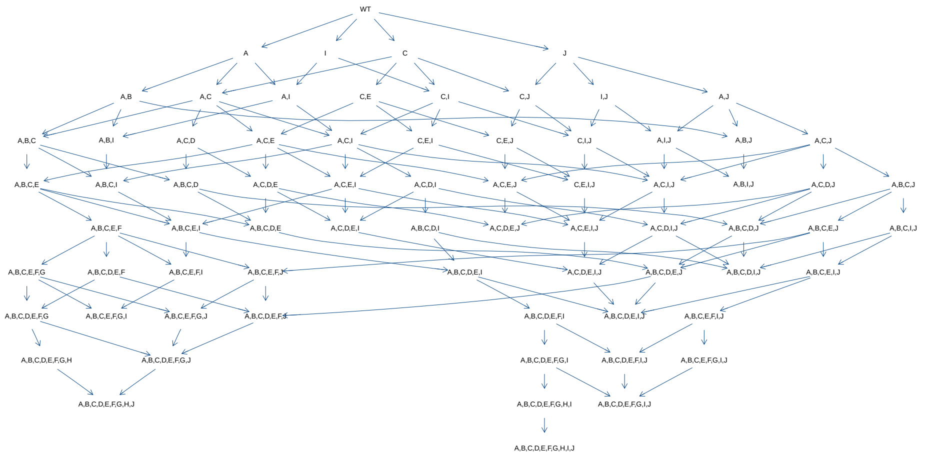

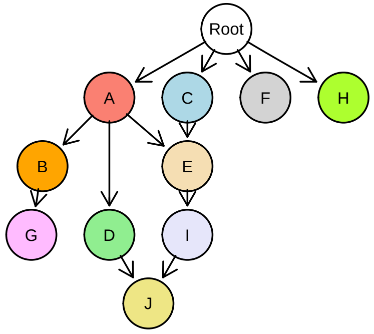


| | |
|---|---|
| *ID* | AjFtq103DguBWTFLW |
| *acc genotypes* | 116 |
| *peaks no back* | 10 |
| *removed edges* | 74 |
| *prop rem edges* | 0.223 |

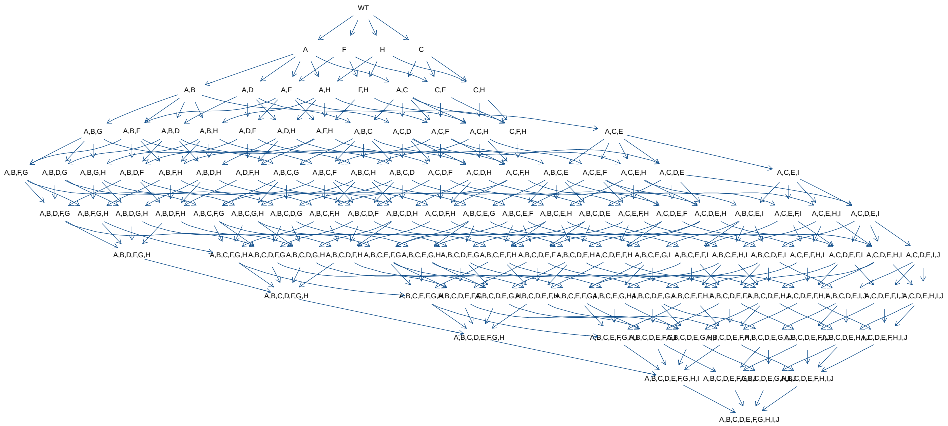

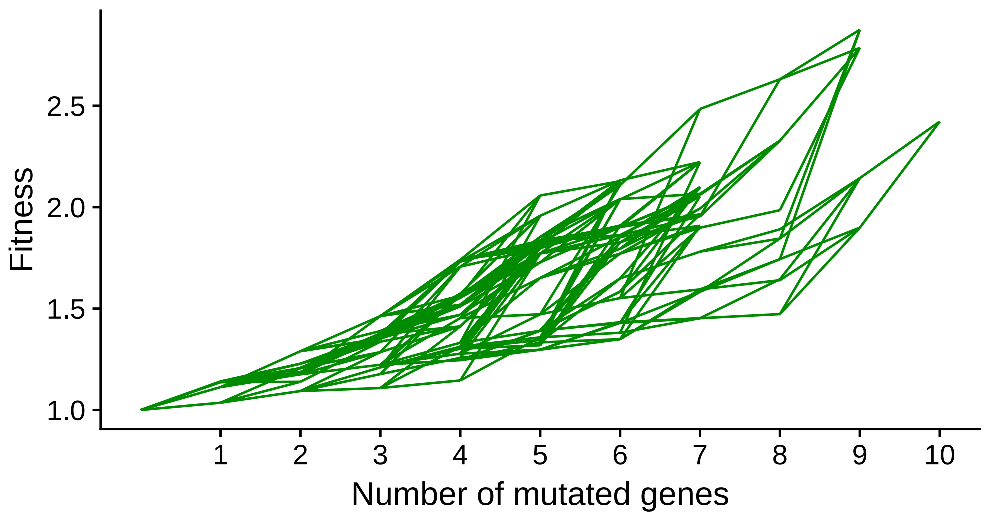


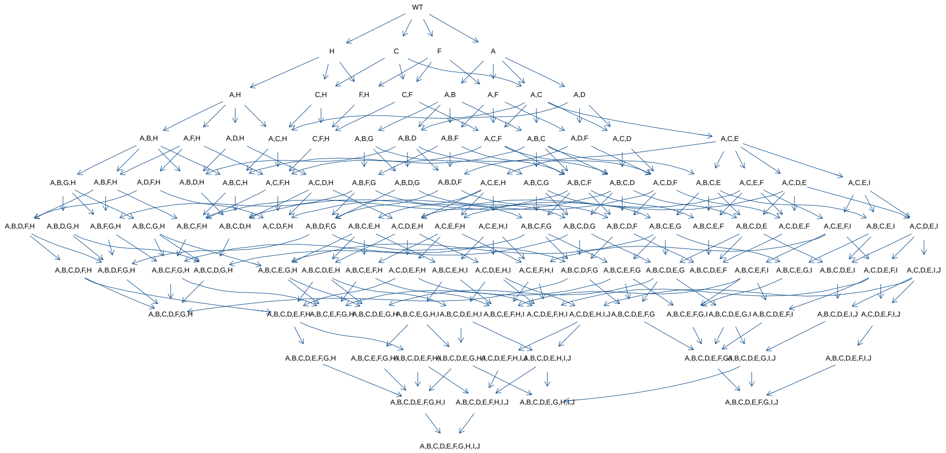

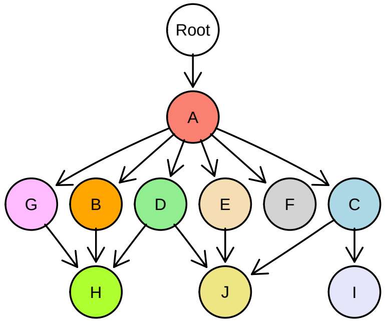


| | |
|---|---|
| *ID* | B10D8cwK5cY9qVLNq |
| *acc genotypes* | 129 |
| *peaks no back* | 13 |
| *removed edges* | 86 |
| *prop rem edges* | 0.217 |

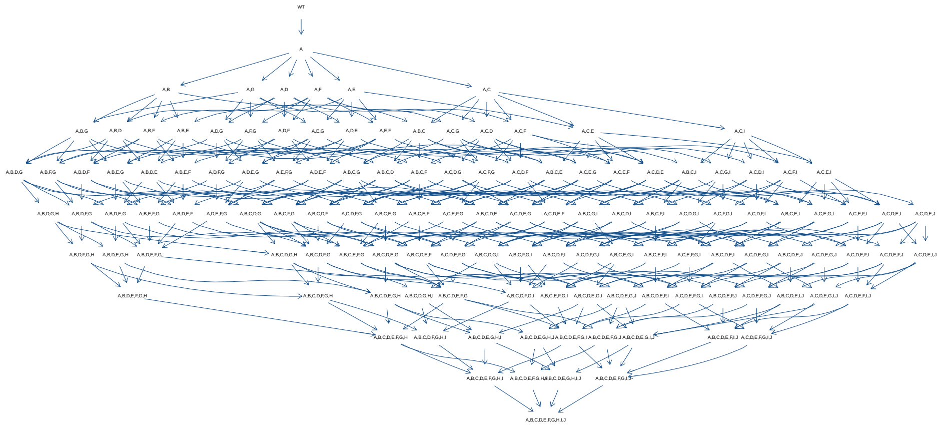

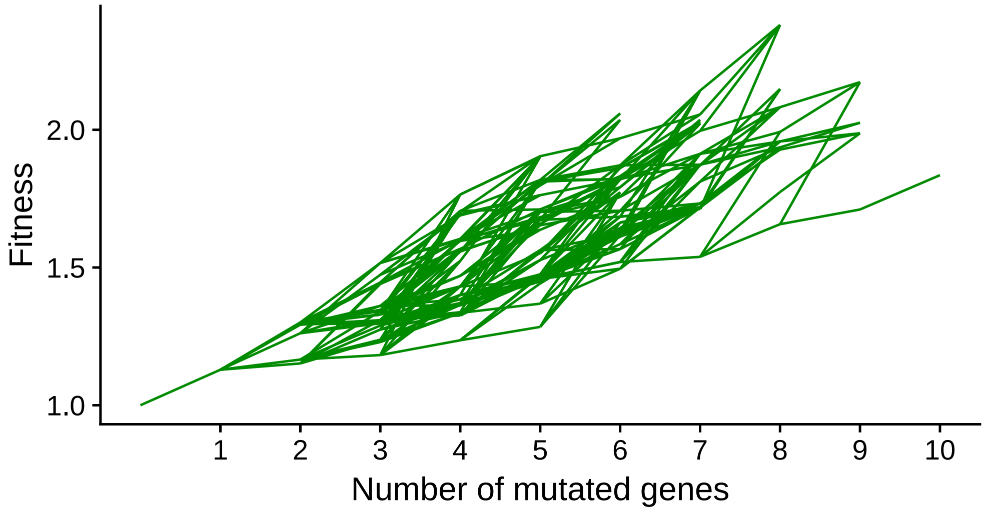


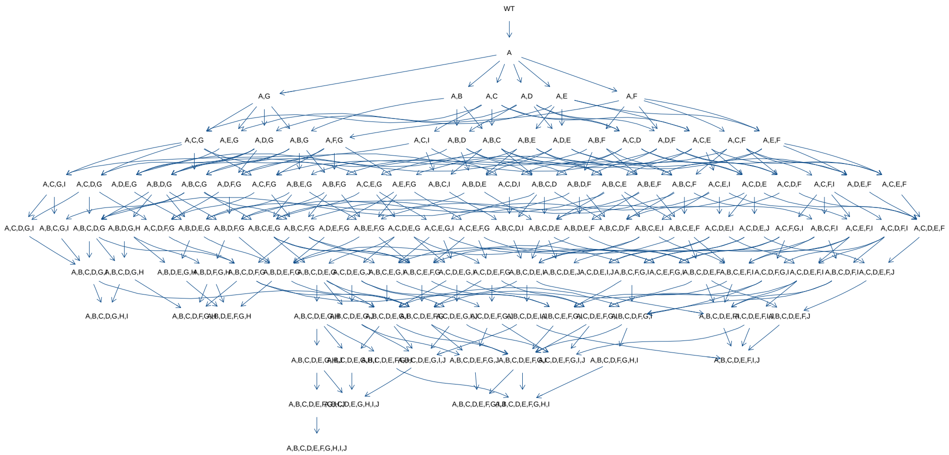

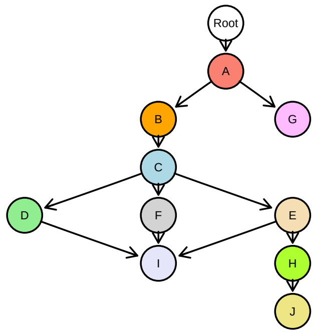


| | |
|---|---|
| *ID* | b2dBTAqsSxgbkkTE |
| *acc genotypes* | 43 |
| *peaks no back* | 7 |
| *removed edges* | 14 |
| *prop rem edges* | 0.152 |

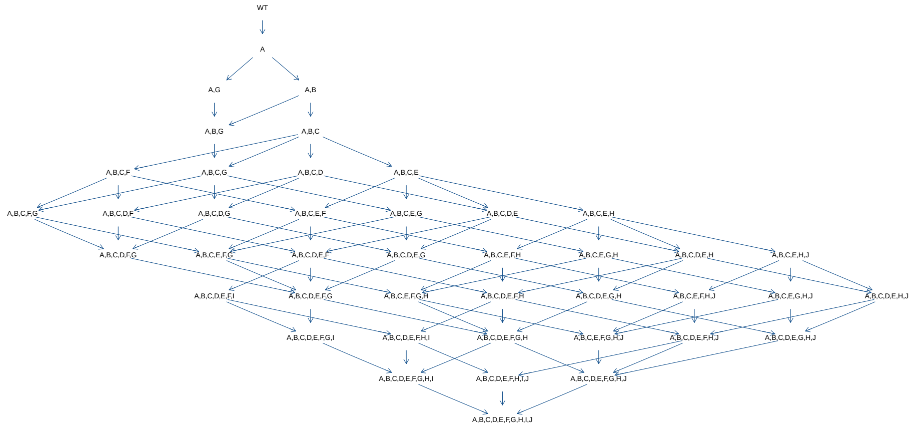

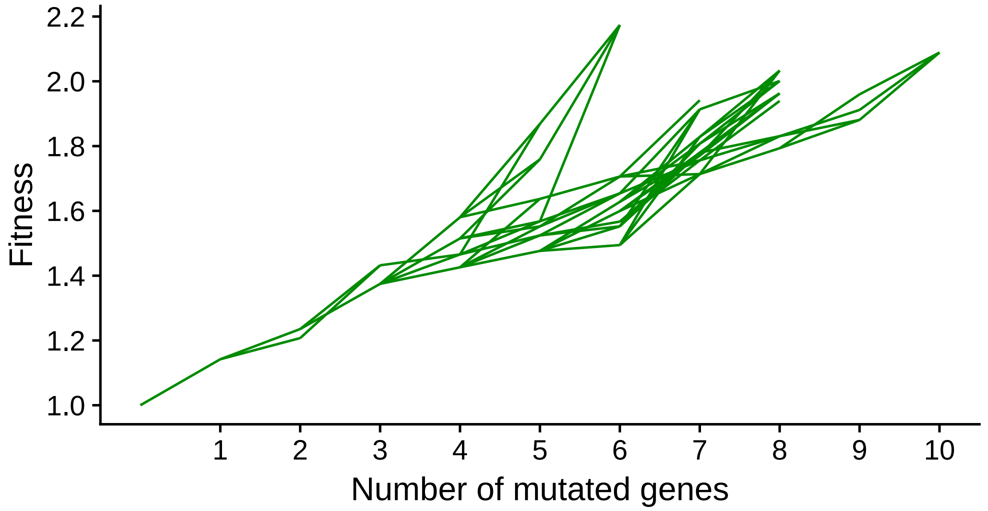


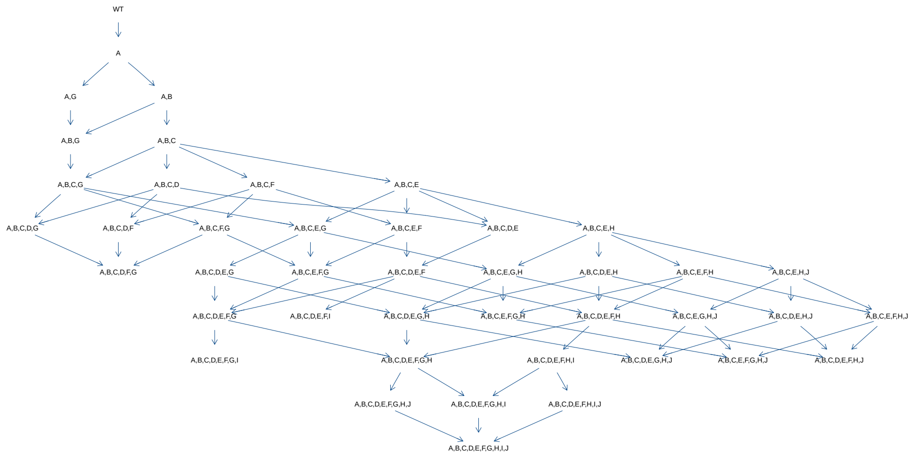

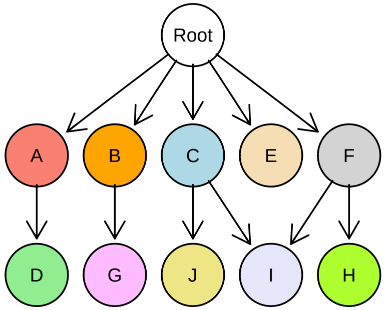


| | |
|---|---|
| *ID* | B8LF4EJa7H2B21J3 |
| *acc genotypes* | 234 |
| *peaks no back* | 14 |
| *removed edges* | 130 |
| *prop rem edges* | 0.165 |

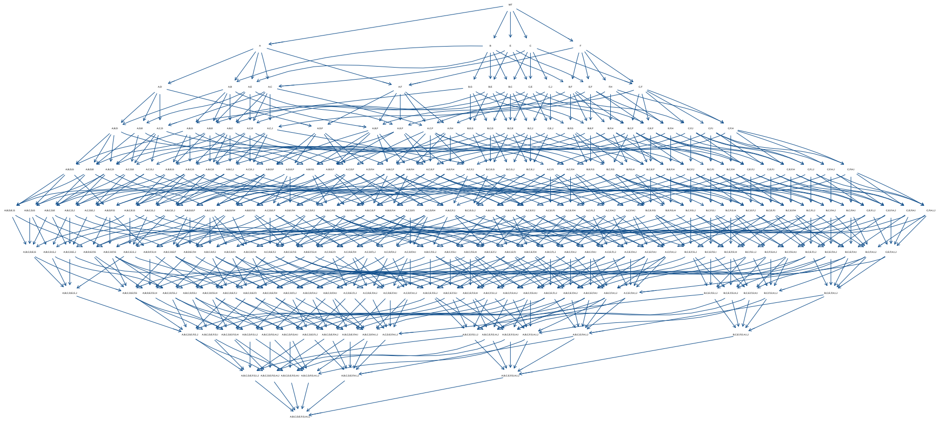

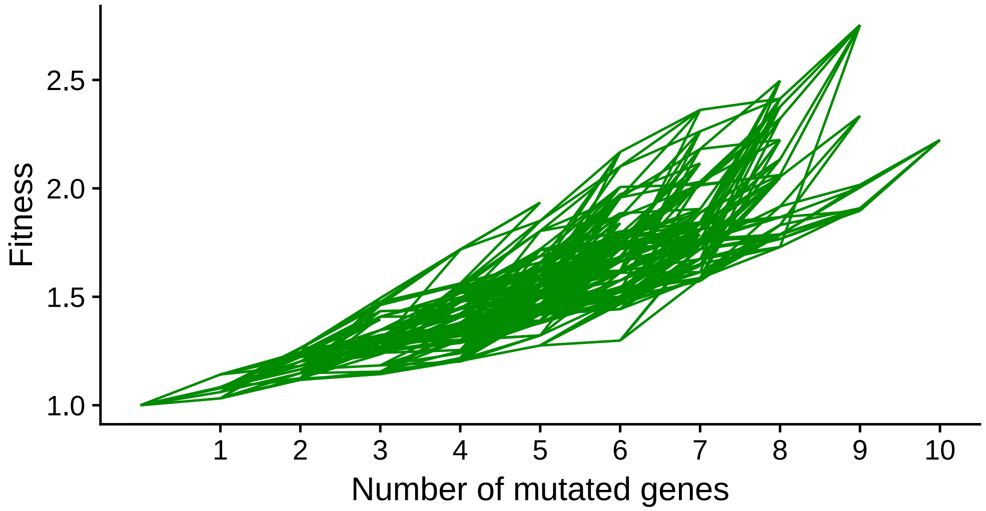


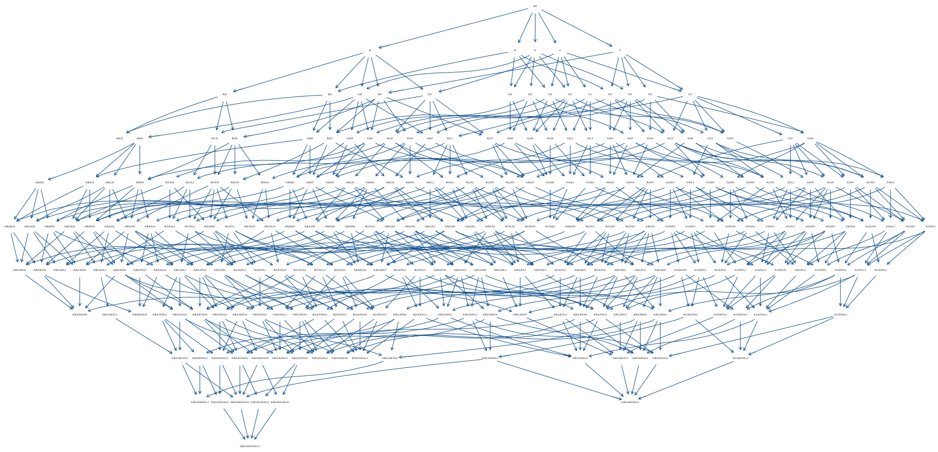

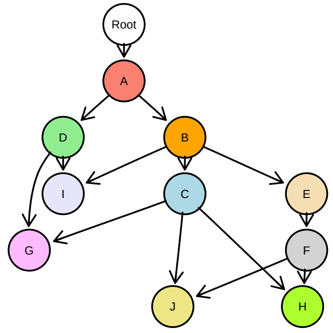


| | |
|---|---|
| *ID* | BoCaqtWu6XR9fw10u |
| *acc genotypes* | 42 |
| *peaks no back* | 9 |
| *removed edges* | 20 |
| *prop rem edges* | 0.235 |

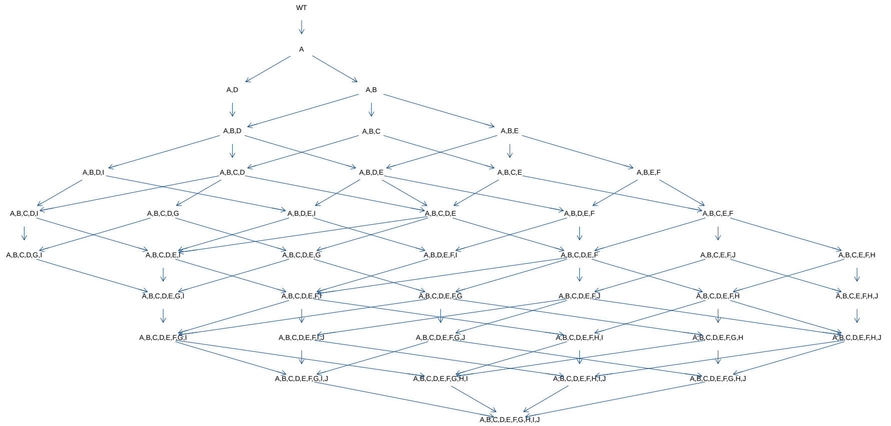

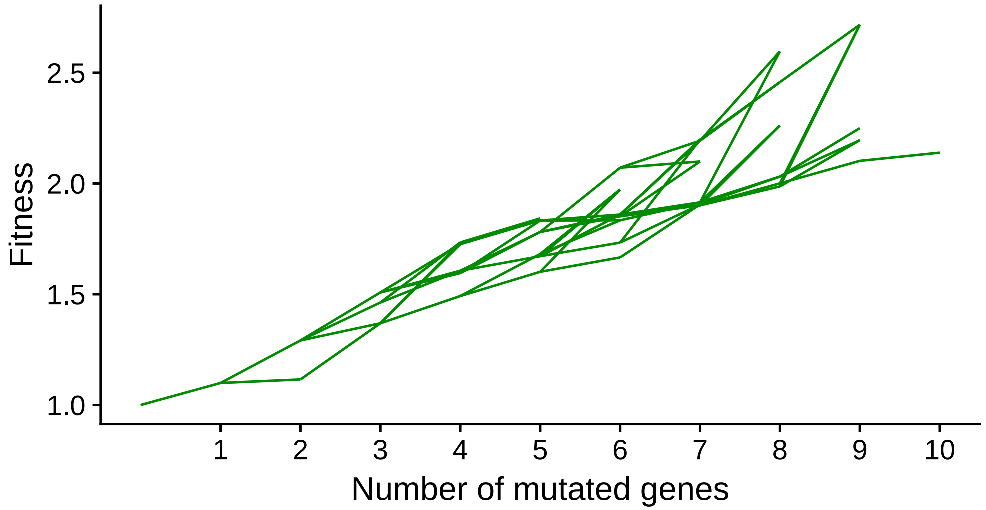


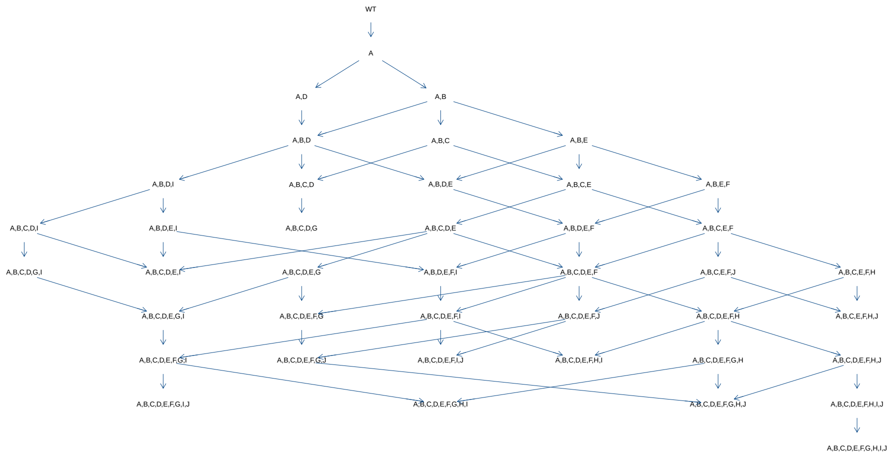

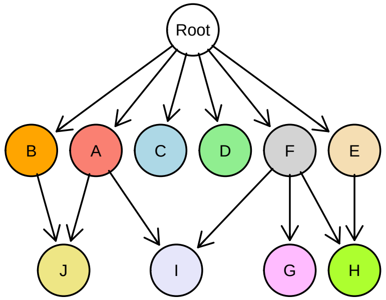


| | |
|---|---|
| *ID* | BpzeoBLsEJ3xTD7v |
| *acc genotypes* | 232 |
| *peaks no back* | 17 |
| *removed edges* | 155 |
| *prop rem edges* | 0.195 |

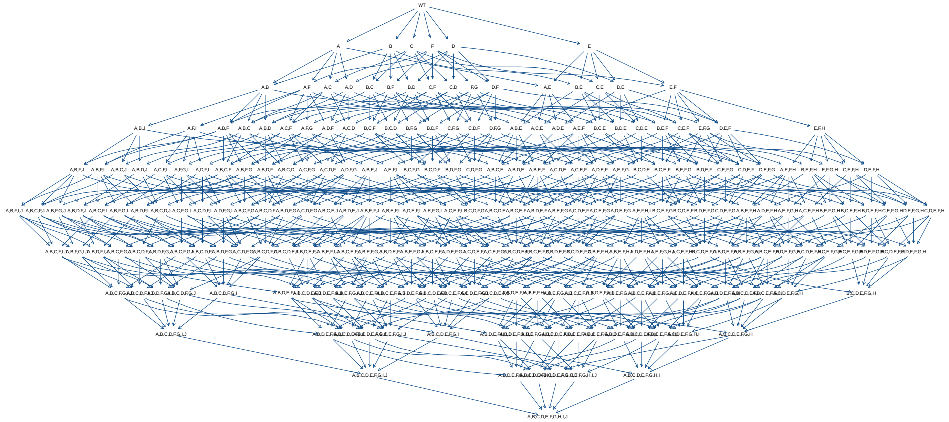

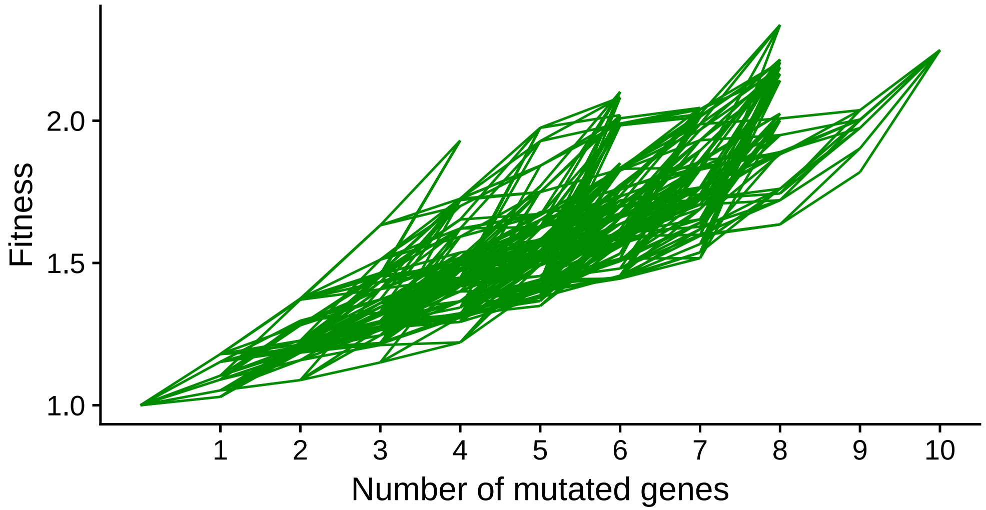


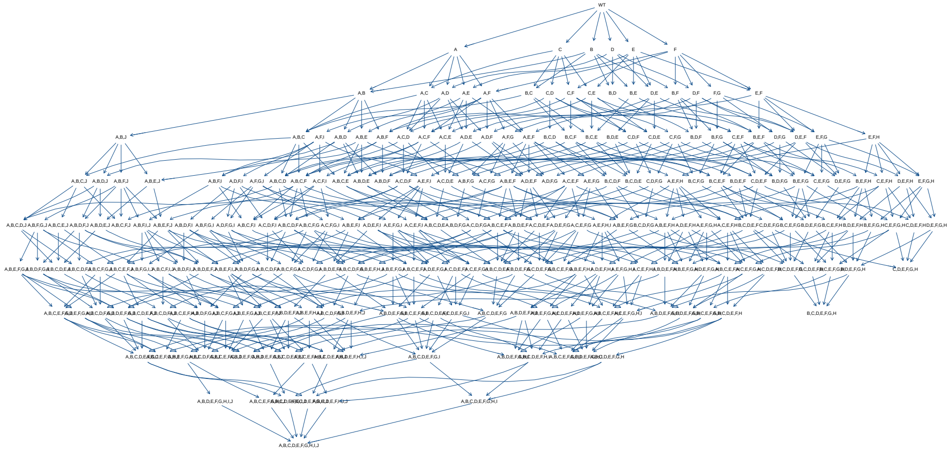

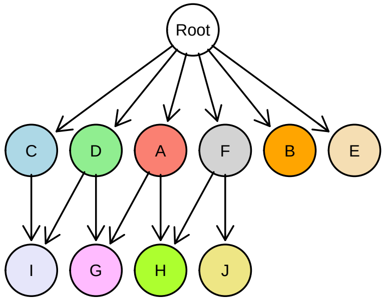


| | |
|---|---|
| *ID* | Bqkt5YbsNE7niJz9 |
| *acc genotypes* | 220 |
| *peaks no back* | 19 |
| *removed edges* | 183 |
| *prop rem edges* | 0.247 |

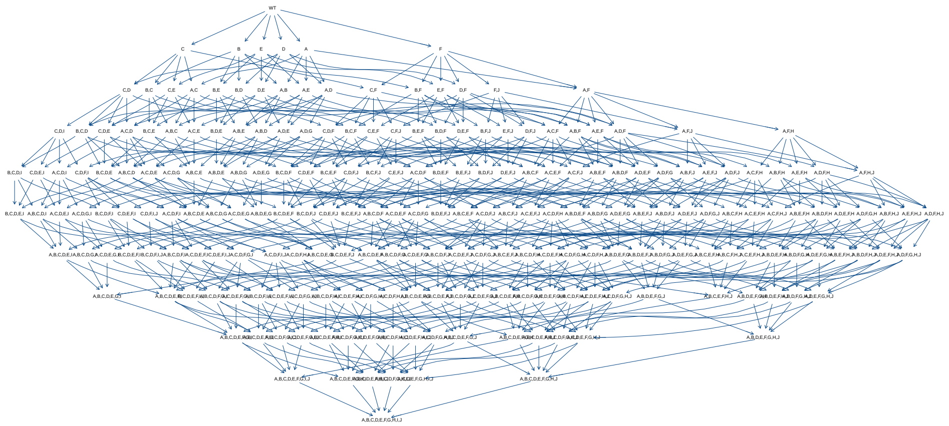

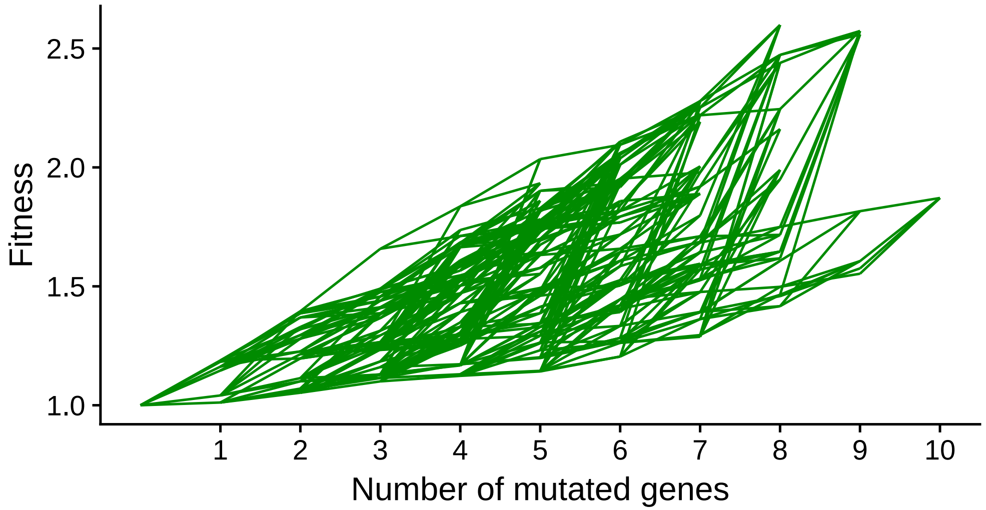


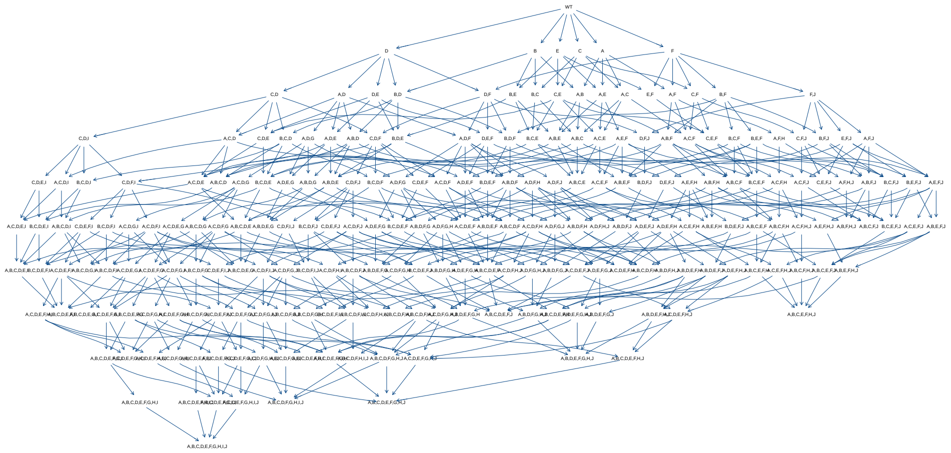

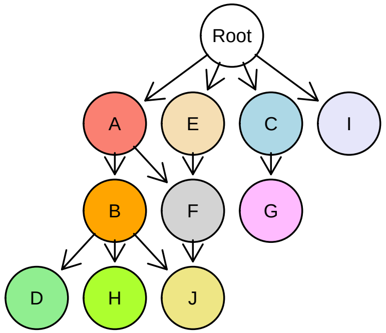


| | |
|---|---|
| *ID* | c10jjy6210kvQdZBVg |
| *acc genotypes* | 126 |
| *peaks no back* | 16 |
| *removed edges* | 70 |
| *prop rem edges* | 0.193 |

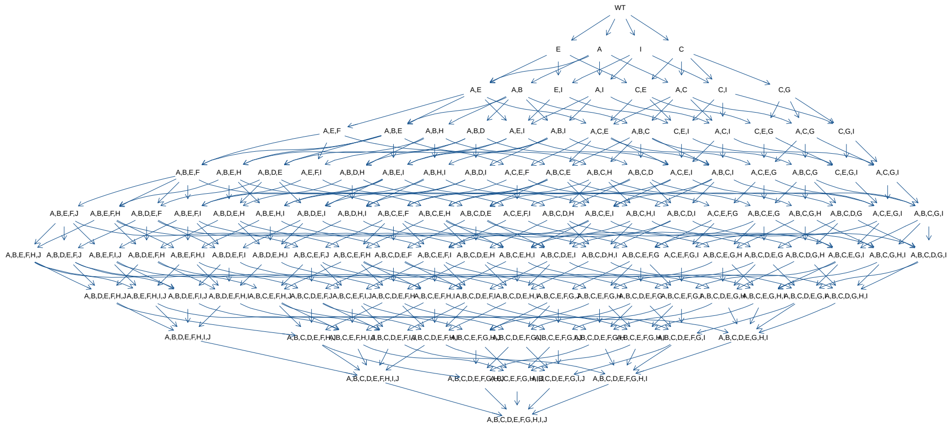

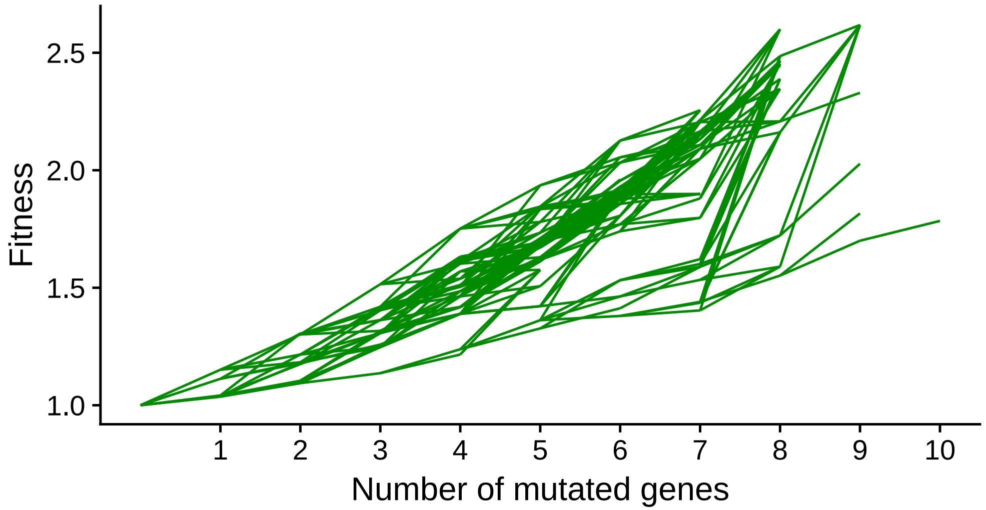


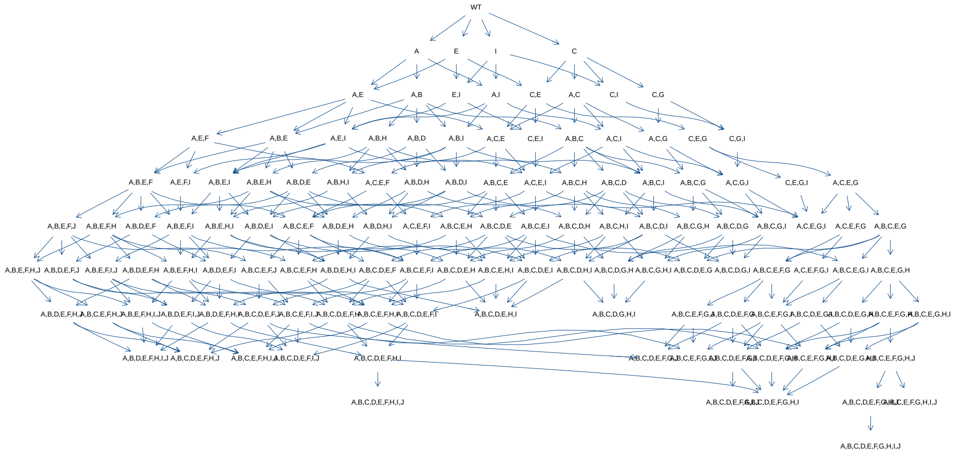

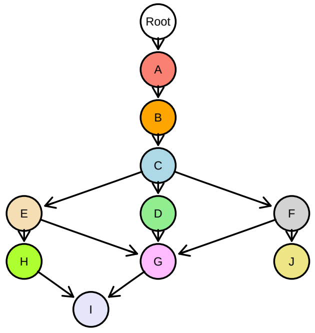


| | |
|---|---|
| *ID* | C4drUyloTXoBU6JC |
| *acc genotypes* | 27 |
| *peaks no back* | 4 |
| *removed edges* | 9 |
| *prop rem edges* | 0.191 |

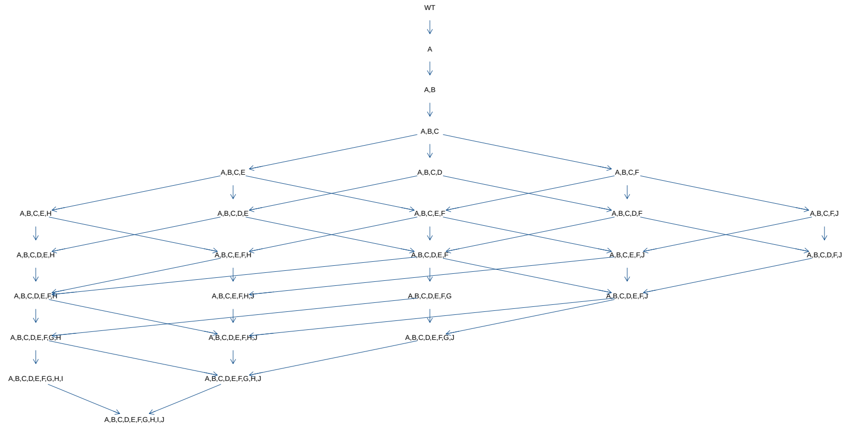

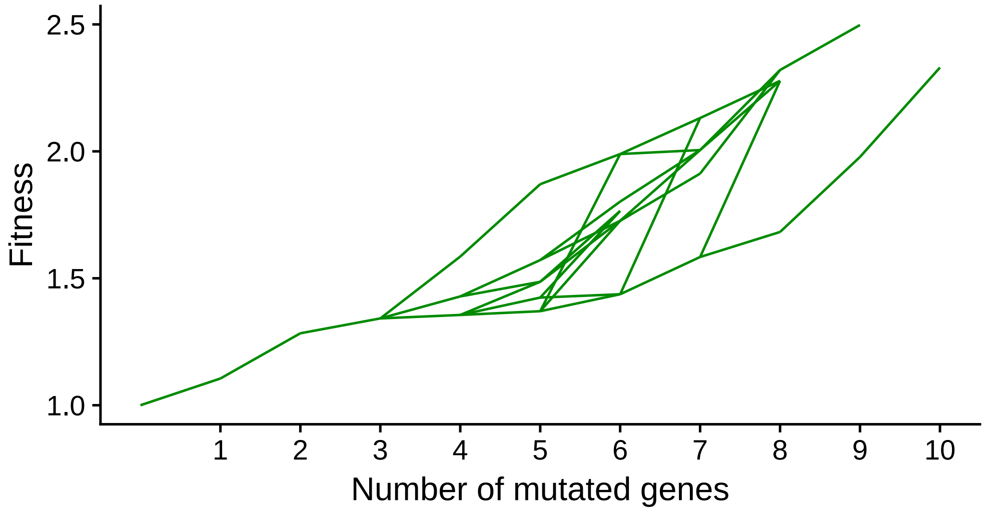


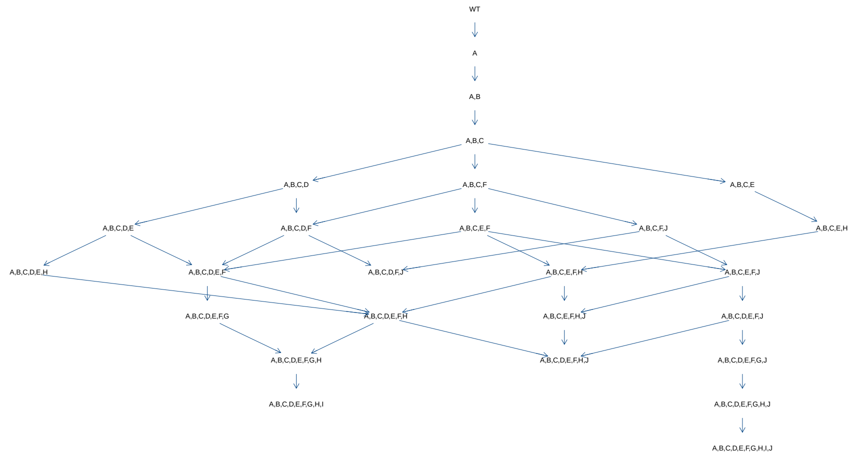

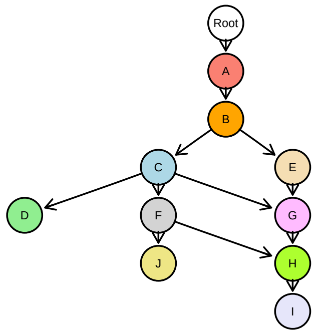


| | |
|---|---|
| *ID* | CDprcI8u4ZcaFn7F |
| *acc genotypes* | 30 |
| *peaks no back* | 4 |
| *removed edges* | 6 |
| *prop rem edges* | 0.111 |

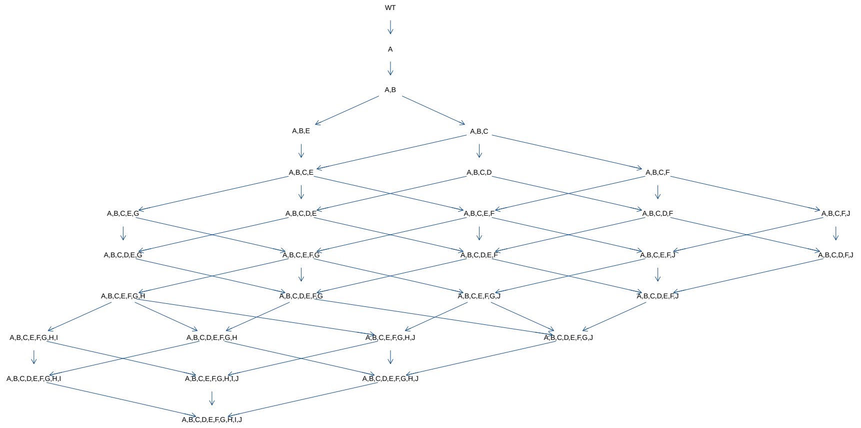

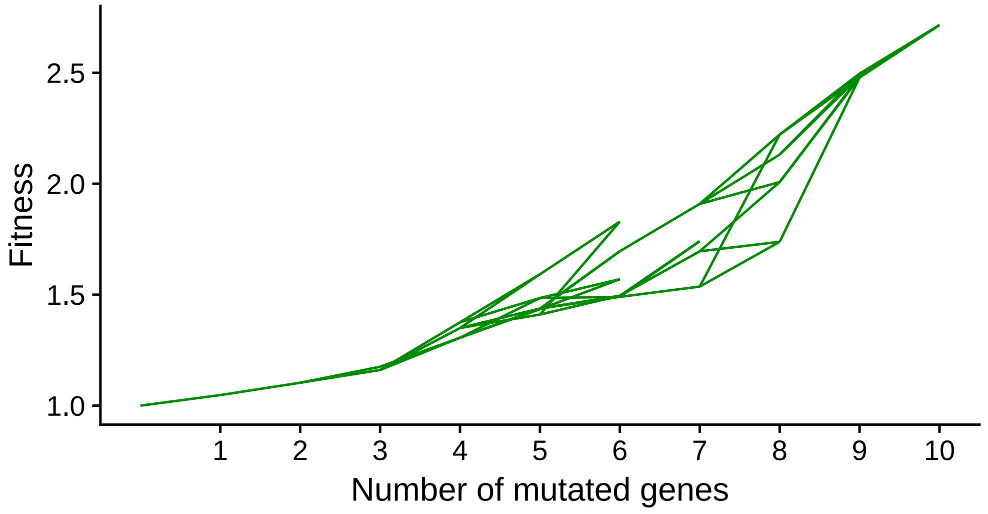


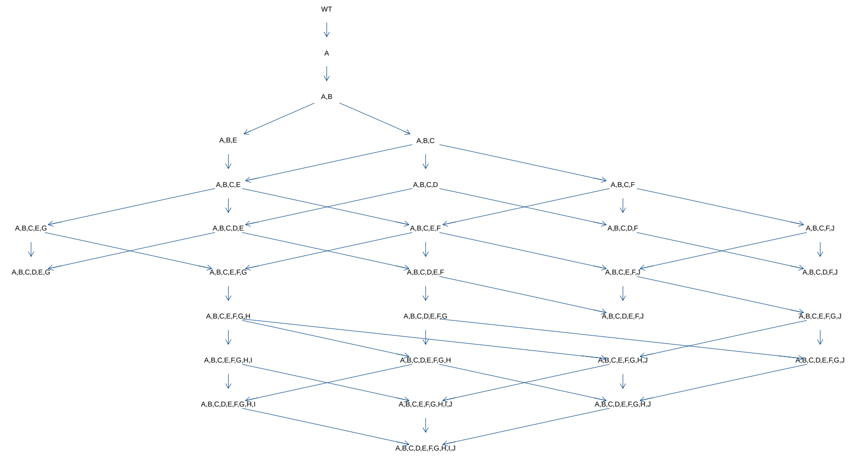

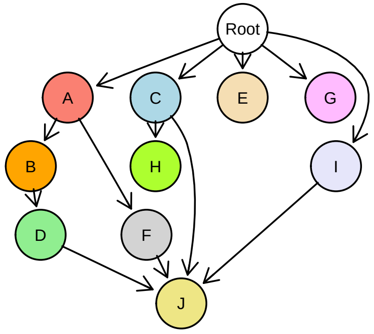


| | |
|---|---|
| *ID* | cGm7gRR3s42NqkyP |
| *acc genotypes* | 176 |
| *peaks no back* | 16 |
| *removed edges* | 116 |
| *prop rem edges* | 0.201 |

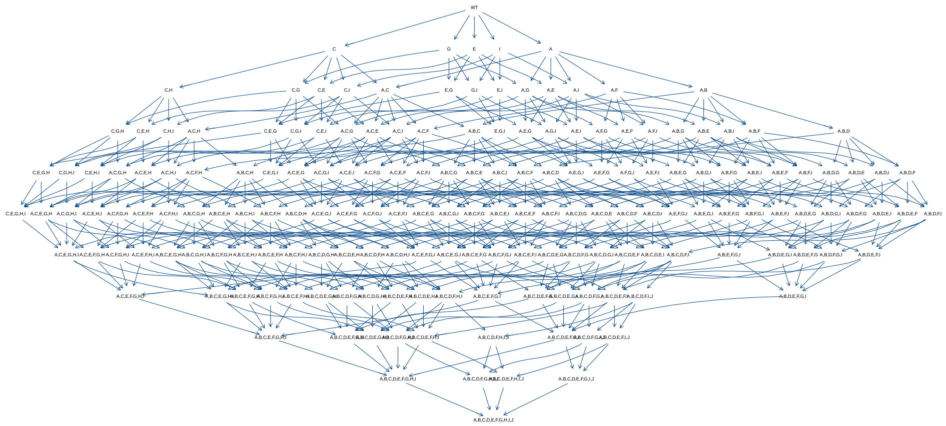

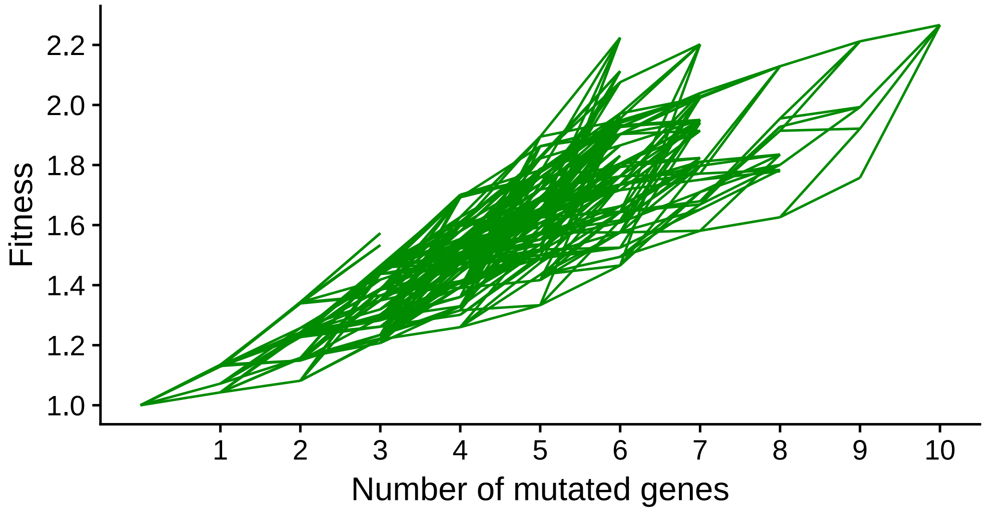


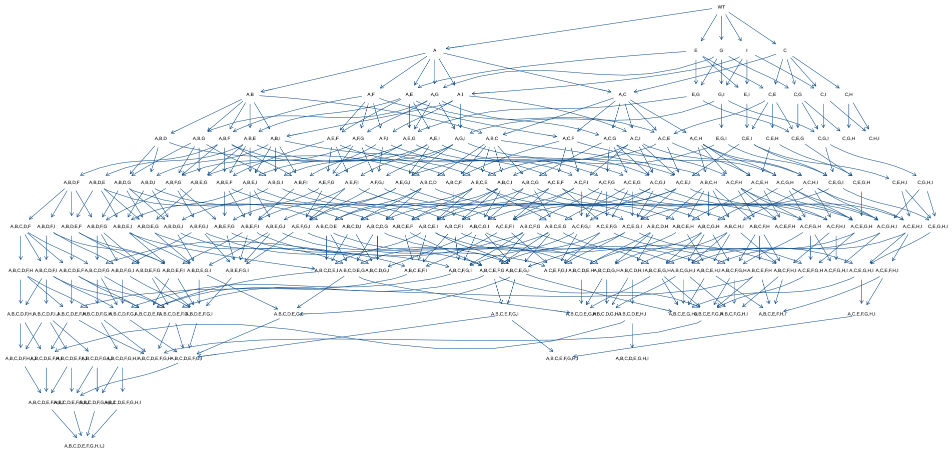

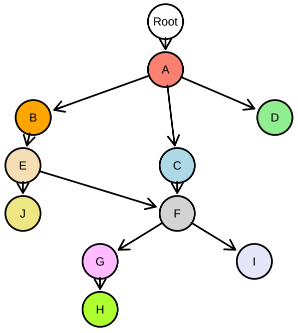


| | |
|---|---|
| *ID* | CjDx5CPhar7wxbYH |
| *acc genotypes* | 41 |
| *peaks no back* | 7 |
| *removed edges* | 22 |
| *prop rem edges* | 0.259 |

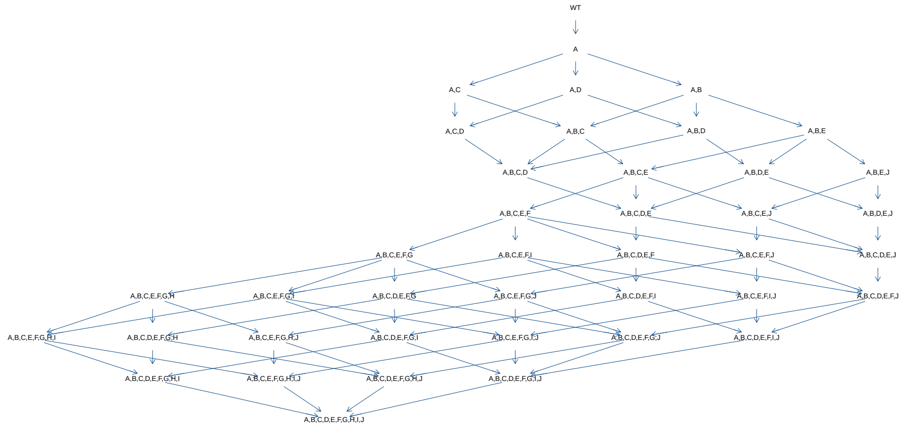

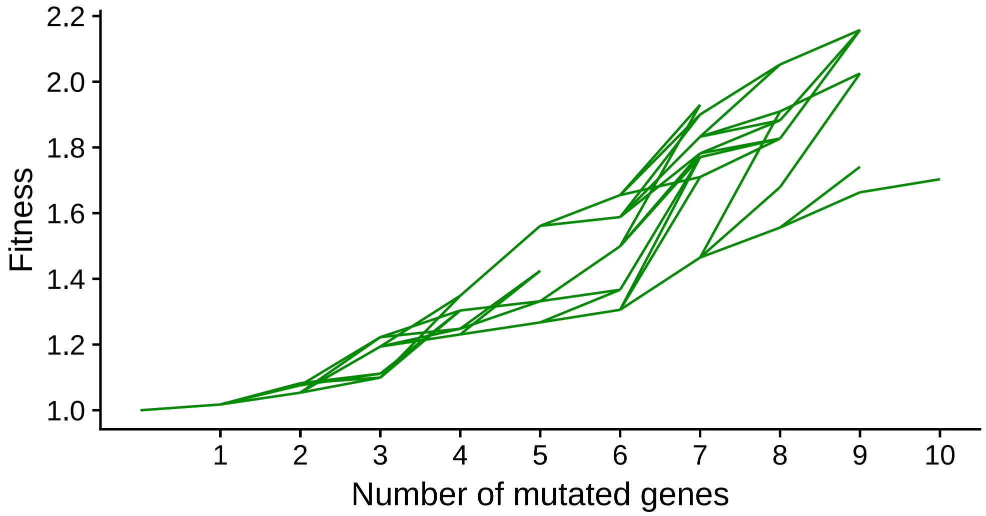


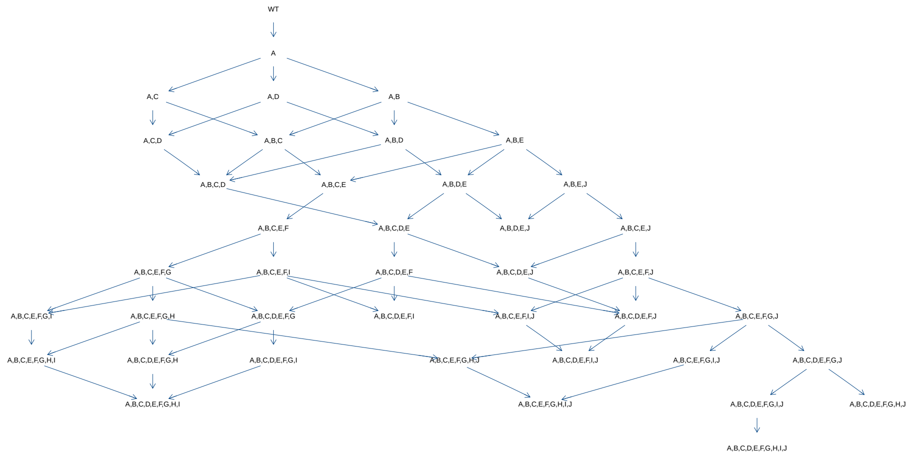

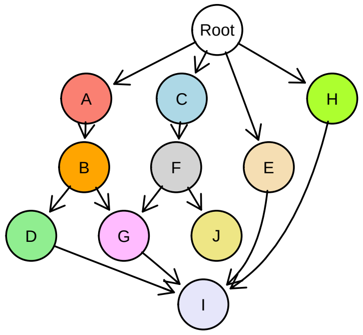


| | |
|---|---|
| *ID* | dC6vMuFmbDjoadOo |
| *acc genotypes* | 82 |
| *peaks no back* | 10 |
| *removed edges* | 39 |
| *prop rem edges* | 0.185 |

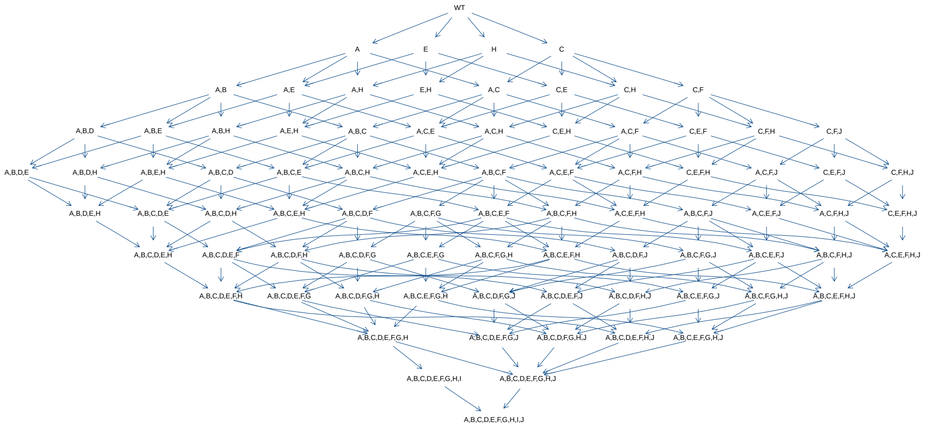

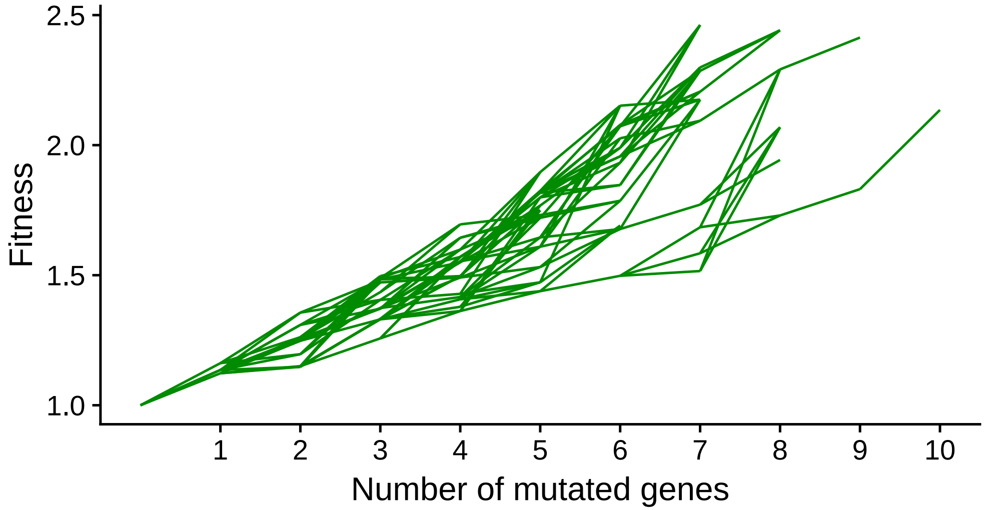


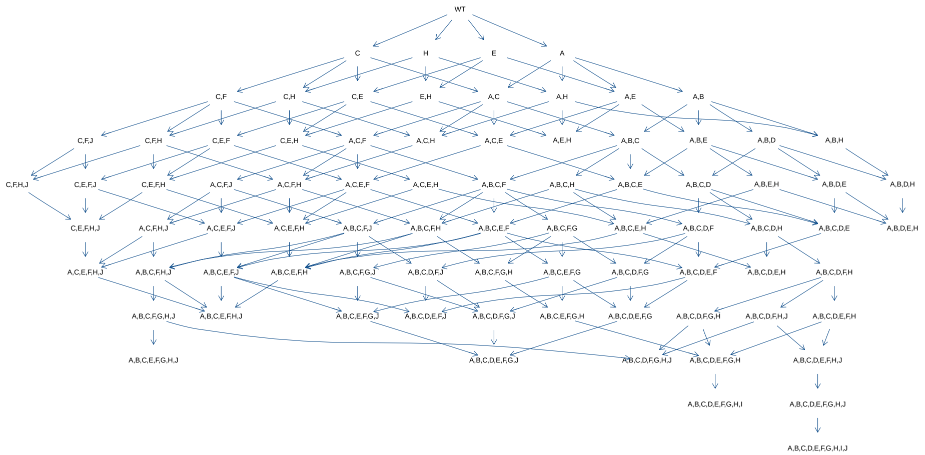

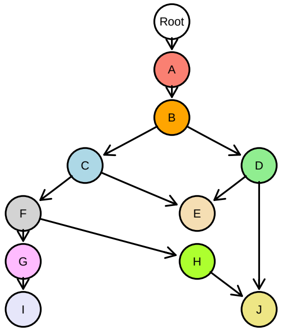


| | |
|---|---|
| *ID* | DcAnAY4JLtfHe55I |
| *acc genotypes* | 31 |
| *peaks no back* | 4 |
| *removed edges* | 8 |
| *prop rem edges* | 0.143 |

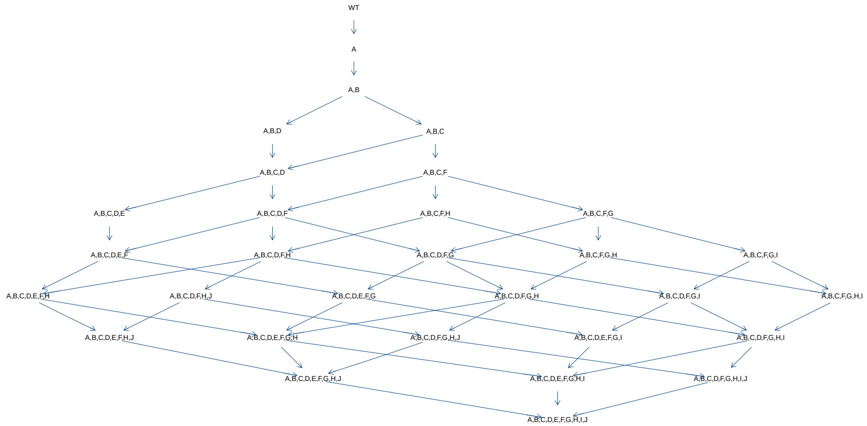

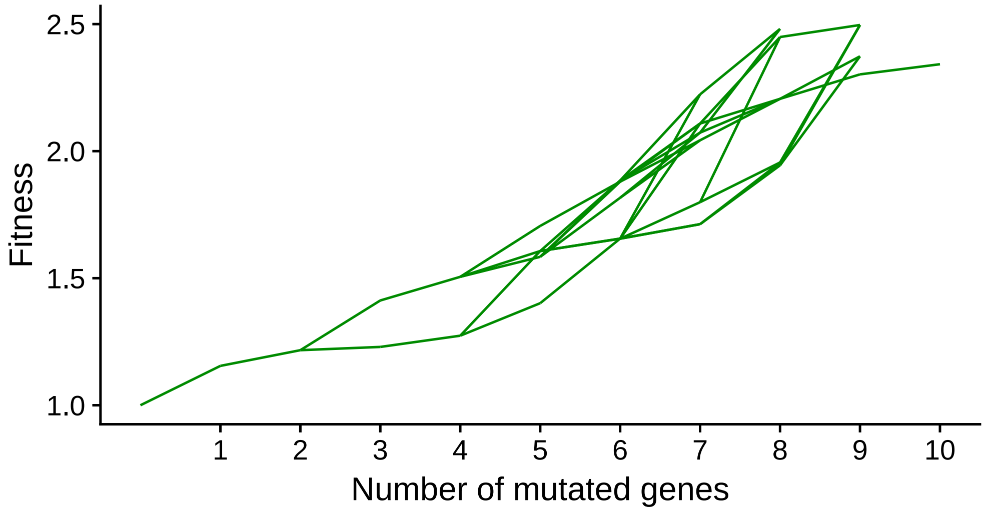


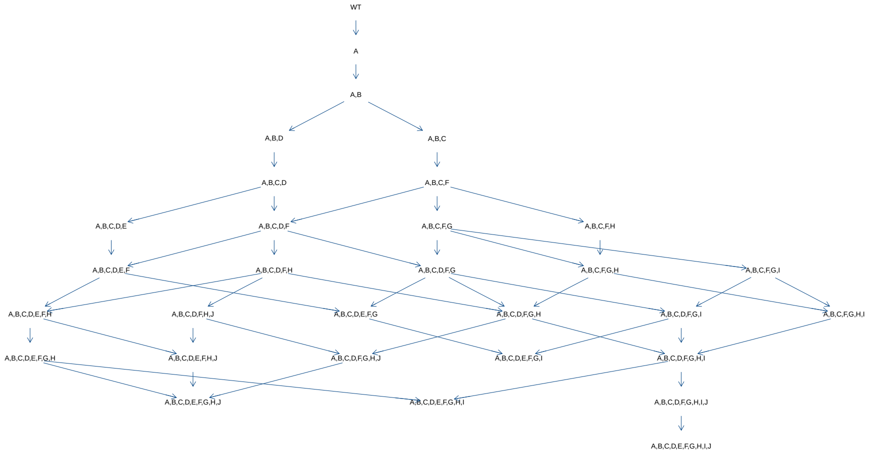

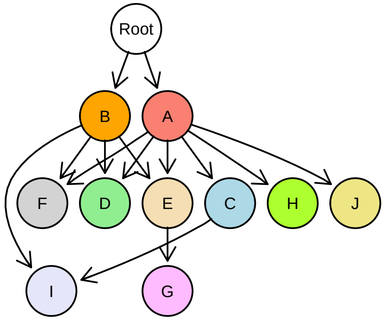


| | |
|---|---|
| *ID* | dDZ4CvpAmrvxLFyS |
| *acc genotypes* | 154 |
| *peaks no back* | 13 |
| *removed edges* | 111 |
| *prop rem edges* | 0.221 |

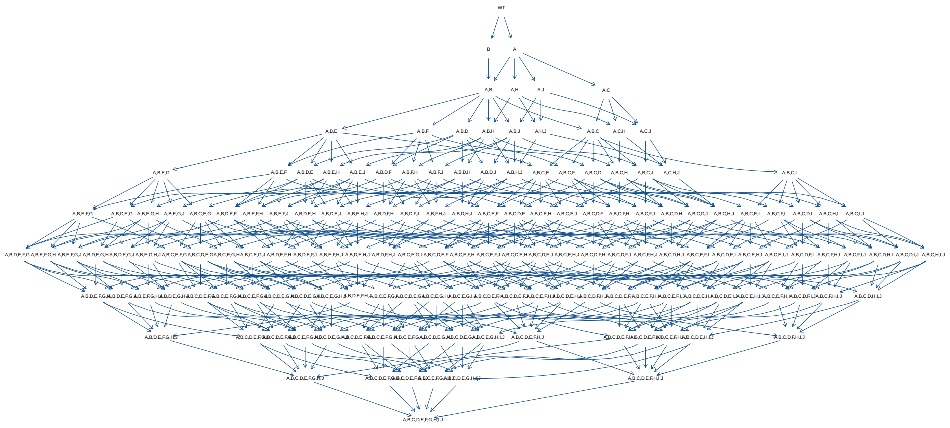

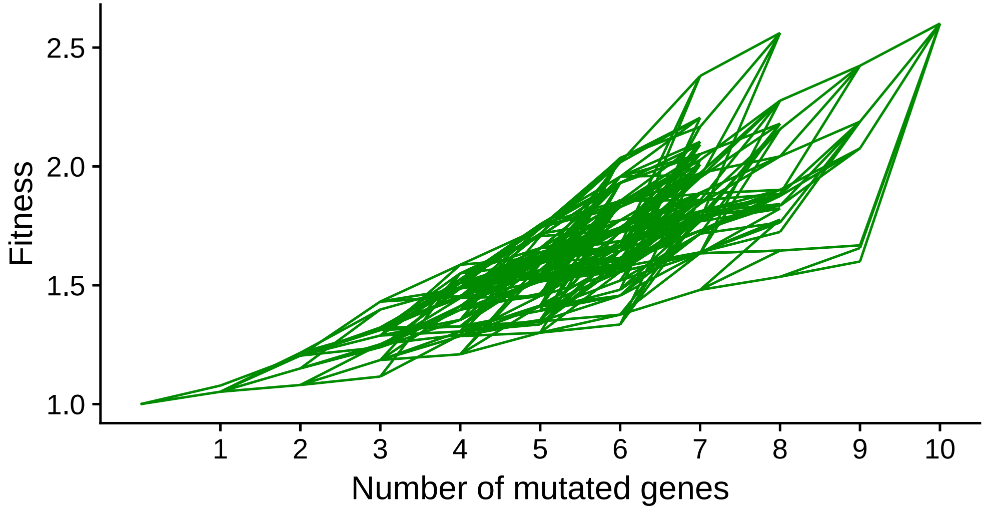


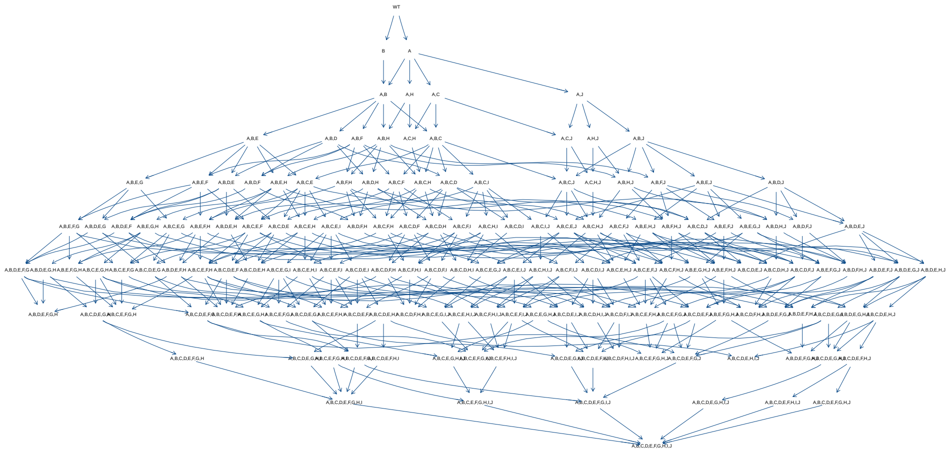

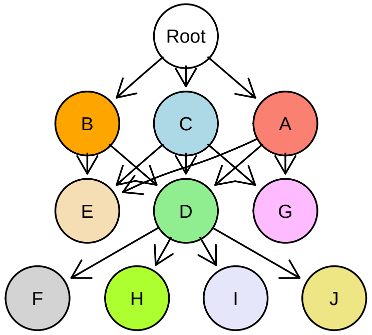


| | |
|---|---|
| *ID* | DFTK3ko7MlzoABxk |
| *acc genotypes* | 76 |
| *peaks no back* | 9 |
| *removed edges* | 40 |
| *prop rem edges* | 0.187 |

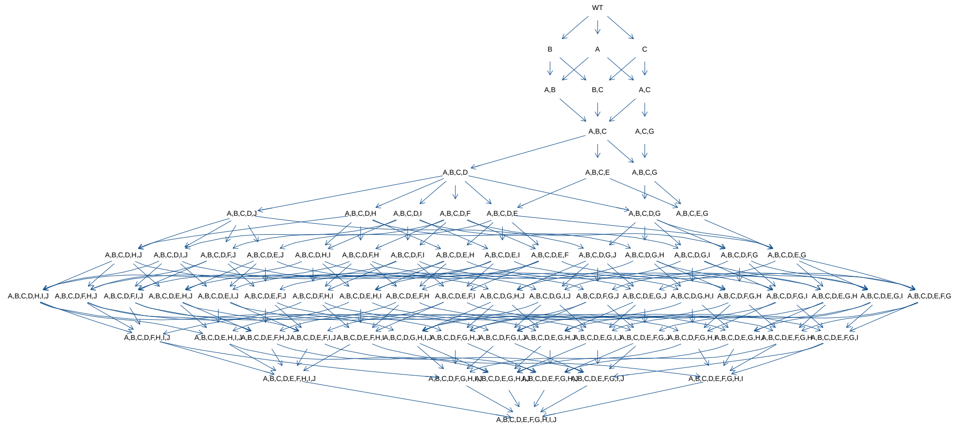

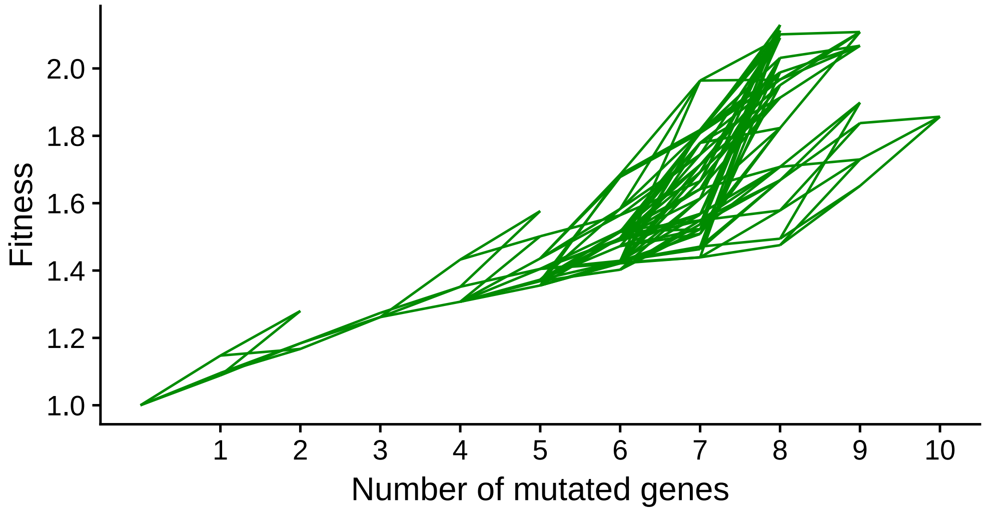


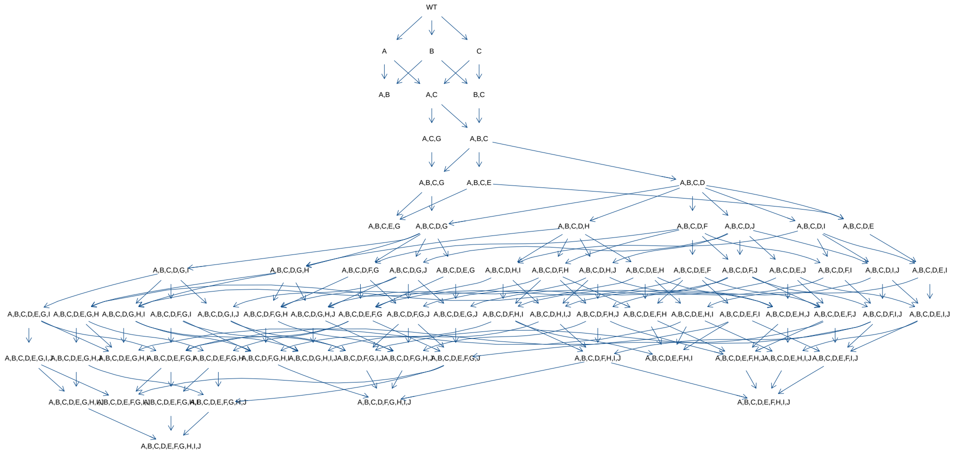

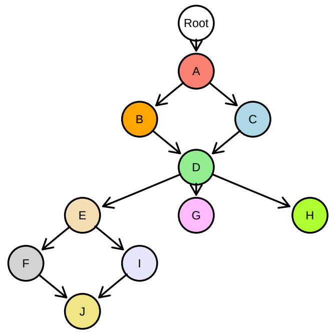


| | |
|---|---|
| *ID* | DXTB5tpZMuV2IF2I |
| *acc genotypes* | 29 |
| *peaks no back* | 5 |
| *removed edges* | 8 |
| *prop rem edges* | 0.148 |

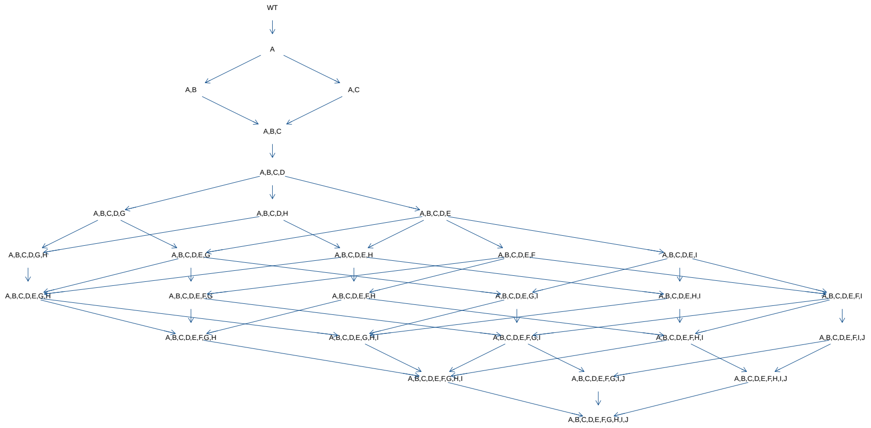

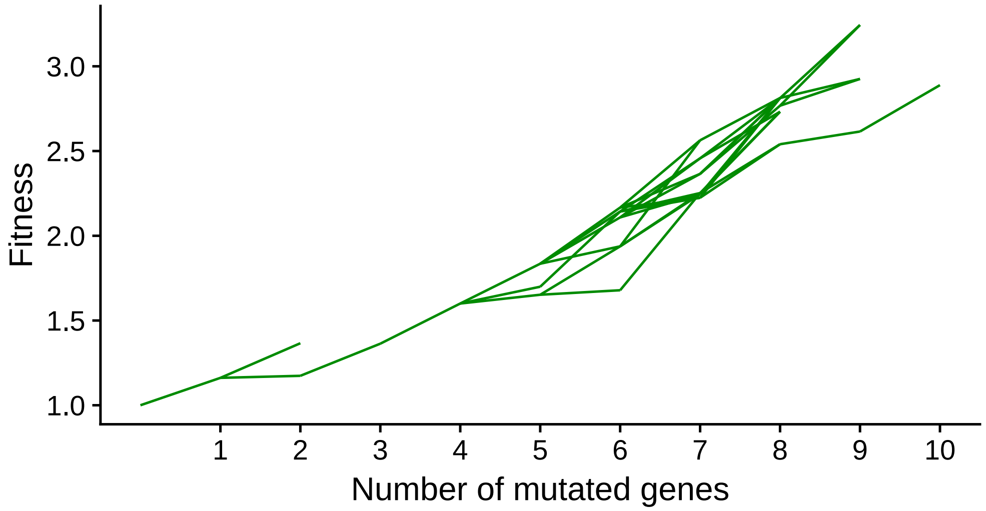


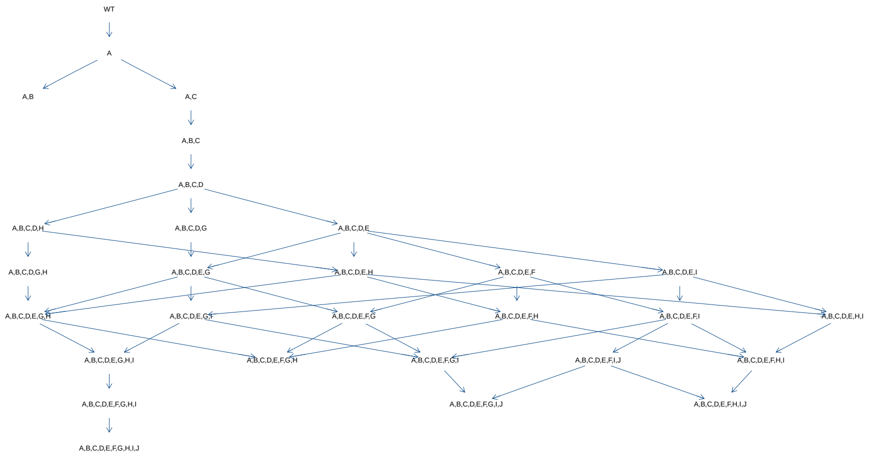

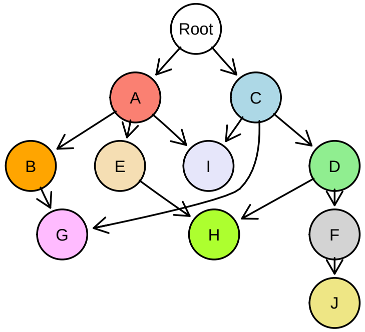


| | |
|---|---|
| *ID* | dYeC4QsDciy4hwa3 |
| *acc genotypes* | 75 |
| *peaks no back* | 10 |
| *removed edges* | 32 |
| *prop rem edges* | 0.174 |

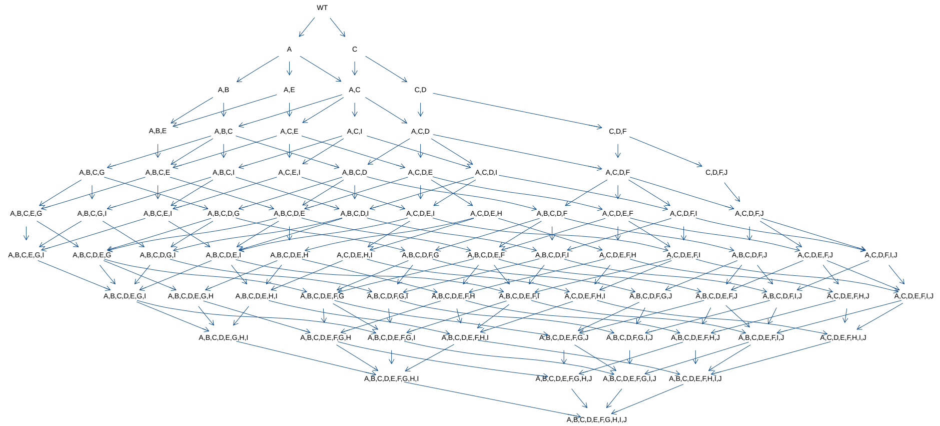

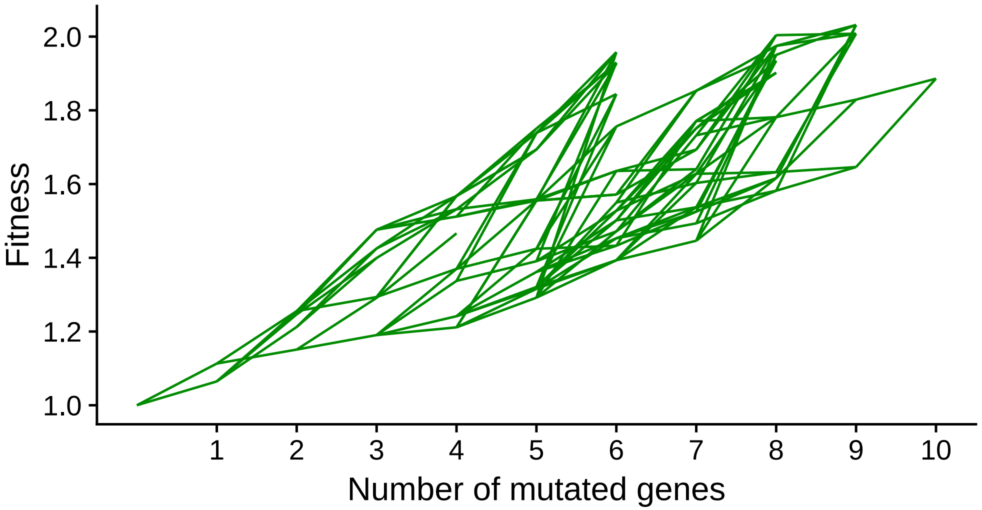


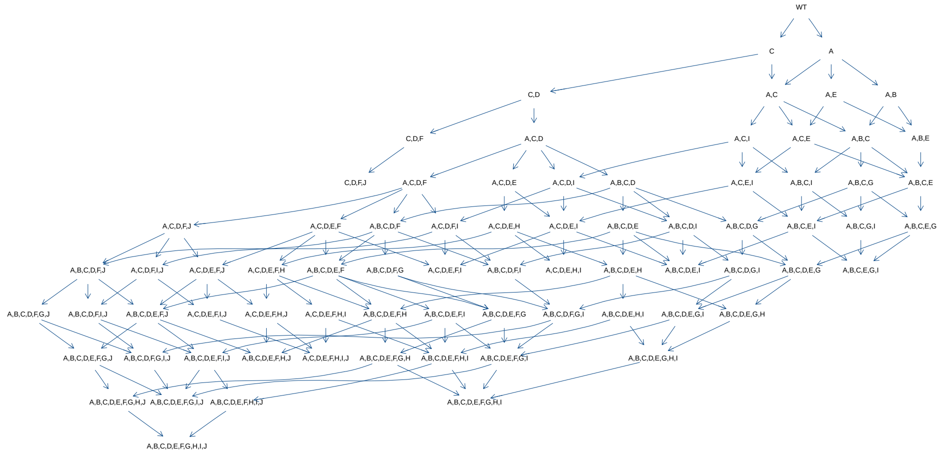

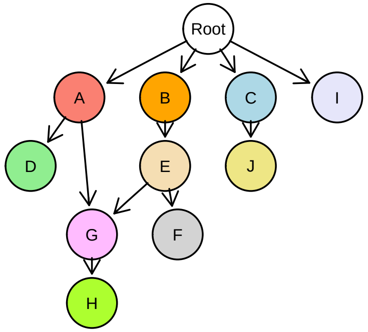


| | |
|---|---|
| *ID* | E8phhmTg4KHAS8DL |
| *acc genotypes* | 120 |
| *peaks no back* | 12 |
| *removed edges* | 61 |
| *prop rem edges* | 0.18 |

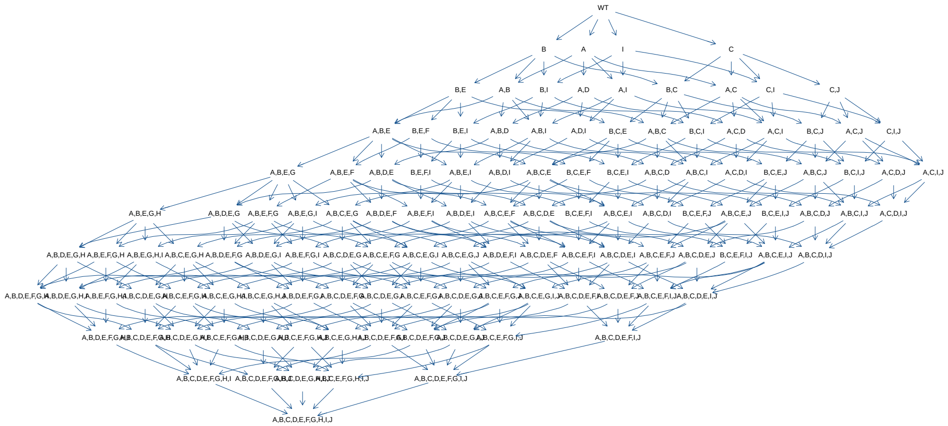

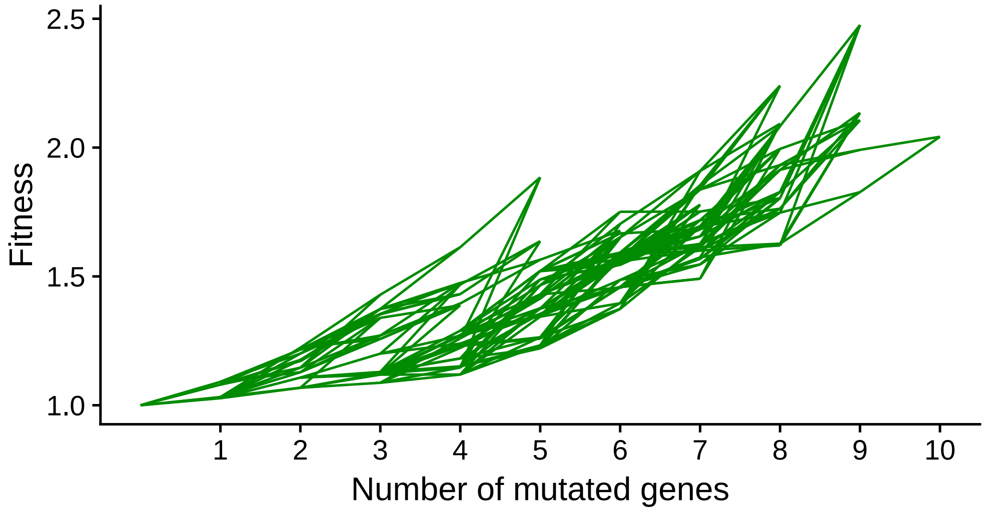


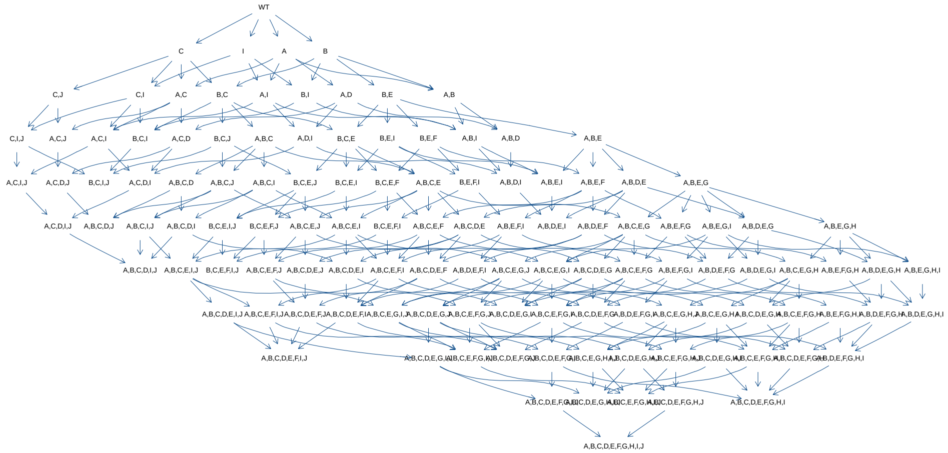

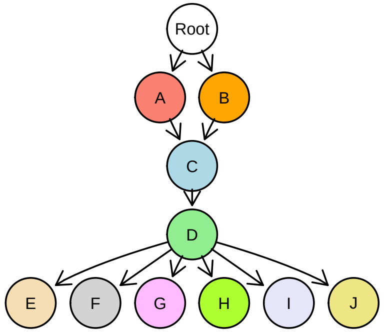


| | |
|---|---|
| *ID* | eBeyysHajtLlNlLT |
| *acc genotypes* | 69 |
| *peaks no back* | 8 |
| *removed edges* | 35 |
| *prop rem edges* | 0.177 |

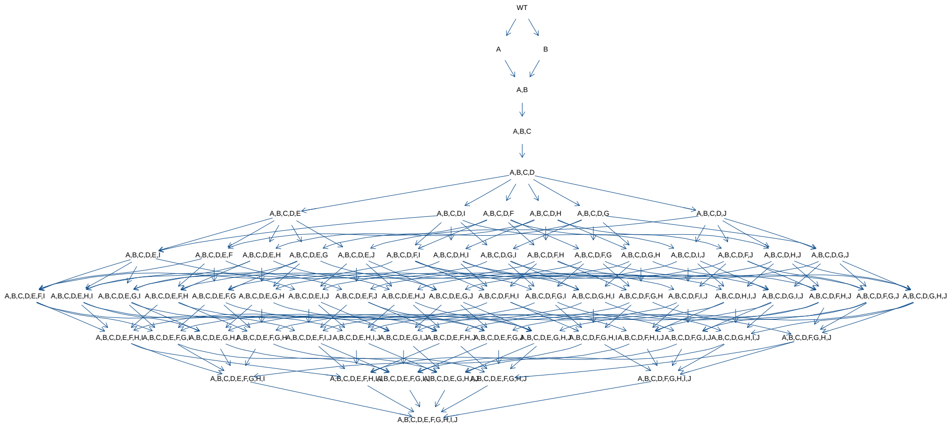

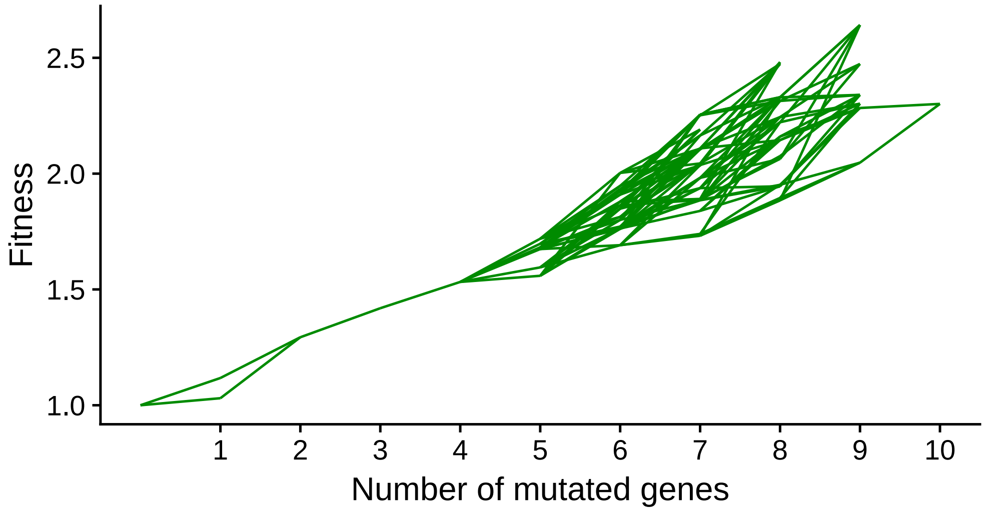


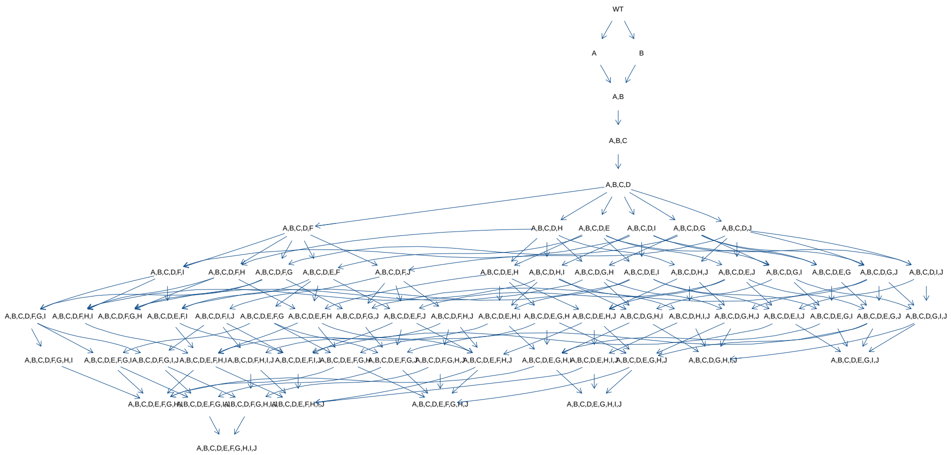

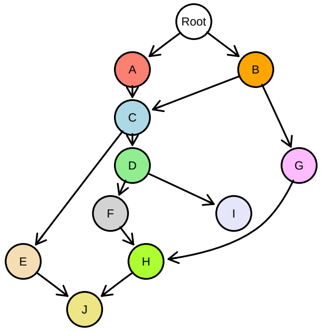


| | |
|---|---|
| *ID* | EflrY474a8aE1bD2 |
| *acc genotypes* | 32 |
| *peaks no back* | 5 |
| *removed edges* | 8 |
| *prop rem edges* | 0.133 |

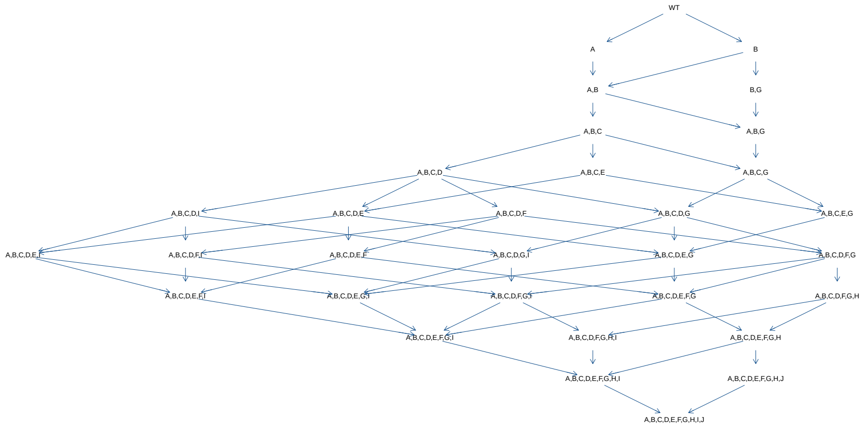

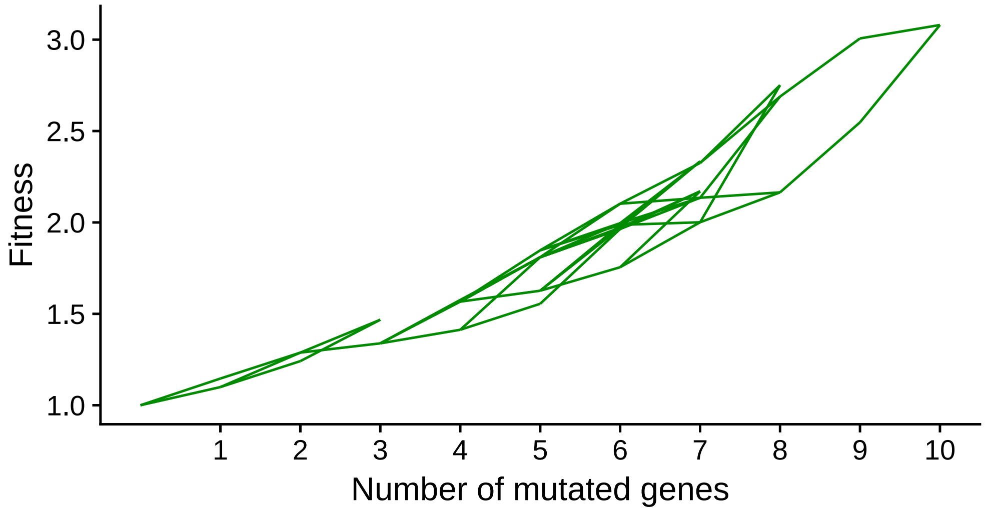


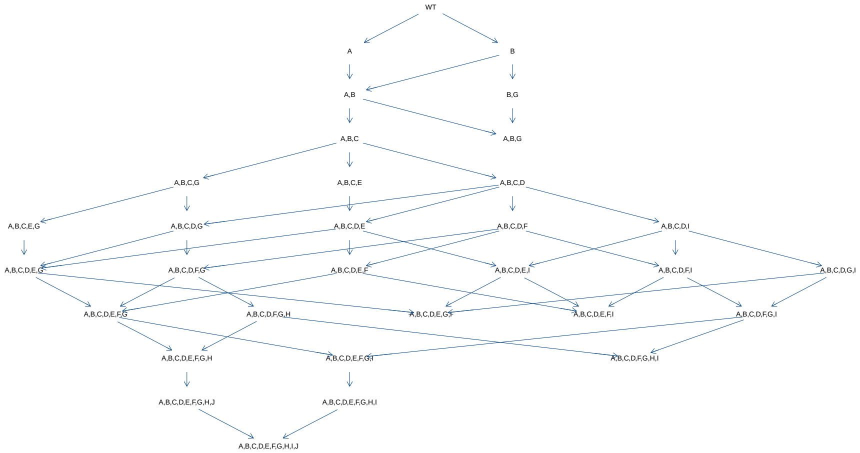

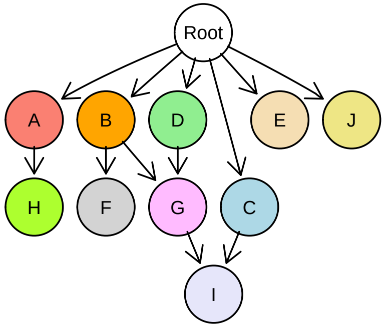


| | |
|---|---|
| *ID* | EjeOkmz5yWV10GMyo |
| *acc genotypes* | 216 |
| *peaks no back* | 14 |
| *removed edges* | 134 |
| *prop rem edges* | 0.183 |

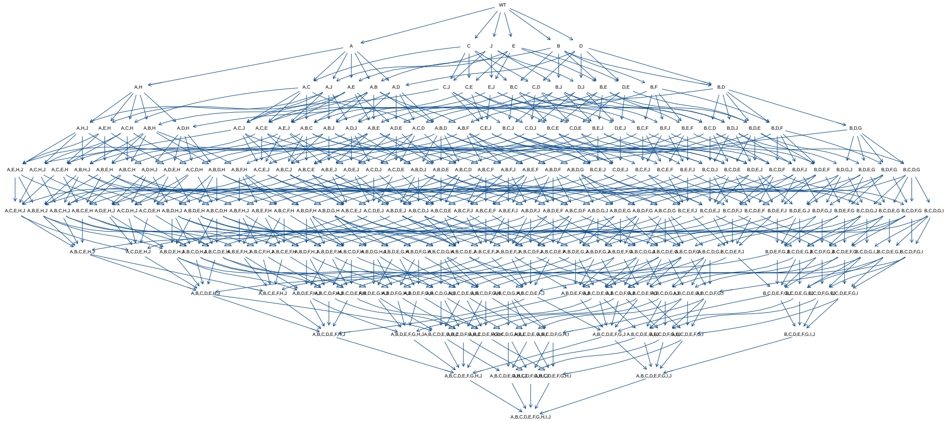

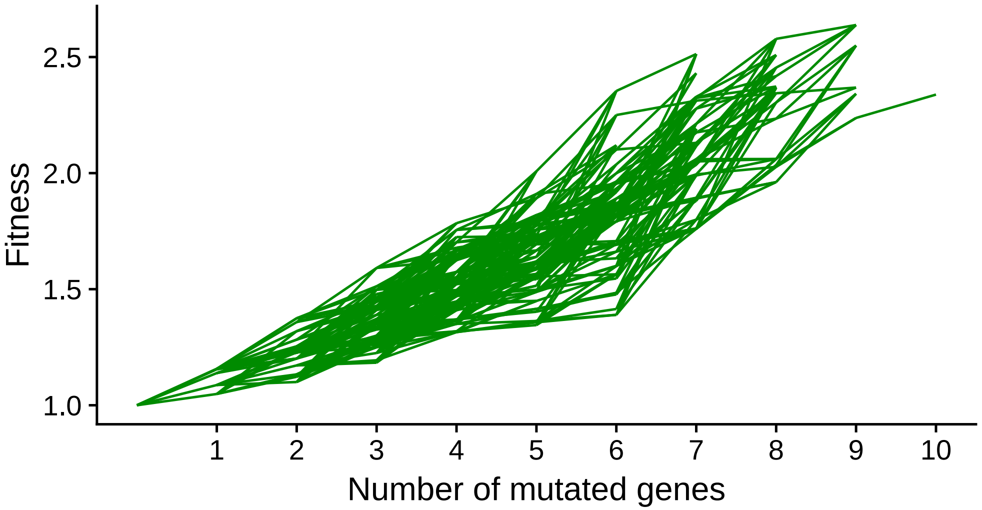


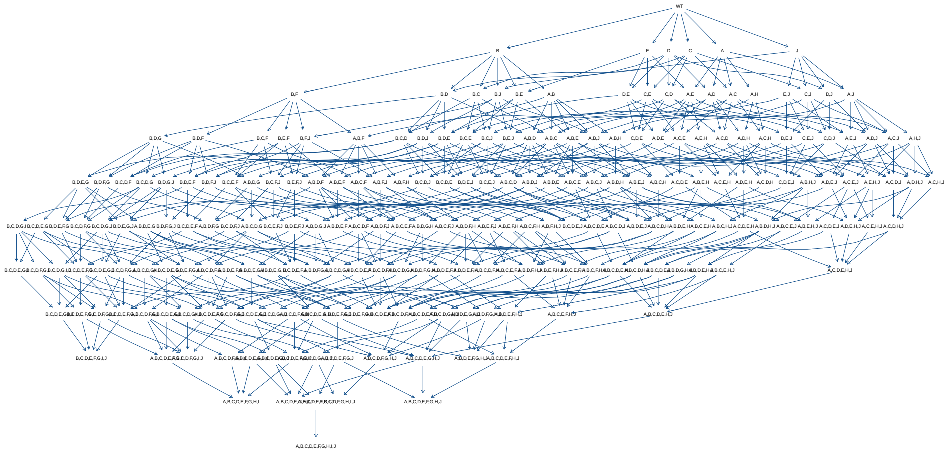

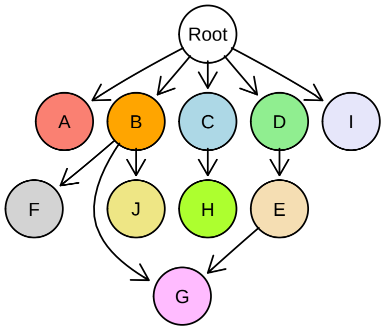


| | |
|---|---|
| *ID* | EJVLcdkFsVSwbuKT |
| *acc genotypes* | 228 |
| *peaks no back* | 18 |
| *removed edges* | 157 |
| *prop rem edges* | 0.202 |

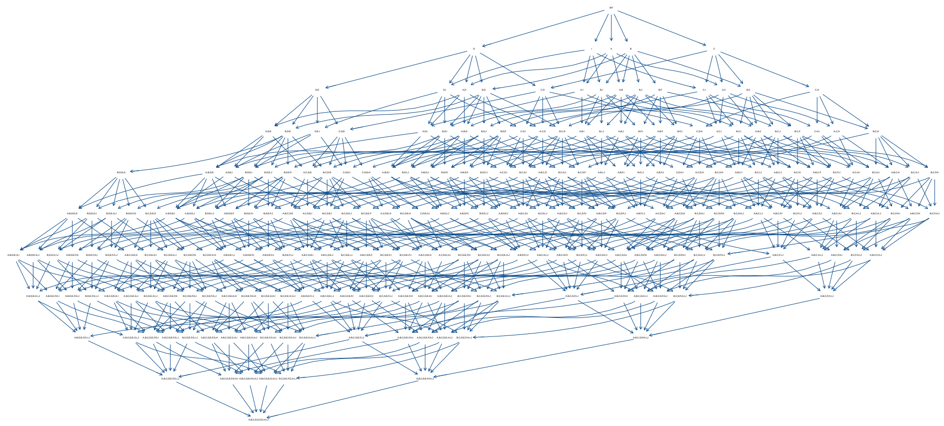

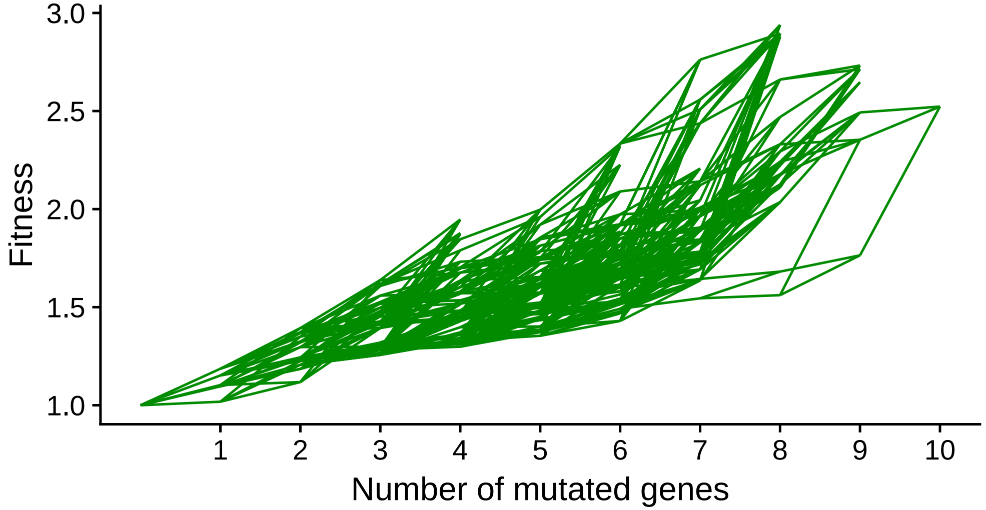


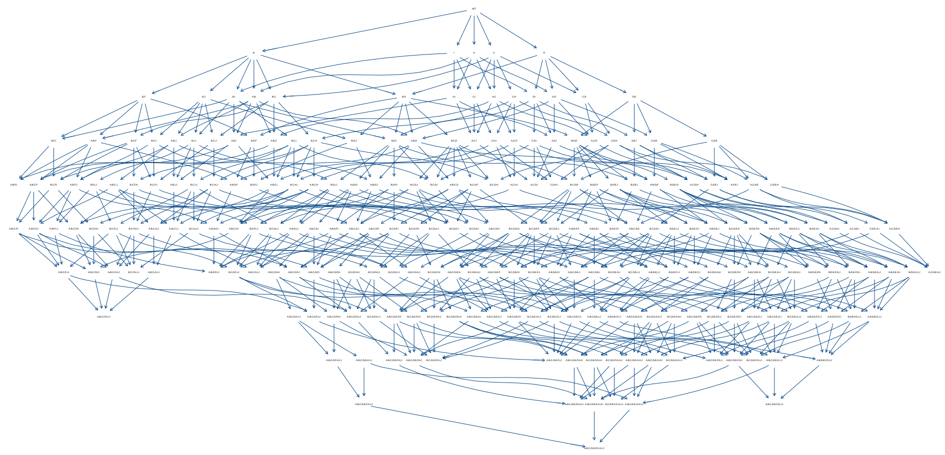

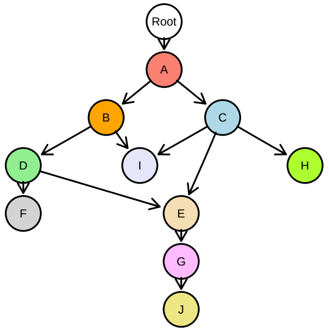


| | |
|---|---|
| *ID* | EkGrTn9N4iF6FhfE |
| *acc genotypes* | 43 |
| *peaks no back* | 5 |
| *removed edges* | 15 |
| *prop rem edges* | 0.165 |

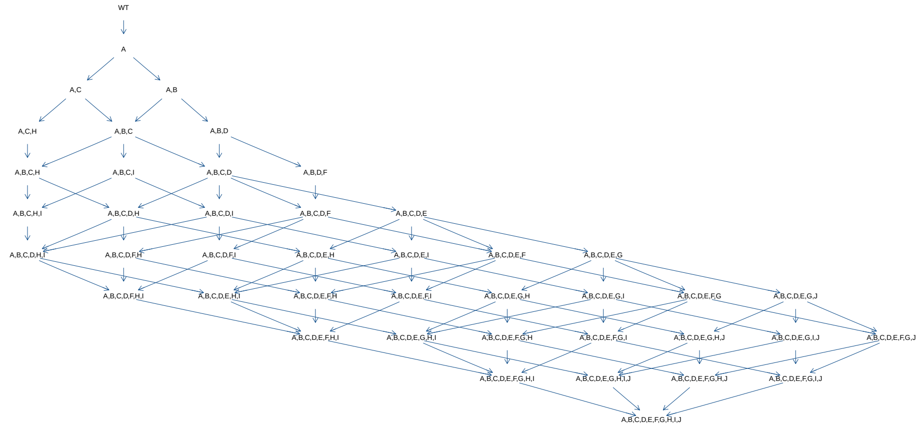

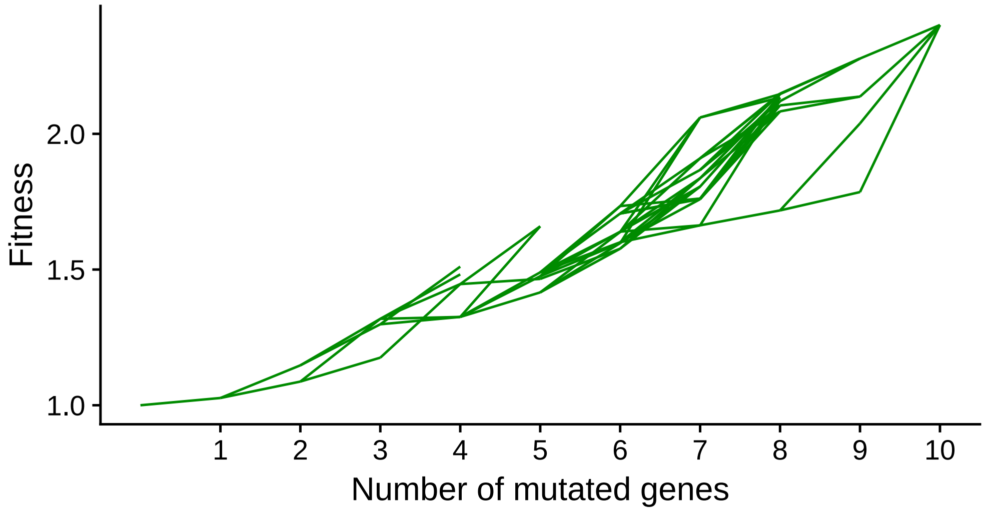


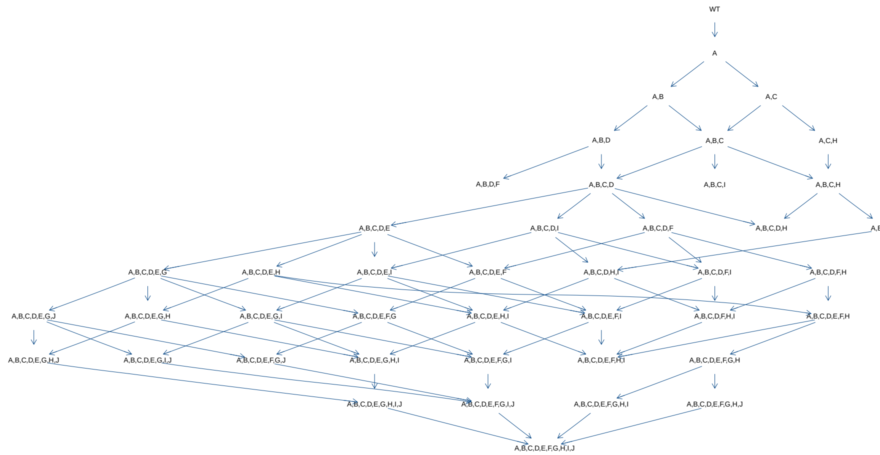

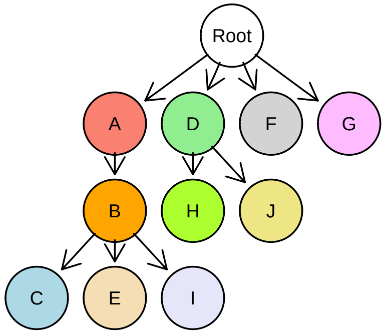


| | |
|---|---|
| *ID* | EMu88ff7108oDY2Nd |
| *acc genotypes* | 200 |
| *peaks no back* | 21 |
| *removed edges* | 136 |
| *prop rem edges* | 0.2 |

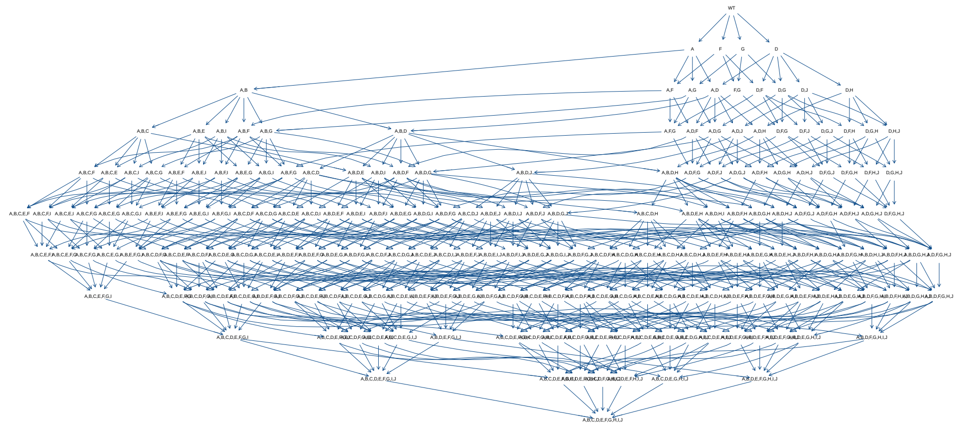

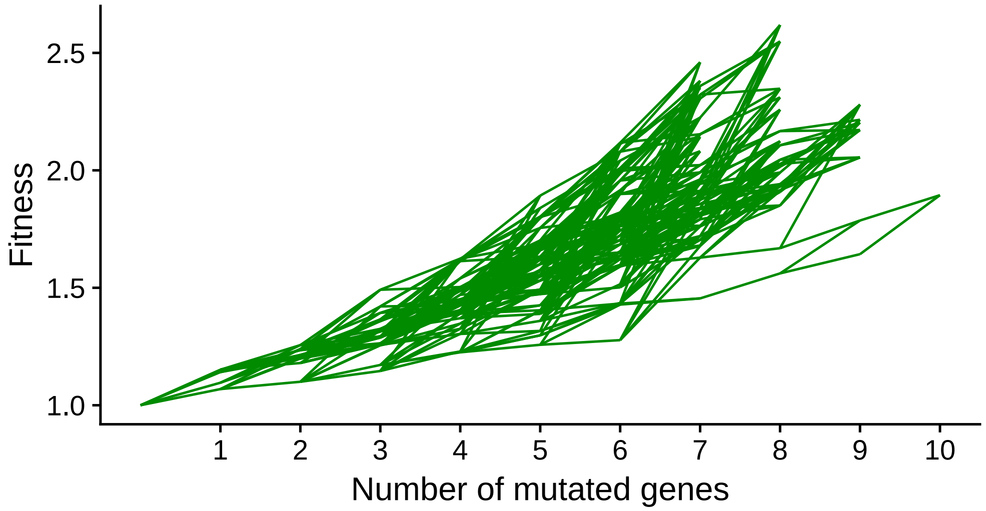


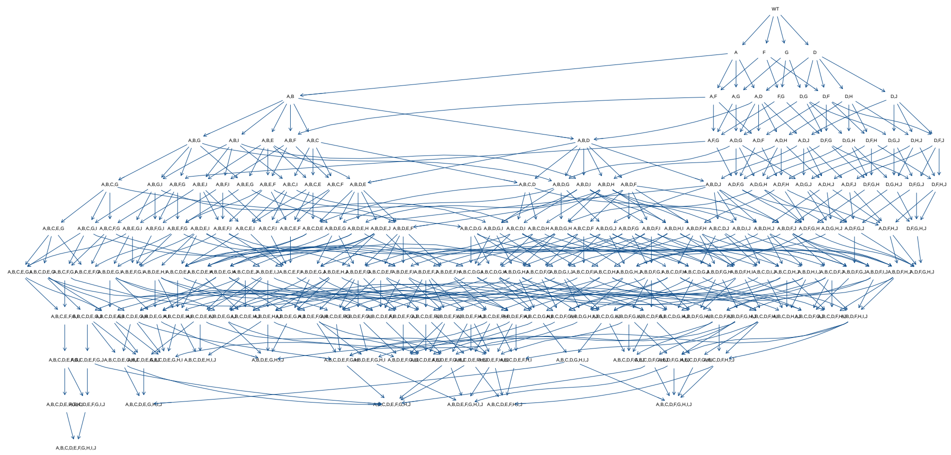

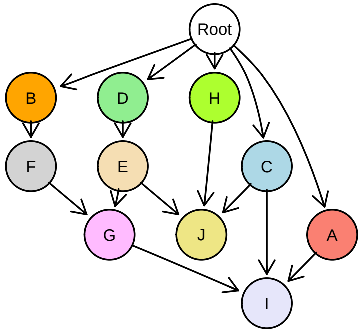


| | |
|---|---|
| *ID* | EnDJ6doxjCtQ9zue |
| *acc genotypes* | 91 |
| *peaks no back* | 11 |
| *removed edges* | 52 |
| *prop rem edges* | 0.211 |

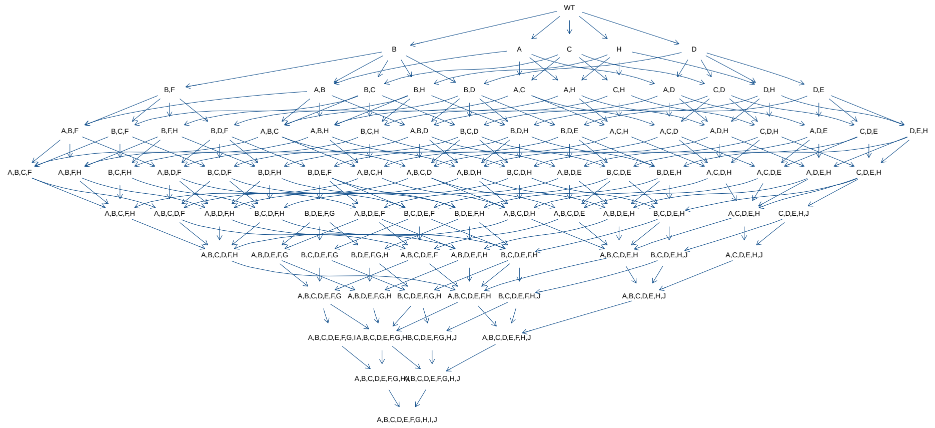

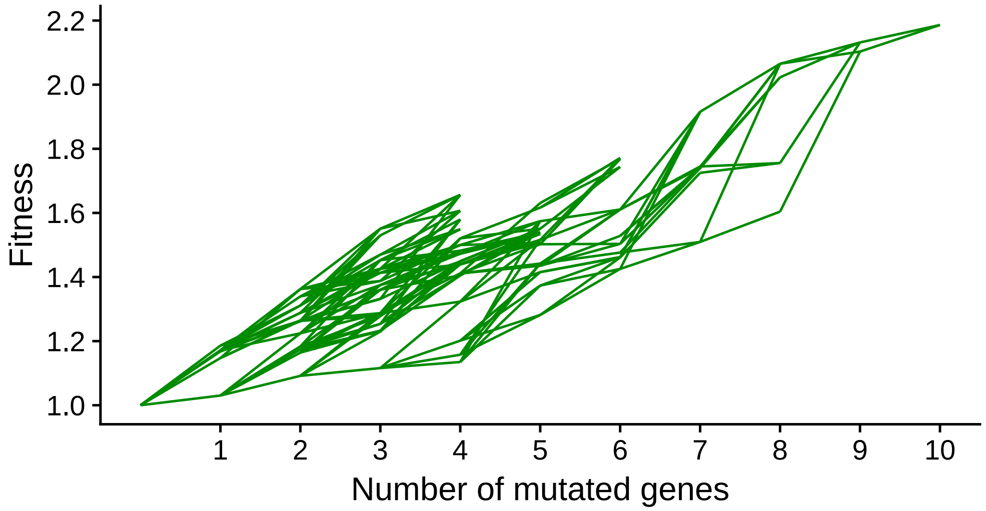


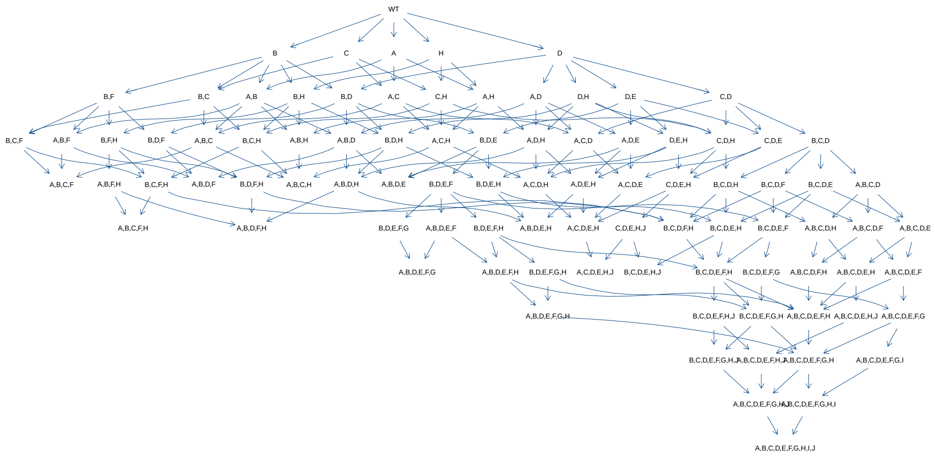

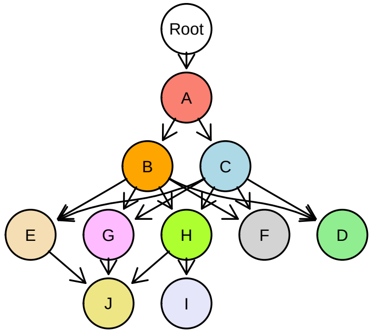


*ID* EPFiT9RotYTgEnMR
*acc genotypes* 60
*peaks no back* 7
*removed edges* 30
*prop rem edges* 0.196

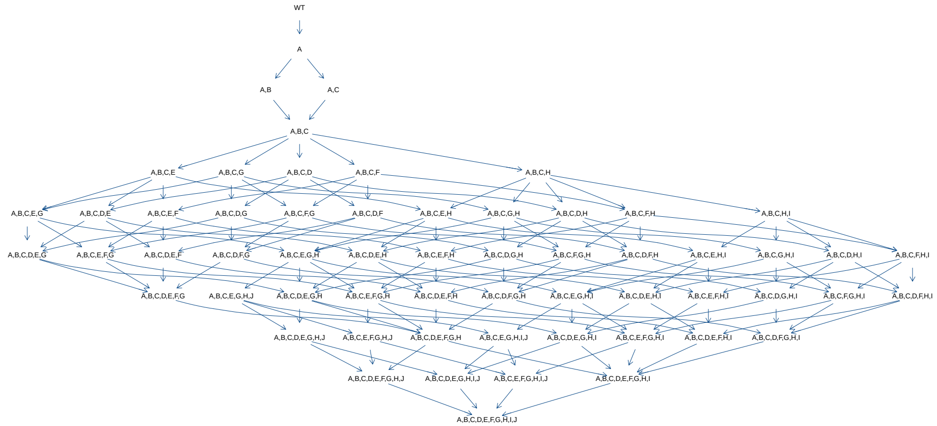

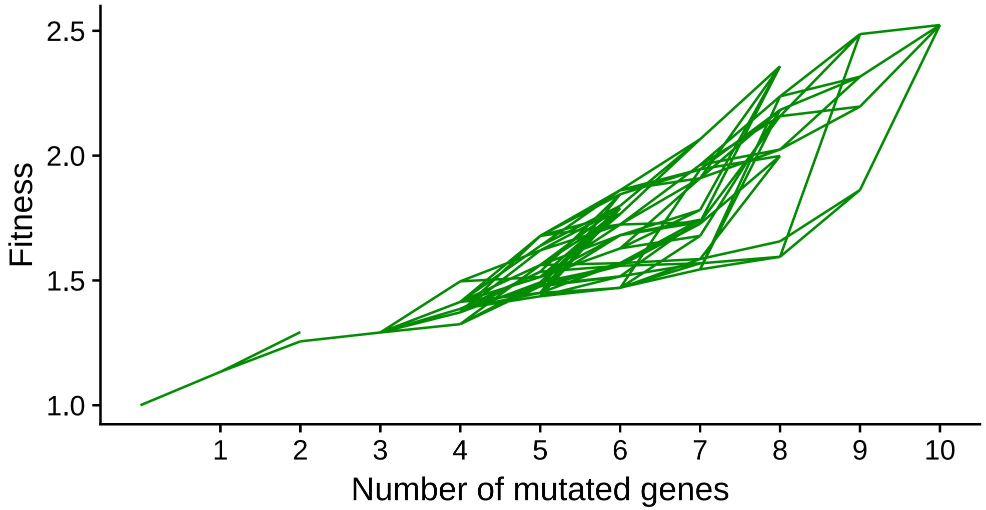


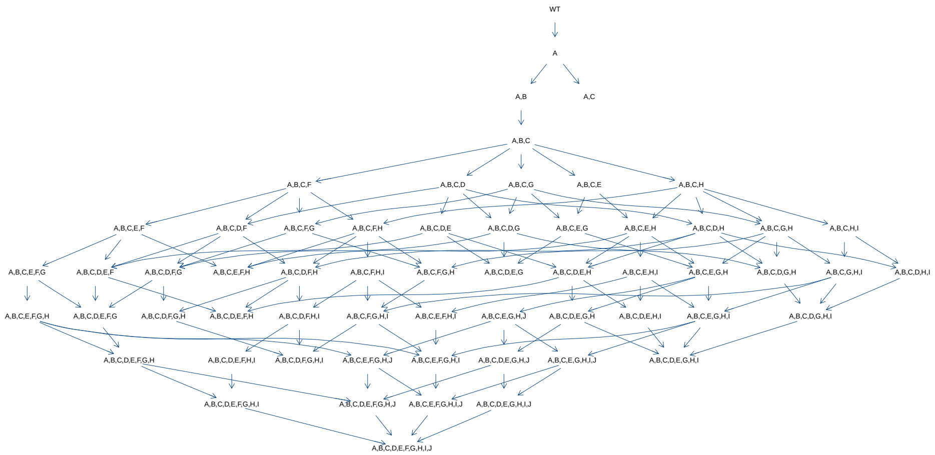

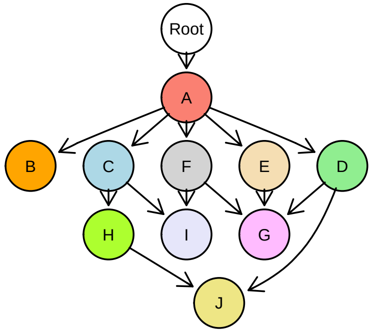


| | |
|---|---|
| *ID* | FcFcGhQtL910uSnWl |
| *acc genotypes* | 91 |
| *peaks no back* | 10 |
| *removed edges* | 51 |
| *prop rem edges* | 0.207 |

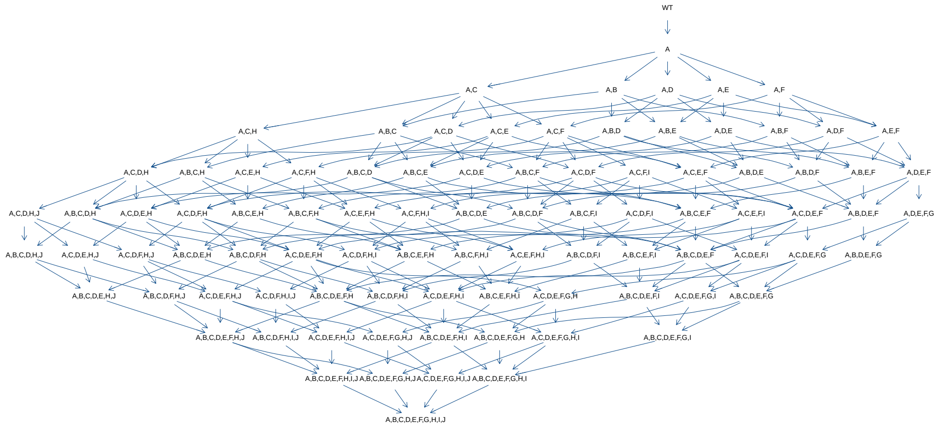

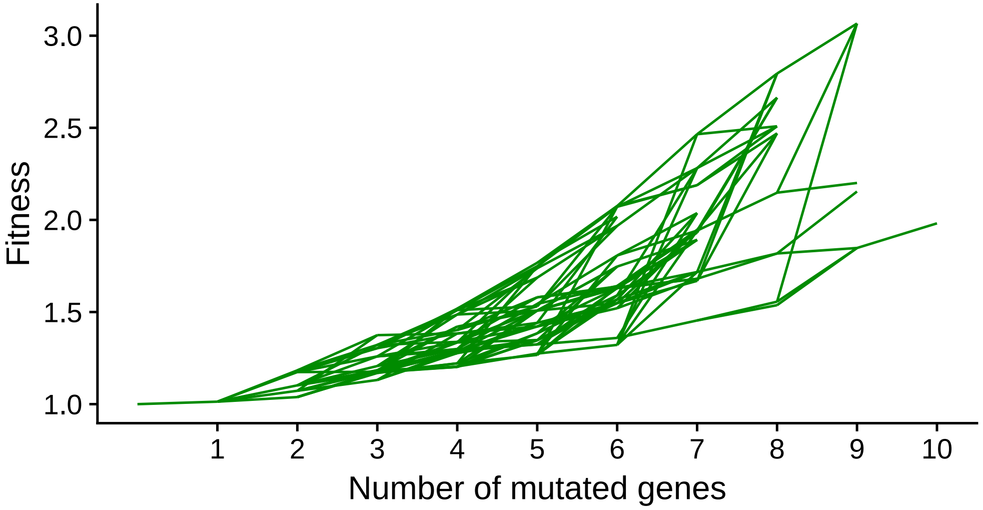


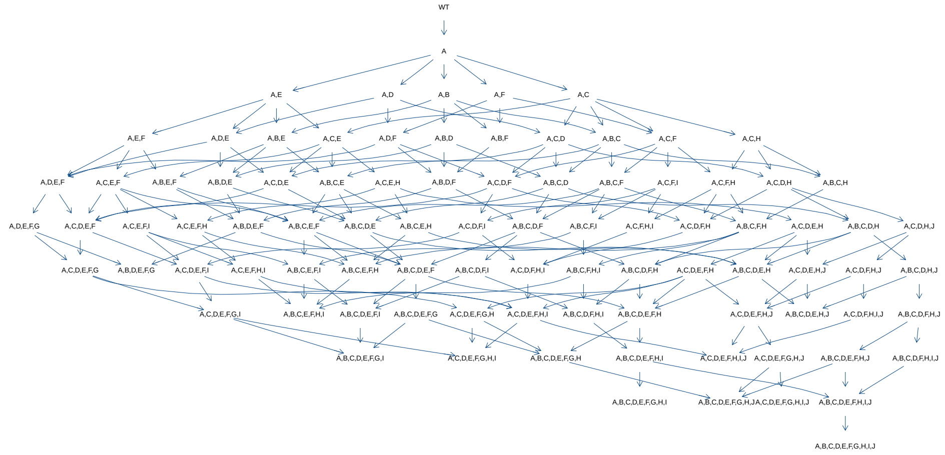

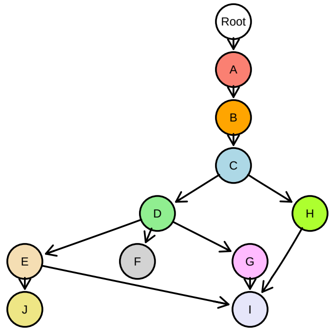


| | |
|---|---|
| *ID* | fCl5UMNFVGXtbOWb |
| *acc genotypes* | 33 |
| *peaks no back* | 2 |
| *removed edges* | 4 |
| *prop rem edges* | 0.061 |

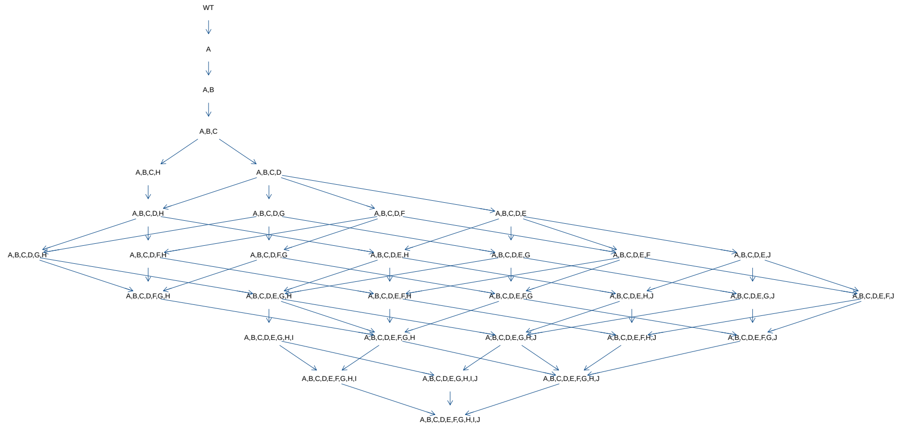

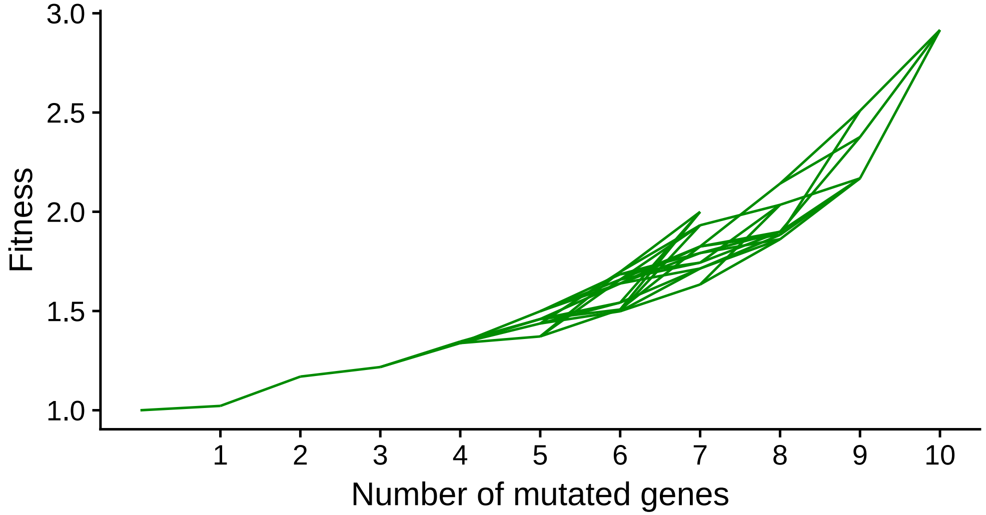


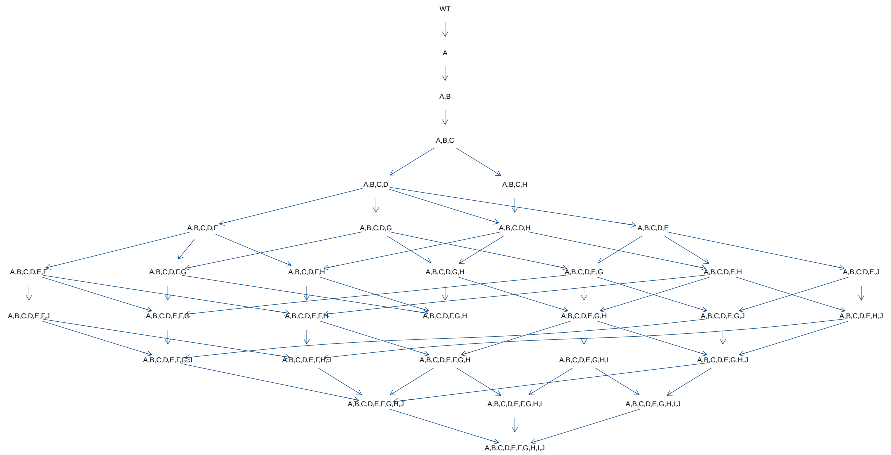

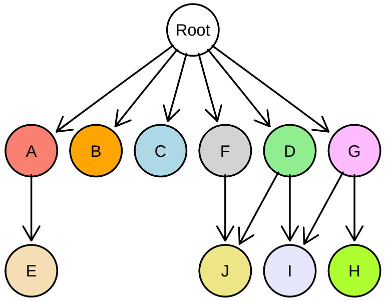


| | |
|---|---|
| *ID* | fjYl5FajfI1l9TqE |
| *acc genotypes* | 252 |
| *peaks no back* | 17 |
| *removed edges* | 166 |
| *prop rem edges* | 0.189 |

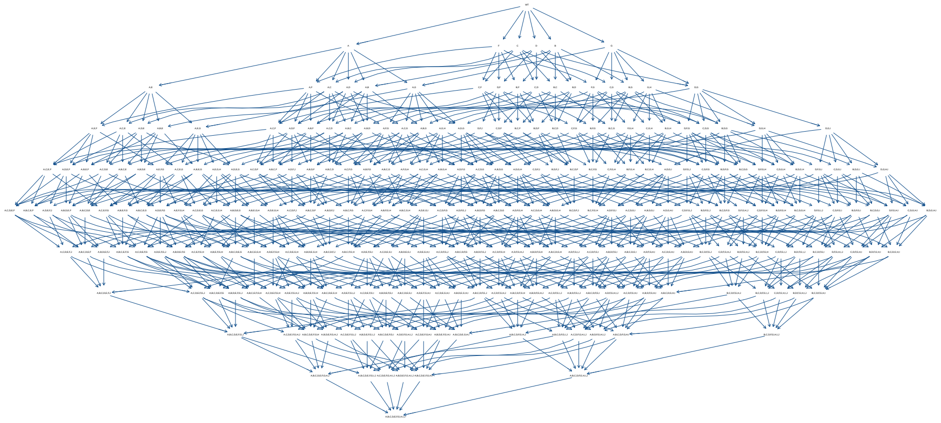

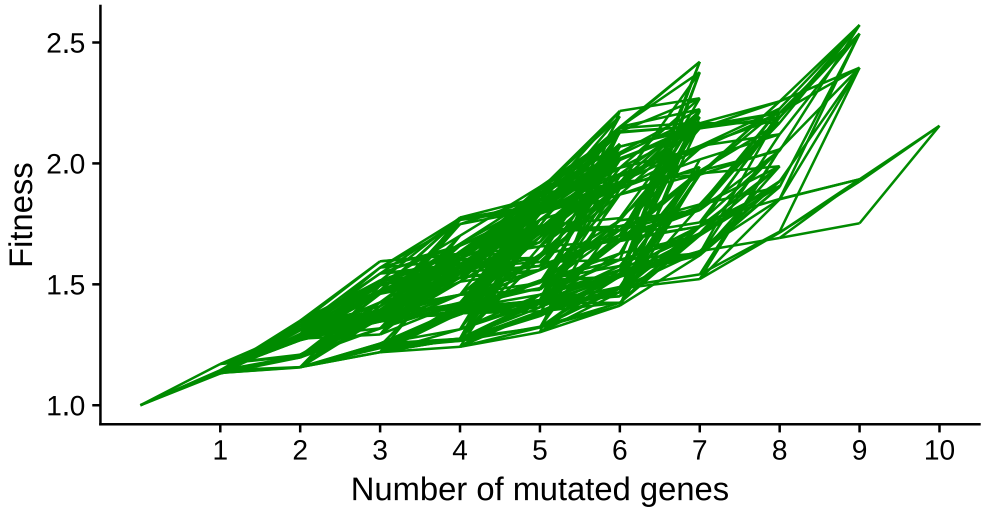


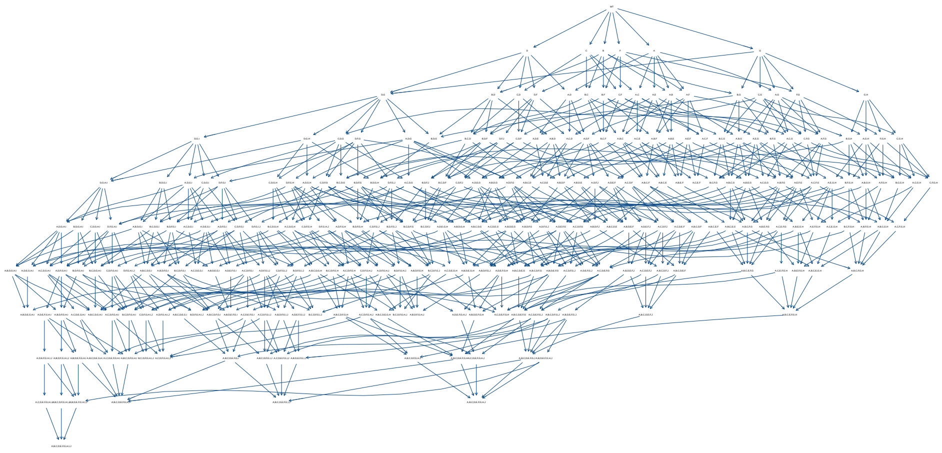

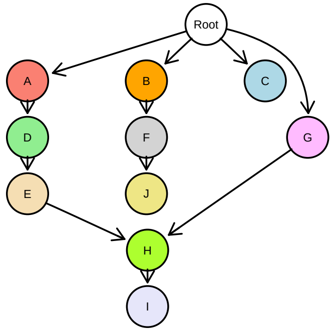


| | |
|---|---|
| *ID* | FNDb716ELrbzr3xJ |
| *acc genotypes* | 80 |
| *peaks no back* | 3 |
| *removed edges* | 22 |
| *prop rem edges* | 0.112 |

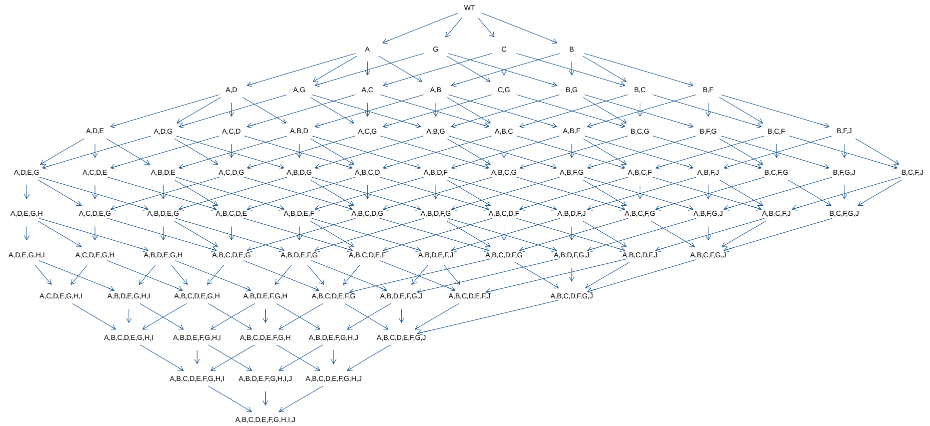

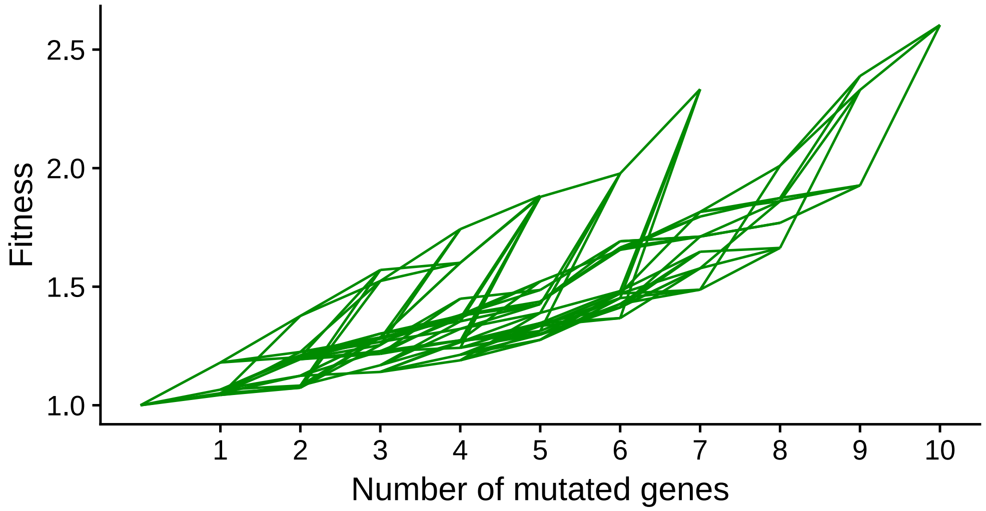


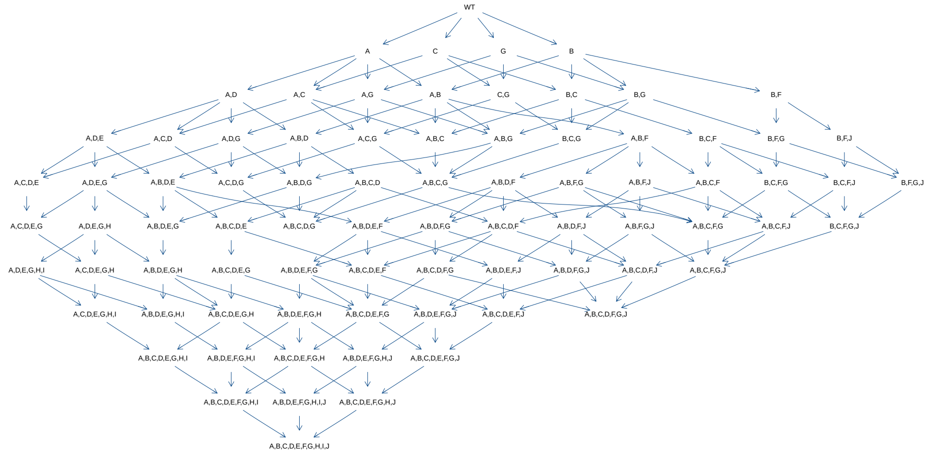

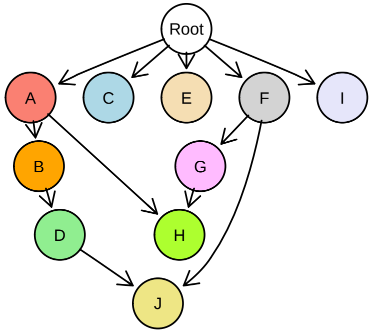


| | |
|---|---|
| *ID* | Fp1fshyDCkcvIW97 |
| *acc genotypes* | 144 |
| *peaks no back* | 13 |
| *removed edges* | 95 |
| *prop rem edges* | 0.22 |

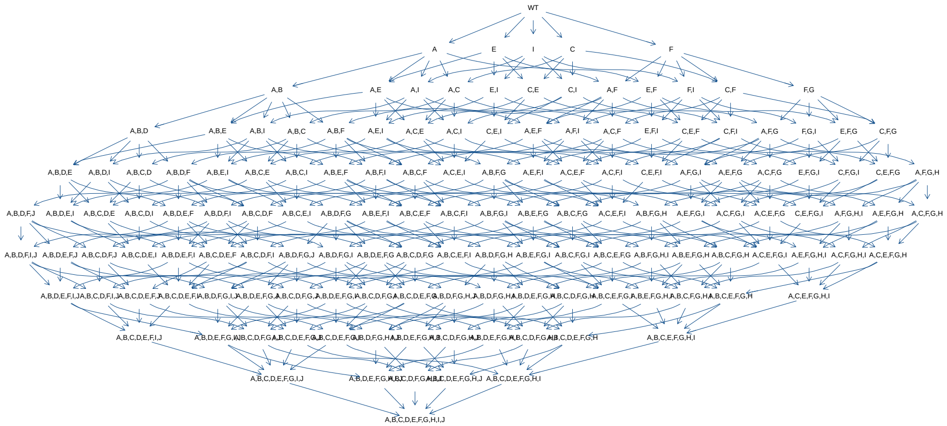

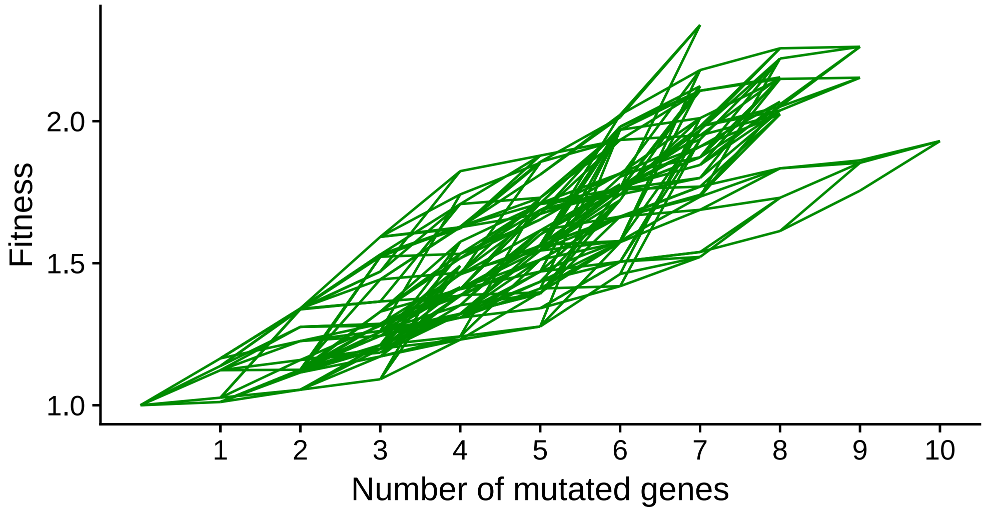


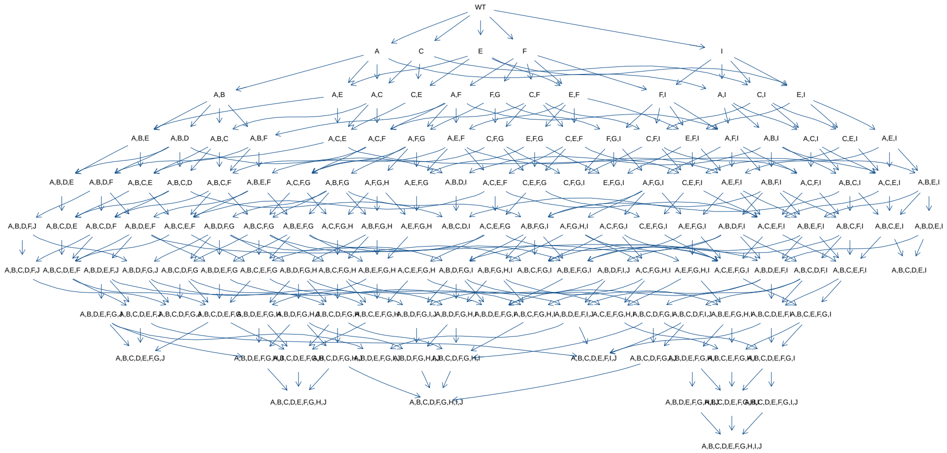

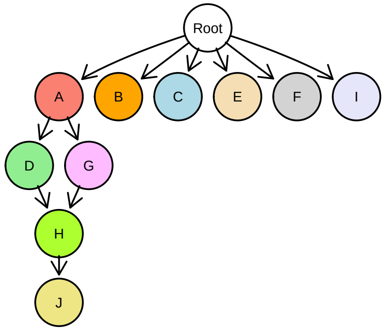


| | |
|---|---|
| *ID* | ftHhRYTFn79eRdaP |
| *acc genotypes* | 224 |
| *peaks no back* | 19 |
| *removed edges* | 143 |
| *prop rem edges* | 0.182 |

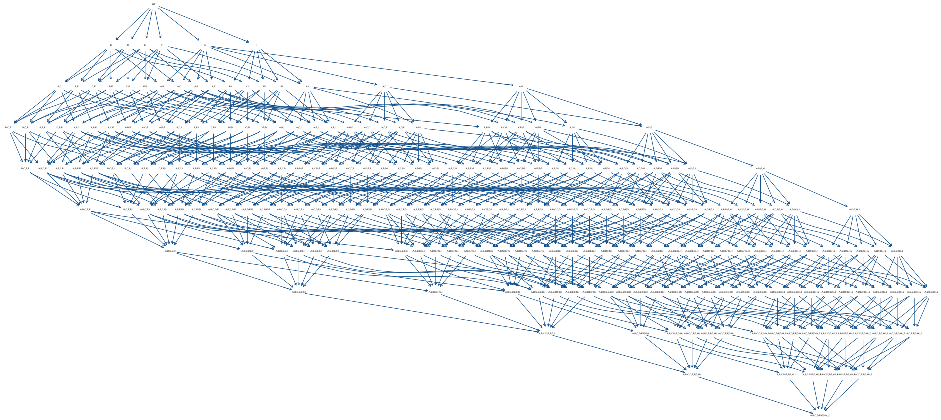

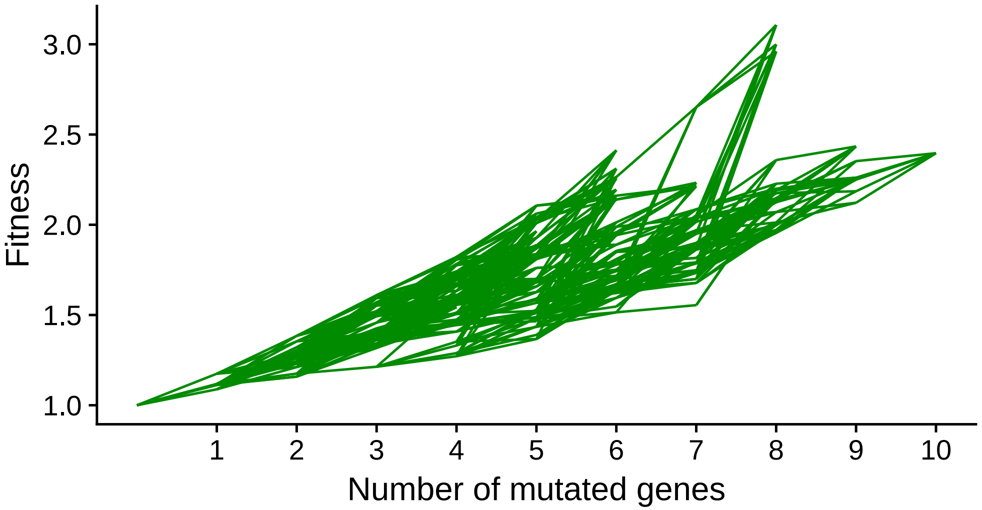


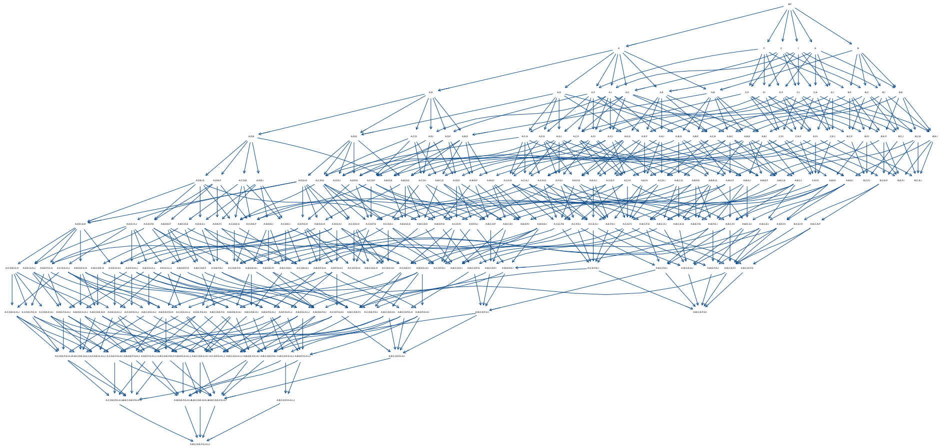

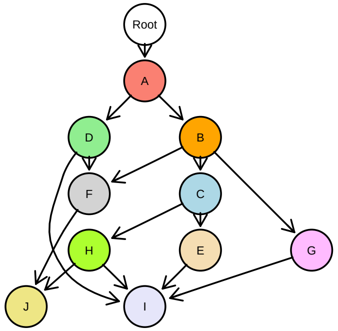


*ID* fV68aMwNJcqHIZQX
*acc genotypes* 40
*peaks no back* 3
*removed edges* 11
*prop rem edges* 0.134

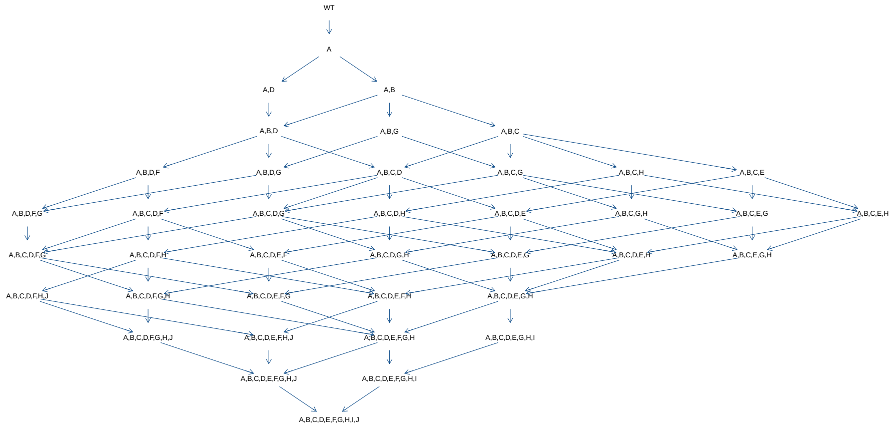

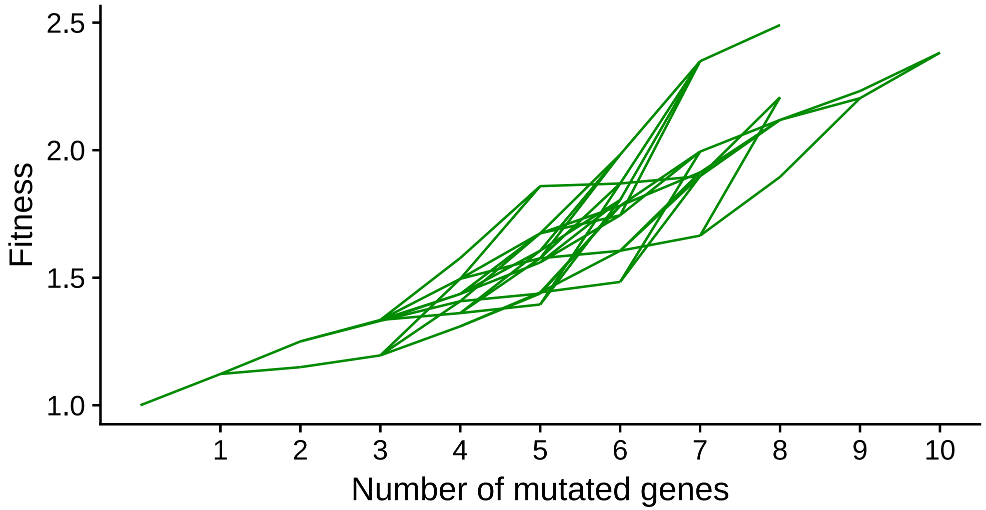


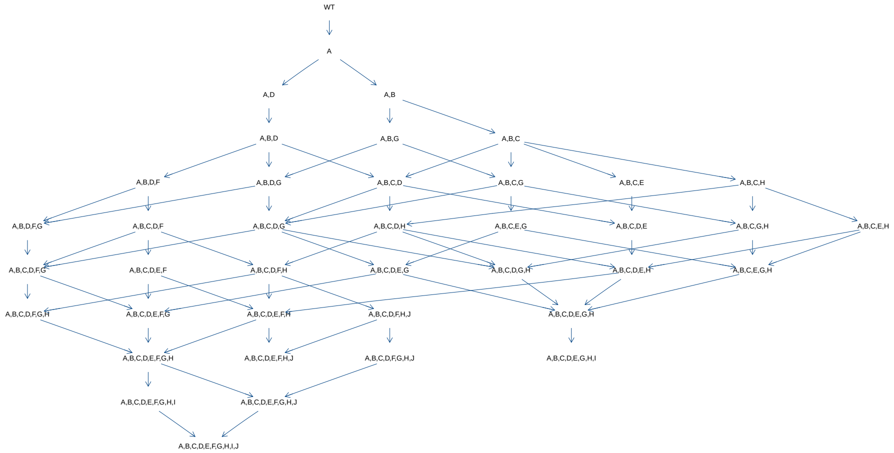

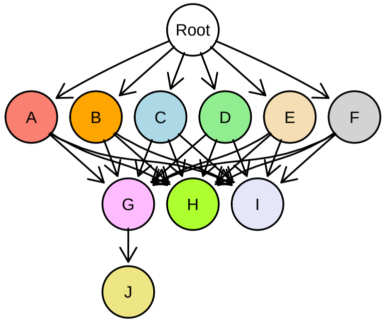


| | |
|---|---|
| *ID* | g632uhVWzFmg59eK |
| *acc genotypes* | 75 |
| *peaks no back* | 10 |
| *removed edges* | 31 |
| *prop rem edges* | 0.146 |

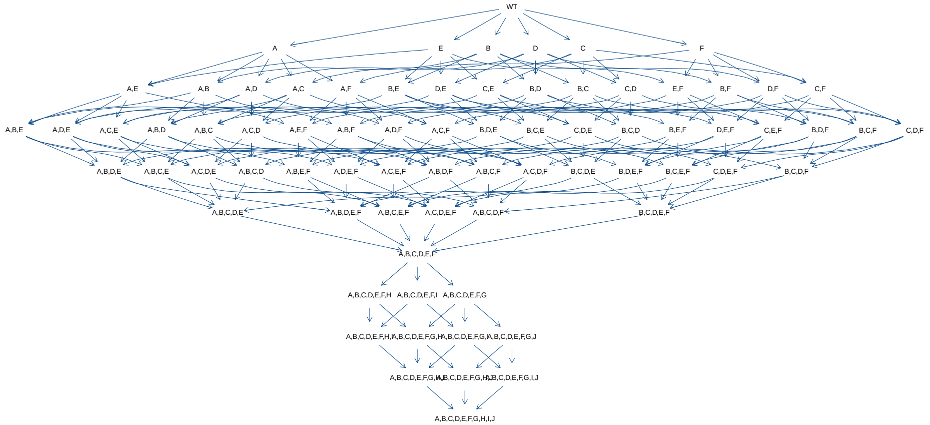

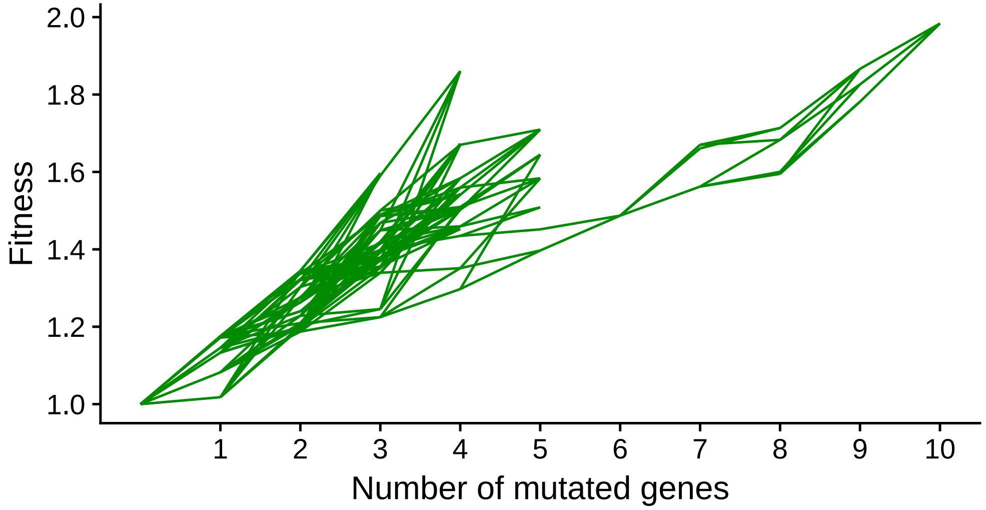


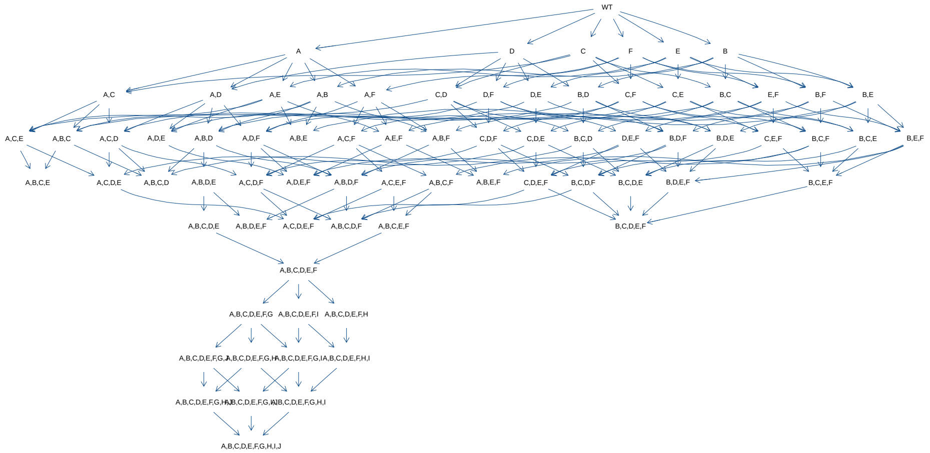

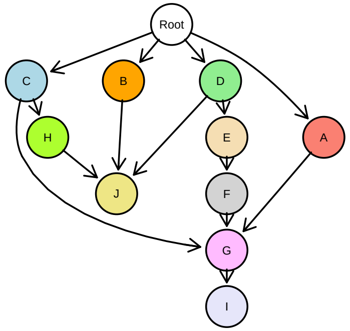


| | |
|---|---|
| *ID* | G9yrL22aEcqd1Y23 |
| *acc genotypes* | 64 |
| *peaks no back* | 8 |
| *removed edges* | 28 |
| *prop rem edges* | 0.188 |

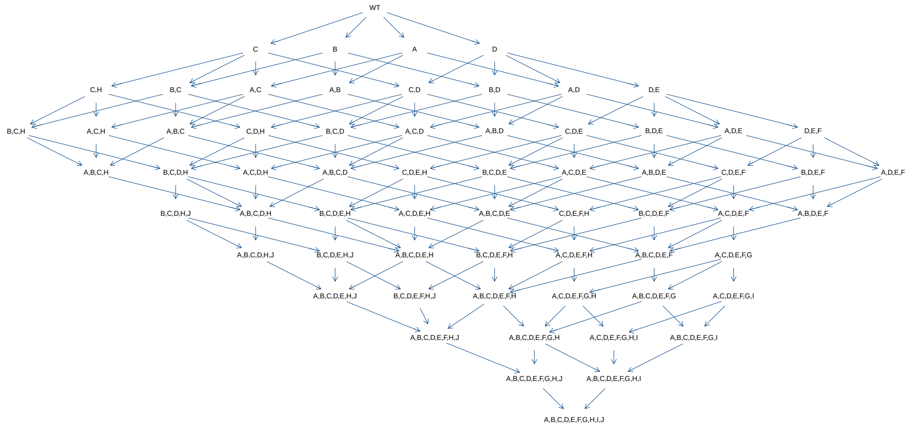

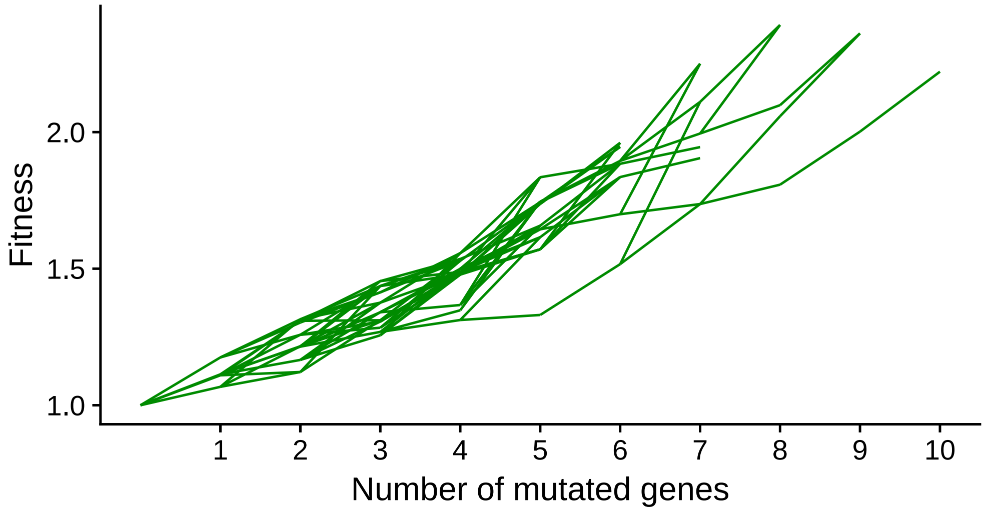


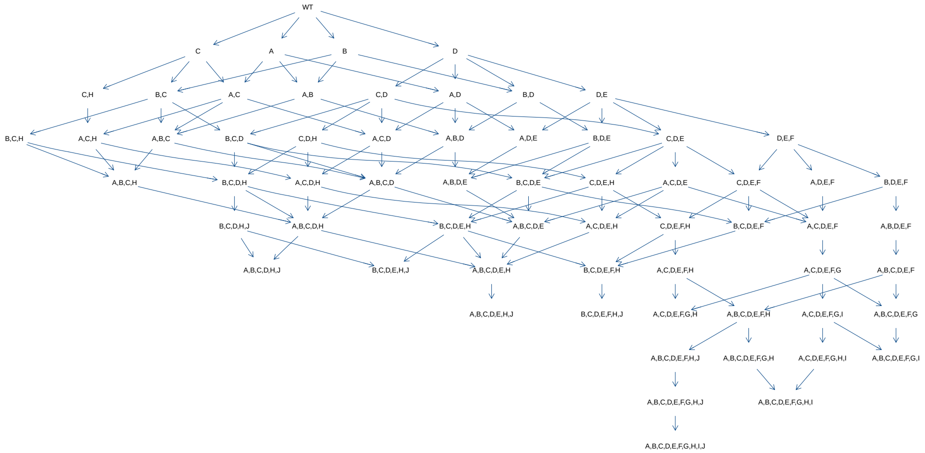

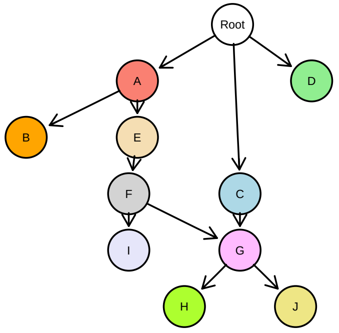


| | |
|---|---|
| *ID* | gA76bffYxtXYMJa2 |
| *acc genotypes* | 68 |
| *peaks no back* | 11 |
| *removed edges* | 30 |
| *prop rem edges* | 0.179 |

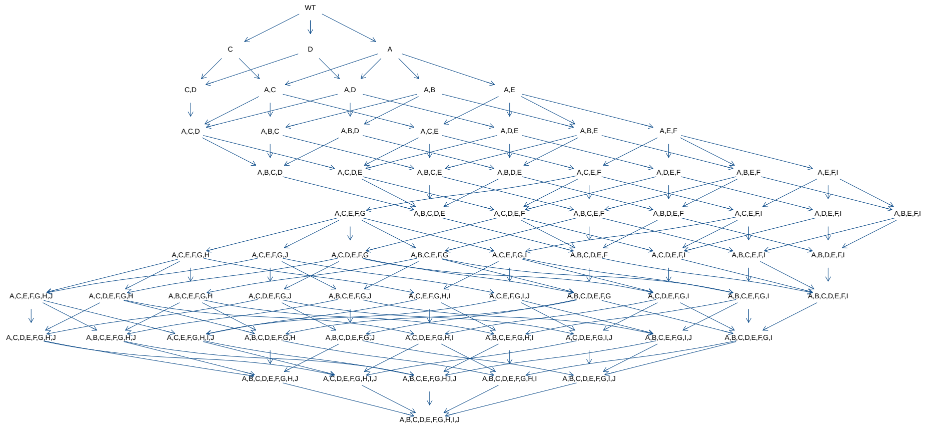

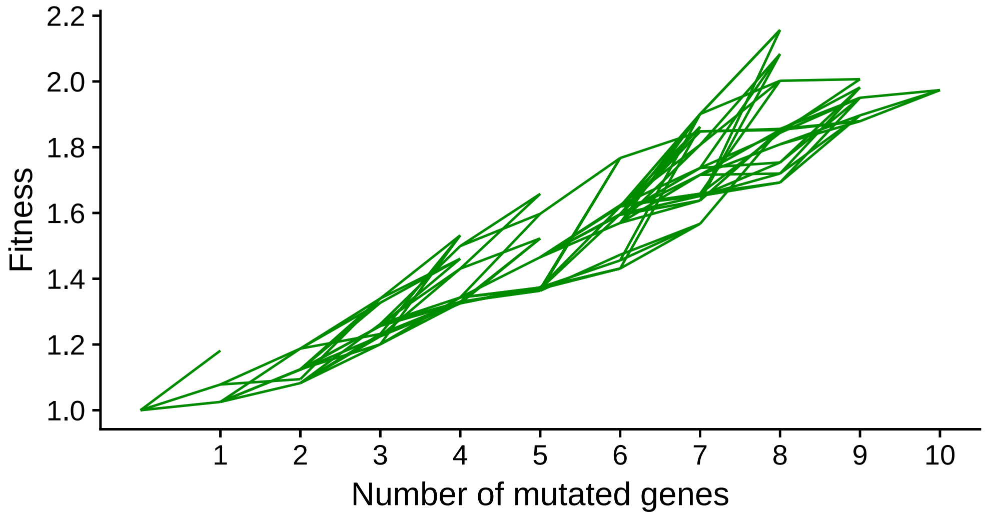


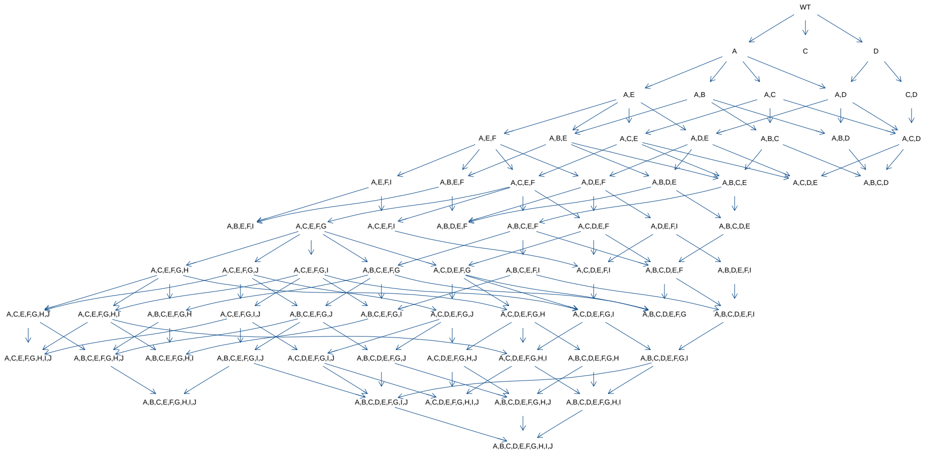

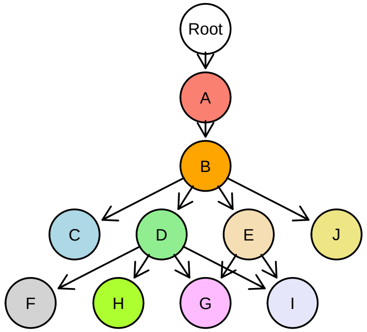


| | |
|---|---|
| *ID* | GeE6ZiWGbvgLEf7V |
| *acc genotypes* | 90 |
| *peaks no back* | 11 |
| *removed edges* | 56 |
| *prop rem edges* | 0.214 |

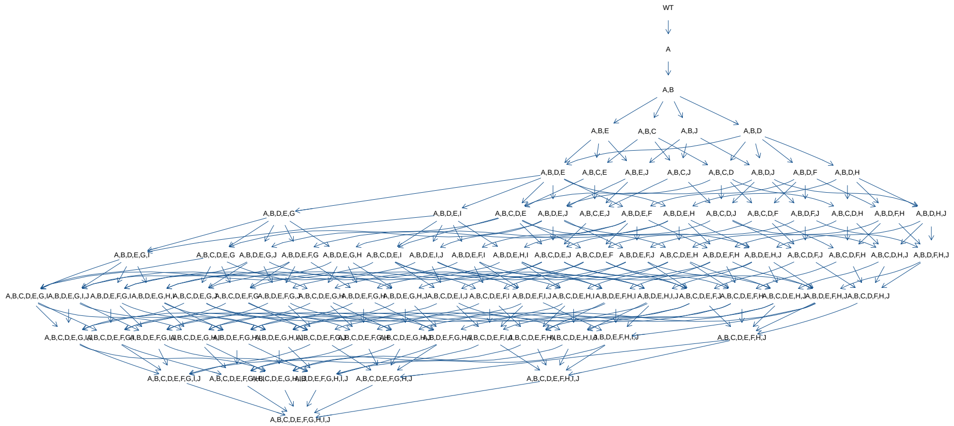

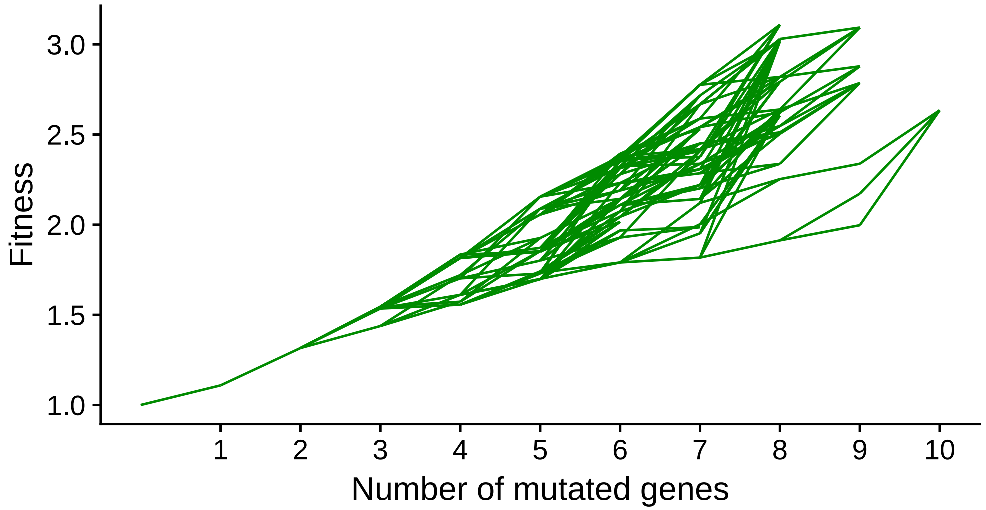


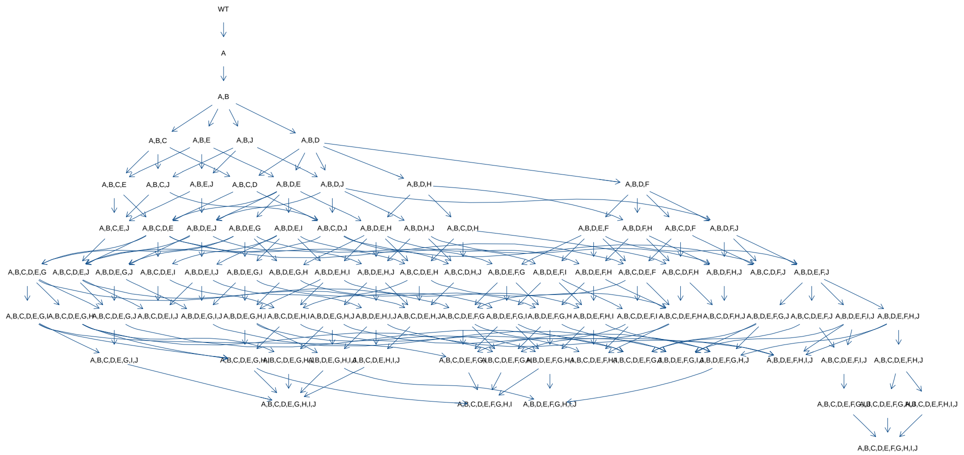

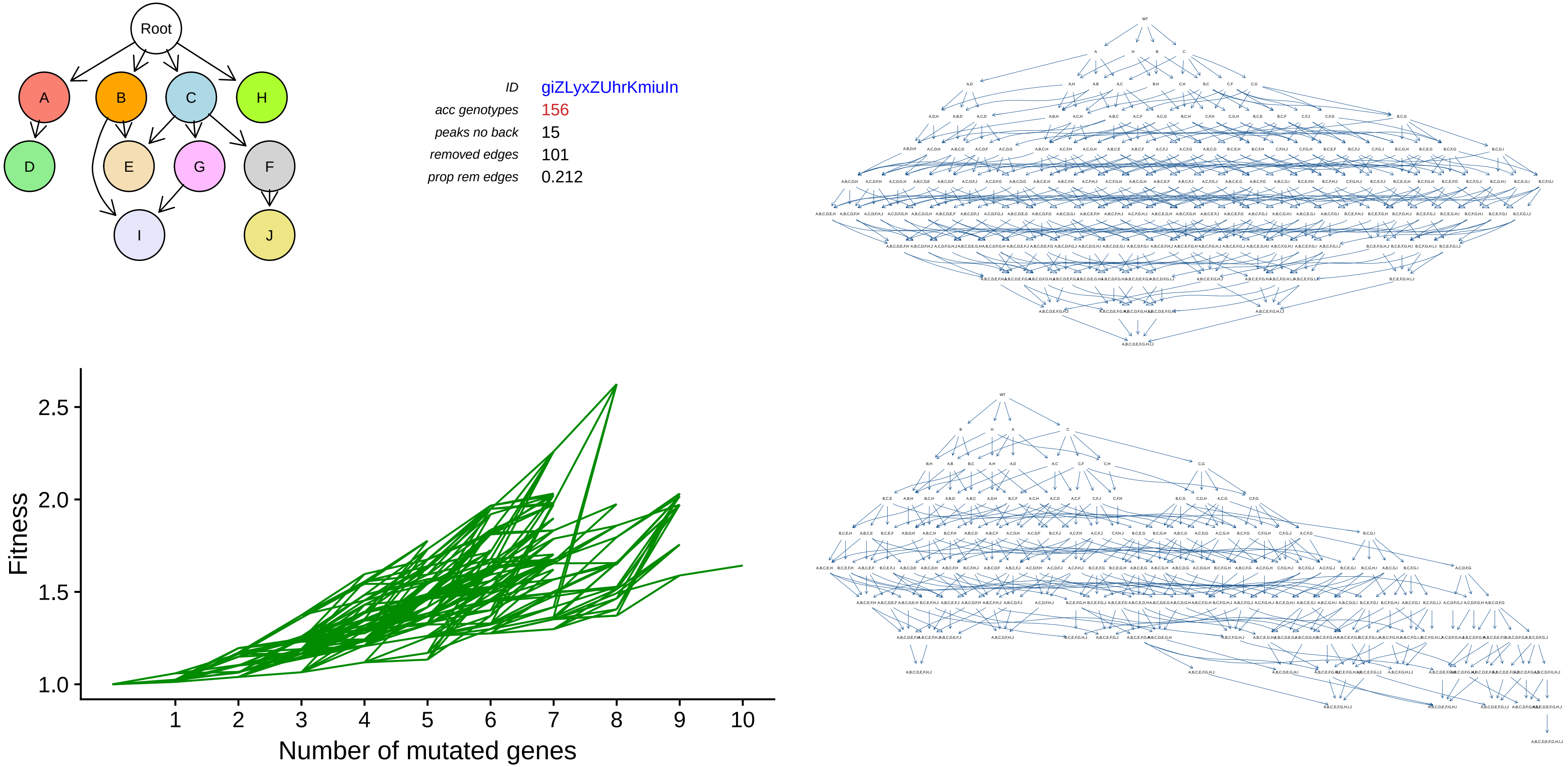
Root
A
B
C
H
D
E
G
F
I
J
ID giZLyxZUhrKmiuIn
acc genotypes 156
peaks no back 15
removed edges 101
prop rem edges 0.212
Fitness
2.5
2.0
1.5
1.0
1
2
3
4
5
6
7
8
9
10
Number of mutated genes

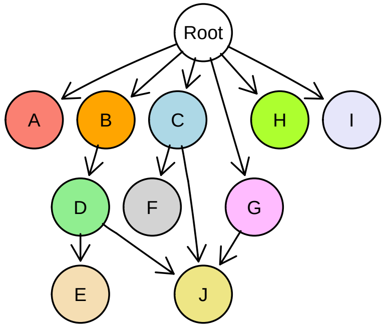


| | |
|---|---|
| *ID* | glDXEi6qi95q3Iss |
| *acc genotypes* | 224 |
| *peaks no back* | 18 |
| *removed edges* | 174 |
| *prop rem edges* | 0.227 |

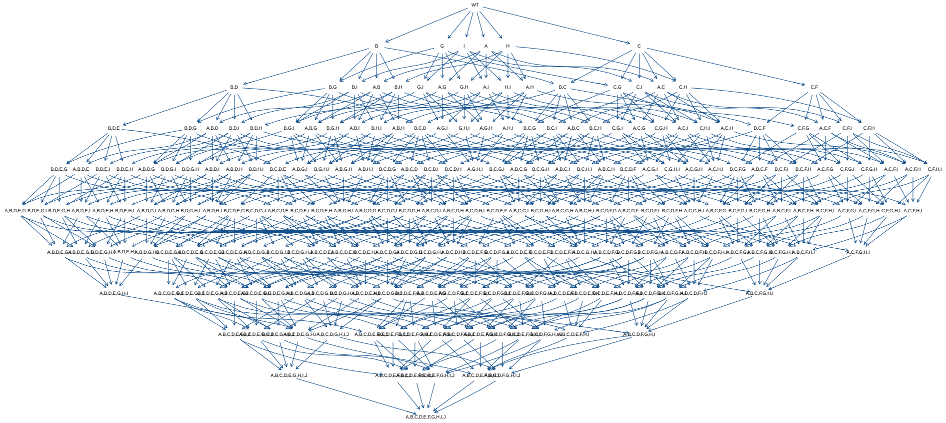

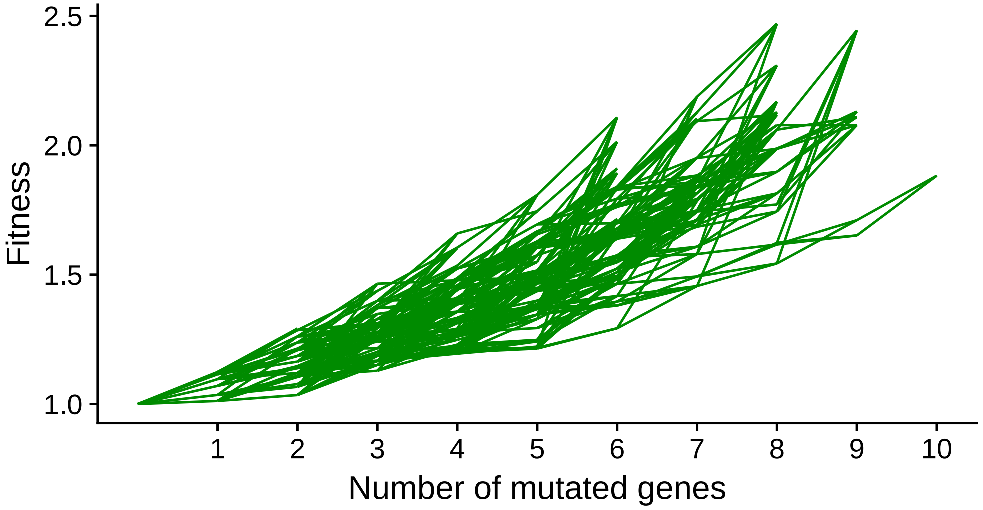


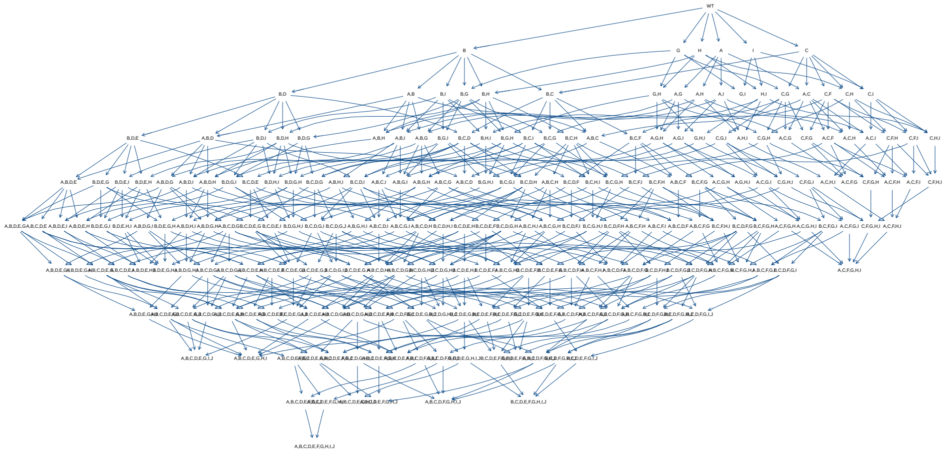

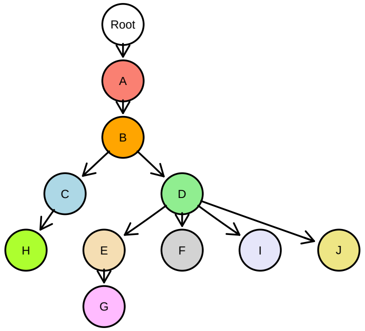


| | |
|---|---|
| *ID* | gnxoxbzJ9TsWlPFr |
| *acc genotypes* | 77 |
| *peaks no back* | 9 |
| *removed edges* | 45 |
| *prop rem edges* | 0.213 |

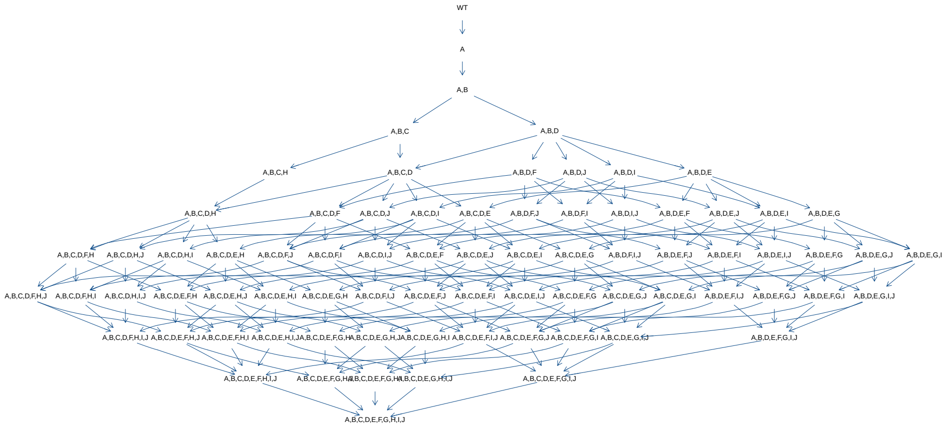

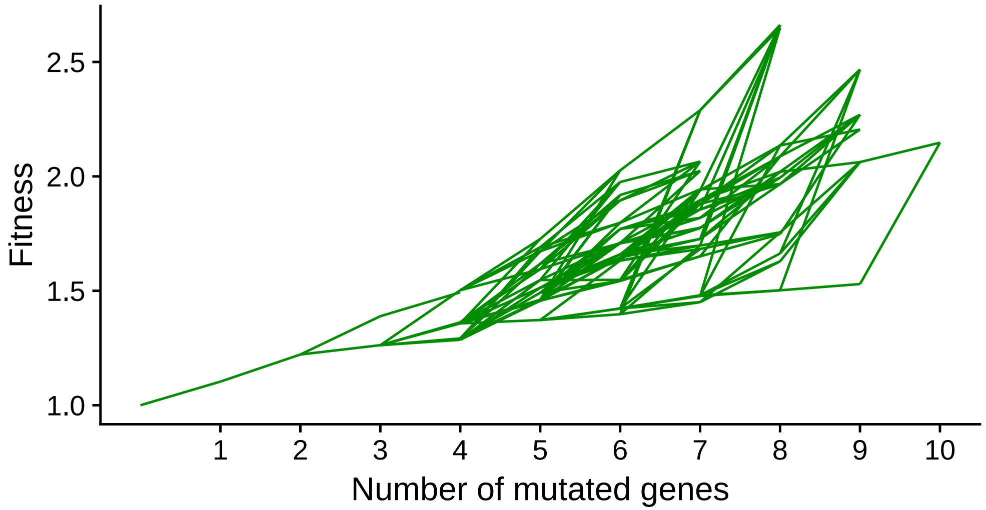


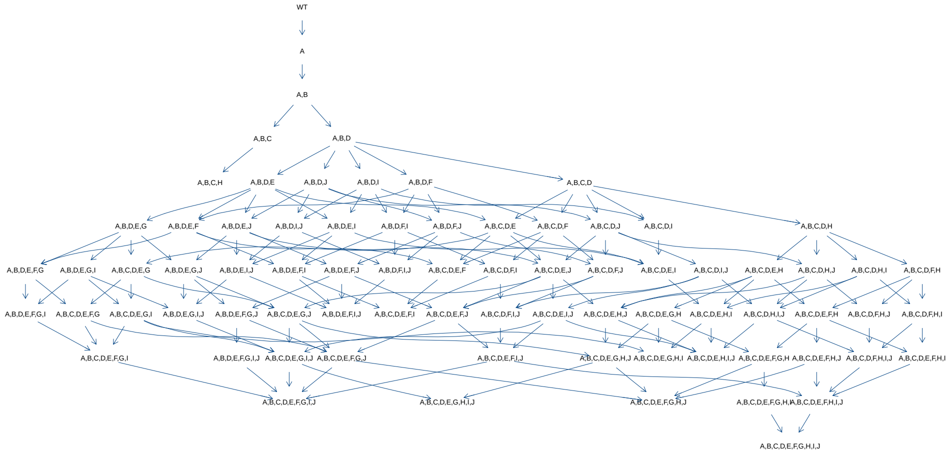

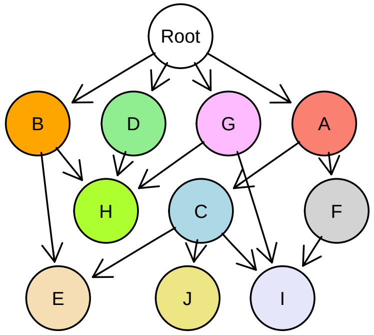


*ID* Gw7Lw136jeS105BFu
*acc genotypes* 99
*peaks no back* 6
*removed edges* 44
*prop rem edges* 0.165

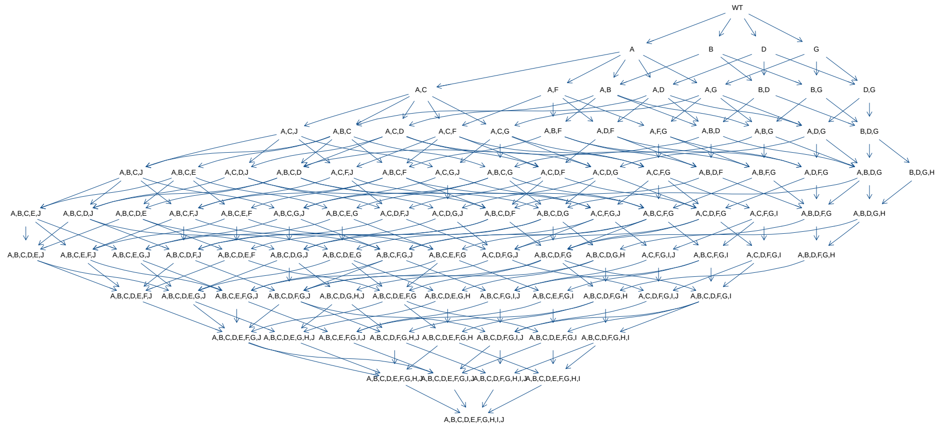

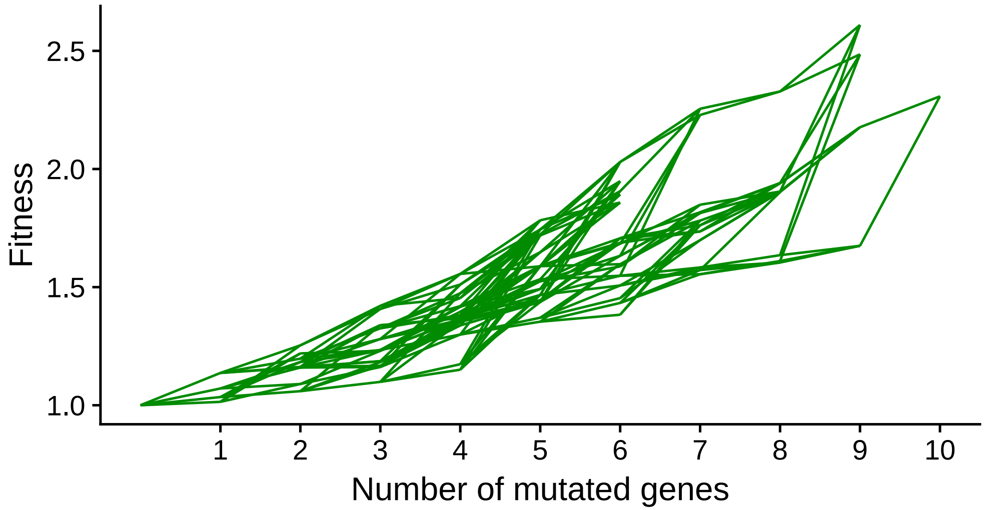


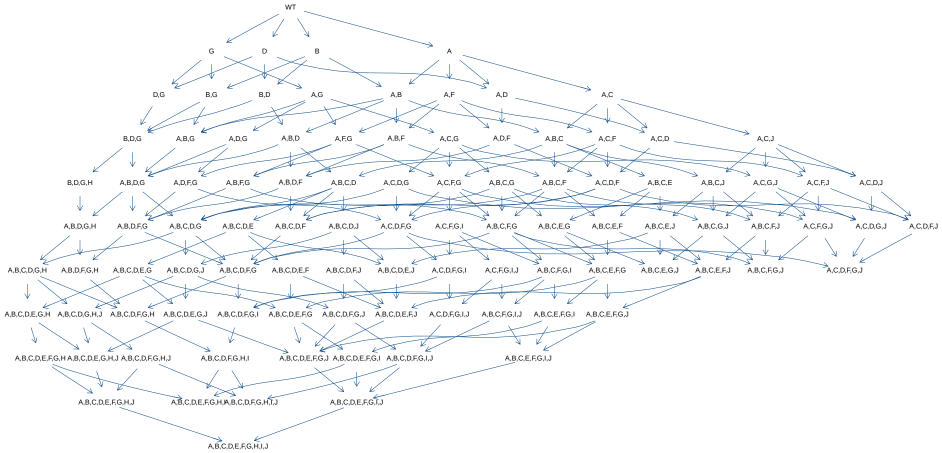

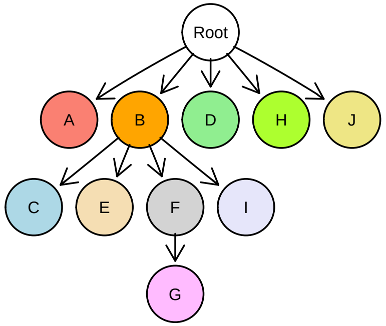


| | |
|---|---|
| *ID* | hEMMzV7G9flteqwr |
| *acc genotypes* | 400 |
| *peaks no back* | 24 |
| *removed edges* | 307 |
| *prop rem edges* | 0.186 |

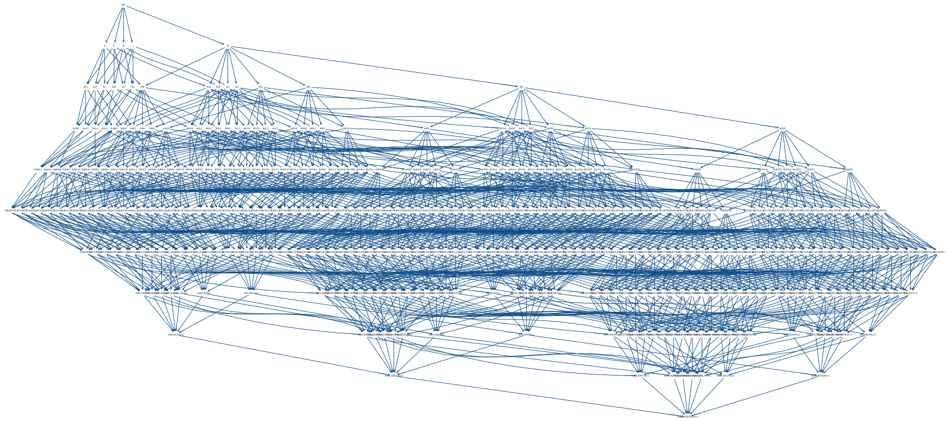

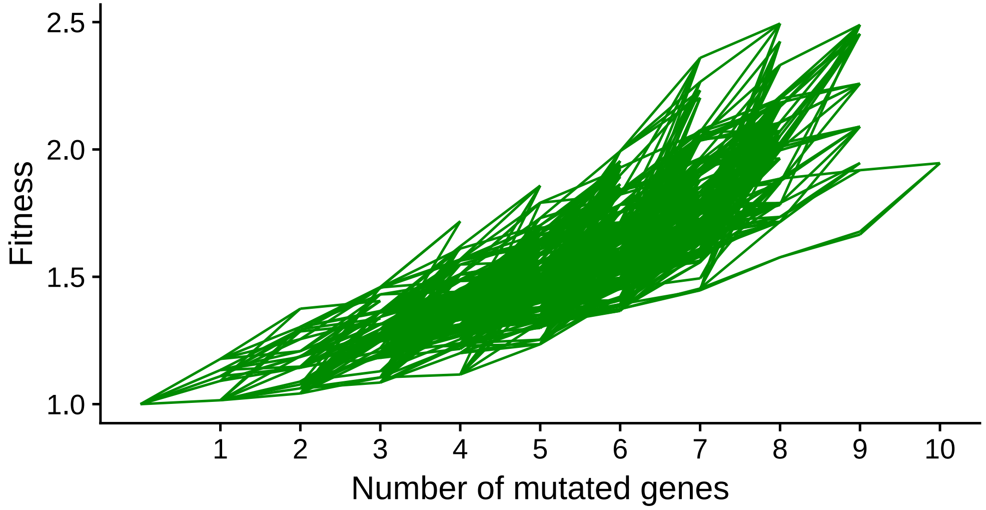


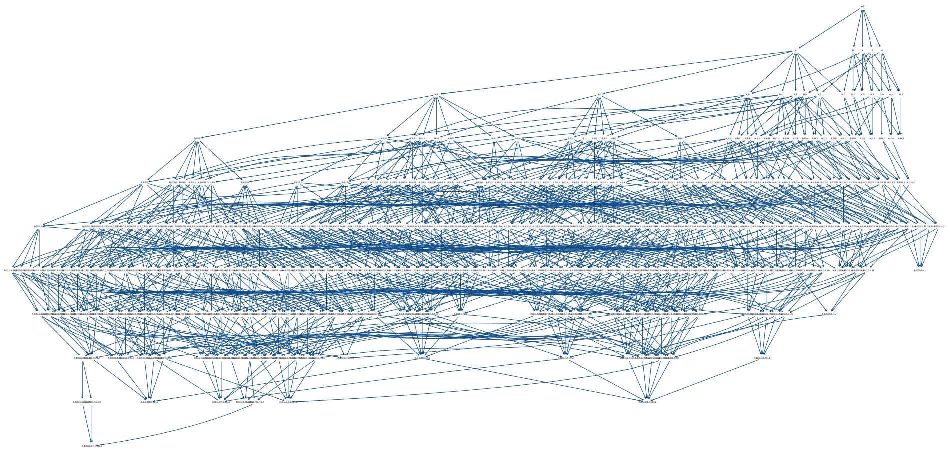

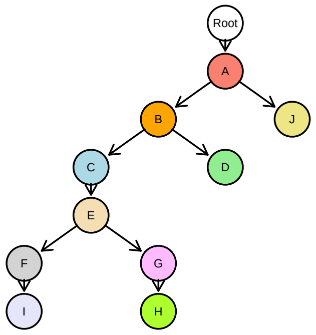


| | |
|---|---|
| *ID* | HJf8BvCZrOXnhdpP |
| *acc genotypes* | 47 |
| *peaks no back* | 5 |
| *removed edges* | 21 |
| *prop rem edges* | 0.202 |

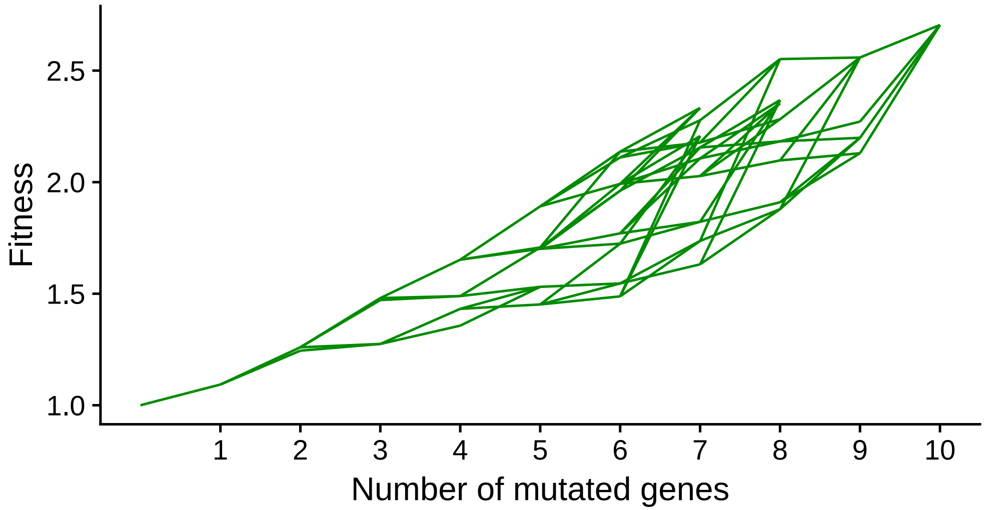


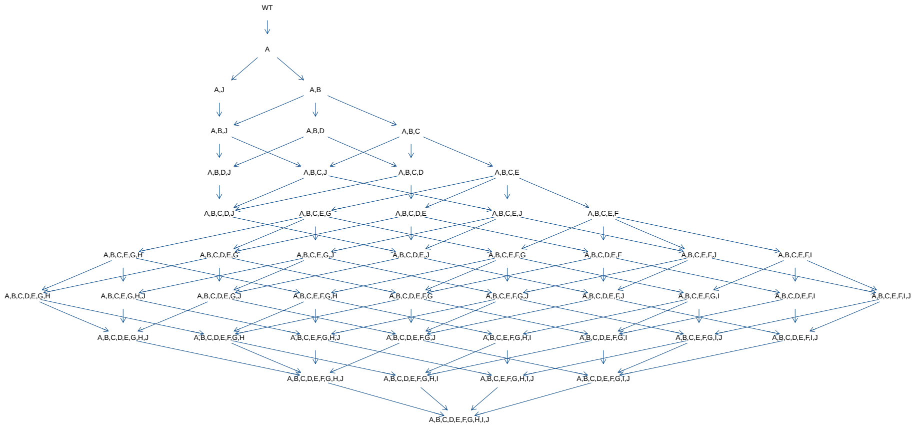

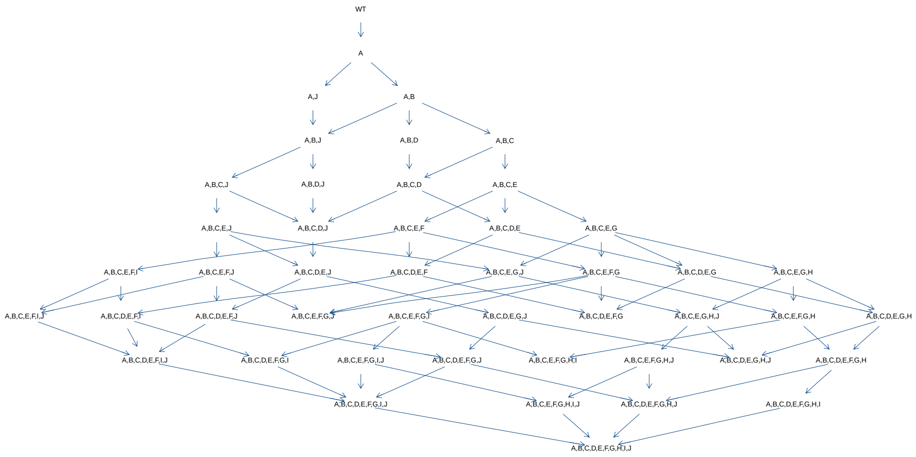

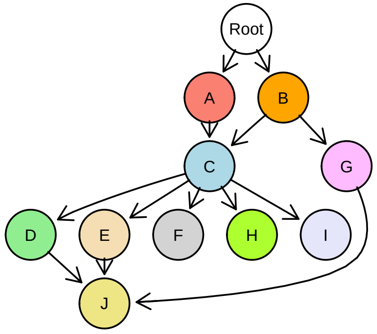


| | |
|---|---|
| *ID* | hjGRWsJhOh106RXPY |
| *acc genotypes* | 78 |
| *peaks no back* | 8 |
| *removed edges* | 48 |
| *prop rem edges* | 0.217 |

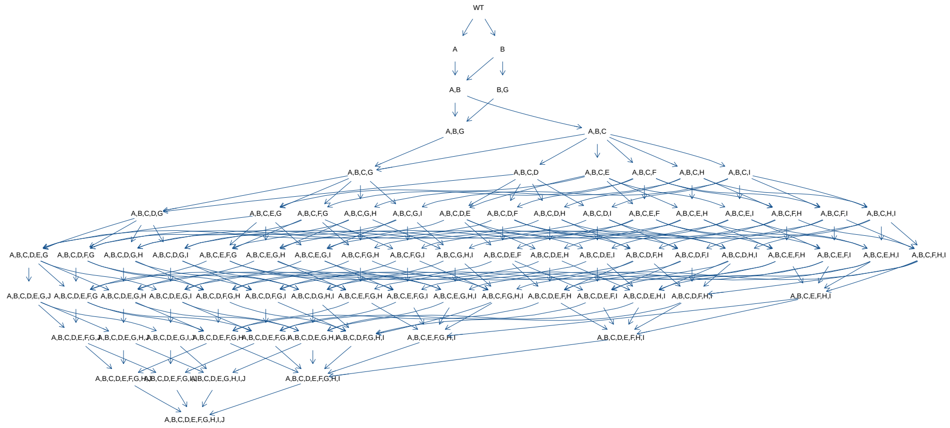

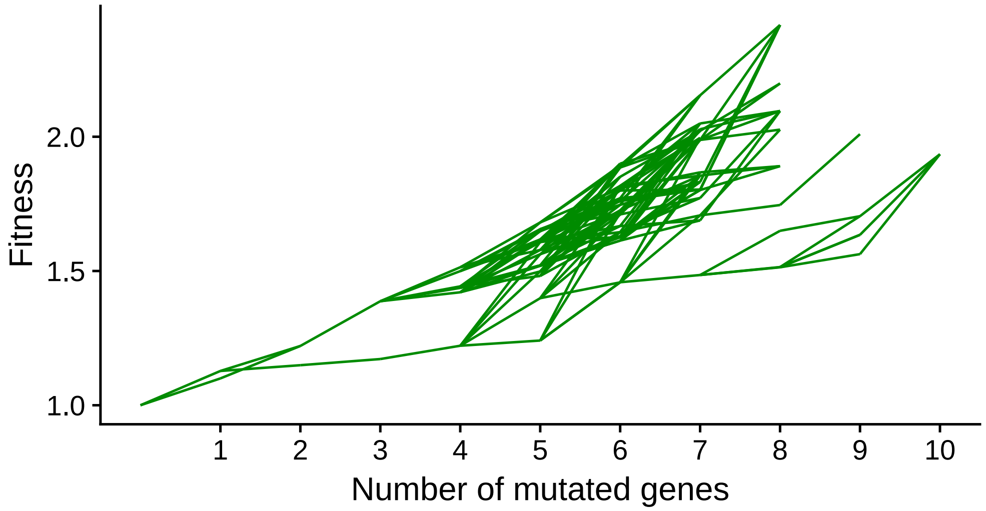


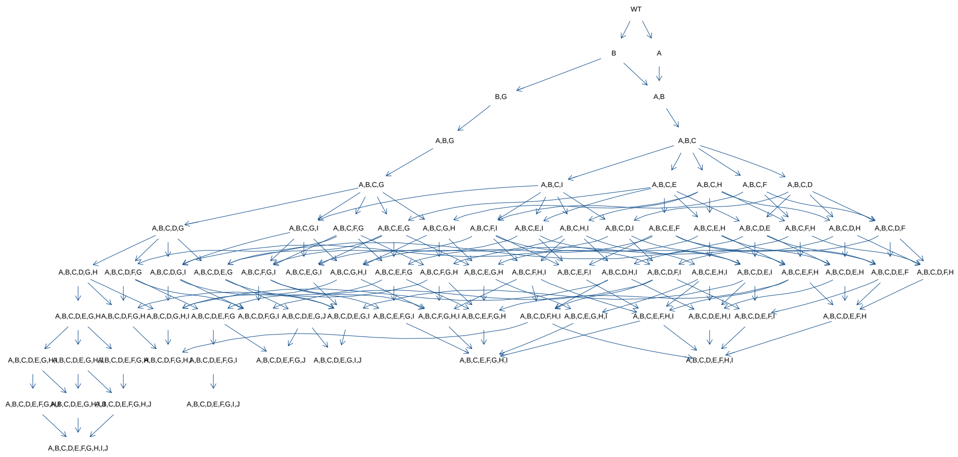

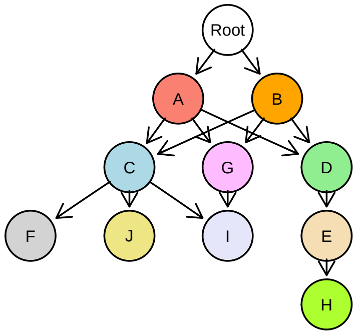


| | |
|---|---|
| *ID* | HjmKaDaJUp3CbAMd |
| *acc genotypes* | 59 |
| *peaks no back* | 9 |
| *removed edges* | 21 |
| *prop rem edges* | 0.152 |

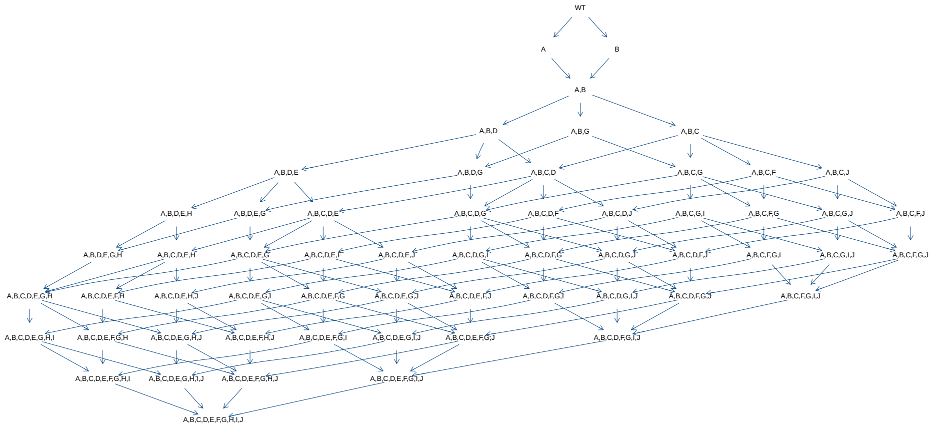

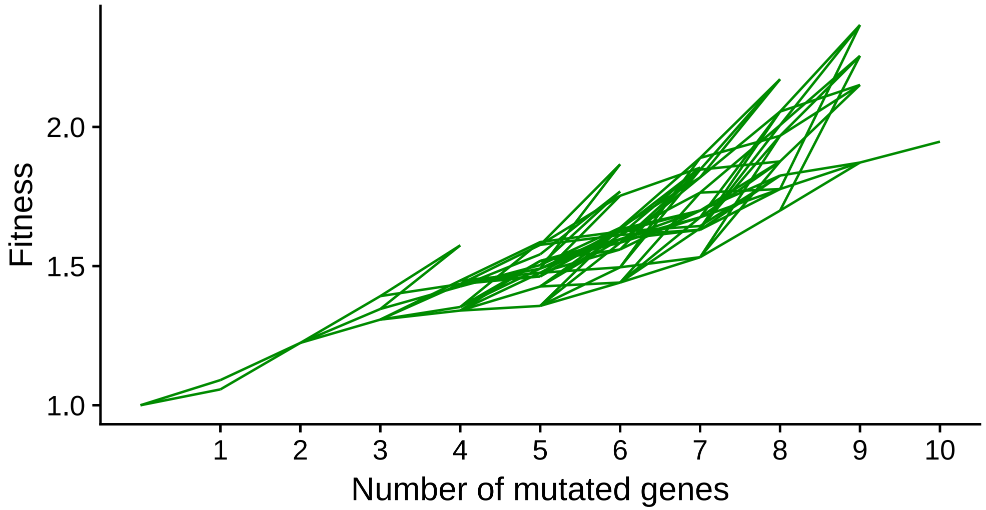


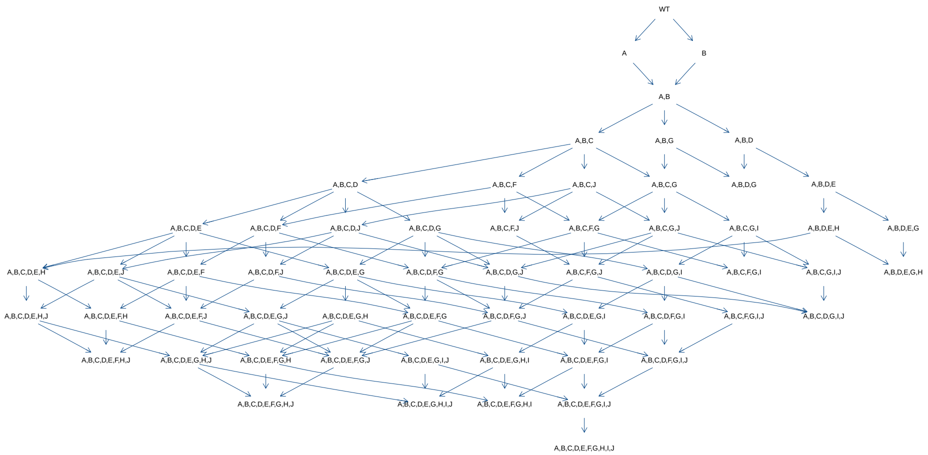

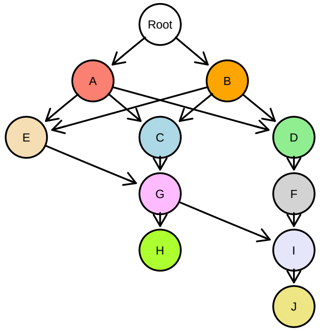


| | |
|---|---|
| *ID* | hootgr4D2eT9wPQw |
| *acc genotypes* | 25 |
| *peaks no back* | 2 |
| *removed edges* | 3 |
| *prop rem edges* | 0.075 |

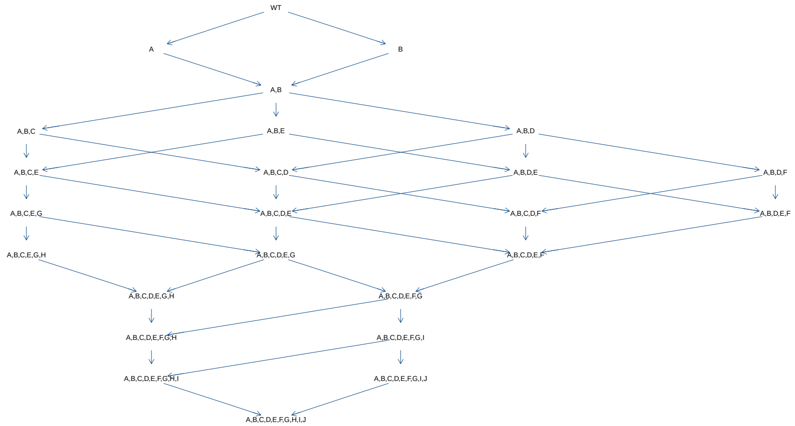


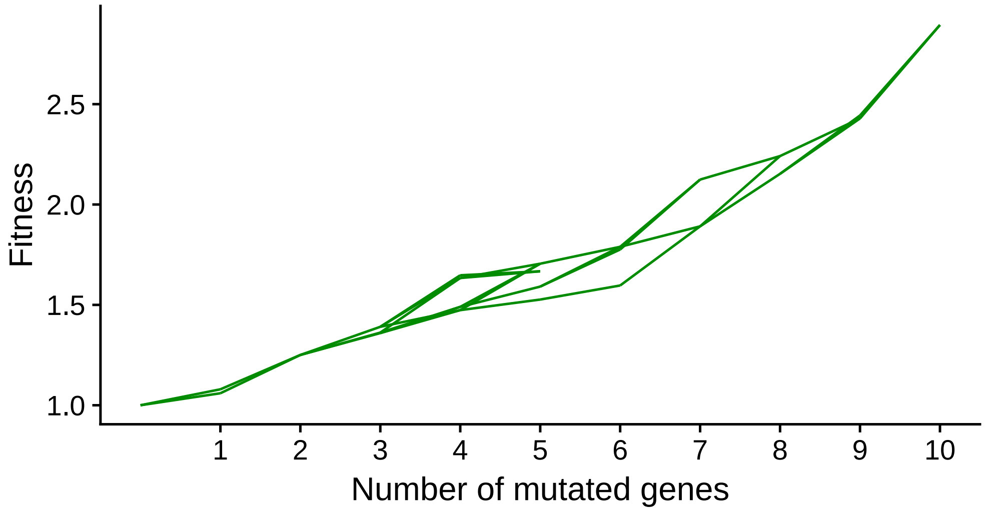


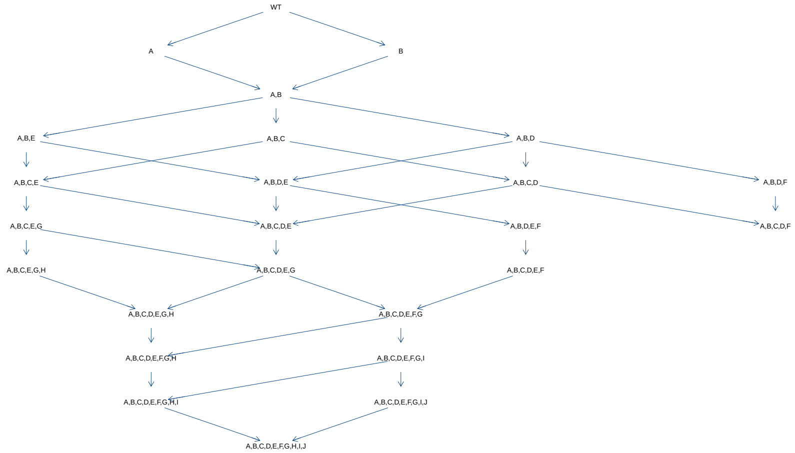

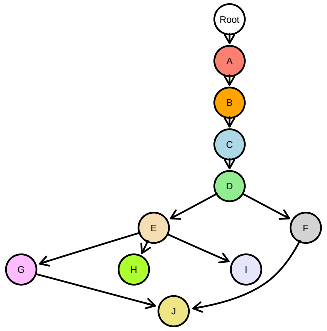


*ID* i210cnkWHYkaWwl3l

*acc genotypes* 26

*peaks no back* 4

*removed edges* 7

*prop rem edges* 0.149

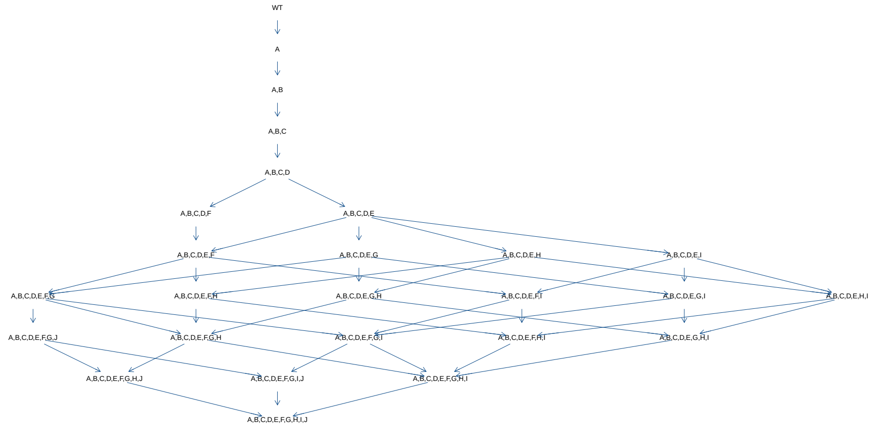

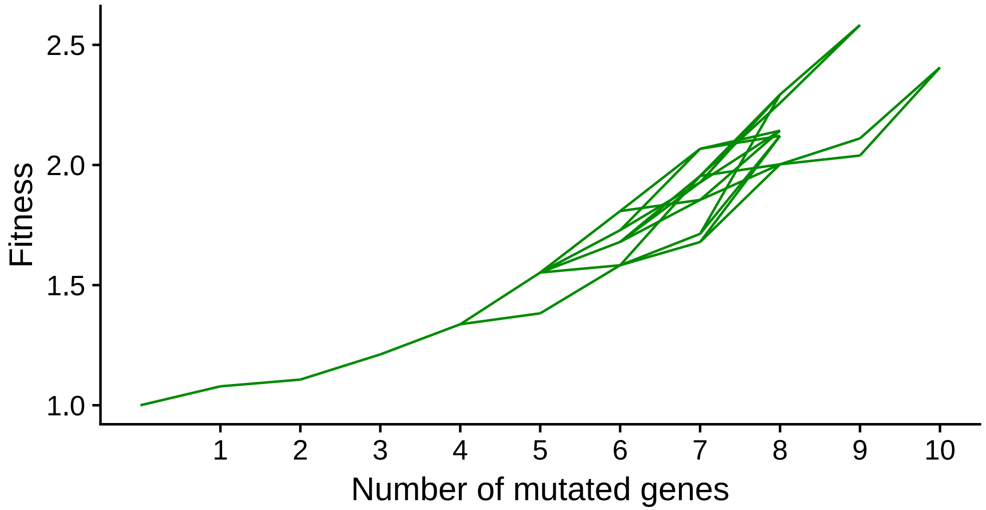


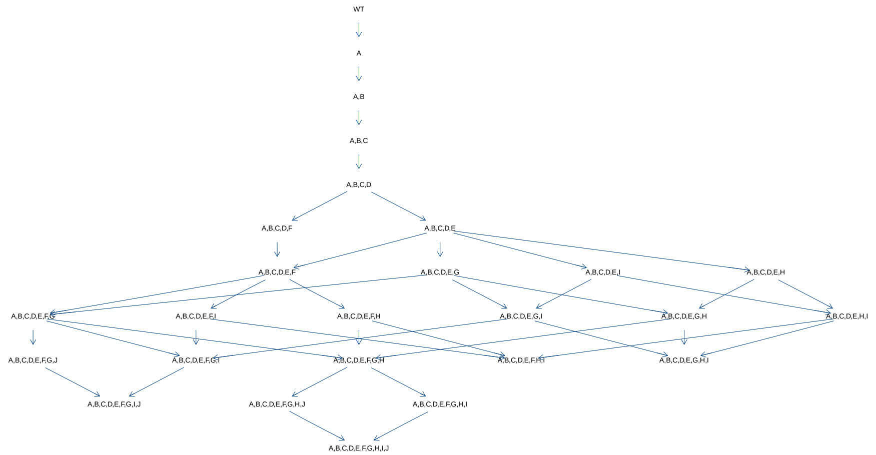

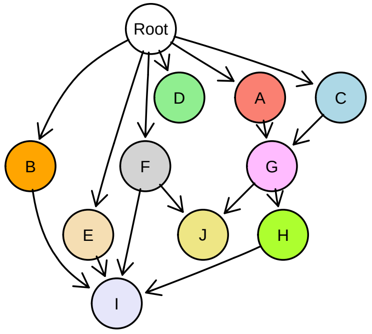


| | |
|---|---|
| *ID* | I9aZnfSBnGVPEGvO |
| *acc genotypes* | 116 |
| *peaks no back* | 14 |
| *removed edges* | 89 |
| *prop rem edges* | 0.259 |

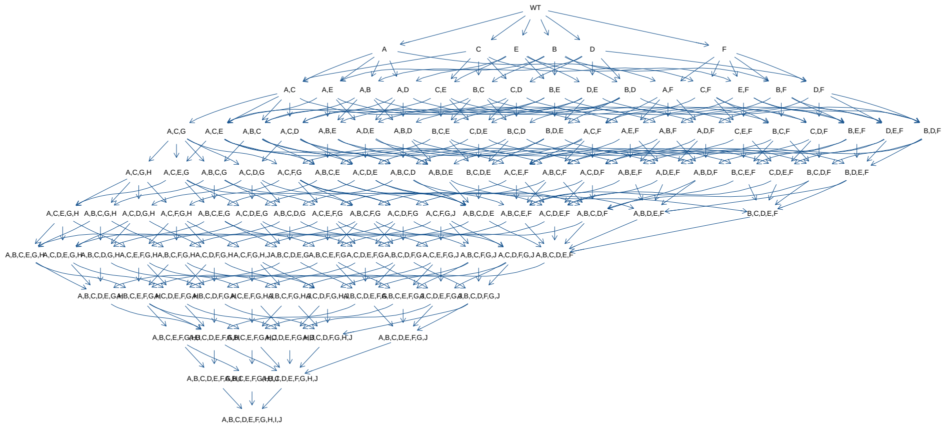

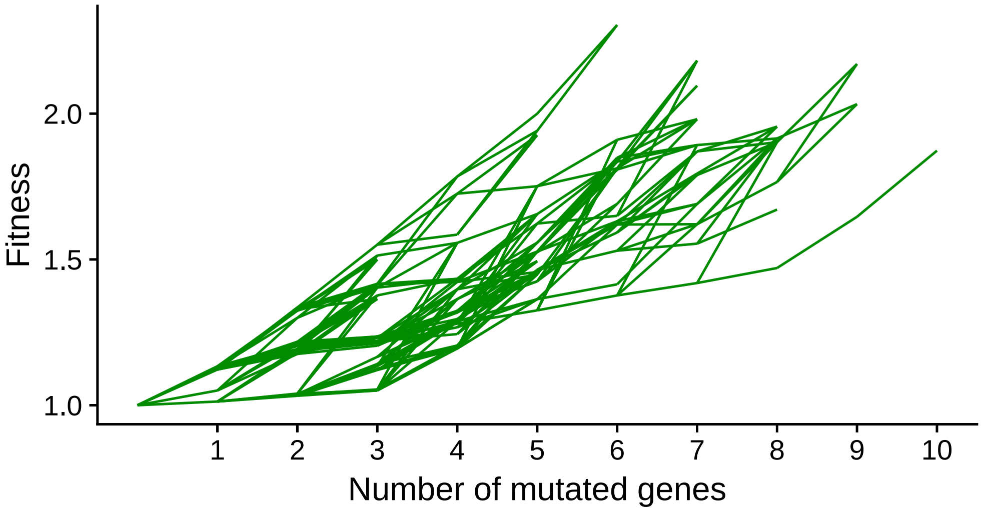


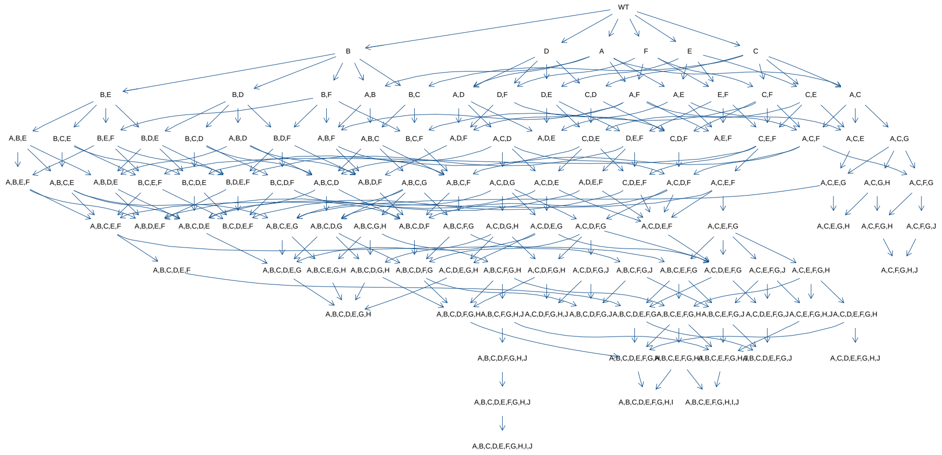

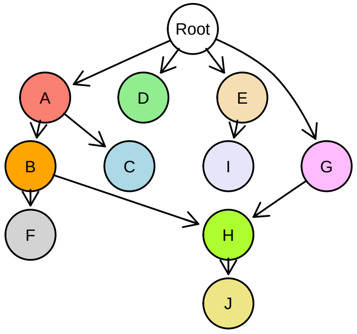


| | |
|---|---|
| *ID* | ibcyCqVRdZgkjfmP |
| *acc genotypes* | 132 |
| *peaks no back* | 13 |
| *removed edges* | 62 |
| *prop rem edges* | 0.16 |

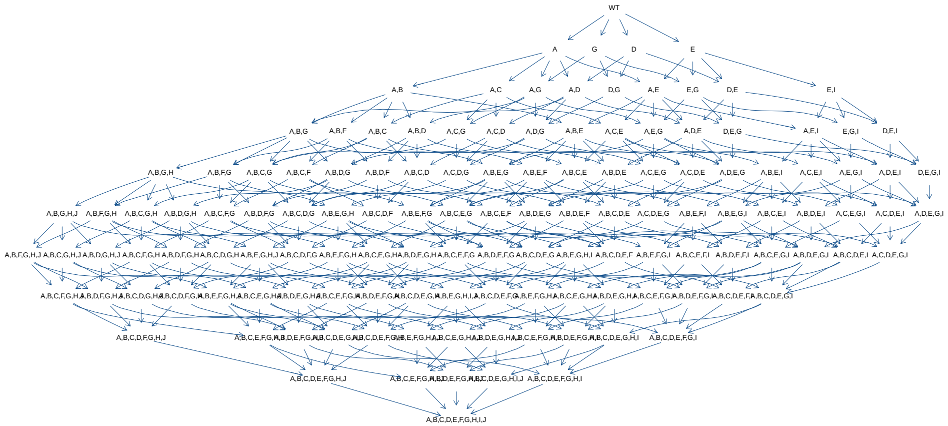

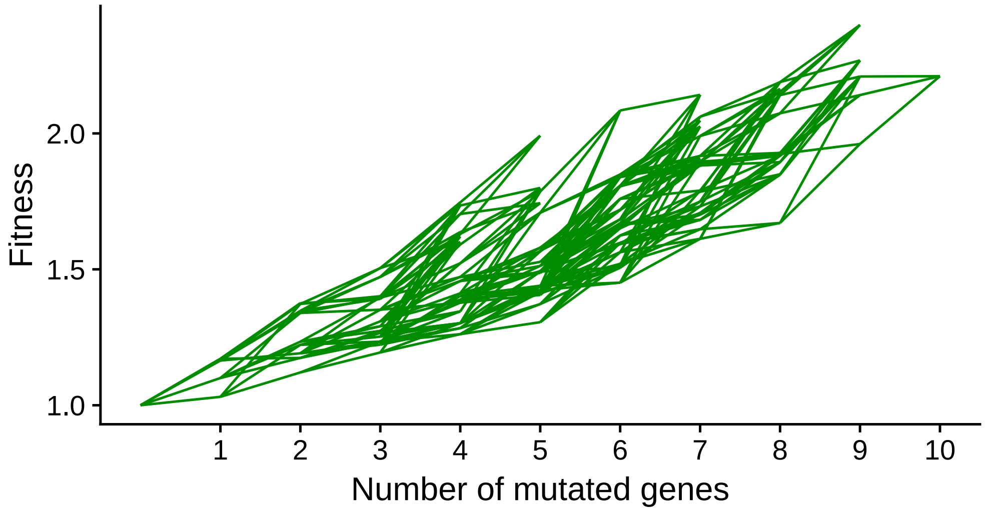


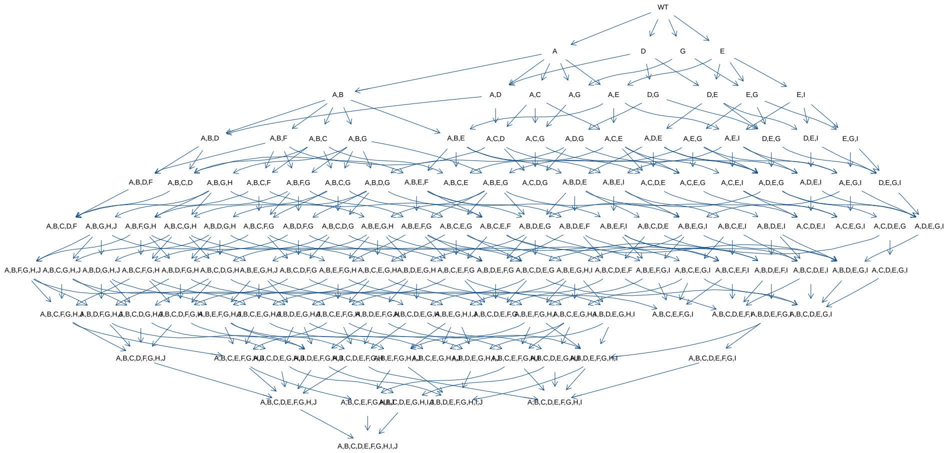

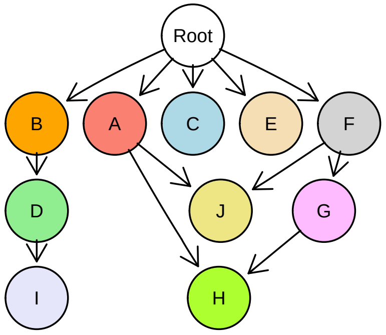


| | |
|---|---|
| *ID* | Ihbe9R5g4X8LVErf |
| *acc genotypes* | 160 |
| *peaks no back* | 12 |
| *removed edges* | 76 |
| *prop rem edges* | 0.156 |

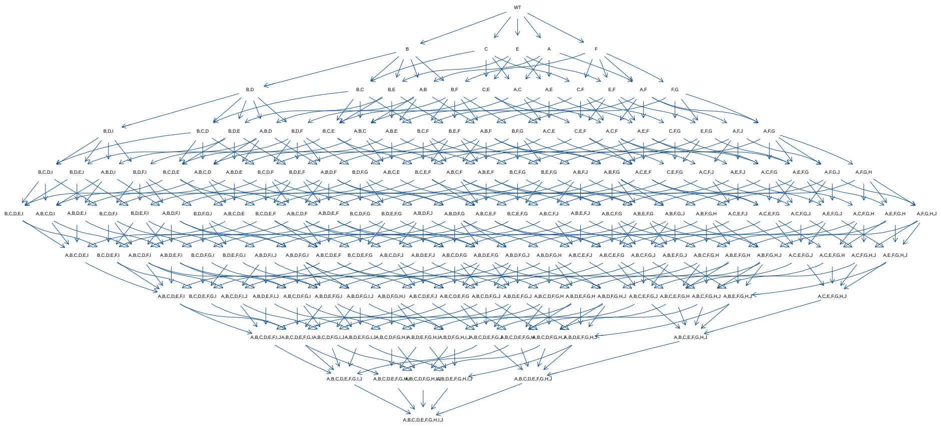

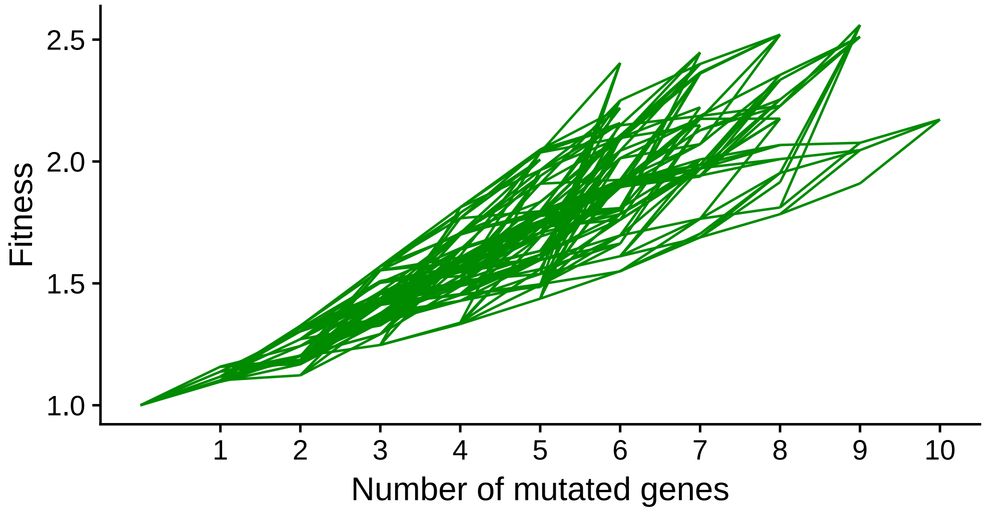


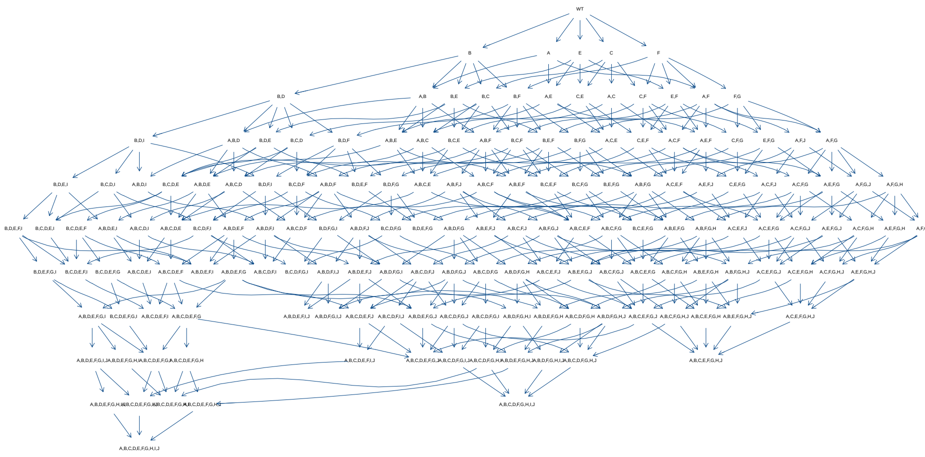

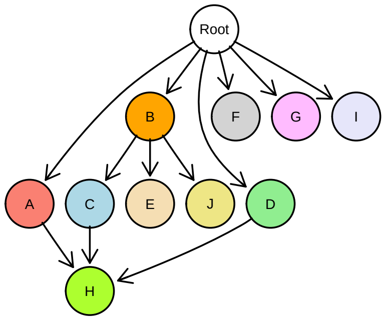


| | |
|---|---|
| *ID* | IJHDmelG9vA10Rtnv |
| *acc genotypes* | 320 |
| *peaks no back* | 21 |
| *removed edges* | 305 |
| *prop rem edges* | 0.244 |

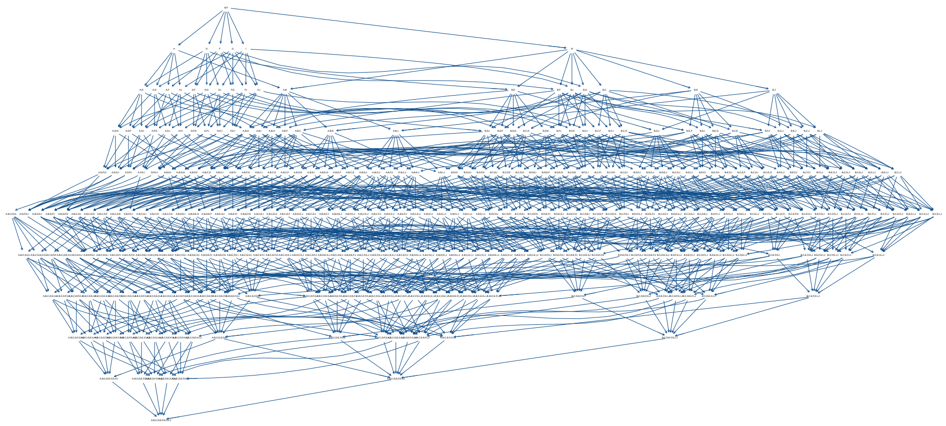

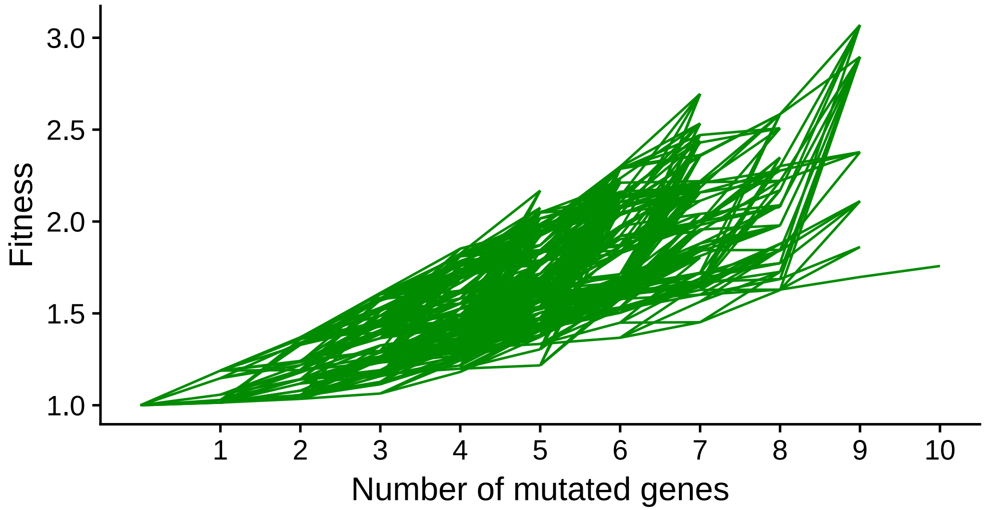


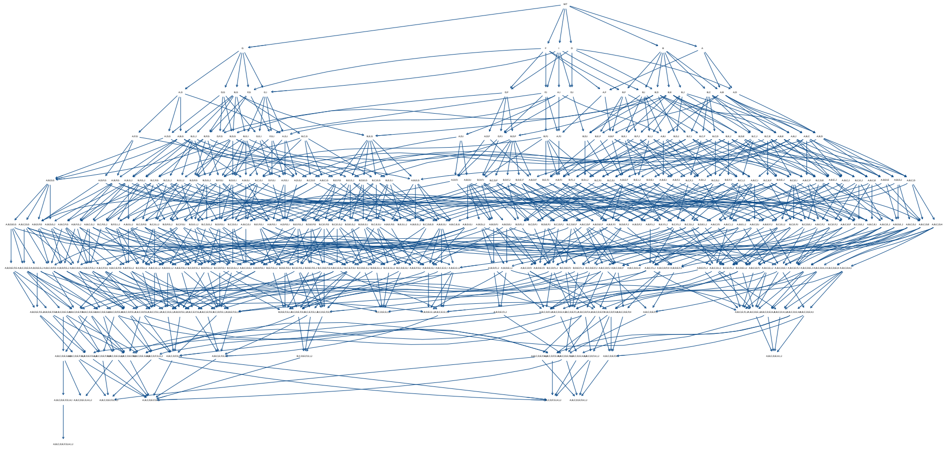

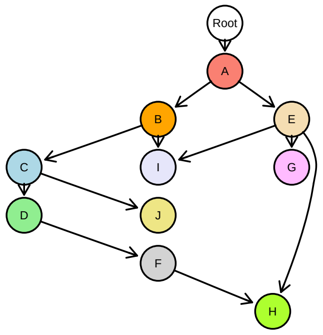


| | |
|---|---|
| *ID* | InQKKevAQbH5g2SM |
| *acc genotypes* | 47 |
| *peaks no back* | 7 |
| *removed edges* | 18 |
| *prop rem edges* | 0.178 |

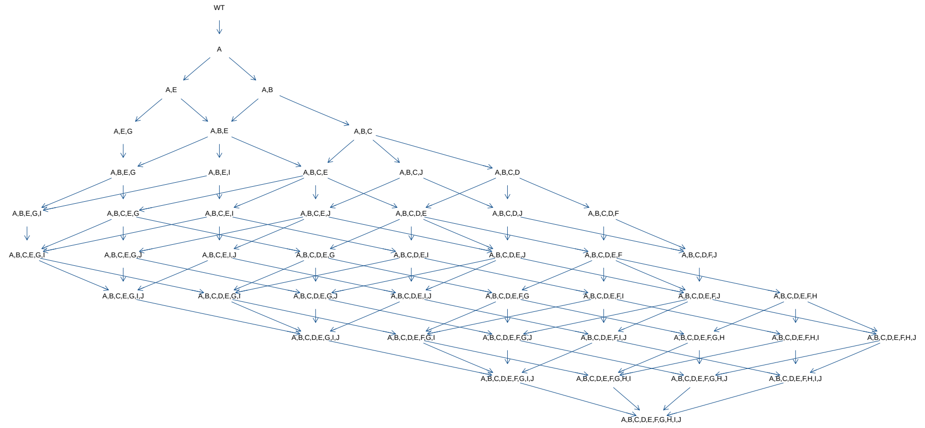

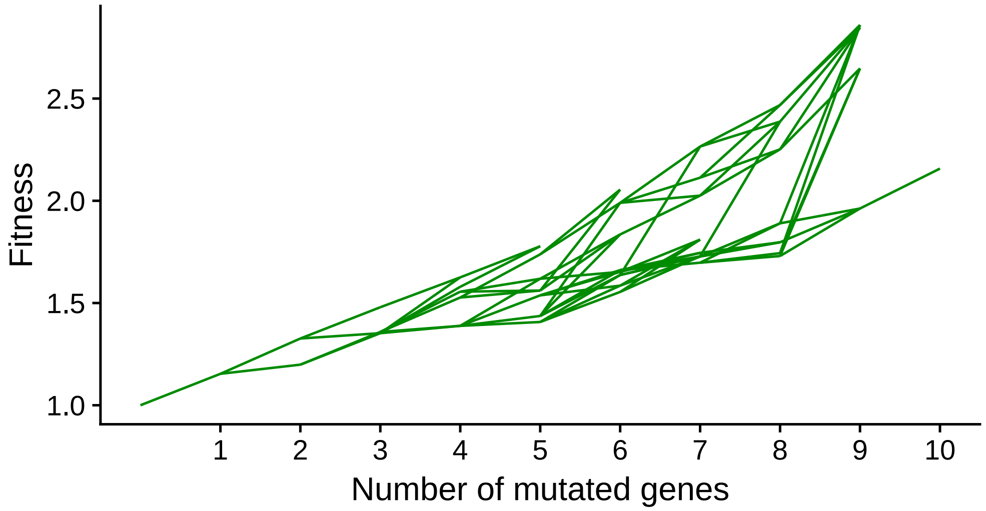


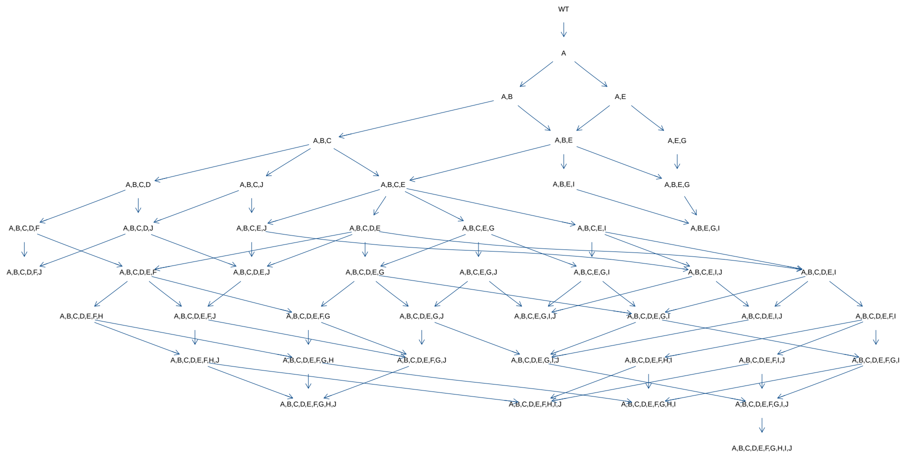

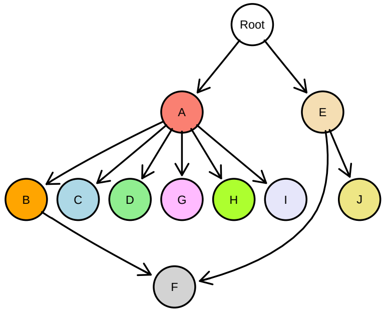


| | |
|---|---|
| *ID* | iudi9ykQn8Kp10Gil |
| *acc genotypes* | 259 |
| *peaks no back* | 19 |
| *removed edges* | 140 |
| *prop rem edges* | 0.145 |

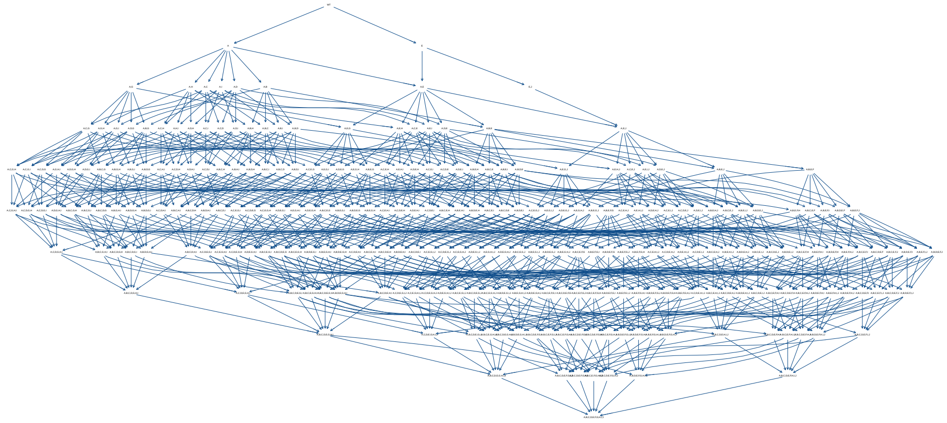

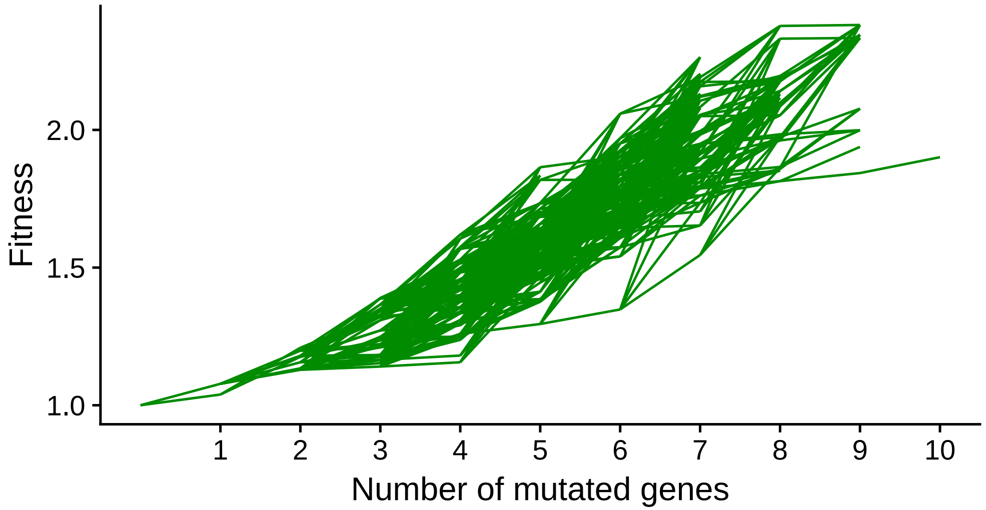


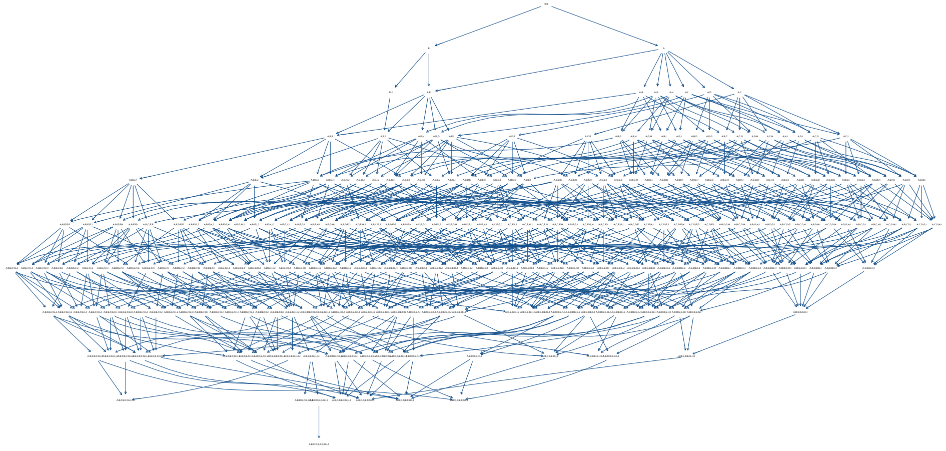

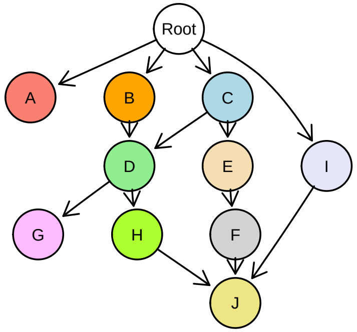


*ID* iyeU83TTZkIieN106
*acc genotypes* 84
*peaks no back* 9
*removed edges* 37
*prop rem edges* 0.168

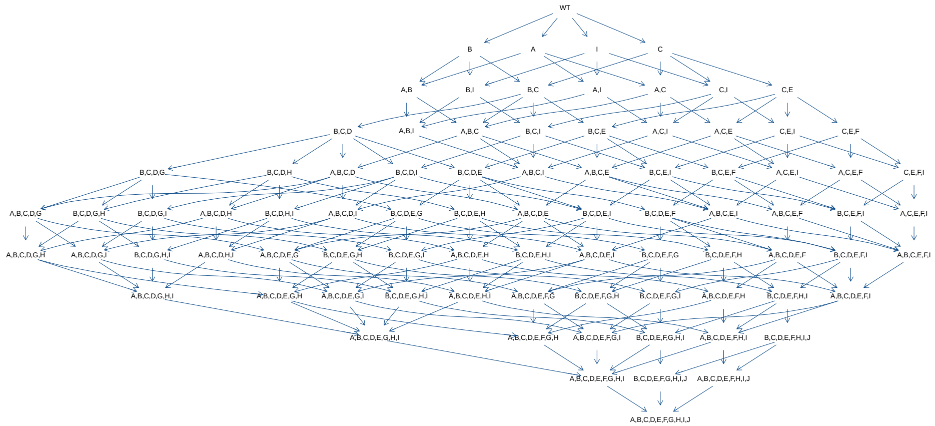

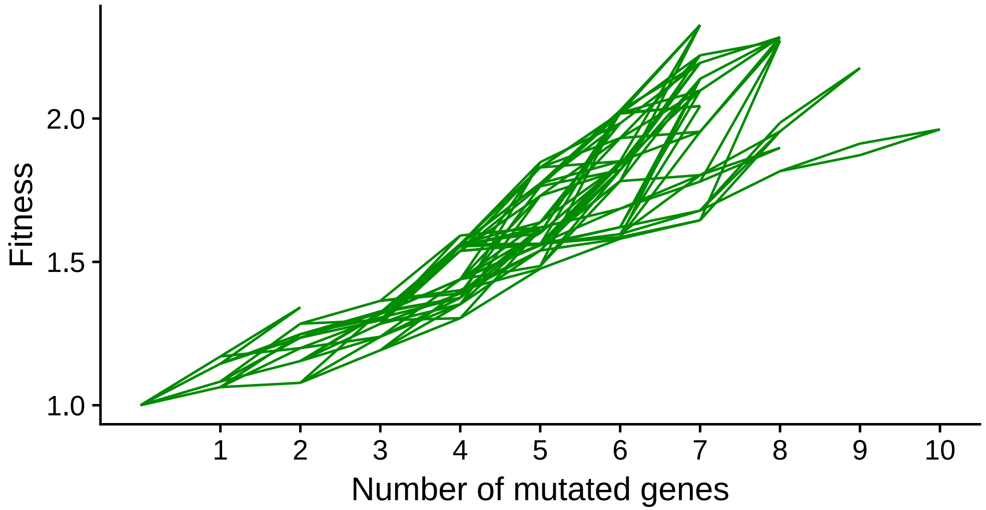


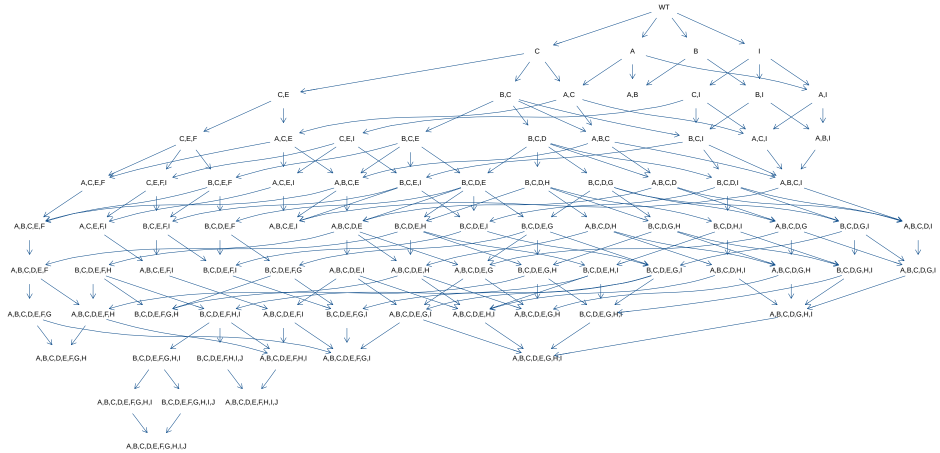

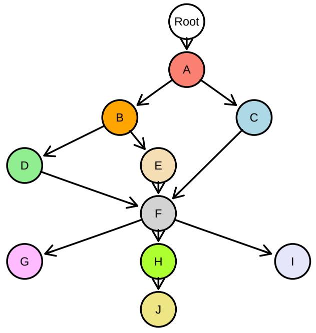


| | |
|---|---|
| *ID* | iyF6c65sIZnguySA |
| *acc genotypes* | 23 |
| *peaks no back* | 2 |
| *removed edges* | 2 |
| *prop rem edges* | 0.054 |

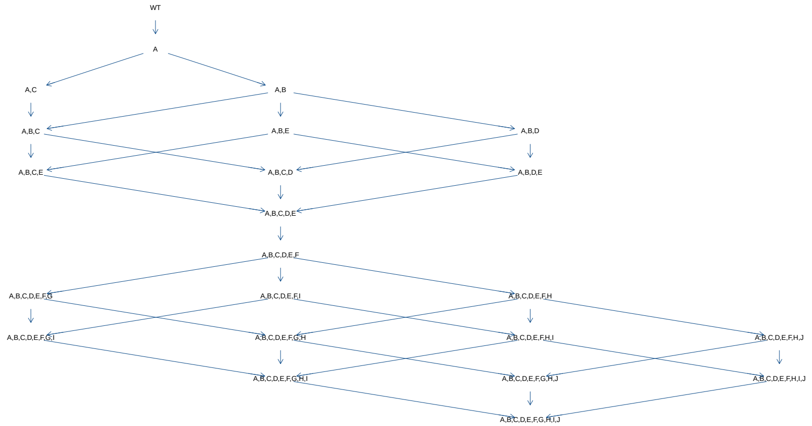

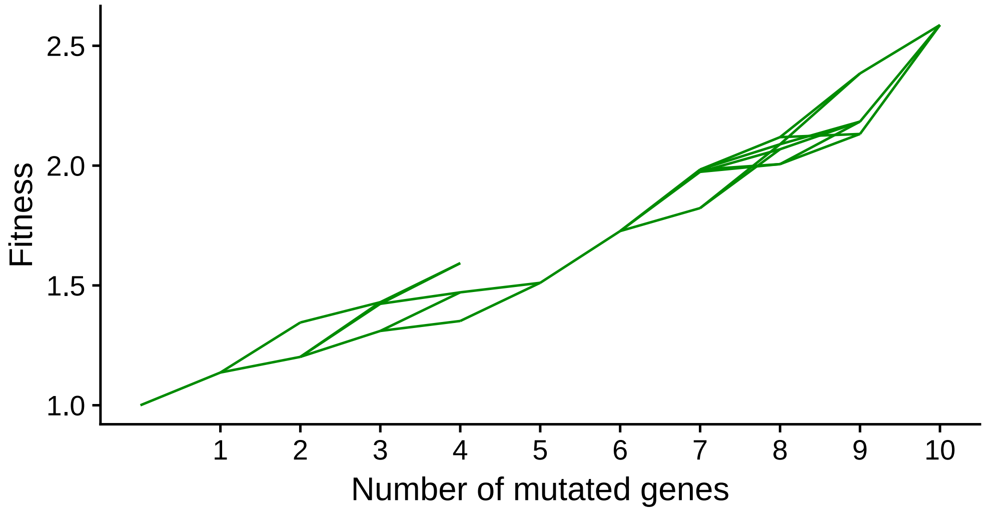


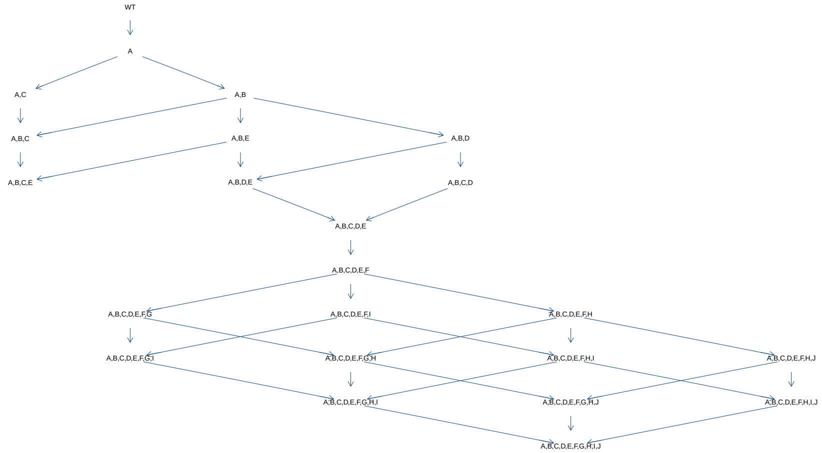

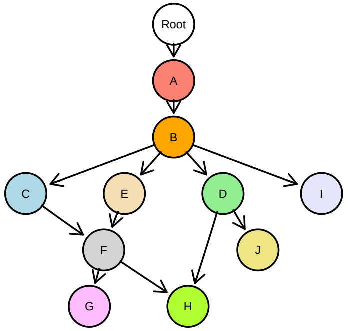


| | |
|---|---|
| *ID* | iYOTCycsSTgt6LB1 |
| *acc genotypes* | 46 |
| *peaks no back* | 4 |
| *removed edges* | 12 |
| *prop rem edges* | 0.12 |

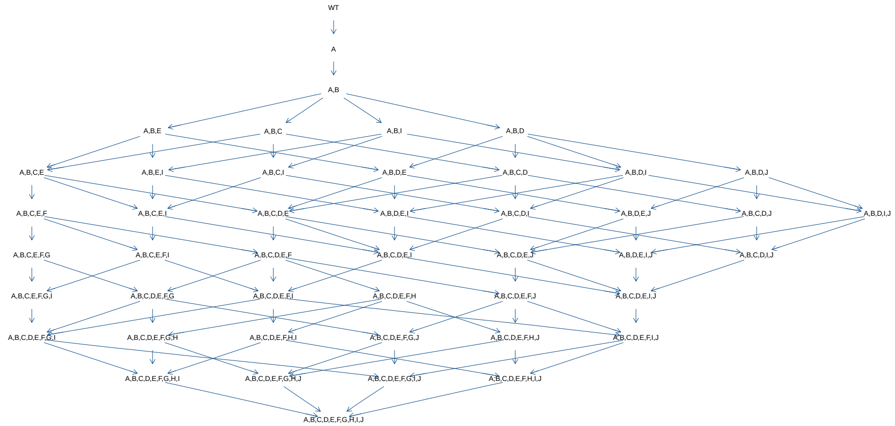

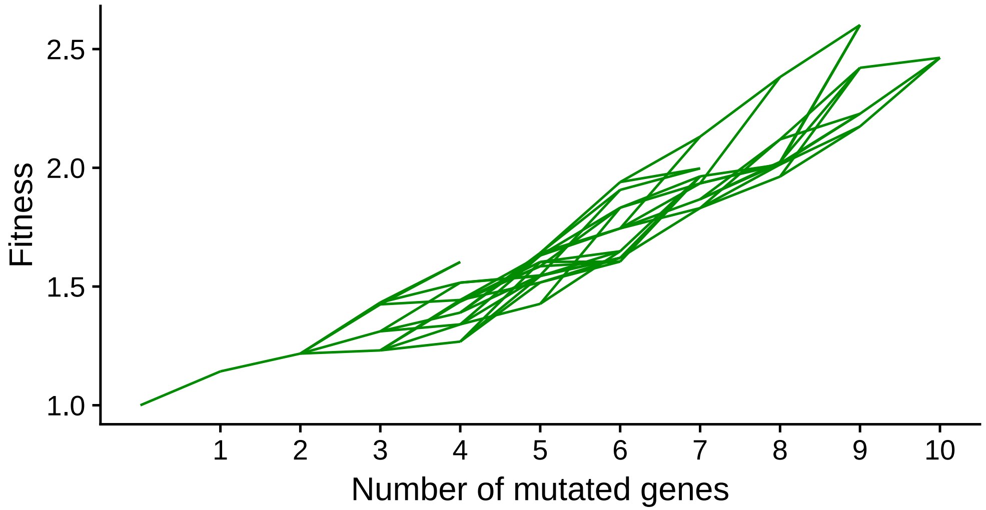


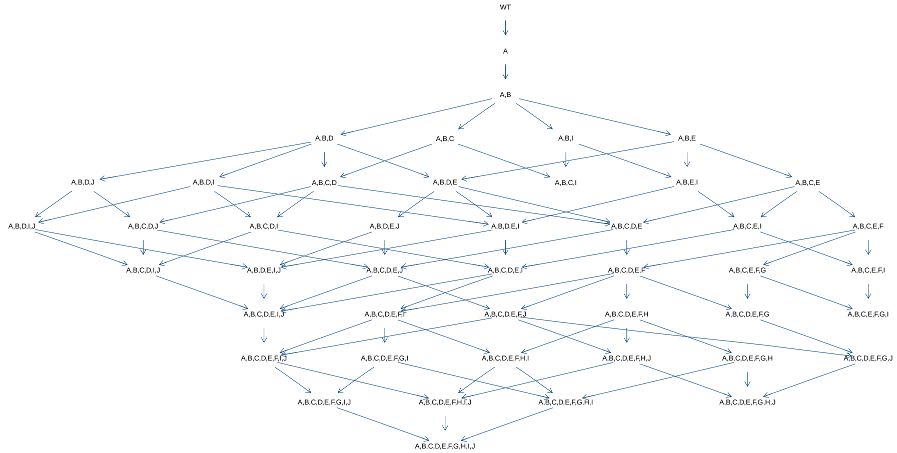

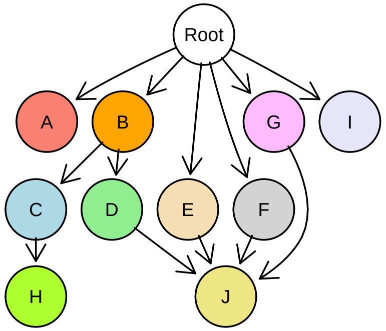


| | |
|---|---|
| *ID* | ja5K99Pb6N9URztp |
| *acc genotypes* | 236 |
| *peaks no back* | 17 |
| *removed edges* | 192 |
| *prop rem edges* | 0.226 |

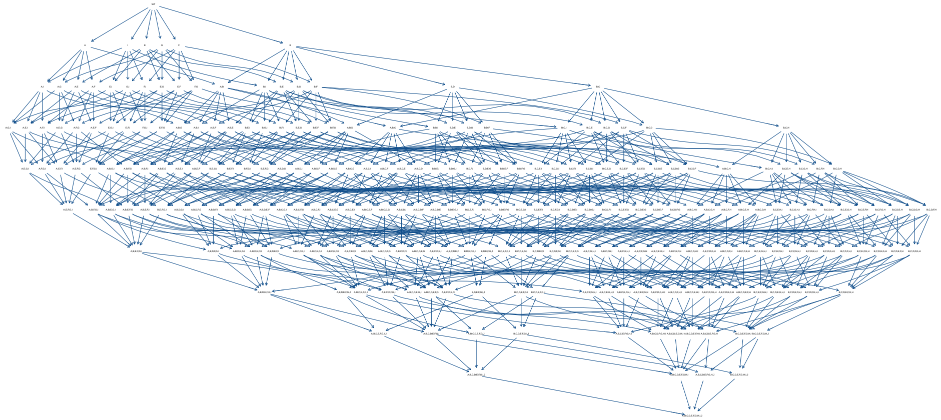

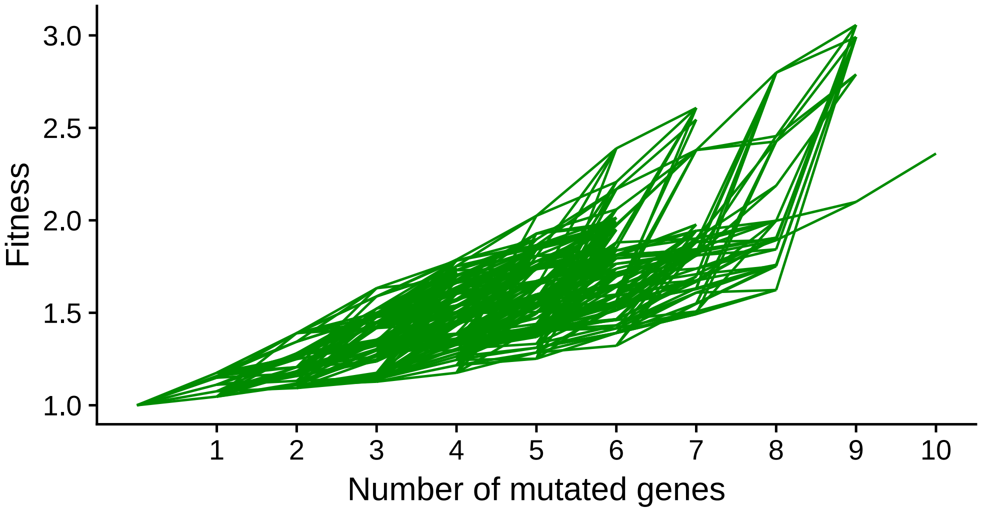


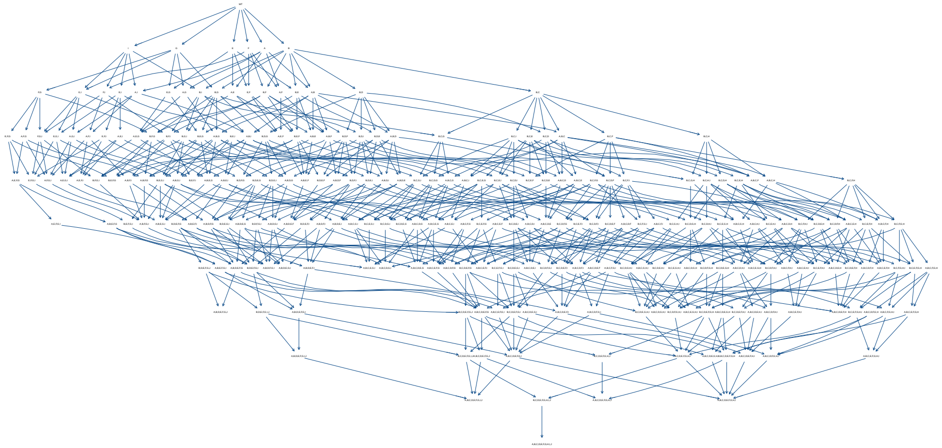

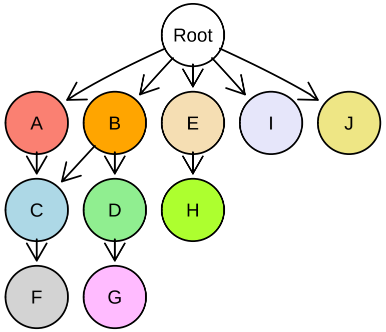


| | |
|---|---|
| *ID* | jBiKG5Scuo10UsyUf |
| *acc genotypes* | 168 |
| *peaks no back* | 14 |
| *removed edges* | 111 |
| *prop rem edges* | 0.213 |

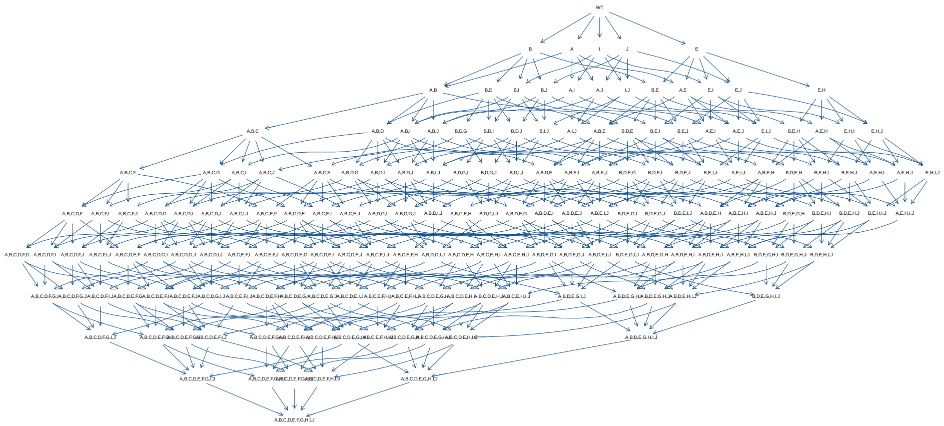

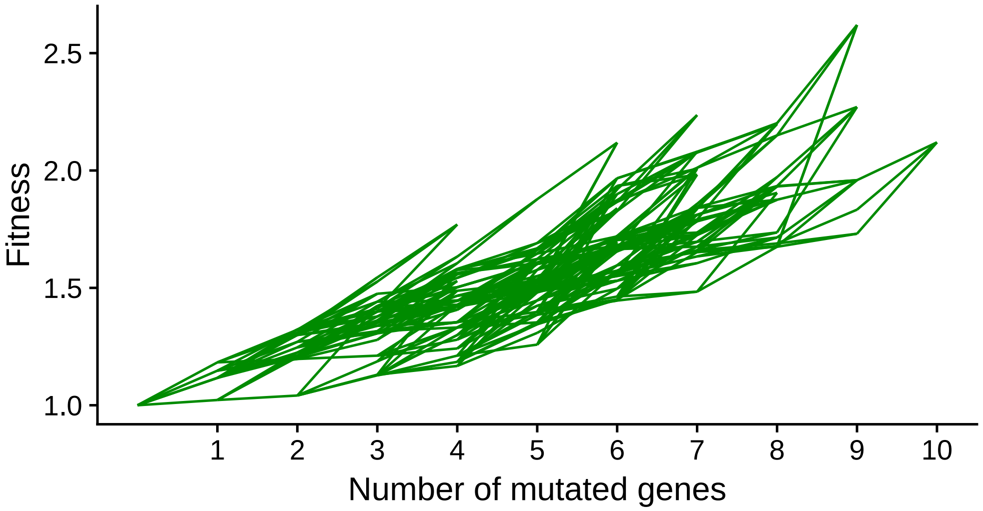


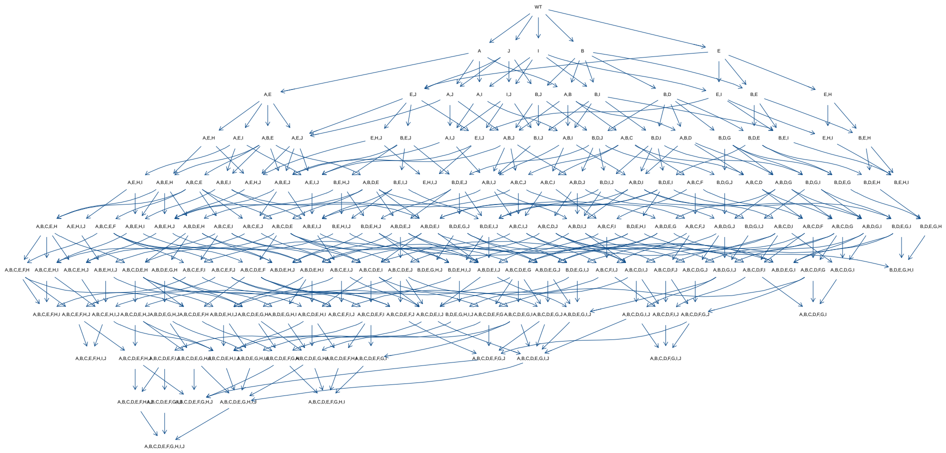

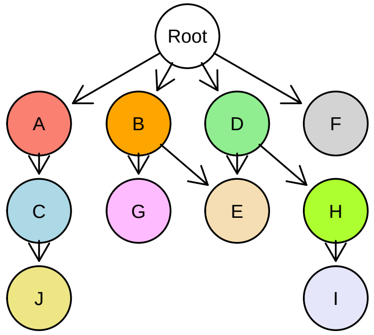


*ID* jeIgNCENfpAlgfTL
*acc genotypes* 144
*peaks no back* 15
*removed edges* 89
*prop rem edges* 0.212

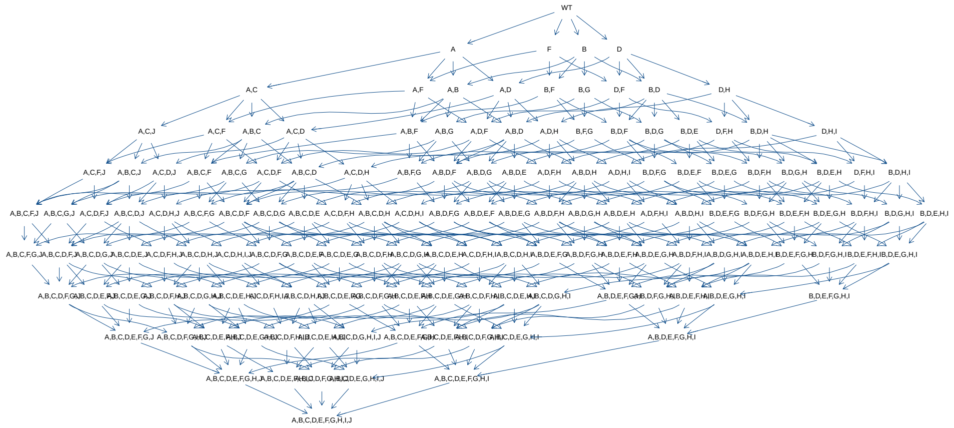

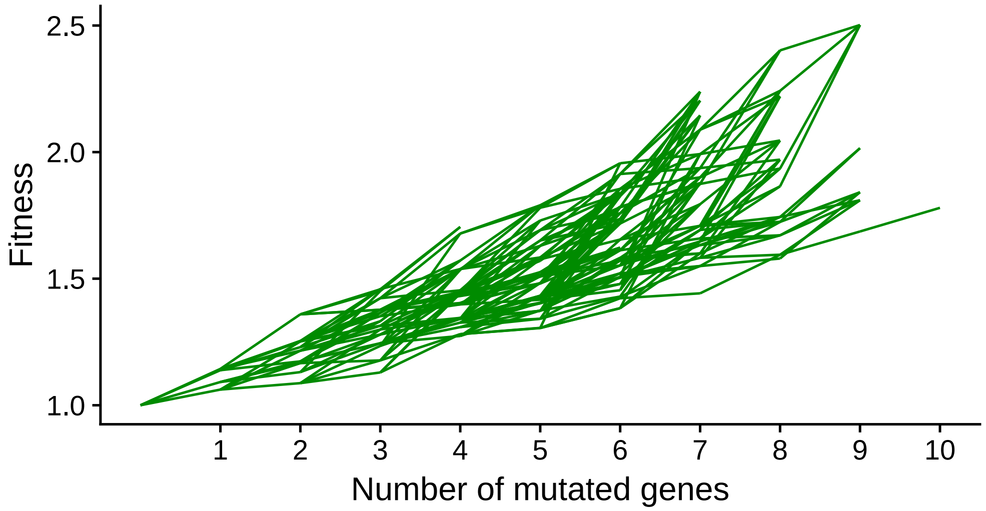


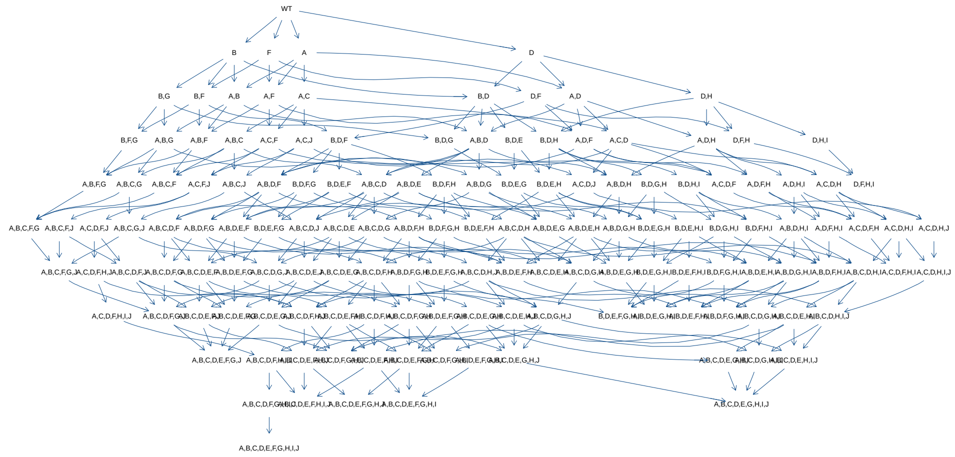

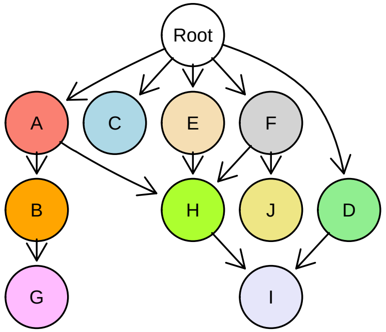


| | |
|---|---|
| *ID* | k9iq8ysBwZDbv105Y |
| *acc genotypes* | 132 |
| *peaks no back* | 12 |
| *removed edges* | 78 |
| *prop rem edges* | 0.201 |

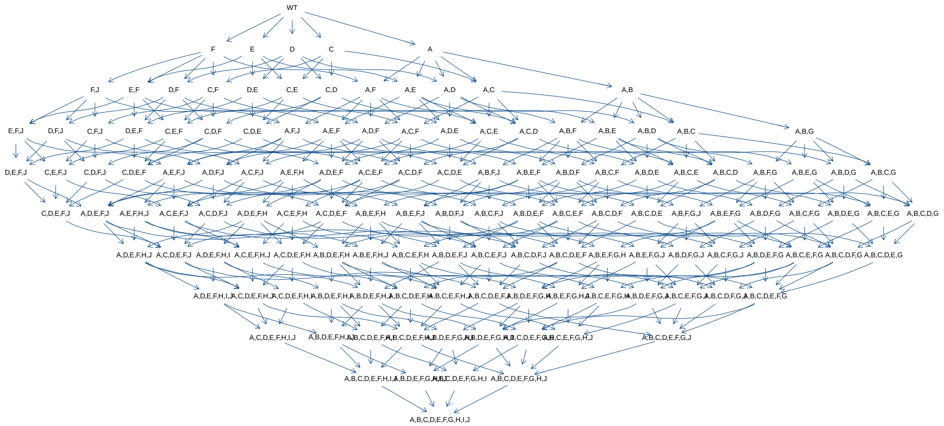

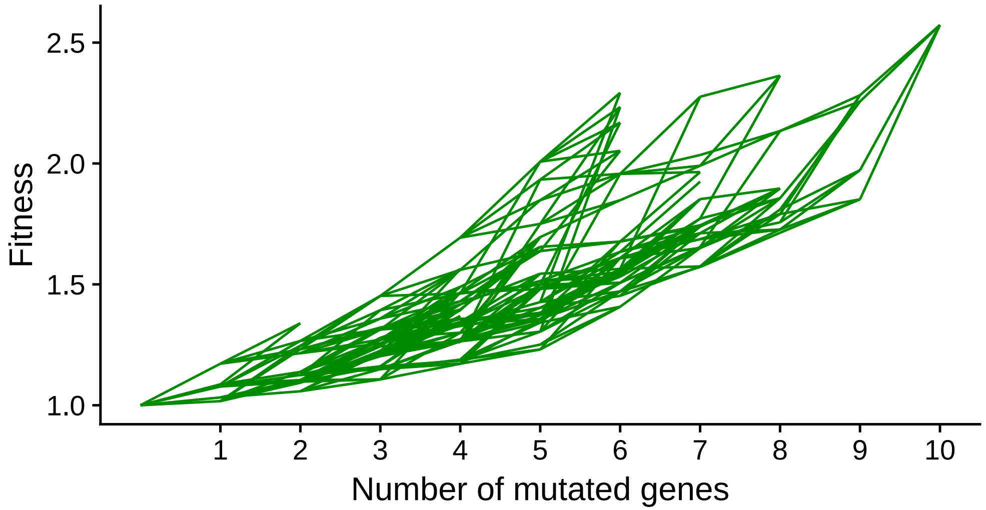


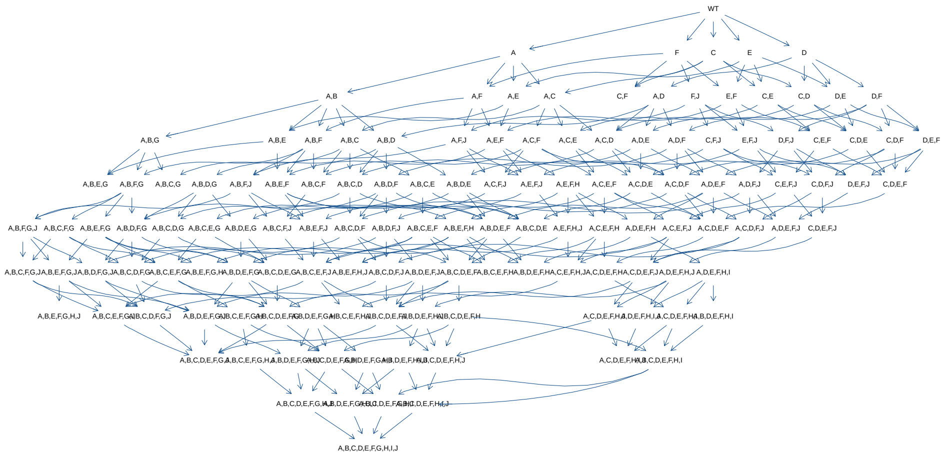

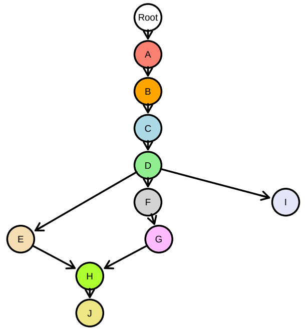


| | |
|---|---|
| *ID* | KbbfhCf9UoSKy10IE |
| *acc genotypes* | 20 |
| *peaks no back* | 2 |
| *removed edges* | 1 |
| *prop rem edges* | 0.033 |

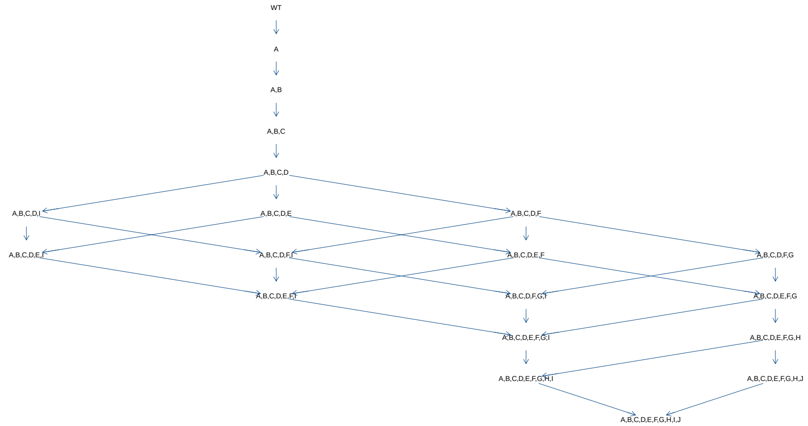

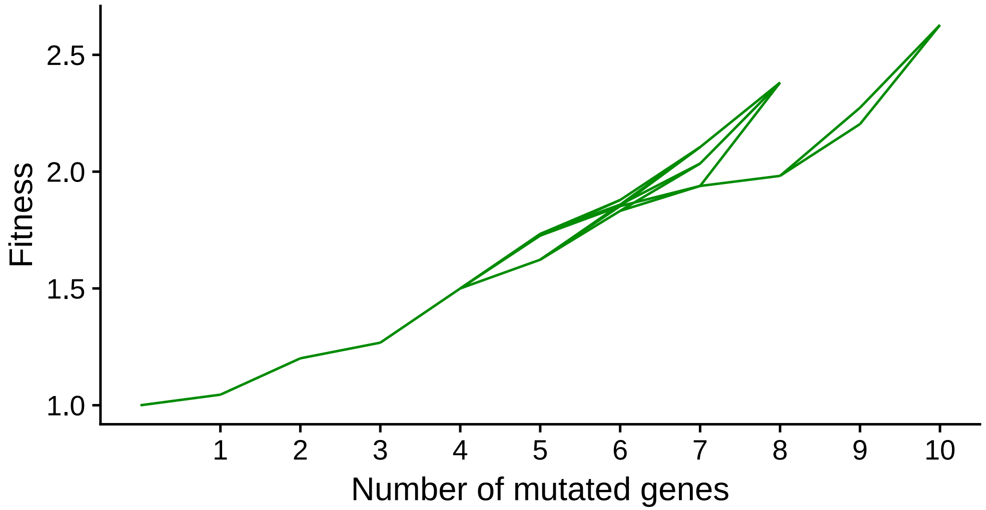


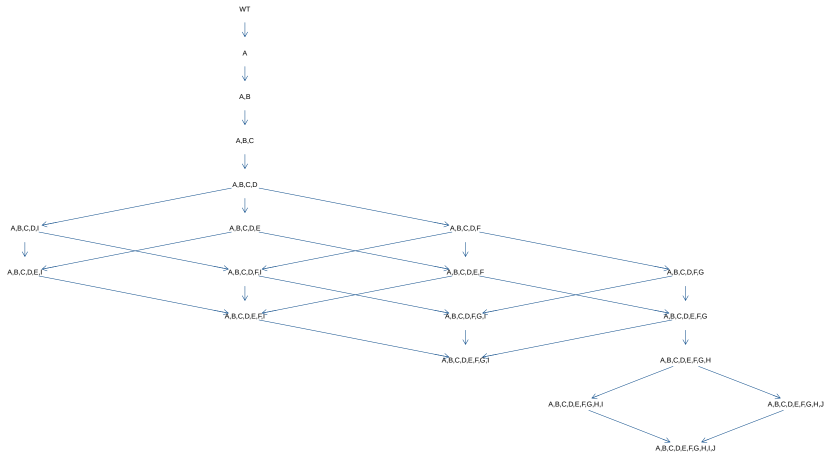

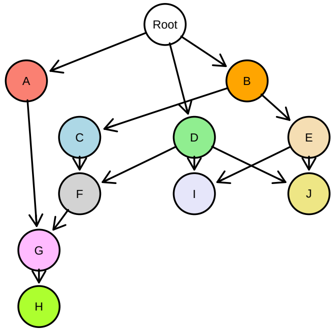


| | |
|---|---|
| *ID* | KEEVYiND4AYuzrNr |
| *acc genotypes* | 52 |
| *peaks no back* | 7 |
| *removed edges* | 19 |
| *prop rem edges* | 0.168 |

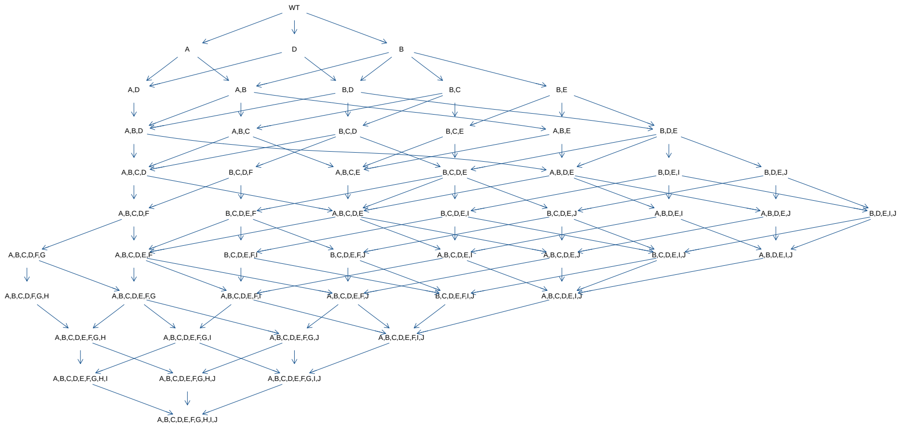

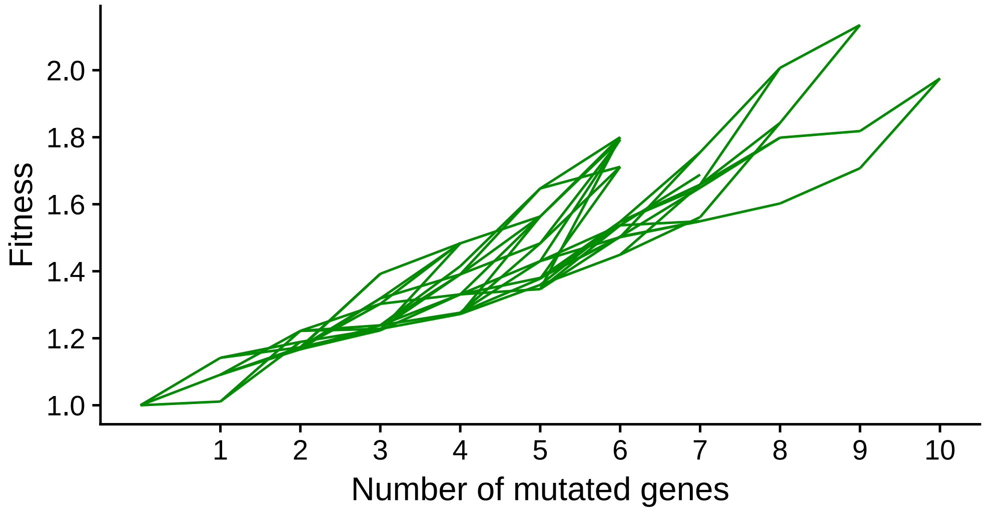


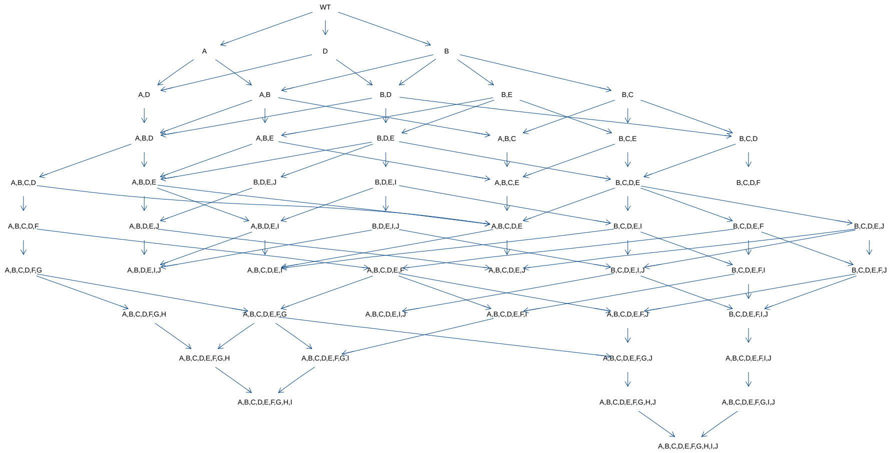

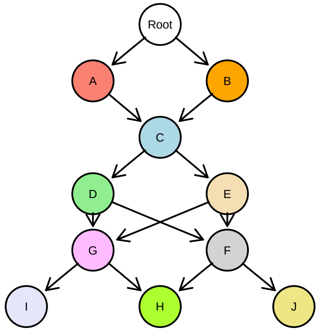


| | |
|---|---|
| *ID* | KpASsdOSpsoQXAqF |
| *acc genotypes* | 20 |
| *peaks no back* | 5 |
| *removed edges* | 5 |
| *prop rem edges* | 0.172 |

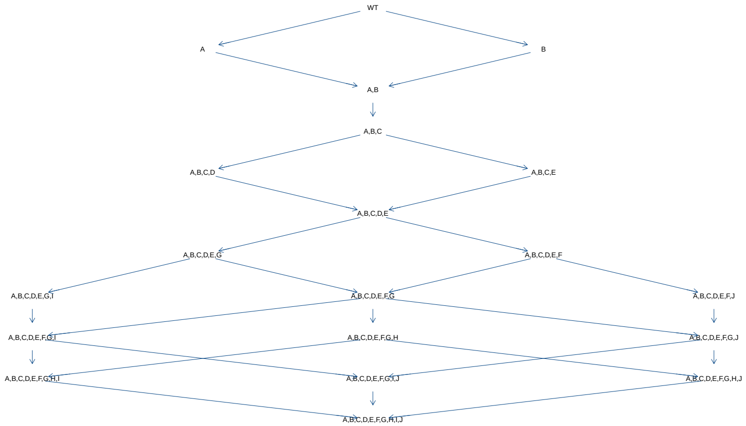

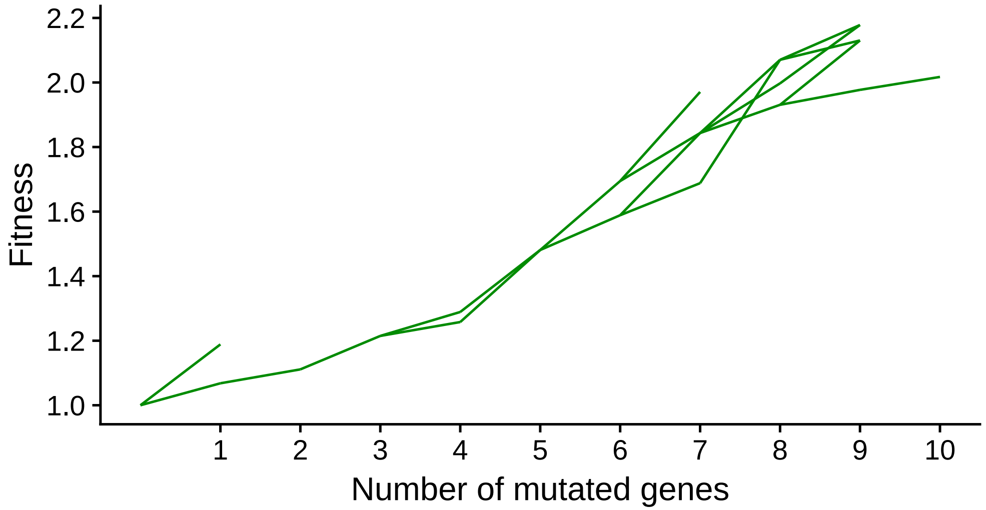


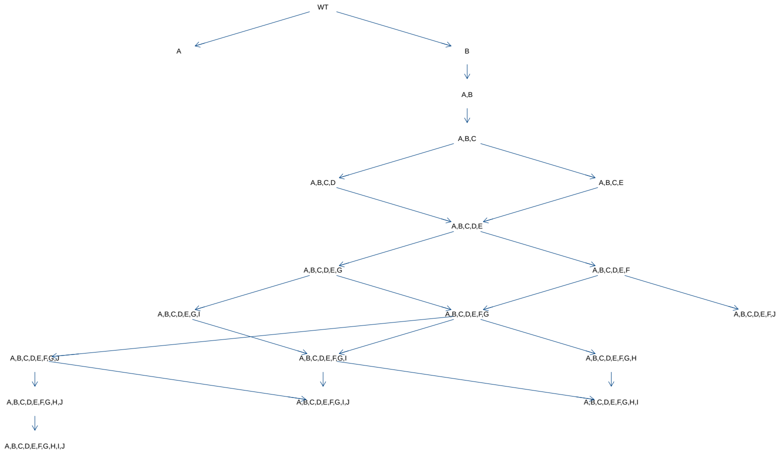

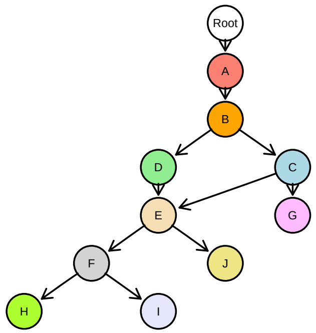


| | |
|---|---|
| *ID* | KTPDesITE10I68VaZ |
| *acc genotypes* | 28 |
| *peaks no back* | 5 |
| *removed edges* | 6 |
| *prop rem edges* | 0.118 |

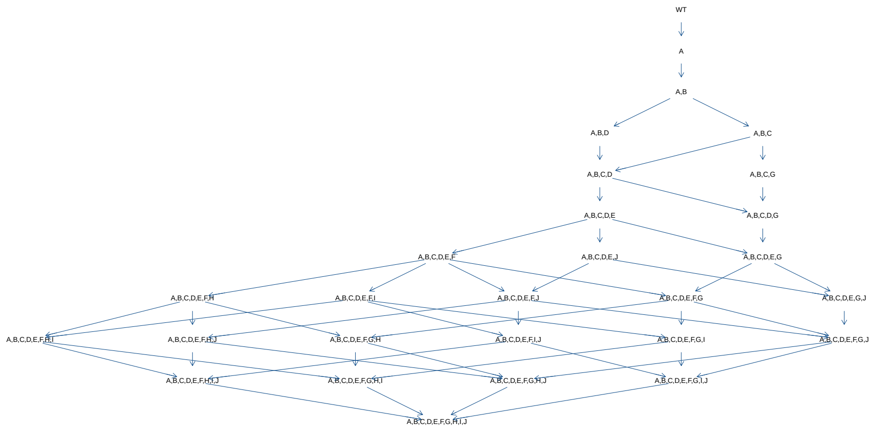

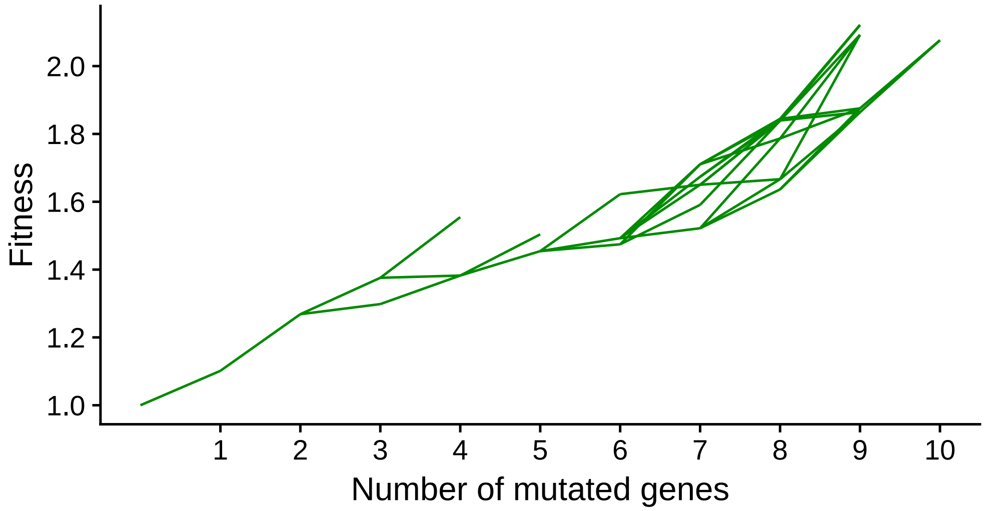


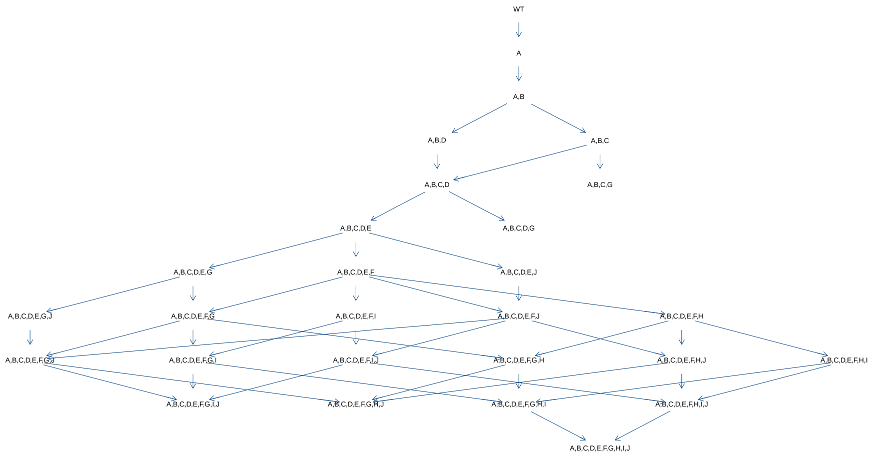

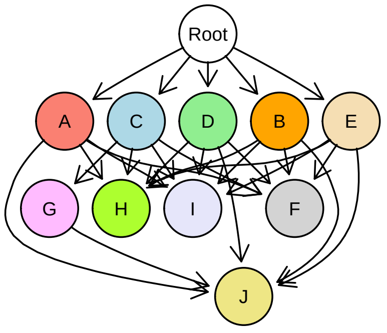


*ID* ku3OjGvU29I3Hlrg
*acc genotypes* 70
*peaks no back* 9
*removed edges* 32
*prop rem edges* 0.179

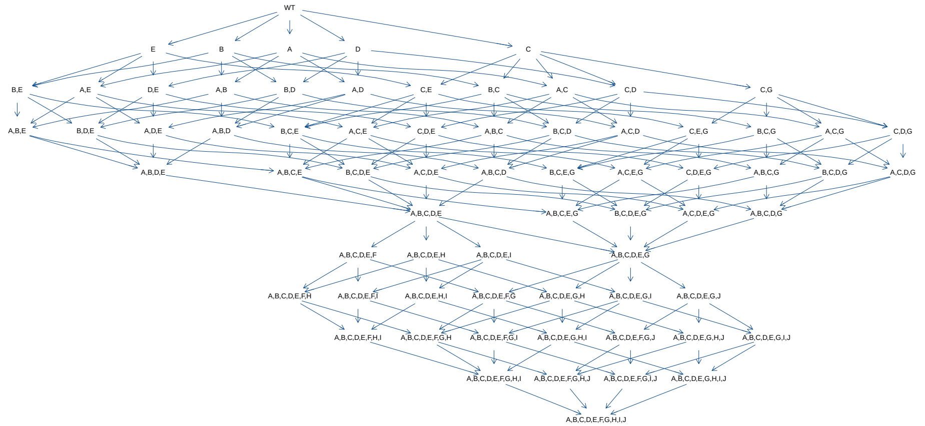

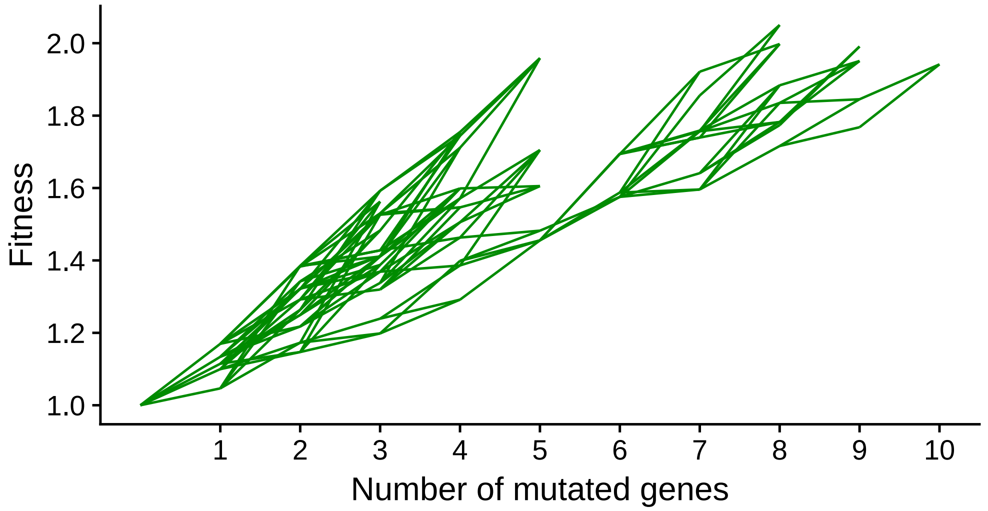


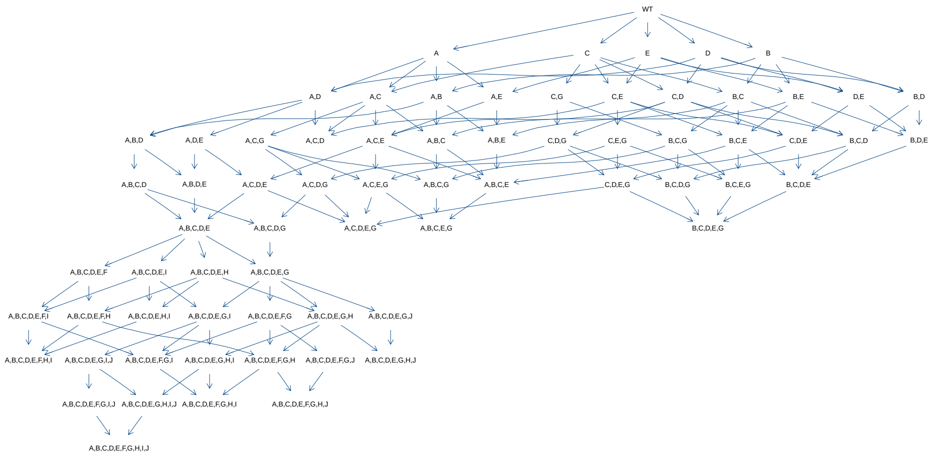

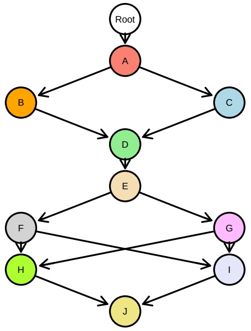


| | |
|---|---|
| *ID* | kuiBBxD31RFF9NEj |
| *acc genotypes* | 14 |
| *peaks no back* | 2 |
| *removed edges* | 1 |
| *prop rem edges* | 0.062 |

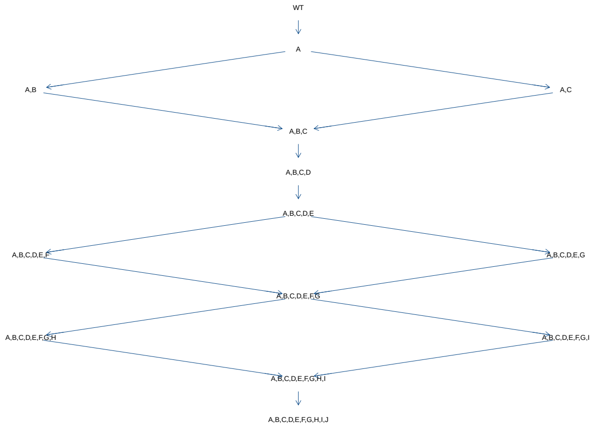


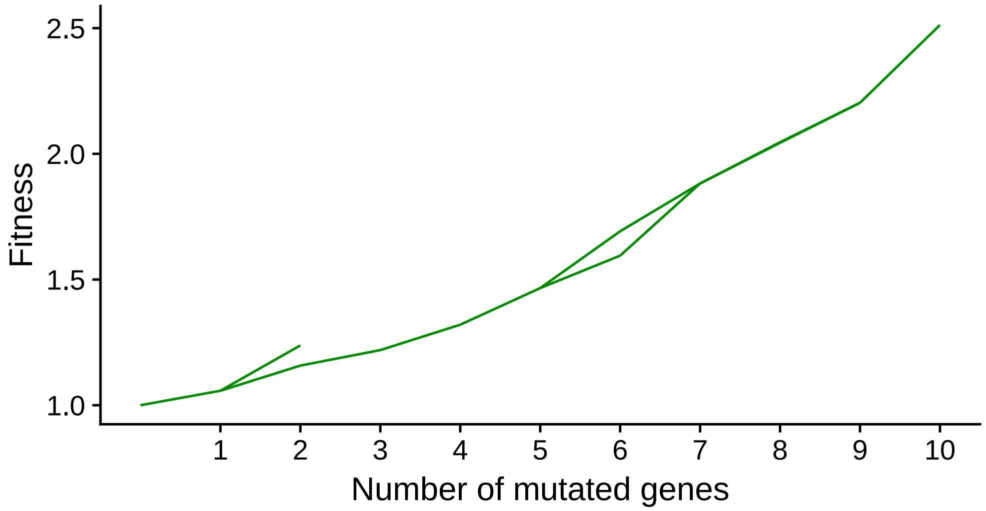


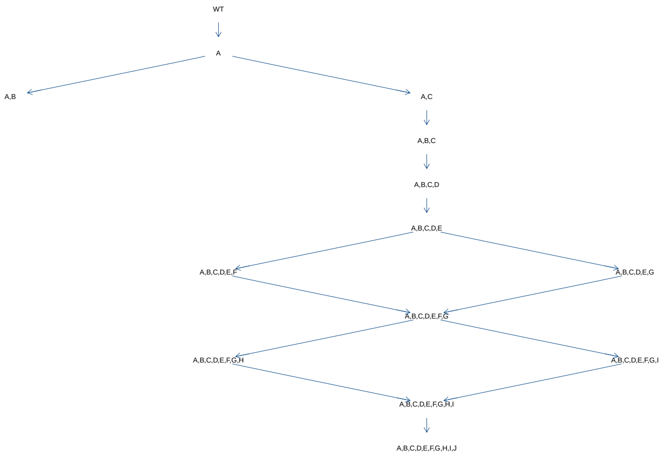

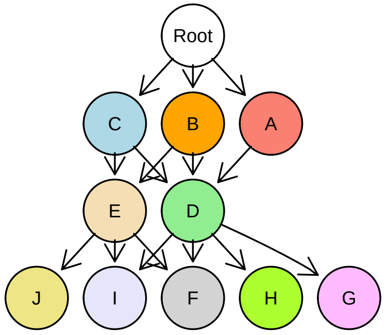


| | |
|---|---|
| *ID* | kvFyL8XlKXwZvz1L |
| *acc genotypes* | 48 |
| *peaks no back* | 9 |
| *removed edges* | 30 |
| *prop rem edges* | 0.275 |

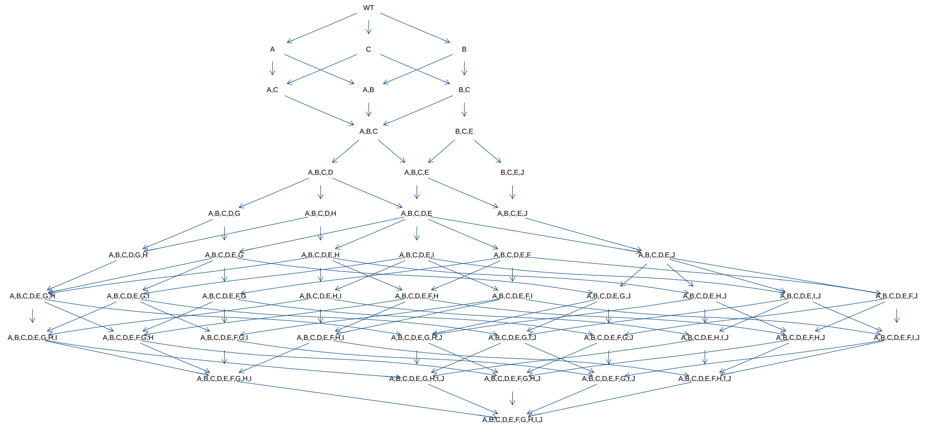

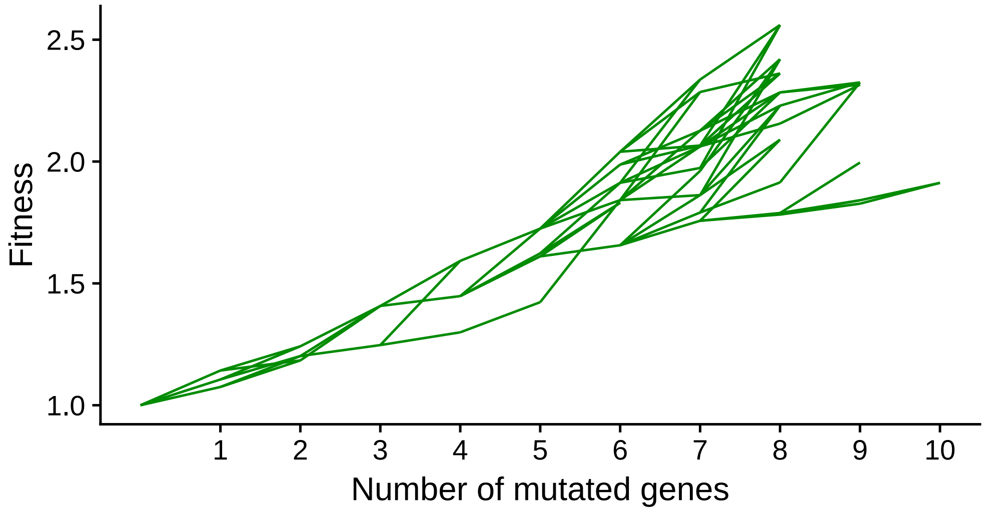


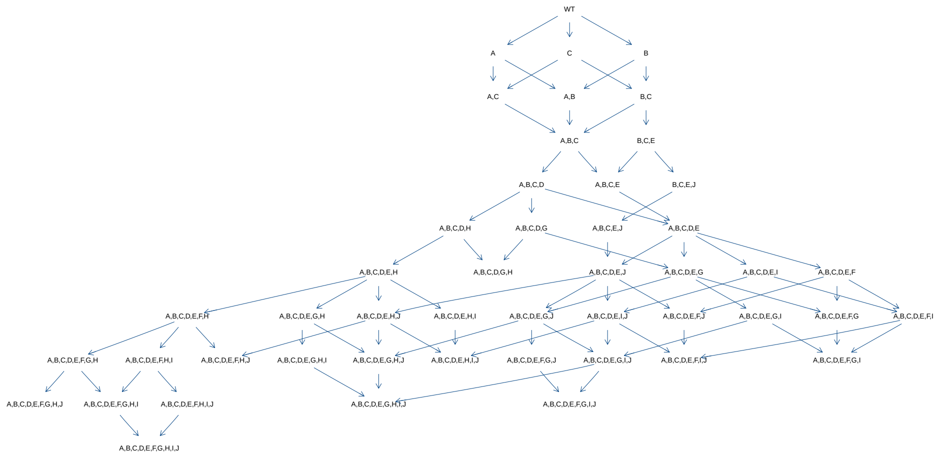

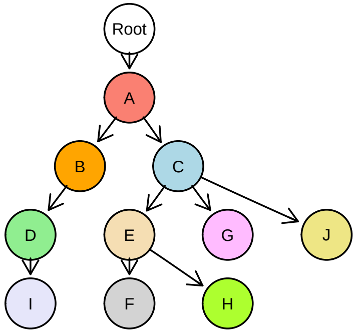


| | |
|---|---|
| *ID* | l37sHoyzCWlv8at2 |
| *acc genotypes* | 85 |
| *peaks no back* | 10 |
| *removed edges* | 42 |
| *prop rem edges* | 0.184 |

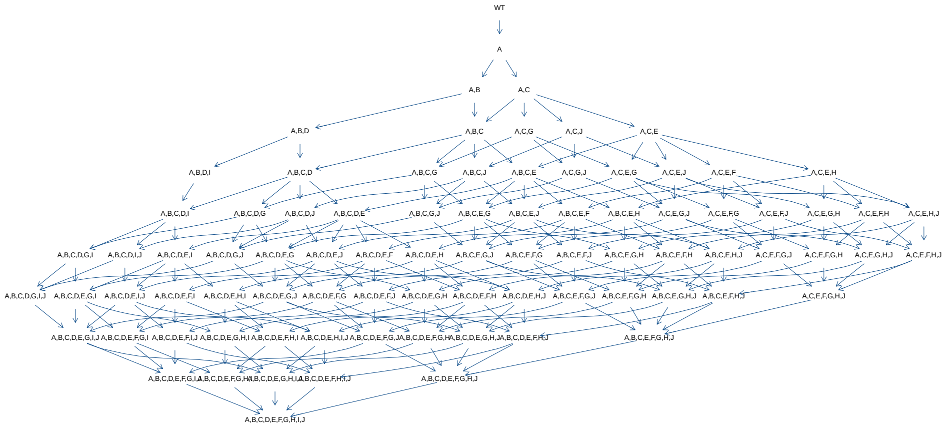

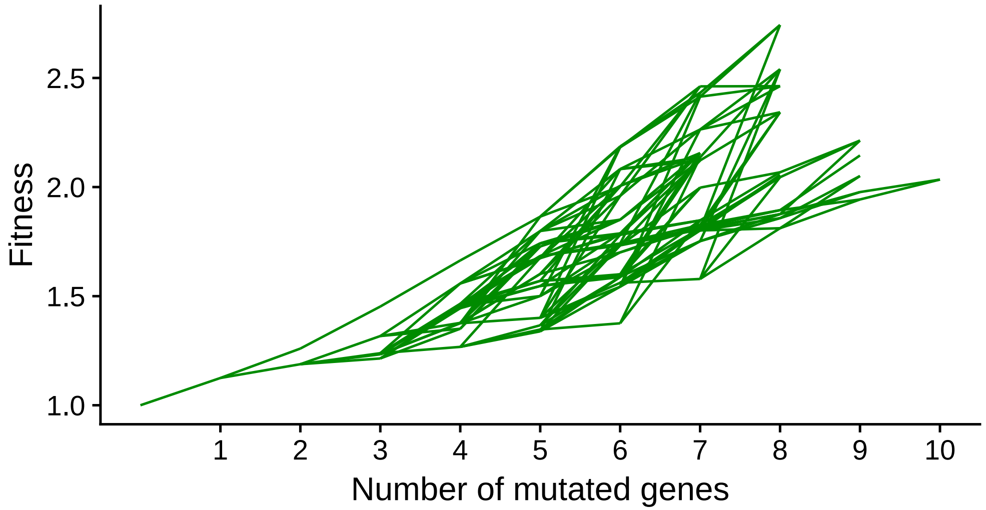


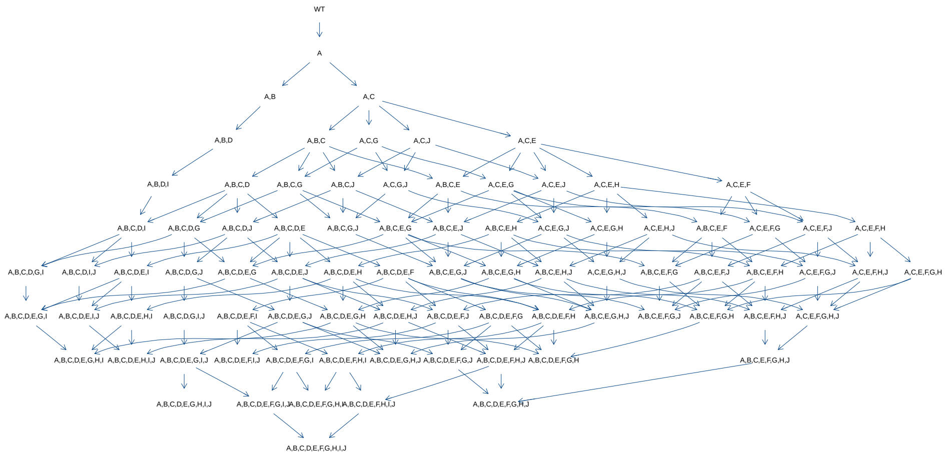

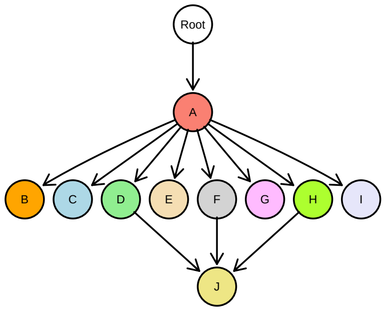


| | |
|---|---|
| *ID* | lMrqdoPtFXSjearn |
| *acc genotypes* | 289 |
| *peaks no back* | 21 |
| *removed edges* | 252 |
| *prop rem edges* | 0.222 |

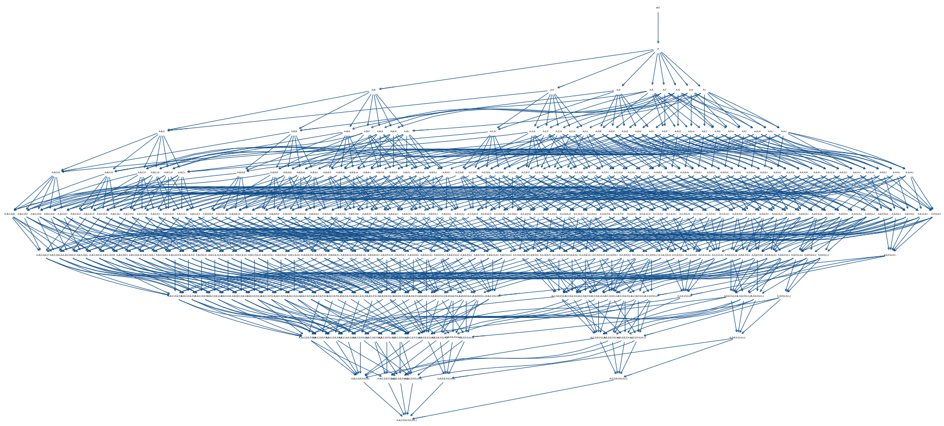

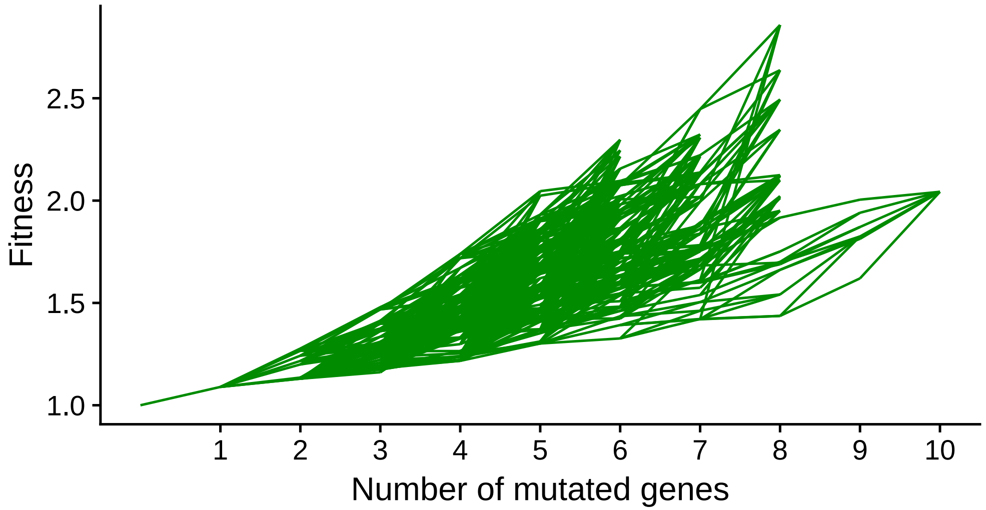


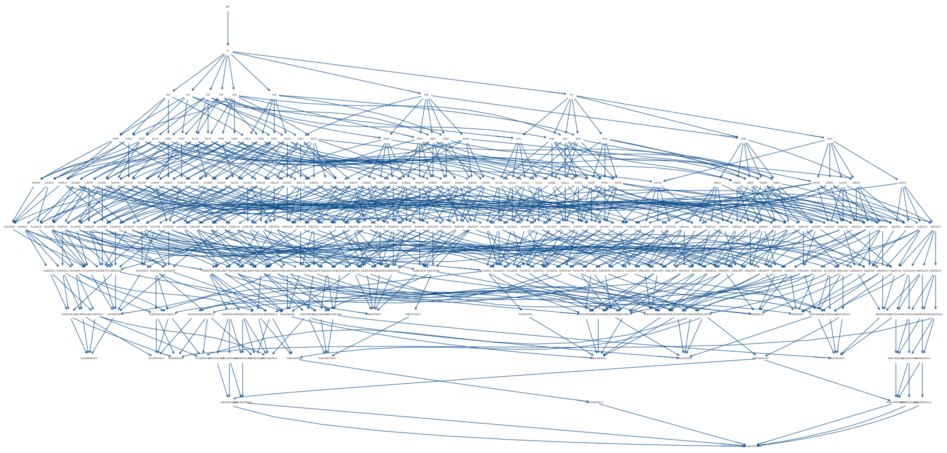

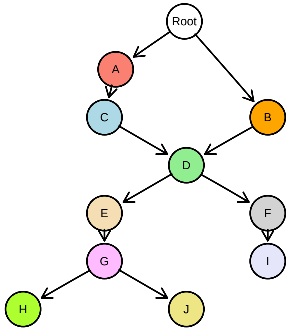


| | |
|---|---|
| *ID* | LsTIGYZJWFqEXMCQ |
| *acc genotypes* | 24 |
| *peaks no back* | 5 |
| *removed edges* | 6 |
| *prop rem edges* | 0.158 |

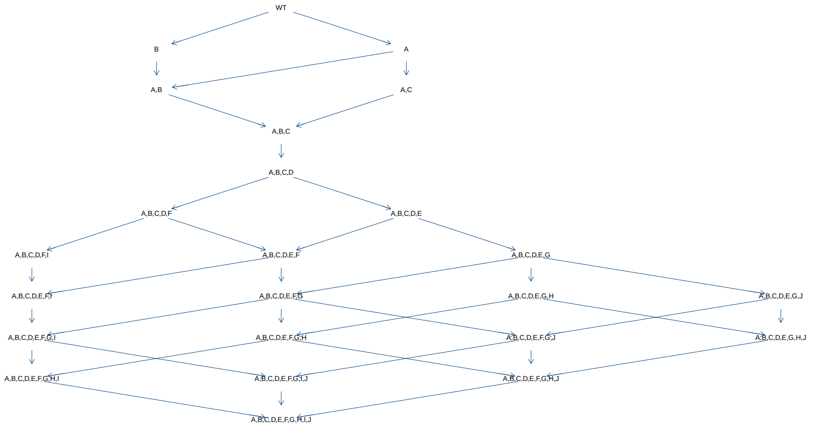

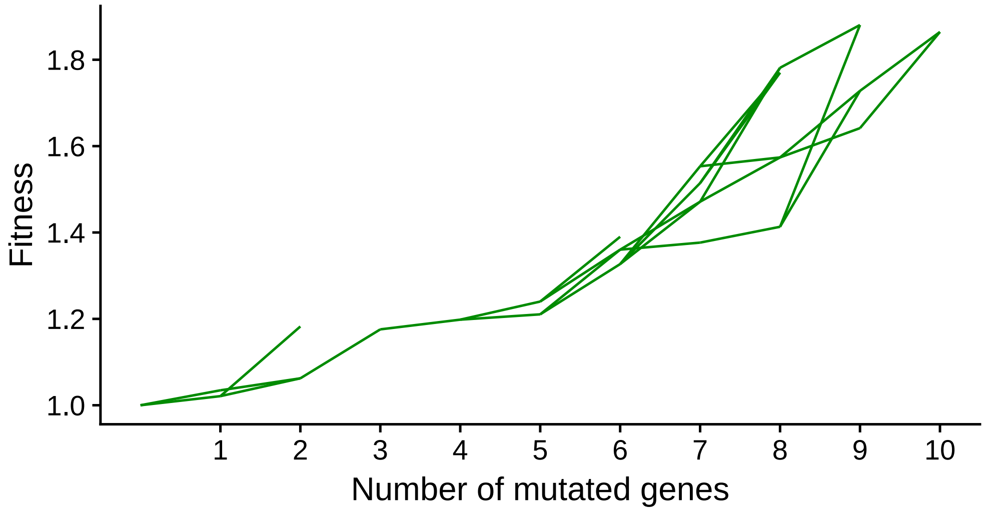


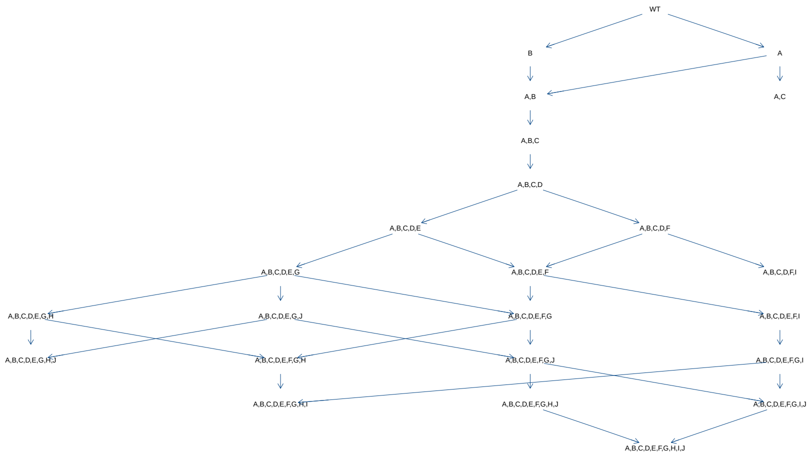

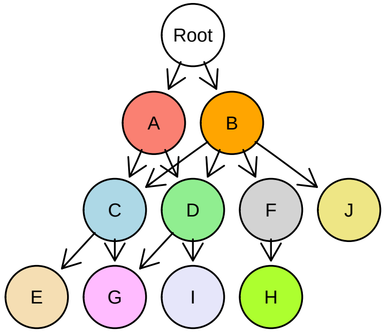


| | |
|---|---|
| *ID* | LZ8ykaYM2CcUTFrI |
| *acc genotypes* | 86 |
| *peaks no back* | 10 |
| *removed edges* | 44 |
| *prop rem edges* | 0.194 |

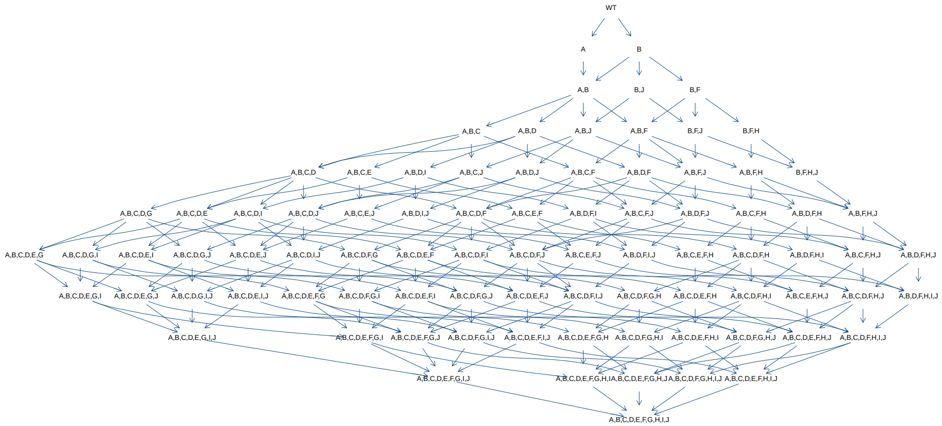

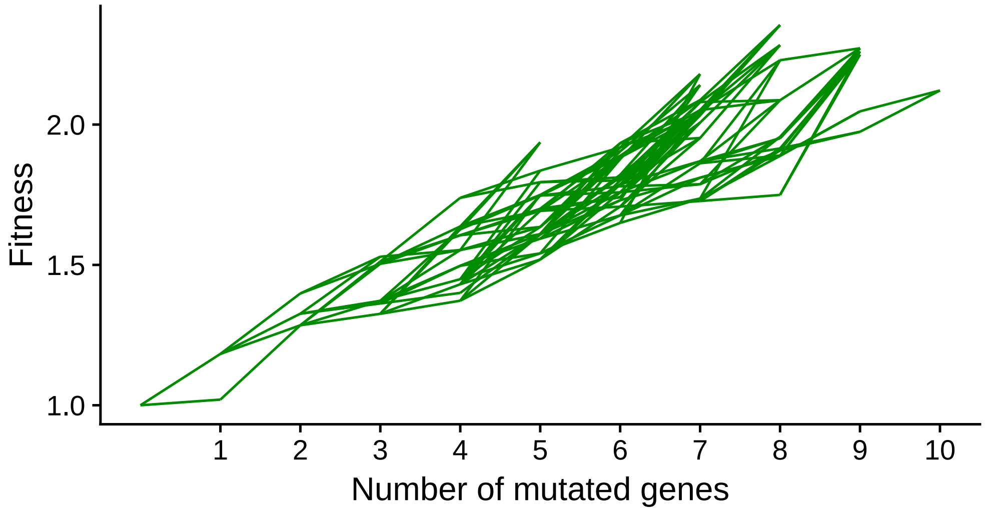


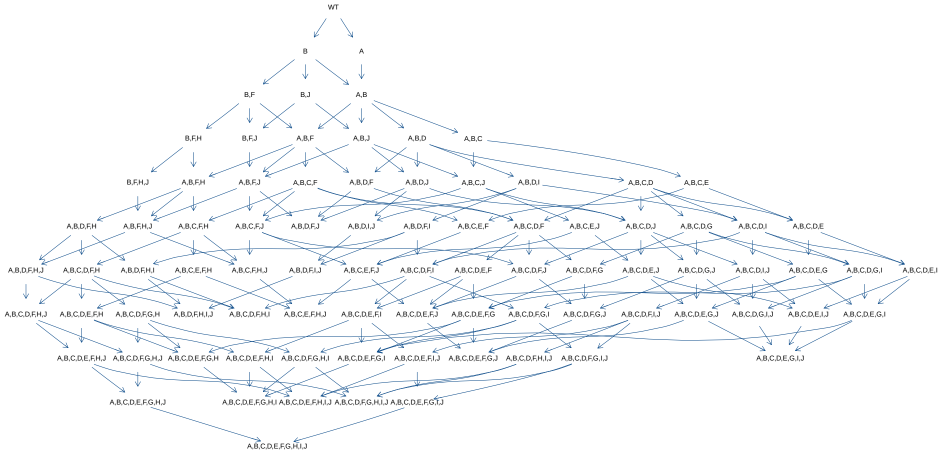

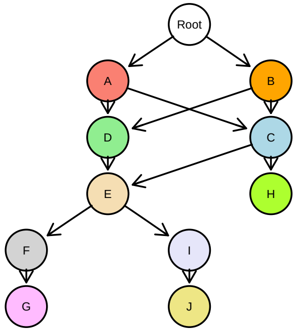


| | |
|---|---|
| *ID* | m39LVsPQFf4Car10C |
| *acc genotypes* | 27 |
| *peaks no back* | 4 |
| *removed edges* | 6 |
| *prop rem edges* | 0.13 |

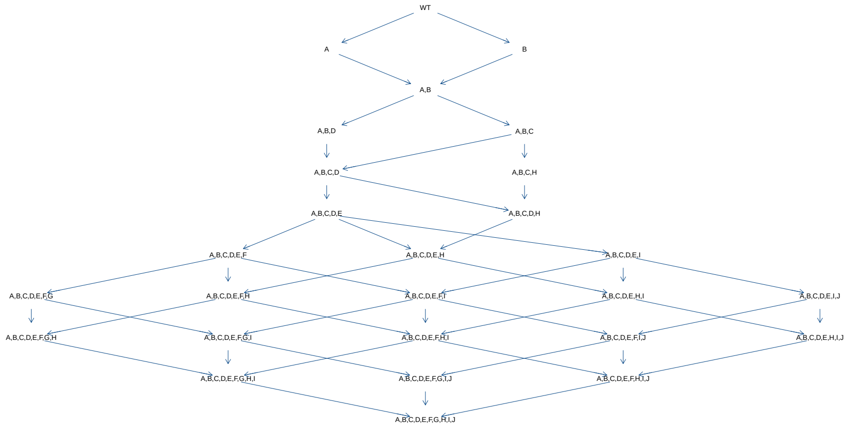

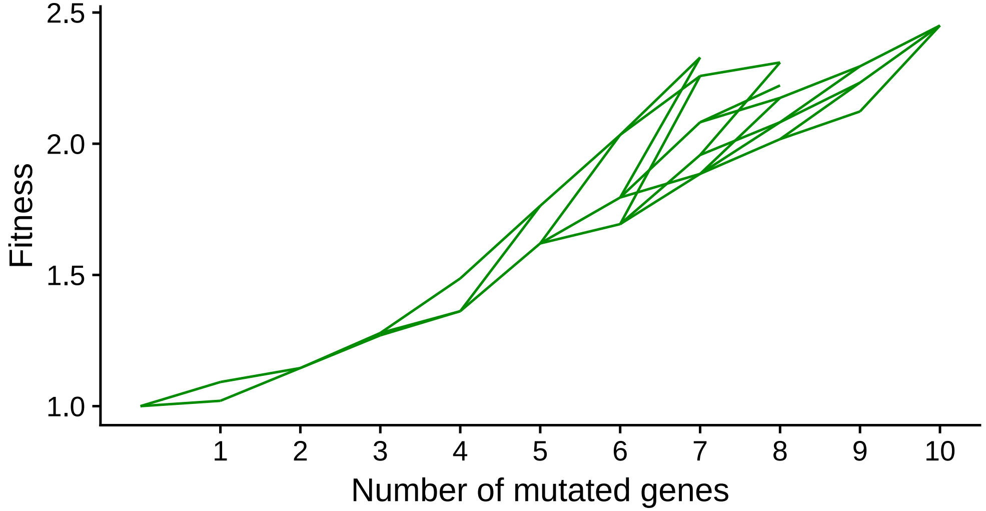


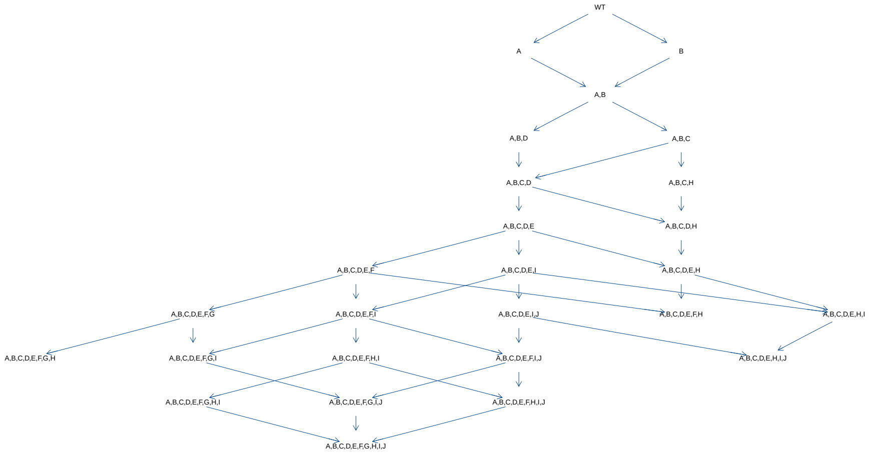

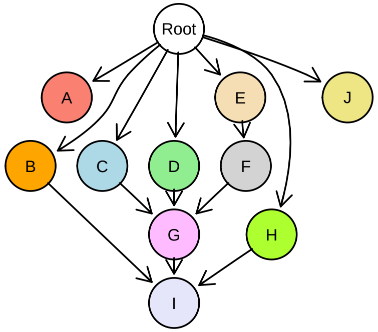


| | |
|---|---|
| *ID* | M7TqwZkMb4FqiZxL |
| *acc genotypes* | 212 |
| *peaks no back* | 18 |
| *removed edges* | 178 |
| *prop rem edges* | 0.234 |

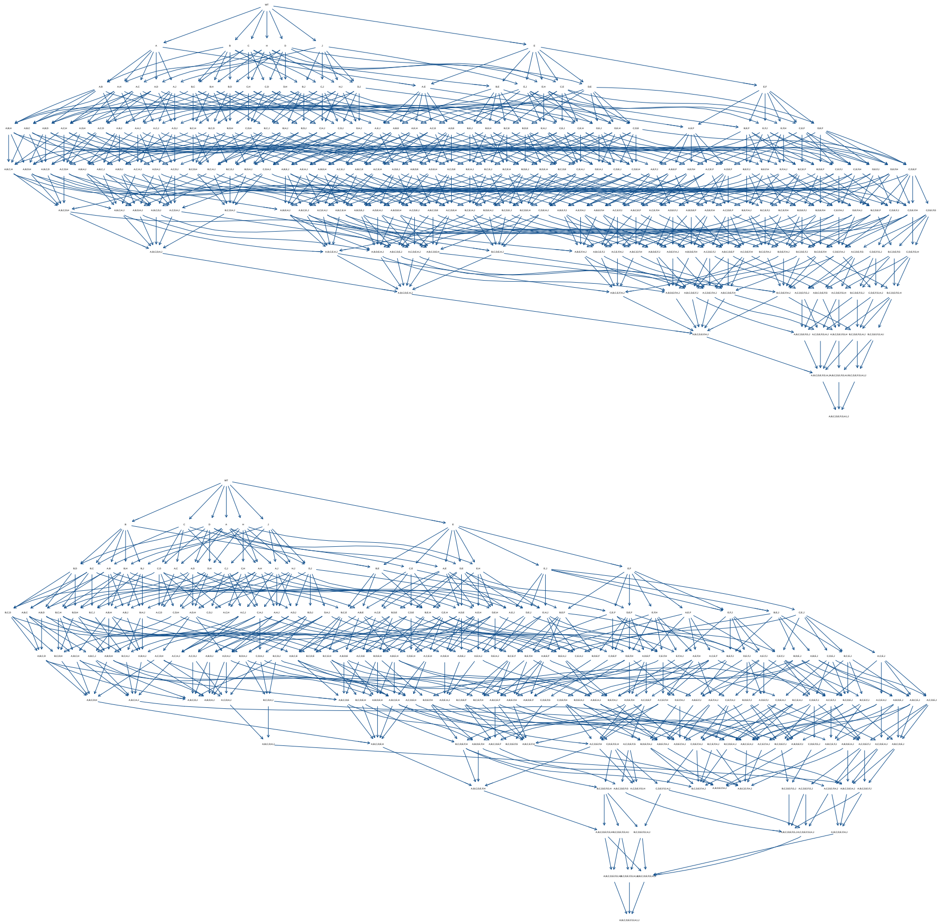

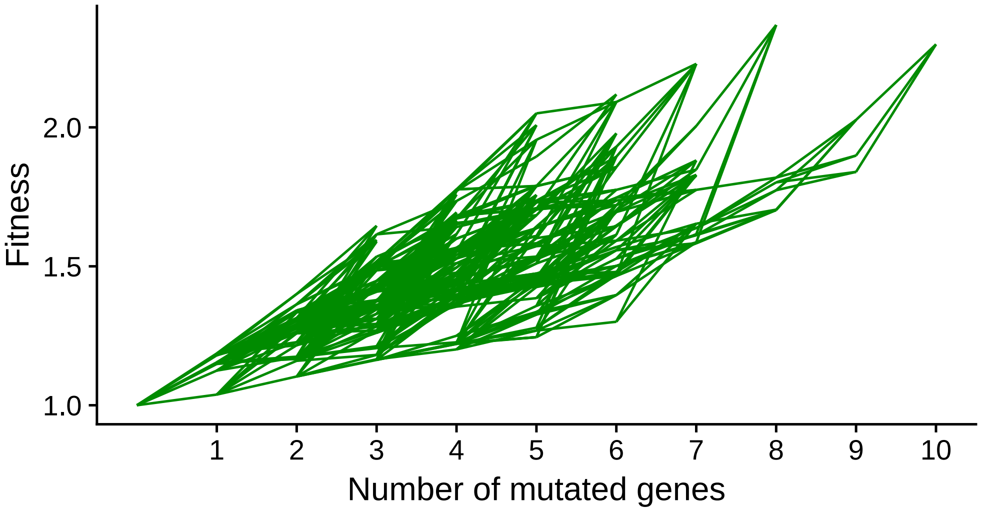

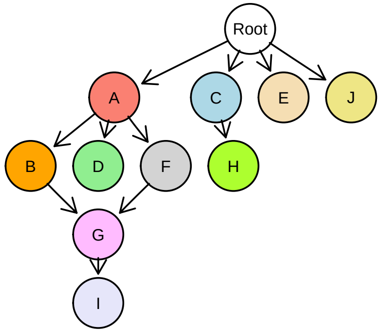


| | |
|---|---|
| *ID* | MbPTZwyHjCzW5F3T |
| *acc genotypes* | 156 |
| *peaks no back* | 12 |
| *removed edges* | 87 |
| *prop rem edges* | 0.178 |

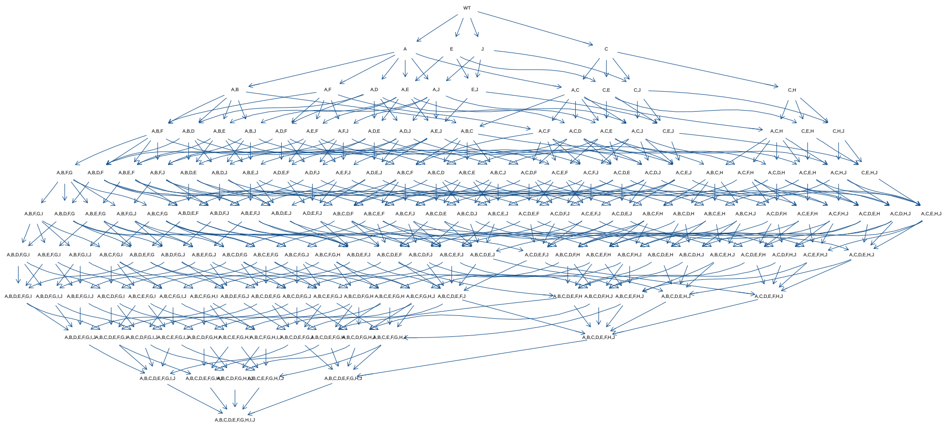

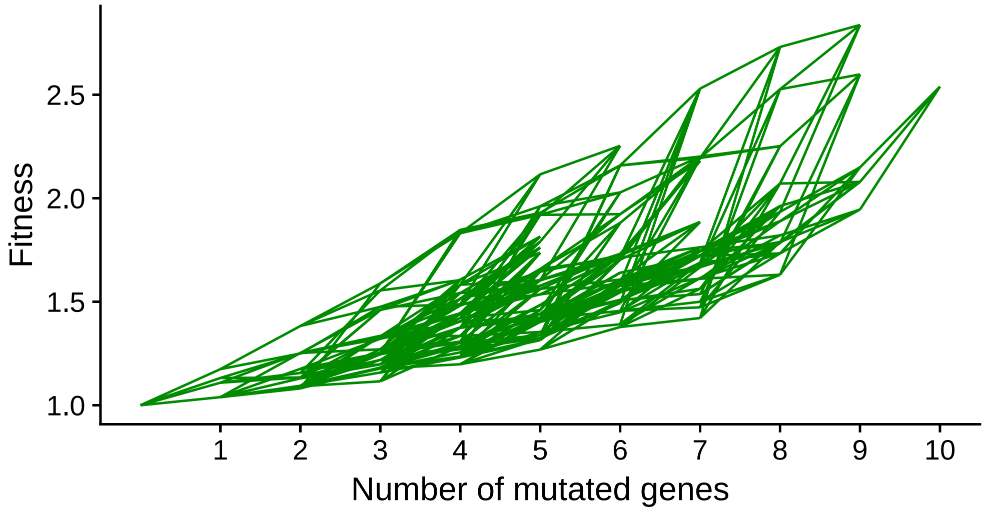


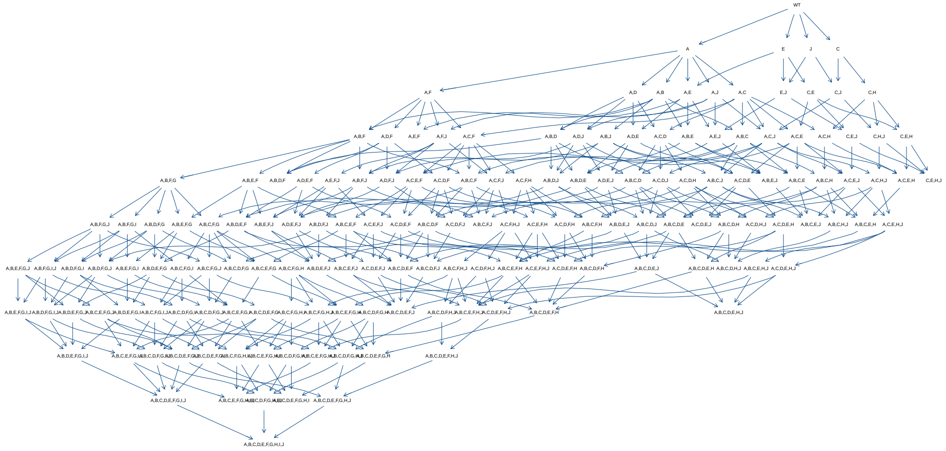

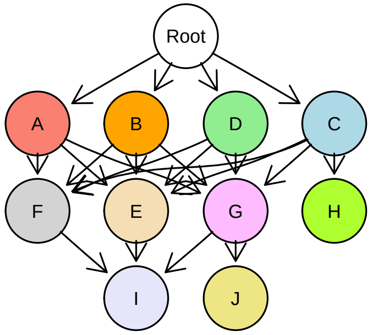


| | |
|---|---|
| *ID* | MEBeweDPZbszojRQ |
| *acc genotypes* | 50 |
| *peaks no back* | 9 |
| *removed edges* | 20 |
| *prop rem edges* | 0.18 |

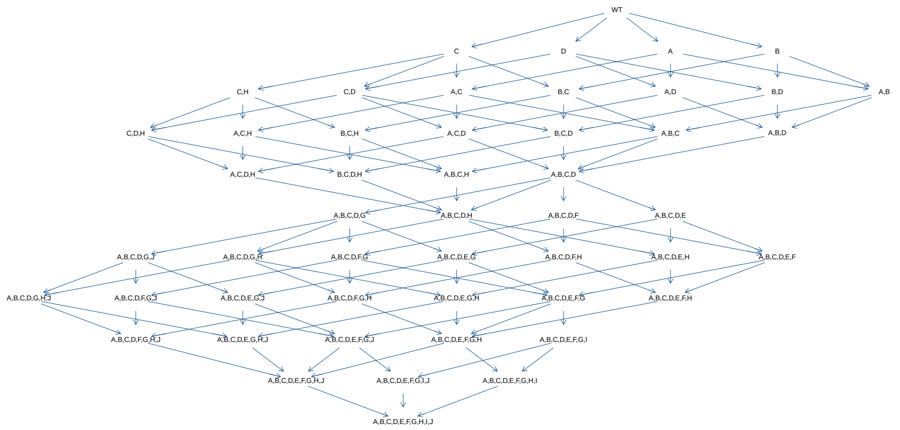

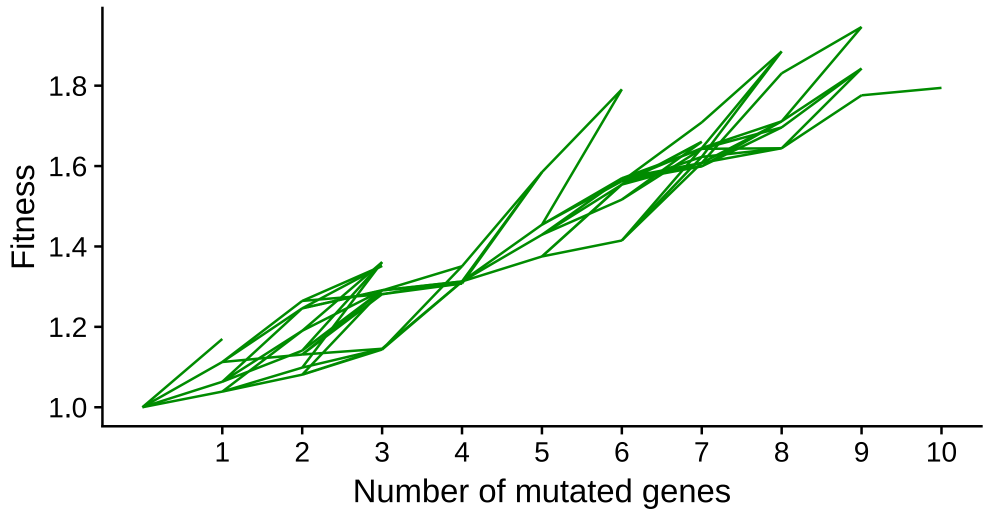


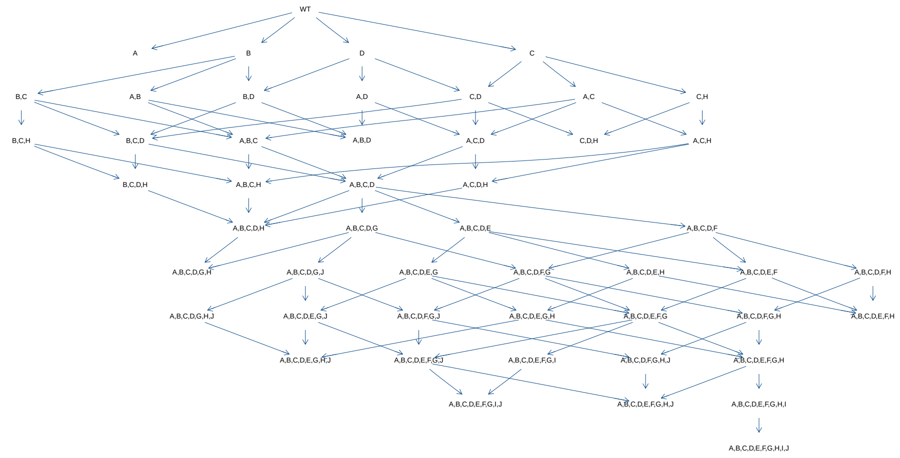

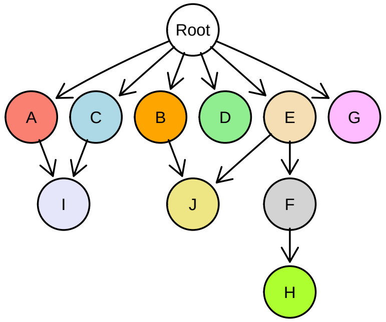


| | |
|---|---|
| *ID* | MeQOZBLaWrCfm8Ns |
| *acc genotypes* | 220 |
| *peaks no back* | 19 |
| *removed edges* | 135 |
| *prop rem edges* | 0.182 |

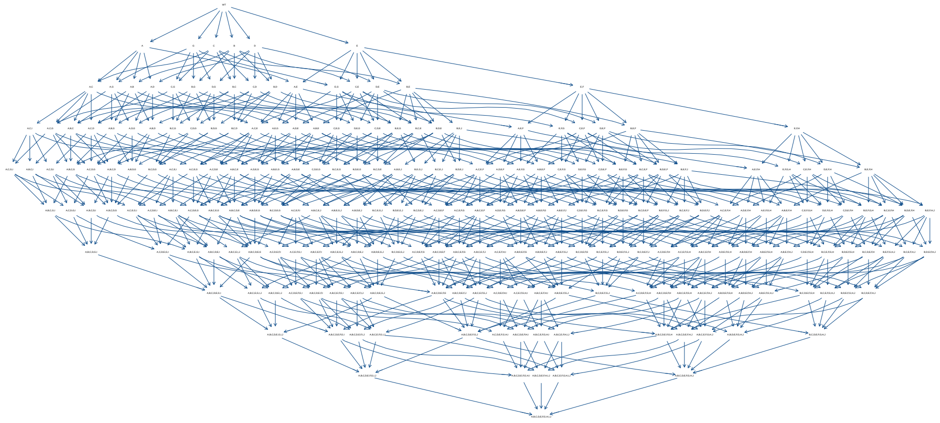

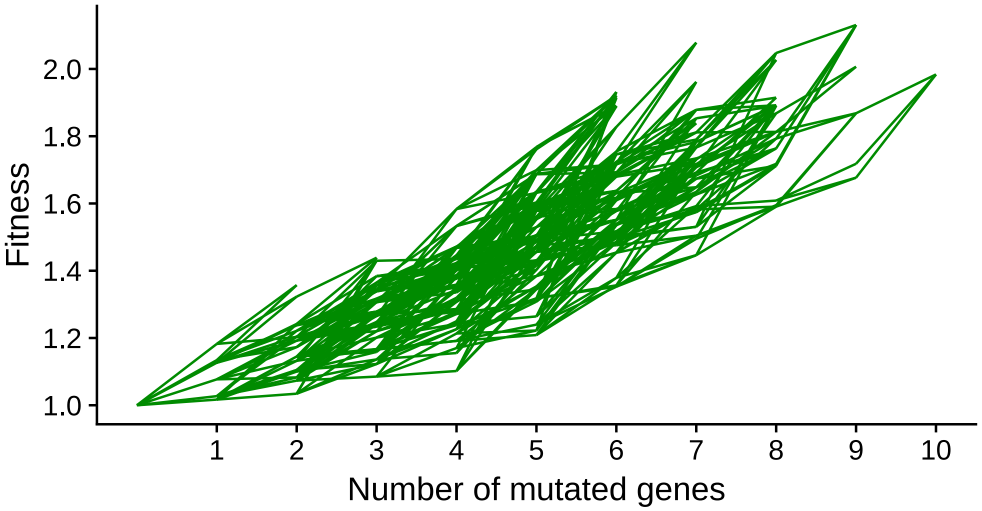


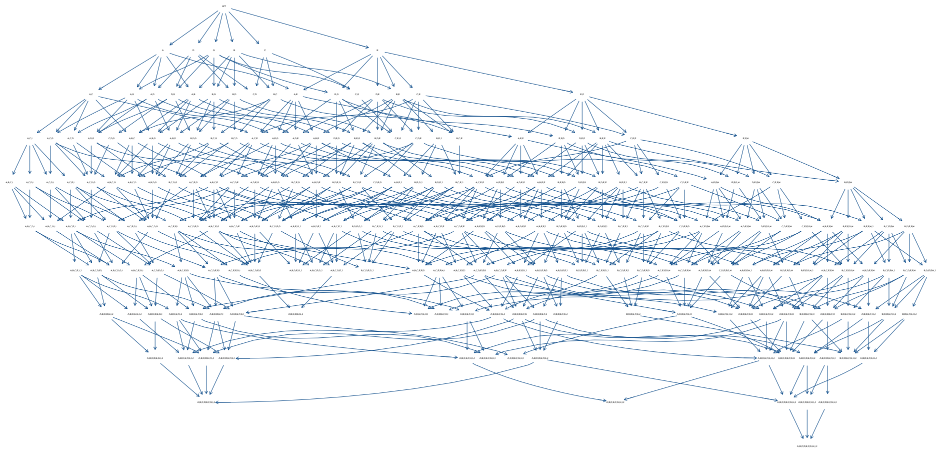

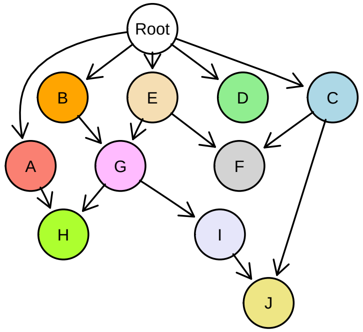


| | |
|---|---|
| *ID* | mHOgzGLqndvBGgxt |
| *acc genotypes* | 88 |
| *peaks no back* | 10 |
| *removed edges* | 35 |
| *prop rem edges* | 0.154 |

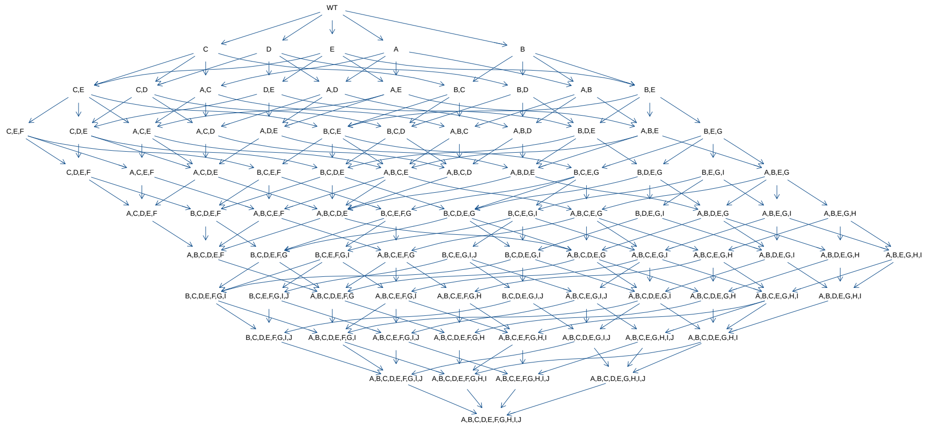

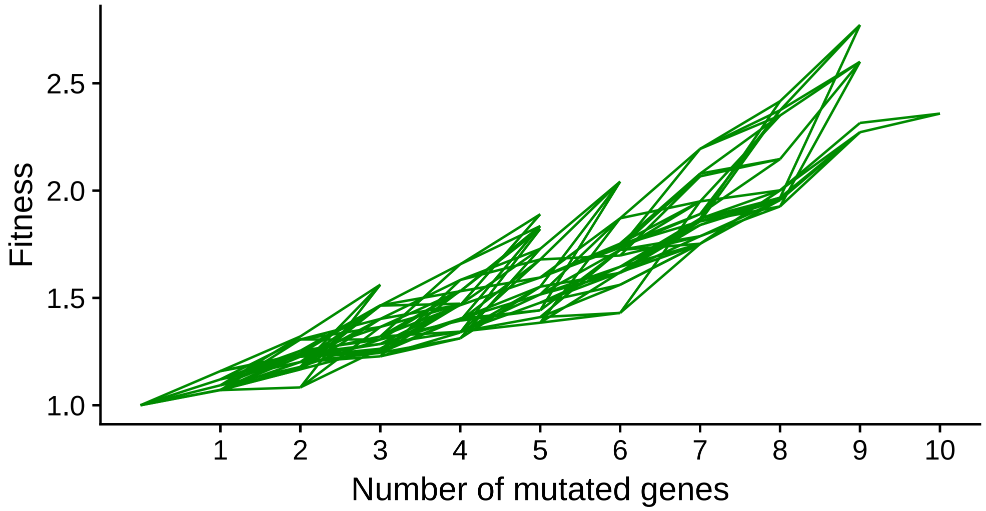


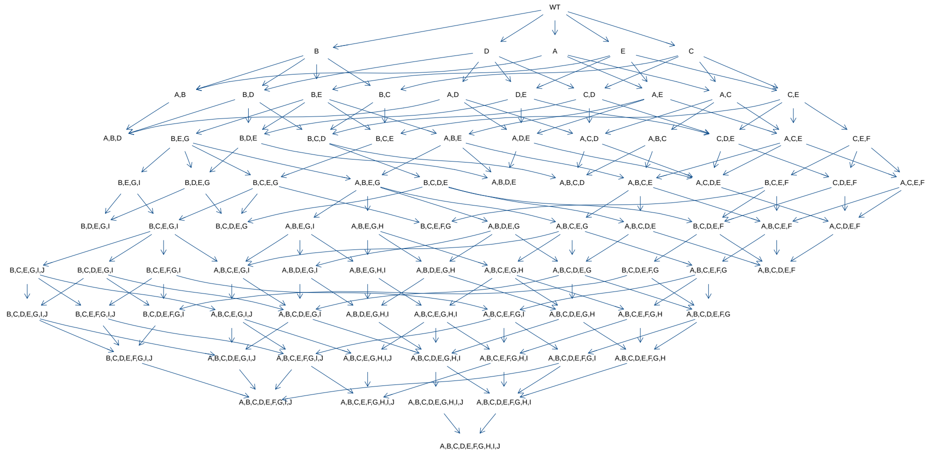

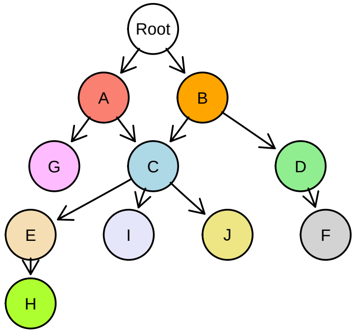


| | |
|---|---|
| *ID* | mHXf10tSNigHlcwWO |
| *acc genotypes* | 84 |
| *peaks no back* | 9 |
| *removed edges* | 44 |
| *prop rem edges* | 0.194 |

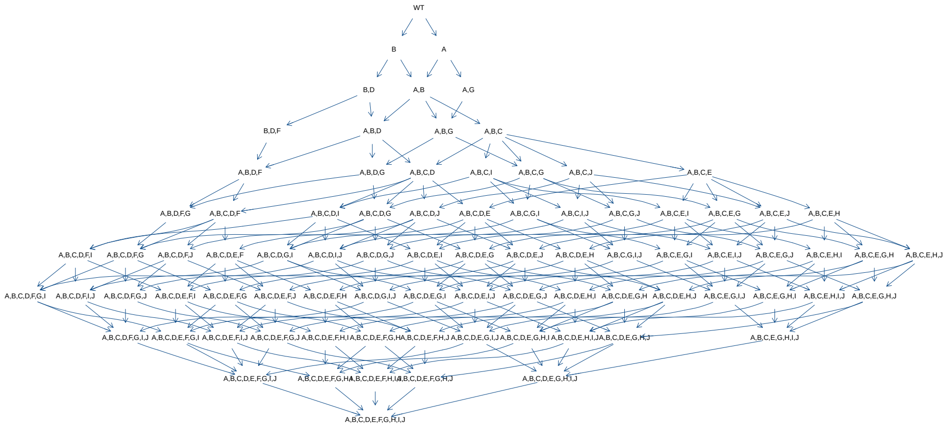

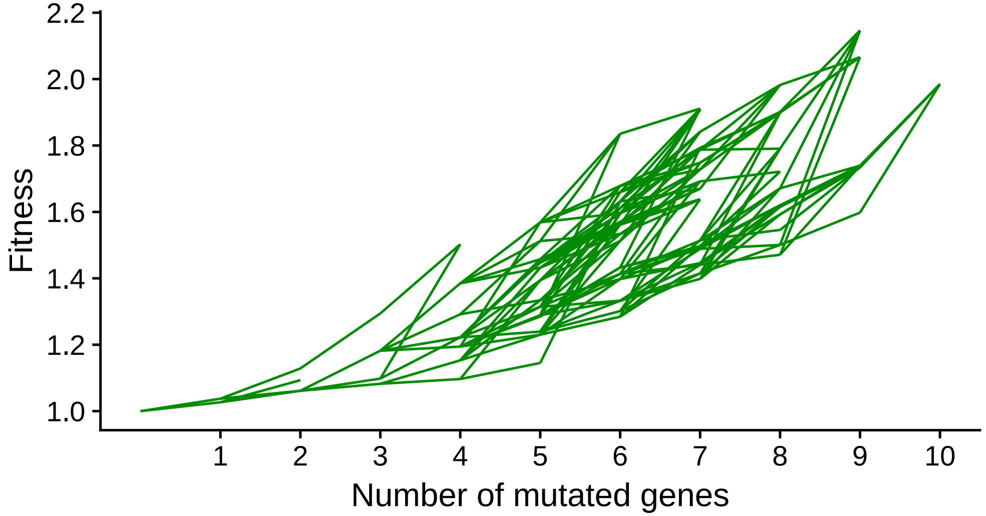


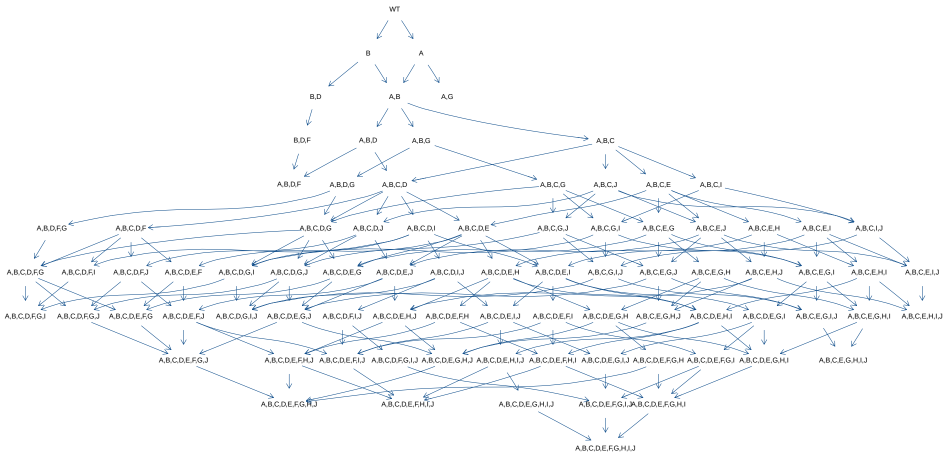

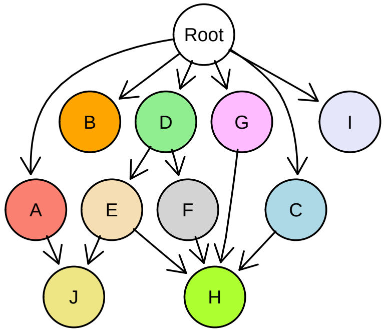


| | |
|---|---|
| *ID* | MW1p5m710opyweI6d |
| *acc genotypes* | 204 |
| *peaks no back* | 15 |
| *removed edges* | 126 |
| *prop rem edges* | 0.179 |

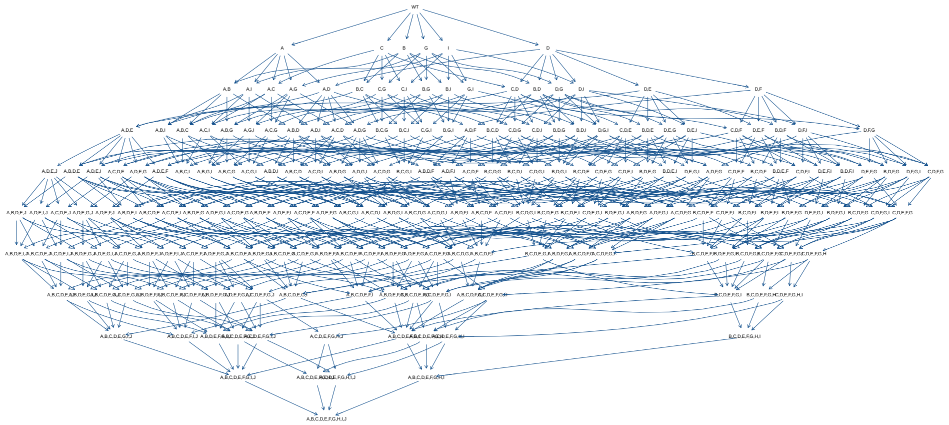

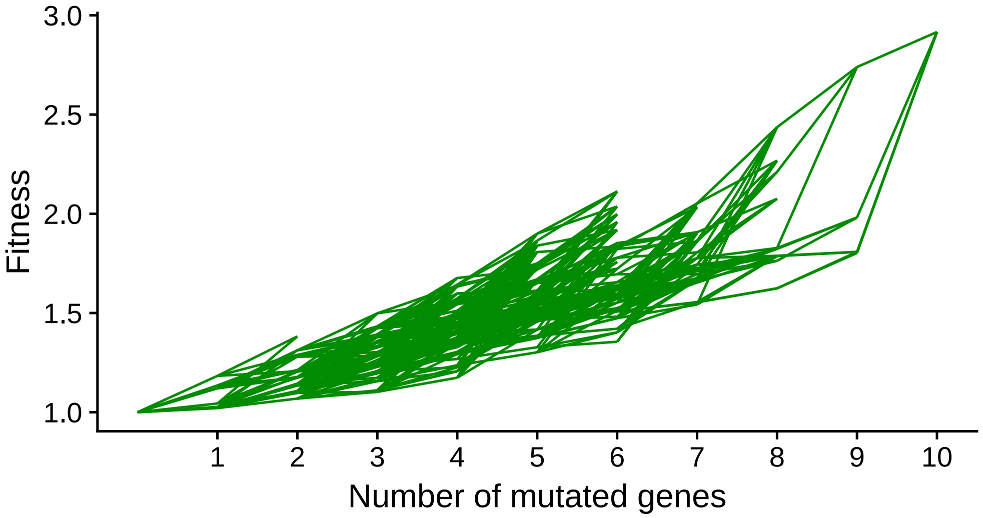


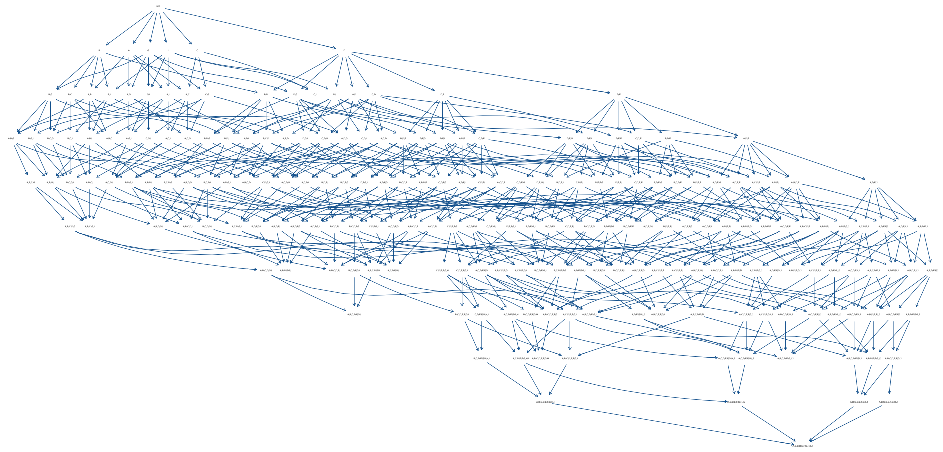

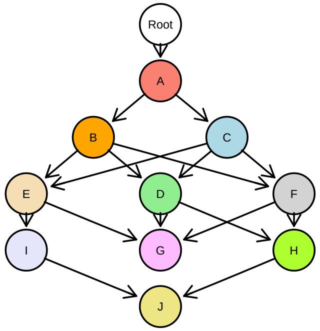


| | |
|---|---|
| *ID* | N3AGZj5WtBq10w7rq |
| *acc genotypes* | 25 |
| *peaks no back* | 5 |
| *removed edges* | 6 |
| *prop rem edges* | 0.146 |

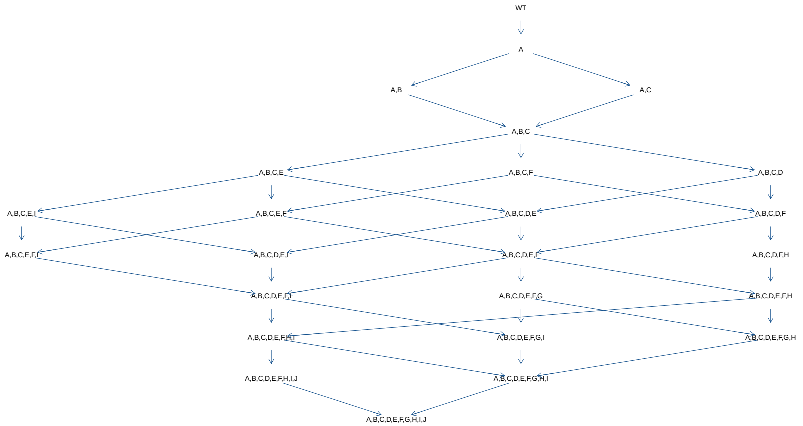

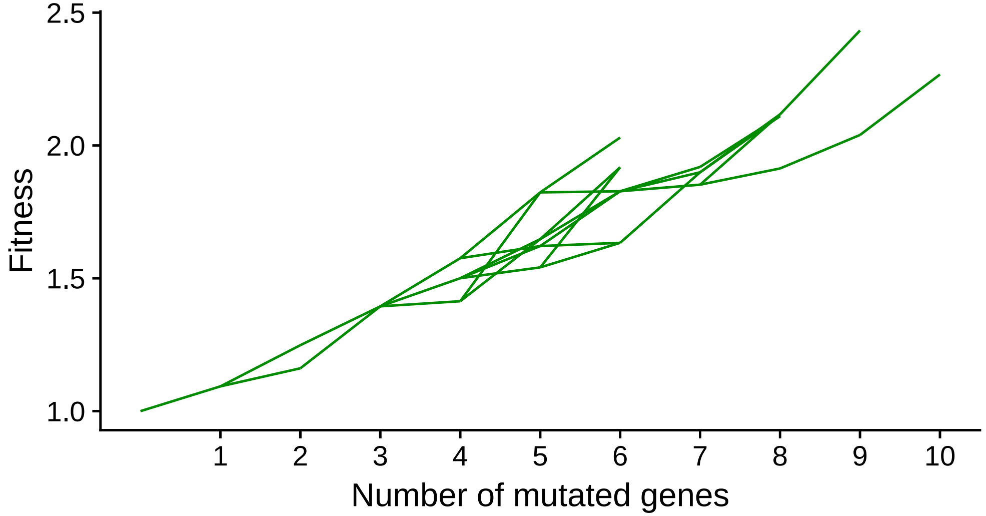


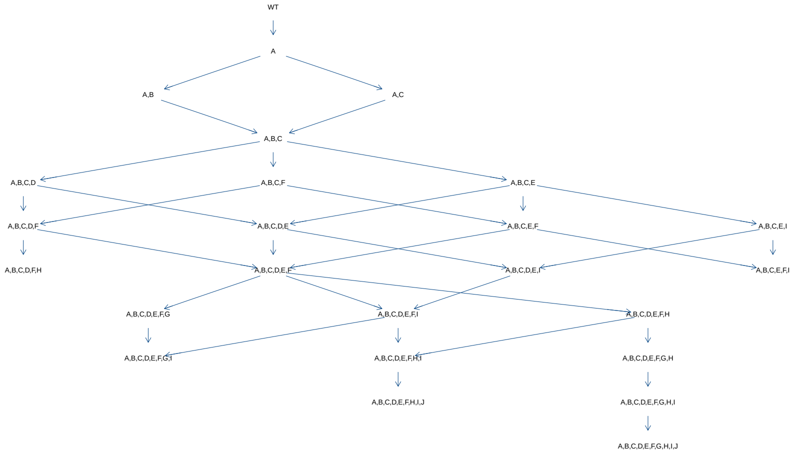

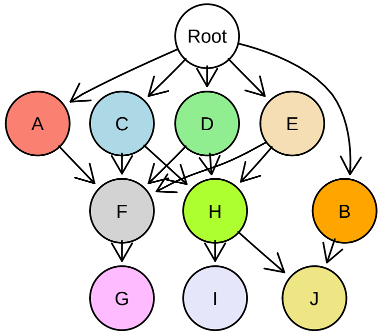


| | |
|---|---|
| *ID* | n3mpftu4orkSLBLy |
| *acc genotypes* | 60 |
| *peaks no back* | 10 |
| *removed edges* | 25 |
| *prop rem edges* | 0.179 |

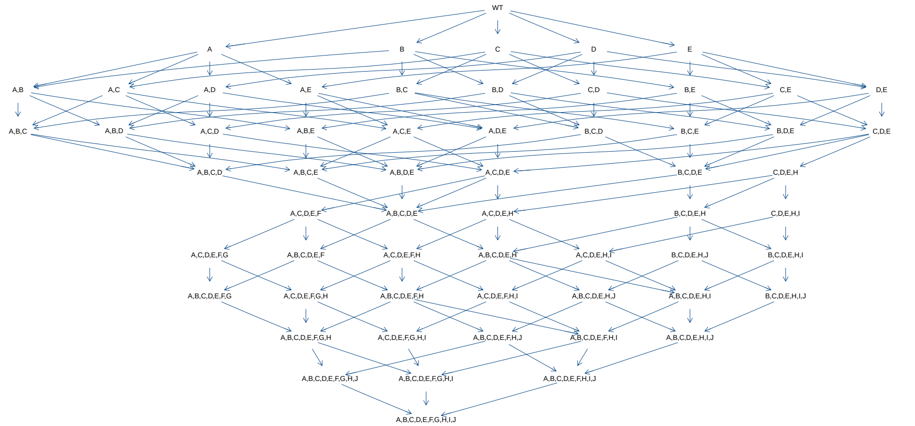

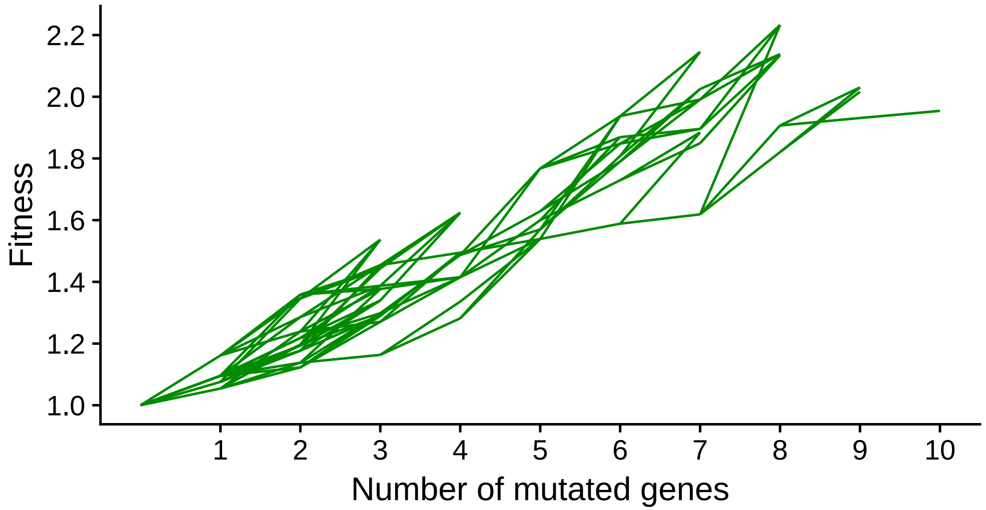


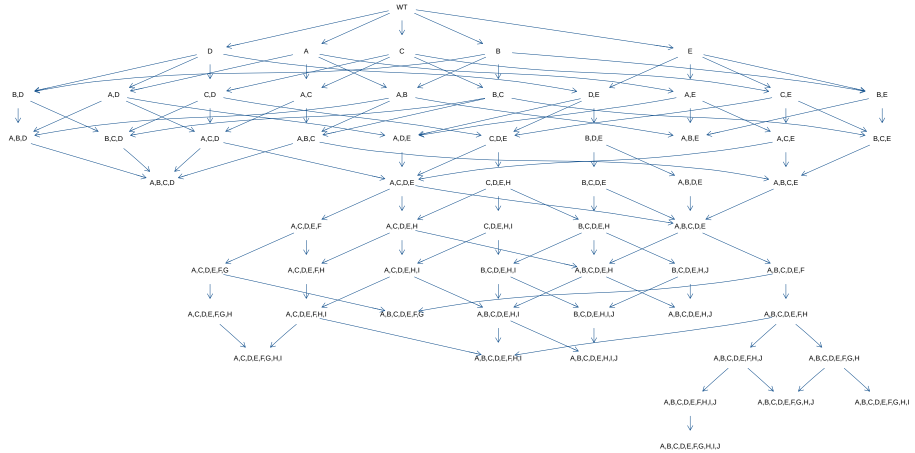

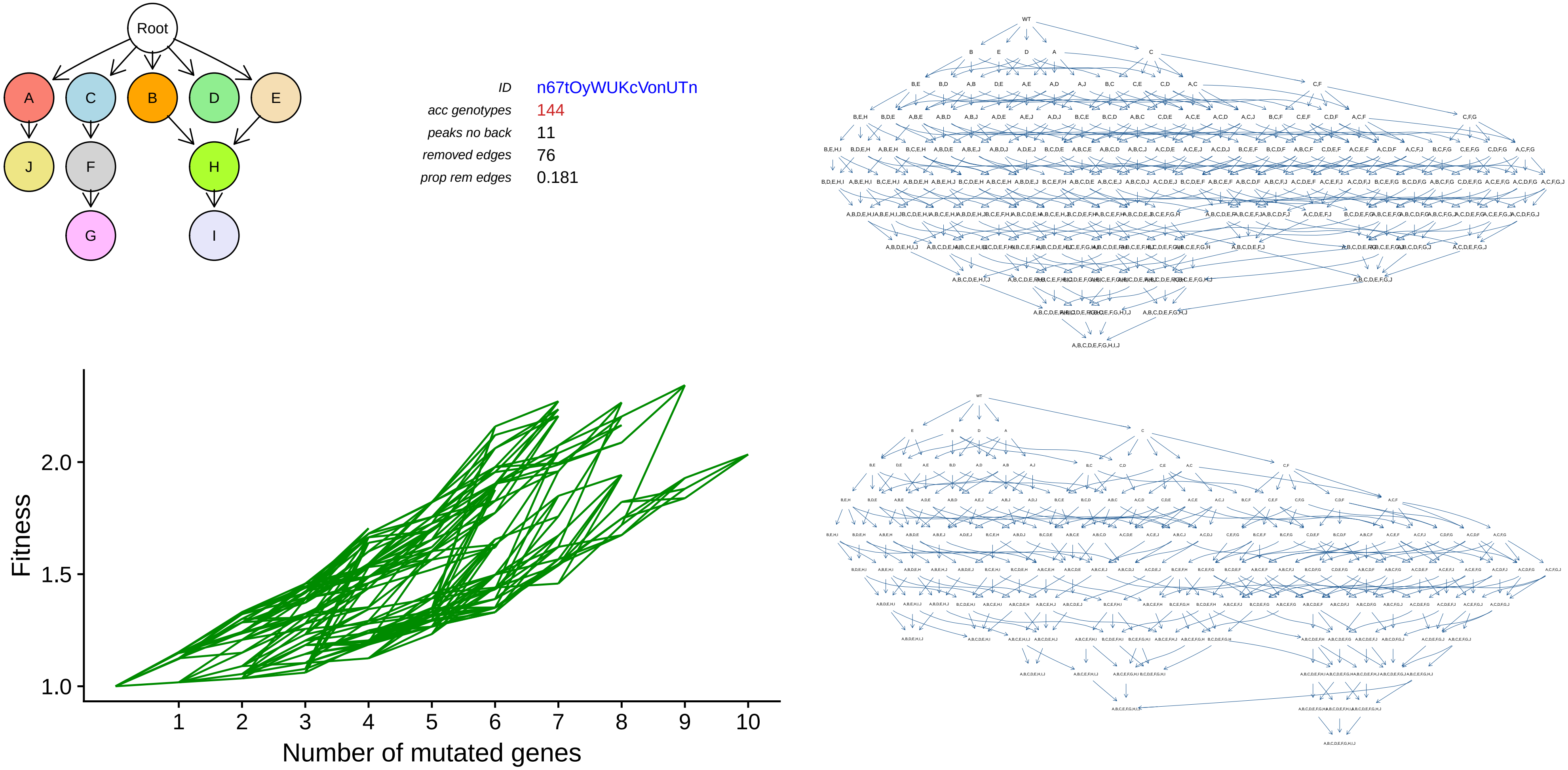
Root
A
C
B
D
E
J
F
H
G
I
ID
n67tOyWUKcVonUTn
acc genotypes
144
peaks no back
11
removed edges
76
prop rem edges
0.181
Fitness
2.0
1.5
1.0
1
2
3
4
5
6
7
8
9
10
Number of mutated genes

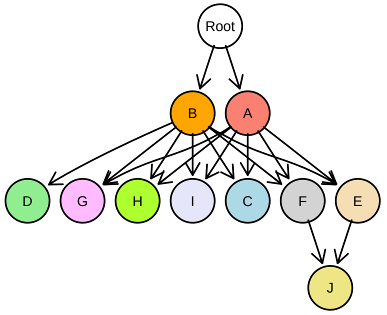


| | |
|---|---|
| *ID* | n6N43lpBlanX2OCS |
| *acc genotypes* | 164 |
| *peaks no back* | 15 |
| *removed edges* | 95 |
| *prop rem edges* | 0.168 |

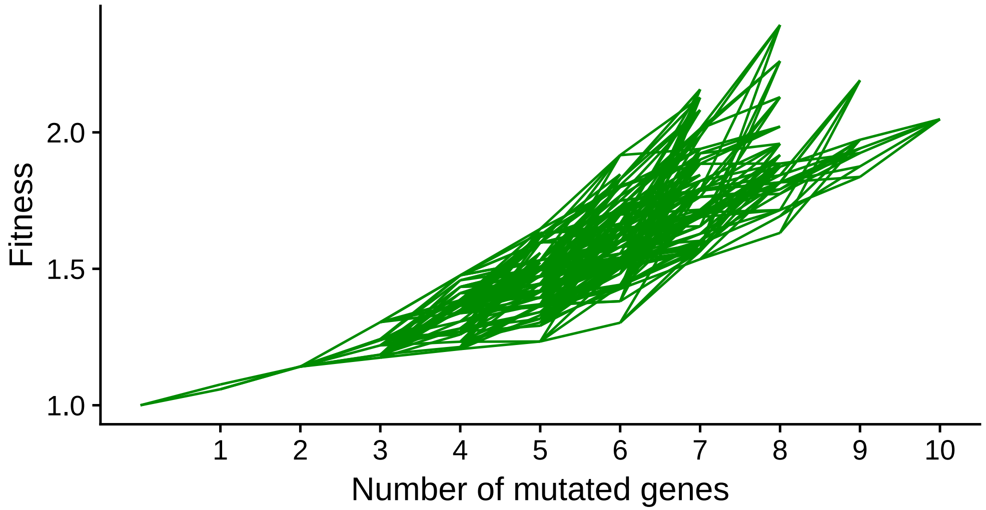


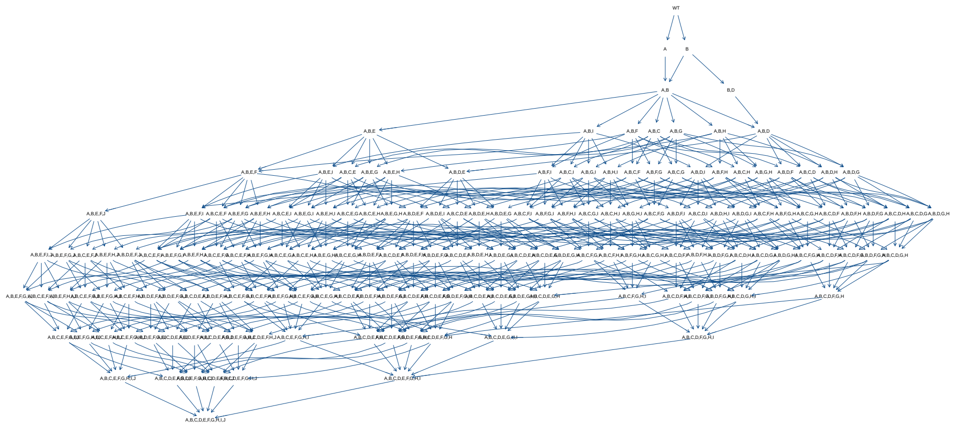

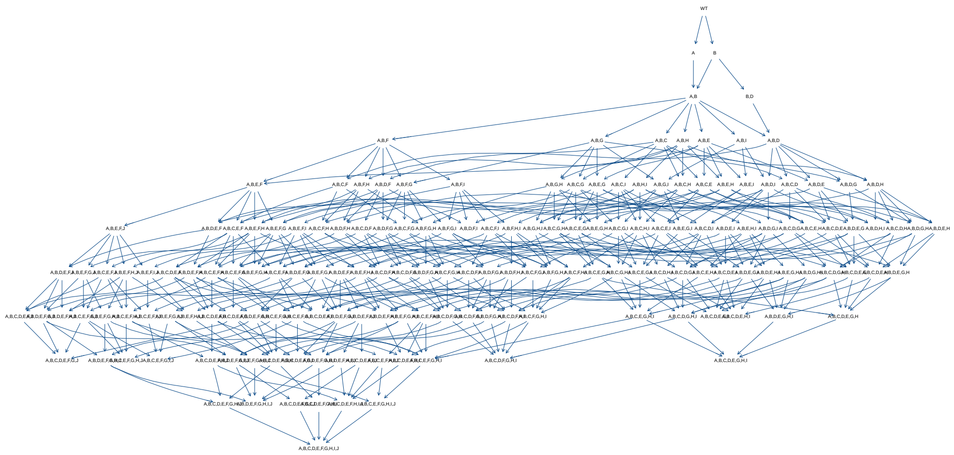

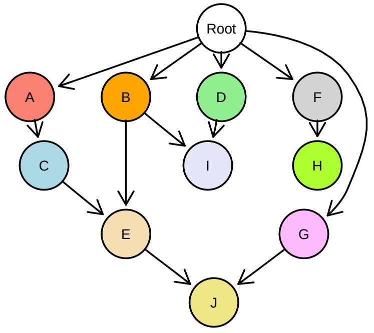


| | |
|---|---|
| *ID* | NHvOV5oluQFLza23 |
| *acc genotypes* | 117 |
| *peaks no back* | 11 |
| *removed edges* | 59 |
| *prop rem edges* | 0.18 |

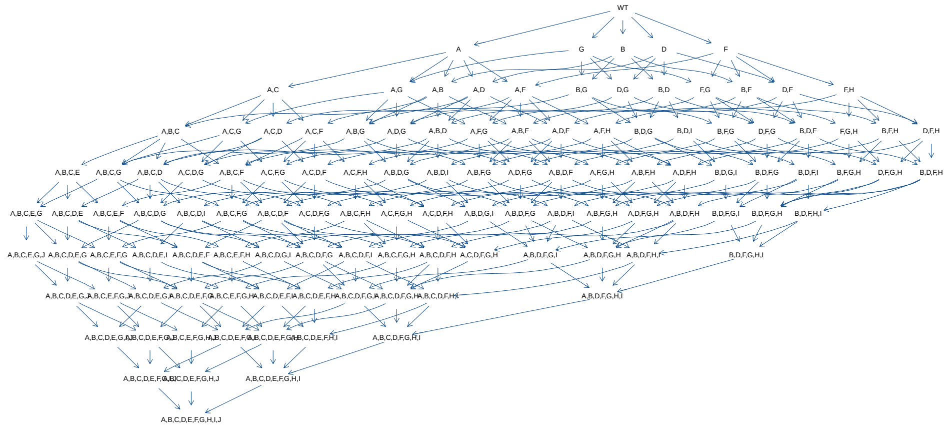

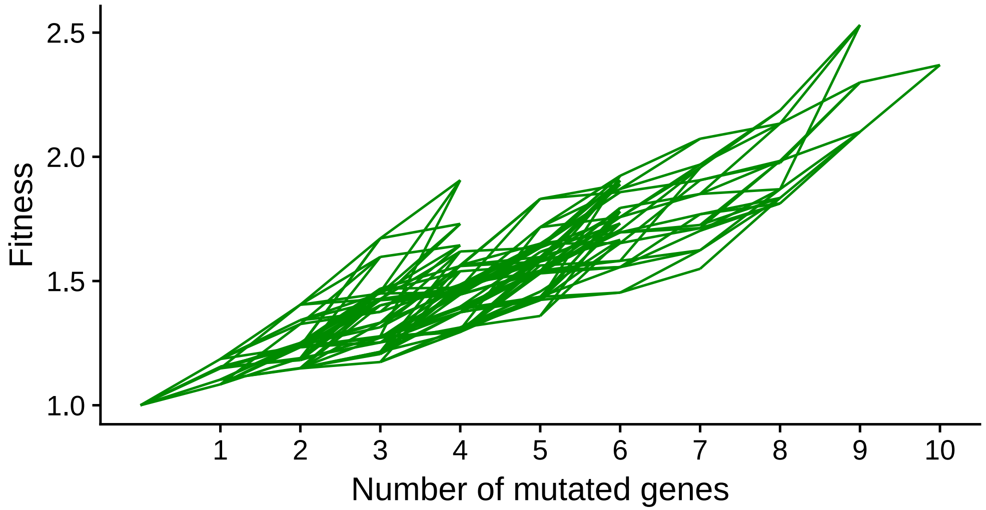


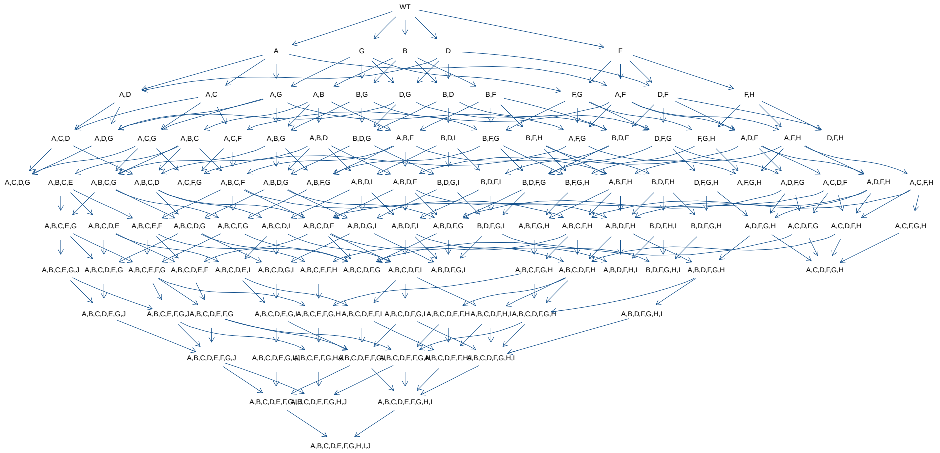

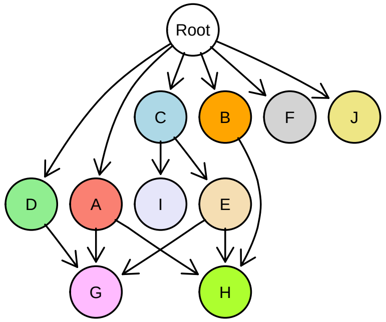


*ID* nLayuUMxBrwIiw43
*acc genotypes* 200
*peaks no back* 17
*removed edges* 135
*prop rem edges* 0.197

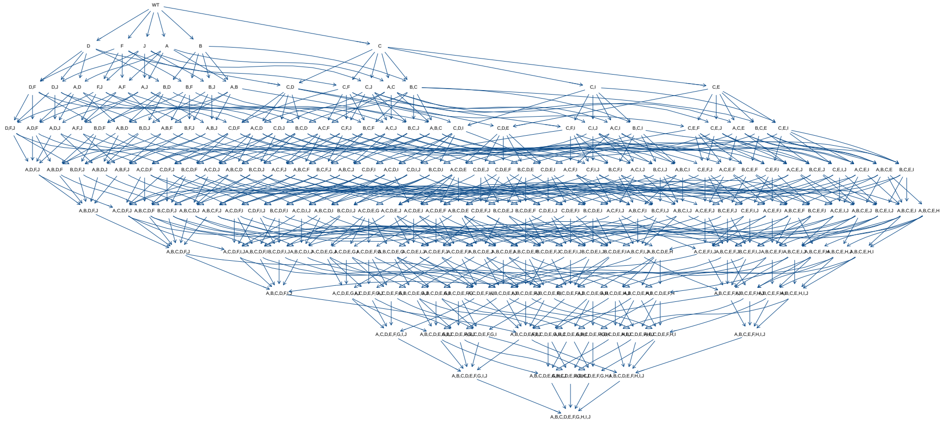

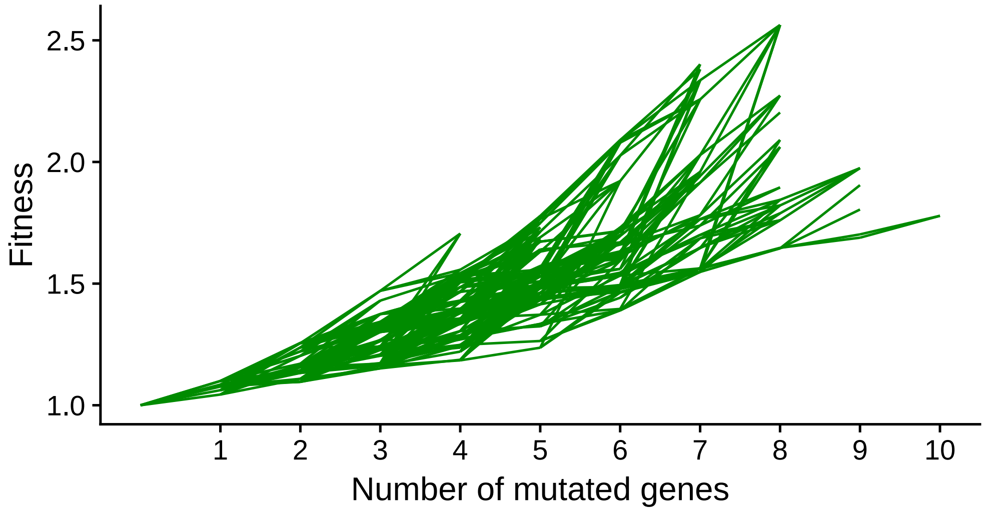


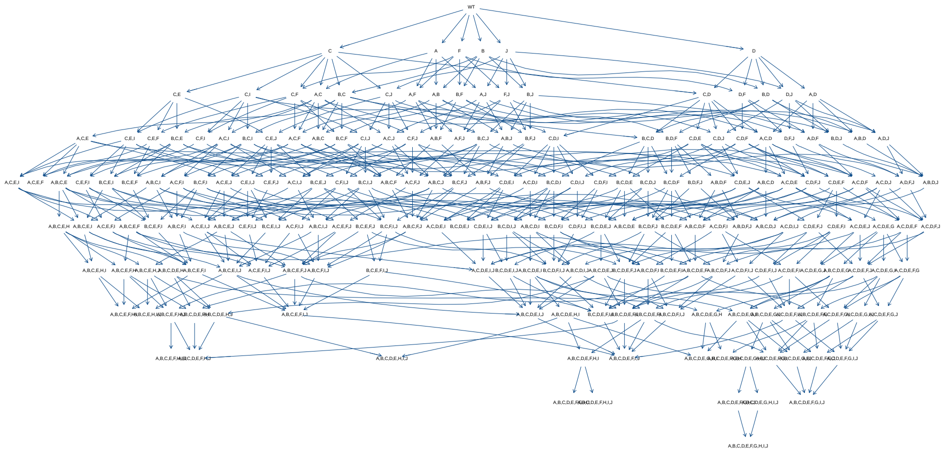

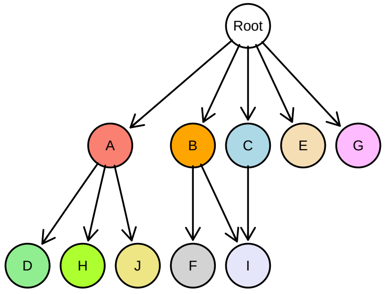


| | |
|---|---|
| *ID* | nOAHFfiFZ4dc4UPV |
| *acc genotypes* | 288 |
| *peaks no back* | 24 |
| *removed edges* | 255 |
| *prop rem edges* | 0.24 |

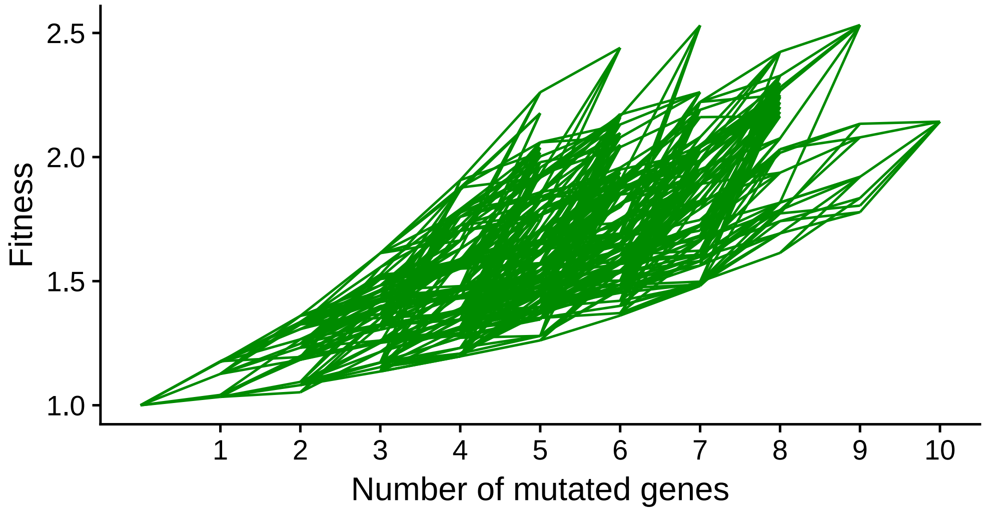


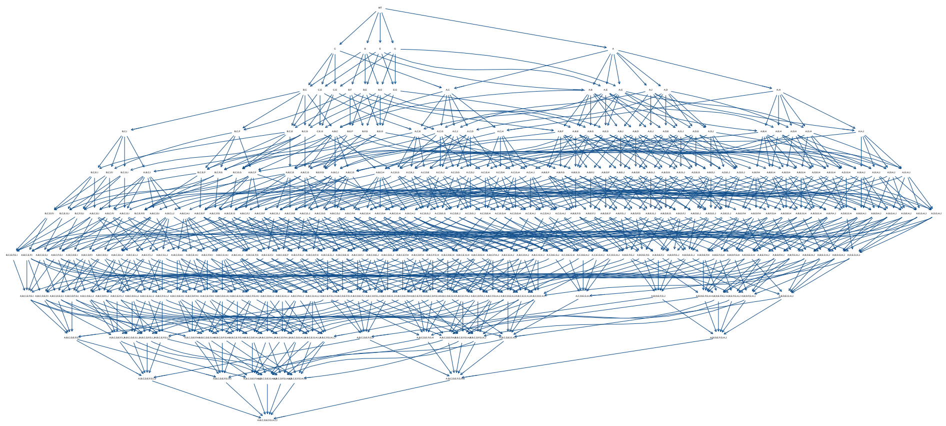

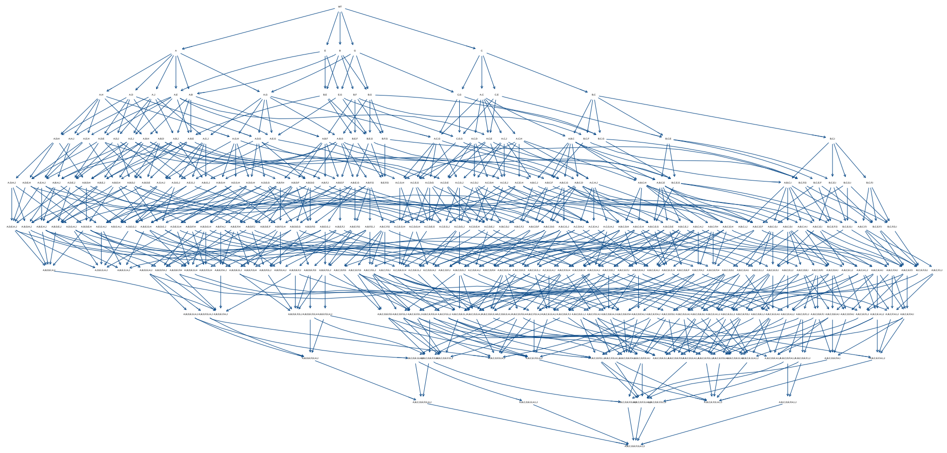

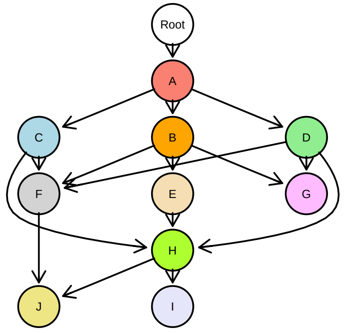


| | |
|---|---|
| *ID* | noHAVtQI4KdH8hEq |
| *acc genotypes* | 33 |
| *peaks no back* | 5 |
| *removed edges* | 12 |
| *prop rem edges* | 0.197 |

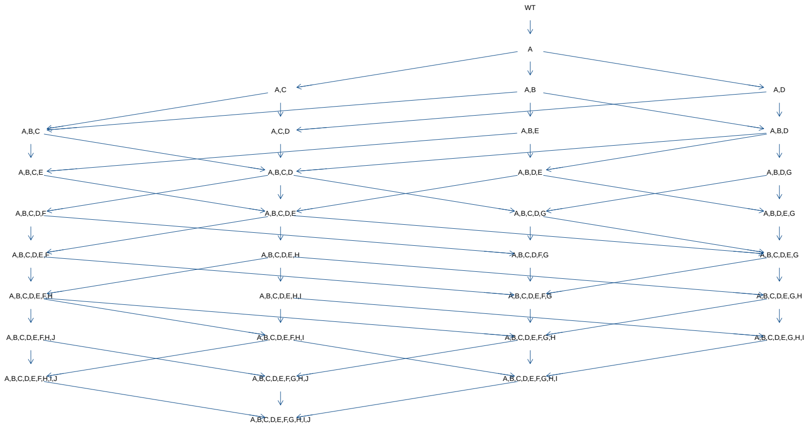

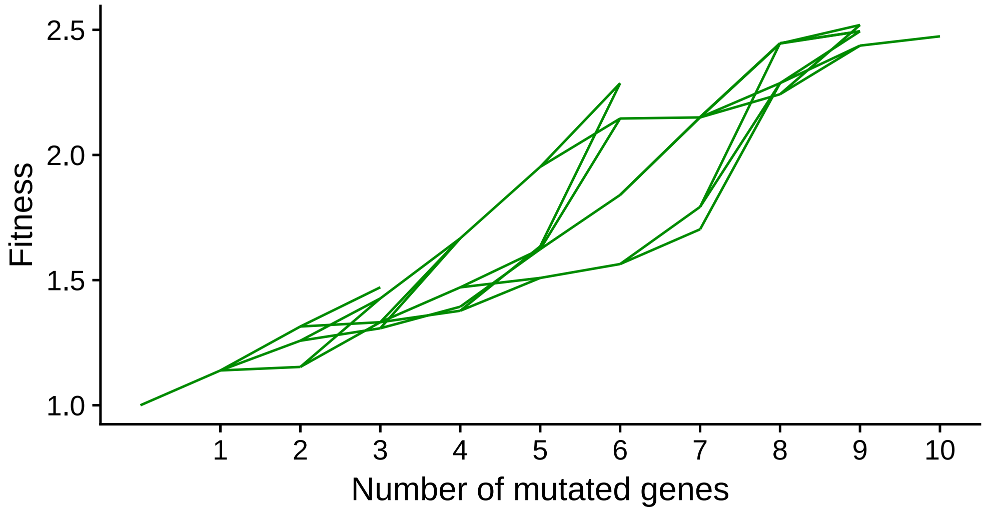


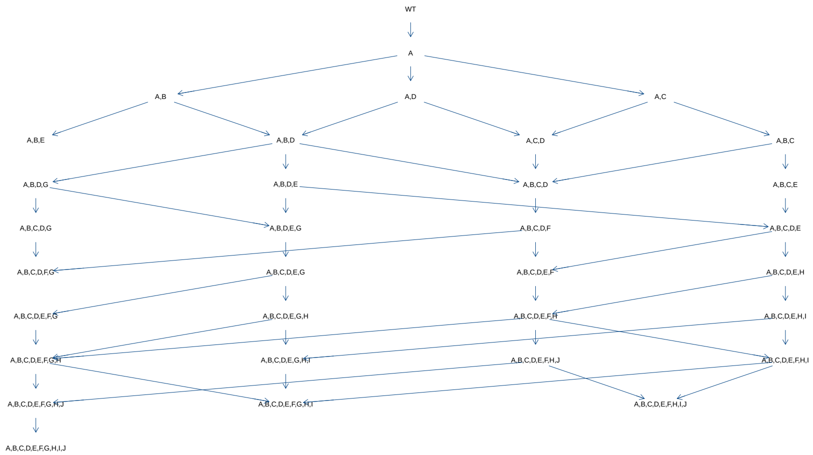

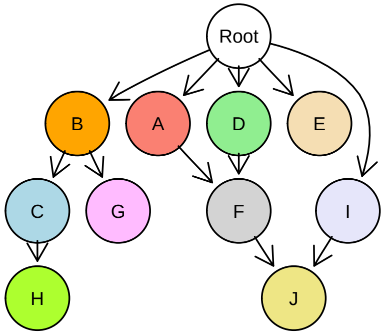


| | |
|---|---|
| *ID* | NSqT8kGDJ9iU108hE |
| *acc genotypes* | 154 |
| *peaks no back* | 11 |
| *removed edges* | 75 |
| *prop rem edges* | 0.157 |

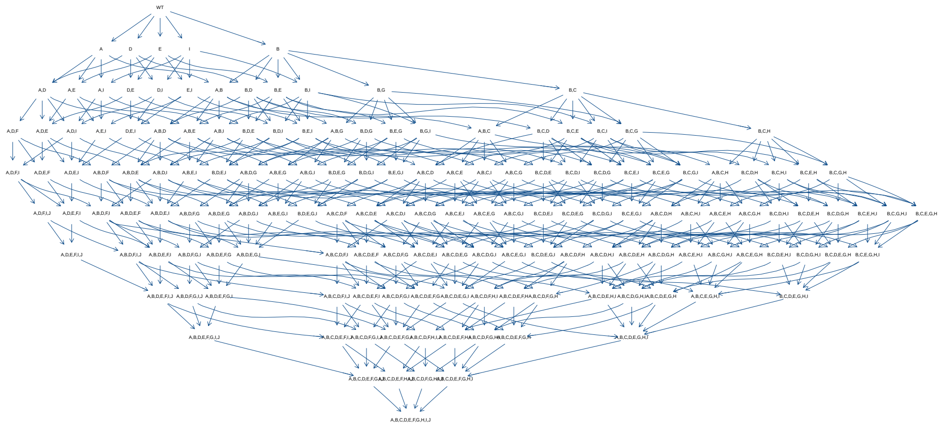

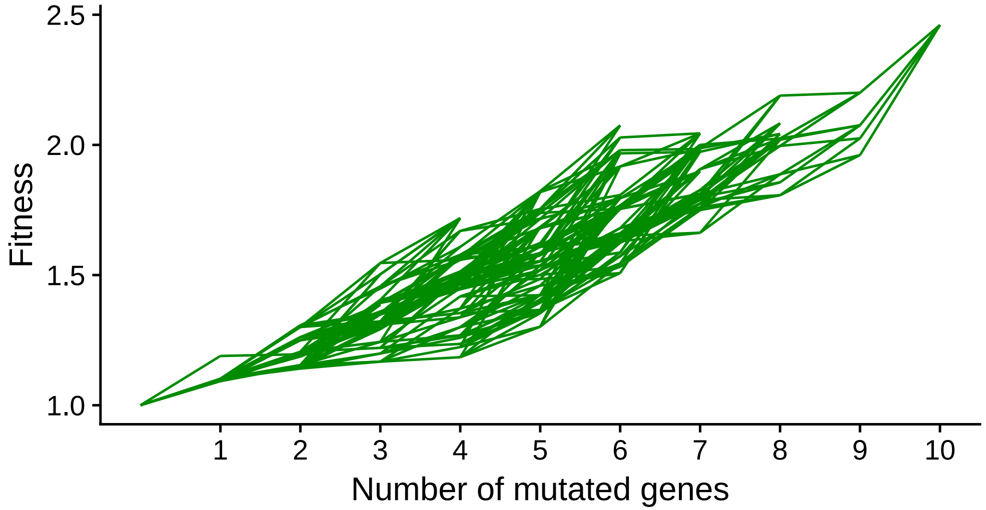


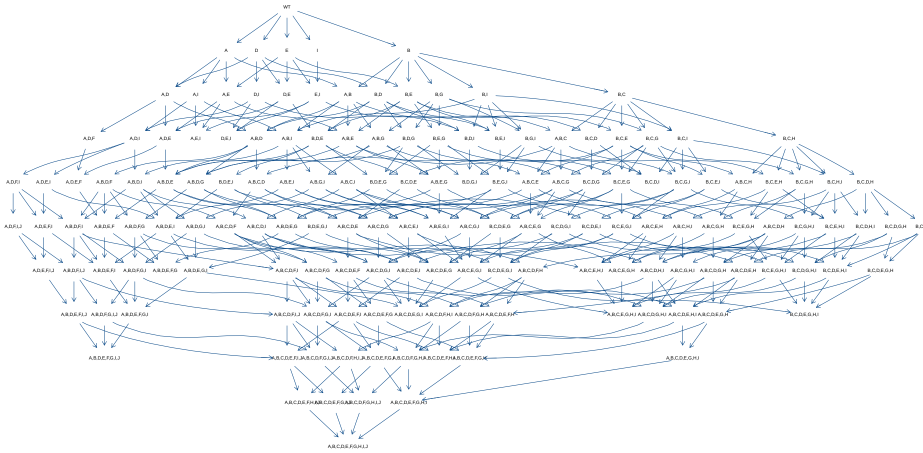

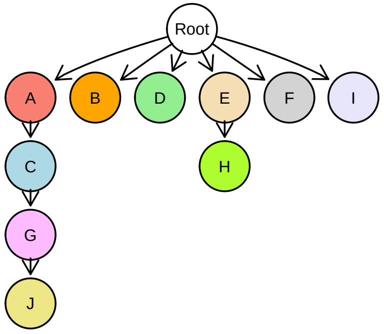


| | |
|---|---|
| *ID* | nTjCF9qeG8JYDoN8 |
| *acc genotypes* | 240 |
| *peaks no back* | 19 |
| *removed edges* | 158 |
| *prop rem edges* | 0.19 |

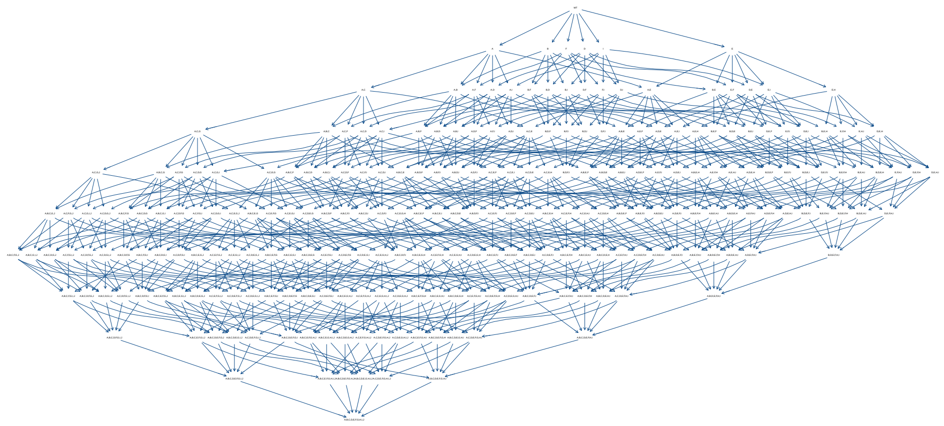

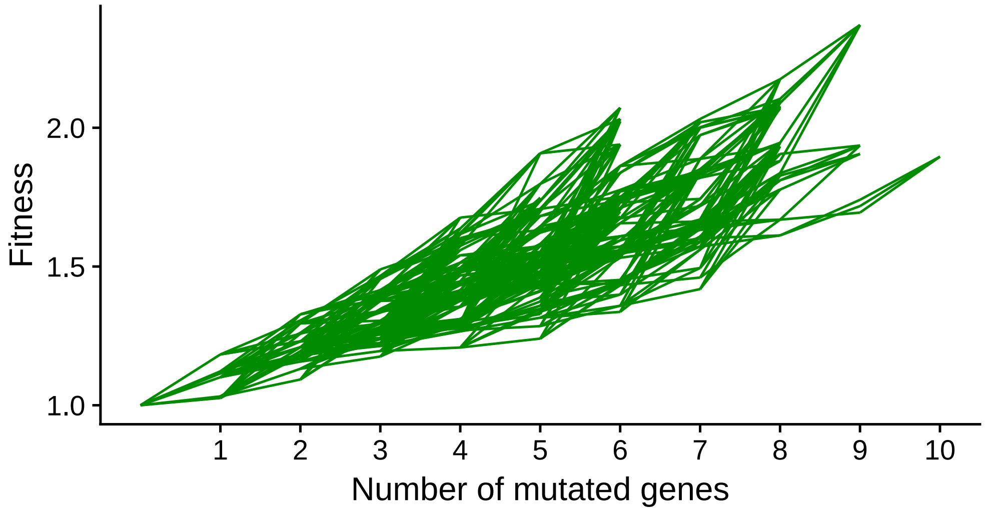


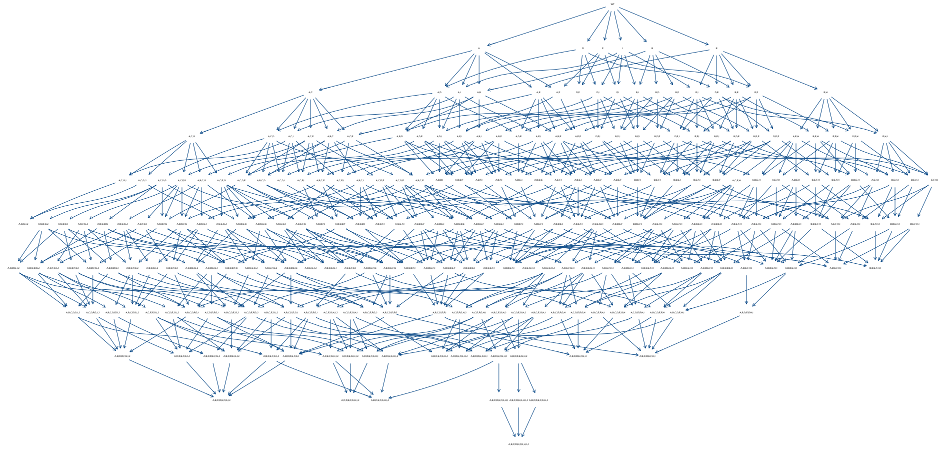

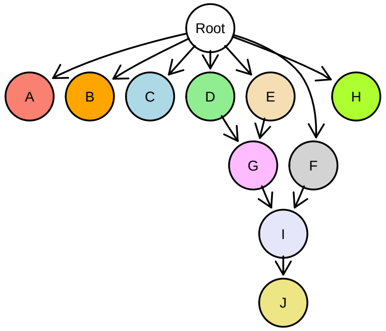


| | |
|---|---|
| *ID* | nVi510c5OJbnQRcl5 |
| *acc genotypes* | 192 |
| *peaks no back* | 17 |
| *removed edges* | 126 |
| *prop rem edges* | 0.192 |

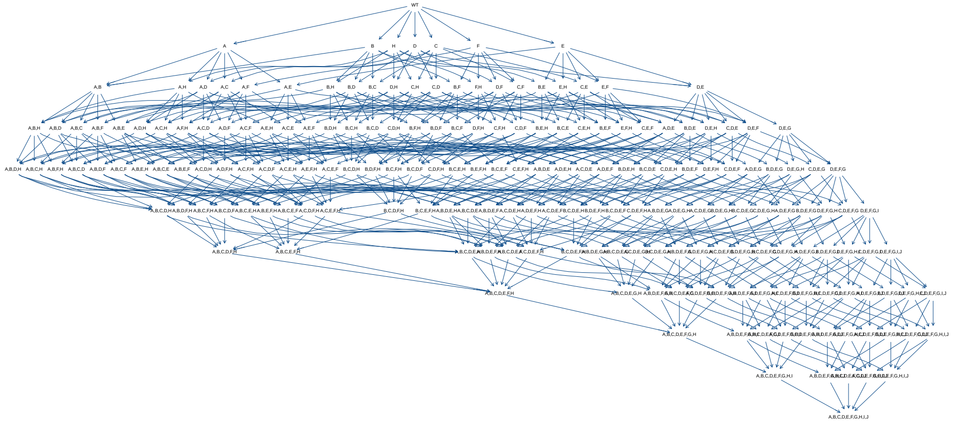


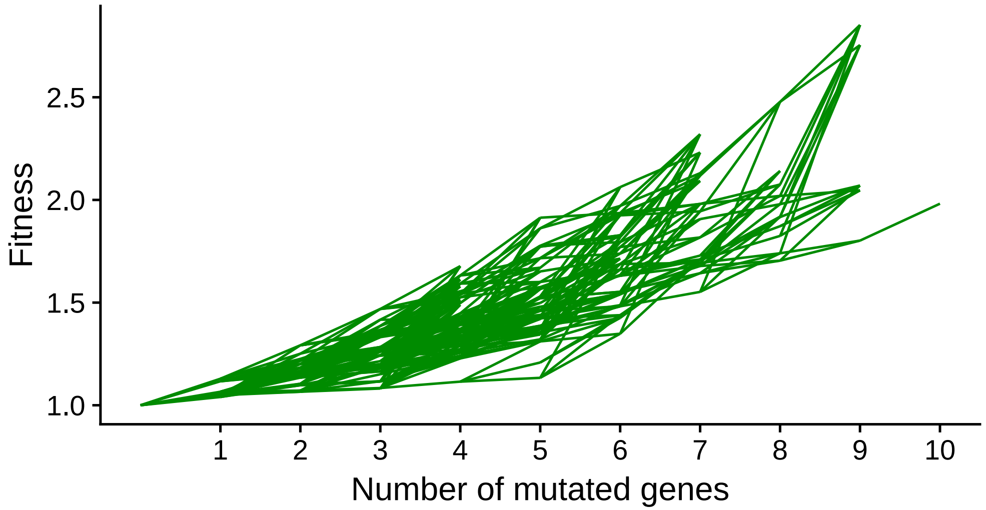


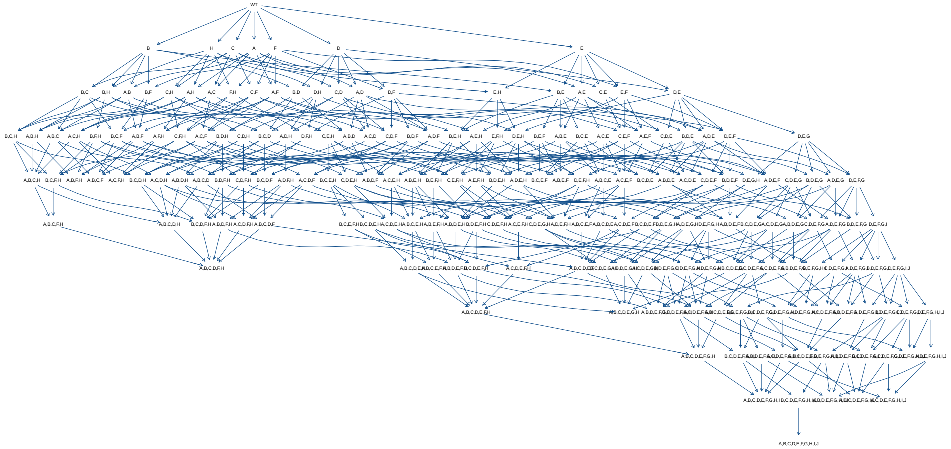

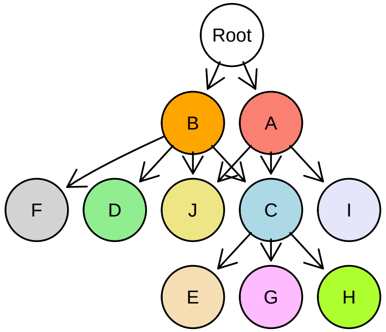


| | |
|---|---|
| *ID* | NXDMDVPfzyoQDyuP |
| *acc genotypes* | 151 |
| *peaks no back* | 11 |
| *removed edges* | 90 |
| *prop rem edges* | 0.177 |

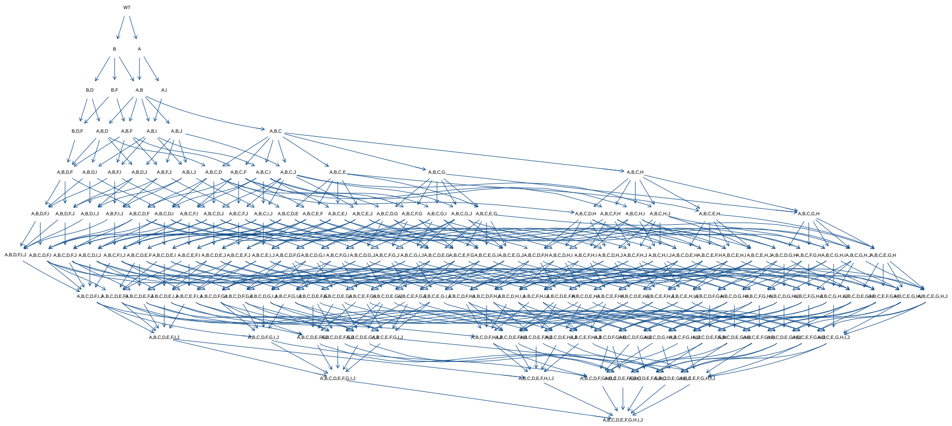

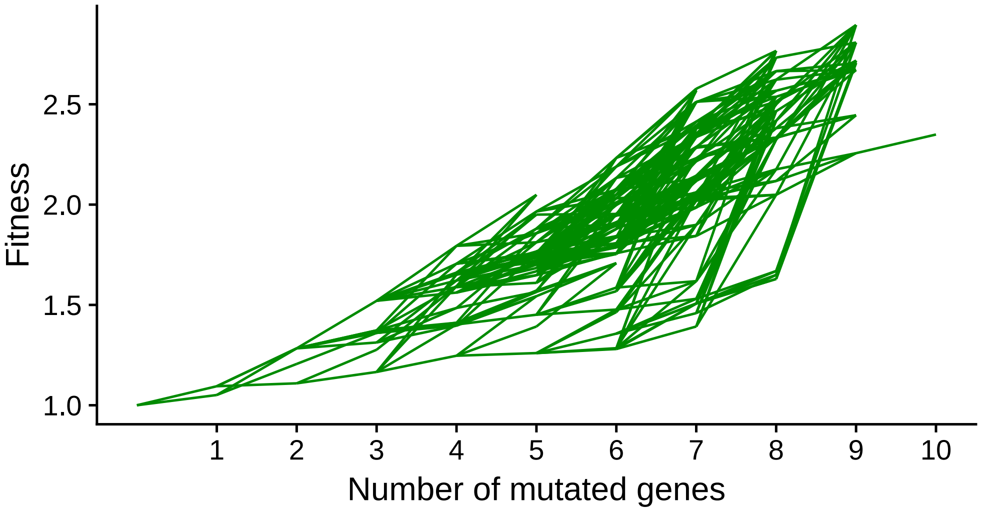


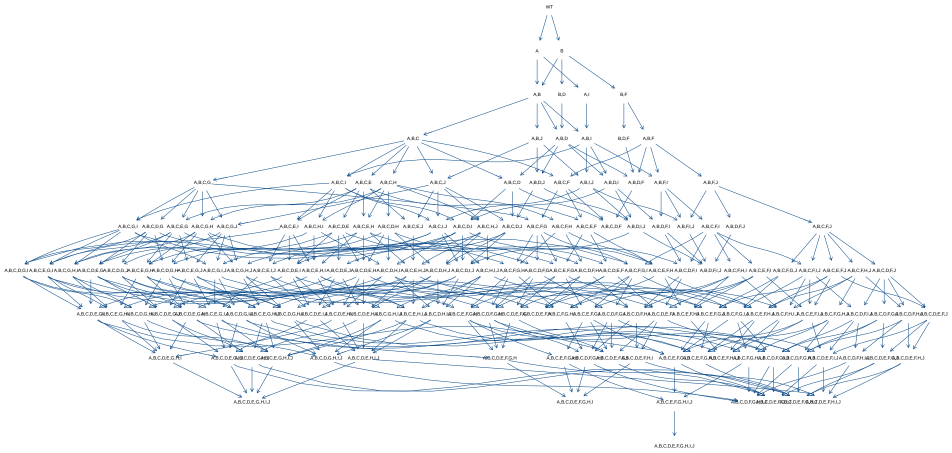

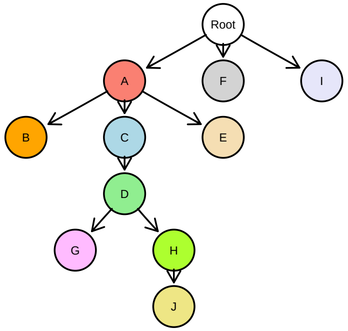


| | |
|---|---|
| *ID* | o8OvpWIeTYvSAYxY |
| *acc genotypes* | 132 |
| *peaks no back* | 13 |
| *removed edges* | 67 |
| *prop rem edges* | 0.164 |

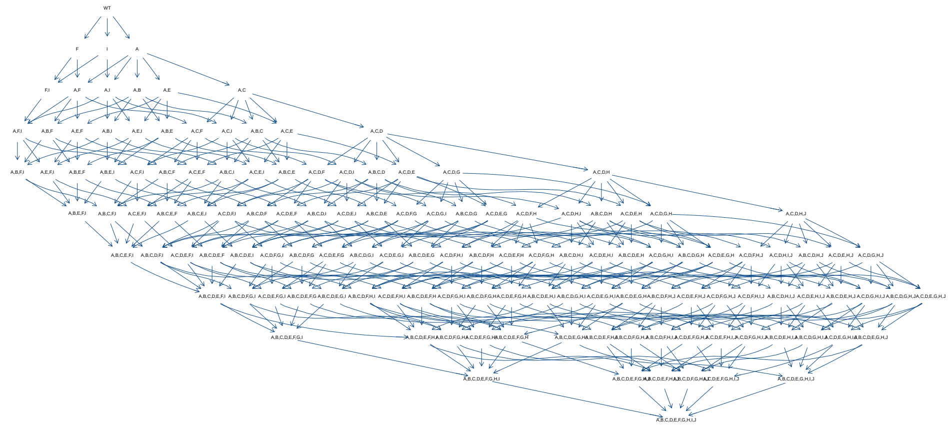

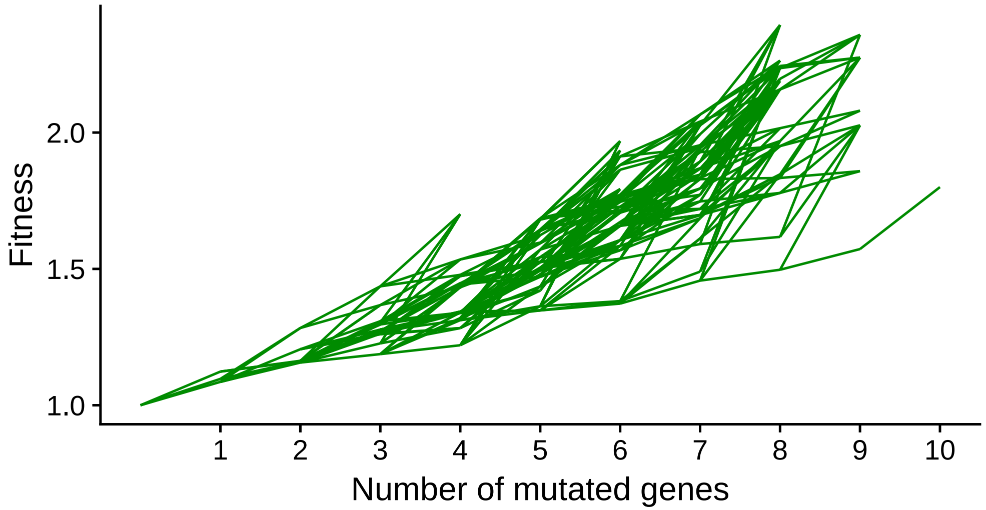


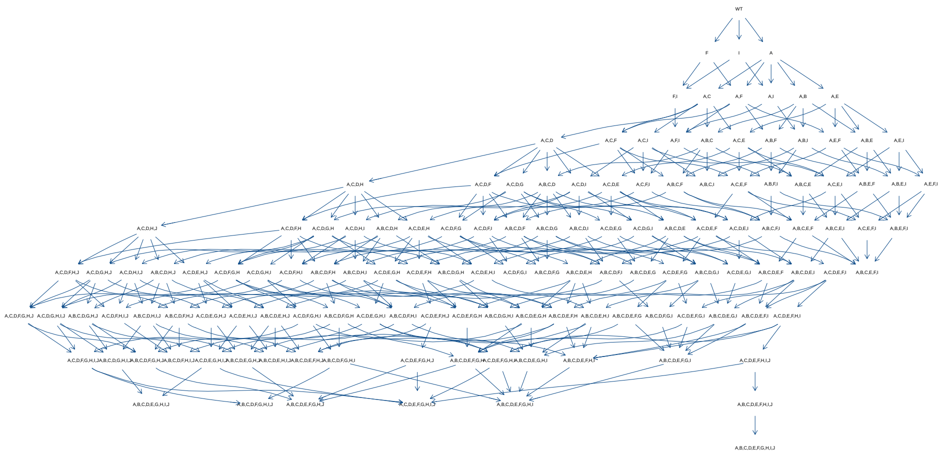

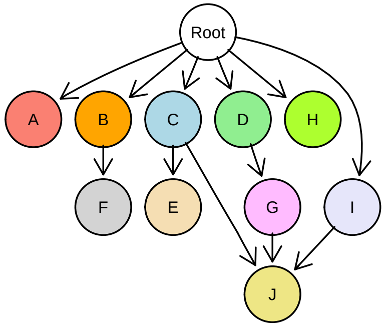


| | |
|---|---|
| *ID* | ochEhkNsBx3ftyMD |
| *acc genotypes* | 240 |
| *peaks no back* | 19 |
| *removed edges* | 173 |
| *prop rem edges* | 0.208 |

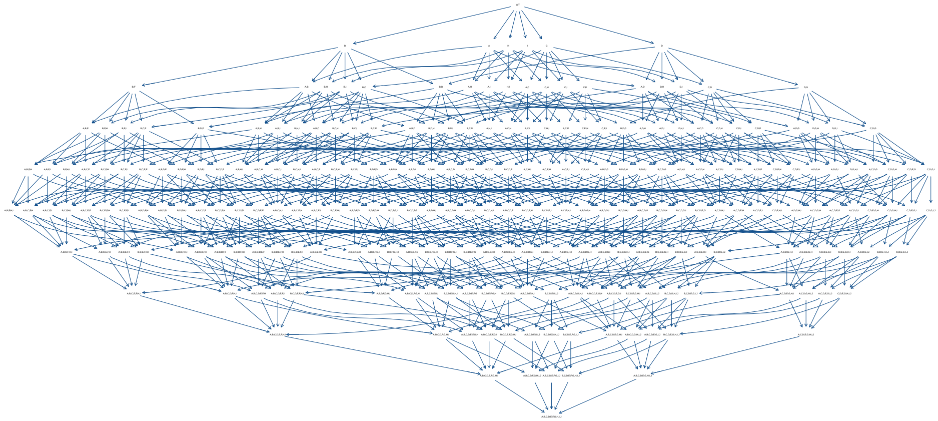

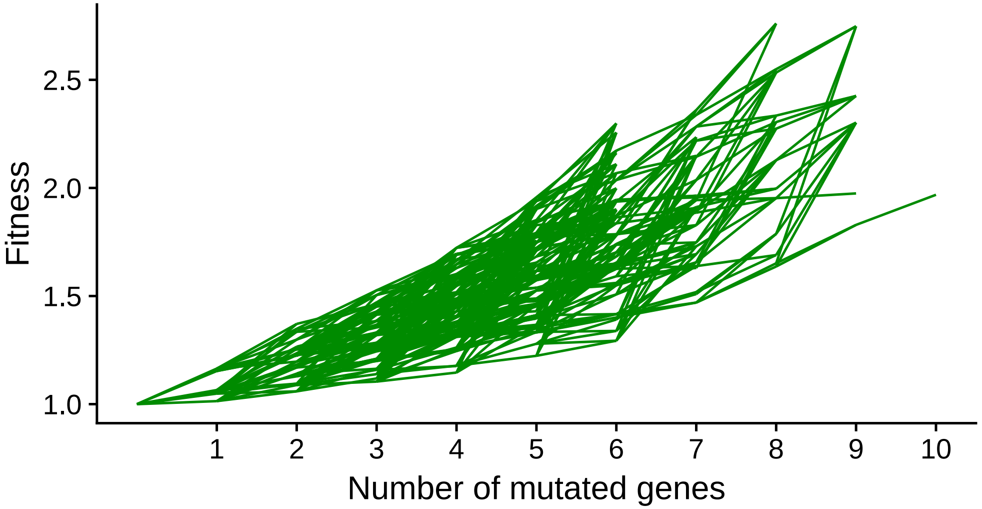


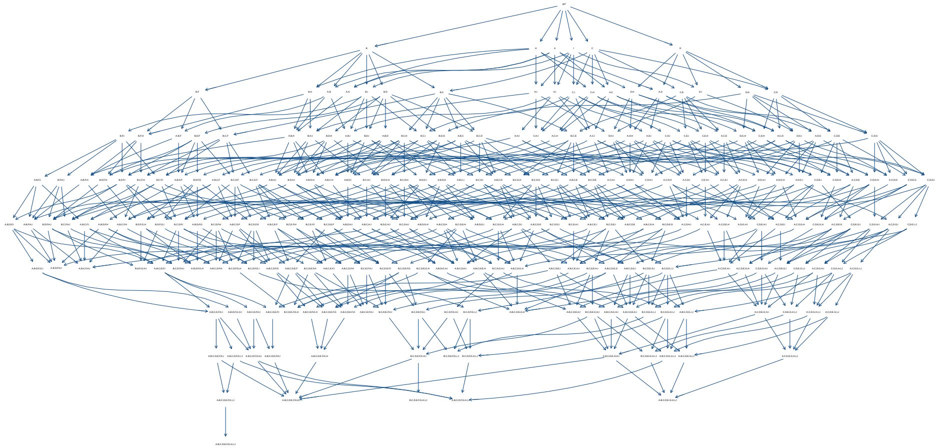

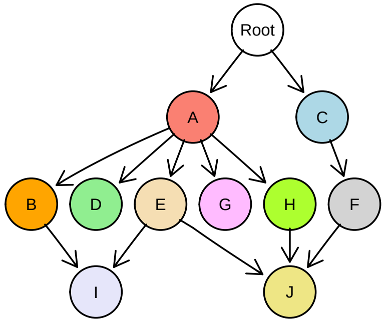


| | |
|---|---|
| *ID* | OCRTvAy6n6Rv7KAn |
| *acc genotypes* | 135 |
| *peaks no back* | 11 |
| *removed edges* | 80 |
| *prop rem edges* | 0.192 |

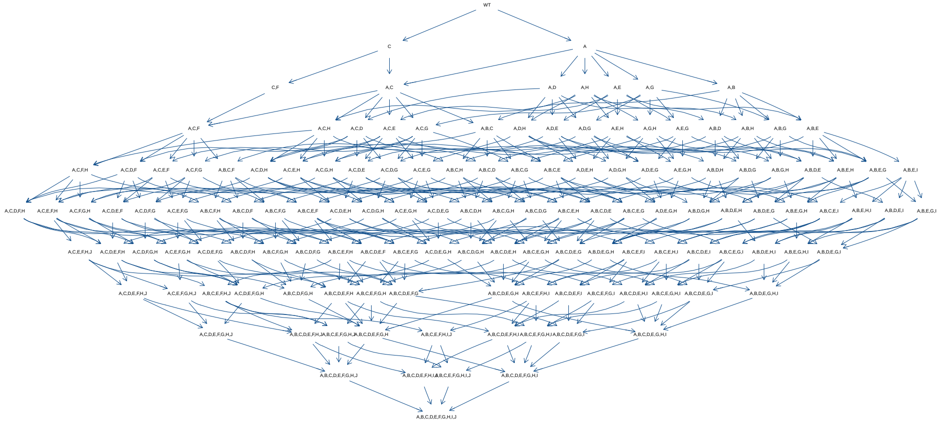

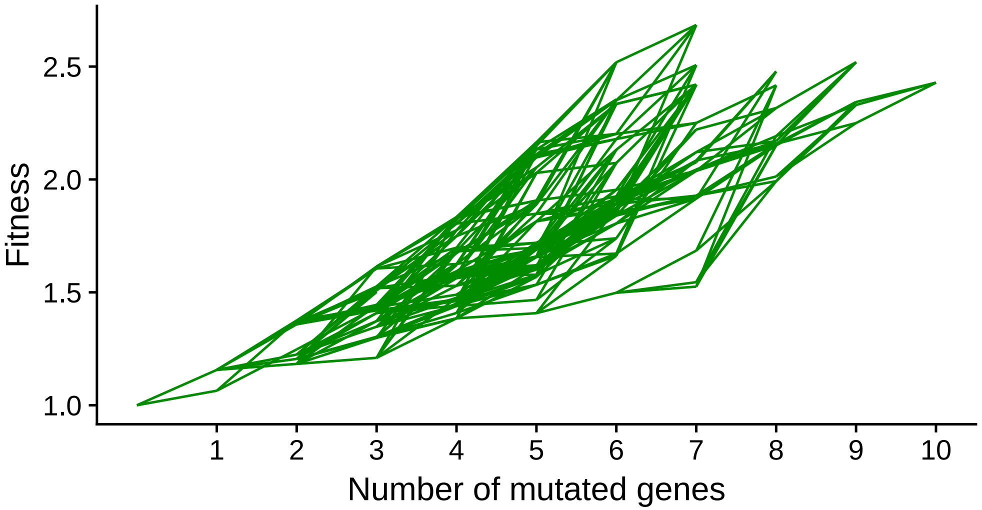


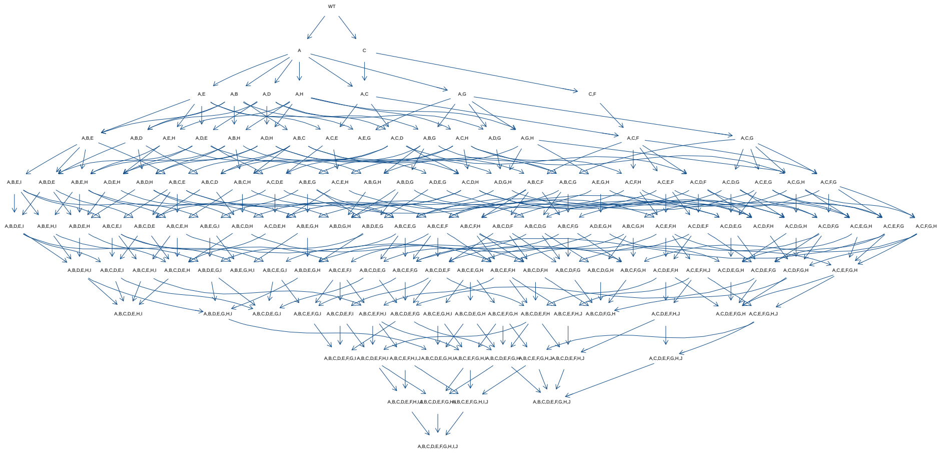

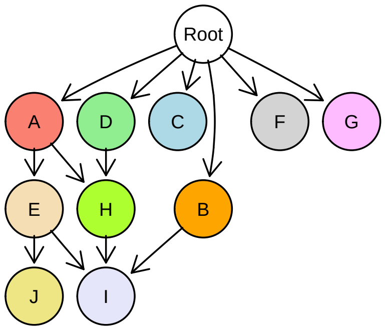


| | |
|---|---|
| *ID* | Oo5hmM8b710XcCc1x |
| *acc genotypes* | 192 |
| *peaks no back* | 15 |
| *removed edges* | 118 |
| *prop rem edges* | 0.184 |

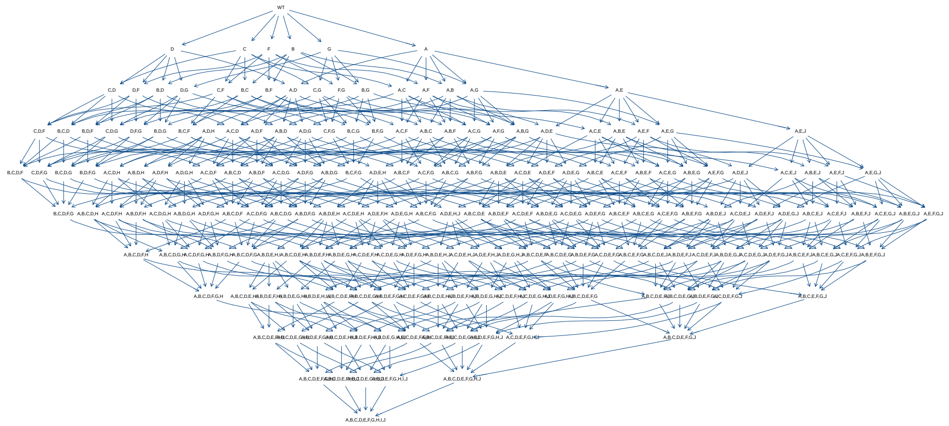

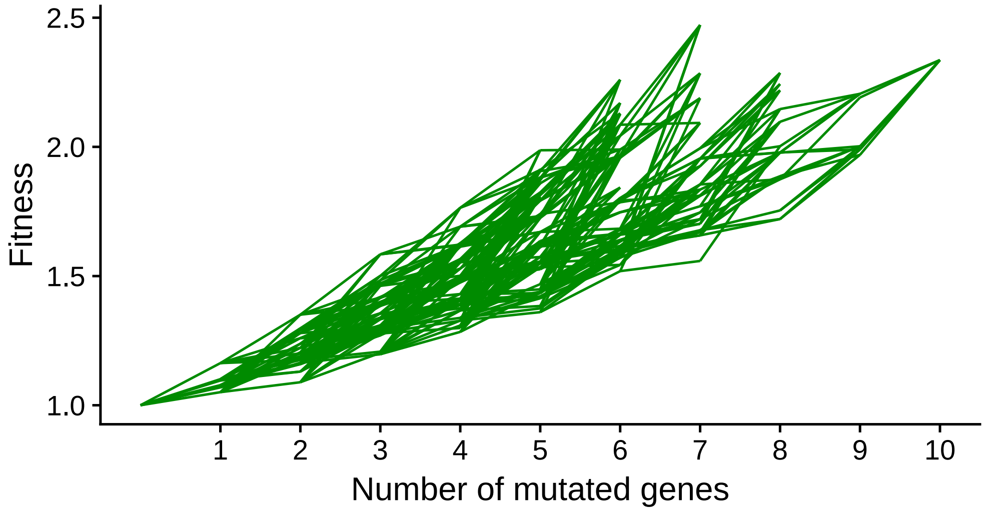


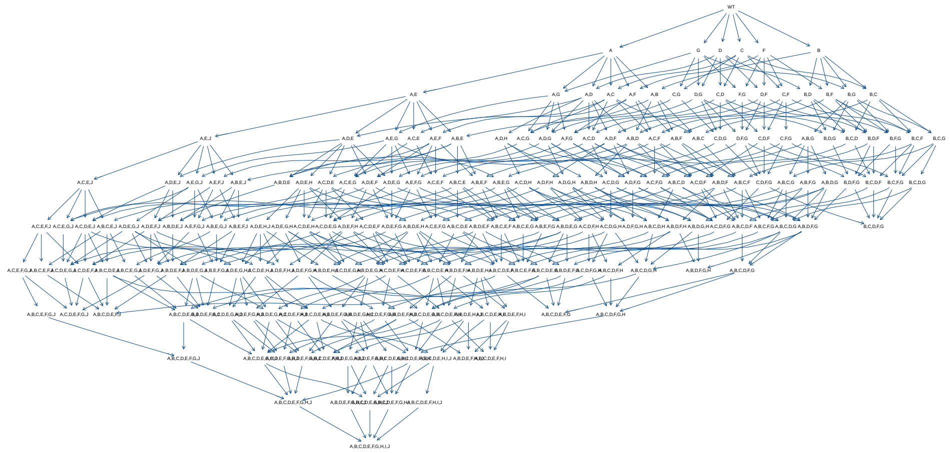

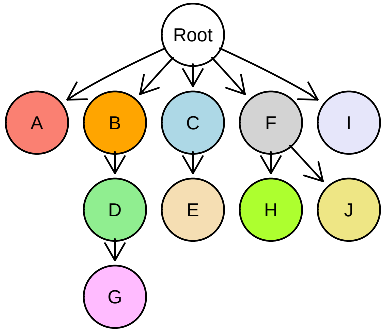


| | |
|---|---|
| *ID* | OrLY4wrCbAM10e8mH |
| *acc genotypes* | 240 |
| *peaks no back* | 16 |
| *removed edges* | 141 |
| *prop rem edges* | 0.172 |

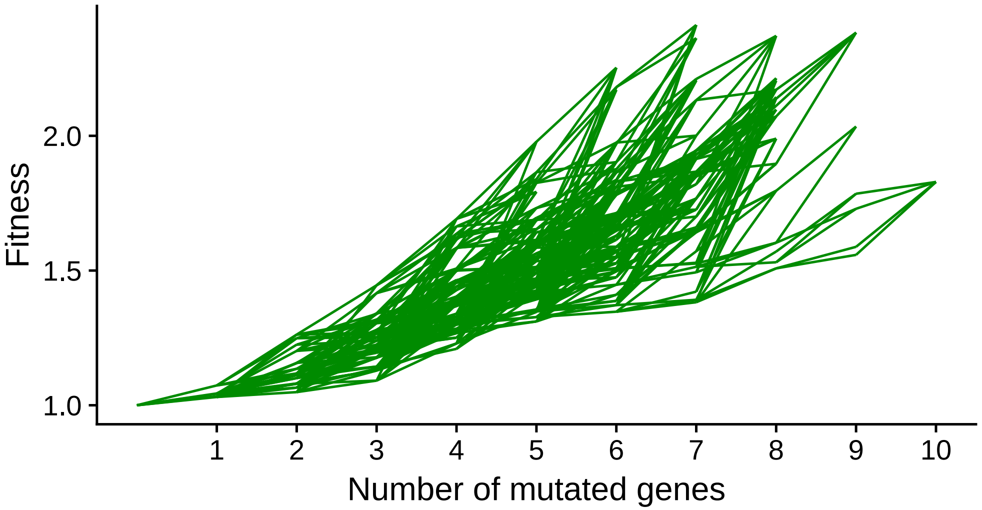


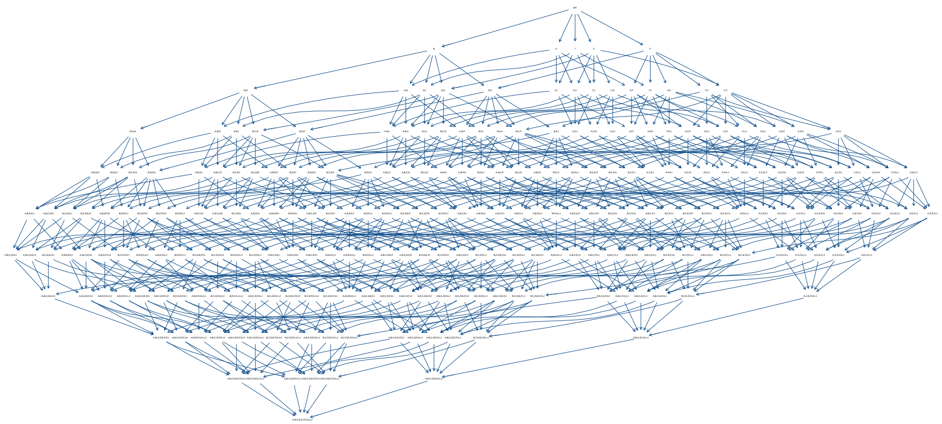

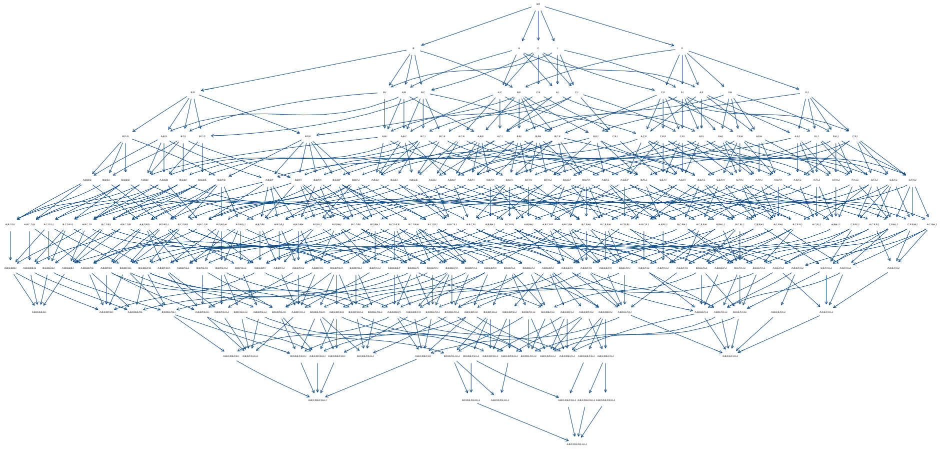

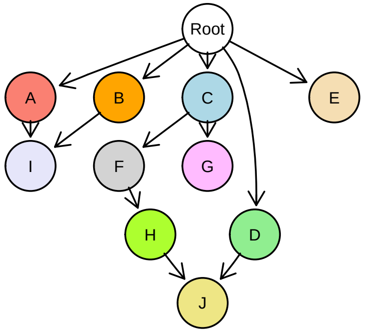


| | |
|---|---|
| *ID* | OSiI7bIkeOUdWfpZ |
| *acc genotypes* | 160 |
| *peaks no back* | 16 |
| *removed edges* | 100 |
| *prop rem edges* | 0.2 |

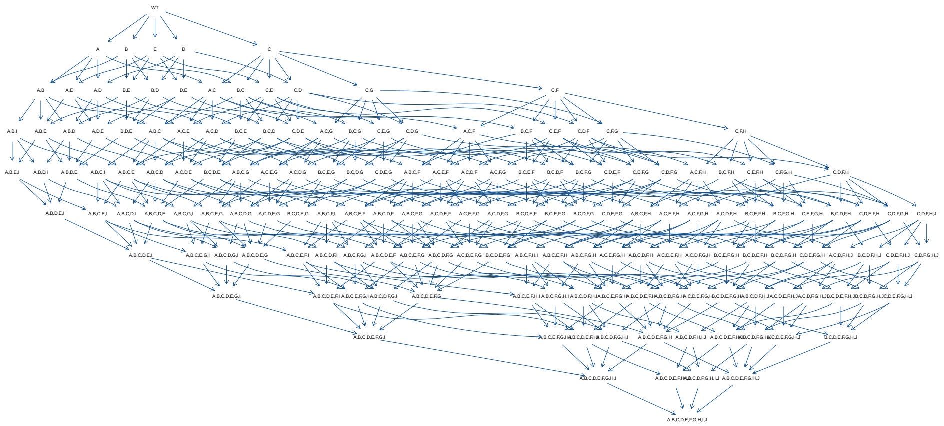

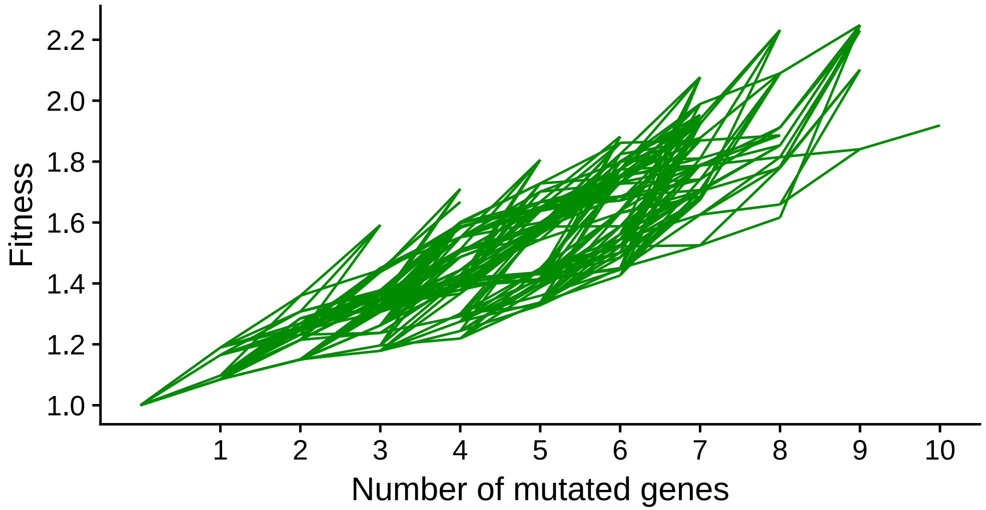


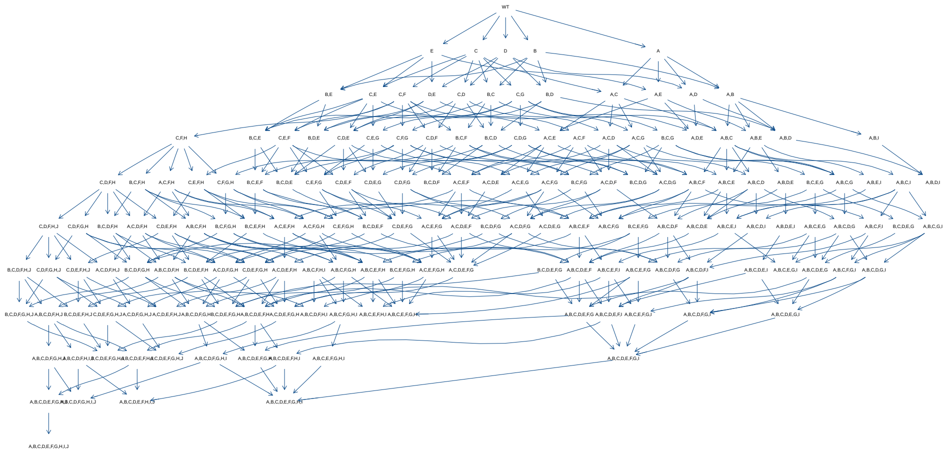

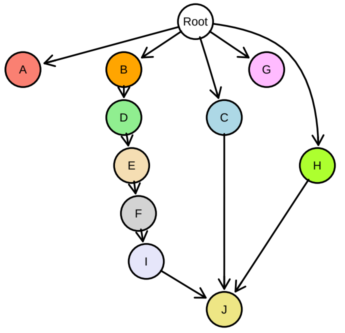


| | |
|---|---|
| *ID* | OXbm8SnMAxb9VCsc |
| *acc genotypes* | 100 |
| *peaks no back* | 10 |
| *removed edges* | 49 |
| *prop rem edges* | 0.175 |

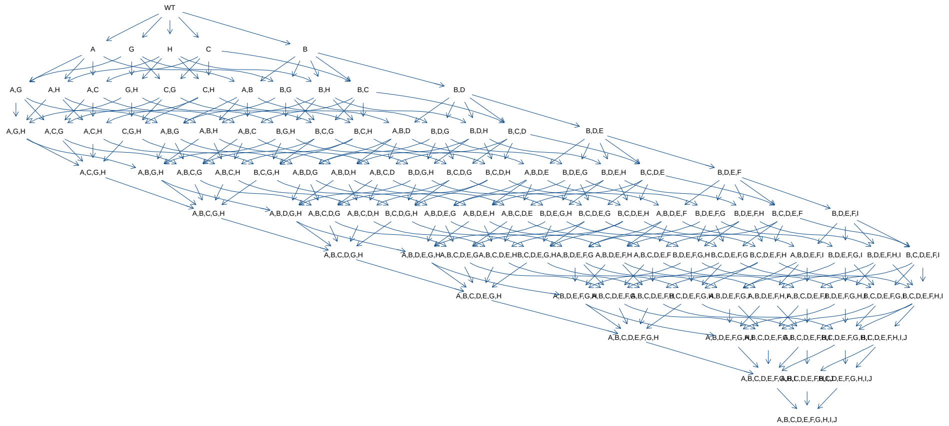

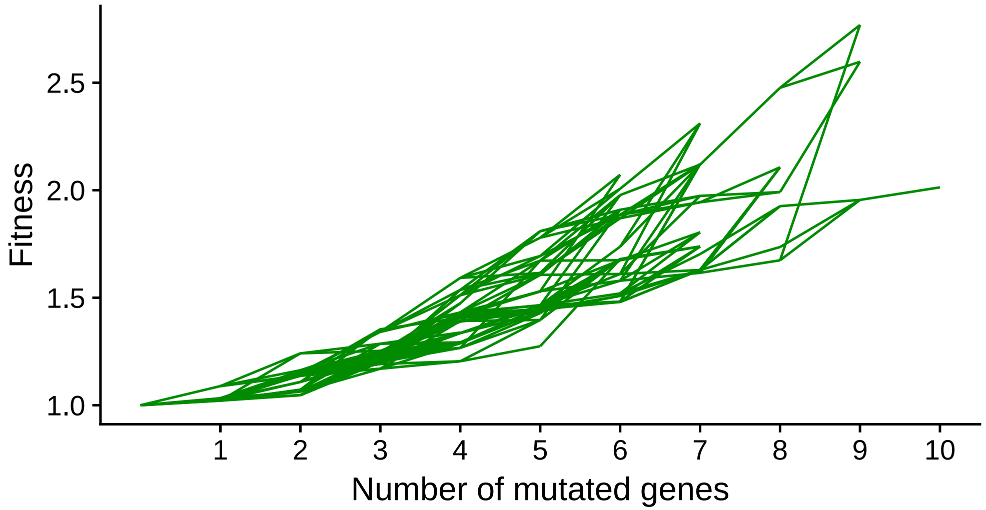


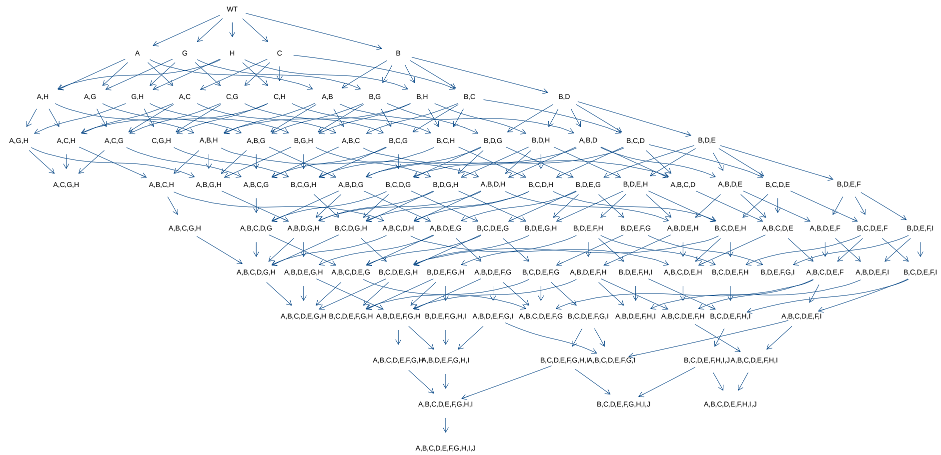

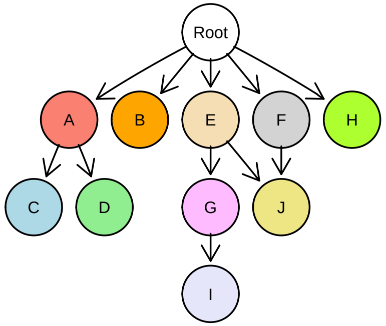


| | |
|---|---|
| *ID* | PG8DkzMLvLzGnrF10 |
| *acc genotypes* | 220 |
| *peaks no back* | 18 |
| *removed edges* | 161 |
| *prop rem edges* | 0.218 |

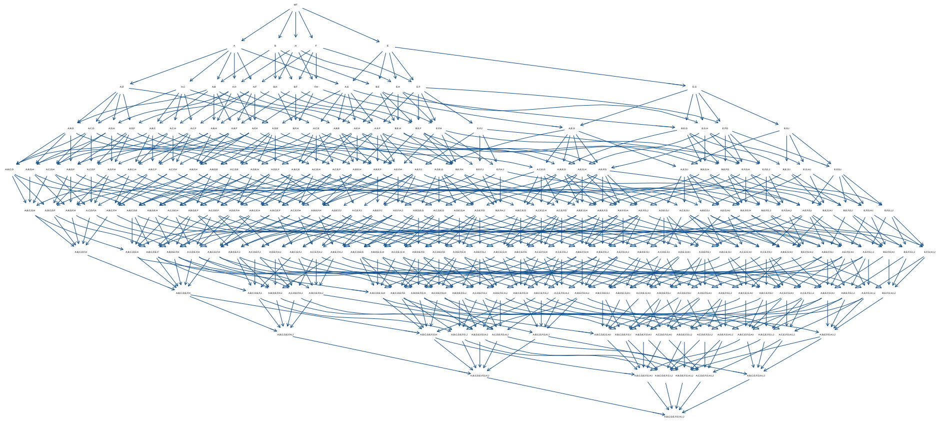

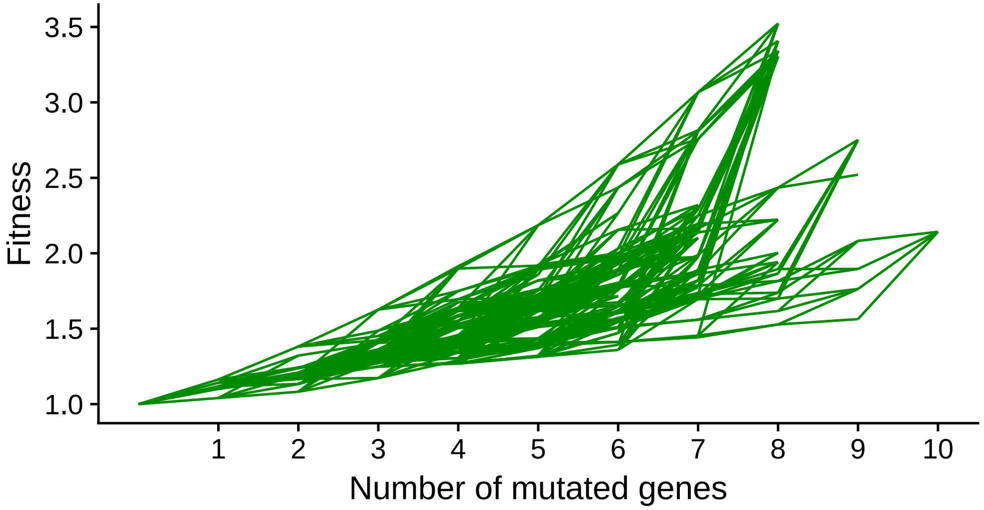


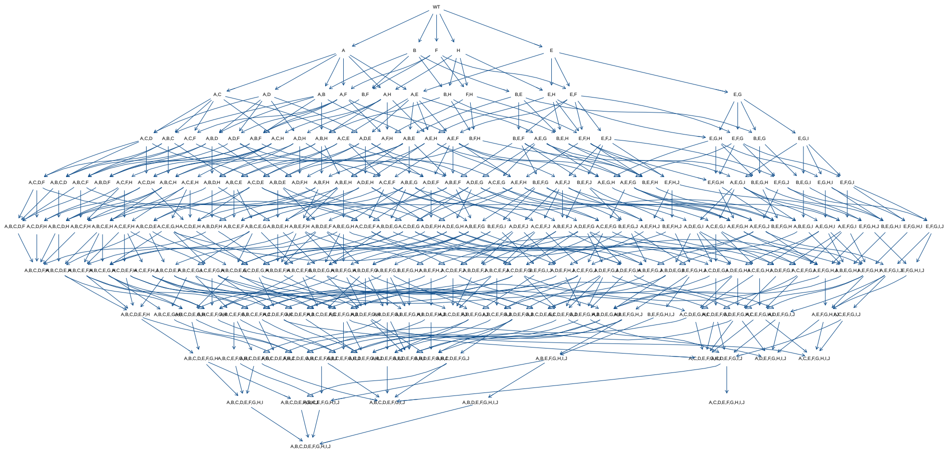

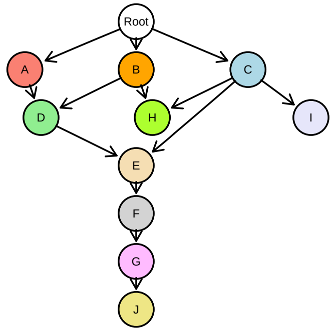


| | |
|---|---|
| *ID* | pLto5Tz84T9OOoht |
| *acc genotypes* | 37 |
| *peaks no back* | 6 |
| *removed edges* | 13 |
| *prop rem edges* | 0.186 |

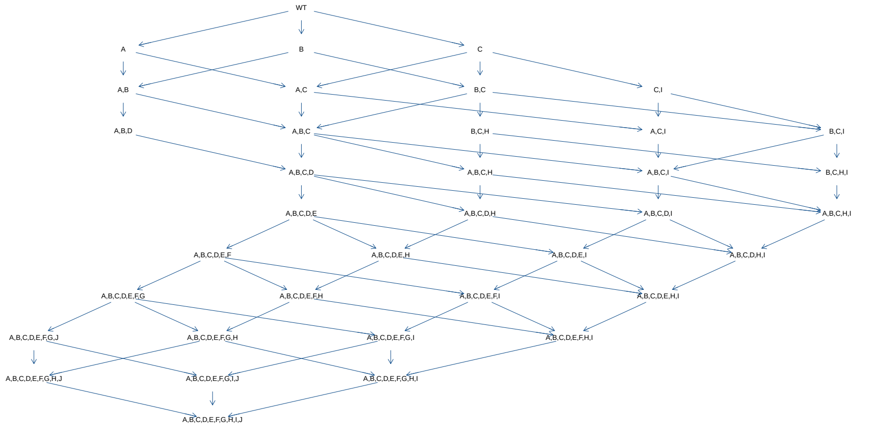

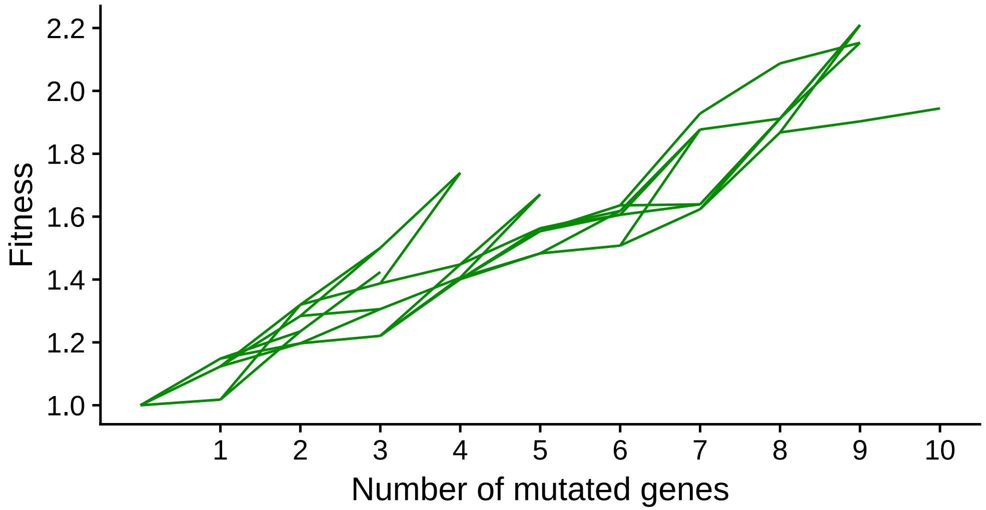


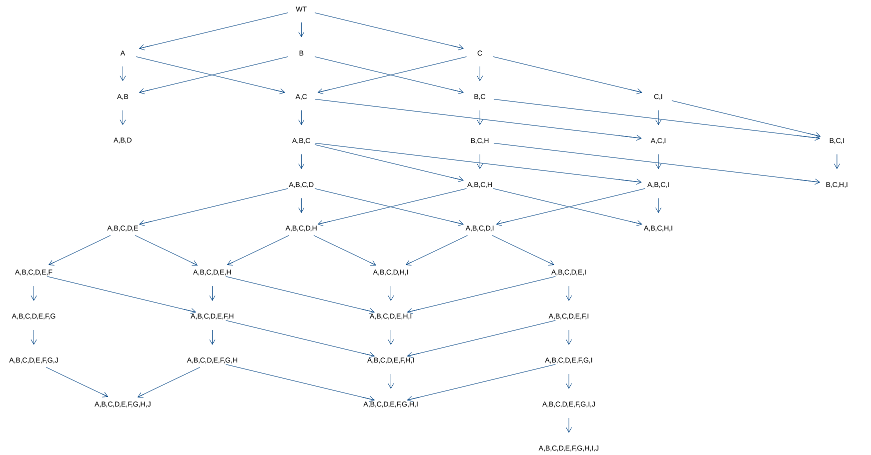

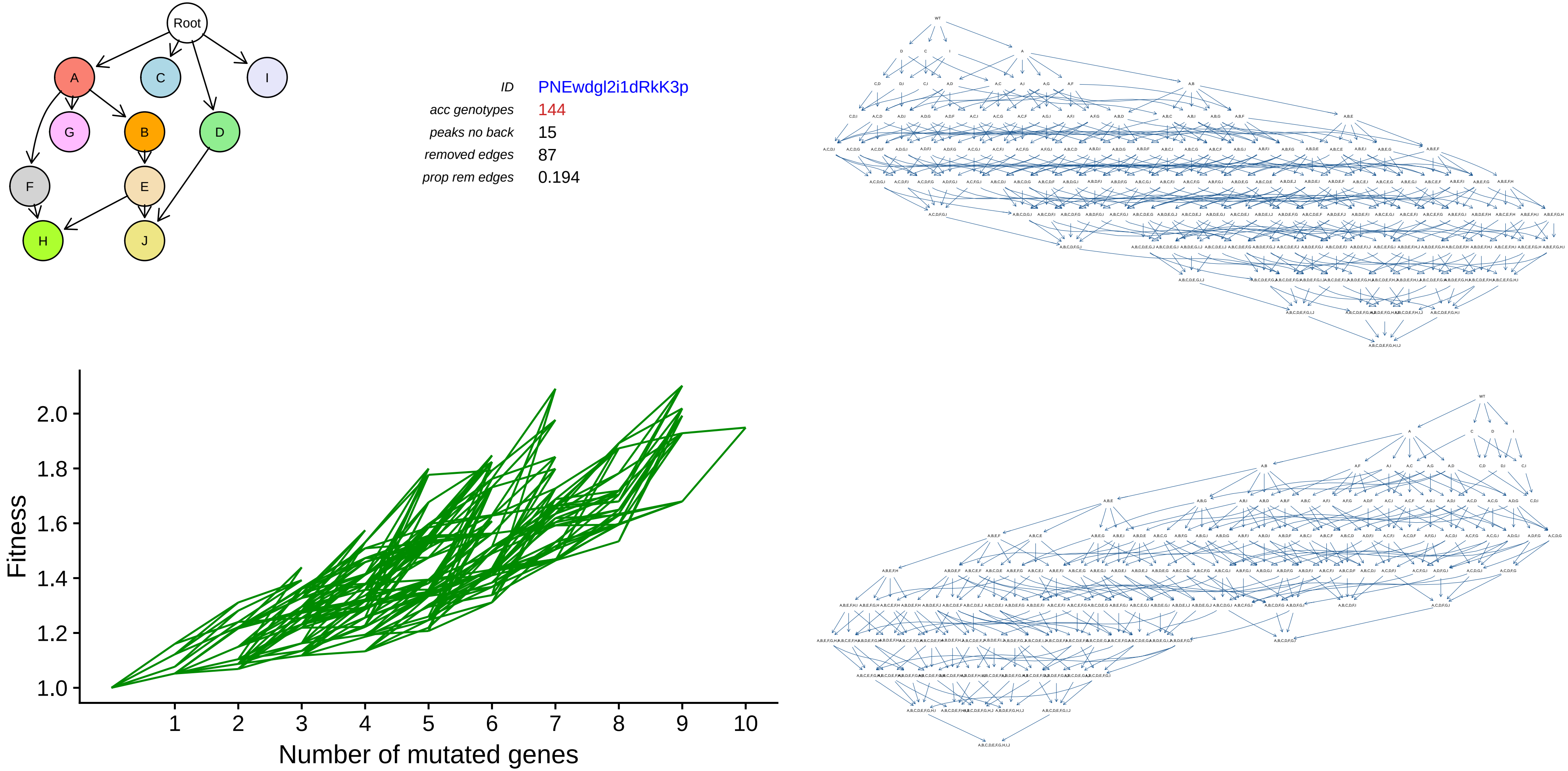

Root
A
C
I
G
B
D
F
E
H
J
ID PNEwdgl2i1dRkK3p
acc genotypes 144
peaks no back 15
removed edges 87
prop rem edges 0.194
Fitness
Number of mutated genes

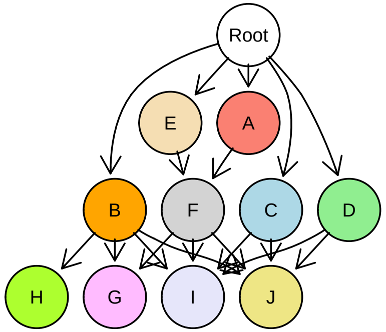


| | |
|---|---|
| *ID* | PpAWVK9tbBJENITR |
| *acc genotypes* | 80 |
| *peaks no back* | 12 |
| *removed edges* | 43 |
| *prop rem edges* | 0.207 |

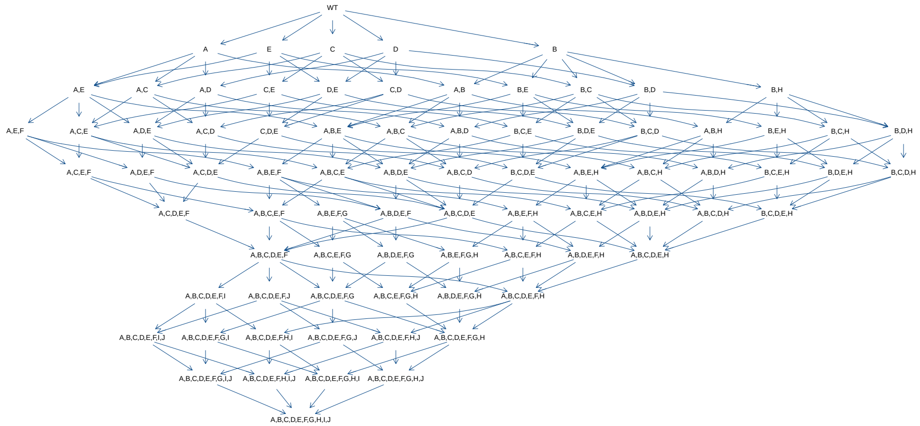

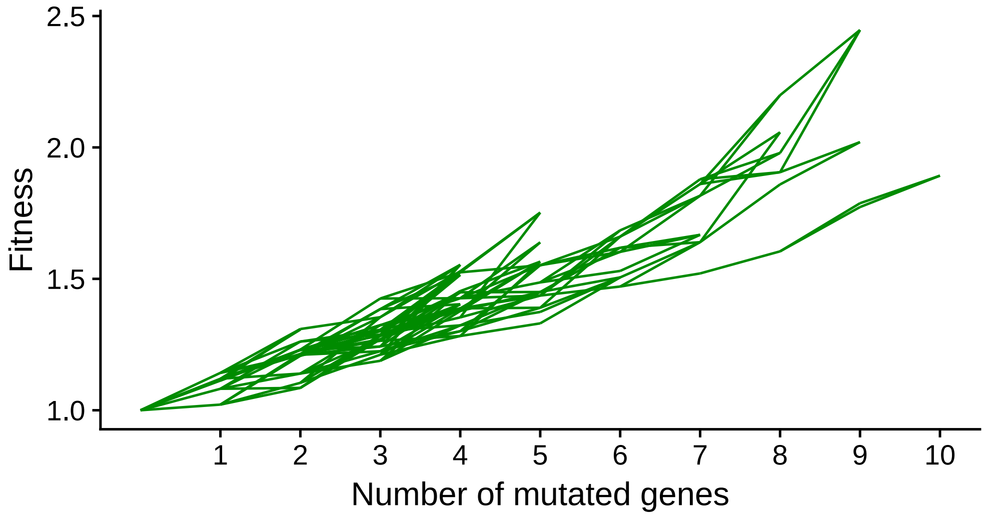


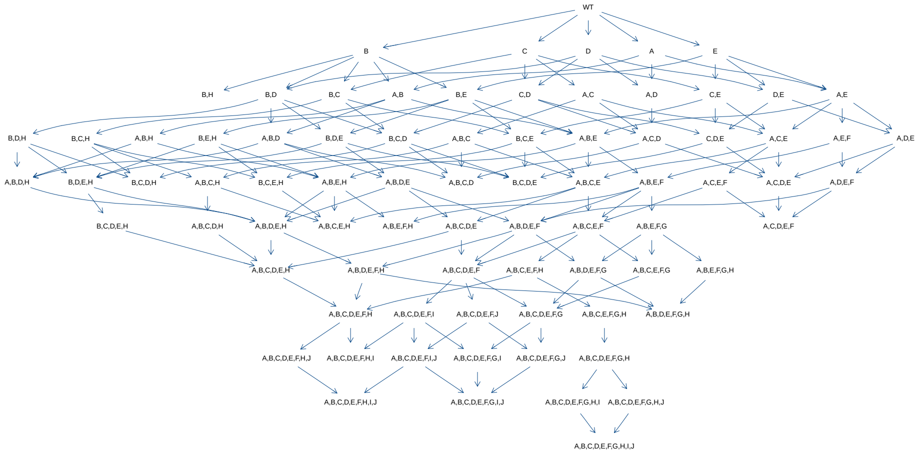

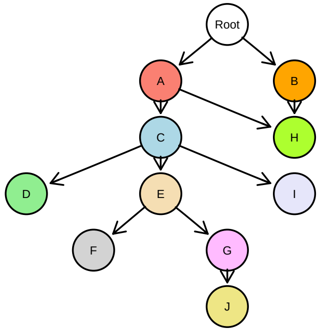


| | |
|---|---|
| *ID* | Pqr5tjYF1SZIZUV10 |
| *acc genotypes* | 89 |
| *peaks no back* | 9 |
| *removed edges* | 46 |
| *prop rem edges* | 0.189 |

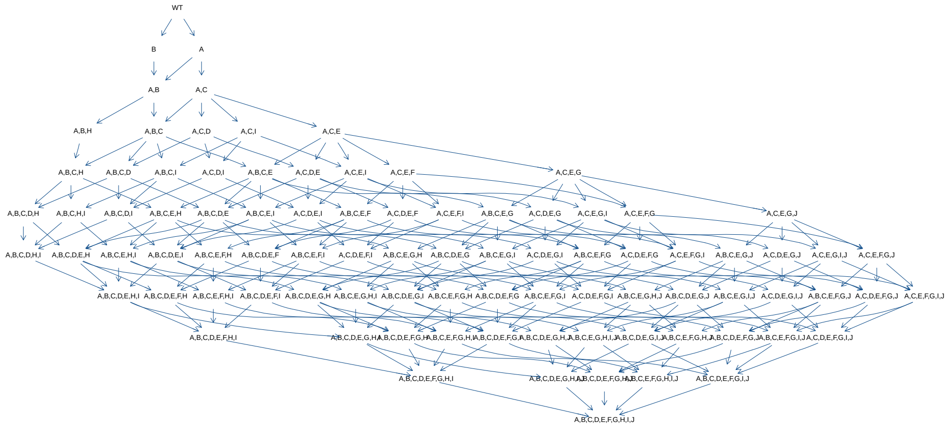

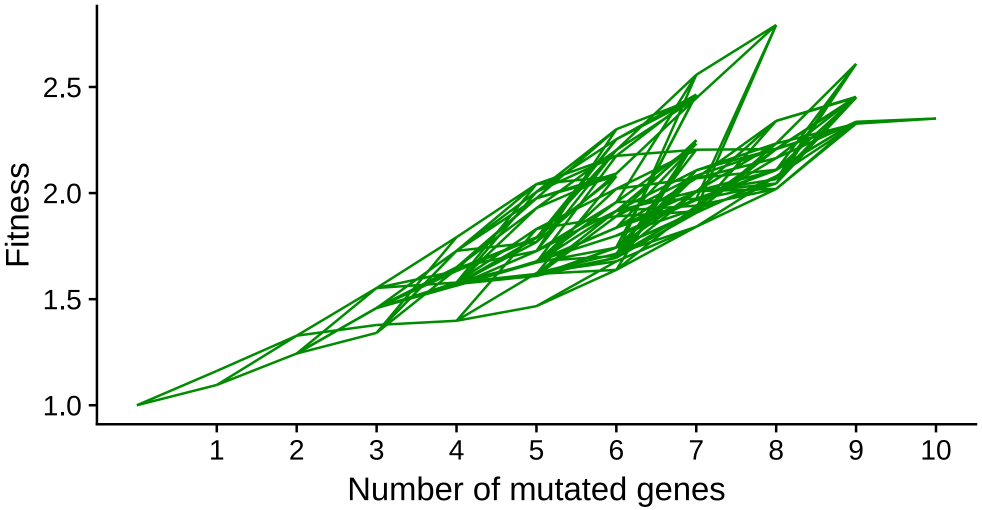


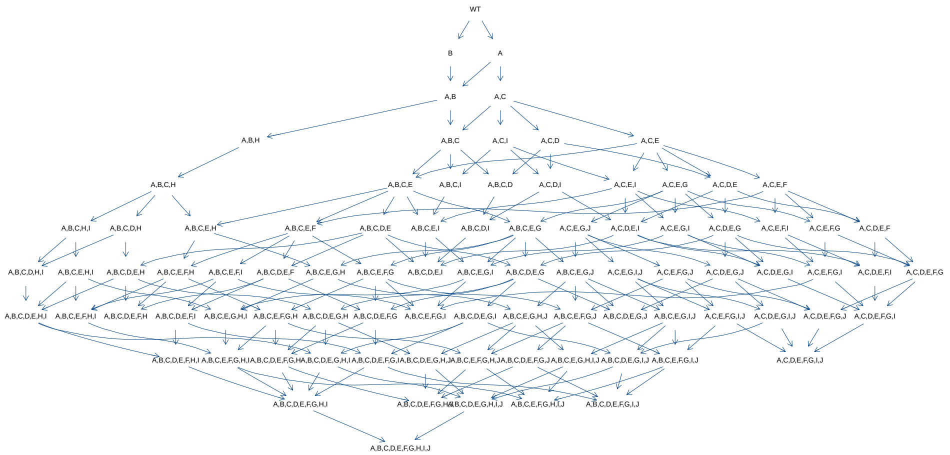

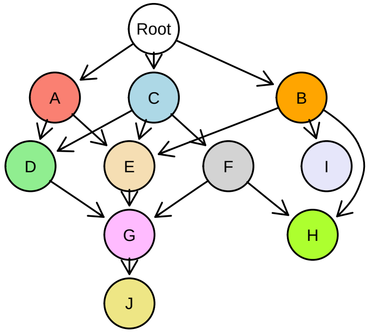


| | |
|---|---|
| *ID* | q4xh5ljukplhq9H9 |
| *acc genotypes* | 50 |
| *peaks no back* | 7 |
| *removed edges* | 20 |
| *prop rem edges* | 0.187 |

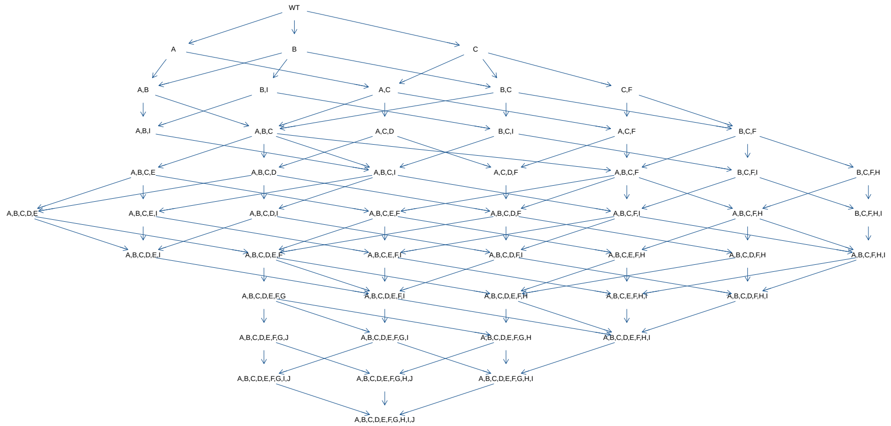

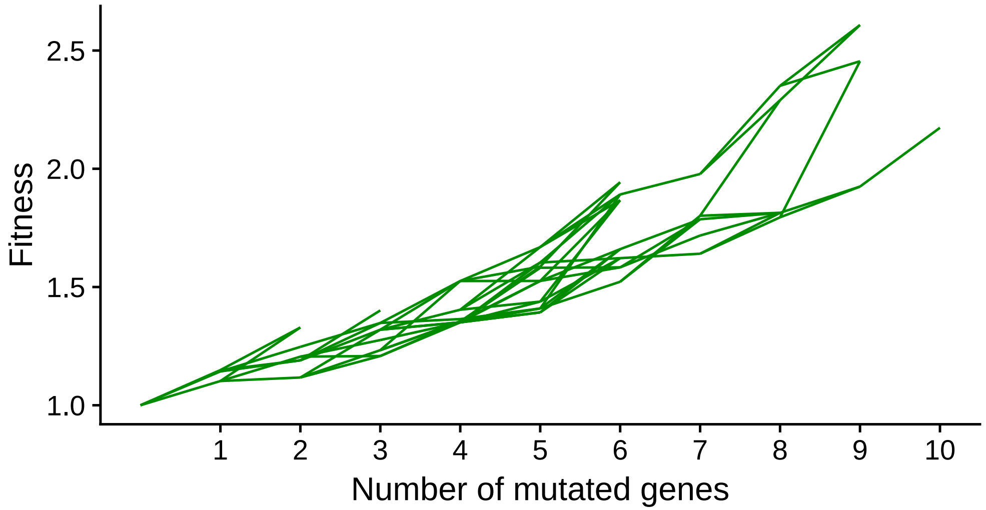


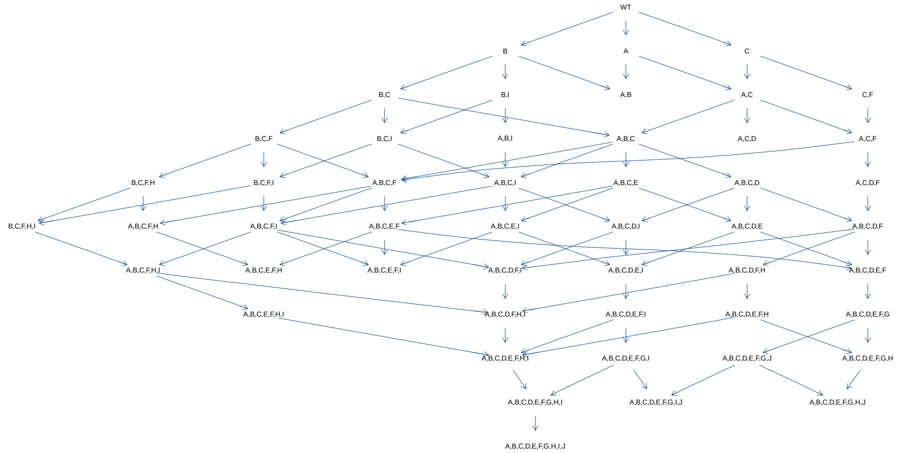

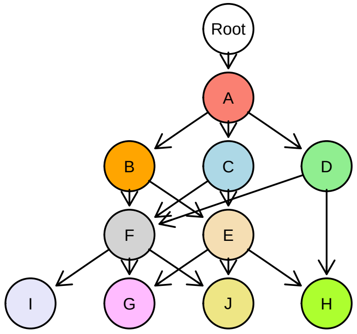


| | |
|---|---|
| *ID* | q7yEvxoMttQHd9KY |
| *acc genotypes* | 30 |
| *peaks no back* | 7 |
| *removed edges* | 10 |
| *prop rem edges* | 0.182 |

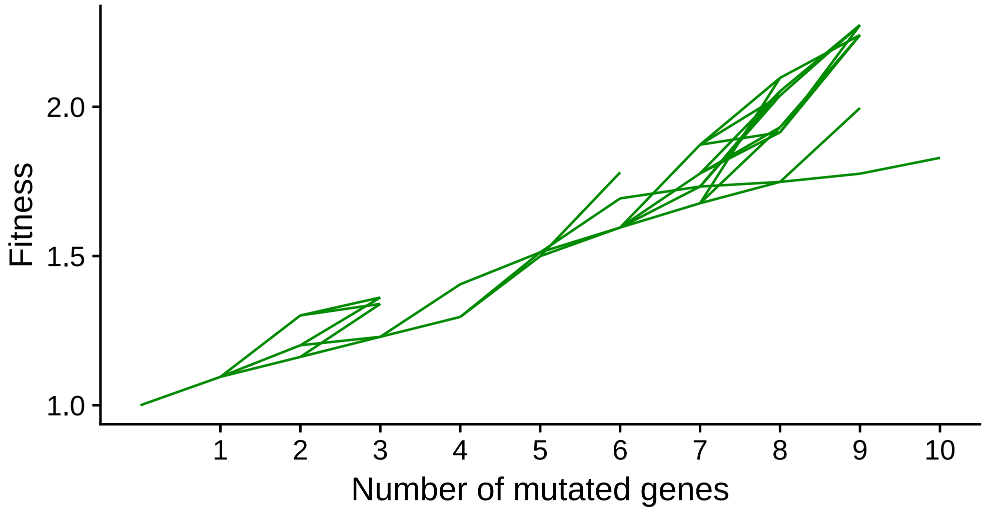


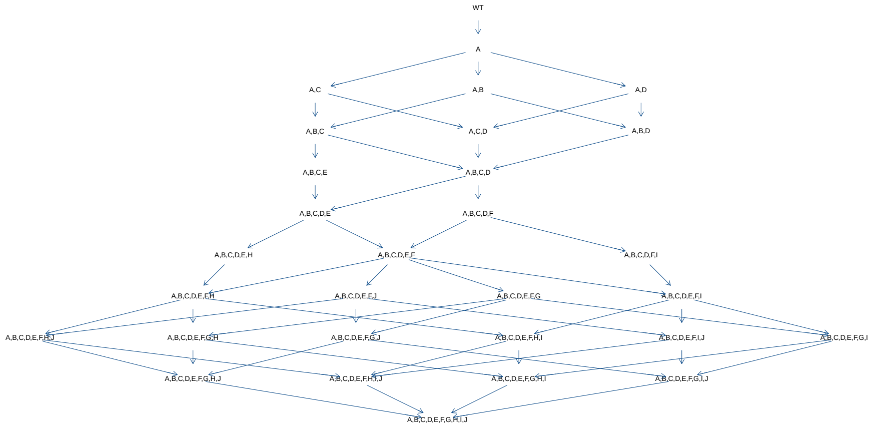

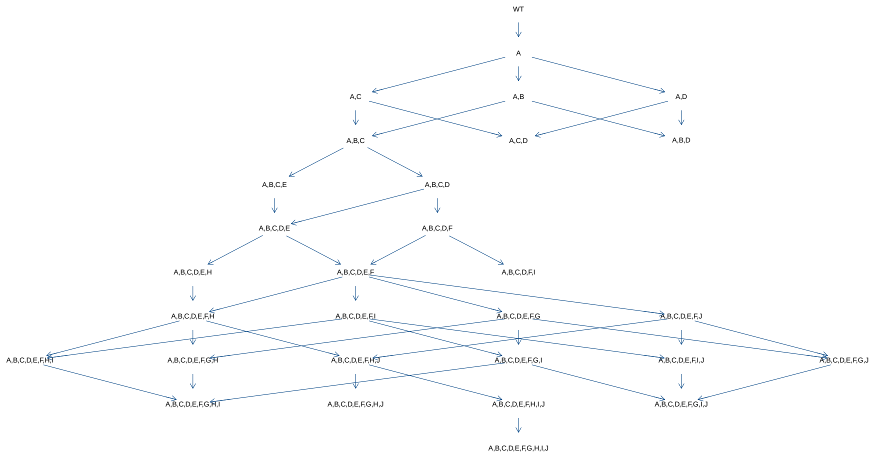

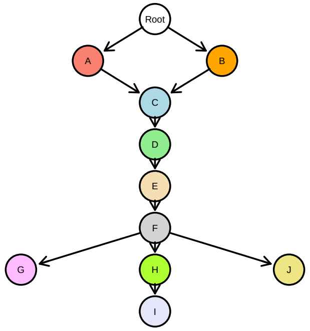


*ID* qA5UjjBYFJWi5Ddc
*acc genotypes* 19
*peaks no back* 3
*removed edges* 2
*prop rem edges* 0.071

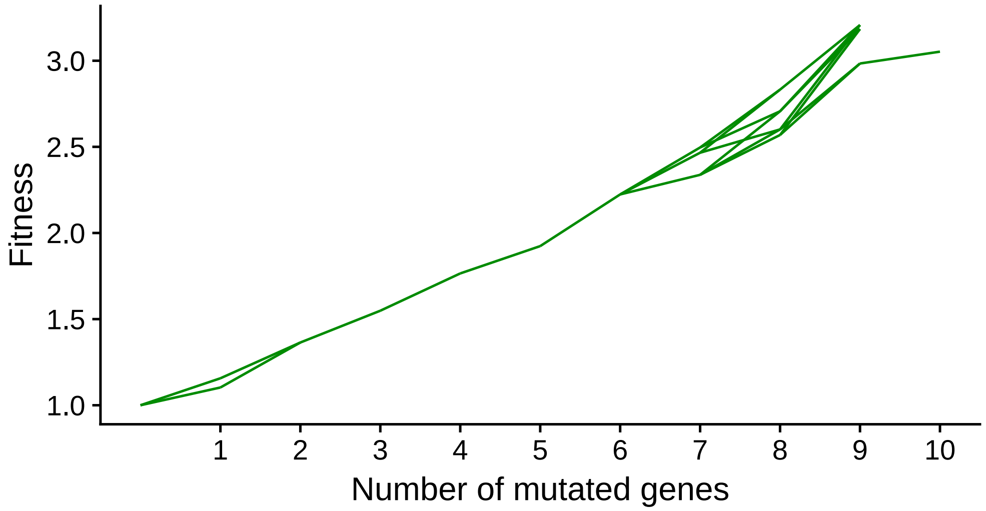


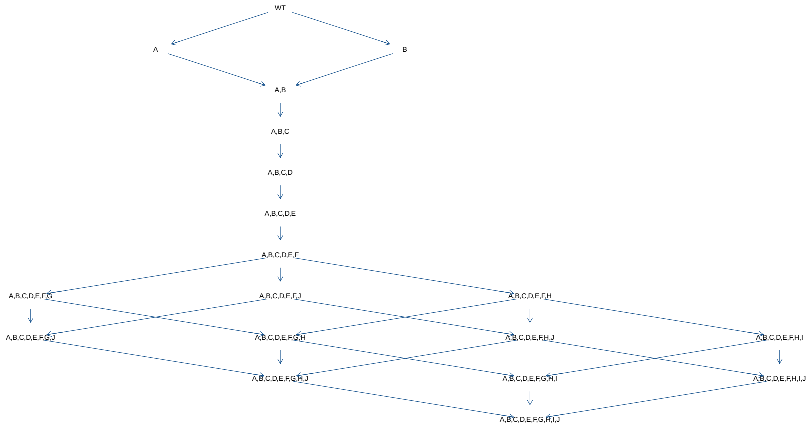


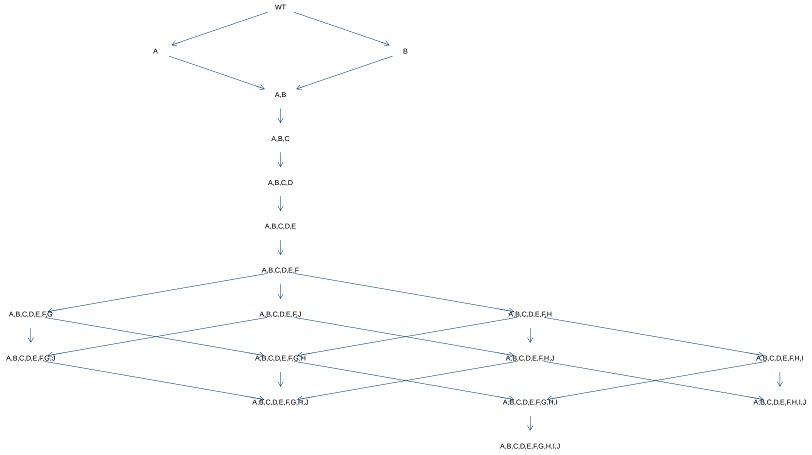

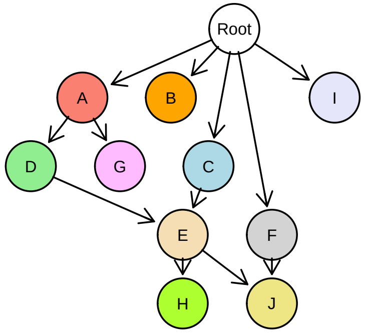


| | |
|---|---|
| *ID* | QpSOnT7M3hBKpMNs |
| *acc genotypes* | 128 |
| *peaks no back* | 13 |
| *removed edges* | 84 |
| *prop rem edges* | 0.219 |

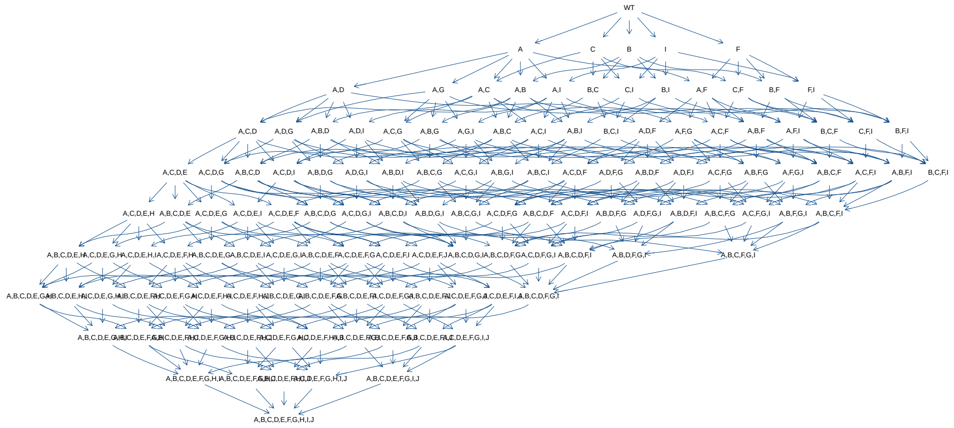

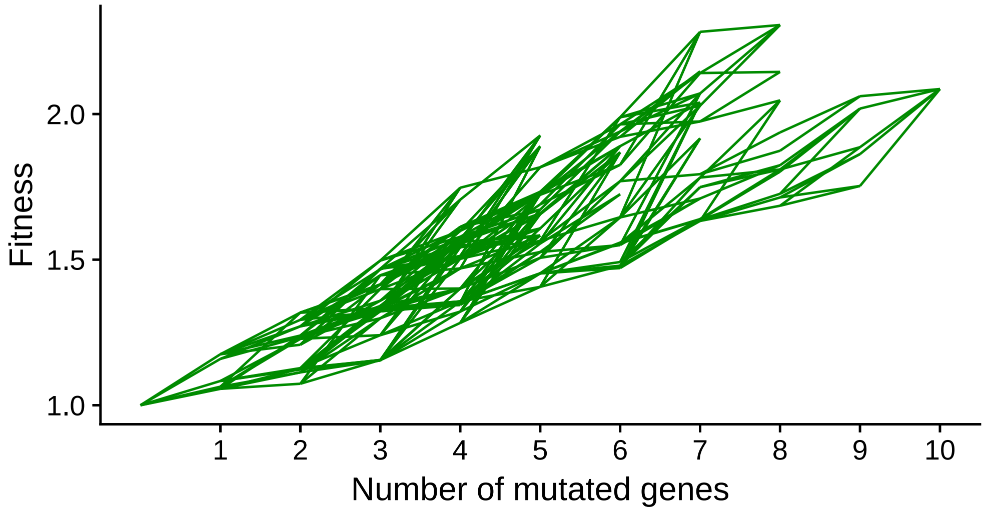


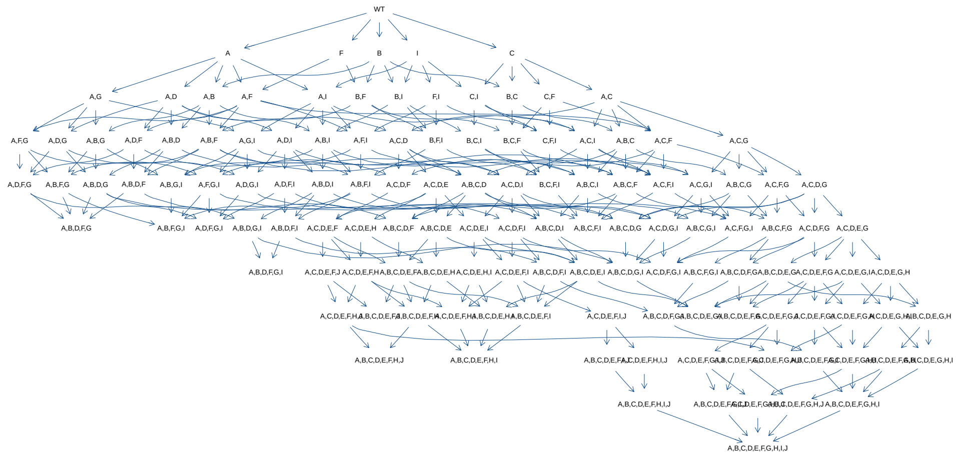

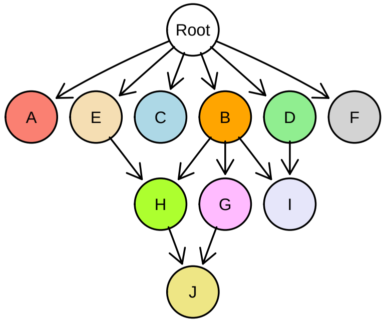


| | |
|---|---|
| *ID* | QTeBb3tzhv6dF9Iy |
| *acc genotypes* | 200 |
| *peaks no back* | 16 |
| *removed edges* | 103 |
| *prop rem edges* | 0.154 |

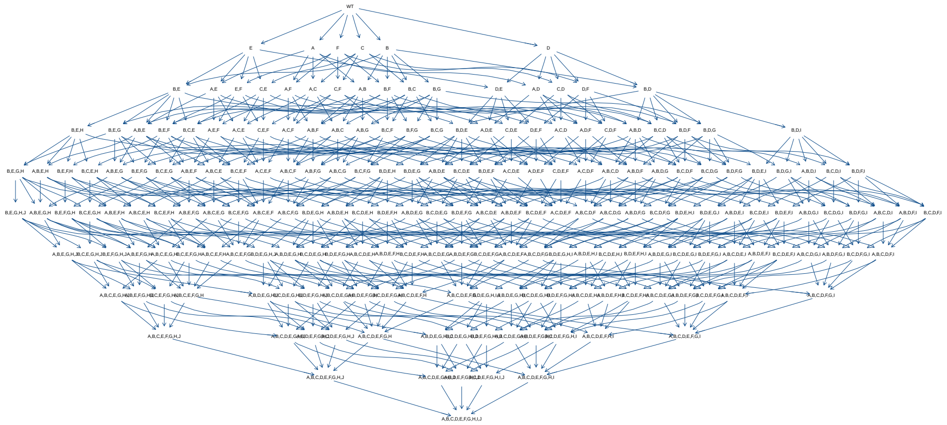

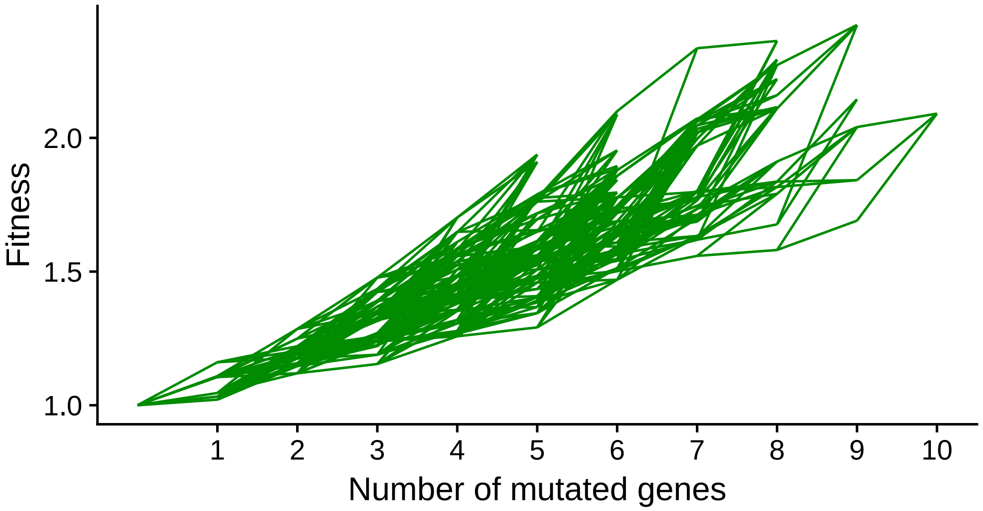


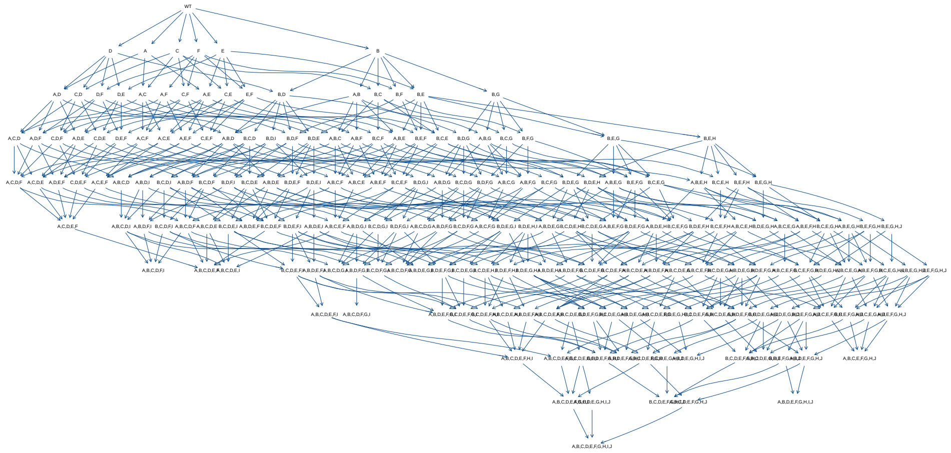

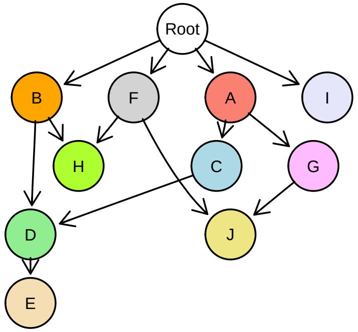


*ID* QVEfHMuWmdeBZFrT
*acc genotypes* 94
*peaks no back* 11
*removed edges* 42
*prop rem edges* 0.171

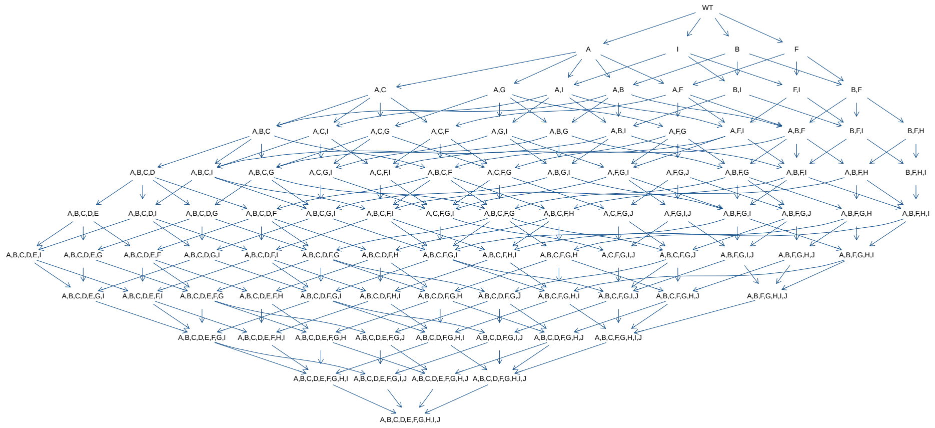

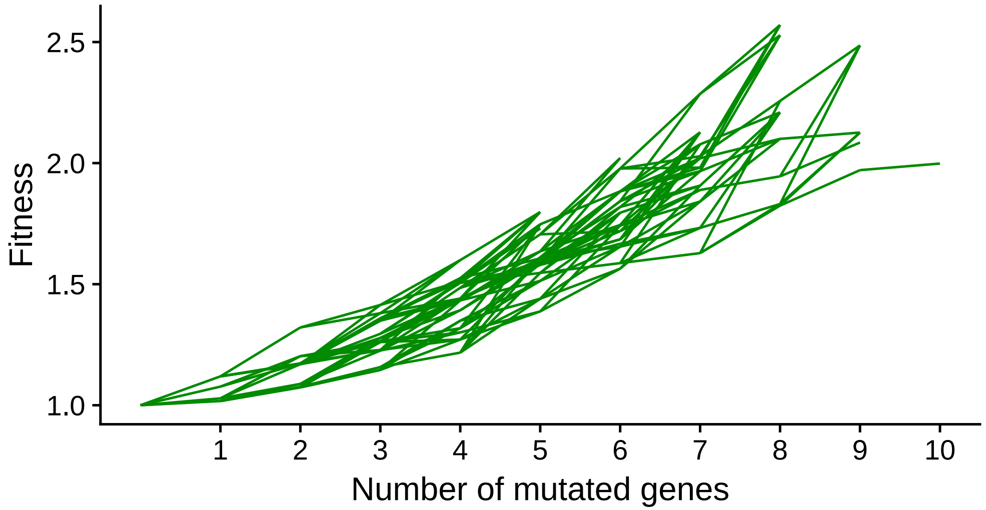


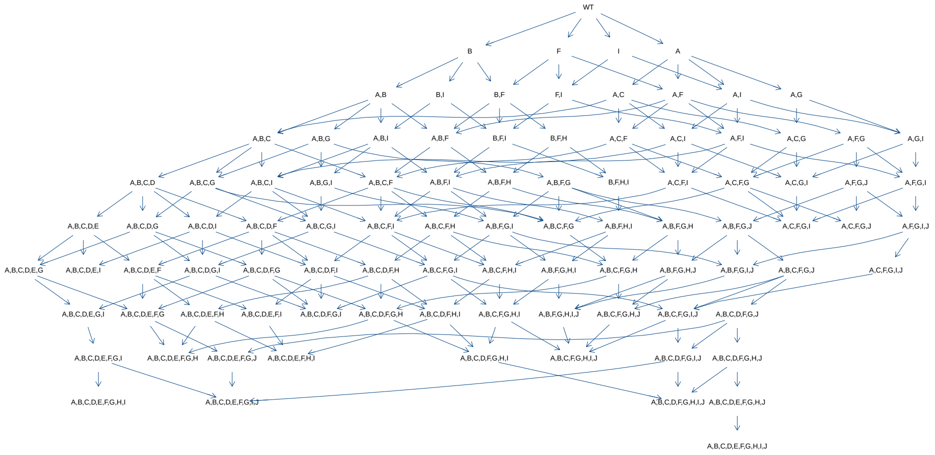

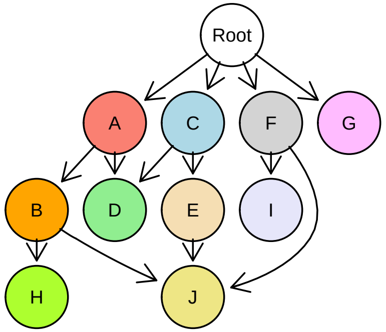


| | |
|---|---|
| *ID* | qzPma2ONgy7W10OcP |
| *acc genotypes* | 124 |
| *peaks no back* | 12 |
| *removed edges* | 68 |
| *prop rem edges* | 0.192 |

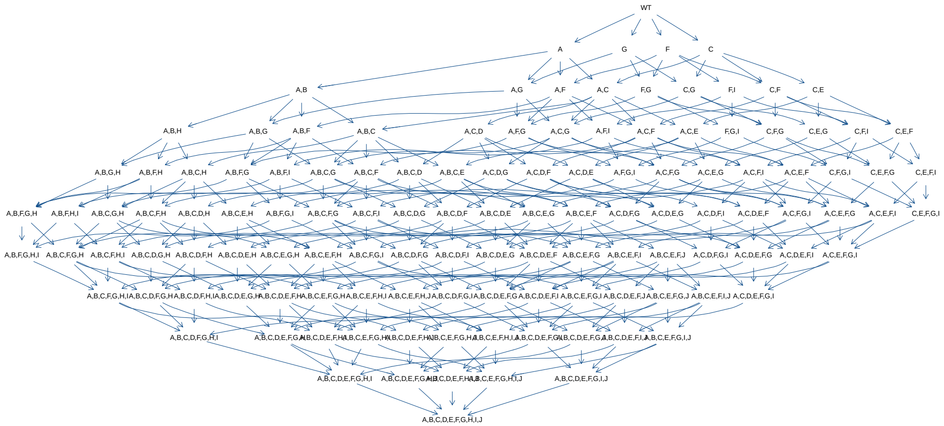

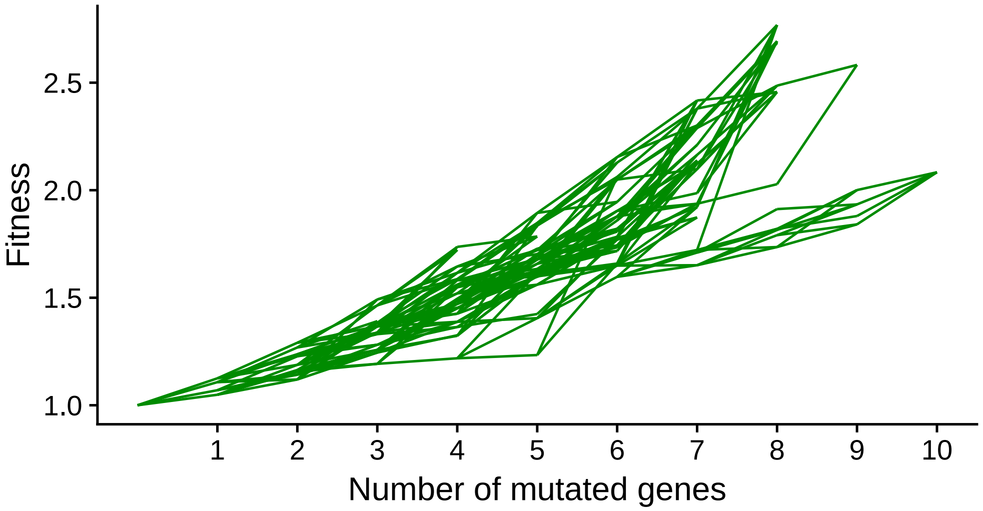


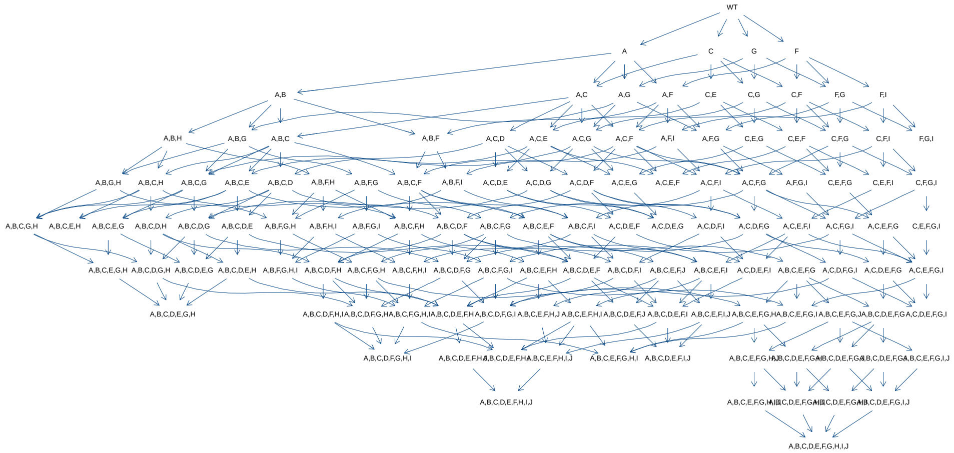

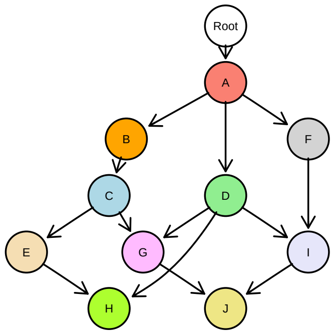


| | |
|---|---|
| *ID* | r10ojqpU7Q5rU9ub3 |
| *acc genotypes* | 36 |
| *peaks no back* | 6 |
| *removed edges* | 12 |
| *prop rem edges* | 0.179 |

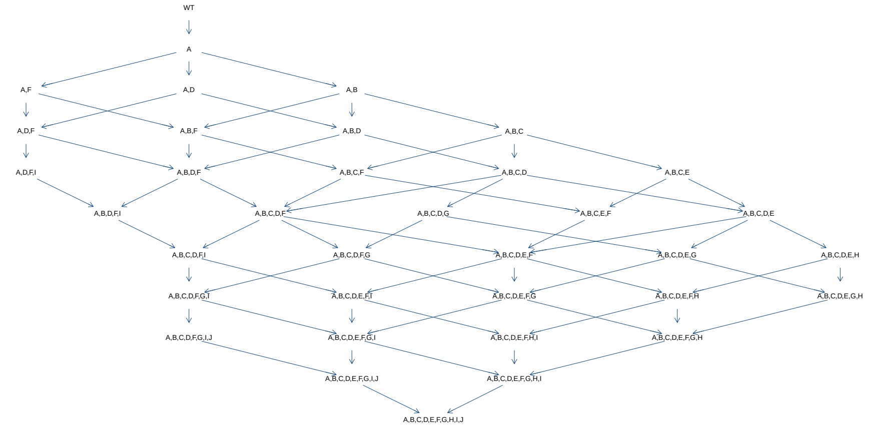

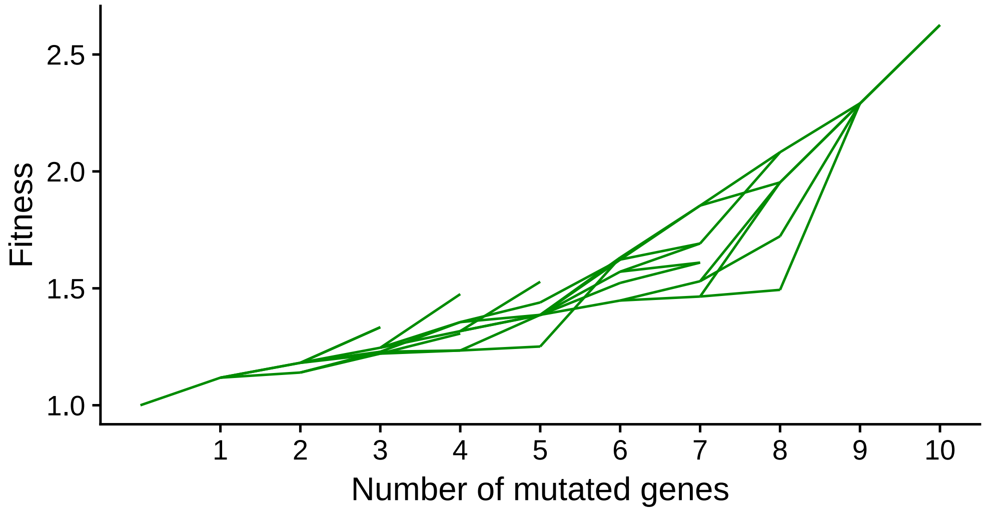


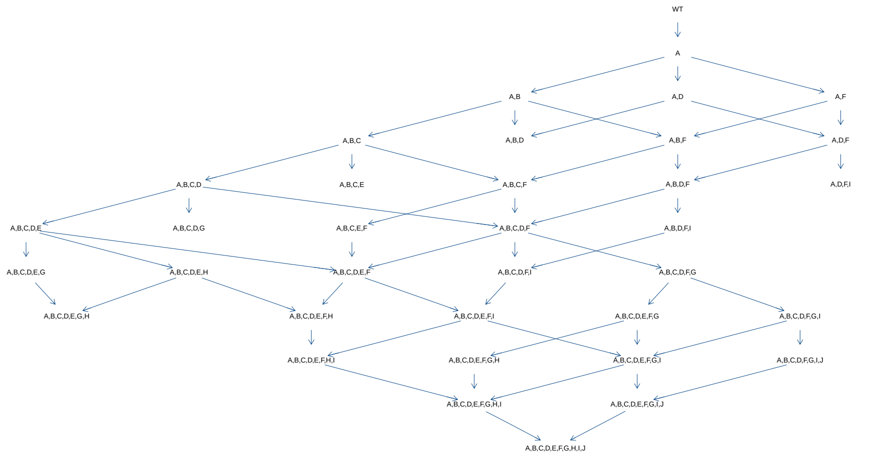

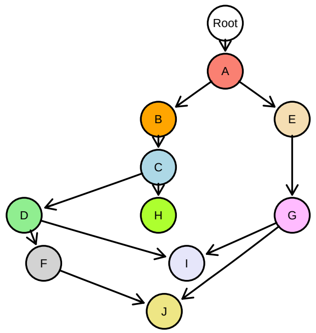


| | |
|---|---|
| *ID* | R4ZN7nnB4ujxYl9L |
| *acc genotypes* | 33 |
| *peaks no back* | 5 |
| *removed edges* | 8 |
| *prop rem edges* | 0.133 |

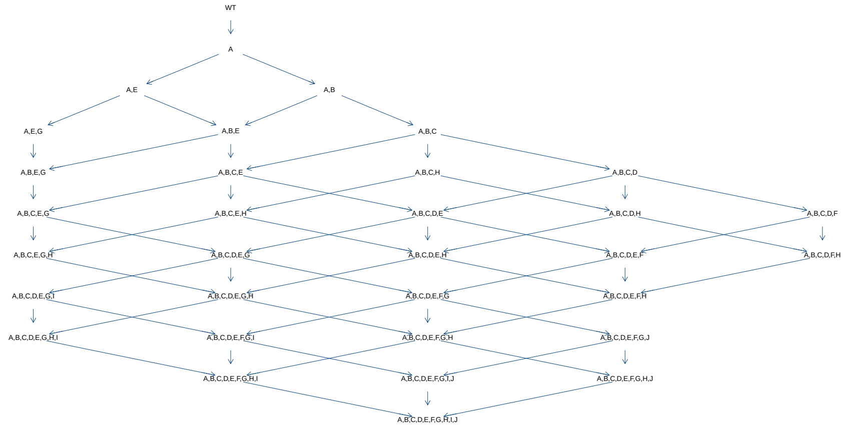

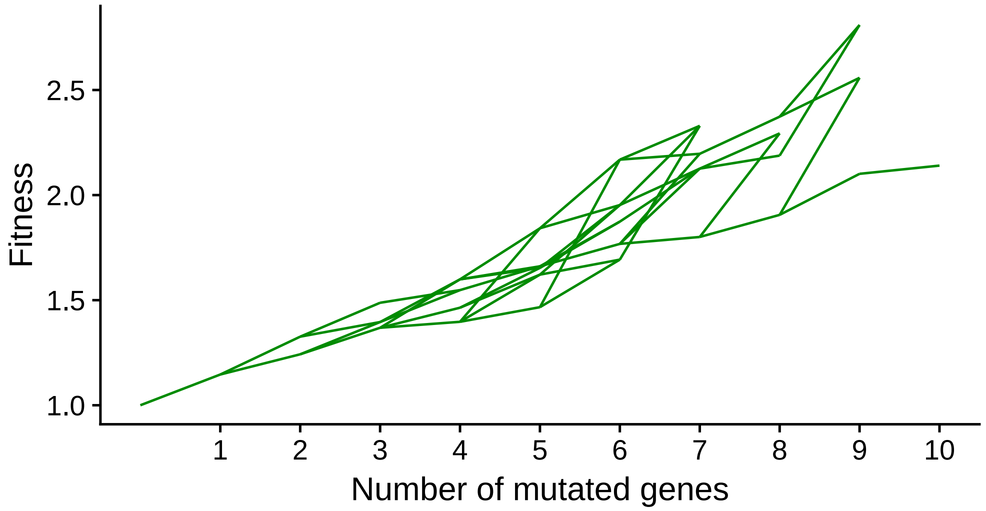


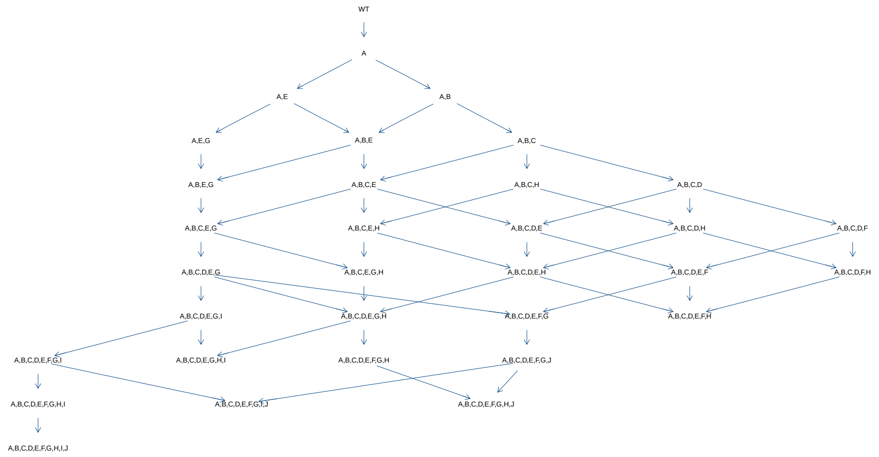

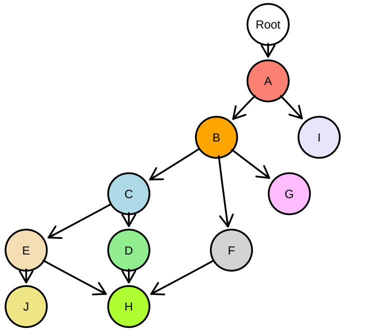


| | |
|---|---|
| *ID* | R5UIg5w7GGy7xUcV |
| *acc genotypes* | 67 |
| *peaks no back* | 8 |
| *removed edges* | 36 |
| *prop rem edges* | 0.209 |

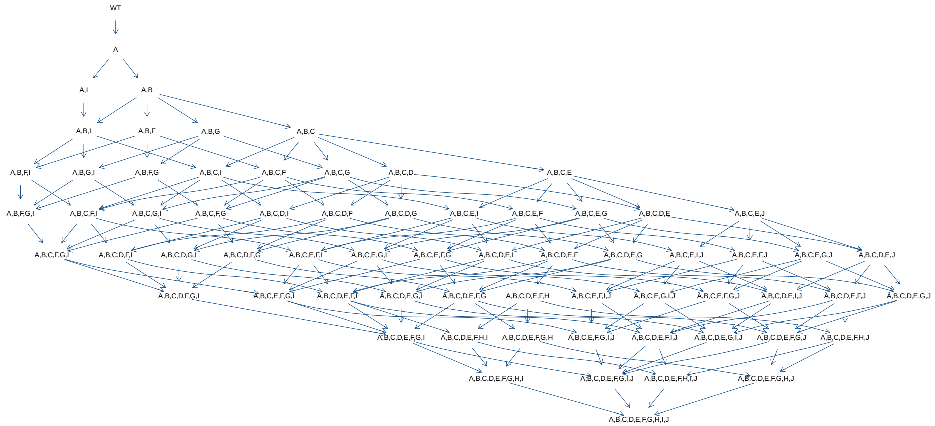

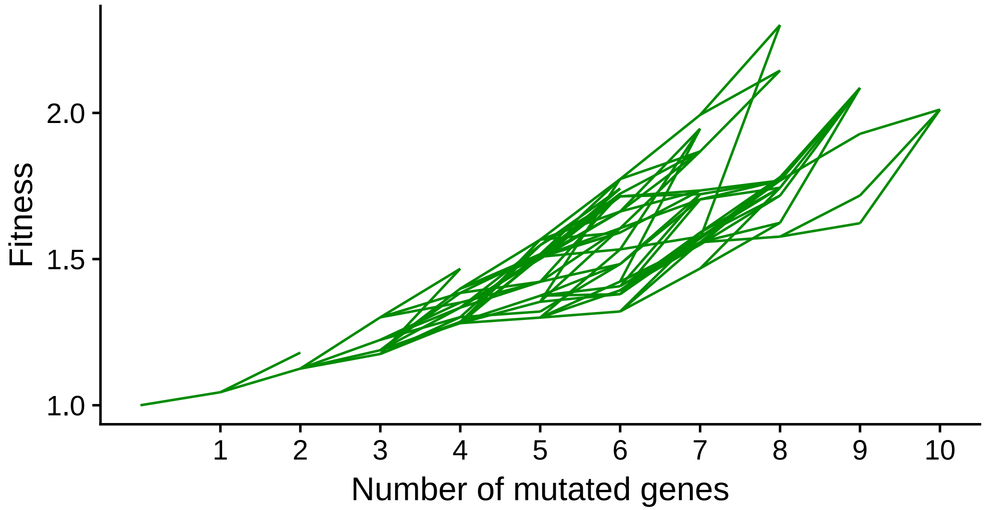


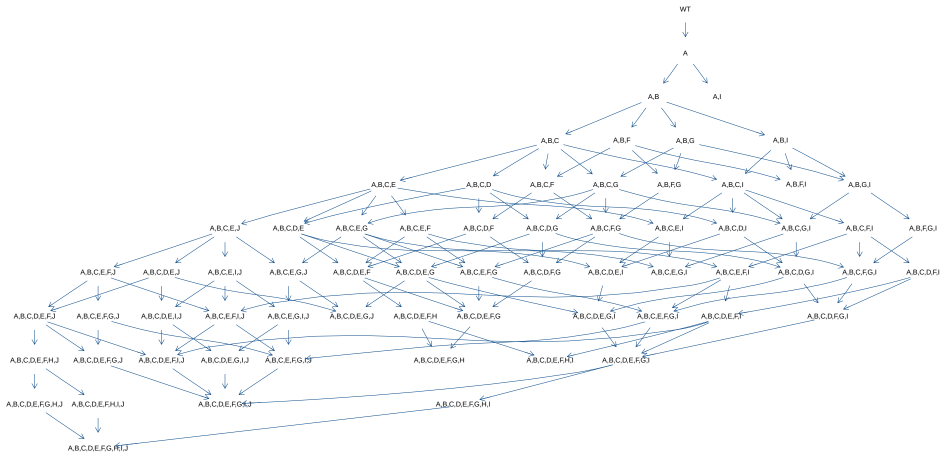

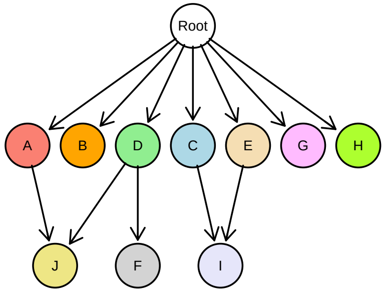


| | |
|---|---|
| *ID* | RcLGnMwFAV7yHcPX |
| *acc genotypes* | 320 |
| *peaks no back* | 19 |
| *removed edges* | 237 |
| *prop rem edges* | 0.198 |

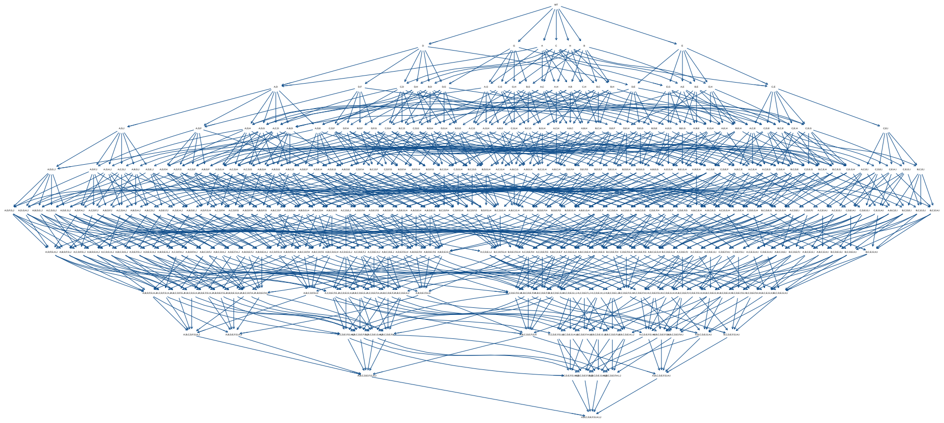

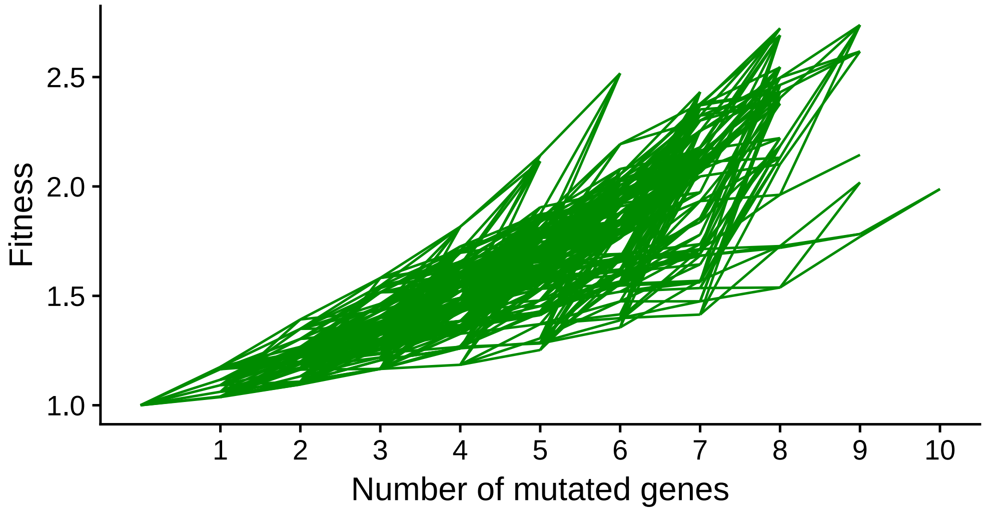


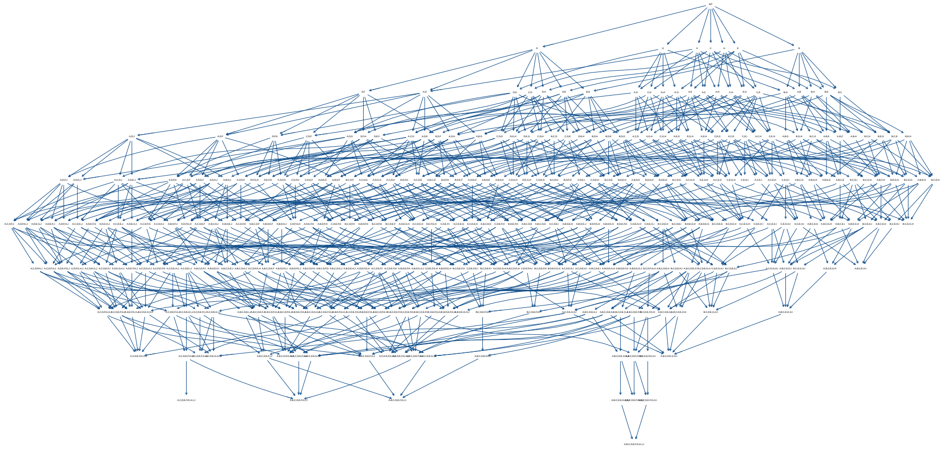

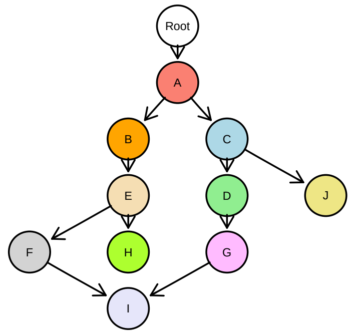


| | |
|---|---|
| *ID* | RMPsjUicmruxvFMT |
| *acc genotypes* | 47 |
| *peaks no back* | 8 |
| *removed edges* | 22 |
| *prop rem edges* | 0.222 |

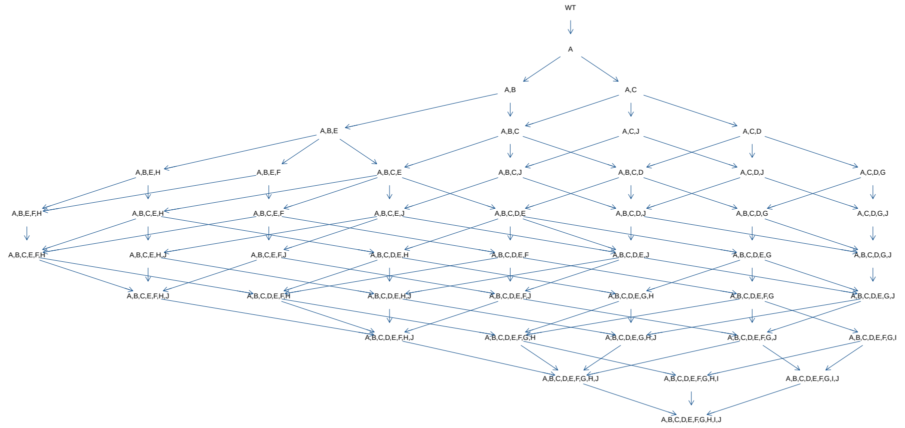

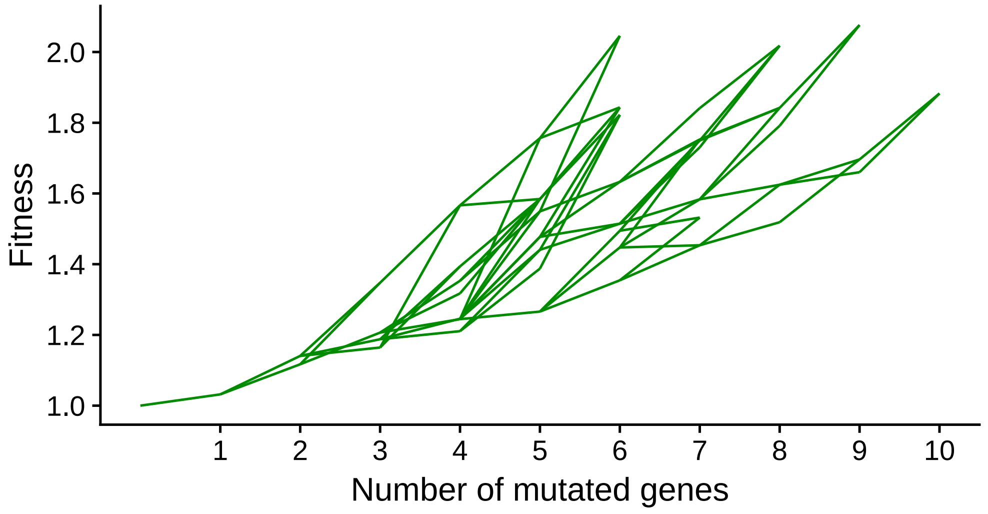


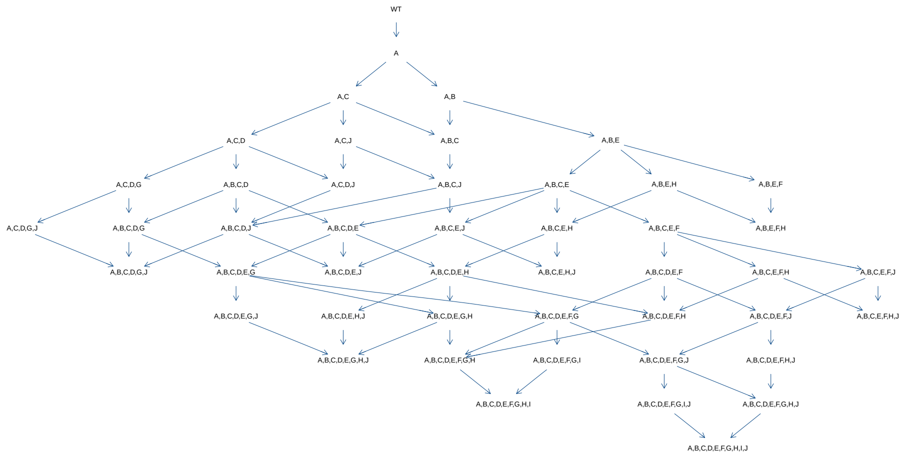

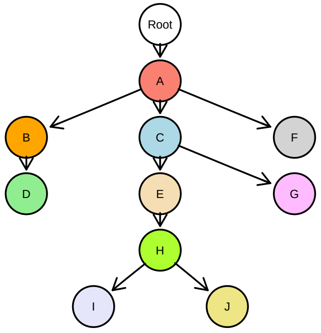


| | |
|---|---|
| *ID* | rRQ3Bws3107fO7mHM |
| *acc genotypes* | 79 |
| *peaks no back* | 11 |
| *removed edges* | 41 |
| *prop rem edges* | 0.199 |

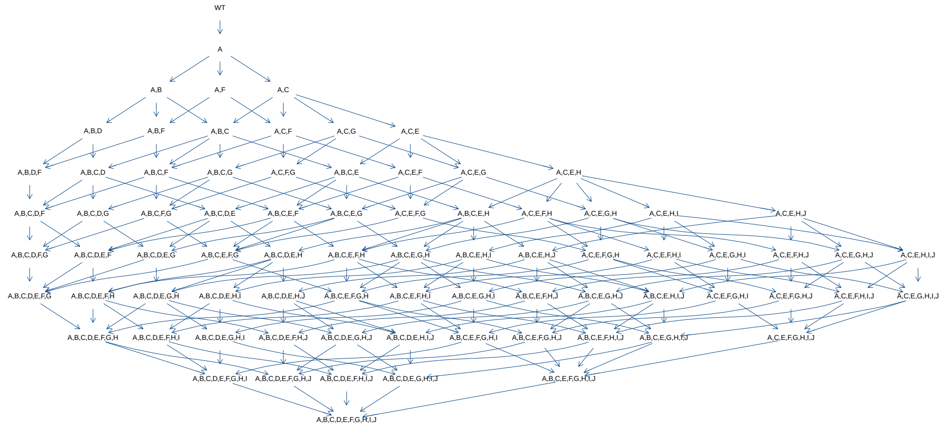

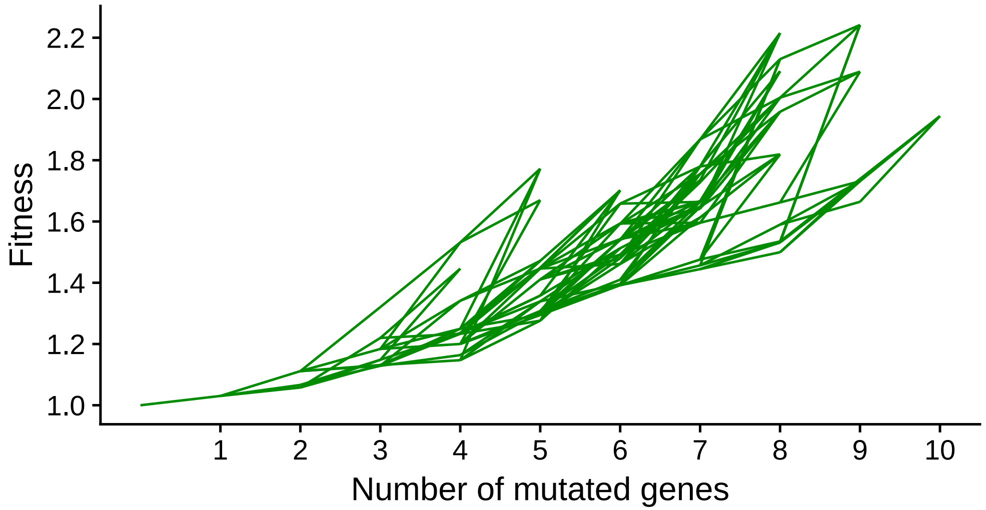


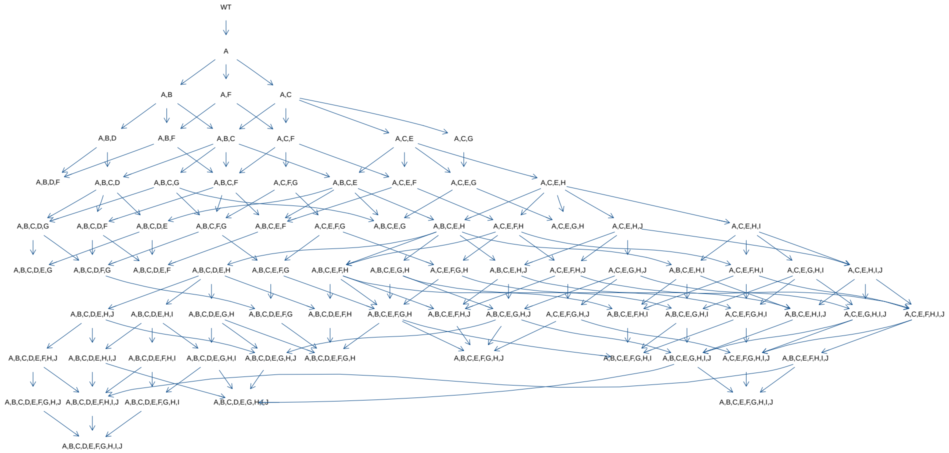

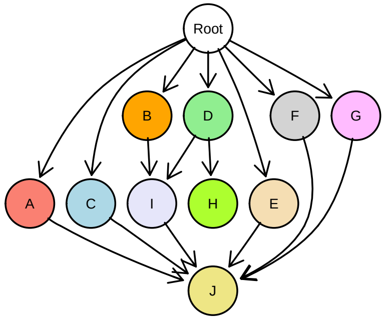


| | |
|---|---|
| *ID* | SArTeHv6MV7tVCry |
| *acc genotypes* | 258 |
| *peaks no back* | 16 |
| *removed edges* | 161 |
| *prop rem edges* | 0.167 |

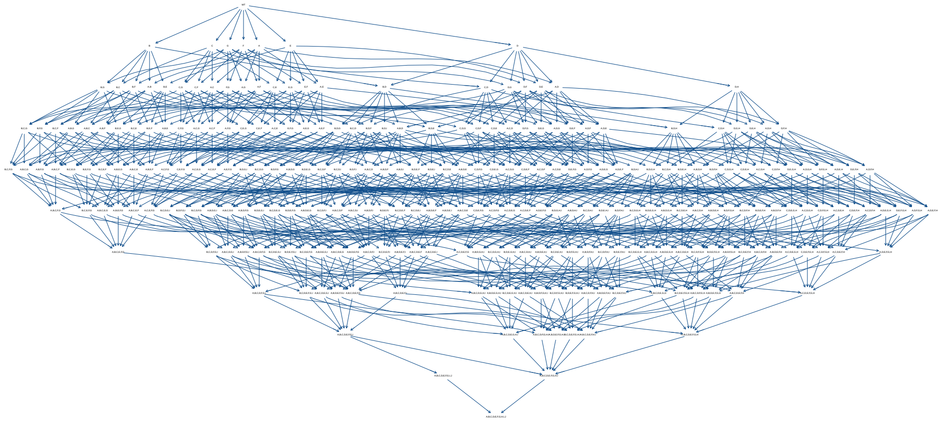

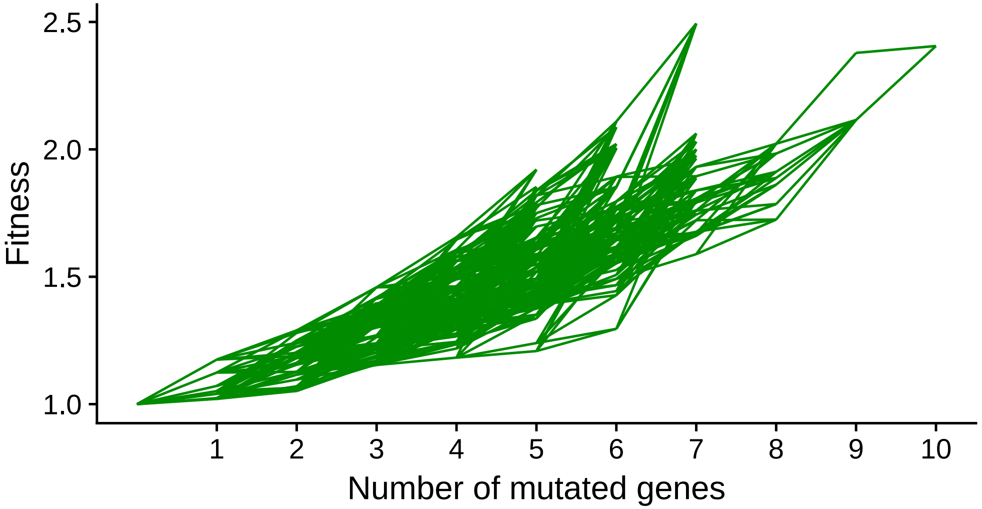


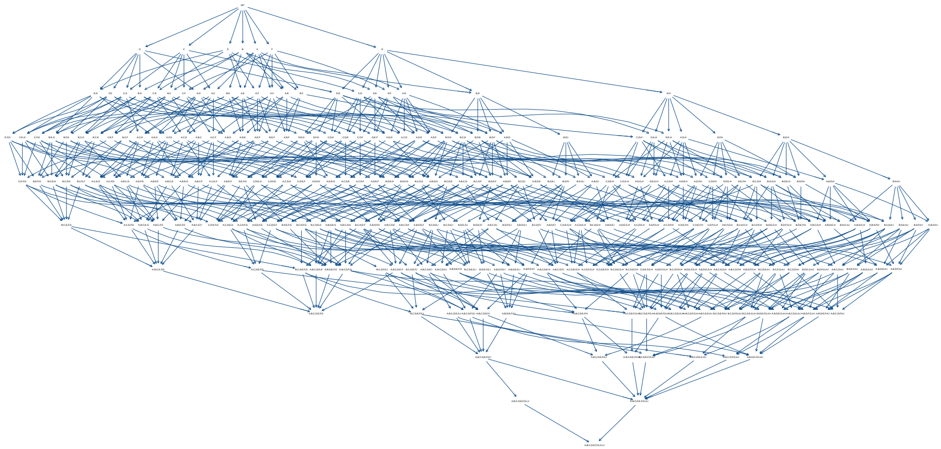

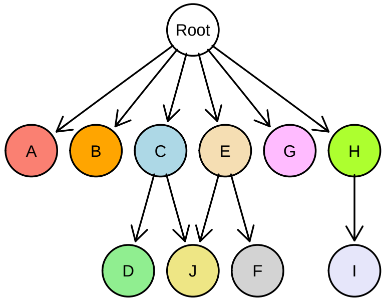


| | |
|---|---|
| *ID* | SlYbZPVjDjsPd3OJ |
| *acc genotypes* | 312 |
| *peaks no back* | 21 |
| *removed edges* | 259 |
| *prop rem edges* | 0.224 |

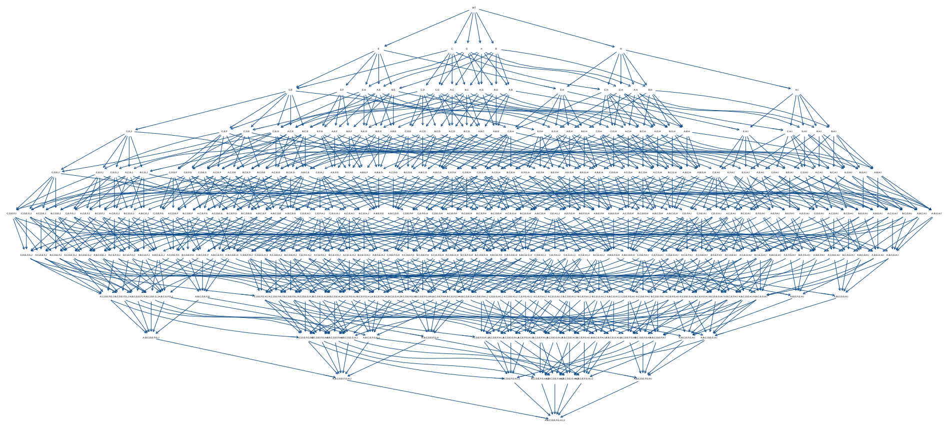

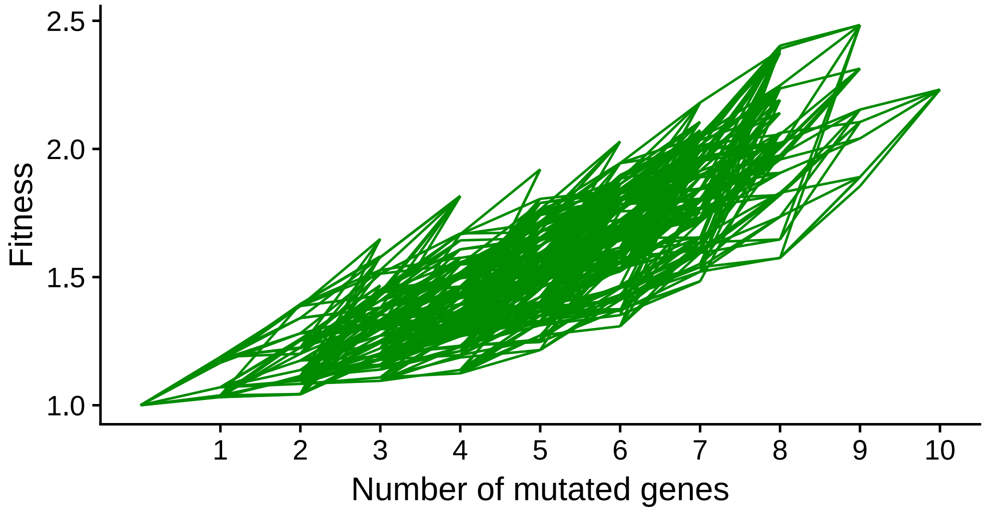


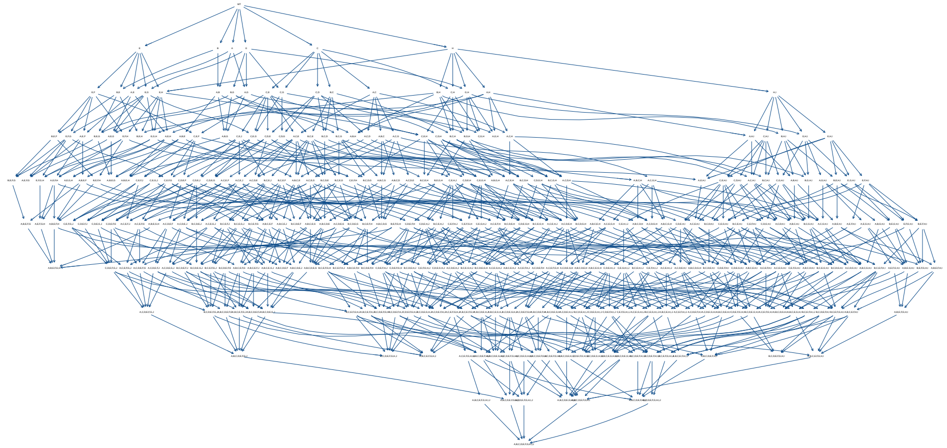

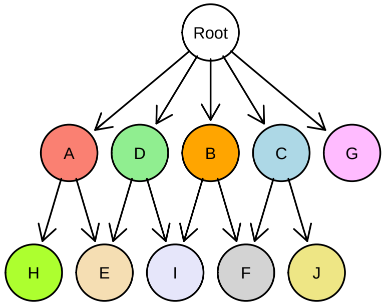


| | |
|---|---|
| *ID* | smwY8F2IPkzSNFOb |
| *acc genotypes* | 178 |
| *peaks no back* | 17 |
| *removed edges* | 101 |
| *prop rem edges* | 0.181 |

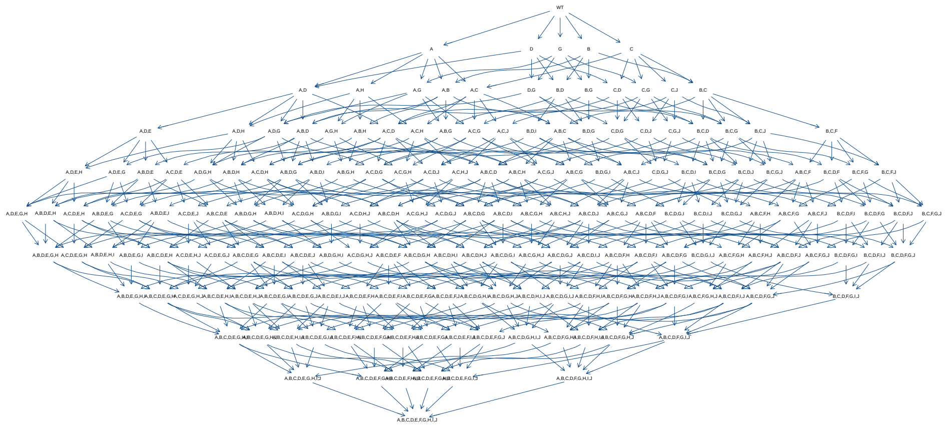

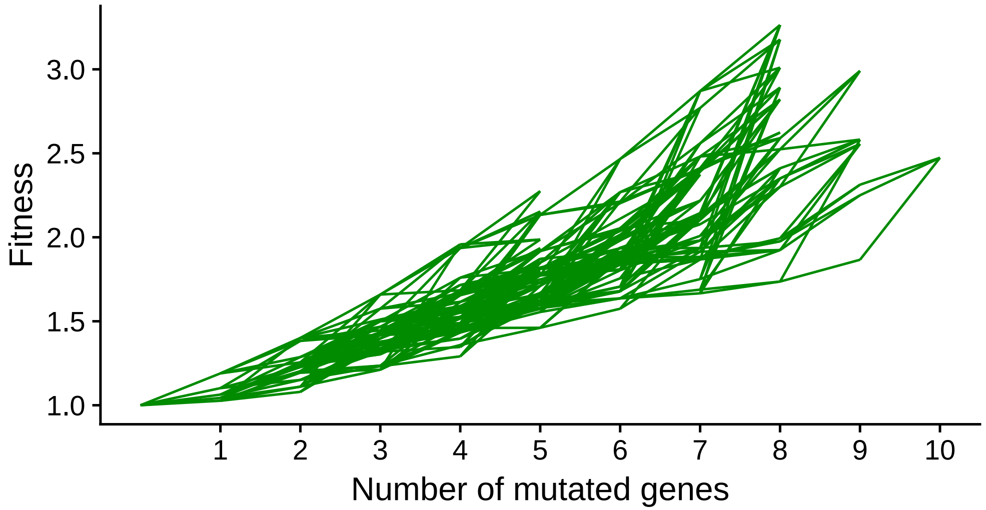


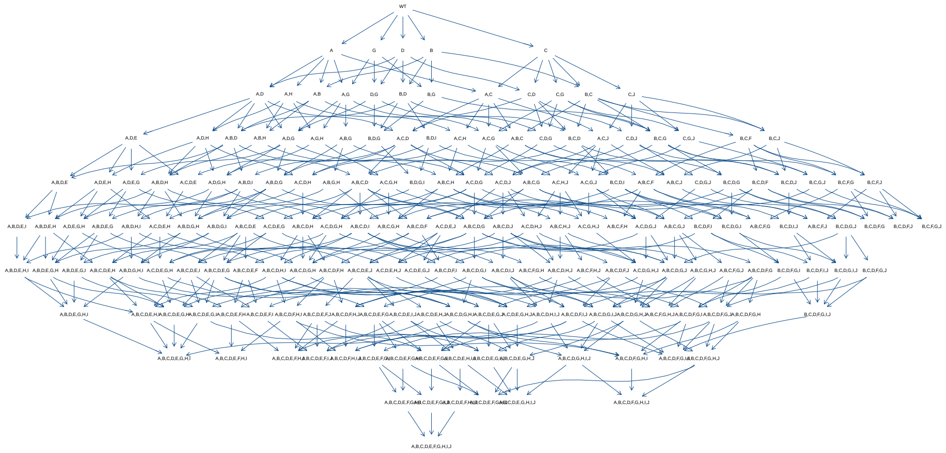

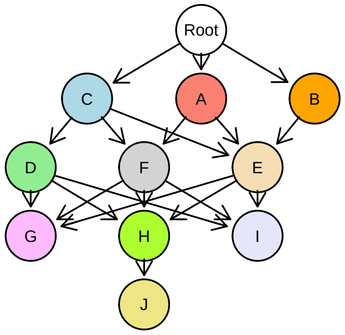


| | |
|---|---|
| *ID* | sO7ye5yMrhvX10WZQ |
| *acc genotypes* | 31 |
| *peaks no back* | 2 |
| *removed edges* | 2 |
| *prop rem edges* | 0.036 |

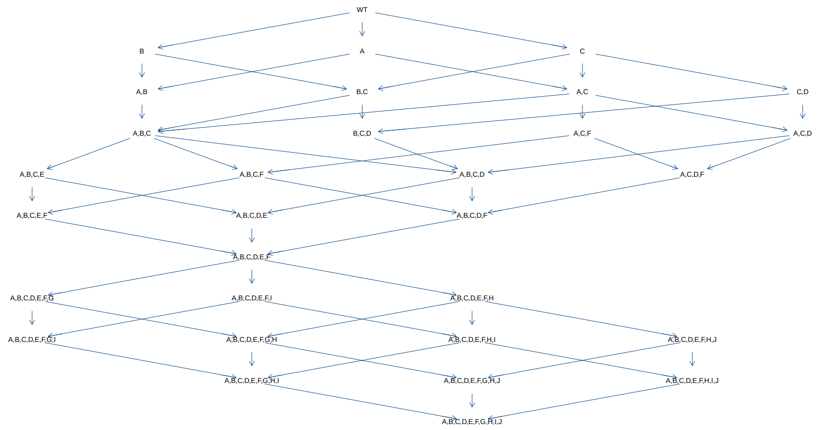

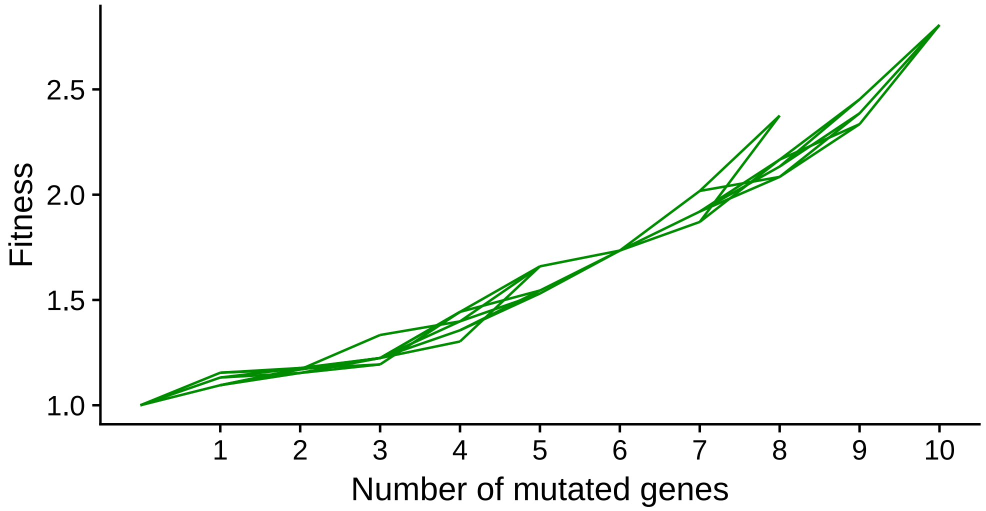


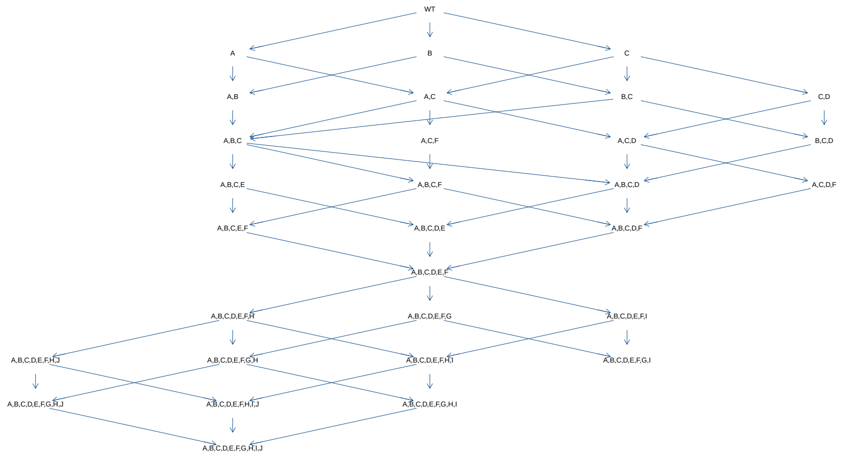

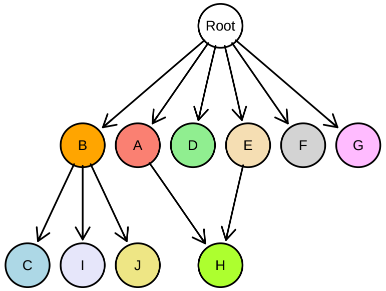


| | |
|---|---|
| *ID* | sUe5FH9IUIMDiI10B |
| *acc genotypes* | 360 |
| *peaks no back* | 23 |
| *removed edges* | 252 |
| *prop rem edges* | 0.177 |

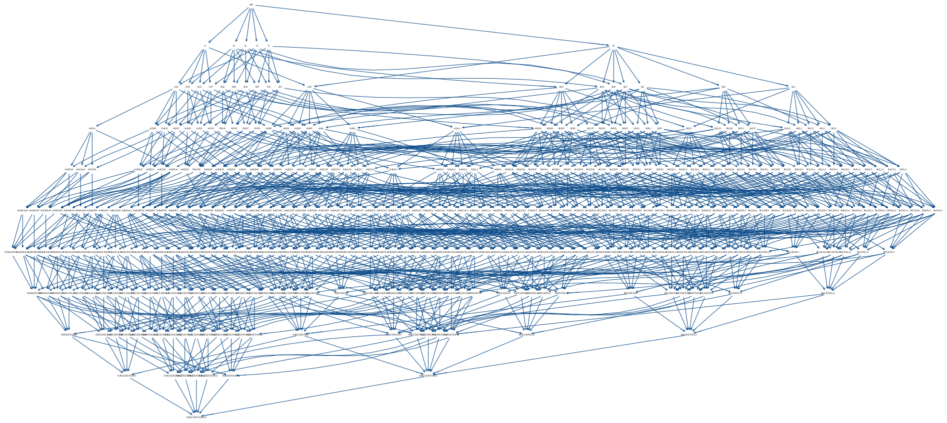

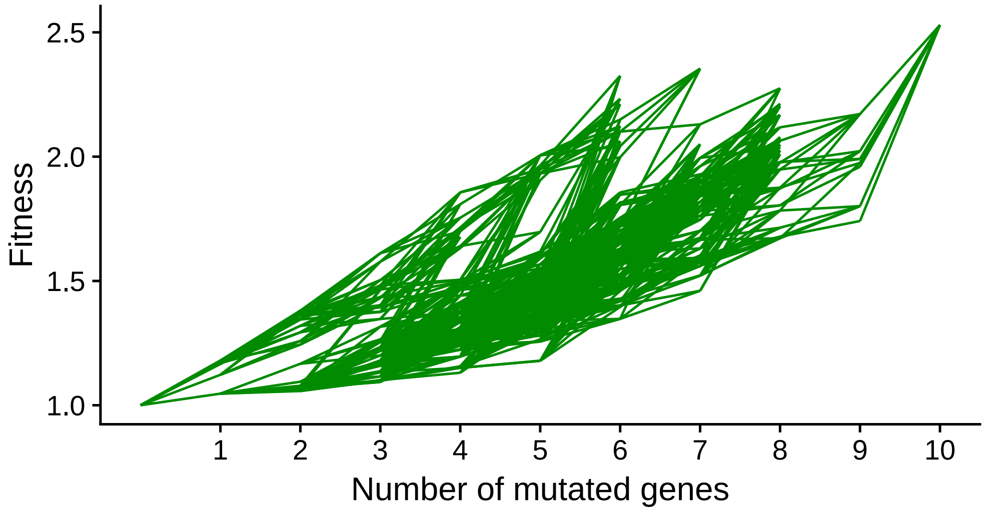


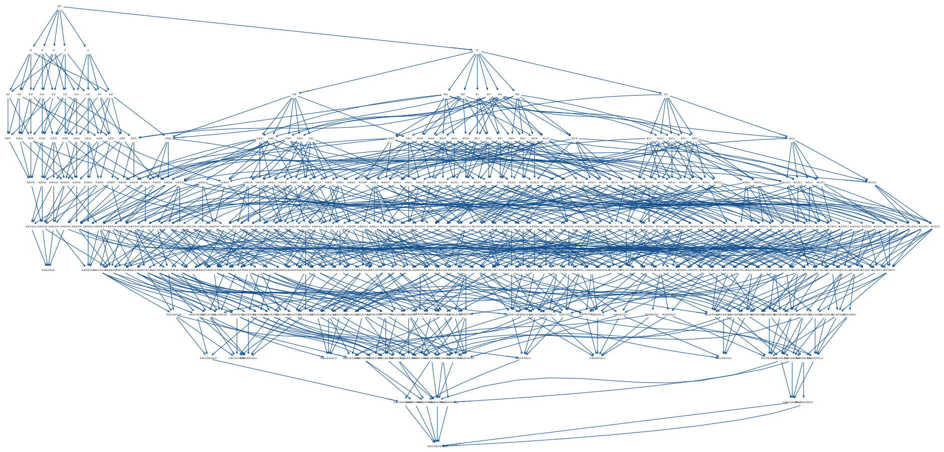

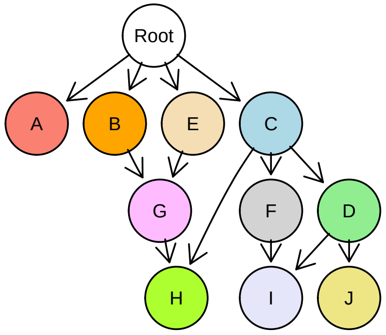


| | |
|---|---|
| *ID* | SWgUDoSPxMcdnnQM |
| *acc genotypes* | 106 |
| *peaks no back* | 12 |
| *removed edges* | 52 |
| *prop rem edges* | 0.18 |

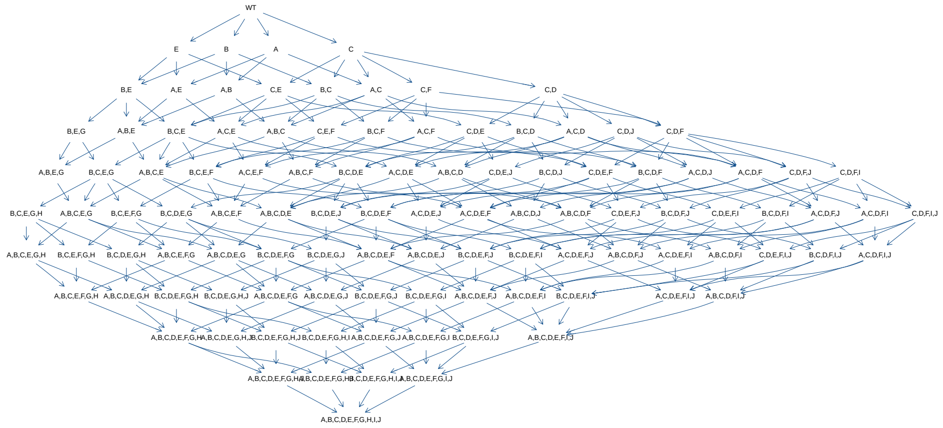

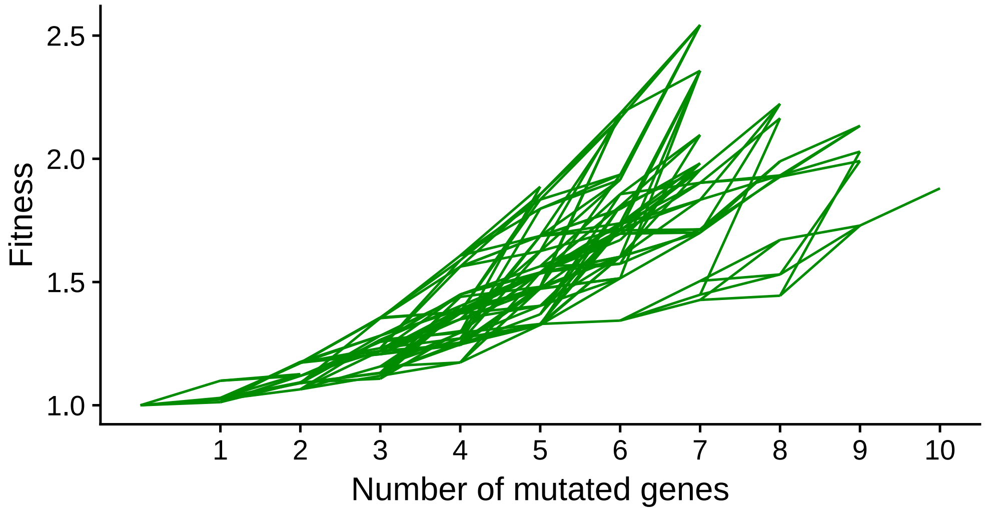


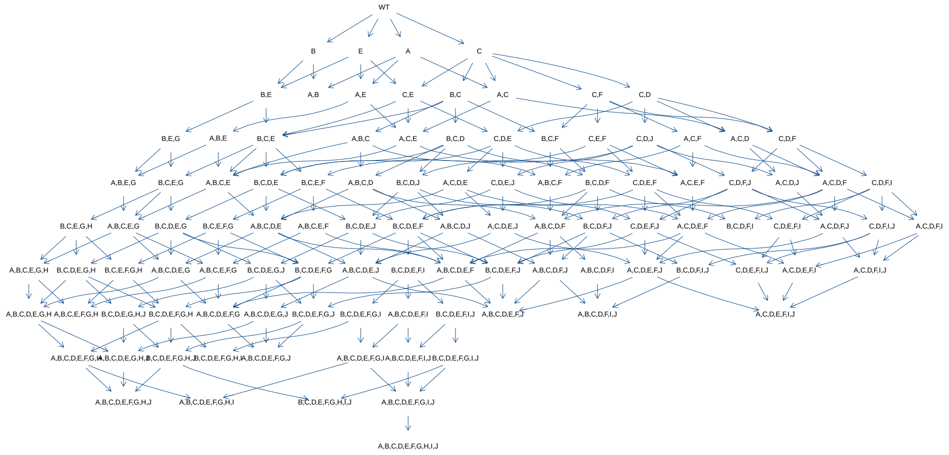

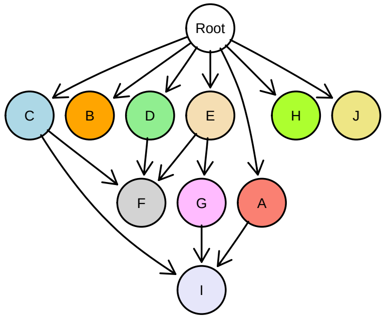


*ID* sxE6qJHh512sjV9a
*acc genotypes* 248
*peaks no back* 16
*removed edges* 163
*prop rem edges* 0.183

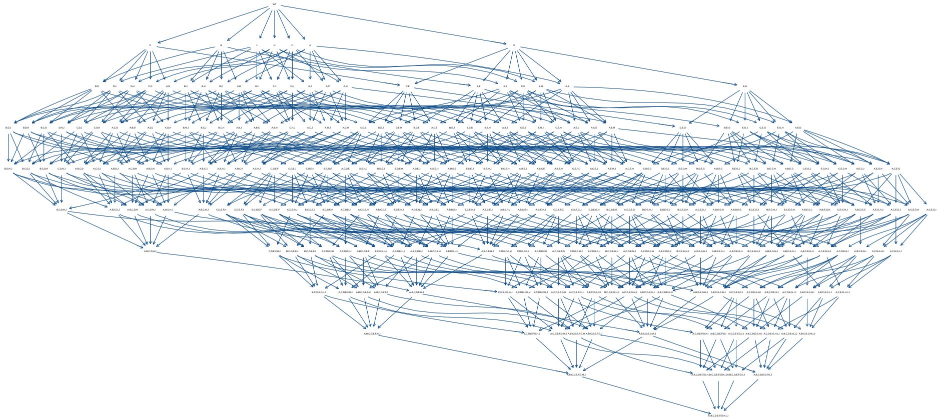

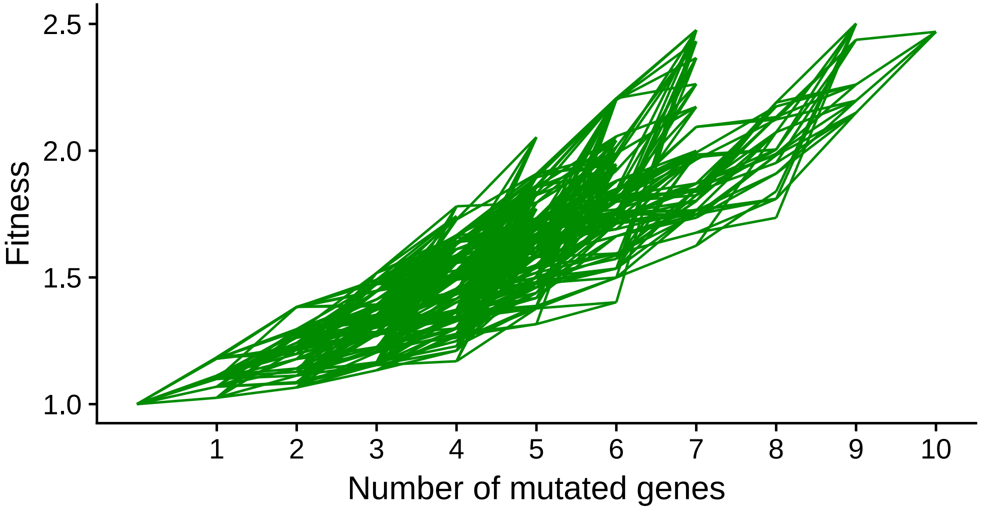


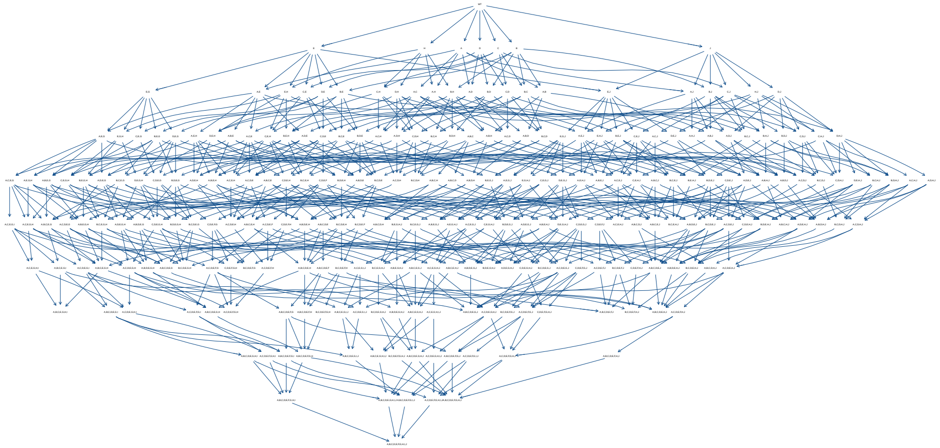

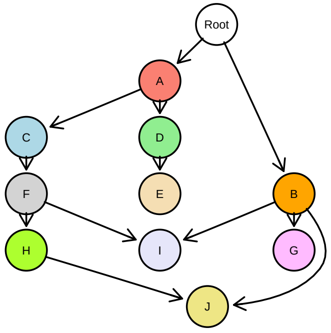


*ID* t10jgW9R9zZrcAqjl
*acc genotypes* 63
*peaks no back* 12
*removed edges* 30
*prop rem edges* 0.208

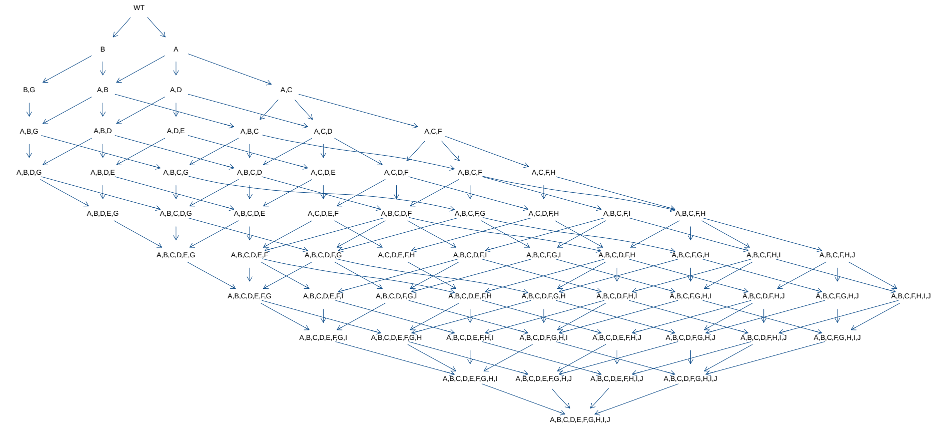

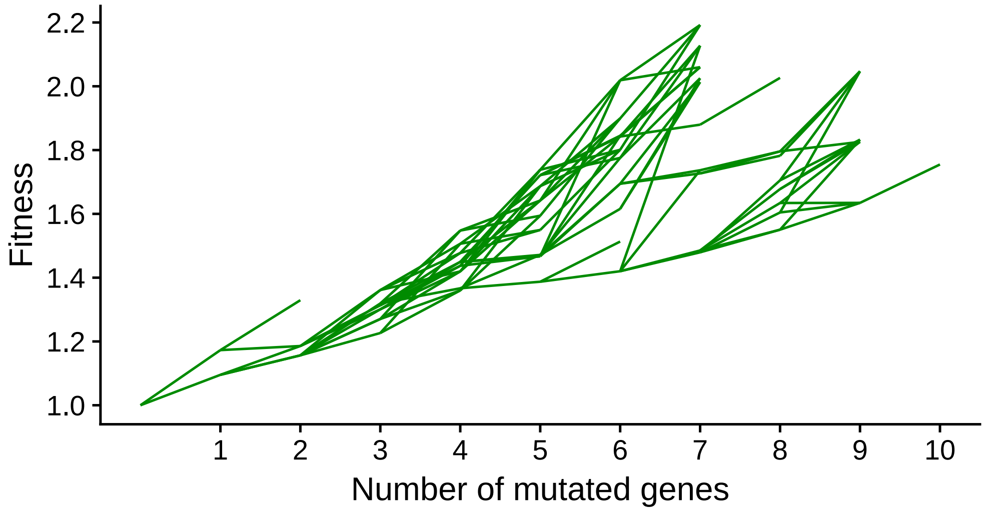


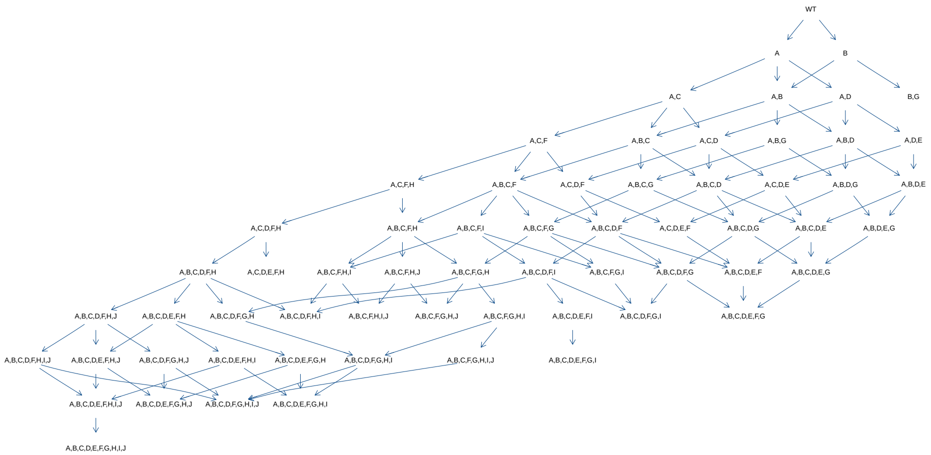

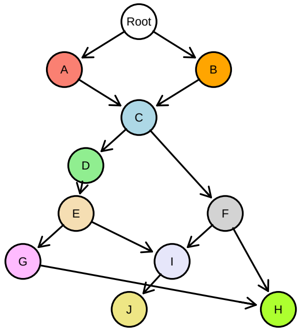


*ID* t1yUXsv5fVuo10GRi

*acc genotypes* 19

*peaks no back* 2

*removed edges* 2

*prop rem edges* 0.077

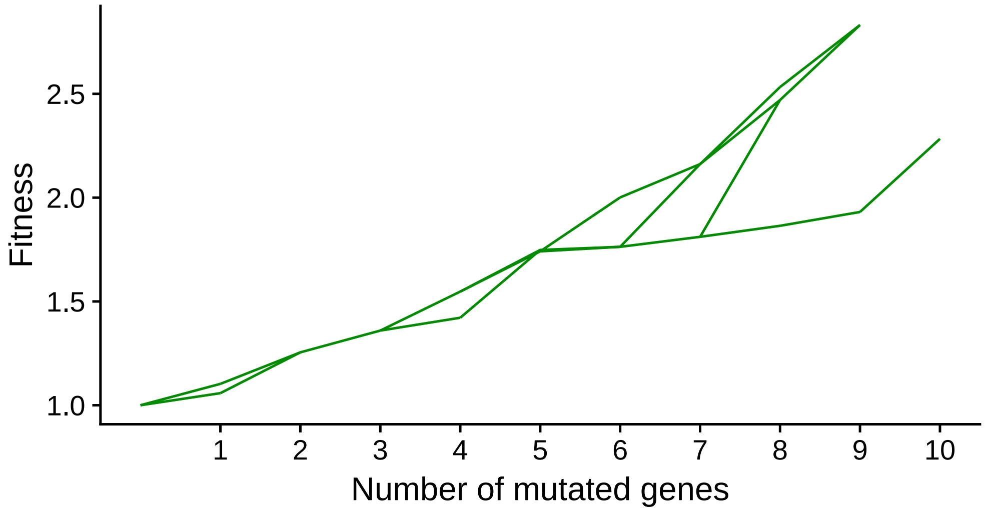


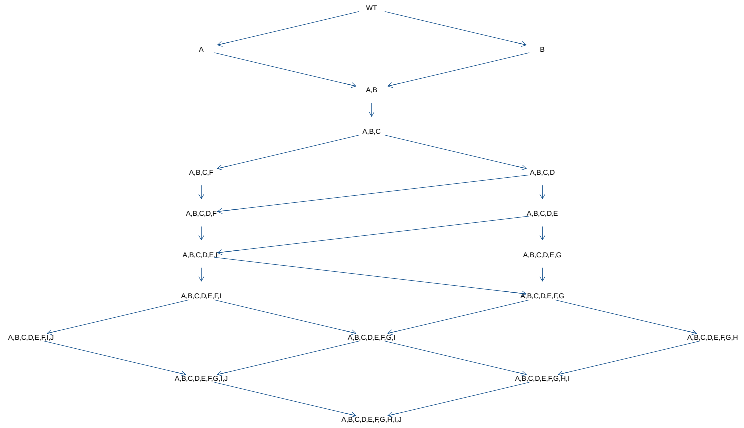


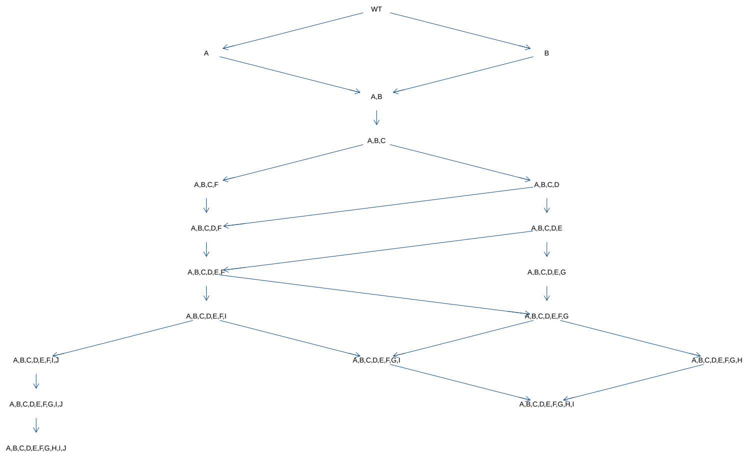

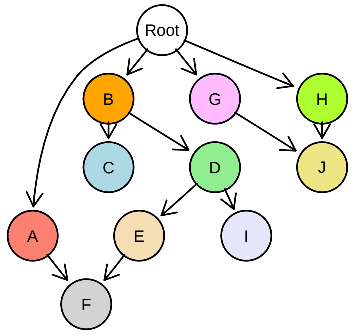


| | |
|---|---|
| *ID* | tBuq3Z94BsQjawuL |
| *acc genotypes* | 130 |
| *peaks no back* | 14 |
| *removed edges* | 92 |
| *prop rem edges* | 0.239 |

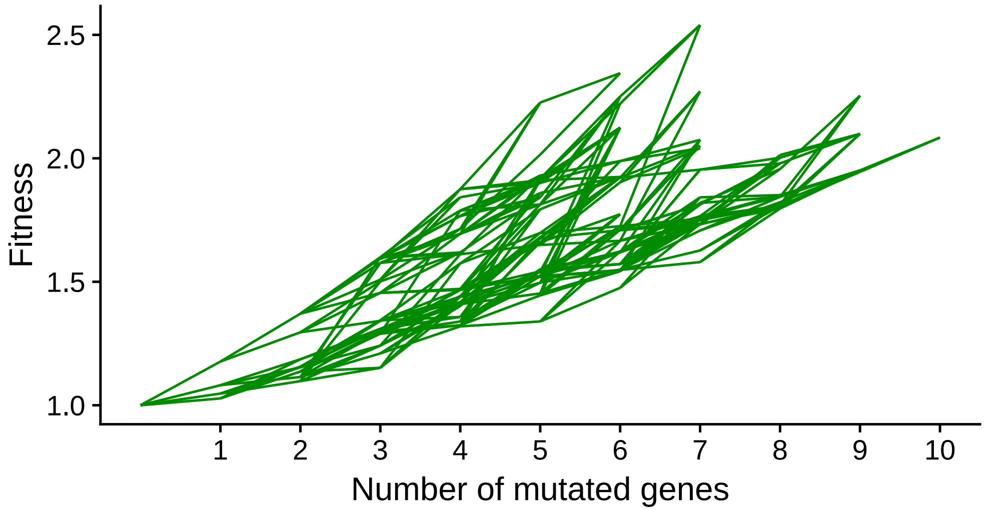


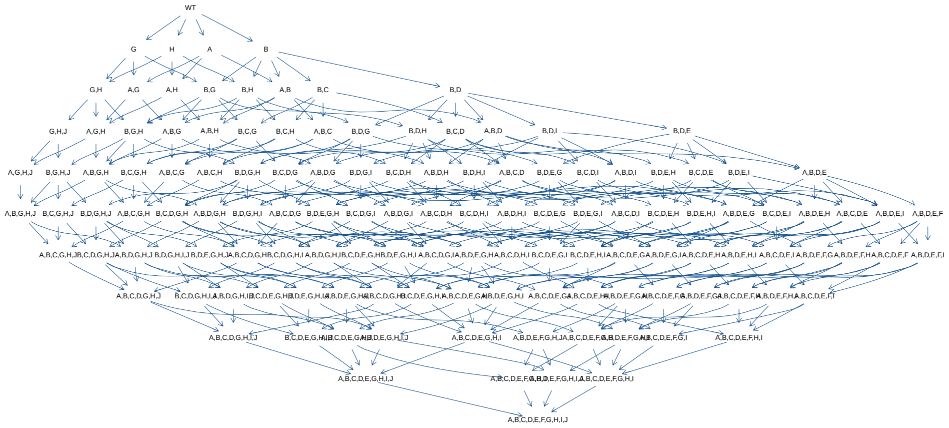

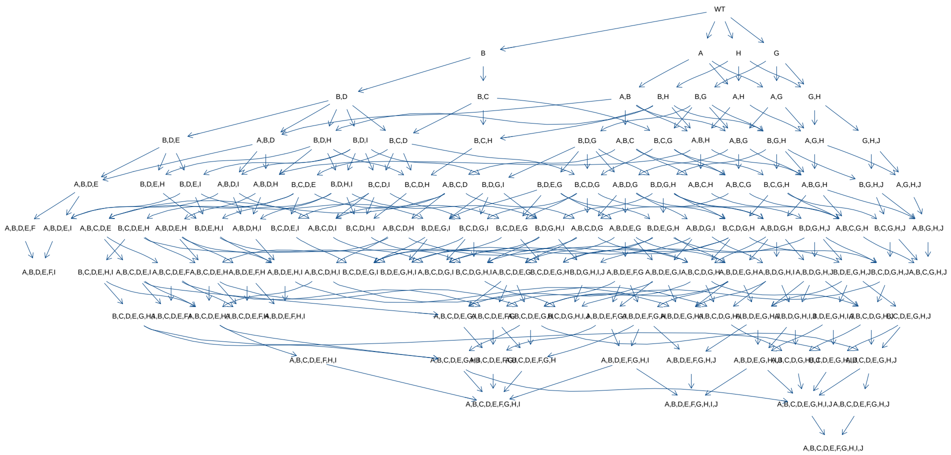

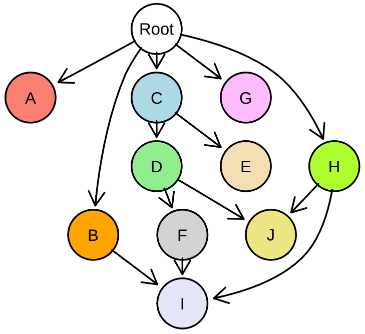


| | |
|---|---|
| *ID* | tGIKPZvUbRWHgDHV |
| *acc genotypes* | 160 |
| *peaks no back* | 16 |
| *removed edges* | 89 |
| *prop rem edges* | 0.174 |

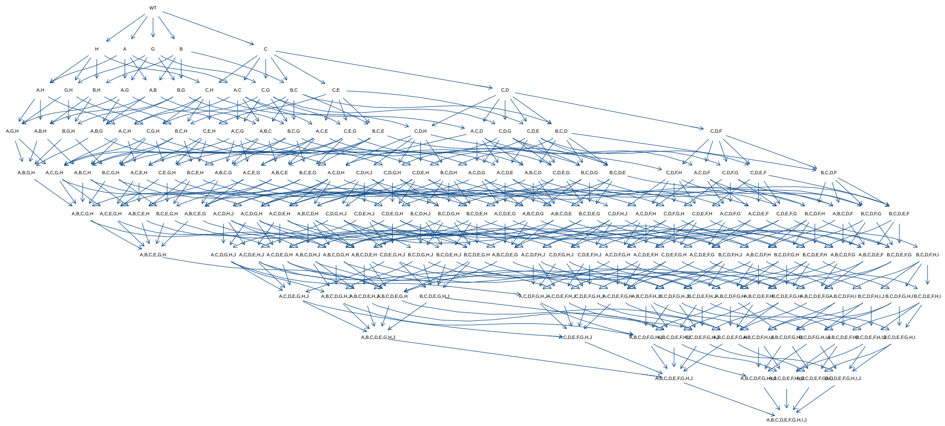

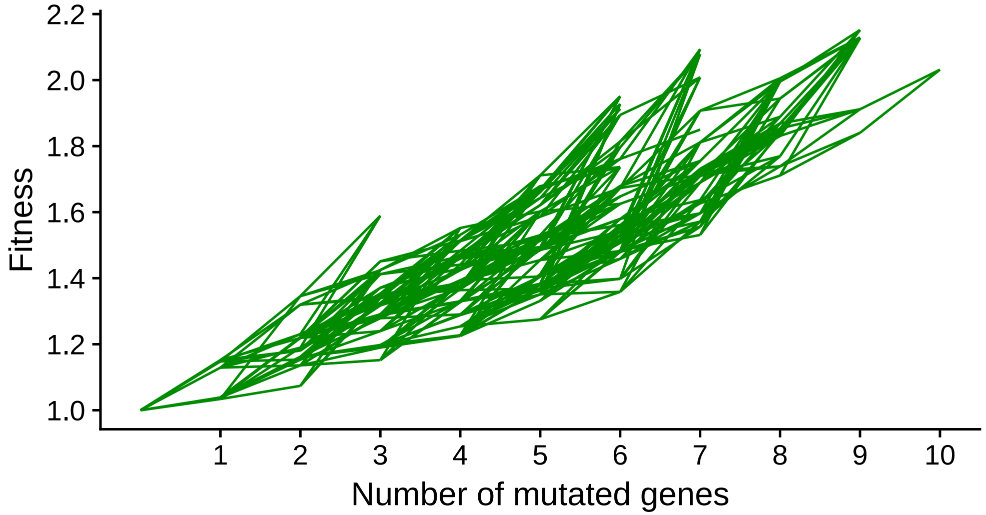


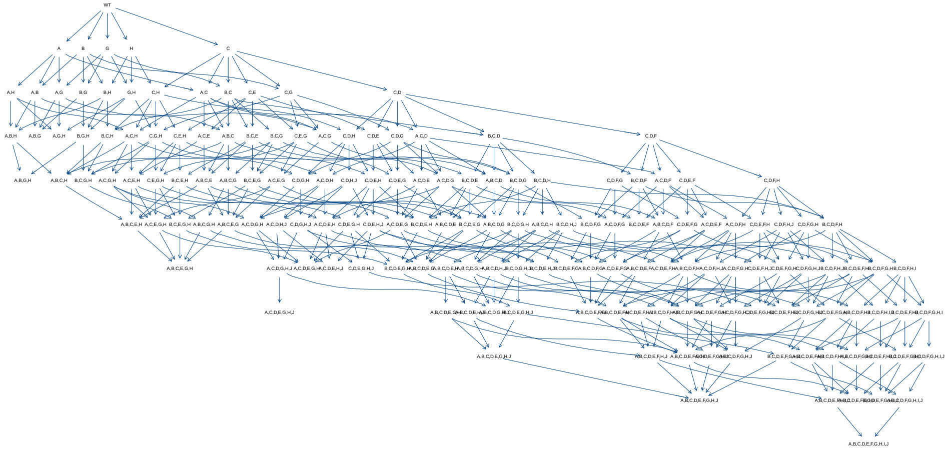

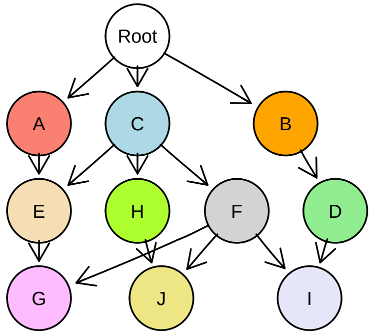


| | |
|---|---|
| *ID* | TGzeGSf7DK4OTwG2 |
| *acc genotypes* | 72 |
| *peaks no back* | 8 |
| *removed edges* | 32 |
| *prop rem edges* | 0.19 |

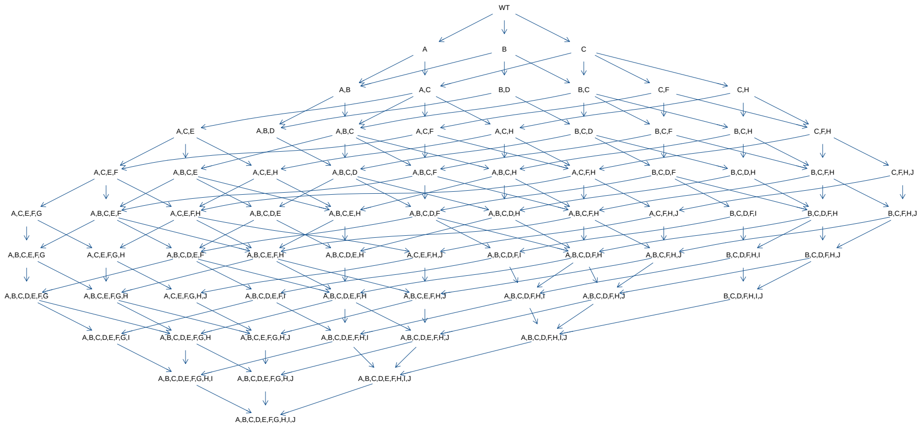

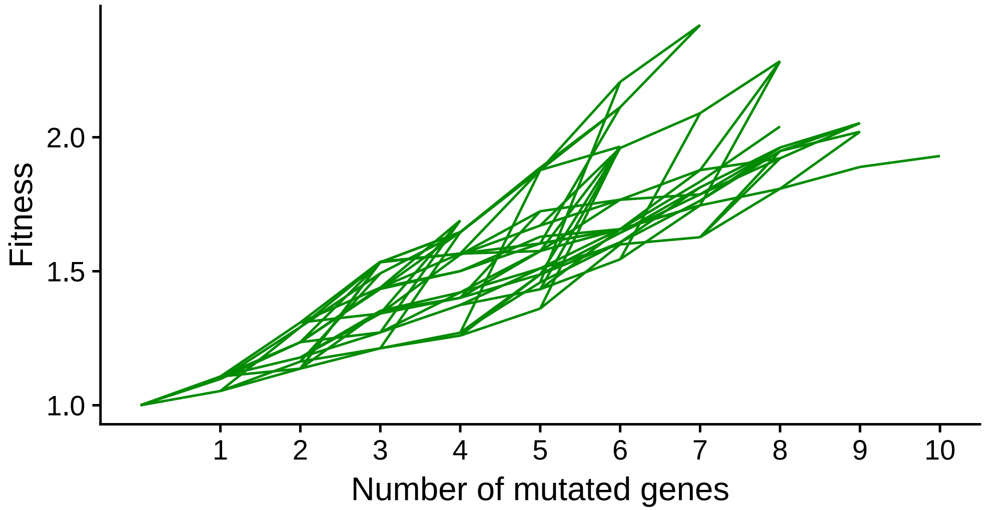


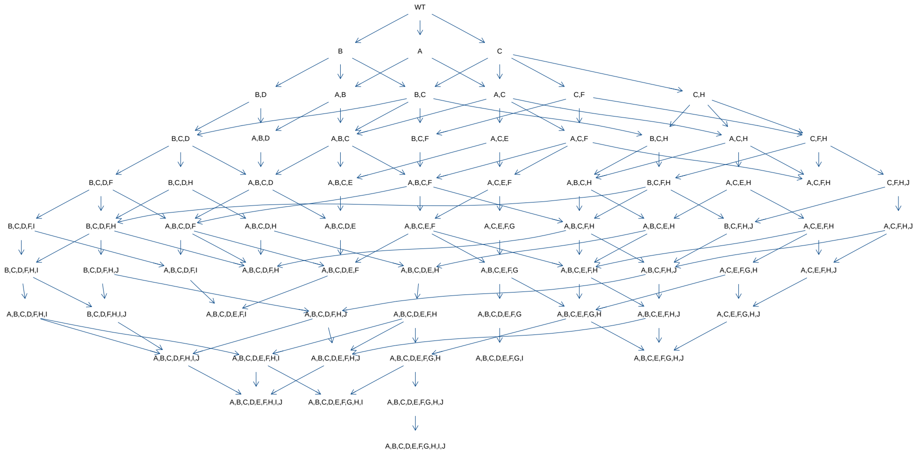

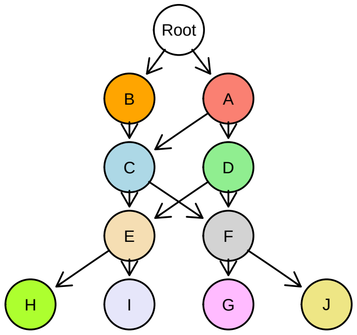


| | |
|---|---|
| *ID* | tI8GpDHXTWuBAEB4 |
| *acc genotypes* | 32 |
| *peaks no back* | 6 |
| *removed edges* | 11 |
| *prop rem edges* | 0.183 |

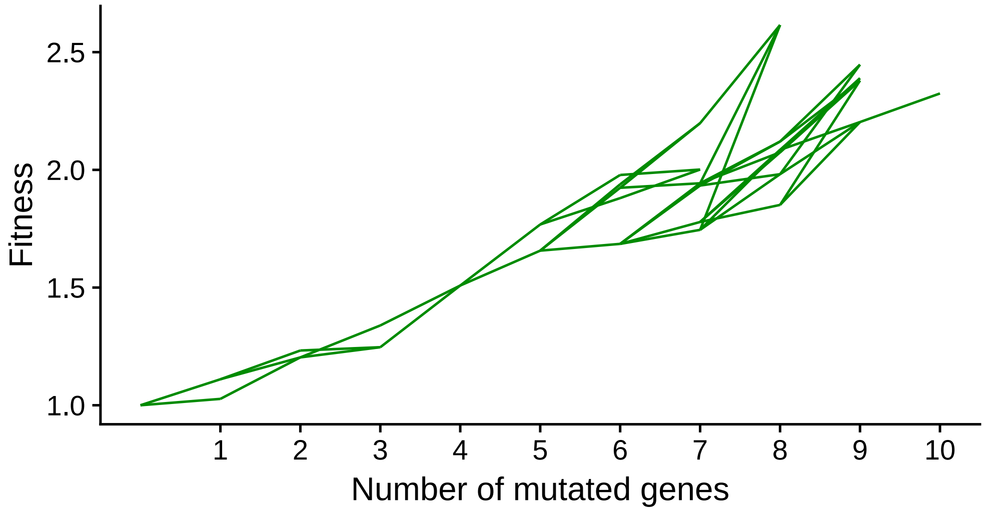


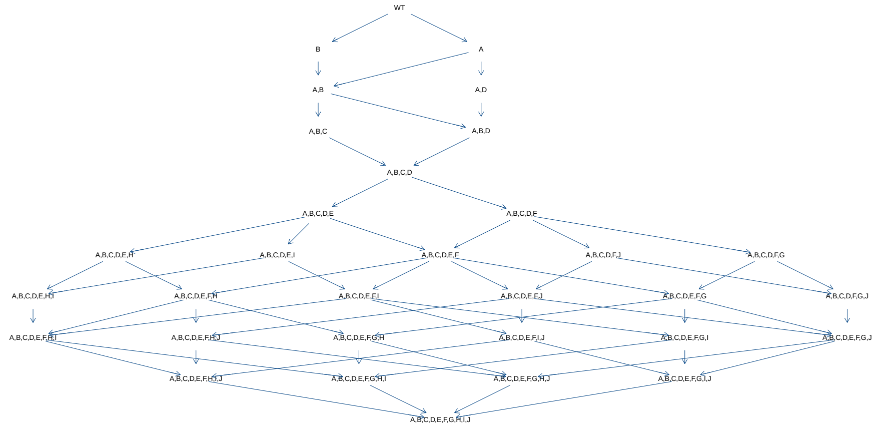

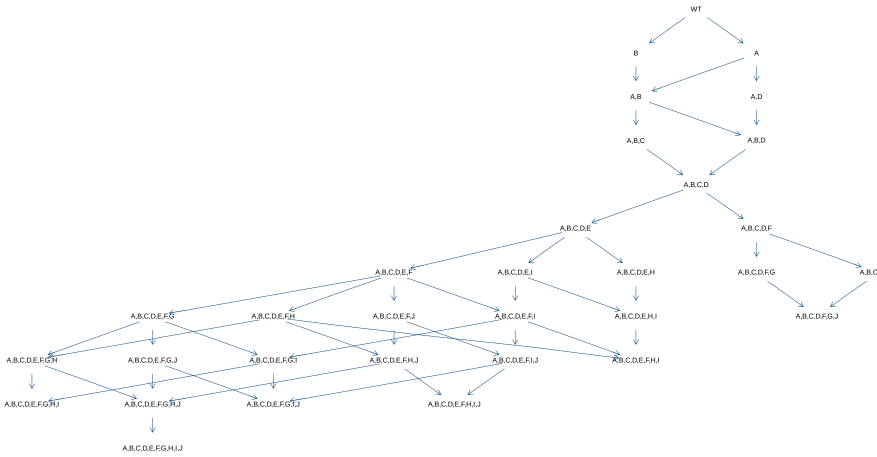

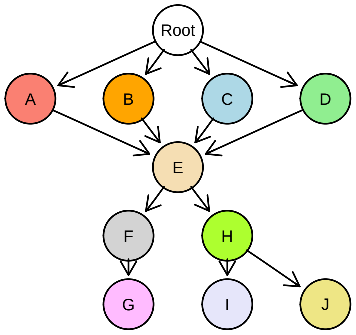


| | |
|---|---|
| *ID* | TnFE10L2B9qFy8hMT |
| *acc genotypes* | 31 |
| *peaks no back* | 5 |
| *removed edges* | 8 |
| *prop rem edges* | 0.138 |

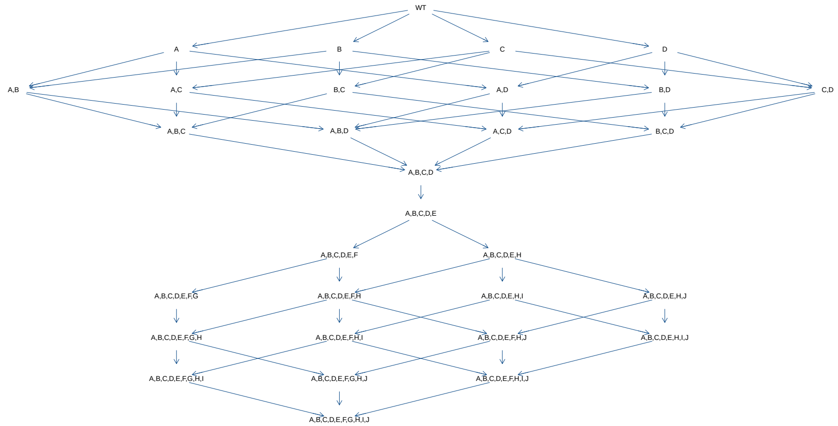

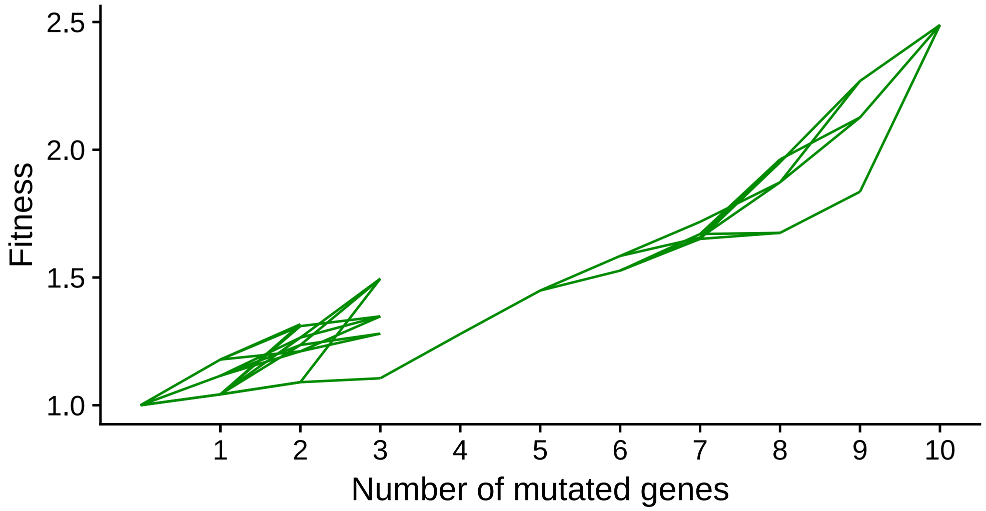


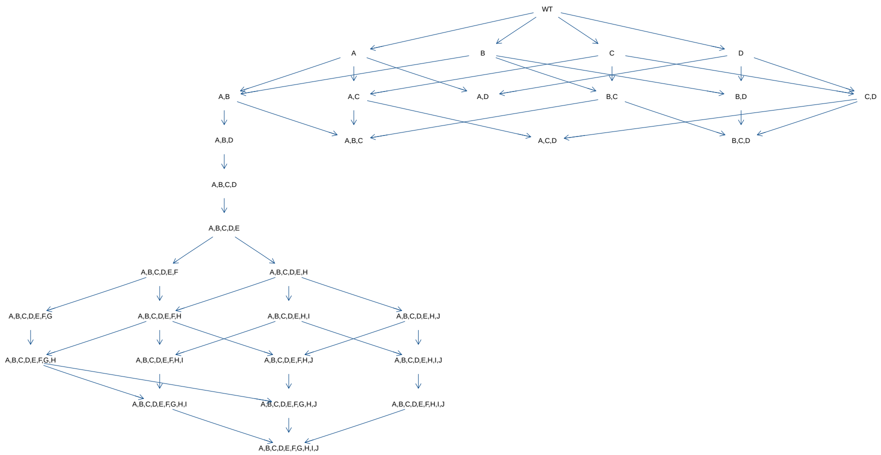

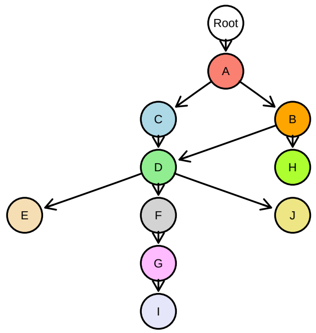


*ID* ToRaV610sJqVtMv2b
*acc genotypes* 39
*peaks no back* 7
*removed edges* 14
*prop rem edges* 0.171

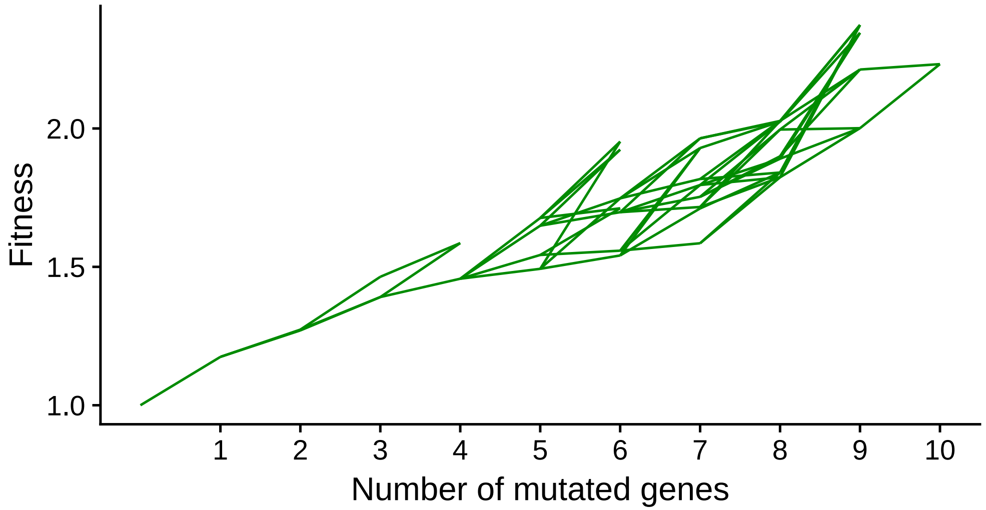


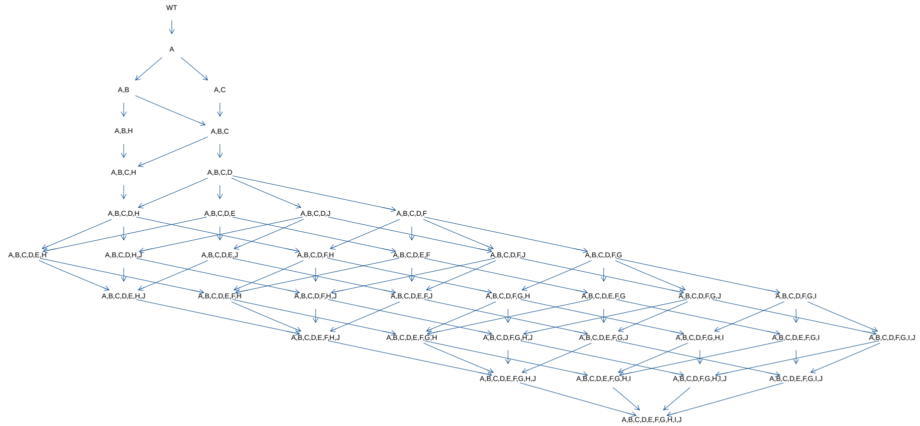

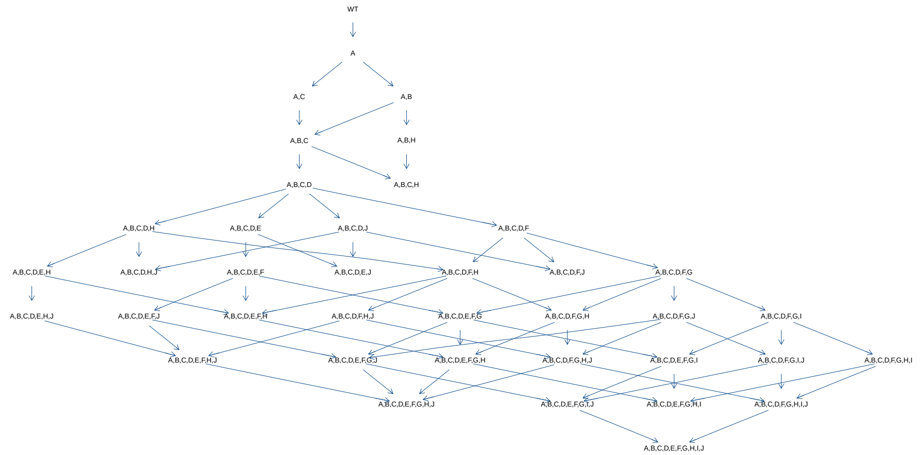

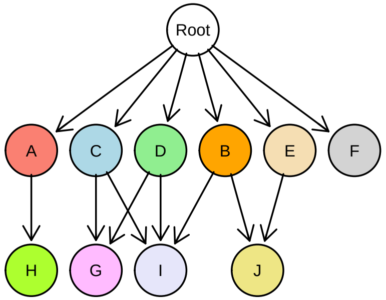


| | |
|---|---|
| *ID* | TQnQiOlfxYRUpqrU |
| *acc genotypes* | 186 |
| *peaks no back* | 13 |
| *removed edges* | 112 |
| *prop rem edges* | 0.188 |

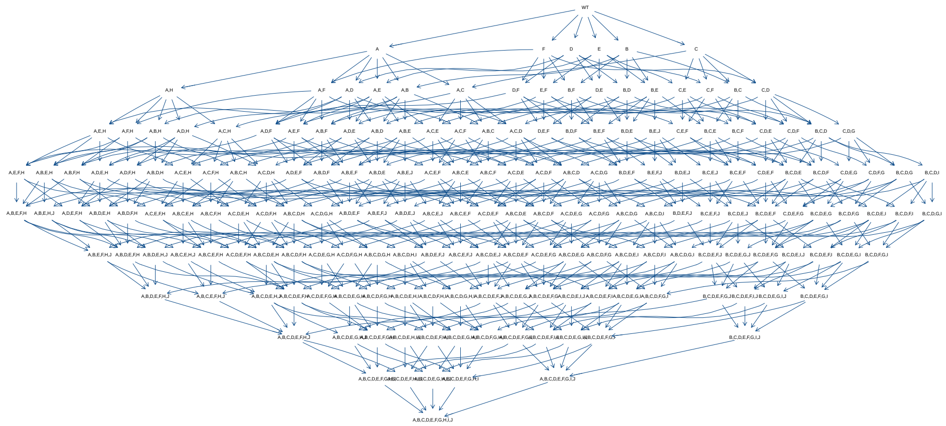

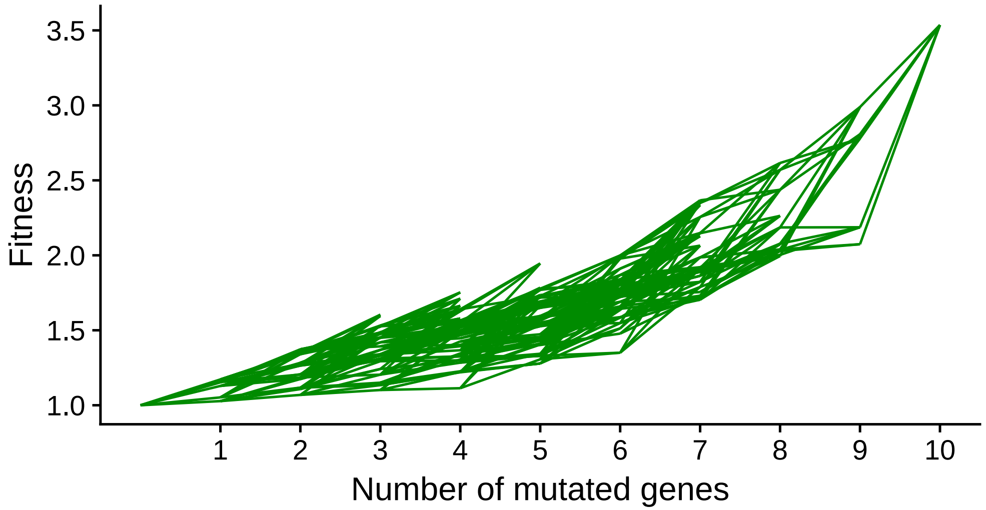


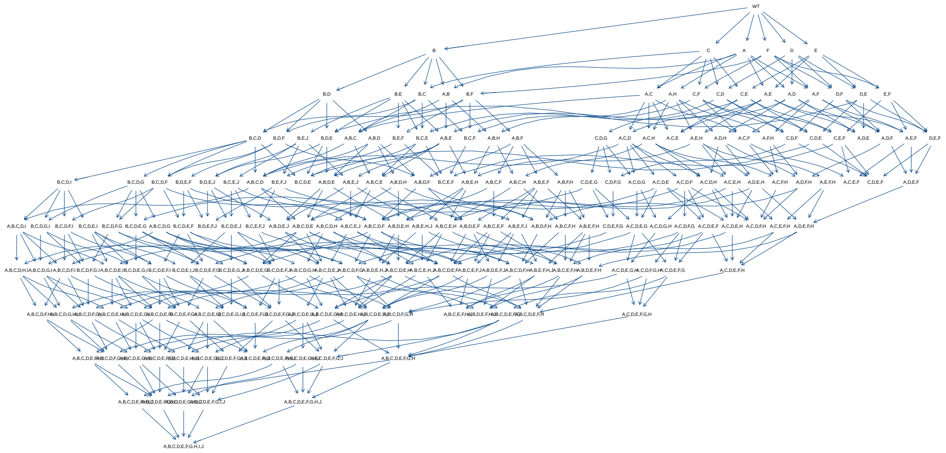

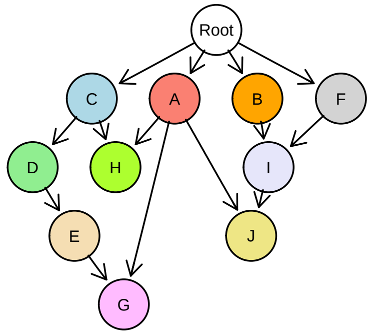


*ID* TVG98obaysdZVe7G
*acc genotypes* 74
*peaks no back* 8
*removed edges* 33
*prop rem edges* 0.189

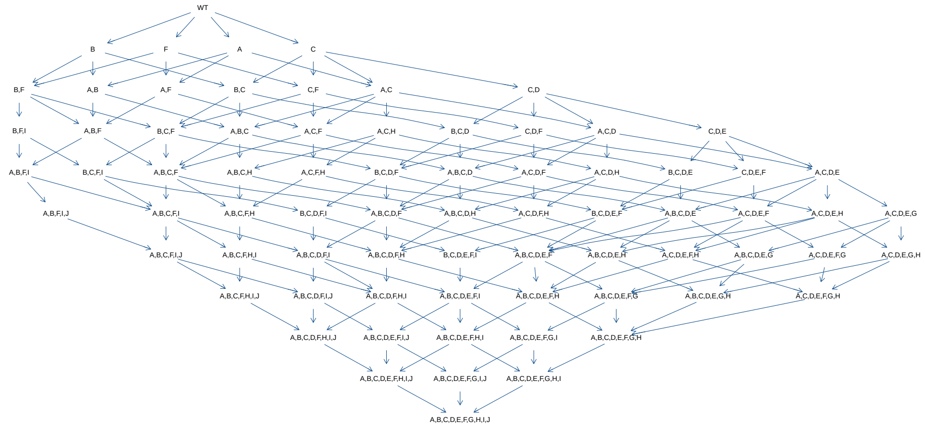

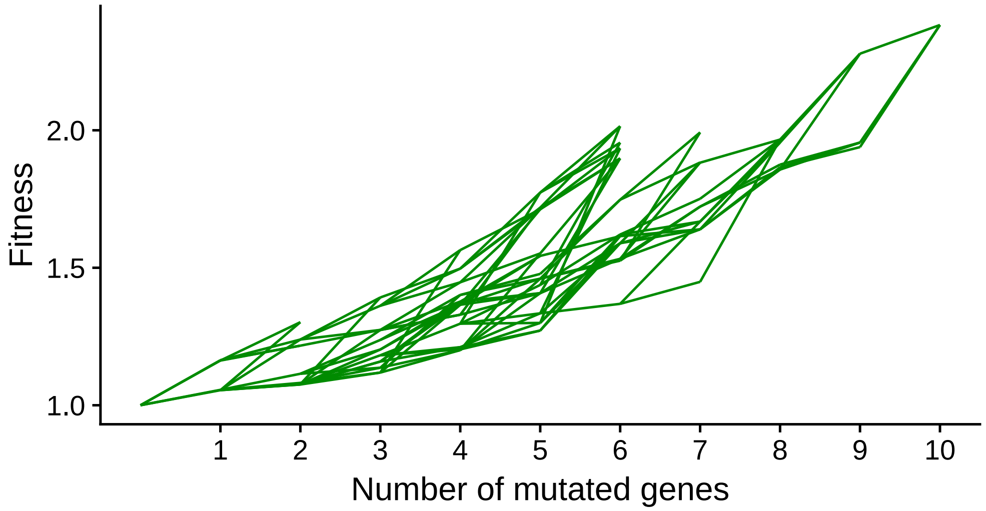


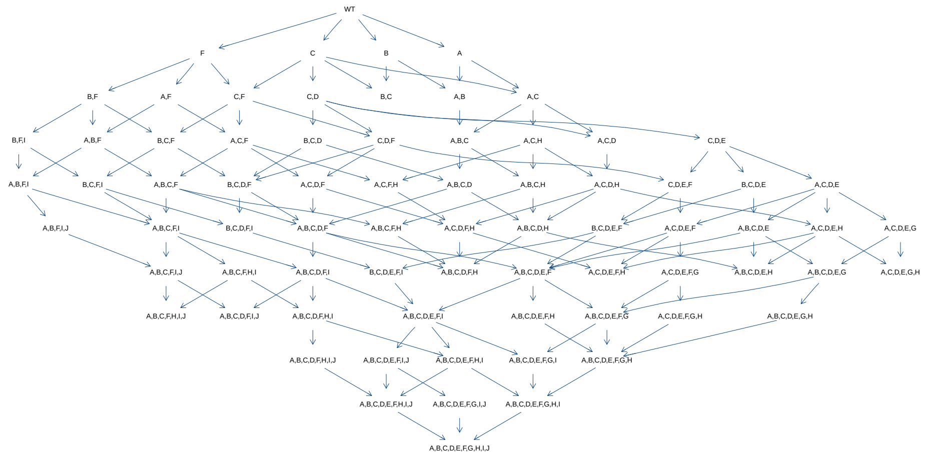

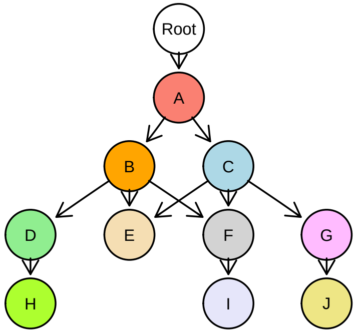


*ID* tytbSnHIzi26I77A
*acc genotypes* 62
*peaks no back* 7
*removed edges* 18
*prop rem edges* 0.122

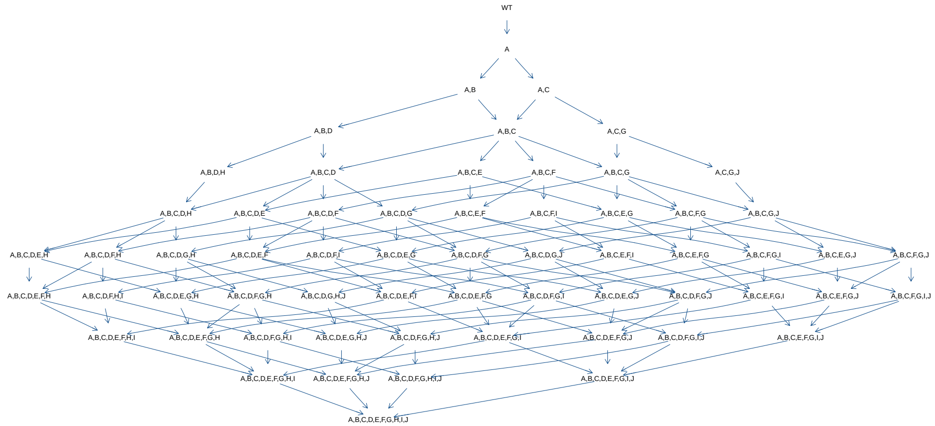

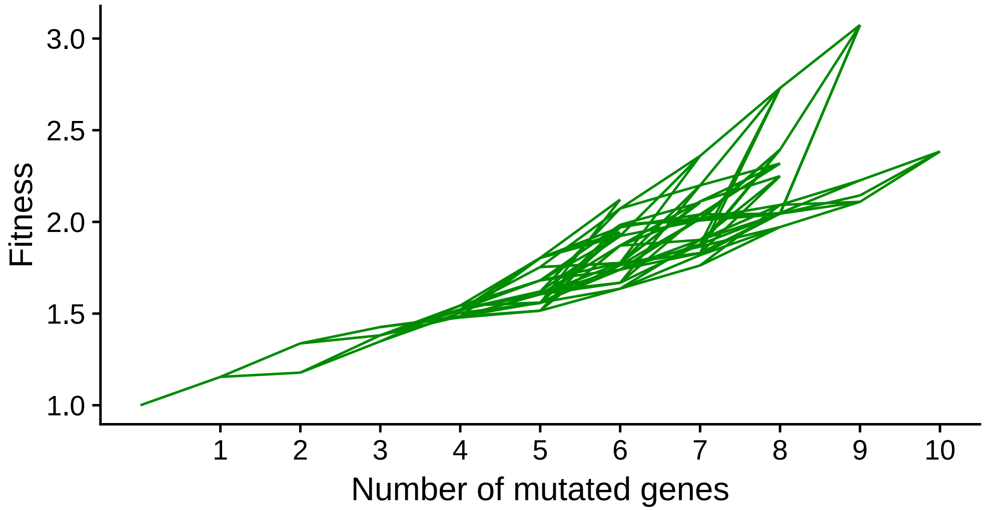


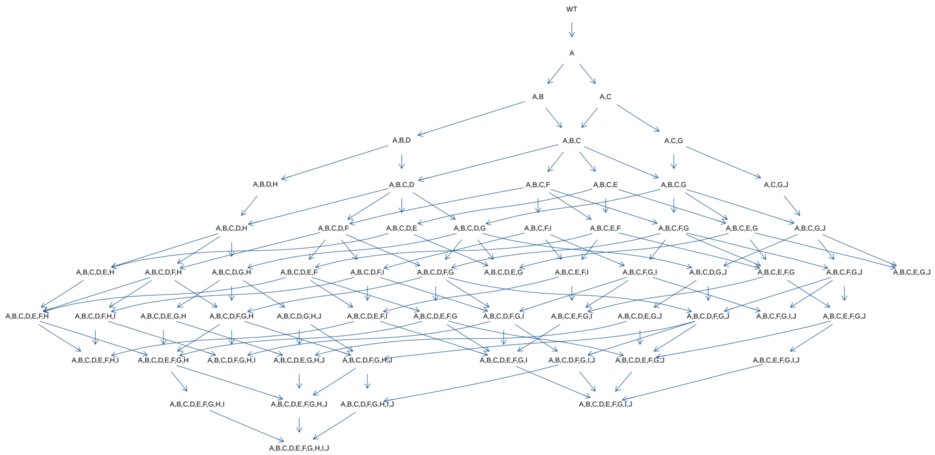

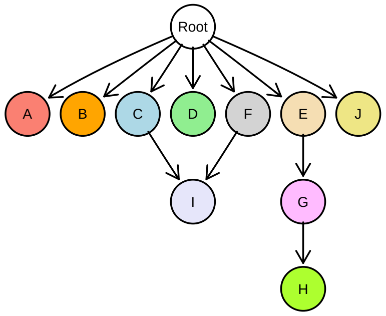


| | |
|---|---|
| *ID* | TZFzcsBmlMqGXCih |
| *acc genotypes* | 320 |
| *peaks no back* | 22 |
| *removed edges* | 231 |
| *prop rem edges* | 0.192 |

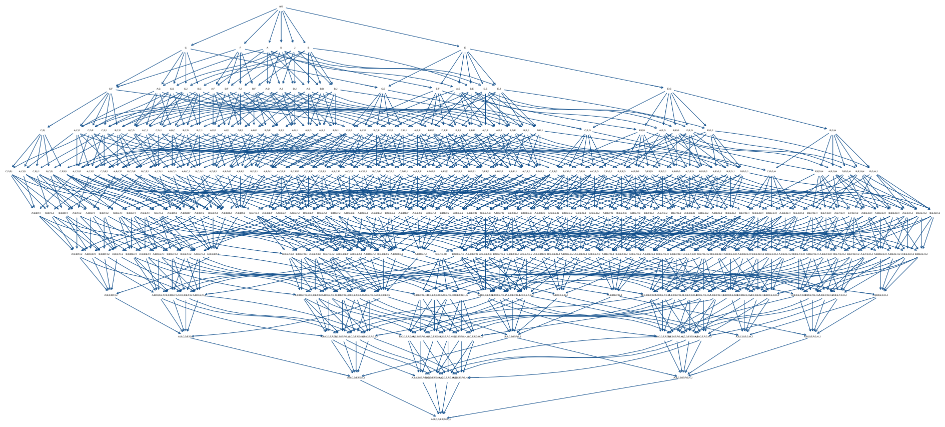

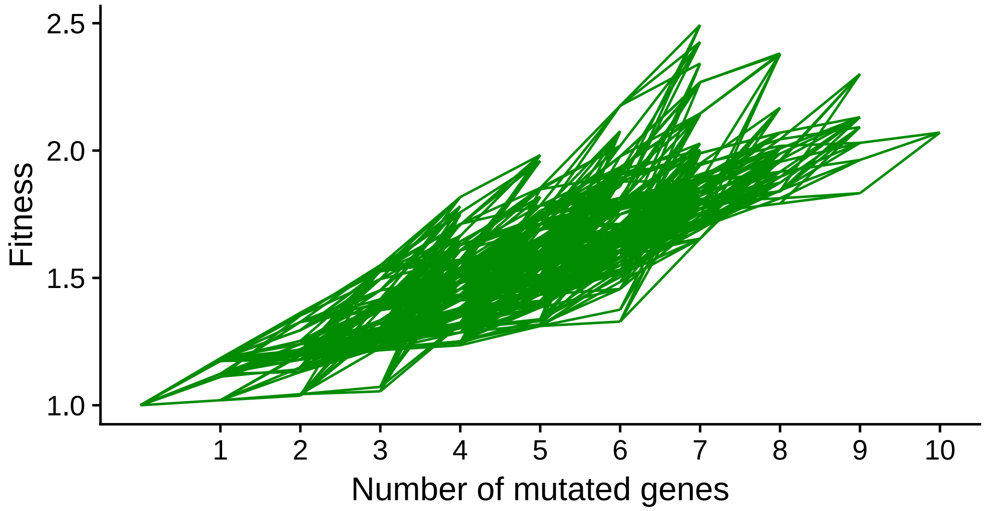


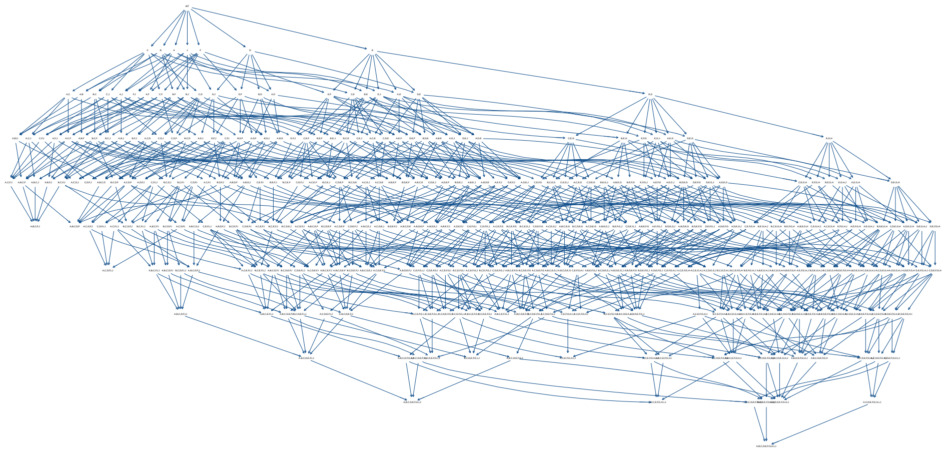

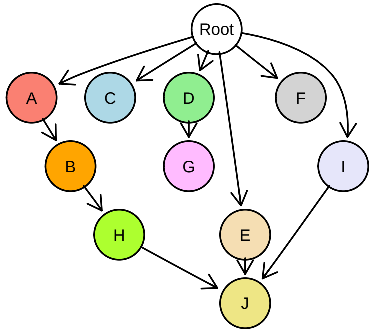


| | |
|---|---|
| *ID* | UBKJVjkElEDOKL7t |
| *acc genotypes* | 204 |
| *peaks no back* | 14 |
| *removed edges* | 117 |
| *prop rem edges* | 0.17 |

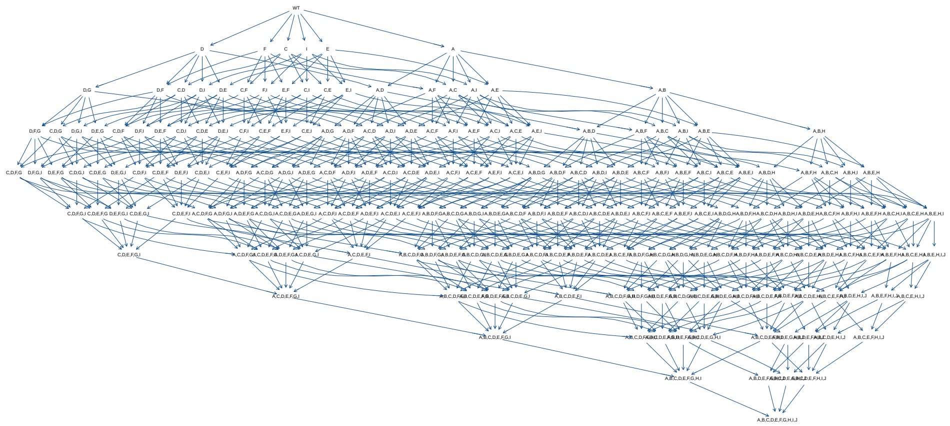

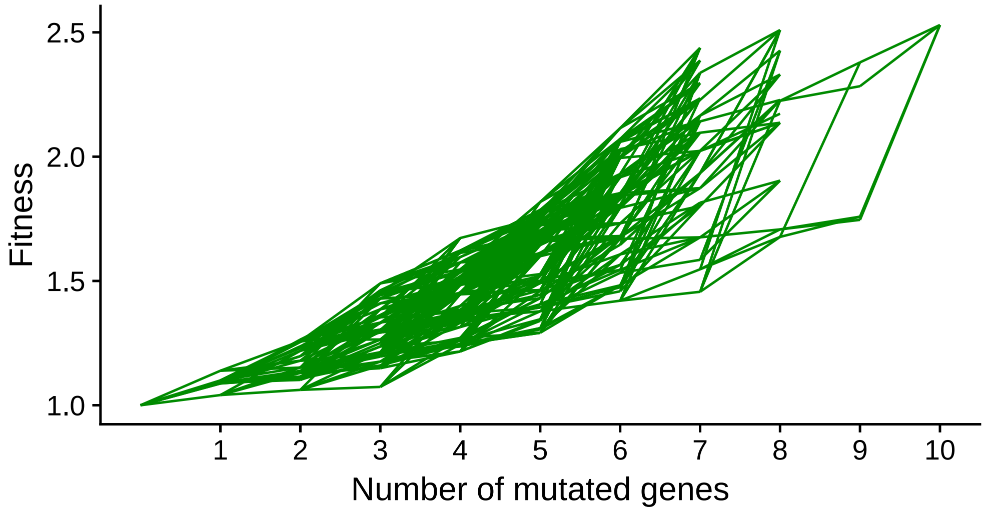


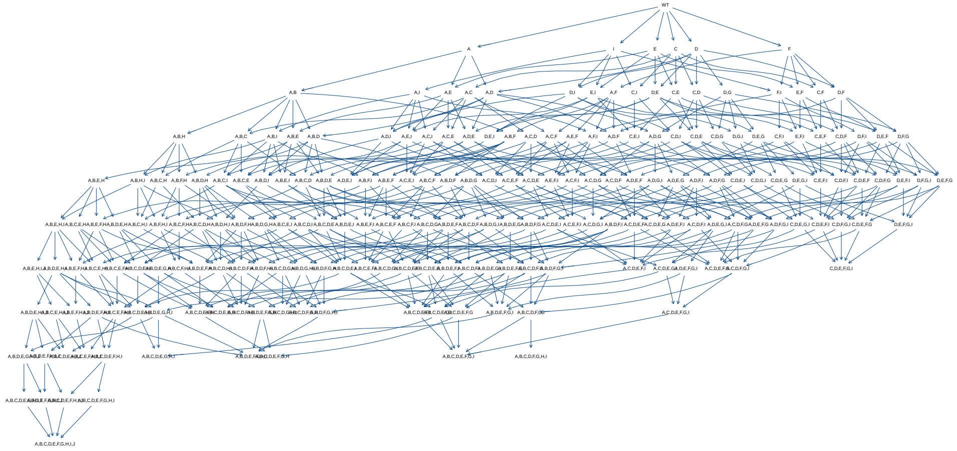

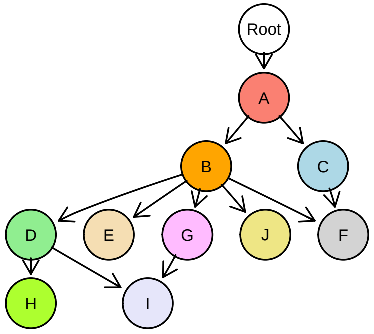


*ID* Ue10Y3AbLFXoQDQkX
*acc genotypes* 99
*peaks no back* 13
*removed edges* 62
*prop rem edges* 0.218

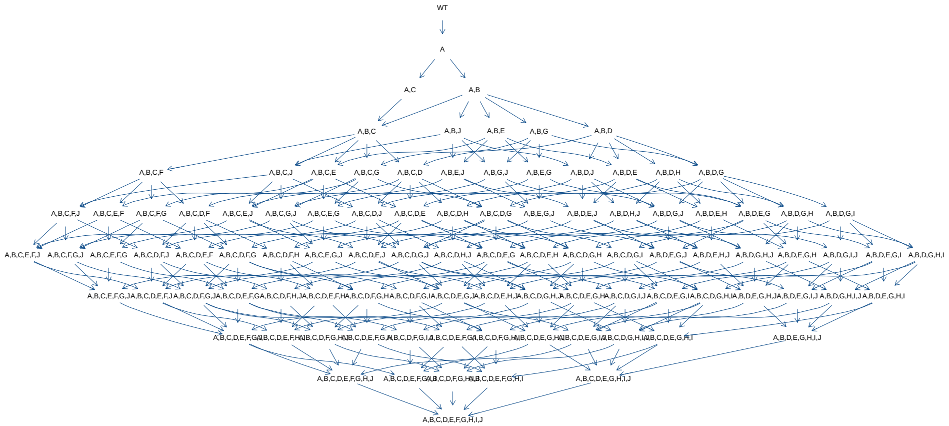

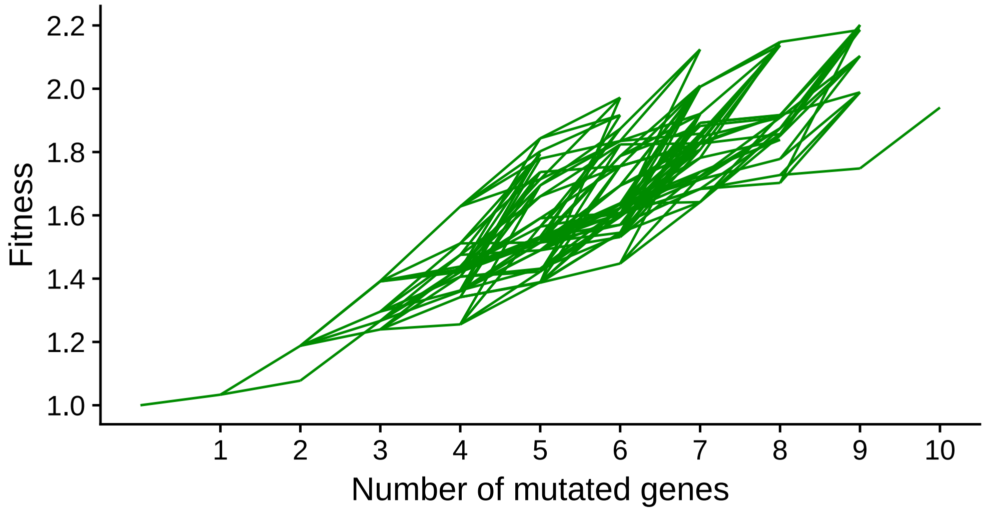


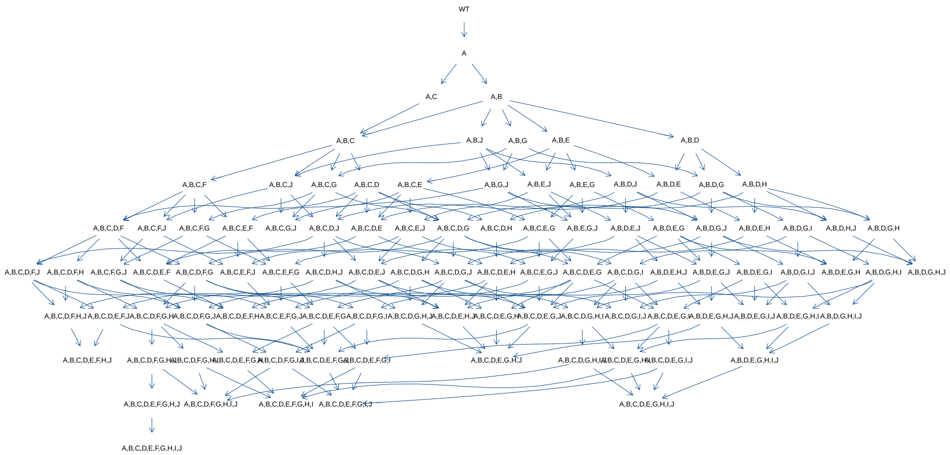

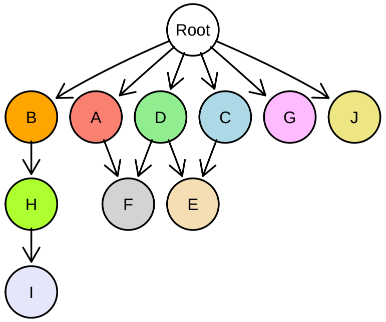


| | |
|---|---|
| *ID* | UEcQih9YNrZ2uptX |
| *acc genotypes* | 208 |
| *peaks no back* | 18 |
| *removed edges* | 127 |
| *prop rem edges* | 0.186 |

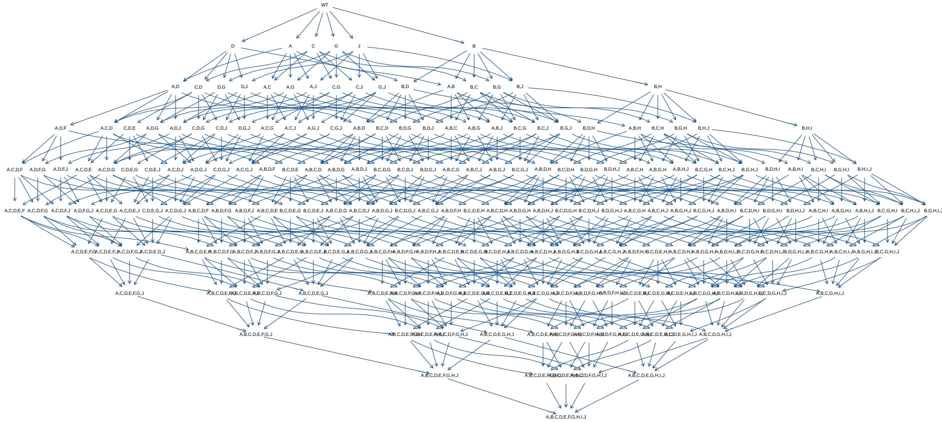

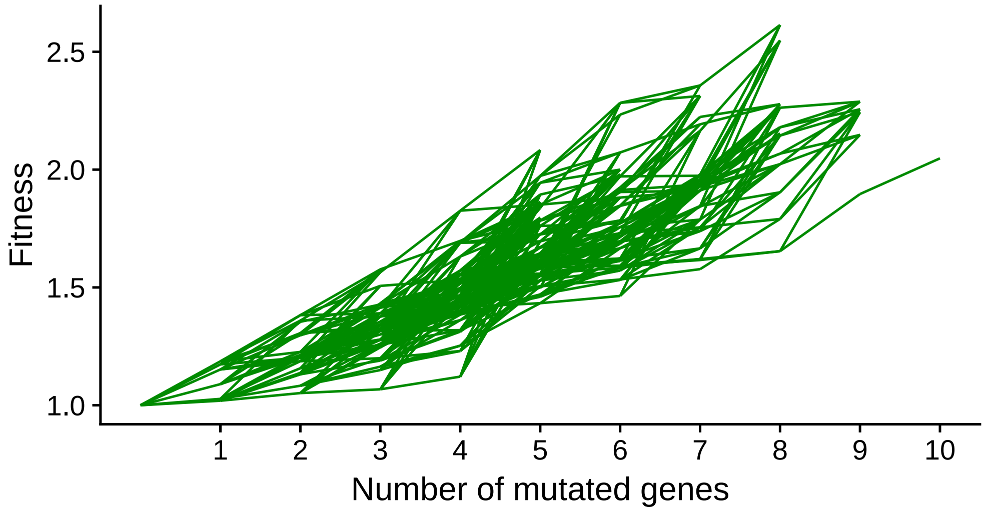


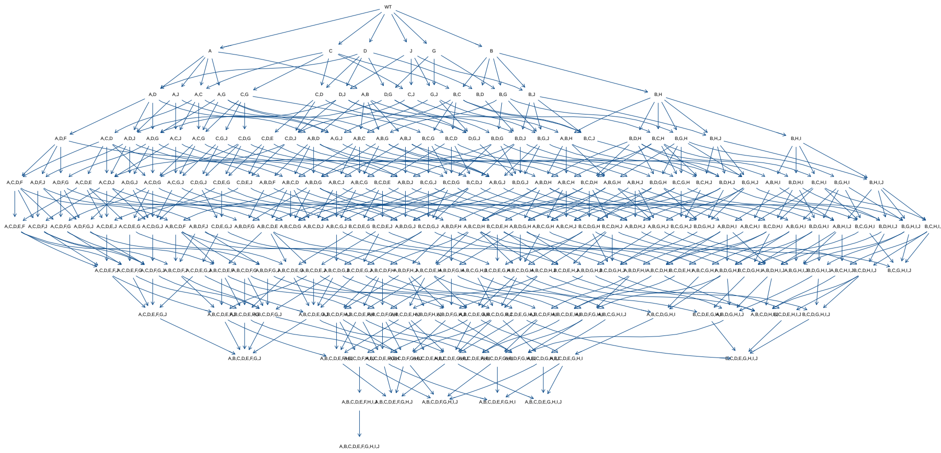

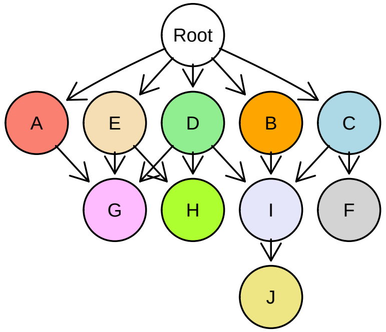


*ID* UHMIcOOkS2hFJbTg
*acc genotypes* 104
*peaks no back* 10
*removed edges* 51
*prop rem edges* 0.182

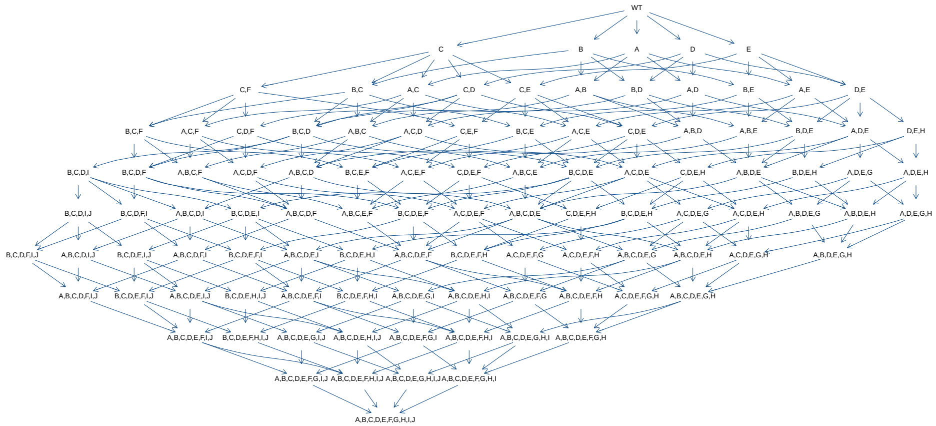

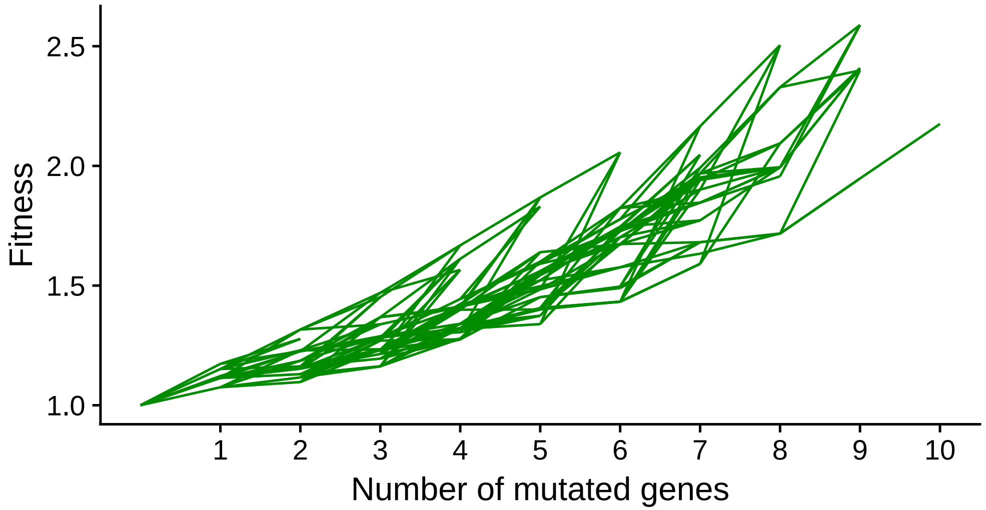


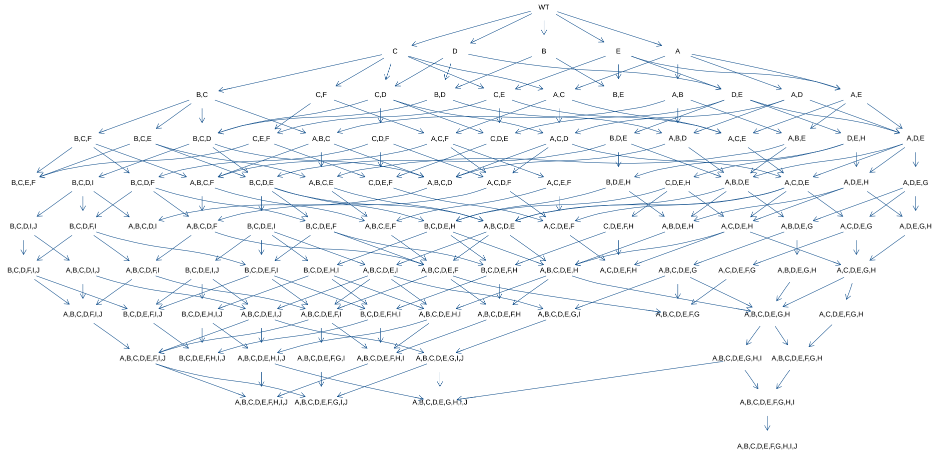

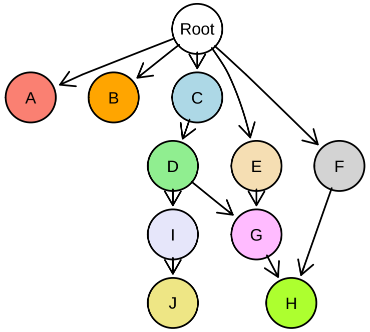


| | |
|---|---|
| *ID* | uKmhMNcyKeLl3N10d |
| *acc genotypes* | 116 |
| *peaks no back* | 12 |
| *removed edges* | 59 |
| *prop rem edges* | 0.178 |

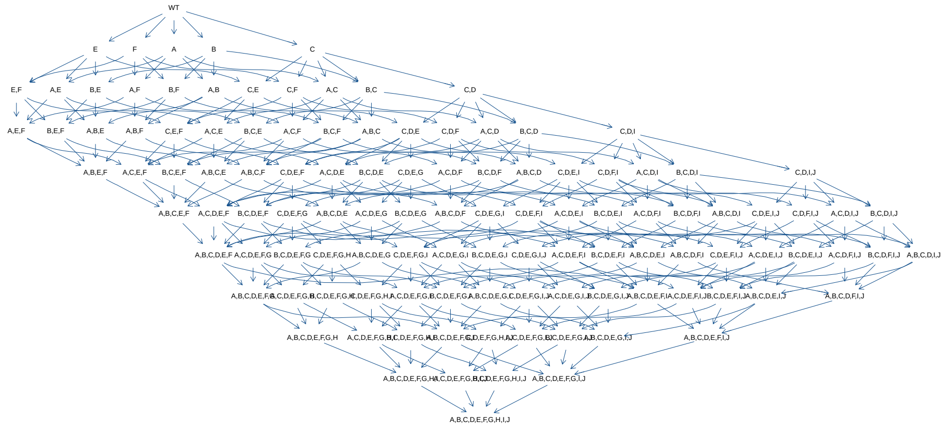


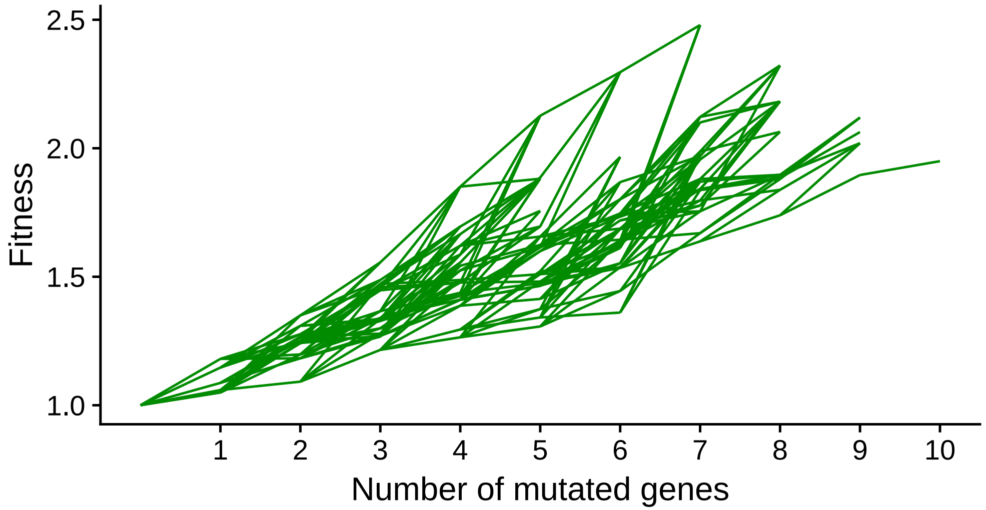


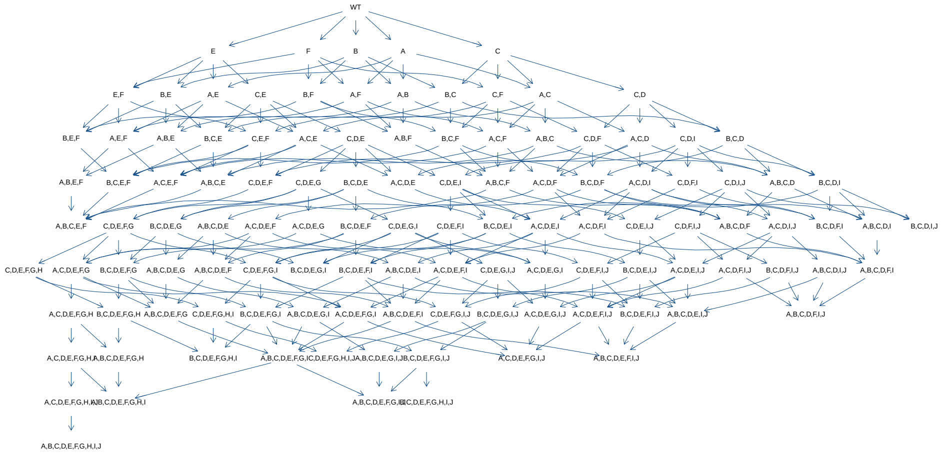

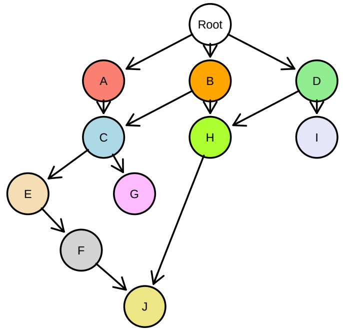


| | |
|---|---|
| *ID* | UoHyw1GDeCOAnn4H |
| *acc genotypes* | 50 |
| *peaks no back* | 6 |
| *removed edges* | 23 |
| *prop rem edges* | 0.217 |

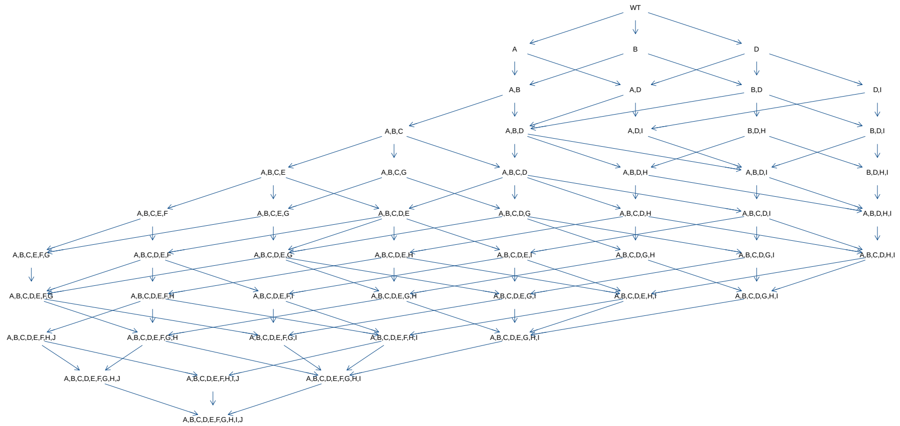

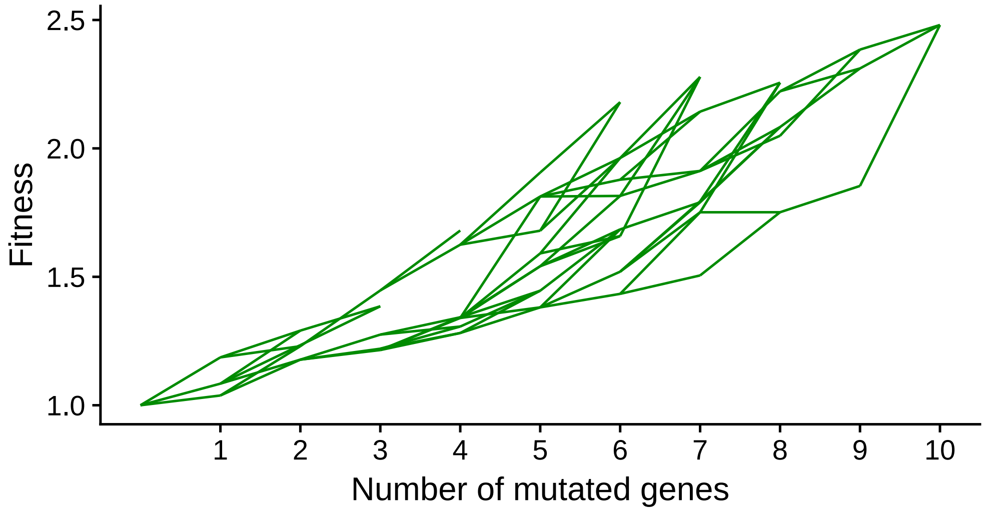


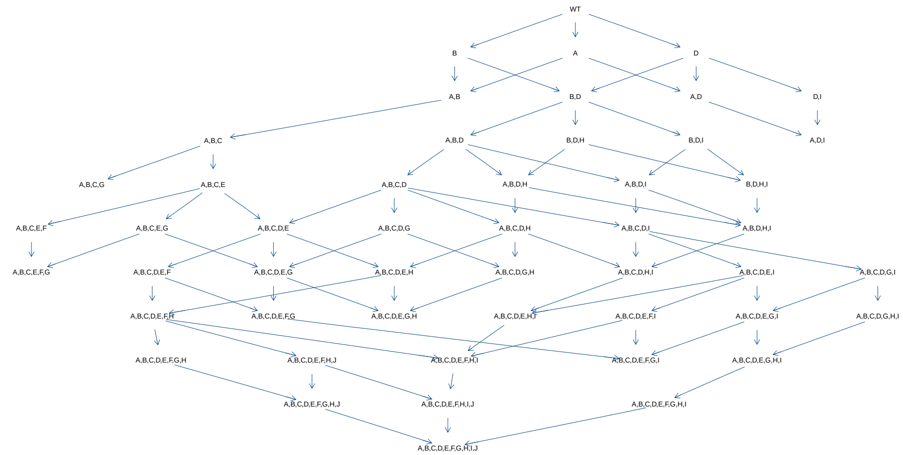

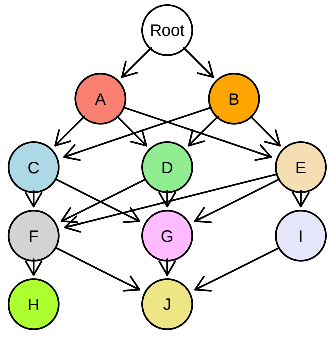


| | |
|---|---|
| *ID* | utFV4f4ZwaTtF6F7 |
| *acc genotypes* | 27 |
| *peaks no back* | 5 |
| *removed edges* | 7 |
| *prop rem edges* | 0.152 |

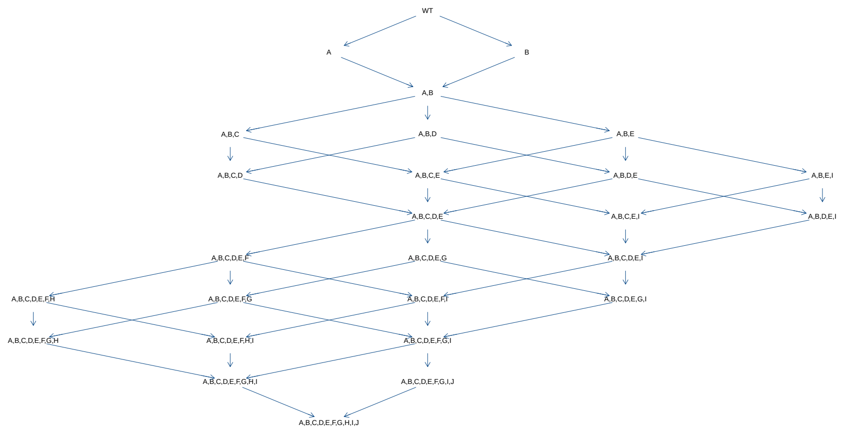

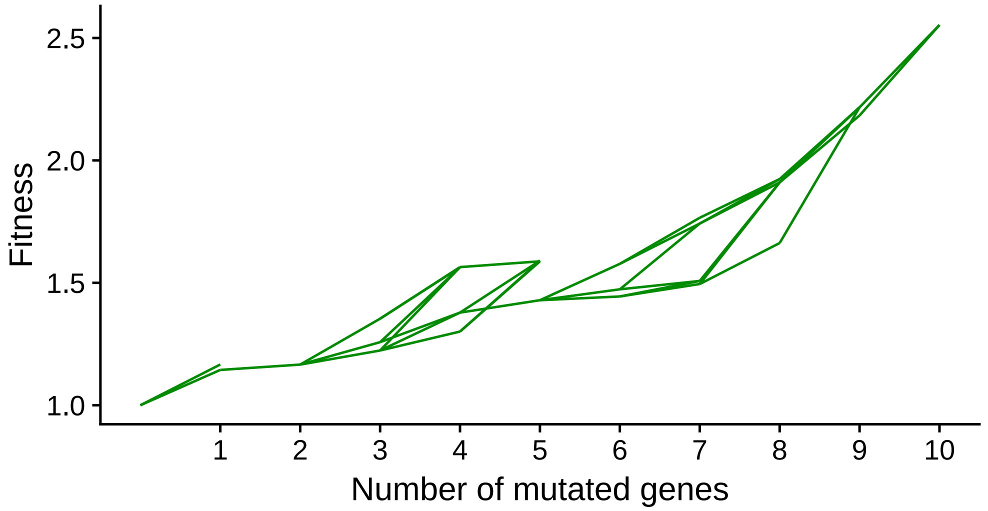


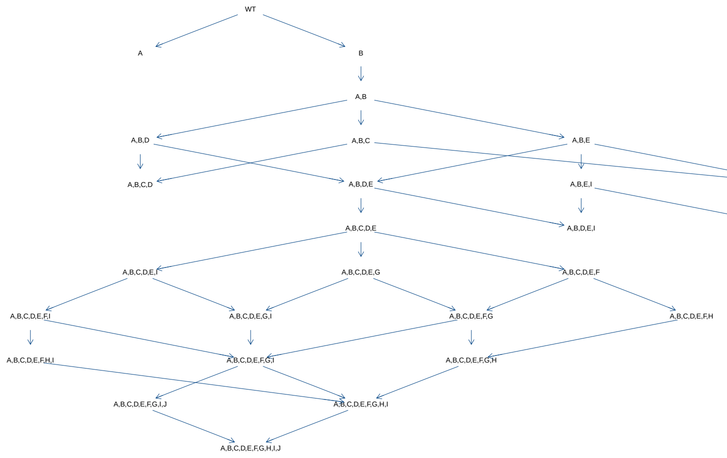

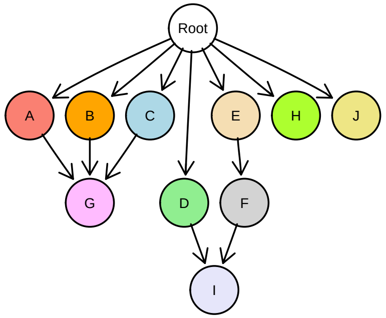


| | |
|---|---|
| *ID* | uzycnOVWW3JNvHXq |
| *acc genotypes* | 252 |
| *peaks no back* | 17 |
| *removed edges* | 162 |
| *prop rem edges* | 0.179 |

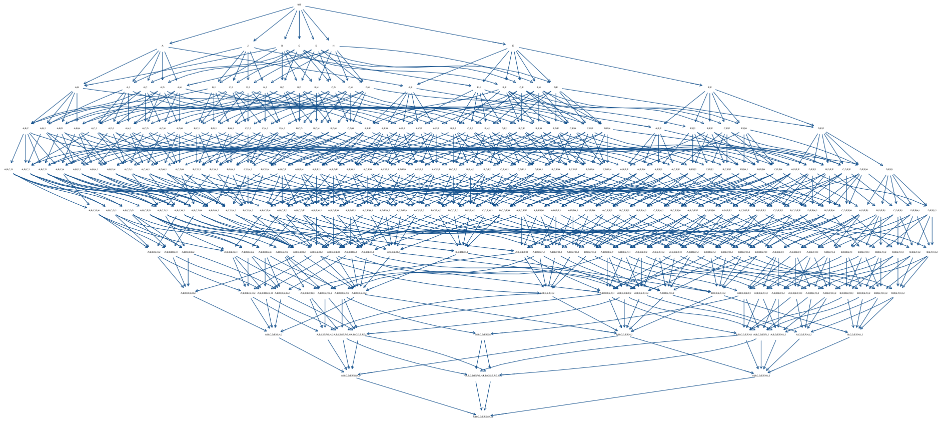

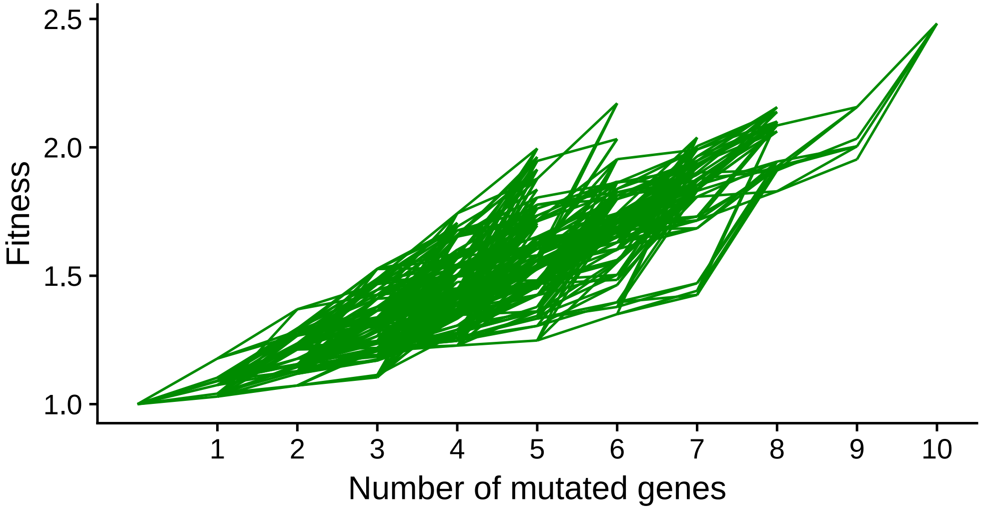


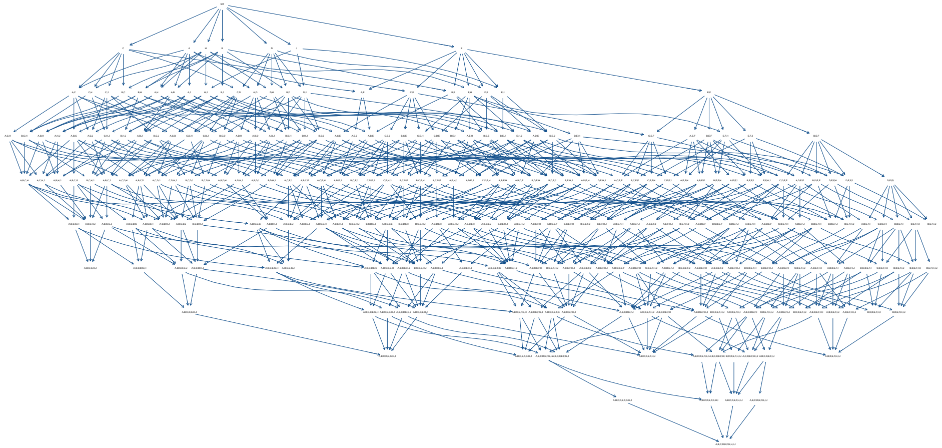

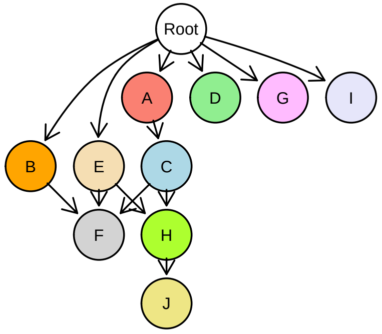


| | |
|---|---|
| *ID* | V3Myq6Ap7yTWkyC1 |
| *acc genotypes* | 152 |
| *peaks no back* | 19 |
| *removed edges* | 116 |
| *prop rem edges* | 0.244 |

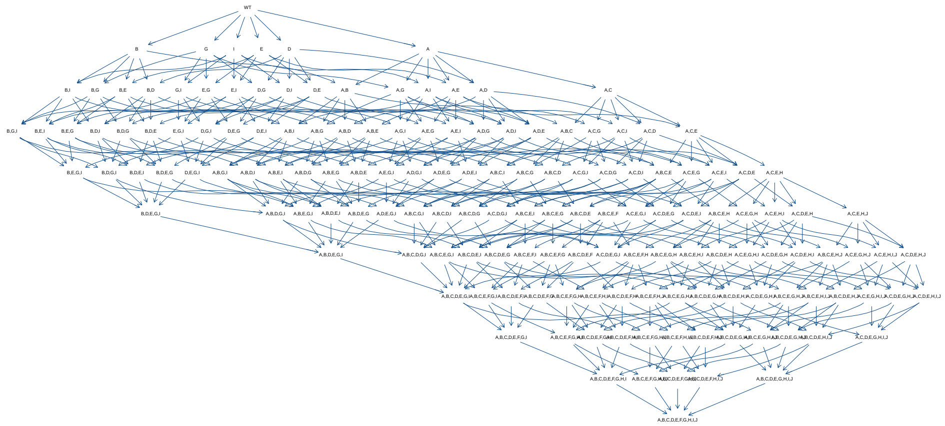

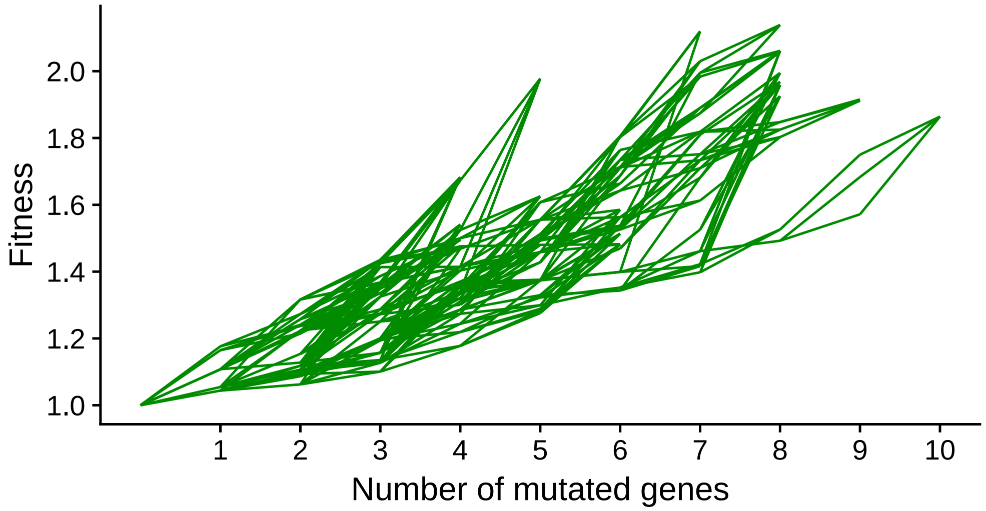


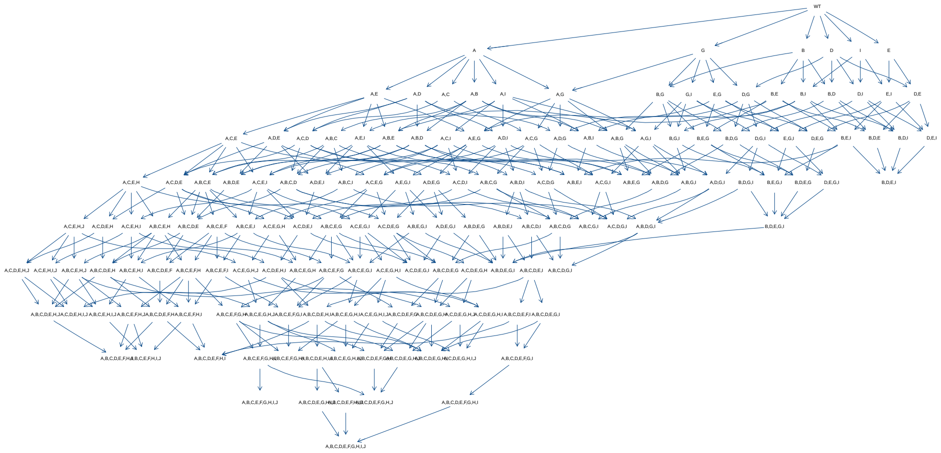

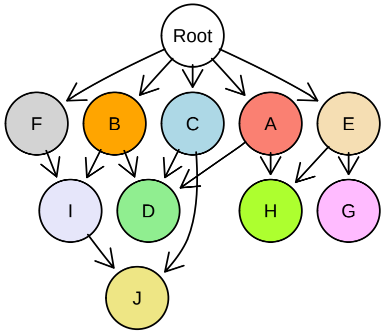


*ID* v3T1IG10jXOD5TPVs
*acc genotypes* 108
*peaks no back* 11
*removed edges* 62
*prop rem edges* 0.212

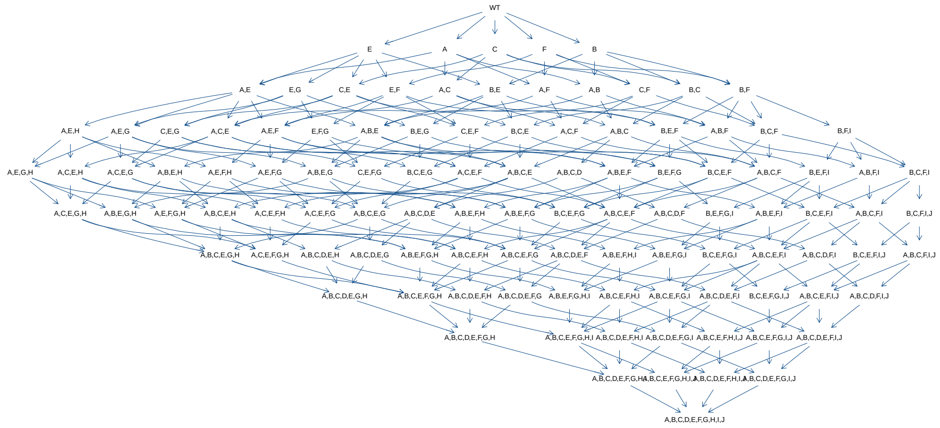

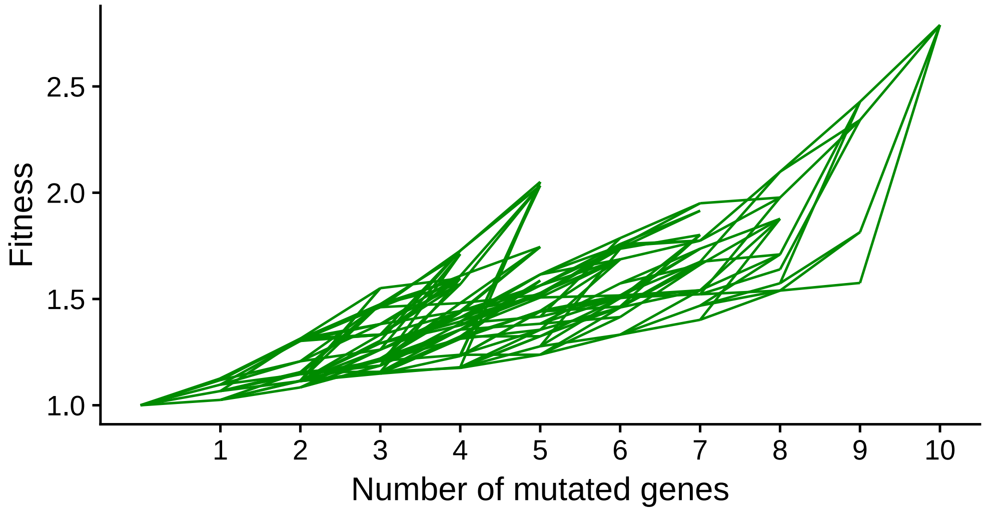


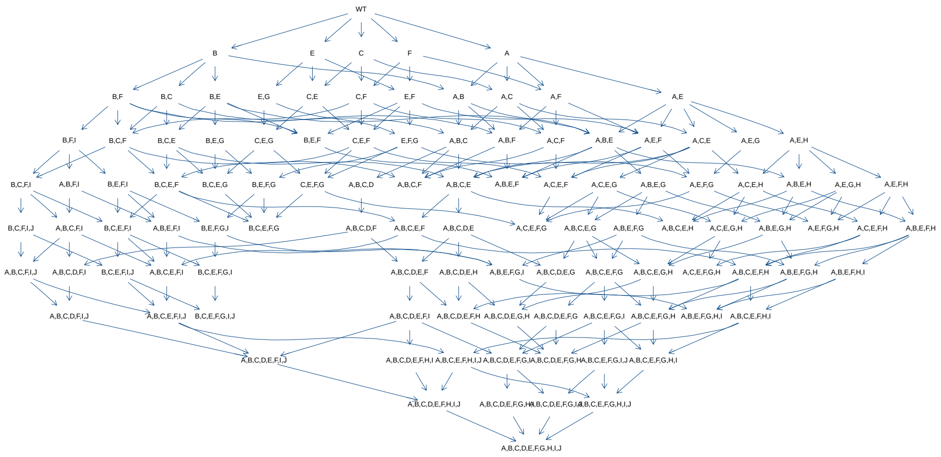

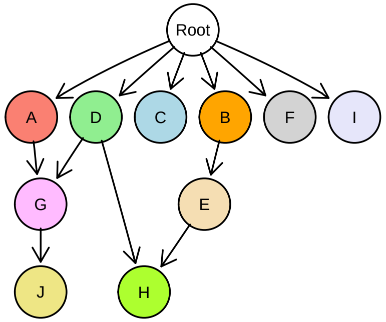


| | |
|---|---|
| *ID* | vDscwX1MSzAXnUjn |
| *acc genotypes* | 176 |
| *peaks no back* | 14 |
| *removed edges* | 95 |
| *prop rem edges* | 0.17 |

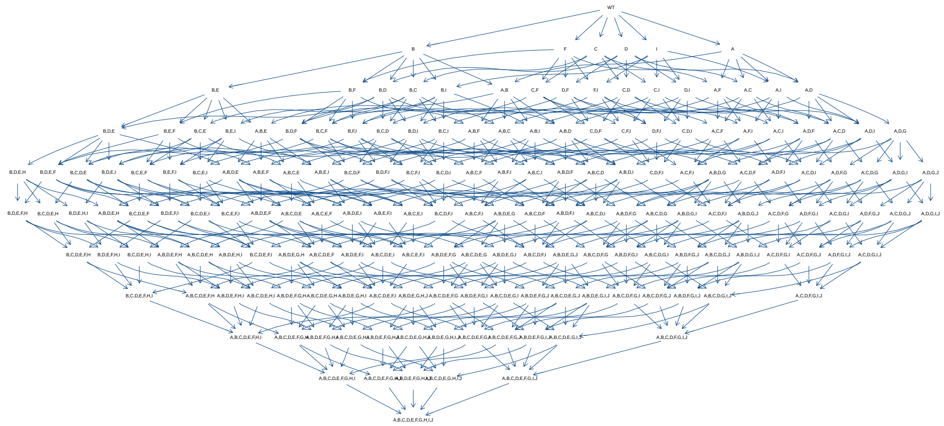

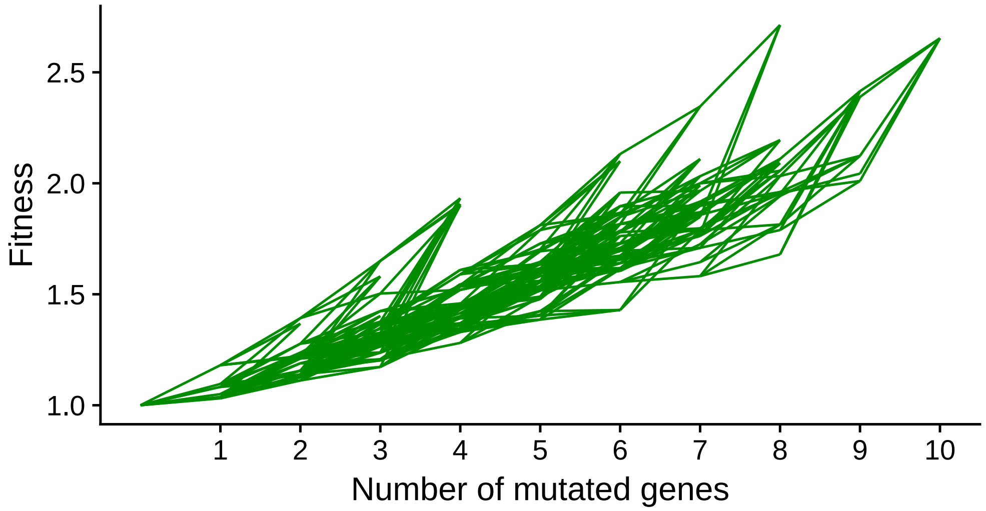


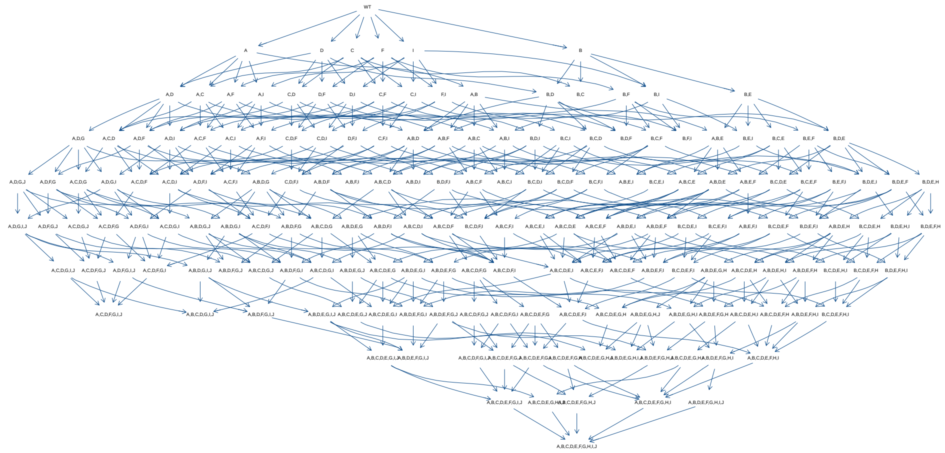

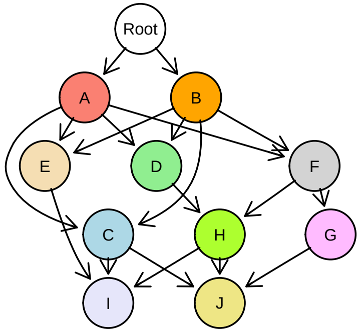


*ID* veSOPRnIsirW3tXS

*acc genotypes* 40

*peaks no back* 7

*removed edges* 18

*prop rem edges* 0.214

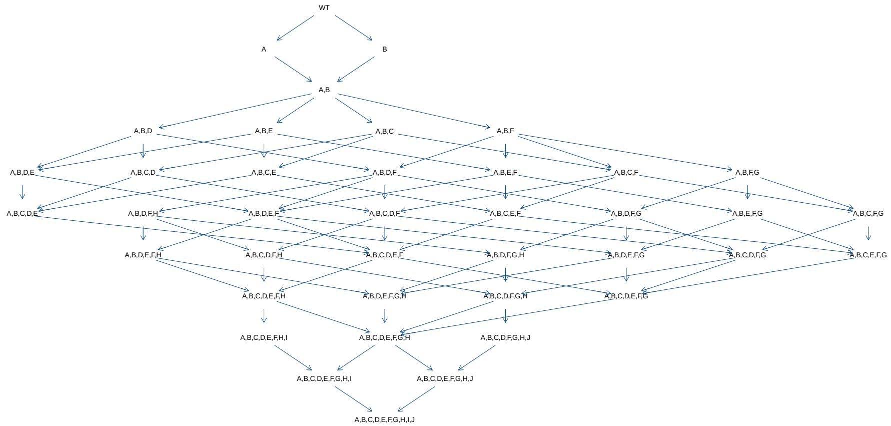

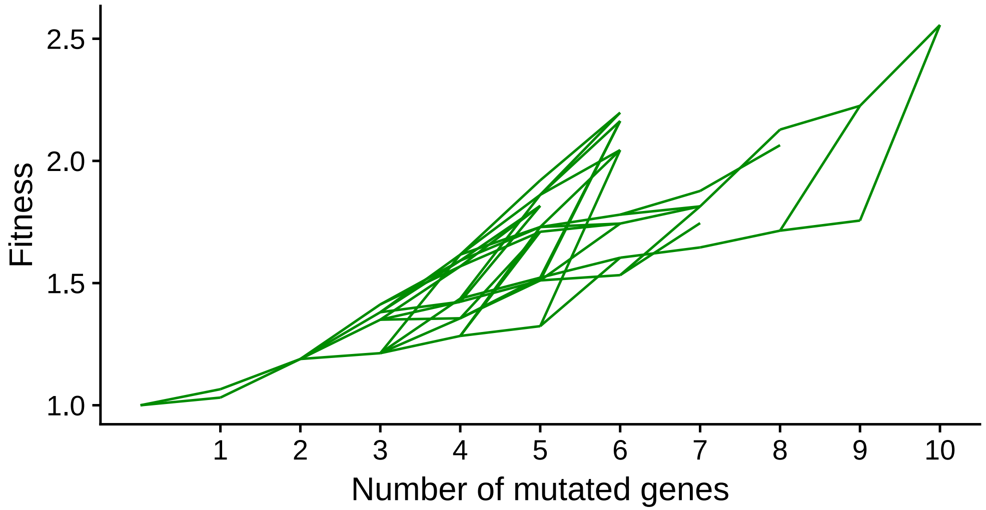


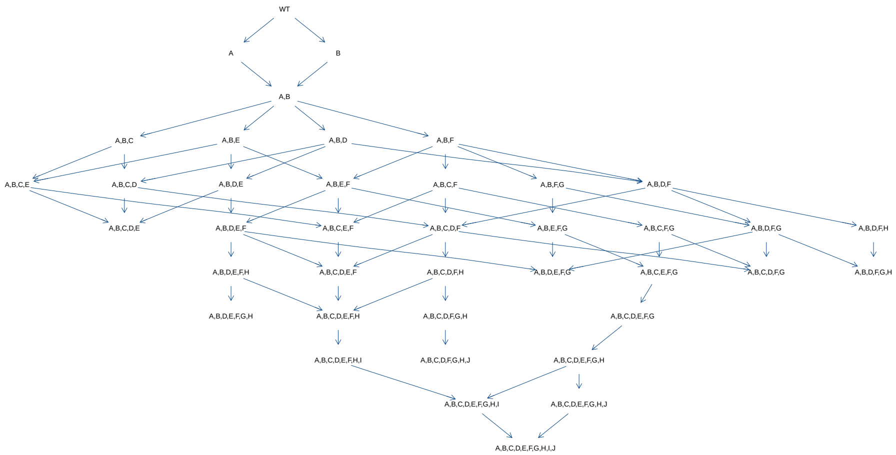

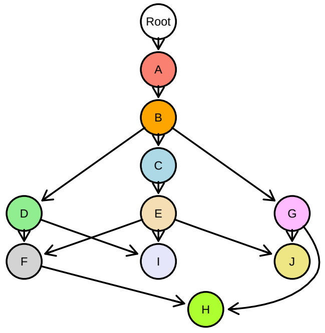


| | |
|---|---|
| *ID* | vm585SzZroFfErOI |
| *acc genotypes* | 29 |
| *peaks no back* | 2 |
| *removed edges* | 3 |
| *prop rem edges* | 0.059 |

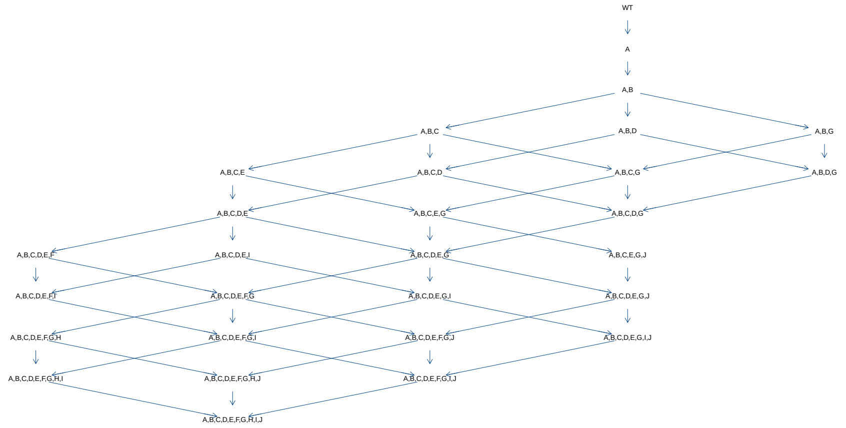

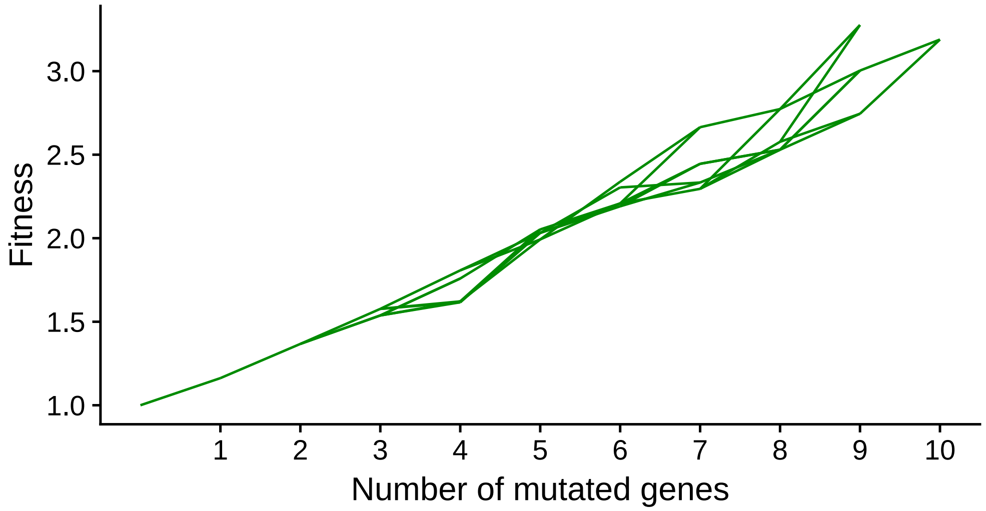


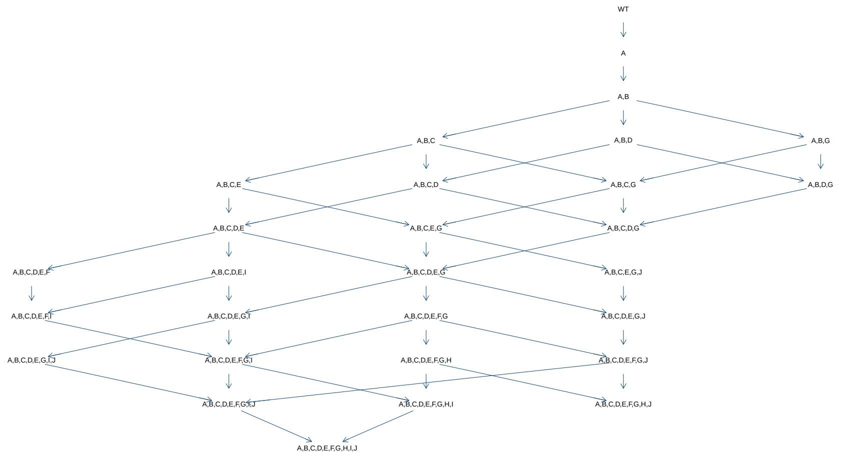

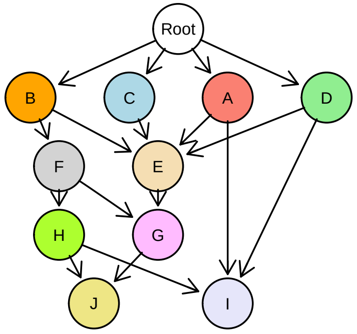


| | |
|---|---|
| *ID* | VNq9x3A103vjj6H6C |
| *acc genotypes* | 43 |
| *peaks no back* | 6 |
| *removed edges* | 18 |
| *prop rem edges* | 0.2 |

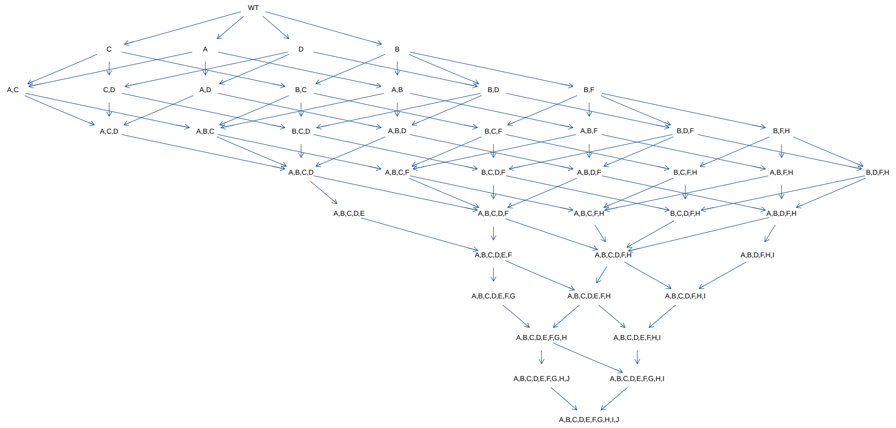

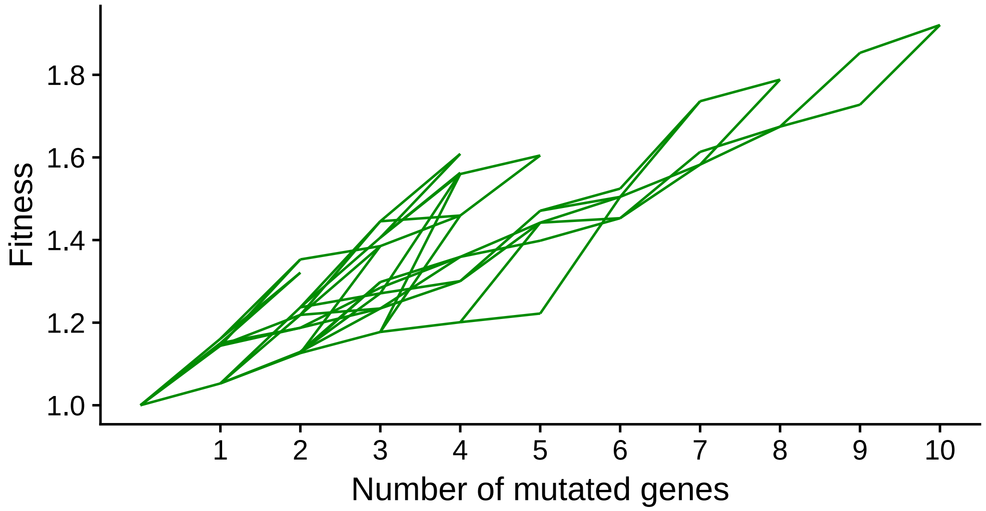


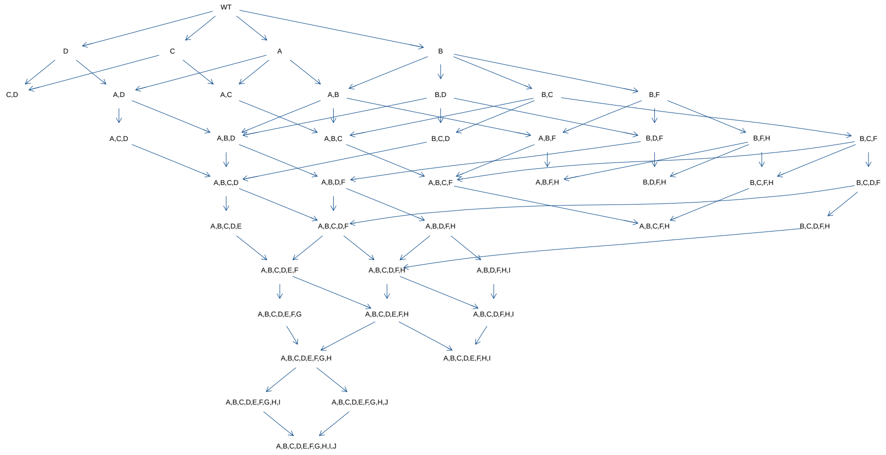

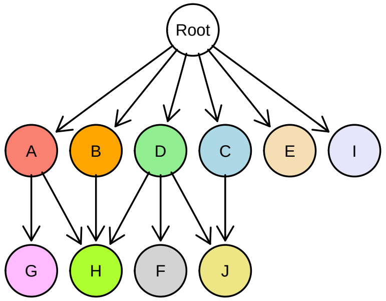


| | |
|---|---|
| *ID* | w2gKASMJutkLe4I3 |
| *acc genotypes* | 240 |
| *peaks no back* | 20 |
| *removed edges* | 158 |
| *prop rem edges* | 0.19 |

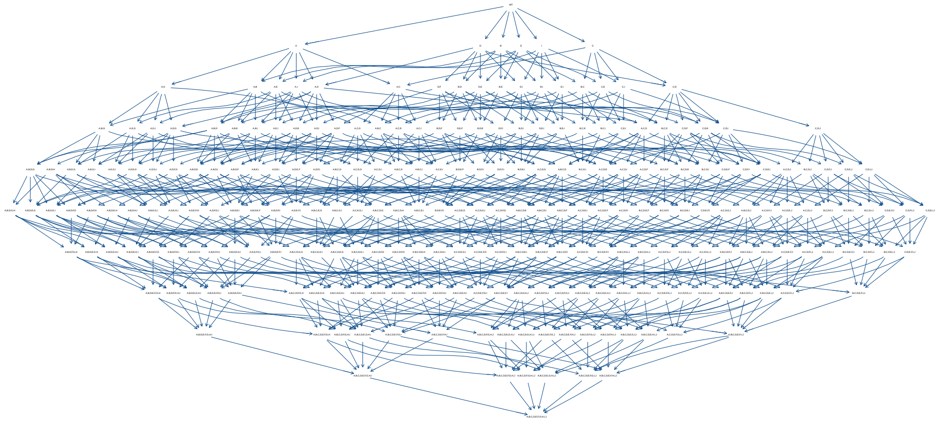

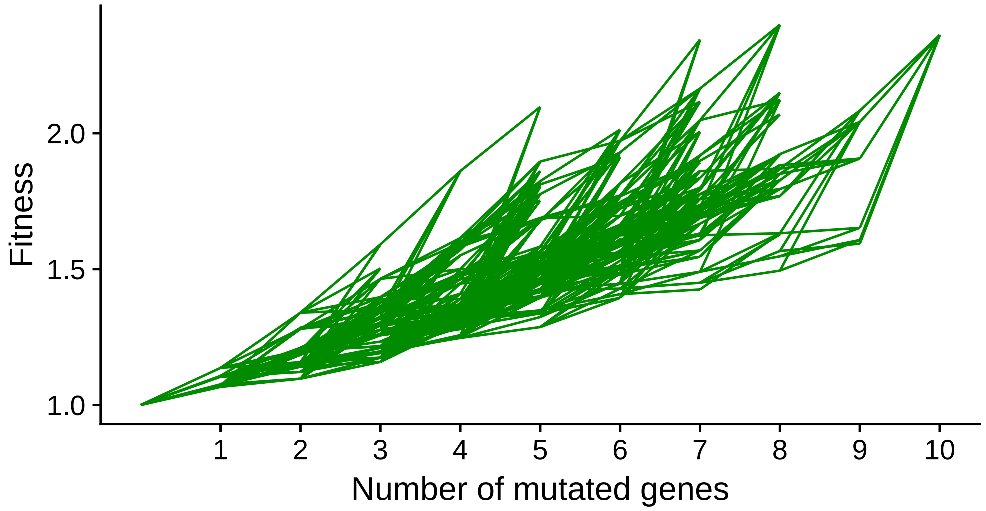


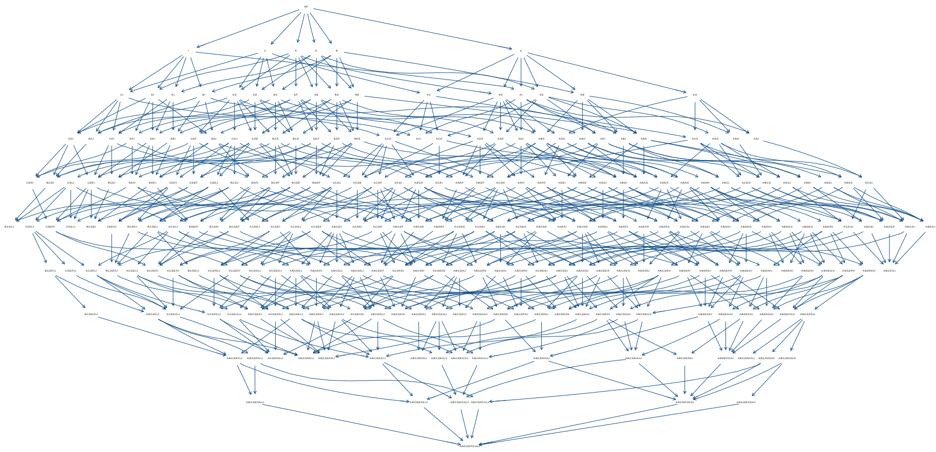

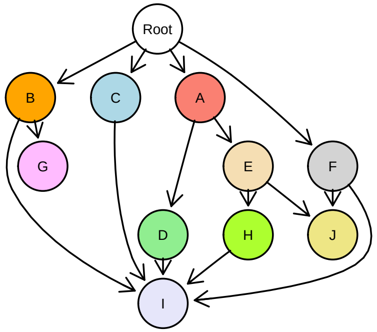


| | |
|---|---|
| *ID* | wavyEmnct10zXrBsJ |
| *acc genotypes* | 112 |
| *peaks no back* | 11 |
| *removed edges* | 68 |
| *prop rem edges* | 0.212 |

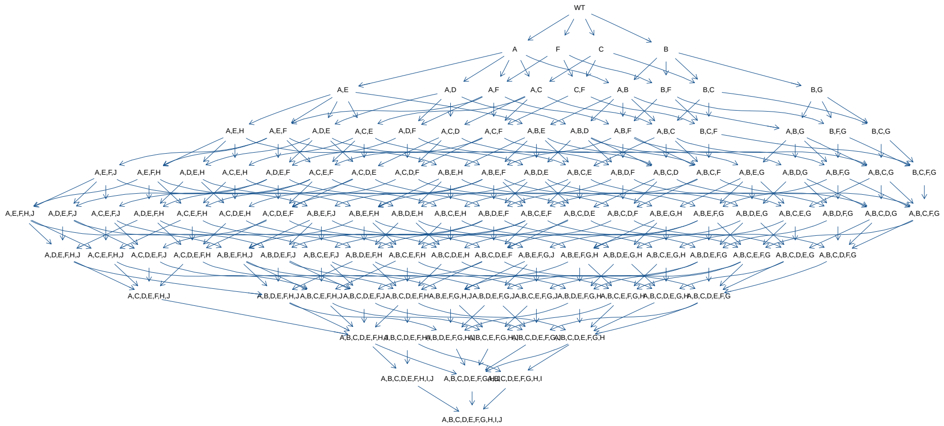

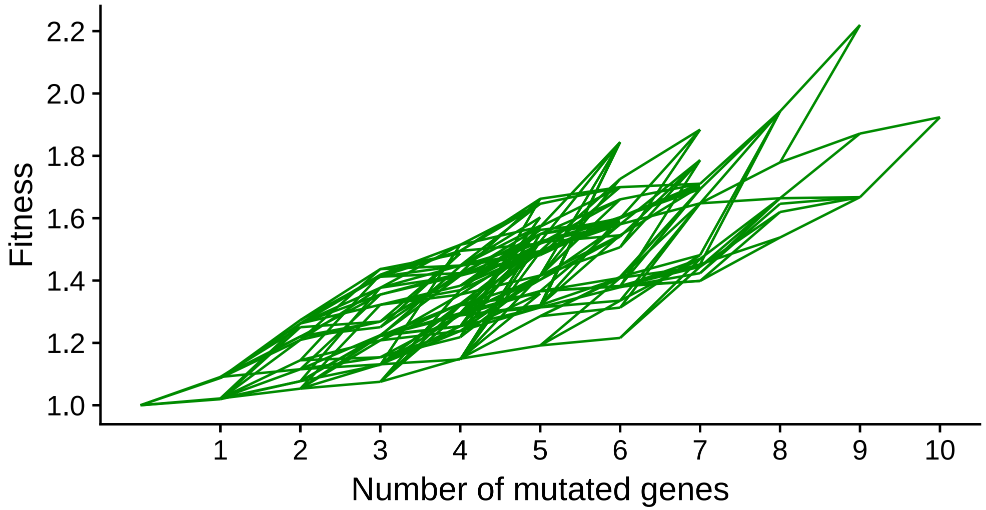


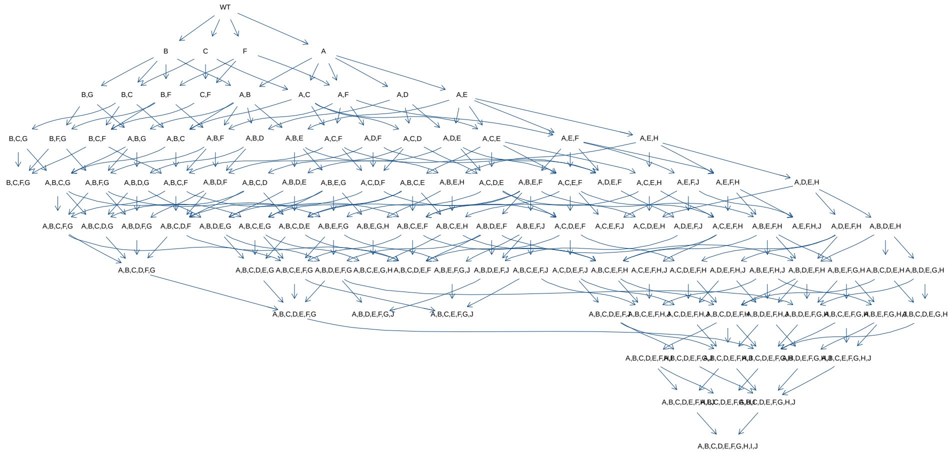

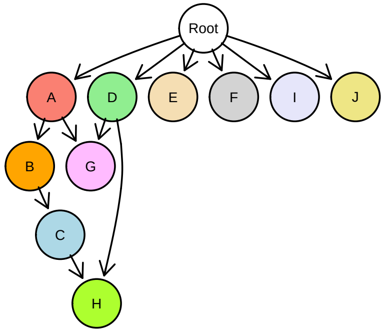


| | |
|---|---|
| *ID* | wcH6gWqLOjM6KUvj |
| *acc genotypes* | 208 |
| *peaks no back* | 21 |
| *removed edges* | 145 |
| *prop rem edges* | 0.206 |

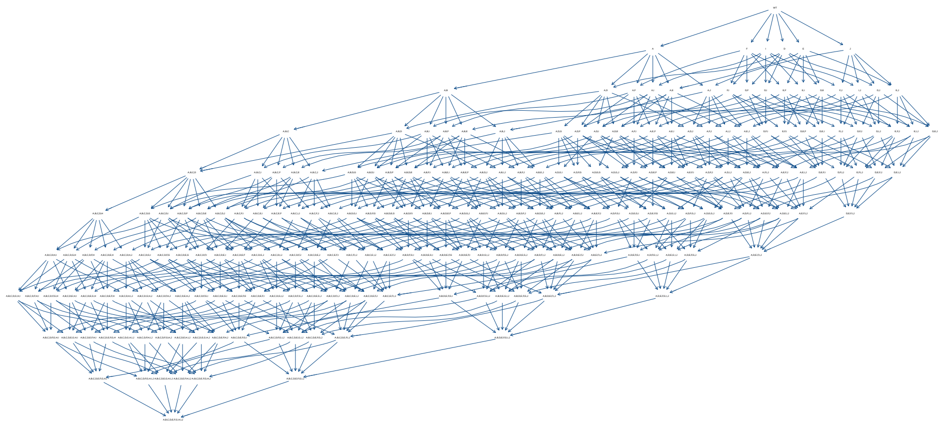

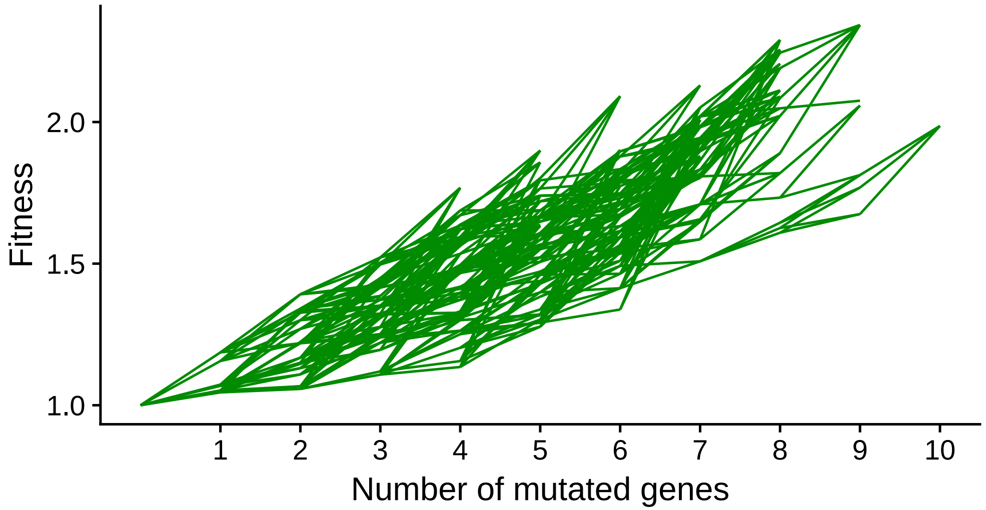


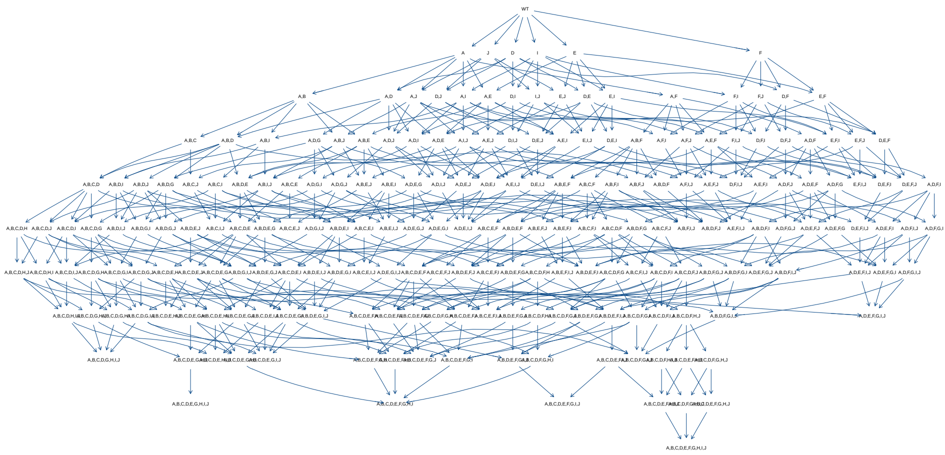

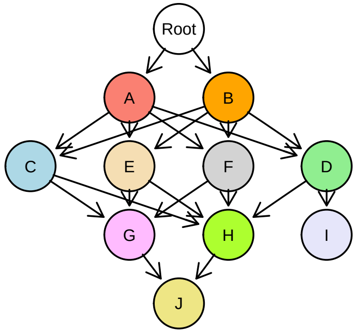


| | |
|---|---|
| *ID* | wCi29Nr110GfPJ9ih |
| *acc genotypes* | 36 |
| *peaks no back* | 6 |
| *removed edges* | 10 |
| *prop rem edges* | 0.139 |

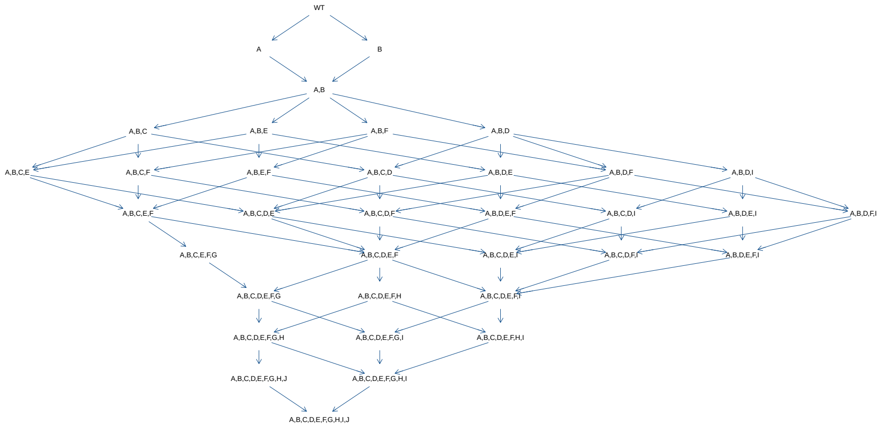

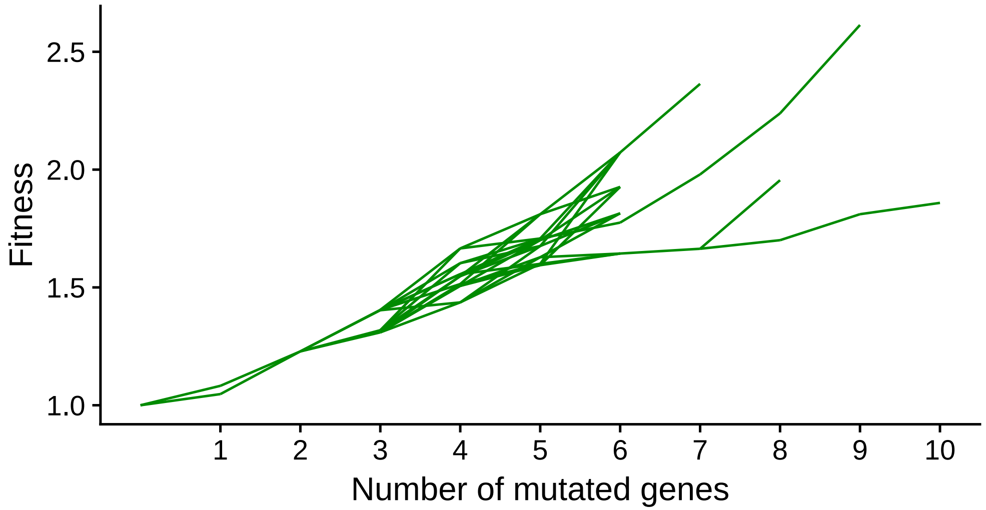


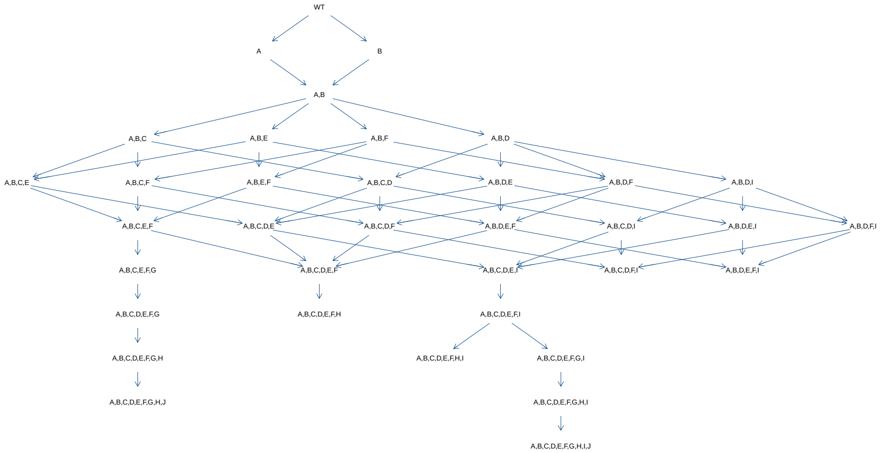

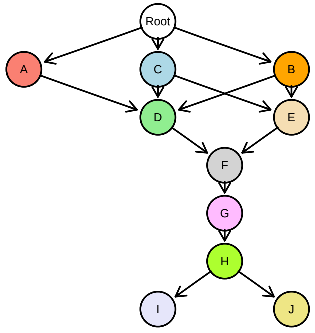


| | |
|---|---|
| *ID* | WMUVgmnyiqxYFBg6 |
| *acc genotypes* | 18 |
| *peaks no back* | 2 |
| *removed edges* | 2 |
| *prop rem edges* | 0.08 |

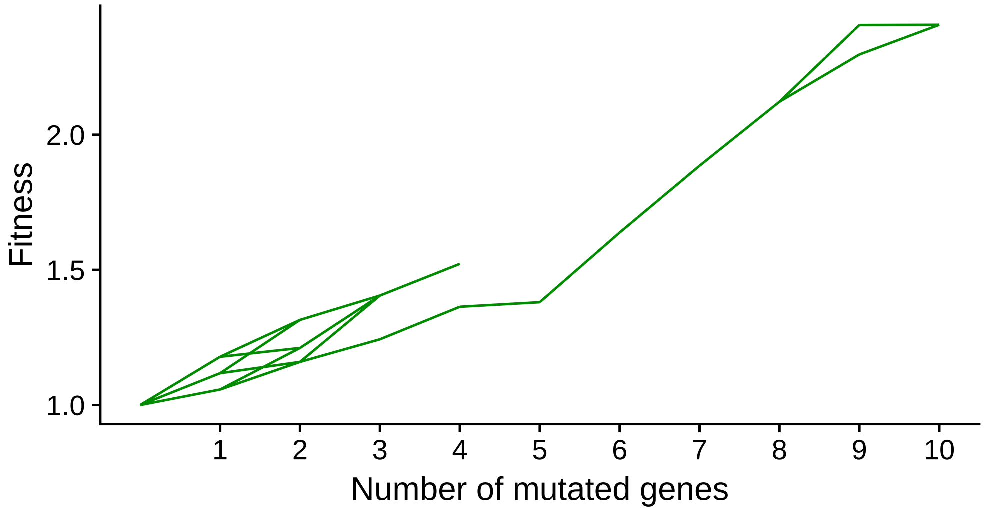


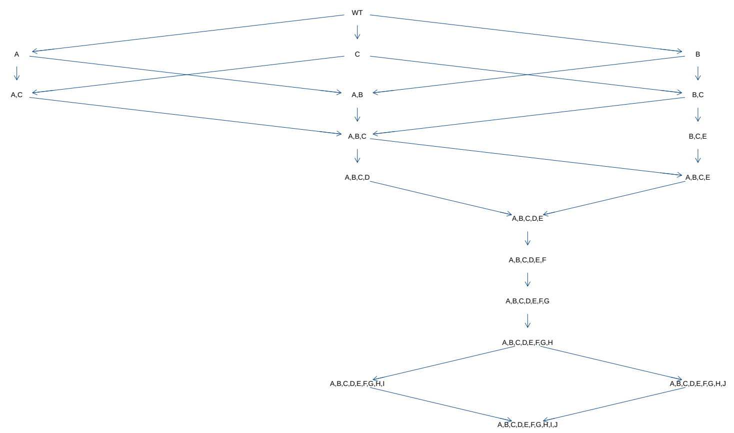


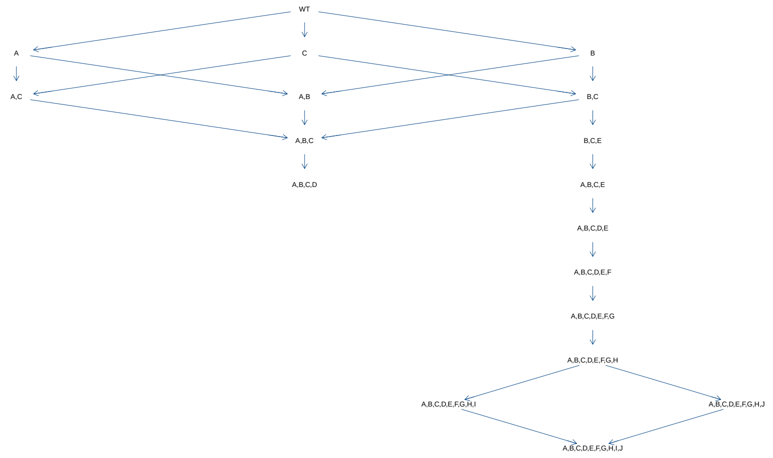

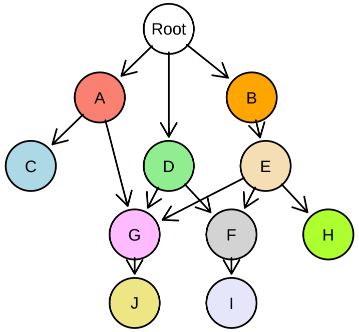


| | |
|---|---|
| *ID* | WNDLFwNiHctCAAKK |
| *acc genotypes* | 60 |
| *peaks no back* | 7 |
| *removed edges* | 24 |
| *prop rem edges* | 0.176 |

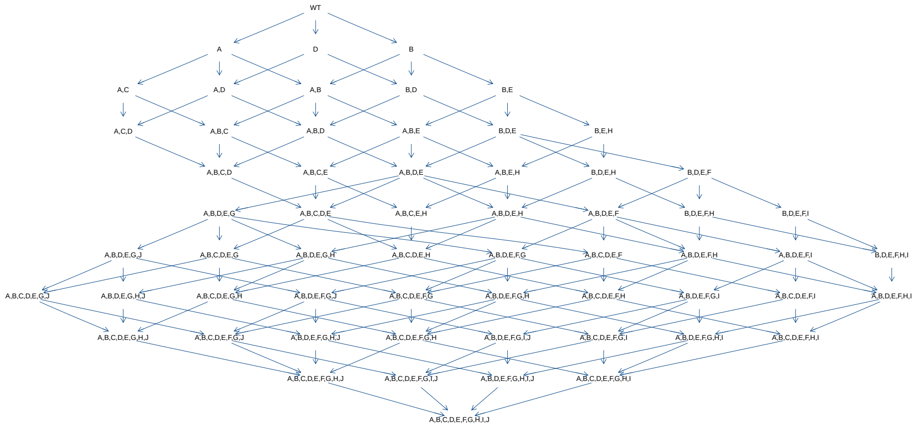

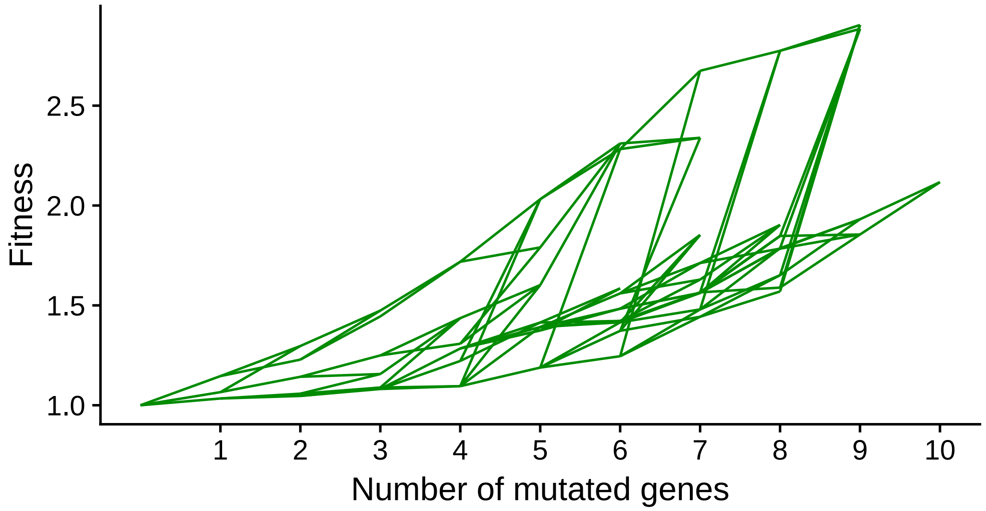


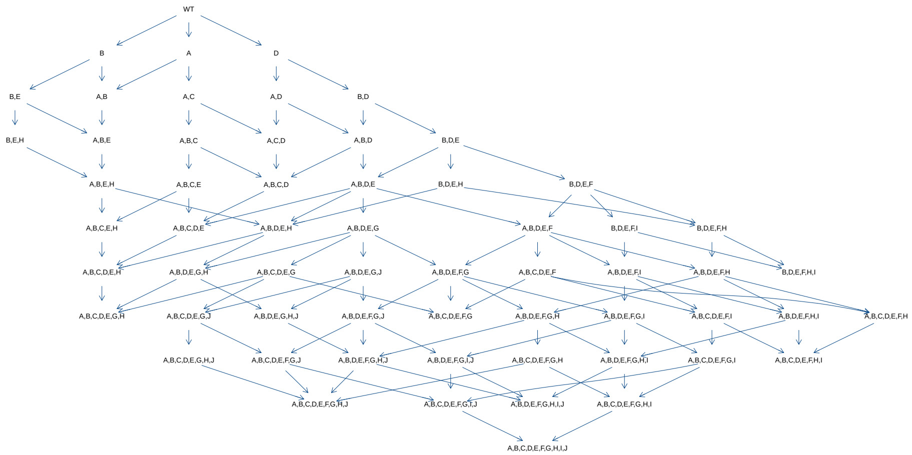

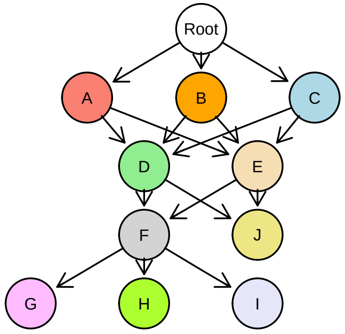


| | |
|---|---|
| *ID* | wrYzolpOzxMG9bWu |
| *acc genotypes* | 28 |
| *peaks no back* | 4 |
| *removed edges* | 6 |
| *prop rem edges* | 0.118 |

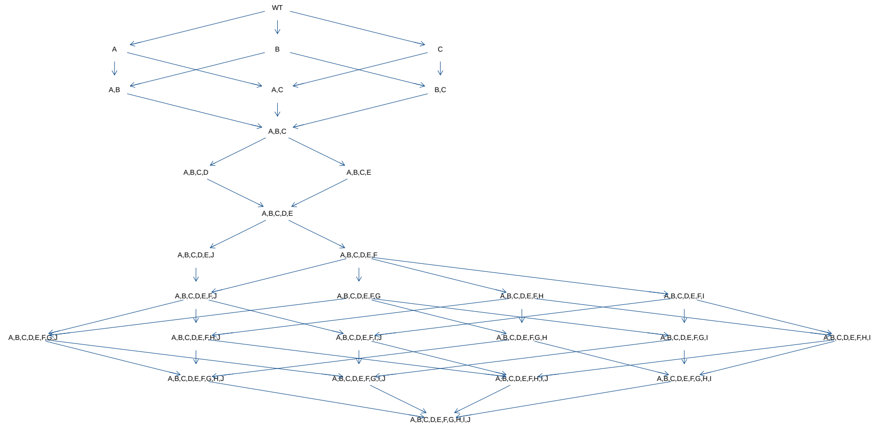


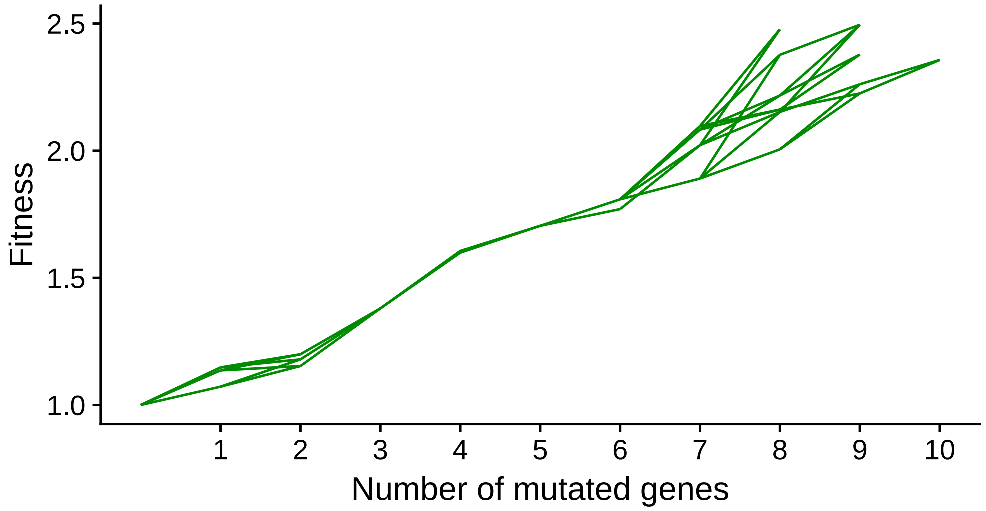


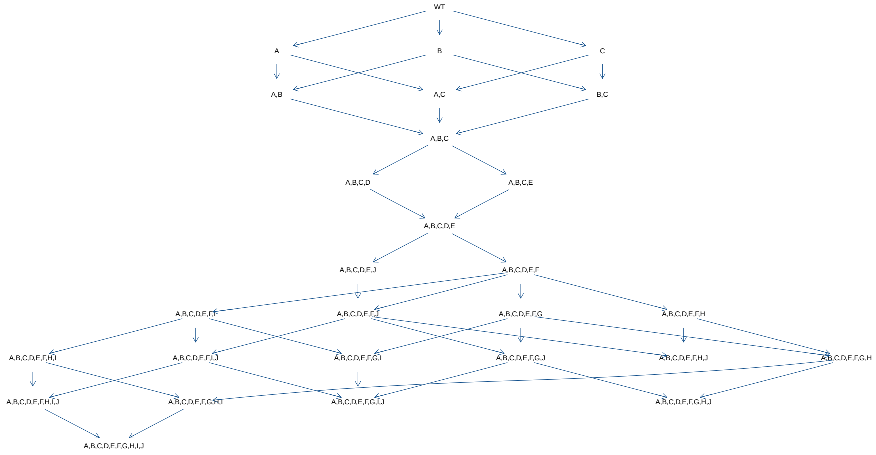

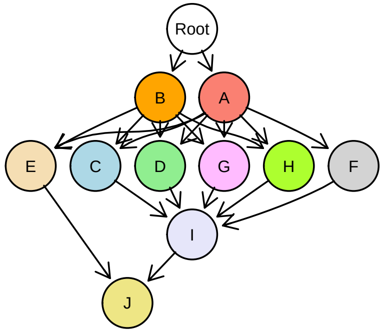


| | |
|---|---|
| *ID* | ws10WbhFUWR9mXKoe |
| *acc genotypes* | 71 |
| *peaks no back* | 8 |
| *removed edges* | 29 |
| *prop rem edges* | 0.144 |

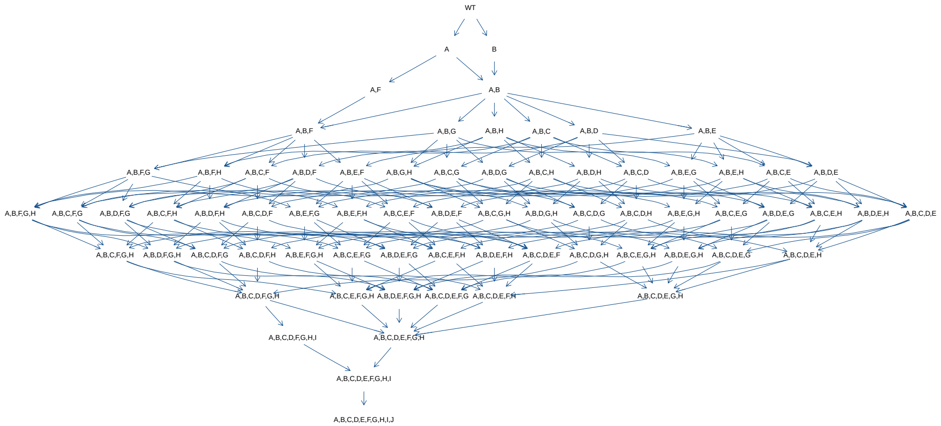

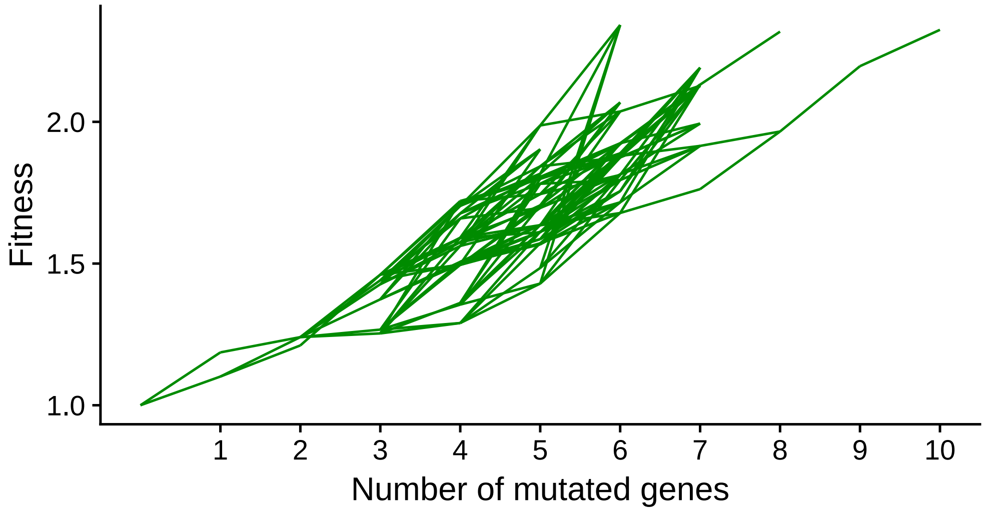


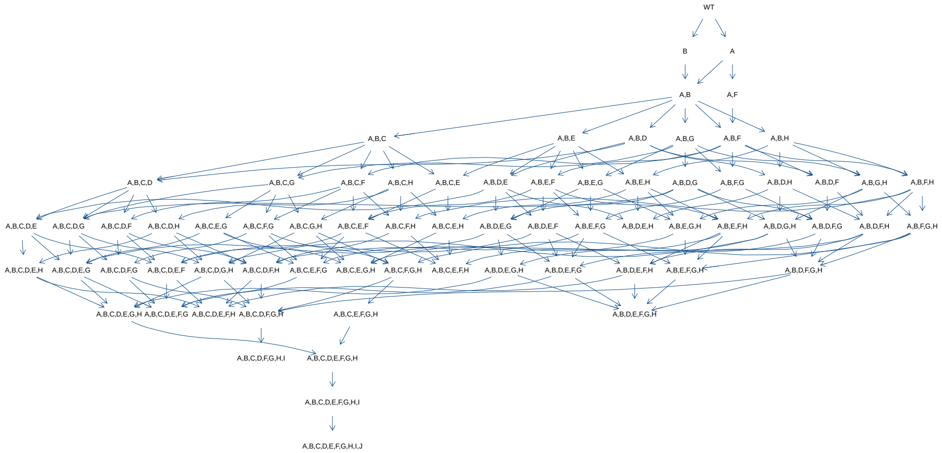

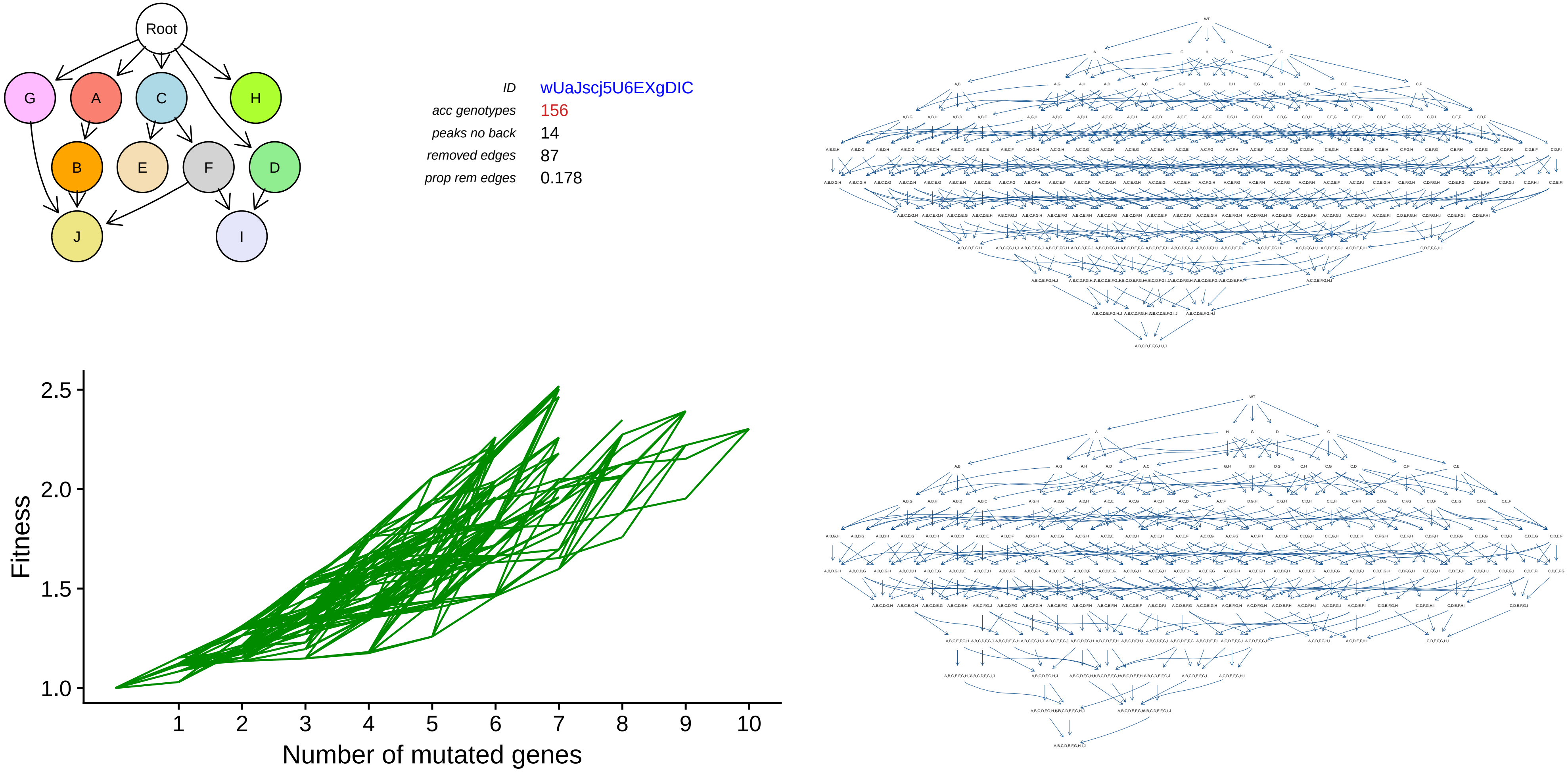
Root
G
A
C
H
B
E
F
D
J
I
ID wUaJscj5U6EXgDIC
acc genotypes 156
peaks no back 14
removed edges 87
prop rem edges 0.178
Fitness
2.5
2.0
1.5
1.0
1 2 3 4 5 6 7 8 9 10
Number of mutated genes

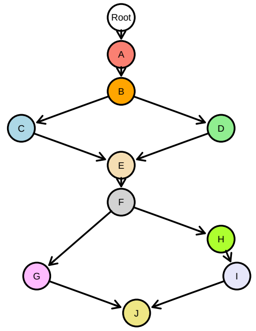


| | |
|---|---|
| *ID* | XajIk5zcAsLjK9cr |
| *acc genotypes* | 14 |
| *peaks no back* | 3 |
| *removed edges* | 2 |
| *prop rem edges* | 0.125 |

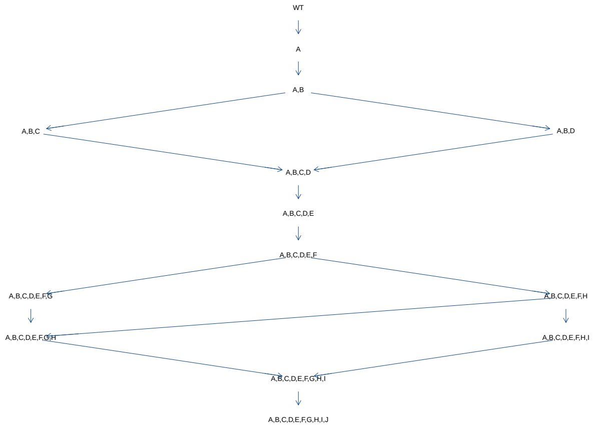


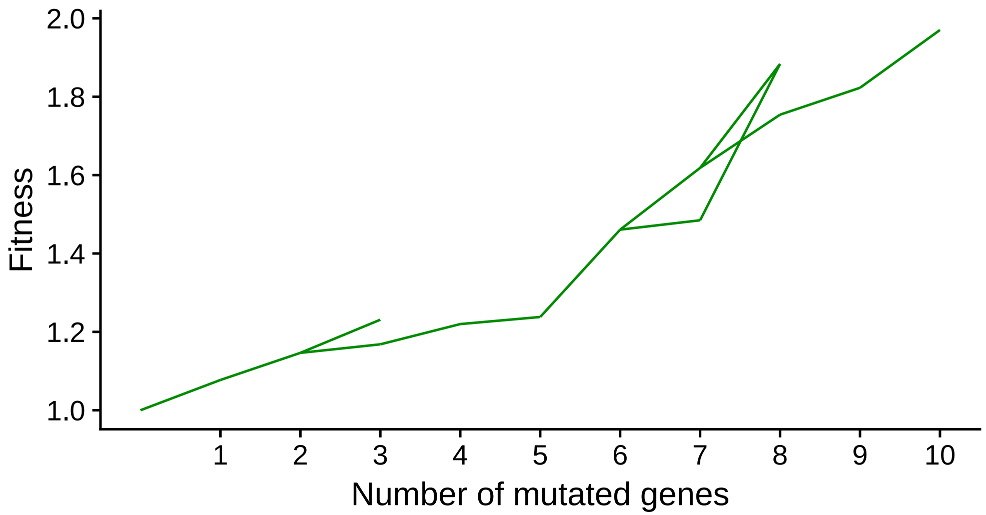


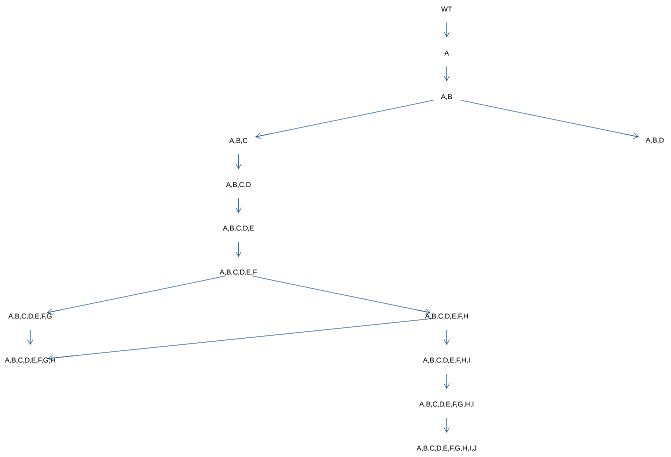

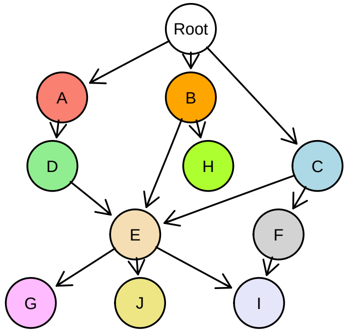


| | |
|---|---|
| *ID* | xAYN10rolXAsrpbMd |
| *acc genotypes* | 51 |
| *peaks no back* | 9 |
| *removed edges* | 16 |
| *prop rem edges* | 0.145 |

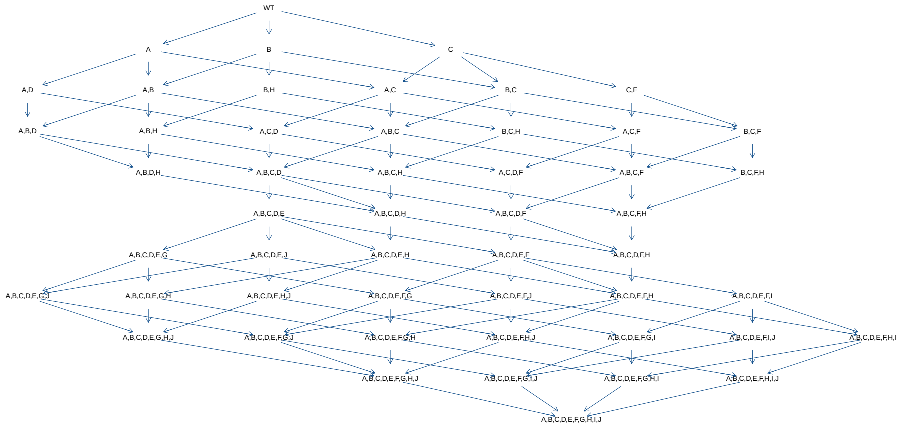

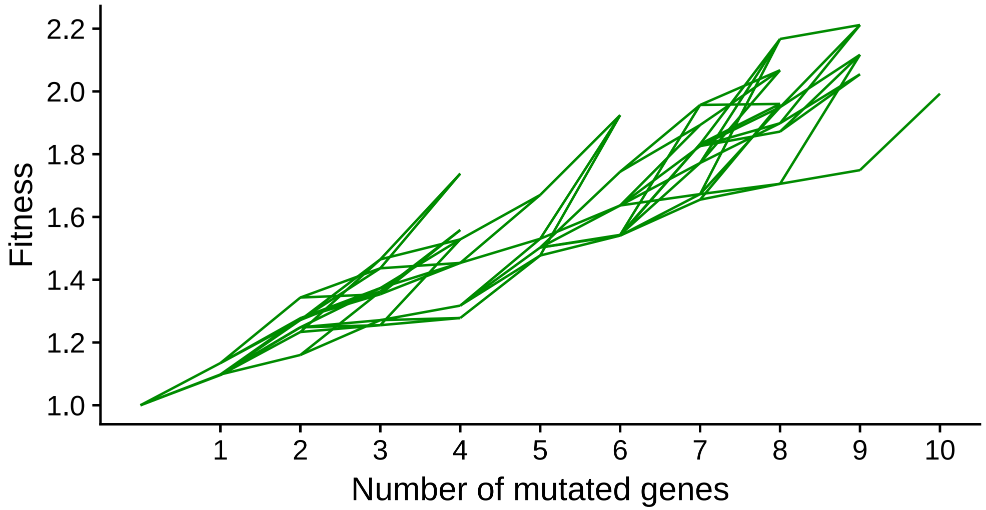


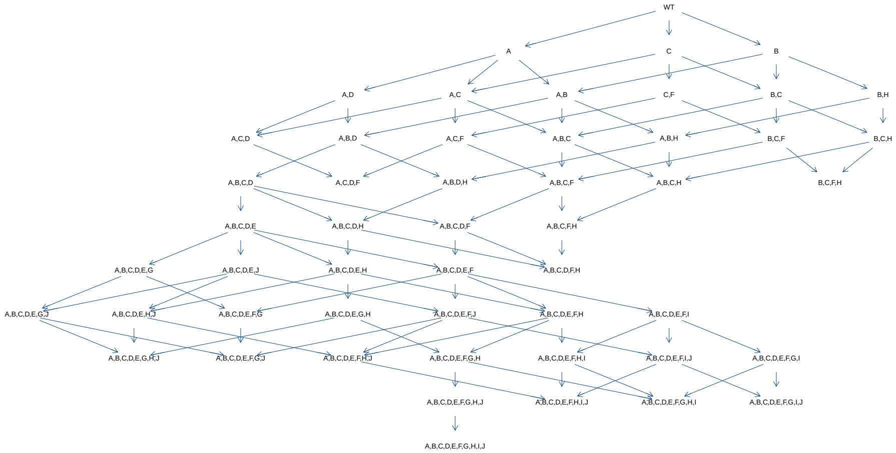

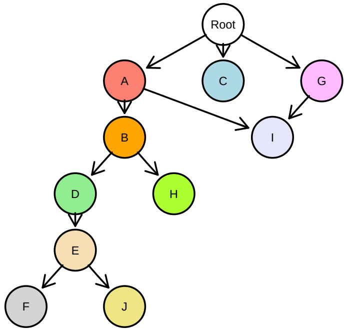


| | |
|---|---|
| *ID* | XmnGrSjLNk1UkGT5 |
| *acc genotypes* | 82 |
| *peaks no back* | 12 |
| *removed edges* | 57 |
| *prop rem edges* | 0.268 |

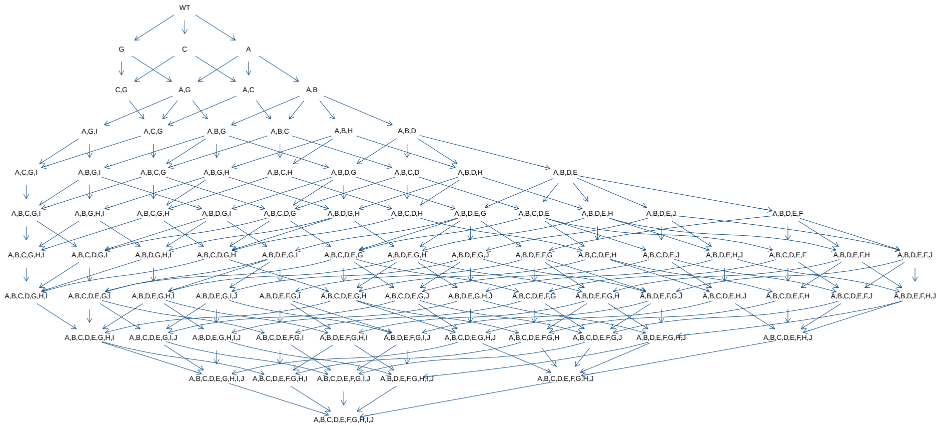

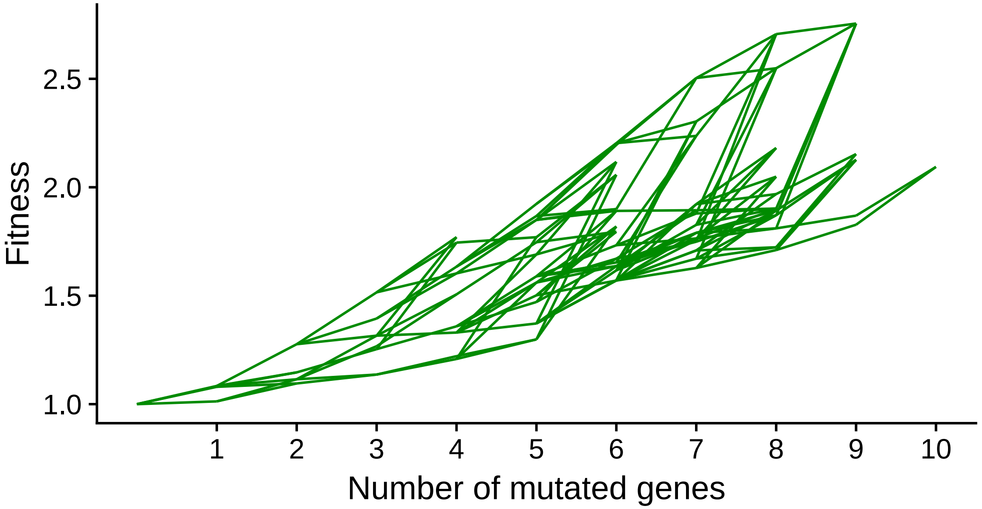


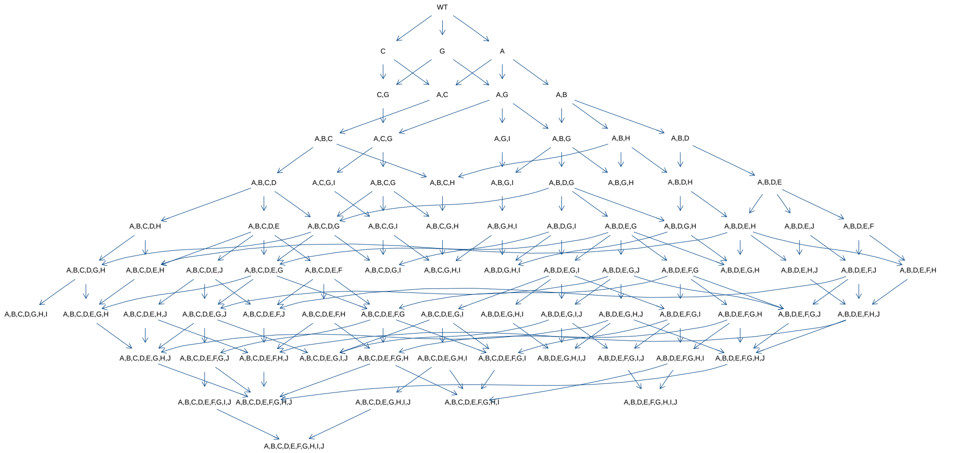

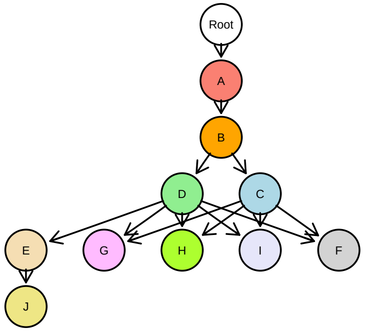


| | |
|---|---|
| *ID* | y8uUVSfWKcZQiY4y |
| *acc genotypes* | 55 |
| *peaks no back* | 8 |
| *removed edges* | 22 |
| *prop rem edges* | 0.159 |

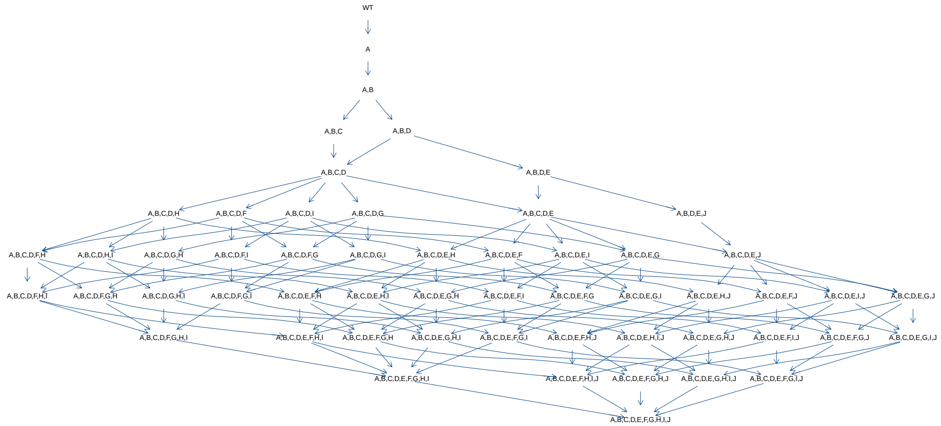

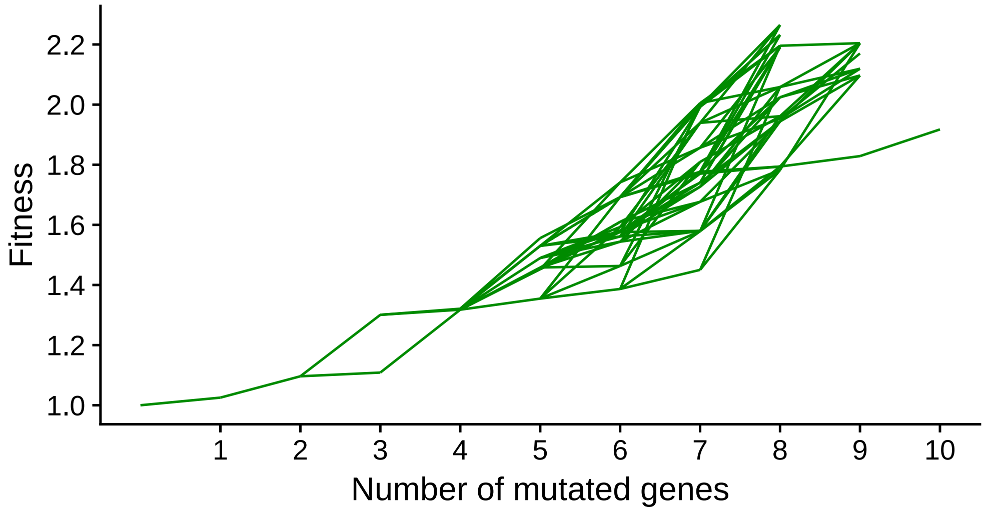


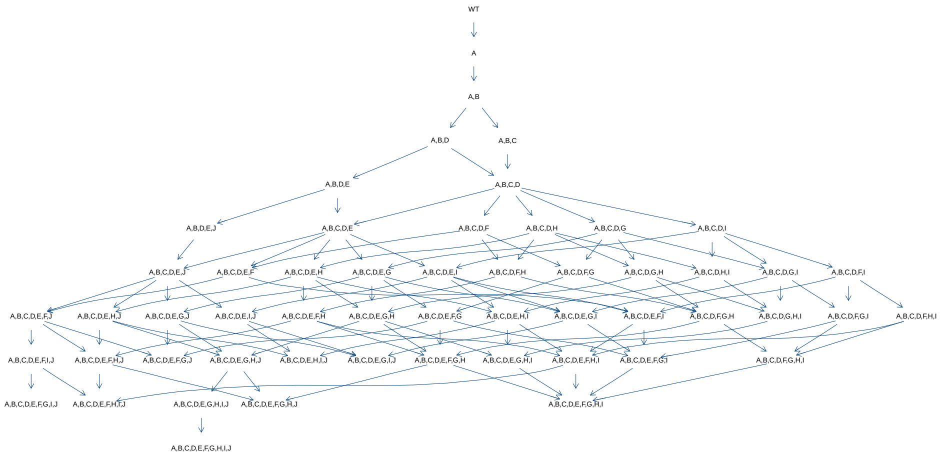

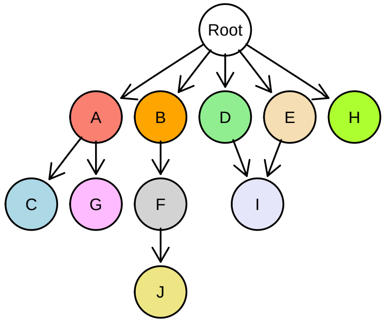


*ID* ypRiC10Qof6jAjsVJ
*acc genotypes* 200
*peaks no back* 18
*removed edges* 112
*prop rem edges* 0.172

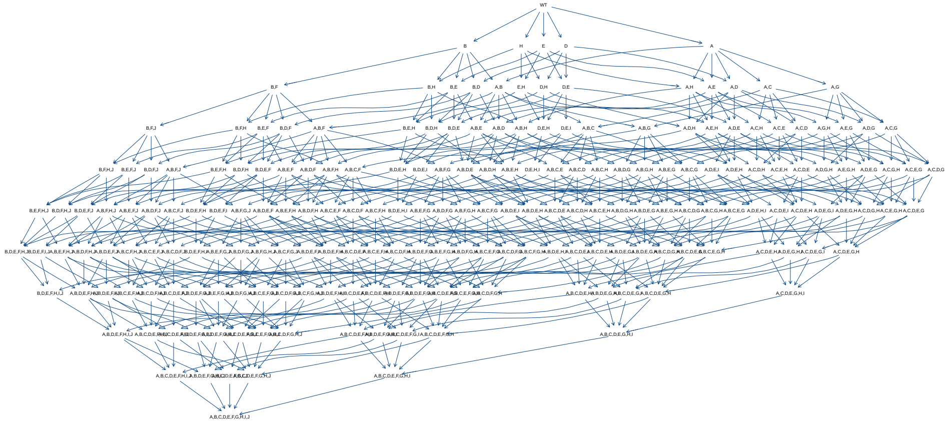

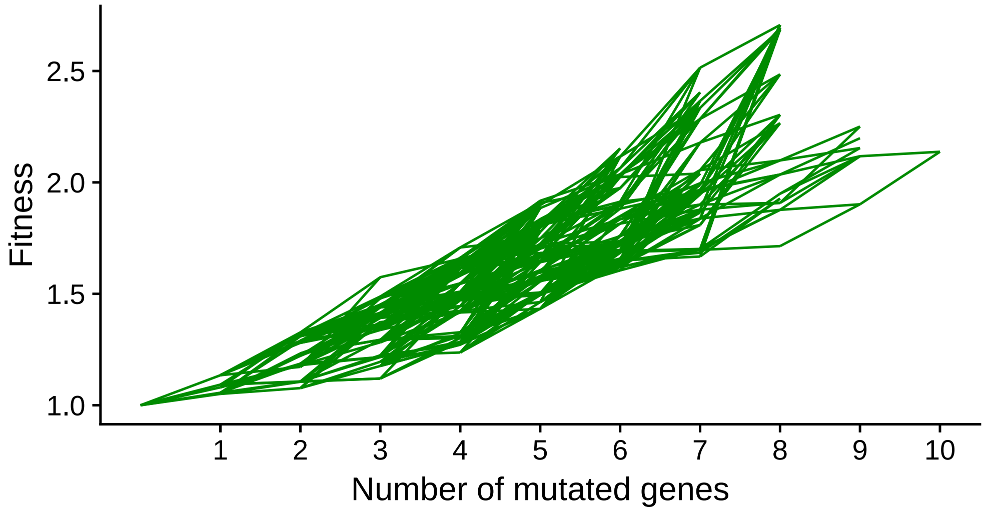


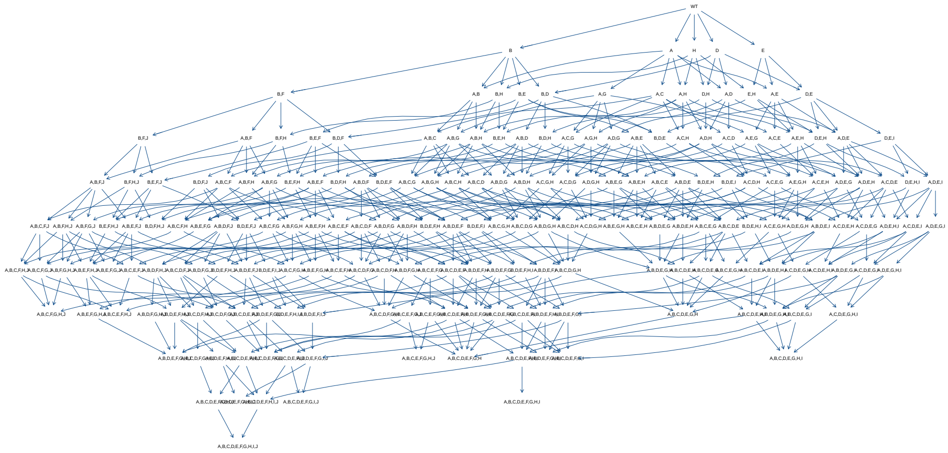

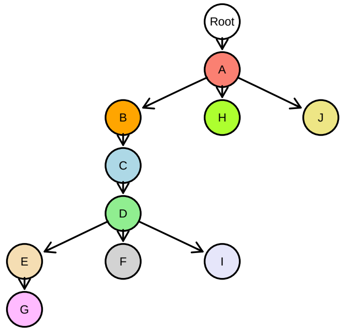


| | |
|---|---|
| *ID* | YW7KQxwKMIe2i7ar |
| *acc genotypes* | 61 |
| *peaks no back* | 9 |
| *removed edges* | 28 |
| *prop rem edges* | 0.183 |

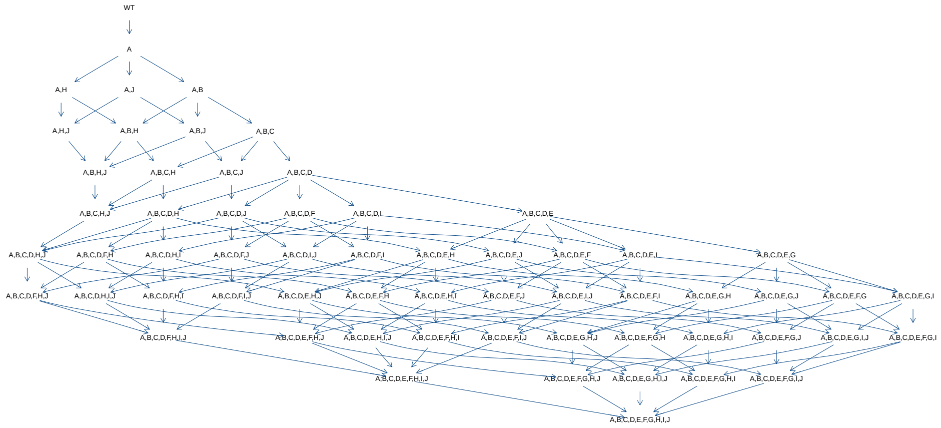

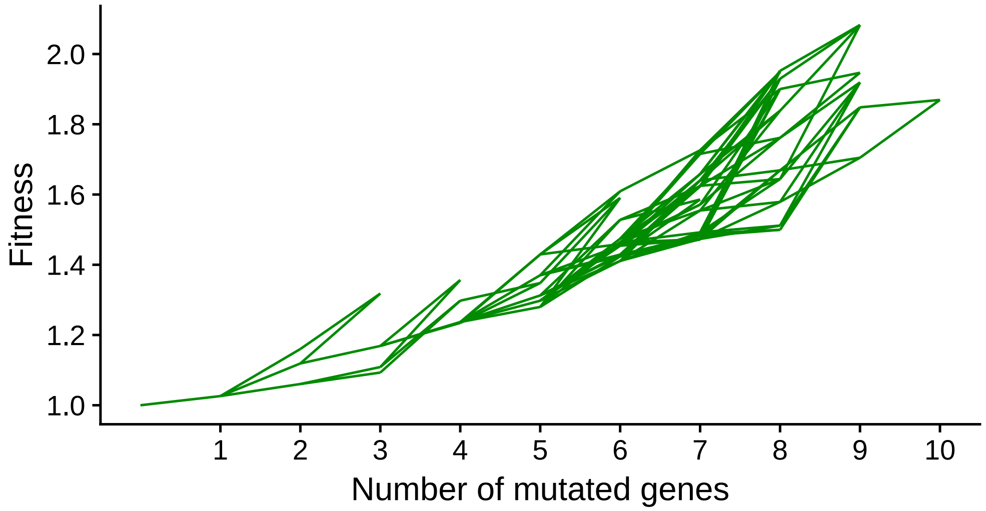


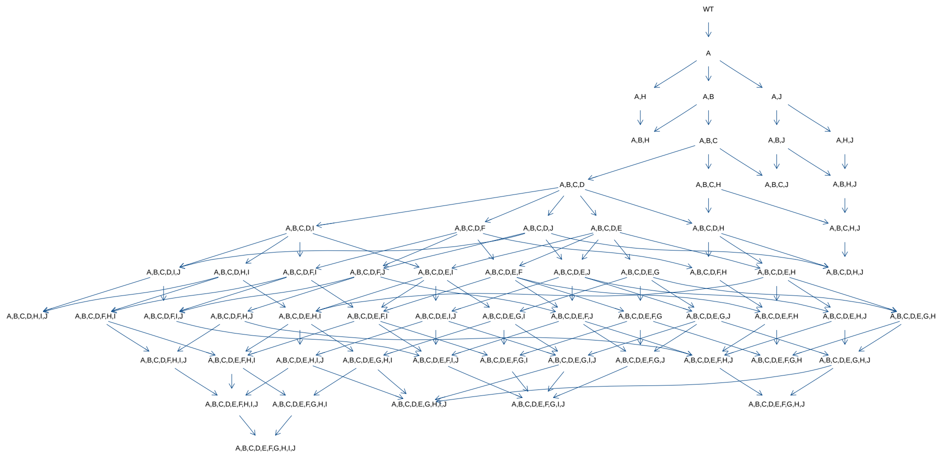

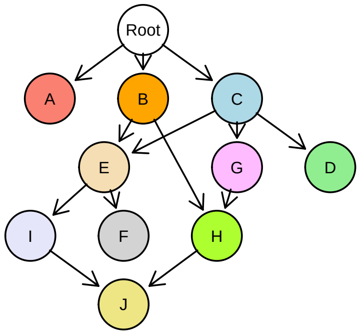


| | |
|---|---|
| *ID* | z4RVEoFHLg5dwFWl |
| *acc genotypes* | 80 |
| *peaks no back* | 9 |
| *removed edges* | 44 |
| *prop rem edges* | 0.212 |

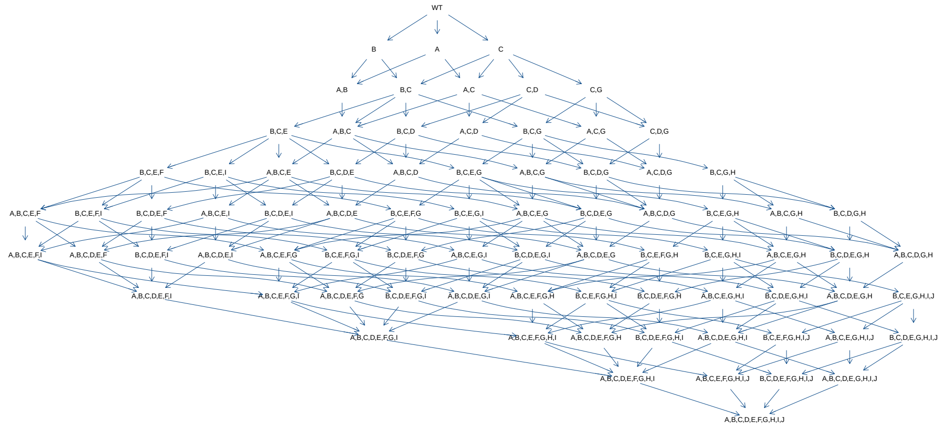

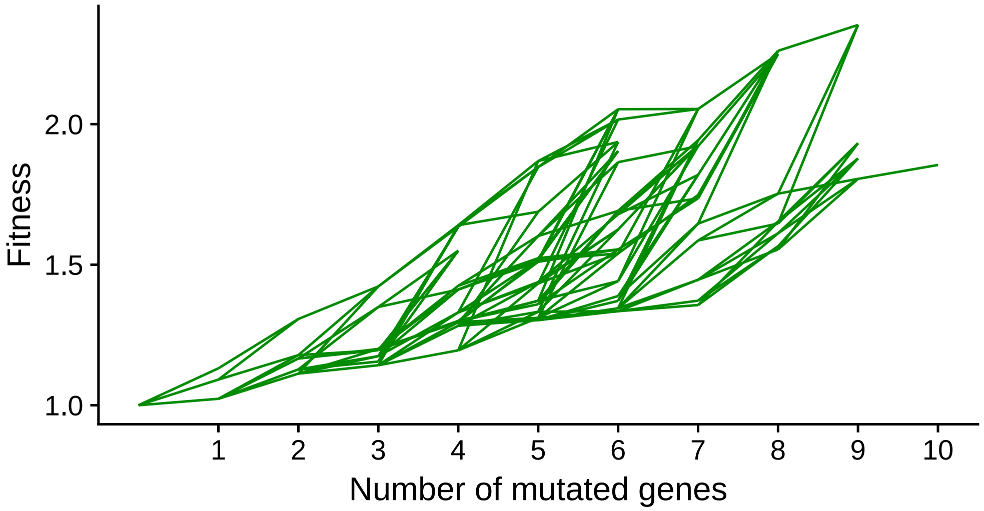


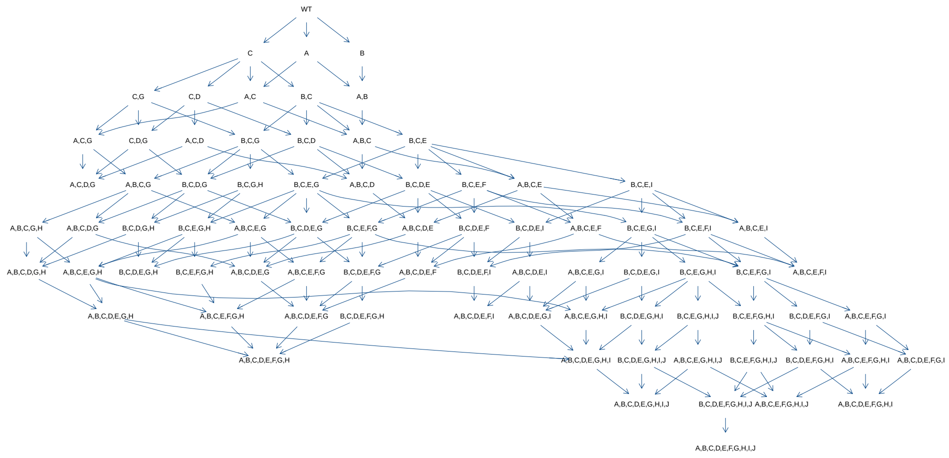

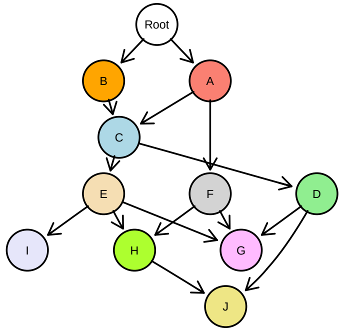


| | |
|---|---|
| *ID* | zAMK2D5LDmq9X37r |
| *acc genotypes* | 30 |
| *peaks no back* | 2 |
| *removed edges* | 3 |
| *prop rem edges* | 0.057 |

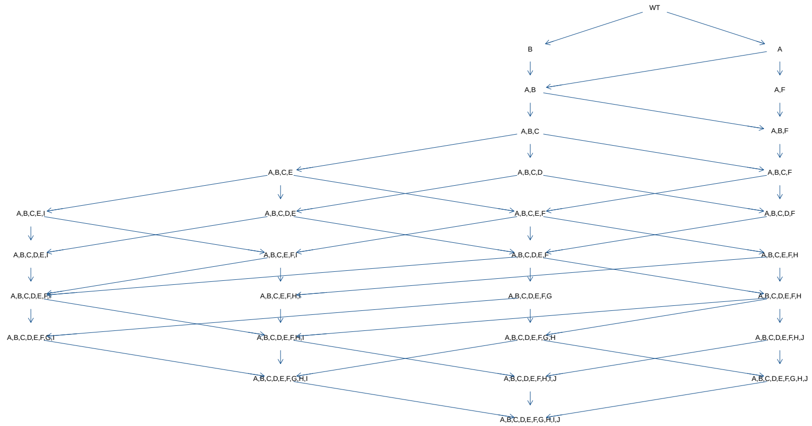

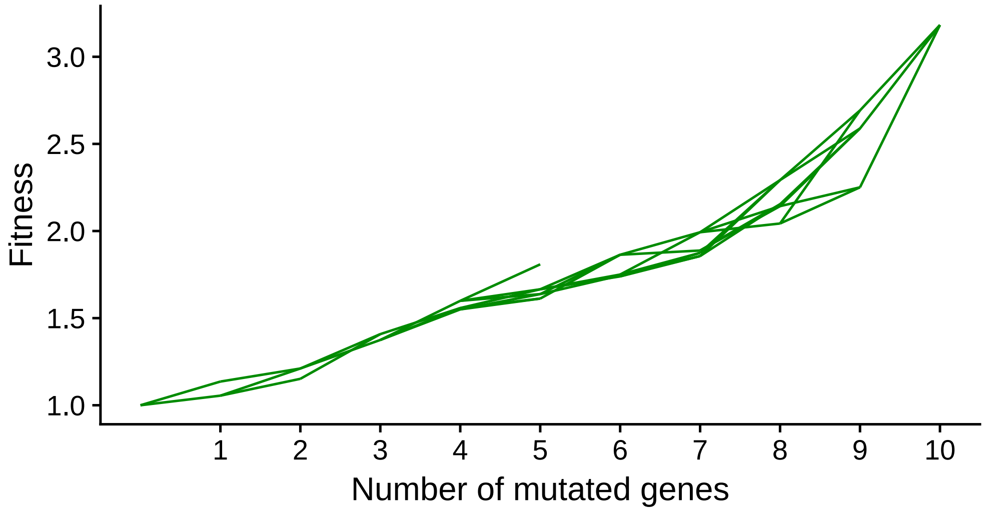


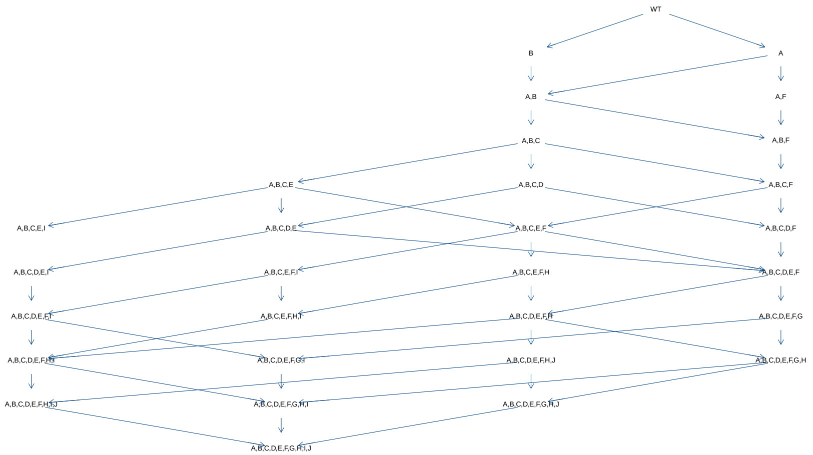

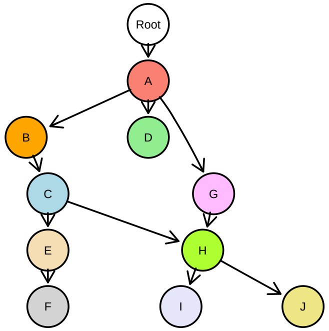


| | |
|---|---|
| *ID* | ZIBdVWiOnxT3sPfI |
| *acc genotypes* | 45 |
| *peaks no back* | 7 |
| *removed edges* | 11 |
| *prop rem edges* | 0.116 |

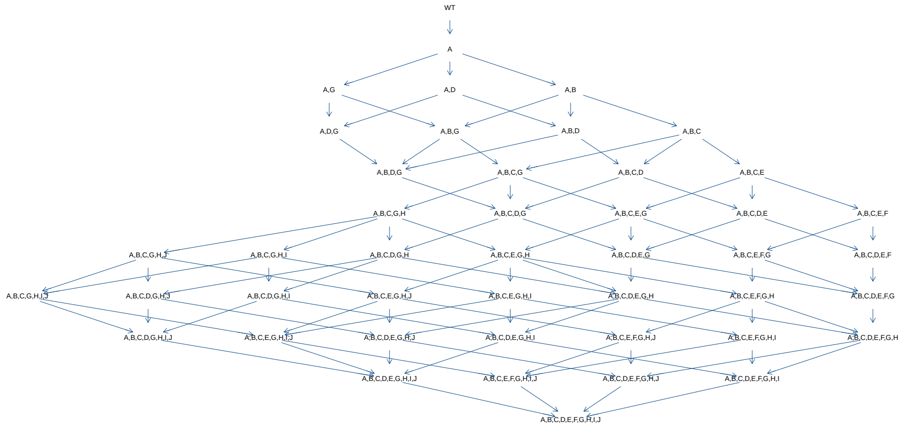

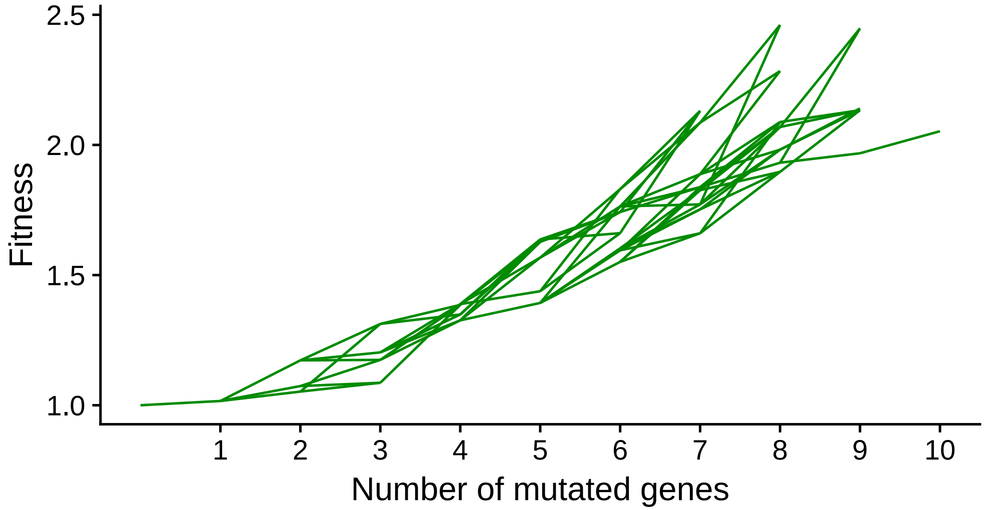


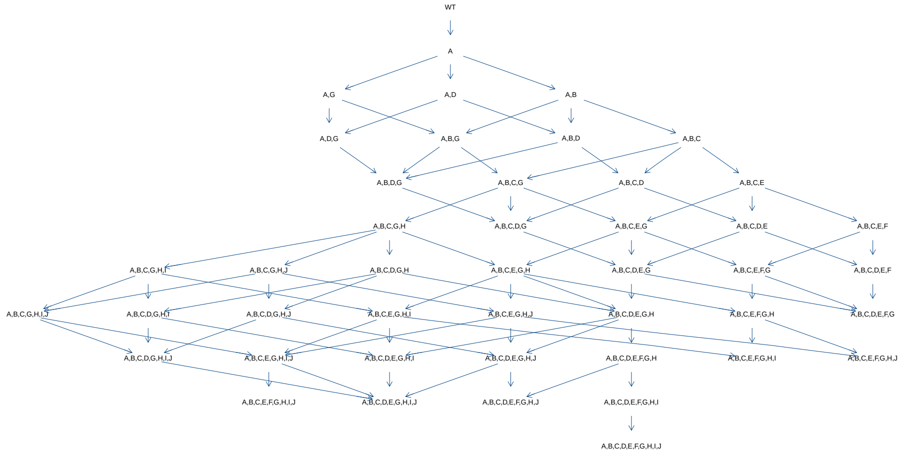

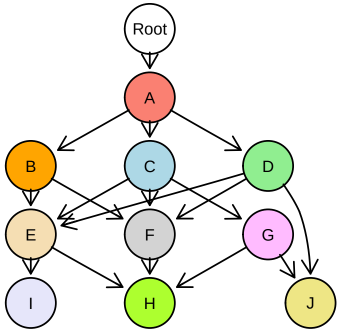


*ID* zJtfsaG9cK5j5Hnm
*acc genotypes* 34
*peaks no back* 5
*removed edges* 12
*prop rem edges* 0.19

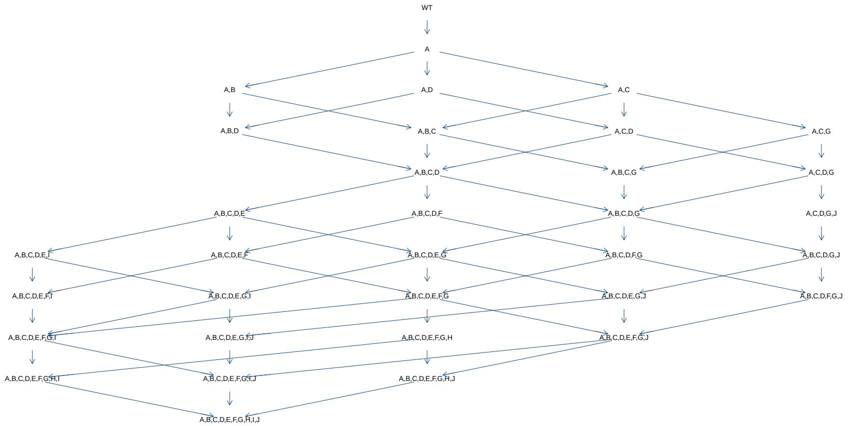

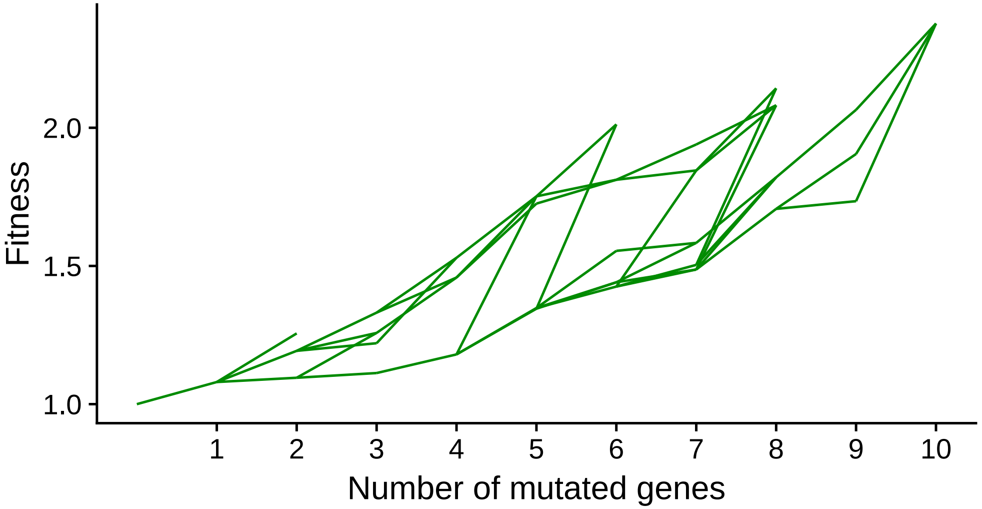


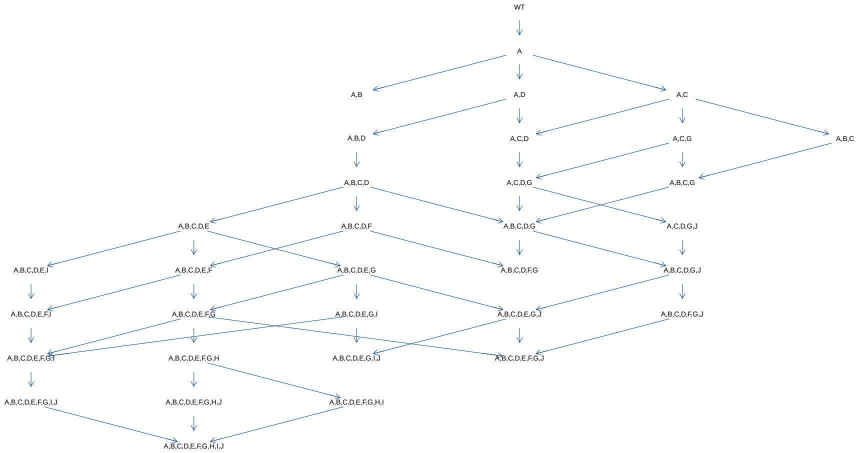

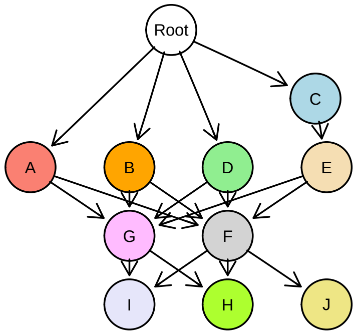


| | |
|---|---|
| *ID* | ZKH10YvDZGVfsdHAY |
| *acc genotypes* | 35 |
| *peaks no back* | 6 |
| *removed edges* | 9 |
| *prop rem edges* | 0.129 |

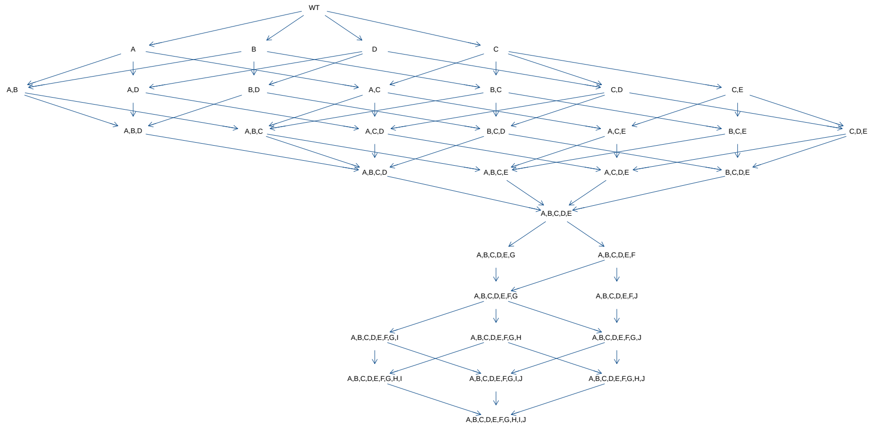

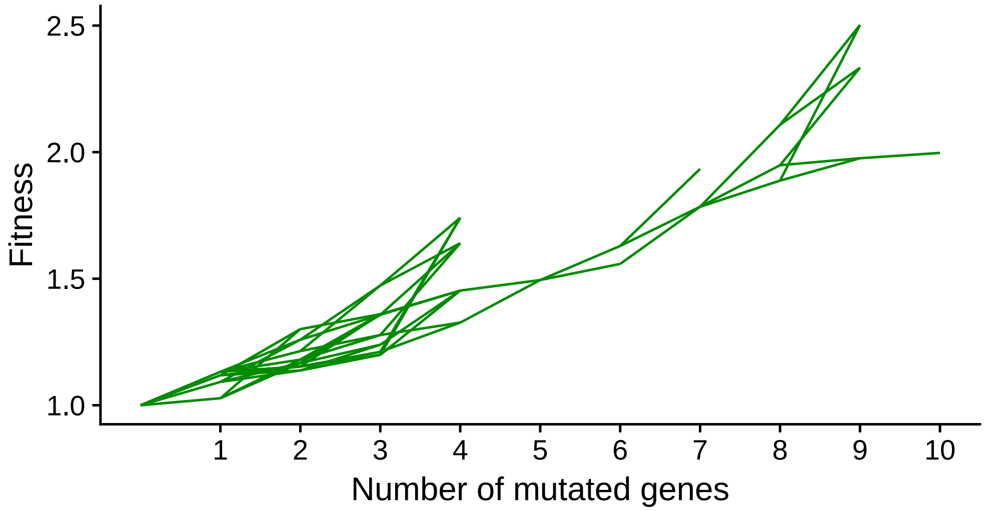


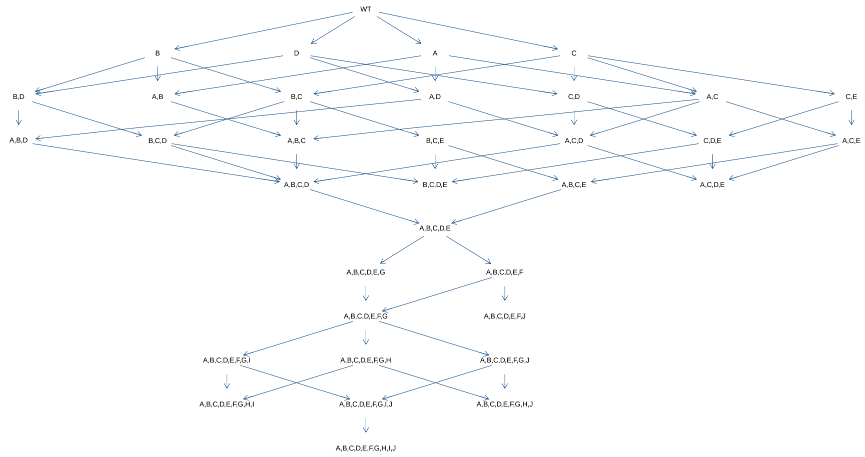

| | |
|---|---|
| *ID* | 103oxrJDcbfc2YpKa |
| *acc genotypes* | 162 |
| *peaks no back* | 57 |
| *removed edges* | NA |
| *prop rem edges* | NA |

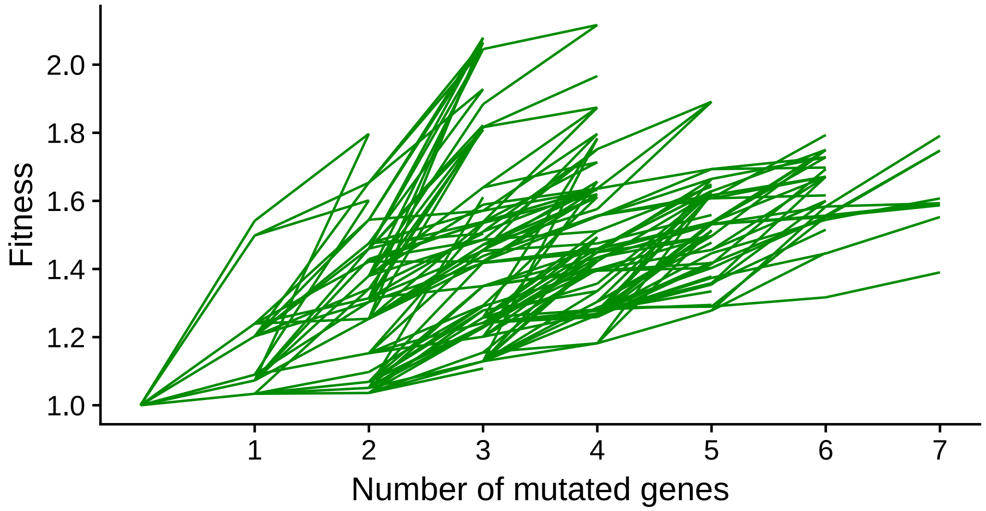


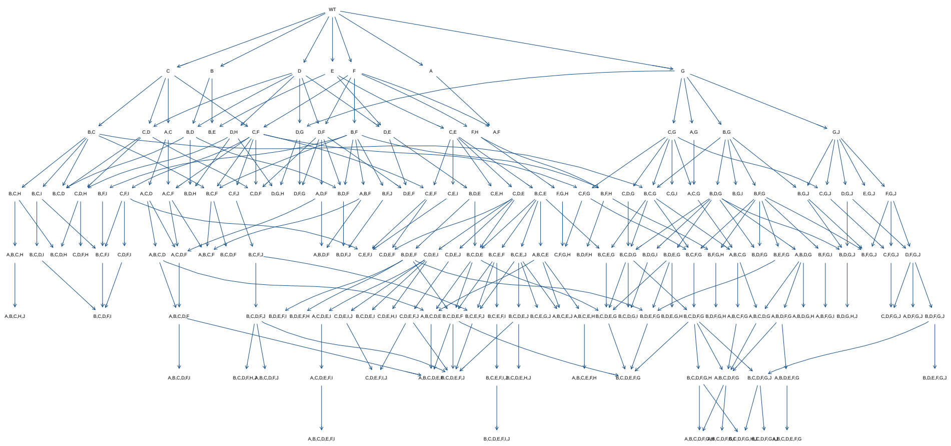

| | |
|---|---|
| *ID* | 10aJ10EAqKH7VA3ccp |
| *acc genotypes* | 79 |
| *peaks no back* | 26 |
| *removed edges* | NA |
| *prop rem edges* | NA |

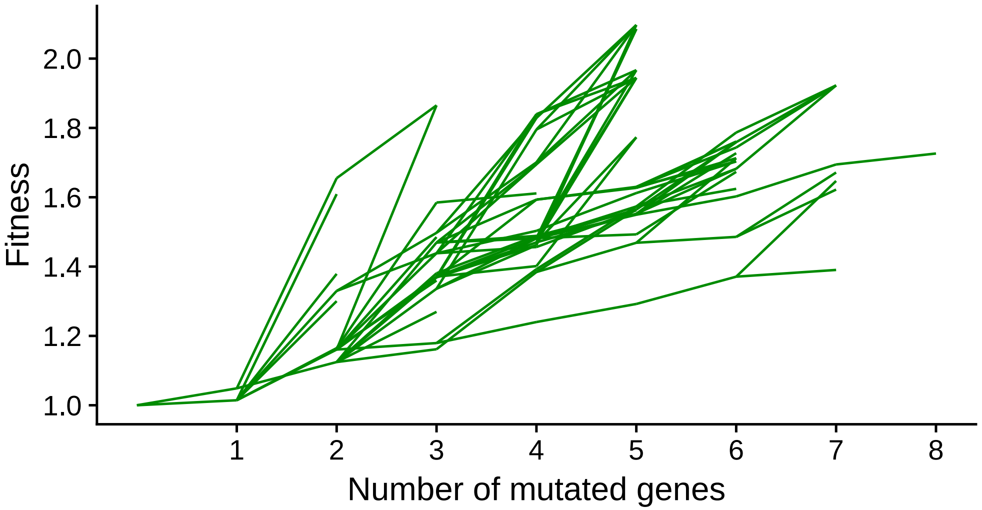


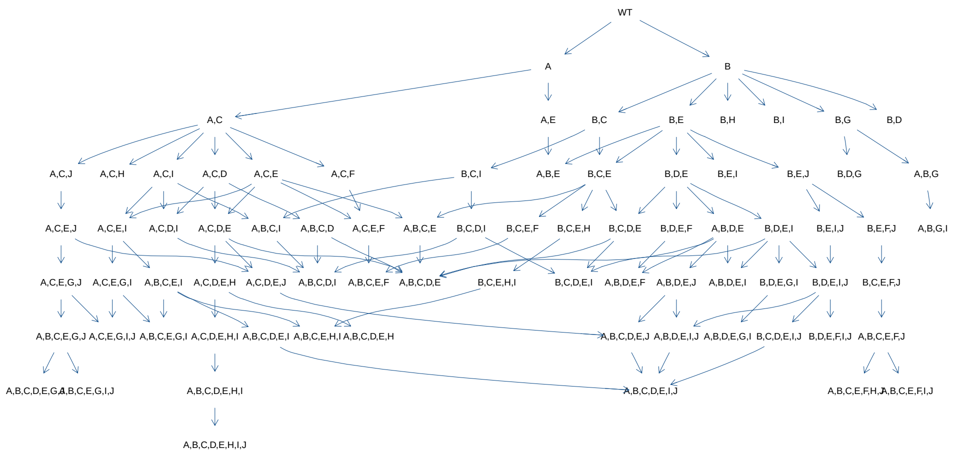

| | |
|---|---|
| *ID* | 10fyev4yPOAHqV3G4 |
| *acc genotypes* | 72 |
| *peaks no back* | 33 |
| *removed edges* | NA |
| *prop rem edges* | NA |

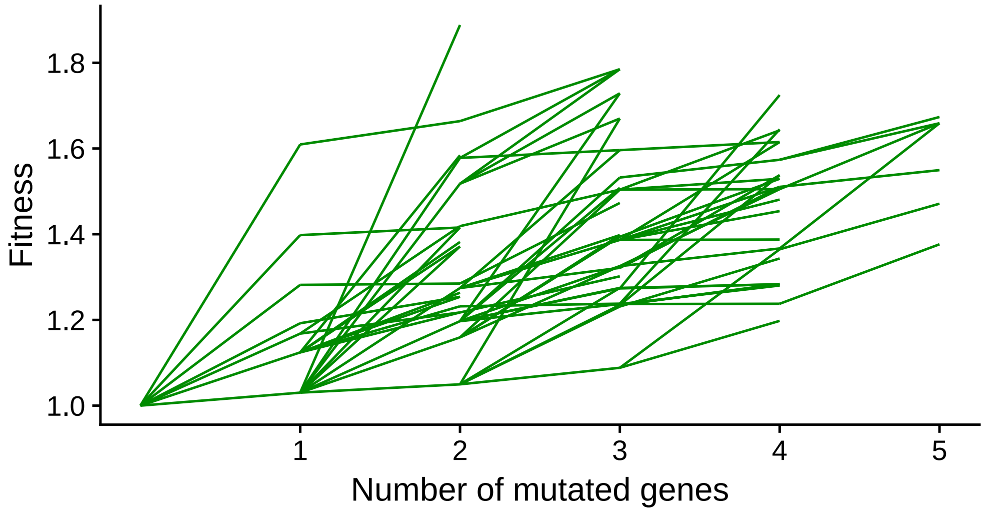


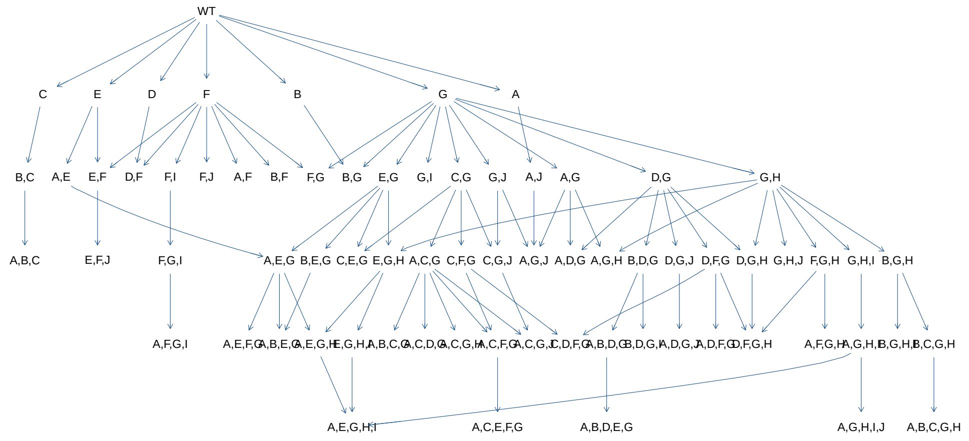

| | |
|---|---|
| *ID* | 10NjHwdZ4eQZuBASX |
| *acc genotypes* | 326 |
| *peaks no back* | 116 |
| *removed edges* | NA |
| *prop rem edges* | NA |

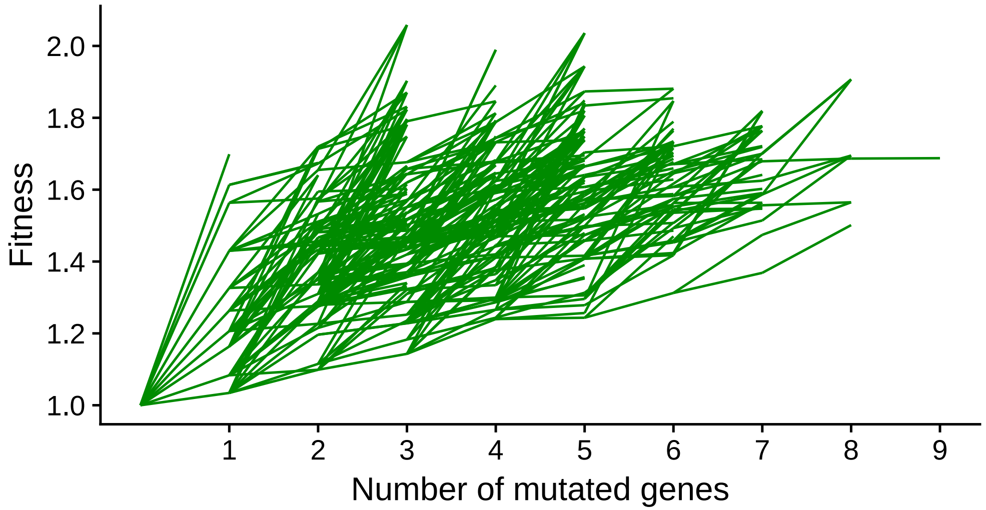


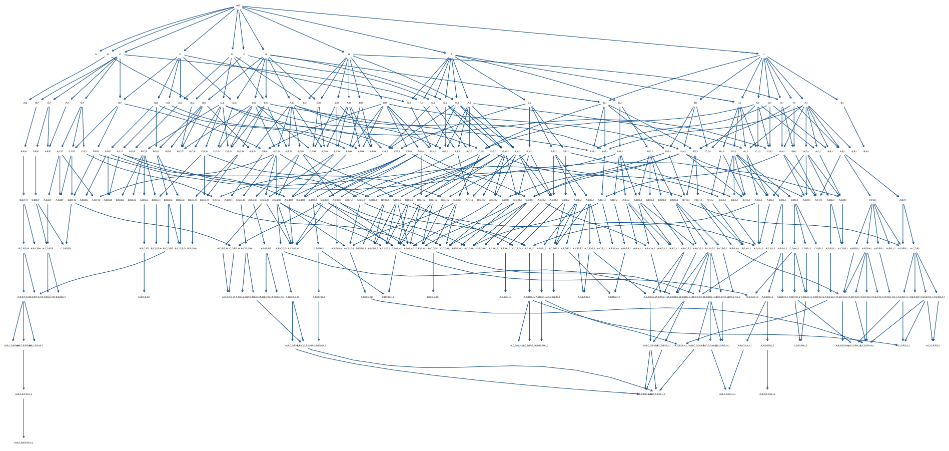

| | |
|---|---|
| *ID* | 10rm10ycggF6sUPqCV |
| *acc genotypes* | 330 |
| *peaks no back* | 87 |
| *removed edges* | NA |
| *prop rem edges* | NA |

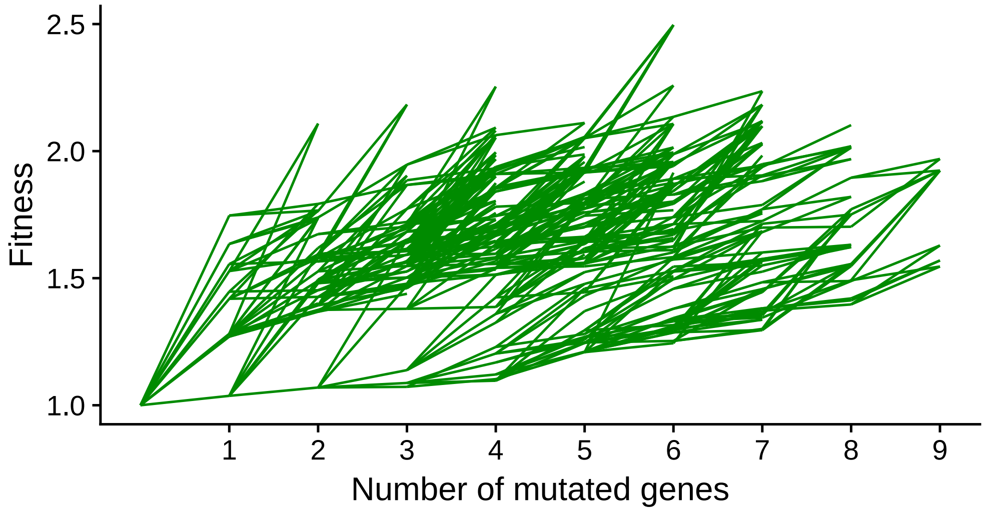


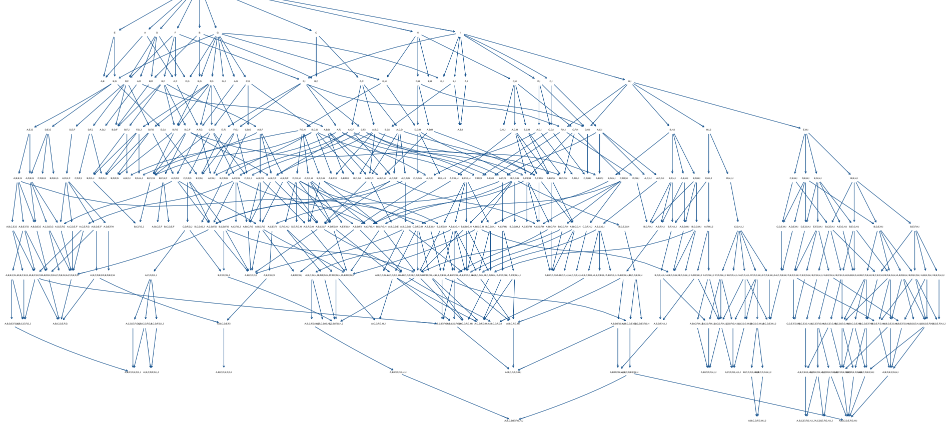

| | |
|---|---|
| *ID* | 10yJX7cQ5FF31Qv6f |
| *acc genotypes* | 95 |
| *peaks no back* | 36 |
| *removed edges* | NA |
| *prop rem edges* | NA |

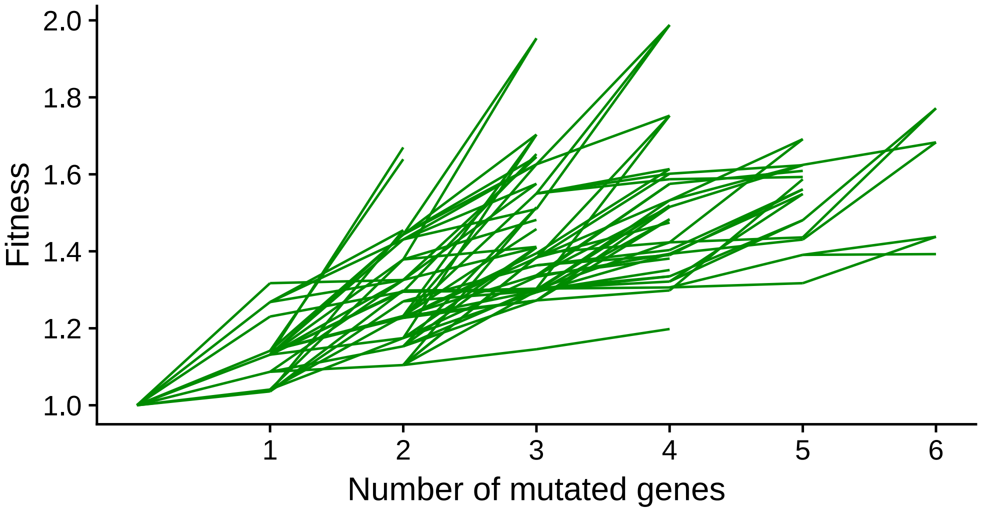


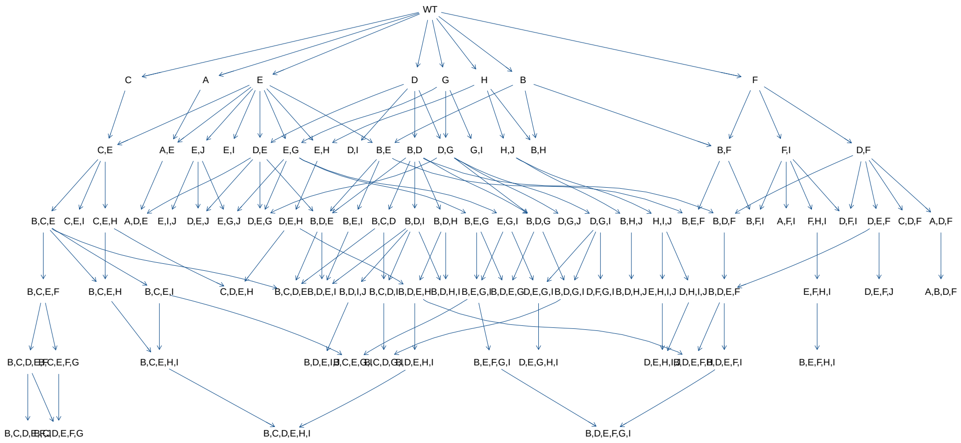

| | |
|---|---|
| *ID* | 1aUzp7cAshgfTlCQ |
| *acc genotypes* | 44 |
| *peaks no back* | 16 |
| *removed edges* | NA |
| *prop rem edges* | NA |

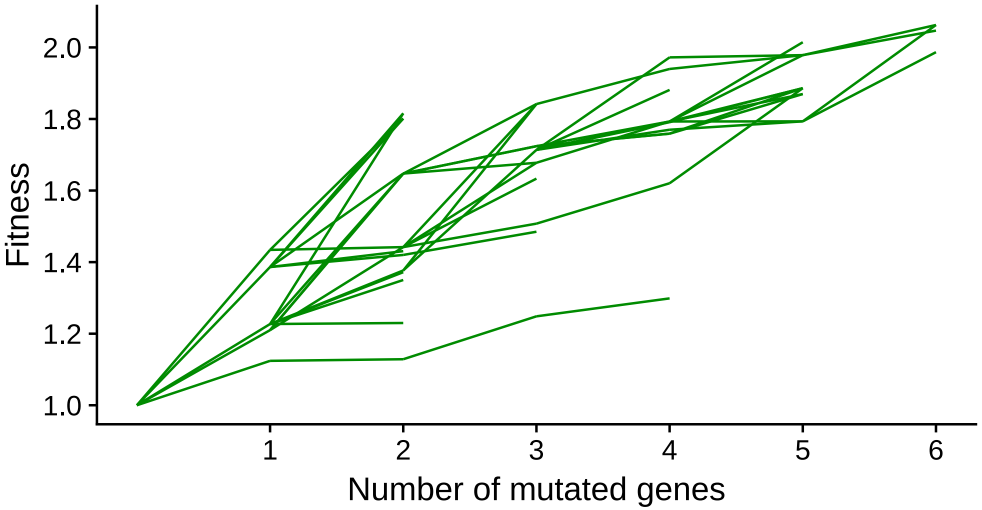


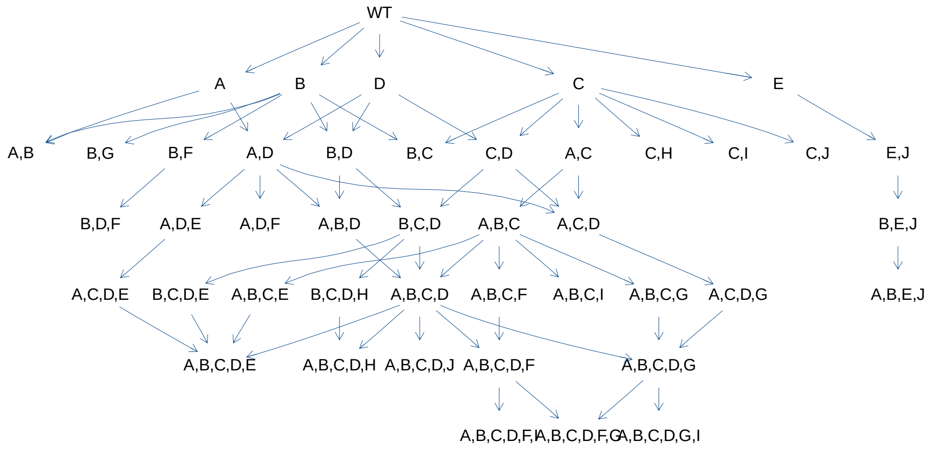

| | |
|---|---|
| *ID* | 1QFuLHW10HcmHRTX10 |
| *acc genotypes* | 217 |
| *peaks no back* | 81 |
| *removed edges* | NA |
| *prop rem edges* | NA |

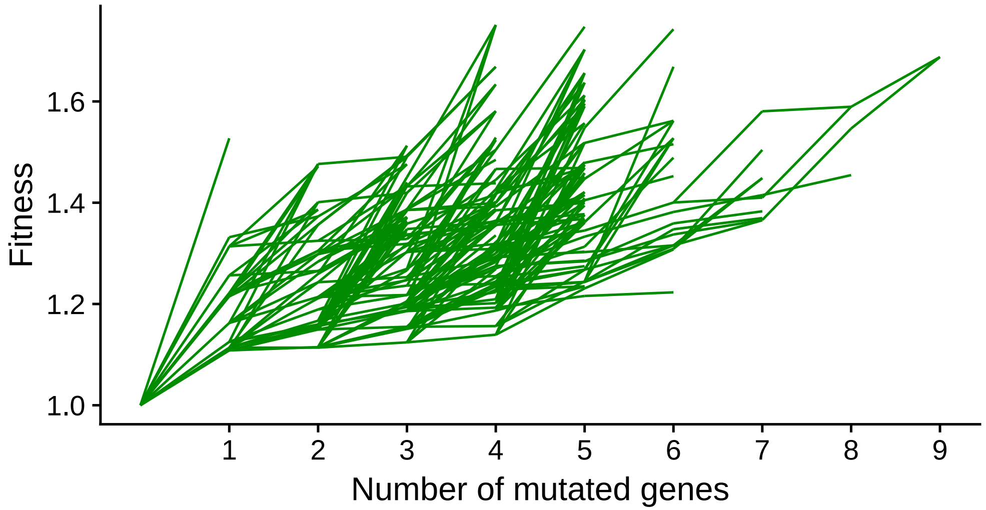


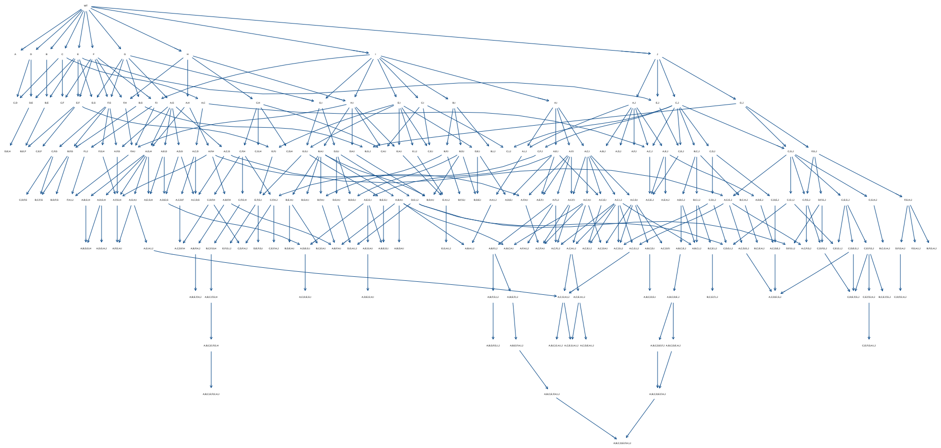

| | |
|---|---|
| *ID* | 2HoRF1oKBcGNaQYL |
| *acc genotypes* | 597 |
| *peaks no back* | 113 |
| *removed edges* | NA |
| *prop rem edges* | NA |

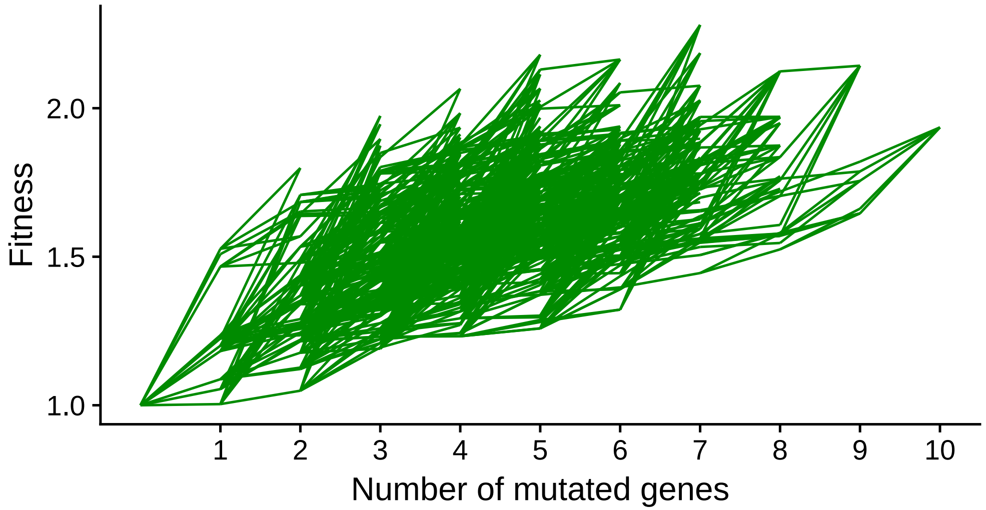


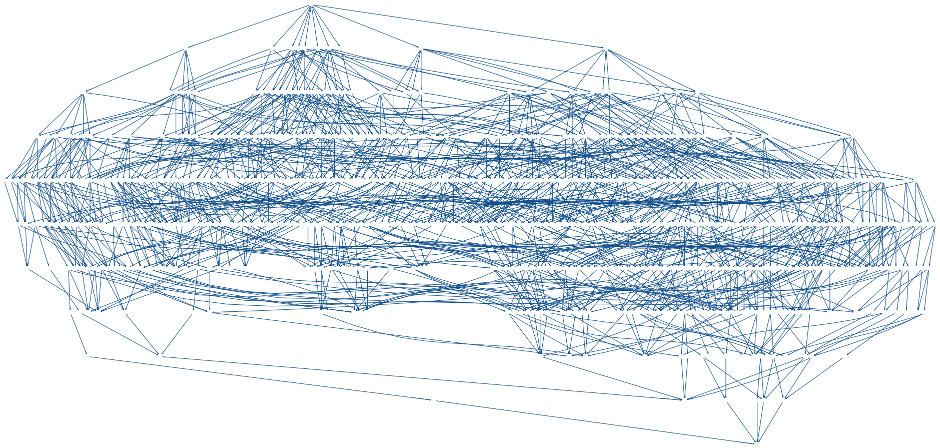

| | |
|---|---|
| *ID* | 2Ytm5OoPJh10HnzIe |
| *acc genotypes* | 52 |
| *peaks no back* | 13 |
| *removed edges* | NA |
| *prop rem edges* | NA |

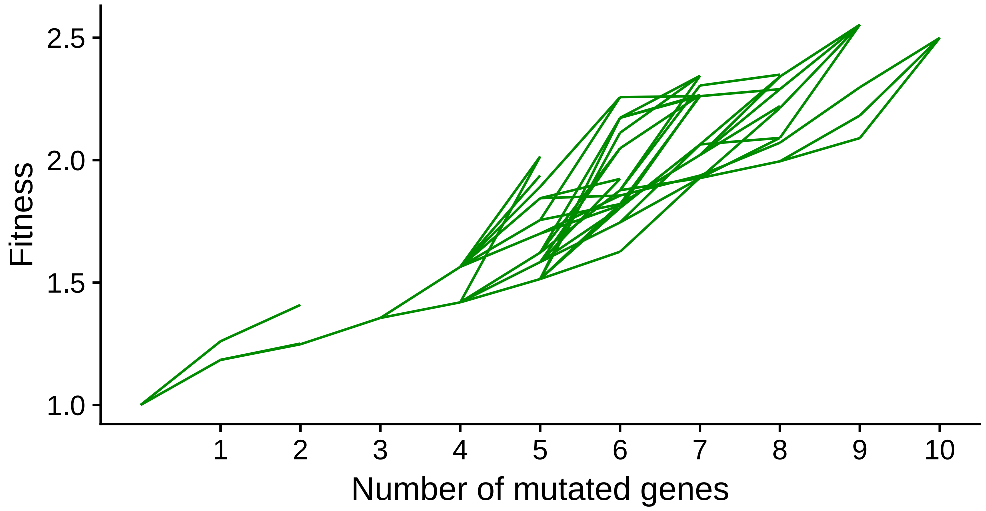


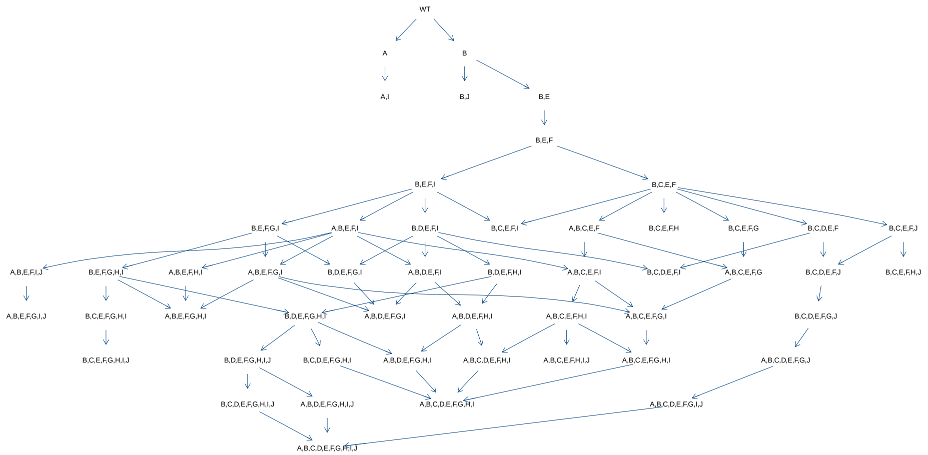

| | |
|---|---|
| *ID* | 2yuFuKxbV3XC3194 |
| *acc genotypes* | 282 |
| *peaks no back* | 63 |
| *removed edges* | NA |
| *prop rem edges* | NA |

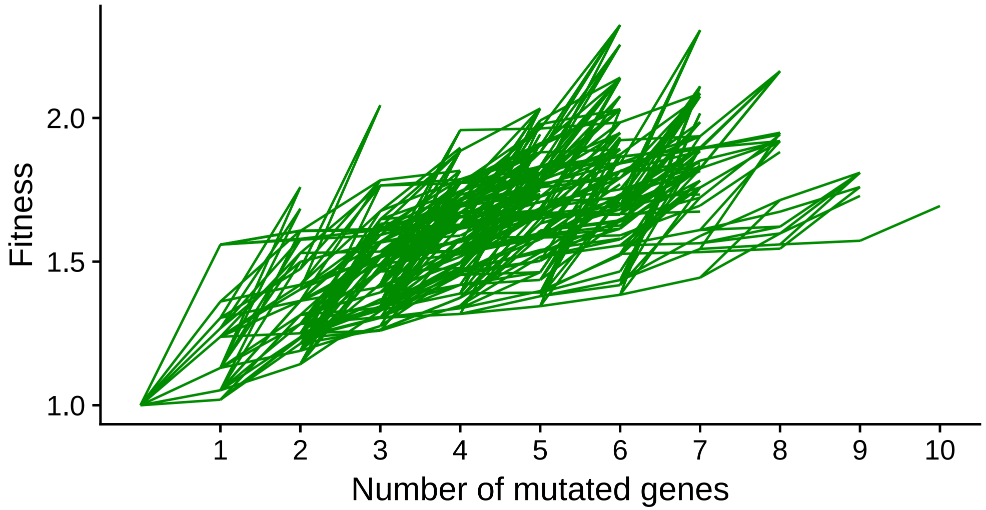


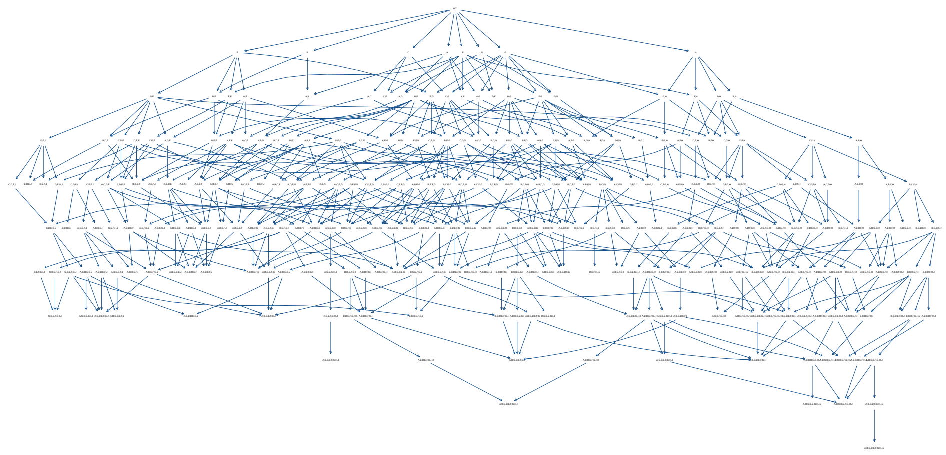

| | |
|---|---|
| *ID* | 341bQ6tMBQ3wbqIX |
| *acc genotypes* | 204 |
| *peaks no back* | 68 |
| *removed edges* | NA |
| *prop rem edges* | NA |

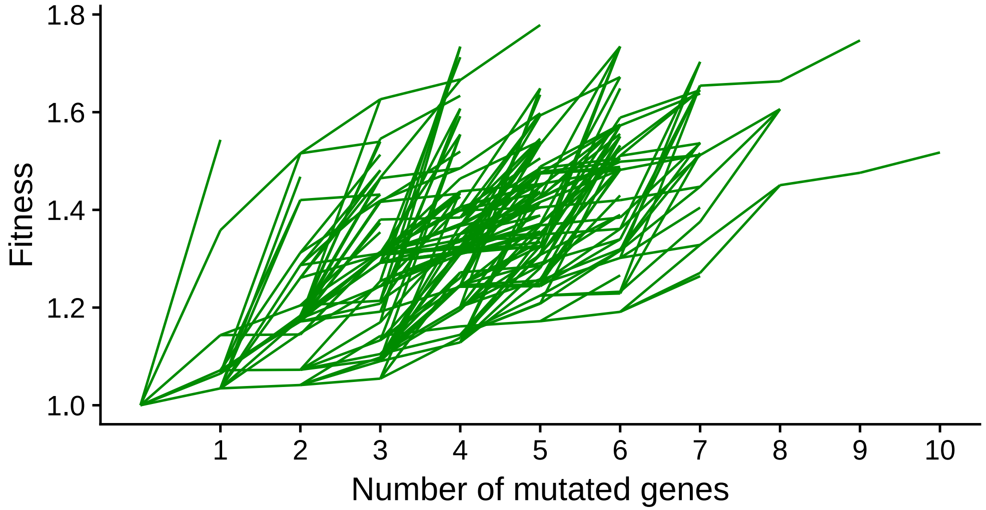


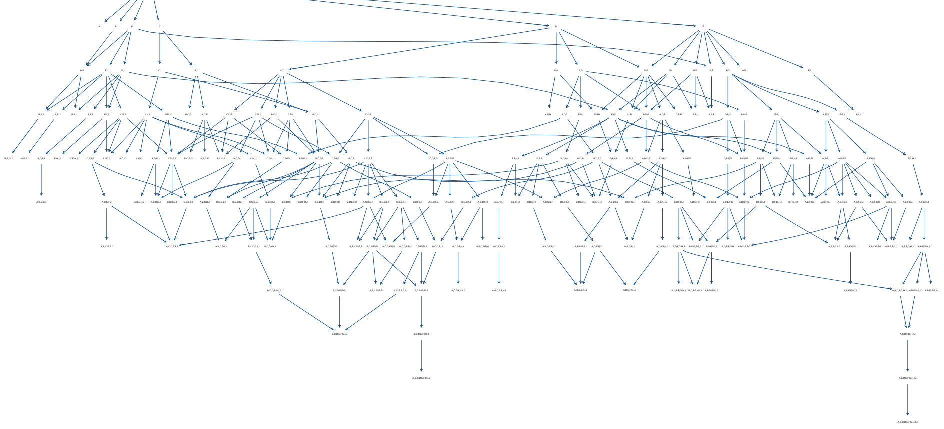

| | |
|---|---|
| *ID* | 3ageR5lDF1410rU5i |
| *acc genotypes* | 51 |
| *peaks no back* | 25 |
| *removed edges* | NA |
| *prop rem edges* | NA |

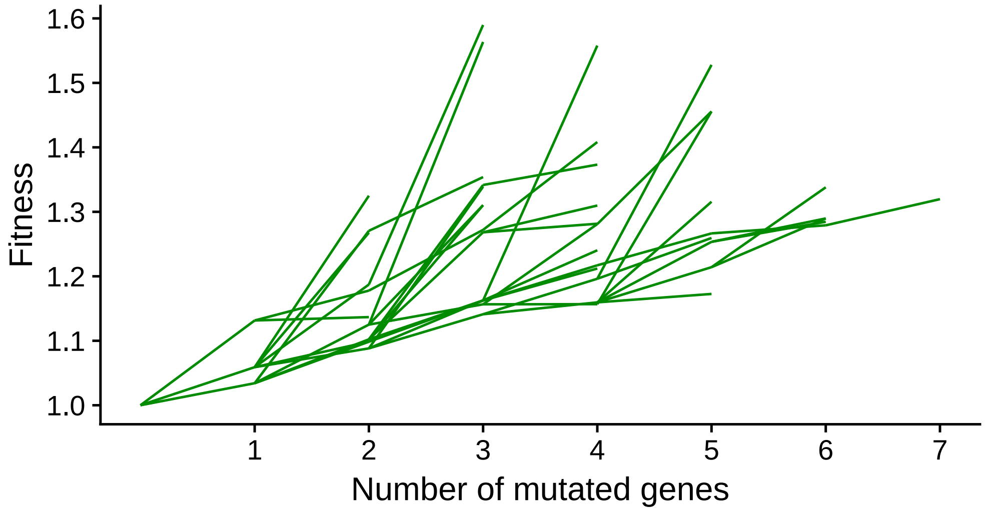


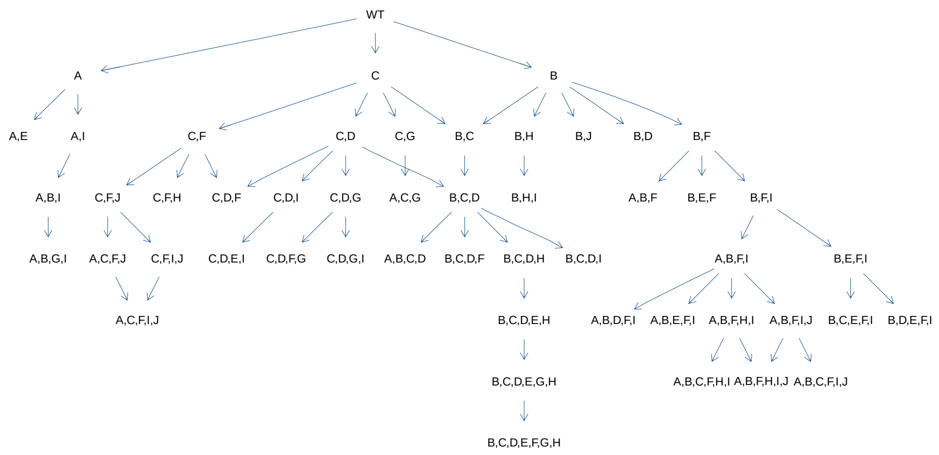

| | |
|---|---|
| *ID* | 3PJ410K7y10k7imair |
| *acc genotypes* | 921 |
| *peaks no back* | 50 |
| *removed edges* | NA |
| *prop rem edges* | NA |

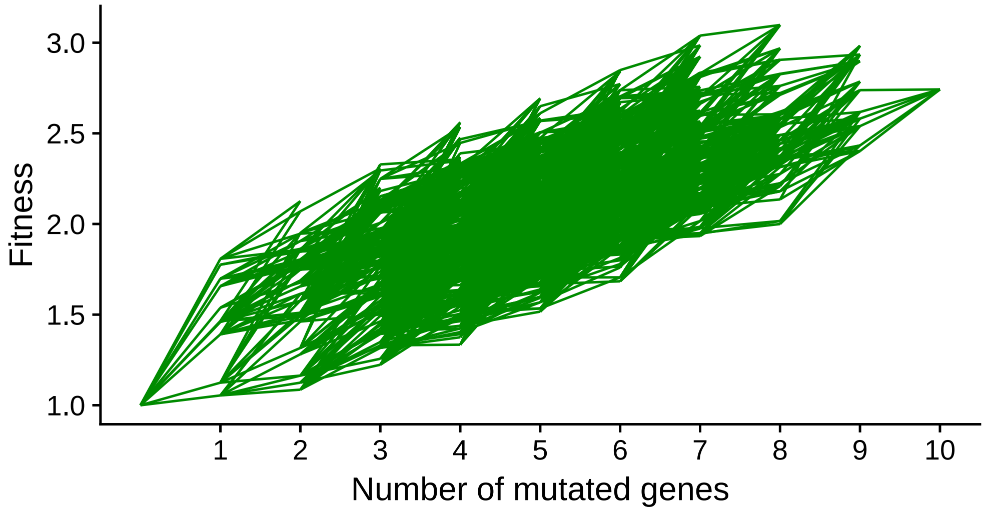


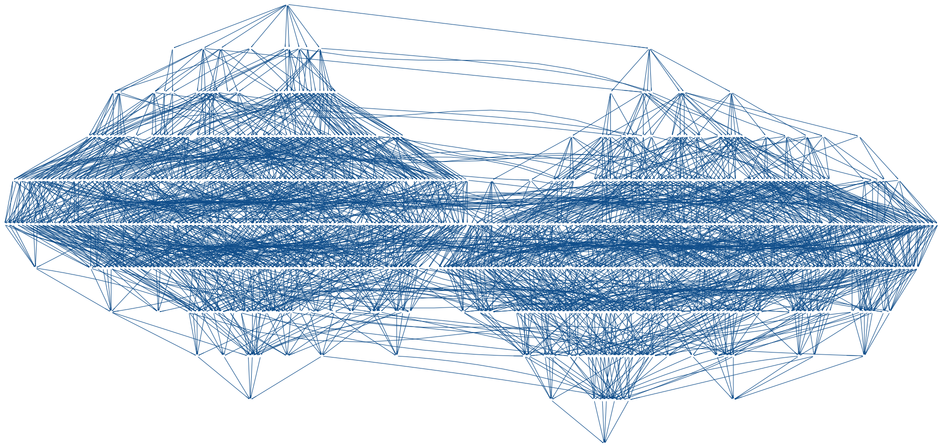

| | |
|---|---|
| *ID* | 3r101LFQ84LQ4QX2E |
| *acc genotypes* | 66 |
| *peaks no back* | 29 |
| *removed edges* | NA |
| *prop rem edges* | NA |

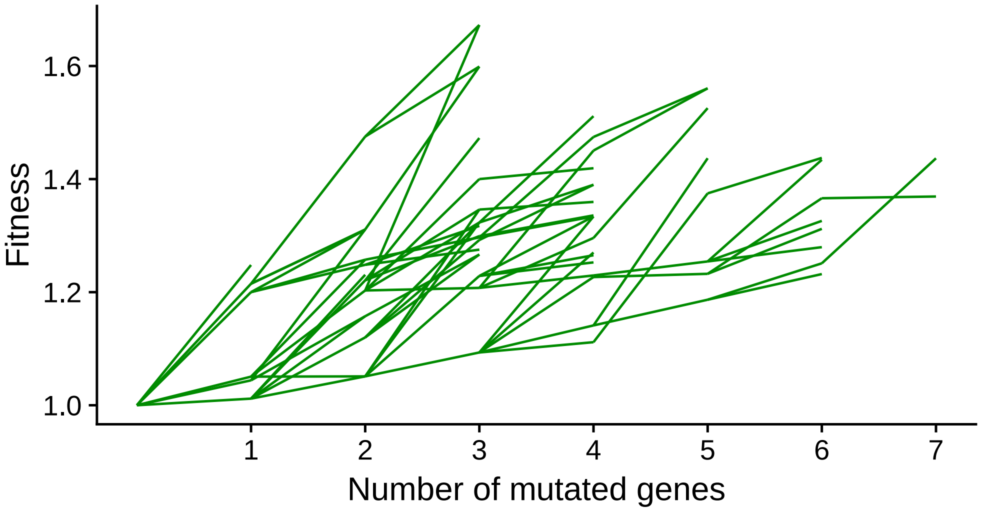


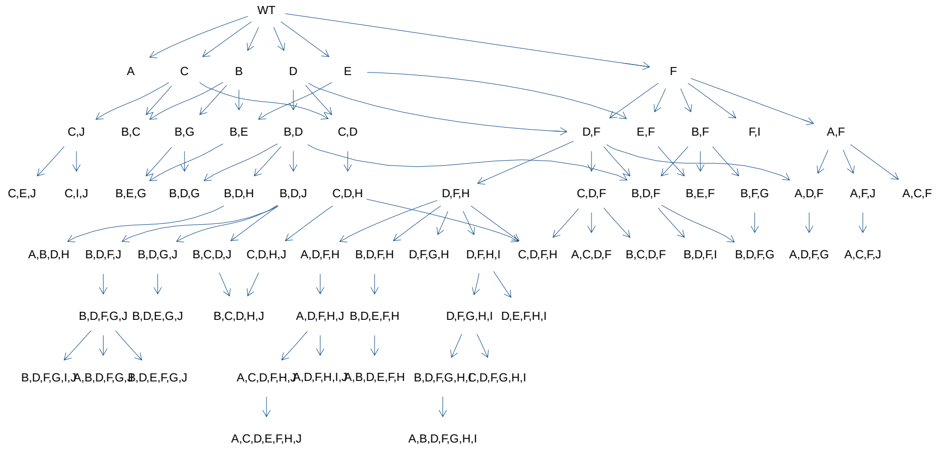

| | |
|---|---|
| *ID* | 3uTPGDsbMFbp45JR |
| *acc genotypes* | 131 |
| *peaks no back* | 38 |
| *removed edges* | NA |
| *prop rem edges* | NA |

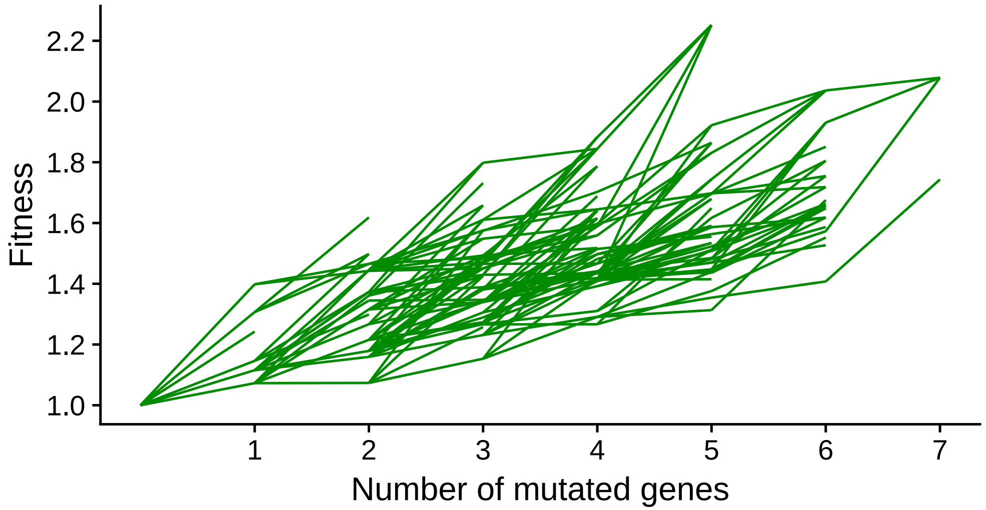


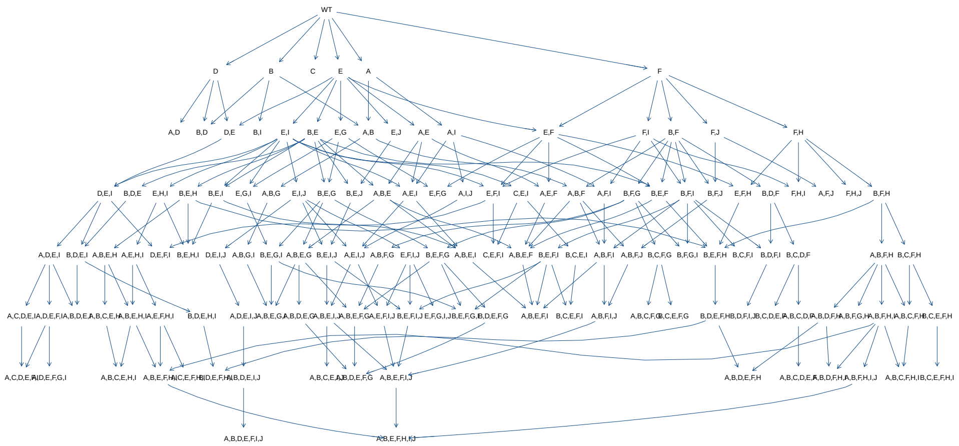

*ID* 3VdNxc10SG10x5vQBJ
*acc genotypes* 82
*peaks no back* 35
*removed edges* NA
*prop rem edges* NA

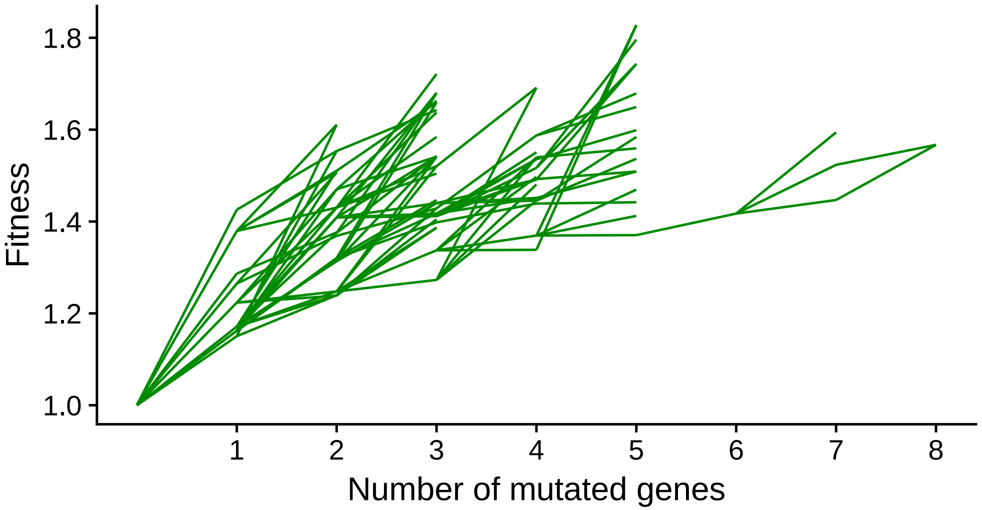


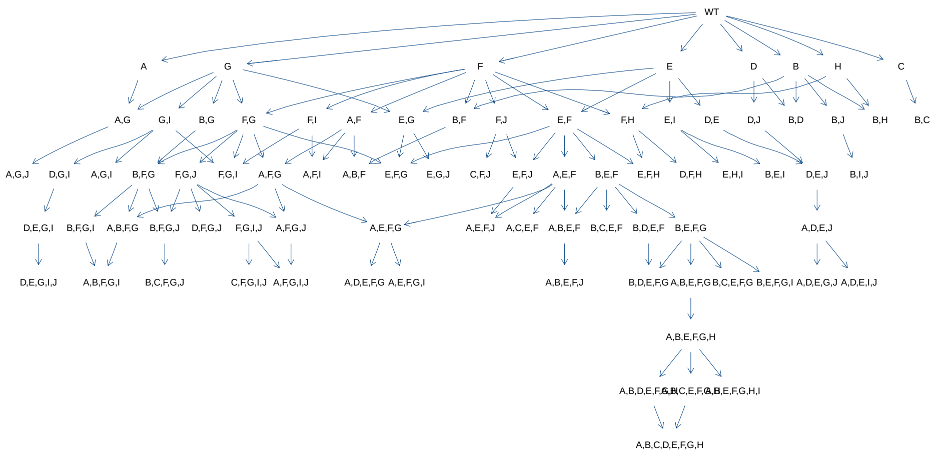

| | |
|---|---|
| *ID* | 4Ckjy1UeVlkV4sax |
| *acc genotypes* | 191 |
| *peaks no back* | 38 |
| *removed edges* | NA |
| *prop rem edges* | NA |

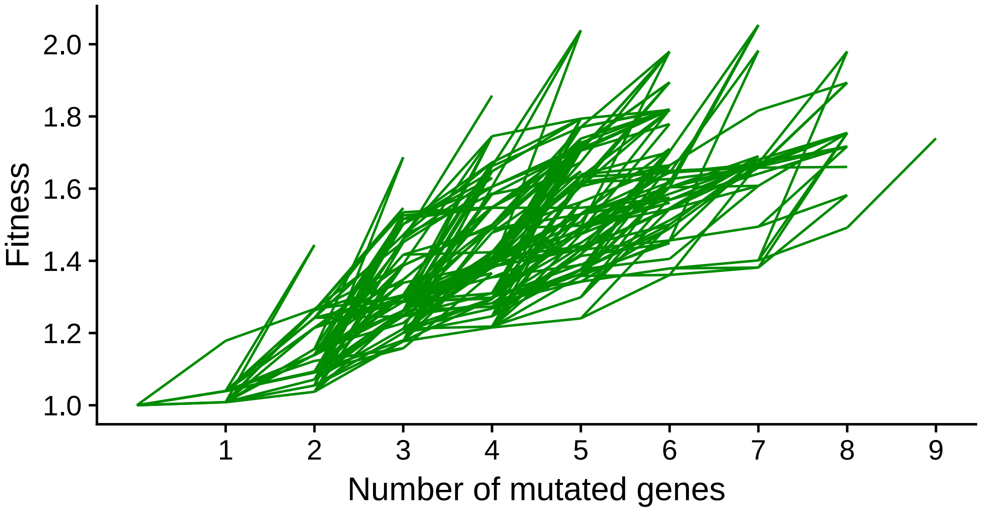


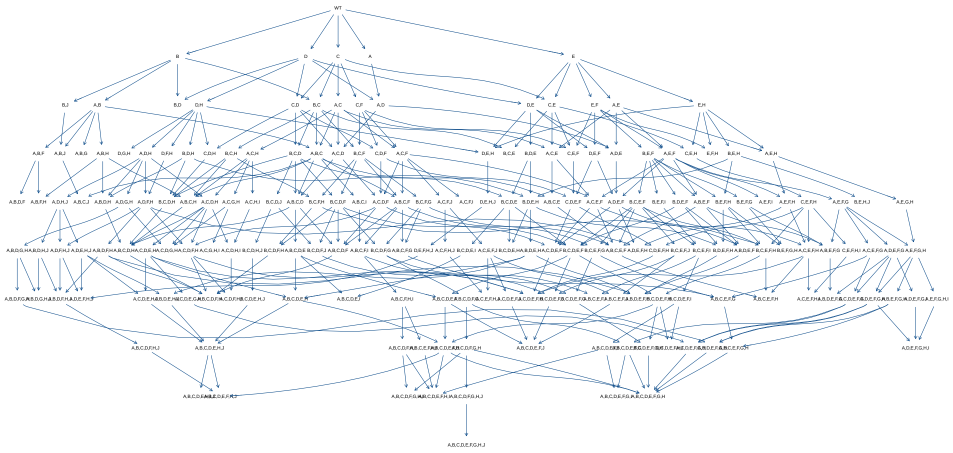

| | |
|---|---|
| *ID* | 4RVESRiVwzxmDTB9 |
| *acc genotypes* | 82 |
| *peaks no back* | 23 |
| *removed edges* | NA |
| *prop rem edges* | NA |

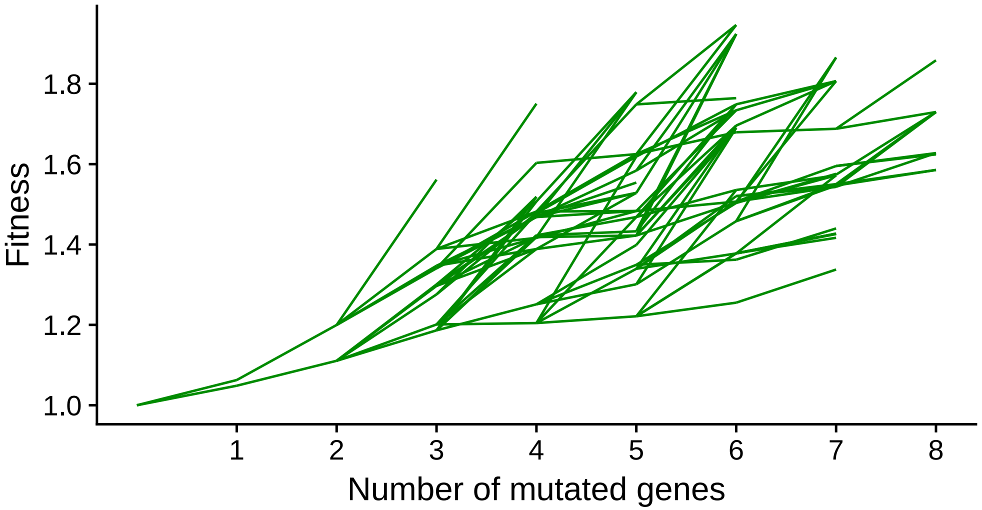


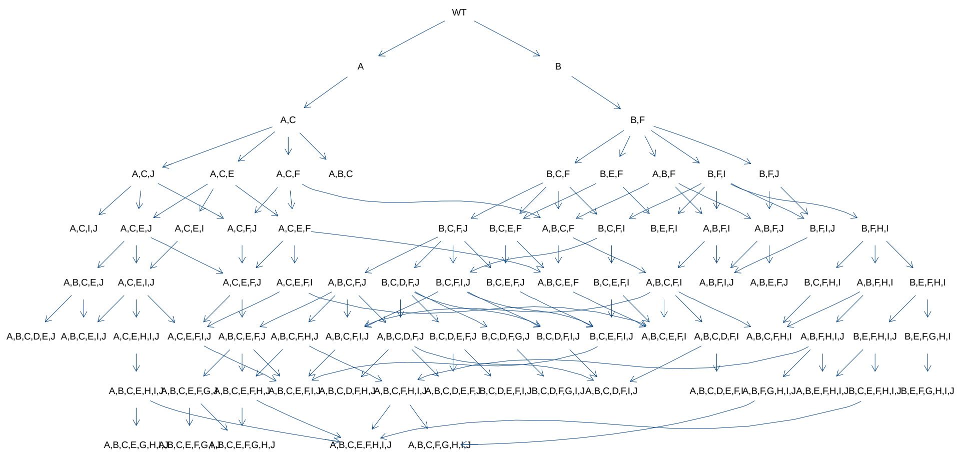

| | |
|---|---|
| *ID* | 5h10KTbvYD9O10OXnK |
| *acc genotypes* | 59 |
| *peaks no back* | 28 |
| *removed edges* | NA |
| *prop rem edges* | NA |

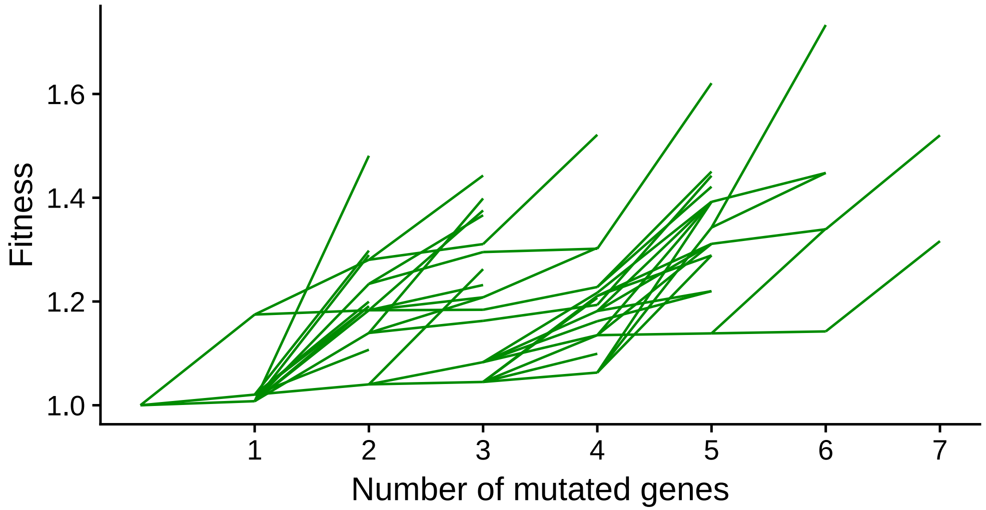


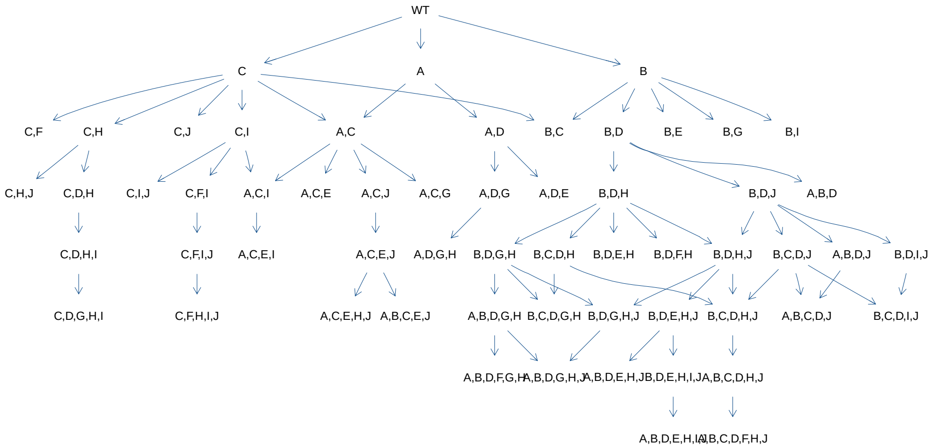

| | |
|---|---|
| *ID* | 5w3nMqdUbSKjUyuj |
| *acc genotypes* | 336 |
| *peaks no back* | 69 |
| *removed edges* | NA |
| *prop rem edges* | NA |

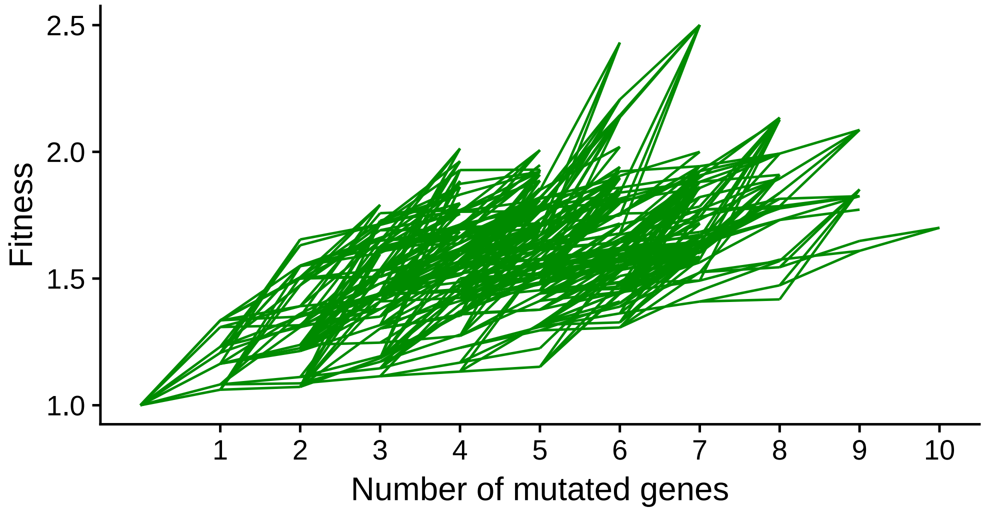


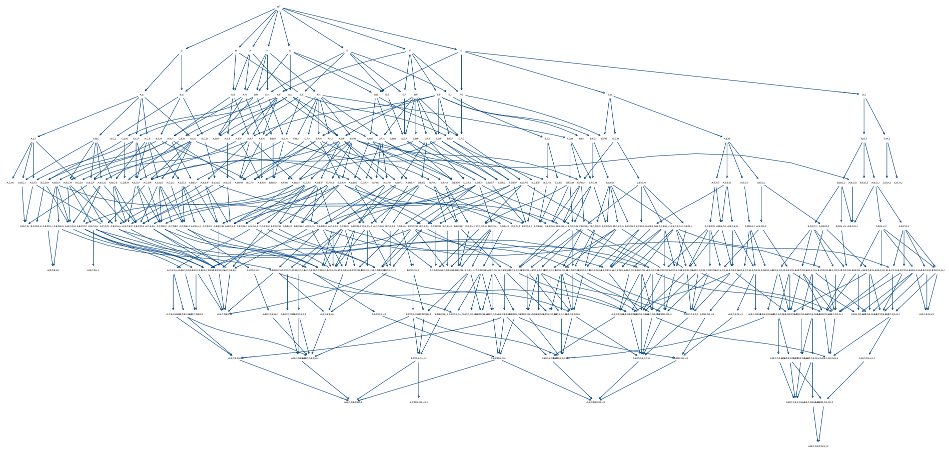

| | |
|---|---|
| *ID* | 5XTuzOyhioi24t2s |
| *acc genotypes* | 48 |
| *peaks no back* | 21 |
| *removed edges* | NA |
| *prop rem edges* | NA |

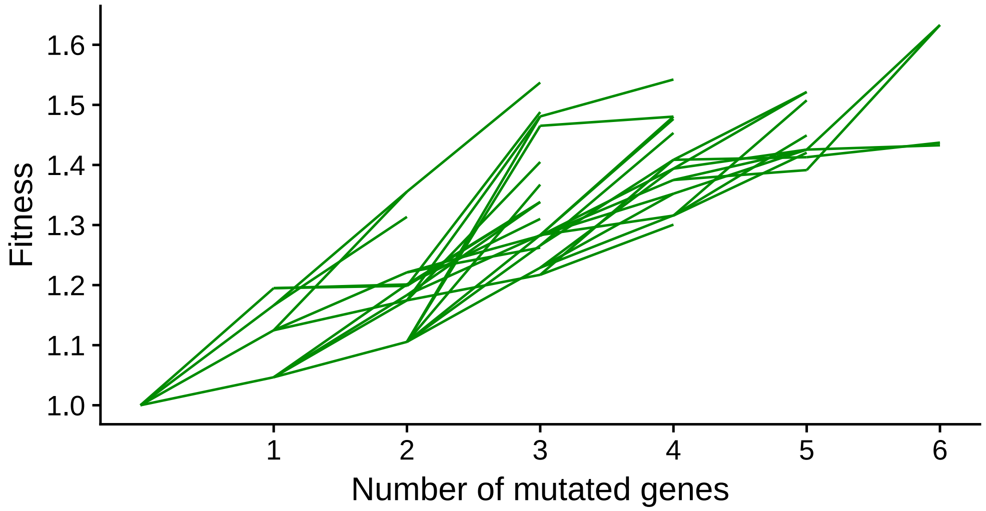


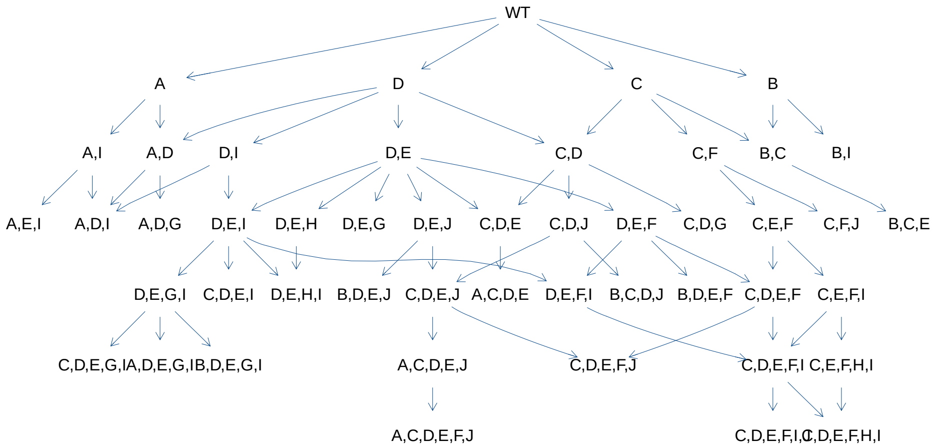

| | |
|---|---|
| *ID* | 75tsJre1YoxpOsyh |
| *acc genotypes* | 187 |
| *peaks no back* | 70 |
| *removed edges* | NA |
| *prop rem edges* | NA |

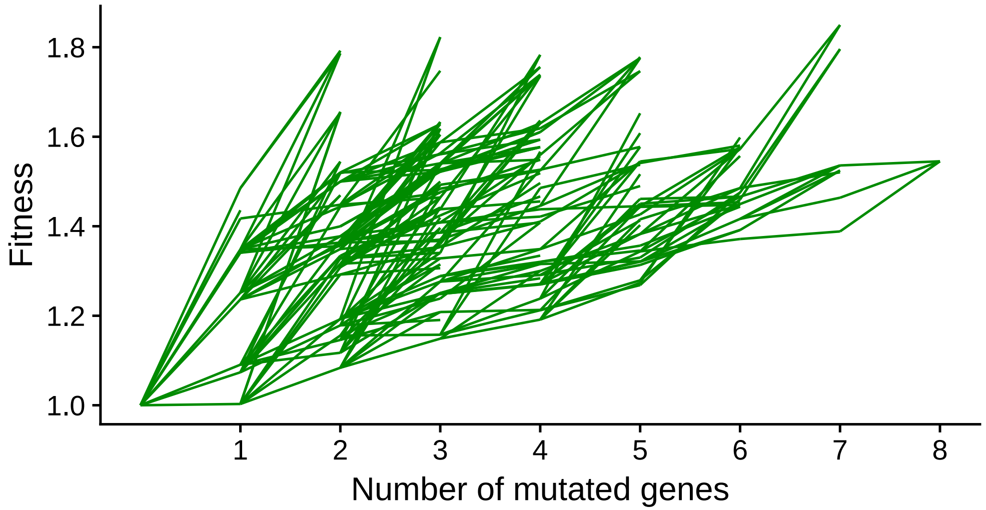


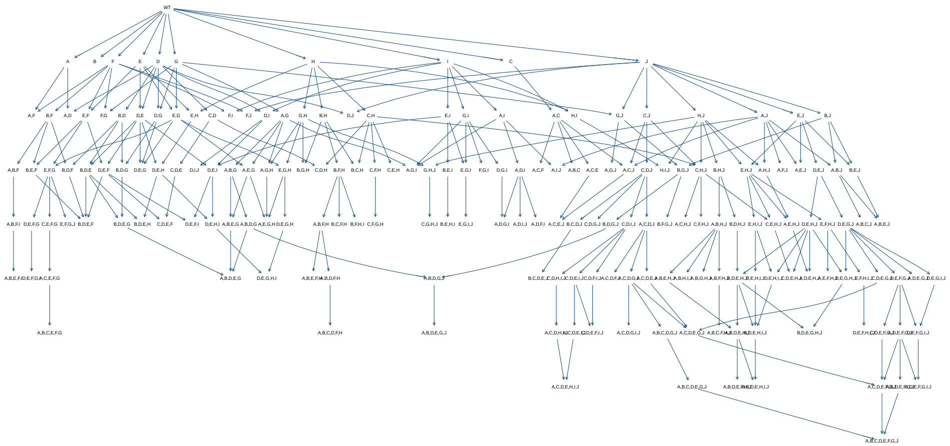

| | |
|---|---|
| *ID* | 7DaZyu10znssTnBfx |
| *acc genotypes* | 674 |
| *peaks no back* | 56 |
| *removed edges* | NA |
| *prop rem edges* | NA |

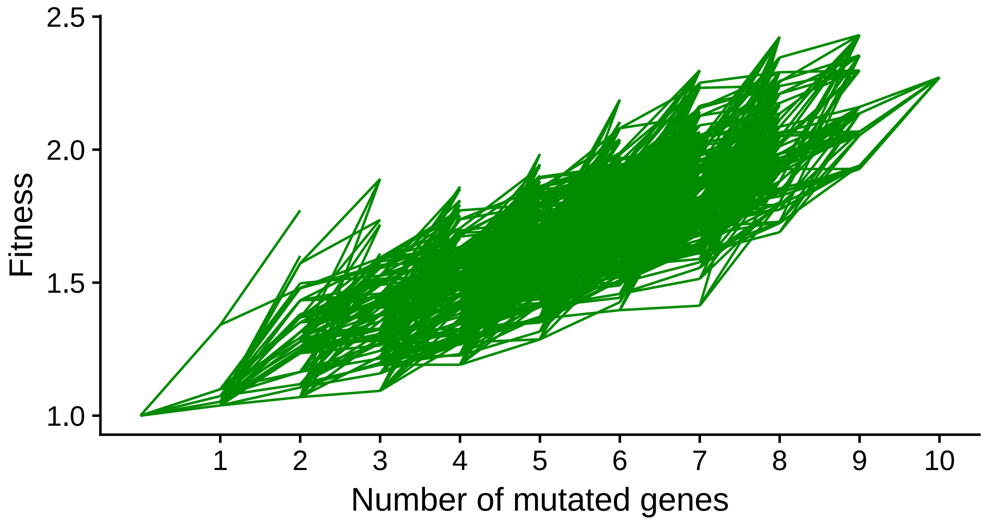


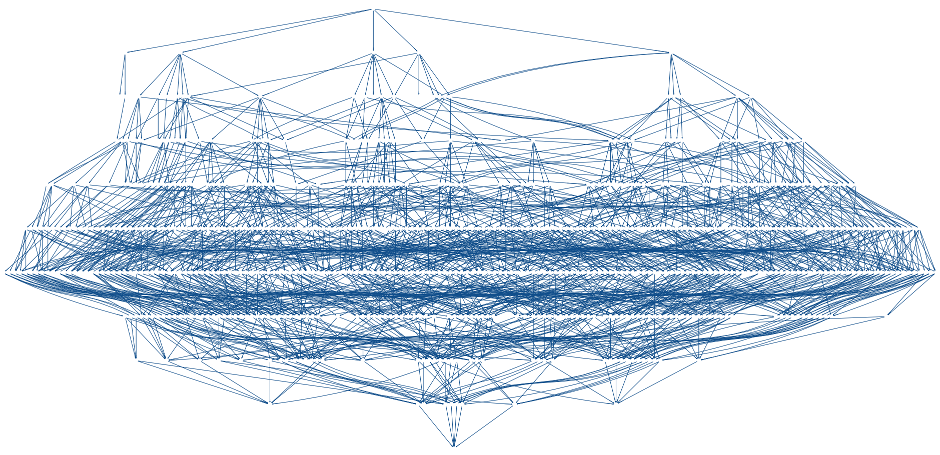

| | |
|---|---|
| *ID* | 7GapUNhPAnaZAI1M |
| *acc genotypes* | 427 |
| *peaks no back* | 53 |
| *removed edges* | NA |
| *prop rem edges* | NA |

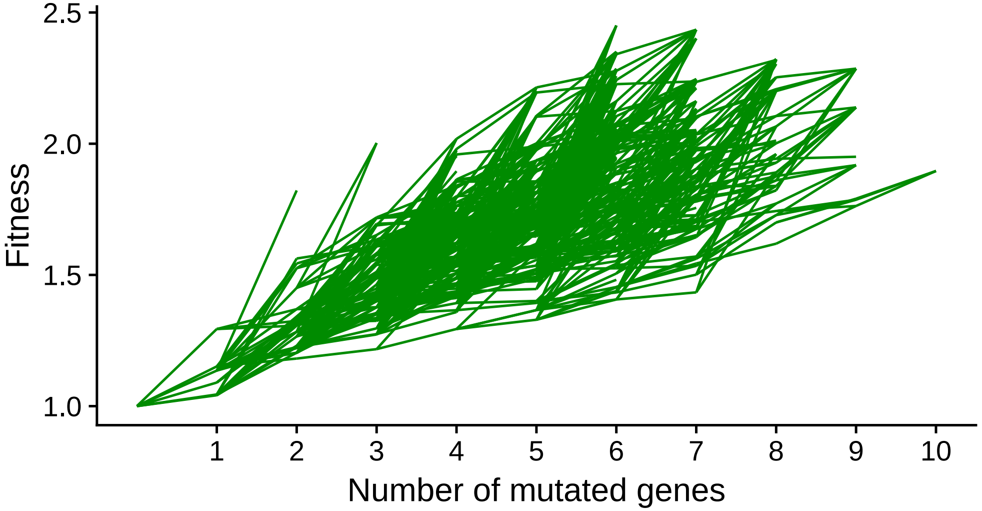


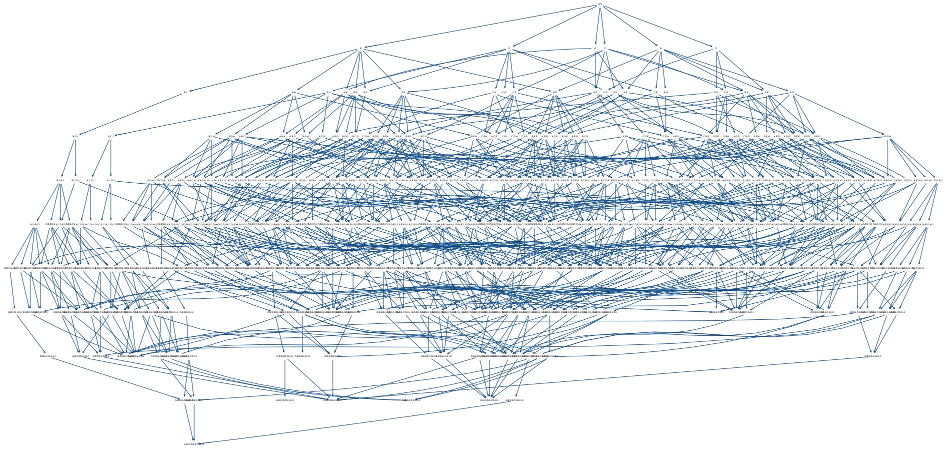

| | |
|---|---|
| *ID* | 7jxvBRIz6DAUbzBL |
| *acc genotypes* | 24 |
| *peaks no back* | 12 |
| *removed edges* | NA |
| *prop rem edges* | NA |

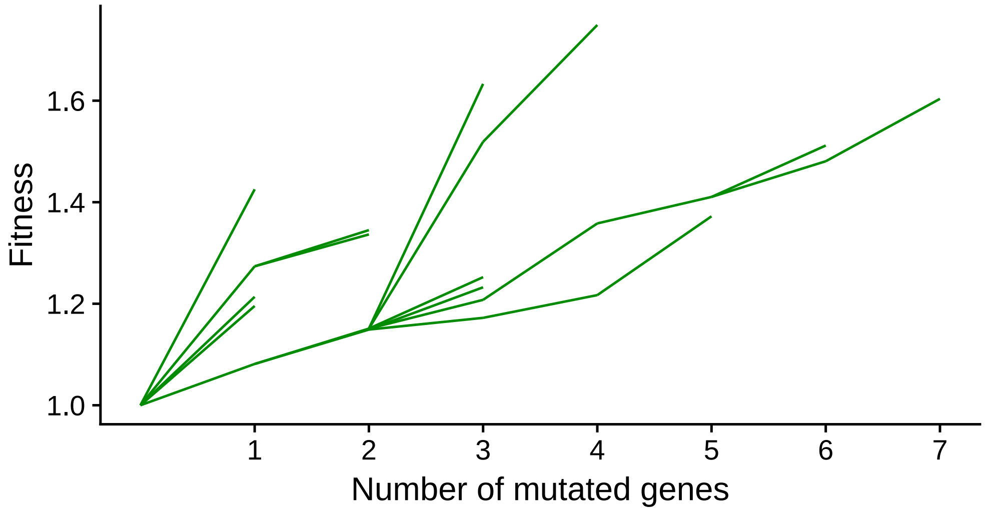


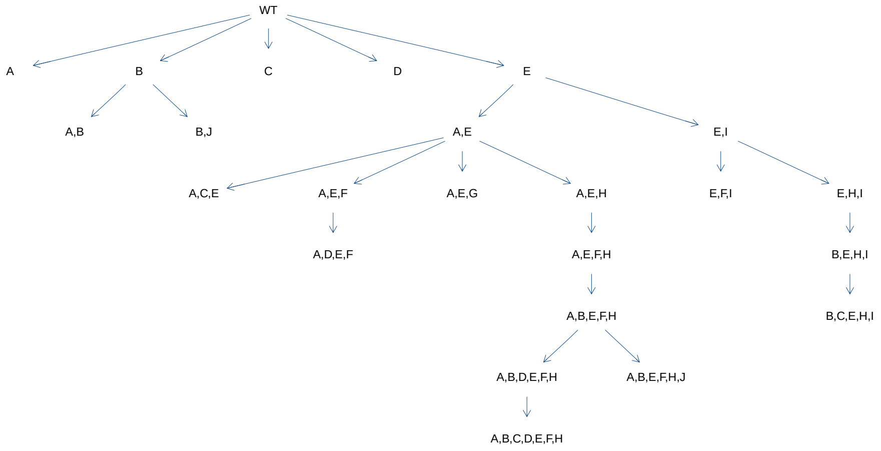

| | |
|---|---|
| *ID* | 7KQQodSTMHz6tzpM |
| *acc genotypes* | 291 |
| *peaks no back* | 55 |
| *removed edges* | NA |
| *prop rem edges* | NA |

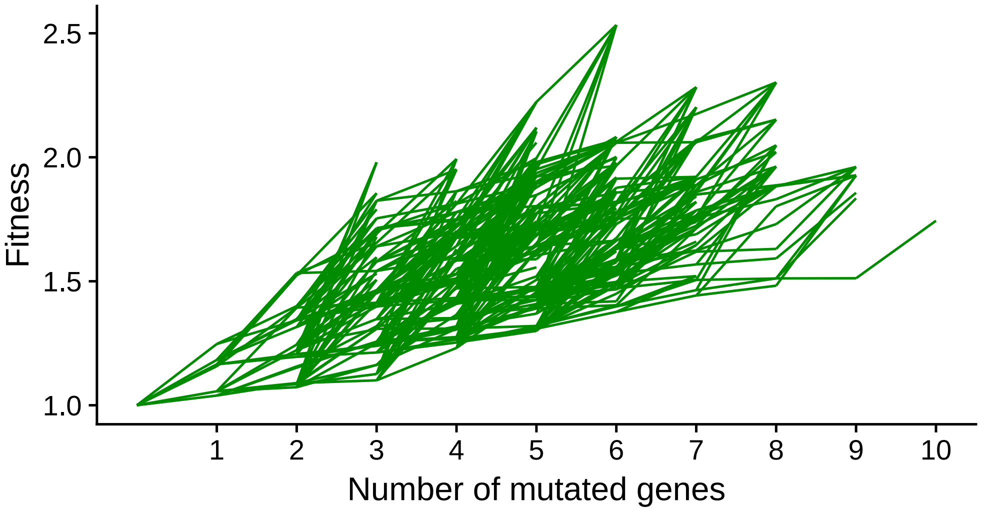


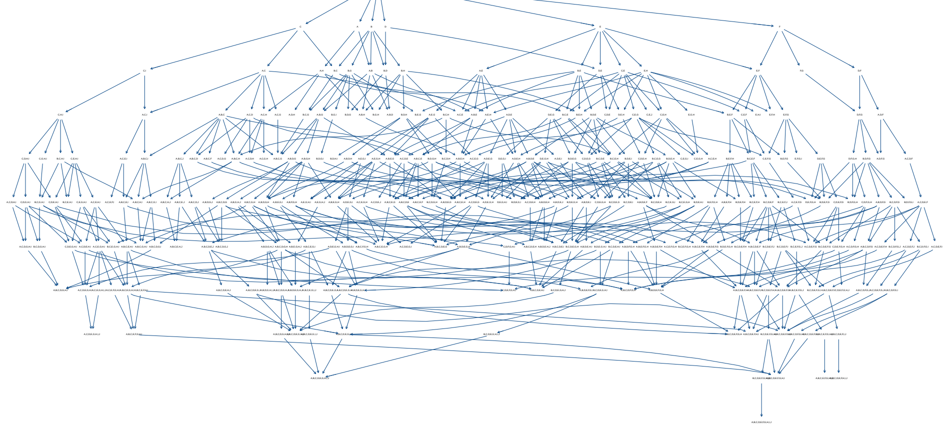

| | |
|---|---|
| *ID* | 7yMvs88xHRDYKybO |
| *acc genotypes* | 182 |
| *peaks no back* | 62 |
| *removed edges* | NA |
| *prop rem edges* | NA |

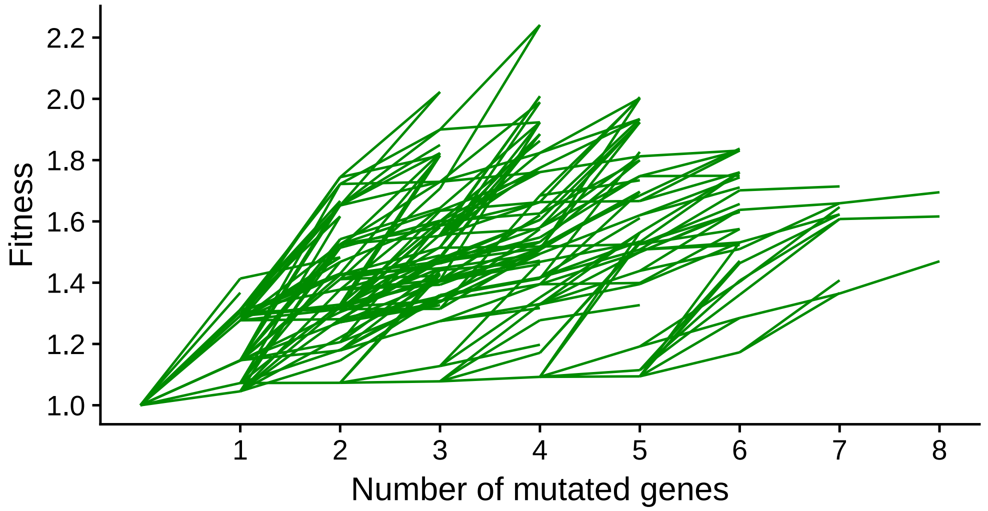


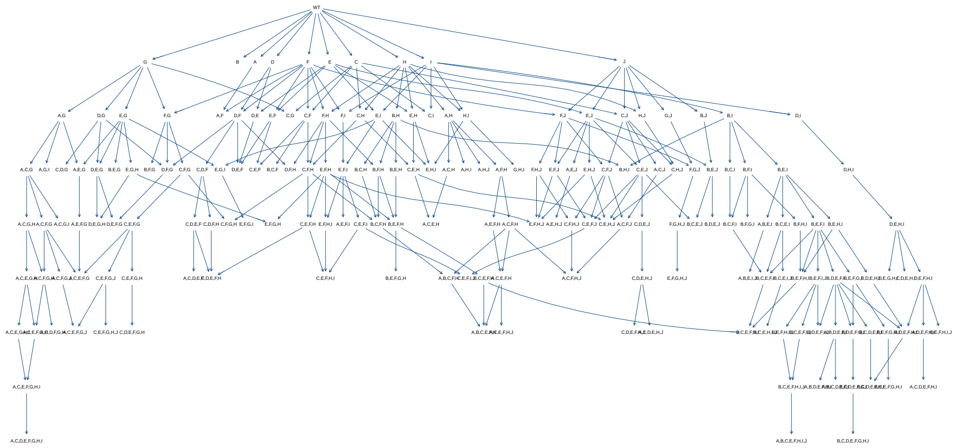

| | |
|---|---|
| *ID* | 7YYdBu6jr3vr7yKp |
| *acc genotypes* | 197 |
| *peaks no back* | 75 |
| *removed edges* | NA |
| *prop rem edges* | NA |

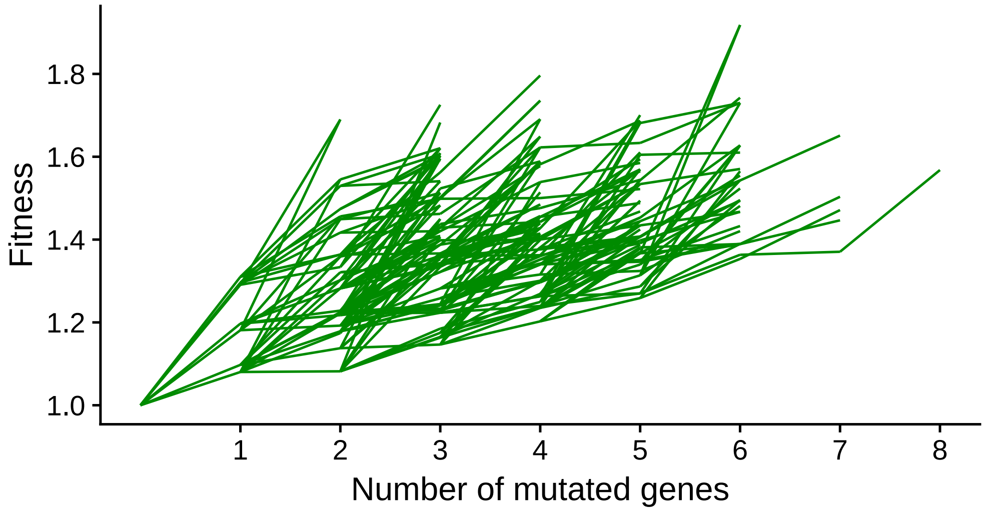


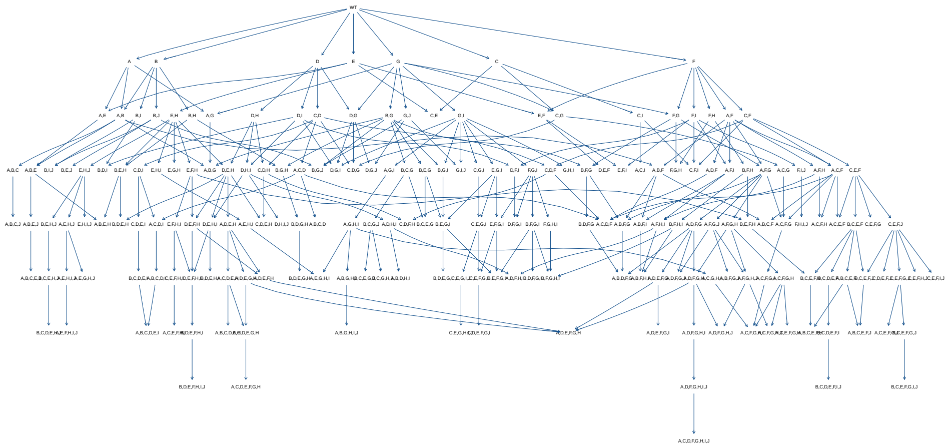

| | |
|---|---|
| *ID* | 8OiLROdLmPboaFyZ |
| *acc genotypes* | 145 |
| *peaks no back* | 62 |
| *removed edges* | NA |
| *prop rem edges* | NA |

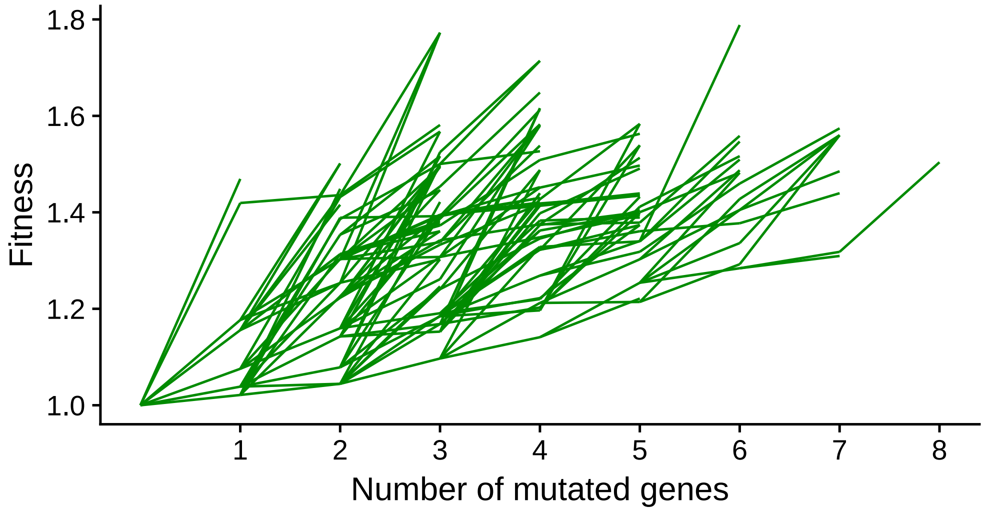


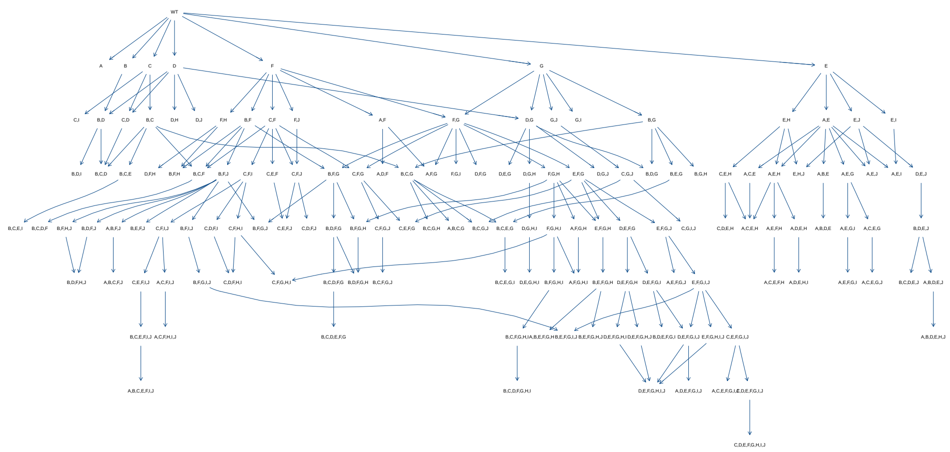

| | |
|---|---|
| *ID* | 8RIqFjyqv5Fe2F7q |
| *acc genotypes* | 130 |
| *peaks no back* | 41 |
| *removed edges* | NA |
| *prop rem edges* | NA |

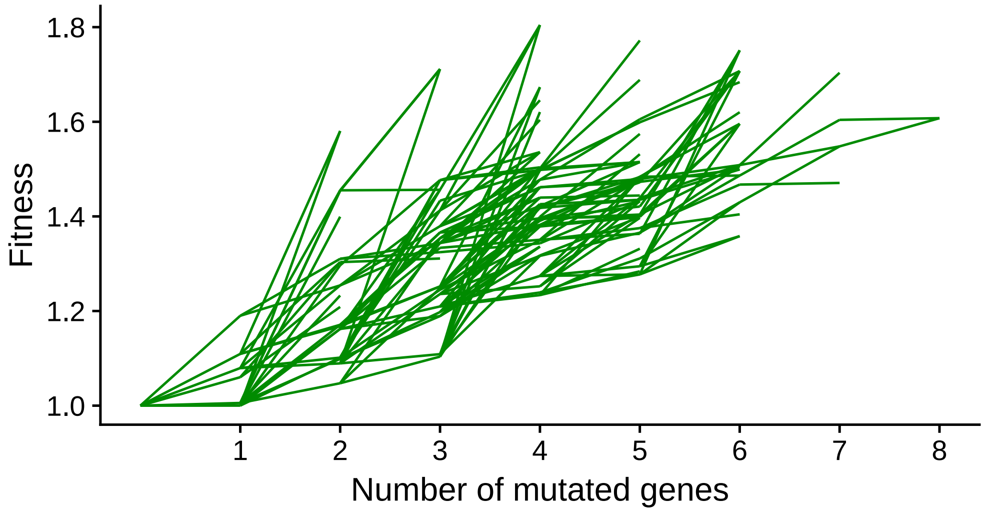


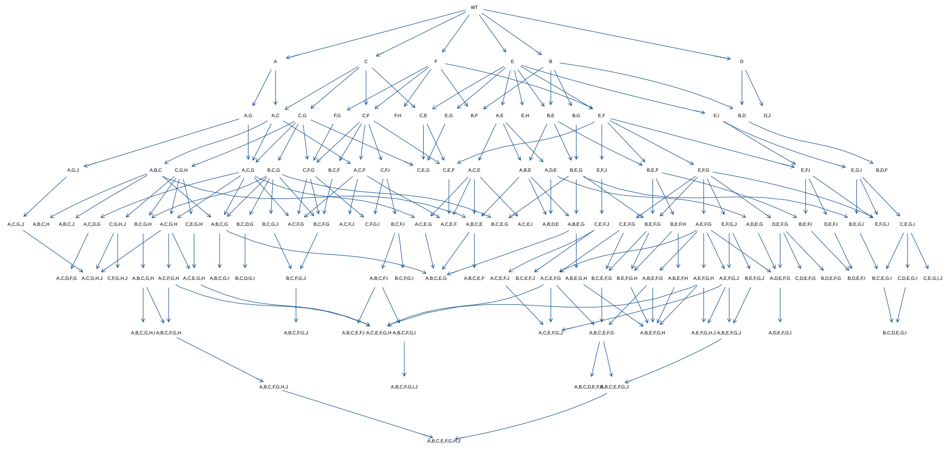

*ID* 8Wuwdsphcy5tFFeb
*acc genotypes* 117
*peaks no back* 46
*removed edges* NA
*prop rem edges* NA

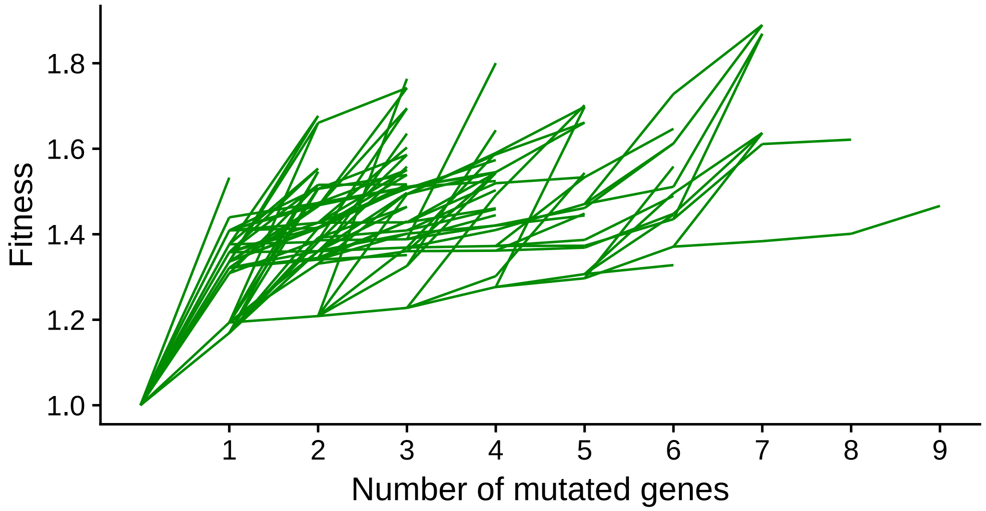


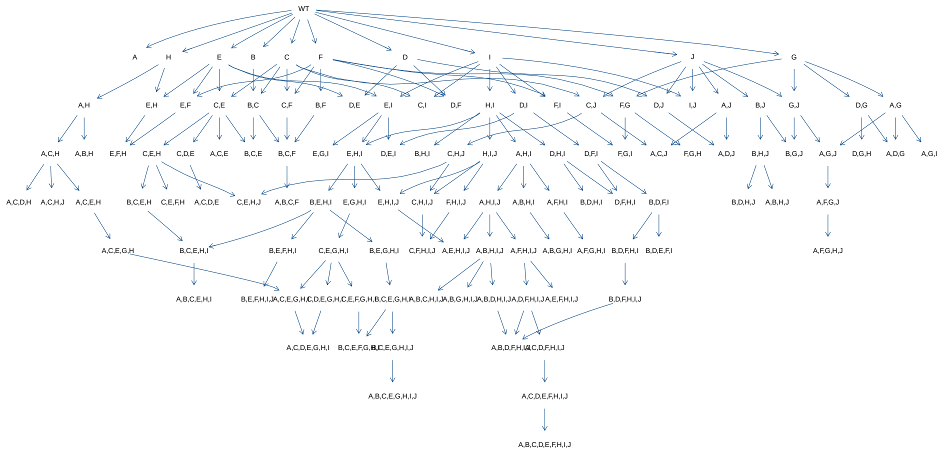

| | |
|---|---|
| *ID* | 913oZplLmQJl3z3r |
| *acc genotypes* | 96 |
| *peaks no back* | 39 |
| *removed edges* | NA |
| *prop rem edges* | NA |

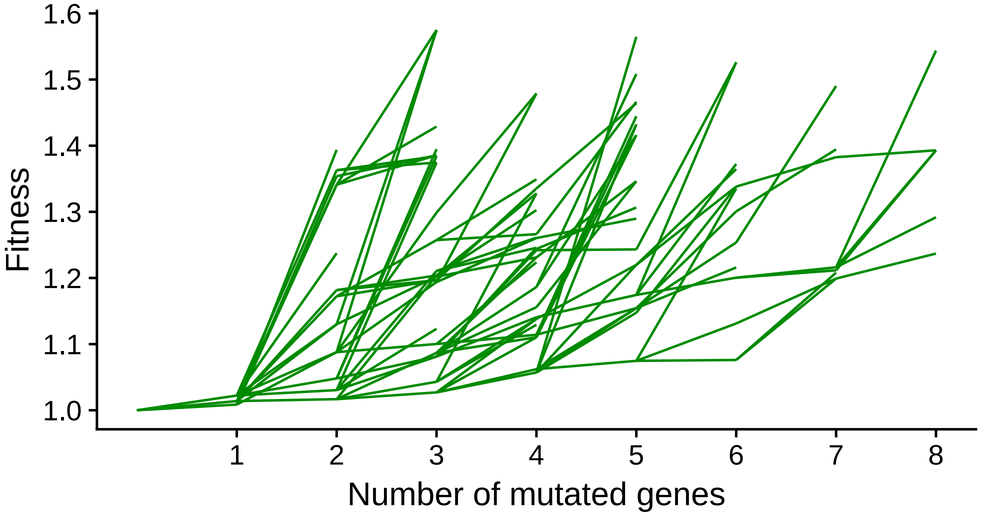


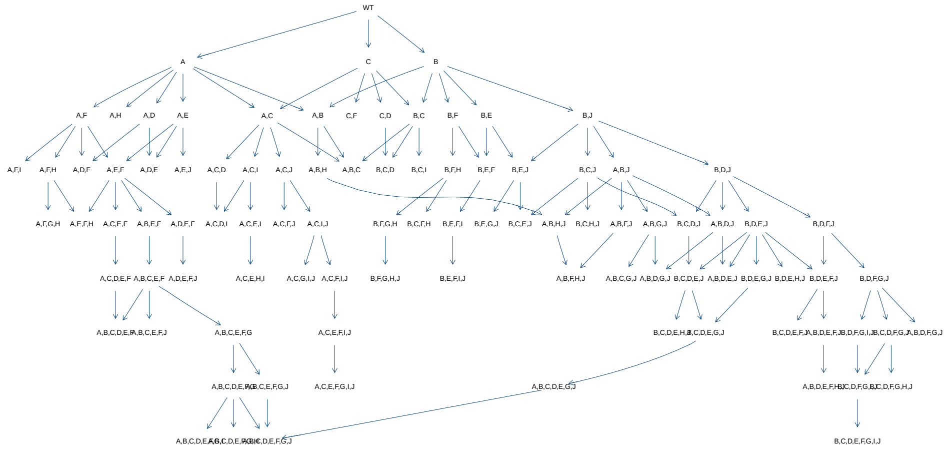

| | |
|---|---|
| *ID* | 9cep5DgsrTOPpWos |
| *acc genotypes* | 163 |
| *peaks no back* | 43 |
| *removed edges* | NA |
| *prop rem edges* | NA |

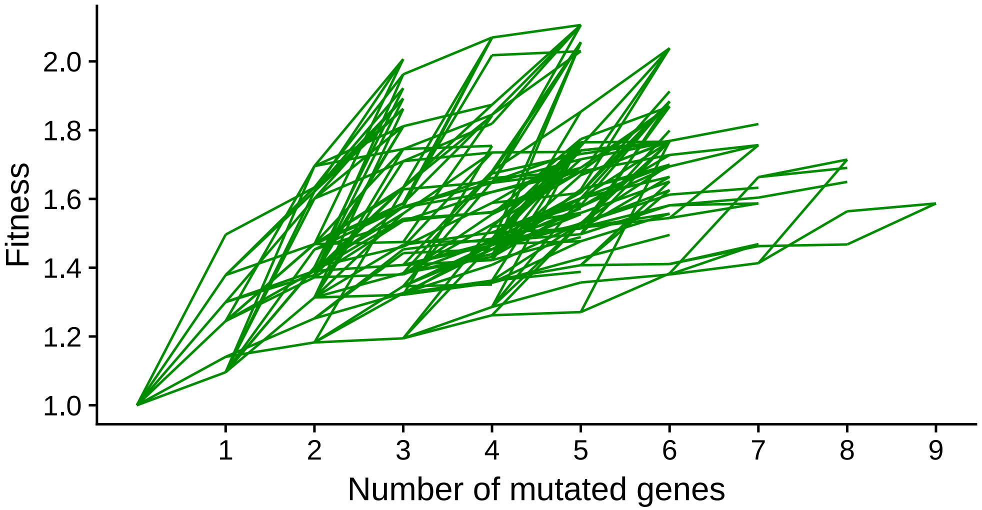


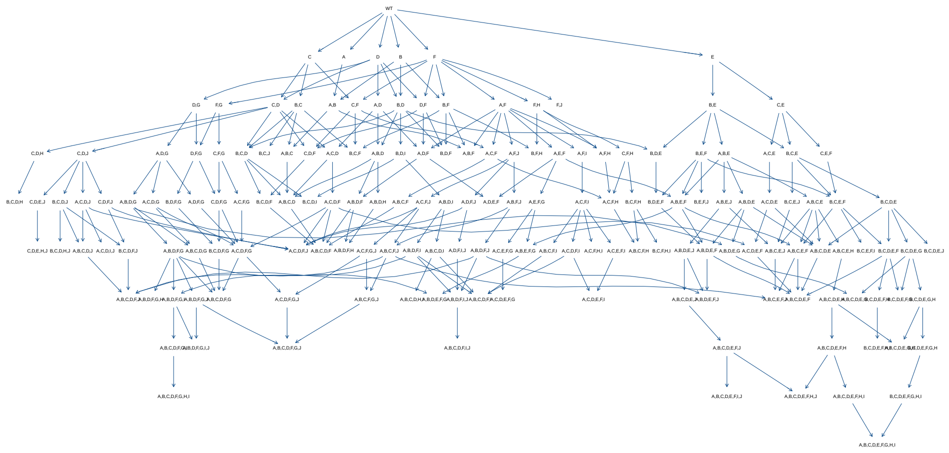

| | |
|---|---|
| *ID* | 9FoiB310FGDvgWcIE |
| *acc genotypes* | 231 |
| *peaks no back* | 69 |
| *removed edges* | NA |
| *prop rem edges* | NA |

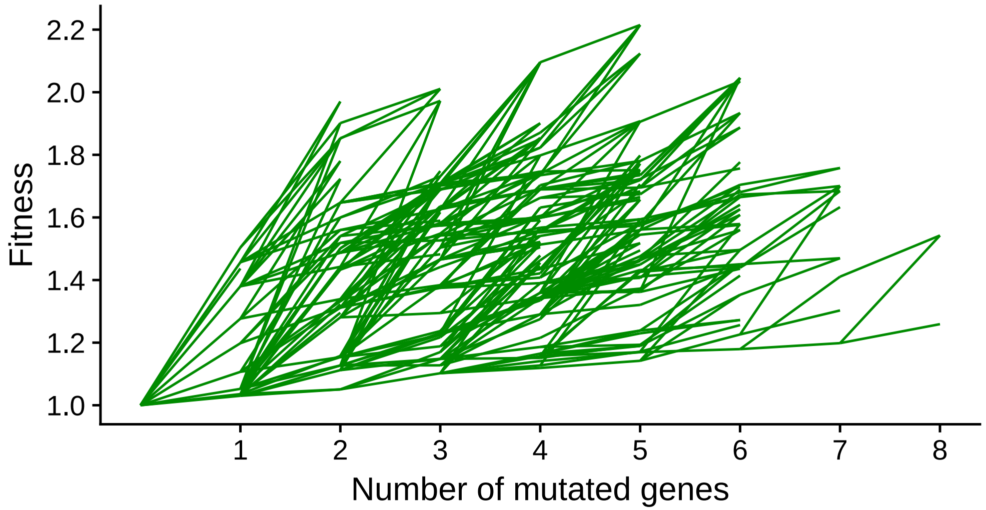


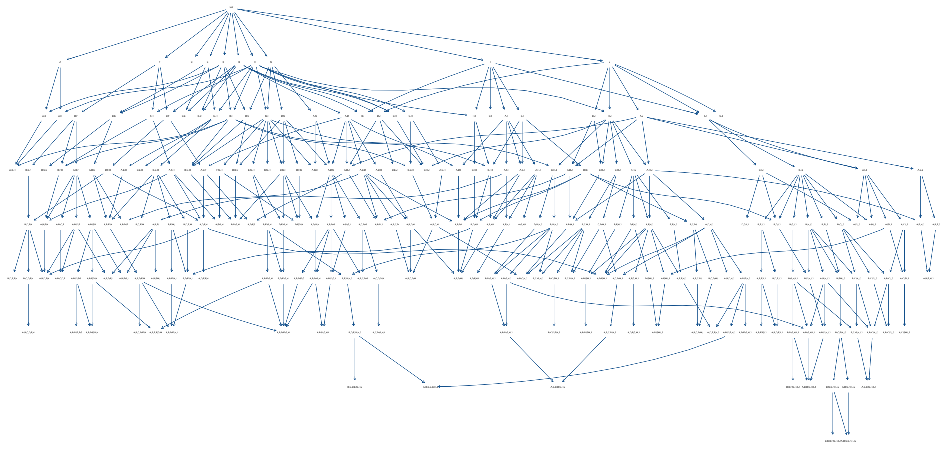

| | |
|---|---|
| *ID* | A1JsKu9z21G7TWcc |
| *acc genotypes* | 261 |
| *peaks no back* | 105 |
| *removed edges* | NA |
| *prop rem edges* | NA |

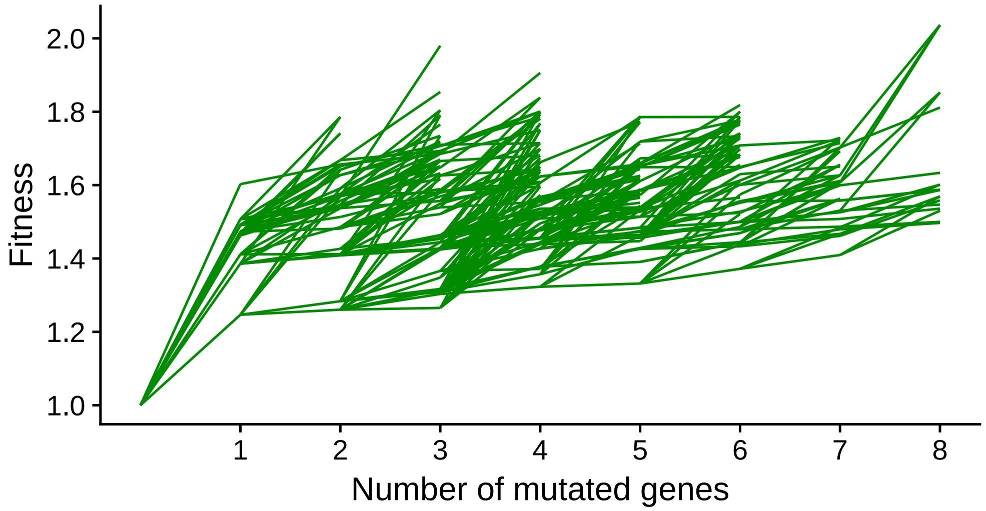


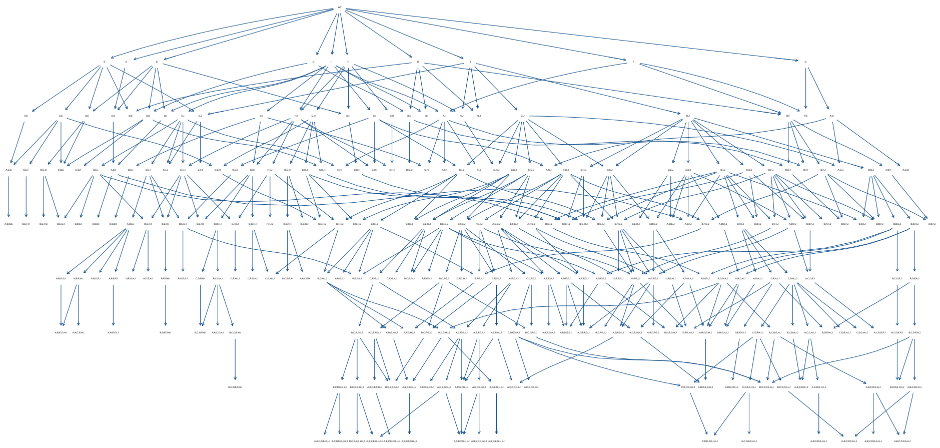

| | |
|---|---|
| *ID* | a3mOmqYxncb9Ad9p |
| *acc genotypes* | 284 |
| *peaks no back* | 99 |
| *removed edges* | NA |
| *prop rem edges* | NA |

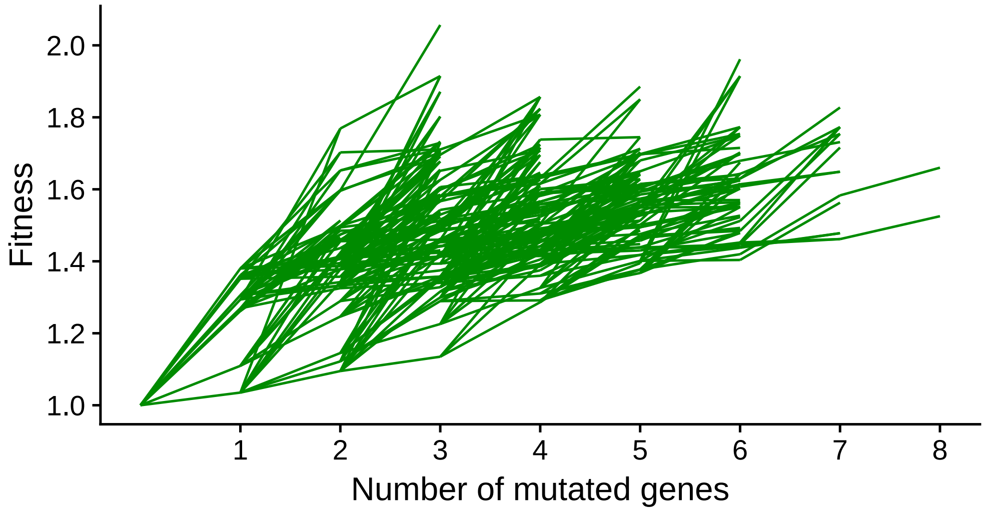


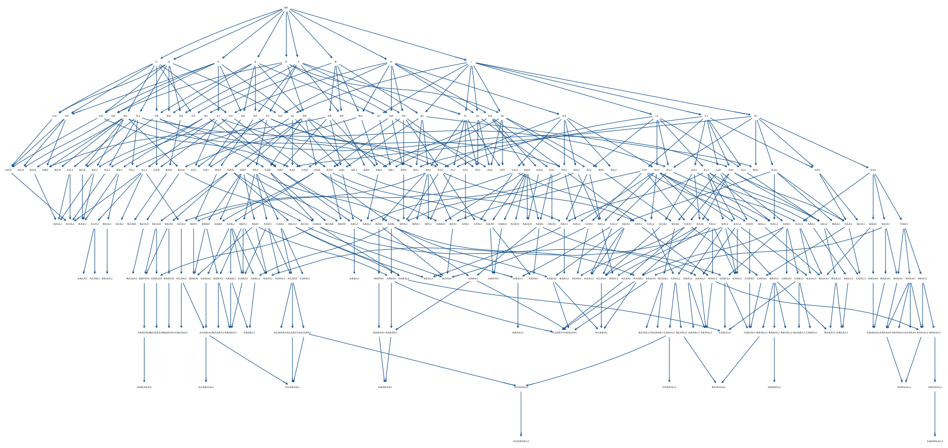

| | |
|---|---|
| *ID* | AbVs2SaYTXrETNTU |
| *acc genotypes* | 47 |
| *peaks no back* | 21 |
| *removed edges* | NA |
| *prop rem edges* | NA |

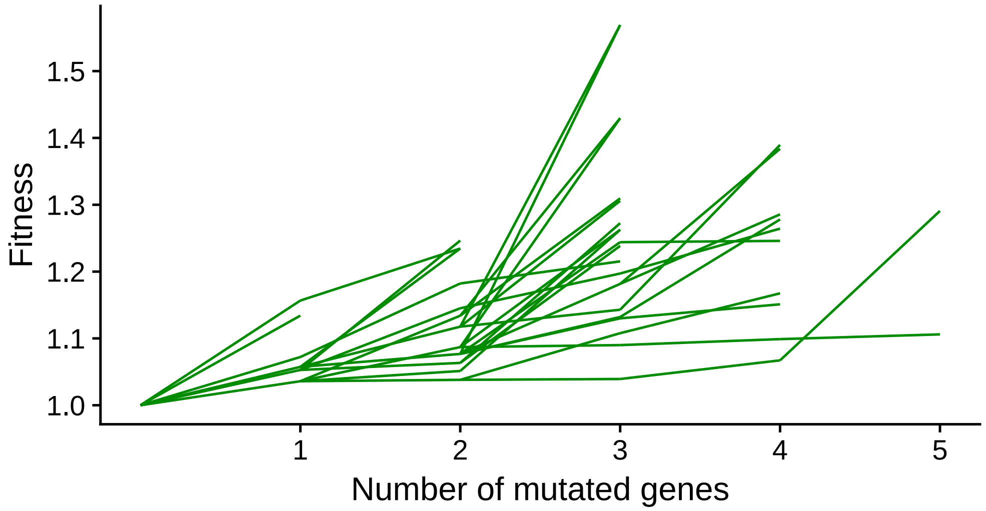


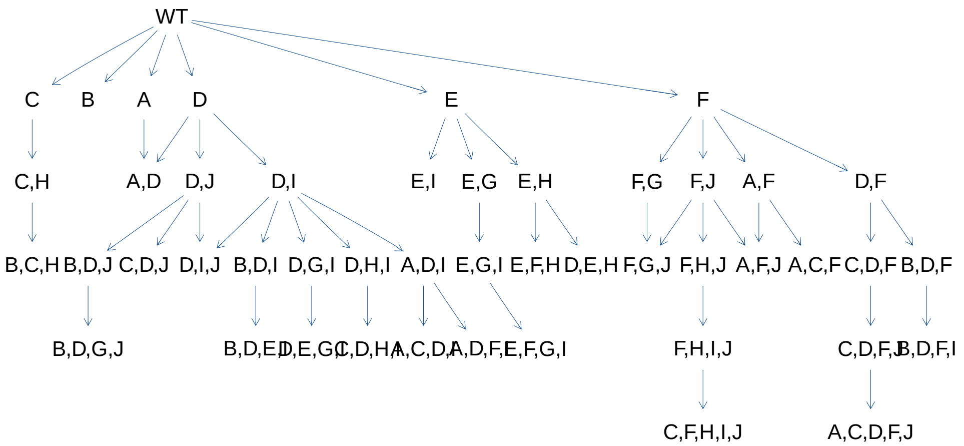

| | |
|---|---|
| *ID* | aCr6IaKy6HuXLp5m |
| *acc genotypes* | 180 |
| *peaks no back* | 53 |
| *removed edges* | NA |
| *prop rem edges* | NA |

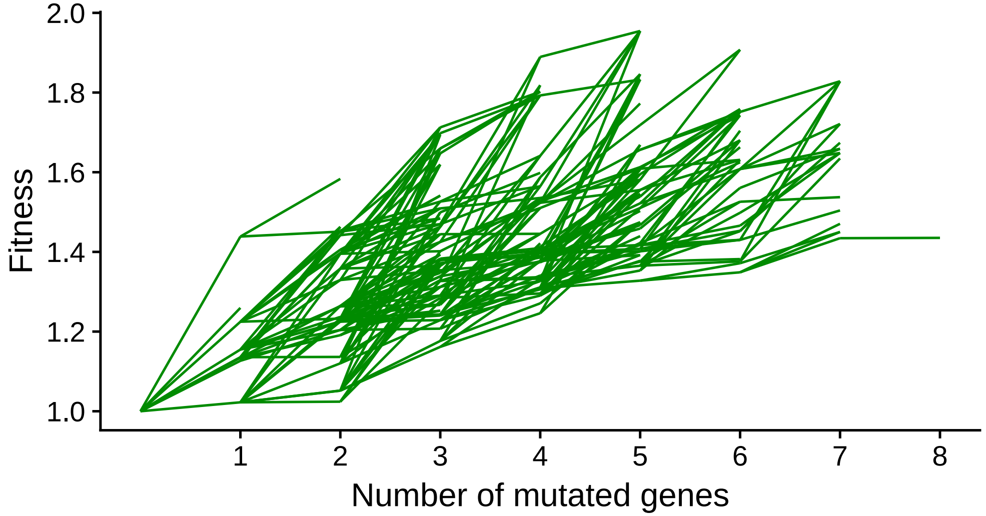


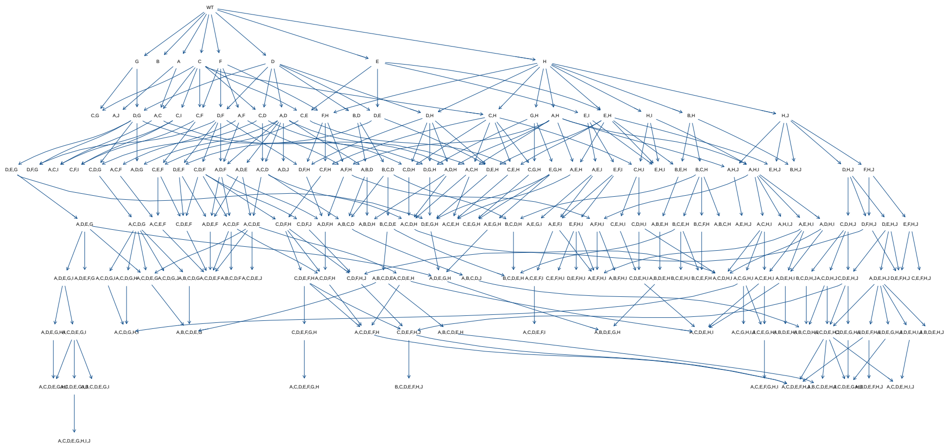

| | |
|---|---|
| *ID* | aEeRemo3Lxmgcy8w |
| *acc genotypes* | 311 |
| *peaks no back* | 94 |
| *removed edges* | NA |
| *prop rem edges* | NA |

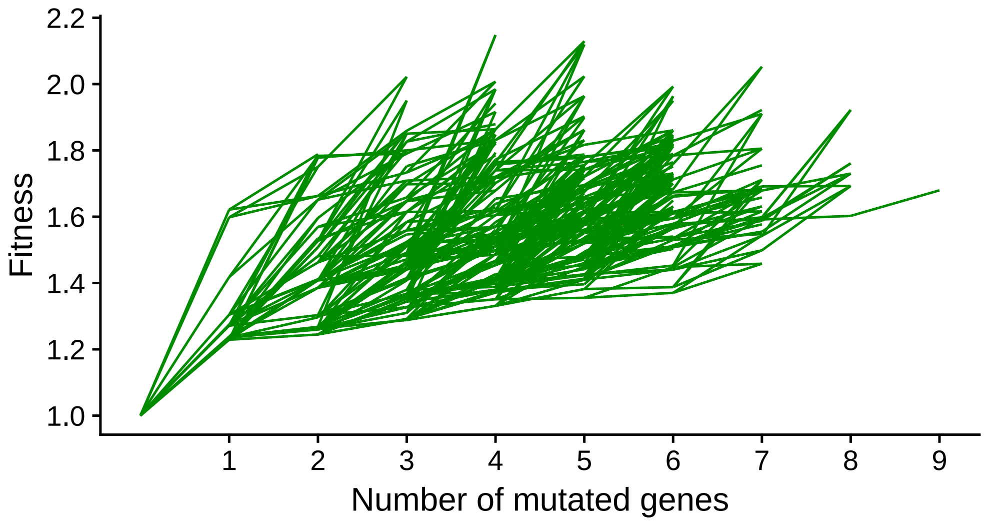


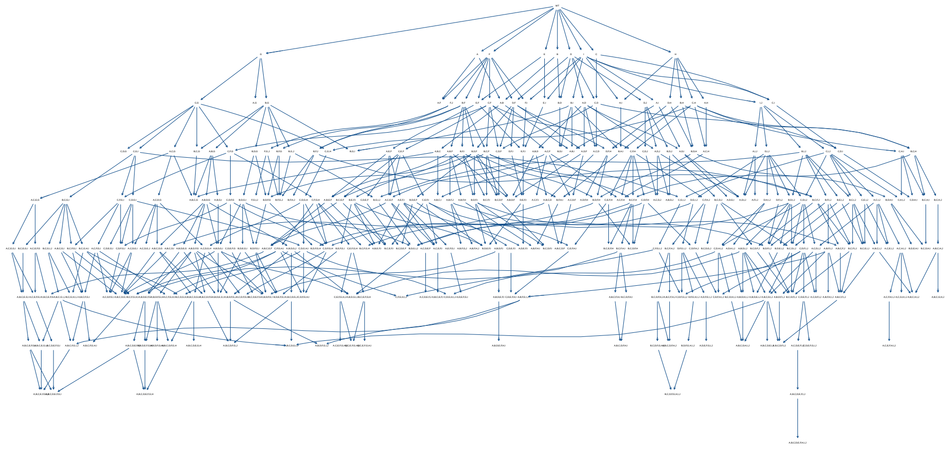

| | |
|---|---|
| *ID* | Ag710LDVSvAEgCIJ3 |
| *acc genotypes* | 50 |
| *peaks no back* | 19 |
| *removed edges* | NA |
| *prop rem edges* | NA |

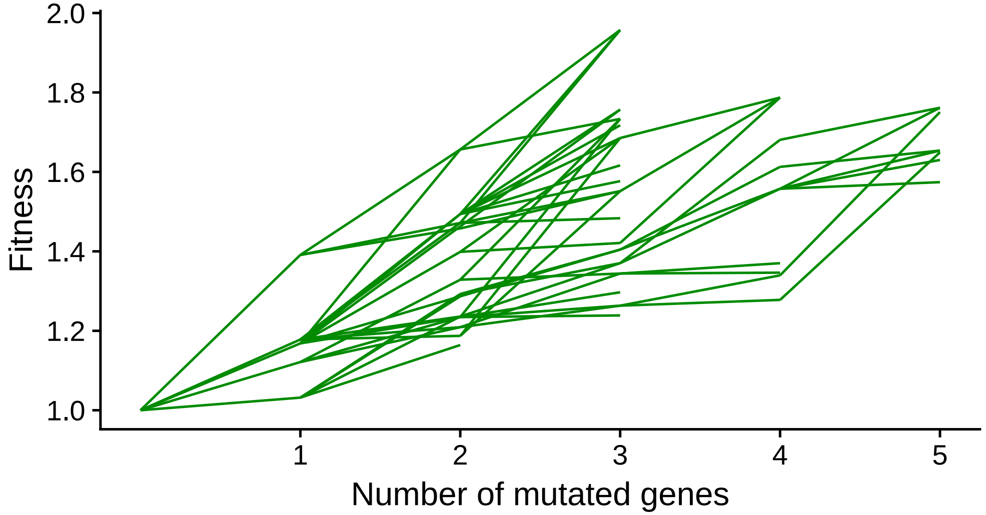


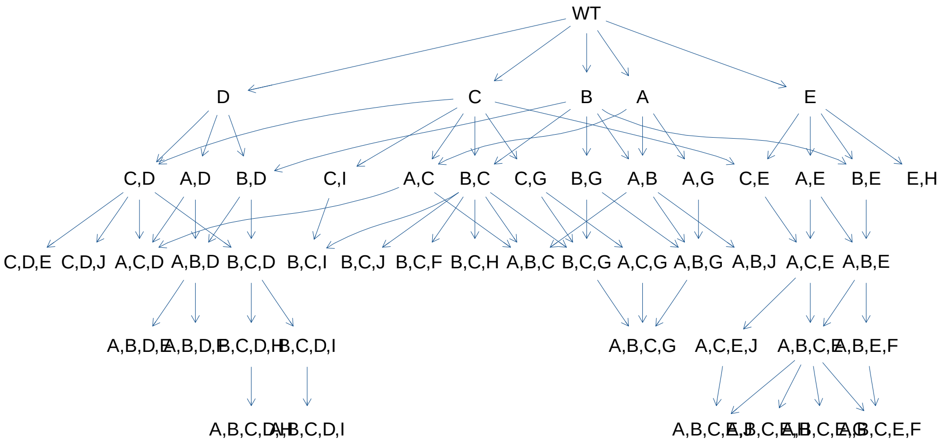

| | |
|---|---|
| *ID* | ahVo16NivbfzL4Zy |
| *acc genotypes* | 442 |
| *peaks no back* | 56 |
| *removed edges* | NA |
| *prop rem edges* | NA |

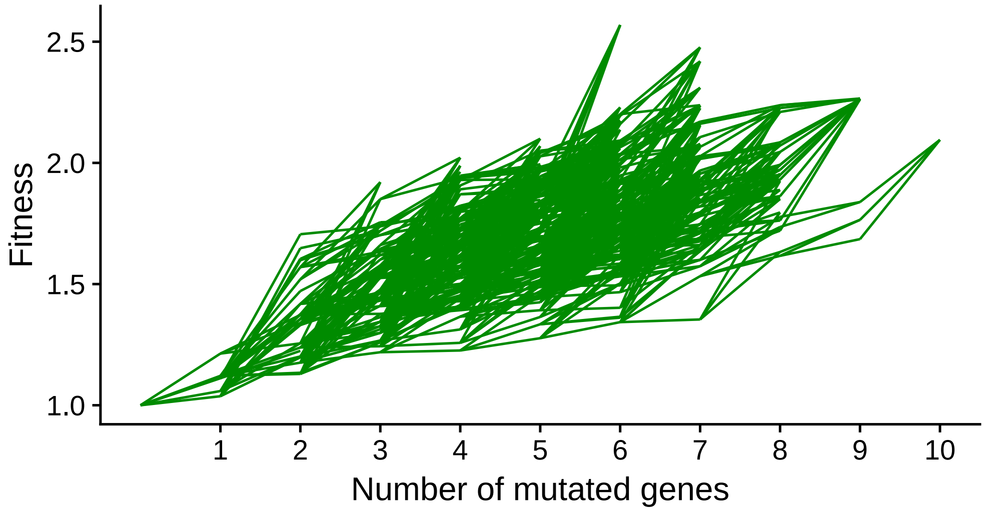


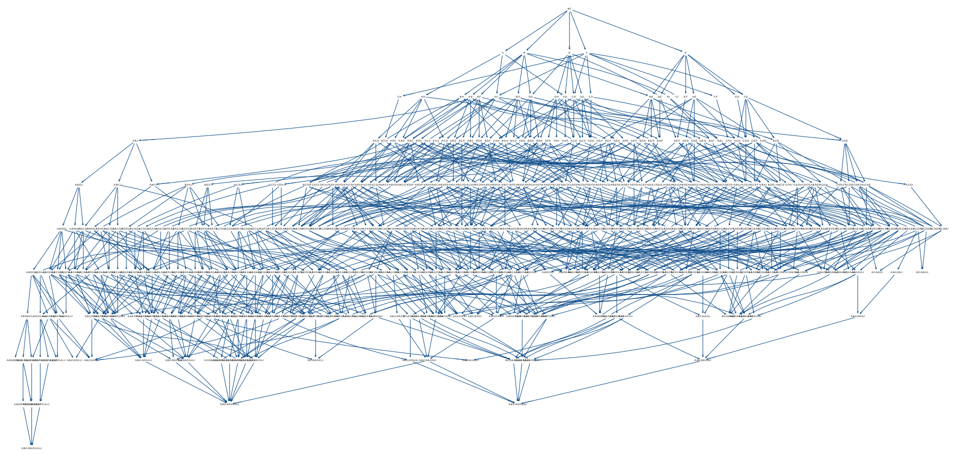

| | |
|---|---|
| *ID* | AOgjTmG9Hs4vtlBq |
| *acc genotypes* | 213 |
| *peaks no back* | 62 |
| *removed edges* | NA |
| *prop rem edges* | NA |

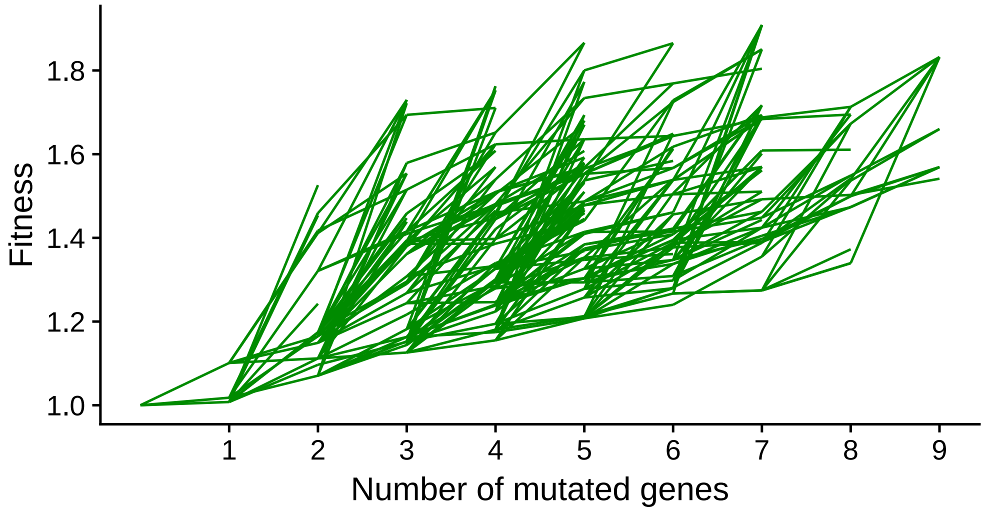


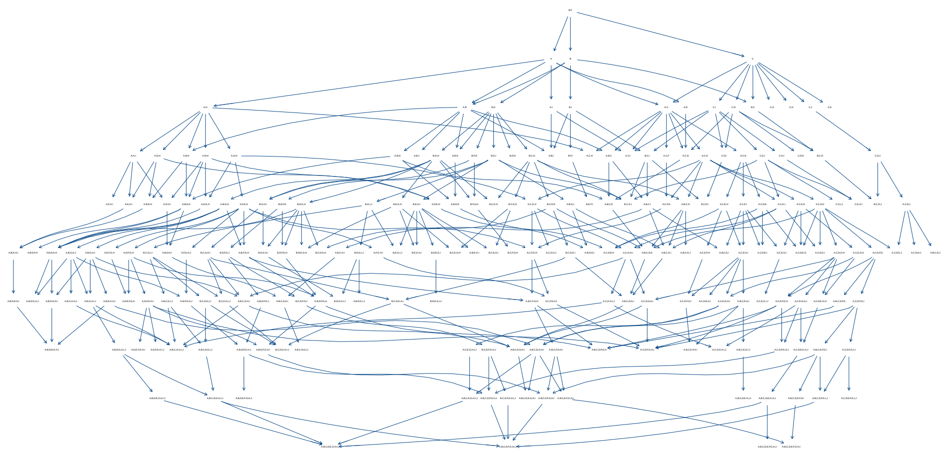

| | |
|---|---|
| *ID* | AXZ6AsdBRokekeDx |
| *acc genotypes* | 327 |
| *peaks no back* | 60 |
| *removed edges* | NA |
| *prop rem edges* | NA |

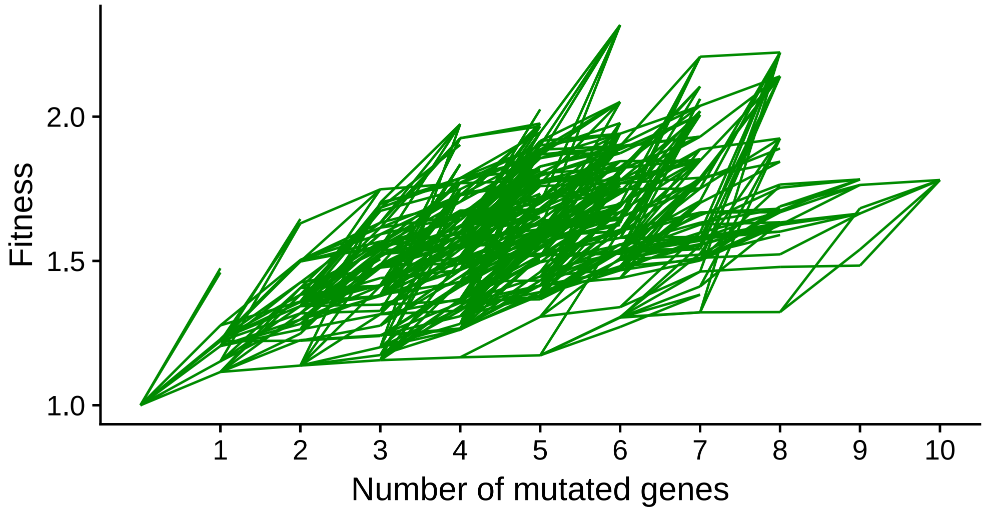


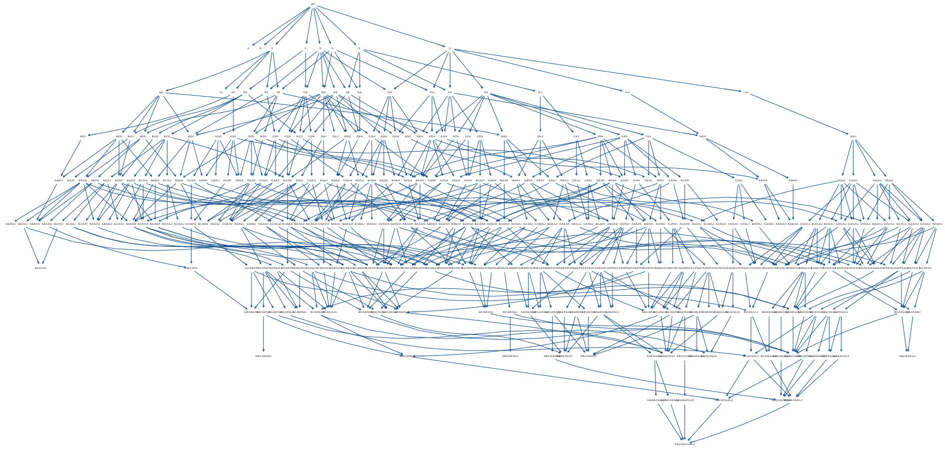

| | |
|---|---|
| *ID* | bFdmuJ1681ocZTYL |
| *acc genotypes* | 359 |
| *peaks no back* | 88 |
| *removed edges* | NA |
| *prop rem edges* | NA |

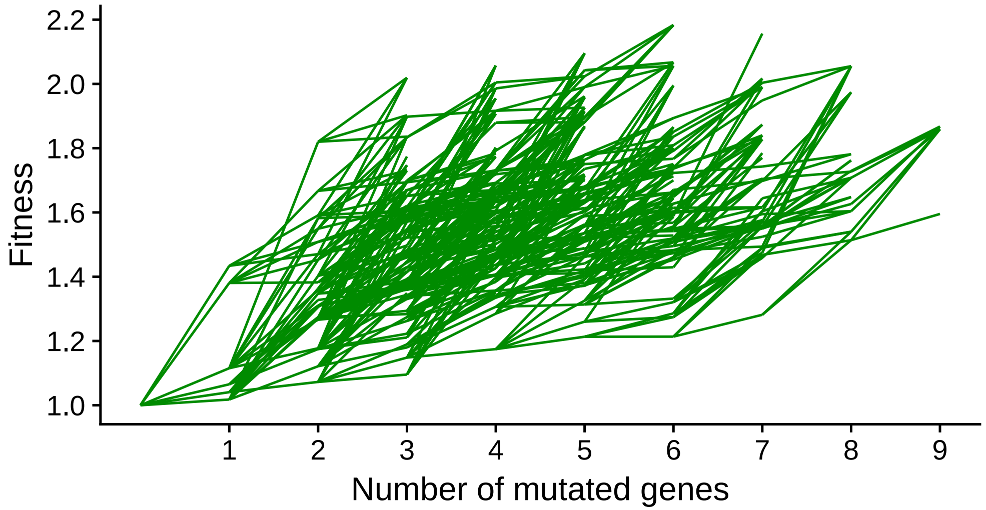


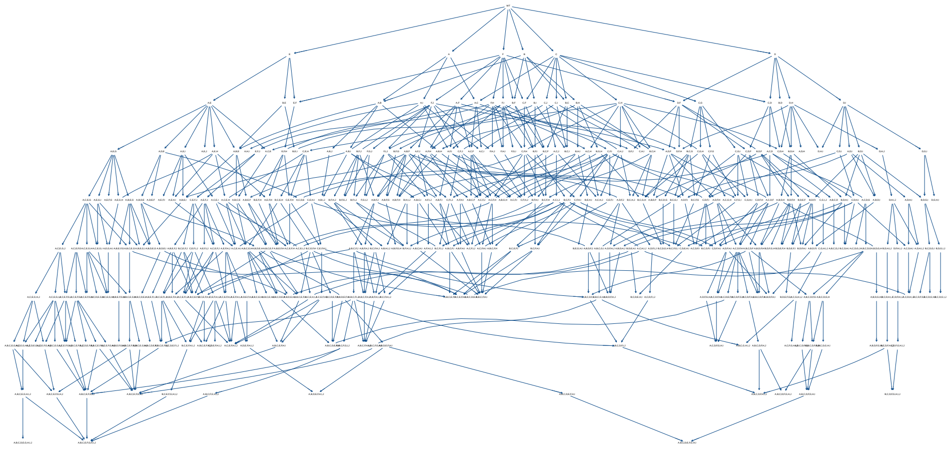

| | |
|---|---|
| *ID* | bGijXeZCPrfEOBkI |
| *acc genotypes* | 183 |
| *peaks no back* | 53 |
| *removed edges* | NA |
| *prop rem edges* | NA |

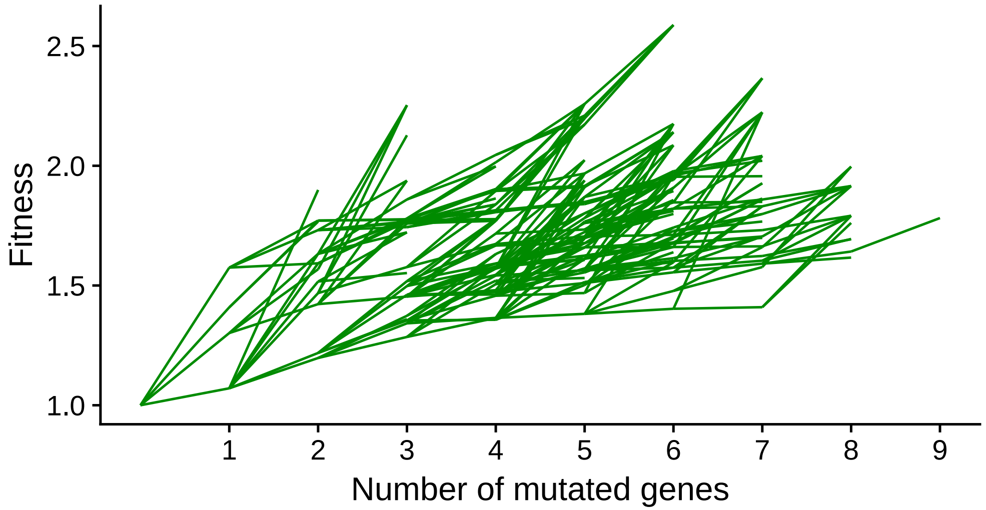


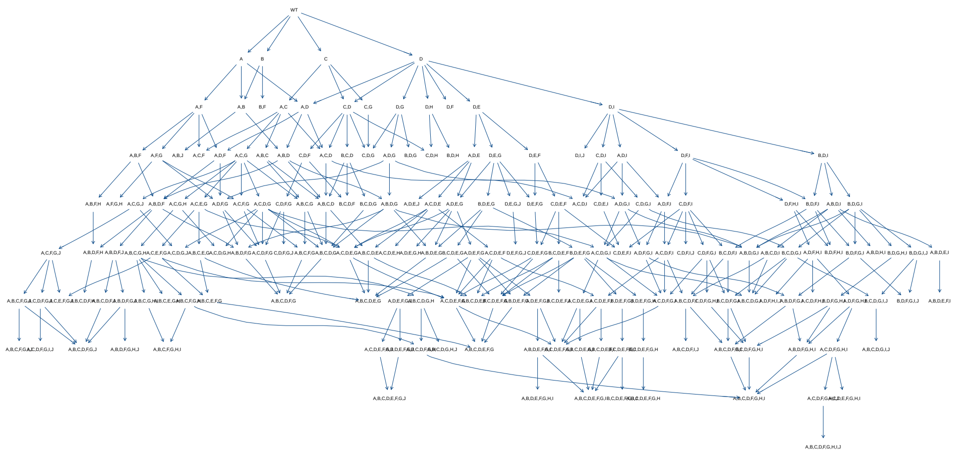

| | |
|---|---|
| *ID* | BnCqeBo4erAk6AAn |
| *acc genotypes* | 616 |
| *peaks no back* | 76 |
| *removed edges* | NA |
| *prop rem edges* | NA |

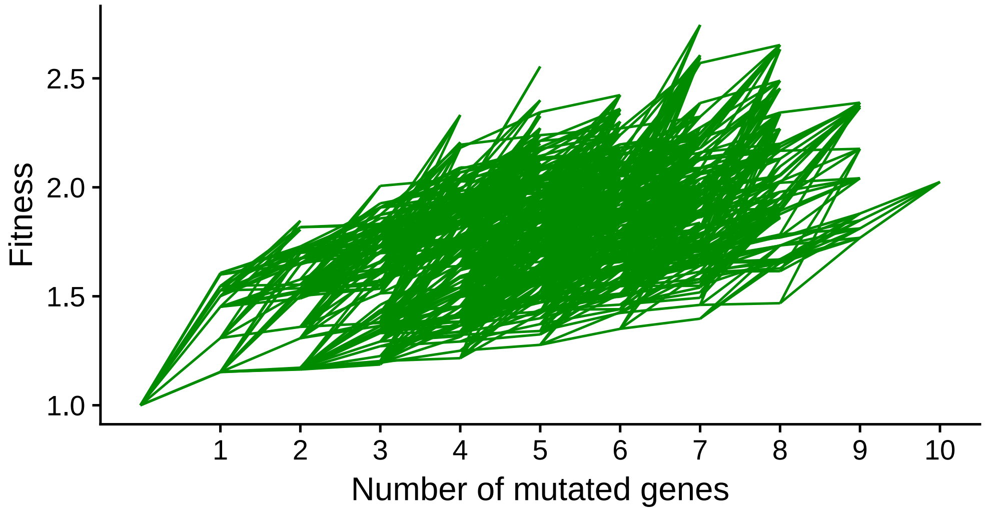


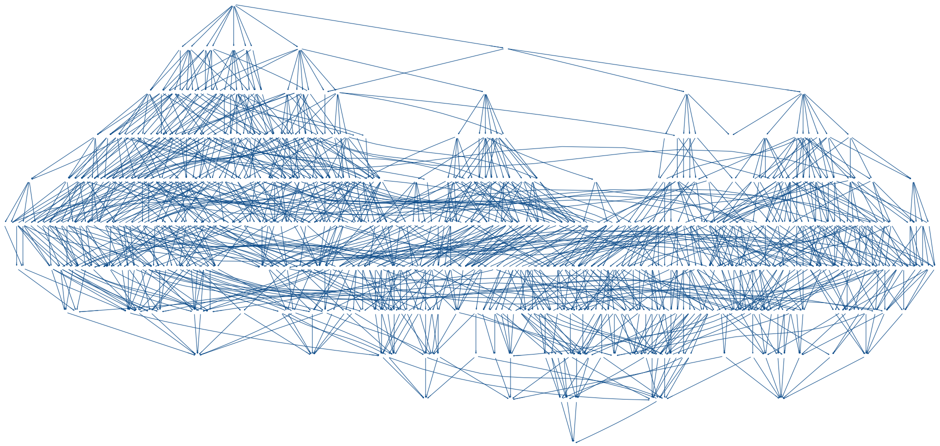

| | |
|---|---|
| *ID* | bo1swth9Yj610ESJy |
| *acc genotypes* | 133 |
| *peaks no back* | 40 |
| *removed edges* | NA |
| *prop rem edges* | NA |

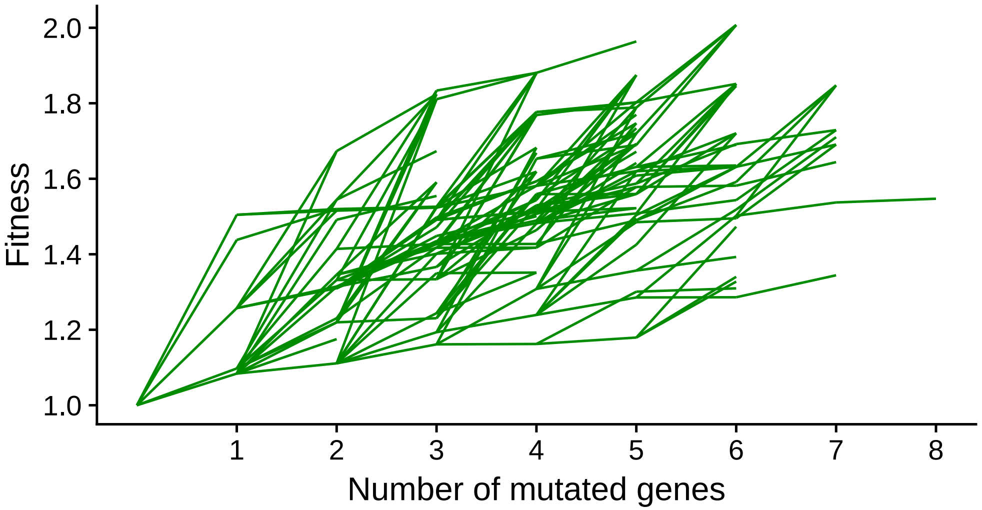


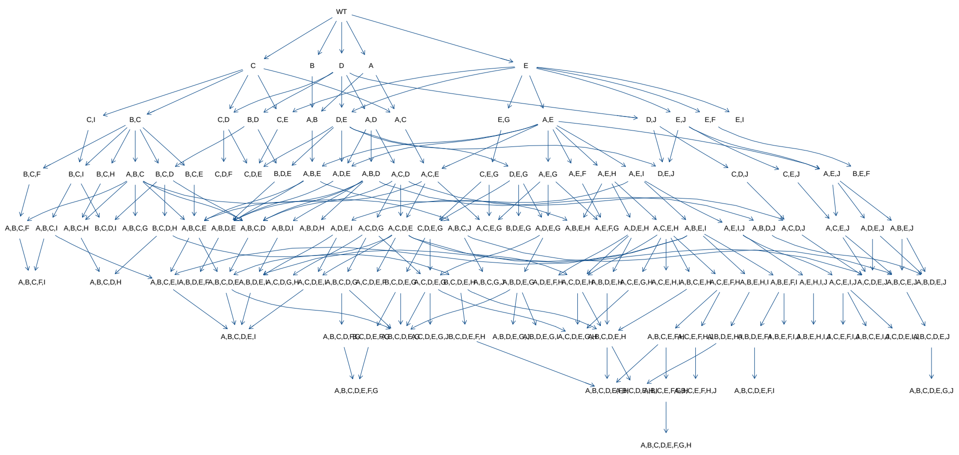

| | |
|---|---|
| *ID* | bOw7SJCwZBwfZRmf |
| *acc genotypes* | 187 |
| *peaks no back* | 71 |
| *removed edges* | NA |
| *prop rem edges* | NA |

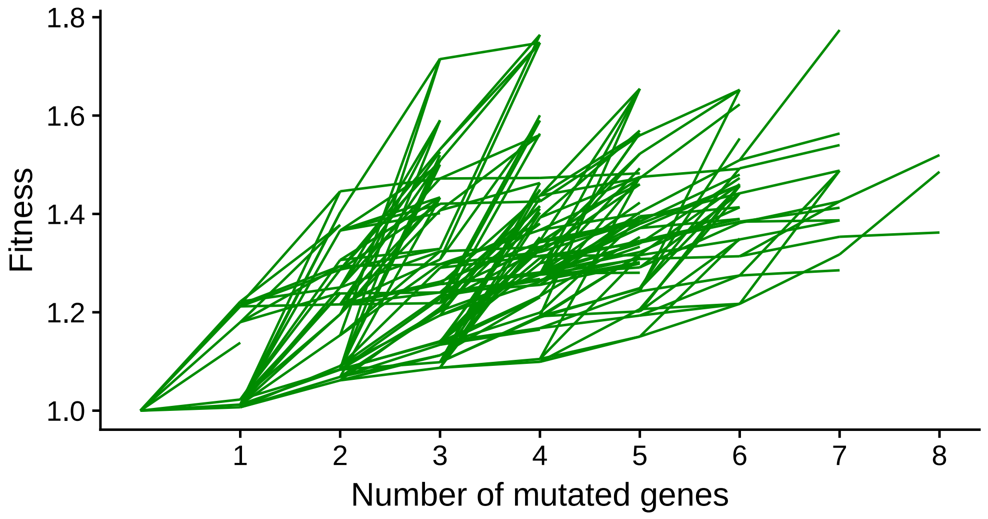


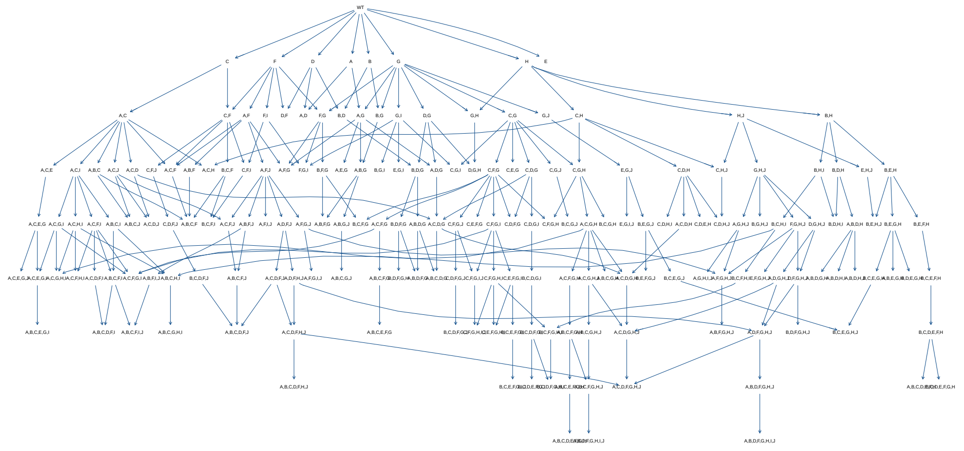

| | |
|---|---|
| *ID* | BtsRtS3YmmkCe2Gg |
| *acc genotypes* | 50 |
| *peaks no back* | 18 |
| *removed edges* | NA |
| *prop rem edges* | NA |

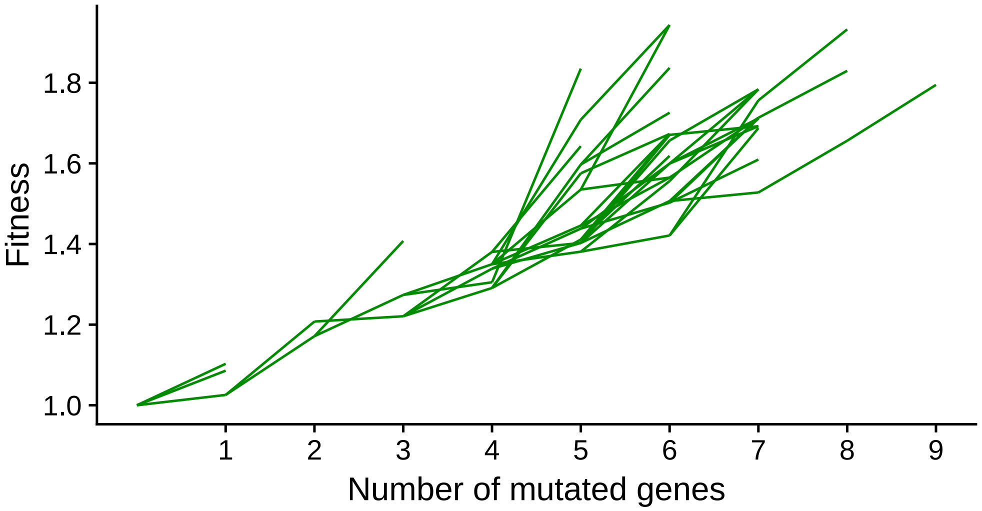


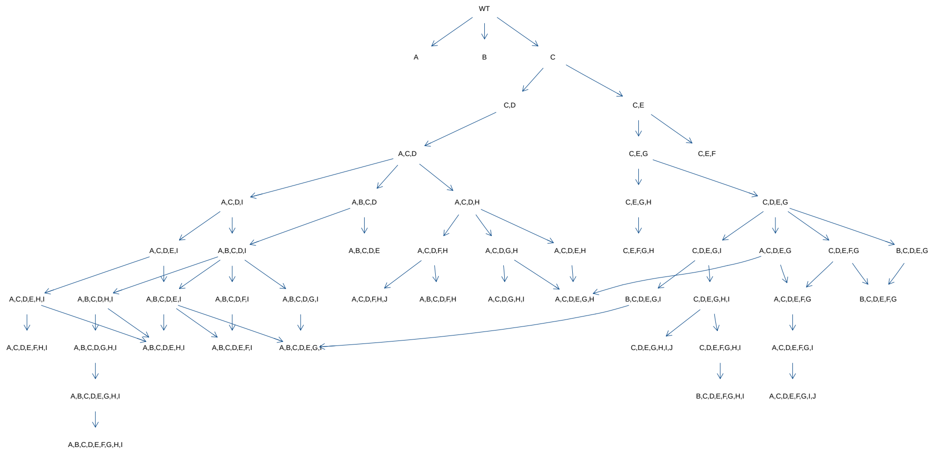

| | |
|---|---|
| *ID* | BWbL4g3bDhB87PnG |
| *acc genotypes* | 405 |
| *peaks no back* | 99 |
| *removed edges* | NA |
| *prop rem edges* | NA |

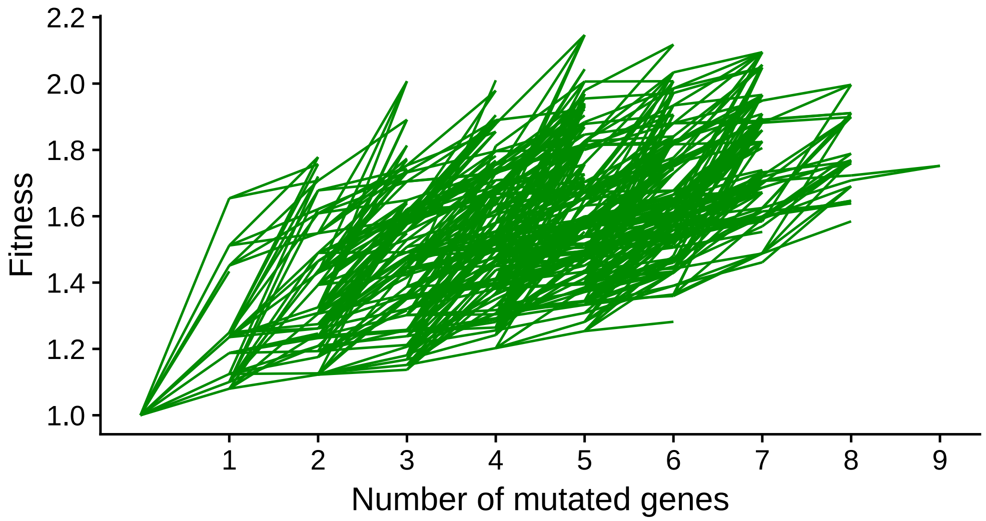


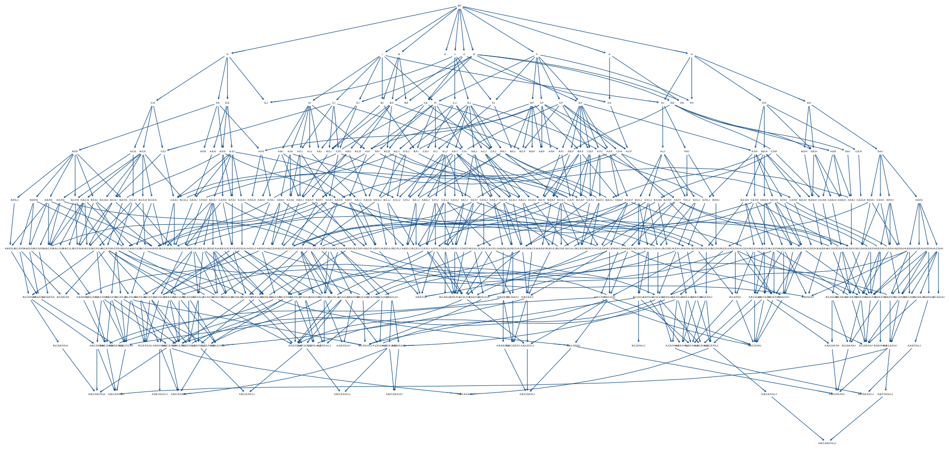

| | |
|---|---|
| *ID* | C1zvP9CZFCg9avLi |
| *acc genotypes* | 102 |
| *peaks no back* | 44 |
| *removed edges* | NA |
| *prop rem edges* | NA |

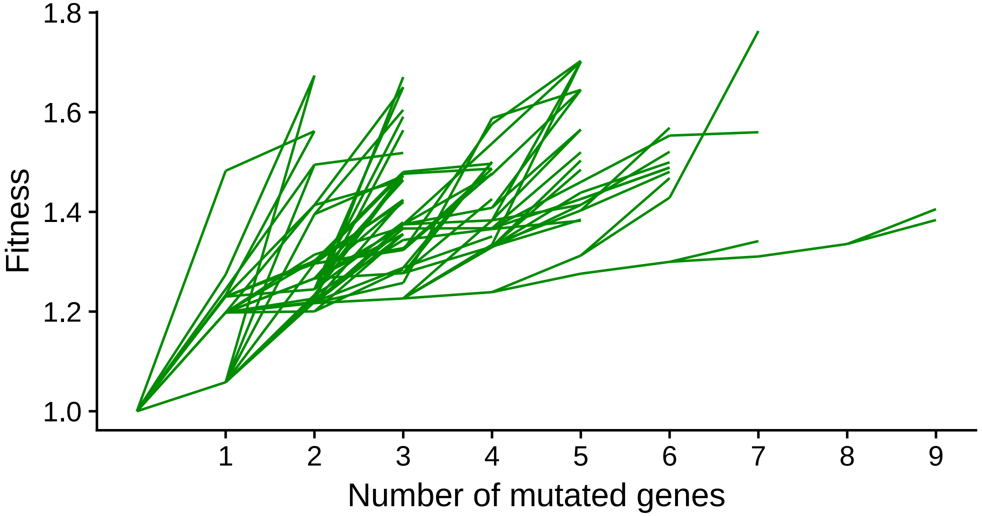


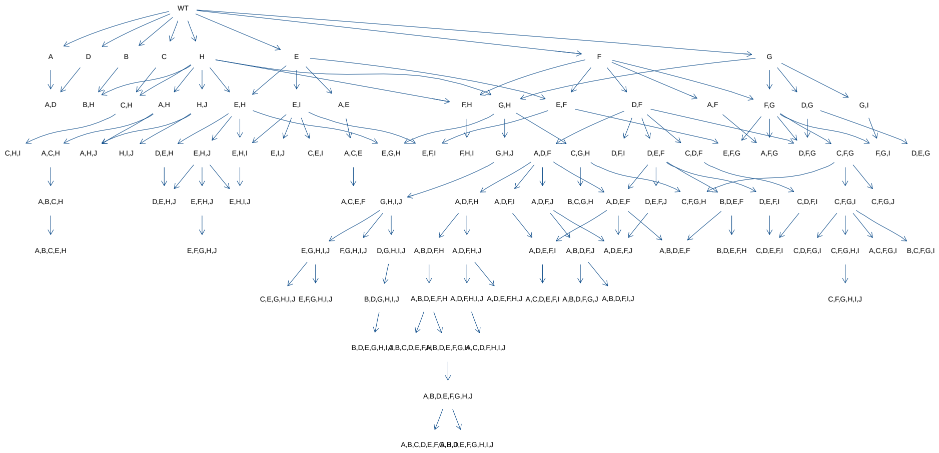

| | |
|---|---|
| *ID* | c21LhpXlCZKWnkC5 |
| *acc genotypes* | 493 |
| *peaks no back* | 93 |
| *removed edges* | NA |
| *prop rem edges* | NA |

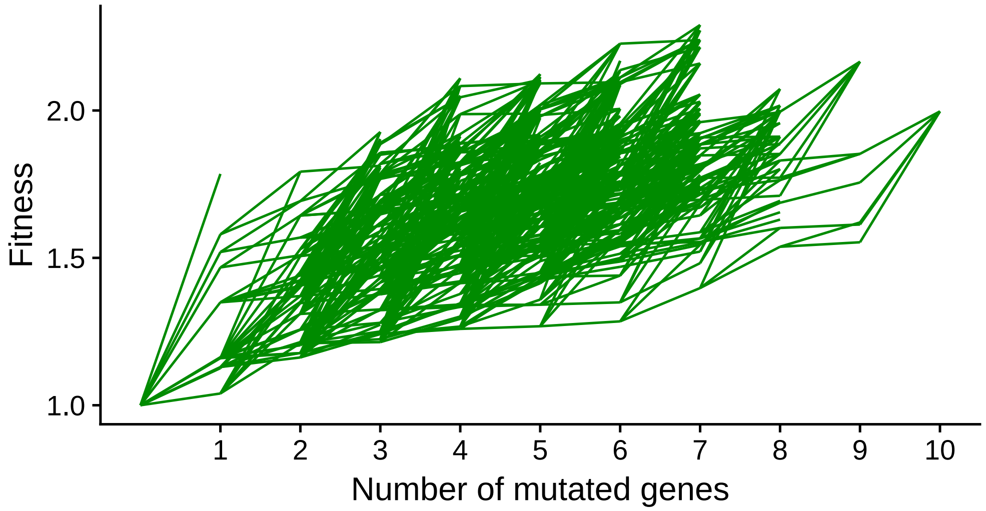


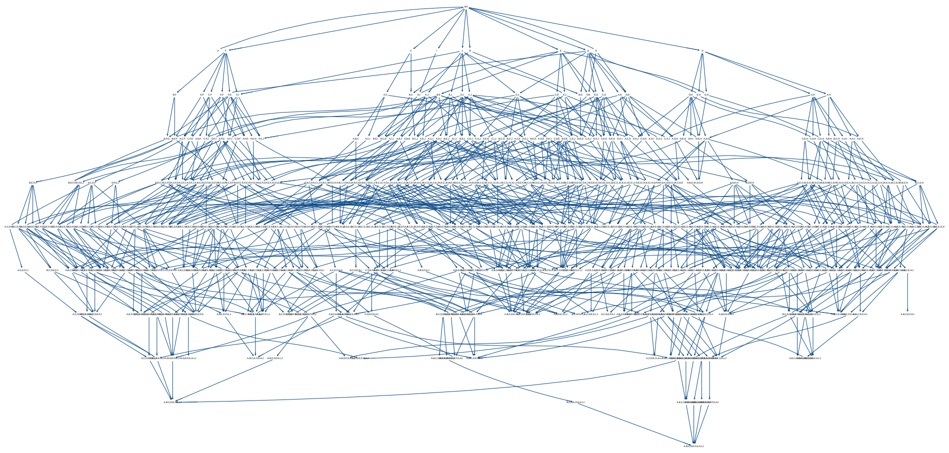

*ID* C46MDiN7E3KDR5Fu
*acc genotypes* 143
*peaks no back* 61
*removed edges* NA
*prop rem edges* NA

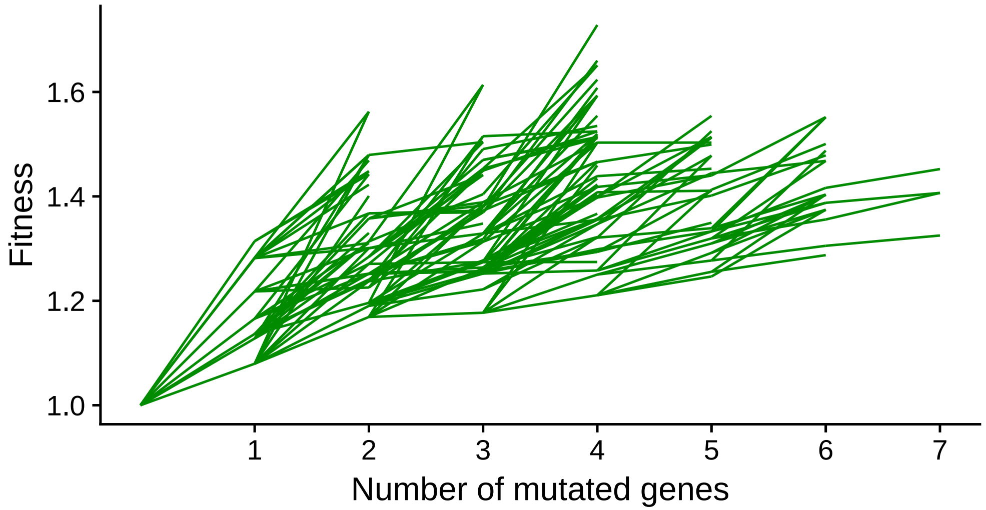


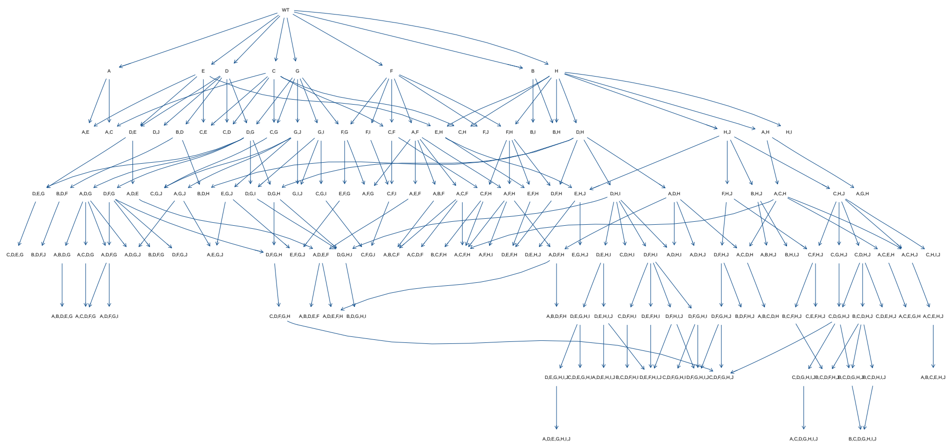

| | |
|---|---|
| *ID* | COoTZ3S17BxN8HQx |
| *acc genotypes* | 209 |
| *peaks no back* | 75 |
| *removed edges* | NA |
| *prop rem edges* | NA |

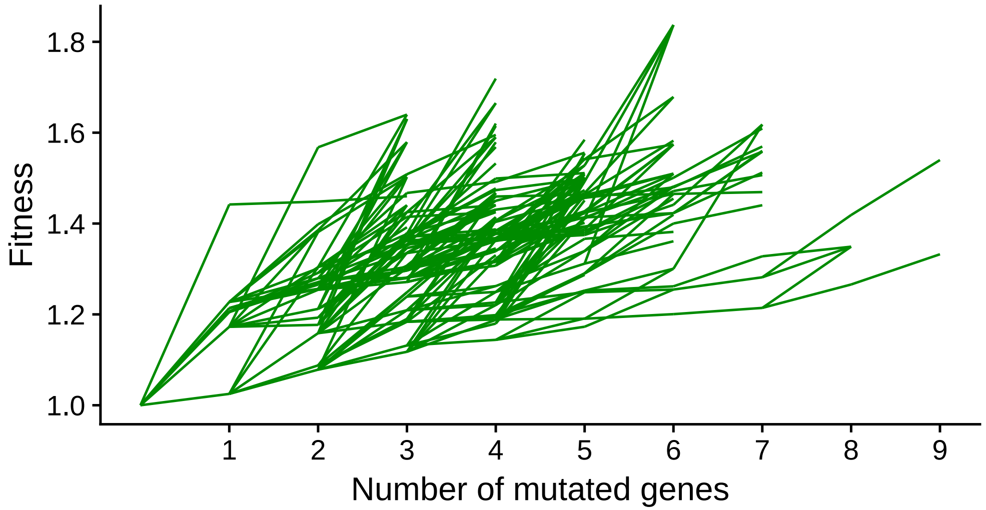


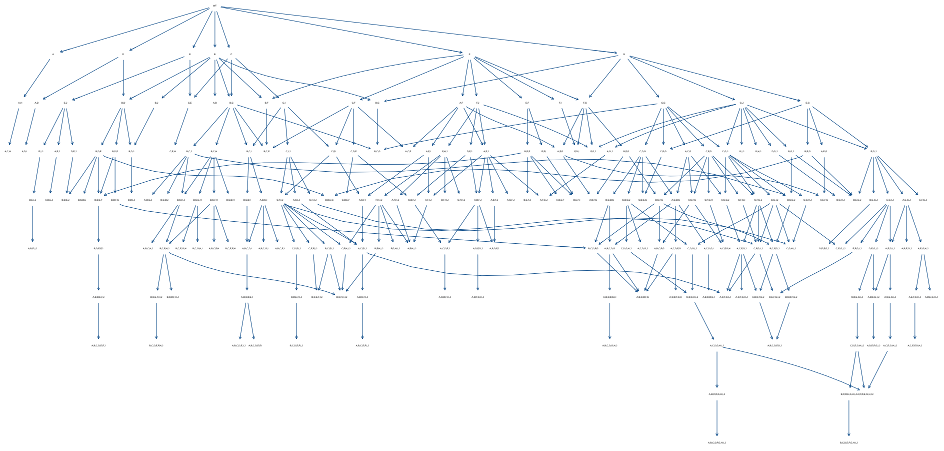

| | |
|---|---|
| *ID* | CscphnR9HTju10jKi |
| *acc genotypes* | 54 |
| *peaks no back* | 14 |
| *removed edges* | NA |
| *prop rem edges* | NA |

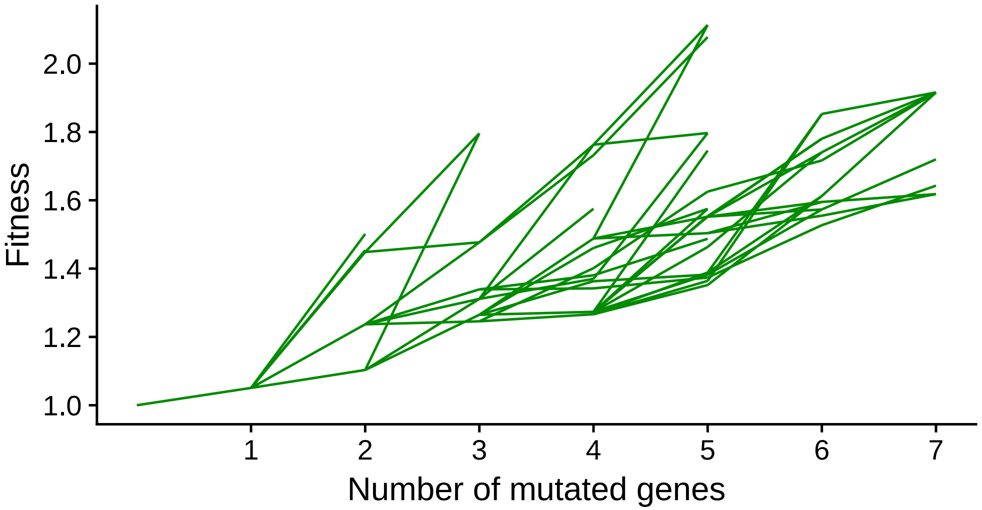


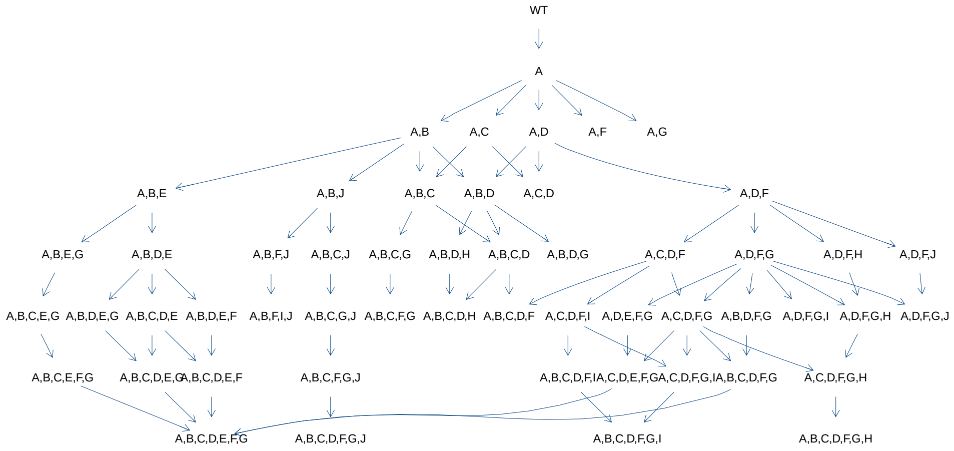

| | |
|---|---|
| *ID* | CxyKl8ZkyGMlHEBy |
| *acc genotypes* | 101 |
| *peaks no back* | 37 |
| *removed edges* | NA |
| *prop rem edges* | NA |

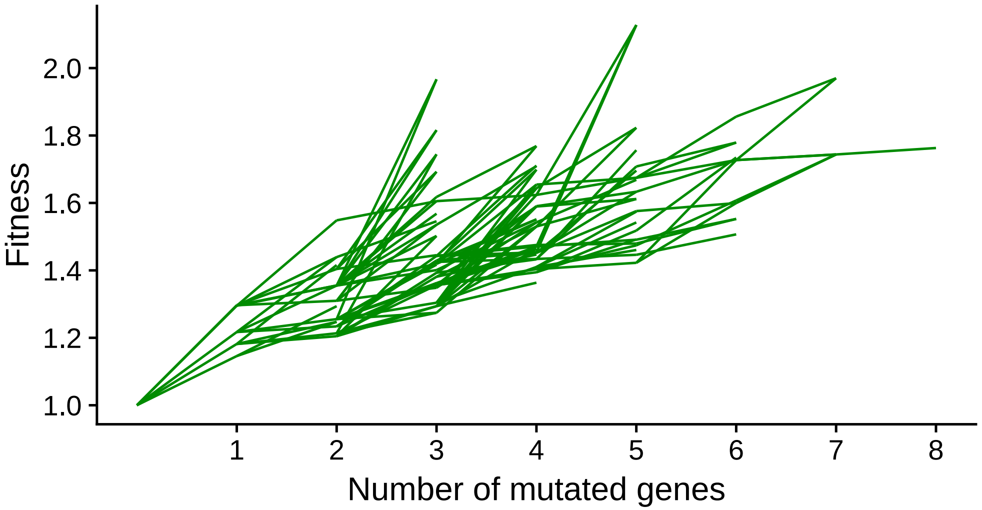


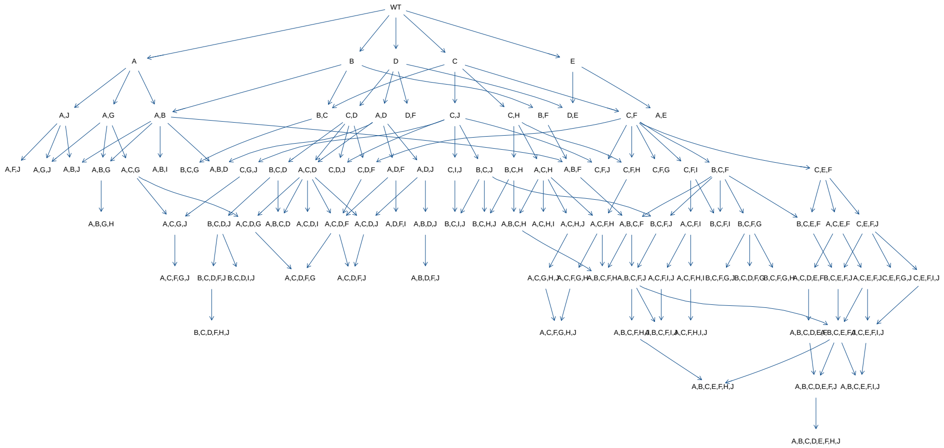

| | |
|---|---|
| *ID* | d107ojrARayuzrHeN |
| *acc genotypes* | 482 |
| *peaks no back* | 107 |
| *removed edges* | NA |
| *prop rem edges* | NA |

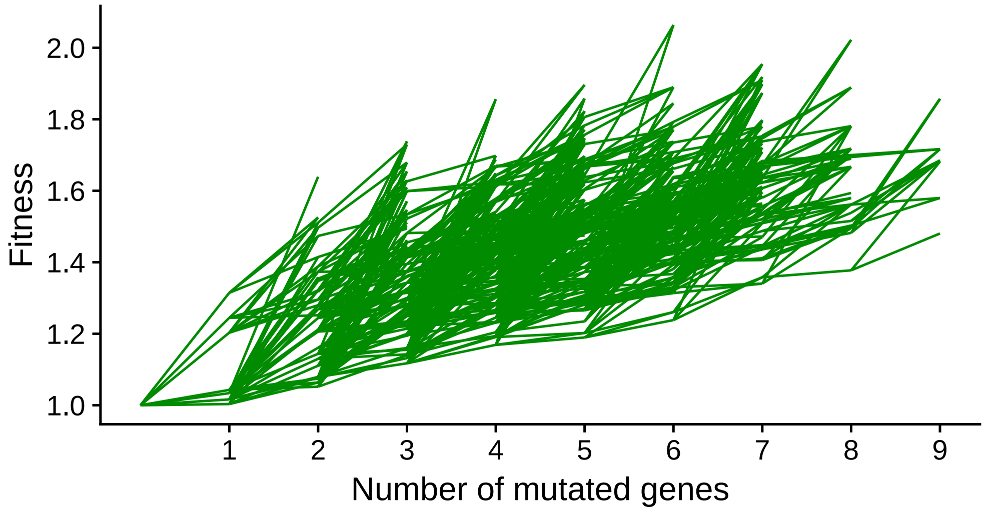


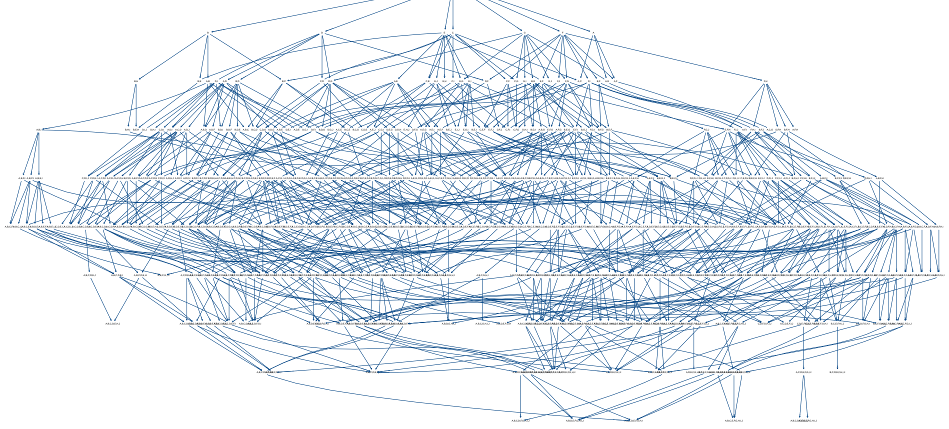

| | |
|---|---|
| *ID* | DF1iteAjggjrOq9G |
| *acc genotypes* | 240 |
| *peaks no back* | 84 |
| *removed edges* | NA |
| *prop rem edges* | NA |

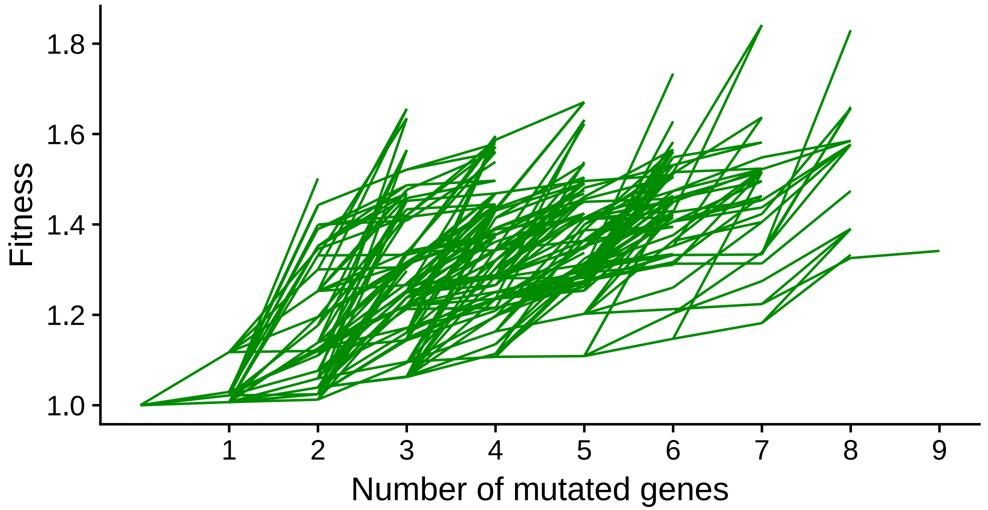


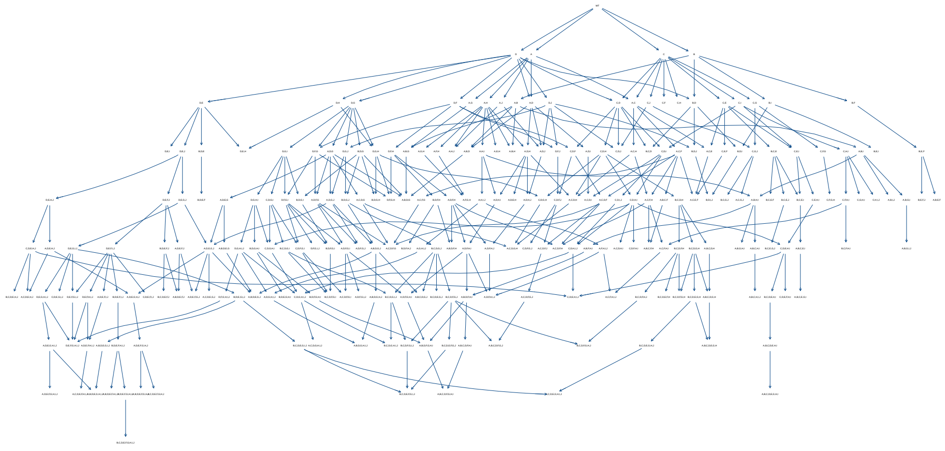

| | |
|---|---|
| *ID* | DvGWOEzyGFaVOowT |
| *acc genotypes* | 577 |
| *peaks no back* | 93 |
| *removed edges* | NA |
| *prop rem edges* | NA |

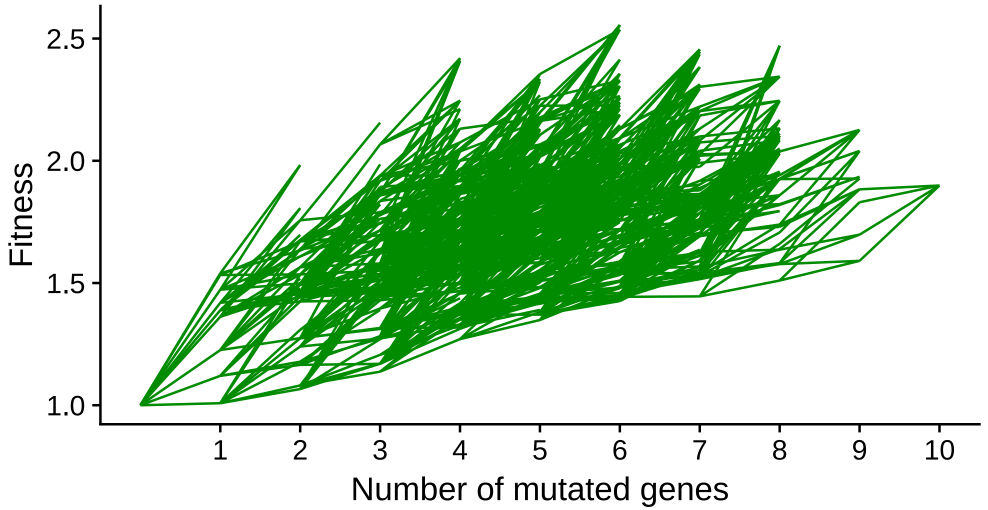


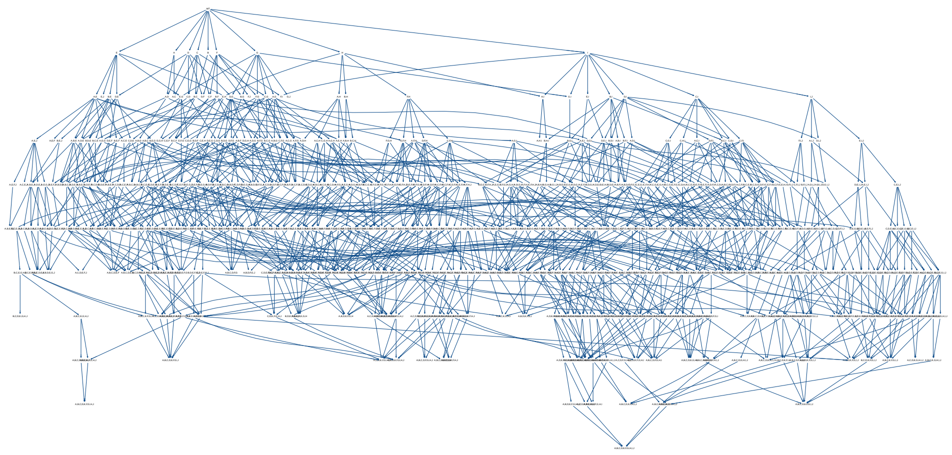

| | |
|---|---|
| *ID* | dzb4OOvYv11WGeeP |
| *acc genotypes* | 394 |
| *peaks no back* | 83 |
| *removed edges* | NA |
| *prop rem edges* | NA |

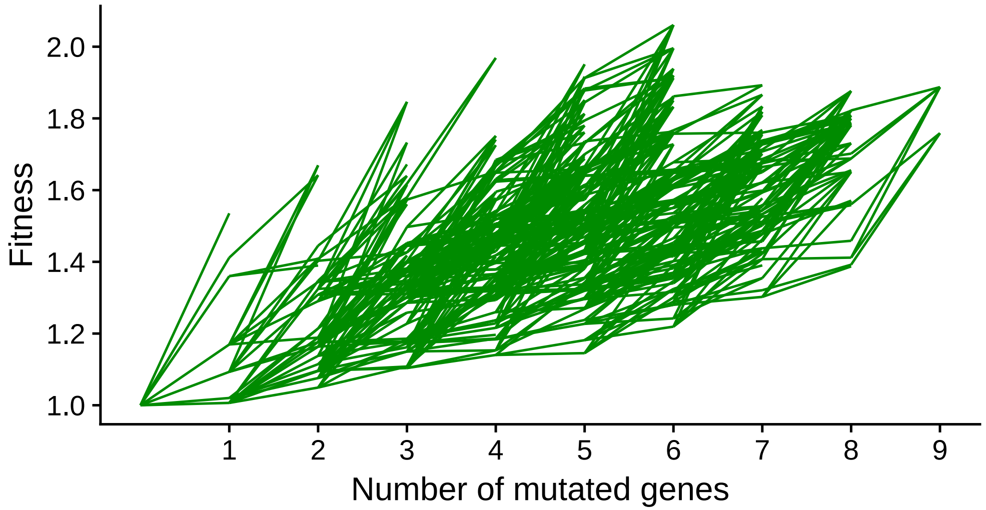


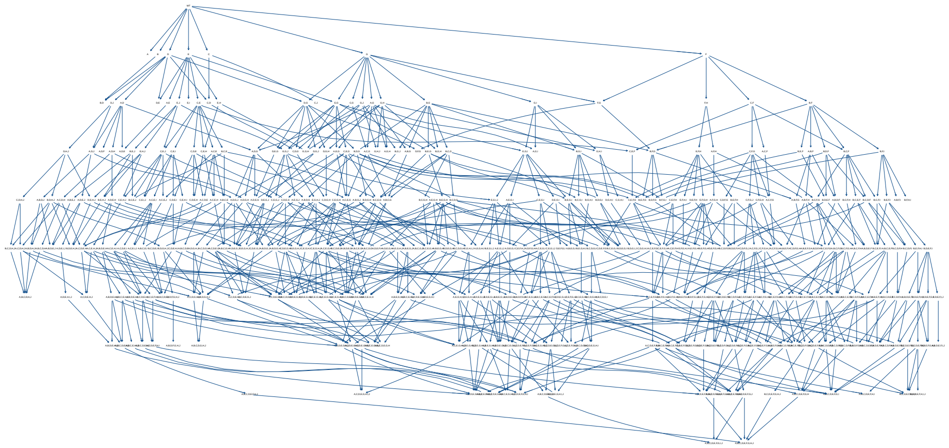

| | |
|---|---|
| *ID* | EgkeUuvhLZsyQQ110 |
| *acc genotypes* | 283 |
| *peaks no back* | 50 |
| *removed edges* | NA |
| *prop rem edges* | NA |

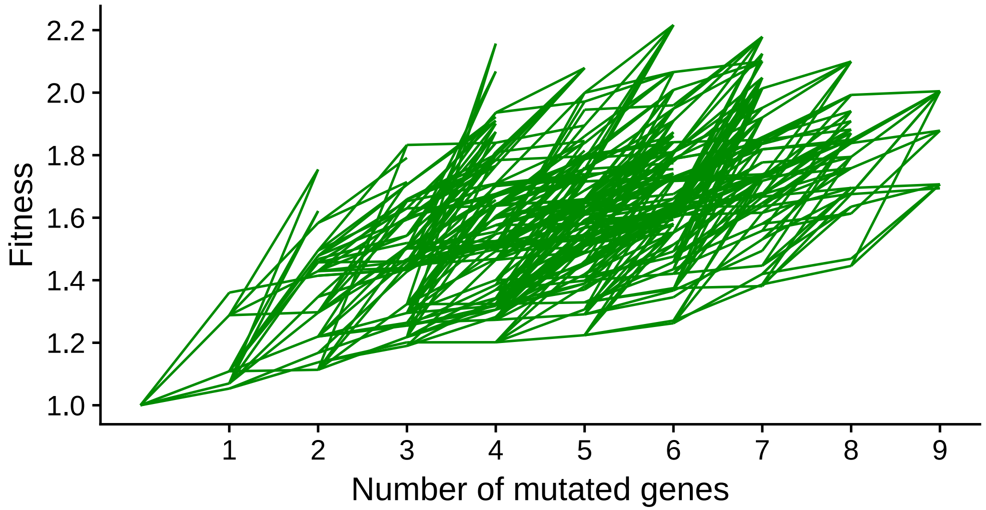


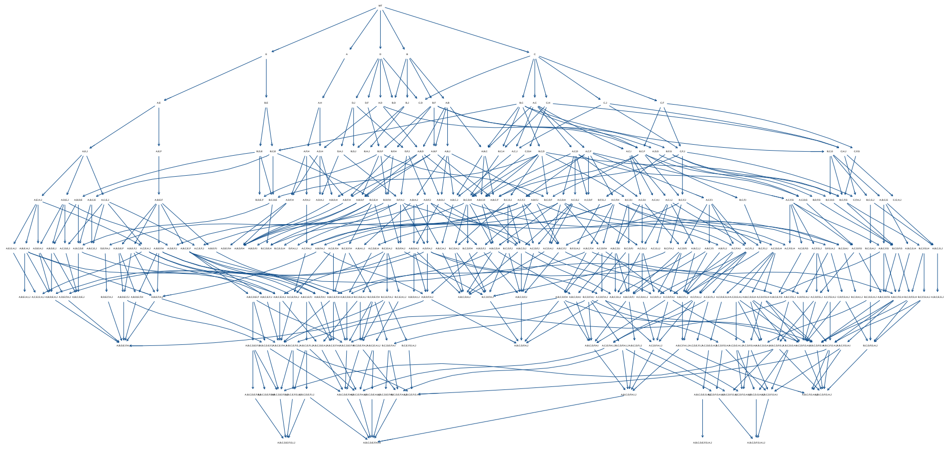

| | |
|---|---|
| *ID* | eIl49J4HyU8o710iq |
| *acc genotypes* | 576 |
| *peaks no back* | 83 |
| *removed edges* | NA |
| *prop rem edges* | NA |

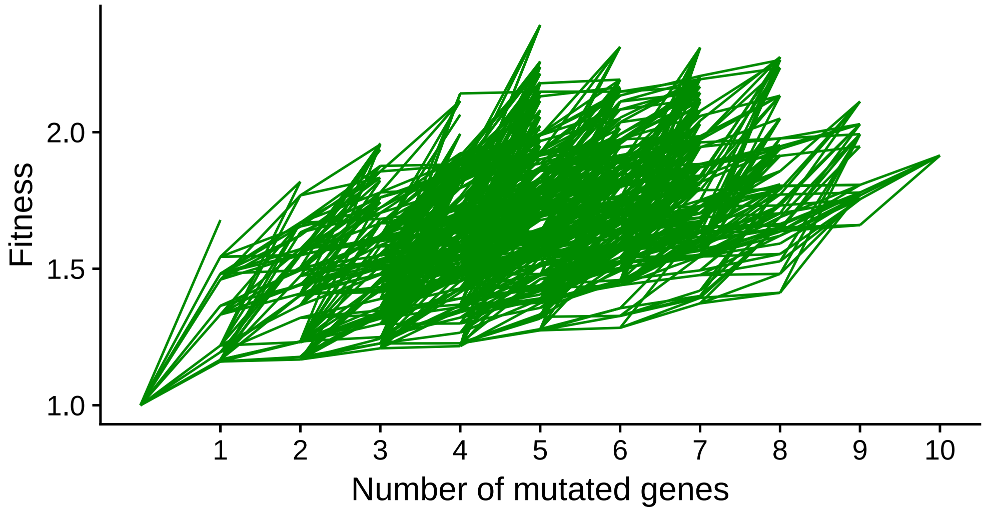


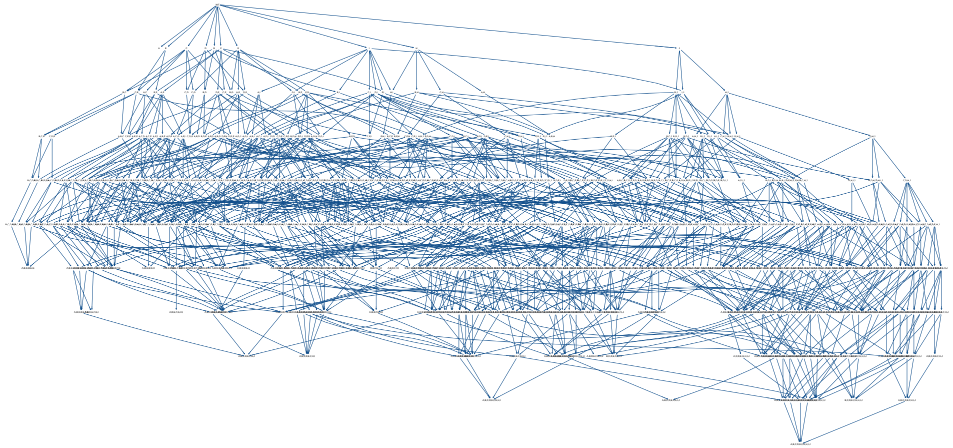

| | |
|---|---|
| *ID* | EMbpnzl5vOYpYkCY |
| *acc genotypes* | 262 |
| *peaks no back* | 64 |
| *removed edges* | NA |
| *prop rem edges* | NA |

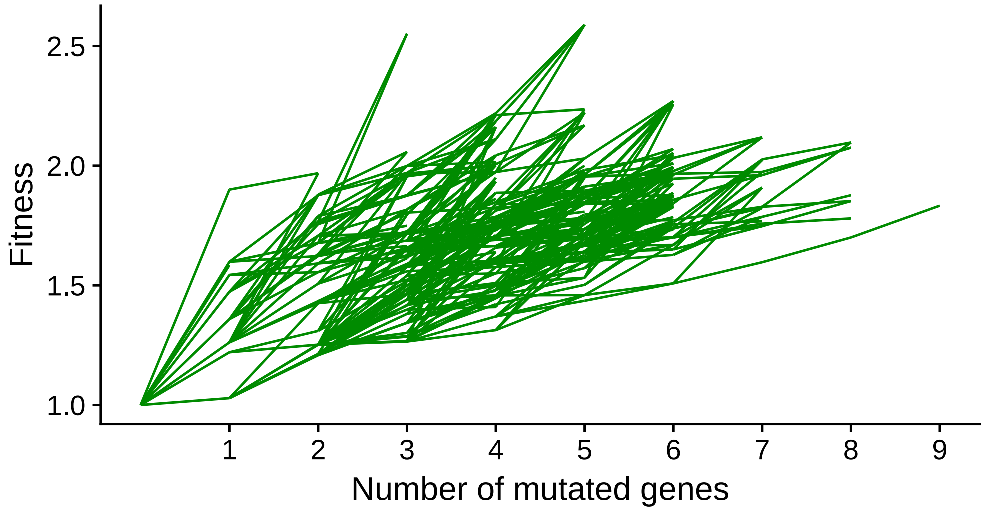


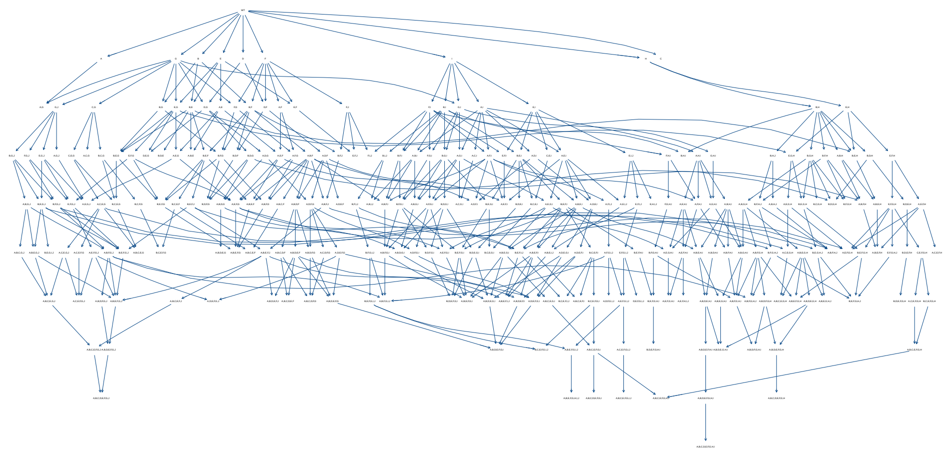

| | |
|---|---|
| *ID* | ERn91KUoSwG1g1w9 |
| *acc genotypes* | 263 |
| *peaks no back* | 93 |
| *removed edges* | NA |
| *prop rem edges* | NA |

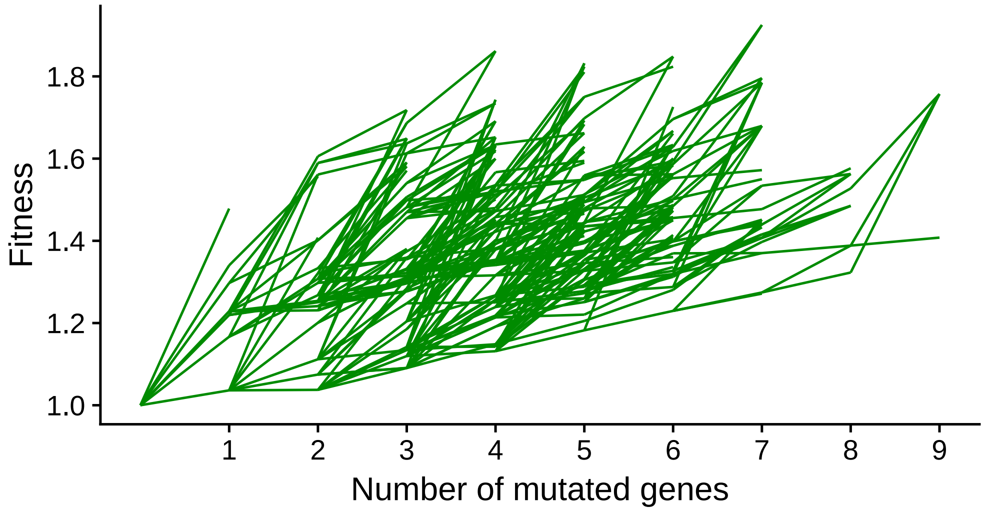


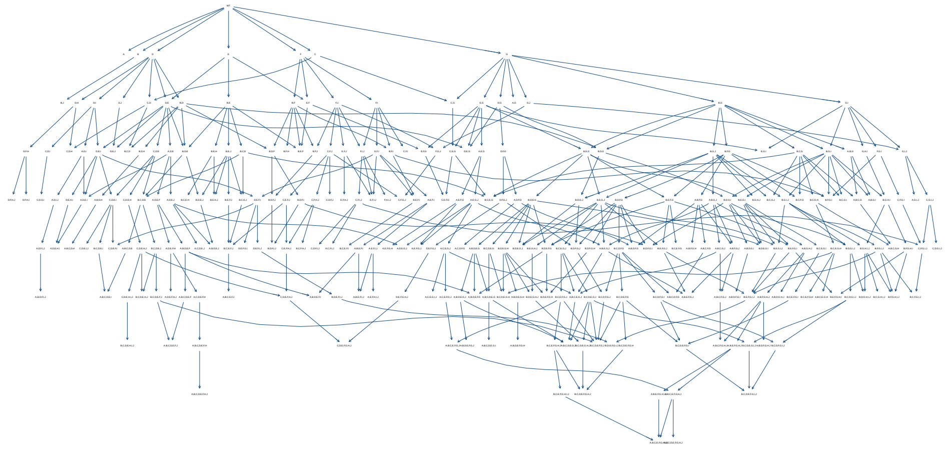

| | |
|---|---|
| *ID* | erQFJpuljZzglgwp |
| *acc genotypes* | 377 |
| *peaks no back* | 75 |
| *removed edges* | NA |
| *prop rem edges* | NA |

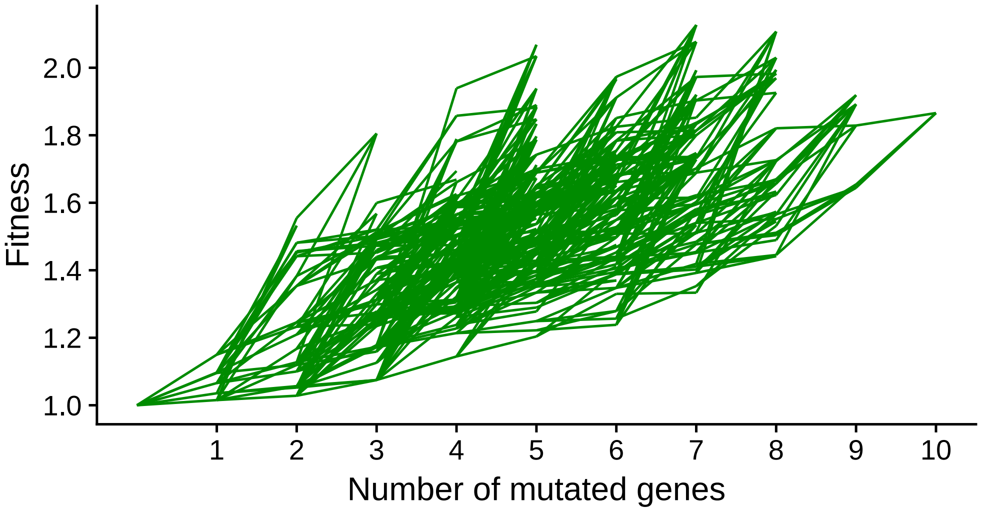


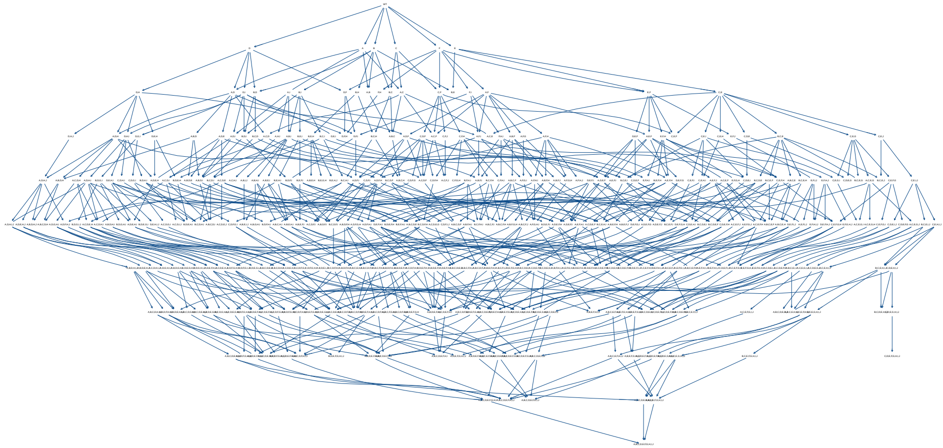

| | |
|---|---|
| *ID* | EV5PzPVJ8yEpuh5C |
| *acc genotypes* | 33 |
| *peaks no back* | 13 |
| *removed edges* | NA |
| *prop rem edges* | NA |

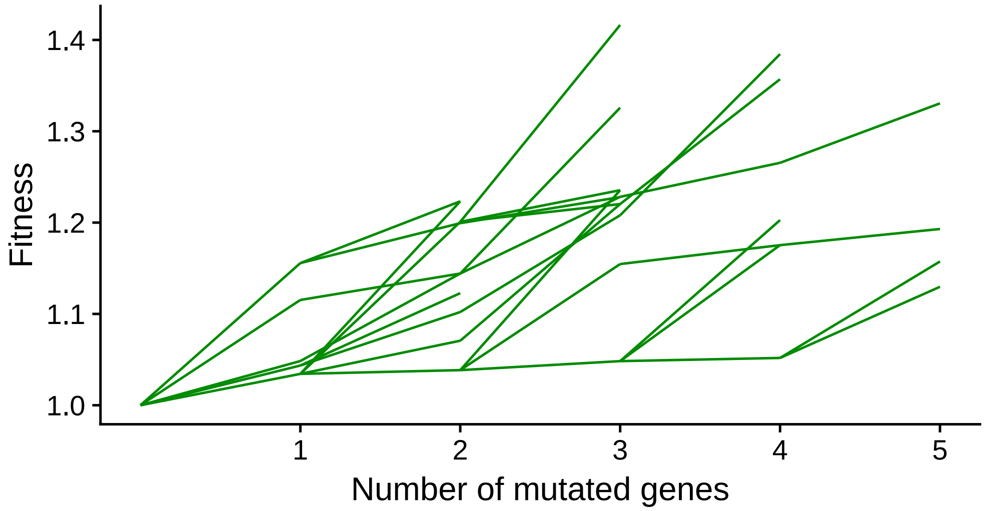


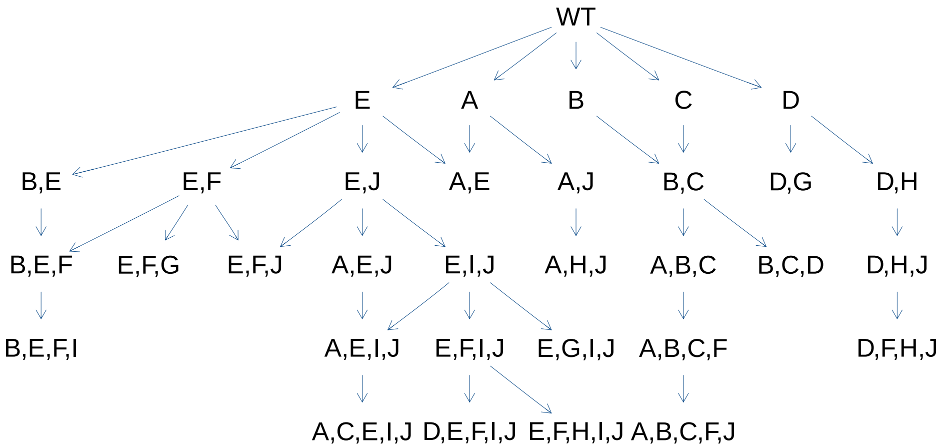

| | |
|---|---|
| *ID* | f4lIMrTliuCuX6MS |
| *acc genotypes* | 540 |
| *peaks no back* | 122 |
| *removed edges* | NA |
| *prop rem edges* | NA |

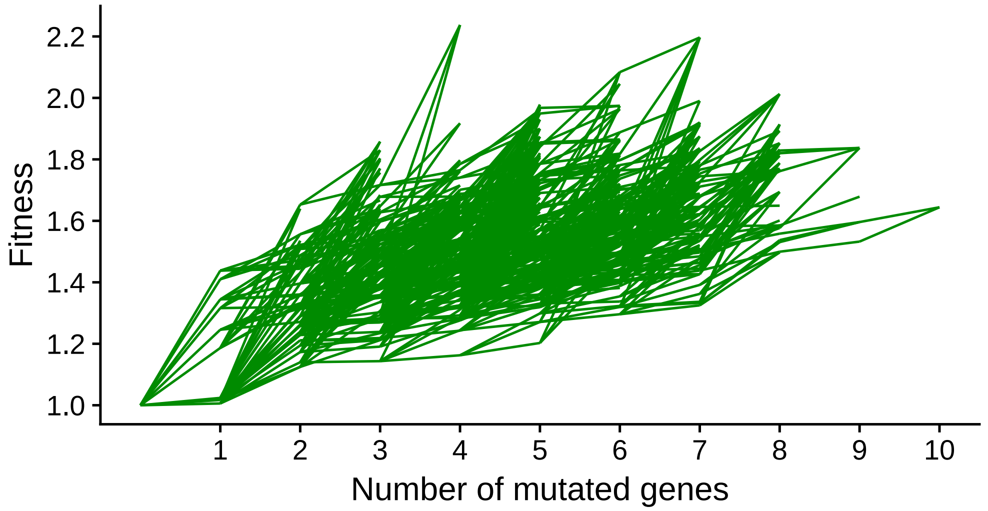


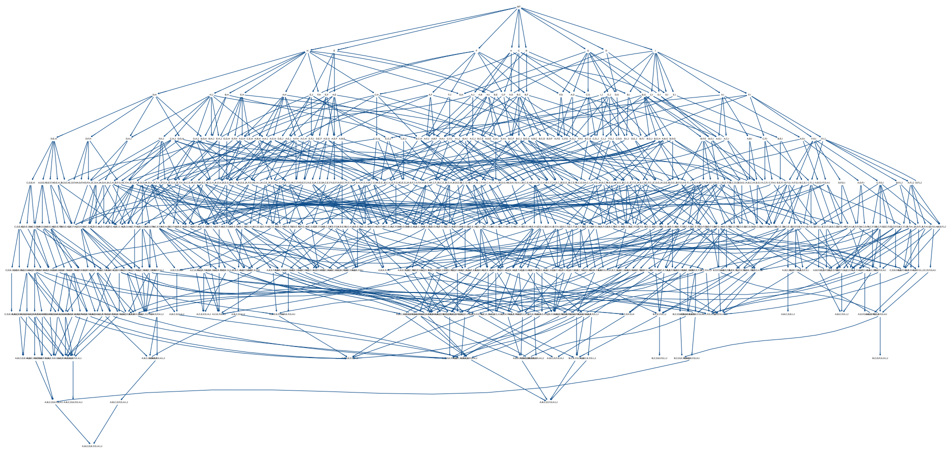

| | |
|---|---|
| *ID* | FCcL7VfUNU6Igd6e |
| *acc genotypes* | 400 |
| *peaks no back* | 62 |
| *removed edges* | NA |
| *prop rem edges* | NA |

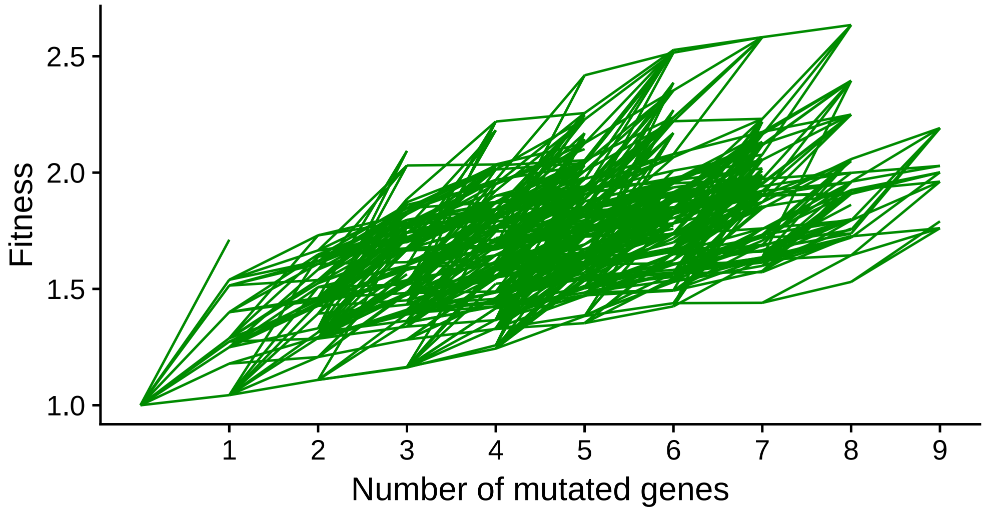


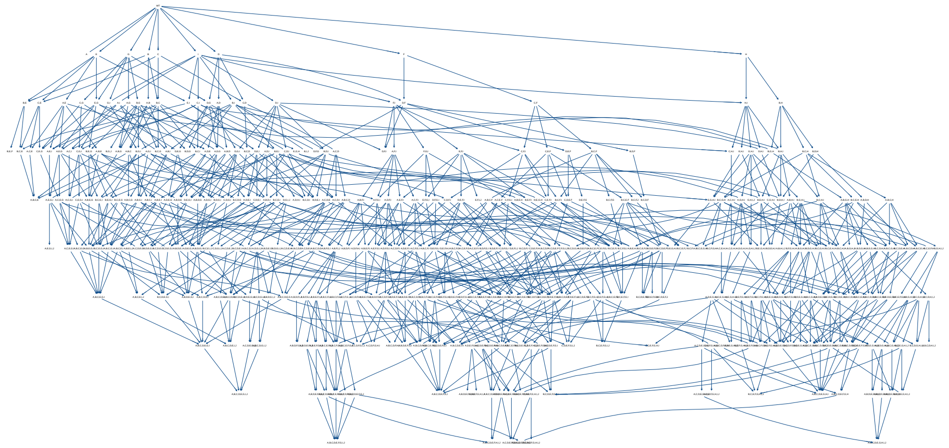

| | |
|---|---|
| *ID* | fhP10WlHL7710GBMFE |
| *acc genotypes* | 412 |
| *peaks no back* | 90 |
| *removed edges* | NA |
| *prop rem edges* | NA |

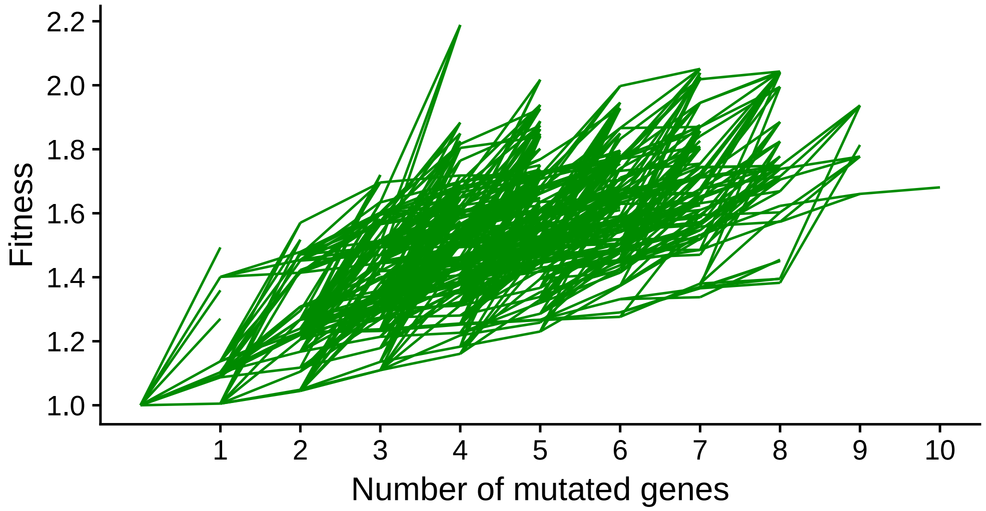


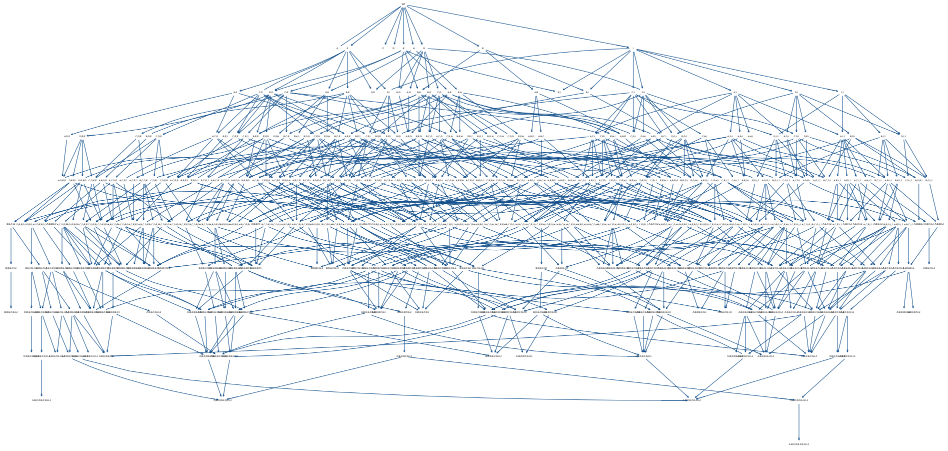

| | |
|---|---|
| *ID* | fIL7P19LmiLt1csg |
| *acc genotypes* | 338 |
| *peaks no back* | 63 |
| *removed edges* | NA |
| *prop rem edges* | NA |

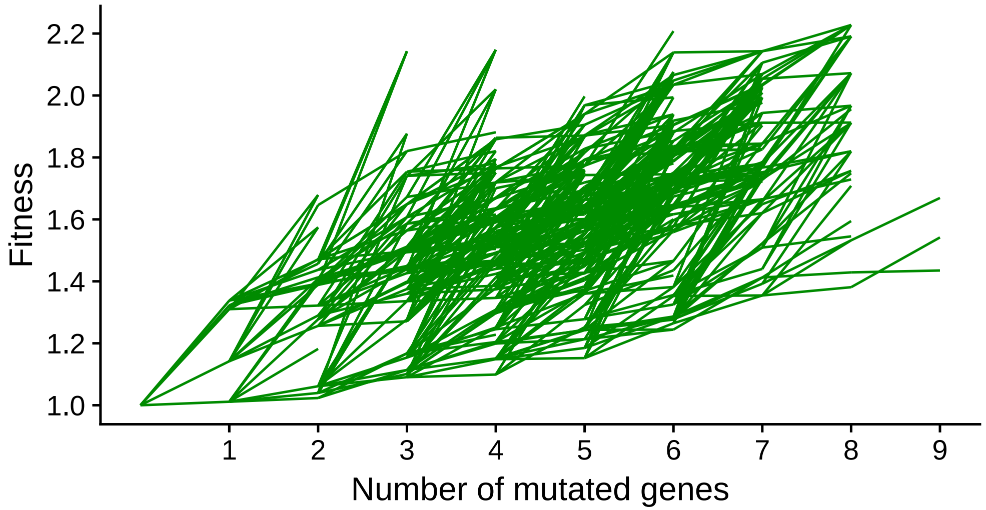


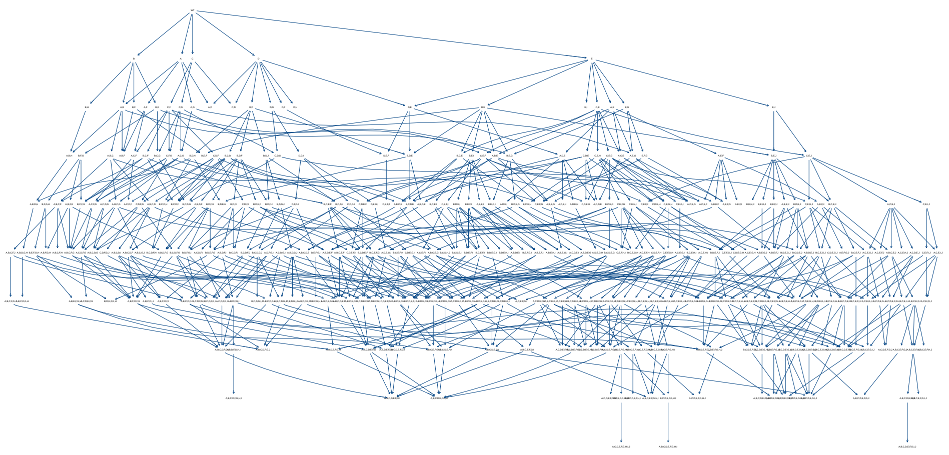

| | |
|---|---|
| *ID* | FJPvUaTzze1FXsAz |
| *acc genotypes* | 459 |
| *peaks no back* | 81 |
| *removed edges* | NA |
| *prop rem edges* | NA |

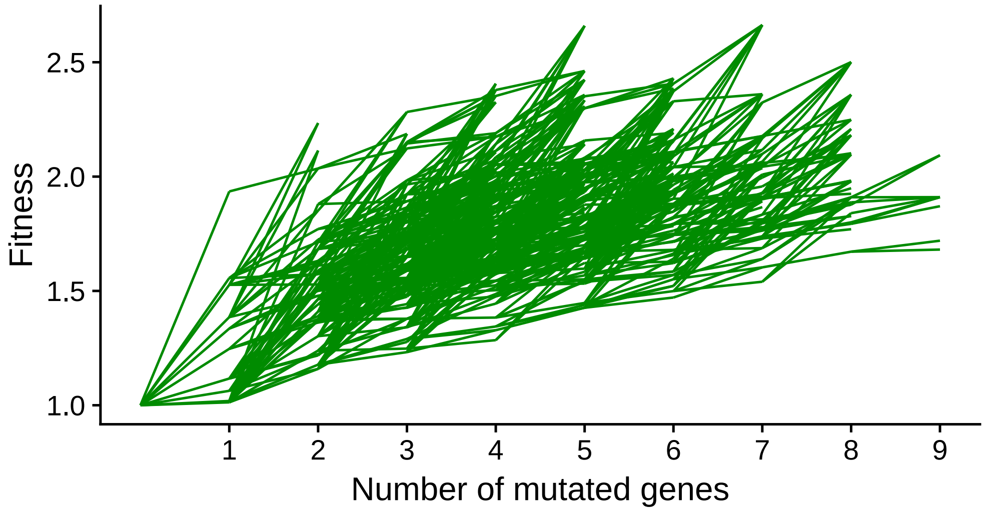


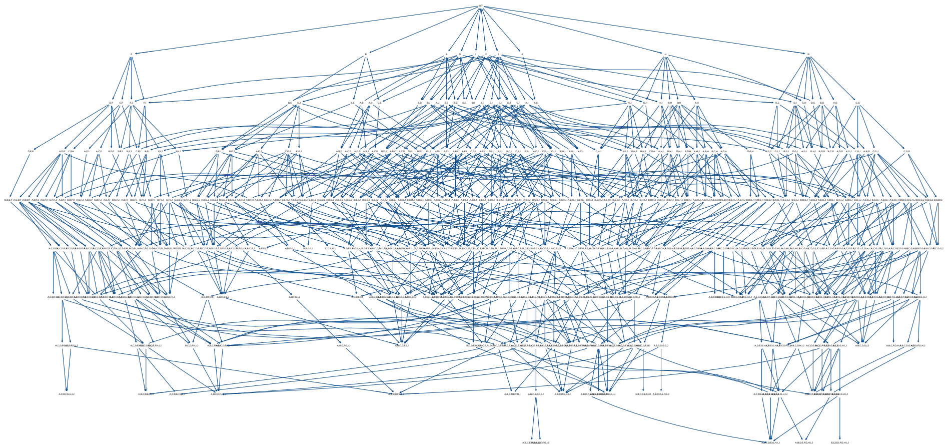

| | |
|---|---|
| *ID* | fpSFxSYrkAEuTBdj |
| *acc genotypes* | 355 |
| *peaks no back* | 123 |
| *removed edges* | NA |
| *prop rem edges* | NA |

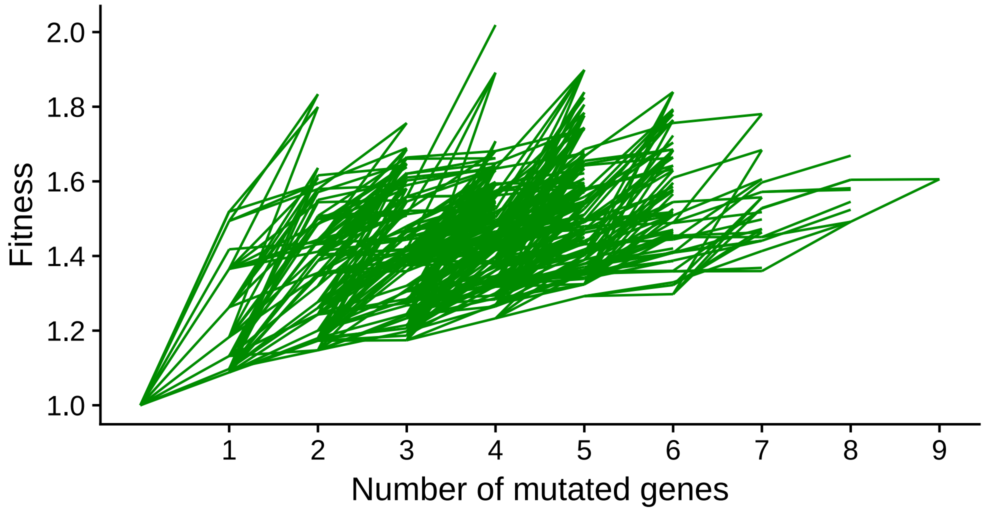


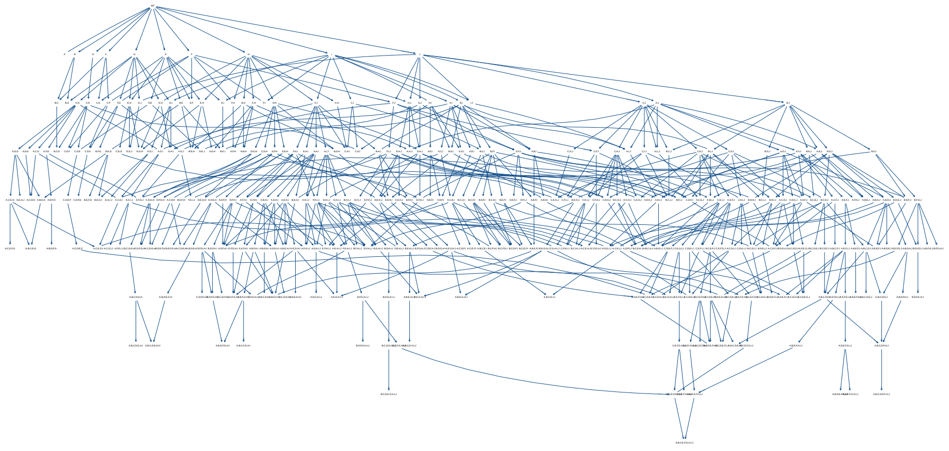

| | |
|---|---|
| *ID* | FSw8uHKnSnOw6Xrh |
| *acc genotypes* | 124 |
| *peaks no back* | 32 |
| *removed edges* | NA |
| *prop rem edges* | NA |

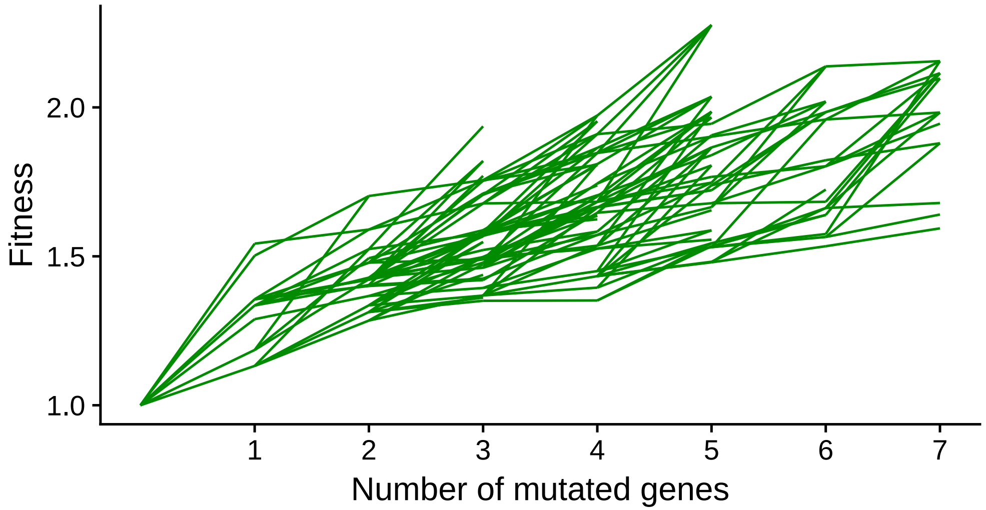


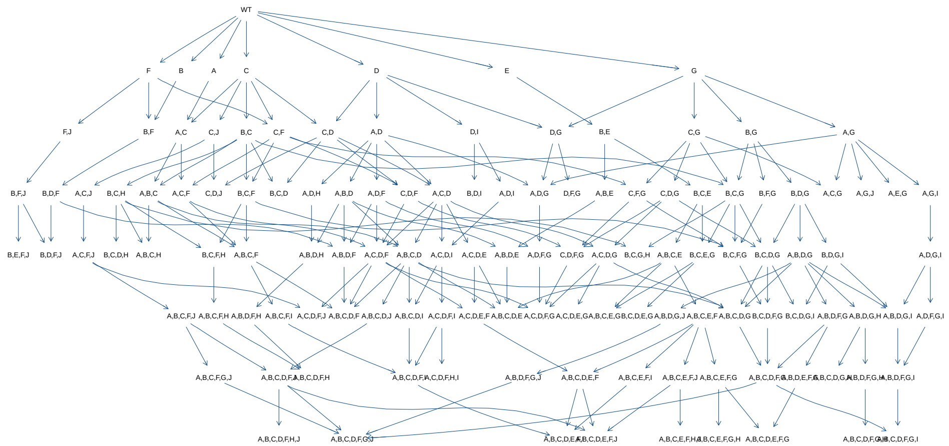

| | |
|---|---|
| *ID* | fWEJplAFKvHJAViu |
| *acc genotypes* | 208 |
| *peaks no back* | 52 |
| *removed edges* | NA |
| *prop rem edges* | NA |

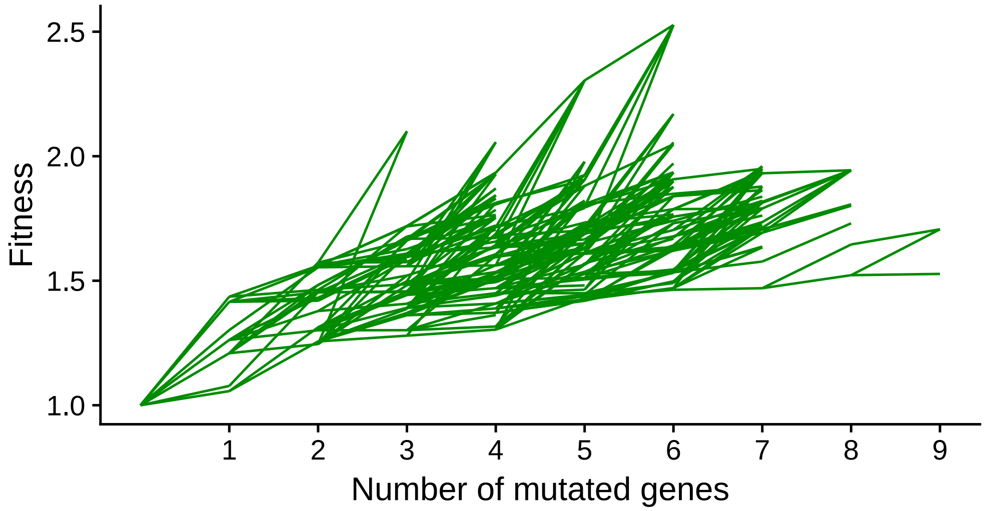


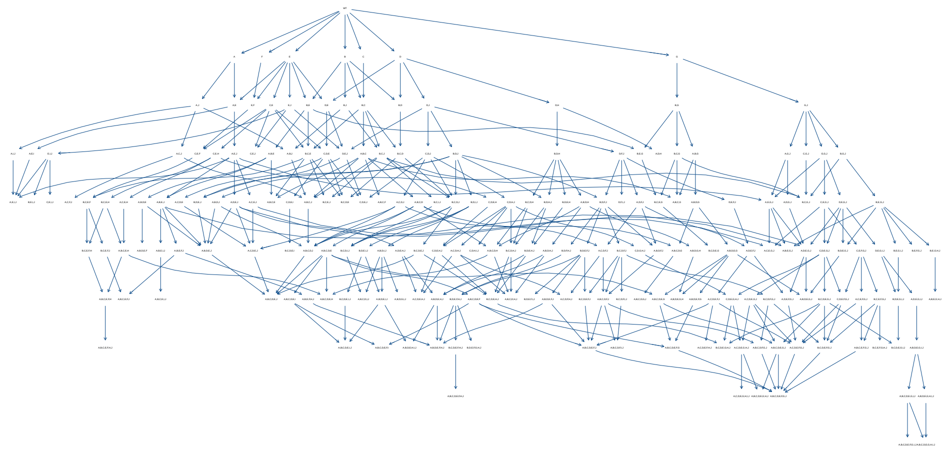

| | |
|---|---|
| *ID* | g4on7kMYrN7IvT9p |
| *acc genotypes* | 106 |
| *peaks no back* | 48 |
| *removed edges* | NA |
| *prop rem edges* | NA |

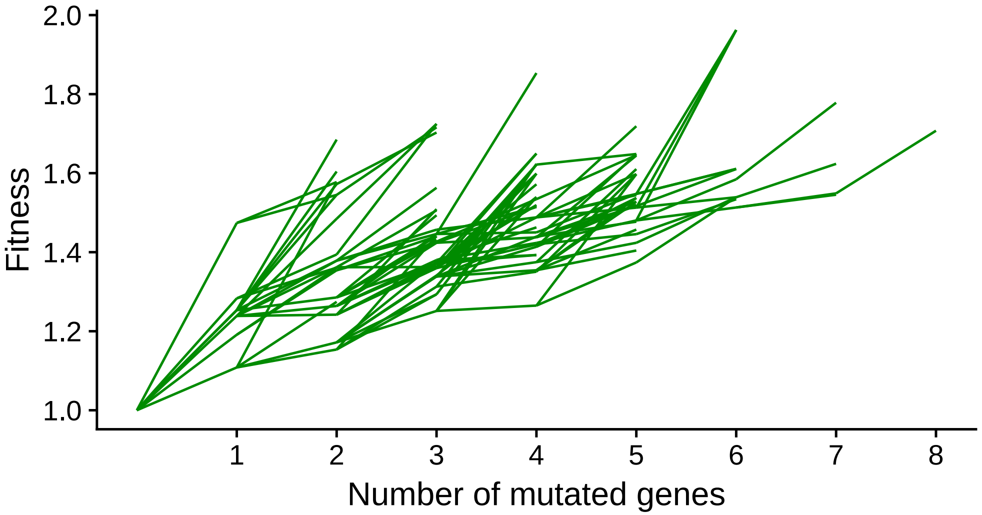


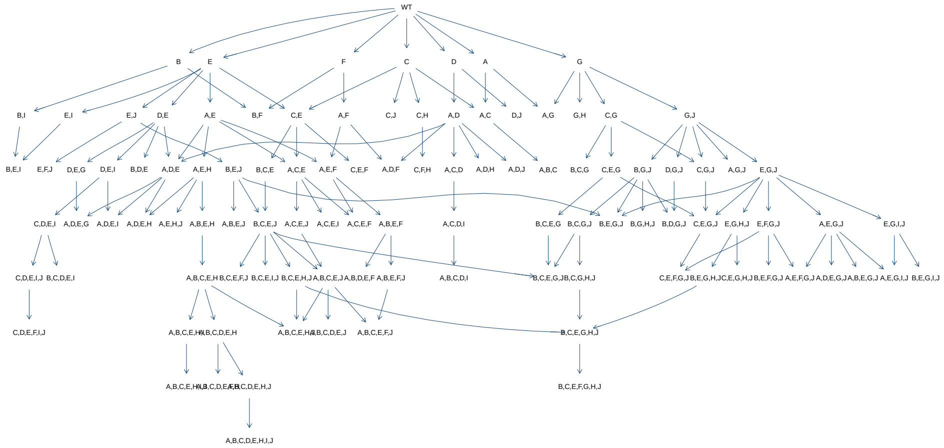

| | |
|---|---|
| *ID* | g5rN9RqrUiS9arMn |
| *acc genotypes* | 39 |
| *peaks no back* | 16 |
| *removed edges* | NA |
| *prop rem edges* | NA |

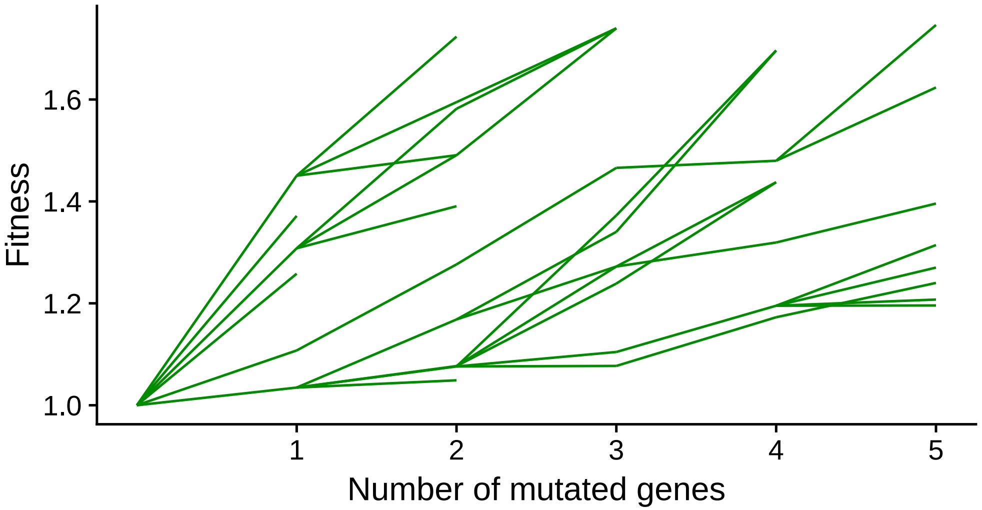


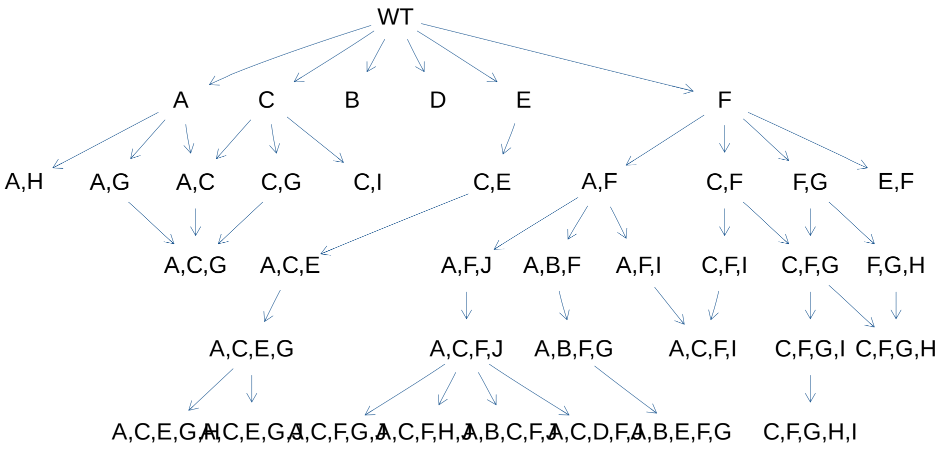

| | |
|---|---|
| *ID* | g7EPSPvUEgZmblyv |
| *acc genotypes* | 282 |
| *peaks no back* | 69 |
| *removed edges* | NA |
| *prop rem edges* | NA |

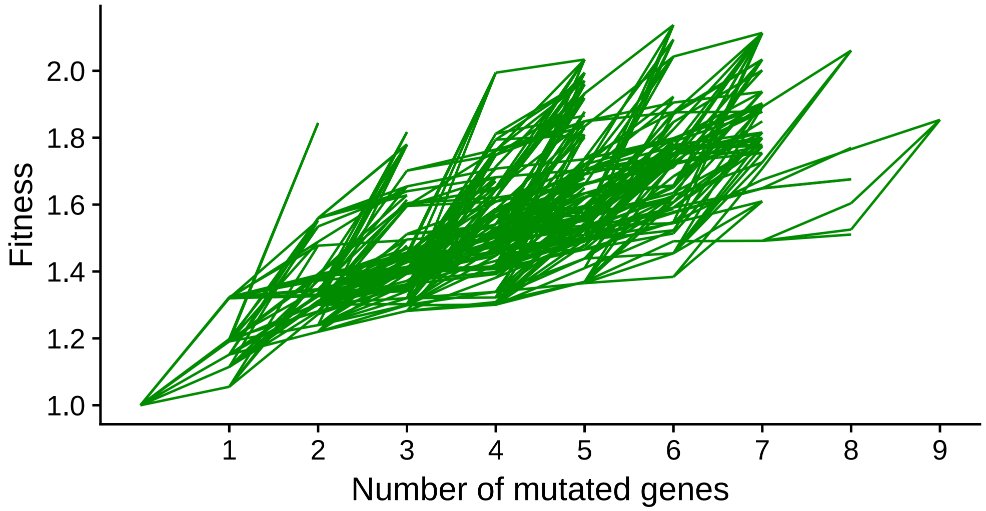


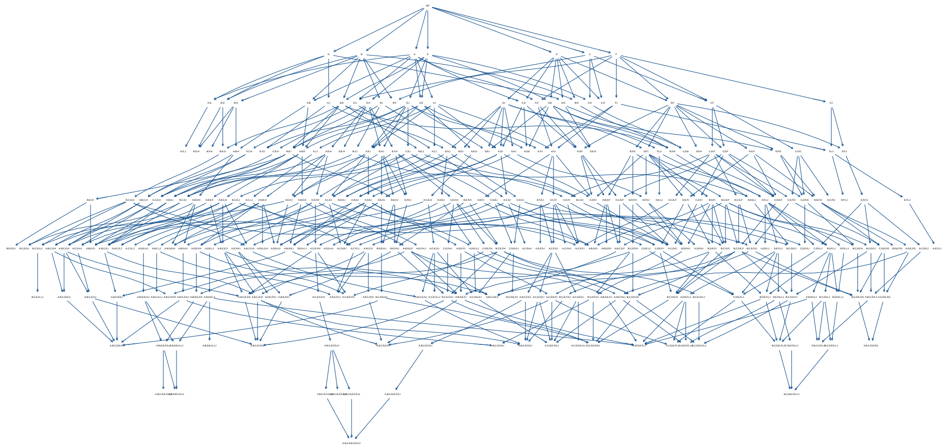

| | |
|---|---|
| *ID* | GDPhLnRCnDMjG2D7 |
| *acc genotypes* | 185 |
| *peaks no back* | 64 |
| *removed edges* | NA |
| *prop rem edges* | NA |

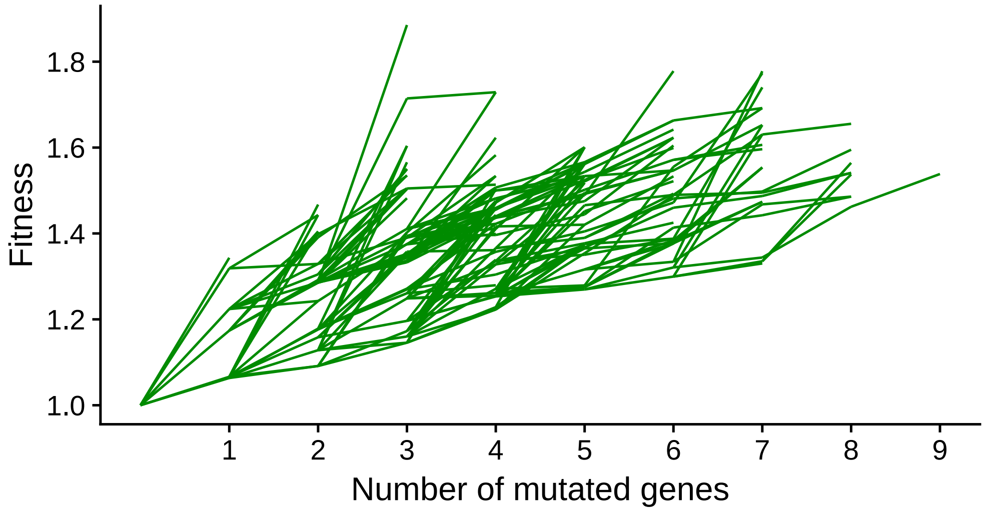


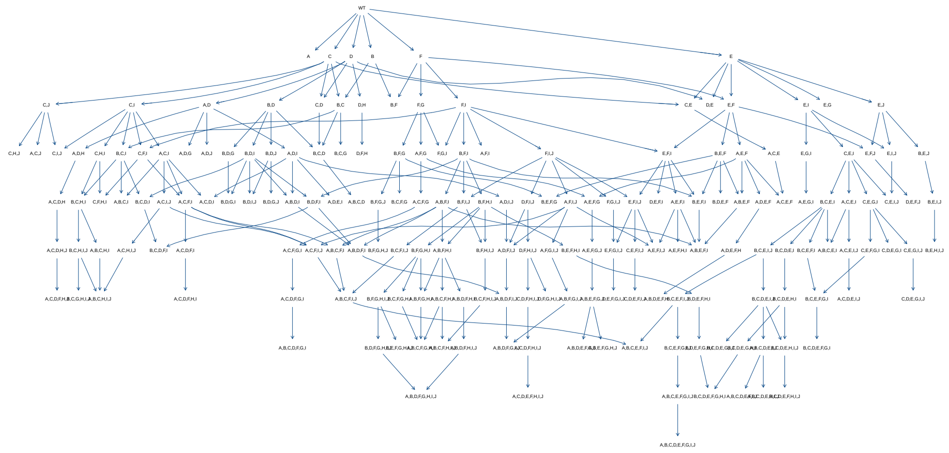

| | |
|---|---|
| *ID* | Ggye5iq3jqoH3lvU |
| *acc genotypes* | 314 |
| *peaks no back* | 112 |
| *removed edges* | NA |
| *prop rem edges* | NA |

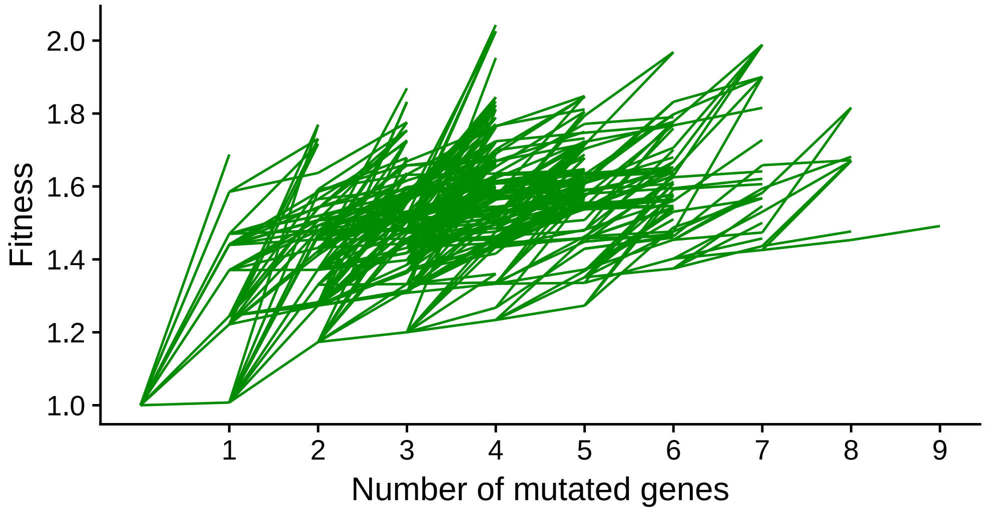


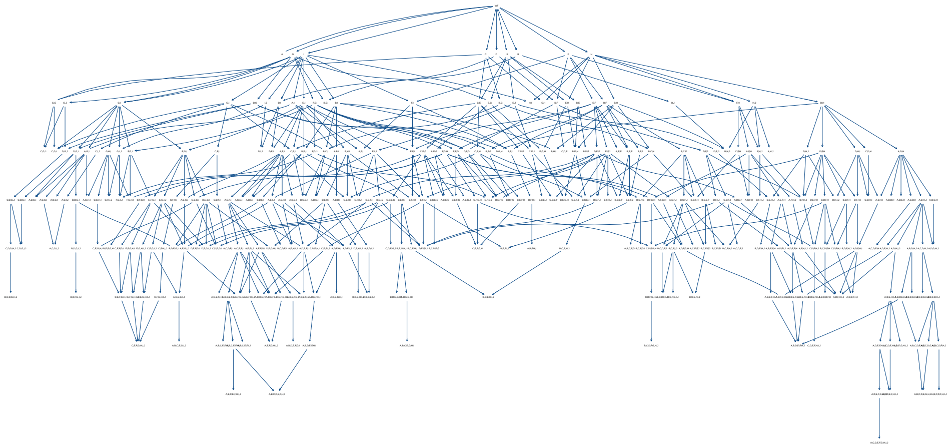

| | |
|---|---|
| *ID* | gQ91ynxxLJpJfC10R |
| *acc genotypes* | 53 |
| *peaks no back* | 23 |
| *removed edges* | NA |
| *prop rem edges* | NA |

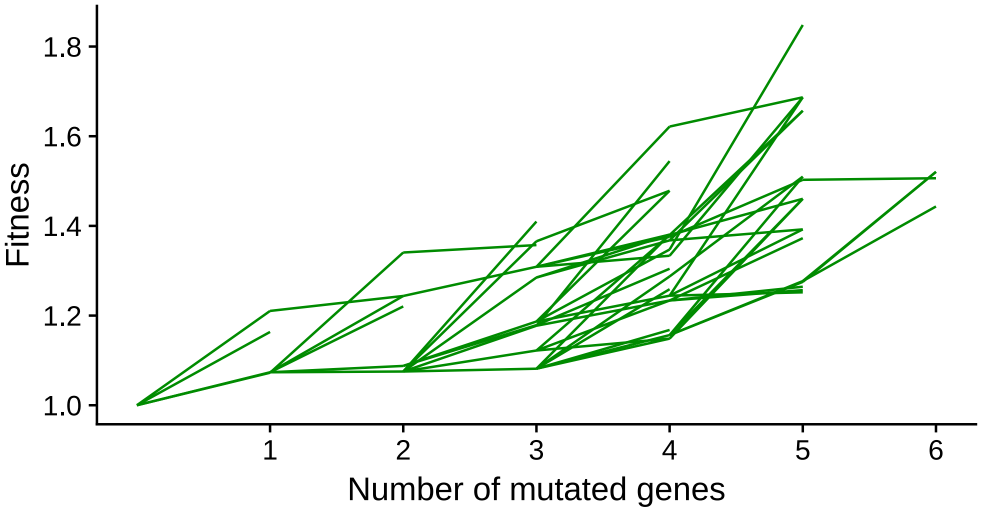


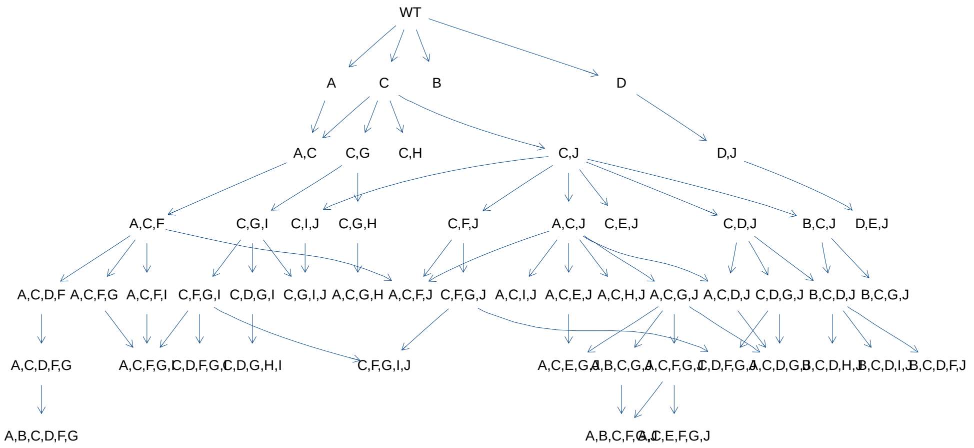

| | |
|---|---|
| *ID* | GtsX2ANXEuqX4ryX |
| *acc genotypes* | 48 |
| *peaks no back* | 21 |
| *removed edges* | NA |
| *prop rem edges* | NA |

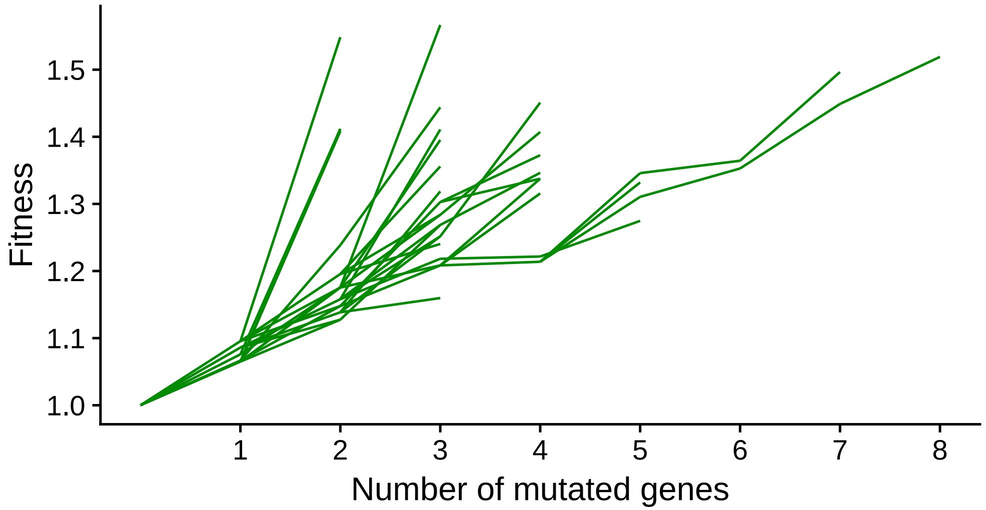


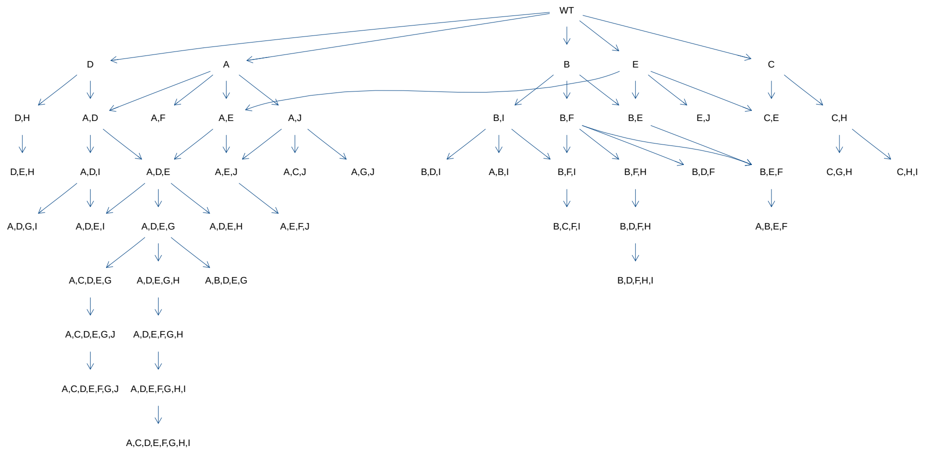

*ID* gU6jF3rPQlacxvDT
*acc genotypes* 198
*peaks no back* 76
*removed edges* NA
*prop rem edges* NA

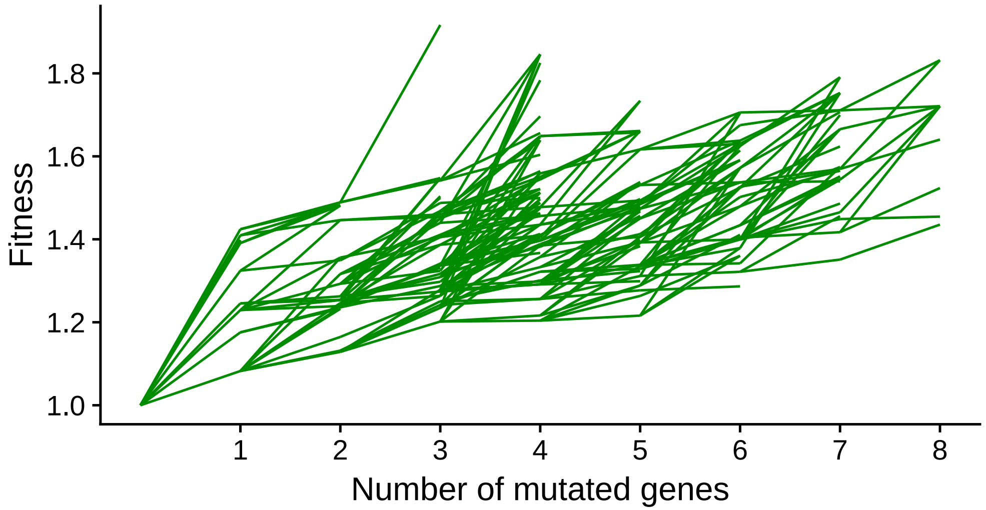


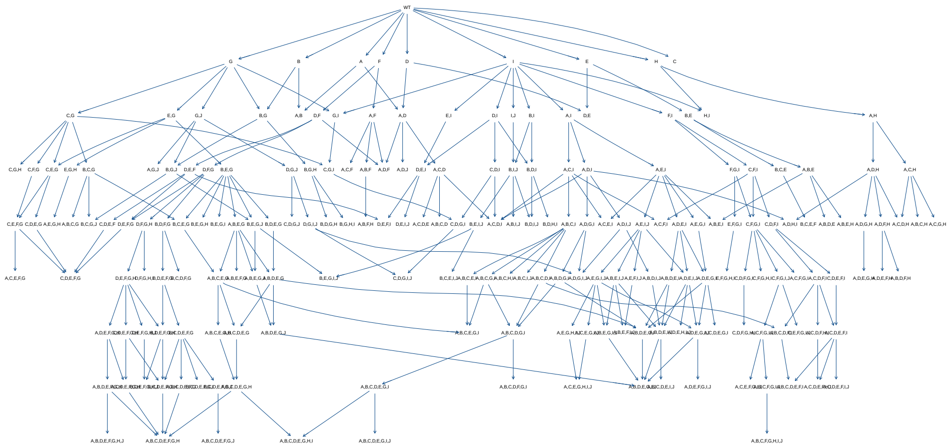

| | |
|---|---|
| *ID* | h10Ra4qDED5CSQYCn |
| *acc genotypes* | 472 |
| *peaks no back* | 79 |
| *removed edges* | NA |
| *prop rem edges* | NA |

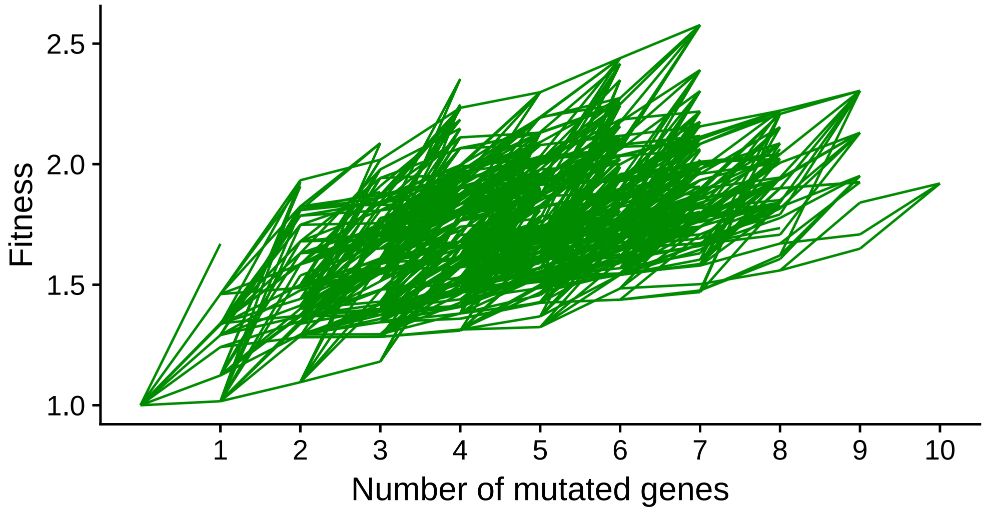


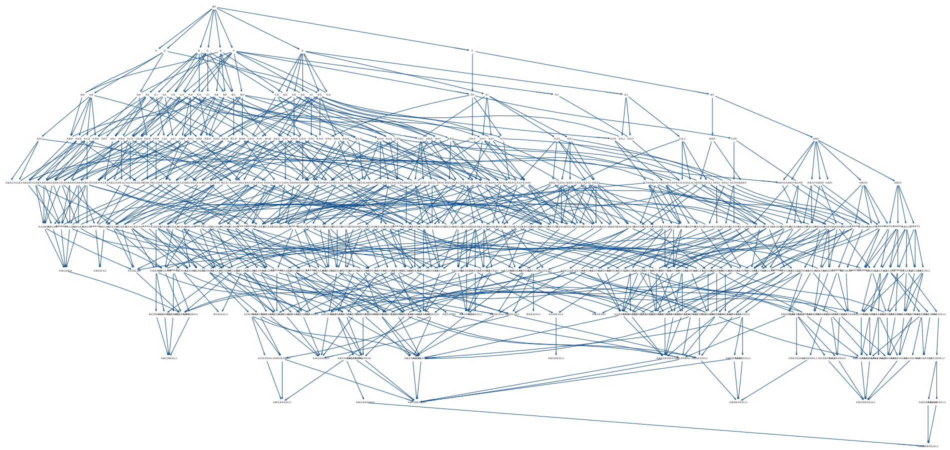

| | |
|---|---|
| *ID* | H5F3U9TPZOWztRjG |
| *acc genotypes* | 501 |
| *peaks no back* | 101 |
| *removed edges* | NA |
| *prop rem edges* | NA |

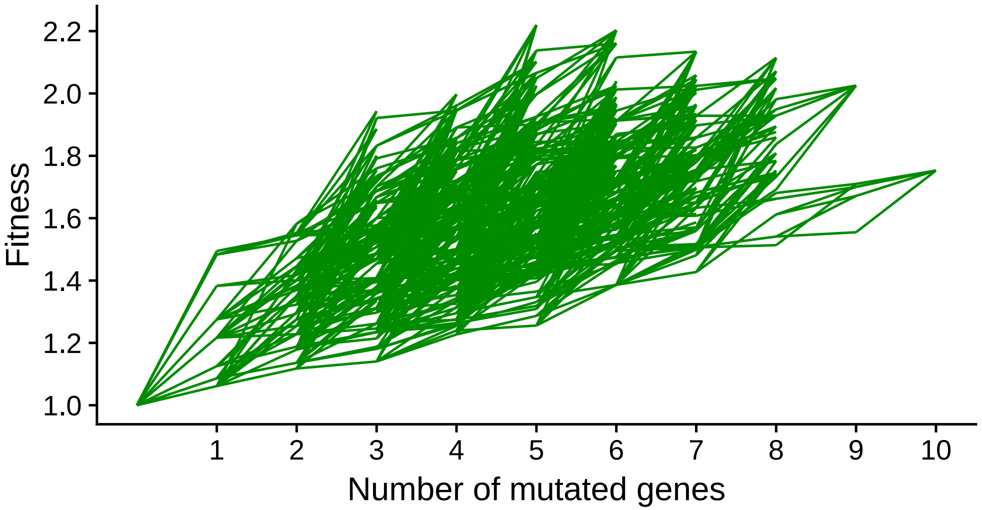


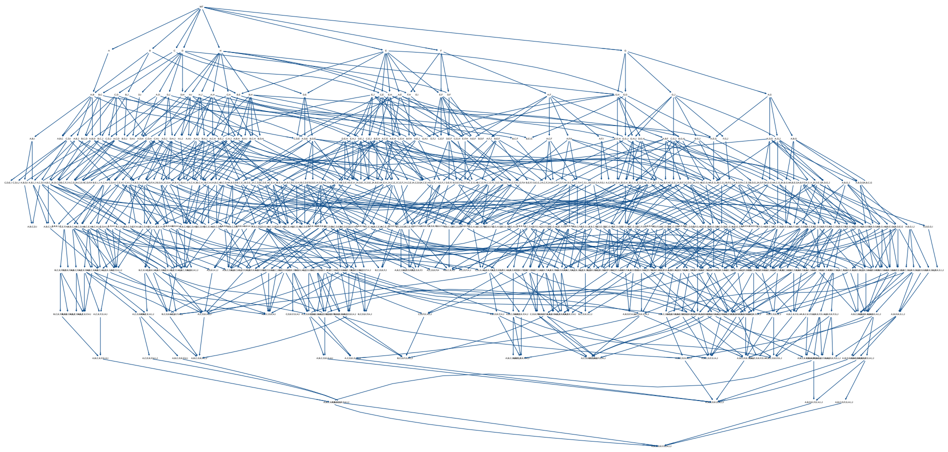

| | |
|---|---|
| *ID* | HdswhOaHWIP1kOLZ |
| *acc genotypes* | 389 |
| *peaks no back* | 95 |
| *removed edges* | NA |
| *prop rem edges* | NA |

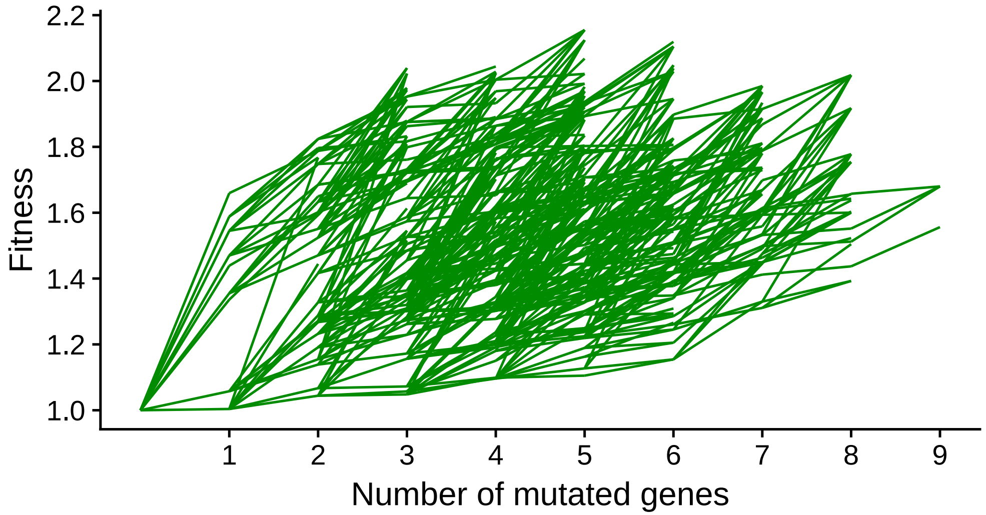


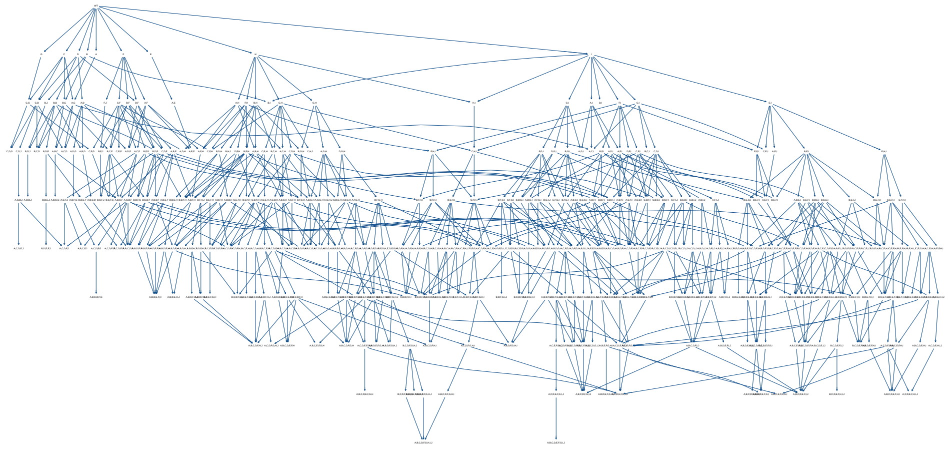

| | |
|---|---|
| *ID* | heIHa8Gt8XkXkA6v |
| *acc genotypes* | 19 |
| *peaks no back* | 8 |
| *removed edges* | NA |
| *prop rem edges* | NA |

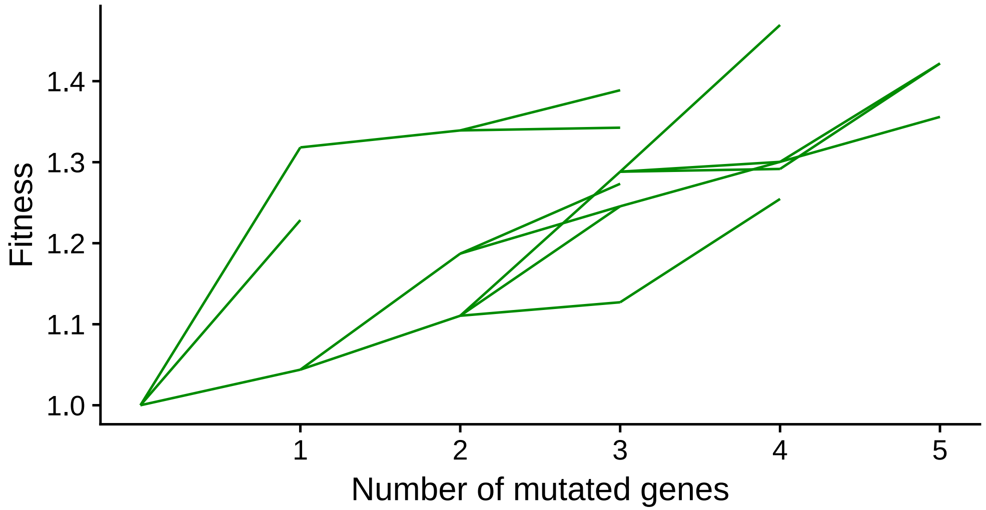


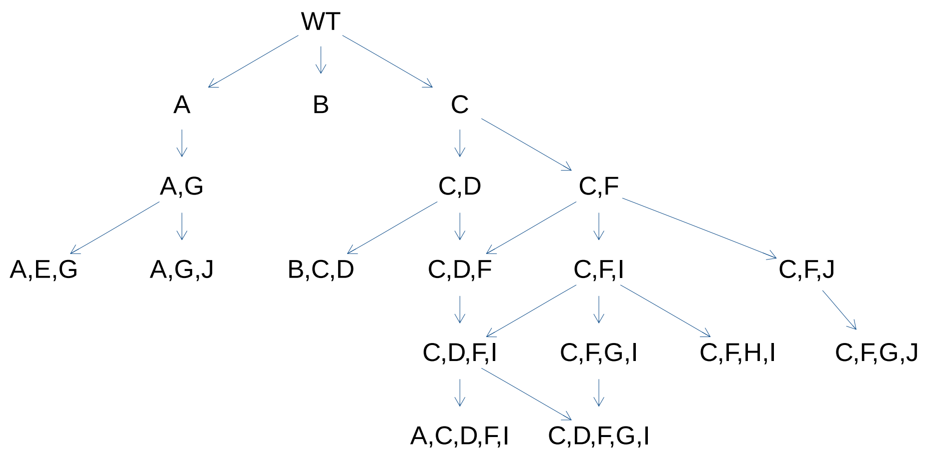

| | |
|---|---|
| *ID* | hm10KP8sr2CYeQUji |
| *acc genotypes* | 309 |
| *peaks no back* | 107 |
| *removed edges* | NA |
| *prop rem edges* | NA |

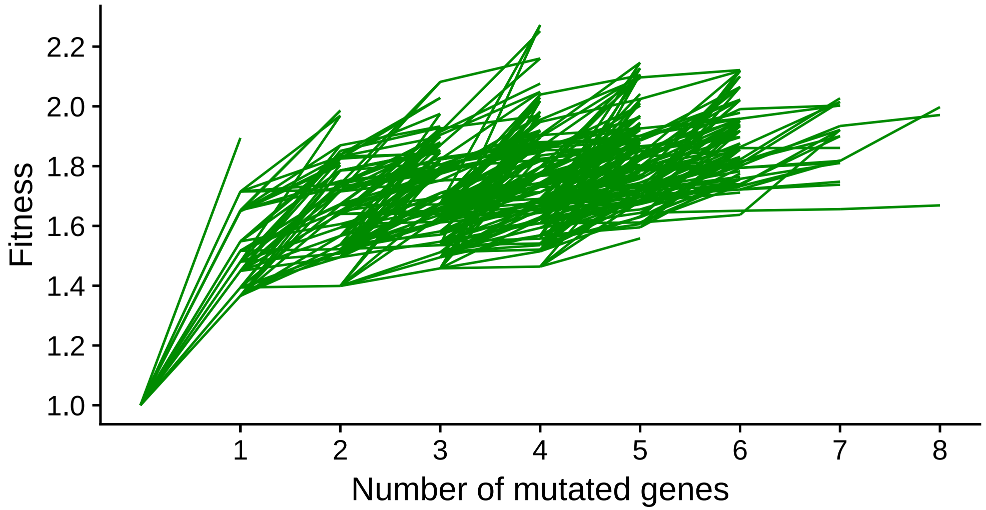


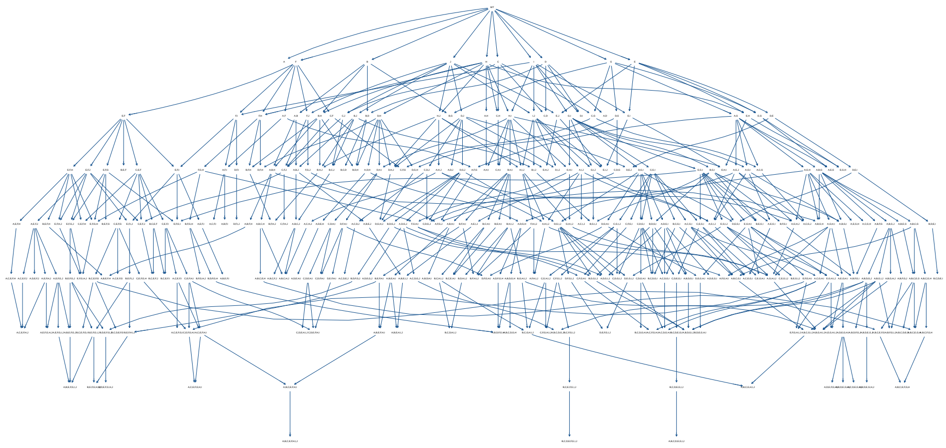

| | |
|---|---|
| *ID* | hQhSt6SyCYjTd10y8 |
| *acc genotypes* | 593 |
| *peaks no back* | 84 |
| *removed edges* | NA |
| *prop rem edges* | NA |

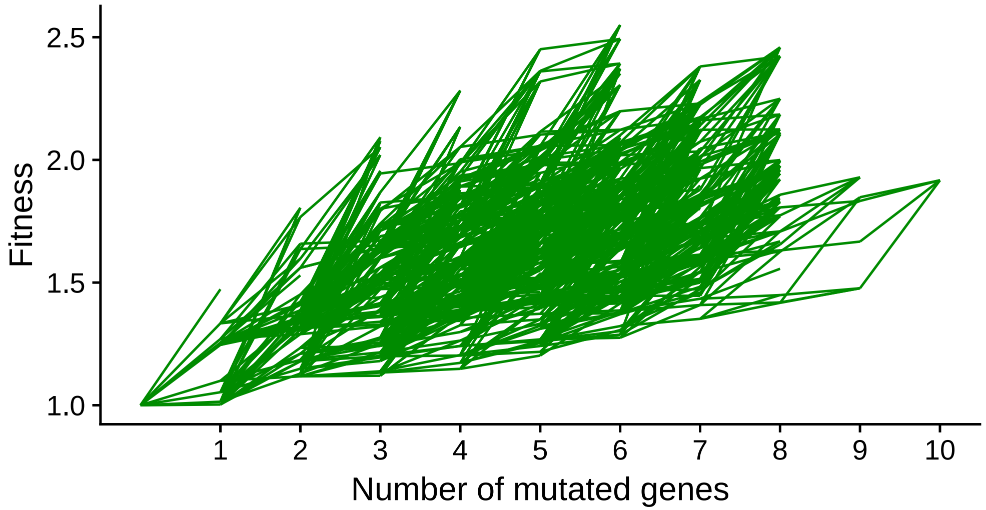


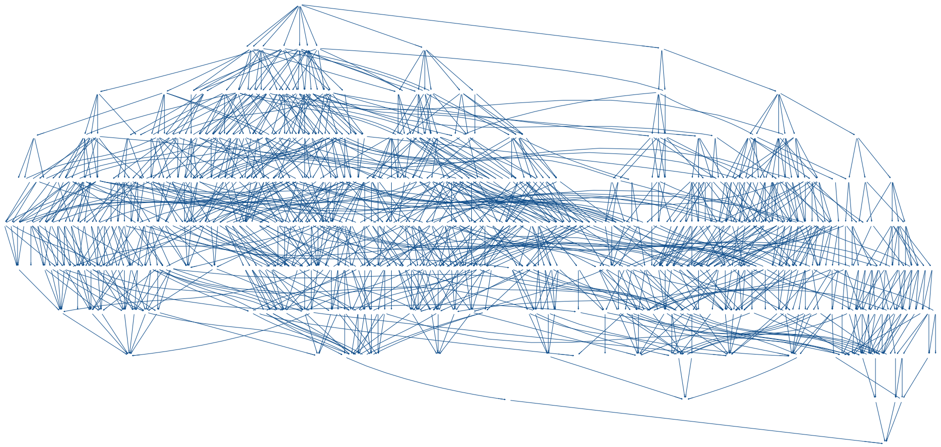

| | |
|---|---|
| *ID* | hrKHhSGSCA4NDMJP |
| *acc genotypes* | 157 |
| *peaks no back* | 43 |
| *removed edges* | NA |
| *prop rem edges* | NA |

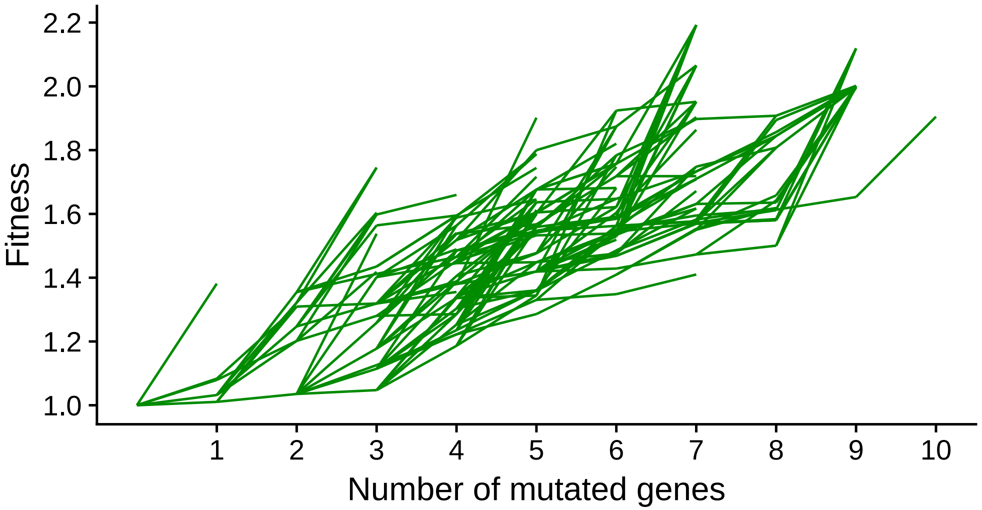


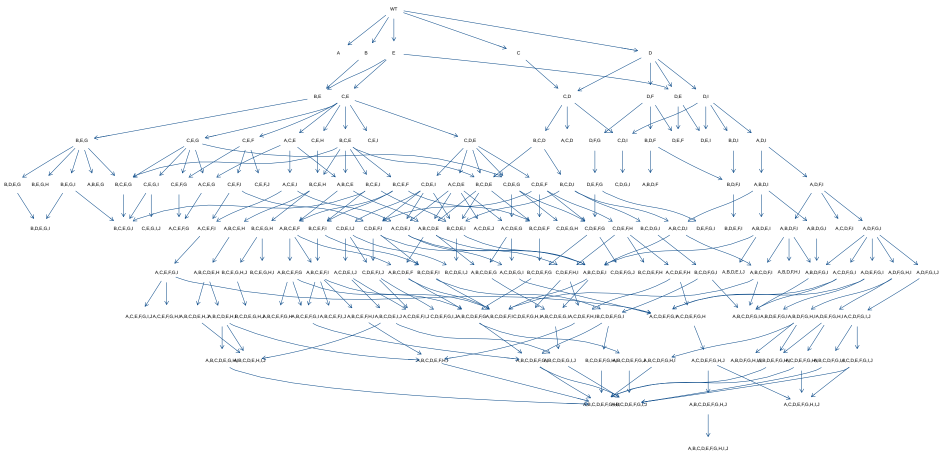

| | |
|---|---|
| *ID* | I9q6GI3hisYql2on |
| *acc genotypes* | 298 |
| *peaks no back* | 79 |
| *removed edges* | NA |
| *prop rem edges* | NA |

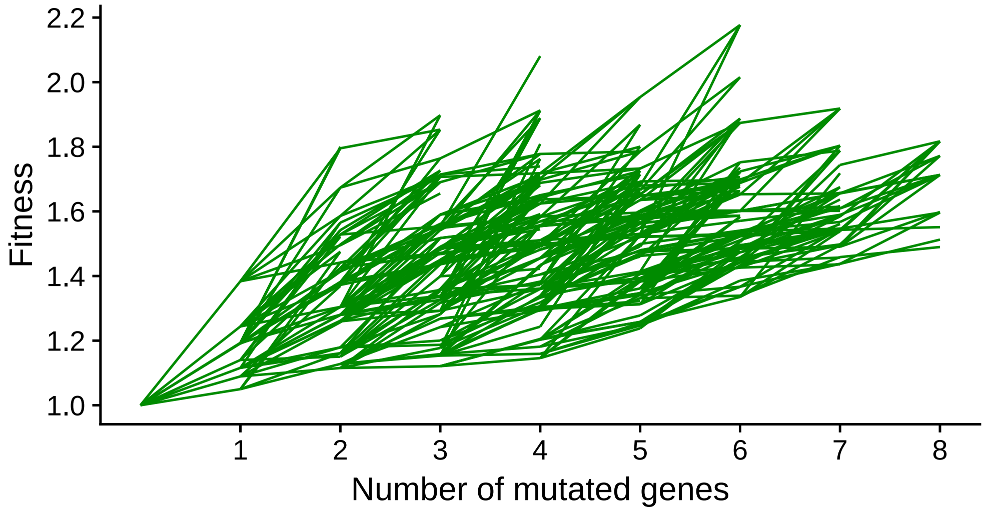


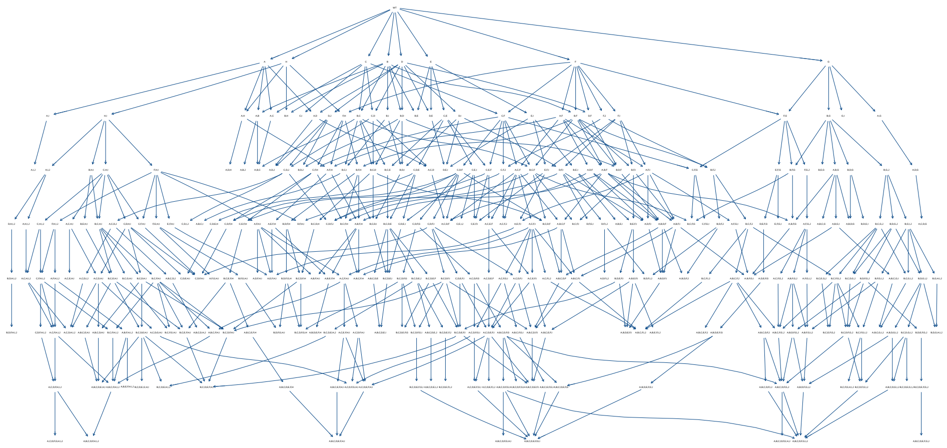

| | |
|---|---|
| *ID* | ikuJV711Pr9P5gMR |
| *acc genotypes* | 54 |
| *peaks no back* | 25 |
| *removed edges* | NA |
| *prop rem edges* | NA |

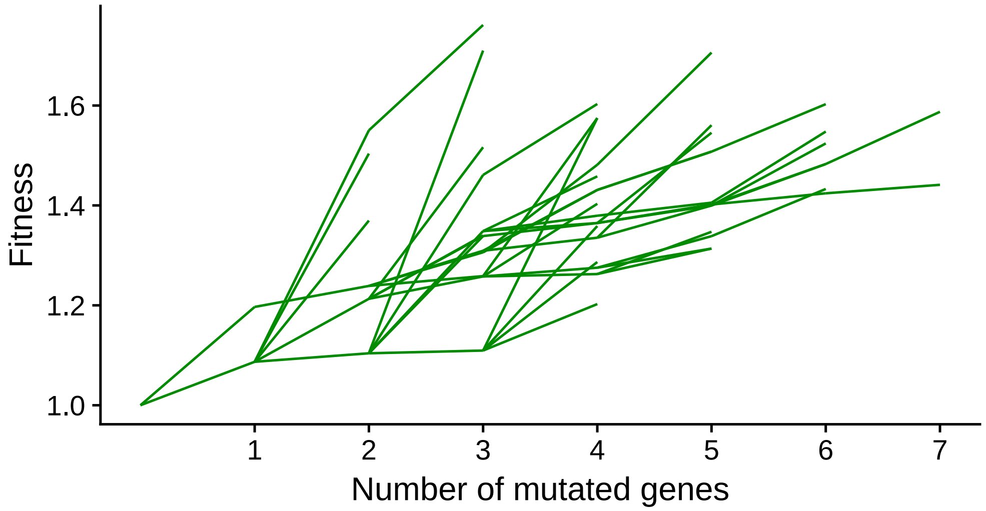


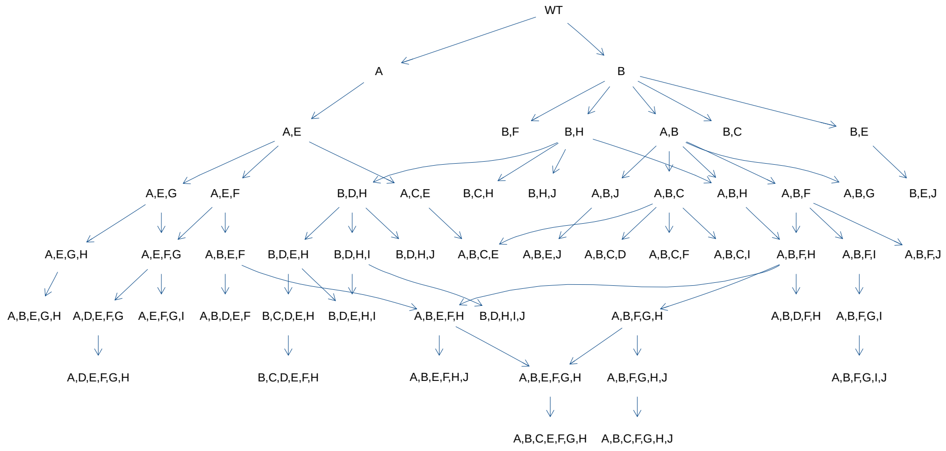

*ID* irlmocvl1nEJmK32
*acc genotypes* 95
*peaks no back* 46
*removed edges* NA
*prop rem edges* NA

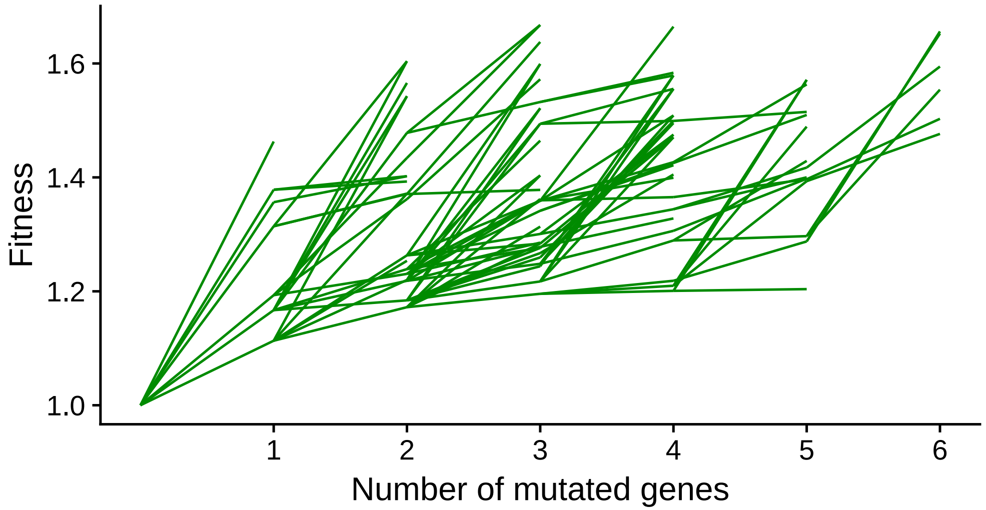


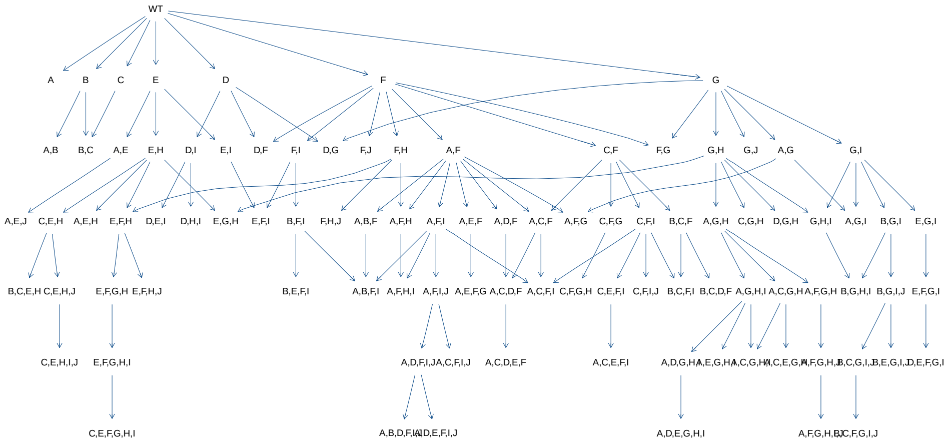

| | |
|---|---|
| *ID* | itu7sgxZ42Rqmcbw |
| *acc genotypes* | 140 |
| *peaks no back* | 36 |
| *removed edges* | NA |
| *prop rem edges* | NA |

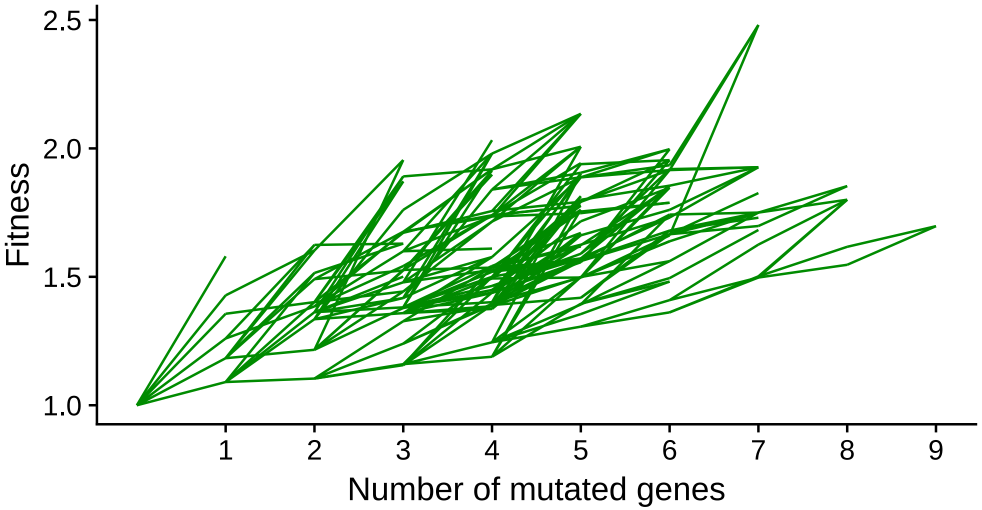


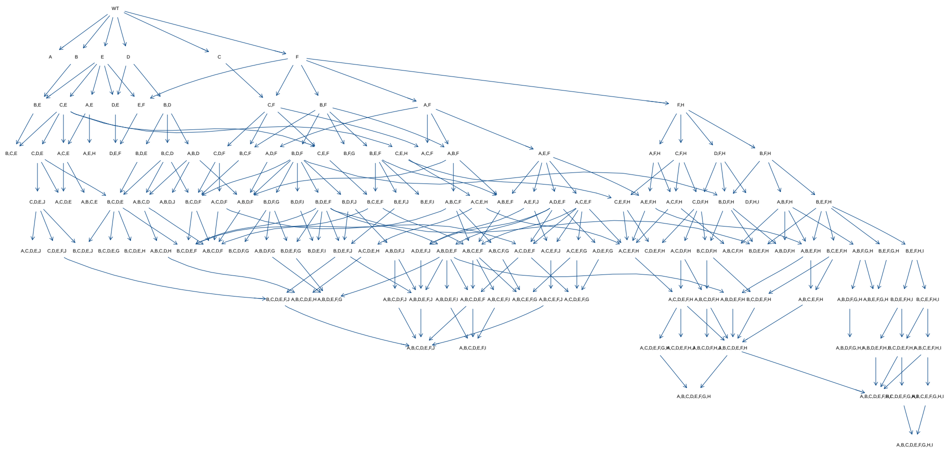

| | |
|---|---|
| *ID* | IxTaa10yvF9m8psni |
| *acc genotypes* | 188 |
| *peaks no back* | 61 |
| *removed edges* | NA |
| *prop rem edges* | NA |

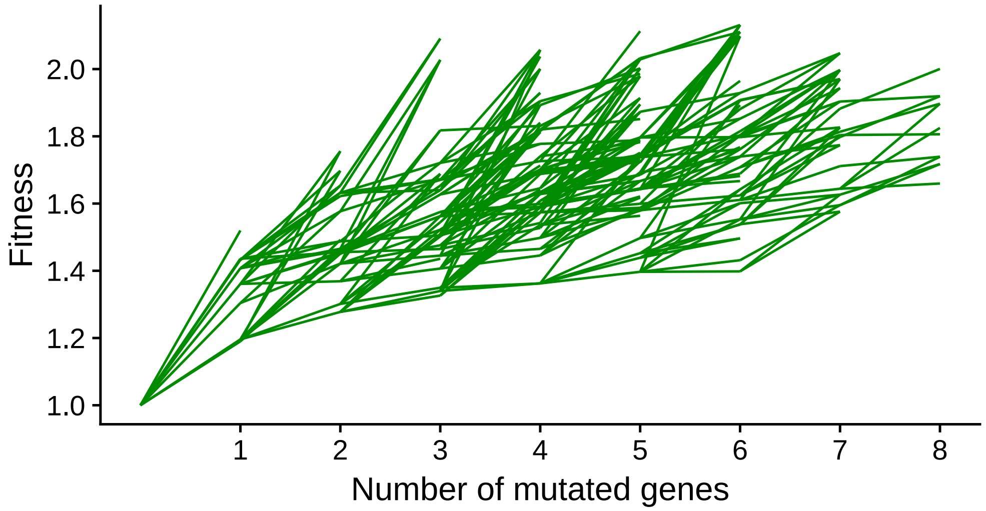


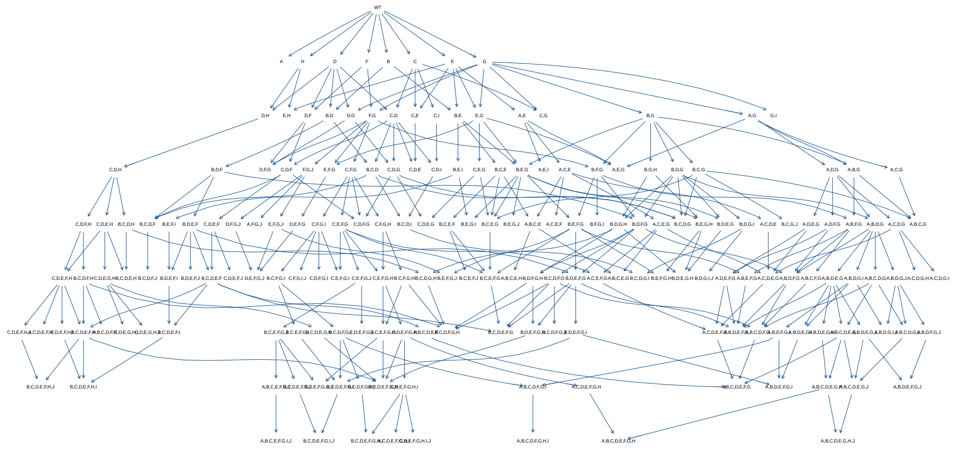

*ID* j5Wlsy8xtbP8JPCd

*acc genotypes* 115

*peaks no back* 54

*removed edges* NA

*prop rem edges* NA

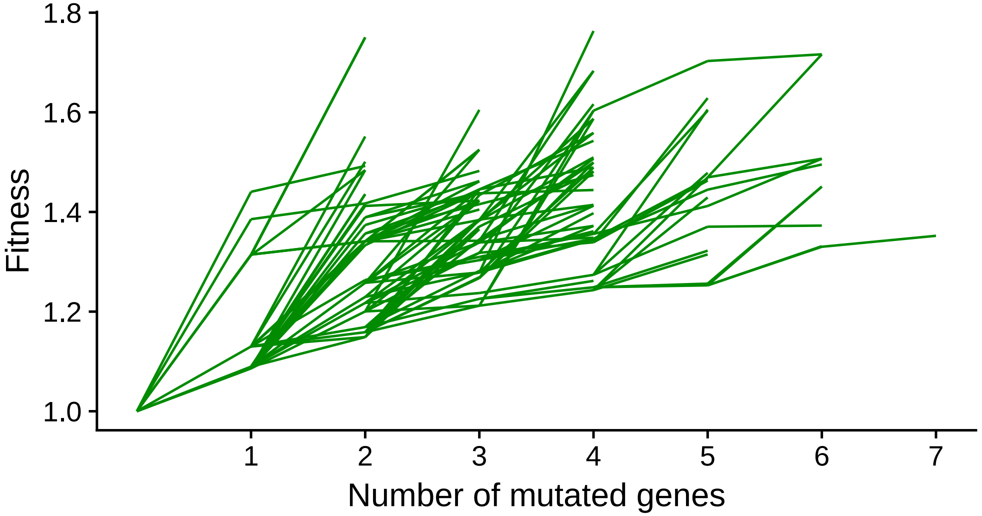


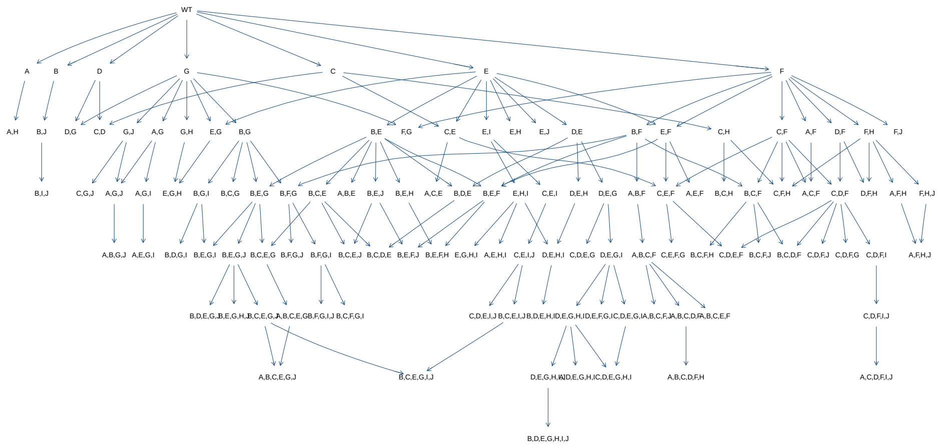

| | |
|---|---|
| *ID* | J6cFc1nMWhianSvF |
| *acc genotypes* | 189 |
| *peaks no back* | 73 |
| *removed edges* | NA |
| *prop rem edges* | NA |

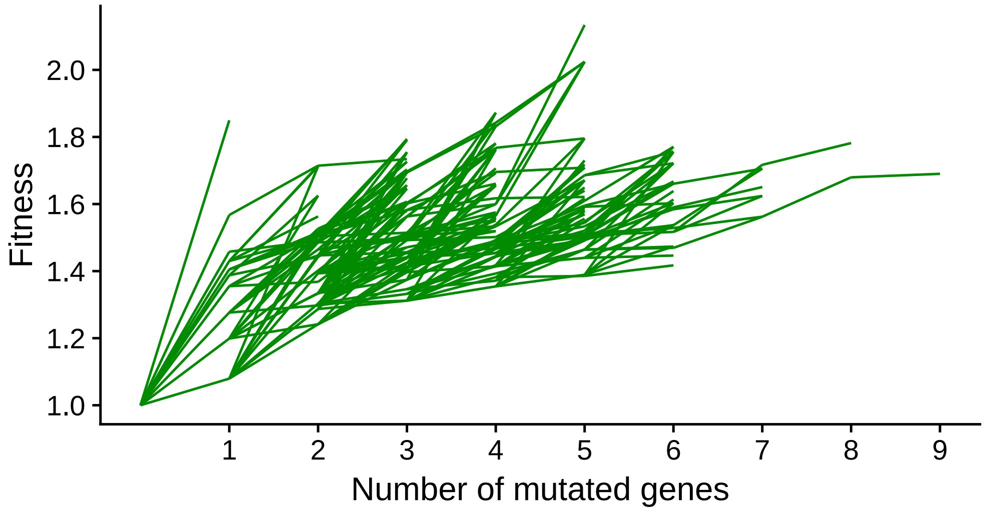


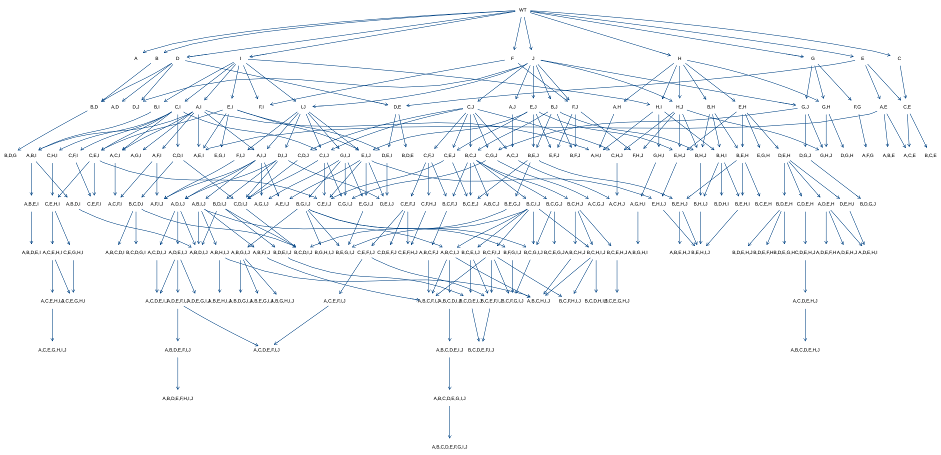

| | |
|---|---|
| *ID* | jANbFMNMm7HJj8B10 |
| *acc genotypes* | 735 |
| *peaks no back* | 93 |
| *removed edges* | NA |
| *prop rem edges* | NA |

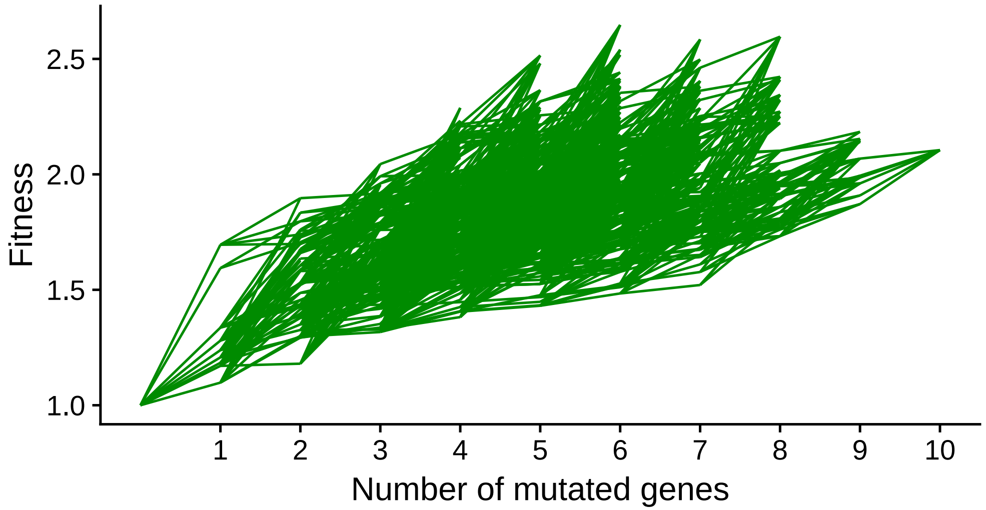


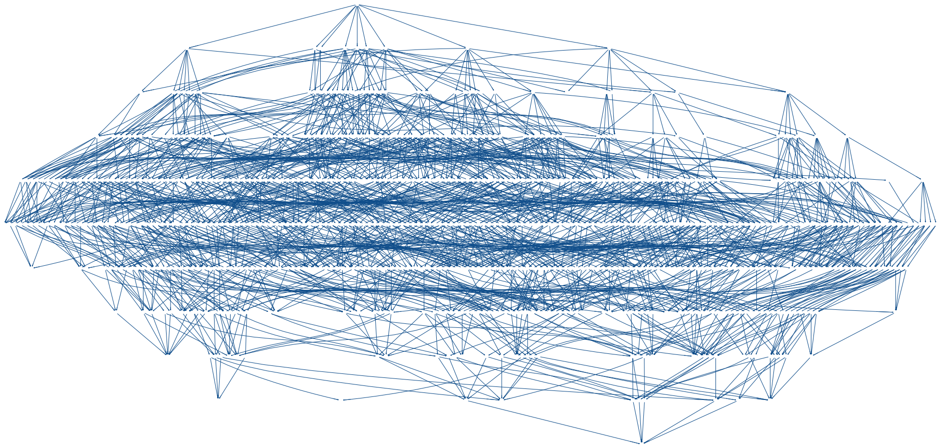

| | |
|---|---|
| *ID* | jQBPZk8ZlI14NJWW |
| *acc genotypes* | 88 |
| *peaks no back* | 31 |
| *removed edges* | NA |
| *prop rem edges* | NA |

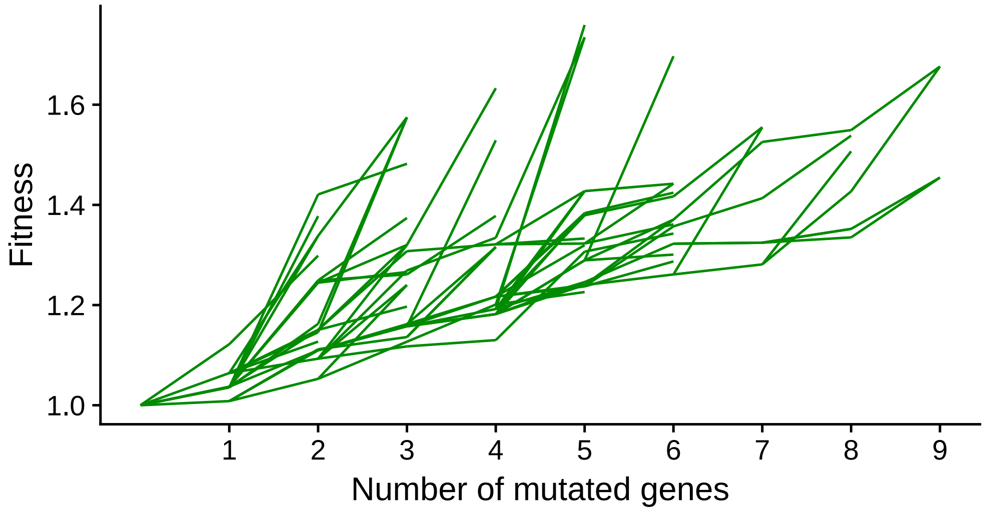


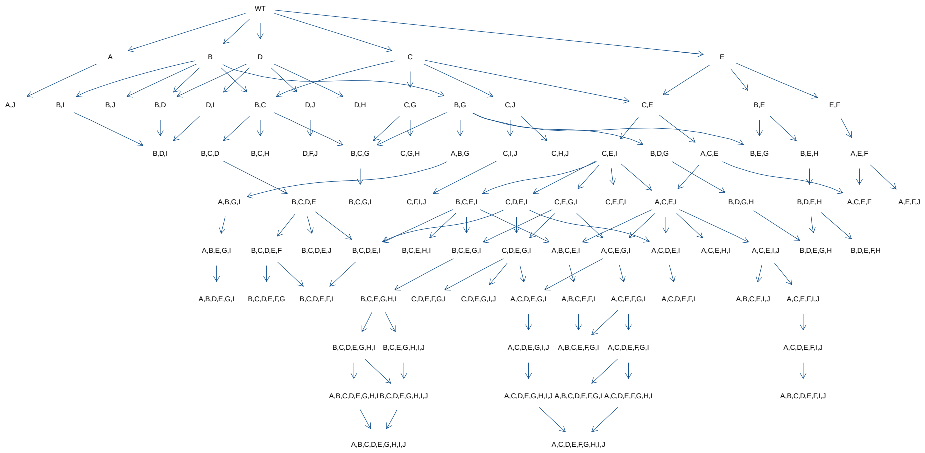

| | |
|---|---|
| *ID* | K4fDyrolukDwi9Ma |
| *acc genotypes* | 145 |
| *peaks no back* | 53 |
| *removed edges* | NA |
| *prop rem edges* | NA |

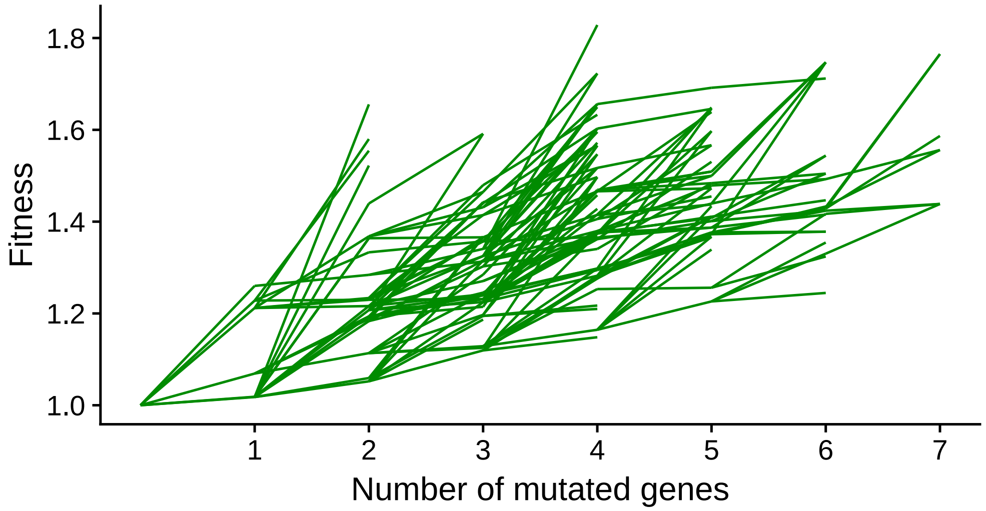


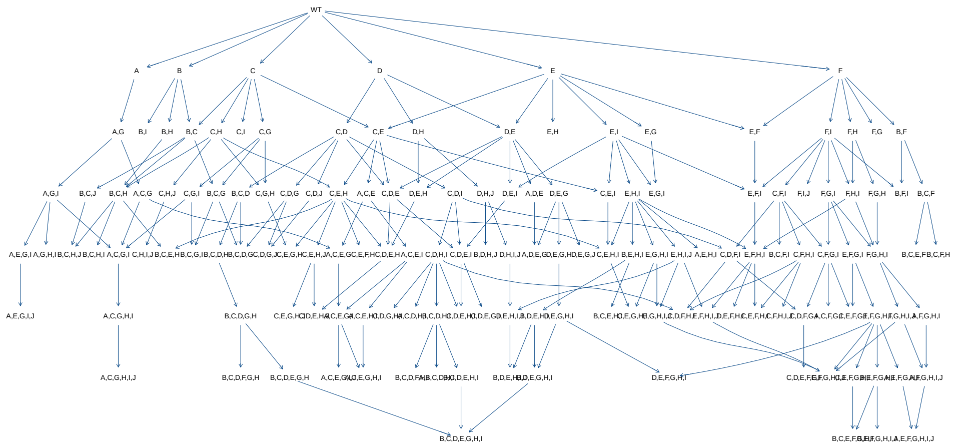

| | |
|---|---|
| *ID* | k4Otptn3TVpfZN7m |
| *acc genotypes* | 166 |
| *peaks no back* | 64 |
| *removed edges* | NA |
| *prop rem edges* | NA |

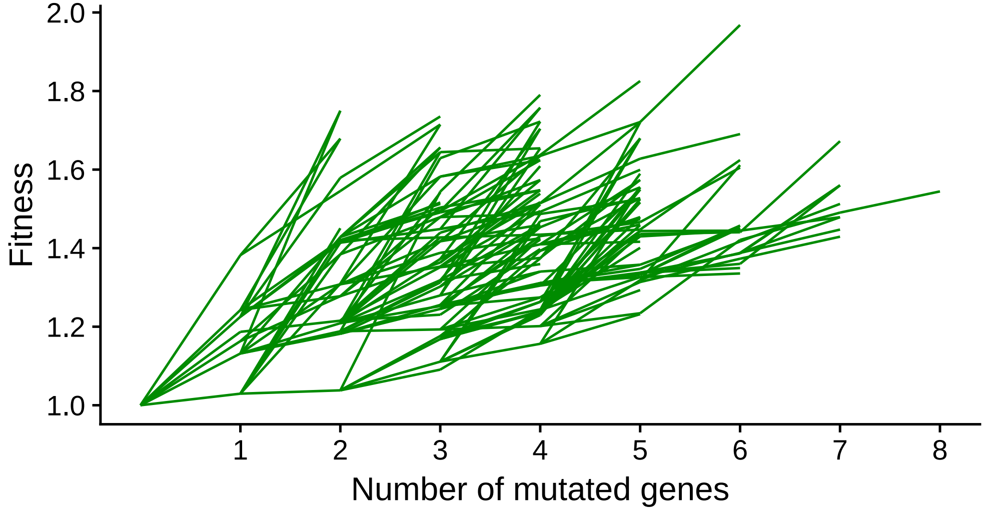


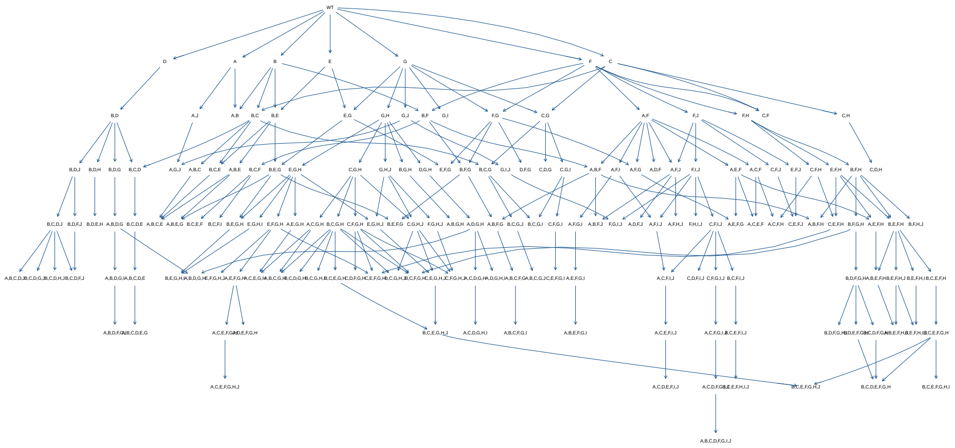

| | |
|---|---|
| *ID* | k9EeYCy4KVgVd19I |
| *acc genotypes* | 82 |
| *peaks no back* | 35 |
| *removed edges* | NA |
| *prop rem edges* | NA |

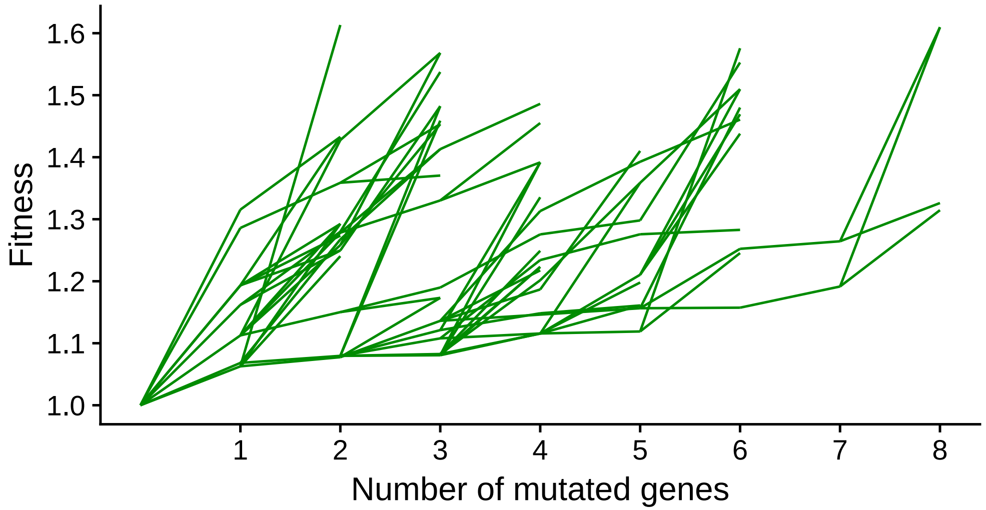


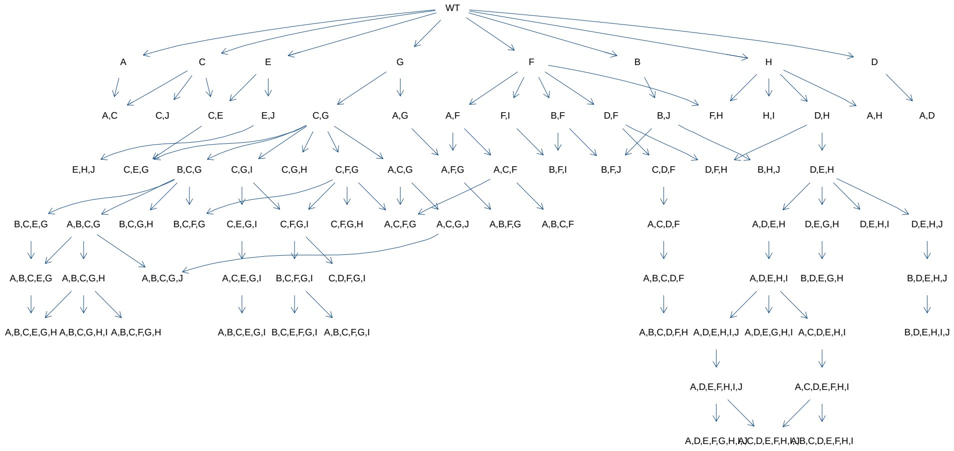

| | |
|---|---|
| *ID* | KB10V10fR8eSnvo1Wn |
| *acc genotypes* | 255 |
| *peaks no back* | 59 |
| *removed edges* | NA |
| *prop rem edges* | NA |

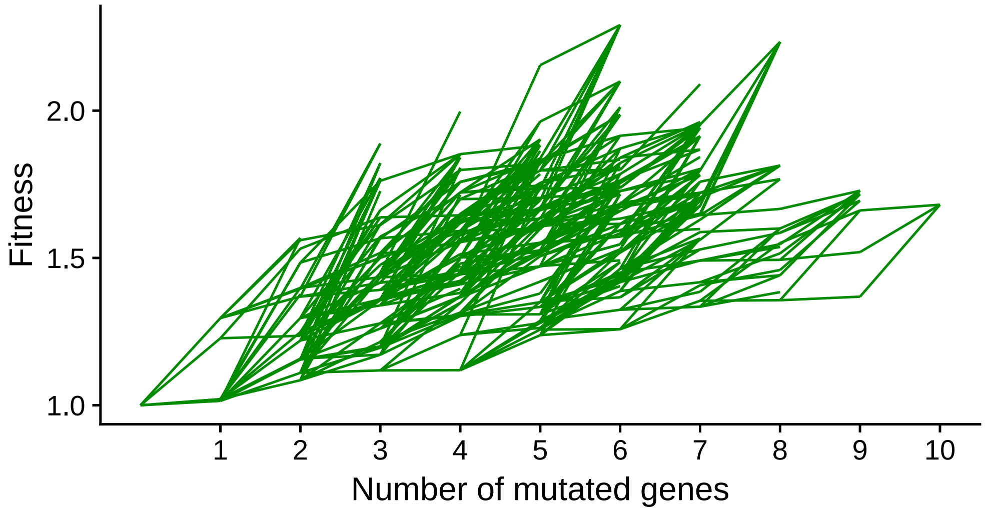


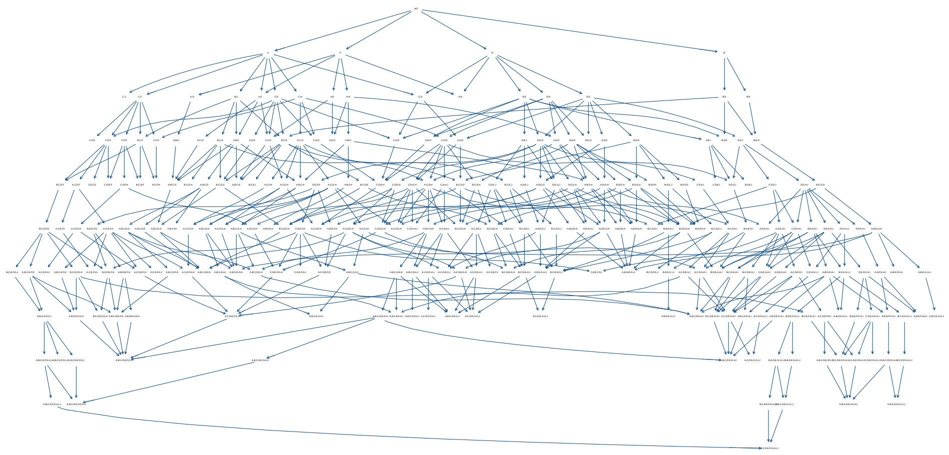

| | |
|---|---|
| *ID* | kEEjEwfuinUR9Smg |
| *acc genotypes* | 707 |
| *peaks no back* | 92 |
| *removed edges* | NA |
| *prop rem edges* | NA |

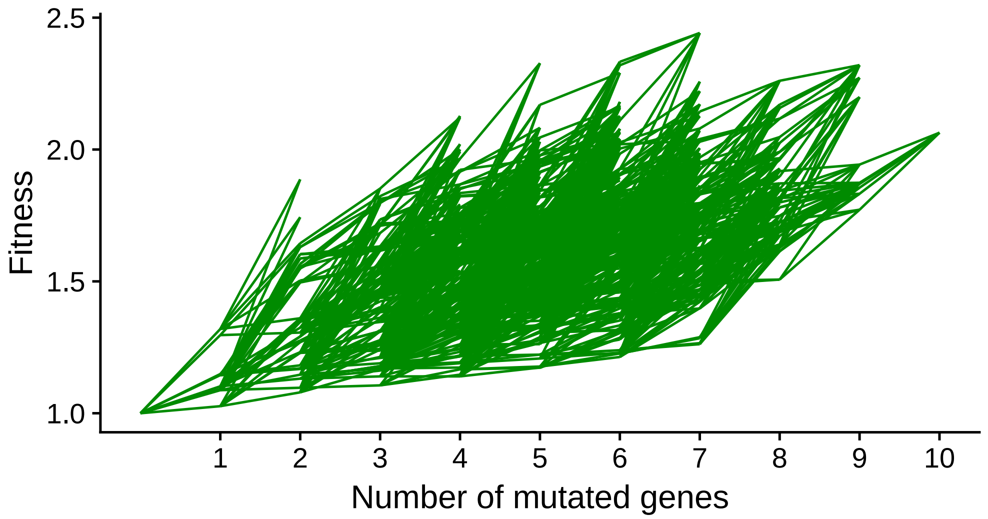


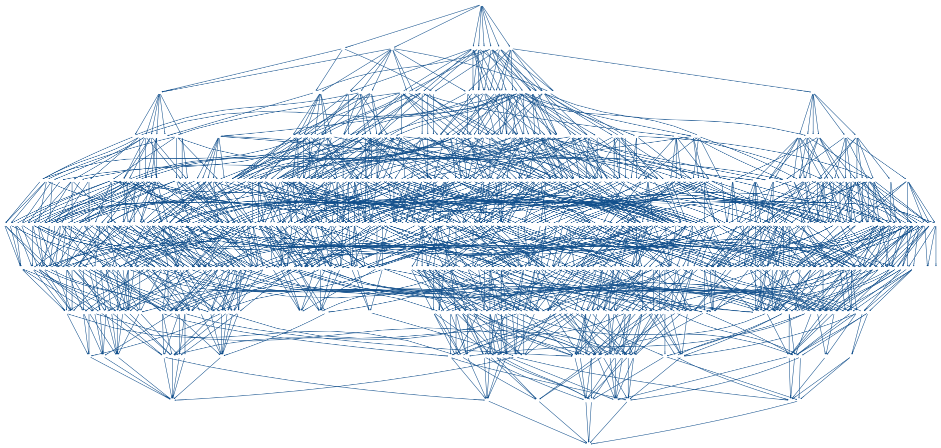

| | |
|---|---|
| *ID* | kfHftKfTcY9lyTX3 |
| *acc genotypes* | 207 |
| *peaks no back* | 79 |
| *removed edges* | NA |
| *prop rem edges* | NA |

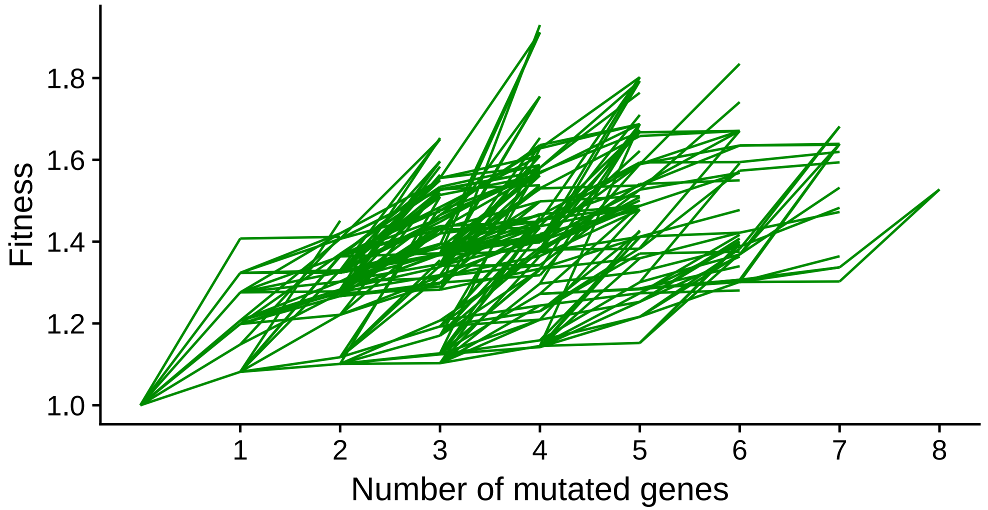


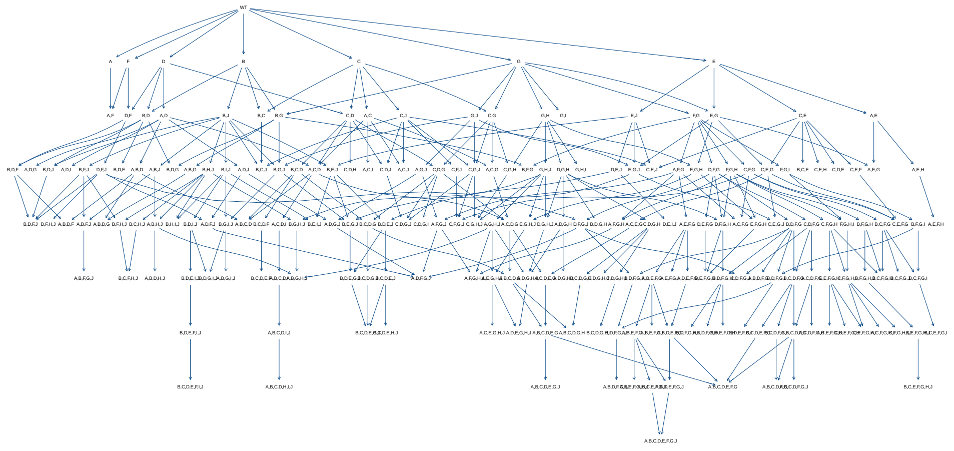

| | |
|---|---|
| *ID* | KJpT1Yp64Tw10qzk10 |
| *acc genotypes* | 35 |
| *peaks no back* | 16 |
| *removed edges* | NA |
| *prop rem edges* | NA |

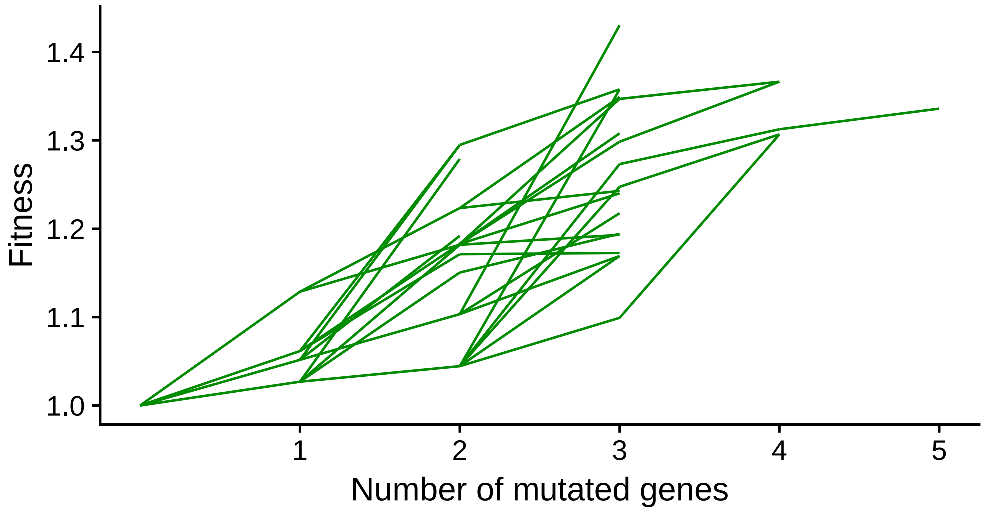


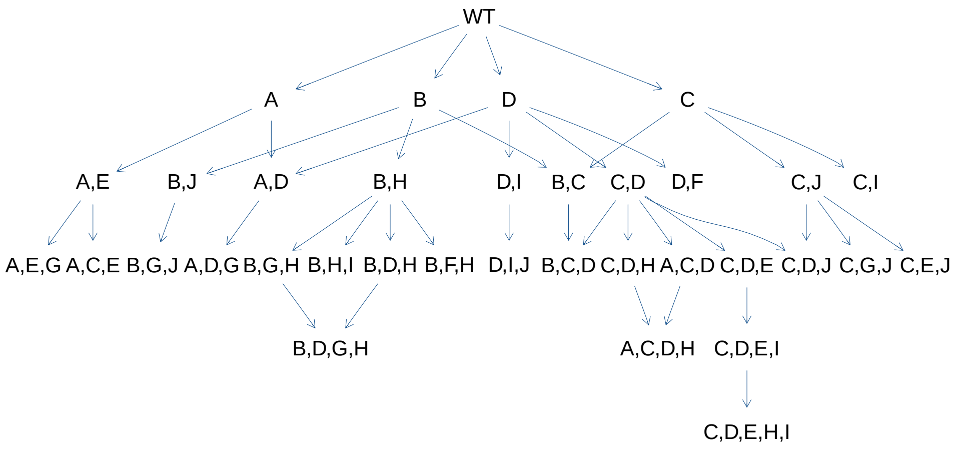

| | |
|---|---|
| *ID* | KVDBUlaSExVhtPy5 |
| *acc genotypes* | 187 |
| *peaks no back* | 57 |
| *removed edges* | NA |
| *prop rem edges* | NA |

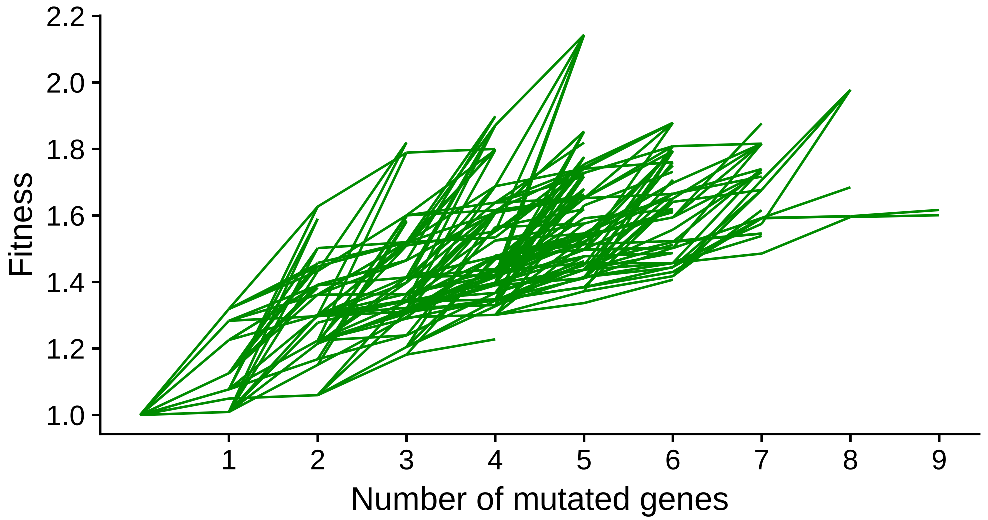


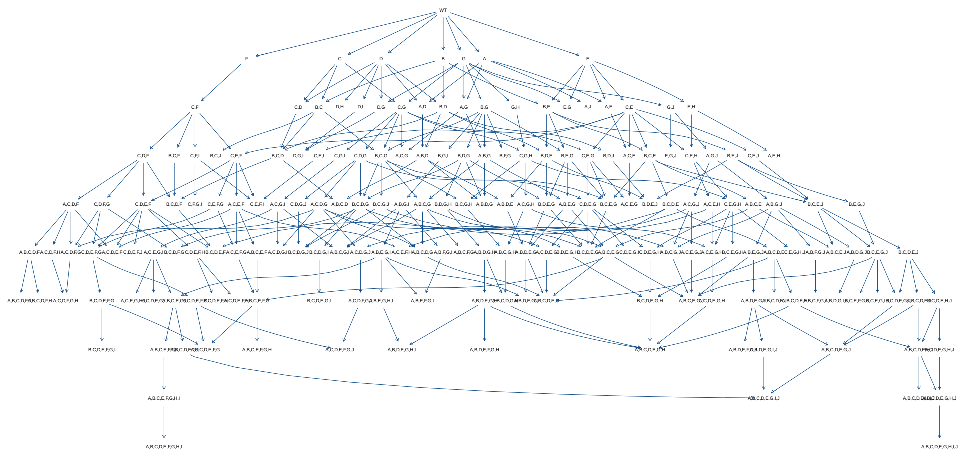

| | |
|---|---|
| *ID* | l10y29Tq5MlxxOTsw |
| *acc genotypes* | 465 |
| *peaks no back* | 98 |
| *removed edges* | NA |
| *prop rem edges* | NA |

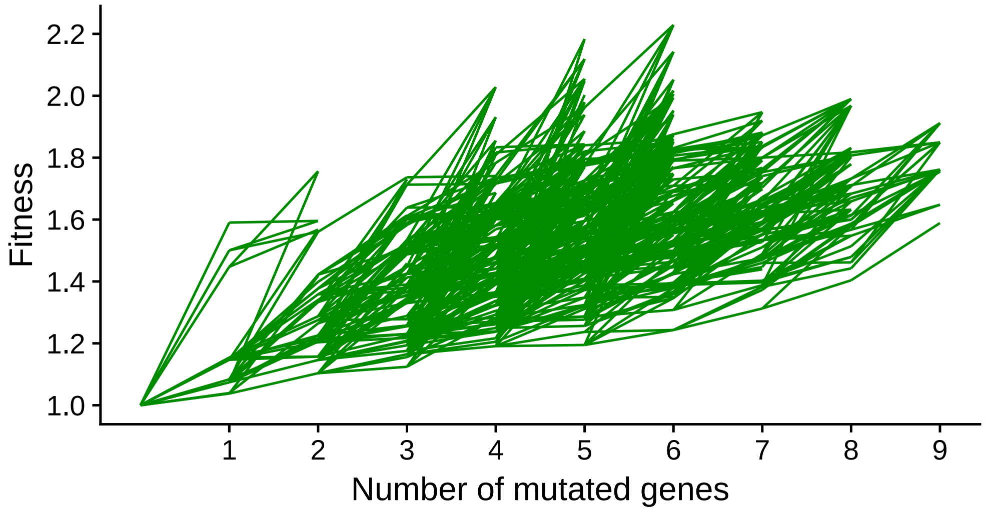


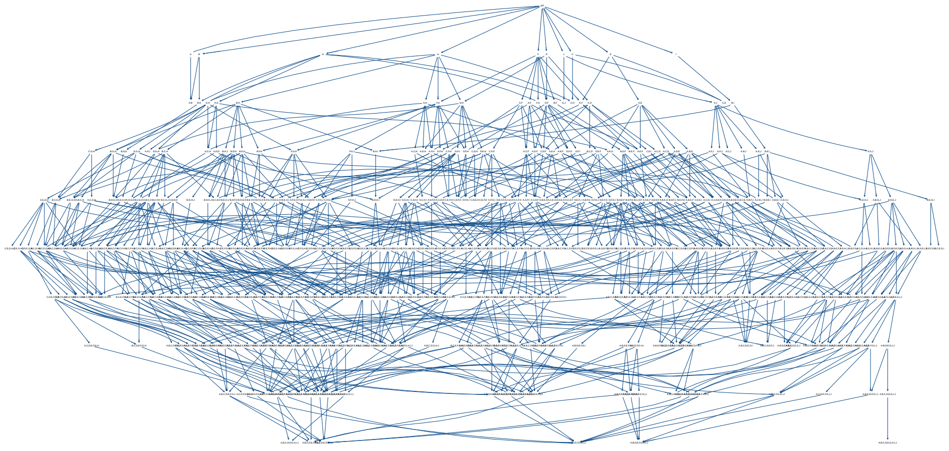

| | |
|---|---|
| *ID* | La9fqNjWh3MIhyOc |
| *acc genotypes* | 286 |
| *peaks no back* | 51 |
| *removed edges* | NA |
| *prop rem edges* | NA |

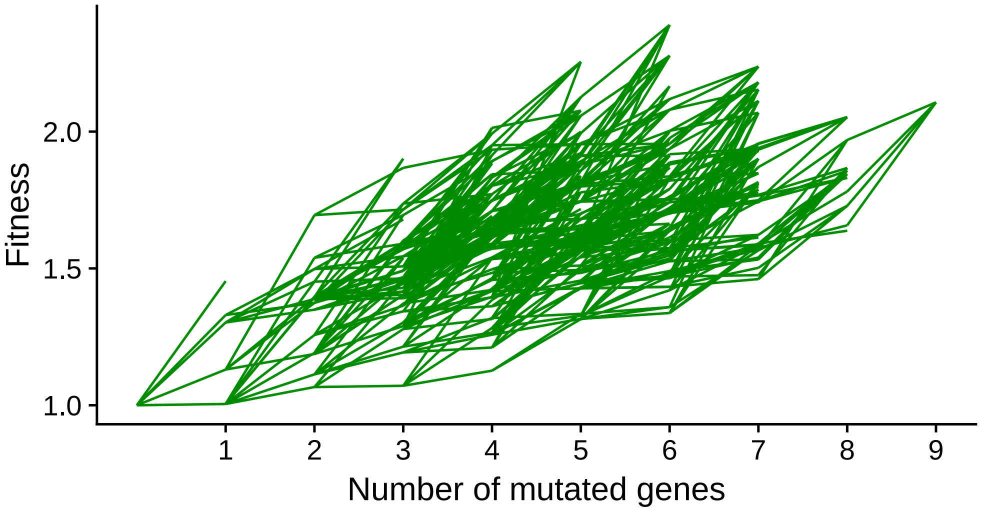


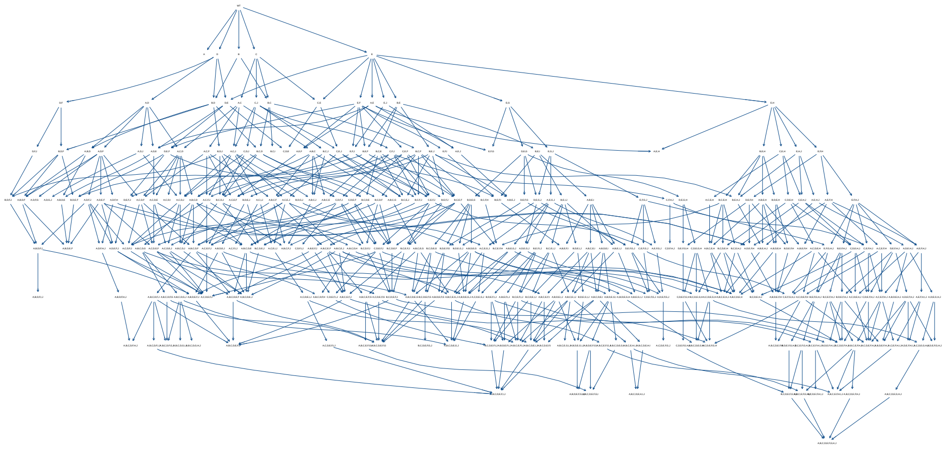

| | |
|---|---|
| *ID* | LarH6pnhHub1WCoT |
| *acc genotypes* | 119 |
| *peaks no back* | 34 |
| *removed edges* | NA |
| *prop rem edges* | NA |

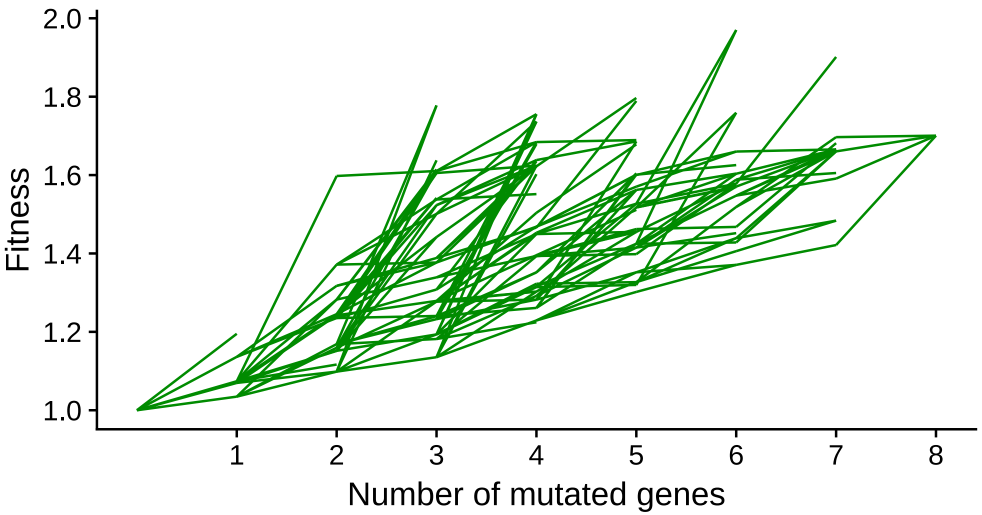


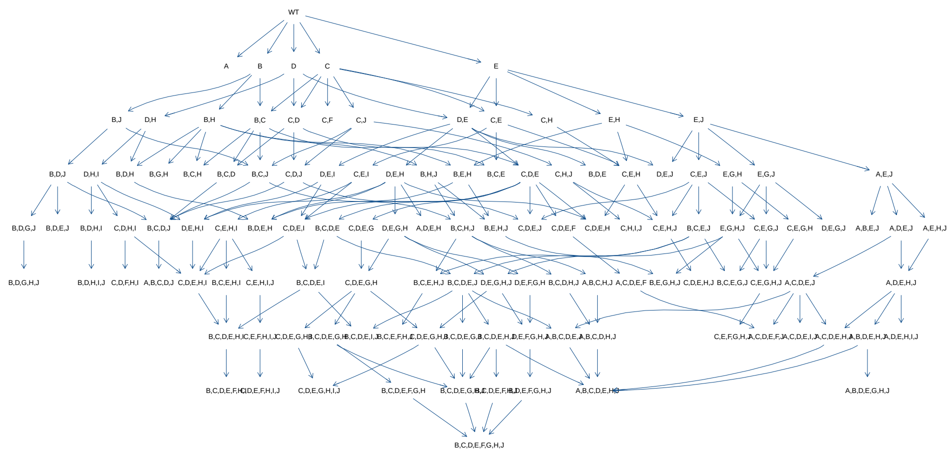

| | |
|---|---|
| *ID* | ligIbsdWsdaVrYD4 |
| *acc genotypes* | 183 |
| *peaks no back* | 56 |
| *removed edges* | NA |
| *prop rem edges* | NA |

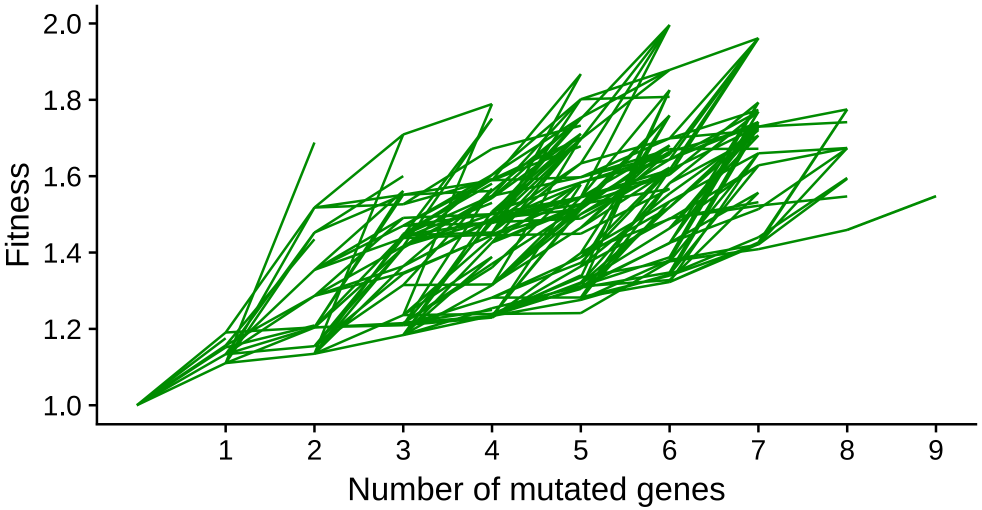


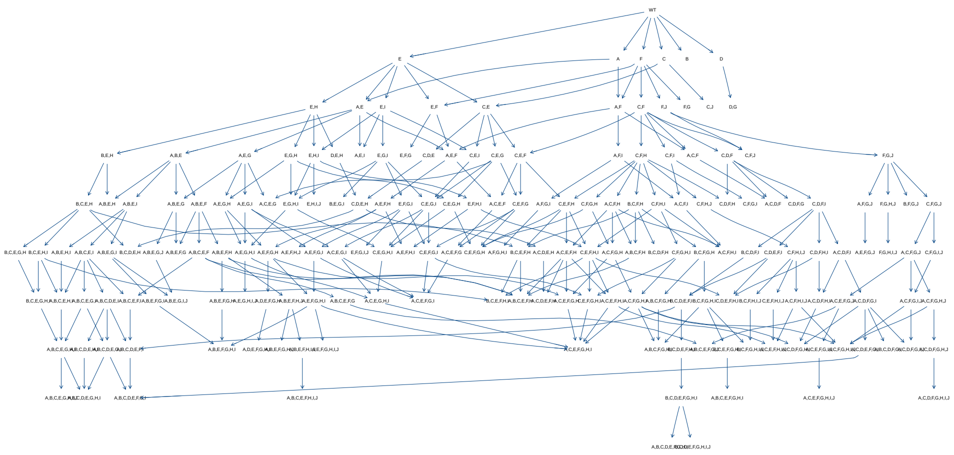

| | |
|---|---|
| *ID* | LjALbpwaXmX67y1J |
| *acc genotypes* | 200 |
| *peaks no back* | 60 |
| *removed edges* | NA |
| *prop rem edges* | NA |

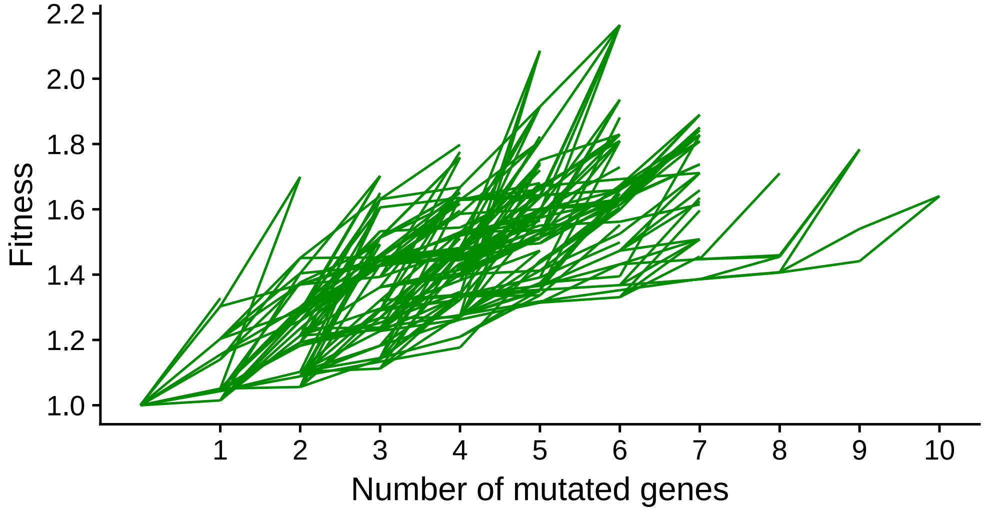


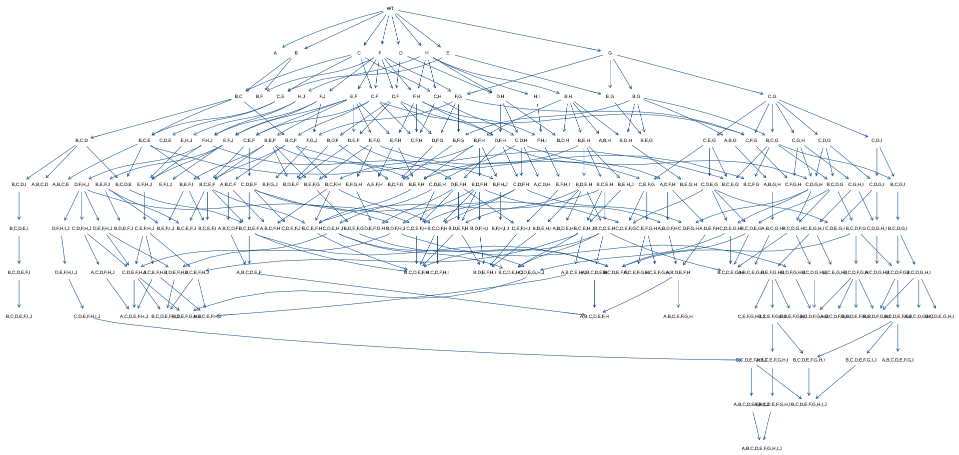

*ID* LOsXXnHBlR9Q10wUS
*acc genotypes* 135
*peaks no back* 40
*removed edges* NA
*prop rem edges* NA

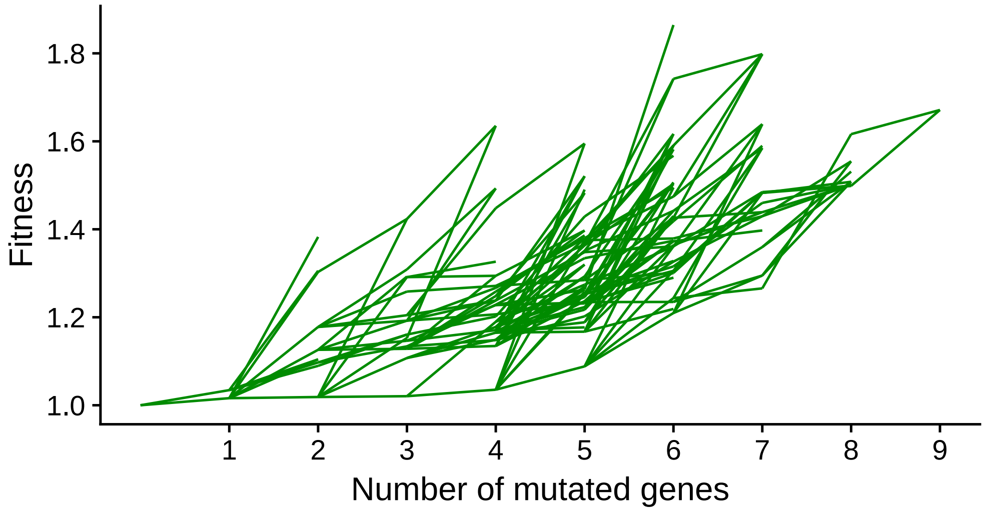


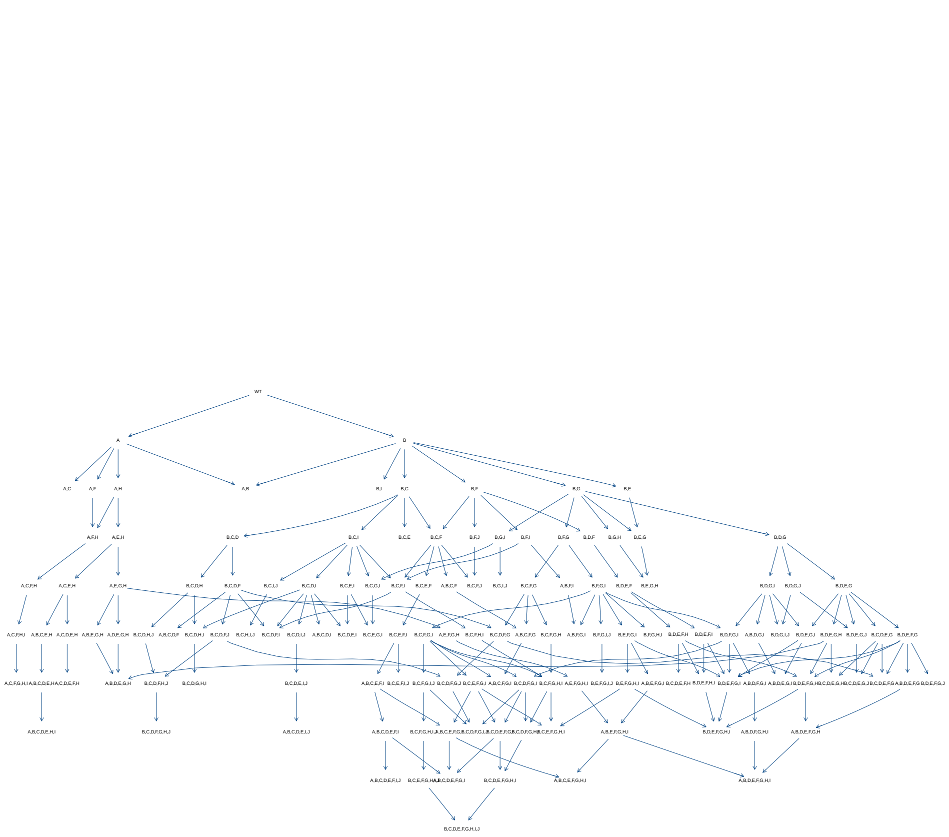

| | |
|---|---|
| *ID* | LPrNa9xFzuJVcvTs |
| *acc genotypes* | 132 |
| *peaks no back* | 51 |
| *removed edges* | NA |
| *prop rem edges* | NA |

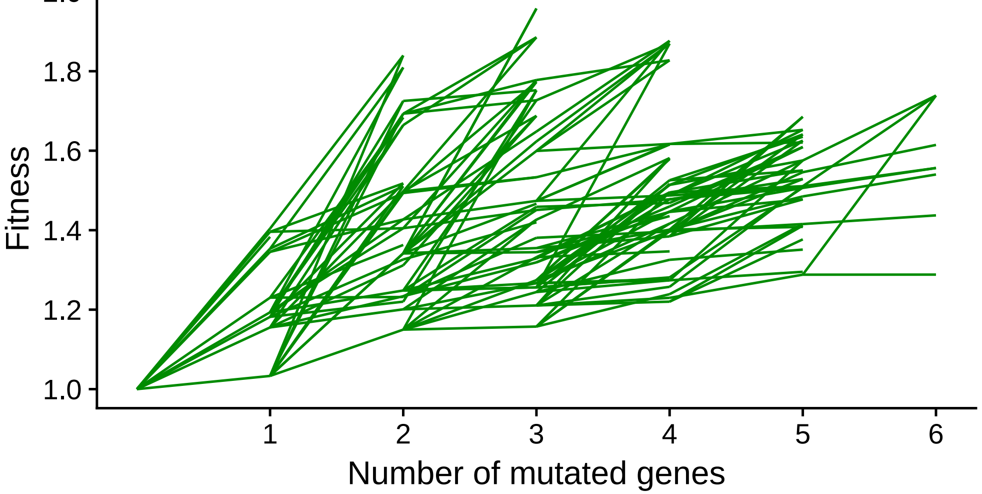


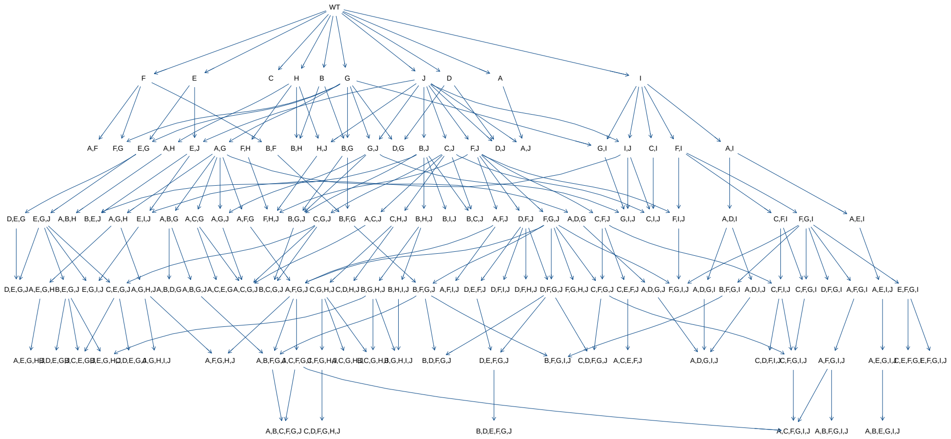

| | |
|---|---|
| *ID* | ltssjb6NCBgdBABA |
| *acc genotypes* | 232 |
| *peaks no back* | 86 |
| *removed edges* | NA |
| *prop rem edges* | NA |

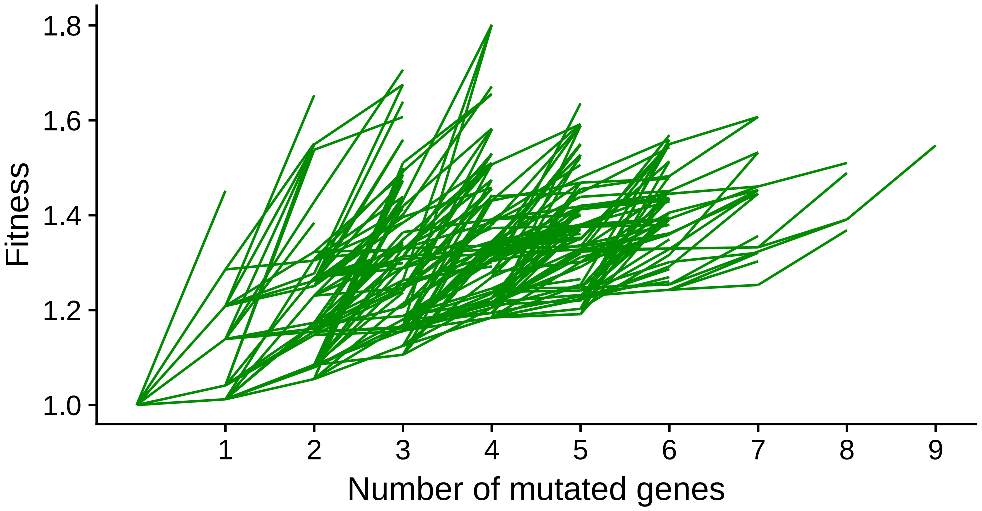


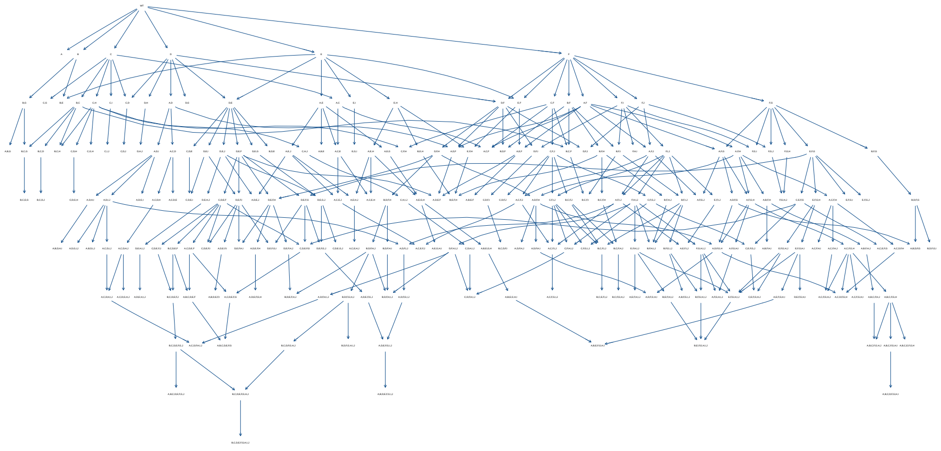

| | |
|---|---|
| *ID* | LYJKKjl6A3xWjKwy |
| *acc genotypes* | 289 |
| *peaks no back* | 75 |
| *removed edges* | NA |
| *prop rem edges* | NA |

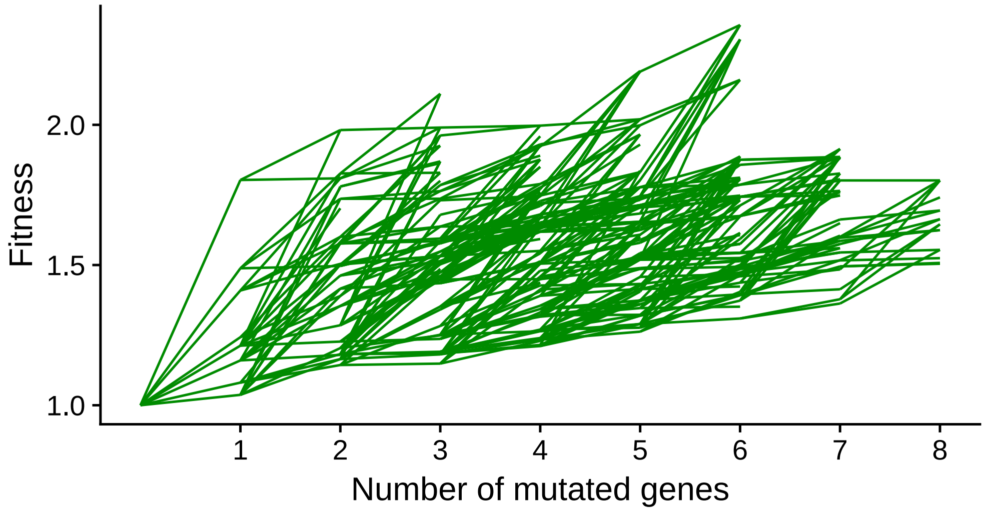


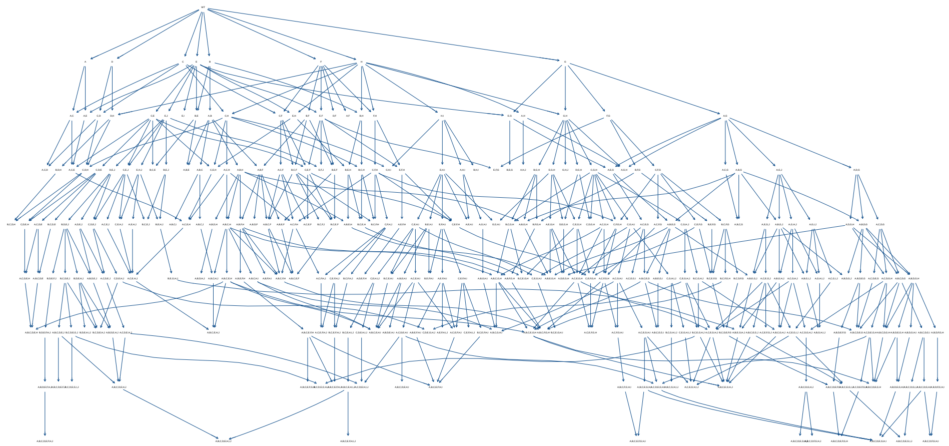

| | |
|---|---|
| *ID* | MkpqetXw6W2muojH |
| *acc genotypes* | 264 |
| *peaks no back* | 74 |
| *removed edges* | NA |
| *prop rem edges* | NA |

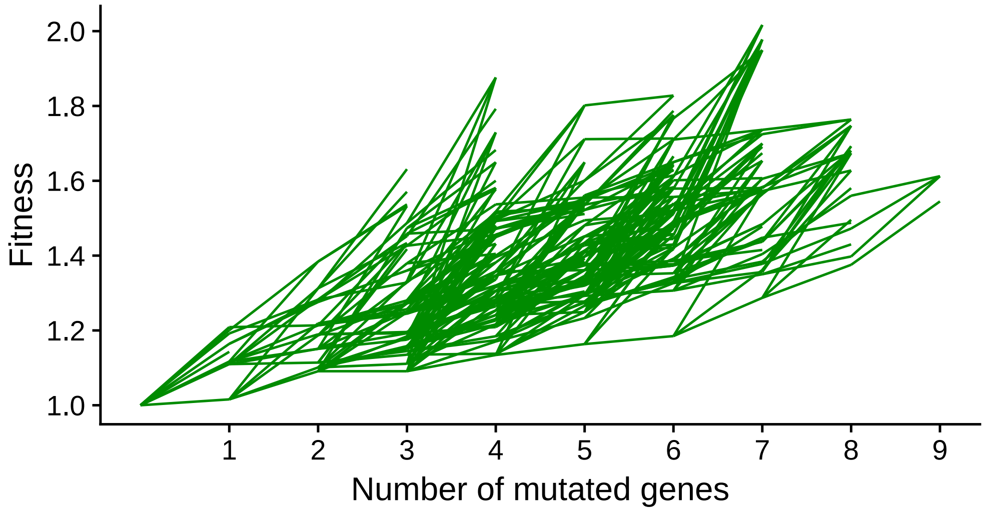


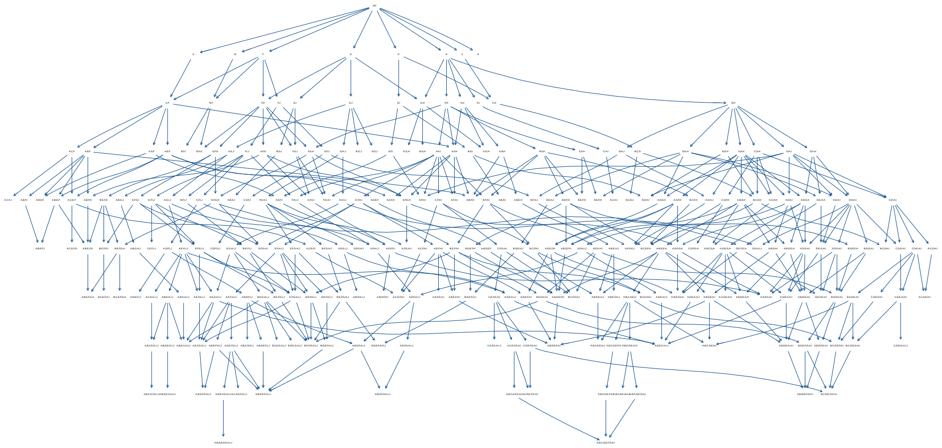

| | |
|---|---|
| *ID* | mLM7OEw142104scrX |
| *acc genotypes* | 107 |
| *peaks no back* | 42 |
| *removed edges* | NA |
| *prop rem edges* | NA |

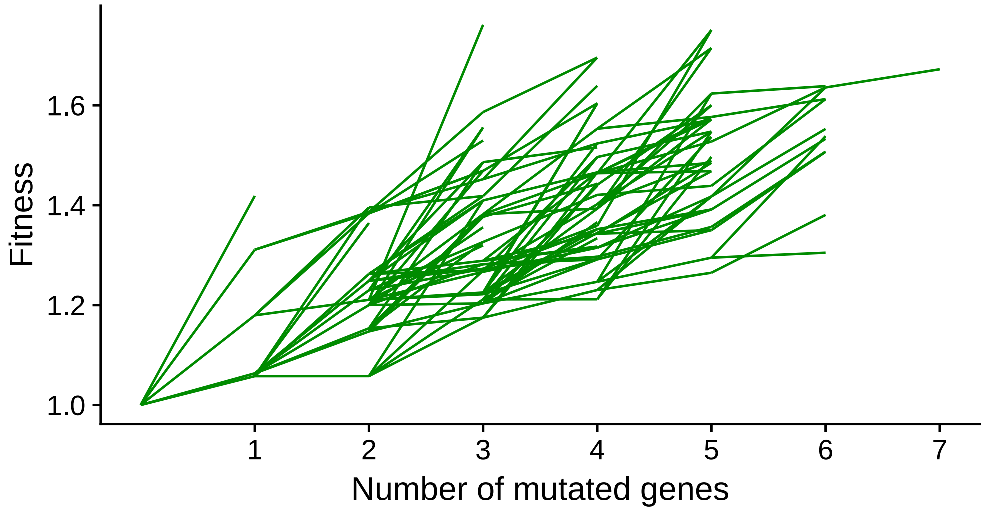


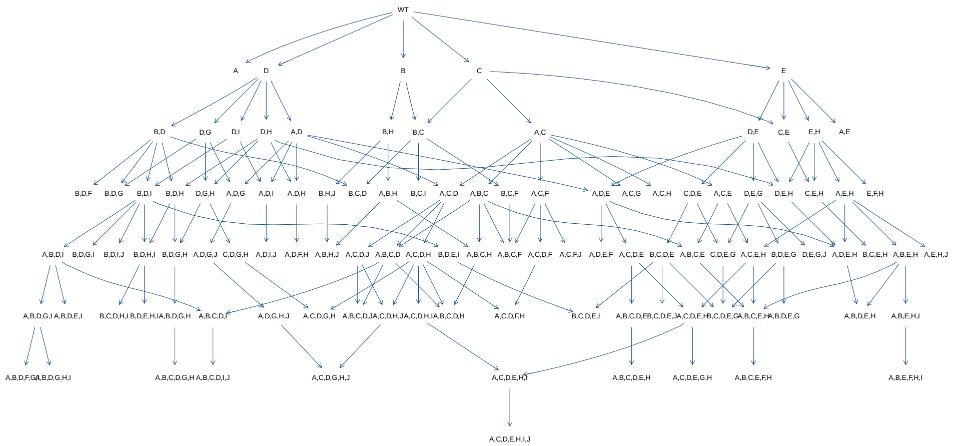

| | |
|---|---|
| *ID* | mqC38cuoz4YoXqrC |
| *acc genotypes* | 214 |
| *peaks no back* | 74 |
| *removed edges* | NA |
| *prop rem edges* | NA |

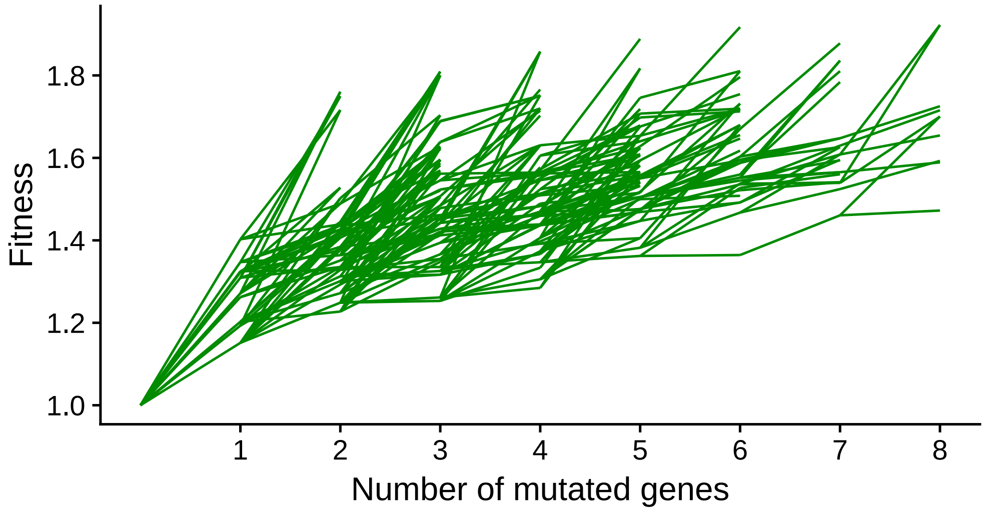


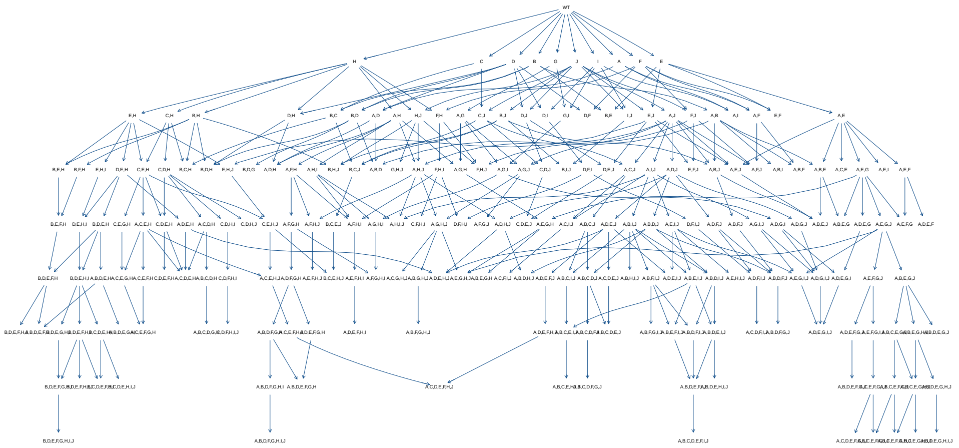

| | |
|---|---|
| *ID* | MtpPxzteweNeL6dl |
| *acc genotypes* | 295 |
| *peaks no back* | 63 |
| *removed edges* | NA |
| *prop rem edges* | NA |

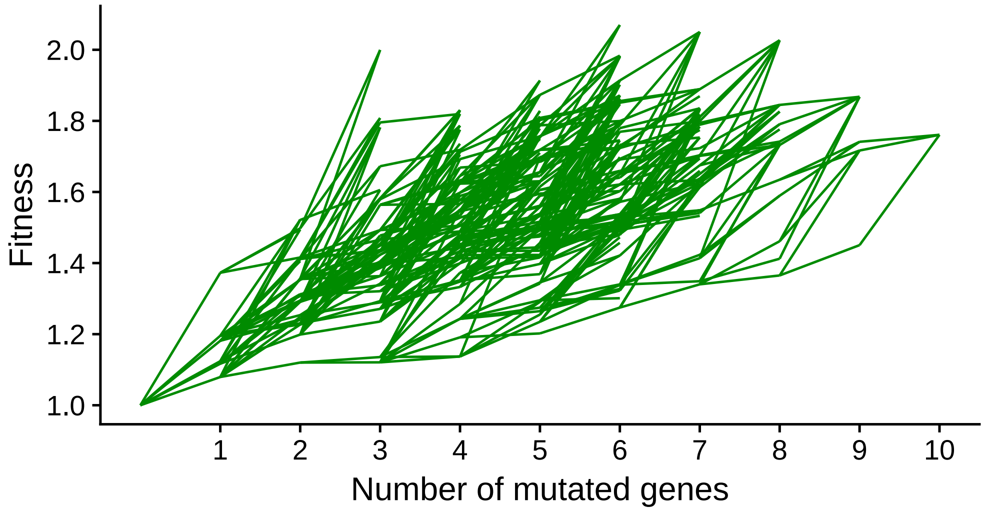


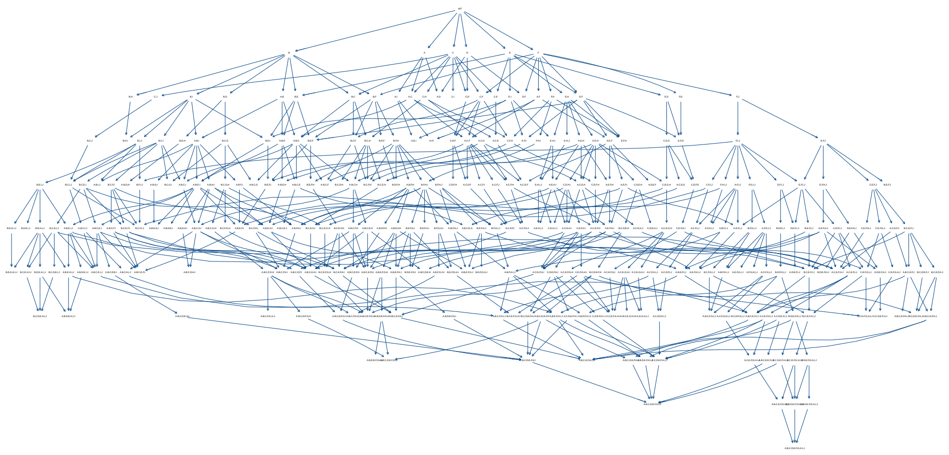

| | |
|---|---|
| *ID* | mYuH5XdoLHUkC10ay |
| *acc genotypes* | 440 |
| *peaks no back* | 88 |
| *removed edges* | NA |
| *prop rem edges* | NA |

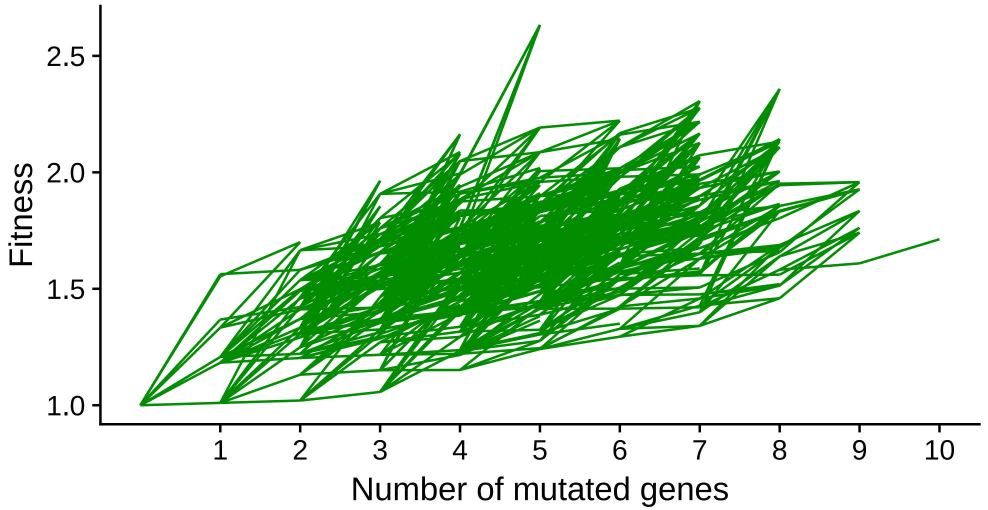


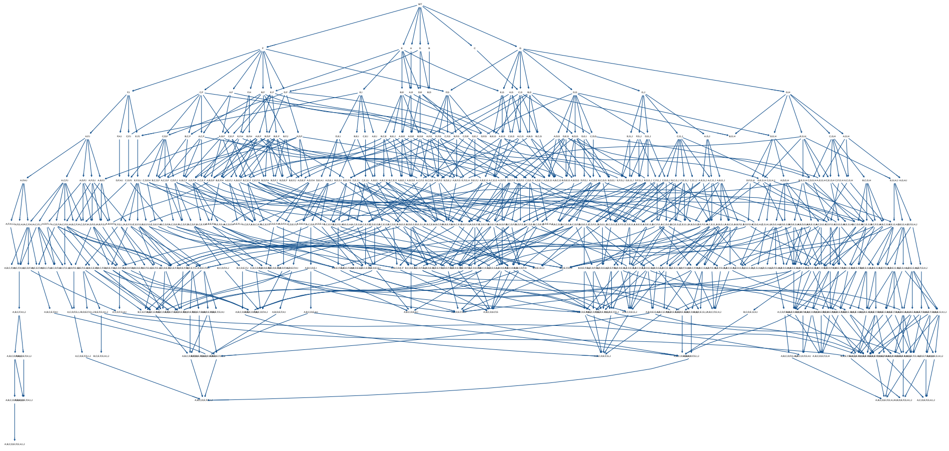

| | |
|---|---|
| *ID* | n10oyDkkU10F10uiAs3 |
| *acc genotypes* | 228 |
| *peaks no back* | 73 |
| *removed edges* | NA |
| *prop rem edges* | NA |

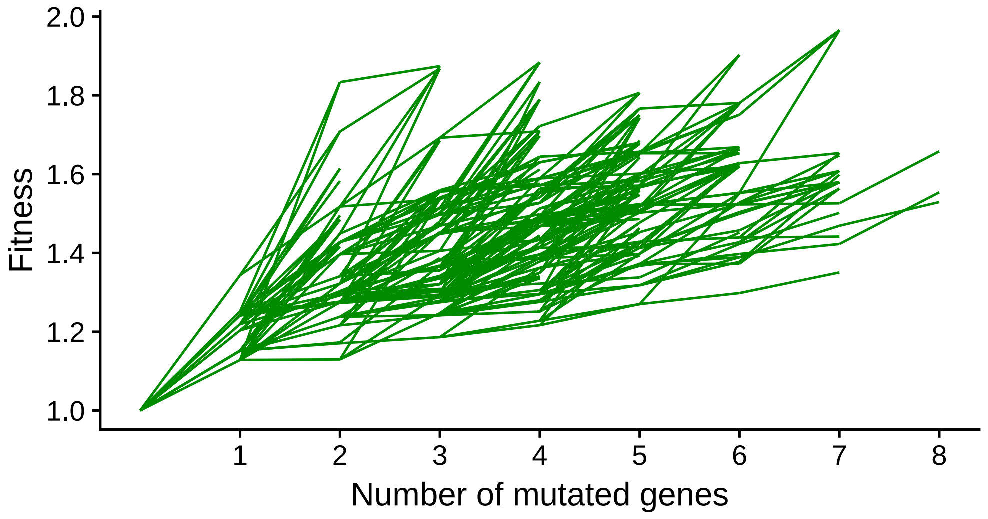


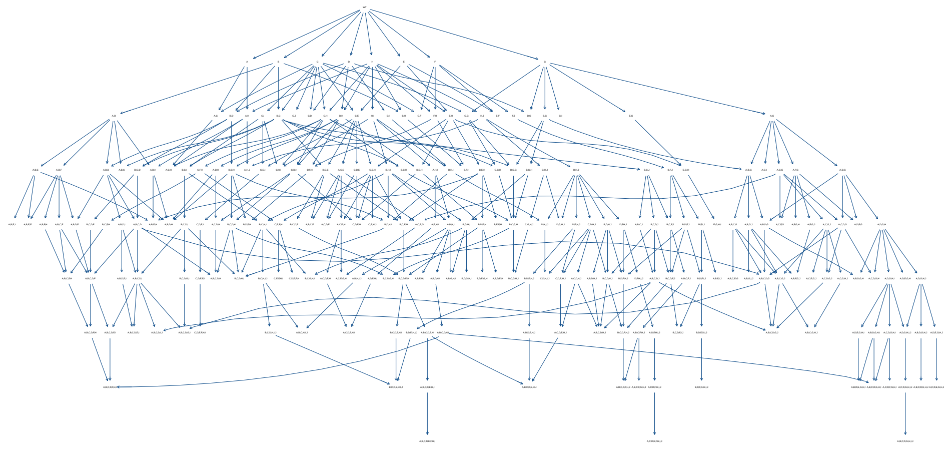

| | |
|---|---|
| *ID* | Na1044PJp2RIHKSl6 |
| *acc genotypes* | 30 |
| *peaks no back* | 13 |
| *removed edges* | NA |
| *prop rem edges* | NA |

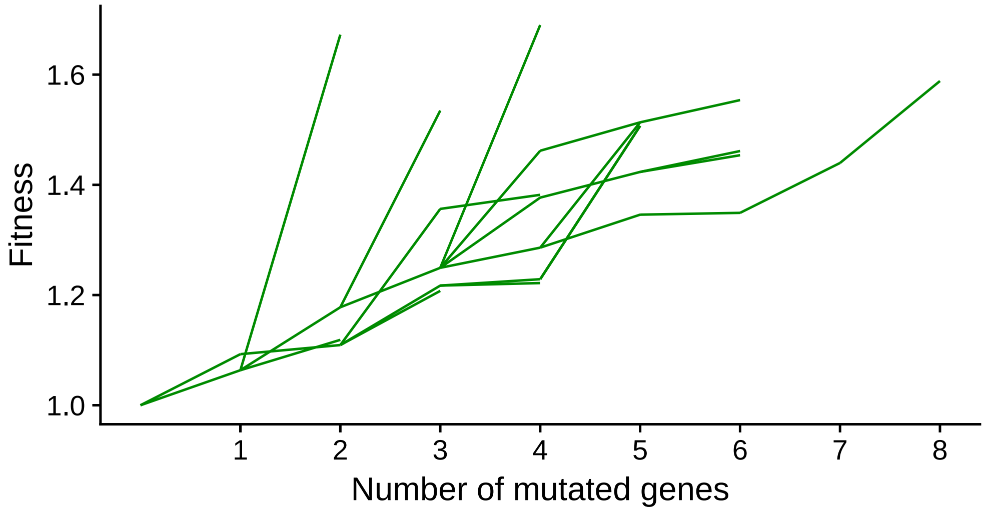


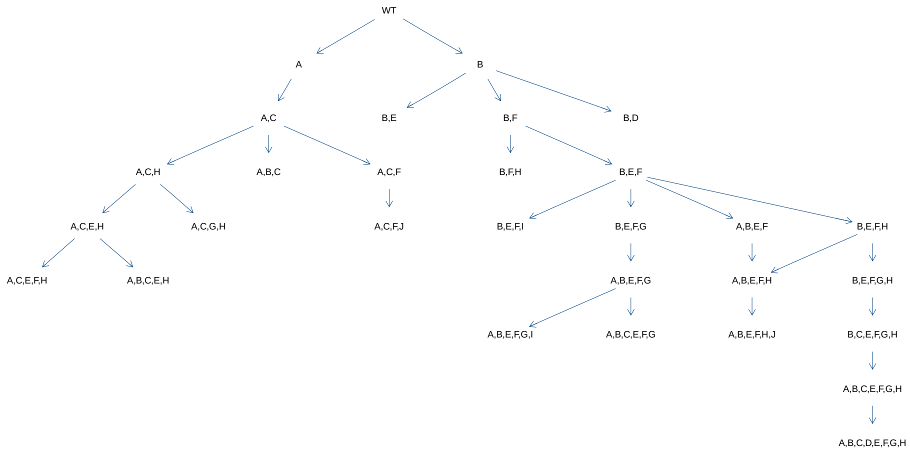

| | |
|---|---|
| *ID* | Nckkjp8YxmOBJiLn |
| *acc genotypes* | 176 |
| *peaks no back* | 45 |
| *removed edges* | NA |
| *prop rem edges* | NA |

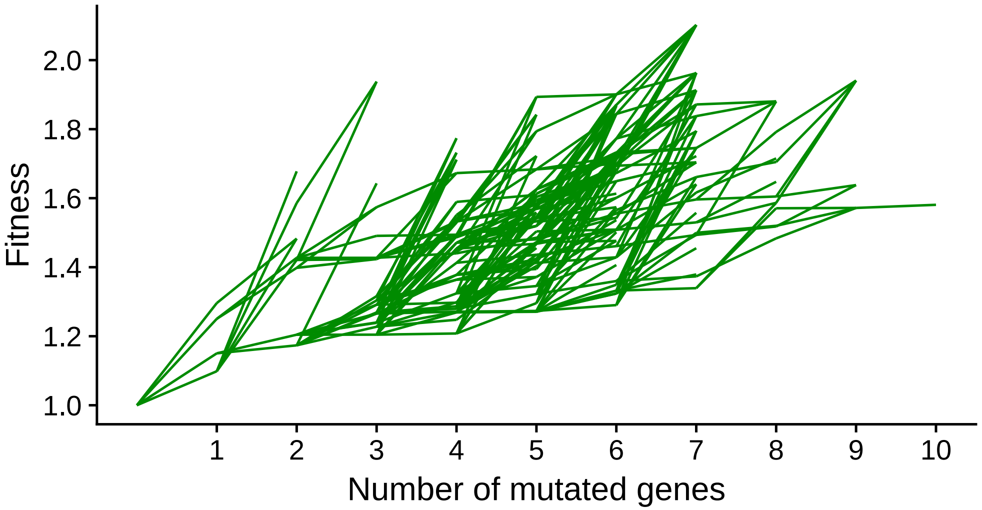


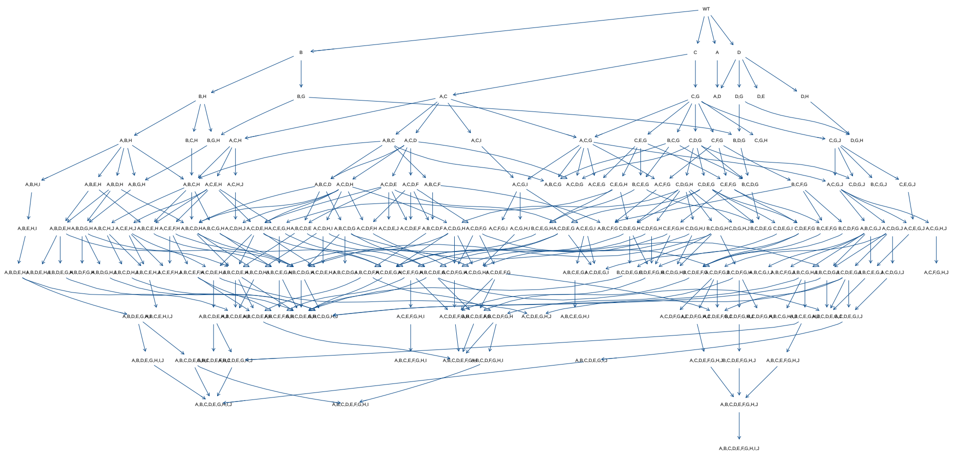

| | |
|---|---|
| *ID* | nFFAJ57LcDAfooEJ |
| *acc genotypes* | 537 |
| *peaks no back* | 73 |
| *removed edges* | NA |
| *prop rem edges* | NA |

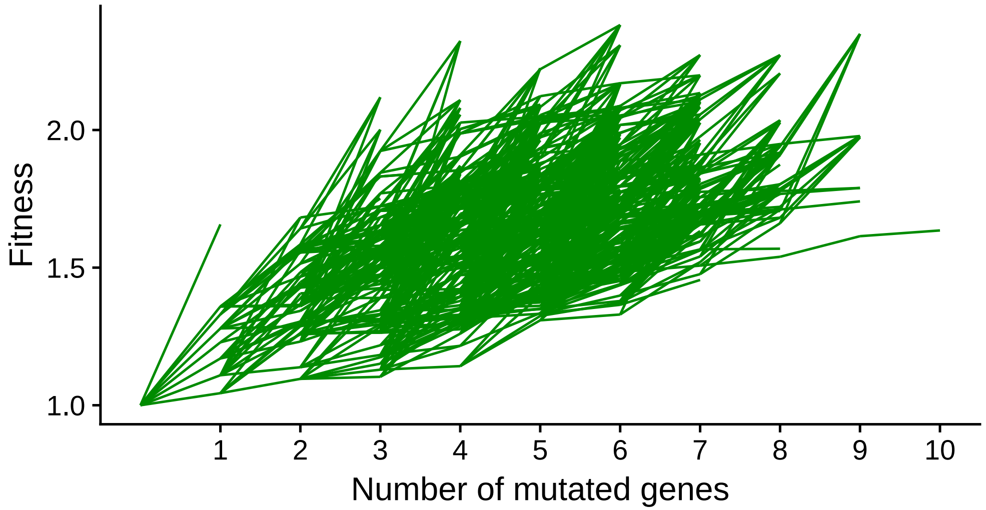


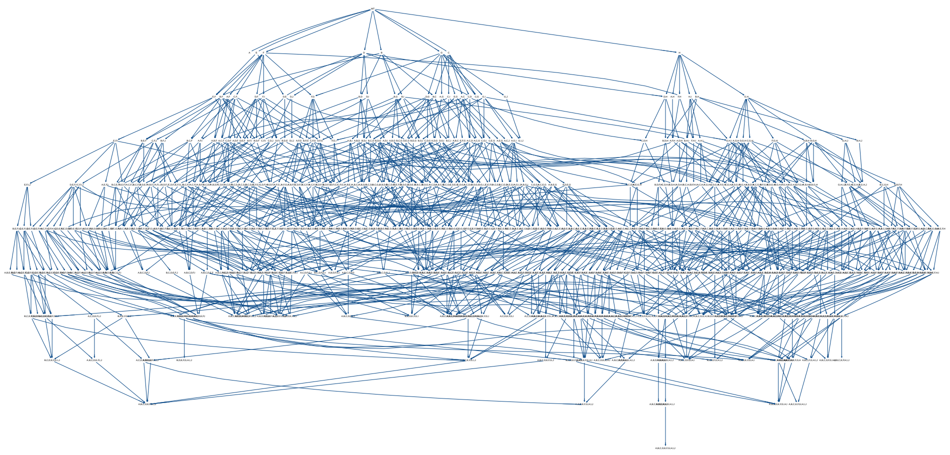

| | |
|---|---|
| *ID* | nFGvQLr121eGqkyD |
| *acc genotypes* | 138 |
| *peaks no back* | 51 |
| *removed edges* | NA |
| *prop rem edges* | NA |

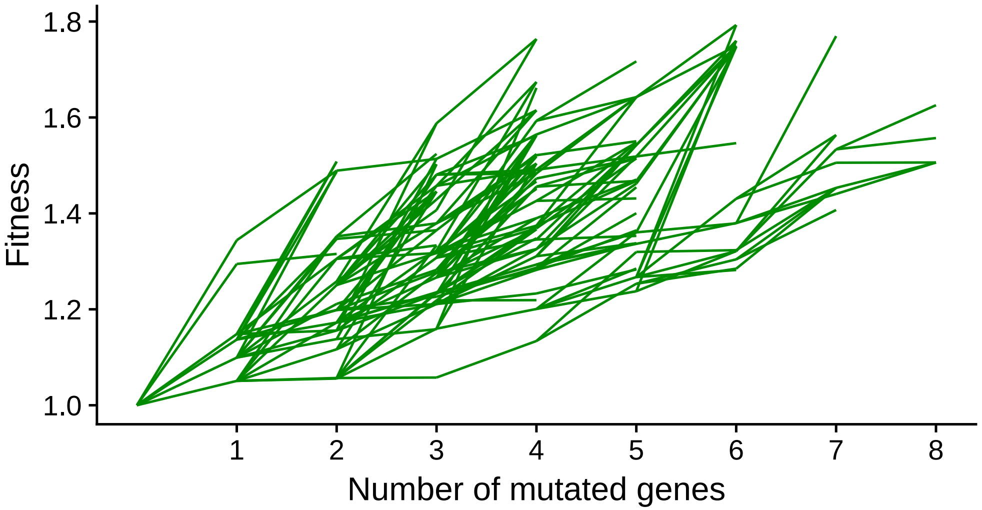


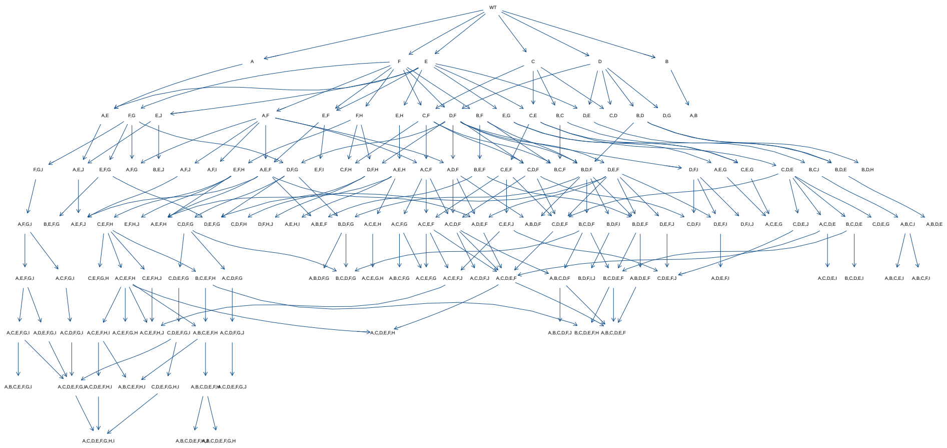

| | |
|---|---|
| *ID* | OcPH3CcYp51kr5sW |
| *acc genotypes* | 541 |
| *peaks no back* | 62 |
| *removed edges* | NA |
| *prop rem edges* | NA |

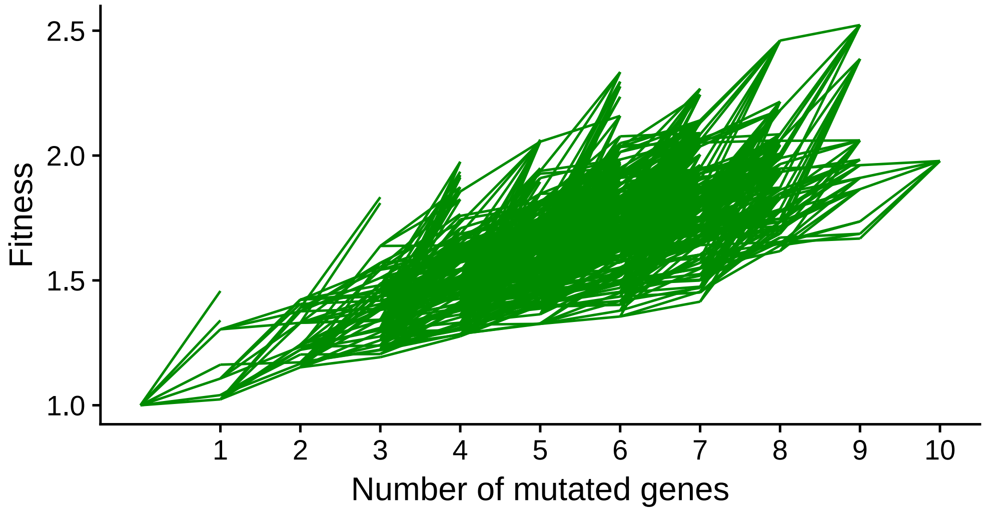


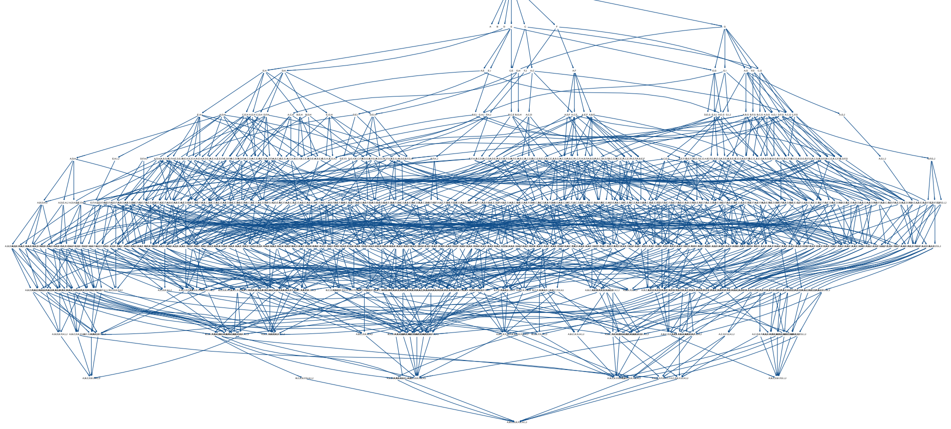

| | |
|---|---|
| *ID* | OGxF8Oyv15NLRVHt |
| *acc genotypes* | 147 |
| *peaks no back* | 63 |
| *removed edges* | NA |
| *prop rem edges* | NA |

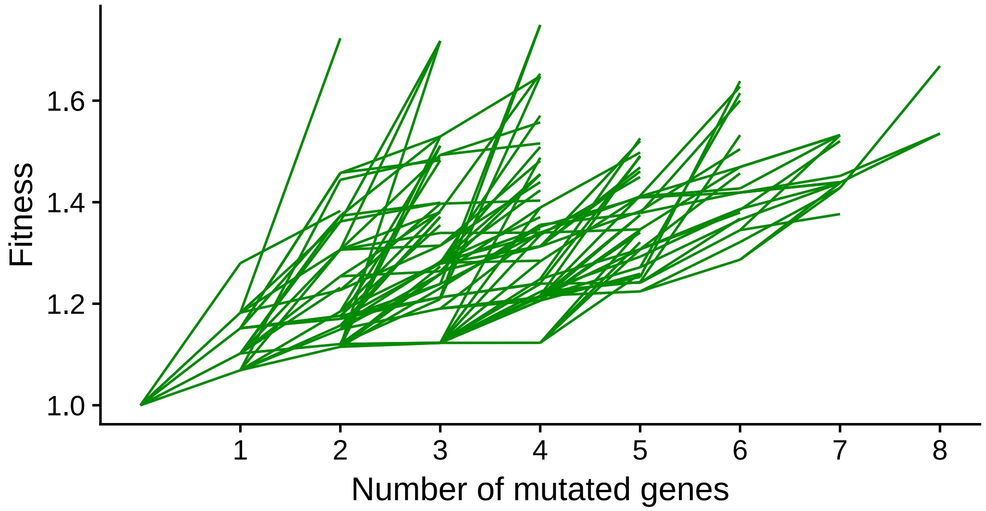


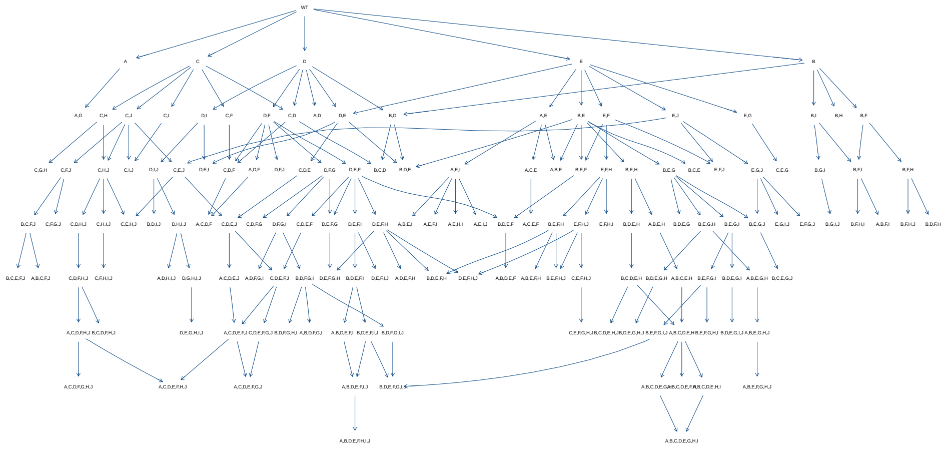

| | |
|---|---|
| *ID* | Oh105Ra4cDXvsiqOA |
| *acc genotypes* | 468 |
| *peaks no back* | 63 |
| *removed edges* | NA |
| *prop rem edges* | NA |

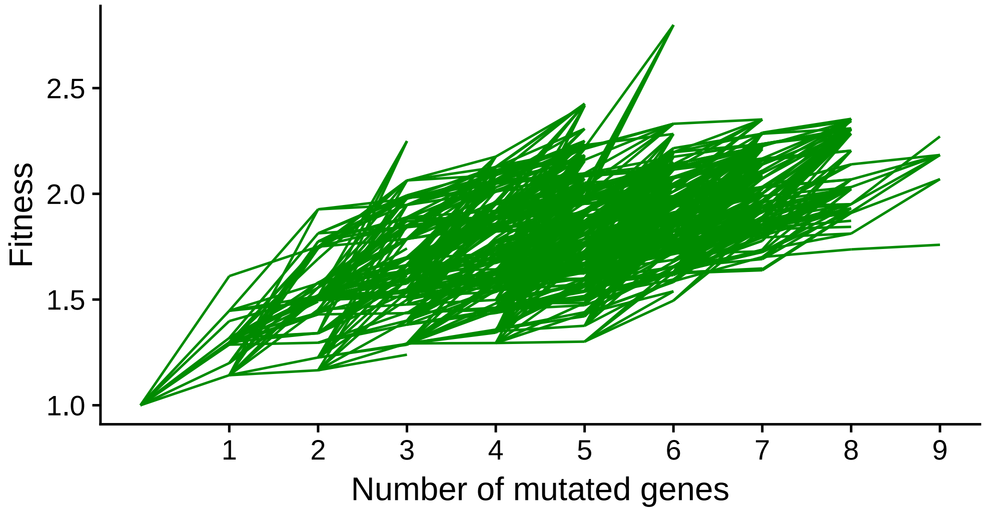


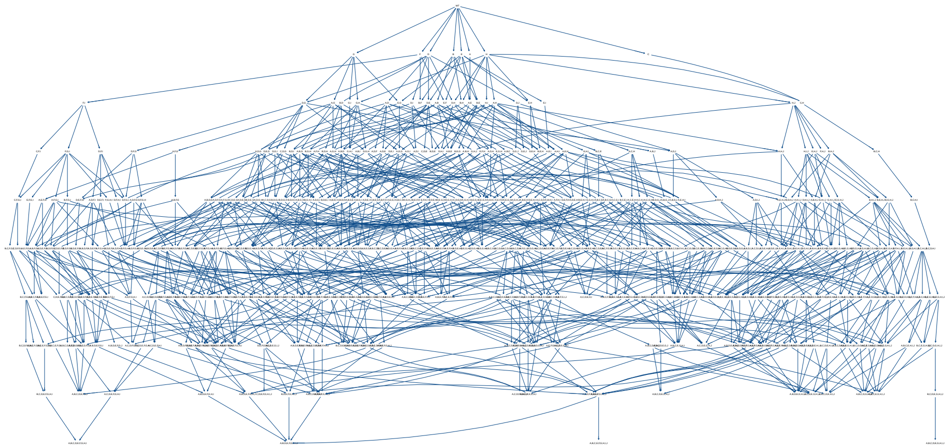

| | |
|---|---|
| *ID* | Oj6oyBsD3bxROStD |
| *acc genotypes* | 134 |
| *peaks no back* | 54 |
| *removed edges* | NA |
| *prop rem edges* | NA |

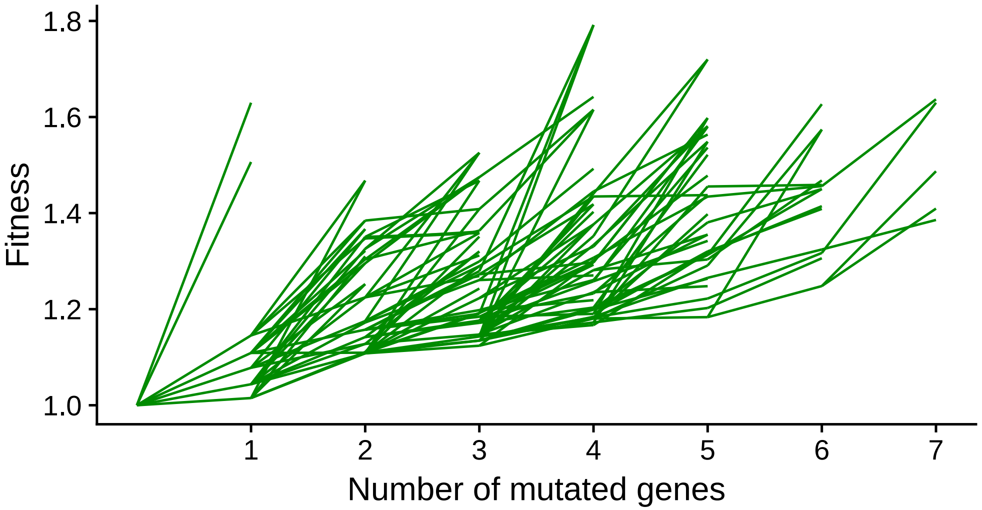


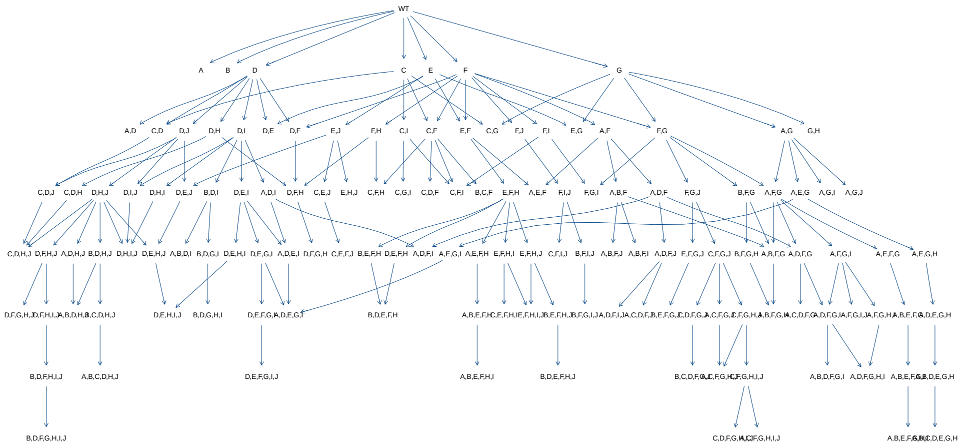

| | |
|---|---|
| *ID* | Oyrh4S6yqbCq9BE1 |
| *acc genotypes* | 84 |
| *peaks no back* | 37 |
| *removed edges* | NA |
| *prop rem edges* | NA |

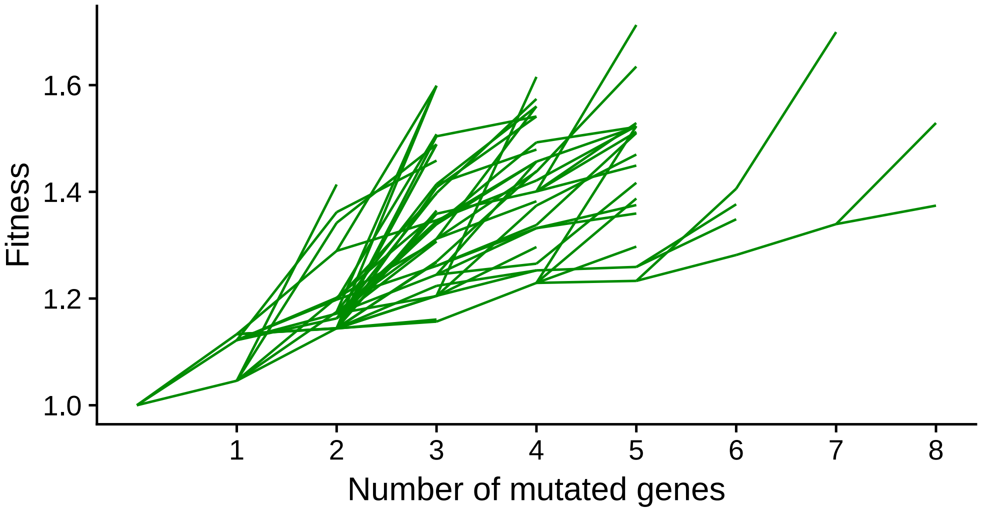


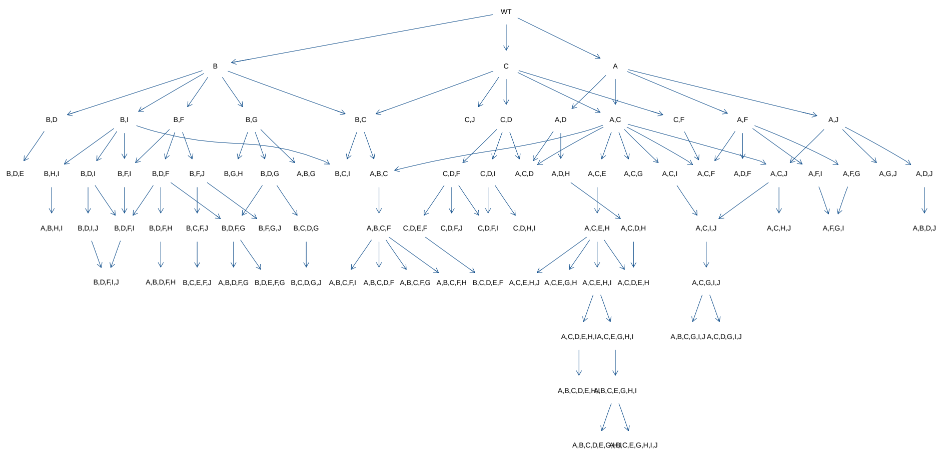

| | |
|---|---|
| *ID* | p2Hy10RKJmoyhCKd10 |
| *acc genotypes* | 140 |
| *peaks no back* | 50 |
| *removed edges* | NA |
| *prop rem edges* | NA |

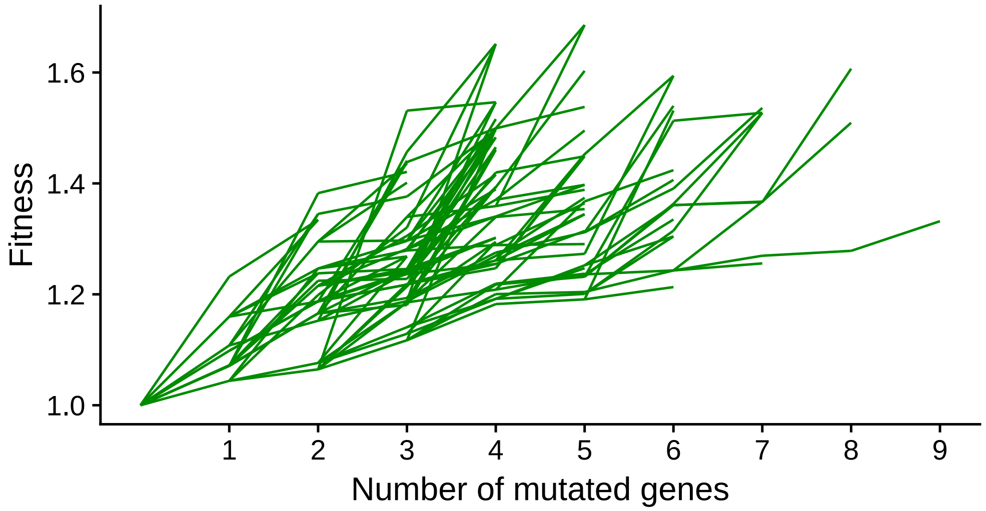


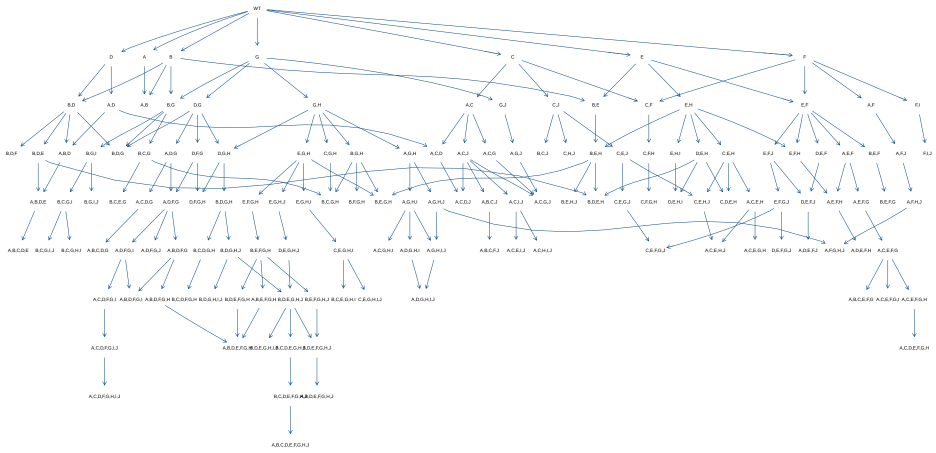

| | |
|---|---|
| *ID* | pcFmFiSb1dmNWs7Q |
| *acc genotypes* | 333 |
| *peaks no back* | 105 |
| *removed edges* | NA |
| *prop rem edges* | NA |

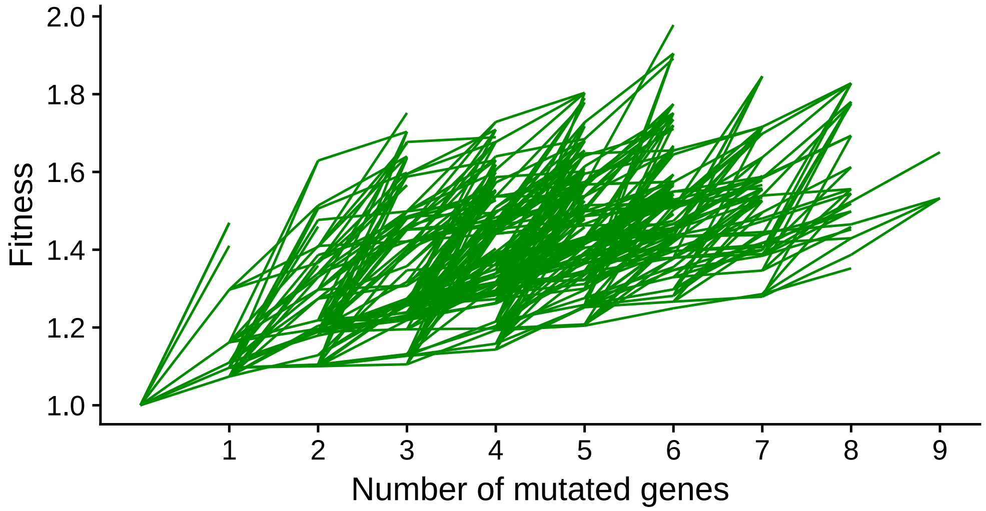


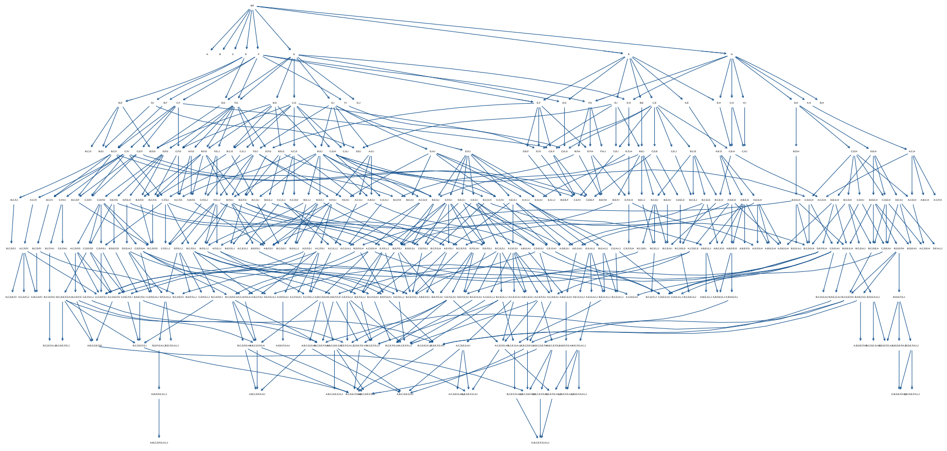

| | |
|---|---|
| *ID* | pF515adN8ZsnzdgE |
| *acc genotypes* | 276 |
| *peaks no back* | 100 |
| *removed edges* | NA |
| *prop rem edges* | NA |

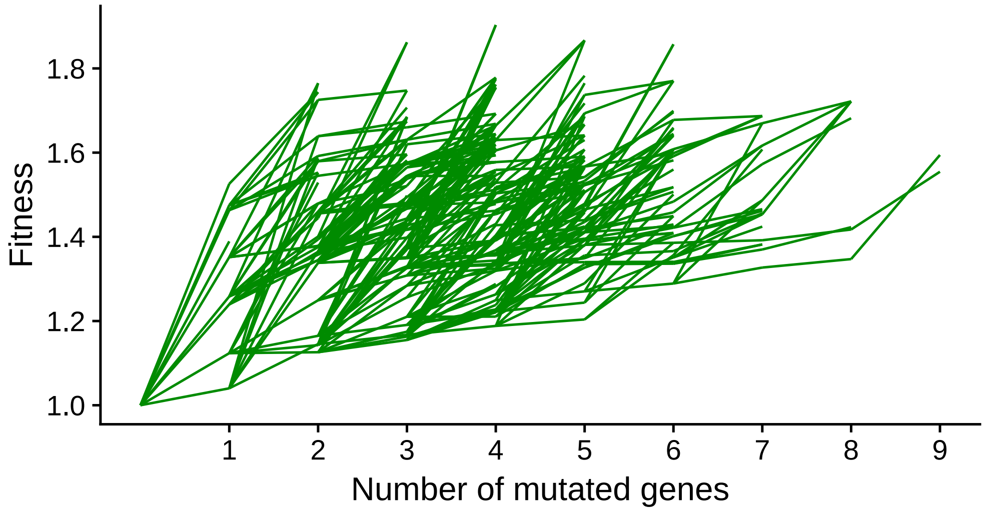


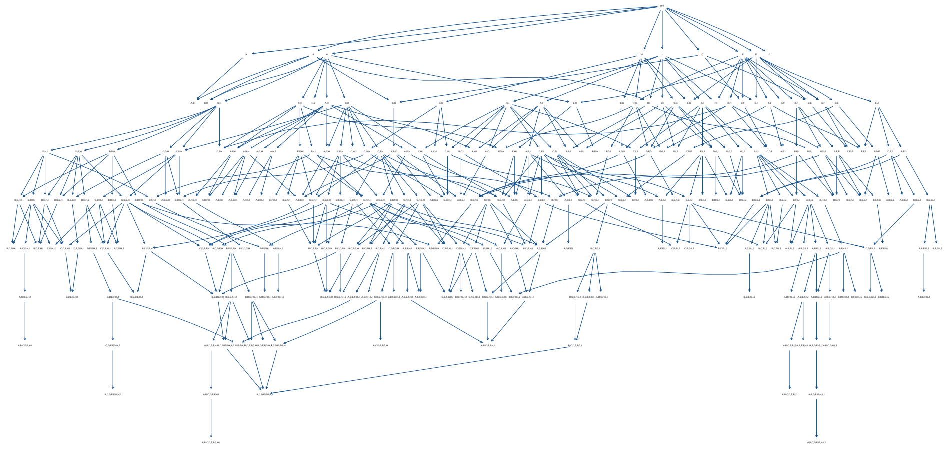

| | |
|---|---|
| *ID* | Pje92ruf6BDq1QiQ |
| *acc genotypes* | 111 |
| *peaks no back* | 40 |
| *removed edges* | NA |
| *prop rem edges* | NA |

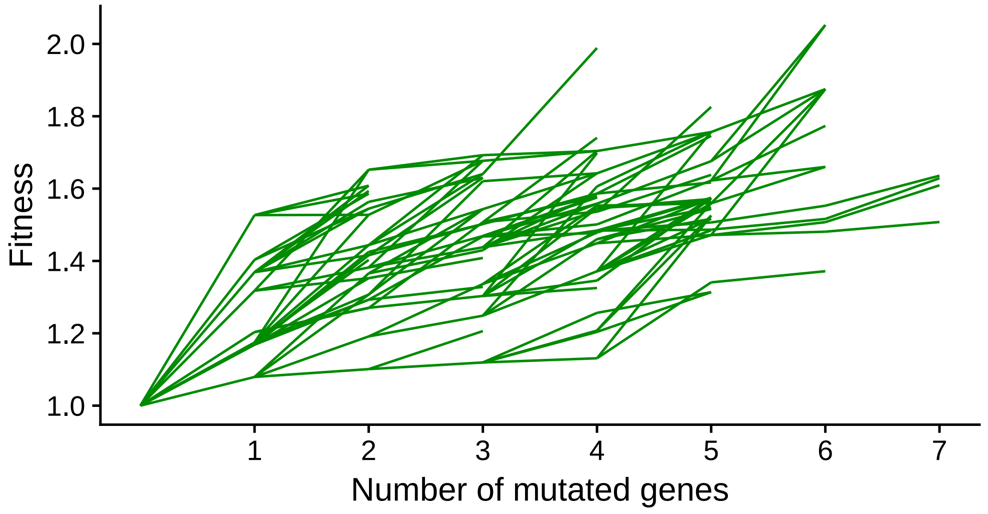


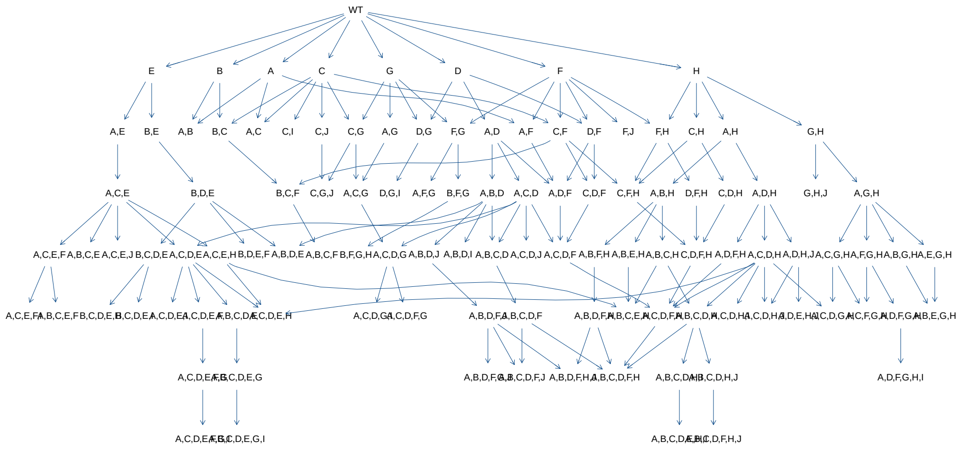

| | |
|---|---|
| *ID* | PmQfeqFvnFM6z8Uz |
| *acc genotypes* | 276 |
| *peaks no back* | 75 |
| *removed edges* | NA |
| *prop rem edges* | NA |

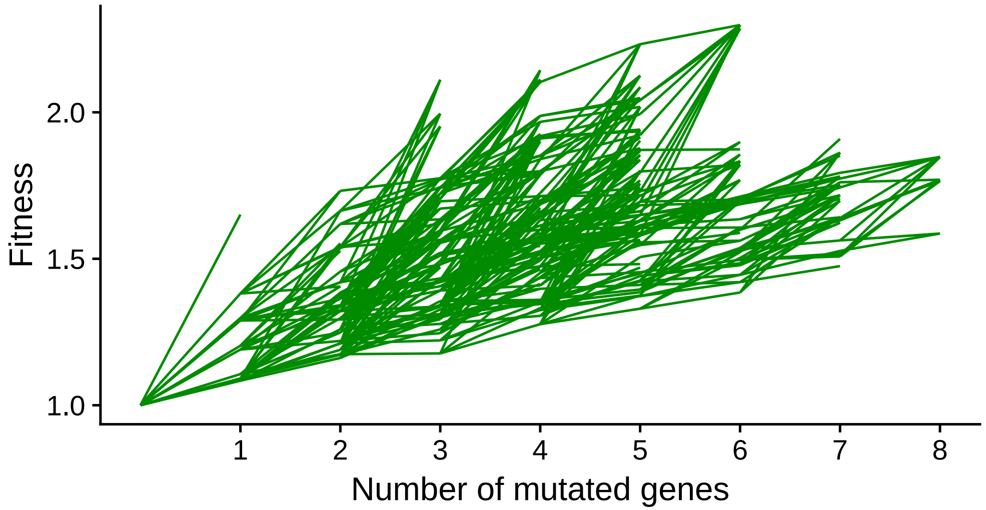


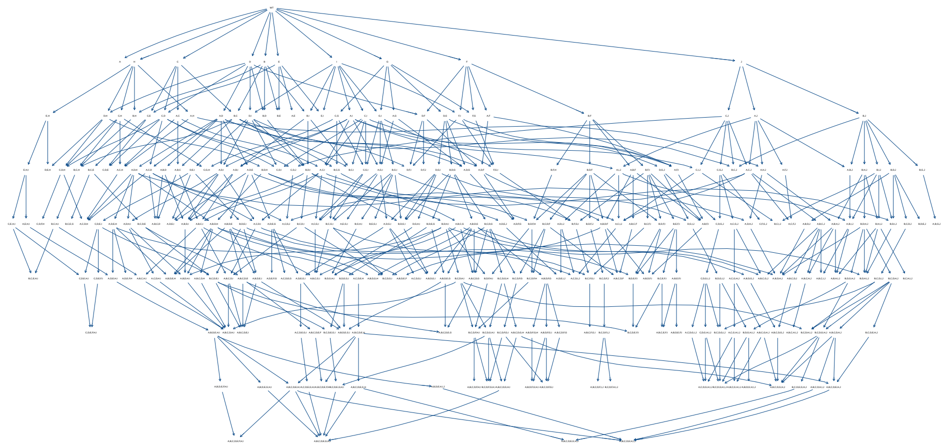

| | |
|---|---|
| *ID* | ppgKWX3qfEYrvpoe |
| *acc genotypes* | 98 |
| *peaks no back* | 22 |
| *removed edges* | NA |
| *prop rem edges* | NA |

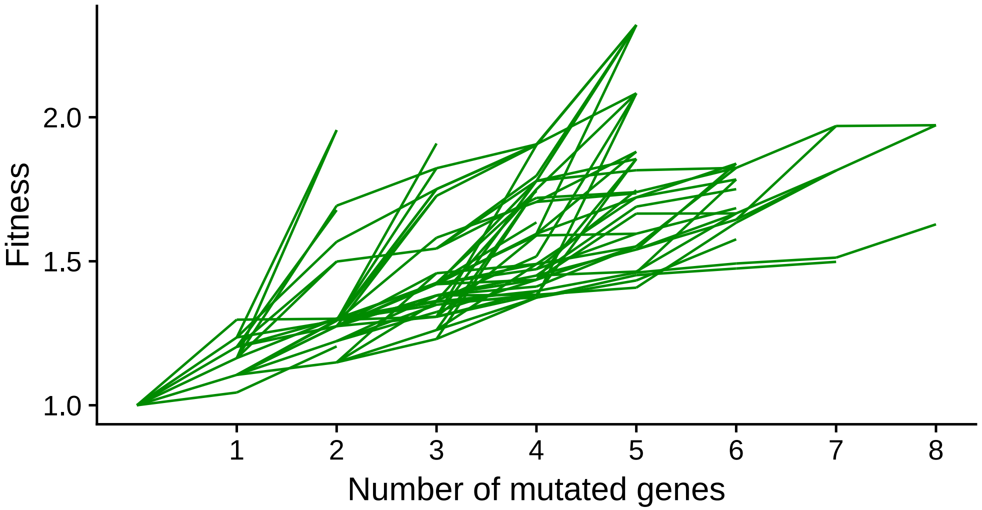


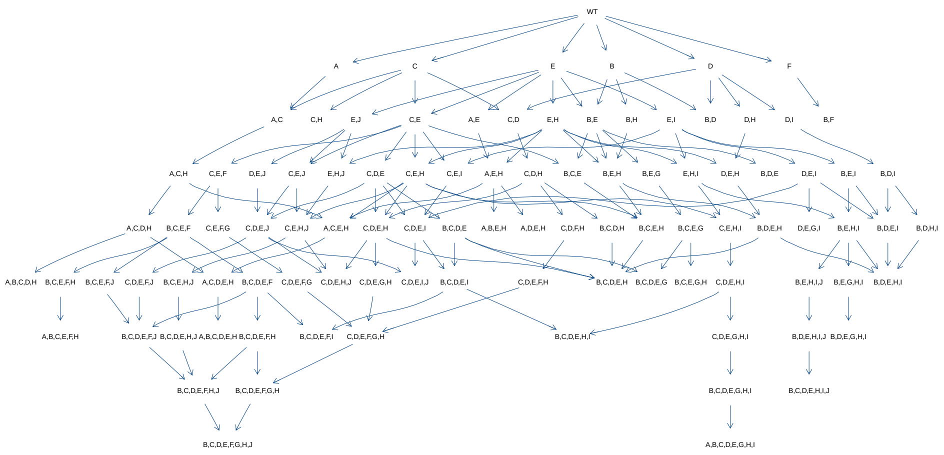

| | |
|---|---|
| *ID* | q6ESfEnjdJLQJKzP |
| *acc genotypes* | 110 |
| *peaks no back* | 41 |
| *removed edges* | NA |
| *prop rem edges* | NA |

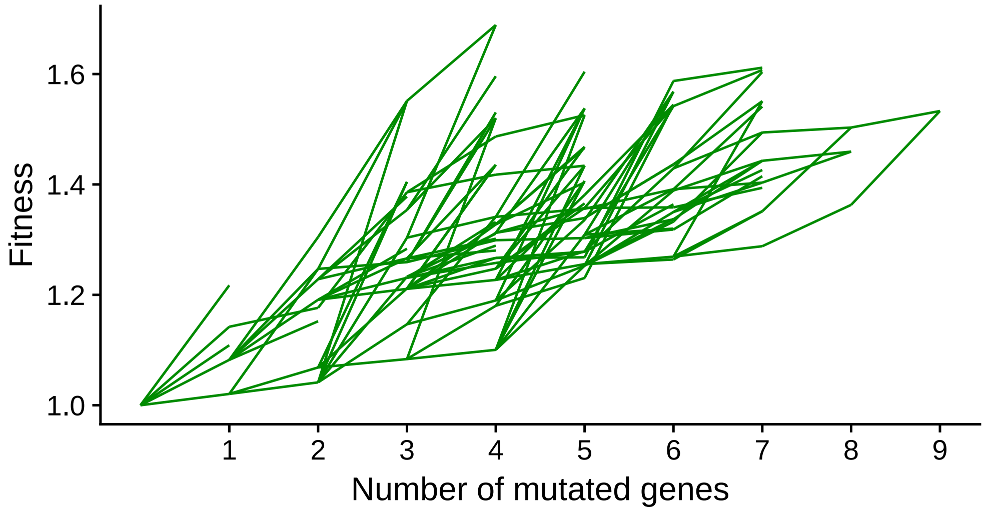


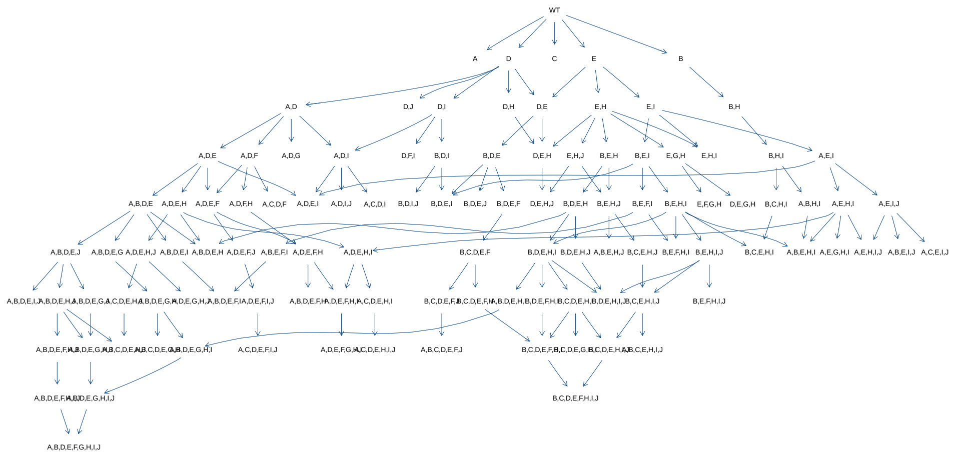

| | |
|---|---|
| *ID* | QDlfwYRc6C9mw910Q |
| *acc genotypes* | 251 |
| *peaks no back* | 69 |
| *removed edges* | NA |
| *prop rem edges* | NA |

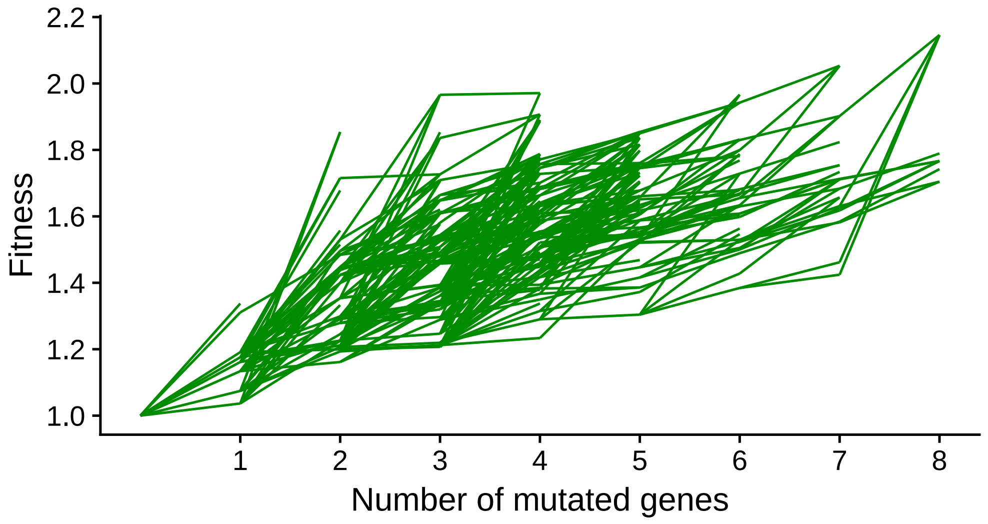


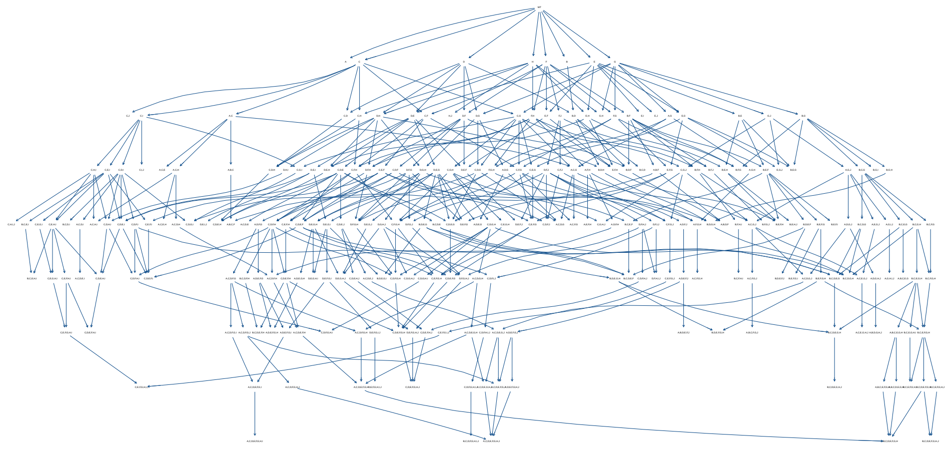

| | |
|---|---|
| *ID* | QDnBQv95LeMoDZkz |
| *acc genotypes* | 397 |
| *peaks no back* | 113 |
| *removed edges* | NA |
| *prop rem edges* | NA |

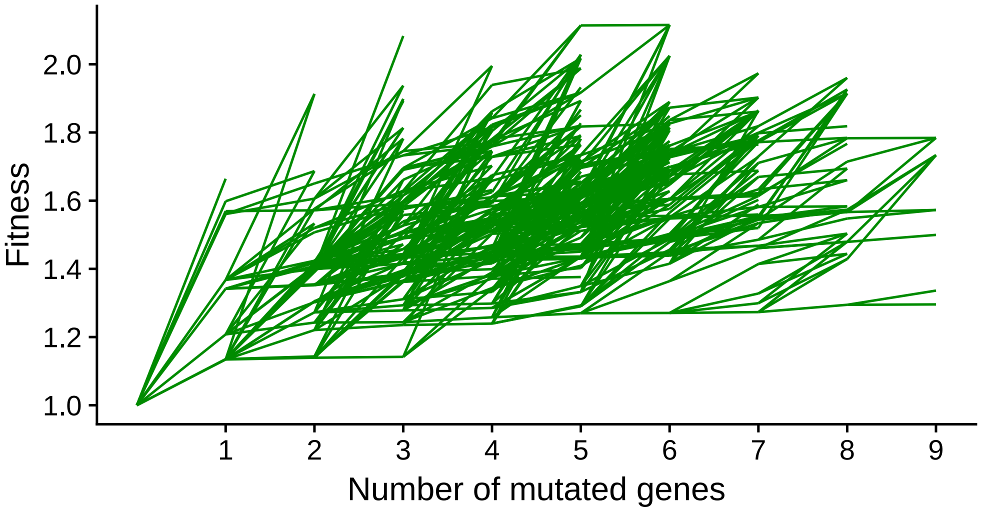


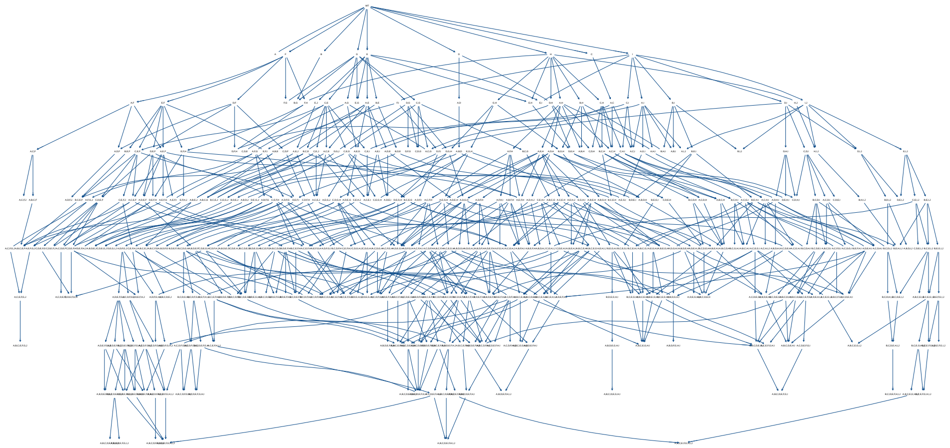

| | |
|---|---|
| *ID* | qhoM9EMECJiKAEai |
| *acc genotypes* | 301 |
| *peaks no back* | 43 |
| *removed edges* | NA |
| *prop rem edges* | NA |

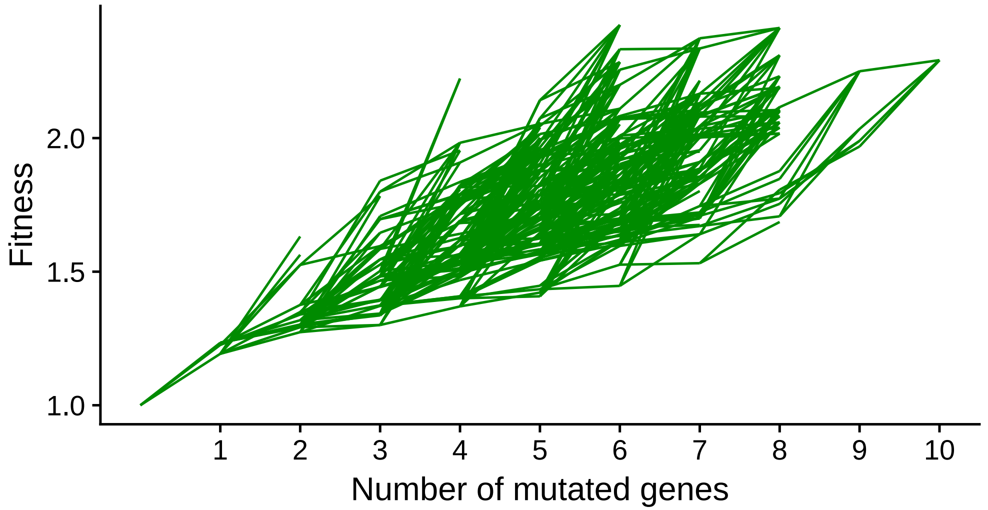


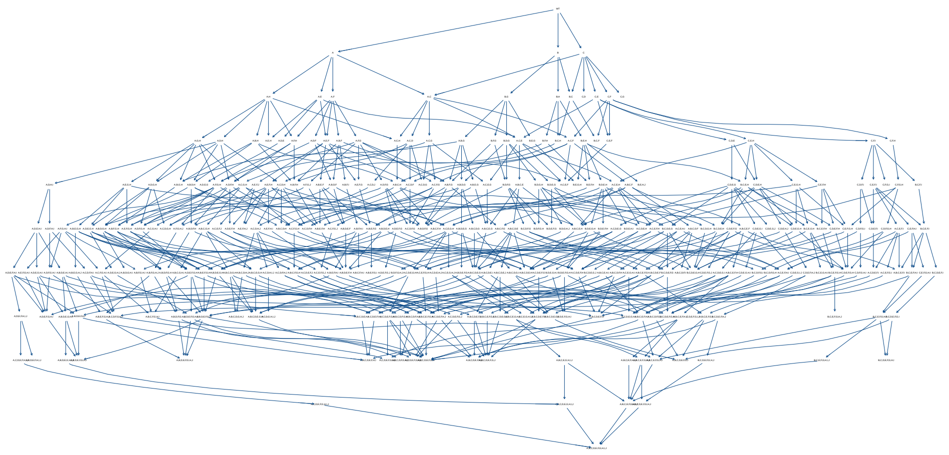

| | |
|---|---|
| *ID* | qIG3omWesEf8erJY |
| *acc genotypes* | 330 |
| *peaks no back* | 79 |
| *removed edges* | NA |
| *prop rem edges* | NA |

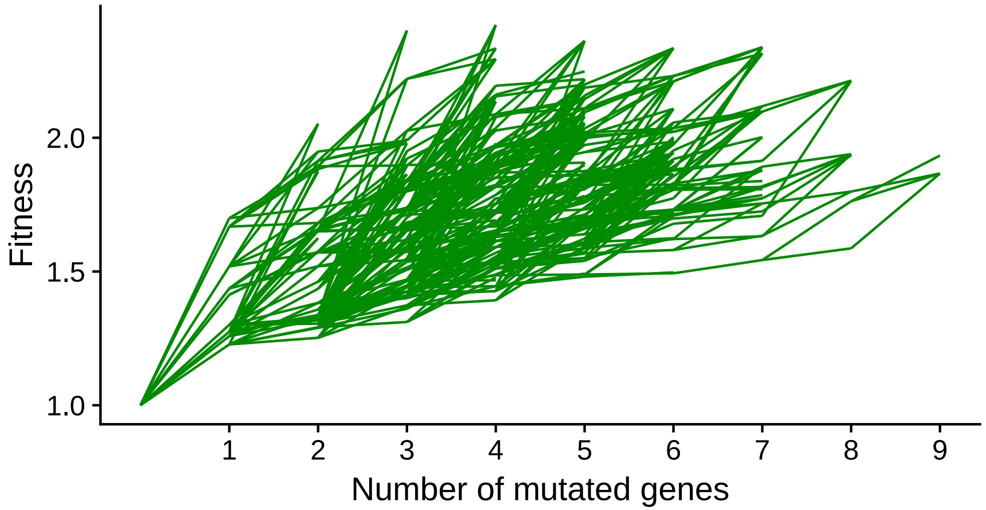


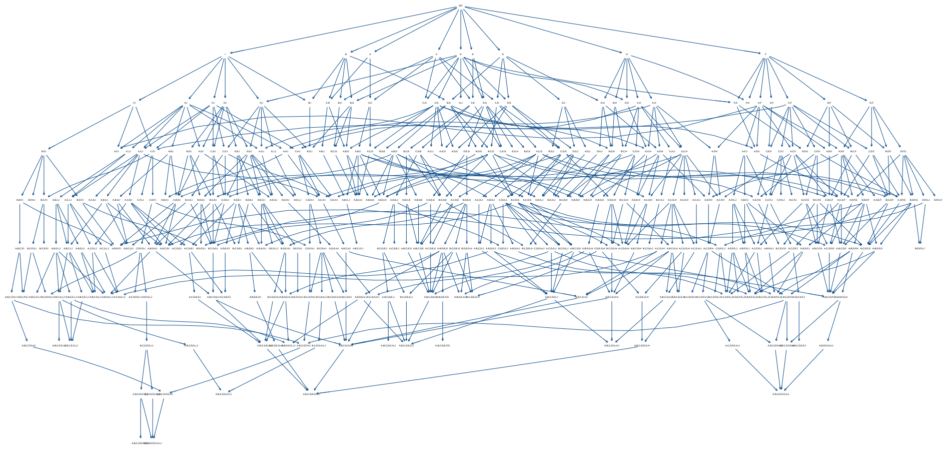

| | |
|---|---|
| *ID* | qiw3nSX8uOEX3EE2 |
| *acc genotypes* | 123 |
| *peaks no back* | 37 |
| *removed edges* | NA |
| *prop rem edges* | NA |

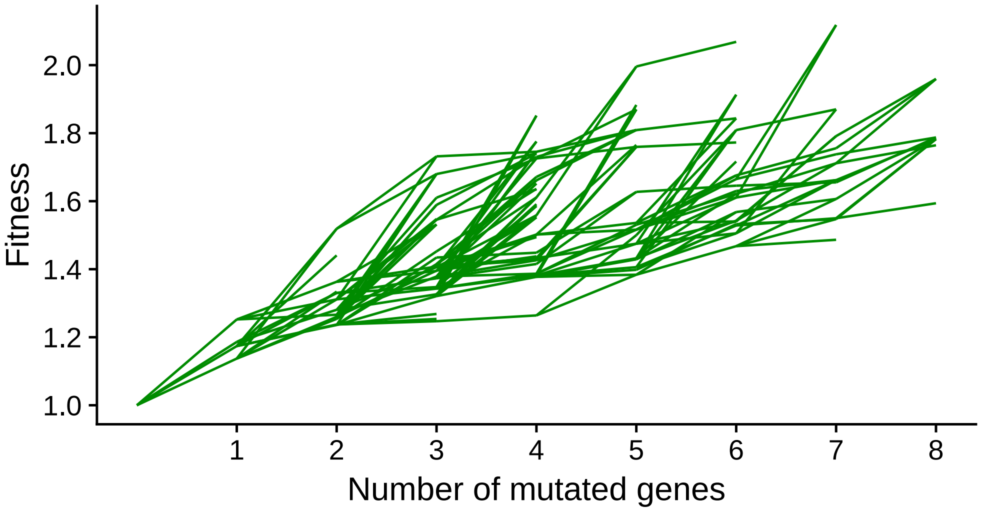


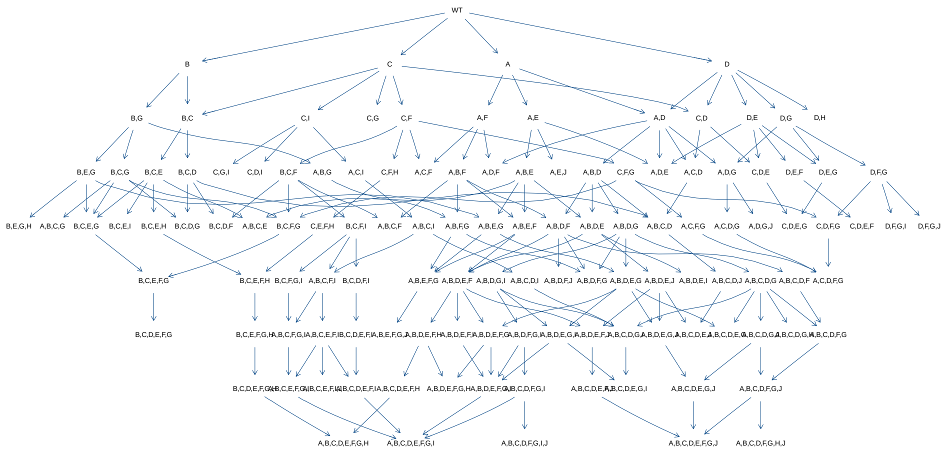

| | |
|---|---|
| *ID* | QLvp54fgddHiWuDQ |
| *acc genotypes* | 365 |
| *peaks no back* | 92 |
| *removed edges* | NA |
| *prop rem edges* | NA |

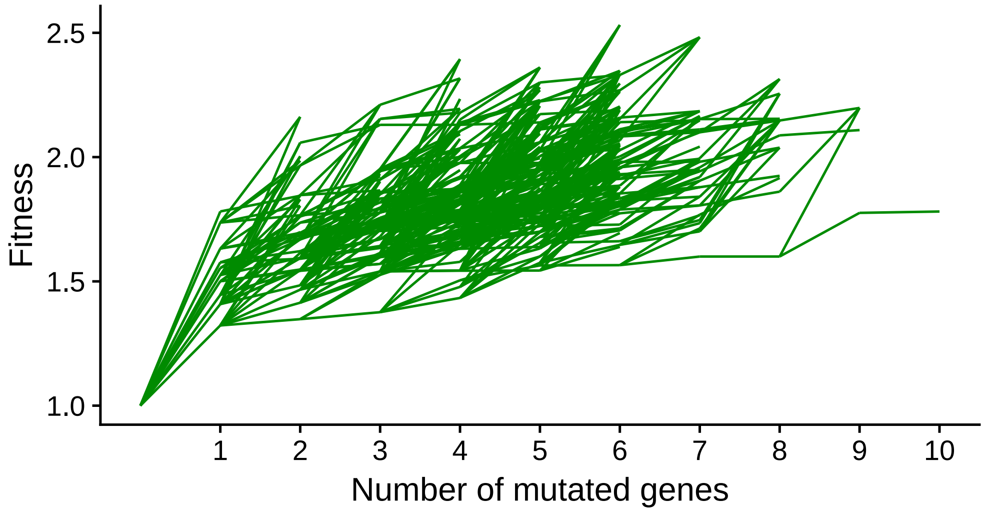


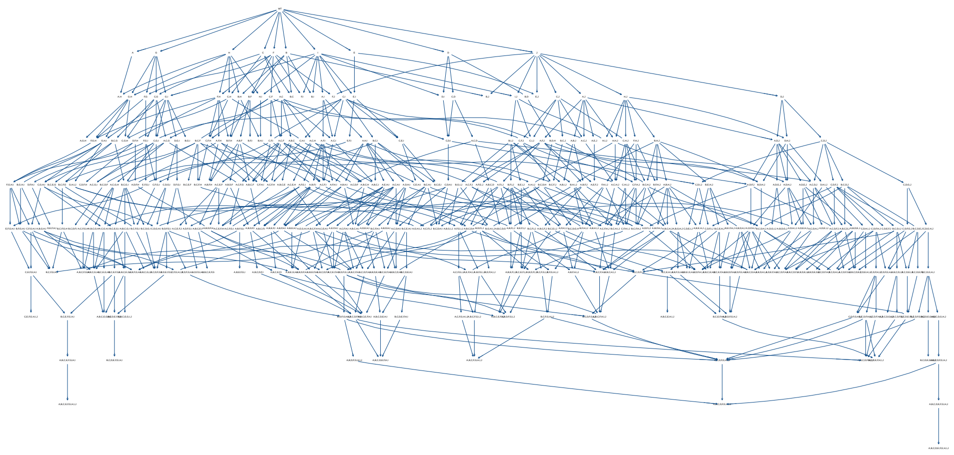

| | |
|---|---|
| *ID* | R1rRFK7kvGTL7JOu |
| *acc genotypes* | 95 |
| *peaks no back* | 39 |
| *removed edges* | NA |
| *prop rem edges* | NA |

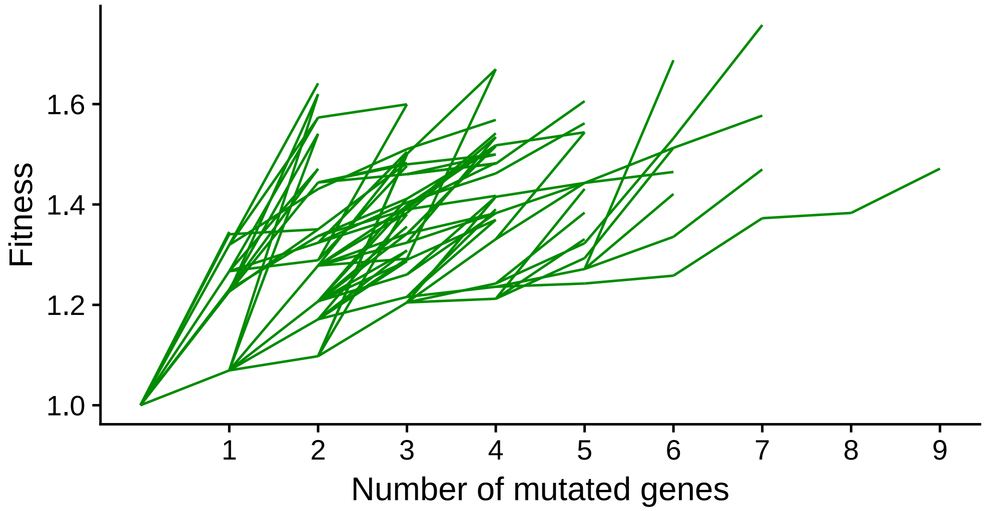


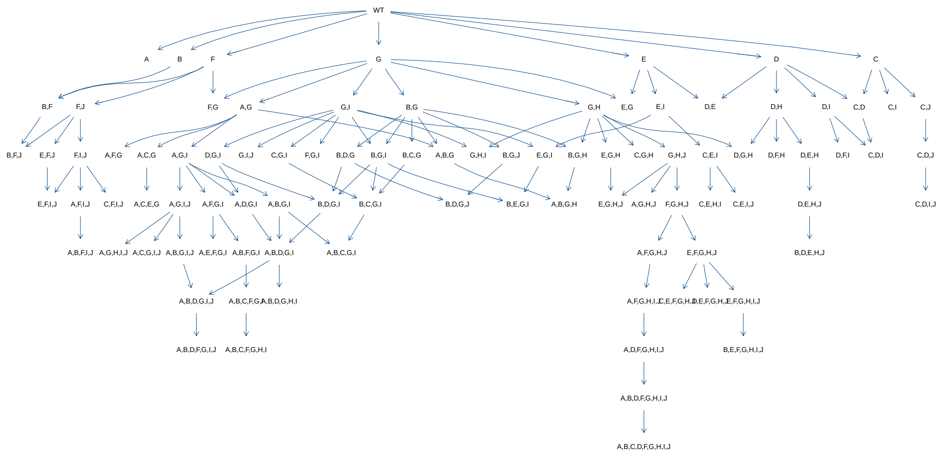

| | |
|---|---|
| *ID* | r8ycAIoMMwveOtjh |
| *acc genotypes* | 474 |
| *peaks no back* | 56 |
| *removed edges* | NA |
| *prop rem edges* | NA |

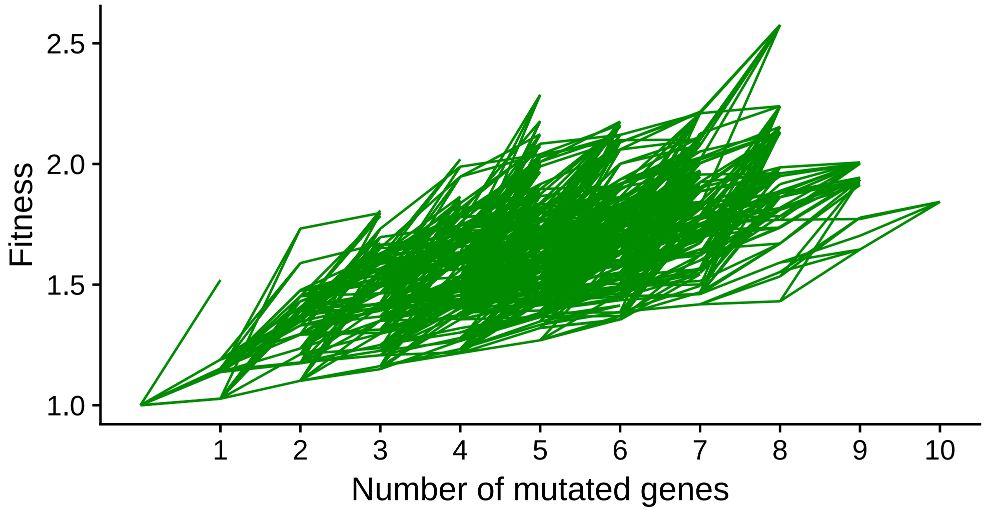


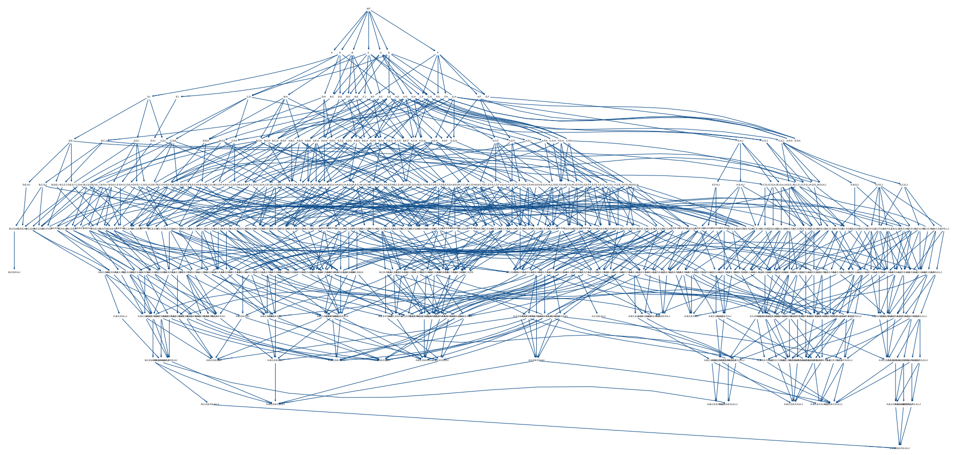

| | |
|---|---|
| *ID* | rFXDokDyxb54xE8I |
| *acc genotypes* | 267 |
| *peaks no back* | 95 |
| *removed edges* | NA |
| *prop rem edges* | NA |

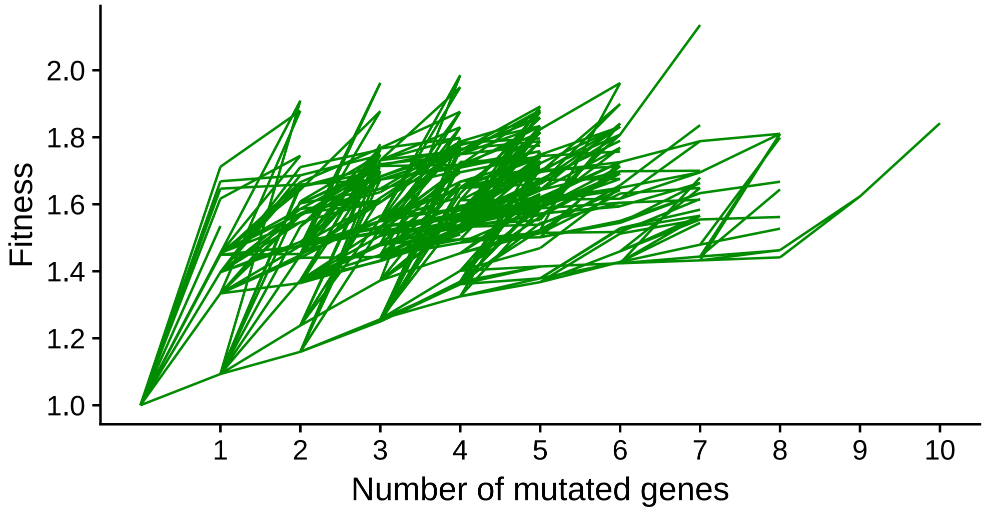


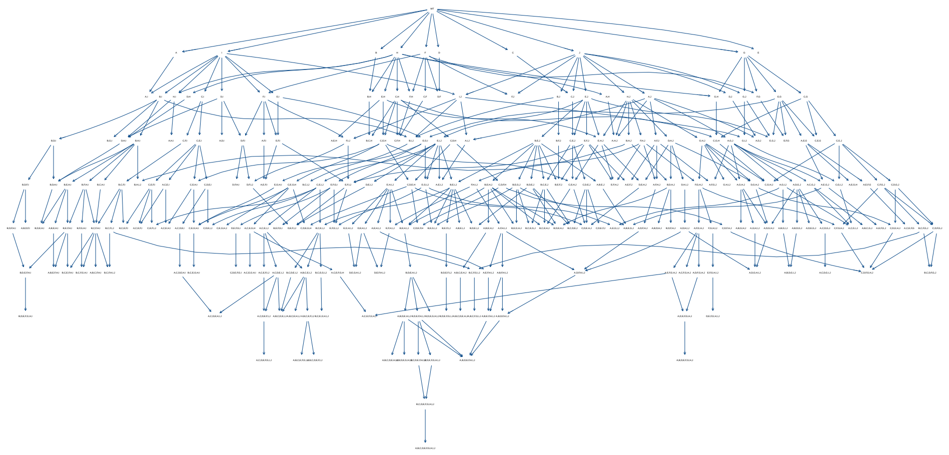

| | |
|---|---|
| *ID* | Rgbd18KZtZQm310aF |
| *acc genotypes* | 305 |
| *peaks no back* | 59 |
| *removed edges* | NA |
| *prop rem edges* | NA |

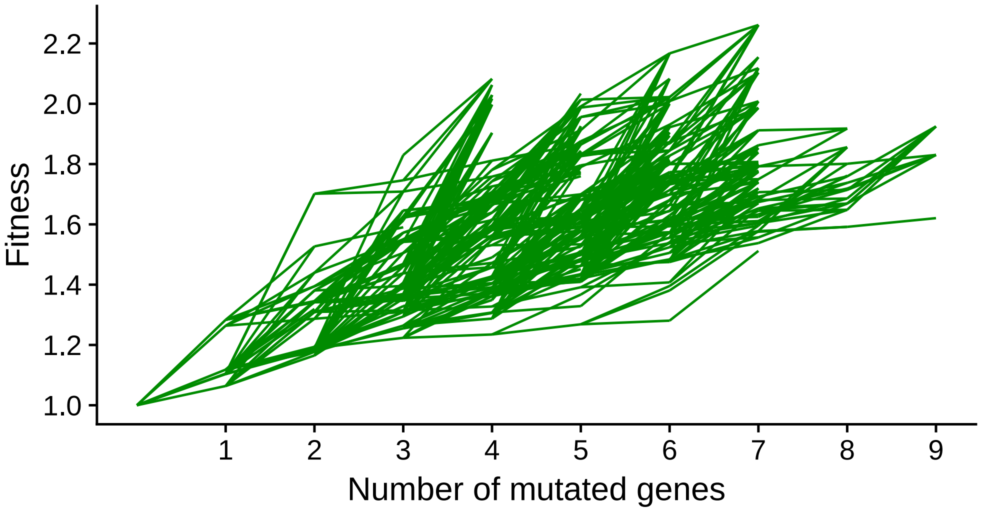


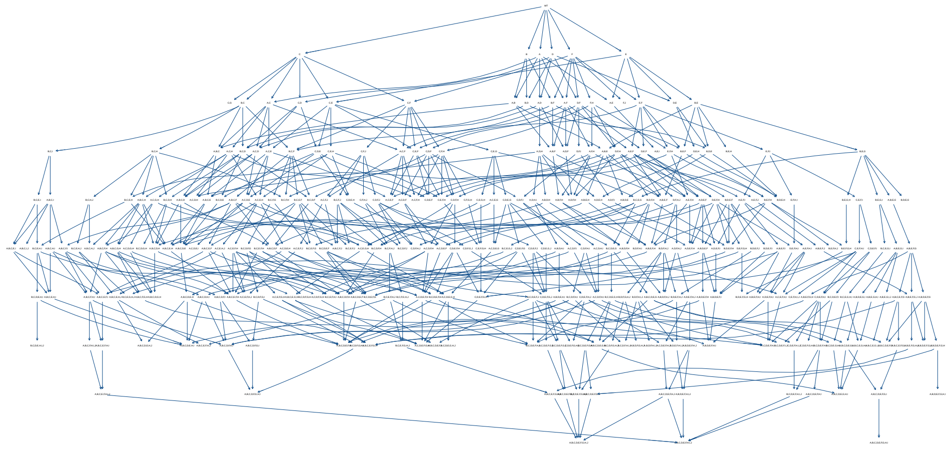

| | |
|---|---|
| *ID* | RgQW10wScrRmd9eE8 |
| *acc genotypes* | 65 |
| *peaks no back* | 28 |
| *removed edges* | NA |
| *prop rem edges* | NA |

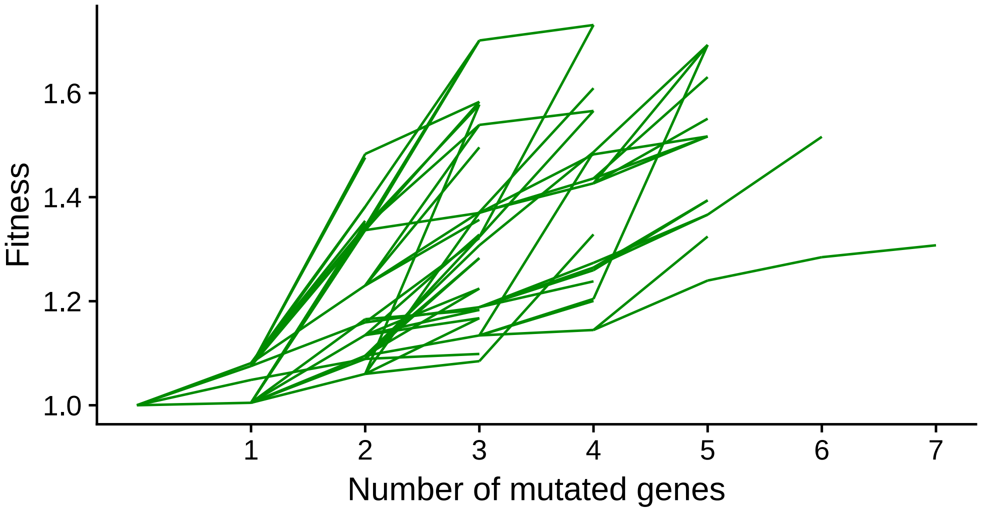


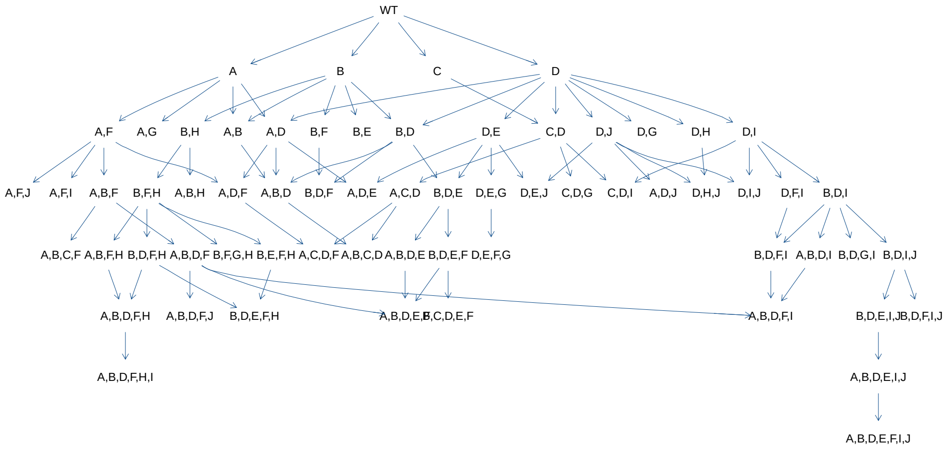

| | |
|---|---|
| *ID* | RM1oRnvkQ74p10MYi |
| *acc genotypes* | 211 |
| *peaks no back* | 85 |
| *removed edges* | NA |
| *prop rem edges* | NA |

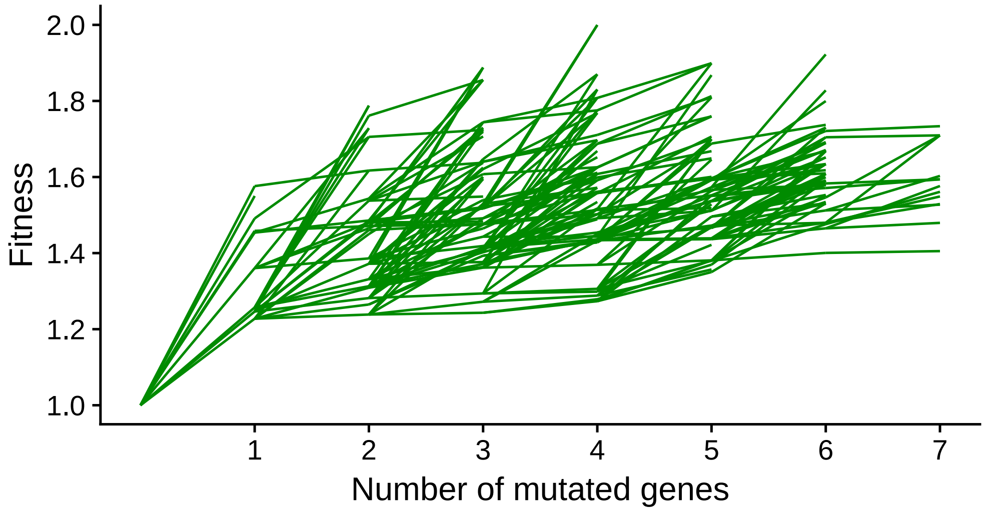


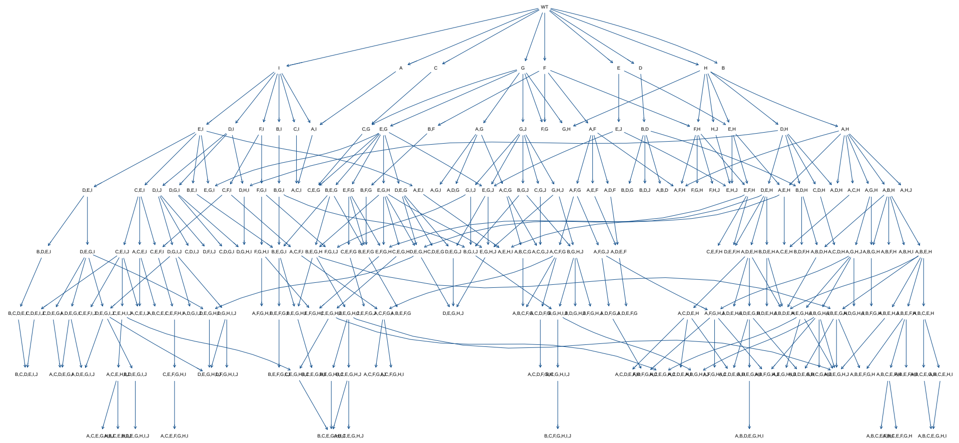

| | |
|---|---|
| *ID* | rwEpMgO6p9XY5Fe3 |
| *acc genotypes* | 59 |
| *peaks no back* | 25 |
| *removed edges* | NA |
| *prop rem edges* | NA |

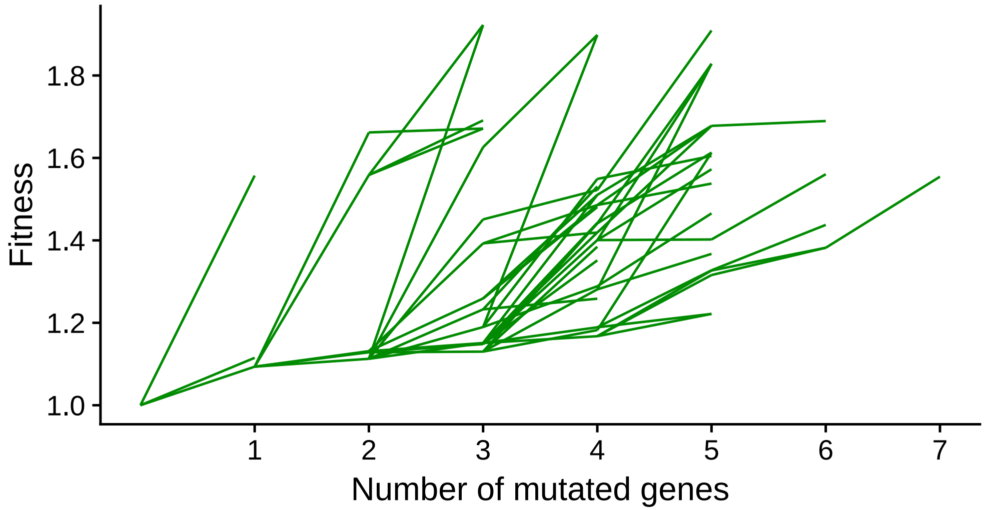


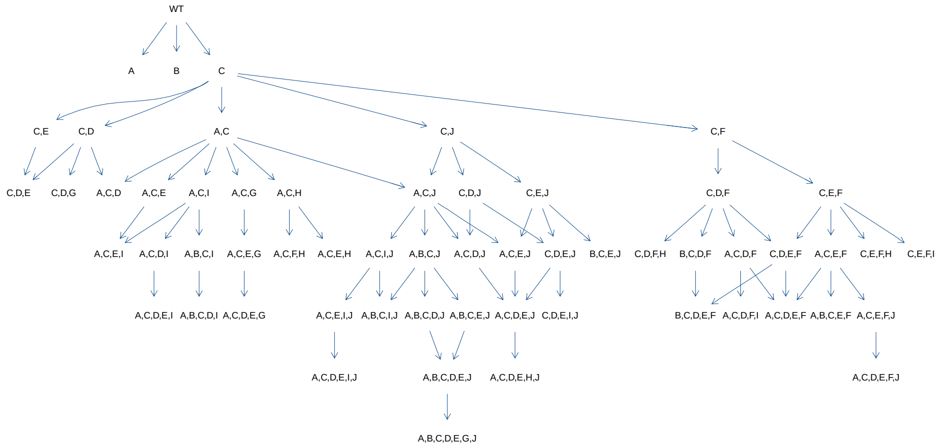

| | |
|---|---|
| *ID* | siB10Dc3bBqHdWLsu |
| *acc genotypes* | 152 |
| *peaks no back* | 59 |
| *removed edges* | NA |
| *prop rem edges* | NA |

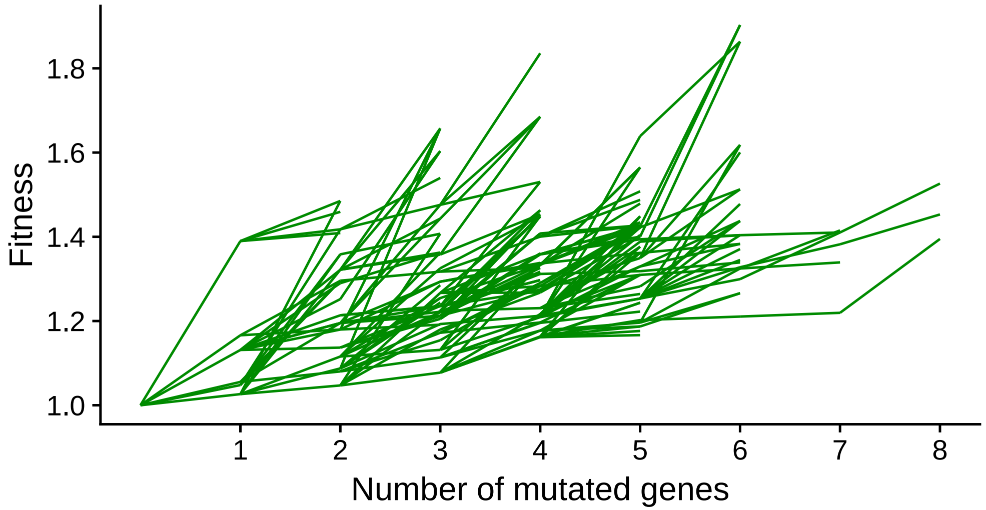


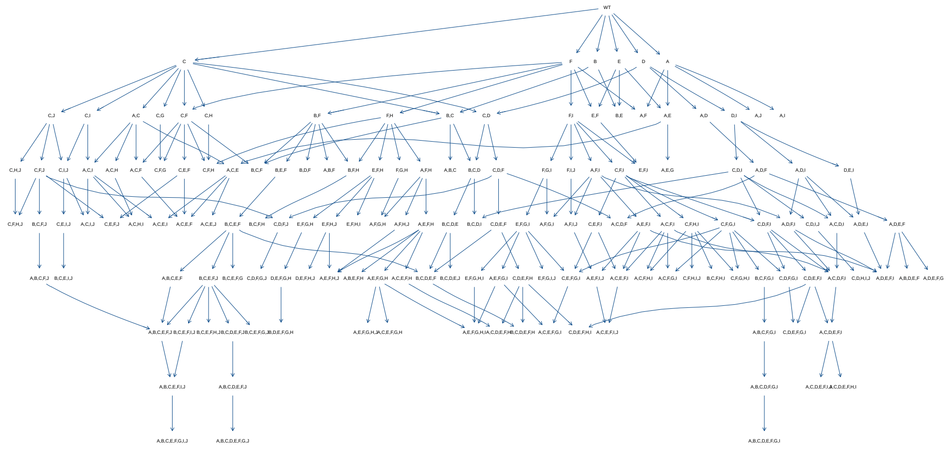

| | |
|---|---|
| *ID* | SkrmpMeOx2A5XIjs |
| *acc genotypes* | 41 |
| *peaks no back* | 14 |
| *removed edges* | NA |
| *prop rem edges* | NA |

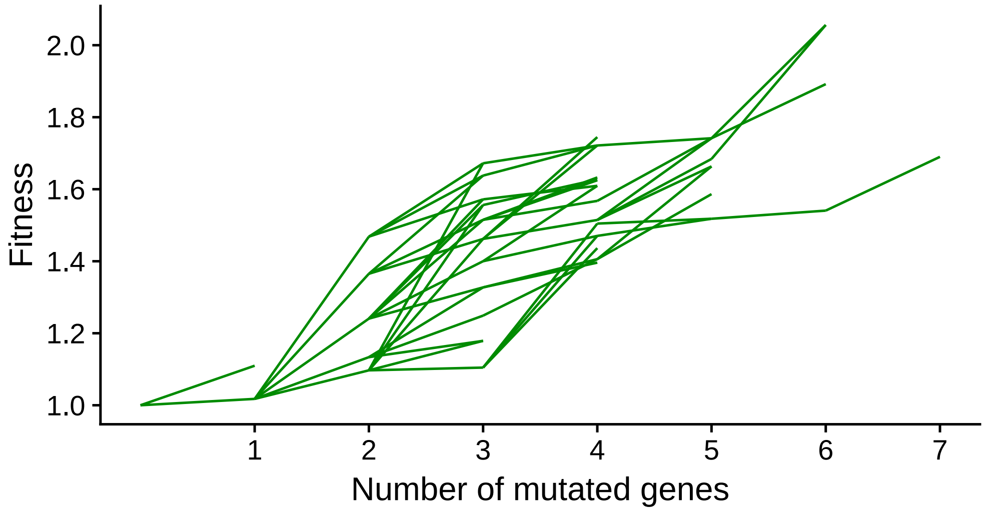


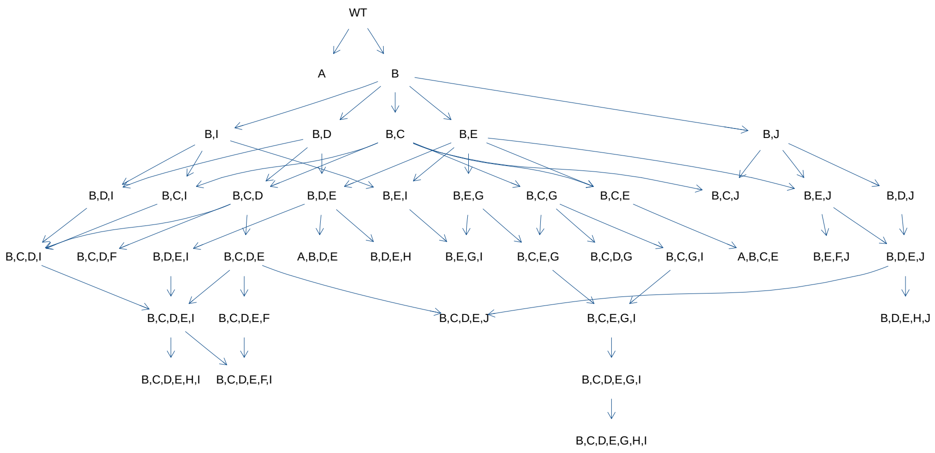

| | |
|---|---|
| *ID* | sKV2JZzX6hoi6ZXJ |
| *acc genotypes* | 302 |
| *peaks no back* | 68 |
| *removed edges* | NA |
| *prop rem edges* | NA |

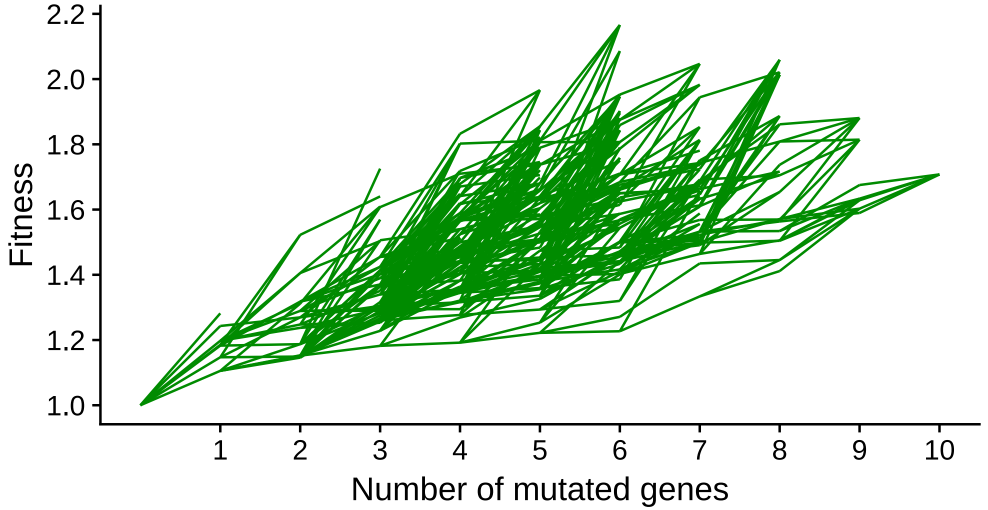


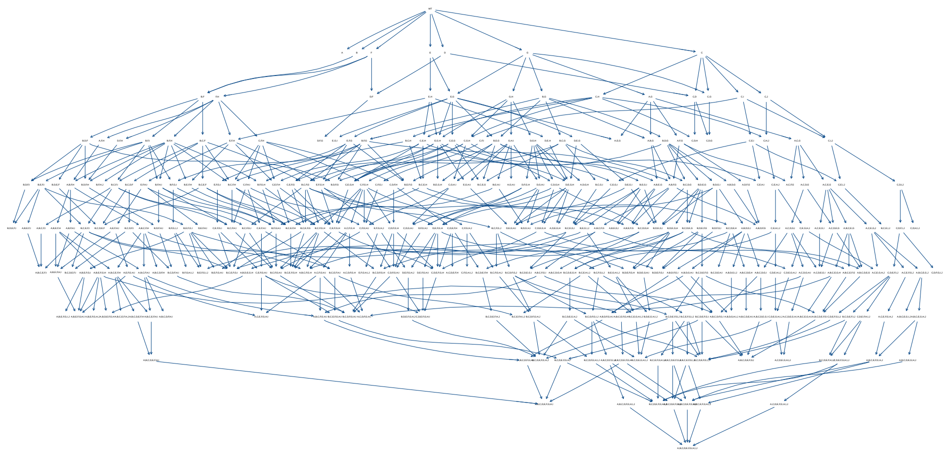

| | |
|---|---|
| *ID* | SRQPFRnma8vPJxWK |
| *acc genotypes* | 327 |
| *peaks no back* | 103 |
| *removed edges* | NA |
| *prop rem edges* | NA |

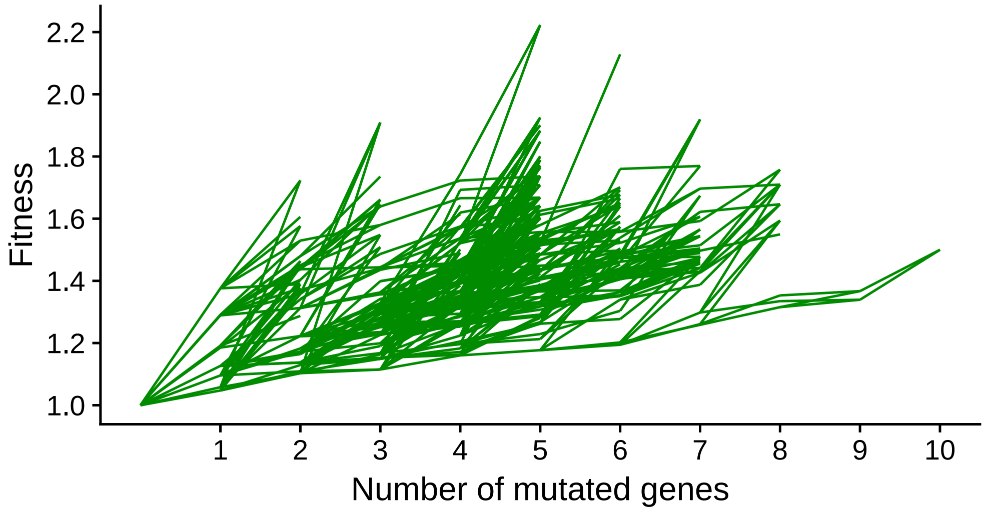


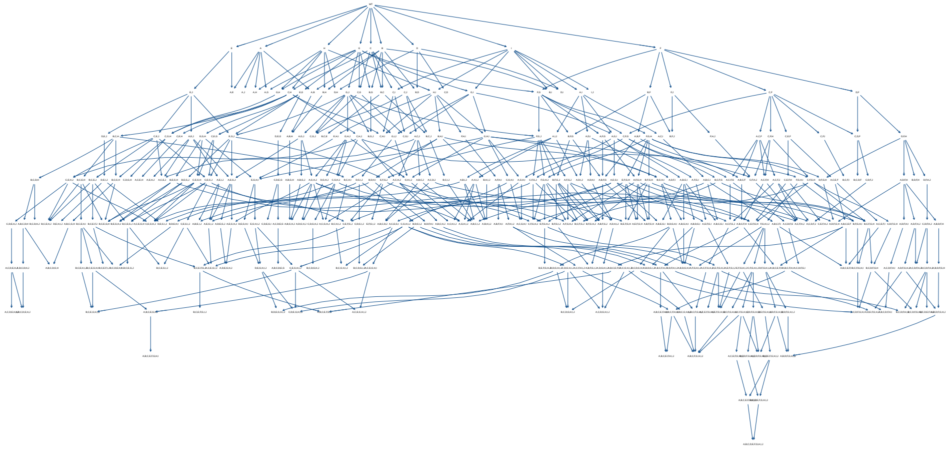

| | |
|---|---|
| *ID* | suRv8SDkQ3Fspdke |
| *acc genotypes* | 330 |
| *peaks no back* | 113 |
| *removed edges* | NA |
| *prop rem edges* | NA |

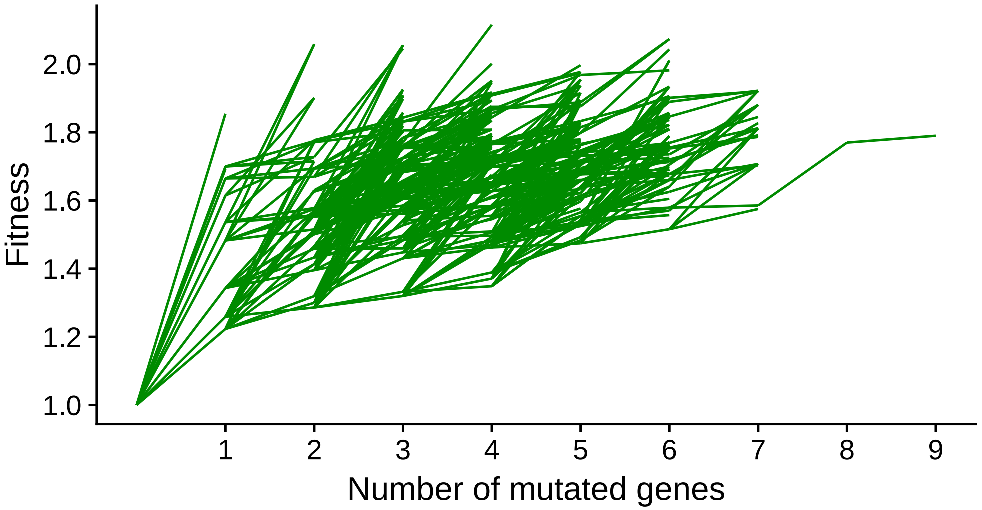


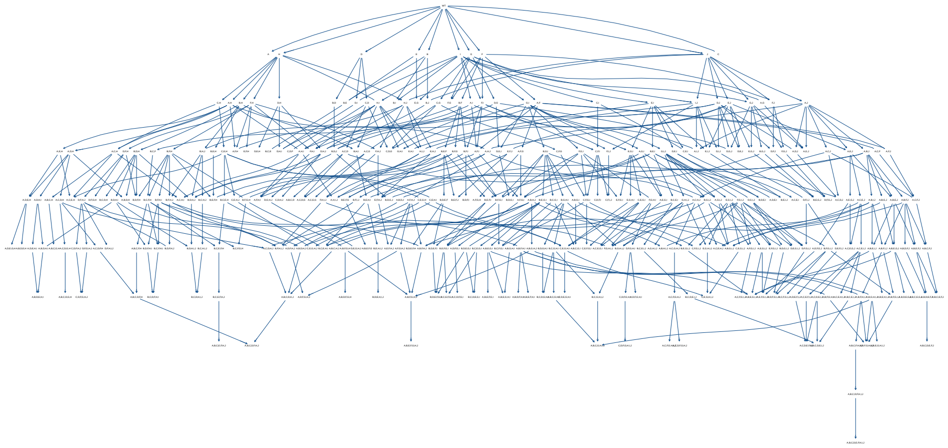

| | |
|---|---|
| *ID* | SUX11JboEBbU7Sly |
| *acc genotypes* | 159 |
| *peaks no back* | 49 |
| *removed edges* | NA |
| *prop rem edges* | NA |

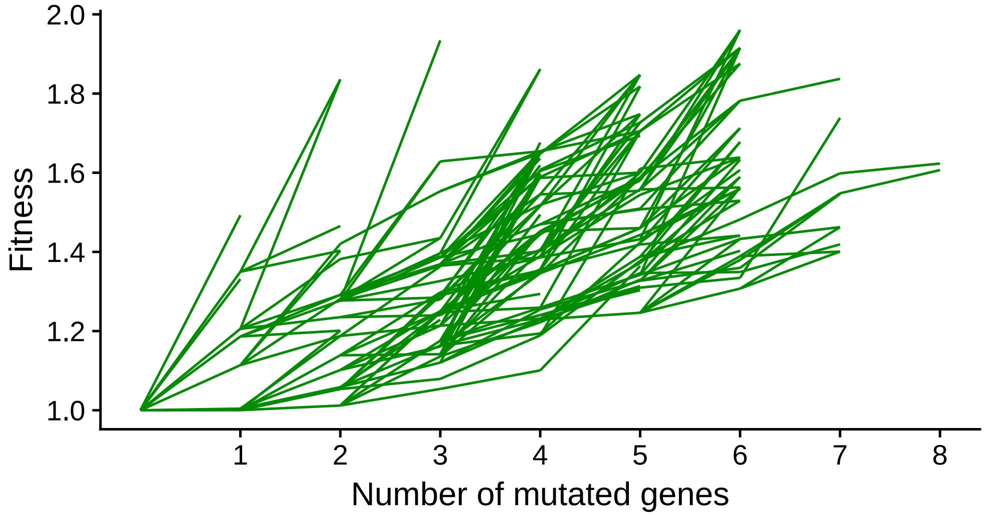


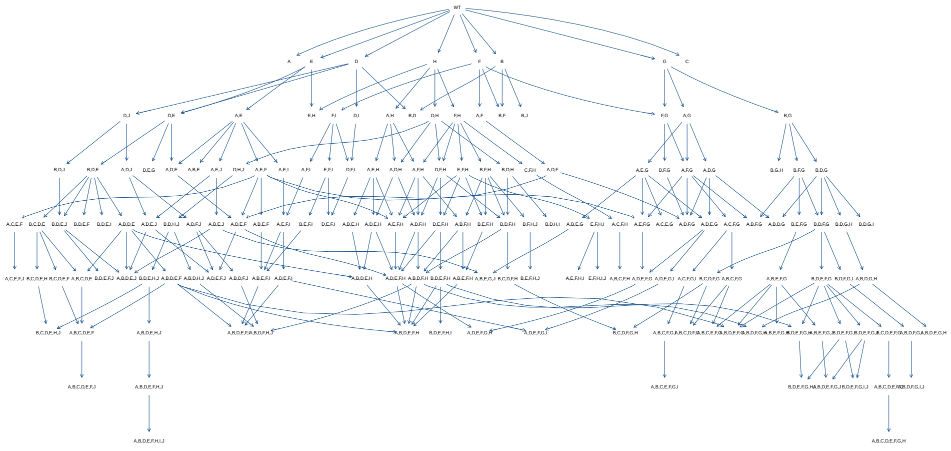

| | |
|---|---|
| *ID* | T1lg10UC4ysvdG1vj |
| *acc genotypes* | 75 |
| *peaks no back* | 31 |
| *removed edges* | NA |
| *prop rem edges* | NA |

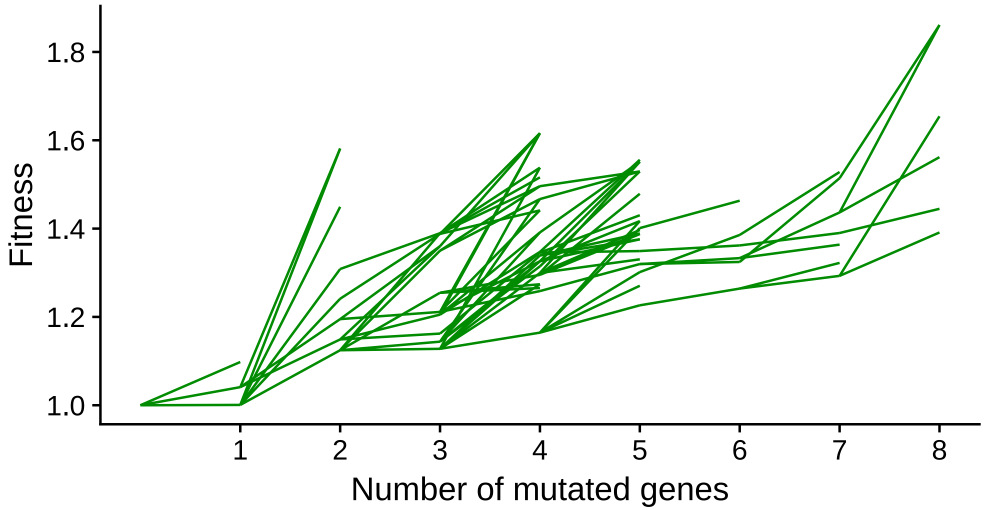


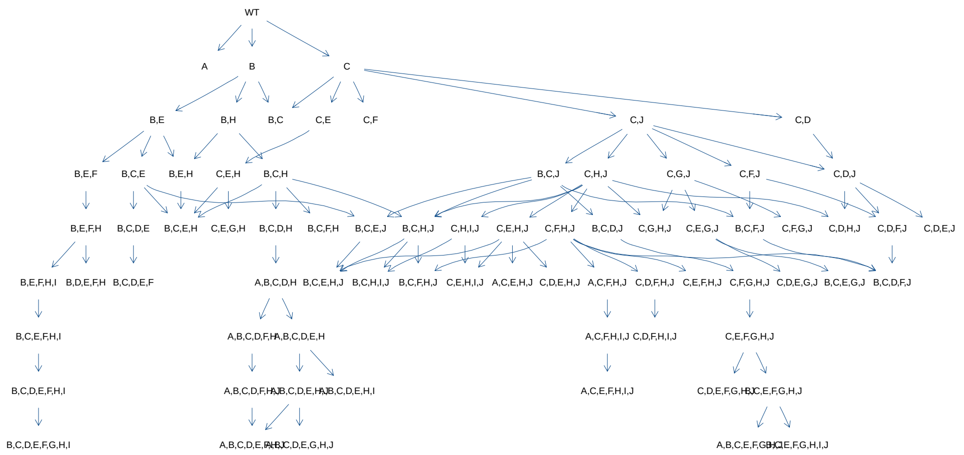

| | |
|---|---|
| *ID* | T5oZEwzVo6JyuGLW |
| *acc genotypes* | 125 |
| *peaks no back* | 52 |
| *removed edges* | NA |
| *prop rem edges* | NA |

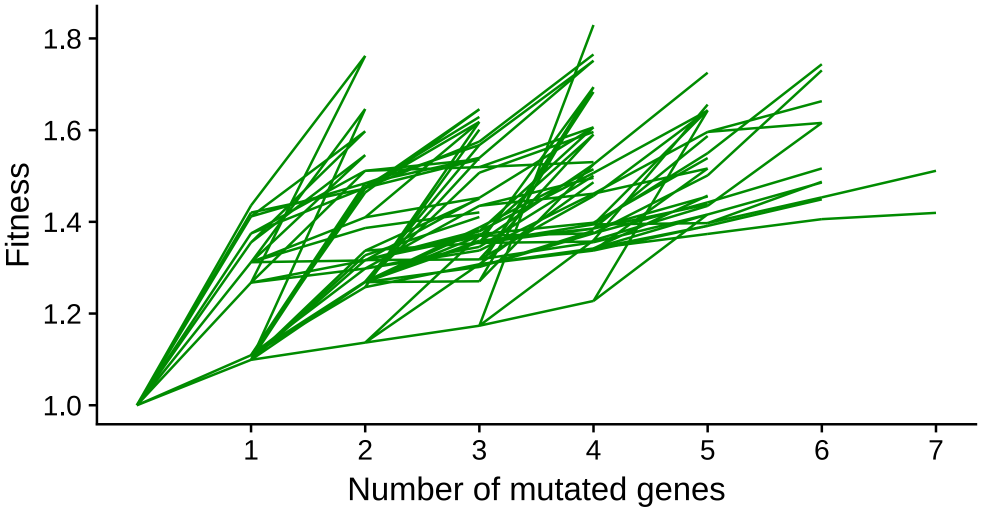


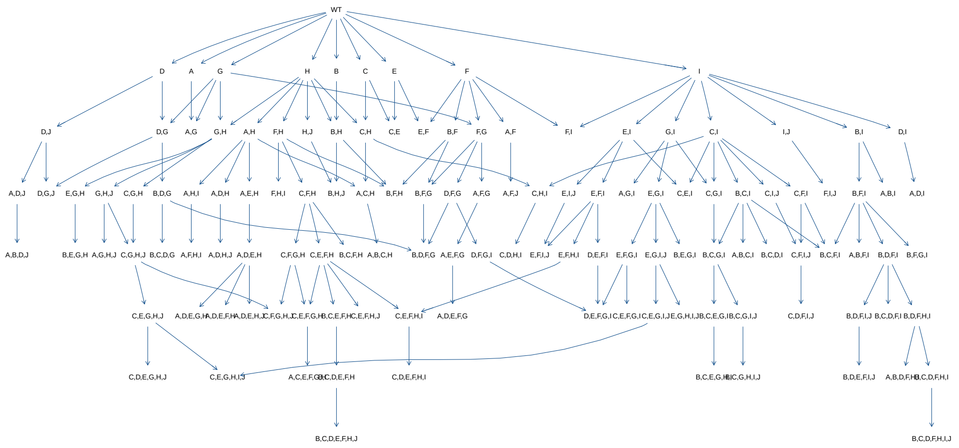

| | |
|---|---|
| *ID* | TkIr7flyal9UtU86 |
| *acc genotypes* | 142 |
| *peaks no back* | 60 |
| *removed edges* | NA |
| *prop rem edges* | NA |

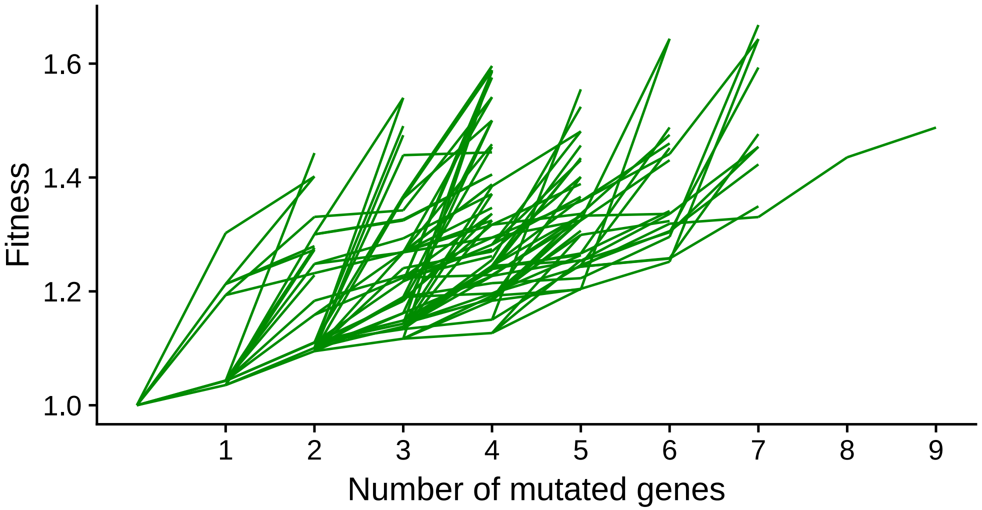


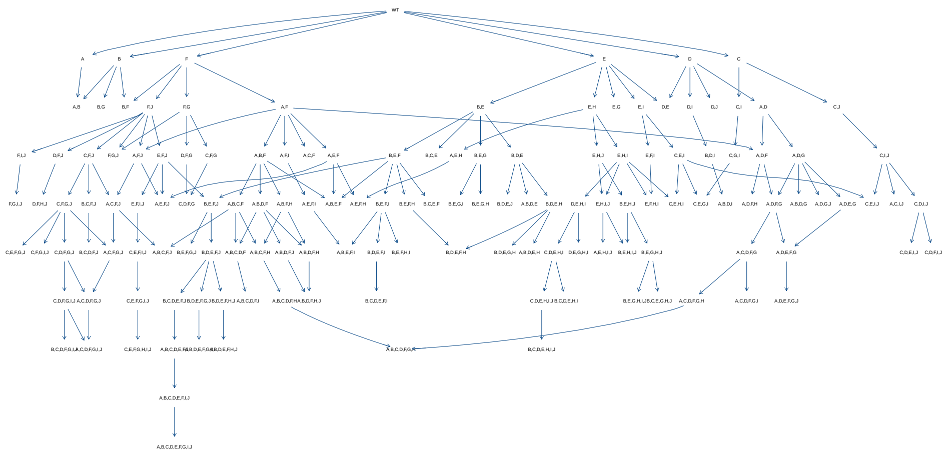

| | |
|---|---|
| *ID* | tKkGKUAR1sBTETkR |
| *acc genotypes* | 209 |
| *peaks no back* | 70 |
| *removed edges* | NA |
| *prop rem edges* | NA |

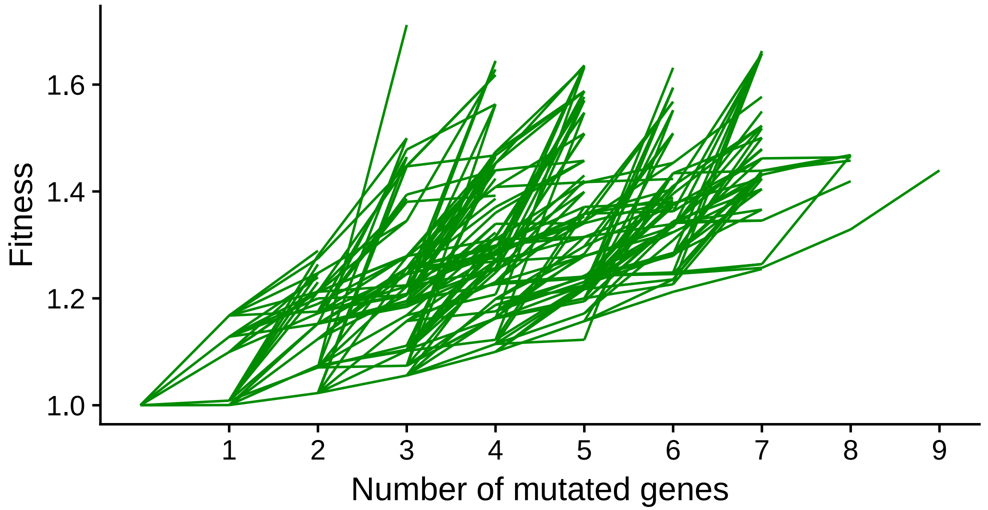


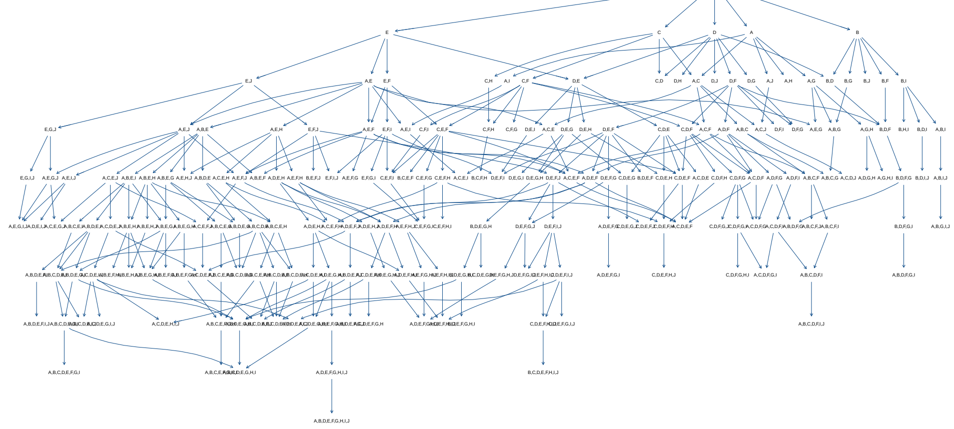

| | |
|---|---|
| *ID* | tyHTZO351lxuXhpp |
| *acc genotypes* | 173 |
| *peaks no back* | 40 |
| *removed edges* | NA |
| *prop rem edges* | NA |

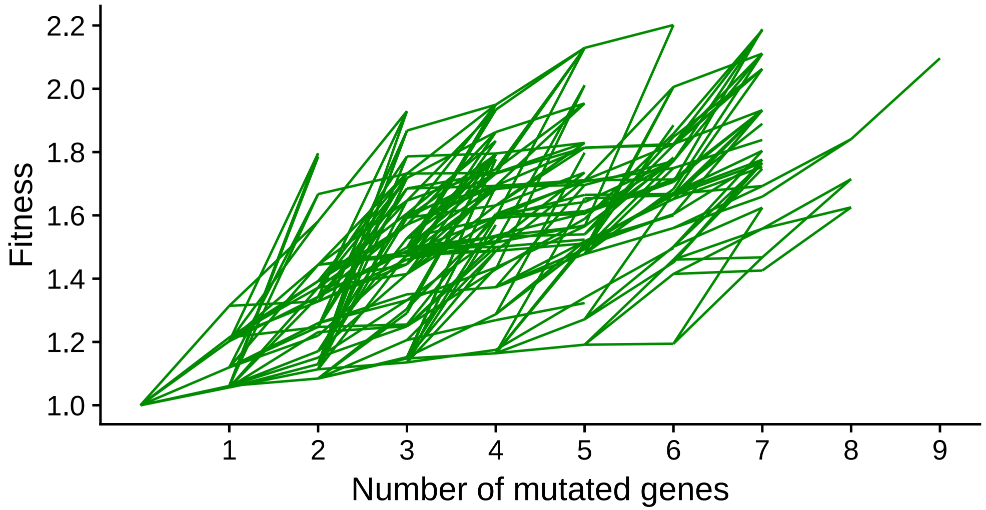


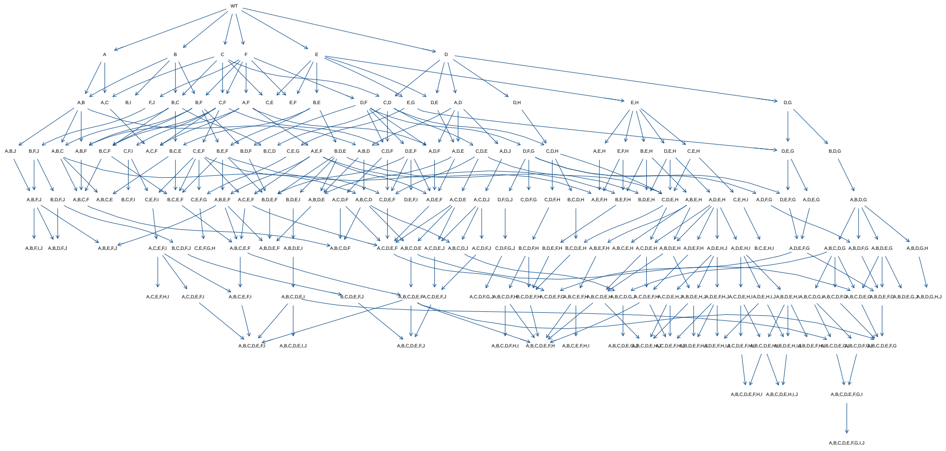

| | |
|---|---|
| *ID* | u4QnEyMMh8u2Dmoq |
| *acc genotypes* | 302 |
| *peaks no back* | 82 |
| *removed edges* | NA |
| *prop rem edges* | NA |

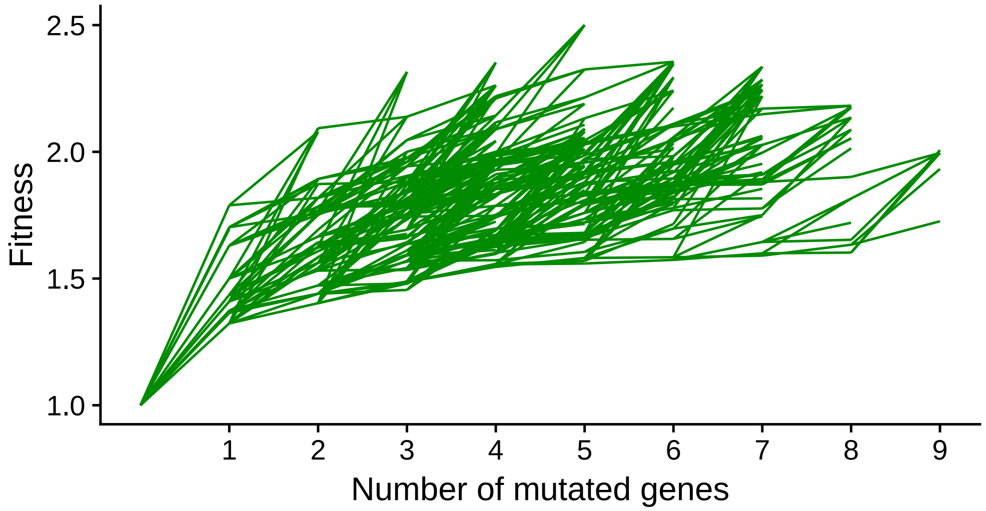


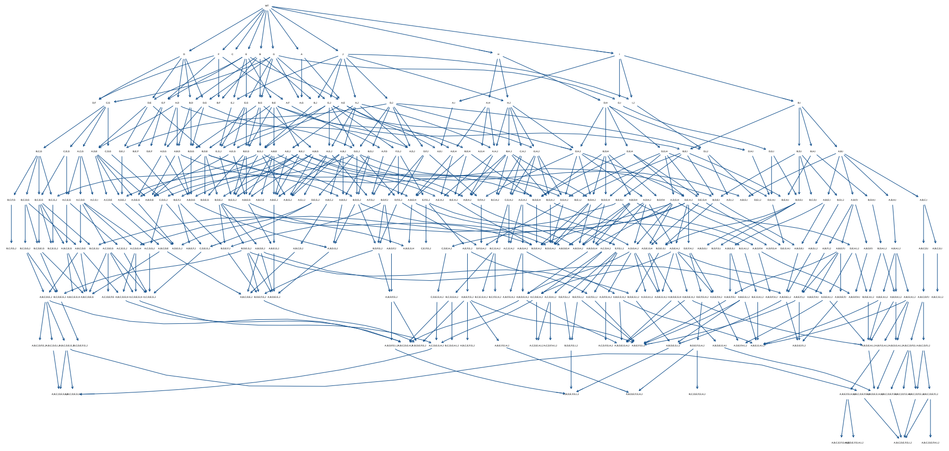

| | |
|---|---|
| *ID* | u6D56PFLKLEytlrF |
| *acc genotypes* | 243 |
| *peaks no back* | 53 |
| *removed edges* | NA |
| *prop rem edges* | NA |

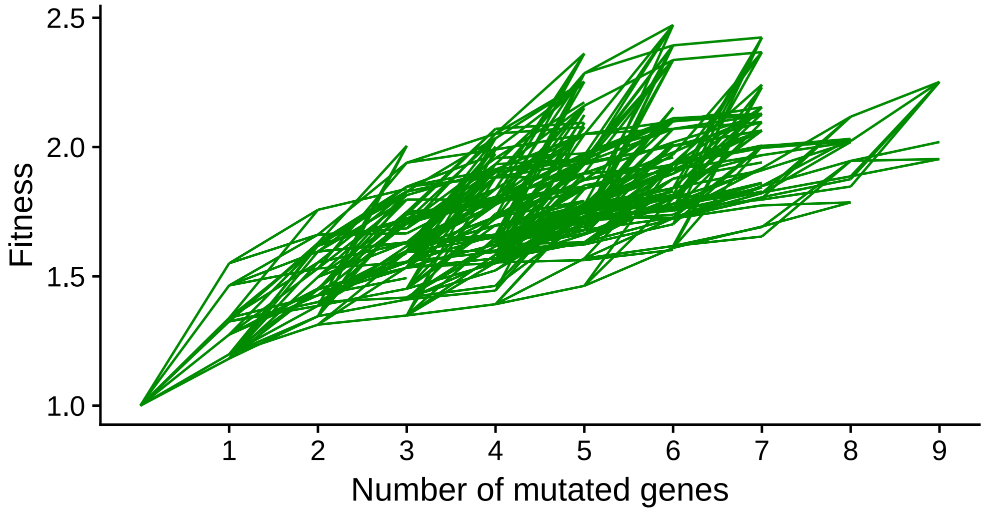


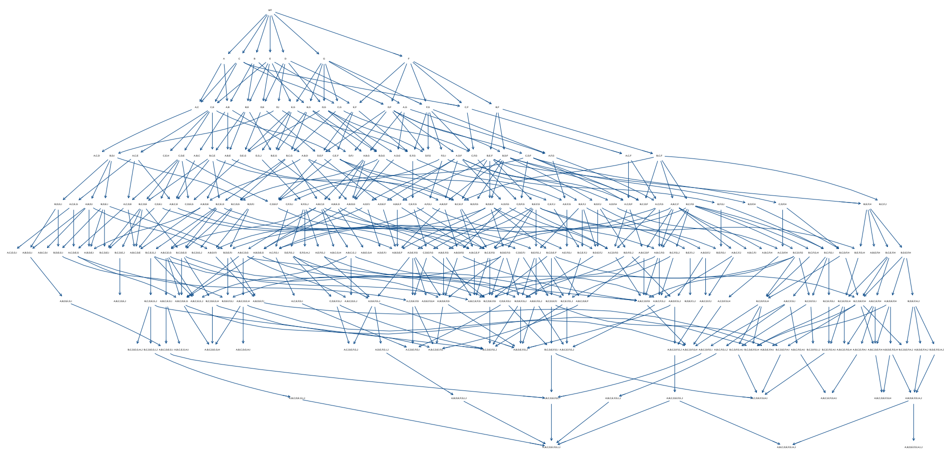

| | |
|---|---|
| *ID* | u8aziQ10mxkwi2VPc |
| *acc genotypes* | 57 |
| *peaks no back* | 19 |
| *removed edges* | NA |
| *prop rem edges* | NA |

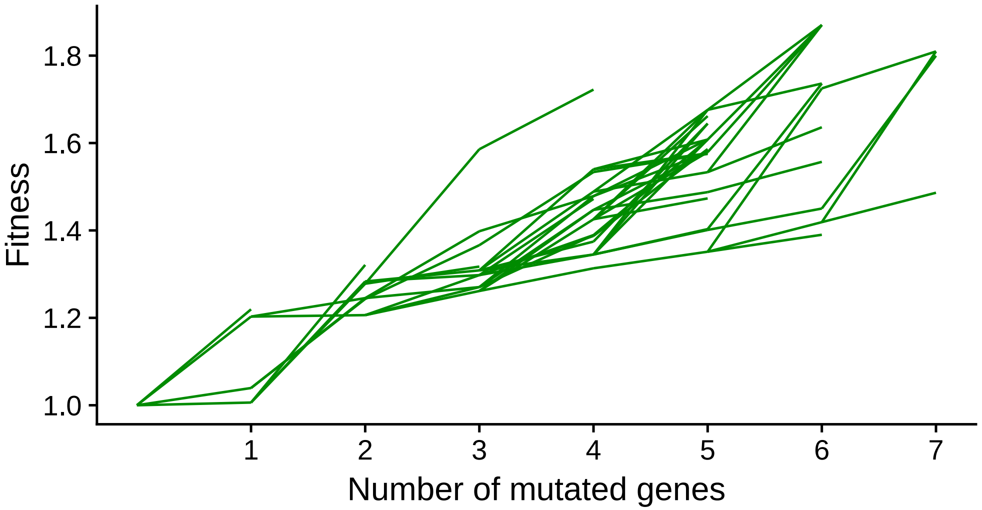


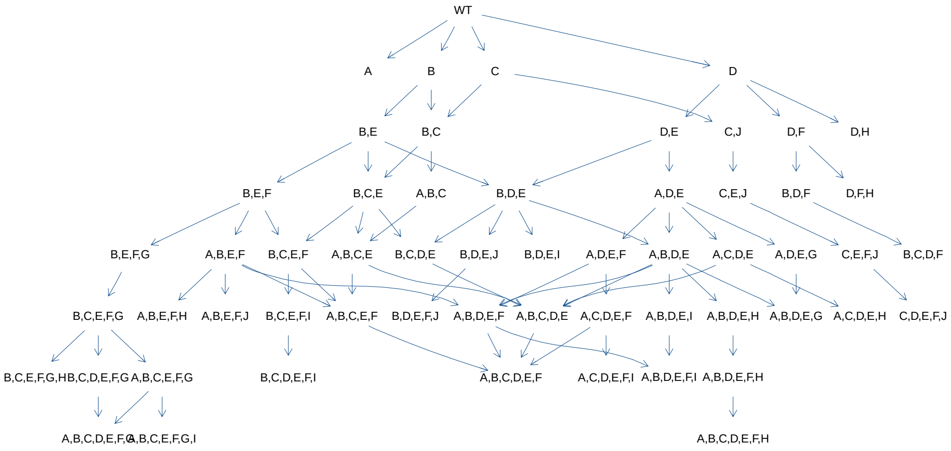

| | |
|---|---|
| *ID* | U9tKbeEaFO915pcB |
| *acc genotypes* | 151 |
| *peaks no back* | 66 |
| *removed edges* | NA |
| *prop rem edges* | NA |

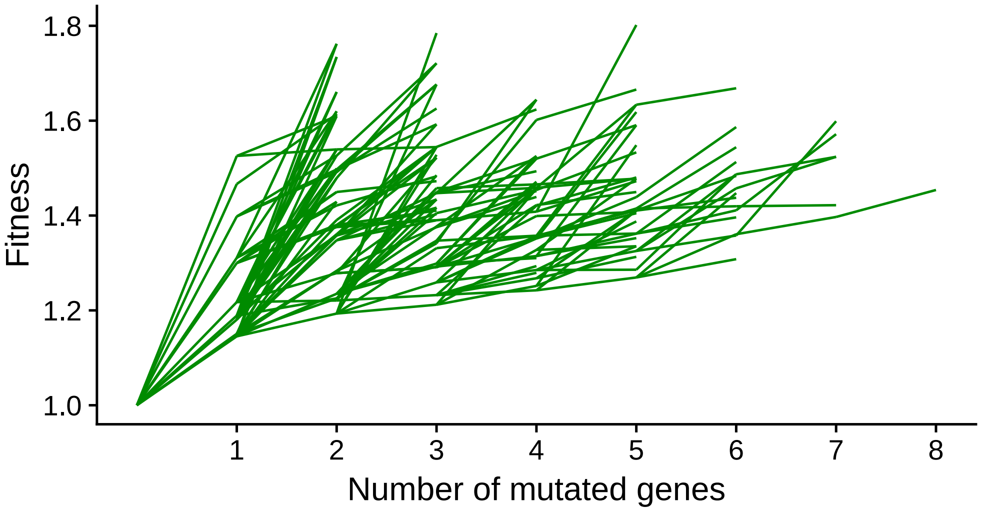


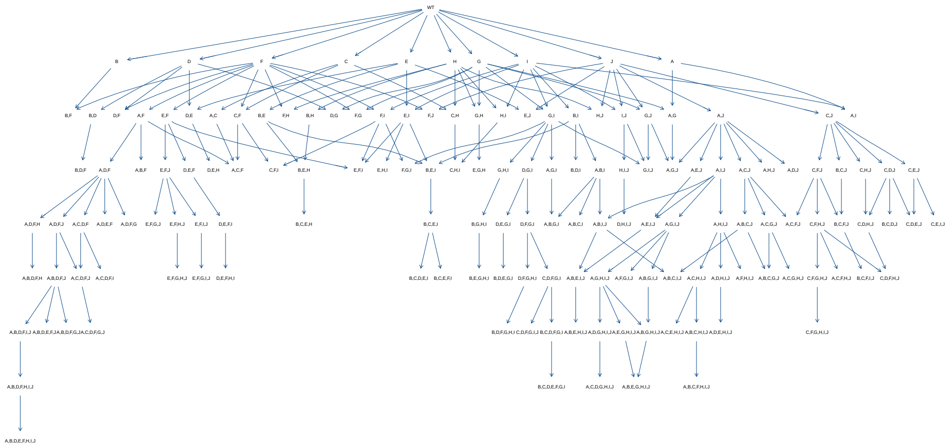

| | |
|---|---|
| *ID* | uaHRvH1XC9BScLS5 |
| *acc genotypes* | 333 |
| *peaks no back* | 60 |
| *removed edges* | NA |
| *prop rem edges* | NA |

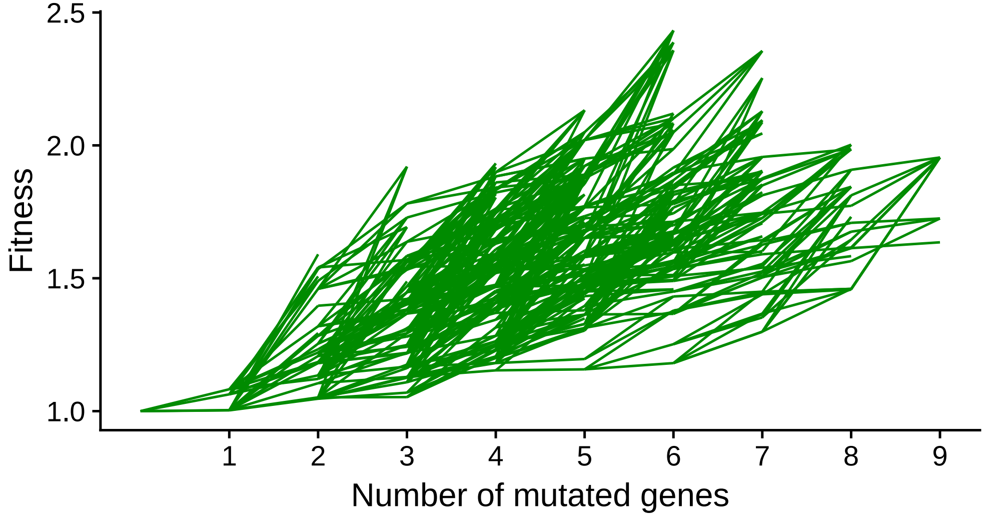


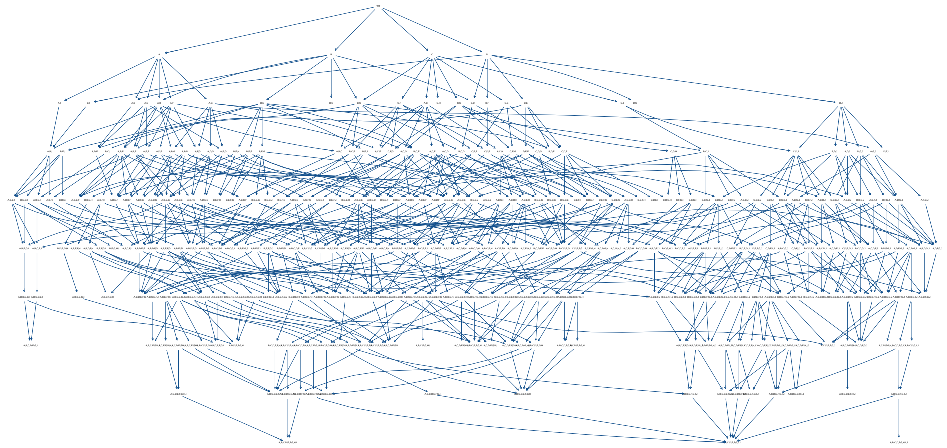

| | |
|---|---|
| *ID* | UdIzVbWbljfForTa |
| *acc genotypes* | 206 |
| *peaks no back* | 78 |
| *removed edges* | NA |
| *prop rem edges* | NA |

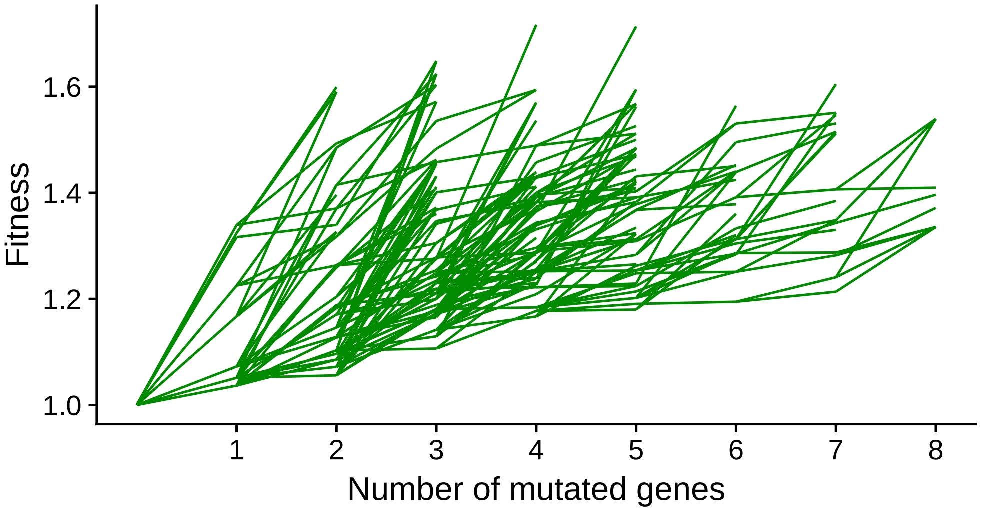


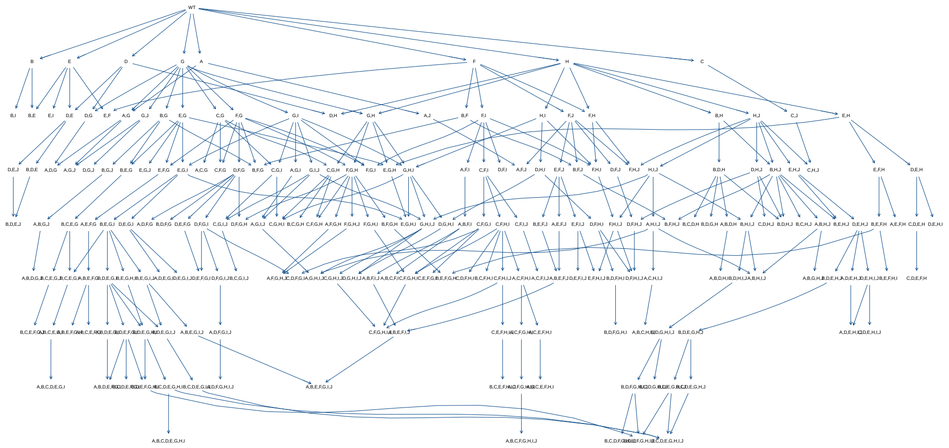

*ID* Ug5ffs10lfz6uMbgO

*acc genotypes* 225

*peaks no back* 65

*removed edges* NA

*prop rem edges* NA

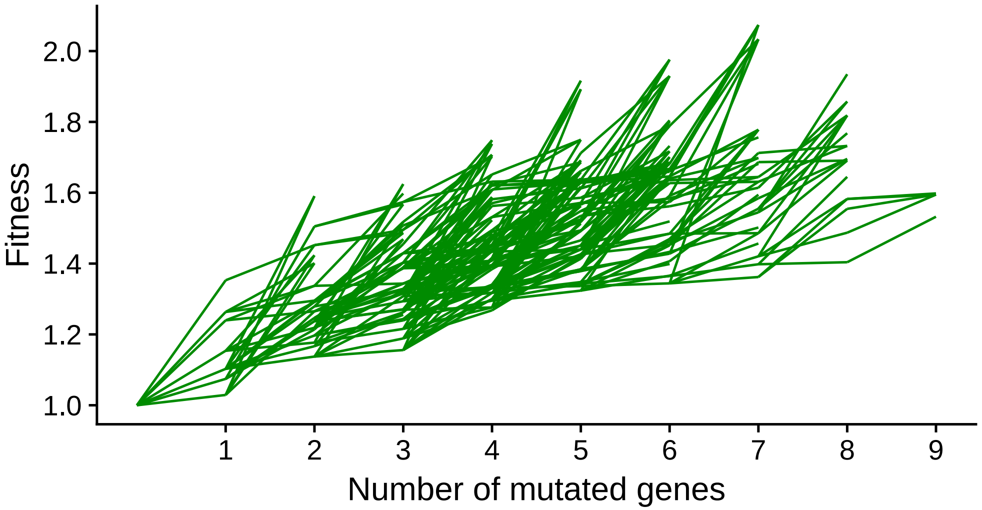


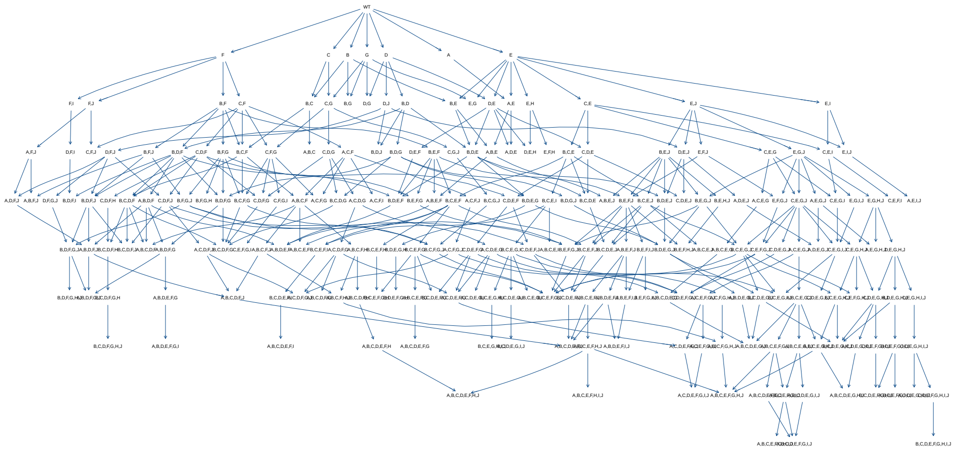

| | |
|---|---|
| *ID* | UmunqSzUuUKy8o5I |
| *acc genotypes* | 172 |
| *peaks no back* | 56 |
| *removed edges* | NA |
| *prop rem edges* | NA |

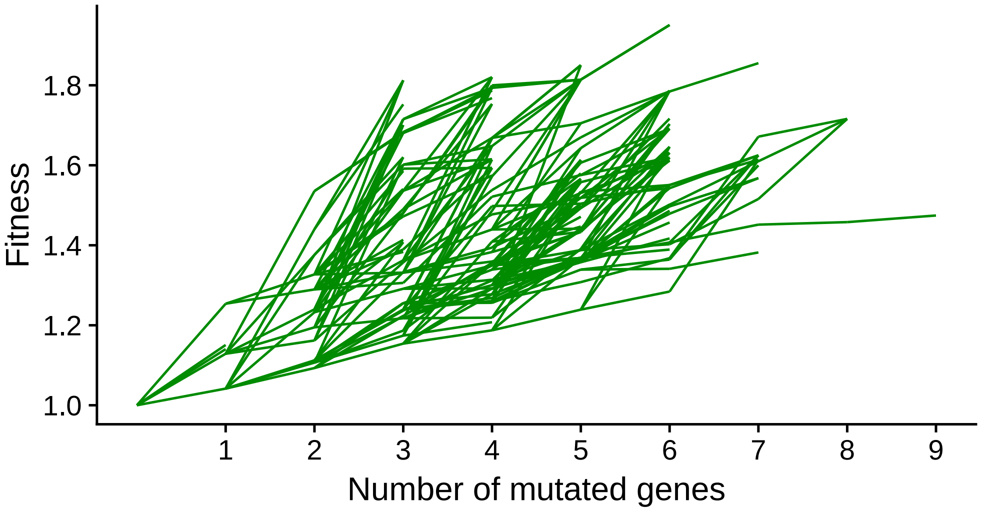


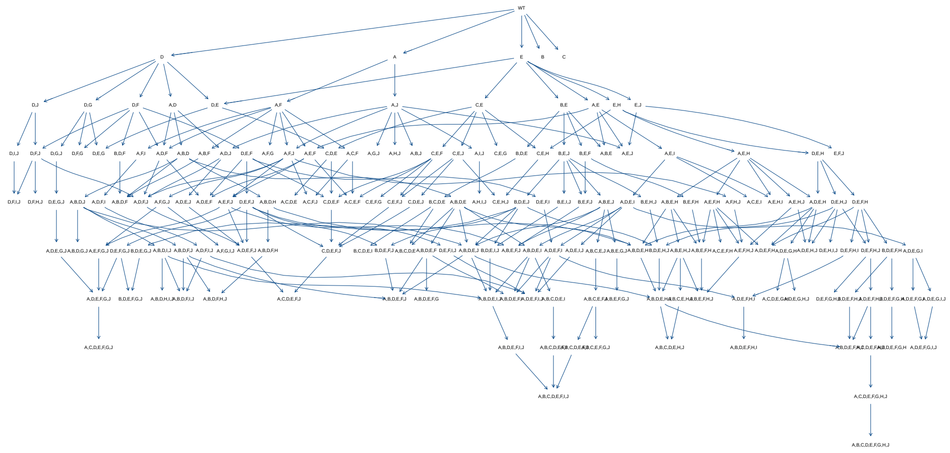

| | |
|---|---|
| *ID* | UOqyH3eyYqb108SQa |
| *acc genotypes* | 360 |
| *peaks no back* | 71 |
| *removed edges* | NA |
| *prop rem edges* | NA |

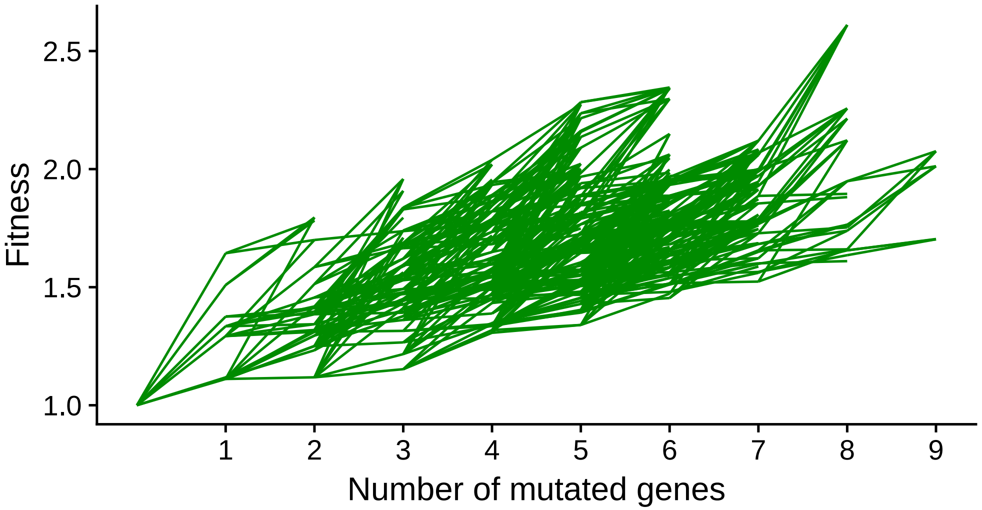


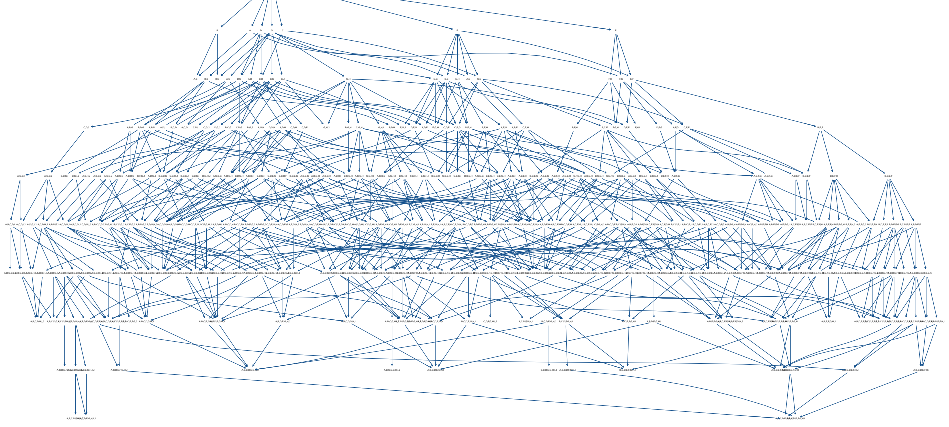

| | |
|---|---|
| *ID* | UpYY82ecse62Y3kB |
| *acc genotypes* | 102 |
| *peaks no back* | 28 |
| *removed edges* | NA |
| *prop rem edges* | NA |

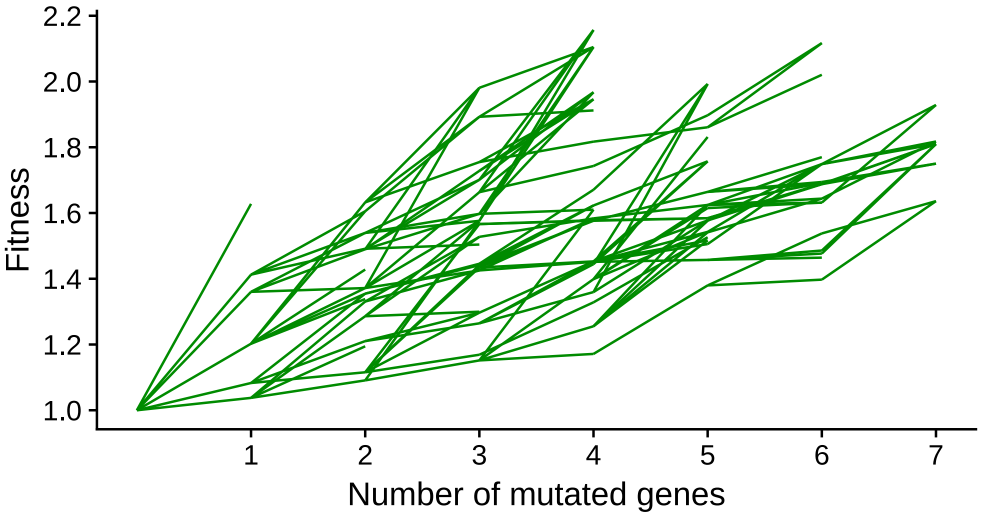


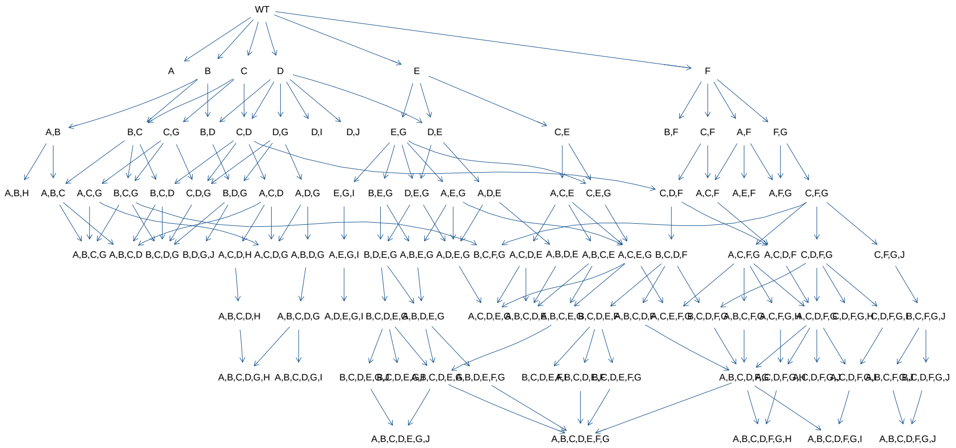

| | |
|---|---|
| *ID* | UR8nbozIYY5kAyVm |
| *acc genotypes* | 238 |
| *peaks no back* | 54 |
| *removed edges* | NA |
| *prop rem edges* | NA |

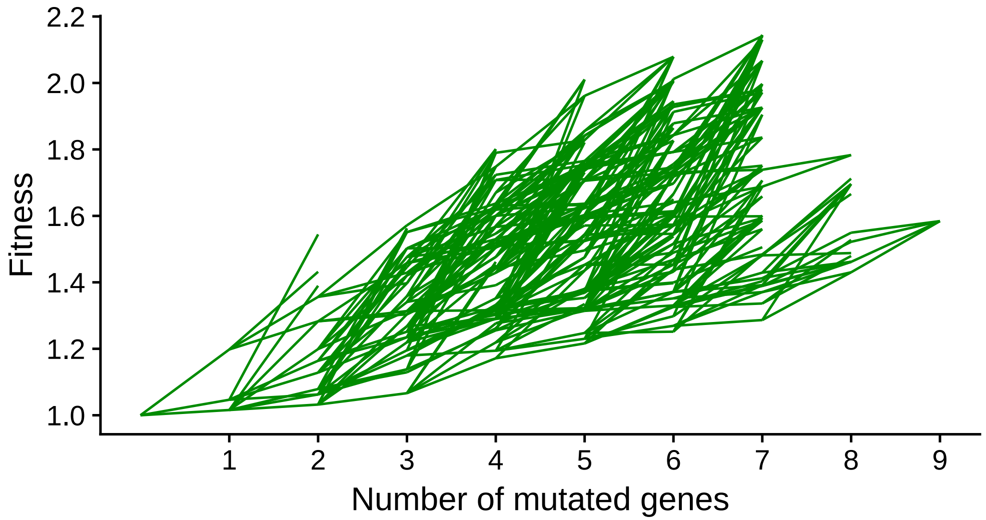


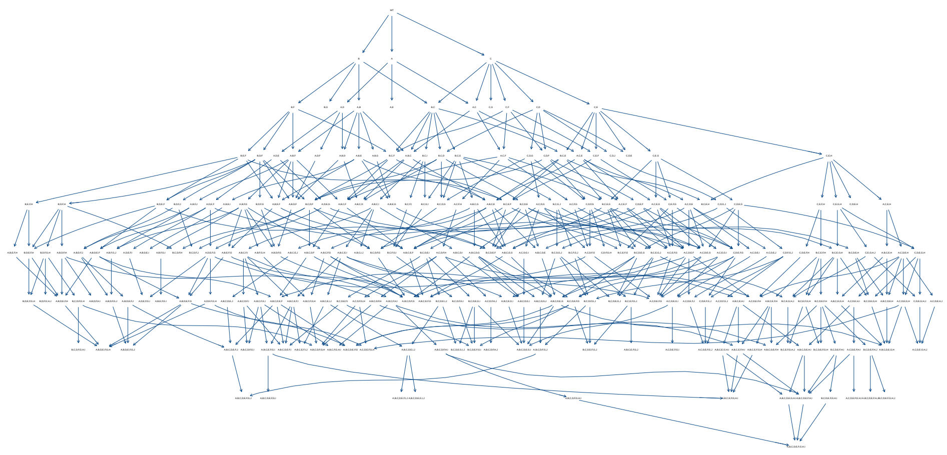

| | |
|---|---|
| *ID* | vhfO10CYjbTCUd5Oc |
| *acc genotypes* | 189 |
| *peaks no back* | 49 |
| *removed edges* | NA |
| *prop rem edges* | NA |

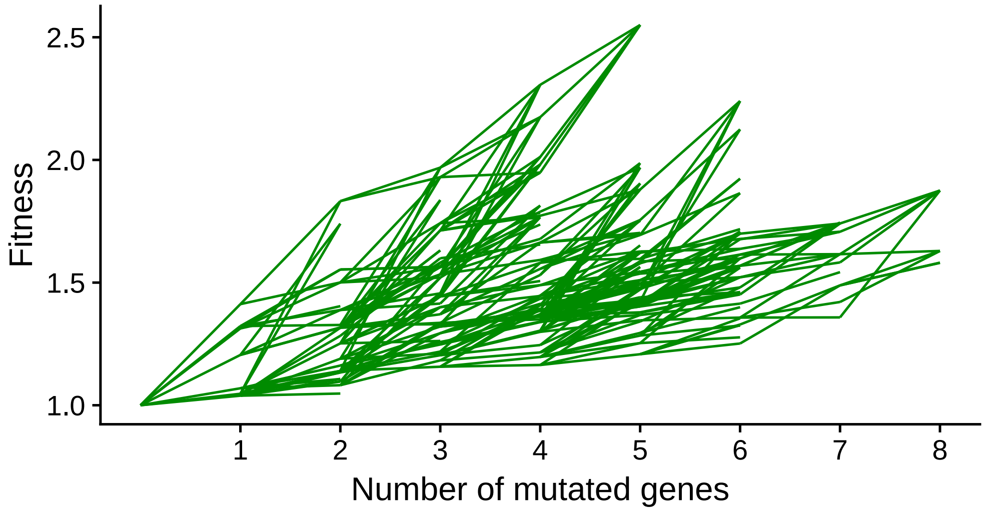


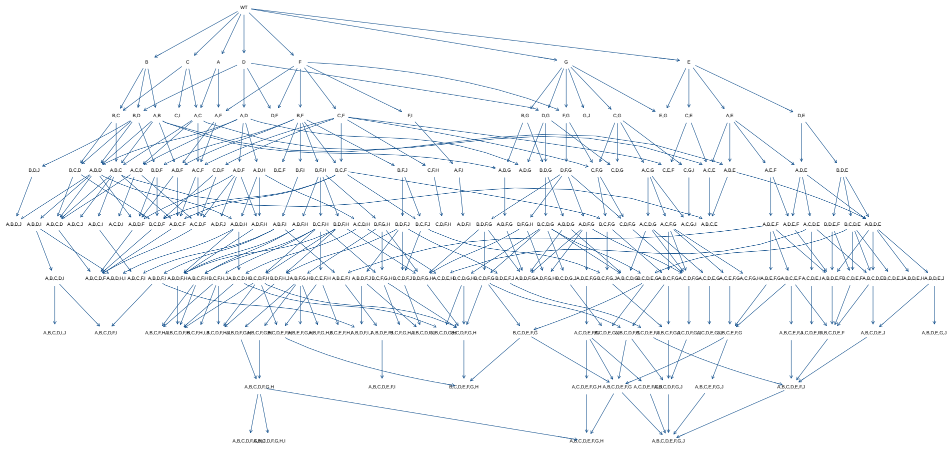

| | |
|---|---|
| *ID* | VLbJrKnG7tDZJCX7 |
| *acc genotypes* | 738 |
| *peaks no back* | 105 |
| *removed edges* | NA |
| *prop rem edges* | NA |

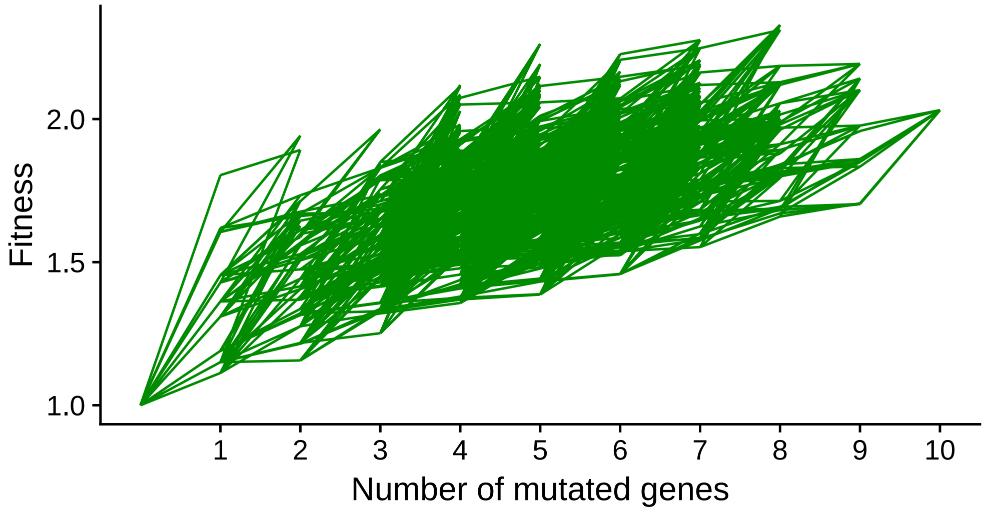


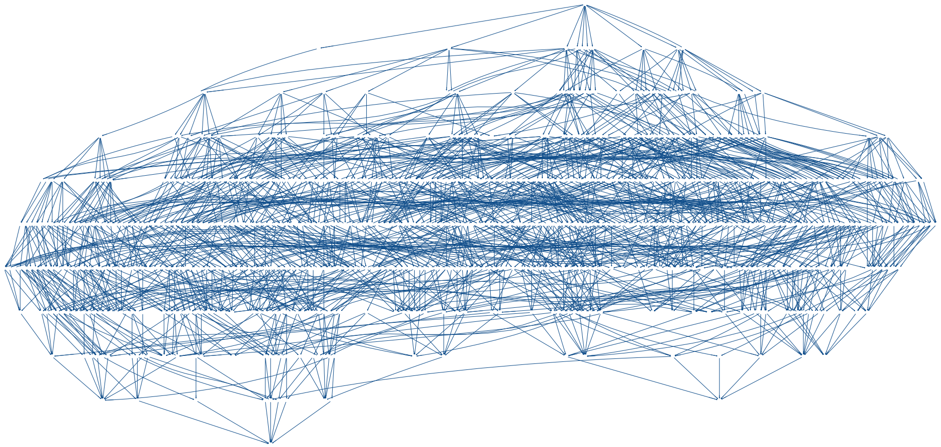

| | |
|---|---|
| *ID* | vlOhJ52j3mkgx4bO |
| *acc genotypes* | 179 |
| *peaks no back* | 73 |
| *removed edges* | NA |
| *prop rem edges* | NA |

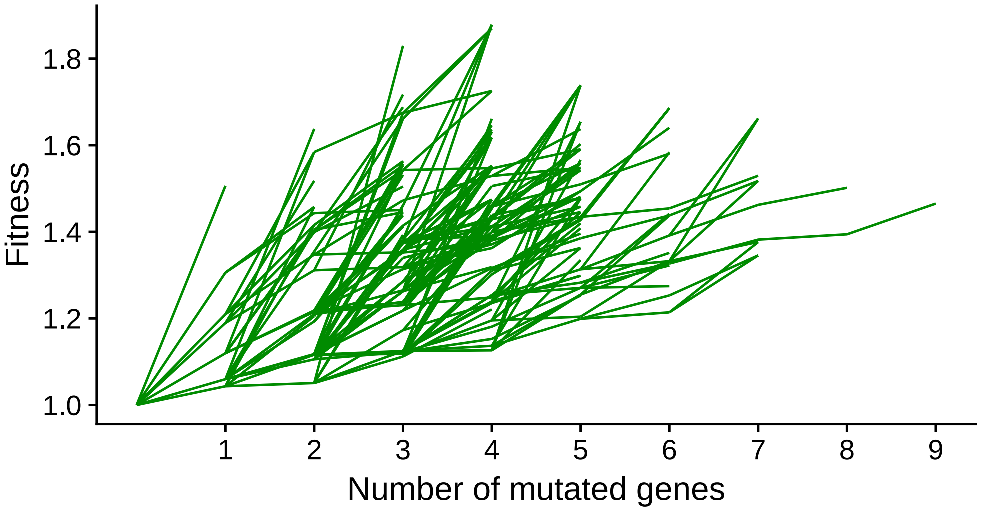


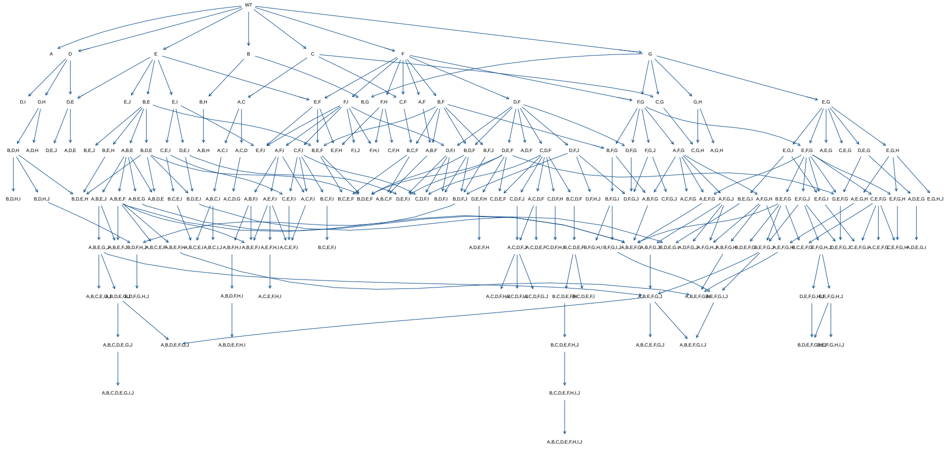

| | |
|---|---|
| *ID* | W10IjZVnZ1043hQUxi |
| *acc genotypes* | 388 |
| *peaks no back* | 70 |
| *removed edges* | NA |
| *prop rem edges* | NA |

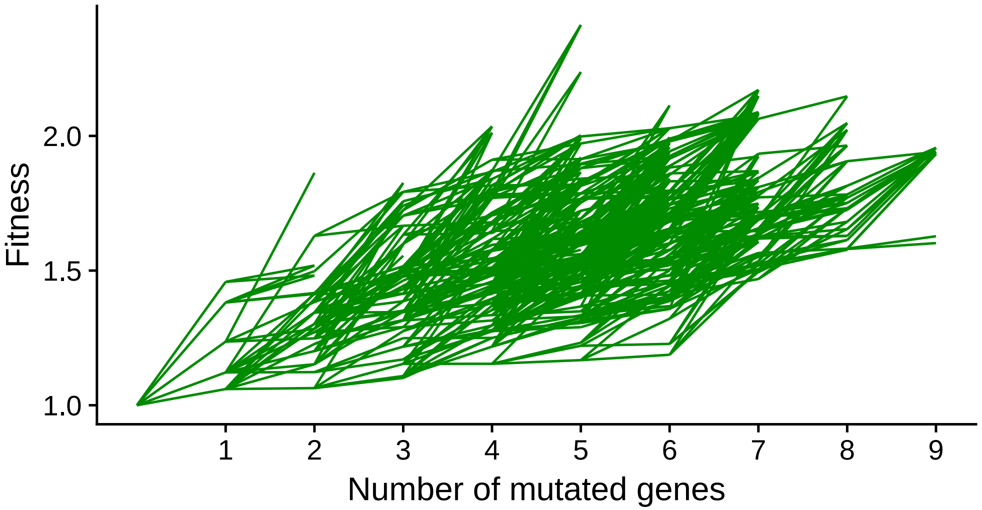


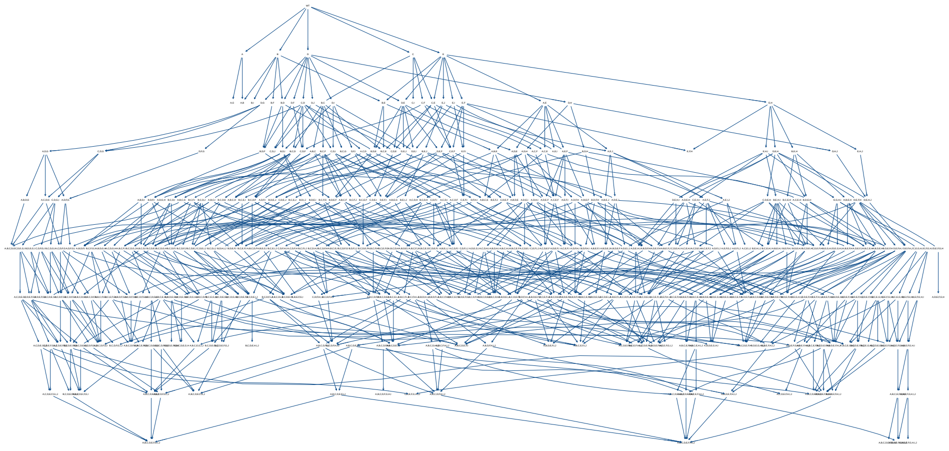

| | |
|---|---|
| *ID* | wM4mouEZ5iUOoAtX |
| *acc genotypes* | 143 |
| *peaks no back* | 52 |
| *removed edges* | NA |
| *prop rem edges* | NA |

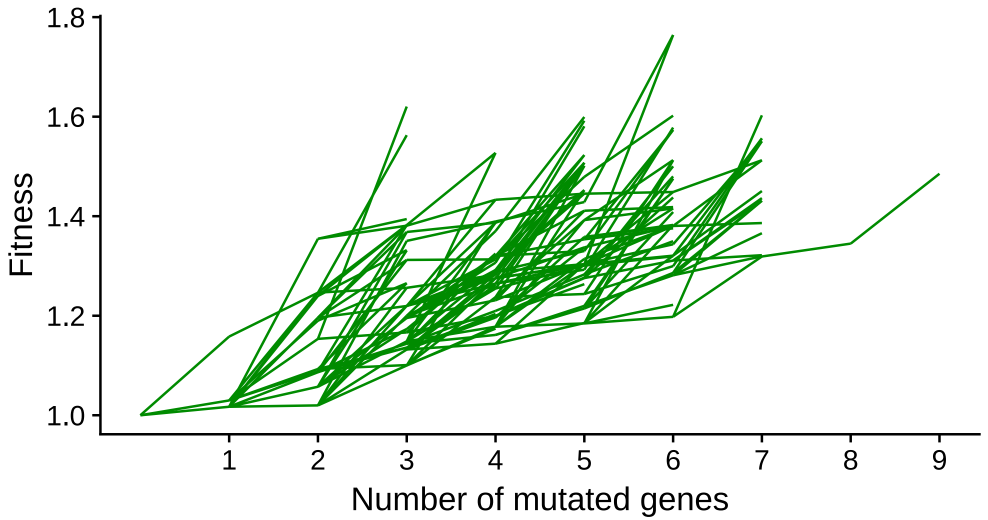


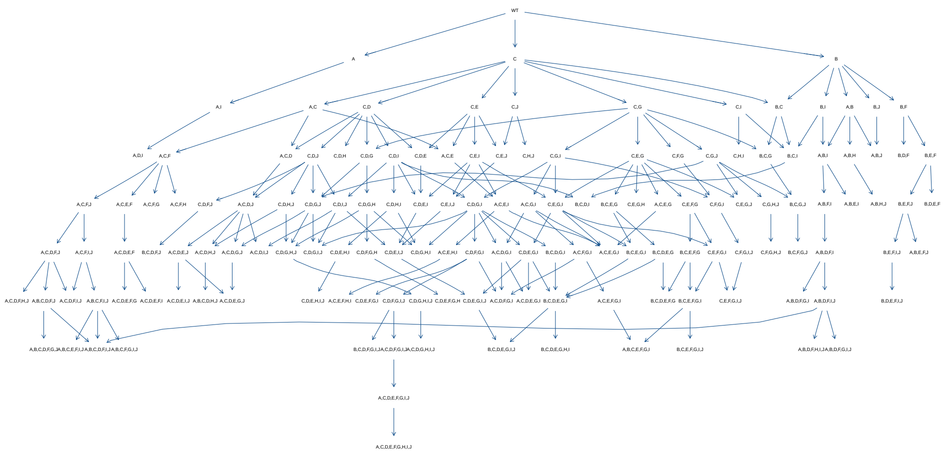

| | |
|---|---|
| *ID* | wOuBIsYKAAzBpTkz |
| *acc genotypes* | 191 |
| *peaks no back* | 69 |
| *removed edges* | NA |
| *prop rem edges* | NA |

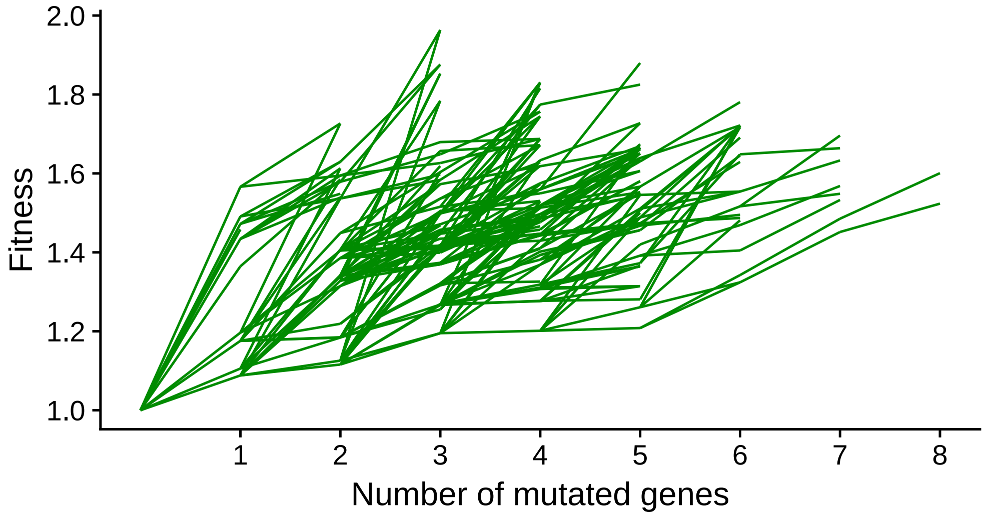


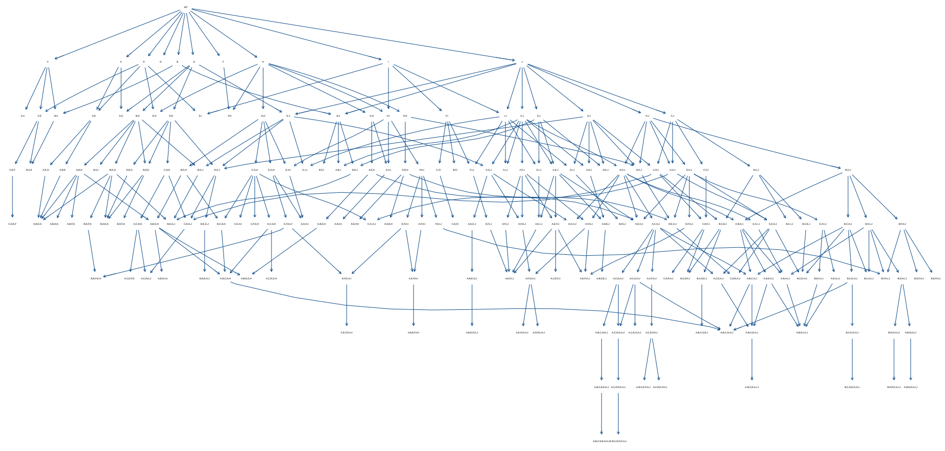

| | |
|---|---|
| *ID* | wuF5q9kOxSCRE7RX |
| *acc genotypes* | 728 |
| *peaks no back* | 82 |
| *removed edges* | NA |
| *prop rem edges* | NA |

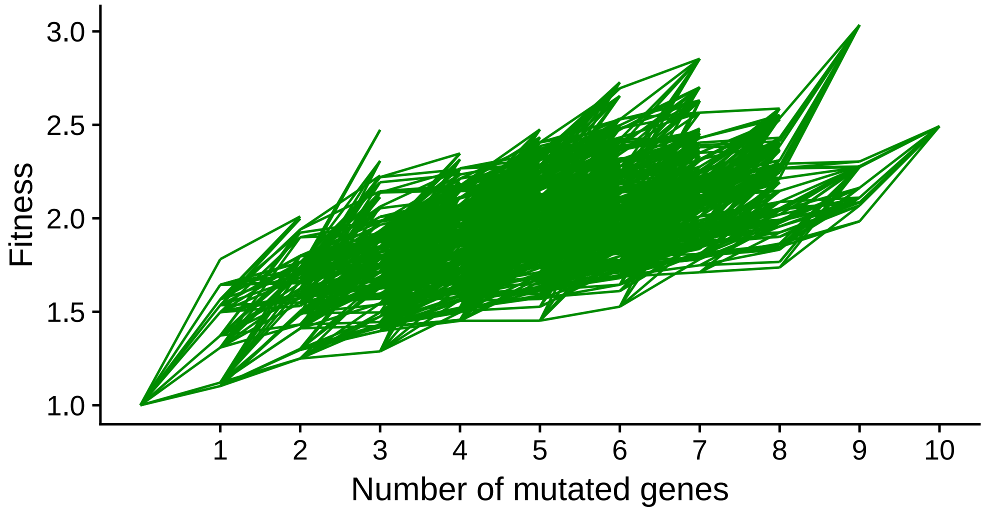


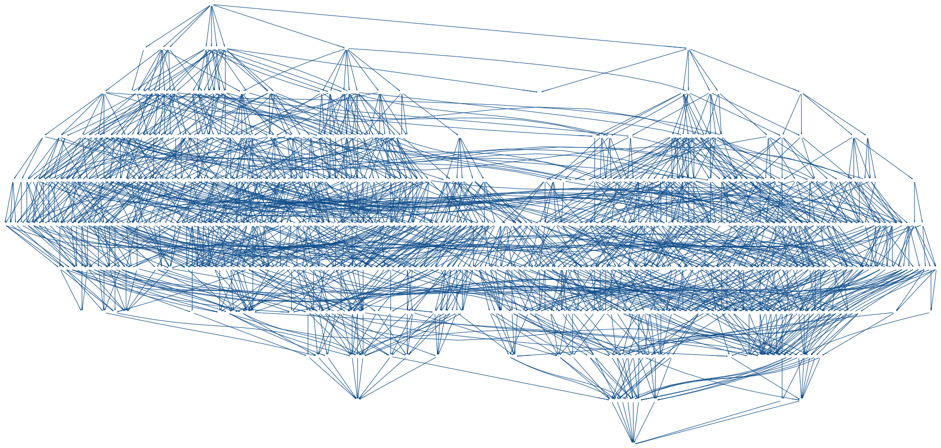

| | |
|---|---|
| *ID* | wwBtyDIZSVYRmVqi |
| *acc genotypes* | 222 |
| *peaks no back* | 83 |
| *removed edges* | NA |
| *prop rem edges* | NA |

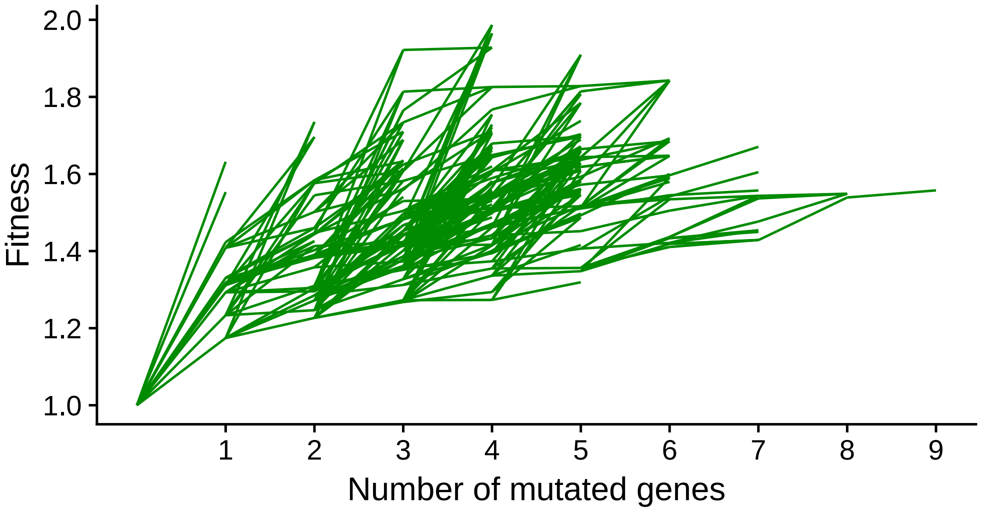


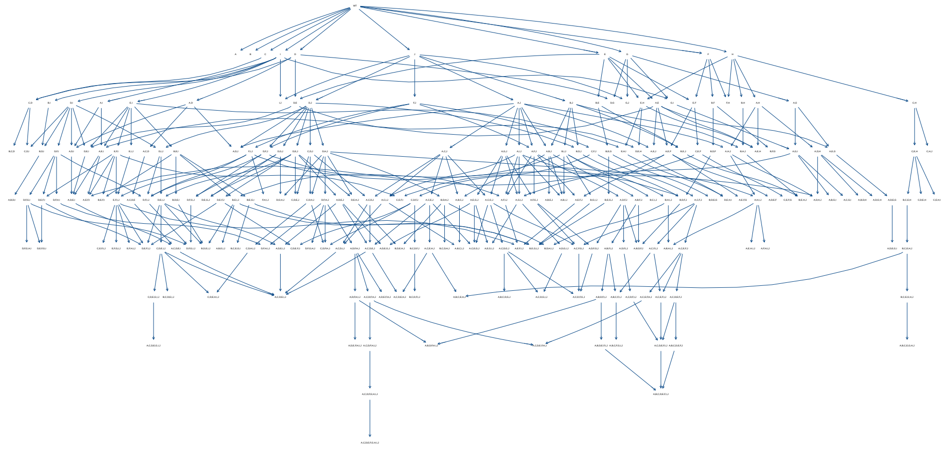

| | |
|---|---|
| *ID* | wXZJGUyFMo10Nl4dV |
| *acc genotypes* | 89 |
| *peaks no back* | 42 |
| *removed edges* | NA |
| *prop rem edges* | NA |

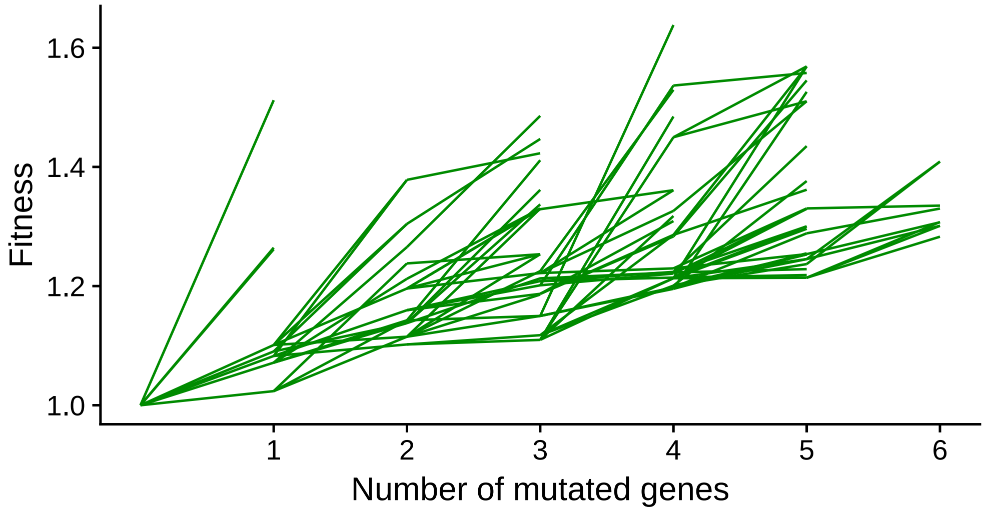


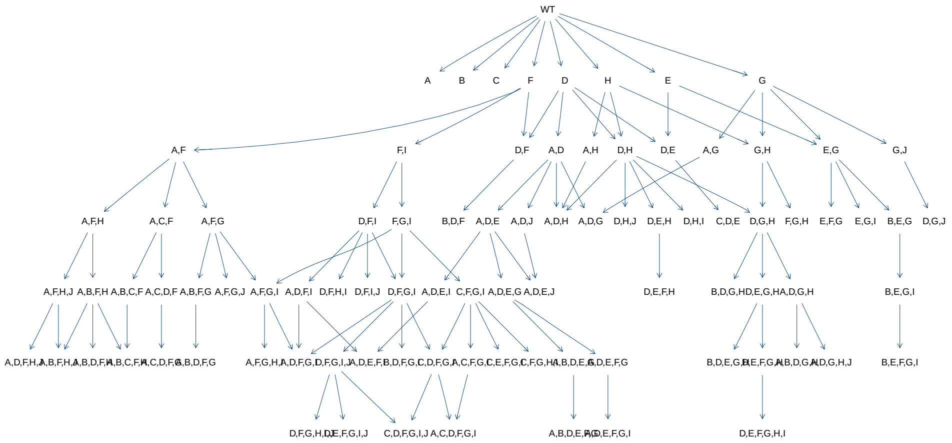

| | |
|---|---|
| *ID* | WYBRgQ10JLk3RhGLo |
| *acc genotypes* | 433 |
| *peaks no back* | 115 |
| *removed edges* | NA |
| *prop rem edges* | NA |

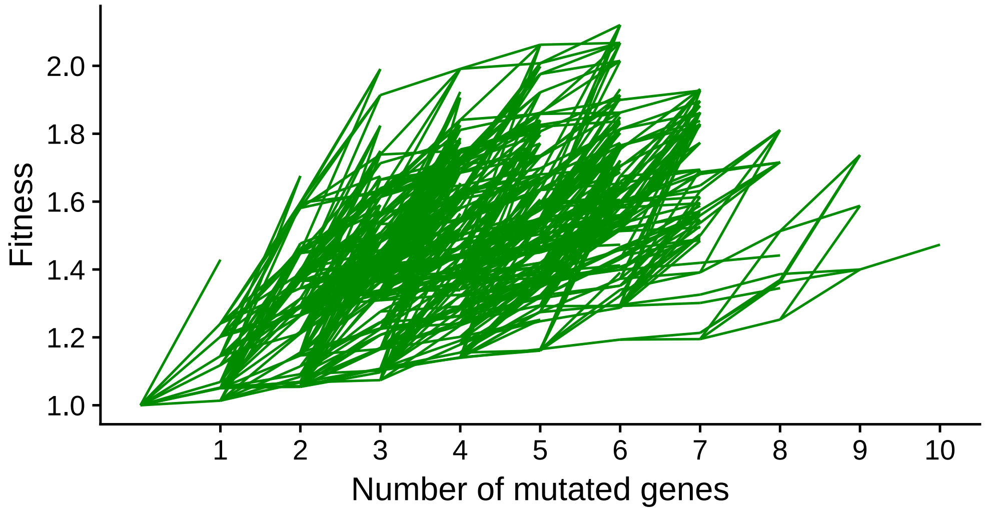


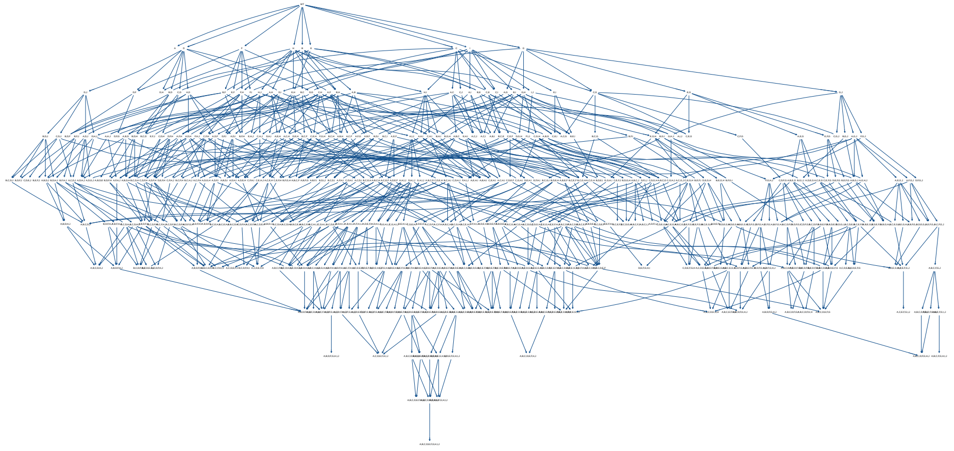

| | |
|---|---|
| *ID* | WZ96wrbzeKdxcOJ3 |
| *acc genotypes* | 141 |
| *peaks no back* | 57 |
| *removed edges* | NA |
| *prop rem edges* | NA |

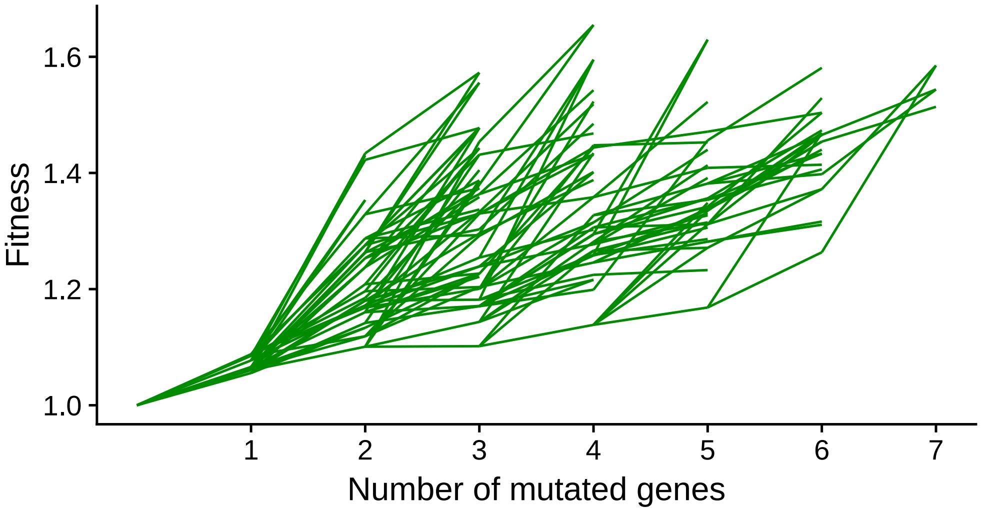


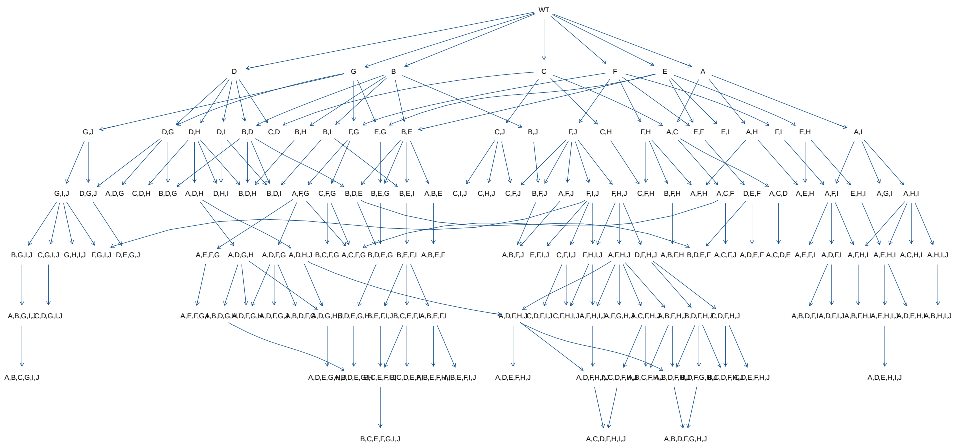

| | |
|---|---|
| *ID* | X6Z5TgU29GZVilQy |
| *acc genotypes* | 620 |
| *peaks no back* | 75 |
| *removed edges* | NA |
| *prop rem edges* | NA |

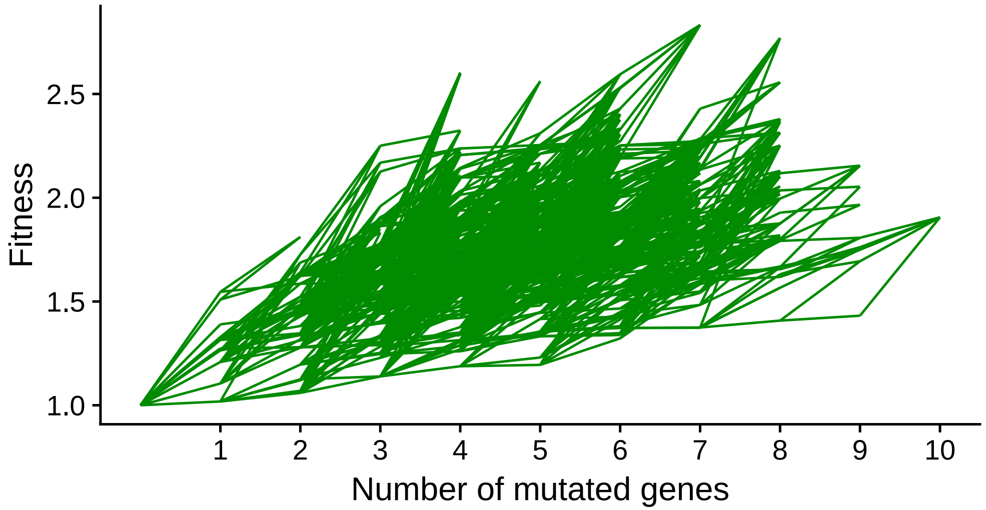


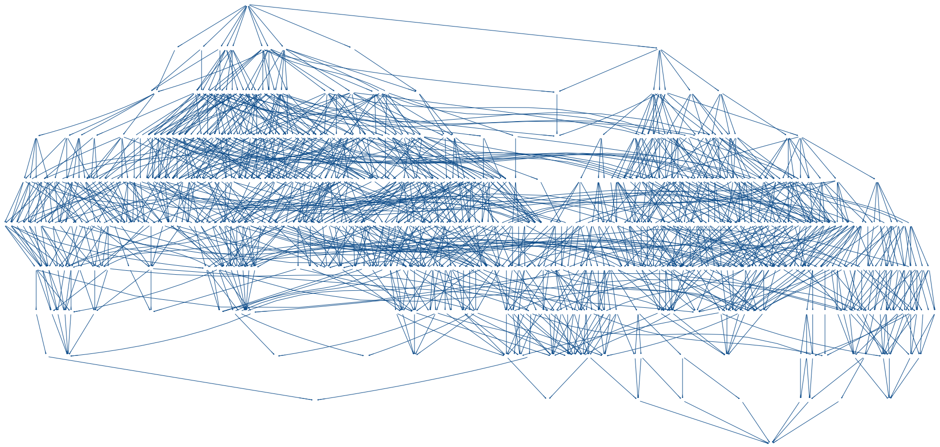

| | |
|---|---|
| *ID* | x8IUOobt10WI39iG2 |
| *acc genotypes* | 209 |
| *peaks no back* | 81 |
| *removed edges* | NA |
| *prop rem edges* | NA |

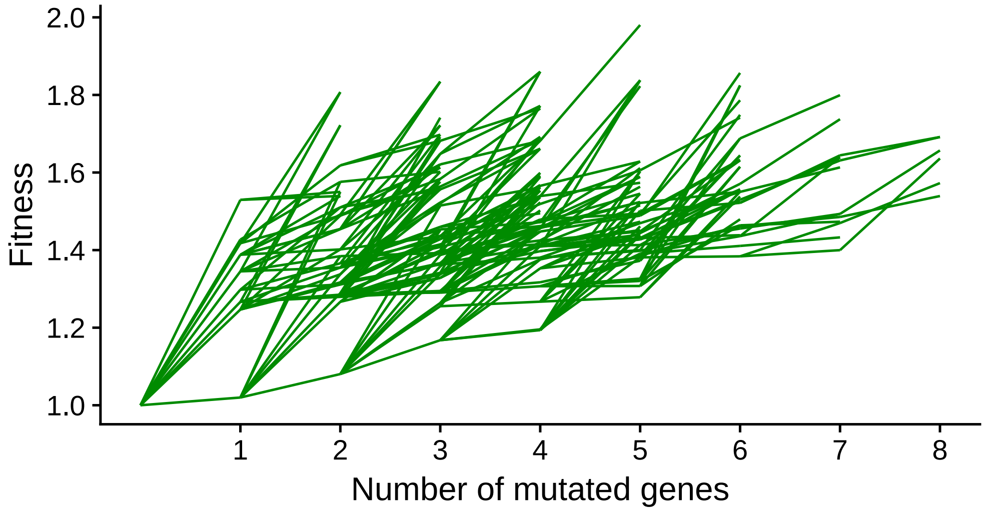


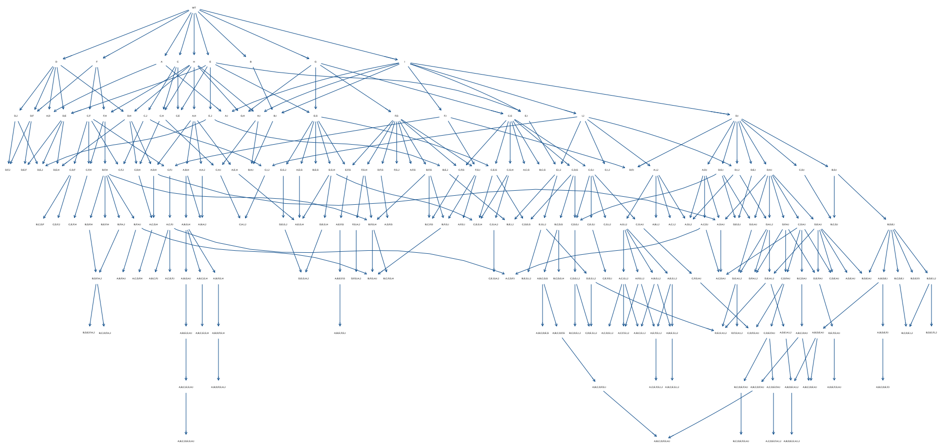

| | |
|---|---|
| *ID* | xcdQXsmxI7kh5NDZ |
| *acc genotypes* | 184 |
| *peaks no back* | 73 |
| *removed edges* | NA |
| *prop rem edges* | NA |

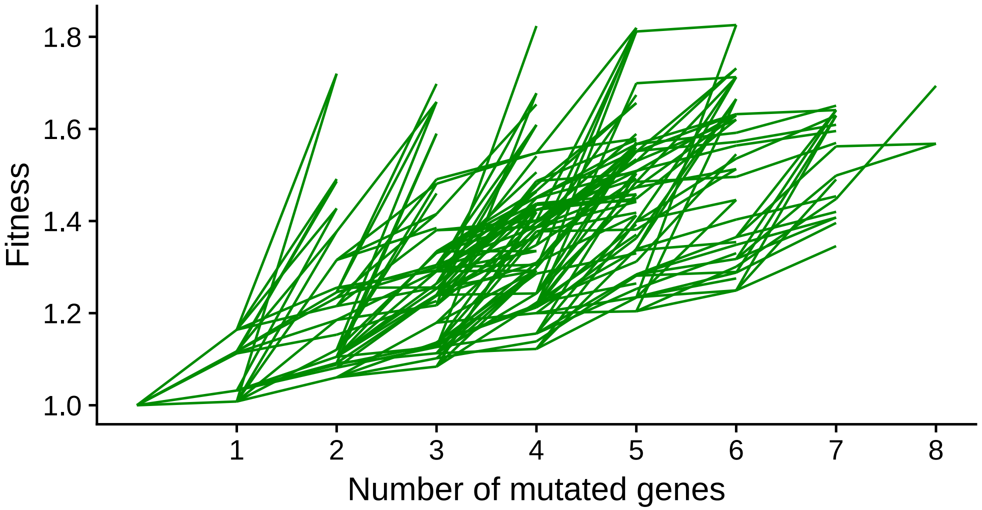


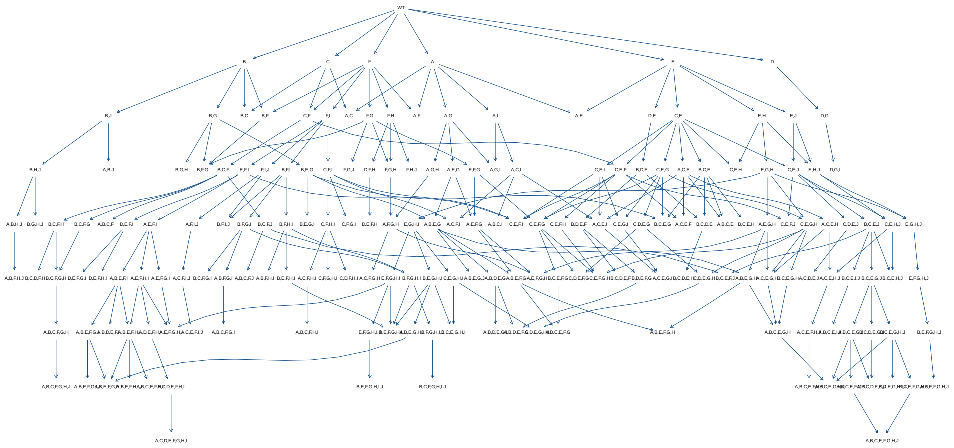

| | |
|---|---|
| *ID* | Xh10HKozlpe9ybd7i |
| *acc genotypes* | 200 |
| *peaks no back* | 89 |
| *removed edges* | NA |
| *prop rem edges* | NA |

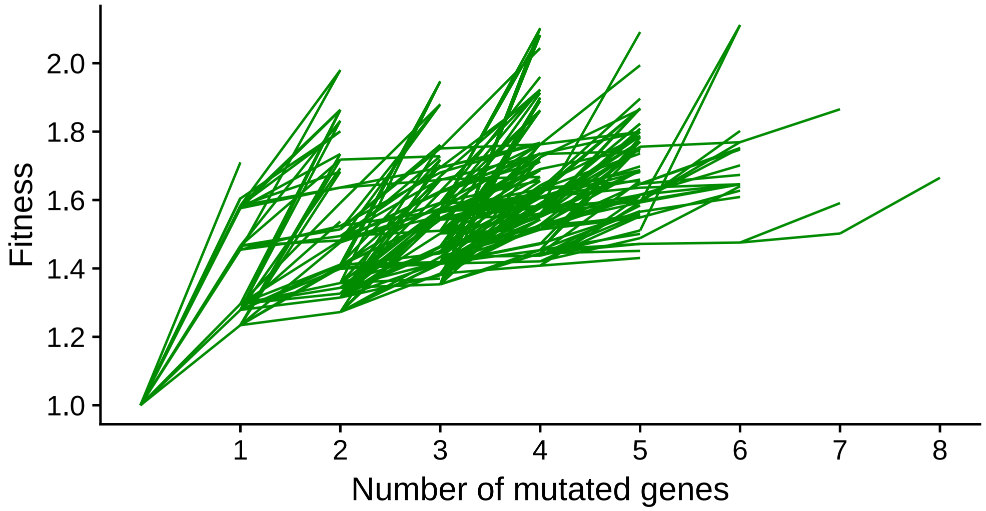


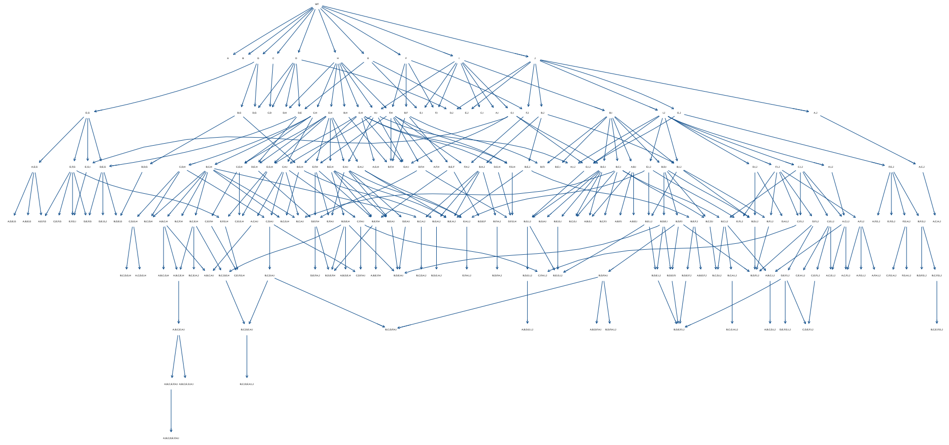

| | |
|---|---|
| *ID* | XKTMaTNASYWPlHNl |
| *acc genotypes* | 352 |
| *peaks no back* | 105 |
| *removed edges* | NA |
| *prop rem edges* | NA |

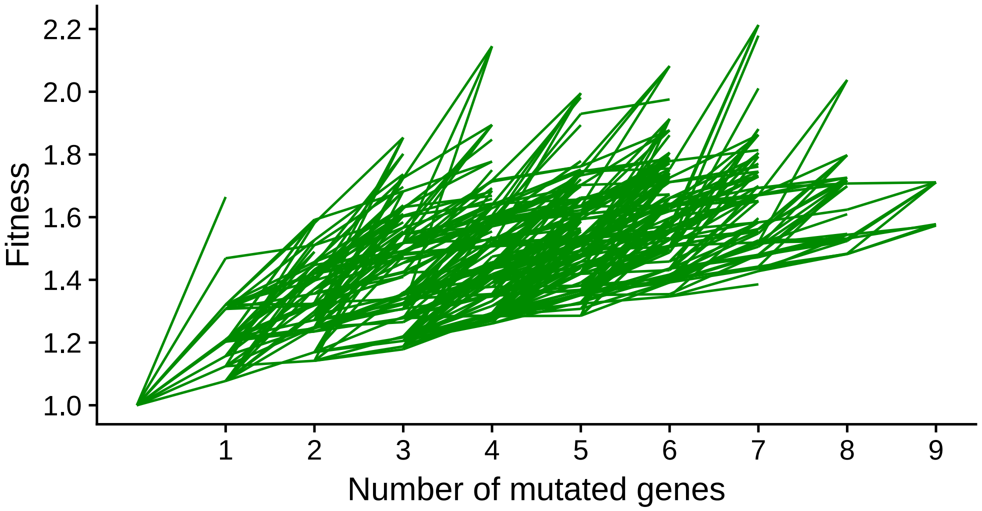


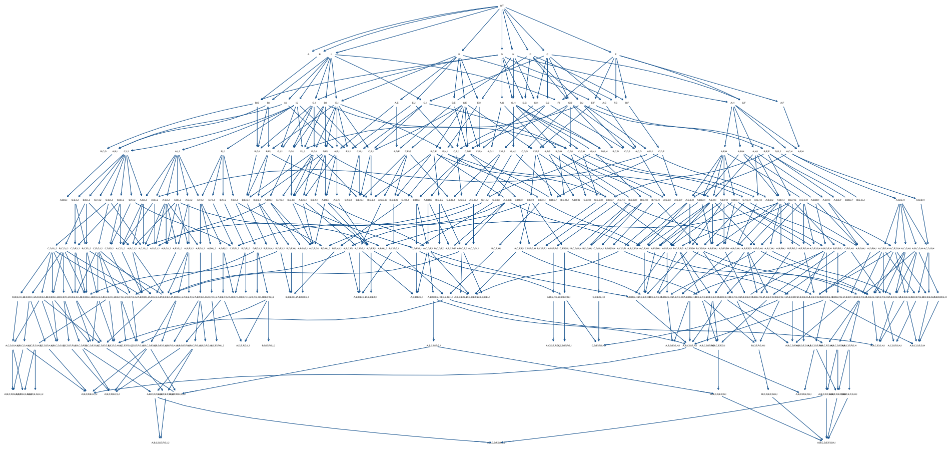

| | |
|---|---|
| *ID* | XNRZEeV3l3oHG9g1 |
| *acc genotypes* | 424 |
| *peaks no back* | 121 |
| *removed edges* | NA |
| *prop rem edges* | NA |

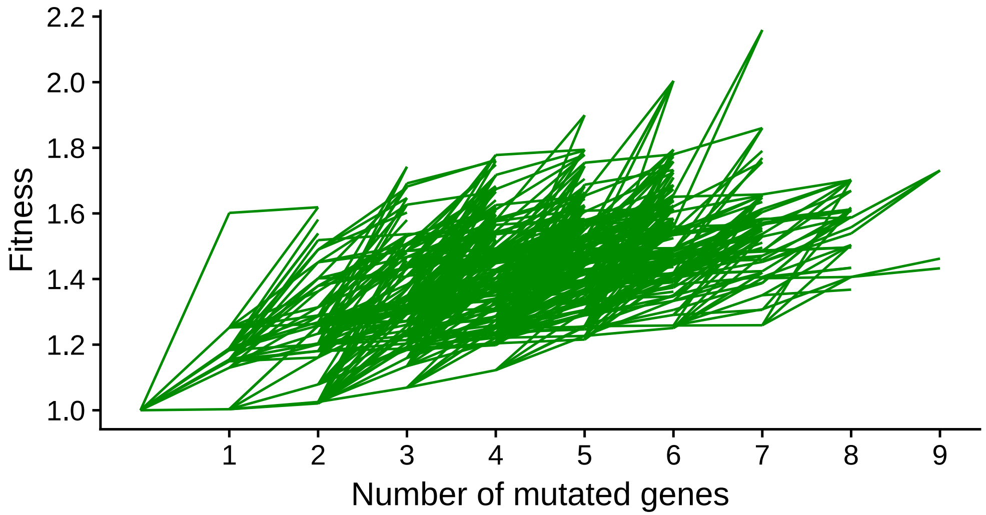


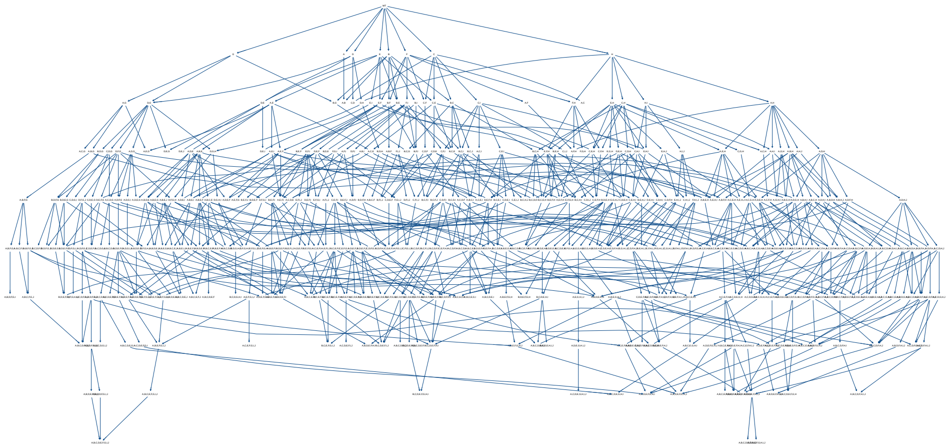

| | |
|---|---|
| *ID* | XrjsetS7KL5noi92 |
| *acc genotypes* | 89 |
| *peaks no back* | 26 |
| *removed edges* | NA |
| *prop rem edges* | NA |

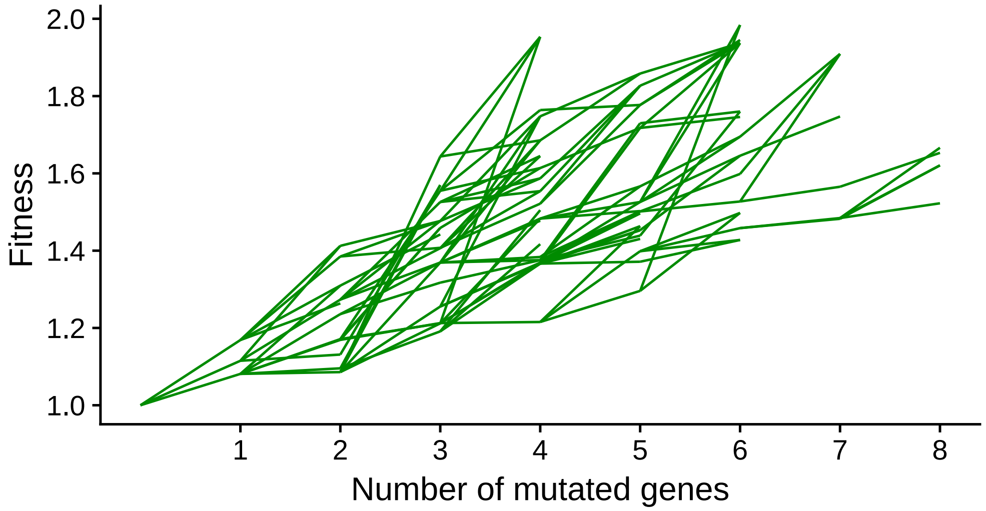


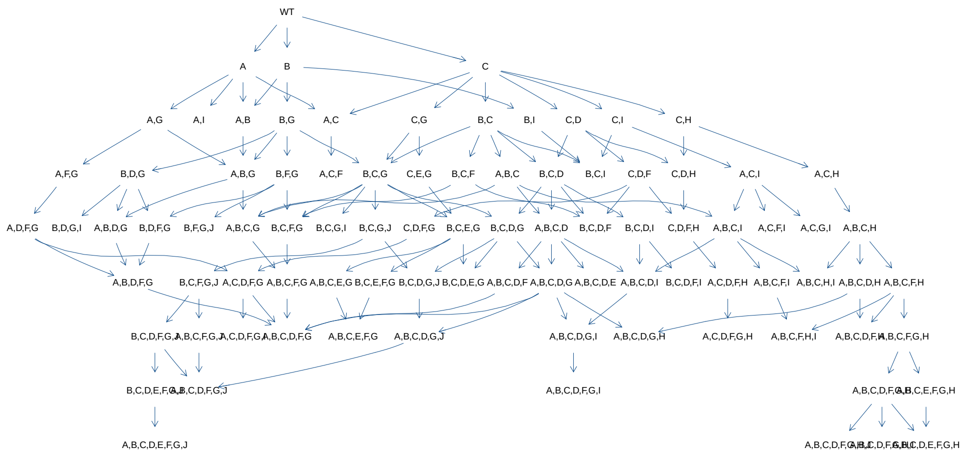

| | |
|---|---|
| *ID* | xvHgdwCzOGa7Ev7e |
| *acc genotypes* | 55 |
| *peaks no back* | 19 |
| *removed edges* | NA |
| *prop rem edges* | NA |

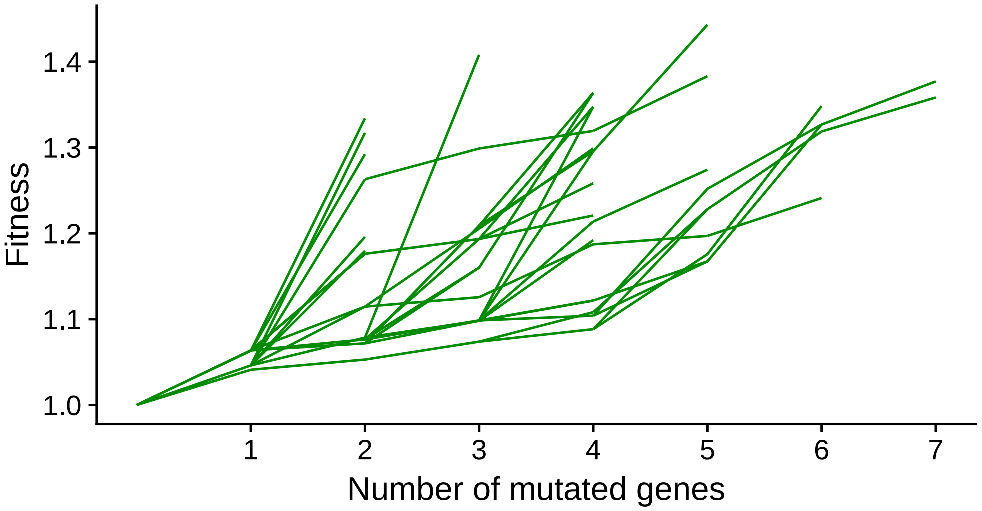


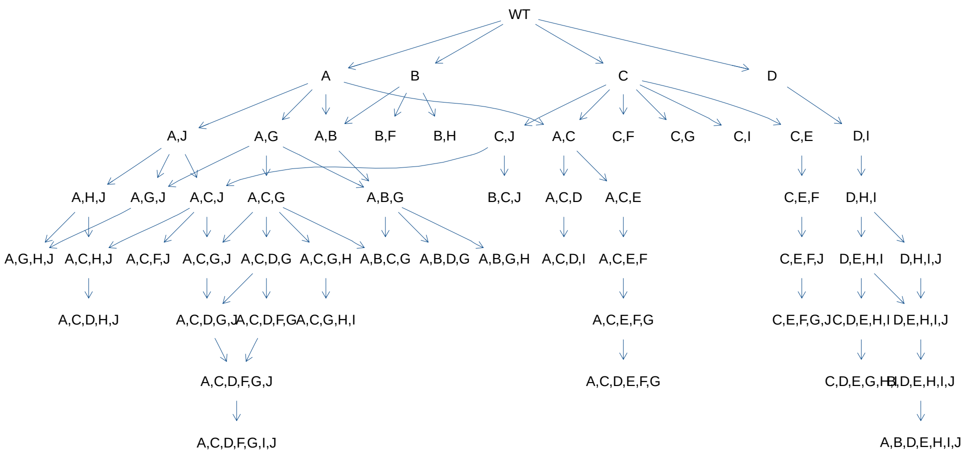

| | |
|---|---|
| *ID* | xwEXGxQHVPacEqsW |
| *acc genotypes* | 67 |
| *peaks no back* | 26 |
| *removed edges* | NA |
| *prop rem edges* | NA |

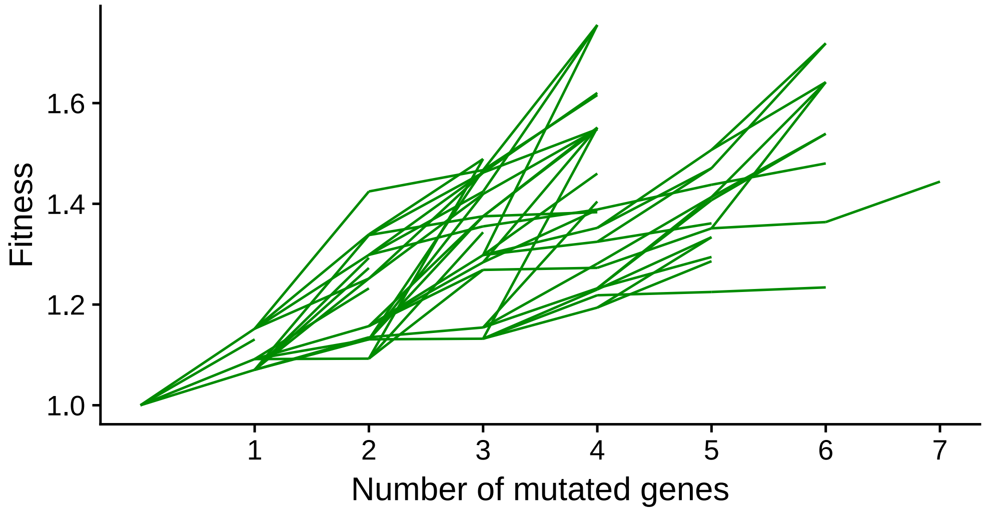


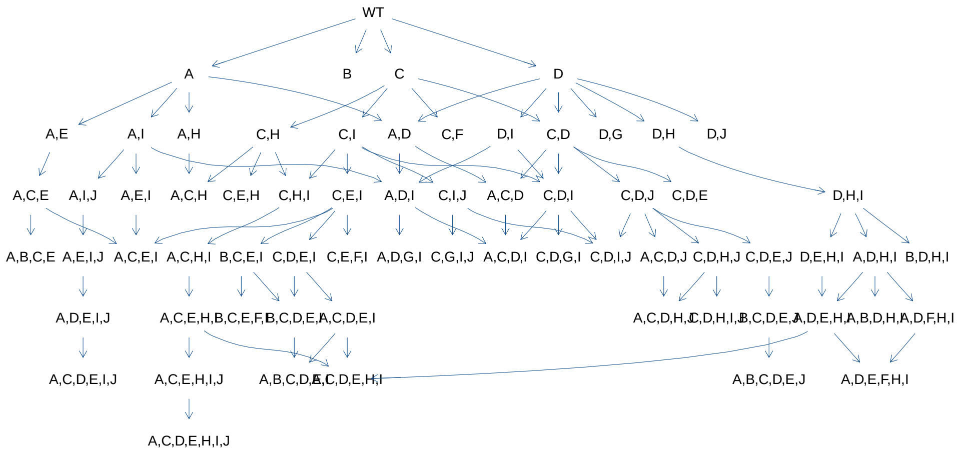

| | |
|---|---|
| *ID* | xZ632I1FnE2fDF10q |
| *acc genotypes* | 663 |
| *peaks no back* | 72 |
| *removed edges* | NA |
| *prop rem edges* | NA |

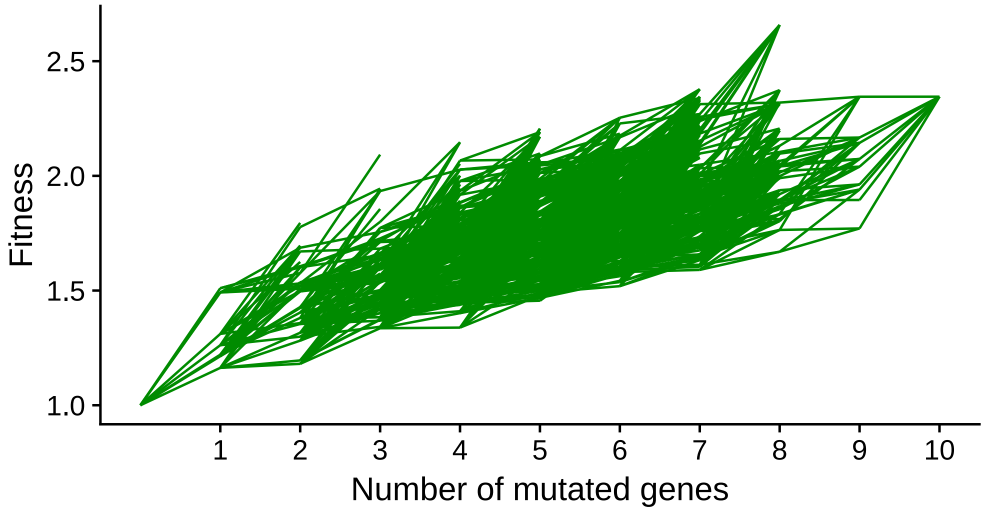


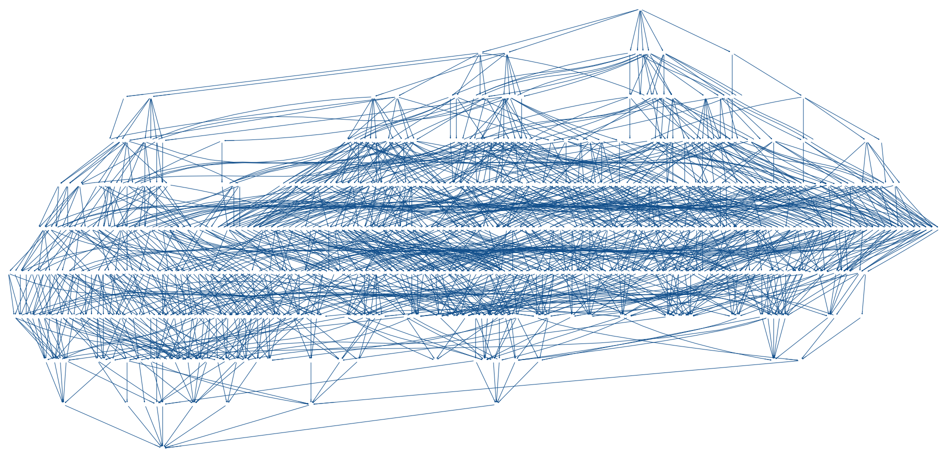

| | |
|---|---|
| *ID* | y5oNzDs3V3q7y3ih |
| *acc genotypes* | 297 |
| *peaks no back* | 91 |
| *removed edges* | NA |
| *prop rem edges* | NA |

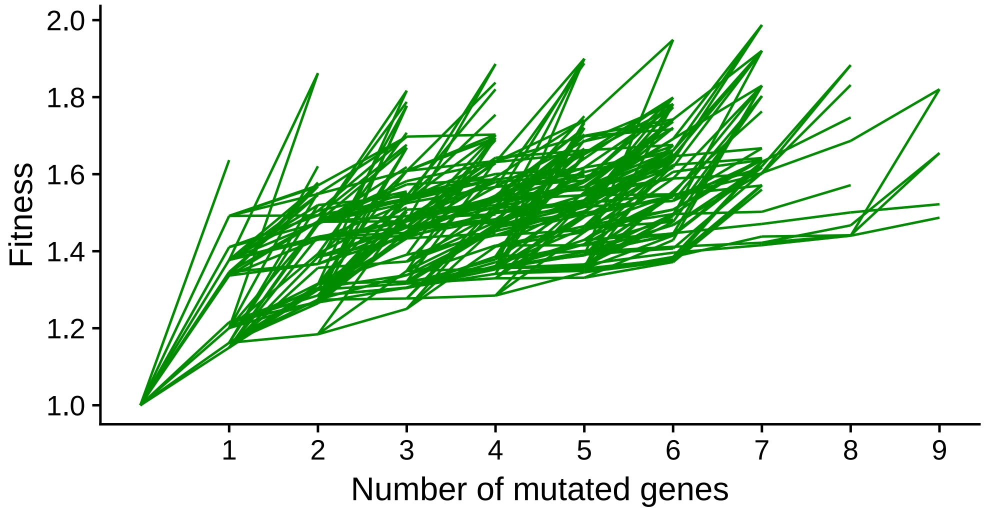


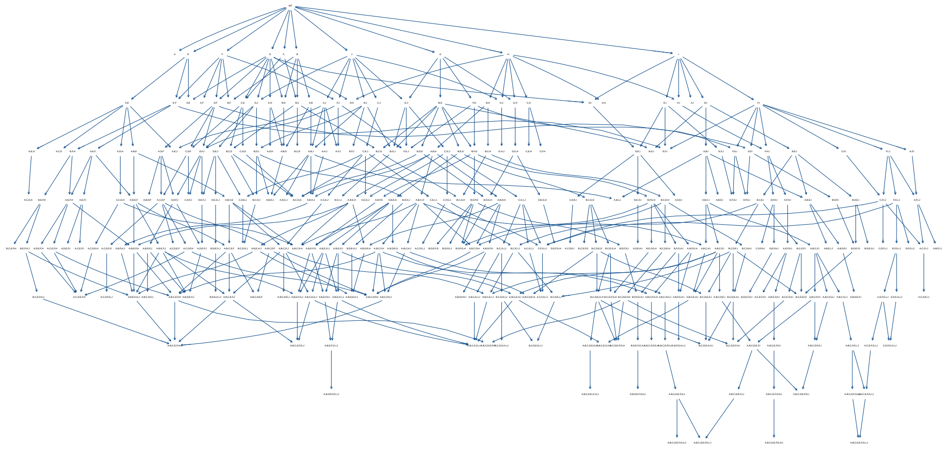

| | |
|---|---|
| *ID* | YeUw4gB10IT8FRdhX |
| *acc genotypes* | 110 |
| *peaks no back* | 46 |
| *removed edges* | NA |
| *prop rem edges* | NA |

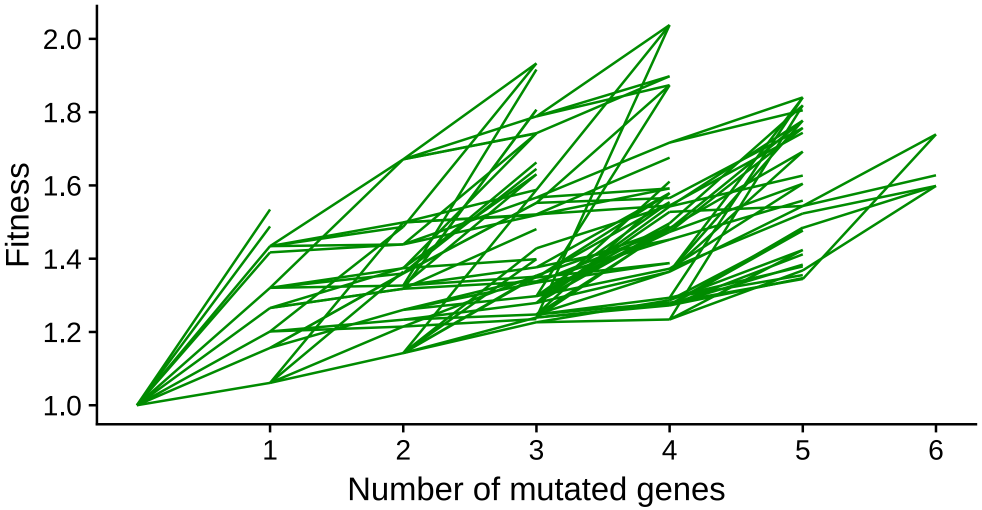


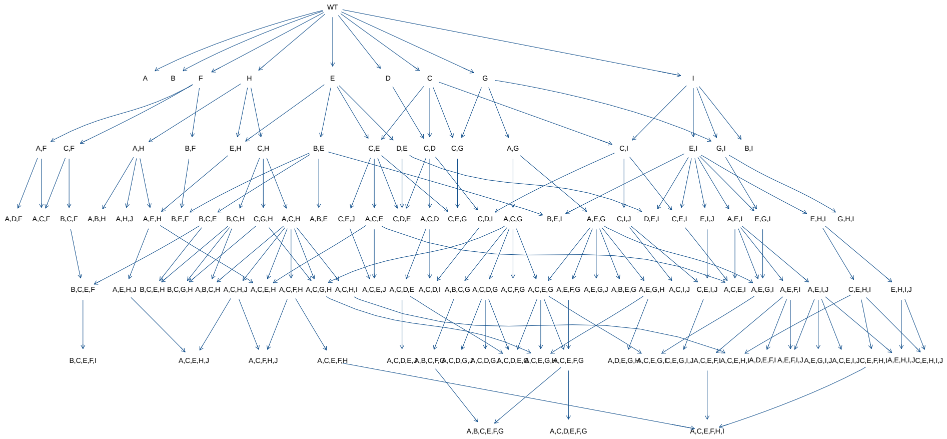

| | |
|---|---|
| *ID* | Yg8zLevZ7wA4DC3B |
| *acc genotypes* | 147 |
| *peaks no back* | 48 |
| *removed edges* | NA |
| *prop rem edges* | NA |

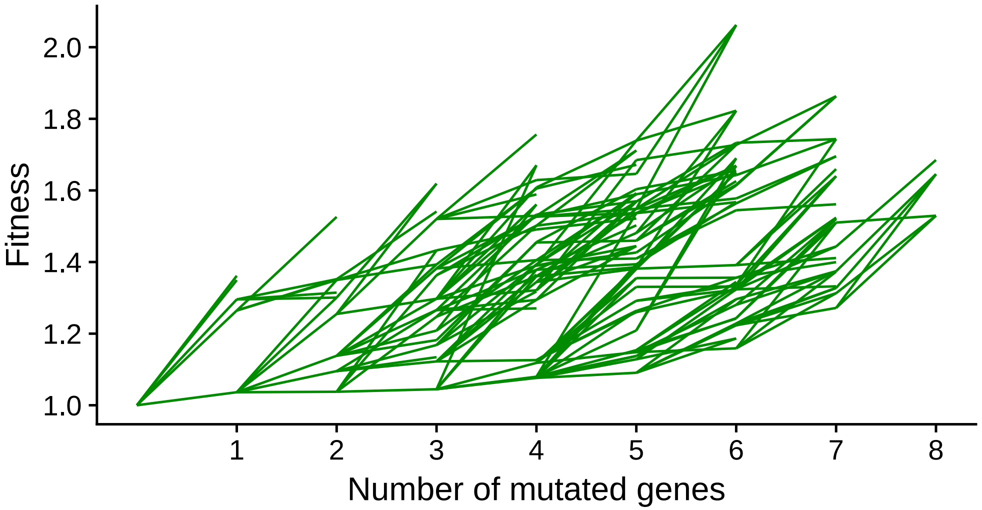


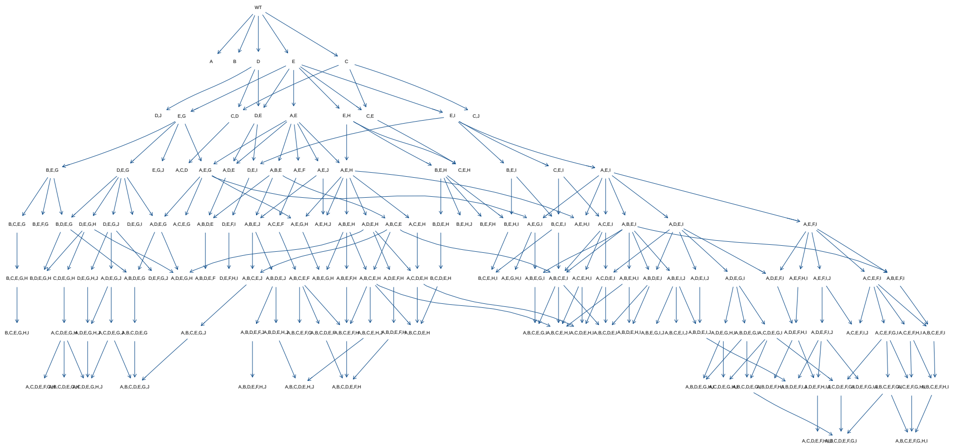

| | |
|---|---|
| *ID* | YqlzMXE6SnRlTBq5 |
| *acc genotypes* | 73 |
| *peaks no back* | 37 |
| *removed edges* | NA |
| *prop rem edges* | NA |

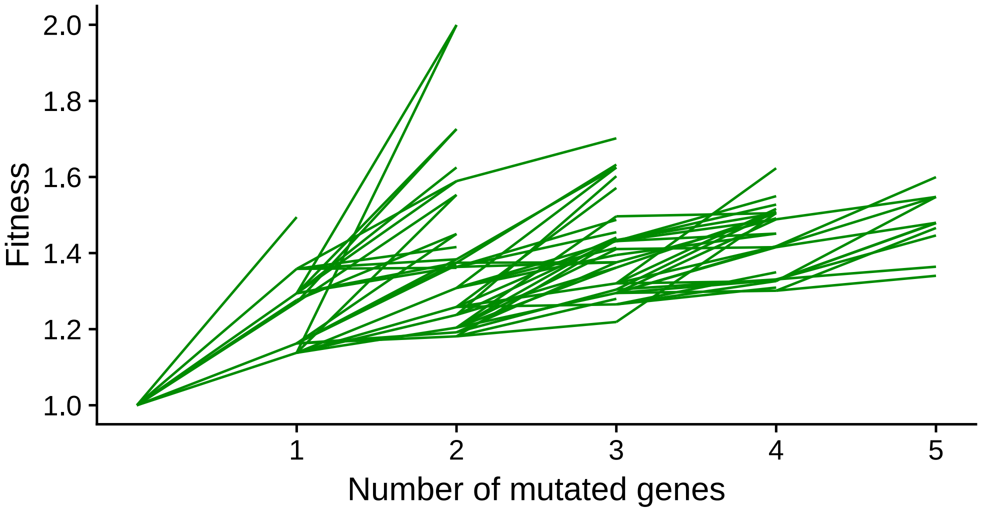


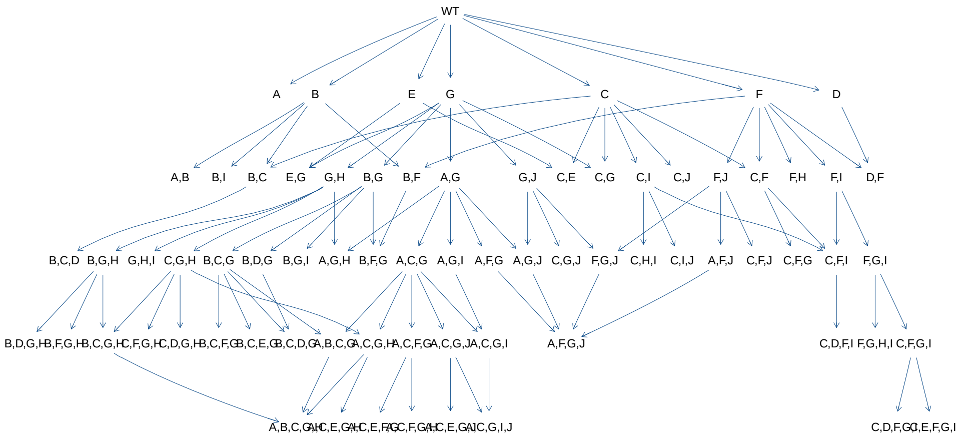

| | |
|---|---|
| *ID* | YSwbWvqbUeR10s1p8 |
| *acc genotypes* | 279 |
| *peaks no back* | 73 |
| *removed edges* | NA |
| *prop rem edges* | NA |

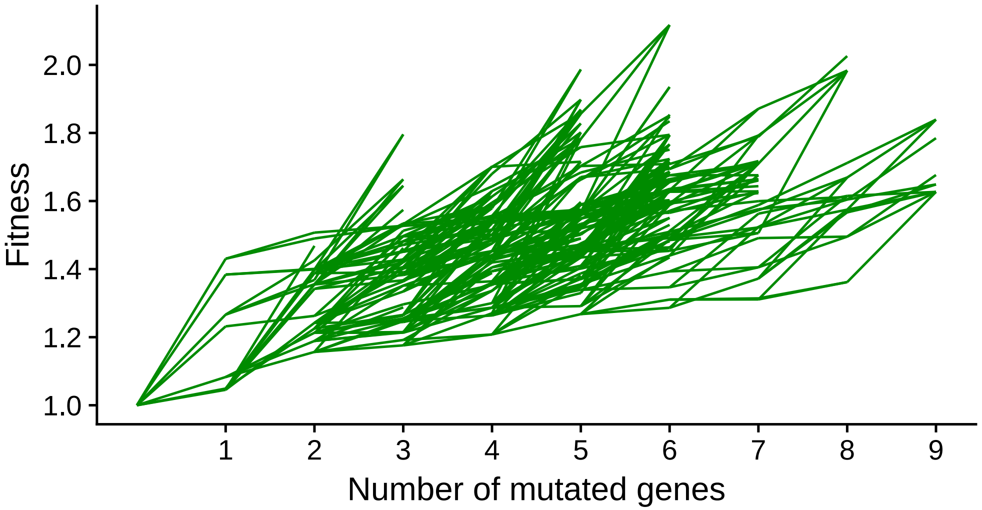


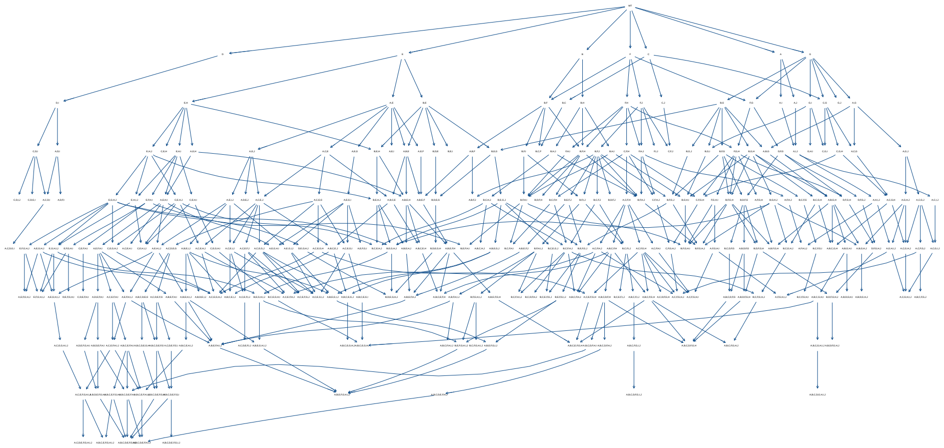

| | |
|---|---|
| *ID* | zaGCXdhS1lAYBVFr |
| *acc genotypes* | 242 |
| *peaks no back* | 91 |
| *removed edges* | NA |
| *prop rem edges* | NA |

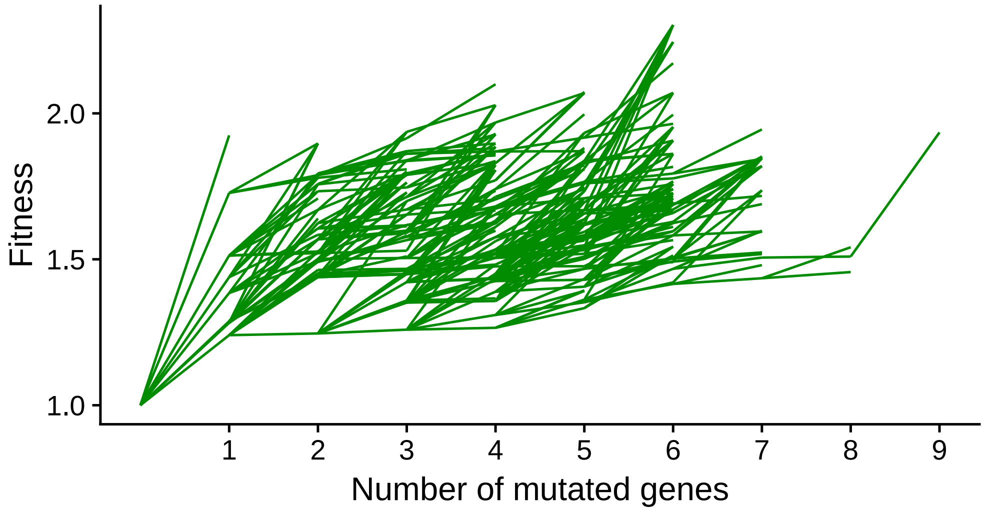


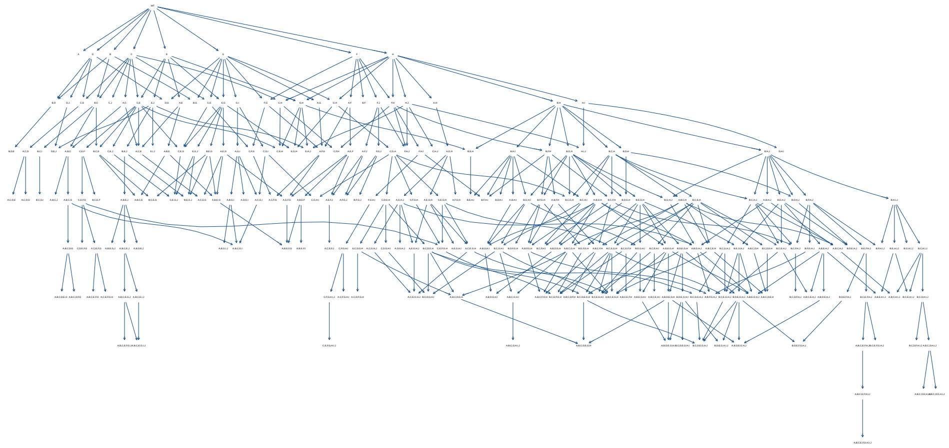

| | |
|---|---|
| *ID* | ZARoxejcvdMx5gej |
| *acc genotypes* | 335 |
| *peaks no back* | 87 |
| *removed edges* | NA |
| *prop rem edges* | NA |

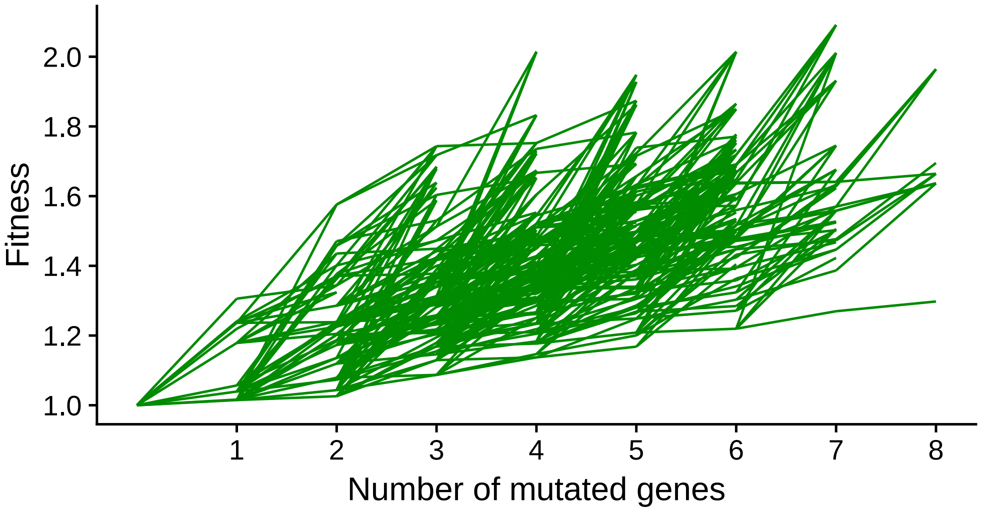


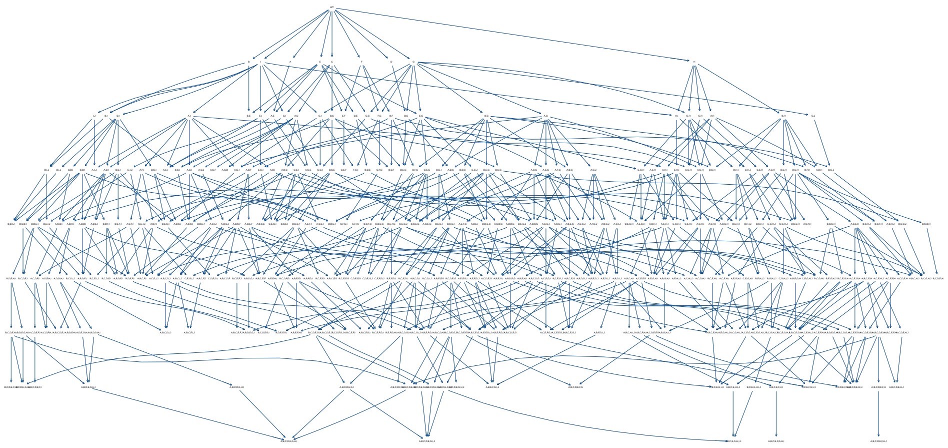

| | |
|---|---|
| *ID* | zdCbK9OwSeu3jN2k |
| *acc genotypes* | 26 |
| *peaks no back* | 12 |
| *removed edges* | NA |
| *prop rem edges* | NA |

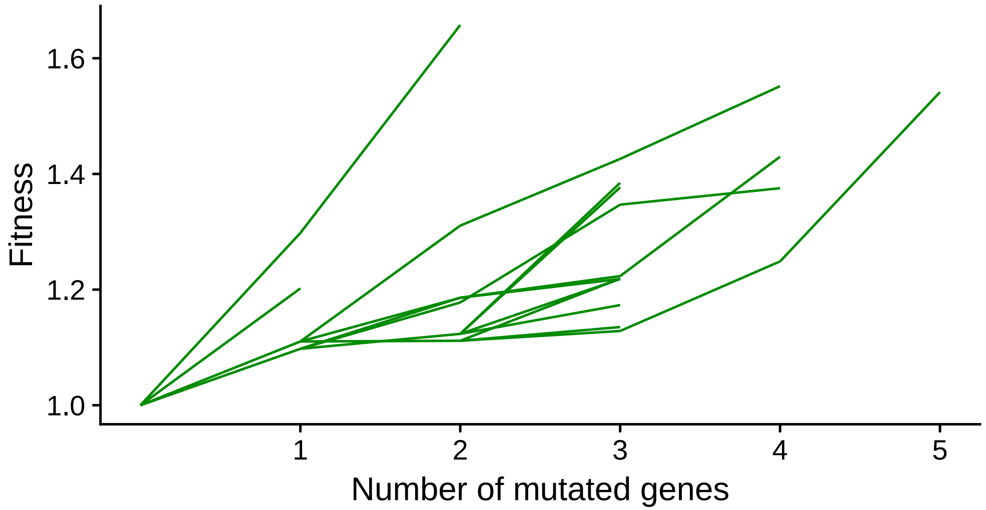


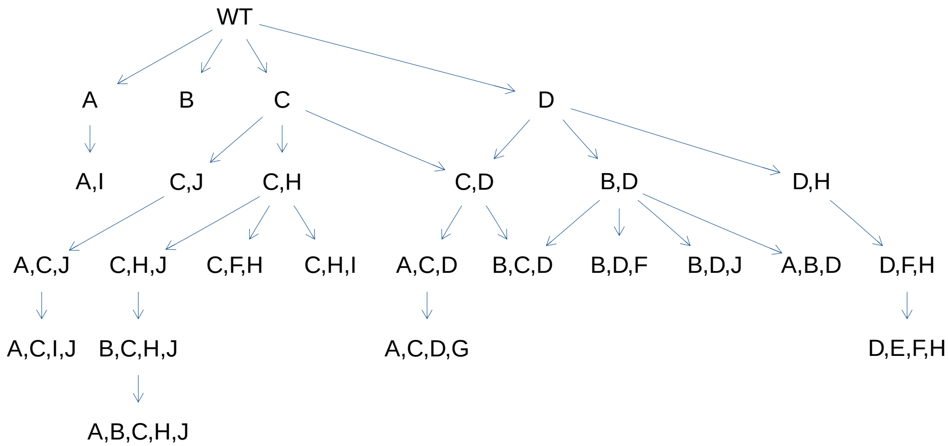

| | |
|---|---|
| *ID* | Zi10rjNfw5yh1JEKK |
| *acc genotypes* | 37 |
| *peaks no back* | 15 |
| *removed edges* | NA |
| *prop rem edges* | NA |

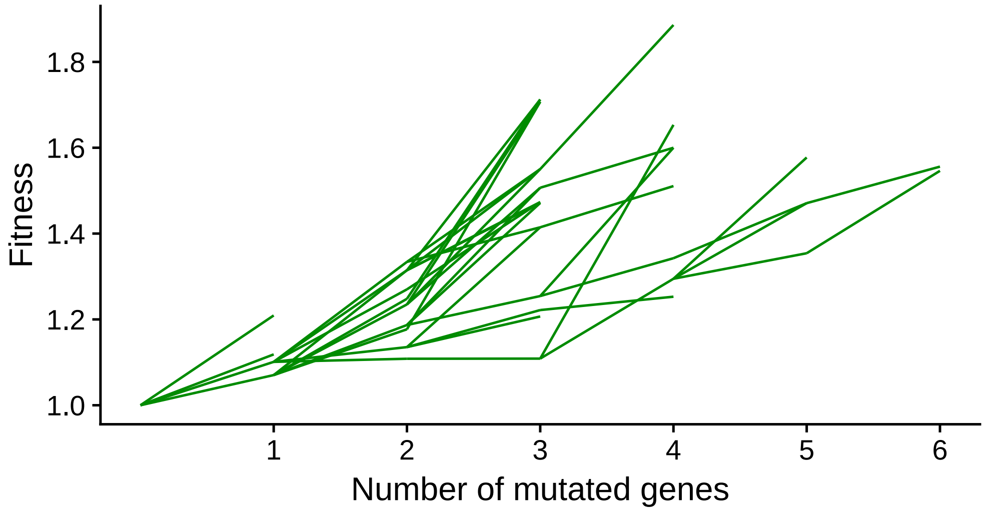


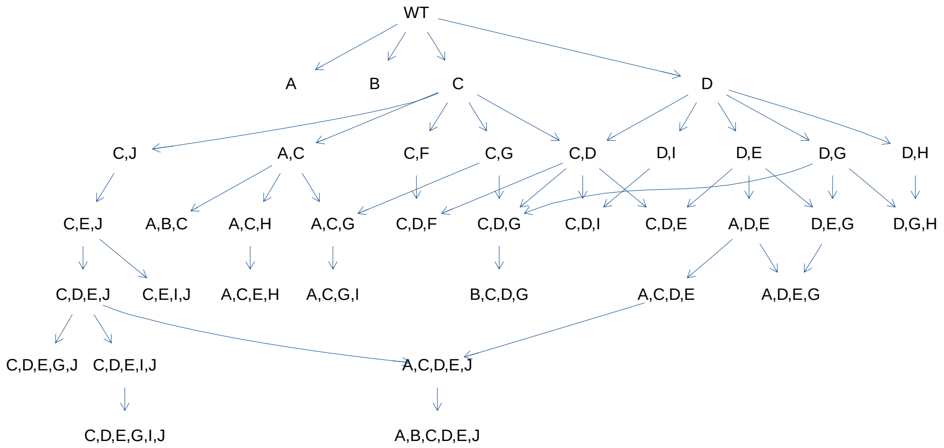

| | |
|---|---|
| *ID* | ZIwKQjyTXgfkZAaS |
| *acc genotypes* | 367 |
| *peaks no back* | 83 |
| *removed edges* | NA |
| *prop rem edges* | NA |

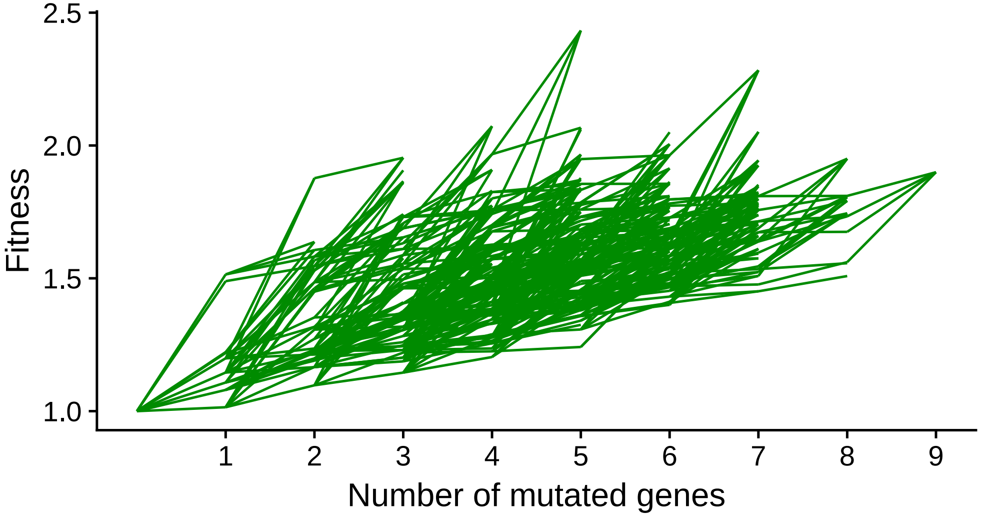


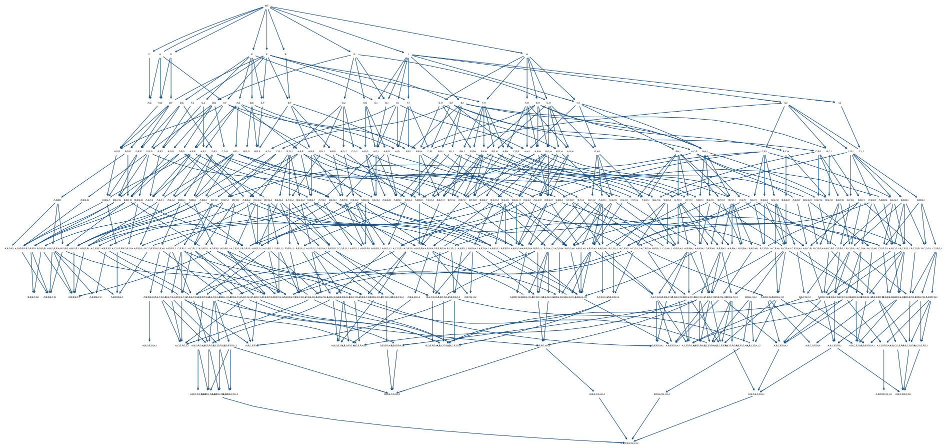

| | |
|---|---|
| *ID* | zja2YeEj3wZTfoRu |
| *acc genotypes* | 121 |
| *peaks no back* | 36 |
| *removed edges* | NA |
| *prop rem edges* | NA |

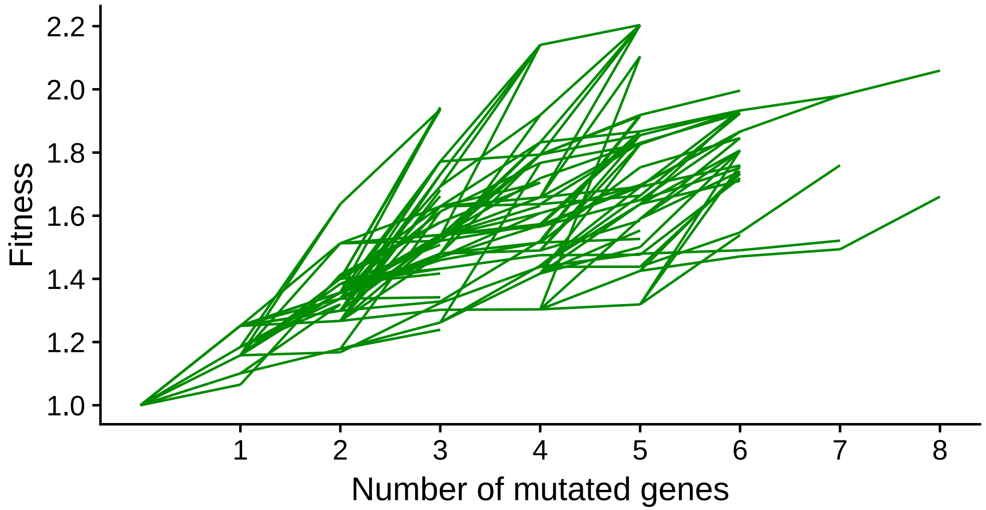


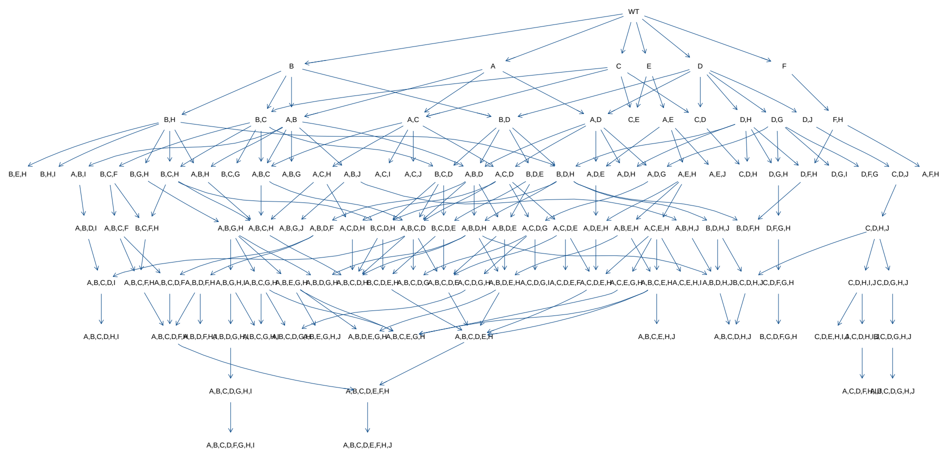

| | |
|---|---|
| *ID* | zk5E1osYtIlv7Cob |
| *acc genotypes* | 307 |
| *peaks no back* | 54 |
| *removed edges* | NA |
| *prop rem edges* | NA |

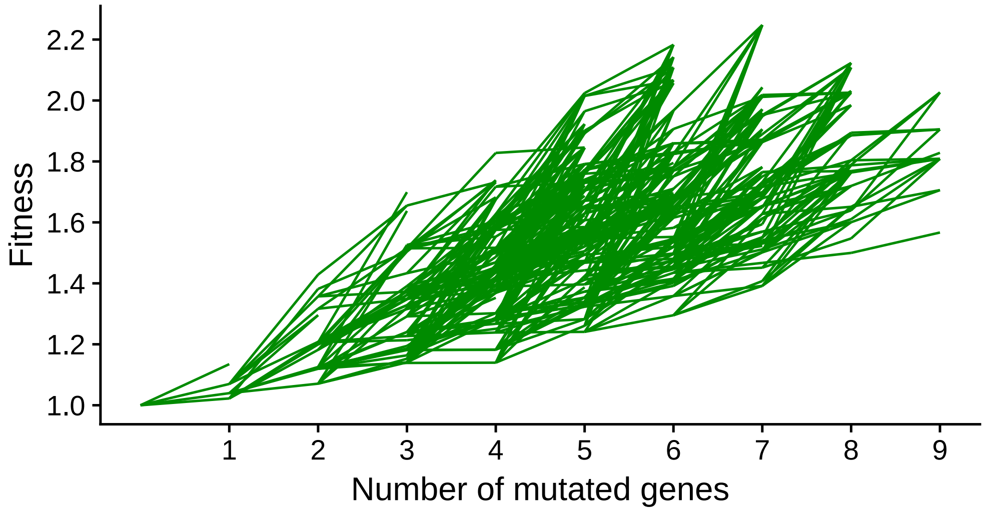


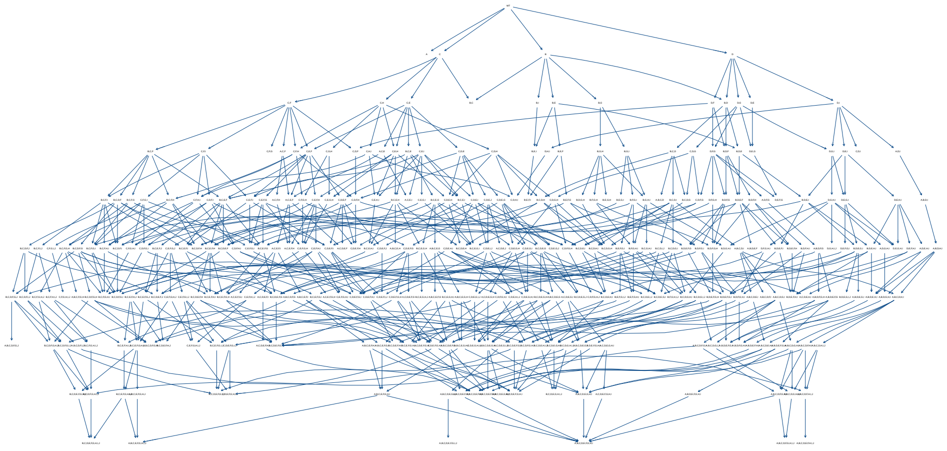

| | |
|---|---|
| *ID* | zSLtW8y3ke10umntK |
| *acc genotypes* | 98 |
| *peaks no back* | 26 |
| *removed edges* | NA |
| *prop rem edges* | NA |

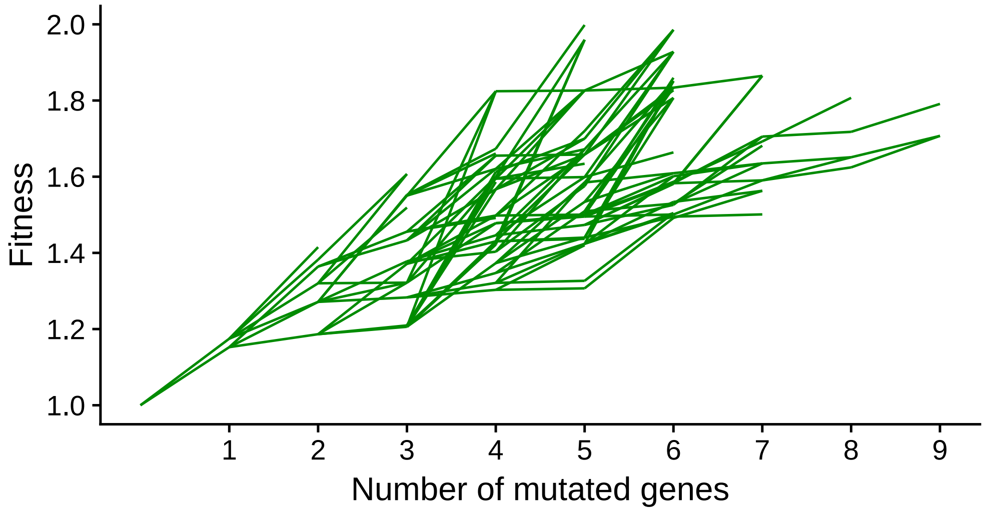


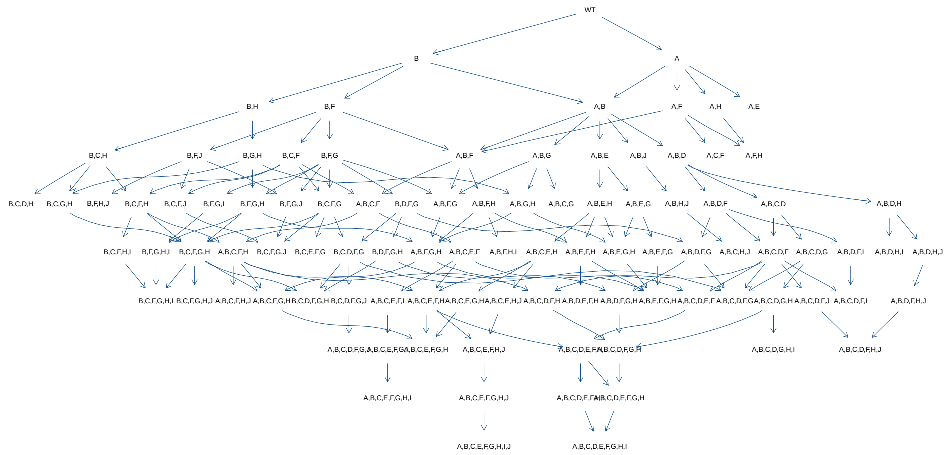

| | |
|---|---|
| *ID* | ZTNMfNi10xGkfS7ED |
| *acc genotypes* | 534 |
| *peaks no back* | 52 |
| *removed edges* | NA |
| *prop rem edges* | NA |

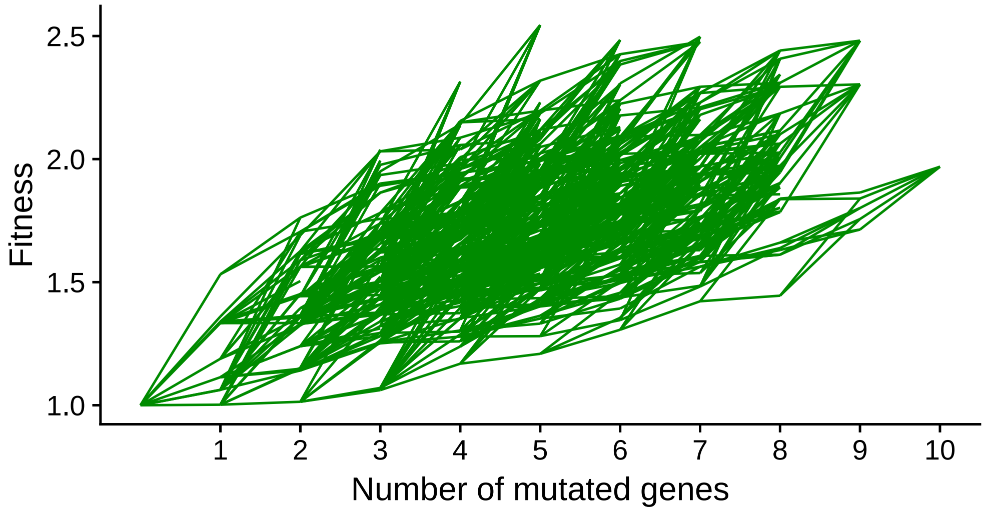


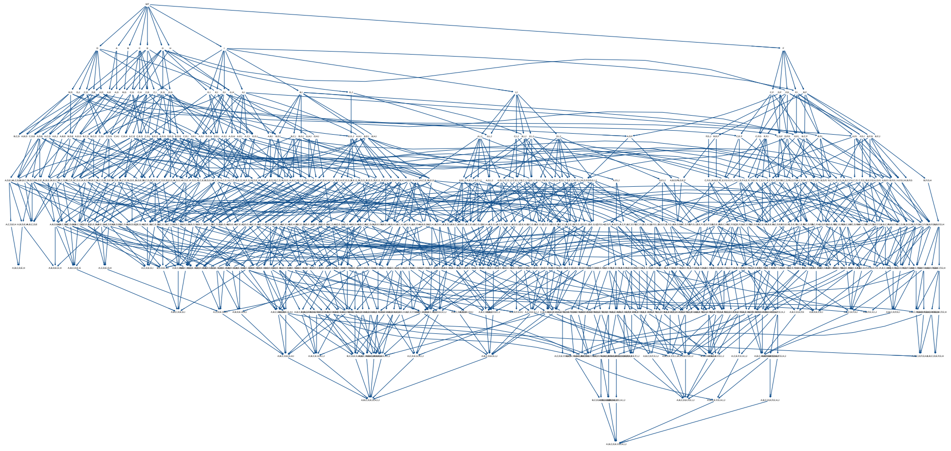

| | |
|---|---|
| *ID* | ZVzsqZQydHbT4TYZ |
| *acc genotypes* | 42 |
| *peaks no back* | 20 |
| *removed edges* | NA |
| *prop rem edges* | NA |

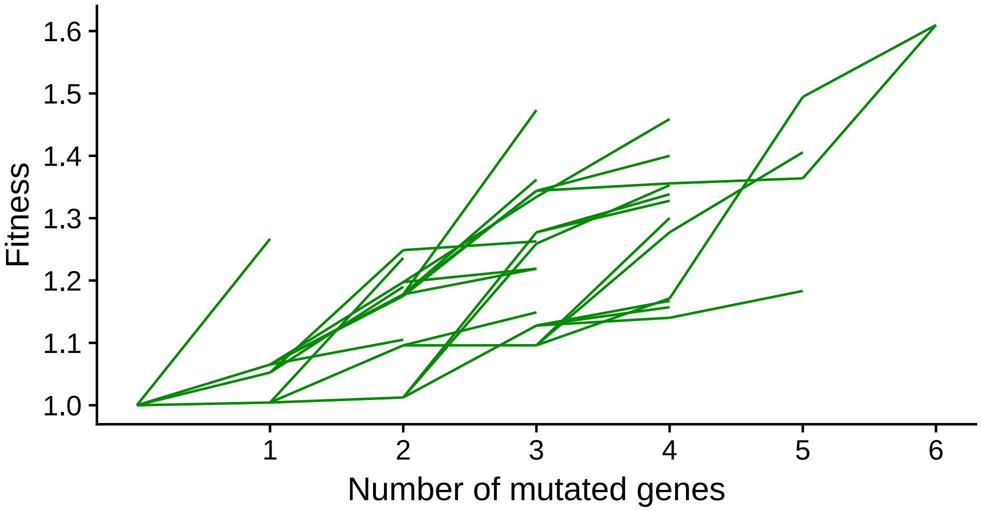


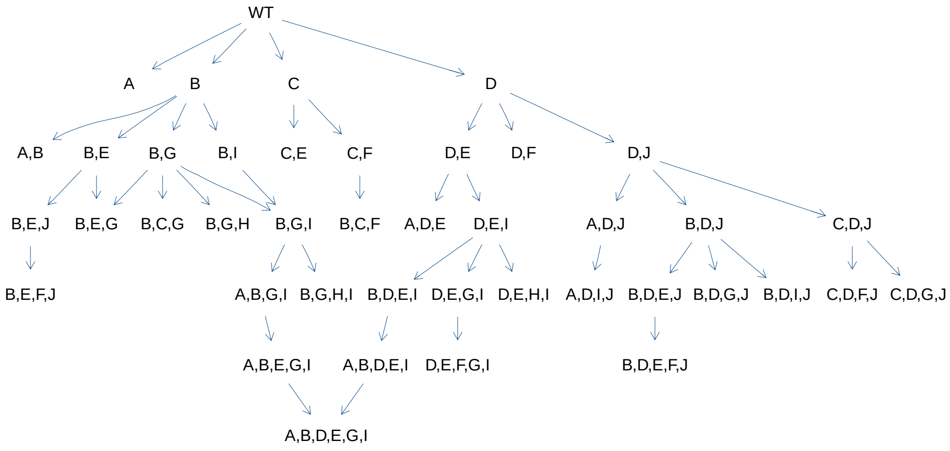

| | |
|---|---|
| *ID* | zWjhPCHtGUy14jOB |
| *acc genotypes* | 154 |
| *peaks no back* | 34 |
| *removed edges* | NA |
| *prop rem edges* | NA |

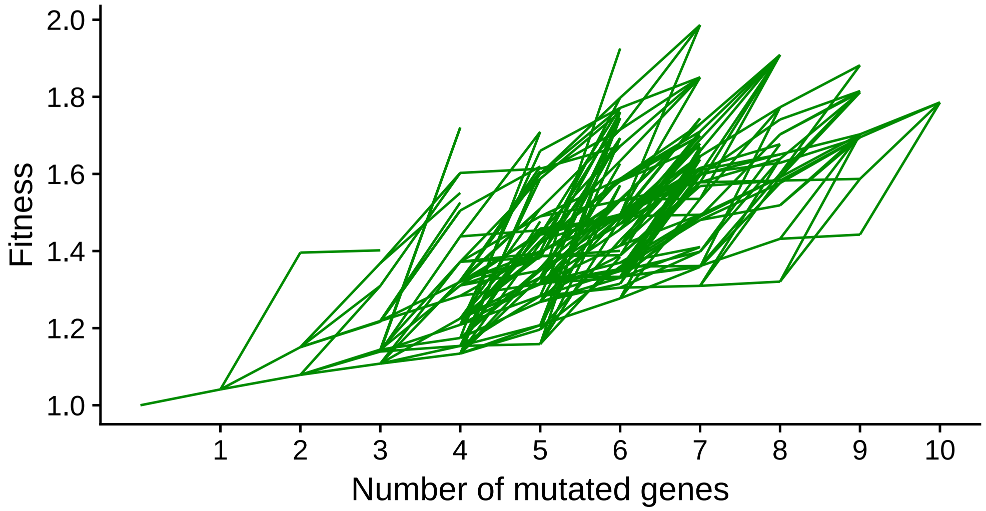


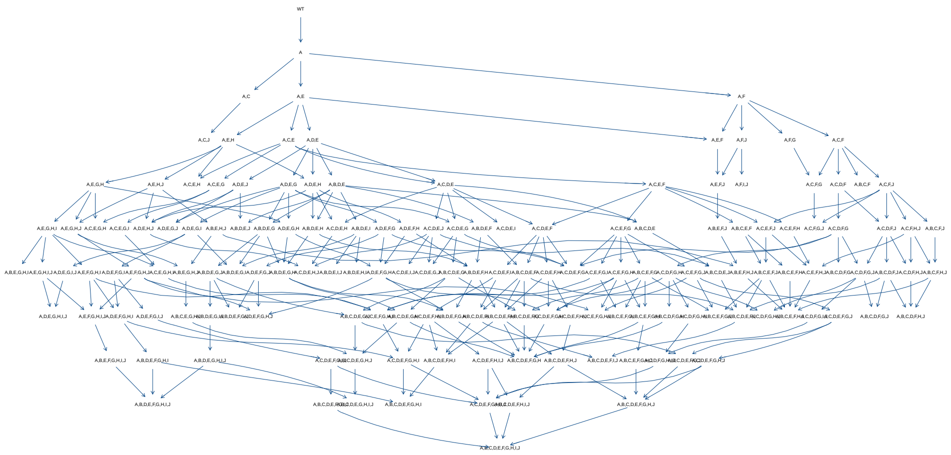
