## Supplementary material for "Every which way? On predicting tumor evolution using cancer progression models": S2_Figure

### Supporting Information for “Every which way? On predicting tumor evolution using cancer progression models”

2019-04-17

#### S2 Figure. Simulated fitness landscapes and fitness graphs: characteristics, evolutionary unpredictability, clonal interference, and sampled genotypes

---

### 1. Description

The following figures show the main fitness landscape characteristics and the resulting variation in evolutionary predictability, clonal interference, and sampling characteristics, between types of fitness landscapes and simulation conditions (initial population size and mutation rate). Note that the "static fitness landscape" characteristics do not depend on the simulations.

#### 1.1. Terminology

The **number of local (fitness) maxima** is a static feature of the landscape (number of genotypes such that all genotypes within a distance of one mutation have lower fitness). The number of **observed local (fitness) maxima** can be smaller, since some peaks (local maxima) might never be visited. For representable fitness landscapes, both numbers are 1. For the other two landscapes, those numbers were $\geq 2$.

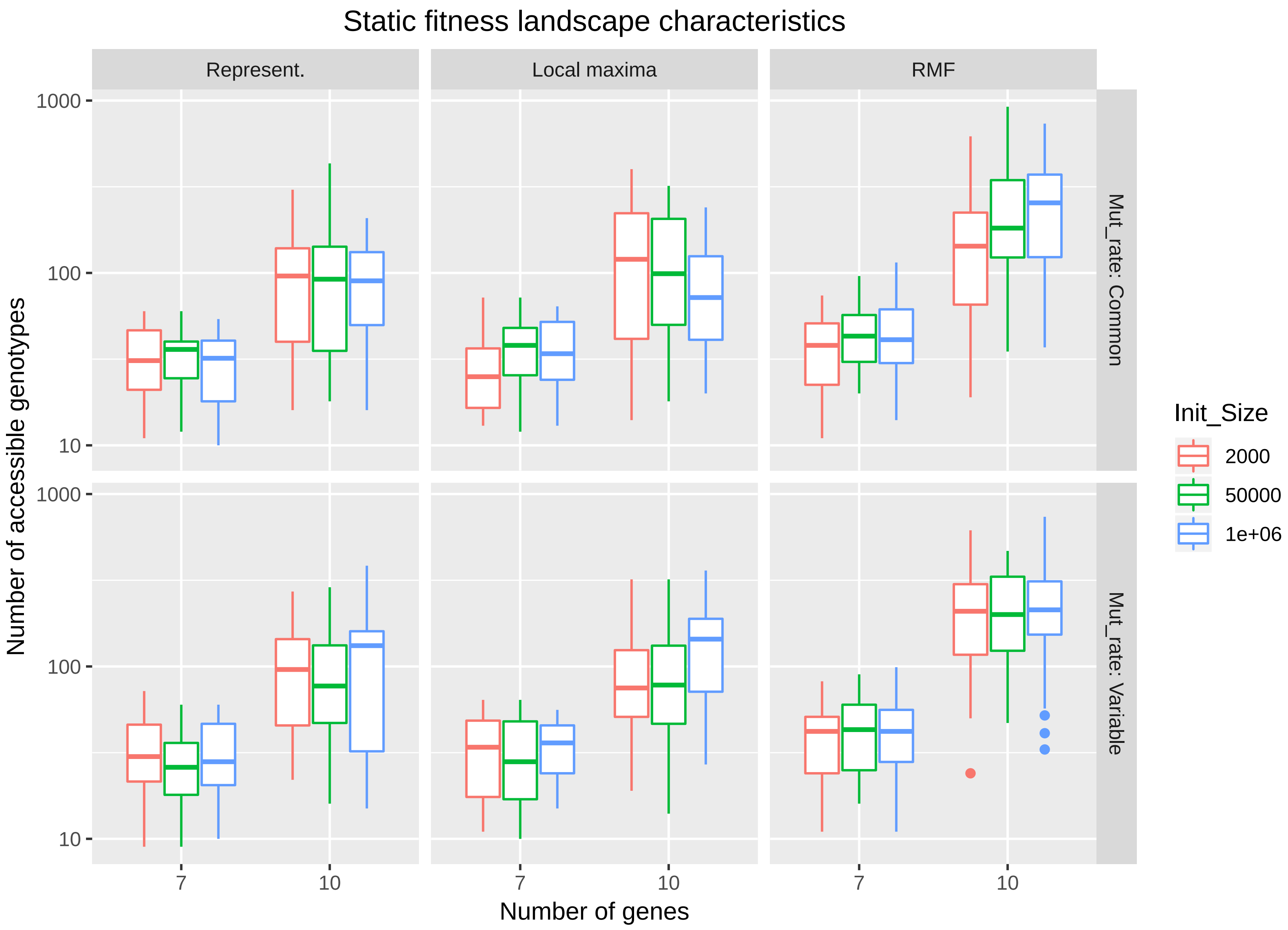

Figure 1: Simulated fitness landscapes: Number of accessible genotypes

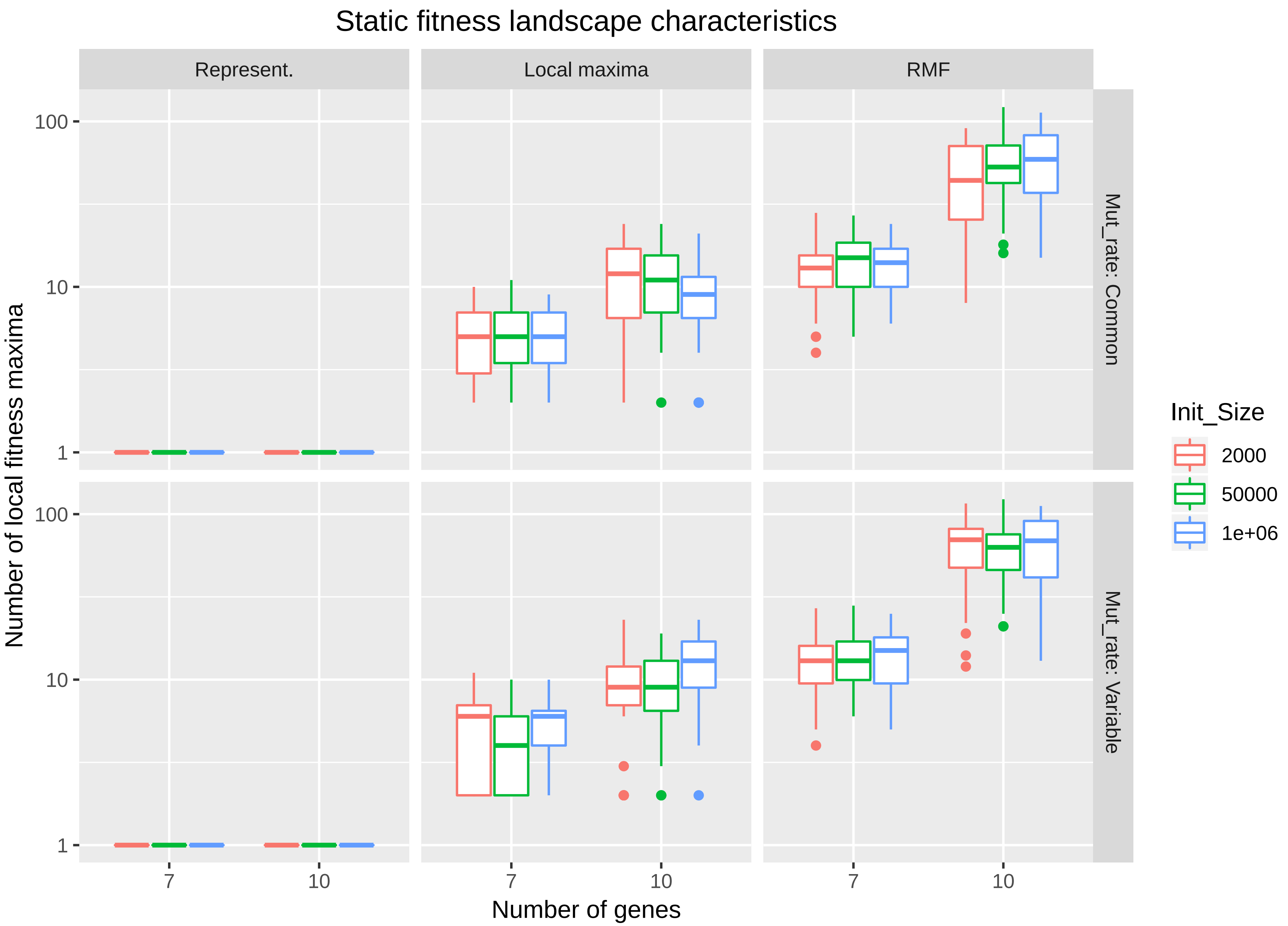

Figure 2: Simulated fitness landscapes: Number of local fitness maxima

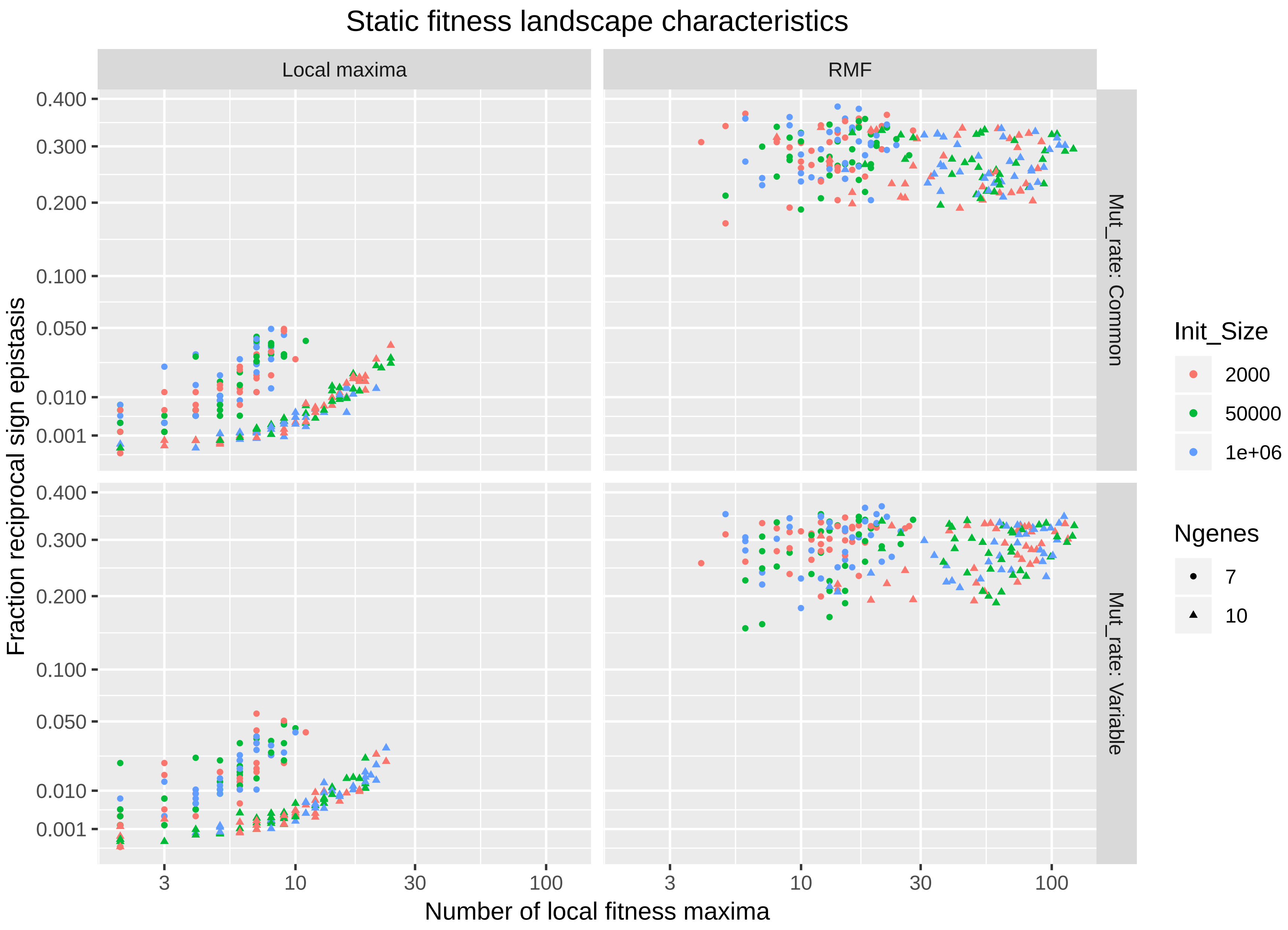

Figure 3: Simulated fitness landscapes: reciprocal sign epistasis

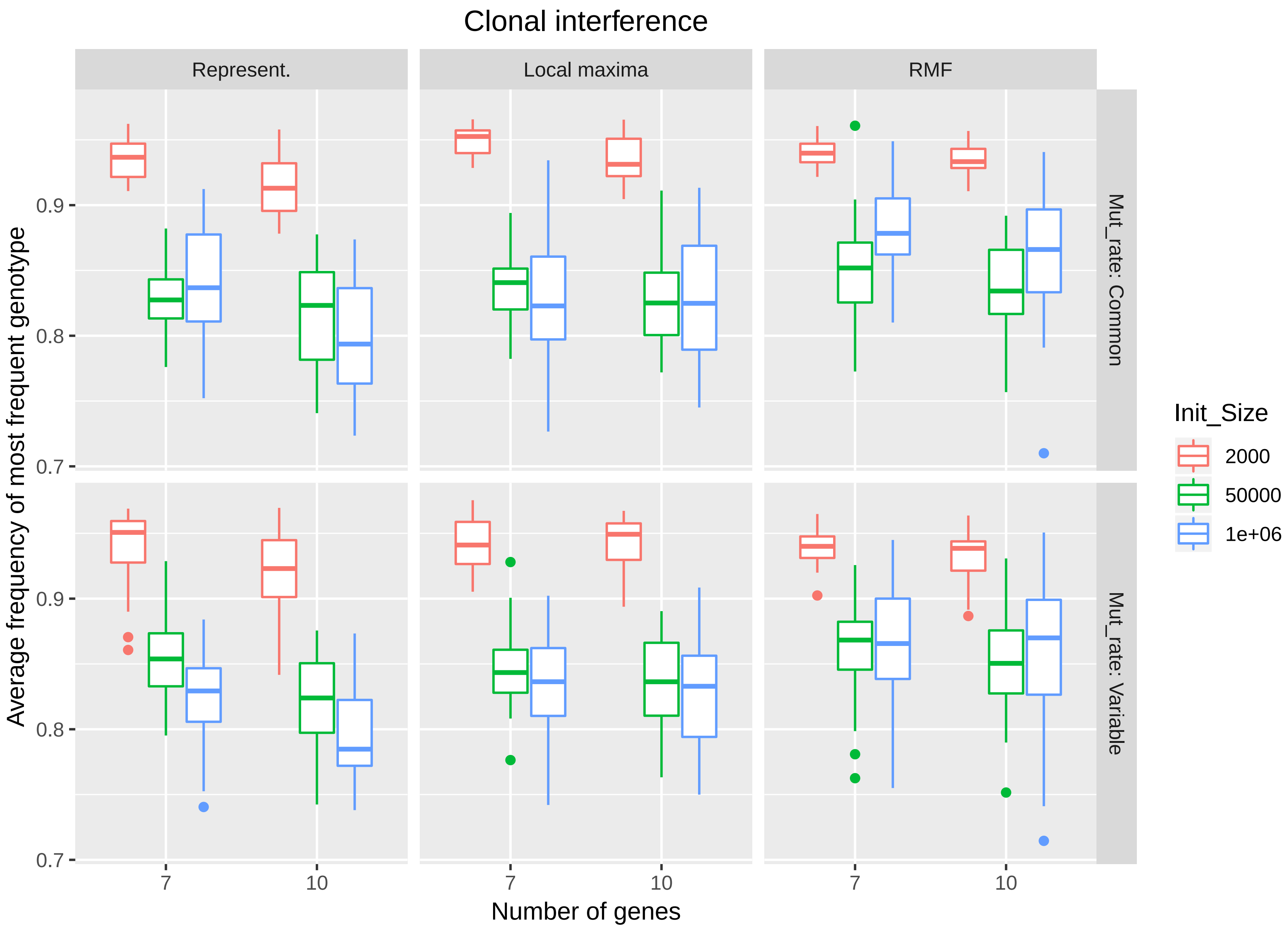

Figure 4: Simulated fitness landscapes: clonal interference (frequency of most frequent genotype)

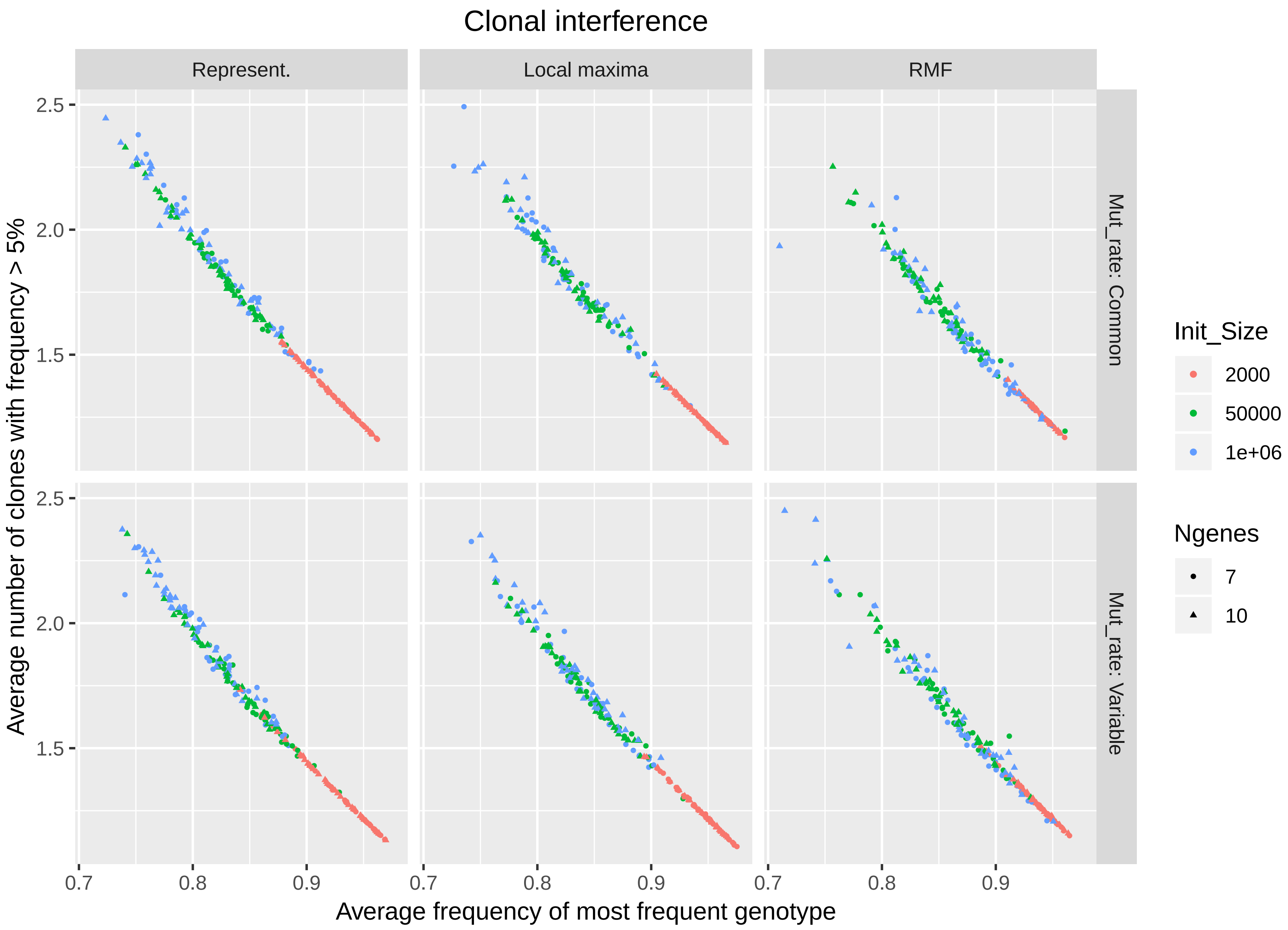

Figure 5: Simulated fitness landscapes: clonal interference (average number of clones with frequency $>$ 5%)

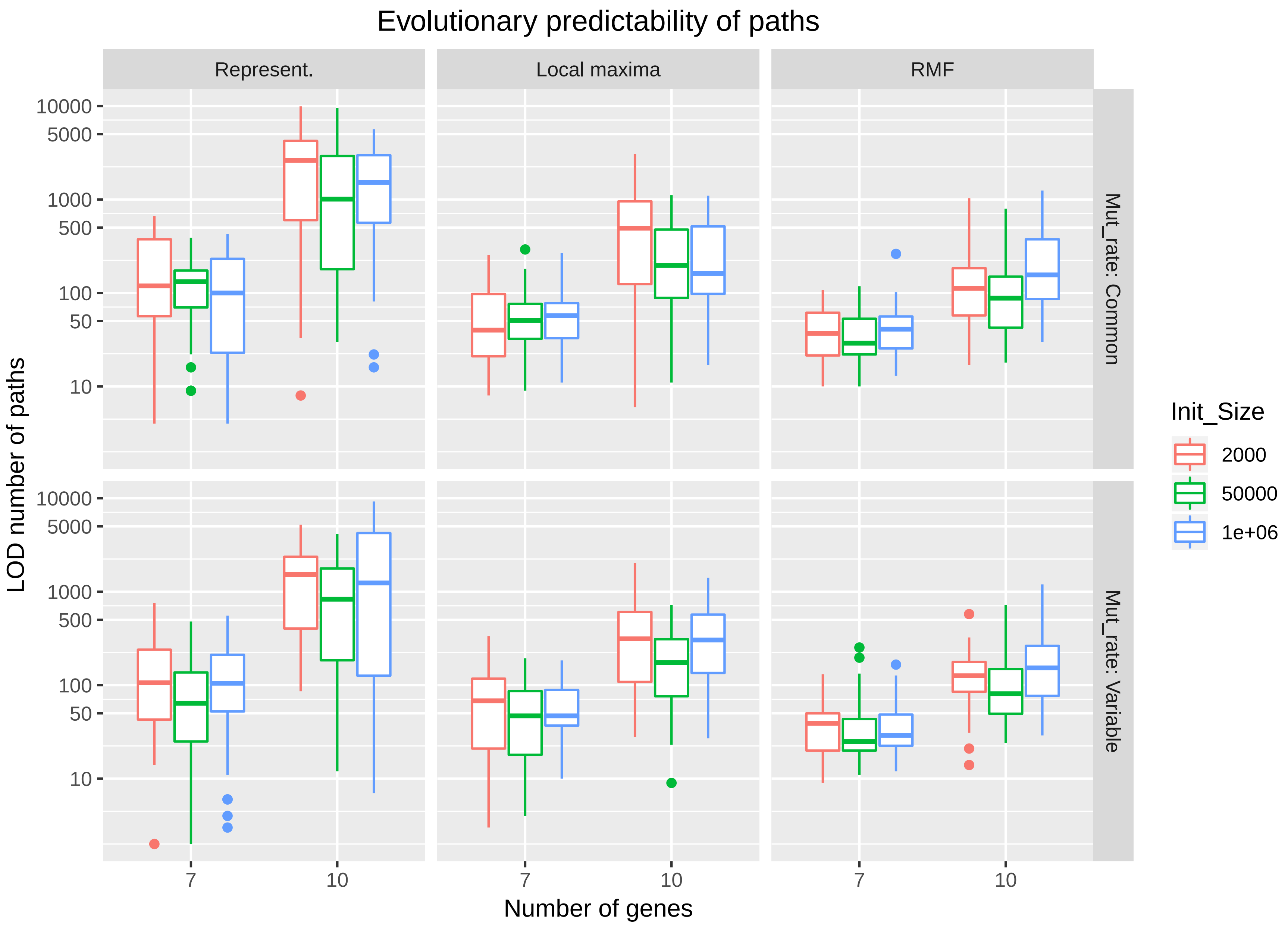

Figure 6: Simulated fitness landscapes: LOD

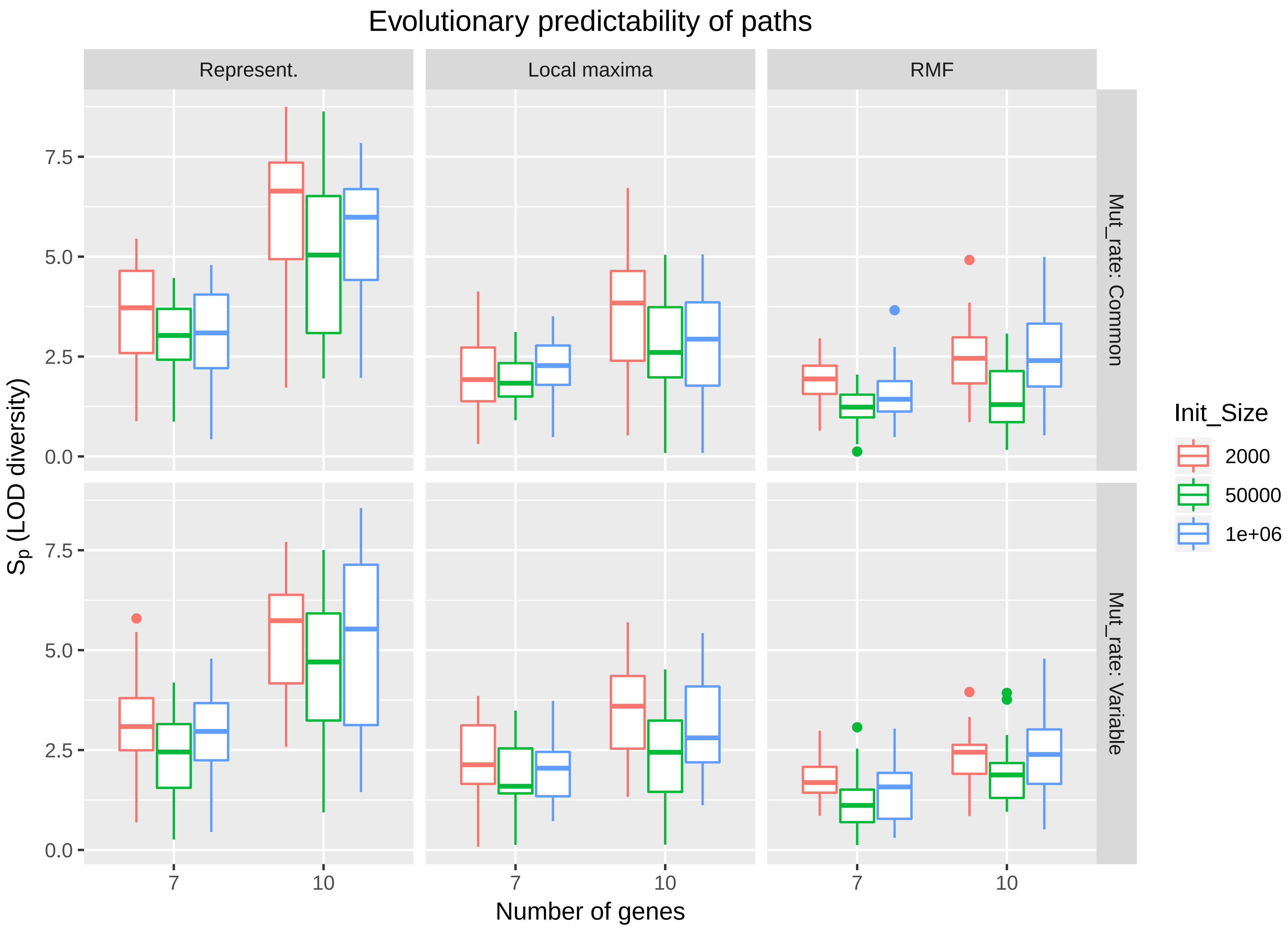

Figure 7: Simulated fitness landscapes: $S_p$

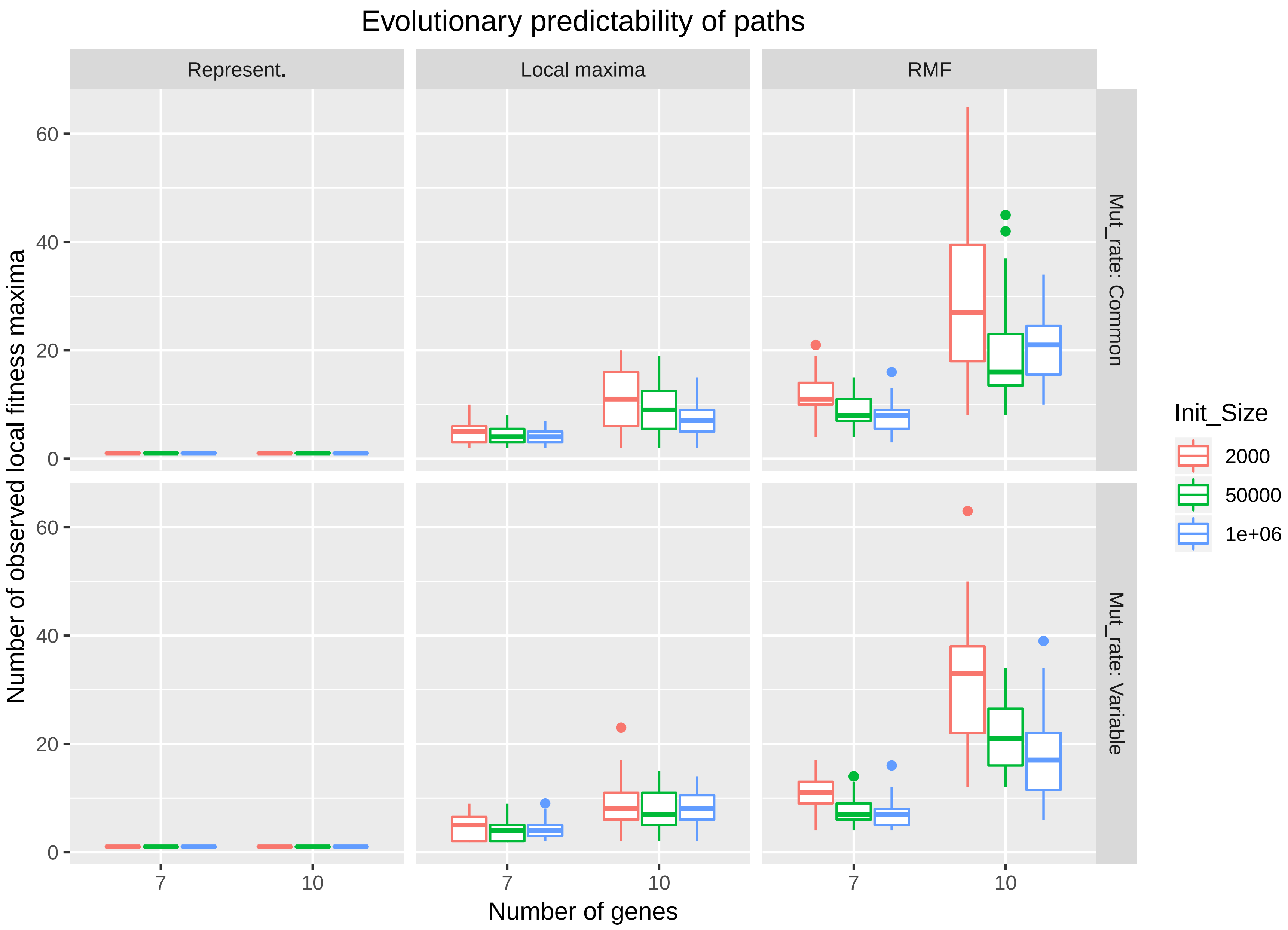

Figure 8: Simulated fitness landscapes: number of observed local fitness maxima

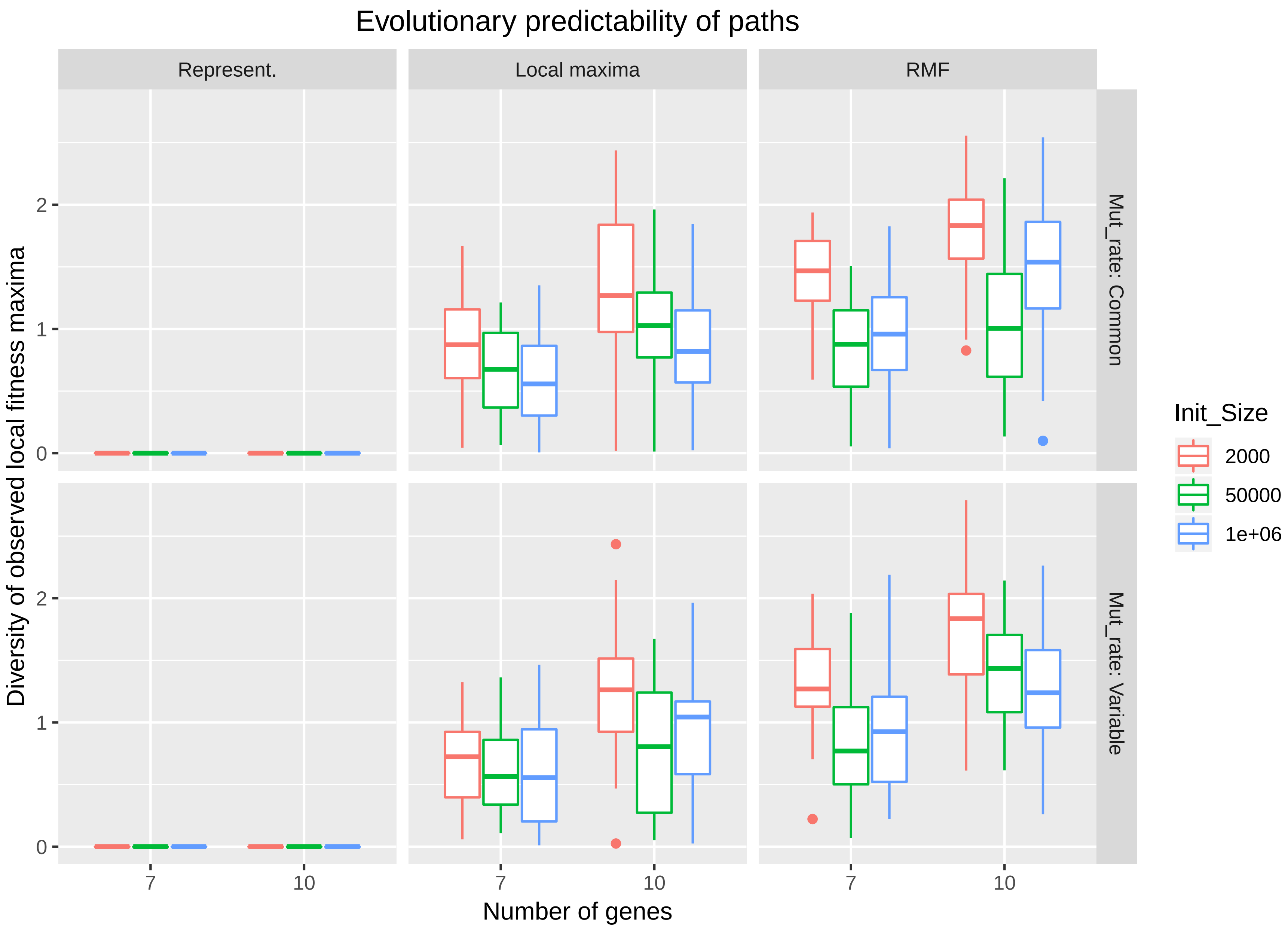

Figure 9: Simulated fitness landscapes: diversity of observed fitness maxima

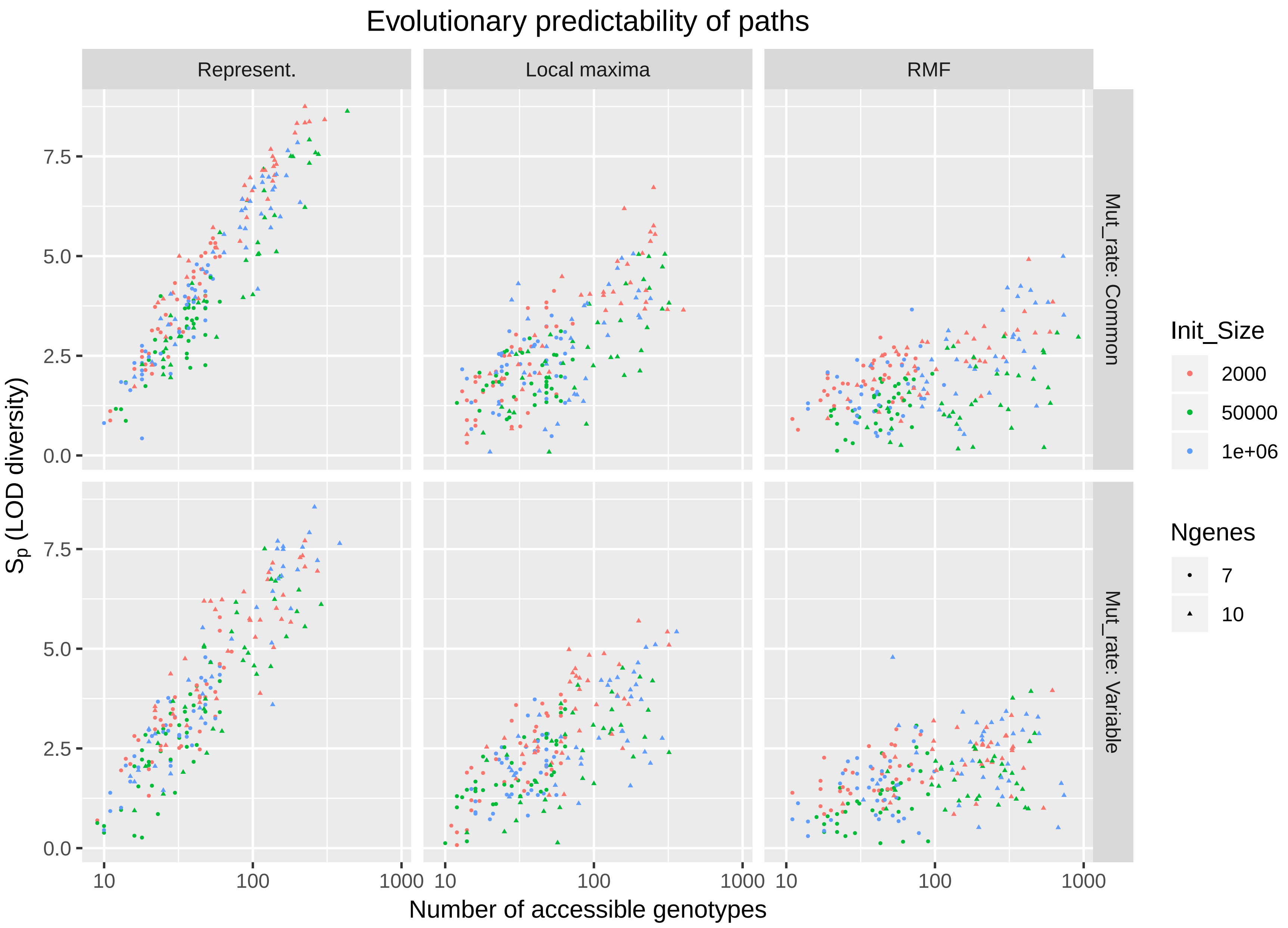

Figure 10: Simulated fitness landscapes: $S_p$ vs. accessible genotypes

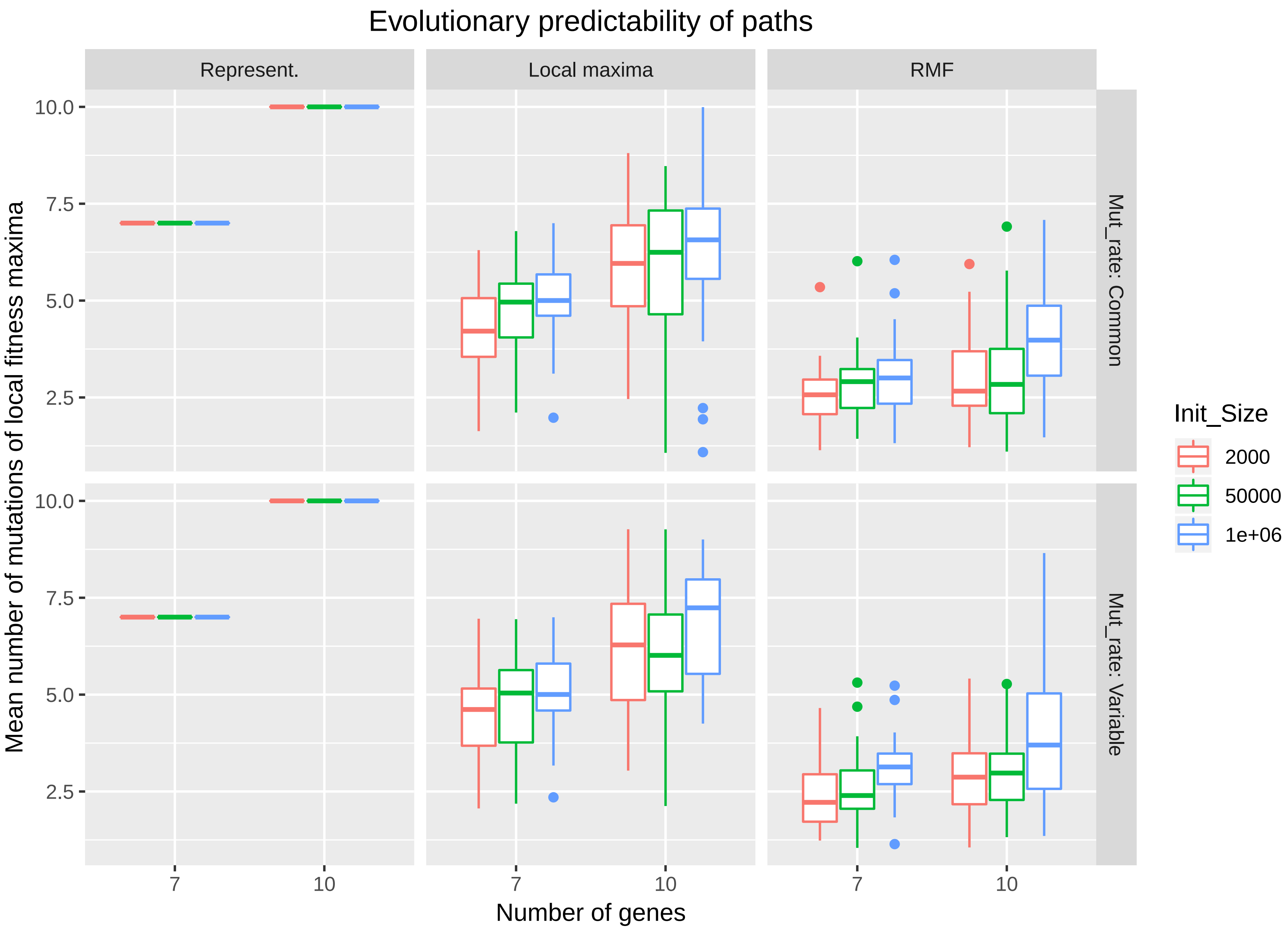

Figure 11: Simulated fitness landscapes: number of mutations of fitness maxima

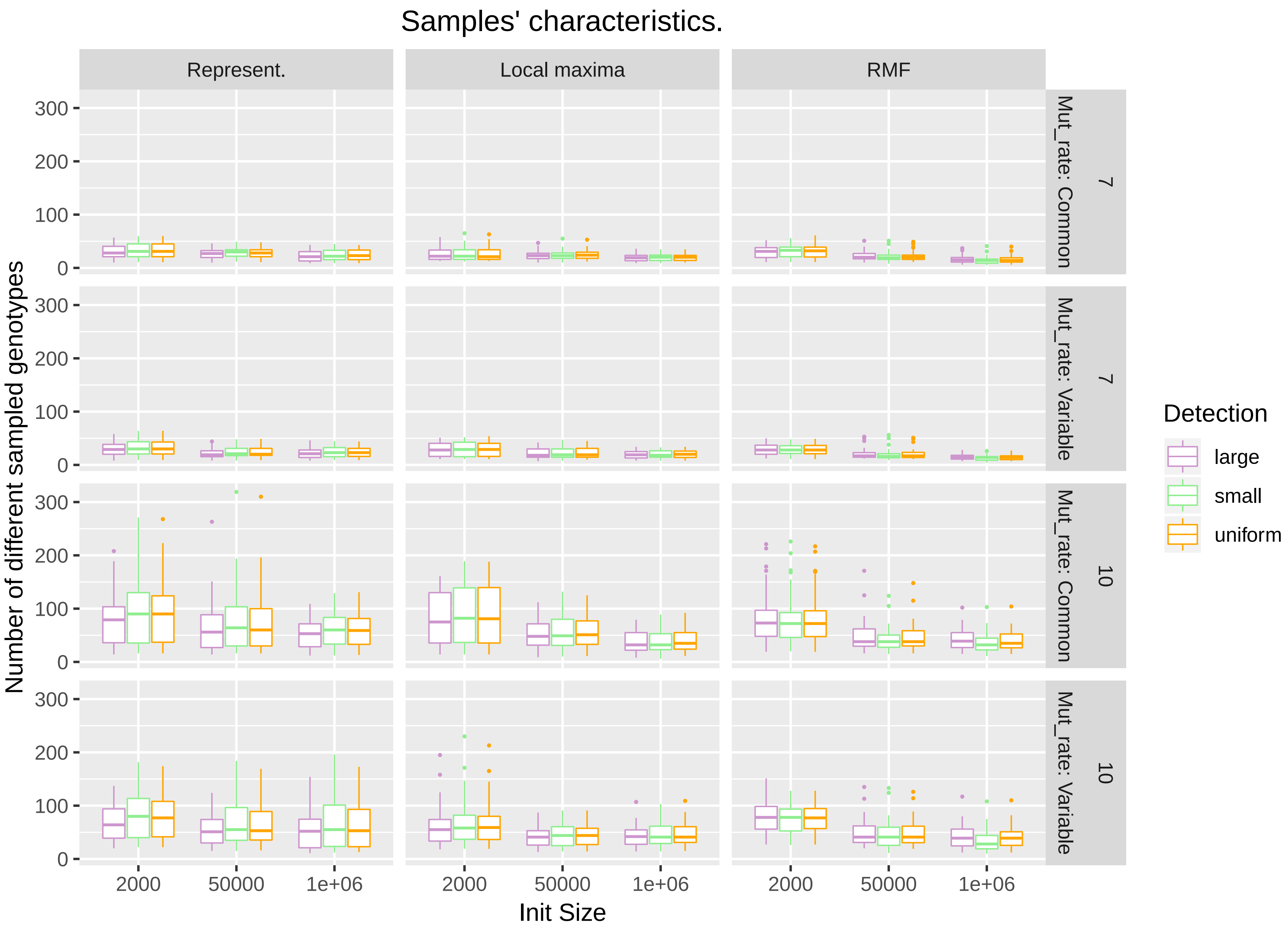

Figure 12: Simulated samples' characteristics: number of genotypes

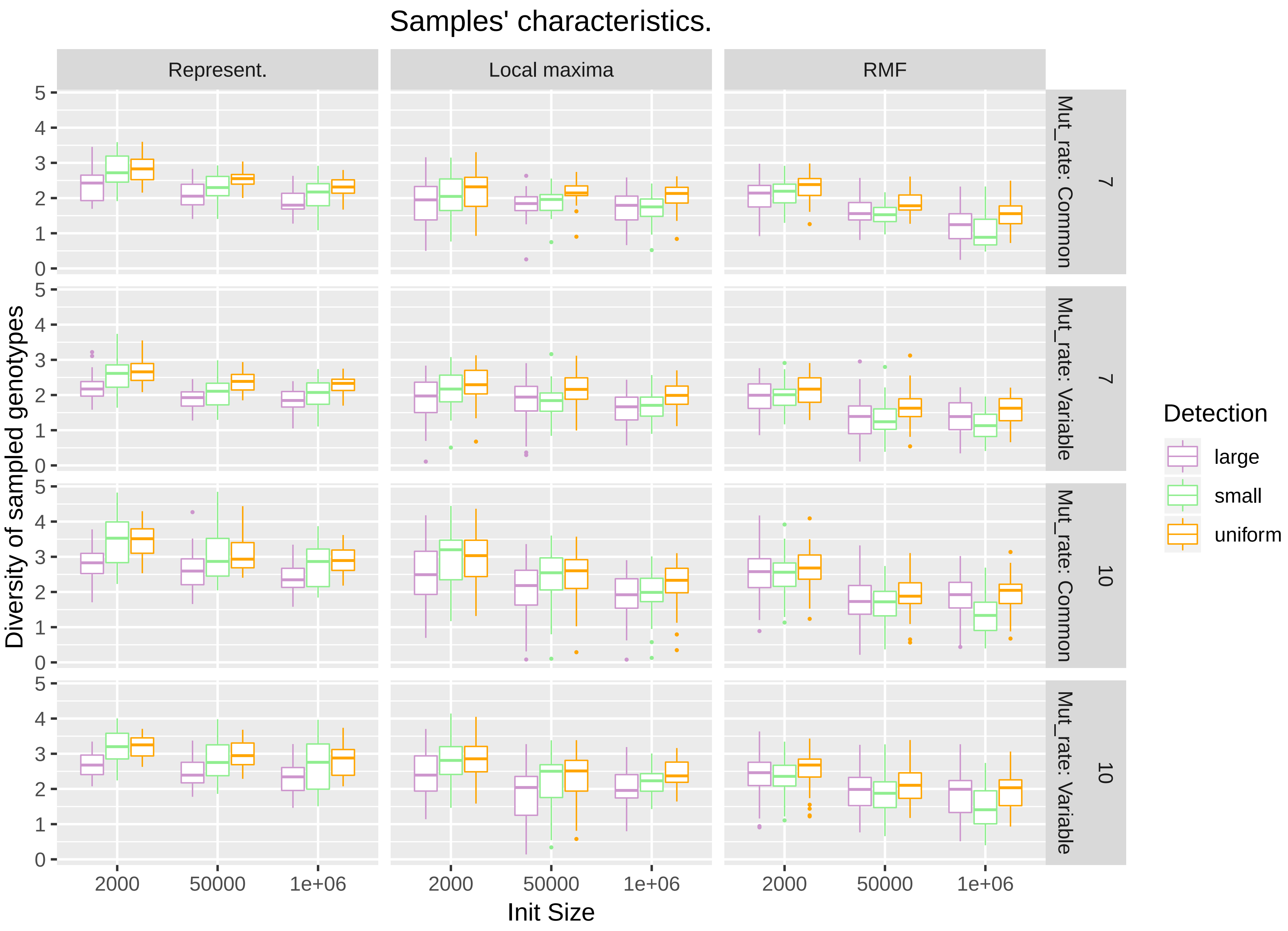

Figure 13: Simulated samples' characteristics: diversity of genotypes

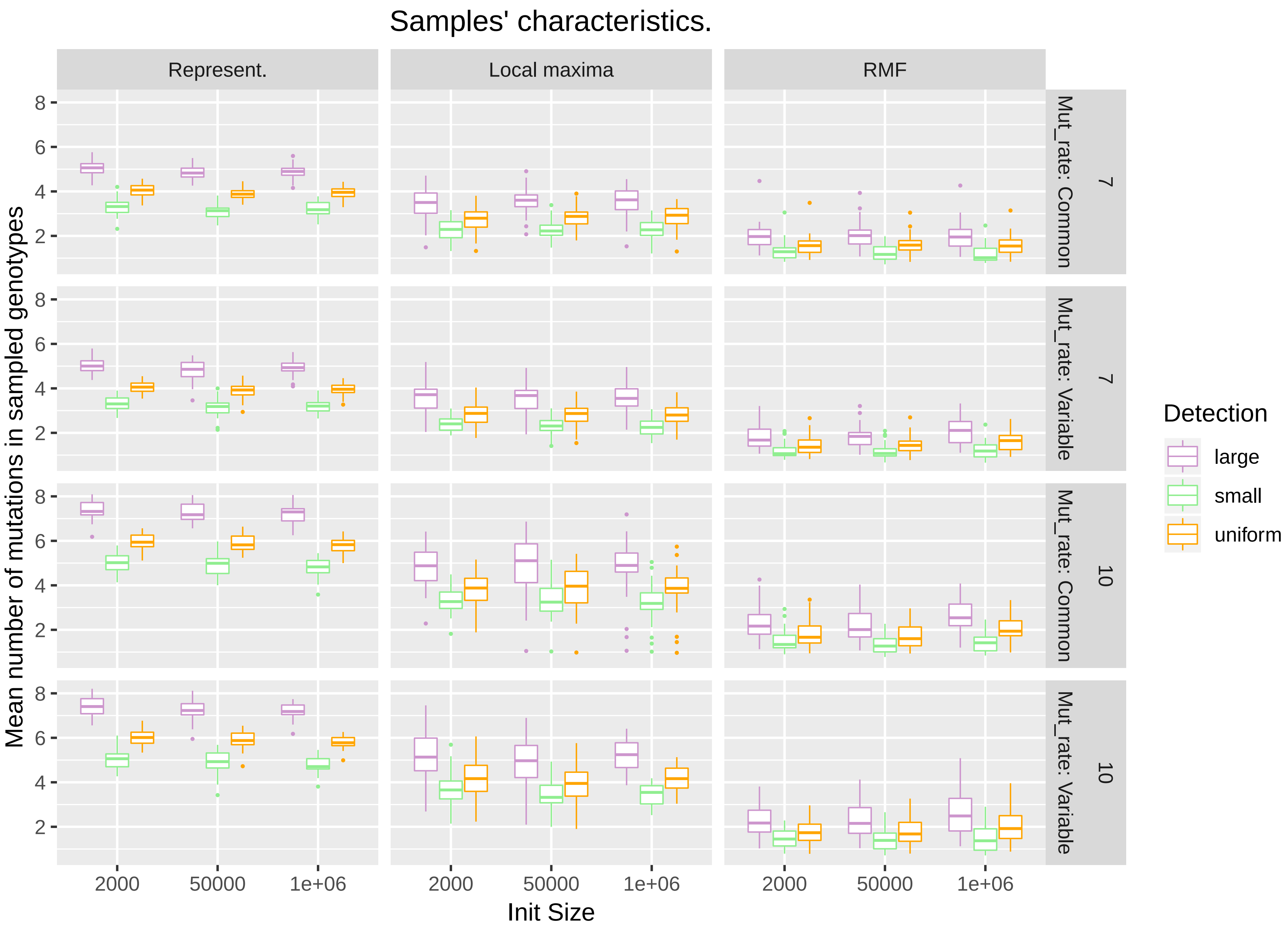

Figure 14: Simulated samples' characteristics: mean number of mutations in genotypes

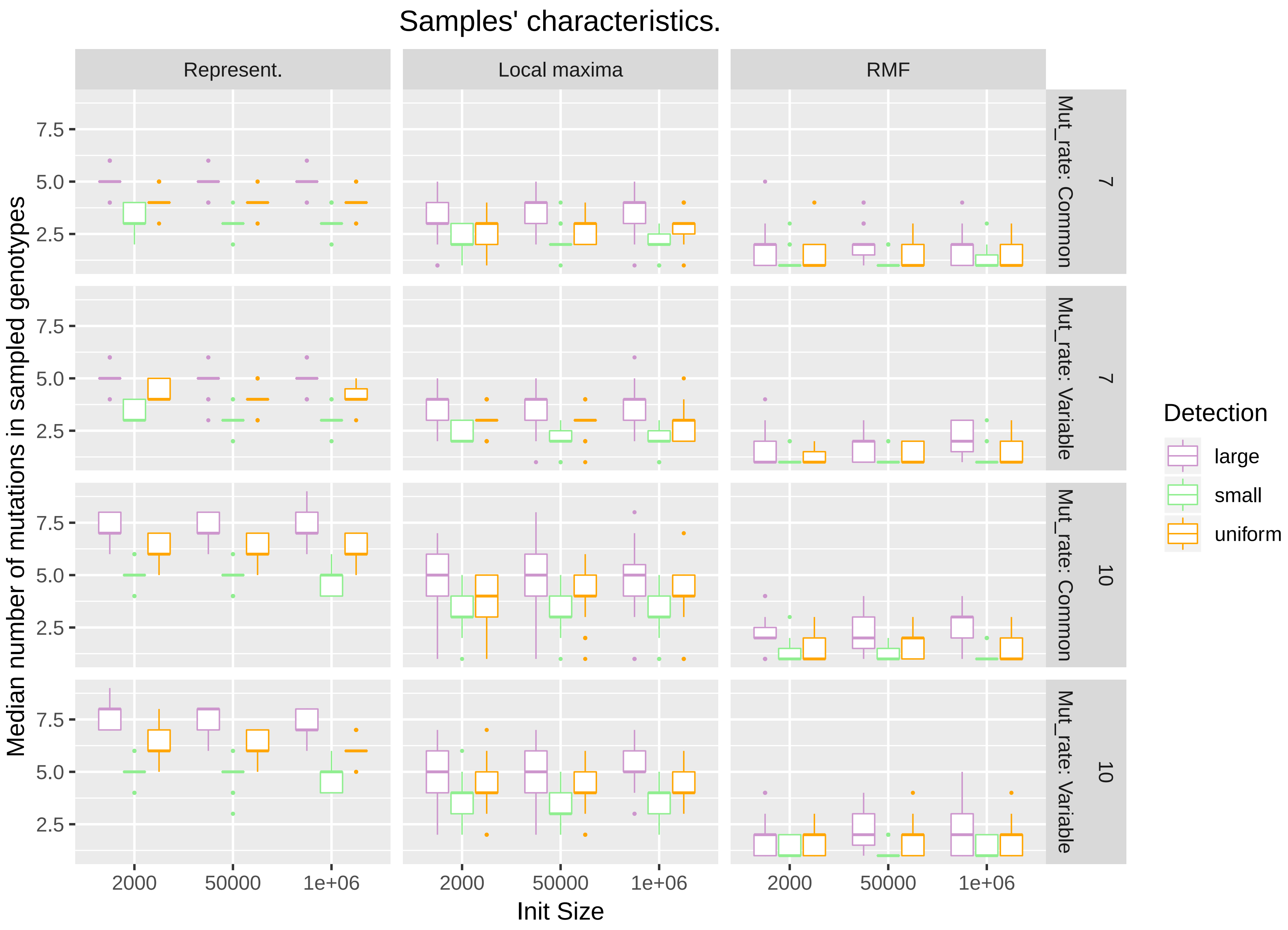

Figure 15: Simulated samples' characteristics: median number of mutations in genotypes

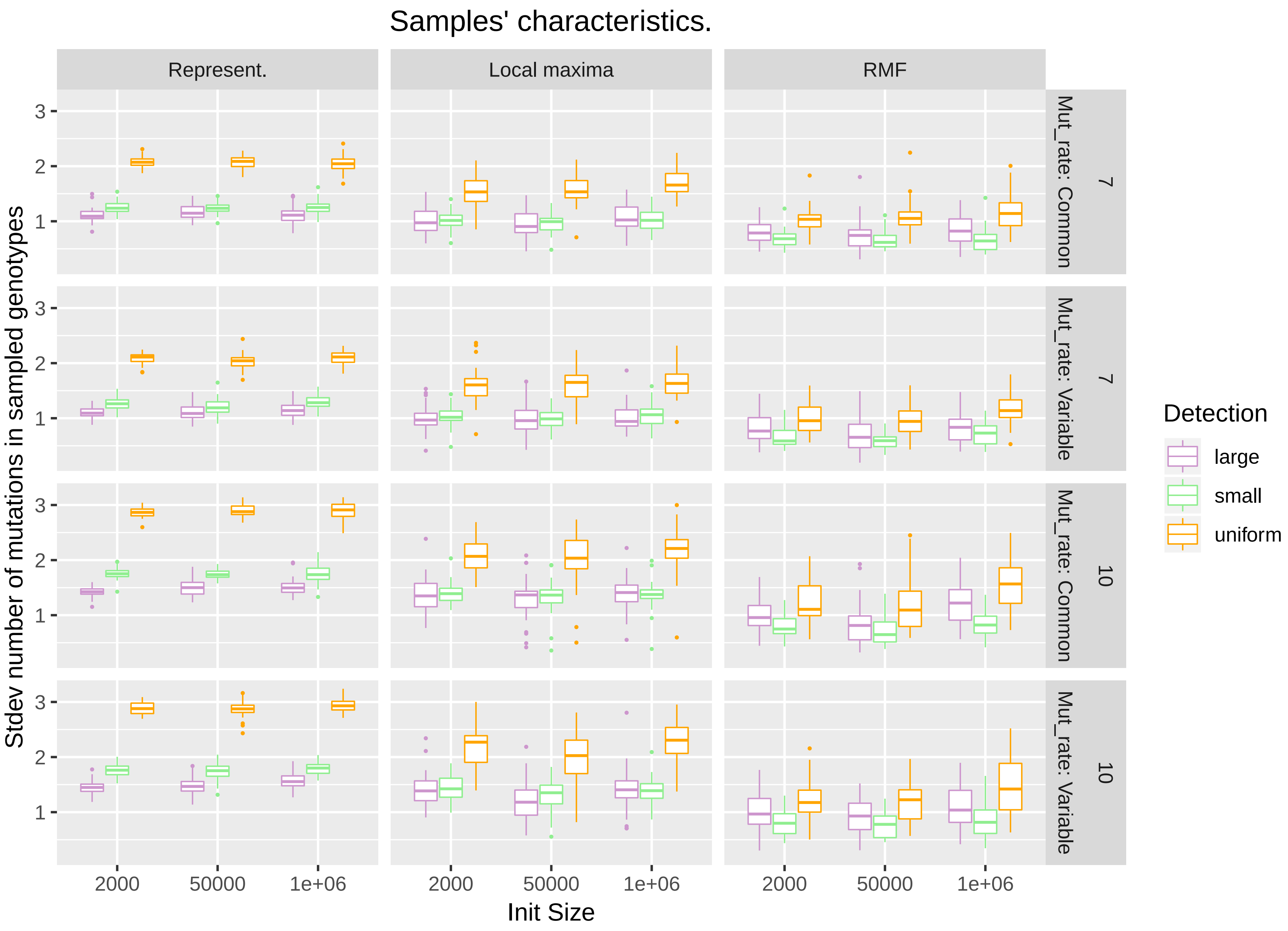

Figure 16: Simulated samples' characteristics: standard deviation number of mutations in genotypes

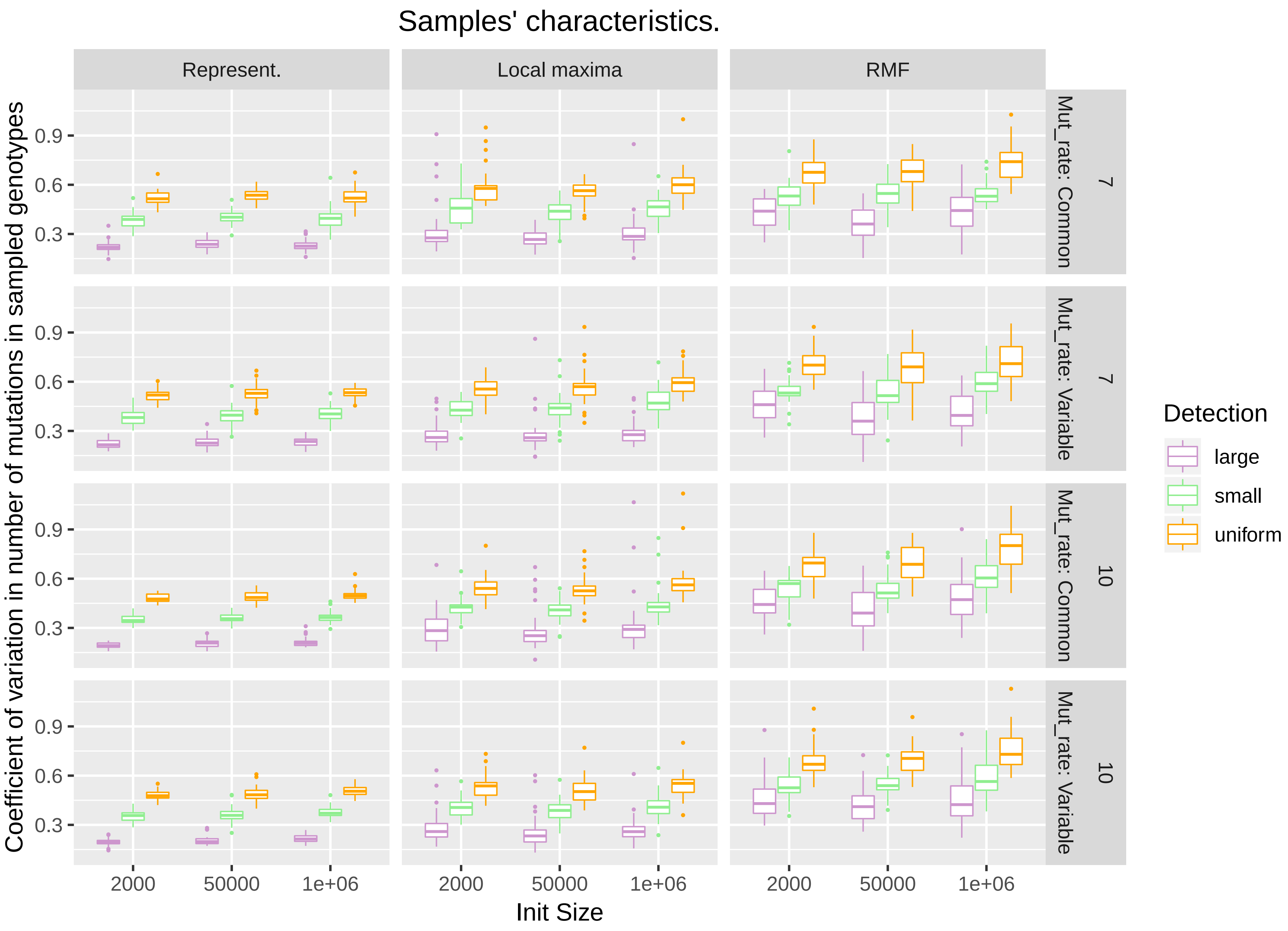

Figure 17: Simulated samples' characteristics: coefficient of variation in number of mutations in genotypes
